# Non-invasive Vagal Nerve Stimulation as a Potential Treatment for Repetitive Blast Trauma

**DOI:** 10.64898/2026.07.13.737563

**Authors:** Britahny M Baskin, Alyssa Easton, Monica Tschang, Suhjung Janet Lee, Emma Skillen, Katrina Wong, Bryan Schuessler, James Meabon, Tami Wolden-Hanson, David G. Cook, Sean M. Gibbons, Abigail G. Schindler

## Abstract

**Background:** Polytrauma caused by exposure to high explosives (blast) is increasingly common among military personnel and civilians yet treatment for related post-concussive symptoms and chronic behavioral dysfunction is limited. Therapeutic targets following these injuries are typically focused on the central nervous system, with less attention placed on potentially more accessible peripheral targets. Vagus nerve stimulation (VNS) has recently gained traction as a potential therapeutic modality but has yet to be examined in a blast trauma setting.

**Methods:** Our well-established blast overpressure model was utilized to induce repetitive (3x) blast trauma, followed by treatment with non-invasive transcutaneous VNS one hour following each blast exposure in male mice. Acutely following repetitive blast exposure, we measured serum and brain cytokine levels, fecal microbial abundance, and locomotion and anxiety-like behavior in the open field assay. Chronically (1-3 months post blast), mice were assessed for behavioral outcomes related to mild traumatic brain injury (mTBI) and posttraumatic stress disorder (PTSD), including acoustic startle (hyperreactivity), probabilistic discounting (risky decision making), and two-bottle choice test (voluntary alcohol consumption).

**Results:** VNS treatment following blast exposure decreased acute blast-induced inflammatory response in the blood and brain, especially serum IL-9 and IP-10, and brain MPC-1. Chronic behavior tests demonstrated a VNS-dependent reduction in blast-induced risky decision-making and a decreased intake and preference for ethanol. Conversely, VNS was not effective in preventing acute blast effects on the microbiome or chronic hyperreactivity behaviors measured with acoustic startle.

**Discussion:** This study identifies the vagus nerve as a novel peripheral target for treating acute and chronic blast-induced dysfunction.

## Introduction

Traumatic brain injury (TBI) is currently a leading cause of death and disability^1,2^, affecting every segment of the population and significantly affecting quality of life and financial burden^3–5^. The majority of TBIs (70-90%) are classified as mild^6,7^ using the Glasgow Coma Scale, yet mild TBI (mTBI) often results in a constellation of physical, sensorimotor, affective, and cognitive disturbances including but not limited to executive dysfunction, balance difficulties, post-traumatic stress disorder, and increased risky substance use^8–10^.

Blast overpressure waves, such as those caused by improvised explosive devices (IEDs), are an increasingly common cause of TBI^11–14^. Blast exposure is the primary source of mTBI in Servicemembers, a significant driver of PTSD in this group, and a major source of morbidity among Veterans enrolled in the VA health care system^9,13,15^. Though awareness of mTBI and the resulting adverse health outcomes has increased in the past decade, the mechanisms mediating such outcomes are still not well understood and treatment options remain limited.

Post-concussive symptoms (PCS; e.g., headache, dizziness, fatigue, sensorimotor impairment) and persistent behavioral dysfunction (e.g., anxiety, depression, impulsivity) following mTBI are increasingly recognized as arising from complex interactions between central and peripheral processes^16^. mTBI frequently induces autonomic dysregulation—manifested as altered heart rate variability, blood-pressure instability, pupillary abnormalities, and gastrointestinal (GI) complaints^17–26^ —alongside acute and sometimes persistent systemic inflammation and peripheral immune activation^17,20,27,28^. While much work has focused on developing and evaluating central nervous system targets for therapeutic efficacy following blast exposure and mTBI, less attention has been placed on understanding and developing peripheral targets for potential therapeutic application.

Among the peripheral systems affected by mTBI, the GI tract and its resident microbiota have emerged as key modulators of immune and autonomic function. mTBI can alter microbial community structure and compromise intestinal barrier integrity^17,26,29,30^, with similar effects documented in moderate TBI^31,32^. Growing preclinical evidence further indicates that gut microbiota–derived metabolites can causally modulate neuroinflammatory tone – including microglial activation, inflammatory signaling, barrier function, and neuroimmune cell balance – and influence functional recovery following mild/repetitive mTBI^33^, moderate TBI^34–37^, and other central nervous system (CNS) injuries such as stroke^38,39^ and spinal cord injury^40,41^. Together, these findings suggest a model wherein central-peripheral interactions sustain chronic low-grade inflammation, barrier dysfunction, and aberrant immune signaling, contributing to ongoing neuroinflammation^42^ and behavioral disturbances^43^ long after the initial injury^27^.

These effects are anatomically and functionally linked in part through the vagus nerve (VN) – the principal parasympathetic conduit between the gut and brain – whose afferent fibers detect microbially-derived metabolites and relay visceral information to brainstem autonomic nuclei^44–46^. Conversely, efferent vagal signaling engages the cholinergic anti-inflammatory reflex, shaping gut physiology^47–49^, systemic immune tone^50,51^, and potentially microbial composition^49,52,53^. Together, these findings highlight a bidirectional gut–vagus–brain axis through which peripheral inflammation and dysbiosis may reinforce central pathology after mTBI, and suggest that modulation of vagal activity could represent a tractable therapeutic entry point. Indeed, vagus nerve stimulation (VNS) has anti-inflammatory properties and can improve dysautonomia^47,54,55^, and is currently approved for therapeutic use in drug-resistant epilepsy, migraine, and treatment-resistant depression^56–60^. Likewise, a recent observational study demonstrated that non-invasive VNS was associated with a reduction in persistent post-concussion symptoms in patients with mTBI^61^. As such, VNS treatment offers a potential early intervention that could be used shortly following blast exposure to temper autonomic dysfunction caused by blast-mTBI and prevent long-term adverse PCS and behavioral outcomes. Critically, VNS has not been examined in a blast exposure setting and was the focus of the current study.

In order to establish a potential therapeutic effect of VNS on adverse outcomes following blast exposure, here we utilized an established mouse model of blast-mTBI^17,62–66^ and non-invasive, transcutaneous vagus nerve stimulation^67–70^. We first demonstrate that blast dose-dependently induces neuroinflammation in the solitary nucleus, or nucleus tractus solitarii (NTS), the main brainstem region receiving afferents from the vagus nerve^71,72^, providing additional support for the potential therapeutic relevance of VNS acutely following blast exposure. Next, we collected brain, blood, and fecal samples acutely following blast and VNS to investigate the potential for VNS to ameliorate blast-induced inflammatory response and gut microbiome changes. We then conducted a series of behavioral tests to assess the potential for VNS to ameliorate chronic behavioral dysfunction related to impulsivity and alcohol use. Finally, we used causal mediation analysis to identify specific blast-altered microbial species that explained within-treatment-group variation in inflammatory and behavioral outcomes. Together, these results highlight the potential of VNS treatment acutely following blast exposure to ameliorate acute blast-induced inflammatory response and chronic behavior dysfunction related to increased impulsivity and substance use, and suggest that the gut microbiota may represent a complementary axis to VNS for therapeutic intervention.

## Methods

### Animals

All animal experiments used male C57Bl/6J mice (Jackson Laboratory) aged 9-11 weeks old at the time of arrival to the VA Puget Sound. Mice were 11-13 weeks old at the time of blast or sham injury. Mice were housed one to three per cage on a 12:12 light:dark cycle (lights on at 06:00), and were given ad libitum food and water. Less than 5% of mice in the study were single housed (resulting from animals needing to be separated due to fighting or animal loss from blast exposure) and single housed mice did not significantly differ from group housed mice on experimental outcomes. Room temperature was set to 21°C and room humidity was set to 35%. All animal experiments were carried out in accordance with Association for Assessment and Accreditation of Laboratory Animal Care guidelines and were approved by the VA Puget Sound Institutional Animal Care and Use Committees. Mice were acclimated to the housing room for a week following arrival and subsequently handled for an additional three days prior to sham or blast exposure. To increase rigor and reproducibility, each experimental timeline included at least two cohorts of mice each run at separate times.

### Blast Trauma Model

We use a shock tube (Baker Engineering and Risk Consultants, San Antonio, TX) designed to generate blast overpressures that mimic open field high explosive detonations encountered by military service members in combat. Descriptions of the design and modeling characteristics have been described in detail elsewhere^17,62,63,73,74,65,66,64^. Briefly, mice were weighed and then anesthetized with isoflurane (induced at 5% and maintained at 2–3% for the duration of the blast), secured against a gurney, and placed into the shock tube oriented perpendicular to the oncoming blast wave (ventral body surface towards the blast). Pressurized helium is built up on one end of the tube and released to generate a blast overpressure wave that induces neuropathological and behavioral changes in line with blast mild traumatic brain injuries (mTBI). Sham (control) animals received anesthesia for a duration matched to blast animals but did not enter the shock tube. Repeated blast/sham exposures occurred successively over three days (one per day). The blast overpressure peak intensity (psi), initial pulse duration (ms), and impulse (psi▪ms) used were in keeping with mild blast injury (19.1 +/− 0.09 psi). Under these experimental conditions, the overall survival rate exceeded 95%, with blast-exposed mice presenting comparable to sham-exposed mice by inspection 2–4 hours-post blast exposure as previously reported. Following sham/blast exposure and removal from isoflurane, loss of righting reflex (LORR) was recorded as the amount of time it took for animals to right themselves. Once animals were able to regain full sternal recumbency, they were weighed and then returned to their home cages to recover.

### Immunohistochemistry

A subset of mice were sacrificed at either 3 days or 3 months post sham/blast exposure and transcardially perfused with 4% PFA, and brains removed, hemisectioned, and post-fixed for 24 hours prior to cryoprotection and shipment to NeuroScience Associates (NSA; Knoxville, TN) for histological and immunohistochemical staining). Brains were coronally sectioned at 40 microns and immunostained by NSA for ionized calcium binding adaptor molecule (Iba1; 1° Ab: Abcam, ab178846, 1:14,000; 2° Ab: Vector: BA-1000, 1:1,000); or glial fibrillary acidic protein (GFAP; 1° Ab: Dako, Z0334, 1:75,000; 2° Ab: Vector, BA-1000). Immunostaining was then visualized using 3,3’-Diaminobenzidine (DAB), wet-mounted on 2%-gelatin-subbed slides, coverslipped, and returned to VA Puget Sound for subsequent slide digitization and analysis (see next section).

Quantitative Analysis: Immunostained sections were digitally scanned at 20x using an AT2 scanner (Leica Biosystems, Nussloch Germany). Images were analyzed using Visiopharm (Hørsholm, Denmark), an AI-based pathology software suite. Briefly, a Visiopharm deep learning APP was trained to identify the NTS and then a second APP was used to measure expression level of DAB staining within the NTS using a threshold method (threshold levels set identically across sections and animals). All images were reviewed for ROI selection accuracy. Expression was calculated as percent of pixels above threshold within the NTS region divided by the total pixel area within the NTS region.

### Vagus Nerve Stimulation (VNS)

#### General VNS methods

Transcutaneous vagus nerve stimulation was delivered via a custom-made gammaCore stimulation device from electroCore (Rockaway, NJ) that was modified for mice^67–70^. Briefly, the leads of a standard gammaCore device are connected via alligator clips to a hand-held applicator with two smaller electrodes spaced 6 mm apart. For optimized conductance, hair around the stimulation area is shaved before stimulation and conductive gel (Signa gel, Parker Laboratories, Fairfield NJ) is applied to both the shaved skin and electrodes. Electrical stimulation (1 msec duration, 5 kHz, 12 V sine waves repeated at 25 Hz; impedance: 350 ohms) was delivered in the form of 2-minute trains.

#### VNS + MouseOx^Ⓡ^ Plus

To assess the acute physiological effects of cervical VNS, the MouseOx^Ⓡ^ Plus (STARR Life Sciences, Alleghany, PA) was used to measure oxygen saturation, heart rate, breath rate, pulse distention, and breath distention before and after VNS in naïve mice. Mice were first anesthetized with isoflurane (5% for induction, 2% for maintenance) and the inner thigh and area around the cervical vagus nerve shaved. One day later, each mouse was recorded for a 35-minute baseline in which they were anesthetized with isoflurane and monitored through a MouseOx clip sensor attached to the thigh. On the following day, each mouse was anesthetized and recorded for 10 minutes before and after two separate 2-minute VNS stimulations to the right cervical vagus nerve for a total of 35 minutes of recorded data. MouseOX data was processed using custom Python scripts; briefly, data points that did not pass MouseOx^Ⓡ^ Plus a priori quality control metrics were removed and then data was downsampled and smoothed. Baseline signal was computed as the average signal across the first 10 minutes of recording and then all individual data points were normalized to baseline for final analysis.

## Sham/Blast +VNS

In a separate group of mice, one hour following each day of sham or blast exposure, mice were anesthetized with isoflurane (5% for induction and 2% for maintenance) and given either sham or transcutaneous VNS. In the VNS group, electrodes were placed on the shaved skin over the right cervical vagus nerve without applying additional pressure and stimulation was delivered in the form of 2-minute trains, every 15 minutes, for 1 hour. In the sham group, mice remained under anesthesia for 1 hour but no stimulation was delivered. Animals were returned to their home cages following stimulation.

### Cytokines

Blood and brain samples were collected at the time of euthanasia for cytokine comparison as previously described^17,75^. Following cervical decapitation at 4 hours post final sham/blast exposure, trunk blood was collected in serum-separator tubes, allowed to clot at room temperature for 30-50 minutes, and then centrifuged at 3,000 x g for 10 minutes. Separated serum was then aliquoted and stored at -80 until analyzed. Whole brains were hemisectioned and flash-frozen at −80°C. One hemisphere of brain tissue was then lysed in a 0.02% Triton-X homogenization buffer of 10mM HEPES, 1.5mM MgCl2, and 10mM KCl, with fresh protease/phosphatase inhibitor cocktail (Sigma) in a bead homogenizer. The samples were then centrifuged at 4°C at 18,000 × g for 10 minutes and the supernatant was aliquoted and stored at −80°C. Serum and brain samples were shipped to IDEXX Bioanalytics (Columbia, MO) for pro- and anti-inflammatory cytokine level analysis using their Cytokine Mouse 25-Plex Panel. Values were normalized by protein concentration quantified with a BCA Protein Assay Kit (Thermo Fisher Scientific, Rockford, IL) using a 50-fold dilution to account for variable protein amounts in brain lysate samples.

### Fecal Microbiome

#### Metagenomic Sequencing

Fecal samples were collected and characterized for microbiome analysis as previously described^17^. In the acute cohort, the lowest fecal pellet was retrieved from the colon during necropsy 4 hours post-final blast or sham. In the chronic cohort, mice were briefly housed individually in sterile cages, and fresh fecal pellets collected and flash-frozen in liquid nitrogen 24 hours post-final blast or sham. Fecal samples were stored at -80 until shipment to Diversigen Inc. (New Brighton, MN) for DNA extraction, library preparation, and metagenomic sequencing via their BoosterShot Shallow Shotgun Sequencing. Libraries were sequenced at a target depth of 2M reads per sample on Illumina NovaSeq 6000 using a 2 x 100 bp flow cell.

#### Species- and SGB-level metagenomic taxonomic profiling

Reverse reads were discarded due to low quality. Forward reads were filtered and classified in a custom nextflow pipeline. The first 5 bp of each read was trimmed, and reads were filtered to meet a quality threshold of 30 and a minimum length of 50 bp. Taxonomic profiling was performed using MetaPhlAn 4.1.1 using reference database vJun23_CHOCOPhlAnSGB_202403 and the additional argument ‘-t rel_ab_w_read_stats’ to obtain estimated read counts^76^. NCBI species-level taxonomic assignments were used for subsequent analyses.

### Behavioral Testing

#### Open Field

72 hours post sham/blast exposure, mice were tested for locomotor deficits and anxiety-like behaviors in an open field assay. For 5 minutes, mice were placed in a large (1 m in diameter), open, circular arena split into center and outer sections. Locomotor behavior measurements like speed, total distance traveled, time spent in the center of the field, time delay to first entry to the center, and the number of entries into the center, were recorded from above and analyzed with ANYmaze (Wood Dale, IL). Less time spent in the center of the arena indicates an anxiety-like phenotype.

#### Acoustic Startle

Similarly to previous publications^17,62^, SR-LAB acoustic startle boxes (San Diego Instruments, San Diego, CA) were used to measure acoustic startle habituation and prepulse inhibition at 1-month post sham/blast exposure. After acclimating to the acoustic startle boxes for 5 minutes, mice were exposed to startle habituation testing which consisted of 50 trials of 120-dB pulses with inter-trial intervals of 7-23 seconds. Prepulse inhibition (PPI) was assessed after a 2-minute break and consisted of 40 trials of 81-dB prepulse followed by a 120-dB pulse with varying interstimulus intervals of 2-1,000 ms.

#### Impulsivity

To assess risky decision-making, mice were subject to a probabilistic discounting task. Mice were food-restricted to 85-90% of starting body weight. Initial lever press training consisted of 30 trials of fixed ratio 1 schedule on two separate levers to a criterion of ≥24 nose-pokes out of 30 trials. On each trial, both levers were extended and mice had unlimited time to lever press; lever press on either lever resulted in pellet delivery (12 mg sucrose), lever retraction, and a 30-45s intertrial interval (ITI). Mice were then assigned a high-reward lever, with lever assignment balanced across sham and blast groups for left vs. right lever preference on the final training day. Next, mice were trained on the probabilistic discounting task over four days. Training sessions consisted of two sets of 24 “forced” trials, with an intertrial interval that progressed from 10s to 45s by day 4. At the start of each trial, a cue light was illuminated over the food tray to indicate the requirement of a head entry into the food port. Mice were initially given 60s to make a head entry, which then pseudo-randomly presented either lever (12 presentations of each) and had 30s to make a lever response thereafter. Head port entry and lever response time limits progressively decreased to 30s and 10s, respectively, by day four. Trials in which the mice failed to make a response in time were not repeated. During training, both levers rewarded one sucrose pellet at FR1. After training, mice were given one probabilistic discounting test session per day for five days. Each session began with 24 forced trials, with either lever presented pseudo randomly as in the training sessions. The previously assigned “risky” lever probabilistically delivered a large reward (1.00, 0.75, 0.50, 0.25 or 0.00 chance; 2 sucrose pellets, FR1) and the “safe” lever always delivered a small reward (1.00 chance; 1 sucrose pellet, FR1). Animals experienced only one probability per test session, beginning with 1.00 and ending with 0.00 chance. The ITI was 45s, head entry time limit 30s and lever response time limit 10s. Missed trials were not repeated. After 24 forced trials, which served to inform the animal of the probability of the risky lever for the given session, 24 “choice” trials were given. Choice trials were identical to forced trials, except both levers were presented simultaneously. Animals were free to choose either the certain but low reward or risky but high reward lever. Animals were given 3 chances to choose the risky lever at least 51% of choice trials when the probability was 1.00 before exclusion from testing.

#### Alcohol intake

As previously described^64^, mice were single-housed 4 days prior to the intermittent 2-bottle choice (I2BC) procedure with two 50mL centrifuge tubes modified with a #6.5 neoprene rubber stopper and 2.5” ball sipper tube (Atrin) replacing their normal water pouches. The two 50 mL bottles were both filled with water during the 4-day baseline period prior to the start of I2BC. Throughout the duration of the study, bottles were weighed immediately before placement in the cage and then again at 24 hour intervals (at ∼5:30 pm). Bottles reversed position each day when they were weighed to prevent any side bias. Every Wednesday night, bottles were also weighed at 8:00 pm (2 hours into the dark cycle) to assess drinking patterns during acute phases of access. At the start of the I2BC procedure, one of the two bottles each mouse had access to contained ethanol every Monday, Wednesday, and Friday. Both bottles contained water every Tuesday, Thursday, Saturday, and Sunday. The concentration of alcohol gradually increased from 3% (Monday), to 6% (Wednesday) and 9% (Friday) during the first week of EtOH exposure. For the remainder of the study (21 more days), 20% EtOH was used. Every Sunday, body weight was recorded for each mouse.

### Statistical analysis

Data are expressed as mean ± SEM. Differences between groups were determined using a two-tailed Student’s t-test, one-way analysis of variance (ANOVA), or two-way (repeated measures (RM) when appropriate) ANOVA followed by post hoc testing using Šidák’s multiple comparisons or Bonferroni’s Multiple Comparison. Reported p values denote two-tailed probabilities of p ≤ 0.05 and non-significance (n.s.) indicates p > 0.05. Statistical analysis and visualization were conducted using Graph Pad Prism 9.0 (GraphPad Software, Inc., La Jolla, CA) and with custom R scripts available on GitHub (https://github.com/Gibbons-Lab/2026_Easton_VNS).

#### Cytokine dimensionality reduction

To reduce the total number of mediation analyses, variation in brain and serum cytokine measurements was collapsed into a few major axes using principal component analysis (PCA). Variance explained by each principal component (PC) was calculated manually as before. Resulting PCs explaining >5% of variance across all cytokines were retained for downstream mediation analysis.

#### Microbiome Statistical Analyses

Statistical analysis for the acute and chronic cohorts was performed separately due to differences in fecal microbiome sampling method.

For alpha and beta diversity metric calculations, species-level read counts were rarefied to the lowest count number per cohort (575,229 in acute cohort, 154,195 in chronic cohort). Treatment effects on alpha diversity metrics (Shannon Index, Richness, and Berger-Parker Index) were assessed using 2-way ANOVA followed by Šidák’s-adjusted post hoc comparisons. Beta diversity was assessed by Bray-Curtis dissimilarity using vegan::vegdist(method = "bray"). Principal coordinate analysis (PCoA) was performed on the Bray-Curtis distance matrix using weighted classical multidimensional scaling (‘wcmdscale(eig=TRUE)), and variance explained by each axis was calculated as the eigenvalue divided by the sum of the absolute eigenvalues. Because batch effects were observed in unconstrained PCoA, batch-conditioned (residualized) ordination was performed using distance-based redundancy analysis (‘capscale(bc_mat∼ batch)’), and variance explained calculated as above. Multivariate effects of Blast, VNS, and their interaction on Bray-Curtis distance were tested using PERMANOVA with batch included as a covariate (vegan::adonis2(BC ∼ Blast + VNS + Blast:VNS + batch, by = "terms", n_perm = 999).

To model differential abundance of microbial species across treatment, batch effects were handled using cohort-specific approaches due to differences in batch structure. In the acute cohort, differential abundance was tested using linear models with batch included as a fixed effect (lm(microbe ∼ Blast + VNS + Blast:VNS + batch)). In the chronic cohort, batch was modeled as a random intercept using linear mixed-effect models (‘lmerTest::lmer(microbe ∼ Blast + VNS + Blast:VNS + (1|batch))’). Overall Blast and VNS effects were tested using likelihood ratio tests (LRTs) comparing the full model to a reduced model in which both the fixed effect in question and interaction effect were removed. For the acute cohort, partial F-tests were used; for the chronic cohort, χ²-based LRTs were used. Resulting p-values were corrected for multiple testing using the Benjamini–Hochberg false discovery rate (FDR) procedure (FDR < 0.20 defining statistical significance).

A similar approach was used to test associations between microbial species and outcomes independent of treatment. For a given outcome, the full acute and chronic models were specified as lm(outcome ∼ microbe + Blast + VNS + Blast:VNS + batch) and lmer(outcome ∼ microbe + Blast + VNS + Blast:VNS + (1|batch), respectively. Significance of the microbe term was assessed using either partial F-test LRTs (acute cohort) or χ² LRTs (chronic cohort), as above.

High-dimensional mediation screening was performed using custom R wrappers implementing M-DACT^77^, which extends the DACT framework^78^ through empirical null estimation^79^ to improve Type I error control. For each exposure (Blast and VNS) and each outcome, M-DACT was applied using raw LRT p-values from the mediator and outcome models outlined above, which controlled for batch and the alternate exposure. Joint DACT statistics were evaluated across FDR thresholds ranging from 0.10 to 0.20, with FDR < 0.20 defining statistical significance.

To contextualize mediation screening results, we estimated nonparametric bootstrap 95% confidence intervals (999 resamples) for the average causal mediated effect (ACME), average direct effect (ADE), and total effect of Blast or VNS exposure for selected exposure–mediator– outcome combinations (M-DACT FDR < 0.2). Because exposure groups were unbalanced and samples were distributed across batches, we implemented a stratified nonparametric bootstrap procedure. Specifically, observations were resampled with replacement within each batch– exposure group combination, preserving the original batch and treatment structure in every bootstrap replicate. For each resampled dataset, mediator and outcome models were refit, and ACME, ADE, and total effects were re-estimated. Effects were calculated at both levels of the secondary exposure variable when applicable (e.g., for the Blast–Richness–PC3 mediation model, effects were estimated separately at VNS(–) and VNS(+)). Bootstrap percentile intervals were then derived from the empirical distribution of effect estimates across resamples.

## Results

### Blast-induced astrocyte and microglia expression in the Nucleus Tractus Solitarii

The nucleus tractus solitarii (NTS), also known as the solitary nucleus, is the main relay of the vagus nerve into the brain. Blast-induced changes in expression levels of IBA1 and GFAP markers were found 3 days and 3 months following blast injury, indicative of activated astrocytes and microglia (Figure 1). Specifically, at the 3-day time point, blast mice who received either 1x or 3x blast exposures had higher percent expression IBA1 compared to sham exposure animals (Figure 1b; One-Way ANOVA: F(2,66) = 19.9, p < 0.0001). Conversely, at three months post-blast, only 3x blast mice had significantly elevated IBA1 expression levels (Figure 1c; One-Way ANOVA: F(2,57) = 3.79, p < 0.05). Likewise, at the 3-day time point, blast mice who received either 1x or 3x blast exposures had higher percent expression GFAP compared to sham exposure animals (Figure 1e; One-Way ANOVA: F(2,58) = 16.53, p < 0.0001). Conversely, at three months post-blast, 1x mice had increased and 3x blast mice had significantly decreased GFAP expression levels (Figure 1f; One-Way ANOVA: F(2,48) = 14.14, p < 0.0001).

**Figure 1:**
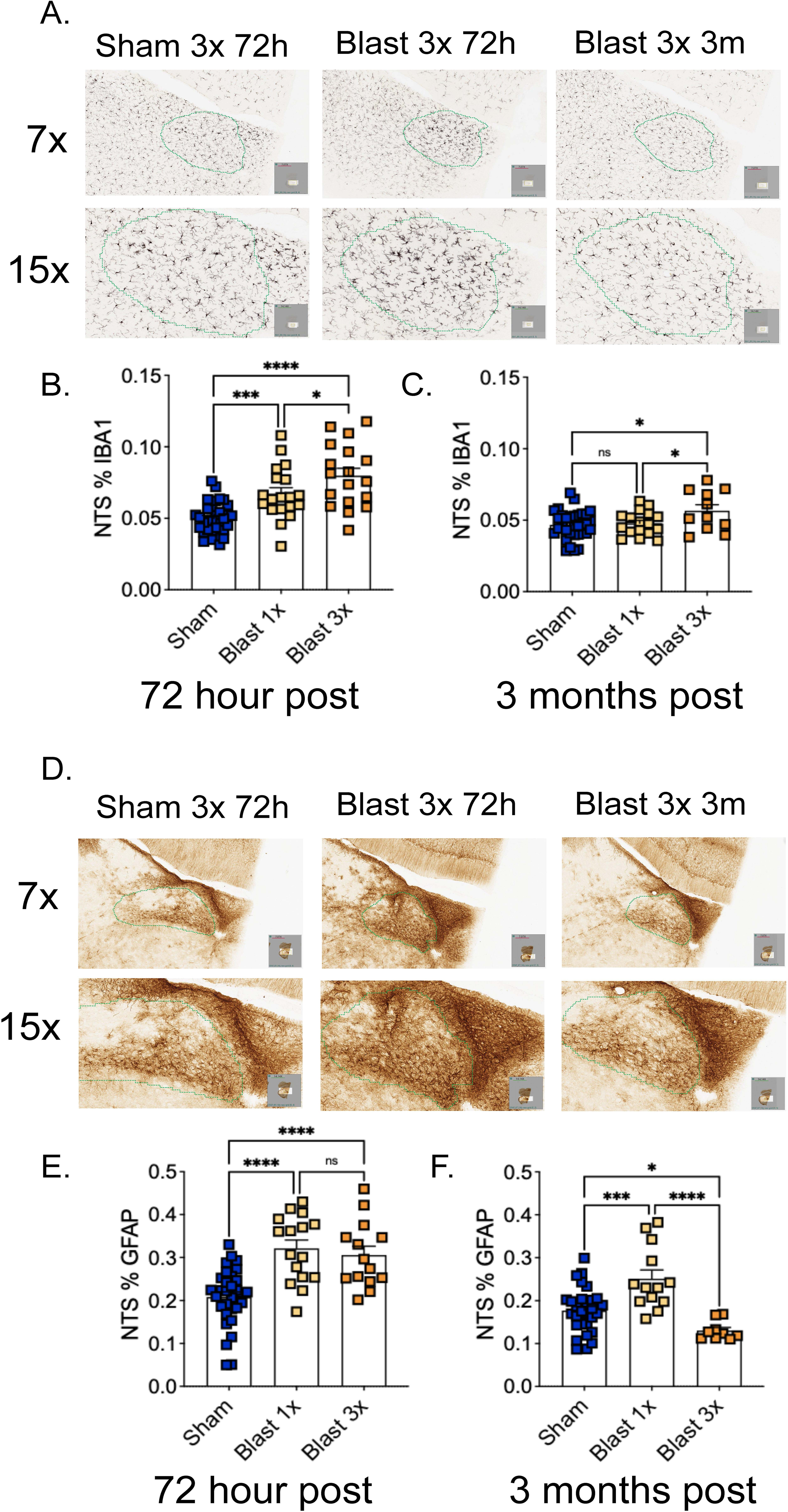
Blast exposure increases microglia and astrocyte expression in the NTS. A: Representative images of IBA1 expression in the NTS at acute and chronic timepoints following sham/blast exposure. B: Single and repetitive blast exposure increases IBA1 expression in the NTS 72-hours following exposure. C: Repetitive, but not single, blast exposure increases IBA1 expression in the NTS 3 months following exposure. D: Representative images of GFAP expression in the NTS at acute and chronic timepoints following sham/blast exposure. E: Single and repetitive blast exposure increases GFAP expression in the NTS 72-hours following exposure. F: Repetitive blast exposure decreases and single blast exposure increases GFAP expression in the NTS 3 months following exposure. One-way ANOVA *post hoc* Bonferroni Multiple Comparison Test (b,c,e,f). *p ≤ 0.05, **p ≤ 0.01, ***p ≤ 0.001, ****p ≤ 0.0001. Values represent mean ± SEM.

### Acute effects of VNS on oxygen saturation, heart rate, and breath rate

To confirm the acute physiological effects of our VNS paradigm, MouseOx was used to collect oxygen saturation, heart rate, and breath rate data on naive mice (n=4) before and after repeated VNS administration (Figure 2a-c). There was no significant effect of VNS on O_2_ saturation (Figure 2a; Two-Way RM ANOVA: main effect of treatment F(1,6) = 0.774, p > 0.05; main effect of time F(2.687,16.12) = 1.217, p > 0.05; interaction effect of treatment by time F(199,1194) = 1.02, p > 0.05). There was a significant effect of VNS on breath rate (Figure 2b; Two-Way RM ANOVA: main effect of treatment F(1,6) = 0.817, p > 0.05; main effect of time F(3.6,21.6) = 1.52, p > 0.05; interaction effect of treatment by time F(199,1194) = 1.993, p < 0.0001). Likewise, there was a significant effect of VNS on heart rate (Figure 2c; Two-Way RM ANOVA: main effect of treatment F(1,6) = 9.12, p < 0.05; main effect of time F(2.489,14.9) = 4.19, p < 0.05; interaction effect of treatment by time F(199,1194) = 5.785, p < 0.0001).

**Figure 2:**
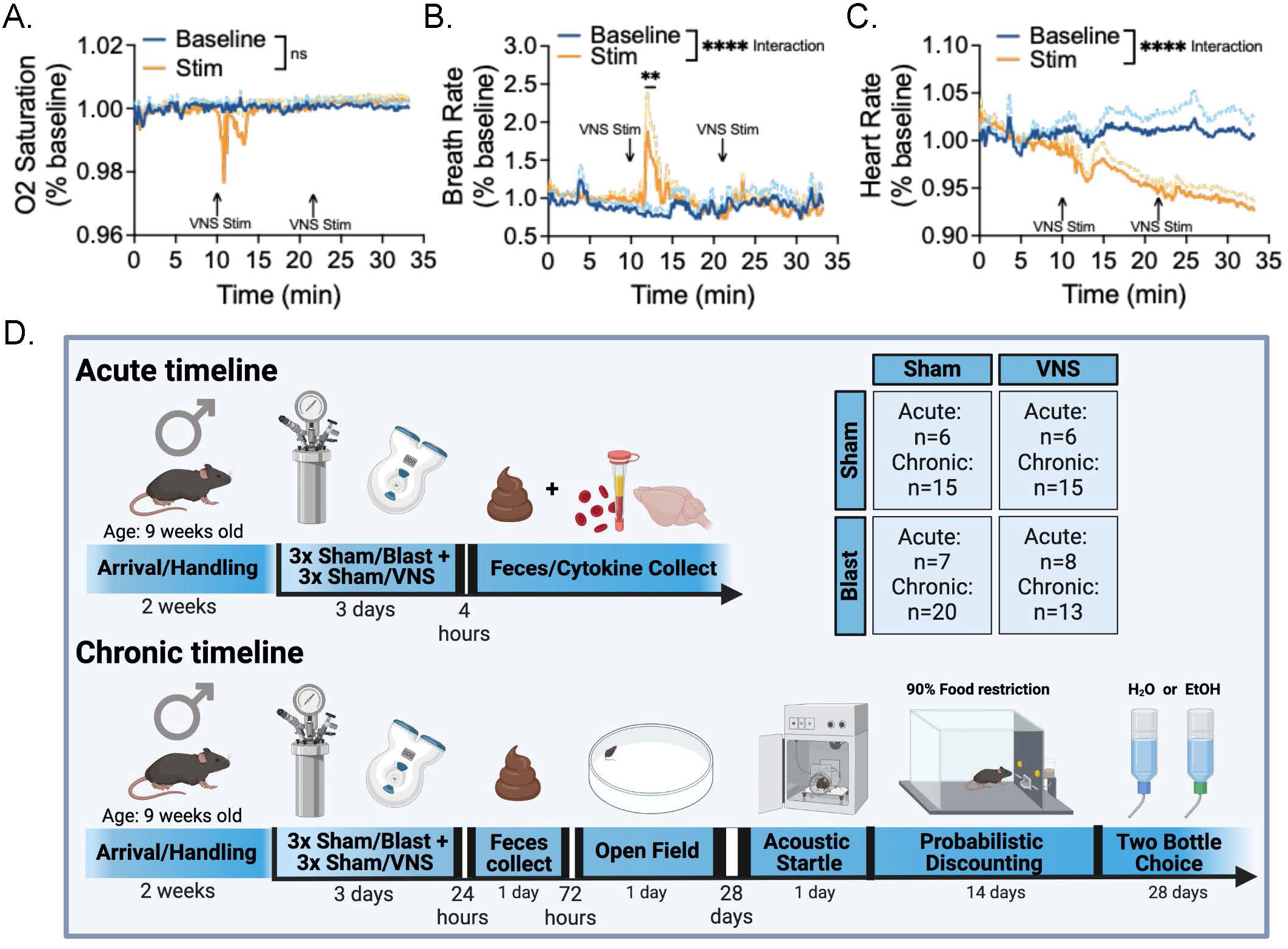
Acute physiological effects of VNS and experimental design. A: VNS has minimal acute effect on O^2^ saturation. B: VNS acutely increases breath rate. C: VNS acutely decreases heart rate. D. Experimental timeline and n per group. Two-way RM ANOVA *post hoc* Bonferroni Multiple Comparison Test (a-c). *p ≤ 0.05, **p ≤ 0.01, ***p ≤ 0.001, ****p ≤ 0.0001. Values represent mean ± SEM.

### Repetitive blast followed by VNS experimental timeline

To understand the potential therapeutic benefit of VNS acutely following repetitive blast exposure, we administered non-invasive VNS 1-hour following either sham or blast exposure for three consecutive days (Figure 2d). Animals were then split into two groups; the first group was sacrificed 4 hours after final sham/blast exposure (acute cohort) and the second group was sacrificed 3 months after final sham/blast exposure (chronic cohort).

### VNS effects on blast-induced LORR, weight loss, and open field

As previously reported^17,75^, blast exposure resulted in significantly greater loss of righting reflex latencies relative to sham controls, which was not ameliorated by VNS stimulation on any single day of blast exposure (Figure 3a; Three-Way RM ANOVA: main effect of injury F(1,86) = 129.2, p < 0.0001; main effect of treatment F(1,86) = 1.115, p > 0.05; main effect of time F(2,172) = 3.70, p < 0.05; two way interaction injury by treatment F (1,86) = 0.3591, p > 0.05; two way interaction injury by time F (2,172) = 0.936, p > 0.05; two way interaction treatment by time F (2,172) = 0.339, p > 0.05; three way interaction F (2,172) = 0.487, p > 0.05). VNS did not ameliorate the average LORR increased induced by blast (Figure 3b; Two-Way ANOVA: main effect of injury F (1,86) = 129.2, p < 0.0001; main effect of treatment F (1,86) = 1.115, p > 0.05; interaction effect injury by treatment F(1,86) = 0.359, p > 0.05).

**Figure 3:**
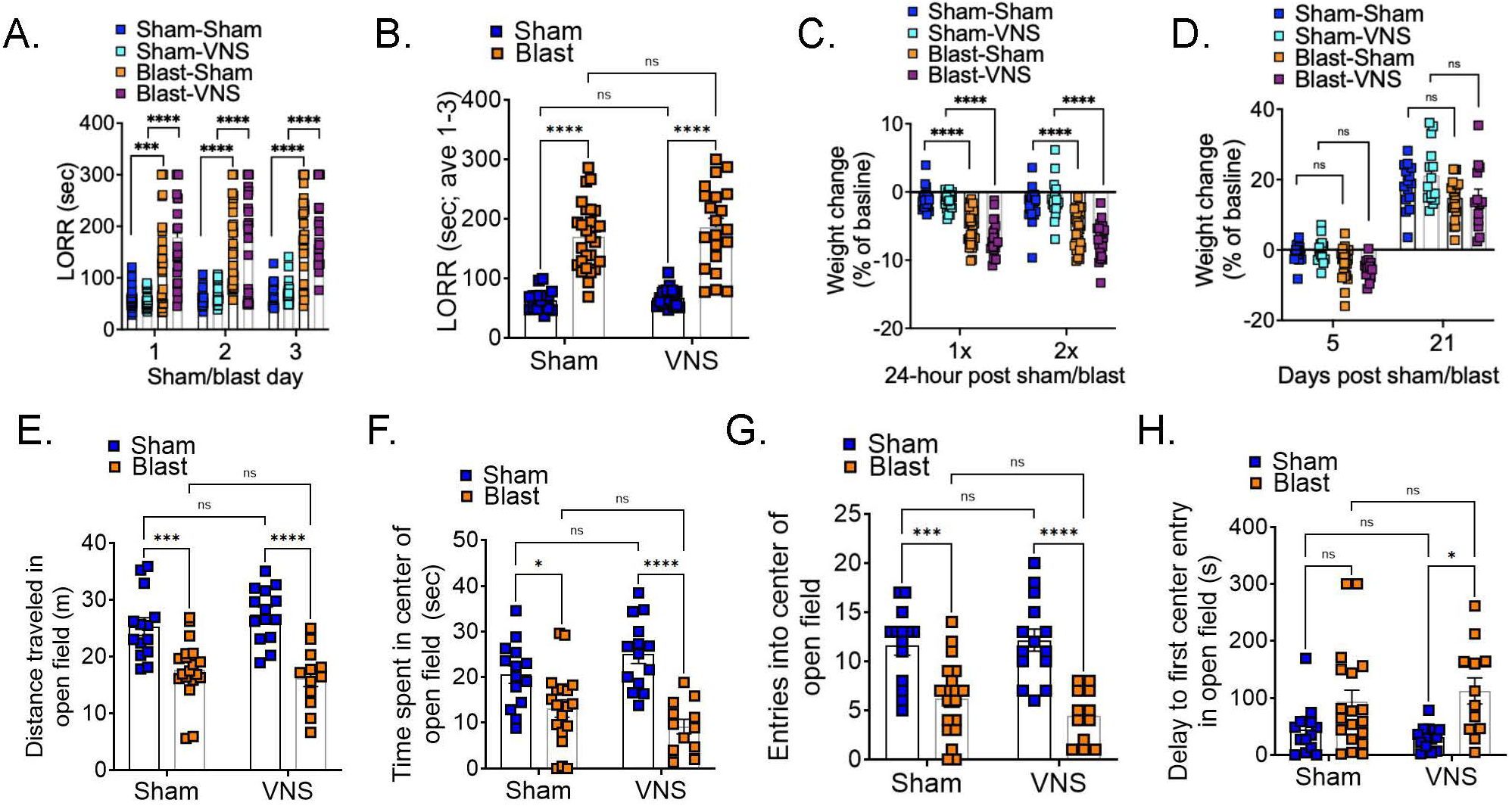
VNS is not effective in ameliorating acute blast behavioral outcomes. A-B: Blast exposure increases LORR which is not ameliorated by VNS treatment. C-D: Blast exposure acutely decreases weight which is not ameliorated by VNS treatment. E-H. Blast exposure results in decreased locomotion and anxiety-like behavior that is not ameliorated by VNS. Three-way RM ANOVA *post hoc* Bonferroni Multiple Comparison Test (a,c,d); Two-way ANOVA *post hoc* Bonferroni Multiple Comparison Test (b,e-h).*p ≤ 0.05, **p ≤ 0.01, ***p ≤ 0.001, ****p ≤ 0.0001. Values represent mean ± SEM.

VNS stimulation also had no effect on the subsequent reduction in weight 24-hours following single or repetitive blast exposure (Figure 3c; Three-Way RM ANOVA: main effect of injury F(1,86) = 110.6, p < 0.0001; main effect of treatment F(1,86) = 1.435, p > 0.05; main effect of time F(1,86) = 1.538, p > 0.05; two way interaction injury by treatment F (1, 86) = 3.548, p > 0.05; two way interaction injury by time F (1, 86) = 0.001, p > 0.05; two way interaction treatment by time F (1,86) = 1.927, p > 0.05; three way interaction F (1,86) = 1.305, p > 0.05). Likewise, VNS stimulation also had no effect on further changes in weight at either 5 days or 1 month post blast exposure (Figure 3d; Three-Way RM ANOVA: main effect of injury F(1,59) = 14.46, p < 0.001; main effect of treatment F(1,59) = 0.2, p > 0.05; main effect of time F(1,59) = 478.9, p < 0.0001; two way interaction injury by treatment F (1,59) = 1.177, p > 0.05; two way interaction injury by time F (1,59) = 0.195, p > 0.05; two way interaction treatment by time F (1,59) = 0.932, p > 0.05; three way interaction F (1,59) = 0.09, p > 0.05).

In keeping with previous reports^17,75^, blast exposure increased anxiety-like behaviors in the open field test (Figure 3e-h). VNS did not have a significant effect on blast-induced decrease in the total distance traveled in the open field (Figure 3e; Two-Way ANOVA: main effect of injury F(1,55) = 43.66, p < 0.0001; main effect of treatment F(1,55) = 0.063, p > 0.05; interaction effect injury by treatment F(1,55) = 1.624, p > 0.05). VNS did not have a significant effect on blast-induced decrease in the time spent in the center of the open field (Figure 3f; Two-Way ANOVA: main effect of injury F(1,55) = 44.07, p < 0.0001; main effect of treatment F(1,55) = 0.426 p > 0.05; interaction effect injury by treatment F(1,55) = 1.339, p > 0.05). VNS did not have a significant effect on blast-induced decrease in entries to the center of the open field (Figure 3g; Two-Way ANOVA: main effect of injury F(1,55) = 35.60, p < 0.0001; main effect of treatment F(1,55) = 0.016, p > 0.05; interaction effect injury by treatment F(1,55) = 4.635, p < 0.05). VNS did not have a significant effect on blast-induced increase in the delay to enter the center of the open field (Figure 3h; Two-Way ANOVA: main effect of injury F(1,55) = 13.75, p < 0.001; main effect of treatment F(1,55) = 0.0278, p > 0.05, interaction injury by treatment F(1,55) = 0.855, p > 0.05).

### Selective VNS effects on blast-induced gut microbiome and cytokines

Four hours following the final sham/blast exposure, a subset of animals was sacrificed by rapid decapitation and brain, blood, and feces were collected. Shannon diversity was increased 4 hours post-blast and was not affected by VNS (Figure 4b; Supplementary Figure 1a-b; Table 1). In batch-adjusted PERMANOVA of Bray–Curtis dissimilarity, blast injury explained 6% of the variance in community composition (p = 0.021; Table 2), whereas VNS and the Blast×VNS interaction each explained ∼3.5% of the variance but were not statistically significant (p = 0.091 and p = 0.089, respectively; Table 2). Batch-adjusted (constrained) PCoA was performed on the Bray–Curtis matrix for visualization (Figure 4c).

**Figure 4:**
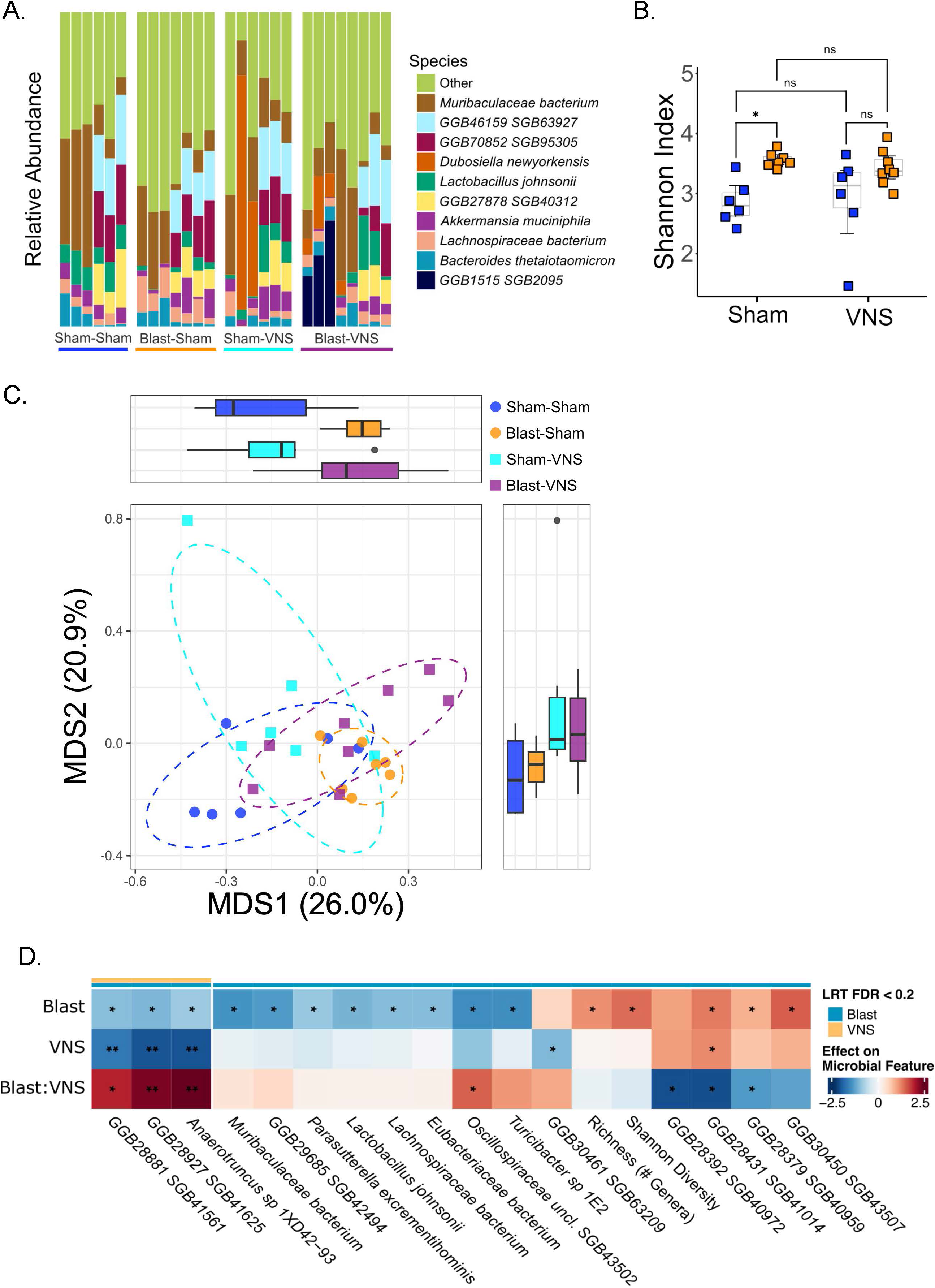
VNS has limited effects on 4-hour blast-induced changes to the gut microbiome. A: Relative abundance of bacterial species across individual samples by experimental group. B: Shannon alpha diversity is increased by blast. C: Blast-induced changes in beta diversity are not affected by VNS. D: Blast results in species-level relative abundance changes that are not ameliorated by VNS. Two-way ANOVA *post hoc* Bonferroni Multiple Comparison Test (b); Batch-adjusted PERMANOVA (c); Joint Likelihood Ratio Test of main and interaction effects from batch-adjusted linear model, FDR-corrected using Benjamini-Hochberg Procedure. Results represent species with Blast and/or VNS effect adjusted p-value < 0.2. Stars denote nominal significance of coefficients in the full linear model (d). *p ≤ 0.05, **p ≤ 0.01, ***p ≤ 0.001, ****p ≤ 0.0001.

**Table 1.**
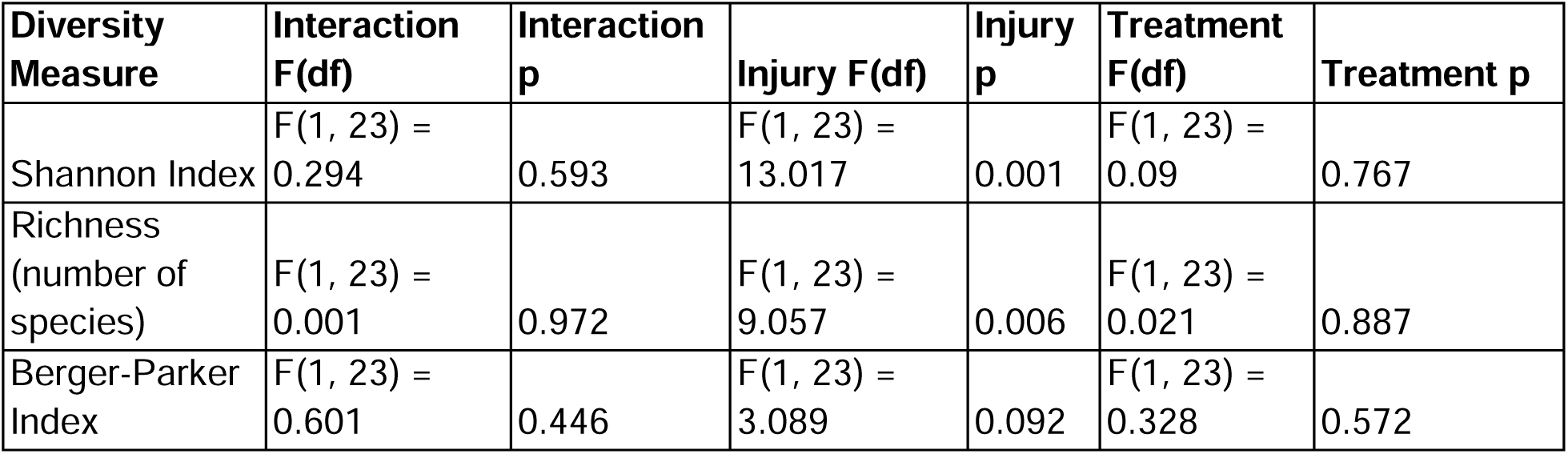
Alpha diversity, acute cohort.

| Diversity Measure | Interaction F(df) | Interaction p | Injury F(df) | Injury p | Treatment F(df) | Treatment p |
| --- | --- | --- | --- | --- | --- | --- |
| Shannon Index | F(1, 23) = 0.294 | 0.593 | F(1, 23) = 13.017 | 0.001 | F(1, 23) = 0.09 | 0.767 |
| Richness (number of species) | F(1, 23) = 0.001 | 0.972 | F(1, 23) = 9.057 | 0.006 | F(1, 23) = 0.021 | 0.887 |
| Berger-Parker Index | F(1, 23) = 0.601 | 0.446 | F(1, 23) = 3.089 | 0.092 | F(1, 23) = 0.328 | 0.572 |

**Table 2.**
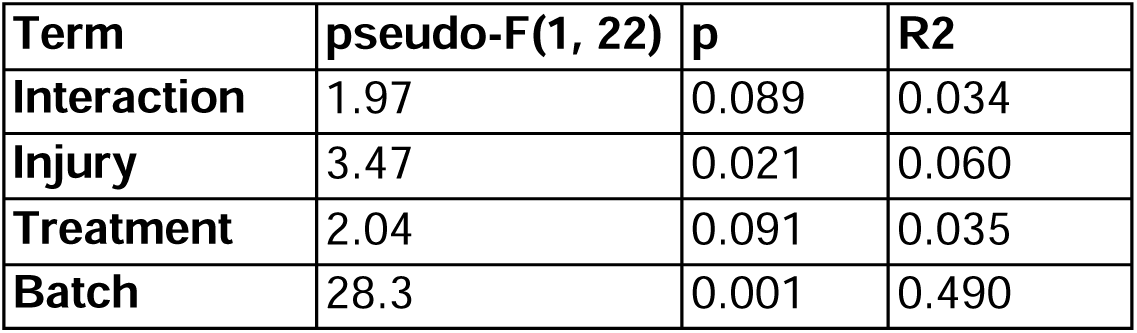
Bray-Curtis beta diversity PERMANOVA, acute cohort.

| Term | pseudo-F(1, 22) | p | R2 |
| --- | --- | --- | --- |
| Interaction | 1.97 | 0.089 | 0.034 |
| Injury | 3.47 | 0.021 | 0.060 |
| Treatment | 2.04 | 0.091 | 0.035 |
| Batch | 28.3 | 0.001 | 0.490 |

The relative abundances of multiple species were associated with blast exposure, with three of these also associated with VNS (FDR-adjusted LRT p < 0.2; Figure 4d; Supplementary Table 1). Blast exposure was associated with both increased and decreased abundance of taxa within the order *Eubacteriales* (including members of Oscillospiraceae, Pumilibacteraceae, Eubacteriaceae, and Lachnospiraceae), as well as decreased abundance of taxa from the orders *Bacteroidales*, *Lactobacillales*, *Burkholderiales*, and *Erysipelotrichales* (Figure 4d; Supplementary Figure 1c-o). Three taxa (Anaerotruncus sp. 1XD42-93, unclassified Bacteria SGB41625, and unclassified Firmicutes SGB41561) were decreased by VNS in blast animals but were not altered in sham animals (Figure 4d; Supplementary Figure 1p-r).

Serum cytokines were significantly elevated following blast trauma (Figure 5a; Table 3), and VNS treatment was successful in blocking (IL-9, IP-10) or significantly reducing (IL-6) select peripheral cytokine effects (Supplementary Figure 2a-k). Blast trauma also induced a significant cytokine response in the brain which was largely not significantly affected by VNS (Figure 5a; Table 4; Supplementary Figure 3a-s). Principal component analysis (PCA) of the cytokine panel was performed to reduce dimensionality (Figure 5b; Table 5; Supplementary Table 2). Together, PCs 1-5 explained 75% of variation in serum and brain cytokine profiles (Figure 5b). PC1 – which explained 30% of variance – was increased by blast injury and trended towards partial attenuation by VNS (Figure 5c; two-Way ANOVA: injury F(1,23) = 161.1, p < 0.00001; treatment F(1,23) = 4.001, p = 0.057; interaction F(1,23) = 0.873, p = 0.36), while the remaining four axes were not significantly affected by either (Table 5; Supplementary Figure 4a-d).

**Figure 5:**
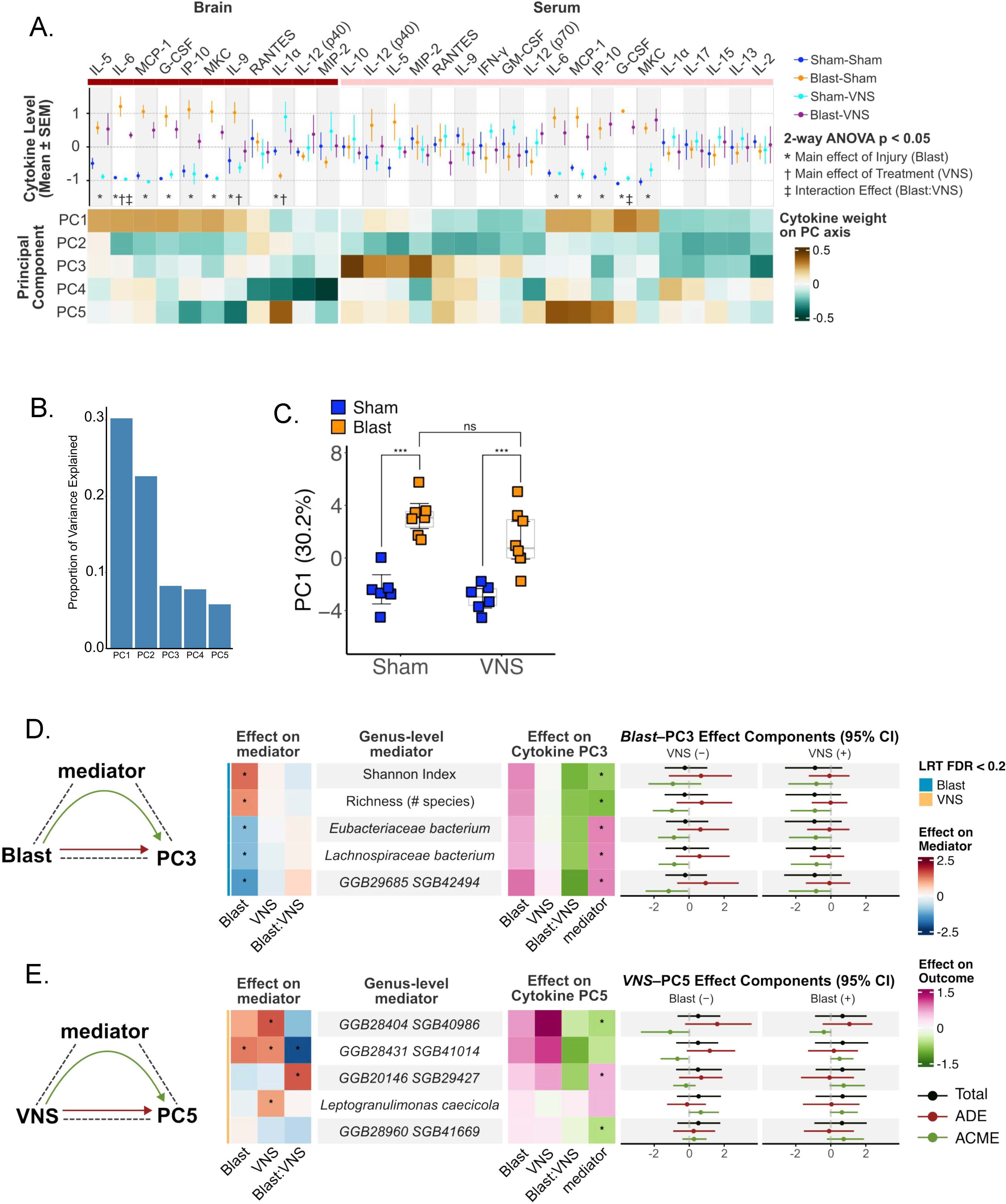
VNS reduces some blast-induced changes in peripheral and brain cytokines. A: VNS ameliorated a select set of blast-induced changes to blood and brain cytokine levels. B: Principal components 1-5 explained a cumulative 75% of variation in blood and brain cytokine levels. C: The blast-induced increase in PC1 is not significantly affected by VNS. D: Changes in microbiome structure (species-level abundance and alpha diversity metrics) mediate the effect of Blast on PC3 in the negative direction. E: Gut microbial species mediate the VNS effect on PC5 in different directions. Two-way ANOVA *post hoc* Bonferroni Multiple Comparison Test (a, c); Causal mediation pathways significant at M-DACT FDR < 0.2 (d). *p ≤ 0.05, **p ≤ 0.01, ***p ≤ 0.001, ****p ≤ 0.0001.

**Table 3.** Serum Cytokine.

| Analyte | Interaction F(df) | Interaction p | Injury F(df) | Injury p | Treatment F(df) | Treatment p |
| --- | --- | --- | --- | --- | --- | --- |
| IL-1α | F(1, 23) = 0.0003 | 0.986 | F(1,23) = 5.202 | 0.0322 | F(1,23) = 10.22 | 0.004 |
| IL-5 | F(1,23) = 0.319 | 0.578 | F(1,23) = 15.33 | < 0.001 | F(1,23) = 0.465 | 0.502 |
| IL-6 | F(1,23) = 5.711 | 0.025 | F(1,23) = 103.7 | <0.0001 | F(1,23) = 7.109 | 0.014 |
| IL-9 | F(1,23) = 2.214 | 0.15 | F(1,23) = 9.183 | 0.006 | F(1,23) = 4.565 | 0.044 |
| IP-10 | F(1, 23) = 2.786 | 0.109 | F(1, 23) = 29.40 | <0.0001 | F(1, 23) = 4.002 | 0.057 |
| MKC | F(1, 23) = 1.611 | 0.217 | F(1, 23) = 59.07 | <0.0001 | F(1, 23) = 2.597 | 0.121 |
| G-CSF | F(1, 23) = 1.168 | 0.291 | F(1, 23) = 45.11 | <0.0001 | F(1, 23) = 0.308 | 0.584 |
| MPC-1 | F(1, 23) = 0.991 | 0.33 | F(1, 23) = 83.86 | <0.0001 | F(1, 23) = 3.748 | 0.065 |
| IL-12(p40) | F(1, 23) = 0.454 | 0.51 | F(1, 23) = 0.118 | 0.73 | F(1, 23) = 0.95 | 0.34 |
| MIP2 | F(1, 23) = 0.002 | 0.96 | F(1, 23) = 1.372 | 0.25 | F(1, 23) = 1.418 | 0.25 |
| RANTES | F(1, 23) = 0.034 | 0.86 | F(1, 23) = 0.048 | 0.94 | F(1, 23) = 0.89 | 0.35 |

**Table 4.** Brain Cytokine.

| Analyte | Interaction F(df) | Interaction p | Injury F(df) | Injury p | Treatment F(df) | Treatment p |
| --- | --- | --- | --- | --- | --- | --- |
| IL-1α | F(1, 23) = 0.023 | 0.881 | F(1, 23) = 0.8948 | 0.354 | F(1, 23) = 0.067 | 0.798 |
| IL-5 | F(1, 23) = 4.261 | 0.051 | F(1, 23) = 3.062 | 0.094 | F(1, 23) = 0.175 | 0.68 |
| IL-6 | F(1, 23) = 0.618 | 0.44 | F(1, 23) = 26.08 | <0.0001 | F(1, 23) = 0.671 | 0.421 |
| IL-9 | F(1, 23) = 0.141 | 0.711 | F(1, 23) = 0.078 | 0.782 | F(1, 23) = 0.948 | 0.34 |
| IP-10 | F(1, 23) = 0.039 | 0.845 | F(1, 23) = 22.90 | <0.0001 | F(1, 23) = 0.417 | 0.525 |
| MKC | F(1, 23) = 0.05 | 0.826 | F(1, 23) = 38.32 | <0.0001 | F(1, 23) = 1.469 | 0.238 |
| G-CSF | F(1, 23) = 5.330 | 0.03 | F(1, 23) = 178.3 | <0.0001 | F(1, 23) = 1.389 | 0.251 |
| MPC-1 | F(1, 23) = 0.437 | 0.515 | F(1, 23) = 18.93 | <0.001 | F(1, 23) = 1.671 | 0.209 |
| IL-2 | F(1, 23) = 0.287 | 0.597 | F(1, 23) = 0.027 | 0.87 | F(1, 23) = 0.004 | 0.95 |
| IL-12(p40) | F(1, 23) = 0.376 | 0.545 | F(1, 23) = 3.79 | 0.064 | F(1, 23) = 1.122 | 0.3 |
| IL-12(p70) | F(1, 23) = 0.535 | 0.472 | F(1, 23) = 0.001 | 0.973 | F(1, 23) = 2.02 | 0.168 |
| IL-13 | F(1, 23) = 0.013 | 0.907 | F(1, 23) = 0.28 | 0.601 | F(1, 23) = 0.272 | 0.607 |
| IL-15 | F(1, 23) = 0.016 | 0.902 | F(1, 23) = 0.055 | 0.816 | F(1, 23) = 1.212 | 0.282 |
| IL-17 | F(1, 23) = 0.033 | 0.856 | F(1, 23) = 0.272 | 0.607 | F(1, 23) = 0.004 | 0.946 |
| GM-CSF | F(1, 23) = 0.343 | 0.563 | F(1, 23) = 2.46 | 0.131 | F(1, 23) = 0.467 | 0.501 |
| IFN-γ | F(1, 23) = 0.055 | 0.815 | F(1, 23) = 1.395 | 0.25 | F(1, 23) = 0.055 | 0.815 |
| IL-10 | F(1, 23) = 0.256 | 0.617 | F(1, 23) = 0.271 | 0.607 | F(1, 23) = 0.004 | 0.951 |
| MIP-2 | F(1, 23) = 0.005 | 0.941 | F(1, 23) = 0.124 | 0.728 | F(1, 23) = 0.767 | 0.39 |
| RANTES | F(1, 23) = 1.386 | 0.251 | F(1, 23) = 0.709 | 0.408 | F(1, 23) = 0.298 | 0.59 |

**Table 5.** Cytokine PCA Two-way ANOVA.

| Principal Component | Interaction F(df) | Interaction p | Injury F(df) | Injury p | Treatment F(df) | Treatment p |
| --- | --- | --- | --- | --- | --- | --- |
| PC1 | F(1,23) = 0.87 | 0.360 | F(1,23) = 60.15 | <0.0001 | F(1,23) = 4.00 | 0.057 |
| PC2 | F(1,23) = 0.73 | 0.402 | F(1,23) = 4.43 | 0.046 | F(1,23) = 0.16 | 0.695 |
| PC3 | F(1,23) = 0.58 | 0.454 | F(1,23) = 1.15 | 0.295 | F(1,23) = 1.07 | 0.311 |
| PC4 | F(1,23) = 0.16 | 0.690 | F(1,23) = 0.00 | 0.983 | F(1,23) = 2.58 | 0.122 |
| PC5 | F(1,23) = 0.00 | 0.963 | F(1,23) = 0.18 | 0.672 | F(1,23) = 1.08 | 0.310 |

Causal mediation analysis (CMA) was used to identify microbial features that statistically mediate the relationship between a given exposure (blast or VNS) and outcome (cytokine PC) (Figure 5d-e). Briefly, CMA tests the joint likelihood of two questions: 1) is the mediator affected by the exposure (Supplementary Table 1), and 2) is the mediator associated with variance in outcome independent of exposure (i.e. mediator explains within-group variance in outcome, across groups; Supplementary Table 3). Significance was tested using M-DACT, a high-throughput screening method for multiple mediators, with an FDR cutoff of 0.2 (Methods; Supplementary Table 4). Effect sizes were then estimated *post hoc* using stratified nonparametric bootstrap of the average causal mediation effect (ACME), average direct effect (ADE), and total effect (Figure 5d-e; Supplementary Table 5).

Although no overall injury or treatment effects were observed for PC3, mediation analysis identified two alpha diversity metrics and multiple taxa (including members of Eubacteriaceae and Lachnospiraceae) as mediators of the blast–PC3 association (Figure 5d; Supplementary Figure 5). In all models, the estimated ACME for blast exposure was negative. Microbial mediators were also identified for the VNS–PC5 association (including taxa from Pumilibacteraceae, Clostridiaceae, Atopobiaceae, and unclassified Firmicutes), with ACME direction varying by blast condition (Figure 5e; Supplementary Figure 6). Additional taxa were associated with within-group variation in PC2 independent of blast or VNS (Supplementary Figure 7).

In a separate cohort, fecal samples were collected 1 day following the final exposure (Figure 6a). In contrast to the 4-hour time point, alpha diversity was not associated with blast exposure (Figure 6b; Table 6; Supplementary Figure 8a-b). However, blast exposure explained 7% of the variance in Bray–Curtis dissimilarity (p = 0.001; Figure 6c; Table 7). At the species level, blast exposure was associated with the relative abundance of *Romboutsia ilealis*, *Clostridiales bacterium*, and an unclassified Firmicutes (*SGB41718*), with no significant associations observed for VNS (FDR < 0.2; Figure 6d; Supplementary Table 6; Supplementary Figure 8c-e). These animals were subsequently assessed in a behavioral battery one month post-exposure, and causal mediation analysis was performed to evaluate microbial mediators of blast and VNS effects on behavioral outcomes (see following sections).

**Figure 6:**
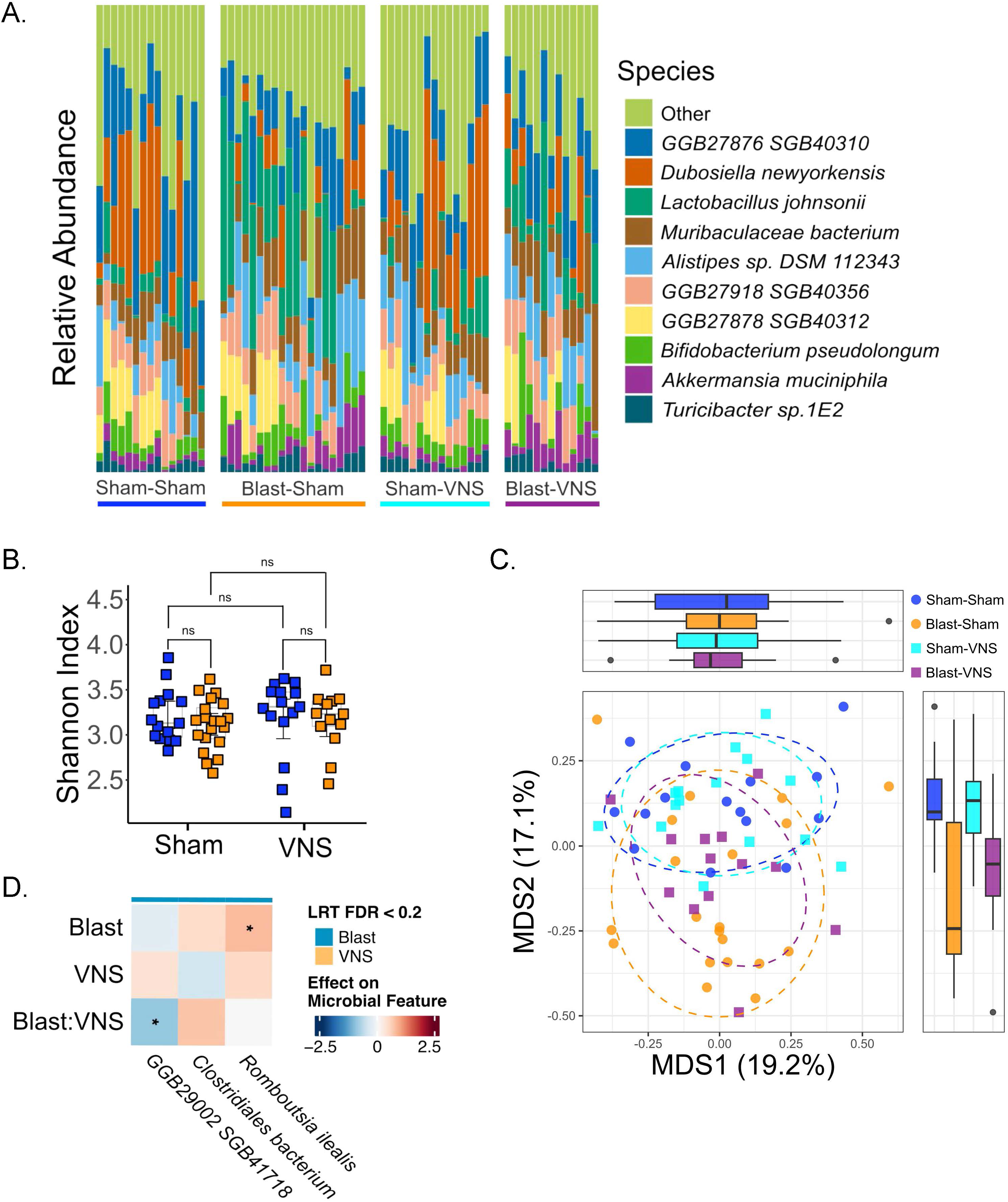
VNS has limited effects on 1-day blast-induced changes to the gut microbiome. A: Relative abundance of bacterial species across individual samples by experimental group. B: Shannon alpha diversity is not changed by blast or by VNS. C: Bray-Curtis Beta Diversity is still altered by Blast, not changed by VNS. D: Blast results in relative abundance changes that are not ameliorated by VNS. Two-way ANOVA *post hoc* Bonferroni Multiple Comparison Test (b); Batch-adjusted PERMANOVA (c); Joint Likelihood Ratio Test of main and interaction effects from batch-adjusted linear mixed effects regression, FDR-corrected using Benjamini-Hochberg Procedure. Results represent species with Blast and/or VNS effect adjusted p-value < 0.2. Stars denote nominal significance of coefficients in the full linear model (d). *p ≤ 0.05, **p ≤ 0.01, ***p ≤ 0.001, ****p ≤ 0.0001.

**Table 6.** Alpha diversity, chronic cohort.

| Diversity Measure | Interaction F(df) | Interaction p | Injury F(df) | Injury p | Treatment F(df) | Treatment p |
| --- | --- | --- | --- | --- | --- | --- |
| Shannon Index | F(1, 59) = 0.14 | 0.71 | F(1, 59) = 0.629 | 0.431 | F(1, 59) = 0.045 | 0.832 |
| Richness (observed species) | F(1, 59) = 0.808 | 0.372 | F(1, 59) = 1.761 | 0.19 | F(1, 59) = 0.007 | 0.932 |
| Berger-Parker Index | F(1, 59) = 0.653 | 0.422 | F(1, 59) = 0.039 | 0.845 | F(1, 59) = 0.983 | 0.326 |

**Table 7.**
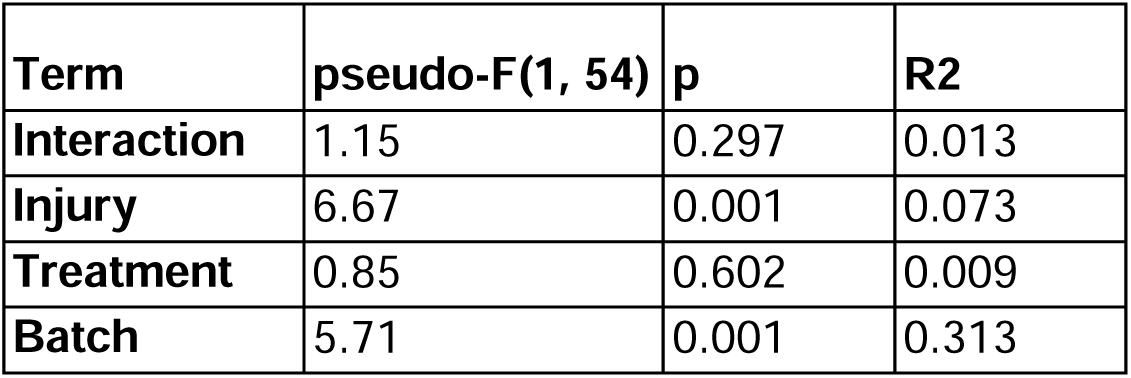
Bray-Curtis beta diversity PERMANOVA, chronic cohort.

| Term | pseudo-F(1, 54) | p | R<sup>2</sup> |
| --- | --- | --- | --- |
| Interaction | 1.15 | 0.297 | 0.013 |
| Injury | 6.67 | 0.001 | 0.073 |
| Treatment | 0.85 | 0.602 | 0.009 |
| Batch | 5.71 | 0.001 | 0.313 |

### VNS effects on chronic behavioral outcomes following repetitive blast

One month following repetitive sham/blast exposure and VNS treatment, mice were tested in a behavioral battery. Blast-exposed mice exhibited deficits in acoustic startle (Figure 7a-c), which were not ameliorated by VNS treatment. Specifically, blast-exposed mice showed decreased startle habituation (Figure 7a) (One-Sample T-Test: Sham/Sham t = 3.185, df = 12, p = 0.008; VNS/Sham t = 2.273, df = 13, p = 0.041; Sham/Blast t = 0.934, df = 19, p = 0.362; VNS/Blast t = 0.613, df = 11, p = 0.553) and impaired prepulse inhibition (Figure 7b,c) (Two-Way ANOVA: main effect of injury F(1,55) = 15.25, p < 0.001; main effect of treatment F(1,55) = 0.0332, p > 0.05; injury by treatment F(1,55) = 0.445, p > 0.05). Conversely, VNS stimulation reduced blast-induced increases in risky decision-making in the probabilistic discounting task (Figure 7d,e) (One-Sample t-test: Sham/Sham t = 0.69, df = 9, p = 0.508; VNS/Sham t = 1.036, df = 7, p = 0.335; Sham/Blast t = 6.511, df = 13, p < 0.0001; VNS/Blast t = 0.847, df = 6, p = 0.429).

**Figure 7:**
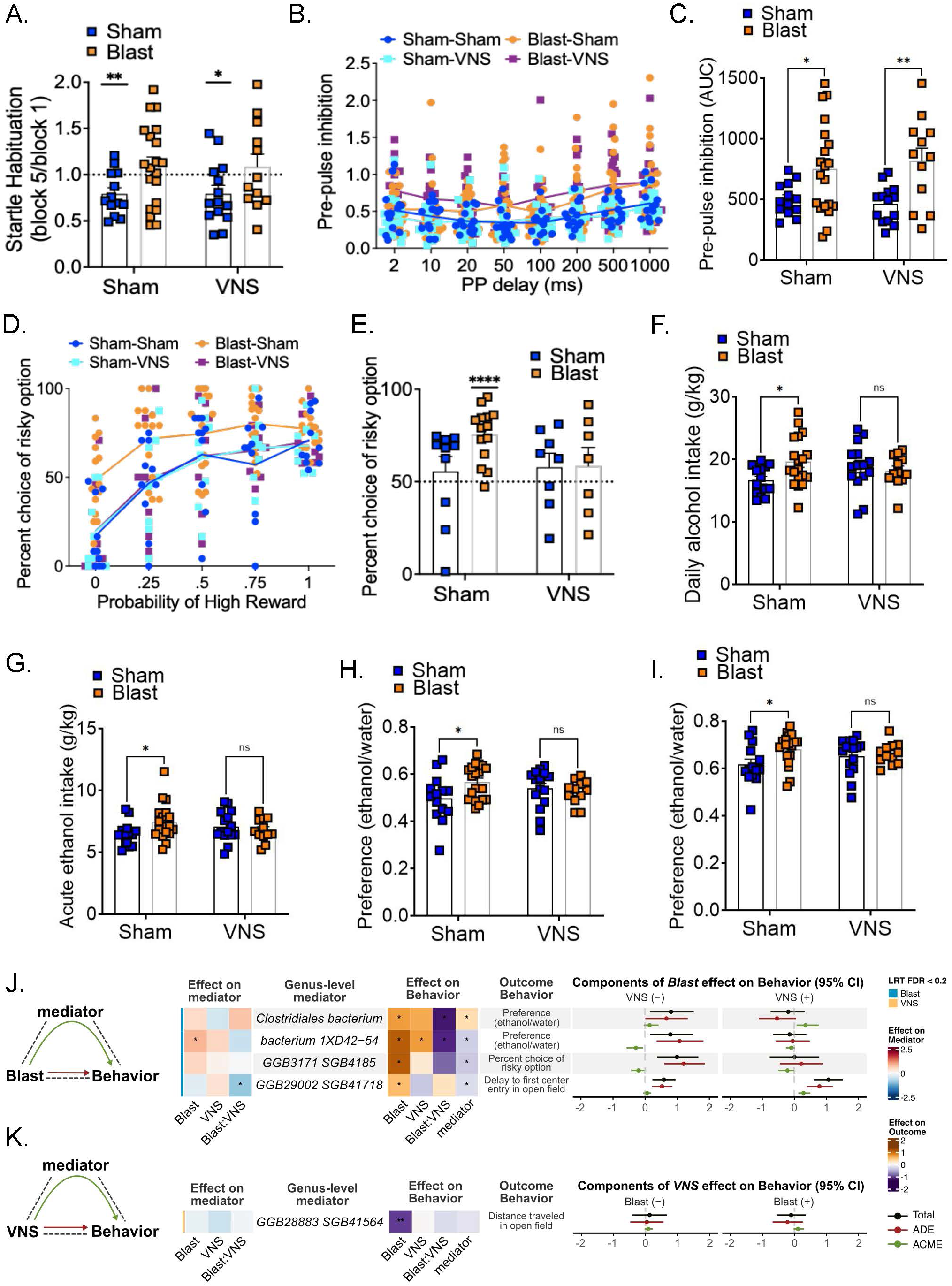
VNS reduces blast-induced risky decision making and alcohol intake independent of microbial mediation, but not hyperreactivity. A. VNS does not ameliorate blast-induced deficits in acoustic startle habituation. B-C: VNS does not ameliorate blast-induced deficits in acoustic startle prepulse inhibition. D-E: Blast-induced increase in risky-decision making is ameliorated by VNS. F-I: Blast-induced increase in alcohol intake is ameliorated by VNS. J: Gut microbial species mediate blast and VNS effects on acute and chronic behavioral outcomes. Two-way ANOVA *post hoc* Bonferroni Multiple Comparison Test (a,c,e-i). *p ≤ 0.05, **p ≤ 0.01, ***p ≤ 0.001, ****p ≤ 0.0001. Values represent mean ± SEM. Causal mediation pathways significant at M-DACT FDR < 0.2 (j,k). *p ≤ 0.05, **p ≤ 0.01, ***p ≤ 0.001, ****p ≤ 0.0001.

VNS stimulation also ameliorated blast-induced increases in alcohol intake behavior (Figure 7g-i). VNS blocked the blast-induced increase in the average daily total alcohol intake (Figure 7f) (Two-Way ANOVA: main effect of injury F(1,56) = 1.735, p > 0.05; main effect of treatment F(1,56) = 0.197, p > 0.05; interaction injury by treatment F(1,56) = 4.307, p = 0.042). VNS blocked the blast-induced increase in the average 8pm acute alcohol intake (Figure 7g) (Two-Way ANOVA: main effect of injury F(1,56) = 1.270, p > 0.05; main effect of treatment F(1,56) = 0.005, p > 0.05; interaction injury by treatment F(1,56) = 4.306, p = 0.043). VNS blocked the blast-induced increase in the average daily preference for alcohol over water (Figure 7h) (Two-Way ANOVA: main effect of injury F(1,56) = 1.706, p = 0.197; main effect of treatment F(1,56) = 0.001, p = 0.975; injury by treatment F(1,56) = 4.143, p = 0.047). VNS blocked the blast-induced increase in the average acute 8pm preference for alcohol over water (Figure 7i) (Two-Way ANOVA: main effect of injury F(1,55) = 4.097, p = 0.048; main effect of treatment F(1,55) = 0.158, p = 0.692; injury by treatment F(1,55) = 2.108, p = 0.152).

Microbial species (24h) were found to mediate injury and treatment effects on later behaviors, including anxiety-like behaviors in the open field (5 days), risky decision-making (1 month), and ethanol preference (2-3 months) (Figure 7j-k ; Supplementary Table 7-9). Blast-induced increase in the delay to enter the center of the open field was mediated by an unclassified Firmicutes species, *SGB41718* (Figure 7j). This species was depleted by blast injury, particularly in Blast-VNS animals (Supplementary Table 6; Supplementary Figure 9a-e), and differences in *SGB41718* abundance within treatment groups were negatively associated with delay to first center entry (Supplementary Table 7; Supplementary Figure 9f-j), resulting in a positive ACME specifically under VNS(+) conditions (Figure 6j; Supplementary Table 9). As noted previously, the effect of VNS on total distance traveled in the open field was not significant; however, this relationship was significantly mediated by a different unclassified species within Firmicutes, *SGB41564* (Figure 7k). *SGB41564* was depleted by VNS, and differences in *SGB41564* abundance within treatment groups were negatively associated with total distance traveled, resulting in a positive ACME on distance traveled regardless of blast condition.

Blast-induced increase in ethanol preference was mediated by two species in opposite directions (Figure 7j). Both *Clostridiales bacterium* and *unclassified bacterium 1XD42-54* were enriched by blast injury; however, *Clostridiales bacterium* relative abundance was associated with increased ethanol preference within treatment groups, whereas *unclassified bacterium 1XD42-54* relative abundance was associated with decreased within-group preference (Supplementary Table 7). For both of these mediation hits, ACME confidence intervals were the same direction regardless of VNS status. Finally, blast-induced increase in risky decision-making was mediated by an unclassified Oscillospiraceae species, *SGB4185* (Figure 7j). *SGB4185* was enriched by blast; but within treatment groups, higher abundance of *SGB4185* was associated with decreased risky decision-making, resulting in a negative ACME independent of the VNS effect.

## Discussion

Repetitive blast exposure produces complex, multi-system pathology involving acute peripheral- and neuroinflammation, autonomic dysregulation, and downstream behavioral dysfunction, yet most therapeutic strategies have focused primarily on central mechanisms. Here, we demonstrate that early transcutaneous vagus nerve stimulation (VNS) selectively modulates components of the acute inflammatory response and improves a subset of chronic behavioral outcomes (risky decision-making and alcohol drinking behaviors). Notably, microbiome features were associated with both inflammatory and behavioral variation beyond what was explained by blast and VNS alone, supporting a model in which gut–immune–brain interactions contribute to heterogeneity in post-blast outcomes. Thus, the gut microbiota may represent a complementary axis to VNS for therapeutic intervention.

### Inflammation/Microbiome

Consistent with prior work, repetitive blast mTBI elicited a robust acute inflammatory response in both serum and brain, characterized by increases in cytokines such as IL-6, IP-10, MCP-1, and G-CSF. Principal component analysis largely captured this response in PC1, aligning with established early recruitment of innate immune pathways following brain injury^80,81^. VNS reduced components of this response, including serum levels of IL-6, IL-9, IP-10, and MCP-1, trending towards partial attenuation of PC1, which is consistent with the known anti-inflammatory effects of vagal stimulation mediated through the cholinergic anti-inflammatory reflex^54,82–85^. Crucially, both peripheral and CNS inflammation are major components of secondary injury cascades after blast/TBI^17,75,86–96^, interacting with blood brain barrier disruption, oxidative stress, and neurovascular dysfunction^97–99^. Therefore, it is possible that the decreasing peripheral inflammation acutely following blast injury may shape the trajectory of recovery at longer timescales.

Beyond this canonical response, PCA identified additional cytokine axes (PC2, PC3, and PC5) that were not strongly shifted in magnitude by blast or VNS, but were consistently associated with variation in gut microbiome composition. These axes appeared to capture regulatory, resolution-related, and coordination aspects of the immune response rather than overall inflammatory load and may be related to the repetitive nature of our blast exposure paradigm. The association of these axes with microbiome variation, and the identification of microbial mediators for blast–PC3 and VNS–PC5 effects, suggests that microbiome-related factors may prime or shape the trajectory of immune response to injury.

### Behavior

VNS stimulation did not prevent acute weight loss following each blast exposure, nor affect loss of righting reflex latencies. Similarly, VNS did not alleviate decreased locomotor movement and increased anxiety-related behaviors in blast-exposed animals during the 72 hours post-blast acute open field assessment. Of note, several rodent studies using a variety of VNS parameters have also failed to observe any significant effects of VNS or vagotomy on locomotor and anxiety-related behavior, both in stress-naive^100,101^ and stressed rodents^102–105^.

At a more chronic timepoint, VNS did not lessen hyperreactivity in blast-exposed mice as measured by the acoustic startle paradigm. As with locomotor and anxiety-related behavior, VNS’ effect on startle hyperreactivity in rodents are mixed. Using a rat model of PTSD, repeated VNS during fear extinction sessions decreased startle amplitude relative to sham stimulated controls^104^. In a follow up study utilizing a more severe rat PTSD model, VNS failed to reduce startle amplitude relative to controls^102^. Thus, differing trauma types and severity may influence VNS effects on hyperreactivity and arousal, in addition to variation in parameters surrounding the VNS procedure itself (i.e., implanted versus transcutaneous stimulation, timing of intervention, duration and intensity of stimulation, etc.).

In contrast to the acoustic startle results, VNS in blast-exposed mice diminished increased risky decision-making in the probabilistic discounting task. This result aligns with non-pathological rodent and human studies that have examined the effects of VNS on risky decision-making^106,107^. Likewise, these data support the idea that VNS both affects and is potentially beneficial for various forms of cognition and behavior^101,108–110^.

Blast exposure significantly increased ethanol preference using the intermittent two-bottle choice paradigm, consistent with a previous report from our lab^64^. Blast exposed animals treated with VNS, however, showed attenuated ethanol preference and intake which were not significantly different from sham controls. Damage or removal of the vagus nerve in rodents can alter ethanol intake^111–113^. Selectively ablating the gastric branch of the vagus nerve specifically increases ethanol preference using the intermittent two-bottle choice paradigm, which is likely mediated, in part, by the altered vagus nerve regulation of the hypothalamic-pituitary-adrenal (HPA) axis^111,114^. Importantly, blast mTBI can disrupt HPA axis functioning in rats^115^, providing a potential mechanism for the increased ethanol intake and preference seen in the present study.

Microbial features measured 24 hours post-blast were also associated with variability in behavioral outcomes assessed weeks later, and mediation analysis identified specific taxa linking blast and VNS to these behaviors. Notably, blast-induced increases in two different taxa exerted opposing effects on ethanol preference, and an Oscillospiraceae species appeared to buffer against blast-induced increases in risky decision-making. Considering that we also demonstrated microbes were related to components of the inflammatory response, it is possible that these microbe–behavior associations are not causal, and rather reflective of inflammation-mediated behavioral differences. Nonetheless, a direct microbiome contribution to reward-related behavior is biologically plausible. There is compelling evidence from preclinical models (including fecal transplant from human patients) that the gut microbiota contribute to alcohol preference and drinking patterns, as well as response to pharmacotherapy^116^, via multiple mechanisms including modulating serum prostaglandin and striatal dopamine, as well as affecting vagal signaling via extracellular vesicle production^117–121^. Risk-taking behavior, like ethanol preference, is also heavily influenced by the brain’s reward system and microbes have been shown to modulate the brain’s reward system^117,122–126^.

### Limitations

The current study has several limitations including the use of male mice only. A practical limitation of the VNS itself is its application during anesthesia, which may be especially important when pairing VNS with motor tasks or learning paradigms^127–129^. The microbiome response exhibited substantial inter-individual variability and batch effects, which may have limited power to detect more subtle VNS-associated changes. The use of different sampling approaches (colon at 4 hours and feces at 24 hours) introduces spatial and temporal variability^130,131^ that complicates direct comparison of taxa across time points. In addition, microbiome and cytokine measurements were collected concurrently, precluding inference of causal direction, and inflammatory measures were not obtained in the behavioral cohort. Future studies incorporating sex differences, longitudinal sampling, baseline controls, and additional interventional approaches (e.g., targeted recolonization or metabolite supplementation) will be necessary to further resolve underlying mechanisms and establish causality^132^.

## Conclusion

Collectively, these findings support a model in which repetitive blast exposure induces acute neuroimmune activation and rapid microbiome perturbation, while early VNS selectively modulates components of the inflammatory response and improves specific chronic behavioral outcomes. Importantly, microbiome variation explained heterogeneity in both inflammatory and behavioral responses beyond what was accounted for by injury or treatment alone, suggesting that gut microbial composition may constrain or shape recovery trajectories. These results highlight the gut–immune–brain axis as a critical and potentially modifiable contributor to post-blast pathology and support the development of combined therapeutic strategies targeting both neural and microbial pathways to improve outcomes following mTBI.

## Abbreviations

ACME: average causal mediated effect
ADE: average direct effect
ANOVA: analysis of variance
CMA: causal mediation analysis
CNS: central nervous system
DAB: diaminobenzidine
FDR: false discovery rate
FR1: fixed ratio 1
GFAP: glial fibrillary acidic protein
GI: gastrointestinal
I2BC: intermittent 2-bottle choice
Iba1: ionized calcium binding adaptor molecule
IEDs: improvised explosive devices
ITI: intertrial interval
LORR: loss of righting reflex
LRT: likelihood ratio test
mTBI: mild traumatic brain injury
NTS: nucleus tractus solitarii
PC: principal component
PCA: principal component analysis
PCoA: principal coordinate analysis
PCS: post-concussive symptoms
PTSD: posttraumatic stress disorder
RM: repeated measures
TBI: traumatic brain injury
VN: vagus nerve
VNS: vagus nerve stimulation

## Disclaimer

The views expressed in this scientific presentation are those of the author(s) and do not reflect the official policy or position of the U.S. government or Department of Veteran Affairs.

## Declarations

All animal experiments were conducted in accordance with Association for Assessment and Accreditation of Laboratory Animal Care guidelines and were approved by the VA Puget Sound Institutional Animal Care and Use Committee.

The generative AI platforms Edison and ChatGPT were used to aid in microbiome-related literature search and computational pipeline construction, respectively. The authors have reviewed and edited this content and take full responsibility for the contents of the published article.

## Availability of data and materials

Raw sequencing data are available under NCBI Bioproject Accession number <u>PRJNA1490884</u>. Corresponding sample metadata are available from the corresponding author upon reasonable request.

## Authors’ contributions

The work presented here was carried out in collaboration among all authors. BMB, JM, DGC, and AGS contributed to the conception and design of the study. BMB, AE, MT, SJL, EK, KW, BS, TWH, and AGS collected and analyzed data. BMB, AE, MT, BS, SMG, and AGS wrote the first draft of the manuscript. All authors contributed to manuscript revision, read, and approved the final manuscript.

## Acknowledgements

We would like to thank Scott Ng Evans, Traci J. Weber, Cindy Pekow, DVM, Kari Koszdin, DVM, and Lena Strait-Bodey for technical assistance and veterinary care.

---

**Supplementary Figure 1: Blast-associated differences in the 4h colonic microbiome are not affected by VNS**

A-B : Species-level alpha diversity metrics capturing community richness and dominance are not significantly affected by blast or VNS. C-O: Species altered by blast injury. P-R: Species altered by both blast and VNS. Two-way ANOVA *post hoc* Bonferroni Multiple Comparison Test (a-r) *p ≤ 0.05, **p ≤ 0.01, ***p ≤ 0.001, ****p ≤ 0.0001.

**Supplementary Figure 2: VNS ameliorates some blast-induced changes to serum cytokine levels**

Two-way ANOVA *post hoc* Bonferroni Multiple Comparison Test (a-k). *p ≤ 0.05, **p ≤ 0.01, ***p ≤ 0.001, ****p ≤ 0.0001.

**Supplementary Figure 3: VNS ameliorates some blast-induced changes to brain cytokine levels**

Two-way ANOVA *post hoc* Bonferroni Multiple Comparison Test (a-s). *p ≤ 0.05, **p ≤ 0.01, ***p ≤ 0.001, ****p ≤ 0.0001.

**Supplementary Figure 4: Principal Component Analysis identifies axes of variation in cytokine profiles not significantly explained by blast or VNS**

Two-way ANOVA *post hoc* Bonferroni Multiple Comparison Test (a-e); Batch-adjusted linear regression (f-j). *p ≤ 0.05, **p ≤ 0.01, ***p ≤ 0.001, ****p ≤ 0.0001.

**Supplementary Figure 5: Gut microbial species and alpha diversity metrics mediate Blast effects on Cytokine PC3**

A-E: Blast affects relative abundance of PC3-associated species and species-level alpha diversity. F-J: Within-group variation in bacterial features explain within-group differences in cytokine PC3 levels, across all groups. Two-way ANOVA *post hoc* Bonferroni Multiple Comparison Test (a-e); Batch-adjusted linear regression (f-j). *p ≤ 0.05, **p ≤ 0.01, ***p ≤ 0.001, ****p ≤ 0.0001.

**Supplementary Figure 6: Gut microbial species mediate VNS effects on Cytokine PC5**

A-E: VNS affects relative abundance of behavior-associated species. F-J: Variation in bacterial relative abundance explains within-experimental-group differences in behavioral assays, across all groups. Two-way ANOVA *post hoc* Bonferroni Multiple Comparison Test (a-e); Batch-adjusted linear mixed effects regression (f-j). *p ≤ 0.05, **p ≤ 0.01, ***p ≤ 0.001, ****p ≤ 0.0001.

**Supplementary Figure 7: Gut microbial species are associated with Cytokine PC2 independent of blast and VNS**

A-C : Variation in bacterial relative abundance explains within-treatment-group differences in Cytokine PC2, across all groups. Batch-adjusted linear mixed effects regression (a-c).

**Supplementary Figure 8: Blast-associated differences in the 24h fecal pellet microbiome are not affected by VNS**

A-B: Species-level alpha diversity metrics capturing community richness and dominance are not significantly affected by blast or VNS. C-E: Species altered by blast injury. Two-way ANOVA *post hoc* Bonferroni Multiple Comparison Test (a-e) *p ≤ 0.05, **p ≤ 0.01, ***p ≤ 0.001, ****p ≤ 0.0001.

**Supplementary Figure 9:**

A: VNS affects relative abundance of behavior-associated species. B-E: Blast affects relative abundance of behavior-associated species. F-J: Variation in bacterial relative abundance explains within-experimental-group differences in behavioral assays, across all groups. Two-way ANOVA *post hoc* Bonferroni Multiple Comparison Test (a-e); Batch-adjusted linear mixed effects regression (f-j). *p ≤ 0.05, **p ≤ 0.01, ***p ≤ 0.001, ****p ≤ 0.0001.

