## Supplementary figures and images for "Non-invasive Vagal Nerve Stimulation as a Potential Treatment for Repetitive Blast Trauma"

### sf1

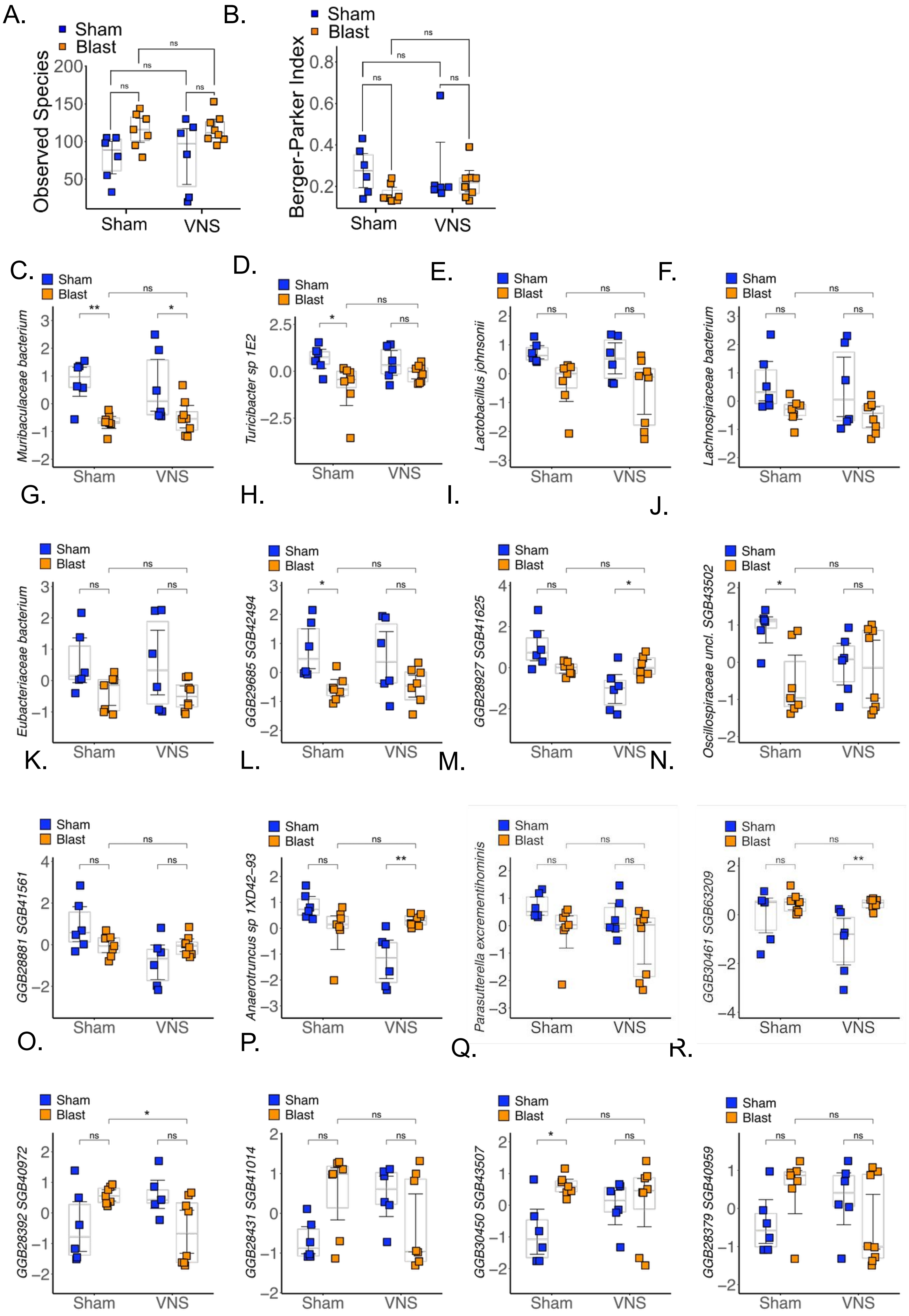

### sf2

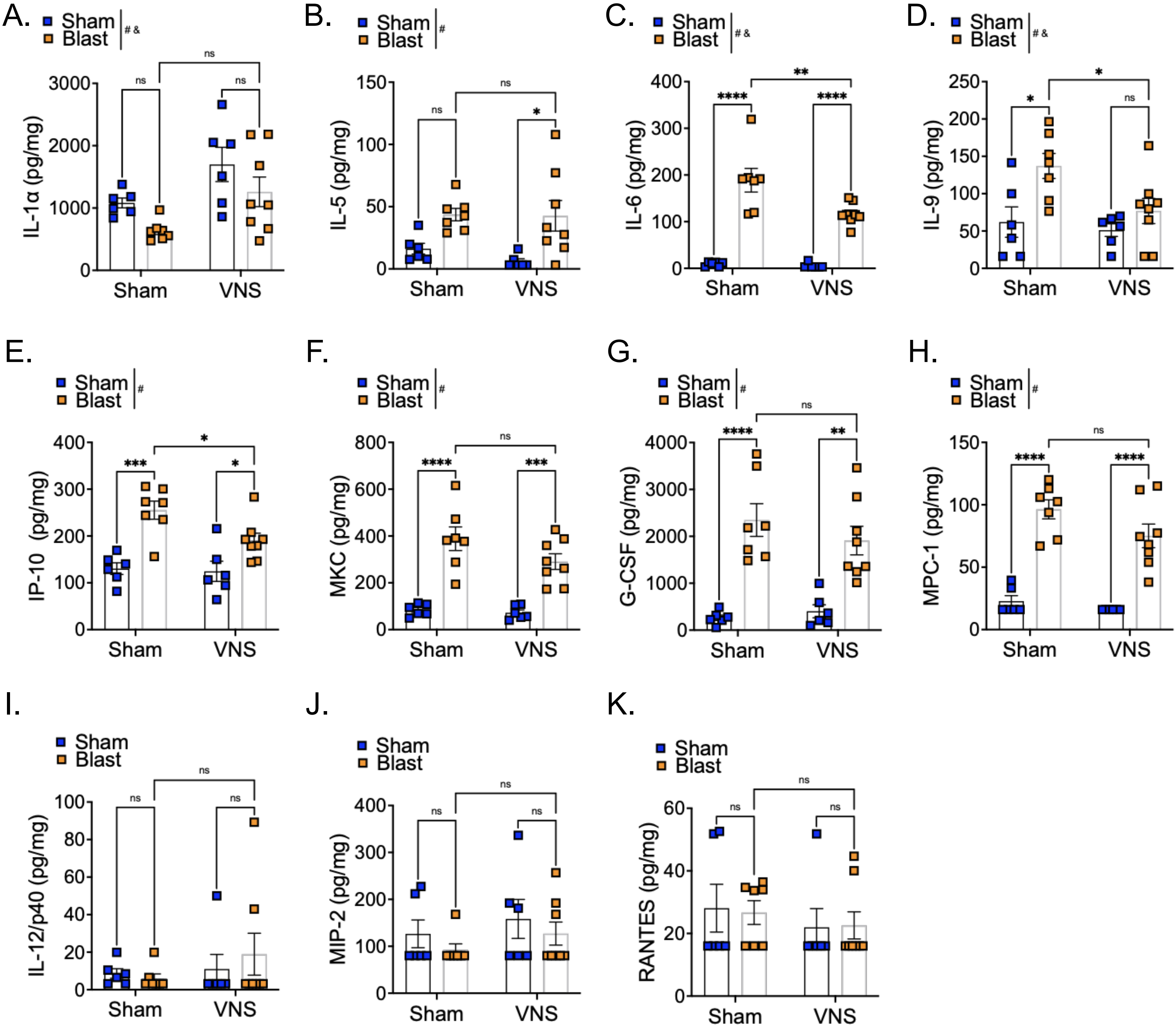

### sf3

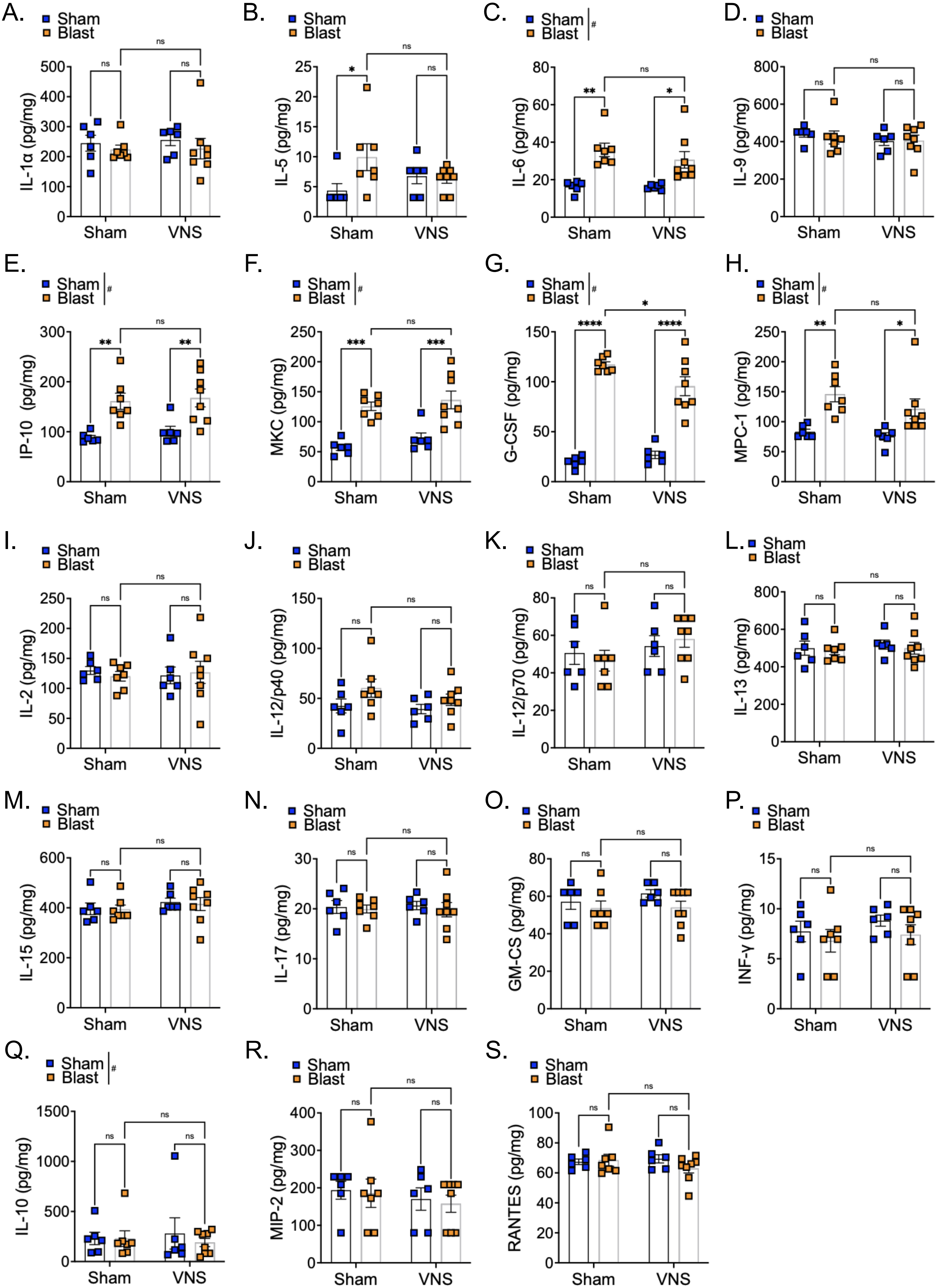

### sf4

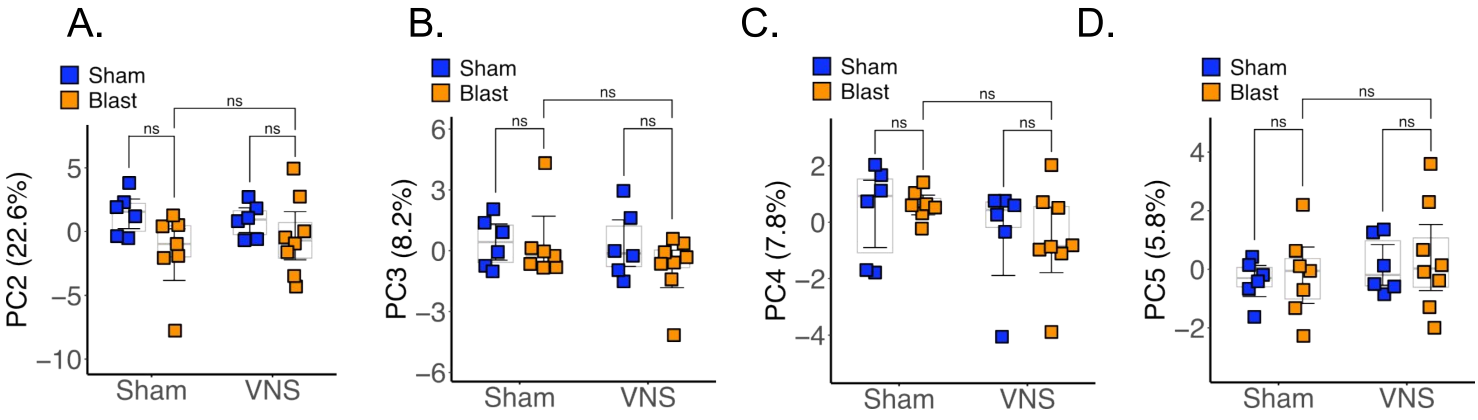

### sf5

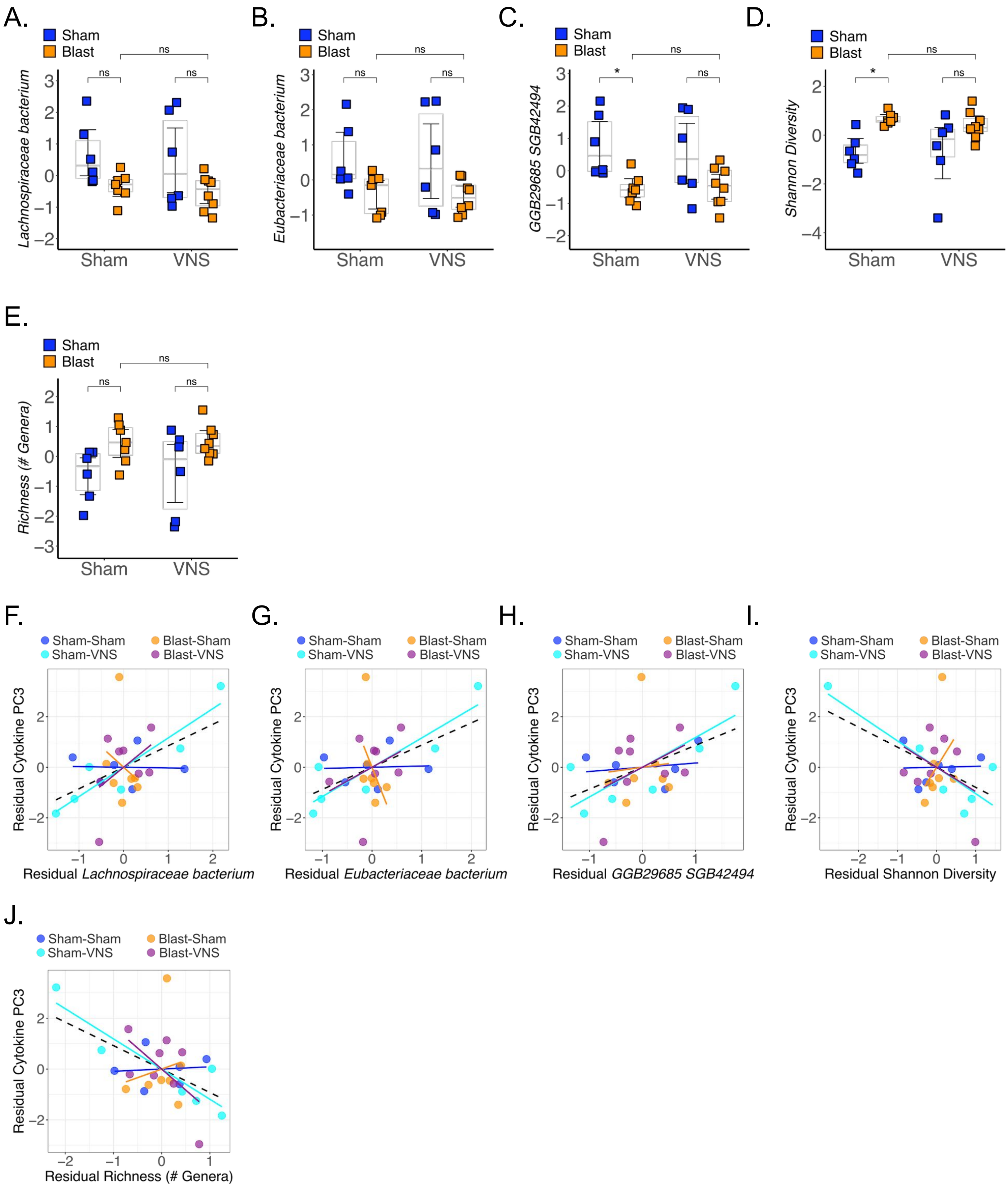

### sf6

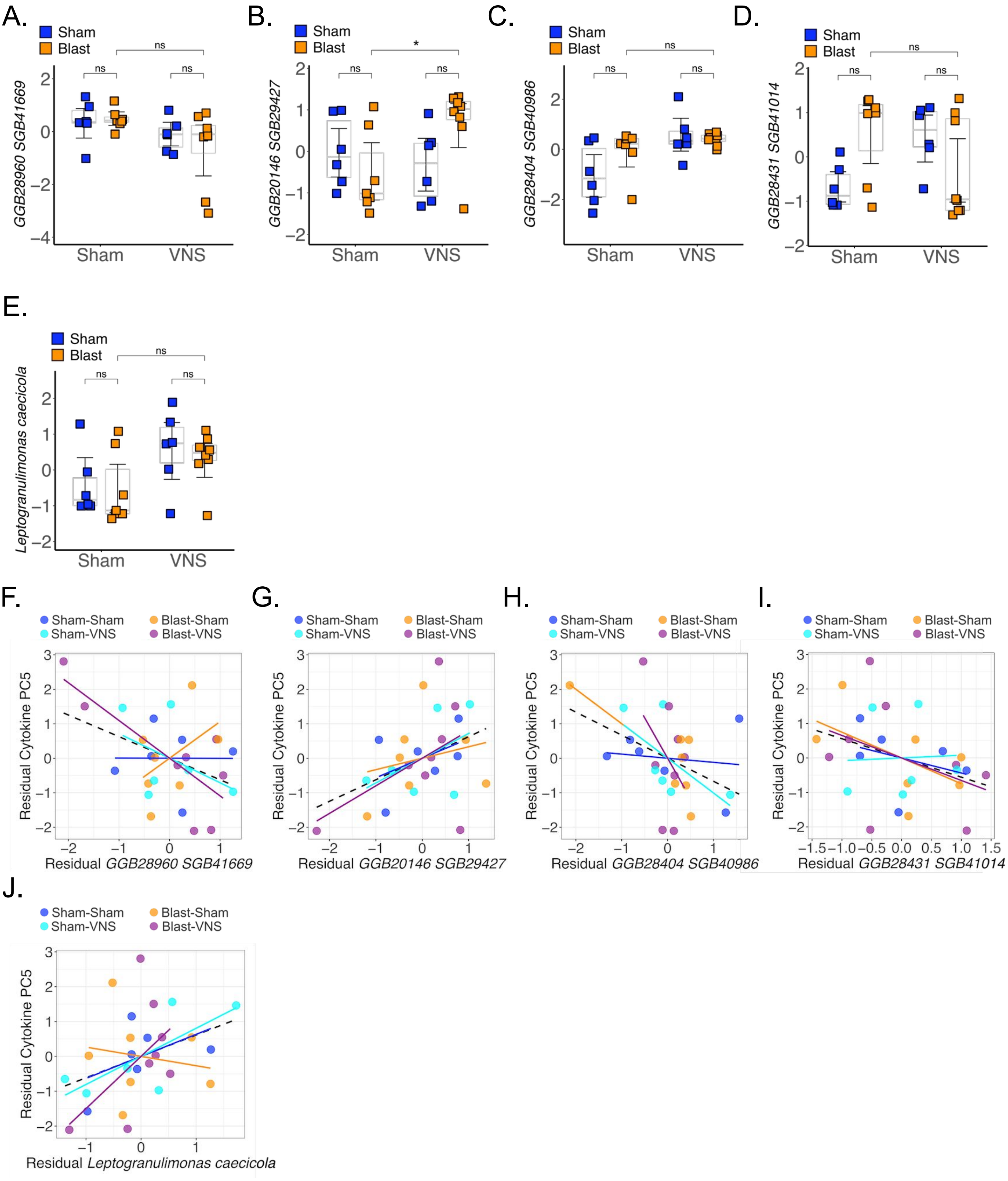

### sf7

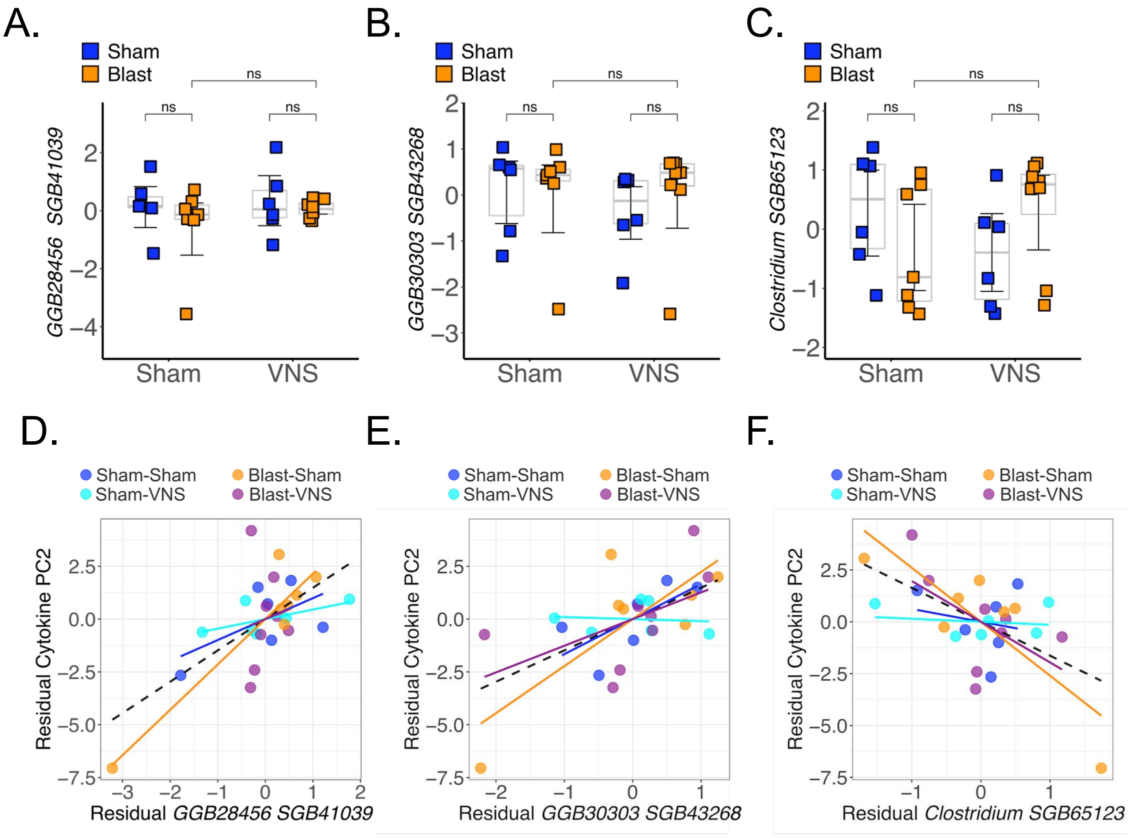

### sf8

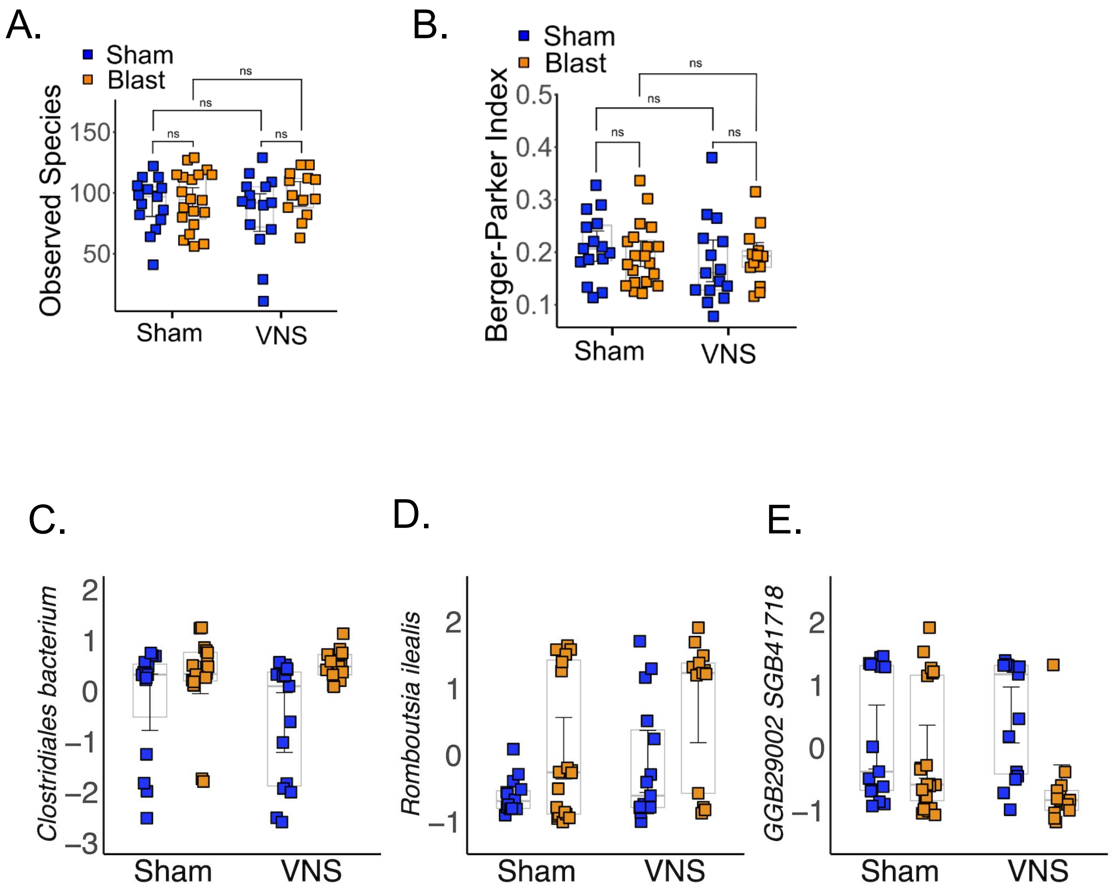

### sf9

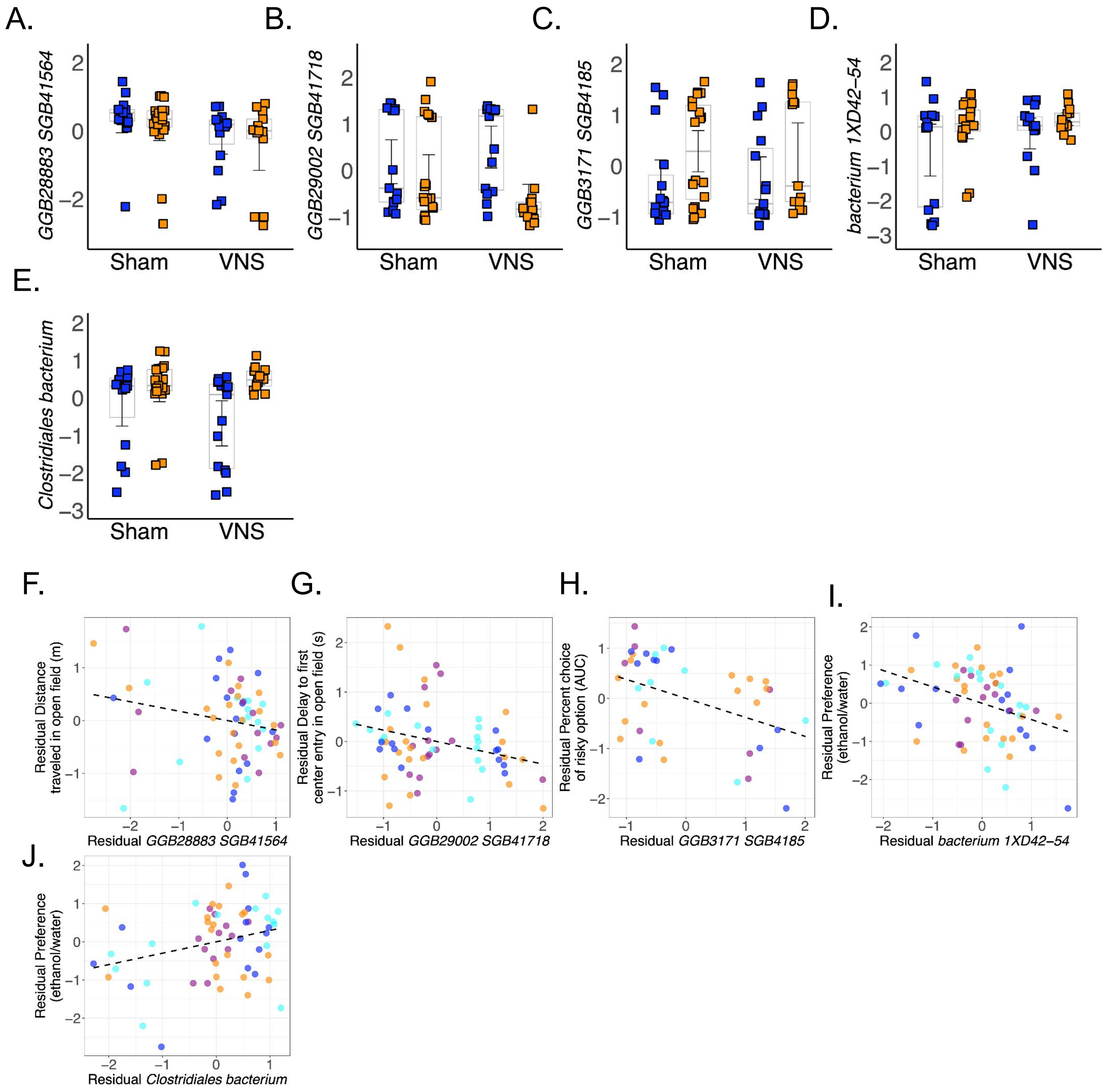
