## Supplementary material for "Non-invasive Vagal Nerve Stimulation as a Potential Treatment for Repetitive Blast Trauma": st1

| Species-level feature | n | LRT_F_blast | LRT_F_vns | LRT_F_interact | LRT_p_blast |
|---|---|---|---|---|---|
| Acetatifactor_SGB41545 | 27 | -1.737559389 | -0.829130954 | 1.647870903 | 0.2103509000 |
| Acutalibacter_muris | 27 | -0.690728962 | -1.081850986 | 0.4367700067 | 0.7869843388 |
| Akkermansia_muciniphila | 27 | -0.746179775 | 0.879050101 | -0.7353449499 | 0.16660254 |
| Anaerotruncus_sp_1XD42_93 | 27 | -2.167500944 | -4.932661639 | 4.238351262 | 0.0009175375 |
| Bacteria_unclassified_SGB41677 | 27 | 0.322609138 | -0.892429325 | 0.2981631831 | 0.714189493 |
| Bacteria_unclassified_SGB43539 | 27 | 2.016693907 | 0.676700601 | -1.237642783 | 0.148902499 |
| Bacteria_unclassified_SGB43546 | 27 | 2.178999982 | 1.536693862 | -1.362268113 | 0.112551908 |
| Bacteria_unclassified_SGB63211 | 27 | -0.562402231 | -0.811577519 | 1.451468956 | 0.295003775 |
| Bacteroides_thetaiotaomicron | 27 | -2.688563861 | -1.611515975 | 1.444773548 | 0.036627754 |
| Clostridia_bacterium | 27 | -1.687109712 | -1.415843682 | 0.9668606116 | 0.2483530870 |
| Clostridiaceae_bacterium | 27 | -1.976156483 | -1.648772466 | 1.265632115 | 0.162346605 |
| Clostridiaceae_unclassified_SGB41663 | 27 | -1.327396058 | -1.997977181 | 0.5329468379 | 0.362409871 |
| Clostridiales_bacterium | 27 | -2.380329549 | -1.055035497 | 2.032232708 | 0.073824740 |
| Clostridium_SGB65123 | 27 | -1.278513916 | -1.546775173 | 1.919649228 | 0.180438982 |
| Clostridium_cocleatum | 27 | 0.136391658 | 1.22710568 | -0.9159713011 | 0.507275085 |
| Eubacteriaceae_bacterium | 27 | -2.349742056 | -0.094316773 | 0.2257123385 | 0.0166040620 |
| Eubacteriaceae_unclassified_SGB94922 | 27 | -1.944640614 | -1.31025256 | 1.450688493 | 0.174224276 |
| GGB20146_SGB29427 | 27 | -1.009664056 | -0.616823623 | 2.1615213 | 0.093844467 |
| GGB20149_SGB29430 | 27 | -1.195873328 | -0.016156629 | 1.304554587 | 0.413225925 |
| GGB22635_SGB63107 | 27 | -0.775565824 | 1.132698371 | -0.8669862322 | 0.114660903 |
| GGB25041_SGB36960 | 27 | -1.9425547 | -1.822629432 | 1.169068899 | 0.167595117 |
| GGB28379_SGB40959 | 27 | 2.46669236 | 1.877464866 | -2.814230903 | 0.029241836 |
| GGB28382_SGB40962 | 27 | -0.447707979 | 1.346358344 | -1.343939082 | 0.072513022 |
| GGB28392_SGB40972 | 27 | 2.025461785 | 2.014129263 | -3.120352237 | 0.0171836690 |
| GGB28404_SGB40986 | 27 | 1.99959786 | 3.16053752 | -1.54135182 | 0.157767928 |
| GGB28418_SGB41001 | 27 | -0.041585992 | 0.209654194 | -0.02293525044 | 0.996265407 |
| GGB28422_SGB41005 | 27 | 1.575141745 | 1.346906441 | -0.9369333963 | 0.298094755 |
| GGB28431_SGB41014 | 27 | 2.612858458 | 2.312986843 | -3.145896704 | 0.015377584 |
| GGB28456_SGB41039 | 27 | -1.133730025 | 0.175605552 | 0.5841506241 | 0.508621549 |
| GGB28782_SGB41435 | 27 | -0.529075003 | -1.325738094 | 1.401029387 | 0.315491599 |
| GGB28792_SGB41445 | 27 | 1.646981447 | 0.106940840 | -0.2260428306 | 0.123770726 |
| GGB28798_SGB41451 | 27 | -0.208945716 | -0.786117249 | -0.06782061124 | 0.931837305 |
| GGB28810_SGB41465 | 27 | 0.4099870410 | -0.002870786 | -0.2812842323 | 0.919521792 |
| GGB28828_SGB41484 | 27 | 2.18871256 | 1.506310347 | -2.764608998 | 0.036453176 |
| GGB28851_SGB41518 | 27 | 0.045952976 | 0.676498427 | 0.09092996276 | 0.983245267 |
| GGB28865_SGB41537 | 27 | 2.467505665 | 2.617363758 | -2.355585473 | 0.051838259 |
| GGB28868_SGB41542 | 27 | 1.423025439 | 0.668299030 | -1.550411877 | 0.292423183 |
| GGB28869_SGB41543 | 27 | -0.196869533 | 0.055224657 | -0.481765068 | 0.6620730770 |
| GGB28875_SGB41555 | 27 | -0.415349022 | -0.564469250 | -0.1068942592 | 0.776644062 |
| GGB28881_SGB41561 | 27 | -2.502061412 | -4.068063868 | 3.298161044 | 0.011894548 |
| GGB28892_SGB41573 | 27 | 0.634742761 | 0.532226659 | -0.08888102133 | 0.715153028 |
| GGB28893_SGB41574 | 27 | -0.690526180 | 0.452694152 | 0.7523813845 | 0.739221992 |
| GGB28898_SGB41580 | 27 | -0.390079089 | -0.993294137 | 0.9355235793 | 0.601924700 |
| GGB28901_SGB41592 | 27 | -2.37580037 | 0.044824382 | 1.32598069 | 0.072063586 |
| GGB28904_SGB41597 | 27 | -0.621785954 | -0.048282861 | 1.611222842 | 0.225752520 |
| GGB28909_SGB41602 | 27 | 0.149052360 | -0.022905988 | -1.360218857 | 0.218283007 |

| | | | | | |
|---|---|---|---|---|---|
| GGB28924_SGB41621 | 27 | 1.001568899 | -0.130095099 | -0.7546637238 | 0.6115781842 |
| GGB28927_SGB41625 | 27 | -2.763553985 | -4.998927254 | 4.089700878 | 0.0019544839 |
| GGB28934_SGB41635 | 27 | -1.483764113 | -1.292771592 | 1.789347276 | 0.2164988531 |
| GGB28949_SGB41655 | 27 | -1.761459223 | -0.770874578 | 0.6077940833 | 0.1595983662 |
| GGB28951_SGB102295 | 27 | -0.817615234 | -1.417425515 | 0.5554203197 | 0.718725508 |
| GGB28951_SGB41658 | 27 | 0.7149686649 | 2.227075952 | -0.3365208725 | 0.7526278431 |
| GGB28954_SGB41662 | 27 | 0.0862181249 | -0.016090960 | 0.4082467004 | 0.7949390323 |
| GGB28960_SGB41669 | 27 | 0.2625165665 | -0.982669415 | -0.954648455 | 0.5366388614 |
| GGB28964_SGB94886 | 27 | -1.57805804 | -1.223257853 | 0.4750808505 | 0.2072130289 |
| GGB28967_SGB41678 | 27 | -0.510056726 | -0.614867762 | 0.5524666292 | 0.8482050661 |
| GGB28996_SGB41712 | 27 | 0.0989955308 | 0.1422517402 | 0.8483129055 | 0.4298847288 |
| GGB29531_SGB42317 | 27 | 1.139878452 | -0.0339946558 | 0.005846641031 | 0.2795934449 |
| GGB29685_SGB42494 | 27 | -3.086960459 | -0.606392337 | 0.7640020298 | 0.0045968812 |
| GGB30141_SGB43066 | 27 | 0.9827381462 | 0.8715180491 | -0.8294030133 | 0.6140281408 |
| GGB30145_SGB43072 | 27 | -0.031208747 | -1.442903132 | 1.140682656 | 0.2949240353 |
| GGB30286_SGB43248 | 27 | -0.230788931 | -0.213833240 | 0.650091804 | 0.7667575811 |
| GGB30300_SGB43264 | 27 | -1.141611788 | -1.493527393 | 1.806092231 | 0.2138469484 |
| GGB30303_SGB43268 | 27 | 0.0612920269 | -0.863776907 | 0.427200127 | 0.7955727513 |
| GGB30450_SGB43507 | 27 | 2.903587772 | 1.664169607 | -1.855239558 | 0.0271094004 |
| GGB30453_SGB43513 | 27 | 1.272461974 | 1.346563515 | -1.540777019 | 0.3157254511 |
| GGB30454_SGB43514 | 27 | -0.885604686 | 0.2665664068 | 0.3976172678 | 0.6428886468 |
| GGB30455_SGB43519 | 27 | 0.8320714881 | 2.025781929 | -0.8603659197 | 0.6637984269 |
| GGB30456_SGB43520 | 27 | 2.094670851 | 1.292716638 | -1.983567641 | 0.1111969489 |
| GGB30457_SGB63218 | 27 | 0.0419297937 | 0.0796109826 | -0.3783903328 | 0.8821640501 |
| GGB30461_SGB43530 | 27 | -0.545894485 | -1.240778268 | -0.206349993 | 0.6024683039 |
| GGB30461_SGB43533 | 27 | -0.649134976 | -0.787774598 | 0.433501048 | 0.8107674231 |
| GGB30461_SGB63209 | 27 | 1.186310369 | -2.388070782 | 1.525246869 | 0.0059141187 |
| GGB30463_SGB43537 | 27 | 1.975195137 | 0.0415191651 | -0.1114196225 | 0.0407681100 |
| GGB30473_SGB43557 | 27 | -0.715707951 | -1.892291501 | 1.45562487 | 0.3284915799 |
| GGB30475_SGB63182 | 27 | -0.054823336 | -0.023471047 | 0.4892695436 | 0.811833041 |
| GGB30861_SGB44083 | 27 | 0.5087882423 | 0.0604976815 | 0.08015704245 | 0.7194016475 |
| GGB31312_SGB44628 | 27 | 0.3705888042 | 0.4802730455 | -0.6041750471 | 0.8310391405 |
| GGB31438_SGB44768 | 27 | -0.975162992 | -0.968230617 | 1.221833074 | 0.4812139003 |
| GGB3171_SGB4185 | 27 | -2.393512711 | -2.507459011 | 2.172233609 | 0.0663422469 |
| GGB31762_SGB45125 | 27 | -1.021684388 | -1.100679979 | 1.293390539 | 0.4428890562 |
| GGB31823_SGB45199 | 27 | -0.415952333 | -1.125005285 | 0.9094533136 | 0.6301752034 |
| GGB31838_SGB45216 | 27 | 0.5715561169 | -0.067387115 | 0.05382542551 | 0.6844031648 |
| GGB31841_SGB65084 | 27 | -0.369087597 | -1.186547181 | 1.233063862 | 0.3717374543 |
| GGB32371_SGB41694 | 27 | -0.143697051 | 1.376444228 | -1.103854608 | 0.2420180555 |
| GGB3793_SGB5158 | 27 | -0.188176521 | -0.096696085 | 0.4542847466 | 0.884993491 |
| GGB42601_SGB59797 | 27 | 0.9883235619 | 0.684691121 | -0.04068618561 | 0.4018920133 |
| GGB45514_SGB63186 | 27 | 0.260326333 | 0.8498537703 | -0.2941763049 | 0.9553647655 |
| GGB45564_SGB63259 | 27 | -0.456513487 | -1.39485483 | 1.213998907 | 0.4152727846 |
| GGB45624_SGB63337 | 27 | 0.9587880791 | 1.387724149 | -0.8540069663 | 0.6203892728 |
| GGB45656_SGB63370 | 27 | 0.9827243777 | 1.978828355 | -1.37318629 | 0.4047054831 |
| GGB47127_SGB65054 | 27 | 1.187950687 | 0.6102248817 | -0.5307932959 | 0.4564936795 |

| | | | | | |
|---|---|---|---|---|---|
| GGB74395_SGB43523 | 27 | 0.666175050 | 0.601580936 | -0.4573035892 | 0.802410438 |
| GGB75053_SGB43494 | 27 | 0.880361518 | 0.099461301 | -1.831835919 | 0.177327401 |
| GGB75109_SGB102238 | 27 | 0.081862972 | -0.029145587 | -0.7684505044 | 0.599792986 |
| Lachnospiraceae_bacterium | 27 | -2.203964115 | -0.387255862 | 0.1852804211 | 0.023925903 |
| Lachnospiraceae_bacterium_A2 | 27 | 0.265356901 | 1.313553919 | 0.3415605907 | 0.724187807 |
| Lachnospiraceae_unclassified_SGB41414 | 27 | -0.550628961 | -0.071448164 | 0.8166452243 | 0.719079500 |
| Lachnospiraceae_unclassified_SGB41424 | 27 | 1.239724625 | -2.049048928 | 0.02832888319 | 0.216930413 |
| Lachnospiraceae_unclassified_SGB94868 | 27 | -1.157161566 | -0.374208568 | -0.1632694205 | 0.207633504 |
| Lactobacillus_johnsonii | 27 | -2.904233498 | -0.545006746 | 0.221966001 | 0.002748365 |
| Leptogranulimonas_caecicola | 27 | -0.424775229 | 2.085478283 | 0.09712389841 | 0.874577980 |
| Muribaculaceae_bacterium | 27 | -3.20434493 | -0.393571486 | 0.4976867034 | 0.001850198 |
| Neglectibacter_sp_X4 | 27 | 1.3707582 | -0.049643228 | -0.8101334702 | 0.394423067 |
| Oscillibacter_SGB43496 | 27 | 0.993141102 | 0.728529128 | 0.3318527618 | 0.220544725 |
| Oscillospiraceae_bacterium | 27 | 1.201927585 | -0.248452492 | -0.9545140101 | 0.492783661 |
| Oscillospiraceae_unclassified_SGB43502 | 27 | -3.128709497 | -1.94852721 | 2.128793537 | 0.017271201 |
| Oscillospiraceae_unclassified_SGB43505 | 27 | -0.010797214 | 0.697591581 | -0.9901098303 | 0.374299908 |
| Oscillospiraceae_unclassified_SGB94989 | 27 | 0.210561514 | 0.654762590 | 0.006552710316 | 0.953368969 |
| Parasutterella_excrementihominis | 27 | -2.246368465 | -0.958588251 | 0.2381937364 | 0.023419378 |
| Schaedlerella_arabinosiphila | 27 | 2.167659768 | 0.713275921 | -1.344943283 | 0.114481983 |
| Turicibacter_sp_1E2 | 27 | -3.247061264 | -0.532201248 | 1.62846818 | 0.009594137 |
| bacterium_0_1xD8_82 | 27 | 0.730564224 | -0.014966699 | -0.4458468647 | 0.763436393 |
| bacterium_1XD42_1 | 27 | 0.660522350 | 0.006280943 | -1.535207489 | 0.272519397 |
| bacterium_1XD42_54 | 27 | -2.731523735 | -2.13093224 | 1.693887731 | 0.038105321 |
| bacterium_1XD42_76 | 27 | -1.502721407 | -0.920803995 | 1.92777027 | 0.177410768 |
| bacterium_1xD8_48 | 27 | 2.076486751 | 1.089861791 | -1.442230665 | 0.139347611 |
| Berger Parker Index | 27 | -2.400004014 | -0.534745917 | 1.115422088 | 0.057475283 |
| Richness (# observed features) | 27 | 2.441438933 | 0.134471858 | -0.2739486566 | 0.013777874 |
| Shannon Index | 27 | 2.839530142 | 0.195304147 | -0.5263857384 | 0.006473070 |

| LRT_p_vns | LRT_p_interact | LRT_q_blast | LRT_q_vns | LRT_q_interact | est_intercept | est_blast |
|---|---|---|---|---|---|---|
| 0.239098669 | 0.1135888756 | 0.499346548 | 0.717296007 | 0.5928157147 | 0.4930595825 | -0.96554993 |
| 0.491902226 | 0.6665350338 | 0.893761146 | 0.817383170 | 0.8978474649 | 0.2279987513 | -0.389763961 |
| 0.678899621 | 0.4698968088 | 0.481170621 | 0.895252247 | 0.7941917894 | -0.1103214233 | -0.399386127 |
| 0.000231687 | 0.000337089034 | 0.078179359 | 0.013901240 | 0.02907734573 | 0.8338932722 | -0.884230098 |
| 0.58777865 | 0.768376414 | 0.886760580 | 0.865929823 | 0.9440157185 | 0.727861165 | 0.137934724 |
| 0.449260123 | 0.2288953073 | 0.481170621 | 0.816836587 | 0.6066894195 | -0.547433953 | 1.117385022 |
| 0.309908808 | 0.1868987777 | 0.416948741 | 0.772603929 | 0.5928157147 | -1.005997881 | 1.143442891 |
| 0.340871629 | 0.1607610545 | 0.576957591 | 0.772603929 | 0.5928157147 | 0.5484779944 | -0.270044308 |
| 0.275405005 | 0.1626148411 | 0.217744694 | 0.738856224 | 0.5928157147 | 0.8184868884 | -1.377170165 |
| 0.380556079 | 0.3441231202 | 0.532185187 | 0.772603929 | 0.757210127 | 0.78033172 | -0.93595003 |
| 0.277071084 | 0.2188848206 | 0.481170621 | 0.738856224 | 0.6066894195 | 0.5655145171 | -1.04826086 |
| 0.070917324 | 0.5994132043 | 0.658927039 | 0.500592877 | 0.8978474649 | 0.6413621793 | -0.671498939 |
| 0.124851167 | 0.05437659762 | 0.316391746 | 0.601953280 | 0.5437080554 | 0.6809451363 | -1.250255975 |
| 0.179389568 | 0.06796350693 | 0.481170621 | 0.632845883 | 0.5437080554 | 0.8492375503 | -0.592547765 |
| 0.482732384 | 0.369613781 | 0.762932324 | 0.817383170 | 0.7585775013 | -0.5009146985 | 0.074631012 |
| 0.969157578 | 0.823508827 | 0.148038868 | 0.999007775 | 0.9535244924 | 0.1354286295 | -1.047054478 |
| 0.346181024 | 0.1609762619 | 0.481170621 | 0.772603929 | 0.5928157147 | 0.7705460849 | -1.074513474 |
| 0.051218919 | 0.04180462925 | 0.388321935 | 0.409751358 | 0.4877585812 | 0.2046435515 | -0.505435513 |
| 0.180434838 | 0.2055274741 | 0.682640193 | 0.632845883 | 0.5979782654 | 0.3051981816 | -0.637854067 |
| 0.535639339 | 0.3953075818 | 0.416948741 | 0.826524236 | 0.7721894584 | 0.1473966249 | -0.421797406 |
| 0.206256260 | 0.2548873782 | 0.481170621 | 0.668939224 | 0.6372184455 | 0.1468927416 | -0.845809113 |
| 0.032794656 | 0.01010347507 | 0.194945578 | 0.357759888 | 0.1938341562 | -1.083284581 | 0.867154966 |
| 0.370430736 | 0.1926651073 | 0.316391746 | 0.772603929 | 0.5928157147 | 0.2289327297 | -0.236246045 |
| 0.016592400 | 0.004982143122 | 0.148038868 | 0.288021378 | 0.1195714349 | -0.6043400677 | 0.989085319 |
| 0.010090214 | 0.1374931891 | 0.481170621 | 0.288021378 | 0.5928157147 | -1.21957616 | 0.941802831 |
| 0.959497945 | 0.9819086792 | 0.996265407 | 0.999007775 | 0.9953877755 | -0.0255073505 | -0.025121584 |
| 0.416676405 | 0.3589651594 | 0.576957591 | 0.793669342 | 0.757210127 | -0.4251724064 | 0.855031643 |
| 0.016801247 | 0.004692592965 | 0.148038868 | 0.288021378 | 0.1195714349 | -0.3931090988 | 1.268171468 |
| 0.567597000 | 0.5650613534 | 0.762932324 | 0.851395500 | 0.8978474649 | 0.04050973727 | -0.651125121 |
| 0.359169919 | 0.1751539938 | 0.591985220 | 0.772603929 | 0.5928157147 | 0.5002608365 | -0.288509786 |
| 0.970858107 | 0.8232549644 | 0.436837858 | 0.999007775 | 0.9535244924 | -0.2445948503 | 0.879403610 |
| 0.471552545 | 0.946541058 | 0.963969626 | 0.817383170 | 0.9953877755 | 0.3090561082 | -0.12167290 |
| 0.915105920 | 0.7811210556 | 0.959501000 | 0.983027427 | 0.9440157185 | -0.7135214484 | 0.172930628 |
| 0.029615316 | 0.01130699244 | 0.217744694 | 0.355383798 | 0.1938341562 | -0.3710884395 | 1.127843444 |
| 0.548244701 | 0.9283710877 | 0.991507832 | 0.832776761 | 0.9948712152 | -0.3534930205 | 0.026967976 |
| 0.043832016 | 0.02782138454 | 0.270460482 | 0.404603226 | 0.417320768 | -0.4150402477 | 1.14205863 |
| 0.254323647 | 0.1353098773 | 0.576957591 | 0.726638993 | 0.5928157147 | -0.1682835008 | 0.804146646 |
| 0.806606148 | 0.6347306827 | 0.875338584 | 0.941377455 | 0.8978474649 | -0.3308795206 | -0.099385824 |
| 0.634070741 | 0.9158416652 | 0.893761146 | 0.874580333 | 0.9948712152 | 0.857760516 | -0.195555641 |
| 0.001988037 | 0.003276334099 | 0.148038868 | 0.079521481 | 0.1195714349 | 0.4963207629 | -1.030353634 |
| 0.784709483 | 0.9299806264 | 0.886760580 | 0.941377455 | 0.9948712152 | -0.6674065486 | 0.352922207 |
| 0.264245168 | 0.4597968228 | 0.893761146 | 0.737428376 | 0.7882231248 | 0.2975744393 | -0.351514322 |
| 0.591718713 | 0.3596748103 | 0.855709341 | 0.865929823 | 0.757210127 | 0.6645917313 | -0.196403008 |
| 0.152665868 | 0.1984495128 | 0.316391746 | 0.631720835 | 0.5953485384 | 0.4827366956 | -1.21629365 |
| 0.085658495 | 0.1213865217 | 0.501672267 | 0.513950974 | 0.5928157147 | 0.3631414204 | -0.271412487 |
| 0.145902665 | 0.1875366336 | 0.499346548 | 0.625297136 | 0.5928157147 | 0.4234922075 | 0.079964967 |

| | | | | | | |
|---|---|---|---|---|---|---|
| 0.456981189 | 0.4584536795 | 0.855709341 | 0.817383170 | 0.7882231248 | 0.5324294399 | 0.455449925 |
| 0.000219425 | 0.000484622428 | 0.078179359 | 0.013901240 | 0.02907734573 | 0.7784783317 | -1.108579812 |
| 0.224102539 | 0.08733647108 | 0.499346548 | 0.689546274 | 0.5515987647 | 0.3808768165 | -0.837820242 |
| 0.744810754 | 0.5495473988 | 0.481170621 | 0.941377455 | 0.8949850654 | 0.657170233 | -0.975862317 |
| 0.298970783 | 0.5842115419 | 0.886760580 | 0.772603929 | 0.8978474649 | 0.7041567913 | -0.460718028 |
| 0.024801176 | 0.7396669805 | 0.893761146 | 0.330682352 | 0.9289265075 | -1.104944222 | 0.331749997 |
| 0.841186745 | 0.6870387291 | 0.893761146 | 0.941377455 | 0.9059851372 | 0.226786524 | 0.047545168 |
| 0.043638773 | 0.3501283184 | 0.795020535 | 0.404603226 | 0.757210127 | 0.6970955679 | 0.132356616 |
| 0.400370147 | 0.6394119482 | 0.499346548 | 0.774909963 | 0.8978474649 | 0.6453720114 | -0.853863173 |
| 0.820661965 | 0.5861985137 | 0.916978449 | 0.941377455 | 0.8978474649 | 0.3497131573 | -0.303270173 |
| 0.377449394 | 0.4053994657 | 0.697110371 | 0.772603929 | 0.7721894584 | -0.01961784964 | 0.053462464 |
| 0.999007775 | 0.9953877755 | 0.576957591 | 0.999007775 | 0.9953877755 | -0.3405672731 | 0.651254204 |
| 0.748416768 | 0.4529827042 | 0.110325149 | 0.941377455 | 0.7882231248 | 0.4695786136 | -1.382159056 |
| 0.664026053 | 0.415785445 | 0.855709341 | 0.885368071 | 0.7755275948 | 0.1741621922 | 0.490346821 |
| 0.367951830 | 0.2662678912 | 0.576957591 | 0.772603929 | 0.6520846316 | 0.3928424426 | -0.017102403 |
| 0.750425732 | 0.5223641345 | 0.893761146 | 0.941377455 | 0.8706068908 | 0.1905115758 | -0.133483697 |
| 0.213511417 | 0.08460524511 | 0.499346548 | 0.674246581 | 0.5515987647 | 0.8297433004 | -0.531185592 |
| 0.660099923 | 0.6733855986 | 0.893761146 | 0.885368071 | 0.8978474649 | 0.5331742804 | 0.032968656 |
| 0.186573367 | 0.07701162963 | 0.191360473 | 0.632845883 | 0.5515987647 | -1.038075753 | 1.461101842 |
| 0.310638137 | 0.1376326802 | 0.591985220 | 0.772603929 | 0.5928157147 | -0.3566686451 | 0.727474260 |
| 0.656930027 | 0.6947440907 | 0.866816153 | 0.885368071 | 0.9061879443 | -0.1407191408 | -0.503273466 |
| 0.104495810 | 0.3988667773 | 0.875338584 | 0.569977148 | 0.7721894584 | -0.7586862338 | 0.451847563 |
| 0.158921661 | 0.05992185166 | 0.416948741 | 0.632845883 | 0.5437080554 | -0.5522432588 | 1.14649791 |
| 0.890757036 | 0.7087673991 | 0.931572095 | 0.971734949 | 0.9145385795 | 0.230945138 | 0.024824711 |
| 0.137401205 | 0.838415279 | 0.855709341 | 0.625297136 | 0.9581888902 | 0.7020068896 | -0.294732404 |
| 0.718693859 | 0.6688718019 | 0.893761146 | 0.937426772 | 0.8978474649 | 0.8113486528 | -0.327268348 |
| 0.075178424 | 0.14144584 | 0.110966921 | 0.501189498 | 0.5928157147 | 0.2479618484 | 0.524097573 |
| 0.991960140 | 0.9122938955 | 0.222371509 | 0.999007775 | 0.9948712152 | -0.878189359 | 0.963975216 |
| 0.189853765 | 0.1596190004 | 0.606445993 | 0.632845883 | 0.5928157147 | 0.3360604264 | -0.397676284 |
| 0.783129024 | 0.6294935129 | 0.893761146 | 0.941377455 | 0.8978474649 | 0.09923559748 | -0.031835993 |
| 0.981045510 | 0.936837061 | 0.886760580 | 0.999007775 | 0.9948712152 | -0.7711297804 | 0.233658212 |
| 0.833457038 | 0.551907457 | 0.906588153 | 0.941377455 | 0.8949850654 | -0.3838494491 | 0.213119124 |
| 0.483694757 | 0.2347017435 | 0.749943740 | 0.817383170 | 0.6066894195 | 0.1064993137 | -0.559220344 |
| 0.057850794 | 0.04089105155 | 0.316391746 | 0.433880958 | 0.4877585812 | 0.2870265978 | -1.057635722 |
| 0.437364142 | 0.2092923929 | 0.708622489 | 0.816836587 | 0.5979782654 | 0.3176836653 | -0.595407027 |
| 0.537240753 | 0.3729672715 | 0.859329822 | 0.826524236 | 0.7585775013 | 0.07883904608 | -0.239844412 |
| 0.997715556 | 0.9575600484 | 0.886760580 | 0.999007775 | 0.9953877755 | -0.7226635987 | 0.270246008 |
| 0.443951934 | 0.2305656554 | 0.660529249 | 0.816836587 | 0.6066894195 | 0.2821383355 | -0.211538409 |
| 0.400159574 | 0.2815841683 | 0.528039393 | 0.774909963 | 0.6625509843 | -0.2436937208 | -0.079653989 |
| 0.847239709 | 0.6540742283 | 0.931572095 | 0.941377455 | 0.8978474649 | -0.3069621728 | -0.108240899 |
| 0.627566595 | 0.9679130587 | 0.682640193 | 0.874580333 | 0.9953877755 | -0.2784167192 | 0.543202690 |
| 0.621527882 | 0.7713808689 | 0.971557388 | 0.874580333 | 0.9440157185 | 0.1626706686 | 0.137427549 |
| 0.380878696 | 0.2376200226 | 0.682640193 | 0.772603929 | 0.6066894195 | 0.09467760706 | -0.258134449 |
| 0.385198794 | 0.4023048237 | 0.855709341 | 0.772603929 | 0.7721894584 | -0.4678310475 | 0.551041127 |
| 0.163643553 | 0.1835292044 | 0.682640193 | 0.632845883 | 0.5928157147 | -0.4271534926 | 0.541175663 |
| 0.82583129 | 0.6008799311 | 0.720779494 | 0.941377455 | 0.8978474649 | -0.6989569267 | 0.664499564 |

| | | | | | | |
|---|---|---|---|---|---|---|
| 0.835585633 | 0.6519367304 | 0.893761146 | 0.9413774552 | 0.8978474649 | -0.6674070068 | 0.3623697714 |
| 0.0504104434 | 0.08055040697 | 0.481170621 | 0.4097513587 | 0.5515987647 | 0.312060403 | 0.4574408088 |
| 0.5108644817 | 0.4503904215 | 0.8557093418 | 0.8173831707 | 0.7882231248 | 0.4154257204 | 0.0463400197 |
| 0.9174922655 | 0.8547066334 | 0.1794442789 | 0.9830274273 | 0.9675924152 | 0.3233001739 | -1.047357365 |
| 0.0838607394 | 0.7359225566 | 0.8867605806 | 0.5139509746 | 0.9289265075 | -0.7607993547 | 0.1392188403 |
| 0.5321281982 | 0.4228865589 | 0.8867605806 | 0.8265242368 | 0.7755275948 | 0.05162971018 | -0.322773627 |
| 0.0220188292 | 0.9776552592 | 0.4993465485 | 0.3302824384 | 0.9953877755 | 0.02875792837 | 0.5888603835 |
| 0.7545629335 | 0.8717968425 | 0.4993465485 | 0.9413774552 | 0.977716085 | 0.8537068209 | -0.597053845 |
| 0.8322162424 | 0.8263878934 | 0.0824509797 | 0.9413774552 | 0.9535244924 | 0.2366433481 | -1.137777585 |
| 0.0144750878 | 0.9235073977 | 0.9315720958 | 0.2880213783 | 0.9948712152 | -0.8363794419 | -0.196059950 |
| 0.8835325068 | 0.6236430334 | 0.0781793592 | 0.9717349493 | 0.8978474649 | 0.8753226863 | -1.455186082 |
| 0.463430059 | 0.4265401771 | 0.6826401939 | 0.8173831707 | 0.7755275948 | -0.3802660815 | 0.7664832764 |
| 0.34795292 | 0.743141206 | 0.4993465485 | 0.7726039291 | 0.9289265075 | -0.4903039331 | 0.5324439746 |
| 0.251973413 | 0.3501948236 | 0.7581287096 | 0.7266389938 | 0.757210127 | -0.09114888039 | 0.6758445324 |
| 0.1123454974 | 0.04471120328 | 0.1480388684 | 0.5861504214 | 0.4877585812 | 0.6864415035 | -1.520440848 |
| 0.6182049908 | 0.332885926 | 0.6605292499 | 0.874580333 | 0.757210127 | -0.3258768721 | -0.005746393 |
| 0.6236152007 | 0.9948307884 | 0.9715573887 | 0.874580333 | 0.9953877755 | -0.4107824406 | 0.1236386855 |
| 0.5041571561 | 0.8139353891 | 0.1794442789 | 0.8173831707 | 0.9535244924 | 0.17216523 | -0.920410932 |
| 0.3863019645 | 0.1923456044 | 0.4169487412 | 0.7726039291 | 0.5928157147 | -0.2400933694 | 1.13422369 |
| 0.183926772 | 0.1176630853 | 0.1439120649 | 0.6328458833 | 0.5928157147 | 0.3678915024 | -1.510200805 |
| 0.7951390939 | 0.6600647536 | 0.893761146 | 0.9413774552 | 0.8978474649 | 0.03040147836 | 0.4298643994 |
| 0.0970919255 | 0.1389903269 | 0.5737250479 | 0.5548110031 | 0.5928157147 | 0.3702698568 | 0.3512119765 |
| 0.1254069334 | 0.1043992389 | 0.2177446949 | 0.6019532803 | 0.5928157147 | 0.3581191356 | -1.178413791 |
| 0.141032802 | 0.06689168433 | 0.481170621 | 0.6252971367 | 0.5437080554 | 0.7698649676 | -0.659807983 |
| 0.3700204072 | 0.1633234022 | 0.4777632406 | 0.7726039291 | 0.5928157147 | -0.7842887733 | 1.128531753 |
| 0.4995059559 | 0.2767059398 | 0.2873764175 | 0.8173831707 | 0.6625509843 | 0.9988602512 | -1.094800423 |
| 0.9582478381 | 0.7866797654 | 0.1480388684 | 0.9990077752 | 0.9440157185 | -0.2325446079 | 1.115083509 |
| 0.8359924459 | 0.6038872084 | 0.1109669215 | 0.9413774552 | 0.8978474649 | -0.7067915721 | 1.366784361 |

| est_vns | est_interact | est_batch | se_intercept | se_blast | se_vns | se_interact |
|---|---|---|---|---|---|---|
| -0.477538093 | 1.280794726 | -0.183860328 | 0.451238152 | 0.555693196 | 0.575950144 | 0.777242151 |
| -0.632719577 | 0.3447212754 | 0.445445924 | 0.4582101943 | 0.564279164 | 0.584849101 | 0.789251253 |
| 0.487655293 | -0.5505057782 | 0.503564804 | 0.4346305347 | 0.535241158 | 0.554752558 | 0.748636103 |
| -2.085629383 | 2.418378155 | 0.046163018 | 0.3312658988 | 0.407949118 | 0.422820281 | 0.570594083 |
| -0.395476418 | 0.1783085747 | -1.354697775 | 0.3471903767 | 0.427559880 | 0.443145923 | 0.598023447 |
| 0.388605760 | -0.9591343373 | 0.019425803 | 0.4499182315 | 0.554067733 | 0.574265428 | 0.774968634 |
| 0.835784708 | -0.9998644076 | 0.485248367 | 0.4261161402 | 0.524755805 | 0.543884978 | 0.733970352 |
| -0.403894383 | 0.9748033435 | -0.992472816 | 0.3899049288 | 0.480162227 | 0.497665809 | 0.671597790 |
| -0.855562372 | 1.035114271 | 0.173489657 | 0.4159469857 | 0.512232640 | 0.530905300 | 0.716454335 |
| -0.814093817 | 0.7502300771 | -0.125710242 | 0.4504847253 | 0.554765362 | 0.574988488 | 0.7759444 |
| -0.906480732 | 0.9390242135 | 0.433350575 | 0.4307435383 | 0.530454378 | 0.549791284 | 0.741940886 |
| -1.047575043 | 0.3770942957 | 0.338846268 | 0.4107859109 | 0.505876852 | 0.524317821 | 0.707564561 |
| -0.574352789 | 1.492988812 | -0.271896192 | 0.4265133047 | 0.525244907 | 0.544391910 | 0.734654455 |
| -0.743010430 | 1.244403001 | -1.045713489 | 0.3763471556 | 0.463466027 | 0.480360975 | 0.648244993 |
| 0.695926386 | -0.7010268573 | 0.636190548 | 0.4443265419 | 0.547181649 | 0.567128322 | 0.765337141 |
| -0.043559996 | 0.1406778281 | 0.887204731 | 0.3618427048 | 0.445604008 | 0.461847823 | 0.623261577 |
| -0.75037325 | 1.121160907 | -0.242392609 | 0.4486867519 | 0.552551184 | 0.572693595 | 0.772847452 |
| -0.320036615 | 1.513455455 | -0.428536478 | 0.4064991019 | 0.500597709 | 0.518846235 | 0.700180680 |
| -0.008931753 | 0.9732393759 | -0.487185411 | 0.4331186472 | 0.533379290 | 0.552822819 | 0.746031929 |
| 0.638482987 | -0.6595059353 | -0.101189393 | 0.4416273629 | 0.543857649 | 0.563683151 | 0.760687898 |
| -0.822521568 | 0.7119680032 | 1.11850697 | 0.3535654634 | 0.435410706 | 0.451282939 | 0.609004314 |
| 0.684074431 | -1.383766207 | 1.364188217 | 0.2854647312 | 0.351545648 | 0.364360709 | 0.491703152 |
| 0.736343047 | -0.9919061008 | -0.385465144 | 0.4284897569 | 0.527678879 | 0.546914609 | 0.738058825 |
| 1.019405222 | -2.131250948 | 0.327633578 | 0.3965340033 | 0.488325835 | 0.50612701 | 0.683016142 |
| 1.542865582 | -1.015407516 | 0.409588910 | 0.3824617966 | 0.470996118 | 0.488165564 | 0.658777251 |
| 0.131266323 | -0.0193787030 | -0.047475129 | 0.4905357764 | 0.604087647 | 0.626108741 | 0.844930954 |
| 0.757791602 | -0.7113650619 | -0.481844739 | 0.4407915685 | 0.542828381 | 0.562616362 | 0.759248271 |
| 1.163550049 | -2.135638877 | -0.585632001 | 0.3941239497 | 0.48535789 | 0.503050872 | 0.678864908 |
| 0.104530459 | 0.4692463621 | 0.265824353 | 0.4663645556 | 0.574321141 | 0.595257143 | 0.803296859 |
| -0.749291582 | 1.068591748 | -0.556772894 | 0.4428065518 | 0.545309803 | 0.565188241 | 0.762719011 |
| 0.059182450 | -0.1688151972 | -0.466540456 | 0.4335810272 | 0.533948704 | 0.553412991 | 0.746828363 |
| -0.474457732 | -0.0552386880 | 0.043453564 | 0.4728584154 | 0.582318235 | 0.603545758 | 0.814482309 |
| -0.001255025 | -0.1659466327 | 1.385866392 | 0.3425094221 | 0.421795352 | 0.437171259 | 0.589960664 |
| 0.804496873 | -1.992576813 | -0.170815809 | 0.4184377147 | 0.515299936 | 0.534084410 | 0.720744529 |
| 0.411482444 | 0.0746385144 | 0.213994582 | 0.4765466713 | 0.586860268 | 0.608253365 | 0.820835203 |
| 1.255579275 | -1.524931637 | -0.869496117 | 0.3758382472 | 0.462839314 | 0.479711416 | 0.647368416 |
| 0.3914202 | -1.225436873 | -0.245764076 | 0.4588738655 | 0.565096465 | 0.585696196 | 0.790394404 |
| 0.028895404 | -0.3401750873 | 0.980107650 | 0.4099365716 | 0.504830903 | 0.523233744 | 0.706101604 |
| -0.275452842 | -0.0703935398 | -1.215900861 | 0.3823207677 | 0.470822443 | 0.487985558 | 0.658534334 |
| -1.736304606 | 1.899684441 | 0.858879450 | 0.3343944607 | 0.411801896 | 0.426813506 | 0.575982923 |
| 0.306709810 | -0.0691212419 | 0.691167406 | 0.4514939788 | 0.556008243 | 0.576276676 | 0.777682804 |
| 0.238845780 | 0.5357006625 | -0.799326249 | 0.4133649112 | 0.509052853 | 0.527609599 | 0.712006800 |
| -0.518350057 | 0.6588271236 | -1.000895984 | 0.4088520683 | 0.503495353 | 0.521849508 | 0.704233584 |
| 0.023784423 | 0.9494824878 | -0.209102143 | 0.4157183799 | 0.511951115 | 0.530613513 | 0.716060569 |
| -0.021843977 | 0.9837070726 | -1.022882456 | 0.3544538111 | 0.436504693 | 0.452416806 | 0.610534462 |
| -0.012736783 | -1.020683191 | -0.329998740 | 0.4356439008 | 0.536489103 | 0.556045996 | 0.750381591 |

| | | | | | | |
|---|---|---|---|---|---|---|
| -0.06131553 | -0.479992602 | -1.269921952 | 0.3692585304 | 0.454736489 | 0.471313214 | 0.636035081 |
| -2.07838328 | 2.294625541 | 0.488472611 | 0.3257389898 | 0.401142810 | 0.415765860 | 0.561074173 |
| -0.75658477 | 1.413194338 | 0.120789367 | 0.4585183478 | 0.564658651 | 0.585242421 | 0.789782037 |
| -0.44263881 | 0.4709704817 | -0.05203707 | 0.4498696418 | 0.554007895 | 0.574203409 | 0.774884939 |
| -0.827820717 | 0.4377531323 | -0.30876907 | 0.4575693912 | 0.563490024 | 0.584031195 | 0.788147492 |
| 1.071047572 | -0.218402292 | 0.893061258 | 0.3767859089 | 0.464006345 | 0.48092099 | 0.649000730 |
| -0.00919686 | 0.3148848474 | -0.709748954 | 0.4477942842 | 0.551452123 | 0.571554470 | 0.771310208 |
| -0.513506791 | -0.673214838 | -0.63323967 | 0.4094111561 | 0.504183861 | 0.522563115 | 0.705196593 |
| -0.68601424 | 0.3595457026 | 0.162364396 | 0.4393757095 | 0.541084771 | 0.560809192 | 0.756809503 |
| -0.37891583 | 0.4594502972 | -0.25107563 | 0.4828163328 | 0.594581264 | 0.616255818 | 0.831634478 |
| 0.079623412 | 0.6407826213 | -0.501018751 | 0.4385348553 | 0.540049271 | 0.559735945 | 0.755361161 |
| -0.020130404 | 0.0046721813 | -0.025311421 | 0.4639409724 | 0.571336534 | 0.592163736 | 0.799122321 |
| -0.281404134 | 0.4784568044 | 0.628135902 | 0.3635780781 | 0.447741095 | 0.464062814 | 0.626250698 |
| 0.450704348 | -0.578831856 | -1.056675964 | 0.4051690683 | 0.498959792 | 0.517148611 | 0.697889744 |
| -0.819535532 | 0.8743129343 | -0.451631994 | 0.4449913119 | 0.548000303 | 0.567976819 | 0.766482184 |
| -0.128185317 | 0.5259073118 | -0.42724778 | 0.4696604446 | 0.578379980 | 0.599463941 | 0.808973915 |
| -0.720262571 | 1.175409056 | -1.058067822 | 0.3778318586 | 0.465294417 | 0.482256016 | 0.650802343 |
| -0.481558074 | 0.3214029854 | -0.82458817 | 0.4367852861 | 0.537894703 | 0.557502834 | 0.752347588 |
| 0.867946443 | -1.305770315 | 0.338956155 | 0.4086168409 | 0.503205674 | 0.521549269 | 0.703828413 |
| 0.797901853 | -1.232065727 | -0.19970466 | 0.4642410622 | 0.571706090 | 0.592546764 | 0.799639215 |
| 0.157007124 | 0.3160462423 | 0.509388394 | 0.4614608492 | 0.568282298 | 0.588998163 | 0.794850394 |
| 1.140181026 | -0.653485219 | 0.228627865 | 0.4409628844 | 0.543039354 | 0.562835026 | 0.759543357 |
| 0.733348873 | -1.518538331 | -0.03119063 | 0.4444554241 | 0.547340365 | 0.567292824 | 0.765559136 |
| 0.048852216 | -0.313345006 | -0.368081567 | 0.4807643017 | 0.592054218 | 0.613636652 | 0.828099925 |
| -0.69432558 | -0.155827787 | -0.27430918 | 0.4384195515 | 0.539907276 | 0.559588773 | 0.755162554 |
| -0.411643042 | 0.3056894229 | -1.052300091 | 0.4093923537 | 0.504160706 | 0.522539116 | 0.705164207 |
| -1.093479964 | 0.9424867003 | -0.522123662 | 0.3587439246 | 0.441787905 | 0.457892610 | 0.617924035 |
| 0.021001692 | -0.0760569331 | 0.735839709 | 0.3963023004 | 0.488040497 | 0.505831269 | 0.682617042 |
| -1.089762104 | 1.131265606 | 0.238312105 | 0.4511953165 | 0.555640445 | 0.575895470 | 0.777168368 |
| -0.014126521 | 0.3973950008 | -0.39870845 | 0.4715455858 | 0.580701504 | 0.601870092 | 0.812221005 |
| 0.028796024 | 0.0514881081 | 1.269275514 | 0.3729191763 | 0.459244521 | 0.475985580 | 0.642340417 |
| 0.286264935 | -0.485975280 | 0.542095493 | 0.4669827743 | 0.575082468 | 0.596046223 | 0.804361720 |
| -0.57548548 | 0.9800289261 | 0.440722239 | 0.4656680955 | 0.573463461 | 0.594368197 | 0.802097231 |
| -1.14837578 | 1.342542885 | 1.034748896 | 0.3588154255 | 0.441875958 | 0.457983872 | 0.618047193 |
| -0.66482612 | 1.054261009 | 0.094394165 | 0.4732252742 | 0.582770016 | 0.604014009 | 0.815114211 |
| -0.672342267 | 0.7334790526 | 0.385690097 | 0.4682273439 | 0.576615138 | 0.597634764 | 0.806505448 |
| -0.03302380 | 0.0355966515 | 1.202752852 | 0.38394688 | 0.472824977 | 0.490061091 | 0.661335256 |
| -0.704846597 | 0.9884771162 | -0.191124882 | 0.4654044337 | 0.573138765 | 0.594031665 | 0.801643083 |
| 0.79080263 | -0.855840460 | 0.273079012 | 0.4501221725 | 0.554318883 | 0.574525733 | 0.775319915 |
| -0.05764805 | 0.3654898957 | 0.599595827 | 0.4670858241 | 0.575209373 | 0.596177754 | 0.804539220 |
| 0.390038314 | -0.031277425 | -0.44931584 | 0.4463067912 | 0.549620297 | 0.569655867 | 0.768748052 |
| 0.464996537 | -0.217212420 | -0.863521034 | 0.4286732991 | 0.527904909 | 0.547148878 | 0.738374970 |
| -0.817468782 | 0.9601341471 | 0.390708548 | 0.4591589844 | 0.565447585 | 0.586060115 | 0.790885512 |
| 0.826636060 | -0.686505573 | -0.13192606 | 0.4666938962 | 0.574726719 | 0.595677506 | 0.803864137 |
| 1.129443386 | -1.057687212 | -0.30270771 | 0.4471747384 | 0.550689161 | 0.570763696 | 0.770243061 |
| 0.353782227 | -0.4152818801 | 0.55951134 | 0.454220802 | 0.559366286 | 0.579757131 | 0.782379663 |

| | | | | | | |
|---|---|---|---|---|---|---|
| 0.3391622255 | -0.3479280752 | 0.8168920045 | 0.4417070502 | 0.5439557835 | 0.5637848620 | 0.7608251574 |
| 0.0535645920 | -1.331317655 | -0.414354312 | 0.421934063 | 0.5196056385 | 0.5385470702 | 0.726766869 |
| -0.017099811 | -0.6084233491 | -0.5234469668 | 0.4596628855 | 0.5660681320 | 0.5867032837 | 0.7917534644 |
| -0.190738421 | 0.1231519142 | 0.6666444894 | 0.3858878496 | 0.4752152529 | 0.4925385007 | 0.6646785098 |
| 0.7142748221 | 0.2506436155 | 0.4960233491 | 0.4260282098 | 0.5246475205 | 0.5437727463 | 0.7338188957 |
| -0.043409009 | 0.669566452 | -0.1000925566 | 0.4760030392 | 0.5861907933 | 0.6075594853 | 0.8198988154 |
| -1.008763248 | 0.01882078915 | 0.3355961020 | 0.3857072856 | 0.4749928911 | 0.492308033 | 0.6643674947 |
| -0.200116565 | -0.1178272161 | -0.796155910 | 0.4189770478 | 0.5159641174 | 0.534772803 | 0.7216735124 |
| -0.221298022 | 0.1216280004 | 0.984803207 | 0.3181241034 | 0.3917651888 | 0.4060463917 | 0.5479577945 |
| 0.9976660397 | 0.06270137670 | 0.8503237395 | 0.3748007382 | 0.4615616371 | 0.4783871632 | 0.6455813429 |
| -0.185247652 | 0.3161233509 | -0.133956998 | 0.3687652638 | 0.4541290385 | 0.4706836205 | 0.6351854464 |
| -0.028770781 | -0.6336065702 | 0.3262752734 | 0.4540592877 | 0.5591673837 | 0.5795509785 | 0.7821014605 |
| 0.4048178871 | 0.2488454261 | -0.185128250 | 0.4353451324 | 0.5361211747 | 0.555664655 | 0.7498669735 |
| -0.144797703 | -0.7507097823 | 0.0273998375 | 0.4566034982 | 0.5623005418 | 0.5827983511 | 0.7864837753 |
| -0.981432816 | 1.446968568 | 0.495154078 | 0.3946163005 | 0.4859642128 | 0.5036792970 | 0.6797129655 |
| 0.3847996519 | -0.7370352605 | 0.7226121851 | 0.4321697448 | 0.5322107312 | 0.5516116623 | 0.7443974778 |
| 0.398482318 | 0.00538167688 | 0.2780569035 | 0.4768108363 | 0.5871855837 | 0.6085905392 | 0.8212902181 |
| -0.40708274 | 0.1365062001 | 1.058831656 | 0.3327143305 | 0.4097328400 | 0.4246690268 | 0.5730889575 |
| 0.3868253907 | -0.9843121211 | -0.620914837 | 0.4248917803 | 0.5232480242 | 0.5423222334 | 0.731861435 |
| -0.256548758 | 1.059362895 | 0.6028246897 | 0.3776721384 | 0.4650977246 | 0.4820521533 | 0.6505272308 |
| -0.009127439 | -0.3669273299 | -0.323507932 | 0.477797431 | 0.5884005607 | 0.6098498064 | 0.8229895935 |
| 0.0034614377 | -1.141747701 | -0.475380330 | 0.4317700681 | 0.5317185352 | 0.5511015241 | 0.7437090486 |
| -0.952823127 | 1.022112562 | 1.013047192 | 0.3503189013 | 0.4314126124 | 0.4471391017 | 0.6034122233 |
| -0.419040597 | 1.18390168 | -1.114913804 | 0.356541285 | 0.4390753871 | 0.4550812112 | 0.6141300646 |
| 0.613911626 | -1.096328262 | 0.3402833784 | 0.4413217698 | 0.5434813163 | 0.5632930994 | 0.7601615252 |
| -0.252825152 | 0.7116779244 | -0.977007052 | 0.3704193976 | 0.4561660802 | 0.4727949194 | 0.6380346349 |
| 0.0636564987 | -0.1750056088 | -0.764514948 | 0.3708790138 | 0.4567320908 | 0.4733815631 | 0.638826308 |
| 0.0974349567 | -0.3543879103 | 0.0040477629 | 0.3908627202 | 0.4813417335 | 0.4988883129 | 0.673247553 |

| se_batch | t_intercept | t_blast | t_vns | t_interact | t_batch | t_p_intercept |
|---|---|---|---|---|---|---|
| 0.38862107 | 1.092681504 | -1.73755938 | -0.829130954 | 1.647870903 | -0.47310951 | 0.2863550335 |
| 0.39462562 | 0.49758550 | -0.690728962 | -1.08185098 | 0.43677000 | 1.128781038 | 0.6237132229 |
| 0.37431805 | -0.25382805 | -0.74617977 | 0.87905010 | -0.73534494 | 1.345285919 | 0.801985587 |
| 0.28529704 | 2.517292831 | -2.167500944 | -4.93266163 | 4.238351262 | 0.16180686 | 0.01961137463 |
| 0.29901172 | 2.09643243 | 0.32260913 | -0.89242932 | 0.29816318 | -4.53058414 | 0.04776309527 |
| 0.387484317 | -1.216740987 | 2.016693907 | 0.67670060 | -1.23764278 | 0.05013313 | 0.236595477 |
| 0.36698517 | -2.36085373 | 2.178999982 | 1.536693862 | -1.36226811 | 1.322256044 | 0.02751013538 |
| 0.33579889 | 1.406696745 | -0.562402231 | -0.81157751 | 1.451468956 | -2.955557122 | 0.1734871571 |
| 0.35822716 | 1.967767327 | -2.688563861 | -1.61151597 | 1.444773548 | 0.48430066 | 0.06182717177 |
| 0.3879722 | 1.732204615 | -1.687109712 | -1.415843682 | 0.96686061 | -0.32401868 | 0.09723628567 |
| 0.37097044 | 1.312879862 | -1.97615648 | -1.64877246 | 1.265632115 | 1.168153914 | 0.2027543022 |
| 0.35378228 | 1.561305201 | -1.32739605 | -1.997977181 | 0.53294683 | 0.957781909 | 0.1327227759 |
| 0.36732722 | 1.596539026 | -2.38032954 | -1.05503549 | 2.032232708 | -0.74020157 | 0.1246340893 |
| 0.32412249 | 2.256527086 | -1.27851391 | -1.54677517 | 1.919649228 | -3.22629098 | 0.03431061361 |
| 0.38266857 | -1.12735713 | 0.13639165 | 1.22710568 | -0.915971301 | 1.662510581 | 0.2717375623 |
| 0.31163078 | 0.37427486 | -2.34974205 | -0.09431677 | 0.22571233 | 2.84697393 | 0.7117830213 |
| 0.38642372 | 1.717336386 | -1.944640614 | -1.31025256 | 1.450688493 | -0.62727155 | 0.09996434665 |
| 0.35009034 | 0.50342928 | -1.00966405 | -0.61682362 | 2.1615213 | -1.22407398 | 0.6196660522 |
| 0.37301596 | 0.70465260 | -1.19587332 | -0.01615662 | 1.304554587 | -1.30607120 | 0.4884197074 |
| 0.38034394 | 0.33375790 | -0.77556582 | 1.132698371 | -0.86698623 | -0.26604707 | 0.7417226266 |
| 0.30450215 | 0.415461228 | -1.9425547 | -1.82262943 | 1.169068899 | 3.673231679 | 0.6818285358 |
| 0.24585157 | -3.79481057 | 2.46669236 | 1.877464866 | -2.81423090 | 5.548828435 | 0.00099351908 |
| 0.36902941 | 0.53427818 | -0.44770797 | 1.346358344 | -1.34393908 | -1.04453772 | 0.5985073233 |
| 0.34150807 | -1.52405610 | 2.025461785 | 2.014129263 | -3.12035223 | 0.95937286 | 0.1417417613 |
| 0.32938862 | -3.18875289 | 1.99959786 | 3.16053752 | -1.54135182 | 1.243482252 | 0.00424298328 |
| 0.422465477 | -0.05199896 | -0.04158599 | 0.20965419 | -0.02293525 | -0.11237635 | 0.9589987883 |
| 0.37962413 | -0.96456565 | 1.575141745 | 1.346906441 | -0.93693339 | -1.26926792 | 0.3452462483 |
| 0.33943245 | -0.99742504 | 2.612858458 | 2.312986843 | -3.145896704 | -1.725327071 | 0.3294031656 |
| 0.40164843 | 0.08686281 | -1.13373002 | 0.17560555 | 0.58415062 | 0.66183341 | 0.9315663221 |
| 0.38135950 | 1.129750304 | -0.52907500 | -1.325738094 | 1.401029387 | -1.45996857 | 0.2707492321 |
| 0.37341418 | -0.56412719 | 1.646981447 | 0.10694084 | -0.22604283 | -1.249391371 | 0.5783739353 |
| 0.40724115 | 0.65359121 | -0.20894571 | -0.78611724 | -0.06782061 | 0.10670229 | 0.5201486819 |
| 0.29498033 | -2.08321699 | 0.40998704 | -0.00287078 | -0.281284232 | 4.6981654 | 0.04906233638 |
| 0.36037226 | -0.88684271 | 2.18871256 | 1.506310347 | -2.76460899 | -0.47399821 | 0.3847559956 |
| 0.41041760 | -0.74178048 | 0.04595297 | 0.67649842 | 0.09092996 | 0.52140693 | 0.4660661916 |
| 0.32368420 | -1.10430551 | 2.467505665 | 2.617363758 | -2.35558547 | -2.68624818 | 0.2813928519 |
| 0.39519720 | -0.36673149 | 1.423025439 | 0.66829903 | -1.55041187 | -0.62187706 | 0.7173229524 |
| 0.353050802 | -0.80714808 | -0.19686953 | 0.05522465 | -0.48176506 | 2.7761094 | 0.4282217778 |
| 0.32926716 | 2.243562444 | -0.415349022 | -0.56446925 | -0.10689425 | -3.69274857 | 0.03525547755 |
| 0.28799146 | 1.484237394 | -2.502061412 | -4.06806386 | 3.298161044 | 2.98230873 | 0.1519333233 |
| 0.38884140 | -1.47821804 | 0.63474276 | 0.53222665 | -0.08888102 | 1.777504664 | 0.1535246957 |
| 0.35600340 | 0.71988316 | -0.69052618 | 0.45269415 | 0.75238138 | -2.24527700 | 0.4791757771 |
| 0.35211679 | 1.62550659 | -0.39007908 | -0.993294137 | 0.93552357 | -2.842511367 | 0.1182956253 |
| 0.35803028 | 1.161210855 | -2.37580037 | 0.04482438 | 1.32598069 | -0.584034792 | 0.2580010177 |
| 0.30526723 | 1.024509849 | -0.621785954 | -0.048282861 | 1.611222842 | -3.35077712 | 0.3167284708 |
| 0.37519079 | 0.97210636 | 0.14905236 | -0.02290598 | -1.360218857 | -0.87954913 | 0.3415652828 |

| | | | | | | |
|---|---|---|---|---|---|---|
| 0.3180175400 | 1.441887989 | 1.001568899 | -0.130095099 | -0.754663723 | -3.993244994 | 0.163419077 |
| 0.2805370869 | 2.389883791 | -2.763553985 | -4.998927254 | 4.089700878 | 1.741205118 | 0.02585188369 |
| 0.3948910189 | 0.8306686489 | -1.483764113 | -1.292771592 | 1.789347276 | 0.3058802600 | 0.4150851059 |
| 0.3874424699 | 1.460801468 | -1.761459223 | -0.770874578 | 0.6077940833 | -0.134309176 | 0.1582056451 |
| 0.3940737463 | 1.538907114 | -0.817615234 | -1.417425515 | 0.5554203197 | -0.783531194 | 0.1380872704 |
| 0.3245003655 | -2.93255187 | 0.7149686649 | 2.227075952 | -0.336520872 | 2.752111718 | 0.00770654939 |
| 0.385655104 | 0.506452476 | 0.0862181249 | -0.016096060 | 0.4082467004 | -1.840372256 | 0.617577124 |
| 0.3525982969 | 1.702678487 | 0.2625165665 | -0.982669415 | -0.954648455 | -1.795923807 | 0.1027174144 |
| 0.3784047517 | 1.468838621 | -1.57805804 | -1.223257853 | 0.4750808505 | 0.429075998 | 0.1560313734 |
| 0.4158172394 | 0.724319236 | -0.510056726 | -0.614867762 | 0.5524666292 | -0.603812481 | 0.476502694 |
| 0.3776805800 | -0.044734983 | 0.0989955308 | 0.1422517402 | 0.8483129055 | -1.326567415 | 0.9647221212 |
| 0.3995611607 | -0.734074577 | 1.139878452 | -0.033994658 | 0.0058466410 | -0.063348053 | 0.470655165 |
| 0.313125349 | 1.291548204 | -3.086960459 | -0.606392337 | 0.7640020298 | 2.006020606 | 0.2099187985 |
| 0.348944872 | 0.4298506619 | 0.9827381462 | 0.8715180491 | -0.829403013 | -3.028203161 | 0.6714852867 |
| 0.3832410925 | 0.8828092417 | -0.031208747 | -1.442903132 | 1.140682656 | -1.178453989 | 0.3868843278 |
| 0.4044869575 | 0.405636834 | -0.230788931 | -0.213833240 | 0.650091804 | -1.056270844 | 0.6889274643 |
| 0.3254011710 | 2.196064947 | -1.141611788 | -1.493527393 | 1.806092231 | -3.251579632 | 0.03892318658 |
| 0.3761737943 | 1.220678208 | 0.0612920269 | -0.863776907 | 0.427200127 | -2.192040451 | 0.2351302258 |
| 0.3519142067 | -2.54046248 | 2.903587772 | 1.664169607 | -1.855239558 | 0.9631783789 | 0.01863974064 |
| 0.3998196079 | -0.768283278 | 1.272461974 | 1.346563515 | -1.540777019 | -0.499486917 | 0.4504877075 |
| 0.3974251974 | -0.304942751 | -0.885604686 | 0.2665664068 | 0.3976172678 | 1.281721436 | 0.7632759849 |
| 0.3797716788 | -1.720521751 | 0.8320714881 | 2.025781929 | -0.860365919 | 0.602013996 | 0.09937445609 |
| 0.3827795683 | -1.242516637 | 2.094670851 | 1.292716638 | -1.983567641 | -0.081484584 | 0.2271275071 |
| 0.4140499629 | 0.4803708121 | 0.0419297937 | 0.0796109826 | -0.378390332 | -0.888978627 | 0.6357058646 |
| 0.3775812777 | 1.601221677 | -0.545894485 | -1.240778268 | -0.206349993 | -0.726490428 | 0.1235906624 |
| 0.3525821037 | 1.981836362 | -0.649134976 | -0.787774598 | 0.433501048 | -2.984553327 | 0.06012805345 |
| 0.3089620177 | 0.6911945581 | 1.186310369 | -2.388070782 | 1.525246869 | -1.689928315 | 0.4966727431 |
| 0.341308521 | -2.215958268 | 1.975195137 | 0.0415191651 | -0.111419622 | 2.155937118 | 0.03734676942 |
| 0.3885841844 | 0.7448225063 | -0.715707951 | -1.892291501 | 1.45562487 | 0.61328308 | 0.4642619635 |
| 0.4061105029 | 0.2104475160 | -0.054823336 | -0.023471047 | 0.4892695430 | -0.981773319 | 0.8352553444 |
| 0.3211702087 | -2.067820132 | 0.5087882423 | 0.0604976815 | 0.0801570424 | 3.95203378 | 0.05061595134 |
| 0.4021808602 | -0.821977747 | 0.3705888042 | 0.4802730455 | -0.604175047 | 1.347889834 | 0.4199092252 |
| 0.4010486157 | 0.2287021909 | -0.975162992 | -0.968230617 | 1.221833074 | 1.098924723 | 0.8212129397 |
| 0.3090235960 | 0.7999282565 | -2.393512711 | -2.507459011 | 2.172233609 | 3.348446226 | 0.4323055733 |
| 0.4075571055 | 0.6713159300 | -1.021684388 | -1.100679979 | 1.293390539 | 0.2316096658 | 0.5090073288 |
| 0.4032527243 | 0.1683777063 | -0.415952333 | -1.125005285 | 0.9094533130 | 0.956447593 | 0.8678245357 |
| 0.3306676283 | -1.882196826 | 0.5715561169 | -0.067387115 | 0.0538254255 | 3.637346838 | 0.07310406651 |
| 0.4008215415 | 0.6062218473 | -0.369087597 | -1.186547181 | 1.233063862 | -0.476832860 | 0.5505720368 |
| 0.3876599570 | -0.541394616 | -0.143697051 | 1.376444228 | -1.103854608 | 0.704429249 | 0.5936764321 |
| 0.4022696101 | -0.657185803 | -0.188176521 | -0.096696085 | 0.4542847460 | 1.490532251 | 0.5178783828 |
| 0.3843740263 | -0.623823622 | 0.9883235619 | 0.684691121 | -0.040686185 | -1.168954742 | 0.5391585193 |
| 0.3691874854 | 0.3794746931 | 0.260326333 | 0.8498537703 | -0.294176304 | -2.338976993 | 0.7079736401 |
| 0.3954427561 | 0.2061978754 | -0.456513487 | -1.39485483 | 1.213998907 | 0.9880280840 | 0.8385326433 |
| 0.4019320689 | -1.00243661 | 0.9587880791 | 1.387724149 | -0.854006966 | -0.328229765 | 0.3270317882 |
| 0.3851215309 | -0.955227243 | 0.9827243772 | 1.978828355 | -1.37318629 | -0.786005703 | 0.3498421095 |
| 0.3911898310 | -1.538804307 | 1.187950687 | 0.6102248817 | -0.530793295 | 1.430280888 | 0.1381122995 |

| | | | | | | |
|---|---|---|---|---|---|---|
| 0.380412578 | -1.51097204 | 0.666175050 | 0.601580936 | -0.45730358 | 2.147384315 | 0.1450269179 |
| 0.363383434 | 0.739595188 | 0.880361518 | 0.099461301 | -1.83183591 | -1.14026747 | 0.4673648597 |
| 0.395876732 | 0.903761720 | 0.081862972 | -0.029145587 | -0.768450504 | -1.322247371 | 0.3759120142 |
| 0.332339254 | 0.837808638 | -2.20396411 | -0.387255862 | 0.185280421 | 2.00591558 | 0.411148176 |
| 0.366909447 | -1.785795722 | 0.265356901 | 1.313553919 | 0.341560590 | 1.351895821 | 0.08792543598 |
| 0.409949407 | 0.108465085 | -0.550628961 | -0.071448164 | 0.816645224 | -0.24415831 | 0.9146099706 |
| 0.332183747 | 0.074558945 | 1.239724625 | -2.04904892 | 0.028328883 | 1.010272493 | 0.9412394468 |
| 0.360836756 | 2.037598063 | -1.15716156 | -0.37420856 | -0.16326942 | -2.206415774 | 0.05379411254 |
| 0.273978897 | 0.743871167 | -2.90423349 | -0.54500674 | 0.221966001 | 3.594449123 | 0.464825757 |
| 0.322790671 | -2.23153093 | -0.42477522 | 2.085478283 | 0.097123898 | 2.634288456 | 0.03615350332 |
| 0.317592723 | 2.373658184 | -3.20434493 | -0.39357148 | 0.497686703 | -0.42178862 | 0.02676698259 |
| 0.391050730 | -0.837481121 | 1.3707582 | -0.04964322 | -0.81013347 | 0.83435536 | 0.411328246 |
| 0.374933486 | -1.12624190 | 0.993141102 | 0.72852912 | 0.33185276 | -0.493762912 | 0.272199031 |
| 0.393241887 | -0.199623701 | 1.201927585 | -0.248452492 | -0.95451401 | 0.06967680 | 0.8436084935 |
| 0.339856482 | 1.739516342 | -3.128709497 | -1.94852721 | 2.128793537 | 1.456950516 | 0.09591820973 |
| 0.372198738 | -0.754048324 | -0.010797214 | 0.69759158 | -0.99010983 | 1.941468655 | 0.4588156064 |
| 0.410645109 | -0.86152077 | 0.210561514 | 0.654762590 | 0.006552710 | 0.677122160 | 0.3982444261 |
| 0.286544478 | 0.517456611 | -2.24636846 | -0.958588251 | 0.238193736 | 3.69517382 | 0.6100016148 |
| 0.365930717 | -0.565069461 | 2.167659768 | 0.713275921 | -1.34494328 | -1.69680982 | 0.577743949 |
| 0.325263615 | 0.974102839 | -3.247061264 | -0.53220124 | 1.62846818 | 1.853341908 | 0.340595223 |
| 0.411494796 | 0.063628383 | 0.730564224 | -0.01496669 | -0.445846864 | -0.78617745 | 0.9498407282 |
| 0.371854524 | 0.857562587 | 0.660522350 | 0.006280943 | -1.53520748 | -1.278404053 | 0.4003800966 |
| 0.301706111 | 1.022266096 | -2.73152373 | -2.13093224 | 1.693887731 | 3.35772844 | 0.3177652991 |
| 0.307065032 | 2.15925897 | -1.502721407 | -0.92080399 | 1.92777027 | -3.630871922 | 0.04199991356 |
| 0.380080762 | -1.77713592 | 2.076486751 | 1.089861791 | -1.44223066 | 0.895292295 | 0.08937588441 |
| 0.319017317 | 2.696565724 | -2.400004014 | -0.534745917 | 1.115422088 | -3.06255177 | 0.01317976274 |
| 0.319413154 | -0.62700934 | 2.441438933 | 0.134471858 | -0.27394865 | -2.39349863 | 0.5371064229 |
| 0.336623776 | -1.808285967 | 2.839530142 | 0.195304147 | -0.52638573 | 0.012024590 | 0.08425295881 |

| t_p_blast | t_p_vns | t_p_interact | t_p_batch | t_q_blast | t_q_vns | t_q_interact |
|---|---|---|---|---|---|---|
| 0.096269480 | 0.415936086 | 0.1135888756 | 0.640795515 | 0.312225342 | 0.805037586 | 0.5928157147 |
| 0.496959682 | 0.291035077 | 0.6665350338 | 0.271149202 | 0.774677428 | 0.698484187 | 0.8978474649 |
| 0.463458308 | 0.388874804 | 0.4698968088 | 0.192236684 | 0.771435749 | 0.794060801 | 0.7941917894 |
| 0.041292415 | 0.000062022 | 0.0003370890 | 0.872934801 | 0.190580377 | 0.003721366 | 0.0290773457 |
| 0.750037207 | 0.381820795 | 0.768376414 | 0.000165072 | 0.909136008 | 0.794060801 | 0.9440157185 |
| 0.056095402 | 0.505649298 | 0.2288953073 | 0.960468681 | 0.221545061 | 0.839826083 | 0.6066894195 |
| 0.040323350 | 0.138626929 | 0.1868987777 | 0.199666061 | 0.190580377 | 0.613233235 | 0.5928157147 |
| 0.579528123 | 0.425728254 | 0.1607610545 | 0.007308493 | 0.824137168 | 0.810910961 | 0.5928157147 |
| 0.013418341 | 0.121322417 | 0.1626148411 | 0.632958948 | 0.161020099 | 0.613233235 | 0.5928157147 |
| 0.105711902 | 0.170823696 | 0.3441231202 | 0.748984247 | 0.333827061 | 0.613233235 | 0.757210127 |
| 0.060808970 | 0.113402486 | 0.2188848206 | 0.255248484 | 0.221545061 | 0.613233235 | 0.6066894195 |
| 0.197988745 | 0.058229805 | 0.5994132043 | 0.348580733 | 0.516492380 | 0.453314142 | 0.8978474649 |
| 0.026387167 | 0.302858373 | 0.0543765976 | 0.467004283 | 0.168280905 | 0.712607937 | 0.5437080554 |
| 0.214392049 | 0.136182794 | 0.0679635069 | 0.003883678 | 0.541234625 | 0.613233235 | 0.5437080554 |
| 0.892752081 | 0.232752993 | 0.369613781 | 0.110594004 | 0.973911362 | 0.638656179 | 0.7585775013 |
| 0.028170406 | 0.925711280 | 0.823508827 | 0.009377156 | 0.169022440 | 0.997735319 | 0.9535244924 |
| 0.064712568 | 0.203626299 | 0.1609762619 | 0.536937710 | 0.222783661 | 0.613233235 | 0.5928157147 |
| 0.323632817 | 0.543682310 | 0.0418046292 | 0.233872000 | 0.627841620 | 0.840915049 | 0.4877585812 |
| 0.244476819 | 0.987255065 | 0.2055274741 | 0.205020099 | 0.561857608 | 0.997735319 | 0.5979782654 |
| 0.446262805 | 0.269535363 | 0.3953075818 | 0.792680408 | 0.754246995 | 0.696284431 | 0.7721894584 |
| 0.064978568 | 0.081980713 | 0.2548873782 | 0.001333661 | 0.222783661 | 0.491884281 | 0.6372184455 |
| 0.021899879 | 0.073777180 | 0.0101034750 | 0.000014104 | 0.168280905 | 0.465961140 | 0.1938341562 |
| 0.658741420 | 0.191896086 | 0.1926651073 | 0.307578822 | 0.866248951 | 0.613233235 | 0.5928157147 |
| 0.055119699 | 0.056383691 | 0.0049821431 | 0.347796753 | 0.221545061 | 0.453314142 | 0.1195714349 |
| 0.058042175 | 0.004534043 | 0.1374931891 | 0.226778506 | 0.221545061 | 0.136021294 | 0.5928157147 |
| 0.967203853 | 0.835866919 | 0.9819086792 | 0.911544080 | 0.983580975 | 0.997735319 | 0.9953877755 |
| 0.129495852 | 0.191722196 | 0.3589651594 | 0.217609497 | 0.379012252 | 0.613233235 | 0.757210127 |
| 0.015886142 | 0.030459760 | 0.0046925929 | 0.098490188 | 0.168280905 | 0.453314142 | 0.1195714349 |
| 0.269111523 | 0.862210131 | 0.5650613534 | 0.514951146 | 0.566550576 | 0.997735319 | 0.8978474649 |
| 0.602051461 | 0.198528572 | 0.1751539938 | 0.158432361 | 0.830415809 | 0.613233235 | 0.5928157147 |
| 0.113773013 | 0.915805137 | 0.8232549644 | 0.224651686 | 0.350070812 | 0.997735319 | 0.9535244924 |
| 0.836413177 | 0.440184469 | 0.946541058 | 0.915992202 | 0.955900774 | 0.812648251 | 0.9953877755 |
| 0.68578042 | 0.997735319 | 0.7811210556 | 0.000109686 | 0.866248951 | 0.997735319 | 0.9440157185 |
| 0.039520943 | 0.146212347 | 0.0113069924 | 0.640171623 | 0.190580377 | 0.613233235 | 0.1938341562 |
| 0.963762313 | 0.505775153 | 0.9283710877 | 0.607292922 | 0.983580975 | 0.839826083 | 0.9948712152 |
| 0.021861208 | 0.015728071 | 0.0278213845 | 0.013488141 | 0.168280905 | 0.377473711 | 0.417320768 |
| 0.168755498 | 0.510894201 | 0.1353098773 | 0.540414466 | 0.460242268 | 0.839826083 | 0.5928157147 |
| 0.845737037 | 0.956457947 | 0.6347306827 | 0.011016512 | 0.956033509 | 0.997735319 | 0.8978474649 |
| 0.681909445 | 0.578145203 | 0.9158416652 | 0.001272195 | 0.866248951 | 0.857029730 | 0.9948712152 |
| 0.020275649 | 0.000510897 | 0.0032763340 | 0.006870384 | 0.168280905 | 0.020435886 | 0.1195714349 |
| 0.532142433 | 0.599903507 | 0.9299806264 | 0.089313706 | 0.788359160 | 0.857029730 | 0.9948712152 |
| 0.497084683 | 0.655201675 | 0.4597968228 | 0.035129172 | 0.774677428 | 0.914234895 | 0.7882231248 |
| 0.700229152 | 0.33136677 | 0.3596748103 | 0.009473134 | 0.875286441 | 0.759672525 | 0.757210127 |
| 0.026644476 | 0.964651670 | 0.1984495128 | 0.565137909 | 0.168280905 | 0.997735319 | 0.5953485384 |
| 0.540473291 | 0.961926464 | 0.1213865217 | 0.002891340 | 0.790936523 | 0.997735319 | 0.5928157147 |
| 0.882870207 | 0.981931756 | 0.1875366336 | 0.388610181 | 0.973911362 | 0.997735319 | 0.5928157147 |

| | | | | | | |
|---|---|---|---|---|---|---|
| 0.327441522 | 0.897673387 | 0.4584536795 | 0.000613156 | 0.627841620 | 0.997735319 | 0.7882231248 |
| 0.011334001 | 0.000052821 | 0.0004846224 | 0.095615958 | 0.161020099 | 0.003721366 | 0.0290773457 |
| 0.152057959 | 0.209502677 | 0.0873364710 | 0.762571532 | 0.424347793 | 0.613233235 | 0.5515987647 |
| 0.092054034 | 0.448981618 | 0.5495473988 | 0.894379243 | 0.306846782 | 0.816330215 | 0.8949850654 |
| 0.422343986 | 0.170366426 | 0.5842115419 | 0.441669498 | 0.724018263 | 0.613233235 | 0.8978474649 |
| 0.482147343 | 0.036491274 | 0.7396669805 | 0.011630882 | 0.771435749 | 0.453314142 | 0.9289265075 |
| 0.932072912 | 0.987306863 | 0.6870387291 | 0.079243620 | 0.983580975 | 0.997735319 | 0.9059851372 |
| 0.795365828 | 0.336454286 | 0.3501283184 | 0.086254846 | 0.937686000 | 0.759672525 | 0.757210127 |
| 0.128824096 | 0.234173932 | 0.6394119482 | 0.672040453 | 0.379012252 | 0.638656179 | 0.8978474649 |
| 0.615091020 | 0.544949898 | 0.5861985137 | 0.552144187 | 0.830515207 | 0.840915049 | 0.8978474649 |
| 0.922038320 | 0.888175881 | 0.4053994657 | 0.198258405 | 0.983580975 | 0.997735319 | 0.7721894584 |
| 0.266595676 | 0.973187961 | 0.9953877755 | 0.950061408 | 0.566550576 | 0.997735319 | 0.9953877755 |
| 0.005386755 | 0.550460878 | 0.4529827042 | 0.057303868 | 0.161020099 | 0.840915049 | 0.7882231248 |
| 0.336421203 | 0.392883068 | 0.415785445 | 0.006176858 | 0.627841620 | 0.794060801 | 0.7755275948 |
| 0.975384467 | 0.163135781 | 0.2662678912 | 0.251205387 | 0.983580975 | 0.613233235 | 0.6520846316 |
| 0.819611514 | 0.832646478 | 0.5223641345 | 0.302306288 | 0.954887201 | 0.997735319 | 0.8706068908 |
| 0.265889556 | 0.149503647 | 0.0846052451 | 0.003658451 | 0.566550576 | 0.613233235 | 0.5515987647 |
| 0.951680060 | 0.397030400 | 0.6733855986 | 0.039249358 | 0.983580975 | 0.794060801 | 0.8978474649 |
| 0.008237275 | 0.110258855 | 0.0770116296 | 0.345926374 | 0.161020099 | 0.613233235 | 0.5515987647 |
| 0.21649385 | 0.191830979 | 0.1376326802 | 0.622395037 | 0.541234625 | 0.613233235 | 0.5928157147 |
| 0.385408447 | 0.792285605 | 0.6947440907 | 0.213284494 | 0.685022910 | 0.997735319 | 0.9061879443 |
| 0.414309716 | 0.055084361 | 0.3988667773 | 0.553319263 | 0.720538637 | 0.453314142 | 0.7721894584 |
| 0.047934476 | 0.209521355 | 0.0599218516 | 0.935793369 | 0.213042118 | 0.613233235 | 0.5437080554 |
| 0.966932885 | 0.937266398 | 0.7087673991 | 0.383632051 | 0.983580975 | 0.997735319 | 0.9145385795 |
| 0.590631637 | 0.227756834 | 0.838415279 | 0.475197570 | 0.824137168 | 0.638656179 | 0.9581888902 |
| 0.522970796 | 0.439234358 | 0.6688718019 | 0.006834792 | 0.784456194 | 0.812648251 | 0.8978474649 |
| 0.248153776 | 0.025952681 | 0.14144584 | 0.105164347 | 0.561857608 | 0.444903106 | 0.5928157147 |
| 0.060924891 | 0.967256524 | 0.9122938955 | 0.042288150 | 0.221545061 | 0.997735319 | 0.9948712152 |
| 0.481699647 | 0.071686041 | 0.1596190004 | 0.545978077 | 0.771435749 | 0.465961140 | 0.5928157147 |
| 0.956774038 | 0.981486122 | 0.6294935129 | 0.336885815 | 0.983580975 | 0.997735319 | 0.8978474649 |
| 0.615965445 | 0.952305483 | 0.936837061 | 0.000677915 | 0.830515207 | 0.997735319 | 0.9948712152 |
| 0.714488095 | 0.635774270 | 0.551907457 | 0.191410514 | 0.876233733 | 0.897563676 | 0.8949850654 |
| 0.340080877 | 0.343453844 | 0.2347017435 | 0.283682054 | 0.627841620 | 0.759672525 | 0.6066894195 |
| 0.025651213 | 0.020037884 | 0.0408910515 | 0.002907420 | 0.168280905 | 0.400757689 | 0.4877585812 |
| 0.318034491 | 0.282933820 | 0.2092923929 | 0.818981879 | 0.627841620 | 0.698484187 | 0.5979782654 |
| 0.681474454 | 0.272711402 | 0.3729672715 | 0.349239171 | 0.866248951 | 0.696284431 | 0.7585775013 |
| 0.573416440 | 0.946882214 | 0.9575600484 | 0.001454402 | 0.824137168 | 0.997735319 | 0.9953877755 |
| 0.715590882 | 0.248062225 | 0.2305656554 | 0.638183437 | 0.876233733 | 0.661499268 | 0.6066894195 |
| 0.887047828 | 0.182533110 | 0.2815841683 | 0.488556028 | 0.973911362 | 0.613233235 | 0.6625509843 |
| 0.852463212 | 0.923843234 | 0.6540742283 | 0.150283502 | 0.956033509 | 0.997735319 | 0.8978474649 |
| 0.333740213 | 0.500689253 | 0.9679130587 | 0.254932410 | 0.627841620 | 0.839826083 | 0.9953877755 |
| 0.797033100 | 0.404560529 | 0.7713808689 | 0.028823970 | 0.937686000 | 0.795856779 | 0.9440157185 |
| 0.652495869 | 0.176984519 | 0.2376200226 | 0.333881672 | 0.866248951 | 0.613233235 | 0.6066894195 |
| 0.348084781 | 0.179117425 | 0.4023048237 | 0.745841474 | 0.632881421 | 0.613233235 | 0.7721894584 |
| 0.336427830 | 0.060487813 | 0.1835292044 | 0.440248461 | 0.627841620 | 0.453314142 | 0.5928157147 |
| 0.247520150 | 0.547965210 | 0.6008799311 | 0.166686525 | 0.561857608 | 0.840915049 | 0.8978474649 |

| | | | | | | |
|---|---|---|---|---|---|---|
| 0.512224951 | 0.553602407 | 0.6519367304 | 0.043038525 | 0.783457173 | 0.840915049 | 0.8978474649 |
| 0.388179649 | 0.921672772 | 0.0805504069 | 0.266437075 | 0.685022910 | 0.997735319 | 0.5515987647 |
| 0.935495908 | 0.977011262 | 0.4503904215 | 0.199668900 | 0.983580975 | 0.997735319 | 0.7882231248 |
| 0.038290131 | 0.702287770 | 0.8547066334 | 0.057315873 | 0.190580377 | 0.957665141 | 0.9675924152 |
| 0.793205169 | 0.202531053 | 0.7359225566 | 0.190144952 | 0.937686000 | 0.613233235 | 0.9289265075 |
| 0.587436417 | 0.943686627 | 0.4228865589 | 0.809370833 | 0.824137168 | 0.997735319 | 0.7755275948 |
| 0.228138923 | 0.052569545 | 0.9776552592 | 0.323347805 | 0.558707567 | 0.453314142 | 0.9953877755 |
| 0.259616462 | 0.711831640 | 0.8717968425 | 0.038095561 | 0.566550576 | 0.959772998 | 0.977716085 |
| 0.008225071 | 0.591231708 | 0.8263878934 | 0.001612909 | 0.161020099 | 0.857029730 | 0.9535244924 |
| 0.675126094 | 0.048837799 | 0.9235073977 | 0.015147368 | 0.866248951 | 0.453314142 | 0.9948712152 |
| 0.004089990 | 0.697685851 | 0.6236430334 | 0.677272316 | 0.161020099 | 0.957665141 | 0.8978474649 |
| 0.184274373 | 0.960854652 | 0.4265401771 | 0.413049316 | 0.491398329 | 0.997735319 | 0.7755275948 |
| 0.331439669 | 0.473974000 | 0.743141206 | 0.626367199 | 0.627841620 | 0.839826083 | 0.9289265075 |
| 0.242170188 | 0.806088876 | 0.3501948236 | 0.945080373 | 0.561857608 | 0.997735319 | 0.757210127 |
| 0.004885562 | 0.064219503 | 0.0447112032 | 0.159256092 | 0.161020099 | 0.453314142 | 0.4877585812 |
| 0.991482551 | 0.492739934 | 0.332885926 | 0.065117443 | 0.991482551 | 0.839826083 | 0.757210127 |
| 0.835167472 | 0.519408254 | 0.9948307884 | 0.505386929 | 0.955900774 | 0.840915049 | 0.9953877755 |
| 0.035048984 | 0.348183240 | 0.8139353891 | 0.001264754 | 0.190580377 | 0.759672525 | 0.9535244924 |
| 0.041278888 | 0.483173342 | 0.1923456044 | 0.103837614 | 0.190580377 | 0.839826083 | 0.5928157147 |
| 0.003697740 | 0.599920811 | 0.1176630853 | 0.077293436 | 0.161020099 | 0.857029730 | 0.5928157147 |
| 0.472754442 | 0.988193648 | 0.6600647536 | 0.440149933 | 0.771435749 | 0.997735319 | 0.8978474649 |
| 0.515775972 | 0.995045172 | 0.1389903269 | 0.214430063 | 0.783457173 | 0.997735319 | 0.5928157147 |
| 0.012183706 | 0.044515836 | 0.1043992389 | 0.002843895 | 0.161020099 | 0.453314142 | 0.5928157147 |
| 0.147130374 | 0.367140347 | 0.0668916843 | 0.001477305 | 0.420372498 | 0.786729315 | 0.5437080554 |
| 0.049736111 | 0.287568214 | 0.1633234022 | 0.380322300 | 0.213154765 | 0.698484187 | 0.5928157147 |
| 0.025295846 | 0.598189224 | 0.2767059398 | 0.005702380 | 0.168280905 | 0.857029730 | 0.6625509843 |
| 0.023132731 | 0.894252113 | 0.7866797654 | 0.025651988 | 0.168280905 | 0.997735319 | 0.9440157185 |
| 0.009537773 | 0.846947375 | 0.6038872084 | 0.990514375 | 0.161020099 | 0.997735319 | 0.8978474649 |

| t_q_batch | Genus | Family | Order |
|---|---|---|---|
| 0.746557882! | Acetatifactor | Lachnospiraceae | Eubacteriales |
| 0.478498591! | Acutalibacter | Oscillospiraceae | Eubacteriales |
| 0.427861930: | Akkermansia | Akkermansiaceae | Verrucomicrobiales |
| 0.935287286! | Anaerotruncus | Oscillospiraceae | Eubacteriales |
| 0.006602889( | Bacteria_unclassified | Bacteria_unclassified | Bacteria_unclassified |
| 0.968539846! | Bacteria_unclassified | Bacteria_unclassified | Bacteria_unclassified |
| 0.427861930: | Bacteria_unclassified | Bacteria_unclassified | Bacteria_unclassified |
| 0.041762821! | Bacteria_unclassified | Bacteria_unclassified | Bacteria_unclassified |
| 0.746557882! | Bacteroides | Bacteroidaceae | Bacteroidales |
| 0.839982334 | Clostridia_unclassified | Clostridia_unclassified | Clostridia_unclassified |
| 0.464088153! | Clostridiaceae_unclassified | Clostridiaceae | Eubacteriales |
| 0.537291032 | Clostridiaceae_unclassified | Clostridiaceae | Eubacteriales |
| 0.651633884: | Eubacteriales_unclassified | Eubacteriales_unclassified | Eubacteriales |
| 0.029127591! | Clostridium | Clostridiaceae | Eubacteriales |
| 0.288506097 | Erysipelatoclostridium | Erysipelotrichaceae | Erysipelotrichales |
| 0.049425050( | Eubacteriaceae_unclassified | Eubacteriaceae | Eubacteriales |
| 0.698929596 | Eubacteriaceae_unclassified | Eubacteriaceae | Eubacteriales |
| 0.445470477: | GGB20146 | FGB77306 | OFGB77306 |
| 0.431621261! | GGB20149 | Lachnospiraceae | Eubacteriales |
| 0.872675679 | GGB22635 | Eggerthellaceae | Eggerthellales |
| 0.017595378: | GGB25041 | Lachnospiraceae | Eubacteriales |
| 0.001692491: | GGB28379 | FGB9506 | OFGB9506 |
| 0.519851531 | GGB28382 | FGB9508 | OFGB9508 |
| 0.537291032 | GGB28392 | FGB9512 | OFGB9512 |
| 0.438926141 | GGB28404 | FGB2838 | OFGB2838 |
| 0.955817950: | GGB28418 | FGB2838 | OFGB2838 |
| 0.435218994: | GGB28422 | FGB2838 | OFGB2838 |
| 0.274856340( | GGB28431 | Pumilibacteraceae | Eubacteriales |
| 0.686601527! | GGB28456 | FGB2833 | OFGB2833 |
| 0.390014921: | GGB28782 | Eubacteriaceae | Eubacteriales |
| 0.438926141 | GGB28792 | Lachnospiraceae | Eubacteriales |
| 0.955817950: | GGB28798 | Lachnospiraceae | Eubacteriales |
| 0.006581191! | GGB28810 | FGB9622 | OFGB9622 |
| 0.746557882! | GGB28828 | FGB77305 | OFGB77305 |
| 0.746557882! | GGB28851 | Clostridiaceae | Eubacteriales |
| 0.062252961: | GGB28865 | Lachnospiraceae | Eubacteriales |
| 0.698929596 | GGB28868 | Lachnospiraceae | Eubacteriales |
| 0.055082564( | GGB28869 | Lachnospiraceae | Eubacteriales |
| 0.017595378: | GGB28875 | Lachnospiraceae | Eubacteriales |
| 0.041222306 | GGB28881 | FGB9633 | OFGB9633 |
| 0.261405970! | GGB28892 | Bacteria_unclassified | Bacteria_unclassified |
| 0.140516689 | GGB28893 | Bacteria_unclassified | Bacteria_unclassified |
| 0.049425050( | GGB28898 | Bacteria_unclassified | Bacteria_unclassified |
| 0.706422386! | GGB28901 | Bacteria_unclassified | Bacteria_unclassified |
| 0.024920750 | GGB28904 | Bacteria_unclassified | Bacteria_unclassified |
| 0.575718787: | GGB28909 | Bacteria_unclassified | Bacteria_unclassified |

| | | | |
|---|---|---|---|
| 0.016269970 | GGB28924 | Lachnospiraceae | Eubacteriales |
| 0.273188452 | GGB28927 | FGB77359 | OFGB77359 |
| 0.847301702 | GGB28934 | FGB9639 | OFGB9639 |
| 0.949783267 | GGB28949 | Lachnospiraceae | Eubacteriales |
| 0.623533408 | GGB28951 | Clostridiaceae | Eubacteriales |
| 0.055828235 | GGB28951 | Clostridiaceae | Eubacteriales |
| 0.243826523 | GGB28954 | Clostridiaceae | Eubacteriales |
| 0.258764540 | GGB28960 | Clostridiaceae | Eubacteriales |
| 0.774025504 | GGB28964 | Clostridiaceae | Eubacteriales |
| 0.698929596 | GGB28967 | Clostridiaceae | Eubacteriales |
| 0.427861930 | GGB28996 | FGB9656 | OFGB9656 |
| 0.966164143 | GGB29531 | FGB9827 | OFGB9827 |
| 0.191052912 | GGB29685 | Eubacteriaceae | Eubacteriales |
| 0.041179056 | GGB30141 | FGB77303 | OFGB77303 |
| 0.464088153 | GGB30145 | Clostridiaceae | Eubacteriales |
| 0.518239352 | GGB30286 | Eubacteriales_unclassified | Eubacteriales |
| 0.029127591 | GGB30300 | FGB72709 | OFGB72709 |
| 0.147185092 | GGB30303 | Oscillospiraceae | Eubacteriales |
| 0.537291032 | GGB30450 | Oscillospiraceae | Eubacteriales |
| 0.746557882 | GGB30453 | Oscillospiraceae | Eubacteriales |
| 0.435218994 | GGB30454 | Oscillospiraceae | Eubacteriales |
| 0.698929596 | GGB30455 | Oscillospiraceae | Eubacteriales |
| 0.966164143 | GGB30456 | Oscillospiraceae | Eubacteriales |
| 0.575448077 | GGB30457 | Oscillospiraceae | Eubacteriales |
| 0.655444924 | GGB30461 | Oscillospiraceae | Eubacteriales |
| 0.041222306 | GGB30461 | Oscillospiraceae | Eubacteriales |
| 0.280438260 | GGB30461 | Oscillospiraceae | Eubacteriales |
| 0.151900678 | GGB30463 | Oscillospiraceae | Eubacteriales |
| 0.698929596 | GGB30473 | Oscillospiraceae | Eubacteriales |
| 0.537291032 | GGB30475 | Oscillospiraceae | Eubacteriales |
| 0.016269970 | GGB30861 | FGB77153 | OFGB77153 |
| 0.427861930 | GGB31312 | FGB1791 | OFGB1791 |
| 0.493360095 | GGB31438 | FGB10290 | OFGB10290 |
| 0.024920750 | GGB3171 | Oscillospiraceae | Eubacteriales |
| 0.885385815 | GGB31762 | FGB10289 | OFGB10289 |
| 0.537291032 | GGB31823 | FGB1765 | OFGB1765 |
| 0.017595378 | GGB31838 | FGB1765 | OFGB1765 |
| 0.746557882 | GGB31841 | FGB1765 | OFGB1765 |
| 0.666212766 | GGB32371 | FGB10667 | OFGB10667 |
| 0.383702558 | GGB3793 | Lachnospiraceae | Eubacteriales |
| 0.464088153 | GGB42601 | Clostridiaceae | Eubacteriales |
| 0.119271600 | GGB45514 | Oscillospiraceae | Eubacteriales |
| 0.537291032 | GGB45564 | FGB75721 | OFGB75721 |
| 0.839982334 | GGB45624 | Oscillospiraceae | Eubacteriales |
| 0.623533408 | GGB45656 | Christensenellaceae | Eubacteriales |
| 0.400047661 | GGB47127 | FGB10299 | OFGB10299 |

| | | | |
|---|---|---|---|
| 0.151900678 | GGB74395 | Oscillospiraceae | Eubacteriales |
| 0.477200732 | GGB75053 | Oscillospiraceae | Eubacteriales |
| 0.427861930 | GGB75109 | Lachnospiraceae | Eubacteriales |
| 0.191052912 | Lachnospiraceae_unclassified | Lachnospiraceae | Eubacteriales |
| 0.427861930 | Lachnospiraceae_unclassified | Lachnospiraceae | Eubacteriales |
| 0.882949999 | Lachnospiraceae_unclassified | Lachnospiraceae | Eubacteriales |
| 0.537291032 | Lachnospiraceae_unclassified | Lachnospiraceae | Eubacteriales |
| 0.147185092 | Lachnospiraceae_unclassified | Lachnospiraceae | Eubacteriales |
| 0.017595378 | Lactobacillus | Lactobacillaceae | Lactobacillales |
| 0.067321635 | Leptogranulimonas | Atopobiaceae | Coriobacteriales |
| 0.774025504 | Muribaculaceae_unclassified | Muribaculaceae | Bacteroidales |
| 0.604462414 | Neglectibacter | Oscillospiraceae | Eubacteriales |
| 0.746557882 | Oscillibacter | Oscillospiraceae | Eubacteriales |
| 0.966164143 | Oscillospiraceae_unclassified | Oscillospiraceae | Eubacteriales |
| 0.390014921 | Oscillospiraceae_unclassified | Oscillospiraceae | Eubacteriales |
| 0.211191709 | Oscillospiraceae_unclassified | Oscillospiraceae | Eubacteriales |
| 0.681420578 | Oscillospiraceae_unclassified | Oscillospiraceae | Eubacteriales |
| 0.017595378 | Parasutterella | Sutterellaceae | Burkholderiales |
| 0.280438260 | Schaedlerella | Lachnospiraceae | Eubacteriales |
| 0.243826523 | Turicibacter | Turicibacteraceae | Erysipelotrichales |
| 0.623533408 | Bacteria_unclassified | Bacteria_unclassified | Bacteria_unclassified |
| 0.435218994 | Bacteria_unclassified | Bacteria_unclassified | Bacteria_unclassified |
| 0.024920750 | Bacteria_unclassified | Bacteria_unclassified | Bacteria_unclassified |
| 0.017595378 | Bacteria_unclassified | Bacteria_unclassified | Bacteria_unclassified |
| 0.575448077 | Bacteria_unclassified | Bacteria_unclassified | Bacteria_unclassified |
| 0.040252096 | | | |
| 0.109937094 | | | |
| 0.990514375 | | | |

| Class | Phylum | Kingdom |
|---|---|---|
| Clostridia | Firmicutes | Bacteria |
| Clostridia | Firmicutes | Bacteria |
| Verrucomicrobiae | Verrucomicrobia | Bacteria |
| Clostridia | Firmicutes | Bacteria |
| Bacteria_unclassified | Bacteria_unclassified | Bacteria |
| Bacteria_unclassified | Bacteria_unclassified | Bacteria |
| Bacteria_unclassified | Bacteria_unclassified | Bacteria |
| Bacteria_unclassified | Bacteria_unclassified | Bacteria |
| Bacteroidia | Bacteroidota | Bacteria |
| Clostridia | Firmicutes | Bacteria |
| Clostridia | Firmicutes | Bacteria |
| Clostridia | Firmicutes | Bacteria |
| Clostridia | Firmicutes | Bacteria |
| Clostridia | Firmicutes | Bacteria |
| Erysipelotrichia | Firmicutes | Bacteria |
| Clostridia | Firmicutes | Bacteria |
| Clostridia | Firmicutes | Bacteria |
| CFGB77306 | Firmicutes | Bacteria |
| Clostridia | Firmicutes | Bacteria |
| Coriobacteriia | Actinobacteria | Bacteria |
| Clostridia | Firmicutes | Bacteria |
| CFGB9506 | Firmicutes | Bacteria |
| CFGB9508 | Firmicutes | Bacteria |
| CFGB9512 | Firmicutes | Bacteria |
| CFGB2838 | Firmicutes | Bacteria |
| CFGB2838 | Firmicutes | Bacteria |
| CFGB2838 | Firmicutes | Bacteria |
| Clostridia | Firmicutes | Bacteria |
| CFGB2833 | Firmicutes | Bacteria |
| Clostridia | Firmicutes | Bacteria |
| Clostridia | Firmicutes | Bacteria |
| Clostridia | Firmicutes | Bacteria |
| CFGB9622 | Firmicutes | Bacteria |
| CFGB77305 | Firmicutes | Bacteria |
| Clostridia | Firmicutes | Bacteria |
| Clostridia | Firmicutes | Bacteria |
| Clostridia | Firmicutes | Bacteria |
| Clostridia | Firmicutes | Bacteria |
| Clostridia | Firmicutes | Bacteria |
| CFGB9633 | Firmicutes | Bacteria |
| Bacteria_unclassified | Bacteria_unclassified | Bacteria |
| Bacteria_unclassified | Bacteria_unclassified | Bacteria |
| Bacteria_unclassified | Bacteria_unclassified | Bacteria |
| Bacteria_unclassified | Bacteria_unclassified | Bacteria |
| Bacteria_unclassified | Bacteria_unclassified | Bacteria |
| Bacteria_unclassified | Bacteria_unclassified | Bacteria |

| | | |
|---|---|---|
| Clostridia | Firmicutes | Bacteria |
| CFGB77359 | Bacteria_unclassified | Bacteria |
| CFGB9639 | Firmicutes | Bacteria |
| Clostridia | Firmicutes | Bacteria |
| Clostridia | Firmicutes | Bacteria |
| Clostridia | Firmicutes | Bacteria |
| Clostridia | Firmicutes | Bacteria |
| Clostridia | Firmicutes | Bacteria |
| Clostridia | Firmicutes | Bacteria |
| Clostridia | Firmicutes | Bacteria |
| CFGB9656 | Firmicutes | Bacteria |
| CFGB9827 | Firmicutes | Bacteria |
| Clostridia | Firmicutes | Bacteria |
| CFGB77303 | Bacteria_unclassified | Bacteria |
| Clostridia | Firmicutes | Bacteria |
| Clostridia | Firmicutes | Bacteria |
| CFGB72709 | Firmicutes | Bacteria |
| Clostridia | Firmicutes | Bacteria |
| Clostridia | Firmicutes | Bacteria |
| Clostridia | Firmicutes | Bacteria |
| Clostridia | Firmicutes | Bacteria |
| Clostridia | Firmicutes | Bacteria |
| Clostridia | Firmicutes | Bacteria |
| Clostridia | Firmicutes | Bacteria |
| Clostridia | Firmicutes | Bacteria |
| Clostridia | Firmicutes | Bacteria |
| Clostridia | Firmicutes | Bacteria |
| Clostridia | Firmicutes | Bacteria |
| Clostridia | Firmicutes | Bacteria |
| Clostridia | Firmicutes | Bacteria |
| CFGB77153 | Actinobacteria | Bacteria |
| CFGB1791 | Tenericutes | Bacteria |
| CFGB10290 | Firmicutes | Bacteria |
| Clostridia | Firmicutes | Bacteria |
| CFGB10289 | Firmicutes | Bacteria |
| CFGB1765 | Firmicutes | Bacteria |
| CFGB1765 | Firmicutes | Bacteria |
| CFGB1765 | Firmicutes | Bacteria |
| CFGB10667 | Firmicutes | Bacteria |
| Clostridia | Firmicutes | Bacteria |
| Clostridia | Firmicutes | Bacteria |
| Clostridia | Firmicutes | Bacteria |
| CFGB75721 | Firmicutes | Bacteria |
| Clostridia | Firmicutes | Bacteria |
| Clostridia | Firmicutes | Bacteria |
| CFGB10299 | Firmicutes | Bacteria |

| | | |
|---|---|---|
| Clostridia | Firmicutes | Bacteria |
| Clostridia | Firmicutes | Bacteria |
| Clostridia | Firmicutes | Bacteria |
| Clostridia | Firmicutes | Bacteria |
| Clostridia | Firmicutes | Bacteria |
| Clostridia | Firmicutes | Bacteria |
| Clostridia | Firmicutes | Bacteria |
| Clostridia | Firmicutes | Bacteria |
| Bacilli | Firmicutes | Bacteria |
| Coriobacteriia | Actinobacteria | Bacteria |
| Bacteroidia | Bacteroidota | Bacteria |
| Clostridia | Firmicutes | Bacteria |
| Clostridia | Firmicutes | Bacteria |
| Clostridia | Firmicutes | Bacteria |
| Clostridia | Firmicutes | Bacteria |
| Clostridia | Firmicutes | Bacteria |
| Clostridia | Firmicutes | Bacteria |
| Betaproteobacteria | Proteobacteria | Bacteria |
| Clostridia | Firmicutes | Bacteria |
| Erysipelotrichia | Firmicutes | Bacteria |
| Bacteria_unclassified | Bacteria_unclassified | Bacteria |
| Bacteria_unclassified | Bacteria_unclassified | Bacteria |
| Bacteria_unclassified | Bacteria_unclassified | Bacteria |
| Bacteria_unclassified | Bacteria_unclassified | Bacteria |
| Bacteria_unclassified | Bacteria_unclassified | Bacteria |

**MetaPhlan Annotation**

k__Bacteria|p__Firmicutes|c__Clostridia|o__Eubacteriales|f__Lachnospiraceae|g__Acetatifactor|s__Acetatifactor_SGB4154

k__Bacteria|p__Firmicutes|c__Clostridia|o__Eubacteriales|f__Oscillospiraceae|g__Acutalibacter|s__Acutalibacter_muris

k__Bacteria|p__Verrucomicrobia|c__Verrucomicrobiae|o__Verrucomicrobiales|f__Akkermansiaceae|g__Akkermansia|s__A

k__Bacteria|p__Firmicutes|c__Clostridia|o__Eubacteriales|f__Oscillospiraceae|g__Anaerotruncus|s__Anaerotruncus_sp_1

k__Bacteria|p__Bacteria_unclassified|c__Bacteria_unclassified|o__Bacteria_unclassified|f__Bacteria_unclassified|g__Bacte

k__Bacteria|p__Bacteria_unclassified|c__Bacteria_unclassified|o__Bacteria_unclassified|f__Bacteria_unclassified|g__Bacte

k__Bacteria|p__Bacteria_unclassified|c__Bacteria_unclassified|o__Bacteria_unclassified|f__Bacteria_unclassified|g__Bacte

k__Bacteria|p__Bacteria_unclassified|c__Bacteria_unclassified|o__Bacteria_unclassified|f__Bacteria_unclassified|g__Bacte

k__Bacteria|p__Bacteroidota|c__Bacteroidia|o__Bacteroidales|f__Bacteroidaceae|g__Bacteroides|s__Bacteroides_thetaio

k__Bacteria|p__Firmicutes|c__Clostridia|o__Clostridia_unclassified|f__Clostridia_unclassified|g__Clostridia_unclassified|s_

k__Bacteria|p__Firmicutes|c__Clostridia|o__Eubacteriales|f__Clostridiaceae|g__Clostridiaceae_unclassified|s__Clostridiac

k__Bacteria|p__Firmicutes|c__Clostridia|o__Eubacteriales|f__Clostridiaceae|g__Clostridiaceae_unclassified|s__Clostridiac

k__Bacteria|p__Firmicutes|c__Clostridia|o__Eubacteriales|f__Eubacteriales_unclassified|g__Eubacteriales_unclassified|s_

k__Bacteria|p__Firmicutes|c__Clostridia|o__Eubacteriales|f__Clostridiaceae|g__Clostridium|s__Clostridium_SGB65123

k__Bacteria|p__Firmicutes|c__Erysipelotrichia|o__Erysipelotrichales|f__Erysipelotrichaceae|g__Erysipelatoclostridium|s_

k__Bacteria|p__Firmicutes|c__Clostridia|o__Eubacteriales|f__Eubacteriaceae|g__Eubacteriaceae_unclassified|s__Eubacte

k__Bacteria|p__Firmicutes|c__Clostridia|o__Eubacteriales|f__Eubacteriaceae|g__Eubacteriaceae_unclassified|s__Eubacte

k__Bacteria|p__Firmicutes|c__CFGB77306|o__OFGB77306|f__FGB77306|g__GGB20146|s__GGB20146_SGB29427

k__Bacteria|p__Firmicutes|c__Clostridia|o__Eubacteriales|f__Lachnospiraceae|g__GGB20149|s__GGB20149_SGB29430

k__Bacteria|p__Actinobacteria|c__Coriobacteriia|o__Eggerthellales|f__Eggerthellaceae|g__GGB22635|s__GGB22635_SGB

k__Bacteria|p__Firmicutes|c__Clostridia|o__Eubacteriales|f__Lachnospiraceae|g__GGB25041|s__GGB25041_SGB36960

k__Bacteria|p__Firmicutes|c__CFGB9506|o__OFGB9506|f__FGB9506|g__GGB28379|s__GGB28379_SGB40959

k__Bacteria|p__Firmicutes|c__CFGB9508|o__OFGB9508|f__FGB9508|g__GGB28382|s__GGB28382_SGB40962

k__Bacteria|p__Firmicutes|c__CFGB9512|o__OFGB9512|f__FGB9512|g__GGB28392|s__GGB28392_SGB40972

k__Bacteria|p__Firmicutes|c__CFGB2838|o__OFGB2838|f__FGB2838|g__GGB28404|s__GGB28404_SGB40986

k__Bacteria|p__Firmicutes|c__CFGB2838|o__OFGB2838|f__FGB2838|g__GGB28418|s__GGB28418_SGB41001

k__Bacteria|p__Firmicutes|c__CFGB2838|o__OFGB2838|f__FGB2838|g__GGB28422|s__GGB28422_SGB41005

k__Bacteria|p__Firmicutes|c__Clostridia|o__Eubacteriales|f__Pumilibacteraceae|g__GGB28431|s__GGB28431_SGB41014

k__Bacteria|p__Firmicutes|c__CFGB2833|o__OFGB2833|f__FGB2833|g__GGB28456|s__GGB28456_SGB41039

k__Bacteria|p__Firmicutes|c__Clostridia|o__Eubacteriales|f__Eubacteriaceae|g__GGB28782|s__GGB28782_SGB41435

k__Bacteria|p__Firmicutes|c__Clostridia|o__Eubacteriales|f__Lachnospiraceae|g__GGB28792|s__GGB28792_SGB41445

k__Bacteria|p__Firmicutes|c__Clostridia|o__Eubacteriales|f__Lachnospiraceae|g__GGB28798|s__GGB28798_SGB41451

k__Bacteria|p__Firmicutes|c__CFGB9622|o__OFGB9622|f__FGB9622|g__GGB28810|s__GGB28810_SGB41465

k__Bacteria|p__Firmicutes|c__CFGB77305|o__OFGB77305|f__FGB77305|g__GGB28828|s__GGB28828_SGB41484

k__Bacteria|p__Firmicutes|c__Clostridia|o__Eubacteriales|f__Clostridiaceae|g__GGB28851|s__GGB28851_SGB41518

k__Bacteria|p__Firmicutes|c__Clostridia|o__Eubacteriales|f__Lachnospiraceae|g__GGB28865|s__GGB28865_SGB41537

k__Bacteria|p__Firmicutes|c__Clostridia|o__Eubacteriales|f__Lachnospiraceae|g__GGB28868|s__GGB28868_SGB41542

k__Bacteria|p__Firmicutes|c__Clostridia|o__Eubacteriales|f__Lachnospiraceae|g__GGB28869|s__GGB28869_SGB41543

k__Bacteria|p__Firmicutes|c__Clostridia|o__Eubacteriales|f__Lachnospiraceae|g__GGB28875|s__GGB28875_SGB41555

k__Bacteria|p__Firmicutes|c__CFGB9633|o__OFGB9633|f__FGB9633|g__GGB28881|s__GGB28881_SGB41561

k__Bacteria|p__Bacteria_unclassified|c__Bacteria_unclassified|o__Bacteria_unclassified|f__Bacteria_unclassified|g__GGB2

k__Bacteria|p__Bacteria_unclassified|c__Bacteria_unclassified|o__Bacteria_unclassified|f__Bacteria_unclassified|g__GGB2

k__Bacteria|p__Bacteria_unclassified|c__Bacteria_unclassified|o__Bacteria_unclassified|f__Bacteria_unclassified|g__GGB2

k__Bacteria|p__Bacteria_unclassified|c__Bacteria_unclassified|o__Bacteria_unclassified|f__Bacteria_unclassified|g__GGB2

k__Bacteria|p__Bacteria_unclassified|c__Bacteria_unclassified|o__Bacteria_unclassified|f__Bacteria_unclassified|g__GGB2

k__Bacteria|p__Bacteria_unclassified|c__Bacteria_unclassified|o__Bacteria_unclassified|f__Bacteria_unclassified|g__GGB2

k__Bacteria|p__Firmicutes|c__Clostridia|o__Eubacteriales|f__Lachnospiraceae|g__GGB28924|s__GGB28924_SGB41621
k__Bacteria|p__Bacteria_unclassified|c__CFGB77359|o__OFGB77359|f__FGB77359|g__GGB28927|s__GGB28927_SGB416[illegible]
k__Bacteria|p__Firmicutes|c__CFGB9639|o__OFGB9639|f__FGB9639|g__GGB28934|s__GGB28934_SGB41635
k__Bacteria|p__Firmicutes|c__Clostridia|o__Eubacteriales|f__Lachnospiraceae|g__GGB28949|s__GGB28949_SGB41655
k__Bacteria|p__Firmicutes|c__Clostridia|o__Eubacteriales|f__Clostridiaceae|g__GGB28951|s__GGB28951_SGB102295
k__Bacteria|p__Firmicutes|c__Clostridia|o__Eubacteriales|f__Clostridiaceae|g__GGB28951|s__GGB28951_SGB41658
k__Bacteria|p__Firmicutes|c__Clostridia|o__Eubacteriales|f__Clostridiaceae|g__GGB28954|s__GGB28954_SGB41662
k__Bacteria|p__Firmicutes|c__Clostridia|o__Eubacteriales|f__Clostridiaceae|g__GGB28960|s__GGB28960_SGB41669
k__Bacteria|p__Firmicutes|c__Clostridia|o__Eubacteriales|f__Clostridiaceae|g__GGB28964|s__GGB28964_SGB94886
k__Bacteria|p__Firmicutes|c__Clostridia|o__Eubacteriales|f__Clostridiaceae|g__GGB28967|s__GGB28967_SGB41678
k__Bacteria|p__Firmicutes|c__CFGB9656|o__OFGB9656|f__FGB9656|g__GGB28996|s__GGB28996_SGB41712
k__Bacteria|p__Firmicutes|c__CFGB9827|o__OFGB9827|f__FGB9827|g__GGB29531|s__GGB29531_SGB42317
k__Bacteria|p__Firmicutes|c__Clostridia|o__Eubacteriales|f__Eubacteriaceae|g__GGB29685|s__GGB29685_SGB42494
k__Bacteria|p__Bacteria_unclassified|c__CFGB77303|o__OFGB77303|f__FGB77303|g__GGB30141|s__GGB30141_SGB430[illegible]
k__Bacteria|p__Firmicutes|c__Clostridia|o__Eubacteriales|f__Clostridiaceae|g__GGB30145|s__GGB30145_SGB43072
k__Bacteria|p__Firmicutes|c__Clostridia|o__Eubacteriales|f__Eubacteriales_unclassified|g__GGB30286|s__GGB30286_SG[illegible]
k__Bacteria|p__Firmicutes|c__CFGB72709|o__OFGB72709|f__FGB72709|g__GGB30300|s__GGB30300_SGB43264
k__Bacteria|p__Firmicutes|c__Clostridia|o__Eubacteriales|f__Oscillospiraceae|g__GGB30303|s__GGB30303_SGB43268
k__Bacteria|p__Firmicutes|c__Clostridia|o__Eubacteriales|f__Oscillospiraceae|g__GGB30450|s__GGB30450_SGB43507
k__Bacteria|p__Firmicutes|c__Clostridia|o__Eubacteriales|f__Oscillospiraceae|g__GGB30453|s__GGB30453_SGB43513
k__Bacteria|p__Firmicutes|c__Clostridia|o__Eubacteriales|f__Oscillospiraceae|g__GGB30454|s__GGB30454_SGB43514
k__Bacteria|p__Firmicutes|c__Clostridia|o__Eubacteriales|f__Oscillospiraceae|g__GGB30455|s__GGB30455_SGB43519
k__Bacteria|p__Firmicutes|c__Clostridia|o__Eubacteriales|f__Oscillospiraceae|g__GGB30456|s__GGB30456_SGB43520
k__Bacteria|p__Firmicutes|c__Clostridia|o__Eubacteriales|f__Oscillospiraceae|g__GGB30457|s__GGB30457_SGB63218
k__Bacteria|p__Firmicutes|c__Clostridia|o__Eubacteriales|f__Oscillospiraceae|g__GGB30461|s__GGB30461_SGB43530
k__Bacteria|p__Firmicutes|c__Clostridia|o__Eubacteriales|f__Oscillospiraceae|g__GGB30461|s__GGB30461_SGB43533
k__Bacteria|p__Firmicutes|c__Clostridia|o__Eubacteriales|f__Oscillospiraceae|g__GGB30461|s__GGB30461_SGB63209
k__Bacteria|p__Firmicutes|c__Clostridia|o__Eubacteriales|f__Oscillospiraceae|g__GGB30463|s__GGB30463_SGB43537
k__Bacteria|p__Firmicutes|c__Clostridia|o__Eubacteriales|f__Oscillospiraceae|g__GGB30473|s__GGB30473_SGB43557
k__Bacteria|p__Firmicutes|c__Clostridia|o__Eubacteriales|f__Oscillospiraceae|g__GGB30475|s__GGB30475_SGB63182
k__Bacteria|p__Actinobacteria|c__CFGB77153|o__OFGB77153|f__FGB77153|g__GGB30861|s__GGB30861_SGB44083
k__Bacteria|p__Tenericutes|c__CFGB1791|o__OFGB1791|f__FGB1791|g__GGB31312|s__GGB31312_SGB44628
k__Bacteria|p__Firmicutes|c__CFGB10290|o__OFGB10290|f__FGB10290|g__GGB31438|s__GGB31438_SGB44768
k__Bacteria|p__Firmicutes|c__Clostridia|o__Eubacteriales|f__Oscillospiraceae|g__GGB3171|s__GGB3171_SGB4185
k__Bacteria|p__Firmicutes|c__CFGB10289|o__OFGB10289|f__FGB10289|g__GGB31762|s__GGB31762_SGB45125
k__Bacteria|p__Firmicutes|c__CFGB1765|o__OFGB1765|f__FGB1765|g__GGB31823|s__GGB31823_SGB45199
k__Bacteria|p__Firmicutes|c__CFGB1765|o__OFGB1765|f__FGB1765|g__GGB31838|s__GGB31838_SGB45216
k__Bacteria|p__Firmicutes|c__CFGB1765|o__OFGB1765|f__FGB1765|g__GGB31841|s__GGB31841_SGB65084
k__Bacteria|p__Firmicutes|c__CFGB10667|o__OFGB10667|f__FGB10667|g__GGB32371|s__GGB32371_SGB41694
k__Bacteria|p__Firmicutes|c__Clostridia|o__Eubacteriales|f__Lachnospiraceae|g__GGB3793|s__GGB3793_SGB5158
k__Bacteria|p__Firmicutes|c__Clostridia|o__Eubacteriales|f__Clostridiaceae|g__GGB42601|s__GGB42601_SGB59797
k__Bacteria|p__Firmicutes|c__Clostridia|o__Eubacteriales|f__Oscillospiraceae|g__GGB45514|s__GGB45514_SGB63186
k__Bacteria|p__Firmicutes|c__CFGB75721|o__OFGB75721|f__FGB75721|g__GGB45564|s__GGB45564_SGB63259
k__Bacteria|p__Firmicutes|c__Clostridia|o__Eubacteriales|f__Oscillospiraceae|g__GGB45624|s__GGB45624_SGB63337
k__Bacteria|p__Firmicutes|c__Clostridia|o__Eubacteriales|f__Christensenellaceae|g__GGB45656|s__GGB45656_SGB6337[illegible]
k__Bacteria|p__Firmicutes|c__CFGB10299|o__OFGB10299|f__FGB10299|g__GGB47127|s__GGB47127_SGB65054

k__Bacteria|p__Firmicutes|c__Clostridia|o__Eubacteriales|f__Oscillospiraceae|g__GGB74395|s__GGB74395_SGB43523
k__Bacteria|p__Firmicutes|c__Clostridia|o__Eubacteriales|f__Oscillospiraceae|g__GGB75053|s__GGB75053_SGB43494
k__Bacteria|p__Firmicutes|c__Clostridia|o__Eubacteriales|f__Lachnospiraceae|g__GGB75109|s__GGB75109_SGB102238
k__Bacteria|p__Firmicutes|c__Clostridia|o__Eubacteriales|f__Lachnospiraceae|g__Lachnospiraceae_unclassified|s__Lachn
k__Bacteria|p__Firmicutes|c__Clostridia|o__Eubacteriales|f__Lachnospiraceae|g__Lachnospiraceae_unclassified|s__Lachn
k__Bacteria|p__Firmicutes|c__Clostridia|o__Eubacteriales|f__Lachnospiraceae|g__Lachnospiraceae_unclassified|s__Lachn
k__Bacteria|p__Firmicutes|c__Clostridia|o__Eubacteriales|f__Lachnospiraceae|g__Lachnospiraceae_unclassified|s__Lachn
k__Bacteria|p__Firmicutes|c__Clostridia|o__Eubacteriales|f__Lachnospiraceae|g__Lachnospiraceae_unclassified|s__Lachn
k__Bacteria|p__Firmicutes|c__Bacilli|o__Lactobacillales|f__Lactobacillaceae|g__Lactobacillus|s__Lactobacillus_johnsonii
k__Bacteria|p__Actinobacteria|c__Coriobacteriia|o__Coriobacteriales|f__Atopobiaceae|g__Leptogranulimonas|s__Leptog
k__Bacteria|p__Bacteroidota|c__Bacteroidia|o__Bacteroidales|f__Muribaculaceae|g__Muribaculaceae_unclassified|s__M
k__Bacteria|p__Firmicutes|c__Clostridia|o__Eubacteriales|f__Oscillospiraceae|g__Neglectibacter|s__Neglectibacter_sp_X
k__Bacteria|p__Firmicutes|c__Clostridia|o__Eubacteriales|f__Oscillospiraceae|g__Oscillibacter|s__Oscillibacter_SGB4349
k__Bacteria|p__Firmicutes|c__Clostridia|o__Eubacteriales|f__Oscillospiraceae|g__Oscillospiraceae_unclassified|s__Oscillo
k__Bacteria|p__Firmicutes|c__Clostridia|o__Eubacteriales|f__Oscillospiraceae|g__Oscillospiraceae_unclassified|s__Oscillo
k__Bacteria|p__Firmicutes|c__Clostridia|o__Eubacteriales|f__Oscillospiraceae|g__Oscillospiraceae_unclassified|s__Oscillo
k__Bacteria|p__Firmicutes|c__Clostridia|o__Eubacteriales|f__Oscillospiraceae|g__Oscillospiraceae_unclassified|s__Oscillo
k__Bacteria|p__Proteobacteria|c__Betaproteobacteria|o__Burkholderiales|f__Sutterellaceae|g__Parasutterella|s__Parasu
k__Bacteria|p__Firmicutes|c__Clostridia|o__Eubacteriales|f__Lachnospiraceae|g__Schaedlerella|s__Schaedlerella_arabinc
k__Bacteria|p__Firmicutes|c__Erysipelotrichia|o__Erysipelotrichales|f__Turicibacteraceae|g__Turicibacter|s__Turicibacter_
k__Bacteria|p__Bacteria_unclassified|c__Bacteria_unclassified|o__Bacteria_unclassified|f__Bacteria_unclassified|g__Bacte
k__Bacteria|p__Bacteria_unclassified|c__Bacteria_unclassified|o__Bacteria_unclassified|f__Bacteria_unclassified|g__Bacte
k__Bacteria|p__Bacteria_unclassified|c__Bacteria_unclassified|o__Bacteria_unclassified|f__Bacteria_unclassified|g__Bacte
k__Bacteria|p__Bacteria_unclassified|c__Bacteria_unclassified|o__Bacteria_unclassified|f__Bacteria_unclassified|g__Bacte
k__Bacteria|p__Bacteria_unclassified|c__Bacteria_unclassified|o__Bacteria_unclassified|f__Bacteria_unclassified|g__Bacte
