## Supplementary material for "Non-invasive Vagal Nerve Stimulation as a Potential Treatment for Repetitive Blast Trauma": st2

| Cytokine | PC1 | PC2 | PC3 | PC4 | PC5 |
|---|---|---|---|---|---|
| serum_IL-1a | -0.13971 | 0.036106 | -0.087596 | -0.357139 | 0.371858 |
| serum_IL-5 | 0.240246 | 0.033876 | 0.024549 | -0.074063 | -0.017311 |
| serum_IL-6 | 0.245772 | -0.242198 | 0.012981 | 0.057687 | 0.011426 |
| serum_IL-9 | 0.174559 | -0.121984 | 0.042901 | -0.061347 | -0.365543 |
| serum_IL-12(p40) | -0.031778 | -0.058041 | -0.163105 | -0.425547 | -0.015543 |
| serum_G-CSF | 0.250196 | -0.166522 | -0.059469 | -0.105981 | -0.129922 |
| serum_IP-10 | 0.22272 | -0.195961 | -0.023552 | 0.027461 | -0.295476 |
| serum_MKC | 0.240829 | -0.196427 | -0.149233 | 0.006243 | -0.205592 |
| serum_MCP-1 | 0.248431 | -0.201177 | -0.121352 | 0.077855 | 0.051629 |
| serum_MIP-2 | -0.094252 | -0.036926 | -0.047439 | -0.490785 | -0.180603 |
| serum_RANTES | 0.080822 | 0.113766 | 0.003994 | -0.329662 | 0.084753 |
| brain_IL-1a | -0.183209 | -0.210215 | -0.2115 | 0.204748 | 0.035073 |
| brain_IL-2 | -0.114026 | -0.176093 | -0.357926 | 0.056552 | -0.112375 |
| brain_IL-5 | 0.039054 | -0.28159 | 0.273279 | -0.044639 | 0.071742 |
| brain_IL-6 | 0.2405 | -0.036503 | -0.019805 | 0.081688 | 0.390307 |
| brain_IL-9 | -0.123845 | -0.268348 | 0.071079 | 0.130707 | 0.069637 |
| brain_IL-10 | -0.054829 | -0.12279 | 0.450084 | 0.002576 | 0.09619 |
| brain_IL-12(p40) | 0.083786 | -0.221444 | 0.293526 | 0.012208 | 0.044444 |
| brain_IL-12(p70) | -0.13614 | -0.217599 | -0.027898 | -0.296259 | 0.128426 |
| brain_IL-13 | -0.162908 | -0.253525 | -0.168449 | -0.004867 | -0.055385 |
| brain_IL-15 | -0.160795 | -0.248007 | -0.17559 | -0.043006 | 0.087001 |
| brain_IL-17 | -0.177583 | -0.264139 | -0.22875 | 0.109316 | -0.006545 |
| brain_G-CSF | 0.306053 | -0.093439 | -0.050826 | -0.010712 | 0.121084 |
| brain_GM-CSF | -0.191808 | -0.194343 | 0.134021 | -0.097393 | -0.074245 |
| brain_INF_gamma | -0.20092 | -0.245934 | 0.047837 | -0.022614 | 0.0235 |
| brain_IP-10 | 0.198248 | -0.106163 | -0.240737 | -0.155292 | 0.324683 |
| brain_MKC | 0.250728 | -0.119331 | -0.001576 | -0.145678 | -0.083827 |
| brain_MCP-1 | 0.241745 | -0.065458 | -0.002953 | 0.034903 | 0.376643 |
| brain_MIP-2 | -0.010219 | -0.149169 | 0.386006 | -0.244862 | -0.10944 |
| brain_RANTES | -0.135069 | -0.259359 | 0.189592 | 0.148015 | 0.192105 |
