## Supplementary material for "Non-invasive Vagal Nerve Stimulation as a Potential Treatment for Repetitive Blast Trauma": st3

| outcome | Species-level feature | n | LRT_F_microbe | LRT_p_microbe |
|---|---|---|---|---|
| acute_cyto_PC1 | Acetatifactor_SGB41545 | 27 | 0.811221818 | 0.4263375897 |
| acute_cyto_PC1 | Acutalibacter_muris | 27 | -0.5653962247 | 0.5777957076 |
| acute_cyto_PC1 | Akkermansia_muciniphila | 27 | -0.5296359608 | 0.6019194146 |
| acute_cyto_PC1 | Anaerotruncus_sp_1XD42_93 | 27 | -0.1053420732 | 0.9171041076 |
| acute_cyto_PC1 | Bacteria_unclassified_SGB41677 | 27 | -0.02999591667 | 0.9763535607 |
| acute_cyto_PC1 | Bacteria_unclassified_SGB43539 | 27 | 1.267139755 | 0.2189780512 |
| acute_cyto_PC1 | Bacteria_unclassified_SGB43546 | 27 | 0.5238735197 | 0.6058519277 |
| acute_cyto_PC1 | Bacteria_unclassified_SGB63211 | 27 | 2.048051107 | 0.05326401688 |
| acute_cyto_PC1 | Bacteroides_thetaiotaomicron | 27 | -0.509179467 | 0.6159351495 |
| acute_cyto_PC1 | Clostridia_bacterium | 27 | -0.5587690197 | 0.5822293971 |
| acute_cyto_PC1 | Clostridiaceae_bacterium | 27 | -0.4431309337 | 0.6622024077 |
| acute_cyto_PC1 | Clostridiaceae_unclassified_SGB41663 | 27 | -1.444867654 | 0.1632521935 |
| acute_cyto_PC1 | Clostridiales_bacterium | 27 | 0.5961201509 | 0.5574665105 |
| acute_cyto_PC1 | Clostridium_SGB65123 | 27 | -3.01130997 | 0.00664767832 |
| acute_cyto_PC1 | Clostridium_cocleatum | 27 | -0.3568938693 | 0.7247324894 |
| acute_cyto_PC1 | Eubacteriaceae_bacterium | 27 | -0.1835035817 | 0.8561631507 |
| acute_cyto_PC1 | Eubacteriaceae_unclassified_SGB94922 | 27 | -0.109767717 | 0.9136358044 |
| acute_cyto_PC1 | GGB20146_SGB29427 | 27 | 0.3045693121 | 0.7636919925 |
| acute_cyto_PC1 | GGB20149_SGB29430 | 27 | 0.4461351552 | 0.6600657249 |
| acute_cyto_PC1 | GGB22635_SGB63107 | 27 | 0.3677705653 | 0.7167250388 |
| acute_cyto_PC1 | GGB25041_SGB36960 | 27 | 0.2196630048 | 0.8282547772 |
| acute_cyto_PC1 | GGB28379_SGB40959 | 27 | 2.176355529 | 0.04108690661 |
| acute_cyto_PC1 | GGB28382_SGB40962 | 27 | -1.573002392 | 0.1306643628 |
| acute_cyto_PC1 | GGB28392_SGB40972 | 27 | -1.591362358 | 0.1264701801 |
| acute_cyto_PC1 | GGB28404_SGB40986 | 27 | 0.3398309152 | 0.7373593019 |
| acute_cyto_PC1 | GGB28418_SGB41001 | 27 | 0.0349175958 | 0.9724752282 |
| acute_cyto_PC1 | GGB28422_SGB41005 | 27 | 0.4177402243 | 0.6803773997 |
| acute_cyto_PC1 | GGB28431_SGB41014 | 27 | -0.1621872213 | 0.872708849 |
| acute_cyto_PC1 | GGB28456_SGB41039 | 27 | 1.16296545 | 0.2578889366 |
| acute_cyto_PC1 | GGB28782_SGB41435 | 27 | -0.7729789869 | 0.4481484807 |
| acute_cyto_PC1 | GGB28792_SGB41445 | 27 | 0.344766815 | 0.7336986581 |
| acute_cyto_PC1 | GGB28798_SGB41451 | 27 | 0.4707769995 | 0.6426532597 |
| acute_cyto_PC1 | GGB28810_SGB41465 | 27 | -0.04987734884 | 0.9606914889 |
| acute_cyto_PC1 | GGB28828_SGB41484 | 27 | -1.554575845 | 0.134989453 |
| acute_cyto_PC1 | GGB28851_SGB41518 | 27 | 0.9308435064 | 0.3625128158 |
| acute_cyto_PC1 | GGB28865_SGB41537 | 27 | -2.075790327 | 0.0503856047 |
| acute_cyto_PC1 | GGB28868_SGB41542 | 27 | 0.3424653491 | 0.7354047075 |
| acute_cyto_PC1 | GGB28869_SGB41543 | 27 | 1.035154156 | 0.3123709457 |
| acute_cyto_PC1 | GGB28875_SGB41555 | 27 | -0.05192365605 | 0.9590802748 |
| acute_cyto_PC1 | GGB28881_SGB41561 | 27 | -0.6436208003 | 0.5267884535 |
| acute_cyto_PC1 | GGB28892_SGB41573 | 27 | -1.053782381 | 0.3039568745 |
| acute_cyto_PC1 | GGB28893_SGB41574 | 27 | 2.21260236 | 0.03813709284 |
| acute_cyto_PC1 | GGB28898_SGB41580 | 27 | -0.4325431875 | 0.6697561454 |
| acute_cyto_PC1 | GGB28901_SGB41592 | 27 | -1.484052936 | 0.1526502905 |
| acute_cyto_PC1 | GGB28904_SGB41597 | 27 | 1.300027778 | 0.2076802308 |
| acute_cyto_PC1 | GGB28909_SGB41602 | 27 | -2.385918078 | 0.02653112285 |

| | | | | |
|---|---|---|---|---|
| acute_cyto_PC1 | GGB28924_SGB41621 | 27 | -2.054891923 | 0.05254079686 |
| acute_cyto_PC1 | GGB28927_SGB41625 | 27 | -0.9235697201 | 0.3662013664 |
| acute_cyto_PC1 | GGB28934_SGB41635 | 27 | 0.6009285837 | 0.5543189854 |
| acute_cyto_PC1 | GGB28949_SGB41655 | 27 | -1.026372543 | 0.3163940609 |
| acute_cyto_PC1 | GGB28951_SGB102295 | 27 | 0.5911817779 | 0.5607087809 |
| acute_cyto_PC1 | GGB28951_SGB41658 | 27 | -1.177054193 | 0.2523431626 |
| acute_cyto_PC1 | GGB28954_SGB41662 | 27 | 0.2678883809 | 0.7913988996 |
| acute_cyto_PC1 | GGB28960_SGB41669 | 27 | -1.191581955 | 0.2467184541 |
| acute_cyto_PC1 | GGB28964_SGB94886 | 27 | -0.2476769268 | 0.8067899815 |
| acute_cyto_PC1 | GGB28967_SGB41678 | 27 | -0.4790088279 | 0.636882447 |
| acute_cyto_PC1 | GGB28996_SGB41712 | 27 | 0.3261232271 | 0.7475587555 |
| acute_cyto_PC1 | GGB29531_SGB42317 | 27 | 0.4017377774 | 0.6919362382 |
| acute_cyto_PC1 | GGB29685_SGB42494 | 27 | 0.2130394145 | 0.8333507625 |
| acute_cyto_PC1 | GGB30141_SGB43066 | 27 | -0.7716703163 | 0.4489067099 |
| acute_cyto_PC1 | GGB30145_SGB43072 | 27 | 1.551965795 | 0.1356115768 |
| acute_cyto_PC1 | GGB30286_SGB43248 | 27 | 0.7790528204 | 0.4446395985 |
| acute_cyto_PC1 | GGB30300_SGB43264 | 27 | -1.26996853 | 0.2179880419 |
| acute_cyto_PC1 | GGB30303_SGB43268 | 27 | 0.8457922497 | 0.4072031284 |
| acute_cyto_PC1 | GGB30450_SGB43507 | 27 | -0.7977695814 | 0.4339330774 |
| acute_cyto_PC1 | GGB30453_SGB43513 | 27 | 0.1317853615 | 0.8964085344 |
| acute_cyto_PC1 | GGB30454_SGB43514 | 27 | 0.1369197874 | 0.892398402 |
| acute_cyto_PC1 | GGB30455_SGB43519 | 27 | 0.4906613313 | 0.6287539192 |
| acute_cyto_PC1 | GGB30456_SGB43520 | 27 | -0.8573757689 | 0.400916417 |
| acute_cyto_PC1 | GGB30457_SGB63218 | 27 | 0.4519449013 | 0.6559421165 |
| acute_cyto_PC1 | GGB30461_SGB43530 | 27 | -0.552541678 | 0.5864110497 |
| acute_cyto_PC1 | GGB30461_SGB43533 | 27 | 0.1921807148 | 0.849446904 |
| acute_cyto_PC1 | GGB30461_SGB63209 | 27 | 1.136870591 | 0.268399194 |
| acute_cyto_PC1 | GGB30463_SGB43537 | 27 | 0.02225685992 | 0.9824531902 |
| acute_cyto_PC1 | GGB30473_SGB43557 | 27 | -0.06014332864 | 0.9526101765 |
| acute_cyto_PC1 | GGB30475_SGB63182 | 27 | 0.699893571 | 0.4916750511 |
| acute_cyto_PC1 | GGB30861_SGB44083 | 27 | 1.825138121 | 0.08223849766 |
| acute_cyto_PC1 | GGB31312_SGB44628 | 27 | 0.5293518882 | 0.6021129872 |
| acute_cyto_PC1 | GGB31438_SGB44768 | 27 | -3.464143267 | 0.00232035856 |
| acute_cyto_PC1 | GGB3171_SGB4185 | 27 | -1.5458549 | 0.1370774432 |
| acute_cyto_PC1 | GGB31762_SGB45125 | 27 | -0.3742329648 | 0.7119831078 |
| acute_cyto_PC1 | GGB31823_SGB45199 | 27 | -0.4338015114 | 0.668856506 |
| acute_cyto_PC1 | GGB31838_SGB45216 | 27 | -0.1235804218 | 0.9028226785 |
| acute_cyto_PC1 | GGB31841_SGB65084 | 27 | -1.997655241 | 0.05887218089 |
| acute_cyto_PC1 | GGB32371_SGB41694 | 27 | -0.2317107619 | 0.8190056127 |
| acute_cyto_PC1 | GGB3793_SGB5158 | 27 | -0.4804734028 | 0.6358581838 |
| acute_cyto_PC1 | GGB42601_SGB59797 | 27 | 0.8867956016 | 0.3852324177 |
| acute_cyto_PC1 | GGB45514_SGB63186 | 27 | 0.4917523126 | 0.6279953191 |
| acute_cyto_PC1 | GGB45564_SGB63259 | 27 | -0.069750953 | 0.9450518329 |
| acute_cyto_PC1 | GGB45624_SGB63337 | 27 | 1.66580504 | 0.1106008108 |
| acute_cyto_PC1 | GGB45656_SGB63370 | 27 | 0.9603113865 | 0.3478258885 |
| acute_cyto_PC1 | GGB47127_SGB65054 | 27 | -0.5504710205 | 0.5878047965 |

| | | | | |
|---|---|---|---|---|
| acute_cyto_PC1 | GGB74395_SGB43523 | 27 | 1.885640797 | 0.07325377808 |
| acute_cyto_PC1 | GGB75053_SGB43494 | 27 | -0.9656016453 | 0.3452327395 |
| acute_cyto_PC1 | GGB75109_SGB102238 | 27 | -1.60283389 | 0.1239070982 |
| acute_cyto_PC1 | Lachnospiraceae_bacterium | 27 | 0.2746479015 | 0.7862705667 |
| acute_cyto_PC1 | Lachnospiraceae_bacterium_A2 | 27 | -0.4828169006 | 0.6342207918 |
| acute_cyto_PC1 | Lachnospiraceae_unclassified_SGB41414 | 27 | 0.5779585099 | 0.5694384296 |
| acute_cyto_PC1 | Lachnospiraceae_unclassified_SGB41424 | 27 | 2.357147748 | 0.02819891813 |
| acute_cyto_PC1 | Lachnospiraceae_unclassified_SGB94868 | 27 | -0.0278115273[illegible] | 0.9780750794 |
| acute_cyto_PC1 | Lactobacillus_johnsonii | 27 | 0.5833192659 | 0.5658910214 |
| acute_cyto_PC1 | Leptogranulimonas_caecicola | 27 | -1.523024296 | 0.1426703397 |
| acute_cyto_PC1 | Muribaculaceae_bacterium | 27 | -0.0076183815[illegible] | 0.99399338 |
| acute_cyto_PC1 | Neglectibacter_sp_X4 | 27 | 0.6379522338 | 0.5304003737 |
| acute_cyto_PC1 | Oscillibacter_SGB43496 | 27 | 1.74444352 | 0.09570201712 |
| acute_cyto_PC1 | Oscillospiraceae_bacterium | 27 | 0.8998424088 | 0.3784074486 |
| acute_cyto_PC1 | Oscillospiraceae_unclassified_SGB43502 | 27 | 0.4149108539 | 0.6824153502 |
| acute_cyto_PC1 | Oscillospiraceae_unclassified_SGB43505 | 27 | 2.885277896 | 0.0088548848[illegible] |
| acute_cyto_PC1 | Oscillospiraceae_unclassified_SGB94989 | 27 | 1.385488261 | 0.1804436782 |
| acute_cyto_PC1 | Parasutterella_excrementihominis | 27 | 1.801264781 | 0.08603923696 |
| acute_cyto_PC1 | Schaedlerella_arabinosiphila | 27 | -0.334776754 | 0.741114287 |
| acute_cyto_PC1 | Turicibacter_sp_1E2 | 27 | -2.276558793 | 0.03339918902 |
| acute_cyto_PC1 | bacterium_0_1xD8_82 | 27 | -0.4427230772 | 0.6624927144 |
| acute_cyto_PC1 | bacterium_1XD42_1 | 27 | -0.2006733119 | 0.8428848194 |
| acute_cyto_PC1 | bacterium_1XD42_54 | 27 | 0.1996301392 | 0.8436902412 |
| acute_cyto_PC1 | bacterium_1XD42_76 | 27 | 3.188114672 | 0.0044237988[illegible] |
| acute_cyto_PC1 | bacterium_1xD8_48 | 27 | -0.4870063818 | 0.6312984042 |
| acute_cyto_PC1 | Berger Parker Index | 27 | 0.5546160728 | 0.5850164374 |
| acute_cyto_PC1 | Richness (# observed features) | 27 | 0.1699098669 | 0.8667072268 |
| acute_cyto_PC1 | Shannon Index | 27 | -0.2057340609 | 0.8389799902 |
| acute_cyto_PC2 | Acetatifactor_SGB41545 | 27 | -0.3501689773 | 0.729699651 |
| acute_cyto_PC2 | Acutalibacter_muris | 27 | 0.6042963891 | 0.5521200392 |
| acute_cyto_PC2 | Akkermansia_muciniphila | 27 | -0.1032435303 | 0.9187492969 |
| acute_cyto_PC2 | Anaerotruncus_sp_1XD42_93 | 27 | -0.5504887108 | 0.5877928823 |
| acute_cyto_PC2 | Bacteria_unclassified_SGB41677 | 27 | -0.6832335355 | 0.5019284422 |
| acute_cyto_PC2 | Bacteria_unclassified_SGB43539 | 27 | 1.06990166 | 0.2968074174 |
| acute_cyto_PC2 | Bacteria_unclassified_SGB43546 | 27 | -0.4402517785 | 0.6642529128 |
| acute_cyto_PC2 | Bacteria_unclassified_SGB63211 | 27 | 0.9424632476 | 0.3566723803 |
| acute_cyto_PC2 | Bacteroides_thetaiotaomicron | 27 | -0.0397891165[illegible] | 0.9686371082 |
| acute_cyto_PC2 | Clostridia_bacterium | 27 | -0.5722310411 | 0.573241047 |
| acute_cyto_PC2 | Clostridiaceae_bacterium | 27 | -0.5236410097 | 0.6060108602 |
| acute_cyto_PC2 | Clostridiaceae_unclassified_SGB41663 | 27 | -1.758758371 | 0.09318366282 |
| acute_cyto_PC2 | Clostridiales_bacterium | 27 | -0.2950912952 | 0.7708218478 |
| acute_cyto_PC2 | Clostridium_SGB65123 | 27 | -3.25528208 | 0.00378456795 |
| acute_cyto_PC2 | Clostridium_cocleatum | 27 | 0.2372376377 | 0.8147715187 |
| acute_cyto_PC2 | Eubacteriaceae_bacterium | 27 | -0.0841345468[illegible] | 0.9337463589 |
| acute_cyto_PC2 | Eubacteriaceae_unclassified_SGB94922 | 27 | 0.9369028527 | 0.3594592285 |
| acute_cyto_PC2 | GGB20146_SGB29427 | 27 | 1.266319437 | 0.2192657912 |

| | | | | |
|---|---|---|---|---|
| acute_cyto_PC2 | GGB20149_SGB29430 | 27 | 1.232347741 | 0.2314390826 |
| acute_cyto_PC2 | GGB22635_SGB63107 | 27 | 1.077431154 | 0.2935094684 |
| acute_cyto_PC2 | GGB25041_SGB36960 | 27 | 0.7369272722 | 0.4693203776 |
| acute_cyto_PC2 | GGB28379_SGB40959 | 27 | 1.432099219 | 0.1668328279 |
| acute_cyto_PC2 | GGB28382_SGB40962 | 27 | -0.338896761 | 0.7380528278 |
| acute_cyto_PC2 | GGB28392_SGB40972 | 27 | -1.067270996 | 0.2979659032 |
| acute_cyto_PC2 | GGB28404_SGB40986 | 27 | -0.3247224273 | 0.7486037584 |
| acute_cyto_PC2 | GGB28418_SGB41001 | 27 | 0.6949720229 | 0.4946914071 |
| acute_cyto_PC2 | GGB28422_SGB41005 | 27 | 0.4839059165 | 0.6334605536 |
| acute_cyto_PC2 | GGB28431_SGB41014 | 27 | -0.3898383231 | 0.7005818042 |
| acute_cyto_PC2 | GGB28456_SGB41039 | 27 | 3.92989586 | 0.00076812473 |
| acute_cyto_PC2 | GGB28782_SGB41435 | 27 | -0.5782861153 | 0.5692213149 |
| acute_cyto_PC2 | GGB28792_SGB41445 | 27 | -0.1055764555 | 0.9169203831 |
| acute_cyto_PC2 | GGB28798_SGB41451 | 27 | -0.5477440335 | 0.5896428186 |
| acute_cyto_PC2 | GGB28810_SGB41465 | 27 | 0.125401195 | 0.9013987003 |
| acute_cyto_PC2 | GGB28828_SGB41484 | 27 | -1.386346406 | 0.1801853188 |
| acute_cyto_PC2 | GGB28851_SGB41518 | 27 | 0.3025941857 | 0.7651760368 |
| acute_cyto_PC2 | GGB28865_SGB41537 | 27 | -2.21165167 | 0.03821193107 |
| acute_cyto_PC2 | GGB28868_SGB41542 | 27 | 1.354382344 | 0.1900092626 |
| acute_cyto_PC2 | GGB28869_SGB41543 | 27 | 0.7548553026 | 0.4587184199 |
| acute_cyto_PC2 | GGB28875_SGB41555 | 27 | 0.5831572802 | 0.5659980463 |
| acute_cyto_PC2 | GGB28881_SGB41561 | 27 | -0.5503074411 | 0.5879149708 |
| acute_cyto_PC2 | GGB28892_SGB41573 | 27 | -1.128786811 | 0.2717183239 |
| acute_cyto_PC2 | GGB28893_SGB41574 | 27 | 2.963520432 | 0.0074139524[illegible] |
| acute_cyto_PC2 | GGB28898_SGB41580 | 27 | -0.5047567358 | 0.6189854942 |
| acute_cyto_PC2 | GGB28901_SGB41592 | 27 | -0.5855905044 | 0.5643914995 |
| acute_cyto_PC2 | GGB28904_SGB41597 | 27 | 1.697022351 | 0.1044657882 |
| acute_cyto_PC2 | GGB28909_SGB41602 | 27 | -1.141357508 | 0.2665698429 |
| acute_cyto_PC2 | GGB28924_SGB41621 | 27 | -2.018021169 | 0.05654533343 |
| acute_cyto_PC2 | GGB28927_SGB41625 | 27 | -0.3588821464 | 0.7232662676 |
| acute_cyto_PC2 | GGB28934_SGB41635 | 27 | 0.2199049322 | 0.828068791 |
| acute_cyto_PC2 | GGB28949_SGB41655 | 27 | -1.648304377 | 0.1141717851 |
| acute_cyto_PC2 | GGB28951_SGB102295 | 27 | 0.3912223347 | 0.6995740795 |
| acute_cyto_PC2 | GGB28951_SGB41658 | 27 | -1.002985021 | 0.3272859349 |
| acute_cyto_PC2 | GGB28954_SGB41662 | 27 | -1.200312306 | 0.2433839379 |
| acute_cyto_PC2 | GGB28960_SGB41669 | 27 | -0.4872633615 | 0.6311193475 |
| acute_cyto_PC2 | GGB28964_SGB94886 | 27 | 0.9154188855 | 0.3703644085 |
| acute_cyto_PC2 | GGB28967_SGB41678 | 27 | -0.5102022612 | 0.6152307406 |
| acute_cyto_PC2 | GGB28996_SGB41712 | 27 | -0.1074183943 | 0.9154767142 |
| acute_cyto_PC2 | GGB29531_SGB42317 | 27 | 1.232461785 | 0.2313973712 |
| acute_cyto_PC2 | GGB29685_SGB42494 | 27 | 0.04658519237 | 0.9632840138 |
| acute_cyto_PC2 | GGB30141_SGB43066 | 27 | -0.2112526391 | 0.8347267496 |
| acute_cyto_PC2 | GGB30145_SGB43072 | 27 | 1.267246398 | 0.2189406657 |
| acute_cyto_PC2 | GGB30286_SGB43248 | 27 | 0.266523701 | 0.7924354436 |
| acute_cyto_PC2 | GGB30300_SGB43264 | 27 | 0.3887679193 | 0.7013615732 |
| acute_cyto_PC2 | GGB30303_SGB43268 | 27 | 3.526249434 | 0.0020042744[illegible] |

| | | | | |
|---|---|---|---|---|
| acute_cyto_PC2 | GGB30450_SGB43507 | 27 | -0.8538722419 | 0.4028112568 |
| acute_cyto_PC2 | GGB30453_SGB43513 | 27 | 0.6299998499 | 0.5354901584 |
| acute_cyto_PC2 | GGB30454_SGB43514 | 27 | 1.699684715 | 0.1039562309 |
| acute_cyto_PC2 | GGB30455_SGB43519 | 27 | 0.8573790641 | 0.4009146375 |
| acute_cyto_PC2 | GGB30456_SGB43520 | 27 | 0.3321798111 | 0.7430462664 |
| acute_cyto_PC2 | GGB30457_SGB63218 | 27 | -0.9419613996 | 0.3569233035 |
| acute_cyto_PC2 | GGB30461_SGB43530 | 27 | -0.1408613113 | 0.8893219555 |
| acute_cyto_PC2 | GGB30461_SGB43533 | 27 | 0.5928693887 | 0.559599685 |
| acute_cyto_PC2 | GGB30461_SGB63209 | 27 | 1.150605812 | 0.2628283018 |
| acute_cyto_PC2 | GGB30463_SGB43537 | 27 | 0.4916179085 | 0.6280887525 |
| acute_cyto_PC2 | GGB30473_SGB43557 | 27 | 1.128753435 | 0.2717320899 |
| acute_cyto_PC2 | GGB30475_SGB63182 | 27 | 0.2843757778 | 0.7789076763 |
| acute_cyto_PC2 | GGB30861_SGB44083 | 27 | 0.00493497603( | 0.9961090555 |
| acute_cyto_PC2 | GGB31312_SGB44628 | 27 | 1.38142302 | 0.1816716013 |
| acute_cyto_PC2 | GGB31438_SGB44768 | 27 | -1.90692963 | 0.07030348686 |
| acute_cyto_PC2 | GGB3171_SGB4185 | 27 | -0.7636350447 | 0.4535793479 |
| acute_cyto_PC2 | GGB31762_SGB45125 | 27 | -0.0857777700! | 0.9324556547 |
| acute_cyto_PC2 | GGB31823_SGB45199 | 27 | 0.6953887028 | 0.4944356186 |
| acute_cyto_PC2 | GGB31838_SGB45216 | 27 | -0.510882786 | 0.6147622668 |
| acute_cyto_PC2 | GGB31841_SGB65084 | 27 | -0.8430903285 | 0.4086785656 |
| acute_cyto_PC2 | GGB32371_SGB41694 | 27 | -0.5098764025 | 0.6154551216 |
| acute_cyto_PC2 | GGB3793_SGB5158 | 27 | -0.125438139 | 0.9013698108 |
| acute_cyto_PC2 | GGB42601_SGB59797 | 27 | -0.9832324678 | 0.3366862375 |
| acute_cyto_PC2 | GGB45514_SGB63186 | 27 | 0.4658515222 | 0.6461173274 |
| acute_cyto_PC2 | GGB45564_SGB63259 | 27 | 1.057529367 | 0.3022840998 |
| acute_cyto_PC2 | GGB45624_SGB63337 | 27 | 0.4250028761 | 0.6751577094 |
| acute_cyto_PC2 | GGB45656_SGB63370 | 27 | -0.6982020031 | 0.4927105963 |
| acute_cyto_PC2 | GGB47127_SGB65054 | 27 | -1.094504391 | 0.2861292433 |
| acute_cyto_PC2 | GGB74395_SGB43523 | 27 | -0.0538059309: | 0.9575983741 |
| acute_cyto_PC2 | GGB75053_SGB43494 | 27 | -1.188495322 | 0.2479055584 |
| acute_cyto_PC2 | GGB75109_SGB102238 | 27 | -1.452133797 | 0.1612424576 |
| acute_cyto_PC2 | Lachnospiraceae_bacterium | 27 | 0.1064494213 | 0.9162361358 |
| acute_cyto_PC2 | Lachnospiraceae_bacterium_A2 | 27 | 0.3156783381 | 0.7553625359 |
| acute_cyto_PC2 | Lachnospiraceae_unclassified_SGB41414 | 27 | 0.3037941729 | 0.7642742957 |
| acute_cyto_PC2 | Lachnospiraceae_unclassified_SGB41424 | 27 | 1.744387803 | 0.09571193232 |
| acute_cyto_PC2 | Lachnospiraceae_unclassified_SGB94868 | 27 | -1.024969862 | 0.3170400346 |
| acute_cyto_PC2 | Lactobacillus_johnsonii | 27 | 0.1715874077 | 0.8654046089 |
| acute_cyto_PC2 | Leptogranulimonas_caecicola | 27 | -1.04517865 | 0.3078227825 |
| acute_cyto_PC2 | Muribaculaceae_bacterium | 27 | 0.1735920162 | 0.8638485361 |
| acute_cyto_PC2 | Neglectibacter_sp_X4 | 27 | 0.3409411768 | 0.7365353302 |
| acute_cyto_PC2 | Oscillibacter_SGB43496 | 27 | 1.250988338 | 0.2246971229 |
| acute_cyto_PC2 | Oscillospiraceae_bacterium | 27 | 1.207185535 | 0.2407827247 |
| acute_cyto_PC2 | Oscillospiraceae_unclassified_SGB43502 | 27 | 0.3100020812 | 0.7596148083 |
| acute_cyto_PC2 | Oscillospiraceae_unclassified_SGB43505 | 27 | 0.8928920425 | 0.3820332914 |
| acute_cyto_PC2 | Oscillospiraceae_unclassified_SGB94989 | 27 | -0.6778944078 | 0.5052399228 |
| acute_cyto_PC2 | Parasutterella_excrementihominis | 27 | 0.5358307928 | 0.5977056413 |

| | | | | |
|---|---|---|---|---|
| acute_cyto_PC2 | Schaedlerella_arabinosiphila | 27 | -0.2442436395 | 0.8094126256 |
| acute_cyto_PC2 | Turicibacter_sp_1E2 | 27 | -0.82848826 | 0.4167112574 |
| acute_cyto_PC2 | bacterium_0_1xD8_82 | 27 | -1.490223907 | 0.1510330056 |
| acute_cyto_PC2 | bacterium_1XD42_1 | 27 | 0.2733842685 | 0.7872285157 |
| acute_cyto_PC2 | bacterium_1XD42_54 | 27 | 0.4546979732 | 0.6539919676 |
| acute_cyto_PC2 | bacterium_1XD42_76 | 27 | 1.992607704 | 0.05946189678 |
| acute_cyto_PC2 | bacterium_1xD8_48 | 27 | 0.1514177317 | 0.8810913739 |
| acute_cyto_PC2 | Berger Parker Index | 27 | 0.3774047316 | 0.7096601025 |
| acute_cyto_PC2 | Richness (# observed features) | 27 | 0.00245786860[illegible] | 0.9980621059 |
| acute_cyto_PC2 | Shannon Index | 27 | -0.1063925193 | 0.9162807346 |
| acute_cyto_PC3 | Acetatifactor_SGB41545 | 27 | 0.1903115271 | 0.8508927151 |
| acute_cyto_PC3 | Acutalibacter_muris | 27 | -0.4067041552 | 0.6883405268 |
| acute_cyto_PC3 | Akkermansia_muciniphila | 27 | 1.216068945 | 0.2374519338 |
| acute_cyto_PC3 | Anaerotruncus_sp_1XD42_93 | 27 | 1.265036412 | 0.2197164171 |
| acute_cyto_PC3 | Bacteria_unclassified_SGB41677 | 27 | 1.046019543 | 0.3074434124 |
| acute_cyto_PC3 | Bacteria_unclassified_SGB43539 | 27 | -1.476311818 | 0.1546990323 |
| acute_cyto_PC3 | Bacteria_unclassified_SGB43546 | 27 | 0.6745329537 | 0.5073311112 |
| acute_cyto_PC3 | Bacteria_unclassified_SGB63211 | 27 | -0.1139195232 | 0.9103837051 |
| acute_cyto_PC3 | Bacteroides_thetaiotaomicron | 27 | -0.4255439057 | 0.6747695356 |
| acute_cyto_PC3 | Clostridia_bacterium | 27 | 1.613678119 | 0.1215242849 |
| acute_cyto_PC3 | Clostridiaceae_bacterium | 27 | 2.088713081 | 0.04909298222 |
| acute_cyto_PC3 | Clostridiaceae_unclassified_SGB41663 | 27 | -0.7662039157 | 0.4520823154 |
| acute_cyto_PC3 | Clostridiales_bacterium | 27 | 0.2339847229 | 0.8172628562 |
| acute_cyto_PC3 | Clostridium_SGB65123 | 27 | 1.111983076 | 0.2787140336 |
| acute_cyto_PC3 | Clostridium_cocleatum | 27 | -0.2385669173 | 0.8137540285 |
| acute_cyto_PC3 | Eubacteriaceae_bacterium | 27 | 2.448423639 | 0.02321785465 |
| acute_cyto_PC3 | Eubacteriaceae_unclassified_SGB94922 | 27 | -1.520004105 | 0.1434240909 |
| acute_cyto_PC3 | GGB20146_SGB29427 | 27 | -0.1365575881 | 0.8926811945 |
| acute_cyto_PC3 | GGB20149_SGB29430 | 27 | -1.345693219 | 0.1927517498 |
| acute_cyto_PC3 | GGB22635_SGB63107 | 27 | 2.118418649 | 0.04623409341 |
| acute_cyto_PC3 | GGB25041_SGB36960 | 27 | -0.5880709274 | 0.5627562164 |
| acute_cyto_PC3 | GGB28379_SGB40959 | 27 | -0.1777867323 | 0.8605942371 |
| acute_cyto_PC3 | GGB28382_SGB40962 | 27 | -1.824176563 | 0.08238871481 |
| acute_cyto_PC3 | GGB28392_SGB40972 | 27 | -0.6570812837 | 0.5182659046 |
| acute_cyto_PC3 | GGB28404_SGB40986 | 27 | 0.2529619094 | 0.8027574125 |
| acute_cyto_PC3 | GGB28418_SGB41001 | 27 | -0.4863103291 | 0.6317835119 |
| acute_cyto_PC3 | GGB28422_SGB41005 | 27 | -0.7236159349 | 0.4772854309 |
| acute_cyto_PC3 | GGB28431_SGB41014 | 27 | -2.087543664 | 0.04920871562 |
| acute_cyto_PC3 | GGB28456_SGB41039 | 27 | -2.201917532 | 0.03898593793 |
| acute_cyto_PC3 | GGB28782_SGB41435 | 27 | 1.413258151 | 0.1722317222 |
| acute_cyto_PC3 | GGB28792_SGB41445 | 27 | 1.219343553 | 0.2362329796 |
| acute_cyto_PC3 | GGB28798_SGB41451 | 27 | -0.2617273938 | 0.7960815948 |
| acute_cyto_PC3 | GGB28810_SGB41465 | 27 | -0.426566864 | 0.674035845 |
| acute_cyto_PC3 | GGB28828_SGB41484 | 27 | 0.07477679813 | 0.9411000538 |
| acute_cyto_PC3 | GGB28851_SGB41518 | 27 | 1.283692982 | 0.2132337182 |
| acute_cyto_PC3 | GGB28865_SGB41537 | 27 | 0.757076298 | 0.4574150989 |

| | | | | |
|---|---|---|---|---|
| acute_cyto_PC3 | GGB28868_SGB41542 | 27 | -1.413940495 | 0.1720337783 |
| acute_cyto_PC3 | GGB28869_SGB41543 | 27 | -1.17740508 | 0.2522061885 |
| acute_cyto_PC3 | GGB28875_SGB41555 | 27 | 0.0801714076 | 0.9368600518 |
| acute_cyto_PC3 | GGB28881_SGB41561 | 27 | -0.81998505 | 0.4214346526 |
| acute_cyto_PC3 | GGB28892_SGB41573 | 27 | 1.331175908 | 0.197403228 |
| acute_cyto_PC3 | GGB28893_SGB41574 | 27 | -3.268564487 | 0.00366926147[illegible] |
| acute_cyto_PC3 | GGB28898_SGB41580 | 27 | -0.2791222852 | 0.7828813756 |
| acute_cyto_PC3 | GGB28901_SGB41592 | 27 | -0.9018604315 | 0.3773589643 |
| acute_cyto_PC3 | GGB28904_SGB41597 | 27 | -1.972317699 | 0.06188576569 |
| acute_cyto_PC3 | GGB28909_SGB41602 | 27 | 0.8372554805 | 0.4118764282 |
| acute_cyto_PC3 | GGB28924_SGB41621 | 27 | -0.6368323151 | 0.5311155611 |
| acute_cyto_PC3 | GGB28927_SGB41625 | 27 | -0.9668539547 | 0.3446208297 |
| acute_cyto_PC3 | GGB28934_SGB41635 | 27 | 0.1732788065 | 0.8640916272 |
| acute_cyto_PC3 | GGB28949_SGB41655 | 27 | -0.2557885878 | 0.8006028914 |
| acute_cyto_PC3 | GGB28951_SGB102295 | 27 | 0.6055960653 | 0.55127267 |
| acute_cyto_PC3 | GGB28951_SGB41658 | 27 | 0.7060390296 | 0.4879234862 |
| acute_cyto_PC3 | GGB28954_SGB41662 | 27 | 0.6178239012 | 0.5433340329 |
| acute_cyto_PC3 | GGB28960_SGB41669 | 27 | 0.5826590041 | 0.5663273256 |
| acute_cyto_PC3 | GGB28964_SGB94886 | 27 | -2.517303115 | 0.02001494513 |
| acute_cyto_PC3 | GGB28967_SGB41678 | 27 | 1.334395624 | 0.1963640665 |
| acute_cyto_PC3 | GGB28996_SGB41712 | 27 | 0.6029555892 | 0.552994939 |
| acute_cyto_PC3 | GGB29531_SGB42317 | 27 | -1.358752407 | 0.1886416677 |
| acute_cyto_PC3 | GGB29685_SGB42494 | 27 | 2.421264353 | 0.02460723615 |
| acute_cyto_PC3 | GGB30141_SGB43066 | 27 | -2.090830133 | 0.04888408709 |
| acute_cyto_PC3 | GGB30145_SGB43072 | 27 | 0.1704252163 | 0.8663070142 |
| acute_cyto_PC3 | GGB30286_SGB43248 | 27 | -0.5906355332 | 0.5610680167 |
| acute_cyto_PC3 | GGB30300_SGB43264 | 27 | 0.02274929599 | 0.982065034 |
| acute_cyto_PC3 | GGB30303_SGB43268 | 27 | -1.885266042 | 0.07330667194 |
| acute_cyto_PC3 | GGB30450_SGB43507 | 27 | -0.1415709371 | 0.8887682654 |
| acute_cyto_PC3 | GGB30453_SGB43513 | 27 | -1.434479821 | 0.1661604772 |
| acute_cyto_PC3 | GGB30454_SGB43514 | 27 | -0.7079392795 | 0.4867668184 |
| acute_cyto_PC3 | GGB30455_SGB43519 | 27 | -1.057668749 | 0.3022220024 |
| acute_cyto_PC3 | GGB30456_SGB43520 | 27 | -1.308199556 | 0.2049445384 |
| acute_cyto_PC3 | GGB30457_SGB63218 | 27 | -0.09911733209 | 0.9219851833 |
| acute_cyto_PC3 | GGB30461_SGB43530 | 27 | 0.6750569582 | 0.5070048031 |
| acute_cyto_PC3 | GGB30461_SGB43533 | 27 | -1.612397941 | 0.1218035645 |
| acute_cyto_PC3 | GGB30461_SGB63209 | 27 | 0.4197563772 | 0.6789267222 |
| acute_cyto_PC3 | GGB30463_SGB43537 | 27 | 0.9602554506 | 0.3478533777 |
| acute_cyto_PC3 | GGB30473_SGB43557 | 27 | -0.6786715152 | 0.5047571718 |
| acute_cyto_PC3 | GGB30475_SGB63182 | 27 | -0.6436797334 | 0.5267509732 |
| acute_cyto_PC3 | GGB30861_SGB44083 | 27 | 1.25507908 | 0.2232379076 |
| acute_cyto_PC3 | GGB31312_SGB44628 | 27 | -0.5463984523 | 0.5905508027 |
| acute_cyto_PC3 | GGB31438_SGB44768 | 27 | 0.25223142 | 0.8033144609 |
| acute_cyto_PC3 | GGB3171_SGB4185 | 27 | -0.9176135124 | 0.3692404109 |
| acute_cyto_PC3 | GGB31762_SGB45125 | 27 | 1.426160789 | 0.1685195711 |
| acute_cyto_PC3 | GGB31823_SGB45199 | 27 | -0.5591139602 | 0.5819982072 |

| | | | | |
|---|---|---|---|---|
| acute_cyto_PC3 | GGB31838_SGB45216 | 27 | 1.563635446 | 0.1328483636 |
| acute_cyto_PC3 | GGB31841_SGB65084 | 27 | 0.6843620681 | 0.5012300742 |
| acute_cyto_PC3 | GGB32371_SGB41694 | 27 | 1.441998506 | 0.1640513269 |
| acute_cyto_PC3 | GGB3793_SGB5158 | 27 | -0.3768358699 | 0.7100765252 |
| acute_cyto_PC3 | GGB42601_SGB59797 | 27 | 2.243849167 | 0.03575052555 |
| acute_cyto_PC3 | GGB45514_SGB63186 | 27 | -0.8883468542 | 0.3844167313 |
| acute_cyto_PC3 | GGB45564_SGB63259 | 27 | -0.736966604 | 0.4692969601 |
| acute_cyto_PC3 | GGB45624_SGB63337 | 27 | -0.5096796176 | 0.6155906432 |
| acute_cyto_PC3 | GGB45656_SGB63370 | 27 | -1.180611779 | 0.2509569817 |
| acute_cyto_PC3 | GGB47127_SGB65054 | 27 | -0.2027366013 | 0.8412922969 |
| acute_cyto_PC3 | GGB74395_SGB43523 | 27 | 1.107172814 | 0.2807406373 |
| acute_cyto_PC3 | GGB75053_SGB43494 | 27 | -1.746321996 | 0.09536824745 |
| acute_cyto_PC3 | GGB75109_SGB102238 | 27 | -0.4173732739 | 0.6806415674 |
| acute_cyto_PC3 | Lachnospiraceae_bacterium | 27 | 2.553869563 | 0.01848733547 |
| acute_cyto_PC3 | Lachnospiraceae_bacterium_A2 | 27 | -0.882398847 | 0.3875504988 |
| acute_cyto_PC3 | Lachnospiraceae_unclassified_SGB41414 | 27 | 0.8951589635 | 0.3808481874 |
| acute_cyto_PC3 | Lachnospiraceae_unclassified_SGB41424 | 27 | 1.305695971 | 0.2057796704 |
| acute_cyto_PC3 | Lachnospiraceae_unclassified_SGB94868 | 27 | 1.561076241 | 0.1334503143 |
| acute_cyto_PC3 | Lactobacillus_johnsonii | 27 | 1.718428996 | 0.1004283297 |
| acute_cyto_PC3 | Leptogranulimonas_caecicola | 27 | 1.430635627 | 0.1672472759 |
| acute_cyto_PC3 | Muribaculaceae_bacterium | 27 | 1.69537538 | 0.1047820704 |
| acute_cyto_PC3 | Neglectibacter_sp_X4 | 27 | -0.172652877 | 0.8645774706 |
| acute_cyto_PC3 | Oscillibacter_SGB43496 | 27 | 0.6495393602 | 0.5230316625 |
| acute_cyto_PC3 | Oscillospiraceae_bacterium | 27 | -2.209192031 | 0.03840617522 |
| acute_cyto_PC3 | Oscillospiraceae_unclassified_SGB43502 | 27 | 1.82747006 | 0.08187518489 |
| acute_cyto_PC3 | Oscillospiraceae_unclassified_SGB43505 | 27 | 0.3485482229 | 0.7308986096 |
| acute_cyto_PC3 | Oscillospiraceae_unclassified_SGB94989 | 27 | 0.3126788071 | 0.7576086032 |
| acute_cyto_PC3 | Parasutterella_excrementihominis | 27 | 0.478950048 | 0.6369235708 |
| acute_cyto_PC3 | Schaedlerella_arabinosiphila | 27 | 0.6965191979 | 0.4937420204 |
| acute_cyto_PC3 | Turicibacter_sp_1E2 | 27 | 1.205639189 | 0.2413661101 |
| acute_cyto_PC3 | bacterium_0_1xD8_82 | 27 | -0.2616725039 | 0.79612335 |
| acute_cyto_PC3 | bacterium_1XD42_1 | 27 | -1.202341667 | 0.2426137194 |
| acute_cyto_PC3 | bacterium_1XD42_54 | 27 | -0.0191220875 | 0.9849242331 |
| acute_cyto_PC3 | bacterium_1XD42_76 | 27 | 0.7868270087 | 0.4401730268 |
| acute_cyto_PC3 | bacterium_1xD8_48 | 27 | -0.6354170628 | 0.5320201004 |
| acute_cyto_PC3 | Berger Parker Index | 27 | 1.916876523 | 0.06896119119 |
| acute_cyto_PC3 | Richness (# observed features) | 27 | -2.701883533 | 0.01335399273 |
| acute_cyto_PC3 | Shannon Index | 27 | -2.379238941 | 0.02690996432 |
| acute_cyto_PC4 | Acetatifactor_SGB41545 | 27 | 0.6022104862 | 0.5534814492 |
| acute_cyto_PC4 | Acutalibacter_muris | 27 | 2.72339589 | 0.01273111231 |
| acute_cyto_PC4 | Akkermansia_muciniphila | 27 | 0.8478703408 | 0.406070667 |
| acute_cyto_PC4 | Anaerotruncus_sp_1XD42_93 | 27 | -1.443746171 | 0.1635641806 |
| acute_cyto_PC4 | Bacteria_unclassified_SGB41677 | 27 | 0.06785558977 | 0.9465425232 |
| acute_cyto_PC4 | Bacteria_unclassified_SGB43539 | 27 | 0.196756952 | 0.8459095008 |
| acute_cyto_PC4 | Bacteria_unclassified_SGB43546 | 27 | 0.8626800771 | 0.3980585952 |
| acute_cyto_PC4 | Bacteria_unclassified_SGB63211 | 27 | -2.004917184 | 0.05803288344 |

| | | | | |
|---|---|---|---|---|
| acute_cyto_PC4 | Bacteroides_thetaiotaomicron | 27 | 0.3379066671 | 0.7387881338 |
| acute_cyto_PC4 | Clostridia_bacterium | 27 | -0.5136877915 | 0.6128330794 |
| acute_cyto_PC4 | Clostridiaceae_bacterium | 27 | -0.2631255119 | 0.7950182484 |
| acute_cyto_PC4 | Clostridiaceae_unclassified_SGB41663 | 27 | -0.7656000031 | 0.4524339813 |
| acute_cyto_PC4 | Clostridiales_bacterium | 27 | 0.2851798641 | 0.7783000156 |
| acute_cyto_PC4 | Clostridium_SGB65123 | 27 | -0.2433449652 | 0.8100994922 |
| acute_cyto_PC4 | Clostridium_cocleatum | 27 | -1.240158691 | 0.2285954642 |
| acute_cyto_PC4 | Eubacteriaceae_bacterium | 27 | 0.5045014686 | 0.6191617671 |
| acute_cyto_PC4 | Eubacteriaceae_unclassified_SGB94922 | 27 | 0.2438670853 | 0.809700411 |
| acute_cyto_PC4 | GGB20146_SGB29427 | 27 | -0.2597158674 | 0.7976121847 |
| acute_cyto_PC4 | GGB20149_SGB29430 | 27 | 2.050527867 | 0.0530011472 |
| acute_cyto_PC4 | GGB22635_SGB63107 | 27 | 1.465279291 | 0.1576575174 |
| acute_cyto_PC4 | GGB25041_SGB36960 | 27 | -0.494745697 | 0.6259160731 |
| acute_cyto_PC4 | GGB28379_SGB40959 | 27 | -1.778356265 | 0.08982850546 |
| acute_cyto_PC4 | GGB28382_SGB40962 | 27 | 1.010667784 | 0.3236794987 |
| acute_cyto_PC4 | GGB28392_SGB40972 | 27 | 0.3129656763 | 0.7573936985 |
| acute_cyto_PC4 | GGB28404_SGB40986 | 27 | -0.8097050351 | 0.4271898241 |
| acute_cyto_PC4 | GGB28418_SGB41001 | 27 | 0.03141968838 | 0.9752315483 |
| acute_cyto_PC4 | GGB28422_SGB41005 | 27 | 0.1454222959 | 0.8857642401 |
| acute_cyto_PC4 | GGB28431_SGB41014 | 27 | -1.183174 | 0.2499621723 |
| acute_cyto_PC4 | GGB28456_SGB41039 | 27 | 0.8729989646 | 0.3925368809 |
| acute_cyto_PC4 | GGB28782_SGB41435 | 27 | 2.913798589 | 0.00830102686[illegible] |
| acute_cyto_PC4 | GGB28792_SGB41445 | 27 | -1.729578537 | 0.09837877017 |
| acute_cyto_PC4 | GGB28798_SGB41451 | 27 | -1.74546042 | 0.09552120907 |
| acute_cyto_PC4 | GGB28810_SGB41465 | 27 | 1.2657596 | 0.2194623305 |
| acute_cyto_PC4 | GGB28828_SGB41484 | 27 | 0.1309810204 | 0.8970370078 |
| acute_cyto_PC4 | GGB28851_SGB41518 | 27 | 0.6315347447 | 0.534505723 |
| acute_cyto_PC4 | GGB28865_SGB41537 | 27 | -1.02579019 | 0.3166621381 |
| acute_cyto_PC4 | GGB28868_SGB41542 | 27 | 1.606382661 | 0.1231230537 |
| acute_cyto_PC4 | GGB28869_SGB41543 | 27 | 0.2434268992 | 0.8100368626 |
| acute_cyto_PC4 | GGB28875_SGB41555 | 27 | -0.1607110803 | 0.873856933 |
| acute_cyto_PC4 | GGB28881_SGB41561 | 27 | -0.9021031857 | 0.3772329684 |
| acute_cyto_PC4 | GGB28892_SGB41573 | 27 | 0.2372756472 | 0.8147424199 |
| acute_cyto_PC4 | GGB28893_SGB41574 | 27 | 0.1532510805 | 0.8796633326 |
| acute_cyto_PC4 | GGB28898_SGB41580 | 27 | -0.746622936 | 0.4635687616 |
| acute_cyto_PC4 | GGB28901_SGB41592 | 27 | 0.8331677553 | 0.4141262171 |
| acute_cyto_PC4 | GGB28904_SGB41597 | 27 | 0.8143131254 | 0.4246039737 |
| acute_cyto_PC4 | GGB28909_SGB41602 | 27 | -0.0108313065 | 0.9914602769 |
| acute_cyto_PC4 | GGB28924_SGB41621 | 27 | 0.4710154836 | 0.6424857454 |
| acute_cyto_PC4 | GGB28927_SGB41625 | 27 | -1.005340608 | 0.3261772135 |
| acute_cyto_PC4 | GGB28934_SGB41635 | 27 | 0.4681026567 | 0.6445330861 |
| acute_cyto_PC4 | GGB28949_SGB41655 | 27 | 0.07515255214 | 0.9408046628 |
| acute_cyto_PC4 | GGB28951_SGB102295 | 27 | -0.3521298613 | 0.7282500278 |
| acute_cyto_PC4 | GGB28951_SGB41658 | 27 | 0.7583156603 | 0.4566887888 |
| acute_cyto_PC4 | GGB28954_SGB41662 | 27 | -1.408452604 | 0.1736309616 |
| acute_cyto_PC4 | GGB28960_SGB41669 | 27 | -0.3082272721 | 0.7609459855 |

| | | | | |
|---|---|---|---|---|
| acute_cyto_PC4 | GGB28964_SGB94886 | 27 | 1.708382911 | 0.1023061965 |
| acute_cyto_PC4 | GGB28967_SGB41678 | 27 | -0.3727434166 | 0.7130750465 |
| acute_cyto_PC4 | GGB28996_SGB41712 | 27 | -1.285459176 | 0.2126277585 |
| acute_cyto_PC4 | GGB29531_SGB42317 | 27 | 0.08047967613 | 0.9366178187 |
| acute_cyto_PC4 | GGB29685_SGB42494 | 27 | 0.02217024415 | 0.9825214644 |
| acute_cyto_PC4 | GGB30141_SGB43066 | 27 | -0.0738448248 | 0.9418327421 |
| acute_cyto_PC4 | GGB30145_SGB43072 | 27 | 0.8437589321 | 0.4083131439 |
| acute_cyto_PC4 | GGB30286_SGB43248 | 27 | 0.03838666721 | 0.9697419788 |
| acute_cyto_PC4 | GGB30300_SGB43264 | 27 | -0.2889320822 | 0.7754663336 |
| acute_cyto_PC4 | GGB30303_SGB43268 | 27 | -0.162425492 | 0.8725235583 |
| acute_cyto_PC4 | GGB30450_SGB43507 | 27 | -0.2060988903 | 0.8386986541 |
| acute_cyto_PC4 | GGB30453_SGB43513 | 27 | -0.1083084215 | 0.9147792384 |
| acute_cyto_PC4 | GGB30454_SGB43514 | 27 | 1.814278493 | 0.08394892999 |
| acute_cyto_PC4 | GGB30455_SGB43519 | 27 | -0.1024729246 | 0.9193535207 |
| acute_cyto_PC4 | GGB30456_SGB43520 | 27 | 1.812951528 | 0.08416003484 |
| acute_cyto_PC4 | GGB30457_SGB63218 | 27 | 0.08957372983 | 0.9294747727 |
| acute_cyto_PC4 | GGB30461_SGB43530 | 27 | -0.920083213 | 0.3679782556 |
| acute_cyto_PC4 | GGB30461_SGB43533 | 27 | -0.3316934091 | 0.7434083157 |
| acute_cyto_PC4 | GGB30461_SGB63209 | 27 | -0.9146294293 | 0.3707692924 |
| acute_cyto_PC4 | GGB30463_SGB43537 | 27 | 0.731243059 | 0.472711939 |
| acute_cyto_PC4 | GGB30473_SGB43557 | 27 | 0.6342815225 | 0.5327464705 |
| acute_cyto_PC4 | GGB30475_SGB63182 | 27 | 0.09688143576 | 0.9237392219 |
| acute_cyto_PC4 | GGB30861_SGB44083 | 27 | -1.034082206 | 0.3128600922 |
| acute_cyto_PC4 | GGB31312_SGB44628 | 27 | 0.6692155784 | 0.5106490299 |
| acute_cyto_PC4 | GGB31438_SGB44768 | 27 | -0.7502155098 | 0.46144833 |
| acute_cyto_PC4 | GGB3171_SGB4185 | 27 | -0.7921267067 | 0.4371440277 |
| acute_cyto_PC4 | GGB31762_SGB45125 | 27 | -0.0233330884 | 0.9816048734 |
| acute_cyto_PC4 | GGB31823_SGB45199 | 27 | 0.09075304814 | 0.9285488945 |
| acute_cyto_PC4 | GGB31838_SGB45216 | 27 | -1.058671834 | 0.3017753752 |
| acute_cyto_PC4 | GGB31841_SGB65084 | 27 | -0.8713747533 | 0.3934026916 |
| acute_cyto_PC4 | GGB32371_SGB41694 | 27 | -0.4284729349 | 0.672669652 |
| acute_cyto_PC4 | GGB3793_SGB5158 | 27 | -0.6156915393 | 0.5447140138 |
| acute_cyto_PC4 | GGB42601_SGB59797 | 27 | -0.8423939252 | 0.409059403 |
| acute_cyto_PC4 | GGB45514_SGB63186 | 27 | 0.6562092896 | 0.518815686 |
| acute_cyto_PC4 | GGB45564_SGB63259 | 27 | 0.6053175763 | 0.5514541828 |
| acute_cyto_PC4 | GGB45624_SGB63337 | 27 | 0.6479772143 | 0.5240217956 |
| acute_cyto_PC4 | GGB45656_SGB63370 | 27 | -1.358415292 | 0.1887468887 |
| acute_cyto_PC4 | GGB47127_SGB65054 | 27 | 2.230691663 | 0.03673833262 |
| acute_cyto_PC4 | GGB74395_SGB43523 | 27 | -1.597338787 | 0.1251293948 |
| acute_cyto_PC4 | GGB75053_SGB43494 | 27 | -0.184662687 | 0.8552653247 |
| acute_cyto_PC4 | GGB75109_SGB102238 | 27 | -0.0238856107 | 0.9811693666 |
| acute_cyto_PC4 | Lachnospiraceae_bacterium | 27 | 0.3213498491 | 0.751121746 |
| acute_cyto_PC4 | Lachnospiraceae_bacterium_A2 | 27 | 0.8717941522 | 0.3931790058 |
| acute_cyto_PC4 | Lachnospiraceae_unclassified_SGB41414 | 27 | -0.9059090322 | 0.375261271 |
| acute_cyto_PC4 | Lachnospiraceae_unclassified_SGB41424 | 27 | -0.594908619 | 0.5582610302 |
| acute_cyto_PC4 | Lachnospiraceae_unclassified_SGB94868 | 27 | -0.0026962215 | 0.9978741781 |

| | | | | |
|---|---|---|---|---|
| acute_cyto_PC4 | Lactobacillus_johnsonii | 27 | -0.493865462 | 0.6265271664 |
| acute_cyto_PC4 | Leptogranulimonas_caecicola | 27 | 0.03811463546 | 0.9699562968 |
| acute_cyto_PC4 | Muribaculaceae_bacterium | 27 | 0.3329868843 | 0.7424456627 |
| acute_cyto_PC4 | Neglectibacter_sp_X4 | 27 | 0.3343993487 | 0.7413949469 |
| acute_cyto_PC4 | Oscillibacter_SGB43496 | 27 | 1.161007398 | 0.258666812 |
| acute_cyto_PC4 | Oscillospiraceae_bacterium | 27 | 0.4284370499 | 0.6726953621 |
| acute_cyto_PC4 | Oscillospiraceae_unclassified_SGB43502 | 27 | 1.119048648 | 0.2757566387 |
| acute_cyto_PC4 | Oscillospiraceae_unclassified_SGB43505 | 27 | -0.3579761213 | 0.723934266 |
| acute_cyto_PC4 | Oscillospiraceae_unclassified_SGB94989 | 27 | 0.1272604688 | 0.899944963 |
| acute_cyto_PC4 | Parasutterella_excrementihominis | 27 | -1.805812581 | 0.08530368235 |
| acute_cyto_PC4 | Schaedlerella_arabinosiphila | 27 | -1.502009509 | 0.1479831614 |
| acute_cyto_PC4 | Turicibacter_sp_1E2 | 27 | 0.601852887 | 0.5537150212 |
| acute_cyto_PC4 | bacterium_0_1xD8_82 | 27 | -0.9656713329 | 0.3451986689 |
| acute_cyto_PC4 | bacterium_1XD42_1 | 27 | -0.2144353541 | 0.8322761366 |
| acute_cyto_PC4 | bacterium_1XD42_54 | 27 | -0.3086900377 | 0.7605988196 |
| acute_cyto_PC4 | bacterium_1XD42_76 | 27 | -0.4054858278 | 0.6892219139 |
| acute_cyto_PC4 | bacterium_1xD8_48 | 27 | 1.142848945 | 0.2659638145 |
| acute_cyto_PC4 | Berger Parker Index | 27 | 0.04234323106 | 0.966625106 |
| acute_cyto_PC4 | Richness (# observed features) | 27 | 0.01267875952 | 0.990003766 |
| acute_cyto_PC4 | Shannon Index | 27 | -0.5582312714 | 0.5825899045 |
| acute_cyto_PC5 | Acetatifactor_SGB41545 | 27 | -0.3096492217 | 0.7598794062 |
| acute_cyto_PC5 | Acutalibacter_muris | 27 | 0.978737021 | 0.3388514184 |
| acute_cyto_PC5 | Akkermansia_muciniphila | 27 | 1.539382489 | 0.1386442973 |
| acute_cyto_PC5 | Anaerotruncus_sp_1XD42_93 | 27 | 1.567419492 | 0.1319624507 |
| acute_cyto_PC5 | Bacteria_unclassified_SGB41677 | 27 | -1.083706231 | 0.2907811712 |
| acute_cyto_PC5 | Bacteria_unclassified_SGB43539 | 27 | -0.9445641512 | 0.3556232263 |
| acute_cyto_PC5 | Bacteria_unclassified_SGB43546 | 27 | -2.652808862 | 0.01488461554 |
| acute_cyto_PC5 | Bacteria_unclassified_SGB63211 | 27 | -0.9325265902 | 0.361662888 |
| acute_cyto_PC5 | Bacteroides_thetaiotaomicron | 27 | -0.486105574 | 0.6319262465 |
| acute_cyto_PC5 | Clostridia_bacterium | 27 | 1.370435118 | 0.1850238283 |
| acute_cyto_PC5 | Clostridiaceae_bacterium | 27 | 1.201930954 | 0.2427694518 |
| acute_cyto_PC5 | Clostridiaceae_unclassified_SGB41663 | 27 | 1.380748843 | 0.1818758808 |
| acute_cyto_PC5 | Clostridiales_bacterium | 27 | -0.147331557 | 0.8842756834 |
| acute_cyto_PC5 | Clostridium_SGB65123 | 27 | 1.341136118 | 0.1942024975 |
| acute_cyto_PC5 | Clostridium_cocleatum | 27 | -0.00094931903 | 0.9992515135 |
| acute_cyto_PC5 | Eubacteriaceae_bacterium | 27 | 0.1638214416 | 0.8714381526 |
| acute_cyto_PC5 | Eubacteriaceae_unclassified_SGB94922 | 27 | 1.749405199 | 0.09482258756 |
| acute_cyto_PC5 | GGB20146_SGB29427 | 27 | 2.09766964 | 0.0482146708 |
| acute_cyto_PC5 | GGB20149_SGB29430 | 27 | 1.422884055 | 0.1694561509 |
| acute_cyto_PC5 | GGB22635_SGB63107 | 27 | 1.101350674 | 0.2832078842 |
| acute_cyto_PC5 | GGB25041_SGB36960 | 27 | -0.8603967561 | 0.3992871712 |
| acute_cyto_PC5 | GGB28379_SGB40959 | 27 | 1.47690336 | 0.1545416901 |
| acute_cyto_PC5 | GGB28382_SGB40962 | 27 | 0.4135737302 | 0.6833793254 |
| acute_cyto_PC5 | GGB28392_SGB40972 | 27 | -0.1896789541 | 0.8513821305 |
| acute_cyto_PC5 | GGB28404_SGB40986 | 27 | -2.115385234 | 0.04651898783 |
| acute_cyto_PC5 | GGB28418_SGB41001 | 27 | -0.8362145597 | 0.4124485877 |

| | | | | |
|---|---|---|---|---|
| acute_cyto_PC5 | GGB28422_SGB41005 | 27 | 1.564880941 | 0.1325562273 |
| acute_cyto_PC5 | GGB28431_SGB41014 | 27 | -1.768670637 | 0.09147343241 |
| acute_cyto_PC5 | GGB28456_SGB41039 | 27 | -0.3766995312 | 0.7101763428 |
| acute_cyto_PC5 | GGB28782_SGB41435 | 27 | -0.7432240988 | 0.4655801956 |
| acute_cyto_PC5 | GGB28792_SGB41445 | 27 | -1.465387927 | 0.1576281629 |
| acute_cyto_PC5 | GGB28798_SGB41451 | 27 | -1.92706551 | 0.06760968239 |
| acute_cyto_PC5 | GGB28810_SGB41465 | 27 | 0.424256454 | 0.6756933987 |
| acute_cyto_PC5 | GGB28828_SGB41484 | 27 | 0.798558248 | 0.4334854718 |
| acute_cyto_PC5 | GGB28851_SGB41518 | 27 | -0.7877056398 | 0.439669961 |
| acute_cyto_PC5 | GGB28865_SGB41537 | 27 | -1.202420784 | 0.2425837289 |
| acute_cyto_PC5 | GGB28868_SGB41542 | 27 | -0.2792224218 | 0.7828055758 |
| acute_cyto_PC5 | GGB28869_SGB41543 | 27 | 0.8392004314 | 0.4108087078 |
| acute_cyto_PC5 | GGB28875_SGB41555 | 27 | -0.07408967694 | 0.9416402419 |
| acute_cyto_PC5 | GGB28881_SGB41561 | 27 | 0.2202244385 | 0.8278231804 |
| acute_cyto_PC5 | GGB28892_SGB41573 | 27 | -0.2281233779 | 0.8217569205 |
| acute_cyto_PC5 | GGB28893_SGB41574 | 27 | -1.702033883 | 0.103508377 |
| acute_cyto_PC5 | GGB28898_SGB41580 | 27 | 0.4805444686 | 0.6358085023 |
| acute_cyto_PC5 | GGB28901_SGB41592 | 27 | 0.3336691597 | 0.7419380615 |
| acute_cyto_PC5 | GGB28904_SGB41597 | 27 | 0.4884155011 | 0.6303168527 |
| acute_cyto_PC5 | GGB28909_SGB41602 | 27 | 0.2822148972 | 0.7805414054 |
| acute_cyto_PC5 | GGB28924_SGB41621 | 27 | 2.380298205 | 0.02684955058 |
| acute_cyto_PC5 | GGB28927_SGB41625 | 27 | -0.2681550885 | 0.7911963677 |
| acute_cyto_PC5 | GGB28934_SGB41635 | 27 | -0.8402299452 | 0.4102442496 |
| acute_cyto_PC5 | GGB28949_SGB41655 | 27 | -0.07220443055 | 0.9431224967 |
| acute_cyto_PC5 | GGB28951_SGB102295 | 27 | -0.2889106881 | 0.7754824814 |
| acute_cyto_PC5 | GGB28951_SGB41658 | 27 | 0.7917506391 | 0.4373585409 |
| acute_cyto_PC5 | GGB28954_SGB41662 | 27 | -0.8286196058 | 0.4166385605 |
| acute_cyto_PC5 | GGB28960_SGB41669 | 27 | -2.12061281 | 0.04602900326 |
| acute_cyto_PC5 | GGB28964_SGB94886 | 27 | 1.198404007 | 0.2441098883 |
| acute_cyto_PC5 | GGB28967_SGB41678 | 27 | -1.204086557 | 0.24195294 |
| acute_cyto_PC5 | GGB28996_SGB41712 | 27 | 0.8495098475 | 0.4051786384 |
| acute_cyto_PC5 | GGB29531_SGB42317 | 27 | 0.3440851382 | 0.7342038304 |
| acute_cyto_PC5 | GGB29685_SGB42494 | 27 | 1.118360637 | 0.2760436028 |
| acute_cyto_PC5 | GGB30141_SGB43066 | 27 | 0.4928579722 | 0.6272269425 |
| acute_cyto_PC5 | GGB30145_SGB43072 | 27 | -1.5363527 | 0.1393828228 |
| acute_cyto_PC5 | GGB30286_SGB43248 | 27 | -1.066598718 | 0.2982624789 |
| acute_cyto_PC5 | GGB30300_SGB43264 | 27 | -1.340409195 | 0.1944347053 |
| acute_cyto_PC5 | GGB30303_SGB43268 | 27 | -0.9979576804 | 0.3296609601 |
| acute_cyto_PC5 | GGB30450_SGB43507 | 27 | 1.652558436 | 0.1132949128 |
| acute_cyto_PC5 | GGB30453_SGB43513 | 27 | 1.242635326 | 0.2276994246 |
| acute_cyto_PC5 | GGB30454_SGB43514 | 27 | -0.8469322003 | 0.4065816592 |
| acute_cyto_PC5 | GGB30455_SGB43519 | 27 | 0.7068452854 | 0.4874325302 |
| acute_cyto_PC5 | GGB30456_SGB43520 | 27 | -0.3935107739 | 0.6979090696 |
| acute_cyto_PC5 | GGB30457_SGB63218 | 27 | -0.6616737976 | 0.5153757219 |
| acute_cyto_PC5 | GGB30461_SGB43530 | 27 | -2.129277161 | 0.04522714171 |
| acute_cyto_PC5 | GGB30461_SGB43533 | 27 | 0.8501040976 | 0.4048556272 |

| | | | | |
|---|---|---|---|---|
| acute_cyto_PC5 | GGB30461_SGB63209 | 27 | 1.261052512 | 0.2211201919 |
| acute_cyto_PC5 | GGB30463_SGB43537 | 27 | 0.2790963436 | 0.7829010128 |
| acute_cyto_PC5 | GGB30473_SGB43557 | 27 | 0.6992152723 | 0.4920901419 |
| acute_cyto_PC5 | GGB30475_SGB63182 | 27 | -0.9755825285 | 0.340376455 |
| acute_cyto_PC5 | GGB30861_SGB44083 | 27 | -1.335965555 | 0.1958589329 |
| acute_cyto_PC5 | GGB31312_SGB44628 | 27 | -0.1581558697 | 0.8758449468 |
| acute_cyto_PC5 | GGB31438_SGB44768 | 27 | 0.8521586764 | 0.4037401113 |
| acute_cyto_PC5 | GGB3171_SGB4185 | 27 | 0.6304434312 | 0.535205558 |
| acute_cyto_PC5 | GGB31762_SGB45125 | 27 | 1.235814588 | 0.2301736483 |
| acute_cyto_PC5 | GGB31823_SGB45199 | 27 | 1.235348085 | 0.2303436188 |
| acute_cyto_PC5 | GGB31838_SGB45216 | 27 | -1.225921432 | 0.233798756 |
| acute_cyto_PC5 | GGB31841_SGB65084 | 27 | 0.1738115173 | 0.8636781831 |
| acute_cyto_PC5 | GGB32371_SGB41694 | 27 | 0.9073681633 | 0.3745071523 |
| acute_cyto_PC5 | GGB3793_SGB5158 | 27 | 0.1454115168 | 0.8857726453 |
| acute_cyto_PC5 | GGB42601_SGB59797 | 27 | -1.814659684 | 0.08388837191 |
| acute_cyto_PC5 | GGB45514_SGB63186 | 27 | -0.1321426422 | 0.8961293948 |
| acute_cyto_PC5 | GGB45564_SGB63259 | 27 | 0.9779238008 | 0.3392441183 |
| acute_cyto_PC5 | GGB45624_SGB63337 | 27 | -1.331030885 | 0.1974501358 |
| acute_cyto_PC5 | GGB45656_SGB63370 | 27 | -0.1027877003 | 0.9191067024 |
| acute_cyto_PC5 | GGB47127_SGB65054 | 27 | 1.287419677 | 0.2119566975 |
| acute_cyto_PC5 | GGB74395_SGB43523 | 27 | -0.06884262351 | 0.9457662025 |
| acute_cyto_PC5 | GGB75053_SGB43494 | 27 | 0.420597881 | 0.6783216138 |
| acute_cyto_PC5 | GGB75109_SGB102238 | 27 | 0.890149429 | 0.3834703182 |
| acute_cyto_PC5 | Lachnospiraceae_bacterium | 27 | 0.6143630085 | 0.5455747303 |
| acute_cyto_PC5 | Lachnospiraceae_bacterium_A2 | 27 | 1.171214928 | 0.2546307738 |
| acute_cyto_PC5 | Lachnospiraceae_unclassified_SGB41414 | 27 | 0.7907994898 | 0.4379013773 |
| acute_cyto_PC5 | Lachnospiraceae_unclassified_SGB41424 | 27 | -1.334221959 | 0.1964200068 |
| acute_cyto_PC5 | Lachnospiraceae_unclassified_SGB94868 | 27 | -0.6014257392 | 0.5539940878 |
| acute_cyto_PC5 | Lactobacillus_johnsonii | 27 | -0.00232328415 | 0.9981682181 |
| acute_cyto_PC5 | Leptogranulimonas_caecicola | 27 | 1.867324302 | 0.07587850089 |
| acute_cyto_PC5 | Muribaculaceae_bacterium | 27 | 0.9501273587 | 0.3528551549 |
| acute_cyto_PC5 | Neglectibacter_sp_X4 | 27 | 0.2423160348 | 0.8108861076 |
| acute_cyto_PC5 | Oscillibacter_SGB43496 | 27 | -1.32725454 | 0.1986746723 |
| acute_cyto_PC5 | Oscillospiraceae_bacterium | 27 | 0.1224919858 | 0.9036740757 |
| acute_cyto_PC5 | Oscillospiraceae_unclassified_SGB43502 | 27 | 1.026278324 | 0.3164374223 |
| acute_cyto_PC5 | Oscillospiraceae_unclassified_SGB43505 | 27 | -0.6574904754 | 0.5180080258 |
| acute_cyto_PC5 | Oscillospiraceae_unclassified_SGB94989 | 27 | -2.01588025 | 0.05678601931 |
| acute_cyto_PC5 | Parasutterella_excrementihominis | 27 | -0.01604049968 | 0.9873535037 |
| acute_cyto_PC5 | Schaedlerella_arabinosiphila | 27 | -0.9128055055 | 0.3717058445 |
| acute_cyto_PC5 | Turicibacter_sp_1E2 | 27 | 0.7452989227 | 0.4643516954 |
| acute_cyto_PC5 | bacterium_0_1xD8_82 | 27 | 0.5112573272 | 0.6145045042 |
| acute_cyto_PC5 | bacterium_1XD42_1 | 27 | -2.33746808 | 0.02939495413 |
| acute_cyto_PC5 | bacterium_1XD42_54 | 27 | -0.6113728507 | 0.5475146111 |
| acute_cyto_PC5 | bacterium_1XD42_76 | 27 | 0.6974615359 | 0.4931642904 |
| acute_cyto_PC5 | bacterium_1xD8_48 | 27 | 0.3740792347 | 0.712095773 |
| acute_cyto_PC5 | Berger Parker Index | 27 | 1.431913881 | 0.1668852645 |

| | | | | |
|---|---|---|---|---|
| acute_cyto_PC5 | Richness (# observed features) | 27 | -1.150101141 | 0.2630314661 |
| acute_cyto_PC5 | Shannon Index | 27 | -1.426489391 | 0.1684258779 |

| LRT_q_microbe | est_intercept | est_microbe | est_blast | est_vns | est_interact | est_batch |
|---|---|---|---|---|---|---|
| 0.9750766376 | -3.234967486 | 0.2621333141 | 5.724219169 | -0.478833464 | -1.255417694 | 1.402141897 |
| 0.9750766376 | -3.064374596 | -0.181341115 | 5.400436133 | -0.718750181 | -0.8571665865 | 1.43472364 |
| 0.9750766376 | -3.125495505 | -0.179252229 | 5.399525511 | -0.516598809 | -1.018358115 | 1.444211093 |
| 0.9907090994 | -3.06646382 | -0.047075957 | 5.429490387 | -0.702195107 | -0.8058312602 | 1.356119148 |
| 0.9907090994 | -3.096408584 | -0.012793044 | 5.47288097 | -0.609071455 | -0.9173976175 | 1.336615271 |
| 0.8476569724 | -2.885674582 | 0.4019581923 | 5.021974301 | -0.760215376 | -0.5341468226 | 1.346137619 |
| 0.9750766376 | -2.923764799 | 0.1808705054 | 5.264301271 | -0.755180910 | -0.7388327463 | 1.266178862 |
| 0.5326401688 | -3.495201505 | 0.7101130135 | 5.662878342 | -0.31720145 | -1.611899267 | 2.058713842 |
| 0.9750766376 | -2.958261949 | -0.180159507 | 5.223006067 | -0.758149803 | -0.7331930502 | 1.385201791 |
| 0.9750766376 | -2.963449423 | -0.182320821 | 5.300473186 | -0.752438361 | -0.782896163 | 1.331026385 |
| 0.9750766376 | -3.019971995 | -0.151628554 | 5.312170086 | -0.741460471 | -0.7772958426 | 1.419654301 |
| 0.7255653047 | -2.787135767 | -0.496730844 | 5.13756213 | -1.124374943 | -0.7323643592 | 1.522261372 |
| 0.9750766376 | -3.245472228 | 0.2052325176 | 5.727709546 | -0.486136238 | -1.22608858 | 1.40974792 |
| 0.2656465465 | -2.264824264 | -0.990177459 | 4.884388924 | -1.339724288 | 0.3125010752 | 0.3185040538 |
| 0.9750766376 | -3.165118749 | -0.118580280 | 5.479966131 | -0.521488961 | -1.002806688 | 1.429385633 |
| 0.9878805585 | -3.095558218 | -0.075035287 | 5.392550331 | -0.607280644 | -0.9091229257 | 1.420517642 |
| 0.9907090994 | -3.077814362 | -0.036215590 | 5.432202224 | -0.631187318 | -0.8790752222 | 1.345167588 |
| 0.9854090226 | -3.128374738 | 0.1107027004 | 5.527069441 | -0.568583190 | -1.087222333 | 1.401386125 |
| 0.9750766376 | -3.152052228 | 0.1518098309 | 5.567948883 | -0.602656179 | -1.067426032 | 1.427905514 |
| 0.9750766376 | -3.123837849 | 0.1229180435 | 5.522962877 | -0.682493187 | -0.8386135478 | 1.366383982 |
| 0.9878805585 | -3.119218404 | 0.0918919488 | 5.548839413 | -0.528428997 | -0.9851028543 | 1.251164194 |
| 0.5326401688 | -2.001022572 | 1.019766727 | 4.586820583 | -1.301608451 | 0.4914400084 | -0.0372077733 |
| 0.6848176303 | -2.988014156 | -0.514150981 | 5.349650229 | -0.225420607 | -1.429668222 | 1.155758697 |
| 0.6848176303 | -3.444981976 | -0.561375706 | 6.026364834 | -0.031742781 | -2.116111233 | 1.537871511 |
| 0.9750766376 | -2.945697437 | 0.1312117371 | 5.347540779 | -0.806454181 | -0.7864453424 | 1.300203107 |
| 0.9907090994 | -3.105451291 | 0.0105402226 | 5.471381152 | -0.605395684 | -0.9194744712 | 1.354446378 |
| 0.9750766376 | -3.046300529 | 0.1397541645 | 5.351622132 | -0.70991664 | -0.8202624972 | 1.421285788 |
| 0.9879722819 | -3.129659575 | -0.060897676 | 5.548345061 | -0.533154613 | -1.049734173 | 1.318282351 |
| 0.9101962467 | -3.120219081 | 0.3579123869 | 5.704162111 | -0.641424854 | -1.087627813 | 1.258804151 |
| 0.9750766376 | -2.978209381 | -0.254888556 | 5.397578522 | -0.794997957 | -0.647306919 | 1.21203094 |
| 0.9750766376 | -3.077001389 | 0.1174135725 | 5.367862445 | -0.610960930 | -0.8998575316 | 1.408724161 |
| 0.9750766376 | -3.151044376 | 0.1466537352 | 5.48896015 | -0.534431109 | -0.9115777671 | 1.347573352 |
| 0.9907090994 | -3.121105264 | -0.021562239 | 5.474845136 | -0.604039168 | -0.923256908 | 1.383828362 |
| 0.6848176303 | -3.299047811 | -0.520974642 | 6.0586942 | -0.184889636 | -1.95776072 | 1.264955274 |
| 0.9750766376 | -3.005521663 | 0.2834524997 | 5.463472225 | -0.720647835 | -0.9408352005 | 1.29328868 |
| 0.5326401688 | -3.414916548 | -0.744979327 | 6.321926435 | 0.3313684964 | -2.055721273 | 0.7061893469 |
| 0.9750766376 | -3.087174401 | 0.1102055332 | 5.3824951 | -0.647148709 | -0.7846290236 | 1.381030496 |
| 0.9750766376 | -2.985038758 | 0.3647290878 | 5.507365266 | -0.614551102 | -0.7956069777 | 0.9964722102 |
| 0.9907090994 | -3.088471129 | -0.020109359 | 5.467183866 | -0.609551288 | -0.921094296 | 1.329494992 |
| 0.9750766376 | -2.96563876 | -0.282239620 | 5.180309747 | -1.09406606 | -0.383512512 | 1.596355789 |
| 0.9750766376 | -3.330516343 | -0.336820488 | 5.589987795 | -0.500705959 | -0.9429601775 | 1.586745323 |
| 0.5326401688 | -3.318118286 | 0.7137647408 | 5.722014894 | -0.774491804 | -1.302042971 | 1.924476873 |
| 0.9750766376 | -3.002067374 | -0.155964579 | 5.440484452 | -0.684856356 | -0.8169250318 | 1.197841658 |
| 0.7045398025 | -2.862951666 | -0.502900401 | 4.8594418 | -0.592050911 | -0.4421836024 | 1.248788428 |
| 0.8476569724 | -3.295455918 | 0.5224845302 | 5.612925191 | -0.592598967 | -1.433650455 | 1.888386239 |
| 0.5326401688 | -2.801090455 | -0.719327730 | 5.528637383 | -0.613174029 | -1.653884451 | 1.116568735 |

| | | | | | | |
|---|---|---|---|---|---|---|
| 0.5326401688 | -2.705384608 | -0.751903455 | 5.813570737 | -0.650115473 | -1.280586824 | 0.399087275 |
| 0.9750766376 | -2.785322348 | -0.411569317 | 5.014858929 | -1.459410899 | 0.0247187396 | 1.554986318 |
| 0.9750766376 | -3.17900886 | 0.192421048 | 5.632330614 | -0.458429271 | -1.191607063 | 1.330703563 |
| 0.9750766376 | -2.889071268 | -0.329669337 | 5.149404481 | -0.749936551 | -0.7644142003 | 1.33679095 |
| 0.9750766376 | -3.239329856 | 0.189744263 | 5.558534968 | -0.446937875 | -1.002739873 | 1.41253314 |
| 0.9101962467 | -3.600779846 | -0.448040446 | 5.619753782 | -0.124139475 | -1.017531788 | 1.754073545 |
| 0.9878805585 | -3.125775968 | 0.088434816 | 5.466911717 | -0.603198785 | -0.947525510 | 1.416712498 |
| 0.9101962467 | -2.814957614 | -0.417105693 | 5.526323063 | -0.818198713 | -1.200480469 | 1.089818106 |
| 0.9878805585 | -3.051928479 | -0.083349857 | 5.399946991 | -0.661191297 | -0.889710644 | 1.367479029 |
| 0.9750766376 | -3.054622301 | -0.146113583 | 5.426804473 | -0.659376858 | -0.8525467977 | 1.317260418 |
| 0.9750766376 | -3.103565275 | 0.109842238 | 5.465243928 | -0.612758121 | -0.9900637244 | 1.408979001 |
| 0.9750766376 | -3.062218154 | 0.127733910 | 5.387929118 | -0.601440772 | -0.920275523 | 1.357179106 |
| 0.9878805585 | -3.14641978 | 0.086672678 | 5.590911792 | -0.579622057 | -0.9611478598 | 1.299503758 |
| 0.9750766376 | -3.057284352 | -0.278107386 | 5.607485438 | -0.478667899 | -1.080656142 | 1.060076588 |
| 0.6848176303 | -3.297878692 | 0.489149154 | 5.479481991 | -0.203136995 | -1.34734816 | 1.574861388 |
| 0.9750766376 | -3.151852562 | 0.242150208 | 5.50343947 | -0.572972006 | -1.047027292 | 1.457404119 |
| 0.8476569724 | -2.707742048 | -0.479640022 | 5.216338495 | -0.949478863 | -0.355905500 | 0.846454305 |
| 0.9750766376 | -3.256061936 | 0.281974953 | 5.461820029 | -0.468224792 | -1.010306319 | 1.586459191 |
| 0.9750766376 | -3.401382332 | -0.284817545 | 5.887263805 | -0.356805732 | -1.291585023 | 1.45048664 |
| 0.9907090994 | -3.090733728 | 0.042017756 | 5.440549529 | -0.637538153 | -0.8679100897 | 1.362337121 |
| 0.9907090994 | -3.099540271 | 0.043916360 | 5.493218304 | -0.610907289 | -0.9335583276 | 1.331575495 |
| 0.9750766376 | -2.981424621 | 0.163829943 | 5.397090204 | -0.790807900 | -0.8126182805 | 1.316489889 |
| 0.9750766376 | -3.26077703 | -0.280776421 | 5.793025946 | -0.398105035 | -1.346048486 | 1.345188385 |
| 0.9750766376 | -3.137712704 | 0.138528831 | 5.467677427 | -0.610779548 | -0.8762714093 | 1.404935889 |
| 0.9750766376 | -2.975651969 | -0.185280482 | 5.416508203 | -0.732657087 | -0.9485505747 | 1.303121842 |
| 0.9878805585 | -3.162068957 | 0.069450800 | 5.493845414 | -0.575423169 | -0.9409091021 | 1.427029063 |
| 0.920225808 | -3.218655674 | 0.455455267 | 5.232413364 | -0.105980897 | -1.348939259 | 1.591749952 |
| 0.9907090994 | -3.098417013 | 0.008316123 | 5.463099828 | -0.604187660 | -0.9190462282 | 1.347826646 |
| 0.9907090994 | -3.099087425 | -0.019736685 | 5.463267553 | -0.625520399 | -0.8973512934 | 1.358649471 |
| 0.9750766376 | -3.127280578 | 0.217265121 | 5.478033216 | -0.600942907 | -1.0060188 | 1.440571421 |
| 0.6452942772 | -2.586530815 | 0.673283981 | 5.313798033 | -0.623400009 | -0.9543448454 | 0.499363108 |
| 0.9750766376 | -3.041714992 | 0.166745456 | 5.435579719 | -0.651745485 | -0.8386445569 | 1.263554019 |
| 0.2654279322 | -3.012135262 | -0.878736945 | 4.979708788 | -1.109712466 | -0.0584911025 | 1.741224893 |
| 0.6848176303 | -2.932217484 | -0.604482862 | 4.831793696 | -1.298185591 | -0.1081345611 | 1.979433954 |
| 0.9750766376 | -3.068642308 | -0.116713070 | 5.401624583 | -0.681606006 | -0.7966326877 | 1.364963012 |
| 0.9750766376 | -3.094952339 | -0.136579593 | 5.438358512 | -0.695840341 | -0.8195004559 | 1.406623376 |
| 0.9907090994 | -3.140150823 | -0.047644131 | 5.483992001 | -0.605585498 | -0.9179827555 | 1.411250095 |
| 0.5434355159 | -2.941335235 | -0.582639393 | 5.347865754 | -1.014683502 | -0.3437530194 | 1.242589094 |
| 0.9878805585 | -3.124272301 | -0.076128992 | 5.465052387 | -0.543809100 | -0.984832999 | 1.37473521 |
| 0.9750766376 | -3.152222202 | -0.15149117 | 5.454718824 | -0.612745279 | -0.8643102351 | 1.444779453 |
| 0.9750766376 | -3.025295168 | 0.288865468 | 5.314203865 | -0.716680708 | -0.9106437587 | 1.48373781 |
| 0.9750766376 | -3.133194768 | 0.168897221 | 5.447905234 | -0.682548731 | -0.8829921526 | 1.499792283 |
| 0.9907090994 | -3.103590668 | -0.022491866 | 5.465310439 | -0.622398506 | -0.8980835176 | 1.362733744 |
| 0.6848176303 | -2.873328731 | 0.496742177 | 5.197390996 | -1.014637104 | -0.5786624537 | 1.419479222 |
| 0.9750766376 | -2.97277419 | 0.311236958 | 5.302682497 | -0.955536632 | -0.5904873757 | 1.44815981 |
| 0.9750766376 | -3.230256363 | -0.178174383 | 5.589513165 | -0.540977177 | -0.9936713198 | 1.453636567 |

| | | | | | | |
|---|---|---|---|---|---|---|
| 0.6278895264 | -2.715564269 | 0.584584625 | 5.259280568 | -0.80228113 | -0.716285323 | 0.876403473 |
| 0.9750766376 | -3.002242264 | -0.33159567 | 5.622801759 | -0.58625032 | -1.361137904 | 1.216547881 |
| 0.6848176303 | -2.903246 | -0.48738952 | 5.493702005 | -0.61234637 | -1.216217892 | 1.098823412 |
| 0.9878805585 | -3.139732074 | 0.105202324 | 5.581300795 | -0.583945982 | -0.932634594 | 1.283813429 |
| 0.9750766376 | -3.232691515 | -0.166892059 | 5.494350884 | -0.484805311 | -0.877848297 | 1.436728338 |
| 0.9750766376 | -3.114929908 | 0.178381092 | 5.528693077 | -0.596268761 | -1.039116722 | 1.371800599 |
| 0.5326401688 | -3.128862226 | 0.804720077 | 4.997248591 | 0.207759931 | -0.934824193 | 1.083885058 |
| 0.9907090994 | -3.097328943 | -0.00982913 | 5.465247841 | -0.60597908 | -0.920836866 | 1.346120455 |
| 0.9750766376 | -3.169458752 | 0.269344599 | 5.777570612 | -0.54440668 | -0.952438572 | 1.088694554 |
| 0.6848176303 | -3.583301716 | -0.57101065 | 5.359164045 | -0.034334169 | -0.883875572 | 1.839489894 |
| 0.99399338 | -3.1030424 | -0.00305915 | 5.466664731 | -0.604578808 | -0.918711658 | 1.353536185 |
| 0.9750766376 | -3.027362052 | 0.206061216 | 5.313173888 | -0.598083565 | -0.7891169861 | 1.2867133 |
| 0.6755436502 | -2.833830972 | 0.554531899 | 5.175859196 | -0.828496539 | -1.057671454 | 1.4566055 |
| 0.9750766376 | -3.079619456 | 0.286352255 | 5.277586759 | -0.562548958 | -0.704711287 | 1.346099974 |
| 0.9750766376 | -3.212158758 | 0.155058535 | 5.706873696 | -0.451832572 | -1.144043554 | 1.277168113 |
| 0.2656465465 | -2.833094012 | 0.836592452 | 5.475923754 | -0.925932592 | -0.303080590 | 0.749414079 |
| 0.7733300493 | -2.936534428 | 0.411862092 | 5.420194277 | -0.768131869 | -0.921895235 | 1.239424881 |
| 0.6452942772 | -3.23417271 | 0.746100511 | 6.157835432 | -0.300287467 | -1.021526073 | 0.563951139 |
| 0.9750766376 | -3.133657806 | -0.116361656 | 5.603096512 | -0.559000464 | -1.034214915 | 1.281695301 |
| 0.5326401688 | -2.811633911 | -0.79938305 | 4.263887434 | -0.809092837 | -0.0728419814 | 1.83583382 |
| 0.9750766376 | -3.101568171 | -0.13657139 | 5.529823546 | -0.605258654 | -0.969790504 | 1.309764049 |
| 0.9878805585 | -3.080261915 | -0.06875587 | 5.495264252 | -0.603774113 | -0.998180591 | 1.321260788 |
| 0.9878805585 | -3.135910422 | 0.084302332 | 5.570459396 | -0.523686895 | -1.0058452 | 1.268543738 |
| 0.2654279322 | -3.942497537 | 1.086914496 | 6.188271227 | -0.148550808 | -2.206478625 | 2.565761955 |
| 0.9750766376 | -3.233160013 | -0.16249100 | 5.654492622 | -0.504256991 | -1.097822206 | 1.409238967 |
| 0.9750766376 | -3.325574258 | 0.220104978 | 5.712087389 | -0.548364032 | -1.076322582 | 1.568990096 |
| 0.9879722819 | -3.08995554 | 0.067791738 | 5.395522915 | -0.608327492 | -0.907814792 | 1.405773777 |
| 0.9878805585 | -3.160753351 | -0.07786341 | 5.577538867 | -0.596425489 | -0.947272581 | 1.354261152 |
| 0.9703291146 | 0.5712867148 | -0.179274501 | -3.201627502 | -0.61969489 | 2.239991968 | 1.79657415 |
| 0.9703291146 | 0.4138244054 | 0.302937179 | -2.910455024 | -0.342410202 | 1.905949241 | 1.694593487 |
| 0.9743440267 | 0.4768235015 | -0.05502287 | -3.050504394 | -0.507252288 | 1.980087719 | 1.857243204 |
| 0.9703291146 | 0.801666998 | -0.382271094 | -3.366544627 | -1.331360314 | 2.934854197 | 1.847182406 |
| 0.9703291146 | 0.8111301872 | -0.450960291 | -2.96632593 | -0.712428647 | 2.090788219 | 1.218620715 |
| 0.9703291146 | 0.7766097982 | 0.536532476 | -3.628042372 | -0.742584097 | 2.524984853 | 1.819113044 |
| 0.9703291146 | 0.2431837289 | -0.23828079 | -2.75606853 | -0.334933042 | 1.772129646 | 1.945160986 |
| 0.9703291146 | 0.1820228917 | 0.548555849 | -2.880394634 | -0.31252586 | 1.475644056 | 2.373962388 |
| 0.9850546863 | 0.501033507 | -0.02216260 | -3.0590507 | -0.55304597 | 2.033318963 | 1.833380602 |
| 0.9703291146 | 0.7107818304 | -0.292040065 | -3.301863927 | -0.771832498 | 2.229475373 | 1.792823191 |
| 0.9703291146 | 0.641146927 | -0.279839364 | -3.321873672 | -0.78775347 | 2.273154071 | 1.950804168 |
| 0.9703291146 | 1.076867116 | -0.926112314 | -3.65041245 | -1.504256634 | 2.359609803 | 2.14334532 |
| 0.9703291146 | 0.5918238961 | -0.159969117 | -3.228531364 | -0.625963195 | 2.249210234 | 1.786040625 |
| 0.151382718 | 1.870376954 | -1.63379875 | -3.99663282 | -1.748014001 | 4.043482204 | 0.121050224 |
| 0.9743440267 | 0.5447771959 | 0.123540978 | -3.03774900 | -0.620059913 | 2.096983676 | 1.750940016 |
| 0.9743440267 | 0.490188325 | -0.053863212 | -3.084926737 | -0.53643076 | 2.017955492 | 1.877323316 |
| 0.9703291146 | 0.1176617372 | 0.473991074 | -2.51921922 | -0.178414263 | 1.478957869 | 1.944427552 |
| 0.9703291146 | 0.340525233 | 0.695689993 | -2.67690259 | -0.31143821 | 0.9574823167 | 2.127664158 |

| | | | | | | |
|---|---|---|---|---|---|---|
| 0.9703291146 | 0.2886005928 | 0.636612938 | -2.622462867 | -0.528398416 | 1.390801353 | 2.139684155 |
| 0.9703291146 | 0.4017885687 | 0.550250966 | -2.796434589 | -0.885410368 | 2.37327191 | 1.88521518 |
| 0.9703291146 | 0.4128576031 | 0.476783945 | -2.625260813 | -0.141919408 | 1.670923219 | 1.296249453 |
| 0.9703291146 | 1.684714552 | 1.109423016 | -3.990570698 | -1.293012406 | 3.545560211 | 0.316073812 |
| 0.9703291146 | 0.5247307866 | -0.182748367 | -3.071702598 | -0.399518997 | 1.829108911 | 1.759092493 |
| 0.9703291146 | 0.1158510435 | -0.607344573 | -2.427813418 | 0.085045742 | 0.7159744348 | 2.028522094 |
| 0.9703291146 | 0.243586132 | -0.196221916 | -2.843726662 | -0.231340444 | 1.811132923 | 1.90990594 |
| 0.9703291146 | 0.4911718904 | 0.324541212 | -3.02037603 | -0.576685818 | 2.01666732 | 1.844943255 |
| 0.9703291146 | 0.5904400537 | 0.252947623 | -3.244807242 | -0.725766071 | 2.190316234 | 1.951417101 |
| 0.9703291146 | 0.3931281134 | -0.228347781 | -2.738944878 | -0.268390414 | 1.522709733 | 1.69580785 |
| 0.09217496846 | 0.4228562908 | 1.48204894 | -2.063529723 | -0.689003743 | 1.314932059 | 1.435570918 |
| 0.9703291146 | 0.6330713897 | -0.300198766 | -3.115139301 | -0.759020895 | 2.331168056 | 1.662393083 |
| 0.9743440267 | 0.4690981296 | -0.056401736 | -2.978929128 | -0.530746493 | 2.000856662 | 1.803221927 |
| 0.9703291146 | 0.5652533301 | -0.266487618 | -3.060953341 | -0.660521597 | 1.995657706 | 1.841115456 |
| 0.9743440267 | 0.5433977663 | 0.084796417 | -3.043192917 | -0.533978065 | 2.024449812 | 1.712019114 |
| 0.9703291146 | 0.2102389545 | -0.734743313 | -2.199853591 | 0.057014211 | 0.5463456434 | 1.704029845 |
| 0.9703291146 | 0.5347859108 | 0.146798391 | -3.032487875 | -0.594489448 | 1.999421318 | 1.798121558 |
| 0.9170863458 | -0.0267289409 | -1.22788729 | -1.626209743 | 1.007625347 | 0.1379339566 | 0.761892388 |
| 0.9703291146 | 0.5932547225 | 0.655804152 | -3.555891729 | -0.790779479 | 2.814024721 | 1.99070872 |
| 0.9703291146 | 0.6221819122 | 0.420963521 | -2.986691213 | -0.546248398 | 2.153579435 | 1.416946051 |
| 0.9703291146 | 0.1821846791 | 0.350574571 | -2.959972184 | -0.437517724 | 2.035056317 | 2.255799541 |
| 0.9703291146 | 0.6707866874 | -0.378571676 | -3.418591722 | -1.191400233 | 2.729544857 | 2.154683053 |
| 0.9703291146 | 0.1075144158 | -0.562444718 | -2.830029788 | -0.361577173 | 1.971501255 | 2.218279076 |
| 0.2224185739 | 0.067834308 | 1.39480863 | -2.538233809 | -0.867228642 | 1.263178225 | 2.94444277 |
| 0.9703291146 | 0.6718507541 | -0.284320495 | -3.08437042 | -0.681462032 | 2.197696186 | 1.544960377 |
| 0.9703291146 | 0.6391720392 | -0.323734111 | -3.422284764 | -0.526384657 | 2.317758002 | 1.761842122 |
| 0.9703291146 | 0.1051405237 | 1.040237106 | -2.746195679 | -0.51136157 | 0.9870895338 | 2.893575905 |
| 0.9703291146 | 0.7323401594 | -0.589022539 | -2.981427851 | -0.541586739 | 1.409172727 | 1.635158923 |
| 0.9703291146 | 1.099886012 | -1.158824553 | -2.500742463 | -0.605138439 | 1.454150919 | 0.35791888 |
| 0.9703291146 | 0.681006169 | -0.254486807 | -3.310647956 | -1.063005614 | 2.594330059 | 1.953845454 |
| 0.9743440267 | 0.4406196185 | 0.110991490 | -2.935538102 | -0.450110015 | 1.853525587 | 1.816129027 |
| 0.9703291146 | 1.007846343 | -0.798807695 | -3.808055349 | -0.887667775 | 2.386592977 | 1.787968 |
| 0.9703291146 | 0.3439080282 | 0.197378875 | -2.937593013 | -0.370690164 | 1.923974911 | 1.890480111 |
| 0.9703291146 | -0.1828243679 | -0.602490206 | -2.828652895 | 0.111211185 | 1.87879289 | 2.36759628 |
| 0.9703291146 | 0.6191431736 | -0.600782918 | -2.999964694 | -0.539609803 | 2.19955557 | 1.403130571 |
| 0.9703291146 | 0.6740400561 | -0.274203941 | -2.992236313 | -0.674890073 | 1.82577997 | 1.655898804 |
| 0.9703291146 | 0.1773958625 | 0.473367044 | -2.624338333 | -0.209347949 | 1.840181046 | 1.752677664 |
| 0.9703291146 | 0.5679895151 | -0.24333031 | -3.102323845 | -0.626286195 | 2.122176316 | 1.768441305 |
| 0.9743440267 | 0.4817806466 | -0.056736971 | -3.025495721 | -0.529566895 | 2.046734198 | 1.801109332 |
| 0.9703291146 | 0.6853167727 | 0.594370289 | -3.415615169 | -0.522119572 | 2.007601126 | 1.844579976 |
| 0.9850546863 | 0.4689542112 | 0.029685109 | -2.987499476 | -0.525730974 | 1.996175089 | 1.810889335 |
| 0.9743440267 | 0.5039106526 | -0.120674575 | -2.969356625 | -0.47969593 | 1.940527843 | 1.702021695 |
| 0.9703291146 | 0.2330678103 | 0.635944252 | -3.017652844 | -0.012905575 | 1.454363846 | 2.11674839 |
| 0.9703291146 | 0.4578873237 | 0.131259111 | -3.011008068 | -0.517258996 | 1.941348005 | 1.885615783 |
| 0.9703291146 | 0.285792513 | 0.237544781 | -2.902348654 | -0.362989871 | 1.731165845 | 2.080874108 |
| 0.1202564645 | -0.3074896218 | 1.482410827 | -3.077402113 | 0.179782416 | 1.533926867 | 3.051914055 |

| | | | | | | |
|---|---|---|---|---|---|---|
| 0.9703291146 | -0.01119551405 | -0.475966437 | -2.333093581 | -0.120971110 | 1.388875287 | 1.990867372 |
| 0.9703291146 | 0.5939918837 | 0.311488495 | -3.255128882 | -0.782621735 | 2.394152432 | 1.891741324 |
| 0.9703291146 | 0.5954860567 | 0.800121092 | -2.625849303 | -0.659709198 | 1.757502867 | 1.42196322 |
| 0.9703291146 | 0.8188422573 | 0.442803017 | -3.228608484 | -1.038960085 | 2.299743359 | 1.72829851 |
| 0.9703291146 | 0.5782717323 | 0.172710172 | -3.226540871 | -0.660741297 | 2.272645149 | 1.834922558 |
| 0.9703291146 | 0.5855851459 | -0.444657302 | -3.01749053 | -0.512361992 | 1.871046987 | 1.665865462 |
| 0.9743440267 | 0.5351271897 | -0.074405944 | -3.050458862 | -0.585746438 | 1.998783618 | 1.809125385 |
| 0.9703291146 | 0.2129126878 | 0.332755856 | -2.91962856 | -0.397107853 | 1.908658186 | 2.179694637 |
| 0.9703291146 | 0.304178112 | 0.720738263 | -3.406266194 | 0.254028363 | 1.331091905 | 2.20585012 |
| 0.9703291146 | 0.7338599422 | 0.285776906 | -3.304010874 | -0.540086285 | 2.032113447 | 1.619249623 |
| 0.9703291146 | 0.2937584486 | 0.562801331 | -2.804716277 | 0.079235075 | 1.373700343 | 1.695413249 |
| 0.9703291146 | 0.4690544496 | 0.139458567 | -3.024089217 | -0.532114422 | 1.954957994 | 1.885138929 |
| 0.9980621059 | 0.4852580359 | 0.003066062 | -3.02924543 | -0.534172777 | 2.010220266 | 1.825643941 |
| 0.9703291146 | 0.7347835041 | 0.656220298 | -3.168382115 | -0.721937348 | 2.329284976 | 1.473801552 |
| 0.9703291146 | 0.5761841314 | -0.875972099 | -3.518390439 | -1.038193717 | 2.868856128 | 2.215596004 |
| 0.9703291146 | 0.6224984341 | -0.486382555 | -3.542944585 | -1.092634438 | 2.663367572 | 2.332819431 |
| 0.9743440267 | 0.4962331112 | -0.041989591 | -3.053529917 | -0.562000264 | 2.054646121 | 1.833499191 |
| 0.9703291146 | 0.4560724188 | 0.340203065 | -2.946933215 | -0.305351586 | 1.76084631 | 1.698322665 |
| 0.9703291146 | 0.2614746487 | -0.306392982 | -2.945727539 | -0.544202748 | 2.021284696 | 2.198050652 |
| 0.9703291146 | 0.5993567938 | -0.412787186 | -3.115849364 | -0.825036131 | 2.41840882 | 1.750641716 |
| 0.9703291146 | 0.4193287078 | -0.260839696 | -3.049305942 | -0.327811769 | 1.787140966 | 1.900765466 |
| 0.9743440267 | 0.4638010619 | -0.062196880 | -3.03526146 | -0.537670119 | 2.033111121 | 1.866829688 |
| 0.9703291146 | 0.3439446143 | -0.499068770 | -2.757433521 | -0.339428544 | 1.994768546 | 1.605296115 |
| 0.9703291146 | 0.4421454774 | 0.250495229 | -3.062953965 | -0.650563901 | 2.064788807 | 2.045843519 |
| 0.9703291146 | 0.4336647483 | 0.519964088 | -2.894308376 | -0.109030076 | 1.511142855 | 1.626381204 |
| 0.9703291146 | 0.5811815734 | 0.210092660 | -3.144298716 | -0.707754656 | 2.154607914 | 1.857252317 |
| 0.9703291146 | 0.3301316355 | -0.357628044 | -2.834989425 | -0.130163857 | 1.632119522 | 1.721278849 |
| 0.9703291146 | 0.1033471001 | -0.543018588 | -2.667693404 | -0.341974161 | 1.784872352 | 2.133360677 |
| 0.9850546863 | 0.4640586208 | -0.028221284 | -3.018302479 | -0.524512893 | 2.000559155 | 1.852589361 |
| 0.9703291146 | 0.6800300676 | -0.631725018 | -2.739552216 | -0.500246393 | 1.169351462 | 1.567777633 |
| 0.9703291146 | 0.7727571374 | -0.697750329 | -2.996195255 | -0.546015886 | 1.58585054 | 1.464300324 |
| 0.9743440267 | 0.4622360073 | 0.063896336 | -2.961606721 | -0.521897000 | 2.002509176 | 1.786939478 |
| 0.9703291146 | 0.6131982143 | 0.171273161 | -3.05237347 | -0.656420593 | 1.967449608 | 1.744580132 |
| 0.9703291146 | 0.4752759302 | 0.147546319 | -2.980904958 | -0.527679647 | 1.911586066 | 1.844303907 |
| 0.9703291146 | 0.454731281 | 0.979292477 | -3.605195563 | 0.453789773 | 1.991947075 | 1.50088888 |
| 0.9703291146 | 0.9551092873 | -0.553135539 | -3.358780721 | -0.644776071 | 1.945203711 | 1.389153489 |
| 0.9743440267 | 0.4533415048 | 0.124880751 | -2.886442499 | -0.506448623 | 1.995189136 | 1.706552654 |
| 0.9703291146 | -0.04396304993 | -0.629925518 | -3.152032185 | 0.094370810 | 2.049875329 | 2.365176241 |
| 0.9743440267 | 0.3874939075 | 0.108988145 | -2.869930987 | -0.513894688 | 1.975924434 | 1.844135344 |
| 0.9703291146 | 0.5488667781 | 0.173491871 | -3.161507637 | -0.529092990 | 2.120303722 | 1.772929511 |
| 0.9703291146 | 0.7978048925 | 0.642277508 | -3.370505809 | -0.794089910 | 1.850550312 | 1.94843933 |
| 0.9703291146 | 0.5368855193 | 0.592347543 | -3.428863868 | -0.448313923 | 2.455059227 | 1.813305392 |
| 0.9703291146 | 0.3582378825 | 0.181597151 | -2.752421293 | -0.355859083 | 1.747612762 | 1.739617049 |
| 0.9703291146 | 0.6360087118 | 0.469855399 | -3.025829045 | -0.714884681 | 2.356678129 | 1.490012382 |
| 0.9703291146 | 0.3490376135 | -0.325856407 | -2.988240561 | -0.404236470 | 2.012131786 | 1.920142242 |
| 0.9703291146 | 0.419087376 | 0.370610999 | -2.687414604 | -0.383215145 | 1.959787433 | 1.43712096 |

| | | | | | | |
|---|---|---|---|---|---|---|
| 0.9743440267 | 0.4509621527 | -0.132996389 | -2.877681364 | -0.482638106 | 1.879468174 | 1.746956187 |
| 0.9703291146 | 0.6668938007 | -0.500147721 | -3.783852512 | -0.662396763 | 2.540216071 | 2.131037014 |
| 0.9703291146 | 0.5037859282 | -0.687210800 | -2.733121561 | -0.540356961 | 1.758221708 | 1.607217474 |
| 0.9703291146 | 0.4286712976 | 0.146440239 | -3.079960585 | -0.534591380 | 2.177575938 | 1.899150428 |
| 0.9703291146 | 0.3757243881 | 0.299256044 | -2.67588157 | -0.248946406 | 1.70450477 | 1.526375124 |
| 0.9703291146 | -0.4312902372 | 1.187460113 | -2.245033357 | -0.036490492 | 0.6045421096 | 3.15345129 |
| 0.9743440267 | 0.5452056525 | 0.079450262 | -3.118191163 | -0.582859926 | 2.0974817 | 1.802500015 |
| 0.9703291146 | 0.2478972013 | 0.235264645 | -2.770961186 | -0.474603667 | 1.842945478 | 2.059390836 |
| 0.9980621059 | 0.4832507658 | 0.001535455 | -3.03024118 | -0.534182228 | 2.010646845 | 1.830709497 |
| 0.9743440267 | 0.4383309434 | -0.063049366 | -2.942354132 | -0.527941274 | 1.988034199 | 1.829790828 |
| 0.9281860866 | -0.1771930248 | 0.061958811 | -0.175160777 | -0.083521029 | -0.789550994 | 1.145948131 |
| 0.8694827707 | -0.1170048373 | -0.129995456 | -0.285652647 | -0.195359392 | -0.665382276 | 1.19246231 |
| 0.6176478085 | -0.1027765652 | 0.397629693 | -0.076177320 | -0.307014947 | -0.4912970317 | 0.934324045 |
| 0.6176478085 | -0.5979870776 | 0.541248447 | 0.243603064 | 1.015734942 | -2.019137897 | 1.109570702 |
| 0.6960982921 | -0.4609908314 | 0.431877956 | -0.294556070 | 0.057688824 | -0.787202018 | 1.71962047 |
| 0.6078766667 | -0.3980358798 | -0.459219307 | 0.278139672 | 0.065346545 | -1.150647482 | 1.143477068 |
| 0.7995209302 | 0.08500816269 | 0.230270665 | -0.498286459 | -0.305565423 | -0.479955032 | 1.022817899 |
| 0.9499656053 | -0.1230886732 | -0.042946054 | -0.246582441 | -0.130454392 | -0.668330518 | 1.091933572 |
| 0.8689041286 | -0.02404895816 | -0.149782094 | -0.441260536 | -0.241256647 | -0.555152891 | 1.160542008 |
| 0.6078766667 | -0.5343039395 | 0.496789109 | 0.229984678 | 0.291324220 | -1.082900608 | 1.197007843 |
| 0.421788991 | -0.5135316113 | 0.648768442 | 0.445093461 | 0.474987369 | -1.319403752 | 0.853412186 |
| 0.7995209302 | 0.02684177803 | -0.270495240 | -0.416622370 | -0.396472786 | -0.608192263 | 1.226212667 |
| 0.9252032334 | -0.2014990715 | 0.080557786 | -0.134267250 | -0.066840133 | -0.830466348 | 1.156459719 |
| 0.6605662054 | -0.5051799003 | 0.422186067 | 0.015180307 | 0.200579928 | -1.235564084 | 1.576042029 |
| 0.9252032334 | -0.1861349916 | -0.078838478 | -0.229101308 | -0.058242945 | -0.765462366 | 1.184712659 |
| 0.421788991 | -0.265484278 | 0.877514891 | 0.683820792 | -0.074884177 | -0.833641364 | 0.356021000 |
| 0.6078766667 | 0.2176510066 | -0.472774636 | -0.742987821 | -0.467866163 | -0.180138034 | 1.019959286 |
| 0.9396644152 | -0.1365399595 | -0.049372087 | -0.259939510 | -0.128909598 | -0.635472019 | 1.113398623 |
| 0.6173390112 | -0.01286781294 | -0.438324453 | -0.514572138 | -0.117023728 | -0.283599858 | 0.921011084 |
| 0.421788991 | -0.2410074969 | 0.640203654 | 0.035051137 | -0.521867865 | -0.287976365 | 1.199338184 |
| 0.7995209302 | -0.1110123652 | -0.242566606 | -0.440150149 | -0.312624988 | -0.537494813 | 1.405868803 |
| 0.9281860866 | -0.2457681013 | -0.091503621 | -0.155637283 | -0.050513435 | -0.836814094 | 1.259384526 |
| 0.5203497778 | -0.01350254053 | -0.581573017 | -0.372379429 | 0.315128525 | -1.2870603 | 0.910380236 |
| 0.7995209302 | -0.2923970777 | -0.241177851 | 0.003560368 | 0.132749237 | -1.224204999 | 1.213574326 |
| 0.9252032334 | -0.02822138742 | 0.097101153 | -0.326435244 | -0.26292275 | -0.611597234 | 1.094784808 |
| 0.8492314277 | -0.1503410211 | -0.144953596 | -0.238626567 | -0.094081197 | -0.713003488 | 1.127674673 |
| 0.7995209302 | -0.2480137427 | -0.238421172 | -0.031127456 | 0.067564839 | -0.879798967 | 1.019674376 |
| 0.421788991 | -0.4252476448 | -0.708719301 | 0.663792493 | 0.711521654 | -2.223762968 | 0.719507660 |
| 0.421788991 | -0.1212955397 | -0.625728552 | -0.642412683 | -0.047701029 | -0.416573628 | 1.300890251 |
| 0.6078766667 | -0.3709766515 | 0.448432090 | -0.105608057 | 0.222897667 | -1.189385307 | 1.384231197 |
| 0.6176478085 | -0.04890485202 | 0.399594622 | -0.586390057 | -0.136757711 | -0.642736830 | 1.320983421 |
| 0.9252032334 | -0.121532512 | -0.081251029 | -0.244871152 | -0.151658902 | -0.714682675 | 1.138087011 |
| 0.8689041286 | -0.2767403136 | -0.182330433 | -0.203454587 | -0.113337552 | -0.740451596 | 1.387241983 |
| 0.9570509022 | -0.1368943342 | 0.026272186 | -0.264616017 | -0.134244614 | -0.657845125 | 1.139044069 |
| 0.6176478085 | -0.01182481164 | 0.381390351 | -0.245270429 | -0.270044157 | -0.738660884 | 1.052940895 |
| 0.7995209302 | -0.02536073233 | 0.292219627 | -0.568717051 | -0.480013631 | -0.264579520 | 1.388640195 |

| | | | | | | |
|---|---|---|---|---|---|---|
| 0.6078766667 | -0.219497128 | -0.432921163 | 0.113146998 | 0.056345365 | -1.240712032 | 1.028159894 |
| 0.6176478085 | -0.2819803977 | -0.409021261 | -0.275636018 | -0.101289888 | -0.849333318 | 1.535441231 |
| 0.9570509022 | -0.1730865464 | 0.030827844 | -0.228956544 | -0.104617105 | -0.708024394 | 1.172039966 |
| 0.7995209302 | 0.02950677989 | -0.354912451 | -0.600670438 | -0.729344847 | -0.035972812 | 1.439383376 |
| 0.6173390112 | 0.131200166 | 0.416303684 | -0.381907919 | -0.240793146 | -0.681419047 | 0.846820826 |
| 0.421788991 | 0.135019515 | -0.946530067 | -0.567703979 | 0.112965989 | -0.203137691 | 0.377970035 |
| 0.9252032334 | -0.08005546331 | -0.100194107 | -0.254663527 | -0.165044344 | -0.644183879 | 1.034272484 |
| 0.7751009975 | 0.00443979554 | -0.312972756 | -0.615651880 | -0.105664846 | -0.413032323 | 1.06911309 |
| 0.4950861255 | 0.1262590862 | -0.751505363 | -0.438953043 | -0.129524589 | 0.029066665 | 0.365854711 |
| 0.7995209302 | -0.2643661666 | 0.277980386 | -0.257213796 | -0.109568146 | -0.426464567 | 1.226289541 |
| 0.7995209302 | -0.01291605397 | -0.251164896 | -0.120592070 | -0.128509033 | -0.830751767 | 0.815596548 |
| 0.758952824 | 0.1857849698 | -0.427023585 | -0.708374830 | -1.000627408 | 0.2696647513 | 1.34314569 |
| 0.9281860866 | -0.1677922087 | 0.055526009 | -0.188464288 | -0.071098589 | -0.788663517 | 1.127849412 |
| 0.9252032334 | -0.09178881141 | -0.083471260 | -0.316441561 | -0.150056342 | -0.670881975 | 1.130212763 |
| 0.7995209302 | -0.2824918386 | 0.192923225 | -0.146101895 | 0.046597120 | -0.794647221 | 1.194125089 |
| 0.7995209302 | 0.1542396426 | 0.272306308 | -0.325322720 | -0.404761732 | -0.650722153 | 0.891370149 |
| 0.7995209302 | -0.1922378661 | 0.201044693 | -0.244543807 | -0.111259742 | -0.773500403 | 1.277247625 |
| 0.7995209302 | -0.2913483641 | 0.207582334 | -0.262459999 | -0.006513784 | -0.570446967 | 1.266005734 |
| 0.421788991 | 0.3298567668 | -0.738334475 | -0.865421722 | -0.619616693 | -0.444729487 | 1.254435595 |
| 0.6173390112 | -0.2830892506 | 0.390164364 | -0.116659889 | 0.034730733 | -0.889455608 | 1.232517131 |
| 0.7995209302 | -0.1427115487 | 0.200434315 | -0.245700816 | -0.129067986 | -0.838629301 | 1.234977714 |
| 0.6173390112 | -0.2872487413 | -0.412855589 | 0.033888834 | -0.121419672 | -0.708265539 | 1.124106402 |
| 0.421788991 | -0.5531871918 | 0.865762496 | 0.961636371 | 0.130520423 | -1.124424433 | 0.590739856 |
| 0.421788991 | -0.02641987838 | -0.690297699 | 0.103500178 | 0.198011452 | -1.109760774 | 0.405135376 |
| 0.9281860866 | -0.1687500676 | 0.056273014 | -0.234022699 | -0.066990987 | -0.759394700 | 1.159971058 |
| 0.7995209302 | -0.111705695 | -0.183390136 | -0.259464697 | -0.136616645 | -0.613748261 | 1.056203335 |
| 0.9849242331 | -0.1539892136 | 0.008852827 | -0.230282609 | -0.106732362 | -0.720600169 | 1.143923256 |
| 0.5174588607 | 0.1662799689 | -0.586906794 | -0.215635575 | -0.395738428 | -0.521560879 | 0.650599961 |
| 0.9396644152 | -0.1995000576 | -0.050917689 | -0.160589174 | -0.068914895 | -0.776681282 | 1.151815228 |
| 0.6078766667 | -0.3012883823 | -0.433580987 | 0.080433904 | 0.232846351 | -1.24439475 | 1.047968219 |
| 0.7995209302 | -0.1780133726 | -0.222924425 | -0.347177052 | -0.078107999 | -0.639740048 | 1.248111479 |
| 0.6960982921 | -0.4073538474 | -0.343633766 | -0.079715023 | 0.278695978 | -0.934754063 | 1.213120619 |
| 0.6173390112 | -0.3764605722 | -0.416151631 | 0.242131872 | 0.192075607 | -1.342136679 | 1.121576331 |
| 0.9537777759 | -0.1396445318 | -0.030306362 | -0.234232757 | -0.111628189 | -0.719690822 | 1.12340115 |
| 0.7995209302 | -0.30387825 | 0.223978729 | -0.168971314 | 0.042405440 | -0.675292365 | 1.195995786 |
| 0.6078766667 | 0.2965701378 | -0.546267964 | -0.413761318 | -0.337976129 | -0.543206136 | 0.559718535 |
| 0.8689041286 | -0.1891252795 | 0.171323293 | -0.324775226 | 0.074229866 | -0.871664401 | 1.224008309 |
| 0.758952824 | 0.1595773887 | 0.348695898 | -0.571119308 | -0.120431926 | -0.683673734 | 0.877972075 |
| 0.7995209302 | -0.07312132481 | -0.218770066 | -0.321987554 | -0.351523678 | -0.462699505 | 1.186693587 |
| 0.7995209302 | -0.1269199069 | -0.198756621 | -0.241312718 | -0.115916462 | -0.631209587 | 1.055310418 |
| 0.6176478085 | 0.2213967285 | 0.477274223 | -0.346504145 | -0.126852322 | -0.734768422 | 0.528763878 |
| 0.8145528314 | -0.2122161967 | -0.170828844 | -0.19857821 | -0.064206414 | -0.793213070 | 1.22716191 |
| 0.9252032334 | -0.1551126097 | 0.079521364 | -0.190515138 | -0.067345331 | -0.788127712 | 1.09950953 |
| 0.7751009975 | -0.04081770085 | -0.368697322 | -0.624932562 | -0.536511801 | -0.215202508 | 1.516065511 |
| 0.6078766667 | -0.2810559398 | 0.423101076 | 0.016932250 | 0.168179927 | -1.156253443 | 1.094618091 |
| 0.8120905217 | -0.1329034009 | -0.174282144 | -0.276785702 | -0.230285975 | -0.582362173 | 1.201775261 |

| | | | | | | |
|---|---|---|---|---|---|---|
| 0.6078766667 | 0.2629030972 | 0.566718369 | -0.38813848 | -0.09439352 | -0.730367751 | 0.452934228 |
| 0.7995209302 | -0.206975372 | 0.213837417 | -0.18975027 | 0.037613853 | -0.9215678691 | 1.175426015 |
| 0.6078766667 | -0.0371485854 | 0.449314217 | -0.199195433 | -0.468427587 | -0.3256531883 | 1.011858081 |
| 0.8875956565 | -0.1829343063 | -0.11822521 | -0.247781906 | -0.11992417 | -0.6669843557 | 1.205443707 |
| 0.421788991 | 0.03820041955 | 0.663911488 | -0.59562361 | -0.37205964 | -0.689429033 | 1.432862313 |
| 0.7751009975 | -0.0979835636 | -0.299132447 | -0.193876064 | 0.025986829 | -0.775169758 | 0.876249203 |
| 0.7995209302 | -0.1245836479 | -0.23300114 | -0.29513072 | -0.30357988 | -0.4864821182 | 1.225591904 |
| 0.8394417861 | -0.2213061352 | -0.159592862 | -0.147042872 | 0.018816492 | -0.8197558653 | 1.113501905 |
| 0.6176478085 | -0.3072183396 | -0.375918032 | -0.031547412 | 0.311469412 | -1.107798171 | 1.020763073 |
| 0.9281860866 | -0.1924693221 | -0.065562957 | -0.191418547 | -0.089913713 | -0.737421583 | 1.171239582 |
| 0.6605662054 | 0.09245134719 | 0.358244644 | -0.364802133 | -0.234611773 | -0.5855511057 | 0.841909177 |
| 0.5715385661 | 0.03081279178 | -0.56866051 | 0.025143424 | -0.082648653 | -1.467262264 | 0.898929425 |
| 0.8689041286 | -0.0914101212 | -0.132956422 | -0.22882390 | -0.115382252 | -0.791088267 | 1.064960727 |
| 0.421788991 | -0.4214515403 | 0.850008516 | 0.655277576 | 0.049020560 | -0.814874651 | 0.567902870 |
| 0.7751009975 | -0.3741581437 | -0.299046658 | -0.193352174 | 0.100492776 | -0.6352403397 | 1.282890489 |
| 0.7751009975 | -0.1606548757 | 0.271379340 | -0.14739100 | -0.101328414 | -0.891900977 | 1.161719416 |
| 0.6173390112 | -0.1604097165 | 0.478688082 | -0.516865551 | 0.369774221 | -0.719203762 | 0.973910509 |
| 0.6078766667 | -0.5893539503 | 0.518574176 | 0.074631602 | -0.00933344 | -0.649092323 | 1.547422259 |
| 0.5715385661 | -0.3226291896 | 0.743674192 | 0.611150722 | 0.051464905 | -0.800646080 | 0.402183634 |
| 0.6078766667 | 0.3014358207 | 0.535737055 | -0.129948523 | -0.647595389 | -0.7437859264 | 0.679006427 |
| 0.5715385661 | -0.701584091 | 0.633983856 | 0.687579380 | 0.004335298 | -0.9106115764 | 1.219482938 |
| 0.9281860866 | -0.1678887407 | -0.055869042 | -0.192162417 | -0.114716118 | -0.7455934677 | 1.152785051 |
| 0.7995209302 | -0.0401470287 | 0.217205294 | -0.350634754 | -0.201037311 | -0.7642450195 | 1.1747672 |
| 0.421788991 | -0.2050535613 | -0.640818867 | 0.198108824 | -0.205897823 | -1.191263468 | 1.152114697 |
| 0.5203497778 | -0.5808031129 | 0.632478473 | 0.726661002 | 0.507626406 | -1.625370946 | 0.821382068 |
| 0.9042044655 | -0.1081111153 | 0.118242584 | -0.234305635 | -0.158608428 | -0.6230455212 | 1.049112831 |
| 0.9252032334 | -0.1071275842 | 0.096197039 | -0.24687877 | -0.151441542 | -0.710712176 | 1.107808113 |
| 0.8492314277 | -0.1828865444 | 0.210512340 | -0.041227244 | -0.027412782 | -0.738930715 | 0.911659234 |
| 0.7995209302 | -0.0894308025 | 0.238294112 | -0.505263931 | -0.20528693 | -0.475638691 | 1.282516714 |
| 0.6176478085 | -0.3136400507 | 0.453928429 | 0.450537975 | 0.003346051 | -1.19106941 | 0.860917099 |
| 0.9252032334 | -0.1441995326 | -0.080394324 | -0.200426445 | -0.113842517 | -0.7396933502 | 1.108548162 |
| 0.6176478085 | -0.0000026088 | -0.39603826 | -0.095891723 | -0.111737861 | -1.162370249 | 0.946287564 |
| 0.9849242331 | -0.1437694622 | -0.008025755 | -0.244442764 | -0.120755848 | -0.7019912497 | 1.142686833 |
| 0.7995209302 | -0.3928463631 | 0.319799879 | -0.023978589 | 0.020900409 | -1.08880609 | 1.491105664 |
| 0.7995209302 | -0.3111043527 | -0.209694081 | 0.001661325 | 0.015624911 | -0.9400880231 | 1.205911774 |
| 0.5172089339 | -0.8477910672 | 0.701947472 | 0.533507285 | 0.064361254 | -1.209754995 | 1.820363995 |
| 0.421788991 | -0.3612159684 | -0.922714705 | 0.793918847 | -0.054371935 | -0.8716747243 | 0.429127178 |
| 0.421788991 | -0.70807465 | -0.794337444 | 0.850702892 | -0.035712488 | -0.9916980624 | 1.137771653 |
| 0.9978741781 | 0.1856168494 | 0.205024892 | 0.454911587 | -0.586168482 | -0.7185191904 | 0.163014117 |
| 0.7638667387 | 0.106204348 | 0.791679727 | 0.565518043 | -0.183164416 | -0.728833234 | -0.227332334 |
| 0.9978741781 | 0.3194963844 | 0.297222843 | 0.375656497 | -0.829017971 | -0.292301496 | -0.02435278 |
| 0.9978741781 | 0.8238086101 | -0.64408994 | -0.31257389 | -2.027408595 | 1.101728666 | 0.155051309 |
| 0.9978741781 | 0.2646668759 | 0.030279759 | 0.252773186 | -0.672100747 | -0.4613235304 | 0.166338096 |
| 0.9978741781 | 0.3237667597 | 0.067698436 | 0.181304598 | -0.71038368 | -0.390992494 | 0.124003077 |
| 0.9978741781 | 0.5968327057 | 0.308277357 | -0.095547735 | -0.94172917 | -0.1476888322 | -0.02427291 |
| 0.9978741781 | 0.6870635553 | -0.729942171 | 0.059833088 | -0.978895221 | 0.2556256796 | -0.59912958 |

| | | | | | | |
|---|---|---|---|---|---|---|
| 0.9978741781 | 0.1839579257 | 0.125534584 | 0.429832301 | -0.576673011 | -0.585867029 | 0.103539221 |
| 0.9978741781 | 0.4237216032 | -0.175585923 | 0.092610166 | -0.827019093 | -0.324194548 | 0.103245224 |
| 0.9978741781 | 0.3401449641 | -0.094495588 | 0.157893790 | -0.769734108 | -0.3671907442 | 0.166267891 |
| 0.9978741781 | 0.4693875701 | -0.284833186 | 0.065684634 | -0.982459815 | -0.348515419 | 0.221832835 |
| 0.9978741781 | 0.2162952341 | 0.103402020 | 0.386228811 | -0.624686643 | -0.6103024497 | 0.153432789 |
| 0.9978741781 | 0.3716700609 | -0.100047064 | 0.197667152 | -0.758411690 | -0.331425522 | 0.020697608 |
| 0.9978741781 | 0.07759726069 | -0.417454463 | 0.288104866 | -0.393558102 | -0.748571179 | 0.390898757 |
| 0.9978741781 | 0.2576246795 | 0.214737886 | 0.481792082 | -0.674721696 | -0.4861332491 | -0.065198295 |
| 0.9978741781 | 0.2219061056 | 0.084096503 | 0.347312643 | -0.620971911 | -0.550210101 | 0.145702544 |
| 0.9978741781 | 0.3069329275 | -0.098838151 | 0.206993505 | -0.715707505 | -0.306337250 | 0.082962420 |
| 0.9978741781 | 0.0823481675 | 0.669591701 | 0.684051607 | -0.678095050 | -1.107597399 | 0.451533482 |
| 0.9978741781 | 0.2145298165 | 0.489675532 | 0.463493686 | -0.996725175 | -0.132980469 | 0.174868143 |
| 0.9978741781 | 0.3183712433 | -0.215564811 | 0.074623134 | -0.861382385 | -0.3024491412 | 0.366428917 |
| 0.9978741781 | -0.6881243884 | -0.899884243 | 1.037288907 | -0.068487876 | -1.701153795 | 1.352929655 |
| 0.9978741781 | 0.2050020879 | 0.356891954 | 0.341264129 | -0.946870587 | -0.101921082 | 0.262887582 |
| 0.9978741781 | 0.3604405911 | 0.122007885 | 0.136273608 | -0.808451154 | -0.195894967 | 0.085344293 |
| 0.9978741781 | -0.1072554522 | -0.323031723 | 0.561182009 | -0.185681149 | -0.783933229 | 0.257628385 |
| 0.9978741781 | 0.2869594817 | 0.009924372 | 0.257199132 | -0.685378414 | -0.4557320681 | 0.125789334 |
| 0.9978741781 | 0.3084296626 | 0.051092980 | 0.213263702 | -0.722793509 | -0.4195786284 | 0.149937057 |
| 0.9978741781 | 0.1096556691 | -0.450385576 | 0.828115954 | -0.160029518 | -1.417785336 | -0.138442033 |
| 0.9978741781 | 0.275164135 | 0.284924142 | 0.442471084 | -0.71385893 | -0.589624006 | 0.049578397 |
| 0.7638667387 | -0.143715902 | 0.860395633 | 0.505182377 | -0.039388473 | -1.375336063 | 0.604363140 |
| 0.9978741781 | 0.1452635255 | -0.578273874 | 0.765485949 | -0.649852013 | -0.5535458077 | -0.144469983 |
| 0.9978741781 | 0.4518952194 | -0.534494798 | 0.191916284 | -0.937670868 | -0.485449181 | 0.148543877 |
| 0.9978741781 | 0.6805307351 | 0.551944722 | 0.161501669 | -0.683382973 | -0.3643310214 | -0.639603467 |
| 0.9978741781 | 0.3046975205 | 0.048482198 | 0.202269487 | -0.723079455 | -0.359319885 | 0.133599699 |
| 0.9978741781 | 0.3586130283 | 0.203417569 | 0.251464056 | -0.767778437 | -0.471107174 | 0.081787915 |
| 0.9978741781 | 0.1154237947 | -0.412688994 | 0.728264844 | -0.165911929 | -1.085246894 | -0.233513304 |
| 0.9978741781 | 0.372847904 | 0.511883615 | -0.154679675 | -0.884437265 | 0.1713566674 | 0.251120777 |
| 0.9978741781 | 0.3171076264 | 0.091880238 | 0.266081410 | -0.686730595 | -0.424669021 | 0.035265649 |
| 0.9978741781 | 0.3425403906 | -0.065092822 | 0.244220548 | -0.702005681 | -0.460506503 | 0.046171754 |
| 0.9978741781 | 0.4902632003 | -0.410131669 | -0.165630839 | -1.396189185 | 0.3231963617 | 0.477571836 |
| 0.9978741781 | 0.3409804046 | 0.081320849 | 0.228249883 | -0.709017580 | -0.450303391 | 0.069111852 |
| 0.9978741781 | 0.2696217328 | 0.057412875 | 0.277131265 | -0.697788501 | -0.486680505 | 0.171209792 |
| 0.9978741781 | 0.4723085457 | -0.279272521 | 0.202099853 | -0.828836606 | -0.271932077 | -0.154204571 |
| 0.9978741781 | 0.139215933 | 0.305529713 | 0.628563667 | -0.691342526 | -0.746019501 | 0.189205091 |
| 0.9978741781 | 0.1594325577 | 0.350479929 | 0.352074446 | -0.676419802 | -0.8006939752 | 0.483817944 |
| 0.9978741781 | 0.2883377937 | -0.003852388 | 0.257257873 | -0.684124745 | -0.459856457 | 0.124046890 |
| 0.9978741781 | 0.182025603 | 0.196609590 | 0.167403993 | -0.672020455 | -0.3615532404 | 0.374997008 |
| 0.9978741781 | 0.6503405313 | -0.467108947 | -0.260877732 | -1.654907109 | 0.6159157314 | 0.353488100 |
| 0.9978741781 | 0.2267690656 | 0.157366552 | 0.388794700 | -0.565015450 | -0.678313910 | 0.106309967 |
| 0.9978741781 | 0.2696981268 | 0.025880981 | 0.282206091 | -0.672619751 | -0.468113567 | 0.126664944 |
| 0.9978741781 | 0.3704243594 | -0.118891166 | 0.202174513 | -0.782496249 | -0.4038794091 | 0.088608258 |
| 0.9978741781 | 0.6266536602 | 0.307660166 | 0.154883557 | -1.013594352 | -0.388730704 | -0.149441201 |
| 0.9978741781 | 0.392354592 | -0.465848909 | 0.279098681 | -0.688360025 | -0.309235626 | -0.205317602 |
| 0.9978741781 | 0.3678408503 | -0.116389368 | 0.27235472 | -0.743842409 | -0.5342794397 | 0.051615807 |

| | | | | | | |
|---|---|---|---|---|---|---|
| 0.9978741781 | -0.07761399214 | 0.564512130 | 0.738965936 | -0.296812313 | -0.6588923003 | 0.033661502 |
| 0.9978741781 | 0.3284019299 | -0.119227978 | 0.220791527 | -0.729253047 | -0.4011450596 | 0.095382932 |
| 0.9978741781 | 0.2781272686 | -0.437309326 | 0.280329451 | -0.649255617 | -0.1757041732 | -0.093781999 |
| 0.9978741781 | 0.2958588748 | 0.026874389 | 0.239447757 | -0.683534686 | -0.4560499516 | 0.125998402 |
| 0.9978741781 | 0.2822696573 | 0.009448215 | 0.270008753 | -0.681416911 | -0.4604449525 | 0.119383410 |
| 0.9978741781 | 0.2916240523 | -0.028236410 | 0.270795451 | -0.671349405 | -0.4722685234 | 0.095481437 |
| 0.9978741781 | 0.1731979959 | 0.288941135 | 0.261891404 | -0.447278151 | -0.7085493612 | 0.255813234 |
| 0.9978741781 | 0.284293741 | 0.012663777 | 0.258640224 | -0.682452368 | -0.4625843627 | 0.130728744 |
| 0.9978741781 | 0.3848272344 | -0.118254521 | 0.194134718 | -0.769249984 | -0.3169269538 | 0.000196869 |
| 0.9978741781 | 0.3174082078 | -0.057583180 | 0.258848257 | -0.711805323 | -0.4374169836 | 0.077835764 |
| 0.9978741781 | 0.2056603083 | -0.078073328 | 0.371022900 | -0.616312211 | -0.5578702238 | 0.151781608 |
| 0.9978741781 | 0.2738165893 | -0.036139279 | 0.283240212 | -0.655240080 | -0.5004503574 | 0.118100990 |
| 0.9978741781 | 0.3664113943 | 0.566412335 | 0.542010116 | -0.773006450 | -0.6349368796 | -0.163205696 |
| 0.9978741781 | 0.2593950086 | -0.035998186 | 0.273215509 | -0.643031229 | -0.4794486724 | 0.133548362 |
| 0.9978741781 | 0.6112663763 | 0.587712088 | -0.416860864 | -1.115073676 | 0.4365389444 | 0.143649286 |
| 0.9978741781 | 0.2800404739 | 0.028863406 | 0.256233291 | -0.685485719 | -0.4468801854 | 0.135942261 |
| 0.9978741781 | 0.5105160011 | -0.318814056 | 0.162984983 | -0.905436435 | -0.5056044786 | 0.037864550 |
| 0.9978741781 | 0.3882965888 | -0.125211586 | 0.215972028 | -0.735618156 | -0.4176485321 | -0.006441989 |
| 0.9978741781 | 0.3827669225 | -0.387400666 | 0.459985566 | -1.107690545 | -0.0908044137 | -0.076952881 |
| 0.9978741781 | 0.5346457024 | 0.282330186 | -0.015209485 | -0.690005090 | -0.4344512215 | -0.082431588 |
| 0.9978741781 | 0.2141966178 | 0.215763933 | 0.342754016 | -0.448944320 | -0.7000107062 | 0.073899016 |
| 0.9978741781 | 0.2835479206 | 0.031827456 | 0.257963075 | -0.683626067 | -0.4685724616 | 0.138008049 |
| 0.9978741781 | -0.03648894958 | -0.419119187 | 0.354880457 | -0.672006712 | -0.4343447355 | 0.657295895 |
| 0.9978741781 | 0.3710448533 | 0.219717694 | 0.210123774 | -0.746973150 | -0.3491470212 | 0.006210201 |
| 0.9978741781 | 0.3129421958 | -0.246347677 | 0.119187184 | -0.825845191 | -0.21449654 | 0.233889073 |
| 0.9978741781 | 0.3834526935 | -0.337064080 | -0.099541195 | -1.071151908 | -0.0034014064 | 0.474094858 |
| 0.9978741781 | 0.2891333699 | -0.007639778 | 0.252401039 | -0.689154802 | -0.4478700693 | 0.126039324 |
| 0.9978741781 | 0.2843390947 | 0.030026269 | 0.26415145 | -0.663887748 | -0.4779480295 | 0.113737338 |
| 0.9978741781 | -0.01412137948 | -0.416276282 | 0.369446820 | -0.697822704 | -0.4411063478 | 0.625995660 |
| 0.9978741781 | 0.367115564 | -0.284999295 | 0.196661519 | -0.884956462 | -0.1742091079 | 0.070847716 |
| 0.9978741781 | 0.2509190083 | -0.146853717 | 0.245252332 | -0.567943372 | -0.5816077432 | 0.165420841 |
| 0.9978741781 | 0.2245695746 | -0.202424820 | 0.235039172 | -0.695745075 | -0.3819401632 | 0.246691251 |
| 0.9978741781 | 0.2066229219 | -0.287638671 | 0.413195917 | -0.571885575 | -0.4649209868 | -0.003922438 |
| 0.9978741781 | 0.248511941 | 0.234795839 | 0.224682400 | -0.793254930 | -0.4049238169 | 0.328069319 |
| 0.9978741781 | 0.2675331891 | 0.202509851 | 0.309224586 | -0.518530196 | -0.650361013 | 0.046195843 |
| 0.9978741781 | 0.3863615275 | 0.213015341 | 0.139569603 | -0.860161841 | -0.3096881705 | 0.15342045 |
| 0.9978741781 | 0.09393936151 | -0.451282686 | 0.501173024 | -0.174377433 | -0.9332403162 | -0.011288579 |
| 0.9978741781 | 0.7649273469 | 0.684192389 | -0.197695727 | -0.926130785 | -0.1717916878 | -0.257495227 |
| 0.9978741781 | -0.06642478324 | -0.529109099 | 0.448682960 | -0.504621858 | -0.6400163002 | 0.557543166 |
| 0.9978741781 | 0.3078511083 | -0.067758584 | 0.287945358 | -0.680446217 | -0.5461325889 | 0.097242111 |
| 0.9978741781 | 0.2900511089 | -0.008051431 | 0.257322920 | -0.684213356 | -0.4608230684 | 0.121103676 |
| 0.9978741781 | 0.245092302 | 0.128716402 | 0.391761888 | -0.659524515 | -0.4717760609 | 0.039510093 |
| 0.9978741781 | 0.5236841921 | 0.311485352 | 0.213585187 | -0.906561822 | -0.5339962044 | -0.029185834 |
| 0.9978741781 | 0.3016422658 | -0.289289413 | 0.163574823 | -0.696633445 | -0.2622259033 | 0.096362456 |
| 0.9978741781 | 0.2935219197 | -0.236998381 | 0.396508774 | -0.923150935 | -0.451463893 | 0.204853906 |
| 0.9978741781 | 0.2875575852 | -0.000997119 | 0.256354482 | -0.684275218 | -0.4560418774 | 0.124524310 |

| | | | | | | |
|---|---|---|---|---|---|---|
| 0.9978741781 | 0.3433017138 | -0.2391589588 | -0.0151598855 | -0.7370010833 | -0.4268359637 | 0.3608426827 |
| 0.9978741781 | 0.2998847086 | 0.0157564506 | 0.2600390259 | -0.699795354 | -0.4569123407 | 0.1119200895 |
| 0.9978741781 | 0.164559039 | 0.13954545 | 0.4600144137 | -0.658225211 | -0.5000379648 | 0.1440112631 |
| 0.9978741781 | 0.3299845493 | 0.1138103402 | 0.1697160945 | -0.6808012666 | -0.3838134102 | 0.0881846736 |
| 0.9978741781 | 0.4831055171 | 0.4005661931 | 0.0436707610 | -0.8462320388 | -0.5556034546 | 0.1994742919 |
| 0.9978741781 | 0.2999007904 | 0.1447571617 | 0.1591164807 | -0.6631151733 | -0.3472537723 | 0.1213518508 |
| 0.9978741781 | -0.0063033982 5 | 0.4268531754 | 0.9059548209 | -0.265147964 | -1.073567517 | -0.086039916 |
| 0.9978741781 | 0.2450084296 | -0.1279560200 | 0.2562145313 | -0.6348382466 | -0.5502324883 | 0.2177807529 |
| 0.9978741781 | 0.3036877553 | 0.0413392015 | 0.2518386924 | -0.7005486199 | -0.4561468638 | 0.1138235231 |
| 0.9978741781 | 0.4214115373 | -0.7824181466 | -0.4631963988 | -1.002584601 | -0.3491194616 | 0.953767275 |
| 0.9978741781 | 0.1617384478 | -0.5204970455 | 0.8473098960 | -0.4827342055 | -0.9682559403 | -0.197866164 |
| 0.9978741781 | 0.196639767 | 0.2448182947 | 0.6266746028 | -0.6212678499 | -0.7152758071 | -0.0222643399 |
| 0.9978741781 | 0.2960222987 | -0.3064311988 | 0.3886736801 | -0.6868726100 | -0.5683623711 | 0.0261852500 |
| 0.9978741781 | 0.3151685946 | -0.0768689566 | 0.283947115 | -0.683809601 | -0.5436893436 | 0.0887761836 |
| 0.9978741781 | 0.3354912525 | -0.13622538 | 0.0964199506 | -0.813874371 | -0.3166867175 | 0.2633209121 |
| 0.9978741781 | 0.4218416865 | -0.1755312366 | 0.1411329061 | -0.757630392 | -0.2481126643 | -0.070384024 |
| 0.9978741781 | 0.5920503086 | 0.3893259496 | -0.1824168799 | -0.923087405 | -0.0290953479 | -0.007162975 |
| 0.9978741781 | 0.2690150861 | 0.0177114377 | 0.2763403065 | -0.679597781 | -0.4685292288 | 0.1426223731 |
| 0.9978741781 | 0.2879381117 | 0.0052969384 | 0.2510432883 | -0.684412863 | -0.4549973956 | 0.1293677621 |
| 0.9978741781 | 0.1314444899 | -0.2196713333 | 0.5571931606 | -0.662672011 | -0.5337732545 | 0.126207351 |
| 0.9301794211 | -0.678170974 | -0.0911506533 | 0.0470577275 | 0.475370443 | 0.2992954962 | 0.658092324 |
| 0.7327832683 | -0.7865199948 | 0.278099408 | 0.2434613610 | 0.694857292 | 0.0866834375 | 0.5509730652 |
| 0.6974568237 | -0.6738015126 | 0.4469862967 | 0.3135883606 | 0.3009231186 | 0.4286187592 | 0.449764746 |
| 0.6974568237 | -1.220139576 | 0.59603059 | 0.6620964219 | 1.761997264 | -1.258877139 | 0.6473367419 |
| 0.7327832683 | -0.4287658946 | -0.4044009999 | 0.1908491751 | 0.3589672935 | 0.2546583859 | 0.1270101792 |
| 0.7327832683 | -0.8729714708 | -0.2737458888 | 0.4409477902 | 0.6252775813 | -0.0800088611 | 0.6801690469 |
| 0.6974568237 | -1.444720562 | -0.7173045766 | 0.9552650535 | 1.118410549 | -0.5346570957 | 1.022922188 |
| 0.7327832683 | -0.5519799202 | -0.312015721 | 0.0508101651 | 0.3928769549 | 0.4867041883 | 0.3651841917 |
| 0.867011594 | -0.5964770339 | -0.1547204299 | -0.0780081255 | 0.3865253741 | 0.3427035451 | 0.7016937074 |
| 0.6974568237 | -1.02590416 | 0.3880279058 | 0.498242965 | 0.8347894715 | -0.1085599856 | 0.723630395 |
| 0.6974568237 | -0.9263225465 | 0.359334488 | 0.511744514 | 0.844628142 | -0.1548735649 | 0.519133506 |
| 0.6974568237 | -0.9979144129 | 0.4284642043 | 0.4227814933 | 0.9677467596 | 0.0209788127 | 0.5296678163 |
| 0.957592049 | -0.6918142932 | -0.0459646199 | 0.0776006949 | 0.4924984449 | 0.2511748823 | 0.6623537081 |
| 0.6974568237 | -1.109795125 | 0.4553277798 | 0.4048716932 | 0.8572116418 | -0.3840610357 | 1.150993714 |
| 0.9992515135 | -0.7232561586 | -0.000284442 | 0.1350894629 | 0.5190963033 | 0.1823508182 | 0.6750322728 |
| 0.957592049 | -0.7312713987 | 0.0602363154 | 0.1981389385 | 0.5215222459 | 0.1740763061 | 0.621409369 |
| 0.6974568237 | -1.096784038 | 0.4849422608 | 0.6561452282 | 0.8827860525 | -0.3611480847 | 0.7923977333 |
| 0.6974568237 | -0.850949126 | 0.6246737215 | 0.450800518 | 0.7188168156 | -0.7628656315 | 0.9425467897 |
| 0.6974568237 | -0.8505939516 | 0.4176967041 | 0.4014977762 | 0.5226291163 | -0.2239686595 | 0.8783470538 |
| 0.7327832683 | -0.7706962909 | 0.32282024 | 0.2712329747 | 0.3127831208 | 0.3954520845 | 0.7075172975 |
| 0.7327832683 | -0.6763411184 | -0.3184130000 | -0.1342483833 | 0.2569967914 | 0.4092500884 | 1.030998474 |
| 0.6974568237 | -0.0129339485 | 0.655580021 | -0.4334212366 | 0.0704328223 | 1.089719699 | -0.2194832277 |
| 0.90115955 | -0.7524120642 | 0.1279781496 | 0.1653025663 | 0.4246625315 | 0.3094925275 | 0.7241824291 |
| 0.957592049 | -0.7615669403 | -0.0636285188 | 0.1980022683 | 0.5837615963 | 0.0469418794 | 0.6956981523 |
| 0.6974568237 | -1.538472795 | -0.6685594099 | 0.7647193792 | 1.550395655 | -0.4963100291 | 0.948685833 |
| 0.7327832683 | -0.7288085552 | -0.2232641966 | 0.1294594843 | 0.5482054225 | 0.1782236496 | 0.6642518165 |

| | | | | | | |
|---|---|---|---|---|---|---|
| 0.6974568237 | -0.5329415986 | 0.447282268 | -0.24737225 | 0.179951605 | 0.5007311988 | 0.890371921 |
| 0.6974568237 | -0.9422221977 | -0.557373312 | 0.841913166 | 1.167430097 | -1.007797895 | 0.348435664 |
| 0.9090584336 | -0.7187720612 | -0.10717462 | 0.065284144 | 0.530101364 | 0.2328415223 | 0.703340938 |
| 0.7549949117 | -0.6127695897 | -0.220573107 | 0.071430734 | 0.353624779 | 0.4182528228 | 0.552042185 |
| 0.6974568237 | -0.8279407247 | -0.428574221 | 0.511957952 | 0.544262424 | 0.1102003784 | 0.474904100 |
| 0.6974568237 | -0.5685422205 | -0.50014043 | 0.074214697 | 0.281602855 | 0.1549231186 | 0.696584197 |
| 0.90115955 | -0.6059495905 | 0.164205416 | 0.106672088 | 0.519104434 | 0.2097995561 | 0.447284545 |
| 0.7327832683 | -0.63022941 | 0.250302238 | -0.147233504 | 0.317530983 | 0.6812966567 | 0.717606892 |
| 0.7327832683 | -0.7997793842 | -0.21688039 | 0.14091706 | 0.608140828 | 0.1987378509 | 0.721262543 |
| 0.6974568237 | -0.8941043703 | -0.411985811 | 0.605580186 | 1.036179199 | -0.445699978 | 0.316631249 |
| 0.9301794211 | -0.7367211857 | -0.080860622 | 0.200092033 | 0.550548833 | 0.0834606317 | 0.654978676 |
| 0.7327832683 | -0.6344101453 | 0.268084079 | 0.161711991 | 0.511151954 | 0.2737457453 | 0.412100055 |
| 0.9700166179 | -0.7009866067 | -0.02579632 | 0.130023617 | 0.511792680 | 0.1807343254 | 0.643485537 |
| 0.9460836347 | -0.766579987 | 0.087577053 | 0.225303569 | 0.670958793 | 0.0161814543 | 0.599633181 |
| 0.9460836347 | -0.7679526498 | -0.06718389 | 0.158778923 | 0.539504312 | 0.1779063857 | 0.721286632 |
| 0.6974568237 | -0.5701974008 | -0.51387570 | -0.045566437 | 0.641635397 | 0.4578337783 | 0.264096969 |
| 0.867011594 | -0.8265402978 | 0.155624296 | 0.165633314 | 0.599566215 | 0.0800207124 | 0.830615046 |
| 0.9274225769 | -0.774561116 | 0.106574535 | 0.264694165 | 0.516363538 | 0.0813595652 | 0.697136276 |
| 0.867011594 | -0.7893563127 | 0.182415532 | 0.184578088 | 0.522883033 | 0.0031067704 | 0.861440961 |
| 0.9301794211 | -0.7595686288 | 0.086081753 | 0.12818471 | 0.519994756 | 0.2704124187 | 0.703258183 |
| 0.6974568237 | -1.128603428 | 0.761584014 | -0.211795147 | 0.565595286 | 0.5481049137 | 1.642003572 |
| 0.9308192561 | -0.6379400072 | -0.10941045 | 0.013778011 | 0.291501487 | 0.4336062486 | 0.728295324 |
| 0.7327832683 | -0.631716945 | -0.23996402 | -0.06597847 | 0.337345227 | 0.5216660148 | 0.703836415 |
| 0.9700166179 | -0.7090730679 | -0.021365254 | 0.114218688 | 0.509441261 | 0.1926126242 | 0.673739527 |
| 0.9301794211 | -0.6640392616 | -0.083893837 | 0.096416831 | 0.449449295 | 0.2192750103 | 0.648947490 |
| 0.7327832683 | -0.4185123752 | 0.275671202 | 0.043614313 | 0.223641380 | 0.2427574426 | 0.428660042 |
| 0.7327832683 | -0.6681352235 | -0.242423811 | 0.146594315 | 0.516668814 | 0.2588858051 | 0.502791266 |
| 0.6974568237 | -0.2868551244 | -0.625823162 | 0.217900070 | 0.197533908 | -0.238763219 | 0.278555256 |
| 0.6974568237 | -0.9498376323 | 0.351307387 | 0.435036675 | 0.759900225 | 0.0562391586 | 0.617811501 |
| 0.6974568237 | -0.6108147239 | -0.32111732 | 0.037682928 | 0.397221912 | 0.33008767 | 0.594226575 |
| 0.7327832683 | -0.718139015 | 0.253578355 | 0.121511310 | 0.498707578 | 0.0200616170 | 0.801898823 |
| 0.9274153647 | -0.689580855 | 0.098461668 | 0.070944658 | 0.520880425 | 0.1820901893 | 0.677343517 |
| 0.7327832683 | -0.9099299963 | 0.397838218 | 0.684943931 | 0.630851672 | -0.0077981827 | 0.424954844 |
| 0.867011594 | -0.7511568304 | 0.161017457 | 0.056113836 | 0.446327083 | 0.2757522539 | 0.844994590 |
| 0.6974568237 | -0.5519103081 | -0.435806701 | 0.127614892 | 0.161739275 | 0.5635816558 | 0.478027063 |
| 0.7327832683 | -0.6670131027 | -0.29447331 | 0.095760848 | 0.481151197 | 0.337415887 | 0.549038245 |
| 0.6974568237 | -0.3469809938 | -0.45331210 | -0.10572462 | 0.192394608 | 0.7153773752 | 0.19521636 |
| 0.7327832683 | -0.5646474526 | -0.297212807 | 0.144866941 | 0.375773125 | 0.2780753036 | 0.429773147 |
| 0.6974568237 | -0.1973307138 | 0.506497682 | -0.604976462 | 0.079285489 | 0.8439198585 | 0.503170806 |
| 0.6974568237 | -0.6004424608 | 0.343936081 | -0.115136412 | 0.244471115 | 0.6063020786 | 0.743536952 |
| 0.7327832683 | -0.7569247193 | -0.240273227 | 0.014145094 | 0.556622960 | 0.2584876709 | 0.797243706 |
| 0.7685677253 | -0.5630976465 | 0.210911999 | 0.039768161 | 0.278420492 | 0.3203780944 | 0.626630952 |
| 0.9090584336 | -0.7879694012 | -0.11744049 | 0.269713521 | 0.605023210 | 0.0042123200 | 0.671188269 |
| 0.7868476341 | -0.6812319317 | -0.181349327 | 0.139570179 | 0.527757668 | 0.125725314 | 0.608099968 |
| 0.6974568237 | -0.3114711784 | -0.58637957 | -0.03775682 | 0.111760010 | 0.0911759887 | 0.514002011 |
| 0.7327832683 | -0.9436487591 | 0.271812963 | 0.224024014 | 0.630788267 | 0.0994598720 | 0.960880119 |

| | | | | | | |
|---|---|---|---|---|---|---|
| 0.6974568237 | -0.8349975228 | 0.451213952 | -0.101411902 | 1.012291769 | -0.2427129292 | 0.910440794 |
| 0.9301794211 | -0.6409279193 | 0.093585463 | 0.044854167 | 0.516932899 | 0.1896680434 | 0.605987413 |
| 0.7685677253 | -0.7916547723 | 0.203954675 | 0.216176172 | 0.741160428 | -0.0481766894 | 0.626246444 |
| 0.7327832683 | -0.6963794527 | -0.269401556 | 0.126491568 | 0.515092645 | 0.2896090518 | 0.567438634 |
| 0.6974568237 | -1.076199132 | -0.457880715 | 0.242055824 | 0.532083496 | 0.2061256319 | 1.256028093 |
| 0.957592049 | -0.7404107132 | -0.045062031 | 0.144671815 | 0.531798031 | 0.1606511866 | 0.699279237 |
| 0.7327832683 | -0.748622688 | 0.239522773 | 0.269014242 | 0.656740231 | -0.0521890259 | 0.569288300 |
| 0.8009553077 | -0.7896272867 | 0.231733261 | 0.380157609 | 0.785015218 | -0.1285616212 | 0.435065576 |
| 0.6974568237 | -0.8297538291 | 0.335680312 | 0.334934651 | 0.742067394 | -0.171344445 | 0.643165050 |
| 0.6974568237 | -0.7498514686 | 0.339144026 | 0.216410034 | 0.746919215 | -0.0662048187 | 0.544046820 |
| 0.6974568237 | -1.019872094 | -0.410645308 | 0.24604349 | 0.505337282 | 0.1971678181 | 1.16875613 |
| 0.957592049 | -0.7371315714 | 0.049684472 | 0.145578408 | 0.553918283 | 0.1334382563 | 0.684347252 |
| 0.7327832683 | -0.6589585241 | 0.263261411 | 0.156038056 | 0.310710535 | 0.4078599882 | 0.602960146 |
| 0.957592049 | -0.710397597 | 0.041425560 | 0.139552174 | 0.521286455 | 0.1674095963 | 0.650012719 |
| 0.6974568237 | -0.8632371778 | -0.503286947 | 0.408455058 | 0.715199545 | 0.1668086999 | 0.448716515 |
| 0.9596538857 | -0.7164405387 | -0.041022382 | 0.140705840 | 0.537973617 | 0.1736396492 | 0.639427623 |
| 0.7327832683 | -0.749368176 | 0.277304208 | 0.206650003 | 0.745585886 | -0.0836990199 | 0.566506188 |
| 0.6974568237 | -0.8936992594 | -0.364630742 | 0.335994770 | 0.820315273 | -0.0677708171 | 0.626747012 |
| 0.9674807393 | -0.7361821077 | -0.030594226 | 0.151625085 | 0.553452798 | 0.1501910982 | 0.665590204 |
| 0.6974568237 | -0.4691962656 | 0.363280485 | -0.106331489 | 0.390376173 | 0.333414023 | 0.471591762 |
| 0.9700166179 | -0.7369604844 | -0.020747171 | 0.142586382 | 0.525935008 | 0.1753316968 | 0.691799511 |
| 0.90115955 | -0.7643542219 | 0.132155648 | 0.074614848 | 0.511819488 | 0.3584913677 | 0.729610575 |
| 0.7327832683 | -0.8282515348 | 0.253084613 | 0.123340288 | 0.523226051 | 0.3365328081 | 0.807327686 |
| 0.8009553077 | -0.791031926 | 0.210077984 | 0.355094959 | 0.558968295 | 0.1566787142 | 0.534803982 |
| 0.7105975083 | -0.4533317152 | 0.354603300 | 0.085700774 | 0.265614143 | 0.0936711669 | 0.498959796 |
| 0.7327832683 | -0.734366697 | 0.217956286 | 0.205418775 | 0.528359618 | 0.0366140029 | 0.696667115 |
| 0.6974568237 | -0.7103978673 | -0.442167099 | 0.395442922 | 0.072856433 | 0.1908721539 | 0.823240868 |
| 0.8009553077 | -0.561351513 | -0.189482103 | 0.021937216 | 0.480979844 | 0.1602240714 | 0.523994016 |
| 0.9992515135 | -0.7228835939 | -0.000972278 | 0.133961998 | 0.518683188 | 0.1826684764 | 0.675808815 |
| 0.6974568237 | -0.2093655963 | 0.614252401 | 0.255498529 | -0.093920408 | 0.1440357489 | 0.152537914 |
| 0.7327832683 | -1.017112245 | 0.335874497 | 0.623828129 | 0.581118314 | 0.0763724483 | 0.719844052 |
| 0.9447216787 | -0.6961340663 | 0.070949296 | 0.080686785 | 0.520939618 | 0.2275041603 | 0.651702312 |
| 0.6974568237 | -0.9142657102 | -0.389864368 | 0.342649168 | 0.676722422 | 0.279566185 | 0.602676404 |
| 0.9596538857 | -0.7198594377 | 0.035702461 | 0.110938921 | 0.524067986 | 0.2093524069 | 0.673873071 |
| 0.7327832683 | -0.9550409664 | 0.337868978 | 0.648778030 | 0.850494055 | -0.3063355713 | 0.507554110 |
| 0.7868476341 | -0.7884491102 | -0.200491163 | 0.133916133 | 0.596047282 | 0.0347811633 | 0.819728670 |
| 0.6974568237 | -0.9347556073 | -0.515216594 | 0.198768937 | 0.724203055 | 0.1853229494 | 0.818110844 |
| 0.9992515135 | -0.7220086473 | -0.006418425 | 0.129160645 | 0.516285521 | 0.183426375 | 0.681647345 |
| 0.7327832683 | -0.7904602823 | -0.280501728 | 0.453219940 | 0.627403543 | -0.0935510314 | 0.500683627 |
| 0.7549949117 | -0.8185142549 | 0.259317155 | 0.526689211 | 0.585425846 | -0.0921607521 | 0.518528529 |
| 0.867011594 | -0.7274178442 | 0.141577560 | 0.074209081 | 0.520190592 | 0.2344988963 | 0.720652776 |
| 0.6974568237 | -0.4853857407 | -0.642039669 | 0.360560255 | 0.521120732 | -0.5504970961 | 0.369638282 |
| 0.8009553077 | -0.6406383726 | -0.230301305 | -0.136321999 | 0.299461941 | 0.4179440776 | 0.908157404 |
| 0.7685677253 | -0.9213289387 | 0.257467568 | 0.304947392 | 0.626787716 | -0.1222660673 | 0.961905459 |
| 0.9090584336 | -0.6349018379 | 0.112473673 | 0.008138123 | 0.449849456 | 0.3058582868 | 0.636578391 |
| 0.6974568237 | -1.213776717 | 0.491222910 | 0.672859284 | 0.643091859 | -0.1670422811 | 1.154779561 |

| | | | | | | |
|---|---|---|---|---|---|---|
| 0.7173585438 | -0.8162310737 | -0.400428104 | 0.581579010 | 0.544388203 | 0.1124730558 | 0.368718041 |
| 0.6974568237 | -1.051010503 | -0.463922942 | 0.769150856 | 0.564100664 | 0.0181415381 | 0.676729163 |

| se_intercept | se_microbe | se_blast | se_vns | se_interact | se_batch | t_intercept |
|---|---|---|---|---|---|---|
| 0.7022242029 | 0.323133955 | 0.898159829 | 0.886462583 | 1.248600354 | 0.591995526 | -4.606744503 |
| 0.6931857473 | 0.320732801 | 0.858040119 | 0.902929797 | 1.192462761 | 0.610612296 | -4.420712064 |
| 0.6909612271 | 0.338444219 | 0.860350063 | 0.895971002 | 1.202934214 | 0.618166823 | -4.523402157 |
| 0.7880416123 | 0.446886566 | 0.941983190 | 1.286144173 | 1.6119701 | 0.598362233 | -3.891246061 |
| 0.7607474654 | 0.426492858 | 0.857323837 | 0.902385158 | 1.198718018 | 0.831626331 | -4.070218732 |
| 0.6915813096 | 0.317216937 | 0.897356898 | 0.863284296 | 1.192525557 | 0.576562834 | -4.172574564 |
| 0.7725316205 | 0.345256056 | 0.937010692 | 0.926829270 | 1.237703397 | 0.617456901 | -3.784653885 |
| 0.662001679 | 0.346726217 | 0.786476469 | 0.821375188 | 1.143313631 | 0.645482985 | -5.279747191 |
| 0.7485834028 | 0.353823197 | 0.979841640 | 0.931631771 | 1.244140749 | 0.597666571 | -3.951813437 |
| 0.7349514519 | 0.326290139 | 0.902286567 | 0.919200934 | 1.212501308 | 0.595181879 | -4.03217031 |
| 0.7178909612 | 0.342175513 | 0.923826454 | 0.935313928 | 1.233363302 | 0.613574439 | -4.206727982 |
| 0.6981335863 | 0.343789856 | 0.847772221 | 0.918981565 | 1.148301738 | 0.582252416 | -3.992267127 |
| 0.7275474076 | 0.344280456 | 0.951145947 | 0.901058797 | 1.292900014 | 0.600507623 | -4.46083952 |
| 0.6441189253 | 0.328819506 | 0.740882927 | 0.780105909 | 1.080281601 | 0.606735286 | -3.516158546 |
| 0.7121681546 | 0.332256422 | 0.853099694 | 0.913570636 | 1.215247537 | 0.632711988 | -4.444341871 |
| 0.6961942602 | 0.408903669 | 0.955882414 | 0.885970037 | 1.196754243 | 0.699169640 | -4.446400085 |
| 0.7394228877 | 0.329929345 | 0.925653869 | 0.920176138 | 1.25188306 | 0.603316806 | -4.162454819 |
| 0.6969959435 | 0.363472930 | 0.872987716 | 0.892165361 | 1.314351142 | 0.616837758 | -4.488368644 |
| 0.6990337595 | 0.340277669 | 0.878530393 | 0.882334637 | 1.235896504 | 0.617999863 | -4.509155939 |
| 0.6940691014 | 0.334224799 | 0.864156831 | 0.909061462 | 1.212695519 | 0.597205824 | -4.500759135 |
| 0.6964641526 | 0.418331475 | 0.924712020 | 0.949988333 | 1.23151529 | 0.758892066 | -4.47864889 |
| 0.8070076457 | 0.468566239 | 0.872944473 | 0.86255012 | 1.260242217 | 0.836985151 | -2.479558382 |
| 0.661168974 | 0.326859631 | 0.812665481 | 0.872338330 | 1.177055795 | 0.579621110 | -4.519289733 |
| 0.689876385 | 0.352764223 | 0.880106542 | 0.911389232 | 1.357362001 | 0.576761903 | -4.99362212 |
| 0.8375502481 | 0.386108890 | 0.927257534 | 1.066049221 | 1.255820381 | 0.617133693 | -3.517039656 |
| 0.6945670338 | 0.301859919 | 0.855330219 | 0.887360262 | 1.196308546 | 0.598318771 | -4.471060588 |
| 0.7061507624 | 0.334548018 | 0.898537527 | 0.918519597 | 1.214925992 | 0.617120320 | -4.31395205 |
| 0.7096304963 | 0.375477649 | 0.978466879 | 0.987811170 | 1.439594227 | 0.636950156 | -4.410266457 |
| 0.6733195944 | 0.307758400 | 0.852915578 | 0.859864529 | 1.168529133 | 0.585528837 | -4.634083289 |
| 0.7044561972 | 0.329748363 | 0.848755833 | 0.908399489 | 1.231166409 | 0.617745312 | -4.227671491 |
| 0.6975784928 | 0.340559379 | 0.903964020 | 0.884232324 | 1.194342076 | 0.617277315 | -4.410975138 |
| 0.6975837629 | 0.311514231 | 0.851687194 | 0.894159521 | 1.190189963 | 0.595186731 | -4.517083889 |
| 0.7599223241 | 0.432305241 | 0.858531873 | 0.886448532 | 1.198407298 | 0.846581310 | -4.107137223 |
| 0.6693822743 | 0.335123335 | 0.893829874 | 0.881739408 | 1.31506604 | 0.569342667 | -4.928495925 |
| 0.6891037699 | 0.304511443 | 0.838244105 | 0.877748899 | 1.17260712 | 0.589804240 | -4.361493573 |
| 0.6499621346 | 0.358889487 | 0.880351085 | 0.924736849 | 1.219448471 | 0.627902340 | -5.254023837 |
| 0.6947269689 | 0.321800011 | 0.891333762 | 0.892964041 | 1.256488262 | 0.601721123 | -4.443723275 |
| 0.6874330431 | 0.352342774 | 0.835035341 | 0.864773863 | 1.173066921 | 0.678001083 | -4.342297461 |
| 0.7698614802 | 0.387287054 | 0.858613305 | 0.892840069 | 1.196562799 | 0.761251692 | -4.011723159 |
| 0.721408458 | 0.438518487 | 0.959985605 | 1.162073051 | 1.448269312 | 0.701949313 | -4.110901012 |
| 0.7096982458 | 0.319630024 | 0.841164437 | 0.869496875 | 1.166109585 | 0.623406418 | -4.692862584 |
| 0.6327796005 | 0.322590608 | 0.778542079 | 0.802027262 | 1.091098355 | 0.597199448 | -5.243718798 |
| 0.7318179738 | 0.360575738 | 0.854476307 | 0.902151476 | 1.214494985 | 0.696339806 | -4.102205031 |
| 0.6807074093 | 0.338869584 | 0.912147103 | 0.843416450 | 1.182740159 | 0.573461937 | -4.205847662 |
| 0.683931086 | 0.401902589 | 0.830049446 | 0.852891691 | 1.216924333 | 0.707214539 | -4.81840347 |
| 0.6291393612 | 0.301488863 | 0.759036561 | 0.786318534 | 1.10484007 | 0.539808052 | -4.452257525 |

| | | | | | | |
|---|---|---|---|---|---|---|
| 0.663014804 | 0.365909003 | 0.798043315 | 0.809209628 | 1.10564397 | 0.716815196 | -4.08042866 |
| 0.7641408474 | 0.445628855 | 0.973175410 | 1.270051979 | 1.555940406 | 0.625474467 | -3.645037899 |
| 0.6993648173 | 0.320206183 | 0.889482370 | 0.911750869 | 1.269557671 | 0.594346689 | -4.545565893 |
| 0.7098626719 | 0.321198515 | 0.891558892 | 0.876674226 | 1.177166923 | 0.583942483 | -4.069901662 |
| 0.7249642977 | 0.320957565 | 0.861083345 | 0.918483919 | 1.194786259 | 0.601468732 | -4.468261218 |
| 0.7933688387 | 0.380645555 | 0.837999607 | 0.950502734 | 1.161695215 | 0.671725539 | -4.538595003 |
| 0.6973909742 | 0.330118147 | 0.854007947 | 0.884995211 | 1.198805319 | 0.641466407 | -4.482099832 |
| 0.7151115945 | 0.350043646 | 0.829089911 | 0.876597486 | 1.181565442 | 0.619899012 | -3.936389279 |
| 0.7267436249 | 0.336526532 | 0.901117416 | 0.914818052 | 1.200697564 | 0.599786739 | -4.199456829 |
| 0.6989690428 | 0.305033174 | 0.855701740 | 0.889240732 | 1.198071703 | 0.599832553 | -4.370182532 |
| 0.692823887 | 0.336812067 | 0.853353702 | 0.884671060 | 1.21267093 | 0.620059714 | -4.479587574 |
| 0.7003131652 | 0.317953446 | 0.876853787 | 0.883137022 | 1.191759098 | 0.595933430 | -4.372641135 |
| 0.7196173343 | 0.406838700 | 1.022837286 | 0.892914356 | 1.210787757 | 0.649873010 | -4.37235129 |
| 0.6877720035 | 0.360396637 | 0.861760980 | 0.889155804 | 1.198021977 | 0.702111443 | -4.445200351 |
| 0.6693933259 | 0.315180370 | 0.810141565 | 0.878487879 | 1.166138058 | 0.584163977 | -4.926668021 |
| 0.6872761957 | 0.310826432 | 0.844243313 | 0.874869342 | 1.190681578 | 0.604471831 | -4.586005715 |
| 0.7390473882 | 0.377678667 | 0.848318232 | 0.896566298 | 1.235393079 | 0.701405093 | -3.663827369 |
| 0.7057596925 | 0.333385595 | 0.841186930 | 0.886436202 | 1.181328304 | 0.649297319 | -4.613556103 |
| 0.7781741637 | 0.357017304 | 0.991041035 | 0.926707935 | 1.267450335 | 0.601598065 | -4.370978234 |
| 0.7035093 | 0.318834776 | 0.885871485 | 0.921929048 | 1.258702566 | 0.601298023 | -4.393308983 |
| 0.6957003646 | 0.320745167 | 0.870045677 | 0.887535298 | 1.200084343 | 0.619819256 | -4.455280504 |
| 0.7355928053 | 0.333896178 | 0.863738785 | 0.960161678 | 1.209376263 | 0.599643915 | -4.053091058 |
| 0.7062465824 | 0.327483504 | 0.920761670 | 0.903870245 | 1.276754665 | 0.588051070 | -4.617051765 |
| 0.6948067197 | 0.306517080 | 0.851225946 | 0.882347910 | 1.194420955 | 0.605873963 | -4.515950429 |
| 0.7286237105 | 0.335323993 | 0.854902868 | 0.910400575 | 1.188875268 | 0.600944492 | -4.083935131 |
| 0.7533364266 | 0.361382776 | 0.862715454 | 0.898127946 | 1.2003725 | 0.708369845 | -4.197419434 |
| 0.6813898652 | 0.400621910 | 0.856297113 | 0.965518964 | 1.22097989 | 0.617098039 | -4.723662382 |
| 0.7681468857 | 0.373643155 | 0.928056626 | 0.886525146 | 1.196652347 | 0.658319925 | -4.033625691 |
| 0.7031863907 | 0.328160846 | 0.865147084 | 0.955844418 | 1.252506885 | 0.603204034 | -4.407206205 |
| 0.6872737098 | 0.310425942 | 0.845574667 | 0.876349915 | 1.18903218 | 0.604122279 | -4.550269468 |
| 0.7051733604 | 0.368894810 | 0.799278335 | 0.823652088 | 1.111584659 | 0.726672870 | -3.667936086 |
| 0.7004711485 | 0.314999267 | 0.852319354 | 0.885249487 | 1.198245083 | 0.618262222 | -4.342384406 |
| 0.5547106668 | 0.253666455 | 0.696897743 | 0.72209006 | 0.986184454 | 0.490089627 | -5.430101569 |
| 0.6676109962 | 0.391034671 | 0.909875526 | 0.952492551 | 1.249233851 | 0.696394008 | -4.392104835 |
| 0.6992943045 | 0.311872767 | 0.872473147 | 0.907561439 | 1.236862081 | 0.596906205 | -4.388198628 |
| 0.6918987133 | 0.314843517 | 0.854856433 | 0.907586612 | 1.213186261 | 0.607757078 | -4.473129201 |
| 0.7481074638 | 0.385531388 | 0.861335191 | 0.886270149 | 1.195972938 | 0.756675276 | -4.197459557 |
| 0.6419757053 | 0.291661635 | 0.786485783 | 0.838243269 | 1.133921883 | 0.551155816 | -4.581692439 |
| 0.6982639074 | 0.328551819 | 0.854630929 | 0.922705764 | 1.227444458 | 0.604101065 | -4.474343107 |
| 0.6975054719 | 0.315295642 | 0.851342852 | 0.881855330 | 1.195374589 | 0.624219571 | -4.519279532 |
| 0.6878986441 | 0.325740754 | 0.858182148 | 0.879578552 | 1.174583255 | 0.605232926 | -4.397879242 |
| 0.692836206 | 0.343459944 | 0.851747795 | 0.895792225 | 1.191837104 | 0.664597686 | -4.522273433 |
| 0.6951348183 | 0.322459634 | 0.859263555 | 0.924763260 | 1.235607729 | 0.611220735 | -4.464732 |
| 0.6674964554 | 0.298199468 | 0.820481299 | 0.868862598 | 1.142833115 | 0.563548951 | -4.30463519 |
| 0.6937327503 | 0.32410004 | 0.855314636 | 0.941710504 | 1.22004353 | 0.593611234 | -4.285186462 |
| 0.7257499952 | 0.323676227 | 0.876030399 | 0.887591107 | 1.19537157 | 0.620893486 | -4.450921646 |

| | | | | | | |
|---|---|---|---|---|---|---|
| 0.6747987211 | 0.310019079 | 0.798914148 | 0.826525355 | 1.111575656 | 0.608381509 | -4.024258174 |
| 0.6880180637 | 0.343408357 | 0.851558043 | 0.867647838 | 1.2567333 | 0.602359692 | -4.363609652 |
| 0.6676584324 | 0.304079870 | 0.807483869 | 0.836808177 | 1.14430217 | 0.586629988 | -4.348400109 |
| 0.7042737196 | 0.383044342 | 0.943347874 | 0.887923968 | 1.195117493 | 0.649404350 | -4.458113353 |
| 0.7390904696 | 0.345663250 | 0.851973481 | 0.915540661 | 1.192894144 | 0.619088103 | -4.373877959 |
| 0.6892698979 | 0.308639961 | 0.854426408 | 0.879635367 | 1.204782275 | 0.594266661 | -4.519172994 |
| 0.6177063659 | 0.341395688 | 0.786719528 | 0.860268475 | 1.063862912 | 0.544120649 | -5.065290564 |
| 0.7572365887 | 0.353419506 | 0.880950184 | 0.889301349 | 1.19703149 | 0.661029958 | -4.090305446 |
| 0.6975960323 | 0.461744733 | 0.997956003 | 0.885321759 | 1.188081291 | 0.747578121 | -4.543401345 |
| 0.7298886656 | 0.374918939 | 0.814989671 | 0.920663183 | 1.135514746 | 0.651034021 | -4.909381231 |
| 0.7784164629 | 0.401548622 | 1.035862879 | 0.889615023 | 1.203043032 | 0.600577350 | -3.986352483 |
| 0.6987900927 | 0.323004146 | 0.882587549 | 0.878082668 | 1.202446724 | 0.601751642 | -4.332291032 |
| 0.6675545636 | 0.317884696 | 0.817085292 | 0.838436668 | 1.120855065 | 0.562119123 | -4.245092652 |
| 0.6821465681 | 0.318224894 | 0.866411769 | 0.871108465 | 1.197972532 | 0.587020376 | -4.514600821 |
| 0.7377528623 | 0.373715302 | 1.023958593 | 0.956043271 | 1.308426952 | 0.623805847 | -4.35397668 |
| 0.595295851 | 0.289952123 | 0.723806388 | 0.758441273 | 1.034687491 | 0.547838445 | -4.759136163 |
| 0.6759452141 | 0.297268554 | 0.819545118 | 0.856793878 | 1.145136837 | 0.578503438 | -4.344337924 |
| 0.6503236768 | 0.414209237 | 0.882619364 | 0.842106792 | 1.11483975 | 0.708708823 | -4.97317386 |
| 0.6977072692 | 0.347579856 | 0.939739867 | 0.894309447 | 1.241231751 | 0.634411949 | -4.491364709 |
| 0.6352895718 | 0.351136573 | 0.931647856 | 0.799022933 | 1.134140401 | 0.576005149 | -4.425751714 |
| 0.6913893695 | 0.308480410 | 0.861622743 | 0.882397017 | 1.196152305 | 0.603698028 | -4.485993433 |
| 0.7053816723 | 0.342625916 | 0.862933898 | 0.885652905 | 1.257575139 | 0.619390880 | -4.366801741 |
| 0.7101754765 | 0.42229261 | 0.988852912 | 0.972777970 | 1.270746664 | 0.734940729 | -4.415683906 |
| 0.6276548461 | 0.340927039 | 0.737275108 | 0.741606122 | 1.061759199 | 0.620954343 | -6.281314582 |
| 0.7385670118 | 0.333652718 | 0.930153325 | 0.905021422 | 1.244597882 | 0.605553788 | -4.37761227 |
| 0.7953410138 | 0.396860079 | 0.953828479 | 0.885779904 | 1.22078352 | 0.709206601 | -4.181318705 |
| 0.700241647 | 0.398986473 | 0.963591876 | 0.886256148 | 1.197544864 | 0.671083849 | -4.412698893 |
| 0.7436237672 | 0.378466345 | 0.998841198 | 0.886376231 | 1.202627695 | 0.597564544 | -4.250473816 |
| 1.112587184 | 0.511965688 | 1.423022891 | 1.404490054 | 1.978252452 | 0.937943513 | 0.513475908 |
| 1.083449876 | 0.501305625 | 1.341117391 | 1.411279994 | 1.863820247 | 0.954387507 | 0.381950669 |
| 1.088045475 | 0.532942641 | 1.354779338 | 1.410871055 | 1.894241063 | 0.973417304 | 0.438238577 |
| 1.22454541 | 0.694421315 | 1.463756703 | 1.99855175 | 2.504855779 | 0.929800806 | 0.654664981 |
| 1.177329098 | 0.660038285 | 1.326790225 | 1.396526907 | 1.855130208 | 1.287020889 | 0.688957903 |
| 1.093299207 | 0.501478309 | 1.418603383 | 1.364739074 | 1.88522626 | 0.911470105 | 0.710336011 |
| 1.211052249 | 0.541237552 | 1.468896388 | 1.452935573 | 1.940274603 | 0.96795076 | 0.200803663 |
| 1.11129366 | 0.582044818 | 1.320247881 | 1.378831909 | 1.919265811 | 1.083563943 | 0.163793692 |
| 1.178448033 | 0.557001730 | 1.542503413 | 1.466609632 | 1.958572971 | 0.940869105 | 0.425163853 |
| 1.149544337 | 0.510353413 | 1.411274733 | 1.43773065 | 1.896484467 | 0.930929460 | 0.618316151 |
| 1.12120419 | 0.534410711 | 1.442834842 | 1.460776011 | 1.926270389 | 0.958282342 | 0.571837791 |
| 1.06930834 | 0.526571659 | 1.298504934 | 1.407573953 | 1.758816148 | 0.891816948 | 1.007068846 |
| 1.145588583 | 0.542100426 | 1.497664518 | 1.418797813 | 2.035786918 | 0.945553059 | 0.516611203 |
| 0.9831469106 | 0.501891606 | 1.130842043 | 1.190709797 | 1.648881094 | 0.926086625 | 1.902438928 |
| 1.116186102 | 0.520747802 | 1.337069085 | 1.431845612 | 1.904665918 | 0.9916539 | 0.488070219 |
| 1.090002069 | 0.640203275 | 1.496584889 | 1.387126021 | 1.873707779 | 1.094660497 | 0.449713205 |
| 1.133828963 | 0.505912724 | 1.419395023 | 1.410995487 | 1.91963394 | 0.925124282 | 0.103773797 |
| 1.053490592 | 0.549379542 | 1.319497416 | 1.348483909 | 1.986606343 | 0.932333654 | 0.323235191 |

| | | | | | | |
|---|---|---|---|---|---|---|
| 1.061223572 | 0.516585471 | 1.333722656 | 1.33949799 | 1.876250589 | 0.938203646 | 0.271950793 |
| 1.060560197 | 0.510706381 | 1.32045979 | 1.389075528 | 1.853038258 | 0.912549954 | 0.378845604 |
| 1.077147604 | 0.646989144 | 1.430154493 | 1.469246699 | 1.904654731 | 1.173698269 | 0.383287863 |
| 1.334230053 | 0.774683067 | 1.443243763 | 1.426058722 | 2.083565191 | 1.383792023 | 1.262686707 |
| 1.090780255 | 0.539244951 | 1.340715453 | 1.439162247 | 1.941877599 | 0.956244601 | 0.481060034 |
| 1.112877069 | 0.569063129 | 1.419747669 | 1.470211474 | 2.189634372 | 0.930405954 | 0.104100485 |
| 1.310799597 | 0.604275838 | 1.451195083 | 1.668409617 | 1.965409064 | 0.965838884 | 0.185830185 |
| 1.074511942 | 0.466984571 | 1.323216466 | 1.372767713 | 1.850718155 | 0.925613561 | 0.457111616 |
| 1.103338167 | 0.522720667 | 1.403936383 | 1.435157735 | 1.898283324 | 0.964230925 | 0.535139698 |
| 1.107032692 | 0.585749957 | 1.526420903 | 1.540998118 | 2.24578549 | 0.993650427 | 0.355118793 |
| 0.8250738823 | 0.377121682 | 1.045147614 | 1.053662735 | 1.431894863 | 0.717496646 | 0.512507182 |
| 1.109015158 | 0.519118059 | 1.336183979 | 1.43008012 | 1.938207393 | 0.972507471 | 0.570841061 |
| 1.09427288 | 0.534226465 | 1.418024383 | 1.387071794 | 1.873532737 | 0.968306553 | 0.428684780 |
| 1.089476459 | 0.486518522 | 1.33015302 | 1.3964857 | 1.858821858 | 0.929554223 | 0.518830237 |
| 1.188651463 | 0.676201028 | 1.34289405 | 1.386560589 | 1.874518676 | 1.324201279 | 0.457154837 |
| 1.058603728 | 0.529985369 | 1.413559447 | 1.394438814 | 2.079729126 | 0.900394727 | 0.198600240 |
| 1.097846765 | 0.485132889 | 1.335449927 | 1.398387053 | 1.868140895 | 0.939647561 | 0.487122545 |
| 1.005469948 | 0.555190180 | 1.361874043 | 1.430537354 | 1.886446496 | 0.971344175 | -0.02658353 |
| 1.045348214 | 0.484209023 | 1.341180346 | 1.343633409 | 1.890624403 | 0.905403316 | 0.567518760 |
| 1.088042319 | 0.557674458 | 1.321661504 | 1.368730481 | 1.856684757 | 1.073113779 | 0.571836133 |
| 1.195017649 | 0.601166414 | 1.33278269 | 1.385911191 | 1.857364864 | 1.181653105 | 0.152453546 |
| 1.131712351 | 0.687927599 | 1.505981187 | 1.823006661 | 2.271978169 | 1.101185741 | 0.592718359 |
| 1.106353922 | 0.498273644 | 1.311297554 | 1.355465205 | 1.817856984 | 0.971832943 | 0.097179043 |
| 0.9232247757 | 0.47065936 | 1.13589208 | 1.170156938 | 1.591911359 | 0.871313371 | 0.073475398 |
| 1.14322737 | 0.563282221 | 1.33484109 | 1.409318024 | 1.897253085 | 1.087804283 | 0.587679031 |
| 1.11050962 | 0.552833608 | 1.488081544 | 1.375953998 | 1.929528468 | 0.935548797 | 0.575566413 |
| 1.043124914 | 0.612977846 | 1.265983189 | 1.300821955 | 1.856040931 | 1.078636607 | 0.100793799 |
| 1.076925835 | 0.516071901 | 1.299276652 | 1.345976417 | 1.891203902 | 0.924013459 | 0.680028406 |
| 1.040500039 | 0.574238056 | 1.252406575 | 1.269930391 | 1.735138623 | 1.124931502 | 1.057074455 |
| 1.215943878 | 0.709109688 | 1.548571427 | 2.020978114 | 2.475899853 | 0.995290138 | 0.560063816 |
| 1.102373534 | 0.504724878 | 1.402046257 | 1.43714697 | 2.001139808 | 0.936838749 | 0.399700831 |
| 1.07103987 | 0.4846239 | 1.34518289 | 1.322724924 | 1.776107911 | 0.881051653 | 0.940997970 |
| 1.139583187 | 0.504518423 | 1.353550935 | 1.443779832 | 1.878103979 | 0.945458497 | 0.301784049 |
| 1.252016118 | 0.600697112 | 1.322447977 | 1.499989268 | 1.833272322 | 1.060050712 | -0.146023973 |
| 1.057377928 | 0.500522168 | 1.294839175 | 1.341821787 | 1.817617851 | 0.972585602 | 0.585545770 |
| 1.149639178 | 0.562742785 | 1.332874829 | 1.409249719 | 1.899527199 | 0.996571998 | 0.586305746 |
| 1.116709141 | 0.517104302 | 1.384650682 | 1.405702983 | 1.844983429 | 0.921628084 | 0.158855924 |
| 1.092860421 | 0.476929110 | 1.337916999 | 1.390356284 | 1.87322337 | 0.937857353 | 0.519727409 |
| 1.086482428 | 0.528186743 | 1.338224361 | 1.38733606 | 1.901703566 | 0.972374071 | 0.443431604 |
| 1.062214889 | 0.482262652 | 1.329986634 | 1.339516864 | 1.807625961 | 0.903894706 | 0.645177147 |
| 1.127119919 | 0.637222008 | 1.602046288 | 1.398551021 | 1.896428745 | 1.017881005 | 0.416064167 |
| 1.090127765 | 0.571233458 | 1.365902606 | 1.409323766 | 1.898880754 | 1.112855968 | 0.462249168 |
| 1.065810999 | 0.501831572 | 1.289910965 | 1.398732268 | 1.856730147 | 0.930108455 | 0.218676491 |
| 1.088947604 | 0.492485701 | 1.337652518 | 1.38617761 | 1.886563014 | 0.957749091 | 0.420486093 |
| 1.195652303 | 0.611019504 | 1.372433844 | 1.450490966 | 1.998654759 | 1.134753506 | 0.239026439 |
| 0.8899499197 | 0.420393070 | 1.060721162 | 1.117779657 | 1.489633144 | 0.818751911 | -0.345513399 |

| | | | | | | |
|---|---|---|---|---|---|---|
| 1.214985165 | 0.557421138 | 1.547340187 | 1.446895112 | 1.978905785 | 0.939291945 | -0.009214527 |
| 1.090952204 | 0.494426301 | 1.373746515 | 1.429662021 | 1.951906447 | 0.932450223 | 0.544471042 |
| 1.021055762 | 0.470746771 | 1.276936447 | 1.302605369 | 1.76132297 | 0.909687639 | 0.583206205 |
| 1.137794215 | 0.516461196 | 1.336006805 | 1.485151017 | 1.870629111 | 0.927512303 | 0.719675180 |
| 1.121273642 | 0.519929768 | 1.461848903 | 1.435031201 | 2.027041812 | 0.933620327 | 0.515727571 |
| 1.070044039 | 0.472054695 | 1.310939034 | 1.358868725 | 1.839479939 | 0.933082257 | 0.547253313 |
| 1.147769228 | 0.528221297 | 1.346691291 | 1.434114414 | 1.872783482 | 0.946641712 | 0.466232389 |
| 1.170006294 | 0.561263345 | 1.339882789 | 1.394881905 | 1.864297718 | 1.100168728 | 0.181975677 |
| 1.065398271 | 0.626398941 | 1.33887736 | 1.509652973 | 1.909083082 | 0.964873735 | 0.285506481 |
| 1.195051658 | 0.581298812 | 1.443832725 | 1.379219737 | 1.861703013 | 1.024187344 | 0.614082192 |
| 1.068414379 | 0.498604313 | 1.314495839 | 1.452300463 | 1.903046452 | 0.916502186 | 0.274948048 |
| 1.085736211 | 0.490402413 | 1.335815735 | 1.38443363 | 1.878400519 | 0.954375855 | 0.432015111 |
| 1.187652207 | 0.621292238 | 1.346143704 | 1.387193952 | 1.872129674 | 1.223861658 | 0.408585975 |
| 1.05633989 | 0.475032114 | 1.285333358 | 1.33499338 | 1.807003875 | 0.932365379 | 0.695593824 |
| 1.004521145 | 0.459362572 | 1.262006595 | 1.307627161 | 1.785873603 | 0.887499417 | 0.573590843 |
| 1.087427609 | 0.636930637 | 1.482036356 | 1.551452423 | 2.034794795 | 1.134310362 | 0.572450459 |
| 1.097613481 | 0.489516004 | 1.369435275 | 1.424509916 | 1.94138074 | 0.936904953 | 0.452101873 |
| 1.075123509 | 0.489227196 | 1.328339294 | 1.410275355 | 1.885138741 | 0.944377999 | 0.424204675 |
| 1.163755617 | 0.599732445 | 1.339892616 | 1.378681425 | 1.860454937 | 1.177083702 | 0.224681750 |
| 1.077683883 | 0.489612053 | 1.320272785 | 1.407158019 | 1.903513369 | 0.92522464 | 0.556152693 |
| 1.087237669 | 0.511574364 | 1.330710252 | 1.436706743 | 1.91120268 | 0.940620626 | 0.385682652 |
| 1.09693582 | 0.495851427 | 1.338869024 | 1.386854641 | 1.879912428 | 0.981682345 | 0.422815130 |
| 1.071905597 | 0.507579628 | 1.337246783 | 1.370587341 | 1.830273074 | 0.943093241 | 0.320872113 |
| 1.084691961 | 0.537714739 | 1.3334811 | 1.402436271 | 1.865918833 | 1.04048224 | 0.407623079 |
| 1.059923687 | 0.491678155 | 1.310182963 | 1.410055228 | 1.884022877 | 0.931973652 | 0.409147142 |
| 1.106524695 | 0.494332326 | 1.360131298 | 1.440334124 | 1.894501542 | 0.934208454 | 0.525231452 |
| 1.096386277 | 0.512212860 | 1.35175286 | 1.488294265 | 1.928176207 | 0.938152639 | 0.301108872 |
| 1.112431881 | 0.496131941 | 1.342782159 | 1.360502448 | 1.832269311 | 0.951707493 | 0.092901958 |
| 1.141648627 | 0.524501374 | 1.351631548 | 1.398345177 | 1.880603478 | 1.029281611 | 0.406481127 |
| 1.064926367 | 0.531533449 | 1.318056402 | 1.342960467 | 1.945193736 | 0.932342844 | 0.638570035 |
| 1.05501852 | 0.480500027 | 1.275967464 | 1.322305063 | 1.808200007 | 0.926979234 | 0.732458362 |
| 1.103634028 | 0.600250668 | 1.478275826 | 1.391423644 | 1.872812084 | 1.017650835 | 0.418830876 |
| 1.16008263 | 0.542555951 | 1.337264757 | 1.437040338 | 1.872376703 | 0.971725904 | 0.528581498 |
| 1.084641202 | 0.485678570 | 1.344532947 | 1.384201987 | 1.895856034 | 0.935143269 | 0.438187235 |
| 1.015765417 | 0.561396081 | 1.293693143 | 1.414638111 | 1.749431791 | 0.894759983 | 0.447673521 |
| 1.156276081 | 0.539660296 | 1.345182791 | 1.357934751 | 1.827828846 | 1.009371631 | 0.826021832 |
| 1.099542806 | 0.727796714 | 1.572966722 | 1.395433929 | 1.872640004 | 1.178323998 | 0.412300005 |
| 1.173323887 | 0.602696504 | 1.310127001 | 1.480001205 | 1.825383293 | 1.046562037 | -0.037468810 |
| 1.217091943 | 0.627840773 | 1.619621917 | 1.390956294 | 1.881016206 | 0.939031854 | 0.318376856 |
| 1.100875781 | 0.508861596 | 1.390430787 | 1.383333784 | 1.894337785 | 0.948001147 | 0.498572852 |
| 1.078168404 | 0.513416063 | 1.319675715 | 1.354160355 | 1.810294743 | 0.907879462 | 0.739963153 |
| 1.051831363 | 0.490684758 | 1.335957866 | 1.343199904 | 1.847205778 | 0.905152165 | 0.510429274 |
| 1.156416936 | 0.585793329 | 1.605040276 | 1.498583994 | 2.050940313 | 0.977806638 | 0.309782632 |
| 1.080368297 | 0.526217478 | 1.3135947 | 1.376451568 | 1.877794983 | 0.994240573 | 0.588696200 |
| 1.093016556 | 0.480689033 | 1.325220395 | 1.385452363 | 1.851708536 | 0.935451310 | 0.319334242 |
| 1.085926479 | 0.691656777 | 1.473819534 | 1.406170645 | 1.861586849 | 1.183419434 | 0.385926104 |

| | | | | | | |
|---|---|---|---|---|---|---|
| 1.09303794 | 0.544523451 | 1.472209583 | 1.401037654 | 1.944530973 | 0.993878608 | 0.412576852 |
| 1.092213642 | 0.603687156 | 1.601723911 | 1.373710173 | 1.949856684 | 0.990289639 | 0.610589151 |
| 1.033554918 | 0.461146004 | 1.288036037 | 1.319091408 | 1.788122804 | 0.902465519 | 0.487430246 |
| 1.102785053 | 0.535657153 | 1.34910027 | 1.384618886 | 1.966077544 | 0.968348104 | 0.388717000 |
| 1.106807523 | 0.658142463 | 1.541125931 | 1.516073156 | 1.980456963 | 1.145404136 | 0.339466781 |
| 1.097126398 | 0.595932711 | 1.288740124 | 1.296310638 | 1.855930934 | 1.085414072 | -0.393108978 |
| 1.161485617 | 0.524709102 | 1.462778179 | 1.423255235 | 1.957280132 | 0.952306295 | 0.469403705 |
| 1.24929583 | 0.623374922 | 1.498242793 | 1.391354302 | 1.9175671 | 1.113998693 | 0.198429543 |
| 1.096398262 | 0.624710167 | 1.508736966 | 1.387649114 | 1.875047154 | 1.050744653 | 0.440762068 |
| 1.164382386 | 0.592610894 | 1.564007431 | 1.387907321 | 1.883100792 | 0.935679655 | 0.376449307 |
| 0.7075077211 | 0.325565207 | 0.904917562 | 0.893132306 | 1.257994793 | 0.596449686 | -0.250446771 |
| 0.6908055355 | 0.319631492 | 0.855093842 | 0.899829381 | 1.188368168 | 0.608515619 | -0.169374492 |
| 0.667555207 | 0.326979563 | 0.831206067 | 0.865620362 | 1.162185326 | 0.597226683 | -0.153959648 |
| 0.7544761032 | 0.427852069 | 0.901860759 | 1.231362695 | 1.543310531 | 0.572875847 | -0.792585842 |
| 0.7364614002 | 0.412877521 | 0.829954673 | 0.873577458 | 1.160450203 | 0.805077533 | -0.625953826 |
| 0.6781549292 | 0.311058478 | 0.879935584 | 0.846524469 | 1.16937383 | 0.565369426 | -0.586939448 |
| 0.7638540818 | 0.341377933 | 0.926485626 | 0.916418568 | 1.22380077 | 0.610521254 | 0.111288483 |
| 0.7197762618 | 0.376985902 | 0.855114286 | 0.893058705 | 1.243093542 | 0.70181594 | -0.171009631 |
| 0.7446795308 | 0.351978004 | 0.974731753 | 0.9267733 | 1.237652539 | 0.594549733 | -0.032294372 |
| 0.6934415481 | 0.307861341 | 0.851325611 | 0.867284658 | 1.144019489 | 0.561566131 | -0.770510421 |
| 0.6516591352 | 0.310606779 | 0.838595248 | 0.849022899 | 1.119574568 | 0.556966739 | -0.788037155 |
| 0.7169035543 | 0.353032965 | 0.870565362 | 0.943689236 | 1.179174894 | 0.597906811 | 0.037441267 |
| 0.7275602218 | 0.344286520 | 0.951162699 | 0.901074667 | 1.292922786 | 0.600518200 | -0.276951742 |
| 0.7437281254 | 0.379669507 | 0.855456111 | 0.900744696 | 1.247340791 | 0.700563326 | -0.679253457 |
| 0.7083325179 | 0.330466936 | 0.848505020 | 0.908650274 | 1.208702386 | 0.629304291 | -0.262779114 |
| 0.6102072475 | 0.358399942 | 0.837821295 | 0.776543802 | 1.048943024 | 0.612815138 | -0.435072312 |
| 0.6970779625 | 0.311035105 | 0.872643955 | 0.867479920 | 1.180190805 | 0.568766340 | 0.312233377 |
| 0.693304205 | 0.361547744 | 0.868363812 | 0.88743988 | 1.307389493 | 0.613570589 | -0.196940907 |
| 0.6691359079 | 0.325723907 | 0.840955425 | 0.844596960 | 1.183036896 | 0.591567851 | -0.019230492 |
| 0.627581882 | 0.302208279 | 0.781376334 | 0.821979399 | 1.096527326 | 0.539997465 | -0.384025581 |
| 0.6867197341 | 0.412478486 | 0.911774124 | 0.936696789 | 1.214284826 | 0.748274202 | -0.161655999 |
| 0.8864323352 | 0.514681946 | 0.958858584 | 0.947441230 | 1.384273689 | 0.919360189 | -0.277255343 |
| 0.6448943276 | 0.318813995 | 0.792661754 | 0.850865759 | 1.148082616 | 0.565353761 | -0.020937601 |
| 0.7178026708 | 0.367044164 | 0.91573337 | 0.948282386 | 1.412308191 | 0.600109300 | -0.407350222 |
| 0.8326650318 | 0.383856816 | 0.921849078 | 1.059831228 | 1.248495502 | 0.613534110 | -0.033892845 |
| 0.6858422163 | 0.298068100 | 0.844585972 | 0.876213668 | 1.181281093 | 0.590802978 | -0.219206426 |
| 0.6954656235 | 0.329485796 | 0.884941282 | 0.904620994 | 1.196542307 | 0.607782348 | -0.356615387 |
| 0.6416332666 | 0.339499150 | 0.884709582 | 0.893158497 | 1.301651424 | 0.575917201 | -0.662758100 |
| 0.621722035 | 0.284174381 | 0.787555291 | 0.793971732 | 1.078982873 | 0.540658824 | -0.195096092 |
| 0.6778701798 | 0.317303735 | 0.816723980 | 0.874116698 | 1.184702467 | 0.594431744 | -0.547267991 |
| 0.6712646633 | 0.327712908 | 0.869864982 | 0.850877599 | 1.149289492 | 0.593992581 | -0.072854798 |
| 0.6951814302 | 0.310441441 | 0.848754161 | 0.891080223 | 1.186091198 | 0.593137033 | -0.174821286 |
| 0.7513646342 | 0.427436935 | 0.848863714 | 0.876465997 | 1.184911711 | 0.837047730 | -0.368316927 |
| 0.7017766154 | 0.351341422 | 0.937086218 | 0.924410641 | 1.378707848 | 0.596895654 | -0.195068247 |
| 0.6723409186 | 0.297104024 | 0.817853328 | 0.856397145 | 1.144082768 | 0.575456907 | -0.017587523 |
| 0.699031991 | 0.385984382 | 0.946814497 | 0.994551231 | 1.311512544 | 0.675306760 | -0.036279787 |

| | | | | | | |
|---|---|---|---|---|---|---|
| 0.6610065876 | 0.306180610 | 0.848070558 | 0.849621707 | 1.195501335 | 0.572515022 | -0.332064962 |
| 0.6777741662 | 0.347392132 | 0.823302557 | 0.852623234 | 1.156584575 | 0.668474731 | -0.41603886 |
| 0.7643693355 | 0.384524172 | 0.852488010 | 0.886470603 | 1.188026593 | 0.755820969 | -0.226443603 |
| 0.7120469393 | 0.432827954 | 0.947528136 | 1.146993149 | 1.42947552 | 0.692840310 | 0.041439374 |
| 0.6943860197 | 0.312733787 | 0.823015737 | 0.850736884 | 1.140949971 | 0.609955997 | 0.188944135 |
| 0.5680389384 | 0.289585862 | 0.698888232 | 0.719970609 | 0.979466390 | 0.536099047 | 0.237694118 |
| 0.7285414073 | 0.358961334 | 0.850650562 | 0.898112276 | 1.20905733 | 0.693222086 | -0.10988457 |
| 0.6971000121 | 0.347030145 | 0.934113170 | 0.863727366 | 1.211222572 | 0.587271885 | 0.006368950 |
| 0.6484055923 | 0.381026527 | 0.786934113 | 0.808589862 | 1.153713523 | 0.670479631 | 0.194722389 |
| 0.6928380613 | 0.332013815 | 0.835887010 | 0.865931210 | 1.216702212 | 0.594462193 | -0.381569924 |
| 0.7146344981 | 0.394397222 | 0.860175791 | 0.872211469 | 1.19172501 | 0.772623574 | -0.01807365 |
| 0.7573403509 | 0.441662966 | 0.964514603 | 1.25874911 | 1.542093264 | 0.619908037 | 0.245312387 |
| 0.6998824159 | 0.320443167 | 0.890140674 | 0.912425654 | 1.270497268 | 0.594786564 | -0.239743426 |
| 0.721201491 | 0.326329102 | 0.905799992 | 0.890677569 | 1.195970113 | 0.593269946 | -0.127272076 |
| 0.7195657329 | 0.318567502 | 0.854671148 | 0.911644279 | 1.185889088 | 0.596989798 | -0.392586563 |
| 0.8038654592 | 0.385681664 | 0.849086712 | 0.963078305 | 1.177064956 | 0.680612765 | 0.191872459 |
| 0.6874400517 | 0.325407762 | 0.841822291 | 0.872367404 | 1.181699822 | 0.632313460 | -0.279643098 |
| 0.7278259713 | 0.356267272 | 0.843830773 | 0.892183012 | 1.202573168 | 0.630920550 | -0.400299488 |
| 0.6334021811 | 0.293303762 | 0.785379764 | 0.797320719 | 1.046482459 | 0.522751375 | 0.520769862 |
| 0.6699985422 | 0.292390320 | 0.820235065 | 0.852383951 | 1.148414658 | 0.574971009 | -0.422522188 |
| 0.6837887653 | 0.332419698 | 0.842225110 | 0.873134058 | 1.196856479 | 0.611973510 | -0.208707068 |
| 0.6692472518 | 0.303849021 | 0.837956527 | 0.843961037 | 1.138892628 | 0.569497805 | -0.429211685 |
| 0.6324641573 | 0.357566283 | 0.898961005 | 0.7847731 | 1.06414871 | 0.5711666 | -0.874653821 |
| 0.6300593184 | 0.330154845 | 0.789448441 | 0.814544496 | 1.097492929 | 0.643195499 | -0.04193236 |
| 0.7012753063 | 0.330191835 | 0.848727127 | 0.920328651 | 1.221679082 | 0.611986639 | -0.240633123 |
| 0.6865461909 | 0.310496281 | 0.843346583 | 0.873940081 | 1.18941687 | 0.603829778 | -0.162706743 |
| 0.7614894527 | 0.389147335 | 0.874078437 | 0.923791614 | 1.272907278 | 0.722704103 | -0.202221072 |
| 0.6590320138 | 0.311312452 | 0.785492743 | 0.827746103 | 1.10311368 | 0.606307961 | 0.252309395 |
| 0.7839387296 | 0.359662020 | 0.998382478 | 0.933572811 | 1.276839365 | 0.606054588 | -0.254484247 |
| 0.6669295263 | 0.302256595 | 0.839809580 | 0.873992289 | 1.193254881 | 0.570032842 | -0.451754451 |
| 0.6830048005 | 0.314892013 | 0.854168553 | 0.871339013 | 1.178184472 | 0.608508416 | -0.260632681 |
| 0.7157678461 | 0.324897343 | 0.840460164 | 0.934284364 | 1.176782367 | 0.583482913 | -0.569114482 |
| 0.6860322525 | 0.318110206 | 0.894407447 | 0.877999491 | 1.240211139 | 0.571219756 | -0.548750544 |
| 0.6930962392 | 0.305762494 | 0.849130392 | 0.880175739 | 1.19148052 | 0.604382418 | -0.20147928 |
| 0.7209498038 | 0.331792342 | 0.845898982 | 0.900812184 | 1.176353966 | 0.594615311 | -0.421497097 |
| 0.7062443952 | 0.338792273 | 0.808785999 | 0.841984810 | 1.125335667 | 0.664088733 | 0.419925651 |
| 0.6941927671 | 0.408149352 | 0.872386416 | 0.98366048 | 1.243921361 | 0.628692937 | -0.272439138 |
| 0.7465300625 | 0.363128268 | 0.901939700 | 0.861576978 | 1.162976728 | 0.639793800 | 0.213758824 |
| 0.6907579085 | 0.322360761 | 0.849856024 | 0.938950326 | 1.230369426 | 0.592542692 | -0.10585665 |
| 0.6836337483 | 0.308781854 | 0.841096307 | 0.871708562 | 1.182734789 | 0.600922707 | -0.18565482 |
| 0.7269260649 | 0.380274224 | 0.823933925 | 0.849059543 | 1.145874061 | 0.749088776 | 0.304565676 |
| 0.695236318 | 0.312645183 | 0.845949717 | 0.878633752 | 1.189290239 | 0.613641763 | -0.305243255 |
| 0.6894267249 | 0.315271445 | 0.866145105 | 0.897455583 | 1.225687478 | 0.609111932 | -0.224987811 |
| 0.6859909702 | 0.401800232 | 0.934925276 | 0.978715589 | 1.283626463 | 0.715566406 | -0.059501804 |
| 0.6652090909 | 0.296671371 | 0.829946798 | 0.863324806 | 1.176574577 | 0.567811623 | -0.422507665 |
| 0.6850153865 | 0.311711309 | 0.846351928 | 0.898557523 | 1.201116925 | 0.601710831 | -0.194015205 |

| | | | | | | |
|---|---|---|---|---|---|---|
| 0.7032925687 | 0.362436379 | 0.8097374619 | 0.8331787070 | 1.124328951 | 0.711347132 | 0.3738175389 |
| 0.6877602279 | 0.3124624045 | 0.8425764970 | 0.8980252329 | 1.214791099 | 0.5904632320 | -0.300941176 |
| 0.6622181327 | 0.3115913198 | 0.8105131778 | 0.8750738534 | 1.164081328 | 0.5729161641 | -0.056097203 |
| 0.6940448094 | 0.3137313070 | 0.8471189281 | 0.8774800192 | 1.189443756 | 0.6211225155 | -0.263577083 |
| 0.6248400803 | 0.2958806220 | 0.7795139694 | 0.7989489991 | 1.066911095 | 0.5497521965 | 0.0611363143 |
| 0.6792590544 | 0.3367293378 | 0.8350565359 | 0.8782378490 | 1.168481291 | 0.6515739105 | -0.144250655 |
| 0.6815597495 | 0.3161624223 | 0.8424832682 | 0.9067038501 | 1.211477935 | 0.599284398 | -0.182791968 |
| 0.7009036124 | 0.3131238868 | 0.8615451094 | 0.9123478171 | 1.20003013 | 0.5917536982 | -0.315744035 |
| 0.6815522568 | 0.3184095224 | 0.8402971034 | 0.925176041 | 1.198622122 | 0.5831886640 | -0.450762706 |
| 0.7251078573 | 0.3233898411 | 0.8752552948 | 0.8868057730 | 1.194313914 | 0.6203441249 | -0.265435438 |
| 0.7042877286 | 0.3235670531 | 0.8338270555 | 0.8626448849 | 1.160152012 | 0.6349680549 | 0.1312692859 |
| 0.652405679 | 0.3256332567 | 0.8074806943 | 0.8227376681 | 1.191683744 | 0.5711810559 | 0.0472294965 |
| 0.6994414283 | 0.3185551905 | 0.8459230699 | 0.8766433233 | 1.19877516 | 0.6145557321 | -0.130690172 |
| 0.6119514882 | 0.3328316094 | 0.8196857553 | 0.7715272892 | 1.038451255 | 0.564274866 | -0.688700899 |
| 0.7246335114 | 0.3389019143 | 0.8353084783 | 0.8976322541 | 1.169560572 | 0.6069784478 | -0.516341209 |
| 0.6770391375 | 0.3031632948 | 0.8392650261 | 0.8640266637 | 1.183403997 | 0.5837216872 | -0.237290382 |
| 0.663337571 | 0.3666152708 | 0.8448360744 | 0.9238182291 | 1.142452594 | 0.5843159302 | -0.241822148 |
| 0.7117506072 | 0.3321901664 | 0.8280329269 | 0.8358824501 | 1.125127738 | 0.6213229544 | -0.828034348 |
| 0.6538124239 | 0.4327639919 | 0.9353207348 | 0.8297558171 | 1.113513083 | 0.7006574590 | -0.493458334 |
| 0.7290241082 | 0.3744748458 | 0.8140243112 | 0.9195726525 | 1.134169723 | 0.6502628671 | 0.4134785350 |
| 0.7249135184 | 0.3739489578 | 0.9646648545 | 0.8284690621 | 1.120354204 | 0.5592978322 | -0.967817640 |
| 0.700061265 | 0.3235917249 | 0.8841930689 | 0.8796799925 | 1.204634101 | 0.602846292 | -0.239820068 |
| 0.7022343706 | 0.3343989723 | 0.8595332988 | 0.8819938899 | 1.179084069 | 0.5913215045 | -0.057170412 |
| 0.6217923438 | 0.290069337 | 0.7897543289 | 0.7940354752 | 1.091979617 | 0.5350826245 | -0.329778202 |
| 0.6832277536 | 0.3460951219 | 0.9482808739 | 0.8853849713 | 1.211725027 | 0.5777022210 | -0.850087119 |
| 0.6964943601 | 0.3392431140 | 0.8468513029 | 0.887374015 | 1.210581261 | 0.6409693370 | -0.155221810 |
| 0.6995613189 | 0.3076544927 | 0.8481783022 | 0.8867284552 | 1.185145511 | 0.5987151328 | -0.153135374 |
| 0.6900763187 | 0.4395288000 | 0.9365716542 | 0.8935826515 | 1.182987085 | 0.7520304021 | -0.265023649 |
| 0.6867503192 | 0.342121385 | 0.9249819835 | 0.880265013 | 1.221739172 | 0.6244490031 | -0.130223168 |
| 0.6811859586 | 0.3765043749 | 0.9989545967 | 0.8567482091 | 1.216076181 | 0.6176185427 | -0.460432348 |
| 0.6885926893 | 0.3072326025 | 0.8581374664 | 0.8788277080 | 1.191313851 | 0.6012560613 | -0.209411942 |
| 0.6781303913 | 0.3293891174 | 0.8295958417 | 0.8514371363 | 1.208990756 | 0.595461715 | -0.000003847 |
| 0.7058344094 | 0.4197112753 | 0.9828083821 | 0.9668317012 | 1.262979011 | 0.7304482804 | -0.203687239 |
| 0.7482702462 | 0.406442428 | 0.8789560547 | 0.8841193523 | 1.265795719 | 0.740282119 | -0.525005992 |
| 0.7305038987 | 0.3300101517 | 0.9199986185 | 0.8951410871 | 1.231010308 | 0.5989428124 | -0.425876375 |
| 0.7338823437 | 0.3661933691 | 0.880122951 | 0.8173327179 | 1.12644948 | 0.6544038268 | -1.155213877 |
| 0.5993639003 | 0.3415079497 | 0.824775544 | 0.7585809040 | 1.025024952 | 0.5744066144 | -0.602665539 |
| 0.6559835847 | 0.3338619880 | 0.8811222268 | 0.7819118795 | 1.060891355 | 0.527138251 | -1.079409099 |
| 0.7398632845 | 0.340453873 | 0.9463010231 | 0.9339768063 | 1.315525091 | 0.6237263725 | 0.2508799305 |
| 0.6282680758 | 0.2906957927 | 0.7776836393 | 0.8183693459 | 1.080787202 | 0.5534277274 | 0.1690430441 |
| 0.7156807217 | 0.3505522357 | 0.8911295309 | 0.9280248270 | 1.245969807 | 0.6402820607 | 0.4464230692 |
| 0.7866970581 | 0.4461240895 | 0.940375982 | 1.283949758 | 1.609219761 | 0.5973413090 | 1.047173879 |
| 0.7959678888 | 0.4462382537 | 0.89701547 | 0.9441630005 | 1.254215221 | 0.8701282431 | 0.3325094889 |
| 0.7501280913 | 0.3440713804 | 0.9733239003 | 0.9363668342 | 1.293480474 | 0.6253725664 | 0.431615298 |
| 0.7995890136 | 0.3573484142 | 0.9698288532 | 0.9592908330 | 1.281053114 | 0.639082908 | 0.7464243449 |
| 0.6951274296 | 0.3640759713 | 0.8258307858 | 0.8624757933 | 1.200523641 | 0.6777821614 | 0.9883994301 |

| | | | | | | |
|---|---|---|---|---|---|---|
| 0.7859963335 | 0.371506681 | 1.028812466 | 0.978193149 | 1.306320797 | 0.627536934 | 0.234044254 |
| 0.7699192089 | 0.341814477 | 0.945215848 | 0.962934972 | 1.270190086 | 0.623499634 | 0.550345540 |
| 0.75345641 | 0.359127427 | 0.969594272 | 0.981650853 | 1.294466062 | 0.643971883 | 0.451446108 |
| 0.7554994175 | 0.372039165 | 0.917433901 | 0.994494536 | 1.242658012 | 0.630096259 | 0.621294416 |
| 0.7662299013 | 0.362585279 | 1.001716806 | 0.948966605 | 1.361641372 | 0.632435621 | 0.282285034 |
| 0.8053607686 | 0.411132665 | 0.926347636 | 0.975389280 | 1.350707744 | 0.758618909 | 0.461495115 |
| 0.72150777 | 0.336613746 | 0.864287534 | 0.925551512 | 1.231184707 | 0.641009588 | 0.107548752 |
| 0.7246956765 | 0.425643729 | 0.995015174 | 0.922240662 | 1.245748026 | 0.727792865 | 0.355493606 |
| 0.772852643 | 0.344845650 | 0.967503239 | 0.961777857 | 1.308481449 | 0.630593123 | 0.287126022 |
| 0.7297671453 | 0.380562620 | 0.914033661 | 0.934112995 | 1.376149014 | 0.645840100 | 0.420590224 |
| 0.6708247653 | 0.326546013 | 0.843077944 | 0.846728670 | 1.186022808 | 0.593060932 | 0.122756600 |
| 0.6939881127 | 0.334185800 | 0.864055995 | 0.908955386 | 1.212554013 | 0.597136138 | 0.309126067 |
| 0.7253941831 | 0.435708309 | 0.963123110 | 0.989449361 | 1.282670507 | 0.790415254 | 0.438894122 |
| 0.8715143312 | 0.506020228 | 0.942721699 | 0.931496492 | 1.360977382 | 0.903888034 | -0.78957323 |
| 0.7142981328 | 0.353124894 | 0.877968354 | 0.942436298 | 1.271639762 | 0.626197376 | 0.286997933 |
| 0.7623913091 | 0.389844302 | 0.972617115 | 1.007188019 | 1.50003829 | 0.637387034 | 0.472776364 |
| 0.8654050096 | 0.398949876 | 0.958095728 | 1.101503269 | 1.297585729 | 0.637657969 | -0.123936712 |
| 0.7267916038 | 0.315864768 | 0.895013428 | 0.928529510 | 1.251811509 | 0.626077885 | 0.394830485 |
| 0.7415991263 | 0.351342137 | 0.943643596 | 0.964628755 | 1.275914581 | 0.648099407 | 0.415898093 |
| 0.7194225339 | 0.380658784 | 0.991968531 | 1.001441763 | 1.459458876 | 0.645739293 | 0.152421788 |
| 0.7140470974 | 0.326373975 | 0.904506416 | 0.911875692 | 1.239210685 | 0.620946086 | 0.385358523 |
| 0.6308266226 | 0.295283152 | 0.760044099 | 0.813453815 | 1.102485223 | 0.553178736 | -0.227821554 |
| 0.6848469424 | 0.334343807 | 0.887465714 | 0.868094142 | 1.172544061 | 0.606011347 | 0.212110935 |
| 0.6857281038 | 0.306219947 | 0.837212498 | 0.878962994 | 1.169962304 | 0.585071342 | 0.659000581 |
| 0.7665192351 | 0.436058097 | 0.865984817 | 0.894143821 | 1.208810712 | 0.853930510 | 0.887819514 |
| 0.7393387543 | 0.370146744 | 0.987243151 | 0.973889122 | 1.452502293 | 0.628844107 | 0.412121667 |
| 0.7289071673 | 0.322100361 | 0.886662013 | 0.928448649 | 1.240338207 | 0.623871986 | 0.491987243 |
| 0.7286041901 | 0.402313259 | 0.986869012 | 1.036625224 | 1.366995427 | 0.703875275 | 0.158417692 |
| 0.6879396231 | 0.318656088 | 0.882625606 | 0.884239958 | 1.244212619 | 0.595842426 | 0.541977655 |
| 0.7364081383 | 0.377444886 | 0.894526132 | 0.926383329 | 1.256640244 | 0.726304212 | 0.430613962 |
| 0.8051316433 | 0.405030087 | 0.897949513 | 0.933744330 | 1.251381707 | 0.796127410 | 0.425446439 |
| 0.7479290663 | 0.454639420 | 0.995276849 | 1.204793487 | 1.501510971 | 0.727754559 | 0.655494247 |
| 0.7609829089 | 0.342727331 | 0.901949193 | 0.932329008 | 1.250375732 | 0.668455406 | 0.448078926 |
| 0.7348632141 | 0.374632764 | 0.904140927 | 0.931414874 | 1.267120564 | 0.693543069 | 0.366900570 |
| 0.7591602298 | 0.374047605 | 0.886401335 | 0.93585775 | 1.259871068 | 0.722356523 | 0.622146059 |
| 0.7366291256 | 0.366708518 | 0.987082135 | 0.912705097 | 1.279905048 | 0.620573185 | 0.188990535 |
| 0.7324251648 | 0.430399460 | 0.888904036 | 0.913365908 | 1.303210255 | 0.757359529 | 0.217677607 |
| 0.7422066014 | 0.355671634 | 0.895448578 | 0.927633594 | 1.303398967 | 0.636820909 | 0.388487239 |
| 0.7563444857 | 0.417416406 | 0.910380367 | 0.923118513 | 1.261280616 | 0.817718122 | 0.240664943 |
| 0.796718813 | 0.464627553 | 1.014665242 | 1.324198685 | 1.622275524 | 0.652140606 | 0.816273597 |
| 0.7342524285 | 0.336179576 | 0.933853940 | 0.957233297 | 1.332889187 | 0.623995501 | 0.308843466 |
| 0.7610931468 | 0.344379270 | 0.955902303 | 0.939943417 | 1.262122538 | 0.626085353 | 0.354356267 |
| 0.7626332481 | 0.337634433 | 0.905825005 | 0.966208097 | 1.256867032 | 0.632720886 | 0.485717558 |
| 0.8456207305 | 0.405715169 | 0.893190916 | 1.013103587 | 1.23820537 | 0.715965908 | 0.741057589 |
| 0.6987306089 | 0.330752279 | 0.855648431 | 0.88669522 | 1.201108132 | 0.642698615 | 0.561524838 |
| 0.7714252857 | 0.377608924 | 0.894379180 | 0.945627887 | 1.274611495 | 0.668714892 | 0.476832762 |

| | | | | | | |
|---|---|---|---|---|---|---|
| 0.713592013 | 0.330436535 | 0.884810225 | 0.898262926 | 1.178968982 | 0.588932619 | -0.10876521 |
| 0.7329579572 | 0.319866087 | 0.897312128 | 0.932482028 | 1.256330587 | 0.629000736 | 0.448050159 |
| 0.6997866612 | 0.340196977 | 0.861929777 | 0.893561869 | 1.224858088 | 0.626291219 | 0.397445798 |
| 0.7354974038 | 0.333927656 | 0.920907554 | 0.927506463 | 1.251633935 | 0.625873555 | 0.402256858 |
| 0.7538043994 | 0.426166502 | 1.071429508 | 0.935334292 | 1.268308995 | 0.680746712 | 0.374460082 |
| 0.729714849 | 0.382374938 | 0.914314308 | 0.943379768 | 1.271081727 | 0.744928760 | 0.399641110 |
| 0.7272993633 | 0.342445127 | 0.880223064 | 0.954481694 | 1.267015123 | 0.634697228 | 0.238138522 |
| 0.7294511599 | 0.329900414 | 0.896050624 | 0.928556031 | 1.263748204 | 0.641565474 | 0.389736498 |
| 0.800888061 | 0.409281381 | 0.919302273 | 0.971587554 | 1.33876607 | 0.760096001 | 0.480500650 |
| 0.7505013425 | 0.354520582 | 0.894513993 | 0.942631843 | 1.256218636 | 0.690459536 | 0.422928234 |
| 0.8256853528 | 0.378814888 | 1.051548748 | 0.983287809 | 1.344834133 | 0.638328453 | 0.249078305 |
| 0.7362434148 | 0.333670079 | 0.927090867 | 0.964826180 | 1.317269687 | 0.629276272 | 0.371910408 |
| 0.6771593198 | 0.312197017 | 0.846858171 | 0.863881679 | 1.168101008 | 0.603300511 | 0.541100718 |
| 0.7739225771 | 0.351294614 | 0.908745901 | 1.010193133 | 1.272393622 | 0.630889754 | 0.335169196 |
| 0.6991097526 | 0.324174187 | 0.911457102 | 0.894736369 | 1.263852683 | 0.582108641 | 0.874349662 |
| 0.7304260438 | 0.322230704 | 0.894864115 | 0.927581548 | 1.255653044 | 0.636934142 | 0.383393330 |
| 0.7529203289 | 0.346505676 | 0.883410379 | 0.940758708 | 1.228519392 | 0.620983532 | 0.678047837 |
| 0.786917479 | 0.377491932 | 0.901172234 | 0.938163288 | 1.253880827 | 0.739946448 | 0.493440035 |
| 0.7204040816 | 0.423560246 | 0.905325962 | 1.02080151 | 1.290889315 | 0.652431110 | 0.531322534 |
| 0.7937482867 | 0.386096227 | 0.958987625 | 0.916071950 | 1.236535314 | 0.680260927 | 0.673570842 |
| 0.7289210364 | 0.340170611 | 0.896809036 | 0.990825637 | 1.298345117 | 0.625279607 | 0.293854350 |
| 0.727332659 | 0.328519660 | 0.894860465 | 0.927429502 | 1.258336998 | 0.639334602 | 0.389846265 |
| 0.7747754129 | 0.405305482 | 0.878168741 | 0.904948234 | 1.221300338 | 0.798396968 | -0.04709616 |
| 0.7300955487 | 0.328321249 | 0.888365735 | 0.922688552 | 1.248921392 | 0.644409833 | 0.508214101 |
| 0.7180685561 | 0.328369214 | 0.902128598 | 0.934739852 | 1.27660795 | 0.634417132 | 0.435811028 |
| 0.7264839662 | 0.425517884 | 0.990112482 | 1.036487671 | 1.359396967 | 0.757805194 | 0.527819898 |
| 0.7341605738 | 0.327422500 | 0.915973978 | 0.952811746 | 1.298531061 | 0.626667483 | 0.393828516 |
| 0.7270895938 | 0.330856874 | 0.898335558 | 0.953747664 | 1.274890513 | 0.638668404 | 0.391064728 |
| 0.7630000293 | 0.393206156 | 0.878481779 | 0.903913117 | 1.219781156 | 0.771738401 | -0.01850770 |
| 0.719909825 | 0.327068571 | 0.881963036 | 0.940003742 | 1.271576942 | 0.618064646 | 0.509946595 |
| 0.7284122977 | 0.342737442 | 0.891530655 | 0.962544703 | 1.280440859 | 0.630183860 | 0.344473877 |
| 0.7273278666 | 0.328776355 | 0.887742685 | 0.919559748 | 1.246483769 | 0.650908569 | 0.308759756 |
| 0.7210815834 | 0.341453876 | 0.899579244 | 0.922007770 | 1.231242946 | 0.634428227 | 0.286545831 |
| 0.7217761099 | 0.357806333 | 0.887325467 | 0.933209641 | 1.241620373 | 0.692358061 | 0.344306132 |
| 0.7212013895 | 0.334551414 | 0.891484721 | 0.959440572 | 1.281941269 | 0.634140647 | 0.370954899 |
| 0.7358566974 | 0.328738938 | 0.904509161 | 0.957845329 | 1.259873958 | 0.621263629 | 0.525049957 |
| 0.7110976166 | 0.332212607 | 0.876724068 | 0.965282516 | 1.250582513 | 0.608469952 | 0.132104734 |
| 0.6877251918 | 0.306717598 | 0.830131834 | 0.841086832 | 1.132741505 | 0.588362514 | 1.112257274 |
| 0.7209979351 | 0.331244131 | 0.853610762 | 0.883112335 | 1.187678234 | 0.650033555 | -0.09212895 |
| 0.7351469362 | 0.366931648 | 0.909889318 | 0.927081255 | 1.342818865 | 0.643621011 | 0.418761329 |
| 0.7401221644 | 0.337082917 | 0.895123434 | 0.927630431 | 1.268498019 | 0.650299368 | 0.391896260 |
| 0.7364584031 | 0.400549128 | 0.986458034 | 0.928501304 | 1.249733301 | 0.679081552 | 0.332798568 |
| 0.7639555131 | 0.357292316 | 0.880636221 | 0.946341976 | 1.233026396 | 0.639915935 | 0.685490428 |
| 0.7131569315 | 0.319336051 | 0.884037033 | 0.910119622 | 1.246534679 | 0.614861304 | 0.422967586 |
| 0.7208072809 | 0.398377791 | 0.918030306 | 1.003855254 | 1.241431488 | 0.634939426 | 0.407212756 |
| 0.7923788023 | 0.369821175 | 0.921833760 | 0.930572489 | 1.252583925 | 0.691707367 | 0.362904187 |

| | | | | | | |
|---|---|---|---|---|---|---|
| 0.7316107004 | 0.484259331 | 1.046616174 | 0.928489903 | 1.24601194 | 0.784029908 | 0.469240968 |
| 0.8047960999 | 0.413396336 | 0.898630900 | 1.015149535 | 1.252051008 | 0.717848715 | 0.372621970 |
| 0.8123859883 | 0.419071911 | 1.081067177 | 0.928437173 | 1.25554295 | 0.626786107 | 0.202562625 |
| 0.7363002366 | 0.340342589 | 0.929963702 | 0.925217004 | 1.266992501 | 0.634052888 | 0.448165752 |
| 0.7245301354 | 0.345016055 | 0.886823265 | 0.909996974 | 1.216519692 | 0.610095813 | 0.666784573 |
| 0.7242634795 | 0.337872650 | 0.919905534 | 0.924892211 | 1.271937432 | 0.623264032 | 0.414076919 |
| 0.7530079669 | 0.381442912 | 1.045131801 | 0.975812141 | 1.33548234 | 0.636704778 | -0.008370958 |
| 0.7338600197 | 0.357442892 | 0.892283339 | 0.934980021 | 1.27552675 | 0.675356180 | 0.333862620 |
| 0.7386370567 | 0.324839299 | 0.895555411 | 0.936258879 | 1.251344762 | 0.632157856 | 0.411146113 |
| 0.6802617092 | 0.433277602 | 0.923251265 | 0.880873673 | 1.166162053 | 0.741334650 | 0.619484430 |
| 0.6956074664 | 0.346533788 | 0.936911649 | 0.891617955 | 1.23749617 | 0.632502638 | 0.232513961 |
| 0.7359514759 | 0.406774313 | 1.079267857 | 0.925628458 | 1.313845432 | 0.667273410 | 0.267191212 |
| 0.7112114647 | 0.317324526 | 0.886325419 | 0.907695291 | 1.230445926 | 0.621006018 | 0.416222619 |
| 0.7380037292 | 0.358471468 | 0.902842333 | 0.926612034 | 1.315734699 | 0.648036088 | 0.427055558 |
| 0.7421430146 | 0.441301510 | 1.033364718 | 1.016566186 | 1.327947516 | 0.76802304 | 0.452057414 |
| 0.796963021 | 0.432891173 | 0.936153049 | 0.941652342 | 1.348165833 | 0.788455076 | 0.529311493 |
| 0.7540840556 | 0.340662649 | 0.949695533 | 0.924035617 | 1.270746464 | 0.618276269 | 0.785125085 |
| 0.8382735928 | 0.418282622 | 1.005316226 | 0.933594383 | 1.286681524 | 0.747489637 | 0.320915615 |
| 0.7332261148 | 0.417780495 | 1.008981301 | 0.928002719 | 1.253954506 | 0.702694857 | 0.392700295 |
| 0.7731882069 | 0.393513128 | 1.038552382 | 0.921616116 | 1.250440871 | 0.621322070 | 0.170003226 |
| 0.6397097801 | 0.294367455 | 0.818202541 | 0.807546623 | 1.13744564 | 0.539294041 | -1.060122879 |
| 0.614101693 | 0.284141094 | 0.760148188 | 0.799916500 | 1.056417278 | 0.540948867 | -1.280765065 |
| 0.5928082578 | 0.290367273 | 0.738134937 | 0.768695822 | 1.032054055 | 0.530354502 | -1.136626394 |
| 0.6705561621 | 0.380262331 | 0.801547308 | 1.094398934 | 1.371648992 | 0.509155197 | -1.819593414 |
| 0.6656246552 | 0.373164782 | 0.750125251 | 0.789552167 | 1.048831976 | 0.727640926 | -0.644155668 |
| 0.6318340484 | 0.289811854 | 0.819832222 | 0.788703228 | 1.089500599 | 0.526752277 | -1.381646768 |
| 0.6050239983 | 0.270394368 | 0.733839160 | 0.725865370 | 0.969332824 | 0.483574048 | -2.387873152 |
| 0.63883347 | 0.334591768 | 0.758951990 | 0.792629351 | 1.103300849 | 0.622892885 | -0.86404352 |
| 0.6733966532 | 0.318285652 | 0.881427612 | 0.838059880 | 1.11918086 | 0.537637713 | -0.885773683 |
| 0.637762799 | 0.283142120 | 0.782969821 | 0.797647462 | 1.052162325 | 0.516476102 | -1.608598309 |
| 0.6272330575 | 0.298964334 | 0.807162261 | 0.817199054 | 1.07760966 | 0.536090008 | -1.476839486 |
| 0.6301520156 | 0.310312919 | 0.765219414 | 0.829494665 | 1.03648452 | 0.525554909 | -1.583609015 |
| 0.6592904957 | 0.311980814 | 0.861911507 | 0.816523425 | 1.171602953 | 0.544169307 | -1.049331513 |
| 0.6650583879 | 0.339508998 | 0.764968061 | 0.805466131 | 1.115400141 | 0.626459455 | -1.668718335 |
| 0.6422316543 | 0.299628101 | 0.769323402 | 0.823855963 | 1.09590752 | 0.570578260 | -1.12616087 |
| 0.6260327973 | 0.367694941 | 0.859549950 | 0.796683242 | 1.07614706 | 0.628708322 | -1.168103975 |
| 0.6212573847 | 0.277204081 | 0.777727213 | 0.773124866 | 1.051822454 | 0.506902108 | -1.765426157 |
| 0.5710502447 | 0.297794137 | 0.715240675 | 0.730952959 | 1.076850659 | 0.505376475 | -1.490147555 |
| 0.603054047 | 0.293556388 | 0.757905182 | 0.761187091 | 1.066203711 | 0.533146380 | -1.410477147 |
| 0.6086942136 | 0.293113036 | 0.757860077 | 0.797241154 | 1.063526302 | 0.523745732 | -1.266146899 |
| 0.6161270172 | 0.370076942 | 0.818046495 | 0.840407185 | 1.089460009 | 0.671353871 | -1.097730013 |
| 0.7645049229 | 0.443888231 | 0.826969052 | 0.817122138 | 1.19386896 | 0.792903601 | -0.01691807 |
| 0.6259419538 | 0.309444580 | 0.769366741 | 0.825860226 | 1.114342374 | 0.548738952 | -1.202047665 |
| 0.6560235225 | 0.335453761 | 0.836918913 | 0.866666532 | 1.290755011 | 0.548459671 | -1.160883588 |
| 0.6855696873 | 0.316046173 | 0.758998829 | 0.872605592 | 1.027941175 | 0.505149576 | -2.244079374 |
| 0.6143418102 | 0.266993911 | 0.756536216 | 0.784866662 | 1.058130205 | 0.529210600 | -1.186324198 |

| | | | | | | |
|---|---|---|---|---|---|---|
| 0.6033083703 | 0.285825111 | 0.767676309 | 0.784748230 | 1.037986588 | 0.527244144 | -0.883365165 |
| 0.5955899833 | 0.315136861 | 0.821223263 | 0.829065889 | 1.208245567 | 0.534589670 | -1.581998059 |
| 0.6224553774 | 0.284509574 | 0.788484239 | 0.794908249 | 1.080255571 | 0.54129655 | -1.154736689 |
| 0.6340217018 | 0.296778734 | 0.763893653 | 0.817573885 | 1.10806921 | 0.555980536 | -0.966480465 |
| 0.5990646218 | 0.292464687 | 0.776303841 | 0.759358707 | 1.025673944 | 0.530103788 | -1.382055783 |
| 0.5811844159 | 0.259534734 | 0.709574033 | 0.744959396 | 0.991593978 | 0.495873429 | -0.978247532 |
| 0.6803583561 | 0.387042825 | 0.768643472 | 0.793637253 | 1.072933895 | 0.757944135 | -0.890632980 |
| 0.6260768876 | 0.313442681 | 0.836003951 | 0.824695672 | 1.229988431 | 0.532509298 | -1.006632608 |
| 0.6230707385 | 0.275331783 | 0.757919774 | 0.793639041 | 1.060242617 | 0.533286537 | -1.283609284 |
| 0.6205161701 | 0.342630314 | 0.840467551 | 0.882842457 | 1.164202428 | 0.599455776 | -1.440904224 |
| 0.6251941332 | 0.289592153 | 0.802123227 | 0.803590338 | 1.130730669 | 0.541496923 | -1.178387874 |
| 0.6232615838 | 0.319451789 | 0.757085297 | 0.784047746 | 1.063561833 | 0.614710091 | -1.017887452 |
| 0.6921174117 | 0.348177069 | 0.771906679 | 0.802677071 | 1.075728516 | 0.684377079 | -1.012814582 |
| 0.6542114478 | 0.397671820 | 0.870565857 | 1.05382947 | 1.313367417 | 0.636564863 | -1.171761805 |
| 0.653915392 | 0.294506847 | 0.775048234 | 0.801153720 | 1.074452432 | 0.574406171 | -1.174391457 |
| 0.5922303217 | 0.301918613 | 0.728652165 | 0.75063239 | 1.021179459 | 0.558930189 | -0.962796702 |
| 0.6572799564 | 0.323849938 | 0.767445143 | 0.810264443 | 1.09079476 | 0.625415354 | -1.257516359 |
| 0.6416013475 | 0.319401815 | 0.859745027 | 0.794962892 | 1.114792743 | 0.540517036 | -1.207231124 |
| 0.6355707705 | 0.373484322 | 0.771357198 | 0.792584283 | 1.130876417 | 0.657207866 | -1.241964466 |
| 0.6365122268 | 0.305022003 | 0.767931688 | 0.795533377 | 1.117787658 | 0.546134047 | -1.193329204 |
| 0.5797444244 | 0.319953194 | 0.697814226 | 0.707578122 | 0.966782224 | 0.626787834 | -1.946725799 |
| 0.6996372109 | 0.408011861 | 0.891026480 | 1.162842724 | 1.424598371 | 0.572676115 | -0.911815434 |
| 0.6237664905 | 0.285593273 | 0.793333154 | 0.813194524 | 1.13232395 | 0.530100370 | -1.012745883 |
| 0.6539507293 | 0.295899491 | 0.821335747 | 0.807623463 | 1.084448018 | 0.537948574 | -1.084291272 |
| 0.6558967136 | 0.290379833 | 0.779047655 | 0.830979657 | 1.080958584 | 0.544166611 | -1.012414375 |
| 0.7257003845 | 0.348179323 | 0.766524481 | 0.869431928 | 1.062611263 | 0.614432352 | -0.576701327 |
| 0.6180548142 | 0.292563451 | 0.756854824 | 0.784316934 | 1.06242757 | 0.568492303 | -1.081029074 |
| 0.6028952634 | 0.295114298 | 0.698987940 | 0.739040558 | 0.996152508 | 0.522623575 | -0.475795949 |
| 0.6330615693 | 0.293146038 | 0.784957426 | 0.796891960 | 1.045919714 | 0.522470265 | -1.500387448 |
| 0.611106488 | 0.266689568 | 0.748137404 | 0.777460442 | 1.047470411 | 0.524431759 | -0.999522564 |
| 0.6140149879 | 0.298499606 | 0.756284495 | 0.784039494 | 1.074729293 | 0.549527759 | -1.169578967 |
| 0.6302749279 | 0.286154959 | 0.789159743 | 0.794814592 | 1.072571411 | 0.536334197 | -1.09409533 |
| 0.6292221274 | 0.355733388 | 0.894352904 | 0.78075033 | 1.058693853 | 0.568238783 | -1.446118877 |
| 0.6234691118 | 0.326701537 | 0.781191078 | 0.806024637 | 1.086013525 | 0.636467893 | -1.204801996 |
| 0.6024558029 | 0.283663185 | 0.729129599 | 0.790641466 | 1.049527405 | 0.525749158 | -0.916100908 |
| 0.6104614431 | 0.276086315 | 0.749884827 | 0.777087879 | 1.057602749 | 0.536911868 | -1.092637562 |
| 0.6617740758 | 0.338189344 | 0.759619779 | 0.802823125 | 1.106222857 | 0.628067583 | -0.524319411 |
| 0.6304714341 | 0.297821052 | 0.751451713 | 0.791873933 | 1.055307859 | 0.580032292 | -0.895595616 |
| 0.6680487845 | 0.306493054 | 0.850791237 | 0.795562406 | 1.088083742 | 0.516461320 | -0.295383688 |
| 0.6107144581 | 0.276779578 | 0.769022561 | 0.800324031 | 1.092676182 | 0.521985134 | -0.983180360 |
| 0.6153452429 | 0.283698302 | 0.769553238 | 0.785022765 | 1.061471618 | 0.548228591 | -1.23008137 |
| 0.6573595063 | 0.298384956 | 0.771876638 | 0.858044562 | 1.080754158 | 0.535869335 | -0.856605314 |
| 0.6436180154 | 0.298442907 | 0.839110324 | 0.823716797 | 1.163534556 | 0.535903850 | -1.224281146 |
| 0.6212713715 | 0.274076634 | 0.761135861 | 0.788964010 | 1.06800859 | 0.541750874 | -1.096512672 |
| 0.5983913476 | 0.275389029 | 0.702099687 | 0.747677874 | 0.976378703 | 0.493533191 | -0.520514174 |
| 0.666529804 | 0.319740799 | 0.763305135 | 0.794637061 | 1.062054109 | 0.626744702 | -1.415763786 |

| | | | | | | |
|---|---|---|---|---|---|---|
| 0.6085696944 | 0.357807425 | 0.764784595 | 0.862333902 | 1.090493705 | 0.551148739 | -1.372065567 |
| 0.6893527136 | 0.335315976 | 0.832859399 | 0.795588092 | 1.073903388 | 0.590791469 | -0.929753240 |
| 0.6250380459 | 0.291690819 | 0.768999301 | 0.849617022 | 1.113310022 | 0.53616719 | -1.266570535 |
| 0.6113751886 | 0.276144302 | 0.752194307 | 0.779570915 | 1.057722364 | 0.537406520 | -1.139037805 |
| 0.6551646297 | 0.342733922 | 0.742595968 | 0.765241210 | 1.032754486 | 0.675139460 | -1.642639244 |
| 0.6335868632 | 0.284921653 | 0.770935887 | 0.800721695 | 1.083830998 | 0.559227632 | -1.168601744 |
| 0.614652733 | 0.281077667 | 0.772204553 | 0.800119153 | 1.092751602 | 0.54304874 | -1.217960399 |
| 0.6275530798 | 0.367571854 | 0.855281282 | 0.895341205 | 1.174277469 | 0.654609056 | -1.258263742 |
| 0.6090530051 | 0.27162676 | 0.759883769 | 0.790444049 | 1.077249682 | 0.519877704 | -1.362367187 |
| 0.6033128728 | 0.274533170 | 0.745406633 | 0.791385611 | 1.057858432 | 0.529944140 | -1.242889888 |
| 0.6499926922 | 0.334968700 | 0.748370530 | 0.770035253 | 1.039120324 | 0.657436830 | -1.569051631 |
| 0.6291894286 | 0.285852589 | 0.770821288 | 0.821547917 | 1.111337478 | 0.540178406 | -1.171557464 |
| 0.6166225267 | 0.290137369 | 0.754707035 | 0.814822524 | 1.083931011 | 0.533469253 | -1.068657883 |
| 0.6302302443 | 0.284885004 | 0.769229827 | 0.796799341 | 1.080079296 | 0.564012855 | -1.127203277 |
| 0.5856967745 | 0.277345086 | 0.730681068 | 0.748898584 | 1.000074109 | 0.515312795 | -1.473863636 |
| 0.6262278387 | 0.310440154 | 0.769861875 | 0.809671931 | 1.077255443 | 0.600704133 | -1.144057313 |
| 0.6112869573 | 0.283564229 | 0.755618321 | 0.813217385 | 1.086567482 | 0.537494675 | -1.225886087 |
| 0.6132072041 | 0.273946117 | 0.753749386 | 0.798195706 | 1.049883476 | 0.517714025 | -1.457418069 |
| 0.6371056228 | 0.297644817 | 0.785498109 | 0.864841766 | 1.120455381 | 0.545156697 | -1.155510297 |
| 0.6327004808 | 0.282177204 | 0.763713205 | 0.773791696 | 1.042111156 | 0.541287785 | -0.741577223 |
| 0.6559750244 | 0.301371012 | 0.776628217 | 0.803469202 | 1.080567947 | 0.591410539 | -1.123458146 |
| 0.6295172408 | 0.314209020 | 0.779151737 | 0.793873449 | 1.149875739 | 0.551142232 | -1.214191086 |
| 0.624265725 | 0.284316997 | 0.755003574 | 0.782422026 | 1.069931254 | 0.548503512 | -1.326761188 |
| 0.6287064478 | 0.341944390 | 0.842128386 | 0.792651363 | 1.066883589 | 0.579724460 | -1.258189619 |
| 0.6473670138 | 0.302765352 | 0.746240888 | 0.801919180 | 1.044852223 | 0.542257319 | -0.700270025 |
| 0.6155171736 | 0.275615107 | 0.763001735 | 0.785513304 | 1.075869094 | 0.530679399 | -1.193088883 |
| 0.5996286647 | 0.331404453 | 0.763695513 | 0.835091988 | 1.032728061 | 0.528196496 | -1.184729665 |
| 0.6750365063 | 0.315054862 | 0.785320656 | 0.792765279 | 1.06709048 | 0.589273366 | -0.831586896 |
| 0.6322520229 | 0.418492979 | 0.904477194 | 0.802393430 | 1.076793395 | 0.677552276 | -1.143347222 |
| 0.6403926717 | 0.328947896 | 0.715058936 | 0.807775190 | 0.996282524 | 0.571206864 | -0.326933154 |
| 0.6852816109 | 0.353504711 | 0.911925448 | 0.783175646 | 1.059103071 | 0.528720336 | -1.484225213 |
| 0.6334386789 | 0.292796538 | 0.800047249 | 0.795963668 | 1.089992935 | 0.545475343 | -1.098976254 |
| 0.6168456026 | 0.293737453 | 0.755017637 | 0.774747114 | 1.035712369 | 0.519419278 | -1.48216297 |
| 0.6247899393 | 0.291467730 | 0.793561651 | 0.797863436 | 1.097243936 | 0.537662201 | -1.152162339 |
| 0.6499099465 | 0.329217689 | 0.902037671 | 0.842208906 | 1.152634891 | 0.549530398 | -1.469497384 |
| 0.6260547745 | 0.304933943 | 0.761205448 | 0.797629917 | 1.088149771 | 0.576145245 | -1.259393175 |
| 0.5811491933 | 0.255578968 | 0.704610336 | 0.736635249 | 0.984540367 | 0.497372863 | -1.608460647 |
| 0.6282325391 | 0.400138777 | 0.852637269 | 0.813500888 | 1.076969257 | 0.684634374 | -1.149269741 |
| 0.6168448452 | 0.307296271 | 0.830826506 | 0.790661351 | 1.097376279 | 0.560885285 | -1.281457223 |
| 0.6295009868 | 0.347937112 | 0.923158935 | 0.791742454 | 1.123806424 | 0.570756747 | -1.300258891 |
| 0.6206546119 | 0.276920354 | 0.773471726 | 0.792120623 | 1.073776193 | 0.541934808 | -1.172017142 |
| 0.5654837573 | 0.274673127 | 0.691788746 | 0.710001906 | 1.008161032 | 0.496547466 | -0.858354876 |
| 0.6334939051 | 0.376695342 | 0.882080997 | 0.867741756 | 1.133537123 | 0.655585116 | -1.011277879 |
| 0.6796130504 | 0.369149488 | 0.798307842 | 0.802997383 | 1.149653209 | 0.672357870 | -1.355666931 |
| 0.665552802 | 0.300668047 | 0.838199028 | 0.815551648 | 1.12155782 | 0.545689171 | -0.953946607 |
| 0.6875078762 | 0.343053389 | 0.824507451 | 0.765684970 | 1.055268458 | 0.613051654 | -1.76547318 |

| | | | | | | |
|---|---|---|---|---|---|---|
| 0.6110521569 | 0.3481677311 | 0.8408595761 | 0.7733740683 | 1.045014068 | 0.5856081765 | -1.335779711 |
| 0.6390035925 | 0.3252200438 | 0.85831457 | 0.7616722608 | 1.033430414 | 0.5134933923 | -1.644764623 |

| t_microbe | t_blast | t_vns | t_interact | t_batch | t_p_intercept | t_p_microbe |
|---|---|---|---|---|---|---|
| 0.811221818 | 6.373274533 | -0.540162070 | -1.005459986 | 2.368500831 | 0.000152420297 | 0.4263375891 |
| -0.565396224 | 6.293920311 | -0.796020004 | -0.718820423 | 2.349647473 | 0.000237611751 | 0.5777957070 |
| -0.529635960 | 6.275963404 | -0.576579831 | -0.846561767 | 2.336280497 | 0.000185938222 | 0.6019194140 |
| -0.105342073 | 5.763893071 | -0.545969202 | -0.499904595 | 2.266384929 | 0.000842338810 | 0.9171041070 |
| -0.029995916 | 6.383679922 | -0.674957305 | -0.765315615 | 1.607230579 | 0.000549338900 | 0.9763535601 |
| 1.267139755 | 5.596406857 | -0.880608369 | -0.447912264 | 2.334763079 | 0.000430072880 | 0.2189780512 |
| 0.523873519 | 5.618186977 | -0.814800453 | -0.596938449 | 2.050635209 | 0.001085946059 | 0.6058519271 |
| 2.048051107 | 7.200315026 | -0.386183384 | -1.409848726 | 3.189416122 | 0.000031054619 | 0.0532640168 |
| -0.509179467 | 5.330459385 | -0.813789776 | -0.589316804 | 2.31768323 | 0.000728965471 | 0.6159351495 |
| -0.558769019 | 5.874489742 | -0.818578760 | -0.645686860 | 2.236335533 | 0.000601641671 | 0.5822293971 |
| -0.443130933 | 5.75018182 | -0.792739687 | -0.630224558 | 2.313744202 | 0.000396338260 | 0.6622024071 |
| -1.444867654 | 6.060073684 | -1.223501086 | -0.637780415 | 2.614435476 | 0.000661829685 | 0.1632521935 |
| 0.596120150 | 6.02190396 | -0.539516666 | -0.948324361 | 2.34759371 | 0.000215892241 | 0.5574665105 |
| -3.01130997 | 6.592659569 | -1.717362055 | 0.289277420 | 0.524947305 | 0.002052588351 | 0.0066476783 |
| -0.356893869 | 6.42359406 | -0.570825003 | -0.825187179 | 2.259141062 | 0.000224569455 | 0.7247324894 |
| -0.183503581 | 5.641436905 | -0.685441515 | -0.759657156 | 2.031721002 | 0.000223468058 | 0.8561631501 |
| -0.109767717 | 5.868502693 | -0.685941845 | -0.702202346 | 2.229620614 | 0.000440609010 | 0.9136358044 |
| 0.304569312 | 6.331211007 | -0.637306955 | -0.827193204 | 2.271887713 | 0.000202156231 | 0.7636919925 |
| 0.446135155 | 6.337798809 | -0.683024505 | -0.863685614 | 2.310527234 | 0.000192369141 | 0.6600657249 |
| 0.367770565 | 6.391158036 | -0.750766824 | -0.691528528 | 2.28796158 | 0.000196263840 | 0.7167250388 |
| 0.219663004 | 6.000613475 | -0.556247881 | -0.799911184 | 1.648672123 | 0.000206902731 | 0.8282547772 |
| 2.176355529 | 5.254424217 | -1.509023558 | 0.389956789 | -0.044454520 | 0.02171505831 | 0.0410869060 |
| -1.573002392 | 6.582844175 | -0.258409609 | -1.214613809 | 1.993989999 | 0.000187772099 | 0.1306643628 |
| -1.591362358 | 6.847312844 | -0.034829006 | -1.558988119 | 2.666388853 | 0.000060847211 | 0.1264701801 |
| 0.339830915 | 5.767050233 | -0.756488693 | -0.626240308 | 2.106841872 | 0.002048324505 | 0.7373593019 |
| 0.034917595 | 6.396805614 | -0.682243402 | -0.768593080 | 2.263753774 | 0.000210686100 | 0.9724752282 |
| 0.417740224 | 5.955925013 | -0.772892208 | -0.675154291 | 2.303093483 | 0.000306688441 | 0.6803773991 |
| -0.162187221 | 5.670447492 | -0.539733330 | -0.729187540 | 2.069678982 | 0.000243617005 | 0.872708849 |
| 1.16296545 | 6.687839048 | -0.745960360 | -0.930766535 | 2.149858504 | 0.000142807407 | 0.2578889360 |
| -0.772978986 | 6.359400796 | -0.875163369 | -0.525767202 | 1.962023695 | 0.000376973837 | 0.4481484801 |
| 0.344766815 | 5.938137277 | -0.690950685 | -0.753433668 | 2.282157671 | 0.000243204804 | 0.7336986581 |
| 0.470776999 | 6.444807655 | -0.597691011 | -0.765909472 | 2.264118604 | 0.000188763248 | 0.6426532597 |
| -0.049877348 | 6.376985301 | -0.681414821 | -0.770403275 | 1.63460774 | 0.000502927821 | 0.9606914889 |
| -1.554575845 | 6.778352762 | -0.209687391 | -1.488716658 | 2.221781971 | 0.000070974671 | 0.134989453 |
| 0.930843506 | 6.517758002 | -0.821018215 | -0.802344778 | 2.192742256 | 0.000273736861 | 0.3625128158 |
| -2.075790327 | 7.181142319 | 0.358338154 | -1.685779531 | 1.124680226 | 0.000032981395 | 0.0503856047 |
| 0.342465349 | 6.038697656 | -0.724719786 | -0.624461880 | 2.295133813 | 0.000224901545 | 0.7354047075 |
| 1.035154156 | 6.595367873 | -0.710649486 | -0.678228124 | 1.469720676 | 0.000286591714 | 0.3123709457 |
| -0.051923656 | 6.367457655 | -0.682710497 | -0.769783497 | 1.746459159 | 0.000631769151 | 0.9590802748 |
| -0.643620800 | 5.396236901 | -0.941477869 | -0.264807456 | 2.274175298 | 0.000498421811 | 0.5267884535 |
| -1.053782381 | 6.645535105 | -0.575857111 | -0.808637704 | 2.545282302 | 0.000124156604 | 0.3039568745 |
| 2.21260236 | 7.34965398 | -0.965667677 | -1.193332357 | 3.222502761 | 0.000033786957 | 0.0381370928 |
| -0.432543187 | 6.367039563 | -0.759136768 | -0.672645866 | 1.720197017 | 0.000508894231 | 0.6697561454 |
| -1.484052936 | 5.327475999 | -0.70196747 | -0.373863691 | 2.177630888 | 0.000397173650 | 0.1526502905 |
| 1.300027778 | 6.762157621 | -0.694811513 | -1.178093343 | 2.670174513 | 0.000092130421 | 0.2076802308 |
| -2.385918078 | 7.283756358 | -0.779803606 | -1.496944667 | 2.068455128 | 0.000220363210 | 0.0265311228 |

| | | | | | | |
|---|---|---|---|---|---|---|
| -2.054891923 | 7.284780949 | -0.803395623 | -1.158227113 | 0.556750578 | 0.000536092390 | 0.0525407968 |
| -0.923569720 | 5.153088411 | -1.149095409 | 0.0158866878 | 2.486090798 | 0.00151314854 | 0.3662013664 |
| 0.6009285837 | 6.33214418 | -0.502801025 | -0.938600183 | 2.238934929 | 0.000176360330 | 0.5543189854 |
| -1.026372543 | 5.775731165 | -0.855433556 | -0.649367719 | 2.289251063 | 0.000549755455 | 0.3163940609 |
| 0.5911817779 | 6.455281009 | -0.486603919 | -0.839262977 | 2.3484731 | 0.000212099301 | 0.5607087809 |
| -1.177054193 | 6.70615324 | -0.130604017 | -0.875902538 | 2.611295004 | 0.000179318008 | 0.2523431626 |
| 0.2678883809 | 6.401476393 | -0.681584235 | -0.790391480 | 2.208552905 | 0.000205204827 | 0.7913988996 |
| -1.191581955 | 6.665529261 | -0.933380173 | -1.016008446 | 1.758057496 | 0.000756310128 | 0.2467184541 |
| -0.247676926 | 5.992500971 | -0.722757160 | -0.740994793 | 2.279942083 | 0.000403291270 | 0.8067899815 |
| -0.479008827 | 6.341934596 | -0.74150546 | -0.711599143 | 2.196046899 | 0.000268109891 | 0.636882447 |
| 0.3261232271 | 6.404429851 | -0.692639500 | -0.816432306 | 2.272327919 | 0.000206439499 | 0.7475587555 |
| 0.4017377774 | 6.144615206 | -0.681027696 | -0.772199284 | 2.277400524 | 0.000266538861 | 0.6919362382 |
| 0.2130394145 | 5.466081327 | -0.649135108 | -0.793820266 | 1.999627214 | 0.000266723591 | 0.8333507625 |
| -0.771670316 | 6.507007817 | -0.538339733 | -0.902033654 | 1.50984092 | 0.000224109401 | 0.4489067099 |
| 1.551965795 | 6.763610489 | -0.231234829 | -1.155393351 | 2.695923488 | 0.000071282321 | 0.1356115768 |
| 0.7790528204 | 6.518783602 | -0.654922945 | -0.879351215 | 2.411037279 | 0.000160145049 | 0.4446395985 |
| -1.26996853 | 6.149034994 | -1.059016901 | -0.288090897 | 1.20679806 | 0.001447191645 | 0.2179880419 |
| 0.8457922497 | 6.492992022 | -0.528210367 | -0.855229079 | 2.443347821 | 0.000149965871 | 0.4072031284 |
| -0.797769581 | 5.940484392 | -0.385025009 | -1.01904192 | 2.411056025 | 0.000267600430 | 0.4339330774 |
| 0.1317853615 | 6.141465911 | -0.691526266 | -0.689527544 | 2.265660401 | 0.000253692121 | 0.8964085344 |
| 0.1369197874 | 6.313712542 | -0.688318864 | -0.777910597 | 2.148328698 | 0.000218777811 | 0.892398402 |
| 0.4906613313 | 6.248521303 | -0.823619519 | -0.671931726 | 2.195452762 | 0.000572297497 | 0.6287539192 |
| -0.857375768 | 6.291558533 | -0.440444895 | -1.054273403 | 2.287536665 | 0.000148721771 | 0.400916417 |
| 0.4519449013 | 6.423297418 | -0.692220768 | -0.733637002 | 2.318858334 | 0.000189274589 | 0.6559421165 |
| -0.552541678 | 6.33581709 | -0.804763428 | -0.797855418 | 2.16845625 | 0.000531616930 | 0.5864110497 |
| 0.1921807148 | 6.368085079 | -0.640691753 | -0.783847598 | 2.014525423 | 0.000405261309 | 0.849446904 |
| 1.136870591 | 6.110511507 | -0.109765734 | -1.104800554 | 2.579411777 | 0.000115384999 | 0.268399194 |
| 0.0222568599 | 5.886601819 | -0.681522417 | -0.768014394 | 2.047373311 | 0.000599552651 | 0.9824531902 |
| -0.060143328 | 6.314842474 | -0.654416543 | -0.716442000 | 2.252387903 | 0.000245405051 | 0.9526101765 |
| 0.699893571 | 6.478473663 | -0.685739700 | -0.846082063 | 2.384569264 | 0.000174392440 | 0.4916750511 |
| 1.825138121 | 6.648244793 | -0.756872978 | -0.858544454 | 0.6871910717 | 0.001433151255 | 0.0822384970 |
| 0.5293518882 | 6.377397964 | -0.736228029 | -0.699894010 | 2.043718625 | 0.000286532147 | 0.6021129872 |
| -3.464143267 | 7.145537252 | -1.53680618 | -0.059310509 | 3.552870327 | 0.000021869400 | 0.0023203585 |
| -1.5458549 | 5.310389781 | -1.362935163 | -0.086560703 | 2.842405206 | 0.000254423259 | 0.1370774432 |
| -0.374232964 | 6.19116428 | -0.751030152 | -0.644075600 | 2.286729474 | 0.000256809618 | 0.7119831078 |
| -0.433801511 | 6.361721452 | -0.766693042 | -0.675494342 | 2.314450009 | 0.000209647899 | 0.668856506 |
| -0.123580421 | 6.366850047 | -0.683296733 | -0.767561477 | 1.865067009 | 0.000405222420 | 0.9028226785 |
| -1.997655241 | 6.799697934 | -1.210488099 | -0.303154057 | 2.254515068 | 0.000161800528 | 0.0588721808 |
| -0.231710761 | 6.39463445 | -0.589363501 | -0.802344246 | 2.275670891 | 0.000209041051 | 0.8190056127 |
| -0.480473402 | 6.407194011 | -0.694836508 | -0.723045515 | 2.314537255 | 0.000187776671 | 0.6358581838 |
| 0.8867956010 | 6.192396188 | -0.814800118 | -0.775290942 | 2.45151535 | 0.000250936287 | 0.3852324177 |
| 0.4917523120 | 6.396148323 | -0.761949827 | -0.740866473 | 2.256692001 | 0.000186439759 | 0.6279953191 |
| -0.069750953 | 6.360458796 | -0.673035503 | -0.726835464 | 2.229528001 | 0.000213894525 | 0.9450518329 |
| 1.66580504 | 6.334563627 | -1.167776247 | -0.506340292 | 2.518821511 | 0.000313597751 | 0.1106008108 |
| 0.9603113865 | 6.199686376 | -1.01468193 | -0.483988776 | 2.439576151 | 0.000328527424 | 0.3478258885 |
| -0.550471020 | 6.38050137 | -0.609489181 | -0.831265645 | 2.341201186 | 0.000221067481 | 0.5878047965 |

| | | | | | | |
|---|---|---|---|---|---|---|
| 1.885640797 | 6.583035963 | -0.970667294 | -0.64438737 | 1.440549162 | 0.00061312608 | 0.073253778 |
| -0.96560164 | 6.60295772 | -0.67567772 | -1.083076181 | 2.019636933 | 0.00027235567 | 0.34523273 |
| -1.60283389 | 6.803482045 | -0.73176433 | -1.062846794 | 1.873111558 | 0.00028244070 | 0.12390709 |
| 0.27464790 | 5.91648208 | -0.65765313 | -0.78037063 | 1.976909191 | 0.00021730252 | 0.78627056 |
| -0.48281690 | 6.448969367 | -0.52952897 | -0.735897901 | 2.320717084 | 0.00026575203 | 0.63422079 |
| 0.57795850 | 6.470648637 | -0.677859012 | -0.862493367 | 2.308392324 | 0.00018782442 | 0.56943842 |
| 2.357147748 | 6.35200781 | 0.24150592 | -0.878707381 | 1.99199398 | 0.00005138206 | 0.02819891 |
| -0.02781152 | 6.203810312 | -0.68141027 | -0.76926703 | 2.036398557 | 0.00052358134 | 0.97807507 |
| 0.58331926 | 5.789404135 | -0.614925222 | -0.80166111 | 1.456295366 | 0.00017727343 | 0.56589102 |
| -1.52302429 | 6.575744741 | -0.037292867 | -0.77839198 | 2.825489657 | 0.00007425936 | 0.14267033 |
| -0.00761838 | 5.277401907 | -0.67959599 | -0.76365652 | 2.253724992 | 0.00067124722 | 0.99399338 |
| 0.63795223 | 6.019996417 | -0.68112443 | -0.656259417 | 2.138279662 | 0.00029353084 | 0.53040037 |
| 1.74444352 | 6.334539666 | -0.98814444 | -0.943629097 | 2.591275477 | 0.00036158922 | 0.09570201 |
| 0.89984240 | 6.09131471 | -0.64578520 | -0.58825329 | 2.293106044 | 0.00018988525 | 0.37840744 |
| 0.41491085 | 5.573344213 | -0.472606822 | -0.87436562 | 2.047380799 | 0.00027870027 | 0.68241535 |
| 2.885277896 | 7.565453753 | -1.22083624 | -0.29291993 | 1.367947223 | 0.00010605368 | 0.00885488 |
| 1.385488261 | 6.61366184 | -0.896518857 | -0.80505246 | 2.142467612 | 0.00028519703 | 0.18044367 |
| 1.801264781 | 6.976773546 | -0.35659071 | -0.91629857 | 0.79574448 | 0.00006385877 | 0.08603923 |
| -0.33477675 | 5.962390984 | -0.62506380 | -0.83321661 | 2.020288713 | 0.00020071532 | 0.741114287 |
| -2.27655879 | 4.576715768 | -1.01260277 | -0.06422659 | 3.187183003 | 0.00023476779 | 0.03339918 |
| -0.44272307 | 6.417917343 | -0.685925544 | -0.81075837 | 2.169568206 | 0.00020330593 | 0.66249271 |
| -0.200673311 | 6.36811726 | -0.68172769 | -0.79373435 | 2.133161512 | 0.00027028537 | 0.84288481 |
| 0.19963013 | 5.633253772 | -0.538341647 | -0.79153872 | 1.726049037 | 0.00024048369 | 0.84369024 |
| 3.188114672 | 8.393435709 | -0.20030957 | -2.07813469 | 4.131965549 | 0.00000314071 | 0.00442379 |
| -0.48700638 | 6.079097357 | -0.55717685 | -0.88206980 | 2.327190407 | 0.00026339050 | 0.63129840 |
| 0.55461607 | 5.988589679 | -0.619074817 | -0.88166539 | 2.212317387 | 0.00042117193 | 0.58501643 |
| 0.16990986 | 5.599386055 | -0.686401435 | -0.75806328 | 2.094781117 | 0.00024220511 | 0.86670722 |
| -0.20573406 | 5.584009628 | -0.672880734 | -0.787669022 | 2.266301045 | 0.00035696534 | 0.83897999 |
| -0.35016897 | -2.24987772 | -0.441224121 | 1.13230845 | 1.915439603 | 0.6129787052 | 0.729699651 |
| 0.60429638 | -2.170171712 | -0.24262386 | 1.022603571 | 1.775582218 | 0.7063357006 | 0.55212003 |
| -0.10324353 | -2.25166144 | -0.35953128 | 1.045319816 | 1.907961976 | 0.6656882987 | 0.91874929 |
| -0.55048871 | -2.29993455 | -0.66616254 | 1.171665939 | 1.986643155 | 0.5197901455 | 0.58779288 |
| -0.68323353 | -2.23571584 | -0.51014315 | 1.127030442 | 0.94685387 | 0.4983917522 | 0.50192844 |
| 1.06990166 | -2.55747477 | -0.54412166 | 1.339353746 | 1.995801105 | 0.4853102145 | 0.29680741 |
| -0.44025177 | -1.87628518 | -0.23052160 | 0.91333960 | 2.00956605 | 0.8427841884 | 0.66425291 |
| 0.94246324 | -2.181707447 | -0.22665986 | 0.76885861 | 2.190883522 | 0.8714597263 | 0.35667238 |
| -0.03978911 | -1.983172727 | -0.37709146 | 1.038163496 | 1.948603256 | 0.6750422029 | 0.96863710 |
| -0.572231041 | -2.339632284 | -0.53684081 | 1.17558325 | 1.925842147 | 0.5430157332 | 0.573241047 |
| -0.52364100 | -2.30232426 | -0.539270547 | 1.180080473 | 2.035730058 | 0.5735026095 | 0.60601086 |
| -1.758758371 | -2.811242652 | -1.068687461 | 1.341589799 | 2.403346701 | 0.3253654382 | 0.09318366 |
| -0.29509129 | -2.15571065 | -0.44119267 | 1.104835783 | 1.888884613 | 0.6108255146 | 0.77082184 |
| -3.25528208 | -3.53420961 | -1.468043687 | 2.452258211 | 0.13071155 | 0.0709169847 | 0.00378456 |
| 0.23723763 | -2.27194618 | -0.433049421 | 1.100971911 | 1.765676529 | 0.6305573011 | 0.81477151 |
| -0.08413454 | -2.06131089 | -0.386721004 | 1.07698517 | 1.714982245 | 0.6575247956 | 0.93374635 |
| 0.93690285 | -1.774854204 | -0.12644566 | 0.77043744 | 2.101801443 | 0.9183335491 | 0.35945922 |
| 1.266319437 | -2.028728937 | -0.23095434 | 0.48196882 | 2.282084474 | 0.7497137854 | 0.21926579 |

| | | | | | | |
|---|---|---|---|---|---|---|
| 1.232347741 | -1.966273014 | -0.39447496 | 0.741266311 | 2.280618032 | 0.7883156384 | 0.231439082 |
| 1.077431154 | -2.117773378 | -0.637409809 | 1.28074631 | 2.065876143 | 0.70860576 | 0.293509468 |
| 0.736927272 | -1.835648404 | -0.096593314 | 0.877284051 | 1.104414556 | 0.7053589626 | 0.469320377 |
| 1.432099219 | -2.765001173 | -0.906703479 | 1.701679519 | 0.228411355 | 0.2205435316 | 0.166832827 |
| -0.338896761 | -2.291092112 | -0.277605251 | 0.941928014 | 1.839584233 | 0.6354481273 | 0.738052827 |
| -1.067270996 | -1.710031628 | 0.057845924 | 0.326983556 | 2.180254849 | 0.9180774262 | 0.297965903 |
| -0.324722427 | -1.959575729 | -0.138659261 | 0.921504309 | 1.977458116 | 0.8543612009 | 0.748603758 |
| 0.694972022 | -2.282601606 | -0.420089876 | 1.089667443 | 1.993211133 | 0.6522843363 | 0.494691407 |
| 0.483905916 | -2.311220992 | -0.505704741 | 1.153840529 | 2.023806797 | 0.598175016 | 0.633460553 |
| -0.389838323 | -1.794357554 | -0.174166607 | 0.678029909 | 1.706644312 | 0.7260424058 | 0.700581804 |
| 3.92989586 | -1.974390693 | -0.653912984 | 0.918316066 | 2.000805056 | 0.6136447125 | 0.000768124 |
| -0.578286115 | -2.331370043 | -0.530754105 | 1.202744383 | 1.709388494 | 0.5741658364 | 0.569221314 |
| -0.105576455 | -2.100760159 | -0.382638084 | 1.067959274 | 1.86224282 | 0.6725178811 | 0.916920383 |
| -0.547744033 | -2.301203918 | -0.472988443 | 1.073614288 | 1.98064342 | 0.6093037515 | 0.589642818 |
| 0.125401195 | -2.266145208 | -0.385109795 | 1.079983805 | 1.29286925 | 0.6522537755 | 0.901398700 |
| -1.386346406 | -1.556251204 | 0.040886850 | 0.262700385 | 1.892536455 | 0.8444855867 | 0.180185318 |
| 0.302594185 | -2.270761198 | -0.425125108 | 1.070273299 | 1.913612755 | 0.6312174614 | 0.765176036 |
| -2.21165167 | -1.194097025 | 0.704368427 | 0.073118403 | 0.784369132 | 0.9790429132 | 0.038211931 |
| 1.354382344 | -2.65131512 | -0.588538118 | 1.48841024 | 2.198698286 | 0.576379314 | 0.190009262 |
| 0.754855302 | -2.259800414 | -0.399091278 | 1.159905809 | 1.320406167 | 0.5735037123 | 0.458718419 |
| 0.583157280 | -2.22089633 | -0.315689582 | 1.095668577 | 1.909020111 | 0.8802845003 | 0.565998046 |
| -0.550307441 | -2.270009581 | -0.653535863 | 1.201395724 | 1.956693564 | 0.559698895 | 0.587914970 |
| -1.128786811 | -2.158190396 | -0.266755039 | 1.084519449 | 2.282572421 | 0.9235057285 | 0.271718323 |
| 2.963520432 | -2.234573031 | -0.741121651 | 0.793497839 | 3.379315486 | 0.9421231884 | 0.007413952 |
| -0.504756735 | -2.310664875 | -0.483540280 | 1.158356892 | 1.420255832 | 0.5630144205 | 0.618985494 |
| -0.585590504 | -2.299796525 | -0.382559779 | 1.201204357 | 1.883217773 | 0.5710250341 | 0.564391499 |
| 1.697022351 | -2.169219704 | -0.393106503 | 0.531825304 | 2.682623496 | 0.9206702775 | 0.104465788 |
| -1.141357508 | -2.29468285 | -0.402374612 | 0.745119405 | 1.769626737 | 0.5039148759 | 0.266569842 |
| -2.018021169 | -1.996749708 | -0.476513077 | 0.838060371 | 0.318169488 | 0.302486836 | 0.056545333 |
| -0.358882146 | -2.137872298 | -0.525985712 | 1.047833197 | 1.963091342 | 0.5813618218 | 0.723266267 |
| 0.219904932 | -2.093752676 | -0.313196927 | 0.926234928 | 1.938571636 | 0.6934131724 | 0.828068791 |
| -1.648304377 | -2.830882981 | -0.671090231 | 1.343720706 | 2.029356612 | 0.3574053511 | 0.114171785 |
| 0.391222334 | -2.170286272 | -0.256749787 | 1.024424064 | 1.999537914 | 0.7657850145 | 0.699574079 |
| -1.002985021 | -2.138952113 | 0.074141320 | 1.024830227 | 2.233474545 | 0.8852950952 | 0.327285934 |
| -1.200312306 | -2.316862782 | -0.402147147 | 1.210130924 | 1.442680796 | 0.5644210138 | 0.243383937 |
| -0.487263361 | -2.244949225 | -0.478900271 | 0.961176008 | 1.661594753 | 0.5639197061 | 0.631119347 |
| 0.915418885 | -1.895307147 | -0.148927584 | 0.997397058 | 1.901719027 | 0.8753002039 | 0.370364408 |
| -0.510202261 | -2.318771528 | -0.450450148 | 1.132900833 | 1.885618639 | 0.6086890055 | 0.615230740 |
| -0.107418394 | -2.260828461 | -0.381714936 | 1.076263532 | 1.852280294 | 0.6619884292 | 0.915476714 |
| 1.232461785 | -2.568157515 | -0.389782007 | 1.11062862 | 2.040702267 | 0.5257991354 | 0.231397371 |
| 0.046585192 | -1.864802221 | -0.375911186 | 1.052596937 | 1.77907764 | 0.6815843354 | 0.963284013 |
| -0.211252639 | -2.173915338 | -0.340373122 | 1.021932441 | 1.529417772 | 0.648656076 | 0.834726749 |
| 1.267246398 | -2.339427236 | -0.009226622 | 0.783293064 | 2.275808137 | 0.8290132842 | 0.218940665 |
| 0.266523701 | -2.250964302 | -0.373154920 | 1.029039576 | 1.968799343 | 0.6784019855 | 0.792435443 |
| 0.388767919 | -2.114745761 | -0.250253107 | 0.866165523 | 1.833767508 | 0.8134023678 | 0.701361573 |
| 3.526249434 | -2.901235709 | 0.160838869 | 1.029734652 | 3.727519913 | 0.7331455264 | 0.002004274 |

| | | | | | | |
|---|---|---|---|---|---|---|
| -0.853872241 | -1.507809078 | -0.083607380 | 0.701840026 | 2.119540556 | 0.9927349524 | 0.4028112568 |
| 0.629999849 | -2.369526581 | -0.547417308 | 1.226571302 | 2.028785319 | 0.5918526001 | 0.5354901584 |
| 1.699684715 | -2.056366478 | -0.506453615 | 0.9978311174 | 1.563133495 | 0.5659657198 | 0.1039562309 |
| 0.857379064 | -2.416610807 | -0.699565278 | 1.229395686 | 1.863369901 | 0.4796585904 | 0.4009146375 |
| 0.332179811 | -2.20716441 | -0.460436885 | 1.121163429 | 1.965384113 | 0.611431992 | 0.7430462664 |
| -0.941961399 | -2.301777926 | -0.377050396 | 1.017160855 | 1.785336125 | 0.5899738712 | 0.3569233035 |
| -0.140861311 | -2.265150805 | -0.408437731 | 1.067279607 | 1.911098319 | 0.6458491699 | 0.8893219555 |
| 0.592869388 | -2.179017885 | -0.284689228 | 1.023794734 | 1.981236679 | 0.857346948 | 0.559599685 |
| 1.150605812 | -2.544121138 | 0.1682693756 | 0.6972414754 | 2.286154175 | 0.778053227 | 0.2628283018 |
| 0.491617908 | -2.288361261 | -0.391588280 | 1.091534704 | 1.581009209 | 0.5457567556 | 0.6280887525 |
| 1.128753435 | -2.133682126 | 0.0545583217 | 0.721842781 | 1.849873655 | 0.7860430788 | 0.2717320899 |
| 0.284375777 | -2.26385207 | -0.384355313 | 1.040756737 | 1.975258403 | 0.6701338455 | 0.7789076763 |
| 0.004934976 | -2.25031356 | -0.385074326 | 1.073761232 | 1.491707767 | 0.6869800342 | 0.9961090555 |
| 1.38142302 | -2.465027532 | -0.540779721 | 1.289031534 | 1.580712439 | 0.4943097277 | 0.1816716013 |
| -1.90692963 | -2.787933481 | -0.793952396 | 1.606416111 | 2.496447841 | 0.5723370694 | 0.0703034868 |
| -0.763635044 | -2.390592222 | -0.704265513 | 1.308912122 | 2.056597126 | 0.5730951318 | 0.4535793479 |
| -0.085777770 | -2.229773085 | -0.394521833 | 1.058342693 | 1.956974595 | 0.6558308568 | 0.9324556547 |
| 0.695388702 | -2.218509403 | -0.216519125 | 0.9340672237 | 1.798350518 | 0.6757305657 | 0.4944356186 |
| -0.510882786 | -2.198480315 | -0.394726975 | 1.086446469 | 1.867369881 | 0.8243986549 | 0.6147622668 |
| -0.843090328 | -2.360004235 | -0.586313775 | 1.270497418 | 1.892126129 | 0.5839844373 | 0.4086785656 |
| -0.509876402 | -2.291487525 | -0.228168880 | 0.9350870973 | 2.020756735 | 0.7036110182 | 0.6154551216 |
| -0.125438139 | -2.267033897 | -0.387690320 | 1.081492463 | 1.901663707 | 0.676728301 | 0.9013698108 |
| -0.983232467 | -2.062022924 | -0.247651889 | 1.089874825 | 1.702160555 | 0.7514786573 | 0.3366862375 |
| 0.465851522 | -2.296960912 | -0.463881258 | 1.106580185 | 1.966245497 | 0.6876760401 | 0.6461173274 |
| 1.057529367 | -2.209087171 | -0.077323266 | 0.8020830712 | 1.745093545 | 0.6865745397 | 0.3022840998 |
| 0.425002876 | -2.311761167 | -0.491382273 | 1.137295413 | 1.988049143 | 0.6049241107 | 0.6751577094 |
| -0.698202003 | -2.097269041 | -0.087458414 | 0.8464576608 | 1.834753511 | 0.7662926617 | 0.4927105963 |
| -1.094504391 | -1.98669113 | -0.251358725 | 0.9741321001 | 2.241613826 | 0.9268620613 | 0.2861292433 |
| -0.053805930 | -2.233080815 | -0.375095435 | 1.063785736 | 1.799885804 | 0.6885018403 | 0.9575983741 |
| -1.188495322 | -2.078478745 | -0.372495249 | 0.6011490994 | 1.681546271 | 0.5300060662 | 0.2479055584 |
| -1.452133797 | -2.348175279 | -0.412927320 | 0.8770327029 | 1.579647386 | 0.4719855977 | 0.1612424576 |
| 0.106449421 | -2.003419571 | -0.375081308 | 1.069252593 | 1.755945572 | 0.6795924874 | 0.9162361358 |
| 0.315678338 | -2.28254985 | -0.456786477 | 1.050776591 | 1.795341798 | 0.6026380976 | 0.7553625359 |
| 0.303794172 | -2.21705609 | -0.381215785 | 1.00829706 | 1.972215347 | 0.6657249219 | 0.7642742957 |
| 1.744387803 | -2.78674706 | 0.3207815268 | 1.138625173 | 1.677420657 | 0.6589727464 | 0.0957119323 |
| -1.024969862 | -2.496895399 | -0.474821099 | 1.064215457 | 1.376255729 | 0.4180778584 | 0.3170400346 |
| 0.171587407 | -1.835030874 | -0.362932714 | 1.065441906 | 1.448288125 | 0.6842981099 | 0.8654046089 |
| -1.04517865 | -2.405898193 | 0.0637640093 | 1.122983506 | 2.259948439 | 0.9704651143 | 0.3078227825 |
| 0.173592016 | -1.771975889 | -0.369454231 | 1.050455827 | 1.963868781 | 0.7533437618 | 0.8638485361 |
| 0.340941176 | -2.273761246 | -0.382476735 | 1.119284923 | 1.870176545 | 0.6232623216 | 0.7365353302 |
| 1.250988338 | -2.554040944 | -0.586407590 | 1.022237024 | 2.146143195 | 0.4675149066 | 0.2246971229 |
| 1.207185535 | -2.566595815 | -0.333765601 | 1.329066451 | 2.003315532 | 0.6150744461 | 0.2407827247 |
| 0.310002081 | -1.714861199 | -0.237463555 | 0.8521031799 | 1.779101287 | 0.7597793623 | 0.7596148083 |
| 0.892892042 | -2.30347233 | -0.519367842 | 1.25502419 | 1.49864371 | 0.5623443763 | 0.3820332914 |
| -0.677894407 | -2.254900825 | -0.291772190 | 1.08663526 | 2.052637289 | 0.7526279714 | 0.5052399228 |
| 0.535830792 | -1.823435326 | -0.272523926 | 1.052751008 | 1.214380057 | 0.7034334174 | 0.5977056413 |

| | | | | | | |
|---|---|---|---|---|---|---|
| -0.244243639 | -1.954668273 | -0.344486178 | 0.966540620 | 1.757715854 | 0.6840983672 | 0.809412625 |
| -0.82848826 | -2.362362505 | -0.482195427 | 1.302770656 | 2.151933059 | 0.5480236448 | 0.416711257 |
| -1.490223907 | -2.12192942 | -0.409643303 | 0.983277940 | 1.780918427 | 0.631003079 | 0.151033005 |
| 0.273384268 | -2.282973811 | -0.386092798 | 1.10757378 | 1.96122698 | 0.7013986752 | 0.787228515 |
| 0.454697973 | -1.736315973 | -0.164204745 | 0.860662363 | 1.332608357 | 0.737629611 | 0.653991967 |
| 1.992607704 | -1.742037292 | -0.028149496 | 0.325735240 | 2.905297962 | 0.6982012933 | 0.059461896 |
| 0.151417731 | -2.131691057 | -0.409525932 | 1.071630813 | 1.892773389 | 0.6436182582 | 0.881091373 |
| 0.377404731 | -1.849474063 | -0.341109138 | 0.961085261 | 1.848647443 | 0.8446174246 | 0.709660102 |
| 0.002457868 | -2.008462209 | -0.38495483 | 1.072318017 | 1.742297229 | 0.6638892921 | 0.998062105 |
| -0.106392519 | -1.881291658 | -0.380386547 | 1.055723734 | 1.955574022 | 0.7103595524 | 0.916280734 |
| 0.190311527 | -0.193565452 | -0.093514733 | -0.627626600 | 1.921282141 | 0.804675828 | 0.850892715 |
| -0.406704155 | -0.334059998 | -0.217107150 | -0.559912571 | 1.959624817 | 0.8671230291 | 0.688340526 |
| 1.216068945 | -0.091646732 | -0.354676207 | -0.422735531 | 1.564437879 | 0.8791115233 | 0.237451933 |
| 1.265036412 | 0.270111612 | 0.824886888 | -1.308316023 | 1.936843223 | 0.4368822198 | 0.219716417 |
| 1.046019543 | -0.354906213 | 0.066037446 | -0.678359154 | 2.135968773 | 0.5380898461 | 0.307443412 |
| -1.476311818 | 0.316090947 | 0.077193924 | -0.983986003 | 2.022530782 | 0.5635018691 | 0.154699032 |
| 0.674532953 | -0.537824273 | -0.333434343 | -0.392183960 | 1.675319067 | 0.9124444083 | 0.507331111 |
| -0.113919523 | -0.288361971 | -0.146075943 | -0.537634937 | 1.555868868 | 0.8658532107 | 0.910383705 |
| -0.425543905 | -0.452699457 | -0.260318944 | -0.448553106 | 1.951967924 | 0.9745422738 | 0.674769535 |
| 1.613678119 | 0.270148901 | 0.335903809 | -0.946575314 | 2.13155277 | 0.4495793926 | 0.121524284 |
| 2.088713081 | 0.530760772 | 0.559451777 | -1.178486713 | 1.532249818 | 0.4394802414 | 0.049092982 |
| -0.766203915 | -0.478565296 | -0.420130664 | -0.515777825 | 2.050842445 | 0.9704868139 | 0.452082315 |
| 0.233984722 | -0.141161181 | -0.074178240 | -0.642317049 | 1.925769642 | 0.784524939 | 0.817262856 |
| 1.111983076 | 0.017745278 | 0.222682331 | -0.990558549 | 2.249678179 | 0.5043958315 | 0.278714033 |
| -0.238566917 | -0.270005836 | -0.064098308 | -0.633292673 | 1.882575211 | 0.7952816652 | 0.813754028 |
| 2.448423639 | 0.816189318 | -0.096432650 | -0.794744180 | 0.580959866 | 0.6679484637 | 0.023217854 |
| -1.520004105 | -0.851421495 | -0.539339473 | -0.152634670 | 1.793283485 | 0.7579423316 | 0.143424090 |
| -0.136557588 | -0.299344015 | -0.145260092 | -0.486061745 | 1.814621891 | 0.8457673731 | 0.892681194 |
| -1.345693219 | -0.611889909 | -0.138555706 | -0.239721904 | 1.556898472 | 0.9848387783 | 0.192751749 |
| 2.118418649 | 0.044858202 | -0.634891659 | -0.262625799 | 2.221007062 | 0.7048203277 | 0.046234093 |
| -0.588070927 | -0.482740338 | -0.333752598 | -0.442643111 | 1.878815011 | 0.873121979 | 0.562756216 |
| -0.177786732 | -0.162315159 | -0.053315639 | -0.604514917 | 1.36984888 | 0.7842949867 | 0.860594237 |
| -1.824176563 | -0.469783519 | 0.370362212 | -1.121051988 | 1.610284213 | 0.9834930997 | 0.082388714 |
| -0.657081283 | 0.003887996 | 0.139989142 | -0.866811512 | 2.022255487 | 0.6878733197 | 0.518265904 |
| 0.252961909 | -0.354109205 | -0.248079829 | -0.489867391 | 1.784391102 | 0.9732826873 | 0.802757412 |
| -0.486310329 | -0.282536740 | -0.107372437 | -0.603584949 | 1.90871528 | 0.8286058087 | 0.631783511 |
| -0.723615934 | -0.035174600 | 0.074688560 | -0.735284462 | 1.677696595 | 0.724937938 | 0.477285430 |
| -2.087543664 | 0.750294228 | 0.796635375 | -1.708416652 | 1.249324831 | 0.5146946553 | 0.049208715 |
| -2.201917532 | -0.815704866 | -0.060079002 | -0.386079926 | 2.406120445 | 0.8471929539 | 0.038985937 |
| 1.413258151 | -0.129306913 | 0.254997608 | -1.003952756 | 2.32866298 | 0.5899639681 | 0.172231722 |
| 1.219343553 | -0.674116178 | -0.160725481 | -0.559247113 | 2.22390559 | 0.9426111274 | 0.236232979 |
| -0.261727393 | -0.288506570 | -0.170196687 | -0.602552887 | 1.918758984 | 0.8628945975 | 0.796081594 |
| -0.426566864 | -0.239678742 | -0.129311978 | -0.624900226 | 1.657303321 | 0.7163236761 | 0.674035845 |
| 0.074776798 | -0.282381718 | -0.145221840 | -0.477146138 | 1.90828005 | 0.8472144756 | 0.941100053 |
| 1.283692982 | -0.299895373 | -0.315325849 | -0.64563588 | 1.829747598 | 0.9861339401 | 0.213233718 |
| 0.757076298 | -0.600663649 | -0.482643443 | -0.201736172 | 2.056310223 | 0.9714019264 | 0.457415098 |

| | | | | | | |
|---|---|---|---|---|---|---|
| -1.413940495 | 0.133416963 | 0.066318803 | -1.037817354 | 1.795865353 | 0.7431317472 | 0.172033778 |
| -1.17740508 | -0.334793104 | -0.118797945 | -0.734346053 | 2.296932344 | 0.6816025597 | 0.252206188 |
| 0.080171407 | -0.268574504 | -0.118015312 | -0.595966789 | 1.550684637 | 0.823046018 | 0.936860051 |
| -0.81998505 | -0.633934144 | -0.635875504 | -0.025165042 | 2.077511013 | 0.967337093 | 0.421434652 |
| 1.331175908 | -0.464034770 | -0.283040680 | -0.597238323 | 1.388331011 | 0.8519507296 | 0.197403228 |
| -3.268564487 | -0.812295804 | 0.156903612 | -0.207396285 | 0.705037692 | 0.8144220696 | 0.003669261 |
| -0.279122285 | -0.299375018 | -0.183768053 | -0.532798456 | 1.49197855 | 0.9135442479 | 0.782881375 |
| -0.901860431 | -0.65907633 | -0.122335878 | -0.341004480 | 1.820473816 | 0.9949784633 | 0.377358964 |
| -1.972317699 | -0.557801518 | -0.160185769 | 0.025194006 | 0.545661187 | 0.8474817985 | 0.061885765 |
| 0.837255480 | -0.307713594 | -0.126532160 | -0.350508582 | 2.062855392 | 0.7066139065 | 0.411876428 |
| -0.636832315 | -0.140194680 | -0.147337014 | -0.697100221 | 1.055619548 | 0.9857507193 | 0.531115561 |
| -0.966853954 | -0.734436604 | -0.794937926 | 0.174869288 | 2.166685394 | 0.8085959748 | 0.344620829 |
| 0.173278806 | -0.211724162 | -0.077922610 | -0.620751840 | 1.896225436 | 0.8128537565 | 0.864091627 |
| -0.255788587 | -0.349350368 | -0.168474369 | -0.560952124 | 1.905056492 | 0.8999358882 | 0.800602891 |
| 0.605596065 | -0.170945159 | 0.051113270 | -0.670085617 | 2.000243712 | 0.698581314 | 0.55127267 |
| 0.706039029 | -0.383144284 | -0.420279151 | -0.552834531 | 1.30965829 | 0.8496853015 | 0.487923486 |
| 0.617823901 | -0.290493385 | -0.127537712 | -0.654565896 | 2.019959567 | 0.7824871631 | 0.543334032 |
| 0.582659004 | -0.311033926 | -0.007300950 | -0.474355309 | 2.006600884 | 0.6929789725 | 0.566327325 |
| -2.517303115 | -1.101914974 | -0.777123531 | -0.424975577 | 2.399679188 | 0.6079750842 | 0.020014945 |
| 1.334395624 | -0.142227385 | 0.040745409 | -0.774507362 | 2.143616131 | 0.6769387196 | 0.196364066 |
| 0.602955589 | -0.291728200 | -0.147821500 | -0.700693287 | 2.018024788 | 0.8366880184 | 0.552994939 |
| -1.358752407 | 0.040442234 | -0.143868813 | -0.621889651 | 1.973855548 | 0.6721404569 | 0.188641667 |
| 2.421264353 | 1.069719783 | 0.166316128 | -1.056642199 | 1.03426891 | 0.3916560099 | 0.024607236 |
| -2.090830133 | 0.131104418 | 0.243094702 | -1.011178063 | 0.629879059 | 0.9669487492 | 0.048884087 |
| 0.170425216 | -0.275733733 | -0.072790287 | -0.621599167 | 1.895418924 | 0.8121731312 | 0.866307014 |
| -0.590635533 | -0.307660815 | -0.156322668 | -0.516007698 | 1.749173976 | 0.8723048535 | 0.561068016 |
| 0.022749295 | -0.263457601 | -0.115537271 | -0.566105781 | 1.582837637 | 0.8416901362 | 0.982065034 |
| -1.885266042 | -0.274522682 | -0.478091563 | -0.472807915 | 1.073051986 | 0.8032549942 | 0.073306671 |
| -0.141570937 | -0.160849351 | -0.073818447 | -0.608284255 | 1.900513995 | 0.801596871 | 0.888768265 |
| -1.434479821 | 0.095776359 | 0.266416939 | -1.042857456 | 1.838434806 | 0.6560771157 | 0.166160477 |
| -0.707939279 | -0.406450285 | -0.089641343 | -0.542988015 | 2.051099779 | 0.7969144681 | 0.486766818 |
| -1.057668749 | -0.094846878 | 0.298298878 | -0.794330446 | 2.079102215 | 0.5753156276 | 0.302222002 |
| -1.308199556 | 0.270717638 | 0.218765055 | -1.082184022 | 1.963476086 | 0.5889640871 | 0.204944538 |
| -0.099117332 | -0.275850162 | -0.126824888 | -0.604030708 | 1.85875882 | 0.8422626605 | 0.921985183 |
| 0.675056958 | -0.199753537 | 0.047074674 | -0.574055416 | 2.01137738 | 0.6776752511 | 0.507004803 |
| -1.612397941 | -0.511583185 | -0.401404070 | -0.482705873 | 0.842836969 | 0.6788049825 | 0.121803564 |
| 0.419756377 | -0.372283680 | 0.075462893 | -0.700739153 | 1.94690959 | 0.7879452362 | 0.678926722 |
| 0.960255450 | -0.633212295 | -0.139780808 | -0.587865361 | 1.372273496 | 0.8327969024 | 0.347853377 |
| -0.678671515 | -0.378873062 | -0.374379420 | -0.376065509 | 2.002714069 | 0.9167007465 | 0.504757171 |
| -0.643679733 | -0.286902600 | -0.132976166 | -0.533686498 | 1.756150008 | 0.8544969855 | 0.526750973 |
| 1.25507908 | -0.420548462 | -0.149403329 | -0.641229649 | 0.705876118 | 0.7636947236 | 0.223237907 |
| -0.546398452 | -0.234739968 | -0.073075288 | -0.666963407 | 1.999801811 | 0.763185827 | 0.590550802 |
| 0.25223142 | -0.219957531 | -0.075040294 | -0.643008700 | 1.805102594 | 0.8241636408 | 0.803314460 |
| -0.917613512 | -0.668430492 | -0.548179478 | -0.167651972 | 2.118692966 | 0.953115036 | 0.369240410 |
| 1.426160789 | 0.020401609 | 0.194804928 | -0.982728563 | 1.927783875 | 0.6769491527 | 0.168519571 |
| -0.559113960 | -0.327033817 | -0.256284065 | -0.484850526 | 1.997263799 | 0.848028459 | 0.581998207 |

| | | | | | | |
|---|---|---|---|---|---|---|
| 1.563635446 | -0.479338673 | -0.113293253 | -0.64960326 | 0.6367274268 | 0.7122875792 | 0.1328483630 |
| 0.6843620681 | -0.225202432 | 0.041885074 | -0.758622507 | 1.990684517 | 0.7664187647 | 0.5012300742 |
| 1.441998506 | -0.245764583 | -0.535300633 | -0.279751234 | 1.766153836 | 0.9557946852 | 0.1640513269 |
| -0.376835869 | -0.292499551 | -0.136668839 | -0.560753168 | 1.940750297 | 0.7946748907 | 0.7100765252 |
| 2.243849167 | -0.764096133 | -0.465686347 | -0.646191642 | 2.606378513 | 0.9518287676 | 0.0357505259 |
| -0.888346854 | -0.232171183 | 0.029589739 | -0.663399375 | 1.34481935 | 0.8866779252 | 0.3844167313 |
| -0.736966604 | -0.350310489 | -0.334817026 | -0.401560857 | 2.045092293 | 0.8567144552 | 0.4692969601 |
| -0.509679617 | -0.170673445 | 0.020624253 | -0.683112735 | 1.881698261 | 0.7553133659 | 0.6155906432 |
| -1.180611779 | -0.037543165 | 0.336659618 | -0.924226368 | 1.750313638 | 0.6567803028 | 0.2509569817 |
| -0.202736601 | -0.218700244 | -0.101390537 | -0.617443684 | 1.888048157 | 0.7932623168 | 0.8412922969 |
| 1.107172814 | -0.437503354 | -0.271967965 | -0.504719294 | 1.325907928 | 0.8968117631 | 0.2807406373 |
| -1.746321996 | 0.0311381110 | -0.100455658 | -1.231251388 | 1.573808193 | 0.9627766 | 0.0953682474 |
| -0.417373273 | -0.270502021 | -0.131618241 | -0.659913796 | 1.732895278 | 0.8972642791 | 0.6806415674 |
| 2.553869563 | 0.7994253550 | 0.063537039 | -0.784701878 | 1.006429498 | 0.4985502318 | 0.0184873354 |
| -0.882398847 | -0.231473976 | 0.1119531690 | -0.543144455 | 2.113568437 | 0.6110107935 | 0.3875504988 |
| 0.8951589635 | -0.175619148 | -0.117274638 | -0.753674129 | 1.990194028 | 0.8147311394 | 0.3808481874 |
| 1.305695971 | -0.611793893 | 0.4002672920 | -0.629526132 | 1.666753307 | 0.8112637557 | 0.2057796704 |
| 1.561076241 | 0.0901312014 | -0.011165971 | -0.576905450 | 2.490528071 | 0.4169625488 | 0.1334503143 |
| 1.718428996 | 0.6534129948 | 0.0620241570 | -0.719027097 | 0.574008924 | 0.6268099033 | 0.1004283297 |
| 1.430635627 | -0.159637152 | -0.704235155 | -0.655797727 | 1.044202985 | 0.6834479754 | 0.1672472759 |
| 1.69537538 | 0.7127650364 | 0.0052329028 | -0.812789002 | 2.180382022 | 0.3441504541 | 0.1047820704 |
| -0.172652877 | -0.217330834 | -0.130406647 | -0.618937706 | 1.91223711 | 0.8127951188 | 0.8645774700 |
| 0.6495393602 | -0.407936207 | -0.227935038 | -0.648168387 | 1.986681003 | 0.9549499385 | 0.5230316625 |
| -2.209192031 | 0.2508486710 | -0.259305571 | -1.090920974 | 2.153152885 | 0.7448344751 | 0.0384061752 |
| 1.82747006 | 0.766293007 | 0.5733397593 | -1.34136946 | 1.421808741 | 0.4048648536 | 0.0818751848 |
| 0.3485482229 | -0.276678602 | -0.178739094 | -0.514666418 | 1.636759779 | 0.8781287503 | 0.7308986090 |
| 0.3126788071 | -0.291069435 | -0.170786830 | -0.599683473 | 1.850309191 | 0.879753447 | 0.7576086032 |
| 0.478950048 | -0.044019316 | -0.030677388 | -0.624631261 | 1.212263801 | 0.7935752634 | 0.6369235708 |
| 0.6965191979 | -0.546241916 | -0.233210377 | -0.389312795 | 2.053837395 | 0.8976292203 | 0.4937420204 |
| 1.205639189 | 0.4510094624 | 0.0039055254 | -0.979436509 | 1.393930136 | 0.6499381407 | 0.2413661101 |
| -0.261672503 | -0.233559835 | -0.129539062 | -0.620905523 | 1.84372056 | 0.8361448292 | 0.79612335 |
| -1.202341667 | -0.115588481 | -0.131234422 | -0.961438492 | 1.589166088 | 0.9999969667 | 0.2426137194 |
| -0.019122087 | -0.248718640 | -0.124898519 | -0.555821786 | 1.56436378 | 0.8405587936 | 0.9849242331 |
| 0.7868270087 | -0.027280760 | 0.0236398058 | -0.860175203 | 2.014239742 | 0.6050781109 | 0.4401730268 |
| -0.635417062 | 0.0018057912 | 0.0174552501 | -0.763671934 | 2.013400527 | 0.6745310437 | 0.5320201004 |
| 1.916876523 | 0.6061735860 | 0.0787454761 | -1.073954063 | 2.781713554 | 0.2609786169 | 0.0689611911 |
| -2.701883533 | 0.9625877653 | -0.071675855 | -0.850393663 | 0.7470791032 | 0.5531842982 | 0.0133539927 |
| -2.379238941 | 0.9654766011 | -0.045673290 | -0.934778153 | 2.15839327 | 0.2926475097 | 0.0269099643 |
| 0.6022104862 | 0.4807260867 | -0.627604966 | -0.546184329 | 0.2613551780 | 0.8043453469 | 0.5534814492 |
| 2.72339589 | 0.7271826423 | -0.223816320 | -0.674354057 | -0.410771494 | 0.8673804711 | 0.0127311123 |
| 0.8478703408 | 0.4215509467 | -0.893314431 | -0.234595776 | -0.038034471 | 0.6598611084 | 0.406070667 |
| -1.443746171 | -0.332392474 | -1.579040444 | 0.6846353077 | 0.2595690389 | 0.3069231726 | 0.1635641800 |
| 0.0678555897 | 0.2817935644 | -0.711848215 | -0.367818475 | 0.1911650359 | 0.7428009081 | 0.9465425232 |
| 0.196756952 | 0.186273653 | -0.758659592 | -0.302279394 | 0.1982867232 | 0.6704198663 | 0.8459095008 |
| 0.8626800771 | -0.098520203 | -0.981693086 | -0.115287048 | -0.037980848 | 0.4636861449 | 0.3980585952 |
| -2.004917184 | 0.0724519955 | -1.134982836 | 0.2129284840 | -0.883955971 | 0.3342094417 | 0.0580328834 |

| | | | | | | |
|---|---|---|---|---|---|---|
| 0.337906667 | 0.417794608 | -0.589528777 | -0.448486337 | 0.164993031 | 0.8172172445 | 0.738788133 |
| -0.513687791 | 0.097977796 | -0.858852484 | -0.255233096 | 0.165589871 | 0.5878893091 | 0.612833079 |
| -0.263125511 | 0.162845217 | -0.784122079 | -0.283661932 | 0.258191227 | 0.6562957082 | 0.795018248 |
| -0.765600003 | 0.071596040 | -0.987898655 | -0.280459641 | 0.352061819 | 0.5410920501 | 0.452433981 |
| 0.285179864 | 0.385566867 | -0.65828074 | -0.448210859 | 0.242606178 | 0.7804883618 | 0.778300015 |
| -0.243344965 | 0.213383339 | -0.777547699 | -0.245371749 | 0.027283275 | 0.6491880488 | 0.810099492 |
| -1.240158691 | 0.333343771 | -0.425214693 | -0.608008835 | 0.609817332 | 0.9153745537 | 0.228595464 |
| 0.504501468 | 0.484205764 | -0.731611307 | -0.390234011 | -0.089583587 | 0.7257657409 | 0.619161767 |
| 0.243867085 | 0.358978274 | -0.645650039 | -0.420495148 | 0.231056348 | 0.7768298754 | 0.809700411 |
| -0.259715867 | 0.226461578 | -0.766189432 | -0.222604708 | 0.128456595 | 0.6783271185 | 0.797612184 |
| 2.050527867 | 0.811374098 | -0.800841017 | -0.933875294 | 0.761361029 | 0.9034670779 | 0.053001147 |
| 1.465279291 | 0.536416261 | -1.096561162 | -0.109669728 | 0.292844683 | 0.7602717578 | 0.157657517 |
| -0.494745697 | 0.077480369 | -0.870567427 | -0.235796441 | 0.463590391 | 0.6652207592 | 0.625916073 |
| -1.778356265 | 1.100312964 | -0.073524567 | -1.249950086 | 1.496788987 | 0.4386018349 | 0.089828505 |
| 1.010667784 | 0.388697529 | -1.004705134 | -0.080149336 | 0.419815847 | 0.7769266087 | 0.323679498 |
| 0.312965676 | 0.140110230 | -0.802681464 | -0.130593311 | 0.133897128 | 0.6412494845 | 0.757393698 |
| -0.809705035 | 0.585726449 | -0.168570675 | -0.604147543 | 0.404022842 | 0.9025440061 | 0.427189824 |
| 0.031419688 | 0.287369021 | -0.738133152 | -0.364058058 | 0.200916431 | 0.6969495893 | 0.975231548 |
| 0.145422295 | 0.226000264 | -0.74929708 | -0.32884539 | 0.231348857 | 0.6817039741 | 0.885764240 |
| -1.183174 | 0.834820791 | -0.159791126 | -0.971445896 | -0.214393075 | 0.8803092375 | 0.249962172 |
| 0.872998964 | 0.489185124 | -0.782846758 | -0.475806103 | 0.079843320 | 0.7038475002 | 0.392536880 |
| 2.913798589 | 0.664675086 | -0.048421277 | -1.247487073 | 1.092527786 | 0.8219885085 | 0.008301026 |
| -1.729578537 | 0.862552702 | -0.748596242 | -0.472089558 | -0.238394848 | 0.8340657118 | 0.098378770 |
| -1.74546042 | 0.229232464 | -1.066792202 | -0.414927198 | 0.253890195 | 0.5170569503 | 0.095521209 |
| 1.2657596 | 0.186494804 | -0.76428753 | -0.301396254 | -0.749011142 | 0.3846938922 | 0.219462330 |
| 0.130981020 | 0.204883150 | -0.742465891 | -0.247379909 | 0.212452812 | 0.6844267913 | 0.897037007 |
| 0.631534744 | 0.283607567 | -0.826947659 | -0.379821545 | 0.131097272 | 0.6278320176 | 0.534505723 |
| -1.02579019 | 0.737954922 | -0.16005006 | -0.793892117 | -0.33175381 | 0.8756412035 | 0.316662138 |
| 1.606382661 | -0.175249476 | -1.000223138 | 0.137722978 | 0.421455012 | 0.5935387603 | 0.123123053 |
| 0.243426899 | 0.297455155 | -0.741302842 | -0.337940013 | 0.048554928 | 0.6711364331 | 0.810036862 |
| -0.16071108 | 0.271975812 | -0.751817878 | -0.36799843 | 0.057995433 | 0.6748394584 | 0.873856933 |
| -0.902103185 | -0.166416851 | -1.15886183 | 0.215247419 | 0.656226512 | 0.5192667523 | 0.377232968 |
| 0.237275647 | 0.253062905 | -0.760480017 | -0.360134461 | 0.103390371 | 0.6586848443 | 0.814742419 |
| 0.153251080 | 0.306513350 | -0.749170451 | -0.384083819 | 0.246862523 | 0.7173643176 | 0.879663332 |
| -0.746622936 | 0.228000393 | -0.885643791 | -0.215841195 | -0.213474325 | 0.54054264 | 0.463568761 |
| 0.833167755 | 0.636789629 | -0.757465394 | -0.582870974 | 0.304887635 | 0.8519148228 | 0.414126217 |
| 0.814313125 | 0.396077003 | -0.740579209 | -0.614401223 | 0.638822020 | 0.8297814776 | 0.424603973 |
| -0.010831306 | 0.287294970 | -0.737494577 | -0.352813274 | 0.194790856 | 0.701566099 | 0.991460276 |
| 0.471015483 | 0.183883571 | -0.727989359 | -0.28665567 | 0.458589577 | 0.8121487916 | 0.642485745 |
| -1.005340608 | -0.257107192 | -1.249742299 | 0.379661606 | 0.542042770 | 0.4235068276 | 0.326177213 |
| 0.468102656 | 0.416333522 | -0.590257925 | -0.508904954 | 0.170369765 | 0.7604837287 | 0.644533086 |
| 0.075152552 | 0.295224826 | -0.715596002 | -0.370893913 | 0.202312581 | 0.7266053753 | 0.940804662 |
| -0.352129861 | 0.223193786 | -0.809863063 | -0.321338215 | 0.140043200 | 0.6321967721 | 0.728250027 |
| 0.758315660 | 0.173404761 | -1.000484418 | -0.313946873 | -0.20872670 | 0.4668650479 | 0.456688788 |
| -1.408452604 | 0.326183828 | -0.776320893 | -0.257458607 | -0.319461715 | 0.5803836476 | 0.173630961 |
| -0.308227272 | 0.304518179 | -0.786612175 | -0.419170423 | 0.077186567 | 0.6384056722 | 0.760945985 |

| | | | | | | |
|---|---|---|---|---|---|---|
| 1.708382911 | 0.835168847 | -0.330429214 | -0.558871616 | 0.057156796 | 0.9144212927 | 0.102306196 |
| -0.372743416 | 0.246058779 | -0.782055873 | -0.319298967 | 0.151642004 | 0.6587052712 | 0.713075046 |
| -1.285459176 | 0.325234675 | -0.726592796 | -0.143448596 | -0.149741839 | 0.6950496942 | 0.212627758 |
| 0.080479676 | 0.260012806 | -0.736959485 | -0.364363684 | 0.201316066 | 0.6915600675 | 0.936617818 |
| 0.022170244 | 0.252007949 | -0.728527668 | -0.363038466 | 0.175371262 | 0.7118166711 | 0.982521464 |
| -0.073844824 | 0.296173261 | -0.711642784 | -0.371548511 | 0.128175259 | 0.6934564938 | 0.941832742 |
| 0.843758932 | 0.297528450 | -0.468608412 | -0.559227232 | 0.403047662 | 0.8140819042 | 0.408313143 |
| 0.038386667 | 0.288644656 | -0.734960891 | -0.366041559 | 0.203765242 | 0.7006559668 | 0.969741978 |
| -0.288932082 | 0.211176154 | -0.791745407 | -0.236730643 | 0.000259005 | 0.6358391348 | 0.775466333 |
| -0.162425492 | 0.289373066 | -0.755125480 | -0.348201317 | 0.112730377 | 0.676647066 | 0.872523558 |
| -0.206098890 | 0.352834712 | -0.626787198 | -0.414824557 | 0.237779794 | 0.805720154 | 0.838698654 |
| -0.108308421 | 0.305515049 | -0.679127591 | -0.379914881 | 0.187677489 | 0.7136859725 | 0.914779238 |
| 1.814278493 | 0.640024663 | -0.894805931 | -0.543563335 | -0.270521395 | 0.5941323519 | 0.083948929 |
| -0.102472924 | 0.300651160 | -0.636542863 | -0.376808453 | 0.211682566 | 0.7408224834 | 0.919353520 |
| 1.812951528 | -0.457356537 | -1.246259472 | 0.345403345 | 0.246774014 | 0.3918178151 | 0.084160034 |
| 0.089573729 | 0.286337653 | -0.739003185 | -0.355894637 | 0.213432209 | 0.7052819469 | 0.929474772 |
| -0.920083213 | 0.18449521 | -0.962453419 | -0.411555960 | 0.060975127 | 0.5051445891 | 0.367978255 |
| -0.331693409 | 0.239656771 | -0.784104607 | -0.333084710 | -0.008706021 | 0.6268226125 | 0.743408315 |
| -0.914629429 | 0.508088340 | -1.085118444 | -0.070342524 | -0.117947902 | 0.6007707708 | 0.370769292 |
| 0.731243059 | -0.015859939 | -0.753221501 | -0.351345583 | -0.121176427 | 0.5079305449 | 0.472711939 |
| 0.634281522 | 0.382192866 | -0.453101235 | -0.539156112 | 0.118185553 | 0.7717538915 | 0.532746470 |
| 0.096881435 | 0.288271843 | -0.737119172 | -0.372374381 | 0.215862005 | 0.7005760199 | 0.923739221 |
| -1.034082206 | 0.404114198 | -0.742591329 | -0.355641214 | 0.823269527 | 0.9628816034 | 0.312860092 |
| 0.669215578 | 0.236528454 | -0.809561523 | -0.279558844 | 0.009637036 | 0.6166003549 | 0.510649029 |
| -0.750215509 | 0.132117731 | -0.883502708 | -0.168020683 | 0.368667649 | 0.6674208583 | 0.46144833 |
| -0.792126706 | -0.100535239 | -1.033443945 | -0.002502143 | 0.625615741 | 0.6031574333 | 0.437144027 |
| -0.023333088 | 0.275554814 | -0.723285376 | -0.344905164 | 0.201126318 | 0.697678011 | 0.981604873 |
| 0.090753048 | 0.294045412 | -0.696083223 | -0.374893392 | 0.178085118 | 0.6996888068 | 0.928548894 |
| -1.058671834 | 0.420551489 | -0.772001966 | -0.361627449 | 0.811150072 | 0.9854085506 | 0.301775375 |
| -0.871374753 | 0.222981588 | -0.941439296 | -0.137002411 | 0.114628327 | 0.6154067848 | 0.393402691 |
| -0.428472934 | 0.275091306 | -0.590043631 | -0.454224604 | 0.262496157 | 0.7339157313 | 0.672669652 |
| -0.615691539 | 0.264760472 | -0.756606709 | -0.306414068 | 0.378995242 | 0.7605465215 | 0.544714013 |
| -0.842393925 | 0.459321310 | -0.620261123 | -0.377602964 | -0.006182634 | 0.777268066 | 0.409059403 |
| 0.656209289 | 0.253213063 | -0.850028649 | -0.326125300 | 0.473843431 | 0.7340400439 | 0.518815686 |
| 0.605317576 | 0.346864706 | -0.540450561 | -0.507325123 | 0.072847945 | 0.7143869831 | 0.551454182 |
| 0.647977214 | 0.154304244 | -0.898017471 | -0.245808851 | 0.246949029 | 0.6050480793 | 0.524021795 |
| -1.358415292 | 0.571642826 | -0.180649115 | -0.746244495 | -0.018552402 | 0.8961590114 | 0.188746888 |
| 2.230691663 | -0.23814979 | -1.101111978 | -0.151660098 | -0.437647234 | 0.278598834 | 0.036738332 |
| -1.597338787 | 0.525629455 | -0.571412988 | -0.538880213 | 0.857714438 | 0.9274688099 | 0.125129394 |
| -0.184662687 | 0.316461961 | -0.733966104 | -0.406706074 | 0.151085981 | 0.6796425275 | 0.855265324 |
| -0.023885610 | 0.287471996 | -0.737592615 | -0.363282450 | 0.186227577 | 0.6990835866 | 0.981169366 |
| 0.321349849 | 0.397139944 | -0.710310811 | -0.377501392 | 0.058181661 | 0.7425857879 | 0.751121746 |
| 0.871794152 | 0.242535092 | -0.957964293 | -0.433077674 | -0.045608856 | 0.5005323652 | 0.393179005 |
| -0.905909032 | 0.185031641 | -0.765430640 | -0.210363905 | 0.156722265 | 0.6766188034 | 0.375261271 |
| -0.594908619 | 0.431912511 | -0.919605622 | -0.363663961 | 0.322635354 | 0.6879727184 | 0.558261030 |
| -0.002696221 | 0.278091879 | -0.735327151 | -0.364080895 | 0.180024554 | 0.7203036153 | 0.997874178 |

| | | | | | | |
|---|---|---|---|---|---|---|
| -0.493865462 | -0.014484665 | -0.793763163 | -0.342561696 | 0.460240965 | 0.6437326543 | 0.6265271664 |
| 0.0381146354 | 0.2893724500 | -0.689351992 | -0.364931091 | 0.1559104120 | 0.7131641025 | 0.9699562968 |
| 0.3329868843 | 0.4255188052 | -0.708960423 | -0.398264324 | 0.2297614153 | 0.8414265508 | 0.7424456627 |
| 0.3343993487 | 0.182497547 | -0.735828743 | -0.302932661 | 0.1390809429 | 0.6586231911 | 0.7413949469 |
| 1.161007398 | 0.0492440408 | -0.929928408 | -0.456715545 | 0.3269556805 | 0.5121699649 | 0.258666812 |
| 0.4284370499 | 0.1729704571 | -0.716964815 | -0.273011677 | 0.1947037604 | 0.6830164952 | 0.6726953621 |
| 1.119048648 | 0.8668330824 | -0.271720296 | -0.803879980 | -0.135133141 | 0.993400034 | 0.2757566387 |
| -0.357976121 | 0.2871448114 | -0.678985894 | -0.431376674 | 0.3224679943 | 0.7417941517 | 0.723934266 |
| 0.1272604688 | 0.2812095033 | -0.748242430 | -0.364525331 | 0.1800555384 | 0.6851308872 | 0.899944963 |
| -1.805812581 | -0.501701341 | -1.138170695 | -0.299374740 | 1.28655429 | 0.5422606982 | 0.0853036823 |
| -1.502009509 | 0.9043647792 | -0.541413732 | -0.782431464 | -0.312830575 | 0.8183899329 | 0.1479831614 |
| 0.601852887 | 0.5806478889 | -0.671184904 | -0.544413969 | -0.033366141 | 0.7919283855 | 0.5537150212 |
| -0.965671332 | 0.4385225465 | -0.756721574 | -0.461915764 | 0.0421658554 | 0.6814701963 | 0.3451986689 |
| -0.214435354 | 0.3145035450 | -0.737967537 | -0.413221103 | 0.136992654 | 0.6736854589 | 0.8322761360 |
| -0.308690037 | 0.0933067950 | -0.800611295 | -0.238478338 | 0.3428554853 | 0.6558623677 | 0.7605988190 |
| -0.405485827 | 0.1507583682 | -0.804575487 | -0.184037199 | -0.089268274 | 0.6021405158 | 0.6892219139 |
| 1.142848945 | -0.192079327 | -0.998973835 | -0.022896265 | -0.011585396 | 0.4411484808 | 0.2659638145 |
| 0.0423432310 | 0.2748789870 | -0.727936879 | -0.364137682 | 0.1908018064 | 0.7514461549 | 0.966625106 |
| 0.0126787595 | 0.2488086628 | -0.737511699 | -0.362850002 | 0.1841023324 | 0.6984985754 | 0.990003766 |
| -0.558231271 | 0.5365094434 | -0.719032577 | -0.426868048 | 0.203127101 | 0.8666347224 | 0.5825899045 |
| -0.309649221 | 0.0575135430 | 0.5886600588 | 0.2631294945 | 1.220284805 | 0.3011301266 | 0.7598794062 |
| 0.978737021 | 0.320281447 | 0.8686622807 | 0.0820541648 | 1.018530767 | 0.2142411944 | 0.3388514184 |
| 1.539382489 | 0.4248387994 | 0.3914722959 | 0.415306502 | 0.8480454944 | 0.2684990196 | 0.1386442973 |
| 1.567419492 | 0.8260228873 | 1.610013688 | -0.917783737 | 1.271393762 | 0.08310798813 | 0.1319624507 |
| -1.083706231 | 0.2544230770 | 0.4546467078 | 0.2428018898 | 0.1745506261 | 0.5264483409 | 0.2907811712 |
| -0.944564151 | 0.5378512554 | 0.7927919634 | -0.073436270 | 1.29125032 | 0.1816038449 | 0.3556232263 |
| -2.652808862 | 1.301736273 | 1.540796123 | -0.551572258 | 2.115337231 | 0.02642116863 | 0.0148846155 |
| -0.932526590 | 0.0669477987 | 0.4956628897 | 0.4411346087 | 0.5862712523 | 0.39732614 | 0.361662888 |
| -0.486105574 | -0.088502021 | 0.4612145064 | 0.3062092619 | 1.305142272 | 0.3857703867 | 0.6319262465 |
| 1.370435118 | 0.6363501523 | 1.046564442 | -0.103177982 | 1.401091729 | 0.1226356512 | 0.1850238283 |
| 1.201930954 | 0.6340045097 | 1.033564757 | -0.143719540 | 0.9683700459 | 0.1545586732 | 0.2427694518 |
| 1.380748843 | 0.5524970810 | 1.166670263 | 0.0202403531 | 1.007825837 | 0.1282274631 | 0.1818758808 |
| -0.147331557 | 0.0900332507 | 0.6031651142 | 0.2143856685 | 1.217183144 | 0.3059524416 | 0.8842756834 |
| 1.341136118 | 0.5292661402 | 1.064242938 | -0.344325791 | 1.837299611 | 0.1100156351 | 0.1942024975 |
| -0.000949319 | 0.1755951559 | 0.6300813814 | 0.1663925238 | 1.183066933 | 0.2728029745 | 0.9992515135 |
| 0.1638214410 | 0.2305147459 | 0.6546168137 | 0.1617588459 | 0.9883905563 | 0.2558558325 | 0.8714381520 |
| 1.749405199 | 0.8436701415 | 1.141841493 | -0.343354606 | 1.563216486 | 0.09203022303 | 0.0948225875 |
| 2.09766964 | 0.6302780779 | 0.9833968192 | -0.708422867 | 1.865038909 | 0.15105293 | 0.0482146708 |
| 1.422884055 | 0.5297467088 | 0.6865974507 | -0.210061789 | 1.647478228 | 0.1730403653 | 0.1694561509 |
| 1.101350674 | 0.3578932085 | 0.3923318797 | 0.3718310338 | 1.350879356 | 0.2193263488 | 0.2832078842 |
| -0.860396756 | -0.1641085 | 0.3058003261 | 0.3756448928 | 1.535700496 | 0.2847501365 | 0.3992871712 |
| 1.47690336 | -0.524108169 | 0.0861961988 | 0.9127632392 | -0.276809471 | 0.9866616848 | 0.1545416901 |
| 0.4135737302 | 0.2148553570 | 0.5142062999 | 0.2777355817 | 1.3197212 | 0.2427251902 | 0.6833793254 |
| -0.189678954 | 0.2365847697 | 0.673571177 | 0.0363677685 | 1.268458171 | 0.2587160568 | 0.8513821305 |
| -2.115385234 | 1.007536968 | 1.776742743 | -0.482819485 | 1.878029551 | 0.03573346079 | 0.0465189878 |
| -0.836214559 | 0.1711213311 | 0.6984694965 | 0.1684326265 | 1.255174812 | 0.2487431277 | 0.4124485877 |

| | | | | | | |
|---|---|---|---|---|---|---|
| 1.564880941 | -0.322235108 | 0.2293112594 | 0.4824062322 | 1.688727945 | 0.3870402502 | 0.1325562273 |
| -1.7686706374 | 1.025194005 | 1.408127041 | -0.834100221 | 0.6517815133 | 0.1285951267 | 0.0914734324 |
| -0.376699531 | 0.0827970194 | 0.6668711329 | 0.2155429961 | 1.299363423 | 0.2611697117 | 0.7101763428 |
| -0.743224098 | 0.0935087419 | 0.4325294458 | 0.3774609196 | 0.9929163871 | 0.344803248 | 0.4655801956 |
| -1.465387927 | 0.6594814107 | 0.7167395585 | 0.1074419205 | 0.8958700365 | 0.1814800369 | 0.1576281629 |
| -1.92706551 | 0.1045904916 | 0.3780110119 | 0.1562364456 | 1.404762095 | 0.3390877527 | 0.0676096823 |
| 0.424256454 | 0.1387796716 | 0.6540827459 | 0.1955381939 | 0.5901286446 | 0.3832166978 | 0.6756933987 |
| 0.798558248 | -0.176115799 | 0.385028071 | 0.5539049306 | 1.347595046 | 0.3255702119 | 0.4334854718 |
| -0.787705639 | 0.1859260899 | 0.7662687914 | 0.1874456353 | 1.352485939 | 0.2132624671 | 0.439669961 |
| -1.202420784 | 0.7205277408 | 1.173685282 | -0.382837183 | 0.5281978461 | 0.1643569447 | 0.2425837289 |
| -0.279222421 | 0.2494529844 | 0.685111315 | 0.0738112391 | 1.20957045 | 0.2518228364 | 0.7828055758 |
| 0.8392004314 | 0.2135981143 | 0.6519398299 | 0.2573858303 | 0.6703974138 | 0.3203158469 | 0.4108087078 |
| -0.074089676 | 0.1684447368 | 0.6376072011 | 0.1680110945 | 0.9402499832 | 0.3226767332 | 0.9416402419 |
| 0.2202244385 | 0.2588012935 | 0.6366863068 | 0.0123205846 | 0.9419828456 | 0.2544158725 | 0.8278231804 |
| -0.228123377 | 0.2048632804 | 0.6734092321 | 0.1655786524 | 1.255708362 | 0.2533844132 | 0.8217569205 |
| -1.702033883 | -0.062535239 | 0.8547931125 | 0.4483382171 | 0.472504392 | 0.3466059998 | 0.103508377 |
| 0.4805444686 | 0.215824305 | 0.7399636256 | 0.0733600081 | 1.328101461 | 0.2223719648 | 0.6358085023 |
| 0.3336691597 | 0.3078751915 | 0.6495442029 | 0.0729817859 | 1.289758194 | 0.2407655416 | 0.7419380615 |
| 0.4884155013 | 0.2392900313 | 0.6597191541 | 0.0027472236 | 1.310758749 | 0.2279418744 | 0.6303168527 |
| 0.2822148972 | 0.1669220217 | 0.653642916 | 0.2419175204 | 1.28770251 | 0.2460483663 | 0.7805414054 |
| 2.380298205 | -0.303512224 | 0.7993397039 | 0.5669373099 | 2.619711937 | 0.06506776697 | 0.0268495505 |
| -0.268155088 | 0.0154630776 | 0.250680063 | 0.3043708722 | 1.271740352 | 0.3722148868 | 0.7911963677 |
| -0.840229945 | -0.083166168 | 0.4148395214 | 0.46070386 | 1.327741792 | 0.3227087887 | 0.4102442496 |
| -0.072204430 | 0.1390645524 | 0.6307905659 | 0.1776135149 | 1.252423669 | 0.2905277405 | 0.9431224967 |
| -0.288910688 | 0.1237624304 | 0.5408667844 | 0.2028523697 | 1.192552937 | 0.3228635038 | 0.7754824814 |
| 0.7917506391 | 0.0568987886 | 0.2572270159 | 0.2284536697 | 0.6976521348 | 0.5702719989 | 0.4373585409 |
| -0.828619605 | 0.1936888169 | 0.6587500426 | 0.2436738393 | 0.8844293288 | 0.291942908 | 0.4166385605 |
| -2.12061281 | 0.3117365239 | 0.2672842589 | -0.239685406 | 0.5329940513 | 0.6391320087 | 0.0460290032 |
| 1.198404007 | 0.5542169049 | 0.9535799875 | 0.0537700532 | 1.182481649 | 0.1483998961 | 0.2441098883 |
| -1.204086557 | 0.0503689934 | 0.5109223458 | 0.3151283954 | 1.133086555 | 0.3289203945 | 0.24195294 |
| 0.8495098475 | 0.1606687849 | 0.636074562 | 0.0186666699 | 1.459250804 | 0.2552744505 | 0.4051786384 |
| 0.3440851382 | 0.0898989836 | 0.6553483418 | 0.1697697585 | 1.262913162 | 0.2863044817 | 0.7342038304 |
| 1.118360637 | 0.7658542036 | 0.8080069232 | -0.007365852 | 0.7478455484 | 0.1629046821 | 0.2760436028 |
| 0.4928579722 | 0.0718311278 | 0.5537387609 | 0.253912357 | 1.327631133 | 0.2416824003 | 0.6272269425 |
| -1.5363527 | 0.1750236073 | 0.2045671549 | 0.5369861264 | 0.9092302965 | 0.3700148607 | 0.1393828228 |
| -1.066598718 | 0.1277007413 | 0.6191721818 | 0.3190383981 | 1.022585414 | 0.2869296097 | 0.2982624789 |
| -1.340409195 | -0.139180979 | 0.2396475673 | 0.6466846811 | 0.3108206269 | 0.605547194 | 0.1944347053 |
| -0.997957680 | 0.1927827683 | 0.474536551 | 0.2635015945 | 0.7409469317 | 0.3806201919 | 0.3296609601 |
| 1.652558436 | -0.711075098 | 0.0996596738 | 0.7756019378 | 0.9742661959 | 0.7706015802 | 0.1132949128 |
| 1.242635326 | -0.149717859 | 0.3054651681 | 0.5548780955 | 1.424440857 | 0.3367112797 | 0.2276994246 |
| -0.846932200 | 0.0183809176 | 0.7090532725 | 0.2435182125 | 1.454217675 | 0.2322691914 | 0.4065816592 |
| 0.7068452854 | 0.0515213954 | 0.3244825552 | 0.2964393816 | 1.169372664 | 0.4013326147 | 0.4874325302 |
| -0.393510773 | 0.3214279623 | 0.7345039117 | 0.0036202793 | 1.252441588 | 0.2344039723 | 0.6979090696 |
| -0.661673797 | 0.1833709149 | 0.668924896 | 0.1177193847 | 1.122471595 | 0.2852700399 | 0.5153757219 |
| -2.129277161 | -0.053777016 | 0.1494761509 | 0.0933817876 | 1.041474051 | 0.6081501543 | 0.0452271417 |
| 0.8501040976 | 0.2934920835 | 0.7938067561 | 0.0936485921 | 1.533128427 | 0.171505748 | 0.4048556272 |

| | | | | | | |
|---|---|---|---|---|---|---|
| 1.261052512 | -0.13260191 | 1.173897682 | -0.22257160 | 1.651896718 | 0.1845233252 | 0.22112019 |
| 0.279096343 | 0.05385563 | 0.649749417 | 0.176615555 | 1.025721333 | 0.3630640965 | 0.782901012 |
| 0.699215272 | 0.281113613 | 0.872346491 | -0.043273381 | 1.168005907 | 0.2191776834 | 0.492090141 |
| -0.97558252 | 0.168163421 | 0.660738664 | 0.273804413 | 1.05588342 | 0.2675144523 | 0.340376455 |
| -1.33596555 | 0.325958979 | 0.695314744 | 0.199588222 | 1.860397987 | 0.1153484049 | 0.195858932 |
| -0.15815586 | 0.187657388 | 0.664148398 | 0.148225310 | 1.250437562 | 0.2556595219 | 0.875844946 |
| 0.852158676 | 0.348371737 | 0.820803038 | -0.04775927 | 1.048318978 | 0.2367472706 | 0.403740111 |
| 0.630443431 | 0.444482555 | 0.876777717 | -0.10948146 | 0.664618938 | 0.2221069439 | 0.535205558 |
| 1.235814588 | 0.440770898 | 0.938798127 | -0.159057317 | 1.237146822 | 0.1875163294 | 0.230173648 |
| 1.235348085 | 0.290324803 | 0.943811973 | -0.062583817 | 1.026611635 | 0.2276074775 | 0.230343618 |
| -1.22592143 | 0.328772285 | 0.656252139 | 0.189744934 | 1.777746659 | 0.1315818558 | 0.233798756 |
| 0.173811517 | 0.188861427 | 0.674237340 | 0.120069968 | 1.266891167 | 0.2544961547 | 0.863678183 |
| 0.907368163 | 0.206753149 | 0.381322958 | 0.376278549 | 1.130262228 | 0.2973547458 | 0.374507152 |
| 0.145411516 | 0.181418049 | 0.654225509 | 0.154997505 | 1.152478554 | 0.2723720261 | 0.885772645 |
| -1.81465968 | 0.559005941 | 0.955001864 | 0.166796338 | 0.870765328 | 0.1553516045 | 0.083888371 |
| -0.13214264 | 0.182767642 | 0.664434071 | 0.161187070 | 1.064463498 | 0.2654735542 | 0.896129394 |
| 0.977923800 | 0.273484639 | 0.916834661 | -0.07703066 | 1.053975442 | 0.2338117846 | 0.339244118 |
| -1.33103088 | 0.445764568 | 1.027711959 | -0.06455079 | 1.210604661 | 0.1597935083 | 0.197450135 |
| -0.10278770 | 0.193030490 | 0.639946889 | 0.134044693 | 1.220915395 | 0.2608599644 | 0.919106702 |
| 1.287419677 | -0.13922960 | 0.504497753 | 0.31994094 | 0.871240354 | 0.466556686 | 0.211956697 |
| -0.06884262 | 0.183596705 | 0.654580172 | 0.162258835 | 1.169744983 | 0.2739226549 | 0.945766202 |
| 0.420597881 | 0.095764206 | 0.644711684 | 0.311765311 | 1.323815402 | 0.2381531031 | 0.678321613 |
| 0.890149429 | 0.163363847 | 0.668726127 | 0.314536851 | 1.471873322 | 0.1988350879 | 0.383470318 |
| 0.614363008 | 0.421663685 | 0.705188083 | 0.146856429 | 0.922514088 | 0.222133217 | 0.545574730 |
| 1.171214928 | 0.114843311 | 0.331223082 | 0.089650158 | 0.92015318 | 0.4914447635 | 0.254630773 |
| 0.790799489 | 0.269224519 | 0.672629751 | 0.034032024 | 1.312783416 | 0.2461404506 | 0.437901377 |
| -1.33422195 | 0.517801813 | 0.087243601 | 0.184823247 | 1.558588279 | 0.2493596118 | 0.196420006 |
| -0.60142573 | 0.027934087 | 0.606711541 | 0.150150408 | 0.8892206 | 0.4149983728 | 0.553994087 |
| -0.00232328 | 0.148109868 | 0.646420034 | 0.169641156 | 0.997426824 | 0.2657615721 | 0.998168218 |
| 1.867324302 | 0.357311148 | -0.11627047 | 0.144573196 | 0.267044959 | 0.7469547729 | 0.075878500 |
| 0.950127358 | 0.684077991 | 0.742002534 | 0.072110496 | 1.361483572 | 0.1526049493 | 0.352855154 |
| 0.242316034 | 0.100852525 | 0.654476629 | 0.208720765 | 1.194742018 | 0.2842186029 | 0.810886107 |
| -1.32725454 | 0.453829356 | 0.873475239 | 0.269926471 | 1.160288865 | 0.1531484299 | 0.198674672 |
| 0.122491985 | 0.139798743 | 0.656839206 | 0.190798417 | 1.253339122 | 0.2622024263 | 0.903674075 |
| 1.026278324 | 0.719236070 | 1.009837403 | -0.26576982 | 0.923614256 | 0.1565210068 | 0.316437422 |
| -0.65749047 | 0.175926399 | 0.747272976 | 0.031963581 | 1.422781281 | 0.2217069092 | 0.518008025 |
| -2.01588025 | 0.282097674 | 0.983122998 | 0.188232961 | 1.644864254 | 0.1226658874 | 0.056786019 |
| -0.01604049 | 0.151483696 | 0.634646537 | 0.170317187 | 0.995637044 | 0.2633664144 | 0.987353503 |
| -0.91280550 | 0.545504912 | 0.793517403 | -0.08524972 | 0.892666720 | 0.2140026959 | 0.371705844 |
| 0.745298922 | 0.570529289 | 0.739414493 | -0.082007675 | 0.908493034 | 0.2076024718 | 0.464351695 |
| 0.511257327 | 0.095942849 | 0.656706285 | 0.218387125 | 1.329777614 | 0.2543155818 | 0.614504504 |
| -2.33746808 | 0.521199943 | 0.733970891 | -0.54604083 | 0.744416813 | 0.4003879066 | 0.029394954 |
| -0.61137285 | -0.15454589 | 0.345104911 | 0.368707887 | 1.385262387 | 0.3233943031 | 0.547514611 |
| 0.697461535 | 0.381992229 | 0.780560097 | -0.10635039 | 1.430645051 | 0.1896064452 | 0.493164290 |
| 0.374079234 | 0.009709058 | 0.551589169 | 0.272708443 | 1.166558593 | 0.3509633057 | 0.712095773 |
| 1.431913881 | 0.816074231 | 0.839890926 | -0.15829363 | 1.883657849 | 0.09202213254 | 0.166885264 |

| | | | | | | |
|---|---|---|---|---|---|---|
| -1.150101141 | 0.691648198 | 0.703913184 | 0.107628269 | 0.629632672 | 0.1959186759 | 0.263031466 |
| -1.426489391 | 0.896117675 | 0.740608123 | 0.017554677 | 1.317892642 | 0.1149057812 | 0.168425877 |

| t_p_blast | t_p_vns | t_p_interact | t_p_batch | t_q_microbe | t_q_blast | t_q_vns |
|---|---|---|---|---|---|---|
| 0.000002559 | 0.594768037 | 0.3261210949 | 0.027529537 | 0.975076637 | 0.000005033 | 0.702397450 |
| 0.000003053 | 0.434927067 | 0.4801742436 | 0.028649348 | 0.975076637 | 0.000005188 | 0.702397450 |
| 0.000003178 | 0.570352589 | 0.4067835421 | 0.029468605 | 0.975076637 | 0.000005297 | 0.702397450 |
| 0.000010120 | 0.590840601 | 0.6223401172 | 0.034114898 | 0.990709099 | 0.000011906 | 0.702397450 |
| 0.000002501 | 0.507066849 | 0.4525996413 | 0.122936335 | 0.990709099 | 0.000005033 | 0.702397450 |
| 0.000014875 | 0.388497092 | 0.6588031943 | 0.029562960 | 0.847656972 | 0.000016377 | 0.702397450 |
| 0.000014146 | 0.424331081 | 0.5569302067 | 0.052989780 | 0.975076637 | 0.000015865 | 0.702397450 |
| 0.000000426 | 0.703245748 | 0.1732235146 | 0.004410470 | 0.532640168 | 0.000005033 | 0.775145902 |
| 0.000027584 | 0.424898727 | 0.5619357663 | 0.030644382 | 0.975076637 | 0.000028984 | 0.702397450 |
| 0.000007860 | 0.422219059 | 0.5254753497 | 0.036311601 | 0.975076637 | 0.000009724 | 0.702397450 |
| 0.000010443 | 0.436794515 | 0.5353459757 | 0.030898899 | 0.975076637 | 0.000012166 | 0.702397450 |
| 0.000005159 | 0.234692208 | 0.5305100643 | 0.016195358 | 0.725565304 | 0.000007371 | 0.702397450 |
| 0.000005624 | 0.595205321 | 0.3537506627 | 0.028773841 | 0.975076637 | 0.000007791 | 0.702397450 |
| 0.000001577 | 0.10062636 | 0.7752056937 | 0.605118199 | 0.265646546 | 0.000005033 | 0.702397450 |
| 0.000002289 | 0.574176524 | 0.4185409641 | 0.034633023 | 0.975076637 | 0.000005033 | 0.702397450 |
| 0.000013408 | 0.500562598 | 0.4559034085 | 0.055026614 | 0.987880558 | 0.000015323 | 0.702397450 |
| 0.000007968 | 0.500253391 | 0.4902636879 | 0.036819829 | 0.990709099 | 0.000009757 | 0.702397450 |
| 0.000002810 | 0.530812388 | 0.4174284704 | 0.033726062 | 0.985409022 | 0.000005033 | 0.702397450 |
| 0.000002769 | 0.502057856 | 0.3975183258 | 0.031108202 | 0.975076637 | 0.000005033 | 0.702397450 |
| 0.000002460 | 0.461123444 | 0.4968081775 | 0.032613410 | 0.975076637 | 0.000005033 | 0.702397450 |
| 0.000005902 | 0.583920538 | 0.4327182837 | 0.114095757 | 0.987880558 | 0.000008048 | 0.702397450 |
| 0.000032950 | 0.146192159 | 0.7004955247 | 0.964962110 | 0.532640168 | 0.000033508 | 0.702397450 |
| 0.000001612 | 0.798606576 | 0.237995127 | 0.059299879 | 0.684817630 | 0.000005033 | 0.855649903 |
| 0.000000906 | 0.972545031 | 0.1339431366 | 0.014445235 | 0.684817630 | 0.000005033 | 0.972545031 |
| 0.000010047 | 0.457759699 | 0.5379055499 | 0.047329901 | 0.975076637 | 0.000011906 | 0.702397450 |
| 0.000002429 | 0.502541619 | 0.4506927005 | 0.034302267 | 0.990709099 | 0.000005033 | 0.702397450 |
| 0.000006531 | 0.448198735 | 0.5069442048 | 0.031596855 | 0.975076637 | 0.000008519 | 0.702397450 |
| 0.000012542 | 0.595058506 | 0.4739419473 | 0.051007488 | 0.987972281 | 0.000014471 | 0.702397450 |
| 0.000001281 | 0.463960466 | 0.362551717 | 0.043372783 | 0.910196246 | 0.000005033 | 0.702397450 |
| 0.000002639 | 0.391385039 | 0.604558244 | 0.063148713 | 0.975076637 | 0.000005033 | 0.702397450 |
| 0.000006800 | 0.497163893 | 0.459553832 | 0.033011219 | 0.975076637 | 0.000008681 | 0.702397450 |
| 0.000002184 | 0.556437223 | 0.4522537525 | 0.034276231 | 0.975076637 | 0.000005033 | 0.702397450 |
| 0.000002538 | 0.503055075 | 0.4496415629 | 0.117034018 | 0.990709099 | 0.000005033 | 0.702397450 |
| 0.000001052 | 0.835932586 | 0.151426727 | 0.037421318 | 0.684817630 | 0.000005033 | 0.872240312 |
| 0.000001859 | 0.420858953 | 0.4313404281 | 0.039728548 | 0.975076637 | 0.000005033 | 0.702397450 |
| 0.000000444 | 0.723667317 | 0.1066410966 | 0.273415956 | 0.532640168 | 0.000005033 | 0.783736364 |
| 0.000005415 | 0.476621918 | 0.5390501766 | 0.032127895 | 0.975076637 | 0.000007644 | 0.702397450 |
| 0.000001568 | 0.485119889 | 0.5050325809 | 0.156461029 | 0.975076637 | 0.000005033 | 0.702397450 |
| 0.000002592 | 0.502252300 | 0.4500012858 | 0.095343915 | 0.990709099 | 0.000005033 | 0.702397450 |
| 0.000023662 | 0.357165180 | 0.793739578 | 0.033565616 | 0.975076637 | 0.000025128 | 0.702397450 |
| 0.000001405 | 0.570832102 | 0.4277901629 | 0.018835933 | 0.975076637 | 0.000005033 | 0.702397450 |
| 0.000000311 | 0.345200456 | 0.2460471583 | 0.004084425 | 0.532640168 | 0.000005033 | 0.702397450 |
| 0.000002595 | 0.456207973 | 0.5085072178 | 0.100100908 | 0.975076637 | 0.000005033 | 0.702397450 |
| 0.000027777 | 0.490407161 | 0.7122537506 | 0.040979692 | 0.704539802 | 0.000028984 | 0.702397450 |
| 0.000001090 | 0.494789959 | 0.2519376767 | 0.014324952 | 0.847656972 | 0.000005033 | 0.702397450 |
| 0.000000358 | 0.444207036 | 0.14928759 | 0.051132851 | 0.532640168 | 0.000005033 | 0.702397450 |

| | | | | | | |
|---|---|---|---|---|---|---|
| 0.000000357 | 0.430746303 | 0.2597743306 | 0.583583141 | 0.532640168 | 0.000005033 | 0.702397450 |
| 0.000041791 | 0.263436687 | 0.9874747598 | 0.021411490 | 0.975076637 | 0.000042142 | 0.702397450 |
| 0.000002804 | 0.620336594 | 0.3586069806 | 0.036116589 | 0.975076637 | 0.000005033 | 0.715772994 |
| 0.000009849 | 0.401966128 | 0.5231404027 | 0.032525627 | 0.975076637 | 0.000011819 | 0.702397450 |
| 0.000002134 | 0.631578875 | 0.4107744012 | 0.028720474 | 0.975076637 | 0.000005033 | 0.721804429 |
| 0.000001231 | 0.897331603 | 0.3909921768 | 0.016307312 | 0.910196246 | 0.000005033 | 0.920340106 |
| 0.000002404 | 0.502950068 | 0.4381343621 | 0.038456795 | 0.987880558 | 0.000005033 | 0.702397450 |
| 0.000001345 | 0.361232356 | 0.3211889101 | 0.093305624 | 0.910196246 | 0.000005033 | 0.702397450 |
| 0.000006011 | 0.477802005 | 0.4669023204 | 0.033164249 | 0.987880558 | 0.000008086 | 0.702397450 |
| 0.000002744 | 0.466599264 | 0.4845435713 | 0.039459613 | 0.975076637 | 0.000005033 | 0.702397450 |
| 0.000002388 | 0.49612468 | 0.4234180864 | 0.033695133 | 0.975076637 | 0.000005033 | 0.702397450 |
| 0.000004265 | 0.503295071 | 0.4486001376 | 0.033340596 | 0.975076637 | 0.000006443 | 0.702397450 |
| 0.000020115 | 0.523287790 | 0.4361788076 | 0.058643208 | 0.987880558 | 0.000021552 | 0.702397450 |
| 0.000001904 | 0.596003139 | 0.3772690542 | 0.145984612 | 0.975076637 | 0.000005033 | 0.702397450 |
| 0.000001086 | 0.819370487 | 0.2609067712 | 0.013531605 | 0.684817630 | 0.000005033 | 0.862495250 |
| 0.000001855 | 0.519627299 | 0.3891626298 | 0.025150147 | 0.975076637 | 0.000005033 | 0.702397450 |
| 0.000004223 | 0.301621841 | 0.7761013206 | 0.240928806 | 0.847656972 | 0.000006443 | 0.702397450 |
| 0.000001964 | 0.602891144 | 0.4020767454 | 0.023471842 | 0.975076637 | 0.000005033 | 0.702397450 |
| 0.000006764 | 0.704090861 | 0.3197802594 | 0.025149141 | 0.975076637 | 0.000008681 | 0.775145902 |
| 0.000004295 | 0.496809569 | 0.4980405897 | 0.034166399 | 0.990709099 | 0.000006443 | 0.702397450 |
| 0.000002921 | 0.498785863 | 0.44529818 | 0.043508211 | 0.990709099 | 0.000005081 | 0.702397450 |
| 0.000003379 | 0.419411653 | 0.5089526975 | 0.039507842 | 0.975076637 | 0.000005554 | 0.702397450 |
| 0.000003069 | 0.664115292 | 0.3037372915 | 0.032642384 | 0.975076637 | 0.000005188 | 0.744802197 |
| 0.000002291 | 0.496382231 | 0.4712817989 | 0.030568829 | 0.975076637 | 0.000005033 | 0.702397450 |
| 0.000002781 | 0.429973740 | 0.433884347 | 0.041756660 | 0.975076637 | 0.000005033 | 0.702397450 |
| 0.000002589 | 0.528653112 | 0.441881542 | 0.056938803 | 0.987880558 | 0.000005033 | 0.702397450 |
| 0.000004605 | 0.913637358 | 0.2817440336 | 0.017485892 | 0.920225808 | 0.000006822 | 0.929122737 |
| 0.000007646 | 0.502988383 | 0.4510290446 | 0.053336157 | 0.990709099 | 0.000009558 | 0.702397450 |
| 0.000002914 | 0.519947004 | 0.4816094594 | 0.035122530 | 0.990709099 | 0.000005081 | 0.702397450 |
| 0.000002028 | 0.500381846 | 0.4070450725 | 0.026607227 | 0.975076637 | 0.000005033 | 0.702397450 |
| 0.000001397 | 0.457534317 | 0.4002856266 | 0.499481836 | 0.645294277 | 0.000005033 | 0.702397450 |
| 0.000002536 | 0.469736810 | 0.4916747822 | 0.053726651 | 0.975076637 | 0.000005033 | 0.702397450 |
| 0.000000479 | 0.139272078 | 0.9532655834 | 0.001882116 | 0.265427932 | 0.000005033 | 0.702397450 |
| 0.000028907 | 0.187339993 | 0.9318407499 | 0.009754459 | 0.684817630 | 0.000029904 | 0.702397450 |
| 0.000003842 | 0.460968316 | 0.5264992468 | 0.032697491 | 0.975076637 | 0.000005987 | 0.702397450 |
| 0.000002626 | 0.451797612 | 0.5067325255 | 0.030853151 | 0.975076637 | 0.000005033 | 0.702397450 |
| 0.000002596 | 0.501889318 | 0.4512923956 | 0.076207596 | 0.990709099 | 0.000005033 | 0.702397450 |
| 0.000001005 | 0.239540344 | 0.7647552737 | 0.034967662 | 0.543435515 | 0.000005033 | 0.702397450 |
| 0.000002441 | 0.561905026 | 0.431340729 | 0.033461098 | 0.987880558 | 0.000005033 | 0.702397450 |
| 0.000002374 | 0.494774612 | 0.4776285161 | 0.030847501 | 0.975076637 | 0.000005033 | 0.702397450 |
| 0.000003831 | 0.42433127 | 0.446810868 | 0.023064404 | 0.975076637 | 0.000005987 | 0.702397450 |
| 0.000002433 | 0.454563051 | 0.4669784906 | 0.034809818 | 0.975076637 | 0.000005033 | 0.702397450 |
| 0.000002633 | 0.508264255 | 0.4753517256 | 0.036826884 | 0.990709099 | 0.000005033 | 0.702397450 |
| 0.000002789 | 0.255985143 | 0.6178925064 | 0.019949228 | 0.684817630 | 0.000005033 | 0.702397450 |
| 0.000003769 | 0.321806266 | 0.633402726 | 0.023662243 | 0.975076637 | 0.000005987 | 0.702397450 |
| 0.000002518 | 0.548738527 | 0.415175748 | 0.029164537 | 0.975076637 | 0.000005033 | 0.702397450 |

| | | | | | | |
|---|---|---|---|---|---|---|
| 0.000001611 | 0.342762109 | 0.5263010378 | 0.164456207 | 0.627889526 | 0.000005033 | 0.702397450 |
| 0.000001542 | 0.506618390 | 0.291054277 | 0.056364289 | 0.975076637 | 0.000005033 | 0.702397450 |
| 0.000000996 | 0.472400312 | 0.2999215237 | 0.075040437 | 0.684817630 | 0.000005033 | 0.702397450 |
| 0.000007143 | 0.517905534 | 0.4438805168 | 0.061329709 | 0.987880558 | 0.000009023 | 0.702397450 |
| 0.000002164 | 0.601992310 | 0.469933493 | 0.030449670 | 0.975076637 | 0.000005033 | 0.702397450 |
| 0.000002063 | 0.505261917 | 0.3981589651 | 0.031247823 | 0.975076637 | 0.000005033 | 0.702397450 |
| 0.000002683 | 0.811505581 | 0.389503763 | 0.059533956 | 0.532640168 | 0.000005033 | 0.861775839 |
| 0.000003734 | 0.503057891 | 0.4503011761 | 0.054516484 | 0.990709099 | 0.000005987 | 0.702397450 |
| 0.000009546 | 0.545210399 | 0.4317272302 | 0.160100467 | 0.975076637 | 0.000011570 | 0.702397450 |
| 0.000001637 | 0.970603734 | 0.4450205518 | 0.010132874 | 0.684817630 | 0.000005033 | 0.972545031 |
| 0.000031225 | 0.504183210 | 0.453566819 | 0.035025110 | 0.99399338 | 0.000032026 | 0.702397450 |
| 0.000005648 | 0.503235090 | 0.5187840725 | 0.044407378 | 0.975076637 | 0.000007791 | 0.702397450 |
| 0.000002789 | 0.334331371 | 0.3560899174 | 0.017038285 | 0.675543650 | 0.000005033 | 0.702397450 |
| 0.000004808 | 0.525412890 | 0.5626360835 | 0.032264485 | 0.975076637 | 0.000007037 | 0.702397450 |
| 0.000015689 | 0.641368468 | 0.3918093207 | 0.053335360 | 0.975076637 | 0.000016962 | 0.726077511 |
| 0.000000199 | 0.235678913 | 0.7724582138 | 0.185789619 | 0.265646546 | 0.000005033 | 0.702397450 |
| 0.000001506 | 0.380138423 | 0.4298105966 | 0.044030625 | 0.773330049 | 0.000005033 | 0.702397450 |
| 0.000000686 | 0.724956137 | 0.3699135951 | 0.435083726 | 0.645294277 | 0.000005033 | 0.783736364 |
| 0.000006436 | 0.538662622 | 0.4140992809 | 0.056291405 | 0.975076637 | 0.000008487 | 0.702397450 |
| 0.000163732 | 0.322775572 | 0.949397255 | 0.004433364 | 0.532640168 | 0.000163732 | 0.702397450 |
| 0.000002318 | 0.500263463 | 0.4265978687 | 0.041661786 | 0.975076637 | 0.000005033 | 0.702397450 |
| 0.000002589 | 0.502861158 | 0.4362277398 | 0.044871771 | 0.987880558 | 0.000005033 | 0.702397450 |
| 0.000013663 | 0.596001841 | 0.4374794452 | 0.099023670 | 0.987880558 | 0.000015468 | 0.702397450 |
| 0.000000037 | 0.843165635 | 0.0501488580 | 0.000473937 | 0.265427932 | 0.000004538 | 0.872240312 |
| 0.000004943 | 0.583297110 | 0.3877243463 | 0.030038015 | 0.975076637 | 0.000007146 | 0.702397450 |
| 0.000006065 | 0.542525358 | 0.3879380808 | 0.038159511 | 0.975076637 | 0.000008086 | 0.702397450 |
| 0.000014773 | 0.499969457 | 0.4568366339 | 0.048496371 | 0.987972281 | 0.000016377 | 0.702397450 |
| 0.000015307 | 0.508360755 | 0.4396909193 | 0.034120857 | 0.987880558 | 0.000016699 | 0.702397450 |
| 0.035306090 | 0.663560117 | 0.2702686803 | 0.069153692 | 0.970329114 | 0.065758938 | 0.977261916 |
| 0.041610375 | 0.810650752 | 0.3181318793 | 0.090296997 | 0.970329114 | 0.065758938 | 0.977261916 |
| 0.035175563 | 0.722787804 | 0.3077590722 | 0.070163114 | 0.974344026 | 0.065758938 | 0.977261916 |
| 0.031806634 | 0.512559539 | 0.2544535346 | 0.060165518 | 0.970329114 | 0.065758938 | 0.977261916 |
| 0.036358233 | 0.615271434 | 0.2724434445 | 0.354482159 | 0.970329114 | 0.065758938 | 0.977261916 |
| 0.018342793 | 0.592088725 | 0.1947722466 | 0.059088196 | 0.970329114 | 0.065758938 | 0.977261916 |
| 0.074584307 | 0.819917360 | 0.3714314294 | 0.057501203 | 0.970329114 | 0.085239208 | 0.977261916 |
| 0.040638694 | 0.822880021 | 0.4505384171 | 0.039880546 | 0.970329114 | 0.065758938 | 0.977261916 |
| 0.060578312 | 0.709889408 | 0.3110006297 | 0.064829503 | 0.985054686 | 0.074177525 | 0.977261916 |
| 0.029261170 | 0.597019979 | 0.2529179719 | 0.067770710 | 0.970329114 | 0.065758938 | 0.977261916 |
| 0.031647820 | 0.595372117 | 0.2511636376 | 0.054589130 | 0.970329114 | 0.065758938 | 0.977261916 |
| 0.010462418 | 0.297341721 | 0.1940576845 | 0.025565705 | 0.970329114 | 0.065758938 | 0.977261916 |
| 0.042858228 | 0.663582521 | 0.2817291137 | 0.072797335 | 0.970329114 | 0.065758938 | 0.977261916 |
| 0.001966952 | 0.156911932 | 0.0230276756 | 0.897247569 | 0.151382718 | 0.065758938 | 0.977261916 |
| 0.033721952 | 0.669394152 | 0.2833689368 | 0.091987152 | 0.974344026 | 0.065758938 | 0.977261916 |
| 0.051870191 | 0.702853651 | 0.2937040734 | 0.101069268 | 0.974344026 | 0.068329646 | 0.977261916 |
| 0.090420296 | 0.900582000 | 0.4496217357 | 0.047814281 | 0.970329114 | 0.096540743 | 0.981544683 |
| 0.055355158 | 0.819585542 | 0.6348131203 | 0.033016264 | 0.970329114 | 0.071426011 | 0.977261916 |

| | | | | | | |
|---|---|---|---|---|---|---|
| 0.062624631 | 0.697208018 | 0.4667411733 | 0.033117493 | 0.970329114 | 0.075149557 | 0.977261916 |
| 0.046294564 | 0.530746703 | 0.2142476599 | 0.051397930 | 0.970329114 | 0.065758938 | 0.977261916 |
| 0.080612022 | 0.923965279 | 0.3902586029 | 0.281907546 | 0.970329114 | 0.088851529 | 0.981544683 |
| 0.011603818 | 0.374850554 | 0.1035758287 | 0.821535971 | 0.970329114 | 0.065758938 | 0.977261916 |
| 0.032400670 | 0.784029985 | 0.3569400004 | 0.080010196 | 0.970329114 | 0.065758938 | 0.977261916 |
| 0.101995969 | 0.954418255 | 0.7469171922 | 0.040759905 | 0.970329114 | 0.104611250 | 0.981544683 |
| 0.063452390 | 0.891040487 | 0.3672533032 | 0.061263528 | 0.970329114 | 0.075370672 | 0.981544683 |
| 0.032980634 | 0.678686883 | 0.2882063292 | 0.059391121 | 0.970329114 | 0.065758938 | 0.977261916 |
| 0.031062954 | 0.618331061 | 0.2615288684 | 0.055899450 | 0.970329114 | 0.065758938 | 0.977261916 |
| 0.087166909 | 0.863402614 | 0.5051557282 | 0.102634207 | 0.970329114 | 0.094234496 | 0.981544683 |
| 0.061634162 | 0.520265022 | 0.3688810729 | 0.058506823 | 0.092174968 | 0.074708075 | 0.977261916 |
| 0.029774952 | 0.601157781 | 0.2424610926 | 0.102116887 | 0.970329114 | 0.065758938 | 0.977261916 |
| 0.047914903 | 0.705833521 | 0.2976624886 | 0.076621099 | 0.974344026 | 0.066268672 | 0.977261916 |
| 0.031722185 | 0.641100661 | 0.2951779623 | 0.060880747 | 0.970329114 | 0.065758938 | 0.977261916 |
| 0.034131930 | 0.704028990 | 0.2923974035 | 0.210099983 | 0.974344026 | 0.065758938 | 0.977261916 |
| 0.134591367 | 0.967772341 | 0.7953415374 | 0.072286466 | 0.970329114 | 0.136872576 | 0.984175262 |
| 0.033805331 | 0.675070002 | 0.2966440168 | 0.069399115 | 0.970329114 | 0.065758938 | 0.977261916 |
| 0.245754332 | 0.488941682 | 0.9424038683 | 0.441582179 | 0.917086345 | 0.245754332 | 0.977261916 |
| 0.014933708 | 0.562448483 | 0.1515068715 | 0.039245039 | 0.970329114 | 0.065758938 | 0.977261916 |
| 0.034585566 | 0.693855386 | 0.2591052064 | 0.200910540 | 0.970329114 | 0.065758938 | 0.977261916 |
| 0.037489837 | 0.755354120 | 0.2856309412 | 0.070019491 | 0.970329114 | 0.065758938 | 0.977261916 |
| 0.033858315 | 0.520503259 | 0.2429725113 | 0.063811588 | 0.970329114 | 0.065758938 | 0.977261916 |
| 0.042641869 | 0.792259702 | 0.2904289409 | 0.032982644 | 0.970329114 | 0.065758938 | 0.977261916 |
| 0.036444375 | 0.466827025 | 0.4363624684 | 0.002832227 | 0.222418573 | 0.065758938 | 0.977261916 |
| 0.031099220 | 0.633715756 | 0.2597225554 | 0.170210390 | 0.970329114 | 0.065758938 | 0.977261916 |
| 0.031815830 | 0.705890718 | 0.2430451445 | 0.073596361 | 0.970329114 | 0.065758938 | 0.977261916 |
| 0.041691500 | 0.698203093 | 0.6004285641 | 0.013936048 | 0.970329114 | 0.065758938 | 0.977261916 |
| 0.032158226 | 0.691474743 | 0.4644579109 | 0.091309909 | 0.970329114 | 0.065758938 | 0.977261916 |
| 0.058977591 | 0.638629587 | 0.411434353 | 0.753498831 | 0.970329114 | 0.073721989 | 0.977261916 |
| 0.044444180 | 0.604409052 | 0.3066263099 | 0.063016672 | 0.970329114 | 0.065758938 | 0.977261916 |
| 0.048597026 | 0.757220474 | 0.3648469283 | 0.066111600 | 0.974344026 | 0.066268672 | 0.977261916 |
| 0.010010725 | 0.509477902 | 0.1933786452 | 0.055286091 | 0.970329114 | 0.065758938 | 0.977261916 |
| 0.041600623 | 0.799870625 | 0.3172916404 | 0.058653560 | 0.970329114 | 0.065758938 | 0.977261916 |
| 0.044346687 | 0.941599640 | 0.3171043913 | 0.036527350 | 0.970329114 | 0.065758938 | 0.981544683 |
| 0.030697235 | 0.691639566 | 0.2396744746 | 0.163861005 | 0.970329114 | 0.065758938 | 0.977261916 |
| 0.035669047 | 0.636958396 | 0.3474011672 | 0.111451161 | 0.970329114 | 0.065758938 | 0.977261916 |
| 0.071900967 | 0.883031675 | 0.3299265504 | 0.071015770 | 0.970329114 | 0.083768117 | 0.981544683 |
| 0.030574404 | 0.657001986 | 0.2700253924 | 0.073256904 | 0.970329114 | 0.065758938 | 0.977261916 |
| 0.034511691 | 0.706507943 | 0.2940191571 | 0.078095457 | 0.974344026 | 0.065758938 | 0.977261916 |
| 0.017920696 | 0.700622820 | 0.2792835948 | 0.054050866 | 0.970329114 | 0.065758938 | 0.977261916 |
| 0.076246282 | 0.710753617 | 0.3044874655 | 0.089707021 | 0.985054686 | 0.086316546 | 0.977261916 |
| 0.041292753 | 0.736956866 | 0.3184420302 | 0.141085472 | 0.974344026 | 0.065758938 | 0.977261916 |
| 0.029273821 | 0.992725416 | 0.4421999838 | 0.033451521 | 0.970329114 | 0.065758938 | 0.992725416 |
| 0.035226525 | 0.712773323 | 0.3151683746 | 0.062314881 | 0.970329114 | 0.065758938 | 0.977261916 |
| 0.046579248 | 0.804823597 | 0.3961879144 | 0.080901018 | 0.970329114 | 0.065758938 | 0.977261916 |
| 0.008540774 | 0.873757532 | 0.3148494898 | 0.001244036 | 0.120256464 | 0.065758938 | 0.981544683 |

| | | | | | | |
|---|---|---|---|---|---|---|
| 0.146500991 | 0.934160472 | 0.490485021 | 0.046129124 | 0.970329114 | 0.147732092 | 0.981544683 |
| 0.027469788 | 0.589863225 | 0.2335593022 | 0.055348951 | 0.970329114 | 0.065758938 | 0.977261916 |
| 0.052386062 | 0.617814324 | 0.3297209053 | 0.132966249 | 0.970329114 | 0.068329646 | 0.977261916 |
| 0.024852909 | 0.491875927 | 0.2325207846 | 0.076455843 | 0.970329114 | 0.065758938 | 0.977261916 |
| 0.038566977 | 0.649934937 | 0.2748759475 | 0.062733942 | 0.970329114 | 0.065758938 | 0.977261916 |
| 0.031684064 | 0.709919474 | 0.3206532561 | 0.088658989 | 0.970329114 | 0.065758938 | 0.977261916 |
| 0.034202665 | 0.687087170 | 0.297962106 | 0.069738164 | 0.974344026 | 0.065758938 | 0.977261916 |
| 0.040863381 | 0.778670779 | 0.3175819277 | 0.060809689 | 0.970329114 | 0.065758938 | 0.977261916 |
| 0.018883541 | 0.867981451 | 0.4932991707 | 0.032736818 | 0.970329114 | 0.065758938 | 0.981544683 |
| 0.032586178 | 0.699307722 | 0.2874031996 | 0.128821243 | 0.970329114 | 0.065758938 | 0.977261916 |
| 0.044824333 | 0.957006066 | 0.4783523872 | 0.078455310 | 0.970329114 | 0.065758938 | 0.981544683 |
| 0.034295251 | 0.704579628 | 0.3098232029 | 0.061529116 | 0.970329114 | 0.065758938 | 0.977261916 |
| 0.035274156 | 0.704054873 | 0.2951136014 | 0.150646211 | 0.998062105 | 0.065758938 | 0.977261916 |
| 0.022404774 | 0.594349704 | 0.2114062048 | 0.12888917 | 0.970329114 | 0.065758938 | 0.977261916 |
| 0.011023707 | 0.436103557 | 0.1231156833 | 0.020938294 | 0.970329114 | 0.065758938 | 0.977261916 |
| 0.026268948 | 0.489004445 | 0.2047073287 | 0.052361896 | 0.970329114 | 0.065758938 | 0.977261916 |
| 0.036808217 | 0.697173944 | 0.3019218744 | 0.063776484 | 0.974344026 | 0.065758938 | 0.977261916 |
| 0.037675077 | 0.830672628 | 0.3608860706 | 0.086513468 | 0.970329114 | 0.065758938 | 0.977261916 |
| 0.039262639 | 0.697024826 | 0.2895955185 | 0.075871868 | 0.970329114 | 0.065758938 | 0.977261916 |
| 0.028029097 | 0.563914410 | 0.2178033262 | 0.072343711 | 0.970329114 | 0.065758938 | 0.977261916 |
| 0.032373890 | 0.821722008 | 0.3603724486 | 0.056239121 | 0.970329114 | 0.065758938 | 0.977261916 |
| 0.034068828 | 0.702146924 | 0.2917415842 | 0.071023366 | 0.974344026 | 0.065758938 | 0.977261916 |
| 0.051796278 | 0.806809098 | 0.2881170519 | 0.103484274 | 0.970329114 | 0.068329646 | 0.977261916 |
| 0.032005278 | 0.647505320 | 0.280991058 | 0.062628012 | 0.970329114 | 0.065758938 | 0.977261916 |
| 0.038414476 | 0.939098374 | 0.431488471 | 0.095586406 | 0.970329114 | 0.065758938 | 0.981544683 |
| 0.031027766 | 0.628252574 | 0.2682255951 | 0.059998994 | 0.970329114 | 0.065758938 | 0.977261916 |
| 0.048253650 | 0.931135754 | 0.406840291 | 0.080749409 | 0.970329114 | 0.066268672 | 0.981544683 |
| 0.060159829 | 0.803980090 | 0.3410792443 | 0.035916614 | 0.970329114 | 0.074177525 | 0.977261916 |
| 0.036557133 | 0.711351148 | 0.2995057195 | 0.086263353 | 0.985054686 | 0.065758938 | 0.977261916 |
| 0.050114197 | 0.713257031 | 0.5541748632 | 0.107470085 | 0.970329114 | 0.067569705 | 0.977261916 |
| 0.028738538 | 0.683845541 | 0.3903920003 | 0.129133189 | 0.970329114 | 0.065758938 | 0.977261916 |
| 0.058205092 | 0.711361498 | 0.2970929517 | 0.093673958 | 0.974344026 | 0.073522221 | 0.977261916 |
| 0.032984198 | 0.652514255 | 0.3053035153 | 0.087005444 | 0.970329114 | 0.065758938 | 0.977261916 |
| 0.037788271 | 0.706872709 | 0.3247893879 | 0.061898211 | 0.970329114 | 0.065758938 | 0.977261916 |
| 0.011053030 | 0.751546339 | 0.267682739 | 0.108283254 | 0.970329114 | 0.065758938 | 0.977261916 |
| 0.020918067 | 0.639815276 | 0.299315558 | 0.183241990 | 0.970329114 | 0.065758938 | 0.977261916 |
| 0.080706805 | 0.720282618 | 0.2987733014 | 0.162303618 | 0.974344026 | 0.088851529 | 0.977261916 |
| 0.025427135 | 0.949761200 | 0.2741196352 | 0.034574920 | 0.970329114 | 0.065758938 | 0.981544683 |
| 0.090909200 | 0.715488472 | 0.3054474726 | 0.062920677 | 0.974344026 | 0.096540743 | 0.977261916 |
| 0.033594605 | 0.705951379 | 0.2756581409 | 0.075464444 | 0.970329114 | 0.065758938 | 0.977261916 |
| 0.018480440 | 0.563852543 | 0.318301246 | 0.043702351 | 0.970329114 | 0.065758938 | 0.977261916 |
| 0.017981828 | 0.741866320 | 0.1980863916 | 0.058217072 | 0.970329114 | 0.065758938 | 0.977261916 |
| 0.101091840 | 0.814598567 | 0.4037702167 | 0.089703040 | 0.970329114 | 0.104577766 | 0.977261916 |
| 0.031571782 | 0.608935346 | 0.2232574391 | 0.148848963 | 0.970329114 | 0.065758938 | 0.977261916 |
| 0.034939644 | 0.773323607 | 0.2895139605 | 0.052778181 | 0.970329114 | 0.065758938 | 0.977261916 |
| 0.082504675 | 0.787880931 | 0.3044184678 | 0.238082472 | 0.970329114 | 0.090005100 | 0.977261916 |

| | | | | | | |
|---|---|---|---|---|---|---|
| 0.064065071 | 0.733906615 | 0.3447738629 | 0.093365124 | 0.974344026 | 0.075370672 | 0.977261916 |
| 0.027889609 | 0.634654826 | 0.2067588377 | 0.043189739 | 0.970329114 | 0.065758938 | 0.977261916 |
| 0.045906333 | 0.686216099 | 0.3366643848 | 0.089397662 | 0.970329114 | 0.065758938 | 0.977261916 |
| 0.032955011 | 0.703311822 | 0.2805712979 | 0.063247406 | 0.970329114 | 0.065758938 | 0.977261916 |
| 0.097157706 | 0.871140164 | 0.3991441316 | 0.196940374 | 0.970329114 | 0.101381954 | 0.981544683 |
| 0.096131022 | 0.977808718 | 0.7478481454 | 0.008462534 | 0.970329114 | 0.101190550 | 0.986025598 |
| 0.045006001 | 0.686300884 | 0.2960476988 | 0.072253430 | 0.974344026 | 0.065758938 | 0.977261916 |
| 0.078515198 | 0.736410707 | 0.3474457273 | 0.078639215 | 0.970329114 | 0.088054428 | 0.977261916 |
| 0.057627052 | 0.704142074 | 0.295746158 | 0.096084599 | 0.998062105 | 0.073522221 | 0.977261916 |
| 0.073869690 | 0.707478854 | 0.3030893707 | 0.063951599 | 0.974344026 | 0.085234258 | 0.977261916 |
| 0.848376163 | 0.926381114 | 0.5370142169 | 0.068373917 | 0.928186086 | 0.998576232 | 0.996920713 |
| 0.741647338 | 0.830220267 | 0.58146313 | 0.063446288 | 0.869482770 | 0.998576232 | 0.996920713 |
| 0.927847335 | 0.726369145 | 0.6767854736 | 0.132660088 | 0.617647808 | 0.998576232 | 0.996920713 |
| 0.789711089 | 0.418707659 | 0.2049057526 | 0.066334749 | 0.617647808 | 0.998576232 | 0.996920713 |
| 0.726199336 | 0.947972669 | 0.5049511837 | 0.044616514 | 0.696098292 | 0.998576232 | 0.996920713 |
| 0.755053744 | 0.939200035 | 0.3363242416 | 0.056041330 | 0.607876666 | 0.998576232 | 0.996920713 |
| 0.596352723 | 0.742112747 | 0.6988742357 | 0.108699486 | 0.799520930 | 0.998576232 | 0.996920713 |
| 0.775896677 | 0.885254574 | 0.5964811565 | 0.134682129 | 0.949965605 | 0.998576232 | 0.996920713 |
| 0.655407372 | 0.797153209 | 0.6583481694 | 0.064404431 | 0.868904128 | 0.998576232 | 0.996920713 |
| 0.78968279 | 0.740276365 | 0.3546208452 | 0.045018644 | 0.607876666 | 0.998576232 | 0.996920713 |
| 0.601153241 | 0.581771836 | 0.2517843073 | 0.140388093 | 0.421788991 | 0.998576232 | 0.996920713 |
| 0.637192781 | 0.678657552 | 0.6113974925 | 0.052967842 | 0.799520930 | 0.998576232 | 0.996920713 |
| 0.889087972 | 0.941570614 | 0.5276179887 | 0.067780265 | 0.925203233 | 0.998576232 | 0.996920713 |
| 0.986009579 | 0.825934357 | 0.3331782046 | 0.035320719 | 0.660566205 | 0.998576232 | 0.996920713 |
| 0.789791367 | 0.949498187 | 0.5333794447 | 0.073687446 | 0.925203233 | 0.998576232 | 0.996920713 |
| 0.423553956 | 0.924091337 | 0.4356527917 | 0.567450919 | 0.421788991 | 0.998576232 | 0.996920713 |
| 0.40414013 | 0.595325403 | 0.8801434231 | 0.087343406 | 0.607876666 | 0.998576232 | 0.996920713 |
| 0.767620118 | 0.885890722 | 0.631956801 | 0.083894374 | 0.939664415 | 0.998576232 | 0.996920713 |
| 0.547178905 | 0.891121317 | 0.8128702228 | 0.134437829 | 0.617339011 | 0.998576232 | 0.996920713 |
| 0.964644161 | 0.532356117 | 0.7953982609 | 0.037481264 | 0.421788991 | 0.998576232 | 0.996920713 |
| 0.634274255 | 0.741875992 | 0.6625496394 | 0.074222450 | 0.799520930 | 0.998576232 | 0.996920713 |
| 0.872609357 | 0.957984362 | 0.5519775143 | 0.185204050 | 0.928186086 | 0.998576232 | 0.996920713 |
| 0.643351302 | 0.714821938 | 0.2749223051 | 0.122265866 | 0.520349777 | 0.998576232 | 0.996920713 |
| 0.996934533 | 0.890002550 | 0.3958418307 | 0.056071982 | 0.799520930 | 0.998576232 | 0.996920713 |
| 0.726787814 | 0.806482361 | 0.6293062399 | 0.088816560 | 0.925203233 | 0.998576232 | 0.996920713 |
| 0.780298010 | 0.915512730 | 0.5525841778 | 0.070060840 | 0.849231427 | 0.998576232 | 0.996920713 |
| 0.972272723 | 0.941169421 | 0.4702990961 | 0.108228703 | 0.799520930 | 0.998576232 | 0.996920713 |
| 0.461401933 | 0.434577295 | 0.1022998395 | 0.225292601 | 0.421788991 | 0.998576232 | 0.996920713 |
| 0.423824925 | 0.952660798 | 0.7033212118 | 0.025415098 | 0.421788991 | 0.998576232 | 0.996920713 |
| 0.898345296 | 0.801205620 | 0.3268301269 | 0.029945089 | 0.607876666 | 0.998576232 | 0.996920713 |
| 0.507590730 | 0.873845730 | 0.5819089763 | 0.037257485 | 0.617647808 | 0.998576232 | 0.996920713 |
| 0.775787520 | 0.866484482 | 0.5532578527 | 0.068709715 | 0.925203233 | 0.998576232 | 0.996920713 |
| 0.812903246 | 0.898341337 | 0.538767928 | 0.112323587 | 0.868904128 | 0.998576232 | 0.996920713 |
| 0.780415243 | 0.885920550 | 0.6381862112 | 0.070119914 | 0.957050902 | 0.998576232 | 0.996920713 |
| 0.767205329 | 0.755626367 | 0.5255077292 | 0.081521695 | 0.617647808 | 0.998576232 | 0.996920713 |
| 0.554492164 | 0.634341919 | 0.8420643784 | 0.052391958 | 0.799520930 | 0.998576232 | 0.996920713 |

| | | | | | | |
|---|---|---|---|---|---|---|
| 0.895133894 | 0.947751833 | 0.3111580299 | 0.086919660 | 0.607876666 | 0.998576232 | 0.996920713 |
| 0.741102129 | 0.906564511 | 0.4708587042 | 0.032007192 | 0.617647808 | 0.998576232 | 0.996920713 |
| 0.790877903 | 0.907177060 | 0.5575670521 | 0.135917817 | 0.957050902 | 0.998576232 | 0.996920713 |
| 0.532968785 | 0.531727001 | 0.9801609216 | 0.050211742 | 0.799520930 | 0.998576232 | 0.996920713 |
| 0.647397127 | 0.779916950 | 0.55673374 | 0.179588948 | 0.617339011 | 0.998576232 | 0.996920713 |
| 0.425734792 | 0.876819540 | 0.8376983541 | 0.488533632 | 0.421788991 | 0.998576232 | 0.996920713 |
| 0.767596792 | 0.855958277 | 0.5997664642 | 0.150575714 | 0.925203233 | 0.998576232 | 0.996920713 |
| 0.517009269 | 0.903796195 | 0.7364883599 | 0.082969397 | 0.775100997 | 0.998576232 | 0.996920713 |
| 0.582878091 | 0.874265567 | 0.9801380931 | 0.591048594 | 0.495086125 | 0.998576232 | 0.996920713 |
| 0.761331406 | 0.900514373 | 0.7294485169 | 0.051709983 | 0.799520930 | 0.998576232 | 0.996920713 |
| 0.889842151 | 0.884271429 | 0.49338576 | 0.303135881 | 0.799520930 | 0.998576232 | 0.996920713 |
| 0.470804688 | 0.435542535 | 0.8628573516 | 0.041908158 | 0.758952824 | 0.998576232 | 0.996920713 |
| 0.834363578 | 0.938627312 | 0.5414422317 | 0.071773600 | 0.928186086 | 0.998576232 | 0.996920713 |
| 0.730305131 | 0.867822205 | 0.5807669886 | 0.070558816 | 0.925203233 | 0.998576232 | 0.996920713 |
| 0.865903271 | 0.959718332 | 0.5101053142 | 0.058571787 | 0.799520930 | 0.998576232 | 0.996920713 |
| 0.705463813 | 0.678550779 | 0.5862140645 | 0.204459162 | 0.799520930 | 0.998576232 | 0.996920713 |
| 0.774288176 | 0.899728221 | 0.519852703 | 0.056328200 | 0.799520930 | 0.998576232 | 0.996920713 |
| 0.758841233 | 0.994243649 | 0.6401418611 | 0.057839819 | 0.799520930 | 0.998576232 | 0.996920713 |
| 0.282968063 | 0.445752332 | 0.6751772978 | 0.025766111 | 0.421788991 | 0.998576232 | 0.996920713 |
| 0.888256118 | 0.967883762 | 0.4472639452 | 0.043927809 | 0.617339011 | 0.998576232 | 0.996920713 |
| 0.773356780 | 0.883893768 | 0.4911859164 | 0.056544927 | 0.799520930 | 0.998576232 | 0.996920713 |
| 0.968122593 | 0.886975733 | 0.5407080223 | 0.061699026 | 0.617339011 | 0.998576232 | 0.996920713 |
| 0.296887408 | 0.869499083 | 0.3026795656 | 0.312774857 | 0.421788991 | 0.998576232 | 0.996920713 |
| 0.896940585 | 0.810290798 | 0.3234409509 | 0.535567671 | 0.421788991 | 0.998576232 | 0.996920713 |
| 0.785447683 | 0.942661849 | 0.5408954158 | 0.071885453 | 0.928186086 | 0.998576232 | 0.996920713 |
| 0.761371010 | 0.877271739 | 0.6112396972 | 0.094863415 | 0.799520930 | 0.998576232 | 0.996920713 |
| 0.794765735 | 0.90911696 | 0.5773220163 | 0.128403402 | 0.984924233 | 0.998576232 | 0.996920713 |
| 0.786365478 | 0.637524323 | 0.6412273429 | 0.29542434 | 0.517458860 | 0.998576232 | 0.996920713 |
| 0.873749379 | 0.94185348 | 0.5495221897 | 0.071181397 | 0.939664415 | 0.998576232 | 0.996920713 |
| 0.924606290 | 0.792516551 | 0.3088717169 | 0.080185549 | 0.607876666 | 0.998576232 | 0.996920713 |
| 0.688524149 | 0.929421686 | 0.5928552172 | 0.052940613 | 0.799520930 | 0.998576232 | 0.996920713 |
| 0.925335657 | 0.768406575 | 0.4358882964 | 0.050051444 | 0.696098292 | 0.998576232 | 0.996920713 |
| 0.789251196 | 0.828945182 | 0.2914413151 | 0.062969149 | 0.617339011 | 0.998576232 | 0.996920713 |
| 0.785359465 | 0.900285504 | 0.5522933433 | 0.077133908 | 0.953777775 | 0.998576232 | 0.996920713 |
| 0.843594957 | 0.962898526 | 0.5720283948 | 0.057295224 | 0.799520930 | 0.998576232 | 0.996920713 |
| 0.614280286 | 0.692178114 | 0.6342983225 | 0.408817092 | 0.607876666 | 0.998576232 | 0.996920713 |
| 0.713412191 | 0.940560701 | 0.4911578722 | 0.065044408 | 0.868904128 | 0.998576232 | 0.996920713 |
| 0.533430913 | 0.890165136 | 0.5628916476 | 0.184459574 | 0.758952824 | 0.998576232 | 0.996920713 |
| 0.708585673 | 0.711875780 | 0.7106405986 | 0.058286374 | 0.799520930 | 0.998576232 | 0.996920713 |
| 0.776998606 | 0.895478225 | 0.59916258 | 0.093638248 | 0.799520930 | 0.998576232 | 0.996920713 |
| 0.678357143 | 0.882660920 | 0.5283104127 | 0.488022723 | 0.617647808 | 0.998576232 | 0.996920713 |
| 0.816684252 | 0.942437767 | 0.5120579924 | 0.058622973 | 0.814552831 | 0.998576232 | 0.996920713 |
| 0.828028356 | 0.940892910 | 0.527177824 | 0.085418154 | 0.925203233 | 0.998576232 | 0.996920713 |
| 0.511139934 | 0.589349132 | 0.8684611047 | 0.046208407 | 0.775100997 | 0.998576232 | 0.996920713 |
| 0.983915605 | 0.847418000 | 0.3369284615 | 0.067515283 | 0.607876666 | 0.998576232 | 0.996920713 |
| 0.746879717 | 0.800225400 | 0.632801461 | 0.058917727 | 0.812090521 | 0.998576232 | 0.996920713 |

| | | | | | | |
|---|---|---|---|---|---|---|
| 0.636651701 | 0.910874158 | 0.5229911828 | 0.531182570 | 0.607876666 | 0.998576232 | 0.996920713 |
| 0.823998850 | 0.966986002 | 0.4565090731 | 0.059687966 | 0.799520930 | 0.998576232 | 0.996920713 |
| 0.808250510 | 0.598065697 | 0.7824053208 | 0.091905091 | 0.607876666 | 0.998576232 | 0.996920713 |
| 0.772775144 | 0.892594331 | 0.5809001877 | 0.065831266 | 0.887595656 | 0.998576232 | 0.996920713 |
| 0.453310423 | 0.646233637 | 0.5251548038 | 0.016484043 | 0.421788991 | 0.998576232 | 0.996920713 |
| 0.818652669 | 0.976673660 | 0.5142920971 | 0.193029281 | 0.775100997 | 0.998576232 | 0.996920713 |
| 0.729595000 | 0.741084340 | 0.6920644685 | 0.053579579 | 0.799520930 | 0.998576232 | 0.996920713 |
| 0.866114256 | 0.983740101 | 0.5020032293 | 0.073811916 | 0.839441786 | 0.998576232 | 0.996920713 |
| 0.970406532 | 0.739714635 | 0.3658673505 | 0.094662325 | 0.617647808 | 0.998576232 | 0.996920713 |
| 0.828995019 | 0.920202293 | 0.5435799574 | 0.072914795 | 0.928186086 | 0.998576232 | 0.996920713 |
| 0.666212831 | 0.788302613 | 0.6190113473 | 0.199112769 | 0.660566205 | 0.998576232 | 0.996920713 |
| 0.975453442 | 0.920935474 | 0.2318403636 | 0.130477885 | 0.571538566 | 0.998576232 | 0.996920713 |
| 0.789414812 | 0.896539108 | 0.5164822722 | 0.097776025 | 0.868904128 | 0.998576232 | 0.996920713 |
| 0.432993677 | 0.949939774 | 0.4413912469 | 0.325665583 | 0.421788991 | 0.998576232 | 0.996920713 |
| 0.81918714 | 0.911923747 | 0.5927494149 | 0.046690376 | 0.775100997 | 0.998576232 | 0.996920713 |
| 0.862275555 | 0.907756825 | 0.4594124627 | 0.059745745 | 0.775100997 | 0.998576232 | 0.996920713 |
| 0.547241236 | 0.693002321 | 0.5357941844 | 0.110410049 | 0.617339011 | 0.998576232 | 0.996920713 |
| 0.929037091 | 0.991196428 | 0.5701366139 | 0.021207549 | 0.607876666 | 0.998576232 | 0.996920713 |
| 0.520580892 | 0.951130140 | 0.4800495325 | 0.572059281 | 0.571538566 | 0.998576232 | 0.996920713 |
| 0.874692370 | 0.489022961 | 0.5190752831 | 0.308263371 | 0.607876666 | 0.998576232 | 0.996920713 |
| 0.483836571 | 0.995874159 | 0.425458154 | 0.040749280 | 0.571538566 | 0.998576232 | 0.996920713 |
| 0.830048205 | 0.897485837 | 0.5426139646 | 0.069584428 | 0.928186086 | 0.998576232 | 0.996920713 |
| 0.687449672 | 0.821901430 | 0.5239005695 | 0.060161030 | 0.799520930 | 0.998576232 | 0.996920713 |
| 0.804369194 | 0.797924485 | 0.2876669926 | 0.043082438 | 0.421788991 | 0.998576232 | 0.996920713 |
| 0.452030450 | 0.572503932 | 0.194128005 | 0.169764415 | 0.520349777 | 0.998576232 | 0.996920713 |
| 0.784731837 | 0.859855737 | 0.6121606889 | 0.116580336 | 0.904204465 | 0.998576232 | 0.996920713 |
| 0.773853630 | 0.866026212 | 0.5551331194 | 0.078390079 | 0.925203233 | 0.998576232 | 0.996920713 |
| 0.965304893 | 0.975816515 | 0.5389411041 | 0.238874356 | 0.849231427 | 0.998576232 | 0.996920713 |
| 0.590656476 | 0.817856203 | 0.7009645991 | 0.052651706 | 0.799520930 | 0.998576232 | 0.996920713 |
| 0.656605312 | 0.996920713 | 0.3385138892 | 0.177914901 | 0.617647808 | 0.998576232 | 0.996920713 |
| 0.817588415 | 0.898163857 | 0.5413430319 | 0.079381936 | 0.925203233 | 0.998576232 | 0.996920713 |
| 0.909076864 | 0.896839004 | 0.3472722993 | 0.126965908 | 0.617647808 | 0.998576232 | 0.996920713 |
| 0.805994688 | 0.901791795 | 0.5842065989 | 0.132677464 | 0.984924233 | 0.998576232 | 0.996920713 |
| 0.978493393 | 0.981363113 | 0.3994065109 | 0.056971066 | 0.799520930 | 0.998576232 | 0.996920713 |
| 0.998576232 | 0.986238213 | 0.453557829 | 0.057065936 | 0.799520930 | 0.998576232 | 0.996920713 |
| 0.550896355 | 0.937980609 | 0.2950291581 | 0.011178263 | 0.517208933 | 0.998576232 | 0.996920713 |
| 0.346708441 | 0.943538121 | 0.4046982902 | 0.463299197 | 0.421788991 | 0.998576232 | 0.996920713 |
| 0.345293880 | 0.964002198 | 0.3605279848 | 0.042624212 | 0.421788991 | 0.998576232 | 0.996920713 |
| 0.635681542 | 0.537028120 | 0.590695354 | 0.796364753 | 0.997874178 | 0.967795776 | 0.637363585 |
| 0.475143480 | 0.825063280 | 0.5074425407 | 0.685401342 | 0.763866738 | 0.967795776 | 0.868487663 |
| 0.677636552 | 0.381812290 | 0.8167933327 | 0.970019454 | 0.997874178 | 0.967795776 | 0.636773498 |
| 0.742887991 | 0.129272421 | 0.5010610689 | 0.797723940 | 0.997874178 | 0.967795776 | 0.636773498 |
| 0.780860074 | 0.484392482 | 0.7166898402 | 0.850232461 | 0.997874178 | 0.967795776 | 0.636773498 |
| 0.854017828 | 0.456487355 | 0.7654126458 | 0.844727735 | 0.997874178 | 0.967795776 | 0.636773498 |
| 0.922453585 | 0.337426585 | 0.9093128759 | 0.970061700 | 0.997874178 | 0.967795776 | 0.636773498 |
| 0.942927839 | 0.269171605 | 0.8334361731 | 0.386728501 | 0.997874178 | 0.967795776 | 0.636773498 |

| | | | | | | |
|---|---|---|---|---|---|---|
| 0.680338252 | 0.561796237 | 0.6583955721 | 0.870527395 | 0.997874178 | 0.967795776 | 0.642052842 |
| 0.922879087 | 0.400119476 | 0.8010261646 | 0.870063501 | 0.997874178 | 0.967795776 | 0.636773498 |
| 0.872197177 | 0.441723973 | 0.7794472623 | 0.798772854 | 0.997874178 | 0.967795776 | 0.636773498 |
| 0.943600882 | 0.334448936 | 0.7818692289 | 0.728300311 | 0.997874178 | 0.967795776 | 0.636773498 |
| 0.703695490 | 0.517510190 | 0.6585911623 | 0.810664272 | 0.997874178 | 0.967795776 | 0.637363585 |
| 0.833085969 | 0.445507544 | 0.8085506216 | 0.978491411 | 0.997874178 | 0.967795776 | 0.636773498 |
| 0.742180129 | 0.675005725 | 0.5497014008 | 0.548525207 | 0.997874178 | 0.967795776 | 0.729735919 |
| 0.633251303 | 0.472491780 | 0.7002936386 | 0.929467033 | 0.997874178 | 0.967795776 | 0.636773498 |
| 0.723195407 | 0.525498736 | 0.6783954745 | 0.819507331 | 0.997874178 | 0.967795776 | 0.637363585 |
| 0.823032220 | 0.452090747 | 0.8259939915 | 0.899009920 | 0.997874178 | 0.967795776 | 0.636773498 |
| 0.426252086 | 0.432191507 | 0.3609827837 | 0.454907050 | 0.997874178 | 0.967795776 | 0.636773498 |
| 0.597308145 | 0.285249317 | 0.9137125774 | 0.772514942 | 0.997874178 | 0.967795776 | 0.636773498 |
| 0.938974895 | 0.393833510 | 0.8158750536 | 0.647710338 | 0.997874178 | 0.967795776 | 0.636773498 |
| 0.283649284 | 0.942084530 | 0.2250686404 | 0.149327833 | 0.997874178 | 0.967795776 | 0.950001207 |
| 0.701412863 | 0.326476058 | 0.9368773956 | 0.678883950 | 0.997874178 | 0.967795776 | 0.636773498 |
| 0.889908054 | 0.431150018 | 0.8973399697 | 0.894758834 | 0.997874178 | 0.967795776 | 0.636773498 |
| 0.564301810 | 0.867747394 | 0.5522171283 | 0.690280894 | 0.997874178 | 0.967795776 | 0.889389515 |
| 0.776646372 | 0.468602733 | 0.7194544971 | 0.842697135 | 0.997874178 | 0.967795776 | 0.636773498 |
| 0.823386332 | 0.461989857 | 0.7455294341 | 0.819283064 | 0.997874178 | 0.967795776 | 0.636773498 |
| 0.413215486 | 0.874566357 | 0.3423834452 | 0.832308678 | 0.997874178 | 0.967795776 | 0.889389515 |
| 0.629781050 | 0.442456377 | 0.6391248939 | 0.937117864 | 0.997874178 | 0.967795776 | 0.636773498 |
| 0.513491796 | 0.961838070 | 0.2259518615 | 0.286976724 | 0.763866738 | 0.967795776 | 0.961838070 |
| 0.398127066 | 0.462403343 | 0.6417315435 | 0.813885718 | 0.997874178 | 0.967795776 | 0.636773498 |
| 0.820906064 | 0.298177102 | 0.6824035704 | 0.802049685 | 0.997874178 | 0.967795776 | 0.636773498 |
| 0.853846604 | 0.453198821 | 0.7660765733 | 0.462158531 | 0.997874178 | 0.967795776 | 0.636773498 |
| 0.839636250 | 0.466029612 | 0.8070167777 | 0.833802441 | 0.997874178 | 0.967795776 | 0.636773498 |
| 0.779488360 | 0.417564543 | 0.7078919656 | 0.89694617 | 0.997874178 | 0.967795776 | 0.636773498 |
| 0.468708760 | 0.874371140 | 0.4361378865 | 0.743363353 | 0.997874178 | 0.967795776 | 0.889389515 |
| 0.862562363 | 0.328589230 | 0.8917713512 | 0.677705496 | 0.997874178 | 0.967795776 | 0.636773498 |
| 0.769041655 | 0.466719494 | 0.7387633643 | 0.961732823 | 0.997874178 | 0.967795776 | 0.636773498 |
| 0.788296661 | 0.460504447 | 0.7165576369 | 0.954300583 | 0.997874178 | 0.967795776 | 0.636773498 |
| 0.869420811 | 0.259521183 | 0.831651145 | 0.518804824 | 0.997874178 | 0.967795776 | 0.636773498 |
| 0.802680404 | 0.455422067 | 0.722343327 | 0.918634165 | 0.997874178 | 0.967795776 | 0.636773498 |
| 0.762232215 | 0.462064551 | 0.7047778127 | 0.807411884 | 0.997874178 | 0.967795776 | 0.636773498 |
| 0.821851283 | 0.385838801 | 0.8311942279 | 0.833015922 | 0.997874178 | 0.967795776 | 0.636773498 |
| 0.531142830 | 0.457186999 | 0.5661872359 | 0.763452901 | 0.997874178 | 0.967795776 | 0.636773498 |
| 0.696043801 | 0.467149036 | 0.5455499617 | 0.529845284 | 0.997874178 | 0.967795776 | 0.636773498 |
| 0.776702291 | 0.468982681 | 0.7277450452 | 0.847428877 | 0.997874178 | 0.967795776 | 0.636773498 |
| 0.855868793 | 0.474659799 | 0.7771851042 | 0.651239654 | 0.997874178 | 0.967795776 | 0.636773498 |
| 0.799598393 | 0.225143049 | 0.7080089248 | 0.593494696 | 0.997874178 | 0.967795776 | 0.636773498 |
| 0.681390313 | 0.561316418 | 0.6161242739 | 0.866350075 | 0.997874178 | 0.967795776 | 0.642052842 |
| 0.770721252 | 0.482122368 | 0.7144317344 | 0.841619514 | 0.997874178 | 0.967795776 | 0.636773498 |
| 0.825541452 | 0.427100983 | 0.7511304368 | 0.889960363 | 0.997874178 | 0.967795776 | 0.636773498 |
| 0.863993868 | 0.328465781 | 0.7566587983 | 0.836672888 | 0.997874178 | 0.967795776 | 0.636773498 |
| 0.747513557 | 0.446215761 | 0.7993307499 | 0.752532682 | 0.997874178 | 0.967795776 | 0.636773498 |
| 0.763730399 | 0.440296084 | 0.6793482012 | 0.939205817 | 0.997874178 | 0.967795776 | 0.636773498 |

| | | | | | | |
|---|---|---|---|---|---|---|
| 0.413023889 | 0.744349593 | 0.582160629 | 0.954960655 | 0.997874178 | 0.967795776 | 0.797517421 |
| 0.808025775 | 0.442910948 | 0.7526543402 | 0.880916659 | 0.997874178 | 0.967795776 | 0.636773498 |
| 0.748221561 | 0.475497315 | 0.8873034911 | 0.882397131 | 0.997874178 | 0.967795776 | 0.636773498 |
| 0.797386188 | 0.469301198 | 0.7192296539 | 0.842388643 | 0.997874178 | 0.967795776 | 0.636773498 |
| 0.803484893 | 0.474337209 | 0.7202047819 | 0.862467874 | 0.997874178 | 0.967795776 | 0.636773498 |
| 0.770006871 | 0.484517096 | 0.7139514487 | 0.899229834 | 0.997874178 | 0.967795776 | 0.636773498 |
| 0.768986479 | 0.644177396 | 0.5819222985 | 0.690987139 | 0.997874178 | 0.967795776 | 0.709186125 |
| 0.775683285 | 0.470492009 | 0.7179957349 | 0.840498613 | 0.997874178 | 0.967795776 | 0.636773498 |
| 0.834785662 | 0.437361525 | 0.8151596829 | 0.999795788 | 0.997874178 | 0.967795776 | 0.636773498 |
| 0.775133511 | 0.458559755 | 0.7311553262 | 0.911314997 | 0.997874178 | 0.967795776 | 0.636773498 |
| 0.727729206 | 0.537553825 | 0.6824775475 | 0.814356486 | 0.997874178 | 0.967795776 | 0.637363585 |
| 0.762981725 | 0.504473971 | 0.7078237148 | 0.852931047 | 0.997874178 | 0.967795776 | 0.636773498 |
| 0.529078291 | 0.381032586 | 0.5924661667 | 0.789400109 | 0.997874178 | 0.967795776 | 0.636773498 |
| 0.766636864 | 0.531300492 | 0.7100965975 | 0.834395615 | 0.997874178 | 0.967795776 | 0.637363585 |
| 0.652111165 | 0.226393061 | 0.7332270542 | 0.807479480 | 0.997874178 | 0.967795776 | 0.636773498 |
| 0.777425311 | 0.468085362 | 0.7254697656 | 0.833048346 | 0.997874178 | 0.967795776 | 0.636773498 |
| 0.855395037 | 0.346774322 | 0.6848350487 | 0.951955606 | 0.997874178 | 0.967795776 | 0.636773498 |
| 0.812920056 | 0.441734002 | 0.7423728745 | 0.993135864 | 0.997874178 | 0.967795776 | 0.636773498 |
| 0.616687037 | 0.290169694 | 0.9445866076 | 0.907229824 | 0.997874178 | 0.967795776 | 0.636773498 |
| 0.987495846 | 0.459678588 | 0.7288296962 | 0.904703291 | 0.997874178 | 0.988580044 | 0.636773498 |
| 0.706158751 | 0.655122715 | 0.5954496776 | 0.907043812 | 0.997874178 | 0.967795776 | 0.714679325 |
| 0.775964716 | 0.46920613 | 0.7133456715 | 0.831178215 | 0.997874178 | 0.967795776 | 0.636773498 |
| 0.690214747 | 0.465955242 | 0.7256567957 | 0.419606196 | 0.997874178 | 0.967795776 | 0.636773498 |
| 0.815314496 | 0.427270514 | 0.7825509323 | 0.992401842 | 0.997874178 | 0.967795776 | 0.636773498 |
| 0.896148856 | 0.386967658 | 0.8681746513 | 0.716066076 | 0.997874178 | 0.967795776 | 0.636773498 |
| 0.920873059 | 0.313151596 | 0.9980271975 | 0.538307382 | 0.997874178 | 0.967795776 | 0.636773498 |
| 0.785583258 | 0.477484231 | 0.7335961459 | 0.842535112 | 0.997874178 | 0.967795776 | 0.636773498 |
| 0.771609902 | 0.49400944 | 0.7114991724 | 0.860362842 | 0.997874178 | 0.967795776 | 0.636773498 |
| 0.678354967 | 0.448714481 | 0.7212435807 | 0.426377877 | 0.997874178 | 0.967795776 | 0.636773498 |
| 0.825704459 | 0.357184480 | 0.8923338938 | 0.909828659 | 0.997874178 | 0.967795776 | 0.636773498 |
| 0.785934507 | 0.561457413 | 0.6543271002 | 0.795496857 | 0.997874178 | 0.967795776 | 0.642052842 |
| 0.793775288 | 0.457690475 | 0.7623067444 | 0.708496298 | 0.997874178 | 0.967795776 | 0.636773498 |
| 0.650722709 | 0.541759047 | 0.7095150122 | 0.995125360 | 0.997874178 | 0.967795776 | 0.637363585 |
| 0.802565914 | 0.404896629 | 0.7475572093 | 0.640500847 | 0.997874178 | 0.967795776 | 0.636773498 |
| 0.732144744 | 0.594572625 | 0.6172132226 | 0.942616515 | 0.997874178 | 0.967795776 | 0.660636250 |
| 0.878843186 | 0.379357268 | 0.8082166935 | 0.807345819 | 0.997874178 | 0.967795776 | 0.636773498 |
| 0.573632309 | 0.858375027 | 0.4637924659 | 0.985373315 | 0.997874178 | 0.967795776 | 0.889389515 |
| 0.814073275 | 0.283309371 | 0.8809025648 | 0.666110169 | 0.997874178 | 0.967795776 | 0.636773498 |
| 0.604652302 | 0.573785226 | 0.5956366945 | 0.400733556 | 0.997874178 | 0.967795776 | 0.649568181 |
| 0.754776118 | 0.471085394 | 0.688339139 | 0.881349827 | 0.997874178 | 0.967795776 | 0.636773498 |
| 0.776568614 | 0.468924337 | 0.7200252159 | 0.854053502 | 0.997874178 | 0.967795776 | 0.636773498 |
| 0.695271776 | 0.485325516 | 0.7095893535 | 0.954154011 | 0.997874178 | 0.967795776 | 0.636773498 |
| 0.810718621 | 0.348980617 | 0.6693739513 | 0.964052945 | 0.997874178 | 0.967795776 | 0.636773498 |
| 0.854979580 | 0.452532633 | 0.8354113617 | 0.876960693 | 0.997874178 | 0.967795776 | 0.636773498 |
| 0.670207239 | 0.368222106 | 0.719744466 | 0.750161642 | 0.997874178 | 0.967795776 | 0.636773498 |
| 0.783661483 | 0.470273648 | 0.7194376954 | 0.858859148 | 0.997874178 | 0.967795776 | 0.636773498 |

| | | | | | | |
|---|---|---|---|---|---|---|
| 0.988580044 | 0.436211331 | 0.7353332583 | 0.650073257 | 0.997874178 | 0.988580044 | 0.636773498 |
| 0.775133975 | 0.498148796 | 0.7188122912 | 0.877592659 | 0.997874178 | 0.967795776 | 0.636773498 |
| 0.674787542 | 0.486145912 | 0.6944554973 | 0.820500350 | 0.997874178 | 0.967795776 | 0.636773498 |
| 0.856942572 | 0.469974702 | 0.7649216516 | 0.890711354 | 0.997874178 | 0.967795776 | 0.636773498 |
| 0.961190176 | 0.362975486 | 0.6525644192 | 0.746937977 | 0.997874178 | 0.977481534 | 0.636773498 |
| 0.864330959 | 0.481294798 | 0.7875110395 | 0.847496197 | 0.997874178 | 0.967795776 | 0.636773498 |
| 0.395830278 | 0.788490484 | 0.4304726307 | 0.893793494 | 0.997874178 | 0.967795776 | 0.837335028 |
| 0.776815686 | 0.504561948 | 0.6705905985 | 0.750286615 | 0.997874178 | 0.967795776 | 0.636773498 |
| 0.781301886 | 0.462612173 | 0.7191107434 | 0.858835130 | 0.997874178 | 0.967795776 | 0.636773498 |
| 0.621096914 | 0.267868182 | 0.7675970022 | 0.212252708 | 0.997874178 | 0.967795776 | 0.636773498 |
| 0.376060478 | 0.593920441 | 0.4426950372 | 0.757494904 | 0.997874178 | 0.967795776 | 0.660636250 |
| 0.567657346 | 0.509418798 | 0.5918911696 | 0.973697719 | 0.997874178 | 0.967795776 | 0.636773498 |
| 0.665485752 | 0.457623107 | 0.6488912633 | 0.966764826 | 0.997874178 | 0.967795776 | 0.636773498 |
| 0.756241964 | 0.468701255 | 0.6836336381 | 0.892341512 | 0.997874178 | 0.967795776 | 0.636773498 |
| 0.926544314 | 0.432321614 | 0.8138218201 | 0.735115405 | 0.997874178 | 0.967795776 | 0.636773498 |
| 0.881605070 | 0.430079842 | 0.8557497933 | 0.929714600 | 0.997874178 | 0.967795776 | 0.636773498 |
| 0.849525312 | 0.329179943 | 0.9819491885 | 0.990865756 | 0.997874178 | 0.967795776 | 0.636773498 |
| 0.786095419 | 0.474691256 | 0.7193959165 | 0.850513433 | 0.997874178 | 0.967795776 | 0.636773498 |
| 0.805925971 | 0.468972491 | 0.7203434981 | 0.855699342 | 0.997874178 | 0.967795776 | 0.636773498 |
| 0.597244892 | 0.480046226 | 0.6738198914 | 0.840990973 | 0.997874178 | 0.967795776 | 0.636773498 |
| 0.954679863 | 0.562368177 | 0.79501522 | 0.235883483 | 0.930179421 | 0.992345060 | 0.775680244 |
| 0.751920016 | 0.394851380 | 0.9353807063 | 0.320017319 | 0.732783268 | 0.992345060 | 0.771430006 |
| 0.675275448 | 0.699392138 | 0.6821302212 | 0.405975308 | 0.697456823 | 0.992345060 | 0.818326604 |
| 0.418077273 | 0.122325140 | 0.3691533239 | 0.217490551 | 0.697456823 | 0.992345060 | 0.771430006 |
| 0.801643494 | 0.654028258 | 0.8105146467 | 0.863104617 | 0.732783268 | 0.992345060 | 0.809107124 |
| 0.596334422 | 0.436764717 | 0.9421539511 | 0.210650243 | 0.732783268 | 0.992345060 | 0.771430006 |
| 0.207105935 | 0.138300824 | 0.5870633551 | 0.046523508 | 0.697456823 | 0.992345060 | 0.771430006 |
| 0.947256566 | 0.625279615 | 0.6636238825 | 0.563942454 | 0.732783268 | 0.992345060 | 0.798229296 |
| 0.930316255 | 0.649386063 | 0.7624604959 | 0.205964727 | 0.867011594 | 0.992345060 | 0.809107124 |
| 0.531423634 | 0.307197757 | 0.9188006905 | 0.175791886 | 0.697456823 | 0.992345060 | 0.771430006 |
| 0.532923748 | 0.313096405 | 0.8870921599 | 0.343881023 | 0.697456823 | 0.992345060 | 0.771430006 |
| 0.586441049 | 0.256421888 | 0.9840427202 | 0.325010314 | 0.697456823 | 0.992345060 | 0.771430006 |
| 0.929113992 | 0.552858171 | 0.83231438 | 0.237036645 | 0.957592049 | 0.992345060 | 0.771430006 |
| 0.602171423 | 0.299303399 | 0.734025475 | 0.080359058 | 0.697456823 | 0.992345060 | 0.771430006 |
| 0.862294169 | 0.535437842 | 0.8694397163 | 0.250003682 | 0.999251513 | 0.992345060 | 0.771430006 |
| 0.819922621 | 0.519820555 | 0.8730419931 | 0.334213684 | 0.957592049 | 0.992345060 | 0.771430006 |
| 0.408361659 | 0.266373069 | 0.7347453448 | 0.132946752 | 0.697456823 | 0.992345060 | 0.771430006 |
| 0.535311638 | 0.336607260 | 0.486472716 | 0.076211701 | 0.697456823 | 0.992345060 | 0.771430006 |
| 0.601843956 | 0.499848387 | 0.8356441191 | 0.114342738 | 0.697456823 | 0.992345060 | 0.771430006 |
| 0.723995407 | 0.698766608 | 0.7137441961 | 0.191111155 | 0.732783268 | 0.992345060 | 0.818326604 |
| 0.871214985 | 0.762767520 | 0.7109486545 | 0.139542224 | 0.732783268 | 0.992345060 | 0.840021118 |
| 0.605691552 | 0.932127021 | 0.3717275661 | 0.784632704 | 0.697456823 | 0.992345060 | 0.932127021 |
| 0.831952874 | 0.612476781 | 0.7839312865 | 0.201135248 | 0.90115955 | 0.992345060 | 0.798229296 |
| 0.815271376 | 0.507930336 | 0.9713326063 | 0.218516204 | 0.957592049 | 0.992345060 | 0.771430006 |
| 0.325145798 | 0.090100747 | 0.6342189869 | 0.074334635 | 0.697456823 | 0.992345060 | 0.771430006 |
| 0.865766480 | 0.492546758 | 0.8678546324 | 0.223203846 | 0.732783268 | 0.992345060 | 0.771430006 |

| | | | | | | |
|---|---|---|---|---|---|---|
| 0.750460529 | 0.820845624 | 0.6345075857 | 0.106066931 | 0.697456823 | 0.992345060 | 0.864048025 |
| 0.316936748 | 0.173726085 | 0.4136123237 | 0.521612323 | 0.697456823 | 0.992345060 | 0.771430006 |
| 0.934797086 | 0.512115766 | 0.831423688 | 0.207903882 | 0.909058433 | 0.992345060 | 0.771430006 |
| 0.926385816 | 0.669765972 | 0.7096189764 | 0.332054574 | 0.754994911 | 0.992345060 | 0.818326604 |
| 0.516754324 | 0.481430929 | 0.9154582769 | 0.380476951 | 0.697456823 | 0.992345060 | 0.771430006 |
| 0.917693279 | 0.709216390 | 0.8773388576 | 0.174711706 | 0.697456823 | 0.992345060 | 0.818326604 |
| 0.890946501 | 0.520157799 | 0.846851271 | 0.561401477 | 0.90115955 | 0.992345060 | 0.771430006 |
| 0.861890261 | 0.704088627 | 0.5854943508 | 0.192148837 | 0.732783268 | 0.992345060 | 0.818326604 |
| 0.854286940 | 0.452044547 | 0.8531105168 | 0.190605167 | 0.732783268 | 0.992345060 | 0.771430006 |
| 0.479144586 | 0.253661097 | 0.7056880986 | 0.602899683 | 0.697456823 | 0.992345060 | 0.771430006 |
| 0.805434185 | 0.500766724 | 0.9418591471 | 0.239885064 | 0.930179421 | 0.992345060 | 0.771430006 |
| 0.832920622 | 0.521512185 | 0.7993861761 | 0.509910541 | 0.732783268 | 0.992345060 | 0.771430006 |
| 0.867845224 | 0.530620657 | 0.8681821006 | 0.357779906 | 0.970016617 | 0.992345060 | 0.771430006 |
| 0.798308369 | 0.531208841 | 0.9902861444 | 0.356912578 | 0.946083634 | 0.992345060 | 0.771430006 |
| 0.839651576 | 0.508031273 | 0.8700722216 | 0.223014081 | 0.946083634 | 0.992345060 | 0.771430006 |
| 0.950727998 | 0.402312658 | 0.6585007344 | 0.641440358 | 0.697456823 | 0.992345060 | 0.771430006 |
| 0.831207224 | 0.467514626 | 0.9422139108 | 0.198399528 | 0.867011594 | 0.992345060 | 0.771430006 |
| 0.761210150 | 0.523028594 | 0.9425112834 | 0.211158393 | 0.927422576 | 0.992345060 | 0.771430006 |
| 0.813200666 | 0.516604729 | 0.9978339659 | 0.204093591 | 0.867011594 | 0.992345060 | 0.771430006 |
| 0.869028261 | 0.520435626 | 0.8111908262 | 0.211860022 | 0.930179421 | 0.992345060 | 0.771430006 |
| 0.764486137 | 0.433042241 | 0.5767671481 | 0.016008888 | 0.697456823 | 0.992345060 | 0.771430006 |
| 0.987808711 | 0.804497832 | 0.7638410517 | 0.217369702 | 0.930819256 | 0.992345060 | 0.861961963 |
| 0.934507079 | 0.682466762 | 0.649746474 | 0.198516339 | 0.732783268 | 0.992345060 | 0.818326604 |
| 0.890724146 | 0.534982894 | 0.860728571 | 0.224184293 | 0.970016617 | 0.992345060 | 0.771430006 |
| 0.902680319 | 0.594290749 | 0.8412029632 | 0.246345903 | 0.930179421 | 0.992345060 | 0.792387665 |
| 0.955163735 | 0.799507131 | 0.8215035076 | 0.493047485 | 0.732783268 | 0.992345060 | 0.861961963 |
| 0.848280787 | 0.517214674 | 0.8098481119 | 0.386478845 | 0.732783268 | 0.992345060 | 0.771430006 |
| 0.758314645 | 0.791857712 | 0.8128981476 | 0.599633430 | 0.697456823 | 0.992345060 | 0.861961963 |
| 0.585284668 | 0.351144609 | 0.9576266189 | 0.250230693 | 0.697456823 | 0.992345060 | 0.771430006 |
| 0.960304363 | 0.614735039 | 0.755774171 | 0.269949150 | 0.697456823 | 0.992345060 | 0.798229296 |
| 0.873889832 | 0.531599764 | 0.985283238 | 0.159293444 | 0.732783268 | 0.992345060 | 0.771430006 |
| 0.929219406 | 0.519358819 | 0.8668160393 | 0.220463713 | 0.927415364 | 0.992345060 | 0.771430006 |
| 0.452285936 | 0.428145204 | 0.9941924797 | 0.462846490 | 0.732783268 | 0.992345060 | 0.771430006 |
| 0.943416027 | 0.585606051 | 0.8020327919 | 0.198552289 | 0.867011594 | 0.992345060 | 0.791530750 |
| 0.862737613 | 0.839879991 | 0.5969213676 | 0.373546212 | 0.697456823 | 0.992345060 | 0.876396512 |
| 0.899600773 | 0.542462443 | 0.7528491352 | 0.318140267 | 0.732783268 | 0.992345060 | 0.771430006 |
| 0.890633275 | 0.812927098 | 0.5248418192 | 0.759001123 | 0.697456823 | 0.992345060 | 0.863285414 |
| 0.848981332 | 0.640014776 | 0.7947322864 | 0.466930730 | 0.732783268 | 0.992345060 | 0.808439717 |
| 0.484861549 | 0.921559782 | 0.4466311235 | 0.341014227 | 0.697456823 | 0.992345060 | 0.932127021 |
| 0.882415817 | 0.763019182 | 0.5848403976 | 0.169010653 | 0.697456823 | 0.992345060 | 0.840021118 |
| 0.985508498 | 0.486089478 | 0.8099670652 | 0.160669791 | 0.732783268 | 0.992345060 | 0.771430006 |
| 0.959396991 | 0.748782753 | 0.7698064624 | 0.255355707 | 0.768567725 | 0.992345060 | 0.839756359 |
| 0.751063393 | 0.470764540 | 0.9971456126 | 0.224177896 | 0.909058433 | 0.992345060 | 0.771430006 |
| 0.856265925 | 0.510830759 | 0.9074086929 | 0.274332199 | 0.786847634 | 0.992345060 | 0.771430006 |
| 0.957621137 | 0.882604172 | 0.9264854567 | 0.309498074 | 0.697456823 | 0.992345060 | 0.913038798 |
| 0.772026928 | 0.436186502 | 0.926276057 | 0.140172318 | 0.732783268 | 0.992345060 | 0.771430006 |

| | | | | | | |
|---|---|---|---|---|---|---|
| 0.895770589 | 0.253577853 | 0.8260194276 | 0.113430935 | 0.697456823 | 0.992345060 | 0.771430006 |
| 0.957559248 | 0.522898600 | 0.8615025939 | 0.316693846 | 0.930179421 | 0.992345060 | 0.771430006 |
| 0.781374429 | 0.392884542 | 0.9658924364 | 0.255894521 | 0.768567725 | 0.992345060 | 0.771430006 |
| 0.868063762 | 0.515963495 | 0.7869099688 | 0.303018092 | 0.732783268 | 0.992345060 | 0.771430006 |
| 0.747681259 | 0.494481014 | 0.8437226086 | 0.076892267 | 0.697456823 | 0.992345060 | 0.771430006 |
| 0.852946606 | 0.513822122 | 0.8835790178 | 0.224894148 | 0.957592049 | 0.992345060 | 0.771430006 |
| 0.731029207 | 0.420978813 | 0.9623593905 | 0.306407714 | 0.732783268 | 0.992345060 | 0.771430006 |
| 0.661240731 | 0.390527358 | 0.9138600803 | 0.513527005 | 0.800955307 | 0.992345060 | 0.771430006 |
| 0.663883001 | 0.358507679 | 0.8751435024 | 0.229688775 | 0.697456823 | 0.992345060 | 0.771430006 |
| 0.774415361 | 0.355998610 | 0.9506897753 | 0.316284045 | 0.697456823 | 0.992345060 | 0.771430006 |
| 0.745583907 | 0.518788662 | 0.851331079 | 0.089931277 | 0.697456823 | 0.992345060 | 0.771430006 |
| 0.852014733 | 0.507515247 | 0.9055690528 | 0.219065216 | 0.957592049 | 0.992345060 | 0.771430006 |
| 0.838194181 | 0.706794384 | 0.7104845896 | 0.271110291 | 0.732783268 | 0.992345060 | 0.818326604 |
| 0.857779075 | 0.520067637 | 0.8783033891 | 0.262075412 | 0.957592049 | 0.992345060 | 0.771430006 |
| 0.5820706 | 0.350441806 | 0.8691259223 | 0.393727874 | 0.697456823 | 0.992345060 | 0.771430006 |
| 0.856733302 | 0.513642940 | 0.8734866952 | 0.299205833 | 0.959653885 | 0.992345060 | 0.771430006 |
| 0.787152412 | 0.369639045 | 0.9393283533 | 0.303870524 | 0.732783268 | 0.992345060 | 0.771430006 |
| 0.660329135 | 0.315778087 | 0.9491421959 | 0.239496584 | 0.697456823 | 0.992345060 | 0.771430006 |
| 0.848789784 | 0.529127873 | 0.8946435755 | 0.235649559 | 0.967480739 | 0.992345060 | 0.771430006 |
| 0.890595322 | 0.619164332 | 0.7521744907 | 0.393474390 | 0.697456823 | 0.992345060 | 0.798229296 |
| 0.856091010 | 0.519843689 | 0.8726531575 | 0.255209075 | 0.970016617 | 0.992345060 | 0.771430006 |
| 0.924615826 | 0.526094908 | 0.758293072 | 0.199795037 | 0.90115955 | 0.992345060 | 0.771430006 |
| 0.871793921 | 0.510955046 | 0.7562170278 | 0.155883780 | 0.732783268 | 0.992345060 | 0.771430006 |
| 0.677555534 | 0.488441966 | 0.8846460764 | 0.366738759 | 0.800955307 | 0.992345060 | 0.771430006 |
| 0.909660321 | 0.743758457 | 0.929414766 | 0.367942540 | 0.710597508 | 0.992345060 | 0.839756359 |
| 0.790384417 | 0.508517267 | 0.9731730189 | 0.203422332 | 0.732783268 | 0.992345060 | 0.771430006 |
| 0.610008796 | 0.931304446 | 0.8551409728 | 0.134037675 | 0.697456823 | 0.992345060 | 0.932127021 |
| 0.977978486 | 0.550545943 | 0.8820787666 | 0.383957793 | 0.800955307 | 0.992345060 | 0.771430006 |
| 0.883668997 | 0.525009806 | 0.8669159182 | 0.329912445 | 0.999251513 | 0.992345060 | 0.771430006 |
| 0.724424683 | 0.908542917 | 0.8864263806 | 0.792039474 | 0.697456823 | 0.992345060 | 0.931838889 |
| 0.501405817 | 0.466304387 | 0.9431963569 | 0.187790920 | 0.732783268 | 0.992345060 | 0.771430006 |
| 0.920624221 | 0.519909064 | 0.8366774622 | 0.245507539 | 0.944721678 | 0.992345060 | 0.771430006 |
| 0.654606983 | 0.392283230 | 0.789851603 | 0.258952700 | 0.697456823 | 0.992345060 | 0.771430006 |
| 0.890151139 | 0.518418498 | 0.8505160546 | 0.223857680 | 0.959653885 | 0.992345060 | 0.771430006 |
| 0.479923453 | 0.324067951 | 0.7930082212 | 0.366178705 | 0.732783268 | 0.992345060 | 0.771430006 |
| 0.862037190 | 0.463184659 | 0.974802944 | 0.169485594 | 0.786847634 | 0.992345060 | 0.771430006 |
| 0.780630061 | 0.336738848 | 0.8525011106 | 0.114885067 | 0.697456823 | 0.992345060 | 0.771430006 |
| 0.881039985 | 0.532512923 | 0.8663909046 | 0.330761309 | 0.999251513 | 0.992345060 | 0.771430006 |
| 0.591154135 | 0.436351323 | 0.9328703942 | 0.382151218 | 0.732783268 | 0.992345060 | 0.771430006 |
| 0.574373370 | 0.467840894 | 0.9354172319 | 0.373926474 | 0.754994911 | 0.992345060 | 0.771430006 |
| 0.924475652 | 0.518502296 | 0.8292358038 | 0.197855869 | 0.867011594 | 0.992345060 | 0.771430006 |
| 0.607680661 | 0.471082537 | 0.5907922323 | 0.464873754 | 0.697456823 | 0.992345060 | 0.771430006 |
| 0.878655019 | 0.733448149 | 0.7160365255 | 0.180511730 | 0.800955307 | 0.992345060 | 0.838226456 |
| 0.706305335 | 0.443771447 | 0.9163137549 | 0.167244604 | 0.768567725 | 0.992345060 | 0.771430006 |
| 0.992345060 | 0.587051973 | 0.7877409949 | 0.256466017 | 0.909058433 | 0.992345060 | 0.791530750 |
| 0.423618317 | 0.410430071 | 0.8757377399 | 0.073534036 | 0.697456823 | 0.992345060 | 0.771430006 |

| | | | | | | |
|---|---|---|---|---|---|---|
| 0.496734528 | 0.489219355 | 0.9153122378 | 0.535725800 | 0.717358543 | 0.992345060 | 0.771430006 |
| 0.380347720 | 0.467131868 | 0.9861598328 | 0.201736083 | 0.697456823 | 0.992345060 | 0.771430006 |

| t_q_interact | t_q_batch | Genus | Family |
|---|---|---|---|
| 0.6404556554 | 0.065307203 | Acetatifactor | Lachnospiraceae |
| 0.6404556554 | 0.065307203 | Acutalibacter | Oscillospiraceae |
| 0.6404556554 | 0.065307203 | Akkermansia | Akkermansiaceae |
| 0.6979515333 | 0.065307203 | Anaerotruncus | Oscillospiraceae |
| 0.6404556554 | 0.136595928 | Bacteria_unclassified | Bacteria_unclassified |
| 0.7252879204 | 0.065307203 | Bacteria_unclassified | Bacteria_unclassified |
| 0.649195481 | 0.072440428 | Bacteria_unclassified | Bacteria_unclassified |
| 0.6404556554 | 0.065307203 | Bacteria_unclassified | Bacteria_unclassified |
| 0.649195481 | 0.065307203 | Bacteroides | Bacteroidaceae |
| 0.6404556554 | 0.065307203 | Clostridia_unclassified | Clostridia_unclassified |
| 0.6404556554 | 0.065307203 | Clostridiaceae_unclassified | Clostridiaceae |
| 0.6404556554 | 0.065307203 | Clostridiaceae_unclassified | Clostridiaceae |
| 0.6404556554 | 0.065307203 | Eubacteriales_unclassified | Eubacteriales_unclassified |
| 0.8098448563 | 0.610203226 | Clostridium | Clostridiaceae |
| 0.6404556554 | 0.065307203 | Erysipelatoclostridium | Erysipelotrichaceae |
| 0.6404556554 | 0.072562568 | Eubacteriaceae_unclassified | Eubacteriaceae |
| 0.6404556554 | 0.065307203 | Eubacteriaceae_unclassified | Eubacteriaceae |
| 0.6404556554 | 0.065307203 | GGB20146 | FGB77306 |
| 0.6404556554 | 0.065307203 | GGB20149 | Lachnospiraceae |
| 0.6404556554 | 0.065307203 | GGB22635 | Eggerthellaceae |
| 0.6404556554 | 0.129165008 | GGB25041 | Lachnospiraceae |
| 0.764176936 | 0.964962110 | GGB28379 | FGB9506 |
| 0.6404556554 | 0.073650255 | GGB28382 | FGB9508 |
| 0.6404556554 | 0.065307203 | GGB28392 | FGB9512 |
| 0.6404556554 | 0.069263270 | GGB28404 | FGB2838 |
| 0.6404556554 | 0.065307203 | GGB28418 | FGB2838 |
| 0.6404556554 | 0.065307203 | GGB28422 | FGB2838 |
| 0.6404556554 | 0.072187554 | GGB28431 | Pumilibacteraceae |
| 0.6404556554 | 0.066476697 | GGB28456 | FGB2833 |
| 0.6909237074 | 0.076543895 | GGB28782 | Eubacteriaceae |
| 0.6404556554 | 0.065307203 | GGB28792 | Lachnospiraceae |
| 0.6404556554 | 0.065307203 | GGB28798 | Lachnospiraceae |
| 0.6404556554 | 0.131253105 | GGB28810 | FGB9622 |
| 0.6404556554 | 0.065307203 | GGB28828 | FGB77305 |
| 0.6404556554 | 0.065307203 | GGB28851 | Clostridiaceae |
| 0.6404556554 | 0.285303606 | GGB28865 | Lachnospiraceae |
| 0.6404556554 | 0.065307203 | GGB28868 | Lachnospiraceae |
| 0.6404556554 | 0.170684759 | GGB28869 | Lachnospiraceae |
| 0.6404556554 | 0.111080289 | GGB28875 | Lachnospiraceae |
| 0.8211099083 | 0.065307203 | GGB28881 | FGB9633 |
| 0.6404556554 | 0.065307203 | GGB28892 | Bacteria_unclassified |
| 0.6404556554 | 0.065307203 | GGB28893 | Bacteria_unclassified |
| 0.6404556554 | 0.114401038 | GGB28898 | Bacteria_unclassified |
| 0.7700040547 | 0.065931569 | GGB28901 | Bacteria_unclassified |
| 0.6404556554 | 0.065307203 | GGB28904 | Bacteria_unclassified |
| 0.6404556554 | 0.072187554 | GGB28909 | Bacteria_unclassified |

| | | | |
|---|---|---|---|
| 0.6404556554 | 0.5934743807 | GGB28924 | Lachnospiraceae |
| 0.9874747598 | 0.0653072035 | GGB28927 | FGB77359 |
| 0.6404556554 | 0.0653072035 | GGB28934 | FGB9639 |
| 0.6404556554 | 0.0653072035 | GGB28949 | Lachnospiraceae |
| 0.6404556554 | 0.0653072035 | GGB28951 | Clostridiaceae |
| 0.6404556554 | 0.0653072035 | GGB28951 | Clostridiaceae |
| 0.6404556554 | 0.0653072035 | GGB28954 | Clostridiaceae |
| 0.6404556554 | 0.1097713229 | GGB28960 | Clostridiaceae |
| 0.6404556554 | 0.0653072035 | GGB28964 | Clostridiaceae |
| 0.6404556554 | 0.0653072035 | GGB28967 | Clostridiaceae |
| 0.6404556554 | 0.0653072035 | GGB28996 | FGB9656 |
| 0.6404556554 | 0.0653072035 | GGB29531 | FGB9827 |
| 0.6404556554 | 0.0736502558 | GGB29685 | Eubacteriaceae |
| 0.6404556554 | 0.1607170045 | GGB30141 | FGB77303 |
| 0.6404556554 | 0.0653072035 | GGB30145 | Clostridiaceae |
| 0.6404556554 | 0.0653072035 | GGB30286 | Eubacteriales_unclassified |
| 0.8098448563 | 0.2536092697 | GGB30300 | FGB72709 |
| 0.6404556554 | 0.0653072035 | GGB30303 | Oscillospiraceae |
| 0.6404556554 | 0.0653072035 | GGB30450 | Oscillospiraceae |
| 0.6404556554 | 0.0653072035 | GGB30453 | Oscillospiraceae |
| 0.6404556554 | 0.0664766979 | GGB30454 | Oscillospiraceae |
| 0.6404556554 | 0.0653072035 | GGB30455 | Oscillospiraceae |
| 0.6404556554 | 0.0653072035 | GGB30456 | Oscillospiraceae |
| 0.6404556554 | 0.0653072035 | GGB30457 | Oscillospiraceae |
| 0.6404556554 | 0.065931569 | GGB30461 | Oscillospiraceae |
| 0.6404556554 | 0.0726878339 | GGB30461 | Oscillospiraceae |
| 0.6404556554 | 0.0653072035 | GGB30461 | Oscillospiraceae |
| 0.6404556554 | 0.0724404287 | GGB30463 | Oscillospiraceae |
| 0.6404556554 | 0.0653072035 | GGB30473 | Oscillospiraceae |
| 0.6404556554 | 0.0653072035 | GGB30475 | Oscillospiraceae |
| 0.6404556554 | 0.512289062 | GGB30861 | FGB77153 |
| 0.6404556554 | 0.0724404287 | GGB31312 | FGB1791 |
| 0.9612762186 | 0.0653072035 | GGB31438 | FGB10290 |
| 0.9557341024 | 0.0653072035 | GGB3171 | Oscillospiraceae |
| 0.6404556554 | 0.0653072035 | GGB31762 | FGB10289 |
| 0.6404556554 | 0.0653072035 | GGB31823 | FGB1765 |
| 0.6404556554 | 0.0905436793 | GGB31838 | FGB1765 |
| 0.8098448563 | 0.0653072035 | GGB31841 | FGB1765 |
| 0.6404556554 | 0.0653072035 | GGB32371 | FGB10667 |
| 0.6404556554 | 0.0653072035 | GGB3793 | Lachnospiraceae |
| 0.6404556554 | 0.0653072035 | GGB42601 | Clostridiaceae |
| 0.6404556554 | 0.0653072035 | GGB45514 | Oscillospiraceae |
| 0.6404556554 | 0.0653072035 | GGB45564 | FGB75721 |
| 0.6979515333 | 0.0653072035 | GGB45624 | Oscillospiraceae |
| 0.7037808067 | 0.0653072035 | GGB45656 | Christensenellaceae |
| 0.6404556554 | 0.0653072035 | GGB47127 | FGB10299 |

| | | | |
|---|---|---|---|
| 0.6404556554 | 0.176203079 | GGB74395 | Oscillospiraceae |
| 0.6404556554 | 0.072687833 | GGB75053 | Oscillospiraceae |
| 0.6404556554 | 0.090048525 | GGB75109 | Lachnospiraceae |
| 0.6404556554 | 0.075097602 | Lachnospiraceae_unclassified | Lachnospiraceae |
| 0.6404556554 | 0.065307203 | Lachnospiraceae_unclassified | Lachnospiraceae |
| 0.6404556554 | 0.065307203 | Lachnospiraceae_unclassified | Lachnospiraceae |
| 0.6404556554 | 0.073650255 | Lachnospiraceae_unclassified | Lachnospiraceae |
| 0.6404556554 | 0.072562568 | Lachnospiraceae_unclassified | Lachnospiraceae |
| 0.6404556554 | 0.173081586 | Lactobacillus | Lactobacillaceae |
| 0.6404556554 | 0.065307203 | Leptogranulimonas | Atopobiaceae |
| 0.6404556554 | 0.065307203 | Muribaculaceae_unclassified | Muribaculaceae |
| 0.6404556554 | 0.066476697 | Neglectibacter | Oscillospiraceae |
| 0.6404556554 | 0.065307203 | Oscillibacter | Oscillospiraceae |
| 0.649195481 | 0.065307203 | Oscillospiraceae_unclassified | Oscillospiraceae |
| 0.6404556554 | 0.072440428 | Oscillospiraceae_unclassified | Oscillospiraceae |
| 0.8098448563 | 0.197298710 | Oscillospiraceae_unclassified | Oscillospiraceae |
| 0.6404556554 | 0.066476697 | Oscillospiraceae_unclassified | Oscillospiraceae |
| 0.6404556554 | 0.450086613 | Parasutterella | Sutterellaceae |
| 0.6404556554 | 0.072687833 | Schaedlerella | Lachnospiraceae |
| 0.9612762186 | 0.065307203 | Turicibacter | Turicibacteraceae |
| 0.6404556554 | 0.065931569 | Bacteria_unclassified | Bacteria_unclassified |
| 0.6404556554 | 0.066476697 | Bacteria_unclassified | Bacteria_unclassified |
| 0.6404556554 | 0.114258081 | Bacteria_unclassified | Bacteria_unclassified |
| 0.6404556554 | 0.056872449 | Bacteria_unclassified | Bacteria_unclassified |
| 0.6404556554 | 0.065307203 | Bacteria_unclassified | Bacteria_unclassified |
| 0.6404556554 | 0.065307203 | | |
| 0.6404556554 | 0.070115235 | | |
| 0.6404556554 | 0.065307203 | | |
| 0.4691765424 | 0.124463106 | Acetatifactor | Lachnospiraceae |
| 0.4691765424 | 0.124898611 | Acutalibacter | Oscillospiraceae |
| 0.4691765424 | 0.124463106 | Akkermansia | Akkermansiaceae |
| 0.4691765424 | 0.124463106 | Anaerotruncus | Oscillospiraceae |
| 0.4691765424 | 0.366705682 | Bacteria_unclassified | Bacteria_unclassified |
| 0.4691765424 | 0.124463106 | Bacteria_unclassified | Bacteria_unclassified |
| 0.4691765424 | 0.124463106 | Bacteria_unclassified | Bacteria_unclassified |
| 0.5052767294 | 0.124463106 | Bacteria_unclassified | Bacteria_unclassified |
| 0.4691765424 | 0.124463106 | Bacteroides | Bacteroidaceae |
| 0.4691765424 | 0.124463106 | Clostridia_unclassified | Clostridia_unclassified |
| 0.4691765424 | 0.124463106 | Clostridiaceae_unclassified | Clostridiaceae |
| 0.4691765424 | 0.124463106 | Clostridiaceae_unclassified | Clostridiaceae |
| 0.4691765424 | 0.124463106 | Eubacteriales_unclassified | Eubacteriales_unclassified |
| 0.4691765424 | 0.897247569 | Clostridium | Clostridiaceae |
| 0.4691765424 | 0.124898611 | Erysipelatoclostridium | Erysipelotrichaceae |
| 0.4691765424 | 0.129355343 | Eubacteriaceae_unclassified | Eubacteriaceae |
| 0.5052767294 | 0.124463106 | Eubacteriaceae_unclassified | Eubacteriaceae |
| 0.6567032279 | 0.124463106 | GGB20146 | FGB77306 |

| | | | |
|---|---|---|---|
| 0.5138434935 | 0.124463106 | GGB20149 | Lachnospiraceae |
| 0.4691765424 | 0.124463106 | GGB22635 | Eggerthellaceae |
| 0.4829591757 | 0.294164396 | GGB25041 | Lachnospiraceae |
| 0.4691765424 | 0.828439635 | GGB28379 | FGB9506 |
| 0.4691765424 | 0.124463106 | GGB28382 | FGB9508 |
| 0.7605235377 | 0.124463106 | GGB28392 | FGB9512 |
| 0.4691765424 | 0.124463106 | GGB28404 | FGB2838 |
| 0.4691765424 | 0.124463106 | GGB28418 | FGB2838 |
| 0.4691765424 | 0.124463106 | GGB28422 | FGB2838 |
| 0.5364485609 | 0.129355343 | GGB28431 | Pumilibacteraceae |
| 0.4691765424 | 0.124463106 | GGB28456 | FGB2833 |
| 0.4691765424 | 0.129355343 | GGB28782 | Eubacteriaceae |
| 0.4691765424 | 0.124463106 | GGB28792 | Lachnospiraceae |
| 0.4691765424 | 0.124463106 | GGB28798 | Lachnospiraceae |
| 0.4691765424 | 0.223115026 | GGB28810 | FGB9622 |
| 0.8020250797 | 0.124463106 | GGB28828 | FGB77305 |
| 0.4691765424 | 0.124463106 | GGB28851 | Clostridiaceae |
| 0.9424038683 | 0.452904799 | GGB28865 | Lachnospiraceae |
| 0.4691765424 | 0.124463106 | GGB28868 | Lachnospiraceae |
| 0.4691765424 | 0.215261293 | GGB28869 | Lachnospiraceae |
| 0.4691765424 | 0.124463106 | GGB28875 | Lachnospiraceae |
| 0.4691765424 | 0.124463106 | GGB28881 | FGB9633 |
| 0.4691765424 | 0.124463106 | GGB28892 | Bacteria_unclassified |
| 0.5034951558 | 0.124463106 | GGB28893 | Bacteria_unclassified |
| 0.4691765424 | 0.187387585 | GGB28898 | Bacteria_unclassified |
| 0.4691765424 | 0.124463106 | GGB28901 | Bacteria_unclassified |
| 0.6265341538 | 0.124463106 | GGB28904 | Bacteria_unclassified |
| 0.5138434935 | 0.124898611 | GGB28909 | Bacteria_unclassified |
| 0.4840404152 | 0.766269998 | GGB28924 | Lachnospiraceae |
| 0.4691765424 | 0.124463106 | GGB28927 | FGB77359 |
| 0.4691765424 | 0.124463106 | GGB28934 | FGB9639 |
| 0.4691765424 | 0.124463106 | GGB28949 | Lachnospiraceae |
| 0.4691765424 | 0.124463106 | GGB28951 | Clostridiaceae |
| 0.4691765424 | 0.124463106 | GGB28951 | Clostridiaceae |
| 0.4691765424 | 0.182067783 | GGB28954 | Clostridiaceae |
| 0.4691765424 | 0.135092316 | GGB28960 | Clostridiaceae |
| 0.4691765424 | 0.124463106 | GGB28964 | Clostridiaceae |
| 0.4691765424 | 0.124463106 | GGB28967 | Clostridiaceae |
| 0.4691765424 | 0.124463106 | GGB28996 | FGB9656 |
| 0.4691765424 | 0.124463106 | GGB29531 | FGB9827 |
| 0.4691765424 | 0.124898611 | GGB29685 | Eubacteriaceae |
| 0.4691765424 | 0.162790929 | GGB30141 | FGB77303 |
| 0.5052767294 | 0.124463106 | GGB30145 | Clostridiaceae |
| 0.4691765424 | 0.124463106 | GGB30286 | Eubacteriales_unclassified |
| 0.4833746032 | 0.124463106 | GGB30300 | FGB72709 |
| 0.4691765424 | 0.124463106 | GGB30303 | Oscillospiraceae |

| | | | |
|---|---|---|---|
| 0.5285348257 | 0.124463106 | GGB30450 | Oscillospiraceae |
| 0.4691765424 | 0.124463106 | GGB30453 | Oscillospiraceae |
| 0.4691765424 | 0.154912135 | GGB30454 | Oscillospiraceae |
| 0.4691765424 | 0.124463106 | GGB30455 | Oscillospiraceae |
| 0.4691765424 | 0.124463106 | GGB30456 | Oscillospiraceae |
| 0.4691765424 | 0.124898611 | GGB30457 | Oscillospiraceae |
| 0.4691765424 | 0.124463106 | GGB30461 | Oscillospiraceae |
| 0.4691765424 | 0.124463106 | GGB30461 | Oscillospiraceae |
| 0.5285348257 | 0.124463106 | GGB30461 | Oscillospiraceae |
| 0.4691765424 | 0.151921399 | GGB30463 | Oscillospiraceae |
| 0.5218389679 | 0.124463106 | GGB30473 | Oscillospiraceae |
| 0.4691765424 | 0.124463106 | GGB30475 | Oscillospiraceae |
| 0.4691765424 | 0.170542880 | GGB30861 | FGB77153 |
| 0.4691765424 | 0.151921399 | GGB31312 | FGB1791 |
| 0.4691765424 | 0.124463106 | GGB31438 | FGB10290 |
| 0.4691765424 | 0.124463106 | GGB3171 | Oscillospiraceae |
| 0.4691765424 | 0.124463106 | GGB31762 | FGB10289 |
| 0.4691765424 | 0.124898611 | GGB31823 | FGB1765 |
| 0.4691765424 | 0.124463106 | GGB31838 | FGB1765 |
| 0.4691765424 | 0.124463106 | GGB31841 | FGB1765 |
| 0.4691765424 | 0.124463106 | GGB32371 | FGB10667 |
| 0.4691765424 | 0.124463106 | GGB3793 | Lachnospiraceae |
| 0.4691765424 | 0.129355343 | GGB42601 | Clostridiaceae |
| 0.4691765424 | 0.124463106 | GGB45514 | Oscillospiraceae |
| 0.5027050147 | 0.125327738 | GGB45564 | FGB75721 |
| 0.4691765424 | 0.124463106 | GGB45624 | Oscillospiraceae |
| 0.4833746032 | 0.124463106 | GGB45656 | Christensenellaceae |
| 0.4691765424 | 0.124463106 | GGB47127 | FGB10299 |
| 0.4691765424 | 0.124898611 | GGB74395 | Oscillospiraceae |
| 0.5833419612 | 0.13259174 | GGB75053 | Oscillospiraceae |
| 0.4829591757 | 0.151921399 | GGB75109 | Lachnospiraceae |
| 0.4691765424 | 0.124898611 | Lachnospiraceae_unclassified | Lachnospiraceae |
| 0.4691765424 | 0.124898611 | Lachnospiraceae_unclassified | Lachnospiraceae |
| 0.4691765424 | 0.124463106 | Lachnospiraceae_unclassified | Lachnospiraceae |
| 0.4691765424 | 0.13259174 | Lachnospiraceae_unclassified | Lachnospiraceae |
| 0.4691765424 | 0.199900353 | Lachnospiraceae_unclassified | Lachnospiraceae |
| 0.4691765424 | 0.182022749 | Lactobacillus | Lactobacillaceae |
| 0.4691765424 | 0.124463106 | Leptogranulimonas | Atopobiaceae |
| 0.4691765424 | 0.124463106 | Muribaculaceae_unclassified | Muribaculaceae |
| 0.4691765424 | 0.124463106 | Neglectibacter | Oscillospiraceae |
| 0.4691765424 | 0.124463106 | Oscillibacter | Oscillospiraceae |
| 0.4691765424 | 0.124463106 | Oscillospiraceae_unclassified | Oscillospiraceae |
| 0.4833746032 | 0.124898611 | Oscillospiraceae_unclassified | Oscillospiraceae |
| 0.4691765424 | 0.170113100 | Oscillospiraceae_unclassified | Oscillospiraceae |
| 0.4691765424 | 0.124463106 | Oscillospiraceae_unclassified | Oscillospiraceae |
| 0.4691765424 | 0.250613129 | Parasutterella | Sutterellaceae |

| | | | |
|---|---|---|---|
| 0.4691765424 | 0.124898611 | Schaedlerella | Lachnospiraceae |
| 0.4691765424 | 0.124463106 | Turicibacter | Turicibacteraceae |
| 0.4691765424 | 0.124898611 | Bacteria_unclassified | Bacteria_unclassified |
| 0.4691765424 | 0.124463106 | Bacteria_unclassified | Bacteria_unclassified |
| 0.4833746032 | 0.212908512 | Bacteria_unclassified | Bacteria_unclassified |
| 0.7605235377 | 0.124463106 | Bacteria_unclassified | Bacteria_unclassified |
| 0.4691765424 | 0.124463106 | Bacteria_unclassified | Bacteria_unclassified |
| 0.4691765424 | 0.124463106 | | |
| 0.4691765424 | 0.125327738 | | |
| 0.4691765424 | 0.124463106 | | |
| 0.7772452641 | 0.139167094 | Acetatifactor | Lachnospiraceae |
| 0.7772452641 | 0.139167094 | Acutalibacter | Oscillospiraceae |
| 0.7884879304 | 0.173512107 | Akkermansia | Akkermansiaceae |
| 0.7772452641 | 0.139167094 | Anaerotruncus | Oscillospiraceae |
| 0.7772452641 | 0.139167094 | Bacteria_unclassified | Bacteria_unclassified |
| 0.7772452641 | 0.139167094 | Bacteria_unclassified | Bacteria_unclassified |
| 0.7887714525 | 0.159072419 | Bacteria_unclassified | Bacteria_unclassified |
| 0.7772452641 | 0.173512107 | Bacteria_unclassified | Bacteria_unclassified |
| 0.7871876904 | 0.139167094 | Bacteroides | Bacteroidaceae |
| 0.7772452641 | 0.139167094 | Clostridia_unclassified | Clostridia_unclassified |
| 0.7772452641 | 0.177332328 | Clostridiaceae_unclassified | Clostridiaceae |
| 0.7772452641 | 0.139167094 | Clostridiaceae_unclassified | Clostridiaceae |
| 0.7772452641 | 0.139167094 | Eubacteriales_unclassified | Eubacteriales_unclassified |
| 0.7772452641 | 0.139167094 | Clostridium | Clostridiaceae |
| 0.7772452641 | 0.139167094 | Erysipelatoclostridium | Erysipelotrichaceae |
| 0.7772452641 | 0.576866502 | Eubacteriaceae_unclassified | Eubacteriaceae |
| 0.8950611082 | 0.141637955 | Eubacteriaceae_unclassified | Eubacteriaceae |
| 0.7772452641 | 0.141637955 | GGB20146 | FGB77306 |
| 0.8632250154 | 0.173512107 | GGB20149 | Lachnospiraceae |
| 0.8522124224 | 0.139167094 | GGB22635 | Eggerthellaceae |
| 0.7871876904 | 0.139167094 | GGB25041 | Lachnospiraceae |
| 0.7772452641 | 0.220044416 | GGB28379 | FGB9506 |
| 0.7772452641 | 0.170603535 | GGB28382 | FGB9508 |
| 0.7772452641 | 0.139167094 | GGB28392 | FGB9512 |
| 0.7772452641 | 0.142106496 | GGB28404 | FGB2838 |
| 0.7772452641 | 0.139167094 | GGB28418 | FGB2838 |
| 0.7772452641 | 0.159072419 | GGB28422 | FGB2838 |
| 0.7772452641 | 0.257477259 | GGB28431 | Pumilibacteraceae |
| 0.7887714525 | 0.139167094 | GGB28456 | FGB2833 |
| 0.7772452641 | 0.139167094 | GGB28782 | Eubacteriaceae |
| 0.7772452641 | 0.139167094 | GGB28792 | Lachnospiraceae |
| 0.7772452641 | 0.139167094 | GGB28798 | Lachnospiraceae |
| 0.7772452641 | 0.160462267 | GGB28810 | FGB9622 |
| 0.7772452641 | 0.139167094 | GGB28828 | FGB77305 |
| 0.7772452641 | 0.141637955 | GGB28851 | Clostridiaceae |
| 0.8786758731 | 0.139167094 | GGB28865 | Lachnospiraceae |

| | | | |
|---|---|---|---|
| 0.7772452641 | 0.141637955 | GGB28868 | Lachnospiraceae |
| 0.7772452641 | 0.139167094 | GGB28869 | Lachnospiraceae |
| 0.7772452641 | 0.173512107 | GGB28875 | Lachnospiraceae |
| 0.9801609216 | 0.139167094 | GGB28881 | FGB9633 |
| 0.7772452641 | 0.217683573 | GGB28892 | Bacteria_unclassified |
| 0.8786758731 | 0.509774224 | GGB28893 | Bacteria_unclassified |
| 0.7772452641 | 0.188219643 | GGB28898 | Bacteria_unclassified |
| 0.8034418472 | 0.141637955 | GGB28901 | Bacteria_unclassified |
| 0.9801609216 | 0.591048594 | GGB28904 | Bacteria_unclassified |
| 0.8030625874 | 0.139167094 | GGB28909 | Bacteria_unclassified |
| 0.7772452641 | 0.336817646 | GGB28924 | Lachnospiraceae |
| 0.8907293381 | 0.139167094 | GGB28927 | FGB77359 |
| 0.7772452641 | 0.139167094 | GGB28934 | FGB9639 |
| 0.7772452641 | 0.139167094 | GGB28949 | Lachnospiraceae |
| 0.7772452641 | 0.139167094 | GGB28951 | Clostridiaceae |
| 0.7772452641 | 0.235914418 | GGB28951 | Clostridiaceae |
| 0.7772452641 | 0.139167094 | GGB28954 | Clostridiaceae |
| 0.7772452641 | 0.139167094 | GGB28960 | Clostridiaceae |
| 0.7884879304 | 0.139167094 | GGB28964 | Clostridiaceae |
| 0.7772452641 | 0.139167094 | GGB28967 | Clostridiaceae |
| 0.7772452641 | 0.139167094 | GGB28996 | FGB9656 |
| 0.7772452641 | 0.139167094 | GGB29531 | FGB9827 |
| 0.7772452641 | 0.341208935 | GGB29685 | Eubacteriaceae |
| 0.7772452641 | 0.549300175 | GGB30141 | FGB77303 |
| 0.7772452641 | 0.139167094 | GGB30145 | Clostridiaceae |
| 0.7772452641 | 0.144096327 | GGB30286 | Eubacteriales_unclassified |
| 0.7772452641 | 0.173512107 | GGB30300 | FGB72709 |
| 0.7772452641 | 0.331317016 | GGB30303 | Oscillospiraceae |
| 0.7772452641 | 0.139167094 | GGB30450 | Oscillospiraceae |
| 0.7772452641 | 0.141503910 | GGB30453 | Oscillospiraceae |
| 0.7772452641 | 0.139167094 | GGB30454 | Oscillospiraceae |
| 0.7772452641 | 0.139167094 | GGB30455 | Oscillospiraceae |
| 0.7772452641 | 0.139167094 | GGB30456 | Oscillospiraceae |
| 0.7772452641 | 0.141503910 | GGB30457 | Oscillospiraceae |
| 0.7772452641 | 0.139167094 | GGB30461 | Oscillospiraceae |
| 0.7772452641 | 0.438018312 | GGB30461 | Oscillospiraceae |
| 0.7772452641 | 0.139167094 | GGB30461 | Oscillospiraceae |
| 0.7772452641 | 0.220044416 | GGB30463 | Oscillospiraceae |
| 0.7896006651 | 0.139167094 | GGB30473 | Oscillospiraceae |
| 0.7772452641 | 0.144096327 | GGB30475 | Oscillospiraceae |
| 0.7772452641 | 0.509774224 | GGB30861 | FGB77153 |
| 0.7772452641 | 0.139167094 | GGB31312 | FGB1791 |
| 0.7772452641 | 0.141637955 | GGB31438 | FGB10290 |
| 0.8907293381 | 0.139167094 | GGB3171 | Oscillospiraceae |
| 0.7772452641 | 0.139167094 | GGB31762 | FGB10289 |
| 0.7772452641 | 0.139167094 | GGB31823 | FGB1765 |

| | | | |
|---|---|---|---|
| 0.7772452641 | 0.549300175 | GGB31838 | FGB1765 |
| 0.7772452641 | 0.139167094 | GGB31841 | FGB1765 |
| 0.8458435901 | 0.144096327 | GGB32371 | FGB10667 |
| 0.7772452641 | 0.139167094 | GGB3793 | Lachnospiraceae |
| 0.7772452641 | 0.139167094 | GGB42601 | Clostridiaceae |
| 0.7772452641 | 0.227093271 | GGB45514 | Oscillospiraceae |
| 0.7887714525 | 0.139167094 | GGB45564 | FGB75721 |
| 0.7772452641 | 0.139167094 | GGB45624 | Oscillospiraceae |
| 0.7772452641 | 0.144096327 | GGB45656 | Christensenellaceae |
| 0.7772452641 | 0.139167094 | GGB47127 | FGB10299 |
| 0.7772452641 | 0.231976041 | GGB74395 | Oscillospiraceae |
| 0.7772452641 | 0.173512107 | GGB75053 | Oscillospiraceae |
| 0.7772452641 | 0.146664038 | GGB75109 | Lachnospiraceae |
| 0.7772452641 | 0.352070901 | Lachnospiraceae_unclassified | Lachnospiraceae |
| 0.7772452641 | 0.139167094 | Lachnospiraceae_unclassified | Lachnospiraceae |
| 0.7772452641 | 0.139167094 | Lachnospiraceae_unclassified | Lachnospiraceae |
| 0.7772452641 | 0.159628987 | Lachnospiraceae_unclassified | Lachnospiraceae |
| 0.7772452641 | 0.139167094 | Lachnospiraceae_unclassified | Lachnospiraceae |
| 0.7772452641 | 0.576866502 | Lactobacillus | Lactobacillaceae |
| 0.7772452641 | 0.339372519 | Leptogranulimonas | Atopobiaceae |
| 0.7772452641 | 0.139167094 | Muribaculaceae_unclassified | Muribaculaceae |
| 0.7772452641 | 0.139167094 | Neglectibacter | Oscillospiraceae |
| 0.7772452641 | 0.139167094 | Oscillibacter | Oscillospiraceae |
| 0.7772452641 | 0.139167094 | Oscillospiraceae_unclassified | Oscillospiraceae |
| 0.7772452641 | 0.210017833 | Oscillospiraceae_unclassified | Oscillospiraceae |
| 0.7772452641 | 0.164584004 | Oscillospiraceae_unclassified | Oscillospiraceae |
| 0.7772452641 | 0.141503910 | Oscillospiraceae_unclassified | Oscillospiraceae |
| 0.7772452641 | 0.270423799 | Parasutterella | Sutterellaceae |
| 0.7887714525 | 0.139167094 | Schaedlerella | Lachnospiraceae |
| 0.7772452641 | 0.217683573 | Turicibacter | Turicibacteraceae |
| 0.7772452641 | 0.141503910 | Bacteria_unclassified | Bacteria_unclassified |
| 0.7772452641 | 0.173512107 | Bacteria_unclassified | Bacteria_unclassified |
| 0.7772452641 | 0.173512107 | Bacteria_unclassified | Bacteria_unclassified |
| 0.7772452641 | 0.139167094 | Bacteria_unclassified | Bacteria_unclassified |
| 0.7772452641 | 0.139167094 | Bacteria_unclassified | Bacteria_unclassified |
| 0.7772452641 | 0.139167094 | | |
| 0.7772452641 | 0.491999147 | | |
| 0.7772452641 | 0.139167094 | | |
| 0.9369099384 | 0.999795788 | Acetatifactor | Lachnospiraceae |
| 0.9369099384 | 0.999795788 | Acutalibacter | Oscillospiraceae |
| 0.9369099384 | 0.999795788 | Akkermansia | Akkermansiaceae |
| 0.9369099384 | 0.999795788 | Anaerotruncus | Oscillospiraceae |
| 0.9369099384 | 0.999795788 | Bacteria_unclassified | Bacteria_unclassified |
| 0.9369099384 | 0.999795788 | Bacteria_unclassified | Bacteria_unclassified |
| 0.9452199076 | 0.999795788 | Bacteria_unclassified | Bacteria_unclassified |
| 0.9369099384 | 0.999795788 | Bacteria_unclassified | Bacteria_unclassified |

| | | | |
|---|---|---|---|
| 0.9369099384 | 0.999795788 | Bacteroides | Bacteroidaceae |
| 0.9369099384 | 0.999795788 | Clostridia_unclassified | Clostridia_unclassified |
| 0.9369099384 | 0.999795788 | Clostridiaceae_unclassified | Clostridiaceae |
| 0.9369099384 | 0.999795788 | Clostridiaceae_unclassified | Clostridiaceae |
| 0.9369099384 | 0.999795788 | Eubacteriales_unclassified | Eubacteriales_unclassified |
| 0.9369099384 | 0.999795788 | Clostridium | Clostridiaceae |
| 0.9369099384 | 0.999795788 | Erysipelatoclostridium | Erysipelotrichaceae |
| 0.9369099384 | 0.999795788 | Eubacteriaceae_unclassified | Eubacteriaceae |
| 0.9369099384 | 0.999795788 | Eubacteriaceae_unclassified | Eubacteriaceae |
| 0.9369099384 | 0.999795788 | GGB20146 | FGB77306 |
| 0.9369099384 | 0.999795788 | GGB20149 | Lachnospiraceae |
| 0.9452199076 | 0.999795788 | GGB22635 | Eggerthellaceae |
| 0.9369099384 | 0.999795788 | GGB25041 | Lachnospiraceae |
| 0.9369099384 | 0.999795788 | GGB28379 | FGB9506 |
| 0.9605965501 | 0.999795788 | GGB28382 | FGB9508 |
| 0.9445683892 | 0.999795788 | GGB28392 | FGB9512 |
| 0.9369099384 | 0.999795788 | GGB28404 | FGB2838 |
| 0.9369099384 | 0.999795788 | GGB28418 | FGB2838 |
| 0.9369099384 | 0.999795788 | GGB28422 | FGB2838 |
| 0.9369099384 | 0.999795788 | GGB28431 | Pumilibacteraceae |
| 0.9369099384 | 0.999795788 | GGB28456 | FGB2833 |
| 0.9369099384 | 0.999795788 | GGB28782 | Eubacteriaceae |
| 0.9369099384 | 0.999795788 | GGB28792 | Lachnospiraceae |
| 0.9369099384 | 0.999795788 | GGB28798 | Lachnospiraceae |
| 0.9369099384 | 0.999795788 | GGB28810 | FGB9622 |
| 0.9369099384 | 0.999795788 | GGB28828 | FGB77305 |
| 0.9369099384 | 0.999795788 | GGB28851 | Clostridiaceae |
| 0.9369099384 | 0.999795788 | GGB28865 | Lachnospiraceae |
| 0.9445683892 | 0.999795788 | GGB28868 | Lachnospiraceae |
| 0.9369099384 | 0.999795788 | GGB28869 | Lachnospiraceae |
| 0.9369099384 | 0.999795788 | GGB28875 | Lachnospiraceae |
| 0.9369099384 | 0.999795788 | GGB28881 | FGB9633 |
| 0.9369099384 | 0.999795788 | GGB28892 | Bacteria_unclassified |
| 0.9369099384 | 0.999795788 | GGB28893 | Bacteria_unclassified |
| 0.9369099384 | 0.999795788 | GGB28898 | Bacteria_unclassified |
| 0.9369099384 | 0.999795788 | GGB28901 | Bacteria_unclassified |
| 0.9369099384 | 0.999795788 | GGB28904 | Bacteria_unclassified |
| 0.9369099384 | 0.999795788 | GGB28909 | Bacteria_unclassified |
| 0.9369099384 | 0.999795788 | GGB28924 | Lachnospiraceae |
| 0.9369099384 | 0.999795788 | GGB28927 | FGB77359 |
| 0.9369099384 | 0.999795788 | GGB28934 | FGB9639 |
| 0.9369099384 | 0.999795788 | GGB28949 | Lachnospiraceae |
| 0.9369099384 | 0.999795788 | GGB28951 | Clostridiaceae |
| 0.9369099384 | 0.999795788 | GGB28951 | Clostridiaceae |
| 0.9369099384 | 0.999795788 | GGB28954 | Clostridiaceae |
| 0.9369099384 | 0.999795788 | GGB28960 | Clostridiaceae |

| | | | |
|---|---|---|---|
| 0.9369099384 | 0.999795788 | GGB28964 | Clostridiaceae |
| 0.9369099384 | 0.999795788 | GGB28967 | Clostridiaceae |
| 0.9445683892 | 0.999795788 | GGB28996 | FGB9656 |
| 0.9369099384 | 0.999795788 | GGB29531 | FGB9827 |
| 0.9369099384 | 0.999795788 | GGB29685 | Eubacteriaceae |
| 0.9369099384 | 0.999795788 | GGB30141 | FGB77303 |
| 0.9369099384 | 0.999795788 | GGB30145 | Clostridiaceae |
| 0.9369099384 | 0.999795788 | GGB30286 | Eubacteriales_unclassified |
| 0.9369099384 | 0.999795788 | GGB30300 | FGB72709 |
| 0.9369099384 | 0.999795788 | GGB30303 | Oscillospiraceae |
| 0.9369099384 | 0.999795788 | GGB30450 | Oscillospiraceae |
| 0.9369099384 | 0.999795788 | GGB30453 | Oscillospiraceae |
| 0.9369099384 | 0.999795788 | GGB30454 | Oscillospiraceae |
| 0.9369099384 | 0.999795788 | GGB30455 | Oscillospiraceae |
| 0.9369099384 | 0.999795788 | GGB30456 | Oscillospiraceae |
| 0.9369099384 | 0.999795788 | GGB30457 | Oscillospiraceae |
| 0.9369099384 | 0.999795788 | GGB30461 | Oscillospiraceae |
| 0.9369099384 | 0.999795788 | GGB30461 | Oscillospiraceae |
| 0.9605965501 | 0.999795788 | GGB30461 | Oscillospiraceae |
| 0.9369099384 | 0.999795788 | GGB30463 | Oscillospiraceae |
| 0.9369099384 | 0.999795788 | GGB30473 | Oscillospiraceae |
| 0.9369099384 | 0.999795788 | GGB30475 | Oscillospiraceae |
| 0.9369099384 | 0.999795788 | GGB30861 | FGB77153 |
| 0.9369099384 | 0.999795788 | GGB31312 | FGB1791 |
| 0.9445683892 | 0.999795788 | GGB31438 | FGB10290 |
| 0.9980271975 | 0.999795788 | GGB3171 | Oscillospiraceae |
| 0.9369099384 | 0.999795788 | GGB31762 | FGB10289 |
| 0.9369099384 | 0.999795788 | GGB31823 | FGB1765 |
| 0.9369099384 | 0.999795788 | GGB31838 | FGB1765 |
| 0.9445683892 | 0.999795788 | GGB31841 | FGB1765 |
| 0.9369099384 | 0.999795788 | GGB32371 | FGB10667 |
| 0.9369099384 | 0.999795788 | GGB3793 | Lachnospiraceae |
| 0.9369099384 | 0.999795788 | GGB42601 | Clostridiaceae |
| 0.9369099384 | 0.999795788 | GGB45514 | Oscillospiraceae |
| 0.9369099384 | 0.999795788 | GGB45564 | FGB75721 |
| 0.9369099384 | 0.999795788 | GGB45624 | Oscillospiraceae |
| 0.9369099384 | 0.999795788 | GGB45656 | Christensenellaceae |
| 0.9445683892 | 0.999795788 | GGB47127 | FGB10299 |
| 0.9369099384 | 0.999795788 | GGB74395 | Oscillospiraceae |
| 0.9369099384 | 0.999795788 | GGB75053 | Oscillospiraceae |
| 0.9369099384 | 0.999795788 | GGB75109 | Lachnospiraceae |
| 0.9369099384 | 0.999795788 | Lachnospiraceae_unclassified | Lachnospiraceae |
| 0.9369099384 | 0.999795788 | Lachnospiraceae_unclassified | Lachnospiraceae |
| 0.9369099384 | 0.999795788 | Lachnospiraceae_unclassified | Lachnospiraceae |
| 0.9369099384 | 0.999795788 | Lachnospiraceae_unclassified | Lachnospiraceae |
| 0.9369099384 | 0.999795788 | Lachnospiraceae_unclassified | Lachnospiraceae |

| | | | |
|---|---|---|---|
| 0.9369099384 | 0.999795788 | Lactobacillus | Lactobacillaceae |
| 0.9369099384 | 0.999795788 | Leptogranulimonas | Atopobiaceae |
| 0.9369099384 | 0.999795788 | Muribaculaceae_unclassified | Muribaculaceae |
| 0.9369099384 | 0.999795788 | Neglectibacter | Oscillospiraceae |
| 0.9369099384 | 0.999795788 | Oscillibacter | Oscillospiraceae |
| 0.9369099384 | 0.999795788 | Oscillospiraceae_unclassified | Oscillospiraceae |
| 0.9369099384 | 0.999795788 | Oscillospiraceae_unclassified | Oscillospiraceae |
| 0.9369099384 | 0.999795788 | Oscillospiraceae_unclassified | Oscillospiraceae |
| 0.9369099384 | 0.999795788 | Oscillospiraceae_unclassified | Oscillospiraceae |
| 0.9369099384 | 0.999795788 | Parasutterella | Sutterellaceae |
| 0.9369099384 | 0.999795788 | Schaedlerella | Lachnospiraceae |
| 0.9369099384 | 0.999795788 | Turicibacter | Turicibacteraceae |
| 0.9369099384 | 0.999795788 | Bacteria_unclassified | Bacteria_unclassified |
| 0.9369099384 | 0.999795788 | Bacteria_unclassified | Bacteria_unclassified |
| 0.9369099384 | 0.999795788 | Bacteria_unclassified | Bacteria_unclassified |
| 0.9445683892 | 0.999795788 | Bacteria_unclassified | Bacteria_unclassified |
| 0.9902008623 | 0.999795788 | Bacteria_unclassified | Bacteria_unclassified |
| 0.9369099384 | 0.999795788 | | |
| 0.9369099384 | 0.999795788 | | |
| 0.9369099384 | 0.999795788 | | |
| 0.9978339659 | 0.449272135 | Acetatifactor | Lachnospiraceae |
| 0.9978339659 | 0.460984392 | Acutalibacter | Oscillospiraceae |
| 0.9978339659 | 0.472980942 | Akkermansia | Akkermansiaceae |
| 0.9978339659 | 0.449272135 | Anaerotruncus | Oscillospiraceae |
| 0.9978339659 | 0.863104617 | Bacteria_unclassified | Bacteria_unclassified |
| 0.9978339659 | 0.449272135 | Bacteria_unclassified | Bacteria_unclassified |
| 0.9978339659 | 0.449272135 | Bacteria_unclassified | Bacteria_unclassified |
| 0.9978339659 | 0.598876942 | Bacteria_unclassified | Bacteria_unclassified |
| 0.9978339659 | 0.449272135 | Bacteroides | Bacteroidaceae |
| 0.9978339659 | 0.449272135 | Clostridia_unclassified | Clostridia_unclassified |
| 0.9978339659 | 0.463209264 | Clostridiaceae_unclassified | Clostridiaceae |
| 0.9978339659 | 0.460984392 | Clostridiaceae_unclassified | Clostridiaceae |
| 0.9978339659 | 0.449272135 | Eubacteriales_unclassified | Eubacteriales_unclassified |
| 0.9978339659 | 0.449272135 | Clostridium | Clostridiaceae |
| 0.9978339659 | 0.449272135 | Erysipelatoclostridium | Erysipelotrichaceae |
| 0.9978339659 | 0.460984392 | Eubacteriaceae_unclassified | Eubacteriaceae |
| 0.9978339659 | 0.449272135 | Eubacteriaceae_unclassified | Eubacteriaceae |
| 0.9978339659 | 0.449272135 | GGB20146 | FGB77306 |
| 0.9978339659 | 0.449272135 | GGB20149 | Lachnospiraceae |
| 0.9978339659 | 0.449272135 | GGB22635 | Eggerthellaceae |
| 0.9978339659 | 0.449272135 | GGB25041 | Lachnospiraceae |
| 0.9978339659 | 0.797931564 | GGB28379 | FGB9506 |
| 0.9978339659 | 0.449272135 | GGB28382 | FGB9508 |
| 0.9978339659 | 0.449272135 | GGB28392 | FGB9512 |
| 0.9978339659 | 0.449272135 | GGB28404 | FGB2838 |
| 0.9978339659 | 0.449272135 | GGB28418 | FGB2838 |

| | | | |
|---|---|---|---|
| 0.9978339659 | 0.449272135: | GGB28422 | FGB2838 |
| 0.9978339659 | 0.569031625: | GGB28431 | Pumilibacteraceae |
| 0.9978339659 | 0.449272135: | GGB28456 | FGB2833 |
| 0.9978339659 | 0.460984392! | GGB28782 | Eubacteriaceae |
| 0.9978339659 | 0.463209264! | GGB28792 | Lachnospiraceae |
| 0.9978339659 | 0.449272135: | GGB28798 | Lachnospiraceae |
| 0.9978339659 | 0.598876942! | GGB28810 | FGB9622 |
| 0.9978339659 | 0.449272135: | GGB28828 | FGB77305 |
| 0.9978339659 | 0.449272135: | GGB28851 | Clostridiaceae |
| 0.9978339659 | 0.629112713: | GGB28865 | Lachnospiraceae |
| 0.9978339659 | 0.449272135: | GGB28868 | Lachnospiraceae |
| 0.9978339659 | 0.565350831! | GGB28869 | Lachnospiraceae |
| 0.9978339659 | 0.463209264! | GGB28875 | Lachnospiraceae |
| 0.9978339659 | 0.463209264! | GGB28881 | FGB9633 |
| 0.9978339659 | 0.449272135: | GGB28892 | Bacteria_unclassified |
| 0.9978339659 | 0.663558991: | GGB28893 | Bacteria_unclassified |
| 0.9978339659 | 0.449272135: | GGB28898 | Bacteria_unclassified |
| 0.9978339659 | 0.449272135: | GGB28901 | Bacteria_unclassified |
| 0.9978339659 | 0.449272135: | GGB28904 | Bacteria_unclassified |
| 0.9978339659 | 0.449272135: | GGB28909 | Bacteria_unclassified |
| 0.9978339659 | 0.449272135: | GGB28924 | Lachnospiraceae |
| 0.9978339659 | 0.449272135: | GGB28927 | FGB77359 |
| 0.9978339659 | 0.449272135: | GGB28934 | FGB9639 |
| 0.9978339659 | 0.449272135: | GGB28949 | Lachnospiraceae |
| 0.9978339659 | 0.449272135: | GGB28951 | Clostridiaceae |
| 0.9978339659 | 0.552950450 | GGB28951 | Clostridiaceae |
| 0.9978339659 | 0.463209264! | GGB28954 | Clostridiaceae |
| 0.9978339659 | 0.629112713: | GGB28960 | Clostridiaceae |
| 0.9978339659 | 0.449272135: | GGB28964 | Clostridiaceae |
| 0.9978339659 | 0.450957040 | GGB28967 | Clostridiaceae |
| 0.9978339659 | 0.449272135: | GGB28996 | FGB9656 |
| 0.9978339659 | 0.449272135: | GGB29531 | FGB9827 |
| 0.9978339659 | 0.528600826: | GGB29685 | Eubacteriaceae |
| 0.9978339659 | 0.449272135: | GGB30141 | FGB77303 |
| 0.9978339659 | 0.463209264! | GGB30145 | Clostridiaceae |
| 0.9978339659 | 0.460984392! | GGB30286 | Eubacteriales_unclassified |
| 0.9978339659 | 0.778462690! | GGB30300 | FGB72709 |
| 0.9978339659 | 0.528600826: | GGB30303 | Oscillospiraceae |
| 0.9978339659 | 0.463209264! | GGB30450 | Oscillospiraceae |
| 0.9978339659 | 0.449272135: | GGB30453 | Oscillospiraceae |
| 0.9978339659 | 0.449272135: | GGB30454 | Oscillospiraceae |
| 0.9978339659 | 0.449272135: | GGB30455 | Oscillospiraceae |
| 0.9978339659 | 0.449272135: | GGB30456 | Oscillospiraceae |
| 0.9978339659 | 0.450957040 | GGB30457 | Oscillospiraceae |
| 0.9978339659 | 0.460984392! | GGB30461 | Oscillospiraceae |
| 0.9978339659 | 0.449272135: | GGB30461 | Oscillospiraceae |

| | | | |
|---|---|---|---|
| 0.9978339659 | 0.449272135: | GGB30461 | Oscillospiraceae |
| 0.9978339659 | 0.460984392! | GGB30463 | Oscillospiraceae |
| 0.9978339659 | 0.449272135: | GGB30473 | Oscillospiraceae |
| 0.9978339659 | 0.460984392! | GGB30475 | Oscillospiraceae |
| 0.9978339659 | 0.449272135: | GGB30861 | FGB77153 |
| 0.9978339659 | 0.449272135: | GGB31312 | FGB1791 |
| 0.9978339659 | 0.460984392! | GGB31438 | FGB10290 |
| 0.9978339659 | 0.565350831! | GGB3171 | Oscillospiraceae |
| 0.9978339659 | 0.449272135: | GGB31762 | FGB10289 |
| 0.9978339659 | 0.460984392! | GGB31823 | FGB1765 |
| 0.9978339659 | 0.449272135: | GGB31838 | FGB1765 |
| 0.9978339659 | 0.449272135: | GGB31841 | FGB1765 |
| 0.9978339659 | 0.450957040 | GGB32371 | FGB10667 |
| 0.9978339659 | 0.449272135: | GGB3793 | Lachnospiraceae |
| 0.9978339659 | 0.463209264! | GGB42601 | Clostridiaceae |
| 0.9978339659 | 0.460984392! | GGB45514 | Oscillospiraceae |
| 0.9978339659 | 0.460984392! | GGB45564 | FGB75721 |
| 0.9978339659 | 0.449272135: | GGB45624 | Oscillospiraceae |
| 0.9978339659 | 0.449272135: | GGB45656 | Christensenellaceae |
| 0.9978339659 | 0.463209264! | GGB47127 | FGB10299 |
| 0.9978339659 | 0.449272135: | GGB74395 | Oscillospiraceae |
| 0.9978339659 | 0.449272135: | GGB75053 | Oscillospiraceae |
| 0.9978339659 | 0.449272135: | GGB75109 | Lachnospiraceae |
| 0.9978339659 | 0.463209264! | Lachnospiraceae_unclassified | Lachnospiraceae |
| 0.9978339659 | 0.463209264! | Lachnospiraceae_unclassified | Lachnospiraceae |
| 0.9978339659 | 0.449272135: | Lachnospiraceae_unclassified | Lachnospiraceae |
| 0.9978339659 | 0.449272135: | Lachnospiraceae_unclassified | Lachnospiraceae |
| 0.9978339659 | 0.463209264! | Lachnospiraceae_unclassified | Lachnospiraceae |
| 0.9978339659 | 0.460984392! | Lactobacillus | Lactobacillaceae |
| 0.9978339659 | 0.798695268( | Leptogranulimonas | Atopobiaceae |
| 0.9978339659 | 0.449272135: | Muribaculaceae_unclassified | Muribaculaceae |
| 0.9978339659 | 0.449272135: | Neglectibacter | Oscillospiraceae |
| 0.9978339659 | 0.449272135: | Oscillibacter | Oscillospiraceae |
| 0.9978339659 | 0.449272135: | Oscillospiraceae_unclassified | Oscillospiraceae |
| 0.9978339659 | 0.463209264! | Oscillospiraceae_unclassified | Oscillospiraceae |
| 0.9978339659 | 0.449272135: | Oscillospiraceae_unclassified | Oscillospiraceae |
| 0.9978339659 | 0.449272135: | Oscillospiraceae_unclassified | Oscillospiraceae |
| 0.9978339659 | 0.460984392! | Parasutterella | Sutterellaceae |
| 0.9978339659 | 0.463209264! | Schaedlerella | Lachnospiraceae |
| 0.9978339659 | 0.463209264! | Turicibacter | Turicibacteraceae |
| 0.9978339659 | 0.449272135: | Bacteria_unclassified | Bacteria_unclassified |
| 0.9978339659 | 0.528600826! | Bacteria_unclassified | Bacteria_unclassified |
| 0.9978339659 | 0.449272135: | Bacteria_unclassified | Bacteria_unclassified |
| 0.9978339659 | 0.449272135: | Bacteria_unclassified | Bacteria_unclassified |
| 0.9978339659 | 0.449272135: | Bacteria_unclassified | Bacteria_unclassified |
| 0.9978339659 | 0.449272135: | | |

| | |
|---|---|
| 0.9978339659 | 0.579163027[illegible] |
| 0.9978339659 | 0.449272135[illegible] |

| Order | Class | Phylum | Kingdom |
|---|---|---|---|
| Eubacteriales | Clostridia | Firmicutes | Bacteria |
| Eubacteriales | Clostridia | Firmicutes | Bacteria |
| Verrucomicrobiales | Verrucomicrobiae | Verrucomicrobia | Bacteria |
| Eubacteriales | Clostridia | Firmicutes | Bacteria |
| Bacteria_unclassified | Bacteria_unclassified | Bacteria_unclassified | Bacteria |
| Bacteria_unclassified | Bacteria_unclassified | Bacteria_unclassified | Bacteria |
| Bacteria_unclassified | Bacteria_unclassified | Bacteria_unclassified | Bacteria |
| Bacteria_unclassified | Bacteria_unclassified | Bacteria_unclassified | Bacteria |
| Bacteroidales | Bacteroidia | Bacteroidota | Bacteria |
| Clostridia_unclassified | Clostridia | Firmicutes | Bacteria |
| Eubacteriales | Clostridia | Firmicutes | Bacteria |
| Eubacteriales | Clostridia | Firmicutes | Bacteria |
| Eubacteriales | Clostridia | Firmicutes | Bacteria |
| Eubacteriales | Clostridia | Firmicutes | Bacteria |
| Erysipelotrichales | Erysipelotrichia | Firmicutes | Bacteria |
| Eubacteriales | Clostridia | Firmicutes | Bacteria |
| Eubacteriales | Clostridia | Firmicutes | Bacteria |
| OFGB77306 | CFGB77306 | Firmicutes | Bacteria |
| Eubacteriales | Clostridia | Firmicutes | Bacteria |
| Eggerthellales | Coriobacteriia | Actinobacteria | Bacteria |
| Eubacteriales | Clostridia | Firmicutes | Bacteria |
| OFGB9506 | CFGB9506 | Firmicutes | Bacteria |
| OFGB9508 | CFGB9508 | Firmicutes | Bacteria |
| OFGB9512 | CFGB9512 | Firmicutes | Bacteria |
| OFGB2838 | CFGB2838 | Firmicutes | Bacteria |
| OFGB2838 | CFGB2838 | Firmicutes | Bacteria |
| OFGB2838 | CFGB2838 | Firmicutes | Bacteria |
| Eubacteriales | Clostridia | Firmicutes | Bacteria |
| OFGB2833 | CFGB2833 | Firmicutes | Bacteria |
| Eubacteriales | Clostridia | Firmicutes | Bacteria |
| Eubacteriales | Clostridia | Firmicutes | Bacteria |
| Eubacteriales | Clostridia | Firmicutes | Bacteria |
| OFGB9622 | CFGB9622 | Firmicutes | Bacteria |
| OFGB77305 | CFGB77305 | Firmicutes | Bacteria |
| Eubacteriales | Clostridia | Firmicutes | Bacteria |
| Eubacteriales | Clostridia | Firmicutes | Bacteria |
| Eubacteriales | Clostridia | Firmicutes | Bacteria |
| Eubacteriales | Clostridia | Firmicutes | Bacteria |
| Eubacteriales | Clostridia | Firmicutes | Bacteria |
| OFGB9633 | CFGB9633 | Firmicutes | Bacteria |
| Bacteria_unclassified | Bacteria_unclassified | Bacteria_unclassified | Bacteria |
| Bacteria_unclassified | Bacteria_unclassified | Bacteria_unclassified | Bacteria |
| Bacteria_unclassified | Bacteria_unclassified | Bacteria_unclassified | Bacteria |
| Bacteria_unclassified | Bacteria_unclassified | Bacteria_unclassified | Bacteria |
| Bacteria_unclassified | Bacteria_unclassified | Bacteria_unclassified | Bacteria |
| Bacteria_unclassified | Bacteria_unclassified | Bacteria_unclassified | Bacteria |

| | | | |
|---|---|---|---|
| Eubacteriales | Clostridia | Firmicutes | Bacteria |
| OFGB77359 | CFGB77359 | Bacteria_unclassified | Bacteria |
| OFGB9639 | CFGB9639 | Firmicutes | Bacteria |
| Eubacteriales | Clostridia | Firmicutes | Bacteria |
| Eubacteriales | Clostridia | Firmicutes | Bacteria |
| Eubacteriales | Clostridia | Firmicutes | Bacteria |
| Eubacteriales | Clostridia | Firmicutes | Bacteria |
| Eubacteriales | Clostridia | Firmicutes | Bacteria |
| Eubacteriales | Clostridia | Firmicutes | Bacteria |
| Eubacteriales | Clostridia | Firmicutes | Bacteria |
| OFGB9656 | CFGB9656 | Firmicutes | Bacteria |
| OFGB9827 | CFGB9827 | Firmicutes | Bacteria |
| Eubacteriales | Clostridia | Firmicutes | Bacteria |
| OFGB77303 | CFGB77303 | Bacteria_unclassified | Bacteria |
| Eubacteriales | Clostridia | Firmicutes | Bacteria |
| Eubacteriales | Clostridia | Firmicutes | Bacteria |
| OFGB72709 | CFGB72709 | Firmicutes | Bacteria |
| Eubacteriales | Clostridia | Firmicutes | Bacteria |
| Eubacteriales | Clostridia | Firmicutes | Bacteria |
| Eubacteriales | Clostridia | Firmicutes | Bacteria |
| Eubacteriales | Clostridia | Firmicutes | Bacteria |
| Eubacteriales | Clostridia | Firmicutes | Bacteria |
| Eubacteriales | Clostridia | Firmicutes | Bacteria |
| Eubacteriales | Clostridia | Firmicutes | Bacteria |
| Eubacteriales | Clostridia | Firmicutes | Bacteria |
| Eubacteriales | Clostridia | Firmicutes | Bacteria |
| Eubacteriales | Clostridia | Firmicutes | Bacteria |
| Eubacteriales | Clostridia | Firmicutes | Bacteria |
| Eubacteriales | Clostridia | Firmicutes | Bacteria |
| Eubacteriales | Clostridia | Firmicutes | Bacteria |
| OFGB77153 | CFGB77153 | Actinobacteria | Bacteria |
| OFGB1791 | CFGB1791 | Tenericutes | Bacteria |
| OFGB10290 | CFGB10290 | Firmicutes | Bacteria |
| Eubacteriales | Clostridia | Firmicutes | Bacteria |
| OFGB10289 | CFGB10289 | Firmicutes | Bacteria |
| OFGB1765 | CFGB1765 | Firmicutes | Bacteria |
| OFGB1765 | CFGB1765 | Firmicutes | Bacteria |
| OFGB1765 | CFGB1765 | Firmicutes | Bacteria |
| OFGB10667 | CFGB10667 | Firmicutes | Bacteria |
| Eubacteriales | Clostridia | Firmicutes | Bacteria |
| Eubacteriales | Clostridia | Firmicutes | Bacteria |
| Eubacteriales | Clostridia | Firmicutes | Bacteria |
| OFGB75721 | CFGB75721 | Firmicutes | Bacteria |
| Eubacteriales | Clostridia | Firmicutes | Bacteria |
| Eubacteriales | Clostridia | Firmicutes | Bacteria |
| OFGB10299 | CFGB10299 | Firmicutes | Bacteria |

| | | | |
|---|---|---|---|
| Eubacteriales | Clostridia | Firmicutes | Bacteria |
| Eubacteriales | Clostridia | Firmicutes | Bacteria |
| Eubacteriales | Clostridia | Firmicutes | Bacteria |
| Eubacteriales | Clostridia | Firmicutes | Bacteria |
| Eubacteriales | Clostridia | Firmicutes | Bacteria |
| Eubacteriales | Clostridia | Firmicutes | Bacteria |
| Eubacteriales | Clostridia | Firmicutes | Bacteria |
| Eubacteriales | Clostridia | Firmicutes | Bacteria |
| Lactobacillales | Bacilli | Firmicutes | Bacteria |
| Coriobacteriales | Coriobacteriia | Actinobacteria | Bacteria |
| Bacteroidales | Bacteroidia | Bacteroidota | Bacteria |
| Eubacteriales | Clostridia | Firmicutes | Bacteria |
| Eubacteriales | Clostridia | Firmicutes | Bacteria |
| Eubacteriales | Clostridia | Firmicutes | Bacteria |
| Eubacteriales | Clostridia | Firmicutes | Bacteria |
| Eubacteriales | Clostridia | Firmicutes | Bacteria |
| Eubacteriales | Clostridia | Firmicutes | Bacteria |
| Burkholderiales | Betaproteobacteria | Proteobacteria | Bacteria |
| Eubacteriales | Clostridia | Firmicutes | Bacteria |
| Erysipelotrichales | Erysipelotrichia | Firmicutes | Bacteria |
| Bacteria_unclassified | Bacteria_unclassified | Bacteria_unclassified | Bacteria |
| Bacteria_unclassified | Bacteria_unclassified | Bacteria_unclassified | Bacteria |
| Bacteria_unclassified | Bacteria_unclassified | Bacteria_unclassified | Bacteria |
| Bacteria_unclassified | Bacteria_unclassified | Bacteria_unclassified | Bacteria |
| Bacteria_unclassified | Bacteria_unclassified | Bacteria_unclassified | Bacteria |

| | | | |
|---|---|---|---|
| Eubacteriales | Clostridia | Firmicutes | Bacteria |
| Eubacteriales | Clostridia | Firmicutes | Bacteria |
| Verrucomicrobiales | Verrucomicrobiae | Verrucomicrobia | Bacteria |
| Eubacteriales | Clostridia | Firmicutes | Bacteria |
| Bacteria_unclassified | Bacteria_unclassified | Bacteria_unclassified | Bacteria |
| Bacteria_unclassified | Bacteria_unclassified | Bacteria_unclassified | Bacteria |
| Bacteria_unclassified | Bacteria_unclassified | Bacteria_unclassified | Bacteria |
| Bacteria_unclassified | Bacteria_unclassified | Bacteria_unclassified | Bacteria |
| Bacteroidales | Bacteroidia | Bacteroidota | Bacteria |
| Clostridia_unclassified | Clostridia | Firmicutes | Bacteria |
| Eubacteriales | Clostridia | Firmicutes | Bacteria |
| Eubacteriales | Clostridia | Firmicutes | Bacteria |
| Eubacteriales | Clostridia | Firmicutes | Bacteria |
| Eubacteriales | Clostridia | Firmicutes | Bacteria |
| Erysipelotrichales | Erysipelotrichia | Firmicutes | Bacteria |
| Eubacteriales | Clostridia | Firmicutes | Bacteria |
| Eubacteriales | Clostridia | Firmicutes | Bacteria |
| OFGB77306 | CFGB77306 | Firmicutes | Bacteria |

| | | | |
|---|---|---|---|
| Eubacteriales | Clostridia | Firmicutes | Bacteria |
| Eggerthellales | Coriobacteriia | Actinobacteria | Bacteria |
| Eubacteriales | Clostridia | Firmicutes | Bacteria |
| OFGB9506 | CFGB9506 | Firmicutes | Bacteria |
| OFGB9508 | CFGB9508 | Firmicutes | Bacteria |
| OFGB9512 | CFGB9512 | Firmicutes | Bacteria |
| OFGB2838 | CFGB2838 | Firmicutes | Bacteria |
| OFGB2838 | CFGB2838 | Firmicutes | Bacteria |
| OFGB2838 | CFGB2838 | Firmicutes | Bacteria |
| Eubacteriales | Clostridia | Firmicutes | Bacteria |
| OFGB2833 | CFGB2833 | Firmicutes | Bacteria |
| Eubacteriales | Clostridia | Firmicutes | Bacteria |
| Eubacteriales | Clostridia | Firmicutes | Bacteria |
| Eubacteriales | Clostridia | Firmicutes | Bacteria |
| OFGB9622 | CFGB9622 | Firmicutes | Bacteria |
| OFGB77305 | CFGB77305 | Firmicutes | Bacteria |
| Eubacteriales | Clostridia | Firmicutes | Bacteria |
| Eubacteriales | Clostridia | Firmicutes | Bacteria |
| Eubacteriales | Clostridia | Firmicutes | Bacteria |
| Eubacteriales | Clostridia | Firmicutes | Bacteria |
| Eubacteriales | Clostridia | Firmicutes | Bacteria |
| OFGB9633 | CFGB9633 | Firmicutes | Bacteria |
| Bacteria_unclassified | Bacteria_unclassified | Bacteria_unclassified | Bacteria |
| Bacteria_unclassified | Bacteria_unclassified | Bacteria_unclassified | Bacteria |
| Bacteria_unclassified | Bacteria_unclassified | Bacteria_unclassified | Bacteria |
| Bacteria_unclassified | Bacteria_unclassified | Bacteria_unclassified | Bacteria |
| Bacteria_unclassified | Bacteria_unclassified | Bacteria_unclassified | Bacteria |
| Bacteria_unclassified | Bacteria_unclassified | Bacteria_unclassified | Bacteria |
| Eubacteriales | Clostridia | Firmicutes | Bacteria |
| OFGB77359 | CFGB77359 | Bacteria_unclassified | Bacteria |
| OFGB9639 | CFGB9639 | Firmicutes | Bacteria |
| Eubacteriales | Clostridia | Firmicutes | Bacteria |
| Eubacteriales | Clostridia | Firmicutes | Bacteria |
| Eubacteriales | Clostridia | Firmicutes | Bacteria |
| Eubacteriales | Clostridia | Firmicutes | Bacteria |
| Eubacteriales | Clostridia | Firmicutes | Bacteria |
| Eubacteriales | Clostridia | Firmicutes | Bacteria |
| Eubacteriales | Clostridia | Firmicutes | Bacteria |
| OFGB9656 | CFGB9656 | Firmicutes | Bacteria |
| OFGB9827 | CFGB9827 | Firmicutes | Bacteria |
| Eubacteriales | Clostridia | Firmicutes | Bacteria |
| OFGB77303 | CFGB77303 | Bacteria_unclassified | Bacteria |
| Eubacteriales | Clostridia | Firmicutes | Bacteria |
| Eubacteriales | Clostridia | Firmicutes | Bacteria |
| OFGB72709 | CFGB72709 | Firmicutes | Bacteria |
| Eubacteriales | Clostridia | Firmicutes | Bacteria |

| | | | |
|---|---|---|---|
| Eubacteriales | Clostridia | Firmicutes | Bacteria |
| Eubacteriales | Clostridia | Firmicutes | Bacteria |
| Eubacteriales | Clostridia | Firmicutes | Bacteria |
| Eubacteriales | Clostridia | Firmicutes | Bacteria |
| Eubacteriales | Clostridia | Firmicutes | Bacteria |
| Eubacteriales | Clostridia | Firmicutes | Bacteria |
| Eubacteriales | Clostridia | Firmicutes | Bacteria |
| Eubacteriales | Clostridia | Firmicutes | Bacteria |
| Eubacteriales | Clostridia | Firmicutes | Bacteria |
| Eubacteriales | Clostridia | Firmicutes | Bacteria |
| Eubacteriales | Clostridia | Firmicutes | Bacteria |
| Eubacteriales | Clostridia | Firmicutes | Bacteria |
| OFGB77153 | CFGB77153 | Actinobacteria | Bacteria |
| OFGB1791 | CFGB1791 | Tenericutes | Bacteria |
| OFGB10290 | CFGB10290 | Firmicutes | Bacteria |
| Eubacteriales | Clostridia | Firmicutes | Bacteria |
| OFGB10289 | CFGB10289 | Firmicutes | Bacteria |
| OFGB1765 | CFGB1765 | Firmicutes | Bacteria |
| OFGB1765 | CFGB1765 | Firmicutes | Bacteria |
| OFGB1765 | CFGB1765 | Firmicutes | Bacteria |
| OFGB10667 | CFGB10667 | Firmicutes | Bacteria |
| Eubacteriales | Clostridia | Firmicutes | Bacteria |
| Eubacteriales | Clostridia | Firmicutes | Bacteria |
| Eubacteriales | Clostridia | Firmicutes | Bacteria |
| OFGB75721 | CFGB75721 | Firmicutes | Bacteria |
| Eubacteriales | Clostridia | Firmicutes | Bacteria |
| Eubacteriales | Clostridia | Firmicutes | Bacteria |
| OFGB10299 | CFGB10299 | Firmicutes | Bacteria |
| Eubacteriales | Clostridia | Firmicutes | Bacteria |
| Eubacteriales | Clostridia | Firmicutes | Bacteria |
| Eubacteriales | Clostridia | Firmicutes | Bacteria |
| Eubacteriales | Clostridia | Firmicutes | Bacteria |
| Eubacteriales | Clostridia | Firmicutes | Bacteria |
| Eubacteriales | Clostridia | Firmicutes | Bacteria |
| Eubacteriales | Clostridia | Firmicutes | Bacteria |
| Eubacteriales | Clostridia | Firmicutes | Bacteria |
| Lactobacillales | Bacilli | Firmicutes | Bacteria |
| Coriobacteriales | Coriobacteriia | Actinobacteria | Bacteria |
| Bacteroidales | Bacteroidia | Bacteroidota | Bacteria |
| Eubacteriales | Clostridia | Firmicutes | Bacteria |
| Eubacteriales | Clostridia | Firmicutes | Bacteria |
| Eubacteriales | Clostridia | Firmicutes | Bacteria |
| Eubacteriales | Clostridia | Firmicutes | Bacteria |
| Eubacteriales | Clostridia | Firmicutes | Bacteria |
| Eubacteriales | Clostridia | Firmicutes | Bacteria |
| Burkholderiales | Betaproteobacteria | Proteobacteria | Bacteria |

| | | | |
|---|---|---|---|
| Eubacteriales | Clostridia | Firmicutes | Bacteria |
| Erysipelotrichales | Erysipelotrichia | Firmicutes | Bacteria |
| Bacteria_unclassified | Bacteria_unclassified | Bacteria_unclassified | Bacteria |
| Bacteria_unclassified | Bacteria_unclassified | Bacteria_unclassified | Bacteria |
| Bacteria_unclassified | Bacteria_unclassified | Bacteria_unclassified | Bacteria |
| Bacteria_unclassified | Bacteria_unclassified | Bacteria_unclassified | Bacteria |
| Bacteria_unclassified | Bacteria_unclassified | Bacteria_unclassified | Bacteria |

| | | | |
|---|---|---|---|
| Eubacteriales | Clostridia | Firmicutes | Bacteria |
| Eubacteriales | Clostridia | Firmicutes | Bacteria |
| Verrucomicrobiales | Verrucomicrobiae | Verrucomicrobia | Bacteria |
| Eubacteriales | Clostridia | Firmicutes | Bacteria |
| Bacteria_unclassified | Bacteria_unclassified | Bacteria_unclassified | Bacteria |
| Bacteria_unclassified | Bacteria_unclassified | Bacteria_unclassified | Bacteria |
| Bacteria_unclassified | Bacteria_unclassified | Bacteria_unclassified | Bacteria |
| Bacteria_unclassified | Bacteria_unclassified | Bacteria_unclassified | Bacteria |
| Bacteroidales | Bacteroidia | Bacteroidota | Bacteria |
| Clostridia_unclassified | Clostridia | Firmicutes | Bacteria |
| Eubacteriales | Clostridia | Firmicutes | Bacteria |
| Eubacteriales | Clostridia | Firmicutes | Bacteria |
| Eubacteriales | Clostridia | Firmicutes | Bacteria |
| Eubacteriales | Clostridia | Firmicutes | Bacteria |
| Erysipelotrichales | Erysipelotrichia | Firmicutes | Bacteria |
| Eubacteriales | Clostridia | Firmicutes | Bacteria |
| Eubacteriales | Clostridia | Firmicutes | Bacteria |
| OFGB77306 | CFGB77306 | Firmicutes | Bacteria |
| Eubacteriales | Clostridia | Firmicutes | Bacteria |
| Eggerthellales | Coriobacteriia | Actinobacteria | Bacteria |
| Eubacteriales | Clostridia | Firmicutes | Bacteria |
| OFGB9506 | CFGB9506 | Firmicutes | Bacteria |
| OFGB9508 | CFGB9508 | Firmicutes | Bacteria |
| OFGB9512 | CFGB9512 | Firmicutes | Bacteria |
| OFGB2838 | CFGB2838 | Firmicutes | Bacteria |
| OFGB2838 | CFGB2838 | Firmicutes | Bacteria |
| OFGB2838 | CFGB2838 | Firmicutes | Bacteria |
| Eubacteriales | Clostridia | Firmicutes | Bacteria |
| OFGB2833 | CFGB2833 | Firmicutes | Bacteria |
| Eubacteriales | Clostridia | Firmicutes | Bacteria |
| Eubacteriales | Clostridia | Firmicutes | Bacteria |
| Eubacteriales | Clostridia | Firmicutes | Bacteria |
| OFGB9622 | CFGB9622 | Firmicutes | Bacteria |
| OFGB77305 | CFGB77305 | Firmicutes | Bacteria |
| Eubacteriales | Clostridia | Firmicutes | Bacteria |
| Eubacteriales | Clostridia | Firmicutes | Bacteria |

| | | | |
|---|---|---|---|
| Eubacteriales | Clostridia | Firmicutes | Bacteria |
| Eubacteriales | Clostridia | Firmicutes | Bacteria |
| Eubacteriales | Clostridia | Firmicutes | Bacteria |
| OFGB9633 | CFGB9633 | Firmicutes | Bacteria |
| Bacteria_unclassified | Bacteria_unclassified | Bacteria_unclassified | Bacteria |
| Bacteria_unclassified | Bacteria_unclassified | Bacteria_unclassified | Bacteria |
| Bacteria_unclassified | Bacteria_unclassified | Bacteria_unclassified | Bacteria |
| Bacteria_unclassified | Bacteria_unclassified | Bacteria_unclassified | Bacteria |
| Bacteria_unclassified | Bacteria_unclassified | Bacteria_unclassified | Bacteria |
| Bacteria_unclassified | Bacteria_unclassified | Bacteria_unclassified | Bacteria |
| Eubacteriales | Clostridia | Firmicutes | Bacteria |
| OFGB77359 | CFGB77359 | Bacteria_unclassified | Bacteria |
| OFGB9639 | CFGB9639 | Firmicutes | Bacteria |
| Eubacteriales | Clostridia | Firmicutes | Bacteria |
| Eubacteriales | Clostridia | Firmicutes | Bacteria |
| Eubacteriales | Clostridia | Firmicutes | Bacteria |
| Eubacteriales | Clostridia | Firmicutes | Bacteria |
| Eubacteriales | Clostridia | Firmicutes | Bacteria |
| Eubacteriales | Clostridia | Firmicutes | Bacteria |
| Eubacteriales | Clostridia | Firmicutes | Bacteria |
| OFGB9656 | CFGB9656 | Firmicutes | Bacteria |
| OFGB9827 | CFGB9827 | Firmicutes | Bacteria |
| Eubacteriales | Clostridia | Firmicutes | Bacteria |
| OFGB77303 | CFGB77303 | Bacteria_unclassified | Bacteria |
| Eubacteriales | Clostridia | Firmicutes | Bacteria |
| Eubacteriales | Clostridia | Firmicutes | Bacteria |
| OFGB72709 | CFGB72709 | Firmicutes | Bacteria |
| Eubacteriales | Clostridia | Firmicutes | Bacteria |
| Eubacteriales | Clostridia | Firmicutes | Bacteria |
| Eubacteriales | Clostridia | Firmicutes | Bacteria |
| Eubacteriales | Clostridia | Firmicutes | Bacteria |
| Eubacteriales | Clostridia | Firmicutes | Bacteria |
| Eubacteriales | Clostridia | Firmicutes | Bacteria |
| Eubacteriales | Clostridia | Firmicutes | Bacteria |
| Eubacteriales | Clostridia | Firmicutes | Bacteria |
| Eubacteriales | Clostridia | Firmicutes | Bacteria |
| Eubacteriales | Clostridia | Firmicutes | Bacteria |
| Eubacteriales | Clostridia | Firmicutes | Bacteria |
| Eubacteriales | Clostridia | Firmicutes | Bacteria |
| Eubacteriales | Clostridia | Firmicutes | Bacteria |
| OFGB77153 | CFGB77153 | Actinobacteria | Bacteria |
| OFGB1791 | CFGB1791 | Tenericutes | Bacteria |
| OFGB10290 | CFGB10290 | Firmicutes | Bacteria |
| Eubacteriales | Clostridia | Firmicutes | Bacteria |
| OFGB10289 | CFGB10289 | Firmicutes | Bacteria |
| OFGB1765 | CFGB1765 | Firmicutes | Bacteria |

| | | | |
|---|---|---|---|
| OFGB1765 | CFGB1765 | Firmicutes | Bacteria |
| OFGB1765 | CFGB1765 | Firmicutes | Bacteria |
| OFGB10667 | CFGB10667 | Firmicutes | Bacteria |
| Eubacteriales | Clostridia | Firmicutes | Bacteria |
| Eubacteriales | Clostridia | Firmicutes | Bacteria |
| Eubacteriales | Clostridia | Firmicutes | Bacteria |
| OFGB75721 | CFGB75721 | Firmicutes | Bacteria |
| Eubacteriales | Clostridia | Firmicutes | Bacteria |
| Eubacteriales | Clostridia | Firmicutes | Bacteria |
| OFGB10299 | CFGB10299 | Firmicutes | Bacteria |
| Eubacteriales | Clostridia | Firmicutes | Bacteria |
| Eubacteriales | Clostridia | Firmicutes | Bacteria |
| Eubacteriales | Clostridia | Firmicutes | Bacteria |
| Eubacteriales | Clostridia | Firmicutes | Bacteria |
| Eubacteriales | Clostridia | Firmicutes | Bacteria |
| Eubacteriales | Clostridia | Firmicutes | Bacteria |
| Eubacteriales | Clostridia | Firmicutes | Bacteria |
| Eubacteriales | Clostridia | Firmicutes | Bacteria |
| Lactobacillales | Bacilli | Firmicutes | Bacteria |
| Coriobacteriales | Coriobacteriia | Actinobacteria | Bacteria |
| Bacteroidales | Bacteroidia | Bacteroidota | Bacteria |
| Eubacteriales | Clostridia | Firmicutes | Bacteria |
| Eubacteriales | Clostridia | Firmicutes | Bacteria |
| Eubacteriales | Clostridia | Firmicutes | Bacteria |
| Eubacteriales | Clostridia | Firmicutes | Bacteria |
| Eubacteriales | Clostridia | Firmicutes | Bacteria |
| Eubacteriales | Clostridia | Firmicutes | Bacteria |
| Burkholderiales | Betaproteobacteria | Proteobacteria | Bacteria |
| Eubacteriales | Clostridia | Firmicutes | Bacteria |
| Erysipelotrichales | Erysipelotrichia | Firmicutes | Bacteria |
| Bacteria_unclassified | Bacteria_unclassified | Bacteria_unclassified | Bacteria |
| Bacteria_unclassified | Bacteria_unclassified | Bacteria_unclassified | Bacteria |
| Bacteria_unclassified | Bacteria_unclassified | Bacteria_unclassified | Bacteria |
| Bacteria_unclassified | Bacteria_unclassified | Bacteria_unclassified | Bacteria |
| Bacteria_unclassified | Bacteria_unclassified | Bacteria_unclassified | Bacteria |

| | | | |
|---|---|---|---|
| Eubacteriales | Clostridia | Firmicutes | Bacteria |
| Eubacteriales | Clostridia | Firmicutes | Bacteria |
| Verrucomicrobiales | Verrucomicrobiae | Verrucomicrobia | Bacteria |
| Eubacteriales | Clostridia | Firmicutes | Bacteria |
| Bacteria_unclassified | Bacteria_unclassified | Bacteria_unclassified | Bacteria |
| Bacteria_unclassified | Bacteria_unclassified | Bacteria_unclassified | Bacteria |
| Bacteria_unclassified | Bacteria_unclassified | Bacteria_unclassified | Bacteria |
| Bacteria_unclassified | Bacteria_unclassified | Bacteria_unclassified | Bacteria |

| | | | |
|---|---|---|---|
| Bacteroidales | Bacteroidia | Bacteroidota | Bacteria |
| Clostridia_unclassified | Clostridia | Firmicutes | Bacteria |
| Eubacteriales | Clostridia | Firmicutes | Bacteria |
| Eubacteriales | Clostridia | Firmicutes | Bacteria |
| Eubacteriales | Clostridia | Firmicutes | Bacteria |
| Eubacteriales | Clostridia | Firmicutes | Bacteria |
| Erysipelotrichales | Erysipelotrichia | Firmicutes | Bacteria |
| Eubacteriales | Clostridia | Firmicutes | Bacteria |
| Eubacteriales | Clostridia | Firmicutes | Bacteria |
| OFGB77306 | CFGB77306 | Firmicutes | Bacteria |
| Eubacteriales | Clostridia | Firmicutes | Bacteria |
| Eggerthellales | Coriobacteriia | Actinobacteria | Bacteria |
| Eubacteriales | Clostridia | Firmicutes | Bacteria |
| OFGB9506 | CFGB9506 | Firmicutes | Bacteria |
| OFGB9508 | CFGB9508 | Firmicutes | Bacteria |
| OFGB9512 | CFGB9512 | Firmicutes | Bacteria |
| OFGB2838 | CFGB2838 | Firmicutes | Bacteria |
| OFGB2838 | CFGB2838 | Firmicutes | Bacteria |
| OFGB2838 | CFGB2838 | Firmicutes | Bacteria |
| Eubacteriales | Clostridia | Firmicutes | Bacteria |
| OFGB2833 | CFGB2833 | Firmicutes | Bacteria |
| Eubacteriales | Clostridia | Firmicutes | Bacteria |
| Eubacteriales | Clostridia | Firmicutes | Bacteria |
| Eubacteriales | Clostridia | Firmicutes | Bacteria |
| OFGB9622 | CFGB9622 | Firmicutes | Bacteria |
| OFGB77305 | CFGB77305 | Firmicutes | Bacteria |
| Eubacteriales | Clostridia | Firmicutes | Bacteria |
| Eubacteriales | Clostridia | Firmicutes | Bacteria |
| Eubacteriales | Clostridia | Firmicutes | Bacteria |
| Eubacteriales | Clostridia | Firmicutes | Bacteria |
| Eubacteriales | Clostridia | Firmicutes | Bacteria |
| OFGB9633 | CFGB9633 | Firmicutes | Bacteria |
| Bacteria_unclassified | Bacteria_unclassified | Bacteria_unclassified | Bacteria |
| Bacteria_unclassified | Bacteria_unclassified | Bacteria_unclassified | Bacteria |
| Bacteria_unclassified | Bacteria_unclassified | Bacteria_unclassified | Bacteria |
| Bacteria_unclassified | Bacteria_unclassified | Bacteria_unclassified | Bacteria |
| Bacteria_unclassified | Bacteria_unclassified | Bacteria_unclassified | Bacteria |
| Bacteria_unclassified | Bacteria_unclassified | Bacteria_unclassified | Bacteria |
| Eubacteriales | Clostridia | Firmicutes | Bacteria |
| OFGB77359 | CFGB77359 | Bacteria_unclassified | Bacteria |
| OFGB9639 | CFGB9639 | Firmicutes | Bacteria |
| Eubacteriales | Clostridia | Firmicutes | Bacteria |
| Eubacteriales | Clostridia | Firmicutes | Bacteria |
| Eubacteriales | Clostridia | Firmicutes | Bacteria |
| Eubacteriales | Clostridia | Firmicutes | Bacteria |
| Eubacteriales | Clostridia | Firmicutes | Bacteria |

| | | | |
|---|---|---|---|
| Eubacteriales | Clostridia | Firmicutes | Bacteria |
| Eubacteriales | Clostridia | Firmicutes | Bacteria |
| OFGB9656 | CFGB9656 | Firmicutes | Bacteria |
| OFGB9827 | CFGB9827 | Firmicutes | Bacteria |
| Eubacteriales | Clostridia | Firmicutes | Bacteria |
| OFGB77303 | CFGB77303 | Bacteria_unclassified | Bacteria |
| Eubacteriales | Clostridia | Firmicutes | Bacteria |
| Eubacteriales | Clostridia | Firmicutes | Bacteria |
| OFGB72709 | CFGB72709 | Firmicutes | Bacteria |
| Eubacteriales | Clostridia | Firmicutes | Bacteria |
| Eubacteriales | Clostridia | Firmicutes | Bacteria |
| Eubacteriales | Clostridia | Firmicutes | Bacteria |
| Eubacteriales | Clostridia | Firmicutes | Bacteria |
| Eubacteriales | Clostridia | Firmicutes | Bacteria |
| Eubacteriales | Clostridia | Firmicutes | Bacteria |
| Eubacteriales | Clostridia | Firmicutes | Bacteria |
| Eubacteriales | Clostridia | Firmicutes | Bacteria |
| Eubacteriales | Clostridia | Firmicutes | Bacteria |
| Eubacteriales | Clostridia | Firmicutes | Bacteria |
| Eubacteriales | Clostridia | Firmicutes | Bacteria |
| Eubacteriales | Clostridia | Firmicutes | Bacteria |
| Eubacteriales | Clostridia | Firmicutes | Bacteria |
| OFGB77153 | CFGB77153 | Actinobacteria | Bacteria |
| OFGB1791 | CFGB1791 | Tenericutes | Bacteria |
| OFGB10290 | CFGB10290 | Firmicutes | Bacteria |
| Eubacteriales | Clostridia | Firmicutes | Bacteria |
| OFGB10289 | CFGB10289 | Firmicutes | Bacteria |
| OFGB1765 | CFGB1765 | Firmicutes | Bacteria |
| OFGB1765 | CFGB1765 | Firmicutes | Bacteria |
| OFGB1765 | CFGB1765 | Firmicutes | Bacteria |
| OFGB10667 | CFGB10667 | Firmicutes | Bacteria |
| Eubacteriales | Clostridia | Firmicutes | Bacteria |
| Eubacteriales | Clostridia | Firmicutes | Bacteria |
| Eubacteriales | Clostridia | Firmicutes | Bacteria |
| OFGB75721 | CFGB75721 | Firmicutes | Bacteria |
| Eubacteriales | Clostridia | Firmicutes | Bacteria |
| Eubacteriales | Clostridia | Firmicutes | Bacteria |
| OFGB10299 | CFGB10299 | Firmicutes | Bacteria |
| Eubacteriales | Clostridia | Firmicutes | Bacteria |
| Eubacteriales | Clostridia | Firmicutes | Bacteria |
| Eubacteriales | Clostridia | Firmicutes | Bacteria |
| Eubacteriales | Clostridia | Firmicutes | Bacteria |
| Eubacteriales | Clostridia | Firmicutes | Bacteria |
| Eubacteriales | Clostridia | Firmicutes | Bacteria |
| Eubacteriales | Clostridia | Firmicutes | Bacteria |
| Eubacteriales | Clostridia | Firmicutes | Bacteria |

| | | | |
|---|---|---|---|
| Lactobacillales | Bacilli | Firmicutes | Bacteria |
| Coriobacteriales | Coriobacteriia | Actinobacteria | Bacteria |
| Bacteroidales | Bacteroidia | Bacteroidota | Bacteria |
| Eubacteriales | Clostridia | Firmicutes | Bacteria |
| Eubacteriales | Clostridia | Firmicutes | Bacteria |
| Eubacteriales | Clostridia | Firmicutes | Bacteria |
| Eubacteriales | Clostridia | Firmicutes | Bacteria |
| Eubacteriales | Clostridia | Firmicutes | Bacteria |
| Eubacteriales | Clostridia | Firmicutes | Bacteria |
| Burkholderiales | Betaproteobacteria | Proteobacteria | Bacteria |
| Eubacteriales | Clostridia | Firmicutes | Bacteria |
| Erysipelotrichales | Erysipelotrichia | Firmicutes | Bacteria |
| Bacteria_unclassified | Bacteria_unclassified | Bacteria_unclassified | Bacteria |
| Bacteria_unclassified | Bacteria_unclassified | Bacteria_unclassified | Bacteria |
| Bacteria_unclassified | Bacteria_unclassified | Bacteria_unclassified | Bacteria |
| Bacteria_unclassified | Bacteria_unclassified | Bacteria_unclassified | Bacteria |
| Bacteria_unclassified | Bacteria_unclassified | Bacteria_unclassified | Bacteria |
| Eubacteriales | Clostridia | Firmicutes | Bacteria |
| Eubacteriales | Clostridia | Firmicutes | Bacteria |
| Verrucomicrobiales | Verrucomicrobiae | Verrucomicrobia | Bacteria |
| Eubacteriales | Clostridia | Firmicutes | Bacteria |
| Bacteria_unclassified | Bacteria_unclassified | Bacteria_unclassified | Bacteria |
| Bacteria_unclassified | Bacteria_unclassified | Bacteria_unclassified | Bacteria |
| Bacteria_unclassified | Bacteria_unclassified | Bacteria_unclassified | Bacteria |
| Bacteria_unclassified | Bacteria_unclassified | Bacteria_unclassified | Bacteria |
| Bacteroidales | Bacteroidia | Bacteroidota | Bacteria |
| Clostridia_unclassified | Clostridia | Firmicutes | Bacteria |
| Eubacteriales | Clostridia | Firmicutes | Bacteria |
| Eubacteriales | Clostridia | Firmicutes | Bacteria |
| Eubacteriales | Clostridia | Firmicutes | Bacteria |
| Eubacteriales | Clostridia | Firmicutes | Bacteria |
| Erysipelotrichales | Erysipelotrichia | Firmicutes | Bacteria |
| Eubacteriales | Clostridia | Firmicutes | Bacteria |
| Eubacteriales | Clostridia | Firmicutes | Bacteria |
| OFGB77306 | CFGB77306 | Firmicutes | Bacteria |
| Eubacteriales | Clostridia | Firmicutes | Bacteria |
| Eggerthellales | Coriobacteriia | Actinobacteria | Bacteria |
| Eubacteriales | Clostridia | Firmicutes | Bacteria |
| OFGB9506 | CFGB9506 | Firmicutes | Bacteria |
| OFGB9508 | CFGB9508 | Firmicutes | Bacteria |
| OFGB9512 | CFGB9512 | Firmicutes | Bacteria |
| OFGB2838 | CFGB2838 | Firmicutes | Bacteria |
| OFGB2838 | CFGB2838 | Firmicutes | Bacteria |

| | | | |
|---|---|---|---|
| OFGB2838 | CFGB2838 | Firmicutes | Bacteria |
| Eubacteriales | Clostridia | Firmicutes | Bacteria |
| OFGB2833 | CFGB2833 | Firmicutes | Bacteria |
| Eubacteriales | Clostridia | Firmicutes | Bacteria |
| Eubacteriales | Clostridia | Firmicutes | Bacteria |
| Eubacteriales | Clostridia | Firmicutes | Bacteria |
| OFGB9622 | CFGB9622 | Firmicutes | Bacteria |
| OFGB77305 | CFGB77305 | Firmicutes | Bacteria |
| Eubacteriales | Clostridia | Firmicutes | Bacteria |
| Eubacteriales | Clostridia | Firmicutes | Bacteria |
| Eubacteriales | Clostridia | Firmicutes | Bacteria |
| Eubacteriales | Clostridia | Firmicutes | Bacteria |
| Eubacteriales | Clostridia | Firmicutes | Bacteria |
| OFGB9633 | CFGB9633 | Firmicutes | Bacteria |
| Bacteria_unclassified | Bacteria_unclassified | Bacteria_unclassified | Bacteria |
| Bacteria_unclassified | Bacteria_unclassified | Bacteria_unclassified | Bacteria |
| Bacteria_unclassified | Bacteria_unclassified | Bacteria_unclassified | Bacteria |
| Bacteria_unclassified | Bacteria_unclassified | Bacteria_unclassified | Bacteria |
| Bacteria_unclassified | Bacteria_unclassified | Bacteria_unclassified | Bacteria |
| Bacteria_unclassified | Bacteria_unclassified | Bacteria_unclassified | Bacteria |
| Eubacteriales | Clostridia | Firmicutes | Bacteria |
| OFGB77359 | CFGB77359 | Bacteria_unclassified | Bacteria |
| OFGB9639 | CFGB9639 | Firmicutes | Bacteria |
| Eubacteriales | Clostridia | Firmicutes | Bacteria |
| Eubacteriales | Clostridia | Firmicutes | Bacteria |
| Eubacteriales | Clostridia | Firmicutes | Bacteria |
| Eubacteriales | Clostridia | Firmicutes | Bacteria |
| Eubacteriales | Clostridia | Firmicutes | Bacteria |
| Eubacteriales | Clostridia | Firmicutes | Bacteria |
| Eubacteriales | Clostridia | Firmicutes | Bacteria |
| OFGB9656 | CFGB9656 | Firmicutes | Bacteria |
| OFGB9827 | CFGB9827 | Firmicutes | Bacteria |
| Eubacteriales | Clostridia | Firmicutes | Bacteria |
| OFGB77303 | CFGB77303 | Bacteria_unclassified | Bacteria |
| Eubacteriales | Clostridia | Firmicutes | Bacteria |
| Eubacteriales | Clostridia | Firmicutes | Bacteria |
| OFGB72709 | CFGB72709 | Firmicutes | Bacteria |
| Eubacteriales | Clostridia | Firmicutes | Bacteria |
| Eubacteriales | Clostridia | Firmicutes | Bacteria |
| Eubacteriales | Clostridia | Firmicutes | Bacteria |
| Eubacteriales | Clostridia | Firmicutes | Bacteria |
| Eubacteriales | Clostridia | Firmicutes | Bacteria |
| Eubacteriales | Clostridia | Firmicutes | Bacteria |
| Eubacteriales | Clostridia | Firmicutes | Bacteria |
| Eubacteriales | Clostridia | Firmicutes | Bacteria |
| Eubacteriales | Clostridia | Firmicutes | Bacteria |

| | | | |
|---|---|---|---|
| Eubacteriales | Clostridia | Firmicutes | Bacteria |
| Eubacteriales | Clostridia | Firmicutes | Bacteria |
| Eubacteriales | Clostridia | Firmicutes | Bacteria |
| Eubacteriales | Clostridia | Firmicutes | Bacteria |
| OFGB77153 | CFGB77153 | Actinobacteria | Bacteria |
| OFGB1791 | CFGB1791 | Tenericutes | Bacteria |
| OFGB10290 | CFGB10290 | Firmicutes | Bacteria |
| Eubacteriales | Clostridia | Firmicutes | Bacteria |
| OFGB10289 | CFGB10289 | Firmicutes | Bacteria |
| OFGB1765 | CFGB1765 | Firmicutes | Bacteria |
| OFGB1765 | CFGB1765 | Firmicutes | Bacteria |
| OFGB1765 | CFGB1765 | Firmicutes | Bacteria |
| OFGB10667 | CFGB10667 | Firmicutes | Bacteria |
| Eubacteriales | Clostridia | Firmicutes | Bacteria |
| Eubacteriales | Clostridia | Firmicutes | Bacteria |
| Eubacteriales | Clostridia | Firmicutes | Bacteria |
| OFGB75721 | CFGB75721 | Firmicutes | Bacteria |
| Eubacteriales | Clostridia | Firmicutes | Bacteria |
| Eubacteriales | Clostridia | Firmicutes | Bacteria |
| OFGB10299 | CFGB10299 | Firmicutes | Bacteria |
| Eubacteriales | Clostridia | Firmicutes | Bacteria |
| Eubacteriales | Clostridia | Firmicutes | Bacteria |
| Eubacteriales | Clostridia | Firmicutes | Bacteria |
| Eubacteriales | Clostridia | Firmicutes | Bacteria |
| Eubacteriales | Clostridia | Firmicutes | Bacteria |
| Eubacteriales | Clostridia | Firmicutes | Bacteria |
| Eubacteriales | Clostridia | Firmicutes | Bacteria |
| Eubacteriales | Clostridia | Firmicutes | Bacteria |
| Lactobacillales | Bacilli | Firmicutes | Bacteria |
| Coriobacteriales | Coriobacteriia | Actinobacteria | Bacteria |
| Bacteroidales | Bacteroidia | Bacteroidota | Bacteria |
| Eubacteriales | Clostridia | Firmicutes | Bacteria |
| Eubacteriales | Clostridia | Firmicutes | Bacteria |
| Eubacteriales | Clostridia | Firmicutes | Bacteria |
| Eubacteriales | Clostridia | Firmicutes | Bacteria |
| Eubacteriales | Clostridia | Firmicutes | Bacteria |
| Eubacteriales | Clostridia | Firmicutes | Bacteria |
| Burkholderiales | Betaproteobacteria | Proteobacteria | Bacteria |
| Eubacteriales | Clostridia | Firmicutes | Bacteria |
| Erysipelotrichales | Erysipelotrichia | Firmicutes | Bacteria |
| Bacteria_unclassified | Bacteria_unclassified | Bacteria_unclassified | Bacteria |
| Bacteria_unclassified | Bacteria_unclassified | Bacteria_unclassified | Bacteria |
| Bacteria_unclassified | Bacteria_unclassified | Bacteria_unclassified | Bacteria |
| Bacteria_unclassified | Bacteria_unclassified | Bacteria_unclassified | Bacteria |
| Bacteria_unclassified | Bacteria_unclassified | Bacteria_unclassified | Bacteria |

| **MetaPhlan Annotation** |
|---|
| k__Bacteria\|p__Firmicutes\|c__Clostridia\|o__Eubacteriales\|f__Lachnospiraceae\|g__Acetatifactor\|s__Acetatifactor_SGB415 |
| k__Bacteria\|p__Firmicutes\|c__Clostridia\|o__Eubacteriales\|f__Oscillospiraceae\|g__Acutalibacter\|s__Acutalibacter_muris |
| k__Bacteria\|p__Verrucomicrobia\|c__Verrucomicrobiae\|o__Verrucomicrobiales\|f__Akkermansiaceae\|g__Akkermansia\|s__A |
| k__Bacteria\|p__Firmicutes\|c__Clostridia\|o__Eubacteriales\|f__Oscillospiraceae\|g__Anaerotruncus\|s__Anaerotruncus_sp_1 |
| k__Bacteria\|p__Bacteria_unclassified\|c__Bacteria_unclassified\|o__Bacteria_unclassified\|f__Bacteria_unclassified\|g__Bacte |
| k__Bacteria\|p__Bacteria_unclassified\|c__Bacteria_unclassified\|o__Bacteria_unclassified\|f__Bacteria_unclassified\|g__Bacte |
| k__Bacteria\|p__Bacteria_unclassified\|c__Bacteria_unclassified\|o__Bacteria_unclassified\|f__Bacteria_unclassified\|g__Bacte |
| k__Bacteria\|p__Bacteria_unclassified\|c__Bacteria_unclassified\|o__Bacteria_unclassified\|f__Bacteria_unclassified\|g__Bacte |
| k__Bacteria\|p__Bacteroidota\|c__Bacteroidia\|o__Bacteroidales\|f__Bacteroidaceae\|g__Bacteroides\|s__Bacteroides_thetaio |
| k__Bacteria\|p__Firmicutes\|c__Clostridia\|o__Clostridia_unclassified\|f__Clostridia_unclassified\|g__Clostridia_unclassified\|s_ |
| k__Bacteria\|p__Firmicutes\|c__Clostridia\|o__Eubacteriales\|f__Clostridiaceae\|g__Clostridiaceae_unclassified\|s__Clostridiac |
| k__Bacteria\|p__Firmicutes\|c__Clostridia\|o__Eubacteriales\|f__Clostridiaceae\|g__Clostridiaceae_unclassified\|s__Clostridiac |
| k__Bacteria\|p__Firmicutes\|c__Clostridia\|o__Eubacteriales\|f__Eubacteriales_unclassified\|g__Eubacteriales_unclassified\|s_ |
| k__Bacteria\|p__Firmicutes\|c__Clostridia\|o__Eubacteriales\|f__Clostridiaceae\|g__Clostridium\|s__Clostridium_SGB65123 |
| k__Bacteria\|p__Firmicutes\|c__Erysipelotrichia\|o__Erysipelotrichales\|f__Erysipelotrichaceae\|g__Erysipelatoclostridium\|s__ |
| k__Bacteria\|p__Firmicutes\|c__Clostridia\|o__Eubacteriales\|f__Eubacteriaceae\|g__Eubacteriaceae_unclassified\|s__Eubacte |
| k__Bacteria\|p__Firmicutes\|c__Clostridia\|o__Eubacteriales\|f__Eubacteriaceae\|g__Eubacteriaceae_unclassified\|s__Eubacte |
| k__Bacteria\|p__Firmicutes\|c__CFGB77306\|o__OFGB77306\|f__FGB77306\|g__GGB20146\|s__GGB20146_SGB29427 |
| k__Bacteria\|p__Firmicutes\|c__Clostridia\|o__Eubacteriales\|f__Lachnospiraceae\|g__GGB20149\|s__GGB20149_SGB29430 |
| k__Bacteria\|p__Actinobacteria\|c__Coriobacteriia\|o__Eggerthellales\|f__Eggerthellaceae\|g__GGB22635\|s__GGB22635_SGB |
| k__Bacteria\|p__Firmicutes\|c__Clostridia\|o__Eubacteriales\|f__Lachnospiraceae\|g__GGB25041\|s__GGB25041_SGB36960 |
| k__Bacteria\|p__Firmicutes\|c__CFGB9506\|o__OFGB9506\|f__FGB9506\|g__GGB28379\|s__GGB28379_SGB40959 |
| k__Bacteria\|p__Firmicutes\|c__CFGB9508\|o__OFGB9508\|f__FGB9508\|g__GGB28382\|s__GGB28382_SGB40962 |
| k__Bacteria\|p__Firmicutes\|c__CFGB9512\|o__OFGB9512\|f__FGB9512\|g__GGB28392\|s__GGB28392_SGB40972 |
| k__Bacteria\|p__Firmicutes\|c__CFGB2838\|o__OFGB2838\|f__FGB2838\|g__GGB28404\|s__GGB28404_SGB40986 |
| k__Bacteria\|p__Firmicutes\|c__CFGB2838\|o__OFGB2838\|f__FGB2838\|g__GGB28418\|s__GGB28418_SGB41001 |
| k__Bacteria\|p__Firmicutes\|c__CFGB2838\|o__OFGB2838\|f__FGB2838\|g__GGB28422\|s__GGB28422_SGB41005 |
| k__Bacteria\|p__Firmicutes\|c__Clostridia\|o__Eubacteriales\|f__Pumilibacteraceae\|g__GGB28431\|s__GGB28431_SGB41014 |
| k__Bacteria\|p__Firmicutes\|c__CFGB2833\|o__OFGB2833\|f__FGB2833\|g__GGB28456\|s__GGB28456_SGB41039 |
| k__Bacteria\|p__Firmicutes\|c__Clostridia\|o__Eubacteriales\|f__Eubacteriaceae\|g__GGB28782\|s__GGB28782_SGB41435 |
| k__Bacteria\|p__Firmicutes\|c__Clostridia\|o__Eubacteriales\|f__Lachnospiraceae\|g__GGB28792\|s__GGB28792_SGB41445 |
| k__Bacteria\|p__Firmicutes\|c__Clostridia\|o__Eubacteriales\|f__Lachnospiraceae\|g__GGB28798\|s__GGB28798_SGB41451 |
| k__Bacteria\|p__Firmicutes\|c__CFGB9622\|o__OFGB9622\|f__FGB9622\|g__GGB28810\|s__GGB28810_SGB41465 |
| k__Bacteria\|p__Firmicutes\|c__CFGB77305\|o__OFGB77305\|f__FGB77305\|g__GGB28828\|s__GGB28828_SGB41484 |
| k__Bacteria\|p__Firmicutes\|c__Clostridia\|o__Eubacteriales\|f__Clostridiaceae\|g__GGB28851\|s__GGB28851_SGB41518 |
| k__Bacteria\|p__Firmicutes\|c__Clostridia\|o__Eubacteriales\|f__Lachnospiraceae\|g__GGB28865\|s__GGB28865_SGB41537 |
| k__Bacteria\|p__Firmicutes\|c__Clostridia\|o__Eubacteriales\|f__Lachnospiraceae\|g__GGB28868\|s__GGB28868_SGB41542 |
| k__Bacteria\|p__Firmicutes\|c__Clostridia\|o__Eubacteriales\|f__Lachnospiraceae\|g__GGB28869\|s__GGB28869_SGB41543 |
| k__Bacteria\|p__Firmicutes\|c__Clostridia\|o__Eubacteriales\|f__Lachnospiraceae\|g__GGB28875\|s__GGB28875_SGB41555 |
| k__Bacteria\|p__Firmicutes\|c__CFGB9633\|o__OFGB9633\|f__FGB9633\|g__GGB28881\|s__GGB28881_SGB41561 |
| k__Bacteria\|p__Bacteria_unclassified\|c__Bacteria_unclassified\|o__Bacteria_unclassified\|f__Bacteria_unclassified\|g__GGB2 |
| k__Bacteria\|p__Bacteria_unclassified\|c__Bacteria_unclassified\|o__Bacteria_unclassified\|f__Bacteria_unclassified\|g__GGB2 |
| k__Bacteria\|p__Bacteria_unclassified\|c__Bacteria_unclassified\|o__Bacteria_unclassified\|f__Bacteria_unclassified\|g__GGB2 |
| k__Bacteria\|p__Bacteria_unclassified\|c__Bacteria_unclassified\|o__Bacteria_unclassified\|f__Bacteria_unclassified\|g__GGB2 |
| k__Bacteria\|p__Bacteria_unclassified\|c__Bacteria_unclassified\|o__Bacteria_unclassified\|f__Bacteria_unclassified\|g__GGB2 |
| k__Bacteria\|p__Bacteria_unclassified\|c__Bacteria_unclassified\|o__Bacteria_unclassified\|f__Bacteria_unclassified\|g__GGB2 |

k__Bacteria|p__Firmicutes|c__Clostridia|o__Eubacteriales|f__Lachnospiraceae|g__GGB28924|s__GGB28924_SGB41621
k__Bacteria|p__Bacteria_unclassified|c__CFGB77359|o__OFGB77359|f__FGB77359|g__GGB28927|s__GGB28927_SGB416
k__Bacteria|p__Firmicutes|c__CFGB9639|o__OFGB9639|f__FGB9639|g__GGB28934|s__GGB28934_SGB41635
k__Bacteria|p__Firmicutes|c__Clostridia|o__Eubacteriales|f__Lachnospiraceae|g__GGB28949|s__GGB28949_SGB41655
k__Bacteria|p__Firmicutes|c__Clostridia|o__Eubacteriales|f__Clostridiaceae|g__GGB28951|s__GGB28951_SGB102295
k__Bacteria|p__Firmicutes|c__Clostridia|o__Eubacteriales|f__Clostridiaceae|g__GGB28951|s__GGB28951_SGB41658
k__Bacteria|p__Firmicutes|c__Clostridia|o__Eubacteriales|f__Clostridiaceae|g__GGB28954|s__GGB28954_SGB41662
k__Bacteria|p__Firmicutes|c__Clostridia|o__Eubacteriales|f__Clostridiaceae|g__GGB28960|s__GGB28960_SGB41669
k__Bacteria|p__Firmicutes|c__Clostridia|o__Eubacteriales|f__Clostridiaceae|g__GGB28964|s__GGB28964_SGB94886
k__Bacteria|p__Firmicutes|c__Clostridia|o__Eubacteriales|f__Clostridiaceae|g__GGB28967|s__GGB28967_SGB41678
k__Bacteria|p__Firmicutes|c__CFGB9656|o__OFGB9656|f__FGB9656|g__GGB28996|s__GGB28996_SGB41712
k__Bacteria|p__Firmicutes|c__CFGB9827|o__OFGB9827|f__FGB9827|g__GGB29531|s__GGB29531_SGB42317
k__Bacteria|p__Firmicutes|c__Clostridia|o__Eubacteriales|f__Eubacteriaceae|g__GGB29685|s__GGB29685_SGB42494
k__Bacteria|p__Bacteria_unclassified|c__CFGB77303|o__OFGB77303|f__FGB77303|g__GGB30141|s__GGB30141_SGB430
k__Bacteria|p__Firmicutes|c__Clostridia|o__Eubacteriales|f__Clostridiaceae|g__GGB30145|s__GGB30145_SGB43072
k__Bacteria|p__Firmicutes|c__Clostridia|o__Eubacteriales|f__Eubacteriales_unclassified|g__GGB30286|s__GGB30286_SG
k__Bacteria|p__Firmicutes|c__CFGB72709|o__OFGB72709|f__FGB72709|g__GGB30300|s__GGB30300_SGB43264
k__Bacteria|p__Firmicutes|c__Clostridia|o__Eubacteriales|f__Oscillospiraceae|g__GGB30303|s__GGB30303_SGB43268
k__Bacteria|p__Firmicutes|c__Clostridia|o__Eubacteriales|f__Oscillospiraceae|g__GGB30450|s__GGB30450_SGB43507
k__Bacteria|p__Firmicutes|c__Clostridia|o__Eubacteriales|f__Oscillospiraceae|g__GGB30453|s__GGB30453_SGB43513
k__Bacteria|p__Firmicutes|c__Clostridia|o__Eubacteriales|f__Oscillospiraceae|g__GGB30454|s__GGB30454_SGB43514
k__Bacteria|p__Firmicutes|c__Clostridia|o__Eubacteriales|f__Oscillospiraceae|g__GGB30455|s__GGB30455_SGB43519
k__Bacteria|p__Firmicutes|c__Clostridia|o__Eubacteriales|f__Oscillospiraceae|g__GGB30456|s__GGB30456_SGB43520
k__Bacteria|p__Firmicutes|c__Clostridia|o__Eubacteriales|f__Oscillospiraceae|g__GGB30457|s__GGB30457_SGB63218
k__Bacteria|p__Firmicutes|c__Clostridia|o__Eubacteriales|f__Oscillospiraceae|g__GGB30461|s__GGB30461_SGB43530
k__Bacteria|p__Firmicutes|c__Clostridia|o__Eubacteriales|f__Oscillospiraceae|g__GGB30461|s__GGB30461_SGB43533
k__Bacteria|p__Firmicutes|c__Clostridia|o__Eubacteriales|f__Oscillospiraceae|g__GGB30461|s__GGB30461_SGB63209
k__Bacteria|p__Firmicutes|c__Clostridia|o__Eubacteriales|f__Oscillospiraceae|g__GGB30463|s__GGB30463_SGB43537
k__Bacteria|p__Firmicutes|c__Clostridia|o__Eubacteriales|f__Oscillospiraceae|g__GGB30473|s__GGB30473_SGB43557
k__Bacteria|p__Firmicutes|c__Clostridia|o__Eubacteriales|f__Oscillospiraceae|g__GGB30475|s__GGB30475_SGB63182
k__Bacteria|p__Actinobacteria|c__CFGB77153|o__OFGB77153|f__FGB77153|g__GGB30861|s__GGB30861_SGB44083
k__Bacteria|p__Tenericutes|c__CFGB1791|o__OFGB1791|f__FGB1791|g__GGB31312|s__GGB31312_SGB44628
k__Bacteria|p__Firmicutes|c__CFGB10290|o__OFGB10290|f__FGB10290|g__GGB31438|s__GGB31438_SGB44768
k__Bacteria|p__Firmicutes|c__Clostridia|o__Eubacteriales|f__Oscillospiraceae|g__GGB3171|s__GGB3171_SGB4185
k__Bacteria|p__Firmicutes|c__CFGB10289|o__OFGB10289|f__FGB10289|g__GGB31762|s__GGB31762_SGB45125
k__Bacteria|p__Firmicutes|c__CFGB1765|o__OFGB1765|f__FGB1765|g__GGB31823|s__GGB31823_SGB45199
k__Bacteria|p__Firmicutes|c__CFGB1765|o__OFGB1765|f__FGB1765|g__GGB31838|s__GGB31838_SGB45216
k__Bacteria|p__Firmicutes|c__CFGB1765|o__OFGB1765|f__FGB1765|g__GGB31841|s__GGB31841_SGB65084
k__Bacteria|p__Firmicutes|c__CFGB10667|o__OFGB10667|f__FGB10667|g__GGB32371|s__GGB32371_SGB41694
k__Bacteria|p__Firmicutes|c__Clostridia|o__Eubacteriales|f__Lachnospiraceae|g__GGB3793|s__GGB3793_SGB5158
k__Bacteria|p__Firmicutes|c__Clostridia|o__Eubacteriales|f__Clostridiaceae|g__GGB42601|s__GGB42601_SGB59797
k__Bacteria|p__Firmicutes|c__Clostridia|o__Eubacteriales|f__Oscillospiraceae|g__GGB45514|s__GGB45514_SGB63186
k__Bacteria|p__Firmicutes|c__CFGB75721|o__OFGB75721|f__FGB75721|g__GGB45564|s__GGB45564_SGB63259
k__Bacteria|p__Firmicutes|c__Clostridia|o__Eubacteriales|f__Oscillospiraceae|g__GGB45624|s__GGB45624_SGB63337
k__Bacteria|p__Firmicutes|c__Clostridia|o__Eubacteriales|f__Christensenellaceae|g__GGB45656|s__GGB45656_SGB6337
k__Bacteria|p__Firmicutes|c__CFGB10299|o__OFGB10299|f__FGB10299|g__GGB47127|s__GGB47127_SGB65054

k__Bacteria|p__Firmicutes|c__Clostridia|o__Eubacteriales|f__Oscillospiraceae|g__GGB74395|s__GGB74395_SGB43523
k__Bacteria|p__Firmicutes|c__Clostridia|o__Eubacteriales|f__Oscillospiraceae|g__GGB75053|s__GGB75053_SGB43494
k__Bacteria|p__Firmicutes|c__Clostridia|o__Eubacteriales|f__Lachnospiraceae|g__GGB75109|s__GGB75109_SGB102238
k__Bacteria|p__Firmicutes|c__Clostridia|o__Eubacteriales|f__Lachnospiraceae|g__Lachnospiraceae_unclassified|s__Lachn
k__Bacteria|p__Firmicutes|c__Clostridia|o__Eubacteriales|f__Lachnospiraceae|g__Lachnospiraceae_unclassified|s__Lachn
k__Bacteria|p__Firmicutes|c__Clostridia|o__Eubacteriales|f__Lachnospiraceae|g__Lachnospiraceae_unclassified|s__Lachn
k__Bacteria|p__Firmicutes|c__Clostridia|o__Eubacteriales|f__Lachnospiraceae|g__Lachnospiraceae_unclassified|s__Lachn
k__Bacteria|p__Firmicutes|c__Clostridia|o__Eubacteriales|f__Lachnospiraceae|g__Lachnospiraceae_unclassified|s__Lachn
k__Bacteria|p__Firmicutes|c__Bacilli|o__Lactobacillales|f__Lactobacillaceae|g__Lactobacillus|s__Lactobacillus_johnsonii
k__Bacteria|p__Actinobacteria|c__Coriobacteriia|o__Coriobacteriales|f__Atopobiaceae|g__Leptogranulimonas|s__Leptog
k__Bacteria|p__Bacteroidota|c__Bacteroidia|o__Bacteroidales|f__Muribaculaceae|g__Muribaculaceae_unclassified|s__M
k__Bacteria|p__Firmicutes|c__Clostridia|o__Eubacteriales|f__Oscillospiraceae|g__Neglectibacter|s__Neglectibacter_sp_X
k__Bacteria|p__Firmicutes|c__Clostridia|o__Eubacteriales|f__Oscillospiraceae|g__Oscillibacter|s__Oscillibacter_SGB4349
k__Bacteria|p__Firmicutes|c__Clostridia|o__Eubacteriales|f__Oscillospiraceae|g__Oscillospiraceae_unclassified|s__Oscillo
k__Bacteria|p__Firmicutes|c__Clostridia|o__Eubacteriales|f__Oscillospiraceae|g__Oscillospiraceae_unclassified|s__Oscillo
k__Bacteria|p__Firmicutes|c__Clostridia|o__Eubacteriales|f__Oscillospiraceae|g__Oscillospiraceae_unclassified|s__Oscillo
k__Bacteria|p__Firmicutes|c__Clostridia|o__Eubacteriales|f__Oscillospiraceae|g__Oscillospiraceae_unclassified|s__Oscillo
k__Bacteria|p__Proteobacteria|c__Betaproteobacteria|o__Burkholderiales|f__Sutterellaceae|g__Parasutterella|s__Parasu
k__Bacteria|p__Firmicutes|c__Clostridia|o__Eubacteriales|f__Lachnospiraceae|g__Schaedlerella|s__Schaedlerella_arabino
k__Bacteria|p__Firmicutes|c__Erysipelotrichia|o__Erysipelotrichales|f__Turicibacteraceae|g__Turicibacter|s__Turicibacter_
k__Bacteria|p__Bacteria_unclassified|c__Bacteria_unclassified|o__Bacteria_unclassified|f__Bacteria_unclassified|g__Bacte
k__Bacteria|p__Bacteria_unclassified|c__Bacteria_unclassified|o__Bacteria_unclassified|f__Bacteria_unclassified|g__Bacte
k__Bacteria|p__Bacteria_unclassified|c__Bacteria_unclassified|o__Bacteria_unclassified|f__Bacteria_unclassified|g__Bacte
k__Bacteria|p__Bacteria_unclassified|c__Bacteria_unclassified|o__Bacteria_unclassified|f__Bacteria_unclassified|g__Bacte
k__Bacteria|p__Bacteria_unclassified|c__Bacteria_unclassified|o__Bacteria_unclassified|f__Bacteria_unclassified|g__Bacte

k__Bacteria|p__Firmicutes|c__Clostridia|o__Eubacteriales|f__Lachnospiraceae|g__Acetatifactor|s__Acetatifactor_SGB415
k__Bacteria|p__Firmicutes|c__Clostridia|o__Eubacteriales|f__Oscillospiraceae|g__Acutalibacter|s__Acutalibacter_muris
k__Bacteria|p__Verrucomicrobia|c__Verrucomicrobiae|o__Verrucomicrobiales|f__Akkermansiaceae|g__Akkermansia|s__A
k__Bacteria|p__Firmicutes|c__Clostridia|o__Eubacteriales|f__Oscillospiraceae|g__Anaerotruncus|s__Anaerotruncus_sp_1
k__Bacteria|p__Bacteria_unclassified|c__Bacteria_unclassified|o__Bacteria_unclassified|f__Bacteria_unclassified|g__Bacte
k__Bacteria|p__Bacteria_unclassified|c__Bacteria_unclassified|o__Bacteria_unclassified|f__Bacteria_unclassified|g__Bacte
k__Bacteria|p__Bacteria_unclassified|c__Bacteria_unclassified|o__Bacteria_unclassified|f__Bacteria_unclassified|g__Bacte
k__Bacteria|p__Bacteria_unclassified|c__Bacteria_unclassified|o__Bacteria_unclassified|f__Bacteria_unclassified|g__Bacte
k__Bacteria|p__Bacteroidota|c__Bacteroidia|o__Bacteroidales|f__Bacteroidaceae|g__Bacteroides|s__Bacteroides_thetaio
k__Bacteria|p__Firmicutes|c__Clostridia|o__Clostridia_unclassified|f__Clostridia_unclassified|g__Clostridia_unclassified|s_
k__Bacteria|p__Firmicutes|c__Clostridia|o__Eubacteriales|f__Clostridiaceae|g__Clostridiaceae_unclassified|s__Clostridiac
k__Bacteria|p__Firmicutes|c__Clostridia|o__Eubacteriales|f__Clostridiaceae|g__Clostridiaceae_unclassified|s__Clostridiac
k__Bacteria|p__Firmicutes|c__Clostridia|o__Eubacteriales|f__Eubacteriales_unclassified|g__Eubacteriales_unclassified|s_
k__Bacteria|p__Firmicutes|c__Clostridia|o__Eubacteriales|f__Clostridiaceae|g__Clostridium|s__Clostridium_SGB65123
k__Bacteria|p__Firmicutes|c__Erysipelotrichia|o__Erysipelotrichales|f__Erysipelotrichaceae|g__Erysipelatoclostridium|s_
k__Bacteria|p__Firmicutes|c__Clostridia|o__Eubacteriales|f__Eubacteriaceae|g__Eubacteriaceae_unclassified|s__Eubacte
k__Bacteria|p__Firmicutes|c__Clostridia|o__Eubacteriales|f__Eubacteriaceae|g__Eubacteriaceae_unclassified|s__Eubacte
k__Bacteria|p__Firmicutes|c__CFGB77306|o__OFGB77306|f__FGB77306|g__GGB20146|s__GGB20146_SGB29427

k__Bacteria|p__Firmicutes|c__Clostridia|o__Eubacteriales|f__Lachnospiraceae|g__GGB20149|s__GGB20149_SGB29430
k__Bacteria|p__Actinobacteria|c__Coriobacteriia|o__Eggerthellales|f__Eggerthellaceae|g__GGB22635|s__GGB22635_SGB
k__Bacteria|p__Firmicutes|c__Clostridia|o__Eubacteriales|f__Lachnospiraceae|g__GGB25041|s__GGB25041_SGB36960
k__Bacteria|p__Firmicutes|c__CFGB9506|o__OFGB9506|f__FGB9506|g__GGB28379|s__GGB28379_SGB40959
k__Bacteria|p__Firmicutes|c__CFGB9508|o__OFGB9508|f__FGB9508|g__GGB28382|s__GGB28382_SGB40962
k__Bacteria|p__Firmicutes|c__CFGB9512|o__OFGB9512|f__FGB9512|g__GGB28392|s__GGB28392_SGB40972
k__Bacteria|p__Firmicutes|c__CFGB2838|o__OFGB2838|f__FGB2838|g__GGB28404|s__GGB28404_SGB40986
k__Bacteria|p__Firmicutes|c__CFGB2838|o__OFGB2838|f__FGB2838|g__GGB28418|s__GGB28418_SGB41001
k__Bacteria|p__Firmicutes|c__CFGB2838|o__OFGB2838|f__FGB2838|g__GGB28422|s__GGB28422_SGB41005
k__Bacteria|p__Firmicutes|c__Clostridia|o__Eubacteriales|f__Pumilibacteraceae|g__GGB28431|s__GGB28431_SGB41014
k__Bacteria|p__Firmicutes|c__CFGB2833|o__OFGB2833|f__FGB2833|g__GGB28456|s__GGB28456_SGB41039
k__Bacteria|p__Firmicutes|c__Clostridia|o__Eubacteriales|f__Eubacteriaceae|g__GGB28782|s__GGB28782_SGB41435
k__Bacteria|p__Firmicutes|c__Clostridia|o__Eubacteriales|f__Lachnospiraceae|g__GGB28792|s__GGB28792_SGB41445
k__Bacteria|p__Firmicutes|c__Clostridia|o__Eubacteriales|f__Lachnospiraceae|g__GGB28798|s__GGB28798_SGB41451
k__Bacteria|p__Firmicutes|c__CFGB9622|o__OFGB9622|f__FGB9622|g__GGB28810|s__GGB28810_SGB41465
k__Bacteria|p__Firmicutes|c__CFGB77305|o__OFGB77305|f__FGB77305|g__GGB28828|s__GGB28828_SGB41484
k__Bacteria|p__Firmicutes|c__Clostridia|o__Eubacteriales|f__Clostridiaceae|g__GGB28851|s__GGB28851_SGB41518
k__Bacteria|p__Firmicutes|c__Clostridia|o__Eubacteriales|f__Lachnospiraceae|g__GGB28865|s__GGB28865_SGB41537
k__Bacteria|p__Firmicutes|c__Clostridia|o__Eubacteriales|f__Lachnospiraceae|g__GGB28868|s__GGB28868_SGB41542
k__Bacteria|p__Firmicutes|c__Clostridia|o__Eubacteriales|f__Lachnospiraceae|g__GGB28869|s__GGB28869_SGB41543
k__Bacteria|p__Firmicutes|c__Clostridia|o__Eubacteriales|f__Lachnospiraceae|g__GGB28875|s__GGB28875_SGB41555
k__Bacteria|p__Firmicutes|c__CFGB9633|o__OFGB9633|f__FGB9633|g__GGB28881|s__GGB28881_SGB41561
k__Bacteria|p__Bacteria_unclassified|c__Bacteria_unclassified|o__Bacteria_unclassified|f__Bacteria_unclassified|g__GGB
k__Bacteria|p__Bacteria_unclassified|c__Bacteria_unclassified|o__Bacteria_unclassified|f__Bacteria_unclassified|g__GGB
k__Bacteria|p__Bacteria_unclassified|c__Bacteria_unclassified|o__Bacteria_unclassified|f__Bacteria_unclassified|g__GGB
k__Bacteria|p__Bacteria_unclassified|c__Bacteria_unclassified|o__Bacteria_unclassified|f__Bacteria_unclassified|g__GGB
k__Bacteria|p__Bacteria_unclassified|c__Bacteria_unclassified|o__Bacteria_unclassified|f__Bacteria_unclassified|g__GGB
k__Bacteria|p__Bacteria_unclassified|c__Bacteria_unclassified|o__Bacteria_unclassified|f__Bacteria_unclassified|g__GGB
k__Bacteria|p__Firmicutes|c__Clostridia|o__Eubacteriales|f__Lachnospiraceae|g__GGB28924|s__GGB28924_SGB41621
k__Bacteria|p__Bacteria_unclassified|c__CFGB77359|o__OFGB77359|f__FGB77359|g__GGB28927|s__GGB28927_SGB416
k__Bacteria|p__Firmicutes|c__CFGB9639|o__OFGB9639|f__FGB9639|g__GGB28934|s__GGB28934_SGB41635
k__Bacteria|p__Firmicutes|c__Clostridia|o__Eubacteriales|f__Lachnospiraceae|g__GGB28949|s__GGB28949_SGB41655
k__Bacteria|p__Firmicutes|c__Clostridia|o__Eubacteriales|f__Clostridiaceae|g__GGB28951|s__GGB28951_SGB102295
k__Bacteria|p__Firmicutes|c__Clostridia|o__Eubacteriales|f__Clostridiaceae|g__GGB28951|s__GGB28951_SGB41658
k__Bacteria|p__Firmicutes|c__Clostridia|o__Eubacteriales|f__Clostridiaceae|g__GGB28954|s__GGB28954_SGB41662
k__Bacteria|p__Firmicutes|c__Clostridia|o__Eubacteriales|f__Clostridiaceae|g__GGB28960|s__GGB28960_SGB41669
k__Bacteria|p__Firmicutes|c__Clostridia|o__Eubacteriales|f__Clostridiaceae|g__GGB28964|s__GGB28964_SGB94886
k__Bacteria|p__Firmicutes|c__Clostridia|o__Eubacteriales|f__Clostridiaceae|g__GGB28967|s__GGB28967_SGB41678
k__Bacteria|p__Firmicutes|c__CFGB9656|o__OFGB9656|f__FGB9656|g__GGB28996|s__GGB28996_SGB41712
k__Bacteria|p__Firmicutes|c__CFGB9827|o__OFGB9827|f__FGB9827|g__GGB29531|s__GGB29531_SGB42317
k__Bacteria|p__Firmicutes|c__Clostridia|o__Eubacteriales|f__Eubacteriaceae|g__GGB29685|s__GGB29685_SGB42494
k__Bacteria|p__Bacteria_unclassified|c__CFGB77303|o__OFGB77303|f__FGB77303|g__GGB30141|s__GGB30141_SGB430
k__Bacteria|p__Firmicutes|c__Clostridia|o__Eubacteriales|f__Clostridiaceae|g__GGB30145|s__GGB30145_SGB43072
k__Bacteria|p__Firmicutes|c__Clostridia|o__Eubacteriales|f__Eubacteriales_unclassified|g__GGB30286|s__GGB30286_SG
k__Bacteria|p__Firmicutes|c__CFGB72709|o__OFGB72709|f__FGB72709|g__GGB30300|s__GGB30300_SGB43264
k__Bacteria|p__Firmicutes|c__Clostridia|o__Eubacteriales|f__Oscillospiraceae|g__GGB30303|s__GGB30303_SGB43268

k__Bacteria|p__Firmicutes|c__Clostridia|o__Eubacteriales|f__Oscillospiraceae|g__GGB30450|s__GGB30450_SGB43507
k__Bacteria|p__Firmicutes|c__Clostridia|o__Eubacteriales|f__Oscillospiraceae|g__GGB30453|s__GGB30453_SGB43513
k__Bacteria|p__Firmicutes|c__Clostridia|o__Eubacteriales|f__Oscillospiraceae|g__GGB30454|s__GGB30454_SGB43514
k__Bacteria|p__Firmicutes|c__Clostridia|o__Eubacteriales|f__Oscillospiraceae|g__GGB30455|s__GGB30455_SGB43519
k__Bacteria|p__Firmicutes|c__Clostridia|o__Eubacteriales|f__Oscillospiraceae|g__GGB30456|s__GGB30456_SGB43520
k__Bacteria|p__Firmicutes|c__Clostridia|o__Eubacteriales|f__Oscillospiraceae|g__GGB30457|s__GGB30457_SGB63218
k__Bacteria|p__Firmicutes|c__Clostridia|o__Eubacteriales|f__Oscillospiraceae|g__GGB30461|s__GGB30461_SGB43530
k__Bacteria|p__Firmicutes|c__Clostridia|o__Eubacteriales|f__Oscillospiraceae|g__GGB30461|s__GGB30461_SGB43533
k__Bacteria|p__Firmicutes|c__Clostridia|o__Eubacteriales|f__Oscillospiraceae|g__GGB30461|s__GGB30461_SGB63209
k__Bacteria|p__Firmicutes|c__Clostridia|o__Eubacteriales|f__Oscillospiraceae|g__GGB30463|s__GGB30463_SGB43537
k__Bacteria|p__Firmicutes|c__Clostridia|o__Eubacteriales|f__Oscillospiraceae|g__GGB30473|s__GGB30473_SGB43557
k__Bacteria|p__Firmicutes|c__Clostridia|o__Eubacteriales|f__Oscillospiraceae|g__GGB30475|s__GGB30475_SGB63182
k__Bacteria|p__Actinobacteria|c__CFGB77153|o__OFGB77153|f__FGB77153|g__GGB30861|s__GGB30861_SGB44083
k__Bacteria|p__Tenericutes|c__CFGB1791|o__OFGB1791|f__FGB1791|g__GGB31312|s__GGB31312_SGB44628
k__Bacteria|p__Firmicutes|c__CFGB10290|o__OFGB10290|f__FGB10290|g__GGB31438|s__GGB31438_SGB44768
k__Bacteria|p__Firmicutes|c__Clostridia|o__Eubacteriales|f__Oscillospiraceae|g__GGB3171|s__GGB3171_SGB4185
k__Bacteria|p__Firmicutes|c__CFGB10289|o__OFGB10289|f__FGB10289|g__GGB31762|s__GGB31762_SGB45125
k__Bacteria|p__Firmicutes|c__CFGB1765|o__OFGB1765|f__FGB1765|g__GGB31823|s__GGB31823_SGB45199
k__Bacteria|p__Firmicutes|c__CFGB1765|o__OFGB1765|f__FGB1765|g__GGB31838|s__GGB31838_SGB45216
k__Bacteria|p__Firmicutes|c__CFGB1765|o__OFGB1765|f__FGB1765|g__GGB31841|s__GGB31841_SGB65084
k__Bacteria|p__Firmicutes|c__CFGB10667|o__OFGB10667|f__FGB10667|g__GGB32371|s__GGB32371_SGB41694
k__Bacteria|p__Firmicutes|c__Clostridia|o__Eubacteriales|f__Lachnospiraceae|g__GGB3793|s__GGB3793_SGB5158
k__Bacteria|p__Firmicutes|c__Clostridia|o__Eubacteriales|f__Clostridiaceae|g__GGB42601|s__GGB42601_SGB59797
k__Bacteria|p__Firmicutes|c__Clostridia|o__Eubacteriales|f__Oscillospiraceae|g__GGB45514|s__GGB45514_SGB63186
k__Bacteria|p__Firmicutes|c__CFGB75721|o__OFGB75721|f__FGB75721|g__GGB45564|s__GGB45564_SGB63259
k__Bacteria|p__Firmicutes|c__Clostridia|o__Eubacteriales|f__Oscillospiraceae|g__GGB45624|s__GGB45624_SGB63337
k__Bacteria|p__Firmicutes|c__Clostridia|o__Eubacteriales|f__Christensenellaceae|g__GGB45656|s__GGB45656_SGB6337
k__Bacteria|p__Firmicutes|c__CFGB10299|o__OFGB10299|f__FGB10299|g__GGB47127|s__GGB47127_SGB65054
k__Bacteria|p__Firmicutes|c__Clostridia|o__Eubacteriales|f__Oscillospiraceae|g__GGB74395|s__GGB74395_SGB43523
k__Bacteria|p__Firmicutes|c__Clostridia|o__Eubacteriales|f__Oscillospiraceae|g__GGB75053|s__GGB75053_SGB43494
k__Bacteria|p__Firmicutes|c__Clostridia|o__Eubacteriales|f__Lachnospiraceae|g__GGB75109|s__GGB75109_SGB102238
k__Bacteria|p__Firmicutes|c__Clostridia|o__Eubacteriales|f__Lachnospiraceae|g__Lachnospiraceae_unclassified|s__Lachn
k__Bacteria|p__Firmicutes|c__Clostridia|o__Eubacteriales|f__Lachnospiraceae|g__Lachnospiraceae_unclassified|s__Lachn
k__Bacteria|p__Firmicutes|c__Clostridia|o__Eubacteriales|f__Lachnospiraceae|g__Lachnospiraceae_unclassified|s__Lachn
k__Bacteria|p__Firmicutes|c__Clostridia|o__Eubacteriales|f__Lachnospiraceae|g__Lachnospiraceae_unclassified|s__Lachn
k__Bacteria|p__Firmicutes|c__Clostridia|o__Eubacteriales|f__Lachnospiraceae|g__Lachnospiraceae_unclassified|s__Lachn
k__Bacteria|p__Firmicutes|c__Bacilli|o__Lactobacillales|f__Lactobacillaceae|g__Lactobacillus|s__Lactobacillus_johnsonii
k__Bacteria|p__Actinobacteria|c__Coriobacteriia|o__Coriobacteriales|f__Atopobiaceae|g__Leptogranulimonas|s__Leptog
k__Bacteria|p__Bacteroidota|c__Bacteroidia|o__Bacteroidales|f__Muribaculaceae|g__Muribaculaceae_unclassified|s__M
k__Bacteria|p__Firmicutes|c__Clostridia|o__Eubacteriales|f__Oscillospiraceae|g__Neglectibacter|s__Neglectibacter_sp_X
k__Bacteria|p__Firmicutes|c__Clostridia|o__Eubacteriales|f__Oscillospiraceae|g__Oscillibacter|s__Oscillibacter_SGB43496
k__Bacteria|p__Firmicutes|c__Clostridia|o__Eubacteriales|f__Oscillospiraceae|g__Oscillospiraceae_unclassified|s__Oscillo
k__Bacteria|p__Firmicutes|c__Clostridia|o__Eubacteriales|f__Oscillospiraceae|g__Oscillospiraceae_unclassified|s__Oscillo
k__Bacteria|p__Firmicutes|c__Clostridia|o__Eubacteriales|f__Oscillospiraceae|g__Oscillospiraceae_unclassified|s__Oscillo
k__Bacteria|p__Firmicutes|c__Clostridia|o__Eubacteriales|f__Oscillospiraceae|g__Oscillospiraceae_unclassified|s__Oscillo
k__Bacteria|p__Proteobacteria|c__Betaproteobacteria|o__Burkholderiales|f__Sutterellaceae|g__Parasutterella|s__Parasu

k__Bacteria|p__Firmicutes|c__Clostridia|o__Eubacteriales|f__Lachnospiraceae|g__Schaedlerella|s__Schaedlerella_arabino
k__Bacteria|p__Firmicutes|c__Erysipelotrichia|o__Erysipelotrichales|f__Turicibacteraceae|g__Turicibacter|s__Turicibacter_
k__Bacteria|p__Bacteria_unclassified|c__Bacteria_unclassified|o__Bacteria_unclassified|f__Bacteria_unclassified|g__Bacte
k__Bacteria|p__Bacteria_unclassified|c__Bacteria_unclassified|o__Bacteria_unclassified|f__Bacteria_unclassified|g__Bacte
k__Bacteria|p__Bacteria_unclassified|c__Bacteria_unclassified|o__Bacteria_unclassified|f__Bacteria_unclassified|g__Bacte
k__Bacteria|p__Bacteria_unclassified|c__Bacteria_unclassified|o__Bacteria_unclassified|f__Bacteria_unclassified|g__Bacte
k__Bacteria|p__Bacteria_unclassified|c__Bacteria_unclassified|o__Bacteria_unclassified|f__Bacteria_unclassified|g__Bacte

k__Bacteria|p__Firmicutes|c__Clostridia|o__Eubacteriales|f__Lachnospiraceae|g__Acetatifactor|s__Acetatifactor_SGB415
k__Bacteria|p__Firmicutes|c__Clostridia|o__Eubacteriales|f__Oscillospiraceae|g__Acutalibacter|s__Acutalibacter_muris
k__Bacteria|p__Verrucomicrobia|c__Verrucomicrobiae|o__Verrucomicrobiales|f__Akkermansiaceae|g__Akkermansia|s__A
k__Bacteria|p__Firmicutes|c__Clostridia|o__Eubacteriales|f__Oscillospiraceae|g__Anaerotruncus|s__Anaerotruncus_sp_1
k__Bacteria|p__Bacteria_unclassified|c__Bacteria_unclassified|o__Bacteria_unclassified|f__Bacteria_unclassified|g__Bacte
k__Bacteria|p__Bacteria_unclassified|c__Bacteria_unclassified|o__Bacteria_unclassified|f__Bacteria_unclassified|g__Bacte
k__Bacteria|p__Bacteria_unclassified|c__Bacteria_unclassified|o__Bacteria_unclassified|f__Bacteria_unclassified|g__Bacte
k__Bacteria|p__Bacteria_unclassified|c__Bacteria_unclassified|o__Bacteria_unclassified|f__Bacteria_unclassified|g__Bacte
k__Bacteria|p__Bacteroidota|c__Bacteroidia|o__Bacteroidales|f__Bacteroidaceae|g__Bacteroides|s__Bacteroides_thetaio
k__Bacteria|p__Firmicutes|c__Clostridia|o__Clostridia_unclassified|f__Clostridia_unclassified|g__Clostridia_unclassified|s_
k__Bacteria|p__Firmicutes|c__Clostridia|o__Eubacteriales|f__Clostridiaceae|g__Clostridiaceae_unclassified|s__Clostridiac
k__Bacteria|p__Firmicutes|c__Clostridia|o__Eubacteriales|f__Clostridiaceae|g__Clostridiaceae_unclassified|s__Clostridiac
k__Bacteria|p__Firmicutes|c__Clostridia|o__Eubacteriales|f__Eubacteriales_unclassified|g__Eubacteriales_unclassified|s_
k__Bacteria|p__Firmicutes|c__Clostridia|o__Eubacteriales|f__Clostridiaceae|g__Clostridium|s__Clostridium_SGB65123
k__Bacteria|p__Firmicutes|c__Erysipelotrichia|o__Erysipelotrichales|f__Erysipelotrichaceae|g__Erysipelatoclostridium|s__
k__Bacteria|p__Firmicutes|c__Clostridia|o__Eubacteriales|f__Eubacteriaceae|g__Eubacteriaceae_unclassified|s__Eubacte
k__Bacteria|p__Firmicutes|c__Clostridia|o__Eubacteriales|f__Eubacteriaceae|g__Eubacteriaceae_unclassified|s__Eubacte
k__Bacteria|p__Firmicutes|c__CFGB77306|o__OFGB77306|f__FGB77306|g__GGB20146|s__GGB20146_SGB29427
k__Bacteria|p__Firmicutes|c__Clostridia|o__Eubacteriales|f__Lachnospiraceae|g__GGB20149|s__GGB20149_SGB29430
k__Bacteria|p__Actinobacteria|c__Coriobacteriia|o__Eggerthellales|f__Eggerthellaceae|g__GGB22635|s__GGB22635_SGB
k__Bacteria|p__Firmicutes|c__Clostridia|o__Eubacteriales|f__Lachnospiraceae|g__GGB25041|s__GGB25041_SGB36960
k__Bacteria|p__Firmicutes|c__CFGB9506|o__OFGB9506|f__FGB9506|g__GGB28379|s__GGB28379_SGB40959
k__Bacteria|p__Firmicutes|c__CFGB9508|o__OFGB9508|f__FGB9508|g__GGB28382|s__GGB28382_SGB40962
k__Bacteria|p__Firmicutes|c__CFGB9512|o__OFGB9512|f__FGB9512|g__GGB28392|s__GGB28392_SGB40972
k__Bacteria|p__Firmicutes|c__CFGB2838|o__OFGB2838|f__FGB2838|g__GGB28404|s__GGB28404_SGB40986
k__Bacteria|p__Firmicutes|c__CFGB2838|o__OFGB2838|f__FGB2838|g__GGB28418|s__GGB28418_SGB41001
k__Bacteria|p__Firmicutes|c__CFGB2838|o__OFGB2838|f__FGB2838|g__GGB28422|s__GGB28422_SGB41005
k__Bacteria|p__Firmicutes|c__Clostridia|o__Eubacteriales|f__Pumilibacteraceae|g__GGB28431|s__GGB28431_SGB41014
k__Bacteria|p__Firmicutes|c__CFGB2833|o__OFGB2833|f__FGB2833|g__GGB28456|s__GGB28456_SGB41039
k__Bacteria|p__Firmicutes|c__Clostridia|o__Eubacteriales|f__Eubacteriaceae|g__GGB28782|s__GGB28782_SGB41435
k__Bacteria|p__Firmicutes|c__Clostridia|o__Eubacteriales|f__Lachnospiraceae|g__GGB28792|s__GGB28792_SGB41445
k__Bacteria|p__Firmicutes|c__Clostridia|o__Eubacteriales|f__Lachnospiraceae|g__GGB28798|s__GGB28798_SGB41451
k__Bacteria|p__Firmicutes|c__CFGB9622|o__OFGB9622|f__FGB9622|g__GGB28810|s__GGB28810_SGB41465
k__Bacteria|p__Firmicutes|c__CFGB77305|o__OFGB77305|f__FGB77305|g__GGB28828|s__GGB28828_SGB41484
k__Bacteria|p__Firmicutes|c__Clostridia|o__Eubacteriales|f__Clostridiaceae|g__GGB28851|s__GGB28851_SGB41518
k__Bacteria|p__Firmicutes|c__Clostridia|o__Eubacteriales|f__Lachnospiraceae|g__GGB28865|s__GGB28865_SGB41537

k__Bacteria|p__Firmicutes|c__Clostridia|o__Eubacteriales|f__Lachnospiraceae|g__GGB28868|s__GGB28868_SGB41542
k__Bacteria|p__Firmicutes|c__Clostridia|o__Eubacteriales|f__Lachnospiraceae|g__GGB28869|s__GGB28869_SGB41543
k__Bacteria|p__Firmicutes|c__Clostridia|o__Eubacteriales|f__Lachnospiraceae|g__GGB28875|s__GGB28875_SGB41555
k__Bacteria|p__Firmicutes|c__CFGB9633|o__OFGB9633|f__FGB9633|g__GGB28881|s__GGB28881_SGB41561
k__Bacteria|p__Bacteria_unclassified|c__Bacteria_unclassified|o__Bacteria_unclassified|f__Bacteria_unclassified|g__GGB2
k__Bacteria|p__Bacteria_unclassified|c__Bacteria_unclassified|o__Bacteria_unclassified|f__Bacteria_unclassified|g__GGB2
k__Bacteria|p__Bacteria_unclassified|c__Bacteria_unclassified|o__Bacteria_unclassified|f__Bacteria_unclassified|g__GGB2
k__Bacteria|p__Bacteria_unclassified|c__Bacteria_unclassified|o__Bacteria_unclassified|f__Bacteria_unclassified|g__GGB2
k__Bacteria|p__Bacteria_unclassified|c__Bacteria_unclassified|o__Bacteria_unclassified|f__Bacteria_unclassified|g__GGB2
k__Bacteria|p__Bacteria_unclassified|c__Bacteria_unclassified|o__Bacteria_unclassified|f__Bacteria_unclassified|g__GGB2
k__Bacteria|p__Firmicutes|c__Clostridia|o__Eubacteriales|f__Lachnospiraceae|g__GGB28924|s__GGB28924_SGB41621
k__Bacteria|p__Bacteria_unclassified|c__CFGB77359|o__OFGB77359|f__FGB77359|g__GGB28927|s__GGB28927_SGB416
k__Bacteria|p__Firmicutes|c__CFGB9639|o__OFGB9639|f__FGB9639|g__GGB28934|s__GGB28934_SGB41635
k__Bacteria|p__Firmicutes|c__Clostridia|o__Eubacteriales|f__Lachnospiraceae|g__GGB28949|s__GGB28949_SGB41655
k__Bacteria|p__Firmicutes|c__Clostridia|o__Eubacteriales|f__Clostridiaceae|g__GGB28951|s__GGB28951_SGB102295
k__Bacteria|p__Firmicutes|c__Clostridia|o__Eubacteriales|f__Clostridiaceae|g__GGB28951|s__GGB28951_SGB41658
k__Bacteria|p__Firmicutes|c__Clostridia|o__Eubacteriales|f__Clostridiaceae|g__GGB28954|s__GGB28954_SGB41662
k__Bacteria|p__Firmicutes|c__Clostridia|o__Eubacteriales|f__Clostridiaceae|g__GGB28960|s__GGB28960_SGB41669
k__Bacteria|p__Firmicutes|c__Clostridia|o__Eubacteriales|f__Clostridiaceae|g__GGB28964|s__GGB28964_SGB94886
k__Bacteria|p__Firmicutes|c__Clostridia|o__Eubacteriales|f__Clostridiaceae|g__GGB28967|s__GGB28967_SGB41678
k__Bacteria|p__Firmicutes|c__CFGB9656|o__OFGB9656|f__FGB9656|g__GGB28996|s__GGB28996_SGB41712
k__Bacteria|p__Firmicutes|c__CFGB9827|o__OFGB9827|f__FGB9827|g__GGB29531|s__GGB29531_SGB42317
k__Bacteria|p__Firmicutes|c__Clostridia|o__Eubacteriales|f__Eubacteriaceae|g__GGB29685|s__GGB29685_SGB42494
k__Bacteria|p__Bacteria_unclassified|c__CFGB77303|o__OFGB77303|f__FGB77303|g__GGB30141|s__GGB30141_SGB430
k__Bacteria|p__Firmicutes|c__Clostridia|o__Eubacteriales|f__Clostridiaceae|g__GGB30145|s__GGB30145_SGB43072
k__Bacteria|p__Firmicutes|c__Clostridia|o__Eubacteriales|f__Eubacteriales_unclassified|g__GGB30286|s__GGB30286_SG
k__Bacteria|p__Firmicutes|c__CFGB72709|o__OFGB72709|f__FGB72709|g__GGB30300|s__GGB30300_SGB43264
k__Bacteria|p__Firmicutes|c__Clostridia|o__Eubacteriales|f__Oscillospiraceae|g__GGB30303|s__GGB30303_SGB43268
k__Bacteria|p__Firmicutes|c__Clostridia|o__Eubacteriales|f__Oscillospiraceae|g__GGB30450|s__GGB30450_SGB43507
k__Bacteria|p__Firmicutes|c__Clostridia|o__Eubacteriales|f__Oscillospiraceae|g__GGB30453|s__GGB30453_SGB43513
k__Bacteria|p__Firmicutes|c__Clostridia|o__Eubacteriales|f__Oscillospiraceae|g__GGB30454|s__GGB30454_SGB43514
k__Bacteria|p__Firmicutes|c__Clostridia|o__Eubacteriales|f__Oscillospiraceae|g__GGB30455|s__GGB30455_SGB43519
k__Bacteria|p__Firmicutes|c__Clostridia|o__Eubacteriales|f__Oscillospiraceae|g__GGB30456|s__GGB30456_SGB43520
k__Bacteria|p__Firmicutes|c__Clostridia|o__Eubacteriales|f__Oscillospiraceae|g__GGB30457|s__GGB30457_SGB63218
k__Bacteria|p__Firmicutes|c__Clostridia|o__Eubacteriales|f__Oscillospiraceae|g__GGB30461|s__GGB30461_SGB43530
k__Bacteria|p__Firmicutes|c__Clostridia|o__Eubacteriales|f__Oscillospiraceae|g__GGB30461|s__GGB30461_SGB43533
k__Bacteria|p__Firmicutes|c__Clostridia|o__Eubacteriales|f__Oscillospiraceae|g__GGB30461|s__GGB30461_SGB63209
k__Bacteria|p__Firmicutes|c__Clostridia|o__Eubacteriales|f__Oscillospiraceae|g__GGB30463|s__GGB30463_SGB43537
k__Bacteria|p__Firmicutes|c__Clostridia|o__Eubacteriales|f__Oscillospiraceae|g__GGB30473|s__GGB30473_SGB43557
k__Bacteria|p__Firmicutes|c__Clostridia|o__Eubacteriales|f__Oscillospiraceae|g__GGB30475|s__GGB30475_SGB63182
k__Bacteria|p__Actinobacteria|c__CFGB77153|o__OFGB77153|f__FGB77153|g__GGB30861|s__GGB30861_SGB44083
k__Bacteria|p__Tenericutes|c__CFGB1791|o__OFGB1791|f__FGB1791|g__GGB31312|s__GGB31312_SGB44628
k__Bacteria|p__Firmicutes|c__CFGB10290|o__OFGB10290|f__FGB10290|g__GGB31438|s__GGB31438_SGB44768
k__Bacteria|p__Firmicutes|c__Clostridia|o__Eubacteriales|f__Oscillospiraceae|g__GGB3171|s__GGB3171_SGB4185
k__Bacteria|p__Firmicutes|c__CFGB10289|o__OFGB10289|f__FGB10289|g__GGB31762|s__GGB31762_SGB45125
k__Bacteria|p__Firmicutes|c__CFGB1765|o__OFGB1765|f__FGB1765|g__GGB31823|s__GGB31823_SGB45199

k__Bacteria|p__Firmicutes|c__CFGB1765|o__OFGB1765|f__FGB1765|g__GGB31838|s__GGB31838_SGB45216
k__Bacteria|p__Firmicutes|c__CFGB1765|o__OFGB1765|f__FGB1765|g__GGB31841|s__GGB31841_SGB65084
k__Bacteria|p__Firmicutes|c__CFGB10667|o__OFGB10667|f__FGB10667|g__GGB32371|s__GGB32371_SGB41694
k__Bacteria|p__Firmicutes|c__Clostridia|o__Eubacteriales|f__Lachnospiraceae|g__GGB3793|s__GGB3793_SGB5158
k__Bacteria|p__Firmicutes|c__Clostridia|o__Eubacteriales|f__Clostridiaceae|g__GGB42601|s__GGB42601_SGB59797
k__Bacteria|p__Firmicutes|c__Clostridia|o__Eubacteriales|f__Oscillospiraceae|g__GGB45514|s__GGB45514_SGB63186
k__Bacteria|p__Firmicutes|c__CFGB75721|o__OFGB75721|f__FGB75721|g__GGB45564|s__GGB45564_SGB63259
k__Bacteria|p__Firmicutes|c__Clostridia|o__Eubacteriales|f__Oscillospiraceae|g__GGB45624|s__GGB45624_SGB63337
k__Bacteria|p__Firmicutes|c__Clostridia|o__Eubacteriales|f__Christensenellaceae|g__GGB45656|s__GGB45656_SGB6337
k__Bacteria|p__Firmicutes|c__CFGB10299|o__OFGB10299|f__FGB10299|g__GGB47127|s__GGB47127_SGB65054
k__Bacteria|p__Firmicutes|c__Clostridia|o__Eubacteriales|f__Oscillospiraceae|g__GGB74395|s__GGB74395_SGB43523
k__Bacteria|p__Firmicutes|c__Clostridia|o__Eubacteriales|f__Oscillospiraceae|g__GGB75053|s__GGB75053_SGB43494
k__Bacteria|p__Firmicutes|c__Clostridia|o__Eubacteriales|f__Lachnospiraceae|g__GGB75109|s__GGB75109_SGB102238
k__Bacteria|p__Firmicutes|c__Clostridia|o__Eubacteriales|f__Lachnospiraceae|g__Lachnospiraceae_unclassified|s__Lachn
k__Bacteria|p__Firmicutes|c__Clostridia|o__Eubacteriales|f__Lachnospiraceae|g__Lachnospiraceae_unclassified|s__Lachn
k__Bacteria|p__Firmicutes|c__Clostridia|o__Eubacteriales|f__Lachnospiraceae|g__Lachnospiraceae_unclassified|s__Lachn
k__Bacteria|p__Firmicutes|c__Clostridia|o__Eubacteriales|f__Lachnospiraceae|g__Lachnospiraceae_unclassified|s__Lachn
k__Bacteria|p__Firmicutes|c__Clostridia|o__Eubacteriales|f__Lachnospiraceae|g__Lachnospiraceae_unclassified|s__Lachn
k__Bacteria|p__Firmicutes|c__Bacilli|o__Lactobacillales|f__Lactobacillaceae|g__Lactobacillus|s__Lactobacillus_johnsonii
k__Bacteria|p__Actinobacteria|c__Coriobacteriia|o__Coriobacteriales|f__Atopobiaceae|g__Leptogranulimonas|s__Leptog
k__Bacteria|p__Bacteroidota|c__Bacteroidia|o__Bacteroidales|f__Muribaculaceae|g__Muribaculaceae_unclassified|s__M
k__Bacteria|p__Firmicutes|c__Clostridia|o__Eubacteriales|f__Oscillospiraceae|g__Neglectibacter|s__Neglectibacter_sp_X
k__Bacteria|p__Firmicutes|c__Clostridia|o__Eubacteriales|f__Oscillospiraceae|g__Oscillibacter|s__Oscillibacter_SGB43496
k__Bacteria|p__Firmicutes|c__Clostridia|o__Eubacteriales|f__Oscillospiraceae|g__Oscillospiraceae_unclassified|s__Oscillo
k__Bacteria|p__Firmicutes|c__Clostridia|o__Eubacteriales|f__Oscillospiraceae|g__Oscillospiraceae_unclassified|s__Oscillo
k__Bacteria|p__Firmicutes|c__Clostridia|o__Eubacteriales|f__Oscillospiraceae|g__Oscillospiraceae_unclassified|s__Oscillo
k__Bacteria|p__Firmicutes|c__Clostridia|o__Eubacteriales|f__Oscillospiraceae|g__Oscillospiraceae_unclassified|s__Oscillo
k__Bacteria|p__Proteobacteria|c__Betaproteobacteria|o__Burkholderiales|f__Sutterellaceae|g__Parasutterella|s__Parasu
k__Bacteria|p__Firmicutes|c__Clostridia|o__Eubacteriales|f__Lachnospiraceae|g__Schaedlerella|s__Schaedlerella_arabino
k__Bacteria|p__Firmicutes|c__Erysipelotrichia|o__Erysipelotrichales|f__Turicibacteraceae|g__Turicibacter|s__Turicibacter_
k__Bacteria|p__Bacteria_unclassified|c__Bacteria_unclassified|o__Bacteria_unclassified|f__Bacteria_unclassified|g__Bacte
k__Bacteria|p__Bacteria_unclassified|c__Bacteria_unclassified|o__Bacteria_unclassified|f__Bacteria_unclassified|g__Bacte
k__Bacteria|p__Bacteria_unclassified|c__Bacteria_unclassified|o__Bacteria_unclassified|f__Bacteria_unclassified|g__Bacte
k__Bacteria|p__Bacteria_unclassified|c__Bacteria_unclassified|o__Bacteria_unclassified|f__Bacteria_unclassified|g__Bacte
k__Bacteria|p__Bacteria_unclassified|c__Bacteria_unclassified|o__Bacteria_unclassified|f__Bacteria_unclassified|g__Bacte

k__Bacteria|p__Firmicutes|c__Clostridia|o__Eubacteriales|f__Lachnospiraceae|g__Acetatifactor|s__Acetatifactor_SGB415
k__Bacteria|p__Firmicutes|c__Clostridia|o__Eubacteriales|f__Oscillospiraceae|g__Acutalibacter|s__Acutalibacter_muris
k__Bacteria|p__Verrucomicrobia|c__Verrucomicrobiae|o__Verrucomicrobiales|f__Akkermansiaceae|g__Akkermansia|s__A
k__Bacteria|p__Firmicutes|c__Clostridia|o__Eubacteriales|f__Oscillospiraceae|g__Anaerotruncus|s__Anaerotruncus_sp_1
k__Bacteria|p__Bacteria_unclassified|c__Bacteria_unclassified|o__Bacteria_unclassified|f__Bacteria_unclassified|g__Bacte
k__Bacteria|p__Bacteria_unclassified|c__Bacteria_unclassified|o__Bacteria_unclassified|f__Bacteria_unclassified|g__Bacte
k__Bacteria|p__Bacteria_unclassified|c__Bacteria_unclassified|o__Bacteria_unclassified|f__Bacteria_unclassified|g__Bacte
k__Bacteria|p__Bacteria_unclassified|c__Bacteria_unclassified|o__Bacteria_unclassified|f__Bacteria_unclassified|g__Bacte

k__Bacteria|p__Bacteroidota|c__Bacteroidia|o__Bacteroidales|f__Bacteroidaceae|g__Bacteroides|s__Bacteroides_thetaio
k__Bacteria|p__Firmicutes|c__Clostridia|o__Clostridia_unclassified|f__Clostridia_unclassified|g__Clostridia_unclassified|s_
k__Bacteria|p__Firmicutes|c__Clostridia|o__Eubacteriales|f__Clostridiaceae|g__Clostridiaceae_unclassified|s__Clostridiac
k__Bacteria|p__Firmicutes|c__Clostridia|o__Eubacteriales|f__Clostridiaceae|g__Clostridiaceae_unclassified|s__Clostridiac
k__Bacteria|p__Firmicutes|c__Clostridia|o__Eubacteriales|f__Eubacteriales_unclassified|g__Eubacteriales_unclassified|s_
k__Bacteria|p__Firmicutes|c__Clostridia|o__Eubacteriales|f__Clostridiaceae|g__Clostridium|s__Clostridium_SGB65123
k__Bacteria|p__Firmicutes|c__Erysipelotrichia|o__Erysipelotrichales|f__Erysipelotrichaceae|g__Erysipelatoclostridium|s__
k__Bacteria|p__Firmicutes|c__Clostridia|o__Eubacteriales|f__Eubacteriaceae|g__Eubacteriaceae_unclassified|s__Eubacte
k__Bacteria|p__Firmicutes|c__Clostridia|o__Eubacteriales|f__Eubacteriaceae|g__Eubacteriaceae_unclassified|s__Eubacte
k__Bacteria|p__Firmicutes|c__CFGB77306|o__OFGB77306|f__FGB77306|g__GGB20146|s__GGB20146_SGB29427
k__Bacteria|p__Firmicutes|c__Clostridia|o__Eubacteriales|f__Lachnospiraceae|g__GGB20149|s__GGB20149_SGB29430
k__Bacteria|p__Actinobacteria|c__Coriobacteriia|o__Eggerthellales|f__Eggerthellaceae|g__GGB22635|s__GGB22635_SGB
k__Bacteria|p__Firmicutes|c__Clostridia|o__Eubacteriales|f__Lachnospiraceae|g__GGB25041|s__GGB25041_SGB36960
k__Bacteria|p__Firmicutes|c__CFGB9506|o__OFGB9506|f__FGB9506|g__GGB28379|s__GGB28379_SGB40959
k__Bacteria|p__Firmicutes|c__CFGB9508|o__OFGB9508|f__FGB9508|g__GGB28382|s__GGB28382_SGB40962
k__Bacteria|p__Firmicutes|c__CFGB9512|o__OFGB9512|f__FGB9512|g__GGB28392|s__GGB28392_SGB40972
k__Bacteria|p__Firmicutes|c__CFGB2838|o__OFGB2838|f__FGB2838|g__GGB28404|s__GGB28404_SGB40986
k__Bacteria|p__Firmicutes|c__CFGB2838|o__OFGB2838|f__FGB2838|g__GGB28418|s__GGB28418_SGB41001
k__Bacteria|p__Firmicutes|c__CFGB2838|o__OFGB2838|f__FGB2838|g__GGB28422|s__GGB28422_SGB41005
k__Bacteria|p__Firmicutes|c__Clostridia|o__Eubacteriales|f__Pumilibacteraceae|g__GGB28431|s__GGB28431_SGB41014
k__Bacteria|p__Firmicutes|c__CFGB2833|o__OFGB2833|f__FGB2833|g__GGB28456|s__GGB28456_SGB41039
k__Bacteria|p__Firmicutes|c__Clostridia|o__Eubacteriales|f__Eubacteriaceae|g__GGB28782|s__GGB28782_SGB41435
k__Bacteria|p__Firmicutes|c__Clostridia|o__Eubacteriales|f__Lachnospiraceae|g__GGB28792|s__GGB28792_SGB41445
k__Bacteria|p__Firmicutes|c__Clostridia|o__Eubacteriales|f__Lachnospiraceae|g__GGB28798|s__GGB28798_SGB41451
k__Bacteria|p__Firmicutes|c__CFGB9622|o__OFGB9622|f__FGB9622|g__GGB28810|s__GGB28810_SGB41465
k__Bacteria|p__Firmicutes|c__CFGB77305|o__OFGB77305|f__FGB77305|g__GGB28828|s__GGB28828_SGB41484
k__Bacteria|p__Firmicutes|c__Clostridia|o__Eubacteriales|f__Clostridiaceae|g__GGB28851|s__GGB28851_SGB41518
k__Bacteria|p__Firmicutes|c__Clostridia|o__Eubacteriales|f__Lachnospiraceae|g__GGB28865|s__GGB28865_SGB41537
k__Bacteria|p__Firmicutes|c__Clostridia|o__Eubacteriales|f__Lachnospiraceae|g__GGB28868|s__GGB28868_SGB41542
k__Bacteria|p__Firmicutes|c__Clostridia|o__Eubacteriales|f__Lachnospiraceae|g__GGB28869|s__GGB28869_SGB41543
k__Bacteria|p__Firmicutes|c__Clostridia|o__Eubacteriales|f__Lachnospiraceae|g__GGB28875|s__GGB28875_SGB41555
k__Bacteria|p__Firmicutes|c__CFGB9633|o__OFGB9633|f__FGB9633|g__GGB28881|s__GGB28881_SGB41561
k__Bacteria|p__Bacteria_unclassified|c__Bacteria_unclassified|o__Bacteria_unclassified|f__Bacteria_unclassified|g__GGB2
k__Bacteria|p__Bacteria_unclassified|c__Bacteria_unclassified|o__Bacteria_unclassified|f__Bacteria_unclassified|g__GGB2
k__Bacteria|p__Bacteria_unclassified|c__Bacteria_unclassified|o__Bacteria_unclassified|f__Bacteria_unclassified|g__GGB2
k__Bacteria|p__Bacteria_unclassified|c__Bacteria_unclassified|o__Bacteria_unclassified|f__Bacteria_unclassified|g__GGB2
k__Bacteria|p__Bacteria_unclassified|c__Bacteria_unclassified|o__Bacteria_unclassified|f__Bacteria_unclassified|g__GGB2
k__Bacteria|p__Bacteria_unclassified|c__Bacteria_unclassified|o__Bacteria_unclassified|f__Bacteria_unclassified|g__GGB2
k__Bacteria|p__Firmicutes|c__Clostridia|o__Eubacteriales|f__Lachnospiraceae|g__GGB28924|s__GGB28924_SGB41621
k__Bacteria|p__Bacteria_unclassified|c__CFGB77359|o__OFGB77359|f__FGB77359|g__GGB28927|s__GGB28927_SGB416
k__Bacteria|p__Firmicutes|c__CFGB9639|o__OFGB9639|f__FGB9639|g__GGB28934|s__GGB28934_SGB41635
k__Bacteria|p__Firmicutes|c__Clostridia|o__Eubacteriales|f__Lachnospiraceae|g__GGB28949|s__GGB28949_SGB41655
k__Bacteria|p__Firmicutes|c__Clostridia|o__Eubacteriales|f__Clostridiaceae|g__GGB28951|s__GGB28951_SGB102295
k__Bacteria|p__Firmicutes|c__Clostridia|o__Eubacteriales|f__Clostridiaceae|g__GGB28951|s__GGB28951_SGB41658
k__Bacteria|p__Firmicutes|c__Clostridia|o__Eubacteriales|f__Clostridiaceae|g__GGB28954|s__GGB28954_SGB41662
k__Bacteria|p__Firmicutes|c__Clostridia|o__Eubacteriales|f__Clostridiaceae|g__GGB28960|s__GGB28960_SGB41669

k__Bacteria|p__Firmicutes|c__Clostridia|o__Eubacteriales|f__Clostridiaceae|g__GGB28964|s__GGB28964_SGB94886
k__Bacteria|p__Firmicutes|c__Clostridia|o__Eubacteriales|f__Clostridiaceae|g__GGB28967|s__GGB28967_SGB41678
k__Bacteria|p__Firmicutes|c__CFGB9656|o__OFGB9656|f__FGB9656|g__GGB28996|s__GGB28996_SGB41712
k__Bacteria|p__Firmicutes|c__CFGB9827|o__OFGB9827|f__FGB9827|g__GGB29531|s__GGB29531_SGB42317
k__Bacteria|p__Firmicutes|c__Clostridia|o__Eubacteriales|f__Eubacteriaceae|g__GGB29685|s__GGB29685_SGB42494
k__Bacteria|p__Bacteria_unclassified|c__CFGB77303|o__OFGB77303|f__FGB77303|g__GGB30141|s__GGB30141_SGB430
k__Bacteria|p__Firmicutes|c__Clostridia|o__Eubacteriales|f__Clostridiaceae|g__GGB30145|s__GGB30145_SGB43072
k__Bacteria|p__Firmicutes|c__Clostridia|o__Eubacteriales|f__Eubacteriales_unclassified|g__GGB30286|s__GGB30286_SG
k__Bacteria|p__Firmicutes|c__CFGB72709|o__OFGB72709|f__FGB72709|g__GGB30300|s__GGB30300_SGB43264
k__Bacteria|p__Firmicutes|c__Clostridia|o__Eubacteriales|f__Oscillospiraceae|g__GGB30303|s__GGB30303_SGB43268
k__Bacteria|p__Firmicutes|c__Clostridia|o__Eubacteriales|f__Oscillospiraceae|g__GGB30450|s__GGB30450_SGB43507
k__Bacteria|p__Firmicutes|c__Clostridia|o__Eubacteriales|f__Oscillospiraceae|g__GGB30453|s__GGB30453_SGB43513
k__Bacteria|p__Firmicutes|c__Clostridia|o__Eubacteriales|f__Oscillospiraceae|g__GGB30454|s__GGB30454_SGB43514
k__Bacteria|p__Firmicutes|c__Clostridia|o__Eubacteriales|f__Oscillospiraceae|g__GGB30455|s__GGB30455_SGB43519
k__Bacteria|p__Firmicutes|c__Clostridia|o__Eubacteriales|f__Oscillospiraceae|g__GGB30456|s__GGB30456_SGB43520
k__Bacteria|p__Firmicutes|c__Clostridia|o__Eubacteriales|f__Oscillospiraceae|g__GGB30457|s__GGB30457_SGB63218
k__Bacteria|p__Firmicutes|c__Clostridia|o__Eubacteriales|f__Oscillospiraceae|g__GGB30461|s__GGB30461_SGB43530
k__Bacteria|p__Firmicutes|c__Clostridia|o__Eubacteriales|f__Oscillospiraceae|g__GGB30461|s__GGB30461_SGB43533
k__Bacteria|p__Firmicutes|c__Clostridia|o__Eubacteriales|f__Oscillospiraceae|g__GGB30461|s__GGB30461_SGB63209
k__Bacteria|p__Firmicutes|c__Clostridia|o__Eubacteriales|f__Oscillospiraceae|g__GGB30463|s__GGB30463_SGB43537
k__Bacteria|p__Firmicutes|c__Clostridia|o__Eubacteriales|f__Oscillospiraceae|g__GGB30473|s__GGB30473_SGB43557
k__Bacteria|p__Firmicutes|c__Clostridia|o__Eubacteriales|f__Oscillospiraceae|g__GGB30475|s__GGB30475_SGB63182
k__Bacteria|p__Actinobacteria|c__CFGB77153|o__OFGB77153|f__FGB77153|g__GGB30861|s__GGB30861_SGB44083
k__Bacteria|p__Tenericutes|c__CFGB1791|o__OFGB1791|f__FGB1791|g__GGB31312|s__GGB31312_SGB44628
k__Bacteria|p__Firmicutes|c__CFGB10290|o__OFGB10290|f__FGB10290|g__GGB31438|s__GGB31438_SGB44768
k__Bacteria|p__Firmicutes|c__Clostridia|o__Eubacteriales|f__Oscillospiraceae|g__GGB3171|s__GGB3171_SGB4185
k__Bacteria|p__Firmicutes|c__CFGB10289|o__OFGB10289|f__FGB10289|g__GGB31762|s__GGB31762_SGB45125
k__Bacteria|p__Firmicutes|c__CFGB1765|o__OFGB1765|f__FGB1765|g__GGB31823|s__GGB31823_SGB45199
k__Bacteria|p__Firmicutes|c__CFGB1765|o__OFGB1765|f__FGB1765|g__GGB31838|s__GGB31838_SGB45216
k__Bacteria|p__Firmicutes|c__CFGB1765|o__OFGB1765|f__FGB1765|g__GGB31841|s__GGB31841_SGB65084
k__Bacteria|p__Firmicutes|c__CFGB10667|o__OFGB10667|f__FGB10667|g__GGB32371|s__GGB32371_SGB41694
k__Bacteria|p__Firmicutes|c__Clostridia|o__Eubacteriales|f__Lachnospiraceae|g__GGB3793|s__GGB3793_SGB5158
k__Bacteria|p__Firmicutes|c__Clostridia|o__Eubacteriales|f__Clostridiaceae|g__GGB42601|s__GGB42601_SGB59797
k__Bacteria|p__Firmicutes|c__Clostridia|o__Eubacteriales|f__Oscillospiraceae|g__GGB45514|s__GGB45514_SGB63186
k__Bacteria|p__Firmicutes|c__CFGB75721|o__OFGB75721|f__FGB75721|g__GGB45564|s__GGB45564_SGB63259
k__Bacteria|p__Firmicutes|c__Clostridia|o__Eubacteriales|f__Oscillospiraceae|g__GGB45624|s__GGB45624_SGB63337
k__Bacteria|p__Firmicutes|c__Clostridia|o__Eubacteriales|f__Christensenellaceae|g__GGB45656|s__GGB45656_SGB6337
k__Bacteria|p__Firmicutes|c__CFGB10299|o__OFGB10299|f__FGB10299|g__GGB47127|s__GGB47127_SGB65054
k__Bacteria|p__Firmicutes|c__Clostridia|o__Eubacteriales|f__Oscillospiraceae|g__GGB74395|s__GGB74395_SGB43523
k__Bacteria|p__Firmicutes|c__Clostridia|o__Eubacteriales|f__Oscillospiraceae|g__GGB75053|s__GGB75053_SGB43494
k__Bacteria|p__Firmicutes|c__Clostridia|o__Eubacteriales|f__Lachnospiraceae|g__GGB75109|s__GGB75109_SGB102238
k__Bacteria|p__Firmicutes|c__Clostridia|o__Eubacteriales|f__Lachnospiraceae|g__Lachnospiraceae_unclassified|s__Lachn
k__Bacteria|p__Firmicutes|c__Clostridia|o__Eubacteriales|f__Lachnospiraceae|g__Lachnospiraceae_unclassified|s__Lachn
k__Bacteria|p__Firmicutes|c__Clostridia|o__Eubacteriales|f__Lachnospiraceae|g__Lachnospiraceae_unclassified|s__Lachn
k__Bacteria|p__Firmicutes|c__Clostridia|o__Eubacteriales|f__Lachnospiraceae|g__Lachnospiraceae_unclassified|s__Lachn
k__Bacteria|p__Firmicutes|c__Clostridia|o__Eubacteriales|f__Lachnospiraceae|g__Lachnospiraceae_unclassified|s__Lachn

k__Bacteria|p__Firmicutes|c__Bacilli|o__Lactobacillales|f__Lactobacillaceae|g__Lactobacillus|s__Lactobacillus_johnsonii
k__Bacteria|p__Actinobacteria|c__Coriobacteriia|o__Coriobacteriales|f__Atopobiaceae|g__Leptogranulimonas|s__Leptog
k__Bacteria|p__Bacteroidota|c__Bacteroidia|o__Bacteroidales|f__Muribaculaceae|g__Muribaculaceae_unclassified|s__M
k__Bacteria|p__Firmicutes|c__Clostridia|o__Eubacteriales|f__Oscillospiraceae|g__Neglectibacter|s__Neglectibacter_sp_X
k__Bacteria|p__Firmicutes|c__Clostridia|o__Eubacteriales|f__Oscillospiraceae|g__Oscillibacter|s__Oscillibacter_SGB43496
k__Bacteria|p__Firmicutes|c__Clostridia|o__Eubacteriales|f__Oscillospiraceae|g__Oscillospiraceae_unclassified|s__Oscillo
k__Bacteria|p__Firmicutes|c__Clostridia|o__Eubacteriales|f__Oscillospiraceae|g__Oscillospiraceae_unclassified|s__Oscillo
k__Bacteria|p__Firmicutes|c__Clostridia|o__Eubacteriales|f__Oscillospiraceae|g__Oscillospiraceae_unclassified|s__Oscillo
k__Bacteria|p__Firmicutes|c__Clostridia|o__Eubacteriales|f__Oscillospiraceae|g__Oscillospiraceae_unclassified|s__Oscillo
k__Bacteria|p__Proteobacteria|c__Betaproteobacteria|o__Burkholderiales|f__Sutterellaceae|g__Parasutterella|s__Parasu
k__Bacteria|p__Firmicutes|c__Clostridia|o__Eubacteriales|f__Lachnospiraceae|g__Schaedlerella|s__Schaedlerella_arabino
k__Bacteria|p__Firmicutes|c__Erysipelotrichia|o__Erysipelotrichales|f__Turicibacteraceae|g__Turicibacter|s__Turicibacter_
k__Bacteria|p__Bacteria_unclassified|c__Bacteria_unclassified|o__Bacteria_unclassified|f__Bacteria_unclassified|g__Bacte
k__Bacteria|p__Bacteria_unclassified|c__Bacteria_unclassified|o__Bacteria_unclassified|f__Bacteria_unclassified|g__Bacte
k__Bacteria|p__Bacteria_unclassified|c__Bacteria_unclassified|o__Bacteria_unclassified|f__Bacteria_unclassified|g__Bacte
k__Bacteria|p__Bacteria_unclassified|c__Bacteria_unclassified|o__Bacteria_unclassified|f__Bacteria_unclassified|g__Bacte
k__Bacteria|p__Bacteria_unclassified|c__Bacteria_unclassified|o__Bacteria_unclassified|f__Bacteria_unclassified|g__Bacte

k__Bacteria|p__Firmicutes|c__Clostridia|o__Eubacteriales|f__Lachnospiraceae|g__Acetatifactor|s__Acetatifactor_SGB415
k__Bacteria|p__Firmicutes|c__Clostridia|o__Eubacteriales|f__Oscillospiraceae|g__Acutalibacter|s__Acutalibacter_muris
k__Bacteria|p__Verrucomicrobia|c__Verrucomicrobiae|o__Verrucomicrobiales|f__Akkermansiaceae|g__Akkermansia|s__A
k__Bacteria|p__Firmicutes|c__Clostridia|o__Eubacteriales|f__Oscillospiraceae|g__Anaerotruncus|s__Anaerotruncus_sp_1
k__Bacteria|p__Bacteria_unclassified|c__Bacteria_unclassified|o__Bacteria_unclassified|f__Bacteria_unclassified|g__Bacte
k__Bacteria|p__Bacteria_unclassified|c__Bacteria_unclassified|o__Bacteria_unclassified|f__Bacteria_unclassified|g__Bacte
k__Bacteria|p__Bacteria_unclassified|c__Bacteria_unclassified|o__Bacteria_unclassified|f__Bacteria_unclassified|g__Bacte
k__Bacteria|p__Bacteria_unclassified|c__Bacteria_unclassified|o__Bacteria_unclassified|f__Bacteria_unclassified|g__Bacte
k__Bacteria|p__Bacteroidota|c__Bacteroidia|o__Bacteroidales|f__Bacteroidaceae|g__Bacteroides|s__Bacteroides_thetaio
k__Bacteria|p__Firmicutes|c__Clostridia|o__Clostridia_unclassified|f__Clostridia_unclassified|g__Clostridia_unclassified|s_
k__Bacteria|p__Firmicutes|c__Clostridia|o__Eubacteriales|f__Clostridiaceae|g__Clostridiaceae_unclassified|s__Clostridiac
k__Bacteria|p__Firmicutes|c__Clostridia|o__Eubacteriales|f__Clostridiaceae|g__Clostridiaceae_unclassified|s__Clostridiac
k__Bacteria|p__Firmicutes|c__Clostridia|o__Eubacteriales|f__Eubacteriales_unclassified|g__Eubacteriales_unclassified|s_
k__Bacteria|p__Firmicutes|c__Clostridia|o__Eubacteriales|f__Clostridiaceae|g__Clostridium|s__Clostridium_SGB65123
k__Bacteria|p__Firmicutes|c__Erysipelotrichia|o__Erysipelotrichales|f__Erysipelotrichaceae|g__Erysipelatoclostridium|s__
k__Bacteria|p__Firmicutes|c__Clostridia|o__Eubacteriales|f__Eubacteriaceae|g__Eubacteriaceae_unclassified|s__Eubacte
k__Bacteria|p__Firmicutes|c__Clostridia|o__Eubacteriales|f__Eubacteriaceae|g__Eubacteriaceae_unclassified|s__Eubacte
k__Bacteria|p__Firmicutes|c__CFGB77306|o__OFGB77306|f__FGB77306|g__GGB20146|s__GGB20146_SGB29427
k__Bacteria|p__Firmicutes|c__Clostridia|o__Eubacteriales|f__Lachnospiraceae|g__GGB20149|s__GGB20149_SGB29430
k__Bacteria|p__Actinobacteria|c__Coriobacteriia|o__Eggerthellales|f__Eggerthellaceae|g__GGB22635|s__GGB22635_SGB
k__Bacteria|p__Firmicutes|c__Clostridia|o__Eubacteriales|f__Lachnospiraceae|g__GGB25041|s__GGB25041_SGB36960
k__Bacteria|p__Firmicutes|c__CFGB9506|o__OFGB9506|f__FGB9506|g__GGB28379|s__GGB28379_SGB40959
k__Bacteria|p__Firmicutes|c__CFGB9508|o__OFGB9508|f__FGB9508|g__GGB28382|s__GGB28382_SGB40962
k__Bacteria|p__Firmicutes|c__CFGB9512|o__OFGB9512|f__FGB9512|g__GGB28392|s__GGB28392_SGB40972
k__Bacteria|p__Firmicutes|c__CFGB2838|o__OFGB2838|f__FGB2838|g__GGB28404|s__GGB28404_SGB40986
k__Bacteria|p__Firmicutes|c__CFGB2838|o__OFGB2838|f__FGB2838|g__GGB28418|s__GGB28418_SGB41001

k__Bacteria|p__Firmicutes|c__CFGB2838|o__OFGB2838|f__FGB2838|g__GGB28422|s__GGB28422_SGB41005
k__Bacteria|p__Firmicutes|c__Clostridia|o__Eubacteriales|f__Pumilibacteraceae|g__GGB28431|s__GGB28431_SGB41014
k__Bacteria|p__Firmicutes|c__CFGB2833|o__OFGB2833|f__FGB2833|g__GGB28456|s__GGB28456_SGB41039
k__Bacteria|p__Firmicutes|c__Clostridia|o__Eubacteriales|f__Eubacteriaceae|g__GGB28782|s__GGB28782_SGB41435
k__Bacteria|p__Firmicutes|c__Clostridia|o__Eubacteriales|f__Lachnospiraceae|g__GGB28792|s__GGB28792_SGB41445
k__Bacteria|p__Firmicutes|c__Clostridia|o__Eubacteriales|f__Lachnospiraceae|g__GGB28798|s__GGB28798_SGB41451
k__Bacteria|p__Firmicutes|c__CFGB9622|o__OFGB9622|f__FGB9622|g__GGB28810|s__GGB28810_SGB41465
k__Bacteria|p__Firmicutes|c__CFGB77305|o__OFGB77305|f__FGB77305|g__GGB28828|s__GGB28828_SGB41484
k__Bacteria|p__Firmicutes|c__Clostridia|o__Eubacteriales|f__Clostridiaceae|g__GGB28851|s__GGB28851_SGB41518
k__Bacteria|p__Firmicutes|c__Clostridia|o__Eubacteriales|f__Lachnospiraceae|g__GGB28865|s__GGB28865_SGB41537
k__Bacteria|p__Firmicutes|c__Clostridia|o__Eubacteriales|f__Lachnospiraceae|g__GGB28868|s__GGB28868_SGB41542
k__Bacteria|p__Firmicutes|c__Clostridia|o__Eubacteriales|f__Lachnospiraceae|g__GGB28869|s__GGB28869_SGB41543
k__Bacteria|p__Firmicutes|c__Clostridia|o__Eubacteriales|f__Lachnospiraceae|g__GGB28875|s__GGB28875_SGB41555
k__Bacteria|p__Firmicutes|c__CFGB9633|o__OFGB9633|f__FGB9633|g__GGB28881|s__GGB28881_SGB41561
k__Bacteria|p__Bacteria_unclassified|c__Bacteria_unclassified|o__Bacteria_unclassified|f__Bacteria_unclassified|g__GGB
k__Bacteria|p__Bacteria_unclassified|c__Bacteria_unclassified|o__Bacteria_unclassified|f__Bacteria_unclassified|g__GGB
k__Bacteria|p__Bacteria_unclassified|c__Bacteria_unclassified|o__Bacteria_unclassified|f__Bacteria_unclassified|g__GGB
k__Bacteria|p__Bacteria_unclassified|c__Bacteria_unclassified|o__Bacteria_unclassified|f__Bacteria_unclassified|g__GGB
k__Bacteria|p__Bacteria_unclassified|c__Bacteria_unclassified|o__Bacteria_unclassified|f__Bacteria_unclassified|g__GGB
k__Bacteria|p__Bacteria_unclassified|c__Bacteria_unclassified|o__Bacteria_unclassified|f__Bacteria_unclassified|g__GGB
k__Bacteria|p__Firmicutes|c__Clostridia|o__Eubacteriales|f__Lachnospiraceae|g__GGB28924|s__GGB28924_SGB41621
k__Bacteria|p__Bacteria_unclassified|c__CFGB77359|o__OFGB77359|f__FGB77359|g__GGB28927|s__GGB28927_SGB416
k__Bacteria|p__Firmicutes|c__CFGB9639|o__OFGB9639|f__FGB9639|g__GGB28934|s__GGB28934_SGB41635
k__Bacteria|p__Firmicutes|c__Clostridia|o__Eubacteriales|f__Lachnospiraceae|g__GGB28949|s__GGB28949_SGB41655
k__Bacteria|p__Firmicutes|c__Clostridia|o__Eubacteriales|f__Clostridiaceae|g__GGB28951|s__GGB28951_SGB102295
k__Bacteria|p__Firmicutes|c__Clostridia|o__Eubacteriales|f__Clostridiaceae|g__GGB28951|s__GGB28951_SGB41658
k__Bacteria|p__Firmicutes|c__Clostridia|o__Eubacteriales|f__Clostridiaceae|g__GGB28954|s__GGB28954_SGB41662
k__Bacteria|p__Firmicutes|c__Clostridia|o__Eubacteriales|f__Clostridiaceae|g__GGB28960|s__GGB28960_SGB41669
k__Bacteria|p__Firmicutes|c__Clostridia|o__Eubacteriales|f__Clostridiaceae|g__GGB28964|s__GGB28964_SGB94886
k__Bacteria|p__Firmicutes|c__Clostridia|o__Eubacteriales|f__Clostridiaceae|g__GGB28967|s__GGB28967_SGB41678
k__Bacteria|p__Firmicutes|c__CFGB9656|o__OFGB9656|f__FGB9656|g__GGB28996|s__GGB28996_SGB41712
k__Bacteria|p__Firmicutes|c__CFGB9827|o__OFGB9827|f__FGB9827|g__GGB29531|s__GGB29531_SGB42317
k__Bacteria|p__Firmicutes|c__Clostridia|o__Eubacteriales|f__Eubacteriaceae|g__GGB29685|s__GGB29685_SGB42494
k__Bacteria|p__Bacteria_unclassified|c__CFGB77303|o__OFGB77303|f__FGB77303|g__GGB30141|s__GGB30141_SGB430
k__Bacteria|p__Firmicutes|c__Clostridia|o__Eubacteriales|f__Clostridiaceae|g__GGB30145|s__GGB30145_SGB43072
k__Bacteria|p__Firmicutes|c__Clostridia|o__Eubacteriales|f__Eubacteriales_unclassified|g__GGB30286|s__GGB30286_SG
k__Bacteria|p__Firmicutes|c__CFGB72709|o__OFGB72709|f__FGB72709|g__GGB30300|s__GGB30300_SGB43264
k__Bacteria|p__Firmicutes|c__Clostridia|o__Eubacteriales|f__Oscillospiraceae|g__GGB30303|s__GGB30303_SGB43268
k__Bacteria|p__Firmicutes|c__Clostridia|o__Eubacteriales|f__Oscillospiraceae|g__GGB30450|s__GGB30450_SGB43507
k__Bacteria|p__Firmicutes|c__Clostridia|o__Eubacteriales|f__Oscillospiraceae|g__GGB30453|s__GGB30453_SGB43513
k__Bacteria|p__Firmicutes|c__Clostridia|o__Eubacteriales|f__Oscillospiraceae|g__GGB30454|s__GGB30454_SGB43514
k__Bacteria|p__Firmicutes|c__Clostridia|o__Eubacteriales|f__Oscillospiraceae|g__GGB30455|s__GGB30455_SGB43519
k__Bacteria|p__Firmicutes|c__Clostridia|o__Eubacteriales|f__Oscillospiraceae|g__GGB30456|s__GGB30456_SGB43520
k__Bacteria|p__Firmicutes|c__Clostridia|o__Eubacteriales|f__Oscillospiraceae|g__GGB30457|s__GGB30457_SGB63218
k__Bacteria|p__Firmicutes|c__Clostridia|o__Eubacteriales|f__Oscillospiraceae|g__GGB30461|s__GGB30461_SGB43530
k__Bacteria|p__Firmicutes|c__Clostridia|o__Eubacteriales|f__Oscillospiraceae|g__GGB30461|s__GGB30461_SGB43533

k__Bacteria|p__Firmicutes|c__Clostridia|o__Eubacteriales|f__Oscillospiraceae|g__GGB30461|s__GGB30461_SGB63209
k__Bacteria|p__Firmicutes|c__Clostridia|o__Eubacteriales|f__Oscillospiraceae|g__GGB30463|s__GGB30463_SGB43537
k__Bacteria|p__Firmicutes|c__Clostridia|o__Eubacteriales|f__Oscillospiraceae|g__GGB30473|s__GGB30473_SGB43557
k__Bacteria|p__Firmicutes|c__Clostridia|o__Eubacteriales|f__Oscillospiraceae|g__GGB30475|s__GGB30475_SGB63182
k__Bacteria|p__Actinobacteria|c__CFGB77153|o__OFGB77153|f__FGB77153|g__GGB30861|s__GGB30861_SGB44083
k__Bacteria|p__Tenericutes|c__CFGB1791|o__OFGB1791|f__FGB1791|g__GGB31312|s__GGB31312_SGB44628
k__Bacteria|p__Firmicutes|c__CFGB10290|o__OFGB10290|f__FGB10290|g__GGB31438|s__GGB31438_SGB44768
k__Bacteria|p__Firmicutes|c__Clostridia|o__Eubacteriales|f__Oscillospiraceae|g__GGB3171|s__GGB3171_SGB4185
k__Bacteria|p__Firmicutes|c__CFGB10289|o__OFGB10289|f__FGB10289|g__GGB31762|s__GGB31762_SGB45125
k__Bacteria|p__Firmicutes|c__CFGB1765|o__OFGB1765|f__FGB1765|g__GGB31823|s__GGB31823_SGB45199
k__Bacteria|p__Firmicutes|c__CFGB1765|o__OFGB1765|f__FGB1765|g__GGB31838|s__GGB31838_SGB45216
k__Bacteria|p__Firmicutes|c__CFGB1765|o__OFGB1765|f__FGB1765|g__GGB31841|s__GGB31841_SGB65084
k__Bacteria|p__Firmicutes|c__CFGB10667|o__OFGB10667|f__FGB10667|g__GGB32371|s__GGB32371_SGB41694
k__Bacteria|p__Firmicutes|c__Clostridia|o__Eubacteriales|f__Lachnospiraceae|g__GGB3793|s__GGB3793_SGB5158
k__Bacteria|p__Firmicutes|c__Clostridia|o__Eubacteriales|f__Clostridiaceae|g__GGB42601|s__GGB42601_SGB59797
k__Bacteria|p__Firmicutes|c__Clostridia|o__Eubacteriales|f__Oscillospiraceae|g__GGB45514|s__GGB45514_SGB63186
k__Bacteria|p__Firmicutes|c__CFGB75721|o__OFGB75721|f__FGB75721|g__GGB45564|s__GGB45564_SGB63259
k__Bacteria|p__Firmicutes|c__Clostridia|o__Eubacteriales|f__Oscillospiraceae|g__GGB45624|s__GGB45624_SGB63337
k__Bacteria|p__Firmicutes|c__Clostridia|o__Eubacteriales|f__Christensenellaceae|g__GGB45656|s__GGB45656_SGB6337
k__Bacteria|p__Firmicutes|c__CFGB10299|o__OFGB10299|f__FGB10299|g__GGB47127|s__GGB47127_SGB65054
k__Bacteria|p__Firmicutes|c__Clostridia|o__Eubacteriales|f__Oscillospiraceae|g__GGB74395|s__GGB74395_SGB43523
k__Bacteria|p__Firmicutes|c__Clostridia|o__Eubacteriales|f__Oscillospiraceae|g__GGB75053|s__GGB75053_SGB43494
k__Bacteria|p__Firmicutes|c__Clostridia|o__Eubacteriales|f__Lachnospiraceae|g__GGB75109|s__GGB75109_SGB102238
k__Bacteria|p__Firmicutes|c__Clostridia|o__Eubacteriales|f__Lachnospiraceae|g__Lachnospiraceae_unclassified|s__Lachn
k__Bacteria|p__Firmicutes|c__Clostridia|o__Eubacteriales|f__Lachnospiraceae|g__Lachnospiraceae_unclassified|s__Lachn
k__Bacteria|p__Firmicutes|c__Clostridia|o__Eubacteriales|f__Lachnospiraceae|g__Lachnospiraceae_unclassified|s__Lachn
k__Bacteria|p__Firmicutes|c__Clostridia|o__Eubacteriales|f__Lachnospiraceae|g__Lachnospiraceae_unclassified|s__Lachn
k__Bacteria|p__Firmicutes|c__Clostridia|o__Eubacteriales|f__Lachnospiraceae|g__Lachnospiraceae_unclassified|s__Lachn
k__Bacteria|p__Firmicutes|c__Bacilli|o__Lactobacillales|f__Lactobacillaceae|g__Lactobacillus|s__Lactobacillus_johnsonii
k__Bacteria|p__Actinobacteria|c__Coriobacteriia|o__Coriobacteriales|f__Atopobiaceae|g__Leptogranulimonas|s__Leptog
k__Bacteria|p__Bacteroidota|c__Bacteroidia|o__Bacteroidales|f__Muribaculaceae|g__Muribaculaceae_unclassified|s__M
k__Bacteria|p__Firmicutes|c__Clostridia|o__Eubacteriales|f__Oscillospiraceae|g__Neglectibacter|s__Neglectibacter_sp_X
k__Bacteria|p__Firmicutes|c__Clostridia|o__Eubacteriales|f__Oscillospiraceae|g__Oscillibacter|s__Oscillibacter_SGB4349
k__Bacteria|p__Firmicutes|c__Clostridia|o__Eubacteriales|f__Oscillospiraceae|g__Oscillospiraceae_unclassified|s__Oscillo
k__Bacteria|p__Firmicutes|c__Clostridia|o__Eubacteriales|f__Oscillospiraceae|g__Oscillospiraceae_unclassified|s__Oscillo
k__Bacteria|p__Firmicutes|c__Clostridia|o__Eubacteriales|f__Oscillospiraceae|g__Oscillospiraceae_unclassified|s__Oscillo
k__Bacteria|p__Firmicutes|c__Clostridia|o__Eubacteriales|f__Oscillospiraceae|g__Oscillospiraceae_unclassified|s__Oscillo
k__Bacteria|p__Proteobacteria|c__Betaproteobacteria|o__Burkholderiales|f__Sutterellaceae|g__Parasutterella|s__Parasu
k__Bacteria|p__Firmicutes|c__Clostridia|o__Eubacteriales|f__Lachnospiraceae|g__Schaedlerella|s__Schaedlerella_arabino
k__Bacteria|p__Firmicutes|c__Erysipelotrichia|o__Erysipelotrichales|f__Turicibacteraceae|g__Turicibacter|s__Turicibacter_
k__Bacteria|p__Bacteria_unclassified|c__Bacteria_unclassified|o__Bacteria_unclassified|f__Bacteria_unclassified|g__Bacte
k__Bacteria|p__Bacteria_unclassified|c__Bacteria_unclassified|o__Bacteria_unclassified|f__Bacteria_unclassified|g__Bacte
k__Bacteria|p__Bacteria_unclassified|c__Bacteria_unclassified|o__Bacteria_unclassified|f__Bacteria_unclassified|g__Bacte
k__Bacteria|p__Bacteria_unclassified|c__Bacteria_unclassified|o__Bacteria_unclassified|f__Bacteria_unclassified|g__Bacte
k__Bacteria|p__Bacteria_unclassified|c__Bacteria_unclassified|o__Bacteria_unclassified|f__Bacteria_unclassified|g__Bacte
