## Supplementary material for "Non-invasive Vagal Nerve Stimulation as a Potential Treatment for Repetitive Blast Trauma": st4

| Exposure | Mediator (Species-level feature) | Outcome | p_m | p_y |
|---|---|---|---|---|
| blast | Acetatifactor_SGB41545 | acute_cyto_PC1 | 0.210350900 | 0.426337589 |
| vns | Acetatifactor_SGB41545 | acute_cyto_PC1 | 0.239098669 | 0.426337589 |
| blast | Acutalibacter_muris | acute_cyto_PC1 | 0.786984338 | 0.577795707 |
| vns | Acutalibacter_muris | acute_cyto_PC1 | 0.491902226 | 0.577795707 |
| blast | Akkermansia_muciniphila | acute_cyto_PC1 | 0.16660254 | 0.601919414 |
| vns | Akkermansia_muciniphila | acute_cyto_PC1 | 0.678899621 | 0.601919414 |
| blast | Anaerotruncus_sp_1XD42_93 | acute_cyto_PC1 | 0.000917537 | 0.917104107 |
| vns | Anaerotruncus_sp_1XD42_93 | acute_cyto_PC1 | 0.000231687 | 0.917104107 |
| blast | Bacteria_unclassified_SGB41677 | acute_cyto_PC1 | 0.714189493 | 0.976353560 |
| vns | Bacteria_unclassified_SGB41677 | acute_cyto_PC1 | 0.58777865 | 0.976353560 |
| blast | Bacteria_unclassified_SGB43539 | acute_cyto_PC1 | 0.148902499 | 0.218978051 |
| vns | Bacteria_unclassified_SGB43539 | acute_cyto_PC1 | 0.449260123 | 0.218978051 |
| blast | Bacteria_unclassified_SGB43546 | acute_cyto_PC1 | 0.112551908 | 0.605851927 |
| vns | Bacteria_unclassified_SGB43546 | acute_cyto_PC1 | 0.309908808 | 0.605851927 |
| blast | Bacteria_unclassified_SGB63211 | acute_cyto_PC1 | 0.295003775 | 0.053264016 |
| vns | Bacteria_unclassified_SGB63211 | acute_cyto_PC1 | 0.340871629 | 0.053264016 |
| blast | bacterium_0_1xD8_82 | acute_cyto_PC1 | 0.763436393 | 0.662492714 |
| vns | bacterium_0_1xD8_82 | acute_cyto_PC1 | 0.795139093 | 0.662492714 |
| blast | bacterium_1XD42_1 | acute_cyto_PC1 | 0.272519397 | 0.842884819 |
| vns | bacterium_1XD42_1 | acute_cyto_PC1 | 0.097091925 | 0.842884819 |
| blast | bacterium_1XD42_54 | acute_cyto_PC1 | 0.038105321 | 0.843690241 |
| vns | bacterium_1XD42_54 | acute_cyto_PC1 | 0.125406933 | 0.843690241 |
| blast | bacterium_1XD42_76 | acute_cyto_PC1 | 0.177410768 | 0.004423798 |
| vns | bacterium_1XD42_76 | acute_cyto_PC1 | 0.141032802 | 0.004423798 |
| blast | bacterium_1xD8_48 | acute_cyto_PC1 | 0.139347611 | 0.631298404 |
| vns | bacterium_1xD8_48 | acute_cyto_PC1 | 0.370020407 | 0.631298404 |
| blast | Bacteroides_thetaiotaomicron | acute_cyto_PC1 | 0.036627754 | 0.615935149 |
| vns | Bacteroides_thetaiotaomicron | acute_cyto_PC1 | 0.275405005 | 0.615935149 |
| blast | Berger Parker Index | acute_cyto_PC1 | 0.057475283 | 0.585016437 |
| vns | Berger Parker Index | acute_cyto_PC1 | 0.499505955 | 0.585016437 |
| blast | Clostridia_bacterium | acute_cyto_PC1 | 0.248353087 | 0.582229397 |
| vns | Clostridia_bacterium | acute_cyto_PC1 | 0.380556079 | 0.582229397 |
| blast | Clostridiaceae_bacterium | acute_cyto_PC1 | 0.162346605 | 0.662202407 |
| vns | Clostridiaceae_bacterium | acute_cyto_PC1 | 0.277071084 | 0.662202407 |
| blast | Clostridiaceae_unclassified_SGB41663 | acute_cyto_PC1 | 0.362409871 | 0.163252193 |
| vns | Clostridiaceae_unclassified_SGB41663 | acute_cyto_PC1 | 0.070917324 | 0.163252193 |
| blast | Clostridiales_bacterium | acute_cyto_PC1 | 0.073824740 | 0.557466510 |
| vns | Clostridiales_bacterium | acute_cyto_PC1 | 0.124851167 | 0.557466510 |
| blast | Clostridium_cocleatum | acute_cyto_PC1 | 0.507275085 | 0.724732489 |
| vns | Clostridium_cocleatum | acute_cyto_PC1 | 0.482732384 | 0.724732489 |
| blast | Clostridium_SGB65123 | acute_cyto_PC1 | 0.180438982 | 0.006647678 |
| vns | Clostridium_SGB65123 | acute_cyto_PC1 | 0.179389568 | 0.006647678 |
| blast | Eubacteriaceae_bacterium | acute_cyto_PC1 | 0.016604062 | 0.856163150 |
| vns | Eubacteriaceae_bacterium | acute_cyto_PC1 | 0.969157578 | 0.856163150 |
| blast | Eubacteriaceae_unclassified_SGB94922 | acute_cyto_PC1 | 0.174224276 | 0.913635804 |
| vns | Eubacteriaceae_unclassified_SGB94922 | acute_cyto_PC1 | 0.346181024 | 0.913635804 |

| | | | | |
|---|---|---|---|---|
| blast | GGB20146_SGB29427 | acute_cyto_PC1 | 0.093844467 | 0.763691992 |
| vns | GGB20146_SGB29427 | acute_cyto_PC1 | 0.051218919 | 0.763691992 |
| blast | GGB20149_SGB29430 | acute_cyto_PC1 | 0.413225925 | 0.660065724 |
| vns | GGB20149_SGB29430 | acute_cyto_PC1 | 0.180434838 | 0.660065724 |
| blast | GGB22635_SGB63107 | acute_cyto_PC1 | 0.114660903 | 0.716725038 |
| vns | GGB22635_SGB63107 | acute_cyto_PC1 | 0.535639339 | 0.716725038 |
| blast | GGB25041_SGB36960 | acute_cyto_PC1 | 0.167595117 | 0.828254777 |
| vns | GGB25041_SGB36960 | acute_cyto_PC1 | 0.206256260 | 0.828254777 |
| blast | GGB28379_SGB40959 | acute_cyto_PC1 | 0.029241836 | 0.041086906 |
| vns | GGB28379_SGB40959 | acute_cyto_PC1 | 0.032794656 | 0.041086906 |
| blast | GGB28382_SGB40962 | acute_cyto_PC1 | 0.072513022 | 0.130664362 |
| vns | GGB28382_SGB40962 | acute_cyto_PC1 | 0.370430736 | 0.130664362 |
| blast | GGB28392_SGB40972 | acute_cyto_PC1 | 0.017183669 | 0.126470180 |
| vns | GGB28392_SGB40972 | acute_cyto_PC1 | 0.016592400 | 0.126470180 |
| blast | GGB28404_SGB40986 | acute_cyto_PC1 | 0.157767928 | 0.737359301 |
| vns | GGB28404_SGB40986 | acute_cyto_PC1 | 0.010090214 | 0.737359301 |
| blast | GGB28418_SGB41001 | acute_cyto_PC1 | 0.996265407 | 0.972475228 |
| vns | GGB28418_SGB41001 | acute_cyto_PC1 | 0.959497945 | 0.972475228 |
| blast | GGB28422_SGB41005 | acute_cyto_PC1 | 0.298094755 | 0.680377399 |
| vns | GGB28422_SGB41005 | acute_cyto_PC1 | 0.416676405 | 0.680377399 |
| blast | GGB28431_SGB41014 | acute_cyto_PC1 | 0.015377584 | 0.872708849 |
| vns | GGB28431_SGB41014 | acute_cyto_PC1 | 0.016801247 | 0.872708849 |
| blast | GGB28456_SGB41039 | acute_cyto_PC1 | 0.508621549 | 0.257888936 |
| vns | GGB28456_SGB41039 | acute_cyto_PC1 | 0.567597000 | 0.257888936 |
| blast | GGB28782_SGB41435 | acute_cyto_PC1 | 0.315491599 | 0.448148480 |
| vns | GGB28782_SGB41435 | acute_cyto_PC1 | 0.359169919 | 0.448148480 |
| blast | GGB28792_SGB41445 | acute_cyto_PC1 | 0.123770726 | 0.733698658 |
| vns | GGB28792_SGB41445 | acute_cyto_PC1 | 0.970858107 | 0.733698658 |
| blast | GGB28798_SGB41451 | acute_cyto_PC1 | 0.931837305 | 0.642653259 |
| vns | GGB28798_SGB41451 | acute_cyto_PC1 | 0.471552545 | 0.642653259 |
| blast | GGB28810_SGB41465 | acute_cyto_PC1 | 0.919521792 | 0.960691488 |
| vns | GGB28810_SGB41465 | acute_cyto_PC1 | 0.915105920 | 0.960691488 |
| blast | GGB28828_SGB41484 | acute_cyto_PC1 | 0.036453176 | 0.134989453 |
| vns | GGB28828_SGB41484 | acute_cyto_PC1 | 0.029615316 | 0.134989453 |
| blast | GGB28851_SGB41518 | acute_cyto_PC1 | 0.983245267 | 0.362512815 |
| vns | GGB28851_SGB41518 | acute_cyto_PC1 | 0.548244701 | 0.362512815 |
| blast | GGB28865_SGB41537 | acute_cyto_PC1 | 0.051838259 | 0.050385604 |
| vns | GGB28865_SGB41537 | acute_cyto_PC1 | 0.043832016 | 0.050385604 |
| blast | GGB28868_SGB41542 | acute_cyto_PC1 | 0.292423183 | 0.735404707 |
| vns | GGB28868_SGB41542 | acute_cyto_PC1 | 0.254323647 | 0.735404707 |
| blast | GGB28869_SGB41543 | acute_cyto_PC1 | 0.662073077 | 0.312370945 |
| vns | GGB28869_SGB41543 | acute_cyto_PC1 | 0.806606148 | 0.312370945 |
| blast | GGB28875_SGB41555 | acute_cyto_PC1 | 0.776644062 | 0.959080274 |
| vns | GGB28875_SGB41555 | acute_cyto_PC1 | 0.634070741 | 0.959080274 |
| blast | GGB28881_SGB41561 | acute_cyto_PC1 | 0.011894548 | 0.526788453 |
| vns | GGB28881_SGB41561 | acute_cyto_PC1 | 0.001988037 | 0.526788453 |

| | | | | |
|---|---|---|---|---|
| blast | GGB28892_SGB41573 | acute_cyto_PC1 | 0.715153028 | 0.303956874 |
| vns | GGB28892_SGB41573 | acute_cyto_PC1 | 0.784709483 | 0.303956874 |
| blast | GGB28893_SGB41574 | acute_cyto_PC1 | 0.739221992 | 0.038137092 |
| vns | GGB28893_SGB41574 | acute_cyto_PC1 | 0.264245168 | 0.038137092 |
| blast | GGB28898_SGB41580 | acute_cyto_PC1 | 0.601924700 | 0.669756145 |
| vns | GGB28898_SGB41580 | acute_cyto_PC1 | 0.591718713 | 0.669756145 |
| blast | GGB28901_SGB41592 | acute_cyto_PC1 | 0.072063586 | 0.152650290 |
| vns | GGB28901_SGB41592 | acute_cyto_PC1 | 0.152665868 | 0.152650290 |
| blast | GGB28904_SGB41597 | acute_cyto_PC1 | 0.225752520 | 0.207680230 |
| vns | GGB28904_SGB41597 | acute_cyto_PC1 | 0.085658495 | 0.207680230 |
| blast | GGB28909_SGB41602 | acute_cyto_PC1 | 0.218283007 | 0.026531122 |
| vns | GGB28909_SGB41602 | acute_cyto_PC1 | 0.145902665 | 0.026531122 |
| blast | GGB28924_SGB41621 | acute_cyto_PC1 | 0.611578184 | 0.052540796 |
| vns | GGB28924_SGB41621 | acute_cyto_PC1 | 0.456981189 | 0.052540796 |
| blast | GGB28927_SGB41625 | acute_cyto_PC1 | 0.001954483 | 0.366201366 |
| vns | GGB28927_SGB41625 | acute_cyto_PC1 | 0.000219425 | 0.366201366 |
| blast | GGB28934_SGB41635 | acute_cyto_PC1 | 0.216498853 | 0.554318985 |
| vns | GGB28934_SGB41635 | acute_cyto_PC1 | 0.224102539 | 0.554318985 |
| blast | GGB28949_SGB41655 | acute_cyto_PC1 | 0.159598366 | 0.316394060 |
| vns | GGB28949_SGB41655 | acute_cyto_PC1 | 0.744810754 | 0.316394060 |
| blast | GGB28951_SGB102295 | acute_cyto_PC1 | 0.718725508 | 0.560708780 |
| vns | GGB28951_SGB102295 | acute_cyto_PC1 | 0.298970783 | 0.560708780 |
| blast | GGB28951_SGB41658 | acute_cyto_PC1 | 0.752627843 | 0.252343162 |
| vns | GGB28951_SGB41658 | acute_cyto_PC1 | 0.024801176 | 0.252343162 |
| blast | GGB28954_SGB41662 | acute_cyto_PC1 | 0.794939032 | 0.791398899 |
| vns | GGB28954_SGB41662 | acute_cyto_PC1 | 0.841186745 | 0.791398899 |
| blast | GGB28960_SGB41669 | acute_cyto_PC1 | 0.536638861 | 0.246718454 |
| vns | GGB28960_SGB41669 | acute_cyto_PC1 | 0.043638773 | 0.246718454 |
| blast | GGB28964_SGB94886 | acute_cyto_PC1 | 0.207213028 | 0.806789981 |
| vns | GGB28964_SGB94886 | acute_cyto_PC1 | 0.400370147 | 0.806789981 |
| blast | GGB28967_SGB41678 | acute_cyto_PC1 | 0.848205066 | 0.636882447 |
| vns | GGB28967_SGB41678 | acute_cyto_PC1 | 0.820661965 | 0.636882447 |
| blast | GGB28996_SGB41712 | acute_cyto_PC1 | 0.429884728 | 0.747558755 |
| vns | GGB28996_SGB41712 | acute_cyto_PC1 | 0.377449394 | 0.747558755 |
| blast | GGB29531_SGB42317 | acute_cyto_PC1 | 0.279593444 | 0.691936238 |
| vns | GGB29531_SGB42317 | acute_cyto_PC1 | 0.999007775 | 0.691936238 |
| blast | GGB29685_SGB42494 | acute_cyto_PC1 | 0.004596881 | 0.833350762 |
| vns | GGB29685_SGB42494 | acute_cyto_PC1 | 0.748416768 | 0.833350762 |
| blast | GGB30141_SGB43066 | acute_cyto_PC1 | 0.614028140 | 0.448906709 |
| vns | GGB30141_SGB43066 | acute_cyto_PC1 | 0.664026053 | 0.448906709 |
| blast | GGB30145_SGB43072 | acute_cyto_PC1 | 0.294924035 | 0.135611576 |
| vns | GGB30145_SGB43072 | acute_cyto_PC1 | 0.367951830 | 0.135611576 |
| blast | GGB30286_SGB43248 | acute_cyto_PC1 | 0.766757581 | 0.444639598 |
| vns | GGB30286_SGB43248 | acute_cyto_PC1 | 0.750425732 | 0.444639598 |
| blast | GGB30300_SGB43264 | acute_cyto_PC1 | 0.213846948 | 0.217988041 |
| vns | GGB30300_SGB43264 | acute_cyto_PC1 | 0.213511417 | 0.217988041 |

| | | | | |
|---|---|---|---|---|
| blast | GGB30303_SGB43268 | acute_cyto_PC1 | 0.7955727513 | 0.4072031284 |
| vns | GGB30303_SGB43268 | acute_cyto_PC1 | 0.6600999239 | 0.4072031284 |
| blast | GGB30450_SGB43507 | acute_cyto_PC1 | 0.0271094004 | 0.4339330774 |
| vns | GGB30450_SGB43507 | acute_cyto_PC1 | 0.1865733679 | 0.4339330774 |
| blast | GGB30453_SGB43513 | acute_cyto_PC1 | 0.3157254511 | 0.8964085344 |
| vns | GGB30453_SGB43513 | acute_cyto_PC1 | 0.3106381377 | 0.8964085344 |
| blast | GGB30454_SGB43514 | acute_cyto_PC1 | 0.6428886468 | 0.892398402 |
| vns | GGB30454_SGB43514 | acute_cyto_PC1 | 0.6569300273 | 0.892398402 |
| blast | GGB30455_SGB43519 | acute_cyto_PC1 | 0.6637984269 | 0.6287539192 |
| vns | GGB30455_SGB43519 | acute_cyto_PC1 | 0.1044958106 | 0.6287539192 |
| blast | GGB30456_SGB43520 | acute_cyto_PC1 | 0.1111969489 | 0.400916417 |
| vns | GGB30456_SGB43520 | acute_cyto_PC1 | 0.1589216618 | 0.400916417 |
| blast | GGB30457_SGB63218 | acute_cyto_PC1 | 0.8821640501 | 0.6559421165 |
| vns | GGB30457_SGB63218 | acute_cyto_PC1 | 0.8907570369 | 0.6559421165 |
| blast | GGB30461_SGB43530 | acute_cyto_PC1 | 0.6024683039 | 0.5864110497 |
| vns | GGB30461_SGB43530 | acute_cyto_PC1 | 0.1374012053 | 0.5864110497 |
| blast | GGB30461_SGB43533 | acute_cyto_PC1 | 0.8107674231 | 0.849446904 |
| vns | GGB30461_SGB43533 | acute_cyto_PC1 | 0.7186938597 | 0.849446904 |
| blast | GGB30461_SGB63209 | acute_cyto_PC1 | 0.0059141187 | 0.268399194 |
| vns | GGB30461_SGB63209 | acute_cyto_PC1 | 0.0751784248 | 0.268399194 |
| blast | GGB30463_SGB43537 | acute_cyto_PC1 | 0.0407681106 | 0.9824531902 |
| vns | GGB30463_SGB43537 | acute_cyto_PC1 | 0.9919601408 | 0.9824531902 |
| blast | GGB30473_SGB43557 | acute_cyto_PC1 | 0.3284915799 | 0.9526101765 |
| vns | GGB30473_SGB43557 | acute_cyto_PC1 | 0.189853765 | 0.9526101765 |
| blast | GGB30475_SGB63182 | acute_cyto_PC1 | 0.811833041 | 0.4916750511 |
| vns | GGB30475_SGB63182 | acute_cyto_PC1 | 0.7831290245 | 0.4916750511 |
| blast | GGB30861_SGB44083 | acute_cyto_PC1 | 0.7194016475 | 0.0822384976 |
| vns | GGB30861_SGB44083 | acute_cyto_PC1 | 0.9810455107 | 0.0822384976 |
| blast | GGB31312_SGB44628 | acute_cyto_PC1 | 0.8310391405 | 0.6021129872 |
| vns | GGB31312_SGB44628 | acute_cyto_PC1 | 0.8334570383 | 0.6021129872 |
| blast | GGB31438_SGB44768 | acute_cyto_PC1 | 0.4812139003 | 0.0023203585 |
| vns | GGB31438_SGB44768 | acute_cyto_PC1 | 0.4836947579 | 0.0023203585 |
| blast | GGB3171_SGB4185 | acute_cyto_PC1 | 0.0663422469 | 0.1370774432 |
| vns | GGB3171_SGB4185 | acute_cyto_PC1 | 0.0578507944 | 0.1370774432 |
| blast | GGB31762_SGB45125 | acute_cyto_PC1 | 0.4428890562 | 0.7119831078 |
| vns | GGB31762_SGB45125 | acute_cyto_PC1 | 0.437364142 | 0.7119831078 |
| blast | GGB31823_SGB45199 | acute_cyto_PC1 | 0.6301752034 | 0.668856506 |
| vns | GGB31823_SGB45199 | acute_cyto_PC1 | 0.5372407539 | 0.668856506 |
| blast | GGB31838_SGB45216 | acute_cyto_PC1 | 0.6844031648 | 0.9028226785 |
| vns | GGB31838_SGB45216 | acute_cyto_PC1 | 0.997715556 | 0.9028226785 |
| blast | GGB31841_SGB65084 | acute_cyto_PC1 | 0.3717374543 | 0.0588721808 |
| vns | GGB31841_SGB65084 | acute_cyto_PC1 | 0.4439519347 | 0.0588721808 |
| blast | GGB32371_SGB41694 | acute_cyto_PC1 | 0.2420180555 | 0.8190056127 |
| vns | GGB32371_SGB41694 | acute_cyto_PC1 | 0.4001595745 | 0.8190056127 |
| blast | GGB3793_SGB5158 | acute_cyto_PC1 | 0.884993491 | 0.6358581838 |
| vns | GGB3793_SGB5158 | acute_cyto_PC1 | 0.8472397096 | 0.6358581838 |

| | | | | |
|---|---|---|---|---|
| blast | GGB42601_SGB59797 | acute_cyto_PC1 | 0.4018920133 | 0.3852324177 |
| vns | GGB42601_SGB59797 | acute_cyto_PC1 | 0.6275665954 | 0.3852324177 |
| blast | GGB45514_SGB63186 | acute_cyto_PC1 | 0.9553647655 | 0.6279953191 |
| vns | GGB45514_SGB63186 | acute_cyto_PC1 | 0.6215278827 | 0.6279953191 |
| blast | GGB45564_SGB63259 | acute_cyto_PC1 | 0.4152727846 | 0.9450518329 |
| vns | GGB45564_SGB63259 | acute_cyto_PC1 | 0.3808786963 | 0.9450518329 |
| blast | GGB45624_SGB63337 | acute_cyto_PC1 | 0.6203892728 | 0.1106008108 |
| vns | GGB45624_SGB63337 | acute_cyto_PC1 | 0.3851987949 | 0.1106008108 |
| blast | GGB45656_SGB63370 | acute_cyto_PC1 | 0.4047054831 | 0.3478258885 |
| vns | GGB45656_SGB63370 | acute_cyto_PC1 | 0.1636435534 | 0.3478258885 |
| blast | GGB47127_SGB65054 | acute_cyto_PC1 | 0.4564936795 | 0.5878047965 |
| vns | GGB47127_SGB65054 | acute_cyto_PC1 | 0.82583129 | 0.5878047965 |
| blast | GGB74395_SGB43523 | acute_cyto_PC1 | 0.8024104382 | 0.0732537780 |
| vns | GGB74395_SGB43523 | acute_cyto_PC1 | 0.835585633 | 0.0732537780 |
| blast | GGB75053_SGB43494 | acute_cyto_PC1 | 0.1773274016 | 0.3452327395 |
| vns | GGB75053_SGB43494 | acute_cyto_PC1 | 0.0504104434 | 0.3452327395 |
| blast | GGB75109_SGB102238 | acute_cyto_PC1 | 0.5997929865 | 0.1239070982 |
| vns | GGB75109_SGB102238 | acute_cyto_PC1 | 0.5108644817 | 0.1239070982 |
| blast | Lachnospiraceae_bacterium | acute_cyto_PC1 | 0.0239259038 | 0.7862705667 |
| vns | Lachnospiraceae_bacterium | acute_cyto_PC1 | 0.9174922655 | 0.7862705667 |
| blast | Lachnospiraceae_bacterium_A2 | acute_cyto_PC1 | 0.7241878075 | 0.6342207918 |
| vns | Lachnospiraceae_bacterium_A2 | acute_cyto_PC1 | 0.0838607394 | 0.6342207918 |
| blast | Lachnospiraceae_unclassified_SGB41414 | acute_cyto_PC1 | 0.7190795008 | 0.5694384296 |
| vns | Lachnospiraceae_unclassified_SGB41414 | acute_cyto_PC1 | 0.5321281982 | 0.5694384296 |
| blast | Lachnospiraceae_unclassified_SGB41424 | acute_cyto_PC1 | 0.2169304138 | 0.0281989181 |
| vns | Lachnospiraceae_unclassified_SGB41424 | acute_cyto_PC1 | 0.0220188292 | 0.0281989181 |
| blast | Lachnospiraceae_unclassified_SGB94868 | acute_cyto_PC1 | 0.2076335048 | 0.9780750794 |
| vns | Lachnospiraceae_unclassified_SGB94868 | acute_cyto_PC1 | 0.7545629335 | 0.9780750794 |
| blast | Lactobacillus_johnsonii | acute_cyto_PC1 | 0.0027483659 | 0.5658910214 |
| vns | Lactobacillus_johnsonii | acute_cyto_PC1 | 0.8322162424 | 0.5658910214 |
| blast | Leptogranulimonas_caecicola | acute_cyto_PC1 | 0.8745779805 | 0.1426703397 |
| vns | Leptogranulimonas_caecicola | acute_cyto_PC1 | 0.0144750878 | 0.1426703397 |
| blast | Muribaculaceae_bacterium | acute_cyto_PC1 | 0.0018501989 | 0.99399338 |
| vns | Muribaculaceae_bacterium | acute_cyto_PC1 | 0.8835325068 | 0.99399338 |
| blast | Neglectibacter_sp_X4 | acute_cyto_PC1 | 0.3944230674 | 0.5304003737 |
| vns | Neglectibacter_sp_X4 | acute_cyto_PC1 | 0.463430059 | 0.5304003737 |
| blast | Oscillibacter_SGB43496 | acute_cyto_PC1 | 0.2205447256 | 0.0957020171 |
| vns | Oscillibacter_SGB43496 | acute_cyto_PC1 | 0.34795292 | 0.0957020171 |
| blast | Oscillospiraceae_bacterium | acute_cyto_PC1 | 0.4927836612 | 0.3784074486 |
| vns | Oscillospiraceae_bacterium | acute_cyto_PC1 | 0.251973413 | 0.3784074486 |
| blast | Oscillospiraceae_unclassified_SGB43502 | acute_cyto_PC1 | 0.0172712013 | 0.6824153502 |
| vns | Oscillospiraceae_unclassified_SGB43502 | acute_cyto_PC1 | 0.1123454974 | 0.6824153502 |
| blast | Oscillospiraceae_unclassified_SGB43505 | acute_cyto_PC1 | 0.3742999083 | 0.0088548848 |
| vns | Oscillospiraceae_unclassified_SGB43505 | acute_cyto_PC1 | 0.6182049908 | 0.0088548848 |
| blast | Oscillospiraceae_unclassified_SGB94989 | acute_cyto_PC1 | 0.9533689691 | 0.1804436782 |
| vns | Oscillospiraceae_unclassified_SGB94989 | acute_cyto_PC1 | 0.6236152007 | 0.1804436782 |

| | | | | |
|---|---|---|---|---|
| blast | Parasutterella_excrementihominis | acute_cyto_PC1 | 0.0234193782 | 0.0860392369 |
| vns | Parasutterella_excrementihominis | acute_cyto_PC1 | 0.5041571561 | 0.0860392369 |
| blast | Richness (# observed features) | acute_cyto_PC1 | 0.0137778742 | 0.8667072268 |
| vns | Richness (# observed features) | acute_cyto_PC1 | 0.9582478381 | 0.8667072268 |
| blast | Schaedlerella_arabinosiphila | acute_cyto_PC1 | 0.114481983 | 0.741114287 |
| vns | Schaedlerella_arabinosiphila | acute_cyto_PC1 | 0.3863019645 | 0.741114287 |
| blast | Shannon Index | acute_cyto_PC1 | 0.0064730704 | 0.8389799902 |
| vns | Shannon Index | acute_cyto_PC1 | 0.8359924459 | 0.8389799902 |
| blast | Turicibacter_sp_1E2 | acute_cyto_PC1 | 0.0095941370 | 0.0333991890 |
| vns | Turicibacter_sp_1E2 | acute_cyto_PC1 | 0.183926772 | 0.0333991890 |
| blast | Acetatifactor_SGB41545 | acute_cyto_PC2 | 0.2103509000 | 0.729699651 |
| vns | Acetatifactor_SGB41545 | acute_cyto_PC2 | 0.2390986693 | 0.729699651 |
| blast | Acutalibacter_muris | acute_cyto_PC2 | 0.7869843388 | 0.5521200392 |
| vns | Acutalibacter_muris | acute_cyto_PC2 | 0.4919022263 | 0.5521200392 |
| blast | Akkermansia_muciniphila | acute_cyto_PC2 | 0.16660254 | 0.9187492969 |
| vns | Akkermansia_muciniphila | acute_cyto_PC2 | 0.6788996217 | 0.9187492969 |
| blast | Anaerotruncus_sp_1XD42_93 | acute_cyto_PC2 | 0.0009175375 | 0.5877928823 |
| vns | Anaerotruncus_sp_1XD42_93 | acute_cyto_PC2 | 0.0002316873 | 0.5877928823 |
| blast | Bacteria_unclassified_SGB41677 | acute_cyto_PC2 | 0.7141894931 | 0.5019284422 |
| vns | Bacteria_unclassified_SGB41677 | acute_cyto_PC2 | 0.58777865 | 0.5019284422 |
| blast | Bacteria_unclassified_SGB43539 | acute_cyto_PC2 | 0.1489024993 | 0.2968074174 |
| vns | Bacteria_unclassified_SGB43539 | acute_cyto_PC2 | 0.4492601232 | 0.2968074174 |
| blast | Bacteria_unclassified_SGB43546 | acute_cyto_PC2 | 0.1125519082 | 0.6642529128 |
| vns | Bacteria_unclassified_SGB43546 | acute_cyto_PC2 | 0.3099088089 | 0.6642529128 |
| blast | Bacteria_unclassified_SGB63211 | acute_cyto_PC2 | 0.2950037757 | 0.3566723803 |
| vns | Bacteria_unclassified_SGB63211 | acute_cyto_PC2 | 0.3408716294 | 0.3566723803 |
| blast | bacterium_0_1xD8_82 | acute_cyto_PC2 | 0.763436393 | 0.1510330050 |
| vns | bacterium_0_1xD8_82 | acute_cyto_PC2 | 0.7951390939 | 0.1510330050 |
| blast | bacterium_1XD42_1 | acute_cyto_PC2 | 0.2725193977 | 0.7872285157 |
| vns | bacterium_1XD42_1 | acute_cyto_PC2 | 0.0970919255 | 0.7872285157 |
| blast | bacterium_1XD42_54 | acute_cyto_PC2 | 0.0381053210 | 0.6539919670 |
| vns | bacterium_1XD42_54 | acute_cyto_PC2 | 0.1254069334 | 0.6539919670 |
| blast | bacterium_1XD42_76 | acute_cyto_PC2 | 0.1774107688 | 0.0594618967 |
| vns | bacterium_1XD42_76 | acute_cyto_PC2 | 0.141032802 | 0.0594618967 |
| blast | bacterium_1xD8_48 | acute_cyto_PC2 | 0.1393476118 | 0.8810913739 |
| vns | bacterium_1xD8_48 | acute_cyto_PC2 | 0.3700204077 | 0.8810913739 |
| blast | Bacteroides_thetaiotaomicron | acute_cyto_PC2 | 0.0366277544 | 0.9686371082 |
| vns | Bacteroides_thetaiotaomicron | acute_cyto_PC2 | 0.2754050051 | 0.9686371082 |
| blast | Berger Parker Index | acute_cyto_PC2 | 0.0574752835 | 0.7096601025 |
| vns | Berger Parker Index | acute_cyto_PC2 | 0.4995059559 | 0.7096601025 |
| blast | Clostridia_bacterium | acute_cyto_PC2 | 0.2483530870 | 0.573241047 |
| vns | Clostridia_bacterium | acute_cyto_PC2 | 0.3805560795 | 0.573241047 |
| blast | Clostridiaceae_bacterium | acute_cyto_PC2 | 0.1623466057 | 0.6060108602 |
| vns | Clostridiaceae_bacterium | acute_cyto_PC2 | 0.2770710842 | 0.6060108602 |
| blast | Clostridiaceae_unclassified_SGB41663 | acute_cyto_PC2 | 0.3624098717 | 0.0931836628 |
| vns | Clostridiaceae_unclassified_SGB41663 | acute_cyto_PC2 | 0.0709173242 | 0.0931836628 |

| | | | | |
|---|---|---|---|---|
| blast | Clostridiales_bacterium | acute_cyto_PC2 | 0.073824740 | 0.770821847 |
| vns | Clostridiales_bacterium | acute_cyto_PC2 | 0.124851167 | 0.770821847 |
| blast | Clostridium_cocleatum | acute_cyto_PC2 | 0.507275085 | 0.814771518 |
| vns | Clostridium_cocleatum | acute_cyto_PC2 | 0.482732384 | 0.814771518 |
| blast | Clostridium_SGB65123 | acute_cyto_PC2 | 0.180438982 | 0.003784567 |
| vns | Clostridium_SGB65123 | acute_cyto_PC2 | 0.179389568 | 0.003784567 |
| blast | Eubacteriaceae_bacterium | acute_cyto_PC2 | 0.016604062 | 0.933746358 |
| vns | Eubacteriaceae_bacterium | acute_cyto_PC2 | 0.969157578 | 0.933746358 |
| blast | Eubacteriaceae_unclassified_SGB94922 | acute_cyto_PC2 | 0.174224276 | 0.359459228 |
| vns | Eubacteriaceae_unclassified_SGB94922 | acute_cyto_PC2 | 0.346181024 | 0.359459228 |
| blast | GGB20146_SGB29427 | acute_cyto_PC2 | 0.093844467 | 0.219265791 |
| vns | GGB20146_SGB29427 | acute_cyto_PC2 | 0.051218919 | 0.219265791 |
| blast | GGB20149_SGB29430 | acute_cyto_PC2 | 0.413225925 | 0.231439082 |
| vns | GGB20149_SGB29430 | acute_cyto_PC2 | 0.180434838 | 0.231439082 |
| blast | GGB22635_SGB63107 | acute_cyto_PC2 | 0.114660903 | 0.293509468 |
| vns | GGB22635_SGB63107 | acute_cyto_PC2 | 0.535639339 | 0.293509468 |
| blast | GGB25041_SGB36960 | acute_cyto_PC2 | 0.167595117 | 0.469320377 |
| vns | GGB25041_SGB36960 | acute_cyto_PC2 | 0.206256260 | 0.469320377 |
| blast | GGB28379_SGB40959 | acute_cyto_PC2 | 0.029241836 | 0.166832827 |
| vns | GGB28379_SGB40959 | acute_cyto_PC2 | 0.032794656 | 0.166832827 |
| blast | GGB28382_SGB40962 | acute_cyto_PC2 | 0.072513022 | 0.738052827 |
| vns | GGB28382_SGB40962 | acute_cyto_PC2 | 0.370430736 | 0.738052827 |
| blast | GGB28392_SGB40972 | acute_cyto_PC2 | 0.017183669 | 0.297965903 |
| vns | GGB28392_SGB40972 | acute_cyto_PC2 | 0.016592400 | 0.297965903 |
| blast | GGB28404_SGB40986 | acute_cyto_PC2 | 0.157767928 | 0.748603758 |
| vns | GGB28404_SGB40986 | acute_cyto_PC2 | 0.010090214 | 0.748603758 |
| blast | GGB28418_SGB41001 | acute_cyto_PC2 | 0.996265407 | 0.494691407 |
| vns | GGB28418_SGB41001 | acute_cyto_PC2 | 0.959497945 | 0.494691407 |
| blast | GGB28422_SGB41005 | acute_cyto_PC2 | 0.298094755 | 0.633460553 |
| vns | GGB28422_SGB41005 | acute_cyto_PC2 | 0.416676405 | 0.633460553 |
| blast | GGB28431_SGB41014 | acute_cyto_PC2 | 0.015377584 | 0.700581804 |
| vns | GGB28431_SGB41014 | acute_cyto_PC2 | 0.016801247 | 0.700581804 |
| blast | GGB28456_SGB41039 | acute_cyto_PC2 | 0.508621549 | 0.000768124 |
| vns | GGB28456_SGB41039 | acute_cyto_PC2 | 0.567597000 | 0.000768124 |
| blast | GGB28782_SGB41435 | acute_cyto_PC2 | 0.315491599 | 0.569221314 |
| vns | GGB28782_SGB41435 | acute_cyto_PC2 | 0.359169919 | 0.569221314 |
| blast | GGB28792_SGB41445 | acute_cyto_PC2 | 0.123770726 | 0.916920383 |
| vns | GGB28792_SGB41445 | acute_cyto_PC2 | 0.970858107 | 0.916920383 |
| blast | GGB28798_SGB41451 | acute_cyto_PC2 | 0.931837305 | 0.589642818 |
| vns | GGB28798_SGB41451 | acute_cyto_PC2 | 0.471552545 | 0.589642818 |
| blast | GGB28810_SGB41465 | acute_cyto_PC2 | 0.919521792 | 0.901398700 |
| vns | GGB28810_SGB41465 | acute_cyto_PC2 | 0.915105920 | 0.901398700 |
| blast | GGB28828_SGB41484 | acute_cyto_PC2 | 0.036453176 | 0.180185318 |
| vns | GGB28828_SGB41484 | acute_cyto_PC2 | 0.029615316 | 0.180185318 |
| blast | GGB28851_SGB41518 | acute_cyto_PC2 | 0.983245267 | 0.765176036 |
| vns | GGB28851_SGB41518 | acute_cyto_PC2 | 0.548244701 | 0.765176036 |

| | | | | | |
|---|---|---|---|---|---|
| blast | GGB28865_SGB41537 | | acute_cyto_PC2 | 0.051838259 | 0.038211931 |
| vns | GGB28865_SGB41537 | | acute_cyto_PC2 | 0.043832016 | 0.038211931 |
| blast | GGB28868_SGB41542 | | acute_cyto_PC2 | 0.292423183 | 0.190009262 |
| vns | GGB28868_SGB41542 | | acute_cyto_PC2 | 0.254323647 | 0.190009262 |
| blast | GGB28869_SGB41543 | | acute_cyto_PC2 | 0.662073077 | 0.458718419 |
| vns | GGB28869_SGB41543 | | acute_cyto_PC2 | 0.806606148 | 0.458718419 |
| blast | GGB28875_SGB41555 | | acute_cyto_PC2 | 0.776644062 | 0.565998046 |
| vns | GGB28875_SGB41555 | | acute_cyto_PC2 | 0.634070741 | 0.565998046 |
| blast | GGB28881_SGB41561 | | acute_cyto_PC2 | 0.011894548 | 0.587914970 |
| vns | GGB28881_SGB41561 | | acute_cyto_PC2 | 0.001988037 | 0.587914970 |
| blast | GGB28892_SGB41573 | | acute_cyto_PC2 | 0.715153028 | 0.271718323 |
| vns | GGB28892_SGB41573 | | acute_cyto_PC2 | 0.784709483 | 0.271718323 |
| blast | GGB28893_SGB41574 | | acute_cyto_PC2 | 0.739221992 | 0.007413952 |
| vns | GGB28893_SGB41574 | | acute_cyto_PC2 | 0.264245168 | 0.007413952 |
| blast | GGB28898_SGB41580 | | acute_cyto_PC2 | 0.601924700 | 0.618985494 |
| vns | GGB28898_SGB41580 | | acute_cyto_PC2 | 0.591718713 | 0.618985494 |
| blast | GGB28901_SGB41592 | | acute_cyto_PC2 | 0.072063586 | 0.564391499 |
| vns | GGB28901_SGB41592 | | acute_cyto_PC2 | 0.152665868 | 0.564391499 |
| blast | GGB28904_SGB41597 | | acute_cyto_PC2 | 0.225752520 | 0.104465788 |
| vns | GGB28904_SGB41597 | | acute_cyto_PC2 | 0.085658495 | 0.104465788 |
| blast | GGB28909_SGB41602 | | acute_cyto_PC2 | 0.218283007 | 0.266569842 |
| vns | GGB28909_SGB41602 | | acute_cyto_PC2 | 0.145902665 | 0.266569842 |
| blast | GGB28924_SGB41621 | | acute_cyto_PC2 | 0.611578184 | 0.056545333 |
| vns | GGB28924_SGB41621 | | acute_cyto_PC2 | 0.456981189 | 0.056545333 |
| blast | GGB28927_SGB41625 | | acute_cyto_PC2 | 0.001954483 | 0.723266267 |
| vns | GGB28927_SGB41625 | | acute_cyto_PC2 | 0.000219425 | 0.723266267 |
| blast | GGB28934_SGB41635 | | acute_cyto_PC2 | 0.216498853 | 0.828068791 |
| vns | GGB28934_SGB41635 | | acute_cyto_PC2 | 0.224102539 | 0.828068791 |
| blast | GGB28949_SGB41655 | | acute_cyto_PC2 | 0.159598366 | 0.114171785 |
| vns | GGB28949_SGB41655 | | acute_cyto_PC2 | 0.744810754 | 0.114171785 |
| blast | GGB28951_SGB102295 | | acute_cyto_PC2 | 0.718725508 | 0.699574079 |
| vns | GGB28951_SGB102295 | | acute_cyto_PC2 | 0.298970783 | 0.699574079 |
| blast | GGB28951_SGB41658 | | acute_cyto_PC2 | 0.752627843 | 0.327285934 |
| vns | GGB28951_SGB41658 | | acute_cyto_PC2 | 0.024801176 | 0.327285934 |
| blast | GGB28954_SGB41662 | | acute_cyto_PC2 | 0.794939032 | 0.243383937 |
| vns | GGB28954_SGB41662 | | acute_cyto_PC2 | 0.841186745 | 0.243383937 |
| blast | GGB28960_SGB41669 | | acute_cyto_PC2 | 0.536638861 | 0.631119347 |
| vns | GGB28960_SGB41669 | | acute_cyto_PC2 | 0.043638773 | 0.631119347 |
| blast | GGB28964_SGB94886 | | acute_cyto_PC2 | 0.207213028 | 0.370364408 |
| vns | GGB28964_SGB94886 | | acute_cyto_PC2 | 0.400370147 | 0.370364408 |
| blast | GGB28967_SGB41678 | | acute_cyto_PC2 | 0.848205066 | 0.615230740 |
| vns | GGB28967_SGB41678 | | acute_cyto_PC2 | 0.820661965 | 0.615230740 |
| blast | GGB28996_SGB41712 | | acute_cyto_PC2 | 0.429884728 | 0.915476714 |
| vns | GGB28996_SGB41712 | | acute_cyto_PC2 | 0.377449394 | 0.915476714 |
| blast | GGB29531_SGB42317 | | acute_cyto_PC2 | 0.279593444 | 0.231397371 |
| vns | GGB29531_SGB42317 | | acute_cyto_PC2 | 0.999007775 | 0.231397371 |

| | | | | |
|---|---|---|---|---|
| blast | GGB29685_SGB42494 | acute_cyto_PC2 | 0.0045968812 | 0.9632840138 |
| vns | GGB29685_SGB42494 | acute_cyto_PC2 | 0.7484167686 | 0.9632840138 |
| blast | GGB30141_SGB43066 | acute_cyto_PC2 | 0.6140281408 | 0.8347267496 |
| vns | GGB30141_SGB43066 | acute_cyto_PC2 | 0.6640260538 | 0.8347267496 |
| blast | GGB30145_SGB43072 | acute_cyto_PC2 | 0.2949240353 | 0.2189406657 |
| vns | GGB30145_SGB43072 | acute_cyto_PC2 | 0.3679518303 | 0.2189406657 |
| blast | GGB30286_SGB43248 | acute_cyto_PC2 | 0.7667575811 | 0.7924354436 |
| vns | GGB30286_SGB43248 | acute_cyto_PC2 | 0.7504257326 | 0.7924354436 |
| blast | GGB30300_SGB43264 | acute_cyto_PC2 | 0.2138469484 | 0.7013615732 |
| vns | GGB30300_SGB43264 | acute_cyto_PC2 | 0.2135114175 | 0.7013615732 |
| blast | GGB30303_SGB43268 | acute_cyto_PC2 | 0.7955727513 | 0.0020042744 |
| vns | GGB30303_SGB43268 | acute_cyto_PC2 | 0.6600999239 | 0.0020042744 |
| blast | GGB30450_SGB43507 | acute_cyto_PC2 | 0.0271094004 | 0.4028112568 |
| vns | GGB30450_SGB43507 | acute_cyto_PC2 | 0.1865733679 | 0.4028112568 |
| blast | GGB30453_SGB43513 | acute_cyto_PC2 | 0.3157254511 | 0.5354901584 |
| vns | GGB30453_SGB43513 | acute_cyto_PC2 | 0.3106381377 | 0.5354901584 |
| blast | GGB30454_SGB43514 | acute_cyto_PC2 | 0.6428886468 | 0.1039562309 |
| vns | GGB30454_SGB43514 | acute_cyto_PC2 | 0.6569300273 | 0.1039562309 |
| blast | GGB30455_SGB43519 | acute_cyto_PC2 | 0.6637984269 | 0.4009146375 |
| vns | GGB30455_SGB43519 | acute_cyto_PC2 | 0.1044958106 | 0.4009146375 |
| blast | GGB30456_SGB43520 | acute_cyto_PC2 | 0.1111969489 | 0.7430462664 |
| vns | GGB30456_SGB43520 | acute_cyto_PC2 | 0.1589216618 | 0.7430462664 |
| blast | GGB30457_SGB63218 | acute_cyto_PC2 | 0.8821640501 | 0.3569233035 |
| vns | GGB30457_SGB63218 | acute_cyto_PC2 | 0.8907570369 | 0.3569233035 |
| blast | GGB30461_SGB43530 | acute_cyto_PC2 | 0.6024683039 | 0.8893219555 |
| vns | GGB30461_SGB43530 | acute_cyto_PC2 | 0.1374012053 | 0.8893219555 |
| blast | GGB30461_SGB43533 | acute_cyto_PC2 | 0.8107674231 | 0.559599685 |
| vns | GGB30461_SGB43533 | acute_cyto_PC2 | 0.7186938592 | 0.559599685 |
| blast | GGB30461_SGB63209 | acute_cyto_PC2 | 0.0059141187 | 0.2628283018 |
| vns | GGB30461_SGB63209 | acute_cyto_PC2 | 0.0751784248 | 0.2628283018 |
| blast | GGB30463_SGB43537 | acute_cyto_PC2 | 0.0407681106 | 0.6280887525 |
| vns | GGB30463_SGB43537 | acute_cyto_PC2 | 0.9919601408 | 0.6280887525 |
| blast | GGB30473_SGB43557 | acute_cyto_PC2 | 0.3284915799 | 0.2717320899 |
| vns | GGB30473_SGB43557 | acute_cyto_PC2 | 0.189853765 | 0.2717320899 |
| blast | GGB30475_SGB63182 | acute_cyto_PC2 | 0.811833041 | 0.7789076763 |
| vns | GGB30475_SGB63182 | acute_cyto_PC2 | 0.7831290245 | 0.7789076763 |
| blast | GGB30861_SGB44083 | acute_cyto_PC2 | 0.7194016475 | 0.9961090555 |
| vns | GGB30861_SGB44083 | acute_cyto_PC2 | 0.9810455107 | 0.9961090555 |
| blast | GGB31312_SGB44628 | acute_cyto_PC2 | 0.8310391405 | 0.1816716013 |
| vns | GGB31312_SGB44628 | acute_cyto_PC2 | 0.8334570381 | 0.1816716013 |
| blast | GGB31438_SGB44768 | acute_cyto_PC2 | 0.4812139003 | 0.0703034868 |
| vns | GGB31438_SGB44768 | acute_cyto_PC2 | 0.4836947579 | 0.0703034868 |
| blast | GGB3171_SGB4185 | acute_cyto_PC2 | 0.0663422469 | 0.4535793479 |
| vns | GGB3171_SGB4185 | acute_cyto_PC2 | 0.0578507944 | 0.4535793479 |
| blast | GGB31762_SGB45125 | acute_cyto_PC2 | 0.4428890562 | 0.9324556547 |
| vns | GGB31762_SGB45125 | acute_cyto_PC2 | 0.437364142 | 0.9324556547 |

| | | | | |
|---|---|---|---|---|
| blast | GGB31823_SGB45199 | acute_cyto_PC2 | 0.6301752034 | 0.4944356180 |
| vns | GGB31823_SGB45199 | acute_cyto_PC2 | 0.5372407539 | 0.4944356180 |
| blast | GGB31838_SGB45216 | acute_cyto_PC2 | 0.6844031648 | 0.6147622668 |
| vns | GGB31838_SGB45216 | acute_cyto_PC2 | 0.997715556 | 0.6147622668 |
| blast | GGB31841_SGB65084 | acute_cyto_PC2 | 0.3717374543 | 0.4086785650 |
| vns | GGB31841_SGB65084 | acute_cyto_PC2 | 0.4439519347 | 0.4086785650 |
| blast | GGB32371_SGB41694 | acute_cyto_PC2 | 0.2420180559 | 0.6154551210 |
| vns | GGB32371_SGB41694 | acute_cyto_PC2 | 0.4001595749 | 0.6154551210 |
| blast | GGB3793_SGB5158 | acute_cyto_PC2 | 0.884993491 | 0.9013698108 |
| vns | GGB3793_SGB5158 | acute_cyto_PC2 | 0.8472397090 | 0.9013698108 |
| blast | GGB42601_SGB59797 | acute_cyto_PC2 | 0.4018920133 | 0.3366862379 |
| vns | GGB42601_SGB59797 | acute_cyto_PC2 | 0.6275665954 | 0.3366862379 |
| blast | GGB45514_SGB63186 | acute_cyto_PC2 | 0.9553647659 | 0.6461173274 |
| vns | GGB45514_SGB63186 | acute_cyto_PC2 | 0.6215278827 | 0.6461173274 |
| blast | GGB45564_SGB63259 | acute_cyto_PC2 | 0.4152727840 | 0.3022840998 |
| vns | GGB45564_SGB63259 | acute_cyto_PC2 | 0.3808786963 | 0.3022840998 |
| blast | GGB45624_SGB63337 | acute_cyto_PC2 | 0.6203892728 | 0.6751577094 |
| vns | GGB45624_SGB63337 | acute_cyto_PC2 | 0.3851987949 | 0.6751577094 |
| blast | GGB45656_SGB63370 | acute_cyto_PC2 | 0.4047054831 | 0.4927105963 |
| vns | GGB45656_SGB63370 | acute_cyto_PC2 | 0.1636435534 | 0.4927105963 |
| blast | GGB47127_SGB65054 | acute_cyto_PC2 | 0.4564936799 | 0.2861292433 |
| vns | GGB47127_SGB65054 | acute_cyto_PC2 | 0.82583129 | 0.2861292433 |
| blast | GGB74395_SGB43523 | acute_cyto_PC2 | 0.8024104382 | 0.9575983741 |
| vns | GGB74395_SGB43523 | acute_cyto_PC2 | 0.835585633 | 0.9575983741 |
| blast | GGB75053_SGB43494 | acute_cyto_PC2 | 0.1773274010 | 0.2479055584 |
| vns | GGB75053_SGB43494 | acute_cyto_PC2 | 0.0504104434 | 0.2479055584 |
| blast | GGB75109_SGB102238 | acute_cyto_PC2 | 0.5997929865 | 0.1612424570 |
| vns | GGB75109_SGB102238 | acute_cyto_PC2 | 0.5108644817 | 0.1612424570 |
| blast | Lachnospiraceae_bacterium | acute_cyto_PC2 | 0.0239259038 | 0.9162361358 |
| vns | Lachnospiraceae_bacterium | acute_cyto_PC2 | 0.9174922659 | 0.9162361358 |
| blast | Lachnospiraceae_bacterium_A2 | acute_cyto_PC2 | 0.7241878079 | 0.7553625359 |
| vns | Lachnospiraceae_bacterium_A2 | acute_cyto_PC2 | 0.0838607394 | 0.7553625359 |
| blast | Lachnospiraceae_unclassified_SGB41414 | acute_cyto_PC2 | 0.7190795008 | 0.7642742957 |
| vns | Lachnospiraceae_unclassified_SGB41414 | acute_cyto_PC2 | 0.5321281982 | 0.7642742957 |
| blast | Lachnospiraceae_unclassified_SGB41424 | acute_cyto_PC2 | 0.2169304138 | 0.0957119323 |
| vns | Lachnospiraceae_unclassified_SGB41424 | acute_cyto_PC2 | 0.0220188292 | 0.0957119323 |
| blast | Lachnospiraceae_unclassified_SGB94868 | acute_cyto_PC2 | 0.2076335048 | 0.3170400340 |
| vns | Lachnospiraceae_unclassified_SGB94868 | acute_cyto_PC2 | 0.7545629339 | 0.3170400340 |
| blast | Lactobacillus_johnsonii | acute_cyto_PC2 | 0.0027483659 | 0.8654046089 |
| vns | Lactobacillus_johnsonii | acute_cyto_PC2 | 0.8322162424 | 0.8654046089 |
| blast | Leptogranulimonas_caecicola | acute_cyto_PC2 | 0.8745779805 | 0.3078227825 |
| vns | Leptogranulimonas_caecicola | acute_cyto_PC2 | 0.0144750878 | 0.3078227825 |
| blast | Muribaculaceae_bacterium | acute_cyto_PC2 | 0.0018501989 | 0.8638485361 |
| vns | Muribaculaceae_bacterium | acute_cyto_PC2 | 0.8835325068 | 0.8638485361 |
| blast | Neglectibacter_sp_X4 | acute_cyto_PC2 | 0.3944230674 | 0.7365353302 |
| vns | Neglectibacter_sp_X4 | acute_cyto_PC2 | 0.463430059 | 0.7365353302 |

| | | | | |
|---|---|---|---|---|
| blast | Oscillibacter_SGB43496 | acute_cyto_PC2 | 0.220544725 | 0.224697122 |
| vns | Oscillibacter_SGB43496 | acute_cyto_PC2 | 0.34795292 | 0.224697122 |
| blast | Oscillospiraceae_bacterium | acute_cyto_PC2 | 0.492783661 | 0.240782724 |
| vns | Oscillospiraceae_bacterium | acute_cyto_PC2 | 0.251973413 | 0.240782724 |
| blast | Oscillospiraceae_unclassified_SGB43502 | acute_cyto_PC2 | 0.017271201 | 0.759614808 |
| vns | Oscillospiraceae_unclassified_SGB43502 | acute_cyto_PC2 | 0.112345497 | 0.759614808 |
| blast | Oscillospiraceae_unclassified_SGB43505 | acute_cyto_PC2 | 0.374299908 | 0.382033291 |
| vns | Oscillospiraceae_unclassified_SGB43505 | acute_cyto_PC2 | 0.618204990 | 0.382033291 |
| blast | Oscillospiraceae_unclassified_SGB94989 | acute_cyto_PC2 | 0.953368969 | 0.505239922 |
| vns | Oscillospiraceae_unclassified_SGB94989 | acute_cyto_PC2 | 0.623615200 | 0.505239922 |
| blast | Parasutterella_excrementihominis | acute_cyto_PC2 | 0.023419378 | 0.597705641 |
| vns | Parasutterella_excrementihominis | acute_cyto_PC2 | 0.504157156 | 0.597705641 |
| blast | Richness (# observed features) | acute_cyto_PC2 | 0.013777874 | 0.998062105 |
| vns | Richness (# observed features) | acute_cyto_PC2 | 0.958247838 | 0.998062105 |
| blast | Schaedlerella_arabinosiphila | acute_cyto_PC2 | 0.114481983 | 0.809412625 |
| vns | Schaedlerella_arabinosiphila | acute_cyto_PC2 | 0.386301964 | 0.809412625 |
| blast | Shannon Index | acute_cyto_PC2 | 0.006473070 | 0.916280734 |
| vns | Shannon Index | acute_cyto_PC2 | 0.835992445 | 0.916280734 |
| blast | Turicibacter_sp_1E2 | acute_cyto_PC2 | 0.009594137 | 0.416711257 |
| vns | Turicibacter_sp_1E2 | acute_cyto_PC2 | 0.183926772 | 0.416711257 |
| blast | Acetatifactor_SGB41545 | acute_cyto_PC3 | 0.210350900 | 0.850892715 |
| vns | Acetatifactor_SGB41545 | acute_cyto_PC3 | 0.239098669 | 0.850892715 |
| blast | Acutalibacter_muris | acute_cyto_PC3 | 0.786984338 | 0.688340526 |
| vns | Acutalibacter_muris | acute_cyto_PC3 | 0.491902226 | 0.688340526 |
| blast | Akkermansia_muciniphila | acute_cyto_PC3 | 0.16660254 | 0.237451933 |
| vns | Akkermansia_muciniphila | acute_cyto_PC3 | 0.678899621 | 0.237451933 |
| blast | Anaerotruncus_sp_1XD42_93 | acute_cyto_PC3 | 0.000917537 | 0.219716417 |
| vns | Anaerotruncus_sp_1XD42_93 | acute_cyto_PC3 | 0.000231687 | 0.219716417 |
| blast | Bacteria_unclassified_SGB41677 | acute_cyto_PC3 | 0.714189493 | 0.307443412 |
| vns | Bacteria_unclassified_SGB41677 | acute_cyto_PC3 | 0.58777865 | 0.307443412 |
| blast | Bacteria_unclassified_SGB43539 | acute_cyto_PC3 | 0.148902499 | 0.154699032 |
| vns | Bacteria_unclassified_SGB43539 | acute_cyto_PC3 | 0.449260123 | 0.154699032 |
| blast | Bacteria_unclassified_SGB43546 | acute_cyto_PC3 | 0.112551908 | 0.507331111 |
| vns | Bacteria_unclassified_SGB43546 | acute_cyto_PC3 | 0.309908808 | 0.507331111 |
| blast | Bacteria_unclassified_SGB63211 | acute_cyto_PC3 | 0.295003775 | 0.910383705 |
| vns | Bacteria_unclassified_SGB63211 | acute_cyto_PC3 | 0.340871629 | 0.910383705 |
| blast | bacterium_0_1xD8_82 | acute_cyto_PC3 | 0.763436393 | 0.79612335 |
| vns | bacterium_0_1xD8_82 | acute_cyto_PC3 | 0.795139093 | 0.79612335 |
| blast | bacterium_1XD42_1 | acute_cyto_PC3 | 0.272519397 | 0.242613719 |
| vns | bacterium_1XD42_1 | acute_cyto_PC3 | 0.097091925 | 0.242613719 |
| blast | bacterium_1XD42_54 | acute_cyto_PC3 | 0.038105321 | 0.984924233 |
| vns | bacterium_1XD42_54 | acute_cyto_PC3 | 0.125406933 | 0.984924233 |
| blast | bacterium_1XD42_76 | acute_cyto_PC3 | 0.177410768 | 0.440173026 |
| vns | bacterium_1XD42_76 | acute_cyto_PC3 | 0.141032802 | 0.440173026 |
| blast | bacterium_1xD8_48 | acute_cyto_PC3 | 0.139347611 | 0.532020100 |
| vns | bacterium_1xD8_48 | acute_cyto_PC3 | 0.370020407 | 0.532020100 |

| | | | | |
|---|---|---|---|---|
| blast | Bacteroides_thetaiotaomicron | acute_cyto_PC3 | 0.036627754 | 0.674769535 |
| vns | Bacteroides_thetaiotaomicron | acute_cyto_PC3 | 0.275405005 | 0.674769535 |
| blast | Berger Parker Index | acute_cyto_PC3 | 0.057475283 | 0.068961191 |
| vns | Berger Parker Index | acute_cyto_PC3 | 0.499505955 | 0.068961191 |
| blast | Clostridia_bacterium | acute_cyto_PC3 | 0.248353087 | 0.121524284 |
| vns | Clostridia_bacterium | acute_cyto_PC3 | 0.380556079 | 0.121524284 |
| blast | Clostridiaceae_bacterium | acute_cyto_PC3 | 0.162346605 | 0.049092982 |
| vns | Clostridiaceae_bacterium | acute_cyto_PC3 | 0.277071084 | 0.049092982 |
| blast | Clostridiaceae_unclassified_SGB41663 | acute_cyto_PC3 | 0.362409871 | 0.452082315 |
| vns | Clostridiaceae_unclassified_SGB41663 | acute_cyto_PC3 | 0.070917324 | 0.452082315 |
| blast | Clostridiales_bacterium | acute_cyto_PC3 | 0.073824740 | 0.817262856 |
| vns | Clostridiales_bacterium | acute_cyto_PC3 | 0.124851167 | 0.817262856 |
| blast | Clostridium_cocleatum | acute_cyto_PC3 | 0.507275085 | 0.813754028 |
| vns | Clostridium_cocleatum | acute_cyto_PC3 | 0.482732384 | 0.813754028 |
| blast | Clostridium_SGB65123 | acute_cyto_PC3 | 0.180438982 | 0.278714033 |
| vns | Clostridium_SGB65123 | acute_cyto_PC3 | 0.179389568 | 0.278714033 |
| blast | Eubacteriaceae_bacterium | acute_cyto_PC3 | 0.016604062 | 0.023217854 |
| vns | Eubacteriaceae_bacterium | acute_cyto_PC3 | 0.969157578 | 0.023217854 |
| blast | Eubacteriaceae_unclassified_SGB94922 | acute_cyto_PC3 | 0.174224276 | 0.143424090 |
| vns | Eubacteriaceae_unclassified_SGB94922 | acute_cyto_PC3 | 0.346181024 | 0.143424090 |
| blast | GGB20146_SGB29427 | acute_cyto_PC3 | 0.093844467 | 0.892681194 |
| vns | GGB20146_SGB29427 | acute_cyto_PC3 | 0.051218919 | 0.892681194 |
| blast | GGB20149_SGB29430 | acute_cyto_PC3 | 0.413225925 | 0.192751749 |
| vns | GGB20149_SGB29430 | acute_cyto_PC3 | 0.180434838 | 0.192751749 |
| blast | GGB22635_SGB63107 | acute_cyto_PC3 | 0.114660903 | 0.046234093 |
| vns | GGB22635_SGB63107 | acute_cyto_PC3 | 0.535639339 | 0.046234093 |
| blast | GGB25041_SGB36960 | acute_cyto_PC3 | 0.167595117 | 0.562756216 |
| vns | GGB25041_SGB36960 | acute_cyto_PC3 | 0.206256260 | 0.562756216 |
| blast | GGB28379_SGB40959 | acute_cyto_PC3 | 0.029241836 | 0.860594237 |
| vns | GGB28379_SGB40959 | acute_cyto_PC3 | 0.032794656 | 0.860594237 |
| blast | GGB28382_SGB40962 | acute_cyto_PC3 | 0.072513022 | 0.082388714 |
| vns | GGB28382_SGB40962 | acute_cyto_PC3 | 0.370430736 | 0.082388714 |
| blast | GGB28392_SGB40972 | acute_cyto_PC3 | 0.017183669 | 0.518265904 |
| vns | GGB28392_SGB40972 | acute_cyto_PC3 | 0.016592400 | 0.518265904 |
| blast | GGB28404_SGB40986 | acute_cyto_PC3 | 0.157767928 | 0.802757412 |
| vns | GGB28404_SGB40986 | acute_cyto_PC3 | 0.010090214 | 0.802757412 |
| blast | GGB28418_SGB41001 | acute_cyto_PC3 | 0.996265407 | 0.631783511 |
| vns | GGB28418_SGB41001 | acute_cyto_PC3 | 0.959497945 | 0.631783511 |
| blast | GGB28422_SGB41005 | acute_cyto_PC3 | 0.298094755 | 0.477285430 |
| vns | GGB28422_SGB41005 | acute_cyto_PC3 | 0.416676405 | 0.477285430 |
| blast | GGB28431_SGB41014 | acute_cyto_PC3 | 0.015377584 | 0.049208715 |
| vns | GGB28431_SGB41014 | acute_cyto_PC3 | 0.016801247 | 0.049208715 |
| blast | GGB28456_SGB41039 | acute_cyto_PC3 | 0.508621549 | 0.038985937 |
| vns | GGB28456_SGB41039 | acute_cyto_PC3 | 0.567597000 | 0.038985937 |
| blast | GGB28782_SGB41435 | acute_cyto_PC3 | 0.315491599 | 0.172231722 |
| vns | GGB28782_SGB41435 | acute_cyto_PC3 | 0.359169919 | 0.172231722 |

| | | | | |
|---|---|---|---|---|
| blast | GGB28792_SGB41445 | acute_cyto_PC3 | 0.123770726 | 0.236232979 |
| vns | GGB28792_SGB41445 | acute_cyto_PC3 | 0.970858107 | 0.236232979 |
| blast | GGB28798_SGB41451 | acute_cyto_PC3 | 0.931837305 | 0.796081594 |
| vns | GGB28798_SGB41451 | acute_cyto_PC3 | 0.471552545 | 0.796081594 |
| blast | GGB28810_SGB41465 | acute_cyto_PC3 | 0.919521792 | 0.674035845 |
| vns | GGB28810_SGB41465 | acute_cyto_PC3 | 0.915105920 | 0.674035845 |
| blast | GGB28828_SGB41484 | acute_cyto_PC3 | 0.036453176 | 0.941100053 |
| vns | GGB28828_SGB41484 | acute_cyto_PC3 | 0.029615316 | 0.941100053 |
| blast | GGB28851_SGB41518 | acute_cyto_PC3 | 0.983245267 | 0.213233718 |
| vns | GGB28851_SGB41518 | acute_cyto_PC3 | 0.548244701 | 0.213233718 |
| blast | GGB28865_SGB41537 | acute_cyto_PC3 | 0.051838259 | 0.457415098 |
| vns | GGB28865_SGB41537 | acute_cyto_PC3 | 0.043832016 | 0.457415098 |
| blast | GGB28868_SGB41542 | acute_cyto_PC3 | 0.292423183 | 0.172033778 |
| vns | GGB28868_SGB41542 | acute_cyto_PC3 | 0.254323647 | 0.172033778 |
| blast | GGB28869_SGB41543 | acute_cyto_PC3 | 0.662073077 | 0.252206188 |
| vns | GGB28869_SGB41543 | acute_cyto_PC3 | 0.806606148 | 0.252206188 |
| blast | GGB28875_SGB41555 | acute_cyto_PC3 | 0.776644062 | 0.936860051 |
| vns | GGB28875_SGB41555 | acute_cyto_PC3 | 0.634070741 | 0.936860051 |
| blast | GGB28881_SGB41561 | acute_cyto_PC3 | 0.011894548 | 0.421434652 |
| vns | GGB28881_SGB41561 | acute_cyto_PC3 | 0.001988037 | 0.421434652 |
| blast | GGB28892_SGB41573 | acute_cyto_PC3 | 0.715153028 | 0.197403228 |
| vns | GGB28892_SGB41573 | acute_cyto_PC3 | 0.784709483 | 0.197403228 |
| blast | GGB28893_SGB41574 | acute_cyto_PC3 | 0.739221992 | 0.003669261 |
| vns | GGB28893_SGB41574 | acute_cyto_PC3 | 0.264245168 | 0.003669261 |
| blast | GGB28898_SGB41580 | acute_cyto_PC3 | 0.601924700 | 0.782881375 |
| vns | GGB28898_SGB41580 | acute_cyto_PC3 | 0.591718713 | 0.782881375 |
| blast | GGB28901_SGB41592 | acute_cyto_PC3 | 0.072063586 | 0.377358964 |
| vns | GGB28901_SGB41592 | acute_cyto_PC3 | 0.152665868 | 0.377358964 |
| blast | GGB28904_SGB41597 | acute_cyto_PC3 | 0.225752520 | 0.061885765 |
| vns | GGB28904_SGB41597 | acute_cyto_PC3 | 0.085658495 | 0.061885765 |
| blast | GGB28909_SGB41602 | acute_cyto_PC3 | 0.218283007 | 0.411876428 |
| vns | GGB28909_SGB41602 | acute_cyto_PC3 | 0.145902665 | 0.411876428 |
| blast | GGB28924_SGB41621 | acute_cyto_PC3 | 0.611578184 | 0.531115561 |
| vns | GGB28924_SGB41621 | acute_cyto_PC3 | 0.456981189 | 0.531115561 |
| blast | GGB28927_SGB41625 | acute_cyto_PC3 | 0.001954483 | 0.344620829 |
| vns | GGB28927_SGB41625 | acute_cyto_PC3 | 0.000219425 | 0.344620829 |
| blast | GGB28934_SGB41635 | acute_cyto_PC3 | 0.216498853 | 0.864091627 |
| vns | GGB28934_SGB41635 | acute_cyto_PC3 | 0.224102539 | 0.864091627 |
| blast | GGB28949_SGB41655 | acute_cyto_PC3 | 0.159598366 | 0.800602891 |
| vns | GGB28949_SGB41655 | acute_cyto_PC3 | 0.744810754 | 0.800602891 |
| blast | GGB28951_SGB102295 | acute_cyto_PC3 | 0.718725508 | 0.55127267 |
| vns | GGB28951_SGB102295 | acute_cyto_PC3 | 0.298970783 | 0.55127267 |
| blast | GGB28951_SGB41658 | acute_cyto_PC3 | 0.752627843 | 0.487923486 |
| vns | GGB28951_SGB41658 | acute_cyto_PC3 | 0.024801176 | 0.487923486 |
| blast | GGB28954_SGB41662 | acute_cyto_PC3 | 0.794939032 | 0.543334032 |
| vns | GGB28954_SGB41662 | acute_cyto_PC3 | 0.841186745 | 0.543334032 |

| | | | | |
|---|---|---|---|---|
| blast | GGB28960_SGB41669 | acute_cyto_PC3 | 0.536638861 | 0.566327325 |
| vns | GGB28960_SGB41669 | acute_cyto_PC3 | 0.043638773 | 0.566327325 |
| blast | GGB28964_SGB94886 | acute_cyto_PC3 | 0.207213028 | 0.020014945 |
| vns | GGB28964_SGB94886 | acute_cyto_PC3 | 0.400370147 | 0.020014945 |
| blast | GGB28967_SGB41678 | acute_cyto_PC3 | 0.848205066 | 0.196364066 |
| vns | GGB28967_SGB41678 | acute_cyto_PC3 | 0.820661965 | 0.196364066 |
| blast | GGB28996_SGB41712 | acute_cyto_PC3 | 0.429884728 | 0.552994939 |
| vns | GGB28996_SGB41712 | acute_cyto_PC3 | 0.377449394 | 0.552994939 |
| blast | GGB29531_SGB42317 | acute_cyto_PC3 | 0.279593444 | 0.188641667 |
| vns | GGB29531_SGB42317 | acute_cyto_PC3 | 0.999007775 | 0.188641667 |
| blast | GGB29685_SGB42494 | acute_cyto_PC3 | 0.004596881 | 0.024607236 |
| vns | GGB29685_SGB42494 | acute_cyto_PC3 | 0.748416768 | 0.024607236 |
| blast | GGB30141_SGB43066 | acute_cyto_PC3 | 0.614028140 | 0.048884087 |
| vns | GGB30141_SGB43066 | acute_cyto_PC3 | 0.664026053 | 0.048884087 |
| blast | GGB30145_SGB43072 | acute_cyto_PC3 | 0.294924035 | 0.866307014 |
| vns | GGB30145_SGB43072 | acute_cyto_PC3 | 0.367951830 | 0.866307014 |
| blast | GGB30286_SGB43248 | acute_cyto_PC3 | 0.766757581 | 0.561068016 |
| vns | GGB30286_SGB43248 | acute_cyto_PC3 | 0.750425732 | 0.561068016 |
| blast | GGB30300_SGB43264 | acute_cyto_PC3 | 0.213846948 | 0.982065034 |
| vns | GGB30300_SGB43264 | acute_cyto_PC3 | 0.213511417 | 0.982065034 |
| blast | GGB30303_SGB43268 | acute_cyto_PC3 | 0.795572751 | 0.073306671 |
| vns | GGB30303_SGB43268 | acute_cyto_PC3 | 0.660099923 | 0.073306671 |
| blast | GGB30450_SGB43507 | acute_cyto_PC3 | 0.027109400 | 0.888768265 |
| vns | GGB30450_SGB43507 | acute_cyto_PC3 | 0.186573367 | 0.888768265 |
| blast | GGB30453_SGB43513 | acute_cyto_PC3 | 0.315725451 | 0.166160477 |
| vns | GGB30453_SGB43513 | acute_cyto_PC3 | 0.310638137 | 0.166160477 |
| blast | GGB30454_SGB43514 | acute_cyto_PC3 | 0.642888646 | 0.486766818 |
| vns | GGB30454_SGB43514 | acute_cyto_PC3 | 0.656930027 | 0.486766818 |
| blast | GGB30455_SGB43519 | acute_cyto_PC3 | 0.663798426 | 0.302222002 |
| vns | GGB30455_SGB43519 | acute_cyto_PC3 | 0.104495810 | 0.302222002 |
| blast | GGB30456_SGB43520 | acute_cyto_PC3 | 0.111196948 | 0.204944538 |
| vns | GGB30456_SGB43520 | acute_cyto_PC3 | 0.158921661 | 0.204944538 |
| blast | GGB30457_SGB63218 | acute_cyto_PC3 | 0.882164050 | 0.921985183 |
| vns | GGB30457_SGB63218 | acute_cyto_PC3 | 0.890757036 | 0.921985183 |
| blast | GGB30461_SGB43530 | acute_cyto_PC3 | 0.602468303 | 0.507004803 |
| vns | GGB30461_SGB43530 | acute_cyto_PC3 | 0.137401205 | 0.507004803 |
| blast | GGB30461_SGB43533 | acute_cyto_PC3 | 0.810767423 | 0.121803564 |
| vns | GGB30461_SGB43533 | acute_cyto_PC3 | 0.718693859 | 0.121803564 |
| blast | GGB30461_SGB63209 | acute_cyto_PC3 | 0.005914118 | 0.678926722 |
| vns | GGB30461_SGB63209 | acute_cyto_PC3 | 0.075178424 | 0.678926722 |
| blast | GGB30463_SGB43537 | acute_cyto_PC3 | 0.040768110 | 0.347853377 |
| vns | GGB30463_SGB43537 | acute_cyto_PC3 | 0.991960140 | 0.347853377 |
| blast | GGB30473_SGB43557 | acute_cyto_PC3 | 0.328491579 | 0.504757171 |
| vns | GGB30473_SGB43557 | acute_cyto_PC3 | 0.189853765 | 0.504757171 |
| blast | GGB30475_SGB63182 | acute_cyto_PC3 | 0.811833041 | 0.526750973 |
| vns | GGB30475_SGB63182 | acute_cyto_PC3 | 0.783129024 | 0.526750973 |

| | | | | |
|---|---|---|---|---|
| blast | GGB30861_SGB44083 | acute_cyto_PC3 | 0.719401647 | 0.223237907 |
| vns | GGB30861_SGB44083 | acute_cyto_PC3 | 0.981045510 | 0.223237907 |
| blast | GGB31312_SGB44628 | acute_cyto_PC3 | 0.831039140 | 0.590550802 |
| vns | GGB31312_SGB44628 | acute_cyto_PC3 | 0.833457038 | 0.590550802 |
| blast | GGB31438_SGB44768 | acute_cyto_PC3 | 0.481213900 | 0.803314460 |
| vns | GGB31438_SGB44768 | acute_cyto_PC3 | 0.483694757 | 0.803314460 |
| blast | GGB3171_SGB4185 | acute_cyto_PC3 | 0.066342246 | 0.369240410 |
| vns | GGB3171_SGB4185 | acute_cyto_PC3 | 0.057850794 | 0.369240410 |
| blast | GGB31762_SGB45125 | acute_cyto_PC3 | 0.442889056 | 0.168519571 |
| vns | GGB31762_SGB45125 | acute_cyto_PC3 | 0.437364142 | 0.168519571 |
| blast | GGB31823_SGB45199 | acute_cyto_PC3 | 0.630175203 | 0.581998207 |
| vns | GGB31823_SGB45199 | acute_cyto_PC3 | 0.537240753 | 0.581998207 |
| blast | GGB31838_SGB45216 | acute_cyto_PC3 | 0.684403164 | 0.132848363 |
| vns | GGB31838_SGB45216 | acute_cyto_PC3 | 0.997715556 | 0.132848363 |
| blast | GGB31841_SGB65084 | acute_cyto_PC3 | 0.371737454 | 0.501230074 |
| vns | GGB31841_SGB65084 | acute_cyto_PC3 | 0.443951934 | 0.501230074 |
| blast | GGB32371_SGB41694 | acute_cyto_PC3 | 0.242018055 | 0.164051326 |
| vns | GGB32371_SGB41694 | acute_cyto_PC3 | 0.400159574 | 0.164051326 |
| blast | GGB3793_SGB5158 | acute_cyto_PC3 | 0.884993491 | 0.710076525 |
| vns | GGB3793_SGB5158 | acute_cyto_PC3 | 0.847239709 | 0.710076525 |
| blast | GGB42601_SGB59797 | acute_cyto_PC3 | 0.401892013 | 0.035750525 |
| vns | GGB42601_SGB59797 | acute_cyto_PC3 | 0.627566595 | 0.035750525 |
| blast | GGB45514_SGB63186 | acute_cyto_PC3 | 0.955364765 | 0.384416731 |
| vns | GGB45514_SGB63186 | acute_cyto_PC3 | 0.621527882 | 0.384416731 |
| blast | GGB45564_SGB63259 | acute_cyto_PC3 | 0.415272784 | 0.469296960 |
| vns | GGB45564_SGB63259 | acute_cyto_PC3 | 0.380878696 | 0.469296960 |
| blast | GGB45624_SGB63337 | acute_cyto_PC3 | 0.620389272 | 0.615590643 |
| vns | GGB45624_SGB63337 | acute_cyto_PC3 | 0.385198794 | 0.615590643 |
| blast | GGB45656_SGB63370 | acute_cyto_PC3 | 0.404705483 | 0.250956981 |
| vns | GGB45656_SGB63370 | acute_cyto_PC3 | 0.163643553 | 0.250956981 |
| blast | GGB47127_SGB65054 | acute_cyto_PC3 | 0.456493679 | 0.841292296 |
| vns | GGB47127_SGB65054 | acute_cyto_PC3 | 0.82583129 | 0.841292296 |
| blast | GGB74395_SGB43523 | acute_cyto_PC3 | 0.802410438 | 0.280740637 |
| vns | GGB74395_SGB43523 | acute_cyto_PC3 | 0.835585633 | 0.280740637 |
| blast | GGB75053_SGB43494 | acute_cyto_PC3 | 0.177327401 | 0.095368247 |
| vns | GGB75053_SGB43494 | acute_cyto_PC3 | 0.050410443 | 0.095368247 |
| blast | GGB75109_SGB102238 | acute_cyto_PC3 | 0.599792986 | 0.680641567 |
| vns | GGB75109_SGB102238 | acute_cyto_PC3 | 0.510864481 | 0.680641567 |
| blast | Lachnospiraceae_bacterium | acute_cyto_PC3 | 0.023925903 | 0.018487335 |
| vns | Lachnospiraceae_bacterium | acute_cyto_PC3 | 0.917492265 | 0.018487335 |
| blast | Lachnospiraceae_bacterium_A2 | acute_cyto_PC3 | 0.724187807 | 0.387550498 |
| vns | Lachnospiraceae_bacterium_A2 | acute_cyto_PC3 | 0.083860739 | 0.387550498 |
| blast | Lachnospiraceae_unclassified_SGB41414 | acute_cyto_PC3 | 0.719079500 | 0.380848187 |
| vns | Lachnospiraceae_unclassified_SGB41414 | acute_cyto_PC3 | 0.532128198 | 0.380848187 |
| blast | Lachnospiraceae_unclassified_SGB41424 | acute_cyto_PC3 | 0.216930413 | 0.205779670 |
| vns | Lachnospiraceae_unclassified_SGB41424 | acute_cyto_PC3 | 0.022018829 | 0.205779670 |

| | | | | |
|---|---|---|---|---|
| blast | Lachnospiraceae_unclassified_SGB94868 | acute_cyto_PC3 | 0.2076335048 | 0.1334503143 |
| vns | Lachnospiraceae_unclassified_SGB94868 | acute_cyto_PC3 | 0.7545629335 | 0.1334503143 |
| blast | Lactobacillus_johnsonii | acute_cyto_PC3 | 0.0027483659 | 0.1004283291 |
| vns | Lactobacillus_johnsonii | acute_cyto_PC3 | 0.8322162424 | 0.1004283291 |
| blast | Leptogranulimonas_caecicola | acute_cyto_PC3 | 0.8745779805 | 0.1672472759 |
| vns | Leptogranulimonas_caecicola | acute_cyto_PC3 | 0.0144750878 | 0.1672472759 |
| blast | Muribaculaceae_bacterium | acute_cyto_PC3 | 0.0018501989 | 0.1047820704 |
| vns | Muribaculaceae_bacterium | acute_cyto_PC3 | 0.8835325068 | 0.1047820704 |
| blast | Neglectibacter_sp_X4 | acute_cyto_PC3 | 0.3944230674 | 0.8645774706 |
| vns | Neglectibacter_sp_X4 | acute_cyto_PC3 | 0.463430059 | 0.8645774706 |
| blast | Oscillibacter_SGB43496 | acute_cyto_PC3 | 0.2205447256 | 0.5230316625 |
| vns | Oscillibacter_SGB43496 | acute_cyto_PC3 | 0.34795292 | 0.5230316625 |
| blast | Oscillospiraceae_bacterium | acute_cyto_PC3 | 0.4927836612 | 0.0384061752 |
| vns | Oscillospiraceae_bacterium | acute_cyto_PC3 | 0.251973413 | 0.0384061752 |
| blast | Oscillospiraceae_unclassified_SGB43502 | acute_cyto_PC3 | 0.0172712013 | 0.0818751848 |
| vns | Oscillospiraceae_unclassified_SGB43502 | acute_cyto_PC3 | 0.1123454974 | 0.0818751848 |
| blast | Oscillospiraceae_unclassified_SGB43505 | acute_cyto_PC3 | 0.3742999083 | 0.7308986096 |
| vns | Oscillospiraceae_unclassified_SGB43505 | acute_cyto_PC3 | 0.6182049908 | 0.7308986096 |
| blast | Oscillospiraceae_unclassified_SGB94989 | acute_cyto_PC3 | 0.9533689691 | 0.7576086032 |
| vns | Oscillospiraceae_unclassified_SGB94989 | acute_cyto_PC3 | 0.6236152007 | 0.7576086032 |
| blast | Parasutterella_excrementihominis | acute_cyto_PC3 | 0.0234193782 | 0.6369235708 |
| vns | Parasutterella_excrementihominis | acute_cyto_PC3 | 0.5041571561 | 0.6369235708 |
| blast | Richness (# observed features) | acute_cyto_PC3 | 0.0137778742 | 0.0133539927 |
| vns | Richness (# observed features) | acute_cyto_PC3 | 0.9582478381 | 0.0133539927 |
| blast | Schaedlerella_arabinosiphila | acute_cyto_PC3 | 0.114481983 | 0.4937420204 |
| vns | Schaedlerella_arabinosiphila | acute_cyto_PC3 | 0.3863019645 | 0.4937420204 |
| blast | Shannon Index | acute_cyto_PC3 | 0.0064730704 | 0.0269099643 |
| vns | Shannon Index | acute_cyto_PC3 | 0.8359924459 | 0.0269099643 |
| blast | Turicibacter_sp_1E2 | acute_cyto_PC3 | 0.0095941376 | 0.2413661101 |
| vns | Turicibacter_sp_1E2 | acute_cyto_PC3 | 0.183926772 | 0.2413661101 |
| blast | Acetatifactor_SGB41545 | acute_cyto_PC4 | 0.2103509006 | 0.5534814492 |
| vns | Acetatifactor_SGB41545 | acute_cyto_PC4 | 0.2390986693 | 0.5534814492 |
| blast | Acutalibacter_muris | acute_cyto_PC4 | 0.7869843388 | 0.0127311123 |
| vns | Acutalibacter_muris | acute_cyto_PC4 | 0.4919022263 | 0.0127311123 |
| blast | Akkermansia_muciniphila | acute_cyto_PC4 | 0.16660254 | 0.406070667 |
| vns | Akkermansia_muciniphila | acute_cyto_PC4 | 0.6788996212 | 0.406070667 |
| blast | Anaerotruncus_sp_1XD42_93 | acute_cyto_PC4 | 0.0009175375 | 0.1635641806 |
| vns | Anaerotruncus_sp_1XD42_93 | acute_cyto_PC4 | 0.0002316873 | 0.1635641806 |
| blast | Bacteria_unclassified_SGB41677 | acute_cyto_PC4 | 0.7141894931 | 0.9465425232 |
| vns | Bacteria_unclassified_SGB41677 | acute_cyto_PC4 | 0.58777865 | 0.9465425232 |
| blast | Bacteria_unclassified_SGB43539 | acute_cyto_PC4 | 0.1489024993 | 0.8459095008 |
| vns | Bacteria_unclassified_SGB43539 | acute_cyto_PC4 | 0.4492601232 | 0.8459095008 |
| blast | Bacteria_unclassified_SGB43546 | acute_cyto_PC4 | 0.1125519082 | 0.3980585952 |
| vns | Bacteria_unclassified_SGB43546 | acute_cyto_PC4 | 0.3099088089 | 0.3980585952 |
| blast | Bacteria_unclassified_SGB63211 | acute_cyto_PC4 | 0.2950037757 | 0.0580328834 |
| vns | Bacteria_unclassified_SGB63211 | acute_cyto_PC4 | 0.3408716294 | 0.0580328834 |

| | | | | |
|---|---|---|---|---|
| blast | bacterium_0_1xD8_82 | acute_cyto_PC4 | 0.763436393 | 0.3451986689 |
| vns | bacterium_0_1xD8_82 | acute_cyto_PC4 | 0.7951390939 | 0.3451986689 |
| blast | bacterium_1XD42_1 | acute_cyto_PC4 | 0.2725193977 | 0.8322761366 |
| vns | bacterium_1XD42_1 | acute_cyto_PC4 | 0.0970919255 | 0.8322761366 |
| blast | bacterium_1XD42_54 | acute_cyto_PC4 | 0.0381053216 | 0.7605988196 |
| vns | bacterium_1XD42_54 | acute_cyto_PC4 | 0.1254069334 | 0.7605988196 |
| blast | bacterium_1XD42_76 | acute_cyto_PC4 | 0.1774107688 | 0.6892219139 |
| vns | bacterium_1XD42_76 | acute_cyto_PC4 | 0.141032802 | 0.6892219139 |
| blast | bacterium_1xD8_48 | acute_cyto_PC4 | 0.1393476118 | 0.2659638145 |
| vns | bacterium_1xD8_48 | acute_cyto_PC4 | 0.3700204072 | 0.2659638145 |
| blast | Bacteroides_thetaiotaomicron | acute_cyto_PC4 | 0.0366277544 | 0.7387881338 |
| vns | Bacteroides_thetaiotaomicron | acute_cyto_PC4 | 0.2754050051 | 0.7387881338 |
| blast | Berger Parker Index | acute_cyto_PC4 | 0.0574752835 | 0.966625106 |
| vns | Berger Parker Index | acute_cyto_PC4 | 0.4995059559 | 0.966625106 |
| blast | Clostridia_bacterium | acute_cyto_PC4 | 0.2483530876 | 0.6128330794 |
| vns | Clostridia_bacterium | acute_cyto_PC4 | 0.3805560795 | 0.6128330794 |
| blast | Clostridiaceae_bacterium | acute_cyto_PC4 | 0.1623466052 | 0.7950182484 |
| vns | Clostridiaceae_bacterium | acute_cyto_PC4 | 0.2770710842 | 0.7950182484 |
| blast | Clostridiaceae_unclassified_SGB41663 | acute_cyto_PC4 | 0.3624098717 | 0.4524339813 |
| vns | Clostridiaceae_unclassified_SGB41663 | acute_cyto_PC4 | 0.0709173242 | 0.4524339813 |
| blast | Clostridiales_bacterium | acute_cyto_PC4 | 0.0738247407 | 0.7783000156 |
| vns | Clostridiales_bacterium | acute_cyto_PC4 | 0.1248511673 | 0.7783000156 |
| blast | Clostridium_cocleatum | acute_cyto_PC4 | 0.5072750859 | 0.2285954642 |
| vns | Clostridium_cocleatum | acute_cyto_PC4 | 0.4827323844 | 0.2285954642 |
| blast | Clostridium_SGB65123 | acute_cyto_PC4 | 0.1804389829 | 0.8100994922 |
| vns | Clostridium_SGB65123 | acute_cyto_PC4 | 0.1793895684 | 0.8100994922 |
| blast | Eubacteriaceae_bacterium | acute_cyto_PC4 | 0.0166040620 | 0.6191617671 |
| vns | Eubacteriaceae_bacterium | acute_cyto_PC4 | 0.9691575786 | 0.6191617671 |
| blast | Eubacteriaceae_unclassified_SGB94922 | acute_cyto_PC4 | 0.1742242764 | 0.809700411 |
| vns | Eubacteriaceae_unclassified_SGB94922 | acute_cyto_PC4 | 0.3461810244 | 0.809700411 |
| blast | GGB20146_SGB29427 | acute_cyto_PC4 | 0.0938444678 | 0.7976121847 |
| vns | GGB20146_SGB29427 | acute_cyto_PC4 | 0.0512189198 | 0.7976121847 |
| blast | GGB20149_SGB29430 | acute_cyto_PC4 | 0.4132259254 | 0.0530011472 |
| vns | GGB20149_SGB29430 | acute_cyto_PC4 | 0.1804348387 | 0.0530011472 |
| blast | GGB22635_SGB63107 | acute_cyto_PC4 | 0.1146609038 | 0.1576575174 |
| vns | GGB22635_SGB63107 | acute_cyto_PC4 | 0.5356393392 | 0.1576575174 |
| blast | GGB25041_SGB36960 | acute_cyto_PC4 | 0.1675951172 | 0.6259160731 |
| vns | GGB25041_SGB36960 | acute_cyto_PC4 | 0.2062562608 | 0.6259160731 |
| blast | GGB28379_SGB40959 | acute_cyto_PC4 | 0.0292418367 | 0.0898285054 |
| vns | GGB28379_SGB40959 | acute_cyto_PC4 | 0.0327946564 | 0.0898285054 |
| blast | GGB28382_SGB40962 | acute_cyto_PC4 | 0.0725130221 | 0.3236794987 |
| vns | GGB28382_SGB40962 | acute_cyto_PC4 | 0.3704307363 | 0.3236794987 |
| blast | GGB28392_SGB40972 | acute_cyto_PC4 | 0.0171836690 | 0.7573936985 |
| vns | GGB28392_SGB40972 | acute_cyto_PC4 | 0.0165924007 | 0.7573936985 |
| blast | GGB28404_SGB40986 | acute_cyto_PC4 | 0.1577679288 | 0.4271898241 |
| vns | GGB28404_SGB40986 | acute_cyto_PC4 | 0.0100902146 | 0.4271898241 |

| | | | | |
|---|---|---|---|---|
| blast | GGB28418_SGB41001 | acute_cyto_PC4 | 0.996265407 | 0.975231548 |
| vns | GGB28418_SGB41001 | acute_cyto_PC4 | 0.959497945 | 0.975231548 |
| blast | GGB28422_SGB41005 | acute_cyto_PC4 | 0.298094755 | 0.885764240 |
| vns | GGB28422_SGB41005 | acute_cyto_PC4 | 0.416676405 | 0.885764240 |
| blast | GGB28431_SGB41014 | acute_cyto_PC4 | 0.015377584 | 0.249962172 |
| vns | GGB28431_SGB41014 | acute_cyto_PC4 | 0.016801247 | 0.249962172 |
| blast | GGB28456_SGB41039 | acute_cyto_PC4 | 0.508621549 | 0.392536880 |
| vns | GGB28456_SGB41039 | acute_cyto_PC4 | 0.567597000 | 0.392536880 |
| blast | GGB28782_SGB41435 | acute_cyto_PC4 | 0.315491599 | 0.008301026 |
| vns | GGB28782_SGB41435 | acute_cyto_PC4 | 0.359169919 | 0.008301026 |
| blast | GGB28792_SGB41445 | acute_cyto_PC4 | 0.123770726 | 0.098378770 |
| vns | GGB28792_SGB41445 | acute_cyto_PC4 | 0.970858107 | 0.098378770 |
| blast | GGB28798_SGB41451 | acute_cyto_PC4 | 0.931837305 | 0.095521209 |
| vns | GGB28798_SGB41451 | acute_cyto_PC4 | 0.471552545 | 0.095521209 |
| blast | GGB28810_SGB41465 | acute_cyto_PC4 | 0.919521792 | 0.219462330 |
| vns | GGB28810_SGB41465 | acute_cyto_PC4 | 0.915105920 | 0.219462330 |
| blast | GGB28828_SGB41484 | acute_cyto_PC4 | 0.036453176 | 0.897037007 |
| vns | GGB28828_SGB41484 | acute_cyto_PC4 | 0.029615316 | 0.897037007 |
| blast | GGB28851_SGB41518 | acute_cyto_PC4 | 0.983245267 | 0.534505723 |
| vns | GGB28851_SGB41518 | acute_cyto_PC4 | 0.548244701 | 0.534505723 |
| blast | GGB28865_SGB41537 | acute_cyto_PC4 | 0.051838259 | 0.316662138 |
| vns | GGB28865_SGB41537 | acute_cyto_PC4 | 0.043832016 | 0.316662138 |
| blast | GGB28868_SGB41542 | acute_cyto_PC4 | 0.292423183 | 0.123123053 |
| vns | GGB28868_SGB41542 | acute_cyto_PC4 | 0.254323647 | 0.123123053 |
| blast | GGB28869_SGB41543 | acute_cyto_PC4 | 0.662073077 | 0.810036862 |
| vns | GGB28869_SGB41543 | acute_cyto_PC4 | 0.806606148 | 0.810036862 |
| blast | GGB28875_SGB41555 | acute_cyto_PC4 | 0.776644062 | 0.873856933 |
| vns | GGB28875_SGB41555 | acute_cyto_PC4 | 0.634070741 | 0.873856933 |
| blast | GGB28881_SGB41561 | acute_cyto_PC4 | 0.011894548 | 0.377232968 |
| vns | GGB28881_SGB41561 | acute_cyto_PC4 | 0.001988037 | 0.377232968 |
| blast | GGB28892_SGB41573 | acute_cyto_PC4 | 0.715153028 | 0.814742419 |
| vns | GGB28892_SGB41573 | acute_cyto_PC4 | 0.784709483 | 0.814742419 |
| blast | GGB28893_SGB41574 | acute_cyto_PC4 | 0.739221992 | 0.879663332 |
| vns | GGB28893_SGB41574 | acute_cyto_PC4 | 0.264245168 | 0.879663332 |
| blast | GGB28898_SGB41580 | acute_cyto_PC4 | 0.601924700 | 0.463568761 |
| vns | GGB28898_SGB41580 | acute_cyto_PC4 | 0.591718713 | 0.463568761 |
| blast | GGB28901_SGB41592 | acute_cyto_PC4 | 0.072063586 | 0.414126217 |
| vns | GGB28901_SGB41592 | acute_cyto_PC4 | 0.152665868 | 0.414126217 |
| blast | GGB28904_SGB41597 | acute_cyto_PC4 | 0.225752520 | 0.424603973 |
| vns | GGB28904_SGB41597 | acute_cyto_PC4 | 0.085658495 | 0.424603973 |
| blast | GGB28909_SGB41602 | acute_cyto_PC4 | 0.218283007 | 0.991460276 |
| vns | GGB28909_SGB41602 | acute_cyto_PC4 | 0.145902665 | 0.991460276 |
| blast | GGB28924_SGB41621 | acute_cyto_PC4 | 0.611578184 | 0.642485745 |
| vns | GGB28924_SGB41621 | acute_cyto_PC4 | 0.456981189 | 0.642485745 |
| blast | GGB28927_SGB41625 | acute_cyto_PC4 | 0.001954483 | 0.326177213 |
| vns | GGB28927_SGB41625 | acute_cyto_PC4 | 0.000219425 | 0.326177213 |

| | | | | |
|---|---|---|---|---|
| blast | GGB28934_SGB41635 | acute_cyto_PC4 | 0.216498853 | 0.644533086 |
| vns | GGB28934_SGB41635 | acute_cyto_PC4 | 0.224102539 | 0.644533086 |
| blast | GGB28949_SGB41655 | acute_cyto_PC4 | 0.159598366 | 0.940804662 |
| vns | GGB28949_SGB41655 | acute_cyto_PC4 | 0.744810754 | 0.940804662 |
| blast | GGB28951_SGB102295 | acute_cyto_PC4 | 0.718725508 | 0.728250027 |
| vns | GGB28951_SGB102295 | acute_cyto_PC4 | 0.298970783 | 0.728250027 |
| blast | GGB28951_SGB41658 | acute_cyto_PC4 | 0.752627843 | 0.456688788 |
| vns | GGB28951_SGB41658 | acute_cyto_PC4 | 0.024801176 | 0.456688788 |
| blast | GGB28954_SGB41662 | acute_cyto_PC4 | 0.794939032 | 0.173630961 |
| vns | GGB28954_SGB41662 | acute_cyto_PC4 | 0.841186745 | 0.173630961 |
| blast | GGB28960_SGB41669 | acute_cyto_PC4 | 0.536638861 | 0.760945985 |
| vns | GGB28960_SGB41669 | acute_cyto_PC4 | 0.043638773 | 0.760945985 |
| blast | GGB28964_SGB94886 | acute_cyto_PC4 | 0.207213028 | 0.102306196 |
| vns | GGB28964_SGB94886 | acute_cyto_PC4 | 0.400370147 | 0.102306196 |
| blast | GGB28967_SGB41678 | acute_cyto_PC4 | 0.848205066 | 0.713075046 |
| vns | GGB28967_SGB41678 | acute_cyto_PC4 | 0.820661965 | 0.713075046 |
| blast | GGB28996_SGB41712 | acute_cyto_PC4 | 0.429884728 | 0.212627758 |
| vns | GGB28996_SGB41712 | acute_cyto_PC4 | 0.377449394 | 0.212627758 |
| blast | GGB29531_SGB42317 | acute_cyto_PC4 | 0.279593444 | 0.936617818 |
| vns | GGB29531_SGB42317 | acute_cyto_PC4 | 0.999007775 | 0.936617818 |
| blast | GGB29685_SGB42494 | acute_cyto_PC4 | 0.004596881 | 0.982521464 |
| vns | GGB29685_SGB42494 | acute_cyto_PC4 | 0.748416768 | 0.982521464 |
| blast | GGB30141_SGB43066 | acute_cyto_PC4 | 0.614028140 | 0.941832742 |
| vns | GGB30141_SGB43066 | acute_cyto_PC4 | 0.664026053 | 0.941832742 |
| blast | GGB30145_SGB43072 | acute_cyto_PC4 | 0.294924035 | 0.408313143 |
| vns | GGB30145_SGB43072 | acute_cyto_PC4 | 0.367951830 | 0.408313143 |
| blast | GGB30286_SGB43248 | acute_cyto_PC4 | 0.766757581 | 0.969741978 |
| vns | GGB30286_SGB43248 | acute_cyto_PC4 | 0.750425732 | 0.969741978 |
| blast | GGB30300_SGB43264 | acute_cyto_PC4 | 0.213846948 | 0.775466333 |
| vns | GGB30300_SGB43264 | acute_cyto_PC4 | 0.213511417 | 0.775466333 |
| blast | GGB30303_SGB43268 | acute_cyto_PC4 | 0.795572751 | 0.872523558 |
| vns | GGB30303_SGB43268 | acute_cyto_PC4 | 0.660099923 | 0.872523558 |
| blast | GGB30450_SGB43507 | acute_cyto_PC4 | 0.027109400 | 0.838698654 |
| vns | GGB30450_SGB43507 | acute_cyto_PC4 | 0.186573367 | 0.838698654 |
| blast | GGB30453_SGB43513 | acute_cyto_PC4 | 0.315725451 | 0.914779238 |
| vns | GGB30453_SGB43513 | acute_cyto_PC4 | 0.310638137 | 0.914779238 |
| blast | GGB30454_SGB43514 | acute_cyto_PC4 | 0.642888646 | 0.083948929 |
| vns | GGB30454_SGB43514 | acute_cyto_PC4 | 0.656930027 | 0.083948929 |
| blast | GGB30455_SGB43519 | acute_cyto_PC4 | 0.663798426 | 0.919353520 |
| vns | GGB30455_SGB43519 | acute_cyto_PC4 | 0.104495810 | 0.919353520 |
| blast | GGB30456_SGB43520 | acute_cyto_PC4 | 0.111196948 | 0.084160034 |
| vns | GGB30456_SGB43520 | acute_cyto_PC4 | 0.158921661 | 0.084160034 |
| blast | GGB30457_SGB63218 | acute_cyto_PC4 | 0.882164050 | 0.929474772 |
| vns | GGB30457_SGB63218 | acute_cyto_PC4 | 0.890757036 | 0.929474772 |
| blast | GGB30461_SGB43530 | acute_cyto_PC4 | 0.602468303 | 0.367978255 |
| vns | GGB30461_SGB43530 | acute_cyto_PC4 | 0.137401205 | 0.367978255 |

| | | | | |
|---|---|---|---|---|
| blast | GGB30461_SGB43533 | acute_cyto_PC4 | 0.810767423 | 0.743408315 |
| vns | GGB30461_SGB43533 | acute_cyto_PC4 | 0.718693859 | 0.743408315 |
| blast | GGB30461_SGB63209 | acute_cyto_PC4 | 0.005914118 | 0.370769292 |
| vns | GGB30461_SGB63209 | acute_cyto_PC4 | 0.075178424 | 0.370769292 |
| blast | GGB30463_SGB43537 | acute_cyto_PC4 | 0.040768110 | 0.472711939 |
| vns | GGB30463_SGB43537 | acute_cyto_PC4 | 0.991960140 | 0.472711939 |
| blast | GGB30473_SGB43557 | acute_cyto_PC4 | 0.328491579 | 0.532746470 |
| vns | GGB30473_SGB43557 | acute_cyto_PC4 | 0.189853765 | 0.532746470 |
| blast | GGB30475_SGB63182 | acute_cyto_PC4 | 0.811833041 | 0.923739221 |
| vns | GGB30475_SGB63182 | acute_cyto_PC4 | 0.783129024 | 0.923739221 |
| blast | GGB30861_SGB44083 | acute_cyto_PC4 | 0.719401647 | 0.312860092 |
| vns | GGB30861_SGB44083 | acute_cyto_PC4 | 0.981045510 | 0.312860092 |
| blast | GGB31312_SGB44628 | acute_cyto_PC4 | 0.831039140 | 0.510649029 |
| vns | GGB31312_SGB44628 | acute_cyto_PC4 | 0.833457038 | 0.510649029 |
| blast | GGB31438_SGB44768 | acute_cyto_PC4 | 0.481213900 | 0.46144833 |
| vns | GGB31438_SGB44768 | acute_cyto_PC4 | 0.483694757 | 0.46144833 |
| blast | GGB3171_SGB4185 | acute_cyto_PC4 | 0.066342246 | 0.437144027 |
| vns | GGB3171_SGB4185 | acute_cyto_PC4 | 0.057850794 | 0.437144027 |
| blast | GGB31762_SGB45125 | acute_cyto_PC4 | 0.442889056 | 0.981604873 |
| vns | GGB31762_SGB45125 | acute_cyto_PC4 | 0.437364142 | 0.981604873 |
| blast | GGB31823_SGB45199 | acute_cyto_PC4 | 0.630175203 | 0.928548894 |
| vns | GGB31823_SGB45199 | acute_cyto_PC4 | 0.537240753 | 0.928548894 |
| blast | GGB31838_SGB45216 | acute_cyto_PC4 | 0.684403164 | 0.301775375 |
| vns | GGB31838_SGB45216 | acute_cyto_PC4 | 0.997715556 | 0.301775375 |
| blast | GGB31841_SGB65084 | acute_cyto_PC4 | 0.371737454 | 0.393402691 |
| vns | GGB31841_SGB65084 | acute_cyto_PC4 | 0.443951934 | 0.393402691 |
| blast | GGB32371_SGB41694 | acute_cyto_PC4 | 0.242018055 | 0.672669652 |
| vns | GGB32371_SGB41694 | acute_cyto_PC4 | 0.400159574 | 0.672669652 |
| blast | GGB3793_SGB5158 | acute_cyto_PC4 | 0.884993491 | 0.544714013 |
| vns | GGB3793_SGB5158 | acute_cyto_PC4 | 0.847239709 | 0.544714013 |
| blast | GGB42601_SGB59797 | acute_cyto_PC4 | 0.401892013 | 0.409059403 |
| vns | GGB42601_SGB59797 | acute_cyto_PC4 | 0.627566595 | 0.409059403 |
| blast | GGB45514_SGB63186 | acute_cyto_PC4 | 0.955364765 | 0.518815686 |
| vns | GGB45514_SGB63186 | acute_cyto_PC4 | 0.621527882 | 0.518815686 |
| blast | GGB45564_SGB63259 | acute_cyto_PC4 | 0.415272784 | 0.551454182 |
| vns | GGB45564_SGB63259 | acute_cyto_PC4 | 0.380878696 | 0.551454182 |
| blast | GGB45624_SGB63337 | acute_cyto_PC4 | 0.620389272 | 0.524021795 |
| vns | GGB45624_SGB63337 | acute_cyto_PC4 | 0.385198794 | 0.524021795 |
| blast | GGB45656_SGB63370 | acute_cyto_PC4 | 0.404705483 | 0.188746888 |
| vns | GGB45656_SGB63370 | acute_cyto_PC4 | 0.163643553 | 0.188746888 |
| blast | GGB47127_SGB65054 | acute_cyto_PC4 | 0.456493679 | 0.036738332 |
| vns | GGB47127_SGB65054 | acute_cyto_PC4 | 0.82583129 | 0.036738332 |
| blast | GGB74395_SGB43523 | acute_cyto_PC4 | 0.802410438 | 0.125129394 |
| vns | GGB74395_SGB43523 | acute_cyto_PC4 | 0.835585633 | 0.125129394 |
| blast | GGB75053_SGB43494 | acute_cyto_PC4 | 0.177327401 | 0.855265324 |
| vns | GGB75053_SGB43494 | acute_cyto_PC4 | 0.050410443 | 0.855265324 |

| | | | | |
|---|---|---|---|---|
| blast | GGB75109_SGB102238 | acute_cyto_PC4 | 0.599792986 | 0.981169366 |
| vns | GGB75109_SGB102238 | acute_cyto_PC4 | 0.510864481 | 0.981169366 |
| blast | Lachnospiraceae_bacterium | acute_cyto_PC4 | 0.023925903 | 0.751121746 |
| vns | Lachnospiraceae_bacterium | acute_cyto_PC4 | 0.917492265 | 0.751121746 |
| blast | Lachnospiraceae_bacterium_A2 | acute_cyto_PC4 | 0.724187807 | 0.393179005 |
| vns | Lachnospiraceae_bacterium_A2 | acute_cyto_PC4 | 0.083860739 | 0.393179005 |
| blast | Lachnospiraceae_unclassified_SGB41414 | acute_cyto_PC4 | 0.719079500 | 0.375261271 |
| vns | Lachnospiraceae_unclassified_SGB41414 | acute_cyto_PC4 | 0.532128198 | 0.375261271 |
| blast | Lachnospiraceae_unclassified_SGB41424 | acute_cyto_PC4 | 0.216930413 | 0.558261030 |
| vns | Lachnospiraceae_unclassified_SGB41424 | acute_cyto_PC4 | 0.022018829 | 0.558261030 |
| blast | Lachnospiraceae_unclassified_SGB94868 | acute_cyto_PC4 | 0.207633504 | 0.997874178 |
| vns | Lachnospiraceae_unclassified_SGB94868 | acute_cyto_PC4 | 0.754562933 | 0.997874178 |
| blast | Lactobacillus_johnsonii | acute_cyto_PC4 | 0.002748365 | 0.626527166 |
| vns | Lactobacillus_johnsonii | acute_cyto_PC4 | 0.832216242 | 0.626527166 |
| blast | Leptogranulimonas_caecicola | acute_cyto_PC4 | 0.874577980 | 0.969956296 |
| vns | Leptogranulimonas_caecicola | acute_cyto_PC4 | 0.014475087 | 0.969956296 |
| blast | Muribaculaceae_bacterium | acute_cyto_PC4 | 0.001850198 | 0.742445662 |
| vns | Muribaculaceae_bacterium | acute_cyto_PC4 | 0.883532506 | 0.742445662 |
| blast | Neglectibacter_sp_X4 | acute_cyto_PC4 | 0.394423067 | 0.741394946 |
| vns | Neglectibacter_sp_X4 | acute_cyto_PC4 | 0.463430059 | 0.741394946 |
| blast | Oscillibacter_SGB43496 | acute_cyto_PC4 | 0.220544725 | 0.258666812 |
| vns | Oscillibacter_SGB43496 | acute_cyto_PC4 | 0.34795292 | 0.258666812 |
| blast | Oscillospiraceae_bacterium | acute_cyto_PC4 | 0.492783661 | 0.672695362 |
| vns | Oscillospiraceae_bacterium | acute_cyto_PC4 | 0.251973413 | 0.672695362 |
| blast | Oscillospiraceae_unclassified_SGB43502 | acute_cyto_PC4 | 0.017271201 | 0.275756638 |
| vns | Oscillospiraceae_unclassified_SGB43502 | acute_cyto_PC4 | 0.112345497 | 0.275756638 |
| blast | Oscillospiraceae_unclassified_SGB43505 | acute_cyto_PC4 | 0.374299908 | 0.723934266 |
| vns | Oscillospiraceae_unclassified_SGB43505 | acute_cyto_PC4 | 0.618204990 | 0.723934266 |
| blast | Oscillospiraceae_unclassified_SGB94989 | acute_cyto_PC4 | 0.953368969 | 0.899944963 |
| vns | Oscillospiraceae_unclassified_SGB94989 | acute_cyto_PC4 | 0.623615200 | 0.899944963 |
| blast | Parasutterella_excrementihominis | acute_cyto_PC4 | 0.023419378 | 0.085303682 |
| vns | Parasutterella_excrementihominis | acute_cyto_PC4 | 0.504157156 | 0.085303682 |
| blast | Richness (# observed features) | acute_cyto_PC4 | 0.013777874 | 0.990003766 |
| vns | Richness (# observed features) | acute_cyto_PC4 | 0.958247838 | 0.990003766 |
| blast | Schaedlerella_arabinosiphila | acute_cyto_PC4 | 0.114481983 | 0.147983161 |
| vns | Schaedlerella_arabinosiphila | acute_cyto_PC4 | 0.386301964 | 0.147983161 |
| blast | Shannon Index | acute_cyto_PC4 | 0.006473070 | 0.582589904 |
| vns | Shannon Index | acute_cyto_PC4 | 0.835992445 | 0.582589904 |
| blast | Turicibacter_sp_1E2 | acute_cyto_PC4 | 0.009594137 | 0.553715021 |
| vns | Turicibacter_sp_1E2 | acute_cyto_PC4 | 0.183926772 | 0.553715021 |
| blast | Acetatifactor_SGB41545 | acute_cyto_PC5 | 0.210350900 | 0.759879406 |
| vns | Acetatifactor_SGB41545 | acute_cyto_PC5 | 0.239098669 | 0.759879406 |
| blast | Acutalibacter_muris | acute_cyto_PC5 | 0.786984338 | 0.338851418 |
| vns | Acutalibacter_muris | acute_cyto_PC5 | 0.491902226 | 0.338851418 |
| blast | Akkermansia_muciniphila | acute_cyto_PC5 | 0.16660254 | 0.138644297 |
| vns | Akkermansia_muciniphila | acute_cyto_PC5 | 0.678899621 | 0.138644297 |

| | | | | |
|---|---|---|---|---|
| blast | Anaerotruncus_sp_1XD42_93 | acute_cyto_PC5 | 0.000917537 | 0.131962450 |
| vns | Anaerotruncus_sp_1XD42_93 | acute_cyto_PC5 | 0.000231687 | 0.131962450 |
| blast | Bacteria_unclassified_SGB41677 | acute_cyto_PC5 | 0.714189493 | 0.290781171 |
| vns | Bacteria_unclassified_SGB41677 | acute_cyto_PC5 | 0.58777865 | 0.290781171 |
| blast | Bacteria_unclassified_SGB43539 | acute_cyto_PC5 | 0.148902499 | 0.355623226 |
| vns | Bacteria_unclassified_SGB43539 | acute_cyto_PC5 | 0.449260123 | 0.355623226 |
| blast | Bacteria_unclassified_SGB43546 | acute_cyto_PC5 | 0.112551908 | 0.014884615 |
| vns | Bacteria_unclassified_SGB43546 | acute_cyto_PC5 | 0.309908808 | 0.014884615 |
| blast | Bacteria_unclassified_SGB63211 | acute_cyto_PC5 | 0.295003775 | 0.361662888 |
| vns | Bacteria_unclassified_SGB63211 | acute_cyto_PC5 | 0.340871629 | 0.361662888 |
| blast | bacterium_0_1xD8_82 | acute_cyto_PC5 | 0.763436393 | 0.614504504 |
| vns | bacterium_0_1xD8_82 | acute_cyto_PC5 | 0.795139093 | 0.614504504 |
| blast | bacterium_1XD42_1 | acute_cyto_PC5 | 0.272519397 | 0.029394954 |
| vns | bacterium_1XD42_1 | acute_cyto_PC5 | 0.097091925 | 0.029394954 |
| blast | bacterium_1XD42_54 | acute_cyto_PC5 | 0.038105321 | 0.547514611 |
| vns | bacterium_1XD42_54 | acute_cyto_PC5 | 0.125406933 | 0.547514611 |
| blast | bacterium_1XD42_76 | acute_cyto_PC5 | 0.177410768 | 0.493164290 |
| vns | bacterium_1XD42_76 | acute_cyto_PC5 | 0.141032802 | 0.493164290 |
| blast | bacterium_1xD8_48 | acute_cyto_PC5 | 0.139347611 | 0.712095773 |
| vns | bacterium_1xD8_48 | acute_cyto_PC5 | 0.370020407 | 0.712095773 |
| blast | Bacteroides_thetaiotaomicron | acute_cyto_PC5 | 0.036627754 | 0.631926246 |
| vns | Bacteroides_thetaiotaomicron | acute_cyto_PC5 | 0.275405005 | 0.631926246 |
| blast | Berger Parker Index | acute_cyto_PC5 | 0.057475283 | 0.166885264 |
| vns | Berger Parker Index | acute_cyto_PC5 | 0.499505955 | 0.166885264 |
| blast | Clostridia_bacterium | acute_cyto_PC5 | 0.248353087 | 0.185023828 |
| vns | Clostridia_bacterium | acute_cyto_PC5 | 0.380556079 | 0.185023828 |
| blast | Clostridiaceae_bacterium | acute_cyto_PC5 | 0.162346605 | 0.242769451 |
| vns | Clostridiaceae_bacterium | acute_cyto_PC5 | 0.277071084 | 0.242769451 |
| blast | Clostridiaceae_unclassified_SGB41663 | acute_cyto_PC5 | 0.362409871 | 0.181875880 |
| vns | Clostridiaceae_unclassified_SGB41663 | acute_cyto_PC5 | 0.070917324 | 0.181875880 |
| blast | Clostridiales_bacterium | acute_cyto_PC5 | 0.073824740 | 0.884275683 |
| vns | Clostridiales_bacterium | acute_cyto_PC5 | 0.124851167 | 0.884275683 |
| blast | Clostridium_cocleatum | acute_cyto_PC5 | 0.507275085 | 0.999251513 |
| vns | Clostridium_cocleatum | acute_cyto_PC5 | 0.482732384 | 0.999251513 |
| blast | Clostridium_SGB65123 | acute_cyto_PC5 | 0.180438982 | 0.194202497 |
| vns | Clostridium_SGB65123 | acute_cyto_PC5 | 0.179389568 | 0.194202497 |
| blast | Eubacteriaceae_bacterium | acute_cyto_PC5 | 0.016604062 | 0.871438152 |
| vns | Eubacteriaceae_bacterium | acute_cyto_PC5 | 0.969157578 | 0.871438152 |
| blast | Eubacteriaceae_unclassified_SGB94922 | acute_cyto_PC5 | 0.174224276 | 0.094822587 |
| vns | Eubacteriaceae_unclassified_SGB94922 | acute_cyto_PC5 | 0.346181024 | 0.094822587 |
| blast | GGB20146_SGB29427 | acute_cyto_PC5 | 0.093844467 | 0.048214670 |
| vns | GGB20146_SGB29427 | acute_cyto_PC5 | 0.051218919 | 0.048214670 |
| blast | GGB20149_SGB29430 | acute_cyto_PC5 | 0.413225925 | 0.169456150 |
| vns | GGB20149_SGB29430 | acute_cyto_PC5 | 0.180434838 | 0.169456150 |
| blast | GGB22635_SGB63107 | acute_cyto_PC5 | 0.114660903 | 0.283207884 |
| vns | GGB22635_SGB63107 | acute_cyto_PC5 | 0.535639339 | 0.283207884 |

| | | | | |
|---|---|---|---|---|
| blast | GGB25041_SGB36960 | acute_cyto_PC5 | 0.16759511 | 0.39928717 |
| vns | GGB25041_SGB36960 | acute_cyto_PC5 | 0.20625626 | 0.39928717 |
| blast | GGB28379_SGB40959 | acute_cyto_PC5 | 0.02924183 | 0.15454169 |
| vns | GGB28379_SGB40959 | acute_cyto_PC5 | 0.03279465 | 0.15454169 |
| blast | GGB28382_SGB40962 | acute_cyto_PC5 | 0.07251302 | 0.68337932 |
| vns | GGB28382_SGB40962 | acute_cyto_PC5 | 0.37043073 | 0.68337932 |
| blast | GGB28392_SGB40972 | acute_cyto_PC5 | 0.01718366 | 0.85138213 |
| vns | GGB28392_SGB40972 | acute_cyto_PC5 | 0.01659240 | 0.85138213 |
| blast | GGB28404_SGB40986 | acute_cyto_PC5 | 0.15776792 | 0.04651898 |
| vns | GGB28404_SGB40986 | acute_cyto_PC5 | 0.01009021 | 0.04651898 |
| blast | GGB28418_SGB41001 | acute_cyto_PC5 | 0.996265407 | 0.41244858 |
| vns | GGB28418_SGB41001 | acute_cyto_PC5 | 0.959497945 | 0.41244858 |
| blast | GGB28422_SGB41005 | acute_cyto_PC5 | 0.29809475 | 0.13255622 |
| vns | GGB28422_SGB41005 | acute_cyto_PC5 | 0.416676405 | 0.13255622 |
| blast | GGB28431_SGB41014 | acute_cyto_PC5 | 0.01537758 | 0.09147343 |
| vns | GGB28431_SGB41014 | acute_cyto_PC5 | 0.01680124 | 0.09147343 |
| blast | GGB28456_SGB41039 | acute_cyto_PC5 | 0.50862154 | 0.71017634 |
| vns | GGB28456_SGB41039 | acute_cyto_PC5 | 0.56759700 | 0.71017634 |
| blast | GGB28782_SGB41435 | acute_cyto_PC5 | 0.31549159 | 0.46558019 |
| vns | GGB28782_SGB41435 | acute_cyto_PC5 | 0.35916991 | 0.46558019 |
| blast | GGB28792_SGB41445 | acute_cyto_PC5 | 0.12377072 | 0.15762816 |
| vns | GGB28792_SGB41445 | acute_cyto_PC5 | 0.97085810 | 0.15762816 |
| blast | GGB28798_SGB41451 | acute_cyto_PC5 | 0.93183730 | 0.06760968 |
| vns | GGB28798_SGB41451 | acute_cyto_PC5 | 0.47155254 | 0.06760968 |
| blast | GGB28810_SGB41465 | acute_cyto_PC5 | 0.91952179 | 0.67569339 |
| vns | GGB28810_SGB41465 | acute_cyto_PC5 | 0.91510592 | 0.67569339 |
| blast | GGB28828_SGB41484 | acute_cyto_PC5 | 0.03645317 | 0.43348547 |
| vns | GGB28828_SGB41484 | acute_cyto_PC5 | 0.02961531 | 0.43348547 |
| blast | GGB28851_SGB41518 | acute_cyto_PC5 | 0.98324526 | 0.439669961 |
| vns | GGB28851_SGB41518 | acute_cyto_PC5 | 0.54824470 | 0.439669961 |
| blast | GGB28865_SGB41537 | acute_cyto_PC5 | 0.05183825 | 0.24258372 |
| vns | GGB28865_SGB41537 | acute_cyto_PC5 | 0.04383201 | 0.24258372 |
| blast | GGB28868_SGB41542 | acute_cyto_PC5 | 0.29242318 | 0.78280557 |
| vns | GGB28868_SGB41542 | acute_cyto_PC5 | 0.25432364 | 0.78280557 |
| blast | GGB28869_SGB41543 | acute_cyto_PC5 | 0.66207307 | 0.41080870 |
| vns | GGB28869_SGB41543 | acute_cyto_PC5 | 0.80660614 | 0.41080870 |
| blast | GGB28875_SGB41555 | acute_cyto_PC5 | 0.77664406 | 0.94164024 |
| vns | GGB28875_SGB41555 | acute_cyto_PC5 | 0.63407074 | 0.94164024 |
| blast | GGB28881_SGB41561 | acute_cyto_PC5 | 0.01189454 | 0.82782318 |
| vns | GGB28881_SGB41561 | acute_cyto_PC5 | 0.00198803 | 0.82782318 |
| blast | GGB28892_SGB41573 | acute_cyto_PC5 | 0.715153028 | 0.82175692 |
| vns | GGB28892_SGB41573 | acute_cyto_PC5 | 0.78470948 | 0.82175692 |
| blast | GGB28893_SGB41574 | acute_cyto_PC5 | 0.73922199 | 0.103508377 |
| vns | GGB28893_SGB41574 | acute_cyto_PC5 | 0.26424516 | 0.103508377 |
| blast | GGB28898_SGB41580 | acute_cyto_PC5 | 0.60192470 | 0.63580850 |
| vns | GGB28898_SGB41580 | acute_cyto_PC5 | 0.591718713 | 0.63580850 |

| | | | | |
|---|---|---|---|---|
| blast | GGB28901_SGB41592 | acute_cyto_PC5 | 0.072063586 | 0.741938061 |
| vns | GGB28901_SGB41592 | acute_cyto_PC5 | 0.152665868 | 0.741938061 |
| blast | GGB28904_SGB41597 | acute_cyto_PC5 | 0.225752520 | 0.630316852 |
| vns | GGB28904_SGB41597 | acute_cyto_PC5 | 0.085658495 | 0.630316852 |
| blast | GGB28909_SGB41602 | acute_cyto_PC5 | 0.218283007 | 0.780541405 |
| vns | GGB28909_SGB41602 | acute_cyto_PC5 | 0.145902665 | 0.780541405 |
| blast | GGB28924_SGB41621 | acute_cyto_PC5 | 0.611578184 | 0.026849550 |
| vns | GGB28924_SGB41621 | acute_cyto_PC5 | 0.456981189 | 0.026849550 |
| blast | GGB28927_SGB41625 | acute_cyto_PC5 | 0.001954483 | 0.791196367 |
| vns | GGB28927_SGB41625 | acute_cyto_PC5 | 0.000219425 | 0.791196367 |
| blast | GGB28934_SGB41635 | acute_cyto_PC5 | 0.216498853 | 0.410244249 |
| vns | GGB28934_SGB41635 | acute_cyto_PC5 | 0.224102539 | 0.410244249 |
| blast | GGB28949_SGB41655 | acute_cyto_PC5 | 0.159598366 | 0.943122496 |
| vns | GGB28949_SGB41655 | acute_cyto_PC5 | 0.744810754 | 0.943122496 |
| blast | GGB28951_SGB102295 | acute_cyto_PC5 | 0.718725508 | 0.775482481 |
| vns | GGB28951_SGB102295 | acute_cyto_PC5 | 0.298970783 | 0.775482481 |
| blast | GGB28951_SGB41658 | acute_cyto_PC5 | 0.752627843 | 0.437358540 |
| vns | GGB28951_SGB41658 | acute_cyto_PC5 | 0.024801176 | 0.437358540 |
| blast | GGB28954_SGB41662 | acute_cyto_PC5 | 0.794939032 | 0.416638560 |
| vns | GGB28954_SGB41662 | acute_cyto_PC5 | 0.841186745 | 0.416638560 |
| blast | GGB28960_SGB41669 | acute_cyto_PC5 | 0.536638861 | 0.046029003 |
| vns | GGB28960_SGB41669 | acute_cyto_PC5 | 0.043638773 | 0.046029003 |
| blast | GGB28964_SGB94886 | acute_cyto_PC5 | 0.207213028 | 0.244109888 |
| vns | GGB28964_SGB94886 | acute_cyto_PC5 | 0.400370147 | 0.244109888 |
| blast | GGB28967_SGB41678 | acute_cyto_PC5 | 0.848205066 | 0.24195294 |
| vns | GGB28967_SGB41678 | acute_cyto_PC5 | 0.820661965 | 0.24195294 |
| blast | GGB28996_SGB41712 | acute_cyto_PC5 | 0.429884728 | 0.405178638 |
| vns | GGB28996_SGB41712 | acute_cyto_PC5 | 0.377449394 | 0.405178638 |
| blast | GGB29531_SGB42317 | acute_cyto_PC5 | 0.279593444 | 0.734203830 |
| vns | GGB29531_SGB42317 | acute_cyto_PC5 | 0.999007775 | 0.734203830 |
| blast | GGB29685_SGB42494 | acute_cyto_PC5 | 0.004596881 | 0.276043602 |
| vns | GGB29685_SGB42494 | acute_cyto_PC5 | 0.748416768 | 0.276043602 |
| blast | GGB30141_SGB43066 | acute_cyto_PC5 | 0.614028140 | 0.627226942 |
| vns | GGB30141_SGB43066 | acute_cyto_PC5 | 0.664026053 | 0.627226942 |
| blast | GGB30145_SGB43072 | acute_cyto_PC5 | 0.294924035 | 0.139382822 |
| vns | GGB30145_SGB43072 | acute_cyto_PC5 | 0.367951830 | 0.139382822 |
| blast | GGB30286_SGB43248 | acute_cyto_PC5 | 0.766757581 | 0.298262478 |
| vns | GGB30286_SGB43248 | acute_cyto_PC5 | 0.750425732 | 0.298262478 |
| blast | GGB30300_SGB43264 | acute_cyto_PC5 | 0.213846948 | 0.194434705 |
| vns | GGB30300_SGB43264 | acute_cyto_PC5 | 0.213511417 | 0.194434705 |
| blast | GGB30303_SGB43268 | acute_cyto_PC5 | 0.795572751 | 0.329660960 |
| vns | GGB30303_SGB43268 | acute_cyto_PC5 | 0.660099923 | 0.329660960 |
| blast | GGB30450_SGB43507 | acute_cyto_PC5 | 0.027109400 | 0.113294912 |
| vns | GGB30450_SGB43507 | acute_cyto_PC5 | 0.186573367 | 0.113294912 |
| blast | GGB30453_SGB43513 | acute_cyto_PC5 | 0.315725451 | 0.227699424 |
| vns | GGB30453_SGB43513 | acute_cyto_PC5 | 0.310638137 | 0.227699424 |

| | | | | |
|---|---|---|---|---|
| blast | GGB30454_SGB43514 | acute_cyto_PC5 | 0.642888646 | 0.406581659 |
| vns | GGB30454_SGB43514 | acute_cyto_PC5 | 0.656930027 | 0.406581659 |
| blast | GGB30455_SGB43519 | acute_cyto_PC5 | 0.663798426 | 0.487432530 |
| vns | GGB30455_SGB43519 | acute_cyto_PC5 | 0.104495810 | 0.487432530 |
| blast | GGB30456_SGB43520 | acute_cyto_PC5 | 0.111196948 | 0.697909069 |
| vns | GGB30456_SGB43520 | acute_cyto_PC5 | 0.158921661 | 0.697909069 |
| blast | GGB30457_SGB63218 | acute_cyto_PC5 | 0.882164050 | 0.515375721 |
| vns | GGB30457_SGB63218 | acute_cyto_PC5 | 0.890757036 | 0.515375721 |
| blast | GGB30461_SGB43530 | acute_cyto_PC5 | 0.602468303 | 0.045227141 |
| vns | GGB30461_SGB43530 | acute_cyto_PC5 | 0.137401205 | 0.045227141 |
| blast | GGB30461_SGB43533 | acute_cyto_PC5 | 0.810767423 | 0.404855627 |
| vns | GGB30461_SGB43533 | acute_cyto_PC5 | 0.718693859 | 0.404855627 |
| blast | GGB30461_SGB63209 | acute_cyto_PC5 | 0.005914118 | 0.221120191 |
| vns | GGB30461_SGB63209 | acute_cyto_PC5 | 0.075178424 | 0.221120191 |
| blast | GGB30463_SGB43537 | acute_cyto_PC5 | 0.040768110 | 0.782901012 |
| vns | GGB30463_SGB43537 | acute_cyto_PC5 | 0.991960140 | 0.782901012 |
| blast | GGB30473_SGB43557 | acute_cyto_PC5 | 0.328491579 | 0.492090141 |
| vns | GGB30473_SGB43557 | acute_cyto_PC5 | 0.189853765 | 0.492090141 |
| blast | GGB30475_SGB63182 | acute_cyto_PC5 | 0.811833041 | 0.340376455 |
| vns | GGB30475_SGB63182 | acute_cyto_PC5 | 0.783129024 | 0.340376455 |
| blast | GGB30861_SGB44083 | acute_cyto_PC5 | 0.719401647 | 0.195858932 |
| vns | GGB30861_SGB44083 | acute_cyto_PC5 | 0.981045510 | 0.195858932 |
| blast | GGB31312_SGB44628 | acute_cyto_PC5 | 0.831039140 | 0.875844946 |
| vns | GGB31312_SGB44628 | acute_cyto_PC5 | 0.833457038 | 0.875844946 |
| blast | GGB31438_SGB44768 | acute_cyto_PC5 | 0.481213900 | 0.403740111 |
| vns | GGB31438_SGB44768 | acute_cyto_PC5 | 0.483694757 | 0.403740111 |
| blast | GGB3171_SGB4185 | acute_cyto_PC5 | 0.066342246 | 0.535205558 |
| vns | GGB3171_SGB4185 | acute_cyto_PC5 | 0.057850794 | 0.535205558 |
| blast | GGB31762_SGB45125 | acute_cyto_PC5 | 0.442889056 | 0.230173648 |
| vns | GGB31762_SGB45125 | acute_cyto_PC5 | 0.437364142 | 0.230173648 |
| blast | GGB31823_SGB45199 | acute_cyto_PC5 | 0.630175203 | 0.230343618 |
| vns | GGB31823_SGB45199 | acute_cyto_PC5 | 0.537240753 | 0.230343618 |
| blast | GGB31838_SGB45216 | acute_cyto_PC5 | 0.684403164 | 0.233798756 |
| vns | GGB31838_SGB45216 | acute_cyto_PC5 | 0.997715556 | 0.233798756 |
| blast | GGB31841_SGB65084 | acute_cyto_PC5 | 0.371737454 | 0.863678183 |
| vns | GGB31841_SGB65084 | acute_cyto_PC5 | 0.443951934 | 0.863678183 |
| blast | GGB32371_SGB41694 | acute_cyto_PC5 | 0.242018055 | 0.374507152 |
| vns | GGB32371_SGB41694 | acute_cyto_PC5 | 0.400159574 | 0.374507152 |
| blast | GGB3793_SGB5158 | acute_cyto_PC5 | 0.884993491 | 0.885772645 |
| vns | GGB3793_SGB5158 | acute_cyto_PC5 | 0.847239709 | 0.885772645 |
| blast | GGB42601_SGB59797 | acute_cyto_PC5 | 0.401892013 | 0.083888371 |
| vns | GGB42601_SGB59797 | acute_cyto_PC5 | 0.627566595 | 0.083888371 |
| blast | GGB45514_SGB63186 | acute_cyto_PC5 | 0.955364765 | 0.896129394 |
| vns | GGB45514_SGB63186 | acute_cyto_PC5 | 0.621527882 | 0.896129394 |
| blast | GGB45564_SGB63259 | acute_cyto_PC5 | 0.415272784 | 0.339244118 |
| vns | GGB45564_SGB63259 | acute_cyto_PC5 | 0.380878696 | 0.339244118 |

| | | | | |
|---|---|---|---|---|
| blast | GGB45624_SGB63337 | acute_cyto_PC5 | 0.6203892728 | 0.1974501358 |
| vns | GGB45624_SGB63337 | acute_cyto_PC5 | 0.3851987949 | 0.1974501358 |
| blast | GGB45656_SGB63370 | acute_cyto_PC5 | 0.4047054831 | 0.9191067024 |
| vns | GGB45656_SGB63370 | acute_cyto_PC5 | 0.1636435534 | 0.9191067024 |
| blast | GGB47127_SGB65054 | acute_cyto_PC5 | 0.4564936795 | 0.2119566975 |
| vns | GGB47127_SGB65054 | acute_cyto_PC5 | 0.82583129 | 0.2119566975 |
| blast | GGB74395_SGB43523 | acute_cyto_PC5 | 0.8024104382 | 0.9457662025 |
| vns | GGB74395_SGB43523 | acute_cyto_PC5 | 0.835585633 | 0.9457662025 |
| blast | GGB75053_SGB43494 | acute_cyto_PC5 | 0.1773274016 | 0.6783216138 |
| vns | GGB75053_SGB43494 | acute_cyto_PC5 | 0.0504104434 | 0.6783216138 |
| blast | GGB75109_SGB102238 | acute_cyto_PC5 | 0.5997929865 | 0.3834703182 |
| vns | GGB75109_SGB102238 | acute_cyto_PC5 | 0.5108644817 | 0.3834703182 |
| blast | Lachnospiraceae_bacterium | acute_cyto_PC5 | 0.0239259038 | 0.5455747303 |
| vns | Lachnospiraceae_bacterium | acute_cyto_PC5 | 0.9174922655 | 0.5455747303 |
| blast | Lachnospiraceae_bacterium_A2 | acute_cyto_PC5 | 0.7241878075 | 0.2546307738 |
| vns | Lachnospiraceae_bacterium_A2 | acute_cyto_PC5 | 0.0838607394 | 0.2546307738 |
| blast | Lachnospiraceae_unclassified_SGB41414 | acute_cyto_PC5 | 0.7190795008 | 0.4379013773 |
| vns | Lachnospiraceae_unclassified_SGB41414 | acute_cyto_PC5 | 0.5321281982 | 0.4379013773 |
| blast | Lachnospiraceae_unclassified_SGB41424 | acute_cyto_PC5 | 0.2169304138 | 0.1964200068 |
| vns | Lachnospiraceae_unclassified_SGB41424 | acute_cyto_PC5 | 0.0220188292 | 0.1964200068 |
| blast | Lachnospiraceae_unclassified_SGB94868 | acute_cyto_PC5 | 0.2076335048 | 0.5539940878 |
| vns | Lachnospiraceae_unclassified_SGB94868 | acute_cyto_PC5 | 0.7545629335 | 0.5539940878 |
| blast | Lactobacillus_johnsonii | acute_cyto_PC5 | 0.0027483659 | 0.9981682181 |
| vns | Lactobacillus_johnsonii | acute_cyto_PC5 | 0.8322162424 | 0.9981682181 |
| blast | Leptogranulimonas_caecicola | acute_cyto_PC5 | 0.8745779805 | 0.0758785008 |
| vns | Leptogranulimonas_caecicola | acute_cyto_PC5 | 0.0144750878 | 0.0758785008 |
| blast | Muribaculaceae_bacterium | acute_cyto_PC5 | 0.0018501989 | 0.3528551549 |
| vns | Muribaculaceae_bacterium | acute_cyto_PC5 | 0.8835325068 | 0.3528551549 |
| blast | Neglectibacter_sp_X4 | acute_cyto_PC5 | 0.3944230674 | 0.8108861076 |
| vns | Neglectibacter_sp_X4 | acute_cyto_PC5 | 0.463430059 | 0.8108861076 |
| blast | Oscillibacter_SGB43496 | acute_cyto_PC5 | 0.2205447256 | 0.1986746723 |
| vns | Oscillibacter_SGB43496 | acute_cyto_PC5 | 0.34795292 | 0.1986746723 |
| blast | Oscillospiraceae_bacterium | acute_cyto_PC5 | 0.4927836612 | 0.9036740757 |
| vns | Oscillospiraceae_bacterium | acute_cyto_PC5 | 0.251973413 | 0.9036740757 |
| blast | Oscillospiraceae_unclassified_SGB43502 | acute_cyto_PC5 | 0.0172712013 | 0.3164374223 |
| vns | Oscillospiraceae_unclassified_SGB43502 | acute_cyto_PC5 | 0.1123454974 | 0.3164374223 |
| blast | Oscillospiraceae_unclassified_SGB43505 | acute_cyto_PC5 | 0.3742999083 | 0.5180080258 |
| vns | Oscillospiraceae_unclassified_SGB43505 | acute_cyto_PC5 | 0.6182049908 | 0.5180080258 |
| blast | Oscillospiraceae_unclassified_SGB94989 | acute_cyto_PC5 | 0.9533689691 | 0.0567860193 |
| vns | Oscillospiraceae_unclassified_SGB94989 | acute_cyto_PC5 | 0.6236152007 | 0.0567860193 |
| blast | Parasutterella_excrementihominis | acute_cyto_PC5 | 0.0234193782 | 0.9873535037 |
| vns | Parasutterella_excrementihominis | acute_cyto_PC5 | 0.5041571561 | 0.9873535037 |
| blast | Richness (# observed features) | acute_cyto_PC5 | 0.0137778742 | 0.2630314661 |
| vns | Richness (# observed features) | acute_cyto_PC5 | 0.9582478381 | 0.2630314661 |
| blast | Schaedlerella_arabinosiphila | acute_cyto_PC5 | 0.114481983 | 0.3717058445 |
| vns | Schaedlerella_arabinosiphila | acute_cyto_PC5 | 0.3863019645 | 0.3717058445 |

| | | | | |
|---|---|---|---|---|
| blast | Shannon Index | acute_cyto_PC5 | 0.00647307[illegible] | 0.16842587[illegible] |
| vns | Shannon Index | acute_cyto_PC5 | 0.83599244[illegible] | 0.16842587[illegible] |
| blast | Turicibacter_sp_1E2 | acute_cyto_PC5 | 0.00959413[illegible] | 0.46435169[illegible] |
| vns | Turicibacter_sp_1E2 | acute_cyto_PC5 | 0.183926772 | 0.46435169[illegible] |

| p_composite | dact_stat | dact_thr_10 | dact_thr_15 | dact_thr_20 | signif_at |
|---|---|---|---|---|---|
| 0.284979380 | 0.290673579 | 0.001892477 | 0.002328539 | 0.002949229 | |
| 0.207157594 | 0.182429176 | 0.000232278 | 0.000313216 | 0.000521262 | |
| 0.654198757 | 0.601665270 | 0.001892477 | 0.002328539 | 0.002949229 | |
| 0.338901524 | 0.334917577 | 0.000232278 | 0.000313216 | 0.000521262 | |
| 0.464672568 | 0.468967998 | 0.001892477 | 0.002328539 | 0.002949229 | |
| 0.510656370 | 0.462185514 | 0.000232278 | 0.000313216 | 0.000521262 | |
| 0.911112899 | 0.874869378 | 0.001892477 | 0.002328539 | 0.002949229 | |
| 0.846440848 | 0.841997280 | 0.000232278 | 0.000313216 | 0.000521262 | |
| 0.971180381 | 0.964241480 | 0.001892477 | 0.002328539 | 0.002949229 | |
| 0.937376209 | 0.954830407 | 0.000232278 | 0.000313216 | 0.000521262 | |
| 0.138659486 | 0.124112140 | 0.001892477 | 0.002328539 | 0.002949229 | |
| 0.233901430 | 0.202502896 | 0.000232278 | 0.000313216 | 0.000521262 | |
| 0.476823640 | 0.473300385 | 0.001892477 | 0.002328539 | 0.002949229 | |
| 0.382106246 | 0.367972319 | 0.000232278 | 0.000313216 | 0.000521262 | |
| 0.080019612 | 0.072316359 | 0.001892477 | 0.002328539 | 0.002949229 | |
| 0.128347795 | 0.116587603 | 0.000232278 | 0.000313216 | 0.000521262 | |
| 0.674500717 | 0.619001935 | 0.001892477 | 0.002328539 | 0.002949229 | |
| 0.650898816 | 0.633703810 | 0.000232278 | 0.000313216 | 0.000521262 | |
| 0.816116443 | 0.769585116 | 0.001892477 | 0.002328539 | 0.002949229 | |
| 0.742927949 | 0.711394795 | 0.000232278 | 0.000313216 | 0.000521262 | |
| 0.821437986 | 0.770463336 | 0.001892477 | 0.002328539 | 0.002949229 | |
| 0.75021983 | 0.712782320 | 0.000232278 | 0.000313216 | 0.000521262 | |
| 0.015960153 | 0.019629421 | 0.001892477 | 0.002328539 | 0.002949229 | |
| 0.038851975 | 0.020035707 | 0.000232278 | 0.000313216 | 0.000521262 | |
| 0.514351344 | 0.502126235 | 0.001892477 | 0.002328539 | 0.002949229 | |
| 0.424465542 | 0.399538993 | 0.000232278 | 0.000313216 | 0.000521262 | |
| 0.487885383 | 0.484550087 | 0.001892477 | 0.002328539 | 0.002949229 | |
| 0.386981644 | 0.380267448 | 0.000232278 | 0.000313216 | 0.000521262 | |
| 0.437690012 | 0.450200465 | 0.001892477 | 0.002328539 | 0.002949229 | |
| 0.348666333 | 0.343328754 | 0.000232278 | 0.000313216 | 0.000521262 | |
| 0.432723270 | 0.447345346 | 0.001892477 | 0.002328539 | 0.002949229 | |
| 0.342808590 | 0.339953856 | 0.000232278 | 0.000313216 | 0.000521262 | |
| 0.573531849 | 0.538092321 | 0.001892477 | 0.002328539 | 0.002949229 | |
| 0.478985942 | 0.439451302 | 0.000232278 | 0.000313216 | 0.000521262 | |
| 0.157897499 | 0.145886115 | 0.001892477 | 0.002328539 | 0.002949229 | |
| 0.050081281 | 0.026885448 | 0.000232278 | 0.000313216 | 0.000521262 | |
| 0.392382453 | 0.420486112 | 0.001892477 | 0.002328539 | 0.002949229 | |
| 0.312193875 | 0.311451228 | 0.000232278 | 0.000313216 | 0.000521262 | |
| 0.670647244 | 0.614409037 | 0.001892477 | 0.002328539 | 0.002949229 | |
| 0.552782245 | 0.526444646 | 0.000232278 | 0.000313216 | 0.000521262 | |
| 0.017499687 | 0.021222866 | 0.001892477 | 0.002328539 | 0.002949229 | |
| 0.054507717 | 0.032366654 | 0.000232278 | 0.000313216 | 0.000521262 | |
| 0.840550310 | 0.787764304 | 0.001892477 | 0.002328539 | 0.002949229 | |
| 0.915495678 | 0.941091732 | 0.000232278 | 0.000313216 | 0.000521262 | |
| 0.904127486 | 0.869973683 | 0.001892477 | 0.002328539 | 0.002949229 | |
| 0.833902834 | 0.835990199 | 0.000232278 | 0.000313216 | 0.000521262 | |

| | | | | |
|---|---|---|---|---|
| 0.725431196 | 0.66352665 | 0.00189247 | 0.00232853 | 0.00294922 |
| 0.612503649 | 0.58404037 | 0.00023227 | 0.00031321 | 0.00052126 |
| 0.56877551 | 0.535823971 | 0.00189247 | 0.00232853 | 0.00294922 |
| 0.46818311 | 0.43652726 | 0.00023227 | 0.00031321 | 0.00052126 |
| 0.65638112 | 0.60404511 | 0.00189247 | 0.00232853 | 0.00294922 |
| 0.54098394 | 0.51494714 | 0.00023227 | 0.00031321 | 0.00052126 |
| 0.788816833 | 0.74939526 | 0.00189247 | 0.00232853 | 0.00294922 |
| 0.68789830 | 0.687040487 | 0.00023227 | 0.00031321 | 0.00052126 |
| 0.01543921 | 0.01922794 | 0.00189247 | 0.00232853 | 0.00294922 |
| 0.00510555 | 0.00176201 | 0.00023227 | 0.00031321 | 0.00052126 |
| 0.07140144 | 0.06763066 | 0.00189247 | 0.00232853 | 0.00294922 |
| 0.15996793 | 0.13772002 | 0.00023227 | 0.00031321 | 0.00052126 |
| 0.06749067 | 0.06511210 | 0.00189247 | 0.00232853 | 0.00294922 |
| 0.03047970 | 0.01613776 | 0.00023227 | 0.00031321 | 0.00052126 |
| 0.69274956 | 0.62992786 | 0.00189247 | 0.00232853 | 0.00294922 |
| 0.56748153 | 0.54444618 | 0.00023227 | 0.00031321 | 0.00052126 |
| 0.99055610 | 0.98462123 | 0.00189247 | 0.00232853 | 0.00294922 |
| 0.93243941 | 0.94764004 | 0.00023227 | 0.00031321 | 0.00052126 |
| 0.61012718 | 0.55986216 | 0.00189247 | 0.00232853 | 0.00294922 |
| 0.51486373 | 0.46401045 | 0.00023227 | 0.00031321 | 0.00052126 |
| 0.86186860 | 0.81100860 | 0.00189247 | 0.00232853 | 0.00294922 |
| 0.78607088 | 0.76251024 | 0.00023227 | 0.00031321 | 0.00052126 |
| 0.25954003 | 0.25884586 | 0.00189247 | 0.00232853 | 0.00294922 |
| 0.32738342 | 0.32299184 | 0.00023227 | 0.00031321 | 0.00052126 |
| 0.30822772 | 0.311068739 | 0.00189247 | 0.00232853 | 0.00294922 |
| 0.22753663 | 0.20164437 | 0.00023227 | 0.00031321 | 0.00052126 |
| 0.67820935 | 0.62527524 | 0.00189247 | 0.00232853 | 0.00294922 |
| 0.92250664 | 0.94427002 | 0.00023227 | 0.00031321 | 0.00052126 |
| 0.81366715 | 0.76895593 | 0.00189247 | 0.00232853 | 0.00294922 |
| 0.45235593 | 0.414117418 | 0.00023227 | 0.00031321 | 0.00052126 |
| 0.95587238 | 0.94063137 | 0.00189247 | 0.00232853 | 0.00294922 |
| 0.90819769 | 0.92480393 | 0.00023227 | 0.00031321 | 0.00052126 |
| 0.07472239 | 0.07015518 | 0.00189247 | 0.00232853 | 0.00294922 |
| 0.03373590 | 0.01838675 | 0.00023227 | 0.00031321 | 0.00052126 |
| 0.75136183 | 0.69919519 | 0.00189247 | 0.00232853 | 0.00294922 |
| 0.30495815 | 0.30148301 | 0.00023227 | 0.00031321 | 0.00052126 |
| 0.02024514 | 0.02393833 | 0.00189247 | 0.00232853 | 0.00294922 |
| 0.00742684 | 0.00263292 | 0.00023227 | 0.00031321 | 0.00052126 |
| 0.68672225 | 0.62759456 | 0.00189247 | 0.00232853 | 0.00294922 |
| 0.56334668 | 0.54180981 | 0.00023227 | 0.00031321 | 0.00052126 |
| 0.35802088 | 0.383016249 | 0.00189247 | 0.00232853 | 0.00294922 |
| 0.65486385 | 0.65173245 | 0.00023227 | 0.00031321 | 0.00052126 |
| 0.94659455 | 0.93805397 | 0.00189247 | 0.00232853 | 0.00294922 |
| 0.88991355 | 0.92142812 | 0.00023227 | 0.00031321 | 0.00052126 |
| 0.361018871 | 0.38831013 | 0.00189247 | 0.00232853 | 0.00294922 |
| 0.27951615 | 0.27803485 | 0.00023227 | 0.00031321 | 0.00052126 |

| | | | | |
|---|---|---|---|---|
| 0.391481730 | 0.419942571 | 0.001892477 | 0.002328539 | 0.002949229 |
| 0.628022645 | 0.616857639 | 0.000232278 | 0.000313216 | 0.000521262 |
| 0.318987754 | 0.321271904 | 0.001892477 | 0.002328539 | 0.002949229 |
| 0.097722027 | 0.070127891 | 0.000232278 | 0.000313216 | 0.000521262 |
| 0.599487188 | 0.547478708 | 0.001892477 | 0.002328539 | 0.002949229 |
| 0.504312788 | 0.449834769 | 0.000232278 | 0.000313216 | 0.000521262 |
| 0.090793707 | 0.080862254 | 0.001892477 | 0.002328539 | 0.002949229 |
| 0.046667440 | 0.023612183 | 0.000232278 | 0.000313216 | 0.000521262 |
| 0.132977220 | 0.120841522 | 0.001892477 | 0.002328539 | 0.002949229 |
| 0.070007841 | 0.043424416 | 0.000232278 | 0.000313216 | 0.000521262 |
| 0.037470659 | 0.038480710 | 0.001892477 | 0.002328539 | 0.002949229 |
| 0.042106386 | 0.021460021 | 0.000232278 | 0.000313216 | 0.000521262 |
| 0.231519125 | 0.231756307 | 0.001892477 | 0.002328539 | 0.002949229 |
| 0.236765527 | 0.209341329 | 0.000232278 | 0.000313216 | 0.000521262 |
| 0.237738517 | 0.237260028 | 0.001892477 | 0.002328539 | 0.002949229 |
| 0.149467714 | 0.134469861 | 0.000232278 | 0.000313216 | 0.000521262 |
| 0.387269449 | 0.417285791 | 0.001892477 | 0.002328539 | 0.002949229 |
| 0.308398697 | 0.308047959 | 0.000232278 | 0.000313216 | 0.000521262 |
| 0.204232876 | 0.196393182 | 0.001892477 | 0.002328539 | 0.002949229 |
| 0.582646414 | 0.555804265 | 0.000232278 | 0.000313216 | 0.000521262 |
| 0.571472326 | 0.536903936 | 0.001892477 | 0.002328539 | 0.002949229 |
| 0.318305662 | 0.315254016 | 0.000232278 | 0.000313216 | 0.000521262 |
| 0.404919517 | 0.427598850 | 0.001892477 | 0.002328539 | 0.002949229 |
| 0.089590317 | 0.063954216 | 0.000232278 | 0.000313216 | 0.000521262 |
| 0.759575969 | 0.703598930 | 0.001892477 | 0.002328539 | 0.002949229 |
| 0.741653265 | 0.709227725 | 0.000232278 | 0.000313216 | 0.000521262 |
| 0.269172670 | 0.270178878 | 0.001892477 | 0.002328539 | 0.002949229 |
| 0.085361360 | 0.061160352 | 0.000232278 | 0.000313216 | 0.000521262 |
| 0.767012807 | 0.720397246 | 0.001892477 | 0.002328539 | 0.002949229 |
| 0.661278165 | 0.652117234 | 0.000232278 | 0.000313216 | 0.000521262 |
| 0.738966668 | 0.683602533 | 0.001892477 | 0.002328539 | 0.002949229 |
| 0.677760161 | 0.674943606 | 0.000232278 | 0.000313216 | 0.000521262 |
| 0.713157846 | 0.643147161 | 0.001892477 | 0.002328539 | 0.002949229 |
| 0.589467317 | 0.559969101 | 0.000232278 | 0.000313216 | 0.000521262 |
| 0.629598925 | 0.573793342 | 0.001892477 | 0.002328539 | 0.002949229 |
| 0.994576308 | 0.999707478 | 0.000232278 | 0.000313216 | 0.000521262 |
| 0.796049155 | 0.756201320 | 0.001892477 | 0.002328539 | 0.002949229 |
| 0.711209412 | 0.696055260 | 0.000232278 | 0.000313216 | 0.000521262 |
| 0.381417723 | 0.409589542 | 0.001892477 | 0.002328539 | 0.002949229 |
| 0.491591161 | 0.442043533 | 0.000232278 | 0.000313216 | 0.000521262 |
| 0.121103445 | 0.108889061 | 0.001892477 | 0.002328539 | 0.002949229 |
| 0.153734526 | 0.135892112 | 0.000232278 | 0.000313216 | 0.000521262 |
| 0.553478036 | 0.525005017 | 0.001892477 | 0.002328539 | 0.002949229 |
| 0.602615565 | 0.564333845 | 0.000232278 | 0.000313216 | 0.000521262 |
| 0.136878256 | 0.123496746 | 0.001892477 | 0.002328539 | 0.002949229 |
| 0.075929012 | 0.047950285 | 0.000232278 | 0.000313216 | 0.000521262 |

| | | | | |
|---|---|---|---|---|
| 0.563907406 | 0.533405853 | 0.001892477 | 0.002328539 | 0.002949229 |
| 0.472722662 | 0.436799212 | 0.000232278 | 0.000313216 | 0.000521262 |
| 0.290886039 | 0.297496208 | 0.001892477 | 0.002328539 | 0.002949229 |
| 0.210686438 | 0.188918422 | 0.000232278 | 0.000313216 | 0.000521262 |
| 0.886628709 | 0.845135218 | 0.001892477 | 0.002328539 | 0.002949229 |
| 0.818465689 | 0.804755307 | 0.000232278 | 0.000313216 | 0.000521262 |
| 0.881004150 | 0.839694905 | 0.001892477 | 0.002328539 | 0.002949229 |
| 0.813853797 | 0.797924236 | 0.000232278 | 0.000313216 | 0.000521262 |
| 0.553417194 | 0.524903513 | 0.001892477 | 0.002328539 | 0.002949229 |
| 0.416912333 | 0.396064740 | 0.000232278 | 0.000313216 | 0.000521262 |
| 0.265340556 | 0.267592923 | 0.001892477 | 0.002328539 | 0.002949229 |
| 0.194435428 | 0.161293811 | 0.000232278 | 0.000313216 | 0.000521262 |
| 0.773416489 | 0.724752777 | 0.001892477 | 0.002328539 | 0.002949229 |
| 0.806401368 | 0.794994797 | 0.000232278 | 0.000313216 | 0.000521262 |
| 0.459654237 | 0.462878522 | 0.001892477 | 0.002328539 | 0.002949229 |
| 0.350534795 | 0.344601731 | 0.000232278 | 0.000313216 | 0.000521262 |
| 0.837522512 | 0.779209415 | 0.001892477 | 0.002328539 | 0.002949229 |
| 0.769546063 | 0.723128183 | 0.000232278 | 0.000313216 | 0.000521262 |
| 0.172180381 | 0.159315626 | 0.001892477 | 0.002328539 | 0.002949229 |
| 0.100684831 | 0.072381704 | 0.000232278 | 0.000313216 | 0.000521262 |
| 0.982227077 | 0.972916780 | 0.001892477 | 0.002328539 | 0.002949229 |
| 0.973126203 | 0.985959334 | 0.000232278 | 0.000313216 | 0.000521262 |
| 0.934413459 | 0.927858652 | 0.001892477 | 0.002328539 | 0.002949229 |
| 0.870557428 | 0.908608612 | 0.000232278 | 0.000313216 | 0.000521262 |
| 0.639610606 | 0.585485681 | 0.001892477 | 0.002328539 | 0.002949229 |
| 0.623013437 | 0.614565873 | 0.000232278 | 0.000313216 | 0.000521262 |
| 0.320962505 | 0.324791362 | 0.001892477 | 0.002328539 | 0.002949229 |
| 0.949384063 | 0.963513578 | 0.000232278 | 0.000313216 | 0.000521262 |
| 0.719810682 | 0.652117951 | 0.001892477 | 0.002328539 | 0.002949229 |
| 0.711740718 | 0.696086204 | 0.000232278 | 0.000313216 | 0.000521262 |
| 0.144102321 | 0.130160716 | 0.001892477 | 0.002328539 | 0.002949229 |
| 0.253341680 | 0.234446633 | 0.000232278 | 0.000313216 | 0.000521262 |
| 0.078473927 | 0.071428664 | 0.001892477 | 0.002328539 | 0.002949229 |
| 0.036495119 | 0.018985153 | 0.000232278 | 0.000313216 | 0.000521262 |
| 0.650280269 | 0.598502018 | 0.001892477 | 0.002328539 | 0.002949229 |
| 0.534046057 | 0.508069293 | 0.000232278 | 0.000313216 | 0.000521262 |
| 0.593515976 | 0.546438080 | 0.001892477 | 0.002328539 | 0.002949229 |
| 0.498214074 | 0.448575122 | 0.000232278 | 0.000313216 | 0.000521262 |
| 0.898740000 | 0.854766031 | 0.001892477 | 0.002328539 | 0.002949229 |
| 0.988924383 | 0.997336868 | 0.000232278 | 0.000313216 | 0.000521262 |
| 0.116105036 | 0.103308670 | 0.001892477 | 0.002328539 | 0.002949229 |
| 0.217385910 | 0.197596144 | 0.000232278 | 0.000313216 | 0.000521262 |
| 0.779007018 | 0.736894620 | 0.001892477 | 0.002328539 | 0.002949229 |
| 0.668967030 | 0.671989358 | 0.000232278 | 0.000313216 | 0.000521262 |
| 0.764720495 | 0.718607230 | 0.001892477 | 0.002328539 | 0.002949229 |
| 0.761683655 | 0.719298223 | 0.000232278 | 0.000313216 | 0.000521262 |

| | | | | |
|---|---|---|---|---|
| 0.260951683 | 0.261348072 | 0.001892477 | 0.002328539 | 0.002949229 |
| 0.408734417 | 0.394852630 | 0.000232278 | 0.000313216 | 0.000521262 |
| 0.840215773 | 0.787132081 | 0.001892477 | 0.002328539 | 0.002949229 |
| 0.415979852 | 0.395627644 | 0.000232278 | 0.000313216 | 0.000521262 |
| 0.926309491 | 0.916617735 | 0.001892477 | 0.002328539 | 0.002949229 |
| 0.864525231 | 0.894448897 | 0.000232278 | 0.000313216 | 0.000521262 |
| 0.262221349 | 0.263600110 | 0.001892477 | 0.002328539 | 0.002949229 |
| 0.177189504 | 0.148873911 | 0.000232278 | 0.000313216 | 0.000521262 |
| 0.245782773 | 0.245986504 | 0.001892477 | 0.002328539 | 0.002949229 |
| 0.138775690 | 0.121494318 | 0.000232278 | 0.000313216 | 0.000521262 |
| 0.452876495 | 0.453655558 | 0.001892477 | 0.002328539 | 0.002949229 |
| 0.684193146 | 0.683410955 | 0.000232278 | 0.000313216 | 0.000521262 |
| 0.366829693 | 0.391061040 | 0.001892477 | 0.002328539 | 0.002949229 |
| 0.714567100 | 0.699112189 | 0.000232278 | 0.000313216 | 0.000521262 |
| 0.220239212 | 0.219828347 | 0.001892477 | 0.002328539 | 0.002949229 |
| 0.131325834 | 0.119581287 | 0.000232278 | 0.000313216 | 0.000521262 |
| 0.257644954 | 0.255531629 | 0.001892477 | 0.002328539 | 0.002949229 |
| 0.275970317 | 0.261617290 | 0.000232278 | 0.000313216 | 0.000521262 |
| 0.742701027 | 0.692933846 | 0.001892477 | 0.002328539 | 0.002949229 |
| 0.860467105 | 0.843495820 | 0.000232278 | 0.000313216 | 0.000521262 |
| 0.631087016 | 0.573960084 | 0.001892477 | 0.002328539 | 0.002949229 |
| 0.429736793 | 0.402954094 | 0.000232278 | 0.000313216 | 0.000521262 |
| 0.585044281 | 0.541066897 | 0.001892477 | 0.002328539 | 0.002949229 |
| 0.334421081 | 0.325361691 | 0.000232278 | 0.000313216 | 0.000521262 |
| 0.037865227 | 0.038893563 | 0.001892477 | 0.002328539 | 0.002949229 |
| 0.002690811 | 0.000845396 | 0.000232278 | 0.000313216 | 0.000521262 |
| 0.975167082 | 0.966369258 | 0.001892477 | 0.002328539 | 0.002949229 |
| 0.946863753 | 0.958363499 | 0.000232278 | 0.000313216 | 0.000521262 |
| 0.405946274 | 0.429416895 | 0.001892477 | 0.002328539 | 0.002949229 |
| 0.697806034 | 0.693981981 | 0.000232278 | 0.000313216 | 0.000521262 |
| 0.495448499 | 0.489220642 | 0.001892477 | 0.002328539 | 0.002949229 |
| 0.040344583 | 0.020511971 | 0.000232278 | 0.000313216 | 0.000521262 |
| 0.990739122 | 0.990678263 | 0.001892477 | 0.002328539 | 0.002949229 |
| 0.980528694 | 0.989900365 | 0.000232278 | 0.000313216 | 0.000521262 |
| 0.370871838 | 0.392419342 | 0.001892477 | 0.002328539 | 0.002949229 |
| 0.287779566 | 0.282318386 | 0.000232278 | 0.000313216 | 0.000521262 |
| 0.073975006 | 0.069776983 | 0.001892477 | 0.002328539 | 0.002949229 |
| 0.139120623 | 0.121514889 | 0.000232278 | 0.000313216 | 0.000521262 |
| 0.297702627 | 0.303582614 | 0.001892477 | 0.002328539 | 0.002949229 |
| 0.167692584 | 0.143822578 | 0.000232278 | 0.000313216 | 0.000521262 |
| 0.614756459 | 0.562030043 | 0.001892477 | 0.002328539 | 0.002949229 |
| 0.518134628 | 0.466485471 | 0.000232278 | 0.000313216 | 0.000521262 |
| 0.092809015 | 0.082143371 | 0.001892477 | 0.002328539 | 0.002949229 |
| 0.389920018 | 0.382804470 | 0.000232278 | 0.000313216 | 0.000521262 |
| 0.639937021 | 0.586101887 | 0.001892477 | 0.002328539 | 0.002949229 |
| 0.401772259 | 0.389699977 | 0.000232278 | 0.000313216 | 0.000521262 |

| | | | | |
|---|---|---|---|---|
| 0.042092110 | 0.042375719 | 0.001892477 | 0.002328539 | 0.002949229 |
| 0.271228990 | 0.254764634 | 0.000232278 | 0.000313216 | 0.000521262 |
| 0.852150131 | 0.802539999 | 0.001892477 | 0.002328539 | 0.002949229 |
| 0.883936101 | 0.920063874 | 0.000232278 | 0.000313216 | 0.000521262 |
| 0.700607219 | 0.634637713 | 0.001892477 | 0.002328539 | 0.002949229 |
| 0.576923093 | 0.550377802 | 0.000232278 | 0.000313216 | 0.000521262 |
| 0.805319791 | 0.763935037 | 0.001892477 | 0.002328539 | 0.002949229 |
| 0.734960763 | 0.705562396 | 0.000232278 | 0.000313216 | 0.000521262 |
| 0.011311770 | 0.015473403 | 0.001892477 | 0.002328539 | 0.002949229 |
| 0.059873799 | 0.034046383 | 0.000232278 | 0.000313216 | 0.000521262 |
| 0.649950804 | 0.606636208 | 0.001148170 | 0.001551604 | 0.001983832 |
| 0.554829221 | 0.545849884 | 0.000567631 | 0.000840464 | 0.001143763 |
| 0.636189341 | 0.594922217 | 0.001148170 | 0.001551604 | 0.001983832 |
| 0.331050627 | 0.321814006 | 0.000567631 | 0.000840464 | 0.001143763 |
| 0.926373129 | 0.872260258 | 0.001148170 | 0.001551604 | 0.001983832 |
| 0.875061618 | 0.849755771 | 0.000567631 | 0.000840464 | 0.001143763 |
| 0.431519475 | 0.436361043 | 0.001148170 | 0.001551604 | 0.001983832 |
| 0.368335404 | 0.361653531 | 0.000567631 | 0.000840464 | 0.001143763 |
| 0.544102756 | 0.507729000 | 0.001148170 | 0.001551604 | 0.001983832 |
| 0.366184468 | 0.356501166 | 0.000567631 | 0.000840464 | 0.001143763 |
| 0.165906835 | 0.166510835 | 0.001148170 | 0.001551604 | 0.001983832 |
| 0.227991955 | 0.208615435 | 0.000567631 | 0.000840464 | 0.001143763 |
| 0.561207975 | 0.524977351 | 0.001148170 | 0.001551604 | 0.001983832 |
| 0.493177366 | 0.456409906 | 0.000567631 | 0.000840464 | 0.001143763 |
| 0.216069291 | 0.213556638 | 0.001148170 | 0.001551604 | 0.001983832 |
| 0.158934819 | 0.142853204 | 0.000567631 | 0.000840464 | 0.001143763 |
| 0.421824006 | 0.421672767 | 0.001148170 | 0.001551604 | 0.001983832 |
| 0.632657845 | 0.600960439 | 0.000567631 | 0.000840464 | 0.001143763 |
| 0.757974361 | 0.682813672 | 0.001148170 | 0.001551604 | 0.001983832 |
| 0.660447719 | 0.630992479 | 0.000567631 | 0.000840464 | 0.001143763 |
| 0.547479309 | 0.512601026 | 0.001148170 | 0.001551604 | 0.001983832 |
| 0.480547801 | 0.442916665 | 0.000567631 | 0.000840464 | 0.001143763 |
| 0.033891787 | 0.042147235 | 0.001148170 | 0.001551604 | 0.001983832 |
| 0.025190240 | 0.022669393 | 0.000567631 | 0.000840464 | 0.001143763 |
| 0.855005156 | 0.815749868 | 0.001148170 | 0.001551604 | 0.001983832 |
| 0.794123153 | 0.783676653 | 0.000567631 | 0.000840464 | 0.001143763 |
| 0.973068295 | 0.949686698 | 0.001148170 | 0.001551604 | 0.001983832 |
| 0.952628903 | 0.940558536 | 0.000567631 | 0.000840464 | 0.001143763 |
| 0.622534483 | 0.580940927 | 0.001148170 | 0.001551604 | 0.001983832 |
| 0.532591665 | 0.517853143 | 0.000567631 | 0.000840464 | 0.001143763 |
| 0.418395773 | 0.420592057 | 0.001148170 | 0.001551604 | 0.001983832 |
| 0.361508198 | 0.345294904 | 0.000567631 | 0.000840464 | 0.001143763 |
| 0.466415728 | 0.456947145 | 0.001148170 | 0.001551604 | 0.001983832 |
| 0.402691597 | 0.383443680 | 0.000567631 | 0.000840464 | 0.001143763 |
| 0.106798247 | 0.117394355 | 0.001148170 | 0.001551604 | 0.001983832 |
| 0.015100333 | 0.014387476 | 0.000567631 | 0.000840464 | 0.001143763 |

| | | | | |
|---|---|---|---|---|
| 0.732659315 | 0.660485968 | 0.001148170 | 0.001551604 | 0.001983832 |
| 0.635770290 | 0.606068207 | 0.000567631 | 0.000840464 | 0.001143763 |
| 0.795899462 | 0.720954486 | 0.001148170 | 0.001551604 | 0.001983832 |
| 0.714236015 | 0.674396619 | 0.000567631 | 0.000840464 | 0.001143763 |
| 0.015255644 | 0.021948543 | 0.001148170 | 0.001551604 | 0.001983832 |
| 0.030997555 | 0.030466936 | 0.000567631 | 0.000840464 | 0.001143763 |
| 0.944113996 | 0.895097902 | 0.001148170 | 0.001551604 | 0.001983832 |
| 0.949808652 | 0.939867566 | 0.000567631 | 0.000840464 | 0.001143763 |
| 0.221516842 | 0.215728270 | 0.001148170 | 0.001551604 | 0.001983832 |
| 0.166067237 | 0.144907004 | 0.000567631 | 0.000840464 | 0.001143763 |
| 0.100502655 | 0.112366945 | 0.001148170 | 0.001551604 | 0.001983832 |
| 0.059834632 | 0.059541259 | 0.000567631 | 0.000840464 | 0.001143763 |
| 0.198495501 | 0.193925172 | 0.001148170 | 0.001551604 | 0.001983832 |
| 0.069439456 | 0.065602819 | 0.000567631 | 0.000840464 | 0.001143763 |
| 0.161452523 | 0.164023091 | 0.001148170 | 0.001551604 | 0.001983832 |
| 0.309032414 | 0.287885138 | 0.000567631 | 0.000840464 | 0.001143763 |
| 0.305520159 | 0.313826247 | 0.001148170 | 0.001551604 | 0.001983832 |
| 0.256076765 | 0.237071790 | 0.000567631 | 0.000840464 | 0.001143763 |
| 0.069520957 | 0.079987297 | 0.001148170 | 0.001551604 | 0.001983832 |
| 0.039309495 | 0.037132629 | 0.000567631 | 0.000840464 | 0.001143763 |
| 0.666556810 | 0.617293558 | 0.001148170 | 0.001551604 | 0.001983832 |
| 0.574343690 | 0.557981130 | 0.000567631 | 0.000840464 | 0.001143763 |
| 0.167791214 | 0.167244197 | 0.001148170 | 0.001551604 | 0.001983832 |
| 0.112202197 | 0.102745753 | 0.000567631 | 0.000840464 | 0.001143763 |
| 0.686803343 | 0.631138919 | 0.001148170 | 0.001551604 | 0.001983832 |
| 0.590784910 | 0.572964088 | 0.000567631 | 0.000840464 | 0.001143763 |
| 0.851682728 | 0.806846018 | 0.001148170 | 0.001551604 | 0.001983832 |
| 0.905126184 | 0.893199477 | 0.000567631 | 0.000840464 | 0.001143763 |
| 0.523546031 | 0.488640973 | 0.001148170 | 0.001551604 | 0.001983832 |
| 0.459526590 | 0.417168168 | 0.000567631 | 0.000840464 | 0.001143763 |
| 0.603478549 | 0.569492844 | 0.001148170 | 0.001551604 | 0.001983832 |
| 0.514899665 | 0.504816128 | 0.000567631 | 0.000840464 | 0.001143763 |
| 0.159408168 | 0.162481593 | 0.001148170 | 0.001551604 | 0.001983832 |
| 0.316189439 | 0.301307403 | 0.000567631 | 0.000840464 | 0.001143763 |
| 0.411350208 | 0.416281550 | 0.001148170 | 0.001551604 | 0.001983832 |
| 0.355254966 | 0.340719302 | 0.000567631 | 0.000840464 | 0.001143763 |
| 0.917918310 | 0.869433282 | 0.001148170 | 0.001551604 | 0.001983832 |
| 0.959356493 | 0.941826651 | 0.000567631 | 0.000840464 | 0.001143763 |
| 0.826168924 | 0.764748371 | 0.001148170 | 0.001551604 | 0.001983832 |
| 0.383292959 | 0.364281150 | 0.000567631 | 0.000840464 | 0.001143763 |
| 0.895968684 | 0.867394238 | 0.001148170 | 0.001551604 | 0.001983832 |
| 0.838060748 | 0.842599275 | 0.000567631 | 0.000840464 | 0.001143763 |
| 0.076493260 | 0.087897665 | 0.001148170 | 0.001551604 | 0.001983832 |
| 0.044738206 | 0.042344269 | 0.000567631 | 0.000840464 | 0.001143763 |
| 0.935849531 | 0.892156293 | 0.001148170 | 0.001551604 | 0.001983832 |
| 0.629647909 | 0.598021389 | 0.000567631 | 0.000840464 | 0.001143763 |

| | | | | |
|---|---|---|---|---|
| 0.011383237 | 0.016060815 | 0.001148170 | 0.001551604 | 0.001983832 |
| 0.005332443 | 0.004384456 | 0.000567631 | 0.000840464 | 0.001143763 |
| 0.115896489 | 0.124990470 | 0.001148170 | 0.001551604 | 0.001983832 |
| 0.078777633 | 0.073290091 | 0.000567631 | 0.000840464 | 0.001143763 |
| 0.453215143 | 0.446644455 | 0.001148170 | 0.001551604 | 0.001983832 |
| 0.669449935 | 0.638627081 | 0.000567631 | 0.000840464 | 0.001143763 |
| 0.633454897 | 0.590010911 | 0.001148170 | 0.001551604 | 0.001983832 |
| 0.448799240 | 0.413609932 | 0.000567631 | 0.000840464 | 0.001143763 |
| 0.432936770 | 0.436507516 | 0.001148170 | 0.001551604 | 0.001983832 |
| 0.369626131 | 0.361797398 | 0.000567631 | 0.000840464 | 0.001143763 |
| 0.422217520 | 0.422261932 | 0.001148170 | 0.001551604 | 0.001983832 |
| 0.616936317 | 0.593616973 | 0.000567631 | 0.000840464 | 0.001143763 |
| 0.336729818 | 0.345050175 | 0.001148170 | 0.001551604 | 0.001983832 |
| 0.070316855 | 0.065928983 | 0.000567631 | 0.000840464 | 0.001143763 |
| 0.500157123 | 0.472185886 | 0.001148170 | 0.001551604 | 0.001983832 |
| 0.436160514 | 0.399457590 | 0.000567631 | 0.000840464 | 0.001143763 |
| 0.396360897 | 0.410804978 | 0.001148170 | 0.001551604 | 0.001983832 |
| 0.342013593 | 0.335080679 | 0.000567631 | 0.000840464 | 0.001143763 |
| 0.061500764 | 0.071253048 | 0.001148170 | 0.001551604 | 0.001983832 |
| 0.019705068 | 0.017235605 | 0.000567631 | 0.000840464 | 0.001143763 |
| 0.139636559 | 0.144594149 | 0.001148170 | 0.001551604 | 0.001983832 |
| 0.090320369 | 0.084239407 | 0.000567631 | 0.000840464 | 0.001143763 |
| 0.259111589 | 0.255583500 | 0.001148170 | 0.001551604 | 0.001983832 |
| 0.221545652 | 0.199136356 | 0.000567631 | 0.000840464 | 0.001143763 |
| 0.637885597 | 0.598173113 | 0.001148170 | 0.001551604 | 0.001983832 |
| 0.541975844 | 0.536457791 | 0.000567631 | 0.000840464 | 0.001143763 |
| 0.806497393 | 0.739303497 | 0.001148170 | 0.001551604 | 0.001983832 |
| 0.738490855 | 0.695413416 | 0.000567631 | 0.000840464 | 0.001143763 |
| 0.050616163 | 0.058893791 | 0.001148170 | 0.001551604 | 0.001983832 |
| 0.536840404 | 0.526116453 | 0.000567631 | 0.000840464 | 0.001143763 |
| 0.631023601 | 0.585912977 | 0.001148170 | 0.001551604 | 0.001983832 |
| 0.511259654 | 0.503714209 | 0.000567631 | 0.000840464 | 0.001143763 |
| 0.503620510 | 0.477515272 | 0.001148170 | 0.001551604 | 0.001983832 |
| 0.130862926 | 0.121818874 | 0.000567631 | 0.000840464 | 0.001143763 |
| 0.520027187 | 0.487018956 | 0.001148170 | 0.001551604 | 0.001983832 |
| 0.720161772 | 0.677488913 | 0.000567631 | 0.000840464 | 0.001143763 |
| 0.518701039 | 0.486151163 | 0.001148170 | 0.001551604 | 0.001983832 |
| 0.449315052 | 0.413875784 | 0.000567631 | 0.000840464 | 0.001143763 |
| 0.229891761 | 0.224824988 | 0.001148170 | 0.001551604 | 0.001983832 |
| 0.196789474 | 0.174701169 | 0.000567631 | 0.000840464 | 0.001143763 |
| 0.754935162 | 0.681217129 | 0.001148170 | 0.001551604 | 0.001983832 |
| 0.706213524 | 0.670423035 | 0.000567631 | 0.000840464 | 0.001143763 |
| 0.897242285 | 0.867544661 | 0.001148170 | 0.001551604 | 0.001983832 |
| 0.841440973 | 0.843633670 | 0.000567631 | 0.000840464 | 0.001143763 |
| 0.129746399 | 0.135911416 | 0.001148170 | 0.001551604 | 0.001983832 |
| 0.963010646 | 0.947907598 | 0.000567631 | 0.000840464 | 0.001143763 |

| | | | | |
|---|---|---|---|---|
| 0.964231934 | 0.941183659 | 0.001148170 | 0.001551604 | 0.001983832 |
| 0.941604052 | 0.931022369 | 0.000567631 | 0.000840464 | 0.001143763 |
| 0.819715619 | 0.749117025 | 0.001148170 | 0.001551604 | 0.001983832 |
| 0.756971573 | 0.706629972 | 0.000567631 | 0.000840464 | 0.001143763 |
| 0.130290125 | 0.136760290 | 0.001148170 | 0.001551604 | 0.001983832 |
| 0.152973159 | 0.141326642 | 0.000567631 | 0.000840464 | 0.001143763 |
| 0.772185833 | 0.690401256 | 0.001148170 | 0.001551604 | 0.001983832 |
| 0.676250919 | 0.639669792 | 0.000567631 | 0.000840464 | 0.001143763 |
| 0.611662676 | 0.570666972 | 0.001148170 | 0.001551604 | 0.001983832 |
| 0.522568351 | 0.506085135 | 0.000567631 | 0.000840464 | 0.001143763 |
| 0.378332458 | 0.397132177 | 0.001148170 | 0.001551604 | 0.001983832 |
| 0.440426865 | 0.407476833 | 0.000567631 | 0.000840464 | 0.001143763 |
| 0.254171071 | 0.252491898 | 0.001148170 | 0.001551604 | 0.001983832 |
| 0.203028289 | 0.178480438 | 0.000567631 | 0.000840464 | 0.001143763 |
| 0.367221510 | 0.380343103 | 0.001148170 | 0.001551604 | 0.001983832 |
| 0.320024721 | 0.303643044 | 0.000567631 | 0.000840464 | 0.001143763 |
| 0.288935460 | 0.297942607 | 0.001148170 | 0.001551604 | 0.001983832 |
| 0.444530080 | 0.410373935 | 0.000567631 | 0.000840464 | 0.001143763 |
| 0.424962677 | 0.426399507 | 0.001148170 | 0.001551604 | 0.001983832 |
| 0.198610711 | 0.176849181 | 0.000567631 | 0.000840464 | 0.001143763 |
| 0.676430794 | 0.623827143 | 0.001148170 | 0.001551604 | 0.001983832 |
| 0.58233216 | 0.565005243 | 0.000567631 | 0.000840464 | 0.001143763 |
| 0.674078927 | 0.621111785 | 0.001148170 | 0.001551604 | 0.001983832 |
| 0.786125349 | 0.765237202 | 0.000567631 | 0.000840464 | 0.001143763 |
| 0.868951090 | 0.828406664 | 0.001148170 | 0.001551604 | 0.001983832 |
| 0.805040786 | 0.797592836 | 0.000567631 | 0.000840464 | 0.001143763 |
| 0.674283049 | 0.621500533 | 0.001148170 | 0.001551604 | 0.001983832 |
| 0.534884943 | 0.520111478 | 0.000567631 | 0.000840464 | 0.001143763 |
| 0.133757544 | 0.141740724 | 0.001148170 | 0.001551604 | 0.001983832 |
| 0.086593316 | 0.082070533 | 0.000567631 | 0.000840464 | 0.001143763 |
| 0.506809475 | 0.482133726 | 0.001148170 | 0.001551604 | 0.001983832 |
| 0.969817360 | 0.961250469 | 0.000567631 | 0.000840464 | 0.001143763 |
| 0.171822372 | 0.169669724 | 0.001148170 | 0.001551604 | 0.001983832 |
| 0.095990820 | 0.087221099 | 0.000567631 | 0.000840464 | 0.001143763 |
| 0.778693046 | 0.704822765 | 0.001148170 | 0.001551604 | 0.001983832 |
| 0.657433933 | 0.625115305 | 0.000567631 | 0.000840464 | 0.001143763 |
| 0.988425843 | 0.994406078 | 0.001148170 | 0.001551604 | 0.001983832 |
| 0.989556813 | 0.993472682 | 0.000567631 | 0.000840464 | 0.001143763 |
| 0.534198858 | 0.500599172 | 0.001148170 | 0.001551604 | 0.001983832 |
| 0.691201183 | 0.661285489 | 0.000567631 | 0.000840464 | 0.001143763 |
| 0.175663632 | 0.171574282 | 0.001148170 | 0.001551604 | 0.001983832 |
| 0.241506613 | 0.223533838 | 0.000567631 | 0.000840464 | 0.001143763 |
| 0.289805900 | 0.298742488 | 0.001148170 | 0.001551604 | 0.001983832 |
| 0.237373696 | 0.222315083 | 0.000567631 | 0.000840464 | 0.001143763 |
| 0.939733735 | 0.893534727 | 0.001148170 | 0.001551604 | 0.001983832 |
| 0.884777181 | 0.874109719 | 0.000567631 | 0.000840464 | 0.001143763 |

| | | | | |
|---|---|---|---|---|
| 0.430108494 | 0.434244024 | 0.001148170 | 0.001551604 | 0.001983832 |
| 0.317032522 | 0.302885401 | 0.000567631 | 0.000840464 | 0.001143763 |
| 0.559677084 | 0.523975060 | 0.001148170 | 0.001551604 | 0.001983832 |
| 0.974075513 | 0.971055775 | 0.000567631 | 0.000840464 | 0.001143763 |
| 0.263588046 | 0.258012555 | 0.001148170 | 0.001551604 | 0.001983832 |
| 0.230092486 | 0.211642955 | 0.000567631 | 0.000840464 | 0.001143763 |
| 0.488282816 | 0.467778317 | 0.001148170 | 0.001551604 | 0.001983832 |
| 0.426560962 | 0.394963173 | 0.000567631 | 0.000840464 | 0.001143763 |
| 0.889455447 | 0.846690882 | 0.001148170 | 0.001551604 | 0.001983832 |
| 0.830907019 | 0.819241593 | 0.000567631 | 0.000840464 | 0.001143763 |
| 0.232824821 | 0.227607475 | 0.001148170 | 0.001551604 | 0.001983832 |
| 0.410174008 | 0.390657158 | 0.000567631 | 0.000840464 | 0.001143763 |
| 0.853364198 | 0.813700509 | 0.001148170 | 0.001551604 | 0.001983832 |
| 0.474006251 | 0.433332443 | 0.000567631 | 0.000840464 | 0.001143763 |
| 0.226383647 | 0.221553988 | 0.001148170 | 0.001551604 | 0.001983832 |
| 0.181467243 | 0.155930494 | 0.000567631 | 0.000840464 | 0.001143763 |
| 0.579124801 | 0.538703238 | 0.001148170 | 0.001551604 | 0.001983832 |
| 0.501905663 | 0.470844449 | 0.000567631 | 0.000840464 | 0.001143763 |
| 0.323308922 | 0.336898511 | 0.001148170 | 0.001551604 | 0.001983832 |
| 0.274718363 | 0.259590499 | 0.000567631 | 0.000840464 | 0.001143763 |
| 0.242033571 | 0.237996509 | 0.001148170 | 0.001551604 | 0.001983832 |
| 0.686852125 | 0.656431945 | 0.000567631 | 0.000840464 | 0.001143763 |
| 0.962451203 | 0.933023454 | 0.001148170 | 0.001551604 | 0.001983832 |
| 0.928419270 | 0.920537147 | 0.000567631 | 0.000840464 | 0.001143763 |
| 0.126032641 | 0.131552640 | 0.001148170 | 0.001551604 | 0.001983832 |
| 0.079724396 | 0.073937469 | 0.000567631 | 0.000840464 | 0.001143763 |
| 0.282798526 | 0.285910481 | 0.001148170 | 0.001551604 | 0.001983832 |
| 0.270830174 | 0.254844045 | 0.000567631 | 0.000840464 | 0.001143763 |
| 0.901606745 | 0.868292887 | 0.001148170 | 0.001551604 | 0.001983832 |
| 0.869362615 | 0.847672488 | 0.000567631 | 0.000840464 | 0.001143763 |
| 0.702532981 | 0.640592989 | 0.001148170 | 0.001551604 | 0.001983832 |
| 0.601698153 | 0.582975753 | 0.000567631 | 0.000840464 | 0.001143763 |
| 0.721240205 | 0.652393939 | 0.001148170 | 0.001551604 | 0.001983832 |
| 0.622593253 | 0.596657933 | 0.000567631 | 0.000840464 | 0.001143763 |
| 0.055892755 | 0.065520657 | 0.001148170 | 0.001551604 | 0.001983832 |
| 0.015843636 | 0.014952870 | 0.000567631 | 0.000840464 | 0.001143763 |
| 0.187020834 | 0.181919136 | 0.001148170 | 0.001551604 | 0.001983832 |
| 0.562734258 | 0.553298104 | 0.000567631 | 0.000840464 | 0.001143763 |
| 0.840664953 | 0.792607687 | 0.001148170 | 0.001551604 | 0.001983832 |
| 0.783923405 | 0.757522651 | 0.000567631 | 0.000840464 | 0.001143763 |
| 0.635897557 | 0.594362273 | 0.001148170 | 0.001551604 | 0.001983832 |
| 0.119602144 | 0.108973868 | 0.000567631 | 0.000840464 | 0.001143763 |
| 0.832871596 | 0.790341484 | 0.001148170 | 0.001551604 | 0.001983832 |
| 0.799728717 | 0.787061146 | 0.000567631 | 0.000840464 | 0.001143763 |
| 0.662246027 | 0.615647854 | 0.001148170 | 0.001551604 | 0.001983832 |
| 0.567908974 | 0.555884458 | 0.000567631 | 0.000840464 | 0.001143763 |

| | | | | |
|---|---|---|---|---|
| 0.105827117 | 0.116037464 | 0.001148170 | 0.001551604 | 0.001983832 |
| 0.138126468 | 0.12832758 | 0.000567631 | 0.000840464 | 0.001143763 |
| 0.244771201 | 0.242558640 | 0.001148170 | 0.001551604 | 0.001983832 |
| 0.082529045 | 0.075562049 | 0.000567631 | 0.000840464 | 0.001143763 |
| 0.705534273 | 0.645506985 | 0.001148170 | 0.001551604 | 0.001983832 |
| 0.60999513 | 0.589300345 | 0.000567631 | 0.000840464 | 0.001143763 |
| 0.240086465 | 0.234855181 | 0.001148170 | 0.001551604 | 0.001983832 |
| 0.399951414 | 0.382786007 | 0.000567631 | 0.000840464 | 0.001143763 |
| 0.824358495 | 0.758488584 | 0.001148170 | 0.001551604 | 0.001983832 |
| 0.429883266 | 0.397275800 | 0.000567631 | 0.000840464 | 0.001143763 |
| 0.453949355 | 0.447445555 | 0.001148170 | 0.001551604 | 0.001983832 |
| 0.393465087 | 0.373786431 | 0.000567631 | 0.000840464 | 0.001143763 |
| 0.990851573 | 0.996867047 | 0.001148170 | 0.001551604 | 0.001983832 |
| 0.998477099 | 0.997215157 | 0.000567631 | 0.000840464 | 0.001143763 |
| 0.782994712 | 0.713112215 | 0.001148170 | 0.001551604 | 0.001983832 |
| 0.698548185 | 0.665819355 | 0.000567631 | 0.000840464 | 0.001143763 |
| 0.902085755 | 0.868343239 | 0.001148170 | 0.001551604 | 0.001983832 |
| 0.860750259 | 0.845520400 | 0.000567631 | 0.000840464 | 0.001143763 |
| 0.267887326 | 0.264806485 | 0.001148170 | 0.001551604 | 0.001983832 |
| 0.214756792 | 0.190036397 | 0.000567631 | 0.000840464 | 0.001143763 |
| 0.846554648 | 0.669868640 | 0.004929348 | 0.016005343 | 0.019029080 |
| 0.710282337 | 0.600518494 | 0.005375604 | 0.006489046 | 0.007577204 |
| 0.860766524 | 0.683017754 | 0.004929348 | 0.016005343 | 0.019029080 |
| 0.624008042 | 0.521053789 | 0.005375604 | 0.006489046 | 0.007577204 |
| 0.178442024 | 0.154790951 | 0.004929348 | 0.016005343 | 0.019029080 |
| 0.584271891 | 0.484065108 | 0.005375604 | 0.006489046 | 0.007577204 |
| 0.111273561 | 0.109842899 | 0.004929348 | 0.016005343 | 0.019029080 |
| 0.053332373 | 0.066931081 | 0.005375604 | 0.006489046 | 0.007577204 |
| 0.603334476 | 0.468165886 | 0.004929348 | 0.016005343 | 0.019029080 |
| 0.505628678 | 0.412121566 | 0.005375604 | 0.006489046 | 0.007577204 |
| 0.103137585 | 0.104453705 | 0.004929348 | 0.016005343 | 0.019029080 |
| 0.304594164 | 0.268271623 | 0.005375604 | 0.006489046 | 0.007577204 |
| 0.399108775 | 0.331355598 | 0.004929348 | 0.016005343 | 0.019029080 |
| 0.385039563 | 0.322034502 | 0.005375604 | 0.006489046 | 0.007577204 |
| 0.929416286 | 0.751585097 | 0.004929348 | 0.016005343 | 0.019029080 |
| 0.831870935 | 0.695573519 | 0.005375604 | 0.006489046 | 0.007577204 |
| 0.907614928 | 0.728446956 | 0.004929348 | 0.016005343 | 0.019029080 |
| 0.854102735 | 0.714667124 | 0.005375604 | 0.006489046 | 0.007577204 |
| 0.219853514 | 0.185713982 | 0.004929348 | 0.016005343 | 0.019029080 |
| 0.103891299 | 0.106272575 | 0.005375604 | 0.006489046 | 0.007577204 |
| 0.953317266 | 0.782102110 | 0.004929348 | 0.016005343 | 0.019029080 |
| 0.854327176 | 0.714869703 | 0.005375604 | 0.006489046 | 0.007577204 |
| 0.348459700 | 0.293023114 | 0.004929348 | 0.016005343 | 0.019029080 |
| 0.242246012 | 0.225558702 | 0.005375604 | 0.006489046 | 0.007577204 |
| 0.437265707 | 0.356847814 | 0.004929348 | 0.016005343 | 0.019029080 |
| 0.436005676 | 0.358032837 | 0.005375604 | 0.006489046 | 0.007577204 |

| | | | | | |
|---|---|---|---|---|---|
| 0.591415898 | 0.459527477 | 0.004929348 | 0.016005343 | 0.019029080 | |
| 0.541284534 | 0.443013679 | 0.005375604 | 0.006489046 | 0.007577204 | |
| 0.028146785 | 0.042491214 | 0.004929348 | 0.016005343 | 0.019029080 | |
| 0.338113566 | 0.290789040 | 0.005375604 | 0.006489046 | 0.007577204 | |
| 0.134630943 | 0.125505066 | 0.004929348 | 0.016005343 | 0.019029080 | |
| 0.224433999 | 0.212322211 | 0.005375604 | 0.006489046 | 0.007577204 | |
| 0.050661666 | 0.064161276 | 0.004929348 | 0.016005343 | 0.019029080 | |
| 0.134452508 | 0.132590659 | 0.005375604 | 0.006489046 | 0.007577204 | |
| 0.412012145 | 0.340511595 | 0.004929348 | 0.016005343 | 0.019029080 | |
| 0.223723731 | 0.211763287 | 0.005375604 | 0.006489046 | 0.007577204 | |
| 0.775720716 | 0.606375810 | 0.004929348 | 0.016005343 | 0.019029080 | |
| 0.634395741 | 0.531020477 | 0.005375604 | 0.006489046 | 0.007577204 | |
| 0.871122061 | 0.693069845 | 0.004929348 | 0.016005343 | 0.019029080 | |
| 0.758349660 | 0.636829321 | 0.005375604 | 0.006489046 | 0.007577204 | |
| 0.216418486 | 0.182852869 | 0.004929348 | 0.016005343 | 0.019029080 | |
| 0.151007747 | 0.147848630 | 0.005375604 | 0.006489046 | 0.007577204 | |
| 0.004681174 | 0.013335436 | 0.004929348 | 0.016005343 | 0.019029080 | 0.15 |
| 0.904757658 | 0.770279271 | 0.005375604 | 0.006489046 | 0.007577204 | |
| 0.107647218 | 0.107439549 | 0.004929348 | 0.016005343 | 0.019029080 | |
| 0.200425638 | 0.193586203 | 0.005375604 | 0.006489046 | 0.007577204 | |
| 0.868284722 | 0.690331321 | 0.004929348 | 0.016005343 | 0.019029080 | |
| 0.695928148 | 0.587666981 | 0.005375604 | 0.006489046 | 0.007577204 | |
| 0.274776276 | 0.230435338 | 0.004929348 | 0.016005343 | 0.019029080 | |
| 0.109871613 | 0.111188992 | 0.005375604 | 0.006489046 | 0.007577204 | |
| 0.033592130 | 0.048082064 | 0.004929348 | 0.016005343 | 0.019029080 | |
| 0.376066932 | 0.316112289 | 0.005375604 | 0.006489046 | 0.007577204 | |
| 0.487373932 | 0.388157865 | 0.004929348 | 0.016005343 | 0.019029080 | |
| 0.398133260 | 0.330794845 | 0.005375604 | 0.006489046 | 0.007577204 | |
| 0.815989467 | 0.642406396 | 0.004929348 | 0.016005343 | 0.019029080 | |
| 0.652245297 | 0.547669578 | 0.005375604 | 0.006489046 | 0.007577204 | |
| 0.037707004 | 0.051980973 | 0.004929348 | 0.016005343 | 0.019029080 | |
| 0.205897981 | 0.197816662 | 0.005375604 | 0.006489046 | 0.007577204 | |
| 0.383580798 | 0.320248888 | 0.004929348 | 0.016005343 | 0.019029080 | |
| 0.262811101 | 0.2401436 | 0.005375604 | 0.006489046 | 0.007577204 | |
| 0.778699539 | 0.609007872 | 0.004929348 | 0.016005343 | 0.019029080 | |
| 0.581342050 | 0.481384571 | 0.005375604 | 0.006489046 | 0.007577204 | |
| 0.983366837 | 0.843002708 | 0.004929348 | 0.016005343 | 0.019029080 | |
| 0.970432192 | 0.876344874 | 0.005375604 | 0.006489046 | 0.007577204 | |
| 0.420800099 | 0.346397522 | 0.004929348 | 0.016005343 | 0.019029080 | |
| 0.402825417 | 0.334023982 | 0.005375604 | 0.006489046 | 0.007577204 | |
| 0.012691633 | 0.024615356 | 0.004929348 | 0.016005343 | 0.019029080 | |
| 0.006421982 | 0.015912824 | 0.005375604 | 0.006489046 | 0.007577204 | |
| 0.260915510 | 0.219217918 | 0.004929348 | 0.016005343 | 0.019029080 | |
| 0.414380086 | 0.342096193 | 0.005375604 | 0.006489046 | 0.007577204 | |
| 0.206227859 | 0.174816240 | 0.004929348 | 0.016005343 | 0.019029080 | |
| 0.218563476 | 0.207737381 | 0.005375604 | 0.006489046 | 0.007577204 | |

| | | | | |
|---|---|---|---|---|
| 0.164236772 | 0.145143233 | 0.004929348 | 0.016005343 | 0.019029080 |
| 0.937215175 | 0.813868011 | 0.005375604 | 0.006489046 | 0.007577204 |
| 0.985551363 | 0.851453722 | 0.004929348 | 0.016005343 | 0.019029080 |
| 0.729916616 | 0.615752096 | 0.005375604 | 0.006489046 | 0.007577204 |
| 0.959088379 | 0.789485431 | 0.004929348 | 0.016005343 | 0.019029080 |
| 0.947375055 | 0.829387646 | 0.005375604 | 0.006489046 | 0.007577204 |
| 0.911187994 | 0.731845425 | 0.004929348 | 0.016005343 | 0.019029080 |
| 0.755727866 | 0.634875457 | 0.005375604 | 0.006489046 | 0.007577204 |
| 0.831939662 | 0.656229367 | 0.004929348 | 0.016005343 | 0.019029080 |
| 0.437686732 | 0.359267452 | 0.005375604 | 0.006489046 | 0.007577204 |
| 0.332373208 | 0.279849976 | 0.004929348 | 0.016005343 | 0.019029080 |
| 0.217543673 | 0.206949227 | 0.005375604 | 0.006489046 | 0.007577204 |
| 0.192259246 | 0.164668982 | 0.004929348 | 0.016005343 | 0.019029080 |
| 0.146087669 | 0.143501274 | 0.005375604 | 0.006489046 | 0.007577204 |
| 0.517154178 | 0.407395588 | 0.004929348 | 0.016005343 | 0.019029080 |
| 0.736621598 | 0.620809821 | 0.005375604 | 0.006489046 | 0.007577204 |
| 0.991313536 | 0.881299235 | 0.004929348 | 0.016005343 | 0.019029080 |
| 0.937821649 | 0.814764448 | 0.005375604 | 0.006489046 | 0.007577204 |
| 0.291414308 | 0.244678490 | 0.004929348 | 0.016005343 | 0.019029080 |
| 0.175750028 | 0.171356665 | 0.005375604 | 0.006489046 | 0.007577204 |
| 0.540692451 | 0.423033004 | 0.004929348 | 0.016005343 | 0.019029080 |
| 0.694195962 | 0.586040789 | 0.005375604 | 0.006489046 | 0.007577204 |
| 0.443165860 | 0.360451873 | 0.004929348 | 0.016005343 | 0.019029080 |
| 0.115994310 | 0.116367801 | 0.005375604 | 0.006489046 | 0.007577204 |
| 0.859086792 | 0.681440270 | 0.004929348 | 0.016005343 | 0.019029080 |
| 0.761884659 | 0.639481498 | 0.005375604 | 0.006489046 | 0.007577204 |
| 0.269009447 | 0.225646069 | 0.004929348 | 0.016005343 | 0.019029080 |
| 0.197421034 | 0.191223152 | 0.005375604 | 0.006489046 | 0.007577204 |
| 0.085290499 | 0.091929085 | 0.004929348 | 0.016005343 | 0.019029080 |
| 0.027269241 | 0.041875461 | 0.005375604 | 0.006489046 | 0.007577204 |
| 0.333433318 | 0.280706627 | 0.004929348 | 0.016005343 | 0.019029080 |
| 0.220819921 | 0.209489549 | 0.005375604 | 0.006489046 | 0.007577204 |
| 0.631235295 | 0.488970725 | 0.004929348 | 0.016005343 | 0.019029080 |
| 0.469925374 | 0.383947480 | 0.005375604 | 0.006489046 | 0.007577204 |
| 0.223231333 | 0.188535515 | 0.004929348 | 0.016005343 | 0.019029080 |
| 0.127543118 | 0.126458410 | 0.005375604 | 0.006489046 | 0.007577204 |
| 0.862983730 | 0.685137483 | 0.004929348 | 0.016005343 | 0.019029080 |
| 0.722027372 | 0.609820762 | 0.005375604 | 0.006489046 | 0.007577204 |
| 0.776653850 | 0.607196068 | 0.004929348 | 0.016005343 | 0.019029080 |
| 0.841530842 | 0.703736343 | 0.005375604 | 0.006489046 | 0.007577204 |
| 0.733680613 | 0.573143810 | 0.004929348 | 0.016005343 | 0.019029080 |
| 0.425840274 | 0.350494870 | 0.005375604 | 0.006489046 | 0.007577204 |
| 0.732924947 | 0.572517171 | 0.004929348 | 0.016005343 | 0.019029080 |
| 0.236714200 | 0.221486812 | 0.005375604 | 0.006489046 | 0.007577204 |
| 0.800905408 | 0.628974503 | 0.004929348 | 0.016005343 | 0.019029080 |
| 0.856041245 | 0.716475137 | 0.005375604 | 0.006489046 | 0.007577204 |

| | | | | | |
|---|---|---|---|---|---|
| 0.603162288 | 0.468041971 | 0.004929348 | 0.016005343 | 0.019029080 | |
| 0.327539066 | 0.283816591 | 0.005375604 | 0.006489046 | 0.007577204 | |
| 0.054794076 | 0.067610406 | 0.004929348 | 0.016005343 | 0.019029080 | |
| 0.216803459 | 0.206375245 | 0.005375604 | 0.006489046 | 0.007577204 | |
| 0.681234202 | 0.528322187 | 0.004929348 | 0.016005343 | 0.019029080 | |
| 0.743205904 | 0.625682755 | 0.005375604 | 0.006489046 | 0.007577204 | |
| 0.557609510 | 0.434650152 | 0.004929348 | 0.016005343 | 0.019029080 | |
| 0.459715570 | 0.375760254 | 0.005375604 | 0.006489046 | 0.007577204 | |
| 0.194379720 | 0.166164014 | 0.004929348 | 0.016005343 | 0.019029080 | |
| 0.954295327 | 0.840940967 | 0.005375604 | 0.006489046 | 0.007577204 | |
| 0.003574834 | 0.011437767 | 0.004929348 | 0.016005343 | 0.019029080 | 0.15 |
| 0.615954108 | 0.513531471 | 0.005375604 | 0.006489046 | 0.007577204 | |
| 0.344393357 | 0.289677358 | 0.004929348 | 0.016005343 | 0.019029080 | |
| 0.529682053 | 0.432867389 | 0.005375604 | 0.006489046 | 0.007577204 | |
| 0.882118701 | 0.703836704 | 0.004929348 | 0.016005343 | 0.019029080 | |
| 0.783559849 | 0.656122233 | 0.005375604 | 0.006489046 | 0.007577204 | |
| 0.784189695 | 0.613988448 | 0.004929348 | 0.016005343 | 0.019029080 | |
| 0.735494064 | 0.619962700 | 0.005375604 | 0.006489046 | 0.007577204 | |
| 0.974119675 | 0.815414610 | 0.004929348 | 0.016005343 | 0.019029080 | |
| 0.877666110 | 0.738422555 | 0.005375604 | 0.006489046 | 0.007577204 | |
| 0.556191223 | 0.433639770 | 0.004929348 | 0.016005343 | 0.019029080 | |
| 0.530752016 | 0.433817228 | 0.005375604 | 0.006489046 | 0.007577204 | |
| 0.849061926 | 0.672183788 | 0.004929348 | 0.016005343 | 0.019029080 | |
| 0.741986941 | 0.624779095 | 0.005375604 | 0.006489046 | 0.007577204 | |
| 0.203015168 | 0.172390628 | 0.004929348 | 0.016005343 | 0.019029080 | |
| 0.180142368 | 0.175474216 | 0.005375604 | 0.006489046 | 0.007577204 | |
| 0.634937889 | 0.491744322 | 0.004929348 | 0.016005343 | 0.019029080 | |
| 0.613432250 | 0.511156253 | 0.005375604 | 0.006489046 | 0.007577204 | |
| 0.550187505 | 0.429452805 | 0.004929348 | 0.016005343 | 0.019029080 | |
| 0.138515830 | 0.136363734 | 0.005375604 | 0.006489046 | 0.007577204 | |
| 0.132821720 | 0.124310437 | 0.004929348 | 0.016005343 | 0.019029080 | |
| 0.107683612 | 0.109410855 | 0.005375604 | 0.006489046 | 0.007577204 | |
| 0.992182228 | 0.886715924 | 0.004929348 | 0.016005343 | 0.019029080 | |
| 0.970489599 | 0.876478997 | 0.005375604 | 0.006489046 | 0.007577204 | |
| 0.609767243 | 0.472879252 | 0.004929348 | 0.016005343 | 0.019029080 | |
| 0.305782583 | 0.269094904 | 0.005375604 | 0.006489046 | 0.007577204 | |
| 0.600565020 | 0.466165295 | 0.004929348 | 0.016005343 | 0.019029080 | |
| 0.603432815 | 0.501545359 | 0.005375604 | 0.006489046 | 0.007577204 | |
| 0.587795860 | 0.456971302 | 0.004929348 | 0.016005343 | 0.019029080 | |
| 0.471823318 | 0.385455461 | 0.005375604 | 0.006489046 | 0.007577204 | |
| 0.235984395 | 0.198808003 | 0.004929348 | 0.016005343 | 0.019029080 | |
| 0.964344961 | 0.862729549 | 0.005375604 | 0.006489046 | 0.007577204 | |
| 0.465162231 | 0.37429366 | 0.004929348 | 0.016005343 | 0.019029080 | |
| 0.327137976 | 0.283548001 | 0.005375604 | 0.006489046 | 0.007577204 | |
| 0.808488889 | 0.635763788 | 0.004929348 | 0.016005343 | 0.019029080 | |
| 0.773653700 | 0.648358759 | 0.005375604 | 0.006489046 | 0.007577204 | |

| | | | | | |
|---|---|---|---|---|---|
| 0.560885395 | 0.436968162 | 0.004929348 | 0.016005343 | 0.019029080 | |
| 0.944130255 | 0.824396424 | 0.005375604 | 0.006489046 | 0.007577204 | |
| 0.855572543 | 0.678180758 | 0.004929348 | 0.016005343 | 0.019029080 | |
| 0.856407922 | 0.716822068 | 0.005375604 | 0.006489046 | 0.007577204 | |
| 0.854257427 | 0.676960050 | 0.004929348 | 0.016005343 | 0.019029080 | |
| 0.744536859 | 0.626652716 | 0.005375604 | 0.006489046 | 0.007577204 | |
| 0.260381242 | 0.218798403 | 0.004929348 | 0.016005343 | 0.019029080 | |
| 0.161745954 | 0.157642618 | 0.005375604 | 0.006489046 | 0.007577204 | |
| 0.281317859 | 0.236041559 | 0.004929348 | 0.016005343 | 0.019029080 | |
| 0.295553040 | 0.262050434 | 0.005375604 | 0.006489046 | 0.007577204 | |
| 0.674379066 | 0.522706048 | 0.004929348 | 0.016005343 | 0.019029080 | |
| 0.545195143 | 0.446684171 | 0.005375604 | 0.006489046 | 0.007577204 | |
| 0.464035571 | 0.373590362 | 0.004929348 | 0.016005343 | 0.019029080 | |
| 0.946738348 | 0.828407322 | 0.005375604 | 0.006489046 | 0.007577204 | |
| 0.475107570 | 0.380503012 | 0.004929348 | 0.016005343 | 0.019029080 | |
| 0.436956854 | 0.358729177 | 0.005375604 | 0.006489046 | 0.007577204 | |
| 0.157846031 | 0.140739925 | 0.004929348 | 0.016005343 | 0.019029080 | |
| 0.254471895 | 0.234233692 | 0.005375604 | 0.006489046 | 0.007577204 | |
| 0.9467656 | 0.773943917 | 0.004929348 | 0.016005343 | 0.019029080 | |
| 0.892772960 | 0.755856798 | 0.005375604 | 0.006489046 | 0.007577204 | |
| 0.184590582 | 0.159192501 | 0.004929348 | 0.016005343 | 0.019029080 | |
| 0.484890311 | 0.395627866 | 0.005375604 | 0.006489046 | 0.007577204 | |
| 0.879723774 | 0.701478652 | 0.004929348 | 0.016005343 | 0.019029080 | |
| 0.557121200 | 0.457807449 | 0.005375604 | 0.006489046 | 0.007577204 | |
| 0.449881036 | 0.364645427 | 0.004929348 | 0.016005343 | 0.019029080 | |
| 0.378535332 | 0.317751606 | 0.005375604 | 0.006489046 | 0.007577204 | |
| 0.683312853 | 0.530074935 | 0.004929348 | 0.016005343 | 0.019029080 | |
| 0.522827907 | 0.426873732 | 0.005375604 | 0.006489046 | 0.007577204 | |
| 0.297896457 | 0.250298999 | 0.004929348 | 0.016005343 | 0.019029080 | |
| 0.131910306 | 0.130289091 | 0.005375604 | 0.006489046 | 0.007577204 | |
| 0.889631619 | 0.711056080 | 0.004929348 | 0.016005343 | 0.019029080 | |
| 0.904378972 | 0.769808316 | 0.005375604 | 0.006489046 | 0.007577204 | |
| 0.677949567 | 0.525592548 | 0.004929348 | 0.016005343 | 0.019029080 | |
| 0.787048976 | 0.659007964 | 0.005375604 | 0.006489046 | 0.007577204 | |
| 0.080783388 | 0.088471856 | 0.004929348 | 0.016005343 | 0.019029080 | |
| 0.024320777 | 0.038494568 | 0.005375604 | 0.006489046 | 0.007577204 | |
| 0.745413008 | 0.582284879 | 0.004929348 | 0.016005343 | 0.019029080 | |
| 0.622962534 | 0.520080923 | 0.005375604 | 0.006489046 | 0.007577204 | |
| 0.004429370 | 0.012902288 | 0.004929348 | 0.016005343 | 0.019029080 | 0.15 |
| 0.842816499 | 0.704835647 | 0.005375604 | 0.006489046 | 0.007577204 | |
| 0.657142981 | 0.509019827 | 0.004929348 | 0.016005343 | 0.019029080 | |
| 0.180786159 | 0.176078850 | 0.005375604 | 0.006489046 | 0.007577204 | |
| 0.648721429 | 0.502398222 | 0.004929348 | 0.016005343 | 0.019029080 | |
| 0.462842458 | 0.378228751 | 0.005375604 | 0.006489046 | 0.007577204 | |
| 0.169329289 | 0.148582417 | 0.004929348 | 0.016005343 | 0.019029080 | |
| 0.054510935 | 0.067913325 | 0.005375604 | 0.006489046 | 0.007577204 | |

| | | | | | |
|---|---|---|---|---|---|
| 0.119242928 | 0.115028154 | 0.004929348 | 0.016005343 | 0.019029080 | |
| 0.645054865 | 0.541192178 | 0.005375604 | 0.006489046 | 0.007577204 | |
| 0.031697589 | 0.046199907 | 0.004929348 | 0.016005343 | 0.019029080 | |
| 0.735687190 | 0.620107964 | 0.005375604 | 0.006489046 | 0.007577204 | |
| 0.693689124 | 0.538722602 | 0.004929348 | 0.016005343 | 0.019029080 | |
| 0.035678570 | 0.050929072 | 0.005375604 | 0.006489046 | 0.007577204 | |
| 0.033674751 | 0.048161884 | 0.004929348 | 0.016005343 | 0.019029080 | |
| 0.813389884 | 0.680657402 | 0.005375604 | 0.006489046 | 0.007577204 | |
| 0.901576122 | 0.722722411 | 0.004929348 | 0.016005343 | 0.019029080 | |
| 0.816892671 | 0.683463016 | 0.005375604 | 0.006489046 | 0.007577204 | |
| 0.452729505 | 0.366462472 | 0.004929348 | 0.016005343 | 0.019029080 | |
| 0.418154762 | 0.344800608 | 0.005375604 | 0.006489046 | 0.007577204 | |
| 0.249170562 | 0.209729237 | 0.004929348 | 0.016005343 | 0.019029080 | |
| 0.115801597 | 0.116205044 | 0.005375604 | 0.006489046 | 0.007577204 | |
| 0.026112030 | 0.040226657 | 0.004929348 | 0.016005343 | 0.019029080 | |
| 0.041443791 | 0.056558721 | 0.005375604 | 0.006489046 | 0.007577204 | |
| 0.746155206 | 0.582850026 | 0.004929348 | 0.016005343 | 0.019029080 | |
| 0.707561375 | 0.598121532 | 0.005375604 | 0.006489046 | 0.007577204 | |
| 0.986366699 | 0.855130933 | 0.004929348 | 0.016005343 | 0.019029080 | |
| 0.742078883 | 0.624847270 | 0.005375604 | 0.006489046 | 0.007577204 | |
| 0.539725380 | 0.422390721 | 0.004929348 | 0.016005343 | 0.019029080 | |
| 0.580623609 | 0.480730087 | 0.005375604 | 0.006489046 | 0.007577204 | |
| 0.002080287 | 0.008505740 | 0.004929348 | 0.016005343 | 0.019029080 | 0.15 |
| 0.891611096 | 0.754514019 | 0.005375604 | 0.006489046 | 0.007577204 | |
| 0.384612988 | 0.320994198 | 0.004929348 | 0.016005343 | 0.019029080 | |
| 0.405580966 | 0.335932695 | 0.005375604 | 0.006489046 | 0.007577204 | |
| 0.004389035 | 0.012832596 | 0.004929348 | 0.016005343 | 0.019029080 | 0.15 |
| 0.722410960 | 0.610116314 | 0.005375604 | 0.006489046 | 0.007577204 | |
| 0.132978712 | 0.124414590 | 0.004929348 | 0.016005343 | 0.019029080 | |
| 0.134096591 | 0.132260945 | 0.005375604 | 0.006489046 | 0.007577204 | |
| 0.344063258 | 0.339504030 | 0.001162785 | 0.001437343 | 0.001785486 | |
| 0.331547431 | 0.307134294 | 0.000303578 | 0.000516412 | 0.000664109 | |
| 0.513438486 | 0.539249568 | 0.001162785 | 0.001437343 | 0.001785486 | |
| 0.261852339 | 0.242472433 | 0.000303578 | 0.000516412 | 0.000664109 | |
| 0.219621909 | 0.197216959 | 0.001162785 | 0.001437343 | 0.001785486 | |
| 0.452432325 | 0.461989666 | 0.000303578 | 0.000516412 | 0.000664109 | |
| 0.056576056 | 0.044995617 | 0.001162785 | 0.001437343 | 0.001785486 | |
| 0.035889323 | 0.026917037 | 0.000303578 | 0.000516412 | 0.000664109 | |
| 0.900831032 | 0.903403574 | 0.001162785 | 0.001437343 | 0.001785486 | |
| 0.850393253 | 0.897477069 | 0.000303578 | 0.000516412 | 0.000664109 | |
| 0.729629055 | 0.733091334 | 0.001162785 | 0.001437343 | 0.001785486 | |
| 0.696610973 | 0.716858053 | 0.000303578 | 0.000516412 | 0.000664109 | |
| 0.212336110 | 0.190510923 | 0.001162785 | 0.001437343 | 0.001785486 | |
| 0.172226635 | 0.159158612 | 0.000303578 | 0.000516412 | 0.000664109 | |
| 0.100916206 | 0.083456318 | 0.001162785 | 0.001437343 | 0.001785486 | |
| 0.108495069 | 0.116592372 | 0.000303578 | 0.000516412 | 0.000664109 | |

| | | | | |
|---|---|---|---|---|
| 0.526038643 | 0.551913701 | 0.001162785 | 0.001437343 | 0.001785486 |
| 0.586459729 | 0.633386516 | 0.000303578 | 0.000516412 | 0.000664109 |
| 0.711251956 | 0.711568429 | 0.001162785 | 0.001437343 | 0.001785486 |
| 0.646991954 | 0.693612935 | 0.000303578 | 0.000516412 | 0.000664109 |
| 0.598717037 | 0.602827103 | 0.001162785 | 0.001437343 | 0.001785486 |
| 0.543299834 | 0.579396570 | 0.000303578 | 0.000516412 | 0.000664109 |
| 0.481718328 | 0.503763599 | 0.001162785 | 0.001437343 | 0.001785486 |
| 0.455672281 | 0.475857101 | 0.000303578 | 0.000516412 | 0.000664109 |
| 0.116246322 | 0.096906373 | 0.001162785 | 0.001437343 | 0.001785486 |
| 0.130225403 | 0.137551085 | 0.000303578 | 0.000516412 | 0.000664109 |
| 0.542682517 | 0.571575231 | 0.001162785 | 0.001437343 | 0.001785486 |
| 0.483581608 | 0.546822099 | 0.000303578 | 0.000516412 | 0.000664109 |
| 0.910419227 | 0.938723038 | 0.001162785 | 0.001437343 | 0.001785486 |
| 0.872315377 | 0.935830226 | 0.000303578 | 0.000516412 | 0.000664109 |
| 0.398133525 | 0.407448562 | 0.001162785 | 0.001437343 | 0.001785486 |
| 0.367384527 | 0.376557772 | 0.000303578 | 0.000516412 | 0.000664109 |
| 0.651143853 | 0.653944926 | 0.001162785 | 0.001437343 | 0.001785486 |
| 0.582130405 | 0.633126104 | 0.000303578 | 0.000516412 | 0.000664109 |
| 0.264153792 | 0.238090580 | 0.001162785 | 0.001437343 | 0.001785486 |
| 0.221526848 | 0.205219858 | 0.000303578 | 0.000516412 | 0.000664109 |
| 0.632176694 | 0.628831285 | 0.001162785 | 0.001437343 | 0.001785486 |
| 0.561562567 | 0.606654065 | 0.000303578 | 0.000516412 | 0.000664109 |
| 0.282861808 | 0.254004281 | 0.001162785 | 0.001437343 | 0.001785486 |
| 0.255171258 | 0.233741882 | 0.000303578 | 0.000516412 | 0.000664109 |
| 0.681053863 | 0.676953400 | 0.001162785 | 0.001437343 | 0.001785486 |
| 0.615409649 | 0.657250676 | 0.000303578 | 0.000516412 | 0.000664109 |
| 0.404251990 | 0.414817960 | 0.001162785 | 0.001437343 | 0.001785486 |
| 0.881199842 | 0.940854731 | 0.000303578 | 0.000516412 | 0.000664109 |
| 0.671853671 | 0.676333733 | 0.001162785 | 0.001437343 | 0.001785486 |
| 0.607763463 | 0.656770637 | 0.000303578 | 0.000516412 | 0.000664109 |
| 0.658524280 | 0.657802639 | 0.001162785 | 0.001437343 | 0.001785486 |
| 0.592655489 | 0.637034028 | 0.000303578 | 0.000516412 | 0.000664109 |
| 0.168490002 | 0.155468288 | 0.001162785 | 0.001437343 | 0.001785486 |
| 0.043005274 | 0.032790167 | 0.000303578 | 0.000516412 | 0.000664109 |
| 0.053283667 | 0.042677436 | 0.001162785 | 0.001437343 | 0.001785486 |
| 0.307500287 | 0.287602798 | 0.000303578 | 0.000516412 | 0.000664109 |
| 0.413513104 | 0.423157878 | 0.001162785 | 0.001437343 | 0.001785486 |
| 0.391569108 | 0.392603102 | 0.000303578 | 0.000516412 | 0.000664109 |
| 0.021159926 | 0.018999648 | 0.001162785 | 0.001437343 | 0.001785486 |
| 0.013149047 | 0.008191783 | 0.000303578 | 0.000516412 | 0.000664109 |
| 0.149350765 | 0.134029075 | 0.001162785 | 0.001437343 | 0.001785486 |
| 0.134387432 | 0.137913040 | 0.000303578 | 0.000516412 | 0.000664109 |
| 0.589438386 | 0.598162196 | 0.001162785 | 0.001437343 | 0.001785486 |
| 0.536227339 | 0.574419200 | 0.000303578 | 0.000516412 | 0.000664109 |
| 0.244888825 | 0.215275404 | 0.001162785 | 0.001437343 | 0.001785486 |
| 0.20169474 | 0.182928425 | 0.000303578 | 0.000516412 | 0.000664109 |

| | | | | |
|---|---|---|---|---|
| 0.990414667 | 0.991232598 | 0.001162785 | 0.001437343 | 0.001785486 |
| 0.916089261 | 0.953011302 | 0.000303578 | 0.000516412 | 0.000664109 |
| 0.785374181 | 0.798367844 | 0.001162785 | 0.001437343 | 0.001785486 |
| 0.745287008 | 0.785880729 | 0.000303578 | 0.000516412 | 0.000664109 |
| 0.103820670 | 0.087493943 | 0.001162785 | 0.001437343 | 0.001785486 |
| 0.066634097 | 0.062747851 | 0.000303578 | 0.000516412 | 0.000664109 |
| 0.295763037 | 0.277049968 | 0.001162785 | 0.001437343 | 0.001785486 |
| 0.347399147 | 0.323126488 | 0.000303578 | 0.000516412 | 0.000664109 |
| 0.105647730 | 0.087685918 | 0.001162785 | 0.001437343 | 0.001785486 |
| 0.119264362 | 0.129370501 | 0.000303578 | 0.000516412 | 0.000664109 |
| 0.031658559 | 0.026517573 | 0.001162785 | 0.001437343 | 0.001785486 |
| 0.901015190 | 0.943634702 | 0.000303578 | 0.000516412 | 0.000664109 |
| 0.751818797 | 0.766212660 | 0.001162785 | 0.001437343 | 0.001785486 |
| 0.245610647 | 0.222928877 | 0.000303578 | 0.000516412 | 0.000664109 |
| 0.750833889 | 0.762965448 | 0.001162785 | 0.001437343 | 0.001785486 |
| 0.778642694 | 0.838553414 | 0.000303578 | 0.000516412 | 0.000664109 |
| 0.795142611 | 0.817026728 | 0.001162785 | 0.001437343 | 0.001785486 |
| 0.756341252 | 0.805602045 | 0.000303578 | 0.000516412 | 0.000664109 |
| 0.904678627 | 0.910119096 | 0.001162785 | 0.001437343 | 0.001785486 |
| 0.324634263 | 0.301655003 | 0.000303578 | 0.000516412 | 0.000664109 |
| 0.144448424 | 0.129178378 | 0.001162785 | 0.001437343 | 0.001785486 |
| 0.100462934 | 0.100635403 | 0.000303578 | 0.000516412 | 0.000664109 |
| 0.108284930 | 0.090818639 | 0.001162785 | 0.001437343 | 0.001785486 |
| 0.071870560 | 0.065057964 | 0.000303578 | 0.000516412 | 0.000664109 |
| 0.687756817 | 0.677338744 | 0.001162785 | 0.001437343 | 0.001785486 |
| 0.624191313 | 0.657776361 | 0.000303578 | 0.000516412 | 0.000664109 |
| 0.769377475 | 0.779100049 | 0.001162785 | 0.001437343 | 0.001785486 |
| 0.725702390 | 0.765133867 | 0.000303578 | 0.000516412 | 0.000664109 |
| 0.188823065 | 0.173640374 | 0.001162785 | 0.001437343 | 0.001785486 |
| 0.140370199 | 0.142683933 | 0.000303578 | 0.000516412 | 0.000664109 |
| 0.699509691 | 0.684645325 | 0.001162785 | 0.001437343 | 0.001785486 |
| 0.631423187 | 0.665404662 | 0.000303578 | 0.000516412 | 0.000664109 |
| 0.778734655 | 0.788660901 | 0.001162785 | 0.001437343 | 0.001785486 |
| 0.729627744 | 0.774951487 | 0.000303578 | 0.000516412 | 0.000664109 |
| 0.379599048 | 0.376415991 | 0.001162785 | 0.001437343 | 0.001785486 |
| 0.363117169 | 0.351186322 | 0.000303578 | 0.000516412 | 0.000664109 |
| 0.232236597 | 0.203922679 | 0.001162785 | 0.001437343 | 0.001785486 |
| 0.191791714 | 0.172067315 | 0.000303578 | 0.000516412 | 0.000664109 |
| 0.240896613 | 0.213089678 | 0.001162785 | 0.001437343 | 0.001785486 |
| 0.198669906 | 0.180798796 | 0.000303578 | 0.000516412 | 0.000664109 |
| 0.983293924 | 0.984340669 | 0.001162785 | 0.001437343 | 0.001785486 |
| 0.957880463 | 0.984130843 | 0.000303578 | 0.000516412 | 0.000664109 |
| 0.433571225 | 0.444025886 | 0.001162785 | 0.001437343 | 0.001785486 |
| 0.414734524 | 0.4138874 | 0.000303578 | 0.000516412 | 0.000664109 |
| 0.15220159 | 0.135698280 | 0.001162785 | 0.001437343 | 0.001785486 |
| 0.105434494 | 0.106717971 | 0.000303578 | 0.000516412 | 0.000664109 |

| | | | | |
|---|---|---|---|---|
| 0.436366515 | 0.446187422 | 0.001162785 | 0.001437343 | 0.001785486 |
| 0.418151958 | 0.416291534 | 0.000303578 | 0.000516412 | 0.000664109 |
| 0.877614304 | 0.892698511 | 0.001162785 | 0.001437343 | 0.001785486 |
| 0.836273539 | 0.886799029 | 0.000303578 | 0.000516412 | 0.000664109 |
| 0.534692805 | 0.557453751 | 0.001162785 | 0.001437343 | 0.001785486 |
| 0.478529210 | 0.531375323 | 0.000303578 | 0.000516412 | 0.000664109 |
| 0.526354775 | 0.552566647 | 0.001162785 | 0.001437343 | 0.001785486 |
| 0.225077369 | 0.209046139 | 0.000303578 | 0.000516412 | 0.000664109 |
| 0.543053289 | 0.571616723 | 0.001162785 | 0.001437343 | 0.001785486 |
| 0.687421808 | 0.708609958 | 0.000303578 | 0.000516412 | 0.000664109 |
| 0.609827498 | 0.603829724 | 0.001162785 | 0.001437343 | 0.001785486 |
| 0.54736547 | 0.579843377 | 0.000303578 | 0.000516412 | 0.000664109 |
| 0.066377602 | 0.051060313 | 0.001162785 | 0.001437343 | 0.001785486 |
| 0.176140738 | 0.160798931 | 0.000303578 | 0.000516412 | 0.000664109 |
| 0.717934895 | 0.719449800 | 0.001162785 | 0.001437343 | 0.001785486 |
| 0.637339834 | 0.675019799 | 0.000303578 | 0.000516412 | 0.000664109 |
| 0.210535824 | 0.188941015 | 0.001162785 | 0.001437343 | 0.001785486 |
| 0.145257851 | 0.143058122 | 0.000303578 | 0.000516412 | 0.000664109 |
| 0.868949558 | 0.885447849 | 0.001162785 | 0.001437343 | 0.001785486 |
| 0.99671111 | 0.999952160 | 0.000303578 | 0.000516412 | 0.000664109 |
| 0.963254980 | 0.967642763 | 0.001162785 | 0.001437343 | 0.001785486 |
| 0.946691099 | 0.967079366 | 0.000303578 | 0.000516412 | 0.000664109 |
| 0.889831749 | 0.894967452 | 0.001162785 | 0.001437343 | 0.001785486 |
| 0.843414613 | 0.888654772 | 0.000303578 | 0.000516412 | 0.000664109 |
| 0.224962838 | 0.199227016 | 0.001162785 | 0.001437343 | 0.001785486 |
| 0.186497518 | 0.167495888 | 0.000303578 | 0.000516412 | 0.000664109 |
| 0.928432639 | 0.945078592 | 0.001162785 | 0.001437343 | 0.001785486 |
| 0.893120354 | 0.942119673 | 0.000303578 | 0.000516412 | 0.000664109 |
| 0.624722411 | 0.624777654 | 0.001162785 | 0.001437343 | 0.001785486 |
| 0.555998678 | 0.602337012 | 0.000303578 | 0.000516412 | 0.000664109 |
| 0.761630337 | 0.776923092 | 0.001162785 | 0.001437343 | 0.001785486 |
| 0.718713003 | 0.762829983 | 0.000303578 | 0.000516412 | 0.000664109 |
| 0.718682838 | 0.721480304 | 0.001162785 | 0.001437343 | 0.001785486 |
| 0.680603352 | 0.704440704 | 0.000303578 | 0.000516412 | 0.000664109 |
| 0.820182158 | 0.847531205 | 0.001162785 | 0.001437343 | 0.001785486 |
| 0.772609779 | 0.838046472 | 0.000303578 | 0.000516412 | 0.000664109 |
| 0.371405016 | 0.370034449 | 0.001162785 | 0.001437343 | 0.001785486 |
| 0.422310268 | 0.432297939 | 0.000303578 | 0.000516412 | 0.000664109 |
| 0.833103641 | 0.855760377 | 0.001162785 | 0.001437343 | 0.001785486 |
| 0.793513095 | 0.846234745 | 0.000303578 | 0.000516412 | 0.000664109 |
| 0.026019155 | 0.022048661 | 0.001162785 | 0.001437343 | 0.001785486 |
| 0.034999895 | 0.025499176 | 0.000303578 | 0.000516412 | 0.000664109 |
| 0.862999045 | 0.873545706 | 0.001162785 | 0.001437343 | 0.001785486 |
| 0.822578581 | 0.865743584 | 0.000303578 | 0.000516412 | 0.000664109 |
| 0.368340205 | 0.364238552 | 0.001162785 | 0.001437343 | 0.001785486 |
| 0.124386383 | 0.135913376 | 0.000303578 | 0.000516412 | 0.000664109 |

| | | | | |
|---|---|---|---|---|
| 0.668251282 | 0.669629848 | 0.001162785 | 0.001437343 | 0.001785486 |
| 0.505402574 | 0.554118026 | 0.000303578 | 0.000516412 | 0.000664109 |
| 0.179630760 | 0.168582372 | 0.001162785 | 0.001437343 | 0.001785486 |
| 0.134433623 | 0.137915815 | 0.000303578 | 0.000516412 | 0.000664109 |
| 0.284385876 | 0.256731393 | 0.001162785 | 0.001437343 | 0.001785486 |
| 0.968850383 | 0.985449593 | 0.000303578 | 0.000516412 | 0.000664109 |
| 0.319295278 | 0.317337649 | 0.001162785 | 0.001437343 | 0.001785486 |
| 0.298786912 | 0.284541402 | 0.000303578 | 0.000516412 | 0.000664109 |
| 0.843978256 | 0.863498659 | 0.001162785 | 0.001437343 | 0.001785486 |
| 0.806567187 | 0.855001018 | 0.000303578 | 0.000516412 | 0.000664109 |
| 0.473646578 | 0.490967647 | 0.001162785 | 0.001437343 | 0.001785486 |
| 0.919167536 | 0.963744199 | 0.000303578 | 0.000516412 | 0.000664109 |
| 0.667565897 | 0.667460155 | 0.001162785 | 0.001437343 | 0.001785486 |
| 0.662749201 | 0.695994740 | 0.000303578 | 0.000516412 | 0.000664109 |
| 0.290565760 | 0.2626989 | 0.001162785 | 0.001437343 | 0.001785486 |
| 0.261189139 | 0.234905761 | 0.000303578 | 0.000516412 | 0.000664109 |
| 0.251342850 | 0.223967793 | 0.001162785 | 0.001437343 | 0.001785486 |
| 0.206080009 | 0.191589895 | 0.000303578 | 0.000516412 | 0.000664109 |
| 0.958518248 | 0.966398582 | 0.001162785 | 0.001437343 | 0.001785486 |
| 0.935950113 | 0.964967096 | 0.000303578 | 0.000516412 | 0.000664109 |
| 0.852591724 | 0.871679337 | 0.001162785 | 0.001437343 | 0.001785486 |
| 0.811957837 | 0.863668839 | 0.000303578 | 0.000516412 | 0.000664109 |
| 0.440158776 | 0.446874452 | 0.001162785 | 0.001437343 | 0.001785486 |
| 0.975586233 | 0.996735821 | 0.000303578 | 0.000516412 | 0.000664109 |
| 0.209169686 | 0.186955683 | 0.001162785 | 0.001437343 | 0.001785486 |
| 0.216460929 | 0.197930674 | 0.000303578 | 0.000516412 | 0.000664109 |
| 0.461800172 | 0.482084504 | 0.001162785 | 0.001437343 | 0.001785486 |
| 0.444298552 | 0.453557289 | 0.000303578 | 0.000516412 | 0.000664109 |
| 0.747623797 | 0.752298544 | 0.001162785 | 0.001437343 | 0.001785486 |
| 0.704426066 | 0.719207079 | 0.000303578 | 0.000516412 | 0.000664109 |
| 0.229602371 | 0.199962128 | 0.001162785 | 0.001437343 | 0.001785486 |
| 0.403712529 | 0.394876457 | 0.000303578 | 0.000516412 | 0.000664109 |
| 0.836527143 | 0.861156380 | 0.001162785 | 0.001437343 | 0.001785486 |
| 0.383822150 | 0.387437252 | 0.000303578 | 0.000516412 | 0.000664109 |
| 0.340316302 | 0.337497317 | 0.001162785 | 0.001437343 | 0.001785486 |
| 0.328052598 | 0.305034048 | 0.000303578 | 0.000516412 | 0.000664109 |
| 0.394600259 | 0.404055098 | 0.001162785 | 0.001437343 | 0.001785486 |
| 0.291399422 | 0.275508062 | 0.000303578 | 0.000516412 | 0.000664109 |
| 0.178977409 | 0.167519281 | 0.001162785 | 0.001437343 | 0.001785486 |
| 0.049050298 | 0.035977778 | 0.000303578 | 0.000516412 | 0.000664109 |
| 0.199574505 | 0.185956553 | 0.001162785 | 0.001437343 | 0.001785486 |
| 0.638307621 | 0.682859889 | 0.000303578 | 0.000516412 | 0.000664109 |
| 0.553205148 | 0.575500506 | 0.001162785 | 0.001437343 | 0.001785486 |
| 0.666257792 | 0.699164065 | 0.000303578 | 0.000516412 | 0.000664109 |
| 0.740939902 | 0.748160976 | 0.001162785 | 0.001437343 | 0.001785486 |
| 0.707195789 | 0.732384451 | 0.000303578 | 0.000516412 | 0.000664109 |

| | | | | |
|---|---|---|---|---|
| 0.951538330 | 0.965756591 | 0.001162785 | 0.001437343 | 0.001785486 |
| 0.926511090 | 0.964185359 | 0.000303578 | 0.000516412 | 0.000664109 |
| 0.578090881 | 0.589132852 | 0.001162785 | 0.001437343 | 0.001785486 |
| 0.791342219 | 0.843460671 | 0.000303578 | 0.000516412 | 0.000664109 |
| 0.487987196 | 0.507669638 | 0.001162785 | 0.001437343 | 0.001785486 |
| 0.163627177 | 0.155066770 | 0.000303578 | 0.000516412 | 0.000664109 |
| 0.477883454 | 0.498885867 | 0.001162785 | 0.001437343 | 0.001785486 |
| 0.295887482 | 0.284067808 | 0.000303578 | 0.000516412 | 0.000664109 |
| 0.355646825 | 0.344753061 | 0.001162785 | 0.001437343 | 0.001785486 |
| 0.339202958 | 0.312235657 | 0.000303578 | 0.000516412 | 0.000664109 |
| 0.992424866 | 0.996243349 | 0.001162785 | 0.001437343 | 0.001785486 |
| 0.987818020 | 0.997505312 | 0.000303578 | 0.000516412 | 0.000664109 |
| 0.417534238 | 0.423737822 | 0.001162785 | 0.001437343 | 0.001785486 |
| 0.654138091 | 0.694042617 | 0.000303578 | 0.000516412 | 0.000664109 |
| 0.934332301 | 0.945575272 | 0.001162785 | 0.001437343 | 0.001785486 |
| 0.887791995 | 0.941799649 | 0.000303578 | 0.000516412 | 0.000664109 |
| 0.557889684 | 0.576723425 | 0.001162785 | 0.001437343 | 0.001785486 |
| 0.740736992 | 0.782255668 | 0.000303578 | 0.000516412 | 0.000664109 |
| 0.554314989 | 0.575624687 | 0.001162785 | 0.001437343 | 0.001785486 |
| 0.491076242 | 0.550871292 | 0.000303578 | 0.000516412 | 0.000664109 |
| 0.111528061 | 0.092696836 | 0.001162785 | 0.001437343 | 0.001785486 |
| 0.118630405 | 0.121677854 | 0.000303578 | 0.000516412 | 0.000664109 |
| 0.466672285 | 0.482368675 | 0.001162785 | 0.001437343 | 0.001785486 |
| 0.442407584 | 0.453443719 | 0.000303578 | 0.000516412 | 0.000664109 |
| 0.121801335 | 0.102687650 | 0.001162785 | 0.001437343 | 0.001785486 |
| 0.083320091 | 0.076429825 | 0.000303578 | 0.000516412 | 0.000664109 |
| 0.522696012 | 0.551102247 | 0.001162785 | 0.001437343 | 0.001785486 |
| 0.475450883 | 0.525422960 | 0.000303578 | 0.000516412 | 0.000664109 |
| 0.904263389 | 0.908670103 | 0.001162785 | 0.001437343 | 0.001785486 |
| 0.768593399 | 0.811424496 | 0.000303578 | 0.000516412 | 0.000664109 |
| 0.019277169 | 0.017703732 | 0.001162785 | 0.001437343 | 0.001785486 |
| 0.271195983 | 0.254763898 | 0.000303578 | 0.000516412 | 0.000664109 |
| 0.973055900 | 0.981440742 | 0.001162785 | 0.001437343 | 0.001785486 |
| 0.956427356 | 0.982055708 | 0.000303578 | 0.000516412 | 0.000664109 |
| 0.047554320 | 0.038824717 | 0.001162785 | 0.001437343 | 0.001785486 |
| 0.157538614 | 0.149763492 | 0.000303578 | 0.000516412 | 0.000664109 |
| 0.372403511 | 0.371841324 | 0.001162785 | 0.001437343 | 0.001785486 |
| 0.676901736 | 0.700301951 | 0.000303578 | 0.000516412 | 0.000664109 |
| 0.344622597 | 0.339558545 | 0.001162785 | 0.001437343 | 0.001785486 |
| 0.334786202 | 0.307337966 | 0.000303578 | 0.000516412 | 0.000664109 |
| 0.643294874 | 0.508512860 | 0.004605957 | 0.005800995 | 0.006869736 |
| 0.579952597 | 0.467638327 | 0.002626182 | 0.003422277 | 0.012806334 |
| 0.710021700 | 0.557369896 | 0.004605957 | 0.005800995 | 0.006869736 |
| 0.285344294 | 0.221297552 | 0.002626182 | 0.003422277 | 0.012806334 |
| 0.079969752 | 0.086487989 | 0.004605957 | 0.005800995 | 0.006869736 |
| 0.483228053 | 0.384284412 | 0.002626182 | 0.003422277 | 0.012806334 |

| | | | | | |
|---|---|---|---|---|---|
| 0.026556281 | 0.041331279 | 0.004605957 | 0.005800995 | 0.006869736 | |
| 0.015335828 | 0.018210688 | 0.002626182 | 0.003422277 | 0.012806334 | 0.25 |
| 0.600725662 | 0.477370602 | 0.004605957 | 0.005800995 | 0.006869736 | |
| 0.380919898 | 0.300767275 | 0.002626182 | 0.003422277 | 0.012806334 | |
| 0.219522149 | 0.182668066 | 0.004605957 | 0.005800995 | 0.006869736 | |
| 0.244189869 | 0.190067497 | 0.002626182 | 0.003422277 | 0.012806334 | |
| 0.022093592 | 0.035894967 | 0.004605957 | 0.005800995 | 0.006869736 | |
| 0.106500283 | 0.090021189 | 0.002626182 | 0.003422277 | 0.012806334 | |
| 0.278360654 | 0.220289957 | 0.004605957 | 0.005800995 | 0.006869736 | |
| 0.160740504 | 0.131748796 | 0.002626182 | 0.003422277 | 0.012806334 | |
| 0.772324909 | 0.60375101 | 0.004605957 | 0.005800995 | 0.006869736 | |
| 0.667139542 | 0.536313386 | 0.002626182 | 0.003422277 | 0.012806334 | |
| 0.109194757 | 0.105594448 | 0.004605957 | 0.005800995 | 0.006869736 | |
| 0.011378077 | 0.014166774 | 0.002626182 | 0.003422277 | 0.012806334 | 0.225 |
| 0.373345061 | 0.285663610 | 0.004605957 | 0.005800995 | 0.006869736 | |
| 0.321099068 | 0.249234974 | 0.002626182 | 0.003422277 | 0.012806334 | |
| 0.363790535 | 0.278185393 | 0.004605957 | 0.005800995 | 0.006869736 | |
| 0.265914413 | 0.206704429 | 0.002626182 | 0.003422277 | 0.012806334 | |
| 0.561297731 | 0.447175901 | 0.004605957 | 0.005800995 | 0.006869736 | |
| 0.527819612 | 0.422543050 | 0.002626182 | 0.003422277 | 0.012806334 | |
| 0.451217671 | 0.352882402 | 0.004605957 | 0.005800995 | 0.006869736 | |
| 0.421979297 | 0.334721767 | 0.002626182 | 0.003422277 | 0.012806334 | |
| 0.053981771 | 0.068129204 | 0.004605957 | 0.005800995 | 0.006869736 | |
| 0.283704159 | 0.220104279 | 0.002626182 | 0.003422277 | 0.012806334 | |
| 0.150149090 | 0.132988663 | 0.004605957 | 0.005800995 | 0.006869736 | |
| 0.168791990 | 0.137094134 | 0.002626182 | 0.003422277 | 0.012806334 | |
| 0.140221298 | 0.126077174 | 0.004605957 | 0.005800995 | 0.006869736 | |
| 0.098862971 | 0.083409284 | 0.002626182 | 0.003422277 | 0.012806334 | |
| 0.233023749 | 0.191323700 | 0.004605957 | 0.005800995 | 0.006869736 | |
| 0.036134055 | 0.036130479 | 0.002626182 | 0.003422277 | 0.012806334 | |
| 0.771606982 | 0.603201653 | 0.004605957 | 0.005800995 | 0.006869736 | |
| 0.767105498 | 0.614815142 | 0.002626182 | 0.003422277 | 0.012806334 | |
| 0.980407154 | 0.834145405 | 0.004605957 | 0.005800995 | 0.006869736 | |
| 0.982983206 | 0.799881786 | 0.002626182 | 0.003422277 | 0.012806334 | |
| 0.113107634 | 0.108253205 | 0.004605957 | 0.005800995 | 0.006869736 | |
| 0.051103238 | 0.046803855 | 0.002626182 | 0.003422277 | 0.012806334 | |
| 0.735251408 | 0.576122564 | 0.004605957 | 0.005800995 | 0.006869736 | |
| 0.963736611 | 0.781764030 | 0.002626182 | 0.003422277 | 0.012806334 | |
| 0.068258966 | 0.078523224 | 0.004605957 | 0.005800995 | 0.006869736 | |
| 0.134658908 | 0.112971537 | 0.002626182 | 0.003422277 | 0.012806334 | |
| 0.023933749 | 0.038060542 | 0.004605957 | 0.005800995 | 0.006869736 | |
| 0.004677150 | 0.007123006 | 0.002626182 | 0.003422277 | 0.012806334 | 0.175 |
| 0.275593159 | 0.218469397 | 0.004605957 | 0.005800995 | 0.006869736 | |
| 0.044760842 | 0.042077133 | 0.002626182 | 0.003422277 | 0.012806334 | |
| 0.152949478 | 0.134985977 | 0.004605957 | 0.005800995 | 0.006869736 | |
| 0.327958534 | 0.254482632 | 0.002626182 | 0.003422277 | 0.012806334 | |

| | | | | | |
|---|---|---|---|---|---|
| 0.267696315 | 0.213328101 | 0.004605957 | 0.005800995 | 0.006869736 | |
| 0.182779849 | 0.145779202 | 0.002626182 | 0.003422277 | 0.012806334 | |
| 0.040864352 | 0.056603980 | 0.004605957 | 0.005800995 | 0.006869736 | |
| 0.023776642 | 0.025903688 | 0.002626182 | 0.003422277 | 0.012806334 | |
| 0.510222868 | 0.405701277 | 0.004605957 | 0.005800995 | 0.006869736 | |
| 0.492864596 | 0.392152217 | 0.002626182 | 0.003422277 | 0.012806334 | |
| 0.707070561 | 0.555119103 | 0.004605957 | 0.005800995 | 0.006869736 | |
| 0.702285103 | 0.565048855 | 0.002626182 | 0.003422277 | 0.012806334 | |
| 0.044826078 | 0.060057932 | 0.004605957 | 0.005800995 | 0.006869736 | |
| 0.002032777 | 0.004146858 | 0.002626182 | 0.003422277 | 0.012806334 | 0.175 |
| 0.968275417 | 0.798761630 | 0.004605957 | 0.005800995 | 0.006869736 | |
| 0.92442045 | 0.748451286 | 0.002626182 | 0.003422277 | 0.012806334 | |
| 0.165566727 | 0.144162725 | 0.004605957 | 0.005800995 | 0.006869736 | |
| 0.201433091 | 0.158162804 | 0.002626182 | 0.003422277 | 0.012806334 | |
| 0.017585451 | 0.030361242 | 0.004605957 | 0.005800995 | 0.006869736 | |
| 0.008150220 | 0.010962691 | 0.002626182 | 0.003422277 | 0.012806334 | 0.175 |
| 0.675199219 | 0.531585984 | 0.004605957 | 0.005800995 | 0.006869736 | |
| 0.539505767 | 0.432821021 | 0.002626182 | 0.003422277 | 0.012806334 | |
| 0.380199851 | 0.291166717 | 0.004605957 | 0.005800995 | 0.006869736 | |
| 0.256689932 | 0.199905047 | 0.002626182 | 0.003422277 | 0.012806334 | |
| 0.070115494 | 0.079881974 | 0.004605957 | 0.005800995 | 0.006869736 | |
| 0.931476930 | 0.754533790 | 0.002626182 | 0.003422277 | 0.012806334 | |
| 0.818794629 | 0.639547481 | 0.004605957 | 0.005800995 | 0.006869736 | |
| 0.249622156 | 0.194427919 | 0.002626182 | 0.003422277 | 0.012806334 | |
| 0.957820435 | 0.778054587 | 0.004605957 | 0.005800995 | 0.006869736 | |
| 0.865040793 | 0.695965920 | 0.002626182 | 0.003422277 | 0.012806334 | |
| 0.253869208 | 0.204568281 | 0.004605957 | 0.005800995 | 0.006869736 | |
| 0.199531993 | 0.156930308 | 0.002626182 | 0.003422277 | 0.012806334 | |
| 0.964636678 | 0.790501305 | 0.004605957 | 0.005800995 | 0.006869736 | |
| 0.349772191 | 0.271752143 | 0.002626182 | 0.003422277 | 0.012806334 | |
| 0.101143658 | 0.100203396 | 0.004605957 | 0.005800995 | 0.006869736 | |
| 0.063046599 | 0.055756329 | 0.002626182 | 0.003422277 | 0.012806334 | |
| 0.700238700 | 0.549899602 | 0.004605957 | 0.005800995 | 0.006869736 | |
| 0.615072434 | 0.495334208 | 0.002626182 | 0.003422277 | 0.012806334 | |
| 0.580319924 | 0.461744916 | 0.004605957 | 0.005800995 | 0.006869736 | |
| 0.673809369 | 0.541935575 | 0.002626182 | 0.003422277 | 0.012806334 | |
| 0.979229626 | 0.830413969 | 0.004605957 | 0.005800995 | 0.006869736 | |
| 0.898256714 | 0.725406558 | 0.002626182 | 0.003422277 | 0.012806334 | |
| 0.672355669 | 0.529533758 | 0.004605957 | 0.005800995 | 0.006869736 | |
| 0.664711153 | 0.534308398 | 0.002626182 | 0.003422277 | 0.012806334 | |
| 0.877264235 | 0.687441006 | 0.004605957 | 0.005800995 | 0.006869736 | |
| 0.717031805 | 0.575679115 | 0.002626182 | 0.003422277 | 0.012806334 | |
| 0.569380422 | 0.453371831 | 0.004605957 | 0.005800995 | 0.006869736 | |
| 0.084378977 | 0.071742605 | 0.002626182 | 0.003422277 | 0.012806334 | |
| 0.615253259 | 0.488052366 | 0.004605957 | 0.005800995 | 0.006869736 | |
| 0.451926648 | 0.358241853 | 0.002626182 | 0.003422277 | 0.012806334 | |

| | | | | | |
|---|---|---|---|---|---|
| 0.577110582 | 0.459186326 | 0.004605957 | 0.005800995 | 0.006869736 | |
| 0.550079674 | 0.441841599 | 0.002626182 | 0.003422277 | 0.012806334 | |
| 0.499316811 | 0.395661157 | 0.004605957 | 0.005800995 | 0.006869736 | |
| 0.403909550 | 0.321314289 | 0.002626182 | 0.003422277 | 0.012806334 | |
| 0.673534379 | 0.530382332 | 0.004605957 | 0.005800995 | 0.006869736 | |
| 0.603351096 | 0.485882537 | 0.002626182 | 0.003422277 | 0.012806334 | |
| 0.417648326 | 0.323960305 | 0.004605957 | 0.005800995 | 0.006869736 | |
| 0.233522953 | 0.181953248 | 0.002626182 | 0.003422277 | 0.012806334 | |
| 0.618522229 | 0.490384594 | 0.004605957 | 0.005800995 | 0.006869736 | |
| 0.607783267 | 0.489432505 | 0.002626182 | 0.003422277 | 0.012806334 | |
| 0.295596488 | 0.231617622 | 0.004605957 | 0.005800995 | 0.006869736 | |
| 0.194812076 | 0.153818920 | 0.002626182 | 0.003422277 | 0.012806334 | |
| 0.878011746 | 0.688110930 | 0.004605957 | 0.005800995 | 0.006869736 | |
| 0.908246934 | 0.734426641 | 0.002626182 | 0.003422277 | 0.012806334 | |
| 0.822123422 | 0.642214009 | 0.004605957 | 0.005800995 | 0.006869736 | |
| 0.607866055 | 0.489498647 | 0.002626182 | 0.003422277 | 0.012806334 | |
| 0.699458121 | 0.549295511 | 0.004605957 | 0.005800995 | 0.006869736 | |
| 0.203081887 | 0.159250164 | 0.002626182 | 0.003422277 | 0.012806334 | |
| 0.746990977 | 0.584545178 | 0.004605957 | 0.005800995 | 0.006869736 | |
| 0.731161840 | 0.585888901 | 0.002626182 | 0.003422277 | 0.012806334 | |
| 0.354274928 | 0.271114243 | 0.004605957 | 0.005800995 | 0.006869736 | |
| 0.003840325 | 0.006190161 | 0.002626182 | 0.003422277 | 0.012806334 | 0.175 |
| 0.155939772 | 0.137185676 | 0.004605957 | 0.005800995 | 0.006869736 | |
| 0.192234691 | 0.152077065 | 0.002626182 | 0.003422277 | 0.012806334 | |
| 0.759819272 | 0.594146939 | 0.004605957 | 0.005800995 | 0.006869736 | |
| 0.686319893 | 0.552456332 | 0.002626182 | 0.003422277 | 0.012806334 | |
| 0.376044184 | 0.287841507 | 0.004605957 | 0.005800995 | 0.006869736 | |
| 0.204506154 | 0.160180124 | 0.002626182 | 0.003422277 | 0.012806334 | |
| 0.632183892 | 0.500348551 | 0.004605957 | 0.005800995 | 0.006869736 | |
| 0.996801390 | 0.820750202 | 0.002626182 | 0.003422277 | 0.012806334 | |
| 0.109267400 | 0.105643539 | 0.004605957 | 0.005800995 | 0.006869736 | |
| 0.578775538 | 0.466704283 | 0.002626182 | 0.003422277 | 0.012806334 | |
| 0.609139229 | 0.483672999 | 0.004605957 | 0.005800995 | 0.006869736 | |
| 0.489268961 | 0.389147980 | 0.002626182 | 0.003422277 | 0.012806334 | |
| 0.165689188 | 0.144252067 | 0.004605957 | 0.005800995 | 0.006869736 | |
| 0.154730333 | 0.127525424 | 0.002626182 | 0.003422277 | 0.012806334 | |
| 0.670001142 | 0.527837076 | 0.004605957 | 0.005800995 | 0.006869736 | |
| 0.582850338 | 0.469951238 | 0.002626182 | 0.003422277 | 0.012806334 | |
| 0.130886208 | 0.119847205 | 0.004605957 | 0.005800995 | 0.006869736 | |
| 0.061561005 | 0.054686521 | 0.002626182 | 0.003422277 | 0.012806334 | |
| 0.717972232 | 0.563419016 | 0.004605957 | 0.005800995 | 0.006869736 | |
| 0.468819480 | 0.372713302 | 0.002626182 | 0.003422277 | 0.012806334 | |
| 0.026009135 | 0.040639265 | 0.004605957 | 0.005800995 | 0.006869736 | |
| 0.044297800 | 0.041763472 | 0.002626182 | 0.003422277 | 0.012806334 | |
| 0.210990864 | 0.177112656 | 0.004605957 | 0.005800995 | 0.006869736 | |
| 0.117514173 | 0.099264064 | 0.002626182 | 0.003422277 | 0.012806334 | |

| | | | | |
|---|---|---|---|---|
| 0.557874868 | 0.444528811 | 0.004605957 | 0.005800995 | 0.006869736 |
| 0.468773722 | 0.372676052 | 0.002626182 | 0.003422277 | 0.012806334 |
| 0.607304604 | 0.482370996 | 0.004605957 | 0.005800995 | 0.006869736 |
| 0.256921015 | 0.200083729 | 0.002626182 | 0.003422277 | 0.012806334 |
| 0.536729506 | 0.427725854 | 0.004605957 | 0.005800995 | 0.006869736 |
| 0.495239993 | 0.394172511 | 0.002626182 | 0.003422277 | 0.012806334 |
| 0.888727946 | 0.697848467 | 0.004605957 | 0.005800995 | 0.006869736 |
| 0.817823568 | 0.655702208 | 0.002626182 | 0.003422277 | 0.012806334 |
| 0.414574336 | 0.321267816 | 0.004605957 | 0.005800995 | 0.006869736 |
| 0.021821149 | 0.024238043 | 0.002626182 | 0.003422277 | 0.012806334 |
| 0.763711940 | 0.597153418 | 0.004605957 | 0.005800995 | 0.006869736 |
| 0.546102656 | 0.438417146 | 0.002626182 | 0.003422277 | 0.012806334 |
| 0.069626338 | 0.079477273 | 0.004605957 | 0.005800995 | 0.006869736 |
| 0.054722869 | 0.049563969 | 0.002626182 | 0.003422277 | 0.012806334 |
| 0.619750992 | 0.491273676 | 0.004605957 | 0.005800995 | 0.006869736 |
| 0.992295198 | 0.812107389 | 0.002626182 | 0.003422277 | 0.012806334 |
| 0.404847681 | 0.312675167 | 0.004605957 | 0.005800995 | 0.006869736 |
| 0.268803717 | 0.208939298 | 0.002626182 | 0.003422277 | 0.012806334 |
| 0.743643679 | 0.582089170 | 0.004605957 | 0.005800995 | 0.006869736 |
| 0.633874277 | 0.510319320 | 0.002626182 | 0.003422277 | 0.012806334 |
| 0.576003519 | 0.458343191 | 0.004605957 | 0.005800995 | 0.006869736 |
| 0.951512737 | 0.771262806 | 0.002626182 | 0.003422277 | 0.012806334 |
| 0.953420565 | 0.770852409 | 0.004605957 | 0.005800995 | 0.006869736 |
| 0.808292762 | 0.647931541 | 0.002626182 | 0.003422277 | 0.012806334 |
| 0.414591545 | 0.321282786 | 0.004605957 | 0.005800995 | 0.006869736 |
| 0.280194365 | 0.217649711 | 0.002626182 | 0.003422277 | 0.012806334 |
| 0.369930093 | 0.282955575 | 0.004605957 | 0.005800995 | 0.006869736 |
| 0.302738420 | 0.234782044 | 0.002626182 | 0.003422277 | 0.012806334 |
| 0.326943569 | 0.252421192 | 0.004605957 | 0.005800995 | 0.006869736 |
| 0.225876522 | 0.176406218 | 0.002626182 | 0.003422277 | 0.012806334 |
| 0.492907591 | 0.389949819 | 0.004605957 | 0.005800995 | 0.006869736 |
| 0.326817736 | 0.253632843 | 0.002626182 | 0.003422277 | 0.012806334 |
| 0.547924864 | 0.436734017 | 0.004605957 | 0.005800995 | 0.006869736 |
| 0.981143291 | 0.797937828 | 0.002626182 | 0.003422277 | 0.012806334 |
| 0.832722857 | 0.650763620 | 0.004605957 | 0.005800995 | 0.006869736 |
| 0.758362771 | 0.607648173 | 0.002626182 | 0.003422277 | 0.012806334 |
| 0.271103623 | 0.215550284 | 0.004605957 | 0.005800995 | 0.006869736 |
| 0.200212622 | 0.157374208 | 0.002626182 | 0.003422277 | 0.012806334 |
| 0.966327137 | 0.794085126 | 0.004605957 | 0.005800995 | 0.006869736 |
| 0.825611262 | 0.661958868 | 0.002626182 | 0.003422277 | 0.012806334 |
| 0.231162966 | 0.190121584 | 0.004605957 | 0.005800995 | 0.006869736 |
| 0.415317082 | 0.329893138 | 0.002626182 | 0.003422277 | 0.012806334 |
| 0.991206720 | 0.872887650 | 0.004605957 | 0.005800995 | 0.006869736 |
| 0.825346437 | 0.661739972 | 0.002626182 | 0.003422277 | 0.012806334 |
| 0.341323186 | 0.262185372 | 0.004605957 | 0.005800995 | 0.006869736 |
| 0.179377177 | 0.143719264 | 0.002626182 | 0.003422277 | 0.012806334 |

| | | | | | |
|---|---|---|---|---|---|
| 0.475081628 | 0.373732027 | 0.004605957 | 0.005800995 | 0.006869736 | |
| 0.174113333 | 0.140494386 | 0.002626182 | 0.003422277 | 0.012806334 | |
| 0.910618056 | 0.718427949 | 0.004605957 | 0.005800995 | 0.006869736 | |
| 0.828558510 | 0.664492575 | 0.002626182 | 0.003422277 | 0.012806334 | |
| 0.333303953 | 0.25675139 | 0.004605957 | 0.005800995 | 0.006869736 | |
| 0.692750753 | 0.557735696 | 0.002626182 | 0.003422277 | 0.012806334 | |
| 0.982450290 | 0.841091717 | 0.004605957 | 0.005800995 | 0.006869736 | |
| 0.919161640 | 0.743851440 | 0.002626182 | 0.003422277 | 0.012806334 | |
| 0.534223779 | 0.425679562 | 0.004605957 | 0.005800995 | 0.006869736 | |
| 0.461782328 | 0.366918374 | 0.002626182 | 0.003422277 | 0.012806334 | |
| 0.508057707 | 0.403703368 | 0.004605957 | 0.005800995 | 0.006869736 | |
| 0.306845347 | 0.238206713 | 0.002626182 | 0.003422277 | 0.012806334 | |
| 0.367284880 | 0.280880560 | 0.004605957 | 0.005800995 | 0.006869736 | |
| 0.862526552 | 0.693882255 | 0.002626182 | 0.003422277 | 0.012806334 | |
| 0.600734208 | 0.477377239 | 0.004605957 | 0.005800995 | 0.006869736 | |
| 0.073204793 | 0.063127767 | 0.002626182 | 0.003422277 | 0.012806334 | |
| 0.657063829 | 0.518562381 | 0.004605957 | 0.005800995 | 0.006869736 | |
| 0.332635555 | 0.257975042 | 0.002626182 | 0.003422277 | 0.012806334 | |
| 0.133690835 | 0.121682873 | 0.004605957 | 0.005800995 | 0.006869736 | |
| 0.038390579 | 0.037692191 | 0.002626182 | 0.003422277 | 0.012806334 | |
| 0.424638528 | 0.330170749 | 0.004605957 | 0.005800995 | 0.006869736 | |
| 0.602735755 | 0.485405410 | 0.002626182 | 0.003422277 | 0.012806334 | |
| 0.907122775 | 0.715141909 | 0.004605957 | 0.005800995 | 0.006869736 | |
| 0.996509183 | 0.820101622 | 0.002626182 | 0.003422277 | 0.012806334 | |
| 0.740721255 | 0.579984921 | 0.004605957 | 0.005800995 | 0.006869736 | |
| 0.005643742 | 0.008264508 | 0.002626182 | 0.003422277 | 0.012806334 | 0.175 |
| 0.168987344 | 0.146748656 | 0.004605957 | 0.005800995 | 0.006869736 | |
| 0.797023235 | 0.639132229 | 0.002626182 | 0.003422277 | 0.012806334 | |
| 0.769513416 | 0.601603839 | 0.004605957 | 0.005800995 | 0.006869736 | |
| 0.674123577 | 0.542204836 | 0.002626182 | 0.003422277 | 0.012806334 | |
| 0.136917082 | 0.123827759 | 0.004605957 | 0.005800995 | 0.006869736 | |
| 0.141984975 | 0.118306277 | 0.002626182 | 0.003422277 | 0.012806334 | |
| 0.914199107 | 0.721993896 | 0.004605957 | 0.005800995 | 0.006869736 | |
| 0.809422211 | 0.648858492 | 0.002626182 | 0.003422277 | 0.012806334 | |
| 0.145746362 | 0.129867868 | 0.004605957 | 0.005800995 | 0.006869736 | |
| 0.110331529 | 0.093219871 | 0.002626182 | 0.003422277 | 0.012806334 | |
| 0.438238875 | 0.342060398 | 0.004605957 | 0.005800995 | 0.006869736 | |
| 0.427848592 | 0.338892508 | 0.002626182 | 0.003422277 | 0.012806334 | |
| 0.845912154 | 0.660808380 | 0.004605957 | 0.005800995 | 0.006869736 | |
| 0.408593535 | 0.324911974 | 0.002626182 | 0.003422277 | 0.012806334 | |
| 0.898804405 | 0.707240803 | 0.004605957 | 0.005800995 | 0.006869736 | |
| 0.965726383 | 0.783490029 | 0.002626182 | 0.003422277 | 0.012806334 | |
| 0.102726332 | 0.101259294 | 0.004605957 | 0.005800995 | 0.006869736 | |
| 0.915166722 | 0.740399346 | 0.002626182 | 0.003422277 | 0.012806334 | |
| 0.221797599 | 0.184116033 | 0.004605957 | 0.005800995 | 0.006869736 | |
| 0.186953416 | 0.148444580 | 0.002626182 | 0.003422277 | 0.012806334 | |

| | | | | |
|---|---|---|---|---|
| 0.0411484507 | 0.0568549148 | 0.0046059573 | 0.0058009956 | 0.0068697368 |
| 0.7074816312 | 0.5688697297 | 0.0026261825 | 0.0034222778 | 0.0128063345 |
| 0.2763273599 | 0.2189503879 | 0.0046059573 | 0.0058009956 | 0.0068697368 |
| 0.2416772727 | 0.188068148 | 0.0026261825 | 0.0034222778 | 0.0128063345 |

| Genus | Family | Order |
|---|---|---|
| Acetatifactor | Lachnospiraceae | Eubacteriales |
| Acetatifactor | Lachnospiraceae | Eubacteriales |
| Acutalibacter | Oscillospiraceae | Eubacteriales |
| Acutalibacter | Oscillospiraceae | Eubacteriales |
| Akkermansia | Akkermansiaceae | Verrucomicrobiales |
| Akkermansia | Akkermansiaceae | Verrucomicrobiales |
| Anaerotruncus | Oscillospiraceae | Eubacteriales |
| Anaerotruncus | Oscillospiraceae | Eubacteriales |
| Bacteria_unclassified | Bacteria_unclassified | Bacteria_unclassified |
| Bacteria_unclassified | Bacteria_unclassified | Bacteria_unclassified |
| Bacteria_unclassified | Bacteria_unclassified | Bacteria_unclassified |
| Bacteria_unclassified | Bacteria_unclassified | Bacteria_unclassified |
| Bacteria_unclassified | Bacteria_unclassified | Bacteria_unclassified |
| Bacteria_unclassified | Bacteria_unclassified | Bacteria_unclassified |
| Bacteria_unclassified | Bacteria_unclassified | Bacteria_unclassified |
| Bacteria_unclassified | Bacteria_unclassified | Bacteria_unclassified |
| Bacteria_unclassified | Bacteria_unclassified | Bacteria_unclassified |
| Bacteria_unclassified | Bacteria_unclassified | Bacteria_unclassified |
| Bacteria_unclassified | Bacteria_unclassified | Bacteria_unclassified |
| Bacteria_unclassified | Bacteria_unclassified | Bacteria_unclassified |
| Bacteria_unclassified | Bacteria_unclassified | Bacteria_unclassified |
| Bacteria_unclassified | Bacteria_unclassified | Bacteria_unclassified |
| Bacteria_unclassified | Bacteria_unclassified | Bacteria_unclassified |
| Bacteria_unclassified | Bacteria_unclassified | Bacteria_unclassified |
| Bacteria_unclassified | Bacteria_unclassified | Bacteria_unclassified |
| Bacteria_unclassified | Bacteria_unclassified | Bacteria_unclassified |
| Bacteroides | Bacteroidaceae | Bacteroidales |
| Bacteroides | Bacteroidaceae | Bacteroidales |
| Clostridia_unclassified | Clostridia_unclassified | Clostridia_unclassified |
| Clostridia_unclassified | Clostridia_unclassified | Clostridia_unclassified |
| Clostridiaceae_unclassified | Clostridiaceae | Eubacteriales |
| Clostridiaceae_unclassified | Clostridiaceae | Eubacteriales |
| Clostridiaceae_unclassified | Clostridiaceae | Eubacteriales |
| Clostridiaceae_unclassified | Clostridiaceae | Eubacteriales |
| Eubacteriales_unclassified | Eubacteriales_unclassified | Eubacteriales |
| Eubacteriales_unclassified | Eubacteriales_unclassified | Eubacteriales |
| Erysipelatoclostridium | Erysipelotrichaceae | Erysipelotrichales |
| Erysipelatoclostridium | Erysipelotrichaceae | Erysipelotrichales |
| Clostridium | Clostridiaceae | Eubacteriales |
| Clostridium | Clostridiaceae | Eubacteriales |
| Eubacteriaceae_unclassified | Eubacteriaceae | Eubacteriales |
| Eubacteriaceae_unclassified | Eubacteriaceae | Eubacteriales |
| Eubacteriaceae_unclassified | Eubacteriaceae | Eubacteriales |
| Eubacteriaceae_unclassified | Eubacteriaceae | Eubacteriales |

| | | |
|---|---|---|
| GGB20146 | FGB77306 | OFGB77306 |
| GGB20146 | FGB77306 | OFGB77306 |
| GGB20149 | Lachnospiraceae | Eubacteriales |
| GGB20149 | Lachnospiraceae | Eubacteriales |
| GGB22635 | Eggerthellaceae | Eggerthellales |
| GGB22635 | Eggerthellaceae | Eggerthellales |
| GGB25041 | Lachnospiraceae | Eubacteriales |
| GGB25041 | Lachnospiraceae | Eubacteriales |
| GGB28379 | FGB9506 | OFGB9506 |
| GGB28379 | FGB9506 | OFGB9506 |
| GGB28382 | FGB9508 | OFGB9508 |
| GGB28382 | FGB9508 | OFGB9508 |
| GGB28392 | FGB9512 | OFGB9512 |
| GGB28392 | FGB9512 | OFGB9512 |
| GGB28404 | FGB2838 | OFGB2838 |
| GGB28404 | FGB2838 | OFGB2838 |
| GGB28418 | FGB2838 | OFGB2838 |
| GGB28418 | FGB2838 | OFGB2838 |
| GGB28422 | FGB2838 | OFGB2838 |
| GGB28422 | FGB2838 | OFGB2838 |
| GGB28431 | Pumilibacteraceae | Eubacteriales |
| GGB28431 | Pumilibacteraceae | Eubacteriales |
| GGB28456 | FGB2833 | OFGB2833 |
| GGB28456 | FGB2833 | OFGB2833 |
| GGB28782 | Eubacteriaceae | Eubacteriales |
| GGB28782 | Eubacteriaceae | Eubacteriales |
| GGB28792 | Lachnospiraceae | Eubacteriales |
| GGB28792 | Lachnospiraceae | Eubacteriales |
| GGB28798 | Lachnospiraceae | Eubacteriales |
| GGB28798 | Lachnospiraceae | Eubacteriales |
| GGB28810 | FGB9622 | OFGB9622 |
| GGB28810 | FGB9622 | OFGB9622 |
| GGB28828 | FGB77305 | OFGB77305 |
| GGB28828 | FGB77305 | OFGB77305 |
| GGB28851 | Clostridiaceae | Eubacteriales |
| GGB28851 | Clostridiaceae | Eubacteriales |
| GGB28865 | Lachnospiraceae | Eubacteriales |
| GGB28865 | Lachnospiraceae | Eubacteriales |
| GGB28868 | Lachnospiraceae | Eubacteriales |
| GGB28868 | Lachnospiraceae | Eubacteriales |
| GGB28869 | Lachnospiraceae | Eubacteriales |
| GGB28869 | Lachnospiraceae | Eubacteriales |
| GGB28875 | Lachnospiraceae | Eubacteriales |
| GGB28875 | Lachnospiraceae | Eubacteriales |
| GGB28881 | FGB9633 | OFGB9633 |
| GGB28881 | FGB9633 | OFGB9633 |

| | | |
|---|---|---|
| GGB28892 | Bacteria_unclassified | Bacteria_unclassified |
| GGB28892 | Bacteria_unclassified | Bacteria_unclassified |
| GGB28893 | Bacteria_unclassified | Bacteria_unclassified |
| GGB28893 | Bacteria_unclassified | Bacteria_unclassified |
| GGB28898 | Bacteria_unclassified | Bacteria_unclassified |
| GGB28898 | Bacteria_unclassified | Bacteria_unclassified |
| GGB28901 | Bacteria_unclassified | Bacteria_unclassified |
| GGB28901 | Bacteria_unclassified | Bacteria_unclassified |
| GGB28904 | Bacteria_unclassified | Bacteria_unclassified |
| GGB28904 | Bacteria_unclassified | Bacteria_unclassified |
| GGB28909 | Bacteria_unclassified | Bacteria_unclassified |
| GGB28909 | Bacteria_unclassified | Bacteria_unclassified |
| GGB28924 | Lachnospiraceae | Eubacteriales |
| GGB28924 | Lachnospiraceae | Eubacteriales |
| GGB28927 | FGB77359 | OFGB77359 |
| GGB28927 | FGB77359 | OFGB77359 |
| GGB28934 | FGB9639 | OFGB9639 |
| GGB28934 | FGB9639 | OFGB9639 |
| GGB28949 | Lachnospiraceae | Eubacteriales |
| GGB28949 | Lachnospiraceae | Eubacteriales |
| GGB28951 | Clostridiaceae | Eubacteriales |
| GGB28951 | Clostridiaceae | Eubacteriales |
| GGB28951 | Clostridiaceae | Eubacteriales |
| GGB28951 | Clostridiaceae | Eubacteriales |
| GGB28954 | Clostridiaceae | Eubacteriales |
| GGB28954 | Clostridiaceae | Eubacteriales |
| GGB28960 | Clostridiaceae | Eubacteriales |
| GGB28960 | Clostridiaceae | Eubacteriales |
| GGB28964 | Clostridiaceae | Eubacteriales |
| GGB28964 | Clostridiaceae | Eubacteriales |
| GGB28967 | Clostridiaceae | Eubacteriales |
| GGB28967 | Clostridiaceae | Eubacteriales |
| GGB28996 | FGB9656 | OFGB9656 |
| GGB28996 | FGB9656 | OFGB9656 |
| GGB29531 | FGB9827 | OFGB9827 |
| GGB29531 | FGB9827 | OFGB9827 |
| GGB29685 | Eubacteriaceae | Eubacteriales |
| GGB29685 | Eubacteriaceae | Eubacteriales |
| GGB30141 | FGB77303 | OFGB77303 |
| GGB30141 | FGB77303 | OFGB77303 |
| GGB30145 | Clostridiaceae | Eubacteriales |
| GGB30145 | Clostridiaceae | Eubacteriales |
| GGB30286 | Eubacteriales_unclassified | Eubacteriales |
| GGB30286 | Eubacteriales_unclassified | Eubacteriales |
| GGB30300 | FGB72709 | OFGB72709 |
| GGB30300 | FGB72709 | OFGB72709 |

| | | |
|---|---|---|
| GGB30303 | Oscillospiraceae | Eubacteriales |
| GGB30303 | Oscillospiraceae | Eubacteriales |
| GGB30450 | Oscillospiraceae | Eubacteriales |
| GGB30450 | Oscillospiraceae | Eubacteriales |
| GGB30453 | Oscillospiraceae | Eubacteriales |
| GGB30453 | Oscillospiraceae | Eubacteriales |
| GGB30454 | Oscillospiraceae | Eubacteriales |
| GGB30454 | Oscillospiraceae | Eubacteriales |
| GGB30455 | Oscillospiraceae | Eubacteriales |
| GGB30455 | Oscillospiraceae | Eubacteriales |
| GGB30456 | Oscillospiraceae | Eubacteriales |
| GGB30456 | Oscillospiraceae | Eubacteriales |
| GGB30457 | Oscillospiraceae | Eubacteriales |
| GGB30457 | Oscillospiraceae | Eubacteriales |
| GGB30461 | Oscillospiraceae | Eubacteriales |
| GGB30461 | Oscillospiraceae | Eubacteriales |
| GGB30461 | Oscillospiraceae | Eubacteriales |
| GGB30461 | Oscillospiraceae | Eubacteriales |
| GGB30461 | Oscillospiraceae | Eubacteriales |
| GGB30461 | Oscillospiraceae | Eubacteriales |
| GGB30463 | Oscillospiraceae | Eubacteriales |
| GGB30463 | Oscillospiraceae | Eubacteriales |
| GGB30473 | Oscillospiraceae | Eubacteriales |
| GGB30473 | Oscillospiraceae | Eubacteriales |
| GGB30475 | Oscillospiraceae | Eubacteriales |
| GGB30475 | Oscillospiraceae | Eubacteriales |
| GGB30861 | FGB77153 | OFGB77153 |
| GGB30861 | FGB77153 | OFGB77153 |
| GGB31312 | FGB1791 | OFGB1791 |
| GGB31312 | FGB1791 | OFGB1791 |
| GGB31438 | FGB10290 | OFGB10290 |
| GGB31438 | FGB10290 | OFGB10290 |
| GGB3171 | Oscillospiraceae | Eubacteriales |
| GGB3171 | Oscillospiraceae | Eubacteriales |
| GGB31762 | FGB10289 | OFGB10289 |
| GGB31762 | FGB10289 | OFGB10289 |
| GGB31823 | FGB1765 | OFGB1765 |
| GGB31823 | FGB1765 | OFGB1765 |
| GGB31838 | FGB1765 | OFGB1765 |
| GGB31838 | FGB1765 | OFGB1765 |
| GGB31841 | FGB1765 | OFGB1765 |
| GGB31841 | FGB1765 | OFGB1765 |
| GGB32371 | FGB10667 | OFGB10667 |
| GGB32371 | FGB10667 | OFGB10667 |
| GGB3793 | Lachnospiraceae | Eubacteriales |
| GGB3793 | Lachnospiraceae | Eubacteriales |

| | | |
|---|---|---|
| GGB42601 | Clostridiaceae | Eubacteriales |
| GGB42601 | Clostridiaceae | Eubacteriales |
| GGB45514 | Oscillospiraceae | Eubacteriales |
| GGB45514 | Oscillospiraceae | Eubacteriales |
| GGB45564 | FGB75721 | OFGB75721 |
| GGB45564 | FGB75721 | OFGB75721 |
| GGB45624 | Oscillospiraceae | Eubacteriales |
| GGB45624 | Oscillospiraceae | Eubacteriales |
| GGB45656 | Christensenellaceae | Eubacteriales |
| GGB45656 | Christensenellaceae | Eubacteriales |
| GGB47127 | FGB10299 | OFGB10299 |
| GGB47127 | FGB10299 | OFGB10299 |
| GGB74395 | Oscillospiraceae | Eubacteriales |
| GGB74395 | Oscillospiraceae | Eubacteriales |
| GGB75053 | Oscillospiraceae | Eubacteriales |
| GGB75053 | Oscillospiraceae | Eubacteriales |
| GGB75109 | Lachnospiraceae | Eubacteriales |
| GGB75109 | Lachnospiraceae | Eubacteriales |
| Lachnospiraceae_unclassified | Lachnospiraceae | Eubacteriales |
| Lachnospiraceae_unclassified | Lachnospiraceae | Eubacteriales |
| Lachnospiraceae_unclassified | Lachnospiraceae | Eubacteriales |
| Lachnospiraceae_unclassified | Lachnospiraceae | Eubacteriales |
| Lachnospiraceae_unclassified | Lachnospiraceae | Eubacteriales |
| Lachnospiraceae_unclassified | Lachnospiraceae | Eubacteriales |
| Lachnospiraceae_unclassified | Lachnospiraceae | Eubacteriales |
| Lachnospiraceae_unclassified | Lachnospiraceae | Eubacteriales |
| Lachnospiraceae_unclassified | Lachnospiraceae | Eubacteriales |
| Lachnospiraceae_unclassified | Lachnospiraceae | Eubacteriales |
| Lactobacillus | Lactobacillaceae | Lactobacillales |
| Lactobacillus | Lactobacillaceae | Lactobacillales |
| Leptogranulimonas | Atopobiaceae | Coriobacteriales |
| Leptogranulimonas | Atopobiaceae | Coriobacteriales |
| Muribaculaceae_unclassified | Muribaculaceae | Bacteroidales |
| Muribaculaceae_unclassified | Muribaculaceae | Bacteroidales |
| Neglectibacter | Oscillospiraceae | Eubacteriales |
| Neglectibacter | Oscillospiraceae | Eubacteriales |
| Oscillibacter | Oscillospiraceae | Eubacteriales |
| Oscillibacter | Oscillospiraceae | Eubacteriales |
| Oscillospiraceae_unclassified | Oscillospiraceae | Eubacteriales |
| Oscillospiraceae_unclassified | Oscillospiraceae | Eubacteriales |
| Oscillospiraceae_unclassified | Oscillospiraceae | Eubacteriales |
| Oscillospiraceae_unclassified | Oscillospiraceae | Eubacteriales |
| Oscillospiraceae_unclassified | Oscillospiraceae | Eubacteriales |
| Oscillospiraceae_unclassified | Oscillospiraceae | Eubacteriales |
| Oscillospiraceae_unclassified | Oscillospiraceae | Eubacteriales |
| Oscillospiraceae_unclassified | Oscillospiraceae | Eubacteriales |

| | | |
|---|---|---|
| Parasutterella | Sutterellaceae | Burkholderiales |
| Parasutterella | Sutterellaceae | Burkholderiales |
| Schaedlerella | Lachnospiraceae | Eubacteriales |
| Schaedlerella | Lachnospiraceae | Eubacteriales |
| Turicibacter | Turicibacteraceae | Erysipelotrichales |
| Turicibacter | Turicibacteraceae | Erysipelotrichales |
| Acetatifactor | Lachnospiraceae | Eubacteriales |
| Acetatifactor | Lachnospiraceae | Eubacteriales |
| Acutalibacter | Oscillospiraceae | Eubacteriales |
| Acutalibacter | Oscillospiraceae | Eubacteriales |
| Akkermansia | Akkermansiaceae | Verrucomicrobiales |
| Akkermansia | Akkermansiaceae | Verrucomicrobiales |
| Anaerotruncus | Oscillospiraceae | Eubacteriales |
| Anaerotruncus | Oscillospiraceae | Eubacteriales |
| Bacteria_unclassified | Bacteria_unclassified | Bacteria_unclassified |
| Bacteria_unclassified | Bacteria_unclassified | Bacteria_unclassified |
| Bacteria_unclassified | Bacteria_unclassified | Bacteria_unclassified |
| Bacteria_unclassified | Bacteria_unclassified | Bacteria_unclassified |
| Bacteria_unclassified | Bacteria_unclassified | Bacteria_unclassified |
| Bacteria_unclassified | Bacteria_unclassified | Bacteria_unclassified |
| Bacteria_unclassified | Bacteria_unclassified | Bacteria_unclassified |
| Bacteria_unclassified | Bacteria_unclassified | Bacteria_unclassified |
| Bacteria_unclassified | Bacteria_unclassified | Bacteria_unclassified |
| Bacteria_unclassified | Bacteria_unclassified | Bacteria_unclassified |
| Bacteria_unclassified | Bacteria_unclassified | Bacteria_unclassified |
| Bacteria_unclassified | Bacteria_unclassified | Bacteria_unclassified |
| Bacteria_unclassified | Bacteria_unclassified | Bacteria_unclassified |
| Bacteria_unclassified | Bacteria_unclassified | Bacteria_unclassified |
| Bacteria_unclassified | Bacteria_unclassified | Bacteria_unclassified |
| Bacteria_unclassified | Bacteria_unclassified | Bacteria_unclassified |
| Bacteria_unclassified | Bacteria_unclassified | Bacteria_unclassified |
| Bacteria_unclassified | Bacteria_unclassified | Bacteria_unclassified |
| Bacteroides | Bacteroidaceae | Bacteroidales |
| Bacteroides | Bacteroidaceae | Bacteroidales |
| Clostridia_unclassified | Clostridia_unclassified | Clostridia_unclassified |
| Clostridia_unclassified | Clostridia_unclassified | Clostridia_unclassified |
| Clostridiaceae_unclassified | Clostridiaceae | Eubacteriales |
| Clostridiaceae_unclassified | Clostridiaceae | Eubacteriales |
| Clostridiaceae_unclassified | Clostridiaceae | Eubacteriales |
| Clostridiaceae_unclassified | Clostridiaceae | Eubacteriales |

| | | |
|---|---|---|
| Eubacteriales_unclassified | Eubacteriales_unclassified | Eubacteriales |
| Eubacteriales_unclassified | Eubacteriales_unclassified | Eubacteriales |
| Erysipelatoclostridium | Erysipelotrichaceae | Erysipelotrichales |
| Erysipelatoclostridium | Erysipelotrichaceae | Erysipelotrichales |
| Clostridium | Clostridiaceae | Eubacteriales |
| Clostridium | Clostridiaceae | Eubacteriales |
| Eubacteriaceae_unclassified | Eubacteriaceae | Eubacteriales |
| Eubacteriaceae_unclassified | Eubacteriaceae | Eubacteriales |
| Eubacteriaceae_unclassified | Eubacteriaceae | Eubacteriales |
| Eubacteriaceae_unclassified | Eubacteriaceae | Eubacteriales |
| GGB20146 | FGB77306 | OFGB77306 |
| GGB20146 | FGB77306 | OFGB77306 |
| GGB20149 | Lachnospiraceae | Eubacteriales |
| GGB20149 | Lachnospiraceae | Eubacteriales |
| GGB22635 | Eggerthellaceae | Eggerthellales |
| GGB22635 | Eggerthellaceae | Eggerthellales |
| GGB25041 | Lachnospiraceae | Eubacteriales |
| GGB25041 | Lachnospiraceae | Eubacteriales |
| GGB28379 | FGB9506 | OFGB9506 |
| GGB28379 | FGB9506 | OFGB9506 |
| GGB28382 | FGB9508 | OFGB9508 |
| GGB28382 | FGB9508 | OFGB9508 |
| GGB28392 | FGB9512 | OFGB9512 |
| GGB28392 | FGB9512 | OFGB9512 |
| GGB28404 | FGB2838 | OFGB2838 |
| GGB28404 | FGB2838 | OFGB2838 |
| GGB28418 | FGB2838 | OFGB2838 |
| GGB28418 | FGB2838 | OFGB2838 |
| GGB28422 | FGB2838 | OFGB2838 |
| GGB28422 | FGB2838 | OFGB2838 |
| GGB28431 | Pumilibacteraceae | Eubacteriales |
| GGB28431 | Pumilibacteraceae | Eubacteriales |
| GGB28456 | FGB2833 | OFGB2833 |
| GGB28456 | FGB2833 | OFGB2833 |
| GGB28782 | Eubacteriaceae | Eubacteriales |
| GGB28782 | Eubacteriaceae | Eubacteriales |
| GGB28792 | Lachnospiraceae | Eubacteriales |
| GGB28792 | Lachnospiraceae | Eubacteriales |
| GGB28798 | Lachnospiraceae | Eubacteriales |
| GGB28798 | Lachnospiraceae | Eubacteriales |
| GGB28810 | FGB9622 | OFGB9622 |
| GGB28810 | FGB9622 | OFGB9622 |
| GGB28828 | FGB77305 | OFGB77305 |
| GGB28828 | FGB77305 | OFGB77305 |
| GGB28851 | Clostridiaceae | Eubacteriales |
| GGB28851 | Clostridiaceae | Eubacteriales |

| | | |
|---|---|---|
| GGB28865 | Lachnospiraceae | Eubacteriales |
| GGB28865 | Lachnospiraceae | Eubacteriales |
| GGB28868 | Lachnospiraceae | Eubacteriales |
| GGB28868 | Lachnospiraceae | Eubacteriales |
| GGB28869 | Lachnospiraceae | Eubacteriales |
| GGB28869 | Lachnospiraceae | Eubacteriales |
| GGB28875 | Lachnospiraceae | Eubacteriales |
| GGB28875 | Lachnospiraceae | Eubacteriales |
| GGB28881 | FGB9633 | OFGB9633 |
| GGB28881 | FGB9633 | OFGB9633 |
| GGB28892 | Bacteria_unclassified | Bacteria_unclassified |
| GGB28892 | Bacteria_unclassified | Bacteria_unclassified |
| GGB28893 | Bacteria_unclassified | Bacteria_unclassified |
| GGB28893 | Bacteria_unclassified | Bacteria_unclassified |
| GGB28898 | Bacteria_unclassified | Bacteria_unclassified |
| GGB28898 | Bacteria_unclassified | Bacteria_unclassified |
| GGB28901 | Bacteria_unclassified | Bacteria_unclassified |
| GGB28901 | Bacteria_unclassified | Bacteria_unclassified |
| GGB28904 | Bacteria_unclassified | Bacteria_unclassified |
| GGB28904 | Bacteria_unclassified | Bacteria_unclassified |
| GGB28909 | Bacteria_unclassified | Bacteria_unclassified |
| GGB28909 | Bacteria_unclassified | Bacteria_unclassified |
| GGB28924 | Lachnospiraceae | Eubacteriales |
| GGB28924 | Lachnospiraceae | Eubacteriales |
| GGB28927 | FGB77359 | OFGB77359 |
| GGB28927 | FGB77359 | OFGB77359 |
| GGB28934 | FGB9639 | OFGB9639 |
| GGB28934 | FGB9639 | OFGB9639 |
| GGB28949 | Lachnospiraceae | Eubacteriales |
| GGB28949 | Lachnospiraceae | Eubacteriales |
| GGB28951 | Clostridiaceae | Eubacteriales |
| GGB28951 | Clostridiaceae | Eubacteriales |
| GGB28951 | Clostridiaceae | Eubacteriales |
| GGB28951 | Clostridiaceae | Eubacteriales |
| GGB28954 | Clostridiaceae | Eubacteriales |
| GGB28954 | Clostridiaceae | Eubacteriales |
| GGB28960 | Clostridiaceae | Eubacteriales |
| GGB28960 | Clostridiaceae | Eubacteriales |
| GGB28964 | Clostridiaceae | Eubacteriales |
| GGB28964 | Clostridiaceae | Eubacteriales |
| GGB28967 | Clostridiaceae | Eubacteriales |
| GGB28967 | Clostridiaceae | Eubacteriales |
| GGB28996 | FGB9656 | OFGB9656 |
| GGB28996 | FGB9656 | OFGB9656 |
| GGB29531 | FGB9827 | OFGB9827 |
| GGB29531 | FGB9827 | OFGB9827 |

| | | |
|---|---|---|
| GGB29685 | Eubacteriaceae | Eubacteriales |
| GGB29685 | Eubacteriaceae | Eubacteriales |
| GGB30141 | FGB77303 | OFGB77303 |
| GGB30141 | FGB77303 | OFGB77303 |
| GGB30145 | Clostridiaceae | Eubacteriales |
| GGB30145 | Clostridiaceae | Eubacteriales |
| GGB30286 | Eubacteriales_unclassified | Eubacteriales |
| GGB30286 | Eubacteriales_unclassified | Eubacteriales |
| GGB30300 | FGB72709 | OFGB72709 |
| GGB30300 | FGB72709 | OFGB72709 |
| GGB30303 | Oscillospiraceae | Eubacteriales |
| GGB30303 | Oscillospiraceae | Eubacteriales |
| GGB30450 | Oscillospiraceae | Eubacteriales |
| GGB30450 | Oscillospiraceae | Eubacteriales |
| GGB30453 | Oscillospiraceae | Eubacteriales |
| GGB30453 | Oscillospiraceae | Eubacteriales |
| GGB30454 | Oscillospiraceae | Eubacteriales |
| GGB30454 | Oscillospiraceae | Eubacteriales |
| GGB30455 | Oscillospiraceae | Eubacteriales |
| GGB30455 | Oscillospiraceae | Eubacteriales |
| GGB30456 | Oscillospiraceae | Eubacteriales |
| GGB30456 | Oscillospiraceae | Eubacteriales |
| GGB30457 | Oscillospiraceae | Eubacteriales |
| GGB30457 | Oscillospiraceae | Eubacteriales |
| GGB30461 | Oscillospiraceae | Eubacteriales |
| GGB30461 | Oscillospiraceae | Eubacteriales |
| GGB30461 | Oscillospiraceae | Eubacteriales |
| GGB30461 | Oscillospiraceae | Eubacteriales |
| GGB30461 | Oscillospiraceae | Eubacteriales |
| GGB30461 | Oscillospiraceae | Eubacteriales |
| GGB30463 | Oscillospiraceae | Eubacteriales |
| GGB30463 | Oscillospiraceae | Eubacteriales |
| GGB30473 | Oscillospiraceae | Eubacteriales |
| GGB30473 | Oscillospiraceae | Eubacteriales |
| GGB30475 | Oscillospiraceae | Eubacteriales |
| GGB30475 | Oscillospiraceae | Eubacteriales |
| GGB30861 | FGB77153 | OFGB77153 |
| GGB30861 | FGB77153 | OFGB77153 |
| GGB31312 | FGB1791 | OFGB1791 |
| GGB31312 | FGB1791 | OFGB1791 |
| GGB31438 | FGB10290 | OFGB10290 |
| GGB31438 | FGB10290 | OFGB10290 |
| GGB3171 | Oscillospiraceae | Eubacteriales |
| GGB3171 | Oscillospiraceae | Eubacteriales |
| GGB31762 | FGB10289 | OFGB10289 |
| GGB31762 | FGB10289 | OFGB10289 |

| | | |
|---|---|---|
| GGB31823 | FGB1765 | OFGB1765 |
| GGB31823 | FGB1765 | OFGB1765 |
| GGB31838 | FGB1765 | OFGB1765 |
| GGB31838 | FGB1765 | OFGB1765 |
| GGB31841 | FGB1765 | OFGB1765 |
| GGB31841 | FGB1765 | OFGB1765 |
| GGB32371 | FGB10667 | OFGB10667 |
| GGB32371 | FGB10667 | OFGB10667 |
| GGB3793 | Lachnospiraceae | Eubacteriales |
| GGB3793 | Lachnospiraceae | Eubacteriales |
| GGB42601 | Clostridiaceae | Eubacteriales |
| GGB42601 | Clostridiaceae | Eubacteriales |
| GGB45514 | Oscillospiraceae | Eubacteriales |
| GGB45514 | Oscillospiraceae | Eubacteriales |
| GGB45564 | FGB75721 | OFGB75721 |
| GGB45564 | FGB75721 | OFGB75721 |
| GGB45624 | Oscillospiraceae | Eubacteriales |
| GGB45624 | Oscillospiraceae | Eubacteriales |
| GGB45656 | Christensenellaceae | Eubacteriales |
| GGB45656 | Christensenellaceae | Eubacteriales |
| GGB47127 | FGB10299 | OFGB10299 |
| GGB47127 | FGB10299 | OFGB10299 |
| GGB74395 | Oscillospiraceae | Eubacteriales |
| GGB74395 | Oscillospiraceae | Eubacteriales |
| GGB75053 | Oscillospiraceae | Eubacteriales |
| GGB75053 | Oscillospiraceae | Eubacteriales |
| GGB75109 | Lachnospiraceae | Eubacteriales |
| GGB75109 | Lachnospiraceae | Eubacteriales |
| Lachnospiraceae_unclassified | Lachnospiraceae | Eubacteriales |
| Lachnospiraceae_unclassified | Lachnospiraceae | Eubacteriales |
| Lachnospiraceae_unclassified | Lachnospiraceae | Eubacteriales |
| Lachnospiraceae_unclassified | Lachnospiraceae | Eubacteriales |
| Lachnospiraceae_unclassified | Lachnospiraceae | Eubacteriales |
| Lachnospiraceae_unclassified | Lachnospiraceae | Eubacteriales |
| Lachnospiraceae_unclassified | Lachnospiraceae | Eubacteriales |
| Lachnospiraceae_unclassified | Lachnospiraceae | Eubacteriales |
| Lachnospiraceae_unclassified | Lachnospiraceae | Eubacteriales |
| Lachnospiraceae_unclassified | Lachnospiraceae | Eubacteriales |
| Lactobacillus | Lactobacillaceae | Lactobacillales |
| Lactobacillus | Lactobacillaceae | Lactobacillales |
| Leptogranulimonas | Atopobiaceae | Coriobacteriales |
| Leptogranulimonas | Atopobiaceae | Coriobacteriales |
| Muribaculaceae_unclassified | Muribaculaceae | Bacteroidales |
| Muribaculaceae_unclassified | Muribaculaceae | Bacteroidales |
| Neglectibacter | Oscillospiraceae | Eubacteriales |
| Neglectibacter | Oscillospiraceae | Eubacteriales |

| | | |
|---|---|---|
| Oscillibacter | Oscillospiraceae | Eubacteriales |
| Oscillibacter | Oscillospiraceae | Eubacteriales |
| Oscillospiraceae_unclassified | Oscillospiraceae | Eubacteriales |
| Oscillospiraceae_unclassified | Oscillospiraceae | Eubacteriales |
| Oscillospiraceae_unclassified | Oscillospiraceae | Eubacteriales |
| Oscillospiraceae_unclassified | Oscillospiraceae | Eubacteriales |
| Oscillospiraceae_unclassified | Oscillospiraceae | Eubacteriales |
| Oscillospiraceae_unclassified | Oscillospiraceae | Eubacteriales |
| Oscillospiraceae_unclassified | Oscillospiraceae | Eubacteriales |
| Oscillospiraceae_unclassified | Oscillospiraceae | Eubacteriales |
| Parasutterella | Sutterellaceae | Burkholderiales |
| Parasutterella | Sutterellaceae | Burkholderiales |
| Schaedlerella | Lachnospiraceae | Eubacteriales |
| Schaedlerella | Lachnospiraceae | Eubacteriales |
| Turicibacter | Turicibacteraceae | Erysipelotrichales |
| Turicibacter | Turicibacteraceae | Erysipelotrichales |
| Acetatifactor | Lachnospiraceae | Eubacteriales |
| Acetatifactor | Lachnospiraceae | Eubacteriales |
| Acutalibacter | Oscillospiraceae | Eubacteriales |
| Acutalibacter | Oscillospiraceae | Eubacteriales |
| Akkermansia | Akkermansiaceae | Verrucomicrobiales |
| Akkermansia | Akkermansiaceae | Verrucomicrobiales |
| Anaerotruncus | Oscillospiraceae | Eubacteriales |
| Anaerotruncus | Oscillospiraceae | Eubacteriales |
| Bacteria_unclassified | Bacteria_unclassified | Bacteria_unclassified |
| Bacteria_unclassified | Bacteria_unclassified | Bacteria_unclassified |
| Bacteria_unclassified | Bacteria_unclassified | Bacteria_unclassified |
| Bacteria_unclassified | Bacteria_unclassified | Bacteria_unclassified |
| Bacteria_unclassified | Bacteria_unclassified | Bacteria_unclassified |
| Bacteria_unclassified | Bacteria_unclassified | Bacteria_unclassified |
| Bacteria_unclassified | Bacteria_unclassified | Bacteria_unclassified |
| Bacteria_unclassified | Bacteria_unclassified | Bacteria_unclassified |
| Bacteria_unclassified | Bacteria_unclassified | Bacteria_unclassified |
| Bacteria_unclassified | Bacteria_unclassified | Bacteria_unclassified |
| Bacteria_unclassified | Bacteria_unclassified | Bacteria_unclassified |
| Bacteria_unclassified | Bacteria_unclassified | Bacteria_unclassified |
| Bacteria_unclassified | Bacteria_unclassified | Bacteria_unclassified |
| Bacteria_unclassified | Bacteria_unclassified | Bacteria_unclassified |
| Bacteria_unclassified | Bacteria_unclassified | Bacteria_unclassified |
| Bacteria_unclassified | Bacteria_unclassified | Bacteria_unclassified |
| Bacteria_unclassified | Bacteria_unclassified | Bacteria_unclassified |
| Bacteria_unclassified | Bacteria_unclassified | Bacteria_unclassified |

| | | |
|---|---|---|
| Bacteroides | Bacteroidaceae | Bacteroidales |
| Bacteroides | Bacteroidaceae | Bacteroidales |
| Clostridia_unclassified | Clostridia_unclassified | Clostridia_unclassified |
| Clostridia_unclassified | Clostridia_unclassified | Clostridia_unclassified |
| Clostridiaceae_unclassified | Clostridiaceae | Eubacteriales |
| Clostridiaceae_unclassified | Clostridiaceae | Eubacteriales |
| Clostridiaceae_unclassified | Clostridiaceae | Eubacteriales |
| Clostridiaceae_unclassified | Clostridiaceae | Eubacteriales |
| Eubacteriales_unclassified | Eubacteriales_unclassified | Eubacteriales |
| Eubacteriales_unclassified | Eubacteriales_unclassified | Eubacteriales |
| Erysipelatoclostridium | Erysipelotrichaceae | Erysipelotrichales |
| Erysipelatoclostridium | Erysipelotrichaceae | Erysipelotrichales |
| Clostridium | Clostridiaceae | Eubacteriales |
| Clostridium | Clostridiaceae | Eubacteriales |
| Eubacteriaceae_unclassified | Eubacteriaceae | Eubacteriales |
| Eubacteriaceae_unclassified | Eubacteriaceae | Eubacteriales |
| Eubacteriaceae_unclassified | Eubacteriaceae | Eubacteriales |
| Eubacteriaceae_unclassified | Eubacteriaceae | Eubacteriales |
| GGB20146 | FGB77306 | OFGB77306 |
| GGB20146 | FGB77306 | OFGB77306 |
| GGB20149 | Lachnospiraceae | Eubacteriales |
| GGB20149 | Lachnospiraceae | Eubacteriales |
| GGB22635 | Eggerthellaceae | Eggerthellales |
| GGB22635 | Eggerthellaceae | Eggerthellales |
| GGB25041 | Lachnospiraceae | Eubacteriales |
| GGB25041 | Lachnospiraceae | Eubacteriales |
| GGB28379 | FGB9506 | OFGB9506 |
| GGB28379 | FGB9506 | OFGB9506 |
| GGB28382 | FGB9508 | OFGB9508 |
| GGB28382 | FGB9508 | OFGB9508 |
| GGB28392 | FGB9512 | OFGB9512 |
| GGB28392 | FGB9512 | OFGB9512 |
| GGB28404 | FGB2838 | OFGB2838 |
| GGB28404 | FGB2838 | OFGB2838 |
| GGB28418 | FGB2838 | OFGB2838 |
| GGB28418 | FGB2838 | OFGB2838 |
| GGB28422 | FGB2838 | OFGB2838 |
| GGB28422 | FGB2838 | OFGB2838 |
| GGB28431 | Pumilibacteraceae | Eubacteriales |
| GGB28431 | Pumilibacteraceae | Eubacteriales |
| GGB28456 | FGB2833 | OFGB2833 |
| GGB28456 | FGB2833 | OFGB2833 |
| GGB28782 | Eubacteriaceae | Eubacteriales |
| GGB28782 | Eubacteriaceae | Eubacteriales |

| | | |
|---|---|---|
| GGB28792 | Lachnospiraceae | Eubacteriales |
| GGB28792 | Lachnospiraceae | Eubacteriales |
| GGB28798 | Lachnospiraceae | Eubacteriales |
| GGB28798 | Lachnospiraceae | Eubacteriales |
| GGB28810 | FGB9622 | OFGB9622 |
| GGB28810 | FGB9622 | OFGB9622 |
| GGB28828 | FGB77305 | OFGB77305 |
| GGB28828 | FGB77305 | OFGB77305 |
| GGB28851 | Clostridiaceae | Eubacteriales |
| GGB28851 | Clostridiaceae | Eubacteriales |
| GGB28865 | Lachnospiraceae | Eubacteriales |
| GGB28865 | Lachnospiraceae | Eubacteriales |
| GGB28868 | Lachnospiraceae | Eubacteriales |
| GGB28868 | Lachnospiraceae | Eubacteriales |
| GGB28869 | Lachnospiraceae | Eubacteriales |
| GGB28869 | Lachnospiraceae | Eubacteriales |
| GGB28875 | Lachnospiraceae | Eubacteriales |
| GGB28875 | Lachnospiraceae | Eubacteriales |
| GGB28881 | FGB9633 | OFGB9633 |
| GGB28881 | FGB9633 | OFGB9633 |
| GGB28892 | Bacteria_unclassified | Bacteria_unclassified |
| GGB28892 | Bacteria_unclassified | Bacteria_unclassified |
| GGB28893 | Bacteria_unclassified | Bacteria_unclassified |
| GGB28893 | Bacteria_unclassified | Bacteria_unclassified |
| GGB28898 | Bacteria_unclassified | Bacteria_unclassified |
| GGB28898 | Bacteria_unclassified | Bacteria_unclassified |
| GGB28901 | Bacteria_unclassified | Bacteria_unclassified |
| GGB28901 | Bacteria_unclassified | Bacteria_unclassified |
| GGB28904 | Bacteria_unclassified | Bacteria_unclassified |
| GGB28904 | Bacteria_unclassified | Bacteria_unclassified |
| GGB28909 | Bacteria_unclassified | Bacteria_unclassified |
| GGB28909 | Bacteria_unclassified | Bacteria_unclassified |
| GGB28924 | Lachnospiraceae | Eubacteriales |
| GGB28924 | Lachnospiraceae | Eubacteriales |
| GGB28927 | FGB77359 | OFGB77359 |
| GGB28927 | FGB77359 | OFGB77359 |
| GGB28934 | FGB9639 | OFGB9639 |
| GGB28934 | FGB9639 | OFGB9639 |
| GGB28949 | Lachnospiraceae | Eubacteriales |
| GGB28949 | Lachnospiraceae | Eubacteriales |
| GGB28951 | Clostridiaceae | Eubacteriales |
| GGB28951 | Clostridiaceae | Eubacteriales |
| GGB28951 | Clostridiaceae | Eubacteriales |
| GGB28951 | Clostridiaceae | Eubacteriales |
| GGB28954 | Clostridiaceae | Eubacteriales |
| GGB28954 | Clostridiaceae | Eubacteriales |

| | | |
|---|---|---|
| GGB28960 | Clostridiaceae | Eubacteriales |
| GGB28960 | Clostridiaceae | Eubacteriales |
| GGB28964 | Clostridiaceae | Eubacteriales |
| GGB28964 | Clostridiaceae | Eubacteriales |
| GGB28967 | Clostridiaceae | Eubacteriales |
| GGB28967 | Clostridiaceae | Eubacteriales |
| GGB28996 | FGB9656 | OFGB9656 |
| GGB28996 | FGB9656 | OFGB9656 |
| GGB29531 | FGB9827 | OFGB9827 |
| GGB29531 | FGB9827 | OFGB9827 |
| GGB29685 | Eubacteriaceae | Eubacteriales |
| GGB29685 | Eubacteriaceae | Eubacteriales |
| GGB30141 | FGB77303 | OFGB77303 |
| GGB30141 | FGB77303 | OFGB77303 |
| GGB30145 | Clostridiaceae | Eubacteriales |
| GGB30145 | Clostridiaceae | Eubacteriales |
| GGB30286 | Eubacteriales_unclassified | Eubacteriales |
| GGB30286 | Eubacteriales_unclassified | Eubacteriales |
| GGB30300 | FGB72709 | OFGB72709 |
| GGB30300 | FGB72709 | OFGB72709 |
| GGB30303 | Oscillospiraceae | Eubacteriales |
| GGB30303 | Oscillospiraceae | Eubacteriales |
| GGB30450 | Oscillospiraceae | Eubacteriales |
| GGB30450 | Oscillospiraceae | Eubacteriales |
| GGB30453 | Oscillospiraceae | Eubacteriales |
| GGB30453 | Oscillospiraceae | Eubacteriales |
| GGB30454 | Oscillospiraceae | Eubacteriales |
| GGB30454 | Oscillospiraceae | Eubacteriales |
| GGB30455 | Oscillospiraceae | Eubacteriales |
| GGB30455 | Oscillospiraceae | Eubacteriales |
| GGB30456 | Oscillospiraceae | Eubacteriales |
| GGB30456 | Oscillospiraceae | Eubacteriales |
| GGB30457 | Oscillospiraceae | Eubacteriales |
| GGB30457 | Oscillospiraceae | Eubacteriales |
| GGB30461 | Oscillospiraceae | Eubacteriales |
| GGB30461 | Oscillospiraceae | Eubacteriales |
| GGB30461 | Oscillospiraceae | Eubacteriales |
| GGB30461 | Oscillospiraceae | Eubacteriales |
| GGB30461 | Oscillospiraceae | Eubacteriales |
| GGB30461 | Oscillospiraceae | Eubacteriales |
| GGB30463 | Oscillospiraceae | Eubacteriales |
| GGB30463 | Oscillospiraceae | Eubacteriales |
| GGB30473 | Oscillospiraceae | Eubacteriales |
| GGB30473 | Oscillospiraceae | Eubacteriales |
| GGB30475 | Oscillospiraceae | Eubacteriales |
| GGB30475 | Oscillospiraceae | Eubacteriales |

| | | |
|---|---|---|
| GGB30861 | FGB77153 | OFGB77153 |
| GGB30861 | FGB77153 | OFGB77153 |
| GGB31312 | FGB1791 | OFGB1791 |
| GGB31312 | FGB1791 | OFGB1791 |
| GGB31438 | FGB10290 | OFGB10290 |
| GGB31438 | FGB10290 | OFGB10290 |
| GGB3171 | Oscillospiraceae | Eubacteriales |
| GGB3171 | Oscillospiraceae | Eubacteriales |
| GGB31762 | FGB10289 | OFGB10289 |
| GGB31762 | FGB10289 | OFGB10289 |
| GGB31823 | FGB1765 | OFGB1765 |
| GGB31823 | FGB1765 | OFGB1765 |
| GGB31838 | FGB1765 | OFGB1765 |
| GGB31838 | FGB1765 | OFGB1765 |
| GGB31841 | FGB1765 | OFGB1765 |
| GGB31841 | FGB1765 | OFGB1765 |
| GGB32371 | FGB10667 | OFGB10667 |
| GGB32371 | FGB10667 | OFGB10667 |
| GGB3793 | Lachnospiraceae | Eubacteriales |
| GGB3793 | Lachnospiraceae | Eubacteriales |
| GGB42601 | Clostridiaceae | Eubacteriales |
| GGB42601 | Clostridiaceae | Eubacteriales |
| GGB45514 | Oscillospiraceae | Eubacteriales |
| GGB45514 | Oscillospiraceae | Eubacteriales |
| GGB45564 | FGB75721 | OFGB75721 |
| GGB45564 | FGB75721 | OFGB75721 |
| GGB45624 | Oscillospiraceae | Eubacteriales |
| GGB45624 | Oscillospiraceae | Eubacteriales |
| GGB45656 | Christensenellaceae | Eubacteriales |
| GGB45656 | Christensenellaceae | Eubacteriales |
| GGB47127 | FGB10299 | OFGB10299 |
| GGB47127 | FGB10299 | OFGB10299 |
| GGB74395 | Oscillospiraceae | Eubacteriales |
| GGB74395 | Oscillospiraceae | Eubacteriales |
| GGB75053 | Oscillospiraceae | Eubacteriales |
| GGB75053 | Oscillospiraceae | Eubacteriales |
| GGB75109 | Lachnospiraceae | Eubacteriales |
| GGB75109 | Lachnospiraceae | Eubacteriales |
| Lachnospiraceae_unclassified | Lachnospiraceae | Eubacteriales |
| Lachnospiraceae_unclassified | Lachnospiraceae | Eubacteriales |
| Lachnospiraceae_unclassified | Lachnospiraceae | Eubacteriales |
| Lachnospiraceae_unclassified | Lachnospiraceae | Eubacteriales |
| Lachnospiraceae_unclassified | Lachnospiraceae | Eubacteriales |
| Lachnospiraceae_unclassified | Lachnospiraceae | Eubacteriales |
| Lachnospiraceae_unclassified | Lachnospiraceae | Eubacteriales |
| Lachnospiraceae_unclassified | Lachnospiraceae | Eubacteriales |

| | | |
|---|---|---|
| Lachnospiraceae_unclassified | Lachnospiraceae | Eubacteriales |
| Lachnospiraceae_unclassified | Lachnospiraceae | Eubacteriales |
| Lactobacillus | Lactobacillaceae | Lactobacillales |
| Lactobacillus | Lactobacillaceae | Lactobacillales |
| Leptogranulimonas | Atopobiaceae | Coriobacteriales |
| Leptogranulimonas | Atopobiaceae | Coriobacteriales |
| Muribaculaceae_unclassified | Muribaculaceae | Bacteroidales |
| Muribaculaceae_unclassified | Muribaculaceae | Bacteroidales |
| Neglectibacter | Oscillospiraceae | Eubacteriales |
| Neglectibacter | Oscillospiraceae | Eubacteriales |
| Oscillibacter | Oscillospiraceae | Eubacteriales |
| Oscillibacter | Oscillospiraceae | Eubacteriales |
| Oscillospiraceae_unclassified | Oscillospiraceae | Eubacteriales |
| Oscillospiraceae_unclassified | Oscillospiraceae | Eubacteriales |
| Oscillospiraceae_unclassified | Oscillospiraceae | Eubacteriales |
| Oscillospiraceae_unclassified | Oscillospiraceae | Eubacteriales |
| Oscillospiraceae_unclassified | Oscillospiraceae | Eubacteriales |
| Oscillospiraceae_unclassified | Oscillospiraceae | Eubacteriales |
| Oscillospiraceae_unclassified | Oscillospiraceae | Eubacteriales |
| Oscillospiraceae_unclassified | Oscillospiraceae | Eubacteriales |
| Parasutterella | Sutterellaceae | Burkholderiales |
| Parasutterella | Sutterellaceae | Burkholderiales |
| Schaedlerella | Lachnospiraceae | Eubacteriales |
| Schaedlerella | Lachnospiraceae | Eubacteriales |
| Turicibacter | Turicibacteraceae | Erysipelotrichales |
| Turicibacter | Turicibacteraceae | Erysipelotrichales |
| Acetatifactor | Lachnospiraceae | Eubacteriales |
| Acetatifactor | Lachnospiraceae | Eubacteriales |
| Acutalibacter | Oscillospiraceae | Eubacteriales |
| Acutalibacter | Oscillospiraceae | Eubacteriales |
| Akkermansia | Akkermansiaceae | Verrucomicrobiales |
| Akkermansia | Akkermansiaceae | Verrucomicrobiales |
| Anaerotruncus | Oscillospiraceae | Eubacteriales |
| Anaerotruncus | Oscillospiraceae | Eubacteriales |
| Bacteria_unclassified | Bacteria_unclassified | Bacteria_unclassified |
| Bacteria_unclassified | Bacteria_unclassified | Bacteria_unclassified |
| Bacteria_unclassified | Bacteria_unclassified | Bacteria_unclassified |
| Bacteria_unclassified | Bacteria_unclassified | Bacteria_unclassified |
| Bacteria_unclassified | Bacteria_unclassified | Bacteria_unclassified |
| Bacteria_unclassified | Bacteria_unclassified | Bacteria_unclassified |
| Bacteria_unclassified | Bacteria_unclassified | Bacteria_unclassified |
| Bacteria_unclassified | Bacteria_unclassified | Bacteria_unclassified |

| | | |
|---|---|---|
| Bacteria_unclassified | Bacteria_unclassified | Bacteria_unclassified |
| Bacteria_unclassified | Bacteria_unclassified | Bacteria_unclassified |
| Bacteria_unclassified | Bacteria_unclassified | Bacteria_unclassified |
| Bacteria_unclassified | Bacteria_unclassified | Bacteria_unclassified |
| Bacteria_unclassified | Bacteria_unclassified | Bacteria_unclassified |
| Bacteria_unclassified | Bacteria_unclassified | Bacteria_unclassified |
| Bacteria_unclassified | Bacteria_unclassified | Bacteria_unclassified |
| Bacteria_unclassified | Bacteria_unclassified | Bacteria_unclassified |
| Bacteria_unclassified | Bacteria_unclassified | Bacteria_unclassified |
| Bacteria_unclassified | Bacteria_unclassified | Bacteria_unclassified |
| Bacteroides | Bacteroidaceae | Bacteroidales |
| Bacteroides | Bacteroidaceae | Bacteroidales |

| | | |
|---|---|---|
| Clostridia_unclassified | Clostridia_unclassified | Clostridia_unclassified |
| Clostridia_unclassified | Clostridia_unclassified | Clostridia_unclassified |
| Clostridiaceae_unclassified | Clostridiaceae | Eubacteriales |
| Clostridiaceae_unclassified | Clostridiaceae | Eubacteriales |
| Clostridiaceae_unclassified | Clostridiaceae | Eubacteriales |
| Clostridiaceae_unclassified | Clostridiaceae | Eubacteriales |
| Eubacteriales_unclassified | Eubacteriales_unclassified | Eubacteriales |
| Eubacteriales_unclassified | Eubacteriales_unclassified | Eubacteriales |
| Erysipelatoclostridium | Erysipelotrichaceae | Erysipelotrichales |
| Erysipelatoclostridium | Erysipelotrichaceae | Erysipelotrichales |
| Clostridium | Clostridiaceae | Eubacteriales |
| Clostridium | Clostridiaceae | Eubacteriales |
| Eubacteriaceae_unclassified | Eubacteriaceae | Eubacteriales |
| Eubacteriaceae_unclassified | Eubacteriaceae | Eubacteriales |
| Eubacteriaceae_unclassified | Eubacteriaceae | Eubacteriales |
| Eubacteriaceae_unclassified | Eubacteriaceae | Eubacteriales |
| GGB20146 | FGB77306 | OFGB77306 |
| GGB20146 | FGB77306 | OFGB77306 |
| GGB20149 | Lachnospiraceae | Eubacteriales |
| GGB20149 | Lachnospiraceae | Eubacteriales |
| GGB22635 | Eggerthellaceae | Eggerthellales |
| GGB22635 | Eggerthellaceae | Eggerthellales |
| GGB25041 | Lachnospiraceae | Eubacteriales |
| GGB25041 | Lachnospiraceae | Eubacteriales |
| GGB28379 | FGB9506 | OFGB9506 |
| GGB28379 | FGB9506 | OFGB9506 |
| GGB28382 | FGB9508 | OFGB9508 |
| GGB28382 | FGB9508 | OFGB9508 |
| GGB28392 | FGB9512 | OFGB9512 |
| GGB28392 | FGB9512 | OFGB9512 |
| GGB28404 | FGB2838 | OFGB2838 |
| GGB28404 | FGB2838 | OFGB2838 |

| | | |
|---|---|---|
| GGB28418 | FGB2838 | OFGB2838 |
| GGB28418 | FGB2838 | OFGB2838 |
| GGB28422 | FGB2838 | OFGB2838 |
| GGB28422 | FGB2838 | OFGB2838 |
| GGB28431 | Pumilibacteraceae | Eubacteriales |
| GGB28431 | Pumilibacteraceae | Eubacteriales |
| GGB28456 | FGB2833 | OFGB2833 |
| GGB28456 | FGB2833 | OFGB2833 |
| GGB28782 | Eubacteriaceae | Eubacteriales |
| GGB28782 | Eubacteriaceae | Eubacteriales |
| GGB28792 | Lachnospiraceae | Eubacteriales |
| GGB28792 | Lachnospiraceae | Eubacteriales |
| GGB28798 | Lachnospiraceae | Eubacteriales |
| GGB28798 | Lachnospiraceae | Eubacteriales |
| GGB28810 | FGB9622 | OFGB9622 |
| GGB28810 | FGB9622 | OFGB9622 |
| GGB28828 | FGB77305 | OFGB77305 |
| GGB28828 | FGB77305 | OFGB77305 |
| GGB28851 | Clostridiaceae | Eubacteriales |
| GGB28851 | Clostridiaceae | Eubacteriales |
| GGB28865 | Lachnospiraceae | Eubacteriales |
| GGB28865 | Lachnospiraceae | Eubacteriales |
| GGB28868 | Lachnospiraceae | Eubacteriales |
| GGB28868 | Lachnospiraceae | Eubacteriales |
| GGB28869 | Lachnospiraceae | Eubacteriales |
| GGB28869 | Lachnospiraceae | Eubacteriales |
| GGB28875 | Lachnospiraceae | Eubacteriales |
| GGB28875 | Lachnospiraceae | Eubacteriales |
| GGB28881 | FGB9633 | OFGB9633 |
| GGB28881 | FGB9633 | OFGB9633 |
| GGB28892 | Bacteria_unclassified | Bacteria_unclassified |
| GGB28892 | Bacteria_unclassified | Bacteria_unclassified |
| GGB28893 | Bacteria_unclassified | Bacteria_unclassified |
| GGB28893 | Bacteria_unclassified | Bacteria_unclassified |
| GGB28898 | Bacteria_unclassified | Bacteria_unclassified |
| GGB28898 | Bacteria_unclassified | Bacteria_unclassified |
| GGB28901 | Bacteria_unclassified | Bacteria_unclassified |
| GGB28901 | Bacteria_unclassified | Bacteria_unclassified |
| GGB28904 | Bacteria_unclassified | Bacteria_unclassified |
| GGB28904 | Bacteria_unclassified | Bacteria_unclassified |
| GGB28909 | Bacteria_unclassified | Bacteria_unclassified |
| GGB28909 | Bacteria_unclassified | Bacteria_unclassified |
| GGB28924 | Lachnospiraceae | Eubacteriales |
| GGB28924 | Lachnospiraceae | Eubacteriales |
| GGB28927 | FGB77359 | OFGB77359 |
| GGB28927 | FGB77359 | OFGB77359 |

| | | |
|---|---|---|
| GGB28934 | FGB9639 | OFGB9639 |
| GGB28934 | FGB9639 | OFGB9639 |
| GGB28949 | Lachnospiraceae | Eubacteriales |
| GGB28949 | Lachnospiraceae | Eubacteriales |
| GGB28951 | Clostridiaceae | Eubacteriales |
| GGB28951 | Clostridiaceae | Eubacteriales |
| GGB28951 | Clostridiaceae | Eubacteriales |
| GGB28951 | Clostridiaceae | Eubacteriales |
| GGB28954 | Clostridiaceae | Eubacteriales |
| GGB28954 | Clostridiaceae | Eubacteriales |
| GGB28960 | Clostridiaceae | Eubacteriales |
| GGB28960 | Clostridiaceae | Eubacteriales |
| GGB28964 | Clostridiaceae | Eubacteriales |
| GGB28964 | Clostridiaceae | Eubacteriales |
| GGB28967 | Clostridiaceae | Eubacteriales |
| GGB28967 | Clostridiaceae | Eubacteriales |
| GGB28996 | FGB9656 | OFGB9656 |
| GGB28996 | FGB9656 | OFGB9656 |
| GGB29531 | FGB9827 | OFGB9827 |
| GGB29531 | FGB9827 | OFGB9827 |
| GGB29685 | Eubacteriaceae | Eubacteriales |
| GGB29685 | Eubacteriaceae | Eubacteriales |
| GGB30141 | FGB77303 | OFGB77303 |
| GGB30141 | FGB77303 | OFGB77303 |
| GGB30145 | Clostridiaceae | Eubacteriales |
| GGB30145 | Clostridiaceae | Eubacteriales |
| GGB30286 | Eubacteriales_unclassified | Eubacteriales |
| GGB30286 | Eubacteriales_unclassified | Eubacteriales |
| GGB30300 | FGB72709 | OFGB72709 |
| GGB30300 | FGB72709 | OFGB72709 |
| GGB30303 | Oscillospiraceae | Eubacteriales |
| GGB30303 | Oscillospiraceae | Eubacteriales |
| GGB30450 | Oscillospiraceae | Eubacteriales |
| GGB30450 | Oscillospiraceae | Eubacteriales |
| GGB30453 | Oscillospiraceae | Eubacteriales |
| GGB30453 | Oscillospiraceae | Eubacteriales |
| GGB30454 | Oscillospiraceae | Eubacteriales |
| GGB30454 | Oscillospiraceae | Eubacteriales |
| GGB30455 | Oscillospiraceae | Eubacteriales |
| GGB30455 | Oscillospiraceae | Eubacteriales |
| GGB30456 | Oscillospiraceae | Eubacteriales |
| GGB30456 | Oscillospiraceae | Eubacteriales |
| GGB30457 | Oscillospiraceae | Eubacteriales |
| GGB30457 | Oscillospiraceae | Eubacteriales |
| GGB30461 | Oscillospiraceae | Eubacteriales |
| GGB30461 | Oscillospiraceae | Eubacteriales |

| | | |
|---|---|---|
| GGB30461 | Oscillospiraceae | Eubacteriales |
| GGB30461 | Oscillospiraceae | Eubacteriales |
| GGB30461 | Oscillospiraceae | Eubacteriales |
| GGB30461 | Oscillospiraceae | Eubacteriales |
| GGB30463 | Oscillospiraceae | Eubacteriales |
| GGB30463 | Oscillospiraceae | Eubacteriales |
| GGB30473 | Oscillospiraceae | Eubacteriales |
| GGB30473 | Oscillospiraceae | Eubacteriales |
| GGB30475 | Oscillospiraceae | Eubacteriales |
| GGB30475 | Oscillospiraceae | Eubacteriales |
| GGB30861 | FGB77153 | OFGB77153 |
| GGB30861 | FGB77153 | OFGB77153 |
| GGB31312 | FGB1791 | OFGB1791 |
| GGB31312 | FGB1791 | OFGB1791 |
| GGB31438 | FGB10290 | OFGB10290 |
| GGB31438 | FGB10290 | OFGB10290 |
| GGB3171 | Oscillospiraceae | Eubacteriales |
| GGB3171 | Oscillospiraceae | Eubacteriales |
| GGB31762 | FGB10289 | OFGB10289 |
| GGB31762 | FGB10289 | OFGB10289 |
| GGB31823 | FGB1765 | OFGB1765 |
| GGB31823 | FGB1765 | OFGB1765 |
| GGB31838 | FGB1765 | OFGB1765 |
| GGB31838 | FGB1765 | OFGB1765 |
| GGB31841 | FGB1765 | OFGB1765 |
| GGB31841 | FGB1765 | OFGB1765 |
| GGB32371 | FGB10667 | OFGB10667 |
| GGB32371 | FGB10667 | OFGB10667 |
| GGB3793 | Lachnospiraceae | Eubacteriales |
| GGB3793 | Lachnospiraceae | Eubacteriales |
| GGB42601 | Clostridiaceae | Eubacteriales |
| GGB42601 | Clostridiaceae | Eubacteriales |
| GGB45514 | Oscillospiraceae | Eubacteriales |
| GGB45514 | Oscillospiraceae | Eubacteriales |
| GGB45564 | FGB75721 | OFGB75721 |
| GGB45564 | FGB75721 | OFGB75721 |
| GGB45624 | Oscillospiraceae | Eubacteriales |
| GGB45624 | Oscillospiraceae | Eubacteriales |
| GGB45656 | Christensenellaceae | Eubacteriales |
| GGB45656 | Christensenellaceae | Eubacteriales |
| GGB47127 | FGB10299 | OFGB10299 |
| GGB47127 | FGB10299 | OFGB10299 |
| GGB74395 | Oscillospiraceae | Eubacteriales |
| GGB74395 | Oscillospiraceae | Eubacteriales |
| GGB75053 | Oscillospiraceae | Eubacteriales |
| GGB75053 | Oscillospiraceae | Eubacteriales |

| | | |
|---|---|---|
| GGB75109 | Lachnospiraceae | Eubacteriales |
| GGB75109 | Lachnospiraceae | Eubacteriales |
| Lachnospiraceae_unclassified | Lachnospiraceae | Eubacteriales |
| Lachnospiraceae_unclassified | Lachnospiraceae | Eubacteriales |
| Lachnospiraceae_unclassified | Lachnospiraceae | Eubacteriales |
| Lachnospiraceae_unclassified | Lachnospiraceae | Eubacteriales |
| Lachnospiraceae_unclassified | Lachnospiraceae | Eubacteriales |
| Lachnospiraceae_unclassified | Lachnospiraceae | Eubacteriales |
| Lachnospiraceae_unclassified | Lachnospiraceae | Eubacteriales |
| Lachnospiraceae_unclassified | Lachnospiraceae | Eubacteriales |
| Lachnospiraceae_unclassified | Lachnospiraceae | Eubacteriales |
| Lachnospiraceae_unclassified | Lachnospiraceae | Eubacteriales |
| Lactobacillus | Lactobacillaceae | Lactobacillales |
| Lactobacillus | Lactobacillaceae | Lactobacillales |
| Leptogranulimonas | Atopobiaceae | Coriobacteriales |
| Leptogranulimonas | Atopobiaceae | Coriobacteriales |
| Muribaculaceae_unclassified | Muribaculaceae | Bacteroidales |
| Muribaculaceae_unclassified | Muribaculaceae | Bacteroidales |
| Neglectibacter | Oscillospiraceae | Eubacteriales |
| Neglectibacter | Oscillospiraceae | Eubacteriales |
| Oscillibacter | Oscillospiraceae | Eubacteriales |
| Oscillibacter | Oscillospiraceae | Eubacteriales |
| Oscillospiraceae_unclassified | Oscillospiraceae | Eubacteriales |
| Oscillospiraceae_unclassified | Oscillospiraceae | Eubacteriales |
| Oscillospiraceae_unclassified | Oscillospiraceae | Eubacteriales |
| Oscillospiraceae_unclassified | Oscillospiraceae | Eubacteriales |
| Oscillospiraceae_unclassified | Oscillospiraceae | Eubacteriales |
| Oscillospiraceae_unclassified | Oscillospiraceae | Eubacteriales |
| Oscillospiraceae_unclassified | Oscillospiraceae | Eubacteriales |
| Oscillospiraceae_unclassified | Oscillospiraceae | Eubacteriales |
| Parasutterella | Sutterellaceae | Burkholderiales |
| Parasutterella | Sutterellaceae | Burkholderiales |
| Schaedlerella | Lachnospiraceae | Eubacteriales |
| Schaedlerella | Lachnospiraceae | Eubacteriales |
| Turicibacter | Turicibacteraceae | Erysipelotrichales |
| Turicibacter | Turicibacteraceae | Erysipelotrichales |
| Acetatifactor | Lachnospiraceae | Eubacteriales |
| Acetatifactor | Lachnospiraceae | Eubacteriales |
| Acutalibacter | Oscillospiraceae | Eubacteriales |
| Acutalibacter | Oscillospiraceae | Eubacteriales |
| Akkermansia | Akkermansiaceae | Verrucomicrobiales |
| Akkermansia | Akkermansiaceae | Verrucomicrobiales |

| | | |
|---|---|---|
| Anaerotruncus | Oscillospiraceae | Eubacteriales |
| Anaerotruncus | Oscillospiraceae | Eubacteriales |
| Bacteria_unclassified | Bacteria_unclassified | Bacteria_unclassified |
| Bacteria_unclassified | Bacteria_unclassified | Bacteria_unclassified |
| Bacteria_unclassified | Bacteria_unclassified | Bacteria_unclassified |
| Bacteria_unclassified | Bacteria_unclassified | Bacteria_unclassified |
| Bacteria_unclassified | Bacteria_unclassified | Bacteria_unclassified |
| Bacteria_unclassified | Bacteria_unclassified | Bacteria_unclassified |
| Bacteria_unclassified | Bacteria_unclassified | Bacteria_unclassified |
| Bacteria_unclassified | Bacteria_unclassified | Bacteria_unclassified |
| Bacteria_unclassified | Bacteria_unclassified | Bacteria_unclassified |
| Bacteria_unclassified | Bacteria_unclassified | Bacteria_unclassified |
| Bacteria_unclassified | Bacteria_unclassified | Bacteria_unclassified |
| Bacteria_unclassified | Bacteria_unclassified | Bacteria_unclassified |
| Bacteria_unclassified | Bacteria_unclassified | Bacteria_unclassified |
| Bacteria_unclassified | Bacteria_unclassified | Bacteria_unclassified |
| Bacteria_unclassified | Bacteria_unclassified | Bacteria_unclassified |
| Bacteria_unclassified | Bacteria_unclassified | Bacteria_unclassified |
| Bacteria_unclassified | Bacteria_unclassified | Bacteria_unclassified |
| Bacteria_unclassified | Bacteria_unclassified | Bacteria_unclassified |
| Bacteroides | Bacteroidaceae | Bacteroidales |
| Bacteroides | Bacteroidaceae | Bacteroidales |
| Clostridia_unclassified | Clostridia_unclassified | Clostridia_unclassified |
| Clostridia_unclassified | Clostridia_unclassified | Clostridia_unclassified |
| Clostridiaceae_unclassified | Clostridiaceae | Eubacteriales |
| Clostridiaceae_unclassified | Clostridiaceae | Eubacteriales |
| Clostridiaceae_unclassified | Clostridiaceae | Eubacteriales |
| Clostridiaceae_unclassified | Clostridiaceae | Eubacteriales |
| Eubacteriales_unclassified | Eubacteriales_unclassified | Eubacteriales |
| Eubacteriales_unclassified | Eubacteriales_unclassified | Eubacteriales |
| Erysipelatoclostridium | Erysipelotrichaceae | Erysipelotrichales |
| Erysipelatoclostridium | Erysipelotrichaceae | Erysipelotrichales |
| Clostridium | Clostridiaceae | Eubacteriales |
| Clostridium | Clostridiaceae | Eubacteriales |
| Eubacteriaceae_unclassified | Eubacteriaceae | Eubacteriales |
| Eubacteriaceae_unclassified | Eubacteriaceae | Eubacteriales |
| Eubacteriaceae_unclassified | Eubacteriaceae | Eubacteriales |
| Eubacteriaceae_unclassified | Eubacteriaceae | Eubacteriales |
| GGB20146 | FGB77306 | OFGB77306 |
| GGB20146 | FGB77306 | OFGB77306 |
| GGB20149 | Lachnospiraceae | Eubacteriales |
| GGB20149 | Lachnospiraceae | Eubacteriales |
| GGB22635 | Eggerthellaceae | Eggerthellales |
| GGB22635 | Eggerthellaceae | Eggerthellales |

| | | |
|---|---|---|
| GGB25041 | Lachnospiraceae | Eubacteriales |
| GGB25041 | Lachnospiraceae | Eubacteriales |
| GGB28379 | FGB9506 | OFGB9506 |
| GGB28379 | FGB9506 | OFGB9506 |
| GGB28382 | FGB9508 | OFGB9508 |
| GGB28382 | FGB9508 | OFGB9508 |
| GGB28392 | FGB9512 | OFGB9512 |
| GGB28392 | FGB9512 | OFGB9512 |
| GGB28404 | FGB2838 | OFGB2838 |
| GGB28404 | FGB2838 | OFGB2838 |
| GGB28418 | FGB2838 | OFGB2838 |
| GGB28418 | FGB2838 | OFGB2838 |
| GGB28422 | FGB2838 | OFGB2838 |
| GGB28422 | FGB2838 | OFGB2838 |
| GGB28431 | Pumilibacteraceae | Eubacteriales |
| GGB28431 | Pumilibacteraceae | Eubacteriales |
| GGB28456 | FGB2833 | OFGB2833 |
| GGB28456 | FGB2833 | OFGB2833 |
| GGB28782 | Eubacteriaceae | Eubacteriales |
| GGB28782 | Eubacteriaceae | Eubacteriales |
| GGB28792 | Lachnospiraceae | Eubacteriales |
| GGB28792 | Lachnospiraceae | Eubacteriales |
| GGB28798 | Lachnospiraceae | Eubacteriales |
| GGB28798 | Lachnospiraceae | Eubacteriales |
| GGB28810 | FGB9622 | OFGB9622 |
| GGB28810 | FGB9622 | OFGB9622 |
| GGB28828 | FGB77305 | OFGB77305 |
| GGB28828 | FGB77305 | OFGB77305 |
| GGB28851 | Clostridiaceae | Eubacteriales |
| GGB28851 | Clostridiaceae | Eubacteriales |
| GGB28865 | Lachnospiraceae | Eubacteriales |
| GGB28865 | Lachnospiraceae | Eubacteriales |
| GGB28868 | Lachnospiraceae | Eubacteriales |
| GGB28868 | Lachnospiraceae | Eubacteriales |
| GGB28869 | Lachnospiraceae | Eubacteriales |
| GGB28869 | Lachnospiraceae | Eubacteriales |
| GGB28875 | Lachnospiraceae | Eubacteriales |
| GGB28875 | Lachnospiraceae | Eubacteriales |
| GGB28881 | FGB9633 | OFGB9633 |
| GGB28881 | FGB9633 | OFGB9633 |
| GGB28892 | Bacteria_unclassified | Bacteria_unclassified |
| GGB28892 | Bacteria_unclassified | Bacteria_unclassified |
| GGB28893 | Bacteria_unclassified | Bacteria_unclassified |
| GGB28893 | Bacteria_unclassified | Bacteria_unclassified |
| GGB28898 | Bacteria_unclassified | Bacteria_unclassified |
| GGB28898 | Bacteria_unclassified | Bacteria_unclassified |

| | | |
|---|---|---|
| GGB28901 | Bacteria_unclassified | Bacteria_unclassified |
| GGB28901 | Bacteria_unclassified | Bacteria_unclassified |
| GGB28904 | Bacteria_unclassified | Bacteria_unclassified |
| GGB28904 | Bacteria_unclassified | Bacteria_unclassified |
| GGB28909 | Bacteria_unclassified | Bacteria_unclassified |
| GGB28909 | Bacteria_unclassified | Bacteria_unclassified |
| GGB28924 | Lachnospiraceae | Eubacteriales |
| GGB28924 | Lachnospiraceae | Eubacteriales |
| GGB28927 | FGB77359 | OFGB77359 |
| GGB28927 | FGB77359 | OFGB77359 |
| GGB28934 | FGB9639 | OFGB9639 |
| GGB28934 | FGB9639 | OFGB9639 |
| GGB28949 | Lachnospiraceae | Eubacteriales |
| GGB28949 | Lachnospiraceae | Eubacteriales |
| GGB28951 | Clostridiaceae | Eubacteriales |
| GGB28951 | Clostridiaceae | Eubacteriales |
| GGB28951 | Clostridiaceae | Eubacteriales |
| GGB28951 | Clostridiaceae | Eubacteriales |
| GGB28954 | Clostridiaceae | Eubacteriales |
| GGB28954 | Clostridiaceae | Eubacteriales |
| GGB28960 | Clostridiaceae | Eubacteriales |
| GGB28960 | Clostridiaceae | Eubacteriales |
| GGB28964 | Clostridiaceae | Eubacteriales |
| GGB28964 | Clostridiaceae | Eubacteriales |
| GGB28967 | Clostridiaceae | Eubacteriales |
| GGB28967 | Clostridiaceae | Eubacteriales |
| GGB28996 | FGB9656 | OFGB9656 |
| GGB28996 | FGB9656 | OFGB9656 |
| GGB29531 | FGB9827 | OFGB9827 |
| GGB29531 | FGB9827 | OFGB9827 |
| GGB29685 | Eubacteriaceae | Eubacteriales |
| GGB29685 | Eubacteriaceae | Eubacteriales |
| GGB30141 | FGB77303 | OFGB77303 |
| GGB30141 | FGB77303 | OFGB77303 |
| GGB30145 | Clostridiaceae | Eubacteriales |
| GGB30145 | Clostridiaceae | Eubacteriales |
| GGB30286 | Eubacteriales_unclassified | Eubacteriales |
| GGB30286 | Eubacteriales_unclassified | Eubacteriales |
| GGB30300 | FGB72709 | OFGB72709 |
| GGB30300 | FGB72709 | OFGB72709 |
| GGB30303 | Oscillospiraceae | Eubacteriales |
| GGB30303 | Oscillospiraceae | Eubacteriales |
| GGB30450 | Oscillospiraceae | Eubacteriales |
| GGB30450 | Oscillospiraceae | Eubacteriales |
| GGB30453 | Oscillospiraceae | Eubacteriales |
| GGB30453 | Oscillospiraceae | Eubacteriales |

| | | |
|---|---|---|
| GGB30454 | Oscillospiraceae | Eubacteriales |
| GGB30454 | Oscillospiraceae | Eubacteriales |
| GGB30455 | Oscillospiraceae | Eubacteriales |
| GGB30455 | Oscillospiraceae | Eubacteriales |
| GGB30456 | Oscillospiraceae | Eubacteriales |
| GGB30456 | Oscillospiraceae | Eubacteriales |
| GGB30457 | Oscillospiraceae | Eubacteriales |
| GGB30457 | Oscillospiraceae | Eubacteriales |
| GGB30461 | Oscillospiraceae | Eubacteriales |
| GGB30461 | Oscillospiraceae | Eubacteriales |
| GGB30461 | Oscillospiraceae | Eubacteriales |
| GGB30461 | Oscillospiraceae | Eubacteriales |
| GGB30461 | Oscillospiraceae | Eubacteriales |
| GGB30461 | Oscillospiraceae | Eubacteriales |
| GGB30463 | Oscillospiraceae | Eubacteriales |
| GGB30463 | Oscillospiraceae | Eubacteriales |
| GGB30473 | Oscillospiraceae | Eubacteriales |
| GGB30473 | Oscillospiraceae | Eubacteriales |
| GGB30475 | Oscillospiraceae | Eubacteriales |
| GGB30475 | Oscillospiraceae | Eubacteriales |
| GGB30861 | FGB77153 | OFGB77153 |
| GGB30861 | FGB77153 | OFGB77153 |
| GGB31312 | FGB1791 | OFGB1791 |
| GGB31312 | FGB1791 | OFGB1791 |
| GGB31438 | FGB10290 | OFGB10290 |
| GGB31438 | FGB10290 | OFGB10290 |
| GGB3171 | Oscillospiraceae | Eubacteriales |
| GGB3171 | Oscillospiraceae | Eubacteriales |
| GGB31762 | FGB10289 | OFGB10289 |
| GGB31762 | FGB10289 | OFGB10289 |
| GGB31823 | FGB1765 | OFGB1765 |
| GGB31823 | FGB1765 | OFGB1765 |
| GGB31838 | FGB1765 | OFGB1765 |
| GGB31838 | FGB1765 | OFGB1765 |
| GGB31841 | FGB1765 | OFGB1765 |
| GGB31841 | FGB1765 | OFGB1765 |
| GGB32371 | FGB10667 | OFGB10667 |
| GGB32371 | FGB10667 | OFGB10667 |
| GGB3793 | Lachnospiraceae | Eubacteriales |
| GGB3793 | Lachnospiraceae | Eubacteriales |
| GGB42601 | Clostridiaceae | Eubacteriales |
| GGB42601 | Clostridiaceae | Eubacteriales |
| GGB45514 | Oscillospiraceae | Eubacteriales |
| GGB45514 | Oscillospiraceae | Eubacteriales |
| GGB45564 | FGB75721 | OFGB75721 |
| GGB45564 | FGB75721 | OFGB75721 |

| | | |
|---|---|---|
| GGB45624 | Oscillospiraceae | Eubacteriales |
| GGB45624 | Oscillospiraceae | Eubacteriales |
| GGB45656 | Christensenellaceae | Eubacteriales |
| GGB45656 | Christensenellaceae | Eubacteriales |
| GGB47127 | FGB10299 | OFGB10299 |
| GGB47127 | FGB10299 | OFGB10299 |
| GGB74395 | Oscillospiraceae | Eubacteriales |
| GGB74395 | Oscillospiraceae | Eubacteriales |
| GGB75053 | Oscillospiraceae | Eubacteriales |
| GGB75053 | Oscillospiraceae | Eubacteriales |
| GGB75109 | Lachnospiraceae | Eubacteriales |
| GGB75109 | Lachnospiraceae | Eubacteriales |
| Lachnospiraceae_unclassified | Lachnospiraceae | Eubacteriales |
| Lachnospiraceae_unclassified | Lachnospiraceae | Eubacteriales |
| Lachnospiraceae_unclassified | Lachnospiraceae | Eubacteriales |
| Lachnospiraceae_unclassified | Lachnospiraceae | Eubacteriales |
| Lachnospiraceae_unclassified | Lachnospiraceae | Eubacteriales |
| Lachnospiraceae_unclassified | Lachnospiraceae | Eubacteriales |
| Lachnospiraceae_unclassified | Lachnospiraceae | Eubacteriales |
| Lachnospiraceae_unclassified | Lachnospiraceae | Eubacteriales |
| Lachnospiraceae_unclassified | Lachnospiraceae | Eubacteriales |
| Lachnospiraceae_unclassified | Lachnospiraceae | Eubacteriales |
| Lactobacillus | Lactobacillaceae | Lactobacillales |
| Lactobacillus | Lactobacillaceae | Lactobacillales |
| Leptogranulimonas | Atopobiaceae | Coriobacteriales |
| Leptogranulimonas | Atopobiaceae | Coriobacteriales |
| Muribaculaceae_unclassified | Muribaculaceae | Bacteroidales |
| Muribaculaceae_unclassified | Muribaculaceae | Bacteroidales |
| Neglectibacter | Oscillospiraceae | Eubacteriales |
| Neglectibacter | Oscillospiraceae | Eubacteriales |
| Oscillibacter | Oscillospiraceae | Eubacteriales |
| Oscillibacter | Oscillospiraceae | Eubacteriales |
| Oscillospiraceae_unclassified | Oscillospiraceae | Eubacteriales |
| Oscillospiraceae_unclassified | Oscillospiraceae | Eubacteriales |
| Oscillospiraceae_unclassified | Oscillospiraceae | Eubacteriales |
| Oscillospiraceae_unclassified | Oscillospiraceae | Eubacteriales |
| Oscillospiraceae_unclassified | Oscillospiraceae | Eubacteriales |
| Oscillospiraceae_unclassified | Oscillospiraceae | Eubacteriales |
| Oscillospiraceae_unclassified | Oscillospiraceae | Eubacteriales |
| Oscillospiraceae_unclassified | Oscillospiraceae | Eubacteriales |
| Parasutterella | Sutterellaceae | Burkholderiales |
| Parasutterella | Sutterellaceae | Burkholderiales |
| Schaedlerella | Lachnospiraceae | Eubacteriales |
| Schaedlerella | Lachnospiraceae | Eubacteriales |

k__Bacteria|p__Proteobacteria|c__Betaproteobacteria|o__Burkholderiales|f__Sutterellaceae|g__Parasutterella|s__Parasu
k__Bacteria|p__Proteobacteria|c__Betaproteobacteria|o__Burkholderiales|f__Sutterellaceae|g__Parasutterella|s__Parasu

k__Bacteria|p__Firmicutes|c__Clostridia|o__Eubacteriales|f__Lachnospiraceae|g__Schaedlerella|s__Schaedlerella_arabin
k__Bacteria|p__Firmicutes|c__Clostridia|o__Eubacteriales|f__Lachnospiraceae|g__Schaedlerella|s__Schaedlerella_arabin

k__Bacteria|p__Firmicutes|c__Erysipelotrichia|o__Erysipelotrichales|f__Turicibacteraceae|g__Turicibacter|s__Turicibacter_
k__Bacteria|p__Firmicutes|c__Erysipelotrichia|o__Erysipelotrichales|f__Turicibacteraceae|g__Turicibacter|s__Turicibacter_
k__Bacteria|p__Firmicutes|c__Clostridia|o__Eubacteriales|f__Lachnospiraceae|g__Acetatifactor|s__Acetatifactor_SGB415
k__Bacteria|p__Firmicutes|c__Clostridia|o__Eubacteriales|f__Lachnospiraceae|g__Acetatifactor|s__Acetatifactor_SGB415
k__Bacteria|p__Firmicutes|c__Clostridia|o__Eubacteriales|f__Oscillospiraceae|g__Acutalibacter|s__Acutalibacter_muris
k__Bacteria|p__Firmicutes|c__Clostridia|o__Eubacteriales|f__Oscillospiraceae|g__Acutalibacter|s__Acutalibacter_muris
k__Bacteria|p__Verrucomicrobia|c__Verrucomicrobiae|o__Verrucomicrobiales|f__Akkermansiaceae|g__Akkermansia|s__
k__Bacteria|p__Verrucomicrobia|c__Verrucomicrobiae|o__Verrucomicrobiales|f__Akkermansiaceae|g__Akkermansia|s__
k__Bacteria|p__Firmicutes|c__Clostridia|o__Eubacteriales|f__Oscillospiraceae|g__Anaerotruncus|s__Anaerotruncus_sp_1
k__Bacteria|p__Firmicutes|c__Clostridia|o__Eubacteriales|f__Oscillospiraceae|g__Anaerotruncus|s__Anaerotruncus_sp_1
k__Bacteria|p__Bacteria_unclassified|c__Bacteria_unclassified|o__Bacteria_unclassified|f__Bacteria_unclassified|g__Bact
k__Bacteria|p__Bacteria_unclassified|c__Bacteria_unclassified|o__Bacteria_unclassified|f__Bacteria_unclassified|g__Bact
k__Bacteria|p__Bacteria_unclassified|c__Bacteria_unclassified|o__Bacteria_unclassified|f__Bacteria_unclassified|g__Bact
k__Bacteria|p__Bacteria_unclassified|c__Bacteria_unclassified|o__Bacteria_unclassified|f__Bacteria_unclassified|g__Bact
k__Bacteria|p__Bacteria_unclassified|c__Bacteria_unclassified|o__Bacteria_unclassified|f__Bacteria_unclassified|g__Bact
k__Bacteria|p__Bacteria_unclassified|c__Bacteria_unclassified|o__Bacteria_unclassified|f__Bacteria_unclassified|g__Bact
k__Bacteria|p__Bacteria_unclassified|c__Bacteria_unclassified|o__Bacteria_unclassified|f__Bacteria_unclassified|g__Bact
k__Bacteria|p__Bacteria_unclassified|c__Bacteria_unclassified|o__Bacteria_unclassified|f__Bacteria_unclassified|g__Bact
k__Bacteria|p__Bacteria_unclassified|c__Bacteria_unclassified|o__Bacteria_unclassified|f__Bacteria_unclassified|g__Bact
k__Bacteria|p__Bacteria_unclassified|c__Bacteria_unclassified|o__Bacteria_unclassified|f__Bacteria_unclassified|g__Bact
k__Bacteria|p__Bacteria_unclassified|c__Bacteria_unclassified|o__Bacteria_unclassified|f__Bacteria_unclassified|g__Bact
k__Bacteria|p__Bacteria_unclassified|c__Bacteria_unclassified|o__Bacteria_unclassified|f__Bacteria_unclassified|g__Bact
k__Bacteria|p__Bacteria_unclassified|c__Bacteria_unclassified|o__Bacteria_unclassified|f__Bacteria_unclassified|g__Bact
k__Bacteria|p__Bacteria_unclassified|c__Bacteria_unclassified|o__Bacteria_unclassified|f__Bacteria_unclassified|g__Bact
k__Bacteria|p__Bacteria_unclassified|c__Bacteria_unclassified|o__Bacteria_unclassified|f__Bacteria_unclassified|g__Bact
k__Bacteria|p__Bacteria_unclassified|c__Bacteria_unclassified|o__Bacteria_unclassified|f__Bacteria_unclassified|g__Bact
k__Bacteria|p__Bacteria_unclassified|c__Bacteria_unclassified|o__Bacteria_unclassified|f__Bacteria_unclassified|g__Bact
k__Bacteria|p__Bacteria_unclassified|c__Bacteria_unclassified|o__Bacteria_unclassified|f__Bacteria_unclassified|g__Bact
k__Bacteria|p__Bacteroidota|c__Bacteroidia|o__Bacteroidales|f__Bacteroidaceae|g__Bacteroides|s__Bacteroides_thetaio
k__Bacteria|p__Bacteroidota|c__Bacteroidia|o__Bacteroidales|f__Bacteroidaceae|g__Bacteroides|s__Bacteroides_thetaio

k__Bacteria|p__Firmicutes|c__Clostridia|o__Eubacteriales|f__Lachnospiraceae|g__GGB75109|s__GGB75109_SGB102238

k__Bacteria|p__Firmicutes|c__Clostridia|o__Eubacteriales|f__Lachnospiraceae|g__Lachnospiraceae_unclassified|s__Lach

k__Bacteria|p__Firmicutes|c__Clostridia|o__Eubacteriales|f__Lachnospiraceae|g__Lachnospiraceae_unclassified|s__Lach

k__Bacteria|p__Firmicutes|c__Clostridia|o__Eubacteriales|f__Lachnospiraceae|g__Lachnospiraceae_unclassified|s__Lach

k__Bacteria|p__Firmicutes|c__Clostridia|o__Eubacteriales|f__Lachnospiraceae|g__Lachnospiraceae_unclassified|s__Lach

k__Bacteria|p__Firmicutes|c__Clostridia|o__Eubacteriales|f__Lachnospiraceae|g__Lachnospiraceae_unclassified|s__Lach

k__Bacteria|p__Firmicutes|c__Clostridia|o__Eubacteriales|f__Lachnospiraceae|g__Lachnospiraceae_unclassified|s__Lach

k__Bacteria|p__Firmicutes|c__Clostridia|o__Eubacteriales|f__Lachnospiraceae|g__Lachnospiraceae_unclassified|s__Lach

k__Bacteria|p__Firmicutes|c__Clostridia|o__Eubacteriales|f__Lachnospiraceae|g__Lachnospiraceae_unclassified|s__Lach

k__Bacteria|p__Firmicutes|c__Clostridia|o__Eubacteriales|f__Lachnospiraceae|g__Lachnospiraceae_unclassified|s__Lach

k__Bacteria|p__Firmicutes|c__Clostridia|o__Eubacteriales|f__Lachnospiraceae|g__Lachnospiraceae_unclassified|s__Lach

k__Bacteria|p__Firmicutes|c__Bacilli|o__Lactobacillales|f__Lactobacillaceae|g__Lactobacillus|s__Lactobacillus_johnsonii

k__Bacteria|p__Proteobacteria|c__Betaproteobacteria|o__Burkholderiales|f__Sutterellaceae|g__Parasutterella|s__Parasu

k__Bacteria|p__Firmicutes|c__Clostridia|o__Eubacteriales|f__Lachnospiraceae|g__Schaedlerella|s__Schaedlerella_arabin

k__Bacteria|p__Firmicutes|c__Clostridia|o__Eubacteriales|f__Lachnospiraceae|g__Schaedlerella|s__Schaedlerella_arabin

k__Bacteria|p__Firmicutes|c__Erysipelotrichia|o__Erysipelotrichales|f__Turicibacteraceae|g__Turicibacter|s__Turicibacter_
