## Supplementary material for "Non-invasive Vagal Nerve Stimulation as a Potential Treatment for Repetitive Blast Trauma": st5

| | | | 95% Confidence Intervals | |
|---|---|---|---|---|
| | | | VNS (–) | |
| exposure | mediator (Species-level feature) | outcome (Cytokine PC) | ACME | ADE |
| Blast | Lachnospiraceae_bacterium | PC3 | [-1.741, -0.029] | [-0.756, 2.319] |
| Blast | Eubacteriaceae_bacterium | PC3 | [-1.77, 0.027] | [-0.671, 2.28] |
| Blast | GGB29685_SGB42494 | PC3 | [-2.478, -0.062] | [-0.675, 2.835] |
| Blast | Shannon Index | PC3 | [-2.317, 0.697] | [-1.151, 2.439] |
| Blast | Richness (# observed features) | PC3 | [-2.067, -0.046] | [-0.715, 2.463] |

| | | | 95% Confidence Intervals | |
|---|---|---|---|---|
| | | | Blast (–) | |
| exposure | mediator (Species-level feature) | outcome (Cytokine PC) | ACME (Blast-) | ADE (Blast-) |
| VNS | GGB28960_SGB41669 | PC5 | [-0.393, 0.989] | [-0.915, 1.48] |
| VNS | GGB20146_SGB29427 | PC5 | [-0.918, 0.363] | [-0.58, 1.902] |
| VNS | GGB28404_SGB40986 | PC5 | [-2.765, 0.025] | [-0.238, 3.563] |
| VNS | GGB28431_SGB41014 | PC5 | [-1.641, -0.046] | [-0.141, 2.629] |
| VNS | Leptogranulimonas_caecicola | PC5 | [-0.04, 1.699] | [-1.271, 0.956] |

| of Blast effect on outcome (at specified VNS condition) | | | | | |
|---|---|---|---|---|---|
| | VNS (+) | | | Taxonomic Information | |
| Total | ACME | ADE | Total | Genus | Family |
| [-1.347, 1.142] | [-2.284, 0.066] | [-1.203, 0.762] | [-2.575, 0.575] | Lachnospiraceae_unclassifie | Lachnospiraceae |
| [-1.312, 1.099] | [-2.391, 0.062] | [-1.353, 1.059] | [-2.628, 0.62] | Eubacteriaceae_unclassified | Eubacteriaceae |
| [-1.351, 1.022] | [-2.403, 0.01] | [-1.424, 1.1] | [-2.666, 0.621] | GGB29685 | Eubacteriaceae |
| [-1.391, 1.04] | [-2.427, 0.109] | [-1.259, 1.065] | [-2.617, 0.654] | | |
| [-1.397, 1.081] | [-2.301, 0.026] | [-1.096, 0.946] | [-2.593, 0.58] | | |

| of VNS effect on outcome (at specified Blast condition) | | | | | |
|---|---|---|---|---|---|
| | Blast (+) | | | Taxonomic Information | |
| Total (Blast-) | ACME (Blast+) | ADE (Blast+) | Total (Blast+) | Genus | Family |
| [-0.677, 1.777] | [-0.223, 1.896] | [-1.536, 1.323] | [-0.937, 2.053] | GGB28960 | Clostridiaceae |
| [-0.752, 1.859] | [-0.075, 1.95] | [-1.698, 1.327] | [-0.868, 1.968] | GGB20146 | FGB77306 |
| [-0.671, 1.777] | [-1.212, 0.031] | [-0.467, 2.377] | [-0.902, 2.047] | GGB28404 | FGB2838 |
| [-0.747, 1.661] | [-0.03, 1.305] | [-1.298, 1.558] | [-0.907, 2.141] | GGB28431 | Pumilibacteraceae |
| [-0.709, 1.814] | [-0.062, 1.547] | [-1.595, 1.645] | [-0.87, 2.054] | Leptogranulimonas | Atopobiaceae |

| Order | Class | Phylum | Kingdom |
|---|---|---|---|
| Eubacteriales | Clostridia | Firmicutes | Bacteria |
| Eubacteriales | Clostridia | Firmicutes | Bacteria |
| Eubacteriales | Clostridia | Firmicutes | Bacteria |

| Order | Class | Phylum | Kingdom |
|---|---|---|---|
| Eubacteriales | Clostridia | Firmicutes | Bacteria |
| OFGB77306 | CFGB77306 | Firmicutes | Bacteria |
| OFGB2838 | CFGB2838 | Firmicutes | Bacteria |
| Eubacteriales | Clostridia | Firmicutes | Bacteria |
| Coriobacteriales | Coriobacteriia | Actinobacteria | Bacteria |

| MetaPhlan Annotation |
| --- |
| k__Bacteria\|p__Firmicutes\|c__Clostridia\|o__Eubacteriales\|f__Lachnospiraceae\|g__Lachnospiraceae_unclassified\|s__Lachn |
| k__Bacteria\|p__Firmicutes\|c__Clostridia\|o__Eubacteriales\|f__Eubacteriaceae\|g__Eubacteriaceae_unclassified\|s__Eubacte |
| k__Bacteria\|p__Firmicutes\|c__Clostridia\|o__Eubacteriales\|f__Eubacteriaceae\|g__GGB29685\|s__GGB29685_SGB42494 |

| MetaPhlan Annotation |
| --- |
| k__Bacteria\|p__Firmicutes\|c__Clostridia\|o__Eubacteriales\|f__Clostridiaceae\|g__GGB28960\|s__GGB28960_SGB41669 |
| k__Bacteria\|p__Firmicutes\|c__CFGB77306\|o__OFGB77306\|f__FGB77306\|g__GGB20146\|s__GGB20146_SGB29427 |
| k__Bacteria\|p__Firmicutes\|c__CFGB2838\|o__OFGB2838\|f__FGB2838\|g__GGB28404\|s__GGB28404_SGB40986 |
| k__Bacteria\|p__Firmicutes\|c__Clostridia\|o__Eubacteriales\|f__Pumilibacteraceae\|g__GGB28431\|s__GGB28431_SGB41014 |
| k__Bacteria\|p__Actinobacteria\|c__Coriobacteriia\|o__Coriobacteriales\|f__Atopobiaceae\|g__Leptogranulimonas\|s__Leptog |
