## Supplementary material for "Non-invasive Vagal Nerve Stimulation as a Potential Treatment for Repetitive Blast Trauma": st6

| Species-level feature | n | Blast Chi(df=2) | VNS Chi (df=2) | Blast:VNS Chi (df= | Blast Pr(>Chi) |
|---|---|---|---|---|---|
| Acetatifactor_SGB41546 | 63 | -0.4734503914 | -0.6735865311 | 0.05527887593 | 0.8402457971 |
| Acetatifactor_muris | 63 | -0.6551189368 | -0.5673238041 | 0.7556277609 | 0.7395220796 |
| Acutalibacter_muris | 63 | -1.256302585 | -0.1120146742 | 0.5269336414 | 0.4221546334 |
| Acutalibacter_sp_1XD8_36 | 63 | 0.285010825 | 0.6121719594 | -0.8331188827 | 0.6605798863 |
| Adlercreutzia_caecimuris | 63 | 0.0032540798 | 0.6923226625 | -0.109097947 | 0.9896089529 |
| Adlercreutzia_mucosicola | 63 | -1.293145404 | -0.8107578137 | 1.60909155 | 0.2718339856 |
| Adlercreutzia_muris | 63 | 0.1453285309 | -0.9977873679 | 0.5602471455 | 0.6704217743 |
| Akkermansia_muciniphila | 63 | 1.164601827 | 0.0441076170 | 0.3446653734 | 0.1698125404 |
| Alistipes_sp_DSM_112343 | 63 | 1.362207172 | 0.4106786184 | -0.6059096635 | 0.3725159774 |
| Anaerotruncus_sp_1XD42_93 | 63 | 0.162751673 | 0.9054957631 | -0.9596841053 | 0.5202714535 |
| Bacteria_unclassified_SGB102200 | 63 | -1.193792207 | -0.7298338535 | 0.5832040956 | 0.4742692127 |
| Bacteria_unclassified_SGB41677 | 63 | -0.4715195856 | 0.1799007488 | 1.195085048 | 0.4441438231 |
| Bacteria_unclassified_SGB43546 | 63 | 0.1743046341 | -0.6360104272 | 0.6428209411 | 0.5863583659 |
| Bacteroides_thetaiotaomicron | 63 | 0.0121161501 | -0.7991227673 | 0.2034814256 | 0.9602014777 |
| Bifidobacterium_pseudolongum | 63 | -0.0723725935 | -0.1327509122 | -0.9242887686 | 0.4280103985 |
| Clostridia_bacterium | 63 | -1.172211165 | -0.6868310968 | 0.1995598485 | 0.3745590434 |
| Clostridiaceae_bacterium | 63 | 0.1034305113 | -1.201920773 | 0.02066687185 | 0.9871697806 |
| Clostridiaceae_unclassified_SGB41663 | 63 | -0.3486267188 | -0.1904672991 | 0.6891000506 | 0.780382224 |
| Clostridiales_bacterium | 63 | 1.484229444 | -1.424419397 | 1.488073748 | 0.0023319564 |
| Clostridium_cocleatum | 63 | 2.222775114 | 1.218321169 | 0.09951137337 | 0.0115678325 |
| Coriobacteriaceae_bacterium | 63 | -1.041914279 | 0.2056027034 | 0.01023882737 | 0.3724483697 |
| Dorea_sp_5_2 | 63 | 0.1593344522 | -0.0829399946 | -0.8581176899 | 0.5884919176 |
| Dubosiella_newyorkensis | 63 | -1.915269676 | -0.6328732472 | 0.6214032232 | 0.1169987394 |
| Erysipelotrichales_bacterium | 63 | 0.2784071645 | 1.339697821 | -0.4645866706 | 0.8970006656 |
| Eubacteriaceae_bacterium | 63 | 2.751672549 | 1.510462675 | -1.650700676 | 0.0271233687 |
| Eubacteriaceae_unclassified_SGB9492 | 63 | 1.242193062 | 1.358359623 | -0.6975490476 | 0.4594880799 |
| GGB20149_SGB29430 | 63 | 0.9685682607 | -0.0204831086 | -0.6658386857 | 0.6280700484 |
| GGB22635_SGB63107 | 63 | 1.092976537 | 0.4376483775 | -0.1528866375 | 0.4117103362 |
| GGB25041_SGB36960 | 63 | -2.252489312 | -1.39219352 | 1.780688688 | 0.0824966676 |
| GGB27876_SGB40310 | 63 | -1.812623277 | -0.5532913055 | 0.7280341665 | 0.164666135 |
| GGB27878_SGB40312 | 63 | -1.527971618 | -1.626099912 | 0.3766334312 | 0.2133541578 |
| GGB27918_SGB40356 | 63 | 0.2812774952 | -0.1203583756 | -0.2447416231 | 0.9587581408 |
| GGB28382_SGB40962 | 63 | 1.114151113 | 0.4940115299 | -0.5441437287 | 0.5193113079 |
| GGB28399_SGB40980 | 63 | 1.161889363 | 1.030230259 | -1.59173109 | 0.2894655025 |
| GGB28411_SGB40993 | 63 | 0.4381956321 | 0.3093870572 | -1.591442774 | 0.2018022626 |
| GGB28415_SGB40997 | 63 | -2.129244403 | 0.0888312568 | 0.1520170106 | 0.0267267446 |
| GGB28430_SGB41013 | 63 | 0.4099890819 | -0.5583147446 | -0.3724148892 | 0.9120709472 |
| GGB28439_SGB41022 | 63 | 0.5177245932 | -0.7763942741 | 0.6099566625 | 0.3804444035 |
| GGB28778_SGB41431 | 63 | 0.4438702725 | -1.272313732 | 0.05934871732 | 0.8046989411 |
| GGB28784_SGB41437 | 63 | -0.6744740061 | 1.534318282 | -1.589122652 | 0.0207266217 |
| GGB28792_SGB41445 | 63 | -1.397060151 | -0.0925532547 | 0.5796802482 | 0.3385777757 |
| GGB28798_SGB41451 | 63 | 0.3906643009 | 0.9447774686 | -1.657947258 | 0.1640226305 |
| GGB28802_SGB41455 | 63 | 0.4123962239 | 1.468186023 | -1.158799927 | 0.4578986259 |
| GGB28818_SGB41473 | 63 | 0.1651347708 | -0.0781731024 | -0.6862728452 | 0.7292095104 |
| GGB28828_SGB41484 | 63 | -1.176824424 | -0.0692565511 | 1.56387429 | 0.2986969933 |

| | | | | | |
|---|---|---|---|---|---|
| GGB28851_SGB41518 | 63 | -0.2609061865 | 0.23485215 | 0.5473952433 | 0.852552179 |
| GGB28859_SGB41528 | 63 | -0.408183271 | 0.8331838433 | -1.013533439 | 0.2057493298 |
| GGB28864_SGB41535 | 63 | -0.6385726577 | -0.2654400632 | 0.2215820789 | 0.7854652488 |
| GGB28869_SGB41543 | 63 | -0.349754911 | -1.327948185 | 1.297592468 | 0.3479316259 |
| GGB28883_SGB41564 | 63 | -0.6129437991 | -1.554159022 | -0.1933407745 | 0.5948772487 |
| GGB28892_SGB41573 | 63 | 0.8997775489 | 0.8742773198 | -0.783674421 | 0.6512272078 |
| GGB28893_SGB41574 | 63 | -1.686186836 | -1.606896813 | 1.822989034 | 0.165594509 |
| GGB28898_SGB41580 | 63 | -1.432725994 | 0.2309245518 | 1.422712869 | 0.3046549939 |
| GGB28904_SGB41597 | 63 | -0.7529335363 | -1.057389145 | -0.7915484519 | 0.1729063974 |
| GGB28916_SGB41612 | 63 | -0.6453289102 | -0.4630564471 | 0.5573992644 | 0.8014326092 |
| GGB28924_SGB41621 | 63 | 0.5930936828 | -0.5792444894 | -0.6488827416 | 0.7964816629 |
| GGB28926_SGB41624 | 63 | -0.0917956606 | -1.234746851 | 1.285464272 | 0.3067808341 |
| GGB28927_SGB41625 | 63 | 0.1394694394 | -1.227048855 | 1.266111807 | 0.1937208057 |
| GGB28934_SGB41635 | 63 | -1.167727928 | -1.191704148 | 2.087082539 | 0.1158733118 |
| GGB28946_SGB41652 | 63 | 1.180726699 | 1.280762636 | -1.230855982 | 0.4278432256 |
| GGB28949_SGB41655 | 63 | -0.8878623449 | -0.2311637267 | -0.09376521063 | 0.4436081164 |
| GGB28949_SGB41656 | 63 | -1.538392369 | -0.6241183205 | 0.7023127676 | 0.2827282346 |
| GGB28950_SGB41657 | 63 | -2.62149732 | -1.459396162 | 1.26111009 | 0.0305270365 |
| GGB28951_SGB102295 | 63 | 0.3873186134 | 1.566467804 | -0.3192449274 | 0.9253521562 |
| GGB28951_SGB41658 | 63 | -0.2133320024 | -0.0471791042 | -0.2519688292 | 0.8470013515 |
| GGB28954_SGB41662 | 63 | -1.146373063 | -0.5975216265 | 0.6380008981 | 0.5157582975 |
| GGB28956_SGB41665 | 63 | 1.783382845 | 0.6557111785 | -1.761647725 | 0.163329713 |
| GGB28960_SGB41669 | 63 | 1.228625729 | 1.483860342 | -1.964078262 | 0.155727292 |
| GGB28967_SGB41678 | 63 | 1.456567246 | 1.137338312 | -0.0926969036 | 0.1792695292 |
| GGB28991_SGB41705 | 63 | 0.5758405906 | 1.26407252 | -1.603233185 | 0.2288938824 |
| GGB29002_SGB41718 | 63 | -0.8607108206 | 1.090754042 | -2.052520505 | 0.0024308675 |
| GGB29003_SGB41719 | 63 | -0.0411721157 | -0.2727235644 | 0.9024894356 | 0.4993986443 |
| GGB29011_SGB41731 | 63 | -0.3361936314 | -1.277894627 | 0.4112990586 | 0.916276033 |
| GGB29531_SGB42317 | 63 | -1.013385715 | -1.80711559 | 1.975506786 | 0.1450820268 |
| GGB29685_SGB42494 | 63 | 0.4671803769 | 1.112099711 | -0.1138923044 | 0.8645438528 |
| GGB30141_SGB43066 | 63 | 0.1302436989 | -0.4471762074 | -0.09088196888 | 0.9915548039 |
| GGB30286_SGB43248 | 63 | 2.39301438 | 1.182046798 | -1.168443427 | 0.0552616333 |
| GGB30303_SGB43268 | 63 | 0.813311188 | -0.1282774814 | 0.3114034245 | 0.3719368939 |
| GGB30413_SGB43452 | 63 | 0.273835828 | -0.2921852478 | -0.07842307499 | 0.9532769507 |
| GGB30454_SGB43514 | 63 | -0.9314504202 | -1.153409769 | -0.154817533 | 0.3798589507 |
| GGB30455_SGB43519 | 63 | -1.39997214 | -1.557994347 | 1.037789879 | 0.3794677965 |
| GGB30461_SGB43527 | 63 | 1.34676679 | 0.559511253 | -0.7201983713 | 0.3975036627 |
| GGB30461_SGB43530 | 63 | -1.224896578 | -0.7897291579 | 0.3673765275 | 0.395449 |
| GGB30463_SGB43537 | 63 | -1.426341545 | -0.1505261436 | 0.1656987748 | 0.2100542578 |
| GGB30473_SGB43557 | 63 | 0.4322874818 | -2.470434655 | 1.804763781 | 0.0206421669 |
| GGB30475_SGB63182 | 63 | 1.476781394 | -0.1271862005 | -0.5408442462 | 0.2879653492 |
| GGB30861_SGB44083 | 63 | 0.2580569327 | 0.5739850527 | -0.2783889895 | 0.9575523797 |
| GGB31312_SGB44628 | 63 | 0.1805007501 | -0.2225797524 | 0.4695208346 | 0.7117271287 |
| GGB31438_SGB44768 | 63 | -0.3936000439 | -1.146891051 | 0.1185015685 | 0.9063572977 |
| GGB3171_SGB4185 | 63 | 1.710840682 | 0.3357514743 | -0.07198001314 | 0.0853457671 |
| GGB31823_SGB45199 | 63 | 1.489891782 | 0.7051407982 | -0.1831198431 | 0.1889777175 |

| | | | | | |
|---|---|---|---|---|---|
| GGB31853_SGB45233 | 63 | 0.1850728337 | 1.374462655 | -0.9880911527 | 0.4968427437 |
| GGB32371_SGB41694 | 63 | 0.1610064189 | 2.619816891 | -1.191350461 | 0.3468118508 |
| GGB3793_SGB5158 | 63 | -0.1242037497 | -0.0816857986 | -0.08652598792 | 0.9657904747 |
| GGB42598_SGB59794 | 63 | 0.5353666761 | -0.2871673312 | -1.143170137 | 0.5017400018 |
| GGB45656_SGB63370 | 63 | 0.7711268946 | 0.2415073292 | -0.07994670085 | 0.621044138 |
| GGB47127_SGB65054 | 63 | 0.5882221237 | 0.873303962 | 0.3569004709 | 0.4967451202 |
| GGB74395_SGB43521 | 63 | -0.2269015992 | -0.7197230491 | 1.674079714 | 0.1233397821 |
| GGB75053_SGB43494 | 63 | -1.227698761 | -0.982606753 | 0.4220938725 | 0.4105959194 |
| GGB75109_SGB102238 | 63 | -0.9660070503 | -0.3571691551 | 0.5237401709 | 0.6208422921 |
| GGB81440_SGB45230 | 63 | 0.9851844807 | 1.469346382 | -2.189691771 | 0.0796107313 |
| Lachnospiraceae_bacterium | 63 | -0.5775196456 | -0.4403681809 | -0.4033409882 | 0.4848161772 |
| Lachnospiraceae_bacterium_A2 | 63 | -1.30536731 | 0.4121900628 | 0.7190113276 | 0.420834896 |
| Lachnospiraceae_bacterium_MD308 | 63 | -1.253742897 | -1.527073537 | -0.07314364088 | 0.2131194519 |
| Lachnospiraceae_bacterium_MD329 | 63 | 1.569434144 | -0.8172981207 | 0.7446499022 | 0.0209761491 |
| Lachnospiraceae_unclassified_SGB4141 | 63 | -1.427172411 | -1.877633067 | 2.005384557 | 0.142639053 |
| Lachnospiraceae_unclassified_SGB4141 | 63 | 0.0448776106 | 0.6149767571 | -0.04645368359 | 0.9987579817 |
| Lachnospiraceae_unclassified_SGB4142 | 63 | 1.659917229 | 1.593184432 | -2.560059828 | 0.0483863782 |
| Lachnospiraceae_unclassified_SGB4158 | 63 | -1.905534628 | -1.507502957 | 1.520310011 | 0.1654249525 |
| Lactobacillus_johnsonii | 63 | 1.729169167 | 0.0093960181 | -0.6307021833 | 0.1826784534 |
| Muribaculaceae_bacterium | 63 | 0.5862997367 | 0.4519000148 | -0.4279988911 | 0.8415252525 |
| Neglectibacter_sp_X4 | 63 | -2.14515562 | -1.259408626 | 1.998620137 | 0.0847821517 |
| Oscillospiraceae_bacterium | 63 | 0.7143316233 | -0.1585713961 | -0.2605384149 | 0.7416725656 |
| Oscillospiraceae_unclassified_SGB4350 | 63 | -0.9535157468 | -2.459249644 | 2.57855137 | 0.0259412852 |
| Oscillospiraceae_unclassified_SGB4350 | 63 | 0.0443117420 | 0.4774133311 | -0.9098576154 | 0.4959230188 |
| Parasutterella_excrementihominis | 63 | 0.0722400841 | 0.5907446666 | -0.8298439721 | 0.5671048722 |
| Romboutsia_ilealis | 63 | 2.857654195 | 1.685300007 | 0.02578551976 | 0.0010997742 |
| Schaedlerella_arabinosiphila | 63 | -0.1376867522 | -0.4224508602 | 0.3336080387 | 0.9390244043 |
| Turicibacter_sp_1E2 | 63 | 0.1707283912 | 0.1536888746 | -1.455449929 | 0.1991772312 |
| bacterium_1XD42_54 | 63 | 2.903031513 | 1.724849481 | -1.278669218 | 0.0144245959 |
| bacterium_1XD42_76 | 63 | 0.2168984617 | 0.9064029235 | -0.8861147937 | 0.6000316038 |
| bacterium_1xD8_48 | 63 | -0.1464346409 | 0.2151082771 | 0.4220521683 | 0.8997089453 |
| Berger Parker Index | 63 | -0.80005855 | -1.72652844 | 1.105207518 | 0.5457417127 |
| Richness (# observed features) | 63 | 0.3862999043 | -0.7185444522 | 0.9243577639 | 0.2717524423 |
| Shannon Index | 63 | -0.826127489 | -0.1244671288 | 0.3867142166 | 0.6947352216 |

| Likelihood Ratio Test Results | | | | |
|---|---|---|---|---|
| VNS Pr(>Chi) | Blast:VNS Pr(>Ch | Blast Adjusted p-valu | VNS Adjusted p-valu | Blast:VNS Adjusted p-valu |
| 0.6630009771 | 0.9567155372 | 0.9867502072 | 0.9748269319 | 0.9867298836 |
| 0.7528971403 | 0.4518126699 | 0.9364552596 | 0.9748269319 | 0.9634120475 |
| 0.8105784495 | 0.6011691176 | 0.7867838207 | 0.9748269319 | 0.9634120475 |
| 0.7091849336 | 0.407256538 | 0.8878761912 | 0.9748269319 | 0.9634120475 |
| 0.6765426913 | 0.9138311674 | 0.9987579817 | 0.9748269319 | 0.9867298836 |
| 0.2498256574 | 0.1114340708 | 0.7669520852 | 0.8202808144 | 0.6923511791 |
| 0.5958277454 | 0.5757855918 | 0.8915183169 | 0.9748269319 | 0.9634120475 |
| 0.8643625099 | 0.7304679937 | 0.6504699932 | 0.9748269319 | 0.9867298836 |
| 0.8326983074 | 0.5456838431 | 0.7867838207 | 0.9748269319 | 0.9634120475 |
| 0.6174107217 | 0.3492927804 | 0.793096728 | 0.9748269319 | 0.9634120475 |
| 0.7641667885 | 0.56028433 | 0.793096728 | 0.9748269319 | 0.9634120475 |
| 0.1699894243 | 0.2347115362 | 0.793096728 | 0.8202808144 | 0.870124474 |
| 0.7890099125 | 0.5236172725 | 0.8523176191 | 0.9748269319 | 0.9634120475 |
| 0.6343063159 | 0.8395609106 | 0.9977174325 | 0.9748269319 | 0.9867298836 |
| 0.3477990653 | 0.3569574351 | 0.7867838207 | 0.8202808144 | 0.9634120475 |
| 0.7249401071 | 0.8418494906 | 0.7867838207 | 0.9748269319 | 0.9867298836 |
| 0.2419370425 | 0.983839374 | 0.9987579817 | 0.8202808144 | 0.9917735625 |
| 0.719154368 | 0.4915744456 | 0.9671862273 | 0.9748269319 | 0.9634120475 |
| 0.2964785018 | 0.1401477562 | 0.1012861489 | 0.8202808144 | 0.7299362302 |
| 0.1960355146 | 0.9207356632 | 0.308220099 | 0.8202808144 | 0.9867298836 |
| 0.953919137 | 0.9918495863 | 0.7867838207 | 0.9803444972 | 0.9918495863 |
| 0.4214161244 | 0.392260953 | 0.8523176191 | 0.8953154676 | 0.9634120475 |
| 0.799186063 | 0.5419962879 | 0.6504699932 | 0.9748269319 | 0.9634120475 |
| 0.3284081083 | 0.6425119949 | 0.9977174325 | 0.8202808144 | 0.9867298836 |
| 0.2383186135 | 0.102464393 | 0.308220099 | 0.8202808144 | 0.6923511791 |
| 0.3726201484 | 0.4865673277 | 0.793096728 | 0.8625466399 | 0.9634120475 |
| 0.6299827926 | 0.5066260299 | 0.862733583 | 0.9748269319 | 0.9634120475 |
| 0.8871521383 | 0.8796922403 | 0.7867838207 | 0.9748269319 | 0.9867298836 |
| 0.2101871969 | 0.0786399911 | 0.5926789385 | 0.8202808144 | 0.6923511791 |
| 0.7672351949 | 0.4675262866 | 0.6504699932 | 0.9748269319 | 0.9634120475 |
| 0.1474794298 | 0.7066228553 | 0.6504699932 | 0.8202808144 | 0.9867298836 |
| 0.8856989297 | 0.8067114261 | 0.9977174325 | 0.9748269319 | 0.9867298836 |
| 0.8550578348 | 0.5892195811 | 0.793096728 | 0.9748269319 | 0.9634120475 |
| 0.287117034 | 0.1161986568 | 0.7669520852 | 0.8202808144 | 0.6923511791 |
| 0.1468484741 | 0.1150921455 | 0.6504699932 | 0.8202808144 | 0.6923511791 |
| 0.9490516966 | 0.8792510459 | 0.308220099 | 0.9803444972 | 0.9867298836 |
| 0.4662504826 | 0.7102045329 | 0.9977174325 | 0.9491235182 | 0.9867298836 |
| 0.7387095529 | 0.5427880088 | 0.7867838207 | 0.9748269319 | 0.9634120475 |
| 0.2184532434 | 0.9526874177 | 0.9671862273 | 0.8202808144 | 0.9867298836 |
| 0.2544569976 | 0.1175944723 | 0.308220099 | 0.8202808144 | 0.6923511791 |
| 0.7612444262 | 0.5639713269 | 0.7867838207 | 0.9748269319 | 0.9634120475 |
| 0.2489158244 | 0.1026632614 | 0.6504699932 | 0.8202808144 | 0.6923511791 |
| 0.342689298 | 0.2505958485 | 0.793096728 | 0.8202808144 | 0.870124474 |
| 0.5694099484 | 0.4934268922 | 0.9364552596 | 0.9748269319 | 0.9634120475 |
| 0.1067065352 | 0.1218538075 | 0.7669520852 | 0.8202808144 | 0.6923511791 |

| | | | | |
|---|---|---|---|---|
| 0.578308429[illegible] | 0.5845532243 | 0.9867502072 | 0.9748269319 | 0.9634120475 |
| 0.595366745[illegible] | 0.3132677119 | 0.6504699932 | 0.9748269319 | 0.9634120475 |
| 0.964658985[illegible] | 0.8275909328 | 0.9671862273 | 0.9803444972 | 0.9867298836 |
| 0.381477475[illegible] | 0.2033782479 | 0.7867838207 | 0.8669942626 | 0.870124474 |
| 0.060524458[illegible] | 0.8487218661 | 0.8523176191 | 0.8202808144 | 0.9867298836 |
| 0.667618882[illegible] | 0.4344240852 | 0.8848195758 | 0.9748269319 | 0.9634120475 |
| 0.1826993761 | 0.0721706083 | 0.6504699932 | 0.8202808144 | 0.6923511791 |
| 0.082893330[illegible] | 0.1581963478 | 0.7669520852 | 0.8202808144 | 0.7605593646 |
| 0.056067199[illegible] | 0.429766694 | 0.6504699932 | 0.8202808144 | 0.9634120475 |
| 0.853813296[illegible] | 0.5782729426 | 0.9671862273 | 0.9748269319 | 0.9634120475 |
| 0.278658378[illegible] | 0.5208167584 | 0.9671862273 | 0.8202808144 | 0.9634120475 |
| 0.470765265 | 0.2436965868 | 0.7669520852 | 0.9491235182 | 0.870124474 |
| 0.418243373[illegible] | 0.2127798207 | 0.6504699932 | 0.8953154676 | 0.870124474 |
| 0.1113740731 | 0.04018812587 | 0.6504699932 | 0.8202808144 | 0.6923511791 |
| 0.422588900[illegible] | 0.2250536625 | 0.7867838207 | 0.8953154676 | 0.870124474 |
| 0.9089481791 | 0.9252982663 | 0.793096728 | 0.9748269319 | 0.9867298836 |
| 0.770863949[illegible] | 0.4835409848 | 0.7669520852 | 0.9748269319 | 0.9634120475 |
| 0.336458904[illegible] | 0.2102296777 | 0.3179899638 | 0.8202808144 | 0.870124474 |
| 0.158790397[illegible] | 0.7497999296 | 0.9977174325 | 0.8202808144 | 0.9867298836 |
| 0.920236623[illegible] | 0.8012238941 | 0.9867502072 | 0.9748269319 | 0.9867298836 |
| 0.8004849551 | 0.5245336238 | 0.793096728 | 0.9748269319 | 0.9634120475 |
| 0.153416502[illegible] | 0.08251291477 | 0.6504699932 | 0.8202808144 | 0.6923511791 |
| 0.155957479[illegible] | 0.05400847696 | 0.6504699932 | 0.8202808144 | 0.6923511791 |
| 0.315046519[illegible] | 0.9261468679 | 0.6504699932 | 0.8202808144 | 0.9867298836 |
| 0.279982342 | 0.1124600555 | 0.6812317928 | 0.8202808144 | 0.6923511791 |
| 0.113465587[illegible] | 0.0434459902 | 0.1012861489 | 0.8202808144 | 0.6923511791 |
| 0.583967886[illegible] | 0.3713550509 | 0.793096728 | 0.9748269319 | 0.9634120475 |
| 0.345599236[illegible] | 0.6810658224 | 0.9977174325 | 0.8202808144 | 0.9867298836 |
| 0.135569966[illegible] | 0.05447985183 | 0.6504699932 | 0.8202808144 | 0.6923511791 |
| 0.340104263 | 0.9093514035 | 0.9914493724 | 0.8202808144 | 0.9867298836 |
| 0.760178789[illegible] | 0.927588801 | 0.9987579817 | 0.9748269319 | 0.9867298836 |
| 0.454603367[illegible] | 0.2472501155 | 0.4934074404 | 0.9470903485 | 0.870124474 |
| 0.944722174[illegible] | 0.7558806993 | 0.7867838207 | 0.9803444972 | 0.9867298836 |
| 0.879642604[illegible] | 0.9389931831 | 0.9977174325 | 0.9748269319 | 0.9867298836 |
| 0.200125591[illegible] | 0.8769767752 | 0.7867838207 | 0.8202808144 | 0.9867298836 |
| 0.302348921[illegible] | 0.3029893204 | 0.7867838207 | 0.8202808144 | 0.9634120475 |
| 0.771038655[illegible] | 0.4723848681 | 0.7867838207 | 0.9748269319 | 0.9634120475 |
| 0.703863693[illegible] | 0.7143974322 | 0.7867838207 | 0.9748269319 | 0.9867298836 |
| 0.985361479[illegible] | 0.8685968913 | 0.6504699932 | 0.9853614792 | 0.9867298836 |
| 0.0562117431 | 0.08022178104 | 0.308220099 | 0.8202808144 | 0.6923511791 |
| 0.660855930[illegible] | 0.5933201721 | 0.7669520852 | 0.9748269319 | 0.9634120475 |
| 0.8334455291 | 0.7807794431 | 0.9977174325 | 0.9748269319 | 0.9867298836 |
| 0.884193943[illegible] | 0.6393869554 | 0.9267280322 | 0.9748269319 | 0.9867298836 |
| 0.318960086[illegible] | 0.905742012 | 0.9977174325 | 0.8202808144 | 0.9867298836 |
| 0.9185517801 | 0.942738773 | 0.5926789385 | 0.9748269319 | 0.9867298836 |
| 0.702813258[illegible] | 0.8547230999 | 0.6504699932 | 0.9748269319 | 0.9867298836 |

| | | | | |
|---|---|---|---|---|
| 0.3962595415 | 0.3252604076 | 0.793096728 | 0.8845079052 | 0.9634120475 |
| 0.0250540925 | 0.236127701 | 0.7867838207 | 0.8202808144 | 0.870124474 |
| 0.9751451519 | 0.931079469 | 0.9977174325 | 0.9830092257 | 0.9867298836 |
| 0.156668604 | 0.2580707395 | 0.793096728 | 0.8202808144 | 0.8718606065 |
| 0.9633746015 | 0.9363521938 | 0.8625613028 | 0.9803444972 | 0.9867298836 |
| 0.2574173127 | 0.7215181373 | 0.793096728 | 0.8202808144 | 0.9867298836 |
| 0.2040214986 | 0.09887835326 | 0.6504699932 | 0.8202808144 | 0.6923511791 |
| 0.5718574569 | 0.6733052542 | 0.7867838207 | 0.9748269319 | 0.9867298836 |
| 0.8718184088 | 0.6008549705 | 0.8625613028 | 0.9748269319 | 0.9634120475 |
| 0.0981405799 | 0.03158525082 | 0.5926789385 | 0.8202808144 | 0.6923511791 |
| 0.5378613321 | 0.6868886622 | 0.793096728 | 0.9748269319 | 0.9867298836 |
| 0.3236483287 | 0.4731647787 | 0.7867838207 | 0.8202808144 | 0.9634120475 |
| 0.0830669656 | 0.9416934878 | 0.6504699932 | 0.8202808144 | 0.9867298836 |
| 0.6991611194 | 0.4603075655 | 0.308220099 | 0.9748269319 | 0.9634120475 |
| 0.119876468 | 0.04864539177 | 0.6504699932 | 0.8202808144 | 0.6923511791 |
| 0.7122365734 | 0.9630483664 | 0.9987579817 | 0.9748269319 | 0.9867298836 |
| 0.0460673576 | 0.01394245873 | 0.4652536372 | 0.8202808144 | 0.6923511791 |
| 0.2735277008 | 0.1326996547 | 0.6504699932 | 0.8202808144 | 0.7211937753 |
| 0.6729515885 | 0.5408046464 | 0.6504699932 | 0.9748269319 | 0.9634120475 |
| 0.8930529806 | 0.6688780979 | 0.9867502072 | 0.9748269319 | 0.9867298836 |
| 0.140656264 | 0.04955953861 | 0.5926789385 | 0.8202808144 | 0.6923511791 |
| 0.856188039 | 0.7960795889 | 0.9364552596 | 0.9748269319 | 0.9867298836 |
| 0.0305351331 | 0.01200183308 | 0.308220099 | 0.8202808144 | 0.6923511791 |
| 0.6425337316 | 0.3652642266 | 0.793096728 | 0.9748269319 | 0.9634120475 |
| 0.7099929171 | 0.4079042386 | 0.8439060598 | 0.9748269319 | 0.9634120475 |
| 0.0569970896 | 0.9795116329 | 0.1012861489 | 0.8202808144 | 0.9917735625 |
| 0.9139116075 | 0.7388784276 | 0.9977174325 | 0.9748269319 | 0.9867298836 |
| 0.1655749312 | 0.1496824151 | 0.6504699932 | 0.8202808144 | 0.7484120754 |
| 0.2345775748 | 0.2077670838 | 0.308220099 | 0.8202808144 | 0.870124474 |
| 0.6429924539 | 0.3849808138 | 0.8523176191 | 0.9748269319 | 0.9634120475 |
| 0.6968168809 | 0.6732043704 | 0.9977174325 | 0.9748269319 | 0.9867298836 |
| 0.2337330546 | 0.2720400368 | 0.8219001698 | 0.8202808144 | 0.8948685421 |
| 0.6607230685 | 0.3632131264 | 0.7669520852 | 0.9748269319 | 0.9634120475 |
| 0.9062773413 | 0.6991374962 | 0.9141252915 | 0.9748269319 | 0.9867298836 |

| Intercept Estimate | Blast Estimate | VNS Estimate | Blast:VNS Estimat | Intercept Std. Erro | Blast Std. Erro |
|---|---|---|---|---|---|
| 0.1862036252 | -0.1585939777 | -0.2414121727 | 0.02768837327 | 0.2564678696 | 0.3349748581 |
| 0.1320788204 | -0.210843694 | -0.1964330009 | 0.365964301 | 0.2787717747 | 0.3218403288 |
| 0.1796058645 | -0.4146335832 | -0.0396196586 | 0.2605908371 | 0.2599806818 | 0.3300427684 |
| -0.0720460121 | 0.0931080639 | 0.2148541588 | -0.4089721397 | 0.2718391799 | 0.3266825531 |
| -0.08417876058 | 0.0010548158 | 0.24117751 | -0.05315828283 | 0.2719847962 | 0.3241518222 |
| 0.1090122372 | -0.3833676609 | -0.259402485 | 0.7200935783 | 0.2970369774 | 0.2964613721 |
| 0.07534050249 | 0.0486005314 | -0.3567172486 | 0.2798318283 | 0.2527965312 | 0.3344183769 |
| -0.2427809083 | 0.3835303967 | 0.0155285852 | 0.1695302879 | 0.248944937 | 0.3293231967 |
| -0.2528121122 | 0.449920403 | 0.1453639231 | -0.2998672791 | 0.2601333911 | 0.3302877948 |
| -0.1068970553 | 0.0506544077 | 0.3038766818 | -0.4504865524 | 0.2956379602 | 0.3112374014 |
| 0.2650911331 | -0.3994019262 | -0.2610365934 | 0.2914266764 | 0.2529078974 | 0.3345657006 |
| -0.09349046117 | -0.1364455461 | 0.0562094751 | 0.5222624623 | 0.2996464452 | 0.2893740797 |
| 0.00901581808 | 0.0581012266 | -0.2270183327 | 0.3207537912 | 0.2588255314 | 0.3333315086 |
| 0.09883247034 | 0.0040541863 | -0.286195728 | 0.10185882 | 0.2577015618 | 0.3346101072 |
| 0.1279196311 | -0.0239936006 | -0.0470494745 | -0.4576740499 | 0.2506122327 | 0.3315288217 |
| 0.2922843076 | -0.3898508417 | -0.2441960002 | 0.09912726211 | 0.2514048192 | 0.3325773149 |
| 0.1217954104 | 0.0328637308 | -0.4109114176 | 0.009883164737 | 0.276501201 | 0.3177372944 |
| 0.02046925437 | -0.1175028811 | -0.0686284584 | 0.3468945298 | 0.254782047 | 0.3370449674 |
| -0.1712712765 | 0.4545159281 | -0.4663177181 | 0.6806123769 | 0.2314882971 | 0.3062302328 |
| -0.5658004526 | 0.6730665607 | 0.3965381484 | 0.04530300529 | 0.2553879661 | 0.3028046142 |
| -0.05881865091 | -0.1933910963 | 0.0413557631 | 0.00287954607 | 0.4227788288 | 0.1856113311 |
| 0.1433831779 | 0.0450902938 | -0.0253644546 | -0.3670249921 | 0.3158803996 | 0.2829914886 |
| 0.3672031281 | -0.6141939786 | -0.2179055039 | 0.2992381223 | 0.262770854 | 0.3206827667 |
| -0.2126757667 | 0.0926350809 | 0.4765384003 | -0.2308815058 | 0.251522044 | 0.3327323888 |
| -0.5216405645 | 0.8493352243 | 0.5011866638 | -0.7661107563 | 0.2611181654 | 0.3086614447 |
| -0.2688510543 | 0.3298666093 | 0.389967474 | -0.2800665686 | 0.3099935256 | 0.2655518046 |
| -0.1002520089 | 0.3225158349 | -0.0072997648 | -0.3316656356 | 0.2562682992 | 0.3329820395 |
| -0.250187041 | 0.3626941195 | 0.1555119813 | -0.07594340189 | 0.2575501447 | 0.3318407186 |
| 0.416167342 | -0.7092584479 | -0.4711543561 | 0.8429083961 | 0.2649076032 | 0.3148776085 |
| 0.3252687531 | -0.5967649727 | -0.1947357066 | 0.3579932643 | 0.2488724064 | 0.3292272478 |
| 0.1589598841 | -0.2237793896 | -0.2581122692 | 0.08358733208 | 0.3709085894 | 0.1464552004 |
| -0.06455449758 | 0.0280698397 | -0.0130219460 | -0.03702122538 | 0.4211922243 | 0.0997941186 |
| -0.1692060535 | 0.3422611857 | 0.1636424689 | -0.2521240277 | 0.2927215766 | 0.3071945821 |
| -0.1963217056 | 0.326156363 | 0.3124298741 | -0.6751264066 | 0.3043622289 | 0.2807120655 |
| 0.0778694057 | 0.1356511277 | 0.1031175911 | -0.7419492816 | 0.2754196774 | 0.3095675031 |
| 0.1298867135 | -0.4699303057 | 0.0212271521 | 0.05079583395 | 0.3380365577 | 0.2207028489 |
| 0.1025410348 | 0.1246150709 | -0.1830395082 | -0.1707771485 | 0.2941111792 | 0.3039472912 |
| -0.00248189524 | 0.1659698631 | -0.2675588637 | 0.294015138 | 0.271187432 | 0.3205755826 |
| 0.09298163417 | 0.1383495383 | -0.4268497528 | 0.02785119213 | 0.2746038484 | 0.3116891283 |
| 0.03795657332 | -0.1891483353 | 0.4645308279 | -0.6729353303 | 0.2876608864 | 0.2804382876 |
| 0.1873783265 | -0.4181241248 | -0.0298784796 | 0.2617531377 | 0.2899522637 | 0.2992885629 |
| 0.01993959587 | 0.1059594045 | 0.2769962359 | -0.6798179453 | 0.3119108042 | 0.2712287872 |
| -0.1787713052 | 0.1354584702 | 0.5168613722 | -0.5703957475 | 0.2591449273 | 0.3284668055 |
| 0.06161408894 | 0.0528276641 | -0.0269022009 | -0.3303494808 | 0.2763664619 | 0.3199063641 |
| 0.06677732509 | -0.3579862672 | -0.0227122056 | 0.7173734013 | 0.2864071236 | 0.3041968367 |

| | | | | | |
|---|---|---|---|---|---|
| -0.04824489354 | -0.0876396606 | 0.0843348009 | 0.2746278975 | 0.25392022 | 0.3359048775 |
| 0.08431351822 | -0.113477078 | 0.250392416 | -0.4259851133 | 0.3212958158 | 0.278005215 |
| 0.1319192454 | -0.2069427253 | -0.0925118629 | 0.1080209263 | 0.2776162037 | 0.3240707582 |
| 0.1550069923 | -0.1131513606 | -0.461603014 | 0.6308786115 | 0.2699727326 | 0.3235161454 |
| 0.365894064 | -0.1940106873 | -0.5280813505 | -0.09187822025 | 0.2583041955 | 0.3165227996 |
| -0.07978902738 | 0.221631913 | 0.233049718 | -0.2921240939 | 0.3357138957 | 0.2463185632 |
| 0.3347677136 | -0.5373333233 | -0.550278359 | 0.8731785322 | 0.266991347 | 0.3186677253 |
| 0.02982167546 | -0.4035053563 | 0.0702584552 | 0.6054021629 | 0.3044176405 | 0.2816347005 |
| 0.3634739045 | -0.2399058875 | -0.3601762613 | -0.376694337 | 0.2408603096 | 0.31862824 |
| 0.1249573777 | -0.2158771894 | -0.1659538035 | 0.2792908071 | 0.2619631384 | 0.3345227309 |
| 0.08229197167 | 0.1831792359 | -0.1928196932 | -0.3021367566 | 0.2873343203 | 0.308853797 |
| 0.08627837896 | -0.0301751771 | -0.4350782551 | 0.6333391021 | 0.2600368729 | 0.3287211716 |
| -0.007403762026 | 0.0417535634 | -0.3964447291 | 0.5721613326 | 0.299565664 | 0.2993742833 |
| 0.1756310539 | -0.3696989799 | -0.4053998529 | 0.9930611173 | 0.2644733387 | 0.3165968467 |
| -0.2656703559 | 0.3879578944 | 0.4512111358 | -0.606354383 | 0.2612664521 | 0.3285755244 |
| 0.2018382594 | -0.2967935866 | -0.0826084617 | -0.04681423499 | 0.2526910443 | 0.3342788308 |
| 0.2012822146 | -0.4615361389 | -0.2020252587 | 0.3179803862 | 0.2955793747 | 0.3000119788 |
| 0.3389357668 | -0.6320465923 | -0.3807371666 | 0.4600895402 | 0.3214396109 | 0.2411013689 |
| -0.3003867302 | 0.115227992 | 0.5027718024 | -0.1433198595 | 0.2914606571 | 0.2975018189 |
| 0.05080678249 | -0.0701587750 | -0.0166605248 | -0.1244461837 | 0.2700334284 | 0.3288713096 |
| 0.2529519639 | -0.3573601214 | -0.2007380462 | 0.2998100181 | 0.2887515532 | 0.3117310874 |
| -0.236894716 | 0.5335894813 | 0.2115369012 | -0.794942056 | 0.28392315 | 0.2992007482 |
| -0.2373253181 | 0.4016116375 | 0.5194696687 | -0.9612290826 | 0.2544297229 | 0.326878746 |
| -0.4147560344 | 0.4739015801 | 0.3955881373 | -0.04504537092 | 0.2459453369 | 0.3253550988 |
| -0.1350301162 | 0.190523203 | 0.4471091555 | -0.792263441 | 0.2501074193 | 0.3308610163 |
| 0.1740001502 | -0.2649840273 | 0.3589923864 | -0.9437941061 | 0.2327251984 | 0.3078664994 |
| -0.01606083432 | -0.0132584405 | -0.0944379619 | 0.4371253956 | 0.2747778361 | 0.3220247559 |
| 0.2551044445 | -0.1005301176 | -0.4122720003 | 0.1856001833 | 0.2938595898 | 0.2990244556 |
| 0.2417328274 | -0.3270412043 | -0.6256864191 | 0.9565335973 | 0.2598990061 | 0.3227213482 |
| -0.2296424993 | 0.1523793025 | 0.3892167188 | -0.0557444768 | 0.263506357 | 0.3261680285 |
| 0.09867256838 | 0.0384566528 | -0.1425461018 | -0.04052016523 | 0.3028776308 | 0.2952668972 |
| -0.468304578 | 0.7716240233 | 0.4078273497 | -0.5634188497 | 0.2471303504 | 0.3224485527 |
| -0.1449740372 | 0.2671629549 | -0.0451861662 | 0.1533899662 | 0.2619185281 | 0.3284879869 |
| 0.02802320831 | 0.0853702726 | -0.0982233068 | -0.03687584641 | 0.2968704061 | 0.3117571329 |
| 0.3540539676 | -0.3043301692 | -0.4028699395 | -0.07554969753 | 0.2469825329 | 0.3267271801 |
| 0.3745222909 | -0.4540906218 | -0.5422286994 | 0.5051077194 | 0.2618732528 | 0.3243568987 |
| -0.2349796394 | 0.4410470829 | 0.196621351 | -0.3539457954 | 0.2648781061 | 0.3274858618 |
| 0.2832290556 | -0.3582841105 | -0.2493747357 | 0.1622570035 | 0.2992523001 | 0.2925015197 |
| 0.2465460105 | -0.4459389728 | -0.0506702241 | 0.07802123374 | 0.2784901355 | 0.3126452949 |
| 0.09569724572 | 0.1299508273 | -0.7982636354 | 0.8156893506 | 0.2536355586 | 0.3006120527 |
| -0.224081214 | 0.4448873185 | -0.0413470137 | -0.2459256915 | 0.299830494 | 0.3012546884 |
| -0.1088221784 | 0.0871873002 | 0.2073166847 | -0.1404808338 | 0.2553986954 | 0.3378607166 |
| -0.005131767126 | 0.0535660418 | -0.0712808830 | 0.2103146284 | 0.2956464557 | 0.2967635415 |
| 0.2081340033 | -0.0939960435 | -0.2964726301 | 0.04283592046 | 0.3429560812 | 0.2388110596 |
| -0.2631342547 | 0.5170264461 | 0.1093642054 | -0.03279567922 | 0.2816877152 | 0.3022060741 |
| -0.3484375405 | 0.4902040442 | 0.2480241567 | -0.08998806135 | 0.2487156658 | 0.3290198994 |

| | | | | | |
|---|---|---|---|---|---|
| -0.1806549853 | 0.0548984985 | 0.4399747438 | -0.4424039564 | 0.2955213881 | 0.2966318579 |
| -0.3046639223 | 0.0509946691 | 0.8870508155 | -0.5635707548 | 0.2394211783 | 0.3167244482 |
| -0.02598198852 | -0.0358307559 | -0.0254597354 | -0.03771802252 | 0.3147146499 | 0.2884836893 |
| 0.05180008554 | 0.1709260195 | -0.0984699878 | -0.5482580081 | 0.2636167733 | 0.3192690676 |
| -0.1417504503 | 0.235075737 | 0.0794164052 | -0.03677190635 | 0.2961674995 | 0.3048470215 |
| -0.2036200355 | 0.1594077335 | 0.2557767649 | 0.1461931165 | 0.3053567462 | 0.2709992145 |
| 0.02893440529 | -0.0669933525 | -0.2292832369 | 0.745956402 | 0.2914109425 | 0.2952528885 |
| 0.3045652185 | -0.3983775462 | -0.3422571016 | 0.2056184102 | 0.2643745275 | 0.3244912831 |
| 0.172932547 | -0.3248245825 | -0.1283921622 | 0.2630339466 | 0.2541847953 | 0.3362548777 |
| -0.1512685845 | 0.2593575532 | 0.4181473479 | -0.8714941011 | 0.3039636882 | 0.263257855 |
| 0.2116514492 | -0.1921743705 | -0.1566536829 | -0.2004603044 | 0.2515415197 | 0.3327581528 |
| 0.04233502942 | -0.363129333 | 0.1238923341 | 0.3022562315 | 0.3056164715 | 0.2781817272 |
| 0.3384724004 | -0.3206162562 | -0.4222551992 | -0.02828513019 | 0.3048817818 | 0.2557272763 |
| -0.2189550189 | 0.4886043674 | -0.2733594385 | 0.3483547435 | 0.2589814162 | 0.31132518 |
| 0.2199680497 | -0.3902769739 | -0.5550390204 | 0.8290627242 | 0.3151185399 | 0.2734616861 |
| -0.1006650338 | 0.0149722563 | 0.2197980371 | -0.02321226666 | 0.2610402954 | 0.3336241862 |
| -0.2285475353 | 0.4620926613 | 0.4792360556 | -1.077021707 | 0.3073738926 | 0.2783829537 |
| 0.2151878806 | -0.270110837 | -0.2316194046 | 0.3265915091 | 0.3910712273 | 0.1417506841 |
| -0.2348994365 | 0.5674198304 | 0.0032961490 | -0.3091140064 | 0.2480550097 | 0.3281459336 |
| -0.1317291893 | 0.1980771619 | 0.1632123563 | -0.2159656458 | 0.255385174 | 0.3378428294 |
| 0.2715843844 | -0.6161624254 | -0.3905872148 | 0.8669522589 | 0.2973279573 | 0.287234371 |
| -0.06620517248 | 0.2267946157 | -0.0542008088 | -0.1245676065 | 0.2818096345 | 0.317492056 |
| 0.2360393563 | -0.2936868619 | -0.8145040934 | 1.194555638 | 0.2628020617 | 0.3080042075 |
| 0.01227320994 | 0.0141256003 | 0.1637859349 | -0.4366225838 | 0.2793681454 | 0.3187778162 |
| 0.0001352241089 | 0.0171726102 | 0.1520072575 | -0.2985951038 | 0.3410986492 | 0.2377158114 |
| -0.6165574123 | 0.775886241 | 0.4935272053 | 0.01056203603 | 0.2619785709 | 0.2715115924 |
| 0.04896773132 | -0.0407209482 | -0.1348866452 | 0.1489861081 | 0.3036361184 | 0.2957506631 |
| 0.1054452303 | 0.0545402760 | 0.0527225602 | -0.6983128555 | 0.2627343205 | 0.3194563932 |
| -0.5442798437 | 0.8177437636 | 0.5245013235 | -0.5438414756 | 0.2874137686 | 0.2816861477 |
| -0.120128647 | 0.0710905163 | 0.3191635069 | -0.4364130542 | 0.2726102811 | 0.3277594307 |
| -0.05241013265 | -0.0493304516 | 0.0774683596 | 0.2123560915 | 0.2546550192 | 0.3368769255 |
| 0.3195058697 | -0.2601185666 | -0.6025054407 | 0.5393957355 | 0.2644431858 | 0.3251244132 |
| -0.04836798411 | 0.1280607399 | -0.2546876214 | 0.4577778046 | 0.251177294 | 0.3315060099 |
| 0.1254652467 | -0.2781161792 | -0.0447950294 | 0.1944449381 | 0.2544838091 | 0.3366504358 |

| Full model fit coefficient estimates (microbe ~ Blast + VNS + Blast:VNS + (1\|I | | | | | |
|---|---|---|---|---|---|
| VNS Std. Error | Blast:VNS Std. Errc | Intercept t-value | Blast t-value | VNS t-value | Blast:VNS t-valu |
| 0.3583981591 | 0.5008852441 | 0.7260310055 | -0.4734503914 | -0.6735865311 | 0.05527887593 |
| 0.3462449478 | 0.4843182317 | 0.4737883544 | -0.6551189368 | -0.5673238041 | 0.7556277609 |
| 0.35370061 | 0.4945420383 | 0.6908431167 | -1.256302585 | -0.1120146742 | 0.5269336414 |
| 0.3509702716 | 0.4908928944 | -0.2650317446 | 0.285010825 | 0.6121719594 | -0.8331188827 |
| 0.3483599816 | 0.4872528246 | -0.3094980372 | 0.0032540798 | 0.6923226625 | -0.109097947 |
| 0.3199506445 | 0.4475156049 | 0.3669988773 | -1.293145404 | -0.8107578137 | 1.60909155 |
| 0.3575082829 | 0.4994792575 | 0.298028229 | 0.1453285309 | -0.9977873679 | 0.5602471455 |
| 0.3520613062 | 0.4918692188 | -0.9752393891 | 1.164601827 | 0.0441076170 | 0.3446653734 |
| 0.3539602907 | 0.4949042689 | -0.9718556744 | 1.362207172 | 0.4106786184 | -0.6059096635 |
| 0.3355915004 | 0.4694112885 | -0.3615809526 | 0.162751673 | 0.9054957631 | -0.9596841053 |
| 0.3576657785 | 0.4996992966 | 1.048172619 | -1.193792207 | -0.7298338535 | 0.5832040956 |
| 0.3124471439 | 0.4370086155 | -0.3120025706 | -0.4715195856 | 0.1799007488 | 1.195085048 |
| 0.356941212 | 0.4989784413 | 0.03483357314 | 0.1743046341 | -0.6360104272 | 0.6428209411 |
| 0.3581373723 | 0.5005804324 | 0.3835152167 | 0.0121161501 | -0.7991227673 | 0.2034814256 |
| 0.3544192184 | 0.4951634873 | 0.5104285201 | -0.0723725935 | -0.1327509122 | -0.9242887686 |
| 0.3555401049 | 0.4967294916 | 1.162604235 | -1.172211165 | -0.6868310968 | 0.1995598485 |
| 0.3418789548 | 0.4782129009 | 0.4404878168 | 0.1034305113 | -1.201920773 | 0.02066687185 |
| 0.3603162263 | 0.5034022701 | 0.08034025401 | -0.3486267188 | -0.1904672991 | 0.6891000506 |
| 0.3273738893 | 0.4573781224 | -0.7398701301 | 1.484229444 | -1.424419397 | 1.488073748 |
| 0.3254791581 | 0.4552545478 | -2.215454632 | 2.222775114 | 1.218321169 | 0.09951137337 |
| 0.2011440631 | 0.2812378767 | -0.139123927 | -1.041914279 | 0.2056027034 | 0.01023882737 |
| 0.3058169316 | 0.4277093882 | 0.453916033 | 0.1593344522 | -0.0829399946 | -0.8581176899 |
| 0.3443114477 | 0.4815522532 | 1.39742716 | -1.915269676 | -0.6328732472 | 0.6214032232 |
| 0.3557058859 | 0.4969611063 | -0.8455551781 | 0.2784071645 | 1.339697821 | -0.4645866706 |
| 0.3318100289 | 0.464112463 | -1.997718404 | 2.751672549 | 1.510462675 | -1.650700676 |
| 0.2870870625 | 0.4015008975 | -0.867279579 | 1.242193062 | 1.358359623 | -0.6975490476 |
| 0.3563797356 | 0.4981171007 | -0.3911994158 | 0.9685682607 | -0.0204831086 | -0.6658386857 |
| 0.3553354456 | 0.496730147 | -0.9714109899 | 1.092976537 | 0.4376483775 | -0.1528866375 |
| 0.3384259079 | 0.4733608979 | 1.570990553 | -2.252489312 | -1.39219352 | 1.780688688 |
| 0.3519587325 | 0.4917259118 | 1.306969936 | -1.812623277 | -0.5532913055 | 0.7280341665 |
| 0.1587308795 | 0.2219328534 | 0.4285688945 | -1.527971618 | -1.626099912 | 0.3766334312 |
| 0.1081931025 | 0.151266568 | -0.153266119 | 0.2812774952 | -0.1203583756 | -0.2447416231 |
| 0.331252327 | 0.4633408682 | -0.5780443499 | 1.114151113 | 0.4940115299 | -0.5441437287 |
| 0.30326218 | 0.4241460198 | -0.6450265077 | 1.161889363 | 1.030230259 | -1.59173109 |
| 0.3332963959 | 0.4662117254 | 0.2827300011 | 0.4381956321 | 0.3093870572 | -1.591442774 |
| 0.2389603939 | 0.3341457233 | 0.3842386588 | -2.129244403 | 0.0888312568 | 0.1520170106 |
| 0.3278428699 | 0.4585669194 | 0.3486471853 | 0.4099890819 | -0.5583147446 | -0.3724148892 |
| 0.3446172552 | 0.482026275 | -0.00915195524 | 0.5177245932 | -0.7763942741 | 0.6099566625 |
| 0.3354909579 | 0.4692804393 | 0.338602808 | 0.4438702725 | -1.272313732 | 0.05934871732 |
| 0.3027604072 | 0.4234634308 | 0.1319490244 | -0.6744740061 | 1.534318282 | -1.589122652 |
| 0.3228247323 | 0.4515474497 | 0.6462385364 | -1.397060151 | -0.0925532547 | 0.5796802482 |
| 0.2931867504 | 0.4100359297 | 0.06392723689 | 0.3906643009 | 0.9447774686 | -1.657947258 |
| 0.3520407934 | 0.4922297062 | -0.6898506835 | 0.4123962239 | 1.468186023 | -1.158799927 |
| 0.3441362829 | 0.481367554 | 0.2229434372 | 0.1651347708 | -0.0781731024 | -0.6862728452 |
| 0.3279430642 | 0.4587155155 | 0.233155252 | -1.176824424 | -0.0692565511 | 1.56387429 |

| | | | | | |
|---|---|---|---|---|---|
| 0.3590974189 | 0.5016994591 | -0.1900002038 | -0.2609061865 | 0.23485215 | 0.5473952433 |
| 0.3005248097 | 0.4202970487 | 0.2624171062 | -0.408183271 | 0.8331838433 | -1.013533439 |
| 0.3485226073 | 0.4874984784 | 0.4751856831 | -0.6385726577 | -0.2654400632 | 0.2215820789 |
| 0.3476061938 | 0.486191641 | 0.5741579558 | -0.349754911 | -1.327948185 | 1.297592468 |
| 0.3397859183 | 0.4752138833 | 1.41652389 | -0.6129437991 | -1.554159022 | -0.1933407745 |
| 0.2665626944 | 0.3727620631 | -0.2376697194 | 0.8997775489 | 0.8742773198 | -0.783674421 |
| 0.3424478502 | 0.47898178 | 1.253852297 | -1.686186836 | -1.606896813 | 1.822989034 |
| 0.3042485294 | 0.4255265951 | 0.0979630333 | -1.432725994 | 0.2309245518 | 1.422712869 |
| 0.3406279165 | 0.4758954882 | 1.509065172 | -0.7529335363 | -1.057389145 | -0.7915484519 |
| 0.3583878479 | 0.5010605951 | 0.4770036674 | -0.6453289102 | -0.4630564471 | 0.5573992644 |
| 0.3328813596 | 0.4656261251 | 0.2863979896 | 0.5930936828 | -0.5792444894 | -0.6488827416 |
| 0.352362312 | 0.4926928859 | 0.3317928647 | -0.0917956606 | -1.234746851 | 1.285464272 |
| 0.3230879745 | 0.4519042705 | -0.02471498879 | 0.1394694394 | -1.227048855 | 1.266111807 |
| 0.340184981 | 0.4758130542 | 0.6640784843 | -1.167727928 | -1.191704148 | 2.087082539 |
| 0.352298797 | 0.4926282132 | -1.016855987 | 1.180726699 | 1.280762636 | -1.230855982 |
| 0.3573591019 | 0.4992708349 | 0.7987550963 | -0.8878623449 | -0.2311637267 | -0.09376521063 |
| 0.3236970493 | 0.4527617906 | 0.6809751689 | -1.538392369 | -0.6241183205 | 0.7023127676 |
| 0.2608867808 | 0.3648290058 | 1.054430615 | -2.62149732 | -1.459396162 | 1.26111009 |
| 0.3209589123 | 0.4489338662 | -1.03062531 | 0.3873186134 | 1.566467804 | -0.3192449274 |
| 0.3531335556 | 0.4938951541 | 0.1881499738 | -0.2133320024 | -0.0471791042 | -0.2519688292 |
| 0.3359510975 | 0.469920997 | 0.8760194053 | -1.146373063 | -0.5975216265 | 0.6380008981 |
| 0.3226068247 | 0.4512491599 | -0.8343621012 | 1.783382845 | 0.6557111785 | -1.761647725 |
| 0.3500798922 | 0.4894046745 | -0.9327735589 | 1.228625729 | 1.483860342 | -1.964078262 |
| 0.3478192311 | 0.485942563 | -1.686374865 | 1.456567246 | 1.137338312 | -0.0926969036 |
| 0.3537053044 | 0.4941660692 | -0.539888487 | 0.5758405906 | 1.26407252 | -1.603233185 |
| 0.3291231319 | 0.4598220109 | 0.7476635593 | -0.8607108206 | 1.090754042 | -2.052520505 |
| 0.3462772354 | 0.4843551386 | -0.05845025403 | -0.0411721157 | -0.2727235644 | 0.9024894356 |
| 0.3226181498 | 0.4512536058 | 0.8681167922 | -0.3361936314 | -1.277894627 | 0.4112990586 |
| 0.3462348631 | 0.4841965638 | 0.9301029314 | -1.013385715 | -1.80711559 | 1.975506786 |
| 0.3499836524 | 0.4894490203 | -0.8714875115 | 0.4671803769 | 1.112099711 | -0.1138923044 |
| 0.3187694234 | 0.4458548349 | 0.3257836114 | 0.1302436989 | -0.4471762074 | -0.09088196888 |
| 0.3450179387 | 0.4821960881 | -1.89496991 | 2.39301438 | 1.182046798 | -1.168443427 |
| 0.3522533005 | 0.4925763629 | -0.5535081394 | 0.813311188 | -0.1282774814 | 0.3114034245 |
| 0.3361679195 | 0.4702167878 | 0.09439542553 | 0.273835828 | -0.2921852478 | -0.07842307499 |
| 0.3492860477 | 0.4879918707 | 1.433518247 | -0.9314504202 | -1.153409769 | -0.154817533 |
| 0.348029953 | 0.4867148252 | 1.430166261 | -1.39997214 | -1.557994347 | 1.037789879 |
| 0.3514162582 | 0.4914559787 | -0.8871236768 | 1.34676679 | 0.559511253 | -0.7201983713 |
| 0.3157724812 | 0.4416640459 | 0.9464557349 | -1.224896578 | -0.7897291579 | 0.3673765275 |
| 0.3366207551 | 0.4708618626 | 0.8852953087 | -1.426341545 | -0.1505261436 | 0.1656987748 |
| 0.3231267963 | 0.4519646056 | 0.3773021664 | 0.4322874818 | -2.470434655 | 1.804763781 |
| 0.3250904071 | 0.4547070497 | -0.747359653 | 1.476781394 | -0.1271862005 | -0.5408442462 |
| 0.3611882988 | 0.5046206535 | -0.4260874483 | 0.2580569327 | 0.5739850527 | -0.2783889895 |
| 0.3202487302 | 0.4479346024 | -0.01735778335 | 0.1805007501 | -0.2225797524 | 0.4695208346 |
| 0.2585011278 | 0.3614797761 | 0.6068823813 | -0.3936000439 | -1.146891051 | 0.1185015685 |
| 0.3257296358 | 0.4556220232 | -0.934134648 | 1.710840682 | 0.3357514743 | -0.07198001314 |
| 0.3517370677 | 0.4914162212 | -1.4009473 | 1.489891782 | 0.7051407982 | -0.1831198431 |

| | | | | | |
|---|---|---|---|---|---|
| 0.3201067284 | 0.4477359758 | -0.611309342 | 0.1850728337 | 1.374462655 | -0.9880911527 |
| 0.3385926775 | 0.4730520305 | -1.272501975 | 0.1610064189 | 2.619816891 | -1.191350461 |
| 0.3116788459 | 0.4359155374 | -0.08255728967 | -0.1242037497 | -0.0816857986 | -0.08652598792 |
| 0.3429010794 | 0.4795944105 | 0.1964976845 | 0.5353666761 | -0.2871673312 | -1.143170137 |
| 0.3288364189 | 0.4599552696 | -0.4786158188 | 0.7711268946 | 0.2415073292 | -0.07994670085 |
| 0.2928840083 | 0.4096187267 | -0.6668267134 | 0.5882221237 | 0.873303962 | 0.3569004709 |
| 0.3185714799 | 0.4455919247 | 0.09929073026 | -0.2269015992 | -0.7197230491 | 1.674079714 |
| 0.3483154381 | 0.487139055 | 1.152021798 | -1.227698761 | -0.982606753 | 0.4220938725 |
| 0.3594715848 | 0.502222211 | 0.680341823 | -0.9660070503 | -0.3571691551 | 0.5237401709 |
| 0.2845805134 | 0.3979985277 | -0.4976534709 | 0.9851844807 | 1.469346382 | -2.189691771 |
| 0.3557334287 | 0.4969995867 | 0.84141755 | -0.5775196456 | -0.4403681809 | -0.4033409882 |
| 0.3005708902 | 0.4203775656 | 0.1385233892 | -1.30536731 | 0.4121900628 | 0.7190113276 |
| 0.276512682 | 0.3867066207 | 1.11017588 | -1.253742897 | -1.527073537 | -0.07314364088 |
| 0.33446723 | 0.4678100977 | -0.8454468362 | 1.569434144 | -0.8172981207 | 0.7446499022 |
| 0.2956056911 | 0.4134183249 | 0.6980485813 | -1.427172411 | -1.877633067 | 2.005384557 |
| 0.3574086899 | 0.4996862436 | -0.3856302478 | 0.0448776106 | 0.6149767571 | -0.04645368359 |
| 0.3008038781 | 0.4207017723 | -0.7435489509 | 1.659917229 | 1.593184432 | -2.560059828 |
| 0.1536444114 | 0.2148190215 | 0.5502523979 | -1.905534628 | -1.507502957 | 1.520310011 |
| 0.3508027589 | 0.4901108868 | -0.9469650976 | 1.729169167 | 0.0093960181 | -0.6307021833 |
| 0.3611691767 | 0.5045939376 | -0.5158059383 | 0.5862997367 | 0.4519000148 | -0.4279988911 |
| 0.3101354133 | 0.4337754048 | 0.9134169113 | -2.14515562 | -1.259408626 | 1.998620137 |
| 0.3418069729 | 0.4781160834 | -0.234928705 | 0.7143316233 | -0.1585713961 | -0.2605384149 |
| 0.3312002486 | 0.463266178 | 0.8981640204 | -0.9535157468 | -2.459249644 | 2.57855137 |
| 0.3430694624 | 0.4798801224 | 0.04393203069 | 0.0443117420 | 0.4774133311 | -0.9098576154 |
| 0.2573146507 | 0.3598207782 | 0.000396436952 | 0.0722400841 | 0.5907446666 | -0.8298439721 |
| 0.2928423445 | 0.4096111356 | -2.353465057 | 2.857654195 | 1.685300007 | 0.02578551976 |
| 0.3192954681 | 0.446590282 | 0.1612711017 | -0.1376867522 | -0.4224508602 | 0.3336080387 |
| 0.3430473444 | 0.4797917411 | 0.4013378616 | 0.1707283912 | 0.1536888746 | -1.455449929 |
| 0.3040852719 | 0.425318345 | -1.893715275 | 2.903031513 | 1.724849481 | -1.278669218 |
| 0.3521210034 | 0.4925017135 | -0.4406607355 | 0.2168984617 | 0.9064029235 | -0.8861147937 |
| 0.3601365819 | 0.5031512867 | -0.2058083631 | -0.1464346409 | 0.2151082771 | 0.4220521683 |
| 0.3489693114 | 0.488049282 | 1.208221224 | -0.80005855 | -1.72652844 | 1.105207518 |
| 0.3544493603 | 0.4952387728 | -0.1925651134 | 0.3862999043 | -0.7185444522 | 0.9243577639 |
| 0.3598944542 | 0.5028130071 | 0.4930185819 | -0.826127489 | -0.1244671288 | 0.3867142166 |

Batch))

| Intercept Pr(>t | Blast Pr(>t) | VNS Pr(>t) | Blast:VNS Pr(>t | Blast Adjusted p-valu | VNS Adjusted p-valu |
|---|---|---|---|---|---|
| 0.4717721113 | 0.6375874105 | 0.5030830517 | 0.9560935902 | 0.9715330331 | 0.9094096952 |
| 0.6393388214 | 0.5149119155 | 0.5725884718 | 0.4527962518 | 0.9446238668 | 0.9094096952 |
| 0.4937732241 | 0.213837116 | 0.9111760475 | 0.6001265862 | 0.6794305346 | 0.9762566711 |
| 0.7927261947 | 0.7766207887 | 0.5427016509 | 0.4080271324 | 0.9715330331 | 0.9094096952 |
| 0.7589245883 | 0.997414402 | 0.4913573178 | 0.9134819371 | 0.997414402 | 0.9094096952 |
| 0.7181529924 | 0.2010588494 | 0.4207528074 | 0.1129215333 | 0.6794305346 | 0.9094096952 |
| 0.7666622971 | 0.8849154585 | 0.3221995489 | 0.5772977573 | 0.9715330331 | 0.856913694 |
| 0.3331714454 | 0.2485713156 | 0.9649581275 | 0.7314939168 | 0.6794305346 | 0.9806485036 |
| 0.3374445177 | 0.1782112372 | 0.6827377942 | 0.5468000845 | 0.6794305346 | 0.9482469364 |
| 0.7217371056 | 0.8712768598 | 0.3688659403 | 0.3411117346 | 0.9715330331 | 0.9040831871 |
| 0.2985637064 | 0.2370351399 | 0.468196845 | 0.5618379035 | 0.6794305346 | 0.9094096952 |
| 0.7589794566 | 0.6390255376 | 0.8578449308 | 0.2368236948 | 0.9715330331 | 0.9762566711 |
| 0.9723826395 | 0.8622068001 | 0.5271209461 | 0.5227118487 | 0.9715330331 | 0.9094096952 |
| 0.7032443352 | 0.990372594 | 0.4272767101 | 0.839423832 | 0.997414402 | 0.9094096952 |
| 0.6115351518 | 0.9425346728 | 0.8948134811 | 0.3588627933 | 0.9819388528 | 0.9762566711 |
| 0.2493752298 | 0.2455260069 | 0.4947096444 | 0.8424675507 | 0.6794305346 | 0.9094096952 |
| 0.6638456378 | 0.9179767377 | 0.2341448852 | 0.9835803476 | 0.9807443779 | 0.772928993 |
| 0.9362213161 | 0.7285319709 | 0.849555396 | 0.4932905013 | 0.9715330331 | 0.9762566711 |
| 0.4621270535 | 0.1427343843 | 0.1592599022 | 0.1417200842 | 0.6794305346 | 0.772928993 |
| 0.0344858250€ | 0.0300302745 | 0.2278073795 | 0.92105925 | 0.5212109185 | 0.772928993 |
| 0.893270234 | 0.3018461465 | 0.8378313054 | 0.9918663807 | 0.7546153662 | 0.9762566711 |
| 0.657078761 | 0.8739590166 | 0.9341816646 | 0.3943235963 | 0.9715330331 | 0.9762566711 |
| 0.1717931078 | 0.0602427903 | 0.5291811331 | 0.5366448212 | 0.6794305346 | 0.9094096952 |
| 0.4010009889 | 0.781611578 | 0.1851562804 | 0.6438291186 | 0.9715330331 | 0.772928993 |
| 0.0548588263! | 0.0078373247 | 0.1361012583 | 0.1039478603 | 0.3265551964 | 0.772928993 |
| 0.4014423538 | 0.2191540565 | 0.1795479942 | 0.4882155992 | 0.6794305346 | 0.772928993 |
| 0.6975480993 | 0.3365854845 | 0.9837238137 | 0.5079763023 | 0.7817925236 | 0.9916570702 |
| 0.3374482993 | 0.2787581979 | 0.6631745033 | 0.8789859123 | 0.7111178519 | 0.9482469364 |
| 0.1265270507 | 0.0279765337 | 0.1689292277 | 0.0799482369ɕ | 0.5212109185 | 0.772928993 |
| 0.1959734672 | 0.0746550233 | 0.5820222049 | 0.4692900302 | 0.6794305346 | 0.9094096952 |
| 0.6812228107 | 0.1320333044 | 0.1094178193 | 0.7078378771 | 0.6794305346 | 0.772928993 |
| 0.8829838185 | 0.7795154295 | 0.9046219323 | 0.8075342898 | 0.9715330331 | 0.9762566711 |
| 0.5697713331 | 0.2697570467 | 0.6231132529 | 0.5883729548 | 0.702492309 | 0.9482469364 |
| 0.52904787 | 0.2500304367 | 0.3071200405 | 0.1168106813 | 0.6794305346 | 0.8345653273 |
| 0.7798117931 | 0.6628440166 | 0.7580967742 | 0.1167540956 | 0.9715330331 | 0.9762566711 |
| 0.7096363272 | 0.0375271861 | 0.9295229466 | 0.8797029514 | 0.5212109185 | 0.9762566711 |
| 0.7311385894 | 0.6833064194 | 0.5787213272 | 0.7109060082 | 0.9715330331 | 0.9094096952 |
| 0.9927589906 | 0.6065637629 | 0.4405259563 | 0.5441613632 | 0.9715330331 | 0.9094096952 |
| 0.7375782808 | 0.6587465335 | 0.2081295336 | 0.9528699051 | 0.9715330331 | 0.772928993 |
| 0.8967316058 | 0.5026900618 | 0.1303110547 | 0.1173963871 | 0.9446238668 | 0.772928993 |
| 0.5256324441 | 0.167645075 | 0.9265685161 | 0.5643138282 | 0.6794305346 | 0.9762566711 |
| 0.9499663114 | 0.6974718529 | 0.348639919 | 0.1026603036 | 0.9715330331 | 0.8893875485 |
| 0.4943371964 | 0.6815047686 | 0.1471384327 | 0.2509944134 | 0.9715330331 | 0.772928993 |
| 0.8252205201 | 0.8693975968 | 0.9379479524 | 0.4951577094 | 0.9715330331 | 0.9762566711 |
| 0.8179144568 | 0.2440021445 | 0.9450164577 | 0.1231259036 | 0.6794305346 | 0.9762566711 |

| | | | | | |
|---|---|---|---|---|---|
| 0.849919849 | 0.7950158331 | 0.8150851459 | 0.5860412737 | 0.9715330331 | 0.9762566711 |
| 0.7972177278 | 0.6846453742 | 0.4081306707 | 0.3149835856 | 0.9715330331 | 0.9094096952 |
| 0.6383808149 | 0.5255545575 | 0.7915719073 | 0.825384153 | 0.9446238668 | 0.9762566711 |
| 0.5702028779 | 0.7277578393 | 0.1891652927 | 0.1993286301 | 0.9715330331 | 0.772928993 |
| 0.1656247697 | 0.5422248538 | 0.1252999104 | 0.8473308128 | 0.9446238668 | 0.772928993 |
| 0.8168135946 | 0.371979503 | 0.385567217 | 0.4364168505 | 0.8146380077 | 0.9094096952 |
| 0.2191770264 | 0.0969636223 | 0.1132441231 | 0.0732034259 | 0.6794305346 | 0.772928993 |
| 0.9232631499 | 0.1572770807 | 0.8181725753 | 0.1600912349 | 0.6794305346 | 0.9762566711 |
| 0.1362812815 | 0.4542942779 | 0.2943717656 | 0.431594186 | 0.8872935116 | 0.8176993489 |
| 0.636031698 | 0.5211572211 | 0.6449563288 | 0.5792680689 | 0.9446238668 | 0.9482469364 |
| 0.7772372143 | 0.5553821875 | 0.5645924176 | 0.5188907467 | 0.9446238668 | 0.9094096952 |
| 0.7425437131 | 0.9271759556 | 0.2217228225 | 0.2035256414 | 0.9819388528 | 0.772928993 |
| 0.9805662211 | 0.8895590367 | 0.2246573106 | 0.2104318127 | 0.9715330331 | 0.772928993 |
| 0.5116183206 | 0.2475585865 | 0.2380034586 | 0.0410707095 | 0.6794305346 | 0.772928993 |
| 0.3176173642 | 0.2424807414 | 0.2051697968 | 0.2231230037 | 0.6794305346 | 0.772928993 |
| 0.4274334461 | 0.3779920356 | 0.8179367273 | 0.9255932377 | 0.8146380077 | 0.9762566711 |
| 0.5049307657 | 0.1293681941 | 0.5349490552 | 0.4852400673 | 0.6794305346 | 0.9094096952 |
| 0.3151984334 | 0.0111729076 | 0.1498359531 | 0.2123075888 | 0.3491533638 | 0.772928993 |
| 0.3159149841 | 0.6999225808 | 0.1225421309 | 0.7506595597 | 0.9715330331 | 0.772928993 |
| 0.8518969626 | 0.8317908003 | 0.9625242264 | 0.8019087222 | 0.9715330331 | 0.9806485036 |
| 0.3906028886 | 0.2562748461 | 0.552411327 | 0.5259053849 | 0.6815820375 | 0.9094096952 |
| 0.4136454563 | 0.0796777038 | 0.5145250893 | 0.0832431009 | 0.6794305346 | 0.9094096952 |
| 0.3572991307 | 0.2240367471 | 0.1429800259 | 0.0540446994 | 0.6794305346 | 0.772928993 |
| 0.0966702528 | 0.1502019837 | 0.2597041417 | 0.9264385079 | 0.6794305346 | 0.772928993 |
| 0.5911772736 | 0.5667743201 | 0.2108610377 | 0.1138854662 | 0.9446238668 | 0.772928993 |
| 0.4574448822 | 0.3926612168 | 0.2795353704 | 0.0442786897 | 0.8319093576 | 0.7941345749 |
| 0.953794351 | 0.9672963029 | 0.7859913587 | 0.3703616722 | 0.9830246981 | 0.9762566711 |
| 0.3962876991 | 0.7379233157 | 0.2062447178 | 0.6823304281 | 0.9715330331 | 0.772928993 |
| 0.3583486918 | 0.3149165639 | 0.0756384753 | 0.0526953114 | 0.7718543233 | 0.772928993 |
| 0.3893786197 | 0.6420580518 | 0.270437334 | 0.9096945314 | 0.9715330331 | 0.7861550406 |
| 0.7486460982 | 0.8968208456 | 0.6563789407 | 0.9278932819 | 0.9715330331 | 0.9482469364 |
| 0.0648437405 | 0.0198126012 | 0.2417096599 | 0.2470896107 | 0.4953150309 | 0.772928993 |
| 0.5833229669 | 0.419246275 | 0.8983489076 | 0.7565465861 | 0.8591112192 | 0.9762566711 |
| 0.9258550357 | 0.7851835108 | 0.7711684939 | 0.9377566636 | 0.9715330331 | 0.9762566711 |
| 0.1566542989 | 0.3551764382 | 0.2530994548 | 0.877460103 | 0.7928045496 | 0.772928993 |
| 0.1615064687 | 0.1665934566 | 0.1243839941 | 0.3034362016 | 0.6794305346 | 0.772928993 |
| 0.3808141221 | 0.1830870948 | 0.5778469958 | 0.4741287596 | 0.6794305346 | 0.9094096952 |
| 0.3572429263 | 0.2255224618 | 0.432833165 | 0.7146473241 | 0.6794305346 | 0.9094096952 |
| 0.3843698182 | 0.159010472 | 0.8808515258 | 0.8689478398 | 0.6794305346 | 0.9762566711 |
| 0.7090090348 | 0.667113208 | 0.0163434312 | 0.0761059050 | 0.9715330331 | 0.6999447309 |
| 0.4652968507 | 0.145131803 | 0.8992252243 | 0.5906513112 | 0.6794305346 | 0.9762566711 |
| 0.6714964041 | 0.7972041132 | 0.5680216041 | 0.7816254649 | 0.9715330331 | 0.9094096952 |
| 0.9863359987 | 0.8573801603 | 0.8246237427 | 0.6404120668 | 0.9715330331 | 0.9762566711 |
| 0.5579545306 | 0.6953302106 | 0.2561407544 | 0.9060804012 | 0.9715330331 | 0.772928993 |
| 0.3613020759 | 0.0924048404 | 0.7382376279 | 0.9428589714 | 0.6794305346 | 0.9762566711 |
| 0.1661365026 | 0.1412423841 | 0.4833216116 | 0.8552920411 | 0.6794305346 | 0.9094096952 |

| | | | | | |
|---|---|---|---|---|---|
| 0.548587347 | 0.8538104184 | 0.1744556574 | 0.3271140384 | 0.9715330331 | 0.772928993 |
| 0.2078712917 | 0.872603542 | 0.0110104044 | 0.2379850259 | 0.9715330331 | 0.6999447309 |
| 0.9353513634 | 0.9015826547 | 0.9351740161 | 0.931342549 | 0.9715330331 | 0.9762566711 |
| 0.8454244841 | 0.5943741128 | 0.7749554593 | 0.2574192036 | 0.9715330331 | 0.9762566711 |
| 0.6377884532 | 0.4437328753 | 0.8099929762 | 0.9365483203 | 0.8804223717 | 0.9762566711 |
| 0.5157560599 | 0.5586574741 | 0.3860467632 | 0.7224438426 | 0.9446238668 | 0.9094096952 |
| 0.921999877 | 0.821290696 | 0.474517601 | 0.09937615863 | 0.9715330331 | 0.9094096952 |
| 0.2568248324 | 0.2243190756 | 0.3296544646 | 0.6744274021 | 0.6794305346 | 0.8584751683 |
| 0.4987807408 | 0.3377343702 | 0.7221588474 | 0.6022969057 | 0.7817925236 | 0.9762566711 |
| 0.6268627924 | 0.3286155404 | 0.1470758093 | 0.03253525753 | 0.7817925236 | 0.772928993 |
| 0.4032966186 | 0.5656468272 | 0.6611777445 | 0.6880630648 | 0.9446238668 | 0.9482469364 |
| 0.8917118074 | 0.1968943778 | 0.6816947144 | 0.4749758721 | 0.6794305346 | 0.9482469364 |
| 0.2875587774 | 0.2149573465 | 0.1321194864 | 0.941940959 | 0.6794305346 | 0.772928993 |
| 0.4046598148 | 0.1218555255 | 0.4169542348 | 0.4593522963 | 0.6794305346 | 0.9094096952 |
| 0.4974626098 | 0.1588925488 | 0.0654111136 | 0.04955127073 | 0.6794305346 | 0.772928993 |
| 0.7018444115 | 0.9643526587 | 0.5408343293 | 0.9630983287 | 0.9830246981 | 0.9094096952 |
| 0.4692064571 | 0.1023094199 | 0.1164747553 | 0.0130547426 | 0.6794305346 | 0.772928993 |
| 0.5998803301 | 0.0617482817 | 0.1371845111 | 0.1339422117 | 0.6794305346 | 0.772928993 |
| 0.3472747416 | 0.0886764636 | 0.9925328634 | 0.5305166572 | 0.6794305346 | 0.9925328634 |
| 0.6077955002 | 0.5597689896 | 0.6528930531 | 0.6701115396 | 0.9446238668 | 0.9482469364 |
| 0.3746909509 | 0.0361211458 | 0.2128387804 | 0.0502641679 | 0.5212109185 | 0.772928993 |
| 0.8161258243 | 0.4778195828 | 0.8745354341 | 0.7953343793 | 0.9188838131 | 0.9762566711 |
| 0.3769316034 | 0.3441994847 | 0.0167986735 | 0.01236954631 | 0.7822715562 | 0.6999447309 |
| 0.9652973506 | 0.9648048601 | 0.6347919209 | 0.3665153345 | 0.9830246981 | 0.9482469364 |
| 0.9996917953 | 0.9426612987 | 0.556989603 | 0.4100363086 | 0.9819388528 | 0.9094096952 |
| 0.02905014552 | 0.0058922377 | 0.0971511679 | 0.9795144529 | 0.3265551964 | 0.772928993 |
| 0.8737985282 | 0.8909606352 | 0.6742241699 | 0.7398533071 | 0.9715330331 | 0.9482469364 |
| 0.6906174469 | 0.8650087975 | 0.8783597328 | 0.1506382462 | 0.9715330331 | 0.9762566711 |
| 0.07516864414 | 0.0052018825 | 0.0897579617 | 0.2059935391 | 0.3265551964 | 0.772928993 |
| 0.6630611911 | 0.8290400029 | 0.3683366297 | 0.3790757126 | 0.9715330331 | 0.9040831871 |
| 0.8376042222 | 0.884045855 | 0.8303777408 | 0.6744237641 | 0.9715330331 | 0.9762566711 |
| 0.236048894 | 0.4268609362 | 0.0893272907 | 0.2734109383 | 0.8606067263 | 0.772928993 |
| 0.848277261 | 0.7006369664 | 0.475139329 | 0.3588816154 | 0.9715330331 | 0.9094096952 |
| 0.6237135778 | 0.4118495084 | 0.9013417198 | 0.7002709377 | 0.8580198093 | 0.9762566711 |

| | Taxonomic Information | | |
|---|---|---|---|
| Blast:VNS Adjusted p-valu | Genus | Family | Order |
| 0.9867810745 | Acetatifactor | Lachnospiraceae | Eubacteriales |
| 0.9652194001 | Acetatifactor | Lachnospiraceae | Eubacteriales |
| 0.9652194001 | Acutalibacter | Oscillospiraceae | Eubacteriales |
| 0.9652194001 | Acutalibacter | Oscillospiraceae | Eubacteriales |
| 0.9867810745 | Adlercreutzia | Eggerthellaceae | Eggerthellales |
| 0.6987880186 | Adlercreutzia | Eggerthellaceae | Eggerthellales |
| 0.9652194001 | Adlercreutzia | Eggerthellaceae | Eggerthellales |
| 0.9867810745 | Akkermansia | Akkermansiaceae | Verrucomicrobiales |
| 0.9652194001 | Alistipes | Rikenellaceae | Bacteroidales |
| 0.9652194001 | Anaerotruncus | Oscillospiraceae | Eubacteriales |
| 0.9652194001 | Bacteria_unclassified | Bacteria_unclassified | Bacteria_unclassified |
| 0.8696594716 | Bacteria_unclassified | Bacteria_unclassified | Bacteria_unclassified |
| 0.9652194001 | Bacteria_unclassified | Bacteria_unclassified | Bacteria_unclassified |
| 0.9867810745 | Bacteroides | Bacteroidaceae | Bacteroidales |
| 0.9652194001 | Bifidobacterium | Bifidobacteriaceae | Bifidobacteriales |
| 0.9867810745 | Clostridia_unclassified | Clostridia_unclassified | Clostridia_unclassifie |
| 0.9915124472 | Clostridiaceae_unclassified | Clostridiaceae | Eubacteriales |
| 0.9652194001 | Clostridiaceae_unclassified | Clostridiaceae | Eubacteriales |
| 0.7381254386 | Eubacteriales_unclassified | Eubacteriales_unclassified | Eubacteriales |
| 0.9867810745 | Erysipelatoclostridium | Erysipelotrichaceae | Erysipelotrichales |
| 0.9918663807 | Coriobacteriaceae_unclassifie | Coriobacteriaceae | Coriobacteriales |
| 0.9652194001 | Dorea | Lachnospiraceae | Eubacteriales |
| 0.9652194001 | Dubosiella | Erysipelotrichaceae | Erysipelotrichales |
| 0.9867810745 | Erysipelotrichales_unclassifie | Erysipelotrichales_unclassifie | Erysipelotrichales |
| 0.6987880186 | Eubacteriaceae_unclassified | Eubacteriaceae | Eubacteriales |
| 0.9652194001 | Eubacteriaceae_unclassified | Eubacteriaceae | Eubacteriales |
| 0.9652194001 | GGB20149 | Lachnospiraceae | Eubacteriales |
| 0.9867810745 | GGB22635 | Eggerthellaceae | Eggerthellales |
| 0.6987880186 | GGB25041 | Lachnospiraceae | Eubacteriales |
| 0.9652194001 | GGB27876 | Muribaculaceae | Bacteroidales |
| 0.9867810745 | GGB27878 | Muribaculaceae | Bacteroidales |
| 0.9867810745 | GGB27918 | Muribaculaceae | Bacteroidales |
| 0.9652194001 | GGB28382 | FGB9508 | OFGB9508 |
| 0.6987880186 | GGB28399 | FGB2838 | OFGB2838 |
| 0.6987880186 | GGB28411 | FGB2838 | OFGB2838 |
| 0.9867810745 | GGB28415 | FGB2838 | OFGB2838 |
| 0.9867810745 | GGB28430 | Pumilibacteraceae | Eubacteriales |
| 0.9652194001 | GGB28439 | FGB28439 | OFGB28439 |
| 0.9867810745 | GGB28778 | Clostridia_unclassified | Clostridia_unclassifie |
| 0.6987880186 | GGB28784 | Eubacteriaceae | Eubacteriales |
| 0.9652194001 | GGB28792 | Lachnospiraceae | Eubacteriales |
| 0.6987880186 | GGB28798 | Lachnospiraceae | Eubacteriales |
| 0.8696594716 | GGB28802 | Lachnospiraceae | Eubacteriales |
| 0.9652194001 | GGB28818 | Lachnospiraceae | Eubacteriales |
| 0.6995789976 | GGB28828 | FGB77305 | OFGB77305 |

| | | | |
|---|---|---|---|
| 0.9652194001 | GGB28851 | Clostridiaceae | Eubacteriales |
| 0.9652194001 | GGB28859 | Lachnospiraceae | Eubacteriales |
| 0.9867810745 | GGB28864 | Lachnospiraceae | Eubacteriales |
| 0.8560789869 | GGB28869 | Lachnospiraceae | Eubacteriales |
| 0.9867810745 | GGB28883 | FGB9633 | OFGB9633 |
| 0.9652194001 | GGB28892 | Bacteria_unclassified | Bacteria_unclassified |
| 0.6987880186 | GGB28893 | Bacteria_unclassified | Bacteria_unclassified |
| 0.7696693983 | GGB28898 | Bacteria_unclassified | Bacteria_unclassified |
| 0.9652194001 | GGB28904 | Bacteria_unclassified | Bacteria_unclassified |
| 0.9652194001 | GGB28916 | Lachnospiraceae | Eubacteriales |
| 0.9652194001 | GGB28924 | Lachnospiraceae | Eubacteriales |
| 0.8560789869 | GGB28926 | Lachnospiraceae | Eubacteriales |
| 0.8560789869 | GGB28927 | FGB77359 | OFGB77359 |
| 0.6987880186 | GGB28934 | FGB9639 | OFGB9639 |
| 0.8696594716 | GGB28946 | Lachnospiraceae | Eubacteriales |
| 0.9867810745 | GGB28949 | Lachnospiraceae | Eubacteriales |
| 0.9652194001 | GGB28949 | Lachnospiraceae | Eubacteriales |
| 0.8560789869 | GGB28950 | Clostridiaceae | Eubacteriales |
| 0.9867810745 | GGB28951 | Clostridiaceae | Eubacteriales |
| 0.9867810745 | GGB28951 | Clostridiaceae | Eubacteriales |
| 0.9652194001 | GGB28954 | Clostridiaceae | Eubacteriales |
| 0.6987880186 | GGB28956 | Clostridiaceae | Eubacteriales |
| 0.6987880186 | GGB28960 | Clostridiaceae | Eubacteriales |
| 0.9867810745 | GGB28967 | Clostridiaceae | Eubacteriales |
| 0.6987880186 | GGB28991 | Eubacteriaceae | Eubacteriales |
| 0.6987880186 | GGB29002 | FGB9658 | OFGB9658 |
| 0.9652194001 | GGB29003 | FGB9659 | OFGB9659 |
| 0.9867810745 | GGB29011 | Bacteria_unclassified | Bacteria_unclassified |
| 0.6987880186 | GGB29531 | FGB9827 | OFGB9827 |
| 0.9867810745 | GGB29685 | Eubacteriaceae | Eubacteriales |
| 0.9867810745 | GGB30141 | FGB77303 | OFGB77303 |
| 0.8696594716 | GGB30286 | Eubacteriales_unclassified | Eubacteriales |
| 0.9867810745 | GGB30303 | Oscillospiraceae | Eubacteriales |
| 0.9867810745 | GGB30413 | FGB30328 | OFGB30328 |
| 0.9867810745 | GGB30454 | Oscillospiraceae | Eubacteriales |
| 0.9652194001 | GGB30455 | Oscillospiraceae | Eubacteriales |
| 0.9652194001 | GGB30461 | Oscillospiraceae | Eubacteriales |
| 0.9867810745 | GGB30461 | Oscillospiraceae | Eubacteriales |
| 0.9867810745 | GGB30463 | Oscillospiraceae | Eubacteriales |
| 0.6987880186 | GGB30473 | Oscillospiraceae | Eubacteriales |
| 0.9652194001 | GGB30475 | Oscillospiraceae | Eubacteriales |
| 0.9867810745 | GGB30861 | FGB77153 | OFGB77153 |
| 0.9867810745 | GGB31312 | FGB1791 | OFGB1791 |
| 0.9867810745 | GGB31438 | FGB10290 | OFGB10290 |
| 0.9867810745 | GGB3171 | Oscillospiraceae | Eubacteriales |
| 0.9867810745 | GGB31823 | FGB1765 | OFGB1765 |

| | | | |
|---|---|---|---|
| 0.9652194001 | GGB31853 | FGB10349 | OFGB10349 |
| 0.8696594716 | GGB32371 | FGB10667 | OFGB10667 |
| 0.9867810745 | GGB3793 | Lachnospiraceae | Eubacteriales |
| 0.8696594716 | GGB42598 | Lachnospiraceae | Eubacteriales |
| 0.9867810745 | GGB45656 | Christensenellaceae | Eubacteriales |
| 0.9867810745 | GGB47127 | FGB10299 | OFGB10299 |
| 0.6987880186 | GGB74395 | Oscillospiraceae | Eubacteriales |
| 0.9867810745 | GGB75053 | Oscillospiraceae | Eubacteriales |
| 0.9652194001 | GGB75109 | Lachnospiraceae | Eubacteriales |
| 0.6987880186 | GGB81440 | Clostridia_unclassified | Clostridia_unclassifie |
| 0.9867810745 | Lachnospiraceae_unclassified | Lachnospiraceae | Eubacteriales |
| 0.9652194001 | Lachnospiraceae_unclassified | Lachnospiraceae | Eubacteriales |
| 0.9867810745 | Lachnospiraceae_unclassified | Lachnospiraceae | Eubacteriales |
| 0.9652194001 | Lachnospiraceae_unclassified | Lachnospiraceae | Eubacteriales |
| 0.6987880186 | Lachnospiraceae_unclassified | Lachnospiraceae | Eubacteriales |
| 0.9867810745 | Lachnospiraceae_unclassified | Lachnospiraceae | Eubacteriales |
| 0.6987880186 | Lachnospiraceae_unclassified | Lachnospiraceae | Eubacteriales |
| 0.7279468025 | Lachnospiraceae_unclassified | Lachnospiraceae | Eubacteriales |
| 0.9652194001 | Lactobacillus | Lactobacillaceae | Lactobacillales |
| 0.9867810745 | Muribaculaceae_unclassified | Muribaculaceae | Bacteroidales |
| 0.6987880186 | Neglectibacter | Oscillospiraceae | Eubacteriales |
| 0.9867810745 | Oscillospiraceae_unclassified | Oscillospiraceae | Eubacteriales |
| 0.6987880186 | Oscillospiraceae_unclassified | Oscillospiraceae | Eubacteriales |
| 0.9652194001 | Oscillospiraceae_unclassified | Oscillospiraceae | Eubacteriales |
| 0.9652194001 | Parasutterella | Sutterellaceae | Burkholderiales |
| 0.9915124472 | Romboutsia | Peptostreptococcaceae | Eubacteriales |
| 0.9867810745 | Schaedlerella | Lachnospiraceae | Eubacteriales |
| 0.7531912309 | Turicibacter | Turicibacteraceae | Erysipelotrichales |
| 0.8560789869 | Bacteria_unclassified | Bacteria_unclassified | Bacteria_unclassified |
| 0.9652194001 | Bacteria_unclassified | Bacteria_unclassified | Bacteria_unclassified |
| 0.9867810745 | Bacteria_unclassified | Bacteria_unclassified | Bacteria_unclassified |
| 0.8993780864 | | | |
| 0.9652194001 | | | |
| 0.9867810745 | | | |

| Class | Phylum | Kingdom |
|---|---|---|
| Clostridia | Firmicutes | Bacteria |
| Clostridia | Firmicutes | Bacteria |
| Clostridia | Firmicutes | Bacteria |
| Clostridia | Firmicutes | Bacteria |
| Coriobacteriia | Actinobacteria | Bacteria |
| Coriobacteriia | Actinobacteria | Bacteria |
| Coriobacteriia | Actinobacteria | Bacteria |
| Verrucomicrobiae | Verrucomicrobia | Bacteria |
| Bacteroidia | Bacteroidota | Bacteria |
| Clostridia | Firmicutes | Bacteria |
| Bacteria_unclassifie | Bacteria_unclassifie | Bacteria |
| Bacteria_unclassifie | Bacteria_unclassifie | Bacteria |
| Bacteria_unclassifie | Bacteria_unclassifie | Bacteria |
| Bacteroidia | Bacteroidota | Bacteria |
| Actinomycetia | Actinobacteria | Bacteria |
| Clostridia | Firmicutes | Bacteria |
| Clostridia | Firmicutes | Bacteria |
| Clostridia | Firmicutes | Bacteria |
| Clostridia | Firmicutes | Bacteria |
| Erysipelotrichia | Firmicutes | Bacteria |
| Coriobacteriia | Actinobacteria | Bacteria |
| Clostridia | Firmicutes | Bacteria |
| Erysipelotrichia | Firmicutes | Bacteria |
| Erysipelotrichia | Firmicutes | Bacteria |
| Clostridia | Firmicutes | Bacteria |
| Clostridia | Firmicutes | Bacteria |
| Clostridia | Firmicutes | Bacteria |
| Coriobacteriia | Actinobacteria | Bacteria |
| Clostridia | Firmicutes | Bacteria |
| Bacteroidia | Bacteroidota | Bacteria |
| Bacteroidia | Bacteroidota | Bacteria |
| Bacteroidia | Bacteroidota | Bacteria |
| CFGB9508 | Firmicutes | Bacteria |
| CFGB2838 | Firmicutes | Bacteria |
| CFGB2838 | Firmicutes | Bacteria |
| CFGB2838 | Firmicutes | Bacteria |
| Clostridia | Firmicutes | Bacteria |
| CFGB28439 | Firmicutes | Bacteria |
| Clostridia | Firmicutes | Bacteria |
| Clostridia | Firmicutes | Bacteria |
| Clostridia | Firmicutes | Bacteria |
| Clostridia | Firmicutes | Bacteria |
| Clostridia | Firmicutes | Bacteria |
| Clostridia | Firmicutes | Bacteria |
| CFGB77305 | Firmicutes | Bacteria |

| | | |
|---|---|---|
| Clostridia | Firmicutes | Bacteria |
| Clostridia | Firmicutes | Bacteria |
| Clostridia | Firmicutes | Bacteria |
| Clostridia | Firmicutes | Bacteria |
| CFGB9633 | Firmicutes | Bacteria |
| Bacteria_unclassifie | Bacteria_unclassifie | Bacteria |
| Bacteria_unclassifie | Bacteria_unclassifie | Bacteria |
| Bacteria_unclassifie | Bacteria_unclassifie | Bacteria |
| Bacteria_unclassifie | Bacteria_unclassifie | Bacteria |
| Clostridia | Firmicutes | Bacteria |
| Clostridia | Firmicutes | Bacteria |
| Clostridia | Firmicutes | Bacteria |
| CFGB77359 | Bacteria_unclassifie | Bacteria |
| CFGB9639 | Firmicutes | Bacteria |
| Clostridia | Firmicutes | Bacteria |
| Clostridia | Firmicutes | Bacteria |
| Clostridia | Firmicutes | Bacteria |
| Clostridia | Firmicutes | Bacteria |
| Clostridia | Firmicutes | Bacteria |
| Clostridia | Firmicutes | Bacteria |
| Clostridia | Firmicutes | Bacteria |
| Clostridia | Firmicutes | Bacteria |
| Clostridia | Firmicutes | Bacteria |
| Clostridia | Firmicutes | Bacteria |
| Clostridia | Firmicutes | Bacteria |
| CFGB9658 | Firmicutes | Bacteria |
| CFGB9659 | Firmicutes | Bacteria |
| Bacteria_unclassifie | Bacteria_unclassifie | Bacteria |
| CFGB9827 | Firmicutes | Bacteria |
| Clostridia | Firmicutes | Bacteria |
| CFGB77303 | Bacteria_unclassifie | Bacteria |
| Clostridia | Firmicutes | Bacteria |
| Clostridia | Firmicutes | Bacteria |
| CFGB30328 | Firmicutes | Bacteria |
| Clostridia | Firmicutes | Bacteria |
| Clostridia | Firmicutes | Bacteria |
| Clostridia | Firmicutes | Bacteria |
| Clostridia | Firmicutes | Bacteria |
| Clostridia | Firmicutes | Bacteria |
| Clostridia | Firmicutes | Bacteria |
| Clostridia | Firmicutes | Bacteria |
| CFGB77153 | Actinobacteria | Bacteria |
| CFGB1791 | Tenericutes | Bacteria |
| CFGB10290 | Firmicutes | Bacteria |
| Clostridia | Firmicutes | Bacteria |
| CFGB1765 | Firmicutes | Bacteria |

| | | |
|---|---|---|
| CFGB10349 | Firmicutes | Bacteria |
| CFGB10667 | Firmicutes | Bacteria |
| Clostridia | Firmicutes | Bacteria |
| Clostridia | Firmicutes | Bacteria |
| Clostridia | Firmicutes | Bacteria |
| CFGB10299 | Firmicutes | Bacteria |
| Clostridia | Firmicutes | Bacteria |
| Clostridia | Firmicutes | Bacteria |
| Clostridia | Firmicutes | Bacteria |
| Clostridia | Firmicutes | Bacteria |
| Clostridia | Firmicutes | Bacteria |
| Clostridia | Firmicutes | Bacteria |
| Clostridia | Firmicutes | Bacteria |
| Clostridia | Firmicutes | Bacteria |
| Clostridia | Firmicutes | Bacteria |
| Clostridia | Firmicutes | Bacteria |
| Clostridia | Firmicutes | Bacteria |
| Clostridia | Firmicutes | Bacteria |
| Bacilli | Firmicutes | Bacteria |
| Bacteroidia | Bacteroidota | Bacteria |
| Clostridia | Firmicutes | Bacteria |
| Clostridia | Firmicutes | Bacteria |
| Clostridia | Firmicutes | Bacteria |
| Clostridia | Firmicutes | Bacteria |
| Betaproteobacteria | Proteobacteria | Bacteria |
| Clostridia | Firmicutes | Bacteria |
| Clostridia | Firmicutes | Bacteria |
| Erysipelotrichia | Firmicutes | Bacteria |
| Bacteria_unclassifie | Bacteria_unclassifie | Bacteria |
| Bacteria_unclassifie | Bacteria_unclassifie | Bacteria |
| Bacteria_unclassifie | Bacteria_unclassifie | Bacteria |

| MetaPhlan Annotation |
|---|
| k__Bacteria\|p__Firmicutes\|c__Clostridia\|o__Eubacteriales\|f__Lachnospiraceae\|g__Acetatifactor\|s__Acetatifactor_SGB415 |
| k__Bacteria\|p__Firmicutes\|c__Clostridia\|o__Eubacteriales\|f__Lachnospiraceae\|g__Acetatifactor\|s__Acetatifactor_muris |
| k__Bacteria\|p__Firmicutes\|c__Clostridia\|o__Eubacteriales\|f__Oscillospiraceae\|g__Acutalibacter\|s__Acutalibacter_muris |
| k__Bacteria\|p__Firmicutes\|c__Clostridia\|o__Eubacteriales\|f__Oscillospiraceae\|g__Acutalibacter\|s__Acutalibacter_sp_1XD |
| k__Bacteria\|p__Actinobacteria\|c__Coriobacteriia\|o__Eggerthellales\|f__Eggerthellaceae\|g__Adlercreutzia\|s__Adlercreutzia |
| k__Bacteria\|p__Actinobacteria\|c__Coriobacteriia\|o__Eggerthellales\|f__Eggerthellaceae\|g__Adlercreutzia\|s__Adlercreutzia |
| k__Bacteria\|p__Actinobacteria\|c__Coriobacteriia\|o__Eggerthellales\|f__Eggerthellaceae\|g__Adlercreutzia\|s__Adlercreutzia |
| k__Bacteria\|p__Verrucomicrobia\|c__Verrucomicrobiae\|o__Verrucomicrobiales\|f__Akkermansiaceae\|g__Akkermansia\|s__ |
| k__Bacteria\|p__Bacteroidota\|c__Bacteroidia\|o__Bacteroidales\|f__Rikenellaceae\|g__Alistipes\|s__Alistipes_sp_DSM_11234 |
| k__Bacteria\|p__Firmicutes\|c__Clostridia\|o__Eubacteriales\|f__Oscillospiraceae\|g__Anaerotruncus\|s__Anaerotruncus_sp_1 |
| k__Bacteria\|p__Bacteria_unclassified\|c__Bacteria_unclassified\|o__Bacteria_unclassified\|f__Bacteria_unclassified\|g__Bact |
| k__Bacteria\|p__Bacteria_unclassified\|c__Bacteria_unclassified\|o__Bacteria_unclassified\|f__Bacteria_unclassified\|g__Bact |
| k__Bacteria\|p__Bacteria_unclassified\|c__Bacteria_unclassified\|o__Bacteria_unclassified\|f__Bacteria_unclassified\|g__Bact |
| k__Bacteria\|p__Bacteroidota\|c__Bacteroidia\|o__Bacteroidales\|f__Bacteroidaceae\|g__Bacteroides\|s__Bacteroides_thetaio |
| k__Bacteria\|p__Actinobacteria\|c__Actinomycetia\|o__Bifidobacteriales\|f__Bifidobacteriaceae\|g__Bifidobacterium\|s__Bifid |
| k__Bacteria\|p__Firmicutes\|c__Clostridia\|o__Clostridia_unclassified\|f__Clostridia_unclassified\|g__Clostridia_unclassified\|s_ |
| k__Bacteria\|p__Firmicutes\|c__Clostridia\|o__Eubacteriales\|f__Clostridiaceae\|g__Clostridiaceae_unclassified\|s__Clostridiac |
| k__Bacteria\|p__Firmicutes\|c__Clostridia\|o__Eubacteriales\|f__Clostridiaceae\|g__Clostridiaceae_unclassified\|s__Clostridiac |
| k__Bacteria\|p__Firmicutes\|c__Clostridia\|o__Eubacteriales\|f__Eubacteriales_unclassified\|g__Eubacteriales_unclassified\|s_ |
| k__Bacteria\|p__Firmicutes\|c__Erysipelotrichia\|o__Erysipelotrichales\|f__Erysipelotrichaceae\|g__Erysipelatoclostridium\|s_ |
| k__Bacteria\|p__Actinobacteria\|c__Coriobacteriia\|o__Coriobacteriales\|f__Coriobacteriaceae\|g__Coriobacteriaceae_unclass |
| k__Bacteria\|p__Firmicutes\|c__Clostridia\|o__Eubacteriales\|f__Lachnospiraceae\|g__Dorea\|s__Dorea_sp_5_2 |
| k__Bacteria\|p__Firmicutes\|c__Erysipelotrichia\|o__Erysipelotrichales\|f__Erysipelotrichaceae\|g__Dubosiella\|s__Dubosiella_ |
| k__Bacteria\|p__Firmicutes\|c__Erysipelotrichia\|o__Erysipelotrichales\|f__Erysipelotrichales_unclassified\|g__Erysipelotrichal |
| k__Bacteria\|p__Firmicutes\|c__Clostridia\|o__Eubacteriales\|f__Eubacteriaceae\|g__Eubacteriaceae_unclassified\|s__Eubacte |
| k__Bacteria\|p__Firmicutes\|c__Clostridia\|o__Eubacteriales\|f__Eubacteriaceae\|g__Eubacteriaceae_unclassified\|s__Eubacte |
| k__Bacteria\|p__Firmicutes\|c__Clostridia\|o__Eubacteriales\|f__Lachnospiraceae\|g__GGB20149\|s__GGB20149_SGB29430 |
| k__Bacteria\|p__Actinobacteria\|c__Coriobacteriia\|o__Eggerthellales\|f__Eggerthellaceae\|g__GGB22635\|s__GGB22635_SGB |
| k__Bacteria\|p__Firmicutes\|c__Clostridia\|o__Eubacteriales\|f__Lachnospiraceae\|g__GGB25041\|s__GGB25041_SGB36960 |
| k__Bacteria\|p__Bacteroidota\|c__Bacteroidia\|o__Bacteroidales\|f__Muribaculaceae\|g__GGB27876\|s__GGB27876_SGB403 |
| k__Bacteria\|p__Bacteroidota\|c__Bacteroidia\|o__Bacteroidales\|f__Muribaculaceae\|g__GGB27878\|s__GGB27878_SGB403 |
| k__Bacteria\|p__Bacteroidota\|c__Bacteroidia\|o__Bacteroidales\|f__Muribaculaceae\|g__GGB27918\|s__GGB27918_SGB403 |
| k__Bacteria\|p__Firmicutes\|c__CFGB9508\|o__OFGB9508\|f__FGB9508\|g__GGB28382\|s__GGB28382_SGB40962 |
| k__Bacteria\|p__Firmicutes\|c__CFGB2838\|o__OFGB2838\|f__FGB2838\|g__GGB28399\|s__GGB28399_SGB40980 |
| k__Bacteria\|p__Firmicutes\|c__CFGB2838\|o__OFGB2838\|f__FGB2838\|g__GGB28411\|s__GGB28411_SGB40993 |
| k__Bacteria\|p__Firmicutes\|c__CFGB2838\|o__OFGB2838\|f__FGB2838\|g__GGB28415\|s__GGB28415_SGB40997 |
| k__Bacteria\|p__Firmicutes\|c__Clostridia\|o__Eubacteriales\|f__Pumilibacteraceae\|g__GGB28430\|s__GGB28430_SGB41013 |
| k__Bacteria\|p__Firmicutes\|c__CFGB28439\|o__OFGB28439\|f__FGB28439\|g__GGB28439\|s__GGB28439_SGB41022 |
| k__Bacteria\|p__Firmicutes\|c__Clostridia\|o__Clostridia_unclassified\|f__Clostridia_unclassified\|g__GGB28778\|s__GGB2877 |
| k__Bacteria\|p__Firmicutes\|c__Clostridia\|o__Eubacteriales\|f__Eubacteriaceae\|g__GGB28784\|s__GGB28784_SGB41437 |
| k__Bacteria\|p__Firmicutes\|c__Clostridia\|o__Eubacteriales\|f__Lachnospiraceae\|g__GGB28792\|s__GGB28792_SGB41445 |
| k__Bacteria\|p__Firmicutes\|c__Clostridia\|o__Eubacteriales\|f__Lachnospiraceae\|g__GGB28798\|s__GGB28798_SGB41451 |
| k__Bacteria\|p__Firmicutes\|c__Clostridia\|o__Eubacteriales\|f__Lachnospiraceae\|g__GGB28802\|s__GGB28802_SGB41455 |
| k__Bacteria\|p__Firmicutes\|c__Clostridia\|o__Eubacteriales\|f__Lachnospiraceae\|g__GGB28818\|s__GGB28818_SGB41473 |
| k__Bacteria\|p__Firmicutes\|c__CFGB77305\|o__OFGB77305\|f__FGB77305\|g__GGB28828\|s__GGB28828_SGB41484 |

k__Bacteria|p__Firmicutes|c__Clostridia|o__Eubacteriales|f__Clostridiaceae|g__GGB28851|s__GGB28851_SGB41518
k__Bacteria|p__Firmicutes|c__Clostridia|o__Eubacteriales|f__Lachnospiraceae|g__GGB28859|s__GGB28859_SGB41528
k__Bacteria|p__Firmicutes|c__Clostridia|o__Eubacteriales|f__Lachnospiraceae|g__GGB28864|s__GGB28864_SGB41535
k__Bacteria|p__Firmicutes|c__Clostridia|o__Eubacteriales|f__Lachnospiraceae|g__GGB28869|s__GGB28869_SGB41543
k__Bacteria|p__Firmicutes|c__CFGB9633|o__OFGB9633|f__FGB9633|g__GGB28883|s__GGB28883_SGB41564
k__Bacteria|p__Bacteria_unclassified|c__Bacteria_unclassified|o__Bacteria_unclassified|f__Bacteria_unclassified|g__GGB[illegible]
k__Bacteria|p__Bacteria_unclassified|c__Bacteria_unclassified|o__Bacteria_unclassified|f__Bacteria_unclassified|g__GGB[illegible]
k__Bacteria|p__Bacteria_unclassified|c__Bacteria_unclassified|o__Bacteria_unclassified|f__Bacteria_unclassified|g__GGB[illegible]
k__Bacteria|p__Bacteria_unclassified|c__Bacteria_unclassified|o__Bacteria_unclassified|f__Bacteria_unclassified|g__GGB[illegible]
k__Bacteria|p__Firmicutes|c__Clostridia|o__Eubacteriales|f__Lachnospiraceae|g__GGB28916|s__GGB28916_SGB41612
k__Bacteria|p__Firmicutes|c__Clostridia|o__Eubacteriales|f__Lachnospiraceae|g__GGB28924|s__GGB28924_SGB41621
k__Bacteria|p__Firmicutes|c__Clostridia|o__Eubacteriales|f__Lachnospiraceae|g__GGB28926|s__GGB28926_SGB41624
k__Bacteria|p__Bacteria_unclassified|c__CFGB77359|o__OFGB77359|f__FGB77359|g__GGB28927|s__GGB28927_SGB416[illegible]
k__Bacteria|p__Firmicutes|c__CFGB9639|o__OFGB9639|f__FGB9639|g__GGB28934|s__GGB28934_SGB41635
k__Bacteria|p__Firmicutes|c__Clostridia|o__Eubacteriales|f__Lachnospiraceae|g__GGB28946|s__GGB28946_SGB41652
k__Bacteria|p__Firmicutes|c__Clostridia|o__Eubacteriales|f__Lachnospiraceae|g__GGB28949|s__GGB28949_SGB41655
k__Bacteria|p__Firmicutes|c__Clostridia|o__Eubacteriales|f__Lachnospiraceae|g__GGB28949|s__GGB28949_SGB41656
k__Bacteria|p__Firmicutes|c__Clostridia|o__Eubacteriales|f__Clostridiaceae|g__GGB28950|s__GGB28950_SGB41657
k__Bacteria|p__Firmicutes|c__Clostridia|o__Eubacteriales|f__Clostridiaceae|g__GGB28951|s__GGB28951_SGB102295
k__Bacteria|p__Firmicutes|c__Clostridia|o__Eubacteriales|f__Clostridiaceae|g__GGB28951|s__GGB28951_SGB41658
k__Bacteria|p__Firmicutes|c__Clostridia|o__Eubacteriales|f__Clostridiaceae|g__GGB28954|s__GGB28954_SGB41662
k__Bacteria|p__Firmicutes|c__Clostridia|o__Eubacteriales|f__Clostridiaceae|g__GGB28956|s__GGB28956_SGB41665
k__Bacteria|p__Firmicutes|c__Clostridia|o__Eubacteriales|f__Clostridiaceae|g__GGB28960|s__GGB28960_SGB41669
k__Bacteria|p__Firmicutes|c__Clostridia|o__Eubacteriales|f__Clostridiaceae|g__GGB28967|s__GGB28967_SGB41678
k__Bacteria|p__Firmicutes|c__Clostridia|o__Eubacteriales|f__Eubacteriaceae|g__GGB28991|s__GGB28991_SGB41705
k__Bacteria|p__Firmicutes|c__CFGB9658|o__OFGB9658|f__FGB9658|g__GGB29002|s__GGB29002_SGB41718
k__Bacteria|p__Firmicutes|c__CFGB9659|o__OFGB9659|f__FGB9659|g__GGB29003|s__GGB29003_SGB41719
k__Bacteria|p__Bacteria_unclassified|c__Bacteria_unclassified|o__Bacteria_unclassified|f__Bacteria_unclassified|g__GGB[illegible]
k__Bacteria|p__Firmicutes|c__CFGB9827|o__OFGB9827|f__FGB9827|g__GGB29531|s__GGB29531_SGB42317
k__Bacteria|p__Firmicutes|c__Clostridia|o__Eubacteriales|f__Eubacteriaceae|g__GGB29685|s__GGB29685_SGB42494
k__Bacteria|p__Bacteria_unclassified|c__CFGB77303|o__OFGB77303|f__FGB77303|g__GGB30141|s__GGB30141_SGB43[illegible]
k__Bacteria|p__Firmicutes|c__Clostridia|o__Eubacteriales|f__Eubacteriales_unclassified|g__GGB30286|s__GGB30286_SG[illegible]
k__Bacteria|p__Firmicutes|c__Clostridia|o__Eubacteriales|f__Oscillospiraceae|g__GGB30303|s__GGB30303_SGB43268
k__Bacteria|p__Firmicutes|c__CFGB30328|o__OFGB30328|f__FGB30328|g__GGB30413|s__GGB30413_SGB43452
k__Bacteria|p__Firmicutes|c__Clostridia|o__Eubacteriales|f__Oscillospiraceae|g__GGB30454|s__GGB30454_SGB43514
k__Bacteria|p__Firmicutes|c__Clostridia|o__Eubacteriales|f__Oscillospiraceae|g__GGB30455|s__GGB30455_SGB43519
k__Bacteria|p__Firmicutes|c__Clostridia|o__Eubacteriales|f__Oscillospiraceae|g__GGB30461|s__GGB30461_SGB43527
k__Bacteria|p__Firmicutes|c__Clostridia|o__Eubacteriales|f__Oscillospiraceae|g__GGB30461|s__GGB30461_SGB43530
k__Bacteria|p__Firmicutes|c__Clostridia|o__Eubacteriales|f__Oscillospiraceae|g__GGB30463|s__GGB30463_SGB43537
k__Bacteria|p__Firmicutes|c__Clostridia|o__Eubacteriales|f__Oscillospiraceae|g__GGB30473|s__GGB30473_SGB43557
k__Bacteria|p__Firmicutes|c__Clostridia|o__Eubacteriales|f__Oscillospiraceae|g__GGB30475|s__GGB30475_SGB63182
k__Bacteria|p__Actinobacteria|c__CFGB77153|o__OFGB77153|f__FGB77153|g__GGB30861|s__GGB30861_SGB44083
k__Bacteria|p__Tenericutes|c__CFGB1791|o__OFGB1791|f__FGB1791|g__GGB31312|s__GGB31312_SGB44628
k__Bacteria|p__Firmicutes|c__CFGB10290|o__OFGB10290|f__FGB10290|g__GGB31438|s__GGB31438_SGB44768
k__Bacteria|p__Firmicutes|c__Clostridia|o__Eubacteriales|f__Oscillospiraceae|g__GGB3171|s__GGB3171_SGB4185
k__Bacteria|p__Firmicutes|c__CFGB1765|o__OFGB1765|f__FGB1765|g__GGB31823|s__GGB31823_SGB45199

k__Bacteria|p__Firmicutes|c__CFGB10349|o__OFGB10349|f__FGB10349|g__GGB31853|s__GGB31853_SGB45233
k__Bacteria|p__Firmicutes|c__CFGB10667|o__OFGB10667|f__FGB10667|g__GGB32371|s__GGB32371_SGB41694
k__Bacteria|p__Firmicutes|c__Clostridia|o__Eubacteriales|f__Lachnospiraceae|g__GGB3793|s__GGB3793_SGB5158
k__Bacteria|p__Firmicutes|c__Clostridia|o__Eubacteriales|f__Lachnospiraceae|g__GGB42598|s__GGB42598_SGB59794
k__Bacteria|p__Firmicutes|c__Clostridia|o__Eubacteriales|f__Christensenellaceae|g__GGB45656|s__GGB45656_SGB6337
k__Bacteria|p__Firmicutes|c__CFGB10299|o__OFGB10299|f__FGB10299|g__GGB47127|s__GGB47127_SGB65054
k__Bacteria|p__Firmicutes|c__Clostridia|o__Eubacteriales|f__Oscillospiraceae|g__GGB74395|s__GGB74395_SGB43521
k__Bacteria|p__Firmicutes|c__Clostridia|o__Eubacteriales|f__Oscillospiraceae|g__GGB75053|s__GGB75053_SGB43494
k__Bacteria|p__Firmicutes|c__Clostridia|o__Eubacteriales|f__Lachnospiraceae|g__GGB75109|s__GGB75109_SGB102238
k__Bacteria|p__Firmicutes|c__Clostridia|o__Clostridia_unclassified|f__Clostridia_unclassified|g__GGB81440|s__GGB8144
k__Bacteria|p__Firmicutes|c__Clostridia|o__Eubacteriales|f__Lachnospiraceae|g__Lachnospiraceae_unclassified|s__Lachn
k__Bacteria|p__Firmicutes|c__Clostridia|o__Eubacteriales|f__Lachnospiraceae|g__Lachnospiraceae_unclassified|s__Lachn
k__Bacteria|p__Firmicutes|c__Clostridia|o__Eubacteriales|f__Lachnospiraceae|g__Lachnospiraceae_unclassified|s__Lachn
k__Bacteria|p__Firmicutes|c__Clostridia|o__Eubacteriales|f__Lachnospiraceae|g__Lachnospiraceae_unclassified|s__Lachn
k__Bacteria|p__Firmicutes|c__Clostridia|o__Eubacteriales|f__Lachnospiraceae|g__Lachnospiraceae_unclassified|s__Lachn
k__Bacteria|p__Firmicutes|c__Clostridia|o__Eubacteriales|f__Lachnospiraceae|g__Lachnospiraceae_unclassified|s__Lachn
k__Bacteria|p__Firmicutes|c__Clostridia|o__Eubacteriales|f__Lachnospiraceae|g__Lachnospiraceae_unclassified|s__Lachn
k__Bacteria|p__Firmicutes|c__Clostridia|o__Eubacteriales|f__Lachnospiraceae|g__Lachnospiraceae_unclassified|s__Lachn
k__Bacteria|p__Firmicutes|c__Bacilli|o__Lactobacillales|f__Lactobacillaceae|g__Lactobacillus|s__Lactobacillus_johnsonii
k__Bacteria|p__Bacteroidota|c__Bacteroidia|o__Bacteroidales|f__Muribaculaceae|g__Muribaculaceae_unclassified|s__M
k__Bacteria|p__Firmicutes|c__Clostridia|o__Eubacteriales|f__Oscillospiraceae|g__Neglectibacter|s__Neglectibacter_sp_X
k__Bacteria|p__Firmicutes|c__Clostridia|o__Eubacteriales|f__Oscillospiraceae|g__Oscillospiraceae_unclassified|s__Oscillo
k__Bacteria|p__Firmicutes|c__Clostridia|o__Eubacteriales|f__Oscillospiraceae|g__Oscillospiraceae_unclassified|s__Oscillo
k__Bacteria|p__Firmicutes|c__Clostridia|o__Eubacteriales|f__Oscillospiraceae|g__Oscillospiraceae_unclassified|s__Oscillo
k__Bacteria|p__Proteobacteria|c__Betaproteobacteria|o__Burkholderiales|f__Sutterellaceae|g__Parasutterella|s__Parasu
k__Bacteria|p__Firmicutes|c__Clostridia|o__Eubacteriales|f__Peptostreptococcaceae|g__Romboutsia|s__Romboutsia_ilea
k__Bacteria|p__Firmicutes|c__Clostridia|o__Eubacteriales|f__Lachnospiraceae|g__Schaedlerella|s__Schaedlerella_arabinc
k__Bacteria|p__Firmicutes|c__Erysipelotrichia|o__Erysipelotrichales|f__Turicibacteraceae|g__Turicibacter|s__Turicibacter_
k__Bacteria|p__Bacteria_unclassified|c__Bacteria_unclassified|o__Bacteria_unclassified|f__Bacteria_unclassified|g__Bacte
k__Bacteria|p__Bacteria_unclassified|c__Bacteria_unclassified|o__Bacteria_unclassified|f__Bacteria_unclassified|g__Bacte
k__Bacteria|p__Bacteria_unclassified|c__Bacteria_unclassified|o__Bacteria_unclassified|f__Bacteria_unclassified|g__Bacte
