## Supplementary material for "Non-invasive Vagal Nerve Stimulation as a Potential Treatment for Repetitive Blast Trauma": st7

| | | | Likelihood Ratio Tes | |
|---|---|---|---|---|
| outcome | Species-level feature | n | Microbe Chi (df=1 | Microbe Pr(>Chi |
| EtOH_24h_intakeave_final | Acetatifactor_SGB41546 | 63 | 2.643234361 | 0.01014668495 |
| EtOH_24h_intakeave_final | Acetatifactor_muris | 63 | 0.07887962006 | 0.93713 |
| EtOH_24h_intakeave_final | Acutalibacter_muris | 63 | 1.653866013 | 0.1019508547 |
| EtOH_24h_intakeave_final | Acutalibacter_sp_1XD8_36 | 63 | 0.8762493561 | 0.3824095081 |
| EtOH_24h_intakeave_final | Adlercreutzia_caecimuris | 63 | -2.506293877 | 0.01451844488 |
| EtOH_24h_intakeave_final | Adlercreutzia_mucosicola | 63 | 0.8430779108 | 0.4005737629 |
| EtOH_24h_intakeave_final | Adlercreutzia_muris | 63 | 0.4048584518 | 0.6857845285 |
| EtOH_24h_intakeave_final | Akkermansia_muciniphila | 63 | -1.481223827 | 0.14211927 |
| EtOH_24h_intakeave_final | Alistipes_sp_DSM_112343 | 63 | -1.164067037 | 0.247037984 |
| EtOH_24h_intakeave_final | Anaerotruncus_sp_1XD42_93 | 63 | 0.6921483781 | 0.4897083888 |
| EtOH_24h_intakeave_final | Bacteria_unclassified_SGB102200 | 63 | -0.2215233595 | 0.8247201987 |
| EtOH_24h_intakeave_final | Bacteria_unclassified_SGB41677 | 63 | -0.4106691382 | 0.6815264835 |
| EtOH_24h_intakeave_final | Bacteria_unclassified_SGB43546 | 63 | 0.320152722 | 0.748956111 |
| EtOH_24h_intakeave_final | Bacteroides_thetaiotaomicron | 63 | -0.9335646025 | 0.3522671042 |
| EtOH_24h_intakeave_final | Bifidobacterium_pseudolongum | 63 | -1.643744416 | 0.1040182897 |
| EtOH_24h_intakeave_final | Clostridia_bacterium | 63 | -1.436975182 | 0.1542042991 |
| EtOH_24h_intakeave_final | Clostridiaceae_bacterium | 63 | -2.705183306 | 0.008595498925 |
| EtOH_24h_intakeave_final | Clostridiaceae_unclassified_SGB41663 | 63 | 1.022121776 | 0.3088139377 |
| EtOH_24h_intakeave_final | Clostridiales_bacterium | 63 | 0.4240784591 | 0.6717399822 |
| EtOH_24h_intakeave_final | Clostridium_cocleatum | 63 | 0.2905248716 | 0.7714928216 |
| EtOH_24h_intakeave_final | Coriobacteriaceae_bacterium | 63 | -0.5364630185 | 0.5920819959 |
| EtOH_24h_intakeave_final | Dorea_sp_5_2 | 63 | -1.350288739 | 0.1801816088 |
| EtOH_24h_intakeave_final | Dubosiella_newyorkensis | 63 | -1.853825593 | 0.06753860213 |
| EtOH_24h_intakeave_final | Erysipelotrichales_bacterium | 63 | 0.3876428347 | 0.6984597667 |
| EtOH_24h_intakeave_final | Eubacteriaceae_bacterium | 63 | -0.5223830263 | 0.6018161862 |
| EtOH_24h_intakeave_final | Eubacteriaceae_unclassified_SGB9492 | 63 | -0.8018527157 | 0.4238748354 |
| EtOH_24h_intakeave_final | GGB20149_SGB29430 | 63 | 1.013870335 | 0.3127024455 |
| EtOH_24h_intakeave_final | GGB22635_SGB63107 | 63 | -1.393313529 | 0.1668990797 |
| EtOH_24h_intakeave_final | GGB25041_SGB36960 | 63 | -0.8826170116 | 0.3789827761 |
| EtOH_24h_intakeave_final | GGB27876_SGB40310 | 63 | -1.102343481 | 0.2727189781 |
| EtOH_24h_intakeave_final | GGB27878_SGB40312 | 63 | -0.1299359205 | 0.8966243555 |
| EtOH_24h_intakeave_final | GGB27918_SGB40356 | 63 | 0.6806167409 | 0.4969424086 |
| EtOH_24h_intakeave_final | GGB28382_SGB40962 | 63 | 1.136960937 | 0.2580940175 |
| EtOH_24h_intakeave_final | GGB28399_SGB40980 | 63 | 0.2706503609 | 0.7867234573 |
| EtOH_24h_intakeave_final | GGB28411_SGB40993 | 63 | -0.462376347 | 0.6441062366 |
| EtOH_24h_intakeave_final | GGB28415_SGB40997 | 63 | -0.3019997873 | 0.7627396417 |
| EtOH_24h_intakeave_final | GGB28430_SGB41013 | 63 | 0.7423300279 | 0.4589153734 |
| EtOH_24h_intakeave_final | GGB28439_SGB41022 | 63 | 1.608962401 | 0.1113843636 |
| EtOH_24h_intakeave_final | GGB28778_SGB41431 | 63 | 0.2801404161 | 0.7794400064 |
| EtOH_24h_intakeave_final | GGB28784_SGB41437 | 63 | -0.5707917168 | 0.5686647231 |
| EtOH_24h_intakeave_final | GGB28792_SGB41445 | 63 | 0.01198879917 | 0.9904345571 |
| EtOH_24h_intakeave_final | GGB28798_SGB41451 | 63 | -0.7996461555 | 0.4251445275 |
| EtOH_24h_intakeave_final | GGB28802_SGB41455 | 63 | -0.06059249991 | 0.951684485 |
| EtOH_24h_intakeave_final | GGB28818_SGB41473 | 63 | -0.4990469888 | 0.6181103259 |
| EtOH_24h_intakeave_final | GGB28828_SGB41484 | 63 | -0.4505927261 | 0.6525574162 |

| | | | | |
|---|---|---|---|---|
| EtOH_24h_intakeave_final | GGB28851_SGB41518 | 63 | 0.3738803283 | 0.7086551764 |
| EtOH_24h_intakeave_final | GGB28859_SGB41528 | 63 | -0.5768605856 | 0.5645725029 |
| EtOH_24h_intakeave_final | GGB28864_SGB41535 | 63 | 0.8492169195 | 0.397172611 |
| EtOH_24h_intakeave_final | GGB28869_SGB41543 | 63 | 1.519880099 | 0.132183137 |
| EtOH_24h_intakeave_final | GGB28883_SGB41564 | 63 | -1.341714031 | 0.1829224984 |
| EtOH_24h_intakeave_final | GGB28892_SGB41573 | 63 | -1.407206108 | 0.1627752179 |
| EtOH_24h_intakeave_final | GGB28893_SGB41574 | 63 | 1.409591058 | 0.1620752609 |
| EtOH_24h_intakeave_final | GGB28898_SGB41580 | 63 | -1.944815675 | 0.05547079326 |
| EtOH_24h_intakeave_final | GGB28904_SGB41597 | 63 | 0.9283630238 | 0.354937354 |
| EtOH_24h_intakeave_final | GGB28916_SGB41612 | 63 | 0.3686543291 | 0.7125408334 |
| EtOH_24h_intakeave_final | GGB28924_SGB41621 | 63 | 0.1690292605 | 0.8657894494 |
| EtOH_24h_intakeave_final | GGB28926_SGB41624 | 63 | -0.7209932954 | 0.4718705137 |
| EtOH_24h_intakeave_final | GGB28927_SGB41625 | 63 | 2.508346013 | 0.01444196717 |
| EtOH_24h_intakeave_final | GGB28934_SGB41635 | 63 | 0.7106768549 | 0.4782077753 |
| EtOH_24h_intakeave_final | GGB28946_SGB41652 | 63 | 0.6858701256 | 0.4936396379 |
| EtOH_24h_intakeave_final | GGB28949_SGB41655 | 63 | -1.184618857 | 0.2388848902 |
| EtOH_24h_intakeave_final | GGB28949_SGB41656 | 63 | 0.8054138121 | 0.4218305221 |
| EtOH_24h_intakeave_final | GGB28950_SGB41657 | 63 | -0.4173026666 | 0.6766781508 |
| EtOH_24h_intakeave_final | GGB28951_SGB102295 | 63 | 0.4000679325 | 0.6893027149 |
| EtOH_24h_intakeave_final | GGB28951_SGB41658 | 63 | 2.902957924 | 0.004983215651 |
| EtOH_24h_intakeave_final | GGB28954_SGB41662 | 63 | -0.2145016379 | 0.8301879415 |
| EtOH_24h_intakeave_final | GGB28956_SGB41665 | 63 | 0.2839554365 | 0.7765175887 |
| EtOH_24h_intakeave_final | GGB28960_SGB41669 | 63 | -0.02057285725 | 0.9835864216 |
| EtOH_24h_intakeave_final | GGB28967_SGB41678 | 63 | 1.106891304 | 0.2707651359 |
| EtOH_24h_intakeave_final | GGB28991_SGB41705 | 63 | -0.756447692 | 0.4504569676 |
| EtOH_24h_intakeave_final | GGB29002_SGB41718 | 63 | 1.748620771 | 0.08417889984 |
| EtOH_24h_intakeave_final | GGB29003_SGB41719 | 63 | -0.526868663 | 0.5987070189 |
| EtOH_24h_intakeave_final | GGB29011_SGB41731 | 63 | -0.916490799 | 0.3610809192 |
| EtOH_24h_intakeave_final | GGB29531_SGB42317 | 63 | 3.424776826 | 0.00106378805€ |
| EtOH_24h_intakeave_final | GGB29685_SGB42494 | 63 | -2.719328316 | 0.00827324495 |
| EtOH_24h_intakeave_final | GGB30141_SGB43066 | 63 | 0.6620577055 | 0.5087063797 |
| EtOH_24h_intakeave_final | GGB30286_SGB43248 | 63 | -0.1384540699 | 0.8898903048 |
| EtOH_24h_intakeave_final | GGB30303_SGB43268 | 63 | 1.459516725 | 0.1479513531 |
| EtOH_24h_intakeave_final | GGB30413_SGB43452 | 63 | -0.6351439783 | 0.5260284467 |
| EtOH_24h_intakeave_final | GGB30454_SGB43514 | 63 | -1.396380619 | 0.1659817921 |
| EtOH_24h_intakeave_final | GGB30455_SGB43519 | 63 | 1.763591265 | 0.08162112993 |
| EtOH_24h_intakeave_final | GGB30461_SGB43527 | 63 | -1.679195656 | 0.09692369261 |
| EtOH_24h_intakeave_final | GGB30461_SGB43530 | 63 | 1.180151732 | 0.2406402822 |
| EtOH_24h_intakeave_final | GGB30463_SGB43537 | 63 | 0.9891352589 | 0.3245562107 |
| EtOH_24h_intakeave_final | GGB30473_SGB43557 | 63 | -0.3354831787 | 0.7373791736 |
| EtOH_24h_intakeave_final | GGB30475_SGB63182 | 63 | -1.275636235 | 0.2051152624 |
| EtOH_24h_intakeave_final | GGB30861_SGB44083 | 63 | -2.632621552 | 0.01043674126 |
| EtOH_24h_intakeave_final | GGB31312_SGB44628 | 63 | -1.922243558 | 0.05827852977 |
| EtOH_24h_intakeave_final | GGB31438_SGB44768 | 63 | -1.53035277 | 0.1295884264 |
| EtOH_24h_intakeave_final | GGB3171_SGB4185 | 63 | 0.2751050637 | 0.7833021252 |
| EtOH_24h_intakeave_final | GGB31823_SGB45199 | 63 | -1.003778775 | 0.317502865 |

| | | | | |
|---|---|---|---|---|
| EtOH_24h_intakeave_final | GGB31853_SGB45233 | 63 | 2.06763554 | 0.04213730763 |
| EtOH_24h_intakeave_final | GGB32371_SGB41694 | 63 | -0.688934247 | 0.4917188186 |
| EtOH_24h_intakeave_final | GGB3793_SGB5158 | 63 | 0.4185992032 | 0.675732129 |
| EtOH_24h_intakeave_final | GGB42598_SGB59794 | 63 | -0.6006072016 | 0.5487012594 |
| EtOH_24h_intakeave_final | GGB45656_SGB63370 | 63 | 1.323001411 | 0.189013771 |
| EtOH_24h_intakeave_final | GGB47127_SGB65054 | 63 | -1.998784947 | 0.04922229004 |
| EtOH_24h_intakeave_final | GGB74395_SGB43521 | 63 | -1.637328832 | 0.105346324 |
| EtOH_24h_intakeave_final | GGB75053_SGB43494 | 63 | 1.223982884 | 0.2238157793 |
| EtOH_24h_intakeave_final | GGB75109_SGB102238 | 63 | -0.8925170102 | 0.3736937255 |
| EtOH_24h_intakeave_final | GGB81440_SGB45230 | 63 | 1.780035132 | 0.07888645053 |
| EtOH_24h_intakeave_final | Lachnospiraceae_bacterium | 63 | -1.365260355 | 0.1754710972 |
| EtOH_24h_intakeave_final | Lachnospiraceae_bacterium_A2 | 63 | -1.156848284 | 0.2499486122 |
| EtOH_24h_intakeave_final | Lachnospiraceae_bacterium_MD308 | 63 | -1.477078829 | 0.1432187154 |
| EtOH_24h_intakeave_final | Lachnospiraceae_bacterium_MD329 | 63 | -0.8388782919 | 0.4029107489 |
| EtOH_24h_intakeave_final | Lachnospiraceae_unclassified_SGB4141 | 63 | 0.9097484048 | 0.3646001881 |
| EtOH_24h_intakeave_final | Lachnospiraceae_unclassified_SGB4141 | 63 | 0.2510776692 | 0.8018050194 |
| EtOH_24h_intakeave_final | Lachnospiraceae_unclassified_SGB4142 | 63 | 0.663070971 | 0.5080602637 |
| EtOH_24h_intakeave_final | Lachnospiraceae_unclassified_SGB4158 | 63 | -0.3894526247 | 0.6971231456 |
| EtOH_24h_intakeave_final | Lactobacillus_johnsonii | 63 | -0.05579705059 | 0.9555040612 |
| EtOH_24h_intakeave_final | Muribaculaceae_bacterium | 63 | -2.107493665 | 0.03845406567 |
| EtOH_24h_intakeave_final | Neglectibacter_sp_X4 | 63 | 2.044202327 | 0.04444256131 |
| EtOH_24h_intakeave_final | Oscillospiraceae_bacterium | 63 | -0.04317403543 | 0.9655630698 |
| EtOH_24h_intakeave_final | Oscillospiraceae_unclassified_SGB4350 | 63 | 0.934605306 | 0.3517344238 |
| EtOH_24h_intakeave_final | Oscillospiraceae_unclassified_SGB4350 | 63 | 0.6632027772 | 0.5079762489 |
| EtOH_24h_intakeave_final | Parasutterella_excrementihominis | 63 | -1.275085543 | 0.2053083017 |
| EtOH_24h_intakeave_final | Romboutsia_ilealis | 63 | -0.1016791819 | 0.9190148034 |
| EtOH_24h_intakeave_final | Schaedlerella_arabinosiphila | 63 | -1.283136531 | 0.2024995636 |
| EtOH_24h_intakeave_final | Turicibacter_sp_1E2 | 63 | -1.393310511 | 0.1668999841 |
| EtOH_24h_intakeave_final | bacterium_1XD42_54 | 63 | -2.480615243 | 0.01550683875 |
| EtOH_24h_intakeave_final | bacterium_1XD42_76 | 63 | -0.3947983951 | 0.6931806615 |
| EtOH_24h_intakeave_final | bacterium_1xD8_48 | 63 | 0.5664291063 | 0.5716153682 |
| EtOH_24h_intakeave_final | Berger Parker Index | 63 | -1.824870881 | 0.07181587317 |
| EtOH_24h_intakeave_final | Richness (# observed features) | 63 | 1.328132768 | 0.1873283705 |
| EtOH_24h_intakeave_final | Shannon Index | 63 | 0.9345445312 | 0.3517655168 |
| EtOH_8pm_intakeave_fina | Acetatifactor_SGB41546 | 63 | 0.1269479285 | 0.9033652436 |
| EtOH_8pm_intakeave_fina | Acetatifactor_muris | 63 | -1.554120016 | 0.1418367732 |
| EtOH_8pm_intakeave_fina | Acutalibacter_muris | 63 | -0.07888209967 | 0.9371657906 |
| EtOH_8pm_intakeave_fina | Acutalibacter_sp_1XD8_36 | 63 | 0.5309571262 | 0.598230628 |
| EtOH_8pm_intakeave_fina | Adlercreutzia_caecimuris | 63 | -1.539337551 | 0.1278423439 |
| EtOH_8pm_intakeave_fina | Adlercreutzia_mucosicola | 63 | 0.5737974271 | 0.6003383763 |
| EtOH_8pm_intakeave_fina | Adlercreutzia_muris | 63 | -0.07316337356 | 0.942729343 |
| EtOH_8pm_intakeave_fina | Akkermansia_muciniphila | 63 | -0.1278160409 | 0.8984966586 |
| EtOH_8pm_intakeave_fina | Alistipes_sp_DSM_112343 | 63 | -0.09100153497 | 0.9303283236 |
| EtOH_8pm_intakeave_fina | Anaerotruncus_sp_1XD42_93 | 63 | 1.616745063 | 0.1351392145 |
| EtOH_8pm_intakeave_fina | Bacteria_unclassified_SGB102200 | 63 | -0.5846899654 | 0.5610027826 |
| EtOH_8pm_intakeave_fina | Bacteria_unclassified_SGB41677 | 63 | 1.835199181 | 0.07990658851 |

| | | | | |
|---|---|---|---|---|
| EtOH_8pm_intakeave_fina | Bacteria_unclassified_SGB43546 | 63 | -0.289474308 | 0.7771851965 |
| EtOH_8pm_intakeave_fina | Bacteroides_thetaiotaomicron | 63 | -0.9579341796 | 0.3413404388 |
| EtOH_8pm_intakeave_fina | Bifidobacterium_pseudolongum | 63 | -1.212773179 | 0.2294182698 |
| EtOH_8pm_intakeave_fina | Clostridia_bacterium | 63 | -0.7777415287 | 0.4390603691 |
| EtOH_8pm_intakeave_fina | Clostridiaceae_bacterium | 63 | -0.6929243781 | 0.5055735079 |
| EtOH_8pm_intakeave_fina | Clostridiaceae_unclassified_SGB41663 | 63 | 0.8239171355 | 0.4182340115 |
| EtOH_8pm_intakeave_fina | Clostridiales_bacterium | 63 | -0.3205314504 | 0.7567058736 |
| EtOH_8pm_intakeave_fina | Clostridium_cocleatum | 63 | -0.808616513 | 0.4213957656 |
| EtOH_8pm_intakeave_fina | Coriobacteriaceae_bacterium | 63 | -1.115254127 | 0.2940522676 |
| EtOH_8pm_intakeave_fina | Dorea_sp_5_2 | 63 | -0.5971319561 | 0.5943998363 |
| EtOH_8pm_intakeave_fina | Dubosiella_newyorkensis | 63 | -1.893271667 | 0.06212066834 |
| EtOH_8pm_intakeave_fina | Erysipelotrichales_bacterium | 63 | -0.9622145803 | 0.3381752078 |
| EtOH_8pm_intakeave_fina | Eubacteriaceae_bacterium | 63 | 1.378596061 | 0.1804642666 |
| EtOH_8pm_intakeave_fina | Eubacteriaceae_unclassified_SGB9492 | 63 | -1.086264458 | 0.2836808213 |
| EtOH_8pm_intakeave_fina | GGB20149_SGB29430 | 63 | -0.05203064248 | 0.9586771904 |
| EtOH_8pm_intakeave_fina | GGB22635_SGB63107 | 63 | -1.200575448 | 0.2555200888 |
| EtOH_8pm_intakeave_fina | GGB25041_SGB36960 | 63 | 0.9141274723 | 0.3772106307 |
| EtOH_8pm_intakeave_fina | GGB27876_SGB40310 | 63 | -1.033948322 | 0.3035189707 |
| EtOH_8pm_intakeave_fina | GGB27878_SGB40312 | 63 | 0.120040608 | 0.9101681622 |
| EtOH_8pm_intakeave_fina | GGB27918_SGB40356 | 63 | -0.1192929489 | 0.90902181 |
| EtOH_8pm_intakeave_fina | GGB28382_SGB40962 | 63 | 1.200277712 | 0.2578916345 |
| EtOH_8pm_intakeave_fina | GGB28399_SGB40980 | 63 | 2.045511993 | 0.05020343618 |
| EtOH_8pm_intakeave_fina | GGB28411_SGB40993 | 63 | -0.4784546379 | 0.6336251375 |
| EtOH_8pm_intakeave_fina | GGB28415_SGB40997 | 63 | -0.7385223469 | 0.552000583 |
| EtOH_8pm_intakeave_fina | GGB28430_SGB41013 | 63 | 1.900187735 | 0.1115038421 |
| EtOH_8pm_intakeave_fina | GGB28439_SGB41022 | 63 | 1.63437783 | 0.1412674171 |
| EtOH_8pm_intakeave_fina | GGB28778_SGB41431 | 63 | 1.944177325 | 0.06304099233 |
| EtOH_8pm_intakeave_fina | GGB28784_SGB41437 | 63 | 0.374272144 | 0.7098434603 |
| EtOH_8pm_intakeave_fina | GGB28792_SGB41445 | 63 | 0.7842906854 | 0.4599351279 |
| EtOH_8pm_intakeave_fina | GGB28798_SGB41451 | 63 | -0.2352053671 | 0.8146765248 |
| EtOH_8pm_intakeave_fina | GGB28802_SGB41455 | 63 | -0.1410211783 | 0.8901194529 |
| EtOH_8pm_intakeave_fina | GGB28818_SGB41473 | 63 | -2.130352083 | 0.04126921894 |
| EtOH_8pm_intakeave_fina | GGB28828_SGB41484 | 63 | 0.2458133347 | 0.8100278231 |
| EtOH_8pm_intakeave_fina | GGB28851_SGB41518 | 63 | -0.7611262568 | 0.4705578506 |
| EtOH_8pm_intakeave_fina | GGB28859_SGB41528 | 63 | -0.5408020479 | 0.6011606568 |
| EtOH_8pm_intakeave_fina | GGB28864_SGB41535 | 63 | 0.3465868932 | 0.7307320276 |
| EtOH_8pm_intakeave_fina | GGB28869_SGB41543 | 63 | 0.9403416306 | 0.3767491917 |
| EtOH_8pm_intakeave_fina | GGB28883_SGB41564 | 63 | -1.369408093 | 0.2008001019 |
| EtOH_8pm_intakeave_fina | GGB28892_SGB41573 | 63 | -1.295178873 | 0.2558959723 |
| EtOH_8pm_intakeave_fina | GGB28893_SGB41574 | 63 | -0.9739814581 | 0.3409365904 |
| EtOH_8pm_intakeave_fina | GGB28898_SGB41580 | 63 | -1.23000593 | 0.221682008 |
| EtOH_8pm_intakeave_fina | GGB28904_SGB41597 | 63 | 0.8354969696 | 0.4060494568 |
| EtOH_8pm_intakeave_fina | GGB28916_SGB41612 | 63 | 0.5822121123 | 0.5612309939 |
| EtOH_8pm_intakeave_fina | GGB28924_SGB41621 | 63 | 0.9410525387 | 0.4221711435 |
| EtOH_8pm_intakeave_fina | GGB28926_SGB41624 | 63 | -0.7530036385 | 0.4525161498 |
| EtOH_8pm_intakeave_fina | GGB28927_SGB41625 | 63 | 1.798080347 | 0.0793857384 |

| | | | | |
|---|---|---|---|---|
| EtOH_8pm_intakeave_fina | GGB28934_SGB41635 | 63 | 0.3874243369 | 0.6994375182 |
| EtOH_8pm_intakeave_fina | GGB28946_SGB41652 | 63 | 1.624309639 | 0.1125450451 |
| EtOH_8pm_intakeave_fina | GGB28949_SGB41655 | 63 | -1.433323615 | 0.1580025882 |
| EtOH_8pm_intakeave_fina | GGB28949_SGB41656 | 63 | 0.4917894739 | 0.6242686649 |
| EtOH_8pm_intakeave_fina | GGB28950_SGB41657 | 63 | -1.489340947 | 0.1506172199 |
| EtOH_8pm_intakeave_fina | GGB28951_SGB102295 | 63 | 1.179943358 | 0.2801556966 |
| EtOH_8pm_intakeave_fina | GGB28951_SGB41658 | 63 | 2.725497796 | 0.00858195644 |
| EtOH_8pm_intakeave_fina | GGB28954_SGB41662 | 63 | 0.6012426565 | 0.6043333808 |
| EtOH_8pm_intakeave_fina | GGB28956_SGB41665 | 63 | 1.468584273 | 0.1670902269 |
| EtOH_8pm_intakeave_fina | GGB28960_SGB41669 | 63 | 1.892301327 | 0.06254288138 |
| EtOH_8pm_intakeave_fina | GGB28967_SGB41678 | 63 | 1.370579351 | 0.1885998892 |
| EtOH_8pm_intakeave_fina | GGB28991_SGB41705 | 63 | -1.200696279 | 0.236512847 |
| EtOH_8pm_intakeave_fina | GGB29002_SGB41718 | 63 | 0.8462623819 | 0.3996561123 |
| EtOH_8pm_intakeave_fina | GGB29003_SGB41719 | 63 | -0.4535049257 | 0.6573454362 |
| EtOH_8pm_intakeave_fina | GGB29011_SGB41731 | 63 | -1.286809445 | 0.2041102158 |
| EtOH_8pm_intakeave_fina | GGB29531_SGB42317 | 63 | 3.347517278 | 0.00192898841 |
| EtOH_8pm_intakeave_fina | GGB29685_SGB42494 | 63 | -2.785511015 | 0.00984696801 |
| EtOH_8pm_intakeave_fina | GGB30141_SGB43066 | 63 | -0.399380372 | 0.7409062324 |
| EtOH_8pm_intakeave_fina | GGB30286_SGB43248 | 63 | -1.355290016 | 0.206012049 |
| EtOH_8pm_intakeave_fina | GGB30303_SGB43268 | 63 | -0.7084879117 | 0.4894786549 |
| EtOH_8pm_intakeave_fina | GGB30413_SGB43452 | 63 | 0.582121701 | 0.5721068156 |
| EtOH_8pm_intakeave_fina | GGB30454_SGB43514 | 63 | -0.8233265133 | 0.4128011557 |
| EtOH_8pm_intakeave_fina | GGB30455_SGB43519 | 63 | -0.3009936403 | 0.7663632045 |
| EtOH_8pm_intakeave_fina | GGB30461_SGB43527 | 63 | -1.423712348 | 0.1628818294 |
| EtOH_8pm_intakeave_fina | GGB30461_SGB43530 | 63 | -0.4396963662 | 0.6906625349 |
| EtOH_8pm_intakeave_fina | GGB30463_SGB43537 | 63 | 0.1110881217 | 0.9176070924 |
| EtOH_8pm_intakeave_fina | GGB30473_SGB43557 | 63 | -0.0180628908 | 0.9857401854 |
| EtOH_8pm_intakeave_fina | GGB30475_SGB63182 | 63 | -0.7060612878 | 0.5105386436 |
| EtOH_8pm_intakeave_fina | GGB30861_SGB44083 | 63 | -1.379488896 | 0.1755293023 |
| EtOH_8pm_intakeave_fina | GGB31312_SGB44628 | 63 | -0.5215756988 | 0.6067032805 |
| EtOH_8pm_intakeave_fina | GGB31438_SGB44768 | 63 | -1.107392309 | 0.4079368044 |
| EtOH_8pm_intakeave_fina | GGB3171_SGB4185 | 63 | 0.113938486 | 0.9093433478 |
| EtOH_8pm_intakeave_fina | GGB31823_SGB45199 | 63 | -0.2801433176 | 0.7794655299 |
| EtOH_8pm_intakeave_fina | GGB31853_SGB45233 | 63 | 1.571170099 | 0.1451817329 |
| EtOH_8pm_intakeave_fina | GGB32371_SGB41694 | 63 | -0.7542431616 | 0.4555430904 |
| EtOH_8pm_intakeave_fina | GGB3793_SGB5158 | 63 | -1.369548524 | 0.1761896911 |
| EtOH_8pm_intakeave_fina | GGB42598_SGB59794 | 63 | 0.7613088249 | 0.4825346275 |
| EtOH_8pm_intakeave_fina | GGB45656_SGB63370 | 63 | 0.8363380493 | 0.4088445512 |
| EtOH_8pm_intakeave_fina | GGB47127_SGB65054 | 63 | -1.914754799 | 0.09870239831 |
| EtOH_8pm_intakeave_fina | GGB74395_SGB43521 | 63 | -0.4276743401 | 0.6878371447 |
| EtOH_8pm_intakeave_fina | GGB75053_SGB43494 | 63 | -0.3808041501 | 0.7071725039 |
| EtOH_8pm_intakeave_fina | GGB75109_SGB102238 | 63 | -2.144240706 | 0.03976246471 |
| EtOH_8pm_intakeave_fina | GGB81440_SGB45230 | 63 | 0.9337425203 | 0.3859456233 |
| EtOH_8pm_intakeave_fina | Lachnospiraceae_bacterium | 63 | -0.9162059812 | 0.3632274278 |
| EtOH_8pm_intakeave_fina | Lachnospiraceae_bacterium_A2 | 63 | 0.1481153175 | 0.8824398251 |
| EtOH_8pm_intakeave_fina | Lachnospiraceae_bacterium_MD308 | 63 | -1.592137369 | 0.1316625572 |

| | | | | |
|---|---|---|---|---|
| EtOH_8pm_intakeave_fina | Lachnospiraceae_bacterium_MD329 | 63 | -1.295684678 | 0.2006763306 |
| EtOH_8pm_intakeave_fina | Lachnospiraceae_unclassified_SGB4141 | 63 | 1.78788985 | 0.08834442656 |
| EtOH_8pm_intakeave_fina | Lachnospiraceae_unclassified_SGB4141 | 63 | 0.8302654844 | 0.4102784028 |
| EtOH_8pm_intakeave_fina | Lachnospiraceae_unclassified_SGB4142 | 63 | 0.9993086844 | 0.3225004134 |
| EtOH_8pm_intakeave_fina | Lachnospiraceae_unclassified_SGB4158 | 63 | 1.151379959 | 0.2939987945 |
| EtOH_8pm_intakeave_fina | Lactobacillus_johnsonii | 63 | -0.3332841997 | 0.7413871316 |
| EtOH_8pm_intakeave_fina | Muribaculaceae_bacterium | 63 | -1.857285589 | 0.06723337548 |
| EtOH_8pm_intakeave_fina | Neglectibacter_sp_X4 | 63 | 2.476351109 | 0.01766820147 |
| EtOH_8pm_intakeave_fina | Oscillospiraceae_bacterium | 63 | 0.6188171623 | 0.6150398682 |
| EtOH_8pm_intakeave_fina | Oscillospiraceae_unclassified_SGB4350 | 63 | 1.069605757 | 0.2984063192 |
| EtOH_8pm_intakeave_fina | Oscillospiraceae_unclassified_SGB4350 | 63 | 0.99480151 | 0.3547664904 |
| EtOH_8pm_intakeave_fina | Parasutterella_excrementihominis | 63 | 0.2652419208 | 0.8230381162 |
| EtOH_8pm_intakeave_fina | Romboutsia_ilealis | 63 | 1.093903318 | 0.2858626563 |
| EtOH_8pm_intakeave_fina | Schaedlerella_arabinosiphila | 63 | -1.247471065 | 0.275353565 |
| EtOH_8pm_intakeave_fina | Turicibacter_sp_1E2 | 63 | -0.7201506595 | 0.4801366629 |
| EtOH_8pm_intakeave_fina | bacterium_1XD42_54 | 63 | -0.8551736252 | 0.3941583251 |
| EtOH_8pm_intakeave_fina | bacterium_1XD42_76 | 63 | -1.383149714 | 0.1710571147 |
| EtOH_8pm_intakeave_fina | bacterium_1xD8_48 | 63 | 0.3618565104 | 0.7196012766 |
| EtOH_8pm_intakeave_fina | Berger Parker Index | 63 | -1.366413049 | 0.1761792057 |
| EtOH_8pm_intakeave_fina | Richness (# observed features) | 63 | 1.206798294 | 0.2461953866 |
| EtOH_8pm_intakeave_fina | Shannon Index | 63 | 1.154150513 | 0.2511436448 |
| EtOH_pref_24h_final | Acetatifactor_SGB41546 | 63 | 1.751176327 | 0.08373762376 |
| EtOH_pref_24h_final | Acetatifactor_muris | 63 | 1.017169183 | 0.3111439081 |
| EtOH_pref_24h_final | Acutalibacter_muris | 63 | 1.433737522 | 0.1551190807 |
| EtOH_pref_24h_final | Acutalibacter_sp_1XD8_36 | 63 | -0.09993490134 | 0.920399309 |
| EtOH_pref_24h_final | Adlercreutzia_caecimuris | 63 | -2.92050843 | 0.004742590723 |
| EtOH_pref_24h_final | Adlercreutzia_mucosicola | 63 | 0.6495023356 | 0.5167488902 |
| EtOH_pref_24h_final | Adlercreutzia_muris | 63 | 0.7655451132 | 0.4450546575 |
| EtOH_pref_24h_final | Akkermansia_muciniphila | 63 | -1.392862884 | 0.1670341845 |
| EtOH_pref_24h_final | Alistipes_sp_DSM_112343 | 63 | -2.788241066 | 0.006856048457 |
| EtOH_pref_24h_final | Anaerotruncus_sp_1XD42_93 | 63 | 0.9318008048 | 0.3531710923 |
| EtOH_pref_24h_final | Bacteria_unclassified_SGB102200 | 63 | 0.2451008255 | 0.8064258411 |
| EtOH_pref_24h_final | Bacteria_unclassified_SGB41677 | 63 | -0.6806634993 | 0.4969129587 |
| EtOH_pref_24h_final | Bacteria_unclassified_SGB43546 | 63 | 0.3483179185 | 0.727733522 |
| EtOH_pref_24h_final | Bacteroides_thetaiotaomicron | 63 | -1.616880923 | 0.1096713177 |
| EtOH_pref_24h_final | Bifidobacterium_pseudolongum | 63 | -1.621208574 | 0.1087441566 |
| EtOH_pref_24h_final | Clostridia_bacterium | 63 | -2.085255142 | 0.04047295056 |
| EtOH_pref_24h_final | Clostridiaceae_bacterium | 63 | -3.998267895 | 0.000196028189 |
| EtOH_pref_24h_final | Clostridiaceae_unclassified_SGB41663 | 63 | 0.2986012269 | 0.7653289155 |
| EtOH_pref_24h_final | Clostridiales_bacterium | 63 | 2.338330966 | 0.02216452052 |
| EtOH_pref_24h_final | Clostridium_cocleatum | 63 | 0.5828261859 | 0.5605641232 |
| EtOH_pref_24h_final | Coriobacteriaceae_bacterium | 63 | -1.804773612 | 0.07491658018 |
| EtOH_pref_24h_final | Dorea_sp_5_2 | 63 | 0.5962303693 | 0.5516095073 |
| EtOH_pref_24h_final | Dubosiella_newyorkensis | 63 | -1.71361528 | 0.09041991597 |
| EtOH_pref_24h_final | Erysipelotrichales_bacterium | 63 | -0.1223171371 | 0.9026538983 |
| EtOH_pref_24h_final | Eubacteriaceae_bacterium | 63 | -0.9536021644 | 0.3421029335 |

| | | | | |
|---|---|---|---|---|
| EtOH_pref_24h_final | Eubacteriaceae_unclassified_SGB9492 | 63 | -1.256134705 | 0.2120340334 |
| EtOH_pref_24h_final | GGB20149_SGB29430 | 63 | 0.7716789504 | 0.4414336778 |
| EtOH_pref_24h_final | GGB22635_SGB63107 | 63 | -0.2443051781 | 0.8070414992 |
| EtOH_pref_24h_final | GGB25041_SGB36960 | 63 | -0.1953985918 | 0.845105244 |
| EtOH_pref_24h_final | GGB27876_SGB40310 | 63 | -1.094733476 | 0.2760104692 |
| EtOH_pref_24h_final | GGB27878_SGB40312 | 63 | -1.543672341 | 0.1263471145 |
| EtOH_pref_24h_final | GGB27918_SGB40356 | 63 | -0.8529118028 | 0.395134193 |
| EtOH_pref_24h_final | GGB28382_SGB40962 | 63 | 1.805631742 | 0.07478192925 |
| EtOH_pref_24h_final | GGB28399_SGB40980 | 63 | 0.4272480062 | 0.6694349891 |
| EtOH_pref_24h_final | GGB28411_SGB40993 | 63 | 0.3684061778 | 0.7127255313 |
| EtOH_pref_24h_final | GGB28415_SGB40997 | 63 | -0.8772940625 | 0.3818459716 |
| EtOH_pref_24h_final | GGB28430_SGB41013 | 63 | 2.527416394 | 0.01374849 |
| EtOH_pref_24h_final | GGB28439_SGB41022 | 63 | 1.721569809 | 0.08896919664 |
| EtOH_pref_24h_final | GGB28778_SGB41431 | 63 | 0.5400843377 | 0.5895904247 |
| EtOH_pref_24h_final | GGB28784_SGB41437 | 63 | 0.2713496811 | 0.7861860792 |
| EtOH_pref_24h_final | GGB28792_SGB41445 | 63 | -0.04046783737 | 0.9677203682 |
| EtOH_pref_24h_final | GGB28798_SGB41451 | 63 | 0.0302135507 | 0.9758968332 |
| EtOH_pref_24h_final | GGB28802_SGB41455 | 63 | -0.6968714774 | 0.4867623492 |
| EtOH_pref_24h_final | GGB28818_SGB41473 | 63 | 0.8969495152 | 0.3713409178 |
| EtOH_pref_24h_final | GGB28828_SGB41484 | 63 | -0.7849891372 | 0.4336360458 |
| EtOH_pref_24h_final | GGB28851_SGB41518 | 63 | -0.3020771661 | 0.7626807204 |
| EtOH_pref_24h_final | GGB28859_SGB41528 | 63 | -0.07498696339 | 0.9402264822 |
| EtOH_pref_24h_final | GGB28864_SGB41535 | 63 | 1.69283122 | 0.094302699 |
| EtOH_pref_24h_final | GGB28869_SGB41543 | 63 | 1.81819045 | 0.07283438948 |
| EtOH_pref_24h_final | GGB28883_SGB41564 | 63 | -0.8127464924 | 0.4176397691 |
| EtOH_pref_24h_final | GGB28892_SGB41573 | 63 | 0.03155994124 | 0.9748230942 |
| EtOH_pref_24h_final | GGB28893_SGB41574 | 63 | 1.128033666 | 0.2618111392 |
| EtOH_pref_24h_final | GGB28898_SGB41580 | 63 | -1.650832592 | 0.1025669129 |
| EtOH_pref_24h_final | GGB28904_SGB41597 | 63 | 1.491448342 | 0.1394356592 |
| EtOH_pref_24h_final | GGB28916_SGB41612 | 63 | 1.272509265 | 0.2062131865 |
| EtOH_pref_24h_final | GGB28924_SGB41621 | 63 | 0.9509860055 | 0.3434189772 |
| EtOH_pref_24h_final | GGB28926_SGB41624 | 63 | -1.087953313 | 0.278966344 |
| EtOH_pref_24h_final | GGB28927_SGB41625 | 63 | 1.063151096 | 0.2899667358 |
| EtOH_pref_24h_final | GGB28934_SGB41635 | 63 | 0.4712342775 | 0.6377841813 |
| EtOH_pref_24h_final | GGB28946_SGB41652 | 63 | -0.5736161742 | 0.5667583928 |
| EtOH_pref_24h_final | GGB28949_SGB41655 | 63 | -1.909184008 | 0.05995760799 |
| EtOH_pref_24h_final | GGB28949_SGB41656 | 63 | -1.600186346 | 0.1133081415 |
| EtOH_pref_24h_final | GGB28950_SGB41657 | 63 | -0.9194252273 | 0.3595561041 |
| EtOH_pref_24h_final | GGB28951_SGB102295 | 63 | -0.616696723 | 0.5380774745 |
| EtOH_pref_24h_final | GGB28951_SGB41658 | 63 | 1.250205144 | 0.214171622 |
| EtOH_pref_24h_final | GGB28954_SGB41662 | 63 | -0.8980882296 | 0.3707380036 |
| EtOH_pref_24h_final | GGB28956_SGB41665 | 63 | -0.08155608127 | 0.9350015243 |
| EtOH_pref_24h_final | GGB28960_SGB41669 | 63 | 0.2071230936 | 0.8359426294 |
| EtOH_pref_24h_final | GGB28967_SGB41678 | 63 | -0.6456072908 | 0.5192575703 |
| EtOH_pref_24h_final | GGB28991_SGB41705 | 63 | -1.305261613 | 0.1949287407 |
| EtOH_pref_24h_final | GGB29002_SGB41718 | 63 | 1.448232299 | 0.1510563449 |

| | | | | |
|---|---|---|---|---|
| EtOH_pref_24h_final | GGB29003_SGB41719 | 63 | 1.17162297 | 0.244017552 |
| EtOH_pref_24h_final | GGB29011_SGB41731 | 63 | -0.879551068 | 0.3806302791 |
| EtOH_pref_24h_final | GGB29531_SGB42317 | 63 | 2.13702054 | 0.03590944366 |
| EtOH_pref_24h_final | GGB29685_SGB42494 | 63 | -2.265910606 | 0.02941465967 |
| EtOH_pref_24h_final | GGB30141_SGB43066 | 63 | 1.177231413 | 0.2417928679 |
| EtOH_pref_24h_final | GGB30286_SGB43248 | 63 | 0.3996824091 | 0.6895861466 |
| EtOH_pref_24h_final | GGB30303_SGB43268 | 63 | 1.703385313 | 0.09231423979 |
| EtOH_pref_24h_final | GGB30413_SGB43452 | 63 | 1.088673914 | 0.2786511469 |
| EtOH_pref_24h_final | GGB30454_SGB43514 | 63 | -1.021874853 | 0.3089298232 |
| EtOH_pref_24h_final | GGB30455_SGB43519 | 63 | 1.21352438 | 0.2277497993 |
| EtOH_pref_24h_final | GGB30461_SGB43527 | 63 | -0.9262237369 | 0.3560393559 |
| EtOH_pref_24h_final | GGB30461_SGB43530 | 63 | 2.18445958 | 0.03212892139 |
| EtOH_pref_24h_final | GGB30463_SGB43537 | 63 | 0.7234963703 | 0.4703400877 |
| EtOH_pref_24h_final | GGB30473_SGB43557 | 63 | -0.1067668188 | 0.9149779944 |
| EtOH_pref_24h_final | GGB30475_SGB63182 | 63 | -0.6750104288 | 0.5004803345 |
| EtOH_pref_24h_final | GGB30861_SGB44083 | 63 | -3.676387516 | 0.000480626704 |
| EtOH_pref_24h_final | GGB31312_SGB44628 | 63 | 0.01966625909 | 0.9843096322 |
| EtOH_pref_24h_final | GGB31438_SGB44768 | 63 | -0.3579983632 | 0.7204874458 |
| EtOH_pref_24h_final | GGB3171_SGB4185 | 63 | 0.5896049393 | 0.5560265887 |
| EtOH_pref_24h_final | GGB31823_SGB45199 | 63 | -0.4499742918 | 0.6530022331 |
| EtOH_pref_24h_final | GGB31853_SGB45233 | 63 | 0.3895341772 | 0.6970629376 |
| EtOH_pref_24h_final | GGB32371_SGB41694 | 63 | -0.5457471836 | 0.5857041875 |
| EtOH_pref_24h_final | GGB3793_SGB5158 | 63 | -1.015149119 | 0.3120976606 |
| EtOH_pref_24h_final | GGB42598_SGB59794 | 63 | -1.218994423 | 0.2256859588 |
| EtOH_pref_24h_final | GGB45656_SGB63370 | 63 | 0.8574914469 | 0.3926166791 |
| EtOH_pref_24h_final | GGB47127_SGB65054 | 63 | -0.8316530624 | 0.4069509619 |
| EtOH_pref_24h_final | GGB74395_SGB43521 | 63 | 0.1841123181 | 0.8539457364 |
| EtOH_pref_24h_final | GGB75053_SGB43494 | 63 | 0.7492182065 | 0.4547770632 |
| EtOH_pref_24h_final | GGB75109_SGB102238 | 63 | -1.166353808 | 0.2461210438 |
| EtOH_pref_24h_final | GGB81440_SGB45230 | 63 | 1.105586093 | 0.2713248738 |
| EtOH_pref_24h_final | Lachnospiraceae_bacterium | 63 | -2.92523088 | 0.004679709201 |
| EtOH_pref_24h_final | Lachnospiraceae_bacterium_A2 | 63 | -1.219159133 | 0.2256240268 |
| EtOH_pref_24h_final | Lachnospiraceae_bacterium_MD308 | 63 | -1.850212251 | 0.06806036313 |
| EtOH_pref_24h_final | Lachnospiraceae_bacterium_MD329 | 63 | -1.673316022 | 0.09807214061 |
| EtOH_pref_24h_final | Lachnospiraceae_unclassified_SGB4141 | 63 | -0.3769841354 | 0.7063510859 |
| EtOH_pref_24h_final | Lachnospiraceae_unclassified_SGB4141 | 63 | 0.216592729 | 0.8285587358 |
| EtOH_pref_24h_final | Lachnospiraceae_unclassified_SGB414[illegible] | 63 | 0.1241997086 | 0.9011634724 |
| EtOH_pref_24h_final | Lachnospiraceae_unclassified_SGB415[illegible] | 63 | -0.2127910847 | 0.8315212206 |
| EtOH_pref_24h_final | Lactobacillus_johnsonii | 63 | -0.5711781684 | 0.5684037073 |
| EtOH_pref_24h_final | Muribaculaceae_bacterium | 63 | -2.067582716 | 0.04214238537 |
| EtOH_pref_24h_final | Neglectibacter_sp_X4 | 63 | 1.227912478 | 0.2223505874 |
| EtOH_pref_24h_final | Oscillospiraceae_bacterium | 63 | -0.1211104707 | 0.9036093976 |
| EtOH_pref_24h_final | Oscillospiraceae_unclassified_SGB4350 | 63 | -1.375765188 | 0.1722225853 |
| EtOH_pref_24h_final | Oscillospiraceae_unclassified_SGB4350 | 63 | 0.944530789 | 0.3466803967 |
| EtOH_pref_24h_final | Parasutterella_excrementihominis | 63 | -0.6590260988 | 0.510642139 |
| EtOH_pref_24h_final | Romboutsia_ilealis | 63 | 0.5636881863 | 0.5734729949 |

| | | | | |
|---|---|---|---|---|
| EtOH_pref_24h_final | Schaedlerella_arabinosiphila | 63 | -0.246612252 | 0.8052566644 |
| EtOH_pref_24h_final | Turicibacter_sp_1E2 | 63 | -1.002417467 | 0.3181541915 |
| EtOH_pref_24h_final | bacterium_1XD42_54 | 63 | -2.870313725 | 0.00546112408 |
| EtOH_pref_24h_final | bacterium_1XD42_76 | 63 | -0.6066067291 | 0.5447274353 |
| EtOH_pref_24h_final | bacterium_1xD8_48 | 63 | 1.59729399 | 0.1139480152 |
| EtOH_pref_24h_final | Berger Parker Index | 63 | -1.114973078 | 0.2673173421 |
| EtOH_pref_24h_final | Richness (# observed features) | 63 | 1.670838133 | 0.0985594638 |
| EtOH_pref_24h_final | Shannon Index | 63 | 0.9570894165 | 0.3403538395 |
| OFB_Cdistance | Acetatifactor_SGB41546 | 63 | -1.097024985 | 0.2749286589 |
| OFB_Cdistance | Acetatifactor_muris | 63 | -0.364502659 | 0.7167870711 |
| OFB_Cdistance | Acutalibacter_muris | 63 | -0.935597239 | 0.3538994529 |
| OFB_Cdistance | Acutalibacter_sp_1XD8_36 | 63 | 1.788433794 | 0.07734809396 |
| OFB_Cdistance | Adlercreutzia_caecimuris | 63 | -1.439504952 | 0.1538346139 |
| OFB_Cdistance | Adlercreutzia_mucosicola | 63 | 1.2148755 | 0.2302147754 |
| OFB_Cdistance | Adlercreutzia_muris | 63 | 0.3180812294 | 0.7505211343 |
| OFB_Cdistance | Akkermansia_muciniphila | 63 | 0.04891594121 | 0.9611995529 |
| OFB_Cdistance | Alistipes_sp_DSM_112343 | 63 | -0.3400559013 | 0.7340145132 |
| OFB_Cdistance | Anaerotruncus_sp_1XD42_93 | 63 | 0.4097437377 | 0.6855134038 |
| OFB_Cdistance | Bacteria_unclassified_SGB102200 | 63 | -0.1333212018 | 0.8957179311 |
| OFB_Cdistance | Bacteria_unclassified_SGB41677 | 63 | -0.87194838 | 0.4041342254 |
| OFB_Cdistance | Bacteria_unclassified_SGB43546 | 63 | -0.3201214809 | 0.7587180738 |
| OFB_Cdistance | Bacteroides_thetaiotaomicron | 63 | -1.569014562 | 0.1201889326 |
| OFB_Cdistance | Bifidobacterium_pseudolongum | 63 | -0.7597564102 | 0.449135987 |
| OFB_Cdistance | Clostridia_bacterium | 63 | -1.460504214 | 0.149662105 |
| OFB_Cdistance | Clostridiaceae_bacterium | 63 | -0.782278713 | 0.4367078534 |
| OFB_Cdistance | Clostridiaceae_unclassified_SGB41663 | 63 | 0.02834144652 | 0.9777956471 |
| OFB_Cdistance | Clostridiales_bacterium | 63 | 0.8637944696 | 0.3921130335 |
| OFB_Cdistance | Clostridium_cocleatum | 63 | 0.3728638779 | 0.7126632451 |
| OFB_Cdistance | Coriobacteriaceae_bacterium | 63 | 0.5826937517 | 0.5609699392 |
| OFB_Cdistance | Dorea_sp_5_2 | 63 | 1.676551846 | 0.1020232262 |
| OFB_Cdistance | Dubosiella_newyorkensis | 63 | -0.985712763 | 0.3408195587 |
| OFB_Cdistance | Erysipelotrichales_bacterium | 63 | 1.14208663 | 0.2666311715 |
| OFB_Cdistance | Eubacteriaceae_bacterium | 63 | 0.2694487275 | 0.7908067584 |
| OFB_Cdistance | Eubacteriaceae_unclassified_SGB9492 | 63 | -2.34252918 | 0.02193379013 |
| OFB_Cdistance | GGB20149_SGB29430 | 63 | -0.6234549376 | 0.538601681 |
| OFB_Cdistance | GGB22635_SGB63107 | 63 | 0.2264119339 | 0.8219763488 |
| OFB_Cdistance | GGB25041_SGB36960 | 63 | -0.6457937805 | 0.5212561595 |
| OFB_Cdistance | GGB27876_SGB40310 | 63 | 0.2923213017 | 0.7765272052 |
| OFB_Cdistance | GGB27878_SGB40312 | 63 | -0.1100815996 | 0.9174226806 |
| OFB_Cdistance | GGB27918_SGB40356 | 63 | -0.9514525948 | 0.3637891233 |
| OFB_Cdistance | GGB28382_SGB40962 | 63 | 1.634818187 | 0.1187017497 |
| OFB_Cdistance | GGB28399_SGB40980 | 63 | -0.1065674057 | 0.9158616859 |
| OFB_Cdistance | GGB28411_SGB40993 | 63 | -0.006325946417 | 0.9950381856 |
| OFB_Cdistance | GGB28415_SGB40997 | 63 | 0.3096532356 | 0.786140247 |
| OFB_Cdistance | GGB28430_SGB41013 | 63 | -0.2912927275 | 0.7747870695 |
| OFB_Cdistance | GGB28439_SGB41022 | 63 | -0.04583883718 | 0.9642736767 |

| | | | | |
|---|---|---|---|---|
| OFB_Cdistance | GGB28778_SGB41431 | 63 | 0.6909227964 | 0.4909007063 |
| OFB_Cdistance | GGB28784_SGB41437 | 63 | -1.392513287 | 0.170848058 |
| OFB_Cdistance | GGB28792_SGB41445 | 63 | -0.6698390459 | 0.5418389817 |
| OFB_Cdistance | GGB28798_SGB41451 | 63 | -1.121923625 | 0.2661967843 |
| OFB_Cdistance | GGB28802_SGB41455 | 63 | -1.60169579 | 0.1272459527 |
| OFB_Cdistance | GGB28818_SGB41473 | 63 | 0.05919818903 | 0.9530790465 |
| OFB_Cdistance | GGB28828_SGB41484 | 63 | 1.061425841 | 0.3148378525 |
| OFB_Cdistance | GGB28851_SGB41518 | 63 | -0.03709672199 | 0.9709621768 |
| OFB_Cdistance | GGB28859_SGB41528 | 63 | 0.4475457881 | 0.6748791925 |
| OFB_Cdistance | GGB28864_SGB41535 | 63 | 0.7123777113 | 0.4975642384 |
| OFB_Cdistance | GGB28869_SGB41543 | 63 | -0.8044450329 | 0.4223945817 |
| OFB_Cdistance | GGB28883_SGB41564 | 63 | -1.809189771 | 0.07598617036 |
| OFB_Cdistance | GGB28892_SGB41573 | 63 | -1.288939892 | 0.2098522383 |
| OFB_Cdistance | GGB28893_SGB41574 | 63 | 0.7416334263 | 0.4741756048 |
| OFB_Cdistance | GGB28898_SGB41580 | 63 | 0.895830871 | 0.3811568824 |
| OFB_Cdistance | GGB28904_SGB41597 | 63 | 0.933545938 | 0.3599627258 |
| OFB_Cdistance | GGB28916_SGB41612 | 63 | 0.8387661182 | 0.4046763302 |
| OFB_Cdistance | GGB28924_SGB41621 | 63 | -0.08336053142 | 0.9380755815 |
| OFB_Cdistance | GGB28926_SGB41624 | 63 | -0.4103162265 | 0.6817815824 |
| OFB_Cdistance | GGB28927_SGB41625 | 63 | 1.662381646 | 0.1022900959 |
| OFB_Cdistance | GGB28934_SGB41635 | 63 | -0.208288771 | 0.8441291442 |
| OFB_Cdistance | GGB28946_SGB41652 | 63 | -0.3405982232 | 0.7423370841 |
| OFB_Cdistance | GGB28949_SGB41655 | 63 | -0.1914454719 | 0.8483914157 |
| OFB_Cdistance | GGB28949_SGB41656 | 63 | 0.718665579 | 0.4815249552 |
| OFB_Cdistance | GGB28950_SGB41657 | 63 | -0.3757757144 | 0.7173287235 |
| OFB_Cdistance | GGB28951_SGB102295 | 63 | -0.2157280117 | 0.8309215067 |
| OFB_Cdistance | GGB28951_SGB41658 | 63 | 0.1298599249 | 0.9001420322 |
| OFB_Cdistance | GGB28954_SGB41662 | 63 | 0.6960902843 | 0.4902709633 |
| OFB_Cdistance | GGB28956_SGB41665 | 63 | 0.7116834037 | 0.4803977213 |
| OFB_Cdistance | GGB28960_SGB41669 | 63 | 0.8544705931 | 0.3969190854 |
| OFB_Cdistance | GGB28967_SGB41678 | 63 | 0.1844274566 | 0.8542504906 |
| OFB_Cdistance | GGB28991_SGB41705 | 63 | 0.1851001282 | 0.8546185524 |
| OFB_Cdistance | GGB29002_SGB41718 | 63 | 1.666375204 | 0.09949646307 |
| OFB_Cdistance | GGB29003_SGB41719 | 63 | 0.5636723936 | 0.5798437058 |
| OFB_Cdistance | GGB29011_SGB41731 | 63 | -1.429450119 | 0.1590573472 |
| OFB_Cdistance | GGB29531_SGB42317 | 63 | 2.115769879 | 0.03829558788 |
| OFB_Cdistance | GGB29685_SGB42494 | 63 | -1.201453321 | 0.2324815169 |
| OFB_Cdistance | GGB30141_SGB43066 | 63 | -0.1349001277 | 0.8933973747 |
| OFB_Cdistance | GGB30286_SGB43248 | 63 | 0.7868163845 | 0.4332611423 |
| OFB_Cdistance | GGB30303_SGB43268 | 63 | -0.1877435905 | 0.8530452683 |
| OFB_Cdistance | GGB30413_SGB43452 | 63 | 0.6120615268 | 0.5417115591 |
| OFB_Cdistance | GGB30454_SGB43514 | 63 | -0.2763535252 | 0.7825333942 |
| OFB_Cdistance | GGB30455_SGB43519 | 63 | -0.3252580793 | 0.7503977535 |
| OFB_Cdistance | GGB30461_SGB43527 | 63 | 0.4218615548 | 0.6742251546 |
| OFB_Cdistance | GGB30461_SGB43530 | 63 | -0.8836066664 | 0.3951533267 |
| OFB_Cdistance | GGB30463_SGB43537 | 63 | 0.3073879663 | 0.7613290681 |

| | | | | |
|---|---|---|---|---|
| OFB_Cdistance | GGB30473_SGB43557 | 63 | -0.1114972326 | 0.9117231045 |
| OFB_Cdistance | GGB30475_SGB63182 | 63 | -0.460464426 | 0.6456022808 |
| OFB_Cdistance | GGB30861_SGB44083 | 63 | -1.710939725 | 0.09153320834 |
| OFB_Cdistance | GGB31312_SGB44628 | 63 | -1.362975586 | 0.1783220474 |
| OFB_Cdistance | GGB31438_SGB44768 | 63 | -0.3391168461 | 0.7388357455 |
| OFB_Cdistance | GGB3171_SGB4185 | 63 | 0.1299123722 | 0.8997336364 |
| OFB_Cdistance | GGB31823_SGB45199 | 63 | 1.161624854 | 0.247931715 |
| OFB_Cdistance | GGB31853_SGB45233 | 63 | 0.1161639934 | 0.9077587857 |
| OFB_Cdistance | GGB32371_SGB41694 | 63 | 0.8636173458 | 0.3998182216 |
| OFB_Cdistance | GGB3793_SGB5158 | 63 | 0.5095594259 | 0.611444888 |
| OFB_Cdistance | GGB42598_SGB59794 | 63 | -0.08895521968 | 0.9294081678 |
| OFB_Cdistance | GGB45656_SGB63370 | 63 | 1.899893571 | 0.07652807929 |
| OFB_Cdistance | GGB47127_SGB65054 | 63 | -0.8302255858 | 0.4076980441 |
| OFB_Cdistance | GGB74395_SGB43521 | 63 | -1.24102366 | 0.2253548287 |
| OFB_Cdistance | GGB75053_SGB43494 | 63 | -0.03923096588 | 0.9687153911 |
| OFB_Cdistance | GGB75109_SGB102238 | 63 | -0.3749413091 | 0.7080024591 |
| OFB_Cdistance | GGB81440_SGB45230 | 63 | 0.6193845561 | 0.5554564783 |
| OFB_Cdistance | Lachnospiraceae_bacterium | 63 | -1.814260349 | 0.07330615937 |
| OFB_Cdistance | Lachnospiraceae_bacterium_A2 | 63 | 0.4854578724 | 0.6283248353 |
| OFB_Cdistance | Lachnospiraceae_bacterium_MD308 | 63 | 0.2566028034 | 0.7975450374 |
| OFB_Cdistance | Lachnospiraceae_bacterium_MD329 | 63 | -0.8831216906 | 0.3887480883 |
| OFB_Cdistance | Lachnospiraceae_unclassified_SGB4141 | 63 | -0.7373807906 | 0.4637614235 |
| OFB_Cdistance | Lachnospiraceae_unclassified_SGB4141 | 63 | 0.1288308141 | 0.8993058601 |
| OFB_Cdistance | Lachnospiraceae_unclassified_SGB414 | 63 | -0.1056773252 | 0.9160896425 |
| OFB_Cdistance | Lachnospiraceae_unclassified_SGB415 | 63 | -1.158351269 | 0.2598364335 |
| OFB_Cdistance | Lactobacillus_johnsonii | 63 | 0.253416087 | 0.8005875404 |
| OFB_Cdistance | Muribaculaceae_bacterium | 63 | 0.1811217116 | 0.8576935066 |
| OFB_Cdistance | Neglectibacter_sp_X4 | 63 | -0.1542307716 | 0.8793372923 |
| OFB_Cdistance | Oscillospiraceae_bacterium | 63 | -1.005684608 | 0.330391997 |
| OFB_Cdistance | Oscillospiraceae_unclassified_SGB4350 | 63 | -0.4527310602 | 0.6516276948 |
| OFB_Cdistance | Oscillospiraceae_unclassified_SGB4350 | 63 | -0.07755818909 | 0.9388809505 |
| OFB_Cdistance | Parasutterella_excrementihominis | 63 | -0.2802371981 | 0.7927857257 |
| OFB_Cdistance | Romboutsia_ilealis | 63 | 0.8278531487 | 0.4287836008 |
| OFB_Cdistance | Schaedlerella_arabinosiphila | 63 | -1.465153727 | 0.1468669442 |
| OFB_Cdistance | Turicibacter_sp_1E2 | 63 | 0.7353280131 | 0.4645353458 |
| OFB_Cdistance | bacterium_1XD42_54 | 63 | -1.183455045 | 0.2478069564 |
| OFB_Cdistance | bacterium_1XD42_76 | 63 | -0.06120582285 | 0.9515012508 |
| OFB_Cdistance | bacterium_1xD8_48 | 63 | 0.675516001 | 0.5001274219 |
| OFB_Cdistance | Berger Parker Index | 63 | -0.272970109 | 0.786498134 |
| OFB_Cdistance | Richness (# observed features) | 63 | -0.0231978421 | 0.9816188679 |
| OFB_Cdistance | Shannon Index | 63 | -0.4578632832 | 0.652558339 |
| OFB_Centries | Acetatifactor_SGB41546 | 63 | -0.9883790317 | 0.3254504833 |
| OFB_Centries | Acetatifactor_muris | 63 | -0.850005856 | 0.3980103038 |
| OFB_Centries | Acutalibacter_muris | 63 | -1.658185522 | 0.1009210554 |
| OFB_Centries | Acutalibacter_sp_1XD8_36 | 63 | 2.006358204 | 0.04842804857 |
| OFB_Centries | Adlercreutzia_caecimuris | 63 | -1.234769352 | 0.2218181338 |

| | | | | |
|---|---|---|---|---|
| OFB_Centries | Adlercreutzia_mucosicola | 63 | 1.322443525 | 0.1941212822 |
| OFB_Centries | Adlercreutzia_muris | 63 | 0.001948676553 | 0.9984451821 |
| OFB_Centries | Akkermansia_muciniphila | 63 | 0.2279890475 | 0.820673657 |
| OFB_Centries | Alistipes_sp_DSM_112343 | 63 | -0.3843455034 | 0.7013828692 |
| OFB_Centries | Anaerotruncus_sp_1XD42_93 | 63 | 0.1934301147 | 0.8474461831 |
| OFB_Centries | Bacteria_unclassified_SGB102200 | 63 | -0.1697715071 | 0.8664072849 |
| OFB_Centries | Bacteria_unclassified_SGB41677 | 63 | -0.0627670092 | 0.9514339953 |
| OFB_Centries | Bacteria_unclassified_SGB43546 | 63 | -0.3091027308 | 0.7637333326 |
| OFB_Centries | Bacteroides_thetaiotaomicron | 63 | -1.792404086 | 0.07690854458 |
| OFB_Centries | Bifidobacterium_pseudolongum | 63 | -0.7573400789 | 0.4503396884 |
| OFB_Centries | Clostridia_bacterium | 63 | -1.29569679 | 0.1988688031 |
| OFB_Centries | Clostridiaceae_bacterium | 63 | -0.7696567185 | 0.4442892504 |
| OFB_Centries | Clostridiaceae_unclassified_SGB41663 | 63 | 0.7089827236 | 0.4849575143 |
| OFB_Centries | Clostridiales_bacterium | 63 | 0.4769450318 | 0.6353706772 |
| OFB_Centries | Clostridium_cocleatum | 63 | 0.271260193 | 0.7889158902 |
| OFB_Centries | Coriobacteriaceae_bacterium | 63 | 0.682719051 | 0.4955796435 |
| OFB_Centries | Dorea_sp_5_2 | 63 | 2.438962422 | 0.02013932312 |
| OFB_Centries | Dubosiella_newyorkensis | 63 | -0.9692207351 | 0.3425541493 |
| OFB_Centries | Erysipelotrichales_bacterium | 63 | 0.797716524 | 0.4318375118 |
| OFB_Centries | Eubacteriaceae_bacterium | 63 | 0.5150091036 | 0.6102973425 |
| OFB_Centries | Eubacteriaceae_unclassified_SGB9492 | 63 | -2.008183076 | 0.05058860908 |
| OFB_Centries | GGB20149_SGB29430 | 63 | -0.2984746279 | 0.7672991753 |
| OFB_Centries | GGB22635_SGB63107 | 63 | 0.02971023131 | 0.9764327095 |
| OFB_Centries | GGB25041_SGB36960 | 63 | -0.2745972593 | 0.7844394328 |
| OFB_Centries | GGB27876_SGB40310 | 63 | 0.6314079839 | 0.5359221948 |
| OFB_Centries | GGB27878_SGB40312 | 63 | 0.00859048758 | 0.9933382066 |
| OFB_Centries | GGB27918_SGB40356 | 63 | -0.5969911749 | 0.5623736048 |
| OFB_Centries | GGB28382_SGB40962 | 63 | 1.667978653 | 0.1092141505 |
| OFB_Centries | GGB28399_SGB40980 | 63 | 0.6708169457 | 0.5073940756 |
| OFB_Centries | GGB28411_SGB40993 | 63 | -0.7264798488 | 0.4727817822 |
| OFB_Centries | GGB28415_SGB40997 | 63 | 0.7548516694 | 0.506814287 |
| OFB_Centries | GGB28430_SGB41013 | 63 | -0.01024536602 | 0.9920240824 |
| OFB_Centries | GGB28439_SGB41022 | 63 | 0.3351785519 | 0.7437019052 |
| OFB_Centries | GGB28778_SGB41431 | 63 | 0.5796343191 | 0.5644169504 |
| OFB_Centries | GGB28784_SGB41437 | 63 | -0.7540780355 | 0.4536086043 |
| OFB_Centries | GGB28792_SGB41445 | 63 | 0.07757868879 | 0.9417744252 |
| OFB_Centries | GGB28798_SGB41451 | 63 | -0.9944653234 | 0.3296945566 |
| OFB_Centries | GGB28802_SGB41455 | 63 | -2.062695059 | 0.04800735004 |
| OFB_Centries | GGB28818_SGB41473 | 63 | -0.5657844778 | 0.5735988937 |
| OFB_Centries | GGB28828_SGB41484 | 63 | 0.9211969477 | 0.3675503245 |
| OFB_Centries | GGB28851_SGB41518 | 63 | -0.08803070152 | 0.9309415629 |
| OFB_Centries | GGB28859_SGB41528 | 63 | -0.7515163584 | 0.4722968931 |
| OFB_Centries | GGB28864_SGB41535 | 63 | 0.8282250566 | 0.4216269442 |
| OFB_Centries | GGB28869_SGB41543 | 63 | -0.6606146823 | 0.5096367801 |
| OFB_Centries | GGB28883_SGB41564 | 63 | -1.377186928 | 0.1769056149 |
| OFB_Centries | GGB28892_SGB41573 | 63 | -1.339383048 | 0.187668077 |

| | | | | |
|---|---|---|---|---|
| OFB_Centries | GGB28893_SGB41574 | 63 | 0.1426920073 | 0.8883058839 |
| OFB_Centries | GGB28898_SGB41580 | 63 | 1.454401013 | 0.1539663919 |
| OFB_Centries | GGB28904_SGB41597 | 63 | 1.266229412 | 0.2150584201 |
| OFB_Centries | GGB28916_SGB41612 | 63 | 0.9037163782 | 0.3693595747 |
| OFB_Centries | GGB28924_SGB41621 | 63 | 0.5884687761 | 0.5827750578 |
| OFB_Centries | GGB28926_SGB41624 | 63 | -0.3357040357 | 0.7372073553 |
| OFB_Centries | GGB28927_SGB41625 | 63 | 0.8538417734 | 0.3947577393 |
| OFB_Centries | GGB28934_SGB41635 | 63 | 0.1885633556 | 0.8551878709 |
| OFB_Centries | GGB28946_SGB41652 | 63 | -0.4295024735 | 0.6745186828 |
| OFB_Centries | GGB28949_SGB41655 | 63 | -0.01067081827 | 0.9914871894 |
| OFB_Centries | GGB28949_SGB41656 | 63 | 0.6356967484 | 0.528757987 |
| OFB_Centries | GGB28950_SGB41657 | 63 | 0.1993954689 | 0.8463085967 |
| OFB_Centries | GGB28951_SGB102295 | 63 | -0.5537939694 | 0.5856450536 |
| OFB_Centries | GGB28951_SGB41658 | 63 | -0.150879981 | 0.8839388217 |
| OFB_Centries | GGB28954_SGB41662 | 63 | 0.6476116797 | 0.5210324892 |
| OFB_Centries | GGB28956_SGB41665 | 63 | 0.7496270444 | 0.4594036035 |
| OFB_Centries | GGB28960_SGB41669 | 63 | 1.029542676 | 0.3062587285 |
| OFB_Centries | GGB28967_SGB41678 | 63 | 0.4265515206 | 0.6713487096 |
| OFB_Centries | GGB28991_SGB41705 | 63 | -0.5372577136 | 0.5940433579 |
| OFB_Centries | GGB29002_SGB41718 | 63 | 1.041976841 | 0.2995524591 |
| OFB_Centries | GGB29003_SGB41719 | 63 | 0.02314461116 | 0.9817261421 |
| OFB_Centries | GGB29011_SGB41731 | 63 | -1.440214665 | 0.1535311437 |
| OFB_Centries | GGB29531_SGB42317 | 63 | 1.401595052 | 0.1643484275 |
| OFB_Centries | GGB29685_SGB42494 | 63 | -1.095563086 | 0.2756902126 |
| OFB_Centries | GGB30141_SGB43066 | 63 | -0.7756900866 | 0.4400937379 |
| OFB_Centries | GGB30286_SGB43248 | 63 | 1.173585658 | 0.2444696704 |
| OFB_Centries | GGB30303_SGB43268 | 63 | -0.541487251 | 0.5915626065 |
| OFB_Centries | GGB30413_SGB43452 | 63 | 0.5475905362 | 0.5863474001 |
| OFB_Centries | GGB30454_SGB43514 | 63 | -0.07497532514 | 0.9402468598 |
| OFB_Centries | GGB30455_SGB43519 | 63 | -0.6833048884 | 0.5006058143 |
| OFB_Centries | GGB30461_SGB43527 | 63 | 0.3754059925 | 0.7086003264 |
| OFB_Centries | GGB30461_SGB43530 | 63 | -1.285991696 | 0.2145052302 |
| OFB_Centries | GGB30463_SGB43537 | 63 | 0.03536406532 | 0.9719788907 |
| OFB_Centries | GGB30473_SGB43557 | 63 | -0.06097043946 | 0.9515138409 |
| OFB_Centries | GGB30475_SGB63182 | 63 | -0.5681751356 | 0.5704957029 |
| OFB_Centries | GGB30861_SGB44083 | 63 | -1.35364785 | 0.1796722657 |
| OFB_Centries | GGB31312_SGB44628 | 63 | -1.986923894 | 0.05142250813 |
| OFB_Centries | GGB31438_SGB44768 | 63 | -0.7209421804 | 0.4739687758 |
| OFB_Centries | GGB3171_SGB4185 | 63 | 0.1664032987 | 0.8694792801 |
| OFB_Centries | GGB31823_SGB45199 | 63 | 0.9638927758 | 0.3369162068 |
| OFB_Centries | GGB31853_SGB45233 | 63 | 0.300435837 | 0.7645504474 |
| OFB_Centries | GGB32371_SGB41694 | 63 | 0.6787498895 | 0.5027900364 |
| OFB_Centries | GGB3793_SGB5158 | 63 | 0.6361050879 | 0.5255081202 |
| OFB_Centries | GGB42598_SGB59794 | 63 | -0.1226115521 | 0.903235538 |
| OFB_Centries | GGB45656_SGB63370 | 63 | 1.123910576 | 0.275829535 |
| OFB_Centries | GGB47127_SGB65054 | 63 | -0.7890300239 | 0.4313710889 |

| | | | | |
|---|---|---|---|---|
| OFB_Centries | GGB74395_SGB43521 | 63 | -1.435326145 | 0.1595204482 |
| OFB_Centries | GGB75053_SGB43494 | 63 | -0.3038188499 | 0.7617779238 |
| OFB_Centries | GGB75109_SGB102238 | 63 | -0.1320512324 | 0.8950041349 |
| OFB_Centries | GGB81440_SGB45230 | 63 | -0.1397754322 | 0.8945518338 |
| OFB_Centries | Lachnospiraceae_bacterium | 63 | -1.8212301 | 0.07242942705 |
| OFB_Centries | Lachnospiraceae_bacterium_A2 | 63 | 0.2275608306 | 0.8200845241 |
| OFB_Centries | Lachnospiraceae_bacterium_MD308 | 63 | 0.1983349124 | 0.8428074744 |
| OFB_Centries | Lachnospiraceae_bacterium_MD329 | 63 | -0.525190769 | 0.6059232348 |
| OFB_Centries | Lachnospiraceae_unclassified_SGB4141 | 63 | -0.3706176629 | 0.7111933046 |
| OFB_Centries | Lachnospiraceae_unclassified_SGB4141 | 63 | 1.487776622 | 0.146464048 |
| OFB_Centries | Lachnospiraceae_unclassified_SGB4142 | 63 | -0.2722164231 | 0.7865459707 |
| OFB_Centries | Lachnospiraceae_unclassified_SGB4158 | 63 | -0.6413915636 | 0.535281593 |
| OFB_Centries | Lactobacillus_johnsonii | 63 | 0.4453642785 | 0.6565358738 |
| OFB_Centries | Muribaculaceae_bacterium | 63 | 0.198701691 | 0.8435811933 |
| OFB_Centries | Neglectibacter_sp_X4 | 63 | -0.1933456013 | 0.8471861896 |
| OFB_Centries | Oscillospiraceae_bacterium | 63 | -1.139855181 | 0.266935165 |
| OFB_Centries | Oscillospiraceae_unclassified_SGB4350 | 63 | -0.03936293769 | 0.9686085963 |
| OFB_Centries | Oscillospiraceae_unclassified_SGB4350 | 63 | 0.09744949691 | 0.9227291823 |
| OFB_Centries | Parasutterella_excrementihominis | 63 | 0.05597449461 | 0.9573671167 |
| OFB_Centries | Romboutsia_ilealis | 63 | 1.311081375 | 0.2079905343 |
| OFB_Centries | Schaedlerella_arabinosiphila | 63 | -1.820760435 | 0.07229956347 |
| OFB_Centries | Turicibacter_sp_1E2 | 63 | 0.9261386811 | 0.3582024716 |
| OFB_Centries | bacterium_1XD42_54 | 63 | -0.6775256837 | 0.5129654016 |
| OFB_Centries | bacterium_1XD42_76 | 63 | -0.7843611841 | 0.4382414261 |
| OFB_Centries | bacterium_1xD8_48 | 63 | 0.3577498674 | 0.7207145956 |
| OFB_Centries | Berger Parker Index | 63 | -0.04757386599 | 0.9623959157 |
| OFB_Centries | Richness (# observed features) | 63 | -0.2733877822 | 0.7852430771 |
| OFB_Centries | Shannon Index | 63 | -0.7925520116 | 0.4354798526 |
| OFB_Clatency | Acetatifactor_SGB41546 | 63 | -0.5871770132 | 0.5577634466 |
| OFB_Clatency | Acetatifactor_muris | 63 | 0.7820176093 | 0.4384333551 |
| OFB_Clatency | Acutalibacter_muris | 63 | -0.0319673238 | 0.9747726878 |
| OFB_Clatency | Acutalibacter_sp_1XD8_36 | 63 | -0.2332956461 | 0.8159901459 |
| OFB_Clatency | Adlercreutzia_caecimuris | 63 | 2.383655218 | 0.01994207195 |
| OFB_Clatency | Adlercreutzia_mucosicola | 63 | -1.29815407 | 0.1973879111 |
| OFB_Clatency | Adlercreutzia_muris | 63 | -0.2320294056 | 0.8167445486 |
| OFB_Clatency | Akkermansia_muciniphila | 63 | 1.518940182 | 0.1323482992 |
| OFB_Clatency | Alistipes_sp_DSM_112343 | 63 | 0.7023318329 | 0.4833264089 |
| OFB_Clatency | Anaerotruncus_sp_1XD42_93 | 63 | 0.5762525909 | 0.5651045803 |
| OFB_Clatency | Bacteria_unclassified_SGB102200 | 63 | -1.213833262 | 0.2295781845 |
| OFB_Clatency | Bacteria_unclassified_SGB41677 | 63 | 1.048903697 | 0.3003578556 |
| OFB_Clatency | Bacteria_unclassified_SGB43546 | 63 | -0.2481120454 | 0.8062739026 |
| OFB_Clatency | Bacteroides_thetaiotaomicron | 63 | 1.449211734 | 0.1506322813 |
| OFB_Clatency | Bifidobacterium_pseudolongum | 63 | 2.092765442 | 0.03990900409 |
| OFB_Clatency | Clostridia_bacterium | 63 | 0.8002325458 | 0.4257618515 |
| OFB_Clatency | Clostridiaceae_bacterium | 63 | 0.9597075747 | 0.3406392689 |
| OFB_Clatency | Clostridiaceae_unclassified_SGB41663 | 63 | 0.9283632691 | 0.3561105312 |

| | | | | |
|---|---|---|---|---|
| OFB_Clatency | Clostridiales_bacterium | 63 | -0.3708215471 | 0.7110596776 |
| OFB_Clatency | Clostridium_cocleatum | 63 | 0.716867379 | 0.4777248632 |
| OFB_Clatency | Coriobacteriaceae_bacterium | 63 | 1.41668082 | 0.200816851 |
| OFB_Clatency | Dorea_sp_5_2 | 63 | -0.5809971325 | 0.5619919552 |
| OFB_Clatency | Dubosiella_newyorkensis | 63 | 1.2813051 | 0.2087840786 |
| OFB_Clatency | Erysipelotrichales_bacterium | 63 | -0.1236611966 | 0.9020454802 |
| OFB_Clatency | Eubacteriaceae_bacterium | 63 | 1.73206169 | 0.09267304275 |
| OFB_Clatency | Eubacteriaceae_unclassified_SGB9492 | 63 | -0.2646553482 | 0.7940970098 |
| OFB_Clatency | GGB20149_SGB29430 | 63 | -1.531010944 | 0.1295908698 |
| OFB_Clatency | GGB22635_SGB63107 | 63 | 1.160488843 | 0.2512308597 |
| OFB_Clatency | GGB25041_SGB36960 | 63 | 0.1516738413 | 0.8801421205 |
| OFB_Clatency | GGB27876_SGB40310 | 63 | 1.343066132 | 0.1868124822 |
| OFB_Clatency | GGB27878_SGB40312 | 63 | 0.8101573137 | 0.4463471024 |
| OFB_Clatency | GGB27918_SGB40356 | 63 | 2.230561589 | 0.03929396915 |
| OFB_Clatency | GGB28382_SGB40962 | 63 | -0.7158319111 | 0.4768925702 |
| OFB_Clatency | GGB28399_SGB40980 | 63 | -0.3317036103 | 0.7418778416 |
| OFB_Clatency | GGB28411_SGB40993 | 63 | -0.4645364722 | 0.6449385712 |
| OFB_Clatency | GGB28415_SGB40997 | 63 | 1.09102538 | 0.2970570808 |
| OFB_Clatency | GGB28430_SGB41013 | 63 | -0.4992635676 | 0.6184612147 |
| OFB_Clatency | GGB28439_SGB41022 | 63 | -0.4079857976 | 0.6844989969 |
| OFB_Clatency | GGB28778_SGB41431 | 63 | 0.5307248102 | 0.5960282521 |
| OFB_Clatency | GGB28784_SGB41437 | 63 | 0.7158341916 | 0.4871626584 |
| OFB_Clatency | GGB28792_SGB41445 | 63 | 1.509991165 | 0.1395653814 |
| OFB_Clatency | GGB28798_SGB41451 | 63 | 0.6076478696 | 0.544019112 |
| OFB_Clatency | GGB28802_SGB41455 | 63 | 1.433187319 | 0.1620186419 |
| OFB_Clatency | GGB28818_SGB41473 | 63 | -0.311961609 | 0.7569947566 |
| OFB_Clatency | GGB28828_SGB41484 | 63 | -0.8303257543 | 0.4234293901 |
| OFB_Clatency | GGB28851_SGB41518 | 63 | -1.332070342 | 0.1859487227 |
| OFB_Clatency | GGB28859_SGB41528 | 63 | 0.3978629004 | 0.7099541387 |
| OFB_Clatency | GGB28864_SGB41535 | 63 | -1.828087442 | 0.07284523737 |
| OFB_Clatency | GGB28869_SGB41543 | 63 | 0.4142748843 | 0.6788988597 |
| OFB_Clatency | GGB28883_SGB41564 | 63 | 0.2239623791 | 0.8228432893 |
| OFB_Clatency | GGB28892_SGB41573 | 63 | 0.8533818196 | 0.3962781804 |
| OFB_Clatency | GGB28893_SGB41574 | 63 | -0.8171015995 | 0.4210962483 |
| OFB_Clatency | GGB28898_SGB41580 | 63 | 0.9251849863 | 0.3699014347 |
| OFB_Clatency | GGB28904_SGB41597 | 63 | -1.3455841 | 0.1834300824 |
| OFB_Clatency | GGB28916_SGB41612 | 63 | 0.4252112359 | 0.6724506841 |
| OFB_Clatency | GGB28924_SGB41621 | 63 | -0.8370974557 | 0.4092655574 |
| OFB_Clatency | GGB28926_SGB41624 | 63 | -1.315879568 | 0.1947947255 |
| OFB_Clatency | GGB28927_SGB41625 | 63 | -0.6296653181 | 0.5334976101 |
| OFB_Clatency | GGB28934_SGB41635 | 63 | 0.6831973744 | 0.5048072739 |
| OFB_Clatency | GGB28946_SGB41652 | 63 | 0.576787747 | 0.5714516999 |
| OFB_Clatency | GGB28949_SGB41655 | 63 | 1.295509354 | 0.200507977 |
| OFB_Clatency | GGB28949_SGB41656 | 63 | 0.2357945096 | 0.8143513808 |
| OFB_Clatency | GGB28950_SGB41657 | 63 | 0.7515355462 | 0.457939587 |
| OFB_Clatency | GGB28951_SGB102295 | 63 | 0.9631626719 | 0.3381176097 |

| | | | | |
|---|---|---|---|---|
| OFB_Clatency | GGB28951_SGB41658 | 63 | 1.117198331 | 0.271926529 |
| OFB_Clatency | GGB28954_SGB41662 | 63 | -0.991118512 | 0.3238602546 |
| OFB_Clatency | GGB28956_SGB41665 | 63 | 0.1265842372 | 0.899379732 |
| OFB_Clatency | GGB28960_SGB41669 | 63 | 0.575215314 | 0.568835471 |
| OFB_Clatency | GGB28967_SGB41678 | 63 | -2.028226142 | 0.04764490547 |
| OFB_Clatency | GGB28991_SGB41705 | 63 | 0.5168061398 | 0.6073293258 |
| OFB_Clatency | GGB29002_SGB41718 | 63 | -2.39389046 | 0.01953826403 |
| OFB_Clatency | GGB29003_SGB41719 | 63 | -0.08821759487 | 0.9300284713 |
| OFB_Clatency | GGB29011_SGB41731 | 63 | -1.367066828 | 0.1749113362 |
| OFB_Clatency | GGB29531_SGB42317 | 63 | -1.222084293 | 0.2254028831 |
| OFB_Clatency | GGB29685_SGB42494 | 63 | 2.329232445 | 0.02265484287 |
| OFB_Clatency | GGB30141_SGB43066 | 63 | -0.2771065112 | 0.7842829391 |
| OFB_Clatency | GGB30286_SGB43248 | 63 | -0.6159752297 | 0.538610316 |
| OFB_Clatency | GGB30303_SGB43268 | 63 | -1.789058597 | 0.07805440486 |
| OFB_Clatency | GGB30413_SGB43452 | 63 | -0.5105372175 | 0.6119214745 |
| OFB_Clatency | GGB30454_SGB43514 | 63 | -0.9345927976 | 0.3516943082 |
| OFB_Clatency | GGB30455_SGB43519 | 63 | -1.601555969 | 0.1147123289 |
| OFB_Clatency | GGB30461_SGB43527 | 63 | -0.2099249898 | 0.8338967271 |
| OFB_Clatency | GGB30461_SGB43530 | 63 | -0.6148466614 | 0.5417380418 |
| OFB_Clatency | GGB30463_SGB43537 | 63 | -1.712042572 | 0.09165626244 |
| OFB_Clatency | GGB30473_SGB43557 | 63 | 1.160689901 | 0.2482865408 |
| OFB_Clatency | GGB30475_SGB63182 | 63 | 0.6194173889 | 0.5362850771 |
| OFB_Clatency | GGB30861_SGB44083 | 63 | 1.700156149 | 0.09304196639 |
| OFB_Clatency | GGB31312_SGB44628 | 63 | -0.02404280957 | 0.9809167288 |
| OFB_Clatency | GGB31438_SGB44768 | 63 | 1.280533723 | 0.2121177129 |
| OFB_Clatency | GGB3171_SGB4185 | 63 | -0.8057109879 | 0.4261768125 |
| OFB_Clatency | GGB31823_SGB45199 | 63 | -0.1680652524 | 0.8666122301 |
| OFB_Clatency | GGB31853_SGB45233 | 63 | -0.24060765 | 0.8102208765 |
| OFB_Clatency | GGB32371_SGB41694 | 63 | -0.6929897797 | 0.4938531599 |
| OFB_Clatency | GGB3793_SGB5158 | 63 | -1.014762923 | 0.3144336536 |
| OFB_Clatency | GGB42598_SGB59794 | 63 | 1.184016087 | 0.238998106 |
| OFB_Clatency | GGB45656_SGB63370 | 63 | -2.973657192 | 0.00569724008 |
| OFB_Clatency | GGB47127_SGB65054 | 63 | 0.2779117985 | 0.7814550249 |
| OFB_Clatency | GGB74395_SGB43521 | 63 | -0.5184839165 | 0.6045148679 |
| OFB_Clatency | GGB75053_SGB43494 | 63 | -0.76048432 | 0.448105827 |
| OFB_Clatency | GGB75109_SGB102238 | 63 | -1.419924132 | 0.1590504823 |
| OFB_Clatency | GGB81440_SGB45230 | 63 | -0.903335435 | 0.3767188198 |
| OFB_Clatency | Lachnospiraceae_bacterium | 63 | 1.586272633 | 0.1163223214 |
| OFB_Clatency | Lachnospiraceae_bacterium_A2 | 63 | 2.544659452 | 0.01367016647 |
| OFB_Clatency | Lachnospiraceae_bacterium_MD308 | 63 | 2.266545899 | 0.02916222563 |
| OFB_Clatency | Lachnospiraceae_bacterium_MD329 | 63 | 1.409183426 | 0.1633950825 |
| OFB_Clatency | Lachnospiraceae_unclassified_SGB414 | 63 | 0.8689367629 | 0.3947305553 |
| OFB_Clatency | Lachnospiraceae_unclassified_SGB414 | 63 | -2.630223252 | 0.01204365581 |
| OFB_Clatency | Lachnospiraceae_unclassified_SGB414 | 63 | 0.8597158049 | 0.3922327518 |
| OFB_Clatency | Lachnospiraceae_unclassified_SGB415 | 63 | 0.4354516249 | 0.7266077813 |
| OFB_Clatency | Lactobacillus_johnsonii | 63 | 1.495736424 | 0.1392004983 |

| | | | | |
|---|---|---|---|---|
| OFB_Clatency | Muribaculaceae_bacterium | 63 | 1.878796296 | 0.06395893975 |
| OFB_Clatency | Neglectibacter_sp_X4 | 63 | 0.2133545898 | 0.8349025754 |
| OFB_Clatency | Oscillospiraceae_bacterium | 63 | -0.1040487123 | 0.9174293874 |
| OFB_Clatency | Oscillospiraceae_unclassified_SGB4350 | 63 | 0.1200178405 | 0.9049456263 |
| OFB_Clatency | Oscillospiraceae_unclassified_SGB4350 | 63 | -0.3447151153 | 0.7331569679 |
| OFB_Clatency | Parasutterella_excrementihominis | 63 | 1.369400326 | 0.1982449334 |
| OFB_Clatency | Romboutsia_ilealis | 63 | -0.003276127923 | 0.9974164499 |
| OFB_Clatency | Schaedlerella_arabinosiphila | 63 | 1.541685179 | 0.1286996449 |
| OFB_Clatency | Turicibacter_sp_1E2 | 63 | 0.3323608815 | 0.7397542963 |
| OFB_Clatency | bacterium_1XD42_54 | 63 | 1.343463026 | 0.1823548779 |
| OFB_Clatency | bacterium_1XD42_76 | 63 | 0.7493200565 | 0.4585147406 |
| OFB_Clatency | bacterium_1xD8_48 | 63 | -0.4959451592 | 0.6208522922 |
| OFB_Clatency | Berger Parker Index | 63 | 0.7739073552 | 0.4405178516 |
| OFB_Clatency | Richness (# observed features) | 63 | -1.268534043 | 0.2100343789 |
| OFB_Clatency | Shannon Index | 63 | -0.8861642271 | 0.3790876367 |
| OFB_Ctime | Acetatifactor_SGB41546 | 63 | -1.001760818 | 0.3193417225 |
| OFB_Ctime | Acetatifactor_muris | 63 | 0.4477151478 | 0.6602640786 |
| OFB_Ctime | Acutalibacter_muris | 63 | -1.660484903 | 0.101422053 |
| OFB_Ctime | Acutalibacter_sp_1XD8_36 | 63 | 0.3638798653 | 0.716207136 |
| OFB_Ctime | Adlercreutzia_caecimuris | 63 | -1.612339213 | 0.1118266371 |
| OFB_Ctime | Adlercreutzia_mucosicola | 63 | 1.350991436 | 0.1801681257 |
| OFB_Ctime | Adlercreutzia_muris | 63 | 0.07106372678 | 0.9434711917 |
| OFB_Ctime | Akkermansia_muciniphila | 63 | 0.3988173307 | 0.6938934558 |
| OFB_Ctime | Alistipes_sp_DSM_112343 | 63 | 0.05037345174 | 0.9598609979 |
| OFB_Ctime | Anaerotruncus_sp_1XD42_93 | 63 | 0.3255636525 | 0.7451234374 |
| OFB_Ctime | Bacteria_unclassified_SGB102200 | 63 | 0.2695344255 | 0.7921614298 |
| OFB_Ctime | Bacteria_unclassified_SGB41677 | 63 | -1.817904715 | 0.08029184583 |
| OFB_Ctime | Bacteria_unclassified_SGB43546 | 63 | -0.2068843739 | 0.8410551671 |
| OFB_Ctime | Bacteroides_thetaiotaomicron | 63 | -1.119749097 | 0.2651775294 |
| OFB_Ctime | Bifidobacterium_pseudolongum | 63 | -0.9052578274 | 0.3670782778 |
| OFB_Ctime | Clostridia_bacterium | 63 | -1.346551294 | 0.1844492297 |
| OFB_Ctime | Clostridiaceae_bacterium | 63 | -1.012568545 | 0.314345718 |
| OFB_Ctime | Clostridiaceae_unclassified_SGB41663 | 63 | 0.57368602 | 0.5716007348 |
| OFB_Ctime | Clostridiales_bacterium | 63 | 1.138784294 | 0.2588960202 |
| OFB_Ctime | Clostridium_cocleatum | 63 | -0.02650859512 | 0.9793102559 |
| OFB_Ctime | Coriobacteriaceae_bacterium | 63 | 0.5821497656 | 0.5619201788 |
| OFB_Ctime | Dorea_sp_5_2 | 63 | 1.548544852 | 0.1257943335 |
| OFB_Ctime | Dubosiella_newyorkensis | 63 | -1.573974963 | 0.133746852 |
| OFB_Ctime | Erysipelotrichales_bacterium | 63 | 1.881307025 | 0.07032699776 |
| OFB_Ctime | Eubacteriaceae_bacterium | 63 | 0.9675778147 | 0.3541574226 |
| OFB_Ctime | Eubacteriaceae_unclassified_SGB9492: | 63 | -2.059581497 | 0.04688856043 |
| OFB_Ctime | GGB20149_SGB29430 | 63 | -0.479787926 | 0.6403416468 |
| OFB_Ctime | GGB22635_SGB63107 | 63 | 0.3823185651 | 0.702589549 |
| OFB_Ctime | GGB25041_SGB36960 | 63 | -0.4807616019 | 0.639257774 |
| OFB_Ctime | GGB27876_SGB40310 | 63 | 0.1556192288 | 0.8810173092 |
| OFB_Ctime | GGB27878_SGB40312 | 63 | -0.1846908066 | 0.856462496 |

| | | | | |
|---|---|---|---|---|
| OFB_Ctime | GGB27918_SGB40356 | 63 | -1.292633915 | 0.2260075677 |
| OFB_Ctime | GGB28382_SGB40962 | 63 | 1.246862136 | 0.2269447104 |
| OFB_Ctime | GGB28399_SGB40980 | 63 | -0.5769039609 | 0.5647218801 |
| OFB_Ctime | GGB28411_SGB40993 | 63 | 0.05657515714 | 0.9555048821 |
| OFB_Ctime | GGB28415_SGB40997 | 63 | 0.9000596204 | 0.4019558059 |
| OFB_Ctime | GGB28430_SGB41013 | 63 | -0.0100178978 | 0.9920445365 |
| OFB_Ctime | GGB28439_SGB41022 | 63 | -0.2099090243 | 0.8348193272 |
| OFB_Ctime | GGB28778_SGB41431 | 63 | -0.3595872399 | 0.7193194066 |
| OFB_Ctime | GGB28784_SGB41437 | 63 | -0.4392770858 | 0.6661751766 |
| OFB_Ctime | GGB28792_SGB41445 | 63 | -0.7037991589 | 0.5062502222 |
| OFB_Ctime | GGB28798_SGB41451 | 63 | -1.837139921 | 0.07089736455 |
| OFB_Ctime | GGB28802_SGB41455 | 63 | -2.359648162 | 0.02606055131 |
| OFB_Ctime | GGB28818_SGB41473 | 63 | 0.4401779233 | 0.6690540598 |
| OFB_Ctime | GGB28828_SGB41484 | 63 | 0.6716506475 | 0.5357387749 |
| OFB_Ctime | GGB28851_SGB41518 | 63 | 0.3612968917 | 0.7210217902 |
| OFB_Ctime | GGB28859_SGB41528 | 63 | 1.129578446 | 0.3079858972 |
| OFB_Ctime | GGB28864_SGB41535 | 63 | 1.232148985 | 0.2523148241 |
| OFB_Ctime | GGB28869_SGB41543 | 63 | 0.1361985457 | 0.8917935461 |
| OFB_Ctime | GGB28883_SGB41564 | 63 | -1.215546098 | 0.2272496702 |
| OFB_Ctime | GGB28892_SGB41573 | 63 | -2.450152346 | 0.01870374293 |
| OFB_Ctime | GGB28893_SGB41574 | 63 | 1.06043783 | 0.3002372181 |
| OFB_Ctime | GGB28898_SGB41580 | 63 | 2.432495823 | 0.0216470335 |
| OFB_Ctime | GGB28904_SGB41597 | 63 | 0.6621732247 | 0.5161259991 |
| OFB_Ctime | GGB28916_SGB41612 | 63 | 0.4190232414 | 0.6787733904 |
| OFB_Ctime | GGB28924_SGB41621 | 63 | 0.08135429766 | 0.9382739565 |
| OFB_Ctime | GGB28926_SGB41624 | 63 | 0.1341996634 | 0.8933389516 |
| OFB_Ctime | GGB28927_SGB41625 | 63 | 0.5716701404 | 0.5694284994 |
| OFB_Ctime | GGB28934_SGB41635 | 63 | -0.6879575389 | 0.5145470996 |
| OFB_Ctime | GGB28946_SGB41652 | 63 | -0.5617529521 | 0.5908008036 |
| OFB_Ctime | GGB28949_SGB41655 | 63 | 0.006392052134 | 0.9949208784 |
| OFB_Ctime | GGB28949_SGB41656 | 63 | 0.04424090139 | 0.9649965971 |
| OFB_Ctime | GGB28950_SGB41657 | 63 | -0.3333135006 | 0.7401390197 |
| OFB_Ctime | GGB28951_SGB102295 | 63 | -0.8316014239 | 0.4073388043 |
| OFB_Ctime | GGB28951_SGB41658 | 63 | -1.153381556 | 0.2701976199 |
| OFB_Ctime | GGB28954_SGB41662 | 63 | 0.2077344078 | 0.8356300278 |
| OFB_Ctime | GGB28956_SGB41665 | 63 | -0.0668209499 | 0.9468577916 |
| OFB_Ctime | GGB28960_SGB41669 | 63 | -0.2316307572 | 0.8194139887 |
| OFB_Ctime | GGB28967_SGB41678 | 63 | 0.192476023 | 0.8476955891 |
| OFB_Ctime | GGB28991_SGB41705 | 63 | -1.648311765 | 0.1078737586 |
| OFB_Ctime | GGB29002_SGB41718 | 63 | 1.213957913 | 0.227454206 |
| OFB_Ctime | GGB29003_SGB41719 | 63 | 0.02559825177 | 0.9797286821 |
| OFB_Ctime | GGB29011_SGB41731 | 63 | -1.513716934 | 0.1342716892 |
| OFB_Ctime | GGB29531_SGB42317 | 63 | 1.219114188 | 0.2255453721 |
| OFB_Ctime | GGB29685_SGB42494 | 63 | -1.166067863 | 0.247080295 |
| OFB_Ctime | GGB30141_SGB43066 | 63 | -0.1393419567 | 0.8927792095 |
| OFB_Ctime | GGB30286_SGB43248 | 63 | 1.8217218 | 0.07226187933 |

| | | | | |
|---|---|---|---|---|
| OFB_Ctime | GGB30303_SGB43268 | 63 | 0.1255244816 | 0.9032898883 |
| OFB_Ctime | GGB30413_SGB43452 | 63 | 0.6502198157 | 0.5165175508 |
| OFB_Ctime | GGB30454_SGB43514 | 63 | -0.1270317208 | 0.898953702 |
| OFB_Ctime | GGB30455_SGB43519 | 63 | 0.2813380192 | 0.7860094411 |
| OFB_Ctime | GGB30461_SGB43527 | 63 | 0.6641591589 | 0.5117525911 |
| OFB_Ctime | GGB30461_SGB43530 | 63 | -0.2957179066 | 0.7837552975 |
| OFB_Ctime | GGB30463_SGB43537 | 63 | 0.03835966411 | 0.970873424 |
| OFB_Ctime | GGB30473_SGB43557 | 63 | 0.2583025429 | 0.7986153167 |
| OFB_Ctime | GGB30475_SGB63182 | 63 | -0.01110860444 | 0.9912118266 |
| OFB_Ctime | GGB30861_SGB44083 | 63 | -0.6643558352 | 0.513163785 |
| OFB_Ctime | GGB31312_SGB44628 | 63 | -0.5808716281 | 0.5793112021 |
| OFB_Ctime | GGB31438_SGB44768 | 63 | -1.245364748 | 0.2492368402 |
| OFB_Ctime | GGB3171_SGB4185 | 63 | 0.1325171161 | 0.8976909592 |
| OFB_Ctime | GGB31823_SGB45199 | 63 | 2.341650944 | 0.02185577779 |
| OFB_Ctime | GGB31853_SGB45233 | 63 | 0.2980318512 | 0.7704272105 |
| OFB_Ctime | GGB32371_SGB41694 | 63 | 1.161954569 | 0.2580898841 |
| OFB_Ctime | GGB3793_SGB5158 | 63 | 1.449852332 | 0.1522085237 |
| OFB_Ctime | GGB42598_SGB59794 | 63 | -0.2608897144 | 0.7943110635 |
| OFB_Ctime | GGB45656_SGB63370 | 63 | 2.092797895 | 0.05152755498 |
| OFB_Ctime | GGB47127_SGB65054 | 63 | -0.7602106192 | 0.4581615937 |
| OFB_Ctime | GGB74395_SGB43521 | 63 | -0.7654856399 | 0.4451799014 |
| OFB_Ctime | GGB75053_SGB43494 | 63 | -0.3329274017 | 0.7409435884 |
| OFB_Ctime | GGB75109_SGB102238 | 63 | -0.05361550195 | 0.9572422678 |
| OFB_Ctime | GGB81440_SGB45230 | 63 | 0.8049956404 | 0.4892961624 |
| OFB_Ctime | Lachnospiraceae_bacterium | 63 | -1.653485755 | 0.1019292993 |
| OFB_Ctime | Lachnospiraceae_bacterium_A2 | 63 | 0.9522526233 | 0.3428142858 |
| OFB_Ctime | Lachnospiraceae_bacterium_MD308 | 63 | 1.182899903 | 0.2432820103 |
| OFB_Ctime | Lachnospiraceae_bacterium_MD329 | 63 | -1.956854619 | 0.06035222718 |
| OFB_Ctime | Lachnospiraceae_unclassified_SGB4141 | 63 | -1.193732258 | 0.2459974185 |
| OFB_Ctime | Lachnospiraceae_unclassified_SGB4141 | 63 | 0.6958537923 | 0.5043903837 |
| OFB_Ctime | Lachnospiraceae_unclassified_SGB4142 | 63 | -0.6008466616 | 0.5534594918 |
| OFB_Ctime | Lachnospiraceae_unclassified_SGB4158 | 63 | -2.007438672 | 0.07861071271 |
| OFB_Ctime | Lactobacillus_johnsonii | 63 | -0.3934027254 | 0.6954689555 |
| OFB_Ctime | Muribaculaceae_bacterium | 63 | 0.08772806887 | 0.9307165999 |
| OFB_Ctime | Neglectibacter_sp_X4 | 63 | -1.392996729 | 0.1775887113 |
| OFB_Ctime | Oscillospiraceae_bacterium | 63 | -0.4559615879 | 0.6506742156 |
| OFB_Ctime | Oscillospiraceae_unclassified_SGB4350 | 63 | -1.196565999 | 0.2350165484 |
| OFB_Ctime | Oscillospiraceae_unclassified_SGB4350 | 63 | -0.1385365455 | 0.8946303958 |
| OFB_Ctime | Parasutterella_excrementihominis | 63 | 0.3621151045 | 0.7820241105 |
| OFB_Ctime | Romboutsia_ilealis | 63 | 1.376224343 | 0.206879804 |
| OFB_Ctime | Schaedlerella_arabinosiphila | 63 | -1.257618536 | 0.2252702023 |
| OFB_Ctime | Turicibacter_sp_1E2 | 63 | 0.5050256621 | 0.6185016576 |
| OFB_Ctime | bacterium_1XD42_54 | 63 | -1.250981897 | 0.2290999082 |
| OFB_Ctime | bacterium_1XD42_76 | 63 | -0.09502429275 | 0.9253678534 |
| OFB_Ctime | bacterium_1xD8_48 | 63 | 2.02010801 | 0.04675699948 |
| OFB_Ctime | Berger Parker Index | 63 | 0.6866522834 | 0.4994122149 |

| | | | | |
|---|---|---|---|---|
| OFB_Ctime | Richness (# observed features) | 63 | 0.4736440753 | 0.6421160642 |
| OFB_Ctime | Shannon Index | 63 | -0.7385991882 | 0.4722876446 |
| PD_choice_filt | Acetatifactor_SGB41546 | 63 | -0.9299684024 | 0.3550315504 |
| PD_choice_filt | Acetatifactor_muris | 63 | -0.4411615223 | 0.6594934665 |
| PD_choice_filt | Acutalibacter_muris | 63 | 0.4823639393 | 0.63005687 |
| PD_choice_filt | Acutalibacter_sp_1XD8_36 | 63 | -0.07198233514 | 0.9426178684 |
| PD_choice_filt | Adlercreutzia_caecimuris | 63 | 2.034380962 | 0.04733331088 |
| PD_choice_filt | Adlercreutzia_mucosicola | 63 | -1.099647316 | 0.2751568637 |
| PD_choice_filt | Adlercreutzia_muris | 63 | -0.6209325262 | 0.5356489015 |
| PD_choice_filt | Akkermansia_muciniphila | 63 | -0.02315555161 | 0.9815262573 |
| PD_choice_filt | Alistipes_sp_DSM_112343 | 63 | 0.05167121379 | 0.9587913793 |
| PD_choice_filt | Anaerotruncus_sp_1XD42_93 | 63 | 0.738505318 | 0.4617655016 |
| PD_choice_filt | Bacteria_unclassified_SGB102200 | 63 | -1.464755662 | 0.1484046823 |
| PD_choice_filt | Bacteria_unclassified_SGB41677 | 63 | 0.2487427912 | 0.8036360005 |
| PD_choice_filt | Bacteria_unclassified_SGB43546 | 63 | 0.6943609051 | 0.4887934355 |
| PD_choice_filt | Bacteroides_thetaiotaomicron | 63 | 0.4829969153 | 0.6296092112 |
| PD_choice_filt | Bifidobacterium_pseudolongum | 63 | 0.4189999744 | 0.6755599627 |
| PD_choice_filt | Clostridia_bacterium | 63 | 0.8727357119 | 0.3851102284 |
| PD_choice_filt | Clostridiaceae_bacterium | 63 | 1.78887195 | 0.07947807304 |
| PD_choice_filt | Clostridiaceae_unclassified_SGB41663 | 63 | 0.8309914826 | 0.4080399042 |
| PD_choice_filt | Clostridiales_bacterium | 63 | 0.4477372343 | 0.6547570394 |
| PD_choice_filt | Clostridium_cocleatum | 63 | -1.704789454 | 0.0940814706 |
| PD_choice_filt | Coriobacteriaceae_bacterium | 63 | 0.6410065216 | 0.5226097475 |
| PD_choice_filt | Dorea_sp_5_2 | 63 | 0.8243960996 | 0.4117385228 |
| PD_choice_filt | Dubosiella_newyorkensis | 63 | 0.8003786812 | 0.4253808248 |
| PD_choice_filt | Erysipelotrichales_bacterium | 63 | -0.4931256114 | 0.6224652159 |
| PD_choice_filt | Eubacteriaceae_bacterium | 63 | 1.030551016 | 0.3060068864 |
| PD_choice_filt | Eubacteriaceae_unclassified_SGB9492 | 63 | 1.852668169 | 0.06971773566 |
| PD_choice_filt | GGB20149_SGB29430 | 63 | -0.4545983744 | 0.6498303588 |
| PD_choice_filt | GGB22635_SGB63107 | 63 | -0.7280039543 | 0.4681154819 |
| PD_choice_filt | GGB25041_SGB36960 | 63 | 0.5243488902 | 0.6006762588 |
| PD_choice_filt | GGB27876_SGB40310 | 63 | -1.603433621 | 0.1145859386 |
| PD_choice_filt | GGB27878_SGB40312 | 63 | 1.027937131 | 0.3072189877 |
| PD_choice_filt | GGB27918_SGB40356 | 63 | -0.5847120853 | 0.5595993683 |
| PD_choice_filt | GGB28382_SGB40962 | 63 | -1.3324099 | 0.1876343814 |
| PD_choice_filt | GGB28399_SGB40980 | 63 | -1.880030286 | 0.06585553097 |
| PD_choice_filt | GGB28411_SGB40993 | 63 | 1.605358087 | 0.1141650319 |
| PD_choice_filt | GGB28415_SGB40997 | 63 | -0.2158361988 | 0.8291656412 |
| PD_choice_filt | GGB28430_SGB41013 | 63 | -0.6584458932 | 0.5114208509 |
| PD_choice_filt | GGB28439_SGB41022 | 63 | 0.6934090082 | 0.4893858472 |
| PD_choice_filt | GGB28778_SGB41431 | 63 | 0.5216670125 | 0.602533688 |
| PD_choice_filt | GGB28784_SGB41437 | 63 | 0.05699135757 | 0.9545530248 |
| PD_choice_filt | GGB28792_SGB41445 | 63 | 0.6552531471 | 0.513459525 |
| PD_choice_filt | GGB28798_SGB41451 | 63 | -0.1875880123 | 0.8512327815 |
| PD_choice_filt | GGB28802_SGB41455 | 63 | 0.9038180773 | 0.3685790396 |
| PD_choice_filt | GGB28818_SGB41473 | 63 | -0.9926472027 | 0.3239066091 |

| | | | | |
|---|---|---|---|---|
| PD_choice_filt | GGB28828_SGB41484 | 63 | 0.997597252 | 0.3215295573 |
| PD_choice_filt | GGB28851_SGB41518 | 63 | -1.393966388 | 0.1684947691 |
| PD_choice_filt | GGB28859_SGB41528 | 63 | 0.5811854537 | 0.5619598544 |
| PD_choice_filt | GGB28864_SGB41535 | 63 | -1.465376005 | 0.1482374067 |
| PD_choice_filt | GGB28869_SGB41543 | 63 | 0.2566585913 | 0.7975259367 |
| PD_choice_filt | GGB28883_SGB41564 | 63 | -0.624533493 | 0.5332974063 |
| PD_choice_filt | GGB28892_SGB41573 | 63 | 0.2925576931 | 0.7699828183 |
| PD_choice_filt | GGB28893_SGB41574 | 63 | 0.7153493106 | 0.4758338194 |
| PD_choice_filt | GGB28898_SGB41580 | 63 | -0.429045619 | 0.6682575557 |
| PD_choice_filt | GGB28904_SGB41597 | 63 | 0.6938937079 | 0.4890841449 |
| PD_choice_filt | GGB28916_SGB41612 | 63 | -0.5750131331 | 0.5661031726 |
| PD_choice_filt | GGB28924_SGB41621 | 63 | -0.0921512248 | 0.926581875 |
| PD_choice_filt | GGB28926_SGB41624 | 63 | -0.5498103115 | 0.5831774684 |
| PD_choice_filt | GGB28927_SGB41625 | 63 | 0.1592404318 | 0.8734998356 |
| PD_choice_filt | GGB28934_SGB41635 | 63 | -0.4354912355 | 0.663589147 |
| PD_choice_filt | GGB28946_SGB41652 | 63 | -0.1226542089 | 0.9023902909 |
| PD_choice_filt | GGB28949_SGB41655 | 63 | -0.6250677247 | 0.5329490057 |
| PD_choice_filt | GGB28949_SGB41656 | 63 | -0.9073319649 | 0.3667394325 |
| PD_choice_filt | GGB28950_SGB41657 | 63 | -0.2357701883 | 0.8136761473 |
| PD_choice_filt | GGB28951_SGB102295 | 63 | 0.1093504814 | 0.912931155 |
| PD_choice_filt | GGB28951_SGB41658 | 63 | 0.5813540513 | 0.5618468931 |
| PD_choice_filt | GGB28954_SGB41662 | 63 | 0.3826045991 | 0.7022786813 |
| PD_choice_filt | GGB28956_SGB41665 | 63 | -0.08260283118 | 0.9341702226 |
| PD_choice_filt | GGB28960_SGB41669 | 63 | 1.097035377 | 0.2762814748 |
| PD_choice_filt | GGB28967_SGB41678 | 63 | -1.007667311 | 0.3167304264 |
| PD_choice_filt | GGB28991_SGB41705 | 63 | -0.2796854511 | 0.7798263333 |
| PD_choice_filt | GGB29002_SGB41718 | 63 | 0.0127572117 | 0.9898215044 |
| PD_choice_filt | GGB29003_SGB41719 | 63 | 0.4314631963 | 0.6665049865 |
| PD_choice_filt | GGB29011_SGB41731 | 63 | 0.8708460185 | 0.3861302272 |
| PD_choice_filt | GGB29531_SGB42317 | 63 | 0.880433409 | 0.3809729436 |
| PD_choice_filt | GGB29685_SGB42494 | 63 | -0.6820761225 | 0.4964696024 |
| PD_choice_filt | GGB30141_SGB43066 | 63 | -0.6512106289 | 0.5160470968 |
| PD_choice_filt | GGB30286_SGB43248 | 63 | 1.123482505 | 0.2650440549 |
| PD_choice_filt | GGB30303_SGB43268 | 63 | -0.574605138 | 0.5663775848 |
| PD_choice_filt | GGB30413_SGB43452 | 63 | -0.5239985072 | 0.6009187763 |
| PD_choice_filt | GGB30454_SGB43514 | 63 | -1.974774425 | 0.05386573274 |
| PD_choice_filt | GGB30455_SGB43519 | 63 | -1.728793673 | 0.08970037911 |
| PD_choice_filt | GGB30461_SGB43527 | 63 | -0.1495668955 | 0.8811232452 |
| PD_choice_filt | GGB30461_SGB43530 | 63 | -0.1628938939 | 0.8706237862 |
| PD_choice_filt | GGB30463_SGB43537 | 63 | -0.8765312623 | 0.3830666792 |
| PD_choice_filt | GGB30473_SGB43557 | 63 | -0.9252778571 | 0.3574372358 |
| PD_choice_filt | GGB30475_SGB63182 | 63 | 0.586947926 | 0.5581054407 |
| PD_choice_filt | GGB30861_SGB44083 | 63 | 1.241740935 | 0.2188006691 |
| PD_choice_filt | GGB31312_SGB44628 | 63 | 0.4824116231 | 0.6300231417 |
| PD_choice_filt | GGB31438_SGB44768 | 63 | 1.037711346 | 0.3027034028 |
| PD_choice_filt | GGB3171_SGB4185 | 63 | -2.852822815 | 0.00655258972 |

| | | | | |
|---|---|---|---|---|
| PD_choice_filt | GGB31823_SGB45199 | 63 | -0.4881959599 | 0.6259376556 |
| PD_choice_filt | GGB31853_SGB45233 | 63 | 0.1605656257 | 0.8724564242 |
| PD_choice_filt | GGB32371_SGB41694 | 63 | 0.7226751772 | 0.4713568109 |
| PD_choice_filt | GGB3793_SGB5158 | 63 | -1.229972424 | 0.2231142241 |
| PD_choice_filt | GGB42598_SGB59794 | 63 | -0.5159515122 | 0.6065010953 |
| PD_choice_filt | GGB45656_SGB63370 | 63 | 1.026591114 | 0.3078444439 |
| PD_choice_filt | GGB47127_SGB65054 | 63 | 0.7700168018 | 0.4430137541 |
| PD_choice_filt | GGB74395_SGB43521 | 63 | 0.04174055169 | 0.9667058982 |
| PD_choice_filt | GGB75053_SGB43494 | 63 | -0.228321214 | 0.8194558021 |
| PD_choice_filt | GGB75109_SGB102238 | 63 | -0.04559219972 | 0.9636357706 |
| PD_choice_filt | GGB81440_SGB45230 | 63 | -0.7259713709 | 0.4693503245 |
| PD_choice_filt | Lachnospiraceae_bacterium | 63 | 0.9503846866 | 0.3446844102 |
| PD_choice_filt | Lachnospiraceae_bacterium_A2 | 63 | -0.1557110429 | 0.876279863 |
| PD_choice_filt | Lachnospiraceae_bacterium_MD308 | 63 | 0.1371259708 | 0.8909442857 |
| PD_choice_filt | Lachnospiraceae_bacterium_MD329 | 63 | -1.42204684 | 0.160285117 |
| PD_choice_filt | Lachnospiraceae_unclassified_SGB4141 | 63 | 0.9735293507 | 0.3331985176 |
| PD_choice_filt | Lachnospiraceae_unclassified_SGB4141 | 63 | -0.2768398987 | 0.7820073724 |
| PD_choice_filt | Lachnospiraceae_unclassified_SGB4142 | 63 | 0.8451585042 | 0.4001649836 |
| PD_choice_filt | Lachnospiraceae_unclassified_SGB4158 | 63 | -0.1489895009 | 0.8815786412 |
| PD_choice_filt | Lactobacillus_johnsonii | 63 | -1.084530242 | 0.2817108294 |
| PD_choice_filt | Muribaculaceae_bacterium | 63 | -0.4708766077 | 0.6382054204 |
| PD_choice_filt | Neglectibacter_sp_X4 | 63 | 1.379909006 | 0.1727259155 |
| PD_choice_filt | Oscillospiraceae_bacterium | 63 | 0.4808014533 | 0.6311625085 |
| PD_choice_filt | Oscillospiraceae_unclassified_SGB4350 | 63 | -0.6154631427 | 0.539230854 |
| PD_choice_filt | Oscillospiraceae_unclassified_SGB4350 | 63 | -1.030886997 | 0.3058513257 |
| PD_choice_filt | Parasutterella_excrementihominis | 63 | 1.222116469 | 0.2260286639 |
| PD_choice_filt | Romboutsia_ilealis | 63 | 0.3747614607 | 0.7080883536 |
| PD_choice_filt | Schaedlerella_arabinosiphila | 63 | 1.167324661 | 0.2471429805 |
| PD_choice_filt | Turicibacter_sp_1E2 | 63 | -0.06473011595 | 0.9483902682 |
| PD_choice_filt | bacterium_1XD42_54 | 63 | 2.227966153 | 0.0306506842 |
| PD_choice_filt | bacterium_1XD42_76 | 63 | 0.8479971731 | 0.3985985537 |
| PD_choice_filt | bacterium_1xD8_48 | 63 | 0.7690861123 | 0.4435610321 |
| PD_choice_filt | Berger Parker Index | 63 | -0.4749033908 | 0.6353437792 |
| PD_choice_filt | Richness (# observed features) | 63 | 1.318695898 | 0.1921173807 |
| PD_choice_filt | Shannon Index | 63 | 1.035721637 | 0.3036188932 |
| startle_PPI | Acetatifactor_SGB41546 | 63 | 1.6440789 | 0.1037705338 |
| startle_PPI | Acetatifactor_muris | 63 | 1.937298752 | 0.05621809439 |
| startle_PPI | Acutalibacter_muris | 63 | -0.5699347208 | 0.5692189787 |
| startle_PPI | Acutalibacter_sp_1XD8_36 | 63 | -0.01615654923 | 0.987109513 |
| startle_PPI | Adlercreutzia_caecimuris | 63 | -0.196016968 | 0.84462027 |
| startle_PPI | Adlercreutzia_mucosicola | 63 | -0.8827561233 | 0.3788351928 |
| startle_PPI | Adlercreutzia_muris | 63 | 1.88661053 | 0.06277957972 |
| startle_PPI | Akkermansia_muciniphila | 63 | -1.976350727 | 0.05157007534 |
| startle_PPI | Alistipes_sp_DSM_112343 | 63 | -3.703468412 | 0.000427224672 |
| startle_PPI | Anaerotruncus_sp_1XD42_93 | 63 | -0.6549777646 | 0.5131975944 |
| startle_PPI | Bacteria_unclassified_SGB102200 | 63 | 0.9091213447 | 0.3648508324 |

| | | | | |
|---|---|---|---|---|
| startle_PPI | Bacteria_unclassified_SGB41677 | 63 | -1.424884891 | 0.1574790509 |
| startle_PPI | Bacteria_unclassified_SGB43546 | 63 | 0.4389551622 | 0.6609368392 |
| startle_PPI | Bacteroides_thetaiotaomicron | 63 | -1.276407436 | 0.2047019389 |
| startle_PPI | Bifidobacterium_pseudolongum | 63 | -0.430953788 | 0.6667325618 |
| startle_PPI | Clostridia_bacterium | 63 | -1.57834826 | 0.1180359372 |
| startle_PPI | Clostridiaceae_bacterium | 63 | -1.005260237 | 0.3166993457 |
| startle_PPI | Clostridiaceae_unclassified_SGB41663 | 63 | 0.7536219159 | 0.4520921446 |
| startle_PPI | Clostridiales_bacterium | 63 | 0.2192278093 | 0.8265051483 |
| startle_PPI | Clostridium_cocleatum | 63 | -0.8895826445 | 0.3751823172 |
| startle_PPI | Coriobacteriaceae_bacterium | 63 | -1.657243049 | 0.1010892466 |
| startle_PPI | Dorea_sp_5_2 | 63 | 0.05940311119 | 0.9526316976 |
| startle_PPI | Dubosiella_newyorkensis | 63 | -0.5309411171 | 0.595870244 |
| startle_PPI | Erysipelotrichales_bacterium | 63 | 0.3282983607 | 0.7427921429 |
| startle_PPI | Eubacteriaceae_bacterium | 63 | -0.3553081007 | 0.7224919812 |
| startle_PPI | Eubacteriaceae_unclassified_SGB9492[illegible] | 63 | 1.137810661 | 0.2576220309 |
| startle_PPI | GGB20149_SGB29430 | 63 | 0.4669418647 | 0.6408300146 |
| startle_PPI | GGB22635_SGB63107 | 63 | -1.029926979 | 0.3050656536 |
| startle_PPI | GGB25041_SGB36960 | 63 | -1.762848399 | 0.08156595798 |
| startle_PPI | GGB27876_SGB40310 | 63 | -2.146066572 | 0.03500286241 |
| startle_PPI | GGB27878_SGB40312 | 63 | -0.4091281106 | 0.6826447931 |
| startle_PPI | GGB27918_SGB40356 | 63 | -1.28554402 | 0.2015206697 |
| startle_PPI | GGB28382_SGB40962 | 63 | -0.5622322861 | 0.5744370071 |
| startle_PPI | GGB28399_SGB40980 | 63 | -1.697477146 | 0.0932428364 |
| startle_PPI | GGB28411_SGB40993 | 63 | 2.033137472 | 0.04540073618 |
| startle_PPI | GGB28415_SGB40997 | 63 | -0.2528755989 | 0.800413907 |
| startle_PPI | GGB28430_SGB41013 | 63 | 1.665147555 | 0.09950657948 |
| startle_PPI | GGB28439_SGB41022 | 63 | 0.4933418739 | 0.622106997 |
| startle_PPI | GGB28778_SGB41431 | 63 | -1.870249331 | 0.06503136775 |
| startle_PPI | GGB28784_SGB41437 | 63 | 0.6434009634 | 0.5206473959 |
| startle_PPI | GGB28792_SGB41445 | 63 | 0.5622813167 | 0.574403718 |
| startle_PPI | GGB28798_SGB41451 | 63 | 0.1016702388 | 0.9190217358 |
| startle_PPI | GGB28802_SGB41455 | 63 | 2.028788991 | 0.04584955231 |
| startle_PPI | GGB28818_SGB41473 | 63 | -0.839310119 | 0.4026049268 |
| startle_PPI | GGB28828_SGB41484 | 63 | 1.322011668 | 0.1891902696 |
| startle_PPI | GGB28851_SGB41518 | 63 | 1.904845481 | 0.06034802766 |
| startle_PPI | GGB28859_SGB41528 | 63 | 2.055342821 | 0.04316752283 |
| startle_PPI | GGB28864_SGB41535 | 63 | 1.578467362 | 0.1180087309 |
| startle_PPI | GGB28869_SGB41543 | 63 | 1.81652062 | 0.07291112826 |
| startle_PPI | GGB28883_SGB41564 | 63 | 0.6538744385 | 0.5139051391 |
| startle_PPI | GGB28892_SGB41573 | 63 | -0.7119930441 | 0.4773526128 |
| startle_PPI | GGB28893_SGB41574 | 63 | 1.471592217 | 0.1445166404 |
| startle_PPI | GGB28898_SGB41580 | 63 | 0.04047099663 | 0.9677178391 |
| startle_PPI | GGB28904_SGB41597 | 63 | -0.4605066333 | 0.6454302279 |
| startle_PPI | GGB28916_SGB41612 | 63 | -0.5631537466 | 0.5738115429 |
| startle_PPI | GGB28924_SGB41621 | 63 | -0.6038298925 | 0.546535992 |
| startle_PPI | GGB28926_SGB41624 | 63 | -0.1233476026 | 0.9018377434 |

| | | | | |
|---|---|---|---|---|
| startle_PPI | GGB28927_SGB41625 | 63 | 0.50391594 | 0.614676557 |
| startle_PPI | GGB28934_SGB41635 | 63 | 1.665642092 | 0.09940823818 |
| startle_PPI | GGB28946_SGB41652 | 63 | -1.45398536 | 0.1493011204 |
| startle_PPI | GGB28949_SGB41655 | 63 | -0.8349564377 | 0.4050363445 |
| startle_PPI | GGB28949_SGB41656 | 63 | -1.174386666 | 0.2427928833 |
| startle_PPI | GGB28950_SGB41657 | 63 | -0.7198817171 | 0.4725058394 |
| startle_PPI | GGB28951_SGB102295 | 63 | -1.150305092 | 0.2524856339 |
| startle_PPI | GGB28951_SGB41658 | 63 | 0.9758827896 | 0.3309387896 |
| startle_PPI | GGB28954_SGB41662 | 63 | 0.5362549511 | 0.5922042586 |
| startle_PPI | GGB28956_SGB41665 | 63 | -3.116796939 | 0.002649290467 |
| startle_PPI | GGB28960_SGB41669 | 63 | 0.6987156176 | 0.4855727492 |
| startle_PPI | GGB28967_SGB41678 | 63 | 1.541769537 | 0.1266326788 |
| startle_PPI | GGB28991_SGB41705 | 63 | 0.3407167593 | 0.7334348773 |
| startle_PPI | GGB29002_SGB41718 | 63 | 0.6548653977 | 0.5132696297 |
| startle_PPI | GGB29003_SGB41719 | 63 | 0.8654099446 | 0.3882175196 |
| startle_PPI | GGB29011_SGB41731 | 63 | 1.559926342 | 0.1223050242 |
| startle_PPI | GGB29531_SGB42317 | 63 | 1.071157597 | 0.2862753734 |
| startle_PPI | GGB29685_SGB42494 | 63 | -0.7888227827 | 0.4313489761 |
| startle_PPI | GGB30141_SGB43066 | 63 | 2.804809512 | 0.006474774353 |
| startle_PPI | GGB30286_SGB43248 | 63 | 0.3297258444 | 0.7417145215 |
| startle_PPI | GGB30303_SGB43268 | 63 | 1.349449692 | 0.1802947656 |
| startle_PPI | GGB30413_SGB43452 | 63 | -0.7918535576 | 0.4295897917 |
| startle_PPI | GGB30454_SGB43514 | 63 | -1.387579325 | 0.1684659819 |
| startle_PPI | GGB30455_SGB43519 | 63 | 0.9364128893 | 0.3507276339 |
| startle_PPI | GGB30461_SGB43527 | 63 | -0.01977082596 | 0.9842262153 |
| startle_PPI | GGB30461_SGB43530 | 63 | 1.21485511 | 0.2271130332 |
| startle_PPI | GGB30463_SGB43537 | 63 | 1.581083924 | 0.1174123042 |
| startle_PPI | GGB30473_SGB43557 | 63 | -0.6489409337 | 0.5170752589 |
| startle_PPI | GGB30475_SGB63182 | 63 | -1.513248833 | 0.1336760022 |
| startle_PPI | GGB30861_SGB44083 | 63 | -1.148945255 | 0.2530410868 |
| startle_PPI | GGB31312_SGB44628 | 63 | 0.6380030004 | 0.5241404474 |
| startle_PPI | GGB31438_SGB44768 | 63 | -0.9316449454 | 0.3531691049 |
| startle_PPI | GGB3171_SGB4185 | 63 | -0.5701516877 | 0.5690723302 |
| startle_PPI | GGB31823_SGB45199 | 63 | -0.3962144238 | 0.6921286863 |
| startle_PPI | GGB31853_SGB45233 | 63 | 1.22301701 | 0.2240422336 |
| startle_PPI | GGB32371_SGB41694 | 63 | -0.4565392283 | 0.6482732717 |
| startle_PPI | GGB3793_SGB5158 | 63 | -0.1986566543 | 0.8425556625 |
| startle_PPI | GGB42598_SGB59794 | 63 | -3.897454639 | 0.000225155777 |
| startle_PPI | GGB45656_SGB63370 | 63 | 0.5173284956 | 0.6053095031 |
| startle_PPI | GGB47127_SGB65054 | 63 | -0.9505159679 | 0.3435703186 |
| startle_PPI | GGB74395_SGB43521 | 63 | -1.835755189 | 0.07000283681 |
| startle_PPI | GGB75053_SGB43494 | 63 | -0.8687585859 | 0.3863950791 |
| startle_PPI | GGB75109_SGB102238 | 63 | -1.374813351 | 0.17235808 |
| startle_PPI | GGB81440_SGB45230 | 63 | 0.7680021306 | 0.4435492081 |
| startle_PPI | Lachnospiraceae_bacterium | 63 | -1.658731762 | 0.1007896121 |
| startle_PPI | Lachnospiraceae_bacterium_A2 | 63 | -1.125841229 | 0.2626117913 |

| | | | | |
|---|---|---|---|---|
| startle_PPI | Lachnospiraceae_bacterium_MD308 | 63 | 0.1076432914 | 0.9142825809 |
| startle_PPI | Lachnospiraceae_bacterium_MD329 | 63 | -1.303280663 | 0.1954505754 |
| startle_PPI | Lachnospiraceae_unclassified_SGB4141 | 63 | -2.529553663 | 0.01356300109 |
| startle_PPI | Lachnospiraceae_unclassified_SGB4141 | 63 | 0.1554302228 | 0.8764939032 |
| startle_PPI | Lachnospiraceae_unclassified_SGB4142 | 63 | -0.1765920164 | 0.8598460274 |
| startle_PPI | Lachnospiraceae_unclassified_SGB4158 | 63 | -1.52859163 | 0.1298493933 |
| startle_PPI | Lactobacillus_johnsonii | 63 | -0.5242744214 | 0.6004844834 |
| startle_PPI | Muribaculaceae_bacterium | 63 | -1.604007108 | 0.1122901132 |
| startle_PPI | Neglectibacter_sp_X4 | 63 | 0.8297914037 | 0.4079325088 |
| startle_PPI | Oscillospiraceae_bacterium | 63 | -0.4454896748 | 0.6562189958 |
| startle_PPI | Oscillospiraceae_unclassified_SGB4350 | 63 | -0.1506959121 | 0.880226309 |
| startle_PPI | Oscillospiraceae_unclassified_SGB4350 | 63 | 1.461684378 | 0.1471939771 |
| startle_PPI | Parasutterella_excrementihominis | 63 | -0.4930242961 | 0.6223307765 |
| startle_PPI | Romboutsia_ilealis | 63 | -0.4808612753 | 0.6309281171 |
| startle_PPI | Schaedlerella_arabinosiphila | 63 | -1.539827024 | 0.1271028145 |
| startle_PPI | Turicibacter_sp_1E2 | 63 | -1.00297174 | 0.3177935944 |
| startle_PPI | bacterium_1XD42_54 | 63 | -0.2743462059 | 0.783881503 |
| startle_PPI | bacterium_1XD42_76 | 63 | -1.779478608 | 0.0787973386 |
| startle_PPI | bacterium_1xD8_48 | 63 | 0.351081592 | 0.7256558296 |
| startle_PPI | Berger Parker Index | 63 | 0.1665168508 | 0.8677646375 |
| startle_PPI | Richness (# observed features) | 63 | 2.56279297 | 0.01243599885 |
| startle_PPI | Shannon Index | 63 | 0.936667175 | 0.3505977329 |
| startle_habituation | Acetatifactor_SGB41546 | 63 | -0.5401624641 | 0.5902333958 |
| startle_habituation | Acetatifactor_muris | 63 | 1.808727735 | 0.08225583038 |
| startle_habituation | Acutalibacter_muris | 63 | -0.3876582122 | 0.7026534509 |
| startle_habituation | Acutalibacter_sp_1XD8_36 | 63 | -0.2416752419 | 0.8127144751 |
| startle_habituation | Adlercreutzia_caecimuris | 63 | 0.641458477 | 0.5238231388 |
| startle_habituation | Adlercreutzia_mucosicola | 63 | -1.024300524 | 0.3210901906 |
| startle_habituation | Adlercreutzia_muris | 63 | 0.4653815406 | 0.6436409553 |
| startle_habituation | Akkermansia_muciniphila | 63 | -2.242843781 | 0.0287079963 |
| startle_habituation | Alistipes_sp_DSM_112343 | 63 | -2.366626763 | 0.02090288402 |
| startle_habituation | Anaerotruncus_sp_1XD42_93 | 63 | -0.1316934952 | 0.9036972807 |
| startle_habituation | Bacteria_unclassified_SGB102200 | 63 | 1.561964048 | 0.1264589225 |
| startle_habituation | Bacteria_unclassified_SGB41677 | 63 | -0.3133117349 | 0.7647585713 |
| startle_habituation | Bacteria_unclassified_SGB43546 | 63 | 0.2573198942 | 0.8037535788 |
| startle_habituation | Bacteroides_thetaiotaomicron | 63 | -2.248879749 | 0.03104050107 |
| startle_habituation | Bifidobacterium_pseudolongum | 63 | -0.01004608443 | 0.9920945847 |
| startle_habituation | Clostridia_bacterium | 63 | -0.7068517119 | 0.4938757309 |
| startle_habituation | Clostridiaceae_bacterium | 63 | -0.4538058262 | 0.6612819635 |
| startle_habituation | Clostridiaceae_unclassified_SGB41663 | 63 | 0.124543435 | 0.9015736236 |
| startle_habituation | Clostridiales_bacterium | 63 | -1.354875669 | 0.1785752658 |
| startle_habituation | Clostridium_cocleatum | 63 | -2.224893343 | 0.03101051245 |
| startle_habituation | Coriobacteriaceae_bacterium | 63 | -1.545400822 | 0.1270910547 |
| startle_habituation | Dorea_sp_5_2 | 63 | 0.1441318787 | 0.8858584489 |
| startle_habituation | Dubosiella_newyorkensis | 63 | -0.794503983 | 0.4820055547 |
| startle_habituation | Erysipelotrichales_bacterium | 63 | -0.4662338962 | 0.6459403269 |

| | | | | |
|---|---|---|---|---|
| startle_habituation | Eubacteriaceae_bacterium | 63 | 0.2808204486 | 0.7813572425 |
| startle_habituation | Eubacteriaceae_unclassified_SGB9492; | 63 | 0.6243009244 | 0.5332259582 |
| startle_habituation | GGB20149_SGB29430 | 63 | -0.4013111543 | 0.6901424246 |
| startle_habituation | GGB22635_SGB63107 | 63 | -0.5214829839 | 0.6055076337 |
| startle_habituation | GGB25041_SGB36960 | 63 | -0.8727631186 | 0.4122800413 |
| startle_habituation | GGB27876_SGB40310 | 63 | -1.949971715 | 0.05619267284 |
| startle_habituation | GGB27878_SGB40312 | 63 | 0.6668896778 | 0.5780508952 |
| startle_habituation | GGB27918_SGB40356 | 63 | -0.5344178135 | 0.5992079719 |
| startle_habituation | GGB28382_SGB40962 | 63 | -1.668541197 | 0.1229408996 |
| startle_habituation | GGB28399_SGB40980 | 63 | -0.07390656215 | 0.9411447807 |
| startle_habituation | GGB28411_SGB40993 | 63 | 1.117143564 | 0.2742682362 |
| startle_habituation | GGB28415_SGB40997 | 63 | 1.828484353 | 0.07291142053 |
| startle_habituation | GGB28430_SGB41013 | 63 | 1.507088518 | 0.1421543296 |
| startle_habituation | GGB28439_SGB41022 | 63 | 0.165414857 | 0.8688443442 |
| startle_habituation | GGB28778_SGB41431 | 63 | 0.9327789226 | 0.372565146 |
| startle_habituation | GGB28784_SGB41437 | 63 | 1.87786062 | 0.06799480322 |
| startle_habituation | GGB28792_SGB41445 | 63 | 1.398132655 | 0.2328749066 |
| startle_habituation | GGB28798_SGB41451 | 63 | 0.6666185269 | 0.5182868483 |
| startle_habituation | GGB28802_SGB41455 | 63 | 3.656195775 | 0.00067248293 |
| startle_habituation | GGB28818_SGB41473 | 63 | 0.3404863832 | 0.7402622426 |
| startle_habituation | GGB28828_SGB41484 | 63 | 2.032449421 | 0.06355762909 |
| startle_habituation | GGB28851_SGB41518 | 63 | 0.2170864461 | 0.8313974973 |
| startle_habituation | GGB28859_SGB41528 | 63 | 0.6632371833 | 0.5375356189 |
| startle_habituation | GGB28864_SGB41535 | 63 | 1.644097719 | 0.1484954752 |
| startle_habituation | GGB28869_SGB41543 | 63 | 0.9604120529 | 0.3603570436 |
| startle_habituation | GGB28883_SGB41564 | 63 | 1.580470949 | 0.121457029 |
| startle_habituation | GGB28892_SGB41573 | 63 | 0.2618297295 | 0.7981719079 |
| startle_habituation | GGB28893_SGB41574 | 63 | -0.875663561 | 0.3826751741 |
| startle_habituation | GGB28898_SGB41580 | 63 | 1.406539825 | 0.1636985756 |
| startle_habituation | GGB28904_SGB41597 | 63 | -0.6765055124 | 0.5163528432 |
| startle_habituation | GGB28916_SGB41612 | 63 | 0.3769245685 | 0.7102913843 |
| startle_habituation | GGB28924_SGB41621 | 63 | -1.235667939 | 0.2217153737 |
| startle_habituation | GGB28926_SGB41624 | 63 | -0.9131113514 | 0.388613361 |
| startle_habituation | GGB28927_SGB41625 | 63 | 0.3481535104 | 0.7279684386 |
| startle_habituation | GGB28934_SGB41635 | 63 | 1.039331576 | 0.3356180562 |
| startle_habituation | GGB28946_SGB41652 | 63 | 0.2691517524 | 0.7879935173 |
| startle_habituation | GGB28949_SGB41655 | 63 | 0.1387099384 | 0.890101361 |
| startle_habituation | GGB28949_SGB41656 | 63 | 0.1500271537 | 0.8816912043 |
| startle_habituation | GGB28950_SGB41657 | 63 | -0.4843148302 | 0.6284995968 |
| startle_habituation | GGB28951_SGB102295 | 63 | -0.9056373387 | 0.3674906697 |
| startle_habituation | GGB28951_SGB41658 | 63 | 0.8224447449 | 0.4131656052 |
| startle_habituation | GGB28954_SGB41662 | 63 | 0.3133645916 | 0.7548624389 |
| startle_habituation | GGB28956_SGB41665 | 63 | -1.692840493 | 0.1028361255 |
| startle_habituation | GGB28960_SGB41669 | 63 | 2.432826594 | 0.0190216052 |
| startle_habituation | GGB28967_SGB41678 | 63 | 0.597842929 | 0.5580513302 |
| startle_habituation | GGB28991_SGB41705 | 63 | 0.4213548195 | 0.679717763 |

| | | | | |
|---|---|---|---|---|
| startle_habituation | GGB29002_SGB41718 | 63 | 0.3452530271 | 0.7308785659 |
| startle_habituation | GGB29003_SGB41719 | 63 | -0.6768787397 | 0.5169977742 |
| startle_habituation | GGB29011_SGB41731 | 63 | -0.03890322002 | 0.9690804104 |
| startle_habituation | GGB29531_SGB42317 | 63 | 0.6522237111 | 0.516081854 |
| startle_habituation | GGB29685_SGB42494 | 63 | -0.9638482915 | 0.3413918289 |
| startle_habituation | GGB30141_SGB43066 | 63 | 2.096088371 | 0.0454876065 |
| startle_habituation | GGB30286_SGB43248 | 63 | -0.3674270695 | 0.7232240899 |
| startle_habituation | GGB30303_SGB43268 | 63 | 0.637626261 | 0.5337296769 |
| startle_habituation | GGB30413_SGB43452 | 63 | 0.05276728069 | 0.9581674088 |
| startle_habituation | GGB30454_SGB43514 | 63 | -0.6301698374 | 0.5293752132 |
| startle_habituation | GGB30455_SGB43519 | 63 | 2.160843036 | 0.04700316645 |
| startle_habituation | GGB30461_SGB43527 | 63 | -0.8567294905 | 0.4026797198 |
| startle_habituation | GGB30461_SGB43530 | 63 | 1.000696775 | 0.3299897225 |
| startle_habituation | GGB30463_SGB43537 | 63 | 1.431319727 | 0.176235745 |
| startle_habituation | GGB30473_SGB43557 | 63 | -0.712496141 | 0.5233375857 |
| startle_habituation | GGB30475_SGB63182 | 63 | -1.539758644 | 0.1711268487 |
| startle_habituation | GGB30861_SGB44083 | 63 | -1.01837543 | 0.3105070205 |
| startle_habituation | GGB31312_SGB44628 | 63 | 0.7371474034 | 0.4800862301 |
| startle_habituation | GGB31438_SGB44768 | 63 | 0.3075609414 | 0.7683507837 |
| startle_habituation | GGB3171_SGB4185 | 63 | -1.475346438 | 0.1707888844 |
| startle_habituation | GGB31823_SGB45199 | 63 | -1.796154335 | 0.07696819616 |
| startle_habituation | GGB31853_SGB45233 | 63 | -0.1247854507 | 0.9013501256 |
| startle_habituation | GGB32371_SGB41694 | 63 | -1.326569893 | 0.2035605897 |
| startle_habituation | GGB3793_SGB5158 | 63 | -1.286234305 | 0.2253065374 |
| startle_habituation | GGB42598_SGB59794 | 63 | -2.255202518 | 0.02730478375 |
| startle_habituation | GGB45656_SGB63370 | 63 | -0.03540134442 | 0.9741590441 |
| startle_habituation | GGB47127_SGB65054 | 63 | -0.5729971993 | 0.591535064 |
| startle_habituation | GGB74395_SGB43521 | 63 | -0.8307619765 | 0.40771711 |
| startle_habituation | GGB75053_SGB43494 | 63 | 1.059888543 | 0.2945289628 |
| startle_habituation | GGB75109_SGB102238 | 63 | -0.4027008636 | 0.6916114781 |
| startle_habituation | GGB81440_SGB45230 | 63 | 0.7663566438 | 0.4674294258 |
| startle_habituation | Lachnospiraceae_bacterium | 63 | -0.5263951162 | 0.600797878 |
| startle_habituation | Lachnospiraceae_bacterium_A2 | 63 | -1.042444246 | 0.3277029546 |
| startle_habituation | Lachnospiraceae_bacterium_MD308 | 63 | 0.6750159694 | 0.5036106916 |
| startle_habituation | Lachnospiraceae_bacterium_MD329 | 63 | -1.451095273 | 0.1514752768 |
| startle_habituation | Lachnospiraceae_unclassified_SGB4141 | 63 | -1.287250905 | 0.2193774212 |
| startle_habituation | Lachnospiraceae_unclassified_SGB4141 | 63 | -0.5590409134 | 0.5977285077 |
| startle_habituation | Lachnospiraceae_unclassified_SGB4142 | 63 | -1.523665245 | 0.1338604077 |
| startle_habituation | Lachnospiraceae_unclassified_SGB4158 | 63 | -1.043847695 | 0.3070195884 |
| startle_habituation | Lactobacillus_johnsonii | 63 | -0.2304323133 | 0.8183075298 |
| startle_habituation | Muribaculaceae_bacterium | 63 | -2.373016447 | 0.02023268208 |
| startle_habituation | Neglectibacter_sp_X4 | 63 | 1.003187477 | 0.3489764725 |
| startle_habituation | Oscillospiraceae_bacterium | 63 | -0.2934549106 | 0.7736363921 |
| startle_habituation | Oscillospiraceae_unclassified_SGB4350 | 63 | -0.9707920929 | 0.3379221071 |
| startle_habituation | Oscillospiraceae_unclassified_SGB4350 | 63 | 1.603743614 | 0.1124639303 |
| startle_habituation | Parasutterella_excrementihominis | 63 | -1.472335112 | 0.1512104697 |

| | | | | |
|---|---|---|---|---|
| startle_habituation | Romboutsia_ilealis | 63 | -2.046965725 | 0.06145866059 |
| startle_habituation | Schaedlerella_arabinosiphila | 63 | -1.142780164 | 0.3538269807 |
| startle_habituation | Turicibacter_sp_1E2 | 63 | -0.8888290442 | 0.3755851484 |
| startle_habituation | bacterium_1XD42_54 | 63 | -0.2301421334 | 0.8186205229 |
| startle_habituation | bacterium_1XD42_76 | 63 | -1.288693985 | 0.2006233 |
| startle_habituation | bacterium_1xD8_48 | 63 | 0.3768332785 | 0.7066193127 |
| startle_habituation | Berger Parker Index | 63 | -1.815426166 | 0.0747301899 |
| startle_habituation | Richness (# observed features) | 63 | 1.699161377 | 0.09294646481 |
| startle_habituation | Shannon Index | 63 | 1.944141934 | 0.05777189944 |

| st Results | | | | | |
|---|---|---|---|---|---|
| Microbe Adjusted p-valu | Microbe Estimate | Blast Estimate | VNS Estimate | Blast:VNS Estimate | Microbe Std. Erro |
| 0.2153727604 | 0.3148338033 | 0.960831046 | 0.738839469 | -1.080249443 | 0.1191093034 |
| 0.9761770834 | 0.009552748091 | 0.8632601954 | 0.6412276672 | -1.056740335 | 0.1211054019 |
| 0.5486787709 | 0.1971153136 | 0.9450869445 | 0.6388254149 | -1.113242142 | 0.1191845724 |
| 0.7354029002 | 0.1069322161 | 0.8571653925 | 0.62389001 | -1.004091917 | 0.1220340025 |
| 0.2153727604 | -0.2889847642 | 0.8436943497 | 0.7123352018 | -1.02128563 | 0.115303623 |
| 0.7406447589 | 0.1053084105 | 0.8913269603 | 0.6754258438 | -1.12830884 | 0.1249094646 |
| 0.8559555964 | 0.04926325013 | 0.8581097481 | 0.6546882057 | -1.065014855 | 0.1216801821 |
| 0.5762614969 | -0.1816894201 | 0.9489421913 | 0.6559377954 | -1.027357119 | 0.1226616914 |
| 0.6376240108 | -0.146913028 | 0.9294962612 | 0.6656230587 | -1.071785185 | 0.1262066731 |
| 0.7754670422 | 0.08272856977 | 0.8649098453 | 0.624648569 | -1.037345685 | 0.119524328 |
| 0.9023781973 | -0.02750883994 | 0.8540821184 | 0.6336459213 | -1.050857953 | 0.1241803122 |
| 0.8559555964 | -0.05046694283 | 0.8552383502 | 0.640548536 | -1.025580025 | 0.1228895433 |
| 0.8780395729 | 0.03936799383 | 0.8643837284 | 0.6508737499 | -1.069937465 | 0.1229662943 |
| 0.7350810244 | -0.1118724238 | 0.8603211853 | 0.6032485863 | -1.03572227 | 0.1198336178 |
| 0.5486787709 | -0.1973658474 | 0.8681949129 | 0.6409037025 | -1.145331575 | 0.1200708854 |
| 0.5762614969 | -0.1733875235 | 0.791061364 | 0.5971830415 | -1.018255325 | 0.12066146 |
| 0.2153727604 | -0.3121794085 | 0.8472068822 | 0.4837655036 | -0.9945557746 | 0.1154004639 |
| 0.7215974205 | 0.125116097 | 0.8922532146 | 0.6543055951 | -1.104504796 | 0.12240821 |
| 0.8559555964 | 0.05636183019 | 0.8333196035 | 0.6626145002 | -1.091830339 | 0.1329042515 |
| 0.8780395729 | 0.03772743626 | 0.8346079949 | 0.6265013063 | -1.052014239 | 0.1298595747 |
| 0.8358558141 | -0.06583152002 | 0.8210502944 | 0.6109155811 | -1.003364398 | 0.1227139947 |
| 0.5762614969 | -0.1656362709 | 0.860006263 | 0.6136187177 | -1.10511407 | 0.1226672979 |
| 0.526118124 | -0.2234383286 | 0.7412088396 | 0.6031679639 | -0.9990083334 | 0.120528236 |
| 0.8559555964 | 0.04752179282 | 0.8572611271 | 0.6191777731 | -1.040846115 | 0.1225916967 |
| 0.8358558141 | -0.06965992239 | 0.9094374885 | 0.6597582528 | -1.092405389 | 0.1333502792 |
| 0.7484938865 | -0.09981296499 | 0.8849785618 | 0.6977619976 | -1.078627063 | 0.1244779285 |
| 0.7215974205 | 0.1269640464 | 0.8199896351 | 0.6531570093 | -1.00388475 | 0.1252271045 |
| 0.5762614969 | -0.1688613034 | 0.9323965159 | 0.6775691499 | -1.064473677 | 0.1211940456 |
| 0.7354029002 | -0.1088152566 | 0.7799514571 | 0.5880363616 | -0.9511147857 | 0.1232870601 |
| 0.6555744666 | -0.1370380564 | 0.7800700722 | 0.619928237 | -0.9923810497 | 0.1243152056 |
| 0.9498139359 | -0.0160706726 | 0.857926176 | 0.637978923 | -1.054503602 | 0.1236815234 |
| 0.7754670422 | 0.08889284655 | 0.8569837846 | 0.5879082249 | -1.031284821 | 0.1306063181 |
| 0.6452350437 | 0.1380984383 | 0.8167838387 | 0.6369607383 | -1.034282585 | 0.1214627818 |
| 0.8780395729 | 0.03390381431 | 0.8455859341 | 0.623406832 | -1.021893746 | 0.1252679442 |
| 0.8559555964 | -0.05993392106 | 0.8824553144 | 0.6500993561 | -1.117410627 | 0.1296215117 |
| 0.8780395729 | -0.03800434462 | 0.8495809016 | 0.6448410998 | -1.063860388 | 0.1258422894 |
| 0.7754670422 | 0.09153628343 | 0.8518137996 | 0.6677457193 | -1.043476745 | 0.1233094176 |
| 0.5569218181 | 0.1956233859 | 0.856811949 | 0.7130614677 | -1.141240596 | 0.1215835657 |
| 0.8780395729 | 0.03736663209 | 0.8546339038 | 0.6537171052 | -1.061879176 | 0.1333853666 |
| 0.8212864485 | -0.07398707388 | 0.85532467 | 0.6881570918 | -1.118862535 | 0.1296218423 |
| 0.9904345571 | 0.001509548649 | 0.8625944091 | 0.6400092582 | -1.055049258 | 0.1259132485 |
| 0.7484938865 | -0.09646500085 | 0.8652136927 | 0.6654531915 | -1.100898738 | 0.1206346084 |
| 0.9790000627 | -0.007521474665 | 0.8617476165 | 0.6429361354 | -1.057984599 | 0.1241321067 |
| 0.8490526455 | -0.06146551168 | 0.8656589344 | 0.630948579 | -1.069132524 | 0.1231657801 |
| 0.8559555964 | -0.05565387852 | 0.8495905186 | 0.6378603027 | -1.02424777 | 0.1235125986 |

| | | | | | |
|---|---|---|---|---|---|
| 0.8564192709 | 0.04575326194 | 0.8715595177 | 0.6394112918 | -1.071573164 | 0.122374082 |
| 0.8212864485 | -0.07168086638 | 0.8566595234 | 0.6391527334 | -1.089444248 | 0.1242602947 |
| 0.7406447589 | 0.1046833777 | 0.8595415952 | 0.6421518308 | -1.028387172 | 0.1232704805 |
| 0.5762614969 | 0.1855107635 | 0.9023412088 | 0.7350347694 | -1.17604909 | 0.122056183 |
| 0.5762614969 | -0.1650650198 | 0.8223980688 | 0.5495378623 | -1.060047233 | 0.1230254852 |
| 0.5762614969 | -0.1689312638 | 0.9140060392 | 0.6865545867 | -1.109564967 | 0.1200472787 |
| 0.5762614969 | 0.1759215617 | 0.9878131893 | 0.7489446297 | -1.226871267 | 0.1248032617 |
| 0.4856544148 | -0.238349872 | 0.7451342327 | 0.6102041593 | -0.8996297758 | 0.1225565359 |
| 0.7350810244 | 0.1176884173 | 0.8900469265 | 0.6802353973 | -1.00193287 | 0.1267698242 |
| 0.8564192709 | 0.04561396206 | 0.8706919362 | 0.6452374747 | -1.062593925 | 0.1237309817 |
| 0.9329627687 | 0.02038475489 | 0.8593201943 | 0.6436547574 | -1.051482074 | 0.120598971 |
| 0.7754670422 | -0.08947607683 | 0.8615245505 | 0.6017052498 | -0.9947984157 | 0.1241011219 |
| 0.2153727604 | 0.2893182618 | 0.8263340282 | 0.6928203638 | -1.15225712 | 0.1153422456 |
| 0.7754670422 | 0.08997955231 | 0.9088995281 | 0.680509543 | -1.15743858 | 0.1266110634 |
| 0.7754670422 | 0.0853007414 | 0.835928716 | 0.5984602125 | -1.015725428 | 0.1243686497 |
| 0.6376240108 | -0.1420788651 | 0.814593004 | 0.6290022321 | -1.054946142 | 0.1199363527 |
| 0.7484938865 | 0.1010913231 | 0.9075153048 | 0.65629966 | -1.081269865 | 0.1255147622 |
| 0.8559555964 | -0.05286562498 | 0.8209506298 | 0.6192360586 | -1.022175582 | 0.1266841293 |
| 0.8559555964 | 0.050213126 | 0.8638336157 | 0.6163281732 | -1.050453264 | 0.1255114992 |
| 0.2153727604 | 0.3311156562 | 0.8681677116 | 0.6338912592 | -0.9740978422 | 0.1140614728 |
| 0.9023781973 | -0.02661414541 | 0.8525949487 | 0.6315567973 | -1.047236398 | 0.1240743225 |
| 0.8780395729 | 0.03509057993 | 0.8407760896 | 0.633041025 | -1.025970925 | 0.1235777711 |
| 0.9904345571 | -0.00258276364 | 0.8624424785 | 0.6409628262 | -1.05664012 | 0.1255422915 |
| 0.6555744666 | 0.1376303263 | 0.7927373429 | 0.5877641392 | -1.035393658 | 0.1243395135 |
| 0.7754670422 | -0.09183386283 | 0.8803126506 | 0.6823097781 | -1.128559963 | 0.1214014714 |
| 0.526118124 | 0.2267853812 | 0.9270903238 | 0.5719772736 | -0.8540687005 | 0.1296938621 |
| 0.8358558141 | -0.06470454942 | 0.8725556872 | 0.6401692489 | -1.042559897 | 0.1228096373 |
| 0.7350810244 | -0.1131017006 | 0.8658493872 | 0.5983020905 | -1.052257103 | 0.1234073498 |
| 0.132973507 | 0.3870917018 | 0.9785021771 | 0.8555324073 | -1.398368442 | 0.1130268399 |
| 0.2153727604 | -0.3143962736 | 0.9215908136 | 0.7806834698 | -1.064357016 | 0.1156154157 |
| 0.7754670422 | 0.08140428983 | 0.8562103157 | 0.6621309578 | -1.044118084 | 0.1229564872 |
| 0.9498139359 | -0.01764704365 | 0.8766030672 | 0.6488364616 | -1.066176548 | 0.1274577458 |
| 0.5762614969 | 0.1766713335 | 0.8304609941 | 0.6625911009 | -1.090053773 | 0.1210478307 |
| 0.7922115161 | -0.07886231977 | 0.8948730875 | 0.6551855071 | -1.089539999 | 0.1241644768 |
| 0.5762614969 | -0.1702827074 | 0.8136233285 | 0.5696542519 | -1.07317079 | 0.1219457683 |
| 0.526118124 | 0.2151448042 | 0.9648961091 | 0.7752844129 | -1.182139697 | 0.1219924415 |
| 0.5486787709 | -0.2020796255 | 0.9522679441 | 0.6770502282 | -1.143144797 | 0.1203431088 |
| 0.6376240108 | 0.1455861415 | 0.875431205 | 0.672565755 | -1.025392225 | 0.1233622233 |
| 0.7244558275 | 0.123512134 | 0.9160278738 | 0.6616183375 | -1.050593841 | 0.1248688012 |
| 0.8778323496 | -0.04374457325 | 0.8644204635 | 0.6030329682 | -1.010953299 | 0.130392747 |
| 0.5832622209 | -0.154070515 | 0.916091029 | 0.6235919688 | -1.051541467 | 0.1207793498 |
| 0.2153727604 | -0.3000911327 | 0.8918360227 | 0.7119204347 | -1.081081458 | 0.113989469 |
| 0.4856544148 | -0.2259052137 | 0.8626682614 | 0.6361555011 | -0.9736263988 | 0.1175216391 |
| 0.5762614969 | -0.1853764931 | 0.8511550506 | 0.6436877615 | -1.053335081 | 0.1211331771 |
| 0.8780395729 | 0.03514017344 | 0.8390662922 | 0.6380134605 | -1.045305676 | 0.1277336482 |
| 0.7215974205 | -0.1240137515 | 0.9085221355 | 0.660061831 | -1.05383816 | 0.1235468956 |

| | | | | | |
|---|---|---|---|---|---|
| 0.4629433469 | 0.2465644919 | 0.8519521383 | 0.5524752574 | -0.9150754448 | 0.119249494 |
| 0.7754670422 | -0.08939978913 | 0.8654147595 | 0.7131042784 | -1.106755314 | 0.1297653434 |
| 0.8559555964 | 0.05131439435 | 0.8658240093 | 0.6391789638 | -1.051716671 | 0.1225859819 |
| 0.8165197312 | -0.08022140017 | 0.8817681008 | 0.6378927679 | -1.089215708 | 0.1335671633 |
| 0.5762614969 | 0.162784033 | 0.7977432398 | 0.6336954419 | -1.018301227 | 0.1230414659 |
| 0.4732912503 | -0.245893686 | 0.8969864249 | 0.7181350554 | -0.9925581097 | 0.1230215819 |
| 0.5486787709 | -0.2064994227 | 0.8928431105 | 0.6184416687 | -0.9146442629 | 0.126119701 |
| 0.621710498 | 0.1514010048 | 0.9034313247 | 0.6849542018 | -1.081838717 | 0.1236953611 |
| 0.7354029002 | -0.1068558537 | 0.8269190264 | 0.6264254995 | -1.019723281 | 0.1197241649 |
| 0.526118124 | 0.2261662831 | 0.7722621276 | 0.5771882387 | -0.7948219643 | 0.1270572019 |
| 0.5762614969 | -0.1653387341 | 0.8218040317 | 0.6120718586 | -1.065264242 | 0.1211041787 |
| 0.6376240108 | -0.1403263637 | 0.8082300845 | 0.6315678153 | -1.00158222 | 0.1213005765 |
| 0.5762614969 | -0.1879824063 | 0.7749325174 | 0.5306699018 | -1.019017149 | 0.1272663331 |
| 0.7406447589 | -0.1062476102 | 0.906789223 | 0.6175348814 | -1.004655293 | 0.1266543803 |
| 0.7350810244 | 0.1183497308 | 0.9129721279 | 0.686536243 | -1.174342079 | 0.1300906164 |
| 0.8869524551 | 0.03081577877 | 0.8571503937 | 0.6303446268 | -1.048645674 | 0.1227340483 |
| 0.7754670422 | 0.08136361726 | 0.8367133141 | 0.6076753483 | -0.9905210197 | 0.1227072528 |
| 0.8559555964 | -0.04828329297 | 0.856134228 | 0.6516887302 | -1.048700594 | 0.1239773207 |
| 0.9790000627 | -0.006973926818 | 0.865799015 | 0.6403033384 | -1.056308466 | 0.1249873738 |
| 0.4629433469 | -0.2436695149 | 0.9003438889 | 0.6745166552 | -1.088165946 | 0.1156205207 |
| 0.4629433469 | 0.2462954241 | 1.023957447 | 0.6945041243 | -1.266883032 | 0.1204848565 |
| 0.9812632823 | -0.005207160291 | 0.8626933188 | 0.6395141262 | -1.054104782 | 0.1206086075 |
| 0.7350810244 | 0.1173022534 | 0.9032573836 | 0.7346658369 | -1.201005887 | 0.1255099373 |
| 0.7754670422 | 0.08180741236 | 0.869552359 | 0.6396916965 | -1.024015486 | 0.1233520353 |
| 0.5832622209 | -0.1575016988 | 0.8660904844 | 0.7158144808 | -1.096600454 | 0.1235224567 |
| 0.9653516843 | -0.01406900233 | 0.8704989005 | 0.6465110356 | -1.051176061 | 0.1383665965 |
| 0.5832622209 | -0.1526381984 | 0.8500960818 | 0.6252964579 | -1.006686154 | 0.1189570982 |
| 0.5762614969 | -0.1700128169 | 0.8791209152 | 0.6617390744 | -1.177005471 | 0.1220207667 |
| 0.2153727604 | -0.300126169 | 1.068798832 | 0.811961813 | -1.144625707 | 0.1209886014 |
| 0.8559555964 | -0.05094062145 | 0.8637848434 | 0.6539211174 | -1.061698275 | 0.1290294542 |
| 0.8212864485 | 0.07208972602 | 0.8802005926 | 0.6474399519 | -1.0850347 | 0.1272705184 |
| 0.526118124 | -0.218830784 | 0.7974578533 | 0.5091642783 | -0.9440667475 | 0.1199157629 |
| 0.5762614969 | 0.1610379964 | 0.847243299 | 0.6874876144 | -1.121317256 | 0.1212514292 |
| 0.7350810244 | 0.1124264197 | 0.8970005124 | 0.6510641739 | -1.073349245 | 0.1203007625 |
| 0.9558407212 | 0.01596086659 | 0.7957626027 | 0.4993691844 | -1.023258924 | 0.1257276647 |
| 0.6959866521 | -0.1853323559 | 0.7848514702 | 0.4835542986 | -1.009099735 | 0.1192522804 |
| 0.9580582754 | -0.009654314869 | 0.7884592527 | 0.4956641813 | -1.021618307 | 0.1223891721 |
| 0.8243251094 | 0.06593492154 | 0.7919305192 | 0.4886619035 | -0.9971942161 | 0.1241812536 |
| 0.6959866521 | -0.1853463399 | 0.7836903635 | 0.5422556264 | -1.008953828 | 0.1204065605 |
| 0.8243251094 | 0.07302564364 | 0.8247432854 | 0.5273318966 | -1.091741709 | 0.1272672901 |
| 0.9580582754 | -0.008884113283 | 0.7942057236 | 0.4939280719 | -1.024120869 | 0.1214284259 |
| 0.9558407212 | -0.01596294395 | 0.8008682113 | 0.4973848679 | -1.023142074 | 0.1248899891 |
| 0.9580582754 | -0.01163848207 | 0.7970553087 | 0.4972507539 | -1.024758452 | 0.1278932501 |
| 0.6959866521 | 0.1975959001 | 0.7558505421 | 0.4190374883 | -0.9128392785 | 0.1222183414 |
| 0.8243251094 | -0.07202594767 | 0.7755050476 | 0.4825960132 | -1.019668636 | 0.1231865637 |
| 0.6959866521 | 0.2206982788 | 0.844551739 | 0.5082627847 | -1.183623888 | 0.120258488 |

| | | | | | |
|---|---|---|---|---|---|
| 0.8938824884 | -0.03561235078 | 0.7938358848 | 0.4877520755 | -1.015836706 | 0.1230242194 |
| 0.732936013 | -0.1150298957 | 0.7905677671 | 0.4583373845 | -1.005809367 | 0.1200812103 |
| 0.7158466291 | -0.1458273198 | 0.7906082134 | 0.4905587726 | -1.080918617 | 0.1202428634 |
| 0.7518157004 | -0.09442433273 | 0.7538796587 | 0.4715161819 | -1.005284701 | 0.1214083719 |
| 0.7782601275 | -0.08534462623 | 0.7838793193 | 0.4509096658 | -1.002921476 | 0.123165859 |
| 0.732936013 | 0.1010195991 | 0.8089481991 | 0.5044932844 | -1.054383486 | 0.1226089309 |
| 0.892341832 | -0.04253622099 | 0.8185252789 | 0.4818630279 | -1.002883741 | 0.1327052959 |
| 0.732936013 | -0.1057430494 | 0.8655284197 | 0.5329547728 | -1.026358282 | 0.1307703315 |
| 0.7158466291 | -0.1505095783 | 0.7030755494 | 0.4412744563 | -0.9187735762 | 0.1349554103 |
| 0.8243251094 | -0.07524324672 | 0.8052455663 | 0.4937865664 | -1.066943152 | 0.1260077374 |
| 0.6959866521 | -0.2301105221 | 0.6586861351 | 0.4551674152 | -0.9501044073 | 0.1215412063 |
| 0.732936013 | -0.1178408308 | 0.806779015 | 0.545019288 | -1.062146237 | 0.1224683487 |
| 0.6959866521 | 0.1845548056 | 0.6585414467 | 0.4266057486 | -0.9136109231 | 0.1338715602 |
| 0.7158466291 | -0.143004054 | 0.8262208341 | 0.56855557 | -1.058241682 | 0.1316475494 |
| 0.9664084581 | -0.006567543474 | 0.795191529 | 0.4954517559 | -1.027818179 | 0.1262245316 |
| 0.7158466291 | -0.1461108568 | 0.8400627662 | 0.5235830187 | -1.016726495 | 0.1217006869 |
| 0.732936013 | 0.1135249953 | 0.8890757673 | 0.5577547767 | -1.147189699 | 0.1241894579 |
| 0.7158466291 | -0.1282943041 | 0.7147818324 | 0.4734238813 | -0.9647673751 | 0.1240819307 |
| 0.9558407212 | 0.01624221776 | 0.7984623878 | 0.4995330806 | -1.02770908 | 0.1353060271 |
| 0.9558407212 | -0.0170819964 | 0.7920805956 | 0.5029353288 | -1.026285521 | 0.1431936803 |
| 0.7158466291 | 0.1511115162 | 0.7184666384 | 0.472519274 | -0.9600777245 | 0.1258971275 |
| 0.6959866521 | 0.2489541012 | 0.6988841196 | 0.3913832547 | -0.8213046264 | 0.1217074757 |
| 0.8337172861 | -0.0626813045 | 0.8156070719 | 0.5082220491 | -1.092717848 | 0.1310078313 |
| 0.8243251094 | -0.1038566144 | 0.7332963584 | 0.4926836966 | -1.008702513 | 0.1406275854 |
| 0.6959866521 | 0.2426353928 | 0.7089838659 | 0.5269469513 | -0.905344463 | 0.1276902215 |
| 0.6959866521 | 0.2007723077 | 0.7536598642 | 0.5475436957 | -1.063509459 | 0.1228432643 |
| 0.6959866521 | 0.2527623523 | 0.7693659587 | 0.6058872654 | -1.110469731 | 0.1300099271 |
| 0.878519134 | 0.04977310965 | 0.7963545043 | 0.4648444641 | -0.9813138218 | 0.1329864123 |
| 0.7564722499 | 0.1009645504 | 0.8407425197 | 0.500713935 | -1.055231014 | 0.1287335834 |
| 0.9174285189 | -0.02982487093 | 0.7958341558 | 0.5048411317 | -1.042601049 | 0.1268035305 |
| 0.9558407212 | -0.01769246271 | 0.7911500416 | 0.5022517547 | -1.030860578 | 0.1254596148 |
| 0.6959866521 | -0.2545997496 | 0.8333575434 | 0.475033088 | -1.120363614 | 0.1195106441 |
| 0.9174285189 | 0.03104335502 | 0.7975867397 | 0.4945911665 | -1.039067959 | 0.1262883279 |
| 0.7635041575 | -0.09308536677 | 0.7552999462 | 0.4861263382 | -0.9633419546 | 0.122299508 |
| 0.8243251094 | -0.06865103692 | 0.7940456758 | 0.502036045 | -1.067502341 | 0.1269430047 |
| 0.8826037281 | 0.04323503967 | 0.7916859493 | 0.4960368755 | -1.014653689 | 0.1247451664 |
| 0.732936013 | 0.1175599155 | 0.8091535994 | 0.5504699802 | -1.091761456 | 0.1250183037 |
| 0.6959866521 | -0.169088803 | 0.7764555852 | 0.419175716 | -1.064483833 | 0.1234758315 |
| 0.7158466291 | -0.1758048438 | 0.814625155 | 0.5266498534 | -1.043743484 | 0.1357378872 |
| 0.732936013 | -0.1235022636 | 0.7142379374 | 0.4241885958 | -0.9169023211 | 0.1268014525 |
| 0.7158466291 | -0.1604593047 | 0.7112565324 | 0.482977603 | -0.9161734302 | 0.1304540904 |
| 0.732936013 | 0.1059585154 | 0.812085716 | 0.5286752329 | -0.9672081252 | 0.1268209452 |
| 0.8243251094 | 0.07234436808 | 0.8068372485 | 0.5015218705 | -1.036967076 | 0.1242577517 |
| 0.732936013 | 0.1175275166 | 0.746186985 | 0.4990741763 | -0.9589686718 | 0.1248894315 |
| 0.7564722499 | -0.09357188283 | 0.7899344205 | 0.4521995207 | -0.9571331929 | 0.1242648482 |
| 0.6959866521 | 0.2151085209 | 0.7827174007 | 0.5487888877 | -1.122733622 | 0.1196323185 |

| | | | | | |
|---|---|---|---|---|---|
| 0.878519134 | 0.04952449939 | 0.8170724119 | 0.51829722 | -1.079017685 | 0.1278301198 |
| 0.6959866521 | 0.198744285 | 0.7252541984 | 0.4038315347 | -0.9268881441 | 0.1223561569 |
| 0.6959866521 | -0.1706118131 | 0.7448846652 | 0.4904975392 | -1.037932021 | 0.1190323046 |
| 0.8301445012 | 0.0637684134 | 0.8170167682 | 0.5058657664 | -1.035951114 | 0.1296660803 |
| 0.6959866521 | -0.1889119665 | 0.6662436395 | 0.433645951 | -0.936168778 | 0.1268426594 |
| 0.7158466291 | 0.1472287017 | 0.8231204573 | 0.4428530211 | -1.04751857 | 0.1247760756 |
| 0.410290334 | 0.3151893622 | 0.8177575211 | 0.499441974 | -0.9750535302 | 0.1156446953 |
| 0.8243251094 | 0.07609057022 | 0.8047216602 | 0.5084835149 | -1.024783923 | 0.1265555087 |
| 0.6959866521 | 0.1791896359 | 0.7101866117 | 0.4772006431 | -0.9143644927 | 0.1220152219 |
| 0.6959866521 | 0.2311362675 | 0.7202124998 | 0.3800754663 | -0.8036642025 | 0.1221455929 |
| 0.6959866521 | 0.16947743 | 0.6953311169 | 0.4299962052 | -0.9872258524 | 0.123653862 |
| 0.7158466291 | -0.1444758023 | 0.8105415138 | 0.5571133814 | -1.125037535 | 0.1203266844 |
| 0.732936013 | 0.1118372314 | 0.8273723855 | 0.463905144 | -0.9296243802 | 0.1321543221 |
| 0.8559185367 | -0.05627319915 | 0.7962042431 | 0.4923406702 | -1.00602477 | 0.1240850892 |
| 0.6959866521 | -0.1635338194 | 0.7934657924 | 0.4308765302 | -1.016424931 | 0.1270847212 |
| 0.2411235518 | 0.3861009211 | 0.8905867875 | 0.7004923754 | -1.354618966 | 0.1153394857 |
| 0.410290334 | -0.324337361 | 0.8144999062 | 0.6170159587 | -0.9877415345 | 0.1164372926 |
| 0.8826037281 | -0.0499844832 | 0.807389725 | 0.4902243538 | -1.046932781 | 0.125155082 |
| 0.6959866521 | -0.1709111707 | 0.9617276384 | 0.5993196465 | -1.17453303 | 0.1261067143 |
| 0.7648103982 | -0.08738133567 | 0.8142517789 | 0.4907488592 | -1.015494873 | 0.1233349705 |
| 0.8243251094 | 0.07308705886 | 0.7716311857 | 0.488475649 | -1.006304715 | 0.1255528848 |
| 0.732936013 | -0.1005365797 | 0.7630045433 | 0.4540780296 | -1.034815188 | 0.1221102176 |
| 0.8938824884 | -0.03785948463 | 0.7765352999 | 0.4740237084 | -1.004748149 | 0.1257816763 |
| 0.6959866521 | -0.1750115516 | 0.8634885775 | 0.5137974816 | -1.088713792 | 0.1229262019 |
| 0.878519134 | -0.05591798046 | 0.7959197115 | 0.4900757365 | -1.04766601 | 0.1271740791 |
| 0.9558407212 | 0.014158954 | 0.7973026765 | 0.4969048228 | -1.022177999 | 0.1274569574 |
| 0.9857401854 | -0.00237562483 | 0.7929298689 | 0.4938851689 | -1.022553939 | 0.1315196364 |
| 0.7782601275 | -0.08860141853 | 0.8173868829 | 0.4823249752 | -1.016119372 | 0.1254868665 |
| 0.6959866521 | -0.1625518138 | 0.8034972789 | 0.5244065361 | -1.030117184 | 0.1178348113 |
| 0.8243251094 | -0.06521726604 | 0.7912707978 | 0.4920291943 | -1.000131447 | 0.1250389276 |
| 0.732936013 | -0.1554597848 | 0.7480628797 | 0.4560674384 | -0.9728303452 | 0.1403836596 |
| 0.9558407212 | 0.01473699931 | 0.7836088165 | 0.4949558086 | -1.021487968 | 0.1293417161 |
| 0.8938824884 | -0.03459029948 | 0.8054310273 | 0.5016342071 | -1.024104018 | 0.1234735841 |
| 0.6959866521 | 0.2017857413 | 0.7573067641 | 0.4033190626 | -0.8764073052 | 0.1284302326 |
| 0.7564722499 | -0.09774827143 | 0.7922044714 | 0.5743291024 | -1.075300549 | 0.1295978226 |
| 0.6959866521 | -0.1718432146 | 0.7740241263 | 0.4888426886 | -1.021041899 | 0.1254743527 |
| 0.7635041575 | 0.1021123142 | 0.7829115678 | 0.5084162515 | -1.002590082 | 0.1341273224 |
| 0.732936013 | 0.1062148378 | 0.7461464128 | 0.4834408213 | -0.993938374 | 0.1269998871 |
| 0.6959866521 | -0.2608016095 | 0.8000721051 | 0.5549014354 | -0.93040176 | 0.1362062702 |
| 0.878519134 | -0.05719817616 | 0.7929145359 | 0.4852642768 | -0.9752717172 | 0.1337423614 |
| 0.878519134 | -0.0481381433 | 0.7826458313 | 0.4843646654 | -1.020622487 | 0.1264118138 |
| 0.6959866521 | -0.2476748338 | 0.7026488783 | 0.4618222123 | -0.9347595704 | 0.1155070105 |
| 0.732936013 | 0.1287654371 | 0.7334719785 | 0.4460601116 | -0.8674037468 | 0.1379025099 |
| 0.732936013 | -0.1114132788 | 0.7638736776 | 0.4755193915 | -1.029234967 | 0.1216028721 |
| 0.9558407212 | 0.01895382956 | 0.7995357583 | 0.4961175871 | -1.031014805 | 0.1279667078 |
| 0.6959866521 | -0.2029153362 | 0.7230597669 | 0.3938400446 | -1.020516655 | 0.1274483849 |

| | | | | | |
|---|---|---|---|---|---|
| 0.6959866521 | -0.163727847 | 0.8638121777 | 0.4551778166 | -0.9501993209 | 0.1263639602 |
| 0.6959866521 | 0.2291611876 | 0.9160652655 | 0.6021510197 | -1.291130669 | 0.1281741085 |
| 0.732936013 | 0.102109749 | 0.7802876782 | 0.4686661651 | -1.008916544 | 0.1229844561 |
| 0.732936013 | 0.125762546 | 0.7558029764 | 0.4477103604 | -0.9264932164 | 0.1258495477 |
| 0.7158466291 | 0.141975735 | 0.8327720005 | 0.4766047914 | -1.074643676 | 0.1233091943 |
| 0.8826037281 | -0.04162294409 | 0.814388741 | 0.4956001874 | -1.032053009 | 0.1248872408 |
| 0.6959866521 | -0.2153193791 | 0.8302496104 | 0.5272994683 | -1.059719338 | 0.115932294 |
| 0.5521312959 | 0.2951780256 | 1.011190084 | 0.5772027875 | -1.31338953 | 0.1191987778 |
| 0.8266664895 | 0.07700155132 | 0.7550810788 | 0.4839900674 | -0.9937012812 | 0.124433445 |
| 0.7158466291 | 0.134948218 | 0.8515859464 | 0.613955255 | -1.209592358 | 0.1261663161 |
| 0.732936013 | 0.1245986899 | 0.790898347 | 0.4768462106 | -0.9586451244 | 0.1252497997 |
| 0.9185693261 | 0.03448117875 | 0.7987074315 | 0.4853446443 | -1.025283348 | 0.1299989785 |
| 0.7158466291 | 0.151966993 | 0.7060297333 | 0.4305050377 | -1.068916963 | 0.1389217771 |
| 0.7158466291 | -0.1575350141 | 0.7572497296 | 0.4616781346 | -0.9487447832 | 0.1262835015 |
| 0.7635041575 | -0.08974973939 | 0.7964646587 | 0.5023990015 | -1.082573961 | 0.1246263378 |
| 0.732936013 | -0.1112598196 | 0.8764624786 | 0.561044191 | -1.068695255 | 0.1301020242 |
| 0.6959866521 | -0.1784687627 | 0.7993506178 | 0.553749094 | -1.051995757 | 0.1290306906 |
| 0.8818643096 | 0.04598480911 | 0.8026515628 | 0.4982876031 | -1.041385692 | 0.1270802315 |
| 0.6959866521 | -0.1665870408 | 0.7477236935 | 0.3976611743 | -0.9466709835 | 0.121915581 |
| 0.7158466291 | 0.1462071626 | 0.7627377037 | 0.5205959566 | -1.060183133 | 0.1211529411 |
| 0.7158466291 | 0.1381391312 | 0.8371665855 | 0.5067289098 | -1.048795261 | 0.119689009 |
| 0.4747833966 | 0.2153146553 | 0.9178246629 | 0.5773444613 | -1.05331381 | 0.1229542976 |
| 0.6844246943 | 0.1223404944 | 0.8709567447 | 0.5281242959 | -1.068097215 | 0.120275463 |
| 0.570290738 | 0.1720911899 | 0.9228445976 | 0.5087641386 | -1.087046266 | 0.1200297734 |
| 0.9668059968 | -0.01229211541 | 0.8504878867 | 0.5114252295 | -1.041254376 | 0.1230012263 |
| 0.1365281022 | -0.3317132292 | 0.8293973102 | 0.5928802392 | -0.997720102 | 0.1135806443 |
| 0.7727047178 | 0.08145457832 | 0.8729520315 | 0.5371692085 | -1.092821239 | 0.1254107551 |
| 0.7319977919 | 0.09297707155 | 0.843346898 | 0.5377334338 | -1.055916493 | 0.1214521129 |
| 0.5965506588 | -0.1714783741 | 0.9323884234 | 0.5248471771 | -1.010165708 | 0.1231121714 |
| 0.1428343429 | -0.3353581527 | 1.00491226 | 0.5685878808 | -1.075648853 | 0.1202758817 |
| 0.6914540463 | 0.1111962326 | 0.854394931 | 0.4892587175 | -1.012848109 | 0.1193347677 |
| 0.9255063064 | 0.03048325153 | 0.8583306442 | 0.5163970723 | -1.039192418 | 0.1243702524 |
| 0.7723461953 | -0.08357764668 | 0.8394038539 | 0.5108694285 | -0.9881066529 | 0.122788495 |
| 0.88317175 | 0.04289408797 | 0.8529847767 | 0.5216846863 | -1.052647816 | 0.123146372 |
| 0.4747833966 | -0.1913509546 | 0.8477507736 | 0.4471118515 | -1.003899082 | 0.1183457309 |
| 0.4747833966 | -0.1950900492 | 0.8564728004 | 0.5107081542 | -1.125580993 | 0.1203361815 |
| 0.405215244 | -0.2475060652 | 0.7492522887 | 0.4487908984 | -0.9841925307 | 0.1186934204 |
| 0.02450352368 | -0.4373640483 | 0.8281897171 | 0.293778809 | -0.9540433789 | 0.1093883801 |
| 0.9111058518 | 0.03690045378 | 0.8590096775 | 0.5138822103 | -1.050332906 | 0.1235777031 |
| 0.3463206331 | 0.298442096 | 0.7001117247 | 0.6305038092 | -1.234780107 | 0.1276303912 |
| 0.7791752649 | 0.07564773658 | 0.795809655 | 0.4829710091 | -1.031122994 | 0.1297946771 |
| 0.4747833966 | -0.216560905 | 0.7165102366 | 0.4146417822 | -0.8682825882 | 0.1199933906 |
| 0.7791752649 | 0.07414196104 | 0.8506987966 | 0.5213092972 | -1.012702438 | 0.1243511986 |
| 0.4747833966 | -0.2076943328 | 0.7380471111 | 0.4755652096 | -0.984197197 | 0.1212024281 |
| 0.9653946556 | -0.01503629806 | 0.8513553896 | 0.5161175804 | -1.039718534 | 0.1229287933 |
| 0.6914540463 | -0.1267020626 | 0.9369457117 | 0.5459342492 | -1.104995175 | 0.1328667943 |

| | | | | | |
|---|---|---|---|---|---|
| 0.6470164754 | -0.1554211051 | 0.8863447745 | 0.5998819025 | -1.073539242 | 0.1237296482 |
| 0.7319977919 | 0.09713656181 | 0.8181239286 | 0.519832025 | -0.9970042868 | 0.12587691 |
| 0.9255063064 | -0.03011731255 | 0.8625986246 | 0.5163382066 | -1.037328268 | 0.1232774221 |
| 0.9348509336 | -0.02427758786 | 0.8317542398 | 0.4980538251 | -1.012496975 | 0.1242464833 |
| 0.6579394906 | -0.1363312818 | 0.7688443561 | 0.4898480246 | -0.9740001588 | 0.1245337653 |
| 0.5094641713 | -0.1875717164 | 0.8068400822 | 0.488563929 | -1.039101031 | 0.1215100585 |
| 0.6956587904 | -0.1113332819 | 0.8557845997 | 0.5745663786 | -1.064188208 | 0.1305331706 |
| 0.4747833966 | 0.2162293363 | 0.7797698207 | 0.5051824727 | -1.004317398 | 0.1197527332 |
| 0.862673955 | 0.05355855534 | 0.8246432226 | 0.4837317196 | -0.9844685552 | 0.1253570633 |
| 0.8820860536 | 0.04786182506 | 0.8333385364 | 0.5013586017 | -0.9850121433 | 0.1299159133 |
| 0.6917499485 | -0.1099606891 | 0.8151364627 | 0.5242389312 | -1.06346154 | 0.1253407424 |
| 0.24550875 | 0.2981214319 | 0.818031867 | 0.600675492 | -1.000588191 | 0.1179550123 |
| 0.4747833966 | 0.2090286243 | 0.844836022 | 0.5879004549 | -1.12850579 | 0.1214174547 |
| 0.7840298201 | 0.07202844671 | 0.8365054026 | 0.5364617244 | -1.050307762 | 0.1333651833 |
| 0.9255063064 | 0.03530373631 | 0.8529803988 | 0.486515385 | -1.00463796 | 0.1301042115 |
| 0.983766969 | -0.00510363453 | 0.8466877038 | 0.5088271722 | -1.032605575 | 0.1261158209 |
| 0.983766969 | 0.003670082907 | 0.8498389465 | 0.5086217956 | -1.033718055 | 0.12147142 |
| 0.7701935905 | -0.08629848529 | 0.851417809 | 0.5457980151 | -1.078979444 | 0.1238370174 |
| 0.6917499485 | 0.1101460121 | 0.8427412779 | 0.525426383 | -1.008726308 | 0.1228006819 |
| 0.7319977919 | -0.09678170339 | 0.8290530785 | 0.5062582337 | -0.9834175466 | 0.1232905002 |
| 0.9111058518 | -0.03704129946 | 0.8419302355 | 0.5098979118 | -1.021425501 | 0.1226219775 |
| 0.9713083494 | -0.009358416003 | 0.8493277244 | 0.5095164699 | -1.040097019 | 0.1248005731 |
| 0.4747833966 | 0.2054185034 | 0.8458934134 | 0.5142505307 | -0.9848533647 | 0.1213461218 |
| 0.4747833966 | 0.2205260496 | 0.8983806831 | 0.6228313598 | -1.180349508 | 0.1212887515 |
| 0.7151365909 | -0.101087038 | 0.8259546228 | 0.4543328755 | -1.039079117 | 0.1243770831 |
| 0.983766969 | 0.003856909185 | 0.8487796042 | 0.5085306129 | -1.034230775 | 0.1222090103 |
| 0.6579394906 | 0.1418302965 | 0.9517126746 | 0.5976075843 | -1.174709133 | 0.1257323259 |
| 0.4747833966 | -0.2043487481 | 0.7501051325 | 0.4842407718 | -0.9029790929 | 0.1237852639 |
| 0.5446705437 | 0.1872922614 | 0.8952118599 | 0.573978459 | -0.952324269 | 0.1255774378 |
| 0.6470164754 | 0.1557943061 | 0.8809542079 | 0.5282346915 | -1.064182665 | 0.1224307833 |
| 0.6914540463 | 0.114044944 | 0.8370983277 | 0.531270076 | -1.020319159 | 0.1199228415 |
| 0.6579394906 | -0.1344995112 | 0.8498292358 | 0.4523632664 | -0.9462111553 | 0.1236261792 |
| 0.6712192959 | 0.127905323 | 0.8343751001 | 0.5330467438 | -1.078847596 | 0.1203077563 |
| 0.8391897122 | 0.05990045517 | 0.8814488637 | 0.5367119081 | -1.104225661 | 0.1271139602 |
| 0.7791752649 | -0.07153921171 | 0.8715237282 | 0.5442531274 | -1.067757941 | 0.124716169 |
| 0.4747833966 | -0.2252765666 | 0.7754077702 | 0.4925077951 | -1.036686506 | 0.1179962569 |
| 0.4747833966 | -0.1980747856 | 0.7600537102 | 0.4772326483 | -0.9824447066 | 0.1237823245 |
| 0.6914540463 | -0.116020159 | 0.76071742 | 0.464509844 | -0.9652251553 | 0.1261877046 |
| 0.7791752649 | -0.07738648668 | 0.8465670179 | 0.5457432533 | -1.041261011 | 0.1254854839 |
| 0.6470164754 | 0.1505837628 | 0.852952474 | 0.5069199394 | -0.9990687702 | 0.120447243 |
| 0.6917499485 | -0.1109101571 | 0.8123571582 | 0.4753250009 | -1.006497589 | 0.1234958364 |
| 0.9713083494 | -0.0101010798 | 0.8559762224 | 0.5115386925 | -1.043619351 | 0.1238544035 |
| 0.9329716846 | 0.02603565161 | 0.8417039109 | 0.4976862008 | -1.010843207 | 0.1257013458 |
| 0.7727047178 | -0.08094077973 | 0.8904868098 | 0.5401898367 | -1.046551859 | 0.1253715391 |
| 0.6470164754 | -0.1572556126 | 0.8819814667 | 0.582424274 | -1.16283756 | 0.120478233 |
| 0.570290738 | 0.1895804499 | 0.9047036838 | 0.4529181257 | -0.868200363 | 0.1309047244 |

| | | | | | |
|---|---|---|---|---|---|
| 0.654577244 | 0.1428282783 | 0.8258404891 | 0.5087417406 | -1.061177385 | 0.1219063487 |
| 0.6917499485 | -0.1087783736 | 0.8540414714 | 0.4697050276 | -1.033631471 | 0.1236748809 |
| 0.405215244 | 0.2549979493 | 0.9269708583 | 0.6517253936 | -1.262220585 | 0.1193240517 |
| 0.4016115174 | -0.2676726775 | 0.8962423607 | 0.628014 | -1.039667027 | 0.1181302902 |
| 0.654577244 | 0.1438595875 | 0.8404120858 | 0.549095684 | -1.017687415 | 0.1222016215 |
| 0.8795741666 | 0.05096556119 | 0.8067092776 | 0.4834462003 | -1.000890592 | 0.1275151471 |
| 0.4747833966 | 0.2052552246 | 0.8137731063 | 0.5360989556 | -1.077155775 | 0.1204984116 |
| 0.6579394906 | 0.1345259969 | 0.793254636 | 0.4833211754 | -0.9752016493 | 0.1235686785 |
| 0.6844246943 | -0.1257378187 | 0.8145331385 | 0.4578163953 | -1.049507054 | 0.1230462012 |
| 0.6470164754 | 0.1502429163 | 0.9220954695 | 0.6042250609 | -1.12484332 | 0.1238070852 |
| 0.6914540463 | -0.1134305812 | 0.9008566871 | 0.5305182698 | -1.085424116 | 0.1224656384 |
| 0.4016115174 | 0.2627817049 | 0.8749012389 | 0.5687744998 | -0.9835068374 | 0.1202959795 |
| 0.7537501405 | 0.09082805039 | 0.8899845163 | 0.5256571138 | -1.032846999 | 0.1255404369 |
| 0.9668059968 | -0.01395593643 | 0.8508684248 | 0.49787466 | -1.021699558 | 0.130714173 |
| 0.7723461953 | -0.08244696957 | 0.8791233758 | 0.5009352949 | -1.034075167 | 0.1221417715 |
| 0.03003916901 | -0.4005085717 | 0.8903000411 | 0.6058773177 | -1.071378714 | 0.1089407931 |
| 0.9843096322 | 0.002385167378 | 0.8499645507 | 0.5096367892 | -1.036345608 | 0.1212822107 |
| 0.8829503013 | -0.04422817558 | 0.8474783535 | 0.5105305794 | -1.035289868 | 0.1235429548 |
| 0.7791752649 | 0.07526433217 | 0.8016652463 | 0.5058124955 | -1.016456278 | 0.1276521399 |
| 0.8502633244 | -0.05605415091 | 0.8711744719 | 0.518765383 | -1.035333817 | 0.1245718965 |
| 0.8801299717 | 0.04809534153 | 0.8480893924 | 0.4925684618 | -1.008358091 | 0.1234688619 |
| 0.7840298201 | -0.07103780053 | 0.8529894207 | 0.5678616936 | -1.077260085 | 0.1301661331 |
| 0.6844246943 | -0.1237683215 | 0.839840319 | 0.511050869 | -1.041471718 | 0.1219213208 |
| 0.6470164754 | -0.1615824816 | 0.8905541736 | 0.505794992 | -1.106034612 | 0.1325539137 |
| 0.6956587904 | 0.106557478 | 0.808160988 | 0.5056148643 | -1.011999327 | 0.12426652 |
| 0.7065120866 | -0.1052291561 | 0.8651096883 | 0.5431297452 | -1.009117856 | 0.1265301131 |
| 0.9363440092 | 0.0237650363 | 0.8463824998 | 0.5120543641 | -1.051555137 | 0.129079013 |
| 0.7382744533 | 0.09354025705 | 0.8758067519 | 0.5375078506 | -1.052574318 | 0.1248504858 |
| 0.654577244 | -0.1392230843 | 0.8047610077 | 0.4921974064 | -0.9905820199 | 0.119366082 |
| 0.6579394906 | 0.1429192991 | 0.7935072401 | 0.4700447366 | -0.871591819 | 0.1292701672 |
| 0.1365281022 | -0.3370666424 | 0.7688011446 | 0.4531092336 | -1.058061908 | 0.1152273637 |
| 0.6470164754 | -0.1479460307 | 0.7936846084 | 0.5009392326 | -0.9800256914 | 0.1213508776 |
| 0.4747833966 | -0.2335316655 | 0.7422808378 | 0.3740490002 | -0.991793738 | 0.1262188515 |
| 0.4747833966 | -0.208703436 | 0.9386982682 | 0.4658999738 | -0.9381842179 | 0.1247244593 |
| 0.8820860536 | -0.04940077136 | 0.8285413294 | 0.4900822881 | -0.9853437894 | 0.1310420432 |
| 0.9329716846 | 0.02662997727 | 0.8461114544 | 0.5014438098 | -1.030699712 | 0.122949544 |
| 0.9653946556 | 0.01531877127 | 0.8452859197 | 0.5035537819 | -1.023506768 | 0.1233398326 |
| 0.9329716846 | -0.02644744017 | 0.8469696855 | 0.5161209721 | -1.032485552 | 0.1242882906 |
| 0.7791752649 | -0.07131403973 | 0.8926897866 | 0.5149386875 | -1.057110505 | 0.1248542813 |
| 0.405215244 | -0.2397527427 | 0.8880751472 | 0.5437757083 | -1.068920193 | 0.1159579933 |
| 0.6470164754 | 0.1513680132 | 0.9497436169 | 0.5432301266 | -1.166208781 | 0.1232726403 |
| 0.9653946556 | -0.01462908665 | 0.8529856089 | 0.5088484322 | -1.035242492 | 0.1207912624 |
| 0.5979950877 | -0.1715221647 | 0.789095224 | 0.3708557927 | -0.8208240287 | 0.1246740113 |
| 0.6914540463 | 0.1162644711 | 0.8612463187 | 0.5094714307 | -0.9926044126 | 0.1230923041 |
| 0.7727047178 | -0.08233637021 | 0.852311608 | 0.5493460901 | -1.05766322 | 0.1249364333 |
| 0.7791752649 | 0.07792274322 | 0.8008094405 | 0.4723240788 | -1.052213698 | 0.1382373183 |

| | | | | | |
|---|---|---|---|---|---|
| 0.9255063064 | -0.02976918454 | 0.8477277025 | 0.5067735519 | -1.026229318 | 0.1207125125 |
| 0.6856771369 | -0.1234508212 | 0.8626820845 | 0.525542768 | -1.124671151 | 0.1231531025 |
| 0.1365281022 | -0.3424823965 | 1.086390991 | 0.7060787394 | -1.138688373 | 0.1193188025 |
| 0.7791752649 | -0.07825918694 | 0.853298462 | 0.531321551 | -1.047022648 | 0.1290114059 |
| 0.4747833966 | 0.1999546832 | 0.9015071147 | 0.5308417683 | -1.121035762 | 0.1251833941 |
| 0.6579394906 | -0.1361822995 | 0.8100451562 | 0.4283133235 | -0.9669603944 | 0.1221395405 |
| 0.4747833966 | 0.2012524414 | 0.8320063466 | 0.569218289 | -1.119379435 | 0.120449993 |
| 0.6914540463 | 0.1152848841 | 0.8862537 | 0.5211684442 | -1.05513656 | 0.1204536192 |
| 0.9895351491 | -0.1119390071 | -1.025329847 | 0.09218840023 | -0.2429689496 | 0.1020387034 |
| 0.9895351491 | -0.03845475724 | -1.021211375 | 0.1073008809 | -0.220809011 | 0.1054992503 |
| 0.9895351491 | -0.09705512073 | -1.052050811 | 0.1035474353 | -0.2099743342 | 0.1037360059 |
| 0.9895351491 | 0.1827187371 | -1.031471552 | 0.07430403972 | -0.1591722607 | 0.1021669003 |
| 0.9895351491 | -0.1493483789 | -1.009738512 | 0.1480142301 | -0.2472714191 | 0.1037498195 |
| 0.9895351491 | 0.1328366404 | -0.9552966244 | 0.1510076975 | -0.3420810776 | 0.1093417724 |
| 0.9895351491 | 0.03240258738 | -1.014410472 | 0.1231753262 | -0.2425110025 | 0.1018689076 |
| 0.9895351491 | 0.005105109347 | -1.014320654 | 0.1141681588 | -0.2366801754 | 0.1043649416 |
| 0.9895351491 | -0.03543354703 | -0.9958933179 | 0.1194941546 | -0.2468710823 | 0.1041991828 |
| 0.9895351491 | 0.04387252328 | -1.01471687 | 0.1040472839 | -0.2170267834 | 0.1070730782 |
| 0.9895351491 | -0.01357219405 | -1.018630336 | 0.1115909198 | -0.230923245 | 0.1018007178 |
| 0.9895351491 | -0.09813768688 | -1.031424266 | 0.1238274704 | -0.1760052305 | 0.1125498815 |
| 0.9895351491 | -0.033116974 | -1.007824485 | 0.1032817269 | -0.229249567 | 0.1034512708 |
| 0.9895351491 | -0.1586404206 | -1.007519622 | 0.07264456783 | -0.2281150366 | 0.1011083163 |
| 0.9895351491 | -0.07741951329 | -1.017552807 | 0.1073022016 | -0.2639342092 | 0.1019004411 |
| 0.9895351491 | -0.1480091113 | -1.060631889 | 0.07334640465 | -0.2344580807 | 0.1013411052 |
| 0.9895351491 | -0.08321429084 | -1.011145743 | 0.08147817595 | -0.2330530095 | 0.1063742237 |
| 0.9895351491 | 0.002895276329 | -1.012227566 | 0.1142705367 | -0.2365942983 | 0.102156971 |
| 0.9895351491 | 0.09505309756 | -1.05840072 | 0.15965313 | -0.2941118309 | 0.1100413361 |
| 0.9895351491 | 0.04200405796 | -1.039280732 | 0.0953189367 | -0.2394162835 | 0.1126525267 |
| 0.9895351491 | 0.07234386587 | -0.9820667279 | 0.1299065823 | -0.2653429389 | 0.1241541816 |
| 0.9895351491 | 0.1864771645 | -1.031323447 | 0.1291911718 | -0.1597765195 | 0.1112266018 |
| 0.9895351491 | -0.1044516626 | -1.084101555 | 0.09656068205 | -0.1930726018 | 0.1059656185 |
| 0.9895351491 | 0.1172278338 | -1.037668866 | 0.06512512329 | -0.19205213 | 0.1026435568 |
| 0.9895351491 | 0.02977779405 | -1.03649989 | 0.09739937455 | -0.2144215863 | 0.1105137676 |
| 0.9895351491 | -0.263000158 | -0.9350727691 | 0.2254619489 | -0.3035983926 | 0.1122718813 |
| 0.9895351491 | -0.06424395261 | -0.9946766504 | 0.1184257759 | -0.2534580049 | 0.1030450619 |
| 0.9895351491 | 0.02353917472 | -1.021007958 | 0.1083307582 | -0.2334179537 | 0.1039661396 |
| 0.9895351491 | -0.06983801804 | -1.060304821 | 0.0788902618 | -0.1791052601 | 0.108142909 |
| 0.9895351491 | 0.03049327862 | -0.9953493388 | 0.1242912182 | -0.2456937553 | 0.1043142544 |
| 0.9895351491 | -0.01538708559 | -1.017505774 | 0.1114095275 | -0.2337637381 | 0.1397789063 |
| 0.9895351491 | -0.1246992627 | -1.001241717 | 0.1436121667 | -0.2795933543 | 0.1310619818 |
| 0.9895351491 | 0.1745472807 | -1.080629148 | 0.0987946946 | -0.1834636881 | 0.1067686194 |
| 0.9895351491 | -0.01220485067 | -1.008140802 | 0.1190113105 | -0.2450848958 | 0.1145270507 |
| 0.9950381856 | -0.000691019988 | -1.01226428 | 0.1139516995 | -0.2363194952 | 0.109235827 |
| 0.9895351491 | 0.03854549382 | -0.9911750766 | 0.1077313916 | -0.2380414941 | 0.1244795448 |
| 0.9895351491 | -0.03174031063 | -1.007111828 | 0.1059675329 | -0.2419665403 | 0.1089636219 |
| 0.9895351491 | -0.004879890518 | -1.011298035 | 0.1122474934 | -0.2347023057 | 0.1064575547 |

| | | | | | |
|---|---|---|---|---|---|
| 0.9895351491 | 0.0748104055 | -1.022038596 | 0.1447367351 | -0.2386784 | 0.1082760706 |
| 0.9895351491 | -0.1593332623 | -1.030376686 | 0.1978562073 | -0.367121261 | 0.1144213587 |
| 0.9895351491 | -0.07356278495 | -1.038000682 | 0.1021129211 | -0.2202542549 | 0.1098215839 |
| 0.9895351491 | -0.1280057056 | -1.000589873 | 0.1428854361 | -0.3141146202 | 0.1140948482 |
| 0.9895351491 | -0.1637668209 | -1.002981032 | 0.2114153792 | -0.3105285462 | 0.1022458958 |
| 0.9895351491 | 0.006292384658 | -1.012518492 | 0.113827037 | -0.2339486023 | 0.106293533 |
| 0.9895351491 | 0.1149919909 | -0.964151205 | 0.1094217796 | -0.3274421283 | 0.1083372822 |
| 0.9895351491 | -0.003784199066 | -1.013079488 | 0.1142418062 | -0.2341008378 | 0.1020089879 |
| 0.9895351491 | 0.05031023915 | -1.004005913 | 0.1011302113 | -0.2205030722 | 0.1124136133 |
| 0.9895351491 | 0.07457212735 | -0.9922531509 | 0.1139360262 | -0.2493661 | 0.1046806016 |
| 0.9895351491 | -0.0846073917 | -1.022989076 | 0.0743170629 | -0.1805597513 | 0.1051748575 |
| 0.9895351491 | -0.1914618018 | -1.042770165 | 0.0077722436 | -0.26358144 | 0.1058273736 |
| 0.9895351491 | -0.1549777095 | -0.9794753918 | 0.1568006965 | -0.2735418238 | 0.1202365684 |
| 0.9895351491 | 0.0791439055 | -0.9742868171 | 0.1638128958 | -0.2999988983 | 0.1067156667 |
| 0.9895351491 | 0.1008710184 | -0.9690081562 | 0.1079337874 | -0.3027215388 | 0.1126005161 |
| 0.9895351491 | 0.1000883364 | -0.9977011887 | 0.1498723944 | -0.179623957 | 0.1072130812 |
| 0.9895351491 | 0.08599654417 | -0.9917074386 | 0.1219111534 | -0.2598490522 | 0.1025274416 |
| 0.9895351491 | -0.009033893316 | -1.010141963 | 0.1111239337 | -0.2390803797 | 0.1083713499 |
| 0.9895351491 | -0.04298730705 | -1.01536255 | 0.0920981713 | -0.2044402449 | 0.1047662858 |
| 0.9895351491 | 0.1794486558 | -1.026140032 | 0.1832821823 | -0.3239603756 | 0.1079467258 |
| 0.9895351491 | -0.02249534847 | -1.019030553 | 0.1024958313 | -0.2155595687 | 0.1080007739 |
| 0.9895351491 | -0.03570290951 | -0.9968912239 | 0.1253493312 | -0.2588033464 | 0.1048241214 |
| 0.9895351491 | -0.01941506847 | -1.017980308 | 0.1133326245 | -0.237238255 | 0.1014130461 |
| 0.9895351491 | 0.07902319565 | -0.9768745147 | 0.1336113269 | -0.2586008858 | 0.1099582309 |
| 0.9895351491 | -0.04590857035 | -1.040922259 | 0.0946321102 | -0.217087325 | 0.1221701366 |
| 0.9895351491 | -0.02398111122 | -1.010663266 | 0.1266933639 | -0.2380836996 | 0.1111636409 |
| 0.9895351491 | 0.01353624641 | -1.010531881 | 0.1128993947 | -0.2352117395 | 0.1042372881 |
| 0.9895351491 | 0.07463175688 | -0.9882908583 | 0.131260151 | -0.2553612121 | 0.1072156279 |
| 0.9895351491 | 0.07886764944 | -1.052507353 | 0.0963733488 | -0.1766386131 | 0.1108184468 |
| 0.9895351491 | 0.08978906063 | -1.044688307 | 0.0602933603 | -0.1504174816 | 0.1050815105 |
| 0.9895351491 | 0.01958731688 | -1.021060936 | 0.108119436 | -0.2364132214 | 0.1062060782 |
| 0.9895351491 | 0.01925063505 | -1.01463125 | 0.1045443376 | -0.222382083 | 0.1040011978 |
| 0.9895351491 | 0.1820691789 | -0.9665469971 | 0.0493037486 | -0.06406125007 | 0.1092606146 |
| 0.9895351491 | 0.05950851468 | -1.009157221 | 0.1163164803 | -0.2648442818 | 0.1055728742 |
| 0.9895351491 | -0.1558853657 | -1.030631505 | 0.0544923591 | -0.2020192026 | 0.1090526795 |
| 0.9895351491 | 0.2172702105 | -0.9423385149 | 0.2596347598 | -0.4477485557 | 0.1026908515 |
| 0.9895351491 | -0.1250551013 | -0.9949289773 | 0.1634355204 | -0.2421753776 | 0.1040865251 |
| 0.9895351491 | -0.01492012161 | -1.011934344 | 0.1118747172 | -0.2360165551 | 0.1106012416 |
| 0.9895351491 | 0.08370084532 | -1.076744217 | 0.0837962954 | -0.1917290834 | 0.1063791336 |
| 0.9895351491 | -0.01965070383 | -1.00812815 | 0.1142668942 | -0.231690138 | 0.1046677747 |
| 0.9895351491 | 0.0652323743 | -1.02018713 | 0.1158678194 | -0.2286125654 | 0.1065781322 |
| 0.9895351491 | -0.02855119569 | -1.020655808 | 0.1038508788 | -0.2392579391 | 0.1033140275 |
| 0.9895351491 | -0.03439868336 | -1.029685007 | 0.0974811424 | -0.2159761229 | 0.1057581212 |
| 0.9895351491 | 0.04415800757 | -1.032642767 | 0.1069915356 | -0.2196964347 | 0.1046741687 |
| 0.9895351491 | -0.09863076706 | -1.049728714 | 0.0956957541 | -0.2175262161 | 0.1116229322 |
| 0.9895351491 | 0.03320488489 | -0.9966255352 | 0.1142587448 | -0.2395239014 | 0.1080227222 |

| | | | | | |
|---|---|---|---|---|---|
| 0.9895351491 | -0.01266293623 | -1.010315482 | 0.103475537 | -0.2260454584 | 0.1135717536 |
| 0.9895351491 | -0.05029541315 | -0.991854913 | 0.1093678105 | -0.2447090154 | 0.1092275761 |
| 0.9895351491 | -0.170529379 | -0.988210412 | 0.1379361603 | -0.2657352335 | 0.09967000973 |
| 0.9895351491 | -0.1483779647 | -1.004147626 | 0.0991549762 | -0.2018287127 | 0.1088632594 |
| 0.9895351491 | -0.04062674639 | -1.013829463 | 0.1047585698 | -0.2388029733 | 0.1198016166 |
| 0.9895351491 | 0.01438840015 | -1.01935535 | 0.111397902 | -0.2360025444 | 0.1107546565 |
| 0.9895351491 | 0.1180010473 | -1.069411101 | 0.0817232174 | -0.2256316079 | 0.1015827501 |
| 0.9895351491 | 0.01288271041 | -1.012737157 | 0.1081597355 | -0.2304351771 | 0.1109010635 |
| 0.9895351491 | 0.09375919685 | -1.013465482 | 0.0208867326 | -0.1879336199 | 0.1085656712 |
| 0.9895351491 | 0.05660371155 | -1.010241828 | 0.114378986 | -0.2308030463 | 0.1110836316 |
| 0.9895351491 | -0.009542147516 | -1.010973238 | 0.1130436394 | -0.2406339295 | 0.1072691131 |
| 0.9895351491 | 0.1987201356 | -1.041137307 | 0.0951497308 | -0.2576084816 | 0.104595404 |
| 0.9895351491 | -0.09641015383 | -0.998641960 | 0.1366840479 | -0.2160806545 | 0.1161252501 |
| 0.9895351491 | -0.1376432185 | -1.024137558 | 0.0899236077 | -0.1293246224 | 0.1109110349 |
| 0.9895351491 | -0.004147402689 | -1.014020917 | 0.1124761202 | -0.234938788 | 0.1057175779 |
| 0.9895351491 | -0.03829999592 | -1.022805435 | 0.1114708695 | -0.2290157539 | 0.1021493097 |
| 0.9895351491 | 0.07293796884 | -1.030525071 | 0.0795281478 | -0.1725356713 | 0.1177587787 |
| 0.9895351491 | -0.1824029165 | -1.039738265 | 0.0881916504 | -0.279793235 | 0.1005384462 |
| 0.9895351491 | 0.0556698383 | -0.992739375 | 0.1103152604 | -0.253881775 | 0.114674911 |
| 0.9895351491 | 0.03108071063 | -1.002424328 | 0.1277713623 | -0.2348127562 | 0.1211238156 |
| 0.9895351491 | -0.09654479726 | -0.960216029 | 0.0817574073 | -0.2083611437 | 0.10932219 |
| 0.9895351491 | -0.08428585711 | -1.045527459 | 0.0691468782 | -0.1686672087 | 0.1143043841 |
| 0.9895351491 | 0.01329933622 | -1.013258049 | 0.1118887891 | -0.2345034644 | 0.103231019 |
| 0.9895351491 | -0.01210437429 | -1.007266025 | 0.118705841 | -0.2476131396 | 0.1145408843 |
| 0.9895351491 | -0.1499988316 | -1.040807395 | 0.1072518267 | -0.2164204522 | 0.1294933891 |
| 0.9895351491 | 0.0267054011 | -1.026576656 | 0.117453784 | -0.2308885283 | 0.1053816331 |
| 0.9895351491 | 0.01817978582 | -1.016282292 | 0.1123204946 | -0.2314164308 | 0.1003733106 |
| 0.9895351491 | -0.0174838995 | -1.022719931 | 0.1070057518 | -0.2217558306 | 0.1133619402 |
| 0.9895351491 | -0.1061884555 | -0.983834848 | 0.0993882721 | -0.2531196505 | 0.1055882278 |
| 0.9895351491 | -0.05001251278 | -1.026324265 | 0.0722258913 | -0.1769191094 | 0.1104684816 |
| 0.9895351491 | -0.008251285325 | -1.012485264 | 0.115499394 | -0.2390807056 | 0.1063883185 |
| 0.9895351491 | -0.03370203865 | -1.009482607 | 0.1202929912 | -0.2503767899 | 0.120262545 |
| 0.9895351491 | 0.1010244262 | -1.094828617 | 0.0711982092 | -0.233128292 | 0.1220318198 |
| 0.9895351491 | -0.1586087817 | -1.018085669 | 0.0906029340 | -0.2097921369 | 0.1082540206 |
| 0.9895351491 | 0.07850756492 | -1.018220796 | 0.1116861985 | -0.1787078931 | 0.1067653666 |
| 0.9895351491 | -0.1349467908 | -0.904796527 | 0.1763579682 | -0.3038964769 | 0.1140278132 |
| 0.9895351491 | -0.006390482825 | -1.01196695 | 0.1162142625 | -0.2385595632 | 0.10440972 |
| 0.9895351491 | 0.06851722693 | -1.009409584 | 0.1045851004 | -0.2464613033 | 0.1014294655 |
| 0.9895351491 | -0.02881269398 | -1.01936424 | 0.0951194375 | -0.2210213092 | 0.1055525606 |
| 0.9895351491 | -0.002420323823 | -1.012024686 | 0.1135461614 | -0.2348543623 | 0.1043340071 |
| 0.9895351491 | -0.04666794142 | -1.026515095 | 0.1184459544 | -0.2270590197 | 0.1019254942 |
| 0.9984451821 | -0.09983792187 | -0.9456276181 | 0.1207231185 | -0.2873828813 | 0.1010117765 |
| 0.9984451821 | -0.08876308784 | -0.9569311561 | 0.1229296983 | -0.2414484902 | 0.1044264427 |
| 0.9984451821 | -0.1675387226 | -1.010643101 | 0.1218597245 | -0.2232929715 | 0.1010373811 |
| 0.9984451821 | 0.2019911604 | -0.9622380211 | 0.0948579513 | -0.1870091514 | 0.1006755224 |
| 0.9984451821 | -0.1278311358 | -0.9307629804 | 0.1685545645 | -0.2922777476 | 0.103526327 |

| | | | | | |
|---|---|---|---|---|---|
| 0.9984451821 | 0.1453095432 | -0.87460638842 | 0.1762118782 | -0.3910819583 | 0.1098795831 |
| 0.9984451821 | 0.0001954718104 | -0.9361665487 | 0.1390033851 | -0.2770020805 | 0.1003100335 |
| 0.9984451821 | 0.02344326723 | -0.9450277048 | 0.1405845249 | -0.2811357767 | 0.102826287 |
| 0.9984451821 | -0.03960572049 | -0.9175589835 | 0.1456178098 | -0.2894218291 | 0.1030471806 |
| 0.9984451821 | 0.02074387607 | -0.9375586074 | 0.133122141 | -0.2671737873 | 0.107242226 |
| 0.9984451821 | -0.01699943122 | -0.9438989013 | 0.1358536987 | -0.2709758441 | 0.1001312382 |
| 0.9984451821 | -0.007126622963 | -0.9374596569 | 0.139540797 | -0.272717115 | 0.113540904 |
| 0.9984451821 | -0.03160050327 | -0.9322043227 | 0.1298930832 | -0.2707465504 | 0.1022330123 |
| 0.9984451821 | -0.177730402 | -0.9301608609 | 0.0967980477 | -0.2704821351 | 0.09915755232 |
| 0.9984451821 | -0.07586935311 | -0.9417690037 | 0.130721707 | -0.3028940006 | 0.1001787113 |
| 0.9984451821 | -0.1295279319 | -0.9779325127 | 0.1062200519 | -0.2770720689 | 0.09996778023 |
| 0.9984451821 | -0.08150663564 | -0.9327343025 | 0.109061826 | -0.277881393 | 0.1058999859 |
| 0.9984451821 | 0.07098882936 | -0.9336617225 | 0.1480132998 | -0.2953275895 | 0.1001277281 |
| 0.9984451821 | 0.05184095264 | -0.9611070043 | 0.1631155593 | -0.3084831344 | 0.108693768 |
| 0.9984451821 | 0.0303774059 | -0.9557946689 | 0.1254646443 | -0.2795049041 | 0.1119862283 |
| 0.9984451821 | 0.09083551471 | -0.9048297516 | 0.1526754429 | -0.3023841666 | 0.1330496265 |
| 0.9984451821 | 0.2689043507 | -0.9599647355 | 0.1562686376 | -0.1644295485 | 0.110253585 |
| 0.9984451821 | -0.1017881103 | -1.005993049 | 0.1178134611 | -0.2336505187 | 0.1050205661 |
| 0.9984451821 | 0.08142240546 | -0.953884871 | 0.1025416393 | -0.2460652197 | 0.102069348 |
| 0.9984451821 | 0.05648741862 | -0.9824190164 | 0.1080760661 | -0.2358410116 | 0.1096823692 |
| 0.9984451821 | -0.2319453107 | -0.8599385609 | 0.2307731457 | -0.3462842728 | 0.1155000824 |
| 0.9984451821 | -0.03045362725 | -0.9276014804 | 0.1408425082 | -0.2854769341 | 0.1020308743 |
| 0.9984451821 | 0.003055456125 | -0.9373340142 | 0.1381993565 | -0.2765613 | 0.1028418827 |
| 0.9984451821 | -0.0295700069 | -0.9566155302 | 0.12441433 | -0.2531104834 | 0.1076850038 |
| 0.9984451821 | 0.06475251805 | -0.8987421851 | 0.1606786305 | -0.2996550054 | 0.1025525804 |
| 0.9984451821 | 0.001319058968 | -0.9357616307 | 0.139227474 | -0.2771220475 | 0.1535487894 |
| 0.9984451821 | -0.09148782549 | -0.9298914424 | 0.1544315266 | -0.2996851815 | 0.1532482042 |
| 0.9984451821 | 0.1774386655 | -1.004756804 | 0.1159087598 | -0.2195376292 | 0.1063794583 |
| 0.9984451821 | 0.0773435736 | -0.9612755303 | 0.1095058177 | -0.2225236961 | 0.1152975847 |
| 0.9984451821 | -0.0787493056 | -0.9283641846 | 0.1492916696 | -0.3318918269 | 0.1083984721 |
| 0.9984451821 | 0.09709764759 | -0.8832565283 | 0.1282588045 | -0.2856089072 | 0.1286314272 |
| 0.9984451821 | -0.001121823828 | -0.9359656517 | 0.1386985911 | -0.2772178903 | 0.1094957297 |
| 0.9984451821 | 0.03538958376 | -0.9435196559 | 0.1498500307 | -0.2847625186 | 0.1055842731 |
| 0.9984451821 | 0.0626217386 | -0.9435317209 | 0.1646904327 | -0.2805528131 | 0.1080366302 |
| 0.9984451821 | -0.088086687 | -0.9497642568 | 0.1819124809 | -0.3421609181 | 0.1168137551 |
| 0.9984451821 | 0.008625365622 | -0.9330649265 | 0.1397362283 | -0.2785413329 | 0.1111821527 |
| 0.9984451821 | -0.1160054975 | -0.9221829601 | 0.1656957891 | -0.3550334253 | 0.1166511237 |
| 0.9984451821 | -0.2054064271 | -0.9231276484 | 0.2544243862 | -0.3698127534 | 0.09958157712 |
| 0.9984451821 | -0.05965609161 | -0.9343144674 | 0.1393475668 | -0.2953403973 | 0.1054396046 |
| 0.9984451821 | 0.1005631859 | -0.8964754835 | 0.1381873078 | -0.3543278578 | 0.1091657828 |
| 0.9984451821 | -0.00884330546 | -0.9379859761 | 0.1395663395 | -0.272626279 | 0.1004570599 |
| 0.9984451821 | -0.08668278282 | -0.9495645716 | 0.161712121 | -0.3070431353 | 0.1153438403 |
| 0.9984451821 | 0.08622958596 | -0.9125105107 | 0.143591263 | -0.2952981933 | 0.1041137131 |
| 0.9984451821 | -0.06911164487 | -0.9454989684 | 0.1048250796 | -0.2302414642 | 0.1046171796 |
| 0.9984451821 | -0.1461713589 | -0.9562675368 | 0.0605723084 | -0.305038021 | 0.1061376317 |
| 0.9984451821 | -0.1649787327 | -0.9002496942 | 0.1805125033 | -0.3199375742 | 0.1231751686 |

| | | | | | |
|---|---|---|---|---|---|
| 0.9984451821 | 0.01518568075 | -0.9286271335 | 0.1481158545 | -0.2895690699 | 0.1064227846 |
| 0.9984451821 | 0.1646544224 | -0.8661094007 | 0.1289618674 | -0.3842403427 | 0.1132111577 |
| 0.9984451821 | 0.1329791728 | -0.9179295764 | 0.1825845348 | -0.1979142805 | 0.1050198104 |
| 0.9984451821 | 0.09158394349 | -0.9144034896 | 0.1460412714 | -0.3009867858 | 0.101341467 |
| 0.9984451821 | 0.06364040571 | -0.9523708239 | 0.155033412 | -0.251093022 | 0.1081457646 |
| 0.9984451821 | -0.03489339617 | -0.9395399175 | 0.1199319079 | -0.2493741583 | 0.1039409494 |
| 0.9984451821 | 0.09378681774 | -0.9400723999 | 0.1738412285 | -0.3290399054 | 0.1098409807 |
| 0.9984451821 | 0.02020184653 | -0.9296272008 | 0.1485976745 | -0.295637246 | 0.1071355909 |
| 0.9984451821 | -0.04463921248 | -0.9181423785 | 0.153149878 | -0.3036271485 | 0.1039323758 |
| 0.9984451821 | -0.001063656766 | -0.9364474727 | 0.1389235562 | -0.2770669375 | 0.09967902538 |
| 0.9984451821 | 0.07010159819 | -0.904900056 | 0.154572296 | -0.2973940211 | 0.1102752191 |
| 0.9984451821 | 0.02544745831 | -0.9204214818 | 0.1491684969 | -0.2875568444 | 0.1276230521 |
| 0.9984451821 | -0.06169455553 | -0.9311616713 | 0.1717086406 | -0.2829632523 | 0.1114034441 |
| 0.9984451821 | -0.01558063513 | -0.9382025138 | 0.1398936916 | -0.2776424521 | 0.1032650921 |
| 0.9984451821 | 0.06940117236 | -0.9127724988 | 0.1544188163 | -0.2956639927 | 0.1071647942 |
| 0.9984451821 | 0.08327606084 | -0.9781982071 | 0.1195283499 | -0.2143661123 | 0.11108999 |
| 0.9984451821 | 0.1067873888 | -0.974938527 | 0.0747610538 | -0.1729698034 | 0.10372313 |
| 0.9984451821 | 0.04469915076 | -0.955306958 | 0.126757868 | -0.2797015014 | 0.1047919152 |
| 0.9984451821 | -0.05488574477 | -0.9299484577 | 0.1642440549 | -0.3146119314 | 0.1021590633 |
| 0.9984451821 | 0.1138580745 | -0.9066530097 | 0.0968092427 | -0.1709989264 | 0.1092712141 |
| 0.9984451821 | 0.002434767639 | -0.9360037822 | 0.1391349807 | -0.2781977048 | 0.105198036 |
| 0.9984451821 | -0.1573345456 | -0.952938207 | 0.0753412134 | -0.2459107077 | 0.1092438158 |
| 0.9984451821 | 0.1453801347 | -0.8864130997 | 0.234757621 | -0.4237367404 | 0.1037247773 |
| 0.9984451821 | -0.1131477879 | -0.9207171445 | 0.1830052871 | -0.28305938 | 0.103278204 |
| 0.9984451821 | -0.08610928785 | -0.9335384283 | 0.1270655661 | -0.2789886243 | 0.1110099115 |
| 0.9984451821 | 0.1224125794 | -1.029972269 | 0.0962850920 | -0.2140968854 | 0.1043064718 |
| 0.9984451821 | -0.05594628536 | -0.9241544551 | 0.1395623403 | -0.2652948317 | 0.1033196724 |
| 0.9984451821 | 0.05847514682 | -0.9404915066 | 0.1430107871 | -0.2754608157 | 0.1067862627 |
| 0.9984451821 | -0.007608496115 | -0.9382771451 | 0.1363152311 | -0.2780519017 | 0.1014800016 |
| 0.9984451821 | -0.07129962487 | -0.971385501 | 0.1033928275 | -0.2359471007 | 0.104345258 |
| 0.9984451821 | 0.03891115898 | -0.9540995631 | 0.1333369417 | -0.2629523966 | 0.1036508733 |
| 0.9984451821 | -0.143343817 | -0.9925108433 | 0.1087989214 | -0.2467171496 | 0.1114655853 |
| 0.9984451821 | 0.003812594269 | -0.9343461636 | 0.1390485081 | -0.2774130143 | 0.1078098413 |
| 0.9984451821 | -0.006883482172 | -0.9349312094 | 0.1335927281 | -0.2719353228 | 0.1128986806 |
| 0.9984451821 | -0.06230260793 | -0.9095609735 | 0.1358078383 | -0.2903940443 | 0.1096538796 |
| 0.9984451821 | -0.1340850026 | -0.9162064858 | 0.1571874334 | -0.3003924183 | 0.09905456774 |
| 0.9984451821 | -0.2146791752 | -0.9223759013 | 0.1200133147 | -0.2334183373 | 0.1080459981 |
| 0.9984451821 | -0.0897284591 | -0.9416513267 | 0.1180005267 | -0.2796998465 | 0.1244599935 |
| 0.9984451821 | 0.0184356643 | -0.9451300744 | 0.1364185431 | -0.277135493 | 0.1107890555 |
| 0.9984451821 | 0.09638066895 | -0.9827706551 | 0.1121562889 | -0.2684750688 | 0.09999106889 |
| 0.9984451821 | 0.03349900122 | -0.9374068235 | 0.1239940832 | -0.2628804651 | 0.1115013494 |
| 0.9984451821 | 0.07280521742 | -0.9386661601 | 0.0676849783 | -0.2379036098 | 0.1072636895 |
| 0.9984451821 | 0.07142508008 | -0.9333955671 | 0.1399230762 | -0.2726214734 | 0.1122850319 |
| 0.9984451821 | -0.01304034166 | -0.9345091071 | 0.1376401412 | -0.283025673 | 0.1063549187 |
| 0.9984451821 | 0.1212826713 | -0.9587905832 | 0.1269655149 | -0.281731502 | 0.1079113178 |
| 0.9984451821 | -0.09328662333 | -0.9217703461 | 0.1615569275 | -0.2609539852 | 0.1182294976 |

| | | | | | |
|---|---|---|---|---|---|
| 0.9984451821 | -0.1588410966 | -0.9507996575 | 0.1068744665 | -0.1527707147 | 0.1106655077 |
| 0.9984451821 | -0.03177829723 | -0.94933244 | 0.1287589917 | -0.2696771848 | 0.1045962001 |
| 0.9984451821 | -0.01332695751 | -0.9395356802 | 0.138340089 | -0.2750084252 | 0.1009226288 |
| 0.9984451821 | -0.01701566743 | -0.9322813284 | 0.1467250363 | -0.2910488258 | 0.1217357525 |
| 0.9984451821 | -0.1803420231 | -0.9612632695 | 0.1150860325 | -0.3223666746 | 0.09902209672 |
| 0.9984451821 | 0.02650087059 | -0.9263673151 | 0.1361808103 | -0.2856980892 | 0.1164562043 |
| 0.9984451821 | 0.02460129562 | -0.9282413121 | 0.1496746678 | -0.2762781514 | 0.1240391585 |
| 0.9984451821 | -0.05724248983 | -0.9041504912 | 0.1209463176 | -0.2620567722 | 0.108993709 |
| 0.9984451821 | -0.04338576221 | -0.9533741347 | 0.1157606977 | -0.2414889548 | 0.1170633959 |
| 0.9984451821 | 0.1486669431 | -0.9462079307 | 0.1154696908 | -0.2611073686 | 0.09992558082 |
| 0.9984451821 | -0.03162960222 | -0.9219427904 | 0.1525965164 | -0.3097749306 | 0.1161928508 |
| 0.9984451821 | -0.09443219443 | -0.9551848537 | 0.1292448343 | -0.2611091175 | 0.1472301786 |
| 0.9984451821 | 0.04635456317 | -0.9597098343 | 0.1461472795 | -0.2708664699 | 0.1040823555 |
| 0.9984451821 | 0.01959925562 | -0.9401500988 | 0.137177135 | -0.2724690297 | 0.09863658188 |
| 0.9984451821 | -0.02210823291 | -0.9495828663 | 0.1304445918 | -0.2583994378 | 0.114345673 |
| 0.9984451821 | -0.1199966104 | -0.9033470624 | 0.1281322267 | -0.3001690347 | 0.1052735579 |
| 0.9984451821 | -0.004330226247 | -0.937355768 | 0.1353648168 | -0.2718276738 | 0.1100077002 |
| 0.9984451821 | 0.01032527941 | -0.9360882689 | 0.1368771632 | -0.2726441934 | 0.1059551844 |
| 0.9984451821 | 0.007068843606 | -0.9366307274 | 0.1376581622 | -0.2741495818 | 0.1262868679 |
| 0.9984451821 | 0.1588030021 | -1.066154592 | 0.0666469885[illegible] | -0.2688834679 | 0.1211236809 |
| 0.9984451821 | -0.1975268101 | -0.9453046268 | 0.1131499894 | -0.2446867082 | 0.1084858866 |
| 0.9984451821 | 0.09767494472 | -0.9438588877 | 0.136117971 | -0.2047930647 | 0.1054647071 |
| 0.9984451821 | -0.07881692912 | -0.8699690248 | 0.1764674863 | -0.3220959338 | 0.1163305407 |
| 0.9984451821 | -0.08088792296 | -0.9297922945 | 0.1713694457 | -0.3142943971 | 0.1031258617 |
| 0.9984451821 | 0.03583656757 | -0.9348750043 | 0.1336938852 | -0.2816894638 | 0.1001721338 |
| 0.9984451821 | -0.004981976777 | -0.9374341698 | 0.1356881932 | -0.2743584426 | 0.104720873 |
| 0.9984451821 | -0.02816590084 | -0.9319604453 | 0.1358059297 | -0.2666589477 | 0.1030254557 |
| 0.9984451821 | -0.07935677231 | -0.9591611677 | 0.1474499673 | -0.264061031 | 0.1001281571 |
| 0.8210512929 | -0.05861490428 | 0.5799123017 | -0.4886058046 | 0.4620084624 | 0.09982493008 |
| 0.8072442617 | 0.08101468549 | 0.6039003884 | -0.4676503397 | 0.4382282093 | 0.1035970092 |
| 0.9888273476 | -0.003251149668 | 0.5836476368 | -0.4809821234 | 0.4707784297 | 0.1017022785 |
| 0.9075027994 | -0.02399696094 | 0.5884146779 | -0.4742698959 | 0.4586695784 | 0.1028607321 |
| 0.4719758931 | 0.2355316878 | 0.5748598798 | -0.5465701024 | 0.5022320594 | 0.09881113929 |
| 0.6628678528 | -0.1438797548 | 0.5311662463 | -0.5136107606 | 0.5700746217 | 0.1108341129 |
| 0.9075027994 | -0.02280742449 | 0.5868821122 | -0.485599823 | 0.4730393107 | 0.0982954054 |
| 0.6628678528 | 0.1509897178 | 0.5327591855 | -0.4687203569 | 0.4352588096 | 0.09940465042 |
| 0.811937764 | 0.07122290069 | 0.5498038548 | -0.4921149031 | 0.49439165 | 0.1014091877 |
| 0.8210512929 | 0.06174020775 | 0.5800981525 | -0.5024188487 | 0.5017313285 | 0.1071408766 |
| 0.6832684063 | -0.1181598093 | 0.5336634382 | -0.4997970287 | 0.5061234264 | 0.09734434948 |
| 0.7821819156 | 0.1190886032 | 0.6014272401 | -0.4927618312 | 0.4095341113 | 0.1135362604 |
| 0.9075027994 | -0.02492521402 | 0.5872479654 | -0.4878020552 | 0.4760570141 | 0.100459508 |
| 0.6628678528 | 0.142750752 | 0.5773179213 | -0.4480501821 | 0.4698031853 | 0.09850234349 |
| 0.5542917235 | 0.1989268353 | 0.6008995797 | -0.4561194048 | 0.5346511972 | 0.09505452992 |
| 0.8072442617 | 0.07903332379 | 0.6102243655 | -0.45996754 | 0.4698159099 | 0.0987629461 |
| 0.8072442617 | 0.1008199269 | 0.575927947 | -0.4482176334 | 0.479688903 | 0.1050527573 |
| 0.8072442617 | 0.09142411927 | 0.5901902478 | -0.4689807395 | 0.4421936563 | 0.09847882 |

| | | | | | |
|---|---|---|---|---|---|
| 0.8978026233 | -0.03949606293 | 0.6034667346 | -0.4983453582 | 0.4943801568 | 0.1065096223 |
| 0.811937764 | 0.07908457833 | 0.5312967256 | -0.5178078539 | 0.4676950771 | 0.1103196779 |
| 0.6628678528 | 0.2174753154 | 0.6381553016 | -0.4932225142 | 0.4547238322 | 0.1535104537 |
| 0.8210512929 | -0.0673678931 | 0.5880249668 | -0.4814507673 | 0.4447547404 | 0.1159521955 |
| 0.6628678528 | 0.1320821262 | 0.6741369938 | -0.4529691918 | 0.4159250704 | 0.1030840556 |
| 0.9426516941 | -0.0125219992 | 0.5877814901 | -0.4747904957 | 0.4651369121 | 0.1012605372 |
| 0.6628678528 | 0.1830073923 | 0.4314470191 | -0.5906605278 | 0.6125525777 | 0.1056587033 |
| 0.9075027994 | -0.03258334311 | 0.595974438 | -0.469061075 | 0.4607826067 | 0.1231161332 |
| 0.6628678528 | -0.1512869412 | 0.6311851118 | -0.4704546889 | 0.4211967444 | 0.09881506193 |
| 0.6978634991 | 0.1161286406 | 0.5356179477 | -0.5141793099 | 0.4940236008 | 0.1000687264 |
| 0.9403227783 | 0.01618126375 | 0.5965669839 | -0.4721109767 | 0.4562045972 | 0.1066846043 |
| 0.6628678528 | 0.1349231198 | 0.6685322507 | -0.4305779243 | 0.4119171378 | 0.1004590293 |
| 0.8072442617 | 0.1487186887 | 0.6213698934 | -0.4511218408 | 0.4594228401 | 0.1835676679 |
| 0.5542917235 | 0.3754698486 | 0.5697571329 | -0.5057219398 | 0.5227832146 | 0.1683297384 |
| 0.811937764 | -0.0774855406 | 0.6122194854 | -0.4680721957 | 0.4475510595 | 0.1082454406 |
| 0.9003371864 | -0.03896973178 | 0.5977090846 | -0.4698559002 | 0.4441560017 | 0.1174835925 |
| 0.8576310788 | -0.05032141553 | 0.5904713733 | -0.473052226 | 0.4329445271 | 0.108326081 |
| 0.7821819156 | 0.1538074617 | 0.6579404906 | -0.4930378349 | 0.465890106 | 0.1409751455 |
| 0.8344788873 | -0.05469324974 | 0.5919714549 | -0.4908384038 | 0.4595509172 | 0.1095478487 |
| 0.8820863362 | -0.0426619564 | 0.5922016211 | -0.4929615802 | 0.4814097242 | 0.1045672586 |
| 0.8344788873 | 0.05698482827 | 0.5775598197 | -0.4557360918 | 0.4674161699 | 0.107371706 |
| 0.811937764 | 0.08454561101 | 0.6014922622 | -0.5125568323 | 0.5233831348 | 0.1181078132 |
| 0.6628678528 | 0.1658820951 | 0.6523542275 | -0.4683545333 | 0.4286958702 | 0.1098563348 |
| 0.8210512929 | 0.07321434614 | 0.5773801101 | -0.5005581462 | 0.5193080168 | 0.1204881146 |
| 0.6628678528 | 0.14314897 | 0.5716464482 | -0.5629688355 | 0.5423998288 | 0.09988154943 |
| 0.9075027994 | -0.03273925249 | 0.5864985891 | -0.478902763 | 0.4583156422 | 0.104946415 |
| 0.8072442617 | -0.09105479999 | 0.5514340805 | -0.4739636027 | 0.5339560213 | 0.1096615389 |
| 0.6628678528 | -0.1296108582 | 0.5602973881 | -0.4770763641 | 0.5317211468 | 0.09730031069 |
| 0.8978026233 | 0.04682050716 | 0.5908091321 | -0.496616415 | 0.4905465506 | 0.1176800026 |
| 0.6628678528 | -0.1854002525 | 0.5373155177 | -0.4942952535 | 0.5056722884 | 0.1014176063 |
| 0.8820863362 | 0.04301546165 | 0.5920216849 | -0.4583305058 | 0.4385454561 | 0.103833139 |
| 0.9075027994 | 0.02375090364 | 0.588679988 | -0.4687421663 | 0.4741611565 | 0.1060486307 |
| 0.8072442617 | 0.1100210766 | 0.5602907384 | -0.508124607 | 0.5022435679 | 0.1289236237 |
| 0.8072442617 | -0.08542849973 | 0.5406415494 | -0.5347321619 | 0.5459008086 | 0.1045506456 |
| 0.8072442617 | 0.107650087 | 0.6292551144 | -0.4958123655 | 0.4061690998 | 0.1163552031 |
| 0.6628678528 | -0.1388272664 | 0.5638721431 | -0.5235364907 | 0.3894017562 | 0.1031724932 |
| 0.8820863362 | 0.04259616103 | 0.5945244004 | -0.4792390205 | 0.4608009267 | 0.100176471 |
| 0.8072442617 | -0.09001117839 | 0.6040389427 | -0.5008119101 | 0.4385891263 | 0.1075277171 |
| 0.6628678528 | -0.1333371977 | 0.5679895482 | -0.5658564874 | 0.5867780162 | 0.1013293321 |
| 0.8210512929 | -0.06972839014 | 0.5877383538 | -0.5125559288 | 0.5123557292 | 0.1107388134 |
| 0.8194923277 | 0.07247245023 | 0.6118807573 | -0.4431502453 | 0.3965323607 | 0.10607835 |
| 0.8210512929 | 0.05934420663 | 0.5644975287 | -0.4951492772 | 0.4982213076 | 0.102887426 |
| 0.6628678528 | 0.1242203891 | 0.6184850502 | -0.4830939633 | 0.4850259471 | 0.09588536645 |
| 0.9075027994 | 0.02619434667 | 0.5967002205 | -0.4744313897 | 0.4614919339 | 0.1110897226 |
| 0.8072442617 | 0.09925834877 | 0.6483947478 | -0.4396254612 | 0.4228528691 | 0.1320740573 |
| 0.8072442617 | 0.1070854212 | 0.5717187281 | -0.5365801736 | 0.4864685908 | 0.1111810334 |

| | | | | | |
|---|---|---|---|---|---|
| 0.7389307852 | 0.112593333 | 0.5950143946 | -0.4894858927 | 0.4835850967 | 0.1007818665 |
| 0.8072442617 | -0.1055538062 | 0.5455219195 | -0.5030152554 | 0.5030648972 | 0.1064996818 |
| 0.9426516941 | 0.0141353918 | 0.5775867544 | -0.4838613546 | 0.4810751723 | 0.1116678673 |
| 0.8210512929 | 0.05905926737 | 0.5636275986 | -0.5191972539 | 0.5295200722 | 0.1026733224 |
| 0.5955613183 | -0.2022989023 | 0.6637548872 | -0.4366866896 | 0.4987852608 | 0.09974178824 |
| 0.8344788873 | 0.05186914792 | 0.5789518924 | -0.5044826395 | 0.5061169162 | 0.1003648059 |
| 0.4719758931 | -0.2468409757 | 0.5239176597 | -0.3939701521 | 0.2389428046 | 0.103112895 |
| 0.9528980239 | -0.009202587627 | 0.5846213515 | -0.4811838278 | 0.47411719 | 0.1043169182 |
| 0.6628678528 | -0.1501023199 | 0.5691836949 | -0.5436081778 | 0.497823885 | 0.1097988166 |
| 0.6832684063 | -0.1253175591 | 0.5408674859 | -0.5670307232 | 0.6008665096 | 0.1025441206 |
| 0.4719758931 | 0.2296440822 | 0.5545675473 | -0.5708636763 | 0.48405877 | 0.09859217041 |
| 0.9075027994 | -0.03115493961 | 0.5861057307 | -0.4833134458 | 0.4680667453 | 0.1124294751 |
| 0.8210512929 | -0.06378047466 | 0.6326927577 | -0.4585784964 | 0.4395459373 | 0.1035438952 |
| 0.6628678528 | -0.1790529053 | 0.6259763055 | -0.4762757832 | 0.500915103 | 0.1000821916 |
| 0.8344788873 | -0.05462028217 | 0.5875726524 | -0.4901115153 | 0.4724249785 | 0.106985897 |
| 0.8072442617 | -0.09207251245 | 0.560542735 | -0.5108519577 | 0.4539658071 | 0.09851618018 |
| 0.6628678528 | -0.1631608291 | 0.5086508839 | -0.560269045 | 0.5567169409 | 0.1018764453 |
| 0.9075027994 | -0.02149290849 | 0.5946664815 | -0.4776648793 | 0.4627332198 | 0.1023837539 |
| 0.8210512929 | -0.06964621223 | 0.5591982111 | -0.4955560091 | 0.4814948596 | 0.1132741163 |
| 0.6628678528 | -0.1799460853 | 0.501645245 | -0.4827192491 | 0.4866892461 | 0.105106081 |
| 0.6978634991 | 0.1286821468 | 0.5599247765 | -0.386366824 | 0.3812069231 | 0.1108669479 |
| 0.8210512929 | 0.0682812245 | 0.5532648231 | -0.4787699901 | 0.488960777 | 0.1102345942 |
| 0.6628678528 | 0.1641561867 | 0.5605053171 | -0.4995204357 | 0.4954510971 | 0.09655359406 |
| 0.9888273476 | -0.00269679101 | 0.5853060173 | -0.4806902457 | 0.4701378597 | 0.1121662176 |
| 0.6628678528 | 0.1662755294 | 0.598825984 | -0.4283296416 | 0.4644527372 | 0.1298486142 |
| 0.8072442617 | -0.0889436267 | 0.628522343 | -0.466601953 | 0.4680872734 | 0.1103914779 |
| 0.9338493859 | -0.01652659387 | 0.593175995 | -0.4755589599 | 0.4679777958 | 0.09833438877 |
| 0.9075027994 | -0.026995723 | 0.5867244593 | -0.4678440507 | 0.4574364744 | 0.1121981076 |
| 0.8122584867 | -0.07311521644 | 0.590025303 | -0.4059784655 | 0.425851929 | 0.1055069188 |
| 0.8021266674 | -0.1150649318 | 0.5815062614 | -0.4868959374 | 0.4664179903 | 0.11339095 |
| 0.694761936 | 0.1231497293 | 0.5679832459 | -0.4631003121 | 0.5274633387 | 0.1040101825 |
| 0.4719758931 | -0.3080227448 | 0.6549042464 | -0.4246221975 | 0.4518504794 | 0.1035838111 |
| 0.9075027994 | 0.03367967646 | 0.579714747 | -0.488570937 | 0.464620791 | 0.121188365 |
| 0.8344788873 | -0.05838964801 | 0.580414319 | -0.4943092351 | 0.5140252474 | 0.112616122 |
| 0.8072442617 | -0.07818182571 | 0.5536618424 | -0.5063544914 | 0.4863620642 | 0.1028053093 |
| 0.6628678528 | -0.1387113291 | 0.5544013811 | -0.4851336703 | 0.4831039649 | 0.09768925393 |
| 0.8072442617 | -0.111670727 | 0.613965362 | -0.4282709701 | 0.3706260482 | 0.1236204434 |
| 0.6628678528 | 0.1553782336 | 0.6028289584 | -0.4608793982 | 0.5140154424 | 0.09795178356 |
| 0.4719758931 | 0.2868399863 | 0.6896805252 | -0.5271791124 | 0.3871621608 | 0.1127223472 |
| 0.5207540292 | 0.278288939 | 0.6751931409 | -0.3757836693 | 0.4803753411 | 0.1227810736 |
| 0.6628678528 | 0.1502957701 | 0.5018721762 | -0.4336585188 | 0.4268980147 | 0.1066545116 |
| 0.8072442617 | 0.1034916724 | 0.6252369067 | -0.4179635991 | 0.3827165216 | 0.1191015006 |
| 0.4719758931 | -0.2483873231 | 0.5955884351 | -0.4503226811 | 0.4562322694 | 0.09443583275 |
| 0.8072442617 | 0.1011282054 | 0.5379903317 | -0.531112471 | 0.5800275969 | 0.1176298084 |
| 0.9003371864 | 0.0754818601 | 0.6034003632 | -0.460795889 | 0.4476392721 | 0.1733415511 |
| 0.6628678528 | 0.1515307298 | 0.5117335359 | -0.4482895431 | 0.4797411292 | 0.101308444 |

| | | | | | |
|---|---|---|---|---|---|
| 0.6628678528 | 0.1769052032 | 0.5545980479 | -0.4961122736 | 0.5017320976 | 0.09415879921 |
| 0.9075027994 | 0.02464364947 | 0.6001214198 | -0.4690590777 | 0.4478237021 | 0.1155055979 |
| 0.9477576316 | -0.01098713159 | 0.5879466122 | -0.4810071354 | 0.4676048384 | 0.1055960363 |
| 0.9426516941 | 0.01310091824 | 0.5886293257 | -0.469120106 | 0.4539462456 | 0.1091580901 |
| 0.9003371864 | -0.0363013333 | 0.5854412137 | -0.4710230103 | 0.4525159149 | 0.1053082145 |
| 0.6628678528 | 0.1775710026 | 0.5804954392 | -0.4996092142 | 0.5227958104 | 0.1296706297 |
| 0.9974164499 | -0.000402724322 | 0.5854543977 | -0.4802341152 | 0.4695949982 | 0.1229269222 |
| 0.6628678528 | 0.1701259452 | 0.5913910045 | -0.4521457957 | 0.4441642543 | 0.1103506394 |
| 0.9003371864 | 0.03486647894 | 0.5829608765 | -0.4817364961 | 0.4946124228 | 0.1049054834 |
| 0.6628678528 | 0.1562628613 | 0.4549479671 | -0.5617376448 | 0.5577845653 | 0.1163134812 |
| 0.8072442617 | 0.07664616439 | 0.5753285063 | -0.5152873264 | 0.512097207 | 0.1022876189 |
| 0.8344788873 | -0.04875591793 | 0.5832398876 | -0.4712280047 | 0.4751944744 | 0.09830909128 |
| 0.8072442617 | 0.07986021053 | 0.6090631496 | -0.4275226655 | 0.4230578117 | 0.1031909181 |
| 0.6628678528 | -0.1277041516 | 0.6072570539 | -0.4877904566 | 0.5084344299 | 0.1006706539 |
| 0.8072442617 | -0.08755466166 | 0.5632059963 | -0.4686344399 | 0.4771212026 | 0.09880184618 |
| 0.8677764199 | -0.10749473 | -0.5718154454 | 0.4728064328 | -0.7780620408 | 0.1073057841 |
| 0.9949208784 | 0.04957521283 | -0.5457177013 | 0.4969355139 | -0.7924472426 | 0.1107293624 |
| 0.804159583 | -0.1784587885 | -0.6321783173 | 0.4726333415 | -0.7230567695 | 0.1074739 |
| 0.9949208784 | 0.03999834017 | -0.561650036 | 0.4814636024 | -0.7562867732 | 0.1099218286 |
| 0.804159583 | -0.175052581 | -0.5543936899 | 0.527105468 | -0.7854928619 | 0.108570566 |
| 0.804159583 | 0.1548537885 | -0.4941934197 | 0.5363537663 | -0.8925377551 | 0.1146223316 |
| 0.9949208784 | 0.007617889605 | -0.5582364224 | 0.4926352703 | -0.7743543096 | 0.1071980031 |
| 0.9949208784 | 0.04365721251 | -0.5751165291 | 0.494608217 | -0.7798390039 | 0.1094666885 |
| 0.9949208784 | 0.005521280813 | -0.5604803117 | 0.4898626134 | -0.7708752006 | 0.1096069581 |
| 0.9949208784 | 0.03659411844 | -0.5588233293 | 0.4817813048 | -0.7584812941 | 0.1124023464 |
| 0.9949208784 | 0.02888916835 | -0.5444651963 | 0.4947315047 | -0.7828590722 | 0.1071817386 |
| 0.7720369792 | -0.21143013 | -0.5949918854 | 0.5131746817 | -0.6512483174 | 0.116304297 |
| 0.9949208784 | -0.0224928846 | -0.5551388854 | 0.4832083635 | -0.7674881982 | 0.1087220082 |
| 0.804159583 | -0.1200955099 | -0.554444645 | 0.4595095855 | -0.7664529962 | 0.1072521606 |
| 0.9364241781 | -0.09684978107 | -0.5636013273 | 0.4820978628 | -0.8090521179 | 0.1069858532 |
| 0.804159583 | -0.1438714885 | -0.6048961637 | 0.4488786068 | -0.7703211535 | 0.1068444174 |
| 0.8677764199 | -0.1127710306 | -0.5559187013 | 0.4466896228 | -0.7695331813 | 0.111371256 |
| 0.9949208784 | 0.06136013823 | -0.5548604731 | 0.4998635638 | -0.7900758134 | 0.1069577018 |
| 0.804159583 | 0.1312090175 | -0.6203569651 | 0.5537405442 | -0.8550140939 | 0.1152184994 |
| 0.9949208784 | -0.003139993328 | -0.5558497821 | 0.4920940621 | -0.7724134439 | 0.1184518951 |
| 0.9949208784 | 0.07533842203 | -0.5254856659 | 0.5071519312 | -0.8045554996 | 0.1294141585 |
| 0.804159583 | 0.1800922557 | -0.5735223719 | 0.5024481297 | -0.7043867621 | 0.1162977329 |
| 0.804159583 | -0.1730621459 | -0.6768842254 | 0.4662840134 | -0.7031603103 | 0.1099522864 |
| 0.7720369792 | 0.1991478434 | -0.6007093552 | 0.4116927169 | -0.7006553823 | 0.10585611 |
| 0.9222849546 | 0.1115990691 | -0.6466283659 | 0.424023693 | -0.6935357243 | 0.1153385985 |
| 0.7720369792 | -0.2402769767 | -0.4837915093 | 0.5815622441 | -0.8386261772 | 0.11666301 |
| 0.9949208784 | -0.05202400088 | -0.5439750758 | 0.4962664378 | -0.7869438162 | 0.1084312424 |
| 0.9949208784 | 0.04173616047 | -0.5739049158 | 0.4816370478 | -0.7674859062 | 0.1091659267 |
| 0.9949208784 | -0.05470037797 | -0.5943266086 | 0.4604618215 | -0.7289904379 | 0.1137785916 |
| 0.9949208784 | 0.01707940432 | -0.5483539444 | 0.4969744415 | -0.7783669542 | 0.1097512464 |
| 0.9949208784 | -0.02688436401 | -0.566218087 | 0.4859542785 | -0.7701372193 | 0.1455641702 |

| | | | | | |
|---|---|---|---|---|---|
| 0.804159583 | -0.1694750468 | -0.5417236412 | 0.5302169961 | -0.8361277727 | 0.1311083091 |
| 0.804159583 | 0.1407290966 | -0.6109122797 | 0.480495921 | -0.7355998335 | 0.1128666054 |
| 0.9949208784 | -0.06912924233 | -0.5325274213 | 0.5184583838 | -0.8272697248 | 0.1198279905 |
| 0.9949208784 | 0.006489545572 | -0.5585823745 | 0.4895803411 | -0.7680224129 | 0.1147066292 |
| 0.9949208784 | 0.1143660899 | -0.4986855426 | 0.471920175 | -0.7712511799 | 0.1270650158 |
| 0.9949208784 | -0.001147466391 | -0.557684455 | 0.490347423 | -0.7728581023 | 0.1145416347 |
| 0.9949208784 | -0.0234671172 | -0.553440907 | 0.483358258 | -0.7663347041 | 0.1117966094 |
| 0.9949208784 | -0.04100547425 | -0.5520053172 | 0.4729496489 | -0.7717500193 | 0.1140348425 |
| 0.9949208784 | -0.05358133412 | -0.5638005749 | 0.5171556261 | -0.8163825347 | 0.1219761646 |
| 0.9949208784 | -0.08113007217 | -0.5885259814 | 0.4775147519 | -0.751258958 | 0.1152744659 |
| 0.7720369792 | -0.21536702 | -0.5392940614 | 0.5373678987 | -0.900958968 | 0.1172295139 |
| 0.7720369792 | -0.2475179926 | -0.5415702714 | 0.6425704178 | -0.8908580204 | 0.1048961436 |
| 0.9949208784 | 0.04906265753 | -0.5582309509 | 0.4879030464 | -0.758844172 | 0.1114609682 |
| 0.9949208784 | 0.07683549082 | -0.5252292361 | 0.4835701377 | -0.8330626212 | 0.1143980001 |
| 0.9949208784 | 0.0387524993 | -0.5509144342 | 0.486464596 | -0.7892232199 | 0.1072594318 |
| 0.8677764199 | 0.1300166542 | -0.5353833489 | 0.4532182234 | -0.7343312688 | 0.1151019256 |
| 0.804159583 | 0.1343802021 | -0.5223830378 | 0.4831395069 | -0.7927003181 | 0.1090616505 |
| 0.9949208784 | 0.01512481282 | -0.5558935416 | 0.4974505371 | -0.7825616774 | 0.111049738 |
| 0.804159583 | -0.1368724063 | -0.5817067684 | 0.417660692 | -0.7900090594 | 0.1126015759 |
| 0.7720369792 | -0.3029229265 | -0.4919090292 | 0.5773916812 | -0.8522696518 | 0.123634323 |
| 0.8677764199 | 0.1183594953 | -0.4987740187 | 0.565086779 | -0.872276783 | 0.1116137994 |
| 0.7720369792 | 0.2708966256 | -0.4401736205 | 0.4697325778 | -0.9540419166 | 0.1113657105 |
| 0.9949208784 | 0.07492780522 | -0.5465030733 | 0.5190650522 | -0.7318885383 | 0.1131543868 |
| 0.9949208784 | 0.04533141921 | -0.5465357256 | 0.4935518759 | -0.7856186213 | 0.1081835439 |
| 0.9949208784 | 0.009261737616 | -0.5599013579 | 0.4934814365 | -0.7697179866 | 0.1138444788 |
| 0.9949208784 | 0.01478942595 | -0.556926476 | 0.4982444262 | -0.7833233453 | 0.1102046426 |
| 0.9949208784 | 0.06581231482 | -0.5626392194 | 0.5152518742 | -0.8048253287 | 0.1151228832 |
| 0.9949208784 | -0.07782582809 | -0.5816836982 | 0.4499518602 | -0.700523995 | 0.1131259179 |
| 0.9949208784 | -0.06180528489 | -0.5309969489 | 0.5090485804 | -0.8119853773 | 0.1100221808 |
| 0.9949208784 | 0.0006820376366 | -0.5576446585 | 0.4906179211 | -0.7726275375 | 0.1067008877 |
| 0.9949208784 | 0.005115377357 | -0.5554334335 | 0.4918503484 | -0.774298788 | 0.1156255229 |
| 0.9949208784 | -0.04284765039 | -0.5860526116 | 0.4742139821 | -0.7530653227 | 0.1285505997 |
| 0.9949208784 | -0.09653685401 | -0.549110319 | 0.5400453887 | -0.7844436519 | 0.1160854843 |
| 0.804159583 | -0.1247720985 | -0.5737688742 | 0.5034911599 | -0.7811589159 | 0.1081793773 |
| 0.9949208784 | 0.02344332417 | -0.5495877121 | 0.4949283403 | -0.7796894731 | 0.1128523889 |
| 0.9949208784 | -0.007813335882 | -0.5537789932 | 0.492302073 | -0.7786806326 | 0.1169294345 |
| 0.9949208784 | -0.02568054794 | -0.548826086 | 0.5068068467 | -0.7970985455 | 0.1108684712 |
| 0.9949208784 | 0.02148559405 | -0.5673016945 | 0.4842451 | -0.773486718 | 0.1116273794 |
| 0.804159583 | -0.1756207858 | -0.5377642089 | 0.5787737277 | -0.8956735147 | 0.1065458547 |
| 0.804159583 | 0.1409767951 | -0.521284015 | 0.4394294399 | -0.6409625259 | 0.1161298869 |
| 0.9949208784 | 0.002845556923 | -0.5577414608 | 0.4907546272 | -0.7739876516 | 0.1111621586 |
| 0.804159583 | -0.1726795036 | -0.57568123 | 0.4230763048 | -0.7382714523 | 0.1140764827 |
| 0.804159583 | 0.1348073614 | -0.5120773516 | 0.5763448243 | -0.9060650839 | 0.1105781253 |
| 0.804159583 | -0.1277340485 | -0.5373129812 | 0.5370494913 | -0.7818874235 | 0.1095425511 |
| 0.9949208784 | -0.01617719512 | -0.5577650555 | 0.4893978223 | -0.7725993448 | 0.1160970859 |
| 0.7720369792 | 0.1994680345 | -0.7101688284 | 0.4180110028 | -0.6691741362 | 0.1094942348 |

| | | | | | |
|---|---|---|---|---|---|
| 0.9949208784 | 0.0138163659 | -0.5606513559 | 0.4897001531 | -0.7756413679 | 0.1100690935 |
| 0.9949208784 | 0.07277192984 | -0.5679202446 | 0.4950093682 | -0.7631229181 | 0.1119189666 |
| 0.9949208784 | -0.01381194307 | -0.5618187717 | 0.4857180183 | -0.7743994415 | 0.1087283001 |
| 0.9949208784 | 0.03130654265 | -0.5418952456 | 0.5043606913 | -0.7905193743 | 0.1112773266 |
| 0.9949208784 | 0.07286334247 | -0.5921212717 | 0.4816112176 | -0.7455973018 | 0.1097076529 |
| 0.9949208784 | -0.03472378075 | -0.5712484393 | 0.4862595986 | -0.7664858571 | 0.1174219754 |
| 0.9949208784 | 0.004355342554 | -0.5556798994 | 0.4903698247 | -0.7732308888 | 0.1135396426 |
| 0.9949208784 | 0.03080073933 | -0.5629415671 | 0.5165865225 | -0.7964627172 | 0.1192428808 |
| 0.9949208784 | -0.001275839056 | -0.5573078314 | 0.4904478668 | -0.7729152914 | 0.1148514256 |
| 0.9949208784 | -0.07094766438 | -0.5466197877 | 0.4976299239 | -0.7861067596 | 0.1067916629 |
| 0.9949208784 | -0.06706191 | -0.5519709745 | 0.4787318839 | -0.7590339356 | 0.115450483 |
| 0.804159583 | -0.1487984158 | -0.5570372712 | 0.4497117845 | -0.7911691922 | 0.1194817952 |
| 0.9949208784 | 0.01540847409 | -0.5655005166 | 0.4877064437 | -0.7725297838 | 0.1162753503 |
| 0.7720369792 | 0.2425511748 | -0.6753894375 | 0.4247420294 | -0.7513543143 | 0.1035812683 |
| 0.9949208784 | 0.03461638984 | -0.5581057673 | 0.473588858 | -0.7588473245 | 0.1161499675 |
| 0.804159583 | 0.1321025013 | -0.5600594924 | 0.357863971 | -0.7032009368 | 0.1136899022 |
| 0.804159583 | 0.1670927882 | -0.5523253715 | 0.4950007232 | -0.7581965704 | 0.1152481425 |
| 0.9949208784 | -0.02938901493 | -0.5532034032 | 0.4877307079 | -0.7880800369 | 0.1126491897 |
| 0.7720369792 | 0.2278892371 | -0.5925027521 | 0.4642827931 | -0.7927333595 | 0.1088921379 |
| 0.9949208784 | -0.09193122563 | -0.5417307507 | 0.505618734 | -0.7562086926 | 0.120928626 |
| 0.9949208784 | -0.08942135295 | -0.5619823619 | 0.471807004 | -0.7076924352 | 0.1168164996 |
| 0.9949208784 | -0.03694063159 | -0.5734645553 | 0.4796925282 | -0.7641069413 | 0.1109570177 |
| 0.9949208784 | -0.005764284697 | -0.5593978586 | 0.4901958718 | -0.7716747844 | 0.1075115309 |
| 0.9949208784 | 0.09784417161 | -0.579819518 | 0.4352016518 | -0.6885921318 | 0.121546213 |
| 0.804159583 | -0.1754844793 | -0.5844488446 | 0.4663732337 | -0.8148741632 | 0.1061300219 |
| 0.9117401219 | 0.1138487192 | -0.5168200248 | 0.4826733096 | -0.8110855649 | 0.119557265 |
| 0.804159583 | 0.1471767873 | -0.5073959962 | 0.5504898408 | -0.7713678421 | 0.1244203224 |
| 0.7720369792 | -0.2197559038 | -0.4421396471 | 0.4121309014 | -0.7050075204 | 0.1123005775 |
| 0.804159583 | -0.1407750242 | -0.6108002692 | 0.4101949201 | -0.6634869864 | 0.1179284746 |
| 0.9949208784 | 0.0750734143 | -0.5638453007 | 0.4832239595 | -0.7650971242 | 0.1078867646 |
| 0.9949208784 | -0.07169383828 | -0.5274280451 | 0.5164324851 | -0.8419826412 | 0.1193213558 |
| 0.7720369792 | -0.2344239342 | -0.5850026432 | 0.472919568 | -0.768930499 | 0.1167776319 |
| 0.9949208784 | -0.04356953516 | -0.534428546 | 0.4844908029 | -0.7810036002 | 0.110750466 |
| 0.9949208784 | 0.009264233832 | -0.5598141927 | 0.4899106253 | -0.7705345916 | 0.1056017071 |
| 0.804159583 | -0.1619164031 | -0.6536286834 | 0.4246793713 | -0.6434503417 | 0.1162360253 |
| 0.9949208784 | -0.05092258819 | -0.545689218 | 0.4853498794 | -0.7790123809 | 0.1116817503 |
| 0.804159583 | -0.1375198984 | -0.5967461108 | 0.3756502919 | -0.6100454293 | 0.1149288033 |
| 0.9949208784 | -0.01547929888 | -0.5584762379 | 0.4946929889 | -0.7785228637 | 0.1117344079 |
| 0.9949208784 | 0.0468217943 | -0.5632986649 | 0.4877998464 | -0.7522444291 | 0.1293008596 |
| 0.804159583 | 0.1754371135 | -0.7026251143 | 0.424010041 | -0.76761113 | 0.1274771184 |
| 0.804159583 | -0.1424356881 | -0.5579945724 | 0.459133942 | -0.7531302602 | 0.1132582608 |
| 0.9949208784 | 0.05677458931 | -0.5628096688 | 0.4910467239 | -0.7308977319 | 0.1124192166 |
| 0.804159583 | -0.1489362697 | -0.4392545895 | 0.5535544765 | -0.8457881173 | 0.1190554956 |
| 0.9949208784 | -0.01042221318 | -0.5573403936 | 0.4947644137 | -0.7770683735 | 0.1096794607 |
| 0.7720369792 | 0.2094970418 | -0.5490908919 | 0.4628673551 | -0.8051621571 | 0.1037058616 |
| 0.9949208784 | 0.07584617485 | -0.5395118444 | 0.5421493575 | -0.8120147314 | 0.1104579081 |

| | | | | | |
|---|---|---|---|---|---|
| 0.9949208784 | 0.05190748147 | -0.5637786353 | 0.4955538058 | -0.7944667373 | 0.1095917466 |
| 0.9949208784 | -0.07882627338 | -0.5819447429 | 0.5009587142 | -0.75856604 | 0.1067240184 |
| 0.8886403667 | -0.135294183 | 1.015254398 | 0.1769695848 | -0.9678863909 | 0.145482559 |
| 0.8886403667 | -0.06271159722 | 1.005197719 | 0.2317137512 | -0.9640794046 | 0.1421511035 |
| 0.8886403667 | 0.08201883058 | 1.051331066 | 0.2998129067 | -1.05523865 | 0.170035162 |
| 0.9824246933 | -0.01085485698 | 1.01717441 | 0.257173985 | -0.9945967516 | 0.1507989003 |
| 0.8886403667 | 0.2780610459 | 0.963988131 | 0.1768897216 | -0.9608681144 | 0.1366809124 |
| 0.8886403667 | -0.1878437925 | 0.8751522265 | 0.1572015989 | -0.7687504509 | 0.1708218534 |
| 0.8886403667 | -0.1013430959 | 1.020398652 | 0.1688772906 | -0.9393297999 | 0.1632111246 |
| 0.9894417916 | -0.003329172291 | 1.016423004 | 0.2489478045 | -0.9860203428 | 0.1437742597 |
| 0.9824246933 | 0.007151425249 | 1.012957634 | 0.2492771876 | -0.9873369103 | 0.1384025016 |
| 0.8886403667 | 0.1172432812 | 1.031720716 | 0.2377358272 | -1.001097256 | 0.1587575314 |
| 0.8886403667 | -0.2056808883 | 0.9295266667 | 0.2573355986 | -1.034865383 | 0.1404199306 |
| 0.9824246933 | 0.03691866958 | 1.026444272 | 0.2589177198 | -1.017936691 | 0.1484210634 |
| 0.8886403667 | 0.1003551393 | 1.022762969 | 0.2898705062 | -1.008817958 | 0.144528787 |
| 0.8886403667 | 0.07274704688 | 1.023986727 | 0.3135175931 | -1.06466766 | 0.1506159658 |
| 0.8888946878 | 0.05734486266 | 1.023088633 | 0.2625161917 | -0.9504578241 | 0.1368612558 |
| 0.8886403667 | 0.1326483792 | 1.074380043 | 0.3366844994 | -1.064239649 | 0.1519914648 |
| 0.8886403667 | 0.284608699 | 1.04529923 | 0.3569552362 | -1.050715289 | 0.159099537 |
| 0.8886403667 | 0.1184344209 | 1.034603306 | 0.3049968452 | -1.092084339 | 0.1425218229 |
| 0.8886403667 | 0.08468421382 | 0.9706955512 | 0.3071449271 | -1.043081299 | 0.1891381983 |
| 0.8886403667 | -0.2670277655 | 1.187891955 | 0.2957959337 | -0.7909363046 | 0.1566338675 |
| 0.8886403667 | 0.1160638261 | 1.072570697 | 0.2685644954 | -1.054844457 | 0.1810649693 |
| 0.8886403667 | 0.1235729294 | 1.00566264 | 0.2351039928 | -0.8554629578 | 0.1498950923 |
| 0.8886403667 | 0.1171386908 | 1.07065967 | 0.240484282 | -0.971345879 | 0.1463540866 |
| 0.8886403667 | -0.07234043976 | 1.015058534 | 0.2734476799 | -0.9695265784 | 0.1466977948 |
| 0.8886403667 | 0.1586884533 | 0.8139198341 | 0.06901174838 | -0.7915668076 | 0.1539840831 |
| 0.8886403667 | 0.2674195702 | 0.9782598503 | 0.1366553722 | -0.8718530201 | 0.14434294 |
| 0.8886403667 | -0.06905156511 | 1.044020518 | 0.248666384 | -0.9963540236 | 0.151895759 |
| 0.8886403667 | -0.121232799 | 1.057588182 | 0.2658435719 | -1.01013718 | 0.1665276656 |
| 0.8886403667 | 0.07892343358 | 1.088759506 | 0.2713698459 | -1.056243153 | 0.1505170223 |
| 0.8886403667 | -0.2315181608 | 0.8848025517 | 0.208652101 | -0.9117233355 | 0.1443889899 |
| 0.8886403667 | 0.1479982937 | 1.056952744 | 0.1989022376 | -0.9121742702 | 0.1439760169 |
| 0.8886403667 | -0.08778160854 | 1.031313139 | 0.2967733171 | -1.036589449 | 0.1501279189 |
| 0.8886403667 | -0.1898841856 | 1.012186148 | 0.20902465 | -0.9507027043 | 0.1425118394 |
| 0.8886403667 | -0.2683032194 | 1.115415202 | 0.413521646 | -1.207372334 | 0.1427121794 |
| 0.8886403667 | 0.2268009211 | 0.9775773871 | 0.1663716893 | -0.7889278635 | 0.1412774651 |
| 0.9824246933 | -0.03323196 | 0.9978455829 | 0.2640932299 | -1.008658886 | 0.1539684269 |
| 0.8886403667 | -0.1046901718 | 1.060794165 | 0.1977479658 | -1.022635828 | 0.1589958611 |
| 0.8886403667 | 0.1010078015 | 0.9951180066 | 0.2915516538 | -1.020171823 | 0.1456684299 |
| 0.8886403667 | 0.07812491255 | 1.005687822 | 0.2916507859 | -0.9946784382 | 0.1497601165 |
| 0.9824246933 | 0.008827927856 | 1.014425015 | 0.2461612477 | -0.9792920459 | 0.1548994134 |
| 0.8886403667 | 0.09199427991 | 1.036888257 | 0.2376963886 | -0.9995376225 | 0.1403950219 |
| 0.9824246933 | -0.02685470679 | 1.01496851 | 0.2610353789 | -1.018925612 | 0.1431579047 |
| 0.8886403667 | 0.1451067718 | 0.9346689356 | 0.1529659532 | -0.9118107035 | 0.1605486496 |
| 0.8886403667 | -0.1459014274 | 1.015778665 | 0.2158275781 | -1.007499116 | 0.1469821574 |

| | | | | | |
|---|---|---|---|---|---|
| 0.8886403667 | 0.1462559089 | 1.044654986 | 0.2030638294 | -1.002371879 | 0.1466081714 |
| 0.8886403667 | -0.203726362 | 1.02536181 | 0.2349371866 | -0.880707308 | 0.1461486904 |
| 0.8886403667 | 0.08251082429 | 1.033478297 | 0.2457985948 | -0.9434487696 | 0.1419698717 |
| 0.8886403667 | -0.2081541162 | 1.015670563 | 0.3157475339 | -1.065866179 | 0.1420482631 |
| 0.9824246933 | 0.03779698553 | 1.008585258 | 0.2647656369 | -0.9880450609 | 0.1472656159 |
| 0.8886403667 | -0.08696822981 | 0.9917772551 | 0.2218489669 | -1.048145394 | 0.1392531078 |
| 0.9775092155 | 0.04440505842 | 1.014045688 | 0.2315509012 | -0.9745036846 | 0.1517822278 |
| 0.8886403667 | 0.1085398618 | 1.037303979 | 0.3471345098 | -1.13640705 | 0.1517298755 |
| 0.8886403667 | -0.06330458806 | 1.01268153 | 0.2637903917 | -0.9723797946 | 0.1475474524 |
| 0.8886403667 | 0.1075217381 | 1.05861809 | 0.3199975061 | -0.9305316337 | 0.1549541909 |
| 0.8886403667 | -0.08550622032 | 0.9999645962 | 0.256040862 | -0.953024033 | 0.1487030737 |
| 0.9824246933 | -0.01389101106 | 1.016065858 | 0.2503657801 | -0.9992766475 | 0.1507414697 |
| 0.8886403667 | -0.08205512788 | 1.021651752 | 0.2268663574 | -0.9808940184 | 0.1492426136 |
| 0.9824246933 | 0.03387918195 | 1.024624803 | 0.2795701899 | -1.022528504 | 0.2127548988 |
| 0.8886403667 | -0.06619351325 | 1.007642683 | 0.2253073779 | -0.9227836998 | 0.1519973489 |
| 0.9824246933 | -0.01782552639 | 1.01659367 | 0.2491673479 | -0.9904810676 | 0.1453315508 |
| 0.8886403667 | -0.08514411875 | 1.001351461 | 0.253271229 | -0.9917967414 | 0.1362158297 |
| 0.8886403667 | -0.1531015507 | 0.9089501911 | 0.2279456583 | -0.9273871844 | 0.1687381869 |
| 0.9824246933 | -0.03718038548 | 0.9923608656 | 0.2357584087 | -0.9703787873 | 0.1576975687 |
| 0.9824246933 | 0.01682659945 | 1.013118531 | 0.2448007624 | -0.9971034806 | 0.1538776897 |
| 0.8886403667 | 0.08911134249 | 1.045564192 | 0.2962775078 | -1.085213378 | 0.153282397 |
| 0.9124849918 | 0.05387590073 | 1.034076984 | 0.2800650641 | -1.01581967 | 0.140813521 |
| 0.9824246933 | -0.01435774013 | 1.018178101 | 0.2448874469 | -0.9835200489 | 0.173816562 |
| 0.8886403667 | 0.1619534872 | 0.9835864458 | 0.1574616475 | -0.8382110979 | 0.1476283178 |
| 0.8886403667 | -0.1565875358 | 1.12822807 | 0.279448573 | -1.025802012 | 0.1553960659 |
| 0.9775092155 | -0.04002079668 | 1.022075128 | 0.2746234454 | -1.006667984 | 0.1430921648 |
| 0.9898215044 | 0.001972977232 | 1.015939305 | 0.2486513065 | -0.9851816311 | 0.1546558352 |
| 0.8886403667 | 0.06482682975 | 1.051358846 | 0.2723767476 | -1.054276636 | 0.1502488052 |
| 0.8886403667 | 0.1313298986 | 1.036605139 | 0.2751382634 | -0.969823808 | 0.1508072561 |
| 0.8886403667 | 0.1389376243 | 1.06792653 | 0.3723040147 | -1.13469035 | 0.157805943 |
| 0.8886403667 | -0.1575247939 | 1.023749529 | 0.2795017726 | -0.9839611462 | 0.2309489934 |
| 0.8886403667 | -0.09539040779 | 1.015540012 | 0.2303996588 | -1.004476999 | 0.1464816506 |
| 0.8886403667 | 0.1604995379 | 0.9120740878 | 0.1745526769 | -0.8772396172 | 0.1428589561 |
| 0.8886403667 | -0.08788916091 | 1.023170116 | 0.1869251818 | -0.9565607987 | 0.1529557519 |
| 0.8886403667 | -0.07485423485 | 1.033074567 | 0.251477966 | -1.016011248 | 0.1428520002 |
| 0.8886403667 | -0.2697305303 | 0.8854132595 | 0.1156965403 | -1.075637714 | 0.1365880208 |
| 0.8886403667 | -0.2470403693 | 0.9159736692 | 0.1369929426 | -1.014029095 | 0.1428975437 |
| 0.9824246933 | -0.0231276118 | 1.028467661 | 0.2663542099 | -0.9954126229 | 0.1546305533 |
| 0.9824246933 | -0.02264712892 | 1.0120841 | 0.2413189139 | -0.9947926425 | 0.1390299439 |
| 0.8886403667 | -0.1308334394 | 0.9747460861 | 0.2340008736 | -1.04252777 | 0.1492627189 |
| 0.8886403667 | -0.1806466259 | 1.006555601 | 0.03564975969 | -0.769159915 | 0.1952350038 |
| 0.8886403667 | 0.1051922801 | 1.034627524 | 0.2828389892 | -1.017665267 | 0.1792191017 |
| 0.8886403667 | 0.1823290689 | 0.9375599168 | 0.2183098815 | -0.9428492531 | 0.1468334205 |
| 0.8886403667 | 0.07887204872 | 1.003696139 | 0.242601427 | -0.9943479629 | 0.1634953325 |
| 0.8886403667 | 0.151669941 | 1.054919572 | 0.2561809753 | -0.9894811312 | 0.1461581214 |
| 0.8190737159 | -0.3784135434 | 1.103819493 | 0.1636669737 | -0.8751712821 | 0.1326453019 |

| | | | | | |
|---|---|---|---|---|---|
| 0.8886403667 | -0.07253233389 | 1.061849372 | 0.3009062859 | -1.024988692 | 0.1485721715 |
| 0.9824246933 | 0.02543682465 | 1.005957736 | 0.2366534598 | -0.9814066364 | 0.1584201135 |
| 0.8886403667 | 0.1214946683 | 1.003119808 | 0.1230588211 | -0.9317740359 | 0.1681179487 |
| 0.8886403667 | -0.1950569718 | 1.084297552 | 0.4476280926 | -1.116749274 | 0.1585864594 |
| 0.8886403667 | -0.06999238955 | 1.030782099 | 0.2418306043 | -1.013679784 | 0.1356569133 |
| 0.8886403667 | 0.1444945139 | 0.9813658858 | 0.2458067766 | -1.025079081 | 0.1407517676 |
| 0.8886403667 | 0.1183720649 | 1.024385248 | 0.1860685679 | -1.025362327 | 0.1537266 |
| 0.9824246933 | 0.006157394664 | 1.014688559 | 0.2486658945 | -0.9886687873 | 0.1475158908 |
| 0.9824246933 | -0.03243938496 | 0.9999617006 | 0.2302693026 | -0.9747066418 | 0.1420778402 |
| 0.9824246933 | -0.005995387161 | 1.012391426 | 0.2472019117 | -0.9829098373 | 0.1315002829 |
| 0.8886403667 | -0.1035812012 | 1.07111957 | 0.2938191607 | -1.115575811 | 0.1426794573 |
| 0.8886403667 | 0.133160399 | 1.050372978 | 0.3041453972 | -1.026221348 | 0.140112105 |
| 0.9824246933 | -0.02175297174 | 1.016484234 | 0.2541106682 | -0.9839620093 | 0.139700893 |
| 0.9824246933 | 0.0201940531 | 1.0152958 | 0.2546883711 | -0.9734799235 | 0.1472664367 |
| 0.8886403667 | -0.227176032 | 1.127725454 | 0.2588383297 | -0.9424201523 | 0.1597528475 |
| 0.8886403667 | 0.1310426114 | 1.088386873 | 0.2991101132 | -1.089970474 | 0.134605712 |
| 0.9775092155 | -0.03993718014 | 1.025239797 | 0.2652011874 | -0.9980712984 | 0.144260926 |
| 0.8886403667 | 0.1179421626 | 0.9994325998 | 0.229864893 | -0.9251902785 | 0.1395503471 |
| 0.9824246933 | -0.02259663182 | 1.008535472 | 0.2500679711 | -0.9804716549 | 0.1516659341 |
| 0.8886403667 | -0.1509377769 | 1.10298093 | 0.2221010193 | -0.9776237291 | 0.1391734145 |
| 0.8886403667 | -0.06987508145 | 1.028945888 | 0.2523171782 | -0.995410395 | 0.1483936138 |
| 0.8886403667 | 0.2246764266 | 1.175355291 | 0.3800335867 | -1.214823289 | 0.1628197408 |
| 0.8886403667 | 0.06748286573 | 0.9952314859 | 0.2737562695 | -1.015502869 | 0.1403549537 |
| 0.8886403667 | -0.0843696175 | 0.9713644186 | 0.1692079135 | -0.8606394734 | 0.1370831357 |
| 0.8886403667 | -0.1502619635 | 1.016881519 | 0.3010450048 | -1.092553323 | 0.1457598786 |
| 0.8886403667 | 0.1871705495 | 1.036268626 | 0.1790497484 | -0.9636088096 | 0.1531527921 |
| 0.9124849918 | 0.06183703294 | 0.9707403248 | 0.2259395281 | -1.009291565 | 0.1650037142 |
| 0.8886403667 | 0.1913281297 | 1.106167238 | 0.3019940842 | -1.081685895 | 0.1639030992 |
| 0.9824246933 | -0.009010888178 | 1.015077822 | 0.2517004542 | -0.9972856963 | 0.1392070452 |
| 0.8886403667 | 0.348208305 | 0.7045306052 | -0.0984768841 | -0.7036862607 | 0.1562897644 |
| 0.8886403667 | 0.1263191439 | 1.044432525 | 0.2423963833 | -0.976539593 | 0.1489617512 |
| 0.8886403667 | 0.1052610443 | 1.026494685 | 0.2508741098 | -1.050340648 | 0.1368650956 |
| 0.8886403667 | -0.07510081097 | 0.9847795951 | 0.2178451818 | -0.9190701634 | 0.158139134 |
| 0.8886403667 | 0.1908763379 | 0.9685742921 | 0.3239855185 | -1.055368314 | 0.1447462893 |
| 0.8886403667 | 0.147752013 | 1.050239967 | 0.2641762017 | -1.01587481 | 0.1426561035 |
| 0.4914558906 | 0.1908960625 | 0.2764841154 | -0.3019499935 | 0.7835683561 | 0.1161112538 |
| 0.4914558906 | 0.2220572234 | 0.2846028767 | -0.3041100705 | 0.7222536461 | 0.1146220856 |
| 0.8159616578 | -0.06786190196 | 0.2177097855 | -0.3504747533 | 0.8077048259 | 0.1190696048 |
| 0.987109513 | -0.001911645511 | 0.2448007227 | -0.3493785087 | 0.7913910178 | 0.1183201613 |
| 0.9261187171 | -0.02323018698 | 0.2433817019 | -0.3446320223 | 0.7933142287 | 0.1185111025 |
| 0.7352604538 | -0.105436092 | 0.2001193568 | -0.3908621446 | 0.8785639058 | 0.1194396608 |
| 0.4914558906 | 0.2186674759 | 0.2340823521 | -0.2717616522 | 0.7309184493 | 0.1159049377 |
| 0.4914558906 | -0.2320094786 | 0.333692395 | -0.3461613336 | 0.8314412026 | 0.1173928673 |
| 0.026701542 | -0.4005007638 | 0.4198325957 | -0.2925144251 | 0.6755845335 | 0.1081420763 |
| 0.7989945845 | -0.07709693981 | 0.2397446547 | -0.3386627418 | 0.7772040854 | 0.1177092475 |
| 0.7352604538 | 0.1075564258 | 0.2876679512 | -0.3216879496 | 0.7607637572 | 0.1183081075 |

| | | | | | |
|---|---|---|---|---|---|
| 0.5180231938 | -0.1682504504 | 0.2216151809 | -0.3569114016 | 0.8898869374 | 0.1180800298 |
| 0.8170742179 | 0.05202974114 | 0.2421181173 | -0.3381382855 | 0.7741755883 | 0.1185308788 |
| 0.5686164971 | -0.1493613286 | 0.244442716 | -0.3935278197 | 0.8092271804 | 0.1170169683 |
| 0.8170742179 | -0.05171260101 | 0.2434689361 | -0.3521971633 | 0.7684410534 | 0.1199956989 |
| 0.4914558906 | -0.1854450468 | 0.1724138 | -0.3950490512 | 0.8104912287 | 0.1174931106 |
| 0.7093607019 | -0.1200898202 | 0.2380516774 | -0.4100946513 | 0.8109946551 | 0.1194614248 |
| 0.7741303846 | 0.0886835412 | 0.2551302792 | -0.3436778978 | 0.7613447336 | 0.1176764361 |
| 0.9224387816 | 0.02851071809 | 0.2317511321 | -0.3364690596 | 0.7727038213 | 0.1300506454 |
| 0.7352604538 | -0.1122126235 | 0.3205928913 | -0.3096147032 | 0.7955785008 | 0.1261407517 |
| 0.4914558906 | -0.195999485 | 0.1265793736 | -0.4399154375 | 0.9361963864 | 0.1182684007 |
| 0.9760570672 | 0.00707144377 | 0.2438636128 | -0.3491348175 | 0.7945217372 | 0.1190416399 |
| 0.8170742179 | -0.06426303269 | 0.2079651287 | -0.3623169827 | 0.805399354 | 0.1210360822 |
| 0.851825852 | 0.03927612454 | 0.2410713606 | -0.3684806941 | 0.8011766997 | 0.1196354574 |
| 0.851825852 | -0.04411975362 | 0.2830546772 | -0.3298596029 | 0.7573753148 | 0.1241732275 |
| 0.6312783444 | 0.1354886973 | 0.2170269265 | -0.4167175805 | 0.8152149643 | 0.1190784213 |
| 0.8170742179 | 0.05552490204 | 0.2265106612 | -0.3489227416 | 0.8109193808 | 0.1189118094 |
| 0.7061704944 | -0.1218279633 | 0.2874707059 | -0.3330700223 | 0.7855245469 | 0.118287962 |
| 0.4914558906 | -0.2102954507 | 0.0941988613 | -0.4470803736 | 0.9783023362 | 0.1192929867 |
| 0.4914558906 | -0.2506989529 | 0.0951013538 | -0.3985841503 | 0.8818571054 | 0.1168178826 |
| 0.8284524188 | -0.04884903034 | 0.2279414111 | -0.3589331458 | 0.7931993799 | 0.1193978831 |
| 0.5686164971 | -0.152761122 | 0.2486258486 | -0.2685073746 | 0.7238923664 | 0.1188299425 |
| 0.8159616578 | -0.06663167945 | 0.2656684431 | -0.3452777397 | 0.7877125836 | 0.1185127235 |
| 0.4914558906 | -0.2015845858 | 0.3358805542 | -0.2456165193 | 0.5895108484 | 0.1187554049 |
| 0.4914558906 | 0.2397084574 | 0.1993603126 | -0.3675192283 | 0.9775359768 | 0.1179007621 |
| 0.9013670124 | -0.03082885137 | 0.2318428071 | -0.3476056658 | 0.7850998524 | 0.1219131127 |
| 0.4914558906 | 0.1936144007 | 0.2227052546 | -0.304363927 | 0.8019561146 | 0.1162746209 |
| 0.8170742179 | 0.05897267434 | 0.2353373698 | -0.3337036712 | 0.7717455233 | 0.1195371353 |
| 0.4914558906 | -0.219979004 | 0.2791172372 | -0.4412287739 | 0.7914351402 | 0.1176201485 |
| 0.7989945845 | 0.08113461449 | 0.2508421074 | -0.4064378794 | 0.8690294282 | 0.1261027246 |
| 0.8159616578 | 0.06739273289 | 0.2773186759 | -0.345545513 | 0.7616089507 | 0.1198558993 |
| 0.9494026196 | 0.01207254506 | 0.2445853901 | -0.3523198184 | 0.7972782194 | 0.1187421727 |
| 0.4914558906 | 0.2356266571 | 0.2095396687 | -0.4708070828 | 0.9335237087 | 0.1161415298 |
| 0.7390081682 | -0.09929390539 | 0.2518174174 | -0.3580711144 | 0.759390929 | 0.1183041919 |
| 0.5630662785 | 0.1568595615 | 0.2960399686 | -0.33459295 | 0.6859571325 | 0.1186521764 |
| 0.4914558906 | 0.2196898915 | 0.2639632551 | -0.3682916158 | 0.7317755959 | 0.1153321326 |
| 0.4914558906 | 0.2369494364 | 0.2586110371 | -0.3599962418 | 0.8803633536 | 0.1152846299 |
| 0.4914558906 | 0.1827158492 | 0.2675863228 | -0.3368174026 | 0.8020432945 | 0.1157552279 |
| 0.4914558906 | 0.2116102592 | 0.2597275987 | -0.2621925274 | 0.6763594216 | 0.1164920766 |
| 0.7989945845 | 0.08013328889 | 0.2633782496 | -0.3050384025 | 0.7924673531 | 0.1225514933 |
| 0.7955876879 | -0.08388078089 | 0.2680482306 | -0.3326171006 | 0.7720945995 | 0.1178112365 |
| 0.5043956771 | 0.1748801405 | 0.3412172457 | -0.2608469545 | 0.6410726661 | 0.1188373644 |
| 0.9834530885 | 0.004847935501 | 0.2463911044 | -0.3494007937 | 0.7892789465 | 0.1197878952 |
| 0.8170742179 | -0.05748414486 | 0.2309189228 | -0.3704685369 | 0.7704546171 | 0.1248280496 |
| 0.8159616578 | -0.06628785993 | 0.2307948831 | -0.3627523852 | 0.8114390631 | 0.1177082818 |
| 0.8159616578 | -0.07153464418 | 0.2510063891 | -0.3659058275 | 0.7853103965 | 0.1184682061 |
| 0.947308554 | -0.0147857509 | 0.2447904084 | -0.355332925 | 0.8001984976 | 0.1198705981 |

| | | | | | |
|---|---|---|---|---|---|
| 0.8170742179 | 0.05986001572 | 0.2388186219 | -0.3360833335 | 0.7720298899 | 0.1187896849 |
| 0.4914558906 | 0.198793442 | 0.3185790935 | -0.2779161065 | 0.5912362758 | 0.1193494347 |
| 0.5043956771 | -0.1712626084 | 0.3131833441 | -0.2622640771 | 0.6824330672 | 0.1177883995 |
| 0.7390081682 | -0.09896722156 | 0.215336871 | -0.3579396425 | 0.7874754941 | 0.1185298024 |
| 0.6312783444 | -0.1400661237 | 0.1656865294 | -0.3741477535 | 0.844621577 | 0.1192674677 |
| 0.7955876879 | -0.08777258367 | 0.1818559247 | -0.3769051929 | 0.833525873 | 0.1219263965 |
| 0.6312783444 | -0.1371816683 | 0.2534366986 | -0.2821254594 | 0.776459995 | 0.1192567688 |
| 0.7237071821 | 0.1143727649 | 0.2498714906 | -0.3439141215 | 0.8101794574 | 0.1171992847 |
| 0.8170742179 | 0.06333354942 | 0.2638911771 | -0.3366308952 | 0.7748640957 | 0.1181034306 |
| 0.1103871028 | -0.3526992659 | 0.4367769968 | -0.2996812979 | 0.5233020624 | 0.1131608099 |
| 0.7986393901 | 0.08423200734 | 0.2100511126 | -0.3914346683 | 0.8721608739 | 0.1205526329 |
| 0.4914558906 | 0.1853299529 | 0.1568815501 | -0.4230784434 | 0.8004568254 | 0.1202060025 |
| 0.851825852 | 0.04098965886 | 0.2369002265 | -0.3680909643 | 0.8245831771 | 0.120304205 |
| 0.7989945845 | 0.08445835894 | 0.2670898237 | -0.3800840204 | 0.8718198703 | 0.1289705629 |
| 0.7352604538 | 0.1021849798 | 0.2408963566 | -0.3416801445 | 0.754965943 | 0.1180769651 |
| 0.4914558906 | 0.1824683592 | 0.2600654973 | -0.2727396479 | 0.7514148225 | 0.11697242 |
| 0.6751777676 | 0.1285804521 | 0.2840973767 | -0.2727549557 | 0.6768314925 | 0.1200387809 |
| 0.7594172115 | -0.09402804918 | 0.2596763923 | -0.3130447941 | 0.7882189967 | 0.1192004735 |
| 0.2023366985 | 0.3132945563 | 0.2219423176 | -0.2833442266 | 0.8037122278 | 0.1116990494 |
| 0.851825852 | 0.04057060605 | 0.2133021345 | -0.3665527874 | 0.8153684126 | 0.1230434518 |
| 0.5496791634 | 0.158992701 | 0.1988264639 | -0.338856919 | 0.7705085013 | 0.1178203989 |
| 0.7594172115 | -0.09329986511 | 0.2689990027 | -0.3422756267 | 0.7581456057 | 0.117824646 |
| 0.5386189999 | -0.16667166 | 0.1939864931 | -0.4169111141 | 0.7795165754 | 0.1201168517 |
| 0.7237071821 | 0.1120654883 | 0.2943859289 | -0.2876341889 | 0.7391267088 | 0.1196752945 |
| 0.987109513 | -0.002355873817 | 0.2457267001 | -0.3492254002 | 0.7912560708 | 0.1191590995 |
| 0.6040240245 | 0.1432828737 | 0.2769157858 | -0.3114966515 | 0.7939948187 | 0.1179423558 |
| 0.4914558906 | 0.1860407757 | 0.319266067 | -0.3346893432 | 0.7901965912 | 0.1176666039 |
| 0.7989945845 | -0.08304961981 | 0.2493285575 | -0.421490814 | 0.8719248301 | 0.1279771633 |
| 0.4914558906 | -0.1777207871 | 0.300296096 | -0.3769041171 | 0.7894712048 | 0.1174432012 |
| 0.6312783444 | -0.1340864239 | 0.2564003409 | -0.3219657597 | 0.7732719963 | 0.1167039277 |
| 0.7989945845 | 0.07539917502 | 0.2461770245 | -0.3443517372 | 0.7641808066 | 0.1181799693 |
| 0.7237071821 | -0.1090389113 | 0.2370338088 | -0.3561246214 | 0.7848200432 | 0.1170391272 |
| 0.8159616578 | -0.06970626091 | 0.2872395637 | -0.3444867999 | 0.7843577432 | 0.1222591504 |
| 0.8318854403 | -0.04791746802 | 0.2681990442 | -0.337879423 | 0.7877965688 | 0.1209382222 |
| 0.6040240245 | 0.1441172634 | 0.2387930792 | -0.4062987138 | 0.858165795 | 0.1178374972 |
| 0.8170742179 | -0.05733310471 | 0.2476333903 | -0.2989067353 | 0.7597973078 | 0.1255819898 |
| 0.9261187171 | -0.02342676656 | 0.243134915 | -0.3479890722 | 0.7883289681 | 0.1179259091 |
| 0.026701542 | -0.4245415619 | 0.3318918079 | -0.3750435162 | 0.5245780528 | 0.1089279033 |
| 0.8170742179 | 0.06119125502 | 0.2307854694 | -0.3452554224 | 0.7908099513 | 0.1182831712 |
| 0.7237071821 | -0.1140002756 | 0.2562702056 | -0.3258372386 | 0.8291194724 | 0.1199351504 |
| 0.4914558906 | -0.2166040424 | 0.2463941415 | -0.3944657423 | 0.9409787943 | 0.1179917909 |
| 0.7352604538 | -0.1035109009 | 0.2046678797 | -0.3861263585 | 0.8114100681 | 0.119148061 |
| 0.5386189999 | -0.1605026954 | 0.1925744866 | -0.3703714007 | 0.8343262263 | 0.1167450806 |
| 0.7700507085 | 0.09370373288 | 0.2081668851 | -0.3834961231 | 0.8918703835 | 0.1220097304 |
| 0.4914558906 | -0.1943978361 | 0.2073514258 | -0.3802172495 | 0.7531395195 | 0.1171966683 |
| 0.6312783444 | -0.1330604842 | 0.1981123817 | -0.3485474574 | 0.8432592194 | 0.1181876101 |

| | | | | | |
|---|---|---|---|---|---|
| 0.9494026196 | 0.01321703256 | 0.2487895124 | -0.3434918037 | 0.7928655949 | 0.1227854741 |
| 0.5681702775 | -0.1599659187 | 0.3113935459 | -0.3903327694 | 0.860338738 | 0.1227409592 |
| 0.2825625227 | -0.2894610513 | 0.1168425215 | -0.4746446169 | 1.024253502 | 0.1144316705 |
| 0.932443124 | 0.0183913571 | 0.2445986813 | -0.3539963437 | 0.7921946733 | 0.1183254889 |
| 0.932443124 | -0.02112375464 | 0.2510237257 | -0.3426645317 | 0.7765341489 | 0.1196189673 |
| 0.4914558906 | -0.1786430892 | 0.2043269168 | -0.3193725848 | 0.8134710802 | 0.1168677662 |
| 0.8170742179 | -0.06351444052 | 0.2807490607 | -0.3495547595 | 0.7724753657 | 0.1211473189 |
| 0.4914558906 | -0.18540696 | 0.281434592 | -0.3195034058 | 0.7520670351 | 0.1155898618 |
| 0.7390081682 | 0.1007516792 | 0.3109623021 | -0.3241829587 | 0.7096398675 | 0.1214180801 |
| 0.8170742179 | -0.05258656117 | 0.2534711951 | -0.3552898635 | 0.7930544254 | 0.1180421548 |
| 0.932443124 | -0.01875812288 | 0.2386613349 | -0.3643995474 | 0.8144066777 | 0.1244766538 |
| 0.5043956771 | 0.1707244776 | 0.2396959469 | -0.361853252 | 0.8616467231 | 0.1167998237 |
| 0.8170742179 | -0.05873229452 | 0.2442683212 | -0.3258609316 | 0.7712871249 | 0.1191265724 |
| 0.8170742179 | -0.06370351356 | 0.2898303686 | -0.3195616402 | 0.8033583356 | 0.1324779449 |
| 0.4914558906 | -0.1781160448 | 0.2331625743 | -0.3720679765 | 0.8358584303 | 0.1156727619 |
| 0.7093607019 | -0.1207113359 | 0.2495365919 | -0.3427111505 | 0.7117004752 | 0.120353676 |
| 0.8907744352 | -0.03388592884 | 0.266943068 | -0.3326571793 | 0.7817755521 | 0.1235152086 |
| 0.4914558906 | -0.2055797872 | 0.2503058974 | -0.3002168391 | 0.7182472424 | 0.1155281026 |
| 0.851825852 | 0.04147997951 | 0.2467559337 | -0.3529774985 | 0.7833000426 | 0.1181491154 |
| 0.932443124 | 0.01998930311 | 0.2495322196 | -0.338186949 | 0.7814456756 | 0.1200437254 |
| 0.2825625227 | 0.2929463692 | 0.2073939344 | -0.2743300765 | 0.6573211566 | 0.1143074656 |
| 0.7237071821 | 0.1100852501 | 0.2753261967 | -0.3448328406 | 0.7707030493 | 0.1175286729 |
| 0.8699822324 | -0.06410056781 | 0.4344804538 | -0.1505733855 | 0.444343664 | 0.1186690525 |
| 0.5411567788 | 0.2114545954 | 0.504108316 | -0.0820671561 | 0.3296651613 | 0.1169079189 |
| 0.9135982073 | -0.04657499484 | 0.4281132219 | -0.136520321 | 0.4485453796 | 0.1201444813 |
| 0.9136389765 | -0.02902954186 | 0.4449479419 | -0.1311689938 | 0.4355595564 | 0.12011798 |
| 0.8505310426 | 0.07728174369 | 0.4428625371 | -0.1559157465 | 0.4507898119 | 0.1204781704 |
| 0.8069640727 | -0.125966299 | 0.40308298 | -0.1745003983 | 0.5193067837 | 0.1229778723 |
| 0.8971393429 | 0.05495420092 | 0.4393458327 | -0.1200661269 | 0.4343883886 | 0.1180841871 |
| 0.4850078292 | -0.2600882446 | 0.5310258679 | -0.1475551519 | 0.5130000288 | 0.1159636025 |
| 0.4850078292 | -0.2727205319 | 0.5757579679 | -0.0916156444 | 0.3427500753 | 0.1152359705 |
| 0.9413513341 | -0.01599376912 | 0.4466972951 | -0.1313444019 | 0.4326514705 | 0.121446918 |
| 0.6107874066 | 0.1822049679 | 0.5304997201 | -0.0851215234 | 0.3631126938 | 0.1166511919 |
| 0.9136389765 | -0.03932614418 | 0.4432806806 | -0.1339782877 | 0.455665095 | 0.1255176229 |
| 0.9136389765 | 0.03068034333 | 0.439660419 | -0.1306971721 | 0.4399369101 | 0.1192303589 |
| 0.4850078292 | -0.2584110471 | 0.4679874798 | -0.1972883905 | 0.4196325585 | 0.114906565 |
| 0.9920945847 | -0.001196997586 | 0.4445199591 | -0.1361888659 | 0.4422459085 | 0.1191506596 |
| 0.8505310426 | -0.0841860034 | 0.4214493343 | -0.152000595 | 0.4309411248 | 0.1190999498 |
| 0.9083543454 | -0.05533932926 | 0.4492154347 | -0.1583903933 | 0.4357718959 | 0.121944951 |
| 0.9413513341 | 0.0147210117 | 0.4453805574 | -0.135220099 | 0.4391895584 | 0.1181998208 |
| 0.6200530061 | -0.1723406074 | 0.5239339397 | -0.2148066848 | 0.55539469 | 0.1272003117 |
| 0.4850078292 | -0.2774606269 | 0.6169999413 | -0.0354424859 | 0.4863999035 | 0.1247073833 |
| 0.6107874066 | -0.1983547202 | 0.3500856492 | -0.2015641713 | 0.5423108773 | 0.1283516337 |
| 0.9413513341 | 0.01809339808 | 0.4434885057 | -0.1346174462 | 0.4478500441 | 0.1255336311 |
| 0.8505310426 | -0.09719451451 | 0.3686961458 | -0.167130789 | 0.5070687213 | 0.1223335774 |
| 0.8971393429 | -0.05591295108 | 0.4554574182 | -0.1075067908 | 0.4196728453 | 0.1199246806 |

| | | | | | |
|---|---|---|---|---|---|
| 0.9136389765 | 0.03552382421 | 0.4161790299 | -0.1520545968 | 0.4657773967 | 0.1265001334 |
| 0.8505310426 | 0.08095229453 | 0.4227737579 | -0.1734491782 | 0.4625144205 | 0.1296687084 |
| 0.9135982073 | -0.04791339347 | 0.4577554919 | -0.1367579332 | 0.4314342683 | 0.1193921299 |
| 0.8699822324 | -0.06241578186 | 0.4710263879 | -0.1234871378 | 0.4298249755 | 0.1196890096 |
| 0.8069640727 | -0.1075737969 | 0.3816658283 | -0.1783353328 | 0.5055810846 | 0.1232565797 |
| 0.5345013622 | -0.2295838925 | 0.3167102741 | -0.1825623707 | 0.5050760503 | 0.1177370373 |
| 0.8699822324 | 0.08688851795 | 0.4899613034 | -0.1085742147 | 0.4011295689 | 0.130289193 |
| 0.8699822324 | -0.07665403567 | 0.4430633924 | -0.1116377344 | 0.427593713 | 0.1434346568 |
| 0.6107874066 | -0.1969698019 | 0.5358353013 | -0.1048230056 | 0.3620355632 | 0.1180491092 |
| 0.9722570048 | -0.009463969036 | 0.4482945864 | -0.1319764 | 0.4346413419 | 0.1280531628 |
| 0.7972913844 | 0.1373541198 | 0.4307127653 | -0.1426965947 | 0.5270635099 | 0.1229511803 |
| 0.5345013622 | 0.2356250137 | 0.5576499949 | -0.1431480857 | 0.4540647839 | 0.1288635658 |
| 0.6107874066 | 0.1843097187 | 0.4099577405 | -0.1060175742 | 0.4925231785 | 0.1222952179 |
| 0.9413513341 | 0.02016645204 | 0.4408933002 | -0.1308912244 | 0.4372992614 | 0.1219143939 |
| 0.8069640727 | 0.1153410486 | 0.4169263121 | -0.0939692894 | 0.4648577618 | 0.1236531463 |
| 0.5345013622 | 0.2430958254 | 0.4644888365 | -0.2834303705 | 0.6662576609 | 0.1294536042 |
| 0.6930800793 | 0.1761191692 | 0.4921751123 | -0.1472904732 | 0.4516543215 | 0.1259674242 |
| 0.8505310426 | 0.08604950934 | 0.4324188674 | -0.1610963867 | 0.5056717107 | 0.1290835851 |
| 0.08406036695 | 0.4020307682 | 0.4217209815 | -0.322219176 | 0.6014159289 | 0.1099587639 |
| 0.9136389765 | 0.04129225694 | 0.4456344641 | -0.131818521 | 0.4484802785 | 0.1212743269 |
| 0.5345013622 | 0.2391833662 | 0.5599984439 | -0.0925599481 | 0.198268323 | 0.1176823215 |
| 0.919687497 | 0.02568137742 | 0.4497800795 | -0.1365669042 | 0.4293915897 | 0.1183002342 |
| 0.8505310426 | 0.08211176439 | 0.4619200568 | -0.1375240954 | 0.4494894929 | 0.1238045249 |
| 0.6107874066 | 0.1918900641 | 0.50575404 | -0.1020965468 | 0.370564228 | 0.1167145127 |
| 0.8069640727 | 0.1154134285 | 0.4681207713 | -0.0783061325 | 0.3462176033 | 0.1201707414 |
| 0.6107874066 | 0.1931462194 | 0.4716835484 | -0.0405013074 | 0.4831700242 | 0.1222080163 |
| 0.9136389765 | 0.03419334097 | 0.4331181488 | -0.1450759598 | 0.4574015556 | 0.1305938063 |
| 0.8069640727 | -0.1070075294 | 0.3870443157 | -0.1933751116 | 0.5348974088 | 0.122201647 |
| 0.6200530061 | 0.1771290643 | 0.5078577562 | -0.1357035183 | 0.3438378326 | 0.1259324913 |
| 0.8505310426 | -0.08454720724 | 0.4350983131 | -0.1597750319 | 0.3865688745 | 0.1249763759 |
| 0.9135982073 | 0.04470347228 | 0.4516266825 | -0.1312522663 | 0.4366710031 | 0.1186005796 |
| 0.6869101751 | -0.1495957332 | 0.4698929231 | -0.1644396711 | 0.402051293 | 0.1210646715 |
| 0.8069640727 | -0.1094423789 | 0.4258881181 | -0.196118271 | 0.5462055216 | 0.1198565528 |
| 0.9135982073 | 0.04308438683 | 0.4419922093 | -0.1229205048 | 0.4225896205 | 0.12375112 |
| 0.8069640727 | 0.1279172374 | 0.4758980323 | -0.0937957582 | 0.3505644655 | 0.1230764468 |
| 0.9136389765 | 0.03244201678 | 0.4317409125 | -0.1505499881 | 0.4628974597 | 0.1205342952 |
| 0.9413513341 | 0.01637949645 | 0.4498765641 | -0.1347722804 | 0.4425365696 | 0.1180845197 |
| 0.9413513341 | 0.01870550418 | 0.4535540668 | -0.1329819976 | 0.4377491786 | 0.1246807909 |
| 0.892755109 | -0.06483643975 | 0.400088529 | -0.1578279868 | 0.4729311056 | 0.1338725055 |
| 0.8069640727 | -0.1126842905 | 0.4562838357 | -0.0790063230 | 0.4238635678 | 0.1244254026 |
| 0.8069640727 | 0.09794725062 | 0.4487851818 | -0.1354336842 | 0.4605065334 | 0.1190928038 |
| 0.9136389765 | 0.03821432523 | 0.4582757311 | -0.127670791 | 0.4293512878 | 0.1219484468 |
| 0.6107874066 | -0.2061845136 | 0.5735707949 | -0.0935072778 | 0.2454871071 | 0.121797957 |
| 0.4850078292 | 0.2826484984 | 0.3187929758 | -0.3048579713 | 0.7559748318 | 0.1161811117 |
| 0.8699822324 | 0.07322057248 | 0.4168774538 | -0.1591129528 | 0.4308369228 | 0.1224745982 |
| 0.9135982073 | 0.05070128366 | 0.4398203997 | -0.1564307593 | 0.4730487624 | 0.1203291889 |

| | | | | | |
|---|---|---|---|---|---|
| 0.9135982073 | 0.04461671324 | 0.455131715 | -0.1529141952 | 0.4870118305 | 0.1292290284 |
| 0.8505310426 | -0.08183574241 | 0.4361824886 | -0.1484407462 | 0.4951628674 | 0.1209016292 |
| 0.9820151654 | -0.004845954977 | 0.4440772628 | -0.1381483915 | 0.4438554416 | 0.124564367 |
| 0.8505310426 | 0.07962134763 | 0.4727216875 | -0.0851181396 | 0.3620249112 | 0.1220767449 |
| 0.8069640727 | -0.1158758008 | 0.4684551126 | -0.0874086526 | 0.4232659757 | 0.1202220327 |
| 0.5345013622 | 0.2457400054 | 0.4516317522 | -0.0732234048 | 0.3994602695 | 0.117237426 |
| 0.9135982073 | -0.04532170456 | 0.4749472492 | -0.1210004964 | 0.4278305717 | 0.1233488448 |
| 0.8505310426 | 0.07664708375 | 0.4301991393 | -0.1289010916 | 0.4174309949 | 0.1202069118 |
| 0.9817289025 | 0.006410484091 | 0.4437384111 | -0.135927156 | 0.4434069947 | 0.1214859664 |
| 0.8505310426 | -0.07572925362 | 0.4233490354 | -0.1643461159 | 0.4327024816 | 0.1201727679 |
| 0.5345013622 | 0.2537395992 | 0.5942060917 | 0.0249817687[illegible] | 0.2401687141 | 0.1174262059 |
| 0.8069640727 | -0.1027389993 | 0.4825343579 | -0.1208146434 | 0.4227931777 | 0.1199199986 |
| 0.8069640727 | 0.1232196857 | 0.4874303952 | -0.0983643542 | 0.4156936872 | 0.1231338891 |
| 0.6200530061 | 0.1719986242 | 0.5361933636 | -0.1116894651 | 0.3921856035 | 0.120167857 |
| 0.8505310426 | -0.09221000803 | 0.4671899836 | -0.2049505901 | 0.4934595873 | 0.1294182561 |
| 0.6200530061 | -0.1829651691 | 0.5359632185 | -0.1449299335 | 0.3646556646 | 0.1188271745 |
| 0.8069640727 | -0.119001316 | 0.4584970319 | -0.1145401164 | 0.423600616 | 0.1168540722 |
| 0.8505310426 | 0.09228416536 | 0.4333963146 | -0.1349647284 | 0.4367878443 | 0.1251909251 |
| 0.9136389765 | 0.04045384042 | 0.4435405232 | -0.132700041 | 0.4529286498 | 0.1315311373 |
| 0.6200530061 | -0.1846707563 | 0.5232622079 | -0.1301018505 | 0.4788186161 | 0.1251711134 |
| 0.5345013622 | -0.2088511524 | 0.5403654104 | -0.0848239888 | 0.4370861062 | 0.1162768412 |
| 0.9413513341 | -0.0155921101 | 0.4446523624 | -0.1300479635 | 0.4371113729 | 0.1249513466 |
| 0.6696072029 | -0.1642106435 | 0.4395942458 | 0.0076738840[illegible] | 0.3810301117 | 0.1237858965 |
| 0.6869101751 | -0.161113682 | 0.4191193092 | -0.1439527224 | 0.4665097413 | 0.125259979 |
| 0.4850078292 | -0.267318257 | 0.4849335803 | -0.1651243968 | 0.3076481154 | 0.1185340363 |
| 0.9820151654 | -0.004355132517 | 0.4450031637 | -0.1364595673 | 0.4439769527 | 0.1230216702 |
| 0.8699822324 | -0.07293028889 | 0.4626630822 | -0.114441638 | 0.441826448 | 0.1272786132 |
| 0.8069640727 | -0.1043502895 | 0.4408927648 | -0.1590859985 | 0.5193292307 | 0.1256079268 |
| 0.8069640727 | 0.1277062042 | 0.4901907443 | -0.0955311657 | 0.4279793233 | 0.1204902205 |
| 0.9135982073 | -0.04763324551 | 0.4337316769 | -0.1389524021 | 0.4460135247 | 0.1182844384 |
| 0.8505310426 | 0.09871557771 | 0.421118469 | -0.1666764992 | 0.5173853244 | 0.1288115377 |
| 0.8699822324 | -0.06280800307 | 0.4364714124 | -0.1442324964 | 0.4233930233 | 0.1193172222 |
| 0.8069640727 | -0.128689899 | 0.4148806686 | -0.1226547587 | 0.4546730082 | 0.1234501505 |
| 0.8505310426 | 0.08900874909 | 0.4765898118 | -0.0929654024 | 0.4379057176 | 0.1318616938 |
| 0.6107874066 | -0.1798789471 | 0.5332984178 | -0.1808781072 | 0.4984393263 | 0.1239608111 |
| 0.6869101751 | -0.1585543818 | 0.3929023152 | -0.1990448185 | 0.5339019547 | 0.1231728649 |
| 0.8699822324 | -0.06647323653 | 0.454170498 | -0.1167239465 | 0.4222943564 | 0.1189058527 |
| 0.6107874066 | -0.1885610736 | 0.5189740394 | -0.0604172746 | 0.2653899815 | 0.123754922 |
| 0.8069640727 | -0.138860343 | 0.422259894 | -0.1193029433 | 0.4434340075 | 0.1330273982 |
| 0.9136389765 | -0.02803272807 | 0.4588988981 | -0.1383915124 | 0.4381059852 | 0.1216527651 |
| 0.4850078292 | -0.2654352742 | 0.4933640971 | -0.0998608247 | 0.3909783731 | 0.1118556404 |
| 0.8069640727 | 0.128798104 | 0.511491281 | -0.1022104159 | 0.365768124 | 0.1283888674 |
| 0.9136389765 | -0.03557439063 | 0.4542785952 | -0.1373679214 | 0.4348648581 | 0.1212260874 |
| 0.8069640727 | -0.1219032602 | 0.4130636545 | -0.230033287 | 0.5750066306 | 0.1255709241 |
| 0.6107874066 | 0.1908362152 | 0.4427861211 | -0.1579290571 | 0.518763162 | 0.1189942167 |
| 0.6107874066 | -0.1845998187 | 0.4580625454 | -0.0635720585 | 0.349525842 | 0.125378942 |

| | | | | | |
|---|---|---|---|---|---|
| 0.5345013622 | -0.2656756725 | 0.6699046199 | 0.0102661763[illegible] | 0.407047859 | 0.1297899956 |
| 0.8069640727 | -0.1353037337 | 0.4683475818 | -0.1351674767 | 0.4033792386 | 0.1183987419 |
| 0.8069640727 | -0.1088095423 | 0.4502790155 | -0.1302084295 | 0.3672840872 | 0.122418977 |
| 0.9136389765 | -0.02991914998 | 0.4672272573 | -0.1202598869 | 0.4284831716 | 0.1300029227 |
| 0.6696072029 | -0.1523553835 | 0.4549457434 | -0.0901771284 | 0.3763414457 | 0.1182246408 |
| 0.9135982073 | 0.04441163757 | 0.4471916571 | -0.1406547325 | 0.4334844622 | 0.1178548714 |
| 0.5345013622 | -0.2148041473 | 0.3799834582 | -0.2710149263 | 0.5769670266 | 0.1183216103 |
| 0.5809154051 | 0.2001086754 | 0.4151348371 | -0.0971831352 | 0.3627552328 | 0.1177690819 |
| 0.5345013622 | 0.2226926525 | 0.4931719281 | -0.1482829887 | 0.4330072808 | 0.1145454704 |

| | | | Full model fit coefficient estimates (outcome ~ micr | | |
|---|---|---|---|---|---|
| **Blast Std. Error** | **VNS Std. Error** | **Blast:VNS Std. Error** | **Microbe t-value** | **Blast t-value** | **VNS t-value** |
| 0.3147126229 | 0.331791147 | 0.4645238408 | 2.643234361 | 3.053042605 | 2.226820926 |
| 0.3307937059 | 0.3487978707 | 0.4917491689 | 0.07887962006 | 2.609663303 | 1.838393296 |
| 0.3268006651 | 0.3406556494 | 0.4812238318 | 1.653866013 | 2.891937029 | 1.875282022 |
| 0.3281051482 | 0.3466006504 | 0.4909455065 | 0.8762493561 | 2.612471634 | 1.800025503 |
| 0.3142056869 | 0.3326788288 | 0.4670678256 | -2.506293877 | 2.685165752 | 2.141209901 |
| 0.3301055375 | 0.3488598602 | 0.4956908862 | 0.8430779108 | 2.70012726 | 1.936095037 |
| 0.3298222229 | 0.3498019545 | 0.4907737734 | 0.4048584518 | 2.601734172 | 1.871596763 |
| 0.3295972194 | 0.342308069 | 0.4823216686 | -1.481223827 | 2.879096471 | 1.916220664 |
| 0.3316576853 | 0.3451800853 | 0.485500593 | -1.164067037 | 2.802577183 | 1.928335634 |
| 0.3288822411 | 0.3476393094 | 0.4893728631 | 0.6921483781 | 2.629846605 | 1.796829507 |
| 0.3317737073 | 0.3492908636 | 0.4907450992 | -0.2215233595 | 2.574291149 | 1.814092458 |
| 0.3300613671 | 0.3478499959 | 0.4949552876 | -0.4106691382 | 2.591149512 | 1.841450463 |
| 0.3299892972 | 0.3497567467 | 0.4927554878 | 0.320152722 | 2.619429587 | 1.860932651 |
| 0.3277890448 | 0.3480377881 | 0.4875904634 | -0.9335646025 | 2.624618482 | 1.733284738 |
| 0.3229911455 | 0.3407464993 | 0.483216723 | -1.643744416 | 2.687983634 | 1.880881253 |
| 0.3283123449 | 0.3437704973 | 0.4831301546 | -1.436975182 | 2.409477975 | 1.737156173 |
| 0.3117421058 | 0.3338744472 | 0.4637992004 | -2.705183306 | 2.717653042 | 1.448944379 |
| 0.3286898075 | 0.3456321438 | 0.4889808574 | 1.022121776 | 2.714575245 | 1.893069284 |
| 0.3363520116 | 0.3519554272 | 0.4979528874 | 0.4240784591 | 2.477522283 | 1.882665954 |
| 0.3427785835 | 0.3510771335 | 0.4904270742 | 0.2905248716 | 2.434831215 | 1.784511854 |
| 0.3379403187 | 0.3516421706 | 0.4986269708 | -0.5364630185 | 2.429571877 | 1.737321722 |
| 0.3252554518 | 0.3437051189 | 0.484892834 | -1.350288739 | 2.644094844 | 1.785305728 |
| 0.3275944456 | 0.3393418979 | 0.4781647368 | -1.853825593 | 2.262580607 | 1.777463873 |
| 0.3299373437 | 0.3519347393 | 0.491309342 | 0.3876428347 | 2.598254316 | 1.759353948 |
| 0.3418907549 | 0.3496417512 | 0.4950362834 | -0.5223830263 | 2.660023634 | 1.886955006 |
| 0.3296925133 | 0.3539467438 | 0.489056219 | -0.8018527157 | 2.684254346 | 1.971375666 |
| 0.3299310975 | 0.3456393117 | 0.4890879293 | 1.013870335 | 2.485336003 | 1.889706949 |
| 0.3288894856 | 0.3439025104 | 0.483019746 | -1.393313529 | 2.834984263 | 1.970236126 |
| 0.3408364569 | 0.3510243727 | 0.5013520996 | -0.8826170116 | 2.288345162 | 1.675200947 |
| 0.3351323745 | 0.3453287873 | 0.4890449679 | -1.102343481 | 2.327647615 | 1.79518262 |
| 0.3313347894 | 0.3485605615 | 0.4906511723 | -0.1299359205 | 2.589303036 | 1.830324464 |
| 0.3289615704 | 0.3552678756 | 0.4899885567 | 0.6806167409 | 2.605118232 | 1.654830806 |
| 0.3290299968 | 0.3446496243 | 0.4858279769 | 1.136960937 | 2.482399315 | 1.848139947 |
| 0.3352343021 | 0.3533390614 | 0.5047280214 | 0.2706503609 | 2.522372945 | 1.764330356 |
| 0.3326372525 | 0.3484299147 | 0.5085646254 | -0.462376347 | 2.652905854 | 1.865796617 |
| 0.332308085 | 0.348472086 | 0.4913853406 | -0.3019997873 | 2.556606173 | 1.850481361 |
| 0.3289178858 | 0.348785208 | 0.4886897559 | 0.7423300279 | 2.589746062 | 1.914489789 |
| 0.32327198 | 0.3440813034 | 0.4834952787 | 1.608962401 | 2.65043679 | 2.072363307 |
| 0.3308840607 | 0.3516424857 | 0.4911604303 | 0.2801404161 | 2.582880245 | 1.859039029 |
| 0.3294501873 | 0.3575806127 | 0.5023305036 | -0.5707917168 | 2.596218496 | 1.924480991 |
| 0.3399765506 | 0.3488525064 | 0.4958644647 | 0.01198879917 | 2.537217368 | 1.834612756 |
| 0.3284435312 | 0.3479762046 | 0.4916021828 | -0.7996461555 | 2.634284468 | 1.912352577 |
| 0.3301545807 | 0.3521931811 | 0.4946698015 | -0.0605924999 | 2.610133758 | 1.825521248 |
| 0.3295743733 | 0.3480630877 | 0.4906131355 | -0.4990469888 | 2.626596618 | 1.812742004 |
| 0.3306807459 | 0.3477717544 | 0.4943739832 | -0.4505927261 | 2.569216772 | 1.834134873 |

| | | | | | |
|---|---|---|---|---|---|
| 0.3308436555 | 0.3479298423 | 0.4923428501 | 0.3738803283 | 2.634354636 | 1.837759267 |
| 0.3293587651 | 0.347373253 | 0.4931596566 | -0.5768605856 | 2.600992031 | 1.839959547 |
| 0.3282008495 | 0.34627004 | 0.4887374223 | 0.8492169195 | 2.61894994 | 1.854482793 |
| 0.325086188 | 0.3475135321 | 0.4881610612 | 1.519880099 | 2.77569839 | 2.115125604 |
| 0.3266249005 | 0.3497508403 | 0.4835342087 | -1.341714031 | 2.517867032 | 1.571226711 |
| 0.3269671823 | 0.3443318323 | 0.4844125017 | -1.407206108 | 2.795406049 | 1.993874868 |
| 0.3369343855 | 0.3513464954 | 0.4980848481 | 1.409591058 | 2.931767228 | 2.131641099 |
| 0.3257729503 | 0.3381896529 | 0.4825323231 | -1.944815675 | 2.287280856 | 1.804325336 |
| 0.3292389736 | 0.3485926836 | 0.4904689422 | 0.9283630238 | 2.703346195 | 1.951376002 |
| 0.3307013382 | 0.3482543187 | 0.4906888499 | 0.3686543291 | 2.632864871 | 1.852776664 |
| 0.3303603538 | 0.3490038172 | 0.4908600454 | 0.1690292605 | 2.601160171 | 1.844262801 |
| 0.3287371974 | 0.3508317863 | 0.4954987016 | -0.7209932954 | 2.620709057 | 1.715081909 |
| 0.314414765 | 0.3320651109 | 0.4684810378 | 2.508346013 | 2.628165468 | 2.086399146 |
| 0.3354400857 | 0.3515789276 | 0.5097995335 | 0.7106768549 | 2.709573384 | 1.935581144 |
| 0.3309980994 | 0.3521674166 | 0.4920094327 | 0.6858701256 | 2.5254789 | 1.699362815 |
| 0.328769323 | 0.3444502571 | 0.4850750069 | -1.184618857 | 2.477703809 | 1.826104696 |
| 0.3332947097 | 0.3470721289 | 0.4892393628 | 0.8054138121 | 2.722861415 | 1.890960424 |
| 0.3437846188 | 0.3512961288 | 0.4959748718 | -0.4173026666 | 2.387979523 | 1.76271814 |
| 0.3297650253 | 0.3527747138 | 0.4901504581 | 0.4000679325 | 2.619542854 | 1.747087161 |
| 0.3091682904 | 0.3261857582 | 0.4603325761 | 2.902957924 | 2.808074885 | 1.943344378 |
| 0.3327039293 | 0.3503046124 | 0.4915978424 | -0.2145016379 | 2.562623623 | 1.802878909 |
| 0.3380051693 | 0.3489080188 | 0.5003567096 | 0.2839554365 | 2.487465181 | 1.814349315 |
| 0.3325575176 | 0.3530365001 | 0.5049047795 | -0.0205728572 | 2.593363352 | 1.815570985 |
| 0.3327100107 | 0.3480179856 | 0.486076516 | 1.106891304 | 2.382667541 | 1.688890125 |
| 0.3295226611 | 0.3512133979 | 0.4981871611 | -0.756447692 | 2.671478337 | 1.942721383 |
| 0.3242229651 | 0.341988545 | 0.492161073 | 1.748620771 | 2.859422138 | 1.672504187 |
| 0.330050127 | 0.3475312798 | 0.4900809865 | -0.526868663 | 2.643706564 | 1.842047856 |
| 0.3279036093 | 0.3488687347 | 0.487319852 | -0.916490799 | 2.640560709 | 1.714977672 |
| 0.303882857 | 0.3247522876 | 0.4599178 | 3.424776826 | 3.219997951 | 2.634415337 |
| 0.3122986733 | 0.3327276246 | 0.4630262193 | -2.719328316 | 2.950991766 | 2.346313958 |
| 0.3290603092 | 0.3487059861 | 0.4891686172 | 0.6620577055 | 2.601985994 | 1.898823032 |
| 0.347386661 | 0.3543657148 | 0.4982098296 | -0.1384540699 | 2.523421782 | 1.830979789 |
| 0.3251506254 | 0.3426665338 | 0.4828542611 | 1.459516725 | 2.554080876 | 1.933632367 |
| 0.3331923933 | 0.348014513 | 0.4922289183 | -0.6351439783 | 2.685754853 | 1.882638461 |
| 0.3267338836 | 0.3464668079 | 0.4831212148 | -1.396380619 | 2.490171266 | 1.644181315 |
| 0.3272025723 | 0.3482238525 | 0.4839385575 | 1.763591265 | 2.948925805 | 2.226396634 |
| 0.3271478627 | 0.3411489136 | 0.4824916619 | -1.679195656 | 2.910818173 | 1.984617864 |
| 0.3266013712 | 0.3454783899 | 0.4857298475 | 1.180151732 | 2.680427219 | 1.946766497 |
| 0.3320856002 | 0.3462322964 | 0.4867754127 | 0.9891352589 | 2.758408895 | 1.910908787 |
| 0.3299544062 | 0.3648391608 | 0.5069153213 | -0.3354831787 | 2.619817912 | 1.652873466 |
| 0.3285565999 | 0.3439381396 | 0.4841969847 | -1.275636235 | 2.788228966 | 1.813093394 |
| 0.3128077472 | 0.3309421292 | 0.4647256967 | -2.632621552 | 2.851067567 | 2.15119313 |
| 0.3204391364 | 0.3380842075 | 0.4781090813 | -1.922243558 | 2.692143884 | 1.881648083 |
| 0.3239695374 | 0.3417375402 | 0.4814092843 | -1.53035277 | 2.627268778 | 1.883573461 |
| 0.3399842774 | 0.3481733087 | 0.4914685258 | 0.2751050637 | 2.467956161 | 1.832459423 |
| 0.3307374321 | 0.3460354193 | 0.4866454936 | -1.003778775 | 2.746958909 | 1.907497887 |

| | | | | | |
|---|---|---|---|---|---|
| 0.3190236646 | 0.3391883907 | 0.478864804 | 2.06763554 | 2.670498251 | 1.62881535 |
| 0.3289060807 | 0.362920432 | 0.4947032782 | -0.688934247 | 2.631191122 | 1.964905295 |
| 0.329829914 | 0.3478289435 | 0.4900352125 | 0.4185992032 | 2.625062108 | 1.837624429 |
| 0.3308747701 | 0.3473053108 | 0.4927084034 | -0.6006072016 | 2.664960222 | 1.83669166 |
| 0.3290074643 | 0.3433920079 | 0.4844699583 | 1.323001411 | 2.424696478 | 1.845399506 |
| 0.3201756371 | 0.3395556653 | 0.4761498481 | -1.998784947 | 2.801544905 | 2.114925854 |
| 0.3235829442 | 0.3410521425 | 0.487612458 | -1.637328832 | 2.759240332 | 1.813334654 |
| 0.3278960173 | 0.3460382883 | 0.4852264394 | 1.223982884 | 2.755237262 | 1.97941738 |
| 0.3302847228 | 0.3463673485 | 0.4890166447 | -0.8925170102 | 2.503655087 | 1.808558174 |
| 0.325663815 | 0.3413012044 | 0.499954146 | 1.780035132 | 2.371347666 | 1.691140351 |
| 0.3264517634 | 0.3436456591 | 0.4833333951 | -1.365260355 | 2.517382731 | 1.781113314 |
| 0.3297820584 | 0.344585665 | 0.4874578384 | -1.156848284 | 2.450800654 | 1.832832528 |
| 0.329581978 | 0.3500509064 | 0.4826167269 | -1.477078829 | 2.351258774 | 1.515979225 |
| 0.332625583 | 0.3473224384 | 0.4914229837 | -0.8388782919 | 2.726155983 | 1.777987291 |
| 0.3327269636 | 0.3497521576 | 0.5049414266 | 0.9097484048 | 2.743907851 | 1.962922109 |
| 0.3304650942 | 0.350173425 | 0.4909545394 | 0.2510776692 | 2.593769838 | 1.800092702 |
| 0.3310930961 | 0.3504255313 | 0.4982674205 | 0.663070971 | 2.527124014 | 1.734106947 |
| 0.3300425838 | 0.349235052 | 0.4902983648 | -0.3894526247 | 2.59401141 | 1.866046167 |
| 0.3385308317 | 0.3484501802 | 0.4921619381 | -0.05579705059 | 2.557518943 | 1.837574996 |
| 0.319106632 | 0.3365189056 | 0.4737759905 | -2.107493665 | 2.821451511 | 2.004394535 |
| 0.3289579956 | 0.3378646045 | 0.4857438726 | 2.044202327 | 3.112730077 | 2.055569347 |
| 0.3310843599 | 0.3483829757 | 0.4907112287 | -0.04317403543 | 2.605660138 | 1.835664113 |
| 0.3307943062 | 0.3604195286 | 0.5118795178 | 0.934605306 | 2.730571133 | 2.038363015 |
| 0.3291719094 | 0.3470637487 | 0.4910386994 | 0.6632027772 | 2.641635978 | 1.843153308 |
| 0.325792706 | 0.3488421027 | 0.4853388312 | -1.275085543 | 2.658409683 | 2.051972727 |
| 0.3414785391 | 0.3545365264 | 0.4915694351 | -0.1016791819 | 2.549205297 | 1.823538585 |
| 0.325843266 | 0.3438357669 | 0.4855309255 | -1.283136531 | 2.608911003 | 1.818590496 |
| 0.3251858265 | 0.3431934854 | 0.4909408335 | -1.393310511 | 2.703441674 | 1.928180756 |
| 0.3253311748 | 0.3389212383 | 0.4687545974 | -2.480615243 | 3.285264111 | 2.395724202 |
| 0.3297756723 | 0.3497207578 | 0.4904467082 | -0.3947983951 | 2.619310386 | 1.869837872 |
| 0.3309085131 | 0.347668912 | 0.4924269235 | 0.5664291063 | 2.659951491 | 1.862231363 |
| 0.3232782256 | 0.3465241607 | 0.4814356792 | -1.824870881 | 2.466784925 | 1.469347122 |
| 0.3255895065 | 0.3451973195 | 0.486289782 | 1.328132768 | 2.602182447 | 1.991578658 |
| 0.3299599988 | 0.3460363188 | 0.4876124421 | 0.9345445312 | 2.718512897 | 1.881490868 |
| 0.3311726215 | 0.3492117685 | 0.4901402932 | 0.1269479285 | 2.402863495 | 1.429989563 |
| 0.3257319918 | 0.3434606618 | 0.4842245586 | -1.554120016 | 2.409500725 | 1.407888449 |
| 0.3336035529 | 0.3474677615 | 0.4918757147 | -0.07888209967 | 2.363461797 | 1.426504085 |
| 0.3288694617 | 0.3473691641 | 0.493115304 | 0.5309571262 | 2.40803909 | 1.406750955 |
| 0.3226995087 | 0.3419918632 | 0.4806552123 | -1.539337551 | 2.428545264 | 1.585580491 |
| 0.3313799848 | 0.3500799636 | 0.4979735422 | 0.5737974271 | 2.488814422 | 1.506318417 |
| 0.3294968367 | 0.3490970686 | 0.4909433674 | -0.07316337356 | 2.410359175 | 1.414873158 |
| 0.3347184491 | 0.3476132039 | 0.4906323861 | -0.1278160409 | 2.392662291 | 1.430857235 |
| 0.3345431722 | 0.3481667479 | 0.4904743334 | -0.09100153497 | 2.382518536 | 1.428197141 |
| 0.3204115319 | 0.3402970978 | 0.4802969115 | 1.616745063 | 2.358999183 | 1.231387194 |
| 0.3303636031 | 0.3476948523 | 0.4894292943 | -0.5846899654 | 2.347428834 | 1.387987225 |
| 0.3229947798 | 0.3404025555 | 0.4843583348 | 1.835199181 | 2.614753525 | 1.493122705 |

| | | | | | |
|---|---|---|---|---|---|
| 0.3294020411 | 0.3492587875 | 0.492390786 | -0.289474308 | 2.409930073 | 1.39653487 |
| 0.3265079215 | 0.3465261026 | 0.4865271969 | -0.9579341796 | 2.421282043 | 1.322663375 |
| 0.3249565905 | 0.3429374159 | 0.4867299261 | -1.212773179 | 2.432965622 | 1.430461507 |
| 0.3307285689 | 0.3467432938 | 0.4880548079 | -0.7777415287 | 2.279451277 | 1.359842253 |
| 0.327229423 | 0.3499724509 | 0.4877850026 | -0.6929243781 | 2.395503779 | 1.288414744 |
| 0.3280276133 | 0.3453360096 | 0.4892362811 | 0.8239171355 | 2.466097872 | 1.460876568 |
| 0.3363027052 | 0.3514264538 | 0.4978204454 | -0.3205314504 | 2.433894424 | 1.37116322 |
| 0.3403679422 | 0.3487112381 | 0.4874506976 | -0.808616513 | 2.542919918 | 1.528355598 |
| 0.3312200335 | 0.3448604034 | 0.4896197349 | -1.115254127 | 2.122684253 | 1.279574146 |
| 0.3295830444 | 0.3482221904 | 0.4917417028 | -0.5971319561 | 2.443225099 | 1.418021539 |
| 0.3272537227 | 0.3380324338 | 0.4776500521 | -1.893271667 | 2.01276896 | 1.346519948 |
| 0.3270383288 | 0.348628586 | 0.4881734815 | -0.9622145803 | 2.466924956 | 1.563323576 |
| 0.3349670542 | 0.3439074263 | 0.4863549953 | 1.378596061 | 1.96598871 | 1.240466812 |
| 0.3275986183 | 0.3507998424 | 0.4865542035 | -1.086264458 | 2.522052255 | 1.620740665 |
| 0.33189476 | 0.3476947016 | 0.4928449652 | -0.05203064248 | 2.395914684 | 1.424962053 |
| 0.3282898548 | 0.3435646081 | 0.4832656957 | -1.200575448 | 2.558905656 | 1.523972512 |
| 0.3410210341 | 0.3512219826 | 0.5017741656 | 0.9141274723 | 2.6070995 | 1.588040625 |
| 0.3345709858 | 0.3449338317 | 0.4891124134 | -1.033948322 | 2.136413086 | 1.37250637 |
| 0.3308638505 | 0.3481657326 | 0.4904611172 | 0.120040608 | 2.413265718 | 1.434756594 |
| 0.3291987366 | 0.353917129 | 0.4910823838 | -0.1192929489 | 2.406086377 | 1.421053935 |
| 0.3275058533 | 0.3421574677 | 0.4842551943 | 1.200277712 | 2.193752054 | 1.380999448 |
| 0.3257059972 | 0.3432961682 | 0.4903822267 | 2.045511993 | 2.145751462 | 1.140074638 |
| 0.3317644462 | 0.3477716977 | 0.5082553907 | -0.4784546379 | 2.458392035 | 1.461366904 |
| 0.3307491714 | 0.3446071889 | 0.4866553766 | -0.7385223469 | 2.217076933 | 1.429696514 |
| 0.317626205 | 0.3360594982 | 0.4743673818 | 1.900187735 | 2.232132786 | 1.568016837 |
| 0.3199580874 | 0.3406018202 | 0.4798473359 | 1.63437783 | 2.355495591 | 1.607577127 |
| 0.3225107351 | 0.342743849 | 0.4787311636 | 1.944177325 | 2.385551472 | 1.767755328 |
| 0.3290644383 | 0.3559652391 | 0.5016975082 | 0.374272144 | 2.420056414 | 1.305870386 |
| 0.3353135451 | 0.3451126822 | 0.4904319293 | 0.7842906854 | 2.507332412 | 1.450870863 |
| 0.3293542361 | 0.3492188644 | 0.4951295915 | -0.2352053671 | 2.416347107 | 1.445629613 |
| 0.3291336472 | 0.3510993779 | 0.4932689368 | -0.1410211783 | 2.403734922 | 1.430511662 |
| 0.3197937413 | 0.3377337745 | 0.476053428 | -2.130352083 | 2.605921992 | 1.406531191 |
| 0.3302115123 | 0.3471058904 | 0.4950447633 | 0.2458133347 | 2.415381385 | 1.424899952 |
| 0.3286619278 | 0.3449243038 | 0.4907662274 | -0.7611262568 | 2.298105994 | 1.409371079 |
| 0.3292647355 | 0.3472741384 | 0.4936149013 | -0.5408020479 | 2.41157218 | 1.445647659 |
| 0.3288100277 | 0.3470020417 | 0.4901524541 | 0.3465868932 | 2.407730552 | 1.429492671 |
| 0.3272709085 | 0.350456792 | 0.4941794019 | 0.9403416306 | 2.472427516 | 1.570721392 |
| 0.3278205415 | 0.3510311359 | 0.4853042308 | -1.369408093 | 2.368538535 | 1.194126883 |
| 0.3247029654 | 0.3427388358 | 0.4833329713 | -1.295178873 | 2.508831892 | 1.536592293 |
| 0.3397230713 | 0.3547298714 | 0.5029741955 | -0.9739814581 | 2.102412222 | 1.195807374 |
| 0.3322369203 | 0.3434884083 | 0.4926509496 | -1.23000593 | 2.140811238 | 1.406095785 |
| 0.3282158395 | 0.3477245533 | 0.4914429741 | 0.8354969696 | 2.474242917 | 1.520385109 |
| 0.3292963687 | 0.3465947463 | 0.4894019739 | 0.5822121123 | 2.450185684 | 1.446997901 |
| 0.3262381808 | 0.3437483565 | 0.4864083074 | 0.9410525387 | 2.287246034 | 1.451859091 |
| 0.3277945095 | 0.3507328793 | 0.4963276558 | -0.7530036385 | 2.409846405 | 1.289298915 |
| 0.3222563799 | 0.3406051384 | 0.4809497452 | 1.798080347 | 2.428865492 | 1.611217289 |

| | | | | | |
|---|---|---|---|---|---|
| 0.3353591479 | 0.3521205749 | 0.5105270521 | 0.3874243369 | 2.43641009 | 1.471931086 |
| 0.3231502736 | 0.3430871522 | 0.481492106 | 1.624309639 | 2.244324878 | 1.177052338 |
| 0.3268755823 | 0.3425065582 | 0.4829289396 | -1.433323615 | 2.2788018 | 1.432082182 |
| 0.3327053688 | 0.3473779102 | 0.4899533508 | 0.4917894739 | 2.455676538 | 1.456240456 |
| 0.3389451333 | 0.346373288 | 0.4891086662 | -1.489340947 | 1.965638607 | 1.251961297 |
| 0.3278327962 | 0.3507076614 | 0.4872784646 | 1.179943358 | 2.510793511 | 1.262741223 |
| 0.3120315792 | 0.3292031345 | 0.4647771228 | 2.725497796 | 2.620752435 | 1.517123993 |
| 0.3298591821 | 0.3471939449 | 0.488676304 | 0.6012426565 | 2.43959151 | 1.464551794 |
| 0.3337313447 | 0.3444963358 | 0.4940300703 | 1.468584273 | 2.128018908 | 1.385212537 |
| 0.3215173364 | 0.3424048119 | 0.4892566345 | 1.892301327 | 2.240042505 | 1.110017888 |
| 0.3284240485 | 0.3432861663 | 0.4815828098 | 1.370579351 | 2.117174793 | 1.252588212 |
| 0.3254462111 | 0.3471571589 | 0.4925140845 | -1.200696279 | 2.490554464 | 1.604787247 |
| 0.3297658488 | 0.3479316347 | 0.501429087 | 0.8462623819 | 2.508969284 | 1.333322693 |
| 0.328789911 | 0.3465204123 | 0.4898877193 | -0.4535049257 | 2.421620057 | 1.420812895 |
| 0.3243104774 | 0.3454385573 | 0.4830838502 | -1.286809445 | 2.446623984 | 1.247331895 |
| 0.3022821996 | 0.3238402113 | 0.4615231213 | 3.347517278 | 2.946209829 | 2.163080282 |
| 0.3079298099 | 0.3286875295 | 0.4588617746 | -2.785511015 | 2.645083003 | 1.877211343 |
| 0.3300088741 | 0.3496034559 | 0.4910414185 | -0.399380372 | 2.446569739 | 1.402229714 |
| 0.3437044184 | 0.3506094954 | 0.4929288857 | -1.355290016 | 2.798124164 | 1.709365132 |
| 0.3292074282 | 0.347007408 | 0.489804782 | -0.7084879117 | 2.473370006 | 1.41423165 |
| 0.3324571274 | 0.3477301734 | 0.4919825835 | 0.582121701 | 2.320994565 | 1.404754854 |
| 0.3288617687 | 0.348705274 | 0.4876119505 | -0.8233265133 | 2.320137565 | 1.302182856 |
| 0.3349811935 | 0.3560239503 | 0.4963050823 | -0.3009936403 | 2.318145958 | 1.331437697 |
| 0.3279281007 | 0.3413694791 | 0.4844482274 | -1.423712348 | 2.63316433 | 1.505106675 |
| 0.3299109869 | 0.3488450987 | 0.4909525145 | -0.4396963662 | 2.412528661 | 1.404852006 |
| 0.3340965283 | 0.3478966192 | 0.4900963559 | 0.1110881217 | 2.386444063 | 1.428311732 |
| 0.3295521206 | 0.3635326843 | 0.5050087724 | -0.0180628908 | 2.406083346 | 1.358571568 |
| 0.3308852051 | 0.3451837893 | 0.4873734105 | -0.7060612878 | 2.470303508 | 1.397299034 |
| 0.3237271537 | 0.342262408 | 0.4820886388 | -1.379488896 | 2.482020028 | 1.532176844 |
| 0.3282468125 | 0.3463854025 | 0.4902840543 | -0.5215756988 | 2.410597049 | 1.420467464 |
| 0.3237551431 | 0.3421175454 | 0.482945882 | -1.107392309 | 2.310582228 | 1.333072345 |
| 0.3386399565 | 0.3475128107 | 0.4910454539 | 0.113938486 | 2.313988062 | 1.424280755 |
| 0.3322375509 | 0.3478798946 | 0.4899918305 | -0.2801433176 | 2.424262474 | 1.441975276 |
| 0.3208599017 | 0.3422097323 | 0.4841207019 | 1.571170099 | 2.360241215 | 1.178572742 |
| 0.3273870938 | 0.3619522353 | 0.4933820762 | -0.7542431616 | 2.419779174 | 1.586753849 |
| 0.3238623565 | 0.3416627271 | 0.4822855291 | -1.369548524 | 2.389978677 | 1.430775586 |
| 0.3306428856 | 0.3472199949 | 0.4926800147 | 0.7613088249 | 2.367846404 | 1.464248197 |
| 0.330630002 | 0.3451785756 | 0.4877924565 | 0.8363380493 | 2.256741398 | 1.400552802 |
| 0.3175727889 | 0.3375331947 | 0.4736309464 | -1.914754799 | 2.519334569 | 1.643990707 |
| 0.3285874373 | 0.346670121 | 0.4978366294 | -0.4276743401 | 2.413100581 | 1.399786851 |
| 0.33096704 | 0.3491435569 | 0.4905273485 | -0.3808041501 | 2.364724389 | 1.387293724 |
| 0.3174378999 | 0.3334808031 | 0.4717674308 | -2.144240706 | 2.213500274 | 1.384853965 |
| 0.3290621988 | 0.3463517232 | 0.5068338998 | 0.9337425203 | 2.228976714 | 1.287881889 |
| 0.3278149977 | 0.3452897046 | 0.4866872046 | -0.9162059812 | 2.330197468 | 1.377160643 |
| 0.3326843325 | 0.3474092699 | 0.4922549784 | 0.1481153175 | 2.40328648 | 1.428049364 |
| 0.330053438 | 0.3505516467 | 0.4833070998 | -1.592137369 | 2.19073545 | 1.123486506 |

| | | | | | |
|---|---|---|---|---|---|
| 0.3296732495 | 0.3433469039 | 0.4856894227 | -1.295684678 | 2.620207066 | 1.325708231 |
| 0.3278251968 | 0.344599574 | 0.4975025792 | 1.78788985 | 2.794371129 | 1.747393396 |
| 0.3284216208 | 0.3477759119 | 0.4886552918 | 0.8302654844 | 2.375871833 | 1.347609622 |
| 0.3297933628 | 0.3489666443 | 0.4982009379 | 0.9993086844 | 2.291747081 | 1.282960328 |
| 0.3282639506 | 0.3473529886 | 0.4876560968 | 1.151379959 | 2.536897515 | 1.372105055 |
| 0.3364154774 | 0.346997337 | 0.4906249367 | -0.3332841997 | 2.420782621 | 1.428253576 |
| 0.3210445613 | 0.3385719337 | 0.477402821 | -1.857285589 | 2.586088383 | 1.557422266 |
| 0.3254466341 | 0.3342581721 | 0.4805589482 | 2.476351109 | 3.107084167 | 1.72681728 |
| 0.3287592371 | 0.3450612486 | 0.4875280311 | 0.6188171623 | 2.29676004 | 1.40262075 |
| 0.3302402262 | 0.3604077704 | 0.5117329706 | 1.069605757 | 2.578686298 | 1.703501715 |
| 0.3255483907 | 0.3434562183 | 0.4869191684 | 0.99480151 | 2.429434055 | 1.388375534 |
| 0.3297753716 | 0.3526200338 | 0.4920401193 | 0.2652419208 | 2.421974169 | 1.376395547 |
| 0.3389486348 | 0.3513617496 | 0.4872821483 | 1.093903318 | 2.082999195 | 1.225247308 |
| 0.3231156865 | 0.3413721919 | 0.4830388375 | -1.247471065 | 2.343587022 | 1.352418696 |
| 0.3276197498 | 0.3457773425 | 0.496306168 | -0.7201506595 | 2.431064242 | 1.452955239 |
| 0.3412768304 | 0.3535854824 | 0.4899477739 | -0.8551736252 | 2.568186295 | 1.586728582 |
| 0.3238683016 | 0.3445118565 | 0.482879646 | -1.383149714 | 2.468134775 | 1.607344083 |
| 0.3303975674 | 0.3470867311 | 0.4925289847 | 0.3618565104 | 2.429350703 | 1.4356285 |
| 0.3267423368 | 0.3505562051 | 0.4872864772 | -1.366413049 | 2.288419985 | 1.134372088 |
| 0.3244608771 | 0.3433554281 | 0.4853830279 | 1.206798294 | 2.350784817 | 1.516201329 |
| 0.3279520406 | 0.3439100829 | 0.4854674457 | 1.154150513 | 2.552710402 | 1.473434293 |
| 0.3248719319 | 0.3425017716 | 0.4795192395 | 1.751176327 | 2.825189168 | 1.68566854 |
| 0.3285267667 | 0.3464075484 | 0.488379197 | 1.017169183 | 2.651098276 | 1.524575023 |
| 0.3291181818 | 0.3430714191 | 0.484636445 | 1.433737522 | 2.803991541 | 1.48296859 |
| 0.330705662 | 0.349347757 | 0.4948366695 | -0.09993490134 | 2.571736696 | 1.463943075 |
| 0.3095105207 | 0.3277076189 | 0.460088445 | -2.92050843 | 2.679706358 | 1.809174414 |
| 0.3314303269 | 0.3502599151 | 0.4976802078 | 0.6495023356 | 2.63389304 | 1.533630271 |
| 0.3292040264 | 0.3491463093 | 0.4898539001 | 0.7655451132 | 2.56177577 | 1.540137814 |
| 0.3308076783 | 0.343565209 | 0.4840930141 | -1.392862884 | 2.818521106 | 1.527649376 |
| 0.3160721977 | 0.3289591437 | 0.462685613 | -2.788241066 | 3.179375686 | 1.728445285 |
| 0.3283606483 | 0.3470879688 | 0.4885967392 | 0.9318008048 | 2.602001596 | 1.409610132 |
| 0.3322811723 | 0.349825122 | 0.4914957191 | 0.2451008255 | 2.583145588 | 1.476157771 |
| 0.3297899679 | 0.3475639697 | 0.4945483012 | -0.6806634993 | 2.545268006 | 1.469857273 |
| 0.3304725494 | 0.3502689473 | 0.4934771027 | 0.3483179185 | 2.581106292 | 1.489383202 |
| 0.3237191267 | 0.3437164561 | 0.4815364073 | -1.616880923 | 2.618784939 | 1.300815959 |
| 0.3237047931 | 0.3414993773 | 0.4842843883 | -1.621208574 | 2.645845285 | 1.495487805 |
| 0.3229574312 | 0.3381634545 | 0.4752500967 | -2.085255142 | 2.319972283 | 1.327141926 |
| 0.2940842645 | 0.3148229869 | 0.4377799482 | -3.998267895 | 2.816164675 | 0.9331555229 |
| 0.3318301236 | 0.3489343277 | 0.4936526009 | 0.2986012269 | 2.588703123 | 1.472718989 |
| 0.3230050081 | 0.3379892543 | 0.4781932941 | 2.338330966 | 2.167494953 | 1.865455191 |
| 0.3426072791 | 0.350901682 | 0.4901819822 | 0.5828261859 | 2.322804282 | 1.376371314 |
| 0.3304480859 | 0.3438461638 | 0.4875722693 | -1.804773612 | 2.168298947 | 1.205893291 |
| 0.3297203571 | 0.3484232898 | 0.4915491424 | 0.5962303693 | 2.580061493 | 1.496195325 |
| 0.3294268925 | 0.3412400558 | 0.4808394204 | -1.71361528 | 2.240397272 | 1.393638295 |
| 0.3308445892 | 0.3529024722 | 0.4926603203 | -0.1223171371 | 2.573278867 | 1.462493524 |
| 0.34065117 | 0.3483740636 | 0.4932414425 | -0.9536021644 | 2.750454994 | 1.567092118 |

| | | | | | |
|---|---|---|---|---|---|
| 0.3277106165 | 0.3518190463 | 0.4861163313 | -1.256134705 | 2.704656883 | 1.705086489 |
| 0.3316431154 | 0.3474328398 | 0.4916258146 | 0.7716789504 | 2.466880483 | 1.496208664 |
| 0.3345432338 | 0.3498143388 | 0.4913230581 | -0.2443051781 | 2.578436917 | 1.476034998 |
| 0.3434888551 | 0.3537560535 | 0.5052536348 | -0.1953985918 | 2.421488288 | 1.407901915 |
| 0.3357215738 | 0.3459359131 | 0.4899047625 | -1.094733476 | 2.290124961 | 1.416008012 |
| 0.3255175755 | 0.3424409163 | 0.4820368555 | -1.543672341 | 2.478637539 | 1.426710144 |
| 0.3287773321 | 0.3550689041 | 0.4897141335 | -0.8529118028 | 2.602930665 | 1.618182758 |
| 0.3243976536 | 0.3397973758 | 0.4789881084 | 1.805631742 | 2.403746796 | 1.486716816 |
| 0.3354727971 | 0.3535904366 | 0.5050870989 | 0.4272480062 | 2.458152285 | 1.368056569 |
| 0.3333927518 | 0.3492212829 | 0.509719698 | 0.3684061778 | 2.499570047 | 1.435647328 |
| 0.3309836645 | 0.3470832435 | 0.4894269145 | -0.8772940625 | 2.462769466 | 1.510412678 |
| 0.3146354433 | 0.3336400763 | 0.4674696166 | 2.527416394 | 2.599935527 | 1.800369724 |
| 0.3228303163 | 0.3436112094 | 0.4828347132 | 1.721569809 | 2.616966188 | 1.710946671 |
| 0.3308339929 | 0.3515892768 | 0.4910861102 | 0.5400843377 | 2.528474765 | 1.525819357 |
| 0.3306761893 | 0.3589112981 | 0.5041998551 | 0.2713496811 | 2.579503533 | 1.355530984 |
| 0.3405235135 | 0.3494137491 | 0.496662224 | -0.0404678373 | 2.486429484 | 1.456231112 |
| 0.3307218603 | 0.3503900269 | 0.4950123019 | 0.0302135507 | 2.569648543 | 1.451587535 |
| 0.3293697307 | 0.3513559405 | 0.4934938627 | -0.6968714774 | 2.584991059 | 1.55340483 |
| 0.3285974217 | 0.3470313303 | 0.4891588195 | 0.8969495152 | 2.56466187 | 1.514060366 |
| 0.330086121 | 0.3471463967 | 0.4934850077 | -0.7849891372 | 2.51162659 | 1.458342182 |
| 0.3315138518 | 0.3486346504 | 0.4933401983 | -0.3020771661 | 2.539653263 | 1.462556608 |
| 0.3307908025 | 0.3488836167 | 0.4953038932 | -0.0749869633 | 2.567567532 | 1.460419594 |
| 0.3230773507 | 0.3408644655 | 0.4811078089 | 1.69283122 | 2.618238053 | 1.508665709 |
| 0.3230421998 | 0.3453285314 | 0.4850917354 | 1.81819045 | 2.781001007 | 1.803590793 |
| 0.3302133075 | 0.3535933163 | 0.4888464721 | -0.8127464924 | 2.501276006 | 1.284902329 |
| 0.33285499 | 0.3505323311 | 0.4931354801 | 0.03155994124 | 2.549998136 | 1.450738114 |
| 0.3394426027 | 0.3539619996 | 0.5017927064 | 1.128033666 | 2.803751406 | 1.688338254 |
| 0.3290390865 | 0.3415802766 | 0.4873700983 | -1.650832592 | 2.279683974 | 1.417648515 |
| 0.3261421793 | 0.3453138499 | 0.4858556322 | 1.491448342 | 2.74485153 | 1.662193564 |
| 0.3272262395 | 0.3445947686 | 0.4855325593 | 1.272509265 | 2.692186938 | 1.532915586 |
| 0.328508212 | 0.3470471521 | 0.4881080735 | 0.9509860055 | 2.548180828 | 1.530829666 |
| 0.3274790998 | 0.3494891312 | 0.4936023973 | -1.087953313 | 2.59506404 | 1.294355749 |
| 0.3279503943 | 0.3463605916 | 0.4886492562 | 1.063151096 | 2.544211304 | 1.538993629 |
| 0.3367724475 | 0.3529753926 | 0.5118244478 | 0.4712342775 | 2.617342571 | 1.52053633 |
| 0.3319229966 | 0.3531514665 | 0.4933842386 | -0.5736161742 | 2.625680465 | 1.541132287 |
| 0.3234511356 | 0.3388784142 | 0.4772284117 | -1.909184008 | 2.397294938 | 1.453346612 |
| 0.3286943558 | 0.34228161 | 0.4824865578 | -1.600186346 | 2.312341836 | 1.39426903 |
| 0.3424374636 | 0.349919539 | 0.4940313436 | -0.9194252273 | 2.221478374 | 1.327476154 |
| 0.3296966733 | 0.3527015925 | 0.4900488622 | -0.616696723 | 2.56771477 | 1.547322907 |
| 0.326477182 | 0.3444473784 | 0.4861044514 | 1.250205144 | 2.612594451 | 1.471690514 |
| 0.3311527251 | 0.3486713466 | 0.4893058088 | -0.8980882296 | 2.453119351 | 1.363246523 |
| 0.3387618038 | 0.3496890596 | 0.5014767728 | -0.0815560812 | 2.526779031 | 1.462838709 |
| 0.3329788474 | 0.3534837756 | 0.5055444627 | 0.2071230936 | 2.527799941 | 1.407946376 |
| 0.3354715244 | 0.3509065564 | 0.4901109812 | -0.6456072908 | 2.654433372 | 1.539412208 |
| 0.3270166949 | 0.3485424772 | 0.4943985288 | -1.305261613 | 2.697053332 | 1.671028102 |
| 0.3272500115 | 0.3451814564 | 0.4967560418 | 1.448232299 | 2.764564254 | 1.31211604 |

| | | | | | |
|---|---|---|---|---|---|
| 0.3276225446 | 0.3449751202 | 0.4864763463 | 1.17162297 | 2.520707145 | 1.474720091 |
| 0.3286144617 | 0.3496250368 | 0.4883762981 | -0.879551068 | 2.598916271 | 1.343453638 |
| 0.3208134792 | 0.3428456355 | 0.4855418006 | 2.13702054 | 2.889438626 | 1.900929532 |
| 0.3183071907 | 0.3391531859 | 0.4721756448 | -2.265910606 | 2.815652259 | 1.851711929 |
| 0.3270401119 | 0.3465651783 | 0.4861654682 | 1.177231413 | 2.569752319 | 1.584393697 |
| 0.3475431085 | 0.3545253054 | 0.4984342012 | 0.3996824091 | 2.321177598 | 1.363643703 |
| 0.3236748124 | 0.3411112185 | 0.4806626534 | 1.703385313 | 2.514168774 | 1.571625108 |
| 0.3315935828 | 0.3463445791 | 0.4898669773 | 1.088673914 | 2.392249661 | 1.395492248 |
| 0.3296823148 | 0.3495933081 | 0.487480878 | -1.021874853 | 2.4706607 | 1.309568532 |
| 0.3320697272 | 0.3534037 | 0.4911371682 | 1.21352438 | 2.776812801 | 1.709730433 |
| 0.3329178734 | 0.3471658652 | 0.4910015204 | -0.9262237369 | 2.705942694 | 1.528140647 |
| 0.3184834936 | 0.3368913124 | 0.473656734 | 2.18445958 | 2.747085034 | 1.688302662 |
| 0.3338717994 | 0.3480945868 | 0.4893936468 | 0.7234963703 | 2.665647467 | 1.510098501 |
| 0.3307677637 | 0.3657385113 | 0.5081648981 | -0.1067668188 | 2.572404322 | 1.361285849 |
| 0.3322628018 | 0.3478178491 | 0.4896588496 | -0.6750104288 | 2.645867582 | 1.440223083 |
| 0.2989532663 | 0.3162844634 | 0.4441426602 | -3.676387516 | 2.978057581 | 1.915608852 |
| 0.330692859 | 0.3489025542 | 0.4934080799 | 0.01966625909 | 2.570253719 | 1.460685177 |
| 0.3304144651 | 0.3485359378 | 0.4909862588 | -0.3579983632 | 2.564894831 | 1.464786049 |
| 0.3397673293 | 0.3479511351 | 0.4911549138 | 0.5896049393 | 2.359453594 | 1.453688304 |
| 0.3334813795 | 0.3489062857 | 0.4906829248 | -0.4499742918 | 2.612363164 | 1.48683301 |
| 0.3303115801 | 0.3511897884 | 0.4958083289 | 0.3895341772 | 2.567543627 | 1.40257057 |
| 0.329921931 | 0.3640413382 | 0.4962312052 | -0.5457471836 | 2.585428068 | 1.559882447 |
| 0.3280415765 | 0.3459430153 | 0.487378242 | -1.015149119 | 2.560164257 | 1.477268933 |
| 0.3283647317 | 0.3446706291 | 0.4889706841 | -1.218994423 | 2.712088381 | 1.467473435 |
| 0.3322832049 | 0.3468109673 | 0.4892935508 | 0.8574914469 | 2.432145158 | 1.457897564 |
| 0.329306931 | 0.3492396707 | 0.4897294703 | -0.8316530624 | 2.627061889 | 1.555177693 |
| 0.3311755955 | 0.3490546965 | 0.4990539489 | 0.1841123181 | 2.555691033 | 1.466974573 |
| 0.3309580626 | 0.3492697546 | 0.4897577093 | 0.7492182065 | 2.64627713 | 1.5389476 |
| 0.3292968747 | 0.3453313989 | 0.4875540455 | -1.166353808 | 2.443876846 | 1.425290049 |
| 0.3313359274 | 0.3472456744 | 0.5086618868 | 1.105586093 | 2.394872317 | 1.353637414 |
| 0.3106100587 | 0.3269695873 | 0.4598787052 | -2.92523088 | 2.475132801 | 1.385784034 |
| 0.3299188129 | 0.3447285582 | 0.4876599781 | -1.219159133 | 2.405696727 | 1.453141089 |
| 0.326869312 | 0.3471697683 | 0.4786444892 | -1.850212251 | 2.270879555 | 1.077423884 |
| 0.3275571351 | 0.3420300443 | 0.4839348291 | -1.673316022 | 2.865754299 | 1.362160961 |
| 0.3351603855 | 0.3523100944 | 0.508634351 | -0.3769841354 | 2.472074163 | 1.391053778 |
| 0.331045323 | 0.3507882576 | 0.4918165547 | 0.216592729 | 2.555877989 | 1.429477181 |
| 0.3327999455 | 0.3522320432 | 0.5008360861 | 0.1241997086 | 2.53992205 | 1.4296081 |
| 0.3308704231 | 0.3501110313 | 0.4915281705 | -0.2127910847 | 2.559822899 | 1.474163696 |
| 0.3381703481 | 0.348079134 | 0.4916378609 | -0.5711781684 | 2.639763632 | 1.479372468 |
| 0.3200380387 | 0.3375011351 | 0.4751588453 | -2.067582716 | 2.77490498 | 1.611181865 |
| 0.3365694398 | 0.3456821302 | 0.4969830351 | 1.227912478 | 2.82183557 | 1.571472978 |
| 0.3315857686 | 0.3489105822 | 0.4914543834 | -0.1211104707 | 2.572443361 | 1.458392087 |
| 0.3285911375 | 0.358019049 | 0.5084702786 | -1.375765188 | 2.401450112 | 1.035854918 |
| 0.3284788022 | 0.3463329682 | 0.4900047639 | 0.944530789 | 2.621923585 | 1.471045143 |
| 0.3295220947 | 0.3528353406 | 0.4908945637 | -0.6590260988 | 2.586508224 | 1.556947468 |
| 0.3411594899 | 0.354205277 | 0.4911101535 | 0.5636881863 | 2.347316913 | 1.333475556 |

| | | | | | |
|---|---|---|---|---|---|
| 0.3306516374 | 0.3489096483 | 0.4926957599 | -0.246612252 | 2.563809177 | 1.452449235 |
| 0.3282035059 | 0.3463782734 | 0.4954966964 | -1.002417467 | 2.628497468 | 1.51725096 |
| 0.3208411844 | 0.3342436876 | 0.4622851787 | -2.870313725 | 3.386070877 | 2.112466939 |
| 0.329729544 | 0.3496718396 | 0.4903781056 | -0.6066067291 | 2.587873843 | 1.519486246 |
| 0.3254819052 | 0.3419674483 | 0.4843515561 | 1.59729399 | 2.769761084 | 1.552316663 |
| 0.3292732579 | 0.3529502772 | 0.490363662 | -1.114973078 | 2.460100044 | 1.213523125 |
| 0.3234374558 | 0.3429156669 | 0.4830755499 | 1.670838133 | 2.572387124 | 1.659936667 |
| 0.3303792528 | 0.3464759997 | 0.4882320125 | 0.9570894165 | 2.682534368 | 1.504197822 |
| 0.2678625381 | 0.2884700695 | 0.4026322417 | -1.097024985 | -3.827820995 | 0.3195770029 |
| 0.270496856 | 0.2907491427 | 0.4074863318 | -0.364502659 | -3.775316985 | 0.3690496898 |
| 0.2725534394 | 0.2891332408 | 0.4054522531 | -0.935597239 | -3.859979947 | 0.358130511 |
| 0.2635917735 | 0.2841957133 | 0.3985815116 | 1.788433794 | -3.913140151 | 0.2614537667 |
| 0.2661566018 | 0.2873141313 | 0.4003799736 | -1.439504952 | -3.793775937 | 0.5151651587 |
| 0.2714269889 | 0.2899137147 | 0.4113541242 | 1.2148755 | -3.5195344 | 0.5208711758 |
| 0.2699361386 | 0.2917498896 | 0.4065476862 | 0.3180812294 | -3.757964671 | 0.4221949367 |
| 0.2727128184 | 0.2904789707 | 0.4067894214 | 0.04891594121 | -3.719372856 | 0.3930341621 |
| 0.2741712812 | 0.290611616 | 0.4072479743 | -0.3400559013 | -3.632376496 | 0.4111816186 |
| 0.2693600455 | 0.2911720183 | 0.4074869083 | 0.4097437377 | -3.767139512 | 0.3573395702 |
| 0.2732862716 | 0.2911861657 | 0.4073206508 | -0.1333212018 | -3.727338115 | 0.3832287826 |
| 0.2675358966 | 0.2875593829 | 0.4068195963 | -0.87194838 | -3.855274298 | 0.4306153017 |
| 0.2704962437 | 0.2917652513 | 0.4077023435 | -0.3201214809 | -3.725835415 | 0.353989128 |
| 0.26500476 | 0.2862123644 | 0.3987075455 | -1.569014562 | -3.801892548 | 0.2538135205 |
| 0.268647414 | 0.2891328771 | 0.4060120975 | -0.7597564102 | -3.787688824 | 0.3711172618 |
| 0.2688694695 | 0.287549591 | 0.400750366 | -1.460504214 | -3.944783655 | 0.2550739314 |
| 0.2684098267 | 0.292065185 | 0.4039109713 | -0.782278713 | -3.767171102 | 0.2789725723 |
| 0.2701575379 | 0.2906085068 | 0.4075614058 | 0.02834144652 | -3.746804822 | 0.3932112586 |
| 0.2726871594 | 0.292540012 | 0.4093902936 | 0.8637944696 | -3.881373521 | 0.5457480121 |
| 0.2805689224 | 0.2939483949 | 0.4063366854 | 0.3728638779 | -3.704190481 | 0.3242709889 |
| 0.2742654221 | 0.2912709108 | 0.4084180645 | 0.5826937517 | -3.58071652 | 0.4459991626 |
| 0.2628252896 | 0.2829142773 | 0.3978066481 | 1.676551846 | -3.923988625 | 0.4566442281 |
| 0.2750758328 | 0.2881434206 | 0.40327129 | -0.985712763 | -3.94110069 | 0.3351132636 |
| 0.2665505424 | 0.2904675025 | 0.4018728778 | 1.14208663 | -3.892953496 | 0.22420795 |
| 0.2860072833 | 0.2958761772 | 0.4151590612 | 0.2694487275 | -3.6240332 | 0.3291896478 |
| 0.2615793459 | 0.2834079513 | 0.3914237265 | -2.34252918 | -3.574719426 | 0.795538544 |
| 0.2705522072 | 0.2891132285 | 0.4057032631 | -0.6234549376 | -3.676468437 | 0.4096172856 |
| 0.2731808167 | 0.2912667488 | 0.4065849686 | 0.2264119339 | -3.737480439 | 0.3719297127 |
| 0.2802656982 | 0.2943451014 | 0.4155314772 | -0.6457937805 | -3.783212957 | 0.2680196188 |
| 0.276864018 | 0.2913434544 | 0.4075674579 | 0.2923213017 | -3.595083775 | 0.4266140747 |
| 0.2726564628 | 0.2920528043 | 0.4061243781 | -0.1100815996 | -3.731823424 | 0.3814704939 |
| 0.2699001013 | 0.2927317425 | 0.4072233553 | -0.9514525948 | -3.709675218 | 0.4905930784 |
| 0.264730327 | 0.2830020789 | 0.3959925211 | 1.634818187 | -4.081999824 | 0.3490952963 |
| 0.2732681355 | 0.2937848201 | 0.4161652529 | -0.1065674057 | -3.689199986 | 0.405096868 |
| 0.2705075078 | 0.2906927424 | 0.4144301039 | -0.00632594641 | -3.742093107 | 0.3920004969 |
| 0.2766505726 | 0.2911281249 | 0.40722185 | 0.3096532356 | -3.582768932 | 0.3700480385 |
| 0.2705521188 | 0.2913727974 | 0.4067550422 | -0.2912927275 | -3.722431863 | 0.3636836857 |
| 0.2706983566 | 0.2919304934 | 0.4075724274 | -0.04583883718 | -3.735885386 | 0.3845007492 |

| | | | | | |
|---|---|---|---|---|---|
| 0.2696664004 | 0.2932154277 | 0.4050437873 | 0.6909227964 | -3.790010897 | 0.4936190985 |
| 0.2676337825 | 0.2942761808 | 0.4116104604 | -1.392513287 | -3.849950019 | 0.6723486991 |
| 0.2751604268 | 0.291040161 | 0.4086334684 | -0.6698390459 | -3.772347259 | 0.3508550872 |
| 0.2681460947 | 0.2895882003 | 0.4086199922 | -1.121923625 | -3.731510148 | 0.4934090408 |
| 0.2626582369 | 0.2876907955 | 0.3990640619 | -1.60169579 | -3.818578257 | 0.734870154 |
| 0.2701571211 | 0.2905329063 | 0.4078790109 | 0.05919818903 | -3.747887482 | 0.3917870733 |
| 0.2723666831 | 0.2898286485 | 0.4124251382 | 1.061425841 | -3.539901408 | 0.3775395573 |
| 0.2704696123 | 0.2904906181 | 0.408356949 | -0.0370967219 | -3.745631459 | 0.3932719304 |
| 0.2707894669 | 0.2914199943 | 0.4093291701 | 0.4475457881 | -3.707699286 | 0.3470256444 |
| 0.2711160957 | 0.2906609473 | 0.4064506811 | 0.7123777113 | -3.659882857 | 0.3919894546 |
| 0.2689159695 | 0.2931741605 | 0.4098760198 | -0.8044450329 | -3.804121704 | 0.2534911768 |
| 0.2652939776 | 0.2896298467 | 0.3978841931 | -1.809189771 | -3.930621324 | 0.0268350921 |
| 0.2668004221 | 0.286779589 | 0.4007987503 | -1.288939892 | -3.671191312 | 0.5467637953 |
| 0.2738002942 | 0.2944464735 | 0.413937265 | 0.7416334263 | -3.558384844 | 0.5563418501 |
| 0.2729484677 | 0.2896158962 | 0.410867258 | 0.895830871 | -3.550150563 | 0.3726790857 |
| 0.2679857431 | 0.2895922704 | 0.4061649012 | 0.933545938 | -3.722963681 | 0.5175289872 |
| 0.269952492 | 0.2895296529 | 0.4054550532 | 0.8387661182 | -3.67363691 | 0.4210662094 |
| 0.2708200603 | 0.2914441237 | 0.4075104929 | -0.0833605314 | -3.729937739 | 0.3812872679 |
| 0.2698282785 | 0.2948846221 | 0.4129952835 | -0.4103162265 | -3.762995324 | 0.3123193427 |
| 0.2633394003 | 0.2859911135 | 0.400357308 | 1.662381646 | -3.896644525 | 0.640866704 |
| 0.2733276868 | 0.2941391694 | 0.4205842975 | -0.208288771 | -3.728237577 | 0.3484603274 |
| 0.2731785526 | 0.2936779246 | 0.4109694076 | -0.3405982232 | -3.649229467 | 0.4268258549 |
| 0.2714102435 | 0.2903863101 | 0.4062351703 | -0.1914454719 | -3.75070703 | 0.3902822568 |
| 0.2734816852 | 0.2894301397 | 0.4053896613 | 0.718665579 | -3.571992449 | 0.4616358442 |
| 0.2826624174 | 0.2938693361 | 0.4107255834 | -0.3757757144 | -3.682563352 | 0.3220210434 |
| 0.2700104202 | 0.295445407 | 0.4062153525 | -0.2157280117 | -3.743052824 | 0.4288215721 |
| 0.2703460346 | 0.2906311167 | 0.4067109574 | 0.1298599249 | -3.737920116 | 0.3884628597 |
| 0.271120851 | 0.2897540832 | 0.4054025327 | 0.6960902843 | -3.645204176 | 0.4530053539 |
| 0.2758912932 | 0.2905307324 | 0.4144490981 | 0.7116834037 | -3.81493501 | 0.3317148173 |
| 0.2720847475 | 0.295350486 | 0.4176161977 | 0.8544705931 | -3.839569534 | 0.2041417341 |
| 0.2736703468 | 0.2922758741 | 0.4061411653 | 0.1844274566 | -3.73098857 | 0.3699225477 |
| 0.2706065311 | 0.294168758 | 0.4136097443 | 0.1851001282 | -3.749470664 | 0.3553889894 |
| 0.2654650511 | 0.286760778 | 0.4101206125 | 1.666375204 | -3.640957607 | 0.1719333758 |
| 0.2700156356 | 0.2904840592 | 0.4086721775 | 0.5636723936 | -3.737402904 | 0.4004229376 |
| 0.2650399056 | 0.2885808006 | 0.3992594756 | -1.429450119 | -3.888589918 | 0.1888287752 |
| 0.2622901659 | 0.2875726862 | 0.4045868064 | 2.115769879 | -3.592732925 | 0.9028491655 |
| 0.2672265239 | 0.2898256188 | 0.4015142528 | -1.201453321 | -3.723167008 | 0.5639098469 |
| 0.2699942744 | 0.2909294488 | 0.4061843225 | -0.1349001277 | -3.747984459 | 0.3845424302 |
| 0.2803098233 | 0.2915839193 | 0.4076743064 | 0.7868163845 | -3.841264657 | 0.2873831165 |
| 0.2710748984 | 0.2902622803 | 0.4063670063 | -0.1877435905 | -3.719002222 | 0.3936677343 |
| 0.2698395847 | 0.2898531998 | 0.4055972892 | 0.6120615268 | -3.780717092 | 0.3997465597 |
| 0.2713994211 | 0.2926601029 | 0.4062252796 | -0.2763535252 | -3.760714756 | 0.3548515078 |
| 0.2737952187 | 0.2953498609 | 0.4092403127 | -0.3252580793 | -3.760785205 | 0.330053118 |
| 0.2734603132 | 0.2905057376 | 0.4071263741 | 0.4218615548 | -3.776207066 | 0.3682940534 |
| 0.2696025352 | 0.2889346474 | 0.4024712084 | -0.8836066664 | -3.893615885 | 0.3312020728 |
| 0.274297151 | 0.2905829494 | 0.4064395 | 0.3073879663 | -3.633379099 | 0.3932052623 |

| | | | | | |
|---|---|---|---|---|---|
| 0.2705637357 | 0.304093881 | 0.4164217774 | -0.1114972326 | -3.7341127 | 0.3402749724 |
| 0.2732269007 | 0.2902100309 | 0.4061640081 | -0.460464426 | -3.630151025 | 0.3768574442 |
| 0.2650090324 | 0.2850630865 | 0.398379642 | -1.710939725 | -3.728968795 | 0.4838794178 |
| 0.2670392957 | 0.2871511481 | 0.4024841914 | -1.362975586 | -3.760299111 | 0.3453058671 |
| 0.2703308873 | 0.2913351333 | 0.4064284081 | -0.3391168461 | -3.750327878 | 0.3595809698 |
| 0.2767480547 | 0.2908475656 | 0.4065515035 | 0.1299123722 | -3.683333389 | 0.3830112924 |
| 0.2718353872 | 0.2887961697 | 0.4022546622 | 1.161624854 | -3.934039318 | 0.2829788827 |
| 0.2701355049 | 0.2942528302 | 0.4092390732 | 0.1161639934 | -3.748996849 | 0.3675741554 |
| 0.269675966 | 0.3069305383 | 0.4100208558 | 0.8636173458 | -3.758086034 | 0.068050356 |
| 0.2693779061 | 0.2897267972 | 0.4053397297 | 0.5095594259 | -3.75027723 | 0.3947822123 |
| 0.270562687 | 0.2906685035 | 0.4101612642 | -0.0889552196 | -3.736558244 | 0.3889091458 |
| 0.2677691763 | 0.2861378043 | 0.4001540507 | 1.899893571 | -3.888189526 | 0.3325311422 |
| 0.2691296177 | 0.2902675122 | 0.4048740151 | -0.8302255858 | -3.710635675 | 0.4708899278 |
| 0.2656003261 | 0.2869361031 | 0.4079787088 | -1.24102366 | -3.855934865 | 0.313392448 |
| 0.2733658438 | 0.2926649727 | 0.4069236845 | -0.0392309658 | -3.709391424 | 0.384316986 |
| 0.2710604621 | 0.2901964701 | 0.406156021 | -0.3749413091 | -3.773347936 | 0.3841220726 |
| 0.2728934215 | 0.2942684969 | 0.4215442275 | 0.6193845561 | -3.776291365 | 0.2702570907 |
| 0.2634992149 | 0.2834026626 | 0.3966559477 | -1.814260349 | -3.945887524 | 0.3111885033 |
| 0.272607372 | 0.2899366207 | 0.4072204435 | 0.4854578724 | -3.641645376 | 0.3804806034 |
| 0.2726911939 | 0.295305763 | 0.4061175258 | 0.2566028034 | -3.67604217 | 0.4326748012 |
| 0.2748879074 | 0.2912487279 | 0.406924105 | -0.8831216906 | -3.493118482 | 0.2807133544 |
| 0.2734228057 | 0.295324094 | 0.4158155827 | -0.7373807906 | -3.823848769 | 0.2341389668 |
| 0.2699019404 | 0.2910729714 | 0.4061293381 | 0.1288308141 | -3.754171044 | 0.3844011643 |
| 0.2740111392 | 0.2944957815 | 0.4207678023 | -0.1056773252 | -3.67600393 | 0.4030816347 |
| 0.2703809869 | 0.2887830806 | 0.4045905727 | -1.158351269 | -3.849410447 | 0.3713923493 |
| 0.275240873 | 0.2904835984 | 0.4063462272 | 0.253416087 | -3.729739135 | 0.4043387809 |
| 0.2704819351 | 0.2905574812 | 0.4065483562 | 0.1811217116 | -3.757301914 | 0.38656893 |
| 0.279469046 | 0.2931902966 | 0.4177449574 | -0.1542307716 | -3.659510582 | 0.3649703045 |
| 0.2703424697 | 0.2895864577 | 0.4050425023 | -1.005684608 | -3.639216765 | 0.3432075967 |
| 0.2717282454 | 0.303825115 | 0.4267859303 | -0.4527310602 | -3.777024591 | 0.2377219256 |
| 0.2699886915 | 0.2909002824 | 0.4088408915 | -0.0775581890 | -3.750102491 | 0.3970411892 |
| 0.2705002611 | 0.2927753567 | 0.4089121333 | -0.2802371981 | -3.731909918 | 0.4108712994 |
| 0.2831310652 | 0.2939005824 | 0.4025273464 | 0.8278531487 | -3.866861505 | 0.2422526987 |
| 0.2661114727 | 0.2864421421 | 0.4006385529 | -1.465153727 | -3.82578646 | 0.3163044843 |
| 0.268639713 | 0.2889964457 | 0.4111467199 | 0.7353280131 | -3.790283965 | 0.3864621872 |
| 0.2820014604 | 0.2944505158 | 0.4068741778 | -1.183455045 | -3.208481708 | 0.5989392401 |
| 0.2701553958 | 0.2925845077 | 0.4090608172 | -0.0612058228 | -3.745869844 | 0.3971989613 |
| 0.2691469108 | 0.2897802083 | 0.4051899922 | 0.675516001 | -3.750403753 | 0.3609118131 |
| 0.2716181413 | 0.2976621775 | 0.410433605 | -0.272970109 | -3.752931358 | 0.3195550015 |
| 0.2705301956 | 0.2909439771 | 0.4081724467 | -0.0231978421 | -3.740893633 | 0.3902681285 |
| 0.2704842421 | 0.2896271897 | 0.4053163112 | -0.4578632832 | -3.795101286 | 0.4089600653 |
| 0.2641218531 | 0.2849642598 | 0.3979563391 | -0.9883790317 | -3.580270269 | 0.4236430161 |
| 0.2647166177 | 0.2850579675 | 0.3996837463 | -0.850005856 | -3.61492665 | 0.4312445618 |
| 0.2640197587 | 0.2805673541 | 0.3935567897 | -1.658185522 | -3.827907071 | 0.4343332278 |
| 0.257313115 | 0.2780636986 | 0.3900845226 | 2.006358204 | -3.739560734 | 0.3411374869 |
| 0.2632328329 | 0.2848110296 | 0.3968529541 | -1.234769352 | -3.535892427 | 0.5918119278 |

| | | | | | |
|---|---|---|---|---|---|
| 0.2668198555 | 0.2853453439 | 0.4050624235 | 1.322443525 | -3.27789093 | 0.6175389994 |
| 0.265722301 | 0.2875875758 | 0.4009269967 | 0.00194867655 | -3.523101167 | 0.4833428033 |
| 0.2678988967 | 0.2861171004 | 0.4007169386 | 0.2279890475 | -3.527553552 | 0.4913531025 |
| 0.2697769064 | 0.2863207657 | 0.4013086023 | -0.3843455034 | -3.401176905 | 0.508582776 |
| 0.2654747514 | 0.287928788 | 0.403051209 | 0.1934301147 | -3.531630042 | 0.462343977 |
| 0.2688584446 | 0.28689118 | 0.4014595541 | -0.1697715071 | -3.510765312 | 0.4735373834 |
| 0.2659923524 | 0.2862698569 | 0.4049379161 | -0.0627670092 | -3.524385752 | 0.4874449532 |
| 0.2659985955 | 0.2876035645 | 0.4017192664 | -0.3091027308 | -3.504546033 | 0.4516393369 |
| 0.2587345455 | 0.279971498 | 0.3901513086 | -1.792404086 | -3.595039305 | 0.3457425076 |
| 0.2642942335 | 0.2850783826 | 0.4000020416 | -0.7573400789 | -3.56333542 | 0.4585465437 |
| 0.2647883085 | 0.2840749074 | 0.3959348915 | -1.29569679 | -3.693261678 | 0.373915644 |
| 0.2640517067 | 0.2874781745 | 0.3981170726 | -0.7696567185 | -3.532392629 | 0.3793742818 |
| 0.2639668194 | 0.2847073834 | 0.3991703338 | 0.7089827236 | -3.537041983 | 0.519878684 |
| 0.2695855153 | 0.2896698555 | 0.405310261 | 0.4769450318 | -3.565128501 | 0.5631085049 |
| 0.2763450336 | 0.2901800899 | 0.4006634788 | 0.271260193 | -3.458700366 | 0.432368204 |
| 0.2685987195 | 0.2859831936 | 0.4007690915 | 0.682719051 | -3.368704636 | 0.5338615916 |
| 0.251116377 | 0.2710481816 | 0.3812481516 | 2.438962422 | -3.82278825 | 0.576534536 |
| 0.271354171 | 0.2843476852 | 0.3981212902 | -0.9692207351 | -3.70730638 | 0.4143288909 |
| 0.264232555 | 0.2883198439 | 0.3989912863 | 0.797716524 | -3.610020238 | 0.355652382 |
| 0.2812533514 | 0.2918169701 | 0.4089641993 | 0.5150091036 | -3.493003769 | 0.3703556583 |
| 0.2617278995 | 0.2836024183 | 0.39193311 | -2.008183076 | -3.285620533 | 0.8137206556 |
| 0.2670457938 | 0.2859302446 | 0.4012105324 | -0.2984746279 | -3.473567089 | 0.4925764618 |
| 0.268945972 | 0.2872855129 | 0.4008839031 | 0.02971023131 | -3.485213061 | 0.4810522993 |
| 0.2763579441 | 0.2908511486 | 0.41031273 | -0.2745972593 | -3.46150907 | 0.427759459 |
| 0.2715004025 | 0.2864017767 | 0.4004248386 | 0.6314079839 | -3.310279384 | 0.5610252574 |
| 0.2687234107 | 0.2886976523 | 0.4006830804 | 0.00859048758 | -3.482248265 | 0.4822604994 |
| 0.2658422619 | 0.2880130027 | 0.4017652796 | -0.5969911749 | -3.497906751 | 0.5361963701 |
| 0.2602953831 | 0.2787101176 | 0.3900207431 | 1.667978653 | -3.860063872 | 0.4158756803 |
| 0.2683614479 | 0.2888161958 | 0.4089023984 | 0.6708169457 | -3.582017975 | 0.3791540061 |
| 0.2642550633 | 0.284795684 | 0.4059051956 | -0.7264798488 | -3.513136789 | 0.5242062224 |
| 0.2733347545 | 0.287330974 | 0.4018872028 | 0.7548516694 | -3.231409522 | 0.4463800149 |
| 0.2659932456 | 0.2870638162 | 0.4009372557 | -0.0102453660[illegible] | -3.518757214 | 0.483162918 |
| 0.2655153916 | 0.2870478508 | 0.4006485529 | 0.3351785519 | -3.553540343 | 0.5220385045 |
| 0.2656775228 | 0.2896137627 | 0.3999154811 | 0.5796343191 | -3.551417188 | 0.5686554089 |
| 0.2656116941 | 0.2914239079 | 0.4079736389 | -0.7540780355 | -3.575762205 | 0.6242194824 |
| 0.2695546197 | 0.2861848801 | 0.401407547 | 0.07757868879 | -3.461505975 | 0.488272575 |
| 0.2649049634 | 0.2868060906 | 0.405538518 | -0.9944653234 | -3.481184152 | 0.5777275817 |
| 0.2547891222 | 0.2796453287 | 0.387663028 | -2.062695059 | -3.62310463 | 0.9098109644 |
| 0.2646894128 | 0.2853107287 | 0.4006190829 | -0.5657844778 | -3.529852054 | 0.488406333 |
| 0.2678342248 | 0.2855077406 | 0.4069060938 | 0.9211969477 | -3.347128188 | 0.4840054689 |
| 0.2660683577 | 0.2862435884 | 0.4027870063 | -0.0880307015[illegible] | -3.525357108 | 0.4875789193 |
| 0.2634488208 | 0.2849481565 | 0.3999241778 | -0.7515163584 | -3.604360683 | 0.5675141857 |
| 0.2664716647 | 0.2861348794 | 0.4002649595 | 0.8282250566 | -3.424418547 | 0.5018306865 |
| 0.2651687287 | 0.2899864993 | 0.4055031789 | -0.6606146823 | -3.565650343 | 0.3614826201 |
| 0.2637116319 | 0.2884637446 | 0.3967334158 | -1.377186928 | -3.626186414 | 0.2099823968 |
| 0.2625083215 | 0.2827573243 | 0.3951318796 | -1.339383048 | -3.429413929 | 0.6384008043 |

| | | | | | |
|---|---|---|---|---|---|
| 0.2713596152 | 0.2925200889 | 0.4110049112 | 0.1426920073 | -3.422127249 | 0.506344214 |
| 0.2664921921 | 0.2829984194 | 0.4017994219 | 1.454401013 | -3.250036686 | 0.4556981897 |
| 0.2616412943 | 0.2832886013 | 0.3980388682 | 1.266229412 | -3.508351305 | 0.6445177602 |
| 0.2653373265 | 0.2850969522 | 0.399293442 | 0.9037163782 | -3.446192444 | 0.5122512545 |
| 0.2644425704 | 0.2851189942 | 0.3992270955 | 0.5884687761 | -3.601427798 | 0.543749856 |
| 0.2655796396 | 0.2916324541 | 0.408496078 | -0.3357040357 | -3.537695581 | 0.4112433517 |
| 0.2643130245 | 0.2878332248 | 0.4029298262 | 0.8538417734 | -3.556663172 | 0.6039651212 |
| 0.2682323122 | 0.2896912189 | 0.413889946 | 0.1885633556 | -3.465753969 | 0.512951946 |
| 0.2684391465 | 0.2888444584 | 0.4045475697 | -0.4295024735 | -3.420299873 | 0.530215739 |
| 0.26702403 | 0.2863433227 | 0.4006302337 | -0.01067081827 | -3.506978278 | 0.4851642945 |
| 0.2693082539 | 0.2858306324 | 0.400191949 | 0.6356967484 | -3.360090316 | 0.5407828219 |
| 0.2781652576 | 0.2897521324 | 0.4044608211 | 0.1993954689 | -3.308901658 | 0.5148141469 |
| 0.2647040573 | 0.2905219206 | 0.3990893422 | -0.5537939694 | -3.517746123 | 0.5910350592 |
| 0.2654736384 | 0.2860587592 | 0.4002994286 | -0.150879981 | -3.534070348 | 0.4890383081 |
| 0.267069924 | 0.285740907 | 0.3998579781 | 0.6476116797 | -3.41772853 | 0.5404155042 |
| 0.2716973518 | 0.2866444344 | 0.4092250836 | 0.7496270444 | -3.600322935 | 0.4169916997 |
| 0.2667151567 | 0.2908281416 | 0.4108392713 | 1.029542676 | -3.655354795 | 0.2570626538 |
| 0.2684456173 | 0.2872982472 | 0.39970503 | 0.4265515206 | -3.558661034 | 0.4412065485 |
| 0.2650030286 | 0.2888112761 | 0.4057852397 | -0.5372577136 | -3.509199358 | 0.5686898972 |
| 0.2649543651 | 0.2868133282 | 0.4101701649 | 1.041976841 | -3.421921392 | 0.337533975 |
| 0.2656703397 | 0.2865508918 | 0.403519512 | 0.02314461116 | -3.523177572 | 0.4855506811 |
| 0.2611964701 | 0.2849474907 | 0.3940906902 | -1.440214665 | -3.648357907 | 0.2644038496 |
| 0.2638691315 | 0.2898979135 | 0.4080064708 | 1.401595052 | -3.359290625 | 0.8097941034 |
| 0.2633695257 | 0.2862506961 | 0.3965744533 | -1.095563086 | -3.495913744 | 0.6393182254 |
| 0.2640850862 | 0.2851522338 | 0.3982062667 | -0.7756900866 | -3.534991096 | 0.4456060694 |
| 0.2736977969 | 0.2852624115 | 0.3988189637 | 1.173585658 | -3.763173401 | 0.3375316487 |
| 0.2657480566 | 0.285275478 | 0.3994312217 | -0.541487251 | -3.477558658 | 0.4892195475 |
| 0.2655408336 | 0.2861350686 | 0.4000636024 | 0.5475905362 | -3.541796167 | 0.499801677 |
| 0.2670717459 | 0.2885055962 | 0.4007625823 | -0.07497532514 | -3.513202574 | 0.4724873033 |
| 0.2683817798 | 0.2899380501 | 0.4020248719 | -0.6833048884 | -3.61941672 | 0.3566031691 |
| 0.2690813841 | 0.2863032302 | 0.4013002913 | 0.3754059925 | -3.545765777 | 0.465719306 |
| 0.263384034 | 0.2823731992 | 0.3934104583 | -1.285991696 | -3.768302992 | 0.3853018688 |
| 0.2700698463 | 0.2863954906 | 0.4006461242 | 0.03536406532 | -3.459646371 | 0.4855122119 |
| 0.2662918071 | 0.2991110911 | 0.4093865353 | -0.06097043946 | -3.510927428 | 0.4466324791 |
| 0.2690561309 | 0.2855905874 | 0.4001243178 | -0.5681751356 | -3.380562155 | 0.4755333136 |
| 0.2626792528 | 0.2830846366 | 0.3957535477 | -1.35364785 | -3.487928628 | 0.5552665639 |
| 0.2589032992 | 0.2789715945 | 0.3908637664 | -1.986923894 | -3.562627067 | 0.4301990491 |
| 0.2651627837 | 0.2868592849 | 0.399455407 | -0.7209421804 | -3.551219796 | 0.4113533462 |
| 0.271837395 | 0.2866781266 | 0.4006557478 | 0.1664032987 | -3.476821408 | 0.4758596155 |
| 0.2682235802 | 0.2856488123 | 0.3978045009 | 0.9638927758 | -3.663997977 | 0.3926369864 |
| 0.2656401407 | 0.2903072601 | 0.4034019027 | 0.300435837 | -3.528859836 | 0.4271132703 |
| 0.2653906363 | 0.3035576705 | 0.4046710514 | 0.6787498895 | -3.536922678 | 0.2229723869 |
| 0.2647226698 | 0.2853212934 | 0.3991421946 | 0.6361050879 | -3.525937419 | 0.4904053061 |
| 0.265986256 | 0.2865082622 | 0.4038504717 | -0.1226115521 | -3.513373665 | 0.4804054868 |
| 0.2658443332 | 0.2851150665 | 0.3987347044 | 1.123910576 | -3.606586499 | 0.445313243 |
| 0.2650406426 | 0.2865264873 | 0.3992614363 | -0.7890300239 | -3.477845274 | 0.5638463968 |

| | | | | | |
|---|---|---|---|---|---|
| 0.2601879903 | 0.2816788439 | 0.4009427808 | -1.435326145 | -3.654279571 | 0.3794195722 |
| 0.2686394967 | 0.2881139983 | 0.4006814229 | -0.3038188499 | -3.533852809 | 0.4469029357 |
| 0.2667346487 | 0.2863137846 | 0.4006606346 | -0.1320512324 | -3.522360836 | 0.4831764883 |
| 0.2675192828 | 0.290265964 | 0.414917721 | -0.1397754322 | -3.484912634 | 0.5054848123 |
| 0.2587806163 | 0.2790073695 | 0.3906531166 | -1.8212301 | -3.714587604 | 0.4124838447 |
| 0.2689285226 | 0.2864848112 | 0.402156454 | 0.2275608306 | -3.444659964 | 0.4753508911 |
| 0.2685218424 | 0.2913105393 | 0.4004032113 | 0.1983349124 | -3.456855888 | 0.5137976408 |
| 0.2717109329 | 0.2879141659 | 0.4018665106 | -0.525190769 | -3.327619104 | 0.4200776894 |
| 0.2695535192 | 0.292722101 | 0.4116582604 | -0.3706176629 | -3.536863987 | 0.3954627865 |
| 0.2595913136 | 0.2806623057 | 0.3917099092 | 1.487776622 | -3.644990726 | 0.4114185927 |
| 0.2703267622 | 0.2910198041 | 0.4174063452 | -0.2722164231 | -3.410475466 | 0.5243509695 |
| 0.2681164033 | 0.2865520934 | 0.4018829534 | -0.6413915636 | -3.562575217 | 0.4510343398 |
| 0.2701795883 | 0.2861983124 | 0.3999420379 | 0.4453642785 | -3.552118205 | 0.5106503888 |
| 0.2659886151 | 0.2863241293 | 0.400655633 | 0.198701691 | -3.534550148 | 0.479097362 |
| 0.2750053061 | 0.2894629574 | 0.4124200559 | -0.1933456013 | -3.452961981 | 0.4506434707 |
| 0.2655534641 | 0.2848926038 | 0.3986805229 | -1.139855181 | -3.401752131 | 0.4497562414 |
| 0.2674354721 | 0.3003474811 | 0.4214492476 | -0.0393629376 | -3.504979203 | 0.4506940306 |
| 0.2656709312 | 0.2870194755 | 0.4033258454 | 0.09744949691 | -3.52348774 | 0.4768915523 |
| 0.2655282746 | 0.2876818568 | 0.4025249087 | 0.05597449461 | -3.527423695 | 0.4785083207 |
| 0.2766969666 | 0.2870182512 | 0.3926282186 | 1.311081375 | -3.853148827 | 0.2322047056 |
| 0.2587676726 | 0.279289654 | 0.3905168569 | -1.820760435 | -3.653101708 | 0.4051349121 |
| 0.2633755844 | 0.2839371382 | 0.4041585938 | 0.9261386811 | -3.583699263 | 0.4793947417 |
| 0.2812961287 | 0.2926548308 | 0.4048215795 | -0.6775256837 | -3.092715953 | 0.6029884618 |
| 0.2639142453 | 0.286954074 | 0.4010269295 | -0.7843611841 | -3.523084908 | 0.5972016475 |
| 0.2653463422 | 0.2863963393 | 0.4002842016 | 0.3577498674 | -3.523225519 | 0.4668142249 |
| 0.2673242246 | 0.2938597469 | 0.404840596 | -0.0475738659 | -3.506731092 | 0.4617447426 |
| 0.2658801081 | 0.2863327083 | 0.401666703 | -0.2733877822 | -3.505190561 | 0.474294154 |
| 0.2647732528 | 0.2842739687 | 0.3976584433 | -0.7925520116 | -3.62257576 | 0.518689657 |
| 0.2597951194 | 0.2810397262 | 0.3927112941 | -0.5871770132 | 2.232190901 | -1.738564904 |
| 0.2594643698 | 0.2800452764 | 0.3927938846 | 0.7820176093 | 2.327488698 | -1.669909758 |
| 0.2644094031 | 0.2815717618 | 0.3950087711 | -0.0319673238 | 2.207363392 | -1.708204403 |
| 0.2606319804 | 0.2823348668 | 0.3961331972 | -0.2332956461 | 2.257645731 | -1.679813412 |
| 0.2483124067 | 0.269672506 | 0.3756480953 | 2.383655218 | 2.315067086 | -2.026792091 |
| 0.2604245325 | 0.2791460452 | 0.3963096322 | -1.29815407 | 2.039616779 | -1.839935652 |
| 0.2603208028 | 0.2822666427 | 0.3936513213 | -0.2320294056 | 2.254457216 | -1.720358517 |
| 0.2581497897 | 0.2766200838 | 0.3873975846 | 1.518940182 | 2.063759905 | -1.694455263 |
| 0.2640733999 | 0.2807137733 | 0.3934859174 | 0.7023318329 | 2.082011497 | -1.75308428 |
| 0.2596027653 | 0.2829288236 | 0.3961522163 | 0.5762525909 | 2.23456076 | -1.775778241 |
| 0.2614733323 | 0.2795250758 | 0.3912409336 | -1.213833262 | 2.040986106 | -1.788022156 |
| 0.2575548752 | 0.2780871198 | 0.3930821546 | 1.048903697 | 2.335142131 | -1.771969272 |
| 0.2599901569 | 0.2820240181 | 0.3936623676 | -0.2481120454 | 2.258731532 | -1.729647207 |
| 0.2559930519 | 0.2775746815 | 0.3869146984 | 1.449211734 | 2.255209339 | -1.614160844 |
| 0.2510263856 | 0.2716015065 | 0.3806308358 | 2.092765442 | 2.393770592 | -1.679369937 |
| 0.2612092538 | 0.2812369695 | 0.3919598399 | 0.8002325458 | 2.336151405 | -1.635515917 |
| 0.2580111724 | 0.2810650412 | 0.3899412057 | 0.9597075747 | 2.232182202 | -1.594711429 |
| 0.2590101627 | 0.2800501537 | 0.392492949 | 0.9283632691 | 2.278637416 | -1.674631252 |

| | | | | | |
|---|---|---|---|---|---|
| 0.2645258889 | 0.2848798883 | 0.3984860886 | -0.3708215471 | 2.281314457 | -1.749317444 |
| 0.2693623206 | 0.2837105155 | 0.3909978129 | 0.716867379 | 1.972424073 | -1.825127465 |
| 0.2553208378 | 0.2734001527 | 0.3825431455 | 1.41668082 | 2.499425065 | -1.804031597 |
| 0.2597374318 | 0.2807091903 | 0.3948637111 | -0.5809971325 | 2.263920771 | -1.715122924 |
| 0.2647694374 | 0.2776193457 | 0.3888958046 | 1.2813051 | 2.546128437 | -1.631619694 |
| 0.2609902988 | 0.2853541599 | 0.3949711853 | -0.1236611966 | 2.252120071 | -1.663863936 |
| 0.2675134436 | 0.2789097841 | 0.3900527531 | 1.73206169 | 1.612804999 | -2.117747607 |
| 0.2629485909 | 0.2849378066 | 0.394289087 | -0.2646553482 | 2.26650554 | -1.646187568 |
| 0.2577469089 | 0.2767053041 | 0.3881888479 | -1.531010944 | 2.448856184 | -1.700201196 |
| 0.2602264597 | 0.2786826668 | 0.388613969 | 1.160488843 | 2.058276273 | -1.845035128 |
| 0.2710763382 | 0.2860187092 | 0.4031364936 | 0.1516738413 | 2.200734258 | -1.650629702 |
| 0.2654016987 | 0.2806020486 | 0.3920666073 | 1.343066132 | 2.518944882 | -1.534478905 |
| 0.2608737192 | 0.2818920437 | 0.3891751383 | 0.8101573137 | 2.381879997 | -1.600335486 |
| 0.256283508 | 0.2775241567 | 0.3876687294 | 2.230561589 | 2.223151764 | -1.822262774 |
| 0.2614624552 | 0.2804413297 | 0.3924551727 | -0.7158319111 | 2.341519684 | -1.669055685 |
| 0.2624677393 | 0.2830294558 | 0.4002820434 | -0.3317036103 | 2.27726686 | -1.660095409 |
| 0.2604565501 | 0.2814744911 | 0.4009761748 | -0.4645364722 | 2.267062868 | -1.680622014 |
| 0.2642732906 | 0.2767384349 | 0.3870247904 | 1.09102538 | 2.489621592 | -1.78160231 |
| 0.2598997147 | 0.2812175826 | 0.3929176503 | -0.4992635676 | 2.277691823 | -1.745404392 |
| 0.2602353506 | 0.282111539 | 0.3936019861 | -0.4079857976 | 2.275638647 | -1.747399564 |
| 0.2600820399 | 0.2844970016 | 0.3926027909 | 0.5307248102 | 2.220683212 | -1.601901213 |
| 0.261434152 | 0.2862221336 | 0.4009245569 | 0.7158341916 | 2.300740962 | -1.790765885 |
| 0.2612629186 | 0.278082144 | 0.3897747878 | 1.509991165 | 2.496926204 | -1.684230877 |
| 0.2598275559 | 0.2824914646 | 0.4007630014 | 0.6076478696 | 2.222166575 | -1.77194078 |
| 0.2546336057 | 0.2800639874 | 0.3879895951 | 1.433187319 | 2.24497645 | -2.010143613 |
| 0.2603570543 | 0.2813636664 | 0.3950830052 | -0.311961609 | 2.252670244 | -1.70207749 |
| 0.2633950708 | 0.2813165117 | 0.4013554407 | -0.8303257543 | 2.093562643 | -1.684805488 |
| 0.25747591 | 0.2775821589 | 0.3910085877 | -1.332070342 | 2.17611577 | -1.718685257 |
| 0.2593223012 | 0.2815828021 | 0.3947990119 | 0.3978629004 | 2.278281233 | -1.76366032 |
| 0.2562941114 | 0.2757449665 | 0.3858837327 | -1.828087442 | 2.096480152 | -1.792581238 |
| 0.2603884917 | 0.2859753186 | 0.3999438681 | 0.4142748843 | 2.273609256 | -1.602692526 |
| 0.2606500195 | 0.2859204361 | 0.3938342598 | 0.2239623791 | 2.258507362 | -1.639414701 |
| 0.2599563492 | 0.280844002 | 0.3923400972 | 0.8533818196 | 2.155326231 | -1.809277048 |
| 0.2637679494 | 0.2856590407 | 0.4008875858 | -0.8171015995 | 2.049686289 | -1.871924517 |
| 0.2613701969 | 0.2781550765 | 0.3950482964 | 0.9251849863 | 2.407524354 | -1.782503386 |
| 0.2562587845 | 0.2780519202 | 0.3913782776 | -1.3455841 | 2.200401224 | -1.882873135 |
| 0.260499099 | 0.280674208 | 0.3930293582 | 0.4252112359 | 2.282251274 | -1.70745657 |
| 0.2589189147 | 0.2795966512 | 0.3919064424 | -0.8370974557 | 2.332927061 | -1.791194236 |
| 0.2559993454 | 0.2830889207 | 0.3965141603 | -1.315879568 | 2.218714846 | -1.99886483 |
| 0.2588708315 | 0.2837105378 | 0.3970446184 | -0.6296653181 | 2.270392344 | -1.80661576 |
| 0.2633073066 | 0.285311383 | 0.4073060657 | 0.6831973744 | 2.323827489 | -1.553216141 |
| 0.262701534 | 0.2830998329 | 0.3967439165 | 0.576787747 | 2.148817025 | -1.749027091 |
| 0.2572290065 | 0.2767443217 | 0.3871925787 | 1.295509354 | 2.4044141 | -1.745632793 |
| 0.2648978459 | 0.2822330098 | 0.3949014874 | 0.2357945096 | 2.252567281 | -1.680991851 |
| 0.273246457 | 0.2851710092 | 0.3974778692 | 0.7515355462 | 2.37293012 | -1.541620456 |
| 0.2583404548 | 0.2845117152 | 0.3903742351 | 0.9631626719 | 2.213043747 | -1.885968644 |

| | | | | | |
|---|---|---|---|---|---|
| 0.2567196629 | 0.2774483313 | 0.3881559861 | 1.117198331 | 2.317759333 | -1.764241617 |
| 0.2610735558 | 0.2797130795 | 0.3914602048 | -0.991118512 | 2.089533418 | -1.798325828 |
| 0.2669462953 | 0.2824079685 | 0.4035603981 | 0.1265842372 | 2.163681477 | -1.71334172 |
| 0.2617952821 | 0.2871608205 | 0.4050789841 | 0.575215314 | 2.152932604 | -1.808036532 |
| 0.2534892719 | 0.2721224503 | 0.379158685 | -2.028226142 | 2.618473288 | -1.60474334 |
| 0.2596446069 | 0.2837350266 | 0.3983123545 | 0.5168061398 | 2.229785935 | -1.77800621 |
| 0.2493466598 | 0.2709066331 | 0.3873460563 | -2.39389046 | 2.101161732 | -1.454265433 |
| 0.2603484008 | 0.2815932599 | 0.3968875006 | -0.0882175947 | 2.245534636 | -1.708790289 |
| 0.2564891995 | 0.2806668275 | 0.3879014512 | -1.367066828 | 2.219133187 | -1.93684513 |
| 0.2592408563 | 0.2858943059 | 0.4025468883 | -1.222084293 | 2.086351255 | -1.983357876 |
| 0.2494301053 | 0.2718857234 | 0.376648631 | 2.329232445 | 2.223338464 | -2.099645649 |
| 0.2604130674 | 0.2818540644 | 0.3936291821 | -0.2771065112 | 2.250677113 | -1.714764862 |
| 0.2706085054 | 0.2826916691 | 0.3951590782 | -0.6159752297 | 2.33803722 | -1.622186101 |
| 0.2558223917 | 0.2753371507 | 0.3855017092 | -1.789058597 | 2.446917571 | -1.729791211 |
| 0.2593915896 | 0.2808108646 | 0.3921354971 | -0.5105372175 | 2.265195465 | -1.745343849 |
| 0.2598712743 | 0.2813923309 | 0.3911889722 | -0.9345927976 | 2.157001525 | -1.815443783 |
| 0.2606000289 | 0.2819491486 | 0.3911551538 | -1.601555969 | 1.951845078 | -1.98712799 |
| 0.2640089302 | 0.2815187097 | 0.3945813193 | -0.2099249898 | 2.252448358 | -1.696742926 |
| 0.2632977063 | 0.282352644 | 0.393433316 | -0.6148466614 | 2.123824848 | -1.755096046 |
| 0.2599515793 | 0.2760029254 | 0.3861154016 | -1.712042572 | 1.929764175 | -1.748964249 |
| 0.2583623112 | 0.2899110838 | 0.3965869555 | 1.160689901 | 2.167207647 | -1.332708012 |
| 0.2645501853 | 0.2805113126 | 0.393545992 | 0.6194173889 | 2.091341658 | -1.70677605 |
| 0.2553760097 | 0.2758573878 | 0.3857133147 | 1.700156149 | 2.194823694 | -1.810792307 |
| 0.2603168335 | 0.2814052995 | 0.3939973166 | -0.0240428097 | 2.248437065 | -1.708177659 |
| 0.2589562214 | 0.2816014413 | 0.3909439242 | 1.280533723 | 2.31246031 | -1.52104918 |
| 0.2653059501 | 0.280950415 | 0.3925350377 | -0.8057109879 | 2.369047293 | -1.660798234 |
| 0.2645988555 | 0.282662914 | 0.3934995382 | -0.1680652524 | 2.241793503 | -1.682424317 |
| 0.2603113337 | 0.2857560112 | 0.396563548 | -0.24060765 | 2.253933592 | -1.63721508 |
| 0.2600804559 | 0.2991323764 | 0.3978233484 | -0.6929897797 | 2.268626072 | -1.357186642 |
| 0.2575178805 | 0.2783971937 | 0.3893367078 | -1.014762923 | 2.258119942 | -1.74892545 |
| 0.2577681054 | 0.2785829309 | 0.3921174645 | 1.184016087 | 2.20346596 | -1.662342738 |
| 0.2509311951 | 0.2696504881 | 0.3769810018 | -2.973657192 | 2.609895697 | -1.574713254 |
| 0.2609103757 | 0.2829486778 | 0.3937200012 | 0.2779117985 | 2.22189227 | -1.72671221 |
| 0.2598269847 | 0.2819375909 | 0.4017824155 | -0.5184839165 | 2.233849265 | -1.753257639 |
| 0.2622435808 | 0.2819030957 | 0.3920844896 | -0.76048432 | 2.111250314 | -1.796200535 |
| 0.2568480019 | 0.276670838 | 0.387033043 | -1.419924132 | 2.158480413 | -1.753468757 |
| 0.2618139717 | 0.2854156023 | 0.4072783548 | -0.903335435 | 2.3450443 | -1.500517024 |
| 0.2552114009 | 0.2759275647 | 0.3864737986 | 1.586272633 | 2.362076914 | -1.670291254 |
| 0.2493922972 | 0.2664904269 | 0.3735711318 | 2.544659452 | 2.765444374 | -1.978229082 |
| 0.250758235 | 0.2728158521 | 0.3746445044 | 2.266545899 | 2.692606051 | -1.377426079 |
| 0.2636798297 | 0.2794449312 | 0.3896133381 | 1.409183426 | 1.903339276 | -1.551856807 |
| 0.2640070446 | 0.2883938518 | 0.4049305456 | 0.8689367629 | 2.368258421 | -1.449280547 |
| 0.2440503211 | 0.2645399978 | 0.3692361166 | -2.630223252 | 2.44043291 | -1.702285797 |
| 0.2641589849 | 0.2850712795 | 0.4110691067 | 0.8597158049 | 2.036615684 | -1.863086566 |
| 0.2653513512 | 0.2839035244 | 0.3982061116 | 0.4354516249 | 2.273967555 | -1.623072098 |
| 0.2611120538 | 0.2778577259 | 0.3877501859 | 1.495736424 | 1.959823487 | -1.613378004 |

| | | | | | |
|---|---|---|---|---|---|
| 0.2541303594 | 0.2742695337 | 0.3837540229 | 1.878796296 | 2.18233685 | -1.808849372 |
| 0.2699151809 | 0.2853251752 | 0.4063763361 | 0.2133545898 | 2.223370386 | -1.643945639 |
| 0.2615393301 | 0.2812377279 | 0.3937981481 | -0.1040487123 | 2.248023699 | -1.710322221 |
| 0.2619489126 | 0.2957000689 | 0.4143287876 | 0.1200178405 | 2.247114981 | -1.586472765 |
| 0.2603520151 | 0.2823005285 | 0.396530167 | -0.3447151153 | 2.248652516 | -1.668516218 |
| 0.2597409802 | 0.28163 | 0.3944747327 | 1.369400326 | 2.234901242 | -1.773991458 |
| 0.2776463638 | 0.2878745556 | 0.3934235679 | -0.0032761279 | 2.108633406 | -1.668206189 |
| 0.2566164611 | 0.2776674538 | 0.3880971256 | 1.541685179 | 2.304571585 | -1.628371599 |
| 0.2601334274 | 0.2811304772 | 0.4002853508 | 0.3323608815 | 2.241007172 | -1.713569091 |
| 0.274483997 | 0.2843007855 | 0.3937819611 | 1.343463026 | 1.657466271 | -1.975856816 |
| 0.2587831003 | 0.2826108446 | 0.3947165338 | 0.7493200565 | 2.22320741 | -1.823310521 |
| 0.2599632121 | 0.281454881 | 0.3931112678 | -0.4959451592 | 2.243547781 | -1.674257711 |
| 0.2611051978 | 0.288102027 | 0.3964434672 | 0.7739073552 | 2.332635101 | -1.483928003 |
| 0.2585292497 | 0.2788791109 | 0.3910783749 | -1.268534043 | 2.348891101 | -1.749110771 |
| 0.2603706683 | 0.2803397657 | 0.3919163008 | -0.8861642271 | 2.163093101 | -1.671665947 |
| 0.2816205976 | 0.3033182881 | 0.4233713368 | -1.001760818 | -2.030446105 | 1.55877984 |
| 0.2849850903 | 0.3061540926 | 0.4290057284 | 0.4477151478 | -1.91489913 | 1.623154895 |
| 0.2825874729 | 0.2997158843 | 0.4202699779 | -1.660484903 | -2.237106659 | 1.576937915 |
| 0.2839519884 | 0.3060619906 | 0.429230225 | 0.3638798653 | -1.977975358 | 1.573091783 |
| 0.2791737073 | 0.3012023342 | 0.4197384189 | -1.612339213 | -1.985837761 | 1.750004592 |
| 0.2840756555 | 0.3034489237 | 0.4305785093 | 1.350991436 | -1.739654244 | 1.767525683 |
| 0.2840675434 | 0.3069853339 | 0.4277560335 | 0.07106372678 | -1.965153835 | 1.60475181 |
| 0.2860234591 | 0.3046753585 | 0.4266717605 | 0.3988173307 | -2.010732025 | 1.623394223 |
| 0.2885537405 | 0.3058184202 | 0.428547464 | 0.05037345174 | -1.942377564 | 1.601808724 |
| 0.2836384459 | 0.3064216033 | 0.4288040219 | 0.3255636525 | -1.970195992 | 1.572282436 |
| 0.2877230735 | 0.3064627807 | 0.4286390531 | 0.2695344255 | -1.892323718 | 1.614328185 |
| 0.2749859673 | 0.2957038883 | 0.418325397 | -1.817904715 | -2.163717266 | 1.735434338 |
| 0.2843020371 | 0.3066450402 | 0.4284973477 | -0.2068843739 | -1.952637734 | 1.575790572 |
| 0.2811970975 | 0.3036633949 | 0.4230061258 | -1.119749097 | -1.971729616 | 1.513220208 |
| 0.2820315134 | 0.3034585504 | 0.4261659129 | -0.9052578274 | -1.998362951 | 1.588677802 |
| 0.283526939 | 0.3031231884 | 0.4224483125 | -1.346551294 | -2.133469806 | 1.480845491 |
| 0.2812748939 | 0.3060505964 | 0.4232076703 | -1.012568545 | -1.97642489 | 1.459528679 |
| 0.2827885183 | 0.3042495925 | 0.4266860855 | 0.57368602 | -1.962103965 | 1.64293914 |
| 0.2854938434 | 0.3062341281 | 0.4285591786 | 1.138784294 | -2.172925894 | 1.808226103 |
| 0.2949277617 | 0.3090123277 | 0.4271471839 | -0.0265085951 | -1.884698066 | 1.592473885 |
| 0.2885786785 | 0.3063892785 | 0.4296398758 | 0.5821497656 | -1.820944183 | 1.655253518 |
| 0.2782430008 | 0.2992443565 | 0.4206951508 | 1.548544852 | -2.061228387 | 1.679056326 |
| 0.2849827307 | 0.2985536483 | 0.4178982497 | -1.573974963 | -2.375176292 | 1.561809799 |
| 0.2746182478 | 0.2993819151 | 0.4142390083 | 1.881307025 | -2.18743423 | 1.375142239 |
| 0.3001545526 | 0.3100187863 | 0.4352780517 | 0.9675778147 | -2.154318035 | 1.367735479 |
| 0.2792709543 | 0.302524282 | 0.4175742508 | -2.059581497 | -1.732337366 | 1.922365505 |
| 0.2847634479 | 0.3042573478 | 0.4269550979 | -0.479787926 | -1.910270015 | 1.63107462 |
| 0.2869393496 | 0.305899215 | 0.4270183524 | 0.3823185651 | -2.000091366 | 1.574495861 |
| 0.2953635307 | 0.3101008486 | 0.4378192634 | -0.4807616019 | -2.012186837 | 1.484877657 |
| 0.2913957664 | 0.3065385439 | 0.4288511991 | 0.1556192288 | -1.881818501 | 1.621246174 |
| 0.2866108988 | 0.3069215477 | 0.4268614056 | -0.1846908066 | -1.975563697 | 1.583317568 |

| | | | | | |
|---|---|---|---|---|---|
| 0.2832532035 | 0.3073487308 | 0.4271203636 | -1.292633915 | -1.912506671 | 1.725131562 |
| 0.281497413 | 0.3007256523 | 0.4207685805 | 1.246862136 | -2.170223425 | 1.597788274 |
| 0.2865886269 | 0.3080758839 | 0.4364305378 | -0.5769039609 | -1.858159646 | 1.682891816 |
| 0.2844987619 | 0.3056444443 | 0.4357519066 | 0.05657515714 | -1.963391232 | 1.601796958 |
| 0.2900505545 | 0.3054077983 | 0.42721128 | 0.9000596204 | -1.7193056 | 1.545213245 |
| 0.2843283796 | 0.3062163197 | 0.4274793766 | -0.0100178978 | -1.961409747 | 1.601310549 |
| 0.2846242564 | 0.3068726543 | 0.4284423874 | -0.2099090243 | -1.944461494 | 1.575110233 |
| 0.2841545697 | 0.308938928 | 0.4267695012 | -0.3595872399 | -1.942623403 | 1.530883958 |
| 0.28460019 | 0.3128742597 | 0.4376614726 | -0.4392770858 | -1.981026699 | 1.652918417 |
| 0.2889024764 | 0.3055664703 | 0.429032355 | -0.7037991589 | -2.037109508 | 1.56271973 |
| 0.2777998842 | 0.2998423855 | 0.4228821719 | -1.837139921 | -1.941304126 | 1.792167901 |
| 0.2691313732 | 0.2949603152 | 0.4090733405 | -2.359648162 | -2.01228963 | 2.178497868 |
| 0.2842690051 | 0.3055317324 | 0.4289027807 | 0.4401779233 | -1.963741882 | 1.596898111 |
| 0.2874339984 | 0.3058734249 | 0.4352747762 | 0.6716506475 | -1.827303795 | 1.580948518 |
| 0.2844659192 | 0.3054198549 | 0.4292486272 | 0.3612968917 | -1.936662345 | 1.592773319 |
| 0.2845888461 | 0.3056211718 | 0.4292975044 | 1.129578446 | -1.881252046 | 1.482941187 |
| 0.2843243507 | 0.3045798805 | 0.4258297505 | 1.232148985 | -1.837278575 | 1.586248922 |
| 0.284278003 | 0.3097976598 | 0.4331011651 | 0.1361985457 | -1.955457459 | 1.605727226 |
| 0.2817596461 | 0.3077220563 | 0.4228347035 | -1.215546098 | -2.064549614 | 1.357266025 |
| 0.2705387409 | 0.2910056651 | 0.4066933533 | -2.450152346 | -1.818257258 | 1.984125226 |
| 0.2864934778 | 0.3080470698 | 0.4330722581 | 1.06043783 | -1.740961164 | 1.834416991 |
| 0.2789337107 | 0.2958081286 | 0.4190911777 | 2.432495823 | -1.578058168 | 1.587963725 |
| 0.2829957486 | 0.3057194653 | 0.4286567002 | 0.6621732247 | -1.931135277 | 1.697847573 |
| 0.2849637212 | 0.3055927719 | 0.4279423691 | 0.4190232414 | -1.917913351 | 1.615063971 |
| 0.2844112833 | 0.306077768 | 0.4279800388 | 0.08135429766 | -1.968632719 | 1.612274683 |
| 0.2839958044 | 0.3102639527 | 0.4345275815 | 0.1341996634 | -1.961037689 | 1.605872748 |
| 0.2830060671 | 0.3069048706 | 0.4296235536 | 0.5716701404 | -1.988081829 | 1.678865093 |
| 0.2874000688 | 0.3089282634 | 0.4418365453 | -0.6879575389 | -2.023951145 | 1.456493023 |
| 0.2873085773 | 0.308816667 | 0.4320482758 | -0.5617529521 | -1.848176459 | 1.648384413 |
| 0.2855334672 | 0.3054224546 | 0.4272603099 | 0.00639205213 | -1.952992285 | 1.606358386 |
| 0.289644909 | 0.3061988765 | 0.4289298161 | 0.04424090139 | -1.917635754 | 1.606310102 |
| 0.2968495117 | 0.3086365755 | 0.431350671 | -0.3333135006 | -1.974241454 | 1.536480184 |
| 0.2824287831 | 0.3089673524 | 0.4248310114 | -0.8316014239 | -1.944243476 | 1.747904381 |
| 0.2792734245 | 0.3005790989 | 0.4206343023 | -1.153381556 | -2.054505814 | 1.675070428 |
| 0.2865664873 | 0.3061706205 | 0.4283427872 | 0.2077344078 | -1.91783665 | 1.616511537 |
| 0.2907011298 | 0.3061681736 | 0.436783612 | -0.0668209499 | -1.904977093 | 1.607946598 |
| 0.2867648856 | 0.3114943071 | 0.4403866839 | -0.2316307572 | -1.913853869 | 1.627018007 |
| 0.2877962467 | 0.3073049179 | 0.4269755541 | 0.192476023 | -1.971192123 | 1.575780509 |
| 0.2768550801 | 0.3012822932 | 0.423489457 | -1.648311765 | -1.942403255 | 1.92103466 |
| 0.282298433 | 0.304802055 | 0.4359259948 | 1.213957913 | -1.8465707 | 1.441687917 |
| 0.2839883507 | 0.3055822466 | 0.4299499252 | 0.02559825177 | -1.963958942 | 1.605965767 |
| 0.2785803078 | 0.3031612956 | 0.419468305 | -1.513716934 | -2.066482138 | 1.395548545 |
| 0.2832357424 | 0.3101332953 | 0.4362017973 | 1.219114188 | -1.807954559 | 1.858377778 |
| 0.2818924601 | 0.3055250534 | 0.4232514372 | -1.166067863 | -1.906092064 | 1.757791989 |
| 0.2837611892 | 0.3057351377 | 0.4268492956 | -0.1393419567 | -1.965614315 | 1.600724817 |
| 0.2885928751 | 0.3001677633 | 0.419675158 | 1.8217218 | -2.460798203 | 1.392591257 |

| | | | | | |
|---|---|---|---|---|---|
| 0.2854834172 | 0.3055394489 | 0.4277362687 | 0.1255244816 | -1.963866628 | 1.602739531 |
| 0.2832555819 | 0.3042837431 | 0.4257831414 | 0.6502198157 | -2.004974591 | 1.626801889 |
| 0.2855663095 | 0.3078801537 | 0.4273132714 | -0.1270317208 | -1.967384643 | 1.577620423 |
| 0.2888009837 | 0.3113831926 | 0.4313323338 | 0.2813380192 | -1.876362187 | 1.619742822 |
| 0.286539676 | 0.3044167813 | 0.4266257454 | 0.6641591589 | -2.06645474 | 1.582078411 |
| 0.2855764526 | 0.3060176441 | 0.426236716 | -0.2957179066 | -2.000334531 | 1.588992034 |
| 0.2883947657 | 0.3055114316 | 0.4273178327 | 0.03835966411 | -1.926802999 | 1.605078482 |
| 0.2840413159 | 0.3192398087 | 0.4371596186 | 0.2583025429 | -1.981900292 | 1.61817702 |
| 0.287649613 | 0.3055399964 | 0.4275912101 | -0.0111086044 | -1.937453785 | 1.605183847 |
| 0.284002692 | 0.3054547256 | 0.4268627169 | -0.6643558352 | -1.924699319 | 1.629144623 |
| 0.2843549175 | 0.3056697906 | 0.4284665416 | -0.5808716281 | -1.941133916 | 1.566173363 |
| 0.2837846015 | 0.3049445378 | 0.4258510905 | -1.245364748 | -1.962887586 | 1.474733038 |
| 0.2910244656 | 0.3057577462 | 0.4273974614 | 0.1325171161 | -1.943137376 | 1.595074695 |
| 0.2771149988 | 0.2943324315 | 0.4099698046 | 2.341650944 | -2.437217186 | 1.443069074 |
| 0.2842013772 | 0.3093823343 | 0.4303467853 | 0.2980318512 | -1.963768694 | 1.530755979 |
| 0.2826300867 | 0.3213830971 | 0.429487148 | 1.161954569 | -1.981598983 | 1.113512111 |
| 0.2785894029 | 0.299703053 | 0.4192957027 | 1.449852332 | -1.982578539 | 1.651637239 |
| 0.2843618767 | 0.305422708 | 0.4310207584 | -0.2608897144 | -1.945420426 | 1.596903882 |
| 0.2804028226 | 0.2995020894 | 0.4188037929 | 2.092797895 | -2.113041326 | 1.550182151 |
| 0.2841890646 | 0.3062299352 | 0.4272953836 | -0.7602106192 | -1.906233625 | 1.651108125 |
| 0.2825755636 | 0.3049438451 | 0.4333284719 | -0.7654856399 | -1.988786131 | 1.547193071 |
| 0.2869473155 | 0.3071970805 | 0.4271273505 | -0.3329274017 | -1.998501203 | 1.561513955 |
| 0.2854182163 | 0.3054794279 | 0.4275500549 | -0.0536155019 | -1.959923462 | 1.604677196 |
| 0.2880682293 | 0.3098807608 | 0.4442583019 | 0.8049956404 | -2.012785372 | 1.404416494 |
| 0.2782176081 | 0.2991773638 | 0.4187206952 | -1.653485755 | -2.100689631 | 1.558852006 |
| 0.2851572038 | 0.3032350336 | 0.4259161893 | 0.9522526233 | -1.812403888 | 1.59174652 |
| 0.2846373594 | 0.3080279436 | 0.4237004511 | 1.182899903 | -1.782605057 | 1.787142538 |
| 0.283787827 | 0.3006667126 | 0.4202920029 | -1.956854619 | -1.557993702 | 1.37072341 |
| 0.2867346566 | 0.3090352091 | 0.4352897345 | -1.193732258 | -2.130193387 | 1.327340406 |
| 0.2817306732 | 0.3039627316 | 0.4241416777 | 0.6958537923 | -2.001362842 | 1.589747391 |
| 0.287661497 | 0.3090350555 | 0.4410925136 | -0.6008466616 | -1.8335024 | 1.671112956 |
| 0.2832259587 | 0.302325578 | 0.4226490972 | -2.007438672 | -2.065497972 | 1.564272435 |
| 0.2896210718 | 0.3054423119 | 0.4273500292 | -0.3934027254 | -1.845268173 | 1.586194132 |
| 0.2845484413 | 0.3055970173 | 0.4275835939 | 0.08772806887 | -1.967377471 | 1.603126332 |
| 0.291271005 | 0.3049657077 | 0.4344671198 | -1.392996729 | -2.244056814 | 1.392547951 |
| 0.2848571836 | 0.3052458747 | 0.4270021944 | -0.4559615879 | -1.915658967 | 1.590029284 |
| 0.2832988961 | 0.3165184406 | 0.4446910188 | -1.196565999 | -2.106418765 | 1.186819609 |
| 0.2836709487 | 0.305616476 | 0.4295243633 | -0.1385365455 | -1.968746678 | 1.618672512 |
| 0.2822667176 | 0.305643336 | 0.4272466508 | 0.3621151045 | -1.995625519 | 1.595977366 |
| 0.2937860241 | 0.304863414 | 0.4173304531 | 1.376224343 | -2.391621986 | 1.390819696 |
| 0.2825557558 | 0.3037747206 | 0.424915967 | -1.257618536 | -1.974812266 | 1.511429065 |
| 0.2829608619 | 0.3043767692 | 0.4330162049 | 0.5050256621 | -1.989001818 | 1.613285814 |
| 0.2962656177 | 0.3096298027 | 0.4277329538 | -1.250981897 | -1.482637752 | 1.787794559 |
| 0.2839838856 | 0.3074889642 | 0.429906514 | -0.0950242927 | -1.962577533 | 1.60904771 |
| 0.2752307293 | 0.2962715772 | 0.4142789167 | 2.02010801 | -1.995020299 | 1.562307662 |
| 0.283799544 | 0.3111895303 | 0.4290204835 | 0.6866522834 | -1.9010314 | 1.742183797 |

| | | | | | |
|---|---|---|---|---|---|
| 0.2845275727 | 0.3058963739 | 0.4291427723 | 0.4736440753 | -1.981455189 | 1.620005492 |
| 0.2831473624 | 0.3032412617 | 0.4243591538 | -0.7385991882 | -2.055271637 | 1.652013685 |
| 0.3649984522 | 0.4252489645 | 0.5846228207 | -0.9299684024 | 2.781530693 | 0.4161552399 |
| 0.3688014528 | 0.4235110302 | 0.5914312834 | -0.4411615223 | 2.725579607 | 0.5471256583 |
| 0.375480604 | 0.4345344822 | 0.6059987829 | 0.4823639393 | 2.799961048 | 0.6899634414 |
| 0.3700318146 | 0.4381382192 | 0.6014781977 | -0.07198233514 | 2.748883664 | 0.5869699874 |
| 0.3517763846 | 0.4035327213 | 0.5618146667 | 2.034380962 | 2.740343506 | 0.4383528578 |
| 0.3850964403 | 0.4246297227 | 0.6145313144 | -1.099647316 | 2.272553405 | 0.3702086558 |
| 0.3673085329 | 0.4399856622 | 0.5927012704 | -0.6209325262 | 2.77804233 | 0.3838245313 |
| 0.3728707836 | 0.4227745928 | 0.5909887721 | -0.02315555161 | 2.725938982 | 0.5888428697 |
| 0.3715209047 | 0.4228223385 | 0.5909647809 | 0.05167121379 | 2.726515846 | 0.5895553874 |
| 0.3671530716 | 0.4201152358 | 0.5869738014 | 0.738505318 | 2.810056066 | 0.5658824221 |
| 0.3640015494 | 0.4116395655 | 0.5760674833 | -1.464755662 | 2.553633819 | 0.6251478724 |
| 0.3714985632 | 0.424354809 | 0.6036932168 | 0.2487427912 | 2.762983153 | 0.6101444223 |
| 0.3669254805 | 0.4243053619 | 0.5879920418 | 0.6943609051 | 2.787386058 | 0.6831648436 |
| 0.3683754543 | 0.4422406711 | 0.6108346914 | 0.4829969153 | 2.779736584 | 0.7089298058 |
| 0.3686782692 | 0.4230732561 | 0.5956208273 | 0.4189999744 | 2.775017458 | 0.6204981948 |
| 0.3717122323 | 0.4306164779 | 0.5917942031 | 0.8727357119 | 2.890354284 | 0.7818662701 |
| 0.3551549627 | 0.410889124 | 0.5690221672 | 1.78887195 | 2.943220116 | 0.8687385852 |
| 0.366544561 | 0.4244801484 | 0.5992077352 | 0.8309914826 | 2.822585344 | 0.7185185134 |
| 0.3812550053 | 0.4413051959 | 0.6026443564 | 0.4477372343 | 2.546053266 | 0.6959920933 |
| 0.3701309215 | 0.4087729647 | 0.5812965997 | -1.704789454 | 3.209383183 | 0.7236191219 |
| 0.3778529827 | 0.4216785326 | 0.59725118 | 0.6410065216 | 2.838592643 | 0.6368939242 |
| 0.3660316168 | 0.4194663335 | 0.6068098831 | 0.8243960996 | 2.74747479 | 0.5604835811 |
| 0.372533679 | 0.4194706666 | 0.5862359137 | 0.8003786812 | 2.873994301 | 0.5733041691 |
| 0.3678782254 | 0.4243901788 | 0.5898893657 | -0.4931256114 | 2.759224286 | 0.6443308389 |
| 0.4131704544 | 0.4521733096 | 0.6127480438 | 1.030551016 | 1.969937166 | 0.1526223395 |
| 0.3543442915 | 0.4098117467 | 0.569696489 | 1.852668169 | 2.760760858 | 0.3334588951 |
| 0.3734751541 | 0.4216538349 | 0.5895706448 | -0.4545983744 | 2.795421614 | 0.5897405962 |
| 0.3711399746 | 0.4205710805 | 0.5876548766 | -0.7280039543 | 2.849566887 | 0.6321014075 |
| 0.3935901004 | 0.4234633066 | 0.6035226842 | 0.5243488902 | 2.766226857 | 0.6408343808 |
| 0.3665626637 | 0.4102554196 | 0.5740608603 | -1.603433621 | 2.413782524 | 0.5085907244 |
| 0.366383522 | 0.4199803491 | 0.5873414301 | 1.027937131 | 2.884826093 | 0.4735989148 |
| 0.3684501685 | 0.4288214884 | 0.5943757274 | -0.5847120853 | 2.799057313 | 0.6920672708 |
| 0.3609074213 | 0.4145443068 | 0.5783445248 | -1.3324099 | 2.804558976 | 0.5042275254 |
| 0.3573574485 | 0.4141860288 | 0.5777282687 | -1.880030286 | 3.121287122 | 0.9983959314 |
| 0.3581702354 | 0.4126704374 | 0.5852027463 | 1.605358087 | 2.72936523 | 0.4031587296 |
| 0.3774518609 | 0.4283508003 | 0.5992872374 | -0.2158361988 | 2.643636676 | 0.6165349283 |
| 0.3734687365 | 0.4275519992 | 0.590036576 | -0.6584458932 | 2.840382771 | 0.4625120831 |
| 0.3679093061 | 0.4246680974 | 0.5891301824 | 0.6934090082 | 2.704791616 | 0.6865400428 |
| 0.3681928604 | 0.4292031634 | 0.5888873643 | 0.5216670125 | 2.731415871 | 0.6795168599 |
| 0.3692490497 | 0.4254477594 | 0.6038399753 | 0.05699135757 | 2.747265065 | 0.5785933578 |
| 0.3685000618 | 0.4208077984 | 0.587840701 | 0.6552531471 | 2.813807553 | 0.5648573755 |
| 0.3688586933 | 0.4275125104 | 0.6153425794 | -0.1875880123 | 2.751645898 | 0.6105912051 |
| 0.3759260982 | 0.4316597383 | 0.5904352909 | 0.9038180773 | 2.486310315 | 0.3543669693 |
| 0.3644485062 | 0.4188537868 | 0.5837873869 | -0.9926472027 | 2.787166494 | 0.5152814296 |

| | | | | | |
|---|---|---|---|---|---|
| 0.3655985234 | 0.4199955442 | 0.5835484185 | 0.997597252 | 2.857382946 | 0.4834904374 |
| 0.3602339544 | 0.4127364179 | 0.5815046043 | -1.393966388 | 2.846377464 | 0.5692184562 |
| 0.3687811466 | 0.4209837945 | 0.5928152733 | 0.5811854537 | 2.802416302 | 0.583867118 |
| 0.3592654092 | 0.4141113513 | 0.5776540694 | -1.465376005 | 2.827075852 | 0.7624701252 |
| 0.3696076843 | 0.4269193419 | 0.5902606349 | 0.2566585913 | 2.728799483 | 0.6201771877 |
| 0.369099648 | 0.4228927171 | 0.5960416115 | -0.624533493 | 2.687017613 | 0.5245986937 |
| 0.3686395707 | 0.4264430634 | 0.5914852258 | 0.2925576931 | 2.750778182 | 0.5429819853 |
| 0.3679273385 | 0.4419072113 | 0.6232106236 | 0.7153493106 | 2.819317486 | 0.7855371012 |
| 0.3682016798 | 0.4232030632 | 0.5902456251 | -0.429045619 | 2.750344676 | 0.6233187201 |
| 0.3720688949 | 0.4325002472 | 0.5926135977 | 0.6938937079 | 2.845220616 | 0.7398782039 |
| 0.3684210075 | 0.4211722637 | 0.5910994074 | -0.5750131331 | 2.71418995 | 0.6079243198 |
| 0.3691067455 | 0.4230278599 | 0.6068849288 | -0.0921512248 | 2.752769682 | 0.5918422963 |
| 0.3677894403 | 0.4230403871 | 0.5885354772 | -0.5498103115 | 2.777816979 | 0.5362758837 |
| 0.3736354787 | 0.4644846334 | 0.63254517 | 0.1592404318 | 2.742311321 | 0.6018933024 |
| 0.3685364745 | 0.4252063665 | 0.6071477974 | -0.4354912355 | 2.734173557 | 0.5298777149 |
| 0.3691306887 | 0.4226939236 | 0.5915334259 | -0.1226542089 | 2.754021003 | 0.5894746387 |
| 0.3678549405 | 0.4207258116 | 0.5878511346 | -0.6250677247 | 2.722136775 | 0.6019864291 |
| 0.3834996424 | 0.4190126603 | 0.5881984879 | -0.9073319649 | 2.370146124 | 0.5440066133 |
| 0.3812561551 | 0.4261220007 | 0.5942187068 | -0.2357701883 | 2.602871724 | 0.5532650469 |
| 0.3694505398 | 0.4243526657 | 0.5986429611 | 0.1093504814 | 2.74223048 | 0.5768804633 |
| 0.3711317577 | 0.4287696834 | 0.612233578 | 0.5813540513 | 2.817231807 | 0.6909945347 |
| 0.3716274012 | 0.4297744526 | 0.5946018339 | 0.3826045991 | 2.782563881 | 0.651655915 |
| 0.3707652842 | 0.4254956441 | 0.591724761 | -0.08260283118 | 2.746152739 | 0.5755345567 |
| 0.3645975221 | 0.4246512897 | 0.5972983964 | 1.097035377 | 2.697732118 | 0.3708022353 |
| 0.3811922549 | 0.4184714123 | 0.5844897258 | -1.007667311 | 2.959735029 | 0.6677841421 |
| 0.3694758217 | 0.4322543734 | 0.5945527258 | -0.2796854511 | 2.766284199 | 0.6353283213 |
| 0.3737236789 | 0.4231705985 | 0.5987995992 | 0.0127572117 | 2.718423698 | 0.5875911685 |
| 0.3775714191 | 0.4252631662 | 0.6099378626 | 0.4314631963 | 2.784529742 | 0.6404898643 |
| 0.366313535 | 0.4198014858 | 0.5853751453 | 0.8708460185 | 2.829830294 | 0.6554008803 |
| 0.3702872772 | 0.4414752847 | 0.6086968777 | 0.880433409 | 2.884048673 | 0.8433179106 |
| 0.3670565362 | 0.4226600882 | 0.587244457 | -0.6820761225 | 2.789078597 | 0.6612920888 |
| 0.3670335407 | 0.4214464984 | 0.5881927704 | -0.6512106289 | 2.766886127 | 0.5466877995 |
| 0.3746094839 | 0.4213172432 | 0.5894598315 | 1.123482505 | 2.43473304 | 0.4143022385 |
| 0.3677336799 | 0.4345818279 | 0.5905333105 | -0.574605138 | 2.782367164 | 0.4301265488 |
| 0.3693123748 | 0.4213178552 | 0.5913572829 | -0.5239985072 | 2.797292042 | 0.5968841882 |
| 0.3579351182 | 0.4086994454 | 0.5650449055 | -1.974774425 | 2.473669708 | 0.2830846521 |
| 0.3602477807 | 0.4125540297 | 0.5695374279 | -1.728793673 | 2.542621268 | 0.3320606097 |
| 0.3794552854 | 0.4384868845 | 0.5936007279 | -0.1495668955 | 2.710379064 | 0.6074393997 |
| 0.3693887661 | 0.4251713216 | 0.592750898 | -0.1628938939 | 2.739888683 | 0.567580412 |
| 0.3683419015 | 0.41900976 | 0.5884815367 | -0.8765312623 | 2.646307906 | 0.55846163 |
| 0.3651574199 | 0.4774986749 | 0.629762066 | -0.9252778571 | 2.756497736 | 0.07465938979 |
| 0.3688942354 | 0.424870476 | 0.5905358891 | 0.586947926 | 2.804672517 | 0.6657063862 |
| 0.367296699 | 0.4153822839 | 0.5804452019 | 1.241740935 | 2.552595543 | 0.5255637757 |
| 0.3686971645 | 0.421714793 | 0.5891993319 | 0.4824116231 | 2.722277889 | 0.5752736946 |
| 0.366039963 | 0.4171100042 | 0.5827425495 | 1.037711346 | 2.881979233 | 0.6141808462 |
| 0.3370933578 | 0.3857036755 | 0.5387293591 | -2.852822815 | 3.274521635 | 0.4243334562 |

| | | | | | |
|---|---|---|---|---|---|
| 0.380114958 | 0.4347437601 | 0.5942007052 | -0.4881959599 | 2.793495361 | 0.6921463021 |
| 0.3733510586 | 0.4294419244 | 0.5913590672 | 0.1605656257 | 2.694401723 | 0.5510720923 |
| 0.3669569583 | 0.4546291317 | 0.5916634422 | 0.7226751772 | 2.733617077 | 0.270679577 |
| 0.3664018262 | 0.4451609871 | 0.5891956233 | -1.229972424 | 2.95931263 | 1.005542052 |
| 0.3690104634 | 0.4215556084 | 0.5910835465 | -0.5159515122 | 2.793368214 | 0.5736624052 |
| 0.365623232 | 0.4171810498 | 0.5841165043 | 1.026591114 | 2.684090616 | 0.5892088739 |
| 0.3664444833 | 0.4274488417 | 0.5884613916 | 0.7700168018 | 2.795471879 | 0.4353002039 |
| 0.3692069821 | 0.4227928755 | 0.5931380162 | 0.04174055169 | 2.748291901 | 0.5881506262 |
| 0.3747561759 | 0.4302836572 | 0.5925605415 | -0.228321214 | 2.668299457 | 0.5351569801 |
| 0.374067554 | 0.4243698287 | 0.595739812 | -0.04559219972 | 2.706440095 | 0.5825152849 |
| 0.3745646887 | 0.4244778251 | 0.6131467693 | -0.7259713709 | 2.859638409 | 0.6921896582 |
| 0.3666967506 | 0.4219819588 | 0.5855002374 | 0.9503846866 | 2.864418559 | 0.7207545035 |
| 0.3690028752 | 0.4239674036 | 0.5907548906 | -0.1557110429 | 2.754678358 | 0.5993636916 |
| 0.3689351991 | 0.4247797079 | 0.5980884915 | 0.1371259708 | 2.751962409 | 0.5995775372 |
| 0.3684131081 | 0.4122766136 | 0.5768122484 | -1.42204684 | 3.061035097 | 0.6278268551 |
| 0.372291742 | 0.4208976269 | 0.5932863335 | 0.9735293507 | 2.92347842 | 0.7106481342 |
| 0.3704457974 | 0.4264460036 | 0.5916416083 | -0.2768398987 | 2.767583824 | 0.6218869099 |
| 0.3661643045 | 0.4195541884 | 0.5898567477 | 0.8451585042 | 2.729464853 | 0.5478789138 |
| 0.3716085096 | 0.4227233495 | 0.5919108192 | -0.1489895009 | 2.713973028 | 0.5915641314 |
| 0.3724844896 | 0.417266313 | 0.5820707696 | -1.084530242 | 2.961145929 | 0.53227642 |
| 0.3691372542 | 0.4216355174 | 0.5893618686 | -0.4708766077 | 2.78743442 | 0.5984248665 |
| 0.378565379 | 0.4236108663 | 0.6000885515 | 1.379909006 | 3.104761704 | 0.8971289855 |
| 0.3702676086 | 0.4246835981 | 0.592077808 | 0.4808014533 | 2.68787078 | 0.6446122965 |
| 0.3740824536 | 0.4401985909 | 0.6223935706 | -0.6154631427 | 2.596658596 | 0.3843899482 |
| 0.364099605 | 0.4201815685 | 0.5918588654 | -1.030886997 | 2.792866306 | 0.7164640893 |
| 0.3625644221 | 0.4188160206 | 0.5800296082 | 1.222116469 | 2.858164131 | 0.4275140863 |
| 0.3869812428 | 0.4264290713 | 0.5928111934 | 0.3747614607 | 2.508494514 | 0.5298408184 |
| 0.371019515 | 0.4180544928 | 0.5863738744 | 1.167324661 | 2.981426025 | 0.7223797121 |
| 0.3690071758 | 0.4249756257 | 0.6140313847 | -0.06473011595 | 2.750834912 | 0.5922703303 |
| 0.3744924347 | 0.4276232097 | 0.5706683383 | 2.227966153 | 1.881294627 | -0.2302889131 |
| 0.3672903976 | 0.4189946345 | 0.5854698054 | 0.8479971731 | 2.843615111 | 0.5785190629 |
| 0.3665513597 | 0.4196072501 | 0.5921555971 | 0.7690861123 | 2.800411615 | 0.5978783963 |
| 0.3734895324 | 0.426589488 | 0.6058627201 | -0.4749033908 | 2.636699317 | 0.5106670182 |
| 0.3627874153 | 0.4175496833 | 0.5803395381 | 1.318695898 | 2.669812268 | 0.775920882 |
| 0.3656204043 | 0.4173334101 | 0.5834580713 | 1.035721637 | 2.872487296 | 0.6330099514 |
| 0.3102547099 | 0.33230716 | 0.4625182218 | 1.6440789 | 0.8911520327 | -0.9086472693 |
| 0.3078962377 | 0.3292623542 | 0.4602514249 | 1.937298752 | 0.9243467177 | -0.9236102051 |
| 0.3189509169 | 0.3371930848 | 0.471887544 | -0.5699347208 | 0.6825808423 | -1.039388911 |
| 0.3162746939 | 0.3388996743 | 0.4743888764 | -0.01615654923 | 0.7740129937 | -1.030920167 |
| 0.3162013786 | 0.3389685459 | 0.4722024307 | -0.196016968 | 0.7697047464 | -1.01670797 |
| 0.3183207572 | 0.3391976172 | 0.4795201692 | -0.8827561233 | 0.6286720305 | -1.152313945 |
| 0.3077054103 | 0.3314843539 | 0.4606476383 | 1.88661053 | 0.7607352495 | -0.8198325171 |
| 0.3101412677 | 0.3280476919 | 0.4587443225 | -1.976350727 | 1.075936774 | -1.055216489 |
| 0.290441134 | 0.3067416986 | 0.4291629931 | -3.703468412 | 1.445499781 | -0.9536180651 |
| 0.3152451373 | 0.3373398451 | 0.4712561479 | -0.6549777646 | 0.7605023086 | -1.003921555 |
| 0.3177046765 | 0.3372798583 | 0.4705033396 | 0.9091213447 | 0.905457088 | -0.9537715984 |

| | | | | | |
|---|---|---|---|---|---|
| 0.3116713552 | 0.332777661 | 0.4699124571 | -1.424884891 | 0.7110540549 | -1.072522117 |
| 0.3157978988 | 0.3385806332 | 0.47335201 | 0.4389551622 | 0.7666869166 | -0.9986935232 |
| 0.3122139988 | 0.3355271563 | 0.46650805 | -1.276407436 | 0.7829332346 | -1.172864289 |
| 0.3157732211 | 0.3376089502 | 0.4747985673 | -0.430953788 | 0.7710246462 | -1.043210386 |
| 0.3135165988 | 0.3328060325 | 0.46338273 | -1.57834826 | 0.5499351571 | -1.187024911 |
| 0.3137889948 | 0.3407072068 | 0.4689397883 | -1.005260237 | 0.7586361578 | -1.203657108 |
| 0.3151128755 | 0.3366422397 | 0.471960387 | 0.7536219159 | 0.8096472694 | -1.020899511 |
| 0.3215837948 | 0.3433286055 | 0.4803519178 | 0.2192278093 | 0.7206555052 | -0.980020465 |
| 0.3256290443 | 0.3389735692 | 0.4693838205 | -0.8895826445 | 0.9845340792 | -0.9133889225 |
| 0.3176509133 | 0.3353639781 | 0.4704402662 | -1.657243049 | 0.398485785 | -1.311755186 |
| 0.3165369364 | 0.3382154145 | 0.4740369734 | 0.05940311119 | 0.7704112374 | -1.03228535 |
| 0.323020812 | 0.3381327024 | 0.4719175593 | -0.5309411171 | 0.6438134046 | -1.071523044 |
| 0.3161493293 | 0.3425476638 | 0.4727105275 | 0.3282983607 | 0.7625237137 | -1.075706341 |
| 0.3338340574 | 0.3423354757 | 0.4818539758 | -0.3553081007 | 0.8478903543 | -0.9635565881 |
| 0.3139693831 | 0.339772416 | 0.4679678961 | 1.137810661 | 0.6912359555 | -1.226460892 |
| 0.3180762848 | 0.3374802688 | 0.4732090404 | 0.4669418647 | 0.7121268453 | -1.033905605 |
| 0.3163326153 | 0.3356399942 | 0.4684232896 | -1.029926979 | 0.9087608803 | -0.9923430703 |
| 0.3202923683 | 0.3346027298 | 0.4730144537 | -1.762848399 | 0.2941027345 | -1.336152798 |
| 0.3131228116 | 0.3271327031 | 0.4578486812 | -2.146066572 | 0.303719021 | -1.218417317 |
| 0.3184543689 | 0.3383537557 | 0.471687673 | -0.4091281106 | 0.7157741683 | -1.060822112 |
| 0.3121722688 | 0.339643673 | 0.4692408326 | -1.28554402 | 0.7964379718 | -0.7905560913 |
| 0.3176298933 | 0.337308397 | 0.4711907377 | -0.5622322861 | 0.8364088165 | -1.02362628 |
| 0.3138622279 | 0.33622881 | 0.477034345 | -1.697477146 | 1.070152839 | -0.7305040853 |
| 0.307145959 | 0.3276022875 | 0.4665359567 | 2.033137472 | 0.6490735324 | -1.121845733 |
| 0.3201343105 | 0.3379952906 | 0.4728797477 | -0.2528755989 | 0.7242048086 | -1.028433459 |
| 0.3097700068 | 0.3319781134 | 0.4622816899 | 1.665147555 | 0.7189374366 | -0.9168192561 |
| 0.3161873472 | 0.3389746594 | 0.473200252 | 0.4933418739 | 0.7442972398 | -0.9844501999 |
| 0.3083453236 | 0.332662197 | 0.4597168569 | -1.870249331 | 0.9052098923 | -1.32635682 |
| 0.3153354078 | 0.3482767732 | 0.485705775 | 0.6434009634 | 0.795477137 | -1.166996799 |
| 0.3207215984 | 0.3372973048 | 0.4742379649 | 0.5622813167 | 0.8646710334 | -1.024453822 |
| 0.3162016009 | 0.3389645604 | 0.4749969634 | 0.1016702388 | 0.7735109164 | -1.03940016 |
| 0.30686534 | 0.332918691 | 0.462873261 | 2.028788991 | 0.6828391525 | -1.414180386 |
| 0.3145859173 | 0.3363302758 | 0.4713025533 | -0.839310119 | 0.8004726327 | -1.064641337 |
| 0.3143353362 | 0.3336625669 | 0.472757095 | 1.322011668 | 0.9417966566 | -1.002788395 |
| 0.3076602652 | 0.3288689765 | 0.4603568193 | 1.904845481 | 0.8579699264 | -1.119873391 |
| 0.3062027671 | 0.3273025253 | 0.459237359 | 2.055342821 | 0.8445744614 | -1.099888372 |
| 0.3104900511 | 0.3316674611 | 0.4632777661 | 1.578467362 | 0.8618193139 | -1.015527425 |
| 0.3083663084 | 0.3330465012 | 0.4647915159 | 1.81652062 | 0.8422697021 | -0.7872550123 |
| 0.3164481535 | 0.3437909032 | 0.470712099 | 0.6538744385 | 0.8322951064 | -0.8872788653 |
| 0.3166616699 | 0.3375671578 | 0.4712565986 | -0.7119930441 | 0.8464814537 | -0.9853360817 |
| 0.3177672068 | 0.3378414249 | 0.4755984688 | 1.471592217 | 1.073796283 | -0.7720987876 |
| 0.3189385668 | 0.3381737164 | 0.4774472716 | 0.04047099663 | 0.7725346824 | -1.033199142 |
| 0.317111503 | 0.3404729782 | 0.47385226 | -0.4605066333 | 0.728194722 | -1.088099675 |
| 0.3163985045 | 0.3379990264 | 0.4723708051 | -0.5631537466 | 0.7294436599 | -1.073235 |
| 0.3154864434 | 0.3381432325 | 0.4710800573 | -0.6038298925 | 0.7956170362 | -1.082103063 |
| 0.3161876754 | 0.3410198363 | 0.4767819092 | -0.1233476026 | 0.7741933903 | -1.041971426 |

| | | | | | |
|---|---|---|---|---|---|
| 0.3158062564 | 0.3384702604 | 0.4730384706 | 0.50391594 | 0.7562187798 | -0.9929478975 |
| 0.3126455727 | 0.33365252 | 0.4777109958 | 1.665642092 | 1.018978426 | -0.8329507191 |
| 0.3145942062 | 0.3379274015 | 0.4706596724 | -1.45398536 | 0.9955152953 | -0.7760959188 |
| 0.3164513461 | 0.3363464045 | 0.4697473449 | -0.8349564377 | 0.6804738663 | -1.064199402 |
| 0.3199748577 | 0.3350621679 | 0.4693547315 | -1.174386666 | 0.5178110886 | -1.116651742 |
| 0.3268115003 | 0.338781463 | 0.4738811779 | -0.7198817171 | 0.556455096 | -1.112531924 |
| 0.3130476625 | 0.3396916336 | 0.4676210662 | -1.150305092 | 0.8095786327 | -0.8305340242 |
| 0.313906375 | 0.335585932 | 0.4691420548 | 0.9758827896 | 0.7960064225 | -1.024816861 |
| 0.3175270504 | 0.3381779767 | 0.4723280051 | 0.5362549511 | 0.8310825071 | -0.9954252445 |
| 0.3007263187 | 0.3150778874 | 0.4480061113 | -3.116796939 | 1.452406955 | -0.9511340206 |
| 0.318888529 | 0.3419968165 | 0.4842358344 | 0.6987156176 | 0.6586976123 | -1.144556468 |
| 0.315606736 | 0.3352458494 | 0.4636721218 | 1.541769537 | 0.4970792198 | -1.261994576 |
| 0.3167645779 | 0.3420042562 | 0.4814015703 | 0.3407167593 | 0.7478747404 | -1.076275975 |
| 0.317001898 | 0.3400805744 | 0.4861905085 | 0.6548653977 | 0.8425496041 | -1.117629318 |
| 0.3143930327 | 0.3361970738 | 0.4714812964 | 0.8654099446 | 0.7662267655 | -1.016309097 |
| 0.310445605 | 0.3353682786 | 0.464174573 | 1.559926342 | 0.8377167952 | -0.8132541606 |
| 0.3155342243 | 0.3426490232 | 0.4802760968 | 1.071157597 | 0.9003694521 | -0.7960184833 |
| 0.3152464824 | 0.3396070895 | 0.4700168829 | -0.7888227827 | 0.8237249479 | -0.9217852155 |
| 0.2982673497 | 0.3196215637 | 0.4453391648 | 2.804809512 | 0.7441053062 | -0.8864990939 |
| 0.3299990001 | 0.3415838267 | 0.4771428368 | 0.3297258444 | 0.6463720631 | -1.073097608 |
| 0.3136003986 | 0.3333745646 | 0.4658998713 | 1.349449692 | 0.6340121531 | -1.016445029 |
| 0.3161546935 | 0.3365219611 | 0.4719263892 | -0.7918535576 | 0.8508461467 | -1.017097445 |
| 0.3136385974 | 0.3365062614 | 0.4653389451 | -1.387579325 | 0.6185032541 | -1.238940139 |
| 0.3184962846 | 0.3422238779 | 0.4724534721 | 0.9364128893 | 0.9242994131 | -0.8404854467 |
| 0.3203805821 | 0.3391542102 | 0.4742690509 | -0.0197708259 | 0.7669837494 | -1.029695017 |
| 0.3137071246 | 0.3356487371 | 0.4668719983 | 1.21485511 | 0.8827207421 | -0.928043568 |
| 0.3136965099 | 0.3316821682 | 0.4632073545 | 1.581083924 | 1.017754603 | -1.009066435 |
| 0.3152538965 | 0.3546006371 | 0.4865385056 | -0.6489409337 | 0.7908817631 | -1.188635242 |
| 0.3127945495 | 0.3325615907 | 0.4639530455 | -1.513248833 | 0.9600426111 | -1.133336283 |
| 0.3131286875 | 0.3354455488 | 0.4677219286 | -1.148945255 | 0.818833761 | -0.9598152693 |
| 0.3152169125 | 0.3370788518 | 0.4728185343 | 0.6380030004 | 0.7809765743 | -1.021576214 |
| 0.3141771346 | 0.335823378 | 0.4691512749 | -0.9316449454 | 0.7544591338 | -1.060452145 |
| 0.3241130745 | 0.3373171368 | 0.4712886062 | -0.5701516877 | 0.8862325722 | -1.021254963 |
| 0.3213478525 | 0.3389683012 | 0.4718444952 | -0.3962144238 | 0.8346066176 | -0.9967876694 |
| 0.312574195 | 0.3372983806 | 0.4699117687 | 1.22301701 | 0.7639564719 | -1.204567639 |
| 0.3157683193 | 0.3554102153 | 0.4768088693 | -0.4565392283 | 0.7842249369 | -0.8410189758 |
| 0.3162255538 | 0.3380711989 | 0.4725416382 | -0.1986566543 | 0.7688654889 | -1.029336641 |
| 0.2847312024 | 0.3035189754 | 0.4294742914 | -3.897454639 | 1.165632024 | -1.23565097 |
| 0.3167014542 | 0.3374557253 | 0.471313003 | 0.5173284956 | 0.7287161657 | -1.023113246 |
| 0.3142172608 | 0.336603222 | 0.470569397 | -0.9505159679 | 0.8155828387 | -0.9680158041 |
| 0.308093648 | 0.3302634173 | 0.4672503497 | -1.835755189 | 0.7997378172 | -1.194397325 |
| 0.3177088736 | 0.3386484938 | 0.4700277573 | -0.8687585859 | 0.6441994441 | -1.140198068 |
| 0.3138848107 | 0.3334361563 | 0.4663888857 | -1.374813351 | 0.6135196098 | -1.110771564 |
| 0.3183313409 | 0.339341899 | 0.4877269741 | 0.7680021306 | 0.6539314805 | -1.130117219 |
| 0.3103564776 | 0.3314191554 | 0.4629153492 | -1.658731762 | 0.668107292 | -1.147239812 |
| 0.3158151064 | 0.3347103866 | 0.4698275009 | -1.125841229 | 0.627304957 | -1.041340429 |

| | | | | | |
|---|---|---|---|---|---|
| 0.3184595278 | 0.3430133359 | 0.4723152632 | 0.1076432914 | 0.7812280389 | -1.001394896 |
| 0.3162136361 | 0.3350410278 | 0.4689964721 | -1.303280663 | 0.9847568552 | -1.165029764 |
| 0.3055055158 | 0.3258588448 | 0.4592689109 | -2.529553663 | 0.3824563403 | -1.456595776 |
| 0.3161653692 | 0.3390891634 | 0.4722161232 | 0.1554302228 | 0.7736415977 | -1.04396242 |
| 0.3181623734 | 0.3403580543 | 0.4803552317 | -0.1765920164 | 0.7889799258 | -1.00677662 |
| 0.3116409931 | 0.3325538682 | 0.4639945535 | -1.52859163 | 0.6556483945 | -0.9603634639 |
| 0.322938619 | 0.3373241126 | 0.4727648725 | -0.5242744214 | 0.8693573457 | -1.036257849 |
| 0.3108038046 | 0.3318970864 | 0.4636203853 | -1.604007108 | 0.9055056207 | -0.9626580615 |
| 0.3244874006 | 0.3376369065 | 0.4801446815 | 0.8297914037 | 0.9583185709 | -0.960152615 |
| 0.3163402179 | 0.3377555509 | 0.4715689385 | -0.4454896748 | 0.8012613661 | -1.05191421 |
| 0.3187055848 | 0.3516742244 | 0.494860881 | -0.1506959121 | 0.7488457882 | -1.036184975 |
| 0.311014654 | 0.3325712864 | 0.4669256919 | 1.461684378 | 0.7706902033 | -1.088047185 |
| 0.3156181595 | 0.3408741323 | 0.4732857935 | -0.4930242961 | 0.773936207 | -0.9559567615 |
| 0.3292982176 | 0.3432359361 | 0.472022086 | -0.4808612753 | 0.8801455736 | -0.9310261736 |
| 0.3105278834 | 0.332187402 | 0.464531435 | -1.539827024 | 0.7508587369 | -1.120054446 |
| 0.3137672615 | 0.335465565 | 0.4753888232 | -1.00297174 | 0.7952919966 | -1.021598597 |
| 0.326261738 | 0.3435631691 | 0.4735247307 | -0.2743462059 | 0.8181868632 | -0.9682562312 |
| 0.3085816142 | 0.3310435195 | 0.4627315275 | -1.779478608 | 0.8111497442 | -0.9068802783 |
| 0.3159700442 | 0.3378527177 | 0.4725115607 | 0.351081592 | 0.7809472393 | -1.04476738 |
| 0.3174793146 | 0.3450613819 | 0.4765245043 | 0.1665168508 | 0.7859794578 | -0.9800776524 |
| 0.3012798119 | 0.3230490373 | 0.452525182 | 2.56279297 | 0.6883764735 | -0.8491901995 |
| 0.315742483 | 0.3357705892 | 0.46960797 | 0.936667175 | 0.8719960461 | -1.026989414 |
| 0.3132965366 | 0.3366683445 | 0.4695085985 | -0.5401624641 | 1.386802607 | -0.4472454507 |
| 0.3095261827 | 0.3314197708 | 0.463707161 | 1.808727735 | 1.628645149 | -0.2476229949 |
| 0.317880021 | 0.3366622649 | 0.4718251962 | -0.3876582122 | 1.346776122 | -0.4055112059 |
| 0.3135369255 | 0.3372283283 | 0.4727334985 | -0.2416752419 | 1.419124529 | -0.3889619668 |
| 0.3124221261 | 0.3364682146 | 0.4688833397 | 0.641458477 | 1.417513358 | -0.4633892289 |
| 0.3169693027 | 0.3380315092 | 0.4788925085 | -1.024300524 | 1.271678287 | -0.5162252438 |
| 0.3130649302 | 0.3378862651 | 0.4704888837 | 0.4653815406 | 1.403369686 | -0.3553447988 |
| 0.3037159302 | 0.3229077134 | 0.4521312306 | -2.242843781 | 1.748429421 | -0.4569576564 |
| 0.3062790385 | 0.3239775151 | 0.4537216271 | -2.366626763 | 1.879847771 | -0.2827839591 |
| 0.3141125948 | 0.3378374392 | 0.472503735 | -0.1316934952 | 1.422092914 | -0.3887798885 |
| 0.3131513204 | 0.3328546477 | 0.4650083708 | 1.561964048 | 1.694068284 | -0.2557318158 |
| 0.314666904 | 0.3368957681 | 0.476448247 | -0.3133117349 | 1.408729914 | -0.3976846858 |
| 0.3133405802 | 0.3372894301 | 0.4714484751 | 0.2573198942 | 1.403139098 | -0.3874926412 |
| 0.3050322016 | 0.3281668567 | 0.4565582576 | -2.248879749 | 1.534223198 | -0.6011831678 |
| 0.3138461159 | 0.3367877507 | 0.4733497844 | -0.0100460843 | 1.416362786 | -0.4043759479 |
| 0.3167694339 | 0.3375204752 | 0.4702576511 | -0.7068517119 | 1.330460863 | -0.4503448122 |
| 0.3140619094 | 0.3413781947 | 0.4709945865 | -0.4538058262 | 1.430340392 | -0.4639733755 |
| 0.3138860881 | 0.3366936937 | 0.472244881 | 0.124543435 | 1.418924171 | -0.4016116177 |
| 0.3148299151 | 0.3369476555 | 0.4715775361 | -1.354875669 | 1.664180926 | -0.6375075811 |
| 0.3117880082 | 0.3263776045 | 0.4513337255 | -2.224893343 | 1.978908505 | -0.1085934985 |
| 0.3156069417 | 0.3341240993 | 0.4687256004 | -1.545400822 | 1.10924572 | -0.6032613981 |
| 0.3140767901 | 0.3367640406 | 0.4730061501 | 0.1441318787 | 1.412038456 | -0.3997381845 |
| 0.3193493139 | 0.3344136144 | 0.4677899389 | -0.794503983 | 1.154523056 | -0.4997726821 |
| 0.3141618606 | 0.3412567986 | 0.471676216 | -0.4662338962 | 1.449754013 | -0.3150319385 |

| | | | | | |
|---|---|---|---|---|---|
| 0.3319917296 | 0.3421537702 | 0.4807961431 | 0.2808204486 | 1.253582523 | -0.444404271 |
| 0.3145592851 | 0.3407158607 | 0.4701504233 | 0.6243009244 | 1.344019325 | -0.509072803 |
| 0.3153859666 | 0.3360181854 | 0.4714831283 | -0.4013111543 | 1.451413634 | -0.4069956303 |
| 0.3167449982 | 0.3369305143 | 0.4704156606 | -0.5214829839 | 1.487083902 | -0.3665062455 |
| 0.3259175501 | 0.3411367411 | 0.4820589583 | -0.8727631186 | 1.171050249 | -0.5227678854 |
| 0.3141765353 | 0.3291722907 | 0.4607534334 | -1.949971715 | 1.008064698 | -0.5546103845 |
| 0.317994787 | 0.3385948377 | 0.471930514 | 0.6668896778 | 1.540784074 | -0.3206611638 |
| 0.3125709725 | 0.3391897073 | 0.4712760493 | -0.5344178135 | 1.417480929 | -0.3291306665 |
| 0.3134753121 | 0.3330780864 | 0.4654242269 | -1.668541197 | 1.709338122 | -0.314710003 |
| 0.3179644891 | 0.3413151781 | 0.4838517688 | -0.0739065621 | 1.409888845 | -0.3866701759 |
| 0.312756169 | 0.3346407 | 0.4769968897 | 1.117143564 | 1.377151941 | -0.4264173327 |
| 0.3122959443 | 0.3292269175 | 0.4605672776 | 1.828484353 | 1.785645972 | -0.4348006741 |
| 0.3070750372 | 0.3303878852 | 0.4610786502 | 1.507088518 | 1.335040921 | -0.3208882011 |
| 0.3143347005 | 0.3381320352 | 0.4721393909 | 0.165414857 | 1.402623699 | -0.3871009272 |
| 0.3106268826 | 0.3372243981 | 0.4659257673 | 0.9327789226 | 1.342209369 | -0.2786550735 |
| 0.3041047689 | 0.3344838617 | 0.467777752 | 1.87786062 | 1.52739741 | -0.8473663544 |
| 0.3101234901 | 0.328652586 | 0.46120396 | 1.398132655 | 1.587029451 | -0.4481646562 |
| 0.3118513206 | 0.3361622287 | 0.4735735338 | 0.6666185269 | 1.386618683 | -0.479222152 |
| 0.2905294388 | 0.315195846 | 0.4382323163 | 3.656195775 | 1.451560239 | -1.022282432 |
| 0.3141065998 | 0.3368238507 | 0.4726029084 | 0.3404863832 | 1.418736392 | -0.3913574431 |
| 0.3117659805 | 0.3309352318 | 0.4688928106 | 2.032449421 | 1.79621408 | -0.2796920341 |
| 0.3143208555 | 0.3368457556 | 0.4727572532 | 0.2170864461 | 1.430958435 | -0.4054286032 |
| 0.3144556394 | 0.3370354512 | 0.4733313321 | 0.6632371833 | 1.468951416 | -0.4080404447 |
| 0.3130631391 | 0.3344160502 | 0.4671170339 | 1.644097719 | 1.615501721 | -0.3052979865 |
| 0.3135978567 | 0.3398254349 | 0.4746880883 | 0.9604120529 | 1.492742254 | -0.2304304636 |
| 0.3073365773 | 0.335317017 | 0.4604688719 | 1.580470949 | 1.534745888 | -0.1207851238 |
| 0.3151970648 | 0.3374349927 | 0.4715427595 | 0.2618297295 | 1.374118598 | -0.4299375079 |
| 0.3189227816 | 0.3410697311 | 0.4799632114 | -0.875663561 | 1.213598833 | -0.5669664998 |
| 0.3121348307 | 0.331061511 | 0.4692594583 | 1.406539825 | 1.627046091 | -0.4099042438 |
| 0.3150448617 | 0.3391275023 | 0.4736393804 | -0.6765055124 | 1.381067797 | -0.4711355782 |
| 0.3141468478 | 0.3364085691 | 0.4709383552 | 0.3769245685 | 1.437629203 | -0.3901573216 |
| 0.3112788144 | 0.3343006514 | 0.4666339301 | -1.235667939 | 1.509556389 | -0.4918915665 |
| 0.3101342569 | 0.3380118965 | 0.4733207749 | -0.9131113514 | 1.373237908 | -0.5802111494 |
| 0.3137099966 | 0.3384181483 | 0.4735748671 | 0.3481535104 | 1.408919748 | -0.3632207831 |
| 0.3123272454 | 0.3358336365 | 0.4802847717 | 1.039331576 | 1.523716036 | -0.2792923283 |
| 0.3169954737 | 0.3405704983 | 0.4759809514 | 0.2691517524 | 1.361978162 | -0.4420523471 |
| 0.3156630809 | 0.3367496338 | 0.4708980533 | 0.1387099384 | 1.425179539 | -0.4002150763 |
| 0.3203087372 | 0.3373508158 | 0.4726983741 | 0.1500271537 | 1.415990306 | -0.3941949784 |
| 0.32662423 | 0.3390987598 | 0.4742688912 | -0.4843148302 | 1.22491993 | -0.4654336893 |
| 0.3122937176 | 0.3403624568 | 0.4683799954 | -0.9056373387 | 1.461072733 | -0.2321240827 |
| 0.311905103 | 0.3346025099 | 0.4681756354 | 0.8224447449 | 1.438851681 | -0.4047599172 |
| 0.3162583348 | 0.3374348218 | 0.4718693214 | 0.3133645916 | 1.449055031 | -0.3783568937 |
| 0.3163173061 | 0.3319173428 | 0.4725263423 | -1.692840493 | 1.813276681 | -0.2817185659 |
| 0.3012218437 | 0.3267097358 | 0.4620274081 | 2.432826594 | 1.058332862 | -0.9331156616 |
| 0.318500153 | 0.3391586025 | 0.4702983844 | 0.597842929 | 1.308876777 | -0.4691402536 |
| 0.3145643604 | 0.3408694981 | 0.4796146143 | 0.4213548195 | 1.398188909 | -0.4589168587 |

| | | | | | |
|---|---|---|---|---|---|
| 0.3151849013 | 0.3394352065 | 0.4854472305 | 0.3452530271 | 1.444014968 | -0.4504959776 |
| 0.3115359545 | 0.3346980887 | 0.4706098135 | -0.6768787397 | 1.400103206 | -0.4435064053 |
| 0.3139773384 | 0.340585421 | 0.4714249897 | -0.0389032200 | 1.414360874 | -0.4056203905 |
| 0.3155791244 | 0.3443117083 | 0.4837498652 | 0.6522237111 | 1.497949804 | -0.2472124462 |
| 0.3129153625 | 0.3381652715 | 0.4683447697 | -0.9638482915 | 1.497066519 | -0.2584790931 |
| 0.3068906868 | 0.3291921797 | 0.45896296 | 2.096088371 | 1.471637204 | -0.2224336099 |
| 0.3267445276 | 0.3392251401 | 0.474232749 | -0.3674270695 | 1.453573692 | -0.3566967248 |
| 0.3153297278 | 0.3363471581 | 0.4706087992 | 0.637626261 | 1.364283483 | -0.3832382361 |
| 0.3146200274 | 0.3366903881 | 0.4715863535 | 0.05276728069 | 1.410394675 | -0.4037155821 |
| 0.3148866204 | 0.3387805338 | 0.4696338203 | -0.6301698374 | 1.344449106 | -0.4851108593 |
| 0.3125107018 | 0.3357923764 | 0.4635745322 | 2.160843036 | 1.901394379 | 0.0743964739 |
| 0.3154546151 | 0.3346811893 | 0.4688777997 | -0.8567294905 | 1.529647483 | -0.3609842659 |
| 0.3144695395 | 0.3366916763 | 0.4686649247 | 1.000696775 | 1.550008296 | -0.2921496466 |
| 0.3156966039 | 0.3339127337 | 0.4666083367 | 1.431319727 | 1.698445143 | -0.3344869898 |
| 0.3148577397 | 0.3541202188 | 0.4855234579 | -0.712496141 | 1.483812924 | -0.5787599216 |
| 0.3151755028 | 0.3350842083 | 0.4675742569 | -1.539758644 | 1.700523085 | -0.432517946 |
| 0.3116121964 | 0.3346370882 | 0.4673928334 | -1.01837543 | 1.471370624 | -0.3422815952 |
| 0.3112598203 | 0.3343452264 | 0.4687221289 | 0.7371474034 | 1.392394027 | -0.4036687763 |
| 0.3131719167 | 0.3364797803 | 0.4699064013 | 0.3075609414 | 1.416284473 | -0.3943774597 |
| 0.3131173443 | 0.329002458 | 0.4598881531 | -1.475346438 | 1.67113773 | -0.3954433998 |
| 0.3104766102 | 0.3291256685 | 0.458439297 | -1.796154335 | 1.740438386 | -0.2577252307 |
| 0.3137322783 | 0.3404169366 | 0.4738635279 | -0.1247854507 | 1.417298739 | -0.3820255386 |
| 0.3076340274 | 0.3499359884 | 0.4675785093 | -1.326569893 | 1.428951958 | 0.0219293935 |
| 0.3075660554 | 0.3305085203 | 0.4623923531 | -1.286234305 | 1.362696896 | -0.4355492025 |
| 0.301752651 | 0.3233789283 | 0.4567583237 | -2.255202518 | 1.60705657 | -0.5106220053 |
| 0.3149885768 | 0.3365923644 | 0.4707120354 | -0.0354013444 | 1.412759689 | -0.4054149223 |
| 0.3146844826 | 0.3380691449 | 0.4722413011 | -0.5729971993 | 1.470244349 | -0.3385154774 |
| 0.3119330141 | 0.3357475515 | 0.4763830681 | -0.8307619765 | 1.413421295 | -0.4738262358 |
| 0.3140137759 | 0.3356517575 | 0.4665306865 | 1.059888543 | 1.561048534 | -0.2846139299 |
| 0.315763095 | 0.3367940465 | 0.4713628196 | -0.4027008636 | 1.373598384 | -0.4125738076 |
| 0.3171538412 | 0.3398588594 | 0.487804945 | 0.7663566438 | 1.32780504 | -0.490428584 |
| 0.3140944044 | 0.3366902 | 0.4708989818 | -0.5263951162 | 1.389618555 | -0.4283834112 |
| 0.3165660084 | 0.3357665679 | 0.4716221585 | -1.042444246 | 1.310566067 | -0.365297711 |
| 0.3159911848 | 0.3413092382 | 0.4697206051 | 0.6750159694 | 1.508237681 | -0.2723788049 |
| 0.3147259127 | 0.3334390236 | 0.46629797 | -1.451095273 | 1.694485253 | -0.5424623227 |
| 0.3166744746 | 0.3390519134 | 0.4779105818 | -1.287250905 | 1.240713562 | -0.5870629559 |
| 0.314239984 | 0.3378109175 | 0.4709713594 | -0.5590409134 | 1.445298247 | -0.3455304149 |
| 0.3123707589 | 0.334831123 | 0.4752327728 | -1.523665245 | 1.661404036 | -0.1804410359 |
| 0.3138157323 | 0.3350114855 | 0.4685708571 | -1.043847695 | 1.345566365 | -0.3561159794 |
| 0.3202921765 | 0.3365354845 | 0.4712604018 | -0.2304323133 | 1.432750881 | -0.4112241317 |
| 0.3011661854 | 0.3227326237 | 0.4514166101 | -2.373016447 | 1.638178922 | -0.3094227773 |
| 0.3198633508 | 0.3351326034 | 0.4774727085 | 1.003187477 | 1.599093112 | -0.3049849967 |
| 0.3148818515 | 0.3368796722 | 0.4709436653 | -0.2934549106 | 1.442695389 | -0.4077655398 |
| 0.3147250638 | 0.3496019959 | 0.4916806329 | -0.9707920929 | 1.31245872 | -0.6579861947 |
| 0.3078605682 | 0.3305146882 | 0.4644613567 | 1.603743614 | 1.438268381 | -0.477827651 |
| 0.3103054559 | 0.3354222373 | 0.4669801867 | -1.472335112 | 1.476166586 | -0.1895284554 |

| | | | | | |
|---|---|---|---|---|---|
| 0.3226168267 | 0.3362717519 | 0.4624448582 | -2.046965725 | 2.076471419 | 0.0305294045[illegible] |
| 0.3154378253 | 0.3375884792 | 0.4721239666 | -1.142780164 | 1.484754028 | -0.4003912606 |
| 0.3119249557 | 0.3346344041 | 0.4755155114 | -0.8888290442 | 1.443549184 | -0.3891065231 |
| 0.3277088133 | 0.3431181511 | 0.4737385242 | -0.2301421334 | 1.425739066 | -0.3504911837 |
| 0.3100665668 | 0.3343266249 | 0.4675105256 | -1.288693985 | 1.467251849 | -0.2697276309 |
| 0.3135891773 | 0.3366114864 | 0.4708312362 | 0.3768332785 | 1.426043019 | -0.4178548213 |
| 0.3063731533 | 0.3350218722 | 0.4621957828 | -1.815426166 | 1.240263561 | -0.8089469637 |
| 0.3072122435 | 0.3299285392 | 0.4627600795 | 1.699161377 | 1.351296525 | -0.2945581351 |
| 0.3048468888 | 0.3257821369 | 0.4559858115 | 1.944141934 | 1.617769268 | -0.4551599733 |

obe + Blast + VNS + Blast:VNS + (1|Batch))

| Blast:VNS t-value | Microbe Pr(>t | Blast Pr(>t) | VNS Pr(>t) | Blast:VNS Pr(>t | Microbe Adjusted p-valu |
|---|---|---|---|---|---|
| -2.325498388 | 0.0104612000 | 0.0033741515 | 0.0297240381 | 0.0234441675 | 0.2213566899 |
| -2.14894178 | 0.9373908607 | 0.0114265040 | 0.0709555961 | 0.03568337829 | 0.9764488133 |
| -2.313356214 | 0.1033778047 | 0.0053261470 | 0.0656224952 | 0.02414729275 | 0.5562319781 |
| -2.045220709 | 0.3843892261 | 0.0113427499 | 0.0768843534 | 0.04522589771 | 0.7392100501 |
| -2.186589557 | 0.0149290403 | 0.0093601367 | 0.0363285327 | 0.03268317212 | 0.2213566899 |
| -2.276234789 | 0.4025333416 | 0.0089936506 | 0.0575710314 | 0.02641450713 | 0.7442416032 |
| -2.170072879 | 0.6870213735 | 0.0116660243 | 0.0661395955 | 0.03397102758 | 0.8574168621 |
| -2.130024807 | 0.1437796996 | 0.0055199821 | 0.0601044443 | 0.03727985259 | 0.5818748723 |
| -2.207587798 | 0.2490048423 | 0.0068168906 | 0.0585490947 | 0.031107891 | 0.6426517028 |
| -2.119745011 | 0.4915141665 | 0.0108368721 | 0.0773963697 | 0.03817327138 | 0.7781542416 |
| -2.141351905 | 0.8254371636 | 0.0125306718 | 0.0746642780 | 0.03631659247 | 0.903133968 |
| -2.072066005 | 0.6827780987 | 0.0119928767 | 0.0705001785 | 0.04256522111 | 0.8574168621 |
| -2.171335462 | 0.7499648254 | 0.0111376410 | 0.0676553643 | 0.03387103314 | 0.8790053953 |
| -2.12416433 | 0.3542703914 | 0.0109868854 | 0.0881818389 | 0.03778692558 | 0.7391035993 |
| -2.370223382 | 0.1054594933 | 0.0092900849 | 0.0648433756 | 0.02100881605 | 0.5562319781 |
| -2.107621136 | 0.1559195133 | 0.0190602692 | 0.0874909328 | 0.03925089299 | 0.5818748723 |
| -2.144367161 | 0.0088728017 | 0.0085810809 | 0.1525595899 | 0.03606386734 | 0.2213566899 |
| -2.258789437 | 0.3108268522 | 0.0086522554 | 0.0631744598 | 0.02754367837 | 0.7261735405 |
| -2.192637832 | 0.6730252228 | 0.0160632277 | 0.0645966887 | 0.03222240345 | 0.8574168621 |
| -2.145098209 | 0.7724165504 | 0.0178894494 | 0.0793964345 | 0.03600282614 | 0.8790053953 |
| -2.012254564 | 0.5936221525 | 0.0181268795 | 0.0874614887 | 0.04868879185 | 0.8379543986 |
| -2.2790893 | 0.1819949263 | 0.0104374651 | 0.0792662436 | 0.02623370047 | 0.5818748723 |
| -2.089255557 | 0.0686817272 | 0.0272947230 | 0.0805600954 | 0.04093372306 | 0.5341946883 |
| -2.118514806 | 0.6996521595 | 0.0117725769 | 0.0836153947 | 0.0382814291 | 0.8574168621 |
| -2.206717822 | 0.603327167 | 0.0100068758 | 0.0640070878 | 0.0311718057 | 0.8379543986 |
| -2.205527751 | 0.4258025937 | 0.0093828972 | 0.0532977359 | 0.03125942404 | 0.7518844513 |
| -2.052564967 | 0.3147157819 | 0.0157477662 | 0.0636311920 | 0.0444840808 | 0.7261735405 |
| -2.203789153 | 0.1686653941 | 0.0062366794 | 0.0534313949 | 0.03138781673 | 0.5818748723 |
| -1.897099436 | 0.3809658094 | 0.0256549468 | 0.0990995403 | 0.06263067566 | 0.7392100501 |
| -2.029222494 | 0.2747145198 | 0.0233216257 | 0.0776613201 | 0.04687890396 | 0.6603714418 |
| -2.149192057 | 0.8970517631 | 0.0120507407 | 0.0721694762 | 0.03566266342 | 0.9502666982 |
| -2.104712052 | 0.4987321994 | 0.0115632468 | 0.1031811378 | 0.0395133703 | 0.7781542416 |
| -2.128907009 | 0.2600750207 | 0.0158656674 | 0.0695122334 | 0.03737610838 | 0.6501875518 |
| -2.024642387 | 0.7875888342 | 0.0143271819 | 0.0827663959 | 0.04736163303 | 0.8790053953 |
| -2.197185119 | 0.6454838731 | 0.0101972402 | 0.0669604169 | 0.03187975002 | 0.8574168621 |
| -2.16502264 | 0.7636965784 | 0.0131182774 | 0.0691692032 | 0.03437358899 | 0.8790053953 |
| -2.135253978 | 0.4607837117 | 0.0120368342 | 0.0603295004 | 0.03683242564 | 0.7781542416 |
| -2.360396566 | 0.1128741809 | 0.0102640433 | 0.0425365312 | 0.02152365157 | 0.5643709044 |
| -2.161980303 | 0.7803333756 | 0.0122539962 | 0.0679275590 | 0.03461810939 | 0.8790053953 |
| -2.227343406 | 0.5702725284 | 0.0118353228 | 0.0590402045 | 0.02968721411 | 0.8235845603 |
| -2.127696847 | 0.9904743265 | 0.0137911058 | 0.0715221897 | 0.03748056224 | 0.9904743265 |
| -2.239409783 | 0.4270703684 | 0.0107110011 | 0.0606083780 | 0.02884790047 | 0.7518844513 |
| -2.138769329 | 0.9518851263 | 0.0114124343 | 0.0729003186 | 0.03653428121 | 0.9791894213 |
| -2.179176313 | 0.6195711599 | 0.0109299047 | 0.0748750540 | 0.0332558081 | 0.8510592856 |
| -2.071807588 | 0.65390721 | 0.0126967822 | 0.0715940803 | 0.04259017209 | 0.8574168621 |

| | | | | | |
|---|---|---|---|---|---|
| -2.176477559 | 0.7098113057 | 0.0107090216 | 0.0710503546 | 0.0334664481 | 0.8577920996 |
| -2.209110646 | 0.5661917457 | 0.0116886746 | 0.0707219680 | 0.03099628902 | 0.8235845603 |
| -2.104171126 | 0.3991362818 | 0.0111516711 | 0.0685862708 | 0.0395623444 | 0.7442416032 |
| -2.409141539 | 0.1337936252 | 0.0073354134 | 0.0385807850 | 0.01907625727 | 0.5818748723 |
| -2.192290047 | 0.1847447441 | 0.0144935919 | 0.1213890237 | 0.03224874293 | 0.5818748723 |
| -2.290537431 | 0.1645256214 | 0.0069518245 | 0.0507168065 | 0.02551953724 | 0.5818748723 |
| -2.463177251 | 0.1638228992 | 0.0047642565 | 0.0371410529 | 0.0166572335 | 0.5818748723 |
| -1.864392773 | 0.0564884583 | 0.0257209150 | 0.0761999643 | 0.0671603724 | 0.4943906211 |
| -2.042805943 | 0.3569390786 | 0.0089165391 | 0.0556855561 | 0.04547212859 | 0.7391035993 |
| -2.165514716 | 0.7136830269 | 0.0107511194 | 0.0688343132 | 0.03433418206 | 0.8577920996 |
| -2.142121942 | 0.8663421886 | 0.0116835395 | 0.0700834053 | 0.03625190479 | 0.9335583929 |
| -2.007671085 | 0.4737136465 | 0.0111002940 | 0.0914910596 | 0.04918788881 | 0.7781542416 |
| -2.459559783 | 0.0148509757 | 0.0108849051 | 0.0412010094 | 0.01681011574 | 0.2213566899 |
| -2.270379833 | 0.4800379809 | 0.0087690787 | 0.0576353736 | 0.02678882082 | 0.7781542416 |
| -2.064442998 | 0.4954367862 | 0.0142134823 | 0.0944304611 | 0.04330659619 | 0.7781542416 |
| -2.174810343 | 0.24083948 | 0.0160558348 | 0.072811213 | 0.03359715955 | 0.6426517028 |
| -2.210103985 | 0.4237613288 | 0.0084618577 | 0.0634605969 | 0.03092368211 | 0.7518844513 |
| -2.060942278 | 0.677946481 | 0.0201063849 | 0.0830406600 | 0.0436507893 | 0.8574168621 |
| -2.14312411 | 0.6905272937 | 0.0111343302 | 0.0857390759 | 0.03616786745 | 0.8574168621 |
| -2.116074101 | 0.0051648257 | 0.0067150825 | 0.0566698767 | 0.03849680293 | 0.2213566899 |
| -2.130270534 | 0.8308832506 | 0.0129155935 | 0.0764296163 | 0.03725872154 | 0.903133968 |
| -2.050478999 | 0.7774221447 | 0.0156627794 | 0.0746242446 | 0.0446936993 | 0.8790053953 |
| -2.092751274 | 0.9836546565 | 0.0119238332 | 0.0744340829 | 0.0406086647 | 0.9904743265 |
| -2.130104261 | 0.2727590001 | 0.0203726546 | 0.0964315503 | 0.0372730189 | 0.6603714418 |
| -2.265333294 | 0.4523408486 | 0.0097073229 | 0.0567468448 | 0.02711521112 | 0.7781542416 |
| -1.735343869 | 0.0854711501 | 0.0058296684 | 0.0996321893 | 0.08781380184 | 0.5341946883 |
| -2.127321659 | 0.6002273837 | 0.0104481690 | 0.0704114741 | 0.03751299786 | 0.8379543986 |
| -2.159274035 | 0.3630786502 | 0.0105352623 | 0.0915103003 | 0.0348368976 | 0.7391035993 |
| -3.040474714 | 0.0011153631 | 0.0020708341 | 0.0107073095 | 0.0034983056 | 0.1394203961 |
| -2.298697075 | 0.0085425659 | 0.0045131986 | 0.0222810813 | 0.02502109524 | 0.2213566899 |
| -2.134474795 | 0.5104691825 | 0.0116583477 | 0.0623993269 | 0.03689879674 | 0.7781542416 |
| -2.14001508 | 0.890345208 | 0.0142886955 | 0.0720702425 | 0.03642913335 | 0.9502666982 |
| -2.257521287 | 0.1496390042 | 0.0132041919 | 0.0578799285 | 0.02762740115 | 0.5818748723 |
| -2.213482302 | 0.5277493682 | 0.0093454520 | 0.0646004829 | 0.03067786705 | 0.7948032653 |
| -2.221328225 | 0.1677446231 | 0.0155553620 | 0.1053689383 | 0.03011360288 | 0.5818748723 |
| -2.442747491 | 0.0828920337 | 0.0045395772 | 0.0297539715 | 0.017537337 | 0.5341946883 |
| -2.369252958 | 0.0983148113 | 0.0050525471 | 0.0517654092 | 0.02105916338 | 0.5562319781 |
| -2.111034004 | 0.2425976491 | 0.0094790253 | 0.0562486763 | 0.03894489656 | 0.6426517028 |
| -2.158272201 | 0.3265692181 | 0.0076878642 | 0.0607973924 | 0.03491819729 | 0.7289491475 |
| -1.994323818 | 0.7384308639 | 0.0111262939 | 0.1035804478 | 0.0506664182 | 0.8790053953 |
| -2.171722459 | 0.2070008349 | 0.0070893286 | 0.0748201618 | 0.03384043535 | 0.5886203368 |
| -2.326278632 | 0.0107580094 | 0.0059659683 | 0.0354974168 | 0.02339961449 | 0.2213566899 |
| -2.036410595 | 0.0593268745 | 0.0091875385 | 0.0647372844 | 0.04612985738 | 0.4943906211 |
| -2.188024028 | 0.1311850956 | 0.0109106043 | 0.0644715551 | 0.03257337133 | 0.5818748723 |
| -2.12690258 | 0.7841806708 | 0.0164571851 | 0.0718466075 | 0.03754925688 | 0.8790053953 |
| -2.16551509 | 0.3195163578 | 0.0079297678 | 0.0612459213 | 0.03433415217 | 0.7261735405 |

| | | | | | |
|---|---|---|---|---|---|
| -1.910926502 | 0.0429947552 | 0.0097326291 | 0.1085920275 | 0.06079507026 | 0.4721833344 |
| -2.237210389 | 0.4935202008 | 0.0107985960 | 0.0540604771 | 0.0289993073 | 0.7781542416 |
| -2.146206321 | 0.6770037078 | 0.0109740832 | 0.0710705204 | 0.03591047301 | 0.8574168621 |
| -2.210670043 | 0.5503637252 | 0.0098767586 | 0.0712101535 | 0.03088237377 | 0.8189936387 |
| -2.101887247 | 0.1908549581 | 0.0183494993 | 0.0699155409 | 0.03976970361 | 0.5818748723 |
| -2.084549882 | 0.0501680495 | 0.0068361644 | 0.0385984917 | 0.0413748599 | 0.482385092 |
| -1.875760654 | 0.1067965398 | 0.0076705652 | 0.0747824932 | 0.06555558693 | 0.5562319781 |
| -2.229554347 | 0.2257433436 | 0.0077541834 | 0.0523626207 | 0.02953183322 | 0.6270648433 |
| -2.085252705 | 0.3756815645 | 0.0150299663 | 0.0755312214 | 0.04130871215 | 0.7392100501 |
| -1.589789725 | 0.0801339314 | 0.0209506205 | 0.0959986753 | 0.1171380983 | 0.5341946883 |
| -2.203994701 | 0.1772684659 | 0.0145115814 | 0.0799557916 | 0.03137261323 | 0.5818748723 |
| -2.054705334 | 0.2519194675 | 0.0171855383 | 0.0717903078 | 0.04426988024 | 0.6426517028 |
| -2.111441839 | 0.1448843916 | 0.0220124606 | 0.1347757046 | 0.03890846958 | 0.5818748723 |
| -2.044379946 | 0.4048674322 | 0.0083872287 | 0.0804731906 | 0.04531149829 | 0.7442416032 |
| -2.32569961 | 0.3665953852 | 0.0079953962 | 0.0542961211 | 0.02343267024 | 0.7391035993 |
| -2.135932331 | 0.802611948 | 0.0119111954 | 0.0768736181 | 0.03677472823 | 0.8878450753 |
| -1.987930535 | 0.5098245799 | 0.0141535908 | 0.0880347294 | 0.05138804356 | 0.7781542416 |
| -2.1389029 | 0.6983202595 | 0.0119036906 | 0.0669249250 | 0.03652299443 | 0.8574168621 |
| -2.146262001 | 0.9556888751 | 0.0130873485 | 0.0710779147 | 0.03590583796 | 0.9791894213 |
| -2.296794198 | 0.0392623624 | 0.0064731938 | 0.0495473656 | 0.02513655597 | 0.4721833344 |
| -2.608129724 | 0.0453296001 | 0.0028386966 | 0.0441836659 | 0.01147247793 | 0.4721833344 |
| -2.148116285 | 0.9657061631 | 0.0115468653 | 0.0713642414 | 0.03575177729 | 0.9814087024 |
| -2.346266739 | 0.3537380101 | 0.0082881624 | 0.0459281955 | 0.02228366046 | 0.7391035993 |
| -2.085406888 | 0.5097407616 | 0.0105054192 | 0.0702475779 | 0.04129421323 | 0.7781542416 |
| -2.259453361 | 0.2071943585 | 0.0100497541 | 0.0445435082 | 0.02749993587 | 0.5886203368 |
| -2.138408098 | 0.9193503678 | 0.0133715105 | 0.0732037997 | 0.03656482047 | 0.9657041679 |
| -2.073371853 | 0.2043784651 | 0.0114490356 | 0.0739658147 | 0.04243933087 | 0.5886203368 |
| -2.397448716 | 0.1686663019 | 0.0089142611 | 0.0585687600 | 0.01963944182 | 0.5818748723 |
| -2.441844225 | 0.0159376816 | 0.0017041626 | 0.0197237508 | 0.01757719671 | 0.2213566899 |
| -2.164757674 | 0.6943916551 | 0.0111411262 | 0.0663876066 | 0.03439482467 | 0.8574168621 |
| -2.203443086 | 0.573214854 | 0.0100087889 | 0.0674692160 | 0.03141342855 | 0.8235845603 |
| -1.960940554 | 0.0729997520 | 0.0165060127 | 0.1469627859 | 0.05453244844 | 0.5341946883 |
| -2.305862259 | 0.1891644391 | 0.0116523621 | 0.0509751980 | 0.02459054595 | 0.5818748723 |
| -2.201234326 | 0.3537690857 | 0.0085612928 | 0.0647590232 | 0.0315773281 | 0.7391035993 |
| -2.087685787 | 0.8994116466 | 0.0194707101 | 0.1580433647 | 0.04113504581 | 0.9499253008 |
| -2.083949931 | 0.1254151359 | 0.0190591885 | 0.1643243331 | 0.04143139834 | 0.6443053671 |
| -2.076984645 | 0.9373888988 | 0.0214111153 | 0.1589987686 | 0.04212909397 | 0.957242324 |
| -2.022233356 | 0.5975365874 | 0.0191877482 | 0.1647538694 | 0.04764608834 | 0.8118703633 |
| -2.099121786 | 0.1290725025 | 0.0182436778 | 0.1181769677 | 0.04007515906 | 0.6443053671 |
| -2.192368904 | 0.5691777254 | 0.0156638248 | 0.1373560911 | 0.0322812046 | 0.7905246186 |
| -2.086026489 | 0.9419264468 | 0.0190829494 | 0.1624093429 | 0.04128830377 | 0.957242324 |
| -2.085353726 | 0.8987252864 | 0.0199408997 | 0.1577565431 | 0.04134398555 | 0.9499253008 |
| -2.089321259 | 0.9277951185 | 0.0204425890 | 0.1585150385 | 0.04097557642 | 0.957242324 |
| -1.900572868 | 0.1115146049 | 0.0217244689 | 0.2230703685 | 0.06223264998 | 0.6443053671 |
| -2.083382928 | 0.5610737606 | 0.0222861433 | 0.1703782954 | 0.04152601979 | 0.7904682604 |
| -2.443694685 | 0.0714340533 | 0.0112751078 | 0.1406452731 | 0.01749562612 | 0.6159663638 |

| | | | | | |
|---|---|---|---|---|---|
| -2.063070096 | 0.7732243122 | 0.0191061993 | 0.1677872719 | 0.04350060331 | 0.8949355466 |
| -2.067324034 | 0.3419486154 | 0.0185759685 | 0.1910829869 | 0.04308288574 | 0.7118714722 |
| -2.220776982 | 0.2302424509 | 0.0180451628 | 0.1578737212 | 0.03021858779 | 0.6536180385 |
| -2.059778297 | 0.4398967569 | 0.0263196848 | 0.1790704945 | 0.04382457874 | 0.7281989247 |
| -2.0560728 | 0.4910635341 | 0.0198213773 | 0.2027149164 | 0.04421135408 | 0.7395535153 |
| -2.155162091 | 0.4132902064 | 0.0166174964 | 0.1493700299 | 0.03523453981 | 0.7281989247 |
| -2.014549122 | 0.7497093284 | 0.0179780114 | 0.1755559792 | 0.04850993184 | 0.8840911892 |
| -2.105563265 | 0.4219837995 | 0.0136326037 | 0.1317497367 | 0.03948656011 | 0.7281989247 |
| -1.876504378 | 0.2794887157 | 0.0382196636 | 0.2059261997 | 0.06574122442 | 0.6822233369 |
| -2.169722733 | 0.5539961134 | 0.0175734547 | 0.161465159 | 0.03402748798 | 0.7904682604 |
| -1.989122378 | 0.0632069882 | 0.0486601050 | 0.1832926065 | 0.05128115988 | 0.6159663638 |
| -2.175755704 | 0.3398098115 | 0.0165504466 | 0.1233398025 | 0.0335483374 | 0.7118714722 |
| -1.87848574 | 0.1732175009 | 0.0540865581 | 0.219702029 | 0.06527736672 | 0.6443053671 |
| -2.174971821 | 0.2854815212 | 0.0143735151 | 0.1104490806 | 0.03361896071 | 0.6822233369 |
| -2.085479718 | 0.958679205 | 0.0197808018 | 0.1594492747 | 0.04133344958 | 0.9664104889 |
| -2.103866474 | 0.2346781765 | 0.0131104072 | 0.1328797658 | 0.039669362 | 0.6536180385 |
| -2.286266965 | 0.3644781938 | 0.0115331591 | 0.1175914496 | 0.02581580995 | 0.7118714722 |
| -1.972485974 | 0.3053994996 | 0.0368171174 | 0.1750953733 | 0.05321767433 | 0.706943286 |
| -2.095393588 | 0.9071861887 | 0.0189345176 | 0.1566161825 | 0.04041058251 | 0.9499253008 |
| -2.089843894 | 0.9072187268 | 0.0192967881 | 0.1608030157 | 0.04094033468 | 0.9499253008 |
| -1.982586322 | 0.2353024939 | 0.0322134912 | 0.1725365841 | 0.05206294823 | 0.6536180385 |
| -1.674825436 | 0.0451962738 | 0.0359483572 | 0.2587859034 | 0.09917356944 | 0.6159663638 |
| -2.149938532 | 0.6341166691 | 0.0169297081 | 0.1492173333 | 0.03564539612 | 0.8343640383 |
| -2.07272448 | 0.4696719247 | 0.0304755973 | 0.1582333987 | 0.04267376628 | 0.7365881039 |
| -1.908530176 | 0.0623454447 | 0.0295607008 | 0.1224362135 | 0.0613055069 | 0.6159663638 |
| -2.216349618 | 0.1074338676 | 0.0219292666 | 0.1133801196 | 0.03059957199 | 0.6443053671 |
| -2.319610285 | 0.0565671091 | 0.0202277053 | 0.0821862499 | 0.02378282689 | 0.6159663638 |
| -1.955987036 | 0.7097563078 | 0.0186232314 | 0.1967988345 | 0.05522327811 | 0.8697993968 |
| -2.15163604 | 0.4362059266 | 0.0150231458 | 0.1521671808 | 0.03558420185 | 0.7281989247 |
| -2.105713467 | 0.8155982288 | 0.0187924960 | 0.1535469497 | 0.03942551684 | 0.9102658803 |
| -2.089855048 | 0.8883342771 | 0.0194039420 | 0.1578650073 | 0.04096387516 | 0.9499253008 |
| -2.353440912 | 0.0372517111 | 0.0115389571 | 0.1647249066 | 0.02189483994 | 0.6159663638 |
| -2.098937381 | 0.8067616018 | 0.0188401996 | 0.159470736 | 0.04006535813 | 0.9085153174 |
| -1.96293449 | 0.4495697534 | 0.0251155863 | 0.1640116017 | 0.05432212635 | 0.7281989247 |
| -2.162621788 | 0.5914406828 | 0.0190166932 | 0.153562513 | 0.03460031372 | 0.8118703633 |
| -2.070077749 | 0.7301179339 | 0.0192107215 | 0.1581455836 | 0.0428230311 | 0.8775455936 |
| -2.209241121 | 0.3508272053 | 0.0163410892 | 0.1215850561 | 0.03101038683 | 0.7118714722 |
| -2.193436129 | 0.1759757688 | 0.0210962977 | 0.2371279887 | 0.03216201596 | 0.6443053671 |
| -2.159470895 | 0.2056994087 | 0.0149764152 | 0.1298375541 | 0.03495218796 | 0.6536180385 |
| -1.822960958 | 0.3340839867 | 0.0398066817 | 0.2365064016 | 0.07332272916 | 0.7118714722 |
| -1.859680634 | 0.2282348604 | 0.0363703603 | 0.164961605 | 0.06783705132 | 0.6536180385 |
| -1.968098388 | 0.4067902709 | 0.0162820335 | 0.1337908516 | 0.05370418489 | 0.7281989247 |
| -2.118845309 | 0.5626074009 | 0.0172713993 | 0.1532101309 | 0.03829792901 | 0.7904682604 |
| -1.971530209 | 0.3508468321 | 0.0258286941 | 0.1519570582 | 0.0533591798 | 0.7118714722 |
| -1.928430104 | 0.454396129 | 0.0191062595 | 0.2022686709 | 0.05854216819 | 0.7281989247 |
| -2.334409433 | 0.0787691698 | 0.0182130841 | 0.1124176207 | 0.0229442151 | 0.6159663638 |

| | | | | | |
|---|---|---|---|---|---|
| -2.113536746 | 0.6998164247 | 0.0178884899 | 0.1463237617 | 0.03877639522 | 0.8697993968 |
| -1.925032898 | 0.1095695879 | 0.0286292981 | 0.2440040008 | 0.05906754582 | 0.6443053671 |
| -2.149243783 | 0.1571796819 | 0.0263224172 | 0.1573981822 | 0.03569053628 | 0.6443053671 |
| -2.114387241 | 0.6252196916 | 0.0170860416 | 0.1506255075 | 0.03870911228 | 0.8314091643 |
| -1.91403024 | 0.1513286624 | 0.0540156530 | 0.2154779211 | 0.06041453644 | 0.6443053671 |
| -2.149732948 | 0.2426798861 | 0.0147583534 | 0.2115677711 | 0.03561793101 | 0.6594562121 |
| -2.097894845 | 0.0085039130 | 0.0111320625 | 0.1345570338 | 0.04016659679 | 0.3543297105 |
| -2.097060804 | 0.5500394772 | 0.0177816565 | 0.148437872 | 0.0403047418 | 0.7904682604 |
| -1.850827607 | 0.1471691112 | 0.0374527391 | 0.1711160879 | 0.06911859743 | 0.6443053671 |
| -1.642623004 | 0.0632896669 | 0.0289154382 | 0.2714682952 | 0.1057789383 | 0.6159663638 |
| -2.049960739 | 0.1756573319 | 0.0385490382 | 0.2153856502 | 0.04483368249 | 0.6443053671 |
| -2.284274847 | 0.2346536899 | 0.0156228603 | 0.1139105255 | 0.0259938048 | 0.6536180385 |
| -1.853949849 | 0.4008041678 | 0.0148846119 | 0.1875503858 | 0.06870830413 | 0.7281989247 |
| -2.053582342 | 0.6518322766 | 0.0185771351 | 0.1606527125 | 0.04442557401 | 0.8487399435 |
| -2.104034177 | 0.204260505 | 0.0174398106 | 0.2172032487 | 0.03963163221 | 0.6536180385 |
| -2.935105315 | 0.0014150994 | 0.0046292750 | 0.0346423666 | 0.00474845235 | 0.1768874274 |
| -2.152590582 | 0.0071585520 | 0.0105156450 | 0.0655309243 | 0.03551169449 | 0.3543297105 |
| -2.132066138 | 0.6917076187 | 0.0174398787 | 0.1661408852 | 0.03714442827 | 0.8697993968 |
| -2.382763649 | 0.1804055028 | 0.0069003929 | 0.0925512294 | 0.02036780922 | 0.6443053671 |
| -2.073264513 | 0.4814028008 | 0.0162974934 | 0.162558999 | 0.042476583 | 0.7365881039 |
| -2.045407192 | 0.5628134014 | 0.0238492921 | 0.1653712888 | 0.04529949925 | 0.7904682604 |
| -2.122210473 | 0.4138057896 | 0.0238400978 | 0.1979493518 | 0.03800194563 | 0.7281989247 |
| -2.024456701 | 0.764463568 | 0.0239158615 | 0.1881934872 | 0.04740460371 | 0.8930649159 |
| -2.247327434 | 0.159767487 | 0.0107934115 | 0.1376796202 | 0.02836311005 | 0.6443053671 |
| -2.133945705 | 0.6626500001 | 0.0189703093 | 0.1653305898 | 0.03698935023 | 0.8539304125 |
| -2.085667413 | 0.9119282888 | 0.0202385661 | 0.1585083201 | 0.04132547642 | 0.9499253008 |
| -2.024824112 | 0.9856491496 | 0.0192772734 | 0.179545883 | 0.04750996987 | 0.9856491496 |
| -2.084888814 | 0.4832017962 | 0.0163975107 | 0.1676104904 | 0.04141009589 | 0.7365881039 |
| -2.136779632 | 0.1730174759 | 0.0159512798 | 0.1308676111 | 0.0367617425 | 0.6443053671 |
| -2.039902049 | 0.6044479639 | 0.0190773175 | 0.1607497127 | 0.04584208404 | 0.812430059 |
| -2.014367202 | 0.2789299007 | 0.0245475792 | 0.1877768934 | 0.04868748848 | 0.6822233369 |
| -2.080230985 | 0.9096804071 | 0.0242348860 | 0.1596384103 | 0.04184086751 | 0.9499253008 |
| -2.090043047 | 0.780402435 | 0.0184440813 | 0.1545998134 | 0.04090548107 | 0.8949569209 |
| -1.810307433 | 0.1222854218 | 0.0216592267 | 0.2433341946 | 0.07535858536 | 0.6443053671 |
| -2.179447939 | 0.453682115 | 0.0186496266 | 0.117887463 | 0.03328786629 | 0.7281989247 |
| -2.117090057 | 0.1776606498 | 0.0200911788 | 0.1577939794 | 0.03846686381 | 0.6443053671 |
| -2.034972096 | 0.4495531969 | 0.0212195206 | 0.1484306157 | 0.04633822216 | 0.7281989247 |
| -2.037625553 | 0.4066266869 | 0.0277602979 | 0.1665899129 | 0.04606114485 | 0.7281989247 |
| -1.964402384 | 0.0611785149 | 0.0145874446 | 0.1056301924 | 0.05433042731 | 0.6159663638 |
| -1.959019605 | 0.6709409825 | 0.018985447 | 0.1668433162 | 0.05478763371 | 0.8557920696 |
| -2.080663779 | 0.7047315842 | 0.0213632304 | 0.170595407 | 0.04178546139 | 0.8697993968 |
| -1.981399116 | 0.0361703349 | 0.0308271036 | 0.171373315 | 0.05224880477 | 0.6159663638 |
| -1.7114162 | 0.3559479761 | 0.0297295891 | 0.202823632 | 0.09230285367 | 0.7118714722 |
| -2.114777125 | 0.3633239186 | 0.0232667161 | 0.1736841706 | 0.03866072792 | 0.7118714722 |
| -2.09447309 | 0.8830970021 | 0.0194131217 | 0.1585610071 | 0.04050062844 | 0.9499253008 |
| -2.111528375 | 0.1166091267 | 0.0323667114 | 0.2657061294 | 0.03890074419 | 0.6443053671 |

| | | | | | |
|---|---|---|---|---|---|
| -1.956392865 | 0.2000478928 | 0.0111366019 | 0.1900545871 | 0.05517535158 | 0.6536180385 |
| -2.595224071 | 0.0788436945 | 0.0069714996 | 0.0856855203 | 0.01186608235 | 0.6159663638 |
| -2.06467946 | 0.4097185115 | 0.0207701637 | 0.1829527964 | 0.04331261977 | 0.7281989247 |
| -1.859677784 | 0.324282011 | 0.0254662397 | 0.2044717814 | 0.06784577392 | 0.7118714722 |
| -2.203691666 | 0.2541435213 | 0.0138024626 | 0.1751390992 | 0.03139502972 | 0.6618320868 |
| -2.103547806 | 0.7400884532 | 0.0186472436 | 0.1585090188 | 0.0396873466 | 0.8810576824 |
| -2.219759271 | 0.0684452291 | 0.0122021805 | 0.1247249956 | 0.03026181143 | 0.6159663638 |
| -2.733045624 | 0.0161109988 | 0.0028857180 | 0.0893469300 | 0.00823311060 | 0.5034687151 |
| -2.038244404 | 0.5386566508 | 0.0252287440 | 0.1660666855 | 0.04604207669 | 0.7904682604 |
| -2.363717852 | 0.2892626948 | 0.0124342617 | 0.0936608231 | 0.02136279973 | 0.6822233369 |
| -1.96879726 | 0.323900551 | 0.0182322244 | 0.1702863831 | 0.05369891727 | 0.7118714722 |
| -2.08373933 | 0.7936181259 | 0.0185507189 | 0.1740372986 | 0.04148803657 | 0.9018387794 |
| -2.193630459 | 0.2793944084 | 0.0415276211 | 0.2253275953 | 0.03218750443 | 0.6822233369 |
| -1.964116981 | 0.2178254332 | 0.0225831357 | 0.1815008531 | 0.05430624171 | 0.6536180385 |
| -2.181262355 | 0.4742359431 | 0.0181327853 | 0.1515586832 | 0.03313165408 | 0.7365881039 |
| -2.181243211 | 0.3971396584 | 0.0127361788 | 0.1178739824 | 0.03310324091 | 0.7281989247 |
| -2.178587907 | 0.1719380601 | 0.0165149256 | 0.1132609285 | 0.03334175354 | 0.6443053671 |
| -2.114364281 | 0.7187584883 | 0.0182039329 | 0.1564085609 | 0.03869986955 | 0.8722797188 |
| -1.942740108 | 0.1769181676 | 0.0257086915 | 0.2611915749 | 0.05677825997 | 0.6443053671 |
| -2.184219621 | 0.2323285295 | 0.0221331843 | 0.1348950255 | 0.03295186509 | 0.6536180385 |
| -2.160382268 | 0.2530826144 | 0.0133075511 | 0.1459635289 | 0.03479056905 | 0.6618320868 |
| -2.196603856 | 0.0850262291 | 0.0064070565 | 0.0970540706 | 0.03192337091 | 0.4810582845 |
| -2.187024388 | 0.3131571074 | 0.0102461071 | 0.1326191717 | 0.03264985396 | 0.6888398003 |
| -2.243013867 | 0.1568381911 | 0.0067905641 | 0.1433166841 | 0.02860130241 | 0.5766109966 |
| -2.10423851 | 0.9207291752 | 0.0126140441 | 0.1484292979 | 0.0395562407 | 0.967152495 |
| -2.168539795 | 0.0049172416 | 0.0094972320 | 0.0754342729 | 0.03409279283 | 0.1414074649 |
| -2.195830219 | 0.5184925581 | 0.0107220493 | 0.1303770933 | 0.03198150987 | 0.7752904146 |
| -2.15557433 | 0.4469480759 | 0.0129439786 | 0.1287845249 | 0.03513795807 | 0.735111967 |
| -2.08671821 | 0.1688010093 | 0.0065254864 | 0.1318545547 | 0.04117107829 | 0.6028607473 |
| -2.324794251 | 0.0070890947 | 0.0023352397 | 0.0890518353 | 0.0234844394 | 0.1476894731 |
| -2.072973533 | 0.3551738624 | 0.0116578722 | 0.1638172884 | 0.04247769661 | 0.6952978415 |
| -2.114346835 | 0.8072147369 | 0.0122455379 | 0.1451307674 | 0.03864985833 | 0.9264082432 |
| -1.997998275 | 0.4987028156 | 0.0135080286 | 0.1468249337 | 0.05025562434 | 0.7750958893 |
| -2.133123929 | 0.7288206422 | 0.0123106808 | 0.1416244841 | 0.03701411141 | 0.8844910707 |
| -2.084783345 | 0.111150124 | 0.0111565012 | 0.1982952704 | 0.04135287682 | 0.4810582845 |
| -2.324214904 | 0.1102169301 | 0.0103893339 | 0.140028718 | 0.02351762036 | 0.4810582845 |
| -2.07089391 | 0.0413084829 | 0.0237618814 | 0.1894899863 | 0.04267849246 | 0.4134605685 |
| -2.179276102 | 0.0001824908 | 0.0066002339 | 0.3545762691 | 0.03328484707 | 0.02281135096 |
| -2.127676233 | 0.7662760547 | 0.0120695950 | 0.1460535229 | 0.03748234377 | 0.9122333985 |
| -2.582177797 | 0.0227209164 | 0.0341759984 | 0.0670090012 | 0.01227641371 | 0.3550143195 |
| -2.103551398 | 0.5621944543 | 0.0235985853 | 0.1738218103 | 0.03961851785 | 0.7813413209 |
| -1.78082849 | 0.0761289079 | 0.0341119569 | 0.2325925239 | 0.0800028188 | 0.4810582845 |
| -2.060226234 | 0.5532641908 | 0.0123441774 | 0.1398446905 | 0.04372148225 | 0.7813413209 |
| -2.046831344 | 0.0917620856 | 0.0287801487 | 0.1685677124 | 0.04506230439 | 0.4810582845 |
| -2.110416633 | 0.9030566344 | 0.0125636507 | 0.1488246091 | 0.03900009544 | 0.9658207475 |
| -2.24027237 | 0.3441111855 | 0.0078551750 | 0.122352491 | 0.0287887105 | 0.6952978415 |

| | | | | | |
|---|---|---|---|---|---|
| -2.208399867 | 0.2139362871 | 0.0088853145 | 0.0933512981 | 0.03104833455 | 0.6525141531 |
| -2.027973832 | 0.4433333154 | 0.0165020241 | 0.1398412228 | 0.04701008658 | 0.735111967 |
| -2.111295717 | 0.8078279881 | 0.0123964263 | 0.1451636332 | 0.03892151752 | 0.9264082432 |
| -2.003938033 | 0.8457411307 | 0.0184973263 | 0.1643203628 | 0.0495976282 | 0.9355543481 |
| -1.988141846 | 0.278008657 | 0.0255449657 | 0.1619436648 | 0.05136405217 | 0.6626573547 |
| -2.155646438 | 0.1279260483 | 0.0160178561 | 0.1588466623 | 0.03513206867 | 0.5158308398 |
| -2.173080447 | 0.3971002465 | 0.0116295909 | 0.1108687324 | 0.0337332581 | 0.6991201523 |
| -2.096748083 | 0.0759930400 | 0.0193342717 | 0.1423259664 | 0.0402397615 | 0.4810582845 |
| -1.949106515 | 0.6707280829 | 0.0168699401 | 0.1763961953 | 0.05596220357 | 0.8643403131 |
| -1.932458461 | 0.7138670608 | 0.0151874224 | 0.156295781 | 0.0580276701 | 0.8834988376 |
| -2.17287098 | 0.3838262455 | 0.0166744046 | 0.1361870128 | 0.03374977051 | 0.6953374013 |
| -2.140434705 | 0.0141429704 | 0.0117209885 | 0.0768293764 | 0.03639377439 | 0.2525530445 |
| -2.337250738 | 0.0903000283 | 0.0112098686 | 0.0922569279 | 0.02278102352 | 0.4810582845 |
| -2.138744592 | 0.591137952 | 0.0141045866 | 0.1323092724 | 0.03653637181 | 0.7860877021 |
| -1.992539168 | 0.7870535268 | 0.0123620997 | 0.18032919 | 0.05086697516 | 0.9264082432 |
| -2.079090225 | 0.9678545076 | 0.0157040678 | 0.1505419271 | 0.04189184615 | 0.9838679628 |
| -2.088267405 | 0.9759970191 | 0.0126825708 | 0.1518252872 | 0.04102601768 | 0.9838679628 |
| -2.186409042 | 0.4885745013 | 0.0121868565 | 0.1255857532 | 0.03269701251 | 0.7730609198 |
| -2.062165227 | 0.3733307713 | 0.0128475864 | 0.1352608858 | 0.04353027986 | 0.6953374013 |
| -1.992801263 | 0.4355485939 | 0.0147269459 | 0.1499612939 | 0.05083747858 | 0.735111967 |
| -2.070428287 | 0.7636378798 | 0.0137048932 | 0.1488073882 | 0.04272356277 | 0.9122333985 |
| -2.099916906 | 0.9404745444 | 0.0127511990 | 0.1493916291 | 0.03994935495 | 0.971564612 |
| -2.047053377 | 0.0956744536 | 0.0111725241 | 0.1366323317 | 0.04503979254 | 0.4810582845 |
| -2.433250088 | 0.074027712 | 0.0072303332 | 0.0763165168 | 0.01796053339 | 0.4810582845 |
| -2.125573521 | 0.4195766738 | 0.0151214867 | 0.2037646979 | 0.03766445036 | 0.7184532086 |
| -2.097254844 | 0.9749277402 | 0.013344172 | 0.1520609659 | 0.0401931965 | 0.9838679628 |
| -2.341024725 | 0.2637962775 | 0.0067950273 | 0.0965379577 | 0.0225716426 | 0.6626573547 |
| -1.852758501 | 0.1039981391 | 0.0261961716 | 0.1614659234 | 0.06883695776 | 0.4810582845 |
| -1.96009721 | 0.1410830531 | 0.0079750446 | 0.1016903106 | 0.05463329521 | 0.5511056763 |
| -2.191784351 | 0.2081014989 | 0.0091864827 | 0.1305529464 | 0.03228707546 | 0.6525141531 |
| -2.090355015 | 0.3454266754 | 0.0134069112 | 0.1310672796 | 0.04083124429 | 0.6952978415 |
| -1.916950081 | 0.2809667184 | 0.0118710391 | 0.2005021922 | 0.06000981626 | 0.6626573547 |
| -2.207815897 | 0.2919737104 | 0.0135448820 | 0.1290634093 | 0.03109115231 | 0.6758650705 |
| -2.157430474 | 0.6391823968 | 0.0111988054 | 0.1336289721 | 0.03498663247 | 0.8410294695 |
| -2.164150893 | 0.5683715408 | 0.0109562612 | 0.1285425216 | 0.03444349846 | 0.7813413209 |
| -2.172306762 | 0.0610238487 | 0.0196469466 | 0.1513381243 | 0.03379428356 | 0.4810582845 |
| -2.03621156 | 0.1148101179 | 0.0242068728 | 0.1683781272 | 0.04615045822 | 0.4810582845 |
| -1.953773111 | 0.3615548776 | 0.0301028941 | 0.1893801263 | 0.05539462092 | 0.6952978415 |
| -2.124810588 | 0.5397678274 | 0.0127463322 | 0.1270441829 | 0.03773071514 | 0.7813413209 |
| -2.055255341 | 0.2160787526 | 0.0113390998 | 0.1463303941 | 0.04421498165 | 0.6525141531 |
| -2.056990885 | 0.3727283609 | 0.0170854304 | 0.1778987124 | 0.04404213574 | 0.6953374013 |
| -2.081092101 | 0.935271178 | 0.0141661312 | 0.1487303981 | 0.04170163724 | 0.971564612 |
| -1.999513953 | 0.8366150725 | 0.0141290495 | 0.1643072538 | 0.05008701133 | 0.9337221791 |
| -2.135336483 | 0.5209951586 | 0.0101561105 | 0.1289613289 | 0.03682540397 | 0.7752904146 |
| -2.352024718 | 0.1967871682 | 0.0090678577 | 0.0999247276 | 0.021971111 | 0.6525141531 |
| -1.747739917 | 0.152757888 | 0.0075606398 | 0.1944788322 | 0.08562495284 | 0.5766109966 |

| | | | | | |
|---|---|---|---|---|---|
| -2.181354537 | 0.2459800514 | 0.014388500( | 0.1455159931 | 0.03308664497 | 0.6598047917 |
| -2.116465264 | 0.3826117382 | 0.011752240( | 0.1841843367 | 0.03846221486 | 0.6953374013 |
| -2.599612606 | 0.0366823352 | 0.0053633609 | 0.0621175672 | 0.01173088147 | 0.4134605685 |
| -2.201865002 | 0.0271610365 | 0.0066182375 | 0.0690853801 | 0.03155988953 | 0.3772366194 |
| -2.093294323 | 0.2437520173 | 0.0126791572 | 0.1183612538 | 0.04055836967 | 0.6598047917 |
| -2.008069651 | 0.6908097347 | 0.0236922592 | 0.1777742762 | 0.04914431473 | 0.8811348657 |
| -2.240980794 | 0.0936709797 | 0.0146314772 | 0.121296509 | 0.02874017921 | 0.4810582845 |
| -1.990747886 | 0.2806512966 | 0.0198946007 | 0.1680109218 | 0.05106896251 | 0.6626573547 |
| -2.15291943 | 0.3109427542 | 0.0163449323 | 0.1953343562 | 0.03535539909 | 0.6888398003 |
| -2.290283434 | 0.2296849819 | 0.0073132136 | 0.0924831838 | 0.02553519331 | 0.6525141531 |
| -2.210632903 | 0.3580404047 | 0.0088547799 | 0.1317326967 | 0.03088508261 | 0.6952978415 |
| -2.076412657 | 0.0328468094 | 0.0079270655 | 0.0965448234 | 0.04214743268 | 0.4105851175 |
| -2.110462622 | 0.4721862843 | 0.0098587674 | 0.1362670146 | 0.03899598122 | 0.756708789 |
| -2.010566966 | 0.9153301611 | 0.0125922057 | 0.1785139638 | 0.04887204581 | 0.967152495 |
| -2.111827791 | 0.5022621362 | 0.0103887221 | 0.1550021408 | 0.0388740244 | 0.7750958893 |
| -2.412240053 | 0.0005069474 | 0.0041805141 | 0.0601839132 | 0.01892946357 | 0.0316842142 |
| -2.100382321 | 0.9843748613 | 0.0126626761 | 0.1493189236 | 0.03990685585 | 0.9843748613 |
| -2.108592348 | 0.7216009473 | 0.0128398342 | 0.1481997861 | 0.03916360216 | 0.8843148864 |
| -2.069522771 | 0.5576693329 | 0.021573642 | 0.1512436369 | 0.04281133086 | 0.7813413209 |
| -2.10998542 | 0.6543505512 | 0.0113459744 | 0.1422953408 | 0.03903869039 | 0.8520189469 |
| -2.033765937 | 0.6982602639 | 0.0127519893 | 0.1658981308 | 0.04640423795 | 0.8816417474 |
| -2.170883398 | 0.5872631305 | 0.0121729981 | 0.1240471769 | 0.0339068062 | 0.7860877021 |
| -2.136885951 | 0.3141109489 | 0.0129980859 | 0.1448335811 | 0.03669375136 | 0.6888398003 |
| -2.261965078 | 0.2276171955 | 0.0087101591 | 0.1474699622 | 0.02733500721 | 0.6525141531 |
| -2.068286667 | 0.3945856025 | 0.0180103589 | 0.1500834369 | 0.04293139396 | 0.6991201523 |
| -2.060561834 | 0.4089024807 | 0.0109165416 | 0.125163152 | 0.04368833705 | 0.7099001401 |
| -2.107097116 | 0.8545461868 | 0.0131493532 | 0.1476052269 | 0.0392980612 | 0.9370023978 |
| -2.149173557 | 0.4566530978 | 0.0103774908 | 0.1290746384 | 0.03566419429 | 0.741319964 |
| -2.031737874 | 0.2480866017 | 0.0174876146 | 0.1592549485 | 0.04661560038 | 0.6598047917 |
| -1.713499363 | 0.2733192267 | 0.0197655177 | 0.1809295322 | 0.09178353469 | 0.6626573547 |
| -2.300741252 | 0.0048525263 | 0.0161608284 | 0.1709423085 | 0.02489758553 | 0.1414074649 |
| -2.009649624 | 0.2275551433 | 0.0192406538 | 0.1513949792 | 0.04897190756 | 0.6525141531 |
| -2.072088492 | 0.0692085566 | 0.0267566907 | 0.2856038107 | 0.04256305044 | 0.4810582845 |
| -1.938658186 | 0.0994715946 | 0.0057282862 | 0.1782391571 | 0.05725102414 | 0.4810582845 |
| -1.937234061 | 0.707515449 | 0.0162865408 | 0.1693462777 | 0.05742863735 | 0.8834988376 |
| -2.095699509 | 0.8292605048 | 0.0131429992 | 0.1580534555 | 0.04033626132 | 0.9337221791 |
| -2.043596291 | 0.9015723114 | 0.0136954101 | 0.1580160024 | 0.04539141057 | 0.9658207475 |
| -2.100562316 | 0.8322112386 | 0.0130095735 | 0.145665294 | 0.0398904303 | 0.9337221791 |
| -2.150181239 | 0.5700122456 | 0.0105574343 | 0.1442722882 | 0.03558089388 | 0.7813413209 |
| -2.249606007 | 0.0429998991 | 0.0073512564 | 0.1123887837 | 0.02815506748 | 0.4134605685 |
| -2.34657664 | 0.2242752097 | 0.006466369( | 0.1213318295 | 0.02226673829 | 0.6525141531 |
| -2.106487453 | 0.9040082196 | 0.0125909297 | 0.1499475891 | 0.03935300058 | 0.9658207475 |
| -1.614300901 | 0.1740084994 | 0.0194450608 | 0.3044267312 | 0.1117095028 | 0.6041961785 |
| -2.025703597 | 0.3486866076 | 0.0110649481 | 0.1465043418 | 0.04724940511 | 0.6952978415 |
| -2.15456291 | 0.5124003862 | 0.0121388058 | 0.1247424209 | 0.03522065681 | 0.7752904146 |
| -2.142520758 | 0.5750672122 | 0.022226362( | 0.187416315 | 0.03621844148 | 0.7813413209 |

| | | | | | |
|---|---|---|---|---|---|
| -2.082886442 | 0.8060501283 | 0.0128759973 | 0.1515864932 | 0.04153178469 | 0.9264082432 |
| -2.269785368 | 0.3201676749 | 0.0108754039 | 0.1344549084 | 0.02682708695 | 0.6900165407 |
| -2.463173006 | 0.0056562985 | 0.0012557926 | 0.0388170406 | 0.01665741216 | 0.1414074649 |
| -2.135133351 | 0.5464004326 | 0.0120957021 | 0.1338925227 | 0.03684269386 | 0.7813413209 |
| -2.314508436 | 0.1154539883 | 0.0074547365 | 0.1258457062 | 0.02407977451 | 0.4810582845 |
| -1.971925062 | 0.26930805 | 0.0167872031 | 0.2296854576 | 0.0532333984 | 0.6626573547 |
| -2.317193315 | 0.0999624275 | 0.0125927678 | 0.1021454122 | 0.02392309571 | 0.4810582845 |
| -2.1611376 | 0.3423627857 | 0.0094259880 | 0.1377764794 | 0.03468610833 | 0.6952978415 |
| -0.6034512998 | 0.2769133459 | 0.0003145141 | 0.7503952408 | 0.5484629448 | 0.989483196 |
| -0.5418807792 | 0.7167033487 | 0.0003713051 | 0.713381288 | 0.5898861645 | 0.989483196 |
| -0.5178768463 | 0.3530993229 | 0.0002802697 | 0.7214883786 | 0.6064217147 | 0.989483196 |
| -0.3993468239 | 0.0785274036 | 0.0002369731 | 0.7946299287 | 0.6910393252 | 0.989483196 |
| -0.6175918763 | 0.1549622896 | 0.0003501098 | 0.6083090651 | 0.5391561309 | 0.989483196 |
| -0.8315975396 | 0.2292741643 | 0.0008364868 | 0.6043703151 | 0.4089084654 | 0.989483196 |
| -0.5965130555 | 0.7515355661 | 0.0003931123 | 0.6743894809 | 0.5530630019 | 0.989483196 |
| -0.581824804 | 0.9611468914 | 0.0004466459 | 0.6956764044 | 0.5628446879 | 0.989483196 |
| -0.6061935182 | 0.7349613248 | 0.0005840115 | 0.6823919048 | 0.5466491535 | 0.989483196 |
| -0.5325981743 | 0.6834035337 | 0.0003821156 | 0.7220695374 | 0.5962400637 | 0.989483196 |
| -0.5669323286 | 0.8943907575 | 0.0004337167 | 0.7029005269 | 0.5728603611 | 0.989483196 |
| -0.4326370513 | 0.3867393086 | 0.0002889777 | 0.6682917541 | 0.6668323238 | 0.989483196 |
| -0.5622964171 | 0.7499566817 | 0.0004371333 | 0.7245787483 | 0.5759998662 | 0.989483196 |
| -0.5721362415 | 0.1217754493 | 0.0003412657 | 0.8005035379 | 0.5693509957 | 0.989483196 |
| -0.6500648892 | 0.4504289078 | 0.0003572388 | 0.7118458548 | 0.5181252199 | 0.989483196 |
| -0.5850477021 | 0.1493716015 | 0.0002148341 | 0.7995299872 | 0.5606863879 | 0.989483196 |
| -0.5769910352 | 0.4369812637 | 0.0003819768 | 0.7812312387 | 0.566091129 | 0.989483196 |
| -0.5805120282 | 0.9774827193 | 0.0004087564 | 0.695547755 | 0.5637321004 | 0.989483196 |
| -0.7184142748 | 0.3912161602 | 0.0002636484 | 0.5872563441 | 0.4752849144 | 0.989483196 |
| -0.5892066656 | 0.7105018753 | 0.0004643552 | 0.7468419616 | 0.5579154738 | 0.989483196 |
| -0.6496846293 | 0.5660231264 | 0.0007036711 | 0.657226842 | 0.5184239281 | 0.989483196 |
| -0.4016436634 | 0.0990987815 | 0.0002303733 | 0.6495741212 | 0.6893676316 | 0.989483196 |
| -0.4787660479 | 0.3280855759 | 0.0002142904 | 0.738703932 | 0.6338312255 | 0.989483196 |
| -0.4778927383 | 0.2578609584 | 0.0002538700 | 0.8233553185 | 0.6344538257 | 0.989483196 |
| -0.5164805645 | 0.7884652689 | 0.0006009158 | 0.7431362435 | 0.6073974703 | 0.989483196 |
| -0.775625932 | 0.0235080752 | 0.0006958242 | 0.4294030081 | 0.4409646012 | 0.989483196 |
| -0.6247374079 | 0.5353063443 | 0.0005120463 | 0.683537979 | 0.5345004224 | 0.989483196 |
| -0.5740939083 | 0.8216257667 | 0.0004174843 | 0.7112391304 | 0.5680279352 | 0.989483196 |
| -0.4310269376 | 0.5207595769 | 0.0003605996 | 0.7895919305 | 0.6679816642 | 0.989483196 |
| -0.602829668 | 0.7710364933 | 0.0006601039 | 0.671170273 | 0.5488772848 | 0.989483196 |
| -0.5755964199 | 0.9143761289 | 0.0004272959 | 0.7041784091 | 0.5670338134 | 0.989483196 |
| -0.6865847714 | 0.3646432215 | 0.0004568206 | 0.6255279717 | 0.494966086 | 0.989483196 |
| -0.4633008916 | 0.1070892023 | 0.0001360109 | 0.7282391605 | 0.6448219373 | 0.989483196 |
| -0.588912443 | 0.91551651 | 0.0004935388 | 0.6868589838 | 0.5581627988 | 0.989483196 |
| -0.5702276282 | 0.9949726927 | 0.0004155065 | 0.6964367397 | 0.5706395601 | 0.9949726927 |
| -0.5845499059 | 0.7596078641 | 0.0006767336 | 0.7126502664 | 0.5610448429 | 0.989483196 |
| -0.5948704139 | 0.7718244944 | 0.0004411720 | 0.7173654503 | 0.5541443061 | 0.989483196 |
| -0.5758542283 | 0.9635837904 | 0.0004229445 | 0.7019576847 | 0.5668563971 | 0.989483196 |

| | | | | | |
|---|---|---|---|---|---|
| -0.589265673 | 0.4921570332 | 0.0003547901 | 0.6233564332 | 0.5578759241 | 0.989483196 |
| -0.8919143131 | 0.1690667899 | 0.0002894080 | 0.5039888299 | 0.3760347135 | 0.989483196 |
| -0.5390019955 | 0.5056447941 | 0.0003787768 | 0.7269201299 | 0.5918839789 | 0.989483196 |
| -0.768720636 | 0.2676656958 | 0.0004271813 | 0.6234963159 | 0.4449818318 | 0.989483196 |
| -0.7781420977 | 0.1143905112 | 0.0003262312 | 0.465288221 | 0.4395684427 | 0.989483196 |
| -0.5735735255 | 0.9529816402 | 0.0004065628 | 0.6965940098 | 0.5683809237 | 0.989483196 |
| -0.7939431863 | 0.2927808185 | 0.0007814002 | 0.707091595 | 0.4302906602 | 0.989483196 |
| -0.5732750193 | 0.9705307846 | 0.0004087547 | 0.6955048928 | 0.5685708475 | 0.989483196 |
| -0.5386937661 | 0.6564726489 | 0.0004626063 | 0.7297673148 | 0.5920636475 | 0.989483196 |
| -0.6135211764 | 0.4788639542 | 0.0005361680 | 0.6964426497 | 0.5418232741 | 0.989483196 |
| -0.4405228472 | 0.4241712725 | 0.0003380460 | 0.8007395323 | 0.6611097907 | 0.989483196 |
| -0.6624576812 | 0.0752046349 | 0.0002248574 | 0.9786800182 | 0.5101810293 | 0.989483196 |
| -0.6824917085 | 0.2047672778 | 0.0005223929 | 0.5865616971 | 0.4975472757 | 0.989483196 |
| -0.7247448433 | 0.4610808199 | 0.0007447235 | 0.5800220313 | 0.471408727 | 0.989483196 |
| -0.7367867185 | 0.374500196 | 0.0007533318 | 0.7106857769 | 0.4640678466 | 0.989483196 |
| -0.4422439174 | 0.3542363164 | 0.0004425694 | 0.6066919806 | 0.6598810723 | 0.989483196 |
| -0.64088251 | 0.4048174516 | 0.0005146397 | 0.6752019421 | 0.5240195583 | 0.989483196 |
| -0.5866852114 | 0.9338347881 | 0.0004309344 | 0.7043353581 | 0.5595918046 | 0.989483196 |
| -0.4950183527 | 0.6829743424 | 0.0003861274 | 0.7558507874 | 0.6223429929 | 0.989483196 |
| -0.8091781245 | 0.1015748943 | 0.0002522220 | 0.5240079678 | 0.421554613 | 0.989483196 |
| -0.5125240528 | 0.8356762185 | 0.0004337318 | 0.7286970916 | 0.6101495955 | 0.989483196 |
| -0.6297387144 | 0.734545116 | 0.0005597540 | 0.6710508786 | 0.5312589865 | 0.989483196 |
| -0.5839924074 | 0.8488385311 | 0.0004034160 | 0.6977004997 | 0.5613959788 | 0.989483196 |
| -0.6379069584 | 0.4750709199 | 0.0007174297 | 0.6460040491 | 0.5259581498 | 0.989483196 |
| -0.5285459046 | 0.709340501 | 0.0004935327 | 0.7485319394 | 0.5990401795 | 0.989483196 |
| -0.58610217 | 0.829924691 | 0.0004124496 | 0.6695678013 | 0.5599880608 | 0.989483196 |
| -0.5783265368 | 0.8970928835 | 0.0004203937 | 0.699040947 | 0.5651952552 | 0.989483196 |
| -0.6298954533 | 0.4889547616 | 0.0005610546 | 0.6521681205 | 0.5311342512 | 0.989483196 |
| -0.4262009831 | 0.4793741782 | 0.0003249561 | 0.7412449981 | 0.6714563812 | 0.989483196 |
| -0.3601811482 | 0.3961190696 | 0.0003024600 | 0.8389178475 | 0.7199546783 | 0.989483196 |
| -0.5820961813 | 0.8542835808 | 0.0004320574 | 0.7127452626 | 0.5626640235 | 0.989483196 |
| -0.5376616147 | 0.853767301 | 0.0004053092 | 0.7235367555 | 0.5927971767 | 0.989483196 |
| -0.1562010007 | 0.1008053843 | 0.0005726339 | 0.8640640792 | 0.8763951327 | 0.989483196 |
| -0.6480604659 | 0.5749781847 | 0.0004202986 | 0.6902543183 | 0.5193785447 | 0.989483196 |
| -0.5059847417 | 0.1579640977 | 0.0002578770 | 0.8508598534 | 0.6147145765 | 0.989483196 |
| -1.10668106 | 0.0383892346 | 0.0006653626 | 0.3701686548 | 0.2727729905 | 0.989483196 |
| -0.6031551207 | 0.2341157683 | 0.0004403387 | 0.5749026896 | 0.548661561 | 0.989483196 |
| -0.5810577662 | 0.8931561033 | 0.0004066964 | 0.7019294085 | 0.5633619156 | 0.989483196 |
| -0.4702996493 | 0.4344209838 | 0.0003011145 | 0.7748072497 | 0.6398423828 | 0.989483196 |
| -0.5701499739 | 0.8516885786 | 0.0004475249 | 0.6952139642 | 0.5706893679 | 0.989483196 |
| -0.5636442143 | 0.5427371169 | 0.0003678345 | 0.6907487073 | 0.5750908294 | 0.989483196 |
| -0.5889784589 | 0.7832601769 | 0.0003908326 | 0.723942663 | 0.5580666347 | 0.989483196 |
| -0.5277488953 | 0.7460751906 | 0.0003894414 | 0.742506398 | 0.5996022906 | 0.989483196 |
| -0.5396271249 | 0.6745770698 | 0.0003704421 | 0.7139452887 | 0.5914394153 | 0.989483196 |
| -0.5404764654 | 0.3804407507 | 0.0002513694 | 0.7416455478 | 0.5908649081 | 0.989483196 |
| -0.5893223996 | 0.759566193 | 0.0005814190 | 0.6955511632 | 0.557836974 | 0.989483196 |

| | | | | | |
|---|---|---|---|---|---|
| -0.5428281388 | 0.9115767114 | 0.0004229212 | 0.7348503454 | 0.5892913179 | 0.989483196 |
| -0.6024881835 | 0.6468330096 | 0.0005834740 | 0.7076015624 | 0.5490885023 | 0.989483196 |
| -0.6670401934 | 0.0921767758 | 0.0004303648 | 0.6302184404 | 0.5072738793 | 0.989483196 |
| -0.5014574908 | 0.1782096163 | 0.0003894774 | 0.731059157 | 0.6178703605 | 0.989483196 |
| -0.5875646696 | 0.7367377 | 0.0004026679 | 0.720397827 | 0.5590097381 | 0.989483196 |
| -0.5804985158 | 0.8970555469 | 0.0005049785 | 0.7030554973 | 0.5637393527 | 0.989483196 |
| -0.560917322 | 0.250164556 | 0.0002216635 | 0.778157674 | 0.5769243235 | 0.989483196 |
| -0.5630820521 | 0.9079151621 | 0.0004047604 | 0.7144621169 | 0.5754543575 | 0.989483196 |
| -0.4583513674 | 0.3911631669 | 0.0003930429 | 0.9459660638 | 0.6483350984 | 0.989483196 |
| -0.5694064249 | 0.6123846461 | 0.0004037614 | 0.694394718 | 0.5711929786 | 0.989483196 |
| -0.5866812652 | 0.9294008769 | 0.0004222961 | 0.6987073908 | 0.5596179526 | 0.989483196 |
| -0.6437732698 | 0.0626533801 | 0.0002554639 | 0.7406272862 | 0.5221265981 | 0.989483196 |
| -0.5336985 | 0.4103242029 | 0.0004566656 | 0.6394049383 | 0.595511783 | 0.989483196 |
| -0.3169886554 | 0.2193093974 | 0.0002872749 | 0.7550645702 | 0.7523374899 | 0.989483196 |
| -0.5773534375 | 0.9688313107 | 0.0004588095 | 0.702093352 | 0.5658419424 | 0.989483196 |
| -0.5638615262 | 0.7090032934 | 0.0003763150 | 0.7022377654 | 0.574939009 | 0.989483196 |
| -0.4092943516 | 0.538838553 | 0.0003730435 | 0.7878698659 | 0.6837665463 | 0.989483196 |
| -0.7053801579 | 0.0745846056 | 0.0002137063 | 0.756729052 | 0.4832768018 | 0.989483196 |
| -0.6234504652 | 0.6293527009 | 0.0005676728 | 0.7049193217 | 0.5353365865 | 0.989483196 |
| -0.5781891725 | 0.7986461161 | 0.0005083366 | 0.6667827589 | 0.5652821476 | 0.989483196 |
| -0.5120393243 | 0.3805292478 | 0.0008969099 | 0.7798838131 | 0.61049271 | 0.989483196 |
| -0.405629841 | 0.4643406162 | 0.0003164451 | 0.8156503039 | 0.6864220354 | 0.989483196 |
| -0.5774107959 | 0.8979099159 | 0.0003988404 | 0.7020346355 | 0.5658081636 | 0.989483196 |
| -0.5884792948 | 0.9162399433 | 0.0005025633 | 0.6882836449 | 0.5583239431 | 0.989483196 |
| -0.5349122466 | 0.2700010589 | 0.0002874583 | 0.7116349937 | 0.5946430755 | 0.989483196 |
| -0.5682064035 | 0.8007880076 | 0.0004341444 | 0.6873851659 | 0.5720079904 | 0.989483196 |
| -0.5692223995 | 0.8568958413 | 0.0003950190 | 0.7004362321 | 0.5713168528 | 0.989483196 |
| -0.5308402332 | 0.8779639747 | 0.0005365851 | 0.7163899618 | 0.5974465294 | 0.989483196 |
| -0.6249212097 | 0.3184602952 | 0.0005726355 | 0.7326321851 | 0.5343621236 | 0.989483196 |
| -0.4145382893 | 0.6522977827 | 0.0003706348 | 0.8128890386 | 0.6799359341 | 0.989483196 |
| -0.5847768914 | 0.9384254456 | 0.0004039170 | 0.6927298696 | 0.5608665854 | 0.989483196 |
| -0.6122997326 | 0.7811689693 | 0.0004290494 | 0.6826411879 | 0.5426329077 | 0.989483196 |
| -0.5791613765 | 0.4109303178 | 0.0002707781 | 0.8094038839 | 0.5646518691 | 0.989483196 |
| -0.5236444055 | 0.148378217 | 0.0003153887 | 0.752857581 | 0.602437256 | 0.989483196 |
| -0.4346572268 | 0.4648878016 | 0.0003546372 | 0.7005131521 | 0.6653530081 | 0.989483196 |
| -0.7469052929 | 0.2417273883 | 0.0021067620 | 0.5514190627 | 0.4579561597 | 0.989483196 |
| -0.5831884971 | 0.9513893479 | 0.0004084814 | 0.6926039384 | 0.5619155849 | 0.989483196 |
| -0.6082610825 | 0.5019270821 | 0.0004028910 | 0.71941983 | 0.5452929343 | 0.989483196 |
| -0.5385068534 | 0.7857730117 | 0.0003982329 | 0.7503954175 | 0.5921944222 | 0.989483196 |
| -0.5753802448 | 0.9815673304 | 0.0004151183 | 0.6977179349 | 0.5671814781 | 0.989483196 |
| -0.5602020284 | 0.6486817875 | 0.0003503482 | 0.6840162833 | 0.5774199196 | 0.989483196 |
| -0.722146761 | 0.3269068325 | 0.0006966553 | 0.673349925 | 0.473015272 | 0.9984517566 |
| -0.6040988467 | 0.3985733636 | 0.0006234790 | 0.6678435908 | 0.5480609368 | 0.9984517566 |
| -0.5673716661 | 0.1024134451 | 0.0003136906 | 0.6656050708 | 0.572576955 | 0.9984517566 |
| -0.4794067453 | 0.0491988056 | 0.0004194284 | 0.734195605 | 0.6333955325 | 0.9984517566 |
| -0.7364887789 | 0.2215663033 | 0.0007989324 | 0.5561992733 | 0.4643051886 | 0.9984517566 |

| | | | | | |
|---|---|---|---|---|---|
| -0.9654856528 | 0.1908749258 | 0.0017582651 | 0.5392352939 | 0.3381992661 | 0.9984517566 |
| -0.6909040369 | 0.9984517566 | 0.0008316960 | 0.6306258654 | 0.4923047739 | 0.9984517566 |
| -0.7015819637 | 0.8204320567 | 0.0008225800 | 0.6249754936 | 0.4856612186 | 0.9984517566 |
| -0.7211951785 | 0.7020569444 | 0.0012066160 | 0.612914973 | 0.4735957649 | 0.9984517566 |
| -0.6628780197 | 0.8472445139 | 0.0008097072 | 0.6454938625 | 0.509923675 | 0.9984517566 |
| -0.6749766977 | 0.8657728038 | 0.0008639785 | 0.6375599033 | 0.5022955761 | 0.9984517566 |
| -0.6734788325 | 0.9501541927 | 0.0008306444 | 0.6277293122 | 0.5032497621 | 0.9984517566 |
| -0.673969543 | 0.7583034453 | 0.0008825932 | 0.6531629889 | 0.5029367166 | 0.9984517566 |
| -0.6932749657 | 0.0780801630 | 0.0006645213 | 0.7307542213 | 0.4908261671 | 0.9984517566 |
| -0.7572311367 | 0.451884704 | 0.0007336053 | 0.6482199375 | 0.4518991606 | 0.9984517566 |
| -0.6997919982 | 0.2001083533 | 0.0004881362 | 0.7097884622 | 0.486768248 | 0.9984517566 |
| -0.6979891396 | 0.4443957433 | 0.0008079608 | 0.7057694339 | 0.487892696 | 0.9984517566 |
| -0.7398535525 | 0.4810957718 | 0.0007984674 | 0.6050727805 | 0.4622980621 | 0.9984517566 |
| -0.7611036878 | 0.6351839922 | 0.0007300155 | 0.5754730657 | 0.4496001767 | 0.9984517566 |
| -0.6976051447 | 0.7870890522 | 0.0010109749 | 0.6670171841 | 0.4881300344 | 0.9984517566 |
| -0.7545096988 | 0.5009070493 | 0.0013528156 | 0.5954479864 | 0.4535679831 | 0.9984517566 |
| -0.4312927102 | 0.0175835431 | 0.0003236968 | 0.5664456281 | 0.6678247589 | 0.9984517566 |
| -0.5868827529 | 0.3362380903 | 0.0004619169 | 0.6801186924 | 0.5594893012 | 0.9984517566 |
| -0.6167182797 | 0.4281511389 | 0.0006332131 | 0.7233548585 | 0.5397619507 | 0.9984517566 |
| -0.5766788681 | 0.608370111 | 0.0009103211 | 0.7124112266 | 0.5663148412 | 0.9984517566 |
| -0.8835290102 | 0.0496899656 | 0.0017019786 | 0.4190183704 | 0.3804520954 | 0.9984517566 |
| -0.7115389829 | 0.7663694291 | 0.0009710019 | 0.6241173713 | 0.4795218626 | 0.9984517566 |
| -0.6898787851 | 0.9763951351 | 0.0009318313 | 0.6322273339 | 0.4929353101 | 0.9984517566 |
| -0.6168721194 | 0.7845368253 | 0.0010037521 | 0.6703574027 | 0.5396634071 | 0.9984517566 |
| -0.7483427015 | 0.5301750956 | 0.0015940528 | 0.57687138 | 0.4571916034 | 0.9984517566 |
| -0.6916240316 | 0.9932593245 | 0.0009403715 | 0.6313551712 | 0.4918551284 | 0.9984517566 |
| -0.7459210559 | 0.5635361803 | 0.0008979429 | 0.5938493687 | 0.4586471641 | 0.9984517566 |
| -0.5628870594 | 0.1002952369 | 0.0002851744 | 0.678999009 | 0.5756284148 | 0.9984517566 |
| -0.544197581 | 0.5049018451 | 0.0006953900 | 0.7059335281 | 0.5883664522 | 0.9984517566 |
| -0.8176584842 | 0.470249058 | 0.0008607329 | 0.6020774099 | 0.4168142138 | 0.9984517566 |
| -0.7106693251 | 0.456224044 | 0.0019972424 | 0.656943481 | 0.4800682391 | 0.9984517566 |
| -0.6914246217 | 0.9918579565 | 0.0008446801 | 0.6307506217 | 0.4919766011 | 0.9984517566 |
| -0.7107538928 | 0.7386217056 | 0.0007584064 | 0.6035775306 | 0.4800134637 | 0.9984517566 |
| -0.701530264 | 0.564237243 | 0.0007618183 | 0.5717125191 | 0.4856928048 | 0.9984517566 |
| -0.8386838889 | 0.453649202 | 0.0007021762 | 0.5349099768 | 0.4050439391 | 0.9984517566 |
| -0.6939115493 | 0.9384094662 | 0.0010131549 | 0.6271550435 | 0.4904577694 | 0.9984517566 |
| -0.8754616628 | 0.3243106478 | 0.0009458470 | 0.5655995467 | 0.3847477051 | 0.9984517566 |
| -0.9539541475 | 0.0434825500 | 0.0006118771 | 0.3666100844 | 0.3440025826 | 0.9984517566 |
| -0.737210008 | 0.5735700049 | 0.0008154808 | 0.6270512104 | 0.4638803797 | 0.9984517566 |
| -0.8707853316 | 0.3604677728 | 0.0014226194 | 0.6301469397 | 0.3873241991 | 0.9984517566 |
| -0.6768497362 | 0.9301487231 | 0.0008255218 | 0.6276345993 | 0.5010998467 | 0.9984517566 |
| -0.7677533702 | 0.4552780785 | 0.0006486099 | 0.5724883798 | 0.4456690079 | 0.9984517566 |
| -0.7377567941 | 0.4106875364 | 0.0011244880 | 0.6176263526 | 0.4635362101 | 0.9984517566 |
| -0.5677920081 | 0.511301948 | 0.0007270782 | 0.7189997712 | 0.5722781091 | 0.9984517566 |
| -0.7688740318 | 0.1734162113 | 0.0006034351 | 0.8343978319 | 0.4449813319 | 0.9984517566 |
| -0.8096982063 | 0.1864537826 | 0.0011142328 | 0.5256542459 | 0.4213336509 | 0.9984517566 |

| | | | | | |
|---|---|---|---|---|---|
| -0.7045391965 | 0.8869967647 | 0.0011376521 | 0.6144638112 | 0.4838281317 | 0.9984517566 |
| -0.9562988936 | 0.1511269885 | 0.0018979412 | 0.6502482579 | 0.3427346469 | 0.9984517566 |
| -0.4972235032 | 0.2103476088 | 0.0008743920 | 0.5217256786 | 0.6208496431 | 0.9984517566 |
| -0.7537984705 | 0.3696971226 | 0.0010547427 | 0.6103633421 | 0.4539302231 | 0.9984517566 |
| -0.6289478465 | 0.558326264 | 0.0006537957 | 0.5886587307 | 0.5317889937 | 0.9984517566 |
| -0.6104689169 | 0.7382288072 | 0.0007934585 | 0.6823355892 | 0.543818647 | 0.9984517566 |
| -0.8166183886 | 0.3964518035 | 0.0007498744 | 0.5481180417 | 0.4173424379 | 0.9984517566 |
| -0.7142895081 | 0.8510520958 | 0.0009950099 | 0.6098669438 | 0.4778295524 | 0.9984517566 |
| -0.7505350946 | 0.6690477102 | 0.0011463866 | 0.5979528025 | 0.4559004248 | 0.9984517566 |
| -0.6915777048 | 0.9915225386 | 0.0008755858 | 0.6293328063 | 0.4918781977 | 0.9984517566 |
| -0.7431284458 | 0.5272809591 | 0.0013800147 | 0.5906683962 | 0.4603272819 | 0.9984517566 |
| -0.7109634096 | 0.8427719053 | 0.0015910281 | 0.6085628344 | 0.4798668263 | 0.9984517566 |
| -0.7090223225 | 0.5816829751 | 0.0008462786 | 0.5567161595 | 0.4810725908 | 0.9984517566 |
| -0.6935869309 | 0.8805640239 | 0.0008061469 | 0.6266102077 | 0.4906388401 | 0.9984517566 |
| -0.7394225172 | 0.5195908025 | 0.0011474818 | 0.5909199417 | 0.4625438958 | 0.9984517566 |
| -0.5238342439 | 0.4562738074 | 0.0006503307 | 0.6781693547 | 0.602309535 | 0.9984517566 |
| -0.4210157488 | 0.3072688239 | 0.0005495004 | 0.7980012496 | 0.6752421707 | 0.9984517566 |
| -0.6997697811 | 0.6712140238 | 0.0007484998 | 0.6606676003 | 0.4867847444 | 0.9984517566 |
| -0.775316351 | 0.5930861266 | 0.0008704956 | 0.5717026156 | 0.4412318951 | 0.9984517566 |
| -0.4168975245 | 0.3016204052 | 0.0011376523 | 0.7368946259 | 0.6782449936 | 0.9984517566 |
| -0.6894281356 | 0.9816088858 | 0.0008320360 | 0.62906013 | 0.4932069881 | 0.9984517566 |
| -0.6239952219 | 0.1547697076 | 0.0005615617 | 0.7923763778 | 0.5350082426 | 0.9984517566 |
| -1.038553971 | 0.1660923701 | 0.0013737607 | 0.4212446842 | 0.303149733 | 0.9984517566 |
| -0.7137610043 | 0.2775524279 | 0.0009057877 | 0.5250592811 | 0.4781537718 | 0.9984517566 |
| -0.7006133444 | 0.4408664885 | 0.0008027394 | 0.6574959125 | 0.486267322 | 0.9984517566 |
| -0.5368272448 | 0.2451837939 | 0.0003901946 | 0.7369047568 | 0.5933912355 | 0.9984517566 |
| -0.6641815092 | 0.5901413427 | 0.0009604890 | 0.6264798673 | 0.509132105 | 0.9984517566 |
| -0.6885425569 | 0.5859092538 | 0.0007872729 | 0.6190413932 | 0.4937794163 | 0.9984517566 |
| -0.6938070419 | 0.9404932577 | 0.0008591145 | 0.6383061707 | 0.4904876053 | 0.9984517566 |
| -0.5868967747 | 0.4969949575 | 0.0006151383 | 0.722650387 | 0.5594840476 | 0.9984517566 |
| -0.6552509487 | 0.7086562945 | 0.0007750727 | 0.6431136634 | 0.5148258427 | 0.9984517566 |
| -0.6271240238 | 0.203163227 | 0.0003811613 | 0.7013923348 | 0.5329821565 | 0.9984517566 |
| -0.6924140721 | 0.9719016415 | 0.0010080550 | 0.6290885458 | 0.4913579926 | 0.9984517566 |
| -0.6642507735 | 0.9515776377 | 0.0008610135 | 0.6567787405 | 0.5091323191 | 0.9984517566 |
| -0.7257595487 | 0.5719443596 | 0.0012769698 | 0.6361420557 | 0.4708011772 | 0.9984517566 |
| -0.7590391042 | 0.1809265214 | 0.0009263417 | 0.5807836859 | 0.4508061507 | 0.9984517566 |
| -0.5971859185 | 0.0513578194 | 0.0007339269 | 0.6685915982 | 0.5526366663 | 0.9984517566 |
| -0.7002029304 | 0.4748142934 | 0.0007606472 | 0.6822609149 | 0.4865116717 | 0.9984517566 |
| -0.6917047729 | 0.868372489 | 0.0009661603 | 0.6359079091 | 0.4918031287 | 0.9984517566 |
| -0.6748919838 | 0.339126387 | 0.0005343434 | 0.6959813656 | 0.5023452528 | 0.9984517566 |
| -0.6516589618 | 0.7648411691 | 0.0008168911 | 0.6708168381 | 0.5171108974 | 0.9984517566 |
| -0.5878938191 | 0.4998954532 | 0.0007967485 | 0.8243056738 | 0.5588109039 | 0.9984517566 |
| -0.6830184258 | 0.5270965501 | 0.0008251984 | 0.6256425528 | 0.4972369535 | 0.9984517566 |
| -0.700817983 | 0.9028132521 | 0.0008589711 | 0.6326903372 | 0.4861540907 | 0.9984517566 |
| -0.7065637851 | 0.2653549744 | 0.0006390245 | 0.6576926916 | 0.4825724179 | 0.9984517566 |
| -0.6535917608 | 0.4332696252 | 0.0009553462 | 0.5749566789 | 0.5158860755 | 0.9984517566 |

| | | | | | |
|---|---|---|---|---|---|
| -0.3810287205 | 0.1561426986 | 0.0005521651 | 0.7057240198 | 0.7045255625 | 0.9984517566 |
| -0.6730463888 | 0.7622950221 | 0.0008039618 | 0.6565620984 | 0.5035120041 | 0.9984517566 |
| -0.6863874348 | 0.895383396 | 0.0008364988 | 0.6307363612 | 0.4951280314 | 0.9984517566 |
| -0.7014615454 | 0.8893238648 | 0.0009435638 | 0.6150659605 | 0.4857518151 | 0.9984517566 |
| -0.8251992904 | 0.0735740459 | 0.0004556595 | 0.6814626368 | 0.412537087 | 0.9984517566 |
| -0.7104152784 | 0.8207623763 | 0.0010569119 | 0.6362632141 | 0.4802132385 | 0.9984517566 |
| -0.6899998393 | 0.8435041593 | 0.0010173644 | 0.6092798512 | 0.4928639941 | 0.9984517566 |
| -0.6520990559 | 0.6013177965 | 0.0015012519 | 0.6759308304 | 0.516845976 | 0.9984517566 |
| -0.5866248245 | 0.7122632057 | 0.0007955355 | 0.6938800966 | 0.559634622 | 0.9984517566 |
| -0.6665835161 | 0.1420227889 | 0.0005694276 | 0.6822475526 | 0.5076191346 | 0.9984517566 |
| -0.7421423612 | 0.7863945659 | 0.0011627206 | 0.6019370548 | 0.4607988382 | 0.9984517566 |
| -0.6497143392 | 0.5315609252 | 0.0007251219 | 0.6535769735 | 0.5183269226 | 0.9984517566 |
| -0.6772643139 | 0.6576345291 | 0.0007639489 | 0.6114677597 | 0.5008589776 | 0.9984517566 |
| -0.6800579031 | 0.8431866251 | 0.0008039276 | 0.6336205152 | 0.4990978182 | 0.9984517566 |
| -0.6265443063 | 0.8473203139 | 0.0010299777 | 0.6538549494 | 0.5333137217 | 0.9984517566 |
| -0.7529061929 | 0.2586651414 | 0.0012058298 | 0.6545107964 | 0.4544479014 | 0.9984517566 |
| -0.6449831748 | 0.968726724 | 0.0008815449 | 0.65382224 | 0.5213953959 | 0.9984517566 |
| -0.675989889 | 0.9226804776 | 0.0008311328 | 0.6351693592 | 0.5016451516 | 0.9984517566 |
| -0.6810748251 | 0.955615686 | 0.0008240864 | 0.6340603863 | 0.4984655684 | 0.9984517566 |
| -0.6848297071 | 0.1945923947 | 0.0002868562 | 0.8171741247 | 0.496125121 | 0.9984517566 |
| -0.626571437 | 0.0734545266 | 0.0005530304 | 0.6868230073 | 0.5333283012 | 0.9984517566 |
| -0.5067146111 | 0.3580026607 | 0.0006890517 | 0.633409946 | 0.6142172635 | 0.9984517566 |
| -0.7956491208 | 0.5006196769 | 0.0029799131 | 0.548773638 | 0.4293326901 | 0.9984517566 |
| -0.7837239198 | 0.4358084359 | 0.0008320724 | 0.5526083746 | 0.4362777422 | 0.9984517566 |
| -0.7037236609 | 0.721795411 | 0.0008315952 | 0.6423254029 | 0.4843393368 | 0.9984517566 |
| -0.6776949873 | 0.9622092246 | 0.0008732351 | 0.6459243268 | 0.5005699899 | 0.9984517566 |
| -0.6638811376 | 0.7854822254 | 0.0008788857 | 0.6370248219 | 0.5093324077 | 0.9984517566 |
| -0.6640397946 | 0.4311632975 | 0.0006113492 | 0.6058950313 | 0.5092310344 | 0.9984517566 |
| 1.176458302 | 0.5593357919 | 0.0294925099 | 0.0873822162 | 0.244177017 | 0.8151231524 |
| 1.115669634 | 0.4372743286 | 0.0234595695 | 0.1002928154 | 0.2691318144 | 0.8038399796 |
| 1.191817661 | 0.9746060135 | 0.0312589227 | 0.0929023039 | 0.2381470678 | 0.9888060224 |
| 1.157867055 | 0.8163278565 | 0.0277525825 | 0.0983233866 | 0.2516187706 | 0.907007274 |
| 1.33697486 | 0.0203528032 | 0.0241815945 | 0.0472557734 | 0.1864191002 | 0.4797907818 |
| 1.438457649 | 0.1990168261 | 0.0459703064 | 0.0708737335 | 0.1556587345 | 0.654914271 |
| 1.201670832 | 0.8173333639 | 0.0279694020 | 0.0906716982 | 0.2343449489 | 0.907007274 |
| 1.123545491 | 0.1341657621 | 0.0435458605 | 0.0954926736 | 0.2657941887 | 0.654914271 |
| 1.256440518 | 0.4852231269 | 0.0417528510 | 0.0848259699 | 0.2139495688 | 0.8070010522 |
| 1.266511477 | 0.5665386284 | 0.0293195654 | 0.0809245393 | 0.2103018749 | 0.8151231524 |
| 1.293636179 | 0.2297506659 | 0.0458203650 | 0.0789602796 | 0.2008809335 | 0.6837817438 |
| 1.041853736 | 0.2982750352 | 0.0230367760 | 0.0816198091 | 0.3017901845 | 0.7767579042 |
| 1.209302827 | 0.8049136926 | 0.0276933636 | 0.0889754074 | 0.2314310354 | 0.907007274 |
| 1.214229356 | 0.1525996021 | 0.0279210548 | 0.1118904089 | 0.2295494125 | 0.654914271 |
| 1.404644992 | 0.0407824642 | 0.0199393759 | 0.0984144566 | 0.165438272 | 0.5884208072 |
| 1.198632773 | 0.4268449464 | 0.0229826939 | 0.1073163516 | 0.2355106844 | 0.8038399796 |
| 1.230156998 | 0.3410023091 | 0.0294881479 | 0.1162128942 | 0.2235706024 | 0.8038399796 |
| 1.126628281 | 0.3570405032 | 0.0264016079 | 0.0993460204 | 0.2645098478 | 0.8038399796 |

| | | | | | |
|---|---|---|---|---|---|
| 1.240645962 | 0.7121334035 | 0.0262297176 | 0.0854913599 | 0.2197059983 | 0.8901667544 |
| 1.196157783 | 0.47624425 | 0.0533276099 | 0.0730731510 | 0.2364781557 | 0.8054082189 |
| 1.188686394 | 0.1637549057 | 0.0154287919 | 0.0765496894 | 0.2395557615 | 0.654914271 |
| 1.126349999 | 0.5633160331 | 0.0273473639 | 0.0916179826 | 0.2646169841 | 0.8151231524 |
| 1.06950259 | 0.2050682882 | 0.0135592012 | 0.108142165 | 0.289237514 | 0.654914271 |
| 1.17764771 | 0.9020035843 | 0.0281167671 | 0.1014879827 | 0.2437028236 | 0.9425708261 |
| 1.570435211 | 0.0884510219 | 0.112235317 | 0.0384561269 | 0.1217358252 | 0.654914271 |
| 1.168641542 | 0.7921389862 | 0.0271514405 | 0.1051142388 | 0.2472958472 | 0.907007274 |
| 1.085030512 | 0.1311456623 | 0.0173825311 | 0.094401731 | 0.2823619483 | 0.654914271 |
| 1.271245092 | 0.250521666 | 0.044063635 | 0.0700934768 | 0.2086829843 | 0.6958935166 |
| 1.131638054 | 0.8799538267 | 0.0317506179 | 0.1041679466 | 0.2624152874 | 0.9401216098 |
| 1.050630505 | 0.1844185553 | 0.0145472780 | 0.1302745589 | 0.2977428153 | 0.654914271 |
| 1.180504084 | 0.4244260348 | 0.0205441878 | 0.1148766426 | 0.2426550472 | 0.8038399796 |
| 1.348530781 | 0.0470017797 | 0.0300803382 | 0.0735425703 | 0.1826455056 | 0.5884208072 |
| 1.14038772 | 0.4768016656 | 0.0226642982 | 0.1004435776 | 0.258766432 | 0.8054082189 |
| 1.10960761 | 0.7412173087 | 0.0265067026 | 0.1022988909 | 0.2717709137 | 0.8995355688 |
| 1.079726314 | 0.6439194774 | 0.0271500138 | 0.0981728444 | 0.2847105203 | 0.8562759008 |
| 1.203773292 | 0.279645281 | 0.0156532042 | 0.0800613338 | 0.2335788594 | 0.7437374496 |
| 1.169585833 | 0.6193668797 | 0.0264591103 | 0.0861732306 | 0.2469136184 | 0.8357540956 |
| 1.22308764 | 0.6847377084 | 0.0265919078 | 0.0858196501 | 0.226213229 | 0.8733899342 |
| 1.190557431 | 0.5975521447 | 0.0303039553 | 0.1145581574 | 0.2386422916 | 0.8357540956 |
| 1.305440452 | 0.4767504934 | 0.0250194889 | 0.0786019458 | 0.1969376745 | 0.8054082189 |
| 1.09985531 | 0.1361086775 | 0.0154088759 | 0.0974711091 | 0.2759219737 | 0.654914271 |
| 1.295798302 | 0.5456052708 | 0.0301912507 | 0.0815927024 | 0.2000671118 | 0.8151231524 |
| 1.397975192 | 0.1571187127 | 0.0286234474 | 0.0490556075 | 0.1674400451 | 0.654914271 |
| 1.160048992 | 0.756145199 | 0.0280940125 | 0.0940559047 | 0.2507479174 | 0.907007274 |
| 1.330381919 | 0.4095509597 | 0.0406980499 | 0.0973741221 | 0.1885562177 | 0.8038399796 |
| 1.359870764 | 0.1880383131 | 0.0336373034 | 0.0909655471 | 0.1790745883 | 0.654914271 |
| 1.242522235 | 0.6920767545 | 0.0264628200 | 0.0830284325 | 0.2190448208 | 0.8738342859 |
| 1.310426549 | 0.0724659096 | 0.0404070255 | 0.0782093830 | 0.1951645186 | 0.654914271 |
| 1.096517514 | 0.6801489267 | 0.0267094660 | 0.1143512596 | 0.2773136231 | 0.8733899342 |
| 1.203961171 | 0.8235544612 | 0.0277070281 | 0.1065155319 | 0.2334513411 | 0.907007274 |
| 1.280122963 | 0.3967645869 | 0.0353152231 | 0.0755417539 | 0.2055580557 | 0.8038399796 |
| 1.36173039 | 0.4171265698 | 0.0449510313 | 0.0662091678 | 0.178516097 | 0.8038399796 |
| 1.028150491 | 0.3584272 | 0.0192748117 | 0.0798847130 | 0.3081379412 | 0.8038399796 |
| 0.9949498438 | 0.1836370688 | 0.0317951834 | 0.0647146333 | 0.3238472142 | 0.654914271 |
| 1.172433858 | 0.6722292896 | 0.0261756044 | 0.0930540195 | 0.2457933919 | 0.8733899342 |
| 1.119116909 | 0.4058050715 | 0.0231489976 | 0.0784552470 | 0.2676623959 | 0.8038399796 |
| 1.479841264 | 0.1932344686 | 0.0304467367 | 0.0502265047 | 0.144226577 | 0.654914271 |
| 1.290423558 | 0.5312245427 | 0.0269355447 | 0.0759239247 | 0.2019463716 | 0.8151231524 |
| 0.9735488717 | 0.4971126481 | 0.0236751847 | 0.1257448076 | 0.3342897695 | 0.8070010522 |
| 1.255775544 | 0.5662373977 | 0.0358708905 | 0.0855827691 | 0.2142305272 | 0.8151231524 |
| 1.252673666 | 0.2003313494 | 0.0194302657 | 0.0861396959 | 0.2153246689 | 0.654914271 |
| 1.168625464 | 0.8143672225 | 0.0281195069 | 0.0980980244 | 0.247309939 | 0.907007274 |
| 1.063840032 | 0.4552243088 | 0.0209496490 | 0.1285052232 | 0.2917609236 | 0.8038399796 |
| 1.246159575 | 0.3392153345 | 0.0308504398 | 0.0642499447 | 0.2176804571 | 0.8038399796 |

| | | | | | |
|---|---|---|---|---|---|
| 1.245852477 | 0.268416587 | 0.0240308948 | 0.0829313466 | 0.2178089628 | 0.7293928996 |
| 1.285098437 | 0.3255279336 | 0.0410440906 | 0.0772874942 | 0.2038238854 | 0.8038399796 |
| 1.192077257 | 0.8996820288 | 0.0346131704 | 0.0919312950 | 0.2380186197 | 0.9425708261 |
| 1.307202035 | 0.5673257141 | 0.0355193299 | 0.0757151089 | 0.1962566895 | 0.8151231524 |
| 1.315505303 | 0.0470736645 | 0.0112759398 | 0.1139829036 | 0.1934949469 | 0.5884208072 |
| 1.270653322 | 0.6072466917 | 0.0296658593 | 0.0806138296 | 0.2089125828 | 0.8357540956 |
| 0.6168716597 | 0.0199164341 | 0.0400035833 | 0.1512264507 | 0.5397187363 | 0.4797907818 |
| 1.194588364 | 0.9299967395 | 0.0285706649 | 0.0927932478 | 0.2370561059 | 0.9528655118 |
| 1.283377217 | 0.1765473344 | 0.0304129929 | 0.0575925805 | 0.2044292607 | 0.654914271 |
| 1.492662165 | 0.2265306289 | 0.0413605806 | 0.0520113139 | 0.1408751994 | 0.6837817438 |
| 1.285173316 | 0.0232760688 | 0.0301187656 | 0.0400929341 | 0.2038096721 | 0.4797907818 |
| 1.189105805 | 0.7826128 | 0.0282307540 | 0.0916959097 | 0.2392151379 | 0.907007274 |
| 1.112326558 | 0.540295151 | 0.0228733192 | 0.1101595436 | 0.2705676237 | 0.8151231524 |
| 1.299384908 | 0.0787047259 | 0.0174725756 | 0.0889393524 | 0.1989104317 | 0.654914271 |
| 1.204749333 | 0.6114942538 | 0.0272732801 | 0.0861639079 | 0.2331609405 | 0.8357540956 |
| 1.160477006 | 0.3539257833 | 0.0351791433 | 0.0745980954 | 0.2505705702 | 0.8038399796 |
| 1.423263724 | 0.1145507699 | 0.0557970364 | 0.0516046719 | 0.1599662513 | 0.654914271 |
| 1.172719531 | 0.8344466921 | 0.0281042948 | 0.0950722645 | 0.2456745885 | 0.907007274 |
| 1.223828385 | 0.5408877343 | 0.0379297230 | 0.0844780169 | 0.2259192785 | 0.8151231524 |
| 1.260476127 | 0.0919742440 | 0.0585175576 | 0.0855414491 | 0.2125023238 | 0.654914271 |
| 0.961219016 | 0.2503331802 | 0.0343332159 | 0.187856815 | 0.340452552 | 0.6958935166 |
| 1.24244888 | 0.5379014848 | 0.0408348087 | 0.0931739108 | 0.219007831 | 0.8151231524 |
| 1.284506078 | 0.0944179546 | 0.0321997875 | 0.0753134757 | 0.2040319234 | 0.654914271 |
| 1.193251426 | 0.9808955742 | 0.0283751487 | 0.0929112672 | 0.2376045417 | 0.9888060224 |
| 1.188029046 | 0.2055251867 | 0.0243218510 | 0.1335420402 | 0.2396162366 | 0.654914271 |
| 1.192472591 | 0.4235046966 | 0.0212204061 | 0.1021079077 | 0.2378998352 | 0.8038399796 |
| 1.189271525 | 0.8671258194 | 0.0288265214 | 0.0978155642 | 0.2391439658 | 0.9344028227 |
| 1.153501063 | 0.81065099 | 0.0280036549 | 0.1069215948 | 0.253390635 | 0.907007274 |
| 1.070454841 | 0.4910555905 | 0.0270439399 | 0.179889463 | 0.2888118427 | 0.8070010522 |
| 1.19798103 | 0.3141136576 | 0.0277340692 | 0.0855607722 | 0.2357729789 | 0.8013103511 |
| 1.345166656 | 0.2411068466 | 0.0315609778 | 0.1017886271 | 0.18378695 | 0.6958935166 |
| 1.198602787 | 0.0041672126 | 0.0115129095 | 0.1206840185 | 0.2355044342 | 0.4797907818 |
| 1.180079218 | 0.7819902624 | 0.0302093881 | 0.0894828027 | 0.2427583819 | 0.907007274 |
| 1.279362231 | 0.605964118 | 0.0293693708 | 0.0847884056 | 0.205785223 | 0.8357540956 |
| 1.240452191 | 0.4499829649 | 0.0390806291 | 0.0776347664 | 0.2197674192 | 0.8038399796 |
| 1.248224082 | 0.1609112565 | 0.0350649625 | 0.0847643414 | 0.2169335637 | 0.654914271 |
| 0.9100067405 | 0.3697950515 | 0.0224942166 | 0.1388147688 | 0.3665548898 | 0.8038399796 |
| 1.330013688 | 0.1180762383 | 0.0215663944 | 0.1002142623 | 0.1886793858 | 0.654914271 |
| 1.036381368 | 0.0134201954 | 0.0076199585 | 0.0526348208 | 0.3043240807 | 0.4797907818 |
| 1.282216436 | 0.0268682837 | 0.0092648902 | 0.1736643353 | 0.2048710596 | 0.4797907818 |
| 1.095696612 | 0.1639299511 | 0.0619177271 | 0.1260843787 | 0.2777165805 | 0.654914271 |
| 0.9451411501 | 0.388184424 | 0.0212347179 | 0.1525143167 | 0.3484407998 | 0.8038399796 |
| 1.235611168 | 0.0108941715 | 0.0177700482 | 0.0940567368 | 0.22158489 | 0.4797907818 |
| 1.411022107 | 0.3932114497 | 0.0462010528 | 0.0674082857 | 0.1633848651 | 0.8038399796 |
| 1.124139633 | 0.6689182782 | 0.0266680255 | 0.1099331911 | 0.2655458255 | 0.8733899342 |
| 1.237242809 | 0.1400178659 | 0.0548624247 | 0.1120165038 | 0.2209459501 | 0.654914271 |

| | | | | | |
|---|---|---|---|---|---|
| 1.307431499 | 0.0653379778 | 0.0331628114 | 0.0756187776 | 0.1961898055 | 0.654914271 |
| 1.101992568 | 0.8317437891 | 0.0301090633 | 0.1055140739 | 0.2749554996 | 0.907007274 |
| 1.187422645 | 0.9174742615 | 0.0283907297 | 0.0925124956 | 0.2398591567 | 0.9478039892 |
| 1.095618405 | 0.904867993 | 0.0284737609 | 0.1179928241 | 0.2777358927 | 0.9425708261 |
| 1.141189126 | 0.7315015047 | 0.0283649801 | 0.1005440483 | 0.258439099 | 0.8995355688 |
| 1.325296064 | 0.1764691243 | 0.0293126632 | 0.0813105697 | 0.1902297965 | 0.654914271 |
| 1.193611762 | 0.9973965969 | 0.0392469904 | 0.1006050696 | 0.2374617992 | 0.9973965969 |
| 1.144466746 | 0.1281850916 | 0.0248017428 | 0.1088160796 | 0.2570971663 | 0.654914271 |
| 1.235649573 | 0.7407869613 | 0.0288792248 | 0.0919058649 | 0.2215270668 | 0.8995355688 |
| 1.416480744 | 0.1839854945 | 0.1026769576 | 0.0528641302 | 0.1618840384 | 0.654914271 |
| 1.297379672 | 0.4565811084 | 0.0301234422 | 0.0733365065 | 0.199576521 | 0.8038399796 |
| 1.208804004 | 0.6218010472 | 0.0287064219 | 0.0994130410 | 0.2316102266 | 0.8357540956 |
| 1.067132761 | 0.4420294698 | 0.0231543338 | 0.1431573724 | 0.2902845154 | 0.8038399796 |
| 1.300083212 | 0.2095725667 | 0.0222616355 | 0.0855246107 | 0.1986896947 | 0.654914271 |
| 1.217405863 | 0.379142648 | 0.0346867250 | 0.0999223636 | 0.2283460746 | 0.8038399796 |
| -1.837776848 | 0.3203968605 | 0.0468282121 | 0.1243004567 | 0.0710133703 | 0.8706436426 |
| -1.847171704 | 0.6558967896 | 0.0603227881 | 0.1097603641 | 0.06959392964 | 0.9949216742 |
| -1.720457819 | 0.1018415948 | 0.0290084484 | 0.1200040613 | 0.09042430495 | 0.7766647204 |
| -1.761960666 | 0.7171681283 | 0.0525750244 | 0.1209068421 | 0.08310777236 | 0.9949216742 |
| -1.871386622 | 0.1118871444 | 0.0516824335 | 0.0851713477 | 0.06611478629 | 0.7766647204 |
| -2.072880406 | 0.1818576058 | 0.0871066981 | 0.0822237276 | 0.04246050775 | 0.7766647204 |
| -1.810270923 | 0.9435853741 | 0.0540887937 | 0.1137976889 | 0.07523525968 | 0.9949216742 |
| -1.827725845 | 0.6914303912 | 0.0489411007 | 0.1097185786 | 0.07252952144 | 0.9949216742 |
| -1.798809386 | 0.9599862299 | 0.0568046080 | 0.1144081158 | 0.07702773568 | 0.9949216742 |
| -1.768829711 | 0.7458544442 | 0.0534915193 | 0.1210458485 | 0.08189770898 | 0.9949216742 |
| -1.826382982 | 0.7884542828 | 0.063327748 | 0.1116860552 | 0.07272863679 | 0.9949216742 |
| -1.556798421 | 0.0740316806 | 0.0345762524 | 0.0878176311 | 0.1248080055 | 0.7118430836 |
| -1.7911154 | 0.836780664 | 0.0556043493 | 0.1202950992 | 0.07829885216 | 0.9949216742 |
| -1.811919378 | 0.2671726638 | 0.0533152490 | 0.1354539279 | 0.07496787016 | 0.7766647204 |
| -1.898443994 | 0.369019049 | 0.0502692774 | 0.1173428346 | 0.06243820171 | 0.941375125 |
| -1.823468412 | 0.1831969472 | 0.0370645909 | 0.1438335038 | 0.07316391551 | 0.7766647204 |
| -1.818334674 | 0.315139745 | 0.0527741356 | 0.1496775538 | 0.07398031172 | 0.8706436426 |
| -1.851655913 | 0.568302405 | 0.0544734054 | 0.1056005339 | 0.06898127835 | 0.9949216742 |
| -1.995089912 | 0.2594237402 | 0.0337993833 | 0.0755697446 | 0.05057316974 | 0.7766647204 |
| -1.808307471 | 0.9789355206 | 0.0643436973 | 0.1164597551 | 0.07554284459 | 0.9949216742 |
| -1.872627623 | 0.5665374318 | 0.0738115913 | 0.1031766441 | 0.06609436009 | 0.9949216742 |
| -1.674340103 | 0.1272855521 | 0.0436903917 | 0.0983151882 | 0.09922187691 | 0.7766647204 |
| -1.682611283 | 0.1205811334 | 0.0207673266 | 0.1235960526 | 0.09763095661 | 0.7766647204 |
| -1.691427819 | 0.0647046404 | 0.0326848025 | 0.1742083913 | 0.09593829458 | 0.7118430836 |
| -1.593316552 | 0.336960152 | 0.0352641647 | 0.1763805415 | 0.1162765304 | 0.8961706171 |
| -2.008328281 | 0.0462753175 | 0.0882869331 | 0.0592387924 | 0.04901389877 | 0.7118430836 |
| -1.843153578 | 0.6330923249 | 0.0609693384 | 0.1080949755 | 0.07023109614 | 0.9949216742 |
| -1.797313633 | 0.70353157 | 0.0500358756 | 0.1205657602 | 0.0772658055 | 0.9949216742 |
| -1.665048797 | 0.6323514514 | 0.0487269587 | 0.142751368 | 0.101085025 | 0.9949216742 |
| -1.815004729 | 0.8768488224 | 0.0647689570 | 0.1101525554 | 0.07448788001 | 0.9949216742 |
| -1.804185643 | 0.8568634003 | 0.0528395146 | 0.1185052114 | 0.07619154713 | 0.9949216742 |

| | | | | | |
|---|---|---|---|---|---|
| -1.957592857 | 0.2266453748 | 0.0605894475 | 0.0897122520 | 0.05487310033 | 0.7766647204 |
| -1.748228997 | 0.2171091903 | 0.0340111873 | 0.1153207303 | 0.08550941494 | 0.7766647204 |
| -1.895535837 | 0.5663597394 | 0.0681551754 | 0.0976333287 | 0.06290879579 | 0.9949216742 |
| -1.762522209 | 0.9550633178 | 0.0543228586 | 0.1144145669 | 0.08304166105 | 0.9949216742 |
| -1.80531558 | 0.3781060918 | 0.0906099364 | 0.1275087944 | 0.07600465458 | 0.9452652295 |
| -1.807942429 | 0.9920399718 | 0.0545352551 | 0.1145328053 | 0.07557866342 | 0.9949216742 |
| -1.788652866 | 0.8344161797 | 0.0565923149 | 0.1204486247 | 0.07868733347 | 0.9949216742 |
| -1.808353261 | 0.7203613022 | 0.0568206573 | 0.1309954314 | 0.07552181503 | 0.9949216742 |
| -1.865328766 | 0.6620842926 | 0.0521995376 | 0.1036936171 | 0.06709824426 | 0.9949216742 |
| -1.751054318 | 0.4843948778 | 0.0461485180 | 0.1233430112 | 0.08504677955 | 0.9949216742 |
| -2.130520102 | 0.0730616250 | 0.0569682463 | 0.0780678330 | 0.03710434542 | 0.7118430836 |
| -2.177746463 | 0.0215283678 | 0.0488064186 | 0.0333352162 | 0.03341677025 | 0.708748979 |
| -1.769268482 | 0.6613168655 | 0.0542675955 | 0.115508095 | 0.08188386945 | 0.9949216742 |
| -1.913877548 | 0.504413425 | 0.0726700789 | 0.1191205091 | 0.06032378958 | 0.9949216742 |
| -1.838615594 | 0.7191420241 | 0.0575447508 | 0.116432092 | 0.07084453735 | 0.9949216742 |
| -1.710541667 | 0.2650782068 | 0.0648357259 | 0.1432195133 | 0.09222482643 | 0.7766647204 |
| -1.861542828 | 0.2225175186 | 0.0711547404 | 0.1178879431 | 0.06750080982 | 0.7766647204 |
| -1.80687964 | 0.8920991375 | 0.0552356965 | 0.1134885281 | 0.07569299393 | 0.9949216742 |
| -1.868363814 | 0.2287138194 | 0.0433731721 | 0.179774942 | 0.0665433925 | 0.7766647204 |
| -2.095607526 | 0.0184818006 | 0.0741613454 | 0.0518400650 | 0.04036637729 | 0.708748979 |
| -2.014159916 | 0.2930145806 | 0.0869123585 | 0.0714922846 | 0.04845478791 | 0.8324277859 |
| -2.276454307 | 0.0192895080 | 0.1197263117 | 0.1174496699 | 0.02628628918 | 0.708748979 |
| -1.707400206 | 0.5103627356 | 0.0582913864 | 0.0947239569 | 0.09284247078 | 0.9949216742 |
| -1.835804721 | 0.6766445809 | 0.0599431473 | 0.1115206458 | 0.07131004307 | 0.9949216742 |
| -1.798490389 | 0.9354233137 | 0.0536785039 | 0.1121423806 | 0.07707308002 | 0.9949216742 |
| -1.802701091 | 0.8936749059 | 0.054569585 | 0.113409101 | 0.076313227 | 0.9949216742 |
| -1.873326827 | 0.5696939431 | 0.0514462873 | 0.0982705500 | 0.06579013109 | 0.9949216742 |
| -1.585482239 | 0.494004831 | 0.0474922253 | 0.1503791328 | 0.1180499219 | 0.9949216742 |
| -1.87938576 | 0.5762847303 | 0.0696083099 | 0.104566277 | 0.06505395976 | 0.9949216742 |
| -1.808329769 | 0.9949216742 | 0.0555631612 | 0.1134123706 | 0.07552566721 | 0.9949216742 |
| -1.805187606 | 0.9648580204 | 0.0600392383 | 0.113412029 | 0.07603583091 | 0.9949216742 |
| -1.745830883 | 0.7408341924 | 0.0529120092 | 0.1295634173 | 0.08587855167 | 0.9949216742 |
| -1.846483969 | 0.4089188211 | 0.0566108037 | 0.0855247552 | 0.06971560462 | 0.9949216742 |
| -1.857097511 | 0.2531691596 | 0.0443859610 | 0.0991377470 | 0.06822074256 | 0.7766647204 |
| -1.820246532 | 0.8361175404 | 0.0599056176 | 0.1111877722 | 0.07365813795 | 0.9949216742 |
| -1.782760642 | 0.9469421875 | 0.0616025024 | 0.1130360515 | 0.07959268046 | 0.9949216742 |
| -1.809996929 | 0.817586333 | 0.0605019858 | 0.1088734937 | 0.07523919707 | 0.9949216742 |
| -1.811548016 | 0.848001799 | 0.0534273004 | 0.1203491122 | 0.07501950217 | 0.9949216742 |
| -2.11498421 | 0.104497805 | 0.0568851995 | 0.0594844563 | 0.03861787631 | 0.7766647204 |
| -1.470347108 | 0.2294706438 | 0.0698143955 | 0.1545468676 | 0.1466507865 | 0.7766647204 |
| -1.800180919 | 0.979658694 | 0.0542346489 | 0.1134925708 | 0.07678649825 | 0.9949216742 |
| -1.760017249 | 0.1353257733 | 0.0431701742 | 0.1679519953 | 0.0834762293 | 0.7766647204 |
| -2.077169534 | 0.227388129 | 0.0756668607 | 0.0679366231 | 0.0419775732 | 0.7766647204 |
| -1.847335543 | 0.2480085415 | 0.0614956411 | 0.0838409827 | 0.06958457545 | 0.7766647204 |
| -1.810004966 | 0.8896664883 | 0.0540618033 | 0.1146823321 | 0.07528195653 | 0.9949216742 |
| -1.594504996 | 0.0733788068 | 0.0168103150 | 0.1688811144 | 0.1160746743 | 0.7118430836 |

| | | | | | |
|---|---|---|---|---|---|
| -1.813363572 | 0.9005121593 | 0.0542778066 | 0.114218873 | 0.07474322978 | 0.9949216742 |
| -1.792280727 | 0.5179547673 | 0.0495885478 | 0.1089773381 | 0.07812525529 | 0.9949216742 |
| -1.812252259 | 0.8993562777 | 0.0538425872 | 0.1199036145 | 0.07490625663 | 0.9949216742 |
| -1.8327385 | 0.7793790325 | 0.0655222472 | 0.1105130744 | 0.07174818128 | 0.9949216742 |
| -1.747661293 | 0.5090397503 | 0.0431667210 | 0.1188820752 | 0.08562049655 | 0.9949216742 |
| -1.798263332 | 0.7685030226 | 0.0500167644 | 0.1173212315 | 0.07715723675 | 0.9949216742 |
| -1.809498293 | 0.9695228398 | 0.0587717931 | 0.1137127721 | 0.07535849662 | 0.9949216742 |
| -1.821903679 | 0.7970158917 | 0.0521126214 | 0.1109380601 | 0.07354909891 | 0.9949216742 |
| -1.807603321 | 0.9911735326 | 0.0573592059 | 0.1136823781 | 0.07561791508 | 0.9949216742 |
| -1.84159152 | 0.5089713994 | 0.0590618696 | 0.1084915944 | 0.07043082109 | 0.9949216742 |
| -1.771512736 | 0.5636771557 | 0.0570138779 | 0.1225302164 | 0.08153598694 | 0.9949216742 |
| -1.857854095 | 0.2250301489 | 0.0543285613 | 0.1453881611 | 0.06801627639 | 0.7766647204 |
| -1.807520759 | 0.8950049986 | 0.0568337126 | 0.1159152337 | 0.07566612442 | 0.9949216742 |
| -1.832706472 | 0.0226799673 | 0.0178247067 | 0.1541542902 | 0.07176730528 | 0.708748979 |
| -1.763339127 | 0.766744124 | 0.0542600555 | 0.1309970038 | 0.08288216361 | 0.9949216742 |
| -1.637303794 | 0.249745554 | 0.0521631596 | 0.2698102776 | 0.1067299699 | 0.7766647204 |
| -1.808262201 | 0.1527169993 | 0.0520873499 | 0.1038233522 | 0.07556982895 | 0.7766647204 |
| -1.828403903 | 0.7950313618 | 0.0564864824 | 0.1154925087 | 0.0724603296 | 0.9949216742 |
| -1.892851433 | 0.0410875652 | 0.0387625020 | 0.1262582772 | 0.06309507241 | 0.7118430836 |
| -1.769756289 | 0.4510436441 | 0.0614548607 | 0.1038685702 | 0.08181964424 | 0.9949216742 |
| -1.633154711 | 0.4469893072 | 0.0513466246 | 0.1270254689 | 0.107572625 | 0.9949216742 |
| -1.788944071 | 0.7403035908 | 0.0502494353 | 0.1236364872 | 0.07863515043 | 0.9949216742 |
| -1.804875886 | 0.9574164881 | 0.0547491174 | 0.113785984 | 0.07608734558 | 0.9949216742 |
| -1.549981461 | 0.4263172719 | 0.0487898988 | 0.1652445519 | 0.1263721716 | 0.9949216742 |
| -1.946104342 | 0.1034012464 | 0.0399498464 | 0.1242606487 | 0.05628612962 | 0.7766647204 |
| -1.904331381 | 0.3453877136 | 0.0749481958 | 0.1166456526 | 0.06163336324 | 0.8994471709 |
| -1.820549966 | 0.2437320952 | 0.0797116427 | 0.0788861531 | 0.07360619647 | 0.7766647204 |
| -1.677423114 | 0.0548049791 | 0.1244095086 | 0.1754830072 | 0.09861479492 | 0.7118430836 |
| -1.524242209 | 0.2387719631 | 0.0372828421 | 0.18925848 | 0.1325523358 | 0.7766647204 |
| -1.803871594 | 0.4891437992 | 0.0499740522 | 0.117155658 | 0.07626782887 | 0.9949216742 |
| -1.908857247 | 0.5506726702 | 0.0716150169 | 0.0997675378 | 0.06085227572 | 0.9949216742 |
| -1.819311822 | 0.0739883434 | 0.0431339845 | 0.1229138034 | 0.0736931561 | 0.7118430836 |
| -1.827550127 | 0.69537366 | 0.0700493729 | 0.1178873881 | 0.07256866484 | 0.9949216742 |
| -1.802067718 | 0.9303917819 | 0.0538403607 | 0.1141342248 | 0.07652736348 | 0.9949216742 |
| -1.481010443 | 0.1693078947 | 0.0285251255 | 0.1687583278 | 0.1436859221 | 0.7766647204 |
| -1.824375591 | 0.6499938854 | 0.0602081391 | 0.1170495874 | 0.07302197505 | 0.9949216742 |
| -1.371841129 | 0.2359597391 | 0.0394203954 | 0.2398680679 | 0.1751358198 | 0.7766647204 |
| -1.812523177 | 0.890258633 | 0.0536963506 | 0.1107214414 | 0.07487209543 | 0.9949216742 |
| -1.760679522 | 0.719949355 | 0.0508118346 | 0.1160108311 | 0.08352952519 | 0.9949216742 |
| -1.839336488 | 0.1736444608 | 0.0199128938 | 0.1694244897 | 0.0708703566 | 0.7766647204 |
| -1.772421652 | 0.2141077646 | 0.0529484251 | 0.1358648855 | 0.0813448946 | 0.7766647204 |
| -1.687922354 | 0.615314975 | 0.0513357548 | 0.1119060241 | 0.09656586005 | 0.9949216742 |
| -1.977374223 | 0.216557478 | 0.1431898649 | 0.0787458090 | 0.05246367004 | 0.7766647204 |
| -1.807528726 | 0.9245975092 | 0.0543942527 | 0.1127618468 | 0.07561335556 | 0.9949216742 |
| -1.943526751 | 0.0478137925 | 0.0506400071 | 0.1234192381 | 0.0566108524 | 0.7118430836 |
| -1.89271786 | 0.4948386924 | 0.0621537923 | 0.0865246053 | 0.06317990472 | 0.9949216742 |

| | | | | | |
|---|---|---|---|---|---|
| -1.851287703 | 0.6374214491 | 0.0521769873 | 0.1104665224 | 0.06903838904 | 0.9949216742 |
| -1.78755668 | 0.4629981817 | 0.0443064657 | 0.1037397975 | 0.07889587939 | 0.9949216742 |
| -1.655574084 | 0.3581083128 | 0.0082915420 | 0.6795777155 | 0.1058342316 | 0.8912910305 |
| -1.630078475 | 0.6615322152 | 0.0095626019 | 0.58740976 | 0.1111363072 | 0.8912910305 |
| -1.741321401 | 0.6322437985 | 0.0079084745 | 0.4943027056 | 0.08951137551 | 0.8912910305 |
| -1.653587371 | 0.9429838554 | 0.0090127033 | 0.5606066728 | 0.106239765 | 0.9826407449 |
| -1.710293752 | 0.0487534429 | 0.0092107273 | 0.6635491531 | 0.09515697716 | 0.8912910305 |
| -1.250954073 | 0.2782276095 | 0.0286438242 | 0.7132300484 | 0.2184039647 | 0.8912910305 |
| -1.58482839 | 0.5382563276 | 0.0083659593 | 0.7031931488 | 0.1210816467 | 0.8912910305 |
| -1.668424832 | 0.981644266 | 0.0095538909 | 0.5593620778 | 0.1032418083 | 0.989560752 |
| -1.670720392 | 0.95905443 | 0.0095399235 | 0.5588889512 | 0.1027842829 | 0.9826407449 |
| -1.705522892 | 0.4646287096 | 0.0077056954 | 0.5747154742 | 0.0960508018 | 0.8912910305 |
| -1.796430824 | 0.1510030151 | 0.0146830796 | 0.5355149835 | 0.08017500852 | 0.8912910305 |
| -1.686182091 | 0.8048650177 | 0.0086943919 | 0.5453052631 | 0.09974609489 | 0.9826407449 |
| -1.71570002 | 0.4915717897 | 0.0081680107 | 0.4985413152 | 0.09415248389 | 0.8912910305 |
| -1.742971831 | 0.6317983373 | 0.0083297386 | 0.4825846067 | 0.08921908876 | 0.8912910305 |
| -1.595743098 | 0.6775149515 | 0.0084309907 | 0.5385391992 | 0.118618827 | 0.8914670415 |
| -1.798327262 | 0.3881489601 | 0.0062569563 | 0.43901438 | 0.07986891915 | 0.8912910305 |
| -1.846527868 | 0.0814048813 | 0.0054466845 | 0.3903050448 | 0.07241432248 | 0.8912910305 |
| -1.822547131 | 0.4110372141 | 0.0074607492 | 0.4767205427 | 0.07604578538 | 0.8912910305 |
| -1.730840566 | 0.6568201039 | 0.0149582497 | 0.4905609315 | 0.09138625071 | 0.8912910305 |
| -1.360641547 | 0.0961888290 | 0.0026615023 | 0.4736178072 | 0.1814390533 | 0.8912910305 |
| -1.766165547 | 0.5252675705 | 0.0071583552 | 0.527914973 | 0.08519534086 | 0.8912910305 |
| -1.409770971 | 0.4147282122 | 0.0090450971 | 0.5783554845 | 0.16653259 | 0.8912910305 |
| -1.656919776 | 0.4283402572 | 0.0065297049 | 0.5697300297 | 0.1055602703 | 0.8912910305 |
| -1.643573583 | 0.6246891767 | 0.0087782168 | 0.5231327765 | 0.108303331 | 0.8912910305 |
| -1.291830819 | 0.3091004166 | 0.0559764651 | 0.8794829987 | 0.204019025 | 0.8912910305 |
| -1.530381593 | 0.0715085500 | 0.0087438601 | 0.7405741526 | 0.1339940931 | 0.8912910305 |
| -1.689965456 | 0.651918528 | 0.0080012666 | 0.5587660018 | 0.09901410932 | 0.8912910305 |
| -1.718929291 | 0.4709597289 | 0.0069576845 | 0.5310089506 | 0.09355670369 | 0.8912910305 |
| -1.750129996 | 0.6030036159 | 0.0086226531 | 0.5253782411 | 0.08796062358 | 0.8912910305 |
| -1.588199786 | 0.1169081399 | 0.0205827122 | 0.6139043009 | 0.1203165193 | 0.8912910305 |
| -1.553056235 | 0.3103128519 | 0.0063479066 | 0.6384264399 | 0.128488003 | 0.8912910305 |
| -1.743996939 | 0.562108977 | 0.0079268679 | 0.4929951332 | 0.08903794703 | 0.8912910305 |
| -1.643834537 | 0.1904588894 | 0.0078155078 | 0.6169384214 | 0.1082491403 | 0.8912910305 |
| -2.089861963 | 0.0675886695 | 0.0033841446 | 0.3242419884 | 0.04320001858 | 0.8912910305 |
| -1.348127411 | 0.1164832223 | 0.0094712094 | 0.6890332267 | 0.1853957921 | 0.8912910305 |
| -1.683097558 | 0.8302403295 | 0.0117522052 | 0.541123971 | 0.1003461865 | 0.9826407449 |
| -1.73317362 | 0.5141202242 | 0.0071252575 | 0.6462843517 | 0.09096608655 | 0.8912910305 |
| -1.731657711 | 0.4921622229 | 0.0100791838 | 0.4964345294 | 0.09123890549 | 0.8912910305 |
| -1.689080966 | 0.6048524036 | 0.0094220414 | 0.5008239145 | 0.09918483573 | 0.8912910305 |
| -1.621774122 | 0.9548430749 | 0.0090499285 | 0.566190226 | 0.1129096123 | 0.9826407449 |
| -1.70035457 | 0.5161514472 | 0.0076315801 | 0.5754057153 | 0.0970269562 | 0.8912910305 |
| -1.655867229 | 0.8521722061 | 0.0089495040 | 0.5450123934 | 0.1057745022 | 0.9826407449 |
| -1.544302513 | 0.3716410771 | 0.0172974842 | 0.7249728311 | 0.1305916384 | 0.8912910305 |
| -1.725798018 | 0.32700105 | 0.0081726118 | 0.6092649832 | 0.09229991031 | 0.8912910305 |

| | | | | | |
|---|---|---|---|---|---|
| -1.71771844 | 0.3246243643 | 0.0068179758 | 0.6314511142 | 0.09377972887 | 0.8912910305 |
| -1.51453196 | 0.1712199348 | 0.0070154575 | 0.5724718926 | 0.1379542671 | 0.8912910305 |
| -1.591471765 | 0.5644594707 | 0.0078587038 | 0.5626716984 | 0.1195777184 | 0.8912910305 |
| -1.845163455 | 0.1508345772 | 0.0073747396 | 0.4503666231 | 0.07261691037 | 0.8912910305 |
| -1.673913187 | 0.798791508 | 0.0094848161 | 0.5387483147 | 0.1021507169 | 0.9826407449 |
| -1.758510436 | 0.5359140743 | 0.0105412087 | 0.6028315451 | 0.08650620125 | 0.8912910305 |
| -1.647553721 | 0.771411131 | 0.0089693127 | 0.5902322175 | 0.1074792215 | 0.9792063282 |
| -1.823471885 | 0.4786542016 | 0.0075239269 | 0.4368852722 | 0.07590293841 | 0.8912910305 |
| -1.647415505 | 0.6702508549 | 0.0089792242 | 0.5367036285 | 0.1075077531 | 0.8912910305 |
| -1.570216474 | 0.4918615288 | 0.0070365229 | 0.4638046798 | 0.1244435917 | 0.8912910305 |
| -1.612290625 | 0.5685851022 | 0.0098425180 | 0.5467617602 | 0.1149629706 | 0.8912910305 |
| -1.646566919 | 0.9270496256 | 0.0089239099 | 0.5573717614 | 0.1076830618 | 0.9826407449 |
| -1.666669311 | 0.5855845827 | 0.0083707880 | 0.5948137672 | 0.103592833 | 0.8912910305 |
| -1.616530411 | 0.8743012671 | 0.0091647445 | 0.5507284826 | 0.1140412344 | 0.9826407449 |
| -1.519866668 | 0.6656067289 | 0.0093562933 | 0.5992006083 | 0.1366110055 | 0.8912910305 |
| -1.674429583 | 0.9030108141 | 0.0088954910 | 0.5589425599 | 0.1020485493 | 0.9826407449 |
| -1.687156293 | 0.5355670374 | 0.0096464270 | 0.5506671196 | 0.09955718402 | 0.8912910305 |
| -1.576656866 | 0.3698037061 | 0.0228219814 | 0.5895336898 | 0.1229525601 | 0.8912910305 |
| -1.63303305 | 0.8148447686 | 0.0130057197 | 0.5832398931 | 0.1105109229 | 0.9826407449 |
| -1.665606289 | 0.9134852362 | 0.0091666293 | 0.5673353841 | 0.103805868 | 0.9826407449 |
| -1.772547958 | 0.564346989 | 0.0075645096 | 0.4936616197 | 0.08411520812 | 0.8912910305 |
| -1.708403191 | 0.7040902616 | 0.0082696194 | 0.5184451736 | 0.09551034426 | 0.9147784963 |
| -1.662124207 | 0.9345898602 | 0.0090755928 | 0.5682359998 | 0.1045062279 | 0.9826407449 |
| -1.403337265 | 0.2793535563 | 0.0102604031 | 0.7127914006 | 0.1684284739 | 0.8912910305 |
| -1.755038569 | 0.3198255247 | 0.0052144418 | 0.5082041125 | 0.087106277 | 0.8912910305 |
| -1.693151742 | 0.7811968836 | 0.0086213899 | 0.5289246523 | 0.09840110317 | 0.9792063282 |
| -1.645261006 | 0.9898865319 | 0.0097375942 | 0.5601937221 | 0.1079533086 | 0.9898865319 |
| -1.728498427 | 0.668507421 | 0.0082280555 | 0.5255997704 | 0.09180967962 | 0.8912910305 |
| -1.656756041 | 0.3891673362 | 0.0073224399 | 0.5160573641 | 0.105593573 | 0.8912910305 |
| -1.864130393 | 0.3840180408 | 0.0063607954 | 0.4041934693 | 0.06984351797 | 0.8912910305 |
| -1.675556294 | 0.4992219425 | 0.0081326229 | 0.5123131239 | 0.1018259254 | 0.8912910305 |
| -1.707734351 | 0.5187294845 | 0.0086081399 | 0.5877077001 | 0.09563561902 | 0.8912910305 |
| -1.488209324 | 0.2681009649 | 0.0195799596 | 0.6809227002 | 0.1447376829 | 0.8912910305 |
| -1.619825303 | 0.5688583363 | 0.0082737892 | 0.6694711145 | 0.1133291004 | 0.8912910305 |
| -1.718100508 | 0.603245007 | 0.0079629106 | 0.5540342265 | 0.0937093085 | 0.8912910305 |
| -1.903632266 | 0.0554038139 | 0.0178332883 | 0.7786091658 | 0.06435640708 | 0.8912910305 |
| -1.780443296 | 0.0917562130 | 0.0150843743 | 0.7416215853 | 0.08279498693 | 0.8912910305 |
| -1.676906001 | 0.8818771279 | 0.0099378559 | 0.5470801579 | 0.1015597692 | 0.9826407449 |
| -1.678264252 | 0.8714431179 | 0.0092213858 | 0.5735729858 | 0.1012925095 | 0.9826407449 |
| -1.771555614 | 0.3861085987 | 0.0116740816 | 0.5797216186 | 0.08428238858 | 0.8912910305 |
| -1.221350025 | 0.3605116813 | 0.0088394891 | 0.940867365 | 0.2292854943 | 0.8912910305 |
| -1.723291142 | 0.5606213417 | 0.0078132250 | 0.5095172285 | 0.09275695991 | 0.8912910305 |
| -1.624355322 | 0.2217483436 | 0.0147204926 | 0.6021669888 | 0.1123559602 | 0.8912910305 |
| -1.687625747 | 0.6322102358 | 0.0096429779 | 0.5684106383 | 0.09946625652 | 0.8912910305 |
| -1.697973028 | 0.3057958157 | 0.0063952239 | 0.5426623036 | 0.09747952982 | 0.8912910305 |
| -1.624510095 | 0.0068991647 | 0.0022241076 | 0.6736543281 | 0.1123228333 | 0.8623955959 |

| | | | | | |
|---|---|---|---|---|---|
| -1.724987337 | 0.6281447404 | 0.0080409469 | 0.4929460513 | 0.09244750655 | 0.8912910305 |
| -1.659578234 | 0.8732643523 | 0.0103469367 | 0.5847276995 | 0.1050207688 | 0.9826407449 |
| -1.574837939 | 0.474191144 | 0.0093695264 | 0.788064912 | 0.1233721885 | 0.8912910305 |
| -1.895379445 | 0.2260755655 | 0.0052202642 | 0.3208342737 | 0.06547123033 | 0.8912910305 |
| -1.714951787 | 0.6088012432 | 0.0080435726 | 0.5694899333 | 0.09429097814 | 0.8912910305 |
| -1.754922304 | 0.3109384647 | 0.0106191448 | 0.5591190182 | 0.08712643248 | 0.8912910305 |
| -1.742446219 | 0.4459292722 | 0.0080002336 | 0.6657441786 | 0.08931208644 | 0.8912910305 |
| -1.666844411 | 0.966918493 | 0.0090262964 | 0.5598219352 | 0.1035577768 | 0.9826407449 |
| -1.644906425 | 0.820589478 | 0.0110488673 | 0.5955798321 | 0.1080267821 | 0.9826407449 |
| -1.649897854 | 0.9638679433 | 0.0100372954 | 0.5635725636 | 0.1069962744 | 0.9826407449 |
| -1.819427039 | 0.4721908132 | 0.0067781497 | 0.4929191263 | 0.07652942897 | 0.8912910305 |
| -1.752725759 | 0.3477696251 | 0.0066944602 | 0.4753589526 | 0.08750795554 | 0.8912910305 |
| -1.665601123 | 0.8770639717 | 0.0088805954 | 0.5523966279 | 0.1038069041 | 0.9826407449 |
| -1.62765199 | 0.8916367071 | 0.0089422886 | 0.5522555084 | 0.1116520801 | 0.9826407449 |
| -1.633842129 | 0.1629614235 | 0.0039812961 | 0.5337765803 | 0.1103401725 | 0.8912910305 |
| -1.837174417 | 0.3362901483 | 0.0057370232 | 0.4815307607 | 0.07381281672 | 0.8912910305 |
| -1.686952514 | 0.7833650626 | 0.0085928054 | 0.5376350282 | 0.09959667506 | 0.9792063282 |
| -1.568499949 | 0.4031776655 | 0.0094688153 | 0.5868973789 | 0.1248434521 | 0.8912910305 |
| -1.656451653 | 0.882329679 | 0.0098479221 | 0.5575561913 | 0.1056555064 | 0.9826407449 |
| -1.679561628 | 0.284788802 | 0.0051950377 | 0.5975541754 | 0.1010377715 | 0.8912910305 |
| -1.688963009 | 0.6403520464 | 0.0081669975 | 0.5530163886 | 0.0992076225 | 0.8912910305 |
| -2.024406709 | 0.1754747492 | 0.0035380900 | 0.3751549499 | 0.04981531774 | 0.8912910305 |
| -1.715151041 | 0.6333440018 | 0.0105185914 | 0.522952244 | 0.09425408059 | 0.8912910305 |
| -1.382789788 | 0.5418240748 | 0.013207219 | 0.7027774984 | 0.1745961974 | 0.8912910305 |
| -1.845969347 | 0.3089448101 | 0.0080539449 | 0.477973523 | 0.07249719347 | 0.8912910305 |
| -1.661309692 | 0.2289988066 | 0.0068041573 | 0.6713562747 | 0.1046706146 | 0.8912910305 |
| -1.702551464 | 0.7098681132 | 0.0163924801 | 0.5992259501 | 0.09661102083 | 0.9147784963 |
| -1.844703425 | 0.2501669663 | 0.0049235075 | 0.4743706861 | 0.07268532409 | 0.8912910305 |
| -1.624160786 | 0.9487195469 | 0.0089680164 | 0.5570880251 | 0.1123976095 | 0.9826407449 |
| -1.233091471 | 0.0317211330 | 0.0674120665 | 0.8190709865 | 0.2249226552 | 0.8912910305 |
| -1.667958935 | 0.4016141527 | 0.0070658558 | 0.566239872 | 0.1033348708 | 0.8912910305 |
| -1.773757865 | 0.4464751037 | 0.0078993191 | 0.5533773011 | 0.08391175199 | 0.8912910305 |
| -1.516961075 | 0.6375046231 | 0.0119573520 | 0.6124628727 | 0.1373413182 | 0.8912910305 |
| -1.818535951 | 0.1949624084 | 0.0110070149 | 0.4424758519 | 0.07666803329 | 0.8912910305 |
| -1.741127357 | 0.3067116463 | 0.0065553801 | 0.5304216748 | 0.08954579276 | 0.8912910305 |
| 1.694135105 | 0.1051415257 | 0.3762384812 | 0.3669993155 | 0.09517836115 | 0.4971223625 |
| 1.569258903 | 0.0571951581 | 0.3588328663 | 0.3592133468 | 0.1215970309 | 0.4971223625 |
| 1.711646845 | 0.5707488806 | 0.4973740174 | 0.3025966002 | 0.0918810885 | 0.8181148269 |
| 1.668232661 | 0.9871605578 | 0.441818093 | 0.3065198616 | 0.1002328558 | 0.9871605578 |
| 1.680029956 | 0.8452277854 | 0.4443515429 | 0.3131811022 | 0.09790423364 | 0.9267848524 |
| 1.832172998 | 0.380724078 | 0.5318365927 | 0.253545951 | 0.0716530428 | 0.7388208805 |
| 1.586719194 | 0.0638229964 | 0.4496531734 | 0.4154025498 | 0.1175831567 | 0.4971223625 |
| 1.812428322 | 0.0524971028 | 0.286058715 | 0.2953562945 | 0.0746854818 | 0.4971223625 |
| 1.574191029 | 0.0004498818 | 0.1532738308 | 0.3439217072 | 0.1204522394 | 0.02811761644 |
| 1.649217923 | 0.5148664917 | 0.4497913461 | 0.3192568832 | 0.1040810961 | 0.8015000474 |
| 1.616914681 | 0.3667509895 | 0.3686731581 | 0.3438445785 | 0.1108935209 | 0.7388208805 |

| | | | | | |
|---|---|---|---|---|---|
| 1.893729192 | 0.1591257904 | 0.4796751242 | 0.2875768431 | 0.06285465464 | 0.5234401001 |
| 1.635517695 | 0.66219584 | 0.4461312122 | 0.3217637147 | 0.1069276005 | 0.818594287 |
| 1.734647838 | 0.206496827 | 0.4365995748 | 0.245265876 | 0.08769361077 | 0.5736022972 |
| 1.618456976 | 0.6679729382 | 0.4435744797 | 0.3008374958 | 0.1105602652 | 0.818594287 |
| 1.749075173 | 0.1194940315 | 0.5843083175 | 0.2396744983 | 0.08514837903 | 0.4971223625 |
| 1.729421719 | 0.3186170976 | 0.4508991731 | 0.2332254411 | 0.08863095756 | 0.7136413882 |
| 1.613153889 | 0.4538836637 | 0.4211904796 | 0.3112064612 | 0.1117095495 | 0.7771980543 |
| 1.608620248 | 0.8271813153 | 0.4737871338 | 0.3308245667 | 0.1126997144 | 0.9231934322 |
| 1.694942318 | 0.3770743948 | 0.3286190499 | 0.3645204153 | 0.09502426595 | 0.7388208805 |
| 1.990043059 | 0.1024425394 | 0.6916194805 | 0.1943628845 | 0.05093000098 | 0.4971223625 |
| 1.676075458 | 0.9528190666 | 0.4439355122 | 0.305885099 | 0.09867981619 | 0.9762490436 |
| 1.706652652 | 0.5973264518 | 0.522033395 | 0.2880220734 | 0.09281172198 | 0.818594287 |
| 1.694856901 | 0.743774771 | 0.4485931379 | 0.2861609886 | 0.09504056218 | 0.8529527191 |
| 1.571794263 | 0.7235456327 | 0.399708946 | 0.3389523414 | 0.1210074671 | 0.8529527191 |
| 1.742031817 | 0.2595082993 | 0.4919566179 | 0.2245895363 | 0.08638320492 | 0.6358253148 |
| 1.713659951 | 0.6421523594 | 0.4790152245 | 0.3051328955 | 0.09150813736 | 0.818594287 |
| 1.676954508 | 0.3069822203 | 0.366939795 | 0.3248263976 | 0.09850697833 | 0.7106069915 |
| 2.068229266 | 0.0827759108 | 0.7696462239 | 0.186305674 | 0.04272944555 | 0.4971223625 |
| 1.926088556 | 0.0357266229 | 0.762342792 | 0.2276085777 | 0.05860820785 | 0.4971223625 |
| 1.681619905 | 0.6838332374 | 0.4767754673 | 0.2928207287 | 0.09759381088 | 0.8298947056 |
| 1.542688351 | 0.2033077661 | 0.4287686166 | 0.4321690003 | 0.1279149166 | 0.5736022972 |
| 1.671748871 | 0.5759528381 | 0.4060863131 | 0.3099264 | 0.09953412172 | 0.8181148269 |
| 1.23578282 | 0.0945417028 | 0.2886334701 | 0.467790095 | 0.2211274643 | 0.4971223625 |
| 2.095306831 | 0.0462569946 | 0.5186501513 | 0.2661857261 | 0.04016901192 | 0.4971223625 |
| 1.660252646 | 0.8011876486 | 0.4716209025 | 0.307678389 | 0.1018334887 | 0.902238343 |
| 1.73477802 | 0.100849215 | 0.4748377239 | 0.3627337582 | 0.08767036655 | 0.4971223625 |
| 1.630906831 | 0.623486455 | 0.4594639613 | 0.3286599472 | 0.1078996911 | 0.818594287 |
| 1.72157085 | 0.0660964908 | 0.3688030609 | 0.1895098643 | 0.09005460057 | 0.4971223625 |
| 1.789209585 | 0.5222991621 | 0.4293229953 | 0.247609928 | 0.07838806565 | 0.8015000474 |
| 1.605963687 | 0.5759196394 | 0.3904998751 | 0.3095386221 | 0.1132832015 | 0.8181148269 |
| 1.678491192 | 0.9193413267 | 0.4421129013 | 0.3025914119 | 0.09820543149 | 0.9497327755 |
| 2.016801979 | 0.0467111394 | 0.4972118665 | 0.1622320186 | 0.04798297497 | 0.4971223625 |
| 1.611259951 | 0.404468951 | 0.4264453716 | 0.2911017909 | 0.112122335 | 0.7423732369 |
| 1.450971629 | 0.1909443739 | 0.3498942385 | 0.3197991173 | 0.1517490225 | 0.5682868272 |
| 1.589583482 | 0.0613674088 | 0.3941614375 | 0.2670190005 | 0.1169349777 | 0.4971223625 |
| 1.91701162 | 0.0439968188 | 0.4015443802 | 0.2755659302 | 0.05977400707 | 0.4971223625 |
| 1.731236319 | 0.1194666698 | 0.3920554939 | 0.3137387802 | 0.08830456169 | 0.4971223625 |
| 1.455188829 | 0.0740483052 | 0.4028231714 | 0.4340843795 | 0.1505819315 | 0.4971223625 |
| 1.683549997 | 0.5155724283 | 0.4083863116 | 0.3783035944 | 0.09721806123 | 0.8015000474 |
| 1.638374087 | 0.4790975012 | 0.4004881776 | 0.3282281858 | 0.106328971 | 0.7984958353 |
| 1.347928364 | 0.1461088911 | 0.2870097096 | 0.4429426886 | 0.1825084353 | 0.5098455047 |
| 1.65312275 | 0.9678456141 | 0.4426864533 | 0.3054607164 | 0.1032811712 | 0.9835829412 |
| 1.625938467 | 0.6467382588 | 0.4691924452 | 0.2806962872 | 0.1089551576 | 0.818594287 |
| 1.717801046 | 0.57532907 | 0.468433738 | 0.2872594423 | 0.0907448683 | 0.8181148269 |
| 1.66704233 | 0.5481249181 | 0.4292422503 | 0.2833313011 | 0.1004703013 | 0.8181148269 |
| 1.67833234 | 0.9022245245 | 0.4417121964 | 0.3014070493 | 0.09823656831 | 0.9477148366 |

| | | | | | |
|---|---|---|---|---|---|
| 1.632065758 | 0.6160781522 | 0.452336581 | 0.3245338685 | 0.1076546875 | 0.818594287 |
| 1.237644268 | 0.1007502063 | 0.3121104411 | 0.4080192237 | 0.2204408574 | 0.4971223625 |
| 1.44995016 | 0.1509142694 | 0.3232940831 | 0.4405962786 | 0.1520327681 | 0.5098455047 |
| 1.676380937 | 0.4068974404 | 0.498697721 | 0.2913003385 | 0.09861972515 | 0.7423732369 |
| 1.799537791 | 0.2446603018 | 0.6064037237 | 0.2683840291 | 0.07672259677 | 0.6358253148 |
| 1.758934332 | 0.4742601393 | 0.5798710469 | 0.2701367575 | 0.08344447214 | 0.7984958353 |
| 1.660446997 | 0.2543659215 | 0.4212296472 | 0.4093733663 | 0.1017942597 | 0.6358253148 |
| 1.72693846 | 0.3328549812 | 0.4290175575 | 0.3093686089 | 0.08907923957 | 0.7276160897 |
| 1.640521179 | 0.5936708673 | 0.4090657958 | 0.3233375147 | 0.1058807868 | 0.818594287 |
| 1.168069027 | 0.0027535773 | 0.1513510133 | 0.3451711591 | 0.24718038 | 0.1147323911 |
| 1.801107667 | 0.4873011906 | 0.5124902314 | 0.2567228179 | 0.07647205191 | 0.8014822213 |
| 1.726342361 | 0.1281379808 | 0.6208635182 | 0.2116029094 | 0.08918712618 | 0.4971223625 |
| 1.712880115 | 0.7344504268 | 0.4573183885 | 0.2859082121 | 0.09165246289 | 0.8529527191 |
| 1.793165138 | 0.5149383634 | 0.4026677352 | 0.2679693077 | 0.07774668892 | 0.8015000474 |
| 1.601263823 | 0.3900974249 | 0.4464029364 | 0.3133694511 | 0.1143214472 | 0.7388208805 |
| 1.618819441 | 0.1237870297 | 0.4053566774 | 0.4191352853 | 0.1104820623 | 0.4971223625 |
| 1.409255004 | 0.2881850515 | 0.3713526137 | 0.429010599 | 0.1636769106 | 0.6796817253 |
| 1.677001455 | 0.4331740892 | 0.4132033802 | 0.36015725 | 0.09849775447 | 0.7626304388 |
| 1.804719394 | 0.0066870624 | 0.4595792346 | 0.3787202193 | 0.07589822984 | 0.2089707006 |
| 1.708856028 | 0.7427009565 | 0.52038625 | 0.2873205954 | 0.09240018728 | 0.8529527191 |
| 1.653807071 | 0.1820221728 | 0.5283682863 | 0.3133052551 | 0.1031415011 | 0.5549456488 |
| 1.60649123 | 0.431417548 | 0.3980772 | 0.3129972651 | 0.1131671384 | 0.7626304388 |
| 1.675158685 | 0.1701538589 | 0.5384735257 | 0.2199637926 | 0.09886033518 | 0.5439359075 |
| 1.564443384 | 0.3526364838 | 0.358857296 | 0.4038148765 | 0.1227231459 | 0.7276160897 |
| 1.668369608 | 0.9842886749 | 0.4459559804 | 0.3070902739 | 0.1002055672 | 0.9871605578 |
| 1.700669181 | 0.2289550228 | 0.3807430517 | 0.3569270039 | 0.09393688793 | 0.6089229331 |
| 1.705924104 | 0.118866826 | 0.3126872423 | 0.316802732 | 0.09294812721 | 0.4971223625 |
| 1.792098303 | 0.5187352942 | 0.4319803083 | 0.239044527 | 0.07791923947 | 0.8015000474 |
| 1.701618757 | 0.1352172826 | 0.3407039495 | 0.2613676432 | 0.09375757999 | 0.4971223625 |
| 1.653272915 | 0.2549220487 | 0.4159679727 | 0.3408174764 | 0.1032505092 | 0.6358253148 |
| 1.616224304 | 0.5257840311 | 0.4377411369 | 0.3108884554 | 0.1110429589 | 0.8015000474 |
| 1.67285071 | 0.3550766518 | 0.4533845573 | 0.2929876118 | 0.0993159869 | 0.7276160897 |
| 1.664283271 | 0.5706026249 | 0.3788626857 | 0.3110393949 | 0.1010224398 | 0.8181148269 |
| 1.669610596 | 0.6932856435 | 0.4070929564 | 0.3226808321 | 0.09995855972 | 0.8332760139 |
| 1.826227501 | 0.2258785426 | 0.4477449808 | 0.2328760597 | 0.07255510735 | 0.6089229331 |
| 1.59350498 | 0.6495724024 | 0.4358469284 | 0.4035181795 | 0.1160522182 | 0.818594287 |
| 1.668274083 | 0.8431710366 | 0.4448460515 | 0.3072572646 | 0.1002246014 | 0.9267848524 |
| 1.221442269 | 0.0002381904 | 0.2481574476 | 0.2211761579 | 0.2264697486 | 0.02811761644 |
| 1.67788698 | 0.6067385586 | 0.4688755929 | 0.3101669679 | 0.09832390711 | 0.818594287 |
| 1.761949412 | 0.3454824939 | 0.4178116268 | 0.3367380947 | 0.0829290722 | 0.7276160897 |
| 2.013864291 | 0.0711141003 | 0.4268679336 | 0.2368001621 | 0.04829917503 | 0.4971223625 |
| 1.726302448 | 0.3882768154 | 0.5217847051 | 0.2585200499 | 0.08919435375 | 0.7388208805 |
| 1.788906751 | 0.1740594904 | 0.5417417042 | 0.270888128 | 0.07843735014 | 0.5439359075 |
| 1.828626323 | 0.445355096 | 0.5155359187 | 0.2627111571 | 0.07219001333 | 0.7731859305 |
| 1.626948687 | 0.1021408994 | 0.5065055065 | 0.2556207399 | 0.1087398745 | 0.4971223625 |
| 1.794827288 | 0.264503331 | 0.5327263758 | 0.3016974028 | 0.07747848684 | 0.6358253148 |

| | | | | | |
|---|---|---|---|---|---|
| 1.678678748 | 0.9146207347 | 0.4375943281 | 0.3204667731 | 0.09816867869 | 0.9497327755 |
| 1.834424754 | 0.1972219953 | 0.3285104467 | 0.2483993379 | 0.07131386639 | 0.5733197537 |
| 2.23018253 | 0.0139349255 | 0.7034101493 | 0.1501941275 | 0.02930614178 | 0.2903109496 |
| 1.677610387 | 0.8769790973 | 0.4420361571 | 0.300492142 | 0.09837818126 | 0.9329418089 |
| 1.616583099 | 0.8603953355 | 0.4330829101 | 0.3178934238 | 0.1109652744 | 0.9329418089 |
| 1.753191011 | 0.131371418 | 0.5144376567 | 0.3405437685 | 0.08443358895 | 0.4971223625 |
| 1.633952543 | 0.6019274845 | 0.3879518374 | 0.3040431047 | 0.1072567758 | 0.818594287 |
| 1.622161274 | 0.1137145011 | 0.3686476573 | 0.3393996649 | 0.1097631566 | 0.4971223625 |
| 1.477970901 | 0.4097900268 | 0.341565497 | 0.3406490261 | 0.1443978631 | 0.7423732369 |
| 1.681735926 | 0.6574930388 | 0.4259920818 | 0.2968570269 | 0.09757119052 | 0.818594287 |
| 1.645728545 | 0.8806970676 | 0.4567370071 | 0.3040768276 | 0.1048001712 | 0.9329418089 |
| 1.84536156 | 0.1487980386 | 0.4437713007 | 0.2807192781 | 0.06968559141 | 0.5098455047 |
| 1.629643517 | 0.623709563 | 0.4418631729 | 0.3427480698 | 0.1081672774 | 0.818594287 |
| 1.701950734 | 0.6322809 | 0.382125617 | 0.3553941288 | 0.09369495945 | 0.818594287 |
| 1.799358165 | 0.1286105918 | 0.4555331793 | 0.2669424319 | 0.07675130803 | 0.4971223625 |
| 1.497091308 | 0.3197113419 | 0.4294298658 | 0.3108779406 | 0.139363097 | 0.7136413882 |
| 1.650970903 | 0.7847161759 | 0.4163344471 | 0.3366189806 | 0.1037213675 | 0.8917229272 |
| 1.552189984 | 0.0799846887 | 0.4203336398 | 0.3679258304 | 0.1256262568 | 0.4971223625 |
| 1.657737308 | 0.7266985101 | 0.437758265 | 0.3001227833 | 0.1023423133 | 0.8529527191 |
| 1.639885606 | 0.8682835073 | 0.4348258356 | 0.3307965616 | 0.1060132961 | 0.9329418089 |
| 1.452562603 | 0.0127847938 | 0.4937428283 | 0.3989908592 | 0.1513079017 | 0.2903109496 |
| 1.641162626 | 0.3525066497 | 0.386521701 | 0.3083525128 | 0.105747189 | 0.7276160897 |
| 0.946401547 | 0.5910063055 | 0.1706525217 | 0.656284424 | 0.3476569006 | 0.8721887272 |
| 0.710933945 | 0.0756608910 | 0.1085748952 | 0.8052499018 | 0.4797878427 | 0.5394055412 |
| 0.9506600817 | 0.6995767795 | 0.1830732504 | 0.6865143005 | 0.3454923922 | 0.9092262502 |
| 0.9213638504 | 0.8098190913 | 0.1610386562 | 0.6986543111 | 0.3604629346 | 0.9140125553 |
| 0.9614114508 | 0.5235662389 | 0.1615206918 | 0.6447278879 | 0.3401243099 | 0.8382203167 |
| 1.08439112 | 0.3106230187 | 0.2083651442 | 0.6075665337 | 0.2824135342 | 0.7740437187 |
| 0.9232702486 | 0.6433357581 | 0.1656681796 | 0.7235643593 | 0.3594994102 | 0.8935218862 |
| 1.134626396 | 0.0285355540 | 0.0855607812 | 0.6493286691 | 0.2609710535 | 0.4637289028 |
| 0.7554193029 | 0.0210377875 | 0.0649228796 | 0.7782927167 | 0.452870819 | 0.4637289028 |
| 0.9156572497 | 0.8956876649 | 0.1602231742 | 0.6987796883 | 0.3634028065 | 0.9388421843 |
| 0.7808734565 | 0.1235114339 | 0.0954025216 | 0.7990163646 | 0.4378644738 | 0.5716165492 |
| 0.9563789938 | 0.7554642993 | 0.1640828845 | 0.6922476189 | 0.3426278491 | 0.9140125553 |
| 0.9331601083 | 0.7977756378 | 0.1657504325 | 0.6997382484 | 0.3544191369 | 0.9140125553 |
| 0.919121605 | 0.0280255118 | 0.1301984883 | 0.5499304634 | 0.3616030129 | 0.4637289028 |
| 0.9342898699 | 0.99201813 | 0.1618461548 | 0.6873451875 | 0.3538423112 | 0.99201813 |
| 0.9163936489 | 0.4823558287 | 0.1884485966 | 0.654051541 | 0.3630435359 | 0.8382203167 |
| 0.925216358 | 0.6515511713 | 0.1578135081 | 0.6443348013 | 0.358481781 | 0.8949878727 |
| 0.930003852 | 0.901288497 | 0.1611114174 | 0.689368982 | 0.3560227051 | 0.9388421843 |
| 1.1777378 | 0.18057431 | 0.1012869571 | 0.5261802575 | 0.2434750241 | 0.6100483445 |
| 1.077694566 | 0.0296786497 | 0.0524059154 | 0.9138794911 | 0.2854215764 | 0.4637289028 |
| 1.156990096 | 0.1369358019 | 0.2718805264 | 0.5486186015 | 0.2518904814 | 0.5716165492 |
| 0.946816535 | 0.8860413532 | 0.1631159322 | 0.6907421378 | 0.3474321257 | 0.9388421843 |
| 1.08396671 | 0.4298893321 | 0.252842562 | 0.6190488026 | 0.282656187 | 0.826710254 |
| 0.8897477359 | 0.6426655509 | 0.1523085954 | 0.7538075726 | 0.3770599343 | 0.8935218862 |

| | | | | | |
|---|---|---|---|---|---|
| 0.9687627561 | 0.7797804658 | 0.2148291015 | 0.6582991857 | 0.3364624692 | 0.9140125553 |
| 0.9837583836 | 0.5364610027 | 0.183926231 | 0.6125345262 | 0.3290838791 | 0.8382203167 |
| 0.9150577028 | 0.6895616223 | 0.1518912517 | 0.685432523 | 0.3637510736 | 0.9092262502 |
| 0.9137131508 | 0.6038630563 | 0.1421770898 | 0.7152448731 | 0.3644345116 | 0.8721887272 |
| 1.048795123 | 0.3862079837 | 0.2461754839 | 0.6030116652 | 0.2983883225 | 0.7914098027 |
| 1.096195956 | 0.0556967097 | 0.3174518121 | 0.5811667874 | 0.2772583996 | 0.5041041973 |
| 0.8499759117 | 0.5278771591 | 0.1286202436 | 0.7495492622 | 0.3986346227 | 0.8382203167 |
| 0.9073105108 | 0.6070433541 | 0.1615537501 | 0.7432275058 | 0.3678433686 | 0.8721887272 |
| 0.7778614483 | 0.1017374275 | 0.0924897027 | 0.7540432925 | 0.4396041013 | 0.5714001441 |
| 0.8982944157 | 0.9414018242 | 0.1637902421 | 0.700364694 | 0.3726307909 | 0.9725225456 |
| 1.104962153 | 0.2684183983 | 0.1735894913 | 0.6712939004 | 0.2734696813 | 0.762552268 |
| 0.9858815552 | 0.0838504850 | 0.0790558607 | 0.6652339289 | 0.3280693358 | 0.5516479282 |
| 1.068197754 | 0.1371879718 | 0.186939456 | 0.7493975812 | 0.2896391392 | 0.5716165492 |
| 0.9262079584 | 0.8691520828 | 0.1658806261 | 0.7000240295 | 0.3579760825 | 0.9388421843 |
| 0.9977077775 | 0.3545477698 | 0.1846355116 | 0.7814522261 | 0.322373141 | 0.7914012718 |
| 1.424303866 | 0.0654852201 | 0.1319109599 | 0.4002144012 | 0.159577371 | 0.5394055412 |
| 0.97929411 | 0.1671217257 | 0.1179441893 | 0.655660451 | 0.3314227023 | 0.5802837697 |
| 1.067778654 | 0.5090192606 | 0.1707162257 | 0.6334867062 | 0.2897425213 | 0.8382203167 |
| 1.372367821 | 0.0005239178 | 0.1515857053 | 0.3105568136 | 0.1748154429 | 0.06548973439 |
| 0.9489579316 | 0.7346548102 | 0.1611607363 | 0.6968939825 | 0.346360624 | 0.9092262502 |
| 0.4228435978 | 0.0463285985 | 0.0772553071 | 0.7806300357 | 0.6738492284 | 0.4825895682 |
| 0.90827076 | 0.8288621232 | 0.157615708 | 0.6865779704 | 0.3672638292 | 0.9168828796 |
| 0.9496297043 | 0.5112753475 | 0.1470714158 | 0.684659392 | 0.3459993674 | 0.8382203167 |
| 0.7933006102 | 0.1051376265 | 0.1111995425 | 0.7611456388 | 0.4305803755 | 0.5714001441 |
| 0.7293581024 | 0.3405792362 | 0.1407221209 | 0.8185156362 | 0.4685162783 | 0.7740437187 |
| 1.04930008 | 0.119008837 | 0.1301622133 | 0.9042616316 | 0.298173045 | 0.5716165492 |
| 0.9700107706 | 0.7954885504 | 0.1745236682 | 0.6687474714 | 0.3358423572 | 0.9140125553 |
| 1.114455017 | 0.3845636174 | 0.229654878 | 0.572787653 | 0.2694077056 | 0.7914098027 |
| 0.7327243521 | 0.166387421 | 0.1089011259 | 0.6833114019 | 0.4665003435 | 0.5802837697 |
| 0.8161670895 | 0.501219776 | 0.1724202103 | 0.6392276605 | 0.417523241 | 0.8382203167 |
| 0.9272360135 | 0.707500704 | 0.1557431636 | 0.6977832281 | 0.3574521987 | 0.9092262502 |
| 0.861598926 | 0.2217569893 | 0.1363610768 | 0.6245530145 | 0.3922344917 | 0.6599910397 |
| 1.153985945 | 0.3646741791 | 0.1748429333 | 0.5638786714 | 0.2529333001 | 0.7914098027 |
| 0.8923396276 | 0.7290919104 | 0.164027379 | 0.7176747779 | 0.3756502326 | 0.9092262502 |
| 0.7299095997 | 0.3026233288 | 0.1328890459 | 0.7809638511 | 0.4682495189 | 0.7740437187 |
| 0.9725125729 | 0.7886926274 | 0.1783193692 | 0.66004022 | 0.3346434638 | 0.9140125553 |
| 0.9397714994 | 0.8901501179 | 0.1593045434 | 0.690392987 | 0.3510183195 | 0.9388421843 |
| 0.9260644896 | 0.8812774652 | 0.1620151824 | 0.6948054034 | 0.3580486289 | 0.9388421843 |
| 0.9971792675 | 0.6320372725 | 0.225279552 | 0.6432522226 | 0.322566747 | 0.8935218862 |
| 0.9049565992 | 0.3691929666 | 0.1491950252 | 0.8172097529 | 0.3690203315 | 0.7914098027 |
| 0.9836191774 | 0.4139283514 | 0.1553934471 | 0.6870675846 | 0.3291740985 | 0.8084538113 |
| 0.9098944736 | 0.7551252526 | 0.1524746361 | 0.7064736147 | 0.3664304624 | 0.9140125553 |
| 0.5195204693 | 0.0966913166 | 0.0747275025 | 0.7791009736 | 0.6052385284 | 0.5714001441 |
| 1.636212092 | 0.0178604135 | 0.2941999948 | 0.3544090854 | 0.1069312285 | 0.4637289028 |
| 0.9160927128 | 0.5521020249 | 0.195617897 | 0.6406523886 | 0.3631980011 | 0.8520092977 |
| 0.9863101504 | 0.6749545757 | 0.1672266647 | 0.6479198979 | 0.3278800682 | 0.9092262502 |

| | | | | | |
|---|---|---|---|---|---|
| 1.003223007 | 0.7310773145 | 0.1539470131 | 0.6539442154 | 0.3196995493 | 0.9092262502 |
| 1.052172847 | 0.5009737692 | 0.1666767431 | 0.6589726639 | 0.296847269 | 0.8382203167 |
| 0.9415186959 | 0.9691143342 | 0.1624245765 | 0.6864318575 | 0.3501346231 | 0.9797266688 |
| 0.7483721181 | 0.5166299171 | 0.1393512046 | 0.8055614262 | 0.4570500143 | 0.8382203167 |
| 0.9037486978 | 0.3388117732 | 0.1396198456 | 0.7969026014 | 0.3696554758 | 0.7740437187 |
| 0.8703540466 | 0.0423161691 | 0.1463090671 | 0.8247134291 | 0.3874659934 | 0.4825895682 |
| 0.9021531571 | 0.7145460771 | 0.1513164207 | 0.7225574569 | 0.3705397671 | 0.9092262502 |
| 0.8870021036 | 0.5260278499 | 0.1775991064 | 0.702870612 | 0.3785320797 | 0.8382203167 |
| 0.9402456018 | 0.9580944141 | 0.1636543082 | 0.687829582 | 0.3508151317 | 0.9797266688 |
| 0.9213614159 | 0.5310444533 | 0.1838825092 | 0.6293399057 | 0.3604651042 | 0.8382203167 |
| 0.5180800441 | 0.0345167946 | 0.0618259047 | 0.9409306387 | 0.6062171525 | 0.4793999259 |
| 0.9017129368 | 0.3948436499 | 0.1313662161 | 0.7193641692 | 0.3707570116 | 0.7960557457 |
| 0.8869741799 | 0.3222768253 | 0.1262941721 | 0.7711557269 | 0.3785366028 | 0.7740437187 |
| 0.8405027785 | 0.1579641677 | 0.0945055159 | 0.7391503963 | 0.4038640337 | 0.5802837697 |
| 1.016345512 | 0.4788363505 | 0.1430882191 | 0.5649208494 | 0.3135914354 | 0.8382203167 |
| 0.7798882407 | 0.1303867506 | 0.0940814237 | 0.6668792933 | 0.4384187212 | 0.5716165492 |
| 0.9063053296 | 0.3124706744 | 0.146404712 | 0.7333146658 | 0.3683172681 | 0.7740437187 |
| 0.9318694753 | 0.4642780636 | 0.168985843 | 0.6878731107 | 0.3551032872 | 0.8382203167 |
| 0.9638699292 | 0.7611158981 | 0.1618946851 | 0.6946673376 | 0.3389148247 | 0.9140125553 |
| 1.041163189 | 0.1451862082 | 0.1000766841 | 0.6939123138 | 0.3019638414 | 0.5802837697 |
| 0.9534219887 | 0.0776743979 | 0.0869302533 | 0.7974844184 | 0.344128742 | 0.5394055412 |
| 0.9224414778 | 0.9011784834 | 0.1615730202 | 0.7037547447 | 0.3599084128 | 0.9388421843 |
| 0.8149008223 | 0.1895564783 | 0.1582460066 | 0.9825750489 | 0.4183037289 | 0.623541047 |
| 1.008904533 | 0.2040907339 | 0.1781493319 | 0.6647162056 | 0.3170428234 | 0.6303394336 |
| 0.6735468178 | 0.0276052432 | 0.1133345153 | 0.6114561517 | 0.5031627877 | 0.4637289028 |
| 0.9432028911 | 0.9718888555 | 0.1629288741 | 0.6865950504 | 0.349296527 | 0.9797266688 |
| 0.9355946778 | 0.5705573029 | 0.1467045893 | 0.7361263445 | 0.3531612582 | 0.8697519862 |
| 1.090150481 | 0.4097628455 | 0.1627066034 | 0.6373078638 | 0.2798708461 | 0.8084538113 |
| 0.9173658577 | 0.2932419645 | 0.123764093 | 0.776904061 | 0.362550782 | 0.7740437187 |
| 0.9462212677 | 0.6885546229 | 0.1747233506 | 0.68136263 | 0.347756998 | 0.9092262502 |
| 1.060639769 | 0.4492622567 | 0.189314104 | 0.6255593114 | 0.2930010065 | 0.8382203167 |
| 0.8991164553 | 0.6005011718 | 0.1697981485 | 0.6698755358 | 0.3720959591 | 0.8721887272 |
| 0.9640620145 | 0.305116306 | 0.194924597 | 0.7161426572 | 0.3387770246 | 0.7740437187 |
| 0.9322684865 | 0.5046939752 | 0.1366804569 | 0.7862406873 | 0.3548460212 | 0.8382203167 |
| 1.068928793 | 0.1517446407 | 0.0952276544 | 0.5894586205 | 0.2893011202 | 0.5802837697 |
| 1.11715868 | 0.2067513342 | 0.2194972998 | 0.559272587 | 0.2681983428 | 0.6303394336 |
| 0.8966455135 | 0.5781166891 | 0.1535930403 | 0.7308834554 | 0.3734088556 | 0.8706576643 |
| 0.5584420871 | 0.1356381449 | 0.1016968879 | 0.8573911724 | 0.5785220864 | 0.5716165492 |
| 0.9463542188 | 0.3239604976 | 0.1834501369 | 0.7229847848 | 0.3476562172 | 0.7740437187 |
| 0.9296473533 | 0.8185078305 | 0.1571896846 | 0.6823416534 | 0.3562210951 | 0.9140125553 |
| 0.8661142818 | 0.0209104811 | 0.1066457841 | 0.7580554753 | 0.3898120782 | 0.4637289028 |
| 0.7660503261 | 0.3201710687 | 0.115083017 | 0.7614092307 | 0.446576965 | 0.7740437187 |
| 0.923390397 | 0.7701912765 | 0.1542772714 | 0.6848705309 | 0.3594150256 | 0.9140125553 |
| 1.169471791 | 0.3354294137 | 0.1943738562 | 0.5129722201 | 0.2466877904 | 0.7740437187 |
| 1.116913505 | 0.1139284743 | 0.1555560851 | 0.6344679569 | 0.268369725 | 0.5716165492 |
| 0.748481096 | 0.1557507146 | 0.1451125493 | 0.8503111381 | 0.4570055973 | 0.5802837697 |

| | | | | | |
|---|---|---|---|---|---|
| 0.8802084223 | 0.0448380181 | 0.0419355115 | 0.9757413545 | 0.3820918371 | 0.4825895682 |
| 0.8543926323 | 0.26214896 | 0.1428887837 | 0.6902604559 | 0.3962009179 | 0.7620609302 |
| 0.7723913909 | 0.3774842019 | 0.154065811 | 0.6985480219 | 0.4428295683 | 0.7914098027 |
| 0.9044718756 | 0.8189552495 | 0.1588934167 | 0.7271607468 | 0.3692332158 | 0.9140125553 |
| 0.8049903159 | 0.2022279643 | 0.147511372 | 0.788267102 | 0.4239070295 | 0.6303394336 |
| 0.9206790649 | 0.7075969828 | 0.1590413155 | 0.6775125048 | 0.3608295876 | 0.9092262502 |
| 1.248317376 | 0.0742192347 | 0.2197132068 | 0.4216637827 | 0.2166706089 | 0.5394055412 |
| 0.7838948278 | 0.0942610257 | 0.1816785951 | 0.7693377371 | 0.4361430924 | 0.5714001441 |
| 0.9496069175 | 0.0564596701 | 0.1110119618 | 0.6506148552 | 0.3460734485 | 0.5041041973 |

| | | | Taxonomic Information |
|---|---|---|---|
| **Blast Adjusted p-valu** | **VNS Adjusted p-valu** | **Blast:VNS adjusted p-valu** | **Genus** |
| 0.01711752944 | 0.0987256072 | 0.05120735202 | Acetatifactor |
| 0.01711752944 | 0.0987256072 | 0.05120735202 | Acetatifactor |
| 0.01711752944 | 0.0987256072 | 0.05120735202 | Acutalibacter |
| 0.01711752944 | 0.0987256072 | 0.05120735202 | Acutalibacter |
| 0.01711752944 | 0.0987256072 | 0.05120735202 | Adlercreutzia |
| 0.01711752944 | 0.0987256072 | 0.05120735202 | Adlercreutzia |
| 0.01711752944 | 0.0987256072 | 0.05120735202 | Adlercreutzia |
| 0.01711752944 | 0.0987256072 | 0.05120735202 | Akkermansia |
| 0.01711752944 | 0.0987256072 | 0.05120735202 | Alistipes |
| 0.01711752944 | 0.0987256072 | 0.05120735202 | Anaerotruncus |
| 0.01737393672 | 0.0987256072 | 0.05120735202 | Bacteria_unclassified |
| 0.01711752944 | 0.0987256072 | 0.05120735202 | Bacteria_unclassified |
| 0.01711752944 | 0.0987256072 | 0.05120735202 | Bacteria_unclassified |
| 0.01711752944 | 0.1002066352 | 0.05120735202 | Bacteroides |
| 0.01711752944 | 0.0987256072 | 0.05120735202 | Bifidobacterium |
| 0.02036353548 | 0.1002066352 | 0.05120735202 | Clostridia_unclassified |
| 0.01711752944 | 0.1525595899 | 0.05120735202 | Clostridiaceae_unclassified |
| 0.01711752944 | 0.0987256072 | 0.05120735202 | Clostridiaceae_unclassified |
| 0.01825366792 | 0.0987256072 | 0.05120735202 | Eubacteriales_unclassified |
| 0.01961562443 | 0.0987256072 | 0.05120735202 | Erysipelatoclostridium |
| 0.01970312993 | 0.1002066352 | 0.05292259984 | Coriobacteriaceae_unclassifie |
| 0.01711752944 | 0.0987256072 | 0.05120735202 | Dorea |
| 0.02729472302 | 0.0987256072 | 0.05120735202 | Dubosiella |
| 0.01711752944 | 0.09954213661 | 0.05120735202 | Erysipelotrichales_unclassifie |
| 0.01711752944 | 0.0987256072 | 0.05120735202 | Eubacteriaceae_unclassified |
| 0.01711752944 | 0.0987256072 | 0.05120735202 | Eubacteriaceae_unclassified |
| 0.01825366792 | 0.0987256072 | 0.05120735202 | GGB20149 |
| 0.01711752944 | 0.0987256072 | 0.05120735202 | GGB22635 |
| 0.02592834182 | 0.1064446468 | 0.06470111122 | GGB25041 |
| 0.02389510834 | 0.0987256072 | 0.05185719465 | GGB27876 |
| 0.01711752944 | 0.0987256072 | 0.05120735202 | GGB27878 |
| 0.01711752944 | 0.1088029914 | 0.05120735202 | GGB27918 |
| 0.01825366792 | 0.0987256072 | 0.05120735202 | GGB28382 |
| 0.01761114253 | 0.09954213661 | 0.05193161517 | GGB28399 |
| 0.01711752944 | 0.0987256072 | 0.05120735202 | GGB28411 |
| 0.01737393672 | 0.0987256072 | 0.05120735202 | GGB28415 |
| 0.01711752944 | 0.0987256072 | 0.05120735202 | GGB28430 |
| 0.01711752944 | 0.0987256072 | 0.05120735202 | GGB28439 |
| 0.01721066887 | 0.0987256072 | 0.05120735202 | GGB28778 |
| 0.01711752944 | 0.0987256072 | 0.05120735202 | GGB28784 |
| 0.01761114253 | 0.0987256072 | 0.05120735202 | GGB28792 |
| 0.01711752944 | 0.0987256072 | 0.05120735202 | GGB28798 |
| 0.01711752944 | 0.0987256072 | 0.05120735202 | GGB28802 |
| 0.01711752944 | 0.0987256072 | 0.05120735202 | GGB28818 |
| 0.01737393672 | 0.0987256072 | 0.05120735202 | GGB28828 |

| | | | |
|---|---|---|---|
| 0.01711752944 | 0.0987256072 | 0.05120735202 | GGB28851 |
| 0.01711752944 | 0.0987256072 | 0.05120735202 | GGB28859 |
| 0.01711752944 | 0.0987256072 | 0.05120735202 | GGB28864 |
| 0.01711752944 | 0.0987256072 | 0.05120735202 | GGB28869 |
| 0.01761114253 | 0.1243739997 | 0.05120735202 | GGB28883 |
| 0.01711752944 | 0.0987256072 | 0.05120735202 | GGB28892 |
| 0.01711752944 | 0.0987256072 | 0.05120735202 | GGB28893 |
| 0.02592834182 | 0.0987256072 | 0.06825241097 | GGB28898 |
| 0.01711752944 | 0.0987256072 | 0.05120735202 | GGB28904 |
| 0.01711752944 | 0.0987256072 | 0.05120735202 | GGB28916 |
| 0.01711752944 | 0.0987256072 | 0.05120735202 | GGB28924 |
| 0.01711752944 | 0.1021320316 | 0.05300419053 | GGB28926 |
| 0.01711752944 | 0.0987256072 | 0.05120735202 | GGB28927 |
| 0.01711752944 | 0.0987256072 | 0.05120735202 | GGB28934 |
| 0.01761114253 | 0.1044584747 | 0.05120735202 | GGB28946 |
| 0.01825366792 | 0.0987256072 | 0.05120735202 | GGB28949 |
| 0.01711752944 | 0.0987256072 | 0.05120735202 | GGB28949 |
| 0.02129913663 | 0.09954213661 | 0.05120735202 | GGB28950 |
| 0.01711752944 | 0.1002066352 | 0.05120735202 | GGB28951 |
| 0.01711752944 | 0.0987256072 | 0.05120735202 | GGB28951 |
| 0.01737393672 | 0.0987256072 | 0.05120735202 | GGB28954 |
| 0.01825366792 | 0.0987256072 | 0.05120735202 | GGB28956 |
| 0.01711752944 | 0.0987256072 | 0.05120735202 | GGB28960 |
| 0.02139984729 | 0.1048169025 | 0.05120735202 | GGB28967 |
| 0.01711752944 | 0.0987256072 | 0.05120735202 | GGB28991 |
| 0.01711752944 | 0.1064446468 | 0.08852197766 | GGB29002 |
| 0.01711752944 | 0.0987256072 | 0.05120735202 | GGB29003 |
| 0.01711752944 | 0.1021320316 | 0.05120735202 | GGB29011 |
| 0.01711752944 | 0.0987256072 | 0.05120735202 | GGB29531 |
| 0.01711752944 | 0.0987256072 | 0.05120735202 | GGB29685 |
| 0.01711752944 | 0.0987256072 | 0.05120735202 | GGB30141 |
| 0.01761114253 | 0.0987256072 | 0.05120735202 | GGB30286 |
| 0.01737393672 | 0.0987256072 | 0.05120735202 | GGB30303 |
| 0.01711752944 | 0.0987256072 | 0.05120735202 | GGB30413 |
| 0.01825366792 | 0.1097593108 | 0.05120735202 | GGB30454 |
| 0.01711752944 | 0.0987256072 | 0.05120735202 | GGB30455 |
| 0.01711752944 | 0.0987256072 | 0.05120735202 | GGB30461 |
| 0.01711752944 | 0.0987256072 | 0.05120735202 | GGB30461 |
| 0.01711752944 | 0.0987256072 | 0.05120735202 | GGB30463 |
| 0.01711752944 | 0.1088029914 | 0.05413078867 | GGB30473 |
| 0.01711752944 | 0.0987256072 | 0.05120735202 | GGB30475 |
| 0.01711752944 | 0.0987256072 | 0.05120735202 | GGB30861 |
| 0.01711752944 | 0.0987256072 | 0.05148421583 | GGB31312 |
| 0.01711752944 | 0.0987256072 | 0.05120735202 | GGB31438 |
| 0.01842188925 | 0.0987256072 | 0.05120735202 | GGB3171 |
| 0.01711752944 | 0.0987256072 | 0.05120735202 | GGB31823 |

| | | | |
|---|---|---|---|
| 0.01711752944 | 0.1121818466 | 0.06332819819 | GGB31853 |
| 0.01711752944 | 0.0987256072 | 0.05120735202 | GGB32371 |
| 0.01711752944 | 0.0987256072 | 0.05120735202 | GGB3793 |
| 0.01711752944 | 0.0987256072 | 0.05120735202 | GGB42598 |
| 0.01977316736 | 0.0987256072 | 0.05120735202 | GGB45656 |
| 0.01711752944 | 0.0987256072 | 0.05120735202 | GGB47127 |
| 0.01711752944 | 0.0987256072 | 0.06716760956 | GGB74395 |
| 0.01711752944 | 0.0987256072 | 0.05120735202 | GGB75053 |
| 0.01806486337 | 0.0987256072 | 0.05120735202 | GGB75109 |
| 0.02182356312 | 0.1048169025 | 0.1171380983 | GGB81440 |
| 0.01761114253 | 0.0987256072 | 0.05120735202 | Lachnospiraceae_unclassified |
| 0.01901055128 | 0.0987256072 | 0.05120735202 | Lachnospiraceae_unclassified |
| 0.02274014528 | 0.1369671795 | 0.05120735202 | Lachnospiraceae_unclassified |
| 0.01711752944 | 0.0987256072 | 0.05120735202 | Lachnospiraceae_unclassified |
| 0.01711752944 | 0.0987256072 | 0.05120735202 | Lachnospiraceae_unclassified |
| 0.01711752944 | 0.0987256072 | 0.05120735202 | Lachnospiraceae_unclassified |
| 0.01761114253 | 0.1002066352 | 0.05443648682 | Lachnospiraceae_unclassified |
| 0.01711752944 | 0.0987256072 | 0.05120735202 | Lachnospiraceae_unclassified |
| 0.01737393672 | 0.0987256072 | 0.05120735202 | Lactobacillus |
| 0.01711752944 | 0.0987256072 | 0.05120735202 | Muribaculaceae_unclassified |
| 0.01711752944 | 0.0987256072 | 0.05120735202 | Neglectibacter |
| 0.01711752944 | 0.0987256072 | 0.05120735202 | Oscillospiraceae_unclassified |
| 0.01711752944 | 0.0987256072 | 0.05120735202 | Oscillospiraceae_unclassified |
| 0.01711752944 | 0.0987256072 | 0.05120735202 | Oscillospiraceae_unclassified |
| 0.01711752944 | 0.0987256072 | 0.05120735202 | Parasutterella |
| 0.01741082102 | 0.0987256072 | 0.05120735202 | Romboutsia |
| 0.01711752944 | 0.0987256072 | 0.05120735202 | Schaedlerella |
| 0.01711752944 | 0.0987256072 | 0.05120735202 | Turicibacter |
| 0.01711752944 | 0.0987256072 | 0.05120735202 | Bacteria_unclassified |
| 0.01711752944 | 0.0987256072 | 0.05120735202 | Bacteria_unclassified |
| 0.01711752944 | 0.0987256072 | 0.05120735202 | Bacteria_unclassified |
| 0.01842188925 | 0.1481479697 | 0.05728198365 | |
| 0.01711752944 | 0.0987256072 | 0.05120735202 | |
| 0.01711752944 | 0.0987256072 | 0.05120735202 | |
| 0.03061283424 | 0.2318968287 | 0.06391583115 | Acetatifactor |
| 0.03061283424 | 0.2318968287 | 0.06391583115 | Acetatifactor |
| 0.03061283424 | 0.2318968287 | 0.06391583115 | Acutalibacter |
| 0.03061283424 | 0.2318968287 | 0.06391583115 | Acutalibacter |
| 0.03061283424 | 0.2318968287 | 0.06391583115 | Adlercreutzia |
| 0.03061283424 | 0.2318968287 | 0.06391583115 | Adlercreutzia |
| 0.03061283424 | 0.2318968287 | 0.06391583115 | Adlercreutzia |
| 0.03061283424 | 0.2318968287 | 0.06391583115 | Akkermansia |
| 0.03061283424 | 0.2318968287 | 0.06391583115 | Alistipes |
| 0.03061283424 | 0.2403775523 | 0.0682375548 | Anaerotruncus |
| 0.03061283424 | 0.2318968287 | 0.06391583115 | Bacteria_unclassified |
| 0.03061283424 | 0.2318968287 | 0.06391583115 | Bacteria_unclassified |

| | | | |
|---|---|---|---|
| 0.03061283424 | 0.2318968287 | 0.06391583115 | Bacteria_unclassified |
| 0.03061283424 | 0.2318968287 | 0.06391583115 | Bacteroides |
| 0.03061283424 | 0.2318968287 | 0.06391583115 | Bifidobacterium |
| 0.03133621099 | 0.2318968287 | 0.06391583115 | Clostridia_unclassified |
| 0.03061283424 | 0.2340070451 | 0.06391583115 | Clostridiaceae_unclassified |
| 0.03061283424 | 0.2318968287 | 0.06391583115 | Clostridiaceae_unclassified |
| 0.03061283424 | 0.2318968287 | 0.06391583115 | Eubacteriales_unclassified |
| 0.03061283424 | 0.2318968287 | 0.06391583115 | Erysipelatoclostridium |
| 0.04014670546 | 0.2340070451 | 0.07084183666 | Coriobacteriaceae_unclassifie |
| 0.03061283424 | 0.2318968287 | 0.06391583115 | Dorea |
| 0.04945132624 | 0.2318968287 | 0.06391583115 | Dubosiella |
| 0.03061283424 | 0.2318968287 | 0.06391583115 | Erysipelotrichales_unclassifie |
| 0.0540865581 | 0.2388065533 | 0.07084183666 | Eubacteriaceae_unclassified |
| 0.03061283424 | 0.2318968287 | 0.06391583115 | Eubacteriaceae_unclassified |
| 0.03061283424 | 0.2318968287 | 0.06391583115 | GGB20149 |
| 0.03061283424 | 0.2318968287 | 0.06391583115 | GGB22635 |
| 0.03061283424 | 0.2318968287 | 0.06391583115 | GGB25041 |
| 0.03933452723 | 0.2318968287 | 0.06391583115 | GGB27876 |
| 0.03061283424 | 0.2318968287 | 0.06391583115 | GGB27878 |
| 0.03061283424 | 0.2318968287 | 0.06391583115 | GGB27918 |
| 0.03548981522 | 0.2318968287 | 0.06391583115 | GGB28382 |
| 0.0390743013 | 0.2651494912 | 0.09997335629 | GGB28399 |
| 0.03061283424 | 0.2318968287 | 0.06391583115 | GGB28411 |
| 0.03431936643 | 0.2318968287 | 0.06391583115 | GGB28415 |
| 0.03378362403 | 0.2318968287 | 0.06781582621 | GGB28430 |
| 0.03061283424 | 0.2318968287 | 0.06391583115 | GGB28439 |
| 0.03061283424 | 0.2318968287 | 0.06391583115 | GGB28778 |
| 0.03061283424 | 0.2340070451 | 0.06391583115 | GGB28784 |
| 0.03061283424 | 0.2318968287 | 0.06391583115 | GGB28792 |
| 0.03061283424 | 0.2318968287 | 0.06391583115 | GGB28798 |
| 0.03061283424 | 0.2318968287 | 0.06391583115 | GGB28802 |
| 0.03061283424 | 0.2318968287 | 0.06391583115 | GGB28818 |
| 0.03061283424 | 0.2318968287 | 0.06391583115 | GGB28828 |
| 0.03133621099 | 0.2318968287 | 0.06391583115 | GGB28851 |
| 0.03061283424 | 0.2318968287 | 0.06391583115 | GGB28859 |
| 0.03061283424 | 0.2318968287 | 0.06391583115 | GGB28864 |
| 0.03061283424 | 0.2318968287 | 0.06391583115 | GGB28869 |
| 0.03061283424 | 0.2490840217 | 0.06391583115 | GGB28883 |
| 0.03061283424 | 0.2318968287 | 0.06391583115 | GGB28892 |
| 0.04112260514 | 0.2490840217 | 0.07574662104 | GGB28893 |
| 0.03919219869 | 0.2318968287 | 0.07187052322 | GGB28898 |
| 0.03061283424 | 0.2318968287 | 0.06391583115 | GGB28904 |
| 0.03061283424 | 0.2318968287 | 0.06391583115 | GGB28916 |
| 0.03133621099 | 0.2318968287 | 0.06391583115 | GGB28924 |
| 0.03061283424 | 0.2340070451 | 0.06651750655 | GGB28926 |
| 0.03061283424 | 0.2318968287 | 0.06391583115 | GGB28927 |

| | | | |
|---|---|---|---|
| 0.03061283424 | 0.2318968287 | 0.06391583115 | GGB28934 |
| 0.03344544171 | 0.2520702488 | 0.06651750655 | GGB28946 |
| 0.03133621099 | 0.2318968287 | 0.06391583115 | GGB28949 |
| 0.03061283424 | 0.2318968287 | 0.06391583115 | GGB28949 |
| 0.0540865581 | 0.238161457 | 0.06742693799 | GGB28950 |
| 0.03061283424 | 0.238161457 | 0.06391583115 | GGB28951 |
| 0.03061283424 | 0.2318968287 | 0.06391583115 | GGB28951 |
| 0.03061283424 | 0.2318968287 | 0.06391583115 | GGB28954 |
| 0.03967451176 | 0.2318968287 | 0.07199853899 | GGB28956 |
| 0.03346694243 | 0.2714682952 | 0.1057789383 | GGB28960 |
| 0.04015524817 | 0.238161457 | 0.06391583115 | GGB28967 |
| 0.03061283424 | 0.2318968287 | 0.06391583115 | GGB28991 |
| 0.03061283424 | 0.2318968287 | 0.07199853899 | GGB29002 |
| 0.03061283424 | 0.2318968287 | 0.06391583115 | GGB29003 |
| 0.03061283424 | 0.238161457 | 0.06391583115 | GGB29011 |
| 0.03061283424 | 0.2318968287 | 0.06391583115 | GGB29531 |
| 0.03061283424 | 0.2318968287 | 0.06391583115 | GGB29685 |
| 0.03061283424 | 0.2318968287 | 0.06391583115 | GGB30141 |
| 0.03061283424 | 0.2318968287 | 0.06391583115 | GGB30286 |
| 0.03061283424 | 0.2318968287 | 0.06391583115 | GGB30303 |
| 0.0311404447 | 0.2318968287 | 0.06391583115 | GGB30413 |
| 0.0311404447 | 0.2340070451 | 0.06391583115 | GGB30454 |
| 0.0311404447 | 0.2318968287 | 0.06391583115 | GGB30455 |
| 0.03061283424 | 0.2318968287 | 0.06391583115 | GGB30461 |
| 0.03061283424 | 0.2318968287 | 0.06391583115 | GGB30461 |
| 0.03061283424 | 0.2318968287 | 0.06391583115 | GGB30463 |
| 0.03061283424 | 0.2318968287 | 0.06391583115 | GGB30473 |
| 0.03061283424 | 0.2318968287 | 0.06391583115 | GGB30475 |
| 0.03061283424 | 0.2318968287 | 0.06391583115 | GGB30861 |
| 0.03061283424 | 0.2318968287 | 0.06391583115 | GGB31312 |
| 0.03131068776 | 0.2318968287 | 0.06391583115 | GGB31438 |
| 0.03123052325 | 0.2318968287 | 0.06391583115 | GGB3171 |
| 0.03061283424 | 0.2318968287 | 0.06391583115 | GGB31823 |
| 0.03061283424 | 0.2520702488 | 0.07721166532 | GGB31853 |
| 0.03061283424 | 0.2318968287 | 0.06391583115 | GGB32371 |
| 0.03061283424 | 0.2318968287 | 0.06391583115 | GGB3793 |
| 0.03061283424 | 0.2318968287 | 0.06391583115 | GGB42598 |
| 0.03273620042 | 0.2318968287 | 0.06391583115 | GGB45656 |
| 0.03061283424 | 0.2318968287 | 0.06391583115 | GGB47127 |
| 0.03061283424 | 0.2318968287 | 0.06391583115 | GGB74395 |
| 0.03061283424 | 0.2318968287 | 0.06391583115 | GGB75053 |
| 0.03440524964 | 0.2318968287 | 0.06391583115 | GGB75109 |
| 0.03378362403 | 0.2340070451 | 0.09380371308 | GGB81440 |
| 0.0311404447 | 0.2318968287 | 0.06391583115 | Lachnospiraceae_unclassified |
| 0.03061283424 | 0.2318968287 | 0.06391583115 | Lachnospiraceae_unclassified |
| 0.03548981522 | 0.2678489208 | 0.06391583115 | Lachnospiraceae_unclassified |

| | | | |
|---|---|---|---|
| 0.03061283424 | 0.2318968287 | 0.06391583115 | Lachnospiraceae_unclassified |
| 0.03061283424 | 0.2318968287 | 0.06391583115 | Lachnospiraceae_unclassified |
| 0.03061283424 | 0.2318968287 | 0.06391583115 | Lachnospiraceae_unclassified |
| 0.03133621099 | 0.2340070451 | 0.07187052322 | Lachnospiraceae_unclassified |
| 0.03061283424 | 0.2318968287 | 0.06391583115 | Lachnospiraceae_unclassified |
| 0.03061283424 | 0.2318968287 | 0.06391583115 | Lactobacillus |
| 0.03061283424 | 0.2318968287 | 0.06391583115 | Muribaculaceae_unclassified |
| 0.03061283424 | 0.2318968287 | 0.06391583115 | Neglectibacter |
| 0.03133621099 | 0.2318968287 | 0.06391583115 | Oscillospiraceae_unclassified |
| 0.03061283424 | 0.2318968287 | 0.06391583115 | Oscillospiraceae_unclassified |
| 0.03061283424 | 0.2318968287 | 0.06391583115 | Oscillospiraceae_unclassified |
| 0.03061283424 | 0.2318968287 | 0.06391583115 | Parasutterella |
| 0.04254879213 | 0.2407346103 | 0.06391583115 | Romboutsia |
| 0.03068360841 | 0.2318968287 | 0.06391583115 | Schaedlerella |
| 0.03061283424 | 0.2318968287 | 0.06391583115 | Turicibacter |
| 0.03061283424 | 0.2318968287 | 0.06391583115 | Bacteria_unclassified |
| 0.03061283424 | 0.2318968287 | 0.06391583115 | Bacteria_unclassified |
| 0.03061283424 | 0.2318968287 | 0.06391583115 | Bacteria_unclassified |
| 0.03133621099 | 0.2654385924 | 0.06511268346 | |
| 0.03061283424 | 0.2318968287 | 0.06391583115 | |
| 0.03061283424 | 0.2318968287 | 0.06391583115 | |
| 0.02012234551 | 0.2037494731 | 0.05732595108 | Acetatifactor |
| 0.02012234551 | 0.2037494731 | 0.05732595108 | Acetatifactor |
| 0.02012234551 | 0.2037494731 | 0.05732595108 | Acutalibacter |
| 0.02012234551 | 0.2037494731 | 0.05732595108 | Acutalibacter |
| 0.02012234551 | 0.2037494731 | 0.05732595108 | Adlercreutzia |
| 0.02012234551 | 0.2037494731 | 0.05732595108 | Adlercreutzia |
| 0.02012234551 | 0.2037494731 | 0.05732595108 | Adlercreutzia |
| 0.02012234551 | 0.2037494731 | 0.05732595108 | Akkermansia |
| 0.02012234551 | 0.2037494731 | 0.05732595108 | Alistipes |
| 0.02012234551 | 0.2037494731 | 0.05732595108 | Anaerotruncus |
| 0.02012234551 | 0.2037494731 | 0.05732595108 | Bacteria_unclassified |
| 0.02012234551 | 0.2037494731 | 0.05732595108 | Bacteria_unclassified |
| 0.02012234551 | 0.2037494731 | 0.05732595108 | Bacteria_unclassified |
| 0.02012234551 | 0.2100585492 | 0.05732595108 | Bacteroides |
| 0.02012234551 | 0.2037494731 | 0.05732595108 | Bifidobacterium |
| 0.02538662549 | 0.2059673764 | 0.05732595108 | Clostridia_unclassified |
| 0.02012234551 | 0.3545762691 | 0.05732595108 | Clostridiaceae_unclassified |
| 0.02012234551 | 0.2037494731 | 0.05732595108 | Clostridiaceae_unclassified |
| 0.03417599845 | 0.2037494731 | 0.05732595108 | Eubacteriales_unclassified |
| 0.02538662549 | 0.2037494731 | 0.05732595108 | Erysipelatoclostridium |
| 0.03417599845 | 0.2383120122 | 0.08197010123 | Coriobacteriaceae_unclassifie |
| 0.02012234551 | 0.2037494731 | 0.05732595108 | Dorea |
| 0.0294878573 | 0.2037494731 | 0.05732595108 | Dubosiella |
| 0.02012234551 | 0.2037494731 | 0.05732595108 | Erysipelotrichales_unclassifie |
| 0.02012234551 | 0.2037494731 | 0.05732595108 | Eubacteriaceae_unclassified |

| | | | |
|---|---|---|---|
| 0.02012234551 | 0.2037494731 | 0.05732595108 | Eubacteriaceae_unclassified |
| 0.02067394623 | 0.2037494731 | 0.05732595108 | GGB20149 |
| 0.02012234551 | 0.2037494731 | 0.05732595108 | GGB22635 |
| 0.02181288487 | 0.2037494731 | 0.05732595108 | GGB25041 |
| 0.0268329472 | 0.2037494731 | 0.05732595108 | GGB27876 |
| 0.02067394623 | 0.2037494731 | 0.05732595108 | GGB27878 |
| 0.02012234551 | 0.2037494731 | 0.05732595108 | GGB27918 |
| 0.02220379551 | 0.2037494731 | 0.05732595108 | GGB28382 |
| 0.02067394623 | 0.2037494731 | 0.06030409867 | GGB28399 |
| 0.02041320215 | 0.2037494731 | 0.06095343498 | GGB28411 |
| 0.02067394623 | 0.2037494731 | 0.05732595108 | GGB28415 |
| 0.02012234551 | 0.2037494731 | 0.05732595108 | GGB28430 |
| 0.02012234551 | 0.2037494731 | 0.05732595108 | GGB28439 |
| 0.02012234551 | 0.2037494731 | 0.05732595108 | GGB28778 |
| 0.02012234551 | 0.2037494731 | 0.05732595108 | GGB28784 |
| 0.02067394623 | 0.2037494731 | 0.05732595108 | GGB28792 |
| 0.02012234551 | 0.2037494731 | 0.05732595108 | GGB28798 |
| 0.02012234551 | 0.2037494731 | 0.05732595108 | GGB28802 |
| 0.02012234551 | 0.2037494731 | 0.05732595108 | GGB28818 |
| 0.02022932138 | 0.2037494731 | 0.05732595108 | GGB28828 |
| 0.02012234551 | 0.2037494731 | 0.05732595108 | GGB28851 |
| 0.02012234551 | 0.2037494731 | 0.05732595108 | GGB28859 |
| 0.02012234551 | 0.2037494731 | 0.05732595108 | GGB28864 |
| 0.02012234551 | 0.2037494731 | 0.05732595108 | GGB28869 |
| 0.02041320215 | 0.2122548936 | 0.05732595108 | GGB28883 |
| 0.02012234551 | 0.2037494731 | 0.05732595108 | GGB28892 |
| 0.02012234551 | 0.2037494731 | 0.05732595108 | GGB28893 |
| 0.02728767883 | 0.2037494731 | 0.07111255967 | GGB28898 |
| 0.02012234551 | 0.2037494731 | 0.05990492896 | GGB28904 |
| 0.02012234551 | 0.2037494731 | 0.05732595108 | GGB28916 |
| 0.02012234551 | 0.2037494731 | 0.05732595108 | GGB28924 |
| 0.02012234551 | 0.2106115464 | 0.06251022527 | GGB28926 |
| 0.02012234551 | 0.2037494731 | 0.05732595108 | GGB28927 |
| 0.02012234551 | 0.2037494731 | 0.05732595108 | GGB28934 |
| 0.02012234551 | 0.2037494731 | 0.05732595108 | GGB28946 |
| 0.02220379551 | 0.2037494731 | 0.05732595108 | GGB28949 |
| 0.02564287376 | 0.2037494731 | 0.05732595108 | GGB28949 |
| 0.03059237209 | 0.2059673764 | 0.06021154448 | GGB28950 |
| 0.02012234551 | 0.2037494731 | 0.05732595108 | GGB28951 |
| 0.02012234551 | 0.2037494731 | 0.05732595108 | GGB28951 |
| 0.02073474572 | 0.2037494731 | 0.05732595108 | GGB28954 |
| 0.02012234551 | 0.2037494731 | 0.05732595108 | GGB28956 |
| 0.02012234551 | 0.2037494731 | 0.05732595108 | GGB28960 |
| 0.02012234551 | 0.2037494731 | 0.05732595108 | GGB28967 |
| 0.02012234551 | 0.2037494731 | 0.05732595108 | GGB28991 |
| 0.02012234551 | 0.2086905514 | 0.0870172285 | GGB29002 |

| | | | |
|---|---|---|---|
| 0.02020856749 | 0.2037494731 | 0.05732595108 | GGB29003 |
| 0.02012234551 | 0.2055628758 | 0.05732595108 | GGB29011 |
| 0.02012234551 | 0.2037494731 | 0.05732595108 | GGB29531 |
| 0.02012234551 | 0.2037494731 | 0.05732595108 | GGB29685 |
| 0.02012234551 | 0.2037494731 | 0.05732595108 | GGB30141 |
| 0.02538662549 | 0.2037494731 | 0.05732595108 | GGB30286 |
| 0.02022932138 | 0.2037494731 | 0.05732595108 | GGB30303 |
| 0.02220379551 | 0.2037494731 | 0.05732595108 | GGB30413 |
| 0.02067394623 | 0.2086905514 | 0.05732595108 | GGB30454 |
| 0.02012234551 | 0.2037494731 | 0.05732595108 | GGB30455 |
| 0.02012234551 | 0.2037494731 | 0.05732595108 | GGB30461 |
| 0.02012234551 | 0.2037494731 | 0.05732595108 | GGB30461 |
| 0.02012234551 | 0.2037494731 | 0.05732595108 | GGB30463 |
| 0.02012234551 | 0.2037494731 | 0.05732595108 | GGB30473 |
| 0.02012234551 | 0.2037494731 | 0.05732595108 | GGB30475 |
| 0.02012234551 | 0.2037494731 | 0.05732595108 | GGB30861 |
| 0.02012234551 | 0.2037494731 | 0.05732595108 | GGB31312 |
| 0.02012234551 | 0.2037494731 | 0.05732595108 | GGB31438 |
| 0.02386464823 | 0.2037494731 | 0.05732595108 | GGB3171 |
| 0.02012234551 | 0.2037494731 | 0.05732595108 | GGB31823 |
| 0.02012234551 | 0.2037494731 | 0.05732595108 | GGB31853 |
| 0.02012234551 | 0.2037494731 | 0.05732595108 | GGB32371 |
| 0.02012234551 | 0.2037494731 | 0.05732595108 | GGB3793 |
| 0.02012234551 | 0.2037494731 | 0.05732595108 | GGB42598 |
| 0.02144090351 | 0.2037494731 | 0.05732595108 | GGB45656 |
| 0.02012234551 | 0.2037494731 | 0.05732595108 | GGB47127 |
| 0.02012234551 | 0.2037494731 | 0.05732595108 | GGB74395 |
| 0.02012234551 | 0.2037494731 | 0.05732595108 | GGB75053 |
| 0.02101876757 | 0.2037494731 | 0.05732595108 | GGB75109 |
| 0.02220379551 | 0.2037494731 | 0.09252372449 | GGB81440 |
| 0.02067394623 | 0.2037494731 | 0.05732595108 | Lachnospiraceae_unclassified |
| 0.02220379551 | 0.2037494731 | 0.05732595108 | Lachnospiraceae_unclassified |
| 0.02764120945 | 0.2902477751 | 0.05732595108 | Lachnospiraceae_unclassified |
| 0.02012234551 | 0.2037494731 | 0.06083542092 | Lachnospiraceae_unclassified |
| 0.02067394623 | 0.2037494731 | 0.06083542092 | Lachnospiraceae_unclassified |
| 0.02012234551 | 0.2037494731 | 0.05732595108 | Lachnospiraceae_unclassified |
| 0.02012234551 | 0.2037494731 | 0.05732595108 | Lachnospiraceae_unclassified |
| 0.02012234551 | 0.2037494731 | 0.05732595108 | Lachnospiraceae_unclassified |
| 0.02012234551 | 0.2037494731 | 0.05732595108 | Lactobacillus |
| 0.02012234551 | 0.2037494731 | 0.05732595108 | Muribaculaceae_unclassified |
| 0.02012234551 | 0.2037494731 | 0.05732595108 | Neglectibacter |
| 0.02012234551 | 0.2037494731 | 0.05732595108 | Oscillospiraceae_unclassified |
| 0.02220379551 | 0.3068817855 | 0.1117095028 | Oscillospiraceae_unclassified |
| 0.02012234551 | 0.2037494731 | 0.05732595108 | Oscillospiraceae_unclassified |
| 0.02012234551 | 0.2037494731 | 0.05732595108 | Parasutterella |
| 0.02437101104 | 0.2059673764 | 0.05732595108 | Romboutsia |

| | | | |
|---|---|---|---|
| 0.02012234551 | 0.2037494731 | 0.05732595108 | Schaedlerella |
| 0.02012234551 | 0.2037494731 | 0.05732595108 | Turicibacter |
| 0.02012234551 | 0.2037494731 | 0.05732595108 | Bacteria_unclassified |
| 0.02012234551 | 0.2037494731 | 0.05732595108 | Bacteria_unclassified |
| 0.02012234551 | 0.2037494731 | 0.05732595108 | Bacteria_unclassified |
| 0.02067394623 | 0.2372783653 | 0.05888650266 | |
| 0.02012234551 | 0.2037494731 | 0.05732595108 | |
| 0.02012234551 | 0.2037494731 | 0.05732595108 | |
| 0.0006174936392 | 0.8567755293 | 0.7080320955 | Acetatifactor |
| 0.0006174936392 | 0.8567755293 | 0.7080320955 | Acetatifactor |
| 0.0006174936392 | 0.8567755293 | 0.7080320955 | Acutalibacter |
| 0.0006174936392 | 0.8567755293 | 0.7080320955 | Acutalibacter |
| 0.0006174936392 | 0.8567755293 | 0.7080320955 | Adlercreutzia |
| 0.000850088246 | 0.8567755293 | 0.7080320955 | Adlercreutzia |
| 0.0006174936392 | 0.8567755293 | 0.7080320955 | Adlercreutzia |
| 0.0006174936392 | 0.8567755293 | 0.7080320955 | Akkermansia |
| 0.0006517985519 | 0.8567755293 | 0.7080320955 | Alistipes |
| 0.0006174936392 | 0.8567755293 | 0.7080320955 | Anaerotruncus |
| 0.0006174936392 | 0.8567755293 | 0.7080320955 | Bacteria_unclassified |
| 0.0006174936392 | 0.8567755293 | 0.7080320955 | Bacteria_unclassified |
| 0.0006174936392 | 0.8567755293 | 0.7080320955 | Bacteria_unclassified |
| 0.0006174936392 | 0.8567755293 | 0.7080320955 | Bacteroides |
| 0.0006174936392 | 0.8567755293 | 0.7080320955 | Bifidobacterium |
| 0.0006174936392 | 0.8567755293 | 0.7080320955 | Clostridia_unclassified |
| 0.0006174936392 | 0.8567755293 | 0.7080320955 | Clostridiaceae_unclassified |
| 0.0006174936392 | 0.8567755293 | 0.7080320955 | Clostridiaceae_unclassified |
| 0.0006174936392 | 0.8567755293 | 0.7080320955 | Eubacteriales_unclassified |
| 0.0006174936392 | 0.8567755293 | 0.7080320955 | Erysipelatoclostridium |
| 0.0007454143061 | 0.8567755293 | 0.7080320955 | Coriobacteriaceae_unclassifie |
| 0.0006174936392 | 0.8567755293 | 0.7080320955 | Dorea |
| 0.0006174936392 | 0.8567755293 | 0.7080320955 | Dubosiella |
| 0.0006174936392 | 0.8576617901 | 0.7080320955 | Erysipelotrichales_unclassifie |
| 0.0006647299644 | 0.8567755293 | 0.7080320955 | Eubacteriaceae_unclassified |
| 0.0007434019507 | 0.8567755293 | 0.7080320955 | Eubacteriaceae_unclassified |
| 0.0006369303364 | 0.8567755293 | 0.7080320955 | GGB20149 |
| 0.0006174936392 | 0.8567755293 | 0.7080320955 | GGB22635 |
| 0.0006174936392 | 0.8567755293 | 0.7080320955 | GGB25041 |
| 0.0007232202525 | 0.8567755293 | 0.7080320955 | GGB27876 |
| 0.0006174936392 | 0.8567755293 | 0.7080320955 | GGB27878 |
| 0.0006174936392 | 0.8567755293 | 0.7080320955 | GGB27918 |
| 0.0006174936392 | 0.8567755293 | 0.7080320955 | GGB28382 |
| 0.0006369303364 | 0.8567755293 | 0.7080320955 | GGB28399 |
| 0.0006174936392 | 0.8567755293 | 0.7080320955 | GGB28411 |
| 0.0007292388671 | 0.8567755293 | 0.7080320955 | GGB28415 |
| 0.0006174936392 | 0.8567755293 | 0.7080320955 | GGB28430 |
| 0.0006174936392 | 0.8567755293 | 0.7080320955 | GGB28439 |

| | | | |
|---|---|---|---|
| 0.0006174936392 | 0.8567755293 | 0.7080320955 | GGB28778 |
| 0.0006174936392 | 0.8567755293 | 0.7080320955 | GGB28784 |
| 0.0006174936392 | 0.8567755293 | 0.7080320955 | GGB28792 |
| 0.0006174936392 | 0.8567755293 | 0.7080320955 | GGB28798 |
| 0.0006174936392 | 0.8567755293 | 0.7080320955 | GGB28802 |
| 0.0006174936392 | 0.8567755293 | 0.7080320955 | GGB28818 |
| 0.0008006150434 | 0.8567755293 | 0.7080320955 | GGB28828 |
| 0.0006174936392 | 0.8567755293 | 0.7080320955 | GGB28851 |
| 0.0006174936392 | 0.8567755293 | 0.7080320955 | GGB28859 |
| 0.0006449340524 | 0.8567755293 | 0.7080320955 | GGB28864 |
| 0.0006174936392 | 0.8567755293 | 0.7080320955 | GGB28869 |
| 0.0006174936392 | 0.9786800182 | 0.7080320955 | GGB28883 |
| 0.0006401874772 | 0.8567755293 | 0.7080320955 | GGB28892 |
| 0.0007757537257 | 0.8567755293 | 0.7080320955 | GGB28893 |
| 0.0007782354297 | 0.8567755293 | 0.7080320955 | GGB28898 |
| 0.0006174936392 | 0.8567755293 | 0.7080320955 | GGB28904 |
| 0.0006369303364 | 0.8567755293 | 0.7080320955 | GGB28916 |
| 0.0006174936392 | 0.8567755293 | 0.7080320955 | GGB28924 |
| 0.0006174936392 | 0.8567755293 | 0.7080320955 | GGB28926 |
| 0.0006174936392 | 0.8567755293 | 0.7080320955 | GGB28927 |
| 0.0006174936392 | 0.8567755293 | 0.7080320955 | GGB28934 |
| 0.0006517985519 | 0.8567755293 | 0.7080320955 | GGB28946 |
| 0.0006174936392 | 0.8567755293 | 0.7080320955 | GGB28949 |
| 0.0007536026659 | 0.8567755293 | 0.7080320955 | GGB28949 |
| 0.0006369303364 | 0.8567755293 | 0.7080320955 | GGB28950 |
| 0.0006174936392 | 0.8567755293 | 0.7080320955 | GGB28951 |
| 0.0006174936392 | 0.8567755293 | 0.7080320955 | GGB28951 |
| 0.0006517985519 | 0.8567755293 | 0.7080320955 | GGB28954 |
| 0.0006174936392 | 0.8567755293 | 0.7080320955 | GGB28956 |
| 0.0006174936392 | 0.8666506689 | 0.7316612585 | GGB28960 |
| 0.0006174936392 | 0.8567755293 | 0.7080320955 | GGB28967 |
| 0.0006174936392 | 0.8567755293 | 0.7080320955 | GGB28991 |
| 0.0006517985519 | 0.8781139016 | 0.8763951327 | GGB29002 |
| 0.0006174936392 | 0.8567755293 | 0.7080320955 | GGB29003 |
| 0.0006174936392 | 0.8717826367 | 0.7080320955 | GGB29011 |
| 0.0007232202525 | 0.8567755293 | 0.7080320955 | GGB29531 |
| 0.0006174936392 | 0.8567755293 | 0.7080320955 | GGB29685 |
| 0.0006174936392 | 0.8567755293 | 0.7080320955 | GGB30141 |
| 0.0006174936392 | 0.8567755293 | 0.7080320955 | GGB30286 |
| 0.0006174936392 | 0.8567755293 | 0.7080320955 | GGB30303 |
| 0.0006174936392 | 0.8567755293 | 0.7080320955 | GGB30413 |
| 0.0006174936392 | 0.8567755293 | 0.7080320955 | GGB30454 |
| 0.0006174936392 | 0.8567755293 | 0.7080320955 | GGB30455 |
| 0.0006174936392 | 0.8567755293 | 0.7080320955 | GGB30461 |
| 0.0006174936392 | 0.8567755293 | 0.7080320955 | GGB30461 |
| 0.0006517985519 | 0.8567755293 | 0.7080320955 | GGB30463 |

| | | | |
|---|---|---|---|
| 0.0006174936392 | 0.8567755293 | 0.7080320955 | GGB30473 |
| 0.0006517985519 | 0.8567755293 | 0.7080320955 | GGB30475 |
| 0.0006174936392 | 0.8567755293 | 0.7080320955 | GGB30861 |
| 0.0006174936392 | 0.8567755293 | 0.7080320955 | GGB31312 |
| 0.0006174936392 | 0.8567755293 | 0.7080320955 | GGB31438 |
| 0.0006369303364 | 0.8567755293 | 0.7080320955 | GGB3171 |
| 0.0006174936392 | 0.8567755293 | 0.7080320955 | GGB31823 |
| 0.0006174936392 | 0.8567755293 | 0.7080320955 | GGB31853 |
| 0.0006174936392 | 0.9535948224 | 0.7080320955 | GGB32371 |
| 0.0006174936392 | 0.8567755293 | 0.7080320955 | GGB3793 |
| 0.0006174936392 | 0.8567755293 | 0.7080320955 | GGB42598 |
| 0.0006174936392 | 0.8567755293 | 0.7080320955 | GGB45656 |
| 0.0006174936392 | 0.8567755293 | 0.7080320955 | GGB47127 |
| 0.0006174936392 | 0.8567755293 | 0.7584047277 | GGB74395 |
| 0.0006174936392 | 0.8567755293 | 0.7080320955 | GGB75053 |
| 0.0006174936392 | 0.8567755293 | 0.7080320955 | GGB75109 |
| 0.0006174936392 | 0.8567755293 | 0.7080320955 | GGB81440 |
| 0.0006174936392 | 0.8567755293 | 0.7080320955 | Lachnospiraceae_unclassified |
| 0.0006517985519 | 0.8567755293 | 0.7080320955 | Lachnospiraceae_unclassified |
| 0.0006369303364 | 0.8567755293 | 0.7080320955 | Lachnospiraceae_unclassified |
| 0.0009041431421 | 0.8567755293 | 0.7080320955 | Lachnospiraceae_unclassified |
| 0.0006174936392 | 0.8567755293 | 0.7080320955 | Lachnospiraceae_unclassified |
| 0.0006174936392 | 0.8567755293 | 0.7080320955 | Lachnospiraceae_unclassified |
| 0.0006369303364 | 0.8567755293 | 0.7080320955 | Lachnospiraceae_unclassified |
| 0.0006174936392 | 0.8567755293 | 0.7080320955 | Lachnospiraceae_unclassified |
| 0.0006174936392 | 0.8567755293 | 0.7080320955 | Lactobacillus |
| 0.0006174936392 | 0.8567755293 | 0.7080320955 | Muribaculaceae_unclassified |
| 0.0006449340524 | 0.8567755293 | 0.7080320955 | Neglectibacter |
| 0.0006517985519 | 0.8567755293 | 0.7080320955 | Oscillospiraceae_unclassified |
| 0.0006174936392 | 0.8567755293 | 0.7080320955 | Oscillospiraceae_unclassified |
| 0.0006174936392 | 0.8567755293 | 0.7080320955 | Oscillospiraceae_unclassified |
| 0.0006174936392 | 0.8567755293 | 0.7080320955 | Parasutterella |
| 0.0006174936392 | 0.8567755293 | 0.7080320955 | Romboutsia |
| 0.0006174936392 | 0.8567755293 | 0.7080320955 | Schaedlerella |
| 0.0006174936392 | 0.8567755293 | 0.7080320955 | Turicibacter |
| 0.002106762016 | 0.8567755293 | 0.7080320955 | Bacteria_unclassified |
| 0.0006174936392 | 0.8567755293 | 0.7080320955 | Bacteria_unclassified |
| 0.0006174936392 | 0.8567755293 | 0.7080320955 | Bacteria_unclassified |
| 0.0006174936392 | 0.8567755293 | 0.7080320955 | |
| 0.0006174936392 | 0.8567755293 | 0.7080320955 | |
| 0.0006174936392 | 0.8567755293 | 0.7080320955 | |
| 0.001271705588 | 0.7676091217 | 0.6256830596 | Acetatifactor |
| 0.001271705588 | 0.7676091217 | 0.6256830596 | Acetatifactor |
| 0.001271705588 | 0.7676091217 | 0.6256830596 | Acutalibacter |
| 0.001271705588 | 0.7676091217 | 0.6543342278 | Acutalibacter |
| 0.001271705588 | 0.7676091217 | 0.6256830596 | Adlercreutzia |

| | | | |
|---|---|---|---|
| 0.001801501178 | 0.7676091217 | 0.6256830596 | Adlercreutzia |
| 0.001271705588 | 0.7676091217 | 0.6256830596 | Adlercreutzia |
| 0.001271705588 | 0.7676091217 | 0.6256830596 | Akkermansia |
| 0.00134666968 | 0.7676091217 | 0.6256830596 | Alistipes |
| 0.001271705588 | 0.7676091217 | 0.6256830596 | Anaerotruncus |
| 0.001271705588 | 0.7676091217 | 0.6256830596 | Bacteria_unclassified |
| 0.001271705588 | 0.7676091217 | 0.6256830596 | Bacteria_unclassified |
| 0.001271705588 | 0.7676091217 | 0.6256830596 | Bacteria_unclassified |
| 0.001271705588 | 0.7676091217 | 0.6256830596 | Bacteroides |
| 0.001271705588 | 0.7676091217 | 0.6256830596 | Bifidobacterium |
| 0.001271705588 | 0.7676091217 | 0.6256830596 | Clostridia_unclassified |
| 0.001271705588 | 0.7676091217 | 0.6256830596 | Clostridiaceae_unclassified |
| 0.001271705588 | 0.7676091217 | 0.6256830596 | Clostridiaceae_unclassified |
| 0.001271705588 | 0.7676091217 | 0.6256830596 | Eubacteriales_unclassified |
| 0.001271705588 | 0.7676091217 | 0.6256830596 | Erysipelatoclostridium |
| 0.00148335049 | 0.7676091217 | 0.6256830596 | Coriobacteriaceae_unclassifie |
| 0.001271705588 | 0.7676091217 | 0.6837147113 | Dorea |
| 0.001271705588 | 0.7676091217 | 0.6256830596 | Dubosiella |
| 0.001271705588 | 0.7676091217 | 0.6256830596 | Erysipelotrichales_unclassifie |
| 0.001271705588 | 0.7676091217 | 0.6256830596 | Eubacteriaceae_unclassified |
| 0.001758242386 | 0.7676091217 | 0.6256830596 | Eubacteriaceae_unclassified |
| 0.001271705588 | 0.7676091217 | 0.6256830596 | GGB20149 |
| 0.001271705588 | 0.7676091217 | 0.6256830596 | GGB22635 |
| 0.001271705588 | 0.7676091217 | 0.6256830596 | GGB25041 |
| 0.001660471766 | 0.7676091217 | 0.6256830596 | GGB27876 |
| 0.001271705588 | 0.7676091217 | 0.6256830596 | GGB27878 |
| 0.001271705588 | 0.7676091217 | 0.6256830596 | GGB27918 |
| 0.001271705588 | 0.7676091217 | 0.6256830596 | GGB28382 |
| 0.001271705588 | 0.7676091217 | 0.6339649952 | GGB28399 |
| 0.001271705588 | 0.7676091217 | 0.6256830596 | GGB28411 |
| 0.002013349252 | 0.7676091217 | 0.6256830596 | GGB28415 |
| 0.001271705588 | 0.7676091217 | 0.6256830596 | GGB28430 |
| 0.001271705588 | 0.7676091217 | 0.6256830596 | GGB28439 |
| 0.001271705588 | 0.7676091217 | 0.6256830596 | GGB28778 |
| 0.001271705588 | 0.7676091217 | 0.6256830596 | GGB28784 |
| 0.001271705588 | 0.7676091217 | 0.6256830596 | GGB28792 |
| 0.001271705588 | 0.7676091217 | 0.6256830596 | GGB28798 |
| 0.001271705588 | 0.7676091217 | 0.6256830596 | GGB28802 |
| 0.001271705588 | 0.7676091217 | 0.6256830596 | GGB28818 |
| 0.001519892564 | 0.7676091217 | 0.6256830596 | GGB28828 |
| 0.001271705588 | 0.7676091217 | 0.6256830596 | GGB28851 |
| 0.001271705588 | 0.7676091217 | 0.6256830596 | GGB28859 |
| 0.001315919575 | 0.7676091217 | 0.6256830596 | GGB28864 |
| 0.001271705588 | 0.7676091217 | 0.6256830596 | GGB28869 |
| 0.001271705588 | 0.8343978319 | 0.6256830596 | GGB28883 |
| 0.001315919575 | 0.7676091217 | 0.6256830596 | GGB28892 |

| | | | |
|---|---|---|---|
| 0.001315919575 | 0.7676091217 | 0.6256830596 | GGB28893 |
| 0.001928802043 | 0.7676091217 | 0.6256830596 | GGB28898 |
| 0.001271705588 | 0.7676091217 | 0.6467183782 | GGB28904 |
| 0.001282660148 | 0.7676091217 | 0.6256830596 | GGB28916 |
| 0.001271705588 | 0.7676091217 | 0.6256830596 | GGB28924 |
| 0.001271705588 | 0.7676091217 | 0.6256830596 | GGB28926 |
| 0.001271705588 | 0.7676091217 | 0.6256830596 | GGB28927 |
| 0.001271705588 | 0.7676091217 | 0.6256830596 | GGB28934 |
| 0.001315919575 | 0.7676091217 | 0.6256830596 | GGB28946 |
| 0.001271705588 | 0.7676091217 | 0.6256830596 | GGB28949 |
| 0.001487084829 | 0.7676091217 | 0.6256830596 | GGB28949 |
| 0.001660471766 | 0.7676091217 | 0.6256830596 | GGB28950 |
| 0.001271705588 | 0.7676091217 | 0.6256830596 | GGB28951 |
| 0.001271705588 | 0.7676091217 | 0.6256830596 | GGB28951 |
| 0.001315919575 | 0.7676091217 | 0.6256830596 | GGB28954 |
| 0.001271705588 | 0.7676091217 | 0.6380397617 | GGB28956 |
| 0.001271705588 | 0.8176242312 | 0.6837147113 | GGB28960 |
| 0.001271705588 | 0.7676091217 | 0.6256830596 | GGB28967 |
| 0.001271705588 | 0.7676091217 | 0.6256830596 | GGB28991 |
| 0.001315919575 | 0.7676091217 | 0.6837147113 | GGB29002 |
| 0.001271705588 | 0.7676091217 | 0.6256830596 | GGB29003 |
| 0.001271705588 | 0.8176242312 | 0.6256830596 | GGB29011 |
| 0.001487084829 | 0.7676091217 | 0.6256830596 | GGB29531 |
| 0.001271705588 | 0.7676091217 | 0.6256830596 | GGB29685 |
| 0.001271705588 | 0.7676091217 | 0.6256830596 | GGB30141 |
| 0.001271705588 | 0.7676091217 | 0.6339649952 | GGB30286 |
| 0.001271705588 | 0.7676091217 | 0.6256830596 | GGB30303 |
| 0.001271705588 | 0.7676091217 | 0.6256830596 | GGB30413 |
| 0.001271705588 | 0.7676091217 | 0.6256830596 | GGB30454 |
| 0.001271705588 | 0.7676091217 | 0.6256830596 | GGB30455 |
| 0.001271705588 | 0.7676091217 | 0.6256830596 | GGB30461 |
| 0.001271705588 | 0.7676091217 | 0.6256830596 | GGB30461 |
| 0.001271705588 | 0.7676091217 | 0.6256830596 | GGB30463 |
| 0.001271705588 | 0.7676091217 | 0.6256830596 | GGB30473 |
| 0.001412577259 | 0.7676091217 | 0.6256830596 | GGB30475 |
| 0.001271705588 | 0.7676091217 | 0.6256830596 | GGB30861 |
| 0.001271705588 | 0.7676091217 | 0.6256830596 | GGB31312 |
| 0.001271705588 | 0.7676091217 | 0.6256830596 | GGB31438 |
| 0.001271705588 | 0.7676091217 | 0.6256830596 | GGB3171 |
| 0.001271705588 | 0.7676091217 | 0.6256830596 | GGB31823 |
| 0.001271705588 | 0.7676091217 | 0.6256830596 | GGB31853 |
| 0.001271705588 | 0.8309533002 | 0.6256830596 | GGB32371 |
| 0.001271705588 | 0.7676091217 | 0.6256830596 | GGB3793 |
| 0.001271705588 | 0.7676091217 | 0.6256830596 | GGB42598 |
| 0.001271705588 | 0.7676091217 | 0.6256830596 | GGB45656 |
| 0.001271705588 | 0.7676091217 | 0.6256830596 | GGB47127 |

| | | | |
|---|---|---|---|
| 0.001271705588 | 0.7676091217 | 0.7045255625 | GGB74395 |
| 0.001271705588 | 0.7676091217 | 0.6256830596 | GGB75053 |
| 0.001271705588 | 0.7676091217 | 0.6256830596 | GGB75109 |
| 0.001271705588 | 0.7676091217 | 0.6256830596 | GGB81440 |
| 0.001271705588 | 0.7676091217 | 0.6256830596 | Lachnospiraceae_unclassified |
| 0.001282660148 | 0.7676091217 | 0.6256830596 | Lachnospiraceae_unclassified |
| 0.001271705588 | 0.7676091217 | 0.6256830596 | Lachnospiraceae_unclassified |
| 0.001590309304 | 0.7676091217 | 0.6256830596 | Lachnospiraceae_unclassified |
| 0.001271705588 | 0.7676091217 | 0.6256830596 | Lachnospiraceae_unclassified |
| 0.001271705588 | 0.7676091217 | 0.6256830596 | Lachnospiraceae_unclassified |
| 0.001321273517 | 0.7676091217 | 0.6256830596 | Lachnospiraceae_unclassified |
| 0.001271705588 | 0.7676091217 | 0.6256830596 | Lachnospiraceae_unclassified |
| 0.001271705588 | 0.7676091217 | 0.6256830596 | Lactobacillus |
| 0.001271705588 | 0.7676091217 | 0.6256830596 | Muribaculaceae_unclassified |
| 0.001274724956 | 0.7676091217 | 0.6256830596 | Neglectibacter |
| 0.00134666968 | 0.7676091217 | 0.6256830596 | Oscillospiraceae_unclassified |
| 0.001271705588 | 0.7676091217 | 0.6256830596 | Oscillospiraceae_unclassified |
| 0.001271705588 | 0.7676091217 | 0.6256830596 | Oscillospiraceae_unclassified |
| 0.001271705588 | 0.7676091217 | 0.6256830596 | Parasutterella |
| 0.001271705588 | 0.8304615089 | 0.6256830596 | Romboutsia |
| 0.001271705588 | 0.7676091217 | 0.6256830596 | Schaedlerella |
| 0.001271705588 | 0.7676091217 | 0.6451862011 | Turicibacter |
| 0.002979913103 | 0.7676091217 | 0.6256830596 | Bacteria_unclassified |
| 0.001271705588 | 0.7676091217 | 0.6256830596 | Bacteria_unclassified |
| 0.001271705588 | 0.7676091217 | 0.6256830596 | Bacteria_unclassified |
| 0.001271705588 | 0.7676091217 | 0.6256830596 | |
| 0.001271705588 | 0.7676091217 | 0.6256830596 | |
| 0.001271705588 | 0.7676091217 | 0.6256830596 | |
| 0.04367470257 | 0.1305416393 | 0.3171314387 | Acetatifactor |
| 0.04367470257 | 0.1305416393 | 0.3171314387 | Acetatifactor |
| 0.04367470257 | 0.1305416393 | 0.3171314387 | Acutalibacter |
| 0.04367470257 | 0.1305416393 | 0.3171314387 | Acutalibacter |
| 0.04367470257 | 0.1305416393 | 0.3171314387 | Adlercreutzia |
| 0.04894179322 | 0.1305416393 | 0.3171314387 | Adlercreutzia |
| 0.04367470257 | 0.1305416393 | 0.3171314387 | Adlercreutzia |
| 0.0481701997 | 0.1305416393 | 0.3171314387 | Akkermansia |
| 0.04659916416 | 0.1305416393 | 0.3171314387 | Alistipes |
| 0.04367470257 | 0.1305416393 | 0.3171314387 | Anaerotruncus |
| 0.04894179322 | 0.1305416393 | 0.3171314387 | Bacteria_unclassified |
| 0.04367470257 | 0.1305416393 | 0.3223772041 | Bacteria_unclassified |
| 0.04367470257 | 0.1305416393 | 0.3171314387 | Bacteria_unclassified |
| 0.04367470257 | 0.1305416393 | 0.3171314387 | Bacteroides |
| 0.04367470257 | 0.1305416393 | 0.3171314387 | Bifidobacterium |
| 0.04367470257 | 0.1305416393 | 0.3171314387 | Clostridia_unclassified |
| 0.04367470257 | 0.1308703764 | 0.3171314387 | Clostridiaceae_unclassified |
| 0.04367470257 | 0.1305416393 | 0.3171314387 | Clostridiaceae_unclassified |

| | | | |
|---|---|---|---|
| 0.04367470257 | 0.1305416393 | 0.3171314387 | Eubacteriales_unclassified |
| 0.05601639705 | 0.1305416393 | 0.3171314387 | Erysipelatoclostridium |
| 0.04367470257 | 0.1305416393 | 0.3171314387 | Coriobacteriaceae_unclassifie |
| 0.04367470257 | 0.1305416393 | 0.3171314387 | Dorea |
| 0.04367470257 | 0.1305416393 | 0.3171314387 | Dubosiella |
| 0.04367470257 | 0.1305416393 | 0.3171314387 | Erysipelotrichales_unclassifie |
| 0.112235317 | 0.1305416393 | 0.3171314387 | Eubacteriaceae_unclassified |
| 0.04367470257 | 0.1305416393 | 0.3171314387 | Eubacteriaceae_unclassified |
| 0.04367470257 | 0.1305416393 | 0.3171314387 | GGB20149 |
| 0.04831538925 | 0.1305416393 | 0.3171314387 | GGB22635 |
| 0.04367470257 | 0.1305416393 | 0.3171314387 | GGB25041 |
| 0.04367470257 | 0.139182221 | 0.320843551 | GGB27876 |
| 0.04367470257 | 0.1305416393 | 0.3171314387 | GGB27878 |
| 0.04367470257 | 0.1305416393 | 0.3171314387 | GGB27918 |
| 0.04367470257 | 0.1305416393 | 0.3171314387 | GGB28382 |
| 0.04367470257 | 0.1305416393 | 0.3171314387 | GGB28399 |
| 0.04367470257 | 0.1305416393 | 0.3171314387 | GGB28411 |
| 0.04367470257 | 0.1305416393 | 0.3171314387 | GGB28415 |
| 0.04367470257 | 0.1305416393 | 0.3171314387 | GGB28430 |
| 0.04367470257 | 0.1305416393 | 0.3171314387 | GGB28439 |
| 0.04367470257 | 0.1305416393 | 0.3171314387 | GGB28778 |
| 0.04367470257 | 0.1305416393 | 0.3171314387 | GGB28784 |
| 0.04367470257 | 0.1305416393 | 0.3171314387 | GGB28792 |
| 0.04367470257 | 0.1305416393 | 0.3171314387 | GGB28798 |
| 0.04367470257 | 0.1305416393 | 0.3171314387 | GGB28802 |
| 0.04367470257 | 0.1305416393 | 0.3171314387 | GGB28818 |
| 0.04657723048 | 0.1305416393 | 0.3171314387 | GGB28828 |
| 0.04395942468 | 0.1305416393 | 0.3171314387 | GGB28851 |
| 0.04367470257 | 0.1305416393 | 0.3171314387 | GGB28859 |
| 0.04657723048 | 0.1305416393 | 0.3171314387 | GGB28864 |
| 0.04367470257 | 0.1305416393 | 0.3171314387 | GGB28869 |
| 0.04367470257 | 0.1305416393 | 0.3171314387 | GGB28883 |
| 0.04395942468 | 0.1305416393 | 0.3171314387 | GGB28892 |
| 0.04885981673 | 0.1305416393 | 0.3171314387 | GGB28893 |
| 0.04367470257 | 0.1305416393 | 0.3236743079 | GGB28898 |
| 0.04367470257 | 0.1305416393 | 0.3373408481 | GGB28904 |
| 0.04367470257 | 0.1305416393 | 0.3171314387 | GGB28916 |
| 0.04367470257 | 0.1305416393 | 0.3171314387 | GGB28924 |
| 0.04367470257 | 0.1305416393 | 0.3171314387 | GGB28926 |
| 0.04367470257 | 0.1305416393 | 0.3171314387 | GGB28927 |
| 0.04367470257 | 0.1370482377 | 0.345340671 | GGB28934 |
| 0.04395942468 | 0.1305416393 | 0.3171314387 | GGB28946 |
| 0.04367470257 | 0.1305416393 | 0.3171314387 | GGB28949 |
| 0.04367470257 | 0.1305416393 | 0.3171314387 | GGB28949 |
| 0.04367470257 | 0.138475456 | 0.3171314387 | GGB28950 |
| 0.04367470257 | 0.1305416393 | 0.3171314387 | GGB28951 |

| | | | |
|---|---|---|---|
| 0.04367470257 | 0.1305416393 | 0.3171314387 | GGB28951 |
| 0.04657723048 | 0.1305416393 | 0.3171314387 | GGB28954 |
| 0.04395942468 | 0.1305416393 | 0.3171314387 | GGB28956 |
| 0.04395942468 | 0.1305416393 | 0.3171314387 | GGB28960 |
| 0.04367470257 | 0.1305416393 | 0.3171314387 | GGB28967 |
| 0.04367470257 | 0.1305416393 | 0.3171314387 | GGB28991 |
| 0.04657723048 | 0.1562256722 | 0.5397187363 | GGB29002 |
| 0.04367470257 | 0.1305416393 | 0.3171314387 | GGB29003 |
| 0.04367470257 | 0.1305416393 | 0.3171314387 | GGB29011 |
| 0.04657723048 | 0.1305416393 | 0.3171314387 | GGB29531 |
| 0.04367470257 | 0.1305416393 | 0.3171314387 | GGB29685 |
| 0.04367470257 | 0.1305416393 | 0.3171314387 | GGB30141 |
| 0.04367470257 | 0.1305416393 | 0.3171314387 | GGB30286 |
| 0.04367470257 | 0.1305416393 | 0.3171314387 | GGB30303 |
| 0.04367470257 | 0.1305416393 | 0.3171314387 | GGB30413 |
| 0.04395942468 | 0.1305416393 | 0.3171314387 | GGB30454 |
| 0.05764156657 | 0.1305416393 | 0.3171314387 | GGB30455 |
| 0.04367470257 | 0.1305416393 | 0.3171314387 | GGB30461 |
| 0.04603121726 | 0.1305416393 | 0.3171314387 | GGB30461 |
| 0.05995651403 | 0.1305416393 | 0.3171314387 | GGB30463 |
| 0.04395942468 | 0.187856815 | 0.3488243361 | GGB30473 |
| 0.04657723048 | 0.1305416393 | 0.3171314387 | GGB30475 |
| 0.04374971132 | 0.1305416393 | 0.3171314387 | GGB30861 |
| 0.04367470257 | 0.1305416393 | 0.3171314387 | GGB31312 |
| 0.04367470257 | 0.1414640256 | 0.3171314387 | GGB31438 |
| 0.04367470257 | 0.1305416393 | 0.3171314387 | GGB3171 |
| 0.04367470257 | 0.1305416393 | 0.3171314387 | GGB31823 |
| 0.04367470257 | 0.1305416393 | 0.3171314387 | GGB31853 |
| 0.04367470257 | 0.1813401845 | 0.3171314387 | GGB32371 |
| 0.04367470257 | 0.1305416393 | 0.3171314387 | GGB3793 |
| 0.04367470257 | 0.1305416393 | 0.3171314387 | GGB42598 |
| 0.04367470257 | 0.1335000204 | 0.3171314387 | GGB45656 |
| 0.04367470257 | 0.1305416393 | 0.3171314387 | GGB47127 |
| 0.04367470257 | 0.1305416393 | 0.3171314387 | GGB74395 |
| 0.04657723048 | 0.1305416393 | 0.3171314387 | GGB75053 |
| 0.04395942468 | 0.1305416393 | 0.3171314387 | GGB75109 |
| 0.04367470257 | 0.1458138328 | 0.3695109776 | GGB81440 |
| 0.04367470257 | 0.1305416393 | 0.3171314387 | Lachnospiraceae_unclassified |
| 0.04367470257 | 0.1305416393 | 0.3223772041 | Lachnospiraceae_unclassified |
| 0.04367470257 | 0.1764881456 | 0.3171314387 | Lachnospiraceae_unclassified |
| 0.06292451942 | 0.1370482377 | 0.3171314387 | Lachnospiraceae_unclassified |
| 0.04367470257 | 0.1562646687 | 0.3541065039 | Lachnospiraceae_unclassified |
| 0.04367470257 | 0.1305416393 | 0.3171314387 | Lachnospiraceae_unclassified |
| 0.04894179322 | 0.1305416393 | 0.3171314387 | Lachnospiraceae_unclassified |
| 0.04367470257 | 0.1305416393 | 0.3171314387 | Lachnospiraceae_unclassified |
| 0.05714835907 | 0.1305416393 | 0.3171314387 | Lactobacillus |

| | | | |
|---|---|---|---|
| 0.04395942468 | 0.1305416393 | 0.3171314387 | Muribaculaceae_unclassified |
| 0.04367470257 | 0.1305416393 | 0.3171314387 | Neglectibacter |
| 0.04367470257 | 0.1305416393 | 0.3171314387 | Oscillospiraceae_unclassified |
| 0.04367470257 | 0.1316884198 | 0.3171314387 | Oscillospiraceae_unclassified |
| 0.04367470257 | 0.1305416393 | 0.3171314387 | Oscillospiraceae_unclassified |
| 0.04367470257 | 0.1305416393 | 0.3171314387 | Parasutterella |
| 0.04657723048 | 0.1305416393 | 0.3171314387 | Romboutsia |
| 0.04367470257 | 0.1305416393 | 0.3171314387 | Schaedlerella |
| 0.04367470257 | 0.1305416393 | 0.3171314387 | Turicibacter |
| 0.1035049976 | 0.1305416393 | 0.3171314387 | Bacteria_unclassified |
| 0.04367470257 | 0.1305416393 | 0.3171314387 | Bacteria_unclassified |
| 0.04367470257 | 0.1305416393 | 0.3171314387 | Bacteria_unclassified |
| 0.04367470257 | 0.1491222629 | 0.3171314387 | |
| 0.04367470257 | 0.1305416393 | 0.3171314387 | |
| 0.04395942468 | 0.1305416393 | 0.3171314387 | |
| 0.07692301036 | 0.1589769463 | 0.102236371 | Acetatifactor |
| 0.07692301036 | 0.1589769463 | 0.102236371 | Acetatifactor |
| 0.07692301036 | 0.1589769463 | 0.1066324351 | Acutalibacter |
| 0.07692301036 | 0.1589769463 | 0.102236371 | Acutalibacter |
| 0.07692301036 | 0.1589769463 | 0.102236371 | Adlercreutzia |
| 0.09073614391 | 0.1589769463 | 0.102236371 | Adlercreutzia |
| 0.07692301036 | 0.1589769463 | 0.102236371 | Adlercreutzia |
| 0.07692301036 | 0.1589769463 | 0.102236371 | Akkermansia |
| 0.07692301036 | 0.1589769463 | 0.102236371 | Alistipes |
| 0.07692301036 | 0.1589769463 | 0.102236371 | Anaerotruncus |
| 0.07718538808 | 0.1589769463 | 0.102236371 | Bacteria_unclassified |
| 0.07692301036 | 0.1589769463 | 0.130008339 | Bacteria_unclassified |
| 0.07692301036 | 0.1589769463 | 0.102236371 | Bacteria_unclassified |
| 0.07692301036 | 0.1617439113 | 0.102236371 | Bacteroides |
| 0.07692301036 | 0.1589769463 | 0.102236371 | Bifidobacterium |
| 0.07692301036 | 0.1664739627 | 0.102236371 | Clostridia_unclassified |
| 0.07692301036 | 0.1693458703 | 0.102236371 | Clostridiaceae_unclassified |
| 0.07692301036 | 0.1589769463 | 0.102236371 | Clostridiaceae_unclassified |
| 0.07692301036 | 0.1589769463 | 0.102236371 | Eubacteriales_unclassified |
| 0.07718538808 | 0.1589769463 | 0.102236371 | Erysipelatoclostridium |
| 0.08061015811 | 0.1589769463 | 0.102236371 | Coriobacteriaceae_unclassifie |
| 0.07692301036 | 0.1589769463 | 0.1097587134 | Dorea |
| 0.07692301036 | 0.1589769463 | 0.1097587134 | Dubosiella |
| 0.07692301036 | 0.1822113032 | 0.1097339319 | Erysipelotrichales_unclassifie |
| 0.07692301036 | 0.1822113032 | 0.1231742907 | Eubacteriaceae_unclassified |
| 0.09120550943 | 0.1589769463 | 0.102236371 | Eubacteriaceae_unclassified |
| 0.07692301036 | 0.1589769463 | 0.102236371 | GGB20149 |
| 0.07692301036 | 0.1589769463 | 0.102236371 | GGB22635 |
| 0.07692301036 | 0.1664739627 | 0.1108388432 | GGB25041 |
| 0.07718538808 | 0.1589769463 | 0.102236371 | GGB27876 |
| 0.07692301036 | 0.1589769463 | 0.102236371 | GGB27878 |

| | | | |
|---|---|---|---|
| 0.07692301036 | 0.1589769463 | 0.102236371 | GGB27918 |
| 0.07692301036 | 0.1589769463 | 0.102236371 | GGB28382 |
| 0.07960156012 | 0.1589769463 | 0.102236371 | GGB28399 |
| 0.07692301036 | 0.1589769463 | 0.102236371 | GGB28411 |
| 0.09283804963 | 0.1589769463 | 0.102236371 | GGB28415 |
| 0.07692301036 | 0.1589769463 | 0.102236371 | GGB28430 |
| 0.07692301036 | 0.1589769463 | 0.102236371 | GGB28439 |
| 0.07692301036 | 0.1589769463 | 0.102236371 | GGB28778 |
| 0.07692301036 | 0.1589769463 | 0.102236371 | GGB28784 |
| 0.07692301036 | 0.1589769463 | 0.102236371 | GGB28792 |
| 0.07692301036 | 0.1589769463 | 0.102236371 | GGB28798 |
| 0.07692301036 | 0.1589769463 | 0.102236371 | GGB28802 |
| 0.07692301036 | 0.1589769463 | 0.102236371 | GGB28818 |
| 0.08038725545 | 0.1589769463 | 0.102236371 | GGB28828 |
| 0.07692301036 | 0.1589769463 | 0.102236371 | GGB28851 |
| 0.07718538808 | 0.1664739627 | 0.1074565634 | GGB28859 |
| 0.07992747427 | 0.1589769463 | 0.102236371 | GGB28864 |
| 0.07692301036 | 0.1589769463 | 0.102236371 | GGB28869 |
| 0.07692301036 | 0.1841956373 | 0.102236371 | GGB28883 |
| 0.08061015811 | 0.1589769463 | 0.102236371 | GGB28892 |
| 0.09073614391 | 0.1589769463 | 0.102236371 | GGB28893 |
| 0.121673081 | 0.1589769463 | 0.102236371 | GGB28898 |
| 0.07692301036 | 0.1589769463 | 0.1074565634 | GGB28904 |
| 0.07692301036 | 0.1589769463 | 0.102236371 | GGB28916 |
| 0.07692301036 | 0.1589769463 | 0.102236371 | GGB28924 |
| 0.07692301036 | 0.1589769463 | 0.102236371 | GGB28926 |
| 0.07692301036 | 0.1589769463 | 0.102236371 | GGB28927 |
| 0.07692301036 | 0.1693458703 | 0.1240020188 | GGB28934 |
| 0.07960156012 | 0.1589769463 | 0.102236371 | GGB28946 |
| 0.07692301036 | 0.1589769463 | 0.102236371 | GGB28949 |
| 0.07692301036 | 0.1589769463 | 0.102236371 | GGB28949 |
| 0.07692301036 | 0.1589769463 | 0.102236371 | GGB28950 |
| 0.07692301036 | 0.1589769463 | 0.102236371 | GGB28951 |
| 0.07692301036 | 0.1589769463 | 0.102236371 | GGB28951 |
| 0.07692301036 | 0.1589769463 | 0.102236371 | GGB28954 |
| 0.07692301036 | 0.1589769463 | 0.102236371 | GGB28956 |
| 0.07692301036 | 0.1589769463 | 0.102236371 | GGB28960 |
| 0.07692301036 | 0.1589769463 | 0.102236371 | GGB28967 |
| 0.07692301036 | 0.1589769463 | 0.102236371 | GGB28991 |
| 0.07960156012 | 0.1709589243 | 0.1478334542 | GGB29002 |
| 0.07692301036 | 0.1589769463 | 0.102236371 | GGB29003 |
| 0.07692301036 | 0.179475095 | 0.102236371 | GGB29011 |
| 0.08084066323 | 0.1589769463 | 0.102236371 | GGB29531 |
| 0.07692301036 | 0.1589769463 | 0.102236371 | GGB29685 |
| 0.07692301036 | 0.1589769463 | 0.102236371 | GGB30141 |
| 0.07692301036 | 0.179475095 | 0.1231742907 | GGB30286 |

| | | | |
|---|---|---|---|
| 0.07692301036 | 0.1589769463 | 0.102236371 | GGB30303 |
| 0.07692301036 | 0.1589769463 | 0.102236371 | GGB30413 |
| 0.07692301036 | 0.1589769463 | 0.102236371 | GGB30454 |
| 0.077266801 | 0.1589769463 | 0.102236371 | GGB30455 |
| 0.07692301036 | 0.1589769463 | 0.102236371 | GGB30461 |
| 0.07692301036 | 0.1589769463 | 0.102236371 | GGB30461 |
| 0.07692301036 | 0.1589769463 | 0.102236371 | GGB30463 |
| 0.07692301036 | 0.1589769463 | 0.102236371 | GGB30473 |
| 0.07692301036 | 0.1589769463 | 0.102236371 | GGB30475 |
| 0.07692301036 | 0.1589769463 | 0.102236371 | GGB30861 |
| 0.07692301036 | 0.1589769463 | 0.102236371 | GGB31312 |
| 0.07692301036 | 0.1667295426 | 0.102236371 | GGB31438 |
| 0.07692301036 | 0.1589769463 | 0.102236371 | GGB3171 |
| 0.07692301036 | 0.1709589243 | 0.102236371 | GGB31823 |
| 0.07692301036 | 0.1589769463 | 0.102236371 | GGB31853 |
| 0.07692301036 | 0.2698102776 | 0.115918777 | GGB32371 |
| 0.07692301036 | 0.1589769463 | 0.102236371 | GGB3793 |
| 0.07692301036 | 0.1589769463 | 0.102236371 | GGB42598 |
| 0.07692301036 | 0.1589769463 | 0.102236371 | GGB45656 |
| 0.07692301036 | 0.1589769463 | 0.102236371 | GGB47127 |
| 0.07692301036 | 0.1589769463 | 0.115918777 | GGB74395 |
| 0.07692301036 | 0.1589769463 | 0.102236371 | GGB75053 |
| 0.07692301036 | 0.1589769463 | 0.102236371 | GGB75109 |
| 0.07692301036 | 0.179475095 | 0.1305497641 | GGB81440 |
| 0.07692301036 | 0.1589769463 | 0.102236371 | Lachnospiraceae_unclassified |
| 0.08076314209 | 0.1589769463 | 0.102236371 | Lachnospiraceae_unclassified |
| 0.08444029956 | 0.1589769463 | 0.102236371 | Lachnospiraceae_unclassified |
| 0.1254128111 | 0.1822113032 | 0.1097587134 | Lachnospiraceae_unclassified |
| 0.07692301036 | 0.1923358537 | 0.1358118195 | Lachnospiraceae_unclassified |
| 0.07692301036 | 0.1589769463 | 0.102236371 | Lachnospiraceae_unclassified |
| 0.07992747427 | 0.1589769463 | 0.102236371 | Lachnospiraceae_unclassified |
| 0.07692301036 | 0.1589769463 | 0.102236371 | Lachnospiraceae_unclassified |
| 0.07960156012 | 0.1589769463 | 0.102236371 | Lactobacillus |
| 0.07692301036 | 0.1589769463 | 0.102236371 | Muribaculaceae_unclassified |
| 0.07692301036 | 0.179475095 | 0.1460222786 | Neglectibacter |
| 0.07692301036 | 0.1589769463 | 0.102236371 | Oscillospiraceae_unclassified |
| 0.07692301036 | 0.2418024878 | 0.1751358198 | Oscillospiraceae_unclassified |
| 0.07692301036 | 0.1589769463 | 0.102236371 | Oscillospiraceae_unclassified |
| 0.07692301036 | 0.1589769463 | 0.102236371 | Parasutterella |
| 0.07692301036 | 0.179475095 | 0.102236371 | Romboutsia |
| 0.07692301036 | 0.1617439113 | 0.102236371 | Schaedlerella |
| 0.07692301036 | 0.1589769463 | 0.1097339319 | Turicibacter |
| 0.1431898649 | 0.1589769463 | 0.102236371 | Bacteria_unclassified |
| 0.07692301036 | 0.1589769463 | 0.102236371 | Bacteria_unclassified |
| 0.07692301036 | 0.1589769463 | 0.102236371 | Bacteria_unclassified |
| 0.07692301036 | 0.1589769463 | 0.102236371 | |

| | | | |
|---|---|---|---|
| 0.07692301036 | 0.1589769463 | 0.102236371 | |
| 0.07692301036 | 0.1589769463 | 0.102236371 | |
| 0.01259897985 | 0.7599583708 | 0.1422809042 | Acetatifactor |
| 0.01259897985 | 0.756051987 | 0.1422809042 | Acetatifactor |
| 0.01259897985 | 0.756051987 | 0.1422809042 | Acutalibacter |
| 0.01259897985 | 0.756051987 | 0.1422809042 | Acutalibacter |
| 0.01259897985 | 0.7599583708 | 0.1422809042 | Adlercreutzia |
| 0.02910957752 | 0.7619979149 | 0.2219552486 | Adlercreutzia |
| 0.01259897985 | 0.7619979149 | 0.1431690965 | Adlercreutzia |
| 0.01259897985 | 0.756051987 | 0.1422809042 | Akkermansia |
| 0.01259897985 | 0.756051987 | 0.1422809042 | Alistipes |
| 0.01259897985 | 0.756051987 | 0.1422809042 | Anaerotruncus |
| 0.0161408911 | 0.756051987 | 0.1422809042 | Bacteria_unclassified |
| 0.01259897985 | 0.756051987 | 0.1422809042 | Bacteria_unclassified |
| 0.01259897985 | 0.756051987 | 0.1422809042 | Bacteria_unclassified |
| 0.01259897985 | 0.756051987 | 0.1422809042 | Bacteroides |
| 0.01259897985 | 0.756051987 | 0.1431690965 | Bifidobacterium |
| 0.01259897985 | 0.756051987 | 0.1422809042 | Clostridia_unclassified |
| 0.01259897985 | 0.756051987 | 0.1422809042 | Clostridiaceae_unclassified |
| 0.01259897985 | 0.756051987 | 0.1422809042 | Clostridiaceae_unclassified |
| 0.01625471369 | 0.756051987 | 0.1422809042 | Eubacteriales_unclassified |
| 0.01259897985 | 0.756051987 | 0.1889990139 | Erysipelatoclostridium |
| 0.01259897985 | 0.756051987 | 0.1422809042 | Coriobacteriaceae_unclassifie |
| 0.01259897985 | 0.756051987 | 0.1779194338 | Dorea |
| 0.01259897985 | 0.756051987 | 0.1422809042 | Dubosiella |
| 0.01259897985 | 0.756051987 | 0.1422809042 | Erysipelotrichales_unclassifie |
| 0.05642788826 | 0.8865756036 | 0.2090358863 | Eubacteriaceae_unclassified |
| 0.01259897985 | 0.7725224847 | 0.1495469789 | Eubacteriaceae_unclassified |
| 0.01259897985 | 0.756051987 | 0.1422809042 | GGB20149 |
| 0.01259897985 | 0.756051987 | 0.1422809042 | GGB22635 |
| 0.01259897985 | 0.756051987 | 0.1422809042 | GGB25041 |
| 0.02126313253 | 0.756051987 | 0.1431690965 | GGB27876 |
| 0.01259897985 | 0.7599583708 | 0.1460090943 | GGB27878 |
| 0.01259897985 | 0.756051987 | 0.1422809042 | GGB27918 |
| 0.01259897985 | 0.756051987 | 0.1422809042 | GGB28382 |
| 0.01259897985 | 0.756051987 | 0.1422809042 | GGB28399 |
| 0.01259897985 | 0.7619979149 | 0.1915245786 | GGB28411 |
| 0.01347729962 | 0.756051987 | 0.1422809042 | GGB28415 |
| 0.01259897985 | 0.7599583708 | 0.1422809042 | GGB28430 |
| 0.01259897985 | 0.756051987 | 0.1422809042 | GGB28439 |
| 0.01259897985 | 0.756051987 | 0.1422809042 | GGB28778 |
| 0.01259897985 | 0.756051987 | 0.1422809042 | GGB28784 |
| 0.01259897985 | 0.756051987 | 0.1422809042 | GGB28792 |
| 0.01259897985 | 0.756051987 | 0.1422809042 | GGB28798 |
| 0.01832360615 | 0.7679796939 | 0.1470626559 | GGB28802 |
| 0.01259897985 | 0.756051987 | 0.1422809042 | GGB28818 |

| | | | |
|---|---|---|---|
| 0.01259897985 | 0.7599583708 | 0.1422809042 | GGB28828 |
| 0.01259897985 | 0.756051987 | 0.1499502903 | GGB28851 |
| 0.01259897985 | 0.756051987 | 0.1431690965 | GGB28859 |
| 0.01259897985 | 0.756051987 | 0.1422809042 | GGB28864 |
| 0.01259897985 | 0.756051987 | 0.1422809042 | GGB28869 |
| 0.01264183914 | 0.756051987 | 0.1422809042 | GGB28883 |
| 0.01259897985 | 0.756051987 | 0.1422809042 | GGB28892 |
| 0.01259897985 | 0.756051987 | 0.1422809042 | GGB28893 |
| 0.01259897985 | 0.756051987 | 0.1422809042 | GGB28898 |
| 0.01259897985 | 0.756051987 | 0.1431690965 | GGB28904 |
| 0.01259897985 | 0.756051987 | 0.1422809042 | GGB28916 |
| 0.01259897985 | 0.756051987 | 0.1422809042 | GGB28924 |
| 0.01259897985 | 0.756051987 | 0.1422809042 | GGB28926 |
| 0.01259897985 | 0.756051987 | 0.1422809042 | GGB28927 |
| 0.01259897985 | 0.756051987 | 0.1499502903 | GGB28934 |
| 0.01259897985 | 0.756051987 | 0.1422809042 | GGB28946 |
| 0.01259897985 | 0.756051987 | 0.1422809042 | GGB28949 |
| 0.02338317771 | 0.756051987 | 0.1431690965 | GGB28949 |
| 0.01464608076 | 0.756051987 | 0.1422809042 | GGB28950 |
| 0.01259897985 | 0.756051987 | 0.1422809042 | GGB28951 |
| 0.01259897985 | 0.756051987 | 0.1422809042 | GGB28951 |
| 0.01259897985 | 0.756051987 | 0.1422809042 | GGB28954 |
| 0.01259897985 | 0.756051987 | 0.1422809042 | GGB28956 |
| 0.01264183914 | 0.7619979149 | 0.1784199936 | GGB28960 |
| 0.01259897985 | 0.756051987 | 0.1422809042 | GGB28967 |
| 0.01259897985 | 0.756051987 | 0.1422809042 | GGB28991 |
| 0.01259897985 | 0.756051987 | 0.1422809042 | GGB29002 |
| 0.01259897985 | 0.756051987 | 0.1422809042 | GGB29003 |
| 0.01259897985 | 0.756051987 | 0.1422809042 | GGB29011 |
| 0.01259897985 | 0.756051987 | 0.1422809042 | GGB29531 |
| 0.01259897985 | 0.756051987 | 0.1422809042 | GGB29685 |
| 0.01259897985 | 0.756051987 | 0.1422809042 | GGB30141 |
| 0.02039579126 | 0.7599583708 | 0.1559673307 | GGB30286 |
| 0.01259897985 | 0.7599583708 | 0.1422809042 | GGB30303 |
| 0.01259897985 | 0.756051987 | 0.1422809042 | GGB30413 |
| 0.01873244574 | 0.8043483118 | 0.1422809042 | GGB30454 |
| 0.01625471369 | 0.7725224847 | 0.1422809042 | GGB30455 |
| 0.01259897985 | 0.756051987 | 0.1422809042 | GGB30461 |
| 0.01259897985 | 0.756051987 | 0.1422809042 | GGB30461 |
| 0.01347729962 | 0.756051987 | 0.1422809042 | GGB30463 |
| 0.01259897985 | 0.940867365 | 0.2292854943 | GGB30473 |
| 0.01259897985 | 0.756051987 | 0.1422809042 | GGB30475 |
| 0.0161408911 | 0.756051987 | 0.1422809042 | GGB30861 |
| 0.01259897985 | 0.756051987 | 0.1422809042 | GGB31312 |
| 0.01259897985 | 0.756051987 | 0.1422809042 | GGB31438 |
| 0.01259897985 | 0.7599583708 | 0.1422809042 | GGB3171 |

| | | | |
|---|---|---|---|
| 0.01259897985 | 0.756051987 | 0.1422809042 | GGB31823 |
| 0.01264183914 | 0.756051987 | 0.1422809042 | GGB31853 |
| 0.01259897985 | 0.8074435573 | 0.1431690965 | GGB32371 |
| 0.01259897985 | 0.756051987 | 0.1422809042 | GGB3793 |
| 0.01259897985 | 0.756051987 | 0.1422809042 | GGB42598 |
| 0.01264183914 | 0.756051987 | 0.1422809042 | GGB45656 |
| 0.01259897985 | 0.7599583708 | 0.1422809042 | GGB47127 |
| 0.01259897985 | 0.756051987 | 0.1422809042 | GGB74395 |
| 0.01290755531 | 0.756051987 | 0.1422809042 | GGB75053 |
| 0.01259897985 | 0.756051987 | 0.1422809042 | GGB75109 |
| 0.01259897985 | 0.756051987 | 0.1422809042 | GGB81440 |
| 0.01259897985 | 0.756051987 | 0.1422809042 | Lachnospiraceae_unclassified |
| 0.01259897985 | 0.756051987 | 0.1422809042 | Lachnospiraceae_unclassified |
| 0.01259897985 | 0.756051987 | 0.1422809042 | Lachnospiraceae_unclassified |
| 0.01259897985 | 0.756051987 | 0.1422809042 | Lachnospiraceae_unclassified |
| 0.01259897985 | 0.756051987 | 0.1422809042 | Lachnospiraceae_unclassified |
| 0.01259897985 | 0.756051987 | 0.1422809042 | Lachnospiraceae_unclassified |
| 0.01259897985 | 0.756051987 | 0.1431690965 | Lachnospiraceae_unclassified |
| 0.01259897985 | 0.756051987 | 0.1422809042 | Lachnospiraceae_unclassified |
| 0.01259897985 | 0.756051987 | 0.1422809042 | Lactobacillus |
| 0.01259897985 | 0.756051987 | 0.1422809042 | Muribaculaceae_unclassified |
| 0.01259897985 | 0.756051987 | 0.1422809042 | Neglectibacter |
| 0.01264183914 | 0.756051987 | 0.1422809042 | Oscillospiraceae_unclassified |
| 0.01474019978 | 0.7619979149 | 0.183399367 | Oscillospiraceae_unclassified |
| 0.01259897985 | 0.756051987 | 0.1422809042 | Oscillospiraceae_unclassified |
| 0.01259897985 | 0.7599583708 | 0.1422809042 | Parasutterella |
| 0.01751333351 | 0.756051987 | 0.1422809042 | Romboutsia |
| 0.01259897985 | 0.756051987 | 0.1422809042 | Schaedlerella |
| 0.01259897985 | 0.756051987 | 0.1422809042 | Turicibacter |
| 0.06741206655 | 0.8323892139 | 0.2267365476 | Bacteria_unclassified |
| 0.01259897985 | 0.756051987 | 0.1422809042 | Bacteria_unclassified |
| 0.01259897985 | 0.756051987 | 0.1422809042 | Bacteria_unclassified |
| 0.01358790005 | 0.756051987 | 0.1499502903 | |
| 0.01290755531 | 0.756051987 | 0.1422809042 | |
| 0.01259897985 | 0.756051987 | 0.1422809042 | |
| 0.578783872 | 0.4180975346 | 0.1373635008 | Acetatifactor |
| 0.578783872 | 0.4180975346 | 0.1381784442 | Acetatifactor |
| 0.578783872 | 0.4180975346 | 0.1373635008 | Acutalibacter |
| 0.578783872 | 0.4180975346 | 0.1373635008 | Acutalibacter |
| 0.578783872 | 0.4180975346 | 0.1373635008 | Adlercreutzia |
| 0.578783872 | 0.4180975346 | 0.1373635008 | Adlercreutzia |
| 0.578783872 | 0.440045074 | 0.1373635008 | Adlercreutzia |
| 0.578783872 | 0.4180975346 | 0.1373635008 | Akkermansia |
| 0.578783872 | 0.4180975346 | 0.1381784442 | Alistipes |
| 0.578783872 | 0.4180975346 | 0.1373635008 | Anaerotruncus |
| 0.578783872 | 0.4180975346 | 0.1373635008 | Bacteria_unclassified |

| | | | |
|---|---|---|---|
| 0.578783872 | 0.4180975346 | 0.1373635008 | Bacteria_unclassified |
| 0.578783872 | 0.4180975346 | 0.1373635008 | Bacteria_unclassified |
| 0.578783872 | 0.4180975346 | 0.1373635008 | Bacteroides |
| 0.578783872 | 0.4180975346 | 0.1373635008 | Bifidobacterium |
| 0.613769241 | 0.4180975346 | 0.1373635008 | Clostridia_unclassified |
| 0.578783872 | 0.4180975346 | 0.1373635008 | Clostridiaceae_unclassified |
| 0.578783872 | 0.4180975346 | 0.1373635008 | Clostridiaceae_unclassified |
| 0.578783872 | 0.4180975346 | 0.1373635008 | Eubacteriales_unclassified |
| 0.578783872 | 0.4180975346 | 0.1373635008 | Erysipelatoclostridium |
| 0.7086265169 | 0.4180975346 | 0.1373635008 | Coriobacteriaceae_unclassifie |
| 0.578783872 | 0.4180975346 | 0.1373635008 | Dorea |
| 0.578783872 | 0.4180975346 | 0.1373635008 | Dubosiella |
| 0.578783872 | 0.4180975346 | 0.1373635008 | Erysipelotrichales_unclassifie |
| 0.578783872 | 0.4180975346 | 0.1381784442 | Eubacteriaceae_unclassified |
| 0.578783872 | 0.4180975346 | 0.1373635008 | Eubacteriaceae_unclassified |
| 0.578783872 | 0.4180975346 | 0.1373635008 | GGB20149 |
| 0.578783872 | 0.4180975346 | 0.1373635008 | GGB22635 |
| 0.7696462239 | 0.4180975346 | 0.1373635008 | GGB25041 |
| 0.7684907177 | 0.4180975346 | 0.1373635008 | GGB27876 |
| 0.578783872 | 0.4180975346 | 0.1373635008 | GGB27878 |
| 0.578783872 | 0.4447585855 | 0.1414988016 | GGB27918 |
| 0.578783872 | 0.4180975346 | 0.1373635008 | GGB28382 |
| 0.578783872 | 0.467790095 | 0.2247230329 | GGB28399 |
| 0.578783872 | 0.4180975346 | 0.1373635008 | GGB28411 |
| 0.578783872 | 0.4180975346 | 0.1373635008 | GGB28415 |
| 0.578783872 | 0.4180975346 | 0.1373635008 | GGB28430 |
| 0.578783872 | 0.4180975346 | 0.1373635008 | GGB28439 |
| 0.578783872 | 0.4180975346 | 0.1373635008 | GGB28778 |
| 0.578783872 | 0.4180975346 | 0.1373635008 | GGB28784 |
| 0.578783872 | 0.4180975346 | 0.1373635008 | GGB28792 |
| 0.578783872 | 0.4180975346 | 0.1373635008 | GGB28798 |
| 0.578783872 | 0.4180975346 | 0.1373635008 | GGB28802 |
| 0.578783872 | 0.4180975346 | 0.1373635008 | GGB28818 |
| 0.578783872 | 0.4180975346 | 0.1596982858 | GGB28828 |
| 0.578783872 | 0.4180975346 | 0.1373635008 | GGB28851 |
| 0.578783872 | 0.4180975346 | 0.1373635008 | GGB28859 |
| 0.578783872 | 0.4180975346 | 0.1373635008 | GGB28864 |
| 0.578783872 | 0.4447585855 | 0.1596982858 | GGB28869 |
| 0.578783872 | 0.4226788162 | 0.1373635008 | GGB28883 |
| 0.578783872 | 0.4180975346 | 0.1373635008 | GGB28892 |
| 0.578783872 | 0.446514807 | 0.188541772 | GGB28893 |
| 0.578783872 | 0.4180975346 | 0.1373635008 | GGB28898 |
| 0.578783872 | 0.4180975346 | 0.1373635008 | GGB28904 |
| 0.578783872 | 0.4180975346 | 0.1373635008 | GGB28916 |
| 0.578783872 | 0.4180975346 | 0.1373635008 | GGB28924 |
| 0.578783872 | 0.4180975346 | 0.1373635008 | GGB28926 |

| | | | |
|---|---|---|---|
| 0.578783872 | 0.4180975346 | 0.1373635008 | GGB28927 |
| 0.578783872 | 0.4373647075 | 0.2247230329 | GGB28934 |
| 0.578783872 | 0.446514807 | 0.1596982858 | GGB28946 |
| 0.578783872 | 0.4180975346 | 0.1373635008 | GGB28949 |
| 0.6316705456 | 0.4180975346 | 0.1373635008 | GGB28949 |
| 0.613769241 | 0.4180975346 | 0.1373635008 | GGB28950 |
| 0.578783872 | 0.4373647075 | 0.1373635008 | GGB28951 |
| 0.578783872 | 0.4180975346 | 0.1373635008 | GGB28951 |
| 0.578783872 | 0.4180975346 | 0.1373635008 | GGB28954 |
| 0.578783872 | 0.4180975346 | 0.24718038 | GGB28956 |
| 0.578783872 | 0.4180975346 | 0.1373635008 | GGB28960 |
| 0.641387932 | 0.4180975346 | 0.1373635008 | GGB28967 |
| 0.578783872 | 0.4180975346 | 0.1373635008 | GGB28991 |
| 0.578783872 | 0.4180975346 | 0.1373635008 | GGB29002 |
| 0.578783872 | 0.4180975346 | 0.1373635008 | GGB29003 |
| 0.578783872 | 0.4402681568 | 0.1373635008 | GGB29011 |
| 0.578783872 | 0.4447585855 | 0.1704967819 | GGB29531 |
| 0.578783872 | 0.4180975346 | 0.1373635008 | GGB29685 |
| 0.578783872 | 0.4226788162 | 0.1373635008 | GGB30141 |
| 0.578783872 | 0.4180975346 | 0.1373635008 | GGB30286 |
| 0.578783872 | 0.4180975346 | 0.1373635008 | GGB30303 |
| 0.578783872 | 0.4180975346 | 0.1373635008 | GGB30413 |
| 0.578783872 | 0.4180975346 | 0.1373635008 | GGB30454 |
| 0.578783872 | 0.4373647075 | 0.1382017409 | GGB30455 |
| 0.578783872 | 0.4180975346 | 0.1373635008 | GGB30461 |
| 0.578783872 | 0.4180975346 | 0.1373635008 | GGB30461 |
| 0.578783872 | 0.4180975346 | 0.1373635008 | GGB30463 |
| 0.578783872 | 0.4180975346 | 0.1373635008 | GGB30473 |
| 0.578783872 | 0.4180975346 | 0.1373635008 | GGB30475 |
| 0.578783872 | 0.4180975346 | 0.1373635008 | GGB30861 |
| 0.578783872 | 0.4180975346 | 0.1373635008 | GGB31312 |
| 0.578783872 | 0.4180975346 | 0.1373635008 | GGB31438 |
| 0.578783872 | 0.4180975346 | 0.1373635008 | GGB3171 |
| 0.578783872 | 0.4180975346 | 0.1373635008 | GGB31823 |
| 0.578783872 | 0.4180975346 | 0.1373635008 | GGB31853 |
| 0.578783872 | 0.4373647075 | 0.1373635008 | GGB32371 |
| 0.578783872 | 0.4180975346 | 0.1373635008 | GGB3793 |
| 0.578783872 | 0.4180975346 | 0.2282961175 | GGB42598 |
| 0.578783872 | 0.4180975346 | 0.1373635008 | GGB45656 |
| 0.578783872 | 0.4180975346 | 0.1373635008 | GGB47127 |
| 0.578783872 | 0.4180975346 | 0.1373635008 | GGB74395 |
| 0.578783872 | 0.4180975346 | 0.1373635008 | GGB75053 |
| 0.578783872 | 0.4180975346 | 0.1373635008 | GGB75109 |
| 0.578783872 | 0.4180975346 | 0.1373635008 | GGB81440 |
| 0.578783872 | 0.4180975346 | 0.1373635008 | Lachnospiraceae_unclassified |
| 0.578783872 | 0.4180975346 | 0.1373635008 | Lachnospiraceae_unclassified |

| | | | |
|---|---|---|---|
| 0.578783872 | 0.4180975346 | 0.1373635008 | Lachnospiraceae_unclassified |
| 0.578783872 | 0.4180975346 | 0.1373635008 | Lachnospiraceae_unclassified |
| 0.7148477127 | 0.4180975346 | 0.1373635008 | Lachnospiraceae_unclassified |
| 0.578783872 | 0.4180975346 | 0.1373635008 | Lachnospiraceae_unclassified |
| 0.578783872 | 0.4180975346 | 0.1373635008 | Lachnospiraceae_unclassified |
| 0.578783872 | 0.4180975346 | 0.1373635008 | Lachnospiraceae_unclassified |
| 0.578783872 | 0.4180975346 | 0.1373635008 | Lactobacillus |
| 0.578783872 | 0.4180975346 | 0.1373635008 | Muribaculaceae_unclassified |
| 0.578783872 | 0.4180975346 | 0.156954199 | Neglectibacter |
| 0.578783872 | 0.4180975346 | 0.1373635008 | Oscillospiraceae_unclassified |
| 0.578783872 | 0.4180975346 | 0.1373635008 | Oscillospiraceae_unclassified |
| 0.578783872 | 0.4180975346 | 0.1373635008 | Oscillospiraceae_unclassified |
| 0.578783872 | 0.4180975346 | 0.1373635008 | Parasutterella |
| 0.578783872 | 0.4180975346 | 0.1373635008 | Romboutsia |
| 0.578783872 | 0.4180975346 | 0.1373635008 | Schaedlerella |
| 0.578783872 | 0.4180975346 | 0.1528104133 | Turicibacter |
| 0.578783872 | 0.4180975346 | 0.1373635008 | Bacteria_unclassified |
| 0.578783872 | 0.4180975346 | 0.1402078759 | Bacteria_unclassified |
| 0.578783872 | 0.4180975346 | 0.1373635008 | Bacteria_unclassified |
| 0.578783872 | 0.4180975346 | 0.1373635008 | |
| 0.578783872 | 0.4373647075 | 0.1596982858 | |
| 0.578783872 | 0.4180975346 | 0.1373635008 | |
| 0.2131915586 | 0.8736371071 | 0.488522822 | Acetatifactor |
| 0.2131915586 | 0.8736371071 | 0.4997790029 | Acetatifactor |
| 0.2131915586 | 0.8736371071 | 0.488522822 | Acutalibacter |
| 0.2131915586 | 0.8736371071 | 0.488522822 | Acutalibacter |
| 0.2131915586 | 0.8736371071 | 0.488522822 | Adlercreutzia |
| 0.2264838524 | 0.8736371071 | 0.488522822 | Adlercreutzia |
| 0.2131915586 | 0.8736371071 | 0.488522822 | Adlercreutzia |
| 0.2131915586 | 0.8736371071 | 0.488522822 | Akkermansia |
| 0.2131915586 | 0.8736371071 | 0.4921389478 | Alistipes |
| 0.2131915586 | 0.8736371071 | 0.488522822 | Anaerotruncus |
| 0.2131915586 | 0.8736371071 | 0.4921389478 | Bacteria_unclassified |
| 0.2131915586 | 0.8736371071 | 0.488522822 | Bacteria_unclassified |
| 0.2131915586 | 0.8736371071 | 0.488522822 | Bacteria_unclassified |
| 0.2131915586 | 0.8736371071 | 0.488522822 | Bacteroides |
| 0.2131915586 | 0.8736371071 | 0.488522822 | Bifidobacterium |
| 0.2131915586 | 0.8736371071 | 0.488522822 | Clostridia_unclassified |
| 0.2131915586 | 0.8736371071 | 0.488522822 | Clostridiaceae_unclassified |
| 0.2131915586 | 0.8736371071 | 0.488522822 | Clostridiaceae_unclassified |
| 0.2131915586 | 0.8736371071 | 0.488522822 | Eubacteriales_unclassified |
| 0.2131915586 | 0.9363519376 | 0.488522822 | Erysipelatoclostridium |
| 0.276301348 | 0.8736371071 | 0.488522822 | Coriobacteriaceae_unclassifie |
| 0.2131915586 | 0.8736371071 | 0.488522822 | Dorea |
| 0.2590600021 | 0.8736371071 | 0.488522822 | Dubosiella |
| 0.2131915586 | 0.8736371071 | 0.488522822 | Erysipelotrichales_unclassifie |

| | | | |
|---|---|---|---|
| 0.2314968767 | 0.8736371071 | 0.488522822 | Eubacteriaceae_unclassified |
| 0.2131915586 | 0.8736371071 | 0.488522822 | Eubacteriaceae_unclassified |
| 0.2131915586 | 0.8736371071 | 0.488522822 | GGB20149 |
| 0.2131915586 | 0.8736371071 | 0.488522822 | GGB22635 |
| 0.2543135164 | 0.8736371071 | 0.488522822 | GGB25041 |
| 0.3174518121 | 0.8736371071 | 0.488522822 | GGB27876 |
| 0.2131915586 | 0.8736371071 | 0.488522822 | GGB27878 |
| 0.2131915586 | 0.8736371071 | 0.488522822 | GGB27918 |
| 0.2131915586 | 0.8736371071 | 0.4921389478 | GGB28382 |
| 0.2131915586 | 0.8736371071 | 0.488522822 | GGB28399 |
| 0.2131915586 | 0.8736371071 | 0.488522822 | GGB28411 |
| 0.2131915586 | 0.8736371071 | 0.488522822 | GGB28415 |
| 0.2131915586 | 0.8736371071 | 0.488522822 | GGB28430 |
| 0.2131915586 | 0.8736371071 | 0.488522822 | GGB28439 |
| 0.2131915586 | 0.8736371071 | 0.488522822 | GGB28778 |
| 0.2131915586 | 0.8736371071 | 0.488522822 | GGB28784 |
| 0.2131915586 | 0.8736371071 | 0.488522822 | GGB28792 |
| 0.2131915586 | 0.8736371071 | 0.488522822 | GGB28798 |
| 0.2131915586 | 0.8736371071 | 0.488522822 | GGB28802 |
| 0.2131915586 | 0.8736371071 | 0.488522822 | GGB28818 |
| 0.2131915586 | 0.8736371071 | 0.6738492284 | GGB28828 |
| 0.2131915586 | 0.8736371071 | 0.488522822 | GGB28851 |
| 0.2131915586 | 0.8736371071 | 0.488522822 | GGB28859 |
| 0.2131915586 | 0.8736371071 | 0.4921389478 | GGB28864 |
| 0.2131915586 | 0.8736371071 | 0.4921389478 | GGB28869 |
| 0.2131915586 | 0.9341545781 | 0.488522822 | GGB28883 |
| 0.2131915586 | 0.8736371071 | 0.488522822 | GGB28892 |
| 0.2392238313 | 0.8736371071 | 0.488522822 | GGB28893 |
| 0.2131915586 | 0.8736371071 | 0.4921389478 | GGB28898 |
| 0.2131915586 | 0.8736371071 | 0.4921389478 | GGB28904 |
| 0.2131915586 | 0.8736371071 | 0.488522822 | GGB28916 |
| 0.2131915586 | 0.8736371071 | 0.488522822 | GGB28924 |
| 0.2131915586 | 0.8736371071 | 0.488522822 | GGB28926 |
| 0.2131915586 | 0.8736371071 | 0.488522822 | GGB28927 |
| 0.2131915586 | 0.8736371071 | 0.4921389478 | GGB28934 |
| 0.2131915586 | 0.8736371071 | 0.488522822 | GGB28946 |
| 0.2131915586 | 0.8736371071 | 0.488522822 | GGB28949 |
| 0.2131915586 | 0.8736371071 | 0.488522822 | GGB28949 |
| 0.2366381849 | 0.8736371071 | 0.488522822 | GGB28950 |
| 0.2131915586 | 0.8736371071 | 0.488522822 | GGB28951 |
| 0.2131915586 | 0.8736371071 | 0.488522822 | GGB28951 |
| 0.2131915586 | 0.8736371071 | 0.488522822 | GGB28954 |
| 0.2131915586 | 0.8736371071 | 0.6111060006 | GGB28956 |
| 0.2965725754 | 0.8736371071 | 0.488522822 | GGB28960 |
| 0.2144933081 | 0.8736371071 | 0.488522822 | GGB28967 |
| 0.2131915586 | 0.8736371071 | 0.488522822 | GGB28991 |

| | | | |
|---|---|---|---|
| 0.2131915586 | 0.8736371071 | 0.488522822 | GGB29002 |
| 0.2131915586 | 0.8736371071 | 0.488522822 | GGB29003 |
| 0.2131915586 | 0.8736371071 | 0.488522822 | GGB29011 |
| 0.2131915586 | 0.8736371071 | 0.4921389478 | GGB29531 |
| 0.2131915586 | 0.8736371071 | 0.488522822 | GGB29685 |
| 0.2131915586 | 0.8736371071 | 0.488522822 | GGB30141 |
| 0.2131915586 | 0.8736371071 | 0.488522822 | GGB30286 |
| 0.2131915586 | 0.8736371071 | 0.488522822 | GGB30303 |
| 0.2131915586 | 0.8736371071 | 0.488522822 | GGB30413 |
| 0.2131915586 | 0.8736371071 | 0.488522822 | GGB30454 |
| 0.2131915586 | 0.9562303239 | 0.6111060006 | GGB30455 |
| 0.2131915586 | 0.8736371071 | 0.488522822 | GGB30461 |
| 0.2131915586 | 0.8736371071 | 0.488522822 | GGB30461 |
| 0.2131915586 | 0.8736371071 | 0.4901262545 | GGB30463 |
| 0.2131915586 | 0.8736371071 | 0.488522822 | GGB30473 |
| 0.2131915586 | 0.8736371071 | 0.4921389478 | GGB30475 |
| 0.2131915586 | 0.8736371071 | 0.488522822 | GGB30861 |
| 0.2131915586 | 0.8736371071 | 0.488522822 | GGB31312 |
| 0.2131915586 | 0.8736371071 | 0.488522822 | GGB31438 |
| 0.2131915586 | 0.8736371071 | 0.488522822 | GGB3171 |
| 0.2131915586 | 0.8736371071 | 0.488522822 | GGB31823 |
| 0.2131915586 | 0.8736371071 | 0.488522822 | GGB31853 |
| 0.2131915586 | 0.9825750489 | 0.4921389478 | GGB32371 |
| 0.2131915586 | 0.8736371071 | 0.488522822 | GGB3793 |
| 0.2131915586 | 0.8736371071 | 0.5197962683 | GGB42598 |
| 0.2131915586 | 0.8736371071 | 0.488522822 | GGB45656 |
| 0.2131915586 | 0.8736371071 | 0.488522822 | GGB47127 |
| 0.2131915586 | 0.8736371071 | 0.488522822 | GGB74395 |
| 0.2131915586 | 0.8736371071 | 0.488522822 | GGB75053 |
| 0.2131915586 | 0.8736371071 | 0.488522822 | GGB75109 |
| 0.2131915586 | 0.8736371071 | 0.488522822 | GGB81440 |
| 0.2131915586 | 0.8736371071 | 0.488522822 | Lachnospiraceae_unclassified |
| 0.2144933081 | 0.8736371071 | 0.488522822 | Lachnospiraceae_unclassified |
| 0.2131915586 | 0.8736371071 | 0.488522822 | Lachnospiraceae_unclassified |
| 0.2131915586 | 0.8736371071 | 0.488522822 | Lachnospiraceae_unclassified |
| 0.2327470411 | 0.8736371071 | 0.488522822 | Lachnospiraceae_unclassified |
| 0.2131915586 | 0.8736371071 | 0.488522822 | Lachnospiraceae_unclassified |
| 0.2131915586 | 0.8931158046 | 0.5927480394 | Lachnospiraceae_unclassified |
| 0.2131915586 | 0.8736371071 | 0.488522822 | Lachnospiraceae_unclassified |
| 0.2131915586 | 0.8736371071 | 0.488522822 | Lactobacillus |
| 0.2131915586 | 0.8736371071 | 0.488522822 | Muribaculaceae_unclassified |
| 0.2131915586 | 0.8736371071 | 0.4921389478 | Neglectibacter |
| 0.2131915586 | 0.8736371071 | 0.488522822 | Oscillospiraceae_unclassified |
| 0.2144933081 | 0.8736371071 | 0.488522822 | Oscillospiraceae_unclassified |
| 0.2131915586 | 0.8736371071 | 0.488522822 | Oscillospiraceae_unclassified |
| 0.2131915586 | 0.8931158046 | 0.4921389478 | Parasutterella |

| | | | |
|---|---|---|---|
| 0.2131915586 | 0.9825750489 | 0.488522822 | Romboutsia |
| 0.2131915586 | 0.8736371071 | 0.488522822 | Schaedlerella |
| 0.2131915586 | 0.8736371071 | 0.4921389478 | Turicibacter |
| 0.2131915586 | 0.8736371071 | 0.488522822 | Bacteria_unclassified |
| 0.2131915586 | 0.8736371071 | 0.4921389478 | Bacteria_unclassified |
| 0.2131915586 | 0.8736371071 | 0.488522822 | Bacteria_unclassified |
| 0.2327470411 | 0.8736371071 | 0.488522822 | |
| 0.2131915586 | 0.8736371071 | 0.4921389478 | |
| 0.2131915586 | 0.8736371071 | 0.488522822 | |

| Family | Order | Class | Phylum | Kingdom |
|---|---|---|---|---|
| Lachnospiraceae | Eubacteriales | Clostridia | Firmicutes | Bacteria |
| Lachnospiraceae | Eubacteriales | Clostridia | Firmicutes | Bacteria |
| Oscillospiraceae | Eubacteriales | Clostridia | Firmicutes | Bacteria |
| Oscillospiraceae | Eubacteriales | Clostridia | Firmicutes | Bacteria |
| Eggerthellaceae | Eggerthellales | Coriobacteriia | Actinobacteria | Bacteria |
| Eggerthellaceae | Eggerthellales | Coriobacteriia | Actinobacteria | Bacteria |
| Eggerthellaceae | Eggerthellales | Coriobacteriia | Actinobacteria | Bacteria |
| Akkermansiaceae | Verrucomicrobiales | Verrucomicrobiae | Verrucomicrobia | Bacteria |
| Rikenellaceae | Bacteroidales | Bacteroidia | Bacteroidota | Bacteria |
| Oscillospiraceae | Eubacteriales | Clostridia | Firmicutes | Bacteria |
| Bacteria_unclassified | Bacteria_unclassified | Bacteria_unclassifie | Bacteria_unclassifie | Bacteria |
| Bacteria_unclassified | Bacteria_unclassified | Bacteria_unclassifie | Bacteria_unclassifie | Bacteria |
| Bacteria_unclassified | Bacteria_unclassified | Bacteria_unclassifie | Bacteria_unclassifie | Bacteria |
| Bacteroidaceae | Bacteroidales | Bacteroidia | Bacteroidota | Bacteria |
| Bifidobacteriaceae | Bifidobacteriales | Actinomycetia | Actinobacteria | Bacteria |
| Clostridia_unclassified | Clostridia_unclassifie | Clostridia | Firmicutes | Bacteria |
| Clostridiaceae | Eubacteriales | Clostridia | Firmicutes | Bacteria |
| Clostridiaceae | Eubacteriales | Clostridia | Firmicutes | Bacteria |
| Eubacteriales_unclassified | Eubacteriales | Clostridia | Firmicutes | Bacteria |
| Erysipelotrichaceae | Erysipelotrichales | Erysipelotrichia | Firmicutes | Bacteria |
| Coriobacteriaceae | Coriobacteriales | Coriobacteriia | Actinobacteria | Bacteria |
| Lachnospiraceae | Eubacteriales | Clostridia | Firmicutes | Bacteria |
| Erysipelotrichaceae | Erysipelotrichales | Erysipelotrichia | Firmicutes | Bacteria |
| Erysipelotrichales_unclassifie | Erysipelotrichales | Erysipelotrichia | Firmicutes | Bacteria |
| Eubacteriaceae | Eubacteriales | Clostridia | Firmicutes | Bacteria |
| Eubacteriaceae | Eubacteriales | Clostridia | Firmicutes | Bacteria |
| Lachnospiraceae | Eubacteriales | Clostridia | Firmicutes | Bacteria |
| Eggerthellaceae | Eggerthellales | Coriobacteriia | Actinobacteria | Bacteria |
| Lachnospiraceae | Eubacteriales | Clostridia | Firmicutes | Bacteria |
| Muribaculaceae | Bacteroidales | Bacteroidia | Bacteroidota | Bacteria |
| Muribaculaceae | Bacteroidales | Bacteroidia | Bacteroidota | Bacteria |
| Muribaculaceae | Bacteroidales | Bacteroidia | Bacteroidota | Bacteria |
| FGB9508 | OFGB9508 | CFGB9508 | Firmicutes | Bacteria |
| FGB2838 | OFGB2838 | CFGB2838 | Firmicutes | Bacteria |
| FGB2838 | OFGB2838 | CFGB2838 | Firmicutes | Bacteria |
| FGB2838 | OFGB2838 | CFGB2838 | Firmicutes | Bacteria |
| Pumilibacteraceae | Eubacteriales | Clostridia | Firmicutes | Bacteria |
| FGB28439 | OFGB28439 | CFGB28439 | Firmicutes | Bacteria |
| Clostridia_unclassified | Clostridia_unclassifie | Clostridia | Firmicutes | Bacteria |
| Eubacteriaceae | Eubacteriales | Clostridia | Firmicutes | Bacteria |
| Lachnospiraceae | Eubacteriales | Clostridia | Firmicutes | Bacteria |
| Lachnospiraceae | Eubacteriales | Clostridia | Firmicutes | Bacteria |
| Lachnospiraceae | Eubacteriales | Clostridia | Firmicutes | Bacteria |
| Lachnospiraceae | Eubacteriales | Clostridia | Firmicutes | Bacteria |
| FGB77305 | OFGB77305 | CFGB77305 | Firmicutes | Bacteria |

| | | | | |
|---|---|---|---|---|
| Clostridiaceae | Eubacteriales | Clostridia | Firmicutes | Bacteria |
| Lachnospiraceae | Eubacteriales | Clostridia | Firmicutes | Bacteria |
| Lachnospiraceae | Eubacteriales | Clostridia | Firmicutes | Bacteria |
| Lachnospiraceae | Eubacteriales | Clostridia | Firmicutes | Bacteria |
| FGB9633 | OFGB9633 | CFGB9633 | Firmicutes | Bacteria |
| Bacteria_unclassified | Bacteria_unclassified | Bacteria_unclassifie | Bacteria_unclassifie | Bacteria |
| Bacteria_unclassified | Bacteria_unclassified | Bacteria_unclassifie | Bacteria_unclassifie | Bacteria |
| Bacteria_unclassified | Bacteria_unclassified | Bacteria_unclassifie | Bacteria_unclassifie | Bacteria |
| Bacteria_unclassified | Bacteria_unclassified | Bacteria_unclassifie | Bacteria_unclassifie | Bacteria |
| Lachnospiraceae | Eubacteriales | Clostridia | Firmicutes | Bacteria |
| Lachnospiraceae | Eubacteriales | Clostridia | Firmicutes | Bacteria |
| Lachnospiraceae | Eubacteriales | Clostridia | Firmicutes | Bacteria |
| FGB77359 | OFGB77359 | CFGB77359 | Bacteria_unclassifie | Bacteria |
| FGB9639 | OFGB9639 | CFGB9639 | Firmicutes | Bacteria |
| Lachnospiraceae | Eubacteriales | Clostridia | Firmicutes | Bacteria |
| Lachnospiraceae | Eubacteriales | Clostridia | Firmicutes | Bacteria |
| Lachnospiraceae | Eubacteriales | Clostridia | Firmicutes | Bacteria |
| Clostridiaceae | Eubacteriales | Clostridia | Firmicutes | Bacteria |
| Clostridiaceae | Eubacteriales | Clostridia | Firmicutes | Bacteria |
| Clostridiaceae | Eubacteriales | Clostridia | Firmicutes | Bacteria |
| Clostridiaceae | Eubacteriales | Clostridia | Firmicutes | Bacteria |
| Clostridiaceae | Eubacteriales | Clostridia | Firmicutes | Bacteria |
| Clostridiaceae | Eubacteriales | Clostridia | Firmicutes | Bacteria |
| Clostridiaceae | Eubacteriales | Clostridia | Firmicutes | Bacteria |
| Eubacteriaceae | Eubacteriales | Clostridia | Firmicutes | Bacteria |
| FGB9658 | OFGB9658 | CFGB9658 | Firmicutes | Bacteria |
| FGB9659 | OFGB9659 | CFGB9659 | Firmicutes | Bacteria |
| Bacteria_unclassified | Bacteria_unclassified | Bacteria_unclassifie | Bacteria_unclassifie | Bacteria |
| FGB9827 | OFGB9827 | CFGB9827 | Firmicutes | Bacteria |
| Eubacteriaceae | Eubacteriales | Clostridia | Firmicutes | Bacteria |
| FGB77303 | OFGB77303 | CFGB77303 | Bacteria_unclassifie | Bacteria |
| Eubacteriales_unclassified | Eubacteriales | Clostridia | Firmicutes | Bacteria |
| Oscillospiraceae | Eubacteriales | Clostridia | Firmicutes | Bacteria |
| FGB30328 | OFGB30328 | CFGB30328 | Firmicutes | Bacteria |
| Oscillospiraceae | Eubacteriales | Clostridia | Firmicutes | Bacteria |
| Oscillospiraceae | Eubacteriales | Clostridia | Firmicutes | Bacteria |
| Oscillospiraceae | Eubacteriales | Clostridia | Firmicutes | Bacteria |
| Oscillospiraceae | Eubacteriales | Clostridia | Firmicutes | Bacteria |
| Oscillospiraceae | Eubacteriales | Clostridia | Firmicutes | Bacteria |
| Oscillospiraceae | Eubacteriales | Clostridia | Firmicutes | Bacteria |
| Oscillospiraceae | Eubacteriales | Clostridia | Firmicutes | Bacteria |
| FGB77153 | OFGB77153 | CFGB77153 | Actinobacteria | Bacteria |
| FGB1791 | OFGB1791 | CFGB1791 | Tenericutes | Bacteria |
| FGB10290 | OFGB10290 | CFGB10290 | Firmicutes | Bacteria |
| Oscillospiraceae | Eubacteriales | Clostridia | Firmicutes | Bacteria |
| FGB1765 | OFGB1765 | CFGB1765 | Firmicutes | Bacteria |

| | | | | |
|---|---|---|---|---|
| FGB10349 | OFGB10349 | CFGB10349 | Firmicutes | Bacteria |
| FGB10667 | OFGB10667 | CFGB10667 | Firmicutes | Bacteria |
| Lachnospiraceae | Eubacteriales | Clostridia | Firmicutes | Bacteria |
| Lachnospiraceae | Eubacteriales | Clostridia | Firmicutes | Bacteria |
| Christensenellaceae | Eubacteriales | Clostridia | Firmicutes | Bacteria |
| FGB10299 | OFGB10299 | CFGB10299 | Firmicutes | Bacteria |
| Oscillospiraceae | Eubacteriales | Clostridia | Firmicutes | Bacteria |
| Oscillospiraceae | Eubacteriales | Clostridia | Firmicutes | Bacteria |
| Lachnospiraceae | Eubacteriales | Clostridia | Firmicutes | Bacteria |
| Clostridia_unclassified | Clostridia_unclassifie | Clostridia | Firmicutes | Bacteria |
| Lachnospiraceae | Eubacteriales | Clostridia | Firmicutes | Bacteria |
| Lachnospiraceae | Eubacteriales | Clostridia | Firmicutes | Bacteria |
| Lachnospiraceae | Eubacteriales | Clostridia | Firmicutes | Bacteria |
| Lachnospiraceae | Eubacteriales | Clostridia | Firmicutes | Bacteria |
| Lachnospiraceae | Eubacteriales | Clostridia | Firmicutes | Bacteria |
| Lachnospiraceae | Eubacteriales | Clostridia | Firmicutes | Bacteria |
| Lachnospiraceae | Eubacteriales | Clostridia | Firmicutes | Bacteria |
| Lachnospiraceae | Eubacteriales | Clostridia | Firmicutes | Bacteria |
| Lactobacillaceae | Lactobacillales | Bacilli | Firmicutes | Bacteria |
| Muribaculaceae | Bacteroidales | Bacteroidia | Bacteroidota | Bacteria |
| Oscillospiraceae | Eubacteriales | Clostridia | Firmicutes | Bacteria |
| Oscillospiraceae | Eubacteriales | Clostridia | Firmicutes | Bacteria |
| Oscillospiraceae | Eubacteriales | Clostridia | Firmicutes | Bacteria |
| Oscillospiraceae | Eubacteriales | Clostridia | Firmicutes | Bacteria |
| Sutterellaceae | Burkholderiales | Betaproteobacteria | Proteobacteria | Bacteria |
| Peptostreptococcaceae | Eubacteriales | Clostridia | Firmicutes | Bacteria |
| Lachnospiraceae | Eubacteriales | Clostridia | Firmicutes | Bacteria |
| Turicibacteraceae | Erysipelotrichales | Erysipelotrichia | Firmicutes | Bacteria |
| Bacteria_unclassified | Bacteria_unclassified | Bacteria_unclassifie | Bacteria_unclassifie | Bacteria |
| Bacteria_unclassified | Bacteria_unclassified | Bacteria_unclassifie | Bacteria_unclassifie | Bacteria |
| Bacteria_unclassified | Bacteria_unclassified | Bacteria_unclassifie | Bacteria_unclassifie | Bacteria |

| | | | | |
|---|---|---|---|---|
| Lachnospiraceae | Eubacteriales | Clostridia | Firmicutes | Bacteria |
| Lachnospiraceae | Eubacteriales | Clostridia | Firmicutes | Bacteria |
| Oscillospiraceae | Eubacteriales | Clostridia | Firmicutes | Bacteria |
| Oscillospiraceae | Eubacteriales | Clostridia | Firmicutes | Bacteria |
| Eggerthellaceae | Eggerthellales | Coriobacteriia | Actinobacteria | Bacteria |
| Eggerthellaceae | Eggerthellales | Coriobacteriia | Actinobacteria | Bacteria |
| Eggerthellaceae | Eggerthellales | Coriobacteriia | Actinobacteria | Bacteria |
| Akkermansiaceae | Verrucomicrobiales | Verrucomicrobiae | Verrucomicrobia | Bacteria |
| Rikenellaceae | Bacteroidales | Bacteroidia | Bacteroidota | Bacteria |
| Oscillospiraceae | Eubacteriales | Clostridia | Firmicutes | Bacteria |
| Bacteria_unclassified | Bacteria_unclassified | Bacteria_unclassifie | Bacteria_unclassifie | Bacteria |
| Bacteria_unclassified | Bacteria_unclassified | Bacteria_unclassifie | Bacteria_unclassifie | Bacteria |

| | | | | |
|---|---|---|---|---|
| Bacteria_unclassified | Bacteria_unclassified | Bacteria_unclassifie | Bacteria_unclassifie | Bacteria |
| Bacteroidaceae | Bacteroidales | Bacteroidia | Bacteroidota | Bacteria |
| Bifidobacteriaceae | Bifidobacteriales | Actinomycetia | Actinobacteria | Bacteria |
| Clostridia_unclassified | Clostridia_unclassifie | Clostridia | Firmicutes | Bacteria |
| Clostridiaceae | Eubacteriales | Clostridia | Firmicutes | Bacteria |
| Clostridiaceae | Eubacteriales | Clostridia | Firmicutes | Bacteria |
| Eubacteriales_unclassified | Eubacteriales | Clostridia | Firmicutes | Bacteria |
| Erysipelotrichaceae | Erysipelotrichales | Erysipelotrichia | Firmicutes | Bacteria |
| Coriobacteriaceae | Coriobacteriales | Coriobacteriia | Actinobacteria | Bacteria |
| Lachnospiraceae | Eubacteriales | Clostridia | Firmicutes | Bacteria |
| Erysipelotrichaceae | Erysipelotrichales | Erysipelotrichia | Firmicutes | Bacteria |
| Erysipelotrichales_unclassifie | Erysipelotrichales | Erysipelotrichia | Firmicutes | Bacteria |
| Eubacteriaceae | Eubacteriales | Clostridia | Firmicutes | Bacteria |
| Eubacteriaceae | Eubacteriales | Clostridia | Firmicutes | Bacteria |
| Lachnospiraceae | Eubacteriales | Clostridia | Firmicutes | Bacteria |
| Eggerthellaceae | Eggerthellales | Coriobacteriia | Actinobacteria | Bacteria |
| Lachnospiraceae | Eubacteriales | Clostridia | Firmicutes | Bacteria |
| Muribaculaceae | Bacteroidales | Bacteroidia | Bacteroidota | Bacteria |
| Muribaculaceae | Bacteroidales | Bacteroidia | Bacteroidota | Bacteria |
| Muribaculaceae | Bacteroidales | Bacteroidia | Bacteroidota | Bacteria |
| FGB9508 | OFGB9508 | CFGB9508 | Firmicutes | Bacteria |
| FGB2838 | OFGB2838 | CFGB2838 | Firmicutes | Bacteria |
| FGB2838 | OFGB2838 | CFGB2838 | Firmicutes | Bacteria |
| FGB2838 | OFGB2838 | CFGB2838 | Firmicutes | Bacteria |
| Pumilibacteraceae | Eubacteriales | Clostridia | Firmicutes | Bacteria |
| FGB28439 | OFGB28439 | CFGB28439 | Firmicutes | Bacteria |
| Clostridia_unclassified | Clostridia_unclassifie | Clostridia | Firmicutes | Bacteria |
| Eubacteriaceae | Eubacteriales | Clostridia | Firmicutes | Bacteria |
| Lachnospiraceae | Eubacteriales | Clostridia | Firmicutes | Bacteria |
| Lachnospiraceae | Eubacteriales | Clostridia | Firmicutes | Bacteria |
| Lachnospiraceae | Eubacteriales | Clostridia | Firmicutes | Bacteria |
| Lachnospiraceae | Eubacteriales | Clostridia | Firmicutes | Bacteria |
| FGB77305 | OFGB77305 | CFGB77305 | Firmicutes | Bacteria |
| Clostridiaceae | Eubacteriales | Clostridia | Firmicutes | Bacteria |
| Lachnospiraceae | Eubacteriales | Clostridia | Firmicutes | Bacteria |
| Lachnospiraceae | Eubacteriales | Clostridia | Firmicutes | Bacteria |
| Lachnospiraceae | Eubacteriales | Clostridia | Firmicutes | Bacteria |
| FGB9633 | OFGB9633 | CFGB9633 | Firmicutes | Bacteria |
| Bacteria_unclassified | Bacteria_unclassified | Bacteria_unclassifie | Bacteria_unclassifie | Bacteria |
| Bacteria_unclassified | Bacteria_unclassified | Bacteria_unclassifie | Bacteria_unclassifie | Bacteria |
| Bacteria_unclassified | Bacteria_unclassified | Bacteria_unclassifie | Bacteria_unclassifie | Bacteria |
| Bacteria_unclassified | Bacteria_unclassified | Bacteria_unclassifie | Bacteria_unclassifie | Bacteria |
| Lachnospiraceae | Eubacteriales | Clostridia | Firmicutes | Bacteria |
| Lachnospiraceae | Eubacteriales | Clostridia | Firmicutes | Bacteria |
| Lachnospiraceae | Eubacteriales | Clostridia | Firmicutes | Bacteria |
| FGB77359 | OFGB77359 | CFGB77359 | Bacteria_unclassifie | Bacteria |

| | | | | |
|---|---|---|---|---|
| FGB9639 | OFGB9639 | CFGB9639 | Firmicutes | Bacteria |
| Lachnospiraceae | Eubacteriales | Clostridia | Firmicutes | Bacteria |
| Lachnospiraceae | Eubacteriales | Clostridia | Firmicutes | Bacteria |
| Lachnospiraceae | Eubacteriales | Clostridia | Firmicutes | Bacteria |
| Clostridiaceae | Eubacteriales | Clostridia | Firmicutes | Bacteria |
| Clostridiaceae | Eubacteriales | Clostridia | Firmicutes | Bacteria |
| Clostridiaceae | Eubacteriales | Clostridia | Firmicutes | Bacteria |
| Clostridiaceae | Eubacteriales | Clostridia | Firmicutes | Bacteria |
| Clostridiaceae | Eubacteriales | Clostridia | Firmicutes | Bacteria |
| Clostridiaceae | Eubacteriales | Clostridia | Firmicutes | Bacteria |
| Clostridiaceae | Eubacteriales | Clostridia | Firmicutes | Bacteria |
| Eubacteriaceae | Eubacteriales | Clostridia | Firmicutes | Bacteria |
| FGB9658 | OFGB9658 | CFGB9658 | Firmicutes | Bacteria |
| FGB9659 | OFGB9659 | CFGB9659 | Firmicutes | Bacteria |
| Bacteria_unclassified | Bacteria_unclassified | Bacteria_unclassifie | Bacteria_unclassifie | Bacteria |
| FGB9827 | OFGB9827 | CFGB9827 | Firmicutes | Bacteria |
| Eubacteriaceae | Eubacteriales | Clostridia | Firmicutes | Bacteria |
| FGB77303 | OFGB77303 | CFGB77303 | Bacteria_unclassifie | Bacteria |
| Eubacteriales_unclassified | Eubacteriales | Clostridia | Firmicutes | Bacteria |
| Oscillospiraceae | Eubacteriales | Clostridia | Firmicutes | Bacteria |
| FGB30328 | OFGB30328 | CFGB30328 | Firmicutes | Bacteria |
| Oscillospiraceae | Eubacteriales | Clostridia | Firmicutes | Bacteria |
| Oscillospiraceae | Eubacteriales | Clostridia | Firmicutes | Bacteria |
| Oscillospiraceae | Eubacteriales | Clostridia | Firmicutes | Bacteria |
| Oscillospiraceae | Eubacteriales | Clostridia | Firmicutes | Bacteria |
| Oscillospiraceae | Eubacteriales | Clostridia | Firmicutes | Bacteria |
| Oscillospiraceae | Eubacteriales | Clostridia | Firmicutes | Bacteria |
| Oscillospiraceae | Eubacteriales | Clostridia | Firmicutes | Bacteria |
| FGB77153 | OFGB77153 | CFGB77153 | Actinobacteria | Bacteria |
| FGB1791 | OFGB1791 | CFGB1791 | Tenericutes | Bacteria |
| FGB10290 | OFGB10290 | CFGB10290 | Firmicutes | Bacteria |
| Oscillospiraceae | Eubacteriales | Clostridia | Firmicutes | Bacteria |
| FGB1765 | OFGB1765 | CFGB1765 | Firmicutes | Bacteria |
| FGB10349 | OFGB10349 | CFGB10349 | Firmicutes | Bacteria |
| FGB10667 | OFGB10667 | CFGB10667 | Firmicutes | Bacteria |
| Lachnospiraceae | Eubacteriales | Clostridia | Firmicutes | Bacteria |
| Lachnospiraceae | Eubacteriales | Clostridia | Firmicutes | Bacteria |
| Christensenellaceae | Eubacteriales | Clostridia | Firmicutes | Bacteria |
| FGB10299 | OFGB10299 | CFGB10299 | Firmicutes | Bacteria |
| Oscillospiraceae | Eubacteriales | Clostridia | Firmicutes | Bacteria |
| Oscillospiraceae | Eubacteriales | Clostridia | Firmicutes | Bacteria |
| Lachnospiraceae | Eubacteriales | Clostridia | Firmicutes | Bacteria |
| Clostridia_unclassified | Clostridia_unclassifie | Clostridia | Firmicutes | Bacteria |
| Lachnospiraceae | Eubacteriales | Clostridia | Firmicutes | Bacteria |
| Lachnospiraceae | Eubacteriales | Clostridia | Firmicutes | Bacteria |
| Lachnospiraceae | Eubacteriales | Clostridia | Firmicutes | Bacteria |

| | | | | |
|---|---|---|---|---|
| Lachnospiraceae | Eubacteriales | Clostridia | Firmicutes | Bacteria |
| Lachnospiraceae | Eubacteriales | Clostridia | Firmicutes | Bacteria |
| Lachnospiraceae | Eubacteriales | Clostridia | Firmicutes | Bacteria |
| Lachnospiraceae | Eubacteriales | Clostridia | Firmicutes | Bacteria |
| Lachnospiraceae | Eubacteriales | Clostridia | Firmicutes | Bacteria |
| Lactobacillaceae | Lactobacillales | Bacilli | Firmicutes | Bacteria |
| Muribaculaceae | Bacteroidales | Bacteroidia | Bacteroidota | Bacteria |
| Oscillospiraceae | Eubacteriales | Clostridia | Firmicutes | Bacteria |
| Oscillospiraceae | Eubacteriales | Clostridia | Firmicutes | Bacteria |
| Oscillospiraceae | Eubacteriales | Clostridia | Firmicutes | Bacteria |
| Oscillospiraceae | Eubacteriales | Clostridia | Firmicutes | Bacteria |
| Sutterellaceae | Burkholderiales | Betaproteobacteria | Proteobacteria | Bacteria |
| Peptostreptococcaceae | Eubacteriales | Clostridia | Firmicutes | Bacteria |
| Lachnospiraceae | Eubacteriales | Clostridia | Firmicutes | Bacteria |
| Turicibacteraceae | Erysipelotrichales | Erysipelotrichia | Firmicutes | Bacteria |
| Bacteria_unclassified | Bacteria_unclassified | Bacteria_unclassifie | Bacteria_unclassifie | Bacteria |
| Bacteria_unclassified | Bacteria_unclassified | Bacteria_unclassifie | Bacteria_unclassifie | Bacteria |
| Bacteria_unclassified | Bacteria_unclassified | Bacteria_unclassifie | Bacteria_unclassifie | Bacteria |

| | | | | |
|---|---|---|---|---|
| Lachnospiraceae | Eubacteriales | Clostridia | Firmicutes | Bacteria |
| Lachnospiraceae | Eubacteriales | Clostridia | Firmicutes | Bacteria |
| Oscillospiraceae | Eubacteriales | Clostridia | Firmicutes | Bacteria |
| Oscillospiraceae | Eubacteriales | Clostridia | Firmicutes | Bacteria |
| Eggerthellaceae | Eggerthellales | Coriobacteriia | Actinobacteria | Bacteria |
| Eggerthellaceae | Eggerthellales | Coriobacteriia | Actinobacteria | Bacteria |
| Eggerthellaceae | Eggerthellales | Coriobacteriia | Actinobacteria | Bacteria |
| Akkermansiaceae | Verrucomicrobiales | Verrucomicrobiae | Verrucomicrobia | Bacteria |
| Rikenellaceae | Bacteroidales | Bacteroidia | Bacteroidota | Bacteria |
| Oscillospiraceae | Eubacteriales | Clostridia | Firmicutes | Bacteria |
| Bacteria_unclassified | Bacteria_unclassified | Bacteria_unclassifie | Bacteria_unclassifie | Bacteria |
| Bacteria_unclassified | Bacteria_unclassified | Bacteria_unclassifie | Bacteria_unclassifie | Bacteria |
| Bacteria_unclassified | Bacteria_unclassified | Bacteria_unclassifie | Bacteria_unclassifie | Bacteria |
| Bacteroidaceae | Bacteroidales | Bacteroidia | Bacteroidota | Bacteria |
| Bifidobacteriaceae | Bifidobacteriales | Actinomycetia | Actinobacteria | Bacteria |
| Clostridia_unclassified | Clostridia_unclassifie | Clostridia | Firmicutes | Bacteria |
| Clostridiaceae | Eubacteriales | Clostridia | Firmicutes | Bacteria |
| Clostridiaceae | Eubacteriales | Clostridia | Firmicutes | Bacteria |
| Eubacteriales_unclassified | Eubacteriales | Clostridia | Firmicutes | Bacteria |
| Erysipelotrichaceae | Erysipelotrichales | Erysipelotrichia | Firmicutes | Bacteria |
| Coriobacteriaceae | Coriobacteriales | Coriobacteriia | Actinobacteria | Bacteria |
| Lachnospiraceae | Eubacteriales | Clostridia | Firmicutes | Bacteria |
| Erysipelotrichaceae | Erysipelotrichales | Erysipelotrichia | Firmicutes | Bacteria |
| Erysipelotrichales_unclassifie | Erysipelotrichales | Erysipelotrichia | Firmicutes | Bacteria |
| Eubacteriaceae | Eubacteriales | Clostridia | Firmicutes | Bacteria |

| | | | | |
|---|---|---|---|---|
| Eubacteriaceae | Eubacteriales | Clostridia | Firmicutes | Bacteria |
| Lachnospiraceae | Eubacteriales | Clostridia | Firmicutes | Bacteria |
| Eggerthellaceae | Eggerthellales | Coriobacteriia | Actinobacteria | Bacteria |
| Lachnospiraceae | Eubacteriales | Clostridia | Firmicutes | Bacteria |
| Muribaculaceae | Bacteroidales | Bacteroidia | Bacteroidota | Bacteria |
| Muribaculaceae | Bacteroidales | Bacteroidia | Bacteroidota | Bacteria |
| Muribaculaceae | Bacteroidales | Bacteroidia | Bacteroidota | Bacteria |
| FGB9508 | OFGB9508 | CFGB9508 | Firmicutes | Bacteria |
| FGB2838 | OFGB2838 | CFGB2838 | Firmicutes | Bacteria |
| FGB2838 | OFGB2838 | CFGB2838 | Firmicutes | Bacteria |
| FGB2838 | OFGB2838 | CFGB2838 | Firmicutes | Bacteria |
| Pumilibacteraceae | Eubacteriales | Clostridia | Firmicutes | Bacteria |
| FGB28439 | OFGB28439 | CFGB28439 | Firmicutes | Bacteria |
| Clostridia_unclassified | Clostridia_unclassifie | Clostridia | Firmicutes | Bacteria |
| Eubacteriaceae | Eubacteriales | Clostridia | Firmicutes | Bacteria |
| Lachnospiraceae | Eubacteriales | Clostridia | Firmicutes | Bacteria |
| Lachnospiraceae | Eubacteriales | Clostridia | Firmicutes | Bacteria |
| Lachnospiraceae | Eubacteriales | Clostridia | Firmicutes | Bacteria |
| Lachnospiraceae | Eubacteriales | Clostridia | Firmicutes | Bacteria |
| FGB77305 | OFGB77305 | CFGB77305 | Firmicutes | Bacteria |
| Clostridiaceae | Eubacteriales | Clostridia | Firmicutes | Bacteria |
| Lachnospiraceae | Eubacteriales | Clostridia | Firmicutes | Bacteria |
| Lachnospiraceae | Eubacteriales | Clostridia | Firmicutes | Bacteria |
| Lachnospiraceae | Eubacteriales | Clostridia | Firmicutes | Bacteria |
| FGB9633 | OFGB9633 | CFGB9633 | Firmicutes | Bacteria |
| Bacteria_unclassified | Bacteria_unclassified | Bacteria_unclassifie | Bacteria_unclassifie | Bacteria |
| Bacteria_unclassified | Bacteria_unclassified | Bacteria_unclassifie | Bacteria_unclassifie | Bacteria |
| Bacteria_unclassified | Bacteria_unclassified | Bacteria_unclassifie | Bacteria_unclassifie | Bacteria |
| Bacteria_unclassified | Bacteria_unclassified | Bacteria_unclassifie | Bacteria_unclassifie | Bacteria |
| Lachnospiraceae | Eubacteriales | Clostridia | Firmicutes | Bacteria |
| Lachnospiraceae | Eubacteriales | Clostridia | Firmicutes | Bacteria |
| Lachnospiraceae | Eubacteriales | Clostridia | Firmicutes | Bacteria |
| FGB77359 | OFGB77359 | CFGB77359 | Bacteria_unclassifie | Bacteria |
| FGB9639 | OFGB9639 | CFGB9639 | Firmicutes | Bacteria |
| Lachnospiraceae | Eubacteriales | Clostridia | Firmicutes | Bacteria |
| Lachnospiraceae | Eubacteriales | Clostridia | Firmicutes | Bacteria |
| Lachnospiraceae | Eubacteriales | Clostridia | Firmicutes | Bacteria |
| Clostridiaceae | Eubacteriales | Clostridia | Firmicutes | Bacteria |
| Clostridiaceae | Eubacteriales | Clostridia | Firmicutes | Bacteria |
| Clostridiaceae | Eubacteriales | Clostridia | Firmicutes | Bacteria |
| Clostridiaceae | Eubacteriales | Clostridia | Firmicutes | Bacteria |
| Clostridiaceae | Eubacteriales | Clostridia | Firmicutes | Bacteria |
| Clostridiaceae | Eubacteriales | Clostridia | Firmicutes | Bacteria |
| Clostridiaceae | Eubacteriales | Clostridia | Firmicutes | Bacteria |
| Eubacteriaceae | Eubacteriales | Clostridia | Firmicutes | Bacteria |
| FGB9658 | OFGB9658 | CFGB9658 | Firmicutes | Bacteria |

| | | | | |
|---|---|---|---|---|
| FGB9659 | OFGB9659 | CFGB9659 | Firmicutes | Bacteria |
| Bacteria_unclassified | Bacteria_unclassified | Bacteria_unclassified | Bacteria_unclassified | Bacteria |
| FGB9827 | OFGB9827 | CFGB9827 | Firmicutes | Bacteria |
| Eubacteriaceae | Eubacteriales | Clostridia | Firmicutes | Bacteria |
| FGB77303 | OFGB77303 | CFGB77303 | Bacteria_unclassified | Bacteria |
| Eubacteriales_unclassified | Eubacteriales | Clostridia | Firmicutes | Bacteria |
| Oscillospiraceae | Eubacteriales | Clostridia | Firmicutes | Bacteria |
| FGB30328 | OFGB30328 | CFGB30328 | Firmicutes | Bacteria |
| Oscillospiraceae | Eubacteriales | Clostridia | Firmicutes | Bacteria |
| Oscillospiraceae | Eubacteriales | Clostridia | Firmicutes | Bacteria |
| Oscillospiraceae | Eubacteriales | Clostridia | Firmicutes | Bacteria |
| Oscillospiraceae | Eubacteriales | Clostridia | Firmicutes | Bacteria |
| Oscillospiraceae | Eubacteriales | Clostridia | Firmicutes | Bacteria |
| Oscillospiraceae | Eubacteriales | Clostridia | Firmicutes | Bacteria |
| Oscillospiraceae | Eubacteriales | Clostridia | Firmicutes | Bacteria |
| FGB77153 | OFGB77153 | CFGB77153 | Actinobacteria | Bacteria |
| FGB1791 | OFGB1791 | CFGB1791 | Tenericutes | Bacteria |
| FGB10290 | OFGB10290 | CFGB10290 | Firmicutes | Bacteria |
| Oscillospiraceae | Eubacteriales | Clostridia | Firmicutes | Bacteria |
| FGB1765 | OFGB1765 | CFGB1765 | Firmicutes | Bacteria |
| FGB10349 | OFGB10349 | CFGB10349 | Firmicutes | Bacteria |
| FGB10667 | OFGB10667 | CFGB10667 | Firmicutes | Bacteria |
| Lachnospiraceae | Eubacteriales | Clostridia | Firmicutes | Bacteria |
| Lachnospiraceae | Eubacteriales | Clostridia | Firmicutes | Bacteria |
| Christensenellaceae | Eubacteriales | Clostridia | Firmicutes | Bacteria |
| FGB10299 | OFGB10299 | CFGB10299 | Firmicutes | Bacteria |
| Oscillospiraceae | Eubacteriales | Clostridia | Firmicutes | Bacteria |
| Oscillospiraceae | Eubacteriales | Clostridia | Firmicutes | Bacteria |
| Lachnospiraceae | Eubacteriales | Clostridia | Firmicutes | Bacteria |
| Clostridia_unclassified | Clostridia_unclassified | Clostridia | Firmicutes | Bacteria |
| Lachnospiraceae | Eubacteriales | Clostridia | Firmicutes | Bacteria |
| Lachnospiraceae | Eubacteriales | Clostridia | Firmicutes | Bacteria |
| Lachnospiraceae | Eubacteriales | Clostridia | Firmicutes | Bacteria |
| Lachnospiraceae | Eubacteriales | Clostridia | Firmicutes | Bacteria |
| Lachnospiraceae | Eubacteriales | Clostridia | Firmicutes | Bacteria |
| Lachnospiraceae | Eubacteriales | Clostridia | Firmicutes | Bacteria |
| Lachnospiraceae | Eubacteriales | Clostridia | Firmicutes | Bacteria |
| Lachnospiraceae | Eubacteriales | Clostridia | Firmicutes | Bacteria |
| Lactobacillaceae | Lactobacillales | Bacilli | Firmicutes | Bacteria |
| Muribaculaceae | Bacteroidales | Bacteroidia | Bacteroidota | Bacteria |
| Oscillospiraceae | Eubacteriales | Clostridia | Firmicutes | Bacteria |
| Oscillospiraceae | Eubacteriales | Clostridia | Firmicutes | Bacteria |
| Oscillospiraceae | Eubacteriales | Clostridia | Firmicutes | Bacteria |
| Oscillospiraceae | Eubacteriales | Clostridia | Firmicutes | Bacteria |
| Sutterellaceae | Burkholderiales | Betaproteobacteria | Proteobacteria | Bacteria |
| Peptostreptococcaceae | Eubacteriales | Clostridia | Firmicutes | Bacteria |

| | | | | |
|---|---|---|---|---|
| Lachnospiraceae | Eubacteriales | Clostridia | Firmicutes | Bacteria |
| Turicibacteraceae | Erysipelotrichales | Erysipelotrichia | Firmicutes | Bacteria |
| Bacteria_unclassified | Bacteria_unclassified | Bacteria_unclassified | Bacteria_unclassified | Bacteria |
| Bacteria_unclassified | Bacteria_unclassified | Bacteria_unclassified | Bacteria_unclassified | Bacteria |
| Bacteria_unclassified | Bacteria_unclassified | Bacteria_unclassified | Bacteria_unclassified | Bacteria |
| Lachnospiraceae | Eubacteriales | Clostridia | Firmicutes | Bacteria |
| Lachnospiraceae | Eubacteriales | Clostridia | Firmicutes | Bacteria |
| Oscillospiraceae | Eubacteriales | Clostridia | Firmicutes | Bacteria |
| Oscillospiraceae | Eubacteriales | Clostridia | Firmicutes | Bacteria |
| Eggerthellaceae | Eggerthellales | Coriobacteriia | Actinobacteria | Bacteria |
| Eggerthellaceae | Eggerthellales | Coriobacteriia | Actinobacteria | Bacteria |
| Eggerthellaceae | Eggerthellales | Coriobacteriia | Actinobacteria | Bacteria |
| Akkermansiaceae | Verrucomicrobiales | Verrucomicrobiae | Verrucomicrobia | Bacteria |
| Rikenellaceae | Bacteroidales | Bacteroidia | Bacteroidota | Bacteria |
| Oscillospiraceae | Eubacteriales | Clostridia | Firmicutes | Bacteria |
| Bacteria_unclassified | Bacteria_unclassified | Bacteria_unclassified | Bacteria_unclassified | Bacteria |
| Bacteria_unclassified | Bacteria_unclassified | Bacteria_unclassified | Bacteria_unclassified | Bacteria |
| Bacteria_unclassified | Bacteria_unclassified | Bacteria_unclassified | Bacteria_unclassified | Bacteria |
| Bacteroidaceae | Bacteroidales | Bacteroidia | Bacteroidota | Bacteria |
| Bifidobacteriaceae | Bifidobacteriales | Actinomycetia | Actinobacteria | Bacteria |
| Clostridia_unclassified | Clostridia_unclassified | Clostridia | Firmicutes | Bacteria |
| Clostridiaceae | Eubacteriales | Clostridia | Firmicutes | Bacteria |
| Clostridiaceae | Eubacteriales | Clostridia | Firmicutes | Bacteria |
| Eubacteriales_unclassified | Eubacteriales | Clostridia | Firmicutes | Bacteria |
| Erysipelotrichaceae | Erysipelotrichales | Erysipelotrichia | Firmicutes | Bacteria |
| Coriobacteriaceae | Coriobacteriales | Coriobacteriia | Actinobacteria | Bacteria |
| Lachnospiraceae | Eubacteriales | Clostridia | Firmicutes | Bacteria |
| Erysipelotrichaceae | Erysipelotrichales | Erysipelotrichia | Firmicutes | Bacteria |
| Erysipelotrichales_unclassified | Erysipelotrichales | Erysipelotrichia | Firmicutes | Bacteria |
| Eubacteriaceae | Eubacteriales | Clostridia | Firmicutes | Bacteria |
| Eubacteriaceae | Eubacteriales | Clostridia | Firmicutes | Bacteria |
| Lachnospiraceae | Eubacteriales | Clostridia | Firmicutes | Bacteria |
| Eggerthellaceae | Eggerthellales | Coriobacteriia | Actinobacteria | Bacteria |
| Lachnospiraceae | Eubacteriales | Clostridia | Firmicutes | Bacteria |
| Muribaculaceae | Bacteroidales | Bacteroidia | Bacteroidota | Bacteria |
| Muribaculaceae | Bacteroidales | Bacteroidia | Bacteroidota | Bacteria |
| Muribaculaceae | Bacteroidales | Bacteroidia | Bacteroidota | Bacteria |
| FGB9508 | OFGB9508 | CFGB9508 | Firmicutes | Bacteria |
| FGB2838 | OFGB2838 | CFGB2838 | Firmicutes | Bacteria |
| FGB2838 | OFGB2838 | CFGB2838 | Firmicutes | Bacteria |
| FGB2838 | OFGB2838 | CFGB2838 | Firmicutes | Bacteria |
| Pumilibacteraceae | Eubacteriales | Clostridia | Firmicutes | Bacteria |
| FGB28439 | OFGB28439 | CFGB28439 | Firmicutes | Bacteria |

| | | | | |
|---|---|---|---|---|
| Clostridia_unclassified | Clostridia_unclassifie | Clostridia | Firmicutes | Bacteria |
| Eubacteriaceae | Eubacteriales | Clostridia | Firmicutes | Bacteria |
| Lachnospiraceae | Eubacteriales | Clostridia | Firmicutes | Bacteria |
| Lachnospiraceae | Eubacteriales | Clostridia | Firmicutes | Bacteria |
| Lachnospiraceae | Eubacteriales | Clostridia | Firmicutes | Bacteria |
| Lachnospiraceae | Eubacteriales | Clostridia | Firmicutes | Bacteria |
| FGB77305 | OFGB77305 | CFGB77305 | Firmicutes | Bacteria |
| Clostridiaceae | Eubacteriales | Clostridia | Firmicutes | Bacteria |
| Lachnospiraceae | Eubacteriales | Clostridia | Firmicutes | Bacteria |
| Lachnospiraceae | Eubacteriales | Clostridia | Firmicutes | Bacteria |
| Lachnospiraceae | Eubacteriales | Clostridia | Firmicutes | Bacteria |
| FGB9633 | OFGB9633 | CFGB9633 | Firmicutes | Bacteria |
| Bacteria_unclassified | Bacteria_unclassified | Bacteria_unclassifie | Bacteria_unclassifie | Bacteria |
| Bacteria_unclassified | Bacteria_unclassified | Bacteria_unclassifie | Bacteria_unclassifie | Bacteria |
| Bacteria_unclassified | Bacteria_unclassified | Bacteria_unclassifie | Bacteria_unclassifie | Bacteria |
| Bacteria_unclassified | Bacteria_unclassified | Bacteria_unclassifie | Bacteria_unclassifie | Bacteria |
| Lachnospiraceae | Eubacteriales | Clostridia | Firmicutes | Bacteria |
| Lachnospiraceae | Eubacteriales | Clostridia | Firmicutes | Bacteria |
| Lachnospiraceae | Eubacteriales | Clostridia | Firmicutes | Bacteria |
| FGB77359 | OFGB77359 | CFGB77359 | Bacteria_unclassifie | Bacteria |
| FGB9639 | OFGB9639 | CFGB9639 | Firmicutes | Bacteria |
| Lachnospiraceae | Eubacteriales | Clostridia | Firmicutes | Bacteria |
| Lachnospiraceae | Eubacteriales | Clostridia | Firmicutes | Bacteria |
| Lachnospiraceae | Eubacteriales | Clostridia | Firmicutes | Bacteria |
| Clostridiaceae | Eubacteriales | Clostridia | Firmicutes | Bacteria |
| Clostridiaceae | Eubacteriales | Clostridia | Firmicutes | Bacteria |
| Clostridiaceae | Eubacteriales | Clostridia | Firmicutes | Bacteria |
| Clostridiaceae | Eubacteriales | Clostridia | Firmicutes | Bacteria |
| Clostridiaceae | Eubacteriales | Clostridia | Firmicutes | Bacteria |
| Clostridiaceae | Eubacteriales | Clostridia | Firmicutes | Bacteria |
| Clostridiaceae | Eubacteriales | Clostridia | Firmicutes | Bacteria |
| Eubacteriaceae | Eubacteriales | Clostridia | Firmicutes | Bacteria |
| FGB9658 | OFGB9658 | CFGB9658 | Firmicutes | Bacteria |
| FGB9659 | OFGB9659 | CFGB9659 | Firmicutes | Bacteria |
| Bacteria_unclassified | Bacteria_unclassified | Bacteria_unclassifie | Bacteria_unclassifie | Bacteria |
| FGB9827 | OFGB9827 | CFGB9827 | Firmicutes | Bacteria |
| Eubacteriaceae | Eubacteriales | Clostridia | Firmicutes | Bacteria |
| FGB77303 | OFGB77303 | CFGB77303 | Bacteria_unclassifie | Bacteria |
| Eubacteriales_unclassified | Eubacteriales | Clostridia | Firmicutes | Bacteria |
| Oscillospiraceae | Eubacteriales | Clostridia | Firmicutes | Bacteria |
| FGB30328 | OFGB30328 | CFGB30328 | Firmicutes | Bacteria |
| Oscillospiraceae | Eubacteriales | Clostridia | Firmicutes | Bacteria |
| Oscillospiraceae | Eubacteriales | Clostridia | Firmicutes | Bacteria |
| Oscillospiraceae | Eubacteriales | Clostridia | Firmicutes | Bacteria |
| Oscillospiraceae | Eubacteriales | Clostridia | Firmicutes | Bacteria |
| Oscillospiraceae | Eubacteriales | Clostridia | Firmicutes | Bacteria |

| | | | | |
|---|---|---|---|---|
| Oscillospiraceae | Eubacteriales | Clostridia | Firmicutes | Bacteria |
| Oscillospiraceae | Eubacteriales | Clostridia | Firmicutes | Bacteria |
| FGB77153 | OFGB77153 | CFGB77153 | Actinobacteria | Bacteria |
| FGB1791 | OFGB1791 | CFGB1791 | Tenericutes | Bacteria |
| FGB10290 | OFGB10290 | CFGB10290 | Firmicutes | Bacteria |
| Oscillospiraceae | Eubacteriales | Clostridia | Firmicutes | Bacteria |
| FGB1765 | OFGB1765 | CFGB1765 | Firmicutes | Bacteria |
| FGB10349 | OFGB10349 | CFGB10349 | Firmicutes | Bacteria |
| FGB10667 | OFGB10667 | CFGB10667 | Firmicutes | Bacteria |
| Lachnospiraceae | Eubacteriales | Clostridia | Firmicutes | Bacteria |
| Lachnospiraceae | Eubacteriales | Clostridia | Firmicutes | Bacteria |
| Christensenellaceae | Eubacteriales | Clostridia | Firmicutes | Bacteria |
| FGB10299 | OFGB10299 | CFGB10299 | Firmicutes | Bacteria |
| Oscillospiraceae | Eubacteriales | Clostridia | Firmicutes | Bacteria |
| Oscillospiraceae | Eubacteriales | Clostridia | Firmicutes | Bacteria |
| Lachnospiraceae | Eubacteriales | Clostridia | Firmicutes | Bacteria |
| Clostridia_unclassified | Clostridia_unclassifie | Clostridia | Firmicutes | Bacteria |
| Lachnospiraceae | Eubacteriales | Clostridia | Firmicutes | Bacteria |
| Lachnospiraceae | Eubacteriales | Clostridia | Firmicutes | Bacteria |
| Lachnospiraceae | Eubacteriales | Clostridia | Firmicutes | Bacteria |
| Lachnospiraceae | Eubacteriales | Clostridia | Firmicutes | Bacteria |
| Lachnospiraceae | Eubacteriales | Clostridia | Firmicutes | Bacteria |
| Lachnospiraceae | Eubacteriales | Clostridia | Firmicutes | Bacteria |
| Lachnospiraceae | Eubacteriales | Clostridia | Firmicutes | Bacteria |
| Lachnospiraceae | Eubacteriales | Clostridia | Firmicutes | Bacteria |
| Lactobacillaceae | Lactobacillales | Bacilli | Firmicutes | Bacteria |
| Muribaculaceae | Bacteroidales | Bacteroidia | Bacteroidota | Bacteria |
| Oscillospiraceae | Eubacteriales | Clostridia | Firmicutes | Bacteria |
| Oscillospiraceae | Eubacteriales | Clostridia | Firmicutes | Bacteria |
| Oscillospiraceae | Eubacteriales | Clostridia | Firmicutes | Bacteria |
| Oscillospiraceae | Eubacteriales | Clostridia | Firmicutes | Bacteria |
| Sutterellaceae | Burkholderiales | Betaproteobacteria | Proteobacteria | Bacteria |
| Peptostreptococcaceae | Eubacteriales | Clostridia | Firmicutes | Bacteria |
| Lachnospiraceae | Eubacteriales | Clostridia | Firmicutes | Bacteria |
| Turicibacteraceae | Erysipelotrichales | Erysipelotrichia | Firmicutes | Bacteria |
| Bacteria_unclassified | Bacteria_unclassified | Bacteria_unclassifie | Bacteria_unclassifie | Bacteria |
| Bacteria_unclassified | Bacteria_unclassified | Bacteria_unclassifie | Bacteria_unclassifie | Bacteria |
| Bacteria_unclassified | Bacteria_unclassified | Bacteria_unclassifie | Bacteria_unclassifie | Bacteria |

| | | | | |
|---|---|---|---|---|
| Lachnospiraceae | Eubacteriales | Clostridia | Firmicutes | Bacteria |
| Lachnospiraceae | Eubacteriales | Clostridia | Firmicutes | Bacteria |
| Oscillospiraceae | Eubacteriales | Clostridia | Firmicutes | Bacteria |
| Oscillospiraceae | Eubacteriales | Clostridia | Firmicutes | Bacteria |
| Eggerthellaceae | Eggerthellales | Coriobacteriia | Actinobacteria | Bacteria |

| | | | | |
|---|---|---|---|---|
| Eggerthellaceae | Eggerthellales | Coriobacteriia | Actinobacteria | Bacteria |
| Eggerthellaceae | Eggerthellales | Coriobacteriia | Actinobacteria | Bacteria |
| Akkermansiaceae | Verrucomicrobiales | Verrucomicrobiae | Verrucomicrobia | Bacteria |
| Rikenellaceae | Bacteroidales | Bacteroidia | Bacteroidota | Bacteria |
| Oscillospiraceae | Eubacteriales | Clostridia | Firmicutes | Bacteria |
| Bacteria_unclassified | Bacteria_unclassified | Bacteria_unclassifiec | Bacteria_unclassifiec | Bacteria |
| Bacteria_unclassified | Bacteria_unclassified | Bacteria_unclassifiec | Bacteria_unclassifiec | Bacteria |
| Bacteria_unclassified | Bacteria_unclassified | Bacteria_unclassifiec | Bacteria_unclassifiec | Bacteria |
| Bacteroidaceae | Bacteroidales | Bacteroidia | Bacteroidota | Bacteria |
| Bifidobacteriaceae | Bifidobacteriales | Actinomycetia | Actinobacteria | Bacteria |
| Clostridia_unclassified | Clostridia_unclassifiec | Clostridia | Firmicutes | Bacteria |
| Clostridiaceae | Eubacteriales | Clostridia | Firmicutes | Bacteria |
| Clostridiaceae | Eubacteriales | Clostridia | Firmicutes | Bacteria |
| Eubacteriales_unclassified | Eubacteriales | Clostridia | Firmicutes | Bacteria |
| Erysipelotrichaceae | Erysipelotrichales | Erysipelotrichia | Firmicutes | Bacteria |
| Coriobacteriaceae | Coriobacteriales | Coriobacteriia | Actinobacteria | Bacteria |
| Lachnospiraceae | Eubacteriales | Clostridia | Firmicutes | Bacteria |
| Erysipelotrichaceae | Erysipelotrichales | Erysipelotrichia | Firmicutes | Bacteria |
| Erysipelotrichales_unclassifie | Erysipelotrichales | Erysipelotrichia | Firmicutes | Bacteria |
| Eubacteriaceae | Eubacteriales | Clostridia | Firmicutes | Bacteria |
| Eubacteriaceae | Eubacteriales | Clostridia | Firmicutes | Bacteria |
| Lachnospiraceae | Eubacteriales | Clostridia | Firmicutes | Bacteria |
| Eggerthellaceae | Eggerthellales | Coriobacteriia | Actinobacteria | Bacteria |
| Lachnospiraceae | Eubacteriales | Clostridia | Firmicutes | Bacteria |
| Muribaculaceae | Bacteroidales | Bacteroidia | Bacteroidota | Bacteria |
| Muribaculaceae | Bacteroidales | Bacteroidia | Bacteroidota | Bacteria |
| Muribaculaceae | Bacteroidales | Bacteroidia | Bacteroidota | Bacteria |
| FGB9508 | OFGB9508 | CFGB9508 | Firmicutes | Bacteria |
| FGB2838 | OFGB2838 | CFGB2838 | Firmicutes | Bacteria |
| FGB2838 | OFGB2838 | CFGB2838 | Firmicutes | Bacteria |
| FGB2838 | OFGB2838 | CFGB2838 | Firmicutes | Bacteria |
| Pumilibacteraceae | Eubacteriales | Clostridia | Firmicutes | Bacteria |
| FGB28439 | OFGB28439 | CFGB28439 | Firmicutes | Bacteria |
| Clostridia_unclassified | Clostridia_unclassifiec | Clostridia | Firmicutes | Bacteria |
| Eubacteriaceae | Eubacteriales | Clostridia | Firmicutes | Bacteria |
| Lachnospiraceae | Eubacteriales | Clostridia | Firmicutes | Bacteria |
| Lachnospiraceae | Eubacteriales | Clostridia | Firmicutes | Bacteria |
| Lachnospiraceae | Eubacteriales | Clostridia | Firmicutes | Bacteria |
| Lachnospiraceae | Eubacteriales | Clostridia | Firmicutes | Bacteria |
| FGB77305 | OFGB77305 | CFGB77305 | Firmicutes | Bacteria |
| Clostridiaceae | Eubacteriales | Clostridia | Firmicutes | Bacteria |
| Lachnospiraceae | Eubacteriales | Clostridia | Firmicutes | Bacteria |
| Lachnospiraceae | Eubacteriales | Clostridia | Firmicutes | Bacteria |
| Lachnospiraceae | Eubacteriales | Clostridia | Firmicutes | Bacteria |
| FGB9633 | OFGB9633 | CFGB9633 | Firmicutes | Bacteria |
| Bacteria_unclassified | Bacteria_unclassified | Bacteria_unclassifiec | Bacteria_unclassifiec | Bacteria |

| | | | | |
|---|---|---|---|---|
| Bacteria_unclassified | Bacteria_unclassified | Bacteria_unclassifie | Bacteria_unclassifie | Bacteria |
| Bacteria_unclassified | Bacteria_unclassified | Bacteria_unclassifie | Bacteria_unclassifie | Bacteria |
| Bacteria_unclassified | Bacteria_unclassified | Bacteria_unclassifie | Bacteria_unclassifie | Bacteria |
| Lachnospiraceae | Eubacteriales | Clostridia | Firmicutes | Bacteria |
| Lachnospiraceae | Eubacteriales | Clostridia | Firmicutes | Bacteria |
| Lachnospiraceae | Eubacteriales | Clostridia | Firmicutes | Bacteria |
| FGB77359 | OFGB77359 | CFGB77359 | Bacteria_unclassifie | Bacteria |
| FGB9639 | OFGB9639 | CFGB9639 | Firmicutes | Bacteria |
| Lachnospiraceae | Eubacteriales | Clostridia | Firmicutes | Bacteria |
| Lachnospiraceae | Eubacteriales | Clostridia | Firmicutes | Bacteria |
| Lachnospiraceae | Eubacteriales | Clostridia | Firmicutes | Bacteria |
| Clostridiaceae | Eubacteriales | Clostridia | Firmicutes | Bacteria |
| Clostridiaceae | Eubacteriales | Clostridia | Firmicutes | Bacteria |
| Clostridiaceae | Eubacteriales | Clostridia | Firmicutes | Bacteria |
| Clostridiaceae | Eubacteriales | Clostridia | Firmicutes | Bacteria |
| Clostridiaceae | Eubacteriales | Clostridia | Firmicutes | Bacteria |
| Clostridiaceae | Eubacteriales | Clostridia | Firmicutes | Bacteria |
| Clostridiaceae | Eubacteriales | Clostridia | Firmicutes | Bacteria |
| Eubacteriaceae | Eubacteriales | Clostridia | Firmicutes | Bacteria |
| FGB9658 | OFGB9658 | CFGB9658 | Firmicutes | Bacteria |
| FGB9659 | OFGB9659 | CFGB9659 | Firmicutes | Bacteria |
| Bacteria_unclassified | Bacteria_unclassified | Bacteria_unclassifie | Bacteria_unclassifie | Bacteria |
| FGB9827 | OFGB9827 | CFGB9827 | Firmicutes | Bacteria |
| Eubacteriaceae | Eubacteriales | Clostridia | Firmicutes | Bacteria |
| FGB77303 | OFGB77303 | CFGB77303 | Bacteria_unclassifie | Bacteria |
| Eubacteriales_unclassified | Eubacteriales | Clostridia | Firmicutes | Bacteria |
| Oscillospiraceae | Eubacteriales | Clostridia | Firmicutes | Bacteria |
| FGB30328 | OFGB30328 | CFGB30328 | Firmicutes | Bacteria |
| Oscillospiraceae | Eubacteriales | Clostridia | Firmicutes | Bacteria |
| Oscillospiraceae | Eubacteriales | Clostridia | Firmicutes | Bacteria |
| Oscillospiraceae | Eubacteriales | Clostridia | Firmicutes | Bacteria |
| Oscillospiraceae | Eubacteriales | Clostridia | Firmicutes | Bacteria |
| Oscillospiraceae | Eubacteriales | Clostridia | Firmicutes | Bacteria |
| Oscillospiraceae | Eubacteriales | Clostridia | Firmicutes | Bacteria |
| Oscillospiraceae | Eubacteriales | Clostridia | Firmicutes | Bacteria |
| FGB77153 | OFGB77153 | CFGB77153 | Actinobacteria | Bacteria |
| FGB1791 | OFGB1791 | CFGB1791 | Tenericutes | Bacteria |
| FGB10290 | OFGB10290 | CFGB10290 | Firmicutes | Bacteria |
| Oscillospiraceae | Eubacteriales | Clostridia | Firmicutes | Bacteria |
| FGB1765 | OFGB1765 | CFGB1765 | Firmicutes | Bacteria |
| FGB10349 | OFGB10349 | CFGB10349 | Firmicutes | Bacteria |
| FGB10667 | OFGB10667 | CFGB10667 | Firmicutes | Bacteria |
| Lachnospiraceae | Eubacteriales | Clostridia | Firmicutes | Bacteria |
| Lachnospiraceae | Eubacteriales | Clostridia | Firmicutes | Bacteria |
| Christensenellaceae | Eubacteriales | Clostridia | Firmicutes | Bacteria |
| FGB10299 | OFGB10299 | CFGB10299 | Firmicutes | Bacteria |

| | | | | |
|---|---|---|---|---|
| Oscillospiraceae | Eubacteriales | Clostridia | Firmicutes | Bacteria |
| Oscillospiraceae | Eubacteriales | Clostridia | Firmicutes | Bacteria |
| Lachnospiraceae | Eubacteriales | Clostridia | Firmicutes | Bacteria |
| Clostridia_unclassified | Clostridia_unclassifie | Clostridia | Firmicutes | Bacteria |
| Lachnospiraceae | Eubacteriales | Clostridia | Firmicutes | Bacteria |
| Lachnospiraceae | Eubacteriales | Clostridia | Firmicutes | Bacteria |
| Lachnospiraceae | Eubacteriales | Clostridia | Firmicutes | Bacteria |
| Lachnospiraceae | Eubacteriales | Clostridia | Firmicutes | Bacteria |
| Lachnospiraceae | Eubacteriales | Clostridia | Firmicutes | Bacteria |
| Lachnospiraceae | Eubacteriales | Clostridia | Firmicutes | Bacteria |
| Lachnospiraceae | Eubacteriales | Clostridia | Firmicutes | Bacteria |
| Lachnospiraceae | Eubacteriales | Clostridia | Firmicutes | Bacteria |
| Lactobacillaceae | Lactobacillales | Bacilli | Firmicutes | Bacteria |
| Muribaculaceae | Bacteroidales | Bacteroidia | Bacteroidota | Bacteria |
| Oscillospiraceae | Eubacteriales | Clostridia | Firmicutes | Bacteria |
| Oscillospiraceae | Eubacteriales | Clostridia | Firmicutes | Bacteria |
| Oscillospiraceae | Eubacteriales | Clostridia | Firmicutes | Bacteria |
| Oscillospiraceae | Eubacteriales | Clostridia | Firmicutes | Bacteria |
| Sutterellaceae | Burkholderiales | Betaproteobacteria | Proteobacteria | Bacteria |
| Peptostreptococcaceae | Eubacteriales | Clostridia | Firmicutes | Bacteria |
| Lachnospiraceae | Eubacteriales | Clostridia | Firmicutes | Bacteria |
| Turicibacteraceae | Erysipelotrichales | Erysipelotrichia | Firmicutes | Bacteria |
| Bacteria_unclassified | Bacteria_unclassified | Bacteria_unclassifie | Bacteria_unclassifie | Bacteria |
| Bacteria_unclassified | Bacteria_unclassified | Bacteria_unclassifie | Bacteria_unclassifie | Bacteria |
| Bacteria_unclassified | Bacteria_unclassified | Bacteria_unclassifie | Bacteria_unclassifie | Bacteria |

| | | | | |
|---|---|---|---|---|
| Lachnospiraceae | Eubacteriales | Clostridia | Firmicutes | Bacteria |
| Lachnospiraceae | Eubacteriales | Clostridia | Firmicutes | Bacteria |
| Oscillospiraceae | Eubacteriales | Clostridia | Firmicutes | Bacteria |
| Oscillospiraceae | Eubacteriales | Clostridia | Firmicutes | Bacteria |
| Eggerthellaceae | Eggerthellales | Coriobacteriia | Actinobacteria | Bacteria |
| Eggerthellaceae | Eggerthellales | Coriobacteriia | Actinobacteria | Bacteria |
| Eggerthellaceae | Eggerthellales | Coriobacteriia | Actinobacteria | Bacteria |
| Akkermansiaceae | Verrucomicrobiales | Verrucomicrobiae | Verrucomicrobia | Bacteria |
| Rikenellaceae | Bacteroidales | Bacteroidia | Bacteroidota | Bacteria |
| Oscillospiraceae | Eubacteriales | Clostridia | Firmicutes | Bacteria |
| Bacteria_unclassified | Bacteria_unclassified | Bacteria_unclassifie | Bacteria_unclassifie | Bacteria |
| Bacteria_unclassified | Bacteria_unclassified | Bacteria_unclassifie | Bacteria_unclassifie | Bacteria |
| Bacteria_unclassified | Bacteria_unclassified | Bacteria_unclassifie | Bacteria_unclassifie | Bacteria |
| Bacteroidaceae | Bacteroidales | Bacteroidia | Bacteroidota | Bacteria |
| Bifidobacteriaceae | Bifidobacteriales | Actinomycetia | Actinobacteria | Bacteria |
| Clostridia_unclassified | Clostridia_unclassifie | Clostridia | Firmicutes | Bacteria |
| Clostridiaceae | Eubacteriales | Clostridia | Firmicutes | Bacteria |
| Clostridiaceae | Eubacteriales | Clostridia | Firmicutes | Bacteria |

| | | | | |
|---|---|---|---|---|
| Eubacteriales_unclassified | Eubacteriales | Clostridia | Firmicutes | Bacteria |
| Erysipelotrichaceae | Erysipelotrichales | Erysipelotrichia | Firmicutes | Bacteria |
| Coriobacteriaceae | Coriobacteriales | Coriobacteriia | Actinobacteria | Bacteria |
| Lachnospiraceae | Eubacteriales | Clostridia | Firmicutes | Bacteria |
| Erysipelotrichaceae | Erysipelotrichales | Erysipelotrichia | Firmicutes | Bacteria |
| Erysipelotrichales_unclassifie | Erysipelotrichales | Erysipelotrichia | Firmicutes | Bacteria |
| Eubacteriaceae | Eubacteriales | Clostridia | Firmicutes | Bacteria |
| Eubacteriaceae | Eubacteriales | Clostridia | Firmicutes | Bacteria |
| Lachnospiraceae | Eubacteriales | Clostridia | Firmicutes | Bacteria |
| Eggerthellaceae | Eggerthellales | Coriobacteriia | Actinobacteria | Bacteria |
| Lachnospiraceae | Eubacteriales | Clostridia | Firmicutes | Bacteria |
| Muribaculaceae | Bacteroidales | Bacteroidia | Bacteroidota | Bacteria |
| Muribaculaceae | Bacteroidales | Bacteroidia | Bacteroidota | Bacteria |
| Muribaculaceae | Bacteroidales | Bacteroidia | Bacteroidota | Bacteria |
| FGB9508 | OFGB9508 | CFGB9508 | Firmicutes | Bacteria |
| FGB2838 | OFGB2838 | CFGB2838 | Firmicutes | Bacteria |
| FGB2838 | OFGB2838 | CFGB2838 | Firmicutes | Bacteria |
| FGB2838 | OFGB2838 | CFGB2838 | Firmicutes | Bacteria |
| Pumilibacteraceae | Eubacteriales | Clostridia | Firmicutes | Bacteria |
| FGB28439 | OFGB28439 | CFGB28439 | Firmicutes | Bacteria |
| Clostridia_unclassified | Clostridia_unclassifie | Clostridia | Firmicutes | Bacteria |
| Eubacteriaceae | Eubacteriales | Clostridia | Firmicutes | Bacteria |
| Lachnospiraceae | Eubacteriales | Clostridia | Firmicutes | Bacteria |
| Lachnospiraceae | Eubacteriales | Clostridia | Firmicutes | Bacteria |
| Lachnospiraceae | Eubacteriales | Clostridia | Firmicutes | Bacteria |
| Lachnospiraceae | Eubacteriales | Clostridia | Firmicutes | Bacteria |
| FGB77305 | OFGB77305 | CFGB77305 | Firmicutes | Bacteria |
| Clostridiaceae | Eubacteriales | Clostridia | Firmicutes | Bacteria |
| Lachnospiraceae | Eubacteriales | Clostridia | Firmicutes | Bacteria |
| Lachnospiraceae | Eubacteriales | Clostridia | Firmicutes | Bacteria |
| Lachnospiraceae | Eubacteriales | Clostridia | Firmicutes | Bacteria |
| FGB9633 | OFGB9633 | CFGB9633 | Firmicutes | Bacteria |
| Bacteria_unclassified | Bacteria_unclassified | Bacteria_unclassifie | Bacteria_unclassifie | Bacteria |
| Bacteria_unclassified | Bacteria_unclassified | Bacteria_unclassifie | Bacteria_unclassifie | Bacteria |
| Bacteria_unclassified | Bacteria_unclassified | Bacteria_unclassifie | Bacteria_unclassifie | Bacteria |
| Bacteria_unclassified | Bacteria_unclassified | Bacteria_unclassifie | Bacteria_unclassifie | Bacteria |
| Lachnospiraceae | Eubacteriales | Clostridia | Firmicutes | Bacteria |
| Lachnospiraceae | Eubacteriales | Clostridia | Firmicutes | Bacteria |
| Lachnospiraceae | Eubacteriales | Clostridia | Firmicutes | Bacteria |
| FGB77359 | OFGB77359 | CFGB77359 | Bacteria_unclassifie | Bacteria |
| FGB9639 | OFGB9639 | CFGB9639 | Firmicutes | Bacteria |
| Lachnospiraceae | Eubacteriales | Clostridia | Firmicutes | Bacteria |
| Lachnospiraceae | Eubacteriales | Clostridia | Firmicutes | Bacteria |
| Lachnospiraceae | Eubacteriales | Clostridia | Firmicutes | Bacteria |
| Clostridiaceae | Eubacteriales | Clostridia | Firmicutes | Bacteria |
| Clostridiaceae | Eubacteriales | Clostridia | Firmicutes | Bacteria |

| | | | | |
|---|---|---|---|---|
| Clostridiaceae | Eubacteriales | Clostridia | Firmicutes | Bacteria |
| Clostridiaceae | Eubacteriales | Clostridia | Firmicutes | Bacteria |
| Clostridiaceae | Eubacteriales | Clostridia | Firmicutes | Bacteria |
| Clostridiaceae | Eubacteriales | Clostridia | Firmicutes | Bacteria |
| Clostridiaceae | Eubacteriales | Clostridia | Firmicutes | Bacteria |
| Eubacteriaceae | Eubacteriales | Clostridia | Firmicutes | Bacteria |
| FGB9658 | OFGB9658 | CFGB9658 | Firmicutes | Bacteria |
| FGB9659 | OFGB9659 | CFGB9659 | Firmicutes | Bacteria |
| Bacteria_unclassified | Bacteria_unclassified | Bacteria_unclassifie | Bacteria_unclassifie | Bacteria |
| FGB9827 | OFGB9827 | CFGB9827 | Firmicutes | Bacteria |
| Eubacteriaceae | Eubacteriales | Clostridia | Firmicutes | Bacteria |
| FGB77303 | OFGB77303 | CFGB77303 | Bacteria_unclassifie | Bacteria |
| Eubacteriales_unclassified | Eubacteriales | Clostridia | Firmicutes | Bacteria |
| Oscillospiraceae | Eubacteriales | Clostridia | Firmicutes | Bacteria |
| FGB30328 | OFGB30328 | CFGB30328 | Firmicutes | Bacteria |
| Oscillospiraceae | Eubacteriales | Clostridia | Firmicutes | Bacteria |
| Oscillospiraceae | Eubacteriales | Clostridia | Firmicutes | Bacteria |
| Oscillospiraceae | Eubacteriales | Clostridia | Firmicutes | Bacteria |
| Oscillospiraceae | Eubacteriales | Clostridia | Firmicutes | Bacteria |
| Oscillospiraceae | Eubacteriales | Clostridia | Firmicutes | Bacteria |
| Oscillospiraceae | Eubacteriales | Clostridia | Firmicutes | Bacteria |
| Oscillospiraceae | Eubacteriales | Clostridia | Firmicutes | Bacteria |
| FGB77153 | OFGB77153 | CFGB77153 | Actinobacteria | Bacteria |
| FGB1791 | OFGB1791 | CFGB1791 | Tenericutes | Bacteria |
| FGB10290 | OFGB10290 | CFGB10290 | Firmicutes | Bacteria |
| Oscillospiraceae | Eubacteriales | Clostridia | Firmicutes | Bacteria |
| FGB1765 | OFGB1765 | CFGB1765 | Firmicutes | Bacteria |
| FGB10349 | OFGB10349 | CFGB10349 | Firmicutes | Bacteria |
| FGB10667 | OFGB10667 | CFGB10667 | Firmicutes | Bacteria |
| Lachnospiraceae | Eubacteriales | Clostridia | Firmicutes | Bacteria |
| Lachnospiraceae | Eubacteriales | Clostridia | Firmicutes | Bacteria |
| Christensenellaceae | Eubacteriales | Clostridia | Firmicutes | Bacteria |
| FGB10299 | OFGB10299 | CFGB10299 | Firmicutes | Bacteria |
| Oscillospiraceae | Eubacteriales | Clostridia | Firmicutes | Bacteria |
| Oscillospiraceae | Eubacteriales | Clostridia | Firmicutes | Bacteria |
| Lachnospiraceae | Eubacteriales | Clostridia | Firmicutes | Bacteria |
| Clostridia_unclassified | Clostridia_unclassifie | Clostridia | Firmicutes | Bacteria |
| Lachnospiraceae | Eubacteriales | Clostridia | Firmicutes | Bacteria |
| Lachnospiraceae | Eubacteriales | Clostridia | Firmicutes | Bacteria |
| Lachnospiraceae | Eubacteriales | Clostridia | Firmicutes | Bacteria |
| Lachnospiraceae | Eubacteriales | Clostridia | Firmicutes | Bacteria |
| Lachnospiraceae | Eubacteriales | Clostridia | Firmicutes | Bacteria |
| Lachnospiraceae | Eubacteriales | Clostridia | Firmicutes | Bacteria |
| Lachnospiraceae | Eubacteriales | Clostridia | Firmicutes | Bacteria |
| Lachnospiraceae | Eubacteriales | Clostridia | Firmicutes | Bacteria |
| Lactobacillaceae | Lactobacillales | Bacilli | Firmicutes | Bacteria |

| | | | | |
|---|---|---|---|---|
| Muribaculaceae | Bacteroidales | Bacteroidia | Bacteroidota | Bacteria |
| Oscillospiraceae | Eubacteriales | Clostridia | Firmicutes | Bacteria |
| Oscillospiraceae | Eubacteriales | Clostridia | Firmicutes | Bacteria |
| Oscillospiraceae | Eubacteriales | Clostridia | Firmicutes | Bacteria |
| Oscillospiraceae | Eubacteriales | Clostridia | Firmicutes | Bacteria |
| Sutterellaceae | Burkholderiales | Betaproteobacteria | Proteobacteria | Bacteria |
| Peptostreptococcaceae | Eubacteriales | Clostridia | Firmicutes | Bacteria |
| Lachnospiraceae | Eubacteriales | Clostridia | Firmicutes | Bacteria |
| Turicibacteraceae | Erysipelotrichales | Erysipelotrichia | Firmicutes | Bacteria |
| Bacteria_unclassified | Bacteria_unclassified | Bacteria_unclassifie | Bacteria_unclassifie | Bacteria |
| Bacteria_unclassified | Bacteria_unclassified | Bacteria_unclassifie | Bacteria_unclassifie | Bacteria |
| Bacteria_unclassified | Bacteria_unclassified | Bacteria_unclassifie | Bacteria_unclassifie | Bacteria |

| | | | | |
|---|---|---|---|---|
| Lachnospiraceae | Eubacteriales | Clostridia | Firmicutes | Bacteria |
| Lachnospiraceae | Eubacteriales | Clostridia | Firmicutes | Bacteria |
| Oscillospiraceae | Eubacteriales | Clostridia | Firmicutes | Bacteria |
| Oscillospiraceae | Eubacteriales | Clostridia | Firmicutes | Bacteria |
| Eggerthellaceae | Eggerthellales | Coriobacteriia | Actinobacteria | Bacteria |
| Eggerthellaceae | Eggerthellales | Coriobacteriia | Actinobacteria | Bacteria |
| Eggerthellaceae | Eggerthellales | Coriobacteriia | Actinobacteria | Bacteria |
| Akkermansiaceae | Verrucomicrobiales | Verrucomicrobiae | Verrucomicrobia | Bacteria |
| Rikenellaceae | Bacteroidales | Bacteroidia | Bacteroidota | Bacteria |
| Oscillospiraceae | Eubacteriales | Clostridia | Firmicutes | Bacteria |
| Bacteria_unclassified | Bacteria_unclassified | Bacteria_unclassifie | Bacteria_unclassifie | Bacteria |
| Bacteria_unclassified | Bacteria_unclassified | Bacteria_unclassifie | Bacteria_unclassifie | Bacteria |
| Bacteria_unclassified | Bacteria_unclassified | Bacteria_unclassifie | Bacteria_unclassifie | Bacteria |
| Bacteroidaceae | Bacteroidales | Bacteroidia | Bacteroidota | Bacteria |
| Bifidobacteriaceae | Bifidobacteriales | Actinomycetia | Actinobacteria | Bacteria |
| Clostridia_unclassified | Clostridia_unclassifie | Clostridia | Firmicutes | Bacteria |
| Clostridiaceae | Eubacteriales | Clostridia | Firmicutes | Bacteria |
| Clostridiaceae | Eubacteriales | Clostridia | Firmicutes | Bacteria |
| Eubacteriales_unclassified | Eubacteriales | Clostridia | Firmicutes | Bacteria |
| Erysipelotrichaceae | Erysipelotrichales | Erysipelotrichia | Firmicutes | Bacteria |
| Coriobacteriaceae | Coriobacteriales | Coriobacteriia | Actinobacteria | Bacteria |
| Lachnospiraceae | Eubacteriales | Clostridia | Firmicutes | Bacteria |
| Erysipelotrichaceae | Erysipelotrichales | Erysipelotrichia | Firmicutes | Bacteria |
| Erysipelotrichales_unclassifie | Erysipelotrichales | Erysipelotrichia | Firmicutes | Bacteria |
| Eubacteriaceae | Eubacteriales | Clostridia | Firmicutes | Bacteria |
| Eubacteriaceae | Eubacteriales | Clostridia | Firmicutes | Bacteria |
| Lachnospiraceae | Eubacteriales | Clostridia | Firmicutes | Bacteria |
| Eggerthellaceae | Eggerthellales | Coriobacteriia | Actinobacteria | Bacteria |
| Lachnospiraceae | Eubacteriales | Clostridia | Firmicutes | Bacteria |
| Muribaculaceae | Bacteroidales | Bacteroidia | Bacteroidota | Bacteria |
| Muribaculaceae | Bacteroidales | Bacteroidia | Bacteroidota | Bacteria |

| | | | | |
|---|---|---|---|---|
| Muribaculaceae | Bacteroidales | Bacteroidia | Bacteroidota | Bacteria |
| FGB9508 | OFGB9508 | CFGB9508 | Firmicutes | Bacteria |
| FGB2838 | OFGB2838 | CFGB2838 | Firmicutes | Bacteria |
| FGB2838 | OFGB2838 | CFGB2838 | Firmicutes | Bacteria |
| FGB2838 | OFGB2838 | CFGB2838 | Firmicutes | Bacteria |
| Pumilibacteraceae | Eubacteriales | Clostridia | Firmicutes | Bacteria |
| FGB28439 | OFGB28439 | CFGB28439 | Firmicutes | Bacteria |
| Clostridia_unclassified | Clostridia_unclassifie | Clostridia | Firmicutes | Bacteria |
| Eubacteriaceae | Eubacteriales | Clostridia | Firmicutes | Bacteria |
| Lachnospiraceae | Eubacteriales | Clostridia | Firmicutes | Bacteria |
| Lachnospiraceae | Eubacteriales | Clostridia | Firmicutes | Bacteria |
| Lachnospiraceae | Eubacteriales | Clostridia | Firmicutes | Bacteria |
| Lachnospiraceae | Eubacteriales | Clostridia | Firmicutes | Bacteria |
| FGB77305 | OFGB77305 | CFGB77305 | Firmicutes | Bacteria |
| Clostridiaceae | Eubacteriales | Clostridia | Firmicutes | Bacteria |
| Lachnospiraceae | Eubacteriales | Clostridia | Firmicutes | Bacteria |
| Lachnospiraceae | Eubacteriales | Clostridia | Firmicutes | Bacteria |
| Lachnospiraceae | Eubacteriales | Clostridia | Firmicutes | Bacteria |
| FGB9633 | OFGB9633 | CFGB9633 | Firmicutes | Bacteria |
| Bacteria_unclassified | Bacteria_unclassified | Bacteria_unclassifie | Bacteria_unclassifie | Bacteria |
| Bacteria_unclassified | Bacteria_unclassified | Bacteria_unclassifie | Bacteria_unclassifie | Bacteria |
| Bacteria_unclassified | Bacteria_unclassified | Bacteria_unclassifie | Bacteria_unclassifie | Bacteria |
| Bacteria_unclassified | Bacteria_unclassified | Bacteria_unclassifie | Bacteria_unclassifie | Bacteria |
| Lachnospiraceae | Eubacteriales | Clostridia | Firmicutes | Bacteria |
| Lachnospiraceae | Eubacteriales | Clostridia | Firmicutes | Bacteria |
| Lachnospiraceae | Eubacteriales | Clostridia | Firmicutes | Bacteria |
| FGB77359 | OFGB77359 | CFGB77359 | Bacteria_unclassifie | Bacteria |
| FGB9639 | OFGB9639 | CFGB9639 | Firmicutes | Bacteria |
| Lachnospiraceae | Eubacteriales | Clostridia | Firmicutes | Bacteria |
| Lachnospiraceae | Eubacteriales | Clostridia | Firmicutes | Bacteria |
| Lachnospiraceae | Eubacteriales | Clostridia | Firmicutes | Bacteria |
| Clostridiaceae | Eubacteriales | Clostridia | Firmicutes | Bacteria |
| Clostridiaceae | Eubacteriales | Clostridia | Firmicutes | Bacteria |
| Clostridiaceae | Eubacteriales | Clostridia | Firmicutes | Bacteria |
| Clostridiaceae | Eubacteriales | Clostridia | Firmicutes | Bacteria |
| Clostridiaceae | Eubacteriales | Clostridia | Firmicutes | Bacteria |
| Clostridiaceae | Eubacteriales | Clostridia | Firmicutes | Bacteria |
| Clostridiaceae | Eubacteriales | Clostridia | Firmicutes | Bacteria |
| Eubacteriaceae | Eubacteriales | Clostridia | Firmicutes | Bacteria |
| FGB9658 | OFGB9658 | CFGB9658 | Firmicutes | Bacteria |
| FGB9659 | OFGB9659 | CFGB9659 | Firmicutes | Bacteria |
| Bacteria_unclassified | Bacteria_unclassified | Bacteria_unclassifie | Bacteria_unclassifie | Bacteria |
| FGB9827 | OFGB9827 | CFGB9827 | Firmicutes | Bacteria |
| Eubacteriaceae | Eubacteriales | Clostridia | Firmicutes | Bacteria |
| FGB77303 | OFGB77303 | CFGB77303 | Bacteria_unclassifie | Bacteria |
| Eubacteriales_unclassified | Eubacteriales | Clostridia | Firmicutes | Bacteria |

| | | | | |
|---|---|---|---|---|
| Oscillospiraceae | Eubacteriales | Clostridia | Firmicutes | Bacteria |
| FGB30328 | OFGB30328 | CFGB30328 | Firmicutes | Bacteria |
| Oscillospiraceae | Eubacteriales | Clostridia | Firmicutes | Bacteria |
| Oscillospiraceae | Eubacteriales | Clostridia | Firmicutes | Bacteria |
| Oscillospiraceae | Eubacteriales | Clostridia | Firmicutes | Bacteria |
| Oscillospiraceae | Eubacteriales | Clostridia | Firmicutes | Bacteria |
| Oscillospiraceae | Eubacteriales | Clostridia | Firmicutes | Bacteria |
| Oscillospiraceae | Eubacteriales | Clostridia | Firmicutes | Bacteria |
| Oscillospiraceae | Eubacteriales | Clostridia | Firmicutes | Bacteria |
| FGB77153 | OFGB77153 | CFGB77153 | Actinobacteria | Bacteria |
| FGB1791 | OFGB1791 | CFGB1791 | Tenericutes | Bacteria |
| FGB10290 | OFGB10290 | CFGB10290 | Firmicutes | Bacteria |
| Oscillospiraceae | Eubacteriales | Clostridia | Firmicutes | Bacteria |
| FGB1765 | OFGB1765 | CFGB1765 | Firmicutes | Bacteria |
| FGB10349 | OFGB10349 | CFGB10349 | Firmicutes | Bacteria |
| FGB10667 | OFGB10667 | CFGB10667 | Firmicutes | Bacteria |
| Lachnospiraceae | Eubacteriales | Clostridia | Firmicutes | Bacteria |
| Lachnospiraceae | Eubacteriales | Clostridia | Firmicutes | Bacteria |
| Christensenellaceae | Eubacteriales | Clostridia | Firmicutes | Bacteria |
| FGB10299 | OFGB10299 | CFGB10299 | Firmicutes | Bacteria |
| Oscillospiraceae | Eubacteriales | Clostridia | Firmicutes | Bacteria |
| Oscillospiraceae | Eubacteriales | Clostridia | Firmicutes | Bacteria |
| Lachnospiraceae | Eubacteriales | Clostridia | Firmicutes | Bacteria |
| Clostridia_unclassified | Clostridia_unclassifie | Clostridia | Firmicutes | Bacteria |
| Lachnospiraceae | Eubacteriales | Clostridia | Firmicutes | Bacteria |
| Lachnospiraceae | Eubacteriales | Clostridia | Firmicutes | Bacteria |
| Lachnospiraceae | Eubacteriales | Clostridia | Firmicutes | Bacteria |
| Lachnospiraceae | Eubacteriales | Clostridia | Firmicutes | Bacteria |
| Lachnospiraceae | Eubacteriales | Clostridia | Firmicutes | Bacteria |
| Lachnospiraceae | Eubacteriales | Clostridia | Firmicutes | Bacteria |
| Lachnospiraceae | Eubacteriales | Clostridia | Firmicutes | Bacteria |
| Lachnospiraceae | Eubacteriales | Clostridia | Firmicutes | Bacteria |
| Lactobacillaceae | Lactobacillales | Bacilli | Firmicutes | Bacteria |
| Muribaculaceae | Bacteroidales | Bacteroidia | Bacteroidota | Bacteria |
| Oscillospiraceae | Eubacteriales | Clostridia | Firmicutes | Bacteria |
| Oscillospiraceae | Eubacteriales | Clostridia | Firmicutes | Bacteria |
| Oscillospiraceae | Eubacteriales | Clostridia | Firmicutes | Bacteria |
| Oscillospiraceae | Eubacteriales | Clostridia | Firmicutes | Bacteria |
| Sutterellaceae | Burkholderiales | Betaproteobacteria | Proteobacteria | Bacteria |
| Peptostreptococcaceae | Eubacteriales | Clostridia | Firmicutes | Bacteria |
| Lachnospiraceae | Eubacteriales | Clostridia | Firmicutes | Bacteria |
| Turicibacteraceae | Erysipelotrichales | Erysipelotrichia | Firmicutes | Bacteria |
| Bacteria_unclassified | Bacteria_unclassified | Bacteria_unclassifie | Bacteria_unclassifie | Bacteria |
| Bacteria_unclassified | Bacteria_unclassified | Bacteria_unclassifie | Bacteria_unclassifie | Bacteria |
| Bacteria_unclassified | Bacteria_unclassified | Bacteria_unclassifie | Bacteria_unclassifie | Bacteria |

| | | | | |
|---|---|---|---|---|
| Lachnospiraceae | Eubacteriales | Clostridia | Firmicutes | Bacteria |
| Lachnospiraceae | Eubacteriales | Clostridia | Firmicutes | Bacteria |
| Oscillospiraceae | Eubacteriales | Clostridia | Firmicutes | Bacteria |
| Oscillospiraceae | Eubacteriales | Clostridia | Firmicutes | Bacteria |
| Eggerthellaceae | Eggerthellales | Coriobacteriia | Actinobacteria | Bacteria |
| Eggerthellaceae | Eggerthellales | Coriobacteriia | Actinobacteria | Bacteria |
| Eggerthellaceae | Eggerthellales | Coriobacteriia | Actinobacteria | Bacteria |
| Akkermansiaceae | Verrucomicrobiales | Verrucomicrobiae | Verrucomicrobia | Bacteria |
| Rikenellaceae | Bacteroidales | Bacteroidia | Bacteroidota | Bacteria |
| Oscillospiraceae | Eubacteriales | Clostridia | Firmicutes | Bacteria |
| Bacteria_unclassified | Bacteria_unclassified | Bacteria_unclassifie | Bacteria_unclassifie | Bacteria |
| Bacteria_unclassified | Bacteria_unclassified | Bacteria_unclassifie | Bacteria_unclassifie | Bacteria |
| Bacteria_unclassified | Bacteria_unclassified | Bacteria_unclassifie | Bacteria_unclassifie | Bacteria |
| Bacteroidaceae | Bacteroidales | Bacteroidia | Bacteroidota | Bacteria |
| Bifidobacteriaceae | Bifidobacteriales | Actinomycetia | Actinobacteria | Bacteria |
| Clostridia_unclassified | Clostridia_unclassifie | Clostridia | Firmicutes | Bacteria |
| Clostridiaceae | Eubacteriales | Clostridia | Firmicutes | Bacteria |
| Clostridiaceae | Eubacteriales | Clostridia | Firmicutes | Bacteria |
| Eubacteriales_unclassified | Eubacteriales | Clostridia | Firmicutes | Bacteria |
| Erysipelotrichaceae | Erysipelotrichales | Erysipelotrichia | Firmicutes | Bacteria |
| Coriobacteriaceae | Coriobacteriales | Coriobacteriia | Actinobacteria | Bacteria |
| Lachnospiraceae | Eubacteriales | Clostridia | Firmicutes | Bacteria |
| Erysipelotrichaceae | Erysipelotrichales | Erysipelotrichia | Firmicutes | Bacteria |
| Erysipelotrichales_unclassifie | Erysipelotrichales | Erysipelotrichia | Firmicutes | Bacteria |
| Eubacteriaceae | Eubacteriales | Clostridia | Firmicutes | Bacteria |
| Eubacteriaceae | Eubacteriales | Clostridia | Firmicutes | Bacteria |
| Lachnospiraceae | Eubacteriales | Clostridia | Firmicutes | Bacteria |
| Eggerthellaceae | Eggerthellales | Coriobacteriia | Actinobacteria | Bacteria |
| Lachnospiraceae | Eubacteriales | Clostridia | Firmicutes | Bacteria |
| Muribaculaceae | Bacteroidales | Bacteroidia | Bacteroidota | Bacteria |
| Muribaculaceae | Bacteroidales | Bacteroidia | Bacteroidota | Bacteria |
| Muribaculaceae | Bacteroidales | Bacteroidia | Bacteroidota | Bacteria |
| FGB9508 | OFGB9508 | CFGB9508 | Firmicutes | Bacteria |
| FGB2838 | OFGB2838 | CFGB2838 | Firmicutes | Bacteria |
| FGB2838 | OFGB2838 | CFGB2838 | Firmicutes | Bacteria |
| FGB2838 | OFGB2838 | CFGB2838 | Firmicutes | Bacteria |
| Pumilibacteraceae | Eubacteriales | Clostridia | Firmicutes | Bacteria |
| FGB28439 | OFGB28439 | CFGB28439 | Firmicutes | Bacteria |
| Clostridia_unclassified | Clostridia_unclassifie | Clostridia | Firmicutes | Bacteria |
| Eubacteriaceae | Eubacteriales | Clostridia | Firmicutes | Bacteria |
| Lachnospiraceae | Eubacteriales | Clostridia | Firmicutes | Bacteria |
| Lachnospiraceae | Eubacteriales | Clostridia | Firmicutes | Bacteria |
| Lachnospiraceae | Eubacteriales | Clostridia | Firmicutes | Bacteria |
| Lachnospiraceae | Eubacteriales | Clostridia | Firmicutes | Bacteria |

| | | | | |
|---|---|---|---|---|
| FGB77305 | OFGB77305 | CFGB77305 | Firmicutes | Bacteria |
| Clostridiaceae | Eubacteriales | Clostridia | Firmicutes | Bacteria |
| Lachnospiraceae | Eubacteriales | Clostridia | Firmicutes | Bacteria |
| Lachnospiraceae | Eubacteriales | Clostridia | Firmicutes | Bacteria |
| Lachnospiraceae | Eubacteriales | Clostridia | Firmicutes | Bacteria |
| FGB9633 | OFGB9633 | CFGB9633 | Firmicutes | Bacteria |
| Bacteria_unclassified | Bacteria_unclassified | Bacteria_unclassifie | Bacteria_unclassifie | Bacteria |
| Bacteria_unclassified | Bacteria_unclassified | Bacteria_unclassifie | Bacteria_unclassifie | Bacteria |
| Bacteria_unclassified | Bacteria_unclassified | Bacteria_unclassifie | Bacteria_unclassifie | Bacteria |
| Bacteria_unclassified | Bacteria_unclassified | Bacteria_unclassifie | Bacteria_unclassifie | Bacteria |
| Lachnospiraceae | Eubacteriales | Clostridia | Firmicutes | Bacteria |
| Lachnospiraceae | Eubacteriales | Clostridia | Firmicutes | Bacteria |
| Lachnospiraceae | Eubacteriales | Clostridia | Firmicutes | Bacteria |
| FGB77359 | OFGB77359 | CFGB77359 | Bacteria_unclassifie | Bacteria |
| FGB9639 | OFGB9639 | CFGB9639 | Firmicutes | Bacteria |
| Lachnospiraceae | Eubacteriales | Clostridia | Firmicutes | Bacteria |
| Lachnospiraceae | Eubacteriales | Clostridia | Firmicutes | Bacteria |
| Lachnospiraceae | Eubacteriales | Clostridia | Firmicutes | Bacteria |
| Clostridiaceae | Eubacteriales | Clostridia | Firmicutes | Bacteria |
| Clostridiaceae | Eubacteriales | Clostridia | Firmicutes | Bacteria |
| Clostridiaceae | Eubacteriales | Clostridia | Firmicutes | Bacteria |
| Clostridiaceae | Eubacteriales | Clostridia | Firmicutes | Bacteria |
| Clostridiaceae | Eubacteriales | Clostridia | Firmicutes | Bacteria |
| Clostridiaceae | Eubacteriales | Clostridia | Firmicutes | Bacteria |
| Clostridiaceae | Eubacteriales | Clostridia | Firmicutes | Bacteria |
| Eubacteriaceae | Eubacteriales | Clostridia | Firmicutes | Bacteria |
| FGB9658 | OFGB9658 | CFGB9658 | Firmicutes | Bacteria |
| FGB9659 | OFGB9659 | CFGB9659 | Firmicutes | Bacteria |
| Bacteria_unclassified | Bacteria_unclassified | Bacteria_unclassifie | Bacteria_unclassifie | Bacteria |
| FGB9827 | OFGB9827 | CFGB9827 | Firmicutes | Bacteria |
| Eubacteriaceae | Eubacteriales | Clostridia | Firmicutes | Bacteria |
| FGB77303 | OFGB77303 | CFGB77303 | Bacteria_unclassifie | Bacteria |
| Eubacteriales_unclassified | Eubacteriales | Clostridia | Firmicutes | Bacteria |
| Oscillospiraceae | Eubacteriales | Clostridia | Firmicutes | Bacteria |
| FGB30328 | OFGB30328 | CFGB30328 | Firmicutes | Bacteria |
| Oscillospiraceae | Eubacteriales | Clostridia | Firmicutes | Bacteria |
| Oscillospiraceae | Eubacteriales | Clostridia | Firmicutes | Bacteria |
| Oscillospiraceae | Eubacteriales | Clostridia | Firmicutes | Bacteria |
| Oscillospiraceae | Eubacteriales | Clostridia | Firmicutes | Bacteria |
| Oscillospiraceae | Eubacteriales | Clostridia | Firmicutes | Bacteria |
| Oscillospiraceae | Eubacteriales | Clostridia | Firmicutes | Bacteria |
| Oscillospiraceae | Eubacteriales | Clostridia | Firmicutes | Bacteria |
| FGB77153 | OFGB77153 | CFGB77153 | Actinobacteria | Bacteria |
| FGB1791 | OFGB1791 | CFGB1791 | Tenericutes | Bacteria |
| FGB10290 | OFGB10290 | CFGB10290 | Firmicutes | Bacteria |
| Oscillospiraceae | Eubacteriales | Clostridia | Firmicutes | Bacteria |

| | | | | |
|---|---|---|---|---|
| FGB1765 | OFGB1765 | CFGB1765 | Firmicutes | Bacteria |
| FGB10349 | OFGB10349 | CFGB10349 | Firmicutes | Bacteria |
| FGB10667 | OFGB10667 | CFGB10667 | Firmicutes | Bacteria |
| Lachnospiraceae | Eubacteriales | Clostridia | Firmicutes | Bacteria |
| Lachnospiraceae | Eubacteriales | Clostridia | Firmicutes | Bacteria |
| Christensenellaceae | Eubacteriales | Clostridia | Firmicutes | Bacteria |
| FGB10299 | OFGB10299 | CFGB10299 | Firmicutes | Bacteria |
| Oscillospiraceae | Eubacteriales | Clostridia | Firmicutes | Bacteria |
| Oscillospiraceae | Eubacteriales | Clostridia | Firmicutes | Bacteria |
| Lachnospiraceae | Eubacteriales | Clostridia | Firmicutes | Bacteria |
| Clostridia_unclassified | Clostridia_unclassifie | Clostridia | Firmicutes | Bacteria |
| Lachnospiraceae | Eubacteriales | Clostridia | Firmicutes | Bacteria |
| Lachnospiraceae | Eubacteriales | Clostridia | Firmicutes | Bacteria |
| Lachnospiraceae | Eubacteriales | Clostridia | Firmicutes | Bacteria |
| Lachnospiraceae | Eubacteriales | Clostridia | Firmicutes | Bacteria |
| Lachnospiraceae | Eubacteriales | Clostridia | Firmicutes | Bacteria |
| Lachnospiraceae | Eubacteriales | Clostridia | Firmicutes | Bacteria |
| Lachnospiraceae | Eubacteriales | Clostridia | Firmicutes | Bacteria |
| Lachnospiraceae | Eubacteriales | Clostridia | Firmicutes | Bacteria |
| Lactobacillaceae | Lactobacillales | Bacilli | Firmicutes | Bacteria |
| Muribaculaceae | Bacteroidales | Bacteroidia | Bacteroidota | Bacteria |
| Oscillospiraceae | Eubacteriales | Clostridia | Firmicutes | Bacteria |
| Oscillospiraceae | Eubacteriales | Clostridia | Firmicutes | Bacteria |
| Oscillospiraceae | Eubacteriales | Clostridia | Firmicutes | Bacteria |
| Oscillospiraceae | Eubacteriales | Clostridia | Firmicutes | Bacteria |
| Sutterellaceae | Burkholderiales | Betaproteobacteria | Proteobacteria | Bacteria |
| Peptostreptococcaceae | Eubacteriales | Clostridia | Firmicutes | Bacteria |
| Lachnospiraceae | Eubacteriales | Clostridia | Firmicutes | Bacteria |
| Turicibacteraceae | Erysipelotrichales | Erysipelotrichia | Firmicutes | Bacteria |
| Bacteria_unclassified | Bacteria_unclassified | Bacteria_unclassifie | Bacteria_unclassifie | Bacteria |
| Bacteria_unclassified | Bacteria_unclassified | Bacteria_unclassifie | Bacteria_unclassifie | Bacteria |
| Bacteria_unclassified | Bacteria_unclassified | Bacteria_unclassifie | Bacteria_unclassifie | Bacteria |

| | | | | |
|---|---|---|---|---|
| Lachnospiraceae | Eubacteriales | Clostridia | Firmicutes | Bacteria |
| Lachnospiraceae | Eubacteriales | Clostridia | Firmicutes | Bacteria |
| Oscillospiraceae | Eubacteriales | Clostridia | Firmicutes | Bacteria |
| Oscillospiraceae | Eubacteriales | Clostridia | Firmicutes | Bacteria |
| Eggerthellaceae | Eggerthellales | Coriobacteriia | Actinobacteria | Bacteria |
| Eggerthellaceae | Eggerthellales | Coriobacteriia | Actinobacteria | Bacteria |
| Eggerthellaceae | Eggerthellales | Coriobacteriia | Actinobacteria | Bacteria |
| Akkermansiaceae | Verrucomicrobiales | Verrucomicrobiae | Verrucomicrobia | Bacteria |
| Rikenellaceae | Bacteroidales | Bacteroidia | Bacteroidota | Bacteria |
| Oscillospiraceae | Eubacteriales | Clostridia | Firmicutes | Bacteria |
| Bacteria_unclassified | Bacteria_unclassified | Bacteria_unclassifie | Bacteria_unclassifie | Bacteria |

| | | | | |
|---|---|---|---|---|
| Bacteria_unclassified | Bacteria_unclassified | Bacteria_unclassifie | Bacteria_unclassifie | Bacteria |
| Bacteria_unclassified | Bacteria_unclassified | Bacteria_unclassifie | Bacteria_unclassifie | Bacteria |
| Bacteroidaceae | Bacteroidales | Bacteroidia | Bacteroidota | Bacteria |
| Bifidobacteriaceae | Bifidobacteriales | Actinomycetia | Actinobacteria | Bacteria |
| Clostridia_unclassified | Clostridia_unclassifie | Clostridia | Firmicutes | Bacteria |
| Clostridiaceae | Eubacteriales | Clostridia | Firmicutes | Bacteria |
| Clostridiaceae | Eubacteriales | Clostridia | Firmicutes | Bacteria |
| Eubacteriales_unclassified | Eubacteriales | Clostridia | Firmicutes | Bacteria |
| Erysipelotrichaceae | Erysipelotrichales | Erysipelotrichia | Firmicutes | Bacteria |
| Coriobacteriaceae | Coriobacteriales | Coriobacteriia | Actinobacteria | Bacteria |
| Lachnospiraceae | Eubacteriales | Clostridia | Firmicutes | Bacteria |
| Erysipelotrichaceae | Erysipelotrichales | Erysipelotrichia | Firmicutes | Bacteria |
| Erysipelotrichales_unclassifie | Erysipelotrichales | Erysipelotrichia | Firmicutes | Bacteria |
| Eubacteriaceae | Eubacteriales | Clostridia | Firmicutes | Bacteria |
| Eubacteriaceae | Eubacteriales | Clostridia | Firmicutes | Bacteria |
| Lachnospiraceae | Eubacteriales | Clostridia | Firmicutes | Bacteria |
| Eggerthellaceae | Eggerthellales | Coriobacteriia | Actinobacteria | Bacteria |
| Lachnospiraceae | Eubacteriales | Clostridia | Firmicutes | Bacteria |
| Muribaculaceae | Bacteroidales | Bacteroidia | Bacteroidota | Bacteria |
| Muribaculaceae | Bacteroidales | Bacteroidia | Bacteroidota | Bacteria |
| Muribaculaceae | Bacteroidales | Bacteroidia | Bacteroidota | Bacteria |
| FGB9508 | OFGB9508 | CFGB9508 | Firmicutes | Bacteria |
| FGB2838 | OFGB2838 | CFGB2838 | Firmicutes | Bacteria |
| FGB2838 | OFGB2838 | CFGB2838 | Firmicutes | Bacteria |
| FGB2838 | OFGB2838 | CFGB2838 | Firmicutes | Bacteria |
| Pumilibacteraceae | Eubacteriales | Clostridia | Firmicutes | Bacteria |
| FGB28439 | OFGB28439 | CFGB28439 | Firmicutes | Bacteria |
| Clostridia_unclassified | Clostridia_unclassifie | Clostridia | Firmicutes | Bacteria |
| Eubacteriaceae | Eubacteriales | Clostridia | Firmicutes | Bacteria |
| Lachnospiraceae | Eubacteriales | Clostridia | Firmicutes | Bacteria |
| Lachnospiraceae | Eubacteriales | Clostridia | Firmicutes | Bacteria |
| Lachnospiraceae | Eubacteriales | Clostridia | Firmicutes | Bacteria |
| Lachnospiraceae | Eubacteriales | Clostridia | Firmicutes | Bacteria |
| FGB77305 | OFGB77305 | CFGB77305 | Firmicutes | Bacteria |
| Clostridiaceae | Eubacteriales | Clostridia | Firmicutes | Bacteria |
| Lachnospiraceae | Eubacteriales | Clostridia | Firmicutes | Bacteria |
| Lachnospiraceae | Eubacteriales | Clostridia | Firmicutes | Bacteria |
| Lachnospiraceae | Eubacteriales | Clostridia | Firmicutes | Bacteria |
| FGB9633 | OFGB9633 | CFGB9633 | Firmicutes | Bacteria |
| Bacteria_unclassified | Bacteria_unclassified | Bacteria_unclassifie | Bacteria_unclassifie | Bacteria |
| Bacteria_unclassified | Bacteria_unclassified | Bacteria_unclassifie | Bacteria_unclassifie | Bacteria |
| Bacteria_unclassified | Bacteria_unclassified | Bacteria_unclassifie | Bacteria_unclassifie | Bacteria |
| Bacteria_unclassified | Bacteria_unclassified | Bacteria_unclassifie | Bacteria_unclassifie | Bacteria |
| Lachnospiraceae | Eubacteriales | Clostridia | Firmicutes | Bacteria |
| Lachnospiraceae | Eubacteriales | Clostridia | Firmicutes | Bacteria |
| Lachnospiraceae | Eubacteriales | Clostridia | Firmicutes | Bacteria |

| | | | | |
|---|---|---|---|---|
| FGB77359 | OFGB77359 | CFGB77359 | Bacteria_unclassifie | Bacteria |
| FGB9639 | OFGB9639 | CFGB9639 | Firmicutes | Bacteria |
| Lachnospiraceae | Eubacteriales | Clostridia | Firmicutes | Bacteria |
| Lachnospiraceae | Eubacteriales | Clostridia | Firmicutes | Bacteria |
| Lachnospiraceae | Eubacteriales | Clostridia | Firmicutes | Bacteria |
| Clostridiaceae | Eubacteriales | Clostridia | Firmicutes | Bacteria |
| Clostridiaceae | Eubacteriales | Clostridia | Firmicutes | Bacteria |
| Clostridiaceae | Eubacteriales | Clostridia | Firmicutes | Bacteria |
| Clostridiaceae | Eubacteriales | Clostridia | Firmicutes | Bacteria |
| Clostridiaceae | Eubacteriales | Clostridia | Firmicutes | Bacteria |
| Clostridiaceae | Eubacteriales | Clostridia | Firmicutes | Bacteria |
| Clostridiaceae | Eubacteriales | Clostridia | Firmicutes | Bacteria |
| Eubacteriaceae | Eubacteriales | Clostridia | Firmicutes | Bacteria |
| FGB9658 | OFGB9658 | CFGB9658 | Firmicutes | Bacteria |
| FGB9659 | OFGB9659 | CFGB9659 | Firmicutes | Bacteria |
| Bacteria_unclassified | Bacteria_unclassified | Bacteria_unclassifie | Bacteria_unclassifie | Bacteria |
| FGB9827 | OFGB9827 | CFGB9827 | Firmicutes | Bacteria |
| Eubacteriaceae | Eubacteriales | Clostridia | Firmicutes | Bacteria |
| FGB77303 | OFGB77303 | CFGB77303 | Bacteria_unclassifie | Bacteria |
| Eubacteriales_unclassified | Eubacteriales | Clostridia | Firmicutes | Bacteria |
| Oscillospiraceae | Eubacteriales | Clostridia | Firmicutes | Bacteria |
| FGB30328 | OFGB30328 | CFGB30328 | Firmicutes | Bacteria |
| Oscillospiraceae | Eubacteriales | Clostridia | Firmicutes | Bacteria |
| Oscillospiraceae | Eubacteriales | Clostridia | Firmicutes | Bacteria |
| Oscillospiraceae | Eubacteriales | Clostridia | Firmicutes | Bacteria |
| Oscillospiraceae | Eubacteriales | Clostridia | Firmicutes | Bacteria |
| Oscillospiraceae | Eubacteriales | Clostridia | Firmicutes | Bacteria |
| Oscillospiraceae | Eubacteriales | Clostridia | Firmicutes | Bacteria |
| Oscillospiraceae | Eubacteriales | Clostridia | Firmicutes | Bacteria |
| FGB77153 | OFGB77153 | CFGB77153 | Actinobacteria | Bacteria |
| FGB1791 | OFGB1791 | CFGB1791 | Tenericutes | Bacteria |
| FGB10290 | OFGB10290 | CFGB10290 | Firmicutes | Bacteria |
| Oscillospiraceae | Eubacteriales | Clostridia | Firmicutes | Bacteria |
| FGB1765 | OFGB1765 | CFGB1765 | Firmicutes | Bacteria |
| FGB10349 | OFGB10349 | CFGB10349 | Firmicutes | Bacteria |
| FGB10667 | OFGB10667 | CFGB10667 | Firmicutes | Bacteria |
| Lachnospiraceae | Eubacteriales | Clostridia | Firmicutes | Bacteria |
| Lachnospiraceae | Eubacteriales | Clostridia | Firmicutes | Bacteria |
| Christensenellaceae | Eubacteriales | Clostridia | Firmicutes | Bacteria |
| FGB10299 | OFGB10299 | CFGB10299 | Firmicutes | Bacteria |
| Oscillospiraceae | Eubacteriales | Clostridia | Firmicutes | Bacteria |
| Oscillospiraceae | Eubacteriales | Clostridia | Firmicutes | Bacteria |
| Lachnospiraceae | Eubacteriales | Clostridia | Firmicutes | Bacteria |
| Clostridia_unclassified | Clostridia_unclassifie | Clostridia | Firmicutes | Bacteria |
| Lachnospiraceae | Eubacteriales | Clostridia | Firmicutes | Bacteria |
| Lachnospiraceae | Eubacteriales | Clostridia | Firmicutes | Bacteria |

| | | | | |
|---|---|---|---|---|
| Lachnospiraceae | Eubacteriales | Clostridia | Firmicutes | Bacteria |
| Lachnospiraceae | Eubacteriales | Clostridia | Firmicutes | Bacteria |
| Lachnospiraceae | Eubacteriales | Clostridia | Firmicutes | Bacteria |
| Lachnospiraceae | Eubacteriales | Clostridia | Firmicutes | Bacteria |
| Lachnospiraceae | Eubacteriales | Clostridia | Firmicutes | Bacteria |
| Lachnospiraceae | Eubacteriales | Clostridia | Firmicutes | Bacteria |
| Lactobacillaceae | Lactobacillales | Bacilli | Firmicutes | Bacteria |
| Muribaculaceae | Bacteroidales | Bacteroidia | Bacteroidota | Bacteria |
| Oscillospiraceae | Eubacteriales | Clostridia | Firmicutes | Bacteria |
| Oscillospiraceae | Eubacteriales | Clostridia | Firmicutes | Bacteria |
| Oscillospiraceae | Eubacteriales | Clostridia | Firmicutes | Bacteria |
| Oscillospiraceae | Eubacteriales | Clostridia | Firmicutes | Bacteria |
| Sutterellaceae | Burkholderiales | Betaproteobacteria | Proteobacteria | Bacteria |
| Peptostreptococcaceae | Eubacteriales | Clostridia | Firmicutes | Bacteria |
| Lachnospiraceae | Eubacteriales | Clostridia | Firmicutes | Bacteria |
| Turicibacteraceae | Erysipelotrichales | Erysipelotrichia | Firmicutes | Bacteria |
| Bacteria_unclassified | Bacteria_unclassified | Bacteria_unclassifie | Bacteria_unclassifie | Bacteria |
| Bacteria_unclassified | Bacteria_unclassified | Bacteria_unclassifie | Bacteria_unclassifie | Bacteria |
| Bacteria_unclassified | Bacteria_unclassified | Bacteria_unclassifie | Bacteria_unclassifie | Bacteria |

| | | | | |
|---|---|---|---|---|
| Lachnospiraceae | Eubacteriales | Clostridia | Firmicutes | Bacteria |
| Lachnospiraceae | Eubacteriales | Clostridia | Firmicutes | Bacteria |
| Oscillospiraceae | Eubacteriales | Clostridia | Firmicutes | Bacteria |
| Oscillospiraceae | Eubacteriales | Clostridia | Firmicutes | Bacteria |
| Eggerthellaceae | Eggerthellales | Coriobacteriia | Actinobacteria | Bacteria |
| Eggerthellaceae | Eggerthellales | Coriobacteriia | Actinobacteria | Bacteria |
| Eggerthellaceae | Eggerthellales | Coriobacteriia | Actinobacteria | Bacteria |
| Akkermansiaceae | Verrucomicrobiales | Verrucomicrobiae | Verrucomicrobia | Bacteria |
| Rikenellaceae | Bacteroidales | Bacteroidia | Bacteroidota | Bacteria |
| Oscillospiraceae | Eubacteriales | Clostridia | Firmicutes | Bacteria |
| Bacteria_unclassified | Bacteria_unclassified | Bacteria_unclassifie | Bacteria_unclassifie | Bacteria |
| Bacteria_unclassified | Bacteria_unclassified | Bacteria_unclassifie | Bacteria_unclassifie | Bacteria |
| Bacteria_unclassified | Bacteria_unclassified | Bacteria_unclassifie | Bacteria_unclassifie | Bacteria |
| Bacteroidaceae | Bacteroidales | Bacteroidia | Bacteroidota | Bacteria |
| Bifidobacteriaceae | Bifidobacteriales | Actinomycetia | Actinobacteria | Bacteria |
| Clostridia_unclassified | Clostridia_unclassifie | Clostridia | Firmicutes | Bacteria |
| Clostridiaceae | Eubacteriales | Clostridia | Firmicutes | Bacteria |
| Clostridiaceae | Eubacteriales | Clostridia | Firmicutes | Bacteria |
| Eubacteriales_unclassified | Eubacteriales | Clostridia | Firmicutes | Bacteria |
| Erysipelotrichaceae | Erysipelotrichales | Erysipelotrichia | Firmicutes | Bacteria |
| Coriobacteriaceae | Coriobacteriales | Coriobacteriia | Actinobacteria | Bacteria |
| Lachnospiraceae | Eubacteriales | Clostridia | Firmicutes | Bacteria |
| Erysipelotrichaceae | Erysipelotrichales | Erysipelotrichia | Firmicutes | Bacteria |
| Erysipelotrichales_unclassifie | Erysipelotrichales | Erysipelotrichia | Firmicutes | Bacteria |

| | | | | |
|---|---|---|---|---|
| Eubacteriaceae | Eubacteriales | Clostridia | Firmicutes | Bacteria |
| Eubacteriaceae | Eubacteriales | Clostridia | Firmicutes | Bacteria |
| Lachnospiraceae | Eubacteriales | Clostridia | Firmicutes | Bacteria |
| Eggerthellaceae | Eggerthellales | Coriobacteriia | Actinobacteria | Bacteria |
| Lachnospiraceae | Eubacteriales | Clostridia | Firmicutes | Bacteria |
| Muribaculaceae | Bacteroidales | Bacteroidia | Bacteroidota | Bacteria |
| Muribaculaceae | Bacteroidales | Bacteroidia | Bacteroidota | Bacteria |
| Muribaculaceae | Bacteroidales | Bacteroidia | Bacteroidota | Bacteria |
| FGB9508 | OFGB9508 | CFGB9508 | Firmicutes | Bacteria |
| FGB2838 | OFGB2838 | CFGB2838 | Firmicutes | Bacteria |
| FGB2838 | OFGB2838 | CFGB2838 | Firmicutes | Bacteria |
| FGB2838 | OFGB2838 | CFGB2838 | Firmicutes | Bacteria |
| Pumilibacteraceae | Eubacteriales | Clostridia | Firmicutes | Bacteria |
| FGB28439 | OFGB28439 | CFGB28439 | Firmicutes | Bacteria |
| Clostridia_unclassified | Clostridia_unclassifie | Clostridia | Firmicutes | Bacteria |
| Eubacteriaceae | Eubacteriales | Clostridia | Firmicutes | Bacteria |
| Lachnospiraceae | Eubacteriales | Clostridia | Firmicutes | Bacteria |
| Lachnospiraceae | Eubacteriales | Clostridia | Firmicutes | Bacteria |
| Lachnospiraceae | Eubacteriales | Clostridia | Firmicutes | Bacteria |
| Lachnospiraceae | Eubacteriales | Clostridia | Firmicutes | Bacteria |
| FGB77305 | OFGB77305 | CFGB77305 | Firmicutes | Bacteria |
| Clostridiaceae | Eubacteriales | Clostridia | Firmicutes | Bacteria |
| Lachnospiraceae | Eubacteriales | Clostridia | Firmicutes | Bacteria |
| Lachnospiraceae | Eubacteriales | Clostridia | Firmicutes | Bacteria |
| Lachnospiraceae | Eubacteriales | Clostridia | Firmicutes | Bacteria |
| FGB9633 | OFGB9633 | CFGB9633 | Firmicutes | Bacteria |
| Bacteria_unclassified | Bacteria_unclassified | Bacteria_unclassifie | Bacteria_unclassifie | Bacteria |
| Bacteria_unclassified | Bacteria_unclassified | Bacteria_unclassifie | Bacteria_unclassifie | Bacteria |
| Bacteria_unclassified | Bacteria_unclassified | Bacteria_unclassifie | Bacteria_unclassifie | Bacteria |
| Bacteria_unclassified | Bacteria_unclassified | Bacteria_unclassifie | Bacteria_unclassifie | Bacteria |
| Lachnospiraceae | Eubacteriales | Clostridia | Firmicutes | Bacteria |
| Lachnospiraceae | Eubacteriales | Clostridia | Firmicutes | Bacteria |
| Lachnospiraceae | Eubacteriales | Clostridia | Firmicutes | Bacteria |
| FGB77359 | OFGB77359 | CFGB77359 | Bacteria_unclassifie | Bacteria |
| FGB9639 | OFGB9639 | CFGB9639 | Firmicutes | Bacteria |
| Lachnospiraceae | Eubacteriales | Clostridia | Firmicutes | Bacteria |
| Lachnospiraceae | Eubacteriales | Clostridia | Firmicutes | Bacteria |
| Lachnospiraceae | Eubacteriales | Clostridia | Firmicutes | Bacteria |
| Clostridiaceae | Eubacteriales | Clostridia | Firmicutes | Bacteria |
| Clostridiaceae | Eubacteriales | Clostridia | Firmicutes | Bacteria |
| Clostridiaceae | Eubacteriales | Clostridia | Firmicutes | Bacteria |
| Clostridiaceae | Eubacteriales | Clostridia | Firmicutes | Bacteria |
| Clostridiaceae | Eubacteriales | Clostridia | Firmicutes | Bacteria |
| Clostridiaceae | Eubacteriales | Clostridia | Firmicutes | Bacteria |
| Clostridiaceae | Eubacteriales | Clostridia | Firmicutes | Bacteria |
| Eubacteriaceae | Eubacteriales | Clostridia | Firmicutes | Bacteria |

| | | | | |
|---|---|---|---|---|
| FGB9658 | OFGB9658 | CFGB9658 | Firmicutes | Bacteria |
| FGB9659 | OFGB9659 | CFGB9659 | Firmicutes | Bacteria |
| Bacteria_unclassified | Bacteria_unclassified | Bacteria_unclassifie | Bacteria_unclassifie | Bacteria |
| FGB9827 | OFGB9827 | CFGB9827 | Firmicutes | Bacteria |
| Eubacteriaceae | Eubacteriales | Clostridia | Firmicutes | Bacteria |
| FGB77303 | OFGB77303 | CFGB77303 | Bacteria_unclassifie | Bacteria |
| Eubacteriales_unclassified | Eubacteriales | Clostridia | Firmicutes | Bacteria |
| Oscillospiraceae | Eubacteriales | Clostridia | Firmicutes | Bacteria |
| FGB30328 | OFGB30328 | CFGB30328 | Firmicutes | Bacteria |
| Oscillospiraceae | Eubacteriales | Clostridia | Firmicutes | Bacteria |
| Oscillospiraceae | Eubacteriales | Clostridia | Firmicutes | Bacteria |
| Oscillospiraceae | Eubacteriales | Clostridia | Firmicutes | Bacteria |
| Oscillospiraceae | Eubacteriales | Clostridia | Firmicutes | Bacteria |
| Oscillospiraceae | Eubacteriales | Clostridia | Firmicutes | Bacteria |
| Oscillospiraceae | Eubacteriales | Clostridia | Firmicutes | Bacteria |
| Oscillospiraceae | Eubacteriales | Clostridia | Firmicutes | Bacteria |
| FGB77153 | OFGB77153 | CFGB77153 | Actinobacteria | Bacteria |
| FGB1791 | OFGB1791 | CFGB1791 | Tenericutes | Bacteria |
| FGB10290 | OFGB10290 | CFGB10290 | Firmicutes | Bacteria |
| Oscillospiraceae | Eubacteriales | Clostridia | Firmicutes | Bacteria |
| FGB1765 | OFGB1765 | CFGB1765 | Firmicutes | Bacteria |
| FGB10349 | OFGB10349 | CFGB10349 | Firmicutes | Bacteria |
| FGB10667 | OFGB10667 | CFGB10667 | Firmicutes | Bacteria |
| Lachnospiraceae | Eubacteriales | Clostridia | Firmicutes | Bacteria |
| Lachnospiraceae | Eubacteriales | Clostridia | Firmicutes | Bacteria |
| Christensenellaceae | Eubacteriales | Clostridia | Firmicutes | Bacteria |
| FGB10299 | OFGB10299 | CFGB10299 | Firmicutes | Bacteria |
| Oscillospiraceae | Eubacteriales | Clostridia | Firmicutes | Bacteria |
| Oscillospiraceae | Eubacteriales | Clostridia | Firmicutes | Bacteria |
| Lachnospiraceae | Eubacteriales | Clostridia | Firmicutes | Bacteria |
| Clostridia_unclassified | Clostridia_unclassifie | Clostridia | Firmicutes | Bacteria |
| Lachnospiraceae | Eubacteriales | Clostridia | Firmicutes | Bacteria |
| Lachnospiraceae | Eubacteriales | Clostridia | Firmicutes | Bacteria |
| Lachnospiraceae | Eubacteriales | Clostridia | Firmicutes | Bacteria |
| Lachnospiraceae | Eubacteriales | Clostridia | Firmicutes | Bacteria |
| Lachnospiraceae | Eubacteriales | Clostridia | Firmicutes | Bacteria |
| Lachnospiraceae | Eubacteriales | Clostridia | Firmicutes | Bacteria |
| Lachnospiraceae | Eubacteriales | Clostridia | Firmicutes | Bacteria |
| Lachnospiraceae | Eubacteriales | Clostridia | Firmicutes | Bacteria |
| Lactobacillaceae | Lactobacillales | Bacilli | Firmicutes | Bacteria |
| Muribaculaceae | Bacteroidales | Bacteroidia | Bacteroidota | Bacteria |
| Oscillospiraceae | Eubacteriales | Clostridia | Firmicutes | Bacteria |
| Oscillospiraceae | Eubacteriales | Clostridia | Firmicutes | Bacteria |
| Oscillospiraceae | Eubacteriales | Clostridia | Firmicutes | Bacteria |
| Oscillospiraceae | Eubacteriales | Clostridia | Firmicutes | Bacteria |
| Sutterellaceae | Burkholderiales | Betaproteobacteria | Proteobacteria | Bacteria |

| | | | | |
|---|---|---|---|---|
| Peptostreptococcaceae | Eubacteriales | Clostridia | Firmicutes | Bacteria |
| Lachnospiraceae | Eubacteriales | Clostridia | Firmicutes | Bacteria |
| Turicibacteraceae | Erysipelotrichales | Erysipelotrichia | Firmicutes | Bacteria |
| Bacteria_unclassified | Bacteria_unclassified | Bacteria_unclassifie | Bacteria_unclassifie | Bacteria |
| Bacteria_unclassified | Bacteria_unclassified | Bacteria_unclassifie | Bacteria_unclassifie | Bacteria |
| Bacteria_unclassified | Bacteria_unclassified | Bacteria_unclassifie | Bacteria_unclassifie | Bacteria |

| MetaPhlan Annotation |
|---|
| k__Bacteria\|p__Firmicutes\|c__Clostridia\|o__Eubacteriales\|f__Lachnospiraceae\|g__Acetatifactor\|s__Acetatifactor_SGB415 |
| k__Bacteria\|p__Firmicutes\|c__Clostridia\|o__Eubacteriales\|f__Lachnospiraceae\|g__Acetatifactor\|s__Acetatifactor_muris |
| k__Bacteria\|p__Firmicutes\|c__Clostridia\|o__Eubacteriales\|f__Oscillospiraceae\|g__Acutalibacter\|s__Acutalibacter_muris |
| k__Bacteria\|p__Firmicutes\|c__Clostridia\|o__Eubacteriales\|f__Oscillospiraceae\|g__Acutalibacter\|s__Acutalibacter_sp_1XD |
| k__Bacteria\|p__Actinobacteria\|c__Coriobacteriia\|o__Eggerthellales\|f__Eggerthellaceae\|g__Adlercreutzia\|s__Adlercreutzia |
| k__Bacteria\|p__Actinobacteria\|c__Coriobacteriia\|o__Eggerthellales\|f__Eggerthellaceae\|g__Adlercreutzia\|s__Adlercreutzia |
| k__Bacteria\|p__Actinobacteria\|c__Coriobacteriia\|o__Eggerthellales\|f__Eggerthellaceae\|g__Adlercreutzia\|s__Adlercreutzia |
| k__Bacteria\|p__Verrucomicrobia\|c__Verrucomicrobiae\|o__Verrucomicrobiales\|f__Akkermansiaceae\|g__Akkermansia\|s__A |
| k__Bacteria\|p__Bacteroidota\|c__Bacteroidia\|o__Bacteroidales\|f__Rikenellaceae\|g__Alistipes\|s__Alistipes_sp_DSM_11234 |
| k__Bacteria\|p__Firmicutes\|c__Clostridia\|o__Eubacteriales\|f__Oscillospiraceae\|g__Anaerotruncus\|s__Anaerotruncus_sp_1 |
| k__Bacteria\|p__Bacteria_unclassified\|c__Bacteria_unclassified\|o__Bacteria_unclassified\|f__Bacteria_unclassified\|g__Bact |
| k__Bacteria\|p__Bacteria_unclassified\|c__Bacteria_unclassified\|o__Bacteria_unclassified\|f__Bacteria_unclassified\|g__Bact |
| k__Bacteria\|p__Bacteria_unclassified\|c__Bacteria_unclassified\|o__Bacteria_unclassified\|f__Bacteria_unclassified\|g__Bact |
| k__Bacteria\|p__Bacteroidota\|c__Bacteroidia\|o__Bacteroidales\|f__Bacteroidaceae\|g__Bacteroides\|s__Bacteroides_thetaio |
| k__Bacteria\|p__Actinobacteria\|c__Actinomycetia\|o__Bifidobacteriales\|f__Bifidobacteriaceae\|g__Bifidobacterium\|s__Bifid |
| k__Bacteria\|p__Firmicutes\|c__Clostridia\|o__Clostridia_unclassified\|f__Clostridia_unclassified\|g__Clostridia_unclassified\|s_ |
| k__Bacteria\|p__Firmicutes\|c__Clostridia\|o__Eubacteriales\|f__Clostridiaceae\|g__Clostridiaceae_unclassified\|s__Clostridiac |
| k__Bacteria\|p__Firmicutes\|c__Clostridia\|o__Eubacteriales\|f__Clostridiaceae\|g__Clostridiaceae_unclassified\|s__Clostridiac |
| k__Bacteria\|p__Firmicutes\|c__Clostridia\|o__Eubacteriales\|f__Eubacteriales_unclassified\|g__Eubacteriales_unclassified\|s_ |
| k__Bacteria\|p__Firmicutes\|c__Erysipelotrichia\|o__Erysipelotrichales\|f__Erysipelotrichaceae\|g__Erysipelatoclostridium\|s_ |
| k__Bacteria\|p__Actinobacteria\|c__Coriobacteriia\|o__Coriobacteriales\|f__Coriobacteriaceae\|g__Coriobacteriaceae_unclass |
| k__Bacteria\|p__Firmicutes\|c__Clostridia\|o__Eubacteriales\|f__Lachnospiraceae\|g__Dorea\|s__Dorea_sp_5_2 |
| k__Bacteria\|p__Firmicutes\|c__Erysipelotrichia\|o__Erysipelotrichales\|f__Erysipelotrichaceae\|g__Dubosiella\|s__Dubosiella_ |
| k__Bacteria\|p__Firmicutes\|c__Erysipelotrichia\|o__Erysipelotrichales\|f__Erysipelotrichales_unclassified\|g__Erysipelotrichal |
| k__Bacteria\|p__Firmicutes\|c__Clostridia\|o__Eubacteriales\|f__Eubacteriaceae\|g__Eubacteriaceae_unclassified\|s__Eubacte |
| k__Bacteria\|p__Firmicutes\|c__Clostridia\|o__Eubacteriales\|f__Eubacteriaceae\|g__Eubacteriaceae_unclassified\|s__Eubacte |
| k__Bacteria\|p__Firmicutes\|c__Clostridia\|o__Eubacteriales\|f__Lachnospiraceae\|g__GGB20149\|s__GGB20149_SGB29430 |
| k__Bacteria\|p__Actinobacteria\|c__Coriobacteriia\|o__Eggerthellales\|f__Eggerthellaceae\|g__GGB22635\|s__GGB22635_SGB |
| k__Bacteria\|p__Firmicutes\|c__Clostridia\|o__Eubacteriales\|f__Lachnospiraceae\|g__GGB25041\|s__GGB25041_SGB36960 |
| k__Bacteria\|p__Bacteroidota\|c__Bacteroidia\|o__Bacteroidales\|f__Muribaculaceae\|g__GGB27876\|s__GGB27876_SGB403 |
| k__Bacteria\|p__Bacteroidota\|c__Bacteroidia\|o__Bacteroidales\|f__Muribaculaceae\|g__GGB27878\|s__GGB27878_SGB403 |
| k__Bacteria\|p__Bacteroidota\|c__Bacteroidia\|o__Bacteroidales\|f__Muribaculaceae\|g__GGB27918\|s__GGB27918_SGB403 |
| k__Bacteria\|p__Firmicutes\|c__CFGB9508\|o__OFGB9508\|f__FGB9508\|g__GGB28382\|s__GGB28382_SGB40962 |
| k__Bacteria\|p__Firmicutes\|c__CFGB2838\|o__OFGB2838\|f__FGB2838\|g__GGB28399\|s__GGB28399_SGB40980 |
| k__Bacteria\|p__Firmicutes\|c__CFGB2838\|o__OFGB2838\|f__FGB2838\|g__GGB28411\|s__GGB28411_SGB40993 |
| k__Bacteria\|p__Firmicutes\|c__CFGB2838\|o__OFGB2838\|f__FGB2838\|g__GGB28415\|s__GGB28415_SGB40997 |
| k__Bacteria\|p__Firmicutes\|c__Clostridia\|o__Eubacteriales\|f__Pumilibacteraceae\|g__GGB28430\|s__GGB28430_SGB41013 |
| k__Bacteria\|p__Firmicutes\|c__CFGB28439\|o__OFGB28439\|f__FGB28439\|g__GGB28439\|s__GGB28439_SGB41022 |
| k__Bacteria\|p__Firmicutes\|c__Clostridia\|o__Clostridia_unclassified\|f__Clostridia_unclassified\|g__GGB28778\|s__GGB2877 |
| k__Bacteria\|p__Firmicutes\|c__Clostridia\|o__Eubacteriales\|f__Eubacteriaceae\|g__GGB28784\|s__GGB28784_SGB41437 |
| k__Bacteria\|p__Firmicutes\|c__Clostridia\|o__Eubacteriales\|f__Lachnospiraceae\|g__GGB28792\|s__GGB28792_SGB41445 |
| k__Bacteria\|p__Firmicutes\|c__Clostridia\|o__Eubacteriales\|f__Lachnospiraceae\|g__GGB28798\|s__GGB28798_SGB41451 |
| k__Bacteria\|p__Firmicutes\|c__Clostridia\|o__Eubacteriales\|f__Lachnospiraceae\|g__GGB28802\|s__GGB28802_SGB41455 |
| k__Bacteria\|p__Firmicutes\|c__Clostridia\|o__Eubacteriales\|f__Lachnospiraceae\|g__GGB28818\|s__GGB28818_SGB41473 |
| k__Bacteria\|p__Firmicutes\|c__CFGB77305\|o__OFGB77305\|f__FGB77305\|g__GGB28828\|s__GGB28828_SGB41484 |

k__Bacteria|p__Firmicutes|c__Clostridia|o__Eubacteriales|f__Clostridiaceae|g__GGB28851|s__GGB28851_SGB41518
k__Bacteria|p__Firmicutes|c__Clostridia|o__Eubacteriales|f__Lachnospiraceae|g__GGB28859|s__GGB28859_SGB41528
k__Bacteria|p__Firmicutes|c__Clostridia|o__Eubacteriales|f__Lachnospiraceae|g__GGB28864|s__GGB28864_SGB41535
k__Bacteria|p__Firmicutes|c__Clostridia|o__Eubacteriales|f__Lachnospiraceae|g__GGB28869|s__GGB28869_SGB41543
k__Bacteria|p__Firmicutes|c__CFGB9633|o__OFGB9633|f__FGB9633|g__GGB28883|s__GGB28883_SGB41564
k__Bacteria|p__Bacteria_unclassified|c__Bacteria_unclassified|o__Bacteria_unclassified|f__Bacteria_unclassified|g__GGB2
k__Bacteria|p__Bacteria_unclassified|c__Bacteria_unclassified|o__Bacteria_unclassified|f__Bacteria_unclassified|g__GGB2
k__Bacteria|p__Bacteria_unclassified|c__Bacteria_unclassified|o__Bacteria_unclassified|f__Bacteria_unclassified|g__GGB2
k__Bacteria|p__Bacteria_unclassified|c__Bacteria_unclassified|o__Bacteria_unclassified|f__Bacteria_unclassified|g__GGB2
k__Bacteria|p__Firmicutes|c__Clostridia|o__Eubacteriales|f__Lachnospiraceae|g__GGB28916|s__GGB28916_SGB41612
k__Bacteria|p__Firmicutes|c__Clostridia|o__Eubacteriales|f__Lachnospiraceae|g__GGB28924|s__GGB28924_SGB41621
k__Bacteria|p__Firmicutes|c__Clostridia|o__Eubacteriales|f__Lachnospiraceae|g__GGB28926|s__GGB28926_SGB41624
k__Bacteria|p__Bacteria_unclassified|c__CFGB77359|o__OFGB77359|f__FGB77359|g__GGB28927|s__GGB28927_SGB416
k__Bacteria|p__Firmicutes|c__CFGB9639|o__OFGB9639|f__FGB9639|g__GGB28934|s__GGB28934_SGB41635
k__Bacteria|p__Firmicutes|c__Clostridia|o__Eubacteriales|f__Lachnospiraceae|g__GGB28946|s__GGB28946_SGB41652
k__Bacteria|p__Firmicutes|c__Clostridia|o__Eubacteriales|f__Lachnospiraceae|g__GGB28949|s__GGB28949_SGB41655
k__Bacteria|p__Firmicutes|c__Clostridia|o__Eubacteriales|f__Lachnospiraceae|g__GGB28949|s__GGB28949_SGB41656
k__Bacteria|p__Firmicutes|c__Clostridia|o__Eubacteriales|f__Clostridiaceae|g__GGB28950|s__GGB28950_SGB41657
k__Bacteria|p__Firmicutes|c__Clostridia|o__Eubacteriales|f__Clostridiaceae|g__GGB28951|s__GGB28951_SGB102295
k__Bacteria|p__Firmicutes|c__Clostridia|o__Eubacteriales|f__Clostridiaceae|g__GGB28951|s__GGB28951_SGB41658
k__Bacteria|p__Firmicutes|c__Clostridia|o__Eubacteriales|f__Clostridiaceae|g__GGB28954|s__GGB28954_SGB41662
k__Bacteria|p__Firmicutes|c__Clostridia|o__Eubacteriales|f__Clostridiaceae|g__GGB28956|s__GGB28956_SGB41665
k__Bacteria|p__Firmicutes|c__Clostridia|o__Eubacteriales|f__Clostridiaceae|g__GGB28960|s__GGB28960_SGB41669
k__Bacteria|p__Firmicutes|c__Clostridia|o__Eubacteriales|f__Clostridiaceae|g__GGB28967|s__GGB28967_SGB41678
k__Bacteria|p__Firmicutes|c__Clostridia|o__Eubacteriales|f__Eubacteriaceae|g__GGB28991|s__GGB28991_SGB41705
k__Bacteria|p__Firmicutes|c__CFGB9658|o__OFGB9658|f__FGB9658|g__GGB29002|s__GGB29002_SGB41718
k__Bacteria|p__Firmicutes|c__CFGB9659|o__OFGB9659|f__FGB9659|g__GGB29003|s__GGB29003_SGB41719
k__Bacteria|p__Bacteria_unclassified|c__Bacteria_unclassified|o__Bacteria_unclassified|f__Bacteria_unclassified|g__GGB2
k__Bacteria|p__Firmicutes|c__CFGB9827|o__OFGB9827|f__FGB9827|g__GGB29531|s__GGB29531_SGB42317
k__Bacteria|p__Firmicutes|c__Clostridia|o__Eubacteriales|f__Eubacteriaceae|g__GGB29685|s__GGB29685_SGB42494
k__Bacteria|p__Bacteria_unclassified|c__CFGB77303|o__OFGB77303|f__FGB77303|g__GGB30141|s__GGB30141_SGB430
k__Bacteria|p__Firmicutes|c__Clostridia|o__Eubacteriales|f__Eubacteriales_unclassified|g__GGB30286|s__GGB30286_SG
k__Bacteria|p__Firmicutes|c__Clostridia|o__Eubacteriales|f__Oscillospiraceae|g__GGB30303|s__GGB30303_SGB43268
k__Bacteria|p__Firmicutes|c__CFGB30328|o__OFGB30328|f__FGB30328|g__GGB30413|s__GGB30413_SGB43452
k__Bacteria|p__Firmicutes|c__Clostridia|o__Eubacteriales|f__Oscillospiraceae|g__GGB30454|s__GGB30454_SGB43514
k__Bacteria|p__Firmicutes|c__Clostridia|o__Eubacteriales|f__Oscillospiraceae|g__GGB30455|s__GGB30455_SGB43519
k__Bacteria|p__Firmicutes|c__Clostridia|o__Eubacteriales|f__Oscillospiraceae|g__GGB30461|s__GGB30461_SGB43527
k__Bacteria|p__Firmicutes|c__Clostridia|o__Eubacteriales|f__Oscillospiraceae|g__GGB30461|s__GGB30461_SGB43530
k__Bacteria|p__Firmicutes|c__Clostridia|o__Eubacteriales|f__Oscillospiraceae|g__GGB30463|s__GGB30463_SGB43537
k__Bacteria|p__Firmicutes|c__Clostridia|o__Eubacteriales|f__Oscillospiraceae|g__GGB30473|s__GGB30473_SGB43557
k__Bacteria|p__Firmicutes|c__Clostridia|o__Eubacteriales|f__Oscillospiraceae|g__GGB30475|s__GGB30475_SGB63182
k__Bacteria|p__Actinobacteria|c__CFGB77153|o__OFGB77153|f__FGB77153|g__GGB30861|s__GGB30861_SGB44083
k__Bacteria|p__Tenericutes|c__CFGB1791|o__OFGB1791|f__FGB1791|g__GGB31312|s__GGB31312_SGB44628
k__Bacteria|p__Firmicutes|c__CFGB10290|o__OFGB10290|f__FGB10290|g__GGB31438|s__GGB31438_SGB44768
k__Bacteria|p__Firmicutes|c__Clostridia|o__Eubacteriales|f__Oscillospiraceae|g__GGB3171|s__GGB3171_SGB4185
k__Bacteria|p__Firmicutes|c__CFGB1765|o__OFGB1765|f__FGB1765|g__GGB31823|s__GGB31823_SGB45199

k__Bacteria|p__Firmicutes|c__CFGB10349|o__OFGB10349|f__FGB10349|g__GGB31853|s__GGB31853_SGB45233
k__Bacteria|p__Firmicutes|c__CFGB10667|o__OFGB10667|f__FGB10667|g__GGB32371|s__GGB32371_SGB41694
k__Bacteria|p__Firmicutes|c__Clostridia|o__Eubacteriales|f__Lachnospiraceae|g__GGB3793|s__GGB3793_SGB5158
k__Bacteria|p__Firmicutes|c__Clostridia|o__Eubacteriales|f__Lachnospiraceae|g__GGB42598|s__GGB42598_SGB59794
k__Bacteria|p__Firmicutes|c__Clostridia|o__Eubacteriales|f__Christensenellaceae|g__GGB45656|s__GGB45656_SGB6337
k__Bacteria|p__Firmicutes|c__CFGB10299|o__OFGB10299|f__FGB10299|g__GGB47127|s__GGB47127_SGB65054
k__Bacteria|p__Firmicutes|c__Clostridia|o__Eubacteriales|f__Oscillospiraceae|g__GGB74395|s__GGB74395_SGB43521
k__Bacteria|p__Firmicutes|c__Clostridia|o__Eubacteriales|f__Oscillospiraceae|g__GGB75053|s__GGB75053_SGB43494
k__Bacteria|p__Firmicutes|c__Clostridia|o__Eubacteriales|f__Lachnospiraceae|g__GGB75109|s__GGB75109_SGB102238
k__Bacteria|p__Firmicutes|c__Clostridia|o__Clostridia_unclassified|f__Clostridia_unclassified|g__GGB81440|s__GGB8144
k__Bacteria|p__Firmicutes|c__Clostridia|o__Eubacteriales|f__Lachnospiraceae|g__Lachnospiraceae_unclassified|s__Lachn
k__Bacteria|p__Firmicutes|c__Clostridia|o__Eubacteriales|f__Lachnospiraceae|g__Lachnospiraceae_unclassified|s__Lachn
k__Bacteria|p__Firmicutes|c__Clostridia|o__Eubacteriales|f__Lachnospiraceae|g__Lachnospiraceae_unclassified|s__Lachn
k__Bacteria|p__Firmicutes|c__Clostridia|o__Eubacteriales|f__Lachnospiraceae|g__Lachnospiraceae_unclassified|s__Lachn
k__Bacteria|p__Firmicutes|c__Clostridia|o__Eubacteriales|f__Lachnospiraceae|g__Lachnospiraceae_unclassified|s__Lachn
k__Bacteria|p__Firmicutes|c__Clostridia|o__Eubacteriales|f__Lachnospiraceae|g__Lachnospiraceae_unclassified|s__Lachn
k__Bacteria|p__Firmicutes|c__Clostridia|o__Eubacteriales|f__Lachnospiraceae|g__Lachnospiraceae_unclassified|s__Lachn
k__Bacteria|p__Firmicutes|c__Clostridia|o__Eubacteriales|f__Lachnospiraceae|g__Lachnospiraceae_unclassified|s__Lachn
k__Bacteria|p__Firmicutes|c__Bacilli|o__Lactobacillales|f__Lactobacillaceae|g__Lactobacillus|s__Lactobacillus_johnsonii
k__Bacteria|p__Bacteroidota|c__Bacteroidia|o__Bacteroidales|f__Muribaculaceae|g__Muribaculaceae_unclassified|s__M
k__Bacteria|p__Firmicutes|c__Clostridia|o__Eubacteriales|f__Oscillospiraceae|g__Neglectibacter|s__Neglectibacter_sp_X
k__Bacteria|p__Firmicutes|c__Clostridia|o__Eubacteriales|f__Oscillospiraceae|g__Oscillospiraceae_unclassified|s__Oscillo
k__Bacteria|p__Firmicutes|c__Clostridia|o__Eubacteriales|f__Oscillospiraceae|g__Oscillospiraceae_unclassified|s__Oscillo
k__Bacteria|p__Firmicutes|c__Clostridia|o__Eubacteriales|f__Oscillospiraceae|g__Oscillospiraceae_unclassified|s__Oscillo
k__Bacteria|p__Proteobacteria|c__Betaproteobacteria|o__Burkholderiales|f__Sutterellaceae|g__Parasutterella|s__Parasu
k__Bacteria|p__Firmicutes|c__Clostridia|o__Eubacteriales|f__Peptostreptococcaceae|g__Romboutsia|s__Romboutsia_ilea
k__Bacteria|p__Firmicutes|c__Clostridia|o__Eubacteriales|f__Lachnospiraceae|g__Schaedlerella|s__Schaedlerella_arabinc
k__Bacteria|p__Firmicutes|c__Erysipelotrichia|o__Erysipelotrichales|f__Turicibacteraceae|g__Turicibacter|s__Turicibacter_
k__Bacteria|p__Bacteria_unclassified|c__Bacteria_unclassified|o__Bacteria_unclassified|f__Bacteria_unclassified|g__Bacte
k__Bacteria|p__Bacteria_unclassified|c__Bacteria_unclassified|o__Bacteria_unclassified|f__Bacteria_unclassified|g__Bacte
k__Bacteria|p__Bacteria_unclassified|c__Bacteria_unclassified|o__Bacteria_unclassified|f__Bacteria_unclassified|g__Bacte

k__Bacteria|p__Firmicutes|c__Clostridia|o__Eubacteriales|f__Lachnospiraceae|g__Acetatifactor|s__Acetatifactor_SGB415
k__Bacteria|p__Firmicutes|c__Clostridia|o__Eubacteriales|f__Lachnospiraceae|g__Acetatifactor|s__Acetatifactor_muris
k__Bacteria|p__Firmicutes|c__Clostridia|o__Eubacteriales|f__Oscillospiraceae|g__Acutalibacter|s__Acutalibacter_muris
k__Bacteria|p__Firmicutes|c__Clostridia|o__Eubacteriales|f__Oscillospiraceae|g__Acutalibacter|s__Acutalibacter_sp_1XD
k__Bacteria|p__Actinobacteria|c__Coriobacteriia|o__Eggerthellales|f__Eggerthellaceae|g__Adlercreutzia|s__Adlercreutzia
k__Bacteria|p__Actinobacteria|c__Coriobacteriia|o__Eggerthellales|f__Eggerthellaceae|g__Adlercreutzia|s__Adlercreutzia
k__Bacteria|p__Actinobacteria|c__Coriobacteriia|o__Eggerthellales|f__Eggerthellaceae|g__Adlercreutzia|s__Adlercreutzia
k__Bacteria|p__Verrucomicrobia|c__Verrucomicrobiae|o__Verrucomicrobiales|f__Akkermansiaceae|g__Akkermansia|s__
k__Bacteria|p__Bacteroidota|c__Bacteroidia|o__Bacteroidales|f__Rikenellaceae|g__Alistipes|s__Alistipes_sp_DSM_11234
k__Bacteria|p__Firmicutes|c__Clostridia|o__Eubacteriales|f__Oscillospiraceae|g__Anaerotruncus|s__Anaerotruncus_sp_1
k__Bacteria|p__Bacteria_unclassified|c__Bacteria_unclassified|o__Bacteria_unclassified|f__Bacteria_unclassified|g__Bacte
k__Bacteria|p__Bacteria_unclassified|c__Bacteria_unclassified|o__Bacteria_unclassified|f__Bacteria_unclassified|g__Bacte

k__Bacteria|p__Bacteria_unclassified|c__Bacteria_unclassified|o__Bacteria_unclassified|f__Bacteria_unclassified|g__Bacte

k__Bacteria|p__Bacteroidota|c__Bacteroidia|o__Bacteroidales|f__Bacteroidaceae|g__Bacteroides|s__Bacteroides_thetaio

k__Bacteria|p__Actinobacteria|c__Actinomycetia|o__Bifidobacteriales|f__Bifidobacteriaceae|g__Bifidobacterium|s__Bifid

k__Bacteria|p__Firmicutes|c__Clostridia|o__Clostridia_unclassified|f__Clostridia_unclassified|g__Clostridia_unclassified|s_

k__Bacteria|p__Firmicutes|c__Clostridia|o__Eubacteriales|f__Clostridiaceae|g__Clostridiaceae_unclassified|s__Clostridiac

k__Bacteria|p__Firmicutes|c__Clostridia|o__Eubacteriales|f__Clostridiaceae|g__Clostridiaceae_unclassified|s__Clostridiac

k__Bacteria|p__Firmicutes|c__Clostridia|o__Eubacteriales|f__Eubacteriales_unclassified|g__Eubacteriales_unclassified|s_

k__Bacteria|p__Firmicutes|c__Erysipelotrichia|o__Erysipelotrichales|f__Erysipelotrichaceae|g__Erysipelatoclostridium|s__

k__Bacteria|p__Actinobacteria|c__Coriobacteriia|o__Coriobacteriales|f__Coriobacteriaceae|g__Coriobacteriaceae_unclass

k__Bacteria|p__Firmicutes|c__Clostridia|o__Eubacteriales|f__Lachnospiraceae|g__Dorea|s__Dorea_sp_5_2

k__Bacteria|p__Firmicutes|c__Erysipelotrichia|o__Erysipelotrichales|f__Erysipelotrichaceae|g__Dubosiella|s__Dubosiella_

k__Bacteria|p__Firmicutes|c__Erysipelotrichia|o__Erysipelotrichales|f__Erysipelotrichales_unclassified|g__Erysipelotrichal

k__Bacteria|p__Firmicutes|c__Clostridia|o__Eubacteriales|f__Eubacteriaceae|g__Eubacteriaceae_unclassified|s__Eubacte

k__Bacteria|p__Firmicutes|c__Clostridia|o__Eubacteriales|f__Eubacteriaceae|g__Eubacteriaceae_unclassified|s__Eubacte

k__Bacteria|p__Firmicutes|c__Clostridia|o__Eubacteriales|f__Lachnospiraceae|g__GGB20149|s__GGB20149_SGB29430

k__Bacteria|p__Actinobacteria|c__Coriobacteriia|o__Eggerthellales|f__Eggerthellaceae|g__GGB22635|s__GGB22635_SGB

k__Bacteria|p__Firmicutes|c__Clostridia|o__Eubacteriales|f__Lachnospiraceae|g__GGB25041|s__GGB25041_SGB36960

k__Bacteria|p__Bacteroidota|c__Bacteroidia|o__Bacteroidales|f__Muribaculaceae|g__GGB27876|s__GGB27876_SGB4031

k__Bacteria|p__Bacteroidota|c__Bacteroidia|o__Bacteroidales|f__Muribaculaceae|g__GGB27878|s__GGB27878_SGB4031

k__Bacteria|p__Bacteroidota|c__Bacteroidia|o__Bacteroidales|f__Muribaculaceae|g__GGB27918|s__GGB27918_SGB4035

k__Bacteria|p__Firmicutes|c__CFGB9508|o__OFGB9508|f__FGB9508|g__GGB28382|s__GGB28382_SGB40962

k__Bacteria|p__Firmicutes|c__CFGB2838|o__OFGB2838|f__FGB2838|g__GGB28399|s__GGB28399_SGB40980

k__Bacteria|p__Firmicutes|c__CFGB2838|o__OFGB2838|f__FGB2838|g__GGB28411|s__GGB28411_SGB40993

k__Bacteria|p__Firmicutes|c__CFGB2838|o__OFGB2838|f__FGB2838|g__GGB28415|s__GGB28415_SGB40997

k__Bacteria|p__Firmicutes|c__Clostridia|o__Eubacteriales|f__Pumilibacteraceae|g__GGB28430|s__GGB28430_SGB41013

k__Bacteria|p__Firmicutes|c__CFGB28439|o__OFGB28439|f__FGB28439|g__GGB28439|s__GGB28439_SGB41022

k__Bacteria|p__Firmicutes|c__Clostridia|o__Clostridia_unclassified|f__Clostridia_unclassified|g__GGB28778|s__GGB2877

k__Bacteria|p__Firmicutes|c__Clostridia|o__Eubacteriales|f__Eubacteriaceae|g__GGB28784|s__GGB28784_SGB41437

k__Bacteria|p__Firmicutes|c__Clostridia|o__Eubacteriales|f__Lachnospiraceae|g__GGB28792|s__GGB28792_SGB41445

k__Bacteria|p__Firmicutes|c__Clostridia|o__Eubacteriales|f__Lachnospiraceae|g__GGB28798|s__GGB28798_SGB41451

k__Bacteria|p__Firmicutes|c__Clostridia|o__Eubacteriales|f__Lachnospiraceae|g__GGB28802|s__GGB28802_SGB41455

k__Bacteria|p__Firmicutes|c__Clostridia|o__Eubacteriales|f__Lachnospiraceae|g__GGB28818|s__GGB28818_SGB41473

k__Bacteria|p__Firmicutes|c__CFGB77305|o__OFGB77305|f__FGB77305|g__GGB28828|s__GGB28828_SGB41484

k__Bacteria|p__Firmicutes|c__Clostridia|o__Eubacteriales|f__Clostridiaceae|g__GGB28851|s__GGB28851_SGB41518

k__Bacteria|p__Firmicutes|c__Clostridia|o__Eubacteriales|f__Lachnospiraceae|g__GGB28859|s__GGB28859_SGB41528

k__Bacteria|p__Firmicutes|c__Clostridia|o__Eubacteriales|f__Lachnospiraceae|g__GGB28864|s__GGB28864_SGB41535

k__Bacteria|p__Firmicutes|c__Clostridia|o__Eubacteriales|f__Lachnospiraceae|g__GGB28869|s__GGB28869_SGB41543

k__Bacteria|p__Firmicutes|c__CFGB9633|o__OFGB9633|f__FGB9633|g__GGB28883|s__GGB28883_SGB41564

k__Bacteria|p__Bacteria_unclassified|c__Bacteria_unclassified|o__Bacteria_unclassified|f__Bacteria_unclassified|g__GGB

k__Bacteria|p__Bacteria_unclassified|c__Bacteria_unclassified|o__Bacteria_unclassified|f__Bacteria_unclassified|g__GGB

k__Bacteria|p__Bacteria_unclassified|c__Bacteria_unclassified|o__Bacteria_unclassified|f__Bacteria_unclassified|g__GGB

k__Bacteria|p__Bacteria_unclassified|c__Bacteria_unclassified|o__Bacteria_unclassified|f__Bacteria_unclassified|g__GGB

k__Bacteria|p__Firmicutes|c__Clostridia|o__Eubacteriales|f__Lachnospiraceae|g__GGB28916|s__GGB28916_SGB41612

k__Bacteria|p__Firmicutes|c__Clostridia|o__Eubacteriales|f__Lachnospiraceae|g__GGB28924|s__GGB28924_SGB41621

k__Bacteria|p__Firmicutes|c__Clostridia|o__Eubacteriales|f__Lachnospiraceae|g__GGB28926|s__GGB28926_SGB41624

k__Bacteria|p__Bacteria_unclassified|c__CFGB77359|o__OFGB77359|f__FGB77359|g__GGB28927|s__GGB28927_SGB416

k__Bacteria|p__Firmicutes|c__CFGB9639|o__OFGB9639|f__FGB9639|g__GGB28934|s__GGB28934_SGB41635
k__Bacteria|p__Firmicutes|c__Clostridia|o__Eubacteriales|f__Lachnospiraceae|g__GGB28946|s__GGB28946_SGB41652
k__Bacteria|p__Firmicutes|c__Clostridia|o__Eubacteriales|f__Lachnospiraceae|g__GGB28949|s__GGB28949_SGB41655
k__Bacteria|p__Firmicutes|c__Clostridia|o__Eubacteriales|f__Lachnospiraceae|g__GGB28949|s__GGB28949_SGB41656
k__Bacteria|p__Firmicutes|c__Clostridia|o__Eubacteriales|f__Clostridiaceae|g__GGB28950|s__GGB28950_SGB41657
k__Bacteria|p__Firmicutes|c__Clostridia|o__Eubacteriales|f__Clostridiaceae|g__GGB28951|s__GGB28951_SGB102295
k__Bacteria|p__Firmicutes|c__Clostridia|o__Eubacteriales|f__Clostridiaceae|g__GGB28951|s__GGB28951_SGB41658
k__Bacteria|p__Firmicutes|c__Clostridia|o__Eubacteriales|f__Clostridiaceae|g__GGB28954|s__GGB28954_SGB41662
k__Bacteria|p__Firmicutes|c__Clostridia|o__Eubacteriales|f__Clostridiaceae|g__GGB28956|s__GGB28956_SGB41665
k__Bacteria|p__Firmicutes|c__Clostridia|o__Eubacteriales|f__Clostridiaceae|g__GGB28960|s__GGB28960_SGB41669
k__Bacteria|p__Firmicutes|c__Clostridia|o__Eubacteriales|f__Clostridiaceae|g__GGB28967|s__GGB28967_SGB41678
k__Bacteria|p__Firmicutes|c__Clostridia|o__Eubacteriales|f__Eubacteriaceae|g__GGB28991|s__GGB28991_SGB41705
k__Bacteria|p__Firmicutes|c__CFGB9658|o__OFGB9658|f__FGB9658|g__GGB29002|s__GGB29002_SGB41718
k__Bacteria|p__Firmicutes|c__CFGB9659|o__OFGB9659|f__FGB9659|g__GGB29003|s__GGB29003_SGB41719
k__Bacteria|p__Bacteria_unclassified|c__Bacteria_unclassified|o__Bacteria_unclassified|f__Bacteria_unclassified|g__GGB[illegible]
k__Bacteria|p__Firmicutes|c__CFGB9827|o__OFGB9827|f__FGB9827|g__GGB29531|s__GGB29531_SGB42317
k__Bacteria|p__Firmicutes|c__Clostridia|o__Eubacteriales|f__Eubacteriaceae|g__GGB29685|s__GGB29685_SGB42494
k__Bacteria|p__Bacteria_unclassified|c__CFGB77303|o__OFGB77303|f__FGB77303|g__GGB30141|s__GGB30141_SGB430[illegible]
k__Bacteria|p__Firmicutes|c__Clostridia|o__Eubacteriales|f__Eubacteriales_unclassified|g__GGB30286|s__GGB30286_SG[illegible]
k__Bacteria|p__Firmicutes|c__Clostridia|o__Eubacteriales|f__Oscillospiraceae|g__GGB30303|s__GGB30303_SGB43268
k__Bacteria|p__Firmicutes|c__CFGB30328|o__OFGB30328|f__FGB30328|g__GGB30413|s__GGB30413_SGB43452
k__Bacteria|p__Firmicutes|c__Clostridia|o__Eubacteriales|f__Oscillospiraceae|g__GGB30454|s__GGB30454_SGB43514
k__Bacteria|p__Firmicutes|c__Clostridia|o__Eubacteriales|f__Oscillospiraceae|g__GGB30455|s__GGB30455_SGB43519
k__Bacteria|p__Firmicutes|c__Clostridia|o__Eubacteriales|f__Oscillospiraceae|g__GGB30461|s__GGB30461_SGB43527
k__Bacteria|p__Firmicutes|c__Clostridia|o__Eubacteriales|f__Oscillospiraceae|g__GGB30461|s__GGB30461_SGB43530
k__Bacteria|p__Firmicutes|c__Clostridia|o__Eubacteriales|f__Oscillospiraceae|g__GGB30463|s__GGB30463_SGB43537
k__Bacteria|p__Firmicutes|c__Clostridia|o__Eubacteriales|f__Oscillospiraceae|g__GGB30473|s__GGB30473_SGB43557
k__Bacteria|p__Firmicutes|c__Clostridia|o__Eubacteriales|f__Oscillospiraceae|g__GGB30475|s__GGB30475_SGB63182
k__Bacteria|p__Actinobacteria|c__CFGB77153|o__OFGB77153|f__FGB77153|g__GGB30861|s__GGB30861_SGB44083
k__Bacteria|p__Tenericutes|c__CFGB1791|o__OFGB1791|f__FGB1791|g__GGB31312|s__GGB31312_SGB44628
k__Bacteria|p__Firmicutes|c__CFGB10290|o__OFGB10290|f__FGB10290|g__GGB31438|s__GGB31438_SGB44768
k__Bacteria|p__Firmicutes|c__Clostridia|o__Eubacteriales|f__Oscillospiraceae|g__GGB3171|s__GGB3171_SGB4185
k__Bacteria|p__Firmicutes|c__CFGB1765|o__OFGB1765|f__FGB1765|g__GGB31823|s__GGB31823_SGB45199
k__Bacteria|p__Firmicutes|c__CFGB10349|o__OFGB10349|f__FGB10349|g__GGB31853|s__GGB31853_SGB45233
k__Bacteria|p__Firmicutes|c__CFGB10667|o__OFGB10667|f__FGB10667|g__GGB32371|s__GGB32371_SGB41694
k__Bacteria|p__Firmicutes|c__Clostridia|o__Eubacteriales|f__Lachnospiraceae|g__GGB3793|s__GGB3793_SGB5158
k__Bacteria|p__Firmicutes|c__Clostridia|o__Eubacteriales|f__Lachnospiraceae|g__GGB42598|s__GGB42598_SGB59794
k__Bacteria|p__Firmicutes|c__Clostridia|o__Eubacteriales|f__Christensenellaceae|g__GGB45656|s__GGB45656_SGB6337[illegible]
k__Bacteria|p__Firmicutes|c__CFGB10299|o__OFGB10299|f__FGB10299|g__GGB47127|s__GGB47127_SGB65054
k__Bacteria|p__Firmicutes|c__Clostridia|o__Eubacteriales|f__Oscillospiraceae|g__GGB74395|s__GGB74395_SGB43521
k__Bacteria|p__Firmicutes|c__Clostridia|o__Eubacteriales|f__Oscillospiraceae|g__GGB75053|s__GGB75053_SGB43494
k__Bacteria|p__Firmicutes|c__Clostridia|o__Eubacteriales|f__Lachnospiraceae|g__GGB75109|s__GGB75109_SGB102238
k__Bacteria|p__Firmicutes|c__Clostridia|o__Clostridia_unclassified|f__Clostridia_unclassified|g__GGB81440|s__GGB8144[illegible]
k__Bacteria|p__Firmicutes|c__Clostridia|o__Eubacteriales|f__Lachnospiraceae|g__Lachnospiraceae_unclassified|s__Lachn[illegible]
k__Bacteria|p__Firmicutes|c__Clostridia|o__Eubacteriales|f__Lachnospiraceae|g__Lachnospiraceae_unclassified|s__Lachn[illegible]
k__Bacteria|p__Firmicutes|c__Clostridia|o__Eubacteriales|f__Lachnospiraceae|g__Lachnospiraceae_unclassified|s__Lachn[illegible]

k__Bacteria|p__Firmicutes|c__Clostridia|o__Eubacteriales|f__Lachnospiraceae|g__Lachnospiraceae_unclassified|s__Lachn
k__Bacteria|p__Firmicutes|c__Clostridia|o__Eubacteriales|f__Lachnospiraceae|g__Lachnospiraceae_unclassified|s__Lachn
k__Bacteria|p__Firmicutes|c__Clostridia|o__Eubacteriales|f__Lachnospiraceae|g__Lachnospiraceae_unclassified|s__Lachn
k__Bacteria|p__Firmicutes|c__Clostridia|o__Eubacteriales|f__Lachnospiraceae|g__Lachnospiraceae_unclassified|s__Lachn
k__Bacteria|p__Firmicutes|c__Clostridia|o__Eubacteriales|f__Lachnospiraceae|g__Lachnospiraceae_unclassified|s__Lachn
k__Bacteria|p__Firmicutes|c__Bacilli|o__Lactobacillales|f__Lactobacillaceae|g__Lactobacillus|s__Lactobacillus_johnsonii
k__Bacteria|p__Bacteroidota|c__Bacteroidia|o__Bacteroidales|f__Muribaculaceae|g__Muribaculaceae_unclassified|s__M
k__Bacteria|p__Firmicutes|c__Clostridia|o__Eubacteriales|f__Oscillospiraceae|g__Neglectibacter|s__Neglectibacter_sp_X
k__Bacteria|p__Firmicutes|c__Clostridia|o__Eubacteriales|f__Oscillospiraceae|g__Oscillospiraceae_unclassified|s__Oscillo
k__Bacteria|p__Firmicutes|c__Clostridia|o__Eubacteriales|f__Oscillospiraceae|g__Oscillospiraceae_unclassified|s__Oscillo
k__Bacteria|p__Firmicutes|c__Clostridia|o__Eubacteriales|f__Oscillospiraceae|g__Oscillospiraceae_unclassified|s__Oscillo
k__Bacteria|p__Proteobacteria|c__Betaproteobacteria|o__Burkholderiales|f__Sutterellaceae|g__Parasutterella|s__Parasu
k__Bacteria|p__Firmicutes|c__Clostridia|o__Eubacteriales|f__Peptostreptococcaceae|g__Romboutsia|s__Romboutsia_ilea
k__Bacteria|p__Firmicutes|c__Clostridia|o__Eubacteriales|f__Lachnospiraceae|g__Schaedlerella|s__Schaedlerella_arabino
k__Bacteria|p__Firmicutes|c__Erysipelotrichia|o__Erysipelotrichales|f__Turicibacteraceae|g__Turicibacter|s__Turicibacter_
k__Bacteria|p__Bacteria_unclassified|c__Bacteria_unclassified|o__Bacteria_unclassified|f__Bacteria_unclassified|g__Bacte
k__Bacteria|p__Bacteria_unclassified|c__Bacteria_unclassified|o__Bacteria_unclassified|f__Bacteria_unclassified|g__Bacte
k__Bacteria|p__Bacteria_unclassified|c__Bacteria_unclassified|o__Bacteria_unclassified|f__Bacteria_unclassified|g__Bacte

k__Bacteria|p__Firmicutes|c__Clostridia|o__Eubacteriales|f__Lachnospiraceae|g__Acetatifactor|s__Acetatifactor_SGB415
k__Bacteria|p__Firmicutes|c__Clostridia|o__Eubacteriales|f__Lachnospiraceae|g__Acetatifactor|s__Acetatifactor_muris
k__Bacteria|p__Firmicutes|c__Clostridia|o__Eubacteriales|f__Oscillospiraceae|g__Acutalibacter|s__Acutalibacter_muris
k__Bacteria|p__Firmicutes|c__Clostridia|o__Eubacteriales|f__Oscillospiraceae|g__Acutalibacter|s__Acutalibacter_sp_1XD
k__Bacteria|p__Actinobacteria|c__Coriobacteriia|o__Eggerthellales|f__Eggerthellaceae|g__Adlercreutzia|s__Adlercreutzia
k__Bacteria|p__Actinobacteria|c__Coriobacteriia|o__Eggerthellales|f__Eggerthellaceae|g__Adlercreutzia|s__Adlercreutzia
k__Bacteria|p__Actinobacteria|c__Coriobacteriia|o__Eggerthellales|f__Eggerthellaceae|g__Adlercreutzia|s__Adlercreutzia
k__Bacteria|p__Verrucomicrobia|c__Verrucomicrobiae|o__Verrucomicrobiales|f__Akkermansiaceae|g__Akkermansia|s__A
k__Bacteria|p__Bacteroidota|c__Bacteroidia|o__Bacteroidales|f__Rikenellaceae|g__Alistipes|s__Alistipes_sp_DSM_11234
k__Bacteria|p__Firmicutes|c__Clostridia|o__Eubacteriales|f__Oscillospiraceae|g__Anaerotruncus|s__Anaerotruncus_sp_1
k__Bacteria|p__Bacteria_unclassified|c__Bacteria_unclassified|o__Bacteria_unclassified|f__Bacteria_unclassified|g__Bacte
k__Bacteria|p__Bacteria_unclassified|c__Bacteria_unclassified|o__Bacteria_unclassified|f__Bacteria_unclassified|g__Bacte
k__Bacteria|p__Bacteria_unclassified|c__Bacteria_unclassified|o__Bacteria_unclassified|f__Bacteria_unclassified|g__Bacte
k__Bacteria|p__Bacteroidota|c__Bacteroidia|o__Bacteroidales|f__Bacteroidaceae|g__Bacteroides|s__Bacteroides_thetaio
k__Bacteria|p__Actinobacteria|c__Actinomycetia|o__Bifidobacteriales|f__Bifidobacteriaceae|g__Bifidobacterium|s__Bifid
k__Bacteria|p__Firmicutes|c__Clostridia|o__Clostridia_unclassified|f__Clostridia_unclassified|g__Clostridia_unclassified|s_
k__Bacteria|p__Firmicutes|c__Clostridia|o__Eubacteriales|f__Clostridiaceae|g__Clostridiaceae_unclassified|s__Clostridiac
k__Bacteria|p__Firmicutes|c__Clostridia|o__Eubacteriales|f__Clostridiaceae|g__Clostridiaceae_unclassified|s__Clostridiac
k__Bacteria|p__Firmicutes|c__Clostridia|o__Eubacteriales|f__Eubacteriales_unclassified|g__Eubacteriales_unclassified|s_
k__Bacteria|p__Firmicutes|c__Erysipelotrichia|o__Erysipelotrichales|f__Erysipelotrichaceae|g__Erysipelatoclostridium|s_
k__Bacteria|p__Actinobacteria|c__Coriobacteriia|o__Coriobacteriales|f__Coriobacteriaceae|g__Coriobacteriaceae_unclass
k__Bacteria|p__Firmicutes|c__Clostridia|o__Eubacteriales|f__Lachnospiraceae|g__Dorea|s__Dorea_sp_5_2
k__Bacteria|p__Firmicutes|c__Erysipelotrichia|o__Erysipelotrichales|f__Erysipelotrichaceae|g__Dubosiella|s__Dubosiella_
k__Bacteria|p__Firmicutes|c__Erysipelotrichia|o__Erysipelotrichales|f__Erysipelotrichales_unclassified|g__Erysipelotrichal
k__Bacteria|p__Firmicutes|c__Clostridia|o__Eubacteriales|f__Eubacteriaceae|g__Eubacteriaceae_unclassified|s__Eubacte

k__Bacteria|p__Firmicutes|c__Clostridia|o__Eubacteriales|f__Eubacteriaceae|g__Eubacteriaceae_unclassified|s__Eubacte
k__Bacteria|p__Firmicutes|c__Clostridia|o__Eubacteriales|f__Lachnospiraceae|g__GGB20149|s__GGB20149_SGB29430
k__Bacteria|p__Actinobacteria|c__Coriobacteriia|o__Eggerthellales|f__Eggerthellaceae|g__GGB22635|s__GGB22635_SGB
k__Bacteria|p__Firmicutes|c__Clostridia|o__Eubacteriales|f__Lachnospiraceae|g__GGB25041|s__GGB25041_SGB36960
k__Bacteria|p__Bacteroidota|c__Bacteroidia|o__Bacteroidales|f__Muribaculaceae|g__GGB27876|s__GGB27876_SGB4031
k__Bacteria|p__Bacteroidota|c__Bacteroidia|o__Bacteroidales|f__Muribaculaceae|g__GGB27878|s__GGB27878_SGB4031
k__Bacteria|p__Bacteroidota|c__Bacteroidia|o__Bacteroidales|f__Muribaculaceae|g__GGB27918|s__GGB27918_SGB4035
k__Bacteria|p__Firmicutes|c__CFGB9508|o__OFGB9508|f__FGB9508|g__GGB28382|s__GGB28382_SGB40962
k__Bacteria|p__Firmicutes|c__CFGB2838|o__OFGB2838|f__FGB2838|g__GGB28399|s__GGB28399_SGB40980
k__Bacteria|p__Firmicutes|c__CFGB2838|o__OFGB2838|f__FGB2838|g__GGB28411|s__GGB28411_SGB40993
k__Bacteria|p__Firmicutes|c__CFGB2838|o__OFGB2838|f__FGB2838|g__GGB28415|s__GGB28415_SGB40997
k__Bacteria|p__Firmicutes|c__Clostridia|o__Eubacteriales|f__Pumilibacteraceae|g__GGB28430|s__GGB28430_SGB41013
k__Bacteria|p__Firmicutes|c__CFGB28439|o__OFGB28439|f__FGB28439|g__GGB28439|s__GGB28439_SGB41022
k__Bacteria|p__Firmicutes|c__Clostridia|o__Clostridia_unclassified|f__Clostridia_unclassified|g__GGB28778|s__GGB2877
k__Bacteria|p__Firmicutes|c__Clostridia|o__Eubacteriales|f__Eubacteriaceae|g__GGB28784|s__GGB28784_SGB41437
k__Bacteria|p__Firmicutes|c__Clostridia|o__Eubacteriales|f__Lachnospiraceae|g__GGB28792|s__GGB28792_SGB41445
k__Bacteria|p__Firmicutes|c__Clostridia|o__Eubacteriales|f__Lachnospiraceae|g__GGB28798|s__GGB28798_SGB41451
k__Bacteria|p__Firmicutes|c__Clostridia|o__Eubacteriales|f__Lachnospiraceae|g__GGB28802|s__GGB28802_SGB41455
k__Bacteria|p__Firmicutes|c__Clostridia|o__Eubacteriales|f__Lachnospiraceae|g__GGB28818|s__GGB28818_SGB41473
k__Bacteria|p__Firmicutes|c__CFGB77305|o__OFGB77305|f__FGB77305|g__GGB28828|s__GGB28828_SGB41484
k__Bacteria|p__Firmicutes|c__Clostridia|o__Eubacteriales|f__Clostridiaceae|g__GGB28851|s__GGB28851_SGB41518
k__Bacteria|p__Firmicutes|c__Clostridia|o__Eubacteriales|f__Lachnospiraceae|g__GGB28859|s__GGB28859_SGB41528
k__Bacteria|p__Firmicutes|c__Clostridia|o__Eubacteriales|f__Lachnospiraceae|g__GGB28864|s__GGB28864_SGB41535
k__Bacteria|p__Firmicutes|c__Clostridia|o__Eubacteriales|f__Lachnospiraceae|g__GGB28869|s__GGB28869_SGB41543
k__Bacteria|p__Firmicutes|c__CFGB9633|o__OFGB9633|f__FGB9633|g__GGB28883|s__GGB28883_SGB41564
k__Bacteria|p__Bacteria_unclassified|c__Bacteria_unclassified|o__Bacteria_unclassified|f__Bacteria_unclassified|g__GGB
k__Bacteria|p__Bacteria_unclassified|c__Bacteria_unclassified|o__Bacteria_unclassified|f__Bacteria_unclassified|g__GGB
k__Bacteria|p__Bacteria_unclassified|c__Bacteria_unclassified|o__Bacteria_unclassified|f__Bacteria_unclassified|g__GGB
k__Bacteria|p__Bacteria_unclassified|c__Bacteria_unclassified|o__Bacteria_unclassified|f__Bacteria_unclassified|g__GGB
k__Bacteria|p__Firmicutes|c__Clostridia|o__Eubacteriales|f__Lachnospiraceae|g__GGB28916|s__GGB28916_SGB41612
k__Bacteria|p__Firmicutes|c__Clostridia|o__Eubacteriales|f__Lachnospiraceae|g__GGB28924|s__GGB28924_SGB41621
k__Bacteria|p__Firmicutes|c__Clostridia|o__Eubacteriales|f__Lachnospiraceae|g__GGB28926|s__GGB28926_SGB41624
k__Bacteria|p__Bacteria_unclassified|c__CFGB77359|o__OFGB77359|f__FGB77359|g__GGB28927|s__GGB28927_SGB416
k__Bacteria|p__Firmicutes|c__CFGB9639|o__OFGB9639|f__FGB9639|g__GGB28934|s__GGB28934_SGB41635
k__Bacteria|p__Firmicutes|c__Clostridia|o__Eubacteriales|f__Lachnospiraceae|g__GGB28946|s__GGB28946_SGB41652
k__Bacteria|p__Firmicutes|c__Clostridia|o__Eubacteriales|f__Lachnospiraceae|g__GGB28949|s__GGB28949_SGB41655
k__Bacteria|p__Firmicutes|c__Clostridia|o__Eubacteriales|f__Lachnospiraceae|g__GGB28949|s__GGB28949_SGB41656
k__Bacteria|p__Firmicutes|c__Clostridia|o__Eubacteriales|f__Clostridiaceae|g__GGB28950|s__GGB28950_SGB41657
k__Bacteria|p__Firmicutes|c__Clostridia|o__Eubacteriales|f__Clostridiaceae|g__GGB28951|s__GGB28951_SGB102295
k__Bacteria|p__Firmicutes|c__Clostridia|o__Eubacteriales|f__Clostridiaceae|g__GGB28951|s__GGB28951_SGB41658
k__Bacteria|p__Firmicutes|c__Clostridia|o__Eubacteriales|f__Clostridiaceae|g__GGB28954|s__GGB28954_SGB41662
k__Bacteria|p__Firmicutes|c__Clostridia|o__Eubacteriales|f__Clostridiaceae|g__GGB28956|s__GGB28956_SGB41665
k__Bacteria|p__Firmicutes|c__Clostridia|o__Eubacteriales|f__Clostridiaceae|g__GGB28960|s__GGB28960_SGB41669
k__Bacteria|p__Firmicutes|c__Clostridia|o__Eubacteriales|f__Clostridiaceae|g__GGB28967|s__GGB28967_SGB41678
k__Bacteria|p__Firmicutes|c__Clostridia|o__Eubacteriales|f__Eubacteriaceae|g__GGB28991|s__GGB28991_SGB41705
k__Bacteria|p__Firmicutes|c__CFGB9658|o__OFGB9658|f__FGB9658|g__GGB29002|s__GGB29002_SGB41718

k__Bacteria|p__Firmicutes|c__CFGB9659|o__OFGB9659|f__FGB9659|g__GGB29003|s__GGB29003_SGB41719
k__Bacteria|p__Bacteria_unclassified|c__Bacteria_unclassified|o__Bacteria_unclassified|f__Bacteria_unclassified|g__GGB2
k__Bacteria|p__Firmicutes|c__CFGB9827|o__OFGB9827|f__FGB9827|g__GGB29531|s__GGB29531_SGB42317
k__Bacteria|p__Firmicutes|c__Clostridia|o__Eubacteriales|f__Eubacteriaceae|g__GGB29685|s__GGB29685_SGB42494
k__Bacteria|p__Bacteria_unclassified|c__CFGB77303|o__OFGB77303|f__FGB77303|g__GGB30141|s__GGB30141_SGB430
k__Bacteria|p__Firmicutes|c__Clostridia|o__Eubacteriales|f__Eubacteriales_unclassified|g__GGB30286|s__GGB30286_SG
k__Bacteria|p__Firmicutes|c__Clostridia|o__Eubacteriales|f__Oscillospiraceae|g__GGB30303|s__GGB30303_SGB43268
k__Bacteria|p__Firmicutes|c__CFGB30328|o__OFGB30328|f__FGB30328|g__GGB30413|s__GGB30413_SGB43452
k__Bacteria|p__Firmicutes|c__Clostridia|o__Eubacteriales|f__Oscillospiraceae|g__GGB30454|s__GGB30454_SGB43514
k__Bacteria|p__Firmicutes|c__Clostridia|o__Eubacteriales|f__Oscillospiraceae|g__GGB30455|s__GGB30455_SGB43519
k__Bacteria|p__Firmicutes|c__Clostridia|o__Eubacteriales|f__Oscillospiraceae|g__GGB30461|s__GGB30461_SGB43527
k__Bacteria|p__Firmicutes|c__Clostridia|o__Eubacteriales|f__Oscillospiraceae|g__GGB30461|s__GGB30461_SGB43530
k__Bacteria|p__Firmicutes|c__Clostridia|o__Eubacteriales|f__Oscillospiraceae|g__GGB30463|s__GGB30463_SGB43537
k__Bacteria|p__Firmicutes|c__Clostridia|o__Eubacteriales|f__Oscillospiraceae|g__GGB30473|s__GGB30473_SGB43557
k__Bacteria|p__Firmicutes|c__Clostridia|o__Eubacteriales|f__Oscillospiraceae|g__GGB30475|s__GGB30475_SGB63182
k__Bacteria|p__Actinobacteria|c__CFGB77153|o__OFGB77153|f__FGB77153|g__GGB30861|s__GGB30861_SGB44083
k__Bacteria|p__Tenericutes|c__CFGB1791|o__OFGB1791|f__FGB1791|g__GGB31312|s__GGB31312_SGB44628
k__Bacteria|p__Firmicutes|c__CFGB10290|o__OFGB10290|f__FGB10290|g__GGB31438|s__GGB31438_SGB44768
k__Bacteria|p__Firmicutes|c__Clostridia|o__Eubacteriales|f__Oscillospiraceae|g__GGB3171|s__GGB3171_SGB4185
k__Bacteria|p__Firmicutes|c__CFGB1765|o__OFGB1765|f__FGB1765|g__GGB31823|s__GGB31823_SGB45199
k__Bacteria|p__Firmicutes|c__CFGB10349|o__OFGB10349|f__FGB10349|g__GGB31853|s__GGB31853_SGB45233
k__Bacteria|p__Firmicutes|c__CFGB10667|o__OFGB10667|f__FGB10667|g__GGB32371|s__GGB32371_SGB41694
k__Bacteria|p__Firmicutes|c__Clostridia|o__Eubacteriales|f__Lachnospiraceae|g__GGB3793|s__GGB3793_SGB5158
k__Bacteria|p__Firmicutes|c__Clostridia|o__Eubacteriales|f__Lachnospiraceae|g__GGB42598|s__GGB42598_SGB59794
k__Bacteria|p__Firmicutes|c__Clostridia|o__Eubacteriales|f__Christensenellaceae|g__GGB45656|s__GGB45656_SGB6337
k__Bacteria|p__Firmicutes|c__CFGB10299|o__OFGB10299|f__FGB10299|g__GGB47127|s__GGB47127_SGB65054
k__Bacteria|p__Firmicutes|c__Clostridia|o__Eubacteriales|f__Oscillospiraceae|g__GGB74395|s__GGB74395_SGB43521
k__Bacteria|p__Firmicutes|c__Clostridia|o__Eubacteriales|f__Oscillospiraceae|g__GGB75053|s__GGB75053_SGB43494
k__Bacteria|p__Firmicutes|c__Clostridia|o__Eubacteriales|f__Lachnospiraceae|g__GGB75109|s__GGB75109_SGB102238
k__Bacteria|p__Firmicutes|c__Clostridia|o__Clostridia_unclassified|f__Clostridia_unclassified|g__GGB81440|s__GGB8144
k__Bacteria|p__Firmicutes|c__Clostridia|o__Eubacteriales|f__Lachnospiraceae|g__Lachnospiraceae_unclassified|s__Lachn
k__Bacteria|p__Firmicutes|c__Clostridia|o__Eubacteriales|f__Lachnospiraceae|g__Lachnospiraceae_unclassified|s__Lachn
k__Bacteria|p__Firmicutes|c__Clostridia|o__Eubacteriales|f__Lachnospiraceae|g__Lachnospiraceae_unclassified|s__Lachn
k__Bacteria|p__Firmicutes|c__Clostridia|o__Eubacteriales|f__Lachnospiraceae|g__Lachnospiraceae_unclassified|s__Lachn
k__Bacteria|p__Firmicutes|c__Clostridia|o__Eubacteriales|f__Lachnospiraceae|g__Lachnospiraceae_unclassified|s__Lachn
k__Bacteria|p__Firmicutes|c__Clostridia|o__Eubacteriales|f__Lachnospiraceae|g__Lachnospiraceae_unclassified|s__Lachn
k__Bacteria|p__Firmicutes|c__Clostridia|o__Eubacteriales|f__Lachnospiraceae|g__Lachnospiraceae_unclassified|s__Lachn
k__Bacteria|p__Firmicutes|c__Clostridia|o__Eubacteriales|f__Lachnospiraceae|g__Lachnospiraceae_unclassified|s__Lachn
k__Bacteria|p__Firmicutes|c__Bacilli|o__Lactobacillales|f__Lactobacillaceae|g__Lactobacillus|s__Lactobacillus_johnsonii
k__Bacteria|p__Bacteroidota|c__Bacteroidia|o__Bacteroidales|f__Muribaculaceae|g__Muribaculaceae_unclassified|s__M
k__Bacteria|p__Firmicutes|c__Clostridia|o__Eubacteriales|f__Oscillospiraceae|g__Neglectibacter|s__Neglectibacter_sp_X
k__Bacteria|p__Firmicutes|c__Clostridia|o__Eubacteriales|f__Oscillospiraceae|g__Oscillospiraceae_unclassified|s__Oscillo
k__Bacteria|p__Firmicutes|c__Clostridia|o__Eubacteriales|f__Oscillospiraceae|g__Oscillospiraceae_unclassified|s__Oscillo
k__Bacteria|p__Firmicutes|c__Clostridia|o__Eubacteriales|f__Oscillospiraceae|g__Oscillospiraceae_unclassified|s__Oscillo
k__Bacteria|p__Proteobacteria|c__Betaproteobacteria|o__Burkholderiales|f__Sutterellaceae|g__Parasutterella|s__Parasu
k__Bacteria|p__Firmicutes|c__Clostridia|o__Eubacteriales|f__Peptostreptococcaceae|g__Romboutsia|s__Romboutsia_ilea

k__Bacteria|p__Firmicutes|c__Clostridia|o__Eubacteriales|f__Lachnospiraceae|g__Schaedlerella|s__Schaedlerella_arabinc
k__Bacteria|p__Firmicutes|c__Erysipelotrichia|o__Erysipelotrichales|f__Turicibacteraceae|g__Turicibacter|s__Turicibacter_
k__Bacteria|p__Bacteria_unclassified|c__Bacteria_unclassified|o__Bacteria_unclassified|f__Bacteria_unclassified|g__Bacte
k__Bacteria|p__Bacteria_unclassified|c__Bacteria_unclassified|o__Bacteria_unclassified|f__Bacteria_unclassified|g__Bacte
k__Bacteria|p__Bacteria_unclassified|c__Bacteria_unclassified|o__Bacteria_unclassified|f__Bacteria_unclassified|g__Bacte

k__Bacteria|p__Firmicutes|c__Clostridia|o__Eubacteriales|f__Lachnospiraceae|g__Acetatifactor|s__Acetatifactor_SGB415
k__Bacteria|p__Firmicutes|c__Clostridia|o__Eubacteriales|f__Lachnospiraceae|g__Acetatifactor|s__Acetatifactor_muris
k__Bacteria|p__Firmicutes|c__Clostridia|o__Eubacteriales|f__Oscillospiraceae|g__Acutalibacter|s__Acutalibacter_muris
k__Bacteria|p__Firmicutes|c__Clostridia|o__Eubacteriales|f__Oscillospiraceae|g__Acutalibacter|s__Acutalibacter_sp_1XD
k__Bacteria|p__Actinobacteria|c__Coriobacteriia|o__Eggerthellales|f__Eggerthellaceae|g__Adlercreutzia|s__Adlercreutzia
k__Bacteria|p__Actinobacteria|c__Coriobacteriia|o__Eggerthellales|f__Eggerthellaceae|g__Adlercreutzia|s__Adlercreutzia
k__Bacteria|p__Actinobacteria|c__Coriobacteriia|o__Eggerthellales|f__Eggerthellaceae|g__Adlercreutzia|s__Adlercreutzia
k__Bacteria|p__Verrucomicrobia|c__Verrucomicrobiae|o__Verrucomicrobiales|f__Akkermansiaceae|g__Akkermansia|s__A
k__Bacteria|p__Bacteroidota|c__Bacteroidia|o__Bacteroidales|f__Rikenellaceae|g__Alistipes|s__Alistipes_sp_DSM_11234
k__Bacteria|p__Firmicutes|c__Clostridia|o__Eubacteriales|f__Oscillospiraceae|g__Anaerotruncus|s__Anaerotruncus_sp_1
k__Bacteria|p__Bacteria_unclassified|c__Bacteria_unclassified|o__Bacteria_unclassified|f__Bacteria_unclassified|g__Bacte
k__Bacteria|p__Bacteria_unclassified|c__Bacteria_unclassified|o__Bacteria_unclassified|f__Bacteria_unclassified|g__Bacte
k__Bacteria|p__Bacteria_unclassified|c__Bacteria_unclassified|o__Bacteria_unclassified|f__Bacteria_unclassified|g__Bacte
k__Bacteria|p__Bacteroidota|c__Bacteroidia|o__Bacteroidales|f__Bacteroidaceae|g__Bacteroides|s__Bacteroides_thetaio
k__Bacteria|p__Actinobacteria|c__Actinomycetia|o__Bifidobacteriales|f__Bifidobacteriaceae|g__Bifidobacterium|s__Bifid
k__Bacteria|p__Firmicutes|c__Clostridia|o__Clostridia_unclassified|f__Clostridia_unclassified|g__Clostridia_unclassified|s_
k__Bacteria|p__Firmicutes|c__Clostridia|o__Eubacteriales|f__Clostridiaceae|g__Clostridiaceae_unclassified|s__Clostridiac
k__Bacteria|p__Firmicutes|c__Clostridia|o__Eubacteriales|f__Clostridiaceae|g__Clostridiaceae_unclassified|s__Clostridiac
k__Bacteria|p__Firmicutes|c__Clostridia|o__Eubacteriales|f__Eubacteriales_unclassified|g__Eubacteriales_unclassified|s_
k__Bacteria|p__Firmicutes|c__Erysipelotrichia|o__Erysipelotrichales|f__Erysipelotrichaceae|g__Erysipelatoclostridium|s_
k__Bacteria|p__Actinobacteria|c__Coriobacteriia|o__Coriobacteriales|f__Coriobacteriaceae|g__Coriobacteriaceae_unclass
k__Bacteria|p__Firmicutes|c__Clostridia|o__Eubacteriales|f__Lachnospiraceae|g__Dorea|s__Dorea_sp_5_2
k__Bacteria|p__Firmicutes|c__Erysipelotrichia|o__Erysipelotrichales|f__Erysipelotrichaceae|g__Dubosiella|s__Dubosiella_
k__Bacteria|p__Firmicutes|c__Erysipelotrichia|o__Erysipelotrichales|f__Erysipelotrichales_unclassified|g__Erysipelotrichal
k__Bacteria|p__Firmicutes|c__Clostridia|o__Eubacteriales|f__Eubacteriaceae|g__Eubacteriaceae_unclassified|s__Eubacte
k__Bacteria|p__Firmicutes|c__Clostridia|o__Eubacteriales|f__Eubacteriaceae|g__Eubacteriaceae_unclassified|s__Eubacte
k__Bacteria|p__Firmicutes|c__Clostridia|o__Eubacteriales|f__Lachnospiraceae|g__GGB20149|s__GGB20149_SGB29430
k__Bacteria|p__Actinobacteria|c__Coriobacteriia|o__Eggerthellales|f__Eggerthellaceae|g__GGB22635|s__GGB22635_SGB
k__Bacteria|p__Firmicutes|c__Clostridia|o__Eubacteriales|f__Lachnospiraceae|g__GGB25041|s__GGB25041_SGB36960
k__Bacteria|p__Bacteroidota|c__Bacteroidia|o__Bacteroidales|f__Muribaculaceae|g__GGB27876|s__GGB27876_SGB4031
k__Bacteria|p__Bacteroidota|c__Bacteroidia|o__Bacteroidales|f__Muribaculaceae|g__GGB27878|s__GGB27878_SGB4031
k__Bacteria|p__Bacteroidota|c__Bacteroidia|o__Bacteroidales|f__Muribaculaceae|g__GGB27918|s__GGB27918_SGB4035
k__Bacteria|p__Firmicutes|c__CFGB9508|o__OFGB9508|f__FGB9508|g__GGB28382|s__GGB28382_SGB40962
k__Bacteria|p__Firmicutes|c__CFGB2838|o__OFGB2838|f__FGB2838|g__GGB28399|s__GGB28399_SGB40980
k__Bacteria|p__Firmicutes|c__CFGB2838|o__OFGB2838|f__FGB2838|g__GGB28411|s__GGB28411_SGB40993
k__Bacteria|p__Firmicutes|c__CFGB2838|o__OFGB2838|f__FGB2838|g__GGB28415|s__GGB28415_SGB40997
k__Bacteria|p__Firmicutes|c__Clostridia|o__Eubacteriales|f__Pumilibacteraceae|g__GGB28430|s__GGB28430_SGB41013
k__Bacteria|p__Firmicutes|c__CFGB28439|o__OFGB28439|f__FGB28439|g__GGB28439|s__GGB28439_SGB41022

k__Bacteria|p__Firmicutes|c__Clostridia|o__Clostridia_unclassified|f__Clostridia_unclassified|g__GGB28778|s__GGB2877
k__Bacteria|p__Firmicutes|c__Clostridia|o__Eubacteriales|f__Eubacteriaceae|g__GGB28784|s__GGB28784_SGB41437
k__Bacteria|p__Firmicutes|c__Clostridia|o__Eubacteriales|f__Lachnospiraceae|g__GGB28792|s__GGB28792_SGB41445
k__Bacteria|p__Firmicutes|c__Clostridia|o__Eubacteriales|f__Lachnospiraceae|g__GGB28798|s__GGB28798_SGB41451
k__Bacteria|p__Firmicutes|c__Clostridia|o__Eubacteriales|f__Lachnospiraceae|g__GGB28802|s__GGB28802_SGB41455
k__Bacteria|p__Firmicutes|c__Clostridia|o__Eubacteriales|f__Lachnospiraceae|g__GGB28818|s__GGB28818_SGB41473
k__Bacteria|p__Firmicutes|c__CFGB77305|o__OFGB77305|f__FGB77305|g__GGB28828|s__GGB28828_SGB41484
k__Bacteria|p__Firmicutes|c__Clostridia|o__Eubacteriales|f__Clostridiaceae|g__GGB28851|s__GGB28851_SGB41518
k__Bacteria|p__Firmicutes|c__Clostridia|o__Eubacteriales|f__Lachnospiraceae|g__GGB28859|s__GGB28859_SGB41528
k__Bacteria|p__Firmicutes|c__Clostridia|o__Eubacteriales|f__Lachnospiraceae|g__GGB28864|s__GGB28864_SGB41535
k__Bacteria|p__Firmicutes|c__Clostridia|o__Eubacteriales|f__Lachnospiraceae|g__GGB28869|s__GGB28869_SGB41543
k__Bacteria|p__Firmicutes|c__CFGB9633|o__OFGB9633|f__FGB9633|g__GGB28883|s__GGB28883_SGB41564
k__Bacteria|p__Bacteria_unclassified|c__Bacteria_unclassified|o__Bacteria_unclassified|f__Bacteria_unclassified|g__GGB
k__Bacteria|p__Bacteria_unclassified|c__Bacteria_unclassified|o__Bacteria_unclassified|f__Bacteria_unclassified|g__GGB
k__Bacteria|p__Bacteria_unclassified|c__Bacteria_unclassified|o__Bacteria_unclassified|f__Bacteria_unclassified|g__GGB
k__Bacteria|p__Bacteria_unclassified|c__Bacteria_unclassified|o__Bacteria_unclassified|f__Bacteria_unclassified|g__GGB
k__Bacteria|p__Firmicutes|c__Clostridia|o__Eubacteriales|f__Lachnospiraceae|g__GGB28916|s__GGB28916_SGB41612
k__Bacteria|p__Firmicutes|c__Clostridia|o__Eubacteriales|f__Lachnospiraceae|g__GGB28924|s__GGB28924_SGB41621
k__Bacteria|p__Firmicutes|c__Clostridia|o__Eubacteriales|f__Lachnospiraceae|g__GGB28926|s__GGB28926_SGB41624
k__Bacteria|p__Bacteria_unclassified|c__CFGB77359|o__OFGB77359|f__FGB77359|g__GGB28927|s__GGB28927_SGB416
k__Bacteria|p__Firmicutes|c__CFGB9639|o__OFGB9639|f__FGB9639|g__GGB28934|s__GGB28934_SGB41635
k__Bacteria|p__Firmicutes|c__Clostridia|o__Eubacteriales|f__Lachnospiraceae|g__GGB28946|s__GGB28946_SGB41652
k__Bacteria|p__Firmicutes|c__Clostridia|o__Eubacteriales|f__Lachnospiraceae|g__GGB28949|s__GGB28949_SGB41655
k__Bacteria|p__Firmicutes|c__Clostridia|o__Eubacteriales|f__Lachnospiraceae|g__GGB28949|s__GGB28949_SGB41656
k__Bacteria|p__Firmicutes|c__Clostridia|o__Eubacteriales|f__Clostridiaceae|g__GGB28950|s__GGB28950_SGB41657
k__Bacteria|p__Firmicutes|c__Clostridia|o__Eubacteriales|f__Clostridiaceae|g__GGB28951|s__GGB28951_SGB102295
k__Bacteria|p__Firmicutes|c__Clostridia|o__Eubacteriales|f__Clostridiaceae|g__GGB28951|s__GGB28951_SGB41658
k__Bacteria|p__Firmicutes|c__Clostridia|o__Eubacteriales|f__Clostridiaceae|g__GGB28954|s__GGB28954_SGB41662
k__Bacteria|p__Firmicutes|c__Clostridia|o__Eubacteriales|f__Clostridiaceae|g__GGB28956|s__GGB28956_SGB41665
k__Bacteria|p__Firmicutes|c__Clostridia|o__Eubacteriales|f__Clostridiaceae|g__GGB28960|s__GGB28960_SGB41669
k__Bacteria|p__Firmicutes|c__Clostridia|o__Eubacteriales|f__Clostridiaceae|g__GGB28967|s__GGB28967_SGB41678
k__Bacteria|p__Firmicutes|c__Clostridia|o__Eubacteriales|f__Eubacteriaceae|g__GGB28991|s__GGB28991_SGB41705
k__Bacteria|p__Firmicutes|c__CFGB9658|o__OFGB9658|f__FGB9658|g__GGB29002|s__GGB29002_SGB41718
k__Bacteria|p__Firmicutes|c__CFGB9659|o__OFGB9659|f__FGB9659|g__GGB29003|s__GGB29003_SGB41719
k__Bacteria|p__Bacteria_unclassified|c__Bacteria_unclassified|o__Bacteria_unclassified|f__Bacteria_unclassified|g__GGB
k__Bacteria|p__Firmicutes|c__CFGB9827|o__OFGB9827|f__FGB9827|g__GGB29531|s__GGB29531_SGB42317
k__Bacteria|p__Firmicutes|c__Clostridia|o__Eubacteriales|f__Eubacteriaceae|g__GGB29685|s__GGB29685_SGB42494
k__Bacteria|p__Bacteria_unclassified|c__CFGB77303|o__OFGB77303|f__FGB77303|g__GGB30141|s__GGB30141_SGB430
k__Bacteria|p__Firmicutes|c__Clostridia|o__Eubacteriales|f__Eubacteriales_unclassified|g__GGB30286|s__GGB30286_SG
k__Bacteria|p__Firmicutes|c__Clostridia|o__Eubacteriales|f__Oscillospiraceae|g__GGB30303|s__GGB30303_SGB43268
k__Bacteria|p__Firmicutes|c__CFGB30328|o__OFGB30328|f__FGB30328|g__GGB30413|s__GGB30413_SGB43452
k__Bacteria|p__Firmicutes|c__Clostridia|o__Eubacteriales|f__Oscillospiraceae|g__GGB30454|s__GGB30454_SGB43514
k__Bacteria|p__Firmicutes|c__Clostridia|o__Eubacteriales|f__Oscillospiraceae|g__GGB30455|s__GGB30455_SGB43519
k__Bacteria|p__Firmicutes|c__Clostridia|o__Eubacteriales|f__Oscillospiraceae|g__GGB30461|s__GGB30461_SGB43527
k__Bacteria|p__Firmicutes|c__Clostridia|o__Eubacteriales|f__Oscillospiraceae|g__GGB30461|s__GGB30461_SGB43530
k__Bacteria|p__Firmicutes|c__Clostridia|o__Eubacteriales|f__Oscillospiraceae|g__GGB30463|s__GGB30463_SGB43537

k__Bacteria|p__Firmicutes|c__Clostridia|o__Eubacteriales|f__Oscillospiraceae|g__GGB30473|s__GGB30473_SGB43557
k__Bacteria|p__Firmicutes|c__Clostridia|o__Eubacteriales|f__Oscillospiraceae|g__GGB30475|s__GGB30475_SGB63182
k__Bacteria|p__Actinobacteria|c__CFGB77153|o__OFGB77153|f__FGB77153|g__GGB30861|s__GGB30861_SGB44083
k__Bacteria|p__Tenericutes|c__CFGB1791|o__OFGB1791|f__FGB1791|g__GGB31312|s__GGB31312_SGB44628
k__Bacteria|p__Firmicutes|c__CFGB10290|o__OFGB10290|f__FGB10290|g__GGB31438|s__GGB31438_SGB44768
k__Bacteria|p__Firmicutes|c__Clostridia|o__Eubacteriales|f__Oscillospiraceae|g__GGB3171|s__GGB3171_SGB4185
k__Bacteria|p__Firmicutes|c__CFGB1765|o__OFGB1765|f__FGB1765|g__GGB31823|s__GGB31823_SGB45199
k__Bacteria|p__Firmicutes|c__CFGB10349|o__OFGB10349|f__FGB10349|g__GGB31853|s__GGB31853_SGB45233
k__Bacteria|p__Firmicutes|c__CFGB10667|o__OFGB10667|f__FGB10667|g__GGB32371|s__GGB32371_SGB41694
k__Bacteria|p__Firmicutes|c__Clostridia|o__Eubacteriales|f__Lachnospiraceae|g__GGB3793|s__GGB3793_SGB5158
k__Bacteria|p__Firmicutes|c__Clostridia|o__Eubacteriales|f__Lachnospiraceae|g__GGB42598|s__GGB42598_SGB59794
k__Bacteria|p__Firmicutes|c__Clostridia|o__Eubacteriales|f__Christensenellaceae|g__GGB45656|s__GGB45656_SGB6337
k__Bacteria|p__Firmicutes|c__CFGB10299|o__OFGB10299|f__FGB10299|g__GGB47127|s__GGB47127_SGB65054
k__Bacteria|p__Firmicutes|c__Clostridia|o__Eubacteriales|f__Oscillospiraceae|g__GGB74395|s__GGB74395_SGB43521
k__Bacteria|p__Firmicutes|c__Clostridia|o__Eubacteriales|f__Oscillospiraceae|g__GGB75053|s__GGB75053_SGB43494
k__Bacteria|p__Firmicutes|c__Clostridia|o__Eubacteriales|f__Lachnospiraceae|g__GGB75109|s__GGB75109_SGB102238
k__Bacteria|p__Firmicutes|c__Clostridia|o__Clostridia_unclassified|f__Clostridia_unclassified|g__GGB81440|s__GGB8144
k__Bacteria|p__Firmicutes|c__Clostridia|o__Eubacteriales|f__Lachnospiraceae|g__Lachnospiraceae_unclassified|s__Lachn
k__Bacteria|p__Firmicutes|c__Clostridia|o__Eubacteriales|f__Lachnospiraceae|g__Lachnospiraceae_unclassified|s__Lachn
k__Bacteria|p__Firmicutes|c__Clostridia|o__Eubacteriales|f__Lachnospiraceae|g__Lachnospiraceae_unclassified|s__Lachn
k__Bacteria|p__Firmicutes|c__Clostridia|o__Eubacteriales|f__Lachnospiraceae|g__Lachnospiraceae_unclassified|s__Lachn
k__Bacteria|p__Firmicutes|c__Clostridia|o__Eubacteriales|f__Lachnospiraceae|g__Lachnospiraceae_unclassified|s__Lachn
k__Bacteria|p__Firmicutes|c__Clostridia|o__Eubacteriales|f__Lachnospiraceae|g__Lachnospiraceae_unclassified|s__Lachn
k__Bacteria|p__Firmicutes|c__Clostridia|o__Eubacteriales|f__Lachnospiraceae|g__Lachnospiraceae_unclassified|s__Lachn
k__Bacteria|p__Firmicutes|c__Clostridia|o__Eubacteriales|f__Lachnospiraceae|g__Lachnospiraceae_unclassified|s__Lachn
k__Bacteria|p__Firmicutes|c__Bacilli|o__Lactobacillales|f__Lactobacillaceae|g__Lactobacillus|s__Lactobacillus_johnsonii
k__Bacteria|p__Bacteroidota|c__Bacteroidia|o__Bacteroidales|f__Muribaculaceae|g__Muribaculaceae_unclassified|s__M
k__Bacteria|p__Firmicutes|c__Clostridia|o__Eubacteriales|f__Oscillospiraceae|g__Neglectibacter|s__Neglectibacter_sp_X
k__Bacteria|p__Firmicutes|c__Clostridia|o__Eubacteriales|f__Oscillospiraceae|g__Oscillospiraceae_unclassified|s__Oscillo
k__Bacteria|p__Firmicutes|c__Clostridia|o__Eubacteriales|f__Oscillospiraceae|g__Oscillospiraceae_unclassified|s__Oscillo
k__Bacteria|p__Firmicutes|c__Clostridia|o__Eubacteriales|f__Oscillospiraceae|g__Oscillospiraceae_unclassified|s__Oscillo
k__Bacteria|p__Proteobacteria|c__Betaproteobacteria|o__Burkholderiales|f__Sutterellaceae|g__Parasutterella|s__Parasu
k__Bacteria|p__Firmicutes|c__Clostridia|o__Eubacteriales|f__Peptostreptococcaceae|g__Romboutsia|s__Romboutsia_ilea
k__Bacteria|p__Firmicutes|c__Clostridia|o__Eubacteriales|f__Lachnospiraceae|g__Schaedlerella|s__Schaedlerella_arabino
k__Bacteria|p__Firmicutes|c__Erysipelotrichia|o__Erysipelotrichales|f__Turicibacteraceae|g__Turicibacter|s__Turicibacter_
k__Bacteria|p__Bacteria_unclassified|c__Bacteria_unclassified|o__Bacteria_unclassified|f__Bacteria_unclassified|g__Bacte
k__Bacteria|p__Bacteria_unclassified|c__Bacteria_unclassified|o__Bacteria_unclassified|f__Bacteria_unclassified|g__Bacte
k__Bacteria|p__Bacteria_unclassified|c__Bacteria_unclassified|o__Bacteria_unclassified|f__Bacteria_unclassified|g__Bacte

k__Bacteria|p__Firmicutes|c__Clostridia|o__Eubacteriales|f__Lachnospiraceae|g__Acetatifactor|s__Acetatifactor_SGB415
k__Bacteria|p__Firmicutes|c__Clostridia|o__Eubacteriales|f__Lachnospiraceae|g__Acetatifactor|s__Acetatifactor_muris
k__Bacteria|p__Firmicutes|c__Clostridia|o__Eubacteriales|f__Oscillospiraceae|g__Acutalibacter|s__Acutalibacter_muris
k__Bacteria|p__Firmicutes|c__Clostridia|o__Eubacteriales|f__Oscillospiraceae|g__Acutalibacter|s__Acutalibacter_sp_1XD
k__Bacteria|p__Actinobacteria|c__Coriobacteriia|o__Eggerthellales|f__Eggerthellaceae|g__Adlercreutzia|s__Adlercreutzia

k__Bacteria|p__Actinobacteria|c__Coriobacteriia|o__Eggerthellales|f__Eggerthellaceae|g__Adlercreutzia|s__Adlercreutzia
k__Bacteria|p__Actinobacteria|c__Coriobacteriia|o__Eggerthellales|f__Eggerthellaceae|g__Adlercreutzia|s__Adlercreutzia
k__Bacteria|p__Verrucomicrobia|c__Verrucomicrobiae|o__Verrucomicrobiales|f__Akkermansiaceae|g__Akkermansia|s__A
k__Bacteria|p__Bacteroidota|c__Bacteroidia|o__Bacteroidales|f__Rikenellaceae|g__Alistipes|s__Alistipes_sp_DSM_11234
k__Bacteria|p__Firmicutes|c__Clostridia|o__Eubacteriales|f__Oscillospiraceae|g__Anaerotruncus|s__Anaerotruncus_sp_1
k__Bacteria|p__Bacteria_unclassified|c__Bacteria_unclassified|o__Bacteria_unclassified|f__Bacteria_unclassified|g__Bacte
k__Bacteria|p__Bacteria_unclassified|c__Bacteria_unclassified|o__Bacteria_unclassified|f__Bacteria_unclassified|g__Bacte
k__Bacteria|p__Bacteria_unclassified|c__Bacteria_unclassified|o__Bacteria_unclassified|f__Bacteria_unclassified|g__Bacte
k__Bacteria|p__Bacteroidota|c__Bacteroidia|o__Bacteroidales|f__Bacteroidaceae|g__Bacteroides|s__Bacteroides_thetaio
k__Bacteria|p__Actinobacteria|c__Actinomycetia|o__Bifidobacteriales|f__Bifidobacteriaceae|g__Bifidobacterium|s__Bifid
k__Bacteria|p__Firmicutes|c__Clostridia|o__Clostridia_unclassified|f__Clostridia_unclassified|g__Clostridia_unclassified|s_
k__Bacteria|p__Firmicutes|c__Clostridia|o__Eubacteriales|f__Clostridiaceae|g__Clostridiaceae_unclassified|s__Clostridiac
k__Bacteria|p__Firmicutes|c__Clostridia|o__Eubacteriales|f__Clostridiaceae|g__Clostridiaceae_unclassified|s__Clostridiac
k__Bacteria|p__Firmicutes|c__Clostridia|o__Eubacteriales|f__Eubacteriales_unclassified|g__Eubacteriales_unclassified|s_
k__Bacteria|p__Firmicutes|c__Erysipelotrichia|o__Erysipelotrichales|f__Erysipelotrichaceae|g__Erysipelatoclostridium|s_
k__Bacteria|p__Actinobacteria|c__Coriobacteriia|o__Coriobacteriales|f__Coriobacteriaceae|g__Coriobacteriaceae_unclass
k__Bacteria|p__Firmicutes|c__Clostridia|o__Eubacteriales|f__Lachnospiraceae|g__Dorea|s__Dorea_sp_5_2
k__Bacteria|p__Firmicutes|c__Erysipelotrichia|o__Erysipelotrichales|f__Erysipelotrichaceae|g__Dubosiella|s__Dubosiella_
k__Bacteria|p__Firmicutes|c__Erysipelotrichia|o__Erysipelotrichales|f__Erysipelotrichales_unclassified|g__Erysipelotrichal
k__Bacteria|p__Firmicutes|c__Clostridia|o__Eubacteriales|f__Eubacteriaceae|g__Eubacteriaceae_unclassified|s__Eubacte
k__Bacteria|p__Firmicutes|c__Clostridia|o__Eubacteriales|f__Eubacteriaceae|g__Eubacteriaceae_unclassified|s__Eubacte
k__Bacteria|p__Firmicutes|c__Clostridia|o__Eubacteriales|f__Lachnospiraceae|g__GGB20149|s__GGB20149_SGB29430
k__Bacteria|p__Actinobacteria|c__Coriobacteriia|o__Eggerthellales|f__Eggerthellaceae|g__GGB22635|s__GGB22635_SGB
k__Bacteria|p__Firmicutes|c__Clostridia|o__Eubacteriales|f__Lachnospiraceae|g__GGB25041|s__GGB25041_SGB36960
k__Bacteria|p__Bacteroidota|c__Bacteroidia|o__Bacteroidales|f__Muribaculaceae|g__GGB27876|s__GGB27876_SGB4031
k__Bacteria|p__Bacteroidota|c__Bacteroidia|o__Bacteroidales|f__Muribaculaceae|g__GGB27878|s__GGB27878_SGB4031
k__Bacteria|p__Bacteroidota|c__Bacteroidia|o__Bacteroidales|f__Muribaculaceae|g__GGB27918|s__GGB27918_SGB4035
k__Bacteria|p__Firmicutes|c__CFGB9508|o__OFGB9508|f__FGB9508|g__GGB28382|s__GGB28382_SGB40962
k__Bacteria|p__Firmicutes|c__CFGB2838|o__OFGB2838|f__FGB2838|g__GGB28399|s__GGB28399_SGB40980
k__Bacteria|p__Firmicutes|c__CFGB2838|o__OFGB2838|f__FGB2838|g__GGB28411|s__GGB28411_SGB40993
k__Bacteria|p__Firmicutes|c__CFGB2838|o__OFGB2838|f__FGB2838|g__GGB28415|s__GGB28415_SGB40997
k__Bacteria|p__Firmicutes|c__Clostridia|o__Eubacteriales|f__Pumilibacteraceae|g__GGB28430|s__GGB28430_SGB41013
k__Bacteria|p__Firmicutes|c__CFGB28439|o__OFGB28439|f__FGB28439|g__GGB28439|s__GGB28439_SGB41022
k__Bacteria|p__Firmicutes|c__Clostridia|o__Clostridia_unclassified|f__Clostridia_unclassified|g__GGB28778|s__GGB2877
k__Bacteria|p__Firmicutes|c__Clostridia|o__Eubacteriales|f__Eubacteriaceae|g__GGB28784|s__GGB28784_SGB41437
k__Bacteria|p__Firmicutes|c__Clostridia|o__Eubacteriales|f__Lachnospiraceae|g__GGB28792|s__GGB28792_SGB41445
k__Bacteria|p__Firmicutes|c__Clostridia|o__Eubacteriales|f__Lachnospiraceae|g__GGB28798|s__GGB28798_SGB41451
k__Bacteria|p__Firmicutes|c__Clostridia|o__Eubacteriales|f__Lachnospiraceae|g__GGB28802|s__GGB28802_SGB41455
k__Bacteria|p__Firmicutes|c__Clostridia|o__Eubacteriales|f__Lachnospiraceae|g__GGB28818|s__GGB28818_SGB41473
k__Bacteria|p__Firmicutes|c__CFGB77305|o__OFGB77305|f__FGB77305|g__GGB28828|s__GGB28828_SGB41484
k__Bacteria|p__Firmicutes|c__Clostridia|o__Eubacteriales|f__Clostridiaceae|g__GGB28851|s__GGB28851_SGB41518
k__Bacteria|p__Firmicutes|c__Clostridia|o__Eubacteriales|f__Lachnospiraceae|g__GGB28859|s__GGB28859_SGB41528
k__Bacteria|p__Firmicutes|c__Clostridia|o__Eubacteriales|f__Lachnospiraceae|g__GGB28864|s__GGB28864_SGB41535
k__Bacteria|p__Firmicutes|c__Clostridia|o__Eubacteriales|f__Lachnospiraceae|g__GGB28869|s__GGB28869_SGB41543
k__Bacteria|p__Firmicutes|c__CFGB9633|o__OFGB9633|f__FGB9633|g__GGB28883|s__GGB28883_SGB41564
k__Bacteria|p__Bacteria_unclassified|c__Bacteria_unclassified|o__Bacteria_unclassified|f__Bacteria_unclassified|g__GGB2

k__Bacteria|p__Bacteria_unclassified|c__Bacteria_unclassified|o__Bacteria_unclassified|f__Bacteria_unclassified|g__GGB2
k__Bacteria|p__Bacteria_unclassified|c__Bacteria_unclassified|o__Bacteria_unclassified|f__Bacteria_unclassified|g__GGB2
k__Bacteria|p__Bacteria_unclassified|c__Bacteria_unclassified|o__Bacteria_unclassified|f__Bacteria_unclassified|g__GGB2
k__Bacteria|p__Firmicutes|c__Clostridia|o__Eubacteriales|f__Lachnospiraceae|g__GGB28916|s__GGB28916_SGB41612
k__Bacteria|p__Firmicutes|c__Clostridia|o__Eubacteriales|f__Lachnospiraceae|g__GGB28924|s__GGB28924_SGB41621
k__Bacteria|p__Firmicutes|c__Clostridia|o__Eubacteriales|f__Lachnospiraceae|g__GGB28926|s__GGB28926_SGB41624
k__Bacteria|p__Bacteria_unclassified|c__CFGB77359|o__OFGB77359|f__FGB77359|g__GGB28927|s__GGB28927_SGB416
k__Bacteria|p__Firmicutes|c__CFGB9639|o__OFGB9639|f__FGB9639|g__GGB28934|s__GGB28934_SGB41635
k__Bacteria|p__Firmicutes|c__Clostridia|o__Eubacteriales|f__Lachnospiraceae|g__GGB28946|s__GGB28946_SGB41652
k__Bacteria|p__Firmicutes|c__Clostridia|o__Eubacteriales|f__Lachnospiraceae|g__GGB28949|s__GGB28949_SGB41655
k__Bacteria|p__Firmicutes|c__Clostridia|o__Eubacteriales|f__Lachnospiraceae|g__GGB28949|s__GGB28949_SGB41656
k__Bacteria|p__Firmicutes|c__Clostridia|o__Eubacteriales|f__Clostridiaceae|g__GGB28950|s__GGB28950_SGB41657
k__Bacteria|p__Firmicutes|c__Clostridia|o__Eubacteriales|f__Clostridiaceae|g__GGB28951|s__GGB28951_SGB102295
k__Bacteria|p__Firmicutes|c__Clostridia|o__Eubacteriales|f__Clostridiaceae|g__GGB28951|s__GGB28951_SGB41658
k__Bacteria|p__Firmicutes|c__Clostridia|o__Eubacteriales|f__Clostridiaceae|g__GGB28954|s__GGB28954_SGB41662
k__Bacteria|p__Firmicutes|c__Clostridia|o__Eubacteriales|f__Clostridiaceae|g__GGB28956|s__GGB28956_SGB41665
k__Bacteria|p__Firmicutes|c__Clostridia|o__Eubacteriales|f__Clostridiaceae|g__GGB28960|s__GGB28960_SGB41669
k__Bacteria|p__Firmicutes|c__Clostridia|o__Eubacteriales|f__Clostridiaceae|g__GGB28967|s__GGB28967_SGB41678
k__Bacteria|p__Firmicutes|c__Clostridia|o__Eubacteriales|f__Eubacteriaceae|g__GGB28991|s__GGB28991_SGB41705
k__Bacteria|p__Firmicutes|c__CFGB9658|o__OFGB9658|f__FGB9658|g__GGB29002|s__GGB29002_SGB41718
k__Bacteria|p__Firmicutes|c__CFGB9659|o__OFGB9659|f__FGB9659|g__GGB29003|s__GGB29003_SGB41719
k__Bacteria|p__Bacteria_unclassified|c__Bacteria_unclassified|o__Bacteria_unclassified|f__Bacteria_unclassified|g__GGB2
k__Bacteria|p__Firmicutes|c__CFGB9827|o__OFGB9827|f__FGB9827|g__GGB29531|s__GGB29531_SGB42317
k__Bacteria|p__Firmicutes|c__Clostridia|o__Eubacteriales|f__Eubacteriaceae|g__GGB29685|s__GGB29685_SGB42494
k__Bacteria|p__Bacteria_unclassified|c__CFGB77303|o__OFGB77303|f__FGB77303|g__GGB30141|s__GGB30141_SGB430
k__Bacteria|p__Firmicutes|c__Clostridia|o__Eubacteriales|f__Eubacteriales_unclassified|g__GGB30286|s__GGB30286_SG
k__Bacteria|p__Firmicutes|c__Clostridia|o__Eubacteriales|f__Oscillospiraceae|g__GGB30303|s__GGB30303_SGB43268
k__Bacteria|p__Firmicutes|c__CFGB30328|o__OFGB30328|f__FGB30328|g__GGB30413|s__GGB30413_SGB43452
k__Bacteria|p__Firmicutes|c__Clostridia|o__Eubacteriales|f__Oscillospiraceae|g__GGB30454|s__GGB30454_SGB43514
k__Bacteria|p__Firmicutes|c__Clostridia|o__Eubacteriales|f__Oscillospiraceae|g__GGB30455|s__GGB30455_SGB43519
k__Bacteria|p__Firmicutes|c__Clostridia|o__Eubacteriales|f__Oscillospiraceae|g__GGB30461|s__GGB30461_SGB43527
k__Bacteria|p__Firmicutes|c__Clostridia|o__Eubacteriales|f__Oscillospiraceae|g__GGB30461|s__GGB30461_SGB43530
k__Bacteria|p__Firmicutes|c__Clostridia|o__Eubacteriales|f__Oscillospiraceae|g__GGB30463|s__GGB30463_SGB43537
k__Bacteria|p__Firmicutes|c__Clostridia|o__Eubacteriales|f__Oscillospiraceae|g__GGB30473|s__GGB30473_SGB43557
k__Bacteria|p__Firmicutes|c__Clostridia|o__Eubacteriales|f__Oscillospiraceae|g__GGB30475|s__GGB30475_SGB63182
k__Bacteria|p__Actinobacteria|c__CFGB77153|o__OFGB77153|f__FGB77153|g__GGB30861|s__GGB30861_SGB44083
k__Bacteria|p__Tenericutes|c__CFGB1791|o__OFGB1791|f__FGB1791|g__GGB31312|s__GGB31312_SGB44628
k__Bacteria|p__Firmicutes|c__CFGB10290|o__OFGB10290|f__FGB10290|g__GGB31438|s__GGB31438_SGB44768
k__Bacteria|p__Firmicutes|c__Clostridia|o__Eubacteriales|f__Oscillospiraceae|g__GGB3171|s__GGB3171_SGB4185
k__Bacteria|p__Firmicutes|c__CFGB1765|o__OFGB1765|f__FGB1765|g__GGB31823|s__GGB31823_SGB45199
k__Bacteria|p__Firmicutes|c__CFGB10349|o__OFGB10349|f__FGB10349|g__GGB31853|s__GGB31853_SGB45233
k__Bacteria|p__Firmicutes|c__CFGB10667|o__OFGB10667|f__FGB10667|g__GGB32371|s__GGB32371_SGB41694
k__Bacteria|p__Firmicutes|c__Clostridia|o__Eubacteriales|f__Lachnospiraceae|g__GGB3793|s__GGB3793_SGB5158
k__Bacteria|p__Firmicutes|c__Clostridia|o__Eubacteriales|f__Lachnospiraceae|g__GGB42598|s__GGB42598_SGB59794
k__Bacteria|p__Firmicutes|c__Clostridia|o__Eubacteriales|f__Christensenellaceae|g__GGB45656|s__GGB45656_SGB6337
k__Bacteria|p__Firmicutes|c__CFGB10299|o__OFGB10299|f__FGB10299|g__GGB47127|s__GGB47127_SGB65054

k__Bacteria|p__Firmicutes|c__Clostridia|o__Eubacteriales|f__Oscillospiraceae|g__GGB74395|s__GGB74395_SGB43521
k__Bacteria|p__Firmicutes|c__Clostridia|o__Eubacteriales|f__Oscillospiraceae|g__GGB75053|s__GGB75053_SGB43494
k__Bacteria|p__Firmicutes|c__Clostridia|o__Eubacteriales|f__Lachnospiraceae|g__GGB75109|s__GGB75109_SGB102238
k__Bacteria|p__Firmicutes|c__Clostridia|o__Clostridia_unclassified|f__Clostridia_unclassified|g__GGB81440|s__GGB8144
k__Bacteria|p__Firmicutes|c__Clostridia|o__Eubacteriales|f__Lachnospiraceae|g__Lachnospiraceae_unclassified|s__Lachn
k__Bacteria|p__Firmicutes|c__Clostridia|o__Eubacteriales|f__Lachnospiraceae|g__Lachnospiraceae_unclassified|s__Lachn
k__Bacteria|p__Firmicutes|c__Clostridia|o__Eubacteriales|f__Lachnospiraceae|g__Lachnospiraceae_unclassified|s__Lachn
k__Bacteria|p__Firmicutes|c__Clostridia|o__Eubacteriales|f__Lachnospiraceae|g__Lachnospiraceae_unclassified|s__Lachn
k__Bacteria|p__Firmicutes|c__Clostridia|o__Eubacteriales|f__Lachnospiraceae|g__Lachnospiraceae_unclassified|s__Lachn
k__Bacteria|p__Firmicutes|c__Clostridia|o__Eubacteriales|f__Lachnospiraceae|g__Lachnospiraceae_unclassified|s__Lachn
k__Bacteria|p__Firmicutes|c__Clostridia|o__Eubacteriales|f__Lachnospiraceae|g__Lachnospiraceae_unclassified|s__Lachn
k__Bacteria|p__Firmicutes|c__Clostridia|o__Eubacteriales|f__Lachnospiraceae|g__Lachnospiraceae_unclassified|s__Lachn
k__Bacteria|p__Firmicutes|c__Bacilli|o__Lactobacillales|f__Lactobacillaceae|g__Lactobacillus|s__Lactobacillus_johnsonii
k__Bacteria|p__Bacteroidota|c__Bacteroidia|o__Bacteroidales|f__Muribaculaceae|g__Muribaculaceae_unclassified|s__M
k__Bacteria|p__Firmicutes|c__Clostridia|o__Eubacteriales|f__Oscillospiraceae|g__Neglectibacter|s__Neglectibacter_sp_X
k__Bacteria|p__Firmicutes|c__Clostridia|o__Eubacteriales|f__Oscillospiraceae|g__Oscillospiraceae_unclassified|s__Oscillo
k__Bacteria|p__Firmicutes|c__Clostridia|o__Eubacteriales|f__Oscillospiraceae|g__Oscillospiraceae_unclassified|s__Oscillo
k__Bacteria|p__Firmicutes|c__Clostridia|o__Eubacteriales|f__Oscillospiraceae|g__Oscillospiraceae_unclassified|s__Oscillo
k__Bacteria|p__Proteobacteria|c__Betaproteobacteria|o__Burkholderiales|f__Sutterellaceae|g__Parasutterella|s__Parasu
k__Bacteria|p__Firmicutes|c__Clostridia|o__Eubacteriales|f__Peptostreptococcaceae|g__Romboutsia|s__Romboutsia_ilea
k__Bacteria|p__Firmicutes|c__Clostridia|o__Eubacteriales|f__Lachnospiraceae|g__Schaedlerella|s__Schaedlerella_arabinc
k__Bacteria|p__Firmicutes|c__Erysipelotrichia|o__Erysipelotrichales|f__Turicibacteraceae|g__Turicibacter|s__Turicibacter_
k__Bacteria|p__Bacteria_unclassified|c__Bacteria_unclassified|o__Bacteria_unclassified|f__Bacteria_unclassified|g__Bacte
k__Bacteria|p__Bacteria_unclassified|c__Bacteria_unclassified|o__Bacteria_unclassified|f__Bacteria_unclassified|g__Bacte
k__Bacteria|p__Bacteria_unclassified|c__Bacteria_unclassified|o__Bacteria_unclassified|f__Bacteria_unclassified|g__Bacte

k__Bacteria|p__Firmicutes|c__Clostridia|o__Eubacteriales|f__Lachnospiraceae|g__Acetatifactor|s__Acetatifactor_SGB415
k__Bacteria|p__Firmicutes|c__Clostridia|o__Eubacteriales|f__Lachnospiraceae|g__Acetatifactor|s__Acetatifactor_muris
k__Bacteria|p__Firmicutes|c__Clostridia|o__Eubacteriales|f__Oscillospiraceae|g__Acutalibacter|s__Acutalibacter_muris
k__Bacteria|p__Firmicutes|c__Clostridia|o__Eubacteriales|f__Oscillospiraceae|g__Acutalibacter|s__Acutalibacter_sp_1XD
k__Bacteria|p__Actinobacteria|c__Coriobacteriia|o__Eggerthellales|f__Eggerthellaceae|g__Adlercreutzia|s__Adlercreutzia
k__Bacteria|p__Actinobacteria|c__Coriobacteriia|o__Eggerthellales|f__Eggerthellaceae|g__Adlercreutzia|s__Adlercreutzia
k__Bacteria|p__Actinobacteria|c__Coriobacteriia|o__Eggerthellales|f__Eggerthellaceae|g__Adlercreutzia|s__Adlercreutzia
k__Bacteria|p__Verrucomicrobia|c__Verrucomicrobiae|o__Verrucomicrobiales|f__Akkermansiaceae|g__Akkermansia|s__
k__Bacteria|p__Bacteroidota|c__Bacteroidia|o__Bacteroidales|f__Rikenellaceae|g__Alistipes|s__Alistipes_sp_DSM_11234
k__Bacteria|p__Firmicutes|c__Clostridia|o__Eubacteriales|f__Oscillospiraceae|g__Anaerotruncus|s__Anaerotruncus_sp_1
k__Bacteria|p__Bacteria_unclassified|c__Bacteria_unclassified|o__Bacteria_unclassified|f__Bacteria_unclassified|g__Bacte
k__Bacteria|p__Bacteria_unclassified|c__Bacteria_unclassified|o__Bacteria_unclassified|f__Bacteria_unclassified|g__Bacte
k__Bacteria|p__Bacteria_unclassified|c__Bacteria_unclassified|o__Bacteria_unclassified|f__Bacteria_unclassified|g__Bacte
k__Bacteria|p__Bacteroidota|c__Bacteroidia|o__Bacteroidales|f__Bacteroidaceae|g__Bacteroides|s__Bacteroides_thetaio
k__Bacteria|p__Actinobacteria|c__Actinomycetia|o__Bifidobacteriales|f__Bifidobacteriaceae|g__Bifidobacterium|s__Bifid
k__Bacteria|p__Firmicutes|c__Clostridia|o__Clostridia_unclassified|f__Clostridia_unclassified|g__Clostridia_unclassified|s_
k__Bacteria|p__Firmicutes|c__Clostridia|o__Eubacteriales|f__Clostridiaceae|g__Clostridiaceae_unclassified|s__Clostridiac
k__Bacteria|p__Firmicutes|c__Clostridia|o__Eubacteriales|f__Clostridiaceae|g__Clostridiaceae_unclassified|s__Clostridiac

k__Bacteria|p__Firmicutes|c__Clostridia|o__Eubacteriales|f__Eubacteriales_unclassified|g__Eubacteriales_unclassified|s_
k__Bacteria|p__Firmicutes|c__Erysipelotrichia|o__Erysipelotrichales|f__Erysipelotrichaceae|g__Erysipelatoclostridium|s__
k__Bacteria|p__Actinobacteria|c__Coriobacteriia|o__Coriobacteriales|f__Coriobacteriaceae|g__Coriobacteriaceae_unclass
k__Bacteria|p__Firmicutes|c__Clostridia|o__Eubacteriales|f__Lachnospiraceae|g__Dorea|s__Dorea_sp_5_2
k__Bacteria|p__Firmicutes|c__Erysipelotrichia|o__Erysipelotrichales|f__Erysipelotrichaceae|g__Dubosiella|s__Dubosiella_
k__Bacteria|p__Firmicutes|c__Erysipelotrichia|o__Erysipelotrichales|f__Erysipelotrichales_unclassified|g__Erysipelotrichal
k__Bacteria|p__Firmicutes|c__Clostridia|o__Eubacteriales|f__Eubacteriaceae|g__Eubacteriaceae_unclassified|s__Eubacte
k__Bacteria|p__Firmicutes|c__Clostridia|o__Eubacteriales|f__Eubacteriaceae|g__Eubacteriaceae_unclassified|s__Eubacte
k__Bacteria|p__Firmicutes|c__Clostridia|o__Eubacteriales|f__Lachnospiraceae|g__GGB20149|s__GGB20149_SGB29430
k__Bacteria|p__Actinobacteria|c__Coriobacteriia|o__Eggerthellales|f__Eggerthellaceae|g__GGB22635|s__GGB22635_SGB
k__Bacteria|p__Firmicutes|c__Clostridia|o__Eubacteriales|f__Lachnospiraceae|g__GGB25041|s__GGB25041_SGB36960
k__Bacteria|p__Bacteroidota|c__Bacteroidia|o__Bacteroidales|f__Muribaculaceae|g__GGB27876|s__GGB27876_SGB4031
k__Bacteria|p__Bacteroidota|c__Bacteroidia|o__Bacteroidales|f__Muribaculaceae|g__GGB27878|s__GGB27878_SGB4031
k__Bacteria|p__Bacteroidota|c__Bacteroidia|o__Bacteroidales|f__Muribaculaceae|g__GGB27918|s__GGB27918_SGB4035
k__Bacteria|p__Firmicutes|c__CFGB9508|o__OFGB9508|f__FGB9508|g__GGB28382|s__GGB28382_SGB40962
k__Bacteria|p__Firmicutes|c__CFGB2838|o__OFGB2838|f__FGB2838|g__GGB28399|s__GGB28399_SGB40980
k__Bacteria|p__Firmicutes|c__CFGB2838|o__OFGB2838|f__FGB2838|g__GGB28411|s__GGB28411_SGB40993
k__Bacteria|p__Firmicutes|c__CFGB2838|o__OFGB2838|f__FGB2838|g__GGB28415|s__GGB28415_SGB40997
k__Bacteria|p__Firmicutes|c__Clostridia|o__Eubacteriales|f__Pumilibacteraceae|g__GGB28430|s__GGB28430_SGB41013
k__Bacteria|p__Firmicutes|c__CFGB28439|o__OFGB28439|f__FGB28439|g__GGB28439|s__GGB28439_SGB41022
k__Bacteria|p__Firmicutes|c__Clostridia|o__Clostridia_unclassified|f__Clostridia_unclassified|g__GGB28778|s__GGB2877
k__Bacteria|p__Firmicutes|c__Clostridia|o__Eubacteriales|f__Eubacteriaceae|g__GGB28784|s__GGB28784_SGB41437
k__Bacteria|p__Firmicutes|c__Clostridia|o__Eubacteriales|f__Lachnospiraceae|g__GGB28792|s__GGB28792_SGB41445
k__Bacteria|p__Firmicutes|c__Clostridia|o__Eubacteriales|f__Lachnospiraceae|g__GGB28798|s__GGB28798_SGB41451
k__Bacteria|p__Firmicutes|c__Clostridia|o__Eubacteriales|f__Lachnospiraceae|g__GGB28802|s__GGB28802_SGB41455
k__Bacteria|p__Firmicutes|c__Clostridia|o__Eubacteriales|f__Lachnospiraceae|g__GGB28818|s__GGB28818_SGB41473
k__Bacteria|p__Firmicutes|c__CFGB77305|o__OFGB77305|f__FGB77305|g__GGB28828|s__GGB28828_SGB41484
k__Bacteria|p__Firmicutes|c__Clostridia|o__Eubacteriales|f__Clostridiaceae|g__GGB28851|s__GGB28851_SGB41518
k__Bacteria|p__Firmicutes|c__Clostridia|o__Eubacteriales|f__Lachnospiraceae|g__GGB28859|s__GGB28859_SGB41528
k__Bacteria|p__Firmicutes|c__Clostridia|o__Eubacteriales|f__Lachnospiraceae|g__GGB28864|s__GGB28864_SGB41535
k__Bacteria|p__Firmicutes|c__Clostridia|o__Eubacteriales|f__Lachnospiraceae|g__GGB28869|s__GGB28869_SGB41543
k__Bacteria|p__Firmicutes|c__CFGB9633|o__OFGB9633|f__FGB9633|g__GGB28883|s__GGB28883_SGB41564
k__Bacteria|p__Bacteria_unclassified|c__Bacteria_unclassified|o__Bacteria_unclassified|f__Bacteria_unclassified|g__GGB
k__Bacteria|p__Bacteria_unclassified|c__Bacteria_unclassified|o__Bacteria_unclassified|f__Bacteria_unclassified|g__GGB
k__Bacteria|p__Bacteria_unclassified|c__Bacteria_unclassified|o__Bacteria_unclassified|f__Bacteria_unclassified|g__GGB
k__Bacteria|p__Bacteria_unclassified|c__Bacteria_unclassified|o__Bacteria_unclassified|f__Bacteria_unclassified|g__GGB
k__Bacteria|p__Firmicutes|c__Clostridia|o__Eubacteriales|f__Lachnospiraceae|g__GGB28916|s__GGB28916_SGB41612
k__Bacteria|p__Firmicutes|c__Clostridia|o__Eubacteriales|f__Lachnospiraceae|g__GGB28924|s__GGB28924_SGB41621
k__Bacteria|p__Firmicutes|c__Clostridia|o__Eubacteriales|f__Lachnospiraceae|g__GGB28926|s__GGB28926_SGB41624
k__Bacteria|p__Bacteria_unclassified|c__CFGB77359|o__OFGB77359|f__FGB77359|g__GGB28927|s__GGB28927_SGB416
k__Bacteria|p__Firmicutes|c__CFGB9639|o__OFGB9639|f__FGB9639|g__GGB28934|s__GGB28934_SGB41635
k__Bacteria|p__Firmicutes|c__Clostridia|o__Eubacteriales|f__Lachnospiraceae|g__GGB28946|s__GGB28946_SGB41652
k__Bacteria|p__Firmicutes|c__Clostridia|o__Eubacteriales|f__Lachnospiraceae|g__GGB28949|s__GGB28949_SGB41655
k__Bacteria|p__Firmicutes|c__Clostridia|o__Eubacteriales|f__Lachnospiraceae|g__GGB28949|s__GGB28949_SGB41656
k__Bacteria|p__Firmicutes|c__Clostridia|o__Eubacteriales|f__Clostridiaceae|g__GGB28950|s__GGB28950_SGB41657
k__Bacteria|p__Firmicutes|c__Clostridia|o__Eubacteriales|f__Clostridiaceae|g__GGB28951|s__GGB28951_SGB102295

k__Bacteria|p__Firmicutes|c__Clostridia|o__Eubacteriales|f__Clostridiaceae|g__GGB28951|s__GGB28951_SGB41658
k__Bacteria|p__Firmicutes|c__Clostridia|o__Eubacteriales|f__Clostridiaceae|g__GGB28954|s__GGB28954_SGB41662
k__Bacteria|p__Firmicutes|c__Clostridia|o__Eubacteriales|f__Clostridiaceae|g__GGB28956|s__GGB28956_SGB41665
k__Bacteria|p__Firmicutes|c__Clostridia|o__Eubacteriales|f__Clostridiaceae|g__GGB28960|s__GGB28960_SGB41669
k__Bacteria|p__Firmicutes|c__Clostridia|o__Eubacteriales|f__Clostridiaceae|g__GGB28967|s__GGB28967_SGB41678
k__Bacteria|p__Firmicutes|c__Clostridia|o__Eubacteriales|f__Eubacteriaceae|g__GGB28991|s__GGB28991_SGB41705
k__Bacteria|p__Firmicutes|c__CFGB9658|o__OFGB9658|f__FGB9658|g__GGB29002|s__GGB29002_SGB41718
k__Bacteria|p__Firmicutes|c__CFGB9659|o__OFGB9659|f__FGB9659|g__GGB29003|s__GGB29003_SGB41719
k__Bacteria|p__Bacteria_unclassified|c__Bacteria_unclassified|o__Bacteria_unclassified|f__Bacteria_unclassified|g__GGB2
k__Bacteria|p__Firmicutes|c__CFGB9827|o__OFGB9827|f__FGB9827|g__GGB29531|s__GGB29531_SGB42317
k__Bacteria|p__Firmicutes|c__Clostridia|o__Eubacteriales|f__Eubacteriaceae|g__GGB29685|s__GGB29685_SGB42494
k__Bacteria|p__Bacteria_unclassified|c__CFGB77303|o__OFGB77303|f__FGB77303|g__GGB30141|s__GGB30141_SGB430
k__Bacteria|p__Firmicutes|c__Clostridia|o__Eubacteriales|f__Eubacteriales_unclassified|g__GGB30286|s__GGB30286_SG
k__Bacteria|p__Firmicutes|c__Clostridia|o__Eubacteriales|f__Oscillospiraceae|g__GGB30303|s__GGB30303_SGB43268
k__Bacteria|p__Firmicutes|c__CFGB30328|o__OFGB30328|f__FGB30328|g__GGB30413|s__GGB30413_SGB43452
k__Bacteria|p__Firmicutes|c__Clostridia|o__Eubacteriales|f__Oscillospiraceae|g__GGB30454|s__GGB30454_SGB43514
k__Bacteria|p__Firmicutes|c__Clostridia|o__Eubacteriales|f__Oscillospiraceae|g__GGB30455|s__GGB30455_SGB43519
k__Bacteria|p__Firmicutes|c__Clostridia|o__Eubacteriales|f__Oscillospiraceae|g__GGB30461|s__GGB30461_SGB43527
k__Bacteria|p__Firmicutes|c__Clostridia|o__Eubacteriales|f__Oscillospiraceae|g__GGB30461|s__GGB30461_SGB43530
k__Bacteria|p__Firmicutes|c__Clostridia|o__Eubacteriales|f__Oscillospiraceae|g__GGB30463|s__GGB30463_SGB43537
k__Bacteria|p__Firmicutes|c__Clostridia|o__Eubacteriales|f__Oscillospiraceae|g__GGB30473|s__GGB30473_SGB43557
k__Bacteria|p__Firmicutes|c__Clostridia|o__Eubacteriales|f__Oscillospiraceae|g__GGB30475|s__GGB30475_SGB63182
k__Bacteria|p__Actinobacteria|c__CFGB77153|o__OFGB77153|f__FGB77153|g__GGB30861|s__GGB30861_SGB44083
k__Bacteria|p__Tenericutes|c__CFGB1791|o__OFGB1791|f__FGB1791|g__GGB31312|s__GGB31312_SGB44628
k__Bacteria|p__Firmicutes|c__CFGB10290|o__OFGB10290|f__FGB10290|g__GGB31438|s__GGB31438_SGB44768
k__Bacteria|p__Firmicutes|c__Clostridia|o__Eubacteriales|f__Oscillospiraceae|g__GGB3171|s__GGB3171_SGB4185
k__Bacteria|p__Firmicutes|c__CFGB1765|o__OFGB1765|f__FGB1765|g__GGB31823|s__GGB31823_SGB45199
k__Bacteria|p__Firmicutes|c__CFGB10349|o__OFGB10349|f__FGB10349|g__GGB31853|s__GGB31853_SGB45233
k__Bacteria|p__Firmicutes|c__CFGB10667|o__OFGB10667|f__FGB10667|g__GGB32371|s__GGB32371_SGB41694
k__Bacteria|p__Firmicutes|c__Clostridia|o__Eubacteriales|f__Lachnospiraceae|g__GGB3793|s__GGB3793_SGB5158
k__Bacteria|p__Firmicutes|c__Clostridia|o__Eubacteriales|f__Lachnospiraceae|g__GGB42598|s__GGB42598_SGB59794
k__Bacteria|p__Firmicutes|c__Clostridia|o__Eubacteriales|f__Christensenellaceae|g__GGB45656|s__GGB45656_SGB6337
k__Bacteria|p__Firmicutes|c__CFGB10299|o__OFGB10299|f__FGB10299|g__GGB47127|s__GGB47127_SGB65054
k__Bacteria|p__Firmicutes|c__Clostridia|o__Eubacteriales|f__Oscillospiraceae|g__GGB74395|s__GGB74395_SGB43521
k__Bacteria|p__Firmicutes|c__Clostridia|o__Eubacteriales|f__Oscillospiraceae|g__GGB75053|s__GGB75053_SGB43494
k__Bacteria|p__Firmicutes|c__Clostridia|o__Eubacteriales|f__Lachnospiraceae|g__GGB75109|s__GGB75109_SGB102238
k__Bacteria|p__Firmicutes|c__Clostridia|o__Clostridia_unclassified|f__Clostridia_unclassified|g__GGB81440|s__GGB8144
k__Bacteria|p__Firmicutes|c__Clostridia|o__Eubacteriales|f__Lachnospiraceae|g__Lachnospiraceae_unclassified|s__Lachn
k__Bacteria|p__Firmicutes|c__Clostridia|o__Eubacteriales|f__Lachnospiraceae|g__Lachnospiraceae_unclassified|s__Lachn
k__Bacteria|p__Firmicutes|c__Clostridia|o__Eubacteriales|f__Lachnospiraceae|g__Lachnospiraceae_unclassified|s__Lachn
k__Bacteria|p__Firmicutes|c__Clostridia|o__Eubacteriales|f__Lachnospiraceae|g__Lachnospiraceae_unclassified|s__Lachn
k__Bacteria|p__Firmicutes|c__Clostridia|o__Eubacteriales|f__Lachnospiraceae|g__Lachnospiraceae_unclassified|s__Lachn
k__Bacteria|p__Firmicutes|c__Clostridia|o__Eubacteriales|f__Lachnospiraceae|g__Lachnospiraceae_unclassified|s__Lachn
k__Bacteria|p__Firmicutes|c__Clostridia|o__Eubacteriales|f__Lachnospiraceae|g__Lachnospiraceae_unclassified|s__Lachn
k__Bacteria|p__Firmicutes|c__Clostridia|o__Eubacteriales|f__Lachnospiraceae|g__Lachnospiraceae_unclassified|s__Lachn
k__Bacteria|p__Firmicutes|c__Bacilli|o__Lactobacillales|f__Lactobacillaceae|g__Lactobacillus|s__Lactobacillus_johnsonii

k__Bacteria|p__Bacteroidota|c__Bacteroidia|o__Bacteroidales|f__Muribaculaceae|g__Muribaculaceae_unclassified|s__M
k__Bacteria|p__Firmicutes|c__Clostridia|o__Eubacteriales|f__Oscillospiraceae|g__Neglectibacter|s__Neglectibacter_sp_X
k__Bacteria|p__Firmicutes|c__Clostridia|o__Eubacteriales|f__Oscillospiraceae|g__Oscillospiraceae_unclassified|s__Oscillo
k__Bacteria|p__Firmicutes|c__Clostridia|o__Eubacteriales|f__Oscillospiraceae|g__Oscillospiraceae_unclassified|s__Oscillo
k__Bacteria|p__Firmicutes|c__Clostridia|o__Eubacteriales|f__Oscillospiraceae|g__Oscillospiraceae_unclassified|s__Oscillo
k__Bacteria|p__Proteobacteria|c__Betaproteobacteria|o__Burkholderiales|f__Sutterellaceae|g__Parasutterella|s__Parasu
k__Bacteria|p__Firmicutes|c__Clostridia|o__Eubacteriales|f__Peptostreptococcaceae|g__Romboutsia|s__Romboutsia_ilea
k__Bacteria|p__Firmicutes|c__Clostridia|o__Eubacteriales|f__Lachnospiraceae|g__Schaedlerella|s__Schaedlerella_arabino
k__Bacteria|p__Firmicutes|c__Erysipelotrichia|o__Erysipelotrichales|f__Turicibacteraceae|g__Turicibacter|s__Turicibacter_
k__Bacteria|p__Bacteria_unclassified|c__Bacteria_unclassified|o__Bacteria_unclassified|f__Bacteria_unclassified|g__Bacte
k__Bacteria|p__Bacteria_unclassified|c__Bacteria_unclassified|o__Bacteria_unclassified|f__Bacteria_unclassified|g__Bacte
k__Bacteria|p__Bacteria_unclassified|c__Bacteria_unclassified|o__Bacteria_unclassified|f__Bacteria_unclassified|g__Bacte

k__Bacteria|p__Firmicutes|c__Clostridia|o__Eubacteriales|f__Lachnospiraceae|g__Acetatifactor|s__Acetatifactor_SGB415
k__Bacteria|p__Firmicutes|c__Clostridia|o__Eubacteriales|f__Lachnospiraceae|g__Acetatifactor|s__Acetatifactor_muris
k__Bacteria|p__Firmicutes|c__Clostridia|o__Eubacteriales|f__Oscillospiraceae|g__Acutalibacter|s__Acutalibacter_muris
k__Bacteria|p__Firmicutes|c__Clostridia|o__Eubacteriales|f__Oscillospiraceae|g__Acutalibacter|s__Acutalibacter_sp_1XD
k__Bacteria|p__Actinobacteria|c__Coriobacteriia|o__Eggerthellales|f__Eggerthellaceae|g__Adlercreutzia|s__Adlercreutzia
k__Bacteria|p__Actinobacteria|c__Coriobacteriia|o__Eggerthellales|f__Eggerthellaceae|g__Adlercreutzia|s__Adlercreutzia
k__Bacteria|p__Actinobacteria|c__Coriobacteriia|o__Eggerthellales|f__Eggerthellaceae|g__Adlercreutzia|s__Adlercreutzia
k__Bacteria|p__Verrucomicrobia|c__Verrucomicrobiae|o__Verrucomicrobiales|f__Akkermansiaceae|g__Akkermansia|s__A
k__Bacteria|p__Bacteroidota|c__Bacteroidia|o__Bacteroidales|f__Rikenellaceae|g__Alistipes|s__Alistipes_sp_DSM_11234
k__Bacteria|p__Firmicutes|c__Clostridia|o__Eubacteriales|f__Oscillospiraceae|g__Anaerotruncus|s__Anaerotruncus_sp_1
k__Bacteria|p__Bacteria_unclassified|c__Bacteria_unclassified|o__Bacteria_unclassified|f__Bacteria_unclassified|g__Bacte
k__Bacteria|p__Bacteria_unclassified|c__Bacteria_unclassified|o__Bacteria_unclassified|f__Bacteria_unclassified|g__Bacte
k__Bacteria|p__Bacteria_unclassified|c__Bacteria_unclassified|o__Bacteria_unclassified|f__Bacteria_unclassified|g__Bacte
k__Bacteria|p__Bacteroidota|c__Bacteroidia|o__Bacteroidales|f__Bacteroidaceae|g__Bacteroides|s__Bacteroides_thetaio
k__Bacteria|p__Actinobacteria|c__Actinomycetia|o__Bifidobacteriales|f__Bifidobacteriaceae|g__Bifidobacterium|s__Bifid
k__Bacteria|p__Firmicutes|c__Clostridia|o__Clostridia_unclassified|f__Clostridia_unclassified|g__Clostridia_unclassified|s_
k__Bacteria|p__Firmicutes|c__Clostridia|o__Eubacteriales|f__Clostridiaceae|g__Clostridiaceae_unclassified|s__Clostridiac
k__Bacteria|p__Firmicutes|c__Clostridia|o__Eubacteriales|f__Clostridiaceae|g__Clostridiaceae_unclassified|s__Clostridiac
k__Bacteria|p__Firmicutes|c__Clostridia|o__Eubacteriales|f__Eubacteriales_unclassified|g__Eubacteriales_unclassified|s_
k__Bacteria|p__Firmicutes|c__Erysipelotrichia|o__Erysipelotrichales|f__Erysipelotrichaceae|g__Erysipelatoclostridium|s_
k__Bacteria|p__Actinobacteria|c__Coriobacteriia|o__Coriobacteriales|f__Coriobacteriaceae|g__Coriobacteriaceae_unclass
k__Bacteria|p__Firmicutes|c__Clostridia|o__Eubacteriales|f__Lachnospiraceae|g__Dorea|s__Dorea_sp_5_2
k__Bacteria|p__Firmicutes|c__Erysipelotrichia|o__Erysipelotrichales|f__Erysipelotrichaceae|g__Dubosiella|s__Dubosiella_
k__Bacteria|p__Firmicutes|c__Erysipelotrichia|o__Erysipelotrichales|f__Erysipelotrichales_unclassified|g__Erysipelotrichal
k__Bacteria|p__Firmicutes|c__Clostridia|o__Eubacteriales|f__Eubacteriaceae|g__Eubacteriaceae_unclassified|s__Eubacte
k__Bacteria|p__Firmicutes|c__Clostridia|o__Eubacteriales|f__Eubacteriaceae|g__Eubacteriaceae_unclassified|s__Eubacte
k__Bacteria|p__Firmicutes|c__Clostridia|o__Eubacteriales|f__Lachnospiraceae|g__GGB20149|s__GGB20149_SGB29430
k__Bacteria|p__Actinobacteria|c__Coriobacteriia|o__Eggerthellales|f__Eggerthellaceae|g__GGB22635|s__GGB22635_SGB
k__Bacteria|p__Firmicutes|c__Clostridia|o__Eubacteriales|f__Lachnospiraceae|g__GGB25041|s__GGB25041_SGB36960
k__Bacteria|p__Bacteroidota|c__Bacteroidia|o__Bacteroidales|f__Muribaculaceae|g__GGB27876|s__GGB27876_SGB4031
k__Bacteria|p__Bacteroidota|c__Bacteroidia|o__Bacteroidales|f__Muribaculaceae|g__GGB27878|s__GGB27878_SGB4031

k__Bacteria|p__Bacteroidota|c__Bacteroidia|o__Bacteroidales|f__Muribaculaceae|g__GGB27918|s__GGB27918_SGB4035
k__Bacteria|p__Firmicutes|c__CFGB9508|o__OFGB9508|f__FGB9508|g__GGB28382|s__GGB28382_SGB40962
k__Bacteria|p__Firmicutes|c__CFGB2838|o__OFGB2838|f__FGB2838|g__GGB28399|s__GGB28399_SGB40980
k__Bacteria|p__Firmicutes|c__CFGB2838|o__OFGB2838|f__FGB2838|g__GGB28411|s__GGB28411_SGB40993
k__Bacteria|p__Firmicutes|c__CFGB2838|o__OFGB2838|f__FGB2838|g__GGB28415|s__GGB28415_SGB40997
k__Bacteria|p__Firmicutes|c__Clostridia|o__Eubacteriales|f__Pumilibacteraceae|g__GGB28430|s__GGB28430_SGB41013
k__Bacteria|p__Firmicutes|c__CFGB28439|o__OFGB28439|f__FGB28439|g__GGB28439|s__GGB28439_SGB41022
k__Bacteria|p__Firmicutes|c__Clostridia|o__Clostridia_unclassified|f__Clostridia_unclassified|g__GGB28778|s__GGB2877
k__Bacteria|p__Firmicutes|c__Clostridia|o__Eubacteriales|f__Eubacteriaceae|g__GGB28784|s__GGB28784_SGB41437
k__Bacteria|p__Firmicutes|c__Clostridia|o__Eubacteriales|f__Lachnospiraceae|g__GGB28792|s__GGB28792_SGB41445
k__Bacteria|p__Firmicutes|c__Clostridia|o__Eubacteriales|f__Lachnospiraceae|g__GGB28798|s__GGB28798_SGB41451
k__Bacteria|p__Firmicutes|c__Clostridia|o__Eubacteriales|f__Lachnospiraceae|g__GGB28802|s__GGB28802_SGB41455
k__Bacteria|p__Firmicutes|c__Clostridia|o__Eubacteriales|f__Lachnospiraceae|g__GGB28818|s__GGB28818_SGB41473
k__Bacteria|p__Firmicutes|c__CFGB77305|o__OFGB77305|f__FGB77305|g__GGB28828|s__GGB28828_SGB41484
k__Bacteria|p__Firmicutes|c__Clostridia|o__Eubacteriales|f__Clostridiaceae|g__GGB28851|s__GGB28851_SGB41518
k__Bacteria|p__Firmicutes|c__Clostridia|o__Eubacteriales|f__Lachnospiraceae|g__GGB28859|s__GGB28859_SGB41528
k__Bacteria|p__Firmicutes|c__Clostridia|o__Eubacteriales|f__Lachnospiraceae|g__GGB28864|s__GGB28864_SGB41535
k__Bacteria|p__Firmicutes|c__Clostridia|o__Eubacteriales|f__Lachnospiraceae|g__GGB28869|s__GGB28869_SGB41543
k__Bacteria|p__Firmicutes|c__CFGB9633|o__OFGB9633|f__FGB9633|g__GGB28883|s__GGB28883_SGB41564
k__Bacteria|p__Bacteria_unclassified|c__Bacteria_unclassified|o__Bacteria_unclassified|f__Bacteria_unclassified|g__GGB
k__Bacteria|p__Bacteria_unclassified|c__Bacteria_unclassified|o__Bacteria_unclassified|f__Bacteria_unclassified|g__GGB
k__Bacteria|p__Bacteria_unclassified|c__Bacteria_unclassified|o__Bacteria_unclassified|f__Bacteria_unclassified|g__GGB
k__Bacteria|p__Bacteria_unclassified|c__Bacteria_unclassified|o__Bacteria_unclassified|f__Bacteria_unclassified|g__GGB
k__Bacteria|p__Firmicutes|c__Clostridia|o__Eubacteriales|f__Lachnospiraceae|g__GGB28916|s__GGB28916_SGB41612
k__Bacteria|p__Firmicutes|c__Clostridia|o__Eubacteriales|f__Lachnospiraceae|g__GGB28924|s__GGB28924_SGB41621
k__Bacteria|p__Firmicutes|c__Clostridia|o__Eubacteriales|f__Lachnospiraceae|g__GGB28926|s__GGB28926_SGB41624
k__Bacteria|p__Bacteria_unclassified|c__CFGB77359|o__OFGB77359|f__FGB77359|g__GGB28927|s__GGB28927_SGB416
k__Bacteria|p__Firmicutes|c__CFGB9639|o__OFGB9639|f__FGB9639|g__GGB28934|s__GGB28934_SGB41635
k__Bacteria|p__Firmicutes|c__Clostridia|o__Eubacteriales|f__Lachnospiraceae|g__GGB28946|s__GGB28946_SGB41652
k__Bacteria|p__Firmicutes|c__Clostridia|o__Eubacteriales|f__Lachnospiraceae|g__GGB28949|s__GGB28949_SGB41655
k__Bacteria|p__Firmicutes|c__Clostridia|o__Eubacteriales|f__Lachnospiraceae|g__GGB28949|s__GGB28949_SGB41656
k__Bacteria|p__Firmicutes|c__Clostridia|o__Eubacteriales|f__Clostridiaceae|g__GGB28950|s__GGB28950_SGB41657
k__Bacteria|p__Firmicutes|c__Clostridia|o__Eubacteriales|f__Clostridiaceae|g__GGB28951|s__GGB28951_SGB102295
k__Bacteria|p__Firmicutes|c__Clostridia|o__Eubacteriales|f__Clostridiaceae|g__GGB28951|s__GGB28951_SGB41658
k__Bacteria|p__Firmicutes|c__Clostridia|o__Eubacteriales|f__Clostridiaceae|g__GGB28954|s__GGB28954_SGB41662
k__Bacteria|p__Firmicutes|c__Clostridia|o__Eubacteriales|f__Clostridiaceae|g__GGB28956|s__GGB28956_SGB41665
k__Bacteria|p__Firmicutes|c__Clostridia|o__Eubacteriales|f__Clostridiaceae|g__GGB28960|s__GGB28960_SGB41669
k__Bacteria|p__Firmicutes|c__Clostridia|o__Eubacteriales|f__Clostridiaceae|g__GGB28967|s__GGB28967_SGB41678
k__Bacteria|p__Firmicutes|c__Clostridia|o__Eubacteriales|f__Eubacteriaceae|g__GGB28991|s__GGB28991_SGB41705
k__Bacteria|p__Firmicutes|c__CFGB9658|o__OFGB9658|f__FGB9658|g__GGB29002|s__GGB29002_SGB41718
k__Bacteria|p__Firmicutes|c__CFGB9659|o__OFGB9659|f__FGB9659|g__GGB29003|s__GGB29003_SGB41719
k__Bacteria|p__Bacteria_unclassified|c__Bacteria_unclassified|o__Bacteria_unclassified|f__Bacteria_unclassified|g__GGB
k__Bacteria|p__Firmicutes|c__CFGB9827|o__OFGB9827|f__FGB9827|g__GGB29531|s__GGB29531_SGB42317
k__Bacteria|p__Firmicutes|c__Clostridia|o__Eubacteriales|f__Eubacteriaceae|g__GGB29685|s__GGB29685_SGB42494
k__Bacteria|p__Bacteria_unclassified|c__CFGB77303|o__OFGB77303|f__FGB77303|g__GGB30141|s__GGB30141_SGB430
k__Bacteria|p__Firmicutes|c__Clostridia|o__Eubacteriales|f__Eubacteriales_unclassified|g__GGB30286|s__GGB30286_SG

k__Bacteria|p__Firmicutes|c__Clostridia|o__Eubacteriales|f__Oscillospiraceae|g__GGB30303|s__GGB30303_SGB43268
k__Bacteria|p__Firmicutes|c__CFGB30328|o__OFGB30328|f__FGB30328|g__GGB30413|s__GGB30413_SGB43452
k__Bacteria|p__Firmicutes|c__Clostridia|o__Eubacteriales|f__Oscillospiraceae|g__GGB30454|s__GGB30454_SGB43514
k__Bacteria|p__Firmicutes|c__Clostridia|o__Eubacteriales|f__Oscillospiraceae|g__GGB30455|s__GGB30455_SGB43519
k__Bacteria|p__Firmicutes|c__Clostridia|o__Eubacteriales|f__Oscillospiraceae|g__GGB30461|s__GGB30461_SGB43527
k__Bacteria|p__Firmicutes|c__Clostridia|o__Eubacteriales|f__Oscillospiraceae|g__GGB30461|s__GGB30461_SGB43530
k__Bacteria|p__Firmicutes|c__Clostridia|o__Eubacteriales|f__Oscillospiraceae|g__GGB30463|s__GGB30463_SGB43537
k__Bacteria|p__Firmicutes|c__Clostridia|o__Eubacteriales|f__Oscillospiraceae|g__GGB30473|s__GGB30473_SGB43557
k__Bacteria|p__Firmicutes|c__Clostridia|o__Eubacteriales|f__Oscillospiraceae|g__GGB30475|s__GGB30475_SGB63182
k__Bacteria|p__Actinobacteria|c__CFGB77153|o__OFGB77153|f__FGB77153|g__GGB30861|s__GGB30861_SGB44083
k__Bacteria|p__Tenericutes|c__CFGB1791|o__OFGB1791|f__FGB1791|g__GGB31312|s__GGB31312_SGB44628
k__Bacteria|p__Firmicutes|c__CFGB10290|o__OFGB10290|f__FGB10290|g__GGB31438|s__GGB31438_SGB44768
k__Bacteria|p__Firmicutes|c__Clostridia|o__Eubacteriales|f__Oscillospiraceae|g__GGB3171|s__GGB3171_SGB4185
k__Bacteria|p__Firmicutes|c__CFGB1765|o__OFGB1765|f__FGB1765|g__GGB31823|s__GGB31823_SGB45199
k__Bacteria|p__Firmicutes|c__CFGB10349|o__OFGB10349|f__FGB10349|g__GGB31853|s__GGB31853_SGB45233
k__Bacteria|p__Firmicutes|c__CFGB10667|o__OFGB10667|f__FGB10667|g__GGB32371|s__GGB32371_SGB41694
k__Bacteria|p__Firmicutes|c__Clostridia|o__Eubacteriales|f__Lachnospiraceae|g__GGB3793|s__GGB3793_SGB5158
k__Bacteria|p__Firmicutes|c__Clostridia|o__Eubacteriales|f__Lachnospiraceae|g__GGB42598|s__GGB42598_SGB59794
k__Bacteria|p__Firmicutes|c__Clostridia|o__Eubacteriales|f__Christensenellaceae|g__GGB45656|s__GGB45656_SGB6337
k__Bacteria|p__Firmicutes|c__CFGB10299|o__OFGB10299|f__FGB10299|g__GGB47127|s__GGB47127_SGB65054
k__Bacteria|p__Firmicutes|c__Clostridia|o__Eubacteriales|f__Oscillospiraceae|g__GGB74395|s__GGB74395_SGB43521
k__Bacteria|p__Firmicutes|c__Clostridia|o__Eubacteriales|f__Oscillospiraceae|g__GGB75053|s__GGB75053_SGB43494
k__Bacteria|p__Firmicutes|c__Clostridia|o__Eubacteriales|f__Lachnospiraceae|g__GGB75109|s__GGB75109_SGB102238
k__Bacteria|p__Firmicutes|c__Clostridia|o__Clostridia_unclassified|f__Clostridia_unclassified|g__GGB81440|s__GGB8144
k__Bacteria|p__Firmicutes|c__Clostridia|o__Eubacteriales|f__Lachnospiraceae|g__Lachnospiraceae_unclassified|s__Lachn
k__Bacteria|p__Firmicutes|c__Clostridia|o__Eubacteriales|f__Lachnospiraceae|g__Lachnospiraceae_unclassified|s__Lachn
k__Bacteria|p__Firmicutes|c__Clostridia|o__Eubacteriales|f__Lachnospiraceae|g__Lachnospiraceae_unclassified|s__Lachn
k__Bacteria|p__Firmicutes|c__Clostridia|o__Eubacteriales|f__Lachnospiraceae|g__Lachnospiraceae_unclassified|s__Lachn
k__Bacteria|p__Firmicutes|c__Clostridia|o__Eubacteriales|f__Lachnospiraceae|g__Lachnospiraceae_unclassified|s__Lachn
k__Bacteria|p__Firmicutes|c__Clostridia|o__Eubacteriales|f__Lachnospiraceae|g__Lachnospiraceae_unclassified|s__Lachn
k__Bacteria|p__Firmicutes|c__Clostridia|o__Eubacteriales|f__Lachnospiraceae|g__Lachnospiraceae_unclassified|s__Lachn
k__Bacteria|p__Firmicutes|c__Clostridia|o__Eubacteriales|f__Lachnospiraceae|g__Lachnospiraceae_unclassified|s__Lachn
k__Bacteria|p__Firmicutes|c__Bacilli|o__Lactobacillales|f__Lactobacillaceae|g__Lactobacillus|s__Lactobacillus_johnsonii
k__Bacteria|p__Bacteroidota|c__Bacteroidia|o__Bacteroidales|f__Muribaculaceae|g__Muribaculaceae_unclassified|s__M
k__Bacteria|p__Firmicutes|c__Clostridia|o__Eubacteriales|f__Oscillospiraceae|g__Neglectibacter|s__Neglectibacter_sp_X
k__Bacteria|p__Firmicutes|c__Clostridia|o__Eubacteriales|f__Oscillospiraceae|g__Oscillospiraceae_unclassified|s__Oscillo
k__Bacteria|p__Firmicutes|c__Clostridia|o__Eubacteriales|f__Oscillospiraceae|g__Oscillospiraceae_unclassified|s__Oscillo
k__Bacteria|p__Firmicutes|c__Clostridia|o__Eubacteriales|f__Oscillospiraceae|g__Oscillospiraceae_unclassified|s__Oscillo
k__Bacteria|p__Proteobacteria|c__Betaproteobacteria|o__Burkholderiales|f__Sutterellaceae|g__Parasutterella|s__Parasu
k__Bacteria|p__Firmicutes|c__Clostridia|o__Eubacteriales|f__Peptostreptococcaceae|g__Romboutsia|s__Romboutsia_ilea
k__Bacteria|p__Firmicutes|c__Clostridia|o__Eubacteriales|f__Lachnospiraceae|g__Schaedlerella|s__Schaedlerella_arabino
k__Bacteria|p__Firmicutes|c__Erysipelotrichia|o__Erysipelotrichales|f__Turicibacteraceae|g__Turicibacter|s__Turicibacter_
k__Bacteria|p__Bacteria_unclassified|c__Bacteria_unclassified|o__Bacteria_unclassified|f__Bacteria_unclassified|g__Bacte
k__Bacteria|p__Bacteria_unclassified|c__Bacteria_unclassified|o__Bacteria_unclassified|f__Bacteria_unclassified|g__Bacte
k__Bacteria|p__Bacteria_unclassified|c__Bacteria_unclassified|o__Bacteria_unclassified|f__Bacteria_unclassified|g__Bacte

k__Bacteria|p__Firmicutes|c__Clostridia|o__Eubacteriales|f__Lachnospiraceae|g__Acetatifactor|s__Acetatifactor_SGB415
k__Bacteria|p__Firmicutes|c__Clostridia|o__Eubacteriales|f__Lachnospiraceae|g__Acetatifactor|s__Acetatifactor_muris
k__Bacteria|p__Firmicutes|c__Clostridia|o__Eubacteriales|f__Oscillospiraceae|g__Acutalibacter|s__Acutalibacter_muris
k__Bacteria|p__Firmicutes|c__Clostridia|o__Eubacteriales|f__Oscillospiraceae|g__Acutalibacter|s__Acutalibacter_sp_1XD
k__Bacteria|p__Actinobacteria|c__Coriobacteriia|o__Eggerthellales|f__Eggerthellaceae|g__Adlercreutzia|s__Adlercreutzia
k__Bacteria|p__Actinobacteria|c__Coriobacteriia|o__Eggerthellales|f__Eggerthellaceae|g__Adlercreutzia|s__Adlercreutzia
k__Bacteria|p__Actinobacteria|c__Coriobacteriia|o__Eggerthellales|f__Eggerthellaceae|g__Adlercreutzia|s__Adlercreutzia
k__Bacteria|p__Verrucomicrobia|c__Verrucomicrobiae|o__Verrucomicrobiales|f__Akkermansiaceae|g__Akkermansia|s__A
k__Bacteria|p__Bacteroidota|c__Bacteroidia|o__Bacteroidales|f__Rikenellaceae|g__Alistipes|s__Alistipes_sp_DSM_11234
k__Bacteria|p__Firmicutes|c__Clostridia|o__Eubacteriales|f__Oscillospiraceae|g__Anaerotruncus|s__Anaerotruncus_sp_1
k__Bacteria|p__Bacteria_unclassified|c__Bacteria_unclassified|o__Bacteria_unclassified|f__Bacteria_unclassified|g__Bacte
k__Bacteria|p__Bacteria_unclassified|c__Bacteria_unclassified|o__Bacteria_unclassified|f__Bacteria_unclassified|g__Bacte
k__Bacteria|p__Bacteria_unclassified|c__Bacteria_unclassified|o__Bacteria_unclassified|f__Bacteria_unclassified|g__Bacte
k__Bacteria|p__Bacteroidota|c__Bacteroidia|o__Bacteroidales|f__Bacteroidaceae|g__Bacteroides|s__Bacteroides_thetaio
k__Bacteria|p__Actinobacteria|c__Actinomycetia|o__Bifidobacteriales|f__Bifidobacteriaceae|g__Bifidobacterium|s__Bifid
k__Bacteria|p__Firmicutes|c__Clostridia|o__Clostridia_unclassified|f__Clostridia_unclassified|g__Clostridia_unclassified|s_
k__Bacteria|p__Firmicutes|c__Clostridia|o__Eubacteriales|f__Clostridiaceae|g__Clostridiaceae_unclassified|s__Clostridiac
k__Bacteria|p__Firmicutes|c__Clostridia|o__Eubacteriales|f__Clostridiaceae|g__Clostridiaceae_unclassified|s__Clostridiac
k__Bacteria|p__Firmicutes|c__Clostridia|o__Eubacteriales|f__Eubacteriales_unclassified|g__Eubacteriales_unclassified|s_
k__Bacteria|p__Firmicutes|c__Erysipelotrichia|o__Erysipelotrichales|f__Erysipelotrichaceae|g__Erysipelatoclostridium|s__
k__Bacteria|p__Actinobacteria|c__Coriobacteriia|o__Coriobacteriales|f__Coriobacteriaceae|g__Coriobacteriaceae_unclass
k__Bacteria|p__Firmicutes|c__Clostridia|o__Eubacteriales|f__Lachnospiraceae|g__Dorea|s__Dorea_sp_5_2
k__Bacteria|p__Firmicutes|c__Erysipelotrichia|o__Erysipelotrichales|f__Erysipelotrichaceae|g__Dubosiella|s__Dubosiella_
k__Bacteria|p__Firmicutes|c__Erysipelotrichia|o__Erysipelotrichales|f__Erysipelotrichales_unclassified|g__Erysipelotrichal
k__Bacteria|p__Firmicutes|c__Clostridia|o__Eubacteriales|f__Eubacteriaceae|g__Eubacteriaceae_unclassified|s__Eubacte
k__Bacteria|p__Firmicutes|c__Clostridia|o__Eubacteriales|f__Eubacteriaceae|g__Eubacteriaceae_unclassified|s__Eubacte
k__Bacteria|p__Firmicutes|c__Clostridia|o__Eubacteriales|f__Lachnospiraceae|g__GGB20149|s__GGB20149_SGB29430
k__Bacteria|p__Actinobacteria|c__Coriobacteriia|o__Eggerthellales|f__Eggerthellaceae|g__GGB22635|s__GGB22635_SGB
k__Bacteria|p__Firmicutes|c__Clostridia|o__Eubacteriales|f__Lachnospiraceae|g__GGB25041|s__GGB25041_SGB36960
k__Bacteria|p__Bacteroidota|c__Bacteroidia|o__Bacteroidales|f__Muribaculaceae|g__GGB27876|s__GGB27876_SGB403
k__Bacteria|p__Bacteroidota|c__Bacteroidia|o__Bacteroidales|f__Muribaculaceae|g__GGB27878|s__GGB27878_SGB403
k__Bacteria|p__Bacteroidota|c__Bacteroidia|o__Bacteroidales|f__Muribaculaceae|g__GGB27918|s__GGB27918_SGB403
k__Bacteria|p__Firmicutes|c__CFGB9508|o__OFGB9508|f__FGB9508|g__GGB28382|s__GGB28382_SGB40962
k__Bacteria|p__Firmicutes|c__CFGB2838|o__OFGB2838|f__FGB2838|g__GGB28399|s__GGB28399_SGB40980
k__Bacteria|p__Firmicutes|c__CFGB2838|o__OFGB2838|f__FGB2838|g__GGB28411|s__GGB28411_SGB40993
k__Bacteria|p__Firmicutes|c__CFGB2838|o__OFGB2838|f__FGB2838|g__GGB28415|s__GGB28415_SGB40997
k__Bacteria|p__Firmicutes|c__Clostridia|o__Eubacteriales|f__Pumilibacteraceae|g__GGB28430|s__GGB28430_SGB41013
k__Bacteria|p__Firmicutes|c__CFGB28439|o__OFGB28439|f__FGB28439|g__GGB28439|s__GGB28439_SGB41022
k__Bacteria|p__Firmicutes|c__Clostridia|o__Clostridia_unclassified|f__Clostridia_unclassified|g__GGB28778|s__GGB2877
k__Bacteria|p__Firmicutes|c__Clostridia|o__Eubacteriales|f__Eubacteriaceae|g__GGB28784|s__GGB28784_SGB41437
k__Bacteria|p__Firmicutes|c__Clostridia|o__Eubacteriales|f__Lachnospiraceae|g__GGB28792|s__GGB28792_SGB41445
k__Bacteria|p__Firmicutes|c__Clostridia|o__Eubacteriales|f__Lachnospiraceae|g__GGB28798|s__GGB28798_SGB41451
k__Bacteria|p__Firmicutes|c__Clostridia|o__Eubacteriales|f__Lachnospiraceae|g__GGB28802|s__GGB28802_SGB41455
k__Bacteria|p__Firmicutes|c__Clostridia|o__Eubacteriales|f__Lachnospiraceae|g__GGB28818|s__GGB28818_SGB41473

k__Bacteria|p__Firmicutes|c__CFGB77305|o__OFGB77305|f__FGB77305|g__GGB28828|s__GGB28828_SGB41484
k__Bacteria|p__Firmicutes|c__Clostridia|o__Eubacteriales|f__Clostridiaceae|g__GGB28851|s__GGB28851_SGB41518
k__Bacteria|p__Firmicutes|c__Clostridia|o__Eubacteriales|f__Lachnospiraceae|g__GGB28859|s__GGB28859_SGB41528
k__Bacteria|p__Firmicutes|c__Clostridia|o__Eubacteriales|f__Lachnospiraceae|g__GGB28864|s__GGB28864_SGB41535
k__Bacteria|p__Firmicutes|c__Clostridia|o__Eubacteriales|f__Lachnospiraceae|g__GGB28869|s__GGB28869_SGB41543
k__Bacteria|p__Firmicutes|c__CFGB9633|o__OFGB9633|f__FGB9633|g__GGB28883|s__GGB28883_SGB41564
k__Bacteria|p__Bacteria_unclassified|c__Bacteria_unclassified|o__Bacteria_unclassified|f__Bacteria_unclassified|g__GGB[illegible]
k__Bacteria|p__Bacteria_unclassified|c__Bacteria_unclassified|o__Bacteria_unclassified|f__Bacteria_unclassified|g__GGB[illegible]
k__Bacteria|p__Bacteria_unclassified|c__Bacteria_unclassified|o__Bacteria_unclassified|f__Bacteria_unclassified|g__GGB[illegible]
k__Bacteria|p__Bacteria_unclassified|c__Bacteria_unclassified|o__Bacteria_unclassified|f__Bacteria_unclassified|g__GGB[illegible]
k__Bacteria|p__Firmicutes|c__Clostridia|o__Eubacteriales|f__Lachnospiraceae|g__GGB28916|s__GGB28916_SGB41612
k__Bacteria|p__Firmicutes|c__Clostridia|o__Eubacteriales|f__Lachnospiraceae|g__GGB28924|s__GGB28924_SGB41621
k__Bacteria|p__Firmicutes|c__Clostridia|o__Eubacteriales|f__Lachnospiraceae|g__GGB28926|s__GGB28926_SGB41624
k__Bacteria|p__Bacteria_unclassified|c__CFGB77359|o__OFGB77359|f__FGB77359|g__GGB28927|s__GGB28927_SGB416[illegible]
k__Bacteria|p__Firmicutes|c__CFGB9639|o__OFGB9639|f__FGB9639|g__GGB28934|s__GGB28934_SGB41635
k__Bacteria|p__Firmicutes|c__Clostridia|o__Eubacteriales|f__Lachnospiraceae|g__GGB28946|s__GGB28946_SGB41652
k__Bacteria|p__Firmicutes|c__Clostridia|o__Eubacteriales|f__Lachnospiraceae|g__GGB28949|s__GGB28949_SGB41655
k__Bacteria|p__Firmicutes|c__Clostridia|o__Eubacteriales|f__Lachnospiraceae|g__GGB28949|s__GGB28949_SGB41656
k__Bacteria|p__Firmicutes|c__Clostridia|o__Eubacteriales|f__Clostridiaceae|g__GGB28950|s__GGB28950_SGB41657
k__Bacteria|p__Firmicutes|c__Clostridia|o__Eubacteriales|f__Clostridiaceae|g__GGB28951|s__GGB28951_SGB102295
k__Bacteria|p__Firmicutes|c__Clostridia|o__Eubacteriales|f__Clostridiaceae|g__GGB28951|s__GGB28951_SGB41658
k__Bacteria|p__Firmicutes|c__Clostridia|o__Eubacteriales|f__Clostridiaceae|g__GGB28954|s__GGB28954_SGB41662
k__Bacteria|p__Firmicutes|c__Clostridia|o__Eubacteriales|f__Clostridiaceae|g__GGB28956|s__GGB28956_SGB41665
k__Bacteria|p__Firmicutes|c__Clostridia|o__Eubacteriales|f__Clostridiaceae|g__GGB28960|s__GGB28960_SGB41669
k__Bacteria|p__Firmicutes|c__Clostridia|o__Eubacteriales|f__Clostridiaceae|g__GGB28967|s__GGB28967_SGB41678
k__Bacteria|p__Firmicutes|c__Clostridia|o__Eubacteriales|f__Eubacteriaceae|g__GGB28991|s__GGB28991_SGB41705
k__Bacteria|p__Firmicutes|c__CFGB9658|o__OFGB9658|f__FGB9658|g__GGB29002|s__GGB29002_SGB41718
k__Bacteria|p__Firmicutes|c__CFGB9659|o__OFGB9659|f__FGB9659|g__GGB29003|s__GGB29003_SGB41719
k__Bacteria|p__Bacteria_unclassified|c__Bacteria_unclassified|o__Bacteria_unclassified|f__Bacteria_unclassified|g__GGB[illegible]
k__Bacteria|p__Firmicutes|c__CFGB9827|o__OFGB9827|f__FGB9827|g__GGB29531|s__GGB29531_SGB42317
k__Bacteria|p__Firmicutes|c__Clostridia|o__Eubacteriales|f__Eubacteriaceae|g__GGB29685|s__GGB29685_SGB42494
k__Bacteria|p__Bacteria_unclassified|c__CFGB77303|o__OFGB77303|f__FGB77303|g__GGB30141|s__GGB30141_SGB430[illegible]
k__Bacteria|p__Firmicutes|c__Clostridia|o__Eubacteriales|f__Eubacteriales_unclassified|g__GGB30286|s__GGB30286_SG[illegible]
k__Bacteria|p__Firmicutes|c__Clostridia|o__Eubacteriales|f__Oscillospiraceae|g__GGB30303|s__GGB30303_SGB43268
k__Bacteria|p__Firmicutes|c__CFGB30328|o__OFGB30328|f__FGB30328|g__GGB30413|s__GGB30413_SGB43452
k__Bacteria|p__Firmicutes|c__Clostridia|o__Eubacteriales|f__Oscillospiraceae|g__GGB30454|s__GGB30454_SGB43514
k__Bacteria|p__Firmicutes|c__Clostridia|o__Eubacteriales|f__Oscillospiraceae|g__GGB30455|s__GGB30455_SGB43519
k__Bacteria|p__Firmicutes|c__Clostridia|o__Eubacteriales|f__Oscillospiraceae|g__GGB30461|s__GGB30461_SGB43527
k__Bacteria|p__Firmicutes|c__Clostridia|o__Eubacteriales|f__Oscillospiraceae|g__GGB30461|s__GGB30461_SGB43530
k__Bacteria|p__Firmicutes|c__Clostridia|o__Eubacteriales|f__Oscillospiraceae|g__GGB30463|s__GGB30463_SGB43537
k__Bacteria|p__Firmicutes|c__Clostridia|o__Eubacteriales|f__Oscillospiraceae|g__GGB30473|s__GGB30473_SGB43557
k__Bacteria|p__Firmicutes|c__Clostridia|o__Eubacteriales|f__Oscillospiraceae|g__GGB30475|s__GGB30475_SGB63182
k__Bacteria|p__Actinobacteria|c__CFGB77153|o__OFGB77153|f__FGB77153|g__GGB30861|s__GGB30861_SGB44083
k__Bacteria|p__Tenericutes|c__CFGB1791|o__OFGB1791|f__FGB1791|g__GGB31312|s__GGB31312_SGB44628
k__Bacteria|p__Firmicutes|c__CFGB10290|o__OFGB10290|f__FGB10290|g__GGB31438|s__GGB31438_SGB44768
k__Bacteria|p__Firmicutes|c__Clostridia|o__Eubacteriales|f__Oscillospiraceae|g__GGB3171|s__GGB3171_SGB4185

k__Bacteria|p__Firmicutes|c__CFGB1765|o__OFGB1765|f__FGB1765|g__GGB31823|s__GGB31823_SGB45199
k__Bacteria|p__Firmicutes|c__CFGB10349|o__OFGB10349|f__FGB10349|g__GGB31853|s__GGB31853_SGB45233
k__Bacteria|p__Firmicutes|c__CFGB10667|o__OFGB10667|f__FGB10667|g__GGB32371|s__GGB32371_SGB41694
k__Bacteria|p__Firmicutes|c__Clostridia|o__Eubacteriales|f__Lachnospiraceae|g__GGB3793|s__GGB3793_SGB5158
k__Bacteria|p__Firmicutes|c__Clostridia|o__Eubacteriales|f__Lachnospiraceae|g__GGB42598|s__GGB42598_SGB59794
k__Bacteria|p__Firmicutes|c__Clostridia|o__Eubacteriales|f__Christensenellaceae|g__GGB45656|s__GGB45656_SGB6337
k__Bacteria|p__Firmicutes|c__CFGB10299|o__OFGB10299|f__FGB10299|g__GGB47127|s__GGB47127_SGB65054
k__Bacteria|p__Firmicutes|c__Clostridia|o__Eubacteriales|f__Oscillospiraceae|g__GGB74395|s__GGB74395_SGB43521
k__Bacteria|p__Firmicutes|c__Clostridia|o__Eubacteriales|f__Oscillospiraceae|g__GGB75053|s__GGB75053_SGB43494
k__Bacteria|p__Firmicutes|c__Clostridia|o__Eubacteriales|f__Lachnospiraceae|g__GGB75109|s__GGB75109_SGB102238
k__Bacteria|p__Firmicutes|c__Clostridia|o__Clostridia_unclassified|f__Clostridia_unclassified|g__GGB81440|s__GGB8144
k__Bacteria|p__Firmicutes|c__Clostridia|o__Eubacteriales|f__Lachnospiraceae|g__Lachnospiraceae_unclassified|s__Lachn
k__Bacteria|p__Firmicutes|c__Clostridia|o__Eubacteriales|f__Lachnospiraceae|g__Lachnospiraceae_unclassified|s__Lachn
k__Bacteria|p__Firmicutes|c__Clostridia|o__Eubacteriales|f__Lachnospiraceae|g__Lachnospiraceae_unclassified|s__Lachn
k__Bacteria|p__Firmicutes|c__Clostridia|o__Eubacteriales|f__Lachnospiraceae|g__Lachnospiraceae_unclassified|s__Lachn
k__Bacteria|p__Firmicutes|c__Clostridia|o__Eubacteriales|f__Lachnospiraceae|g__Lachnospiraceae_unclassified|s__Lachn
k__Bacteria|p__Firmicutes|c__Clostridia|o__Eubacteriales|f__Lachnospiraceae|g__Lachnospiraceae_unclassified|s__Lachn
k__Bacteria|p__Firmicutes|c__Clostridia|o__Eubacteriales|f__Lachnospiraceae|g__Lachnospiraceae_unclassified|s__Lachn
k__Bacteria|p__Firmicutes|c__Clostridia|o__Eubacteriales|f__Lachnospiraceae|g__Lachnospiraceae_unclassified|s__Lachn
k__Bacteria|p__Firmicutes|c__Bacilli|o__Lactobacillales|f__Lactobacillaceae|g__Lactobacillus|s__Lactobacillus_johnsonii
k__Bacteria|p__Bacteroidota|c__Bacteroidia|o__Bacteroidales|f__Muribaculaceae|g__Muribaculaceae_unclassified|s__M
k__Bacteria|p__Firmicutes|c__Clostridia|o__Eubacteriales|f__Oscillospiraceae|g__Neglectibacter|s__Neglectibacter_sp_X
k__Bacteria|p__Firmicutes|c__Clostridia|o__Eubacteriales|f__Oscillospiraceae|g__Oscillospiraceae_unclassified|s__Oscillo
k__Bacteria|p__Firmicutes|c__Clostridia|o__Eubacteriales|f__Oscillospiraceae|g__Oscillospiraceae_unclassified|s__Oscillo
k__Bacteria|p__Firmicutes|c__Clostridia|o__Eubacteriales|f__Oscillospiraceae|g__Oscillospiraceae_unclassified|s__Oscillo
k__Bacteria|p__Proteobacteria|c__Betaproteobacteria|o__Burkholderiales|f__Sutterellaceae|g__Parasutterella|s__Parasu
k__Bacteria|p__Firmicutes|c__Clostridia|o__Eubacteriales|f__Peptostreptococcaceae|g__Romboutsia|s__Romboutsia_ilea
k__Bacteria|p__Firmicutes|c__Clostridia|o__Eubacteriales|f__Lachnospiraceae|g__Schaedlerella|s__Schaedlerella_arabino
k__Bacteria|p__Firmicutes|c__Erysipelotrichia|o__Erysipelotrichales|f__Turicibacteraceae|g__Turicibacter|s__Turicibacter_
k__Bacteria|p__Bacteria_unclassified|c__Bacteria_unclassified|o__Bacteria_unclassified|f__Bacteria_unclassified|g__Bacte
k__Bacteria|p__Bacteria_unclassified|c__Bacteria_unclassified|o__Bacteria_unclassified|f__Bacteria_unclassified|g__Bacte
k__Bacteria|p__Bacteria_unclassified|c__Bacteria_unclassified|o__Bacteria_unclassified|f__Bacteria_unclassified|g__Bacte

k__Bacteria|p__Firmicutes|c__Clostridia|o__Eubacteriales|f__Lachnospiraceae|g__Acetatifactor|s__Acetatifactor_SGB415
k__Bacteria|p__Firmicutes|c__Clostridia|o__Eubacteriales|f__Lachnospiraceae|g__Acetatifactor|s__Acetatifactor_muris
k__Bacteria|p__Firmicutes|c__Clostridia|o__Eubacteriales|f__Oscillospiraceae|g__Acutalibacter|s__Acutalibacter_muris
k__Bacteria|p__Firmicutes|c__Clostridia|o__Eubacteriales|f__Oscillospiraceae|g__Acutalibacter|s__Acutalibacter_sp_1XD
k__Bacteria|p__Actinobacteria|c__Coriobacteriia|o__Eggerthellales|f__Eggerthellaceae|g__Adlercreutzia|s__Adlercreutzia
k__Bacteria|p__Actinobacteria|c__Coriobacteriia|o__Eggerthellales|f__Eggerthellaceae|g__Adlercreutzia|s__Adlercreutzia
k__Bacteria|p__Actinobacteria|c__Coriobacteriia|o__Eggerthellales|f__Eggerthellaceae|g__Adlercreutzia|s__Adlercreutzia
k__Bacteria|p__Verrucomicrobia|c__Verrucomicrobiae|o__Verrucomicrobiales|f__Akkermansiaceae|g__Akkermansia|s__A
k__Bacteria|p__Bacteroidota|c__Bacteroidia|o__Bacteroidales|f__Rikenellaceae|g__Alistipes|s__Alistipes_sp_DSM_11234
k__Bacteria|p__Firmicutes|c__Clostridia|o__Eubacteriales|f__Oscillospiraceae|g__Anaerotruncus|s__Anaerotruncus_sp_1
k__Bacteria|p__Bacteria_unclassified|c__Bacteria_unclassified|o__Bacteria_unclassified|f__Bacteria_unclassified|g__Bacte

k__Bacteria|p__Bacteria_unclassified|c__Bacteria_unclassified|o__Bacteria_unclassified|f__Bacteria_unclassified|g__Bacte
k__Bacteria|p__Bacteria_unclassified|c__Bacteria_unclassified|o__Bacteria_unclassified|f__Bacteria_unclassified|g__Bacte
k__Bacteria|p__Bacteroidota|c__Bacteroidia|o__Bacteroidales|f__Bacteroidaceae|g__Bacteroides|s__Bacteroides_thetaio
k__Bacteria|p__Actinobacteria|c__Actinomycetia|o__Bifidobacteriales|f__Bifidobacteriaceae|g__Bifidobacterium|s__Bifid
k__Bacteria|p__Firmicutes|c__Clostridia|o__Clostridia_unclassified|f__Clostridia_unclassified|g__Clostridia_unclassified|s_
k__Bacteria|p__Firmicutes|c__Clostridia|o__Eubacteriales|f__Clostridiaceae|g__Clostridiaceae_unclassified|s__Clostridiac
k__Bacteria|p__Firmicutes|c__Clostridia|o__Eubacteriales|f__Clostridiaceae|g__Clostridiaceae_unclassified|s__Clostridiac
k__Bacteria|p__Firmicutes|c__Clostridia|o__Eubacteriales|f__Eubacteriales_unclassified|g__Eubacteriales_unclassified|s_
k__Bacteria|p__Firmicutes|c__Erysipelotrichia|o__Erysipelotrichales|f__Erysipelotrichaceae|g__Erysipelatoclostridium|s__
k__Bacteria|p__Actinobacteria|c__Coriobacteriia|o__Coriobacteriales|f__Coriobacteriaceae|g__Coriobacteriaceae_unclass
k__Bacteria|p__Firmicutes|c__Clostridia|o__Eubacteriales|f__Lachnospiraceae|g__Dorea|s__Dorea_sp_5_2
k__Bacteria|p__Firmicutes|c__Erysipelotrichia|o__Erysipelotrichales|f__Erysipelotrichaceae|g__Dubosiella|s__Dubosiella_
k__Bacteria|p__Firmicutes|c__Erysipelotrichia|o__Erysipelotrichales|f__Erysipelotrichales_unclassified|g__Erysipelotrichal
k__Bacteria|p__Firmicutes|c__Clostridia|o__Eubacteriales|f__Eubacteriaceae|g__Eubacteriaceae_unclassified|s__Eubacte
k__Bacteria|p__Firmicutes|c__Clostridia|o__Eubacteriales|f__Eubacteriaceae|g__Eubacteriaceae_unclassified|s__Eubacte
k__Bacteria|p__Firmicutes|c__Clostridia|o__Eubacteriales|f__Lachnospiraceae|g__GGB20149|s__GGB20149_SGB29430
k__Bacteria|p__Actinobacteria|c__Coriobacteriia|o__Eggerthellales|f__Eggerthellaceae|g__GGB22635|s__GGB22635_SGB
k__Bacteria|p__Firmicutes|c__Clostridia|o__Eubacteriales|f__Lachnospiraceae|g__GGB25041|s__GGB25041_SGB36960
k__Bacteria|p__Bacteroidota|c__Bacteroidia|o__Bacteroidales|f__Muribaculaceae|g__GGB27876|s__GGB27876_SGB4031
k__Bacteria|p__Bacteroidota|c__Bacteroidia|o__Bacteroidales|f__Muribaculaceae|g__GGB27878|s__GGB27878_SGB4031
k__Bacteria|p__Bacteroidota|c__Bacteroidia|o__Bacteroidales|f__Muribaculaceae|g__GGB27918|s__GGB27918_SGB4035
k__Bacteria|p__Firmicutes|c__CFGB9508|o__OFGB9508|f__FGB9508|g__GGB28382|s__GGB28382_SGB40962
k__Bacteria|p__Firmicutes|c__CFGB2838|o__OFGB2838|f__FGB2838|g__GGB28399|s__GGB28399_SGB40980
k__Bacteria|p__Firmicutes|c__CFGB2838|o__OFGB2838|f__FGB2838|g__GGB28411|s__GGB28411_SGB40993
k__Bacteria|p__Firmicutes|c__CFGB2838|o__OFGB2838|f__FGB2838|g__GGB28415|s__GGB28415_SGB40997
k__Bacteria|p__Firmicutes|c__Clostridia|o__Eubacteriales|f__Pumilibacteraceae|g__GGB28430|s__GGB28430_SGB41013
k__Bacteria|p__Firmicutes|c__CFGB28439|o__OFGB28439|f__FGB28439|g__GGB28439|s__GGB28439_SGB41022
k__Bacteria|p__Firmicutes|c__Clostridia|o__Clostridia_unclassified|f__Clostridia_unclassified|g__GGB28778|s__GGB2877
k__Bacteria|p__Firmicutes|c__Clostridia|o__Eubacteriales|f__Eubacteriaceae|g__GGB28784|s__GGB28784_SGB41437
k__Bacteria|p__Firmicutes|c__Clostridia|o__Eubacteriales|f__Lachnospiraceae|g__GGB28792|s__GGB28792_SGB41445
k__Bacteria|p__Firmicutes|c__Clostridia|o__Eubacteriales|f__Lachnospiraceae|g__GGB28798|s__GGB28798_SGB41451
k__Bacteria|p__Firmicutes|c__Clostridia|o__Eubacteriales|f__Lachnospiraceae|g__GGB28802|s__GGB28802_SGB41455
k__Bacteria|p__Firmicutes|c__Clostridia|o__Eubacteriales|f__Lachnospiraceae|g__GGB28818|s__GGB28818_SGB41473
k__Bacteria|p__Firmicutes|c__CFGB77305|o__OFGB77305|f__FGB77305|g__GGB28828|s__GGB28828_SGB41484
k__Bacteria|p__Firmicutes|c__Clostridia|o__Eubacteriales|f__Clostridiaceae|g__GGB28851|s__GGB28851_SGB41518
k__Bacteria|p__Firmicutes|c__Clostridia|o__Eubacteriales|f__Lachnospiraceae|g__GGB28859|s__GGB28859_SGB41528
k__Bacteria|p__Firmicutes|c__Clostridia|o__Eubacteriales|f__Lachnospiraceae|g__GGB28864|s__GGB28864_SGB41535
k__Bacteria|p__Firmicutes|c__Clostridia|o__Eubacteriales|f__Lachnospiraceae|g__GGB28869|s__GGB28869_SGB41543
k__Bacteria|p__Firmicutes|c__CFGB9633|o__OFGB9633|f__FGB9633|g__GGB28883|s__GGB28883_SGB41564
k__Bacteria|p__Bacteria_unclassified|c__Bacteria_unclassified|o__Bacteria_unclassified|f__Bacteria_unclassified|g__GGB2
k__Bacteria|p__Bacteria_unclassified|c__Bacteria_unclassified|o__Bacteria_unclassified|f__Bacteria_unclassified|g__GGB2
k__Bacteria|p__Bacteria_unclassified|c__Bacteria_unclassified|o__Bacteria_unclassified|f__Bacteria_unclassified|g__GGB2
k__Bacteria|p__Bacteria_unclassified|c__Bacteria_unclassified|o__Bacteria_unclassified|f__Bacteria_unclassified|g__GGB2
k__Bacteria|p__Firmicutes|c__Clostridia|o__Eubacteriales|f__Lachnospiraceae|g__GGB28916|s__GGB28916_SGB41612
k__Bacteria|p__Firmicutes|c__Clostridia|o__Eubacteriales|f__Lachnospiraceae|g__GGB28924|s__GGB28924_SGB41621
k__Bacteria|p__Firmicutes|c__Clostridia|o__Eubacteriales|f__Lachnospiraceae|g__GGB28926|s__GGB28926_SGB41624

k__Bacteria|p__Bacteria_unclassified|c__CFGB77359|o__OFGB77359|f__FGB77359|g__GGB28927|s__GGB28927_SGB416
k__Bacteria|p__Firmicutes|c__CFGB9639|o__OFGB9639|f__FGB9639|g__GGB28934|s__GGB28934_SGB41635
k__Bacteria|p__Firmicutes|c__Clostridia|o__Eubacteriales|f__Lachnospiraceae|g__GGB28946|s__GGB28946_SGB41652
k__Bacteria|p__Firmicutes|c__Clostridia|o__Eubacteriales|f__Lachnospiraceae|g__GGB28949|s__GGB28949_SGB41655
k__Bacteria|p__Firmicutes|c__Clostridia|o__Eubacteriales|f__Lachnospiraceae|g__GGB28949|s__GGB28949_SGB41656
k__Bacteria|p__Firmicutes|c__Clostridia|o__Eubacteriales|f__Clostridiaceae|g__GGB28950|s__GGB28950_SGB41657
k__Bacteria|p__Firmicutes|c__Clostridia|o__Eubacteriales|f__Clostridiaceae|g__GGB28951|s__GGB28951_SGB102295
k__Bacteria|p__Firmicutes|c__Clostridia|o__Eubacteriales|f__Clostridiaceae|g__GGB28951|s__GGB28951_SGB41658
k__Bacteria|p__Firmicutes|c__Clostridia|o__Eubacteriales|f__Clostridiaceae|g__GGB28954|s__GGB28954_SGB41662
k__Bacteria|p__Firmicutes|c__Clostridia|o__Eubacteriales|f__Clostridiaceae|g__GGB28956|s__GGB28956_SGB41665
k__Bacteria|p__Firmicutes|c__Clostridia|o__Eubacteriales|f__Clostridiaceae|g__GGB28960|s__GGB28960_SGB41669
k__Bacteria|p__Firmicutes|c__Clostridia|o__Eubacteriales|f__Clostridiaceae|g__GGB28967|s__GGB28967_SGB41678
k__Bacteria|p__Firmicutes|c__Clostridia|o__Eubacteriales|f__Eubacteriaceae|g__GGB28991|s__GGB28991_SGB41705
k__Bacteria|p__Firmicutes|c__CFGB9658|o__OFGB9658|f__FGB9658|g__GGB29002|s__GGB29002_SGB41718
k__Bacteria|p__Firmicutes|c__CFGB9659|o__OFGB9659|f__FGB9659|g__GGB29003|s__GGB29003_SGB41719
k__Bacteria|p__Bacteria_unclassified|c__Bacteria_unclassified|o__Bacteria_unclassified|f__Bacteria_unclassified|g__GGB2
k__Bacteria|p__Firmicutes|c__CFGB9827|o__OFGB9827|f__FGB9827|g__GGB29531|s__GGB29531_SGB42317
k__Bacteria|p__Firmicutes|c__Clostridia|o__Eubacteriales|f__Eubacteriaceae|g__GGB29685|s__GGB29685_SGB42494
k__Bacteria|p__Bacteria_unclassified|c__CFGB77303|o__OFGB77303|f__FGB77303|g__GGB30141|s__GGB30141_SGB430
k__Bacteria|p__Firmicutes|c__Clostridia|o__Eubacteriales|f__Eubacteriales_unclassified|g__GGB30286|s__GGB30286_SG
k__Bacteria|p__Firmicutes|c__Clostridia|o__Eubacteriales|f__Oscillospiraceae|g__GGB30303|s__GGB30303_SGB43268
k__Bacteria|p__Firmicutes|c__CFGB30328|o__OFGB30328|f__FGB30328|g__GGB30413|s__GGB30413_SGB43452
k__Bacteria|p__Firmicutes|c__Clostridia|o__Eubacteriales|f__Oscillospiraceae|g__GGB30454|s__GGB30454_SGB43514
k__Bacteria|p__Firmicutes|c__Clostridia|o__Eubacteriales|f__Oscillospiraceae|g__GGB30455|s__GGB30455_SGB43519
k__Bacteria|p__Firmicutes|c__Clostridia|o__Eubacteriales|f__Oscillospiraceae|g__GGB30461|s__GGB30461_SGB43527
k__Bacteria|p__Firmicutes|c__Clostridia|o__Eubacteriales|f__Oscillospiraceae|g__GGB30461|s__GGB30461_SGB43530
k__Bacteria|p__Firmicutes|c__Clostridia|o__Eubacteriales|f__Oscillospiraceae|g__GGB30463|s__GGB30463_SGB43537
k__Bacteria|p__Firmicutes|c__Clostridia|o__Eubacteriales|f__Oscillospiraceae|g__GGB30473|s__GGB30473_SGB43557
k__Bacteria|p__Firmicutes|c__Clostridia|o__Eubacteriales|f__Oscillospiraceae|g__GGB30475|s__GGB30475_SGB63182
k__Bacteria|p__Actinobacteria|c__CFGB77153|o__OFGB77153|f__FGB77153|g__GGB30861|s__GGB30861_SGB44083
k__Bacteria|p__Tenericutes|c__CFGB1791|o__OFGB1791|f__FGB1791|g__GGB31312|s__GGB31312_SGB44628
k__Bacteria|p__Firmicutes|c__CFGB10290|o__OFGB10290|f__FGB10290|g__GGB31438|s__GGB31438_SGB44768
k__Bacteria|p__Firmicutes|c__Clostridia|o__Eubacteriales|f__Oscillospiraceae|g__GGB3171|s__GGB3171_SGB4185
k__Bacteria|p__Firmicutes|c__CFGB1765|o__OFGB1765|f__FGB1765|g__GGB31823|s__GGB31823_SGB45199
k__Bacteria|p__Firmicutes|c__CFGB10349|o__OFGB10349|f__FGB10349|g__GGB31853|s__GGB31853_SGB45233
k__Bacteria|p__Firmicutes|c__CFGB10667|o__OFGB10667|f__FGB10667|g__GGB32371|s__GGB32371_SGB41694
k__Bacteria|p__Firmicutes|c__Clostridia|o__Eubacteriales|f__Lachnospiraceae|g__GGB3793|s__GGB3793_SGB5158
k__Bacteria|p__Firmicutes|c__Clostridia|o__Eubacteriales|f__Lachnospiraceae|g__GGB42598|s__GGB42598_SGB59794
k__Bacteria|p__Firmicutes|c__Clostridia|o__Eubacteriales|f__Christensenellaceae|g__GGB45656|s__GGB45656_SGB6337
k__Bacteria|p__Firmicutes|c__CFGB10299|o__OFGB10299|f__FGB10299|g__GGB47127|s__GGB47127_SGB65054
k__Bacteria|p__Firmicutes|c__Clostridia|o__Eubacteriales|f__Oscillospiraceae|g__GGB74395|s__GGB74395_SGB43521
k__Bacteria|p__Firmicutes|c__Clostridia|o__Eubacteriales|f__Oscillospiraceae|g__GGB75053|s__GGB75053_SGB43494
k__Bacteria|p__Firmicutes|c__Clostridia|o__Eubacteriales|f__Lachnospiraceae|g__GGB75109|s__GGB75109_SGB102238
k__Bacteria|p__Firmicutes|c__Clostridia|o__Clostridia_unclassified|f__Clostridia_unclassified|g__GGB81440|s__GGB8144
k__Bacteria|p__Firmicutes|c__Clostridia|o__Eubacteriales|f__Lachnospiraceae|g__Lachnospiraceae_unclassified|s__Lachn
k__Bacteria|p__Firmicutes|c__Clostridia|o__Eubacteriales|f__Lachnospiraceae|g__Lachnospiraceae_unclassified|s__Lachn

k__Bacteria|p__Firmicutes|c__Clostridia|o__Eubacteriales|f__Lachnospiraceae|g__Lachnospiraceae_unclassified|s__Lachn
k__Bacteria|p__Firmicutes|c__Clostridia|o__Eubacteriales|f__Lachnospiraceae|g__Lachnospiraceae_unclassified|s__Lachn
k__Bacteria|p__Firmicutes|c__Clostridia|o__Eubacteriales|f__Lachnospiraceae|g__Lachnospiraceae_unclassified|s__Lachn
k__Bacteria|p__Firmicutes|c__Clostridia|o__Eubacteriales|f__Lachnospiraceae|g__Lachnospiraceae_unclassified|s__Lachn
k__Bacteria|p__Firmicutes|c__Clostridia|o__Eubacteriales|f__Lachnospiraceae|g__Lachnospiraceae_unclassified|s__Lachn
k__Bacteria|p__Firmicutes|c__Clostridia|o__Eubacteriales|f__Lachnospiraceae|g__Lachnospiraceae_unclassified|s__Lachn
k__Bacteria|p__Firmicutes|c__Bacilli|o__Lactobacillales|f__Lactobacillaceae|g__Lactobacillus|s__Lactobacillus_johnsonii
k__Bacteria|p__Bacteroidota|c__Bacteroidia|o__Bacteroidales|f__Muribaculaceae|g__Muribaculaceae_unclassified|s__M
k__Bacteria|p__Firmicutes|c__Clostridia|o__Eubacteriales|f__Oscillospiraceae|g__Neglectibacter|s__Neglectibacter_sp_X
k__Bacteria|p__Firmicutes|c__Clostridia|o__Eubacteriales|f__Oscillospiraceae|g__Oscillospiraceae_unclassified|s__Oscillo
k__Bacteria|p__Firmicutes|c__Clostridia|o__Eubacteriales|f__Oscillospiraceae|g__Oscillospiraceae_unclassified|s__Oscillo
k__Bacteria|p__Firmicutes|c__Clostridia|o__Eubacteriales|f__Oscillospiraceae|g__Oscillospiraceae_unclassified|s__Oscillo
k__Bacteria|p__Proteobacteria|c__Betaproteobacteria|o__Burkholderiales|f__Sutterellaceae|g__Parasutterella|s__Parasu
k__Bacteria|p__Firmicutes|c__Clostridia|o__Eubacteriales|f__Peptostreptococcaceae|g__Romboutsia|s__Romboutsia_ilea
k__Bacteria|p__Firmicutes|c__Clostridia|o__Eubacteriales|f__Lachnospiraceae|g__Schaedlerella|s__Schaedlerella_arabino
k__Bacteria|p__Firmicutes|c__Erysipelotrichia|o__Erysipelotrichales|f__Turicibacteraceae|g__Turicibacter|s__Turicibacter_
k__Bacteria|p__Bacteria_unclassified|c__Bacteria_unclassified|o__Bacteria_unclassified|f__Bacteria_unclassified|g__Bacte
k__Bacteria|p__Bacteria_unclassified|c__Bacteria_unclassified|o__Bacteria_unclassified|f__Bacteria_unclassified|g__Bacte
k__Bacteria|p__Bacteria_unclassified|c__Bacteria_unclassified|o__Bacteria_unclassified|f__Bacteria_unclassified|g__Bacte

k__Bacteria|p__Firmicutes|c__Clostridia|o__Eubacteriales|f__Lachnospiraceae|g__Acetatifactor|s__Acetatifactor_SGB415
k__Bacteria|p__Firmicutes|c__Clostridia|o__Eubacteriales|f__Lachnospiraceae|g__Acetatifactor|s__Acetatifactor_muris
k__Bacteria|p__Firmicutes|c__Clostridia|o__Eubacteriales|f__Oscillospiraceae|g__Acutalibacter|s__Acutalibacter_muris
k__Bacteria|p__Firmicutes|c__Clostridia|o__Eubacteriales|f__Oscillospiraceae|g__Acutalibacter|s__Acutalibacter_sp_1XD
k__Bacteria|p__Actinobacteria|c__Coriobacteriia|o__Eggerthellales|f__Eggerthellaceae|g__Adlercreutzia|s__Adlercreutzia
k__Bacteria|p__Actinobacteria|c__Coriobacteriia|o__Eggerthellales|f__Eggerthellaceae|g__Adlercreutzia|s__Adlercreutzia
k__Bacteria|p__Actinobacteria|c__Coriobacteriia|o__Eggerthellales|f__Eggerthellaceae|g__Adlercreutzia|s__Adlercreutzia
k__Bacteria|p__Verrucomicrobia|c__Verrucomicrobiae|o__Verrucomicrobiales|f__Akkermansiaceae|g__Akkermansia|s__A
k__Bacteria|p__Bacteroidota|c__Bacteroidia|o__Bacteroidales|f__Rikenellaceae|g__Alistipes|s__Alistipes_sp_DSM_11234
k__Bacteria|p__Firmicutes|c__Clostridia|o__Eubacteriales|f__Oscillospiraceae|g__Anaerotruncus|s__Anaerotruncus_sp_1
k__Bacteria|p__Bacteria_unclassified|c__Bacteria_unclassified|o__Bacteria_unclassified|f__Bacteria_unclassified|g__Bacte
k__Bacteria|p__Bacteria_unclassified|c__Bacteria_unclassified|o__Bacteria_unclassified|f__Bacteria_unclassified|g__Bacte
k__Bacteria|p__Bacteria_unclassified|c__Bacteria_unclassified|o__Bacteria_unclassified|f__Bacteria_unclassified|g__Bacte
k__Bacteria|p__Bacteroidota|c__Bacteroidia|o__Bacteroidales|f__Bacteroidaceae|g__Bacteroides|s__Bacteroides_thetaio
k__Bacteria|p__Actinobacteria|c__Actinomycetia|o__Bifidobacteriales|f__Bifidobacteriaceae|g__Bifidobacterium|s__Bifid
k__Bacteria|p__Firmicutes|c__Clostridia|o__Clostridia_unclassified|f__Clostridia_unclassified|g__Clostridia_unclassified|s_
k__Bacteria|p__Firmicutes|c__Clostridia|o__Eubacteriales|f__Clostridiaceae|g__Clostridiaceae_unclassified|s__Clostridiac
k__Bacteria|p__Firmicutes|c__Clostridia|o__Eubacteriales|f__Clostridiaceae|g__Clostridiaceae_unclassified|s__Clostridiac
k__Bacteria|p__Firmicutes|c__Clostridia|o__Eubacteriales|f__Eubacteriales_unclassified|g__Eubacteriales_unclassified|s_
k__Bacteria|p__Firmicutes|c__Erysipelotrichia|o__Erysipelotrichales|f__Erysipelotrichaceae|g__Erysipelatoclostridium|s_
k__Bacteria|p__Actinobacteria|c__Coriobacteriia|o__Coriobacteriales|f__Coriobacteriaceae|g__Coriobacteriaceae_unclass
k__Bacteria|p__Firmicutes|c__Clostridia|o__Eubacteriales|f__Lachnospiraceae|g__Dorea|s__Dorea_sp_5_2
k__Bacteria|p__Firmicutes|c__Erysipelotrichia|o__Erysipelotrichales|f__Erysipelotrichaceae|g__Dubosiella|s__Dubosiella_
k__Bacteria|p__Firmicutes|c__Erysipelotrichia|o__Erysipelotrichales|f__Erysipelotrichales_unclassified|g__Erysipelotrichal

k__Bacteria|p__Firmicutes|c__Clostridia|o__Eubacteriales|f__Eubacteriaceae|g__Eubacteriaceae_unclassified|s__Eubacte[illegible]

k__Bacteria|p__Firmicutes|c__Clostridia|o__Eubacteriales|f__Eubacteriaceae|g__Eubacteriaceae_unclassified|s__Eubacte[illegible]

k__Bacteria|p__Firmicutes|c__Clostridia|o__Eubacteriales|f__Lachnospiraceae|g__GGB20149|s__GGB20149_SGB29430

k__Bacteria|p__Actinobacteria|c__Coriobacteriia|o__Eggerthellales|f__Eggerthellaceae|g__GGB22635|s__GGB22635_SGB[illegible]

k__Bacteria|p__Firmicutes|c__Clostridia|o__Eubacteriales|f__Lachnospiraceae|g__GGB25041|s__GGB25041_SGB36960

k__Bacteria|p__Bacteroidota|c__Bacteroidia|o__Bacteroidales|f__Muribaculaceae|g__GGB27876|s__GGB27876_SGB4031[illegible]

k__Bacteria|p__Bacteroidota|c__Bacteroidia|o__Bacteroidales|f__Muribaculaceae|g__GGB27878|s__GGB27878_SGB4031[illegible]

k__Bacteria|p__Bacteroidota|c__Bacteroidia|o__Bacteroidales|f__Muribaculaceae|g__GGB27918|s__GGB27918_SGB4035[illegible]

k__Bacteria|p__Firmicutes|c__CFGB9508|o__OFGB9508|f__FGB9508|g__GGB28382|s__GGB28382_SGB40962

k__Bacteria|p__Firmicutes|c__CFGB2838|o__OFGB2838|f__FGB2838|g__GGB28399|s__GGB28399_SGB40980

k__Bacteria|p__Firmicutes|c__CFGB2838|o__OFGB2838|f__FGB2838|g__GGB28411|s__GGB28411_SGB40993

k__Bacteria|p__Firmicutes|c__CFGB2838|o__OFGB2838|f__FGB2838|g__GGB28415|s__GGB28415_SGB40997

k__Bacteria|p__Firmicutes|c__Clostridia|o__Eubacteriales|f__Pumilibacteraceae|g__GGB28430|s__GGB28430_SGB41013

k__Bacteria|p__Firmicutes|c__CFGB28439|o__OFGB28439|f__FGB28439|g__GGB28439|s__GGB28439_SGB41022

k__Bacteria|p__Firmicutes|c__Clostridia|o__Clostridia_unclassified|f__Clostridia_unclassified|g__GGB28778|s__GGB2877[illegible]

k__Bacteria|p__Firmicutes|c__Clostridia|o__Eubacteriales|f__Eubacteriaceae|g__GGB28784|s__GGB28784_SGB41437

k__Bacteria|p__Firmicutes|c__Clostridia|o__Eubacteriales|f__Lachnospiraceae|g__GGB28792|s__GGB28792_SGB41445

k__Bacteria|p__Firmicutes|c__Clostridia|o__Eubacteriales|f__Lachnospiraceae|g__GGB28798|s__GGB28798_SGB41451

k__Bacteria|p__Firmicutes|c__Clostridia|o__Eubacteriales|f__Lachnospiraceae|g__GGB28802|s__GGB28802_SGB41455

k__Bacteria|p__Firmicutes|c__Clostridia|o__Eubacteriales|f__Lachnospiraceae|g__GGB28818|s__GGB28818_SGB41473

k__Bacteria|p__Firmicutes|c__CFGB77305|o__OFGB77305|f__FGB77305|g__GGB28828|s__GGB28828_SGB41484

k__Bacteria|p__Firmicutes|c__Clostridia|o__Eubacteriales|f__Clostridiaceae|g__GGB28851|s__GGB28851_SGB41518

k__Bacteria|p__Firmicutes|c__Clostridia|o__Eubacteriales|f__Lachnospiraceae|g__GGB28859|s__GGB28859_SGB41528

k__Bacteria|p__Firmicutes|c__Clostridia|o__Eubacteriales|f__Lachnospiraceae|g__GGB28864|s__GGB28864_SGB41535

k__Bacteria|p__Firmicutes|c__Clostridia|o__Eubacteriales|f__Lachnospiraceae|g__GGB28869|s__GGB28869_SGB41543

k__Bacteria|p__Firmicutes|c__CFGB9633|o__OFGB9633|f__FGB9633|g__GGB28883|s__GGB28883_SGB41564

k__Bacteria|p__Bacteria_unclassified|c__Bacteria_unclassified|o__Bacteria_unclassified|f__Bacteria_unclassified|g__GGB[illegible]

k__Bacteria|p__Bacteria_unclassified|c__Bacteria_unclassified|o__Bacteria_unclassified|f__Bacteria_unclassified|g__GGB[illegible]

k__Bacteria|p__Bacteria_unclassified|c__Bacteria_unclassified|o__Bacteria_unclassified|f__Bacteria_unclassified|g__GGB[illegible]

k__Bacteria|p__Bacteria_unclassified|c__Bacteria_unclassified|o__Bacteria_unclassified|f__Bacteria_unclassified|g__GGB[illegible]

k__Bacteria|p__Firmicutes|c__Clostridia|o__Eubacteriales|f__Lachnospiraceae|g__GGB28916|s__GGB28916_SGB41612

k__Bacteria|p__Firmicutes|c__Clostridia|o__Eubacteriales|f__Lachnospiraceae|g__GGB28924|s__GGB28924_SGB41621

k__Bacteria|p__Firmicutes|c__Clostridia|o__Eubacteriales|f__Lachnospiraceae|g__GGB28926|s__GGB28926_SGB41624

k__Bacteria|p__Bacteria_unclassified|c__CFGB77359|o__OFGB77359|f__FGB77359|g__GGB28927|s__GGB28927_SGB416[illegible]

k__Bacteria|p__Firmicutes|c__CFGB9639|o__OFGB9639|f__FGB9639|g__GGB28934|s__GGB28934_SGB41635

k__Bacteria|p__Firmicutes|c__Clostridia|o__Eubacteriales|f__Lachnospiraceae|g__GGB28946|s__GGB28946_SGB41652

k__Bacteria|p__Firmicutes|c__Clostridia|o__Eubacteriales|f__Lachnospiraceae|g__GGB28949|s__GGB28949_SGB41655

k__Bacteria|p__Firmicutes|c__Clostridia|o__Eubacteriales|f__Lachnospiraceae|g__GGB28949|s__GGB28949_SGB41656

k__Bacteria|p__Firmicutes|c__Clostridia|o__Eubacteriales|f__Clostridiaceae|g__GGB28950|s__GGB28950_SGB41657

k__Bacteria|p__Firmicutes|c__Clostridia|o__Eubacteriales|f__Clostridiaceae|g__GGB28951|s__GGB28951_SGB102295

k__Bacteria|p__Firmicutes|c__Clostridia|o__Eubacteriales|f__Clostridiaceae|g__GGB28951|s__GGB28951_SGB41658

k__Bacteria|p__Firmicutes|c__Clostridia|o__Eubacteriales|f__Clostridiaceae|g__GGB28954|s__GGB28954_SGB41662

k__Bacteria|p__Firmicutes|c__Clostridia|o__Eubacteriales|f__Clostridiaceae|g__GGB28956|s__GGB28956_SGB41665

k__Bacteria|p__Firmicutes|c__Clostridia|o__Eubacteriales|f__Clostridiaceae|g__GGB28960|s__GGB28960_SGB41669

k__Bacteria|p__Firmicutes|c__Clostridia|o__Eubacteriales|f__Clostridiaceae|g__GGB28967|s__GGB28967_SGB41678

k__Bacteria|p__Firmicutes|c__Clostridia|o__Eubacteriales|f__Eubacteriaceae|g__GGB28991|s__GGB28991_SGB41705

k__Bacteria|p__Firmicutes|c__CFGB9658|o__OFGB9658|f__FGB9658|g__GGB29002|s__GGB29002_SGB41718
k__Bacteria|p__Firmicutes|c__CFGB9659|o__OFGB9659|f__FGB9659|g__GGB29003|s__GGB29003_SGB41719
k__Bacteria|p__Bacteria_unclassified|c__Bacteria_unclassified|o__Bacteria_unclassified|f__Bacteria_unclassified|g__GGB2
k__Bacteria|p__Firmicutes|c__CFGB9827|o__OFGB9827|f__FGB9827|g__GGB29531|s__GGB29531_SGB42317
k__Bacteria|p__Firmicutes|c__Clostridia|o__Eubacteriales|f__Eubacteriaceae|g__GGB29685|s__GGB29685_SGB42494
k__Bacteria|p__Bacteria_unclassified|c__CFGB77303|o__OFGB77303|f__FGB77303|g__GGB30141|s__GGB30141_SGB430
k__Bacteria|p__Firmicutes|c__Clostridia|o__Eubacteriales|f__Eubacteriales_unclassified|g__GGB30286|s__GGB30286_SG
k__Bacteria|p__Firmicutes|c__Clostridia|o__Eubacteriales|f__Oscillospiraceae|g__GGB30303|s__GGB30303_SGB43268
k__Bacteria|p__Firmicutes|c__CFGB30328|o__OFGB30328|f__FGB30328|g__GGB30413|s__GGB30413_SGB43452
k__Bacteria|p__Firmicutes|c__Clostridia|o__Eubacteriales|f__Oscillospiraceae|g__GGB30454|s__GGB30454_SGB43514
k__Bacteria|p__Firmicutes|c__Clostridia|o__Eubacteriales|f__Oscillospiraceae|g__GGB30455|s__GGB30455_SGB43519
k__Bacteria|p__Firmicutes|c__Clostridia|o__Eubacteriales|f__Oscillospiraceae|g__GGB30461|s__GGB30461_SGB43527
k__Bacteria|p__Firmicutes|c__Clostridia|o__Eubacteriales|f__Oscillospiraceae|g__GGB30461|s__GGB30461_SGB43530
k__Bacteria|p__Firmicutes|c__Clostridia|o__Eubacteriales|f__Oscillospiraceae|g__GGB30463|s__GGB30463_SGB43537
k__Bacteria|p__Firmicutes|c__Clostridia|o__Eubacteriales|f__Oscillospiraceae|g__GGB30473|s__GGB30473_SGB43557
k__Bacteria|p__Firmicutes|c__Clostridia|o__Eubacteriales|f__Oscillospiraceae|g__GGB30475|s__GGB30475_SGB63182
k__Bacteria|p__Actinobacteria|c__CFGB77153|o__OFGB77153|f__FGB77153|g__GGB30861|s__GGB30861_SGB44083
k__Bacteria|p__Tenericutes|c__CFGB1791|o__OFGB1791|f__FGB1791|g__GGB31312|s__GGB31312_SGB44628
k__Bacteria|p__Firmicutes|c__CFGB10290|o__OFGB10290|f__FGB10290|g__GGB31438|s__GGB31438_SGB44768
k__Bacteria|p__Firmicutes|c__Clostridia|o__Eubacteriales|f__Oscillospiraceae|g__GGB3171|s__GGB3171_SGB4185
k__Bacteria|p__Firmicutes|c__CFGB1765|o__OFGB1765|f__FGB1765|g__GGB31823|s__GGB31823_SGB45199
k__Bacteria|p__Firmicutes|c__CFGB10349|o__OFGB10349|f__FGB10349|g__GGB31853|s__GGB31853_SGB45233
k__Bacteria|p__Firmicutes|c__CFGB10667|o__OFGB10667|f__FGB10667|g__GGB32371|s__GGB32371_SGB41694
k__Bacteria|p__Firmicutes|c__Clostridia|o__Eubacteriales|f__Lachnospiraceae|g__GGB3793|s__GGB3793_SGB5158
k__Bacteria|p__Firmicutes|c__Clostridia|o__Eubacteriales|f__Lachnospiraceae|g__GGB42598|s__GGB42598_SGB59794
k__Bacteria|p__Firmicutes|c__Clostridia|o__Eubacteriales|f__Christensenellaceae|g__GGB45656|s__GGB45656_SGB6337
k__Bacteria|p__Firmicutes|c__CFGB10299|o__OFGB10299|f__FGB10299|g__GGB47127|s__GGB47127_SGB65054
k__Bacteria|p__Firmicutes|c__Clostridia|o__Eubacteriales|f__Oscillospiraceae|g__GGB74395|s__GGB74395_SGB43521
k__Bacteria|p__Firmicutes|c__Clostridia|o__Eubacteriales|f__Oscillospiraceae|g__GGB75053|s__GGB75053_SGB43494
k__Bacteria|p__Firmicutes|c__Clostridia|o__Eubacteriales|f__Lachnospiraceae|g__GGB75109|s__GGB75109_SGB102238
k__Bacteria|p__Firmicutes|c__Clostridia|o__Clostridia_unclassified|f__Clostridia_unclassified|g__GGB81440|s__GGB8144
k__Bacteria|p__Firmicutes|c__Clostridia|o__Eubacteriales|f__Lachnospiraceae|g__Lachnospiraceae_unclassified|s__Lachn
k__Bacteria|p__Firmicutes|c__Clostridia|o__Eubacteriales|f__Lachnospiraceae|g__Lachnospiraceae_unclassified|s__Lachn
k__Bacteria|p__Firmicutes|c__Clostridia|o__Eubacteriales|f__Lachnospiraceae|g__Lachnospiraceae_unclassified|s__Lachn
k__Bacteria|p__Firmicutes|c__Clostridia|o__Eubacteriales|f__Lachnospiraceae|g__Lachnospiraceae_unclassified|s__Lachn
k__Bacteria|p__Firmicutes|c__Clostridia|o__Eubacteriales|f__Lachnospiraceae|g__Lachnospiraceae_unclassified|s__Lachn
k__Bacteria|p__Firmicutes|c__Clostridia|o__Eubacteriales|f__Lachnospiraceae|g__Lachnospiraceae_unclassified|s__Lachn
k__Bacteria|p__Firmicutes|c__Clostridia|o__Eubacteriales|f__Lachnospiraceae|g__Lachnospiraceae_unclassified|s__Lachn
k__Bacteria|p__Firmicutes|c__Clostridia|o__Eubacteriales|f__Lachnospiraceae|g__Lachnospiraceae_unclassified|s__Lachn
k__Bacteria|p__Firmicutes|c__Bacilli|o__Lactobacillales|f__Lactobacillaceae|g__Lactobacillus|s__Lactobacillus_johnsonii
k__Bacteria|p__Bacteroidota|c__Bacteroidia|o__Bacteroidales|f__Muribaculaceae|g__Muribaculaceae_unclassified|s__M
k__Bacteria|p__Firmicutes|c__Clostridia|o__Eubacteriales|f__Oscillospiraceae|g__Neglectibacter|s__Neglectibacter_sp_X
k__Bacteria|p__Firmicutes|c__Clostridia|o__Eubacteriales|f__Oscillospiraceae|g__Oscillospiraceae_unclassified|s__Oscillo
k__Bacteria|p__Firmicutes|c__Clostridia|o__Eubacteriales|f__Oscillospiraceae|g__Oscillospiraceae_unclassified|s__Oscillo
k__Bacteria|p__Firmicutes|c__Clostridia|o__Eubacteriales|f__Oscillospiraceae|g__Oscillospiraceae_unclassified|s__Oscillo
k__Bacteria|p__Proteobacteria|c__Betaproteobacteria|o__Burkholderiales|f__Sutterellaceae|g__Parasutterella|s__Parasu

k__Bacteria|p__Firmicutes|c__Clostridia|o__Eubacteriales|f__Peptostreptococcaceae|g__Romboutsia|s__Romboutsia_ilea
k__Bacteria|p__Firmicutes|c__Clostridia|o__Eubacteriales|f__Lachnospiraceae|g__Schaedlerella|s__Schaedlerella_arabino
k__Bacteria|p__Firmicutes|c__Erysipelotrichia|o__Erysipelotrichales|f__Turicibacteraceae|g__Turicibacter|s__Turicibacter_
k__Bacteria|p__Bacteria_unclassified|c__Bacteria_unclassified|o__Bacteria_unclassified|f__Bacteria_unclassified|g__Bacte
k__Bacteria|p__Bacteria_unclassified|c__Bacteria_unclassified|o__Bacteria_unclassified|f__Bacteria_unclassified|g__Bacte
k__Bacteria|p__Bacteria_unclassified|c__Bacteria_unclassified|o__Bacteria_unclassified|f__Bacteria_unclassified|g__Bacte

GB102200
GB41677
GB43546

rium

GB102200
GB41677

GB43546

rium

GB102200
GB41677
GB43546

rium

GB102200
GB41677
GB43546

rium

GB102200
GB41677
GB43546

rium

GB102200
GB41677
GB43546

rium

GB102200
GB41677
GB43546

rium

GB102200
GB41677
GB43546

rium

GB41677

GB43546

rium

GB102200
GB41677
GB43546

rium
