## Supplementary material for "Non-invasive Vagal Nerve Stimulation as a Potential Treatment for Repetitive Blast Trauma": st8

| Exposure | Mediator (Species-level feature) | Outcome | p_m |
|---|---|---|---|
| blast | Acetatifactor_muris | EtOH_24h_intakeave_final | 0.739522079 |
| vns | Acetatifactor_muris | EtOH_24h_intakeave_final | 0.752897140 |
| blast | Acetatifactor_SGB41546 | EtOH_24h_intakeave_final | 0.840245797 |
| vns | Acetatifactor_SGB41546 | EtOH_24h_intakeave_final | 0.663000977 |
| blast | Acutalibacter_muris | EtOH_24h_intakeave_final | 0.422154633 |
| vns | Acutalibacter_muris | EtOH_24h_intakeave_final | 0.810578449 |
| blast | Acutalibacter_sp_1XD8_36 | EtOH_24h_intakeave_final | 0.660579886 |
| vns | Acutalibacter_sp_1XD8_36 | EtOH_24h_intakeave_final | 0.709184933 |
| blast | Adlercreutzia_caecimuris | EtOH_24h_intakeave_final | 0.989608952 |
| vns | Adlercreutzia_caecimuris | EtOH_24h_intakeave_final | 0.676542691 |
| blast | Adlercreutzia_mucosicola | EtOH_24h_intakeave_final | 0.271833985 |
| vns | Adlercreutzia_mucosicola | EtOH_24h_intakeave_final | 0.249825657 |
| blast | Adlercreutzia_muris | EtOH_24h_intakeave_final | 0.670421774 |
| vns | Adlercreutzia_muris | EtOH_24h_intakeave_final | 0.595827745 |
| blast | Akkermansia_muciniphila | EtOH_24h_intakeave_final | 0.169812540 |
| vns | Akkermansia_muciniphila | EtOH_24h_intakeave_final | 0.864362509 |
| blast | Alistipes_sp_DSM_112343 | EtOH_24h_intakeave_final | 0.372515977 |
| vns | Alistipes_sp_DSM_112343 | EtOH_24h_intakeave_final | 0.832698307 |
| blast | Anaerotruncus_sp_1XD42_93 | EtOH_24h_intakeave_final | 0.520271453 |
| vns | Anaerotruncus_sp_1XD42_93 | EtOH_24h_intakeave_final | 0.617410721 |
| blast | Bacteria_unclassified_SGB102200 | EtOH_24h_intakeave_final | 0.474269212 |
| vns | Bacteria_unclassified_SGB102200 | EtOH_24h_intakeave_final | 0.764166788 |
| blast | Bacteria_unclassified_SGB41677 | EtOH_24h_intakeave_final | 0.444143823 |
| vns | Bacteria_unclassified_SGB41677 | EtOH_24h_intakeave_final | 0.169989424 |
| blast | Bacteria_unclassified_SGB43546 | EtOH_24h_intakeave_final | 0.586358365 |
| vns | Bacteria_unclassified_SGB43546 | EtOH_24h_intakeave_final | 0.789009912 |
| blast | bacterium_1XD42_54 | EtOH_24h_intakeave_final | 0.014424595 |
| vns | bacterium_1XD42_54 | EtOH_24h_intakeave_final | 0.234577574 |
| blast | bacterium_1XD42_76 | EtOH_24h_intakeave_final | 0.600031603 |
| vns | bacterium_1XD42_76 | EtOH_24h_intakeave_final | 0.642992453 |
| blast | bacterium_1xD8_48 | EtOH_24h_intakeave_final | 0.899708945 |
| vns | bacterium_1xD8_48 | EtOH_24h_intakeave_final | 0.696816880 |
| blast | Bacteroides_thetaiotaomicron | EtOH_24h_intakeave_final | 0.960201477 |
| vns | Bacteroides_thetaiotaomicron | EtOH_24h_intakeave_final | 0.634306315 |
| blast | Berger Parker Index | EtOH_24h_intakeave_final | 0.545741712 |
| vns | Berger Parker Index | EtOH_24h_intakeave_final | 0.233733054 |
| blast | Bifidobacterium_pseudolongum | EtOH_24h_intakeave_final | 0.428010398 |
| vns | Bifidobacterium_pseudolongum | EtOH_24h_intakeave_final | 0.347799065 |
| blast | Clostridia_bacterium | EtOH_24h_intakeave_final | 0.374559043 |
| vns | Clostridia_bacterium | EtOH_24h_intakeave_final | 0.724940107 |
| blast | Clostridiaceae_bacterium | EtOH_24h_intakeave_final | 0.987169780 |
| vns | Clostridiaceae_bacterium | EtOH_24h_intakeave_final | 0.241937042 |
| blast | Clostridiaceae_unclassified_SGB41663 | EtOH_24h_intakeave_final | 0.780382224 |
| vns | Clostridiaceae_unclassified_SGB41663 | EtOH_24h_intakeave_final | 0.719154368 |
| blast | Clostridiales_bacterium | EtOH_24h_intakeave_final | 0.002331956 |
| vns | Clostridiales_bacterium | EtOH_24h_intakeave_final | 0.296478501 |

| | | | |
|---|---|---|---|
| blast | Clostridium_cocleatum | EtOH_24h_intakeave_final | 0.0115678325 |
| vns | Clostridium_cocleatum | EtOH_24h_intakeave_final | 0.1960355146 |
| blast | Coriobacteriaceae_bacterium | EtOH_24h_intakeave_final | 0.3724483697 |
| vns | Coriobacteriaceae_bacterium | EtOH_24h_intakeave_final | 0.953919137 |
| blast | Dorea_sp_5_2 | EtOH_24h_intakeave_final | 0.5884919176 |
| vns | Dorea_sp_5_2 | EtOH_24h_intakeave_final | 0.4214161244 |
| blast | Dubosiella_newyorkensis | EtOH_24h_intakeave_final | 0.1169987394 |
| vns | Dubosiella_newyorkensis | EtOH_24h_intakeave_final | 0.799186063 |
| blast | Erysipelotrichales_bacterium | EtOH_24h_intakeave_final | 0.8970006656 |
| vns | Erysipelotrichales_bacterium | EtOH_24h_intakeave_final | 0.3284081083 |
| blast | Eubacteriaceae_bacterium | EtOH_24h_intakeave_final | 0.0271233687 |
| vns | Eubacteriaceae_bacterium | EtOH_24h_intakeave_final | 0.2383186135 |
| blast | Eubacteriaceae_unclassified_SGB94922 | EtOH_24h_intakeave_final | 0.4594880799 |
| vns | Eubacteriaceae_unclassified_SGB94922 | EtOH_24h_intakeave_final | 0.3726201484 |
| blast | GGB20149_SGB29430 | EtOH_24h_intakeave_final | 0.6280700484 |
| vns | GGB20149_SGB29430 | EtOH_24h_intakeave_final | 0.6299827926 |
| blast | GGB22635_SGB63107 | EtOH_24h_intakeave_final | 0.4117103362 |
| vns | GGB22635_SGB63107 | EtOH_24h_intakeave_final | 0.8871521383 |
| blast | GGB25041_SGB36960 | EtOH_24h_intakeave_final | 0.0824966676 |
| vns | GGB25041_SGB36960 | EtOH_24h_intakeave_final | 0.2101871969 |
| blast | GGB27876_SGB40310 | EtOH_24h_intakeave_final | 0.164666135 |
| vns | GGB27876_SGB40310 | EtOH_24h_intakeave_final | 0.7672351949 |
| blast | GGB27878_SGB40312 | EtOH_24h_intakeave_final | 0.2133541578 |
| vns | GGB27878_SGB40312 | EtOH_24h_intakeave_final | 0.1474794298 |
| blast | GGB27918_SGB40356 | EtOH_24h_intakeave_final | 0.9587581408 |
| vns | GGB27918_SGB40356 | EtOH_24h_intakeave_final | 0.8856989297 |
| blast | GGB28382_SGB40962 | EtOH_24h_intakeave_final | 0.5193113079 |
| vns | GGB28382_SGB40962 | EtOH_24h_intakeave_final | 0.8550578348 |
| blast | GGB28399_SGB40980 | EtOH_24h_intakeave_final | 0.2894655025 |
| vns | GGB28399_SGB40980 | EtOH_24h_intakeave_final | 0.287117034 |
| blast | GGB28411_SGB40993 | EtOH_24h_intakeave_final | 0.2018022626 |
| vns | GGB28411_SGB40993 | EtOH_24h_intakeave_final | 0.1468484741 |
| blast | GGB28415_SGB40997 | EtOH_24h_intakeave_final | 0.0267267446 |
| vns | GGB28415_SGB40997 | EtOH_24h_intakeave_final | 0.9490516966 |
| blast | GGB28430_SGB41013 | EtOH_24h_intakeave_final | 0.9120709472 |
| vns | GGB28430_SGB41013 | EtOH_24h_intakeave_final | 0.4662504826 |
| blast | GGB28439_SGB41022 | EtOH_24h_intakeave_final | 0.3804440395 |
| vns | GGB28439_SGB41022 | EtOH_24h_intakeave_final | 0.7387095529 |
| blast | GGB28778_SGB41431 | EtOH_24h_intakeave_final | 0.8046989411 |
| vns | GGB28778_SGB41431 | EtOH_24h_intakeave_final | 0.2184532434 |
| blast | GGB28784_SGB41437 | EtOH_24h_intakeave_final | 0.0207266217 |
| vns | GGB28784_SGB41437 | EtOH_24h_intakeave_final | 0.2544569976 |
| blast | GGB28792_SGB41445 | EtOH_24h_intakeave_final | 0.3385777757 |
| vns | GGB28792_SGB41445 | EtOH_24h_intakeave_final | 0.7612444262 |
| blast | GGB28798_SGB41451 | EtOH_24h_intakeave_final | 0.1640226305 |
| vns | GGB28798_SGB41451 | EtOH_24h_intakeave_final | 0.2489158244 |

| | | | |
|---|---|---|---|
| blast | GGB28802_SGB41455 | EtOH_24h_intakeave_final | 0.4578986259 |
| vns | GGB28802_SGB41455 | EtOH_24h_intakeave_final | 0.342689298 |
| blast | GGB28818_SGB41473 | EtOH_24h_intakeave_final | 0.7292095104 |
| vns | GGB28818_SGB41473 | EtOH_24h_intakeave_final | 0.5694099484 |
| blast | GGB28828_SGB41484 | EtOH_24h_intakeave_final | 0.2986969933 |
| vns | GGB28828_SGB41484 | EtOH_24h_intakeave_final | 0.1067065352 |
| blast | GGB28851_SGB41518 | EtOH_24h_intakeave_final | 0.852552179 |
| vns | GGB28851_SGB41518 | EtOH_24h_intakeave_final | 0.5783084298 |
| blast | GGB28859_SGB41528 | EtOH_24h_intakeave_final | 0.2057493298 |
| vns | GGB28859_SGB41528 | EtOH_24h_intakeave_final | 0.5953667458 |
| blast | GGB28864_SGB41535 | EtOH_24h_intakeave_final | 0.7854652488 |
| vns | GGB28864_SGB41535 | EtOH_24h_intakeave_final | 0.9646589853 |
| blast | GGB28869_SGB41543 | EtOH_24h_intakeave_final | 0.3479316259 |
| vns | GGB28869_SGB41543 | EtOH_24h_intakeave_final | 0.3814774755 |
| blast | GGB28883_SGB41564 | EtOH_24h_intakeave_final | 0.5948772487 |
| vns | GGB28883_SGB41564 | EtOH_24h_intakeave_final | 0.0605244586 |
| blast | GGB28892_SGB41573 | EtOH_24h_intakeave_final | 0.6512272078 |
| vns | GGB28892_SGB41573 | EtOH_24h_intakeave_final | 0.6676188822 |
| blast | GGB28893_SGB41574 | EtOH_24h_intakeave_final | 0.165594509 |
| vns | GGB28893_SGB41574 | EtOH_24h_intakeave_final | 0.1826993761 |
| blast | GGB28898_SGB41580 | EtOH_24h_intakeave_final | 0.3046549939 |
| vns | GGB28898_SGB41580 | EtOH_24h_intakeave_final | 0.0828933309 |
| blast | GGB28904_SGB41597 | EtOH_24h_intakeave_final | 0.1729063974 |
| vns | GGB28904_SGB41597 | EtOH_24h_intakeave_final | 0.0560671999 |
| blast | GGB28916_SGB41612 | EtOH_24h_intakeave_final | 0.8014326092 |
| vns | GGB28916_SGB41612 | EtOH_24h_intakeave_final | 0.8538132969 |
| blast | GGB28924_SGB41621 | EtOH_24h_intakeave_final | 0.7964816629 |
| vns | GGB28924_SGB41621 | EtOH_24h_intakeave_final | 0.2786583787 |
| blast | GGB28926_SGB41624 | EtOH_24h_intakeave_final | 0.3067808341 |
| vns | GGB28926_SGB41624 | EtOH_24h_intakeave_final | 0.470765265 |
| blast | GGB28927_SGB41625 | EtOH_24h_intakeave_final | 0.1937208057 |
| vns | GGB28927_SGB41625 | EtOH_24h_intakeave_final | 0.4182433734 |
| blast | GGB28934_SGB41635 | EtOH_24h_intakeave_final | 0.1158733118 |
| vns | GGB28934_SGB41635 | EtOH_24h_intakeave_final | 0.1113740731 |
| blast | GGB28946_SGB41652 | EtOH_24h_intakeave_final | 0.4278432256 |
| vns | GGB28946_SGB41652 | EtOH_24h_intakeave_final | 0.4225889007 |
| blast | GGB28949_SGB41655 | EtOH_24h_intakeave_final | 0.4436081164 |
| vns | GGB28949_SGB41655 | EtOH_24h_intakeave_final | 0.9089481791 |
| blast | GGB28949_SGB41656 | EtOH_24h_intakeave_final | 0.2827282346 |
| vns | GGB28949_SGB41656 | EtOH_24h_intakeave_final | 0.7708639493 |
| blast | GGB28950_SGB41657 | EtOH_24h_intakeave_final | 0.0305270365 |
| vns | GGB28950_SGB41657 | EtOH_24h_intakeave_final | 0.3364589048 |
| blast | GGB28951_SGB102295 | EtOH_24h_intakeave_final | 0.9253521562 |
| vns | GGB28951_SGB102295 | EtOH_24h_intakeave_final | 0.1587903977 |
| blast | GGB28951_SGB41658 | EtOH_24h_intakeave_final | 0.8470013515 |
| vns | GGB28951_SGB41658 | EtOH_24h_intakeave_final | 0.9202366237 |

| | | | |
|---|---|---|---|
| blast | GGB28954_SGB41662 | EtOH_24h_intakeave_final | 0.5157582975 |
| vns | GGB28954_SGB41662 | EtOH_24h_intakeave_final | 0.8004849551 |
| blast | GGB28956_SGB41665 | EtOH_24h_intakeave_final | 0.163329713 |
| vns | GGB28956_SGB41665 | EtOH_24h_intakeave_final | 0.1534165023 |
| blast | GGB28960_SGB41669 | EtOH_24h_intakeave_final | 0.155727292 |
| vns | GGB28960_SGB41669 | EtOH_24h_intakeave_final | 0.1559574792 |
| blast | GGB28967_SGB41678 | EtOH_24h_intakeave_final | 0.1792695292 |
| vns | GGB28967_SGB41678 | EtOH_24h_intakeave_final | 0.3150465193 |
| blast | GGB28991_SGB41705 | EtOH_24h_intakeave_final | 0.2288938824 |
| vns | GGB28991_SGB41705 | EtOH_24h_intakeave_final | 0.279982342 |
| blast | GGB29002_SGB41718 | EtOH_24h_intakeave_final | 0.0024308675 |
| vns | GGB29002_SGB41718 | EtOH_24h_intakeave_final | 0.1134655877 |
| blast | GGB29003_SGB41719 | EtOH_24h_intakeave_final | 0.4993986443 |
| vns | GGB29003_SGB41719 | EtOH_24h_intakeave_final | 0.5839678864 |
| blast | GGB29011_SGB41731 | EtOH_24h_intakeave_final | 0.916276033 |
| vns | GGB29011_SGB41731 | EtOH_24h_intakeave_final | 0.3455992360 |
| blast | GGB29531_SGB42317 | EtOH_24h_intakeave_final | 0.1450820268 |
| vns | GGB29531_SGB42317 | EtOH_24h_intakeave_final | 0.1355699662 |
| blast | GGB29685_SGB42494 | EtOH_24h_intakeave_final | 0.8645438528 |
| vns | GGB29685_SGB42494 | EtOH_24h_intakeave_final | 0.340104263 |
| blast | GGB30141_SGB43066 | EtOH_24h_intakeave_final | 0.9915548039 |
| vns | GGB30141_SGB43066 | EtOH_24h_intakeave_final | 0.7601787897 |
| blast | GGB30286_SGB43248 | EtOH_24h_intakeave_final | 0.0552616333 |
| vns | GGB30286_SGB43248 | EtOH_24h_intakeave_final | 0.4546033673 |
| blast | GGB30303_SGB43268 | EtOH_24h_intakeave_final | 0.3719368939 |
| vns | GGB30303_SGB43268 | EtOH_24h_intakeave_final | 0.9447221743 |
| blast | GGB30413_SGB43452 | EtOH_24h_intakeave_final | 0.9532769507 |
| vns | GGB30413_SGB43452 | EtOH_24h_intakeave_final | 0.8796426046 |
| blast | GGB30454_SGB43514 | EtOH_24h_intakeave_final | 0.3798589507 |
| vns | GGB30454_SGB43514 | EtOH_24h_intakeave_final | 0.2001255913 |
| blast | GGB30455_SGB43519 | EtOH_24h_intakeave_final | 0.3794677965 |
| vns | GGB30455_SGB43519 | EtOH_24h_intakeave_final | 0.3023489219 |
| blast | GGB30461_SGB43527 | EtOH_24h_intakeave_final | 0.3975036627 |
| vns | GGB30461_SGB43527 | EtOH_24h_intakeave_final | 0.7710386558 |
| blast | GGB30461_SGB43530 | EtOH_24h_intakeave_final | 0.395449 |
| vns | GGB30461_SGB43530 | EtOH_24h_intakeave_final | 0.7038636932 |
| blast | GGB30463_SGB43537 | EtOH_24h_intakeave_final | 0.2100542578 |
| vns | GGB30463_SGB43537 | EtOH_24h_intakeave_final | 0.9853614792 |
| blast | GGB30473_SGB43557 | EtOH_24h_intakeave_final | 0.0206421669 |
| vns | GGB30473_SGB43557 | EtOH_24h_intakeave_final | 0.0562117431 |
| blast | GGB30475_SGB63182 | EtOH_24h_intakeave_final | 0.2879653492 |
| vns | GGB30475_SGB63182 | EtOH_24h_intakeave_final | 0.6608559306 |
| blast | GGB30861_SGB44083 | EtOH_24h_intakeave_final | 0.9575523797 |
| vns | GGB30861_SGB44083 | EtOH_24h_intakeave_final | 0.8334455291 |
| blast | GGB31312_SGB44628 | EtOH_24h_intakeave_final | 0.7117271287 |
| vns | GGB31312_SGB44628 | EtOH_24h_intakeave_final | 0.8841939433 |

| | | | |
|---|---|---|---|
| blast | GGB31438_SGB44768 | EtOH_24h_intakeave_final | 0.906357297 |
| vns | GGB31438_SGB44768 | EtOH_24h_intakeave_final | 0.318960086 |
| blast | GGB3171_SGB4185 | EtOH_24h_intakeave_final | 0.085345767 |
| vns | GGB3171_SGB4185 | EtOH_24h_intakeave_final | 0.918551780 |
| blast | GGB31823_SGB45199 | EtOH_24h_intakeave_final | 0.188977717 |
| vns | GGB31823_SGB45199 | EtOH_24h_intakeave_final | 0.702813258 |
| blast | GGB31853_SGB45233 | EtOH_24h_intakeave_final | 0.496842743 |
| vns | GGB31853_SGB45233 | EtOH_24h_intakeave_final | 0.396259541 |
| blast | GGB32371_SGB41694 | EtOH_24h_intakeave_final | 0.346811850 |
| vns | GGB32371_SGB41694 | EtOH_24h_intakeave_final | 0.025054092 |
| blast | GGB3793_SGB5158 | EtOH_24h_intakeave_final | 0.965790474 |
| vns | GGB3793_SGB5158 | EtOH_24h_intakeave_final | 0.975145151 |
| blast | GGB42598_SGB59794 | EtOH_24h_intakeave_final | 0.501740001 |
| vns | GGB42598_SGB59794 | EtOH_24h_intakeave_final | 0.156668604 |
| blast | GGB45656_SGB63370 | EtOH_24h_intakeave_final | 0.621044138 |
| vns | GGB45656_SGB63370 | EtOH_24h_intakeave_final | 0.963374601 |
| blast | GGB47127_SGB65054 | EtOH_24h_intakeave_final | 0.496745120 |
| vns | GGB47127_SGB65054 | EtOH_24h_intakeave_final | 0.257417312 |
| blast | GGB74395_SGB43521 | EtOH_24h_intakeave_final | 0.123339782 |
| vns | GGB74395_SGB43521 | EtOH_24h_intakeave_final | 0.204021498 |
| blast | GGB75053_SGB43494 | EtOH_24h_intakeave_final | 0.410595919 |
| vns | GGB75053_SGB43494 | EtOH_24h_intakeave_final | 0.571857456 |
| blast | GGB75109_SGB102238 | EtOH_24h_intakeave_final | 0.620842292 |
| vns | GGB75109_SGB102238 | EtOH_24h_intakeave_final | 0.871818408 |
| blast | GGB81440_SGB45230 | EtOH_24h_intakeave_final | 0.079610731 |
| vns | GGB81440_SGB45230 | EtOH_24h_intakeave_final | 0.098140579 |
| blast | Lachnospiraceae_bacterium | EtOH_24h_intakeave_final | 0.484816177 |
| vns | Lachnospiraceae_bacterium | EtOH_24h_intakeave_final | 0.537861332 |
| blast | Lachnospiraceae_bacterium_A2 | EtOH_24h_intakeave_final | 0.420834896 |
| vns | Lachnospiraceae_bacterium_A2 | EtOH_24h_intakeave_final | 0.323648328 |
| blast | Lachnospiraceae_bacterium_MD308 | EtOH_24h_intakeave_final | 0.213119451 |
| vns | Lachnospiraceae_bacterium_MD308 | EtOH_24h_intakeave_final | 0.083066965 |
| blast | Lachnospiraceae_bacterium_MD329 | EtOH_24h_intakeave_final | 0.020976149 |
| vns | Lachnospiraceae_bacterium_MD329 | EtOH_24h_intakeave_final | 0.699161119 |
| blast | Lachnospiraceae_unclassified_SGB41414 | EtOH_24h_intakeave_final | 0.142639053 |
| vns | Lachnospiraceae_unclassified_SGB41414 | EtOH_24h_intakeave_final | 0.119876468 |
| blast | Lachnospiraceae_unclassified_SGB41418 | EtOH_24h_intakeave_final | 0.998757981 |
| vns | Lachnospiraceae_unclassified_SGB41418 | EtOH_24h_intakeave_final | 0.712236573 |
| blast | Lachnospiraceae_unclassified_SGB41424 | EtOH_24h_intakeave_final | 0.048386378 |
| vns | Lachnospiraceae_unclassified_SGB41424 | EtOH_24h_intakeave_final | 0.046067357 |
| blast | Lachnospiraceae_unclassified_SGB41589 | EtOH_24h_intakeave_final | 0.165424952 |
| vns | Lachnospiraceae_unclassified_SGB41589 | EtOH_24h_intakeave_final | 0.273527700 |
| blast | Lactobacillus_johnsonii | EtOH_24h_intakeave_final | 0.182678453 |
| vns | Lactobacillus_johnsonii | EtOH_24h_intakeave_final | 0.672951588 |
| blast | Muribaculaceae_bacterium | EtOH_24h_intakeave_final | 0.841525252 |
| vns | Muribaculaceae_bacterium | EtOH_24h_intakeave_final | 0.893052980 |

| | | | |
|---|---|---|---|
| blast | Neglectibacter_sp_X4 | EtOH_24h_intakeave_final | 0.084782151 |
| vns | Neglectibacter_sp_X4 | EtOH_24h_intakeave_final | 0.140656264 |
| blast | Oscillospiraceae_bacterium | EtOH_24h_intakeave_final | 0.741672565 |
| vns | Oscillospiraceae_bacterium | EtOH_24h_intakeave_final | 0.856188039 |
| blast | Oscillospiraceae_unclassified_SGB43502 | EtOH_24h_intakeave_final | 0.025941285 |
| vns | Oscillospiraceae_unclassified_SGB43502 | EtOH_24h_intakeave_final | 0.030535133 |
| blast | Oscillospiraceae_unclassified_SGB43505 | EtOH_24h_intakeave_final | 0.495923018 |
| vns | Oscillospiraceae_unclassified_SGB43505 | EtOH_24h_intakeave_final | 0.642533731 |
| blast | Parasutterella_excrementihominis | EtOH_24h_intakeave_final | 0.567104872 |
| vns | Parasutterella_excrementihominis | EtOH_24h_intakeave_final | 0.709992917 |
| blast | Richness (# observed features) | EtOH_24h_intakeave_final | 0.271752442 |
| vns | Richness (# observed features) | EtOH_24h_intakeave_final | 0.660723068 |
| blast | Romboutsia_ilealis | EtOH_24h_intakeave_final | 0.001099774 |
| vns | Romboutsia_ilealis | EtOH_24h_intakeave_final | 0.056997089 |
| blast | Schaedlerella_arabinosiphila | EtOH_24h_intakeave_final | 0.939024404 |
| vns | Schaedlerella_arabinosiphila | EtOH_24h_intakeave_final | 0.913911607 |
| blast | Shannon Index | EtOH_24h_intakeave_final | 0.694735221 |
| vns | Shannon Index | EtOH_24h_intakeave_final | 0.906277341 |
| blast | Turicibacter_sp_1E2 | EtOH_24h_intakeave_final | 0.199177231 |
| vns | Turicibacter_sp_1E2 | EtOH_24h_intakeave_final | 0.165574931 |
| blast | Acetatifactor_muris | EtOH_8pm_intakeave_final | 0.739522079 |
| vns | Acetatifactor_muris | EtOH_8pm_intakeave_final | 0.752897140 |
| blast | Acetatifactor_SGB41546 | EtOH_8pm_intakeave_final | 0.840245797 |
| vns | Acetatifactor_SGB41546 | EtOH_8pm_intakeave_final | 0.663000977 |
| blast | Acutalibacter_muris | EtOH_8pm_intakeave_final | 0.422154633 |
| vns | Acutalibacter_muris | EtOH_8pm_intakeave_final | 0.810578449 |
| blast | Acutalibacter_sp_1XD8_36 | EtOH_8pm_intakeave_final | 0.660579886 |
| vns | Acutalibacter_sp_1XD8_36 | EtOH_8pm_intakeave_final | 0.709184933 |
| blast | Adlercreutzia_caecimuris | EtOH_8pm_intakeave_final | 0.989608952 |
| vns | Adlercreutzia_caecimuris | EtOH_8pm_intakeave_final | 0.676542691 |
| blast | Adlercreutzia_mucosicola | EtOH_8pm_intakeave_final | 0.271833985 |
| vns | Adlercreutzia_mucosicola | EtOH_8pm_intakeave_final | 0.249825657 |
| blast | Adlercreutzia_muris | EtOH_8pm_intakeave_final | 0.670421774 |
| vns | Adlercreutzia_muris | EtOH_8pm_intakeave_final | 0.595827745 |
| blast | Akkermansia_muciniphila | EtOH_8pm_intakeave_final | 0.169812540 |
| vns | Akkermansia_muciniphila | EtOH_8pm_intakeave_final | 0.864362509 |
| blast | Alistipes_sp_DSM_112343 | EtOH_8pm_intakeave_final | 0.372515977 |
| vns | Alistipes_sp_DSM_112343 | EtOH_8pm_intakeave_final | 0.832698307 |
| blast | Anaerotruncus_sp_1XD42_93 | EtOH_8pm_intakeave_final | 0.520271453 |
| vns | Anaerotruncus_sp_1XD42_93 | EtOH_8pm_intakeave_final | 0.617410721 |
| blast | Bacteria_unclassified_SGB102200 | EtOH_8pm_intakeave_final | 0.474269212 |
| vns | Bacteria_unclassified_SGB102200 | EtOH_8pm_intakeave_final | 0.764166788 |
| blast | Bacteria_unclassified_SGB41677 | EtOH_8pm_intakeave_final | 0.444143823 |
| vns | Bacteria_unclassified_SGB41677 | EtOH_8pm_intakeave_final | 0.169989424 |
| blast | Bacteria_unclassified_SGB43546 | EtOH_8pm_intakeave_final | 0.586358365 |
| vns | Bacteria_unclassified_SGB43546 | EtOH_8pm_intakeave_final | 0.789009912 |

| | | | |
|---|---|---|---|
| blast | bacterium_1XD42_54 | EtOH_8pm_intakeave_final | 0.014424595 |
| vns | bacterium_1XD42_54 | EtOH_8pm_intakeave_final | 0.234577574 |
| blast | bacterium_1XD42_76 | EtOH_8pm_intakeave_final | 0.600031603 |
| vns | bacterium_1XD42_76 | EtOH_8pm_intakeave_final | 0.642992453 |
| blast | bacterium_1xD8_48 | EtOH_8pm_intakeave_final | 0.899708945 |
| vns | bacterium_1xD8_48 | EtOH_8pm_intakeave_final | 0.696816880 |
| blast | Bacteroides_thetaiotaomicron | EtOH_8pm_intakeave_final | 0.960201477 |
| vns | Bacteroides_thetaiotaomicron | EtOH_8pm_intakeave_final | 0.634306315 |
| blast | Berger Parker Index | EtOH_8pm_intakeave_final | 0.545741712 |
| vns | Berger Parker Index | EtOH_8pm_intakeave_final | 0.233733054 |
| blast | Bifidobacterium_pseudolongum | EtOH_8pm_intakeave_final | 0.428010398 |
| vns | Bifidobacterium_pseudolongum | EtOH_8pm_intakeave_final | 0.347799065 |
| blast | Clostridia_bacterium | EtOH_8pm_intakeave_final | 0.374559043 |
| vns | Clostridia_bacterium | EtOH_8pm_intakeave_final | 0.724940107 |
| blast | Clostridiaceae_bacterium | EtOH_8pm_intakeave_final | 0.987169780 |
| vns | Clostridiaceae_bacterium | EtOH_8pm_intakeave_final | 0.241937042 |
| blast | Clostridiaceae_unclassified_SGB41663 | EtOH_8pm_intakeave_final | 0.780382224 |
| vns | Clostridiaceae_unclassified_SGB41663 | EtOH_8pm_intakeave_final | 0.719154368 |
| blast | Clostridiales_bacterium | EtOH_8pm_intakeave_final | 0.002331956 |
| vns | Clostridiales_bacterium | EtOH_8pm_intakeave_final | 0.296478501 |
| blast | Clostridium_cocleatum | EtOH_8pm_intakeave_final | 0.011567832 |
| vns | Clostridium_cocleatum | EtOH_8pm_intakeave_final | 0.196035514 |
| blast | Coriobacteriaceae_bacterium | EtOH_8pm_intakeave_final | 0.372448369 |
| vns | Coriobacteriaceae_bacterium | EtOH_8pm_intakeave_final | 0.953919137 |
| blast | Dorea_sp_5_2 | EtOH_8pm_intakeave_final | 0.588491917 |
| vns | Dorea_sp_5_2 | EtOH_8pm_intakeave_final | 0.421416124 |
| blast | Dubosiella_newyorkensis | EtOH_8pm_intakeave_final | 0.116998739 |
| vns | Dubosiella_newyorkensis | EtOH_8pm_intakeave_final | 0.799186063 |
| blast | Erysipelotrichales_bacterium | EtOH_8pm_intakeave_final | 0.897000665 |
| vns | Erysipelotrichales_bacterium | EtOH_8pm_intakeave_final | 0.328408108 |
| blast | Eubacteriaceae_bacterium | EtOH_8pm_intakeave_final | 0.027123368 |
| vns | Eubacteriaceae_bacterium | EtOH_8pm_intakeave_final | 0.238318613 |
| blast | Eubacteriaceae_unclassified_SGB94922 | EtOH_8pm_intakeave_final | 0.459488079 |
| vns | Eubacteriaceae_unclassified_SGB94922 | EtOH_8pm_intakeave_final | 0.372620148 |
| blast | GGB20149_SGB29430 | EtOH_8pm_intakeave_final | 0.628070048 |
| vns | GGB20149_SGB29430 | EtOH_8pm_intakeave_final | 0.629982792 |
| blast | GGB22635_SGB63107 | EtOH_8pm_intakeave_final | 0.411710336 |
| vns | GGB22635_SGB63107 | EtOH_8pm_intakeave_final | 0.887152138 |
| blast | GGB25041_SGB36960 | EtOH_8pm_intakeave_final | 0.082496667 |
| vns | GGB25041_SGB36960 | EtOH_8pm_intakeave_final | 0.210187196 |
| blast | GGB27876_SGB40310 | EtOH_8pm_intakeave_final | 0.164666135 |
| vns | GGB27876_SGB40310 | EtOH_8pm_intakeave_final | 0.767235194 |
| blast | GGB27878_SGB40312 | EtOH_8pm_intakeave_final | 0.213354157 |
| vns | GGB27878_SGB40312 | EtOH_8pm_intakeave_final | 0.147479429 |
| blast | GGB27918_SGB40356 | EtOH_8pm_intakeave_final | 0.958758140 |
| vns | GGB27918_SGB40356 | EtOH_8pm_intakeave_final | 0.885698929 |

| | | | |
|---|---|---|---|
| blast | GGB28382_SGB40962 | EtOH_8pm_intakeave_final | 0.519311307 |
| vns | GGB28382_SGB40962 | EtOH_8pm_intakeave_final | 0.855057834 |
| blast | GGB28399_SGB40980 | EtOH_8pm_intakeave_final | 0.289465502 |
| vns | GGB28399_SGB40980 | EtOH_8pm_intakeave_final | 0.287117034 |
| blast | GGB28411_SGB40993 | EtOH_8pm_intakeave_final | 0.201802262 |
| vns | GGB28411_SGB40993 | EtOH_8pm_intakeave_final | 0.146848474 |
| blast | GGB28415_SGB40997 | EtOH_8pm_intakeave_final | 0.026726744 |
| vns | GGB28415_SGB40997 | EtOH_8pm_intakeave_final | 0.949051696 |
| blast | GGB28430_SGB41013 | EtOH_8pm_intakeave_final | 0.912070947 |
| vns | GGB28430_SGB41013 | EtOH_8pm_intakeave_final | 0.466250482 |
| blast | GGB28439_SGB41022 | EtOH_8pm_intakeave_final | 0.380444403 |
| vns | GGB28439_SGB41022 | EtOH_8pm_intakeave_final | 0.738709552 |
| blast | GGB28778_SGB41431 | EtOH_8pm_intakeave_final | 0.804698941 |
| vns | GGB28778_SGB41431 | EtOH_8pm_intakeave_final | 0.218453243 |
| blast | GGB28784_SGB41437 | EtOH_8pm_intakeave_final | 0.020726621 |
| vns | GGB28784_SGB41437 | EtOH_8pm_intakeave_final | 0.254456997 |
| blast | GGB28792_SGB41445 | EtOH_8pm_intakeave_final | 0.338577775 |
| vns | GGB28792_SGB41445 | EtOH_8pm_intakeave_final | 0.761244426 |
| blast | GGB28798_SGB41451 | EtOH_8pm_intakeave_final | 0.164022630 |
| vns | GGB28798_SGB41451 | EtOH_8pm_intakeave_final | 0.248915824 |
| blast | GGB28802_SGB41455 | EtOH_8pm_intakeave_final | 0.457898625 |
| vns | GGB28802_SGB41455 | EtOH_8pm_intakeave_final | 0.342689298 |
| blast | GGB28818_SGB41473 | EtOH_8pm_intakeave_final | 0.729209510 |
| vns | GGB28818_SGB41473 | EtOH_8pm_intakeave_final | 0.569409948 |
| blast | GGB28828_SGB41484 | EtOH_8pm_intakeave_final | 0.298696993 |
| vns | GGB28828_SGB41484 | EtOH_8pm_intakeave_final | 0.106706535 |
| blast | GGB28851_SGB41518 | EtOH_8pm_intakeave_final | 0.852552179 |
| vns | GGB28851_SGB41518 | EtOH_8pm_intakeave_final | 0.578308429 |
| blast | GGB28859_SGB41528 | EtOH_8pm_intakeave_final | 0.205749329 |
| vns | GGB28859_SGB41528 | EtOH_8pm_intakeave_final | 0.595366745 |
| blast | GGB28864_SGB41535 | EtOH_8pm_intakeave_final | 0.785465248 |
| vns | GGB28864_SGB41535 | EtOH_8pm_intakeave_final | 0.964658985 |
| blast | GGB28869_SGB41543 | EtOH_8pm_intakeave_final | 0.347931625 |
| vns | GGB28869_SGB41543 | EtOH_8pm_intakeave_final | 0.381477475 |
| blast | GGB28883_SGB41564 | EtOH_8pm_intakeave_final | 0.594877248 |
| vns | GGB28883_SGB41564 | EtOH_8pm_intakeave_final | 0.060524458 |
| blast | GGB28892_SGB41573 | EtOH_8pm_intakeave_final | 0.651227207 |
| vns | GGB28892_SGB41573 | EtOH_8pm_intakeave_final | 0.667618882 |
| blast | GGB28893_SGB41574 | EtOH_8pm_intakeave_final | 0.165594509 |
| vns | GGB28893_SGB41574 | EtOH_8pm_intakeave_final | 0.182699376 |
| blast | GGB28898_SGB41580 | EtOH_8pm_intakeave_final | 0.304654993 |
| vns | GGB28898_SGB41580 | EtOH_8pm_intakeave_final | 0.082893330 |
| blast | GGB28904_SGB41597 | EtOH_8pm_intakeave_final | 0.172906397 |
| vns | GGB28904_SGB41597 | EtOH_8pm_intakeave_final | 0.056067199 |
| blast | GGB28916_SGB41612 | EtOH_8pm_intakeave_final | 0.801432609 |
| vns | GGB28916_SGB41612 | EtOH_8pm_intakeave_final | 0.853813296 |

| | | | |
|---|---|---|---|
| blast | GGB28924_SGB41621 | EtOH_8pm_intakeave_final | 0.7964816629 |
| vns | GGB28924_SGB41621 | EtOH_8pm_intakeave_final | 0.2786583787 |
| blast | GGB28926_SGB41624 | EtOH_8pm_intakeave_final | 0.3067808341 |
| vns | GGB28926_SGB41624 | EtOH_8pm_intakeave_final | 0.470765265 |
| blast | GGB28927_SGB41625 | EtOH_8pm_intakeave_final | 0.1937208057 |
| vns | GGB28927_SGB41625 | EtOH_8pm_intakeave_final | 0.4182433734 |
| blast | GGB28934_SGB41635 | EtOH_8pm_intakeave_final | 0.1158733118 |
| vns | GGB28934_SGB41635 | EtOH_8pm_intakeave_final | 0.1113740731 |
| blast | GGB28946_SGB41652 | EtOH_8pm_intakeave_final | 0.4278432256 |
| vns | GGB28946_SGB41652 | EtOH_8pm_intakeave_final | 0.4225889007 |
| blast | GGB28949_SGB41655 | EtOH_8pm_intakeave_final | 0.4436081164 |
| vns | GGB28949_SGB41655 | EtOH_8pm_intakeave_final | 0.9089481791 |
| blast | GGB28949_SGB41656 | EtOH_8pm_intakeave_final | 0.2827282346 |
| vns | GGB28949_SGB41656 | EtOH_8pm_intakeave_final | 0.7708639493 |
| blast | GGB28950_SGB41657 | EtOH_8pm_intakeave_final | 0.0305270365 |
| vns | GGB28950_SGB41657 | EtOH_8pm_intakeave_final | 0.3364589048 |
| blast | GGB28951_SGB102295 | EtOH_8pm_intakeave_final | 0.9253521562 |
| vns | GGB28951_SGB102295 | EtOH_8pm_intakeave_final | 0.1587903977 |
| blast | GGB28951_SGB41658 | EtOH_8pm_intakeave_final | 0.8470013515 |
| vns | GGB28951_SGB41658 | EtOH_8pm_intakeave_final | 0.9202366237 |
| blast | GGB28954_SGB41662 | EtOH_8pm_intakeave_final | 0.5157582975 |
| vns | GGB28954_SGB41662 | EtOH_8pm_intakeave_final | 0.8004849551 |
| blast | GGB28956_SGB41665 | EtOH_8pm_intakeave_final | 0.163329713 |
| vns | GGB28956_SGB41665 | EtOH_8pm_intakeave_final | 0.1534165023 |
| blast | GGB28960_SGB41669 | EtOH_8pm_intakeave_final | 0.155727292 |
| vns | GGB28960_SGB41669 | EtOH_8pm_intakeave_final | 0.1559574792 |
| blast | GGB28967_SGB41678 | EtOH_8pm_intakeave_final | 0.1792695292 |
| vns | GGB28967_SGB41678 | EtOH_8pm_intakeave_final | 0.3150465193 |
| blast | GGB28991_SGB41705 | EtOH_8pm_intakeave_final | 0.2288938824 |
| vns | GGB28991_SGB41705 | EtOH_8pm_intakeave_final | 0.279982342 |
| blast | GGB29002_SGB41718 | EtOH_8pm_intakeave_final | 0.0024308675 |
| vns | GGB29002_SGB41718 | EtOH_8pm_intakeave_final | 0.1134655877 |
| blast | GGB29003_SGB41719 | EtOH_8pm_intakeave_final | 0.4993986443 |
| vns | GGB29003_SGB41719 | EtOH_8pm_intakeave_final | 0.5839678864 |
| blast | GGB29011_SGB41731 | EtOH_8pm_intakeave_final | 0.916276033 |
| vns | GGB29011_SGB41731 | EtOH_8pm_intakeave_final | 0.3455992366 |
| blast | GGB29531_SGB42317 | EtOH_8pm_intakeave_final | 0.1450820268 |
| vns | GGB29531_SGB42317 | EtOH_8pm_intakeave_final | 0.1355699662 |
| blast | GGB29685_SGB42494 | EtOH_8pm_intakeave_final | 0.8645438528 |
| vns | GGB29685_SGB42494 | EtOH_8pm_intakeave_final | 0.340104263 |
| blast | GGB30141_SGB43066 | EtOH_8pm_intakeave_final | 0.9915548039 |
| vns | GGB30141_SGB43066 | EtOH_8pm_intakeave_final | 0.7601787897 |
| blast | GGB30286_SGB43248 | EtOH_8pm_intakeave_final | 0.0552616333 |
| vns | GGB30286_SGB43248 | EtOH_8pm_intakeave_final | 0.4546033673 |
| blast | GGB30303_SGB43268 | EtOH_8pm_intakeave_final | 0.3719368939 |
| vns | GGB30303_SGB43268 | EtOH_8pm_intakeave_final | 0.9447221743 |

| | | | |
|---|---|---|---|
| blast | GGB30413_SGB43452 | EtOH_8pm_intakeave_final | 0.9532769507 |
| vns | GGB30413_SGB43452 | EtOH_8pm_intakeave_final | 0.8796426046 |
| blast | GGB30454_SGB43514 | EtOH_8pm_intakeave_final | 0.3798589507 |
| vns | GGB30454_SGB43514 | EtOH_8pm_intakeave_final | 0.2001255913 |
| blast | GGB30455_SGB43519 | EtOH_8pm_intakeave_final | 0.3794677965 |
| vns | GGB30455_SGB43519 | EtOH_8pm_intakeave_final | 0.3023489219 |
| blast | GGB30461_SGB43527 | EtOH_8pm_intakeave_final | 0.3975036627 |
| vns | GGB30461_SGB43527 | EtOH_8pm_intakeave_final | 0.7710386558 |
| blast | GGB30461_SGB43530 | EtOH_8pm_intakeave_final | 0.395449 |
| vns | GGB30461_SGB43530 | EtOH_8pm_intakeave_final | 0.7038636932 |
| blast | GGB30463_SGB43537 | EtOH_8pm_intakeave_final | 0.2100542578 |
| vns | GGB30463_SGB43537 | EtOH_8pm_intakeave_final | 0.9853614792 |
| blast | GGB30473_SGB43557 | EtOH_8pm_intakeave_final | 0.0206421669 |
| vns | GGB30473_SGB43557 | EtOH_8pm_intakeave_final | 0.0562117431 |
| blast | GGB30475_SGB63182 | EtOH_8pm_intakeave_final | 0.2879653492 |
| vns | GGB30475_SGB63182 | EtOH_8pm_intakeave_final | 0.6608559306 |
| blast | GGB30861_SGB44083 | EtOH_8pm_intakeave_final | 0.9575523797 |
| vns | GGB30861_SGB44083 | EtOH_8pm_intakeave_final | 0.8334455291 |
| blast | GGB31312_SGB44628 | EtOH_8pm_intakeave_final | 0.7117271287 |
| vns | GGB31312_SGB44628 | EtOH_8pm_intakeave_final | 0.8841939433 |
| blast | GGB31438_SGB44768 | EtOH_8pm_intakeave_final | 0.9063572977 |
| vns | GGB31438_SGB44768 | EtOH_8pm_intakeave_final | 0.3189600867 |
| blast | GGB3171_SGB4185 | EtOH_8pm_intakeave_final | 0.0853457671 |
| vns | GGB3171_SGB4185 | EtOH_8pm_intakeave_final | 0.9185517801 |
| blast | GGB31823_SGB45199 | EtOH_8pm_intakeave_final | 0.1889777175 |
| vns | GGB31823_SGB45199 | EtOH_8pm_intakeave_final | 0.7028132582 |
| blast | GGB31853_SGB45233 | EtOH_8pm_intakeave_final | 0.4968427437 |
| vns | GGB31853_SGB45233 | EtOH_8pm_intakeave_final | 0.3962595415 |
| blast | GGB32371_SGB41694 | EtOH_8pm_intakeave_final | 0.3468118508 |
| vns | GGB32371_SGB41694 | EtOH_8pm_intakeave_final | 0.0250540925 |
| blast | GGB3793_SGB5158 | EtOH_8pm_intakeave_final | 0.9657904747 |
| vns | GGB3793_SGB5158 | EtOH_8pm_intakeave_final | 0.9751451519 |
| blast | GGB42598_SGB59794 | EtOH_8pm_intakeave_final | 0.5017400018 |
| vns | GGB42598_SGB59794 | EtOH_8pm_intakeave_final | 0.156668604 |
| blast | GGB45656_SGB63370 | EtOH_8pm_intakeave_final | 0.621044138 |
| vns | GGB45656_SGB63370 | EtOH_8pm_intakeave_final | 0.9633746015 |
| blast | GGB47127_SGB65054 | EtOH_8pm_intakeave_final | 0.4967451202 |
| vns | GGB47127_SGB65054 | EtOH_8pm_intakeave_final | 0.2574173127 |
| blast | GGB74395_SGB43521 | EtOH_8pm_intakeave_final | 0.1233397821 |
| vns | GGB74395_SGB43521 | EtOH_8pm_intakeave_final | 0.2040214986 |
| blast | GGB75053_SGB43494 | EtOH_8pm_intakeave_final | 0.4105959194 |
| vns | GGB75053_SGB43494 | EtOH_8pm_intakeave_final | 0.5718574569 |
| blast | GGB75109_SGB102238 | EtOH_8pm_intakeave_final | 0.6208422921 |
| vns | GGB75109_SGB102238 | EtOH_8pm_intakeave_final | 0.8718184088 |
| blast | GGB81440_SGB45230 | EtOH_8pm_intakeave_final | 0.0796107313 |
| vns | GGB81440_SGB45230 | EtOH_8pm_intakeave_final | 0.0981405799 |

| | | | |
|---|---|---|---|
| blast | Lachnospiraceae_bacterium | EtOH_8pm_intakeave_final | 0.484816177 |
| vns | Lachnospiraceae_bacterium | EtOH_8pm_intakeave_final | 0.537861332 |
| blast | Lachnospiraceae_bacterium_A2 | EtOH_8pm_intakeave_final | 0.420834896 |
| vns | Lachnospiraceae_bacterium_A2 | EtOH_8pm_intakeave_final | 0.323648328 |
| blast | Lachnospiraceae_bacterium_MD308 | EtOH_8pm_intakeave_final | 0.213119451 |
| vns | Lachnospiraceae_bacterium_MD308 | EtOH_8pm_intakeave_final | 0.083066965 |
| blast | Lachnospiraceae_bacterium_MD329 | EtOH_8pm_intakeave_final | 0.020976149 |
| vns | Lachnospiraceae_bacterium_MD329 | EtOH_8pm_intakeave_final | 0.699161119 |
| blast | Lachnospiraceae_unclassified_SGB41414 | EtOH_8pm_intakeave_final | 0.142639053 |
| vns | Lachnospiraceae_unclassified_SGB41414 | EtOH_8pm_intakeave_final | 0.119876468 |
| blast | Lachnospiraceae_unclassified_SGB41418 | EtOH_8pm_intakeave_final | 0.998757981 |
| vns | Lachnospiraceae_unclassified_SGB41418 | EtOH_8pm_intakeave_final | 0.712236573 |
| blast | Lachnospiraceae_unclassified_SGB41424 | EtOH_8pm_intakeave_final | 0.048386378 |
| vns | Lachnospiraceae_unclassified_SGB41424 | EtOH_8pm_intakeave_final | 0.046067357 |
| blast | Lachnospiraceae_unclassified_SGB41589 | EtOH_8pm_intakeave_final | 0.165424952 |
| vns | Lachnospiraceae_unclassified_SGB41589 | EtOH_8pm_intakeave_final | 0.273527700 |
| blast | Lactobacillus_johnsonii | EtOH_8pm_intakeave_final | 0.182678453 |
| vns | Lactobacillus_johnsonii | EtOH_8pm_intakeave_final | 0.672951588 |
| blast | Muribaculaceae_bacterium | EtOH_8pm_intakeave_final | 0.841525252 |
| vns | Muribaculaceae_bacterium | EtOH_8pm_intakeave_final | 0.893052980 |
| blast | Neglectibacter_sp_X4 | EtOH_8pm_intakeave_final | 0.084782151 |
| vns | Neglectibacter_sp_X4 | EtOH_8pm_intakeave_final | 0.140656264 |
| blast | Oscillospiraceae_bacterium | EtOH_8pm_intakeave_final | 0.741672565 |
| vns | Oscillospiraceae_bacterium | EtOH_8pm_intakeave_final | 0.856188039 |
| blast | Oscillospiraceae_unclassified_SGB43502 | EtOH_8pm_intakeave_final | 0.025941285 |
| vns | Oscillospiraceae_unclassified_SGB43502 | EtOH_8pm_intakeave_final | 0.030535133 |
| blast | Oscillospiraceae_unclassified_SGB43505 | EtOH_8pm_intakeave_final | 0.495923018 |
| vns | Oscillospiraceae_unclassified_SGB43505 | EtOH_8pm_intakeave_final | 0.642533731 |
| blast | Parasutterella_excrementihominis | EtOH_8pm_intakeave_final | 0.567104872 |
| vns | Parasutterella_excrementihominis | EtOH_8pm_intakeave_final | 0.709992917 |
| blast | Richness (# observed features) | EtOH_8pm_intakeave_final | 0.271752442 |
| vns | Richness (# observed features) | EtOH_8pm_intakeave_final | 0.660723068 |
| blast | Romboutsia_ilealis | EtOH_8pm_intakeave_final | 0.001099774 |
| vns | Romboutsia_ilealis | EtOH_8pm_intakeave_final | 0.056997089 |
| blast | Schaedlerella_arabinosiphila | EtOH_8pm_intakeave_final | 0.939024404 |
| vns | Schaedlerella_arabinosiphila | EtOH_8pm_intakeave_final | 0.913911607 |
| blast | Shannon Index | EtOH_8pm_intakeave_final | 0.694735221 |
| vns | Shannon Index | EtOH_8pm_intakeave_final | 0.906277341 |
| blast | Turicibacter_sp_1E2 | EtOH_8pm_intakeave_final | 0.199177231 |
| vns | Turicibacter_sp_1E2 | EtOH_8pm_intakeave_final | 0.165574931 |
| blast | Acetatifactor_muris | EtOH_pref_24h_final | 0.739522079 |
| vns | Acetatifactor_muris | EtOH_pref_24h_final | 0.752897140 |
| blast | Acetatifactor_SGB41546 | EtOH_pref_24h_final | 0.840245797 |
| vns | Acetatifactor_SGB41546 | EtOH_pref_24h_final | 0.663000977 |
| blast | Acutalibacter_muris | EtOH_pref_24h_final | 0.422154633 |
| vns | Acutalibacter_muris | EtOH_pref_24h_final | 0.810578449 |

| | | | |
|---|---|---|---|
| blast | Acutalibacter_sp_1XD8_36 | EtOH_pref_24h_final | 0.6605798863 |
| vns | Acutalibacter_sp_1XD8_36 | EtOH_pref_24h_final | 0.7091849336 |
| blast | Adlercreutzia_caecimuris | EtOH_pref_24h_final | 0.9896089529 |
| vns | Adlercreutzia_caecimuris | EtOH_pref_24h_final | 0.6765426913 |
| blast | Adlercreutzia_mucosicola | EtOH_pref_24h_final | 0.2718339856 |
| vns | Adlercreutzia_mucosicola | EtOH_pref_24h_final | 0.2498256574 |
| blast | Adlercreutzia_muris | EtOH_pref_24h_final | 0.6704217743 |
| vns | Adlercreutzia_muris | EtOH_pref_24h_final | 0.5958277454 |
| blast | Akkermansia_muciniphila | EtOH_pref_24h_final | 0.1698125404 |
| vns | Akkermansia_muciniphila | EtOH_pref_24h_final | 0.8643625099 |
| blast | Alistipes_sp_DSM_112343 | EtOH_pref_24h_final | 0.3725159774 |
| vns | Alistipes_sp_DSM_112343 | EtOH_pref_24h_final | 0.8326983074 |
| blast | Anaerotruncus_sp_1XD42_93 | EtOH_pref_24h_final | 0.5202714535 |
| vns | Anaerotruncus_sp_1XD42_93 | EtOH_pref_24h_final | 0.6174107217 |
| blast | Bacteria_unclassified_SGB102200 | EtOH_pref_24h_final | 0.4742692127 |
| vns | Bacteria_unclassified_SGB102200 | EtOH_pref_24h_final | 0.7641667885 |
| blast | Bacteria_unclassified_SGB41677 | EtOH_pref_24h_final | 0.4441438231 |
| vns | Bacteria_unclassified_SGB41677 | EtOH_pref_24h_final | 0.1699894243 |
| blast | Bacteria_unclassified_SGB43546 | EtOH_pref_24h_final | 0.5863583659 |
| vns | Bacteria_unclassified_SGB43546 | EtOH_pref_24h_final | 0.7890099125 |
| blast | bacterium_1XD42_54 | EtOH_pref_24h_final | 0.0144245959 |
| vns | bacterium_1XD42_54 | EtOH_pref_24h_final | 0.2345775748 |
| blast | bacterium_1XD42_76 | EtOH_pref_24h_final | 0.6000316038 |
| vns | bacterium_1XD42_76 | EtOH_pref_24h_final | 0.6429924539 |
| blast | bacterium_1xD8_48 | EtOH_pref_24h_final | 0.8997089453 |
| vns | bacterium_1xD8_48 | EtOH_pref_24h_final | 0.6968168809 |
| blast | Bacteroides_thetaiotaomicron | EtOH_pref_24h_final | 0.9602014777 |
| vns | Bacteroides_thetaiotaomicron | EtOH_pref_24h_final | 0.6343063159 |
| blast | Berger Parker Index | EtOH_pref_24h_final | 0.5457417127 |
| vns | Berger Parker Index | EtOH_pref_24h_final | 0.2337330546 |
| blast | Bifidobacterium_pseudolongum | EtOH_pref_24h_final | 0.4280103985 |
| vns | Bifidobacterium_pseudolongum | EtOH_pref_24h_final | 0.3477990653 |
| blast | Clostridia_bacterium | EtOH_pref_24h_final | 0.3745590434 |
| vns | Clostridia_bacterium | EtOH_pref_24h_final | 0.7249401071 |
| blast | Clostridiaceae_bacterium | EtOH_pref_24h_final | 0.9871697806 |
| vns | Clostridiaceae_bacterium | EtOH_pref_24h_final | 0.2419370425 |
| blast | Clostridiaceae_unclassified_SGB41663 | EtOH_pref_24h_final | 0.780382224 |
| vns | Clostridiaceae_unclassified_SGB41663 | EtOH_pref_24h_final | 0.719154368 |
| blast | Clostridiales_bacterium | EtOH_pref_24h_final | 0.0023319564 |
| vns | Clostridiales_bacterium | EtOH_pref_24h_final | 0.2964785018 |
| blast | Clostridium_cocleatum | EtOH_pref_24h_final | 0.0115678325 |
| vns | Clostridium_cocleatum | EtOH_pref_24h_final | 0.1960355146 |
| blast | Coriobacteriaceae_bacterium | EtOH_pref_24h_final | 0.3724483697 |
| vns | Coriobacteriaceae_bacterium | EtOH_pref_24h_final | 0.953919137 |
| blast | Dorea_sp_5_2 | EtOH_pref_24h_final | 0.5884919176 |
| vns | Dorea_sp_5_2 | EtOH_pref_24h_final | 0.4214161244 |

| | | | |
|---|---|---|---|
| blast | Dubosiella_newyorkensis | EtOH_pref_24h_final | 0.1169987394 |
| vns | Dubosiella_newyorkensis | EtOH_pref_24h_final | 0.799186063 |
| blast | Erysipelotrichales_bacterium | EtOH_pref_24h_final | 0.8970006656 |
| vns | Erysipelotrichales_bacterium | EtOH_pref_24h_final | 0.3284081083 |
| blast | Eubacteriaceae_bacterium | EtOH_pref_24h_final | 0.0271233687 |
| vns | Eubacteriaceae_bacterium | EtOH_pref_24h_final | 0.2383186135 |
| blast | Eubacteriaceae_unclassified_SGB94922 | EtOH_pref_24h_final | 0.4594880799 |
| vns | Eubacteriaceae_unclassified_SGB94922 | EtOH_pref_24h_final | 0.3726201484 |
| blast | GGB20149_SGB29430 | EtOH_pref_24h_final | 0.6280700484 |
| vns | GGB20149_SGB29430 | EtOH_pref_24h_final | 0.6299827926 |
| blast | GGB22635_SGB63107 | EtOH_pref_24h_final | 0.4117103362 |
| vns | GGB22635_SGB63107 | EtOH_pref_24h_final | 0.8871521383 |
| blast | GGB25041_SGB36960 | EtOH_pref_24h_final | 0.0824966676 |
| vns | GGB25041_SGB36960 | EtOH_pref_24h_final | 0.2101871969 |
| blast | GGB27876_SGB40310 | EtOH_pref_24h_final | 0.164666135 |
| vns | GGB27876_SGB40310 | EtOH_pref_24h_final | 0.7672351949 |
| blast | GGB27878_SGB40312 | EtOH_pref_24h_final | 0.2133541578 |
| vns | GGB27878_SGB40312 | EtOH_pref_24h_final | 0.1474794298 |
| blast | GGB27918_SGB40356 | EtOH_pref_24h_final | 0.9587581408 |
| vns | GGB27918_SGB40356 | EtOH_pref_24h_final | 0.8856989297 |
| blast | GGB28382_SGB40962 | EtOH_pref_24h_final | 0.5193113079 |
| vns | GGB28382_SGB40962 | EtOH_pref_24h_final | 0.8550578348 |
| blast | GGB28399_SGB40980 | EtOH_pref_24h_final | 0.2894655025 |
| vns | GGB28399_SGB40980 | EtOH_pref_24h_final | 0.287117034 |
| blast | GGB28411_SGB40993 | EtOH_pref_24h_final | 0.2018022626 |
| vns | GGB28411_SGB40993 | EtOH_pref_24h_final | 0.1468484741 |
| blast | GGB28415_SGB40997 | EtOH_pref_24h_final | 0.0267267446 |
| vns | GGB28415_SGB40997 | EtOH_pref_24h_final | 0.9490516966 |
| blast | GGB28430_SGB41013 | EtOH_pref_24h_final | 0.9120709472 |
| vns | GGB28430_SGB41013 | EtOH_pref_24h_final | 0.4662504826 |
| blast | GGB28439_SGB41022 | EtOH_pref_24h_final | 0.3804444035 |
| vns | GGB28439_SGB41022 | EtOH_pref_24h_final | 0.7387095529 |
| blast | GGB28778_SGB41431 | EtOH_pref_24h_final | 0.8046989411 |
| vns | GGB28778_SGB41431 | EtOH_pref_24h_final | 0.2184532434 |
| blast | GGB28784_SGB41437 | EtOH_pref_24h_final | 0.0207266217 |
| vns | GGB28784_SGB41437 | EtOH_pref_24h_final | 0.2544569976 |
| blast | GGB28792_SGB41445 | EtOH_pref_24h_final | 0.3385777757 |
| vns | GGB28792_SGB41445 | EtOH_pref_24h_final | 0.7612444262 |
| blast | GGB28798_SGB41451 | EtOH_pref_24h_final | 0.1640226305 |
| vns | GGB28798_SGB41451 | EtOH_pref_24h_final | 0.2489158244 |
| blast | GGB28802_SGB41455 | EtOH_pref_24h_final | 0.4578986259 |
| vns | GGB28802_SGB41455 | EtOH_pref_24h_final | 0.342689298 |
| blast | GGB28818_SGB41473 | EtOH_pref_24h_final | 0.7292095104 |
| vns | GGB28818_SGB41473 | EtOH_pref_24h_final | 0.5694099484 |
| blast | GGB28828_SGB41484 | EtOH_pref_24h_final | 0.2986969933 |
| vns | GGB28828_SGB41484 | EtOH_pref_24h_final | 0.1067065352 |

| | | | |
|---|---|---|---|
| blast | GGB28851_SGB41518 | EtOH_pref_24h_final | 0.852552179 |
| vns | GGB28851_SGB41518 | EtOH_pref_24h_final | 0.5783084298 |
| blast | GGB28859_SGB41528 | EtOH_pref_24h_final | 0.2057493298 |
| vns | GGB28859_SGB41528 | EtOH_pref_24h_final | 0.5953667458 |
| blast | GGB28864_SGB41535 | EtOH_pref_24h_final | 0.7854652488 |
| vns | GGB28864_SGB41535 | EtOH_pref_24h_final | 0.9646589853 |
| blast | GGB28869_SGB41543 | EtOH_pref_24h_final | 0.3479316259 |
| vns | GGB28869_SGB41543 | EtOH_pref_24h_final | 0.3814774755 |
| blast | GGB28883_SGB41564 | EtOH_pref_24h_final | 0.5948772487 |
| vns | GGB28883_SGB41564 | EtOH_pref_24h_final | 0.0605244586 |
| blast | GGB28892_SGB41573 | EtOH_pref_24h_final | 0.6512272078 |
| vns | GGB28892_SGB41573 | EtOH_pref_24h_final | 0.6676188822 |
| blast | GGB28893_SGB41574 | EtOH_pref_24h_final | 0.165594509 |
| vns | GGB28893_SGB41574 | EtOH_pref_24h_final | 0.1826993761 |
| blast | GGB28898_SGB41580 | EtOH_pref_24h_final | 0.3046549939 |
| vns | GGB28898_SGB41580 | EtOH_pref_24h_final | 0.0828933309 |
| blast | GGB28904_SGB41597 | EtOH_pref_24h_final | 0.1729063974 |
| vns | GGB28904_SGB41597 | EtOH_pref_24h_final | 0.0560671999 |
| blast | GGB28916_SGB41612 | EtOH_pref_24h_final | 0.8014326092 |
| vns | GGB28916_SGB41612 | EtOH_pref_24h_final | 0.8538132969 |
| blast | GGB28924_SGB41621 | EtOH_pref_24h_final | 0.7964816629 |
| vns | GGB28924_SGB41621 | EtOH_pref_24h_final | 0.2786583787 |
| blast | GGB28926_SGB41624 | EtOH_pref_24h_final | 0.3067808341 |
| vns | GGB28926_SGB41624 | EtOH_pref_24h_final | 0.470765265 |
| blast | GGB28927_SGB41625 | EtOH_pref_24h_final | 0.1937208057 |
| vns | GGB28927_SGB41625 | EtOH_pref_24h_final | 0.4182433734 |
| blast | GGB28934_SGB41635 | EtOH_pref_24h_final | 0.1158733118 |
| vns | GGB28934_SGB41635 | EtOH_pref_24h_final | 0.1113740731 |
| blast | GGB28946_SGB41652 | EtOH_pref_24h_final | 0.4278432256 |
| vns | GGB28946_SGB41652 | EtOH_pref_24h_final | 0.4225889007 |
| blast | GGB28949_SGB41655 | EtOH_pref_24h_final | 0.4436081164 |
| vns | GGB28949_SGB41655 | EtOH_pref_24h_final | 0.9089481791 |
| blast | GGB28949_SGB41656 | EtOH_pref_24h_final | 0.2827282346 |
| vns | GGB28949_SGB41656 | EtOH_pref_24h_final | 0.7708639493 |
| blast | GGB28950_SGB41657 | EtOH_pref_24h_final | 0.0305270365 |
| vns | GGB28950_SGB41657 | EtOH_pref_24h_final | 0.3364589048 |
| blast | GGB28951_SGB102295 | EtOH_pref_24h_final | 0.9253521562 |
| vns | GGB28951_SGB102295 | EtOH_pref_24h_final | 0.1587903977 |
| blast | GGB28951_SGB41658 | EtOH_pref_24h_final | 0.8470013515 |
| vns | GGB28951_SGB41658 | EtOH_pref_24h_final | 0.9202366237 |
| blast | GGB28954_SGB41662 | EtOH_pref_24h_final | 0.5157582975 |
| vns | GGB28954_SGB41662 | EtOH_pref_24h_final | 0.8004849551 |
| blast | GGB28956_SGB41665 | EtOH_pref_24h_final | 0.163329713 |
| vns | GGB28956_SGB41665 | EtOH_pref_24h_final | 0.1534165023 |
| blast | GGB28960_SGB41669 | EtOH_pref_24h_final | 0.155727292 |
| vns | GGB28960_SGB41669 | EtOH_pref_24h_final | 0.1559574792 |

| | | | |
|---|---|---|---|
| blast | GGB28967_SGB41678 | EtOH_pref_24h_final | 0.1792695292 |
| vns | GGB28967_SGB41678 | EtOH_pref_24h_final | 0.3150465193 |
| blast | GGB28991_SGB41705 | EtOH_pref_24h_final | 0.2288938824 |
| vns | GGB28991_SGB41705 | EtOH_pref_24h_final | 0.279982342 |
| blast | GGB29002_SGB41718 | EtOH_pref_24h_final | 0.0024308675 |
| vns | GGB29002_SGB41718 | EtOH_pref_24h_final | 0.1134655877 |
| blast | GGB29003_SGB41719 | EtOH_pref_24h_final | 0.4993986443 |
| vns | GGB29003_SGB41719 | EtOH_pref_24h_final | 0.5839678864 |
| blast | GGB29011_SGB41731 | EtOH_pref_24h_final | 0.916276033 |
| vns | GGB29011_SGB41731 | EtOH_pref_24h_final | 0.3455992366 |
| blast | GGB29531_SGB42317 | EtOH_pref_24h_final | 0.1450820268 |
| vns | GGB29531_SGB42317 | EtOH_pref_24h_final | 0.1355699662 |
| blast | GGB29685_SGB42494 | EtOH_pref_24h_final | 0.8645438528 |
| vns | GGB29685_SGB42494 | EtOH_pref_24h_final | 0.340104263 |
| blast | GGB30141_SGB43066 | EtOH_pref_24h_final | 0.9915548039 |
| vns | GGB30141_SGB43066 | EtOH_pref_24h_final | 0.7601787897 |
| blast | GGB30286_SGB43248 | EtOH_pref_24h_final | 0.0552616333 |
| vns | GGB30286_SGB43248 | EtOH_pref_24h_final | 0.4546033673 |
| blast | GGB30303_SGB43268 | EtOH_pref_24h_final | 0.3719368939 |
| vns | GGB30303_SGB43268 | EtOH_pref_24h_final | 0.9447221743 |
| blast | GGB30413_SGB43452 | EtOH_pref_24h_final | 0.9532769507 |
| vns | GGB30413_SGB43452 | EtOH_pref_24h_final | 0.8796426046 |
| blast | GGB30454_SGB43514 | EtOH_pref_24h_final | 0.3798589507 |
| vns | GGB30454_SGB43514 | EtOH_pref_24h_final | 0.2001255913 |
| blast | GGB30455_SGB43519 | EtOH_pref_24h_final | 0.3794677965 |
| vns | GGB30455_SGB43519 | EtOH_pref_24h_final | 0.3023489219 |
| blast | GGB30461_SGB43527 | EtOH_pref_24h_final | 0.3975036627 |
| vns | GGB30461_SGB43527 | EtOH_pref_24h_final | 0.7710386558 |
| blast | GGB30461_SGB43530 | EtOH_pref_24h_final | 0.395449 |
| vns | GGB30461_SGB43530 | EtOH_pref_24h_final | 0.7038636932 |
| blast | GGB30463_SGB43537 | EtOH_pref_24h_final | 0.2100542578 |
| vns | GGB30463_SGB43537 | EtOH_pref_24h_final | 0.9853614792 |
| blast | GGB30473_SGB43557 | EtOH_pref_24h_final | 0.0206421669 |
| vns | GGB30473_SGB43557 | EtOH_pref_24h_final | 0.0562117431 |
| blast | GGB30475_SGB63182 | EtOH_pref_24h_final | 0.2879653492 |
| vns | GGB30475_SGB63182 | EtOH_pref_24h_final | 0.6608559306 |
| blast | GGB30861_SGB44083 | EtOH_pref_24h_final | 0.9575523797 |
| vns | GGB30861_SGB44083 | EtOH_pref_24h_final | 0.8334455291 |
| blast | GGB31312_SGB44628 | EtOH_pref_24h_final | 0.7117271287 |
| vns | GGB31312_SGB44628 | EtOH_pref_24h_final | 0.8841939433 |
| blast | GGB31438_SGB44768 | EtOH_pref_24h_final | 0.9063572977 |
| vns | GGB31438_SGB44768 | EtOH_pref_24h_final | 0.3189600867 |
| blast | GGB3171_SGB4185 | EtOH_pref_24h_final | 0.0853457671 |
| vns | GGB3171_SGB4185 | EtOH_pref_24h_final | 0.9185517801 |
| blast | GGB31823_SGB45199 | EtOH_pref_24h_final | 0.1889777175 |
| vns | GGB31823_SGB45199 | EtOH_pref_24h_final | 0.7028132582 |

| | | | |
|---|---|---|---|
| blast | GGB31853_SGB45233 | EtOH_pref_24h_final | 0.496842743 |
| vns | GGB31853_SGB45233 | EtOH_pref_24h_final | 0.396259541 |
| blast | GGB32371_SGB41694 | EtOH_pref_24h_final | 0.346811850 |
| vns | GGB32371_SGB41694 | EtOH_pref_24h_final | 0.025054092 |
| blast | GGB3793_SGB5158 | EtOH_pref_24h_final | 0.965790474 |
| vns | GGB3793_SGB5158 | EtOH_pref_24h_final | 0.975145151 |
| blast | GGB42598_SGB59794 | EtOH_pref_24h_final | 0.501740001 |
| vns | GGB42598_SGB59794 | EtOH_pref_24h_final | 0.156668604 |
| blast | GGB45656_SGB63370 | EtOH_pref_24h_final | 0.621044138 |
| vns | GGB45656_SGB63370 | EtOH_pref_24h_final | 0.963374601 |
| blast | GGB47127_SGB65054 | EtOH_pref_24h_final | 0.496745120 |
| vns | GGB47127_SGB65054 | EtOH_pref_24h_final | 0.257417312 |
| blast | GGB74395_SGB43521 | EtOH_pref_24h_final | 0.123339782 |
| vns | GGB74395_SGB43521 | EtOH_pref_24h_final | 0.204021498 |
| blast | GGB75053_SGB43494 | EtOH_pref_24h_final | 0.410595919 |
| vns | GGB75053_SGB43494 | EtOH_pref_24h_final | 0.571857456 |
| blast | GGB75109_SGB102238 | EtOH_pref_24h_final | 0.620842292 |
| vns | GGB75109_SGB102238 | EtOH_pref_24h_final | 0.871818408 |
| blast | GGB81440_SGB45230 | EtOH_pref_24h_final | 0.079610731 |
| vns | GGB81440_SGB45230 | EtOH_pref_24h_final | 0.098140579 |
| blast | Lachnospiraceae_bacterium | EtOH_pref_24h_final | 0.484816177 |
| vns | Lachnospiraceae_bacterium | EtOH_pref_24h_final | 0.537861332 |
| blast | Lachnospiraceae_bacterium_A2 | EtOH_pref_24h_final | 0.420834896 |
| vns | Lachnospiraceae_bacterium_A2 | EtOH_pref_24h_final | 0.323648328 |
| blast | Lachnospiraceae_bacterium_MD308 | EtOH_pref_24h_final | 0.213119451 |
| vns | Lachnospiraceae_bacterium_MD308 | EtOH_pref_24h_final | 0.083066965 |
| blast | Lachnospiraceae_bacterium_MD329 | EtOH_pref_24h_final | 0.020976149 |
| vns | Lachnospiraceae_bacterium_MD329 | EtOH_pref_24h_final | 0.699161119 |
| blast | Lachnospiraceae_unclassified_SGB41414 | EtOH_pref_24h_final | 0.142639053 |
| vns | Lachnospiraceae_unclassified_SGB41414 | EtOH_pref_24h_final | 0.119876468 |
| blast | Lachnospiraceae_unclassified_SGB41418 | EtOH_pref_24h_final | 0.998757981 |
| vns | Lachnospiraceae_unclassified_SGB41418 | EtOH_pref_24h_final | 0.712236573 |
| blast | Lachnospiraceae_unclassified_SGB41424 | EtOH_pref_24h_final | 0.048386378 |
| vns | Lachnospiraceae_unclassified_SGB41424 | EtOH_pref_24h_final | 0.046067357 |
| blast | Lachnospiraceae_unclassified_SGB41589 | EtOH_pref_24h_final | 0.165424952 |
| vns | Lachnospiraceae_unclassified_SGB41589 | EtOH_pref_24h_final | 0.273527700 |
| blast | Lactobacillus_johnsonii | EtOH_pref_24h_final | 0.182678453 |
| vns | Lactobacillus_johnsonii | EtOH_pref_24h_final | 0.672951588 |
| blast | Muribaculaceae_bacterium | EtOH_pref_24h_final | 0.841525252 |
| vns | Muribaculaceae_bacterium | EtOH_pref_24h_final | 0.893052980 |
| blast | Neglectibacter_sp_X4 | EtOH_pref_24h_final | 0.084782151 |
| vns | Neglectibacter_sp_X4 | EtOH_pref_24h_final | 0.140656264 |
| blast | Oscillospiraceae_bacterium | EtOH_pref_24h_final | 0.741672565 |
| vns | Oscillospiraceae_bacterium | EtOH_pref_24h_final | 0.856188039 |
| blast | Oscillospiraceae_unclassified_SGB43502 | EtOH_pref_24h_final | 0.025941285 |
| vns | Oscillospiraceae_unclassified_SGB43502 | EtOH_pref_24h_final | 0.030535133 |

| | | | |
|---|---|---|---|
| blast | Oscillospiraceae_unclassified_SGB43505 | EtOH_pref_24h_final | 0.4959230188 |
| vns | Oscillospiraceae_unclassified_SGB43505 | EtOH_pref_24h_final | 0.6425337316 |
| blast | Parasutterella_excrementihominis | EtOH_pref_24h_final | 0.5671048722 |
| vns | Parasutterella_excrementihominis | EtOH_pref_24h_final | 0.7099929171 |
| blast | Richness (# observed features) | EtOH_pref_24h_final | 0.2717524423 |
| vns | Richness (# observed features) | EtOH_pref_24h_final | 0.6607230685 |
| blast | Romboutsia_ilealis | EtOH_pref_24h_final | 0.0010997742 |
| vns | Romboutsia_ilealis | EtOH_pref_24h_final | 0.0569970896 |
| blast | Schaedlerella_arabinosiphila | EtOH_pref_24h_final | 0.9390244043 |
| vns | Schaedlerella_arabinosiphila | EtOH_pref_24h_final | 0.9139116075 |
| blast | Shannon Index | EtOH_pref_24h_final | 0.6947352216 |
| vns | Shannon Index | EtOH_pref_24h_final | 0.9062773413 |
| blast | Turicibacter_sp_1E2 | EtOH_pref_24h_final | 0.1991772312 |
| vns | Turicibacter_sp_1E2 | EtOH_pref_24h_final | 0.1655749312 |
| blast | Acetatifactor_muris | OFB_Cdistance | 0.7395220796 |
| vns | Acetatifactor_muris | OFB_Cdistance | 0.7528971403 |
| blast | Acetatifactor_SGB41546 | OFB_Cdistance | 0.8402457971 |
| vns | Acetatifactor_SGB41546 | OFB_Cdistance | 0.6630009771 |
| blast | Acutalibacter_muris | OFB_Cdistance | 0.4221546334 |
| vns | Acutalibacter_muris | OFB_Cdistance | 0.8105784495 |
| blast | Acutalibacter_sp_1XD8_36 | OFB_Cdistance | 0.6605798863 |
| vns | Acutalibacter_sp_1XD8_36 | OFB_Cdistance | 0.7091849336 |
| blast | Adlercreutzia_caecimuris | OFB_Cdistance | 0.9896089529 |
| vns | Adlercreutzia_caecimuris | OFB_Cdistance | 0.6765426913 |
| blast | Adlercreutzia_mucosicola | OFB_Cdistance | 0.2718339856 |
| vns | Adlercreutzia_mucosicola | OFB_Cdistance | 0.2498256574 |
| blast | Adlercreutzia_muris | OFB_Cdistance | 0.6704217743 |
| vns | Adlercreutzia_muris | OFB_Cdistance | 0.5958277454 |
| blast | Akkermansia_muciniphila | OFB_Cdistance | 0.1698125404 |
| vns | Akkermansia_muciniphila | OFB_Cdistance | 0.8643625099 |
| blast | Alistipes_sp_DSM_112343 | OFB_Cdistance | 0.3725159774 |
| vns | Alistipes_sp_DSM_112343 | OFB_Cdistance | 0.8326983074 |
| blast | Anaerotruncus_sp_1XD42_93 | OFB_Cdistance | 0.5202714535 |
| vns | Anaerotruncus_sp_1XD42_93 | OFB_Cdistance | 0.6174107217 |
| blast | Bacteria_unclassified_SGB102200 | OFB_Cdistance | 0.4742692127 |
| vns | Bacteria_unclassified_SGB102200 | OFB_Cdistance | 0.7641667885 |
| blast | Bacteria_unclassified_SGB41677 | OFB_Cdistance | 0.4441438231 |
| vns | Bacteria_unclassified_SGB41677 | OFB_Cdistance | 0.1699894243 |
| blast | Bacteria_unclassified_SGB43546 | OFB_Cdistance | 0.5863583659 |
| vns | Bacteria_unclassified_SGB43546 | OFB_Cdistance | 0.7890099125 |
| blast | bacterium_1XD42_54 | OFB_Cdistance | 0.0144245959 |
| vns | bacterium_1XD42_54 | OFB_Cdistance | 0.2345775748 |
| blast | bacterium_1XD42_76 | OFB_Cdistance | 0.6000316038 |
| vns | bacterium_1XD42_76 | OFB_Cdistance | 0.6429924539 |
| blast | bacterium_1xD8_48 | OFB_Cdistance | 0.8997089453 |
| vns | bacterium_1xD8_48 | OFB_Cdistance | 0.6968168809 |

| | | | |
|---|---|---|---|
| blast | Bacteroides_thetaiotaomicron | OFB_Cdistance | 0.960201477 |
| vns | Bacteroides_thetaiotaomicron | OFB_Cdistance | 0.634306315 |
| blast | Berger Parker Index | OFB_Cdistance | 0.545741712 |
| vns | Berger Parker Index | OFB_Cdistance | 0.233733054 |
| blast | Bifidobacterium_pseudolongum | OFB_Cdistance | 0.428010398 |
| vns | Bifidobacterium_pseudolongum | OFB_Cdistance | 0.347799065 |
| blast | Clostridia_bacterium | OFB_Cdistance | 0.374559043 |
| vns | Clostridia_bacterium | OFB_Cdistance | 0.724940107 |
| blast | Clostridiaceae_bacterium | OFB_Cdistance | 0.987169780 |
| vns | Clostridiaceae_bacterium | OFB_Cdistance | 0.241937042 |
| blast | Clostridiaceae_unclassified_SGB41663 | OFB_Cdistance | 0.780382224 |
| vns | Clostridiaceae_unclassified_SGB41663 | OFB_Cdistance | 0.719154368 |
| blast | Clostridiales_bacterium | OFB_Cdistance | 0.002331956 |
| vns | Clostridiales_bacterium | OFB_Cdistance | 0.296478501 |
| blast | Clostridium_cocleatum | OFB_Cdistance | 0.011567832 |
| vns | Clostridium_cocleatum | OFB_Cdistance | 0.196035514 |
| blast | Coriobacteriaceae_bacterium | OFB_Cdistance | 0.372448369 |
| vns | Coriobacteriaceae_bacterium | OFB_Cdistance | 0.953919137 |
| blast | Dorea_sp_5_2 | OFB_Cdistance | 0.588491917 |
| vns | Dorea_sp_5_2 | OFB_Cdistance | 0.421416124 |
| blast | Dubosiella_newyorkensis | OFB_Cdistance | 0.116998739 |
| vns | Dubosiella_newyorkensis | OFB_Cdistance | 0.799186063 |
| blast | Erysipelotrichales_bacterium | OFB_Cdistance | 0.897000665 |
| vns | Erysipelotrichales_bacterium | OFB_Cdistance | 0.328408108 |
| blast | Eubacteriaceae_bacterium | OFB_Cdistance | 0.027123368 |
| vns | Eubacteriaceae_bacterium | OFB_Cdistance | 0.238318613 |
| blast | Eubacteriaceae_unclassified_SGB94922 | OFB_Cdistance | 0.459488079 |
| vns | Eubacteriaceae_unclassified_SGB94922 | OFB_Cdistance | 0.372620148 |
| blast | GGB20149_SGB29430 | OFB_Cdistance | 0.628070048 |
| vns | GGB20149_SGB29430 | OFB_Cdistance | 0.629982792 |
| blast | GGB22635_SGB63107 | OFB_Cdistance | 0.411710336 |
| vns | GGB22635_SGB63107 | OFB_Cdistance | 0.887152138 |
| blast | GGB25041_SGB36960 | OFB_Cdistance | 0.082496667 |
| vns | GGB25041_SGB36960 | OFB_Cdistance | 0.210187196 |
| blast | GGB27876_SGB40310 | OFB_Cdistance | 0.164666135 |
| vns | GGB27876_SGB40310 | OFB_Cdistance | 0.767235194 |
| blast | GGB27878_SGB40312 | OFB_Cdistance | 0.213354157 |
| vns | GGB27878_SGB40312 | OFB_Cdistance | 0.147479429 |
| blast | GGB27918_SGB40356 | OFB_Cdistance | 0.958758140 |
| vns | GGB27918_SGB40356 | OFB_Cdistance | 0.885698929 |
| blast | GGB28382_SGB40962 | OFB_Cdistance | 0.519311307 |
| vns | GGB28382_SGB40962 | OFB_Cdistance | 0.855057834 |
| blast | GGB28399_SGB40980 | OFB_Cdistance | 0.289465502 |
| vns | GGB28399_SGB40980 | OFB_Cdistance | 0.287117034 |
| blast | GGB28411_SGB40993 | OFB_Cdistance | 0.201802262 |
| vns | GGB28411_SGB40993 | OFB_Cdistance | 0.146848474 |

| | | | |
|---|---|---|---|
| blast | GGB28415_SGB40997 | OFB_Cdistance | 0.026726744 |
| vns | GGB28415_SGB40997 | OFB_Cdistance | 0.949051696 |
| blast | GGB28430_SGB41013 | OFB_Cdistance | 0.912070947 |
| vns | GGB28430_SGB41013 | OFB_Cdistance | 0.466250482 |
| blast | GGB28439_SGB41022 | OFB_Cdistance | 0.380444403 |
| vns | GGB28439_SGB41022 | OFB_Cdistance | 0.738709552 |
| blast | GGB28778_SGB41431 | OFB_Cdistance | 0.804698941 |
| vns | GGB28778_SGB41431 | OFB_Cdistance | 0.218453243 |
| blast | GGB28784_SGB41437 | OFB_Cdistance | 0.020726621 |
| vns | GGB28784_SGB41437 | OFB_Cdistance | 0.254456997 |
| blast | GGB28792_SGB41445 | OFB_Cdistance | 0.338577775 |
| vns | GGB28792_SGB41445 | OFB_Cdistance | 0.761244426 |
| blast | GGB28798_SGB41451 | OFB_Cdistance | 0.164022630 |
| vns | GGB28798_SGB41451 | OFB_Cdistance | 0.248915824 |
| blast | GGB28802_SGB41455 | OFB_Cdistance | 0.457898625 |
| vns | GGB28802_SGB41455 | OFB_Cdistance | 0.342689298 |
| blast | GGB28818_SGB41473 | OFB_Cdistance | 0.729209510 |
| vns | GGB28818_SGB41473 | OFB_Cdistance | 0.569409948 |
| blast | GGB28828_SGB41484 | OFB_Cdistance | 0.298696993 |
| vns | GGB28828_SGB41484 | OFB_Cdistance | 0.106706535 |
| blast | GGB28851_SGB41518 | OFB_Cdistance | 0.852552179 |
| vns | GGB28851_SGB41518 | OFB_Cdistance | 0.578308429 |
| blast | GGB28859_SGB41528 | OFB_Cdistance | 0.205749329 |
| vns | GGB28859_SGB41528 | OFB_Cdistance | 0.595366745 |
| blast | GGB28864_SGB41535 | OFB_Cdistance | 0.785465248 |
| vns | GGB28864_SGB41535 | OFB_Cdistance | 0.964658985 |
| blast | GGB28869_SGB41543 | OFB_Cdistance | 0.347931625 |
| vns | GGB28869_SGB41543 | OFB_Cdistance | 0.381477475 |
| blast | GGB28883_SGB41564 | OFB_Cdistance | 0.594877248 |
| vns | GGB28883_SGB41564 | OFB_Cdistance | 0.060524458 |
| blast | GGB28892_SGB41573 | OFB_Cdistance | 0.651227207 |
| vns | GGB28892_SGB41573 | OFB_Cdistance | 0.667618882 |
| blast | GGB28893_SGB41574 | OFB_Cdistance | 0.165594509 |
| vns | GGB28893_SGB41574 | OFB_Cdistance | 0.182699376 |
| blast | GGB28898_SGB41580 | OFB_Cdistance | 0.304654993 |
| vns | GGB28898_SGB41580 | OFB_Cdistance | 0.082893330 |
| blast | GGB28904_SGB41597 | OFB_Cdistance | 0.172906397 |
| vns | GGB28904_SGB41597 | OFB_Cdistance | 0.056067199 |
| blast | GGB28916_SGB41612 | OFB_Cdistance | 0.801432609 |
| vns | GGB28916_SGB41612 | OFB_Cdistance | 0.853813296 |
| blast | GGB28924_SGB41621 | OFB_Cdistance | 0.796481662 |
| vns | GGB28924_SGB41621 | OFB_Cdistance | 0.278658378 |
| blast | GGB28926_SGB41624 | OFB_Cdistance | 0.306780834 |
| vns | GGB28926_SGB41624 | OFB_Cdistance | 0.470765265 |
| blast | GGB28927_SGB41625 | OFB_Cdistance | 0.193720805 |
| vns | GGB28927_SGB41625 | OFB_Cdistance | 0.418243373 |

| | | | |
|---|---|---|---|
| blast | GGB28934_SGB41635 | OFB_Cdistance | 0.1158733118 |
| vns | GGB28934_SGB41635 | OFB_Cdistance | 0.1113740731 |
| blast | GGB28946_SGB41652 | OFB_Cdistance | 0.4278432256 |
| vns | GGB28946_SGB41652 | OFB_Cdistance | 0.4225889007 |
| blast | GGB28949_SGB41655 | OFB_Cdistance | 0.4436081164 |
| vns | GGB28949_SGB41655 | OFB_Cdistance | 0.9089481791 |
| blast | GGB28949_SGB41656 | OFB_Cdistance | 0.2827282346 |
| vns | GGB28949_SGB41656 | OFB_Cdistance | 0.7708639493 |
| blast | GGB28950_SGB41657 | OFB_Cdistance | 0.0305270365 |
| vns | GGB28950_SGB41657 | OFB_Cdistance | 0.3364589048 |
| blast | GGB28951_SGB102295 | OFB_Cdistance | 0.9253521562 |
| vns | GGB28951_SGB102295 | OFB_Cdistance | 0.1587903977 |
| blast | GGB28951_SGB41658 | OFB_Cdistance | 0.8470013515 |
| vns | GGB28951_SGB41658 | OFB_Cdistance | 0.9202366237 |
| blast | GGB28954_SGB41662 | OFB_Cdistance | 0.5157582975 |
| vns | GGB28954_SGB41662 | OFB_Cdistance | 0.8004849551 |
| blast | GGB28956_SGB41665 | OFB_Cdistance | 0.163329713 |
| vns | GGB28956_SGB41665 | OFB_Cdistance | 0.1534165023 |
| blast | GGB28960_SGB41669 | OFB_Cdistance | 0.155727292 |
| vns | GGB28960_SGB41669 | OFB_Cdistance | 0.1559574792 |
| blast | GGB28967_SGB41678 | OFB_Cdistance | 0.1792695292 |
| vns | GGB28967_SGB41678 | OFB_Cdistance | 0.3150465193 |
| blast | GGB28991_SGB41705 | OFB_Cdistance | 0.2288938824 |
| vns | GGB28991_SGB41705 | OFB_Cdistance | 0.279982342 |
| blast | GGB29002_SGB41718 | OFB_Cdistance | 0.0024308675 |
| vns | GGB29002_SGB41718 | OFB_Cdistance | 0.1134655877 |
| blast | GGB29003_SGB41719 | OFB_Cdistance | 0.4993986443 |
| vns | GGB29003_SGB41719 | OFB_Cdistance | 0.5839678864 |
| blast | GGB29011_SGB41731 | OFB_Cdistance | 0.916276033 |
| vns | GGB29011_SGB41731 | OFB_Cdistance | 0.3455992366 |
| blast | GGB29531_SGB42317 | OFB_Cdistance | 0.1450820268 |
| vns | GGB29531_SGB42317 | OFB_Cdistance | 0.1355699662 |
| blast | GGB29685_SGB42494 | OFB_Cdistance | 0.8645438528 |
| vns | GGB29685_SGB42494 | OFB_Cdistance | 0.340104263 |
| blast | GGB30141_SGB43066 | OFB_Cdistance | 0.9915548039 |
| vns | GGB30141_SGB43066 | OFB_Cdistance | 0.7601787897 |
| blast | GGB30286_SGB43248 | OFB_Cdistance | 0.0552616333 |
| vns | GGB30286_SGB43248 | OFB_Cdistance | 0.4546033673 |
| blast | GGB30303_SGB43268 | OFB_Cdistance | 0.3719368939 |
| vns | GGB30303_SGB43268 | OFB_Cdistance | 0.9447221743 |
| blast | GGB30413_SGB43452 | OFB_Cdistance | 0.9532769507 |
| vns | GGB30413_SGB43452 | OFB_Cdistance | 0.8796426046 |
| blast | GGB30454_SGB43514 | OFB_Cdistance | 0.3798589507 |
| vns | GGB30454_SGB43514 | OFB_Cdistance | 0.2001255913 |
| blast | GGB30455_SGB43519 | OFB_Cdistance | 0.3794677965 |
| vns | GGB30455_SGB43519 | OFB_Cdistance | 0.3023489219 |

| | | | |
|---|---|---|---|
| blast | GGB30461_SGB43527 | OFB_Cdistance | 0.397503662 |
| vns | GGB30461_SGB43527 | OFB_Cdistance | 0.771038655 |
| blast | GGB30461_SGB43530 | OFB_Cdistance | 0.395449 |
| vns | GGB30461_SGB43530 | OFB_Cdistance | 0.703863693 |
| blast | GGB30463_SGB43537 | OFB_Cdistance | 0.210054257 |
| vns | GGB30463_SGB43537 | OFB_Cdistance | 0.985361479 |
| blast | GGB30473_SGB43557 | OFB_Cdistance | 0.020642166 |
| vns | GGB30473_SGB43557 | OFB_Cdistance | 0.056211743 |
| blast | GGB30475_SGB63182 | OFB_Cdistance | 0.287965349 |
| vns | GGB30475_SGB63182 | OFB_Cdistance | 0.660855930 |
| blast | GGB30861_SGB44083 | OFB_Cdistance | 0.957552379 |
| vns | GGB30861_SGB44083 | OFB_Cdistance | 0.833445529 |
| blast | GGB31312_SGB44628 | OFB_Cdistance | 0.711727128 |
| vns | GGB31312_SGB44628 | OFB_Cdistance | 0.884193943 |
| blast | GGB31438_SGB44768 | OFB_Cdistance | 0.906357297 |
| vns | GGB31438_SGB44768 | OFB_Cdistance | 0.318960086 |
| blast | GGB3171_SGB4185 | OFB_Cdistance | 0.085345767 |
| vns | GGB3171_SGB4185 | OFB_Cdistance | 0.918551780 |
| blast | GGB31823_SGB45199 | OFB_Cdistance | 0.188977717 |
| vns | GGB31823_SGB45199 | OFB_Cdistance | 0.702813258 |
| blast | GGB31853_SGB45233 | OFB_Cdistance | 0.496842743 |
| vns | GGB31853_SGB45233 | OFB_Cdistance | 0.396259541 |
| blast | GGB32371_SGB41694 | OFB_Cdistance | 0.346811850 |
| vns | GGB32371_SGB41694 | OFB_Cdistance | 0.025054092 |
| blast | GGB3793_SGB5158 | OFB_Cdistance | 0.965790474 |
| vns | GGB3793_SGB5158 | OFB_Cdistance | 0.975145151 |
| blast | GGB42598_SGB59794 | OFB_Cdistance | 0.501740001 |
| vns | GGB42598_SGB59794 | OFB_Cdistance | 0.156668604 |
| blast | GGB45656_SGB63370 | OFB_Cdistance | 0.621044138 |
| vns | GGB45656_SGB63370 | OFB_Cdistance | 0.963374601 |
| blast | GGB47127_SGB65054 | OFB_Cdistance | 0.496745120 |
| vns | GGB47127_SGB65054 | OFB_Cdistance | 0.257417312 |
| blast | GGB74395_SGB43521 | OFB_Cdistance | 0.123339782 |
| vns | GGB74395_SGB43521 | OFB_Cdistance | 0.204021498 |
| blast | GGB75053_SGB43494 | OFB_Cdistance | 0.410595919 |
| vns | GGB75053_SGB43494 | OFB_Cdistance | 0.571857456 |
| blast | GGB75109_SGB102238 | OFB_Cdistance | 0.620842292 |
| vns | GGB75109_SGB102238 | OFB_Cdistance | 0.871818408 |
| blast | GGB81440_SGB45230 | OFB_Cdistance | 0.079610731 |
| vns | GGB81440_SGB45230 | OFB_Cdistance | 0.098140579 |
| blast | Lachnospiraceae_bacterium | OFB_Cdistance | 0.484816177 |
| vns | Lachnospiraceae_bacterium | OFB_Cdistance | 0.537861332 |
| blast | Lachnospiraceae_bacterium_A2 | OFB_Cdistance | 0.420834896 |
| vns | Lachnospiraceae_bacterium_A2 | OFB_Cdistance | 0.323648328 |
| blast | Lachnospiraceae_bacterium_MD308 | OFB_Cdistance | 0.213119451 |
| vns | Lachnospiraceae_bacterium_MD308 | OFB_Cdistance | 0.083066965 |

| | | | |
|---|---|---|---|
| blast | Lachnospiraceae_bacterium_MD329 | OFB_Cdistance | 0.020976149 |
| vns | Lachnospiraceae_bacterium_MD329 | OFB_Cdistance | 0.699161119 |
| blast | Lachnospiraceae_unclassified_SGB41414 | OFB_Cdistance | 0.142639053 |
| vns | Lachnospiraceae_unclassified_SGB41414 | OFB_Cdistance | 0.119876468 |
| blast | Lachnospiraceae_unclassified_SGB41418 | OFB_Cdistance | 0.998757981 |
| vns | Lachnospiraceae_unclassified_SGB41418 | OFB_Cdistance | 0.712236573 |
| blast | Lachnospiraceae_unclassified_SGB41424 | OFB_Cdistance | 0.048386378 |
| vns | Lachnospiraceae_unclassified_SGB41424 | OFB_Cdistance | 0.046067357 |
| blast | Lachnospiraceae_unclassified_SGB41589 | OFB_Cdistance | 0.165424952 |
| vns | Lachnospiraceae_unclassified_SGB41589 | OFB_Cdistance | 0.273527700 |
| blast | Lactobacillus_johnsonii | OFB_Cdistance | 0.182678453 |
| vns | Lactobacillus_johnsonii | OFB_Cdistance | 0.672951588 |
| blast | Muribaculaceae_bacterium | OFB_Cdistance | 0.841525252 |
| vns | Muribaculaceae_bacterium | OFB_Cdistance | 0.893052980 |
| blast | Neglectibacter_sp_X4 | OFB_Cdistance | 0.084782151 |
| vns | Neglectibacter_sp_X4 | OFB_Cdistance | 0.140656264 |
| blast | Oscillospiraceae_bacterium | OFB_Cdistance | 0.741672565 |
| vns | Oscillospiraceae_bacterium | OFB_Cdistance | 0.856188039 |
| blast | Oscillospiraceae_unclassified_SGB43502 | OFB_Cdistance | 0.025941285 |
| vns | Oscillospiraceae_unclassified_SGB43502 | OFB_Cdistance | 0.030535133 |
| blast | Oscillospiraceae_unclassified_SGB43505 | OFB_Cdistance | 0.495923018 |
| vns | Oscillospiraceae_unclassified_SGB43505 | OFB_Cdistance | 0.642533731 |
| blast | Parasutterella_excrementihominis | OFB_Cdistance | 0.567104872 |
| vns | Parasutterella_excrementihominis | OFB_Cdistance | 0.709992917 |
| blast | Richness (# observed features) | OFB_Cdistance | 0.271752442 |
| vns | Richness (# observed features) | OFB_Cdistance | 0.660723068 |
| blast | Romboutsia_ilealis | OFB_Cdistance | 0.001099774 |
| vns | Romboutsia_ilealis | OFB_Cdistance | 0.056997089 |
| blast | Schaedlerella_arabinosiphila | OFB_Cdistance | 0.939024404 |
| vns | Schaedlerella_arabinosiphila | OFB_Cdistance | 0.913911607 |
| blast | Shannon Index | OFB_Cdistance | 0.694735221 |
| vns | Shannon Index | OFB_Cdistance | 0.906277341 |
| blast | Turicibacter_sp_1E2 | OFB_Cdistance | 0.199177231 |
| vns | Turicibacter_sp_1E2 | OFB_Cdistance | 0.165574931 |
| blast | Acetatifactor_muris | OFB_Centries | 0.739522079 |
| vns | Acetatifactor_muris | OFB_Centries | 0.752897140 |
| blast | Acetatifactor_SGB41546 | OFB_Centries | 0.840245797 |
| vns | Acetatifactor_SGB41546 | OFB_Centries | 0.663000977 |
| blast | Acutalibacter_muris | OFB_Centries | 0.422154633 |
| vns | Acutalibacter_muris | OFB_Centries | 0.810578449 |
| blast | Acutalibacter_sp_1XD8_36 | OFB_Centries | 0.660579886 |
| vns | Acutalibacter_sp_1XD8_36 | OFB_Centries | 0.709184933 |
| blast | Adlercreutzia_caecimuris | OFB_Centries | 0.989608952 |
| vns | Adlercreutzia_caecimuris | OFB_Centries | 0.676542691 |
| blast | Adlercreutzia_mucosicola | OFB_Centries | 0.271833985 |
| vns | Adlercreutzia_mucosicola | OFB_Centries | 0.249825657 |

| | | | |
|---|---|---|---|
| blast | Adlercreutzia_muris | OFB_Centries | 0.6704217743 |
| vns | Adlercreutzia_muris | OFB_Centries | 0.5958277454 |
| blast | Akkermansia_muciniphila | OFB_Centries | 0.1698125404 |
| vns | Akkermansia_muciniphila | OFB_Centries | 0.8643625099 |
| blast | Alistipes_sp_DSM_112343 | OFB_Centries | 0.3725159774 |
| vns | Alistipes_sp_DSM_112343 | OFB_Centries | 0.8326983074 |
| blast | Anaerotruncus_sp_1XD42_93 | OFB_Centries | 0.5202714535 |
| vns | Anaerotruncus_sp_1XD42_93 | OFB_Centries | 0.6174107217 |
| blast | Bacteria_unclassified_SGB102200 | OFB_Centries | 0.4742692127 |
| vns | Bacteria_unclassified_SGB102200 | OFB_Centries | 0.7641667885 |
| blast | Bacteria_unclassified_SGB41677 | OFB_Centries | 0.4441438231 |
| vns | Bacteria_unclassified_SGB41677 | OFB_Centries | 0.1699894243 |
| blast | Bacteria_unclassified_SGB43546 | OFB_Centries | 0.5863583659 |
| vns | Bacteria_unclassified_SGB43546 | OFB_Centries | 0.7890099125 |
| blast | bacterium_1XD42_54 | OFB_Centries | 0.0144245959 |
| vns | bacterium_1XD42_54 | OFB_Centries | 0.2345775748 |
| blast | bacterium_1XD42_76 | OFB_Centries | 0.6000316038 |
| vns | bacterium_1XD42_76 | OFB_Centries | 0.6429924539 |
| blast | bacterium_1xD8_48 | OFB_Centries | 0.8997089453 |
| vns | bacterium_1xD8_48 | OFB_Centries | 0.6968168809 |
| blast | Bacteroides_thetaiotaomicron | OFB_Centries | 0.9602014777 |
| vns | Bacteroides_thetaiotaomicron | OFB_Centries | 0.6343063159 |
| blast | Berger Parker Index | OFB_Centries | 0.5457417127 |
| vns | Berger Parker Index | OFB_Centries | 0.2337330546 |
| blast | Bifidobacterium_pseudolongum | OFB_Centries | 0.4280103985 |
| vns | Bifidobacterium_pseudolongum | OFB_Centries | 0.3477990653 |
| blast | Clostridia_bacterium | OFB_Centries | 0.3745590434 |
| vns | Clostridia_bacterium | OFB_Centries | 0.7249401071 |
| blast | Clostridiaceae_bacterium | OFB_Centries | 0.9871697806 |
| vns | Clostridiaceae_bacterium | OFB_Centries | 0.2419370425 |
| blast | Clostridiaceae_unclassified_SGB41663 | OFB_Centries | 0.780382224 |
| vns | Clostridiaceae_unclassified_SGB41663 | OFB_Centries | 0.719154368 |
| blast | Clostridiales_bacterium | OFB_Centries | 0.0023319564 |
| vns | Clostridiales_bacterium | OFB_Centries | 0.2964785018 |
| blast | Clostridium_cocleatum | OFB_Centries | 0.0115678325 |
| vns | Clostridium_cocleatum | OFB_Centries | 0.1960355146 |
| blast | Coriobacteriaceae_bacterium | OFB_Centries | 0.3724483697 |
| vns | Coriobacteriaceae_bacterium | OFB_Centries | 0.953919137 |
| blast | Dorea_sp_5_2 | OFB_Centries | 0.5884919176 |
| vns | Dorea_sp_5_2 | OFB_Centries | 0.4214161244 |
| blast | Dubosiella_newyorkensis | OFB_Centries | 0.1169987394 |
| vns | Dubosiella_newyorkensis | OFB_Centries | 0.799186063 |
| blast | Erysipelotrichales_bacterium | OFB_Centries | 0.8970006656 |
| vns | Erysipelotrichales_bacterium | OFB_Centries | 0.3284081083 |
| blast | Eubacteriaceae_bacterium | OFB_Centries | 0.0271233687 |
| vns | Eubacteriaceae_bacterium | OFB_Centries | 0.2383186135 |

| | | | |
|---|---|---|---|
| blast | Eubacteriaceae_unclassified_SGB94922 | OFB_Centries | 0.459488079 |
| vns | Eubacteriaceae_unclassified_SGB94922 | OFB_Centries | 0.372620148 |
| blast | GGB20149_SGB29430 | OFB_Centries | 0.628070048 |
| vns | GGB20149_SGB29430 | OFB_Centries | 0.629982792 |
| blast | GGB22635_SGB63107 | OFB_Centries | 0.411710336 |
| vns | GGB22635_SGB63107 | OFB_Centries | 0.887152138 |
| blast | GGB25041_SGB36960 | OFB_Centries | 0.082496667 |
| vns | GGB25041_SGB36960 | OFB_Centries | 0.210187196 |
| blast | GGB27876_SGB40310 | OFB_Centries | 0.164666135 |
| vns | GGB27876_SGB40310 | OFB_Centries | 0.767235194 |
| blast | GGB27878_SGB40312 | OFB_Centries | 0.213354157 |
| vns | GGB27878_SGB40312 | OFB_Centries | 0.147479429 |
| blast | GGB27918_SGB40356 | OFB_Centries | 0.958758140 |
| vns | GGB27918_SGB40356 | OFB_Centries | 0.885698929 |
| blast | GGB28382_SGB40962 | OFB_Centries | 0.519311307 |
| vns | GGB28382_SGB40962 | OFB_Centries | 0.855057834 |
| blast | GGB28399_SGB40980 | OFB_Centries | 0.289465502 |
| vns | GGB28399_SGB40980 | OFB_Centries | 0.287117034 |
| blast | GGB28411_SGB40993 | OFB_Centries | 0.201802262 |
| vns | GGB28411_SGB40993 | OFB_Centries | 0.146848474 |
| blast | GGB28415_SGB40997 | OFB_Centries | 0.026726744 |
| vns | GGB28415_SGB40997 | OFB_Centries | 0.949051696 |
| blast | GGB28430_SGB41013 | OFB_Centries | 0.912070947 |
| vns | GGB28430_SGB41013 | OFB_Centries | 0.466250482 |
| blast | GGB28439_SGB41022 | OFB_Centries | 0.380444403 |
| vns | GGB28439_SGB41022 | OFB_Centries | 0.738709552 |
| blast | GGB28778_SGB41431 | OFB_Centries | 0.804698941 |
| vns | GGB28778_SGB41431 | OFB_Centries | 0.218453243 |
| blast | GGB28784_SGB41437 | OFB_Centries | 0.020726621 |
| vns | GGB28784_SGB41437 | OFB_Centries | 0.254456997 |
| blast | GGB28792_SGB41445 | OFB_Centries | 0.338577775 |
| vns | GGB28792_SGB41445 | OFB_Centries | 0.761244426 |
| blast | GGB28798_SGB41451 | OFB_Centries | 0.164022630 |
| vns | GGB28798_SGB41451 | OFB_Centries | 0.248915824 |
| blast | GGB28802_SGB41455 | OFB_Centries | 0.457898625 |
| vns | GGB28802_SGB41455 | OFB_Centries | 0.342689298 |
| blast | GGB28818_SGB41473 | OFB_Centries | 0.729209510 |
| vns | GGB28818_SGB41473 | OFB_Centries | 0.569409948 |
| blast | GGB28828_SGB41484 | OFB_Centries | 0.298696993 |
| vns | GGB28828_SGB41484 | OFB_Centries | 0.106706535 |
| blast | GGB28851_SGB41518 | OFB_Centries | 0.852552179 |
| vns | GGB28851_SGB41518 | OFB_Centries | 0.578308429 |
| blast | GGB28859_SGB41528 | OFB_Centries | 0.205749329 |
| vns | GGB28859_SGB41528 | OFB_Centries | 0.595366745 |
| blast | GGB28864_SGB41535 | OFB_Centries | 0.785465248 |
| vns | GGB28864_SGB41535 | OFB_Centries | 0.964658985 |

| | | | |
|---|---|---|---|
| blast | GGB28869_SGB41543 | OFB_Centries | 0.347931625 |
| vns | GGB28869_SGB41543 | OFB_Centries | 0.381477475 |
| blast | GGB28883_SGB41564 | OFB_Centries | 0.594877248 |
| vns | GGB28883_SGB41564 | OFB_Centries | 0.060524458 |
| blast | GGB28892_SGB41573 | OFB_Centries | 0.651227207 |
| vns | GGB28892_SGB41573 | OFB_Centries | 0.667618882 |
| blast | GGB28893_SGB41574 | OFB_Centries | 0.165594509 |
| vns | GGB28893_SGB41574 | OFB_Centries | 0.182699376 |
| blast | GGB28898_SGB41580 | OFB_Centries | 0.304654993 |
| vns | GGB28898_SGB41580 | OFB_Centries | 0.082893330 |
| blast | GGB28904_SGB41597 | OFB_Centries | 0.172906397 |
| vns | GGB28904_SGB41597 | OFB_Centries | 0.056067199 |
| blast | GGB28916_SGB41612 | OFB_Centries | 0.801432609 |
| vns | GGB28916_SGB41612 | OFB_Centries | 0.853813296 |
| blast | GGB28924_SGB41621 | OFB_Centries | 0.796481662 |
| vns | GGB28924_SGB41621 | OFB_Centries | 0.278658378 |
| blast | GGB28926_SGB41624 | OFB_Centries | 0.306780834 |
| vns | GGB28926_SGB41624 | OFB_Centries | 0.470765265 |
| blast | GGB28927_SGB41625 | OFB_Centries | 0.193720805 |
| vns | GGB28927_SGB41625 | OFB_Centries | 0.418243373 |
| blast | GGB28934_SGB41635 | OFB_Centries | 0.115873311 |
| vns | GGB28934_SGB41635 | OFB_Centries | 0.111374073 |
| blast | GGB28946_SGB41652 | OFB_Centries | 0.427843225 |
| vns | GGB28946_SGB41652 | OFB_Centries | 0.422588900 |
| blast | GGB28949_SGB41655 | OFB_Centries | 0.443608116 |
| vns | GGB28949_SGB41655 | OFB_Centries | 0.908948179 |
| blast | GGB28949_SGB41656 | OFB_Centries | 0.282728234 |
| vns | GGB28949_SGB41656 | OFB_Centries | 0.770863949 |
| blast | GGB28950_SGB41657 | OFB_Centries | 0.030527036 |
| vns | GGB28950_SGB41657 | OFB_Centries | 0.336458904 |
| blast | GGB28951_SGB102295 | OFB_Centries | 0.925352156 |
| vns | GGB28951_SGB102295 | OFB_Centries | 0.158790397 |
| blast | GGB28951_SGB41658 | OFB_Centries | 0.847001351 |
| vns | GGB28951_SGB41658 | OFB_Centries | 0.920236623 |
| blast | GGB28954_SGB41662 | OFB_Centries | 0.515758297 |
| vns | GGB28954_SGB41662 | OFB_Centries | 0.800484955 |
| blast | GGB28956_SGB41665 | OFB_Centries | 0.163329713 |
| vns | GGB28956_SGB41665 | OFB_Centries | 0.153416502 |
| blast | GGB28960_SGB41669 | OFB_Centries | 0.155727292 |
| vns | GGB28960_SGB41669 | OFB_Centries | 0.155957479 |
| blast | GGB28967_SGB41678 | OFB_Centries | 0.179269529 |
| vns | GGB28967_SGB41678 | OFB_Centries | 0.315046519 |
| blast | GGB28991_SGB41705 | OFB_Centries | 0.228893882 |
| vns | GGB28991_SGB41705 | OFB_Centries | 0.279982342 |
| blast | GGB29002_SGB41718 | OFB_Centries | 0.002430867 |
| vns | GGB29002_SGB41718 | OFB_Centries | 0.113465587 |

| | | | |
|---|---|---|---|
| blast | GGB29003_SGB41719 | OFB_Centries | 0.4993986443 |
| vns | GGB29003_SGB41719 | OFB_Centries | 0.5839678864 |
| blast | GGB29011_SGB41731 | OFB_Centries | 0.916276033 |
| vns | GGB29011_SGB41731 | OFB_Centries | 0.3455992366 |
| blast | GGB29531_SGB42317 | OFB_Centries | 0.1450820268 |
| vns | GGB29531_SGB42317 | OFB_Centries | 0.1355699662 |
| blast | GGB29685_SGB42494 | OFB_Centries | 0.8645438528 |
| vns | GGB29685_SGB42494 | OFB_Centries | 0.340104263 |
| blast | GGB30141_SGB43066 | OFB_Centries | 0.9915548039 |
| vns | GGB30141_SGB43066 | OFB_Centries | 0.7601787897 |
| blast | GGB30286_SGB43248 | OFB_Centries | 0.0552616333 |
| vns | GGB30286_SGB43248 | OFB_Centries | 0.4546033673 |
| blast | GGB30303_SGB43268 | OFB_Centries | 0.3719368939 |
| vns | GGB30303_SGB43268 | OFB_Centries | 0.9447221743 |
| blast | GGB30413_SGB43452 | OFB_Centries | 0.9532769507 |
| vns | GGB30413_SGB43452 | OFB_Centries | 0.8796426046 |
| blast | GGB30454_SGB43514 | OFB_Centries | 0.3798589507 |
| vns | GGB30454_SGB43514 | OFB_Centries | 0.2001255913 |
| blast | GGB30455_SGB43519 | OFB_Centries | 0.3794677965 |
| vns | GGB30455_SGB43519 | OFB_Centries | 0.3023489219 |
| blast | GGB30461_SGB43527 | OFB_Centries | 0.3975036627 |
| vns | GGB30461_SGB43527 | OFB_Centries | 0.7710386558 |
| blast | GGB30461_SGB43530 | OFB_Centries | 0.395449 |
| vns | GGB30461_SGB43530 | OFB_Centries | 0.7038636932 |
| blast | GGB30463_SGB43537 | OFB_Centries | 0.2100542578 |
| vns | GGB30463_SGB43537 | OFB_Centries | 0.9853614792 |
| blast | GGB30473_SGB43557 | OFB_Centries | 0.0206421669 |
| vns | GGB30473_SGB43557 | OFB_Centries | 0.0562117431 |
| blast | GGB30475_SGB63182 | OFB_Centries | 0.2879653492 |
| vns | GGB30475_SGB63182 | OFB_Centries | 0.6608559306 |
| blast | GGB30861_SGB44083 | OFB_Centries | 0.9575523797 |
| vns | GGB30861_SGB44083 | OFB_Centries | 0.8334455291 |
| blast | GGB31312_SGB44628 | OFB_Centries | 0.7117271287 |
| vns | GGB31312_SGB44628 | OFB_Centries | 0.8841939433 |
| blast | GGB31438_SGB44768 | OFB_Centries | 0.9063572977 |
| vns | GGB31438_SGB44768 | OFB_Centries | 0.3189600867 |
| blast | GGB3171_SGB4185 | OFB_Centries | 0.0853457671 |
| vns | GGB3171_SGB4185 | OFB_Centries | 0.9185517801 |
| blast | GGB31823_SGB45199 | OFB_Centries | 0.1889777175 |
| vns | GGB31823_SGB45199 | OFB_Centries | 0.7028132582 |
| blast | GGB31853_SGB45233 | OFB_Centries | 0.4968427437 |
| vns | GGB31853_SGB45233 | OFB_Centries | 0.3962595415 |
| blast | GGB32371_SGB41694 | OFB_Centries | 0.3468118508 |
| vns | GGB32371_SGB41694 | OFB_Centries | 0.0250540925 |
| blast | GGB3793_SGB5158 | OFB_Centries | 0.9657904747 |
| vns | GGB3793_SGB5158 | OFB_Centries | 0.9751451519 |

| | | | |
|---|---|---|---|
| blast | GGB42598_SGB59794 | OFB_Centries | 0.5017400018 |
| vns | GGB42598_SGB59794 | OFB_Centries | 0.156668604 |
| blast | GGB45656_SGB63370 | OFB_Centries | 0.621044138 |
| vns | GGB45656_SGB63370 | OFB_Centries | 0.9633746015 |
| blast | GGB47127_SGB65054 | OFB_Centries | 0.4967451202 |
| vns | GGB47127_SGB65054 | OFB_Centries | 0.2574173127 |
| blast | GGB74395_SGB43521 | OFB_Centries | 0.1233397821 |
| vns | GGB74395_SGB43521 | OFB_Centries | 0.2040214986 |
| blast | GGB75053_SGB43494 | OFB_Centries | 0.4105959194 |
| vns | GGB75053_SGB43494 | OFB_Centries | 0.5718574569 |
| blast | GGB75109_SGB102238 | OFB_Centries | 0.6208422921 |
| vns | GGB75109_SGB102238 | OFB_Centries | 0.8718184088 |
| blast | GGB81440_SGB45230 | OFB_Centries | 0.0796107313 |
| vns | GGB81440_SGB45230 | OFB_Centries | 0.0981405799 |
| blast | Lachnospiraceae_bacterium | OFB_Centries | 0.4848161772 |
| vns | Lachnospiraceae_bacterium | OFB_Centries | 0.5378613321 |
| blast | Lachnospiraceae_bacterium_A2 | OFB_Centries | 0.420834896 |
| vns | Lachnospiraceae_bacterium_A2 | OFB_Centries | 0.3236483287 |
| blast | Lachnospiraceae_bacterium_MD308 | OFB_Centries | 0.2131194519 |
| vns | Lachnospiraceae_bacterium_MD308 | OFB_Centries | 0.0830669656 |
| blast | Lachnospiraceae_bacterium_MD329 | OFB_Centries | 0.0209761491 |
| vns | Lachnospiraceae_bacterium_MD329 | OFB_Centries | 0.6991611194 |
| blast | Lachnospiraceae_unclassified_SGB41414 | OFB_Centries | 0.142639053 |
| vns | Lachnospiraceae_unclassified_SGB41414 | OFB_Centries | 0.119876468 |
| blast | Lachnospiraceae_unclassified_SGB41418 | OFB_Centries | 0.9987579817 |
| vns | Lachnospiraceae_unclassified_SGB41418 | OFB_Centries | 0.7122365734 |
| blast | Lachnospiraceae_unclassified_SGB41424 | OFB_Centries | 0.0483863782 |
| vns | Lachnospiraceae_unclassified_SGB41424 | OFB_Centries | 0.0460673576 |
| blast | Lachnospiraceae_unclassified_SGB41589 | OFB_Centries | 0.1654249525 |
| vns | Lachnospiraceae_unclassified_SGB41589 | OFB_Centries | 0.2735277008 |
| blast | Lactobacillus_johnsonii | OFB_Centries | 0.1826784534 |
| vns | Lactobacillus_johnsonii | OFB_Centries | 0.6729515885 |
| blast | Muribaculaceae_bacterium | OFB_Centries | 0.8415252525 |
| vns | Muribaculaceae_bacterium | OFB_Centries | 0.8930529806 |
| blast | Neglectibacter_sp_X4 | OFB_Centries | 0.0847821517 |
| vns | Neglectibacter_sp_X4 | OFB_Centries | 0.140656264 |
| blast | Oscillospiraceae_bacterium | OFB_Centries | 0.7416725656 |
| vns | Oscillospiraceae_bacterium | OFB_Centries | 0.856188039 |
| blast | Oscillospiraceae_unclassified_SGB43502 | OFB_Centries | 0.0259412852 |
| vns | Oscillospiraceae_unclassified_SGB43502 | OFB_Centries | 0.0305351331 |
| blast | Oscillospiraceae_unclassified_SGB43505 | OFB_Centries | 0.4959230188 |
| vns | Oscillospiraceae_unclassified_SGB43505 | OFB_Centries | 0.6425337316 |
| blast | Parasutterella_excrementihominis | OFB_Centries | 0.5671048722 |
| vns | Parasutterella_excrementihominis | OFB_Centries | 0.7099929171 |
| blast | Richness (# observed features) | OFB_Centries | 0.2717524423 |
| vns | Richness (# observed features) | OFB_Centries | 0.6607230685 |

| | | | |
|---|---|---|---|
| blast | Romboutsia_ilealis | OFB_Centries | 0.0010997742 |
| vns | Romboutsia_ilealis | OFB_Centries | 0.0569970896 |
| blast | Schaedlerella_arabinosiphila | OFB_Centries | 0.9390244043 |
| vns | Schaedlerella_arabinosiphila | OFB_Centries | 0.9139116075 |
| blast | Shannon Index | OFB_Centries | 0.6947352216 |
| vns | Shannon Index | OFB_Centries | 0.9062773413 |
| blast | Turicibacter_sp_1E2 | OFB_Centries | 0.1991772312 |
| vns | Turicibacter_sp_1E2 | OFB_Centries | 0.1655749312 |
| blast | Acetatifactor_muris | OFB_Clatency | 0.7395220796 |
| vns | Acetatifactor_muris | OFB_Clatency | 0.7528971403 |
| blast | Acetatifactor_SGB41546 | OFB_Clatency | 0.8402457971 |
| vns | Acetatifactor_SGB41546 | OFB_Clatency | 0.6630009771 |
| blast | Acutalibacter_muris | OFB_Clatency | 0.4221546334 |
| vns | Acutalibacter_muris | OFB_Clatency | 0.8105784495 |
| blast | Acutalibacter_sp_1XD8_36 | OFB_Clatency | 0.6605798863 |
| vns | Acutalibacter_sp_1XD8_36 | OFB_Clatency | 0.7091849336 |
| blast | Adlercreutzia_caecimuris | OFB_Clatency | 0.9896089529 |
| vns | Adlercreutzia_caecimuris | OFB_Clatency | 0.6765426913 |
| blast | Adlercreutzia_mucosicola | OFB_Clatency | 0.2718339856 |
| vns | Adlercreutzia_mucosicola | OFB_Clatency | 0.2498256574 |
| blast | Adlercreutzia_muris | OFB_Clatency | 0.6704217743 |
| vns | Adlercreutzia_muris | OFB_Clatency | 0.5958277454 |
| blast | Akkermansia_muciniphila | OFB_Clatency | 0.1698125404 |
| vns | Akkermansia_muciniphila | OFB_Clatency | 0.8643625099 |
| blast | Alistipes_sp_DSM_112343 | OFB_Clatency | 0.3725159774 |
| vns | Alistipes_sp_DSM_112343 | OFB_Clatency | 0.8326983074 |
| blast | Anaerotruncus_sp_1XD42_93 | OFB_Clatency | 0.5202714535 |
| vns | Anaerotruncus_sp_1XD42_93 | OFB_Clatency | 0.6174107217 |
| blast | Bacteria_unclassified_SGB102200 | OFB_Clatency | 0.4742692127 |
| vns | Bacteria_unclassified_SGB102200 | OFB_Clatency | 0.7641667885 |
| blast | Bacteria_unclassified_SGB41677 | OFB_Clatency | 0.4441438231 |
| vns | Bacteria_unclassified_SGB41677 | OFB_Clatency | 0.1699894243 |
| blast | Bacteria_unclassified_SGB43546 | OFB_Clatency | 0.5863583659 |
| vns | Bacteria_unclassified_SGB43546 | OFB_Clatency | 0.7890099125 |
| blast | bacterium_1XD42_54 | OFB_Clatency | 0.0144245959 |
| vns | bacterium_1XD42_54 | OFB_Clatency | 0.2345775748 |
| blast | bacterium_1XD42_76 | OFB_Clatency | 0.6000316038 |
| vns | bacterium_1XD42_76 | OFB_Clatency | 0.6429924539 |
| blast | bacterium_1xD8_48 | OFB_Clatency | 0.8997089453 |
| vns | bacterium_1xD8_48 | OFB_Clatency | 0.6968168809 |
| blast | Bacteroides_thetaiotaomicron | OFB_Clatency | 0.9602014777 |
| vns | Bacteroides_thetaiotaomicron | OFB_Clatency | 0.6343063159 |
| blast | Berger Parker Index | OFB_Clatency | 0.5457417121 |
| vns | Berger Parker Index | OFB_Clatency | 0.2337330546 |
| blast | Bifidobacterium_pseudolongum | OFB_Clatency | 0.4280103985 |
| vns | Bifidobacterium_pseudolongum | OFB_Clatency | 0.3477990653 |

| | | | |
|---|---|---|---|
| blast | Clostridia_bacterium | OFB_Clatency | 0.3745590434 |
| vns | Clostridia_bacterium | OFB_Clatency | 0.7249401071 |
| blast | Clostridiaceae_bacterium | OFB_Clatency | 0.9871697806 |
| vns | Clostridiaceae_bacterium | OFB_Clatency | 0.2419370425 |
| blast | Clostridiaceae_unclassified_SGB41663 | OFB_Clatency | 0.780382224 |
| vns | Clostridiaceae_unclassified_SGB41663 | OFB_Clatency | 0.719154368 |
| blast | Clostridiales_bacterium | OFB_Clatency | 0.0023319564 |
| vns | Clostridiales_bacterium | OFB_Clatency | 0.2964785018 |
| blast | Clostridium_cocleatum | OFB_Clatency | 0.0115678325 |
| vns | Clostridium_cocleatum | OFB_Clatency | 0.1960355146 |
| blast | Coriobacteriaceae_bacterium | OFB_Clatency | 0.3724483697 |
| vns | Coriobacteriaceae_bacterium | OFB_Clatency | 0.953919137 |
| blast | Dorea_sp_5_2 | OFB_Clatency | 0.5884919176 |
| vns | Dorea_sp_5_2 | OFB_Clatency | 0.4214161244 |
| blast | Dubosiella_newyorkensis | OFB_Clatency | 0.1169987394 |
| vns | Dubosiella_newyorkensis | OFB_Clatency | 0.799186063 |
| blast | Erysipelotrichales_bacterium | OFB_Clatency | 0.8970006656 |
| vns | Erysipelotrichales_bacterium | OFB_Clatency | 0.3284081083 |
| blast | Eubacteriaceae_bacterium | OFB_Clatency | 0.0271233687 |
| vns | Eubacteriaceae_bacterium | OFB_Clatency | 0.2383186135 |
| blast | Eubacteriaceae_unclassified_SGB94922 | OFB_Clatency | 0.4594880799 |
| vns | Eubacteriaceae_unclassified_SGB94922 | OFB_Clatency | 0.3726201484 |
| blast | GGB20149_SGB29430 | OFB_Clatency | 0.6280700484 |
| vns | GGB20149_SGB29430 | OFB_Clatency | 0.6299827926 |
| blast | GGB22635_SGB63107 | OFB_Clatency | 0.4117103362 |
| vns | GGB22635_SGB63107 | OFB_Clatency | 0.8871521383 |
| blast | GGB25041_SGB36960 | OFB_Clatency | 0.0824966676 |
| vns | GGB25041_SGB36960 | OFB_Clatency | 0.2101871969 |
| blast | GGB27876_SGB40310 | OFB_Clatency | 0.164666135 |
| vns | GGB27876_SGB40310 | OFB_Clatency | 0.7672351949 |
| blast | GGB27878_SGB40312 | OFB_Clatency | 0.2133541578 |
| vns | GGB27878_SGB40312 | OFB_Clatency | 0.1474794298 |
| blast | GGB27918_SGB40356 | OFB_Clatency | 0.9587581408 |
| vns | GGB27918_SGB40356 | OFB_Clatency | 0.8856989297 |
| blast | GGB28382_SGB40962 | OFB_Clatency | 0.5193113079 |
| vns | GGB28382_SGB40962 | OFB_Clatency | 0.8550578348 |
| blast | GGB28399_SGB40980 | OFB_Clatency | 0.2894655025 |
| vns | GGB28399_SGB40980 | OFB_Clatency | 0.287117034 |
| blast | GGB28411_SGB40993 | OFB_Clatency | 0.2018022626 |
| vns | GGB28411_SGB40993 | OFB_Clatency | 0.1468484741 |
| blast | GGB28415_SGB40997 | OFB_Clatency | 0.0267267446 |
| vns | GGB28415_SGB40997 | OFB_Clatency | 0.9490516966 |
| blast | GGB28430_SGB41013 | OFB_Clatency | 0.9120709472 |
| vns | GGB28430_SGB41013 | OFB_Clatency | 0.4662504826 |
| blast | GGB28439_SGB41022 | OFB_Clatency | 0.3804444035 |
| vns | GGB28439_SGB41022 | OFB_Clatency | 0.7387095529 |

| | | | |
|---|---|---|---|
| blast | GGB28778_SGB41431 | OFB_Clatency | 0.804698941 |
| vns | GGB28778_SGB41431 | OFB_Clatency | 0.218453243 |
| blast | GGB28784_SGB41437 | OFB_Clatency | 0.020726621 |
| vns | GGB28784_SGB41437 | OFB_Clatency | 0.254456997 |
| blast | GGB28792_SGB41445 | OFB_Clatency | 0.338577775 |
| vns | GGB28792_SGB41445 | OFB_Clatency | 0.761244426 |
| blast | GGB28798_SGB41451 | OFB_Clatency | 0.164022630 |
| vns | GGB28798_SGB41451 | OFB_Clatency | 0.248915824 |
| blast | GGB28802_SGB41455 | OFB_Clatency | 0.457898625 |
| vns | GGB28802_SGB41455 | OFB_Clatency | 0.342689298 |
| blast | GGB28818_SGB41473 | OFB_Clatency | 0.729209510 |
| vns | GGB28818_SGB41473 | OFB_Clatency | 0.569409948 |
| blast | GGB28828_SGB41484 | OFB_Clatency | 0.298696993 |
| vns | GGB28828_SGB41484 | OFB_Clatency | 0.106706535 |
| blast | GGB28851_SGB41518 | OFB_Clatency | 0.852552179 |
| vns | GGB28851_SGB41518 | OFB_Clatency | 0.578308429 |
| blast | GGB28859_SGB41528 | OFB_Clatency | 0.205749329 |
| vns | GGB28859_SGB41528 | OFB_Clatency | 0.595366745 |
| blast | GGB28864_SGB41535 | OFB_Clatency | 0.785465248 |
| vns | GGB28864_SGB41535 | OFB_Clatency | 0.964658985 |
| blast | GGB28869_SGB41543 | OFB_Clatency | 0.347931625 |
| vns | GGB28869_SGB41543 | OFB_Clatency | 0.381477475 |
| blast | GGB28883_SGB41564 | OFB_Clatency | 0.594877248 |
| vns | GGB28883_SGB41564 | OFB_Clatency | 0.060524458 |
| blast | GGB28892_SGB41573 | OFB_Clatency | 0.651227207 |
| vns | GGB28892_SGB41573 | OFB_Clatency | 0.667618882 |
| blast | GGB28893_SGB41574 | OFB_Clatency | 0.165594509 |
| vns | GGB28893_SGB41574 | OFB_Clatency | 0.182699376 |
| blast | GGB28898_SGB41580 | OFB_Clatency | 0.304654993 |
| vns | GGB28898_SGB41580 | OFB_Clatency | 0.082893330 |
| blast | GGB28904_SGB41597 | OFB_Clatency | 0.172906397 |
| vns | GGB28904_SGB41597 | OFB_Clatency | 0.056067199 |
| blast | GGB28916_SGB41612 | OFB_Clatency | 0.801432609 |
| vns | GGB28916_SGB41612 | OFB_Clatency | 0.853813296 |
| blast | GGB28924_SGB41621 | OFB_Clatency | 0.796481662 |
| vns | GGB28924_SGB41621 | OFB_Clatency | 0.278658378 |
| blast | GGB28926_SGB41624 | OFB_Clatency | 0.306780834 |
| vns | GGB28926_SGB41624 | OFB_Clatency | 0.470765265 |
| blast | GGB28927_SGB41625 | OFB_Clatency | 0.193720805 |
| vns | GGB28927_SGB41625 | OFB_Clatency | 0.418243373 |
| blast | GGB28934_SGB41635 | OFB_Clatency | 0.115873311 |
| vns | GGB28934_SGB41635 | OFB_Clatency | 0.111374073 |
| blast | GGB28946_SGB41652 | OFB_Clatency | 0.427843225 |
| vns | GGB28946_SGB41652 | OFB_Clatency | 0.422588900 |
| blast | GGB28949_SGB41655 | OFB_Clatency | 0.443608116 |
| vns | GGB28949_SGB41655 | OFB_Clatency | 0.908948179 |

| | | | |
|---|---|---|---|
| blast | GGB28949_SGB41656 | OFB_Clatency | 0.282728234 |
| vns | GGB28949_SGB41656 | OFB_Clatency | 0.770863949 |
| blast | GGB28950_SGB41657 | OFB_Clatency | 0.030527036 |
| vns | GGB28950_SGB41657 | OFB_Clatency | 0.336458904 |
| blast | GGB28951_SGB102295 | OFB_Clatency | 0.925352156 |
| vns | GGB28951_SGB102295 | OFB_Clatency | 0.158790397 |
| blast | GGB28951_SGB41658 | OFB_Clatency | 0.847001351 |
| vns | GGB28951_SGB41658 | OFB_Clatency | 0.920236623 |
| blast | GGB28954_SGB41662 | OFB_Clatency | 0.515758297 |
| vns | GGB28954_SGB41662 | OFB_Clatency | 0.800484955 |
| blast | GGB28956_SGB41665 | OFB_Clatency | 0.163329713 |
| vns | GGB28956_SGB41665 | OFB_Clatency | 0.153416502 |
| blast | GGB28960_SGB41669 | OFB_Clatency | 0.155727292 |
| vns | GGB28960_SGB41669 | OFB_Clatency | 0.155957479 |
| blast | GGB28967_SGB41678 | OFB_Clatency | 0.179269529 |
| vns | GGB28967_SGB41678 | OFB_Clatency | 0.315046519 |
| blast | GGB28991_SGB41705 | OFB_Clatency | 0.228893882 |
| vns | GGB28991_SGB41705 | OFB_Clatency | 0.279982342 |
| blast | GGB29002_SGB41718 | OFB_Clatency | 0.002430867 |
| vns | GGB29002_SGB41718 | OFB_Clatency | 0.113465587 |
| blast | GGB29003_SGB41719 | OFB_Clatency | 0.499398644 |
| vns | GGB29003_SGB41719 | OFB_Clatency | 0.583967886 |
| blast | GGB29011_SGB41731 | OFB_Clatency | 0.916276033 |
| vns | GGB29011_SGB41731 | OFB_Clatency | 0.345599236 |
| blast | GGB29531_SGB42317 | OFB_Clatency | 0.145082026 |
| vns | GGB29531_SGB42317 | OFB_Clatency | 0.135569966 |
| blast | GGB29685_SGB42494 | OFB_Clatency | 0.864543852 |
| vns | GGB29685_SGB42494 | OFB_Clatency | 0.340104263 |
| blast | GGB30141_SGB43066 | OFB_Clatency | 0.991554803 |
| vns | GGB30141_SGB43066 | OFB_Clatency | 0.760178789 |
| blast | GGB30286_SGB43248 | OFB_Clatency | 0.055261633 |
| vns | GGB30286_SGB43248 | OFB_Clatency | 0.454603367 |
| blast | GGB30303_SGB43268 | OFB_Clatency | 0.371936893 |
| vns | GGB30303_SGB43268 | OFB_Clatency | 0.944722174 |
| blast | GGB30413_SGB43452 | OFB_Clatency | 0.953276950 |
| vns | GGB30413_SGB43452 | OFB_Clatency | 0.879642604 |
| blast | GGB30454_SGB43514 | OFB_Clatency | 0.379858950 |
| vns | GGB30454_SGB43514 | OFB_Clatency | 0.200125591 |
| blast | GGB30455_SGB43519 | OFB_Clatency | 0.379467796 |
| vns | GGB30455_SGB43519 | OFB_Clatency | 0.302348921 |
| blast | GGB30461_SGB43527 | OFB_Clatency | 0.397503662 |
| vns | GGB30461_SGB43527 | OFB_Clatency | 0.771038655 |
| blast | GGB30461_SGB43530 | OFB_Clatency | 0.395449 |
| vns | GGB30461_SGB43530 | OFB_Clatency | 0.703863693 |
| blast | GGB30463_SGB43537 | OFB_Clatency | 0.210054257 |
| vns | GGB30463_SGB43537 | OFB_Clatency | 0.985361479 |

| | | | |
|---|---|---|---|
| blast | GGB30473_SGB43557 | OFB_Clatency | 0.0206421669 |
| vns | GGB30473_SGB43557 | OFB_Clatency | 0.0562117431 |
| blast | GGB30475_SGB63182 | OFB_Clatency | 0.2879653492 |
| vns | GGB30475_SGB63182 | OFB_Clatency | 0.6608559306 |
| blast | GGB30861_SGB44083 | OFB_Clatency | 0.9575523797 |
| vns | GGB30861_SGB44083 | OFB_Clatency | 0.8334455291 |
| blast | GGB31312_SGB44628 | OFB_Clatency | 0.7117271287 |
| vns | GGB31312_SGB44628 | OFB_Clatency | 0.8841939433 |
| blast | GGB31438_SGB44768 | OFB_Clatency | 0.9063572977 |
| vns | GGB31438_SGB44768 | OFB_Clatency | 0.3189600867 |
| blast | GGB3171_SGB4185 | OFB_Clatency | 0.0853457671 |
| vns | GGB3171_SGB4185 | OFB_Clatency | 0.9185517801 |
| blast | GGB31823_SGB45199 | OFB_Clatency | 0.1889777175 |
| vns | GGB31823_SGB45199 | OFB_Clatency | 0.7028132582 |
| blast | GGB31853_SGB45233 | OFB_Clatency | 0.4968427437 |
| vns | GGB31853_SGB45233 | OFB_Clatency | 0.3962595415 |
| blast | GGB32371_SGB41694 | OFB_Clatency | 0.3468118508 |
| vns | GGB32371_SGB41694 | OFB_Clatency | 0.0250540925 |
| blast | GGB3793_SGB5158 | OFB_Clatency | 0.9657904747 |
| vns | GGB3793_SGB5158 | OFB_Clatency | 0.9751451519 |
| blast | GGB42598_SGB59794 | OFB_Clatency | 0.5017400018 |
| vns | GGB42598_SGB59794 | OFB_Clatency | 0.156668604 |
| blast | GGB45656_SGB63370 | OFB_Clatency | 0.621044138 |
| vns | GGB45656_SGB63370 | OFB_Clatency | 0.9633746015 |
| blast | GGB47127_SGB65054 | OFB_Clatency | 0.4967451202 |
| vns | GGB47127_SGB65054 | OFB_Clatency | 0.2574173127 |
| blast | GGB74395_SGB43521 | OFB_Clatency | 0.1233397821 |
| vns | GGB74395_SGB43521 | OFB_Clatency | 0.2040214986 |
| blast | GGB75053_SGB43494 | OFB_Clatency | 0.4105959194 |
| vns | GGB75053_SGB43494 | OFB_Clatency | 0.5718574569 |
| blast | GGB75109_SGB102238 | OFB_Clatency | 0.6208422921 |
| vns | GGB75109_SGB102238 | OFB_Clatency | 0.8718184088 |
| blast | GGB81440_SGB45230 | OFB_Clatency | 0.0796107313 |
| vns | GGB81440_SGB45230 | OFB_Clatency | 0.0981405799 |
| blast | Lachnospiraceae_bacterium | OFB_Clatency | 0.4848161772 |
| vns | Lachnospiraceae_bacterium | OFB_Clatency | 0.5378613321 |
| blast | Lachnospiraceae_bacterium_A2 | OFB_Clatency | 0.420834896 |
| vns | Lachnospiraceae_bacterium_A2 | OFB_Clatency | 0.3236483287 |
| blast | Lachnospiraceae_bacterium_MD308 | OFB_Clatency | 0.2131194519 |
| vns | Lachnospiraceae_bacterium_MD308 | OFB_Clatency | 0.0830669656 |
| blast | Lachnospiraceae_bacterium_MD329 | OFB_Clatency | 0.0209761491 |
| vns | Lachnospiraceae_bacterium_MD329 | OFB_Clatency | 0.6991611194 |
| blast | Lachnospiraceae_unclassified_SGB41414 | OFB_Clatency | 0.142639053 |
| vns | Lachnospiraceae_unclassified_SGB41414 | OFB_Clatency | 0.119876468 |
| blast | Lachnospiraceae_unclassified_SGB41418 | OFB_Clatency | 0.9987579817 |
| vns | Lachnospiraceae_unclassified_SGB41418 | OFB_Clatency | 0.7122365734 |

| | | | |
|---|---|---|---|
| blast | Lachnospiraceae_unclassified_SGB41424 | OFB_Clatency | 0.0483863782 |
| vns | Lachnospiraceae_unclassified_SGB41424 | OFB_Clatency | 0.0460673576 |
| blast | Lachnospiraceae_unclassified_SGB41589 | OFB_Clatency | 0.1654249525 |
| vns | Lachnospiraceae_unclassified_SGB41589 | OFB_Clatency | 0.2735277008 |
| blast | Lactobacillus_johnsonii | OFB_Clatency | 0.1826784534 |
| vns | Lactobacillus_johnsonii | OFB_Clatency | 0.6729515885 |
| blast | Muribaculaceae_bacterium | OFB_Clatency | 0.8415252525 |
| vns | Muribaculaceae_bacterium | OFB_Clatency | 0.8930529806 |
| blast | Neglectibacter_sp_X4 | OFB_Clatency | 0.0847821517 |
| vns | Neglectibacter_sp_X4 | OFB_Clatency | 0.140656264 |
| blast | Oscillospiraceae_bacterium | OFB_Clatency | 0.7416725656 |
| vns | Oscillospiraceae_bacterium | OFB_Clatency | 0.856188039 |
| blast | Oscillospiraceae_unclassified_SGB43502 | OFB_Clatency | 0.0259412852 |
| vns | Oscillospiraceae_unclassified_SGB43502 | OFB_Clatency | 0.0305351331 |
| blast | Oscillospiraceae_unclassified_SGB43505 | OFB_Clatency | 0.4959230188 |
| vns | Oscillospiraceae_unclassified_SGB43505 | OFB_Clatency | 0.6425337316 |
| blast | Parasutterella_excrementihominis | OFB_Clatency | 0.5671048722 |
| vns | Parasutterella_excrementihominis | OFB_Clatency | 0.7099929171 |
| blast | Richness (# observed features) | OFB_Clatency | 0.2717524423 |
| vns | Richness (# observed features) | OFB_Clatency | 0.6607230685 |
| blast | Romboutsia_ilealis | OFB_Clatency | 0.0010997742 |
| vns | Romboutsia_ilealis | OFB_Clatency | 0.0569970896 |
| blast | Schaedlerella_arabinosiphila | OFB_Clatency | 0.9390244043 |
| vns | Schaedlerella_arabinosiphila | OFB_Clatency | 0.9139116075 |
| blast | Shannon Index | OFB_Clatency | 0.6947352216 |
| vns | Shannon Index | OFB_Clatency | 0.9062773413 |
| blast | Turicibacter_sp_1E2 | OFB_Clatency | 0.1991772312 |
| vns | Turicibacter_sp_1E2 | OFB_Clatency | 0.1655749312 |
| blast | Acetatifactor_muris | OFB_Ctime | 0.7395220796 |
| vns | Acetatifactor_muris | OFB_Ctime | 0.7528971403 |
| blast | Acetatifactor_SGB41546 | OFB_Ctime | 0.8402457971 |
| vns | Acetatifactor_SGB41546 | OFB_Ctime | 0.6630009771 |
| blast | Acutalibacter_muris | OFB_Ctime | 0.4221546334 |
| vns | Acutalibacter_muris | OFB_Ctime | 0.8105784495 |
| blast | Acutalibacter_sp_1XD8_36 | OFB_Ctime | 0.6605798863 |
| vns | Acutalibacter_sp_1XD8_36 | OFB_Ctime | 0.7091849336 |
| blast | Adlercreutzia_caecimuris | OFB_Ctime | 0.9896089529 |
| vns | Adlercreutzia_caecimuris | OFB_Ctime | 0.6765426913 |
| blast | Adlercreutzia_mucosicola | OFB_Ctime | 0.2718339856 |
| vns | Adlercreutzia_mucosicola | OFB_Ctime | 0.2498256574 |
| blast | Adlercreutzia_muris | OFB_Ctime | 0.6704217743 |
| vns | Adlercreutzia_muris | OFB_Ctime | 0.5958277454 |
| blast | Akkermansia_muciniphila | OFB_Ctime | 0.1698125404 |
| vns | Akkermansia_muciniphila | OFB_Ctime | 0.8643625099 |
| blast | Alistipes_sp_DSM_112343 | OFB_Ctime | 0.3725159774 |
| vns | Alistipes_sp_DSM_112343 | OFB_Ctime | 0.8326983074 |

| | | | |
|---|---|---|---|
| blast | Anaerotruncus_sp_1XD42_93 | OFB_Ctime | 0.5202714535 |
| vns | Anaerotruncus_sp_1XD42_93 | OFB_Ctime | 0.6174107217 |
| blast | Bacteria_unclassified_SGB102200 | OFB_Ctime | 0.4742692127 |
| vns | Bacteria_unclassified_SGB102200 | OFB_Ctime | 0.7641667885 |
| blast | Bacteria_unclassified_SGB41677 | OFB_Ctime | 0.4441438231 |
| vns | Bacteria_unclassified_SGB41677 | OFB_Ctime | 0.1699894243 |
| blast | Bacteria_unclassified_SGB43546 | OFB_Ctime | 0.5863583659 |
| vns | Bacteria_unclassified_SGB43546 | OFB_Ctime | 0.7890099125 |
| blast | bacterium_1XD42_54 | OFB_Ctime | 0.0144245959 |
| vns | bacterium_1XD42_54 | OFB_Ctime | 0.2345775748 |
| blast | bacterium_1XD42_76 | OFB_Ctime | 0.6000316038 |
| vns | bacterium_1XD42_76 | OFB_Ctime | 0.6429924539 |
| blast | bacterium_1xD8_48 | OFB_Ctime | 0.8997089453 |
| vns | bacterium_1xD8_48 | OFB_Ctime | 0.6968168809 |
| blast | Bacteroides_thetaiotaomicron | OFB_Ctime | 0.9602014777 |
| vns | Bacteroides_thetaiotaomicron | OFB_Ctime | 0.6343063159 |
| blast | Berger Parker Index | OFB_Ctime | 0.5457417127 |
| vns | Berger Parker Index | OFB_Ctime | 0.2337330546 |
| blast | Bifidobacterium_pseudolongum | OFB_Ctime | 0.4280103985 |
| vns | Bifidobacterium_pseudolongum | OFB_Ctime | 0.3477990653 |
| blast | Clostridia_bacterium | OFB_Ctime | 0.3745590434 |
| vns | Clostridia_bacterium | OFB_Ctime | 0.7249401071 |
| blast | Clostridiaceae_bacterium | OFB_Ctime | 0.9871697806 |
| vns | Clostridiaceae_bacterium | OFB_Ctime | 0.2419370425 |
| blast | Clostridiaceae_unclassified_SGB41663 | OFB_Ctime | 0.780382224 |
| vns | Clostridiaceae_unclassified_SGB41663 | OFB_Ctime | 0.719154368 |
| blast | Clostridiales_bacterium | OFB_Ctime | 0.0023319564 |
| vns | Clostridiales_bacterium | OFB_Ctime | 0.2964785018 |
| blast | Clostridium_cocleatum | OFB_Ctime | 0.0115678325 |
| vns | Clostridium_cocleatum | OFB_Ctime | 0.1960355146 |
| blast | Coriobacteriaceae_bacterium | OFB_Ctime | 0.3724483697 |
| vns | Coriobacteriaceae_bacterium | OFB_Ctime | 0.953919137 |
| blast | Dorea_sp_5_2 | OFB_Ctime | 0.5884919176 |
| vns | Dorea_sp_5_2 | OFB_Ctime | 0.4214161244 |
| blast | Dubosiella_newyorkensis | OFB_Ctime | 0.1169987394 |
| vns | Dubosiella_newyorkensis | OFB_Ctime | 0.799186063 |
| blast | Erysipelotrichales_bacterium | OFB_Ctime | 0.8970006656 |
| vns | Erysipelotrichales_bacterium | OFB_Ctime | 0.3284081083 |
| blast | Eubacteriaceae_bacterium | OFB_Ctime | 0.0271233687 |
| vns | Eubacteriaceae_bacterium | OFB_Ctime | 0.2383186135 |
| blast | Eubacteriaceae_unclassified_SGB94922 | OFB_Ctime | 0.4594880799 |
| vns | Eubacteriaceae_unclassified_SGB94922 | OFB_Ctime | 0.3726201484 |
| blast | GGB20149_SGB29430 | OFB_Ctime | 0.6280700484 |
| vns | GGB20149_SGB29430 | OFB_Ctime | 0.6299827926 |
| blast | GGB22635_SGB63107 | OFB_Ctime | 0.4117103362 |
| vns | GGB22635_SGB63107 | OFB_Ctime | 0.8871521383 |

| | | | |
|---|---|---|---|
| blast | GGB25041_SGB36960 | OFB_Ctime | 0.082496667 |
| vns | GGB25041_SGB36960 | OFB_Ctime | 0.210187196 |
| blast | GGB27876_SGB40310 | OFB_Ctime | 0.164666135 |
| vns | GGB27876_SGB40310 | OFB_Ctime | 0.767235194 |
| blast | GGB27878_SGB40312 | OFB_Ctime | 0.213354157 |
| vns | GGB27878_SGB40312 | OFB_Ctime | 0.147479429 |
| blast | GGB27918_SGB40356 | OFB_Ctime | 0.958758140 |
| vns | GGB27918_SGB40356 | OFB_Ctime | 0.885698929 |
| blast | GGB28382_SGB40962 | OFB_Ctime | 0.519311307 |
| vns | GGB28382_SGB40962 | OFB_Ctime | 0.855057834 |
| blast | GGB28399_SGB40980 | OFB_Ctime | 0.289465502 |
| vns | GGB28399_SGB40980 | OFB_Ctime | 0.287117034 |
| blast | GGB28411_SGB40993 | OFB_Ctime | 0.201802262 |
| vns | GGB28411_SGB40993 | OFB_Ctime | 0.146848474 |
| blast | GGB28415_SGB40997 | OFB_Ctime | 0.026726744 |
| vns | GGB28415_SGB40997 | OFB_Ctime | 0.949051696 |
| blast | GGB28430_SGB41013 | OFB_Ctime | 0.912070947 |
| vns | GGB28430_SGB41013 | OFB_Ctime | 0.466250482 |
| blast | GGB28439_SGB41022 | OFB_Ctime | 0.380444403 |
| vns | GGB28439_SGB41022 | OFB_Ctime | 0.738709552 |
| blast | GGB28778_SGB41431 | OFB_Ctime | 0.804698941 |
| vns | GGB28778_SGB41431 | OFB_Ctime | 0.218453243 |
| blast | GGB28784_SGB41437 | OFB_Ctime | 0.020726621 |
| vns | GGB28784_SGB41437 | OFB_Ctime | 0.254456997 |
| blast | GGB28792_SGB41445 | OFB_Ctime | 0.338577775 |
| vns | GGB28792_SGB41445 | OFB_Ctime | 0.761244426 |
| blast | GGB28798_SGB41451 | OFB_Ctime | 0.164022630 |
| vns | GGB28798_SGB41451 | OFB_Ctime | 0.248915824 |
| blast | GGB28802_SGB41455 | OFB_Ctime | 0.457898625 |
| vns | GGB28802_SGB41455 | OFB_Ctime | 0.342689298 |
| blast | GGB28818_SGB41473 | OFB_Ctime | 0.729209510 |
| vns | GGB28818_SGB41473 | OFB_Ctime | 0.569409948 |
| blast | GGB28828_SGB41484 | OFB_Ctime | 0.298696993 |
| vns | GGB28828_SGB41484 | OFB_Ctime | 0.106706535 |
| blast | GGB28851_SGB41518 | OFB_Ctime | 0.852552179 |
| vns | GGB28851_SGB41518 | OFB_Ctime | 0.578308429 |
| blast | GGB28859_SGB41528 | OFB_Ctime | 0.205749329 |
| vns | GGB28859_SGB41528 | OFB_Ctime | 0.595366745 |
| blast | GGB28864_SGB41535 | OFB_Ctime | 0.785465248 |
| vns | GGB28864_SGB41535 | OFB_Ctime | 0.964658985 |
| blast | GGB28869_SGB41543 | OFB_Ctime | 0.347931625 |
| vns | GGB28869_SGB41543 | OFB_Ctime | 0.381477475 |
| blast | GGB28883_SGB41564 | OFB_Ctime | 0.594877248 |
| vns | GGB28883_SGB41564 | OFB_Ctime | 0.060524458 |
| blast | GGB28892_SGB41573 | OFB_Ctime | 0.651227207 |
| vns | GGB28892_SGB41573 | OFB_Ctime | 0.667618882 |

| | | | |
|---|---|---|---|
| blast | GGB28893_SGB41574 | OFB_Ctime | 0.165594509 |
| vns | GGB28893_SGB41574 | OFB_Ctime | 0.182699376 |
| blast | GGB28898_SGB41580 | OFB_Ctime | 0.304654993 |
| vns | GGB28898_SGB41580 | OFB_Ctime | 0.082893330 |
| blast | GGB28904_SGB41597 | OFB_Ctime | 0.172906397 |
| vns | GGB28904_SGB41597 | OFB_Ctime | 0.056067199 |
| blast | GGB28916_SGB41612 | OFB_Ctime | 0.801432609 |
| vns | GGB28916_SGB41612 | OFB_Ctime | 0.853813296 |
| blast | GGB28924_SGB41621 | OFB_Ctime | 0.796481662 |
| vns | GGB28924_SGB41621 | OFB_Ctime | 0.278658378 |
| blast | GGB28926_SGB41624 | OFB_Ctime | 0.306780834 |
| vns | GGB28926_SGB41624 | OFB_Ctime | 0.470765265 |
| blast | GGB28927_SGB41625 | OFB_Ctime | 0.193720805 |
| vns | GGB28927_SGB41625 | OFB_Ctime | 0.418243373 |
| blast | GGB28934_SGB41635 | OFB_Ctime | 0.115873311 |
| vns | GGB28934_SGB41635 | OFB_Ctime | 0.111374073 |
| blast | GGB28946_SGB41652 | OFB_Ctime | 0.427843225 |
| vns | GGB28946_SGB41652 | OFB_Ctime | 0.422588900 |
| blast | GGB28949_SGB41655 | OFB_Ctime | 0.443608116 |
| vns | GGB28949_SGB41655 | OFB_Ctime | 0.908948179 |
| blast | GGB28949_SGB41656 | OFB_Ctime | 0.282728234 |
| vns | GGB28949_SGB41656 | OFB_Ctime | 0.770863949 |
| blast | GGB28950_SGB41657 | OFB_Ctime | 0.030527036 |
| vns | GGB28950_SGB41657 | OFB_Ctime | 0.336458904 |
| blast | GGB28951_SGB102295 | OFB_Ctime | 0.925352156 |
| vns | GGB28951_SGB102295 | OFB_Ctime | 0.158790397 |
| blast | GGB28951_SGB41658 | OFB_Ctime | 0.847001351 |
| vns | GGB28951_SGB41658 | OFB_Ctime | 0.920236623 |
| blast | GGB28954_SGB41662 | OFB_Ctime | 0.515758297 |
| vns | GGB28954_SGB41662 | OFB_Ctime | 0.800484955 |
| blast | GGB28956_SGB41665 | OFB_Ctime | 0.163329713 |
| vns | GGB28956_SGB41665 | OFB_Ctime | 0.153416502 |
| blast | GGB28960_SGB41669 | OFB_Ctime | 0.155727292 |
| vns | GGB28960_SGB41669 | OFB_Ctime | 0.155957479 |
| blast | GGB28967_SGB41678 | OFB_Ctime | 0.179269529 |
| vns | GGB28967_SGB41678 | OFB_Ctime | 0.315046519 |
| blast | GGB28991_SGB41705 | OFB_Ctime | 0.228893882 |
| vns | GGB28991_SGB41705 | OFB_Ctime | 0.279982342 |
| blast | GGB29002_SGB41718 | OFB_Ctime | 0.002430867 |
| vns | GGB29002_SGB41718 | OFB_Ctime | 0.113465587 |
| blast | GGB29003_SGB41719 | OFB_Ctime | 0.499398644 |
| vns | GGB29003_SGB41719 | OFB_Ctime | 0.583967886 |
| blast | GGB29011_SGB41731 | OFB_Ctime | 0.916276033 |
| vns | GGB29011_SGB41731 | OFB_Ctime | 0.345599236 |
| blast | GGB29531_SGB42317 | OFB_Ctime | 0.145082026 |
| vns | GGB29531_SGB42317 | OFB_Ctime | 0.135569966 |

| | | | |
|---|---|---|---|
| blast | GGB29685_SGB42494 | OFB_Ctime | 0.8645438528 |
| vns | GGB29685_SGB42494 | OFB_Ctime | 0.340104263 |
| blast | GGB30141_SGB43066 | OFB_Ctime | 0.9915548039 |
| vns | GGB30141_SGB43066 | OFB_Ctime | 0.7601787897 |
| blast | GGB30286_SGB43248 | OFB_Ctime | 0.0552616333 |
| vns | GGB30286_SGB43248 | OFB_Ctime | 0.4546033673 |
| blast | GGB30303_SGB43268 | OFB_Ctime | 0.3719368939 |
| vns | GGB30303_SGB43268 | OFB_Ctime | 0.9447221743 |
| blast | GGB30413_SGB43452 | OFB_Ctime | 0.9532769507 |
| vns | GGB30413_SGB43452 | OFB_Ctime | 0.8796426046 |
| blast | GGB30454_SGB43514 | OFB_Ctime | 0.3798589507 |
| vns | GGB30454_SGB43514 | OFB_Ctime | 0.2001255913 |
| blast | GGB30455_SGB43519 | OFB_Ctime | 0.3794677965 |
| vns | GGB30455_SGB43519 | OFB_Ctime | 0.3023489219 |
| blast | GGB30461_SGB43527 | OFB_Ctime | 0.3975036627 |
| vns | GGB30461_SGB43527 | OFB_Ctime | 0.7710386558 |
| blast | GGB30461_SGB43530 | OFB_Ctime | 0.395449 |
| vns | GGB30461_SGB43530 | OFB_Ctime | 0.7038636932 |
| blast | GGB30463_SGB43537 | OFB_Ctime | 0.2100542578 |
| vns | GGB30463_SGB43537 | OFB_Ctime | 0.9853614792 |
| blast | GGB30473_SGB43557 | OFB_Ctime | 0.0206421669 |
| vns | GGB30473_SGB43557 | OFB_Ctime | 0.0562117431 |
| blast | GGB30475_SGB63182 | OFB_Ctime | 0.2879653492 |
| vns | GGB30475_SGB63182 | OFB_Ctime | 0.6608559306 |
| blast | GGB30861_SGB44083 | OFB_Ctime | 0.9575523797 |
| vns | GGB30861_SGB44083 | OFB_Ctime | 0.8334455291 |
| blast | GGB31312_SGB44628 | OFB_Ctime | 0.7117271287 |
| vns | GGB31312_SGB44628 | OFB_Ctime | 0.8841939433 |
| blast | GGB31438_SGB44768 | OFB_Ctime | 0.9063572977 |
| vns | GGB31438_SGB44768 | OFB_Ctime | 0.3189600867 |
| blast | GGB3171_SGB4185 | OFB_Ctime | 0.0853457671 |
| vns | GGB3171_SGB4185 | OFB_Ctime | 0.9185517801 |
| blast | GGB31823_SGB45199 | OFB_Ctime | 0.1889777175 |
| vns | GGB31823_SGB45199 | OFB_Ctime | 0.7028132582 |
| blast | GGB31853_SGB45233 | OFB_Ctime | 0.4968427437 |
| vns | GGB31853_SGB45233 | OFB_Ctime | 0.3962595415 |
| blast | GGB32371_SGB41694 | OFB_Ctime | 0.3468118508 |
| vns | GGB32371_SGB41694 | OFB_Ctime | 0.0250540925 |
| blast | GGB3793_SGB5158 | OFB_Ctime | 0.9657904747 |
| vns | GGB3793_SGB5158 | OFB_Ctime | 0.9751451519 |
| blast | GGB42598_SGB59794 | OFB_Ctime | 0.5017400018 |
| vns | GGB42598_SGB59794 | OFB_Ctime | 0.156668604 |
| blast | GGB45656_SGB63370 | OFB_Ctime | 0.621044138 |
| vns | GGB45656_SGB63370 | OFB_Ctime | 0.9633746015 |
| blast | GGB47127_SGB65054 | OFB_Ctime | 0.4967451202 |
| vns | GGB47127_SGB65054 | OFB_Ctime | 0.2574173127 |

| | | | |
|---|---|---|---|
| blast | GGB74395_SGB43521 | OFB_Ctime | 0.1233397821 |
| vns | GGB74395_SGB43521 | OFB_Ctime | 0.2040214986 |
| blast | GGB75053_SGB43494 | OFB_Ctime | 0.4105959194 |
| vns | GGB75053_SGB43494 | OFB_Ctime | 0.5718574569 |
| blast | GGB75109_SGB102238 | OFB_Ctime | 0.6208422921 |
| vns | GGB75109_SGB102238 | OFB_Ctime | 0.8718184088 |
| blast | GGB81440_SGB45230 | OFB_Ctime | 0.0796107313 |
| vns | GGB81440_SGB45230 | OFB_Ctime | 0.0981405799 |
| blast | Lachnospiraceae_bacterium | OFB_Ctime | 0.4848161772 |
| vns | Lachnospiraceae_bacterium | OFB_Ctime | 0.5378613321 |
| blast | Lachnospiraceae_bacterium_A2 | OFB_Ctime | 0.420834896 |
| vns | Lachnospiraceae_bacterium_A2 | OFB_Ctime | 0.3236483287 |
| blast | Lachnospiraceae_bacterium_MD308 | OFB_Ctime | 0.2131194519 |
| vns | Lachnospiraceae_bacterium_MD308 | OFB_Ctime | 0.0830669656 |
| blast | Lachnospiraceae_bacterium_MD329 | OFB_Ctime | 0.0209761491 |
| vns | Lachnospiraceae_bacterium_MD329 | OFB_Ctime | 0.6991611194 |
| blast | Lachnospiraceae_unclassified_SGB41414 | OFB_Ctime | 0.142639053 |
| vns | Lachnospiraceae_unclassified_SGB41414 | OFB_Ctime | 0.119876468 |
| blast | Lachnospiraceae_unclassified_SGB41418 | OFB_Ctime | 0.9987579817 |
| vns | Lachnospiraceae_unclassified_SGB41418 | OFB_Ctime | 0.7122365734 |
| blast | Lachnospiraceae_unclassified_SGB41424 | OFB_Ctime | 0.0483863782 |
| vns | Lachnospiraceae_unclassified_SGB41424 | OFB_Ctime | 0.0460673576 |
| blast | Lachnospiraceae_unclassified_SGB41589 | OFB_Ctime | 0.1654249525 |
| vns | Lachnospiraceae_unclassified_SGB41589 | OFB_Ctime | 0.2735277008 |
| blast | Lactobacillus_johnsonii | OFB_Ctime | 0.1826784534 |
| vns | Lactobacillus_johnsonii | OFB_Ctime | 0.6729515885 |
| blast | Muribaculaceae_bacterium | OFB_Ctime | 0.8415252525 |
| vns | Muribaculaceae_bacterium | OFB_Ctime | 0.8930529806 |
| blast | Neglectibacter_sp_X4 | OFB_Ctime | 0.0847821517 |
| vns | Neglectibacter_sp_X4 | OFB_Ctime | 0.140656264 |
| blast | Oscillospiraceae_bacterium | OFB_Ctime | 0.7416725656 |
| vns | Oscillospiraceae_bacterium | OFB_Ctime | 0.856188039 |
| blast | Oscillospiraceae_unclassified_SGB43502 | OFB_Ctime | 0.0259412852 |
| vns | Oscillospiraceae_unclassified_SGB43502 | OFB_Ctime | 0.0305351331 |
| blast | Oscillospiraceae_unclassified_SGB43505 | OFB_Ctime | 0.4959230188 |
| vns | Oscillospiraceae_unclassified_SGB43505 | OFB_Ctime | 0.6425337316 |
| blast | Parasutterella_excrementihominis | OFB_Ctime | 0.5671048722 |
| vns | Parasutterella_excrementihominis | OFB_Ctime | 0.7099929171 |
| blast | Richness (# observed features) | OFB_Ctime | 0.2717524423 |
| vns | Richness (# observed features) | OFB_Ctime | 0.6607230685 |
| blast | Romboutsia_ilealis | OFB_Ctime | 0.0010997742 |
| vns | Romboutsia_ilealis | OFB_Ctime | 0.0569970896 |
| blast | Schaedlerella_arabinosiphila | OFB_Ctime | 0.9390244043 |
| vns | Schaedlerella_arabinosiphila | OFB_Ctime | 0.9139116075 |
| blast | Shannon Index | OFB_Ctime | 0.6947352216 |
| vns | Shannon Index | OFB_Ctime | 0.9062773413 |

| | | | |
|---|---|---|---|
| blast | Turicibacter_sp_1E2 | OFB_Ctime | 0.1991772312 |
| vns | Turicibacter_sp_1E2 | OFB_Ctime | 0.1655749312 |
| blast | Acetatifactor_muris | PD_choice_filt | 0.7395220796 |
| vns | Acetatifactor_muris | PD_choice_filt | 0.7528971403 |
| blast | Acetatifactor_SGB41546 | PD_choice_filt | 0.8402457971 |
| vns | Acetatifactor_SGB41546 | PD_choice_filt | 0.6630009771 |
| blast | Acutalibacter_muris | PD_choice_filt | 0.4221546334 |
| vns | Acutalibacter_muris | PD_choice_filt | 0.8105784495 |
| blast | Acutalibacter_sp_1XD8_36 | PD_choice_filt | 0.6605798863 |
| vns | Acutalibacter_sp_1XD8_36 | PD_choice_filt | 0.7091849336 |
| blast | Adlercreutzia_caecimuris | PD_choice_filt | 0.9896089529 |
| vns | Adlercreutzia_caecimuris | PD_choice_filt | 0.6765426913 |
| blast | Adlercreutzia_mucosicola | PD_choice_filt | 0.2718339856 |
| vns | Adlercreutzia_mucosicola | PD_choice_filt | 0.2498256574 |
| blast | Adlercreutzia_muris | PD_choice_filt | 0.6704217743 |
| vns | Adlercreutzia_muris | PD_choice_filt | 0.5958277454 |
| blast | Akkermansia_muciniphila | PD_choice_filt | 0.1698125404 |
| vns | Akkermansia_muciniphila | PD_choice_filt | 0.8643625099 |
| blast | Alistipes_sp_DSM_112343 | PD_choice_filt | 0.3725159774 |
| vns | Alistipes_sp_DSM_112343 | PD_choice_filt | 0.8326983074 |
| blast | Anaerotruncus_sp_1XD42_93 | PD_choice_filt | 0.5202714535 |
| vns | Anaerotruncus_sp_1XD42_93 | PD_choice_filt | 0.6174107217 |
| blast | Bacteria_unclassified_SGB102200 | PD_choice_filt | 0.4742692127 |
| vns | Bacteria_unclassified_SGB102200 | PD_choice_filt | 0.7641667885 |
| blast | Bacteria_unclassified_SGB41677 | PD_choice_filt | 0.4441438231 |
| vns | Bacteria_unclassified_SGB41677 | PD_choice_filt | 0.1699894243 |
| blast | Bacteria_unclassified_SGB43546 | PD_choice_filt | 0.5863583659 |
| vns | Bacteria_unclassified_SGB43546 | PD_choice_filt | 0.7890099125 |
| blast | bacterium_1XD42_54 | PD_choice_filt | 0.0144245959 |
| vns | bacterium_1XD42_54 | PD_choice_filt | 0.2345775748 |
| blast | bacterium_1XD42_76 | PD_choice_filt | 0.6000316038 |
| vns | bacterium_1XD42_76 | PD_choice_filt | 0.6429924539 |
| blast | bacterium_1xD8_48 | PD_choice_filt | 0.8997089453 |
| vns | bacterium_1xD8_48 | PD_choice_filt | 0.6968168809 |
| blast | Bacteroides_thetaiotaomicron | PD_choice_filt | 0.9602014777 |
| vns | Bacteroides_thetaiotaomicron | PD_choice_filt | 0.6343063159 |
| blast | Berger Parker Index | PD_choice_filt | 0.5457417127 |
| vns | Berger Parker Index | PD_choice_filt | 0.2337330546 |
| blast | Bifidobacterium_pseudolongum | PD_choice_filt | 0.4280103985 |
| vns | Bifidobacterium_pseudolongum | PD_choice_filt | 0.3477990653 |
| blast | Clostridia_bacterium | PD_choice_filt | 0.3745590434 |
| vns | Clostridia_bacterium | PD_choice_filt | 0.7249401071 |
| blast | Clostridiaceae_bacterium | PD_choice_filt | 0.9871697806 |
| vns | Clostridiaceae_bacterium | PD_choice_filt | 0.2419370425 |
| blast | Clostridiaceae_unclassified_SGB41663 | PD_choice_filt | 0.780382224 |
| vns | Clostridiaceae_unclassified_SGB41663 | PD_choice_filt | 0.719154368 |

| | | | |
|---|---|---|---|
| blast | Clostridiales_bacterium | PD_choice_filt | 0.0023319564 |
| vns | Clostridiales_bacterium | PD_choice_filt | 0.2964785018 |
| blast | Clostridium_cocleatum | PD_choice_filt | 0.0115678325 |
| vns | Clostridium_cocleatum | PD_choice_filt | 0.1960355146 |
| blast | Coriobacteriaceae_bacterium | PD_choice_filt | 0.3724483697 |
| vns | Coriobacteriaceae_bacterium | PD_choice_filt | 0.953919137 |
| blast | Dorea_sp_5_2 | PD_choice_filt | 0.5884919176 |
| vns | Dorea_sp_5_2 | PD_choice_filt | 0.4214161244 |
| blast | Dubosiella_newyorkensis | PD_choice_filt | 0.1169987394 |
| vns | Dubosiella_newyorkensis | PD_choice_filt | 0.799186063 |
| blast | Erysipelotrichales_bacterium | PD_choice_filt | 0.8970006656 |
| vns | Erysipelotrichales_bacterium | PD_choice_filt | 0.3284081083 |
| blast | Eubacteriaceae_bacterium | PD_choice_filt | 0.0271233687 |
| vns | Eubacteriaceae_bacterium | PD_choice_filt | 0.2383186135 |
| blast | Eubacteriaceae_unclassified_SGB94922 | PD_choice_filt | 0.4594880799 |
| vns | Eubacteriaceae_unclassified_SGB94922 | PD_choice_filt | 0.3726201484 |
| blast | GGB20149_SGB29430 | PD_choice_filt | 0.6280700484 |
| vns | GGB20149_SGB29430 | PD_choice_filt | 0.6299827926 |
| blast | GGB22635_SGB63107 | PD_choice_filt | 0.4117103362 |
| vns | GGB22635_SGB63107 | PD_choice_filt | 0.8871521383 |
| blast | GGB25041_SGB36960 | PD_choice_filt | 0.0824966676 |
| vns | GGB25041_SGB36960 | PD_choice_filt | 0.2101871969 |
| blast | GGB27876_SGB40310 | PD_choice_filt | 0.164666135 |
| vns | GGB27876_SGB40310 | PD_choice_filt | 0.7672351949 |
| blast | GGB27878_SGB40312 | PD_choice_filt | 0.2133541578 |
| vns | GGB27878_SGB40312 | PD_choice_filt | 0.1474794298 |
| blast | GGB27918_SGB40356 | PD_choice_filt | 0.9587581408 |
| vns | GGB27918_SGB40356 | PD_choice_filt | 0.8856989297 |
| blast | GGB28382_SGB40962 | PD_choice_filt | 0.5193113079 |
| vns | GGB28382_SGB40962 | PD_choice_filt | 0.8550578348 |
| blast | GGB28399_SGB40980 | PD_choice_filt | 0.2894655025 |
| vns | GGB28399_SGB40980 | PD_choice_filt | 0.287117034 |
| blast | GGB28411_SGB40993 | PD_choice_filt | 0.2018022626 |
| vns | GGB28411_SGB40993 | PD_choice_filt | 0.1468484741 |
| blast | GGB28415_SGB40997 | PD_choice_filt | 0.0267267446 |
| vns | GGB28415_SGB40997 | PD_choice_filt | 0.9490516966 |
| blast | GGB28430_SGB41013 | PD_choice_filt | 0.9120709472 |
| vns | GGB28430_SGB41013 | PD_choice_filt | 0.4662504826 |
| blast | GGB28439_SGB41022 | PD_choice_filt | 0.3804444035 |
| vns | GGB28439_SGB41022 | PD_choice_filt | 0.7387095529 |
| blast | GGB28778_SGB41431 | PD_choice_filt | 0.8046989411 |
| vns | GGB28778_SGB41431 | PD_choice_filt | 0.2184532434 |
| blast | GGB28784_SGB41437 | PD_choice_filt | 0.0207266217 |
| vns | GGB28784_SGB41437 | PD_choice_filt | 0.2544569976 |
| blast | GGB28792_SGB41445 | PD_choice_filt | 0.3385777757 |
| vns | GGB28792_SGB41445 | PD_choice_filt | 0.7612444262 |

| | | | |
|---|---|---|---|
| blast | GGB28798_SGB41451 | PD_choice_filt | 0.164022630 |
| vns | GGB28798_SGB41451 | PD_choice_filt | 0.248915824 |
| blast | GGB28802_SGB41455 | PD_choice_filt | 0.457898625 |
| vns | GGB28802_SGB41455 | PD_choice_filt | 0.342689298 |
| blast | GGB28818_SGB41473 | PD_choice_filt | 0.729209510 |
| vns | GGB28818_SGB41473 | PD_choice_filt | 0.569409948 |
| blast | GGB28828_SGB41484 | PD_choice_filt | 0.298696993 |
| vns | GGB28828_SGB41484 | PD_choice_filt | 0.106706535 |
| blast | GGB28851_SGB41518 | PD_choice_filt | 0.852552179 |
| vns | GGB28851_SGB41518 | PD_choice_filt | 0.578308429 |
| blast | GGB28859_SGB41528 | PD_choice_filt | 0.205749329 |
| vns | GGB28859_SGB41528 | PD_choice_filt | 0.595366745 |
| blast | GGB28864_SGB41535 | PD_choice_filt | 0.785465248 |
| vns | GGB28864_SGB41535 | PD_choice_filt | 0.964658985 |
| blast | GGB28869_SGB41543 | PD_choice_filt | 0.347931625 |
| vns | GGB28869_SGB41543 | PD_choice_filt | 0.381477475 |
| blast | GGB28883_SGB41564 | PD_choice_filt | 0.594877248 |
| vns | GGB28883_SGB41564 | PD_choice_filt | 0.060524458 |
| blast | GGB28892_SGB41573 | PD_choice_filt | 0.651227207 |
| vns | GGB28892_SGB41573 | PD_choice_filt | 0.667618882 |
| blast | GGB28893_SGB41574 | PD_choice_filt | 0.165594509 |
| vns | GGB28893_SGB41574 | PD_choice_filt | 0.182699376 |
| blast | GGB28898_SGB41580 | PD_choice_filt | 0.304654993 |
| vns | GGB28898_SGB41580 | PD_choice_filt | 0.082893330 |
| blast | GGB28904_SGB41597 | PD_choice_filt | 0.172906397 |
| vns | GGB28904_SGB41597 | PD_choice_filt | 0.056067199 |
| blast | GGB28916_SGB41612 | PD_choice_filt | 0.801432609 |
| vns | GGB28916_SGB41612 | PD_choice_filt | 0.853813296 |
| blast | GGB28924_SGB41621 | PD_choice_filt | 0.796481662 |
| vns | GGB28924_SGB41621 | PD_choice_filt | 0.278658378 |
| blast | GGB28926_SGB41624 | PD_choice_filt | 0.306780834 |
| vns | GGB28926_SGB41624 | PD_choice_filt | 0.470765265 |
| blast | GGB28927_SGB41625 | PD_choice_filt | 0.193720805 |
| vns | GGB28927_SGB41625 | PD_choice_filt | 0.418243373 |
| blast | GGB28934_SGB41635 | PD_choice_filt | 0.115873311 |
| vns | GGB28934_SGB41635 | PD_choice_filt | 0.111374073 |
| blast | GGB28946_SGB41652 | PD_choice_filt | 0.427843225 |
| vns | GGB28946_SGB41652 | PD_choice_filt | 0.422588900 |
| blast | GGB28949_SGB41655 | PD_choice_filt | 0.443608116 |
| vns | GGB28949_SGB41655 | PD_choice_filt | 0.908948179 |
| blast | GGB28949_SGB41656 | PD_choice_filt | 0.282728234 |
| vns | GGB28949_SGB41656 | PD_choice_filt | 0.770863949 |
| blast | GGB28950_SGB41657 | PD_choice_filt | 0.030527036 |
| vns | GGB28950_SGB41657 | PD_choice_filt | 0.336458904 |
| blast | GGB28951_SGB102295 | PD_choice_filt | 0.925352156 |
| vns | GGB28951_SGB102295 | PD_choice_filt | 0.158790397 |

| | | | |
|---|---|---|---|
| blast | GGB28951_SGB41658 | PD_choice_filt | 0.847001351 |
| vns | GGB28951_SGB41658 | PD_choice_filt | 0.920236623 |
| blast | GGB28954_SGB41662 | PD_choice_filt | 0.515758297 |
| vns | GGB28954_SGB41662 | PD_choice_filt | 0.800484955 |
| blast | GGB28956_SGB41665 | PD_choice_filt | 0.163329713 |
| vns | GGB28956_SGB41665 | PD_choice_filt | 0.153416502 |
| blast | GGB28960_SGB41669 | PD_choice_filt | 0.155727292 |
| vns | GGB28960_SGB41669 | PD_choice_filt | 0.155957479 |
| blast | GGB28967_SGB41678 | PD_choice_filt | 0.179269529 |
| vns | GGB28967_SGB41678 | PD_choice_filt | 0.315046519 |
| blast | GGB28991_SGB41705 | PD_choice_filt | 0.228893882 |
| vns | GGB28991_SGB41705 | PD_choice_filt | 0.279982342 |
| blast | GGB29002_SGB41718 | PD_choice_filt | 0.002430867 |
| vns | GGB29002_SGB41718 | PD_choice_filt | 0.113465587 |
| blast | GGB29003_SGB41719 | PD_choice_filt | 0.499398644 |
| vns | GGB29003_SGB41719 | PD_choice_filt | 0.583967886 |
| blast | GGB29011_SGB41731 | PD_choice_filt | 0.916276033 |
| vns | GGB29011_SGB41731 | PD_choice_filt | 0.345599236 |
| blast | GGB29531_SGB42317 | PD_choice_filt | 0.145082026 |
| vns | GGB29531_SGB42317 | PD_choice_filt | 0.135569966 |
| blast | GGB29685_SGB42494 | PD_choice_filt | 0.864543852 |
| vns | GGB29685_SGB42494 | PD_choice_filt | 0.340104263 |
| blast | GGB30141_SGB43066 | PD_choice_filt | 0.991554803 |
| vns | GGB30141_SGB43066 | PD_choice_filt | 0.760178789 |
| blast | GGB30286_SGB43248 | PD_choice_filt | 0.055261633 |
| vns | GGB30286_SGB43248 | PD_choice_filt | 0.454603367 |
| blast | GGB30303_SGB43268 | PD_choice_filt | 0.371936893 |
| vns | GGB30303_SGB43268 | PD_choice_filt | 0.944722174 |
| blast | GGB30413_SGB43452 | PD_choice_filt | 0.953276950 |
| vns | GGB30413_SGB43452 | PD_choice_filt | 0.879642604 |
| blast | GGB30454_SGB43514 | PD_choice_filt | 0.379858950 |
| vns | GGB30454_SGB43514 | PD_choice_filt | 0.200125591 |
| blast | GGB30455_SGB43519 | PD_choice_filt | 0.379467796 |
| vns | GGB30455_SGB43519 | PD_choice_filt | 0.302348921 |
| blast | GGB30461_SGB43527 | PD_choice_filt | 0.397503662 |
| vns | GGB30461_SGB43527 | PD_choice_filt | 0.771038655 |
| blast | GGB30461_SGB43530 | PD_choice_filt | 0.395449 |
| vns | GGB30461_SGB43530 | PD_choice_filt | 0.703863693 |
| blast | GGB30463_SGB43537 | PD_choice_filt | 0.210054257 |
| vns | GGB30463_SGB43537 | PD_choice_filt | 0.985361479 |
| blast | GGB30473_SGB43557 | PD_choice_filt | 0.020642166 |
| vns | GGB30473_SGB43557 | PD_choice_filt | 0.056211743 |
| blast | GGB30475_SGB63182 | PD_choice_filt | 0.287965349 |
| vns | GGB30475_SGB63182 | PD_choice_filt | 0.660855930 |
| blast | GGB30861_SGB44083 | PD_choice_filt | 0.957552379 |
| vns | GGB30861_SGB44083 | PD_choice_filt | 0.833445529 |

| | | | |
|---|---|---|---|
| blast | GGB31312_SGB44628 | PD_choice_filt | 0.7117271287 |
| vns | GGB31312_SGB44628 | PD_choice_filt | 0.8841939433 |
| blast | GGB31438_SGB44768 | PD_choice_filt | 0.9063572977 |
| vns | GGB31438_SGB44768 | PD_choice_filt | 0.3189600867 |
| blast | GGB3171_SGB4185 | PD_choice_filt | 0.0853457671 |
| vns | GGB3171_SGB4185 | PD_choice_filt | 0.9185517801 |
| blast | GGB31823_SGB45199 | PD_choice_filt | 0.1889777175 |
| vns | GGB31823_SGB45199 | PD_choice_filt | 0.7028132582 |
| blast | GGB31853_SGB45233 | PD_choice_filt | 0.4968427437 |
| vns | GGB31853_SGB45233 | PD_choice_filt | 0.3962595415 |
| blast | GGB32371_SGB41694 | PD_choice_filt | 0.3468118508 |
| vns | GGB32371_SGB41694 | PD_choice_filt | 0.0250540925 |
| blast | GGB3793_SGB5158 | PD_choice_filt | 0.9657904747 |
| vns | GGB3793_SGB5158 | PD_choice_filt | 0.9751451519 |
| blast | GGB42598_SGB59794 | PD_choice_filt | 0.5017400018 |
| vns | GGB42598_SGB59794 | PD_choice_filt | 0.156668604 |
| blast | GGB45656_SGB63370 | PD_choice_filt | 0.621044138 |
| vns | GGB45656_SGB63370 | PD_choice_filt | 0.9633746015 |
| blast | GGB47127_SGB65054 | PD_choice_filt | 0.4967451202 |
| vns | GGB47127_SGB65054 | PD_choice_filt | 0.2574173127 |
| blast | GGB74395_SGB43521 | PD_choice_filt | 0.1233397821 |
| vns | GGB74395_SGB43521 | PD_choice_filt | 0.2040214986 |
| blast | GGB75053_SGB43494 | PD_choice_filt | 0.4105959194 |
| vns | GGB75053_SGB43494 | PD_choice_filt | 0.5718574569 |
| blast | GGB75109_SGB102238 | PD_choice_filt | 0.6208422921 |
| vns | GGB75109_SGB102238 | PD_choice_filt | 0.8718184088 |
| blast | GGB81440_SGB45230 | PD_choice_filt | 0.0796107313 |
| vns | GGB81440_SGB45230 | PD_choice_filt | 0.0981405799 |
| blast | Lachnospiraceae_bacterium | PD_choice_filt | 0.4848161772 |
| vns | Lachnospiraceae_bacterium | PD_choice_filt | 0.5378613321 |
| blast | Lachnospiraceae_bacterium_A2 | PD_choice_filt | 0.420834896 |
| vns | Lachnospiraceae_bacterium_A2 | PD_choice_filt | 0.3236483287 |
| blast | Lachnospiraceae_bacterium_MD308 | PD_choice_filt | 0.2131194519 |
| vns | Lachnospiraceae_bacterium_MD308 | PD_choice_filt | 0.0830669656 |
| blast | Lachnospiraceae_bacterium_MD329 | PD_choice_filt | 0.0209761491 |
| vns | Lachnospiraceae_bacterium_MD329 | PD_choice_filt | 0.6991611194 |
| blast | Lachnospiraceae_unclassified_SGB41414 | PD_choice_filt | 0.142639053 |
| vns | Lachnospiraceae_unclassified_SGB41414 | PD_choice_filt | 0.119876468 |
| blast | Lachnospiraceae_unclassified_SGB41418 | PD_choice_filt | 0.9987579817 |
| vns | Lachnospiraceae_unclassified_SGB41418 | PD_choice_filt | 0.7122365734 |
| blast | Lachnospiraceae_unclassified_SGB41424 | PD_choice_filt | 0.0483863782 |
| vns | Lachnospiraceae_unclassified_SGB41424 | PD_choice_filt | 0.0460673576 |
| blast | Lachnospiraceae_unclassified_SGB41589 | PD_choice_filt | 0.1654249525 |
| vns | Lachnospiraceae_unclassified_SGB41589 | PD_choice_filt | 0.2735277008 |
| blast | Lactobacillus_johnsonii | PD_choice_filt | 0.1826784534 |
| vns | Lactobacillus_johnsonii | PD_choice_filt | 0.6729515885 |

| | | | |
|---|---|---|---|
| blast | Muribaculaceae_bacterium | PD_choice_filt | 0.841525252 |
| vns | Muribaculaceae_bacterium | PD_choice_filt | 0.893052980 |
| blast | Neglectibacter_sp_X4 | PD_choice_filt | 0.084782151 |
| vns | Neglectibacter_sp_X4 | PD_choice_filt | 0.140656264 |
| blast | Oscillospiraceae_bacterium | PD_choice_filt | 0.741672565 |
| vns | Oscillospiraceae_bacterium | PD_choice_filt | 0.856188039 |
| blast | Oscillospiraceae_unclassified_SGB43502 | PD_choice_filt | 0.025941285 |
| vns | Oscillospiraceae_unclassified_SGB43502 | PD_choice_filt | 0.030535133 |
| blast | Oscillospiraceae_unclassified_SGB43505 | PD_choice_filt | 0.495923018 |
| vns | Oscillospiraceae_unclassified_SGB43505 | PD_choice_filt | 0.642533731 |
| blast | Parasutterella_excrementihominis | PD_choice_filt | 0.567104872 |
| vns | Parasutterella_excrementihominis | PD_choice_filt | 0.709992917 |
| blast | Richness (# observed features) | PD_choice_filt | 0.271752442 |
| vns | Richness (# observed features) | PD_choice_filt | 0.660723068 |
| blast | Romboutsia_ilealis | PD_choice_filt | 0.001099774 |
| vns | Romboutsia_ilealis | PD_choice_filt | 0.056997089 |
| blast | Schaedlerella_arabinosiphila | PD_choice_filt | 0.939024404 |
| vns | Schaedlerella_arabinosiphila | PD_choice_filt | 0.913911607 |
| blast | Shannon Index | PD_choice_filt | 0.694735221 |
| vns | Shannon Index | PD_choice_filt | 0.906277341 |
| blast | Turicibacter_sp_1E2 | PD_choice_filt | 0.199177231 |
| vns | Turicibacter_sp_1E2 | PD_choice_filt | 0.165574931 |
| blast | Acetatifactor_muris | startle_habituation | 0.739522079 |
| vns | Acetatifactor_muris | startle_habituation | 0.752897140 |
| blast | Acetatifactor_SGB41546 | startle_habituation | 0.840245797 |
| vns | Acetatifactor_SGB41546 | startle_habituation | 0.663000977 |
| blast | Acutalibacter_muris | startle_habituation | 0.422154633 |
| vns | Acutalibacter_muris | startle_habituation | 0.810578449 |
| blast | Acutalibacter_sp_1XD8_36 | startle_habituation | 0.660579886 |
| vns | Acutalibacter_sp_1XD8_36 | startle_habituation | 0.709184933 |
| blast | Adlercreutzia_caecimuris | startle_habituation | 0.989608952 |
| vns | Adlercreutzia_caecimuris | startle_habituation | 0.676542691 |
| blast | Adlercreutzia_mucosicola | startle_habituation | 0.271833985 |
| vns | Adlercreutzia_mucosicola | startle_habituation | 0.249825657 |
| blast | Adlercreutzia_muris | startle_habituation | 0.670421774 |
| vns | Adlercreutzia_muris | startle_habituation | 0.595827745 |
| blast | Akkermansia_muciniphila | startle_habituation | 0.169812540 |
| vns | Akkermansia_muciniphila | startle_habituation | 0.864362509 |
| blast | Alistipes_sp_DSM_112343 | startle_habituation | 0.372515977 |
| vns | Alistipes_sp_DSM_112343 | startle_habituation | 0.832698307 |
| blast | Anaerotruncus_sp_1XD42_93 | startle_habituation | 0.520271453 |
| vns | Anaerotruncus_sp_1XD42_93 | startle_habituation | 0.617410721 |
| blast | Bacteria_unclassified_SGB102200 | startle_habituation | 0.474269212 |
| vns | Bacteria_unclassified_SGB102200 | startle_habituation | 0.764166788 |
| blast | Bacteria_unclassified_SGB41677 | startle_habituation | 0.444143823 |
| vns | Bacteria_unclassified_SGB41677 | startle_habituation | 0.169989424 |

| | | | |
|---|---|---|---|
| blast | Bacteria_unclassified_SGB43546 | startle_habituation | 0.586358365 |
| vns | Bacteria_unclassified_SGB43546 | startle_habituation | 0.789009912 |
| blast | bacterium_1XD42_54 | startle_habituation | 0.014424595 |
| vns | bacterium_1XD42_54 | startle_habituation | 0.234577574 |
| blast | bacterium_1XD42_76 | startle_habituation | 0.600031603 |
| vns | bacterium_1XD42_76 | startle_habituation | 0.642992453 |
| blast | bacterium_1xD8_48 | startle_habituation | 0.899708945 |
| vns | bacterium_1xD8_48 | startle_habituation | 0.696816880 |
| blast | Bacteroides_thetaiotaomicron | startle_habituation | 0.960201477 |
| vns | Bacteroides_thetaiotaomicron | startle_habituation | 0.634306315 |
| blast | Berger Parker Index | startle_habituation | 0.545741712 |
| vns | Berger Parker Index | startle_habituation | 0.233733054 |
| blast | Bifidobacterium_pseudolongum | startle_habituation | 0.428010398 |
| vns | Bifidobacterium_pseudolongum | startle_habituation | 0.347799065 |
| blast | Clostridia_bacterium | startle_habituation | 0.374559043 |
| vns | Clostridia_bacterium | startle_habituation | 0.724940107 |
| blast | Clostridiaceae_bacterium | startle_habituation | 0.987169780 |
| vns | Clostridiaceae_bacterium | startle_habituation | 0.241937042 |
| blast | Clostridiaceae_unclassified_SGB41663 | startle_habituation | 0.780382224 |
| vns | Clostridiaceae_unclassified_SGB41663 | startle_habituation | 0.719154368 |
| blast | Clostridiales_bacterium | startle_habituation | 0.002331956 |
| vns | Clostridiales_bacterium | startle_habituation | 0.296478501 |
| blast | Clostridium_cocleatum | startle_habituation | 0.011567832 |
| vns | Clostridium_cocleatum | startle_habituation | 0.196035514 |
| blast | Coriobacteriaceae_bacterium | startle_habituation | 0.372448369 |
| vns | Coriobacteriaceae_bacterium | startle_habituation | 0.953919137 |
| blast | Dorea_sp_5_2 | startle_habituation | 0.588491917 |
| vns | Dorea_sp_5_2 | startle_habituation | 0.421416124 |
| blast | Dubosiella_newyorkensis | startle_habituation | 0.116998739 |
| vns | Dubosiella_newyorkensis | startle_habituation | 0.799186063 |
| blast | Erysipelotrichales_bacterium | startle_habituation | 0.897000665 |
| vns | Erysipelotrichales_bacterium | startle_habituation | 0.328408108 |
| blast | Eubacteriaceae_bacterium | startle_habituation | 0.027123368 |
| vns | Eubacteriaceae_bacterium | startle_habituation | 0.238318613 |
| blast | Eubacteriaceae_unclassified_SGB94922 | startle_habituation | 0.459488079 |
| vns | Eubacteriaceae_unclassified_SGB94922 | startle_habituation | 0.372620148 |
| blast | GGB20149_SGB29430 | startle_habituation | 0.628070048 |
| vns | GGB20149_SGB29430 | startle_habituation | 0.629982792 |
| blast | GGB22635_SGB63107 | startle_habituation | 0.411710336 |
| vns | GGB22635_SGB63107 | startle_habituation | 0.887152138 |
| blast | GGB25041_SGB36960 | startle_habituation | 0.082496667 |
| vns | GGB25041_SGB36960 | startle_habituation | 0.210187196 |
| blast | GGB27876_SGB40310 | startle_habituation | 0.164666135 |
| vns | GGB27876_SGB40310 | startle_habituation | 0.767235194 |
| blast | GGB27878_SGB40312 | startle_habituation | 0.213354157 |
| vns | GGB27878_SGB40312 | startle_habituation | 0.147479429 |

| | | | |
|---|---|---|---|
| blast | GGB27918_SGB40356 | startle_habituation | 0.958758140 |
| vns | GGB27918_SGB40356 | startle_habituation | 0.885698929 |
| blast | GGB28382_SGB40962 | startle_habituation | 0.519311307 |
| vns | GGB28382_SGB40962 | startle_habituation | 0.855057834 |
| blast | GGB28399_SGB40980 | startle_habituation | 0.289465502 |
| vns | GGB28399_SGB40980 | startle_habituation | 0.287117034 |
| blast | GGB28411_SGB40993 | startle_habituation | 0.201802262 |
| vns | GGB28411_SGB40993 | startle_habituation | 0.146848474 |
| blast | GGB28415_SGB40997 | startle_habituation | 0.026726744 |
| vns | GGB28415_SGB40997 | startle_habituation | 0.949051696 |
| blast | GGB28430_SGB41013 | startle_habituation | 0.912070947 |
| vns | GGB28430_SGB41013 | startle_habituation | 0.466250482 |
| blast | GGB28439_SGB41022 | startle_habituation | 0.380444403 |
| vns | GGB28439_SGB41022 | startle_habituation | 0.738709552 |
| blast | GGB28778_SGB41431 | startle_habituation | 0.804698941 |
| vns | GGB28778_SGB41431 | startle_habituation | 0.218453243 |
| blast | GGB28784_SGB41437 | startle_habituation | 0.020726621 |
| vns | GGB28784_SGB41437 | startle_habituation | 0.254456997 |
| blast | GGB28792_SGB41445 | startle_habituation | 0.338577775 |
| vns | GGB28792_SGB41445 | startle_habituation | 0.761244426 |
| blast | GGB28798_SGB41451 | startle_habituation | 0.164022630 |
| vns | GGB28798_SGB41451 | startle_habituation | 0.248915824 |
| blast | GGB28802_SGB41455 | startle_habituation | 0.457898625 |
| vns | GGB28802_SGB41455 | startle_habituation | 0.342689298 |
| blast | GGB28818_SGB41473 | startle_habituation | 0.729209510 |
| vns | GGB28818_SGB41473 | startle_habituation | 0.569409948 |
| blast | GGB28828_SGB41484 | startle_habituation | 0.298696993 |
| vns | GGB28828_SGB41484 | startle_habituation | 0.106706535 |
| blast | GGB28851_SGB41518 | startle_habituation | 0.852552179 |
| vns | GGB28851_SGB41518 | startle_habituation | 0.578308429 |
| blast | GGB28859_SGB41528 | startle_habituation | 0.205749329 |
| vns | GGB28859_SGB41528 | startle_habituation | 0.595366745 |
| blast | GGB28864_SGB41535 | startle_habituation | 0.785465248 |
| vns | GGB28864_SGB41535 | startle_habituation | 0.964658985 |
| blast | GGB28869_SGB41543 | startle_habituation | 0.347931625 |
| vns | GGB28869_SGB41543 | startle_habituation | 0.381477475 |
| blast | GGB28883_SGB41564 | startle_habituation | 0.594877248 |
| vns | GGB28883_SGB41564 | startle_habituation | 0.060524458 |
| blast | GGB28892_SGB41573 | startle_habituation | 0.651227207 |
| vns | GGB28892_SGB41573 | startle_habituation | 0.667618882 |
| blast | GGB28893_SGB41574 | startle_habituation | 0.165594509 |
| vns | GGB28893_SGB41574 | startle_habituation | 0.182699376 |
| blast | GGB28898_SGB41580 | startle_habituation | 0.304654993 |
| vns | GGB28898_SGB41580 | startle_habituation | 0.082893330 |
| blast | GGB28904_SGB41597 | startle_habituation | 0.172906397 |
| vns | GGB28904_SGB41597 | startle_habituation | 0.056067199 |

| | | | |
|---|---|---|---|
| blast | GGB28916_SGB41612 | startle_habituation | 0.8014326092 |
| vns | GGB28916_SGB41612 | startle_habituation | 0.8538132969 |
| blast | GGB28924_SGB41621 | startle_habituation | 0.7964816629 |
| vns | GGB28924_SGB41621 | startle_habituation | 0.2786583787 |
| blast | GGB28926_SGB41624 | startle_habituation | 0.3067808341 |
| vns | GGB28926_SGB41624 | startle_habituation | 0.470765265 |
| blast | GGB28927_SGB41625 | startle_habituation | 0.1937208057 |
| vns | GGB28927_SGB41625 | startle_habituation | 0.4182433734 |
| blast | GGB28934_SGB41635 | startle_habituation | 0.1158733118 |
| vns | GGB28934_SGB41635 | startle_habituation | 0.1113740731 |
| blast | GGB28946_SGB41652 | startle_habituation | 0.4278432256 |
| vns | GGB28946_SGB41652 | startle_habituation | 0.4225889007 |
| blast | GGB28949_SGB41655 | startle_habituation | 0.4436081164 |
| vns | GGB28949_SGB41655 | startle_habituation | 0.9089481791 |
| blast | GGB28949_SGB41656 | startle_habituation | 0.2827282346 |
| vns | GGB28949_SGB41656 | startle_habituation | 0.7708639493 |
| blast | GGB28950_SGB41657 | startle_habituation | 0.0305270365 |
| vns | GGB28950_SGB41657 | startle_habituation | 0.3364589048 |
| blast | GGB28951_SGB102295 | startle_habituation | 0.9253521562 |
| vns | GGB28951_SGB102295 | startle_habituation | 0.1587903977 |
| blast | GGB28951_SGB41658 | startle_habituation | 0.8470013515 |
| vns | GGB28951_SGB41658 | startle_habituation | 0.9202366237 |
| blast | GGB28954_SGB41662 | startle_habituation | 0.5157582975 |
| vns | GGB28954_SGB41662 | startle_habituation | 0.8004849551 |
| blast | GGB28956_SGB41665 | startle_habituation | 0.163329713 |
| vns | GGB28956_SGB41665 | startle_habituation | 0.1534165023 |
| blast | GGB28960_SGB41669 | startle_habituation | 0.155727292 |
| vns | GGB28960_SGB41669 | startle_habituation | 0.1559574792 |
| blast | GGB28967_SGB41678 | startle_habituation | 0.1792695292 |
| vns | GGB28967_SGB41678 | startle_habituation | 0.3150465193 |
| blast | GGB28991_SGB41705 | startle_habituation | 0.2288938824 |
| vns | GGB28991_SGB41705 | startle_habituation | 0.279982342 |
| blast | GGB29002_SGB41718 | startle_habituation | 0.0024308675 |
| vns | GGB29002_SGB41718 | startle_habituation | 0.1134655877 |
| blast | GGB29003_SGB41719 | startle_habituation | 0.4993986443 |
| vns | GGB29003_SGB41719 | startle_habituation | 0.5839678864 |
| blast | GGB29011_SGB41731 | startle_habituation | 0.916276033 |
| vns | GGB29011_SGB41731 | startle_habituation | 0.3455992366 |
| blast | GGB29531_SGB42317 | startle_habituation | 0.1450820268 |
| vns | GGB29531_SGB42317 | startle_habituation | 0.1355699662 |
| blast | GGB29685_SGB42494 | startle_habituation | 0.8645438528 |
| vns | GGB29685_SGB42494 | startle_habituation | 0.340104263 |
| blast | GGB30141_SGB43066 | startle_habituation | 0.9915548039 |
| vns | GGB30141_SGB43066 | startle_habituation | 0.7601787897 |
| blast | GGB30286_SGB43248 | startle_habituation | 0.0552616333 |
| vns | GGB30286_SGB43248 | startle_habituation | 0.4546033673 |

| | | | |
|---|---|---|---|
| blast | GGB30303_SGB43268 | startle_habituation | 0.371936893 |
| vns | GGB30303_SGB43268 | startle_habituation | 0.944722174 |
| blast | GGB30413_SGB43452 | startle_habituation | 0.953276950 |
| vns | GGB30413_SGB43452 | startle_habituation | 0.879642604 |
| blast | GGB30454_SGB43514 | startle_habituation | 0.379858950 |
| vns | GGB30454_SGB43514 | startle_habituation | 0.200125591 |
| blast | GGB30455_SGB43519 | startle_habituation | 0.379467796 |
| vns | GGB30455_SGB43519 | startle_habituation | 0.302348921 |
| blast | GGB30461_SGB43527 | startle_habituation | 0.397503662 |
| vns | GGB30461_SGB43527 | startle_habituation | 0.771038655 |
| blast | GGB30461_SGB43530 | startle_habituation | 0.395449 |
| vns | GGB30461_SGB43530 | startle_habituation | 0.703863693 |
| blast | GGB30463_SGB43537 | startle_habituation | 0.210054257 |
| vns | GGB30463_SGB43537 | startle_habituation | 0.985361479 |
| blast | GGB30473_SGB43557 | startle_habituation | 0.020642166 |
| vns | GGB30473_SGB43557 | startle_habituation | 0.056211743 |
| blast | GGB30475_SGB63182 | startle_habituation | 0.287965349 |
| vns | GGB30475_SGB63182 | startle_habituation | 0.660855930 |
| blast | GGB30861_SGB44083 | startle_habituation | 0.957552379 |
| vns | GGB30861_SGB44083 | startle_habituation | 0.833445529 |
| blast | GGB31312_SGB44628 | startle_habituation | 0.711727128 |
| vns | GGB31312_SGB44628 | startle_habituation | 0.884193943 |
| blast | GGB31438_SGB44768 | startle_habituation | 0.906357297 |
| vns | GGB31438_SGB44768 | startle_habituation | 0.318960086 |
| blast | GGB3171_SGB4185 | startle_habituation | 0.085345767 |
| vns | GGB3171_SGB4185 | startle_habituation | 0.918551780 |
| blast | GGB31823_SGB45199 | startle_habituation | 0.188977717 |
| vns | GGB31823_SGB45199 | startle_habituation | 0.702813258 |
| blast | GGB31853_SGB45233 | startle_habituation | 0.496842743 |
| vns | GGB31853_SGB45233 | startle_habituation | 0.396259541 |
| blast | GGB32371_SGB41694 | startle_habituation | 0.346811850 |
| vns | GGB32371_SGB41694 | startle_habituation | 0.025054092 |
| blast | GGB3793_SGB5158 | startle_habituation | 0.965790474 |
| vns | GGB3793_SGB5158 | startle_habituation | 0.975145151 |
| blast | GGB42598_SGB59794 | startle_habituation | 0.501740001 |
| vns | GGB42598_SGB59794 | startle_habituation | 0.156668604 |
| blast | GGB45656_SGB63370 | startle_habituation | 0.621044138 |
| vns | GGB45656_SGB63370 | startle_habituation | 0.963374601 |
| blast | GGB47127_SGB65054 | startle_habituation | 0.496745120 |
| vns | GGB47127_SGB65054 | startle_habituation | 0.257417312 |
| blast | GGB74395_SGB43521 | startle_habituation | 0.123339782 |
| vns | GGB74395_SGB43521 | startle_habituation | 0.204021498 |
| blast | GGB75053_SGB43494 | startle_habituation | 0.410595919 |
| vns | GGB75053_SGB43494 | startle_habituation | 0.571857456 |
| blast | GGB75109_SGB102238 | startle_habituation | 0.620842292 |
| vns | GGB75109_SGB102238 | startle_habituation | 0.871818408 |

| | | | |
|---|---|---|---|
| blast | GGB81440_SGB45230 | startle_habituation | 0.079610731 |
| vns | GGB81440_SGB45230 | startle_habituation | 0.098140579 |
| blast | Lachnospiraceae_bacterium | startle_habituation | 0.484816177 |
| vns | Lachnospiraceae_bacterium | startle_habituation | 0.537861332 |
| blast | Lachnospiraceae_bacterium_A2 | startle_habituation | 0.420834896 |
| vns | Lachnospiraceae_bacterium_A2 | startle_habituation | 0.323648328 |
| blast | Lachnospiraceae_bacterium_MD308 | startle_habituation | 0.213119451 |
| vns | Lachnospiraceae_bacterium_MD308 | startle_habituation | 0.083066965 |
| blast | Lachnospiraceae_bacterium_MD329 | startle_habituation | 0.020976149 |
| vns | Lachnospiraceae_bacterium_MD329 | startle_habituation | 0.699161119 |
| blast | Lachnospiraceae_unclassified_SGB41414 | startle_habituation | 0.142639053 |
| vns | Lachnospiraceae_unclassified_SGB41414 | startle_habituation | 0.119876468 |
| blast | Lachnospiraceae_unclassified_SGB41418 | startle_habituation | 0.998757981 |
| vns | Lachnospiraceae_unclassified_SGB41418 | startle_habituation | 0.712236573 |
| blast | Lachnospiraceae_unclassified_SGB41424 | startle_habituation | 0.048386378 |
| vns | Lachnospiraceae_unclassified_SGB41424 | startle_habituation | 0.046067357 |
| blast | Lachnospiraceae_unclassified_SGB41589 | startle_habituation | 0.165424952 |
| vns | Lachnospiraceae_unclassified_SGB41589 | startle_habituation | 0.273527700 |
| blast | Lactobacillus_johnsonii | startle_habituation | 0.182678453 |
| vns | Lactobacillus_johnsonii | startle_habituation | 0.672951588 |
| blast | Muribaculaceae_bacterium | startle_habituation | 0.841525252 |
| vns | Muribaculaceae_bacterium | startle_habituation | 0.893052980 |
| blast | Neglectibacter_sp_X4 | startle_habituation | 0.084782151 |
| vns | Neglectibacter_sp_X4 | startle_habituation | 0.140656264 |
| blast | Oscillospiraceae_bacterium | startle_habituation | 0.741672565 |
| vns | Oscillospiraceae_bacterium | startle_habituation | 0.856188039 |
| blast | Oscillospiraceae_unclassified_SGB43502 | startle_habituation | 0.025941285 |
| vns | Oscillospiraceae_unclassified_SGB43502 | startle_habituation | 0.030535133 |
| blast | Oscillospiraceae_unclassified_SGB43505 | startle_habituation | 0.495923018 |
| vns | Oscillospiraceae_unclassified_SGB43505 | startle_habituation | 0.642533731 |
| blast | Parasutterella_excrementihominis | startle_habituation | 0.567104872 |
| vns | Parasutterella_excrementihominis | startle_habituation | 0.709992917 |
| blast | Richness (# observed features) | startle_habituation | 0.271752442 |
| vns | Richness (# observed features) | startle_habituation | 0.660723068 |
| blast | Romboutsia_ilealis | startle_habituation | 0.001099774 |
| vns | Romboutsia_ilealis | startle_habituation | 0.056997089 |
| blast | Schaedlerella_arabinosiphila | startle_habituation | 0.939024404 |
| vns | Schaedlerella_arabinosiphila | startle_habituation | 0.913911607 |
| blast | Shannon Index | startle_habituation | 0.694735221 |
| vns | Shannon Index | startle_habituation | 0.906277341 |
| blast | Turicibacter_sp_1E2 | startle_habituation | 0.199177231 |
| vns | Turicibacter_sp_1E2 | startle_habituation | 0.165574931 |
| blast | Acetatifactor_muris | startle_PPI | 0.739522079 |
| vns | Acetatifactor_muris | startle_PPI | 0.752897140 |
| blast | Acetatifactor_SGB41546 | startle_PPI | 0.840245797 |
| vns | Acetatifactor_SGB41546 | startle_PPI | 0.663000977 |

| | | | |
|---|---|---|---|
| blast | Acutalibacter_muris | startle_PPI | 0.422154633 |
| vns | Acutalibacter_muris | startle_PPI | 0.810578449 |
| blast | Acutalibacter_sp_1XD8_36 | startle_PPI | 0.660579886 |
| vns | Acutalibacter_sp_1XD8_36 | startle_PPI | 0.709184933 |
| blast | Adlercreutzia_caecimuris | startle_PPI | 0.989608952 |
| vns | Adlercreutzia_caecimuris | startle_PPI | 0.676542691 |
| blast | Adlercreutzia_mucosicola | startle_PPI | 0.271833985 |
| vns | Adlercreutzia_mucosicola | startle_PPI | 0.249825657 |
| blast | Adlercreutzia_muris | startle_PPI | 0.670421774 |
| vns | Adlercreutzia_muris | startle_PPI | 0.595827745 |
| blast | Akkermansia_muciniphila | startle_PPI | 0.169812540 |
| vns | Akkermansia_muciniphila | startle_PPI | 0.864362509 |
| blast | Alistipes_sp_DSM_112343 | startle_PPI | 0.372515977 |
| vns | Alistipes_sp_DSM_112343 | startle_PPI | 0.832698307 |
| blast | Anaerotruncus_sp_1XD42_93 | startle_PPI | 0.520271453 |
| vns | Anaerotruncus_sp_1XD42_93 | startle_PPI | 0.617410721 |
| blast | Bacteria_unclassified_SGB102200 | startle_PPI | 0.474269212 |
| vns | Bacteria_unclassified_SGB102200 | startle_PPI | 0.764166788 |
| blast | Bacteria_unclassified_SGB41677 | startle_PPI | 0.444143823 |
| vns | Bacteria_unclassified_SGB41677 | startle_PPI | 0.169989424 |
| blast | Bacteria_unclassified_SGB43546 | startle_PPI | 0.586358365 |
| vns | Bacteria_unclassified_SGB43546 | startle_PPI | 0.789009912 |
| blast | bacterium_1XD42_54 | startle_PPI | 0.014424595 |
| vns | bacterium_1XD42_54 | startle_PPI | 0.234577574 |
| blast | bacterium_1XD42_76 | startle_PPI | 0.600031603 |
| vns | bacterium_1XD42_76 | startle_PPI | 0.642992453 |
| blast | bacterium_1xD8_48 | startle_PPI | 0.899708945 |
| vns | bacterium_1xD8_48 | startle_PPI | 0.696816880 |
| blast | Bacteroides_thetaiotaomicron | startle_PPI | 0.960201477 |
| vns | Bacteroides_thetaiotaomicron | startle_PPI | 0.634306315 |
| blast | Berger Parker Index | startle_PPI | 0.545741712 |
| vns | Berger Parker Index | startle_PPI | 0.233733054 |
| blast | Bifidobacterium_pseudolongum | startle_PPI | 0.428010398 |
| vns | Bifidobacterium_pseudolongum | startle_PPI | 0.347799065 |
| blast | Clostridia_bacterium | startle_PPI | 0.374559043 |
| vns | Clostridia_bacterium | startle_PPI | 0.724940107 |
| blast | Clostridiaceae_bacterium | startle_PPI | 0.987169780 |
| vns | Clostridiaceae_bacterium | startle_PPI | 0.241937042 |
| blast | Clostridiaceae_unclassified_SGB41663 | startle_PPI | 0.780382224 |
| vns | Clostridiaceae_unclassified_SGB41663 | startle_PPI | 0.719154368 |
| blast | Clostridiales_bacterium | startle_PPI | 0.002331956 |
| vns | Clostridiales_bacterium | startle_PPI | 0.296478501 |
| blast | Clostridium_cocleatum | startle_PPI | 0.011567832 |
| vns | Clostridium_cocleatum | startle_PPI | 0.196035514 |
| blast | Coriobacteriaceae_bacterium | startle_PPI | 0.372448369 |
| vns | Coriobacteriaceae_bacterium | startle_PPI | 0.953919137 |

| | | | |
|---|---|---|---|
| blast | Dorea_sp_5_2 | startle_PPI | 0.588491917 |
| vns | Dorea_sp_5_2 | startle_PPI | 0.421416124 |
| blast | Dubosiella_newyorkensis | startle_PPI | 0.116998739 |
| vns | Dubosiella_newyorkensis | startle_PPI | 0.799186063 |
| blast | Erysipelotrichales_bacterium | startle_PPI | 0.897000665 |
| vns | Erysipelotrichales_bacterium | startle_PPI | 0.328408108 |
| blast | Eubacteriaceae_bacterium | startle_PPI | 0.027123368 |
| vns | Eubacteriaceae_bacterium | startle_PPI | 0.238318613 |
| blast | Eubacteriaceae_unclassified_SGB94922 | startle_PPI | 0.459488079 |
| vns | Eubacteriaceae_unclassified_SGB94922 | startle_PPI | 0.372620148 |
| blast | GGB20149_SGB29430 | startle_PPI | 0.628070048 |
| vns | GGB20149_SGB29430 | startle_PPI | 0.629982792 |
| blast | GGB22635_SGB63107 | startle_PPI | 0.411710336 |
| vns | GGB22635_SGB63107 | startle_PPI | 0.887152138 |
| blast | GGB25041_SGB36960 | startle_PPI | 0.082496667 |
| vns | GGB25041_SGB36960 | startle_PPI | 0.210187196 |
| blast | GGB27876_SGB40310 | startle_PPI | 0.164666135 |
| vns | GGB27876_SGB40310 | startle_PPI | 0.767235194 |
| blast | GGB27878_SGB40312 | startle_PPI | 0.213354157 |
| vns | GGB27878_SGB40312 | startle_PPI | 0.147479429 |
| blast | GGB27918_SGB40356 | startle_PPI | 0.958758140 |
| vns | GGB27918_SGB40356 | startle_PPI | 0.885698929 |
| blast | GGB28382_SGB40962 | startle_PPI | 0.519311307 |
| vns | GGB28382_SGB40962 | startle_PPI | 0.855057834 |
| blast | GGB28399_SGB40980 | startle_PPI | 0.289465502 |
| vns | GGB28399_SGB40980 | startle_PPI | 0.287117034 |
| blast | GGB28411_SGB40993 | startle_PPI | 0.201802262 |
| vns | GGB28411_SGB40993 | startle_PPI | 0.146848474 |
| blast | GGB28415_SGB40997 | startle_PPI | 0.026726744 |
| vns | GGB28415_SGB40997 | startle_PPI | 0.949051696 |
| blast | GGB28430_SGB41013 | startle_PPI | 0.912070947 |
| vns | GGB28430_SGB41013 | startle_PPI | 0.466250482 |
| blast | GGB28439_SGB41022 | startle_PPI | 0.380444403 |
| vns | GGB28439_SGB41022 | startle_PPI | 0.738709552 |
| blast | GGB28778_SGB41431 | startle_PPI | 0.804698941 |
| vns | GGB28778_SGB41431 | startle_PPI | 0.218453243 |
| blast | GGB28784_SGB41437 | startle_PPI | 0.020726621 |
| vns | GGB28784_SGB41437 | startle_PPI | 0.254456997 |
| blast | GGB28792_SGB41445 | startle_PPI | 0.338577775 |
| vns | GGB28792_SGB41445 | startle_PPI | 0.761244426 |
| blast | GGB28798_SGB41451 | startle_PPI | 0.164022630 |
| vns | GGB28798_SGB41451 | startle_PPI | 0.248915824 |
| blast | GGB28802_SGB41455 | startle_PPI | 0.457898625 |
| vns | GGB28802_SGB41455 | startle_PPI | 0.342689298 |
| blast | GGB28818_SGB41473 | startle_PPI | 0.729209510 |
| vns | GGB28818_SGB41473 | startle_PPI | 0.569409948 |

| | | | |
|---|---|---|---|
| blast | GGB28828_SGB41484 | startle_PPI | 0.2986969933 |
| vns | GGB28828_SGB41484 | startle_PPI | 0.1067065352 |
| blast | GGB28851_SGB41518 | startle_PPI | 0.852552179 |
| vns | GGB28851_SGB41518 | startle_PPI | 0.5783084298 |
| blast | GGB28859_SGB41528 | startle_PPI | 0.2057493298 |
| vns | GGB28859_SGB41528 | startle_PPI | 0.5953667458 |
| blast | GGB28864_SGB41535 | startle_PPI | 0.7854652488 |
| vns | GGB28864_SGB41535 | startle_PPI | 0.9646589853 |
| blast | GGB28869_SGB41543 | startle_PPI | 0.3479316259 |
| vns | GGB28869_SGB41543 | startle_PPI | 0.3814774755 |
| blast | GGB28883_SGB41564 | startle_PPI | 0.5948772487 |
| vns | GGB28883_SGB41564 | startle_PPI | 0.0605244586 |
| blast | GGB28892_SGB41573 | startle_PPI | 0.6512272078 |
| vns | GGB28892_SGB41573 | startle_PPI | 0.6676188822 |
| blast | GGB28893_SGB41574 | startle_PPI | 0.165594509 |
| vns | GGB28893_SGB41574 | startle_PPI | 0.1826993761 |
| blast | GGB28898_SGB41580 | startle_PPI | 0.3046549939 |
| vns | GGB28898_SGB41580 | startle_PPI | 0.0828933309 |
| blast | GGB28904_SGB41597 | startle_PPI | 0.1729063974 |
| vns | GGB28904_SGB41597 | startle_PPI | 0.0560671999 |
| blast | GGB28916_SGB41612 | startle_PPI | 0.8014326092 |
| vns | GGB28916_SGB41612 | startle_PPI | 0.8538132969 |
| blast | GGB28924_SGB41621 | startle_PPI | 0.7964816629 |
| vns | GGB28924_SGB41621 | startle_PPI | 0.2786583787 |
| blast | GGB28926_SGB41624 | startle_PPI | 0.3067808341 |
| vns | GGB28926_SGB41624 | startle_PPI | 0.470765265 |
| blast | GGB28927_SGB41625 | startle_PPI | 0.1937208057 |
| vns | GGB28927_SGB41625 | startle_PPI | 0.4182433734 |
| blast | GGB28934_SGB41635 | startle_PPI | 0.1158733118 |
| vns | GGB28934_SGB41635 | startle_PPI | 0.1113740731 |
| blast | GGB28946_SGB41652 | startle_PPI | 0.4278432256 |
| vns | GGB28946_SGB41652 | startle_PPI | 0.4225889007 |
| blast | GGB28949_SGB41655 | startle_PPI | 0.4436081164 |
| vns | GGB28949_SGB41655 | startle_PPI | 0.9089481791 |
| blast | GGB28949_SGB41656 | startle_PPI | 0.2827282346 |
| vns | GGB28949_SGB41656 | startle_PPI | 0.7708639493 |
| blast | GGB28950_SGB41657 | startle_PPI | 0.0305270365 |
| vns | GGB28950_SGB41657 | startle_PPI | 0.3364589048 |
| blast | GGB28951_SGB102295 | startle_PPI | 0.9253521562 |
| vns | GGB28951_SGB102295 | startle_PPI | 0.1587903977 |
| blast | GGB28951_SGB41658 | startle_PPI | 0.8470013515 |
| vns | GGB28951_SGB41658 | startle_PPI | 0.9202366237 |
| blast | GGB28954_SGB41662 | startle_PPI | 0.5157582975 |
| vns | GGB28954_SGB41662 | startle_PPI | 0.8004849551 |
| blast | GGB28956_SGB41665 | startle_PPI | 0.163329713 |
| vns | GGB28956_SGB41665 | startle_PPI | 0.1534165023 |

| | | | |
|---|---|---|---|
| blast | GGB28960_SGB41669 | startle_PPI | 0.155727292 |
| vns | GGB28960_SGB41669 | startle_PPI | 0.1559574792 |
| blast | GGB28967_SGB41678 | startle_PPI | 0.1792695292 |
| vns | GGB28967_SGB41678 | startle_PPI | 0.3150465193 |
| blast | GGB28991_SGB41705 | startle_PPI | 0.2288938824 |
| vns | GGB28991_SGB41705 | startle_PPI | 0.279982342 |
| blast | GGB29002_SGB41718 | startle_PPI | 0.0024308675 |
| vns | GGB29002_SGB41718 | startle_PPI | 0.1134655877 |
| blast | GGB29003_SGB41719 | startle_PPI | 0.4993986443 |
| vns | GGB29003_SGB41719 | startle_PPI | 0.5839678864 |
| blast | GGB29011_SGB41731 | startle_PPI | 0.916276033 |
| vns | GGB29011_SGB41731 | startle_PPI | 0.3455992366 |
| blast | GGB29531_SGB42317 | startle_PPI | 0.1450820268 |
| vns | GGB29531_SGB42317 | startle_PPI | 0.1355699662 |
| blast | GGB29685_SGB42494 | startle_PPI | 0.8645438528 |
| vns | GGB29685_SGB42494 | startle_PPI | 0.340104263 |
| blast | GGB30141_SGB43066 | startle_PPI | 0.9915548039 |
| vns | GGB30141_SGB43066 | startle_PPI | 0.7601787897 |
| blast | GGB30286_SGB43248 | startle_PPI | 0.0552616333 |
| vns | GGB30286_SGB43248 | startle_PPI | 0.4546033673 |
| blast | GGB30303_SGB43268 | startle_PPI | 0.3719368939 |
| vns | GGB30303_SGB43268 | startle_PPI | 0.9447221743 |
| blast | GGB30413_SGB43452 | startle_PPI | 0.9532769507 |
| vns | GGB30413_SGB43452 | startle_PPI | 0.8796426046 |
| blast | GGB30454_SGB43514 | startle_PPI | 0.3798589507 |
| vns | GGB30454_SGB43514 | startle_PPI | 0.2001255913 |
| blast | GGB30455_SGB43519 | startle_PPI | 0.3794677965 |
| vns | GGB30455_SGB43519 | startle_PPI | 0.3023489219 |
| blast | GGB30461_SGB43527 | startle_PPI | 0.3975036627 |
| vns | GGB30461_SGB43527 | startle_PPI | 0.7710386558 |
| blast | GGB30461_SGB43530 | startle_PPI | 0.395449 |
| vns | GGB30461_SGB43530 | startle_PPI | 0.7038636932 |
| blast | GGB30463_SGB43537 | startle_PPI | 0.2100542578 |
| vns | GGB30463_SGB43537 | startle_PPI | 0.9853614792 |
| blast | GGB30473_SGB43557 | startle_PPI | 0.0206421669 |
| vns | GGB30473_SGB43557 | startle_PPI | 0.0562117431 |
| blast | GGB30475_SGB63182 | startle_PPI | 0.2879653492 |
| vns | GGB30475_SGB63182 | startle_PPI | 0.6608559306 |
| blast | GGB30861_SGB44083 | startle_PPI | 0.9575523797 |
| vns | GGB30861_SGB44083 | startle_PPI | 0.8334455291 |
| blast | GGB31312_SGB44628 | startle_PPI | 0.7117271287 |
| vns | GGB31312_SGB44628 | startle_PPI | 0.8841939433 |
| blast | GGB31438_SGB44768 | startle_PPI | 0.9063572977 |
| vns | GGB31438_SGB44768 | startle_PPI | 0.3189600867 |
| blast | GGB3171_SGB4185 | startle_PPI | 0.0853457671 |
| vns | GGB3171_SGB4185 | startle_PPI | 0.9185517801 |

| | | | |
|---|---|---|---|
| blast | GGB31823_SGB45199 | startle_PPI | 0.188977717 |
| vns | GGB31823_SGB45199 | startle_PPI | 0.702813258 |
| blast | GGB31853_SGB45233 | startle_PPI | 0.496842743 |
| vns | GGB31853_SGB45233 | startle_PPI | 0.396259541 |
| blast | GGB32371_SGB41694 | startle_PPI | 0.346811850 |
| vns | GGB32371_SGB41694 | startle_PPI | 0.025054092 |
| blast | GGB3793_SGB5158 | startle_PPI | 0.965790474 |
| vns | GGB3793_SGB5158 | startle_PPI | 0.975145151 |
| blast | GGB42598_SGB59794 | startle_PPI | 0.501740001 |
| vns | GGB42598_SGB59794 | startle_PPI | 0.156668604 |
| blast | GGB45656_SGB63370 | startle_PPI | 0.621044138 |
| vns | GGB45656_SGB63370 | startle_PPI | 0.963374601 |
| blast | GGB47127_SGB65054 | startle_PPI | 0.496745120 |
| vns | GGB47127_SGB65054 | startle_PPI | 0.257417312 |
| blast | GGB74395_SGB43521 | startle_PPI | 0.123339782 |
| vns | GGB74395_SGB43521 | startle_PPI | 0.204021498 |
| blast | GGB75053_SGB43494 | startle_PPI | 0.410595919 |
| vns | GGB75053_SGB43494 | startle_PPI | 0.571857456 |
| blast | GGB75109_SGB102238 | startle_PPI | 0.620842292 |
| vns | GGB75109_SGB102238 | startle_PPI | 0.871818408 |
| blast | GGB81440_SGB45230 | startle_PPI | 0.079610731 |
| vns | GGB81440_SGB45230 | startle_PPI | 0.098140579 |
| blast | Lachnospiraceae_bacterium | startle_PPI | 0.484816177 |
| vns | Lachnospiraceae_bacterium | startle_PPI | 0.537861332 |
| blast | Lachnospiraceae_bacterium_A2 | startle_PPI | 0.420834896 |
| vns | Lachnospiraceae_bacterium_A2 | startle_PPI | 0.323648328 |
| blast | Lachnospiraceae_bacterium_MD308 | startle_PPI | 0.213119451 |
| vns | Lachnospiraceae_bacterium_MD308 | startle_PPI | 0.083066965 |
| blast | Lachnospiraceae_bacterium_MD329 | startle_PPI | 0.020976149 |
| vns | Lachnospiraceae_bacterium_MD329 | startle_PPI | 0.699161119 |
| blast | Lachnospiraceae_unclassified_SGB41414 | startle_PPI | 0.142639053 |
| vns | Lachnospiraceae_unclassified_SGB41414 | startle_PPI | 0.119876468 |
| blast | Lachnospiraceae_unclassified_SGB41418 | startle_PPI | 0.998757981 |
| vns | Lachnospiraceae_unclassified_SGB41418 | startle_PPI | 0.712236573 |
| blast | Lachnospiraceae_unclassified_SGB41424 | startle_PPI | 0.048386378 |
| vns | Lachnospiraceae_unclassified_SGB41424 | startle_PPI | 0.046067357 |
| blast | Lachnospiraceae_unclassified_SGB41589 | startle_PPI | 0.165424952 |
| vns | Lachnospiraceae_unclassified_SGB41589 | startle_PPI | 0.273527700 |
| blast | Lactobacillus_johnsonii | startle_PPI | 0.182678453 |
| vns | Lactobacillus_johnsonii | startle_PPI | 0.672951588 |
| blast | Muribaculaceae_bacterium | startle_PPI | 0.841525252 |
| vns | Muribaculaceae_bacterium | startle_PPI | 0.893052980 |
| blast | Neglectibacter_sp_X4 | startle_PPI | 0.084782151 |
| vns | Neglectibacter_sp_X4 | startle_PPI | 0.140656264 |
| blast | Oscillospiraceae_bacterium | startle_PPI | 0.741672565 |
| vns | Oscillospiraceae_bacterium | startle_PPI | 0.856188039 |

| | | | |
|---|---|---|---|
| blast | Oscillospiraceae_unclassified_SGB43502 | startle_PPI | 0.0259412852 |
| vns | Oscillospiraceae_unclassified_SGB43502 | startle_PPI | 0.0305351331 |
| blast | Oscillospiraceae_unclassified_SGB43505 | startle_PPI | 0.4959230188 |
| vns | Oscillospiraceae_unclassified_SGB43505 | startle_PPI | 0.6425337316 |
| blast | Parasutterella_excrementihominis | startle_PPI | 0.5671048722 |
| vns | Parasutterella_excrementihominis | startle_PPI | 0.7099929171 |
| blast | Richness (# observed features) | startle_PPI | 0.2717524423 |
| vns | Richness (# observed features) | startle_PPI | 0.6607230685 |
| blast | Romboutsia_ilealis | startle_PPI | 0.0010997742 |
| vns | Romboutsia_ilealis | startle_PPI | 0.0569970896 |
| blast | Schaedlerella_arabinosiphila | startle_PPI | 0.9390244043 |
| vns | Schaedlerella_arabinosiphila | startle_PPI | 0.9139116075 |
| blast | Shannon Index | startle_PPI | 0.6947352216 |
| vns | Shannon Index | startle_PPI | 0.9062773413 |
| blast | Turicibacter_sp_1E2 | startle_PPI | 0.1991772312 |
| vns | Turicibacter_sp_1E2 | startle_PPI | 0.1655749312 |

| p_y | p_composite | dact_stat | dact_thr_10 | dact_thr_15 | dact_thr_20 | signif_at |
|---|---|---|---|---|---|---|
| 0.93713 | 0.882395964 | 0.816354558 | 0.003097121 | 0.004400596 | 0.005669218 | |
| 0.93713 | 0.817043014 | 0.803960468 | 0.007197031 | 0.010056868 | 0.012376928 | |
| 0.010146684 | 0.819624877 | 0.714775437 | 0.003097121 | 0.004400596 | 0.005669218 | |
| 0.010146684 | 0.588941984 | 0.573638851 | 0.007197031 | 0.010056868 | 0.012376928 | |
| 0.101950854 | 0.341697162 | 0.289204687 | 0.003097121 | 0.004400596 | 0.005669218 | |
| 0.101950854 | 0.783135489 | 0.749263989 | 0.007197031 | 0.010056868 | 0.012376928 | |
| 0.382409508 | 0.678849487 | 0.539672052 | 0.003097121 | 0.004400596 | 0.005669218 | |
| 0.382409508 | 0.652340244 | 0.627070677 | 0.007197031 | 0.010056868 | 0.012376928 | |
| 0.014518444 | 0.951941334 | 0.907077160 | 0.003097121 | 0.004400596 | 0.005669218 | |
| 0.014518444 | 0.610492800 | 0.589024138 | 0.007197031 | 0.010056868 | 0.012376928 | |
| 0.400573762 | 0.256067769 | 0.233128323 | 0.003097121 | 0.004400596 | 0.005669218 | |
| 0.400573762 | 0.213013119 | 0.214479704 | 0.007197031 | 0.010056868 | 0.012376928 | |
| 0.685784528 | 0.719393457 | 0.583597398 | 0.003097121 | 0.004400596 | 0.005669218 | |
| 0.685784528 | 0.558188394 | 0.546302599 | 0.007197031 | 0.010056868 | 0.012376928 | |
| 0.14211927 | 0.095651013 | 0.105567532 | 0.003097121 | 0.004400596 | 0.005669218 | |
| 0.14211927 | 0.841104495 | 0.817608644 | 0.007197031 | 0.010056868 | 0.012376928 | |
| 0.247037984 | 0.293937423 | 0.259628695 | 0.003097121 | 0.004400596 | 0.005669218 | |
| 0.247037984 | 0.798601700 | 0.777220610 | 0.007197031 | 0.010056868 | 0.012376928 | |
| 0.489708388 | 0.522880156 | 0.408007218 | 0.003097121 | 0.004400596 | 0.005669218 | |
| 0.489708388 | 0.537905925 | 0.523414541 | 0.007197031 | 0.010056868 | 0.012376928 | |
| 0.824720198 | 0.727508646 | 0.592898736 | 0.003097121 | 0.004400596 | 0.005669218 | |
| 0.824720198 | 0.763944001 | 0.731390155 | 0.007197031 | 0.010056868 | 0.012376928 | |
| 0.681526483 | 0.598592159 | 0.472081626 | 0.003097121 | 0.004400596 | 0.005669218 | |
| 0.681526483 | 0.318374440 | 0.288466520 | 0.007197031 | 0.010056868 | 0.012376928 | |
| 0.748956111 | 0.723422227 | 0.588180017 | 0.003097121 | 0.004400596 | 0.005669218 | |
| 0.748956111 | 0.761108388 | 0.723169560 | 0.007197031 | 0.010056868 | 0.012376928 | |
| 0.015506838 | 0.002738188 | 0.008270156 | 0.003097121 | 0.004400596 | 0.005669218 | |
| 0.015506838 | 0.139730194 | 0.162772707 | 0.007197031 | 0.010056868 | 0.012376928 | |
| 0.693180661 | 0.694023573 | 0.554889371 | 0.003097121 | 0.004400596 | 0.005669218 | |
| 0.693180661 | 0.595965903 | 0.578688424 | 0.007197031 | 0.010056868 | 0.012376928 | |
| 0.571615368 | 0.893965040 | 0.833759045 | 0.003097121 | 0.004400596 | 0.005669218 | |
| 0.571615368 | 0.627612815 | 0.612883250 | 0.007197031 | 0.010056868 | 0.012376928 | |
| 0.352267104 | 0.942921080 | 0.894752303 | 0.003097121 | 0.004400596 | 0.005669218 | |
| 0.352267104 | 0.554203191 | 0.541873857 | 0.007197031 | 0.010056868 | 0.012376928 | |
| 0.071815873 | 0.511206758 | 0.398748259 | 0.003097121 | 0.004400596 | 0.005669218 | |
| 0.071815873 | 0.137809687 | 0.162164104 | 0.007197031 | 0.010056868 | 0.012376928 | |
| 0.104018289 | 0.350052885 | 0.294371331 | 0.003097121 | 0.004400596 | 0.005669218 | |
| 0.104018289 | 0.279339173 | 0.257169133 | 0.007197031 | 0.010056868 | 0.012376928 | |
| 0.154204299 | 0.284958845 | 0.253854254 | 0.003097121 | 0.004400596 | 0.005669218 | |
| 0.154204299 | 0.678725241 | 0.645333532 | 0.007197031 | 0.010056868 | 0.012376928 | |
| 0.008595498 | 0.948899643 | 0.903310972 | 0.003097121 | 0.004400596 | 0.005669218 | |
| 0.008595498 | 0.148481313 | 0.168584234 | 0.007197031 | 0.010056868 | 0.012376928 | |
| 0.308813937 | 0.783554351 | 0.667247005 | 0.003097121 | 0.004400596 | 0.005669218 | |
| 0.308813937 | 0.672223456 | 0.638674636 | 0.007197031 | 0.010056868 | 0.012376928 | |
| 0.671739982 | 0.284260480 | 0.253401763 | 0.003097121 | 0.004400596 | 0.005669218 | |
| 0.671739982 | 0.395124644 | 0.359052682 | 0.007197031 | 0.010056868 | 0.012376928 | |

| | | | | | |
|---|---|---|---|---|---|
| 0.771492821 | 0.402935990 | 0.329160501 | 0.003097121 | 0.004400596 | 0.00566921 |
| 0.771492821 | 0.393093369 | 0.356473271 | 0.007197031 | 0.010056868 | 0.01237692 |
| 0.592081995 | 0.484590200 | 0.380388656 | 0.003097121 | 0.004400596 | 0.00566921 |
| 0.592081995 | 0.963186636 | 0.936928252 | 0.007197031 | 0.010056868 | 0.01237692 |
| 0.180181608 | 0.569413180 | 0.449272653 | 0.003097121 | 0.004400596 | 0.00566921 |
| 0.180181608 | 0.357599763 | 0.324066476 | 0.007197031 | 0.010056868 | 0.01237692 |
| 0.067538602 | 0.052362868 | 0.067585513 | 0.003097121 | 0.004400596 | 0.00566921 |
| 0.067538602 | 0.765608396 | 0.735058521 | 0.007197031 | 0.010056868 | 0.01237692 |
| 0.698459766 | 0.898731392 | 0.840465586 | 0.003097121 | 0.004400596 | 0.00566921 |
| 0.698459766 | 0.428527369 | 0.392881743 | 0.007197031 | 0.010056868 | 0.01237692 |
| 0.601816186 | 0.241249151 | 0.220524909 | 0.003097121 | 0.004400596 | 0.00566921 |
| 0.601816186 | 0.318374017 | 0.288466073 | 0.007197031 | 0.010056868 | 0.01237692 |
| 0.423874835 | 0.428585883 | 0.347361155 | 0.003097121 | 0.004400596 | 0.00566921 |
| 0.423874835 | 0.325217906 | 0.295863914 | 0.007197031 | 0.010056868 | 0.01237692 |
| 0.312702445 | 0.634388466 | 0.500057492 | 0.003097121 | 0.004400596 | 0.00566921 |
| 0.312702445 | 0.546894472 | 0.537053705 | 0.007197031 | 0.010056868 | 0.01237692 |
| 0.166899079 | 0.335431258 | 0.285555264 | 0.003097121 | 0.004400596 | 0.00566921 |
| 0.166899079 | 0.881779500 | 0.847273748 | 0.007197031 | 0.010056868 | 0.01237692 |
| 0.378982776 | 0.135774870 | 0.133113318 | 0.003097121 | 0.004400596 | 0.00566921 |
| 0.378982776 | 0.172367162 | 0.183942478 | 0.007197031 | 0.010056868 | 0.01237692 |
| 0.272718978 | 0.136426244 | 0.133582545 | 0.003097121 | 0.004400596 | 0.00566921 |
| 0.272718978 | 0.730231870 | 0.696073773 | 0.007197031 | 0.010056868 | 0.01237692 |
| 0.896624355 | 0.667057288 | 0.527871447 | 0.003097121 | 0.004400596 | 0.00566921 |
| 0.896624355 | 0.441245291 | 0.410958376 | 0.007197031 | 0.010056868 | 0.01237692 |
| 0.496942408 | 0.949764064 | 0.904414856 | 0.003097121 | 0.004400596 | 0.00566921 |
| 0.496942408 | 0.877049479 | 0.845701337 | 0.007197031 | 0.010056868 | 0.01237692 |
| 0.258094017 | 0.497880977 | 0.388578012 | 0.003097121 | 0.004400596 | 0.00566921 |
| 0.258094017 | 0.825717791 | 0.805742355 | 0.007197031 | 0.010056868 | 0.01237692 |
| 0.786723457 | 0.601507533 | 0.474212189 | 0.003097121 | 0.004400596 | 0.00566921 |
| 0.786723457 | 0.447653104 | 0.420630463 | 0.007197031 | 0.010056868 | 0.01237692 |
| 0.644106236 | 0.405517542 | 0.330937636 | 0.003097121 | 0.004400596 | 0.00566921 |
| 0.644106236 | 0.273155687 | 0.254702328 | 0.007197031 | 0.010056868 | 0.01237692 |
| 0.762739641 | 0.403913124 | 0.329827583 | 0.003097121 | 0.004400596 | 0.00566921 |
| 0.762739641 | 0.956404292 | 0.930473406 | 0.007197031 | 0.010056868 | 0.01237692 |
| 0.458915373 | 0.898783119 | 0.840531586 | 0.003097121 | 0.004400596 | 0.00566921 |
| 0.458915373 | 0.405005681 | 0.367165009 | 0.007197031 | 0.010056868 | 0.01237692 |
| 0.111384363 | 0.287024317 | 0.255208758 | 0.003097121 | 0.004400596 | 0.00566921 |
| 0.111384363 | 0.689003650 | 0.661613837 | 0.007197031 | 0.010056868 | 0.01237692 |
| 0.779440006 | 0.833108249 | 0.733528462 | 0.003097121 | 0.004400596 | 0.00566921 |
| 0.779440006 | 0.414877779 | 0.374862075 | 0.007197031 | 0.010056868 | 0.01237692 |
| 0.568664723 | 0.215368590 | 0.197728965 | 0.003097121 | 0.004400596 | 0.00566921 |
| 0.568664723 | 0.309771115 | 0.282594690 | 0.007197031 | 0.010056868 | 0.01237692 |
| 0.990434557 | 0.788267037 | 0.673374766 | 0.003097121 | 0.004400596 | 0.00566921 |
| 0.990434557 | 0.888094087 | 0.850121335 | 0.007197031 | 0.010056868 | 0.01237692 |
| 0.425144527 | 0.20765665 | 0.192271487 | 0.003097121 | 0.004400596 | 0.00566921 |
| 0.425144527 | 0.222873896 | 0.222073786 | 0.007197031 | 0.010056868 | 0.01237692 |

| | | | | | |
|---|---|---|---|---|---|
| 0.951684485 | 0.8056020332 | 0.6944355771 | 0.0030971210 | 0.0044005960 | 0.0056692184 |
| 0.951684485 | 0.5766830084 | 0.5688466069 | 0.0071970319 | 0.0100568680 | 0.0123769289 |
| 0.6181103259 | 0.7605738323 | 0.6334378555 | 0.0030971210 | 0.0044005960 | 0.0056692184 |
| 0.6181103259 | 0.5118603261 | 0.4950882294 | 0.0071970319 | 0.0100568680 | 0.0123769289 |
| 0.6525574162 | 0.4889476259 | 0.3829448699 | 0.0030971210 | 0.0044005960 | 0.0056692184 |
| 0.6525574162 | 0.2402179727 | 0.2350089511 | 0.0071970319 | 0.0100568680 | 0.0123769289 |
| 0.7086551764 | 0.8647849558 | 0.7857293559 | 0.0030971210 | 0.0044005960 | 0.0056692184 |
| 0.7086551764 | 0.5590486867 | 0.5485705767 | 0.0071970319 | 0.0100568680 | 0.0123769289 |
| 0.5645725029 | 0.3332846154 | 0.2841720074 | 0.0030971210 | 0.0044005960 | 0.0056692184 |
| 0.5645725029 | 0.5189624659 | 0.4995692448 | 0.0071970319 | 0.0100568680 | 0.0123769289 |
| 0.397172611 | 0.7940237085 | 0.6802576174 | 0.0030971210 | 0.0044005960 | 0.0056692184 |
| 0.397172611 | 0.9777412182 | 0.9514193469 | 0.0071970319 | 0.0100568680 | 0.0123769289 |
| 0.132183137 | 0.253457436 | 0.2308466540 | 0.0030971210 | 0.0044005960 | 0.0056692184 |
| 0.132183137 | 0.3152071842 | 0.2872286942 | 0.0071970319 | 0.0100568680 | 0.0123769289 |
| 0.1829224984 | 0.5781280937 | 0.4558816133 | 0.0030971210 | 0.0044005960 | 0.0056692184 |
| 0.1829224984 | 0.0218738173 | 0.0498818538 | 0.0071970319 | 0.0100568680 | 0.0123769289 |
| 0.1627752179 | 0.6489050890 | 0.5122137027 | 0.0030971210 | 0.0044005960 | 0.0056692184 |
| 0.1627752179 | 0.5973850092 | 0.5790200933 | 0.0071970319 | 0.0100568680 | 0.0123769289 |
| 0.1620752609 | 0.0942101561 | 0.1045168634 | 0.0030971210 | 0.0044005960 | 0.0056692184 |
| 0.1620752609 | 0.0955428572 | 0.1231333258 | 0.0071970319 | 0.0100568680 | 0.0123769289 |
| 0.0554707932 | 0.2065260953 | 0.1915105132 | 0.0030971210 | 0.0044005960 | 0.0056692184 |
| 0.0554707932 | 0.0236337719 | 0.0525399910 | 0.0071970319 | 0.0100568680 | 0.0123769289 |
| 0.354937354 | 0.1738971170 | 0.1668214902 | 0.0030971210 | 0.0044005960 | 0.0056692184 |
| 0.354937354 | 0.0500093663 | 0.0843874674 | 0.0071970319 | 0.0100568680 | 0.0123769289 |
| 0.7125408334 | 0.8264586297 | 0.724300379 | 0.0030971210 | 0.0044005960 | 0.0056692184 |
| 0.7125408334 | 0.8222857914 | 0.8045993773 | 0.0071970319 | 0.0100568680 | 0.0123769289 |
| 0.8657894494 | 0.8627784940 | 0.7813945572 | 0.0030971210 | 0.0044005960 | 0.0056692184 |
| 0.8657894494 | 0.4796417280 | 0.467897365 | 0.0071970319 | 0.0100568680 | 0.0123769289 |
| 0.4718705137 | 0.3314201405 | 0.2829756254 | 0.0030971210 | 0.0044005960 | 0.0056692184 |
| 0.4718705137 | 0.4126161424 | 0.3719957422 | 0.0071970319 | 0.0100568680 | 0.0123769289 |
| 0.0144419671 | 0.1025775180 | 0.1106535544 | 0.0030971210 | 0.0044005960 | 0.0056692184 |
| 0.0144419671 | 0.350276121 | 0.3209314738 | 0.0071970319 | 0.0100568680 | 0.0123769289 |
| 0.4782077753 | 0.2108805911 | 0.1944961893 | 0.0030971210 | 0.0044005960 | 0.0056692184 |
| 0.4782077753 | 0.1345051020 | 0.1587757222 | 0.0071970319 | 0.0100568680 | 0.0123769289 |
| 0.4936396379 | 0.436256612 | 0.3520751598 | 0.0030971210 | 0.0044005960 | 0.0056692184 |
| 0.4936396379 | 0.3856073293 | 0.3515190169 | 0.0071970319 | 0.0100568680 | 0.0123769289 |
| 0.2388848902 | 0.3882474561 | 0.3186294779 | 0.0030971210 | 0.0044005960 | 0.0056692184 |
| 0.2388848902 | 0.908420264 | 0.8760825092 | 0.0071970319 | 0.0100568680 | 0.0123769289 |
| 0.4218305221 | 0.2754974748 | 0.2477500297 | 0.0030971210 | 0.0044005960 | 0.0056692184 |
| 0.4218305221 | 0.7406513802 | 0.7006326915 | 0.0071970319 | 0.0100568680 | 0.0123769289 |
| 0.6766781508 | 0.3121949679 | 0.2702602903 | 0.0030971210 | 0.0044005960 | 0.0056692184 |
| 0.6766781508 | 0.4232118645 | 0.3857093489 | 0.0071970319 | 0.0100568680 | 0.0123769289 |
| 0.6893027149 | 0.9275269132 | 0.8760749619 | 0.0030971210 | 0.0044005960 | 0.0056692184 |
| 0.6893027149 | 0.313021158 | 0.2860188345 | 0.0071970319 | 0.0100568680 | 0.0123769289 |
| 0.0049832150 | 0.8251336594 | 0.7226202734 | 0.0030971210 | 0.0044005960 | 0.0056692184 |
| 0.0049832150 | 0.9305367925 | 0.8908811349 | 0.0071970319 | 0.0100568680 | 0.0123769289 |

| | | | | | |
|---|---|---|---|---|---|
| 0.830187941 | 0.749027859 | 0.617232306 | 0.003097121 | 0.004400596 | 0.005669218 |
| 0.830187941 | 0.791054592 | 0.756805968 | 0.007197031 | 0.010056868 | 0.012376928 |
| 0.776517588 | 0.520301017 | 0.405830678 | 0.003097121 | 0.004400596 | 0.005669218 |
| 0.776517588 | 0.371449991 | 0.334018245 | 0.007197031 | 0.010056868 | 0.012376928 |
| 0.983586421 | 0.715824630 | 0.579110603 | 0.003097121 | 0.004400596 | 0.005669218 |
| 0.983586421 | 0.500319438 | 0.481534973 | 0.007197031 | 0.010056868 | 0.012376928 |
| 0.270765135 | 0.144883328 | 0.139968638 | 0.003097121 | 0.004400596 | 0.005669218 |
| 0.270765135 | 0.231598091 | 0.229000400 | 0.007197031 | 0.010056868 | 0.012376928 |
| 0.450456967 | 0.258418768 | 0.235186672 | 0.003097121 | 0.004400596 | 0.005669218 |
| 0.450456967 | 0.262448593 | 0.249604454 | 0.007197031 | 0.010056868 | 0.012376928 |
| 0.084178899 | 0.004214518 | 0.011019006 | 0.003097121 | 0.004400596 | 0.005669218 |
| 0.084178899 | 0.040760709 | 0.073313307 | 0.007197031 | 0.010056868 | 0.012376928 |
| 0.598707018 | 0.564263354 | 0.445325952 | 0.003097121 | 0.004400596 | 0.005669218 |
| 0.598707018 | 0.511509835 | 0.494359476 | 0.007197031 | 0.010056868 | 0.012376928 |
| 0.361080919 | 0.896945939 | 0.838106147 | 0.003097121 | 0.004400596 | 0.005669218 |
| 0.361080919 | 0.284197302 | 0.259872395 | 0.007197031 | 0.010056868 | 0.012376928 |
| 0.001063788 | 0.062394214 | 0.078985945 | 0.003097121 | 0.004400596 | 0.005669218 |
| 0.001063788 | 0.052835634 | 0.088694729 | 0.007197031 | 0.010056868 | 0.012376928 |
| 0.008273244 | 0.839721727 | 0.744514781 | 0.003097121 | 0.004400596 | 0.005669218 |
| 0.008273244 | 0.263257250 | 0.250339195 | 0.007197031 | 0.010056868 | 0.012376928 |
| 0.508706379 | 0.986024317 | 0.949242425 | 0.003097121 | 0.004400596 | 0.005669218 |
| 0.508706379 | 0.711423790 | 0.687764697 | 0.007197031 | 0.010056868 | 0.012376928 |
| 0.889890304 | 0.565281289 | 0.446154900 | 0.003097121 | 0.004400596 | 0.005669218 |
| 0.889890304 | 0.616282655 | 0.590413812 | 0.007197031 | 0.010056868 | 0.012376928 |
| 0.147951353 | 0.280942179 | 0.251234120 | 0.003097121 | 0.004400596 | 0.005669218 |
| 0.147951353 | 0.943309917 | 0.923981250 | 0.007197031 | 0.010056868 | 0.012376928 |
| 0.526028446 | 0.946212098 | 0.899499467 | 0.003097121 | 0.004400596 | 0.005669218 |
| 0.526028446 | 0.861832001 | 0.837820035 | 0.007197031 | 0.010056868 | 0.012376928 |
| 0.165981792 | 0.293069393 | 0.259099681 | 0.003097121 | 0.004400596 | 0.005669218 |
| 0.165981792 | 0.109678962 | 0.136261437 | 0.007197031 | 0.010056868 | 0.012376928 |
| 0.081621129 | 0.282165017 | 0.252032388 | 0.003097121 | 0.004400596 | 0.005669218 |
| 0.081621129 | 0.217171563 | 0.218056922 | 0.007197031 | 0.010056868 | 0.012376928 |
| 0.096923692 | 0.308575491 | 0.268079684 | 0.003097121 | 0.004400596 | 0.005669218 |
| 0.096923692 | 0.738819014 | 0.700520360 | 0.007197031 | 0.010056868 | 0.012376928 |
| 0.240640282 | 0.32396174 | 0.277873903 | 0.003097121 | 0.004400596 | 0.005669218 |
| 0.240640282 | 0.643789351 | 0.620728495 | 0.007197031 | 0.010056868 | 0.012376928 |
| 0.324556210 | 0.181268593 | 0.173138703 | 0.003097121 | 0.004400596 | 0.005669218 |
| 0.324556210 | 0.993709402 | 0.979916341 | 0.007197031 | 0.010056868 | 0.012376928 |
| 0.737379173 | 0.371953431 | 0.308138914 | 0.003097121 | 0.004400596 | 0.005669218 |
| 0.737379173 | 0.267591818 | 0.251955643 | 0.007197031 | 0.010056868 | 0.012376928 |
| 0.205115262 | 0.205939585 | 0.191119167 | 0.003097121 | 0.004400596 | 0.005669218 |
| 0.205115262 | 0.581996160 | 0.571410898 | 0.007197031 | 0.010056868 | 0.012376928 |
| 0.010436741 | 0.916438712 | 0.863898967 | 0.003097121 | 0.004400596 | 0.005669218 |
| 0.010436741 | 0.803552344 | 0.777930334 | 0.007197031 | 0.010056868 | 0.012376928 |
| 0.058278529 | 0.707173780 | 0.569175726 | 0.003097121 | 0.004400596 | 0.005669218 |
| 0.058278529 | 0.866051226 | 0.843294216 | 0.007197031 | 0.010056868 | 0.012376928 |

| | | | | | |
|---|---|---|---|---|---|
| 0.129588426 | 0.876273999 | 0.806871339 | 0.003097121 | 0.004400596 | 0.005669218 |
| 0.129588426 | 0.236012104 | 0.232199855 | 0.007197031 | 0.010056868 | 0.012376928 |
| 0.783302125 | 0.472706963 | 0.373597514 | 0.003097121 | 0.004400596 | 0.005669218 |
| 0.783302125 | 0.929211607 | 0.889409319 | 0.007197031 | 0.010056868 | 0.012376928 |
| 0.317502865 | 0.166916282 | 0.160465084 | 0.003097121 | 0.004400596 | 0.005669218 |
| 0.317502865 | 0.638465592 | 0.619584048 | 0.007197031 | 0.010056868 | 0.012376928 |
| 0.042137307 | 0.433676178 | 0.350470694 | 0.003097121 | 0.004400596 | 0.005669218 |
| 0.042137307 | 0.329780020 | 0.300606511 | 0.007197031 | 0.010056868 | 0.012376928 |
| 0.491718818 | 0.378153633 | 0.312193648 | 0.003097121 | 0.004400596 | 0.005669218 |
| 0.491718818 | 0.083044936 | 0.112239133 | 0.007197031 | 0.010056868 | 0.012376928 |
| 0.675732129 | 0.971419935 | 0.928048544 | 0.003097121 | 0.004400596 | 0.005669218 |
| 0.675732129 | 0.988257395 | 0.966126050 | 0.007197031 | 0.010056868 | 0.012376928 |
| 0.548701259 | 0.532702970 | 0.417203453 | 0.003097121 | 0.004400596 | 0.005669218 |
| 0.548701259 | 0.213557531 | 0.214979092 | 0.007197031 | 0.010056868 | 0.012376928 |
| 0.189013771 | 0.613922890 | 0.482928449 | 0.003097121 | 0.004400596 | 0.005669218 |
| 0.189013771 | 0.968281273 | 0.949450023 | 0.007197031 | 0.010056868 | 0.012376928 |
| 0.049222290 | 0.434433900 | 0.350947955 | 0.003097121 | 0.004400596 | 0.005669218 |
| 0.049222290 | 0.167935377 | 0.181005079 | 0.007197031 | 0.010056868 | 0.012376928 |
| 0.105346324 | 0.058596574 | 0.074324390 | 0.003097121 | 0.004400596 | 0.005669218 |
| 0.105346324 | 0.113293535 | 0.139168154 | 0.007197031 | 0.010056868 | 0.012376928 |
| 0.223815779 | 0.341637526 | 0.289170467 | 0.003097121 | 0.004400596 | 0.005669218 |
| 0.223815779 | 0.489126614 | 0.474146670 | 0.007197031 | 0.010056868 | 0.012376928 |
| 0.373693725 | 0.631555711 | 0.497495665 | 0.003097121 | 0.004400596 | 0.005669218 |
| 0.373693725 | 0.851898542 | 0.827491674 | 0.007197031 | 0.010056868 | 0.012376928 |
| 0.078886450 | 0.034513930 | 0.047312729 | 0.003097121 | 0.004400596 | 0.005669218 |
| 0.078886450 | 0.031500720 | 0.062815863 | 0.007197031 | 0.010056868 | 0.012376928 |
| 0.175471097 | 0.433107214 | 0.350170012 | 0.003097121 | 0.004400596 | 0.005669218 |
| 0.175471097 | 0.460212039 | 0.438610195 | 0.007197031 | 0.010056868 | 0.012376928 |
| 0.249948612 | 0.358404192 | 0.299921523 | 0.003097121 | 0.004400596 | 0.005669218 |
| 0.249948612 | 0.243484228 | 0.236338242 | 0.007197031 | 0.010056868 | 0.012376928 |
| 0.143218715 | 0.136432599 | 0.133739590 | 0.003097121 | 0.004400596 | 0.005669218 |
| 0.143218715 | 0.028182006 | 0.058188038 | 0.007197031 | 0.010056868 | 0.012376928 |
| 0.402910748 | 0.106755810 | 0.113729723 | 0.003097121 | 0.004400596 | 0.005669218 |
| 0.402910748 | 0.632479102 | 0.615430090 | 0.007197031 | 0.010056868 | 0.012376928 |
| 0.364600188 | 0.162493274 | 0.156125411 | 0.003097121 | 0.004400596 | 0.005669218 |
| 0.364600188 | 0.097755633 | 0.125463799 | 0.007197031 | 0.010056868 | 0.012376928 |
| 0.801805019 | 0.998087864 | 0.982455935 | 0.003097121 | 0.004400596 | 0.005669218 |
| 0.801805019 | 0.706699585 | 0.685300264 | 0.007197031 | 0.010056868 | 0.012376928 |
| 0.508060263 | 0.186849926 | 0.177445384 | 0.003097121 | 0.004400596 | 0.005669218 |
| 0.508060263 | 0.102717300 | 0.131398567 | 0.007197031 | 0.010056868 | 0.012376928 |
| 0.697123145 | 0.431294271 | 0.349005328 | 0.003097121 | 0.004400596 | 0.005669218 |
| 0.697123145 | 0.395244684 | 0.359206015 | 0.007197031 | 0.010056868 | 0.012376928 |
| 0.955504061 | 0.704062595 | 0.565840707 | 0.003097121 | 0.004400596 | 0.005669218 |
| 0.955504061 | 0.794717115 | 0.769921661 | 0.007197031 | 0.010056868 | 0.012376928 |
| 0.038454065 | 0.822273544 | 0.718600936 | 0.003097121 | 0.004400596 | 0.005669218 |
| 0.038454065 | 0.889295835 | 0.854887692 | 0.007197031 | 0.010056868 | 0.012376928 |

| | | | | | |
|---|---|---|---|---|---|
| 0.044442561 | 0.034621596 | 0.047413563 | 0.003097121 | 0.004400596 | 0.005669218 |
| 0.044442561 | 0.056760223 | 0.092351874 | 0.007197031 | 0.010056868 | 0.012376928 |
| 0.965563069 | 0.901068114 | 0.843465175 | 0.003097121 | 0.004400596 | 0.005669218 |
| 0.965563069 | 0.922705952 | 0.887603203 | 0.007197031 | 0.010056868 | 0.012376928 |
| 0.351734423 | 0.079550964 | 0.095026096 | 0.003097121 | 0.004400596 | 0.005669218 |
| 0.351734423 | 0.034973767 | 0.068159656 | 0.007197031 | 0.010056868 | 0.012376928 |
| 0.507976248 | 0.502707861 | 0.392218691 | 0.003097121 | 0.004400596 | 0.005669218 |
| 0.507976248 | 0.564084518 | 0.551168053 | 0.007197031 | 0.010056868 | 0.012376928 |
| 0.205308301 | 0.546549762 | 0.430142494 | 0.003097121 | 0.004400596 | 0.005669218 |
| 0.205308301 | 0.656810828 | 0.627837035 | 0.007197031 | 0.010056868 | 0.012376928 |
| 0.187328370 | 0.187511640 | 0.177921173 | 0.003097121 | 0.004400596 | 0.005669218 |
| 0.187328370 | 0.579257990 | 0.571243158 | 0.007197031 | 0.010056868 | 0.012376928 |
| 0.919014803 | 0.564686553 | 0.445667887 | 0.003097121 | 0.004400596 | 0.005669218 |
| 0.919014803 | 0.413371979 | 0.372952552 | 0.007197031 | 0.010056868 | 0.012376928 |
| 0.202499563 | 0.909916079 | 0.854909085 | 0.003097121 | 0.004400596 | 0.005669218 |
| 0.202499563 | 0.916326199 | 0.882643234 | 0.007197031 | 0.010056868 | 0.012376928 |
| 0.351765516 | 0.711604395 | 0.573983240 | 0.003097121 | 0.004400596 | 0.005669218 |
| 0.351765516 | 0.901603233 | 0.872653618 | 0.007197031 | 0.010056868 | 0.012376928 |
| 0.166899984 | 0.125904183 | 0.126412560 | 0.003097121 | 0.004400596 | 0.005669218 |
| 0.166899984 | 0.081540955 | 0.110654100 | 0.007197031 | 0.010056868 | 0.012376928 |
| 0.141836773 | 0.638409505 | 0.524954626 | 0.005492180 | 0.007205648 | 0.008582503 |
| 0.141836773 | 0.614956504 | 0.574437662 | 0.003110558 | 0.003920326 | 0.004492969 |
| 0.903365243 | 0.948081795 | 0.840942349 | 0.005492180 | 0.007205648 | 0.008582503 |
| 0.903365243 | 0.845610299 | 0.810849417 | 0.003110558 | 0.003920326 | 0.004492969 |
| 0.937165790 | 0.864530919 | 0.750178573 | 0.005492180 | 0.007205648 | 0.008582503 |
| 0.937165790 | 0.923513855 | 0.876508834 | 0.003110558 | 0.003920326 | 0.004492969 |
| 0.598230628 | 0.650385581 | 0.537497745 | 0.005492180 | 0.007205648 | 0.008582503 |
| 0.598230628 | 0.553523499 | 0.511791167 | 0.003110558 | 0.003920326 | 0.004492969 |
| 0.127842343 | 0.923484062 | 0.812182648 | 0.005492180 | 0.007205648 | 0.008582503 |
| 0.127842343 | 0.493976888 | 0.466591162 | 0.003110558 | 0.003920326 | 0.004492969 |
| 0.600338376 | 0.469650024 | 0.381230472 | 0.005492180 | 0.007205648 | 0.008582503 |
| 0.600338376 | 0.385116898 | 0.356583284 | 0.003110558 | 0.003920326 | 0.004492969 |
| 0.942729343 | 0.941381140 | 0.832586262 | 0.005492180 | 0.007205648 | 0.008582503 |
| 0.942729343 | 0.924128711 | 0.877964908 | 0.003110558 | 0.003920326 | 0.004492969 |
| 0.898496658 | 0.743884089 | 0.630041325 | 0.005492180 | 0.007205648 | 0.008582503 |
| 0.898496658 | 0.845157884 | 0.810477392 | 0.003110558 | 0.003920326 | 0.004492969 |
| 0.930328323 | 0.843498814 | 0.727289220 | 0.005492180 | 0.007205648 | 0.008582503 |
| 0.930328323 | 0.917900508 | 0.865128618 | 0.003110558 | 0.003920326 | 0.004492969 |
| 0.135139214 | 0.389333863 | 0.320114398 | 0.005492180 | 0.007205648 | 0.008582503 |
| 0.135139214 | 0.423953030 | 0.390779727 | 0.003110558 | 0.003920326 | 0.004492969 |
| 0.561002782 | 0.508147828 | 0.412907015 | 0.005492180 | 0.007205648 | 0.008582503 |
| 0.561002782 | 0.645651219 | 0.591720519 | 0.003110558 | 0.003920326 | 0.004492969 |
| 0.079906588 | 0.283553599 | 0.249502199 | 0.005492180 | 0.007205648 | 0.008582503 |
| 0.079906588 | 0.044216656 | 0.034620031 | 0.003110558 | 0.003920326 | 0.004492969 |
| 0.777185196 | 0.747445236 | 0.633235240 | 0.005492180 | 0.007205648 | 0.008582503 |
| 0.777185196 | 0.683313632 | 0.629972758 | 0.003110558 | 0.003920326 | 0.004492969 |

| | | | | | |
|---|---|---|---|---|---|
| 0.394158325 | 0.152285312 | 0.159899861 | 0.005492180 | 0.007205648 | 0.008582503 |
| 0.394158325 | 0.215465398 | 0.158923615 | 0.003110558 | 0.003920326 | 0.004492969 |
| 0.171057114 | 0.487814448 | 0.395839824 | 0.005492180 | 0.007205648 | 0.008582503 |
| 0.171057114 | 0.453245829 | 0.422792479 | 0.003110558 | 0.003920326 | 0.004492969 |
| 0.719601276 | 0.929759273 | 0.819172583 | 0.005492180 | 0.007205648 | 0.008582503 |
| 0.719601276 | 0.575486321 | 0.525705233 | 0.003110558 | 0.003920326 | 0.004492969 |
| 0.341340438 | 0.928332783 | 0.817576733 | 0.005492180 | 0.007205648 | 0.008582503 |
| 0.341340438 | 0.442456037 | 0.411964315 | 0.003110558 | 0.003920326 | 0.004492969 |
| 0.176179205 | 0.429259070 | 0.349528081 | 0.005492180 | 0.007205648 | 0.008582503 |
| 0.176179205 | 0.078188742 | 0.061971396 | 0.003110558 | 0.003920326 | 0.004492969 |
| 0.229418269 | 0.310789657 | 0.267515340 | 0.005492180 | 0.007205648 | 0.008582503 |
| 0.229418269 | 0.186687966 | 0.130267003 | 0.003110558 | 0.003920326 | 0.004492969 |
| 0.439060369 | 0.356343225 | 0.297778690 | 0.005492180 | 0.007205648 | 0.008582503 |
| 0.439060369 | 0.585800305 | 0.533953297 | 0.003110558 | 0.003920326 | 0.004492969 |
| 0.505573507 | 0.974382090 | 0.884602161 | 0.005492180 | 0.007205648 | 0.008582503 |
| 0.505573507 | 0.320360924 | 0.255563444 | 0.003110558 | 0.003920326 | 0.004492969 |
| 0.418234011 | 0.736429165 | 0.623402249 | 0.005492180 | 0.007205648 | 0.008582503 |
| 0.418234011 | 0.575274048 | 0.525680093 | 0.003110558 | 0.003920326 | 0.004492969 |
| 0.756705873 | 0.535294334 | 0.434540839 | 0.005492180 | 0.007205648 | 0.008582503 |
| 0.756705873 | 0.608926028 | 0.562315473 | 0.003110558 | 0.003920326 | 0.004492969 |
| 0.421395765 | 0.172519069 | 0.175429988 | 0.005492180 | 0.007205648 | 0.008582503 |
| 0.421395765 | 0.246967654 | 0.178734232 | 0.003110558 | 0.003920326 | 0.004492969 |
| 0.294052267 | 0.270750086 | 0.241462062 | 0.005492180 | 0.007205648 | 0.008582503 |
| 0.294052267 | 0.951598919 | 0.912014068 | 0.003110558 | 0.003920326 | 0.004492969 |
| 0.594399836 | 0.583411930 | 0.473650996 | 0.005492180 | 0.007205648 | 0.008582503 |
| 0.594399836 | 0.385174285 | 0.356629763 | 0.003110558 | 0.003920326 | 0.004492969 |
| 0.062120668 | 0.027990386 | 0.055056841 | 0.005492180 | 0.007205648 | 0.008582503 |
| 0.062120668 | 0.688889800 | 0.645179992 | 0.003110558 | 0.003920326 | 0.004492969 |
| 0.338175207 | 0.854840839 | 0.739656274 | 0.005492180 | 0.007205648 | 0.008582503 |
| 0.338175207 | 0.171809028 | 0.123262471 | 0.003110558 | 0.003920326 | 0.004492969 |
| 0.180464266 | 0.032418387 | 0.060477246 | 0.005492180 | 0.007205648 | 0.008582503 |
| 0.180464266 | 0.084101961 | 0.064237139 | 0.003110558 | 0.003920326 | 0.004492969 |
| 0.283680821 | 0.362532309 | 0.301802961 | 0.005492180 | 0.007205648 | 0.008582503 |
| 0.283680821 | 0.203107076 | 0.148480430 | 0.003110558 | 0.003920326 | 0.004492969 |
| 0.958677190 | 0.945542257 | 0.837747484 | 0.005492180 | 0.007205648 | 0.008582503 |
| 0.958677190 | 0.948465440 | 0.908457466 | 0.003110558 | 0.003920326 | 0.004492969 |
| 0.255520088 | 0.301381549 | 0.260984518 | 0.005492180 | 0.007205648 | 0.008582503 |
| 0.255520088 | 0.814371012 | 0.791298965 | 0.003110558 | 0.003920326 | 0.004492969 |
| 0.377210630 | 0.166356107 | 0.170936448 | 0.005492180 | 0.007205648 | 0.008582503 |
| 0.377210630 | 0.198344802 | 0.145381044 | 0.003110558 | 0.003920326 | 0.004492969 |
| 0.303518970 | 0.148431527 | 0.156649133 | 0.005492180 | 0.007205648 | 0.008582503 |
| 0.303518970 | 0.652360957 | 0.596096777 | 0.003110558 | 0.003920326 | 0.004492969 |
| 0.910168162 | 0.771822327 | 0.656142575 | 0.005492180 | 0.007205648 | 0.008582503 |
| 0.910168162 | 0.834977027 | 0.802079185 | 0.003110558 | 0.003920326 | 0.004492969 |
| 0.90902181 | 0.989740516 | 0.929303997 | 0.005492180 | 0.007205648 | 0.008582503 |
| 0.90902181 | 0.871066780 | 0.829604803 | 0.003110558 | 0.003920326 | 0.004492969 |

| | | | | | |
|---|---|---|---|---|---|
| 0.257891634 | 0.421937627 | 0.343878017 | 0.005492180 | 0.007205648 | 0.008582503 |
| 0.257891634 | 0.747690794 | 0.736339149 | 0.003110558 | 0.003920326 | 0.004492969 |
| 0.050203436 | 0.128561553 | 0.140146644 | 0.005492180 | 0.007205648 | 0.008582503 |
| 0.050203436 | 0.128912590 | 0.090673628 | 0.003110558 | 0.003920326 | 0.004492969 |
| 0.633625137 | 0.476342235 | 0.386674923 | 0.005492180 | 0.007205648 | 0.008582503 |
| 0.633625137 | 0.428333981 | 0.391929146 | 0.003110558 | 0.003920326 | 0.004492969 |
| 0.552000583 | 0.312814225 | 0.268917275 | 0.005492180 | 0.007205648 | 0.008582503 |
| 0.552000583 | 0.944118670 | 0.903185226 | 0.003110558 | 0.003920326 | 0.004492969 |
| 0.111503842 | 0.829004108 | 0.712393409 | 0.005492180 | 0.007205648 | 0.008582503 |
| 0.111503842 | 0.295655561 | 0.227455455 | 0.003110558 | 0.003920326 | 0.004492969 |
| 0.141267417 | 0.230450391 | 0.216327152 | 0.005492180 | 0.007205648 | 0.008582503 |
| 0.141267417 | 0.600336747 | 0.553553781 | 0.003110558 | 0.003920326 | 0.004492969 |
| 0.063040992 | 0.690044384 | 0.578835797 | 0.005492180 | 0.007205648 | 0.008582503 |
| 0.063040992 | 0.070254372 | 0.054614117 | 0.003110558 | 0.003920326 | 0.004492969 |
| 0.709843460 | 0.489094365 | 0.396904540 | 0.005492180 | 0.007205648 | 0.008582503 |
| 0.709843460 | 0.520746529 | 0.494610752 | 0.003110558 | 0.003920326 | 0.004492969 |
| 0.459935127 | 0.360047650 | 0.300177435 | 0.005492180 | 0.007205648 | 0.008582503 |
| 0.459935127 | 0.635859715 | 0.587223065 | 0.003110558 | 0.003920326 | 0.004492969 |
| 0.814676524 | 0.653350977 | 0.540659846 | 0.005492180 | 0.007205648 | 0.008582503 |
| 0.814676524 | 0.702042923 | 0.647921235 | 0.003110558 | 0.003920326 | 0.004492969 |
| 0.890119452 | 0.825930002 | 0.709320371 | 0.005492180 | 0.007205648 | 0.008582503 |
| 0.890119452 | 0.781667964 | 0.775217826 | 0.003110558 | 0.003920326 | 0.004492969 |
| 0.041269218 | 0.605998283 | 0.494206371 | 0.005492180 | 0.007205648 | 0.008582503 |
| 0.041269218 | 0.354064547 | 0.334076249 | 0.003110558 | 0.003920326 | 0.004492969 |
| 0.810027823 | 0.688490386 | 0.577304212 | 0.005492180 | 0.007205648 | 0.008582503 |
| 0.810027823 | 0.684496159 | 0.634977560 | 0.003110558 | 0.003920326 | 0.004492969 |
| 0.470557850 | 0.830749920 | 0.714137879 | 0.005492180 | 0.007205648 | 0.008582503 |
| 0.470557850 | 0.368904116 | 0.344665909 | 0.003110558 | 0.003920326 | 0.004492969 |
| 0.601160656 | 0.445164201 | 0.361616365 | 0.005492180 | 0.007205648 | 0.008582503 |
| 0.601160656 | 0.412083991 | 0.371354200 | 0.003110558 | 0.003920326 | 0.004492969 |
| 0.730732027 | 0.807635533 | 0.691387206 | 0.005492180 | 0.007205648 | 0.008582503 |
| 0.730732027 | 0.972756022 | 0.932661371 | 0.003110558 | 0.003920326 | 0.004492969 |
| 0.376749191 | 0.287633156 | 0.252072580 | 0.005492180 | 0.007205648 | 0.008582503 |
| 0.376749191 | 0.212490727 | 0.15533991 | 0.003110558 | 0.003920326 | 0.004492969 |
| 0.200800101 | 0.489424717 | 0.397169320 | 0.005492180 | 0.007205648 | 0.008582503 |
| 0.200800101 | 0.052749185 | 0.041329632 | 0.003110558 | 0.003920326 | 0.004492969 |
| 0.255895972 | 0.566869202 | 0.460109997 | 0.005492180 | 0.007205648 | 0.008582503 |
| 0.255895972 | 0.479913851 | 0.454847024 | 0.003110558 | 0.003920326 | 0.004492969 |
| 0.340936590 | 0.173699436 | 0.176313595 | 0.005492180 | 0.007205648 | 0.008582503 |
| 0.340936590 | 0.165399724 | 0.119237159 | 0.003110558 | 0.003920326 | 0.004492969 |
| 0.221682008 | 0.183207064 | 0.183552501 | 0.005492180 | 0.007205648 | 0.008582503 |
| 0.221682008 | 0.065498119 | 0.050714611 | 0.003110558 | 0.003920326 | 0.004492969 |
| 0.406049456 | 0.229319385 | 0.215573426 | 0.005492180 | 0.007205648 | 0.008582503 |
| 0.406049456 | 0.217236268 | 0.160929852 | 0.003110558 | 0.003920326 | 0.004492969 |
| 0.561230993 | 0.790278124 | 0.674884017 | 0.005492180 | 0.007205648 | 0.008582503 |
| 0.561230993 | 0.743279507 | 0.734551022 | 0.003110558 | 0.003920326 | 0.004492969 |

| | | | | | |
|---|---|---|---|---|---|
| 0.422171143 | 0.756763558 | 0.641650904 | 0.005492180 | 0.007205648 | 0.008582503 |
| 0.422171143 | 0.250487525 | 0.182667841 | 0.003110558 | 0.003920326 | 0.004492969 |
| 0.452516149 | 0.338111826 | 0.285602979 | 0.005492180 | 0.007205648 | 0.008582503 |
| 0.452516149 | 0.305874876 | 0.232038264 | 0.003110558 | 0.003920326 | 0.004492969 |
| 0.079385738 | 0.072004555 | 0.093798551 | 0.005492180 | 0.007205648 | 0.008582503 |
| 0.079385738 | 0.252778538 | 0.184739539 | 0.003110558 | 0.003920326 | 0.004492969 |
| 0.699437518 | 0.512847254 | 0.416766987 | 0.005492180 | 0.007205648 | 0.008582503 |
| 0.699437518 | 0.502833077 | 0.474798728 | 0.003110558 | 0.003920326 | 0.004492969 |
| 0.112545045 | 0.274733695 | 0.244018845 | 0.005492180 | 0.007205648 | 0.008582503 |
| 0.112545045 | 0.262525338 | 0.188454224 | 0.003110558 | 0.003920326 | 0.004492969 |
| 0.158002588 | 0.306805623 | 0.264722499 | 0.005492180 | 0.007205648 | 0.008582503 |
| 0.158002588 | 0.871509028 | 0.829655250 | 0.003110558 | 0.003920326 | 0.004492969 |
| 0.624268664 | 0.497290066 | 0.403814665 | 0.005492180 | 0.007205648 | 0.008582503 |
| 0.624268664 | 0.666550945 | 0.601920805 | 0.003110558 | 0.003920326 | 0.004492969 |
| 0.150617219 | 0.024643224 | 0.050676605 | 0.005492180 | 0.007205648 | 0.008582503 |
| 0.150617219 | 0.169324229 | 0.122285384 | 0.003110558 | 0.003920326 | 0.004492969 |
| 0.280155696 | 0.876206814 | 0.762235596 | 0.005492180 | 0.007205648 | 0.008582503 |
| 0.280155696 | 0.111099856 | 0.081979497 | 0.003110558 | 0.003920326 | 0.004492969 |
| 0.008581956 | 0.727658705 | 0.615386375 | 0.005492180 | 0.007205648 | 0.008582503 |
| 0.008581956 | 0.901813878 | 0.849780072 | 0.003110558 | 0.003920326 | 0.004492969 |
| 0.604333380 | 0.565748036 | 0.459207179 | 0.005492180 | 0.007205648 | 0.008582503 |
| 0.604333380 | 0.700762675 | 0.647768848 | 0.003110558 | 0.003920326 | 0.004492969 |
| 0.167090226 | 0.076803779 | 0.097279268 | 0.005492180 | 0.007205648 | 0.008582503 |
| 0.167090226 | 0.041615152 | 0.033106128 | 0.003110558 | 0.003920326 | 0.004492969 |
| 0.062542881 | 0.044460222 | 0.072228767 | 0.005492180 | 0.007205648 | 0.008582503 |
| 0.062542881 | 0.032193091 | 0.029650667 | 0.003110558 | 0.003920326 | 0.004492969 |
| 0.188599889 | 0.093709827 | 0.110243793 | 0.005492180 | 0.007205648 | 0.008582503 |
| 0.188599889 | 0.148312461 | 0.108074597 | 0.003110558 | 0.003920326 | 0.004492969 |
| 0.236512847 | 0.134550912 | 0.145092934 | 0.005492180 | 0.007205648 | 0.008582503 |
| 0.236512847 | 0.123240549 | 0.086690313 | 0.003110558 | 0.003920326 | 0.004492969 |
| 0.399656112 | 0.151774677 | 0.159474359 | 0.005492180 | 0.007205648 | 0.008582503 |
| 0.399656112 | 0.214903804 | 0.158274287 | 0.003110558 | 0.003920326 | 0.004492969 |
| 0.657345436 | 0.609885727 | 0.497788829 | 0.005492180 | 0.007205648 | 0.008582503 |
| 0.657345436 | 0.460875903 | 0.438834962 | 0.003110558 | 0.003920326 | 0.004492969 |
| 0.204110215 | 0.851538053 | 0.735997165 | 0.005492180 | 0.007205648 | 0.008582503 |
| 0.204110215 | 0.182700657 | 0.128689358 | 0.003110558 | 0.003920326 | 0.004492969 |
| 0.001928988 | 0.028156081 | 0.055261691 | 0.005492180 | 0.007205648 | 0.008582503 |
| 0.001928988 | 0.020396850 | 0.023068774 | 0.003110558 | 0.003920326 | 0.004492969 |
| 0.009846968 | 0.750319675 | 0.635831304 | 0.005492180 | 0.007205648 | 0.008582503 |
| 0.009846968 | 0.173116096 | 0.124658090 | 0.003110558 | 0.003920326 | 0.004492969 |
| 0.740906232 | 0.992096386 | 0.937293978 | 0.005492180 | 0.007205648 | 0.008582503 |
| 0.740906232 | 0.631182033 | 0.585904978 | 0.003110558 | 0.003920326 | 0.004492969 |
| 0.206012049 | 0.052456844 | 0.079086853 | 0.005492180 | 0.007205648 | 0.008582503 |
| 0.206012049 | 0.285664436 | 0.216787799 | 0.003110558 | 0.003920326 | 0.004492969 |
| 0.489478654 | 0.403532747 | 0.330153792 | 0.005492180 | 0.007205648 | 0.008582503 |
| 0.489478654 | 0.934261131 | 0.895078352 | 0.003110558 | 0.003920326 | 0.004492969 |

| | | | | | |
|---|---|---|---|---|---|
| 0.572106815 | 0.958489870 | 0.855069854 | 0.005492180 | 0.007205648 | 0.008582503 |
| 0.572106815 | 0.788122395 | 0.778578078 | 0.003110558 | 0.003920326 | 0.004492969 |
| 0.412801155 | 0.334315224 | 0.283116674 | 0.005492180 | 0.007205648 | 0.008582503 |
| 0.412801155 | 0.236495725 | 0.172006427 | 0.003110558 | 0.003920326 | 0.004492969 |
| 0.766363204 | 0.670807183 | 0.559383628 | 0.005492180 | 0.007205648 | 0.008582503 |
| 0.766363204 | 0.620827714 | 0.576680378 | 0.003110558 | 0.003920326 | 0.004492969 |
| 0.162881829 | 0.256270563 | 0.232428675 | 0.005492180 | 0.007205648 | 0.008582503 |
| 0.162881829 | 0.663329391 | 0.601725012 | 0.003110558 | 0.003920326 | 0.004492969 |
| 0.690662534 | 0.606561736 | 0.494730804 | 0.005492180 | 0.007205648 | 0.008582503 |
| 0.690662534 | 0.544497806 | 0.504452344 | 0.003110558 | 0.003920326 | 0.004492969 |
| 0.917607092 | 0.778985394 | 0.663326093 | 0.005492180 | 0.007205648 | 0.008582503 |
| 0.917607092 | 0.998089927 | 0.972431821 | 0.003110558 | 0.003920326 | 0.004492969 |
| 0.985740185 | 0.798518659 | 0.682842266 | 0.005492180 | 0.007205648 | 0.008582503 |
| 0.985740185 | 0.975154075 | 0.936050574 | 0.003110558 | 0.003920326 | 0.004492969 |
| 0.510538643 | 0.387709842 | 0.319004291 | 0.005492180 | 0.007205648 | 0.008582503 |
| 0.510538643 | 0.471031690 | 0.446206114 | 0.003110558 | 0.003920326 | 0.004492969 |
| 0.175529302 | 0.894185188 | 0.781095826 | 0.005492180 | 0.007205648 | 0.008582503 |
| 0.175529302 | 0.732873075 | 0.700359542 | 0.003110558 | 0.003920326 | 0.004492969 |
| 0.606703280 | 0.700846887 | 0.589504358 | 0.005492180 | 0.007205648 | 0.008582503 |
| 0.606703280 | 0.800097764 | 0.786501433 | 0.003110558 | 0.003920326 | 0.004492969 |
| 0.407936804 | 0.878694533 | 0.764802150 | 0.005492180 | 0.007205648 | 0.008582503 |
| 0.407936804 | 0.238135857 | 0.172922279 | 0.003110558 | 0.003920326 | 0.004492969 |
| 0.909343347 | 0.728306073 | 0.615981331 | 0.005492180 | 0.007205648 | 0.008582503 |
| 0.909343347 | 0.900744309 | 0.847639292 | 0.003110558 | 0.003920326 | 0.004492969 |
| 0.779465529 | 0.627144965 | 0.513579085 | 0.005492180 | 0.007205648 | 0.008582503 |
| 0.779465529 | 0.675570600 | 0.612155847 | 0.003110558 | 0.003920326 | 0.004492969 |
| 0.145181732 | 0.364587240 | 0.303182125 | 0.005492180 | 0.007205648 | 0.008582503 |
| 0.145181732 | 0.226539286 | 0.166736322 | 0.003110558 | 0.003920326 | 0.004492969 |
| 0.455543090 | 0.359525438 | 0.299840542 | 0.005492180 | 0.007205648 | 0.008582503 |
| 0.455543090 | 0.271059516 | 0.200676433 | 0.003110558 | 0.003920326 | 0.004492969 |
| 0.176189691 | 0.903832877 | 0.791570962 | 0.005492180 | 0.007205648 | 0.008582503 |
| 0.176189691 | 0.980042162 | 0.952053740 | 0.003110558 | 0.003920326 | 0.004492969 |
| 0.482534627 | 0.461759274 | 0.374567087 | 0.005492180 | 0.007205648 | 0.008582503 |
| 0.482534627 | 0.300626131 | 0.230275358 | 0.003110558 | 0.003920326 | 0.004492969 |
| 0.408844551 | 0.569846688 | 0.462499051 | 0.005492180 | 0.007205648 | 0.008582503 |
| 0.408844551 | 0.962418775 | 0.929910826 | 0.003110558 | 0.003920326 | 0.004492969 |
| 0.098702398 | 0.350346041 | 0.293808469 | 0.005492180 | 0.007205648 | 0.008582503 |
| 0.098702398 | 0.103349214 | 0.074008520 | 0.003110558 | 0.003920326 | 0.004492969 |
| 0.687837144 | 0.503133536 | 0.408797464 | 0.005492180 | 0.007205648 | 0.008582503 |
| 0.687837144 | 0.491410405 | 0.463043837 | 0.003110558 | 0.003920326 | 0.004492969 |
| 0.707172503 | 0.627637269 | 0.514063104 | 0.005492180 | 0.007205648 | 0.008582503 |
| 0.707172503 | 0.538477116 | 0.503670703 | 0.003110558 | 0.003920326 | 0.004492969 |
| 0.039762464 | 0.478592261 | 0.388497070 | 0.005492180 | 0.007205648 | 0.008582503 |
| 0.039762464 | 0.770701706 | 0.764577143 | 0.003110558 | 0.003920326 | 0.004492969 |
| 0.385945623 | 0.172060147 | 0.175087067 | 0.005492180 | 0.007205648 | 0.008582503 |
| 0.385945623 | 0.199909161 | 0.147307432 | 0.003110558 | 0.003920326 | 0.004492969 |

| | | | | | |
|---|---|---|---|---|---|
| 0.363227427 | 0.413357234 | 0.337278831 | 0.005492180 | 0.007205648 | 0.008582503 |
| 0.363227427 | 0.334599025 | 0.299600700 | 0.003110558 | 0.003920326 | 0.004492969 |
| 0.882439825 | 0.805796441 | 0.689702688 | 0.005492180 | 0.007205648 | 0.008582503 |
| 0.882439825 | 0.769111604 | 0.761380416 | 0.003110558 | 0.003920326 | 0.004492969 |
| 0.131662557 | 0.098358761 | 0.114096294 | 0.005492180 | 0.007205648 | 0.008582503 |
| 0.131662557 | 0.017318092 | 0.020095969 | 0.003110558 | 0.003920326 | 0.004492969 |
| 0.200676330 | 0.038142740 | 0.066434973 | 0.005492180 | 0.007205648 | 0.008582503 |
| 0.200676330 | 0.524839977 | 0.497440341 | 0.003110558 | 0.003920326 | 0.004492969 |
| 0.088344426 | 0.043589124 | 0.071448838 | 0.005492180 | 0.007205648 | 0.008582503 |
| 0.088344426 | 0.015907781 | 0.018678956 | 0.003110558 | 0.003920326 | 0.004492969 |
| 0.410278402 | 0.972661769 | 0.880450097 | 0.005492180 | 0.007205648 | 0.008582503 |
| 0.410278402 | 0.565102845 | 0.515889440 | 0.003110558 | 0.003920326 | 0.004492969 |
| 0.322500413 | 0.117453438 | 0.130648453 | 0.005492180 | 0.007205648 | 0.008582503 |
| 0.322500413 | 0.144086200 | 0.102011450 | 0.003110558 | 0.003920326 | 0.004492969 |
| 0.293998794 | 0.143097641 | 0.152171488 | 0.005492180 | 0.007205648 | 0.008582503 |
| 0.293998794 | 0.134533488 | 0.094212986 | 0.003110558 | 0.003920326 | 0.004492969 |
| 0.741387131 | 0.585575875 | 0.475461892 | 0.005492180 | 0.007205648 | 0.008582503 |
| 0.741387131 | 0.606284626 | 0.555328134 | 0.003110558 | 0.003920326 | 0.004492969 |
| 0.067233375 | 0.733601727 | 0.620875696 | 0.005492180 | 0.007205648 | 0.008582503 |
| 0.067233375 | 0.829138266 | 0.801431233 | 0.003110558 | 0.003920326 | 0.004492969 |
| 0.017668201 | 0.013444558 | 0.033079586 | 0.005492180 | 0.007205648 | 0.008582503 |
| 0.017668201 | 0.022983759 | 0.024636735 | 0.003110558 | 0.003920326 | 0.004492969 |
| 0.615039868 | 0.734669075 | 0.621826136 | 0.005492180 | 0.007205648 | 0.008582503 |
| 0.615039868 | 0.757417338 | 0.738598201 | 0.003110558 | 0.003920326 | 0.004492969 |
| 0.298406319 | 0.095326846 | 0.111566114 | 0.005492180 | 0.007205648 | 0.008582503 |
| 0.298406319 | 0.124010362 | 0.087004289 | 0.003110558 | 0.003920326 | 0.004492969 |
| 0.354766490 | 0.422577214 | 0.344366579 | 0.005492180 | 0.007205648 | 0.008582503 |
| 0.354766490 | 0.449477928 | 0.422391728 | 0.003110558 | 0.003920326 | 0.004492969 |
| 0.823038116 | 0.788106758 | 0.672699709 | 0.005492180 | 0.007205648 | 0.008582503 |
| 0.823038116 | 0.723304232 | 0.679518825 | 0.003110558 | 0.003920326 | 0.004492969 |
| 0.246195386 | 0.163751845 | 0.169008858 | 0.005492180 | 0.007205648 | 0.008582503 |
| 0.246195386 | 0.464198607 | 0.445767892 | 0.003110558 | 0.003920326 | 0.004492969 |
| 0.285862656 | 0.077543263 | 0.097823741 | 0.005492180 | 0.007205648 | 0.008582503 |
| 0.285862656 | 0.110113410 | 0.081014506 | 0.003110558 | 0.003920326 | 0.004492969 |
| 0.275353565 | 0.891411941 | 0.778043167 | 0.005492180 | 0.007205648 | 0.008582503 |
| 0.275353565 | 0.884167697 | 0.838656867 | 0.003110558 | 0.003920326 | 0.004492969 |
| 0.251143644 | 0.613972783 | 0.501391664 | 0.005492180 | 0.007205648 | 0.008582503 |
| 0.251143644 | 0.861212304 | 0.824987311 | 0.003110558 | 0.003920326 | 0.004492969 |
| 0.480136662 | 0.315578272 | 0.270837082 | 0.005492180 | 0.007205648 | 0.008582503 |
| 0.480136662 | 0.299192887 | 0.228413100 | 0.003110558 | 0.003920326 | 0.004492969 |
| 0.311143908 | 0.639554055 | 0.5222124 | 0.003882898 | 0.005253553 | 0.009464192 |
| 0.311143908 | 0.675398337 | 0.641582462 | 0.005675962 | 0.007312185 | 0.008680667 |
| 0.083737623 | 0.702794202 | 0.580555072 | 0.003882898 | 0.005253553 | 0.009464192 |
| 0.083737623 | 0.537788242 | 0.529026305 | 0.005675962 | 0.007312185 | 0.008680667 |
| 0.155119080 | 0.269213909 | 0.224827722 | 0.003882898 | 0.005253553 | 0.009464192 |
| 0.155119080 | 0.768329624 | 0.718608952 | 0.005675962 | 0.007312185 | 0.008680667 |

| | | | | | | |
|---|---|---|---|---|---|---|
| 0.920399309 | 0.925678019 | 0.783715291 | 0.003882898 | 0.005253553 | 0.009464192 | |
| 0.920399309 | 0.826916717 | 0.792875305 | 0.005675962 | 0.007312185 | 0.008680667 | |
| 0.004742590 | 0.881864675 | 0.739907458 | 0.003882898 | 0.005253553 | 0.009464192 | |
| 0.004742590 | 0.561536058 | 0.545247827 | 0.005675962 | 0.007312185 | 0.008680667 | |
| 0.516748890 | 0.395508908 | 0.309442986 | 0.003882898 | 0.005253553 | 0.009464192 | |
| 0.516748890 | 0.299671720 | 0.260664661 | 0.005675962 | 0.007312185 | 0.008680667 | |
| 0.445054657 | 0.600966247 | 0.483598856 | 0.003882898 | 0.005253553 | 0.009464192 | |
| 0.445054657 | 0.464261032 | 0.451782574 | 0.005675962 | 0.007312185 | 0.008680667 | |
| 0.167034184 | 0.082166135 | 0.090262305 | 0.003882898 | 0.005253553 | 0.009464192 | |
| 0.167034184 | 0.828543614 | 0.794185567 | 0.005675962 | 0.007312185 | 0.008680667 | |
| 0.006856048 | 0.174207826 | 0.161879187 | 0.003882898 | 0.005253553 | 0.009464192 | |
| 0.006856048 | 0.782703891 | 0.749118061 | 0.005675962 | 0.007312185 | 0.008680667 | |
| 0.353171092 | 0.433712301 | 0.336914117 | 0.003882898 | 0.005253553 | 0.009464192 | |
| 0.353171092 | 0.479233483 | 0.476035059 | 0.005675962 | 0.007312185 | 0.008680667 | |
| 0.806425841 | 0.738192188 | 0.614526376 | 0.003882898 | 0.005253553 | 0.009464192 | |
| 0.806425841 | 0.746277096 | 0.696666723 | 0.005675962 | 0.007312185 | 0.008680667 | |
| 0.496912958 | 0.432976507 | 0.336376727 | 0.003882898 | 0.005253553 | 0.009464192 | |
| 0.496912958 | 0.254223460 | 0.216646175 | 0.005675962 | 0.007312185 | 0.008680667 | |
| 0.727733522 | 0.683970298 | 0.563247772 | 0.003882898 | 0.005253553 | 0.009464192 | |
| 0.727733522 | 0.743050255 | 0.689853683 | 0.005675962 | 0.007312185 | 0.008680667 | |
| 0.005461124 | 0.001077198 | 0.004704348 | 0.003882898 | 0.005253553 | 0.009464192 | 0.15 |
| 0.005461124 | 0.120794110 | 0.126852474 | 0.005675962 | 0.007312185 | 0.008680667 | |
| 0.544727435 | 0.558347187 | 0.441650309 | 0.003882898 | 0.005253553 | 0.009464192 | |
| 0.544727435 | 0.514420400 | 0.505805286 | 0.005675962 | 0.007312185 | 0.008680667 | |
| 0.113948015 | 0.781506137 | 0.654083141 | 0.003882898 | 0.005253553 | 0.009464192 | |
| 0.113948015 | 0.580263028 | 0.570172959 | 0.005675962 | 0.007312185 | 0.008680667 | |
| 0.109671317 | 0.865336221 | 0.725317270 | 0.003882898 | 0.005253553 | 0.009464192 | |
| 0.109671317 | 0.495733429 | 0.495238899 | 0.005675962 | 0.007312185 | 0.008680667 | |
| 0.267317342 | 0.436645199 | 0.339071034 | 0.003882898 | 0.005253553 | 0.009464192 | |
| 0.267317342 | 0.135241196 | 0.136635676 | 0.005675962 | 0.007312185 | 0.008680667 | |
| 0.108744156 | 0.260559326 | 0.219136045 | 0.003882898 | 0.005253553 | 0.009464192 | |
| 0.108744156 | 0.247627131 | 0.211806884 | 0.005675962 | 0.007312185 | 0.008680667 | |
| 0.040472950 | 0.186426732 | 0.170143316 | 0.003882898 | 0.005253553 | 0.009464192 | |
| 0.040472950 | 0.643428311 | 0.605339411 | 0.005675962 | 0.007312185 | 0.008680667 | |
| 0.000196028 | 0.877313543 | 0.735907659 | 0.003882898 | 0.005253553 | 0.009464192 | |
| 0.000196028 | 0.129055301 | 0.131895132 | 0.005675962 | 0.007312185 | 0.008680667 | |
| 0.765328915 | 0.786513577 | 0.658286312 | 0.003882898 | 0.005253553 | 0.009464192 | |
| 0.765328915 | 0.672087250 | 0.639864085 | 0.005675962 | 0.007312185 | 0.008680667 | |
| 0.022164520 | 0.001318174 | 0.005421417 | 0.003882898 | 0.005253553 | 0.009464192 | 0.175 |
| 0.022164520 | 0.183805824 | 0.171353266 | 0.005675962 | 0.007312185 | 0.008680667 | |
| 0.560564123 | 0.352373307 | 0.280260471 | 0.003882898 | 0.005253553 | 0.009464192 | |
| 0.560564123 | 0.304659306 | 0.267514051 | 0.005675962 | 0.007312185 | 0.008680667 | |
| 0.074916580 | 0.194927563 | 0.175993764 | 0.003882898 | 0.005253553 | 0.009464192 | |
| 0.074916580 | 0.957170663 | 0.927619603 | 0.005675962 | 0.007312185 | 0.008680667 | |
| 0.551609507 | 0.549288120 | 0.433290079 | 0.003882898 | 0.005253553 | 0.009464192 | |
| 0.551609507 | 0.382344383 | 0.351681888 | 0.005675962 | 0.007312185 | 0.008680667 | |

| | | | | | |
|---|---|---|---|---|---|
| 0.090419915 | 0.038443613 | 0.053895656 | 0.003882898 | 0.005253553 | 0.009464192 |
| 0.090419915 | 0.749422922 | 0.702983863 | 0.005675962 | 0.007312185 | 0.008680667 |
| 0.902653898 | 0.955942527 | 0.820201609 | 0.003882898 | 0.005253553 | 0.009464192 |
| 0.902653898 | 0.657012993 | 0.621136333 | 0.005675962 | 0.007312185 | 0.008680667 |
| 0.342102933 | 0.141592036 | 0.137588640 | 0.003882898 | 0.005253553 | 0.009464192 |
| 0.342102933 | 0.178740907 | 0.165890198 | 0.005675962 | 0.007312185 | 0.008680667 |
| 0.212034033 | 0.325331277 | 0.262478417 | 0.003882898 | 0.005253553 | 0.009464192 |
| 0.212034033 | 0.270049489 | 0.232567558 | 0.005675962 | 0.007312185 | 0.008680667 |
| 0.441433677 | 0.561228809 | 0.444524673 | 0.003882898 | 0.005253553 | 0.009464192 |
| 0.441433677 | 0.491118000 | 0.490561542 | 0.005675962 | 0.007312185 | 0.008680667 |
| 0.807041499 | 0.723283545 | 0.600148895 | 0.003882898 | 0.005253553 | 0.009464192 |
| 0.807041499 | 0.877525848 | 0.827891246 | 0.005675962 | 0.007312185 | 0.008680667 |
| 0.845105244 | 0.681913383 | 0.561252962 | 0.003882898 | 0.005253553 | 0.009464192 |
| 0.845105244 | 0.519007622 | 0.513441708 | 0.005675962 | 0.007312185 | 0.008680667 |
| 0.276010469 | 0.139511395 | 0.135935120 | 0.003882898 | 0.005253553 | 0.009464192 |
| 0.276010469 | 0.708291019 | 0.660359994 | 0.005675962 | 0.007312185 | 0.008680667 |
| 0.126347114 | 0.095168599 | 0.100791436 | 0.003882898 | 0.005253553 | 0.009464192 |
| 0.126347114 | 0.052095125 | 0.072168228 | 0.005675962 | 0.007312185 | 0.008680667 |
| 0.395134193 | 0.924983175 | 0.782929058 | 0.003882898 | 0.005253553 | 0.009464192 |
| 0.395134193 | 0.866410416 | 0.825352262 | 0.005675962 | 0.007312185 | 0.008680667 |
| 0.074781929 | 0.349371624 | 0.278267683 | 0.003882898 | 0.005253553 | 0.009464192 |
| 0.074781929 | 0.811026355 | 0.780772256 | 0.005675962 | 0.007312185 | 0.008680667 |
| 0.669434989 | 0.554682275 | 0.438163519 | 0.003882898 | 0.005253553 | 0.009464192 |
| 0.669434989 | 0.400295486 | 0.384402171 | 0.005675962 | 0.007312185 | 0.008680667 |
| 0.712725531 | 0.573258746 | 0.456764027 | 0.003882898 | 0.005253553 | 0.009464192 |
| 0.712725531 | 0.389664153 | 0.364238724 | 0.005675962 | 0.007312185 | 0.008680667 |
| 0.381845971 | 0.172239927 | 0.16049123 | 0.003882898 | 0.005253553 | 0.009464192 |
| 0.381845971 | 0.951294461 | 0.920421998 | 0.005675962 | 0.007312185 | 0.008680667 |
| 0.01374849 | 0.773914700 | 0.647675900 | 0.003882898 | 0.005253553 | 0.009464192 |
| 0.01374849 | 0.352671743 | 0.316947649 | 0.005675962 | 0.007312185 | 0.008680667 |
| 0.088969196 | 0.206611187 | 0.183918077 | 0.003882898 | 0.005253553 | 0.009464192 |
| 0.088969196 | 0.658657721 | 0.622987872 | 0.005675962 | 0.007312185 | 0.008680667 |
| 0.589590424 | 0.773490347 | 0.647303231 | 0.003882898 | 0.005253553 | 0.009464192 |
| 0.589590424 | 0.335289221 | 0.296541009 | 0.005675962 | 0.007312185 | 0.008680667 |
| 0.786186079 | 0.602296505 | 0.484962431 | 0.003882898 | 0.005253553 | 0.009464192 |
| 0.786186079 | 0.475284537 | 0.473422115 | 0.005675962 | 0.007312185 | 0.008680667 |
| 0.967720368 | 0.905848172 | 0.762023650 | 0.003882898 | 0.005253553 | 0.009464192 |
| 0.967720368 | 0.915436673 | 0.867355117 | 0.005675962 | 0.007312185 | 0.008680667 |
| 0.975896833 | 0.870661630 | 0.729967782 | 0.003882898 | 0.005253553 | 0.009464192 |
| 0.975896833 | 0.729673015 | 0.671967004 | 0.005675962 | 0.007312185 | 0.008680667 |
| 0.486762349 | 0.427536728 | 0.332454282 | 0.003882898 | 0.005253553 | 0.009464192 |
| 0.486762349 | 0.323512458 | 0.279725032 | 0.005675962 | 0.007312185 | 0.008680667 |
| 0.371340917 | 0.641933040 | 0.524503406 | 0.003882898 | 0.005253553 | 0.009464192 |
| 0.371340917 | 0.426633432 | 0.422671933 | 0.005675962 | 0.007312185 | 0.008680667 |
| 0.433636045 | 0.319030216 | 0.258160168 | 0.003882898 | 0.005253553 | 0.009464192 |
| 0.433636045 | 0.163009507 | 0.155940382 | 0.005675962 | 0.007312185 | 0.008680667 |

| | | | | | |
|---|---|---|---|---|---|
| 0.7626807204 | 0.8767436939 | 0.7353948644 | 0.0038828986 | 0.0052535536 | 0.0094641923 |
| 0.7626807204 | 0.6074187224 | 0.5810951814 | 0.0056759627 | 0.0073121852 | 0.0086806679 |
| 0.9402264822 | 0.831678265 | 0.6975681764 | 0.0038828986 | 0.0052535536 | 0.0094641923 |
| 0.9402264822 | 0.8010731977 | 0.7695024275 | 0.0056759627 | 0.0073121852 | 0.0086806679 |
| 0.094302699 | 0.6414098105 | 0.5240079175 | 0.0038828986 | 0.0052535536 | 0.0094641923 |
| 0.094302699 | 0.9744701611 | 0.9442980716 | 0.0056759627 | 0.0073121852 | 0.0086806679 |
| 0.0728343894 | 0.1724409381 | 0.1606340284 | 0.0038828986 | 0.0052535536 | 0.0094641923 |
| 0.0728343894 | 0.2780218497 | 0.2399788632 | 0.0056759627 | 0.0073121852 | 0.0086806679 |
| 0.4176397691 | 0.5240658798 | 0.4108256295 | 0.0038828986 | 0.0052535536 | 0.0094641923 |
| 0.4176397691 | 0.1247983385 | 0.1292812092 | 0.0056759627 | 0.0073121852 | 0.0086806679 |
| 0.9748230942 | 0.9703301608 | 0.8456011267 | 0.0038828986 | 0.0052535536 | 0.0094641923 |
| 0.9748230942 | 0.8882814833 | 0.8381904149 | 0.0056759627 | 0.0073121852 | 0.0086806679 |
| 0.2618111392 | 0.1312848469 | 0.1292944763 | 0.0038828986 | 0.0052535536 | 0.0094641923 |
| 0.2618111392 | 0.1061476084 | 0.1144686052 | 0.0056759627 | 0.0073121852 | 0.0086806679 |
| 0.1025669129 | 0.1472200751 | 0.141972225 | 0.0038828986 | 0.0052535536 | 0.0094641923 |
| 0.1025669129 | 0.0213726164 | 0.0395718822 | 0.0056759627 | 0.0073121852 | 0.0086806679 |
| 0.1394356592 | 0.0764429891 | 0.0858072231 | 0.0038828986 | 0.0052535536 | 0.0094641923 |
| 0.1394356592 | 0.0168049373 | 0.0342316974 | 0.0056759627 | 0.0073121852 | 0.0086806679 |
| 0.2062131865 | 0.6848640248 | 0.5640908133 | 0.0038828986 | 0.0052535536 | 0.0094641923 |
| 0.2062131865 | 0.8048763573 | 0.7791298195 | 0.0056759627 | 0.0073121852 | 0.0086806679 |
| 0.3434189772 | 0.7100177192 | 0.5873908625 | 0.0038828986 | 0.0052535536 | 0.0094641923 |
| 0.3434189772 | 0.1984328339 | 0.1825687268 | 0.0056759627 | 0.0073121852 | 0.0086806679 |
| 0.278966344 | 0.2006552946 | 0.1798390136 | 0.0038828986 | 0.0052535536 | 0.0094641923 |
| 0.278966344 | 0.3616699505 | 0.3215570332 | 0.0056759627 | 0.0073121852 | 0.0086806679 |
| 0.2899667358 | 0.1577210438 | 0.1498553968 | 0.0038828986 | 0.0052535536 | 0.0094641923 |
| 0.2899667358 | 0.3105662315 | 0.2725438278 | 0.0056759627 | 0.0073121852 | 0.0086806679 |
| 0.6377841813 | 0.4750116609 | 0.3687342594 | 0.0038828986 | 0.0052535536 | 0.0094641923 |
| 0.6377841813 | 0.3302540933 | 0.2892486106 | 0.0056759627 | 0.0073121852 | 0.0086806679 |
| 0.5667583928 | 0.4939281265 | 0.3850302386 | 0.0038828986 | 0.0052535536 | 0.0094641923 |
| 0.5667583928 | 0.3886048379 | 0.3623313641 | 0.0056759627 | 0.0073121852 | 0.0086806679 |
| 0.0599576079 | 0.2614104597 | 0.2196926688 | 0.0038828986 | 0.0052535536 | 0.0094641923 |
| 0.0599576079 | 0.9015760008 | 0.8593513046 | 0.0056759627 | 0.0073121852 | 0.0086806679 |
| 0.1133081415 | 0.134992348 | 0.1323497201 | 0.0038828986 | 0.0052535536 | 0.0094641923 |
| 0.1133081415 | 0.7151855022 | 0.6649976249 | 0.0056759627 | 0.0073121852 | 0.0086806679 |
| 0.3595561041 | 0.1555775819 | 0.1483058215 | 0.0038828986 | 0.0052535536 | 0.0094641923 |
| 0.3595561041 | 0.2502258837 | 0.2125114732 | 0.0056759627 | 0.0073121852 | 0.0086806679 |
| 0.5380774745 | 0.9157912652 | 0.772418258 | 0.0038828986 | 0.0052535536 | 0.0094641923 |
| 0.5380774745 | 0.2756495599 | 0.2377706577 | 0.0056759627 | 0.0073121852 | 0.0086806679 |
| 0.214171622 | 0.7388124967 | 0.615142602 | 0.0038828986 | 0.0052535536 | 0.0094641923 |
| 0.214171622 | 0.9285120572 | 0.8764100873 | 0.0056759627 | 0.0073121852 | 0.0086806679 |
| 0.3707380036 | 0.4339594353 | 0.3370908863 | 0.0038828986 | 0.0052535536 | 0.0094641923 |
| 0.3707380036 | 0.7588091585 | 0.7050304181 | 0.0056759627 | 0.0073121852 | 0.0086806679 |
| 0.9350015243 | 0.8131711 | 0.6812841076 | 0.0038828986 | 0.0052535536 | 0.0094641923 |
| 0.9350015243 | 0.6192478377 | 0.5868383127 | 0.0056759627 | 0.0073121852 | 0.0086806679 |
| 0.8359426294 | 0.6902267717 | 0.5690362577 | 0.0038828986 | 0.0052535536 | 0.0094641923 |
| 0.8359426294 | 0.4841427438 | 0.4824989821 | 0.0056759627 | 0.0073121852 | 0.0086806679 |

| | | | | | |
|---|---|---|---|---|---|
| 0.519257570 | 0.365623155 | 0.289080014 | 0.003882898 | 0.005253553 | 0.009464192 |
| 0.519257570 | 0.329595578 | 0.288314919 | 0.005675962 | 0.007312185 | 0.008680667 |
| 0.194928740 | 0.122442922 | 0.122304623 | 0.003882898 | 0.005253553 | 0.009464192 |
| 0.194928740 | 0.169691636 | 0.159221932 | 0.005675962 | 0.007312185 | 0.008680667 |
| 0.151056344 | 0.029246919 | 0.043685953 | 0.003882898 | 0.005253553 | 0.009464192 |
| 0.151056344 | 0.038712858 | 0.059228103 | 0.005675962 | 0.007312185 | 0.008680667 |
| 0.244017552 | 0.379324048 | 0.298303616 | 0.003882898 | 0.005253553 | 0.009464192 |
| 0.244017552 | 0.446884535 | 0.438442267 | 0.005675962 | 0.007312185 | 0.008680667 |
| 0.380630279 | 0.869487761 | 0.728932971 | 0.003882898 | 0.005253553 | 0.009464192 |
| 0.380630279 | 0.262326494 | 0.225547970 | 0.005675962 | 0.007312185 | 0.008680667 |
| 0.035909443 | 0.037628307 | 0.053065833 | 0.003882898 | 0.005253553 | 0.009464192 |
| 0.035909443 | 0.043214215 | 0.065291425 | 0.005675962 | 0.007312185 | 0.008680667 |
| 0.029414659 | 0.719351753 | 0.596296043 | 0.003882898 | 0.005253553 | 0.009464192 |
| 0.029414659 | 0.232677582 | 0.205473665 | 0.005675962 | 0.007312185 | 0.008680667 |
| 0.241792867 | 0.932596591 | 0.791654705 | 0.003882898 | 0.005253553 | 0.009464192 |
| 0.241792867 | 0.685356615 | 0.651036384 | 0.005675962 | 0.007312185 | 0.008680667 |
| 0.689586146 | 0.511795906 | 0.400167585 | 0.003882898 | 0.005253553 | 0.009464192 |
| 0.689586146 | 0.472844747 | 0.467848365 | 0.005675962 | 0.007312185 | 0.008680667 |
| 0.092314239 | 0.199863833 | 0.179294787 | 0.003882898 | 0.005253553 | 0.009464192 |
| 0.092314239 | 0.940688072 | 0.913481175 | 0.005675962 | 0.007312185 | 0.008680667 |
| 0.278651146 | 0.895395041 | 0.752019222 | 0.003882898 | 0.005253553 | 0.009464192 |
| 0.278651146 | 0.848928097 | 0.816398360 | 0.005675962 | 0.007312185 | 0.008680667 |
| 0.308929823 | 0.275861393 | 0.229301476 | 0.003882898 | 0.005253553 | 0.009464192 |
| 0.308929823 | 0.140058376 | 0.137621747 | 0.005675962 | 0.007312185 | 0.008680667 |
| 0.227749799 | 0.249813031 | 0.212170083 | 0.003882898 | 0.005253553 | 0.009464192 |
| 0.227749799 | 0.191954617 | 0.176016240 | 0.005675962 | 0.007312185 | 0.008680667 |
| 0.356039355 | 0.307243034 | 0.250357755 | 0.003882898 | 0.005253553 | 0.009464192 |
| 0.356039355 | 0.722040773 | 0.665471866 | 0.005675962 | 0.007312185 | 0.008680667 |
| 0.032128921 | 0.203323342 | 0.181657090 | 0.003882898 | 0.005253553 | 0.009464192 |
| 0.032128921 | 0.600845359 | 0.578832065 | 0.005675962 | 0.007312185 | 0.008680667 |
| 0.470340087 | 0.323899911 | 0.261508296 | 0.003882898 | 0.005253553 | 0.009464192 |
| 0.470340087 | 0.994884592 | 0.977177278 | 0.005675962 | 0.007312185 | 0.008680667 |
| 0.914977994 | 0.747738573 | 0.623908124 | 0.003882898 | 0.005253553 | 0.009464192 |
| 0.914977994 | 0.524903888 | 0.525710513 | 0.005675962 | 0.007312185 | 0.008680667 |
| 0.500480334 | 0.383913863 | 0.301457642 | 0.003882898 | 0.005253553 | 0.009464192 |
| 0.500480334 | 0.534746543 | 0.526881189 | 0.005675962 | 0.007312185 | 0.008680667 |
| 0.000480626 | 0.833661004 | 0.699368700 | 0.003882898 | 0.005253553 | 0.009464192 |
| 0.000480626 | 0.789578076 | 0.750157562 | 0.005675962 | 0.007312185 | 0.008680667 |
| 0.984309632 | 0.981278321 | 0.871610025 | 0.003882898 | 0.005253553 | 0.009464192 |
| 0.984309632 | 0.965120837 | 0.935981158 | 0.005675962 | 0.007312185 | 0.008680667 |
| 0.720487445 | 0.929417417 | 0.787986718 | 0.003882898 | 0.005253553 | 0.009464192 |
| 0.720487445 | 0.450265101 | 0.439765817 | 0.005675962 | 0.007312185 | 0.008680667 |
| 0.556026588 | 0.373627099 | 0.294429285 | 0.003882898 | 0.005253553 | 0.009464192 |
| 0.556026588 | 0.923438272 | 0.874219162 | 0.005675962 | 0.007312185 | 0.008680667 |
| 0.653002233 | 0.511027175 | 0.399502017 | 0.003882898 | 0.005253553 | 0.009464192 |
| 0.653002233 | 0.599828939 | 0.578146191 | 0.005675962 | 0.007312185 | 0.008680667 |

| | | | | | |
|---|---|---|---|---|---|
| 0.697062937 | 0.630180296 | 0.513010479 | 0.003882898 | 0.005253553 | 0.009464192 |
| 0.697062937 | 0.455461298 | 0.450738922 | 0.005675962 | 0.007312185 | 0.008680667 |
| 0.585704187 | 0.488885211 | 0.380702605 | 0.003882898 | 0.005253553 | 0.009464192 |
| 0.585704187 | 0.254023915 | 0.216436978 | 0.005675962 | 0.007312185 | 0.008680667 |
| 0.312097660 | 0.917429593 | 0.774274662 | 0.003882898 | 0.005253553 | 0.009464192 |
| 0.312097660 | 0.985344081 | 0.960914998 | 0.005675962 | 0.007312185 | 0.008680667 |
| 0.225685958 | 0.376297565 | 0.296252710 | 0.003882898 | 0.005253553 | 0.009464192 |
| 0.225685958 | 0.080443972 | 0.093453618 | 0.005675962 | 0.007312185 | 0.008680667 |
| 0.392616679 | 0.543644771 | 0.428191122 | 0.003882898 | 0.005253553 | 0.009464192 |
| 0.392616679 | 0.968976009 | 0.942596830 | 0.005675962 | 0.007312185 | 0.008680667 |
| 0.406950961 | 0.424417116 | 0.330203554 | 0.003882898 | 0.005253553 | 0.009464192 |
| 0.406950961 | 0.229025259 | 0.202739327 | 0.005675962 | 0.007312185 | 0.008680667 |
| 0.853945736 | 0.702825897 | 0.580584601 | 0.003882898 | 0.005253553 | 0.009464192 |
| 0.853945736 | 0.522212321 | 0.519996537 | 0.005675962 | 0.007312185 | 0.008680667 |
| 0.454777063 | 0.380390107 | 0.299029604 | 0.003882898 | 0.005253553 | 0.009464192 |
| 0.454777063 | 0.433151371 | 0.425410330 | 0.005675962 | 0.007312185 | 0.008680667 |
| 0.246121043 | 0.508758795 | 0.397543244 | 0.003882898 | 0.005253553 | 0.009464192 |
| 0.246121043 | 0.838728258 | 0.805013887 | 0.005675962 | 0.007312185 | 0.008680667 |
| 0.271324873 | 0.110746655 | 0.113234149 | 0.003882898 | 0.005253553 | 0.009464192 |
| 0.271324873 | 0.068601278 | 0.083697869 | 0.005675962 | 0.007312185 | 0.008680667 |
| 0.004679709 | 0.288650041 | 0.237673185 | 0.003882898 | 0.005253553 | 0.009464192 |
| 0.004679709 | 0.402361157 | 0.388726100 | 0.005675962 | 0.007312185 | 0.008680667 |
| 0.225624026 | 0.290017979 | 0.238606401 | 0.003882898 | 0.005253553 | 0.009464192 |
| 0.225624026 | 0.214149524 | 0.192533899 | 0.005675962 | 0.007312185 | 0.008680667 |
| 0.068060363 | 0.080046688 | 0.088560213 | 0.003882898 | 0.005253553 | 0.009464192 |
| 0.068060363 | 0.019795889 | 0.037434919 | 0.005675962 | 0.007312185 | 0.008680667 |
| 0.098072140 | 0.017236500 | 0.030357771 | 0.003882898 | 0.005253553 | 0.009464192 |
| 0.098072140 | 0.587147347 | 0.573058282 | 0.005675962 | 0.007312185 | 0.008680667 |
| 0.706351085 | 0.552968726 | 0.436607509 | 0.003882898 | 0.005253553 | 0.009464192 |
| 0.706351085 | 0.380189134 | 0.348016052 | 0.005675962 | 0.007312185 | 0.008680667 |
| 0.828558735 | 0.996123422 | 0.922771095 | 0.003882898 | 0.005253553 | 0.009464192 |
| 0.828558735 | 0.746658115 | 0.697628935 | 0.005675962 | 0.007312185 | 0.008680667 |
| 0.901163472 | 0.738513821 | 0.614847682 | 0.003882898 | 0.005253553 | 0.009464192 |
| 0.901163472 | 0.515589541 | 0.506585468 | 0.005675962 | 0.007312185 | 0.008680667 |
| 0.831521220 | 0.687659865 | 0.566669303 | 0.003882898 | 0.005253553 | 0.009464192 |
| 0.831521220 | 0.524614003 | 0.525099125 | 0.005675962 | 0.007312185 | 0.008680667 |
| 0.568403707 | 0.420616661 | 0.327489180 | 0.003882898 | 0.005253553 | 0.009464192 |
| 0.568403707 | 0.558724969 | 0.541467343 | 0.005675962 | 0.007312185 | 0.008680667 |
| 0.042142385 | 0.694910426 | 0.573312029 | 0.003882898 | 0.005253553 | 0.009464192 |
| 0.042142385 | 0.880784070 | 0.835789510 | 0.005675962 | 0.007312185 | 0.008680667 |
| 0.222350587 | 0.084198833 | 0.091909807 | 0.003882898 | 0.005253553 | 0.009464192 |
| 0.222350587 | 0.072251020 | 0.086148726 | 0.005675962 | 0.007312185 | 0.008680667 |
| 0.903609397 | 0.925931918 | 0.784005261 | 0.003882898 | 0.005253553 | 0.009464192 |
| 0.903609397 | 0.879740176 | 0.833284791 | 0.005675962 | 0.007312185 | 0.008680667 |
| 0.172222585 | 0.041919151 | 0.057234443 | 0.003882898 | 0.005253553 | 0.009464192 |
| 0.172222585 | 0.014381055 | 0.030182647 | 0.005675962 | 0.007312185 | 0.008680667 |

| | | | | | |
|---|---|---|---|---|---|
| 0.346680396 | 0.406023330 | 0.317052143 | 0.003882898 | 0.005253553 | 0.009464192 |
| 0.346680396 | 0.50734297 | 0.505069930 | 0.005675962 | 0.007312185 | 0.008680667 |
| 0.510642139 | 0.519668456 | 0.406981997 | 0.003882898 | 0.005253553 | 0.009464192 |
| 0.510642139 | 0.620302332 | 0.586961774 | 0.005675962 | 0.007312185 | 0.008680667 |
| 0.098559463 | 0.124041853 | 0.123531842 | 0.003882898 | 0.005253553 | 0.009464192 |
| 0.098559463 | 0.525988408 | 0.526320770 | 0.005675962 | 0.007312185 | 0.008680667 |
| 0.573472994 | 0.36387695 | 0.287928422 | 0.003882898 | 0.005253553 | 0.009464192 |
| 0.573472994 | 0.257989876 | 0.220695074 | 0.005675962 | 0.007312185 | 0.008680667 |
| 0.805256664 | 0.969733719 | 0.844323861 | 0.003882898 | 0.005253553 | 0.009464192 |
| 0.805256664 | 0.916714867 | 0.867510555 | 0.005675962 | 0.007312185 | 0.008680667 |
| 0.340353839 | 0.601992832 | 0.484650450 | 0.003882898 | 0.005253553 | 0.009464192 |
| 0.340353839 | 0.895713334 | 0.855654462 | 0.005675962 | 0.007312185 | 0.008680667 |
| 0.318154191 | 0.179952328 | 0.165804317 | 0.003882898 | 0.005253553 | 0.009464192 |
| 0.318154191 | 0.123009562 | 0.127281380 | 0.005675962 | 0.007312185 | 0.008680667 |
| 0.716787071 | 0.514913366 | 0.581611261 | 0.003303901 | 0.004786220 | 0.005872570 |
| 0.716787071 | 0.479725178 | 0.568323788 | 0.005915167 | 0.005975504 | 0.006080996 |
| 0.274928658 | 0.564573957 | 0.620636377 | 0.003303901 | 0.004786220 | 0.005872570 |
| 0.274928658 | 0.336156677 | 0.440508225 | 0.005915167 | 0.005975504 | 0.006080996 |
| 0.353899452 | 0.200887337 | 0.213773672 | 0.003303901 | 0.004786220 | 0.005872570 |
| 0.353899452 | 0.602483297 | 0.658201900 | 0.005915167 | 0.005975504 | 0.006080996 |
| 0.077348093 | 0.364633667 | 0.365222827 | 0.003303901 | 0.004786220 | 0.005872570 |
| 0.077348093 | 0.398787061 | 0.503729803 | 0.005915167 | 0.005975504 | 0.006080996 |
| 0.153834613 | 0.770077557 | 0.815217235 | 0.003303901 | 0.004786220 | 0.005872570 |
| 0.153834613 | 0.359896338 | 0.458540390 | 0.005915167 | 0.005975504 | 0.006080996 |
| 0.230214775 | 0.081152671 | 0.105429761 | 0.003303901 | 0.004786220 | 0.005872570 |
| 0.230214775 | 0.059209518 | 0.062892899 | 0.005915167 | 0.005975504 | 0.006080996 |
| 0.750521134 | 0.544903967 | 0.601400227 | 0.003303901 | 0.004786220 | 0.005872570 |
| 0.750521134 | 0.473058294 | 0.564628321 | 0.005915167 | 0.005975504 | 0.006080996 |
| 0.961199552 | 0.934288110 | 0.931533387 | 0.003303901 | 0.004786220 | 0.005872570 |
| 0.961199552 | 0.918940978 | 0.925730142 | 0.005915167 | 0.005975504 | 0.006080996 |
| 0.734014513 | 0.510834004 | 0.578197263 | 0.003303901 | 0.004786220 | 0.005872570 |
| 0.734014513 | 0.623402177 | 0.694953184 | 0.005915167 | 0.005975504 | 0.006080996 |
| 0.685513403 | 0.463489385 | 0.513565853 | 0.003303901 | 0.004786220 | 0.005872570 |
| 0.685513403 | 0.373115692 | 0.471231551 | 0.005915167 | 0.005975504 | 0.006080996 |
| 0.895717931 | 0.784830708 | 0.821466345 | 0.003303901 | 0.004786220 | 0.005872570 |
| 0.895717931 | 0.757189213 | 0.803970496 | 0.005915167 | 0.005975504 | 0.006080996 |
| 0.404134225 | 0.232189753 | 0.239081977 | 0.003303901 | 0.004786220 | 0.005872570 |
| 0.404134225 | 0.151308676 | 0.163898595 | 0.005915167 | 0.005975504 | 0.006080996 |
| 0.758718073 | 0.554323994 | 0.612852465 | 0.003303901 | 0.004786220 | 0.005872570 |
| 0.758718073 | 0.562609908 | 0.62408437 | 0.005915167 | 0.005975504 | 0.006080996 |
| 0.247806956 | 0.074125507 | 0.098702446 | 0.003303901 | 0.004786220 | 0.005872570 |
| 0.247806956 | 0.054706918 | 0.061890672 | 0.005915167 | 0.005975504 | 0.006080996 |
| 0.951501250 | 0.916934666 | 0.915183986 | 0.003303901 | 0.004786220 | 0.005872570 |
| 0.951501250 | 0.894382171 | 0.906949124 | 0.005915167 | 0.005975504 | 0.006080996 |
| 0.500127421 | 0.700441850 | 0.748506142 | 0.003303901 | 0.004786220 | 0.005872570 |
| 0.500127421 | 0.375946833 | 0.486750709 | 0.005915167 | 0.005975504 | 0.006080996 |

| | | | | | |
|---|---|---|---|---|---|
| 0.120188932 | 0.734843251 | 0.762587490 | 0.003303901 | 0.004786220 | 0.005872570 |
| 0.120188932 | 0.298127482 | 0.403098997 | 0.005915167 | 0.005975504 | 0.006080996 |
| 0.786498134 | 0.605215350 | 0.652708820 | 0.003303901 | 0.004786220 | 0.005872570 |
| 0.786498134 | 0.552358752 | 0.619599545 | 0.005915167 | 0.005975504 | 0.006080996 |
| 0.449135987 | 0.246886348 | 0.251633715 | 0.003303901 | 0.004786220 | 0.005872570 |
| 0.449135987 | 0.185103680 | 0.202520069 | 0.005915167 | 0.005975504 | 0.006080996 |
| 0.149662105 | 0.113209162 | 0.142542561 | 0.003303901 | 0.004786220 | 0.005872570 |
| 0.149662105 | 0.437342863 | 0.526412761 | 0.005915167 | 0.005975504 | 0.006080996 |
| 0.436707853 | 0.870452901 | 0.867932081 | 0.003303901 | 0.004786220 | 0.005872570 |
| 0.436707853 | 0.181157215 | 0.191392394 | 0.005915167 | 0.005975504 | 0.006080996 |
| 0.977795647 | 0.976838905 | 0.961206973 | 0.003303901 | 0.004786220 | 0.005872570 |
| 0.977795647 | 0.972025076 | 0.957781277 | 0.005915167 | 0.005975504 | 0.006080996 |
| 0.392113033 | 0.172989999 | 0.201427043 | 0.003303901 | 0.004786220 | 0.005872570 |
| 0.392113033 | 0.136890092 | 0.154441222 | 0.005915167 | 0.005975504 | 0.006080996 |
| 0.712663245 | 0.481963160 | 0.548855337 | 0.003303901 | 0.004786220 | 0.005872570 |
| 0.712663245 | 0.416291054 | 0.508797599 | 0.005915167 | 0.005975504 | 0.006080996 |
| 0.560969939 | 0.362691781 | 0.364316254 | 0.003303901 | 0.004786220 | 0.005872570 |
| 0.560969939 | 0.908937482 | 0.911476609 | 0.005915167 | 0.005975504 | 0.006080996 |
| 0.102023226 | 0.303697170 | 0.298051326 | 0.003303901 | 0.004786220 | 0.005872570 |
| 0.102023226 | 0.163370638 | 0.178114989 | 0.005915167 | 0.005975504 | 0.006080996 |
| 0.340819558 | 0.129615307 | 0.161207287 | 0.003303901 | 0.004786220 | 0.005872570 |
| 0.340819558 | 0.583893643 | 0.639838369 | 0.005915167 | 0.005975504 | 0.006080996 |
| 0.266631171 | 0.650903563 | 0.697911390 | 0.003303901 | 0.004786220 | 0.005872570 |
| 0.266631171 | 0.096354472 | 0.108446924 | 0.005915167 | 0.005975504 | 0.006080996 |
| 0.790806758 | 0.609482617 | 0.658488738 | 0.003303901 | 0.004786220 | 0.005872570 |
| 0.790806758 | 0.564642184 | 0.626404454 | 0.005915167 | 0.005975504 | 0.006080996 |
| 0.021933790 | 0.142908334 | 0.173749682 | 0.003303901 | 0.004786220 | 0.005872570 |
| 0.021933790 | 0.123392110 | 0.139240329 | 0.005915167 | 0.005975504 | 0.006080996 |
| 0.538601681 | 0.397361669 | 0.423925994 | 0.003303901 | 0.004786220 | 0.005872570 |
| 0.538601681 | 0.297083702 | 0.398046903 | 0.005915167 | 0.005975504 | 0.006080996 |
| 0.821976348 | 0.657178257 | 0.705323074 | 0.003303901 | 0.004786220 | 0.005872570 |
| 0.821976348 | 0.736734944 | 0.788748045 | 0.005915167 | 0.005975504 | 0.006080996 |
| 0.521256159 | 0.319966250 | 0.321700115 | 0.003303901 | 0.004786220 | 0.005872570 |
| 0.521256159 | 0.229842643 | 0.272439427 | 0.005915167 | 0.005975504 | 0.006080996 |
| 0.776527205 | 0.578099472 | 0.637865707 | 0.003303901 | 0.004786220 | 0.005872570 |
| 0.776527205 | 0.538739228 | 0.604538262 | 0.005915167 | 0.005975504 | 0.006080996 |
| 0.917422680 | 0.859678694 | 0.857029390 | 0.003303901 | 0.004786220 | 0.005872570 |
| 0.917422680 | 0.831182534 | 0.842729277 | 0.005915167 | 0.005975504 | 0.006080996 |
| 0.363789123 | 0.766330661 | 0.809090320 | 0.003303901 | 0.004786220 | 0.005872570 |
| 0.363789123 | 0.726097265 | 0.785712082 | 0.005915167 | 0.005975504 | 0.006080996 |
| 0.118701749 | 0.233549887 | 0.240007048 | 0.003303901 | 0.004786220 | 0.005872570 |
| 0.118701749 | 0.670652105 | 0.732097660 | 0.005915167 | 0.005975504 | 0.006080996 |
| 0.915861685 | 0.844443233 | 0.854503904 | 0.003303901 | 0.004786220 | 0.005872570 |
| 0.915861685 | 0.817668920 | 0.840005606 | 0.005915167 | 0.005975504 | 0.006080996 |
| 0.995038185 | 0.992385725 | 0.991290231 | 0.003303901 | 0.004786220 | 0.005872570 |
| 0.995038185 | 0.991178441 | 0.991242877 | 0.005915167 | 0.005975504 | 0.006080996 |

| | | | | | | |
|---|---|---|---|---|---|---|
| 0.786140247 | 0.5944025594 | 0.6516679660 | 0.0033039012 | 0.0047862201 | 0.0058725700 | |
| 0.786140247 | 0.8879745597 | 0.9024343148 | 0.0059151673 | 0.0059755040 | 0.0060809960 | |
| 0.7747870695 | 0.7839876498 | 0.821368215 | 0.0033039012 | 0.0047862201 | 0.0058725700 | |
| 0.7747870695 | 0.5303091970 | 0.6015360400 | 0.0059151673 | 0.0059755040 | 0.0060809960 | |
| 0.9642736767 | 0.9450901032 | 0.9370941580 | 0.0033039012 | 0.0047862201 | 0.0058725700 | |
| 0.9642736767 | 0.9333133357 | 0.9315267068 | 0.0059151673 | 0.0059755040 | 0.0060809960 | |
| 0.4909007063 | 0.5632747954 | 0.6170171489 | 0.0033039012 | 0.0047862201 | 0.0058725700 | |
| 0.4909007063 | 0.2197603999 | 0.2416928574 | 0.0059151673 | 0.0059755040 | 0.0060809960 | |
| 0.170848058 | 0.0443687220 | 0.0575415853 | 0.0033039012 | 0.0047862201 | 0.0058725700 | |
| 0.170848058 | 0.0606471550 | 0.0651736680 | 0.0059151673 | 0.0059755040 | 0.0060809960 | |
| 0.5418389817 | 0.3437773527 | 0.3435779598 | 0.0033039012 | 0.0047862201 | 0.0058725700 | |
| 0.5418389817 | 0.4988006988 | 0.5807961598 | 0.0059151673 | 0.0059755040 | 0.0060809960 | |
| 0.2661967843 | 0.0866545691 | 0.1100919619 | 0.0033039012 | 0.0047862201 | 0.0058725700 | |
| 0.2661967843 | 0.0673983068 | 0.0713758405 | 0.0059151673 | 0.0059755040 | 0.0060809960 | |
| 0.1272459527 | 0.1651836678 | 0.1936440104 | 0.0033039012 | 0.0047862201 | 0.0058725700 | |
| 0.1272459527 | 0.1081422110 | 0.1179058902 | 0.0059151673 | 0.0059755040 | 0.0060809960 | |
| 0.9530790465 | 0.9266460703 | 0.918032754 | 0.0033039012 | 0.0047862201 | 0.0058725700 | |
| 0.9530790465 | 0.9013860620 | 0.9098821579 | 0.0059151673 | 0.0059755040 | 0.0060809960 | |
| 0.3148378529 | 0.113401538 | 0.1425645662 | 0.0033039012 | 0.0047862201 | 0.0058725700 | |
| 0.3148378529 | 0.0853960581 | 0.0995444177 | 0.0059151673 | 0.0059755040 | 0.0060809960 | |
| 0.9709621768 | 0.9671481515 | 0.9492590265 | 0.0033039012 | 0.0047862201 | 0.0058725700 | |
| 0.9709621768 | 0.9552569693 | 0.9443168193 | 0.0059151673 | 0.0059755040 | 0.0060809960 | |
| 0.6748791929 | 0.4464095434 | 0.4995511274 | 0.0033039012 | 0.0047862201 | 0.0058725700 | |
| 0.6748791929 | 0.3582242144 | 0.4567321704 | 0.0059151673 | 0.0059755040 | 0.0060809960 | |
| 0.4975642384 | 0.5310497728 | 0.5938628380 | 0.0033039012 | 0.0047862201 | 0.0058725700 | |
| 0.4975642384 | 0.93910472 | 0.9320291811 | 0.0059151673 | 0.0059755040 | 0.0060809960 | |
| 0.4223945817 | 0.2173140314 | 0.2275605941 | 0.0033039012 | 0.0047862201 | 0.0058725700 | |
| 0.4223945817 | 0.1706696710 | 0.1792210547 | 0.0059151673 | 0.0059755040 | 0.0060809960 | |
| 0.0759861703 | 0.3042651955 | 0.2988952642 | 0.0033039012 | 0.0047862201 | 0.0058725700 | |
| 0.0759861703 | 0.0007625708 | 0.0059104087 | 0.0059151673 | 0.0059755040 | 0.0060809960 | 0.1 |
| 0.2098522383 | 0.3721576102 | 0.3818991758 | 0.0033039012 | 0.0047862201 | 0.0058725700 | |
| 0.2098522383 | 0.3407931768 | 0.446592443 | 0.0059151673 | 0.0059755040 | 0.0060809960 | |
| 0.4741756048 | 0.2684381631 | 0.2748747188 | 0.0033039012 | 0.0047862201 | 0.0058725700 | |
| 0.4741756048 | 0.2048207673 | 0.2254993791 | 0.0059151673 | 0.0059755040 | 0.0060809960 | |
| 0.3811568824 | 0.1640893530 | 0.1927604867 | 0.0033039012 | 0.0047862201 | 0.0058725700 | |
| 0.3811568824 | 0.1264780698 | 0.1457446192 | 0.0059151673 | 0.0059755040 | 0.0060809960 | |
| 0.3599627258 | 0.1457437599 | 0.1758239828 | 0.0033039012 | 0.0047862201 | 0.0058725700 | |
| 0.3599627258 | 0.1182641299 | 0.1299891939 | 0.0059151673 | 0.0059755040 | 0.0060809960 | |
| 0.4046763302 | 0.5315918052 | 0.5955720803 | 0.0033039012 | 0.0047862201 | 0.0058725700 | |
| 0.4046763302 | 0.652534538 | 0.7302556355 | 0.0059151673 | 0.0059755040 | 0.0060809960 | |
| 0.9380755815 | 0.8948914227 | 0.8924002352 | 0.0033039012 | 0.0047862201 | 0.0058725700 | |
| 0.9380755815 | 0.8615235137 | 0.8812025300 | 0.0059151673 | 0.0059755040 | 0.0060809960 | |
| 0.6817815824 | 0.4548724924 | 0.5085239982 | 0.0033039012 | 0.0047862201 | 0.0058725700 | |
| 0.6817815824 | 0.3670489998 | 0.465978673 | 0.0059151673 | 0.0059755040 | 0.0060809960 | |
| 0.1022900959 | 0.0384878947 | 0.0506739404 | 0.0033039012 | 0.0047862201 | 0.0058725700 | |
| 0.1022900959 | 0.160131986 | 0.1754480529 | 0.0059151673 | 0.0059755040 | 0.0060809960 | |

| | | | | | |
|---|---|---|---|---|---|
| 0.844129144 | 0.683028904 | 0.738984911 | 0.003303901 | 0.004786220 | 0.005872570 |
| 0.844129144 | 0.630770908 | 0.713509515 | 0.005915167 | 0.005975504 | 0.006080996 |
| 0.742337084 | 0.527215577 | 0.589746737 | 0.003303901 | 0.004786220 | 0.005872570 |
| 0.742337084 | 0.460772320 | 0.552229272 | 0.005915167 | 0.005975504 | 0.006080996 |
| 0.848391415 | 0.695915013 | 0.745936286 | 0.003303901 | 0.004786220 | 0.005872570 |
| 0.848391415 | 0.800477366 | 0.827944131 | 0.005915167 | 0.005975504 | 0.006080996 |
| 0.481524955 | 0.280941760 | 0.282080745 | 0.003303901 | 0.004786220 | 0.005872570 |
| 0.481524955 | 0.520200196 | 0.595483617 | 0.005915167 | 0.005975504 | 0.006080996 |
| 0.717328723 | 0.495267392 | 0.555144669 | 0.003303901 | 0.004786220 | 0.005872570 |
| 0.717328723 | 0.428237703 | 0.515614285 | 0.005915167 | 0.005975504 | 0.006080996 |
| 0.830921506 | 0.840971163 | 0.852130943 | 0.003303901 | 0.004786220 | 0.005872570 |
| 0.830921506 | 0.612347030 | 0.691420262 | 0.005915167 | 0.005975504 | 0.006080996 |
| 0.900142032 | 0.813796307 | 0.829079950 | 0.003303901 | 0.004786220 | 0.005872570 |
| 0.900142032 | 0.851873274 | 0.848655822 | 0.005915167 | 0.005975504 | 0.006080996 |
| 0.490270963 | 0.314148539 | 0.311375248 | 0.003303901 | 0.004786220 | 0.005872570 |
| 0.490270963 | 0.593996503 | 0.642066919 | 0.005915167 | 0.005975504 | 0.006080996 |
| 0.480397721 | 0.275462544 | 0.280868450 | 0.003303901 | 0.004786220 | 0.005872570 |
| 0.480397721 | 0.208105121 | 0.231415784 | 0.005915167 | 0.005975504 | 0.006080996 |
| 0.396919085 | 0.183543575 | 0.205575352 | 0.003303901 | 0.004786220 | 0.005872570 |
| 0.396919085 | 0.145042305 | 0.158097636 | 0.005915167 | 0.005975504 | 0.006080996 |
| 0.854250490 | 0.711909098 | 0.754824488 | 0.003303901 | 0.004786220 | 0.005872570 |
| 0.854250490 | 0.658612551 | 0.730913197 | 0.005915167 | 0.005975504 | 0.006080996 |
| 0.854618552 | 0.719055634 | 0.755450900 | 0.003303901 | 0.004786220 | 0.005872570 |
| 0.854618552 | 0.665138025 | 0.731507471 | 0.005915167 | 0.005975504 | 0.006080996 |
| 0.099496463 | 0.012063550 | 0.027821360 | 0.003303901 | 0.004786220 | 0.005872570 |
| 0.099496463 | 0.007844734 | 0.013087401 | 0.005915167 | 0.005975504 | 0.006080996 |
| 0.579843705 | 0.377642465 | 0.385443118 | 0.003303901 | 0.004786220 | 0.005872570 |
| 0.579843705 | 0.274913852 | 0.342182304 | 0.005915167 | 0.005975504 | 0.006080996 |
| 0.159057347 | 0.653507389 | 0.704377160 | 0.003303901 | 0.004786220 | 0.005872570 |
| 0.159057347 | 0.111062583 | 0.119943488 | 0.005915167 | 0.005975504 | 0.006080996 |
| 0.038295587 | 0.009585328 | 0.024643235 | 0.003303901 | 0.004786220 | 0.005872570 |
| 0.038295587 | 0.011169368 | 0.018553081 | 0.005915167 | 0.005975504 | 0.006080996 |
| 0.232481516 | 0.585570753 | 0.645309705 | 0.003303901 | 0.004786220 | 0.005872570 |
| 0.232481516 | 0.103612264 | 0.116243495 | 0.005915167 | 0.005975504 | 0.006080996 |
| 0.893397374 | 0.979390582 | 0.966215773 | 0.003303901 | 0.004786220 | 0.005872570 |
| 0.893397374 | 0.750086787 | 0.799812445 | 0.005915167 | 0.005975504 | 0.006080996 |
| 0.433261142 | 0.227804305 | 0.236879664 | 0.003303901 | 0.004786220 | 0.005872570 |
| 0.433261142 | 0.189149868 | 0.207552086 | 0.005915167 | 0.005975504 | 0.006080996 |
| 0.853045268 | 0.705803975 | 0.753129974 | 0.003303901 | 0.004786220 | 0.005872570 |
| 0.853045268 | 0.881026783 | 0.894297754 | 0.005915167 | 0.005975504 | 0.006080996 |
| 0.541711559 | 0.818171513 | 0.836285144 | 0.003303901 | 0.004786220 | 0.005872570 |
| 0.541711559 | 0.713830044 | 0.775192466 | 0.005915167 | 0.005975504 | 0.006080996 |
| 0.782533394 | 0.589120679 | 0.646773348 | 0.003303901 | 0.004786220 | 0.005872570 |
| 0.782533394 | 0.539918867 | 0.613341171 | 0.005915167 | 0.005975504 | 0.006080996 |
| 0.750397753 | 0.538040230 | 0.600936449 | 0.003303901 | 0.004786220 | 0.005872570 |
| 0.750397753 | 0.465630574 | 0.564149535 | 0.005915167 | 0.005975504 | 0.006080996 |

| | | | | | |
|---|---|---|---|---|---|
| 0.674225154 | 0.442394743 | 0.498906181 | 0.003303901 | 0.004786220 | 0.005872570 |
| 0.674225154 | 0.526140948 | 0.595945872 | 0.005915167 | 0.005975504 | 0.006080996 |
| 0.395153326 | 0.181545925 | 0.204530043 | 0.003303901 | 0.004786220 | 0.005872570 |
| 0.395153326 | 0.390802484 | 0.496523115 | 0.005915167 | 0.005975504 | 0.006080996 |
| 0.761329068 | 0.558560214 | 0.616173427 | 0.003303901 | 0.004786220 | 0.005872570 |
| 0.761329068 | 0.9881362 | 0.972683935 | 0.005915167 | 0.005975504 | 0.006080996 |
| 0.911723104 | 0.831454740 | 0.847356478 | 0.003303901 | 0.004786220 | 0.005872570 |
| 0.911723104 | 0.801338342 | 0.832206954 | 0.005915167 | 0.005975504 | 0.006080996 |
| 0.645602280 | 0.416764678 | 0.462850265 | 0.003303901 | 0.004786220 | 0.005872570 |
| 0.645602280 | 0.334125997 | 0.438037019 | 0.005915167 | 0.005975504 | 0.006080996 |
| 0.091533208 | 0.702912884 | 0.752789442 | 0.003303901 | 0.004786220 | 0.005872570 |
| 0.091533208 | 0.624823797 | 0.695556428 | 0.005915167 | 0.005975504 | 0.006080996 |
| 0.178322047 | 0.403956278 | 0.441620541 | 0.003303901 | 0.004786220 | 0.005872570 |
| 0.178322047 | 0.717652239 | 0.782861445 | 0.005915167 | 0.005975504 | 0.006080996 |
| 0.738835745 | 0.764374938 | 0.805860347 | 0.003303901 | 0.004786220 | 0.005872570 |
| 0.738835745 | 0.452626200 | 0.546936054 | 0.005915167 | 0.005975504 | 0.006080996 |
| 0.899733636 | 0.798813285 | 0.827648566 | 0.003303901 | 0.004786220 | 0.005872570 |
| 0.899733636 | 0.844525044 | 0.845555658 | 0.005915167 | 0.005975504 | 0.006080996 |
| 0.247931715 | 0.077617158 | 0.098951428 | 0.003303901 | 0.004786220 | 0.005872570 |
| 0.247931715 | 0.384180433 | 0.494897220 | 0.005915167 | 0.005975504 | 0.006080996 |
| 0.907758785 | 0.824805205 | 0.841269410 | 0.003303901 | 0.004786220 | 0.005872570 |
| 0.907758785 | 0.789653145 | 0.825330031 | 0.005915167 | 0.005975504 | 0.006080996 |
| 0.399818221 | 0.190075626 | 0.208194144 | 0.003303901 | 0.004786220 | 0.005872570 |
| 0.399818221 | 0.147083554 | 0.160279482 | 0.005915167 | 0.005975504 | 0.006080996 |
| 0.611444888 | 0.871187368 | 0.869455760 | 0.003303901 | 0.004786220 | 0.005872570 |
| 0.611444888 | 0.963332203 | 0.952494657 | 0.005915167 | 0.005975504 | 0.006080996 |
| 0.929408167 | 0.878762686 | 0.877423007 | 0.003303901 | 0.004786220 | 0.005872570 |
| 0.929408167 | 0.853243218 | 0.864885619 | 0.005915167 | 0.005975504 | 0.006080996 |
| 0.076528079 | 0.324672823 | 0.324483317 | 0.003303901 | 0.004786220 | 0.005872570 |
| 0.076528079 | 0.920315835 | 0.929130525 | 0.005915167 | 0.005975504 | 0.006080996 |
| 0.407698044 | 0.273362522 | 0.279440925 | 0.003303901 | 0.004786220 | 0.005872570 |
| 0.407698044 | 0.158391582 | 0.166882810 | 0.005915167 | 0.005975504 | 0.006080996 |
| 0.225354828 | 0.064236718 | 0.085822144 | 0.003303901 | 0.004786220 | 0.005872570 |
| 0.225354828 | 0.040869826 | 0.051214175 | 0.005915167 | 0.005975504 | 0.006080996 |
| 0.968715391 | 0.954885541 | 0.944881281 | 0.003303901 | 0.004786220 | 0.005872570 |
| 0.968715391 | 0.947440829 | 0.939950081 | 0.005915167 | 0.005975504 | 0.006080996 |
| 0.708002459 | 0.479366271 | 0.543235319 | 0.003303901 | 0.004786220 | 0.005872570 |
| 0.708002459 | 0.701730650 | 0.761647158 | 0.005915167 | 0.005975504 | 0.006080996 |
| 0.555456478 | 0.352926703 | 0.357996425 | 0.003303901 | 0.004786220 | 0.005872570 |
| 0.555456478 | 0.247118398 | 0.309185496 | 0.005915167 | 0.005975504 | 0.006080996 |
| 0.073306159 | 0.177388860 | 0.203183428 | 0.003303901 | 0.004786220 | 0.005872570 |
| 0.073306159 | 0.231390866 | 0.289905980 | 0.005915167 | 0.005975504 | 0.006080996 |
| 0.628324835 | 0.406448875 | 0.441919480 | 0.003303901 | 0.004786220 | 0.005872570 |
| 0.628324835 | 0.290640570 | 0.395744071 | 0.005915167 | 0.005975504 | 0.006080996 |
| 0.797545037 | 0.627801698 | 0.668584596 | 0.003303901 | 0.004786220 | 0.005872570 |
| 0.797545037 | 0.575583150 | 0.636958698 | 0.005915167 | 0.005975504 | 0.006080996 |

| | | | | | |
|---|---|---|---|---|---|
| 0.388748088 | 0.168166522 | 0.198670654 | 0.003303901 | 0.004786220 | 0.005872570 |
| 0.388748088 | 0.380269823 | 0.48991418 | 0.005915167 | 0.005975504 | 0.006080996 |
| 0.463761423 | 0.255627382 | 0.264954650 | 0.003303901 | 0.004786220 | 0.005872570 |
| 0.463761423 | 0.189857369 | 0.215658295 | 0.005915167 | 0.005975504 | 0.006080996 |
| 0.899305860 | 0.989635892 | 0.978873934 | 0.003303901 | 0.004786220 | 0.005872570 |
| 0.899305860 | 0.763688139 | 0.810362572 | 0.005915167 | 0.005975504 | 0.006080996 |
| 0.916089642 | 0.846012346 | 0.854642501 | 0.003303901 | 0.004786220 | 0.005872570 |
| 0.916089642 | 0.819449290 | 0.84018239 | 0.005915167 | 0.005975504 | 0.006080996 |
| 0.259836433 | 0.082899465 | 0.106144689 | 0.003303901 | 0.004786220 | 0.005872570 |
| 0.259836433 | 0.071499514 | 0.075350767 | 0.005915167 | 0.005975504 | 0.006080996 |
| 0.800587540 | 0.635806421 | 0.673052514 | 0.003303901 | 0.004786220 | 0.005872570 |
| 0.800587540 | 0.599238460 | 0.642413949 | 0.005915167 | 0.005975504 | 0.006080996 |
| 0.857693506 | 0.733871651 | 0.760890747 | 0.003303901 | 0.004786220 | 0.005872570 |
| 0.857693506 | 0.745677692 | 0.799294372 | 0.005915167 | 0.005975504 | 0.006080996 |
| 0.879337292 | 0.751411868 | 0.794539499 | 0.003303901 | 0.004786220 | 0.005872570 |
| 0.879337292 | 0.703628028 | 0.774254067 | 0.005915167 | 0.005975504 | 0.006080996 |
| 0.330391997 | 0.452005957 | 0.506882627 | 0.003303901 | 0.004786220 | 0.005872570 |
| 0.330391997 | 0.679099166 | 0.734244538 | 0.005915167 | 0.005975504 | 0.006080996 |
| 0.651627694 | 0.421929616 | 0.470046402 | 0.003303901 | 0.004786220 | 0.005872570 |
| 0.651627694 | 0.315078296 | 0.425300815 | 0.005915167 | 0.005975504 | 0.006080996 |
| 0.938880950 | 0.900440488 | 0.893470064 | 0.003303901 | 0.004786220 | 0.005872570 |
| 0.938880950 | 0.871949338 | 0.883078853 | 0.005915167 | 0.005975504 | 0.006080996 |
| 0.792785725 | 0.621523524 | 0.661931615 | 0.003303901 | 0.004786220 | 0.005872570 |
| 0.792785725 | 0.574573682 | 0.630011985 | 0.005915167 | 0.005975504 | 0.006080996 |
| 0.981618867 | 0.981802728 | 0.967456007 | 0.003303901 | 0.004786220 | 0.005872570 |
| 0.981618867 | 0.979426331 | 0.965217943 | 0.005915167 | 0.005975504 | 0.006080996 |
| 0.428783600 | 0.221473603 | 0.232842120 | 0.003303901 | 0.004786220 | 0.005872570 |
| 0.428783600 | 0.172975870 | 0.184341157 | 0.005915167 | 0.005975504 | 0.006080996 |
| 0.146866944 | 0.680573278 | 0.735725878 | 0.003303901 | 0.004786220 | 0.005872570 |
| 0.146866944 | 0.809289115 | 0.836295204 | 0.005915167 | 0.005975504 | 0.006080996 |
| 0.652558339 | 0.466014691 | 0.517332025 | 0.003303901 | 0.004786220 | 0.005872570 |
| 0.652558339 | 0.784512274 | 0.822897455 | 0.005915167 | 0.005975504 | 0.006080996 |
| 0.464535345 | 0.260201940 | 0.265740716 | 0.003303901 | 0.004786220 | 0.005872570 |
| 0.464535345 | 0.193890933 | 0.216423197 | 0.005915167 | 0.005975504 | 0.006080996 |
| 0.398010303 | 0.428452872 | 0.458302866 | 0.011796907 | 0.012671120 | 0.013958825 |
| 0.398010303 | 0.495273626 | 0.568005011 | 0.003230790 | 0.003257201 | 0.003343834 |
| 0.325450483 | 0.460377344 | 0.478515735 | 0.011796907 | 0.012671120 | 0.013958825 |
| 0.325450483 | 0.378973063 | 0.440558747 | 0.003230790 | 0.003257201 | 0.003343834 |
| 0.100921055 | 0.077124323 | 0.132260601 | 0.011796907 | 0.012671120 | 0.013958825 |
| 0.100921055 | 0.600477763 | 0.657948922 | 0.003230790 | 0.003257201 | 0.003343834 |
| 0.048428048 | 0.141929266 | 0.204263723 | 0.011796907 | 0.012671120 | 0.013958825 |
| 0.048428048 | 0.438252875 | 0.503700883 | 0.003230790 | 0.003257201 | 0.003343834 |
| 0.221818133 | 0.525986985 | 0.525810841 | 0.011796907 | 0.012671120 | 0.013958825 |
| 0.221818133 | 0.403721963 | 0.458608373 | 0.003230790 | 0.003257201 | 0.003343834 |
| 0.194121282 | 0.089848510 | 0.146302089 | 0.011796907 | 0.012671120 | 0.013958825 |
| 0.194121282 | 0.058904357 | 0.062856806 | 0.003230790 | 0.003257201 | 0.003343834 |

| | | | | | |
|---|---|---|---|---|---|
| 0.998445182 | 0.996403889 | 0.998494643 | 0.011796907 | 0.012671120 | 0.013958825 |
| 0.998445182 | 0.993251227 | 0.998487054 | 0.003230790 | 0.003257201 | 0.003343834 |
| 0.820673657 | 0.742597297 | 0.761976107 | 0.011796907 | 0.012671120 | 0.013958825 |
| 0.820673657 | 0.696253044 | 0.748807584 | 0.003230790 | 0.003257201 | 0.003343834 |
| 0.701382869 | 0.600217390 | 0.617977409 | 0.011796907 | 0.012671120 | 0.013958825 |
| 0.701382869 | 0.623150335 | 0.694920552 | 0.003230790 | 0.003257201 | 0.003343834 |
| 0.847446183 | 0.787704766 | 0.796253994 | 0.011796907 | 0.012671120 | 0.013958825 |
| 0.847446183 | 0.662799951 | 0.719629890 | 0.003230790 | 0.003257201 | 0.003343834 |
| 0.866407284 | 0.805494778 | 0.820583273 | 0.011796907 | 0.012671120 | 0.013958825 |
| 0.866407284 | 0.702558899 | 0.752292157 | 0.003230790 | 0.003257201 | 0.003343834 |
| 0.951433995 | 0.907446206 | 0.9333952 | 0.011796907 | 0.012671120 | 0.013958825 |
| 0.951433995 | 0.867929638 | 0.906348070 | 0.003230790 | 0.003257201 | 0.003343834 |
| 0.763733332 | 0.674792446 | 0.692141799 | 0.011796907 | 0.012671120 | 0.013958825 |
| 0.763733332 | 0.587905884 | 0.624089385 | 0.003230790 | 0.003257201 | 0.003343834 |
| 0.512965401 | 0.370713331 | 0.413047066 | 0.011796907 | 0.012671120 | 0.013958825 |
| 0.512965401 | 0.234762125 | 0.263881046 | 0.003230790 | 0.003257201 | 0.003343834 |
| 0.438241426 | 0.359610755 | 0.407560057 | 0.011796907 | 0.012671120 | 0.013958825 |
| 0.438241426 | 0.360820377 | 0.414520529 | 0.003230790 | 0.003257201 | 0.003343834 |
| 0.720714595 | 0.733756601 | 0.757118940 | 0.011796907 | 0.012671120 | 0.013958825 |
| 0.720714595 | 0.471013380 | 0.520847059 | 0.003230790 | 0.003257201 | 0.003343834 |
| 0.076908544 | 0.376816573 | 0.415900079 | 0.011796907 | 0.012671120 | 0.013958825 |
| 0.076908544 | 0.343428235 | 0.403055717 | 0.003230790 | 0.003257201 | 0.003343834 |
| 0.962395915 | 0.929836595 | 0.948465650 | 0.011796907 | 0.012671120 | 0.013958825 |
| 0.962395915 | 0.898498486 | 0.927402027 | 0.003230790 | 0.003257201 | 0.003343834 |
| 0.450339688 | 0.283447793 | 0.351754157 | 0.011796907 | 0.012671120 | 0.013958825 |
| 0.450339688 | 0.177844460 | 0.203603973 | 0.003230790 | 0.003257201 | 0.003343834 |
| 0.198868803 | 0.117560420 | 0.175813631 | 0.011796907 | 0.012671120 | 0.013958825 |
| 0.198868803 | 0.472516475 | 0.526461967 | 0.003230790 | 0.003257201 | 0.003343834 |
| 0.444289250 | 0.638501739 | 0.657362390 | 0.011796907 | 0.012671120 | 0.013958825 |
| 0.444289250 | 0.173791552 | 0.198079164 | 0.003230790 | 0.003257201 | 0.003343834 |
| 0.484957514 | 0.531789336 | 0.535353457 | 0.011796907 | 0.012671120 | 0.013958825 |
| 0.484957514 | 0.463698516 | 0.518387117 | 0.003230790 | 0.003257201 | 0.003343834 |
| 0.635370677 | 0.536049467 | 0.542703097 | 0.011796907 | 0.012671120 | 0.013958825 |
| 0.635370677 | 0.350139221 | 0.404627746 | 0.003230790 | 0.003257201 | 0.003343834 |
| 0.788915890 | 0.717721578 | 0.722316414 | 0.011796907 | 0.012671120 | 0.013958825 |
| 0.788915890 | 0.577463711 | 0.623373233 | 0.003230790 | 0.003257201 | 0.003343834 |
| 0.495579643 | 0.337527703 | 0.395959907 | 0.011796907 | 0.012671120 | 0.013958825 |
| 0.495579643 | 0.885530874 | 0.911411218 | 0.003230790 | 0.003257201 | 0.003343834 |
| 0.020139323 | 0.096801934 | 0.151201180 | 0.011796907 | 0.012671120 | 0.013958825 |
| 0.020139323 | 0.145880759 | 0.178033105 | 0.003230790 | 0.003257201 | 0.003343834 |
| 0.342554149 | 0.183562272 | 0.252586826 | 0.011796907 | 0.012671120 | 0.013958825 |
| 0.342554149 | 0.590305705 | 0.639840103 | 0.003230790 | 0.003257201 | 0.003343834 |
| 0.431837511 | 0.568181157 | 0.581843585 | 0.011796907 | 0.012671120 | 0.013958825 |
| 0.431837511 | 0.161232476 | 0.187243882 | 0.003230790 | 0.003257201 | 0.003343834 |
| 0.610297342 | 0.514341792 | 0.515190667 | 0.011796907 | 0.012671120 | 0.013958825 |
| 0.610297342 | 0.331763024 | 0.373311462 | 0.003230790 | 0.003257201 | 0.003343834 |

| | | | | | |
|---|---|---|---|---|---|
| 0.050588609 | 0.060264671 | 0.115264371 | 0.011796907 | 0.012671120 | 0.013958825 |
| 0.050588609 | 0.129571368 | 0.139268983 | 0.003230790 | 0.003257201 | 0.003343834 |
| 0.767299175 | 0.688979707 | 0.696506785 | 0.011796907 | 0.012671120 | 0.013958825 |
| 0.767299175 | 0.544246935 | 0.590145306 | 0.003230790 | 0.003257201 | 0.003343834 |
| 0.976432709 | 0.953488230 | 0.967639670 | 0.011796907 | 0.012671120 | 0.013958825 |
| 0.976432709 | 0.949930684 | 0.955284421 | 0.003230790 | 0.003257201 | 0.003343834 |
| 0.784439432 | 0.697758743 | 0.716884245 | 0.011796907 | 0.012671120 | 0.013958825 |
| 0.784439432 | 0.558627674 | 0.616339850 | 0.003230790 | 0.003257201 | 0.003343834 |
| 0.535922194 | 0.410101779 | 0.436603022 | 0.011796907 | 0.012671120 | 0.013958825 |
| 0.535922194 | 0.541178468 | 0.589953001 | 0.003230790 | 0.003257201 | 0.003343834 |
| 0.993338206 | 0.984623441 | 0.990904595 | 0.011796907 | 0.012671120 | 0.013958825 |
| 0.993338206 | 0.983244334 | 0.987861610 | 0.003230790 | 0.003257201 | 0.003343834 |
| 0.562373604 | 0.693652254 | 0.706069790 | 0.011796907 | 0.012671120 | 0.013958825 |
| 0.562373604 | 0.741853364 | 0.785910666 | 0.003230790 | 0.003257201 | 0.003343834 |
| 0.109214150 | 0.116269958 | 0.173921495 | 0.011796907 | 0.012671120 | 0.013958825 |
| 0.109214150 | 0.670583769 | 0.732088172 | 0.003230790 | 0.003257201 | 0.003343834 |
| 0.507394075 | 0.360876106 | 0.40770541 | 0.011796907 | 0.012671120 | 0.013958825 |
| 0.507394075 | 0.227366278 | 0.258243259 | 0.003230790 | 0.003257201 | 0.003343834 |
| 0.472781782 | 0.309619817 | 0.373279917 | 0.011796907 | 0.012671120 | 0.013958825 |
| 0.472781782 | 0.200816626 | 0.224142243 | 0.003230790 | 0.003257201 | 0.003343834 |
| 0.506814287 | 0.354134013 | 0.406859587 | 0.011796907 | 0.012671120 | 0.013958825 |
| 0.506814287 | 0.862976692 | 0.902154988 | 0.003230790 | 0.003257201 | 0.003343834 |
| 0.992024082 | 0.982252002 | 0.989771232 | 0.011796907 | 0.012671120 | 0.013958825 |
| 0.992024082 | 0.977866096 | 0.985570054 | 0.003230790 | 0.003257201 | 0.003343834 |
| 0.743701905 | 0.651964991 | 0.667838597 | 0.011796907 | 0.012671120 | 0.013958825 |
| 0.743701905 | 0.492867540 | 0.554574935 | 0.003230790 | 0.003257201 | 0.003343834 |
| 0.564416950 | 0.587705117 | 0.598471023 | 0.011796907 | 0.012671120 | 0.013958825 |
| 0.564416950 | 0.265028489 | 0.319349364 | 0.003230790 | 0.003257201 | 0.003343834 |
| 0.453608604 | 0.285452537 | 0.354490195 | 0.011796907 | 0.012671120 | 0.013958825 |
| 0.453608604 | 0.180517239 | 0.206468831 | 0.003230790 | 0.003257201 | 0.003343834 |
| 0.941774425 | 0.894983902 | 0.920178860 | 0.011796907 | 0.012671120 | 0.013958825 |
| 0.941774425 | 0.850218097 | 0.888642086 | 0.003230790 | 0.003257201 | 0.003343834 |
| 0.329694556 | 0.172196160 | 0.241460156 | 0.011796907 | 0.012671120 | 0.013958825 |
| 0.329694556 | 0.101425730 | 0.109277111 | 0.003230790 | 0.003257201 | 0.003343834 |
| 0.048007350 | 0.058904758 | 0.113130769 | 0.011796907 | 0.012671120 | 0.013958825 |
| 0.048007350 | 0.111376805 | 0.117826651 | 0.003230790 | 0.003257201 | 0.003343834 |
| 0.573598893 | 0.549718521 | 0.557587149 | 0.011796907 | 0.012671120 | 0.013958825 |
| 0.573598893 | 0.285169030 | 0.330158699 | 0.003230790 | 0.003257201 | 0.003343834 |
| 0.367550324 | 0.203647565 | 0.274866188 | 0.011796907 | 0.012671120 | 0.013958825 |
| 0.367550324 | 0.124238221 | 0.135567497 | 0.003230790 | 0.003257201 | 0.003343834 |
| 0.930941562 | 0.882397485 | 0.906078367 | 0.011796907 | 0.012671120 | 0.013958825 |
| 0.930941562 | 0.834258609 | 0.868161443 | 0.003230790 | 0.003257201 | 0.003343834 |
| 0.472296893 | 0.305575339 | 0.372809627 | 0.011796907 | 0.012671120 | 0.013958825 |
| 0.472296893 | 0.320166470 | 0.355529225 | 0.003230790 | 0.003257201 | 0.003343834 |
| 0.421626944 | 0.499388795 | 0.500543894 | 0.011796907 | 0.012671120 | 0.013958825 |
| 0.421626944 | 0.915351941 | 0.931953243 | 0.003230790 | 0.003257201 | 0.003343834 |

| | | | | | |
|---|---|---|---|---|---|
| 0.509636780 | 0.367070437 | 0.410021858 | 0.011796907 | 0.012671120 | 0.013958825 |
| 0.509636780 | 0.232041560 | 0.260620761 | 0.003230790 | 0.003257201 | 0.003343834 |
| 0.176905614 | 0.180110828 | 0.248289822 | 0.011796907 | 0.012671120 | 0.013958825 |
| 0.176905614 | 0.024310792 | 0.031533026 | 0.003230790 | 0.003257201 | 0.003343834 |
| 0.187668077 | 0.211847541 | 0.282890823 | 0.011796907 | 0.012671120 | 0.013958825 |
| 0.187668077 | 0.383316283 | 0.446570258 | 0.003230790 | 0.003257201 | 0.003343834 |
| 0.888305883 | 0.830324563 | 0.848784062 | 0.011796907 | 0.012671120 | 0.013958825 |
| 0.888305883 | 0.752885094 | 0.790158348 | 0.003230790 | 0.003257201 | 0.003343834 |
| 0.153966391 | 0.073973172 | 0.129810356 | 0.011796907 | 0.012671120 | 0.013958825 |
| 0.153966391 | 0.015080814 | 0.023942509 | 0.003230790 | 0.003257201 | 0.003343834 |
| 0.215058420 | 0.093904554 | 0.147708008 | 0.011796907 | 0.012671120 | 0.013958825 |
| 0.215058420 | 0.041835579 | 0.046521249 | 0.003230790 | 0.003257201 | 0.003343834 |
| 0.369359574 | 0.460950264 | 0.479334868 | 0.011796907 | 0.012671120 | 0.013958825 |
| 0.369359574 | 0.66591647 | 0.730220318 | 0.003230790 | 0.003257201 | 0.003343834 |
| 0.582775057 | 0.590606430 | 0.604214732 | 0.011796907 | 0.012671120 | 0.013958825 |
| 0.582775057 | 0.291321411 | 0.340488201 | 0.003230790 | 0.003257201 | 0.003343834 |
| 0.737207355 | 0.642160655 | 0.660021067 | 0.011796907 | 0.012671120 | 0.013958825 |
| 0.737207355 | 0.479491207 | 0.544682657 | 0.003230790 | 0.003257201 | 0.003343834 |
| 0.394757739 | 0.222505023 | 0.299381833 | 0.011796907 | 0.012671120 | 0.013958825 |
| 0.394757739 | 0.145731935 | 0.175740520 | 0.003230790 | 0.003257201 | 0.003343834 |
| 0.855187870 | 0.792338969 | 0.805767113 | 0.011796907 | 0.012671120 | 0.013958825 |
| 0.855187870 | 0.673265419 | 0.732312856 | 0.003230790 | 0.003257201 | 0.003343834 |
| 0.674518682 | 0.57972623 | 0.587129234 | 0.011796907 | 0.012671120 | 0.013958825 |
| 0.674518682 | 0.400078602 | 0.456072561 | 0.003230790 | 0.003257201 | 0.003343834 |
| 0.991487189 | 0.970550380 | 0.988554660 | 0.011796907 | 0.012671120 | 0.013958825 |
| 0.991487189 | 0.973480369 | 0.984947282 | 0.003230790 | 0.003257201 | 0.003343834 |
| 0.528757987 | 0.397319512 | 0.429371524 | 0.011796907 | 0.012671120 | 0.013958825 |
| 0.528757987 | 0.551785258 | 0.595530850 | 0.003230790 | 0.003257201 | 0.003343834 |
| 0.846308596 | 0.768599453 | 0.794310981 | 0.011796907 | 0.012671120 | 0.013958825 |
| 0.846308596 | 0.645881959 | 0.717421008 | 0.003230790 | 0.003257201 | 0.003343834 |
| 0.585645053 | 0.683324808 | 0.694823029 | 0.011796907 | 0.012671120 | 0.013958825 |
| 0.585645053 | 0.299480558 | 0.343724564 | 0.003230790 | 0.003257201 | 0.003343834 |
| 0.883938821 | 0.827594807 | 0.843749430 | 0.011796907 | 0.012671120 | 0.013958825 |
| 0.883938821 | 0.822292381 | 0.848639619 | 0.003230790 | 0.003257201 | 0.003343834 |
| 0.521032489 | 0.385183495 | 0.421725193 | 0.011796907 | 0.012671120 | 0.013958825 |
| 0.521032489 | 0.598043122 | 0.642097680 | 0.003230790 | 0.003257201 | 0.003343834 |
| 0.459403603 | 0.293775617 | 0.360226160 | 0.011796907 | 0.012671120 | 0.013958825 |
| 0.459403603 | 0.186682011 | 0.211664491 | 0.003230790 | 0.003257201 | 0.003343834 |
| 0.306258728 | 0.155652154 | 0.221428727 | 0.011796907 | 0.012671120 | 0.013958825 |
| 0.306258728 | 0.086106540 | 0.094256624 | 0.003230790 | 0.003257201 | 0.003343834 |
| 0.671348709 | 0.570357826 | 0.583272131 | 0.011796907 | 0.012671120 | 0.013958825 |
| 0.671348709 | 0.388754520 | 0.451695485 | 0.003230790 | 0.003257201 | 0.003343834 |
| 0.594043357 | 0.494542806 | 0.497809913 | 0.011796907 | 0.012671120 | 0.013958825 |
| 0.594043357 | 0.313970735 | 0.353761536 | 0.003230790 | 0.003257201 | 0.003343834 |
| 0.299552459 | 0.148639393 | 0.215626576 | 0.011796907 | 0.012671120 | 0.013958825 |
| 0.299552459 | 0.081120430 | 0.090144693 | 0.003230790 | 0.003257201 | 0.003343834 |

| | | | | | |
|---|---|---|---|---|---|
| 0.981726142 | 0.962526109 | 0.975049571 | 0.011796907 | 0.012671120 | 0.013958825 |
| 0.981726142 | 0.955263773 | 0.965351912 | 0.003230790 | 0.003257201 | 0.003343834 |
| 0.153531143 | 0.394679718 | 0.428859669 | 0.011796907 | 0.012671120 | 0.013958825 |
| 0.153531143 | 0.116543481 | 0.119937962 | 0.003230790 | 0.003257201 | 0.003343834 |
| 0.164348427 | 0.056335625 | 0.109558300 | 0.011796907 | 0.012671120 | 0.013958825 |
| 0.164348427 | 0.021332247 | 0.027310324 | 0.003230790 | 0.003257201 | 0.003343834 |
| 0.275690212 | 0.437626754 | 0.465253100 | 0.011796907 | 0.012671120 | 0.013958825 |
| 0.275690212 | 0.109395954 | 0.116286704 | 0.003230790 | 0.003257201 | 0.003343834 |
| 0.440093737 | 0.638929407 | 0.658320169 | 0.011796907 | 0.012671120 | 0.013958825 |
| 0.440093737 | 0.500903579 | 0.579072064 | 0.003230790 | 0.003257201 | 0.003343834 |
| 0.244469670 | 0.112606042 | 0.170643231 | 0.011796907 | 0.012671120 | 0.013958825 |
| 0.244469670 | 0.184012285 | 0.207363294 | 0.003230790 | 0.003257201 | 0.003343834 |
| 0.591562606 | 0.489189029 | 0.495288027 | 0.011796907 | 0.012671120 | 0.013958825 |
| 0.591562606 | 0.856275727 | 0.894036271 | 0.003230790 | 0.003257201 | 0.003343834 |
| 0.586347400 | 0.696912172 | 0.716256494 | 0.011796907 | 0.012671120 | 0.013958825 |
| 0.586347400 | 0.721228211 | 0.775237101 | 0.003230790 | 0.003257201 | 0.003343834 |
| 0.940246859 | 0.887529139 | 0.918153637 | 0.011796907 | 0.012671120 | 0.013958825 |
| 0.940246859 | 0.838300329 | 0.885204529 | 0.003230790 | 0.003257201 | 0.003343834 |
| 0.500605814 | 0.345270068 | 0.400985428 | 0.011796907 | 0.012671120 | 0.013958825 |
| 0.500605814 | 0.216198241 | 0.251409136 | 0.003230790 | 0.003257201 | 0.003343834 |
| 0.708600326 | 0.610019610 | 0.626403468 | 0.011796907 | 0.012671120 | 0.013958825 |
| 0.708600326 | 0.557260730 | 0.595980247 | 0.003230790 | 0.003257201 | 0.003343834 |
| 0.214505230 | 0.129609552 | 0.191650551 | 0.011796907 | 0.012671120 | 0.013958825 |
| 0.214505230 | 0.428778558 | 0.496342467 | 0.003230790 | 0.003257201 | 0.003343834 |
| 0.971978890 | 0.943608432 | 0.961294574 | 0.011796907 | 0.012671120 | 0.013958825 |
| 0.971978890 | 0.966175811 | 0.972894585 | 0.003230790 | 0.003257201 | 0.003343834 |
| 0.951513840 | 0.902376495 | 0.933080382 | 0.011796907 | 0.012671120 | 0.013958825 |
| 0.951513840 | 0.868539457 | 0.906386315 | 0.003230790 | 0.003257201 | 0.003343834 |
| 0.570495702 | 0.449515387 | 0.472771525 | 0.011796907 | 0.012671120 | 0.013958825 |
| 0.570495702 | 0.375831064 | 0.437961912 | 0.003230790 | 0.003257201 | 0.003343834 |
| 0.179672265 | 0.453730646 | 0.475523535 | 0.011796907 | 0.012671120 | 0.013958825 |
| 0.179672265 | 0.625575361 | 0.695644567 | 0.003230790 | 0.003257201 | 0.003343834 |
| 0.051422508 | 0.164295849 | 0.234187434 | 0.011796907 | 0.012671120 | 0.013958825 |
| 0.051422508 | 0.727740924 | 0.782734545 | 0.003230790 | 0.003257201 | 0.003343834 |
| 0.473968775 | 0.595302320 | 0.613881043 | 0.011796907 | 0.012671120 | 0.013958825 |
| 0.473968775 | 0.205912085 | 0.225439329 | 0.003230790 | 0.003257201 | 0.003343834 |
| 0.869479280 | 0.810154330 | 0.824170601 | 0.011796907 | 0.012671120 | 0.013958825 |
| 0.869479280 | 0.814895787 | 0.845525403 | 0.003230790 | 0.003257201 | 0.003343834 |
| 0.336916206 | 0.178601182 | 0.247743714 | 0.011796907 | 0.012671120 | 0.013958825 |
| 0.336916206 | 0.424230665 | 0.494986205 | 0.003230790 | 0.003257201 | 0.003343834 |
| 0.764550447 | 0.680512022 | 0.693042065 | 0.011796907 | 0.012671120 | 0.013958825 |
| 0.764550447 | 0.530642053 | 0.585698196 | 0.003230790 | 0.003257201 | 0.003343834 |
| 0.502790036 | 0.350104328 | 0.403139962 | 0.011796907 | 0.012671120 | 0.013958825 |
| 0.502790036 | 0.217522238 | 0.253325664 | 0.003230790 | 0.003257201 | 0.003343834 |
| 0.525508120 | 0.664474426 | 0.689371159 | 0.011796907 | 0.012671120 | 0.013958825 |
| 0.525508120 | 0.939252458 | 0.952408720 | 0.003230790 | 0.003257201 | 0.003343834 |

| | | | | | |
|---|---|---|---|---|---|
| 0.903235538 | 0.858809958 | 0.868776837 | 0.011796907 | 0.012671120 | 0.013958825 |
| 0.903235538 | 0.780696918 | 0.816894341 | 0.003230790 | 0.003257201 | 0.003343834 |
| 0.275829535 | 0.238568917 | 0.320397093 | 0.011796907 | 0.012671120 | 0.013958825 |
| 0.275829535 | 0.906507398 | 0.929329826 | 0.003230790 | 0.003257201 | 0.003343834 |
| 0.431371088 | 0.291118199 | 0.358021684 | 0.011796907 | 0.012671120 | 0.013958825 |
| 0.431371088 | 0.157250663 | 0.186769804 | 0.003230790 | 0.003257201 | 0.003343834 |
| 0.159520448 | 0.052593161 | 0.106014318 | 0.011796907 | 0.012671120 | 0.013958825 |
| 0.159520448 | 0.036260875 | 0.041988313 | 0.003230790 | 0.003257201 | 0.003343834 |
| 0.761777923 | 0.666381592 | 0.689599592 | 0.011796907 | 0.012671120 | 0.013958825 |
| 0.761777923 | 0.513960696 | 0.581639240 | 0.003230790 | 0.003257201 | 0.003343834 |
| 0.895004134 | 0.850331981 | 0.858036283 | 0.011796907 | 0.012671120 | 0.013958825 |
| 0.895004134 | 0.779520045 | 0.802799224 | 0.003230790 | 0.003257201 | 0.003343834 |
| 0.894551833 | 0.838580690 | 0.856899904 | 0.011796907 | 0.012671120 | 0.013958825 |
| 0.894551833 | 0.766208576 | 0.801215675 | 0.003230790 | 0.003257201 | 0.003343834 |
| 0.072429427 | 0.082559861 | 0.137961162 | 0.011796907 | 0.012671120 | 0.013958825 |
| 0.072429427 | 0.257179261 | 0.289905103 | 0.003230790 | 0.003257201 | 0.003343834 |
| 0.820084524 | 0.737964861 | 0.761487 | 0.011796907 | 0.012671120 | 0.013958825 |
| 0.820084524 | 0.608035312 | 0.673682359 | 0.003230790 | 0.003257201 | 0.003343834 |
| 0.842807474 | 0.755635348 | 0.790027379 | 0.011796907 | 0.012671120 | 0.013958825 |
| 0.842807474 | 0.630681016 | 0.711250313 | 0.003230790 | 0.003257201 | 0.003343834 |
| 0.605923234 | 0.506278833 | 0.510432103 | 0.011796907 | 0.012671120 | 0.013958825 |
| 0.605923234 | 0.415543919 | 0.490131355 | 0.003230790 | 0.003257201 | 0.003343834 |
| 0.711193304 | 0.614924729 | 0.629176988 | 0.011796907 | 0.012671120 | 0.013958825 |
| 0.711193304 | 0.450001408 | 0.506626986 | 0.003230790 | 0.003257201 | 0.003343834 |
| 0.146464048 | 0.471588168 | 0.487884189 | 0.011796907 | 0.012671120 | 0.013958825 |
| 0.146464048 | 0.455828339 | 0.508139637 | 0.003230790 | 0.003257201 | 0.003343834 |
| 0.786545970 | 0.710820555 | 0.719437794 | 0.011796907 | 0.012671120 | 0.013958825 |
| 0.786545970 | 0.569736509 | 0.619487177 | 0.003230790 | 0.003257201 | 0.003343834 |
| 0.535281593 | 0.405355510 | 0.435944934 | 0.011796907 | 0.012671120 | 0.013958825 |
| 0.535281593 | 0.25036683 | 0.287335193 | 0.003230790 | 0.003257201 | 0.003343834 |
| 0.656535873 | 0.556173281 | 0.566519944 | 0.011796907 | 0.012671120 | 0.013958825 |
| 0.656535873 | 0.396681645 | 0.454193327 | 0.003230790 | 0.003257201 | 0.003343834 |
| 0.843581193 | 0.767737097 | 0.791641933 | 0.011796907 | 0.012671120 | 0.013958825 |
| 0.843581193 | 0.765067910 | 0.799280260 | 0.003230790 | 0.003257201 | 0.003343834 |
| 0.847186189 | 0.776066983 | 0.795486271 | 0.011796907 | 0.012671120 | 0.013958825 |
| 0.847186189 | 0.651023916 | 0.718712282 | 0.003230790 | 0.003257201 | 0.003343834 |
| 0.266935165 | 0.320763360 | 0.380934049 | 0.011796907 | 0.012671120 | 0.013958825 |
| 0.266935165 | 0.685451613 | 0.734181081 | 0.003230790 | 0.003257201 | 0.003343834 |
| 0.968608596 | 0.934014249 | 0.956472144 | 0.011796907 | 0.012671120 | 0.013958825 |
| 0.968608596 | 0.919875869 | 0.939201756 | 0.003230790 | 0.003257201 | 0.003343834 |
| 0.922729182 | 0.870437528 | 0.894705089 | 0.011796907 | 0.012671120 | 0.013958825 |
| 0.922729182 | 0.827661096 | 0.852994406 | 0.003230790 | 0.003257201 | 0.003343834 |
| 0.957367116 | 0.921799366 | 0.941608093 | 0.011796907 | 0.012671120 | 0.013958825 |
| 0.957367116 | 0.894781149 | 0.918219156 | 0.003230790 | 0.003257201 | 0.003343834 |
| 0.785243077 | 0.705908786 | 0.718060274 | 0.011796907 | 0.012671120 | 0.013958825 |
| 0.785243077 | 0.568427147 | 0.618052656 | 0.003230790 | 0.003257201 | 0.003343834 |

| | | | | | |
|---|---|---|---|---|---|
| 0.207990534 | 0.085197897 | 0.142099445 | 0.011796907 | 0.012671120 | 0.013958825 |
| 0.207990534 | 0.036914856 | 0.043525049 | 0.003230790 | 0.003257201 | 0.003343834 |
| 0.072299563 | 0.339100332 | 0.397025495 | 0.011796907 | 0.012671120 | 0.013958825 |
| 0.072299563 | 0.801394427 | 0.836220637 | 0.003230790 | 0.003257201 | 0.003343834 |
| 0.435479852 | 0.426232279 | 0.455045458 | 0.011796907 | 0.012671120 | 0.013958825 |
| 0.435479852 | 0.790042968 | 0.822680376 | 0.003230790 | 0.003257201 | 0.003343834 |
| 0.358202471 | 0.195126180 | 0.266444264 | 0.011796907 | 0.012671120 | 0.013958825 |
| 0.358202471 | 0.122055738 | 0.128832788 | 0.003230790 | 0.003257201 | 0.003343834 |
| 0.438433355 | 0.676152909 | 0.574000386 | 0.005238192 | 0.006352035 | 0.007313854 |
| 0.438433355 | 0.683264405 | 0.641709751 | 0.010564931 | 0.012312513 | 0.012954696 |
| 0.557763446 | 0.823971936 | 0.710368731 | 0.005238192 | 0.006352035 | 0.007313854 |
| 0.557763446 | 0.545800452 | 0.529500331 | 0.010564931 | 0.012312513 | 0.012954696 |
| 0.974772687 | 0.930695334 | 0.821333091 | 0.005238192 | 0.006352035 | 0.007313854 |
| 0.974772687 | 0.938150381 | 0.895315228 | 0.010564931 | 0.012312513 | 0.012954696 |
| 0.815990145 | 0.806902891 | 0.694537422 | 0.005238192 | 0.006352035 | 0.007313854 |
| 0.815990145 | 0.741832062 | 0.683993914 | 0.010564931 | 0.012312513 | 0.012954696 |
| 0.019942071 | 0.902585776 | 0.790054163 | 0.005238192 | 0.006352035 | 0.007313854 |
| 0.019942071 | 0.566471401 | 0.545263026 | 0.010564931 | 0.012312513 | 0.012954696 |
| 0.197387911 | 0.139935062 | 0.148737423 | 0.005238192 | 0.006352035 | 0.007313854 |
| 0.197387911 | 0.122838937 | 0.137575366 | 0.010564931 | 0.012312513 | 0.012954696 |
| 0.816744548 | 0.810483380 | 0.697854932 | 0.005238192 | 0.006352035 | 0.007313854 |
| 0.816744548 | 0.678645388 | 0.639390837 | 0.010564931 | 0.012312513 | 0.012954696 |
| 0.132348299 | 0.069189037 | 0.085252680 | 0.005238192 | 0.006352035 | 0.007313854 |
| 0.132348299 | 0.832639690 | 0.794150881 | 0.010564931 | 0.012312513 | 0.012954696 |
| 0.483326408 | 0.383277103 | 0.318739744 | 0.005238192 | 0.006352035 | 0.007313854 |
| 0.483326408 | 0.792340013 | 0.749594532 | 0.010564931 | 0.012312513 | 0.012954696 |
| 0.565104580 | 0.512541912 | 0.419397293 | 0.005238192 | 0.006352035 | 0.007313854 |
| 0.565104580 | 0.481097400 | 0.476246992 | 0.010564931 | 0.012312513 | 0.012954696 |
| 0.229578184 | 0.339266155 | 0.289026273 | 0.005238192 | 0.006352035 | 0.007313854 |
| 0.229578184 | 0.706601034 | 0.656266822 | 0.010564931 | 0.012312513 | 0.012954696 |
| 0.300357855 | 0.325223997 | 0.280425515 | 0.005238192 | 0.006352035 | 0.007313854 |
| 0.300357855 | 0.099975240 | 0.122425032 | 0.010564931 | 0.012312513 | 0.012954696 |
| 0.806273902 | 0.774372221 | 0.665174657 | 0.005238192 | 0.006352035 | 0.007313854 |
| 0.806273902 | 0.769596174 | 0.706456802 | 0.010564931 | 0.012312513 | 0.012954696 |
| 0.182354877 | 0.033602943 | 0.058303678 | 0.005238192 | 0.006352035 | 0.007313854 |
| 0.182354877 | 0.106685904 | 0.127029368 | 0.010564931 | 0.012312513 | 0.012954696 |
| 0.458514740 | 0.536299345 | 0.440531687 | 0.005238192 | 0.006352035 | 0.007313854 |
| 0.458514740 | 0.516039888 | 0.505719073 | 0.010564931 | 0.012312513 | 0.012954696 |
| 0.620852292 | 0.906867881 | 0.794610373 | 0.005238192 | 0.006352035 | 0.007313854 |
| 0.620852292 | 0.590027628 | 0.570679863 | 0.010564931 | 0.012312513 | 0.012954696 |
| 0.150632281 | 0.891006031 | 0.777396990 | 0.005238192 | 0.006352035 | 0.007313854 |
| 0.150632281 | 0.49763987 | 0.49527986 | 0.010564931 | 0.012312513 | 0.012954696 |
| 0.440517851 | 0.476818933 | 0.389174066 | 0.005238192 | 0.006352035 | 0.007313854 |
| 0.440517851 | 0.225595099 | 0.210367326 | 0.010564931 | 0.012312513 | 0.012954696 |
| 0.039909004 | 0.226080080 | 0.216556554 | 0.005238192 | 0.006352035 | 0.007313854 |
| 0.039909004 | 0.227189281 | 0.211738049 | 0.010564931 | 0.012312513 | 0.012954696 |

| | | | | | |
|---|---|---|---|---|---|
| 0.425761851 | 0.323166210 | 0.279136652 | 0.005238192 | 0.006352035 | 0.007313854 |
| 0.425761851 | 0.652801414 | 0.6057247 | 0.010564931 | 0.012312513 | 0.012954696 |
| 0.340639268 | 0.954144288 | 0.850939814 | 0.005238192 | 0.006352035 | 0.007313854 |
| 0.340639268 | 0.161012474 | 0.166736523 | 0.010564931 | 0.012312513 | 0.012954696 |
| 0.356110531 | 0.705767861 | 0.601836976 | 0.005238192 | 0.006352035 | 0.007313854 |
| 0.356110531 | 0.642363430 | 0.598327660 | 0.010564931 | 0.012312513 | 0.012954696 |
| 0.711059677 | 0.512301930 | 0.419200570 | 0.005238192 | 0.006352035 | 0.007313854 |
| 0.711059677 | 0.420896868 | 0.422665979 | 0.010564931 | 0.012312513 | 0.012954696 |
| 0.477724863 | 0.235776219 | 0.223236328 | 0.005238192 | 0.006352035 | 0.007313854 |
| 0.477724863 | 0.233499232 | 0.215824557 | 0.010564931 | 0.012312513 | 0.012954696 |
| 0.200816851 | 0.216397274 | 0.210349348 | 0.005238192 | 0.006352035 | 0.007313854 |
| 0.200816851 | 0.958097001 | 0.927745503 | 0.010564931 | 0.012312513 | 0.012954696 |
| 0.561991955 | 0.547750302 | 0.450806106 | 0.005238192 | 0.006352035 | 0.007313854 |
| 0.561991955 | 0.375064044 | 0.358629416 | 0.010564931 | 0.012312513 | 0.012954696 |
| 0.208784078 | 0.082998667 | 0.095226129 | 0.005238192 | 0.006352035 | 0.007313854 |
| 0.208784078 | 0.757263901 | 0.703102227 | 0.010564931 | 0.012312513 | 0.012954696 |
| 0.902045480 | 0.955127882 | 0.852464304 | 0.005238192 | 0.006352035 | 0.007313854 |
| 0.902045480 | 0.663306304 | 0.620476917 | 0.010564931 | 0.012312513 | 0.012954696 |
| 0.092673042 | 0.013614092 | 0.030100795 | 0.005238192 | 0.006352035 | 0.007313854 |
| 0.092673042 | 0.109798536 | 0.129497575 | 0.010564931 | 0.012312513 | 0.012954696 |
| 0.794097009 | 0.725775591 | 0.619945615 | 0.005238192 | 0.006352035 | 0.007313854 |
| 0.794097009 | 0.540599651 | 0.528196192 | 0.010564931 | 0.012312513 | 0.012954696 |
| 0.129590869 | 0.490853813 | 0.400673938 | 0.005238192 | 0.006352035 | 0.007313854 |
| 0.129590869 | 0.489664132 | 0.490249699 | 0.010564931 | 0.012312513 | 0.012954696 |
| 0.251230859 | 0.274773423 | 0.247209076 | 0.005238192 | 0.006352035 | 0.007313854 |
| 0.251230859 | 0.876256832 | 0.827335436 | 0.010564931 | 0.012312513 | 0.012954696 |
| 0.880142120 | 0.726266322 | 0.620402996 | 0.005238192 | 0.006352035 | 0.007313854 |
| 0.880142120 | 0.574221334 | 0.549745112 | 0.010564931 | 0.012312513 | 0.012954696 |
| 0.186812482 | 0.086720651 | 0.098155984 | 0.005238192 | 0.006352035 | 0.007313854 |
| 0.186812482 | 0.714248403 | 0.660270797 | 0.010564931 | 0.012312513 | 0.012954696 |
| 0.446347102 | 0.282241228 | 0.252229209 | 0.005238192 | 0.006352035 | 0.007313854 |
| 0.446347102 | 0.176136031 | 0.178973560 | 0.010564931 | 0.012312513 | 0.012954696 |
| 0.039293969 | 0.867160881 | 0.753249577 | 0.005238192 | 0.006352035 | 0.007313854 |
| 0.039293969 | 0.866903759 | 0.824996422 | 0.010564931 | 0.012312513 | 0.012954696 |
| 0.476892570 | 0.458606168 | 0.374364760 | 0.005238192 | 0.006352035 | 0.007313854 |
| 0.476892570 | 0.818901516 | 0.781174367 | 0.010564931 | 0.012312513 | 0.012954696 |
| 0.741877841 | 0.620320891 | 0.522582722 | 0.005238192 | 0.006352035 | 0.007313854 |
| 0.741877841 | 0.449943454 | 0.445818330 | 0.010564931 | 0.012312513 | 0.012954696 |
| 0.644938571 | 0.498933062 | 0.407494636 | 0.005238192 | 0.006352035 | 0.007313854 |
| 0.644938571 | 0.333902793 | 0.308951784 | 0.010564931 | 0.012312513 | 0.012954696 |
| 0.297057080 | 0.104947587 | 0.114421648 | 0.005238192 | 0.006352035 | 0.007313854 |
| 0.297057080 | 0.950597754 | 0.920337209 | 0.010564931 | 0.012312513 | 0.012954696 |
| 0.618461214 | 0.920337459 | 0.809507681 | 0.005238192 | 0.006352035 | 0.007313854 |
| 0.618461214 | 0.414363352 | 0.416615218 | 0.010564931 | 0.012312513 | 0.012954696 |
| 0.684498996 | 0.588283965 | 0.489413634 | 0.005238192 | 0.006352035 | 0.007313854 |
| 0.684498996 | 0.670382841 | 0.623583402 | 0.010564931 | 0.012312513 | 0.012954696 |

| | | | | | |
|---|---|---|---|---|---|
| 0.596028252 | 0.787264890 | 0.677051459 | 0.005238192 | 0.006352035 | 0.007313854 |
| 0.596028252 | 0.324693327 | 0.301127132 | 0.010564931 | 0.012312513 | 0.012954696 |
| 0.487162658 | 0.250832229 | 0.232422285 | 0.005238192 | 0.006352035 | 0.007313854 |
| 0.487162658 | 0.266036718 | 0.244666435 | 0.010564931 | 0.012312513 | 0.012954696 |
| 0.139565381 | 0.170292681 | 0.176353197 | 0.005238192 | 0.006352035 | 0.007313854 |
| 0.139565381 | 0.697951894 | 0.652333181 | 0.010564931 | 0.012312513 | 0.012954696 |
| 0.544019112 | 0.374187397 | 0.312145936 | 0.005238192 | 0.006352035 | 0.007313854 |
| 0.544019112 | 0.306747228 | 0.277684425 | 0.010564931 | 0.012312513 | 0.012954696 |
| 0.162018641 | 0.297750097 | 0.263024943 | 0.005238192 | 0.006352035 | 0.007313854 |
| 0.162018641 | 0.218929088 | 0.207699310 | 0.010564931 | 0.012312513 | 0.012954696 |
| 0.756994756 | 0.757921563 | 0.649394474 | 0.005238192 | 0.006352035 | 0.007313854 |
| 0.756994756 | 0.591499326 | 0.572345611 | 0.010564931 | 0.012312513 | 0.012954696 |
| 0.423429390 | 0.290943798 | 0.258368988 | 0.005238192 | 0.006352035 | 0.007313854 |
| 0.423429390 | 0.138576243 | 0.150681512 | 0.010564931 | 0.012312513 | 0.012954696 |
| 0.185948722 | 0.759097547 | 0.650511682 | 0.005238192 | 0.006352035 | 0.007313854 |
| 0.185948722 | 0.434186786 | 0.432173704 | 0.010564931 | 0.012312513 | 0.012954696 |
| 0.709954138 | 0.568412396 | 0.469653058 | 0.005238192 | 0.006352035 | 0.007313854 |
| 0.709954138 | 0.561730439 | 0.541277579 | 0.010564931 | 0.012312513 | 0.012954696 |
| 0.072845237 | 0.650116586 | 0.550822669 | 0.005238192 | 0.006352035 | 0.007313854 |
| 0.072845237 | 0.973932365 | 0.944276614 | 0.010564931 | 0.012312513 | 0.012954696 |
| 0.678898859 | 0.574770441 | 0.475883118 | 0.005238192 | 0.006352035 | 0.007313854 |
| 0.678898859 | 0.431323548 | 0.429812086 | 0.010564931 | 0.012312513 | 0.012954696 |
| 0.822843289 | 0.796679979 | 0.685403083 | 0.005238192 | 0.006352035 | 0.007313854 |
| 0.822843289 | 0.432142342 | 0.431275273 | 0.010564931 | 0.012312513 | 0.012954696 |
| 0.396278180 | 0.574967011 | 0.476072821 | 0.005238192 | 0.006352035 | 0.007313854 |
| 0.396278180 | 0.552879623 | 0.534872814 | 0.010564931 | 0.012312513 | 0.012954696 |
| 0.421096248 | 0.235570221 | 0.223105912 | 0.005238192 | 0.006352035 | 0.007313854 |
| 0.421096248 | 0.176949995 | 0.179894076 | 0.010564931 | 0.012312513 | 0.012954696 |
| 0.369901434 | 0.240644567 | 0.225958351 | 0.005238192 | 0.006352035 | 0.007313854 |
| 0.369901434 | 0.093099539 | 0.115623476 | 0.010564931 | 0.012312513 | 0.012954696 |
| 0.183430082 | 0.087629010 | 0.098882442 | 0.005238192 | 0.006352035 | 0.007313854 |
| 0.183430082 | 0.019633000 | 0.042798267 | 0.010564931 | 0.012312513 | 0.012954696 |
| 0.672450684 | 0.800362759 | 0.688640575 | 0.005238192 | 0.006352035 | 0.007313854 |
| 0.672450684 | 0.813309694 | 0.779596057 | 0.010564931 | 0.012312513 | 0.012954696 |
| 0.409265557 | 0.737441737 | 0.630424527 | 0.005238192 | 0.006352035 | 0.007313854 |
| 0.409265557 | 0.230239514 | 0.212371594 | 0.010564931 | 0.012312513 | 0.012954696 |
| 0.194794725 | 0.160618948 | 0.168126005 | 0.005238192 | 0.006352035 | 0.007313854 |
| 0.194794725 | 0.347852160 | 0.321472861 | 0.010564931 | 0.012312513 | 0.012954696 |
| 0.533497610 | 0.373109522 | 0.311367562 | 0.005238192 | 0.006352035 | 0.007313854 |
| 0.533497610 | 0.362098828 | 0.338602667 | 0.010564931 | 0.012312513 | 0.012954696 |
| 0.504807273 | 0.308031043 | 0.269623303 | 0.005238192 | 0.006352035 | 0.007313854 |
| 0.504807273 | 0.203202542 | 0.197952666 | 0.010564931 | 0.012312513 | 0.012954696 |
| 0.571451699 | 0.491478273 | 0.401195141 | 0.005238192 | 0.006352035 | 0.007313854 |
| 0.571451699 | 0.379242142 | 0.365541239 | 0.010564931 | 0.012312513 | 0.012954696 |
| 0.200507977 | 0.293314430 | 0.260059846 | 0.005238192 | 0.006352035 | 0.007313854 |
| 0.200507977 | 0.90464502 | 0.859491855 | 0.010564931 | 0.012312513 | 0.012954696 |

| | | | | | | |
|---|---|---|---|---|---|---|
| 0.814351380 | 0.700712534 | 0.597026696 | 0.005238192 | 0.006352035 | 0.007313854 | |
| 0.814351380 | 0.769817726 | 0.707060833 | 0.010564931 | 0.012312513 | 0.012954696 | |
| 0.457939587 | 0.222060952 | 0.213962170 | 0.005238192 | 0.006352035 | 0.007313854 | |
| 0.457939587 | 0.283144305 | 0.260866700 | 0.010564931 | 0.012312513 | 0.012954696 | |
| 0.338117609 | 0.884261405 | 0.770143949 | 0.005238192 | 0.006352035 | 0.007313854 | |
| 0.338117609 | 0.115229774 | 0.132448387 | 0.010564931 | 0.012312513 | 0.012954696 | |
| 0.271926529 | 0.770173065 | 0.661143819 | 0.005238192 | 0.006352035 | 0.007313854 | |
| 0.271926529 | 0.930205276 | 0.876467842 | 0.010564931 | 0.012312513 | 0.012954696 | |
| 0.323860254 | 0.414101527 | 0.340847772 | 0.005238192 | 0.006352035 | 0.007313854 | |
| 0.323860254 | 0.765249050 | 0.704983540 | 0.010564931 | 0.012312513 | 0.012954696 | |
| 0.899379732 | 0.772570112 | 0.663442673 | 0.005238192 | 0.006352035 | 0.007313854 | |
| 0.899379732 | 0.573031356 | 0.547596322 | 0.010564931 | 0.012312513 | 0.012954696 | |
| 0.568835471 | 0.398955620 | 0.330105015 | 0.005238192 | 0.006352035 | 0.007313854 | |
| 0.568835471 | 0.279987750 | 0.257096103 | 0.010564931 | 0.012312513 | 0.012954696 | |
| 0.047644905 | 0.050943913 | 0.072512463 | 0.005238192 | 0.006352035 | 0.007313854 | |
| 0.047644905 | 0.182279959 | 0.185618838 | 0.010564931 | 0.012312513 | 0.012954696 | |
| 0.607329325 | 0.467047592 | 0.381089719 | 0.005238192 | 0.006352035 | 0.007313854 | |
| 0.607329325 | 0.358868800 | 0.333909612 | 0.010564931 | 0.012312513 | 0.012954696 | |
| 0.019538264 | 0.000658166 | 0.004732159 | 0.005238192 | 0.006352035 | 0.007313854 | 0.1 |
| 0.019538264 | 0.028152759 | 0.053130437 | 0.010564931 | 0.012312513 | 0.012954696 | |
| 0.930028471 | 0.898198958 | 0.785360967 | 0.005238192 | 0.006352035 | 0.007313854 | |
| 0.930028471 | 0.802674414 | 0.753488957 | 0.010564931 | 0.012312513 | 0.012954696 | |
| 0.174911336 | 0.840159223 | 0.726065962 | 0.005238192 | 0.006352035 | 0.007313854 | |
| 0.174911336 | 0.223963189 | 0.210077905 | 0.010564931 | 0.012312513 | 0.012954696 | |
| 0.225402883 | 0.099714966 | 0.109608888 | 0.005238192 | 0.006352035 | 0.007313854 | |
| 0.225402883 | 0.061458515 | 0.084937265 | 0.010564931 | 0.012312513 | 0.012954696 | |
| 0.022654842 | 0.739448764 | 0.632147131 | 0.005238192 | 0.006352035 | 0.007313854 | |
| 0.022654842 | 0.213049856 | 0.205466905 | 0.010564931 | 0.012312513 | 0.012954696 | |
| 0.784282939 | 0.992367924 | 0.945522712 | 0.005238192 | 0.006352035 | 0.007313854 | |
| 0.784282939 | 0.737427438 | 0.673915635 | 0.010564931 | 0.012312513 | 0.012954696 | |
| 0.538610316 | 0.325038829 | 0.280310263 | 0.005238192 | 0.006352035 | 0.007313854 | |
| 0.538610316 | 0.373713768 | 0.356440600 | 0.010564931 | 0.012312513 | 0.012954696 | |
| 0.078054404 | 0.181796726 | 0.185459149 | 0.005238192 | 0.006352035 | 0.007313854 | |
| 0.078054404 | 0.940664992 | 0.913466916 | 0.010564931 | 0.012312513 | 0.012954696 | |
| 0.611921474 | 0.960147579 | 0.860657318 | 0.005238192 | 0.006352035 | 0.007313854 | |
| 0.611921474 | 0.855467184 | 0.816731630 | 0.010564931 | 0.012312513 | 0.012954696 | |
| 0.351694308 | 0.272045620 | 0.245449872 | 0.005238192 | 0.006352035 | 0.007313854 | |
| 0.351694308 | 0.144009049 | 0.154615262 | 0.010564931 | 0.012312513 | 0.012954696 | |
| 0.114712328 | 0.198037321 | 0.197792016 | 0.005238192 | 0.006352035 | 0.007313854 | |
| 0.114712328 | 0.171945651 | 0.175903203 | 0.010564931 | 0.012312513 | 0.012954696 | |
| 0.833896727 | 0.756047094 | 0.647623390 | 0.005238192 | 0.006352035 | 0.007313854 | |
| 0.833896727 | 0.782871632 | 0.726479609 | 0.010564931 | 0.012312513 | 0.012954696 | |
| 0.541738041 | 0.451886181 | 0.368963813 | 0.005238192 | 0.006352035 | 0.007313854 | |
| 0.541738041 | 0.610767150 | 0.579341674 | 0.010564931 | 0.012312513 | 0.012954696 | |
| 0.091656262 | 0.083561185 | 0.095665457 | 0.005238192 | 0.006352035 | 0.007313854 | |
| 0.091656262 | 0.991531059 | 0.976798594 | 0.010564931 | 0.012312513 | 0.012954696 | |

| | | | | | |
|---|---|---|---|---|---|
| 0.248286540 | 0.074120446 | 0.088586583 | 0.005238192 | 0.006352035 | 0.007313854 |
| 0.248286540 | 0.034060394 | 0.059720707 | 0.010564931 | 0.012312513 | 0.012954696 |
| 0.536285077 | 0.409157902 | 0.337430491 | 0.005238192 | 0.006352035 | 0.007313854 |
| 0.536285077 | 0.538876225 | 0.526916993 | 0.010564931 | 0.012312513 | 0.012954696 |
| 0.093041966 | 0.876653219 | 0.762430582 | 0.005238192 | 0.006352035 | 0.007313854 |
| 0.093041966 | 0.796027189 | 0.750250123 | 0.010564931 | 0.012312513 | 0.012954696 |
| 0.980916728 | 0.979093021 | 0.902488922 | 0.005238192 | 0.006352035 | 0.007313854 |
| 0.980916728 | 0.964499300 | 0.931977071 | 0.010564931 | 0.012312513 | 0.012954696 |
| 0.212117712 | 0.835044193 | 0.721111732 | 0.005238192 | 0.006352035 | 0.007313854 |
| 0.212117712 | 0.188880383 | 0.188837474 | 0.010564931 | 0.012312513 | 0.012954696 |
| 0.426176812 | 0.209931951 | 0.206145539 | 0.005238192 | 0.006352035 | 0.007313854 |
| 0.426176812 | 0.923695549 | 0.874089312 | 0.010564931 | 0.012312513 | 0.012954696 |
| 0.866612230 | 0.738989894 | 0.631752628 | 0.005238192 | 0.006352035 | 0.007313854 |
| 0.866612230 | 0.784540522 | 0.732601969 | 0.010564931 | 0.012312513 | 0.012954696 |
| 0.810220876 | 0.755149166 | 0.646774638 | 0.005238192 | 0.006352035 | 0.007313854 |
| 0.810220876 | 0.576088921 | 0.553188758 | 0.010564931 | 0.012312513 | 0.012954696 |
| 0.493853159 | 0.384960556 | 0.319956683 | 0.005238192 | 0.006352035 | 0.007313854 |
| 0.493853159 | 0.145886532 | 0.156850056 | 0.010564931 | 0.012312513 | 0.012954696 |
| 0.314433653 | 0.927325816 | 0.817457662 | 0.005238192 | 0.006352035 | 0.007313854 |
| 0.314433653 | 0.984914827 | 0.960917334 | 0.010564931 | 0.012312513 | 0.012954696 |
| 0.238998106 | 0.374713435 | 0.312526611 | 0.005238192 | 0.006352035 | 0.007313854 |
| 0.238998106 | 0.073421231 | 0.097178496 | 0.010564931 | 0.012312513 | 0.012954696 |
| 0.005697240 | 0.452332398 | 0.369317678 | 0.005238192 | 0.006352035 | 0.007313854 |
| 0.005697240 | 0.965504748 | 0.942209911 | 0.010564931 | 0.012312513 | 0.012954696 |
| 0.781455024 | 0.721304913 | 0.615967104 | 0.005238192 | 0.006352035 | 0.007313854 |
| 0.781455024 | 0.473295833 | 0.470151553 | 0.010564931 | 0.012312513 | 0.012954696 |
| 0.604514867 | 0.429698196 | 0.351921948 | 0.005238192 | 0.006352035 | 0.007313854 |
| 0.604514867 | 0.324934552 | 0.301476049 | 0.010564931 | 0.012312513 | 0.012954696 |
| 0.448105827 | 0.361467873 | 0.303408826 | 0.005238192 | 0.006352035 | 0.007313854 |
| 0.448105827 | 0.427741820 | 0.425403659 | 0.010564931 | 0.012312513 | 0.012954696 |
| 0.159050482 | 0.489787004 | 0.39980016 | 0.005238192 | 0.006352035 | 0.007313854 |
| 0.159050482 | 0.842138254 | 0.804926816 | 0.010564931 | 0.012312513 | 0.012954696 |
| 0.376718819 | 0.166255070 | 0.173093147 | 0.005238192 | 0.006352035 | 0.007313854 |
| 0.376718819 | 0.101710043 | 0.124783192 | 0.010564931 | 0.012312513 | 0.012954696 |
| 0.116322321 | 0.315908539 | 0.274576772 | 0.005238192 | 0.006352035 | 0.007313854 |
| 0.116322321 | 0.394211460 | 0.388837742 | 0.010564931 | 0.012312513 | 0.012954696 |
| 0.013670166 | 0.209954877 | 0.206161305 | 0.005238192 | 0.006352035 | 0.007313854 |
| 0.013670166 | 0.192885598 | 0.192321946 | 0.010564931 | 0.012312513 | 0.012954696 |
| 0.029162225 | 0.068299593 | 0.084652252 | 0.005238192 | 0.006352035 | 0.007313854 |
| 0.029162225 | 0.015833645 | 0.037396020 | 0.010564931 | 0.012312513 | 0.012954696 |
| 0.163395082 | 0.028619714 | 0.052587855 | 0.005238192 | 0.006352035 | 0.007313854 |
| 0.163395082 | 0.593209950 | 0.573123605 | 0.010564931 | 0.012312513 | 0.012954696 |
| 0.394730555 | 0.201496377 | 0.20025868 | 0.005238192 | 0.006352035 | 0.007313854 |
| 0.394730555 | 0.128841765 | 0.141832644 | 0.010564931 | 0.012312513 | 0.012954696 |
| 0.012043655 | 0.912603547 | 0.800736989 | 0.005238192 | 0.006352035 | 0.007313854 |
| 0.012043655 | 0.629021918 | 0.589275234 | 0.010564931 | 0.012312513 | 0.012954696 |

| | | | | | |
|---|---|---|---|---|---|
| 0.392232751 | 0.168312748 | 0.174807203 | 0.005238192 | 0.006352035 | 0.007313854 |
| 0.392232751 | 0.089821795 | 0.111127094 | 0.010564931 | 0.012312513 | 0.012954696 |
| 0.726607781 | 0.574732726 | 0.475846725 | 0.005238192 | 0.006352035 | 0.007313854 |
| 0.726607781 | 0.429623548 | 0.426913008 | 0.010564931 | 0.012312513 | 0.012954696 |
| 0.139200498 | 0.079271356 | 0.092361682 | 0.005238192 | 0.006352035 | 0.007313854 |
| 0.139200498 | 0.559684292 | 0.541038140 | 0.010564931 | 0.012312513 | 0.012954696 |
| 0.063958939 | 0.718063689 | 0.613108248 | 0.005238192 | 0.006352035 | 0.007313854 |
| 0.063958939 | 0.883934004 | 0.835811326 | 0.010564931 | 0.012312513 | 0.012954696 |
| 0.834902575 | 0.671232552 | 0.569519389 | 0.005238192 | 0.006352035 | 0.007313854 |
| 0.834902575 | 0.476094645 | 0.475334794 | 0.010564931 | 0.012312513 | 0.012954696 |
| 0.917429387 | 0.939179896 | 0.831492846 | 0.005238192 | 0.006352035 | 0.007313854 |
| 0.917429387 | 0.893554983 | 0.848398653 | 0.010564931 | 0.012312513 | 0.012954696 |
| 0.904945626 | 0.742988688 | 0.635240784 | 0.005238192 | 0.006352035 | 0.007313854 |
| 0.904945626 | 0.506555698 | 0.504474950 | 0.010564931 | 0.012312513 | 0.012954696 |
| 0.733156967 | 0.667455537 | 0.566109021 | 0.005238192 | 0.006352035 | 0.007313854 |
| 0.733156967 | 0.612575579 | 0.580258133 | 0.010564931 | 0.012312513 | 0.012954696 |
| 0.198244933 | 0.439609277 | 0.359127892 | 0.005238192 | 0.006352035 | 0.007313854 |
| 0.198244933 | 0.623386239 | 0.586649377 | 0.010564931 | 0.012312513 | 0.012954696 |
| 0.210034378 | 0.142440187 | 0.151221827 | 0.005238192 | 0.006352035 | 0.007313854 |
| 0.210034378 | 0.531125768 | 0.526432245 | 0.010564931 | 0.012312513 | 0.012954696 |
| 0.997416449 | 0.857145926 | 0.743607533 | 0.005238192 | 0.006352035 | 0.007313854 |
| 0.997416449 | 0.663424567 | 0.620699997 | 0.010564931 | 0.012312513 | 0.012954696 |
| 0.128699644 | 0.859198000 | 0.745658646 | 0.005238192 | 0.006352035 | 0.007313854 |
| 0.128699644 | 0.912741315 | 0.866833998 | 0.010564931 | 0.012312513 | 0.012954696 |
| 0.379087636 | 0.613697533 | 0.515669625 | 0.005238192 | 0.006352035 | 0.007313854 |
| 0.379087636 | 0.898108661 | 0.855693195 | 0.010564931 | 0.012312513 | 0.012954696 |
| 0.739754296 | 0.596427828 | 0.497565000 | 0.005238192 | 0.006352035 | 0.007313854 |
| 0.739754296 | 0.401484685 | 0.395311578 | 0.010564931 | 0.012312513 | 0.012954696 |
| 0.660264078 | 0.598332212 | 0.600538975 | 0.008643055 | 0.009628871 | 0.010456710 |
| 0.660264078 | 0.515441172 | 0.568267265 | 0.002262659 | 0.002725363 | 0.003224299 |
| 0.319341722 | 0.515450445 | 0.526406649 | 0.008643055 | 0.009628871 | 0.010456710 |
| 0.319341722 | 0.373645199 | 0.440552638 | 0.002262659 | 0.002725363 | 0.003224299 |
| 0.101422053 | 0.123682792 | 0.142800197 | 0.008643055 | 0.009628871 | 0.010456710 |
| 0.101422053 | 0.613463165 | 0.657949423 | 0.002262659 | 0.002725363 | 0.003224299 |
| 0.716207136 | 0.607380393 | 0.608465329 | 0.008643055 | 0.009628871 | 0.010456710 |
| 0.716207136 | 0.465725062 | 0.514378053 | 0.002262659 | 0.002725363 | 0.003224299 |
| 0.111826637 | 0.569120347 | 0.575482508 | 0.008643055 | 0.009628871 | 0.010456710 |
| 0.111826637 | 0.397456941 | 0.458498382 | 0.002262659 | 0.002725363 | 0.003224299 |
| 0.180168125 | 0.094736284 | 0.123760274 | 0.008643055 | 0.009628871 | 0.010456710 |
| 0.180168125 | 0.082776602 | 0.062842852 | 0.002262659 | 0.002725363 | 0.003224299 |
| 0.943471191 | 0.903311304 | 0.915697185 | 0.008643055 | 0.009628871 | 0.010456710 |
| 0.943471191 | 0.856381139 | 0.891677188 | 0.002262659 | 0.002725363 | 0.003224299 |
| 0.693893455 | 0.573170302 | 0.580780426 | 0.008643055 | 0.009628871 | 0.010456710 |
| 0.693893455 | 0.695228245 | 0.748680804 | 0.002262659 | 0.002725363 | 0.003224299 |
| 0.959860997 | 0.934901039 | 0.939685320 | 0.008643055 | 0.009628871 | 0.010456710 |
| 0.959860997 | 0.911051378 | 0.923125694 | 0.002262659 | 0.002725363 | 0.003224299 |

| | | | | | |
|---|---|---|---|---|---|
| 0.745123437 | 0.652279856 | 0.644355975 | 0.008643055 | 0.009628871 | 0.010456710 |
| 0.745123437 | 0.509677573 | 0.556571471 | 0.002262659 | 0.002725363 | 0.003224299 |
| 0.792161429 | 0.705200650 | 0.704826793 | 0.008643055 | 0.009628871 | 0.010456710 |
| 0.792161429 | 0.587754876 | 0.629076059 | 0.002262659 | 0.002725363 | 0.003224299 |
| 0.080291845 | 0.123965234 | 0.143120997 | 0.008643055 | 0.009628871 | 0.010456710 |
| 0.080291845 | 0.031517223 | 0.029146685 | 0.002262659 | 0.002725363 | 0.003224299 |
| 0.841055167 | 0.768356170 | 0.770344793 | 0.008643055 | 0.009628871 | 0.010456710 |
| 0.841055167 | 0.656662369 | 0.709003859 | 0.002262659 | 0.002725363 | 0.003224299 |
| 0.229099908 | 0.113214093 | 0.134920658 | 0.008643055 | 0.009628871 | 0.010456710 |
| 0.229099908 | 0.065104244 | 0.055490316 | 0.002262659 | 0.002725363 | 0.003224299 |
| 0.925367853 | 0.875943742 | 0.889134717 | 0.008643055 | 0.009628871 | 0.010456710 |
| 0.925367853 | 0.834990345 | 0.857874024 | 0.002262659 | 0.002725363 | 0.003224299 |
| 0.046756999 | 0.452219463 | 0.454440274 | 0.008643055 | 0.009628871 | 0.010456710 |
| 0.046756999 | 0.416967138 | 0.486297339 | 0.002262659 | 0.002725363 | 0.003224299 |
| 0.265177529 | 0.623235918 | 0.61643605 | 0.008643055 | 0.009628871 | 0.010456710 |
| 0.265177529 | 0.324496124 | 0.403243986 | 0.002262659 | 0.002725363 | 0.003224299 |
| 0.499412214 | 0.388485013 | 0.392449584 | 0.008643055 | 0.009628871 | 0.010456710 |
| 0.499412214 | 0.219135125 | 0.250145705 | 0.002262659 | 0.002725363 | 0.003224299 |
| 0.367078277 | 0.266771166 | 0.269434087 | 0.008643055 | 0.009628871 | 0.010456710 |
| 0.367078277 | 0.164376472 | 0.135461339 | 0.002262659 | 0.002725363 | 0.003224299 |
| 0.184449229 | 0.155273475 | 0.161274587 | 0.008643055 | 0.009628871 | 0.010456710 |
| 0.184449229 | 0.484016647 | 0.526447548 | 0.002262659 | 0.002725363 | 0.003224299 |
| 0.314345718 | 0.662953959 | 0.667417398 | 0.008643055 | 0.009628871 | 0.010456710 |
| 0.314345718 | 0.124225861 | 0.099369513 | 0.002262659 | 0.002725363 | 0.003224299 |
| 0.571600734 | 0.594313281 | 0.59232548 | 0.008643055 | 0.009628871 | 0.010456710 |
| 0.571600734 | 0.472833073 | 0.518473760 | 0.002262659 | 0.002725363 | 0.003224299 |
| 0.258896020 | 0.148458407 | 0.156568287 | 0.008643055 | 0.009628871 | 0.010456710 |
| 0.258896020 | 0.113230802 | 0.088454876 | 0.002262659 | 0.002725363 | 0.003224299 |
| 0.979310255 | 0.958043642 | 0.968515595 | 0.008643055 | 0.009628871 | 0.010456710 |
| 0.979310255 | 0.952208616 | 0.960223923 | 0.002262659 | 0.002725363 | 0.003224299 |
| 0.561920178 | 0.419831542 | 0.431004151 | 0.008643055 | 0.009628871 | 0.010456710 |
| 0.561920178 | 0.886077721 | 0.911477559 | 0.002262659 | 0.002725363 | 0.003224299 |
| 0.125794333 | 0.248551383 | 0.243997974 | 0.008643055 | 0.009628871 | 0.010456710 |
| 0.125794333 | 0.187171938 | 0.178138760 | 0.002262659 | 0.002725363 | 0.003224299 |
| 0.133746852 | 0.049876172 | 0.072072580 | 0.008643055 | 0.009628871 | 0.010456710 |
| 0.133746852 | 0.601438467 | 0.639631296 | 0.002262659 | 0.002725363 | 0.003224299 |
| 0.070326997 | 0.461583469 | 0.462841703 | 0.008643055 | 0.009628871 | 0.010456710 |
| 0.070326997 | 0.136805470 | 0.108250620 | 0.002262659 | 0.002725363 | 0.003224299 |
| 0.354157422 | 0.233594897 | 0.232195243 | 0.008643055 | 0.009628871 | 0.010456710 |
| 0.354157422 | 0.159552352 | 0.126019956 | 0.002262659 | 0.002725363 | 0.003224299 |
| 0.046888560 | 0.113412870 | 0.134943107 | 0.008643055 | 0.009628871 | 0.010456710 |
| 0.046888560 | 0.165303981 | 0.139265283 | 0.002262659 | 0.002725363 | 0.003224299 |
| 0.640341646 | 0.505718437 | 0.518140798 | 0.008643055 | 0.009628871 | 0.010456710 |
| 0.640341646 | 0.336989735 | 0.411307749 | 0.002262659 | 0.002725363 | 0.003224299 |
| 0.702589549 | 0.591585591 | 0.591557272 | 0.008643055 | 0.009628871 | 0.010456710 |
| 0.702589549 | 0.735808564 | 0.788628658 | 0.002262659 | 0.002725363 | 0.003224299 |

| | | | | | |
|---|---|---|---|---|---|
| 0.639257774 | 0.495318315 | 0.516349725 | 0.008643055 | 0.009628871 | 0.010456710 |
| 0.639257774 | 0.328681136 | 0.409499946 | 0.002262659 | 0.002725363 | 0.003224299 |
| 0.881017309 | 0.800474592 | 0.825274876 | 0.008643055 | 0.009628871 | 0.010456710 |
| 0.881017309 | 0.715545600 | 0.777839751 | 0.002262659 | 0.002725363 | 0.003224299 |
| 0.856462496 | 0.785150278 | 0.791110789 | 0.008643055 | 0.009628871 | 0.010456710 |
| 0.856462496 | 0.682595092 | 0.734531949 | 0.002262659 | 0.002725363 | 0.003224299 |
| 0.226007567 | 0.596464520 | 0.596678115 | 0.008643055 | 0.009628871 | 0.010456710 |
| 0.226007567 | 0.725019273 | 0.785574300 | 0.002262659 | 0.002725363 | 0.003224299 |
| 0.226944710 | 0.253446829 | 0.250258434 | 0.008643055 | 0.009628871 | 0.010456710 |
| 0.226944710 | 0.671542227 | 0.732205903 | 0.002262659 | 0.002725363 | 0.003224299 |
| 0.564721880 | 0.425137028 | 0.433912103 | 0.008643055 | 0.009628871 | 0.010456710 |
| 0.564721880 | 0.265571772 | 0.319762640 | 0.002262659 | 0.002725363 | 0.003224299 |
| 0.955504882 | 0.917332547 | 0.933031856 | 0.008643055 | 0.009628871 | 0.010456710 |
| 0.955504882 | 0.890352811 | 0.914091933 | 0.002262659 | 0.002725363 | 0.003224299 |
| 0.401955805 | 0.269801895 | 0.273775953 | 0.008643055 | 0.009628871 | 0.010456710 |
| 0.401955805 | 0.877338268 | 0.902050130 | 0.002262659 | 0.002725363 | 0.003224299 |
| 0.992044536 | 0.990191460 | 0.988747447 | 0.008643055 | 0.009628871 | 0.010456710 |
| 0.992044536 | 0.985817588 | 0.985610657 | 0.002262659 | 0.002725363 | 0.003224299 |
| 0.834819327 | 0.750222212 | 0.761655228 | 0.008643055 | 0.009628871 | 0.010456710 |
| 0.834819327 | 0.643298583 | 0.698496838 | 0.002262659 | 0.002725363 | 0.003224299 |
| 0.719319406 | 0.674396846 | 0.681841961 | 0.008643055 | 0.009628871 | 0.010456710 |
| 0.719319406 | 0.470989418 | 0.518358181 | 0.002262659 | 0.002725363 | 0.003224299 |
| 0.666175176 | 0.537639667 | 0.547590137 | 0.008643055 | 0.009628871 | 0.010456710 |
| 0.666175176 | 0.377748732 | 0.444709998 | 0.002262659 | 0.002725363 | 0.003224299 |
| 0.506250222 | 0.356636924 | 0.373276301 | 0.008643055 | 0.009628871 | 0.010456710 |
| 0.506250222 | 0.525685989 | 0.580760571 | 0.002262659 | 0.002725363 | 0.003224299 |
| 0.070897364 | 0.027998168 | 0.047597951 | 0.008643055 | 0.009628871 | 0.010456710 |
| 0.070897364 | 0.077854936 | 0.062278900 | 0.002262659 | 0.002725363 | 0.003224299 |
| 0.026060551 | 0.095181342 | 0.124444103 | 0.008643055 | 0.009628871 | 0.010456710 |
| 0.026060551 | 0.145698045 | 0.117804704 | 0.002262659 | 0.002725363 | 0.003224299 |
| 0.669054059 | 0.596401418 | 0.596552576 | 0.008643055 | 0.009628871 | 0.010456710 |
| 0.669054059 | 0.391011224 | 0.448871799 | 0.002262659 | 0.002725363 | 0.003224299 |
| 0.535738774 | 0.396644895 | 0.403385343 | 0.008643055 | 0.009628871 | 0.010456710 |
| 0.535738774 | 0.249040601 | 0.287658480 | 0.002262659 | 0.002725363 | 0.003224299 |
| 0.721021790 | 0.727808857 | 0.724980170 | 0.008643055 | 0.009628871 | 0.010456710 |
| 0.721021790 | 0.481605435 | 0.521171752 | 0.002262659 | 0.002725363 | 0.003224299 |
| 0.307985897 | 0.193313234 | 0.194522001 | 0.008643055 | 0.009628871 | 0.010456710 |
| 0.307985897 | 0.304602462 | 0.355364914 | 0.002262659 | 0.002725363 | 0.003224299 |
| 0.252314824 | 0.443733875 | 0.447575400 | 0.008643055 | 0.009628871 | 0.010456710 |
| 0.252314824 | 0.921729639 | 0.931783931 | 0.002262659 | 0.002725363 | 0.003224299 |
| 0.891793546 | 0.812927918 | 0.840675975 | 0.008643055 | 0.009628871 | 0.010456710 |
| 0.891793546 | 0.740061805 | 0.796568999 | 0.002262659 | 0.002725363 | 0.003224299 |
| 0.227249670 | 0.289822130 | 0.295380158 | 0.008643055 | 0.009628871 | 0.010456710 |
| 0.227249670 | 0.055771524 | 0.051930186 | 0.002262659 | 0.002725363 | 0.003224299 |
| 0.018703742 | 0.238936537 | 0.235564641 | 0.008643055 | 0.009628871 | 0.010456710 |
| 0.018703742 | 0.381914524 | 0.446401294 | 0.002262659 | 0.002725363 | 0.003224299 |

| | | | | | |
|---|---|---|---|---|---|
| 0.300237218 | 0.185305932 | 0.188352236 | 0.008643055 | 0.009628871 | 0.010456710 |
| 0.300237218 | 0.115290185 | 0.090625323 | 0.002262659 | 0.002725363 | 0.003224299 |
| 0.021647033 | 0.039721355 | 0.059907758 | 0.008643055 | 0.009628871 | 0.010456710 |
| 0.021647033 | 0.006543555 | 0.006975844 | 0.002262659 | 0.002725363 | 0.003224299 |
| 0.516125999 | 0.376510060 | 0.383104264 | 0.008643055 | 0.009628871 | 0.010456710 |
| 0.516125999 | 0.241430114 | 0.266958240 | 0.002262659 | 0.002725363 | 0.003224299 |
| 0.678773390 | 0.660301742 | 0.660119269 | 0.008643055 | 0.009628871 | 0.010456710 |
| 0.678773390 | 0.668362530 | 0.730529732 | 0.002262659 | 0.002725363 | 0.003224299 |
| 0.938273956 | 0.895675367 | 0.908181937 | 0.008643055 | 0.009628871 | 0.010456710 |
| 0.938273956 | 0.846541361 | 0.881574949 | 0.002262659 | 0.002725363 | 0.003224299 |
| 0.893338951 | 0.827397170 | 0.842827348 | 0.008643055 | 0.009628871 | 0.010456710 |
| 0.893338951 | 0.762099231 | 0.799418586 | 0.002262659 | 0.002725363 | 0.003224299 |
| 0.569428499 | 0.431447880 | 0.438859722 | 0.008643055 | 0.009628871 | 0.010456710 |
| 0.569428499 | 0.274188393 | 0.325236487 | 0.002262659 | 0.002725363 | 0.003224299 |
| 0.514547099 | 0.371065459 | 0.381442502 | 0.008643055 | 0.009628871 | 0.010456710 |
| 0.514547099 | 0.237833353 | 0.265384638 | 0.002262659 | 0.002725363 | 0.003224299 |
| 0.590800803 | 0.459979699 | 0.462292532 | 0.008643055 | 0.009628871 | 0.010456710 |
| 0.590800803 | 0.301169829 | 0.350058979 | 0.002262659 | 0.002725363 | 0.003224299 |
| 0.994920878 | 0.994482126 | 0.992669380 | 0.008643055 | 0.009628871 | 0.010456710 |
| 0.994920878 | 0.997294881 | 0.991771423 | 0.002262659 | 0.002725363 | 0.003224299 |
| 0.964996597 | 0.942795978 | 0.947264304 | 0.008643055 | 0.009628871 | 0.010456710 |
| 0.964996597 | 0.933589071 | 0.932954293 | 0.002262659 | 0.002725363 | 0.003224299 |
| 0.740139019 | 0.633569363 | 0.637591812 | 0.008643055 | 0.009628871 | 0.010456710 |
| 0.740139019 | 0.495921790 | 0.548882366 | 0.002262659 | 0.002725363 | 0.003224299 |
| 0.407338804 | 0.655497048 | 0.647697654 | 0.008643055 | 0.009628871 | 0.010456710 |
| 0.407338804 | 0.180441633 | 0.166491030 | 0.002262659 | 0.002725363 | 0.003224299 |
| 0.270197619 | 0.491191260 | 0.509558578 | 0.008643055 | 0.009628871 | 0.010456710 |
| 0.270197619 | 0.824781029 | 0.848025877 | 0.002262659 | 0.002725363 | 0.003224299 |
| 0.835630027 | 0.758437466 | 0.762891127 | 0.008643055 | 0.009628871 | 0.010456710 |
| 0.835630027 | 0.650415008 | 0.699913658 | 0.002262659 | 0.002725363 | 0.003224299 |
| 0.946857791 | 0.907498714 | 0.920184793 | 0.008643055 | 0.009628871 | 0.010456710 |
| 0.946857791 | 0.867840480 | 0.897639951 | 0.002262659 | 0.002725363 | 0.003224299 |
| 0.819413988 | 0.734479119 | 0.740649873 | 0.008643055 | 0.009628871 | 0.010456710 |
| 0.819413988 | 0.619882041 | 0.672414656 | 0.002262659 | 0.002725363 | 0.003224299 |
| 0.847695589 | 0.775865053 | 0.779017377 | 0.008643055 | 0.009628871 | 0.010456710 |
| 0.847695589 | 0.659166572 | 0.719750553 | 0.002262659 | 0.002725363 | 0.003224299 |
| 0.107873758 | 0.055796811 | 0.078512599 | 0.008643055 | 0.009628871 | 0.010456710 |
| 0.107873758 | 0.105961119 | 0.078777967 | 0.002262659 | 0.002725363 | 0.003224299 |
| 0.227454206 | 0.109655104 | 0.133739948 | 0.008643055 | 0.009628871 | 0.010456710 |
| 0.227454206 | 0.058074067 | 0.052076335 | 0.002262659 | 0.002725363 | 0.003224299 |
| 0.979728682 | 0.969862321 | 0.969635871 | 0.008643055 | 0.009628871 | 0.010456710 |
| 0.979728682 | 0.963142891 | 0.961431987 | 0.002262659 | 0.002725363 | 0.003224299 |
| 0.134271689 | 0.492111706 | 0.511342674 | 0.008643055 | 0.009628871 | 0.010456710 |
| 0.134271689 | 0.152330140 | 0.119918703 | 0.002262659 | 0.002725363 | 0.003224299 |
| 0.225545372 | 0.103284903 | 0.132530636 | 0.008643055 | 0.009628871 | 0.010456710 |
| 0.225545372 | 0.051155065 | 0.051231830 | 0.002262659 | 0.002725363 | 0.003224299 |

| | | | | | |
|---|---|---|---|---|---|
| 0.247080295 | 0.493876250 | 0.514801254 | 0.008643055 | 0.009628871 | 0.010456710 |
| 0.247080295 | 0.144552753 | 0.116258094 | 0.002262659 | 0.002725363 | 0.003224299 |
| 0.892779209 | 0.939886224 | 0.941985014 | 0.008643055 | 0.009628871 | 0.010456710 |
| 0.892779209 | 0.752316813 | 0.798707675 | 0.002262659 | 0.002725363 | 0.003224299 |
| 0.072261879 | 0.018794385 | 0.036562420 | 0.008643055 | 0.009628871 | 0.010456710 |
| 0.072261879 | 0.197211244 | 0.207191086 | 0.002262659 | 0.002725363 | 0.003224299 |
| 0.903289888 | 0.860205329 | 0.857071283 | 0.008643055 | 0.009628871 | 0.010456710 |
| 0.903289888 | 0.866336953 | 0.894347998 | 0.002262659 | 0.002725363 | 0.003224299 |
| 0.516517550 | 0.728381635 | 0.726654504 | 0.008643055 | 0.009628871 | 0.010456710 |
| 0.516517550 | 0.706620135 | 0.775167272 | 0.002262659 | 0.002725363 | 0.003224299 |
| 0.898953702 | 0.851532617 | 0.850887724 | 0.008643055 | 0.009628871 | 0.010456710 |
| 0.898953702 | 0.781195741 | 0.809216837 | 0.002262659 | 0.002725363 | 0.003224299 |
| 0.786009441 | 0.695236665 | 0.696682989 | 0.008643055 | 0.009628871 | 0.010456710 |
| 0.786009441 | 0.571830270 | 0.618899199 | 0.002262659 | 0.002725363 | 0.003224299 |
| 0.511752591 | 0.363824049 | 0.378890427 | 0.008643055 | 0.009628871 | 0.010456710 |
| 0.511752591 | 0.555713335 | 0.5957834 | 0.002262659 | 0.002725363 | 0.003224299 |
| 0.783755297 | 0.688045852 | 0.693759849 | 0.008643055 | 0.009628871 | 0.010456710 |
| 0.783755297 | 0.568894038 | 0.615759985 | 0.002262659 | 0.002725363 | 0.003224299 |
| 0.970873424 | 0.950823593 | 0.956001761 | 0.008643055 | 0.009628871 | 0.010456710 |
| 0.970873424 | 0.974065977 | 0.972893479 | 0.002262659 | 0.002725363 | 0.003224299 |
| 0.798615316 | 0.717191410 | 0.712860549 | 0.008643055 | 0.009628871 | 0.010456710 |
| 0.798615316 | 0.594467717 | 0.638641251 | 0.002262659 | 0.002725363 | 0.003224299 |
| 0.991211826 | 0.977216655 | 0.986853956 | 0.008643055 | 0.009628871 | 0.010456710 |
| 0.991211826 | 0.979446427 | 0.984152953 | 0.002262659 | 0.002725363 | 0.003224299 |
| 0.513163785 | 0.729342640 | 0.729450817 | 0.008643055 | 0.009628871 | 0.010456710 |
| 0.513163785 | 0.635077891 | 0.695978059 | 0.002262659 | 0.002725363 | 0.003224299 |
| 0.579311202 | 0.533835297 | 0.541219891 | 0.008643055 | 0.009628871 | 0.010456710 |
| 0.579311202 | 0.720868334 | 0.783262434 | 0.002262659 | 0.002725363 | 0.003224299 |
| 0.249236840 | 0.552998881 | 0.555341443 | 0.008643055 | 0.009628871 | 0.010456710 |
| 0.249236840 | 0.129748718 | 0.102303733 | 0.002262659 | 0.002725363 | 0.003224299 |
| 0.897690959 | 0.841470983 | 0.848793957 | 0.008643055 | 0.009628871 | 0.010456710 |
| 0.897690959 | 0.822487318 | 0.845553615 | 0.002262659 | 0.002725363 | 0.003224299 |
| 0.021855777 | 0.012590035 | 0.029435048 | 0.008643055 | 0.009628871 | 0.010456710 |
| 0.021855777 | 0.430065715 | 0.494671145 | 0.002262659 | 0.002725363 | 0.003224299 |
| 0.770427210 | 0.670282513 | 0.676593853 | 0.008643055 | 0.009628871 | 0.010456710 |
| 0.770427210 | 0.542503902 | 0.594724773 | 0.002262659 | 0.002725363 | 0.003224299 |
| 0.258089884 | 0.181088821 | 0.184937269 | 0.008643055 | 0.009628871 | 0.010456710 |
| 0.258089884 | 0.090335731 | 0.066893532 | 0.002262659 | 0.002725363 | 0.003224299 |
| 0.152208523 | 0.566041816 | 0.569463763 | 0.008643055 | 0.009628871 | 0.010456710 |
| 0.152208523 | 0.944083090 | 0.952035420 | 0.002262659 | 0.002725363 | 0.003224299 |
| 0.794311063 | 0.712782841 | 0.707676271 | 0.008643055 | 0.009628871 | 0.010456710 |
| 0.794311063 | 0.589137104 | 0.631881045 | 0.002262659 | 0.002725363 | 0.003224299 |
| 0.051527554 | 0.231973361 | 0.230371674 | 0.008643055 | 0.009628871 | 0.010456710 |
| 0.051527554 | 0.912435328 | 0.929105524 | 0.002262659 | 0.002725363 | 0.003224299 |
| 0.458161593 | 0.328302515 | 0.345908536 | 0.008643055 | 0.009628871 | 0.010456710 |
| 0.458161593 | 0.202185607 | 0.210627624 | 0.002262659 | 0.002725363 | 0.003224299 |

| | | | | | |
|---|---|---|---|---|---|
| 0.445179901 | 0.301336032 | 0.313572704 | 0.008643055 | 0.009628871 | 0.010456710 |
| 0.445179901 | 0.194827904 | 0.198834346 | 0.002262659 | 0.002725363 | 0.003224299 |
| 0.740943588 | 0.643175700 | 0.638982884 | 0.008643055 | 0.009628871 | 0.010456710 |
| 0.740943588 | 0.503660154 | 0.550310202 | 0.002262659 | 0.002725363 | 0.003224299 |
| 0.957242267 | 0.928738442 | 0.936034038 | 0.008643055 | 0.009628871 | 0.010456710 |
| 0.957242267 | 0.903750590 | 0.91814182 | 0.002262659 | 0.002725363 | 0.003224299 |
| 0.489296162 | 0.334296562 | 0.356103544 | 0.008643055 | 0.009628871 | 0.010456710 |
| 0.489296162 | 0.214995575 | 0.239998171 | 0.002262659 | 0.002725363 | 0.003224299 |
| 0.101929299 | 0.170988164 | 0.173410076 | 0.008643055 | 0.009628871 | 0.010456710 |
| 0.101929299 | 0.253141794 | 0.289934603 | 0.002262659 | 0.002725363 | 0.003224299 |
| 0.342814285 | 0.256799504 | 0.254855240 | 0.008643055 | 0.009628871 | 0.010456710 |
| 0.342814285 | 0.150868263 | 0.118188097 | 0.002262659 | 0.002725363 | 0.003224299 |
| 0.243282010 | 0.127569510 | 0.145310663 | 0.008643055 | 0.009628871 | 0.010456710 |
| 0.243282010 | 0.071741662 | 0.059512485 | 0.002262659 | 0.002725363 | 0.003224299 |
| 0.060352227 | 0.013141896 | 0.030127957 | 0.008643055 | 0.009628871 | 0.010456710 |
| 0.060352227 | 0.422426927 | 0.489585784 | 0.002262659 | 0.002725363 | 0.003224299 |
| 0.245997418 | 0.130675404 | 0.147215957 | 0.008643055 | 0.009628871 | 0.010456710 |
| 0.245997418 | 0.074132808 | 0.060880603 | 0.002262659 | 0.002725363 | 0.003224299 |
| 0.504390383 | 0.762987352 | 0.768390273 | 0.008643055 | 0.009628871 | 0.010456710 |
| 0.504390383 | 0.458689297 | 0.508497563 | 0.002262659 | 0.002725363 | 0.003224299 |
| 0.553459491 | 0.408285725 | 0.421698767 | 0.008643055 | 0.009628871 | 0.010456710 |
| 0.553459491 | 0.255802690 | 0.306916935 | 0.002262659 | 0.002725363 | 0.003224299 |
| 0.078610712 | 0.030465085 | 0.051445312 | 0.008643055 | 0.009628871 | 0.010456710 |
| 0.078610712 | 0.100689328 | 0.075169541 | 0.002262659 | 0.002725363 | 0.003224299 |
| 0.695468955 | 0.579792528 | 0.582695960 | 0.008643055 | 0.009628871 | 0.010456710 |
| 0.695468955 | 0.416500670 | 0.485045488 | 0.002262659 | 0.002725363 | 0.003224299 |
| 0.930716599 | 0.886731727 | 0.897167079 | 0.008643055 | 0.009628871 | 0.010456710 |
| 0.930716599 | 0.844241915 | 0.868057158 | 0.002262659 | 0.002725363 | 0.003224299 |
| 0.177588711 | 0.072591236 | 0.099779647 | 0.008643055 | 0.009628871 | 0.010456710 |
| 0.177588711 | 0.033452064 | 0.031855995 | 0.002262659 | 0.002725363 | 0.003224299 |
| 0.650674215 | 0.596991665 | 0.597764677 | 0.008643055 | 0.009628871 | 0.010456710 |
| 0.650674215 | 0.683120736 | 0.734564820 | 0.002262659 | 0.002725363 | 0.003224299 |
| 0.235016548 | 0.118840297 | 0.139157812 | 0.008643055 | 0.009628871 | 0.010456710 |
| 0.235016548 | 0.065209130 | 0.055498329 | 0.002262659 | 0.002725363 | 0.003224299 |
| 0.894630395 | 0.836578465 | 0.844850665 | 0.008643055 | 0.009628871 | 0.010456710 |
| 0.894630395 | 0.770562981 | 0.801900709 | 0.002262659 | 0.002725363 | 0.003224299 |
| 0.782024110 | 0.682417897 | 0.691677934 | 0.008643055 | 0.009628871 | 0.010456710 |
| 0.782024110 | 0.562798958 | 0.613053726 | 0.002262659 | 0.002725363 | 0.003224299 |
| 0.642116064 | 0.508864235 | 0.519826203 | 0.008643055 | 0.009628871 | 0.010456710 |
| 0.642116064 | 0.369591916 | 0.437857812 | 0.002262659 | 0.002725363 | 0.003224299 |
| 0.206879804 | 0.090117302 | 0.119371276 | 0.008643055 | 0.009628871 | 0.010456710 |
| 0.206879804 | 0.043630410 | 0.043063130 | 0.002262659 | 0.002725363 | 0.003224299 |
| 0.225270202 | 0.569553317 | 0.576340762 | 0.008643055 | 0.009628871 | 0.010456710 |
| 0.225270202 | 0.809939539 | 0.836373608 | 0.002262659 | 0.002725363 | 0.003224299 |
| 0.472287644 | 0.469895823 | 0.478512717 | 0.008643055 | 0.009628871 | 0.010456710 |
| 0.472287644 | 0.797599572 | 0.822717184 | 0.002262659 | 0.002725363 | 0.003224299 |

| | | | | | |
|---|---|---|---|---|---|
| 0.618501657 | 0.478841452 | 0.492856911 | 0.008643055 | 0.009628871 | 0.010456710 |
| 0.618501657 | 0.315727350 | 0.383328377 | 0.002262659 | 0.002725363 | 0.003224299 |
| 0.659493466 | 0.727650768 | 0.637712876 | 0.006047564 | 0.007230944 | 0.008662615 |
| 0.659493466 | 0.563611879 | 0.568266494 | 0.003263504 | 0.003726758 | 0.004421434 |
| 0.355031550 | 0.402167313 | 0.426068086 | 0.006047564 | 0.007230944 | 0.008662615 |
| 0.355031550 | 0.448145715 | 0.440588328 | 0.003263504 | 0.003726758 | 0.004421434 |
| 0.63005687 | 0.637434373 | 0.583861982 | 0.006047564 | 0.007230944 | 0.008662615 |
| 0.63005687 | 0.656574612 | 0.658478058 | 0.003263504 | 0.003726758 | 0.004421434 |
| 0.942617868 | 0.939585863 | 0.932460563 | 0.006047564 | 0.007230944 | 0.008662615 |
| 0.942617868 | 0.887480722 | 0.890180248 | 0.003263504 | 0.003726758 | 0.004421434 |
| 0.047333310 | 0.150674116 | 0.234721433 | 0.006047564 | 0.007230944 | 0.008662615 |
| 0.047333310 | 0.483469411 | 0.458433889 | 0.003263504 | 0.003726758 | 0.004421434 |
| 0.275156863 | 0.153337877 | 0.235539584 | 0.006047564 | 0.007230944 | 0.008662615 |
| 0.275156863 | 0.062593864 | 0.076236282 | 0.003263504 | 0.003726758 | 0.004421434 |
| 0.535648901 | 0.549509573 | 0.519082614 | 0.006047564 | 0.007230944 | 0.008662615 |
| 0.535648901 | 0.334952978 | 0.356142178 | 0.003263504 | 0.003726758 | 0.004421434 |
| 0.981526257 | 0.984343490 | 0.978069577 | 0.006047564 | 0.007230944 | 0.008662615 |
| 0.981526257 | 0.981080316 | 0.965239682 | 0.003263504 | 0.003726758 | 0.004421434 |
| 0.958791379 | 0.961413815 | 0.951261801 | 0.006047564 | 0.007230944 | 0.008662615 |
| 0.958791379 | 0.934082544 | 0.921072398 | 0.003263504 | 0.003726758 | 0.004421434 |
| 0.461765501 | 0.399627123 | 0.424069149 | 0.006047564 | 0.007230944 | 0.008662615 |
| 0.461765501 | 0.357679980 | 0.382275175 | 0.003263504 | 0.003726758 | 0.004421434 |
| 0.148404682 | 0.096727003 | 0.164184272 | 0.006047564 | 0.007230944 | 0.008662615 |
| 0.148404682 | 0.577595024 | 0.584863452 | 0.003263504 | 0.003726758 | 0.004421434 |
| 0.803636000 | 0.805648172 | 0.772519108 | 0.006047564 | 0.007230944 | 0.008662615 |
| 0.803636000 | 0.646474606 | 0.646804446 | 0.003263504 | 0.003726758 | 0.004421434 |
| 0.488793435 | 0.447418145 | 0.460384333 | 0.006047564 | 0.007230944 | 0.008662615 |
| 0.488793435 | 0.623595866 | 0.623814445 | 0.003263504 | 0.003726758 | 0.004421434 |
| 0.030650684 | 0.004800626 | 0.024722864 | 0.006047564 | 0.007230944 | 0.008662615 |
| 0.030650684 | 0.042691759 | 0.055291866 | 0.003263504 | 0.003726758 | 0.004421434 |
| 0.398598553 | 0.352247444 | 0.391486459 | 0.006047564 | 0.007230944 | 0.008662615 |
| 0.398598553 | 0.417082373 | 0.414480886 | 0.003263504 | 0.003726758 | 0.004421434 |
| 0.443561032 | 0.546961116 | 0.517643771 | 0.006047564 | 0.007230944 | 0.008662615 |
| 0.443561032 | 0.492553845 | 0.486694143 | 0.003263504 | 0.003726758 | 0.004421434 |
| 0.629609211 | 0.763429187 | 0.689044946 | 0.006047564 | 0.007230944 | 0.008662615 |
| 0.629609211 | 0.398564862 | 0.403608417 | 0.003263504 | 0.003726758 | 0.004421434 |
| 0.635343779 | 0.653783194 | 0.589553108 | 0.006047564 | 0.007230944 | 0.008662615 |
| 0.635343779 | 0.400462812 | 0.404530794 | 0.003263504 | 0.003726758 | 0.004421434 |
| 0.675559962 | 0.722301197 | 0.632152233 | 0.006047564 | 0.007230944 | 0.008662615 |
| 0.675559962 | 0.480442775 | 0.457404622 | 0.003263504 | 0.003726758 | 0.004421434 |
| 0.385110228 | 0.287197601 | 0.338124719 | 0.006047564 | 0.007230944 | 0.008662615 |
| 0.385110228 | 0.549384406 | 0.526648209 | 0.003263504 | 0.003726758 | 0.004421434 |
| 0.079478073 | 0.173854134 | 0.259470463 | 0.006047564 | 0.007230944 | 0.008662615 |
| 0.079478073 | 0.046496964 | 0.058854947 | 0.003263504 | 0.003726758 | 0.004421434 |
| 0.408039904 | 0.439215732 | 0.449011588 | 0.006047564 | 0.007230944 | 0.008662615 |
| 0.408039904 | 0.543624518 | 0.518310199 | 0.003263504 | 0.003726758 | 0.004421434 |

| | | | | | |
|---|---|---|---|---|---|
| 0.654757039 | 0.678051482 | 0.609549319 | 0.006047564 | 0.007230944 | 0.008662615 |
| 0.654757039 | 0.426598138 | 0.429658016 | 0.003263504 | 0.003726758 | 0.004421434 |
| 0.094081470 | 0.033205677 | 0.077047008 | 0.006047564 | 0.007230944 | 0.008662615 |
| 0.094081470 | 0.029671030 | 0.038720039 | 0.003263504 | 0.003726758 | 0.004421434 |
| 0.522609747 | 0.471672888 | 0.473084436 | 0.006047564 | 0.007230944 | 0.008662615 |
| 0.522609747 | 0.916282100 | 0.911438248 | 0.003263504 | 0.003726758 | 0.004421434 |
| 0.411738522 | 0.365520241 | 0.399243857 | 0.006047564 | 0.007230944 | 0.008662615 |
| 0.411738522 | 0.182692870 | 0.178424704 | 0.003263504 | 0.003726758 | 0.004421434 |
| 0.425380824 | 0.337858866 | 0.376611427 | 0.006047564 | 0.007230944 | 0.008662615 |
| 0.425380824 | 0.635722779 | 0.639922930 | 0.003263504 | 0.003726758 | 0.004421434 |
| 0.622465215 | 0.739352498 | 0.659791212 | 0.006047564 | 0.007230944 | 0.008662615 |
| 0.622465215 | 0.361343906 | 0.388413818 | 0.003263504 | 0.003726758 | 0.004421434 |
| 0.306006886 | 0.182916489 | 0.263560675 | 0.006047564 | 0.007230944 | 0.008662615 |
| 0.306006886 | 0.086768047 | 0.09418454 | 0.003263504 | 0.003726758 | 0.004421434 |
| 0.069717735 | 0.049020520 | 0.098459535 | 0.006047564 | 0.007230944 | 0.008662615 |
| 0.069717735 | 0.141622327 | 0.139288112 | 0.003263504 | 0.003726758 | 0.004421434 |
| 0.649830358 | 0.674860460 | 0.604948256 | 0.006047564 | 0.007230944 | 0.008662615 |
| 0.649830358 | 0.424075170 | 0.423559308 | 0.003263504 | 0.003726758 | 0.004421434 |
| 0.468115481 | 0.387368022 | 0.418730516 | 0.006047564 | 0.007230944 | 0.008662615 |
| 0.468115481 | 0.797801796 | 0.788394184 | 0.003263504 | 0.003726758 | 0.004421434 |
| 0.600676258 | 0.577070297 | 0.552785897 | 0.006047564 | 0.007230944 | 0.008662615 |
| 0.600676258 | 0.336839740 | 0.361622831 | 0.003263504 | 0.003726758 | 0.004421434 |
| 0.114585938 | 0.048195778 | 0.097256404 | 0.006047564 | 0.007230944 | 0.008662615 |
| 0.114585938 | 0.583418349 | 0.589531665 | 0.003263504 | 0.003726758 | 0.004421434 |
| 0.307218987 | 0.188957053 | 0.264865245 | 0.006047564 | 0.007230944 | 0.008662615 |
| 0.307218987 | 0.088464890 | 0.094838204 | 0.003263504 | 0.003726758 | 0.004421434 |
| 0.559599368 | 0.724485169 | 0.632481687 | 0.006047564 | 0.007230944 | 0.008662615 |
| 0.559599368 | 0.791437712 | 0.785907892 | 0.003263504 | 0.003726758 | 0.004421434 |
| 0.187634381 | 0.127671903 | 0.204563663 | 0.006047564 | 0.007230944 | 0.008662615 |
| 0.187634381 | 0.705414501 | 0.732166593 | 0.003263504 | 0.003726758 | 0.004421434 |
| 0.065855530 | 0.028076405 | 0.069731945 | 0.006047564 | 0.007230944 | 0.008662615 |
| 0.065855530 | 0.071813271 | 0.082789163 | 0.003263504 | 0.003726758 | 0.004421434 |
| 0.114165031 | 0.049820107 | 0.099678658 | 0.006047564 | 0.007230944 | 0.008662615 |
| 0.114165031 | 0.011011145 | 0.021825487 | 0.003263504 | 0.003726758 | 0.004421434 |
| 0.829165641 | 0.825452861 | 0.800862371 | 0.006047564 | 0.007230944 | 0.008662615 |
| 0.829165641 | 0.911117475 | 0.902477340 | 0.003263504 | 0.003726758 | 0.004421434 |
| 0.511420850 | 0.615960002 | 0.576423434 | 0.006047564 | 0.007230944 | 0.008662615 |
| 0.511420850 | 0.244881838 | 0.262528958 | 0.003263504 | 0.003726758 | 0.004421434 |
| 0.489385847 | 0.426764328 | 0.439788823 | 0.006047564 | 0.007230944 | 0.008662615 |
| 0.489385847 | 0.556318310 | 0.546919899 | 0.003263504 | 0.003726758 | 0.004421434 |
| 0.602533688 | 0.685076249 | 0.612339726 | 0.006047564 | 0.007230944 | 0.008662615 |
| 0.602533688 | 0.346549504 | 0.363867832 | 0.003263504 | 0.003726758 | 0.004421434 |
| 0.954553024 | 0.950562771 | 0.945897441 | 0.006047564 | 0.007230944 | 0.008662615 |
| 0.954553024 | 0.921761194 | 0.912380487 | 0.003263504 | 0.003726758 | 0.004421434 |
| 0.513459525 | 0.456530228 | 0.463834334 | 0.006047564 | 0.007230944 | 0.008662615 |
| 0.513459525 | 0.574448360 | 0.580767780 | 0.003263504 | 0.003726758 | 0.004421434 |

| | | | | | |
|---|---|---|---|---|---|
| 0.851232781 | 0.837272127 | 0.826069697 | 0.006047564 | 0.007230944 | 0.008662615 |
| 0.851232781 | 0.692395371 | 0.725697397 | 0.003263504 | 0.003726758 | 0.004421434 |
| 0.368579039 | 0.283416135 | 0.337255360 | 0.006047564 | 0.007230944 | 0.008662615 |
| 0.368579039 | 0.140823536 | 0.136561776 | 0.003263504 | 0.003726758 | 0.004421434 |
| 0.323906609 | 0.327893755 | 0.366203798 | 0.006047564 | 0.007230944 | 0.008662615 |
| 0.323906609 | 0.306337175 | 0.325121005 | 0.003263504 | 0.003726758 | 0.004421434 |
| 0.321529557 | 0.208418147 | 0.278198594 | 0.006047564 | 0.007230944 | 0.008662615 |
| 0.321529557 | 0.100455979 | 0.103809492 | 0.003263504 | 0.003726758 | 0.004421434 |
| 0.168494769 | 0.213646347 | 0.281017411 | 0.006047564 | 0.007230944 | 0.008662615 |
| 0.168494769 | 0.313939359 | 0.335187443 | 0.003263504 | 0.003726758 | 0.004421434 |
| 0.561959854 | 0.531231939 | 0.512933408 | 0.006047564 | 0.007230944 | 0.008662615 |
| 0.561959854 | 0.330511364 | 0.355618888 | 0.003263504 | 0.003726758 | 0.004421434 |
| 0.148237406 | 0.162929009 | 0.242766522 | 0.006047564 | 0.007230944 | 0.008662615 |
| 0.148237406 | 0.951755851 | 0.931679854 | 0.003263504 | 0.003726758 | 0.004421434 |
| 0.797525936 | 0.797871395 | 0.765578204 | 0.006047564 | 0.007230944 | 0.008662615 |
| 0.797525936 | 0.629006025 | 0.637226623 | 0.003263504 | 0.003726758 | 0.004421434 |
| 0.533297406 | 0.508132918 | 0.498008590 | 0.006047564 | 0.007230944 | 0.008662615 |
| 0.533297406 | 0.260806914 | 0.284999945 | 0.003263504 | 0.003726758 | 0.004421434 |
| 0.769982818 | 0.778886752 | 0.73521219 | 0.006047564 | 0.007230944 | 0.008662615 |
| 0.769982818 | 0.593964900 | 0.594311142 | 0.003263504 | 0.003726758 | 0.004421434 |
| 0.475833819 | 0.402591155 | 0.426116214 | 0.006047564 | 0.007230944 | 0.008662615 |
| 0.475833819 | 0.223766521 | 0.227076356 | 0.003263504 | 0.003726758 | 0.004421434 |
| 0.668257555 | 0.711929541 | 0.624224331 | 0.006047564 | 0.007230944 | 0.008662615 |
| 0.668257555 | 0.468290284 | 0.447319311 | 0.003263504 | 0.003726758 | 0.004421434 |
| 0.489084144 | 0.420558647 | 0.439280882 | 0.006047564 | 0.007230944 | 0.008662615 |
| 0.489084144 | 0.229098252 | 0.239748452 | 0.003263504 | 0.003726758 | 0.004421434 |
| 0.566103172 | 0.624356619 | 0.582142816 | 0.006047564 | 0.007230944 | 0.008662615 |
| 0.566103172 | 0.701646285 | 0.730417062 | 0.003263504 | 0.003726758 | 0.004421434 |
| 0.926581875 | 0.924571092 | 0.913772775 | 0.006047564 | 0.007230944 | 0.008662615 |
| 0.926581875 | 0.869094800 | 0.859759211 | 0.003263504 | 0.003726758 | 0.004421434 |
| 0.583177468 | 0.562782825 | 0.534867947 | 0.006047564 | 0.007230944 | 0.008662615 |
| 0.583177468 | 0.320802092 | 0.341149902 | 0.003263504 | 0.003726758 | 0.004421434 |
| 0.873499835 | 0.8610699 | 0.851593981 | 0.006047564 | 0.007230944 | 0.008662615 |
| 0.873499835 | 0.748222701 | 0.764293706 | 0.003263504 | 0.003726758 | 0.004421434 |
| 0.663589147 | 0.695470246 | 0.619057302 | 0.006047564 | 0.007230944 | 0.008662615 |
| 0.663589147 | 0.451503147 | 0.441125519 | 0.003263504 | 0.003726758 | 0.004421434 |
| 0.902390290 | 0.903453908 | 0.885201723 | 0.006047564 | 0.007230944 | 0.008662615 |
| 0.902390290 | 0.818956011 | 0.815633216 | 0.003263504 | 0.003726758 | 0.004421434 |
| 0.532949005 | 0.484597376 | 0.483609741 | 0.006047564 | 0.007230944 | 0.008662615 |
| 0.532949005 | 0.834311023 | 0.827628689 | 0.003263504 | 0.003726758 | 0.004421434 |
| 0.366739432 | 0.259950661 | 0.320573836 | 0.006047564 | 0.007230944 | 0.008662615 |
| 0.366739432 | 0.598519427 | 0.595368831 | 0.003263504 | 0.003726758 | 0.004421434 |
| 0.813676147 | 0.809928213 | 0.783385219 | 0.006047564 | 0.007230944 | 0.008662615 |
| 0.813676147 | 0.660681752 | 0.663219007 | 0.003263504 | 0.003726758 | 0.004421434 |
| 0.912931155 | 0.917325139 | 0.902525598 | 0.006047564 | 0.007230944 | 0.008662615 |
| 0.912931155 | 0.838638 | 0.834515015 | 0.003263504 | 0.003726758 | 0.004421434 |

| | | | | | |
|---|---|---|---|---|---|
| 0.561846893 | 0.662567712 | 0.593806773 | 0.006047564 | 0.007230944 | 0.008662615 |
| 0.561846893 | 0.863920418 | 0.848317527 | 0.003263504 | 0.003726758 | 0.004421434 |
| 0.702278681 | 0.744002574 | 0.660977772 | 0.006047564 | 0.007230944 | 0.008662615 |
| 0.702278681 | 0.644327881 | 0.642278927 | 0.003263504 | 0.003726758 | 0.004421434 |
| 0.934170222 | 0.927456076 | 0.922034308 | 0.006047564 | 0.007230944 | 0.008662615 |
| 0.934170222 | 0.875565027 | 0.873761591 | 0.003263504 | 0.003726758 | 0.004421434 |
| 0.276281474 | 0.155214380 | 0.236447197 | 0.006047564 | 0.007230944 | 0.008662615 |
| 0.276281474 | 0.063998021 | 0.076763692 | 0.003263504 | 0.003726758 | 0.004421434 |
| 0.316730426 | 0.201111456 | 0.273627243 | 0.006047564 | 0.007230944 | 0.008662615 |
| 0.316730426 | 0.097086253 | 0.100949939 | 0.003263504 | 0.003726758 | 0.004421434 |
| 0.779826333 | 0.782432182 | 0.745715782 | 0.006047564 | 0.007230944 | 0.008662615 |
| 0.779826333 | 0.609412720 | 0.609188918 | 0.003263504 | 0.003726758 | 0.004421434 |
| 0.989821504 | 0.991057564 | 0.987808956 | 0.006047564 | 0.007230944 | 0.008662615 |
| 0.989821504 | 0.990913421 | 0.980849897 | 0.003263504 | 0.003726758 | 0.004421434 |
| 0.666504986 | 0.706034176 | 0.622549167 | 0.006047564 | 0.007230944 | 0.008662615 |
| 0.666504986 | 0.460671542 | 0.445479369 | 0.003263504 | 0.003726758 | 0.004421434 |
| 0.386130227 | 0.476253219 | 0.477732811 | 0.006047564 | 0.007230944 | 0.008662615 |
| 0.386130227 | 0.15916675 | 0.149828281 | 0.003263504 | 0.003726758 | 0.004421434 |
| 0.380972943 | 0.275796108 | 0.333951513 | 0.006047564 | 0.007230944 | 0.008662615 |
| 0.380972943 | 0.145170504 | 0.145656926 | 0.003263504 | 0.003726758 | 0.004421434 |
| 0.496469602 | 0.573211875 | 0.547527440 | 0.006047564 | 0.007230944 | 0.008662615 |
| 0.496469602 | 0.240288584 | 0.24731864 | 0.003263504 | 0.003726758 | 0.004421434 |
| 0.516047096 | 0.683862369 | 0.610465418 | 0.006047564 | 0.007230944 | 0.008662615 |
| 0.516047096 | 0.568400986 | 0.579148018 | 0.003263504 | 0.003726758 | 0.004421434 |
| 0.265044054 | 0.143651007 | 0.226140175 | 0.006047564 | 0.007230944 | 0.008662615 |
| 0.265044054 | 0.198719815 | 0.207383869 | 0.003263504 | 0.003726758 | 0.004421434 |
| 0.566377584 | 0.546843350 | 0.517630718 | 0.006047564 | 0.007230944 | 0.008662615 |
| 0.566377584 | 0.893871055 | 0.894011086 | 0.003263504 | 0.003726758 | 0.004421434 |
| 0.600918776 | 0.746633361 | 0.663435686 | 0.006047564 | 0.007230944 | 0.008662615 |
| 0.600918776 | 0.763577789 | 0.775251673 | 0.003263504 | 0.003726758 | 0.004421434 |
| 0.053865732 | 0.029480123 | 0.072331009 | 0.006047564 | 0.007230944 | 0.008662615 |
| 0.053865732 | 0.030742528 | 0.040304243 | 0.003263504 | 0.003726758 | 0.004421434 |
| 0.089700379 | 0.050647678 | 0.100938932 | 0.006047564 | 0.007230944 | 0.008662615 |
| 0.089700379 | 0.076013006 | 0.091806919 | 0.003263504 | 0.003726758 | 0.004421434 |
| 0.881123245 | 0.877894621 | 0.860571734 | 0.006047564 | 0.007230944 | 0.008662615 |
| 0.881123245 | 0.771955691 | 0.778030335 | 0.003263504 | 0.003726758 | 0.004421434 |
| 0.870623786 | 0.848416406 | 0.848491633 | 0.006047564 | 0.007230944 | 0.008662615 |
| 0.870623786 | 0.729665343 | 0.759560264 | 0.003263504 | 0.003726758 | 0.004421434 |
| 0.383066679 | 0.281043279 | 0.336011413 | 0.006047564 | 0.007230944 | 0.008662615 |
| 0.383066679 | 0.985130154 | 0.972305672 | 0.003263504 | 0.003726758 | 0.004421434 |
| 0.357437235 | 0.247396512 | 0.311522706 | 0.006047564 | 0.007230944 | 0.008662615 |
| 0.357437235 | 0.132622471 | 0.128175026 | 0.003263504 | 0.003726758 | 0.004421434 |
| 0.558105440 | 0.517909740 | 0.509068654 | 0.006047564 | 0.007230944 | 0.008662615 |
| 0.558105440 | 0.444464737 | 0.437949522 | 0.003263504 | 0.003726758 | 0.004421434 |
| 0.218800669 | 0.315299541 | 0.359379399 | 0.006047564 | 0.007230944 | 0.008662615 |
| 0.218800669 | 0.685904908 | 0.695683696 | 0.003263504 | 0.003726758 | 0.004421434 |

| | | | | | | |
|---|---|---|---|---|---|---|
| 0.6300231417 | 0.6757461049 | 0.6060413410 | 0.0060475640 | 0.0072309443 | 0.0086626157 | |
| 0.6300231417 | 0.7849227182 | 0.7833131464 | 0.0032635047 | 0.0037267584 | 0.0044214342 | |
| 0.3027034028 | 0.371827624 | 0.4073657897 | 0.0060475640 | 0.0072309443 | 0.0086626157 | |
| 0.3027034028 | 0.0994916565 | 0.1023572004 | 0.0032635047 | 0.0037267584 | 0.0044214342 | |
| 0.0065525897 | 0.0010601253 | 0.0067841975 | 0.0060475640 | 0.0072309443 | 0.0086626157 | 0.15 |
| 0.0065525897 | 0.852097342 | 0.8446624772 | 0.0032635047 | 0.0037267584 | 0.0044214342 | |
| 0.6259376550 | 0.6191552824 | 0.5792986919 | 0.0060475640 | 0.0072309443 | 0.0086626157 | |
| 0.6259376550 | 0.5108660848 | 0.4952752269 | 0.0032635047 | 0.0037267584 | 0.0044214342 | |
| 0.8724564242 | 0.8564098781 | 0.8506980245 | 0.0060475640 | 0.0072309443 | 0.0086626157 | |
| 0.8724564242 | 0.7410894879 | 0.7624489281 | 0.0032635047 | 0.0037267584 | 0.0044214342 | |
| 0.4713568109 | 0.3968457327 | 0.4218677092 | 0.0060475640 | 0.0072309443 | 0.0086626157 | |
| 0.4713568109 | 0.2187586672 | 0.2226736541 | 0.0032635047 | 0.0037267584 | 0.0044214342 | |
| 0.2231142241 | 0.3277185234 | 0.366007418 | 0.0060475640 | 0.0072309443 | 0.0086626157 | |
| 0.2231142241 | 0.9680461713 | 0.9521063260 | 0.0032635047 | 0.0037267584 | 0.0044214342 | |
| 0.6065010953 | 0.5990682890 | 0.559271332 | 0.0060475640 | 0.0072309443 | 0.0086626157 | |
| 0.6065010953 | 0.3507682334 | 0.3686067483 | 0.0032635047 | 0.0037267584 | 0.0044214342 | |
| 0.3078444439 | 0.2664941899 | 0.3240357635 | 0.0060475640 | 0.0072309443 | 0.0086626157 | |
| 0.3078444439 | 0.9375979892 | 0.9293618418 | 0.0032635047 | 0.0037267584 | 0.0044214342 | |
| 0.4430137541 | 0.3694396483 | 0.4042588913 | 0.0060475640 | 0.0072309443 | 0.0086626157 | |
| 0.4430137541 | 0.1917005813 | 0.1969616174 | 0.0032635047 | 0.0037267584 | 0.0044214342 | |
| 0.9667058982 | 0.9754759152 | 0.960392117 | 0.0060475640 | 0.0072309443 | 0.0086626157 | |
| 0.9667058982 | 0.959969294 | 0.9356910209 | 0.0032635047 | 0.0037267584 | 0.0044214342 | |
| 0.8194558021 | 0.8203566418 | 0.7902767999 | 0.0060475640 | 0.0072309443 | 0.0086626157 | |
| 0.8194558021 | 0.6690669523 | 0.6728991249 | 0.0032635047 | 0.0037267584 | 0.0044214342 | |
| 0.9636357700 | 0.9715107117 | 0.9572482385 | 0.0060475640 | 0.0072309443 | 0.0086626157 | |
| 0.9636357700 | 0.9498234001 | 0.9304293520 | 0.0032635047 | 0.0037267584 | 0.0044214342 | |
| 0.4693503245 | 0.3891482763 | 0.4196178157 | 0.0060475640 | 0.0072309443 | 0.0086626157 | |
| 0.4693503245 | 0.211849678 | 0.220857218 | 0.0032635047 | 0.0037267584 | 0.0044214342 | |
| 0.3446844102 | 0.2658384775 | 0.3232416895 | 0.0060475640 | 0.0072309443 | 0.0086626157 | |
| 0.3446844102 | 0.2722504375 | 0.2901773583 | 0.0032635047 | 0.0037267584 | 0.0044214342 | |
| 0.876279863 | 0.8703133441 | 0.8550180049 | 0.0060475640 | 0.0072309443 | 0.0086626157 | |
| 0.876279863 | 0.7544036240 | 0.7690663264 | 0.0032635047 | 0.0037267584 | 0.0044214342 | |
| 0.8909442857 | 0.8923122810 | 0.8717248921 | 0.0060475640 | 0.0072309443 | 0.0086626157 | |
| 0.8909442857 | 0.8018532290 | 0.7947557315 | 0.0032635047 | 0.0037267584 | 0.0044214342 | |
| 0.160285117 | 0.0704506883 | 0.1333873335 | 0.0060475640 | 0.0072309443 | 0.0086626157 | |
| 0.160285117 | 0.4958819977 | 0.4896857171 | 0.0032635047 | 0.0037267584 | 0.0044214342 | |
| 0.3331985170 | 0.2215758297 | 0.2889057035 | 0.0060475640 | 0.0072309443 | 0.0086626157 | |
| 0.3331985170 | 0.1108713278 | 0.1114743271 | 0.0032635047 | 0.0037267584 | 0.0044214342 | |
| 0.7820073724 | 0.8375855307 | 0.8261081571 | 0.0060475640 | 0.0072309443 | 0.0086626157 | |
| 0.7820073724 | 0.6191146825 | 0.6130297744 | 0.0032635047 | 0.0037267584 | 0.0044214342 | |
| 0.4001649830 | 0.3062408411 | 0.3522067761 | 0.0060475640 | 0.0072309443 | 0.0086626157 | |
| 0.4001649830 | 0.1664274221 | 0.1605782465 | 0.0032635047 | 0.0037267584 | 0.0044214342 | |
| 0.8815786412 | 0.8815990712 | 0.860864518 | 0.0060475640 | 0.0072309443 | 0.0086626157 | |
| 0.8815786412 | 0.7750235461 | 0.7783360069 | 0.0032635047 | 0.0037267584 | 0.0044214342 | |
| 0.2817108294 | 0.1611433672 | 0.2414235403 | 0.0060475640 | 0.0072309443 | 0.0086626157 | |
| 0.2817108294 | 0.4744420841 | 0.4538185029 | 0.0032635047 | 0.0037267584 | 0.0044214342 | |

| | | | | | |
|---|---|---|---|---|---|
| 0.638205420 | 0.736310710 | 0.653038811 | 0.006047564 | 0.007230944 | 0.008662615 |
| 0.638205420 | 0.813454254 | 0.799074884 | 0.003263504 | 0.003726758 | 0.004421434 |
| 0.172725915 | 0.080140154 | 0.144232362 | 0.006047564 | 0.007230944 | 0.008662615 |
| 0.172725915 | 0.025102236 | 0.030147624 | 0.003263504 | 0.003726758 | 0.004421434 |
| 0.631162508 | 0.692425859 | 0.615687318 | 0.006047564 | 0.007230944 | 0.008662615 |
| 0.631162508 | 0.715764284 | 0.734545308 | 0.003263504 | 0.003726758 | 0.004421434 |
| 0.539230854 | 0.497766223 | 0.489564607 | 0.006047564 | 0.007230944 | 0.008662615 |
| 0.539230854 | 0.274004546 | 0.291339679 | 0.003263504 | 0.003726758 | 0.004421434 |
| 0.305851325 | 0.227848101 | 0.294364911 | 0.006047564 | 0.007230944 | 0.008662615 |
| 0.305851325 | 0.411202186 | 0.413797981 | 0.003263504 | 0.003726758 | 0.004421434 |
| 0.226028663 | 0.165033891 | 0.245711623 | 0.006047564 | 0.007230944 | 0.008662615 |
| 0.226028663 | 0.530437508 | 0.505025963 | 0.003263504 | 0.003726758 | 0.004421434 |
| 0.192117380 | 0.099712125 | 0.168735535 | 0.006047564 | 0.007230944 | 0.008662615 |
| 0.192117380 | 0.436379645 | 0.437407813 | 0.003263504 | 0.003726758 | 0.004421434 |
| 0.708088353 | 0.748172736 | 0.666749606 | 0.006047564 | 0.007230944 | 0.008662615 |
| 0.708088353 | 0.517510420 | 0.502154202 | 0.003263504 | 0.003726758 | 0.004421434 |
| 0.247142980 | 0.335589004 | 0.375006775 | 0.006047564 | 0.007230944 | 0.008662615 |
| 0.247142980 | 0.846659246 | 0.836395480 | 0.003263504 | 0.003726758 | 0.004421434 |
| 0.303618893 | 0.292163841 | 0.340121255 | 0.006047564 | 0.007230944 | 0.008662615 |
| 0.303618893 | 0.825227961 | 0.822548515 | 0.003263504 | 0.003726758 | 0.004421434 |
| 0.948390268 | 0.943858399 | 0.938800212 | 0.006047564 | 0.007230944 | 0.008662615 |
| 0.948390268 | 0.898221603 | 0.900558066 | 0.003263504 | 0.003726758 | 0.004421434 |
| 0.082255830 | 0.410392841 | 0.361777598 | 0.002545798 | 0.004602761 | 0.006730984 |
| 0.082255830 | 0.583119300 | 0.567689256 | 0.000966233 | 0.001005035 | 0.002118303 |
| 0.590233395 | 0.756273187 | 0.660541403 | 0.002545798 | 0.004602761 | 0.006730984 |
| 0.590233395 | 0.450074825 | 0.440823530 | 0.000966233 | 0.001005035 | 0.002118303 |
| 0.702653450 | 0.657887816 | 0.577716658 | 0.002545798 | 0.004602761 | 0.006730984 |
| 0.702653450 | 0.702582890 | 0.658550654 | 0.000966233 | 0.001005035 | 0.002118303 |
| 0.812714475 | 0.828276052 | 0.722049260 | 0.002545798 | 0.004602761 | 0.006730984 |
| 0.812714475 | 0.709228275 | 0.662026717 | 0.000966233 | 0.001005035 | 0.002118303 |
| 0.523823138 | 0.878332064 | 0.798114392 | 0.002545798 | 0.004602761 | 0.006730984 |
| 0.523823138 | 0.465695838 | 0.458910378 | 0.000966233 | 0.001005035 | 0.002118303 |
| 0.321090190 | 0.210832057 | 0.190567256 | 0.002545798 | 0.004602761 | 0.006730984 |
| 0.321090190 | 0.135039949 | 0.103669826 | 0.000966233 | 0.001005035 | 0.002118303 |
| 0.643640955 | 0.608637892 | 0.527806017 | 0.002545798 | 0.004602761 | 0.006730984 |
| 0.643640955 | 0.423852922 | 0.415513148 | 0.000966233 | 0.001005035 | 0.002118303 |
| 0.028707996 | 0.020517960 | 0.028954790 | 0.002545798 | 0.004602761 | 0.006730984 |
| 0.028707996 | 0.765996090 | 0.748015619 | 0.000966233 | 0.001005035 | 0.002118303 |
| 0.020902884 | 0.100063006 | 0.091994561 | 0.002545798 | 0.004602761 | 0.006730984 |
| 0.020902884 | 0.730982316 | 0.694240072 | 0.000966233 | 0.001005035 | 0.002118303 |
| 0.903697280 | 0.940096621 | 0.852000448 | 0.002545798 | 0.004602761 | 0.006730984 |
| 0.903697280 | 0.863922016 | 0.818189883 | 0.000966233 | 0.001005035 | 0.002118303 |
| 0.126458922 | 0.204935705 | 0.186016609 | 0.002545798 | 0.004602761 | 0.006730984 |
| 0.126458922 | 0.608011491 | 0.584841506 | 0.000966233 | 0.001005035 | 0.002118303 |
| 0.764758571 | 0.752003039 | 0.657260975 | 0.002545798 | 0.004602761 | 0.006730984 |
| 0.764758571 | 0.614494055 | 0.585790420 | 0.000966233 | 0.001005035 | 0.002118303 |

| | | | | | |
|---|---|---|---|---|---|
| 0.8037535788 | 0.8158006064 | 0.7096996797 | 0.0025457983 | 0.0046027613 | 0.0067309843 |
| 0.8037535788 | 0.6964766557 | 0.6476125789 | 0.0009662331 | 0.0010050358 | 0.0021183039 |
| 0.8186205229 | 0.8365096438 | 0.7295463701 | 0.0025457983 | 0.0046027613 | 0.0067309843 |
| 0.8186205229 | 0.7182197514 | 0.6711927580 | 0.0009662331 | 0.0010050358 | 0.0021183039 |
| 0.2006233 | 0.3497778353 | 0.296872107 | 0.0025457983 | 0.0046027613 | 0.0067309843 |
| 0.2006233 | 0.4155511274 | 0.4142829115 | 0.0009662331 | 0.0010050358 | 0.0021183039 |
| 0.7066193127 | 0.8686733027 | 0.7692331457 | 0.0025457983 | 0.0046027613 | 0.0067309843 |
| 0.7066193127 | 0.5171758728 | 0.5007142897 | 0.0009662331 | 0.0010050358 | 0.0021183039 |
| 0.0310405010 | 0.6476823737 | 0.5665685285 | 0.0025457983 | 0.0046027613 | 0.0067309843 |
| 0.0310405010 | 0.4039711177 | 0.4030098497 | 0.0009662331 | 0.0010050358 | 0.0021183039 |
| 0.0747301899 | 0.2399667260 | 0.2091382279 | 0.0025457983 | 0.0046027613 | 0.0067309843 |
| 0.0747301899 | 0.0804679620 | 0.0549396040 | 0.0009662331 | 0.0010050358 | 0.0021183039 |
| 0.9920945847 | 0.99437673 | 0.9878168433 | 0.0025457983 | 0.0046027613 | 0.0067309843 |
| 0.9920945847 | 0.9928419074 | 0.9855915580 | 0.0009662331 | 0.0010050358 | 0.0021183039 |
| 0.4938757309 | 0.3952703107 | 0.344272794 | 0.0025457983 | 0.0046027613 | 0.0067309843 |
| 0.4938757309 | 0.5560980497 | 0.5267569747 | 0.0009662331 | 0.0010050358 | 0.0021183039 |
| 0.6612819635 | 0.9354861550 | 0.8502024607 | 0.0025457983 | 0.0046027613 | 0.0067309843 |
| 0.6612819635 | 0.441980397 | 0.4381970547 | 0.0009662331 | 0.0010050358 | 0.0021183039 |
| 0.9015736230 | 0.9342457091 | 0.8491108309 | 0.0025457983 | 0.0046027613 | 0.0067309843 |
| 0.9015736230 | 0.8574249267 | 0.8144557267 | 0.0009662331 | 0.0010050358 | 0.0021183039 |
| 0.1785752658 | 0.0972102870 | 0.0905659135 | 0.0025457983 | 0.0046027613 | 0.0067309843 |
| 0.1785752658 | 0.1173449494 | 0.0883745557 | 0.0009662331 | 0.0010050358 | 0.0021183039 |
| 0.0310105124 | 0.0055514681 | 0.0129927639 | 0.0025457983 | 0.0046027613 | 0.0067309843 |
| 0.0310105124 | 0.0573888187 | 0.038656969 | 0.0009662331 | 0.0010050358 | 0.0021183039 |
| 0.1270910547 | 0.1502729235 | 0.1344395431 | 0.0025457983 | 0.0046027613 | 0.0067309843 |
| 0.1270910547 | 0.9284125473 | 0.9110427301 | 0.0009662331 | 0.0010050358 | 0.0021183039 |
| 0.8858584489 | 0.905019366 | 0.8257789864 | 0.0025457983 | 0.0046027613 | 0.0067309843 |
| 0.8858584489 | 0.81725553 | 0.7860524661 | 0.0009662331 | 0.0010050358 | 0.0021183039 |
| 0.4820055547 | 0.3815927759 | 0.3323168335 | 0.0025457983 | 0.0046027613 | 0.0067309843 |
| 0.4820055547 | 0.6809589498 | 0.639979555 | 0.0009662331 | 0.0010050358 | 0.0021183039 |
| 0.6459403269 | 0.8513460428 | 0.7420392479 | 0.0025457983 | 0.0046027613 | 0.0067309843 |
| 0.6459403269 | 0.4269258145 | 0.4182132544 | 0.0009662331 | 0.0010050358 | 0.0021183039 |
| 0.7813572425 | 0.7780354467 | 0.6788815047 | 0.0025457983 | 0.0046027613 | 0.0067309843 |
| 0.7813572425 | 0.6528594585 | 0.6115388163 | 0.0009662331 | 0.0010050358 | 0.0021183039 |
| 0.5332259587 | 0.4561957355 | 0.3843478248 | 0.0025457983 | 0.0046027613 | 0.0067309843 |
| 0.5332259587 | 0.3068085887 | 0.2852357680 | 0.0009662331 | 0.0010050358 | 0.0021183039 |
| 0.6901424240 | 0.6390535888 | 0.5624629797 | 0.0025457983 | 0.0046027613 | 0.0067309843 |
| 0.6901424240 | 0.4774609837 | 0.4776166915 | 0.0009662331 | 0.0010050358 | 0.0021183039 |
| 0.6055076337 | 0.5490404440 | 0.4625984605 | 0.0025457983 | 0.0046027613 | 0.0067309843 |
| 0.6055076337 | 0.8273217085 | 0.7885315763 | 0.0009662331 | 0.0010050358 | 0.0021183039 |
| 0.4122800413 | 0.3206839277 | 0.2669794127 | 0.0025457983 | 0.0046027613 | 0.0067309843 |
| 0.4122800413 | 0.2205685084 | 0.1705972997 | 0.0009662331 | 0.0010050358 | 0.0021183039 |
| 0.0561926728 | 0.0331710359 | 0.0389106968 | 0.0025457983 | 0.0046027613 | 0.0067309843 |
| 0.0561926728 | 0.6197972715 | 0.5894732727 | 0.0009662331 | 0.0010050358 | 0.0021183039 |
| 0.5780508957 | 0.5005497443 | 0.4319194147 | 0.0025457983 | 0.0046027613 | 0.0067309843 |
| 0.5780508957 | 0.3369444369 | 0.3348683678 | 0.0009662331 | 0.0010050358 | 0.0021183039 |

| | | | | | |
|---|---|---|---|---|---|
| 0.599207971 | 0.876420784 | 0.792172250 | 0.002545798 | 0.004602761 | 0.006730984 |
| 0.599207971 | 0.815578949 | 0.785947501 | 0.000966233 | 0.001005035 | 0.002118303 |
| 0.122940899 | 0.242285066 | 0.211506211 | 0.002545798 | 0.004602761 | 0.006730984 |
| 0.122940899 | 0.752794924 | 0.732101899 | 0.000966233 | 0.001005035 | 0.002118303 |
| 0.941144780 | 0.956470525 | 0.908199476 | 0.002545798 | 0.004602761 | 0.006730984 |
| 0.941144780 | 0.906735188 | 0.88698176 | 0.000966233 | 0.001005035 | 0.002118303 |
| 0.274268236 | 0.170274593 | 0.155042936 | 0.002545798 | 0.004602761 | 0.006730984 |
| 0.274268236 | 0.103536185 | 0.075644182 | 0.000966233 | 0.001005035 | 0.002118303 |
| 0.072911420 | 0.024468975 | 0.032380940 | 0.002545798 | 0.004602761 | 0.006730984 |
| 0.072911420 | 0.920284543 | 0.901721086 | 0.000966233 | 0.001005035 | 0.002118303 |
| 0.142154329 | 0.630621371 | 0.556897850 | 0.002545798 | 0.004602761 | 0.006730984 |
| 0.142154329 | 0.245856178 | 0.217997917 | 0.000966233 | 0.001005035 | 0.002118303 |
| 0.868844344 | 0.882224635 | 0.800852478 | 0.002545798 | 0.004602761 | 0.006730984 |
| 0.868844344 | 0.778702162 | 0.756498048 | 0.000966233 | 0.001005035 | 0.002118303 |
| 0.372565146 | 0.614675021 | 0.538354988 | 0.002545798 | 0.004602761 | 0.006730984 |
| 0.372565146 | 0.188973160 | 0.139395806 | 0.000966233 | 0.001005035 | 0.002118303 |
| 0.067994803 | 0.021639117 | 0.029992623 | 0.002545798 | 0.004602761 | 0.006730984 |
| 0.067994803 | 0.094436156 | 0.065070815 | 0.000966233 | 0.001005035 | 0.002118303 |
| 0.232874906 | 0.177317614 | 0.162269486 | 0.002545798 | 0.004602761 | 0.006730984 |
| 0.232874906 | 0.602683443 | 0.580487195 | 0.000966233 | 0.001005035 | 0.002118303 |
| 0.518286848 | 0.427883067 | 0.368651516 | 0.002545798 | 0.004602761 | 0.006730984 |
| 0.518286848 | 0.288468010 | 0.269388459 | 0.000966233 | 0.001005035 | 0.002118303 |
| 0.000672482 | 0.142830455 | 0.126529582 | 0.002545798 | 0.004602761 | 0.006730984 |
| 0.000672482 | 0.161940729 | 0.117779316 | 0.000966233 | 0.001005035 | 0.002118303 |
| 0.740262242 | 0.725123773 | 0.625627019 | 0.002545798 | 0.004602761 | 0.006730984 |
| 0.740262242 | 0.580695962 | 0.54929786 | 0.000966233 | 0.001005035 | 0.002118303 |
| 0.063557629 | 0.081233121 | 0.079253684 | 0.002545798 | 0.004602761 | 0.006730984 |
| 0.063557629 | 0.015563370 | 0.011556548 | 0.000966233 | 0.001005035 | 0.002118303 |
| 0.831397497 | 0.868780618 | 0.769518681 | 0.002545798 | 0.004602761 | 0.006730984 |
| 0.831397497 | 0.728748578 | 0.692631504 | 0.000966233 | 0.001005035 | 0.002118303 |
| 0.537535618 | 0.465363910 | 0.388586721 | 0.002545798 | 0.004602761 | 0.006730984 |
| 0.537535618 | 0.363606393 | 0.355594464 | 0.000966233 | 0.001005035 | 0.002118303 |
| 0.148495475 | 0.497912961 | 0.430357049 | 0.002545798 | 0.004602761 | 0.006730984 |
| 0.148495475 | 0.951795549 | 0.931680112 | 0.000966233 | 0.001005035 | 0.002118303 |
| 0.360357043 | 0.260283098 | 0.222405068 | 0.002545798 | 0.004602761 | 0.006730984 |
| 0.360357043 | 0.198906047 | 0.146266898 | 0.000966233 | 0.001005035 | 0.002118303 |
| 0.121457029 | 0.312231744 | 0.261505053 | 0.002545798 | 0.004602761 | 0.006730984 |
| 0.121457029 | 0.020680629 | 0.014933791 | 0.000966233 | 0.001005035 | 0.002118303 |
| 0.798171907 | 0.806503549 | 0.702167027 | 0.002545798 | 0.004602761 | 0.006730984 |
| 0.798171907 | 0.676018031 | 0.638544185 | 0.000966233 | 0.001005035 | 0.002118303 |
| 0.382675174 | 0.287095935 | 0.241099837 | 0.002545798 | 0.004602761 | 0.006730984 |
| 0.382675174 | 0.200670512 | 0.147005663 | 0.000966233 | 0.001005035 | 0.002118303 |
| 0.163698575 | 0.136213138 | 0.121472884 | 0.002545798 | 0.004602761 | 0.006730984 |
| 0.163698575 | 0.043213785 | 0.027043815 | 0.000966233 | 0.001005035 | 0.002118303 |
| 0.516352843 | 0.418032952 | 0.366686198 | 0.002545798 | 0.004602761 | 0.006730984 |
| 0.516352843 | 0.277155513 | 0.267192678 | 0.000966233 | 0.001005035 | 0.002118303 |

| | | | | | |
|---|---|---|---|---|---|
| 0.710291384 | 0.773568805 | 0.670294522 | 0.002545798 | 0.004602761 | 0.006730984 |
| 0.710291384 | 0.750609690 | 0.730561250 | 0.000966233 | 0.001005035 | 0.002118303 |
| 0.221715373 | 0.555051026 | 0.470112454 | 0.002545798 | 0.004602761 | 0.006730984 |
| 0.221715373 | 0.106790276 | 0.078150865 | 0.000966233 | 0.001005035 | 0.002118303 |
| 0.388613361 | 0.295792734 | 0.246364331 | 0.002545798 | 0.004602761 | 0.006730984 |
| 0.388613361 | 0.255612825 | 0.222479313 | 0.000966233 | 0.001005035 | 0.002118303 |
| 0.727968438 | 0.698999691 | 0.609343924 | 0.002545798 | 0.004602761 | 0.006730984 |
| 0.727968438 | 0.561412668 | 0.531084259 | 0.000966233 | 0.001005035 | 0.002118303 |
| 0.335618056 | 0.226874397 | 0.201946783 | 0.002545798 | 0.004602761 | 0.006730984 |
| 0.335618056 | 0.148468403 | 0.113086471 | 0.000966233 | 0.001005035 | 0.002118303 |
| 0.787993517 | 0.791966355 | 0.688185520 | 0.002545798 | 0.004602761 | 0.006730984 |
| 0.787993517 | 0.660928357 | 0.622144365 | 0.000966233 | 0.001005035 | 0.002118303 |
| 0.890101361 | 0.912641297 | 0.831852412 | 0.002545798 | 0.004602761 | 0.006730984 |
| 0.890101361 | 0.880861297 | 0.827985841 | 0.000966233 | 0.001005035 | 0.002118303 |
| 0.881691204 | 0.893782208 | 0.819386837 | 0.002545798 | 0.004602761 | 0.006730984 |
| 0.881691204 | 0.800197764 | 0.779031934 | 0.000966233 | 0.001005035 | 0.002118303 |
| 0.628499596 | 0.565594257 | 0.488437411 | 0.002545798 | 0.004602761 | 0.006730984 |
| 0.628499596 | 0.396323044 | 0.395976701 | 0.000966233 | 0.001005035 | 0.002118303 |
| 0.367490669 | 0.757894544 | 0.661687587 | 0.002545798 | 0.004602761 | 0.006730984 |
| 0.367490669 | 0.183463218 | 0.135575673 | 0.000966233 | 0.001005035 | 0.002118303 |
| 0.413165605 | 0.683286249 | 0.596560017 | 0.002545798 | 0.004602761 | 0.006730984 |
| 0.413165605 | 0.899560969 | 0.848168845 | 0.000966233 | 0.001005035 | 0.002118303 |
| 0.754862438 | 0.739697606 | 0.644351114 | 0.002545798 | 0.004602761 | 0.006730984 |
| 0.754862438 | 0.689613879 | 0.642331510 | 0.000966233 | 0.001005035 | 0.002118303 |
| 0.102836125 | 0.051340148 | 0.057303737 | 0.002545798 | 0.004602761 | 0.006730984 |
| 0.102836125 | 0.037378366 | 0.023792875 | 0.000966233 | 0.001005035 | 0.002118303 |
| 0.019021605 | 0.013169144 | 0.022314963 | 0.002545798 | 0.004602761 | 0.006730984 |
| 0.019021605 | 0.038282182 | 0.024497714 | 0.000966233 | 0.001005035 | 0.002118303 |
| 0.558051330 | 0.483068575 | 0.410252573 | 0.002545798 | 0.004602761 | 0.006730984 |
| 0.558051330 | 0.324454514 | 0.312294385 | 0.000966233 | 0.001005035 | 0.002118303 |
| 0.679717763 | 0.622540686 | 0.549325741 | 0.002545798 | 0.004602761 | 0.006730984 |
| 0.679717763 | 0.469182482 | 0.462975937 | 0.000966233 | 0.001005035 | 0.002118303 |
| 0.730878565 | 0.705565080 | 0.612863944 | 0.002545798 | 0.004602761 | 0.006730984 |
| 0.730878565 | 0.564910637 | 0.535027822 | 0.000966233 | 0.001005035 | 0.002118303 |
| 0.516997774 | 0.425856249 | 0.367670527 | 0.002545798 | 0.004602761 | 0.006730984 |
| 0.516997774 | 0.349219215 | 0.342119458 | 0.000966233 | 0.001005035 | 0.002118303 |
| 0.969080410 | 0.981013007 | 0.952018545 | 0.002545798 | 0.004602761 | 0.006730984 |
| 0.969080410 | 0.961186635 | 0.940431521 | 0.000966233 | 0.001005035 | 0.002118303 |
| 0.516081854 | 0.414051123 | 0.366382111 | 0.002545798 | 0.004602761 | 0.006730984 |
| 0.516081854 | 0.275100155 | 0.266992131 | 0.000966233 | 0.001005035 | 0.002118303 |
| 0.341391828 | 0.670746471 | 0.585882919 | 0.002545798 | 0.004602761 | 0.006730984 |
| 0.341391828 | 0.161820050 | 0.117229876 | 0.000966233 | 0.001005035 | 0.002118303 |
| 0.045487606 | 0.697290576 | 0.609095154 | 0.002545798 | 0.004602761 | 0.006730984 |
| 0.045487606 | 0.595072786 | 0.578677458 | 0.000966233 | 0.001005035 | 0.002118303 |
| 0.723224089 | 0.688027298 | 0.603176748 | 0.002545798 | 0.004602761 | 0.006730984 |
| 0.723224089 | 0.549935076 | 0.524230911 | 0.000966233 | 0.001005035 | 0.002118303 |

| | | | | | |
|---|---|---|---|---|---|
| 0.533729676 | 0.460026964 | 0.384784228 | 0.002545798 | 0.004602761 | 0.006730984 |
| 0.533729676 | 0.916317919 | 0.893978438 | 0.000966233 | 0.001005035 | 0.002118303 |
| 0.958167408 | 0.972548591 | 0.935071110 | 0.002545798 | 0.004602761 | 0.006730984 |
| 0.958167408 | 0.942205682 | 0.919922593 | 0.000966233 | 0.001005035 | 0.002118303 |
| 0.529375213 | 0.448882669 | 0.380272814 | 0.002545798 | 0.004602761 | 0.006730984 |
| 0.529375213 | 0.301039178 | 0.280967617 | 0.000966233 | 0.001005035 | 0.002118303 |
| 0.047003166 | 0.114497757 | 0.105578219 | 0.002545798 | 0.004602761 | 0.006730984 |
| 0.047003166 | 0.119208807 | 0.091764222 | 0.000966233 | 0.001005035 | 0.002118303 |
| 0.402679719 | 0.309939986 | 0.258759965 | 0.002545798 | 0.004602761 | 0.006730984 |
| 0.402679719 | 0.640442289 | 0.595674327 | 0.000966233 | 0.001005035 | 0.002118303 |
| 0.329989722 | 0.263763367 | 0.226219285 | 0.002545798 | 0.004602761 | 0.006730984 |
| 0.329989722 | 0.508544976 | 0.496457952 | 0.000966233 | 0.001005035 | 0.002118303 |
| 0.176235745 | 0.105136671 | 0.097178026 | 0.002545798 | 0.004602761 | 0.006730984 |
| 0.176235745 | 0.983511779 | 0.972098841 | 0.000966233 | 0.001005035 | 0.002118303 |
| 0.523337585 | 0.433784487 | 0.373685013 | 0.002545798 | 0.004602761 | 0.006730984 |
| 0.523337585 | 0.291250129 | 0.274461777 | 0.000966233 | 0.001005035 | 0.002118303 |
| 0.171126848 | 0.131569925 | 0.118493130 | 0.002545798 | 0.004602761 | 0.006730984 |
| 0.171126848 | 0.433621990 | 0.437562543 | 0.000966233 | 0.001005035 | 0.002118303 |
| 0.310507020 | 0.776057238 | 0.675304296 | 0.002545798 | 0.004602761 | 0.006730984 |
| 0.310507020 | 0.740044626 | 0.695775402 | 0.000966233 | 0.001005035 | 0.002118303 |
| 0.480086230 | 0.574965416 | 0.496679522 | 0.002545798 | 0.004602761 | 0.006730984 |
| 0.480086230 | 0.804978230 | 0.783163209 | 0.000966233 | 0.001005035 | 0.002118303 |
| 0.768350783 | 0.884592421 | 0.801136801 | 0.002545798 | 0.004602761 | 0.006730984 |
| 0.768350783 | 0.628094553 | 0.591450237 | 0.000966233 | 0.001005035 | 0.002118303 |
| 0.170788884 | 0.089145536 | 0.085902205 | 0.002545798 | 0.004602761 | 0.006730984 |
| 0.170788884 | 0.890176773 | 0.844826713 | 0.000966233 | 0.001005035 | 0.002118303 |
| 0.076968196 | 0.046297243 | 0.052403802 | 0.002545798 | 0.004602761 | 0.006730984 |
| 0.076968196 | 0.497688628 | 0.494726257 | 0.000966233 | 0.001005035 | 0.002118303 |
| 0.901350125 | 0.925946848 | 0.848496122 | 0.002545798 | 0.004602761 | 0.006730984 |
| 0.901350125 | 0.847618900 | 0.813729658 | 0.000966233 | 0.001005035 | 0.002118303 |
| 0.203560589 | 0.167677014 | 0.153938123 | 0.002545798 | 0.004602761 | 0.006730984 |
| 0.203560589 | 0.062884208 | 0.041665528 | 0.000966233 | 0.001005035 | 0.002118303 |
| 0.225306537 | 0.745084194 | 0.650739150 | 0.002545798 | 0.004602761 | 0.006730984 |
| 0.225306537 | 0.975899063 | 0.952108518 | 0.000966233 | 0.001005035 | 0.002118303 |
| 0.027304783 | 0.177496030 | 0.162469471 | 0.002545798 | 0.004602761 | 0.006730984 |
| 0.027304783 | 0.039520083 | 0.024729024 | 0.000966233 | 0.001005035 | 0.002118303 |
| 0.974159044 | 0.986825039 | 0.959676167 | 0.002545798 | 0.004602761 | 0.006730984 |
| 0.974159044 | 0.973996521 | 0.950923376 | 0.000966233 | 0.001005035 | 0.002118303 |
| 0.591535064 | 0.521352164 | 0.447059009 | 0.002545798 | 0.004602761 | 0.006730984 |
| 0.591535064 | 0.356589305 | 0.350762684 | 0.000966233 | 0.001005035 | 0.002118303 |
| 0.40771711 | 0.314398521 | 0.262950128 | 0.002545798 | 0.004602761 | 0.006730984 |
| 0.40771711 | 0.216700985 | 0.166844980 | 0.000966233 | 0.001005035 | 0.002118303 |
| 0.294528962 | 0.254823003 | 0.219375586 | 0.002545798 | 0.004602761 | 0.006730984 |
| 0.294528962 | 0.333497650 | 0.327887337 | 0.000966233 | 0.001005035 | 0.002118303 |
| 0.691611478 | 0.645648965 | 0.564261295 | 0.002545798 | 0.004602761 | 0.006730984 |
| 0.691611478 | 0.785350125 | 0.761630767 | 0.000966233 | 0.001005035 | 0.002118303 |

| | | | | | |
|---|---|---|---|---|---|
| 0.467429425 | 0.364632296 | 0.318145542 | 0.002545798 | 0.004602761 | 0.006730984 |
| 0.467429425 | 0.249303779 | 0.219055838 | 0.000966233 | 0.001005035 | 0.002118303 |
| 0.600797878 | 0.541705597 | 0.457378821 | 0.002545798 | 0.004602761 | 0.006730984 |
| 0.600797878 | 0.382840429 | 0.362096749 | 0.000966233 | 0.001005035 | 0.002118303 |
| 0.327702954 | 0.282869718 | 0.237763222 | 0.002545798 | 0.004602761 | 0.006730984 |
| 0.327702954 | 0.141452444 | 0.108040577 | 0.000966233 | 0.001005035 | 0.002118303 |
| 0.503610691 | 0.403634252 | 0.353831633 | 0.002545798 | 0.004602761 | 0.006730984 |
| 0.503610691 | 0.270318079 | 0.254210406 | 0.000966233 | 0.001005035 | 0.002118303 |
| 0.151475276 | 0.075557809 | 0.074377942 | 0.002545798 | 0.004602761 | 0.006730984 |
| 0.151475276 | 0.490266350 | 0.489676907 | 0.000966233 | 0.001005035 | 0.002118303 |
| 0.219377421 | 0.127630102 | 0.116769479 | 0.002545798 | 0.004602761 | 0.006730984 |
| 0.219377421 | 0.072079324 | 0.048465706 | 0.000966233 | 0.001005035 | 0.002118303 |
| 0.597728507 | 0.918149295 | 0.838600664 | 0.002545798 | 0.004602761 | 0.006730984 |
| 0.597728507 | 0.539229821 | 0.508590901 | 0.000966233 | 0.001005035 | 0.002118303 |
| 0.133860407 | 0.062169615 | 0.064343714 | 0.002545798 | 0.004602761 | 0.006730984 |
| 0.133860407 | 0.025243475 | 0.018098536 | 0.000966233 | 0.001005035 | 0.002118303 |
| 0.307019588 | 0.195898813 | 0.179529876 | 0.002545798 | 0.004602761 | 0.006730984 |
| 0.307019588 | 0.123909842 | 0.094841574 | 0.000966233 | 0.001005035 | 0.002118303 |
| 0.818307529 | 0.834162466 | 0.729282018 | 0.002545798 | 0.004602761 | 0.006730984 |
| 0.818307529 | 0.717027433 | 0.671118472 | 0.000966233 | 0.001005035 | 0.002118303 |
| 0.020232682 | 0.5046381 | 0.433833448 | 0.002545798 | 0.004602761 | 0.006730984 |
| 0.020232682 | 0.837103274 | 0.798456911 | 0.000966233 | 0.001005035 | 0.002118303 |
| 0.348976472 | 0.244022015 | 0.212746118 | 0.002545798 | 0.004602761 | 0.006730984 |
| 0.348976472 | 0.172231518 | 0.122274211 | 0.000966233 | 0.001005035 | 0.002118303 |
| 0.773636392 | 0.772816278 | 0.669304189 | 0.002545798 | 0.004602761 | 0.006730984 |
| 0.773636392 | 0.764885555 | 0.734687782 | 0.000966233 | 0.001005035 | 0.002118303 |
| 0.337922107 | 0.230327158 | 0.203709594 | 0.002545798 | 0.004602761 | 0.006730984 |
| 0.337922107 | 0.153441102 | 0.114559807 | 0.000966233 | 0.001005035 | 0.002118303 |
| 0.112463930 | 0.213256600 | 0.193045279 | 0.002545798 | 0.004602761 | 0.006730984 |
| 0.112463930 | 0.409705204 | 0.413604593 | 0.000966233 | 0.001005035 | 0.002118303 |
| 0.151210469 | 0.302899703 | 0.254016054 | 0.002545798 | 0.004602761 | 0.006730984 |
| 0.151210469 | 0.524205534 | 0.504951145 | 0.000966233 | 0.001005035 | 0.002118303 |
| 0.092946464 | 0.084768322 | 0.081759972 | 0.002545798 | 0.004602761 | 0.006730984 |
| 0.092946464 | 0.430222668 | 0.437308642 | 0.000966233 | 0.001005035 | 0.002118303 |
| 0.061458660 | 0.017545607 | 0.026850864 | 0.002545798 | 0.004602761 | 0.006730984 |
| 0.061458660 | 0.006010513 | 0.003895622 | 0.000966233 | 0.001005035 | 0.002118303 |
| 0.353826980 | 0.774191210 | 0.671529915 | 0.002545798 | 0.004602761 | 0.006730984 |
| 0.353826980 | 0.884187340 | 0.836502164 | 0.000966233 | 0.001005035 | 0.002118303 |
| 0.057771899 | 0.360982993 | 0.313397711 | 0.002545798 | 0.004602761 | 0.006730984 |
| 0.057771899 | 0.866789987 | 0.822302668 | 0.000966233 | 0.001005035 | 0.002118303 |
| 0.375585148 | 0.279082892 | 0.235071758 | 0.002545798 | 0.004602761 | 0.006730984 |
| 0.375585148 | 0.193502502 | 0.141605363 | 0.000966233 | 0.001005035 | 0.002118303 |
| 0.056218094 | 0.615549501 | 0.486919703 | 0.004666501 | 0.006313121 | 0.007603344 |
| 0.056218094 | 0.699404478 | 0.678536143 | 0.006886504 | 0.008698764 | 0.010396563 |
| 0.103770533 | 0.735031024 | 0.601744647 | 0.004666501 | 0.006313121 | 0.007603344 |
| 0.103770533 | 0.592277619 | 0.573732475 | 0.006886504 | 0.008698764 | 0.010396563 |

| | | | | | |
|---|---|---|---|---|---|
| 0.569218978 | 0.519424606 | 0.412303658 | 0.004666501 | 0.006313121 | 0.007603344 |
| 0.569218978 | 0.787165015 | 0.749731257 | 0.006886504 | 0.008698764 | 0.010396563 |
| 0.987109513 | 0.977220493 | 0.883297436 | 0.004666501 | 0.006313121 | 0.007603344 |
| 0.987109513 | 0.840686716 | 0.816252145 | 0.006886504 | 0.008698764 | 0.010396563 |
| 0.84462027 | 0.994780391 | 0.950081455 | 0.004666501 | 0.006313121 | 0.007603344 |
| 0.84462027 | 0.720980274 | 0.692123595 | 0.006886504 | 0.008698764 | 0.010396563 |
| 0.378835192 | 0.264789918 | 0.238488049 | 0.004666501 | 0.006313121 | 0.007603344 |
| 0.378835192 | 0.207422844 | 0.207680671 | 0.006886504 | 0.008698764 | 0.010396563 |
| 0.062779579 | 0.533390824 | 0.422947694 | 0.004666501 | 0.006313121 | 0.007603344 |
| 0.062779579 | 0.523310793 | 0.499563707 | 0.006886504 | 0.008698764 | 0.010396563 |
| 0.051570075 | 0.059652013 | 0.076585277 | 0.004666501 | 0.006313121 | 0.007603344 |
| 0.051570075 | 0.841383308 | 0.817518095 | 0.006886504 | 0.008698764 | 0.010396563 |
| 0.000427224 | 0.177772252 | 0.175690677 | 0.004666501 | 0.006313121 | 0.007603344 |
| 0.000427224 | 0.797131623 | 0.776974000 | 0.006886504 | 0.008698764 | 0.010396563 |
| 0.513197594 | 0.508289948 | 0.403762756 | 0.004666501 | 0.006313121 | 0.007603344 |
| 0.513197594 | 0.542278056 | 0.523438030 | 0.006886504 | 0.008698764 | 0.010396563 |
| 0.364850832 | 0.404345635 | 0.333313355 | 0.004666501 | 0.006313121 | 0.007603344 |
| 0.364850832 | 0.723641072 | 0.692445276 | 0.006886504 | 0.008698764 | 0.010396563 |
| 0.157479050 | 0.299035321 | 0.261779437 | 0.004666501 | 0.006313121 | 0.007603344 |
| 0.157479050 | 0.085848672 | 0.113709695 | 0.006886504 | 0.008698764 | 0.010396563 |
| 0.660936839 | 0.667348131 | 0.535477603 | 0.004666501 | 0.006313121 | 0.007603344 |
| 0.660936839 | 0.760177 | 0.723081541 | 0.006886504 | 0.008698764 | 0.010396563 |
| 0.783881503 | 0.601135297 | 0.475115236 | 0.004666501 | 0.006313121 | 0.007603344 |
| 0.783881503 | 0.428135658 | 0.387318510 | 0.006886504 | 0.008698764 | 0.010396563 |
| 0.078797338 | 0.450586740 | 0.364538288 | 0.004666501 | 0.006313121 | 0.007603344 |
| 0.078797338 | 0.568866988 | 0.551249988 | 0.006886504 | 0.008698764 | 0.010396563 |
| 0.725655829 | 0.943427237 | 0.816429128 | 0.004666501 | 0.006313121 | 0.007603344 |
| 0.725655829 | 0.662080193 | 0.629446337 | 0.006886504 | 0.008698764 | 0.010396563 |
| 0.204701938 | 0.898350861 | 0.761302872 | 0.004666501 | 0.006313121 | 0.007603344 |
| 0.204701938 | 0.556724247 | 0.541726292 | 0.006886504 | 0.008698764 | 0.010396563 |
| 0.867764637 | 0.854520481 | 0.718373141 | 0.004666501 | 0.006313121 | 0.007603344 |
| 0.867764637 | 0.469634228 | 0.442313783 | 0.006886504 | 0.008698764 | 0.010396563 |
| 0.666732561 | 0.622481041 | 0.492903796 | 0.004666501 | 0.006313121 | 0.007603344 |
| 0.666732561 | 0.428018430 | 0.387159095 | 0.006886504 | 0.008698764 | 0.010396563 |
| 0.118035937 | 0.217408821 | 0.205231797 | 0.004666501 | 0.006313121 | 0.007603344 |
| 0.118035937 | 0.679154842 | 0.645297363 | 0.006886504 | 0.008698764 | 0.010396563 |
| 0.316699345 | 0.945905777 | 0.820430698 | 0.004666501 | 0.006313121 | 0.007603344 |
| 0.316699345 | 0.176530270 | 0.185598315 | 0.006886504 | 0.008698764 | 0.010396563 |
| 0.452092144 | 0.757214760 | 0.622755133 | 0.004666501 | 0.006313121 | 0.007603344 |
| 0.452092144 | 0.674156144 | 0.638817915 | 0.006886504 | 0.008698764 | 0.010396563 |
| 0.826505148 | 0.644668699 | 0.513291772 | 0.004666501 | 0.006313121 | 0.007603344 |
| 0.826505148 | 0.475300316 | 0.451957910 | 0.006886504 | 0.008698764 | 0.010396563 |
| 0.375182317 | 0.157126093 | 0.158264520 | 0.004666501 | 0.006313121 | 0.007603344 |
| 0.375182317 | 0.158183779 | 0.174301199 | 0.006886504 | 0.008698764 | 0.010396563 |
| 0.101089246 | 0.209961091 | 0.199806112 | 0.004666501 | 0.006313121 | 0.007603344 |
| 0.101089246 | 0.959211786 | 0.936437259 | 0.006886504 | 0.008698764 | 0.010396563 |

| | | | | | |
|---|---|---|---|---|---|
| 0.952631697 | 0.947247687 | 0.822632472 | 0.004666501 | 0.006313121 | 0.007603344 |
| 0.952631697 | 0.636105448 | 0.616805166 | 0.006886504 | 0.008698764 | 0.010396563 |
| 0.595870244 | 0.416070537 | 0.341436700 | 0.004666501 | 0.006313121 | 0.007603344 |
| 0.595870244 | 0.769963385 | 0.735586853 | 0.006886504 | 0.008698764 | 0.010396563 |
| 0.742792142 | 0.944087284 | 0.817491003 | 0.004666501 | 0.006313121 | 0.007603344 |
| 0.742792142 | 0.45046194 | 0.418483724 | 0.006886504 | 0.008698764 | 0.010396563 |
| 0.722491981 | 0.531681755 | 0.421652631 | 0.004666501 | 0.006313121 | 0.007603344 |
| 0.722491981 | 0.389994693 | 0.352511525 | 0.006886504 | 0.008698764 | 0.010396563 |
| 0.257622030 | 0.351627490 | 0.296795187 | 0.004666501 | 0.006313121 | 0.007603344 |
| 0.257622030 | 0.308931024 | 0.279368021 | 0.006886504 | 0.008698764 | 0.010396563 |
| 0.640830014 | 0.662735142 | 0.531125247 | 0.004666501 | 0.006313121 | 0.007603344 |
| 0.640830014 | 0.560587158 | 0.542895748 | 0.006886504 | 0.008698764 | 0.010396563 |
| 0.305065653 | 0.318702028 | 0.274701342 | 0.004666501 | 0.006313121 | 0.007603344 |
| 0.305065653 | 0.883794780 | 0.847411915 | 0.006886504 | 0.008698764 | 0.010396563 |
| 0.081565957 | 0.028455517 | 0.047455452 | 0.004666501 | 0.006313121 | 0.007603344 |
| 0.081565957 | 0.118704149 | 0.143865347 | 0.006886504 | 0.008698764 | 0.010396563 |
| 0.035002862 | 0.052063362 | 0.070273398 | 0.004666501 | 0.006313121 | 0.007603344 |
| 0.035002862 | 0.728077647 | 0.695836057 | 0.006886504 | 0.008698764 | 0.010396563 |
| 0.682644793 | 0.559106720 | 0.442202797 | 0.004666501 | 0.006313121 | 0.007603344 |
| 0.682644793 | 0.303341849 | 0.275571868 | 0.006886504 | 0.008698764 | 0.010396563 |
| 0.201520669 | 0.895819084 | 0.758832302 | 0.004666501 | 0.006313121 | 0.007603344 |
| 0.201520669 | 0.875548347 | 0.845405916 | 0.006886504 | 0.008698764 | 0.010396563 |
| 0.574437007 | 0.563404292 | 0.445448096 | 0.004666501 | 0.006313121 | 0.007603344 |
| 0.574437007 | 0.829291882 | 0.806058698 | 0.006886504 | 0.008698764 | 0.010396563 |
| 0.093242836 | 0.145347703 | 0.147761459 | 0.004666501 | 0.006313121 | 0.007603344 |
| 0.093242836 | 0.203530914 | 0.205337939 | 0.006886504 | 0.008698764 | 0.010396563 |
| 0.045400736 | 0.077176003 | 0.090169965 | 0.004666501 | 0.006313121 | 0.007603344 |
| 0.045400736 | 0.061989491 | 0.096780274 | 0.006886504 | 0.008698764 | 0.010396563 |
| 0.800413907 | 0.624670182 | 0.494822075 | 0.004666501 | 0.006313121 | 0.007603344 |
| 0.800413907 | 0.956777773 | 0.930511081 | 0.006886504 | 0.008698764 | 0.010396563 |
| 0.099506579 | 0.814350166 | 0.680164633 | 0.004666501 | 0.006313121 | 0.007603344 |
| 0.099506579 | 0.405607194 | 0.366805601 | 0.006886504 | 0.008698764 | 0.010396563 |
| 0.622106997 | 0.558130265 | 0.441466873 | 0.004666501 | 0.006313121 | 0.007603344 |
| 0.622106997 | 0.693717092 | 0.662124560 | 0.006886504 | 0.008698764 | 0.010396563 |
| 0.065031367 | 0.687768006 | 0.554885788 | 0.004666501 | 0.006313121 | 0.007603344 |
| 0.065031367 | 0.125957437 | 0.150225705 | 0.006886504 | 0.008698764 | 0.010396563 |
| 0.520647395 | 0.290194464 | 0.255867268 | 0.004666501 | 0.006313121 | 0.007603344 |
| 0.520647395 | 0.289830022 | 0.261624330 | 0.006886504 | 0.008698764 | 0.010396563 |
| 0.574403718 | 0.490537034 | 0.391202455 | 0.004666501 | 0.006313121 | 0.007603344 |
| 0.574403718 | 0.718328132 | 0.68911829 | 0.006886504 | 0.008698764 | 0.010396563 |
| 0.919021735 | 0.792382876 | 0.658288443 | 0.004666501 | 0.006313121 | 0.007603344 |
| 0.919021735 | 0.512881185 | 0.488108896 | 0.006886504 | 0.008698764 | 0.010396563 |
| 0.045849552 | 0.273824360 | 0.24482221 | 0.004666501 | 0.006313121 | 0.007603344 |
| 0.045849552 | 0.272708688 | 0.252633810 | 0.006886504 | 0.008698764 | 0.010396563 |
| 0.402604926 | 0.692948718 | 0.559991430 | 0.004666501 | 0.006313121 | 0.007603344 |
| 0.402604926 | 0.489306130 | 0.471739649 | 0.006886504 | 0.008698764 | 0.010396563 |

| | | | | | |
|---|---|---|---|---|---|
| 0.189190269 | 0.178241139 | 0.176055913 | 0.004666501 | 0.006313121 | 0.007603344 |
| 0.189190269 | 0.046931303 | 0.078530294 | 0.006886504 | 0.008698764 | 0.010396563 |
| 0.060348027 | 0.737970696 | 0.604597980 | 0.004666501 | 0.006313121 | 0.007603344 |
| 0.060348027 | 0.500356622 | 0.480821661 | 0.006886504 | 0.008698764 | 0.010396563 |
| 0.043167522 | 0.079024079 | 0.091558086 | 0.004666501 | 0.006313121 | 0.007603344 |
| 0.043167522 | 0.518452186 | 0.499047839 | 0.006886504 | 0.008698764 | 0.010396563 |
| 0.118008730 | 0.680411397 | 0.547761272 | 0.004666501 | 0.006313121 | 0.007603344 |
| 0.118008730 | 0.975505614 | 0.951140183 | 0.006886504 | 0.008698764 | 0.010396563 |
| 0.072911128 | 0.180188649 | 0.17756411 | 0.004666501 | 0.006313121 | 0.007603344 |
| 0.072911128 | 0.317622285 | 0.287169422 | 0.006886504 | 0.008698764 | 0.010396563 |
| 0.513905139 | 0.588169690 | 0.464584720 | 0.004666501 | 0.006313121 | 0.007603344 |
| 0.513905139 | 0.116829053 | 0.142467977 | 0.006886504 | 0.008698764 | 0.010396563 |
| 0.477352612 | 0.635892912 | 0.505017352 | 0.004666501 | 0.006313121 | 0.007603344 |
| 0.477352612 | 0.602280057 | 0.579334670 | 0.006886504 | 0.008698764 | 0.010396563 |
| 0.144516640 | 0.086373265 | 0.096976255 | 0.004666501 | 0.006313121 | 0.007603344 |
| 0.144516640 | 0.095380684 | 0.123115767 | 0.006886504 | 0.008698764 | 0.010396563 |
| 0.967717839 | 0.891334967 | 0.754428575 | 0.004666501 | 0.006313121 | 0.007603344 |
| 0.967717839 | 0.454902001 | 0.425294842 | 0.006886504 | 0.008698764 | 0.010396563 |
| 0.645430227 | 0.500878137 | 0.398402056 | 0.004666501 | 0.006313121 | 0.007603344 |
| 0.645430227 | 0.198369725 | 0.200917821 | 0.006886504 | 0.008698764 | 0.010396563 |
| 0.573811542 | 0.807648664 | 0.673599897 | 0.004666501 | 0.006313121 | 0.007603344 |
| 0.573811542 | 0.822113759 | 0.804460648 | 0.006886504 | 0.008698764 | 0.010396563 |
| 0.546535992 | 0.795893988 | 0.661929335 | 0.004666501 | 0.006313121 | 0.007603344 |
| 0.546535992 | 0.318092895 | 0.287222199 | 0.006886504 | 0.008698764 | 0.010396563 |
| 0.901837743 | 0.816838879 | 0.682598513 | 0.004666501 | 0.006313121 | 0.007603344 |
| 0.901837743 | 0.623081063 | 0.608685522 | 0.006886504 | 0.008698764 | 0.010396563 |
| 0.614676557 | 0.473451095 | 0.379439159 | 0.004666501 | 0.006313121 | 0.007603344 |
| 0.614676557 | 0.439403693 | 0.402691608 | 0.006886504 | 0.008698764 | 0.010396563 |
| 0.099408238 | 0.045517147 | 0.064796217 | 0.004666501 | 0.006313121 | 0.007603344 |
| 0.099408238 | 0.040153699 | 0.071885525 | 0.006886504 | 0.008698764 | 0.010396563 |
| 0.149301120 | 0.279049655 | 0.248388156 | 0.004666501 | 0.006313121 | 0.007603344 |
| 0.149301120 | 0.364663324 | 0.325135193 | 0.006886504 | 0.008698764 | 0.010396563 |
| 0.405036344 | 0.386542426 | 0.320813711 | 0.004666501 | 0.006313121 | 0.007603344 |
| 0.405036344 | 0.910466696 | 0.876248660 | 0.006886504 | 0.008698764 | 0.010396563 |
| 0.242792883 | 0.183177458 | 0.179858979 | 0.004666501 | 0.006313121 | 0.007603344 |
| 0.242792883 | 0.737486929 | 0.700453653 | 0.006886504 | 0.008698764 | 0.010396563 |
| 0.472505839 | 0.245465911 | 0.225259925 | 0.004666501 | 0.006313121 | 0.007603344 |
| 0.472505839 | 0.324258415 | 0.291652556 | 0.006886504 | 0.008698764 | 0.010396563 |
| 0.252485633 | 0.868980546 | 0.732089440 | 0.004666501 | 0.006313121 | 0.007603344 |
| 0.252485633 | 0.092475469 | 0.121026322 | 0.006886504 | 0.008698764 | 0.010396563 |
| 0.330938789 | 0.797491267 | 0.663534908 | 0.004666501 | 0.006313121 | 0.007603344 |
| 0.330938789 | 0.933724095 | 0.891207090 | 0.006886504 | 0.008698764 | 0.010396563 |
| 0.592204258 | 0.580236663 | 0.458181217 | 0.004666501 | 0.006313121 | 0.007603344 |
| 0.592204258 | 0.776858712 | 0.737193642 | 0.006886504 | 0.008698764 | 0.010396563 |
| 0.002649290 | 0.042269453 | 0.061905977 | 0.004666501 | 0.006313121 | 0.007603344 |
| 0.002649290 | 0.067552497 | 0.1014672 | 0.006886504 | 0.008698764 | 0.010396563 |

| | | | | | |
|---|---|---|---|---|---|
| 0.485572749 | 0.314345012 | 0.271714859 | 0.004666501 | 0.006313121 | 0.007603344 |
| 0.485572749 | 0.179323181 | 0.188372418 | 0.006886504 | 0.008698764 | 0.010396563 |
| 0.126632678 | 0.089015521 | 0.098955981 | 0.004666501 | 0.006313121 | 0.007603344 |
| 0.126632678 | 0.234266768 | 0.228856268 | 0.006886504 | 0.008698764 | 0.010396563 |
| 0.733434877 | 0.621631281 | 0.492138826 | 0.004666501 | 0.006313121 | 0.007603344 |
| 0.733434877 | 0.425580380 | 0.383893527 | 0.006886504 | 0.008698764 | 0.010396563 |
| 0.513269629 | 0.274229914 | 0.245098999 | 0.004666501 | 0.006313121 | 0.007603344 |
| 0.513269629 | 0.156523670 | 0.173970907 | 0.006886504 | 0.008698764 | 0.010396563 |
| 0.388217519 | 0.440386786 | 0.357715340 | 0.004666501 | 0.006313121 | 0.007603344 |
| 0.388217519 | 0.510837875 | 0.487176346 | 0.006886504 | 0.008698764 | 0.010396563 |
| 0.122305024 | 0.825161672 | 0.690434429 | 0.004666501 | 0.006313121 | 0.007603344 |
| 0.122305024 | 0.278699882 | 0.255257379 | 0.006886504 | 0.008698764 | 0.010396563 |
| 0.286275373 | 0.147761494 | 0.149929348 | 0.004666501 | 0.006313121 | 0.007603344 |
| 0.286275373 | 0.086611984 | 0.114409690 | 0.006886504 | 0.008698764 | 0.010396563 |
| 0.431348976 | 0.842430004 | 0.706707503 | 0.004666501 | 0.006313121 | 0.007603344 |
| 0.431348976 | 0.306457988 | 0.278918682 | 0.006886504 | 0.008698764 | 0.010396563 |
| 0.006474774 | 0.888101376 | 0.751283614 | 0.004666501 | 0.006313121 | 0.007603344 |
| 0.006474774 | 0.707231576 | 0.687262465 | 0.006886504 | 0.008698764 | 0.010396563 |
| 0.741714521 | 0.566348315 | 0.447654573 | 0.004666501 | 0.006313121 | 0.007603344 |
| 0.741714521 | 0.515593364 | 0.493559907 | 0.006886504 | 0.008698764 | 0.010396563 |
| 0.180294765 | 0.235848643 | 0.218486856 | 0.004666501 | 0.006313121 | 0.007603344 |
| 0.180294765 | 0.943688772 | 0.924013594 | 0.006886504 | 0.008698764 | 0.010396563 |
| 0.429589791 | 0.937247560 | 0.807103629 | 0.004666501 | 0.006313121 | 0.007603344 |
| 0.429589791 | 0.862060788 | 0.837723597 | 0.006886504 | 0.008698764 | 0.010396563 |
| 0.168465981 | 0.239056341 | 0.220764219 | 0.004666501 | 0.006313121 | 0.007603344 |
| 0.168465981 | 0.109518517 | 0.136263921 | 0.006886504 | 0.008698764 | 0.010396563 |
| 0.350727633 | 0.302866985 | 0.264253043 | 0.004666501 | 0.006313121 | 0.007603344 |
| 0.350727633 | 0.237861046 | 0.230964030 | 0.006886504 | 0.008698764 | 0.010396563 |
| 0.984226215 | 0.932977069 | 0.801067962 | 0.004666501 | 0.006313121 | 0.007603344 |
| 0.984226215 | 0.889113729 | 0.851087916 | 0.006886504 | 0.008698764 | 0.010396563 |
| 0.227113033 | 0.274197269 | 0.245076587 | 0.004666501 | 0.006313121 | 0.007603344 |
| 0.227113033 | 0.644530102 | 0.620714968 | 0.006886504 | 0.008698764 | 0.010396563 |
| 0.117412304 | 0.104213950 | 0.111491714 | 0.004666501 | 0.006313121 | 0.007603344 |
| 0.117412304 | 0.992091531 | 0.979709197 | 0.006886504 | 0.008698764 | 0.010396563 |
| 0.517075258 | 0.286276107 | 0.253279451 | 0.004666501 | 0.006313121 | 0.007603344 |
| 0.517075258 | 0.115681088 | 0.141190850 | 0.006886504 | 0.008698764 | 0.010396563 |
| 0.133676002 | 0.155230164 | 0.156616904 | 0.004666501 | 0.006313121 | 0.007603344 |
| 0.133676002 | 0.583500232 | 0.571339458 | 0.006886504 | 0.008698764 | 0.010396563 |
| 0.253041086 | 0.906637297 | 0.769772592 | 0.004666501 | 0.006313121 | 0.007603344 |
| 0.253041086 | 0.806149561 | 0.778172938 | 0.006886504 | 0.008698764 | 0.010396563 |
| 0.524140447 | 0.705075824 | 0.572327378 | 0.004666501 | 0.006313121 | 0.007603344 |
| 0.524140447 | 0.870794161 | 0.843760078 | 0.006886504 | 0.008698764 | 0.010396563 |
| 0.353169104 | 0.871426610 | 0.734550208 | 0.004666501 | 0.006313121 | 0.007603344 |
| 0.353169104 | 0.256118715 | 0.241620587 | 0.006886504 | 0.008698764 | 0.010396563 |
| 0.569072330 | 0.372669796 | 0.311149015 | 0.004666501 | 0.006313121 | 0.007603344 |
| 0.569072330 | 0.928030028 | 0.889195089 | 0.006886504 | 0.008698764 | 0.010396563 |

| | | | | | |
|---|---|---|---|---|---|
| 0.692128686 | 0.560378921 | 0.443163574 | 0.004666501 | 0.006313121 | 0.007603344 |
| 0.692128686 | 0.642422605 | 0.619958674 | 0.006886504 | 0.008698764 | 0.010396563 |
| 0.224042233 | 0.380257538 | 0.316375206 | 0.004666501 | 0.006313121 | 0.007603344 |
| 0.224042233 | 0.334214592 | 0.300788416 | 0.006886504 | 0.008698764 | 0.010396563 |
| 0.648273271 | 0.573444960 | 0.452947928 | 0.004666501 | 0.006313121 | 0.007603344 |
| 0.648273271 | 0.174671739 | 0.183784022 | 0.006886504 | 0.008698764 | 0.010396563 |
| 0.842555662 | 0.988351410 | 0.921016072 | 0.004666501 | 0.006313121 | 0.007603344 |
| 0.842555662 | 0.989632541 | 0.966292873 | 0.006886504 | 0.008698764 | 0.010396563 |
| 0.000225155 | 0.306155495 | 0.266377831 | 0.004666501 | 0.006313121 | 0.007603344 |
| 0.000225155 | 0.071843451 | 0.103819408 | 0.006886504 | 0.008698764 | 0.010396563 |
| 0.605309503 | 0.640156812 | 0.5090076 | 0.004666501 | 0.006313121 | 0.007603344 |
| 0.605309503 | 0.971638892 | 0.949866319 | 0.006886504 | 0.008698764 | 0.010396563 |
| 0.343570318 | 0.421305877 | 0.344988041 | 0.004666501 | 0.006313121 | 0.007603344 |
| 0.343570318 | 0.199478637 | 0.202010183 | 0.006886504 | 0.008698764 | 0.010396563 |
| 0.070002836 | 0.041185011 | 0.060800458 | 0.004666501 | 0.006313121 | 0.007603344 |
| 0.070002836 | 0.112818907 | 0.139132810 | 0.006886504 | 0.008698764 | 0.010396563 |
| 0.386395079 | 0.346611613 | 0.293464538 | 0.004666501 | 0.006313121 | 0.007603344 |
| 0.386395079 | 0.495047807 | 0.474309249 | 0.006886504 | 0.008698764 | 0.010396563 |
| 0.17235808 | 0.509854908 | 0.404923396 | 0.004666501 | 0.006313121 | 0.007603344 |
| 0.17235808 | 0.851294836 | 0.827290338 | 0.006886504 | 0.008698764 | 0.010396563 |
| 0.443549208 | 0.239154250 | 0.220833543 | 0.004666501 | 0.006313121 | 0.007603344 |
| 0.443549208 | 0.111308026 | 0.138022257 | 0.006886504 | 0.008698764 | 0.010396563 |
| 0.100789612 | 0.323129010 | 0.277755853 | 0.004666501 | 0.006313121 | 0.007603344 |
| 0.100789612 | 0.463890177 | 0.438535513 | 0.006886504 | 0.008698764 | 0.010396563 |
| 0.262611791 | 0.312883040 | 0.270744223 | 0.004666501 | 0.006313121 | 0.007603344 |
| 0.262611791 | 0.247791590 | 0.236350905 | 0.006886504 | 0.008698764 | 0.010396563 |
| 0.914282580 | 0.801871654 | 0.667883468 | 0.004666501 | 0.006313121 | 0.007603344 |
| 0.914282580 | 0.426500879 | 0.385119517 | 0.006886504 | 0.008698764 | 0.010396563 |
| 0.195450575 | 0.052667574 | 0.070773409 | 0.004666501 | 0.006313121 | 0.007603344 |
| 0.195450575 | 0.631691768 | 0.615222630 | 0.006886504 | 0.008698764 | 0.010396563 |
| 0.013563001 | 0.035923325 | 0.055405949 | 0.004666501 | 0.006313121 | 0.007603344 |
| 0.013563001 | 0.045448590 | 0.077687590 | 0.006886504 | 0.008698764 | 0.010396563 |
| 0.876493903 | 0.997485857 | 0.968843984 | 0.004666501 | 0.006313121 | 0.007603344 |
| 0.876493903 | 0.769366195 | 0.735515062 | 0.006886504 | 0.008698764 | 0.010396563 |
| 0.859846027 | 0.693907901 | 0.560973147 | 0.004666501 | 0.006313121 | 0.007603344 |
| 0.859846027 | 0.362103789 | 0.324234337 | 0.006886504 | 0.008698764 | 0.010396563 |
| 0.129849393 | 0.081481184 | 0.093379431 | 0.004666501 | 0.006313121 | 0.007603344 |
| 0.129849393 | 0.186051672 | 0.194173431 | 0.006886504 | 0.008698764 | 0.010396563 |
| 0.600484483 | 0.450974653 | 0.364787354 | 0.004666501 | 0.006313121 | 0.007603344 |
| 0.600484483 | 0.610462480 | 0.585516973 | 0.006886504 | 0.008698764 | 0.010396563 |
| 0.112290113 | 0.738582448 | 0.605162988 | 0.004666501 | 0.006313121 | 0.007603344 |
| 0.112290113 | 0.890699454 | 0.854961528 | 0.006886504 | 0.008698764 | 0.010396563 |
| 0.407932508 | 0.210068454 | 0.199886556 | 0.004666501 | 0.006313121 | 0.007603344 |
| 0.407932508 | 0.128216974 | 0.151365263 | 0.006886504 | 0.008698764 | 0.010396563 |
| 0.656218995 | 0.767664997 | 0.633030298 | 0.004666501 | 0.006313121 | 0.007603344 |
| 0.656218995 | 0.837124930 | 0.807592225 | 0.006886504 | 0.008698764 | 0.010396563 |

| | | | | | |
|---|---|---|---|---|---|
| 0.880226309 | 0.708098281 | 0.575443943 | 0.004666501 | 0.006313121 | 0.007603344 |
| 0.880226309 | 0.370376322 | 0.329120648 | 0.006886504 | 0.008698764 | 0.010396563 |
| 0.147193977 | 0.352281981 | 0.297235694 | 0.004666501 | 0.006313121 | 0.007603344 |
| 0.147193977 | 0.564551833 | 0.550807271 | 0.006886504 | 0.008698764 | 0.010396563 |
| 0.622330776 | 0.627812364 | 0.497646821 | 0.004666501 | 0.006313121 | 0.007603344 |
| 0.622330776 | 0.661030439 | 0.628254058 | 0.006886504 | 0.008698764 | 0.010396563 |
| 0.012435998 | 0.112020533 | 0.118481091 | 0.004666501 | 0.006313121 | 0.007603344 |
| 0.012435998 | 0.580185253 | 0.571068266 | 0.006886504 | 0.008698764 | 0.010396563 |
| 0.630928117 | 0.406525936 | 0.334865013 | 0.004666501 | 0.006313121 | 0.007603344 |
| 0.630928117 | 0.185010431 | 0.194057297 | 0.006886504 | 0.008698764 | 0.010396563 |
| 0.127102814 | 0.853939393 | 0.717824739 | 0.004666501 | 0.006313121 | 0.007603344 |
| 0.127102814 | 0.916361846 | 0.882567837 | 0.006886504 | 0.008698764 | 0.010396563 |
| 0.350597732 | 0.645995098 | 0.514586255 | 0.004666501 | 0.006313121 | 0.007603344 |
| 0.350597732 | 0.902314751 | 0.872652450 | 0.006886504 | 0.008698764 | 0.010396563 |
| 0.317793594 | 0.186703492 | 0.182480305 | 0.004666501 | 0.006313121 | 0.007603344 |
| 0.317793594 | 0.114711525 | 0.140059859 | 0.006886504 | 0.008698764 | 0.010396563 |

| Species | Genus |
|---|---|
| Acetatifactor_muris | Acetatifactor |
| Acetatifactor_muris | Acetatifactor |
| Acetatifactor_SGB41546 | Acetatifactor |
| Acetatifactor_SGB41546 | Acetatifactor |
| Acutalibacter_muris | Acutalibacter |
| Acutalibacter_muris | Acutalibacter |
| Acutalibacter_sp_1XD8_36 | Acutalibacter |
| Acutalibacter_sp_1XD8_36 | Acutalibacter |
| Adlercreutzia_caecimuris | Adlercreutzia |
| Adlercreutzia_caecimuris | Adlercreutzia |
| Adlercreutzia_mucosicola | Adlercreutzia |
| Adlercreutzia_mucosicola | Adlercreutzia |
| Adlercreutzia_muris | Adlercreutzia |
| Adlercreutzia_muris | Adlercreutzia |
| Akkermansia_muciniphila | Akkermansia |
| Akkermansia_muciniphila | Akkermansia |
| Alistipes_sp_DSM_112343 | Alistipes |
| Alistipes_sp_DSM_112343 | Alistipes |
| Anaerotruncus_sp_1XD42_93 | Anaerotruncus |
| Anaerotruncus_sp_1XD42_93 | Anaerotruncus |
| Bacteria_unclassified_SGB102200 | Bacteria_unclassified |
| Bacteria_unclassified_SGB102200 | Bacteria_unclassified |
| Bacteria_unclassified_SGB41677 | Bacteria_unclassified |
| Bacteria_unclassified_SGB41677 | Bacteria_unclassified |
| Bacteria_unclassified_SGB43546 | Bacteria_unclassified |
| Bacteria_unclassified_SGB43546 | Bacteria_unclassified |
| bacterium_1XD42_54 | Bacteria_unclassified |
| bacterium_1XD42_54 | Bacteria_unclassified |
| bacterium_1XD42_76 | Bacteria_unclassified |
| bacterium_1XD42_76 | Bacteria_unclassified |
| bacterium_1xD8_48 | Bacteria_unclassified |
| bacterium_1xD8_48 | Bacteria_unclassified |
| Bacteroides_thetaiotaomicron | Bacteroides |
| Bacteroides_thetaiotaomicron | Bacteroides |
| berger_parker | |
| berger_parker | |
| Bifidobacterium_pseudolongum | Bifidobacterium |
| Bifidobacterium_pseudolongum | Bifidobacterium |
| Clostridia_bacterium | Clostridia_unclassified |
| Clostridia_bacterium | Clostridia_unclassified |
| Clostridiaceae_bacterium | Clostridiaceae_unclassified |
| Clostridiaceae_bacterium | Clostridiaceae_unclassified |
| Clostridiaceae_unclassified_SGB41663 | Clostridiaceae_unclassified |
| Clostridiaceae_unclassified_SGB41663 | Clostridiaceae_unclassified |
| Clostridiales_bacterium | Eubacteriales_unclassified |
| Clostridiales_bacterium | Eubacteriales_unclassified |

| | |
|---|---|
| Clostridium_cocleatum | Erysipelatoclostridium |
| Clostridium_cocleatum | Erysipelatoclostridium |
| Coriobacteriaceae_bacterium | Coriobacteriaceae_unclassified |
| Coriobacteriaceae_bacterium | Coriobacteriaceae_unclassified |
| Dorea_sp_5_2 | Dorea |
| Dorea_sp_5_2 | Dorea |
| Dubosiella_newyorkensis | Dubosiella |
| Dubosiella_newyorkensis | Dubosiella |
| Erysipelotrichales_bacterium | Erysipelotrichales_unclassified |
| Erysipelotrichales_bacterium | Erysipelotrichales_unclassified |
| Eubacteriaceae_bacterium | Eubacteriaceae_unclassified |
| Eubacteriaceae_bacterium | Eubacteriaceae_unclassified |
| Eubacteriaceae_unclassified_SGB94922 | Eubacteriaceae_unclassified |
| Eubacteriaceae_unclassified_SGB94922 | Eubacteriaceae_unclassified |
| GGB20149_SGB29430 | GGB20149 |
| GGB20149_SGB29430 | GGB20149 |
| GGB22635_SGB63107 | GGB22635 |
| GGB22635_SGB63107 | GGB22635 |
| GGB25041_SGB36960 | GGB25041 |
| GGB25041_SGB36960 | GGB25041 |
| GGB27876_SGB40310 | GGB27876 |
| GGB27876_SGB40310 | GGB27876 |
| GGB27878_SGB40312 | GGB27878 |
| GGB27878_SGB40312 | GGB27878 |
| GGB27918_SGB40356 | GGB27918 |
| GGB27918_SGB40356 | GGB27918 |
| GGB28382_SGB40962 | GGB28382 |
| GGB28382_SGB40962 | GGB28382 |
| GGB28399_SGB40980 | GGB28399 |
| GGB28399_SGB40980 | GGB28399 |
| GGB28411_SGB40993 | GGB28411 |
| GGB28411_SGB40993 | GGB28411 |
| GGB28415_SGB40997 | GGB28415 |
| GGB28415_SGB40997 | GGB28415 |
| GGB28430_SGB41013 | GGB28430 |
| GGB28430_SGB41013 | GGB28430 |
| GGB28439_SGB41022 | GGB28439 |
| GGB28439_SGB41022 | GGB28439 |
| GGB28778_SGB41431 | GGB28778 |
| GGB28778_SGB41431 | GGB28778 |
| GGB28784_SGB41437 | GGB28784 |
| GGB28784_SGB41437 | GGB28784 |
| GGB28792_SGB41445 | GGB28792 |
| GGB28792_SGB41445 | GGB28792 |
| GGB28798_SGB41451 | GGB28798 |
| GGB28798_SGB41451 | GGB28798 |

| | |
|---|---|
| GGB28802_SGB41455 | GGB28802 |
| GGB28802_SGB41455 | GGB28802 |
| GGB28818_SGB41473 | GGB28818 |
| GGB28818_SGB41473 | GGB28818 |
| GGB28828_SGB41484 | GGB28828 |
| GGB28828_SGB41484 | GGB28828 |
| GGB28851_SGB41518 | GGB28851 |
| GGB28851_SGB41518 | GGB28851 |
| GGB28859_SGB41528 | GGB28859 |
| GGB28859_SGB41528 | GGB28859 |
| GGB28864_SGB41535 | GGB28864 |
| GGB28864_SGB41535 | GGB28864 |
| GGB28869_SGB41543 | GGB28869 |
| GGB28869_SGB41543 | GGB28869 |
| GGB28883_SGB41564 | GGB28883 |
| GGB28883_SGB41564 | GGB28883 |
| GGB28892_SGB41573 | GGB28892 |
| GGB28892_SGB41573 | GGB28892 |
| GGB28893_SGB41574 | GGB28893 |
| GGB28893_SGB41574 | GGB28893 |
| GGB28898_SGB41580 | GGB28898 |
| GGB28898_SGB41580 | GGB28898 |
| GGB28904_SGB41597 | GGB28904 |
| GGB28904_SGB41597 | GGB28904 |
| GGB28916_SGB41612 | GGB28916 |
| GGB28916_SGB41612 | GGB28916 |
| GGB28924_SGB41621 | GGB28924 |
| GGB28924_SGB41621 | GGB28924 |
| GGB28926_SGB41624 | GGB28926 |
| GGB28926_SGB41624 | GGB28926 |
| GGB28927_SGB41625 | GGB28927 |
| GGB28927_SGB41625 | GGB28927 |
| GGB28934_SGB41635 | GGB28934 |
| GGB28934_SGB41635 | GGB28934 |
| GGB28946_SGB41652 | GGB28946 |
| GGB28946_SGB41652 | GGB28946 |
| GGB28949_SGB41655 | GGB28949 |
| GGB28949_SGB41655 | GGB28949 |
| GGB28949_SGB41656 | GGB28949 |
| GGB28949_SGB41656 | GGB28949 |
| GGB28950_SGB41657 | GGB28950 |
| GGB28950_SGB41657 | GGB28950 |
| GGB28951_SGB102295 | GGB28951 |
| GGB28951_SGB102295 | GGB28951 |
| GGB28951_SGB41658 | GGB28951 |
| GGB28951_SGB41658 | GGB28951 |

| | |
|---|---|
| GGB28954_SGB41662 | GGB28954 |
| GGB28954_SGB41662 | GGB28954 |
| GGB28956_SGB41665 | GGB28956 |
| GGB28956_SGB41665 | GGB28956 |
| GGB28960_SGB41669 | GGB28960 |
| GGB28960_SGB41669 | GGB28960 |
| GGB28967_SGB41678 | GGB28967 |
| GGB28967_SGB41678 | GGB28967 |
| GGB28991_SGB41705 | GGB28991 |
| GGB28991_SGB41705 | GGB28991 |
| GGB29002_SGB41718 | GGB29002 |
| GGB29002_SGB41718 | GGB29002 |
| GGB29003_SGB41719 | GGB29003 |
| GGB29003_SGB41719 | GGB29003 |
| GGB29011_SGB41731 | GGB29011 |
| GGB29011_SGB41731 | GGB29011 |
| GGB29531_SGB42317 | GGB29531 |
| GGB29531_SGB42317 | GGB29531 |
| GGB29685_SGB42494 | GGB29685 |
| GGB29685_SGB42494 | GGB29685 |
| GGB30141_SGB43066 | GGB30141 |
| GGB30141_SGB43066 | GGB30141 |
| GGB30286_SGB43248 | GGB30286 |
| GGB30286_SGB43248 | GGB30286 |
| GGB30303_SGB43268 | GGB30303 |
| GGB30303_SGB43268 | GGB30303 |
| GGB30413_SGB43452 | GGB30413 |
| GGB30413_SGB43452 | GGB30413 |
| GGB30454_SGB43514 | GGB30454 |
| GGB30454_SGB43514 | GGB30454 |
| GGB30455_SGB43519 | GGB30455 |
| GGB30455_SGB43519 | GGB30455 |
| GGB30461_SGB43527 | GGB30461 |
| GGB30461_SGB43527 | GGB30461 |
| GGB30461_SGB43530 | GGB30461 |
| GGB30461_SGB43530 | GGB30461 |
| GGB30463_SGB43537 | GGB30463 |
| GGB30463_SGB43537 | GGB30463 |
| GGB30473_SGB43557 | GGB30473 |
| GGB30473_SGB43557 | GGB30473 |
| GGB30475_SGB63182 | GGB30475 |
| GGB30475_SGB63182 | GGB30475 |
| GGB30861_SGB44083 | GGB30861 |
| GGB30861_SGB44083 | GGB30861 |
| GGB31312_SGB44628 | GGB31312 |
| GGB31312_SGB44628 | GGB31312 |

| | |
|---|---|
| GGB31438_SGB44768 | GGB31438 |
| GGB31438_SGB44768 | GGB31438 |
| GGB3171_SGB4185 | GGB3171 |
| GGB3171_SGB4185 | GGB3171 |
| GGB31823_SGB45199 | GGB31823 |
| GGB31823_SGB45199 | GGB31823 |
| GGB31853_SGB45233 | GGB31853 |
| GGB31853_SGB45233 | GGB31853 |
| GGB32371_SGB41694 | GGB32371 |
| GGB32371_SGB41694 | GGB32371 |
| GGB3793_SGB5158 | GGB3793 |
| GGB3793_SGB5158 | GGB3793 |
| GGB42598_SGB59794 | GGB42598 |
| GGB42598_SGB59794 | GGB42598 |
| GGB45656_SGB63370 | GGB45656 |
| GGB45656_SGB63370 | GGB45656 |
| GGB47127_SGB65054 | GGB47127 |
| GGB47127_SGB65054 | GGB47127 |
| GGB74395_SGB43521 | GGB74395 |
| GGB74395_SGB43521 | GGB74395 |
| GGB75053_SGB43494 | GGB75053 |
| GGB75053_SGB43494 | GGB75053 |
| GGB75109_SGB102238 | GGB75109 |
| GGB75109_SGB102238 | GGB75109 |
| GGB81440_SGB45230 | GGB81440 |
| GGB81440_SGB45230 | GGB81440 |
| Lachnospiraceae_bacterium | Lachnospiraceae_unclassified |
| Lachnospiraceae_bacterium | Lachnospiraceae_unclassified |
| Lachnospiraceae_bacterium_A2 | Lachnospiraceae_unclassified |
| Lachnospiraceae_bacterium_A2 | Lachnospiraceae_unclassified |
| Lachnospiraceae_bacterium_MD308 | Lachnospiraceae_unclassified |
| Lachnospiraceae_bacterium_MD308 | Lachnospiraceae_unclassified |
| Lachnospiraceae_bacterium_MD329 | Lachnospiraceae_unclassified |
| Lachnospiraceae_bacterium_MD329 | Lachnospiraceae_unclassified |
| Lachnospiraceae_unclassified_SGB41414 | Lachnospiraceae_unclassified |
| Lachnospiraceae_unclassified_SGB41414 | Lachnospiraceae_unclassified |
| Lachnospiraceae_unclassified_SGB41418 | Lachnospiraceae_unclassified |
| Lachnospiraceae_unclassified_SGB41418 | Lachnospiraceae_unclassified |
| Lachnospiraceae_unclassified_SGB41424 | Lachnospiraceae_unclassified |
| Lachnospiraceae_unclassified_SGB41424 | Lachnospiraceae_unclassified |
| Lachnospiraceae_unclassified_SGB41589 | Lachnospiraceae_unclassified |
| Lachnospiraceae_unclassified_SGB41589 | Lachnospiraceae_unclassified |
| Lactobacillus_johnsonii | Lactobacillus |
| Lactobacillus_johnsonii | Lactobacillus |
| Muribaculaceae_bacterium | Muribaculaceae_unclassified |
| Muribaculaceae_bacterium | Muribaculaceae_unclassified |

| | |
|---|---|
| Neglectibacter_sp_X4 | Neglectibacter |
| Neglectibacter_sp_X4 | Neglectibacter |
| Oscillospiraceae_bacterium | Oscillospiraceae_unclassified |
| Oscillospiraceae_bacterium | Oscillospiraceae_unclassified |
| Oscillospiraceae_unclassified_SGB43502 | Oscillospiraceae_unclassified |
| Oscillospiraceae_unclassified_SGB43502 | Oscillospiraceae_unclassified |
| Oscillospiraceae_unclassified_SGB43505 | Oscillospiraceae_unclassified |
| Oscillospiraceae_unclassified_SGB43505 | Oscillospiraceae_unclassified |
| Parasutterella_excrementihominis | Parasutterella |
| Parasutterella_excrementihominis | Parasutterella |
| richness | |
| richness | |
| Romboutsia_ilealis | Romboutsia |
| Romboutsia_ilealis | Romboutsia |
| Schaedlerella_arabinosiphila | Schaedlerella |
| Schaedlerella_arabinosiphila | Schaedlerella |
| shannon | |
| shannon | |
| Turicibacter_sp_1E2 | Turicibacter |
| Turicibacter_sp_1E2 | Turicibacter |
| Acetatifactor_muris | Acetatifactor |
| Acetatifactor_muris | Acetatifactor |
| Acetatifactor_SGB41546 | Acetatifactor |
| Acetatifactor_SGB41546 | Acetatifactor |
| Acutalibacter_muris | Acutalibacter |
| Acutalibacter_muris | Acutalibacter |
| Acutalibacter_sp_1XD8_36 | Acutalibacter |
| Acutalibacter_sp_1XD8_36 | Acutalibacter |
| Adlercreutzia_caecimuris | Adlercreutzia |
| Adlercreutzia_caecimuris | Adlercreutzia |
| Adlercreutzia_mucosicola | Adlercreutzia |
| Adlercreutzia_mucosicola | Adlercreutzia |
| Adlercreutzia_muris | Adlercreutzia |
| Adlercreutzia_muris | Adlercreutzia |
| Akkermansia_muciniphila | Akkermansia |
| Akkermansia_muciniphila | Akkermansia |
| Alistipes_sp_DSM_112343 | Alistipes |
| Alistipes_sp_DSM_112343 | Alistipes |
| Anaerotruncus_sp_1XD42_93 | Anaerotruncus |
| Anaerotruncus_sp_1XD42_93 | Anaerotruncus |
| Bacteria_unclassified_SGB102200 | Bacteria_unclassified |
| Bacteria_unclassified_SGB102200 | Bacteria_unclassified |
| Bacteria_unclassified_SGB41677 | Bacteria_unclassified |
| Bacteria_unclassified_SGB41677 | Bacteria_unclassified |
| Bacteria_unclassified_SGB43546 | Bacteria_unclassified |
| Bacteria_unclassified_SGB43546 | Bacteria_unclassified |

| | |
|---|---|
| bacterium_1XD42_54 | Bacteria_unclassified |
| bacterium_1XD42_54 | Bacteria_unclassified |
| bacterium_1XD42_76 | Bacteria_unclassified |
| bacterium_1XD42_76 | Bacteria_unclassified |
| bacterium_1xD8_48 | Bacteria_unclassified |
| bacterium_1xD8_48 | Bacteria_unclassified |
| Bacteroides_thetaiotaomicron | Bacteroides |
| Bacteroides_thetaiotaomicron | Bacteroides |
| berger_parker | |
| berger_parker | |
| Bifidobacterium_pseudolongum | Bifidobacterium |
| Bifidobacterium_pseudolongum | Bifidobacterium |
| Clostridia_bacterium | Clostridia_unclassified |
| Clostridia_bacterium | Clostridia_unclassified |
| Clostridiaceae_bacterium | Clostridiaceae_unclassified |
| Clostridiaceae_bacterium | Clostridiaceae_unclassified |
| Clostridiaceae_unclassified_SGB41663 | Clostridiaceae_unclassified |
| Clostridiaceae_unclassified_SGB41663 | Clostridiaceae_unclassified |
| Clostridiales_bacterium | Eubacteriales_unclassified |
| Clostridiales_bacterium | Eubacteriales_unclassified |
| Clostridium_cocleatum | Erysipelatoclostridium |
| Clostridium_cocleatum | Erysipelatoclostridium |
| Coriobacteriaceae_bacterium | Coriobacteriaceae_unclassified |
| Coriobacteriaceae_bacterium | Coriobacteriaceae_unclassified |
| Dorea_sp_5_2 | Dorea |
| Dorea_sp_5_2 | Dorea |
| Dubosiella_newyorkensis | Dubosiella |
| Dubosiella_newyorkensis | Dubosiella |
| Erysipelotrichales_bacterium | Erysipelotrichales_unclassified |
| Erysipelotrichales_bacterium | Erysipelotrichales_unclassified |
| Eubacteriaceae_bacterium | Eubacteriaceae_unclassified |
| Eubacteriaceae_bacterium | Eubacteriaceae_unclassified |
| Eubacteriaceae_unclassified_SGB94922 | Eubacteriaceae_unclassified |
| Eubacteriaceae_unclassified_SGB94922 | Eubacteriaceae_unclassified |
| GGB20149_SGB29430 | GGB20149 |
| GGB20149_SGB29430 | GGB20149 |
| GGB22635_SGB63107 | GGB22635 |
| GGB22635_SGB63107 | GGB22635 |
| GGB25041_SGB36960 | GGB25041 |
| GGB25041_SGB36960 | GGB25041 |
| GGB27876_SGB40310 | GGB27876 |
| GGB27876_SGB40310 | GGB27876 |
| GGB27878_SGB40312 | GGB27878 |
| GGB27878_SGB40312 | GGB27878 |
| GGB27918_SGB40356 | GGB27918 |
| GGB27918_SGB40356 | GGB27918 |

| | |
|---|---|
| GGB28382_SGB40962 | GGB28382 |
| GGB28382_SGB40962 | GGB28382 |
| GGB28399_SGB40980 | GGB28399 |
| GGB28399_SGB40980 | GGB28399 |
| GGB28411_SGB40993 | GGB28411 |
| GGB28411_SGB40993 | GGB28411 |
| GGB28415_SGB40997 | GGB28415 |
| GGB28415_SGB40997 | GGB28415 |
| GGB28430_SGB41013 | GGB28430 |
| GGB28430_SGB41013 | GGB28430 |
| GGB28439_SGB41022 | GGB28439 |
| GGB28439_SGB41022 | GGB28439 |
| GGB28778_SGB41431 | GGB28778 |
| GGB28778_SGB41431 | GGB28778 |
| GGB28784_SGB41437 | GGB28784 |
| GGB28784_SGB41437 | GGB28784 |
| GGB28792_SGB41445 | GGB28792 |
| GGB28792_SGB41445 | GGB28792 |
| GGB28798_SGB41451 | GGB28798 |
| GGB28798_SGB41451 | GGB28798 |
| GGB28802_SGB41455 | GGB28802 |
| GGB28802_SGB41455 | GGB28802 |
| GGB28818_SGB41473 | GGB28818 |
| GGB28818_SGB41473 | GGB28818 |
| GGB28828_SGB41484 | GGB28828 |
| GGB28828_SGB41484 | GGB28828 |
| GGB28851_SGB41518 | GGB28851 |
| GGB28851_SGB41518 | GGB28851 |
| GGB28859_SGB41528 | GGB28859 |
| GGB28859_SGB41528 | GGB28859 |
| GGB28864_SGB41535 | GGB28864 |
| GGB28864_SGB41535 | GGB28864 |
| GGB28869_SGB41543 | GGB28869 |
| GGB28869_SGB41543 | GGB28869 |
| GGB28883_SGB41564 | GGB28883 |
| GGB28883_SGB41564 | GGB28883 |
| GGB28892_SGB41573 | GGB28892 |
| GGB28892_SGB41573 | GGB28892 |
| GGB28893_SGB41574 | GGB28893 |
| GGB28893_SGB41574 | GGB28893 |
| GGB28898_SGB41580 | GGB28898 |
| GGB28898_SGB41580 | GGB28898 |
| GGB28904_SGB41597 | GGB28904 |
| GGB28904_SGB41597 | GGB28904 |
| GGB28916_SGB41612 | GGB28916 |
| GGB28916_SGB41612 | GGB28916 |

| | |
|---|---|
| GGB28924_SGB41621 | GGB28924 |
| GGB28924_SGB41621 | GGB28924 |
| GGB28926_SGB41624 | GGB28926 |
| GGB28926_SGB41624 | GGB28926 |
| GGB28927_SGB41625 | GGB28927 |
| GGB28927_SGB41625 | GGB28927 |
| GGB28934_SGB41635 | GGB28934 |
| GGB28934_SGB41635 | GGB28934 |
| GGB28946_SGB41652 | GGB28946 |
| GGB28946_SGB41652 | GGB28946 |
| GGB28949_SGB41655 | GGB28949 |
| GGB28949_SGB41655 | GGB28949 |
| GGB28949_SGB41656 | GGB28949 |
| GGB28949_SGB41656 | GGB28949 |
| GGB28950_SGB41657 | GGB28950 |
| GGB28950_SGB41657 | GGB28950 |
| GGB28951_SGB102295 | GGB28951 |
| GGB28951_SGB102295 | GGB28951 |
| GGB28951_SGB41658 | GGB28951 |
| GGB28951_SGB41658 | GGB28951 |
| GGB28954_SGB41662 | GGB28954 |
| GGB28954_SGB41662 | GGB28954 |
| GGB28956_SGB41665 | GGB28956 |
| GGB28956_SGB41665 | GGB28956 |
| GGB28960_SGB41669 | GGB28960 |
| GGB28960_SGB41669 | GGB28960 |
| GGB28967_SGB41678 | GGB28967 |
| GGB28967_SGB41678 | GGB28967 |
| GGB28991_SGB41705 | GGB28991 |
| GGB28991_SGB41705 | GGB28991 |
| GGB29002_SGB41718 | GGB29002 |
| GGB29002_SGB41718 | GGB29002 |
| GGB29003_SGB41719 | GGB29003 |
| GGB29003_SGB41719 | GGB29003 |
| GGB29011_SGB41731 | GGB29011 |
| GGB29011_SGB41731 | GGB29011 |
| GGB29531_SGB42317 | GGB29531 |
| GGB29531_SGB42317 | GGB29531 |
| GGB29685_SGB42494 | GGB29685 |
| GGB29685_SGB42494 | GGB29685 |
| GGB30141_SGB43066 | GGB30141 |
| GGB30141_SGB43066 | GGB30141 |
| GGB30286_SGB43248 | GGB30286 |
| GGB30286_SGB43248 | GGB30286 |
| GGB30303_SGB43268 | GGB30303 |
| GGB30303_SGB43268 | GGB30303 |

| | |
|---|---|
| GGB30413_SGB43452 | GGB30413 |
| GGB30413_SGB43452 | GGB30413 |
| GGB30454_SGB43514 | GGB30454 |
| GGB30454_SGB43514 | GGB30454 |
| GGB30455_SGB43519 | GGB30455 |
| GGB30455_SGB43519 | GGB30455 |
| GGB30461_SGB43527 | GGB30461 |
| GGB30461_SGB43527 | GGB30461 |
| GGB30461_SGB43530 | GGB30461 |
| GGB30461_SGB43530 | GGB30461 |
| GGB30463_SGB43537 | GGB30463 |
| GGB30463_SGB43537 | GGB30463 |
| GGB30473_SGB43557 | GGB30473 |
| GGB30473_SGB43557 | GGB30473 |
| GGB30475_SGB63182 | GGB30475 |
| GGB30475_SGB63182 | GGB30475 |
| GGB30861_SGB44083 | GGB30861 |
| GGB30861_SGB44083 | GGB30861 |
| GGB31312_SGB44628 | GGB31312 |
| GGB31312_SGB44628 | GGB31312 |
| GGB31438_SGB44768 | GGB31438 |
| GGB31438_SGB44768 | GGB31438 |
| GGB3171_SGB4185 | GGB3171 |
| GGB3171_SGB4185 | GGB3171 |
| GGB31823_SGB45199 | GGB31823 |
| GGB31823_SGB45199 | GGB31823 |
| GGB31853_SGB45233 | GGB31853 |
| GGB31853_SGB45233 | GGB31853 |
| GGB32371_SGB41694 | GGB32371 |
| GGB32371_SGB41694 | GGB32371 |
| GGB3793_SGB5158 | GGB3793 |
| GGB3793_SGB5158 | GGB3793 |
| GGB42598_SGB59794 | GGB42598 |
| GGB42598_SGB59794 | GGB42598 |
| GGB45656_SGB63370 | GGB45656 |
| GGB45656_SGB63370 | GGB45656 |
| GGB47127_SGB65054 | GGB47127 |
| GGB47127_SGB65054 | GGB47127 |
| GGB74395_SGB43521 | GGB74395 |
| GGB74395_SGB43521 | GGB74395 |
| GGB75053_SGB43494 | GGB75053 |
| GGB75053_SGB43494 | GGB75053 |
| GGB75109_SGB102238 | GGB75109 |
| GGB75109_SGB102238 | GGB75109 |
| GGB81440_SGB45230 | GGB81440 |
| GGB81440_SGB45230 | GGB81440 |

| | |
|---|---|
| Lachnospiraceae_bacterium | Lachnospiraceae_unclassified |
| Lachnospiraceae_bacterium | Lachnospiraceae_unclassified |
| Lachnospiraceae_bacterium_A2 | Lachnospiraceae_unclassified |
| Lachnospiraceae_bacterium_A2 | Lachnospiraceae_unclassified |
| Lachnospiraceae_bacterium_MD308 | Lachnospiraceae_unclassified |
| Lachnospiraceae_bacterium_MD308 | Lachnospiraceae_unclassified |
| Lachnospiraceae_bacterium_MD329 | Lachnospiraceae_unclassified |
| Lachnospiraceae_bacterium_MD329 | Lachnospiraceae_unclassified |
| Lachnospiraceae_unclassified_SGB41414 | Lachnospiraceae_unclassified |
| Lachnospiraceae_unclassified_SGB41414 | Lachnospiraceae_unclassified |
| Lachnospiraceae_unclassified_SGB41418 | Lachnospiraceae_unclassified |
| Lachnospiraceae_unclassified_SGB41418 | Lachnospiraceae_unclassified |
| Lachnospiraceae_unclassified_SGB41424 | Lachnospiraceae_unclassified |
| Lachnospiraceae_unclassified_SGB41424 | Lachnospiraceae_unclassified |
| Lachnospiraceae_unclassified_SGB41589 | Lachnospiraceae_unclassified |
| Lachnospiraceae_unclassified_SGB41589 | Lachnospiraceae_unclassified |
| Lactobacillus_johnsonii | Lactobacillus |
| Lactobacillus_johnsonii | Lactobacillus |
| Muribaculaceae_bacterium | Muribaculaceae_unclassified |
| Muribaculaceae_bacterium | Muribaculaceae_unclassified |
| Neglectibacter_sp_X4 | Neglectibacter |
| Neglectibacter_sp_X4 | Neglectibacter |
| Oscillospiraceae_bacterium | Oscillospiraceae_unclassified |
| Oscillospiraceae_bacterium | Oscillospiraceae_unclassified |
| Oscillospiraceae_unclassified_SGB43502 | Oscillospiraceae_unclassified |
| Oscillospiraceae_unclassified_SGB43502 | Oscillospiraceae_unclassified |
| Oscillospiraceae_unclassified_SGB43505 | Oscillospiraceae_unclassified |
| Oscillospiraceae_unclassified_SGB43505 | Oscillospiraceae_unclassified |
| Parasutterella_excrementihominis | Parasutterella |
| Parasutterella_excrementihominis | Parasutterella |
| richness | |
| richness | |
| Romboutsia_ilealis | Romboutsia |
| Romboutsia_ilealis | Romboutsia |
| Schaedlerella_arabinosiphila | Schaedlerella |
| Schaedlerella_arabinosiphila | Schaedlerella |
| shannon | |
| shannon | |
| Turicibacter_sp_1E2 | Turicibacter |
| Turicibacter_sp_1E2 | Turicibacter |
| Acetatifactor_muris | Acetatifactor |
| Acetatifactor_muris | Acetatifactor |
| Acetatifactor_SGB41546 | Acetatifactor |
| Acetatifactor_SGB41546 | Acetatifactor |
| Acutalibacter_muris | Acutalibacter |
| Acutalibacter_muris | Acutalibacter |

| | |
|---|---|
| Acutalibacter_sp_1XD8_36 | Acutalibacter |
| Acutalibacter_sp_1XD8_36 | Acutalibacter |
| Adlercreutzia_caecimuris | Adlercreutzia |
| Adlercreutzia_caecimuris | Adlercreutzia |
| Adlercreutzia_mucosicola | Adlercreutzia |
| Adlercreutzia_mucosicola | Adlercreutzia |
| Adlercreutzia_muris | Adlercreutzia |
| Adlercreutzia_muris | Adlercreutzia |
| Akkermansia_muciniphila | Akkermansia |
| Akkermansia_muciniphila | Akkermansia |
| Alistipes_sp_DSM_112343 | Alistipes |
| Alistipes_sp_DSM_112343 | Alistipes |
| Anaerotruncus_sp_1XD42_93 | Anaerotruncus |
| Anaerotruncus_sp_1XD42_93 | Anaerotruncus |
| Bacteria_unclassified_SGB102200 | Bacteria_unclassified |
| Bacteria_unclassified_SGB102200 | Bacteria_unclassified |
| Bacteria_unclassified_SGB41677 | Bacteria_unclassified |
| Bacteria_unclassified_SGB41677 | Bacteria_unclassified |
| Bacteria_unclassified_SGB43546 | Bacteria_unclassified |
| Bacteria_unclassified_SGB43546 | Bacteria_unclassified |
| bacterium_1XD42_54 | Bacteria_unclassified |
| bacterium_1XD42_54 | Bacteria_unclassified |
| bacterium_1XD42_76 | Bacteria_unclassified |
| bacterium_1XD42_76 | Bacteria_unclassified |
| bacterium_1xD8_48 | Bacteria_unclassified |
| bacterium_1xD8_48 | Bacteria_unclassified |
| Bacteroides_thetaiotaomicron | Bacteroides |
| Bacteroides_thetaiotaomicron | Bacteroides |
| berger_parker | |
| berger_parker | |
| Bifidobacterium_pseudolongum | Bifidobacterium |
| Bifidobacterium_pseudolongum | Bifidobacterium |
| Clostridia_bacterium | Clostridia_unclassified |
| Clostridia_bacterium | Clostridia_unclassified |
| Clostridiaceae_bacterium | Clostridiaceae_unclassified |
| Clostridiaceae_bacterium | Clostridiaceae_unclassified |
| Clostridiaceae_unclassified_SGB41663 | Clostridiaceae_unclassified |
| Clostridiaceae_unclassified_SGB41663 | Clostridiaceae_unclassified |
| Clostridiales_bacterium | Eubacteriales_unclassified |
| Clostridiales_bacterium | Eubacteriales_unclassified |
| Clostridium_cocleatum | Erysipelatoclostridium |
| Clostridium_cocleatum | Erysipelatoclostridium |
| Coriobacteriaceae_bacterium | Coriobacteriaceae_unclassified |
| Coriobacteriaceae_bacterium | Coriobacteriaceae_unclassified |
| Dorea_sp_5_2 | Dorea |
| Dorea_sp_5_2 | Dorea |

| | |
|---|---|
| Dubosiella_newyorkensis | Dubosiella |
| Dubosiella_newyorkensis | Dubosiella |
| Erysipelotrichales_bacterium | Erysipelotrichales_unclassified |
| Erysipelotrichales_bacterium | Erysipelotrichales_unclassified |
| Eubacteriaceae_bacterium | Eubacteriaceae_unclassified |
| Eubacteriaceae_bacterium | Eubacteriaceae_unclassified |
| Eubacteriaceae_unclassified_SGB94922 | Eubacteriaceae_unclassified |
| Eubacteriaceae_unclassified_SGB94922 | Eubacteriaceae_unclassified |
| GGB20149_SGB29430 | GGB20149 |
| GGB20149_SGB29430 | GGB20149 |
| GGB22635_SGB63107 | GGB22635 |
| GGB22635_SGB63107 | GGB22635 |
| GGB25041_SGB36960 | GGB25041 |
| GGB25041_SGB36960 | GGB25041 |
| GGB27876_SGB40310 | GGB27876 |
| GGB27876_SGB40310 | GGB27876 |
| GGB27878_SGB40312 | GGB27878 |
| GGB27878_SGB40312 | GGB27878 |
| GGB27918_SGB40356 | GGB27918 |
| GGB27918_SGB40356 | GGB27918 |
| GGB28382_SGB40962 | GGB28382 |
| GGB28382_SGB40962 | GGB28382 |
| GGB28399_SGB40980 | GGB28399 |
| GGB28399_SGB40980 | GGB28399 |
| GGB28411_SGB40993 | GGB28411 |
| GGB28411_SGB40993 | GGB28411 |
| GGB28415_SGB40997 | GGB28415 |
| GGB28415_SGB40997 | GGB28415 |
| GGB28430_SGB41013 | GGB28430 |
| GGB28430_SGB41013 | GGB28430 |
| GGB28439_SGB41022 | GGB28439 |
| GGB28439_SGB41022 | GGB28439 |
| GGB28778_SGB41431 | GGB28778 |
| GGB28778_SGB41431 | GGB28778 |
| GGB28784_SGB41437 | GGB28784 |
| GGB28784_SGB41437 | GGB28784 |
| GGB28792_SGB41445 | GGB28792 |
| GGB28792_SGB41445 | GGB28792 |
| GGB28798_SGB41451 | GGB28798 |
| GGB28798_SGB41451 | GGB28798 |
| GGB28802_SGB41455 | GGB28802 |
| GGB28802_SGB41455 | GGB28802 |
| GGB28818_SGB41473 | GGB28818 |
| GGB28818_SGB41473 | GGB28818 |
| GGB28828_SGB41484 | GGB28828 |
| GGB28828_SGB41484 | GGB28828 |

| | |
|---|---|
| GGB28851_SGB41518 | GGB28851 |
| GGB28851_SGB41518 | GGB28851 |
| GGB28859_SGB41528 | GGB28859 |
| GGB28859_SGB41528 | GGB28859 |
| GGB28864_SGB41535 | GGB28864 |
| GGB28864_SGB41535 | GGB28864 |
| GGB28869_SGB41543 | GGB28869 |
| GGB28869_SGB41543 | GGB28869 |
| GGB28883_SGB41564 | GGB28883 |
| GGB28883_SGB41564 | GGB28883 |
| GGB28892_SGB41573 | GGB28892 |
| GGB28892_SGB41573 | GGB28892 |
| GGB28893_SGB41574 | GGB28893 |
| GGB28893_SGB41574 | GGB28893 |
| GGB28898_SGB41580 | GGB28898 |
| GGB28898_SGB41580 | GGB28898 |
| GGB28904_SGB41597 | GGB28904 |
| GGB28904_SGB41597 | GGB28904 |
| GGB28916_SGB41612 | GGB28916 |
| GGB28916_SGB41612 | GGB28916 |
| GGB28924_SGB41621 | GGB28924 |
| GGB28924_SGB41621 | GGB28924 |
| GGB28926_SGB41624 | GGB28926 |
| GGB28926_SGB41624 | GGB28926 |
| GGB28927_SGB41625 | GGB28927 |
| GGB28927_SGB41625 | GGB28927 |
| GGB28934_SGB41635 | GGB28934 |
| GGB28934_SGB41635 | GGB28934 |
| GGB28946_SGB41652 | GGB28946 |
| GGB28946_SGB41652 | GGB28946 |
| GGB28949_SGB41655 | GGB28949 |
| GGB28949_SGB41655 | GGB28949 |
| GGB28949_SGB41656 | GGB28949 |
| GGB28949_SGB41656 | GGB28949 |
| GGB28950_SGB41657 | GGB28950 |
| GGB28950_SGB41657 | GGB28950 |
| GGB28951_SGB102295 | GGB28951 |
| GGB28951_SGB102295 | GGB28951 |
| GGB28951_SGB41658 | GGB28951 |
| GGB28951_SGB41658 | GGB28951 |
| GGB28954_SGB41662 | GGB28954 |
| GGB28954_SGB41662 | GGB28954 |
| GGB28956_SGB41665 | GGB28956 |
| GGB28956_SGB41665 | GGB28956 |
| GGB28960_SGB41669 | GGB28960 |
| GGB28960_SGB41669 | GGB28960 |

| | |
|---|---|
| GGB28967_SGB41678 | GGB28967 |
| GGB28967_SGB41678 | GGB28967 |
| GGB28991_SGB41705 | GGB28991 |
| GGB28991_SGB41705 | GGB28991 |
| GGB29002_SGB41718 | GGB29002 |
| GGB29002_SGB41718 | GGB29002 |
| GGB29003_SGB41719 | GGB29003 |
| GGB29003_SGB41719 | GGB29003 |
| GGB29011_SGB41731 | GGB29011 |
| GGB29011_SGB41731 | GGB29011 |
| GGB29531_SGB42317 | GGB29531 |
| GGB29531_SGB42317 | GGB29531 |
| GGB29685_SGB42494 | GGB29685 |
| GGB29685_SGB42494 | GGB29685 |
| GGB30141_SGB43066 | GGB30141 |
| GGB30141_SGB43066 | GGB30141 |
| GGB30286_SGB43248 | GGB30286 |
| GGB30286_SGB43248 | GGB30286 |
| GGB30303_SGB43268 | GGB30303 |
| GGB30303_SGB43268 | GGB30303 |
| GGB30413_SGB43452 | GGB30413 |
| GGB30413_SGB43452 | GGB30413 |
| GGB30454_SGB43514 | GGB30454 |
| GGB30454_SGB43514 | GGB30454 |
| GGB30455_SGB43519 | GGB30455 |
| GGB30455_SGB43519 | GGB30455 |
| GGB30461_SGB43527 | GGB30461 |
| GGB30461_SGB43527 | GGB30461 |
| GGB30461_SGB43530 | GGB30461 |
| GGB30461_SGB43530 | GGB30461 |
| GGB30463_SGB43537 | GGB30463 |
| GGB30463_SGB43537 | GGB30463 |
| GGB30473_SGB43557 | GGB30473 |
| GGB30473_SGB43557 | GGB30473 |
| GGB30475_SGB63182 | GGB30475 |
| GGB30475_SGB63182 | GGB30475 |
| GGB30861_SGB44083 | GGB30861 |
| GGB30861_SGB44083 | GGB30861 |
| GGB31312_SGB44628 | GGB31312 |
| GGB31312_SGB44628 | GGB31312 |
| GGB31438_SGB44768 | GGB31438 |
| GGB31438_SGB44768 | GGB31438 |
| GGB3171_SGB4185 | GGB3171 |
| GGB3171_SGB4185 | GGB3171 |
| GGB31823_SGB45199 | GGB31823 |
| GGB31823_SGB45199 | GGB31823 |

| | |
|---|---|
| GGB31853_SGB45233 | GGB31853 |
| GGB31853_SGB45233 | GGB31853 |
| GGB32371_SGB41694 | GGB32371 |
| GGB32371_SGB41694 | GGB32371 |
| GGB3793_SGB5158 | GGB3793 |
| GGB3793_SGB5158 | GGB3793 |
| GGB42598_SGB59794 | GGB42598 |
| GGB42598_SGB59794 | GGB42598 |
| GGB45656_SGB63370 | GGB45656 |
| GGB45656_SGB63370 | GGB45656 |
| GGB47127_SGB65054 | GGB47127 |
| GGB47127_SGB65054 | GGB47127 |
| GGB74395_SGB43521 | GGB74395 |
| GGB74395_SGB43521 | GGB74395 |
| GGB75053_SGB43494 | GGB75053 |
| GGB75053_SGB43494 | GGB75053 |
| GGB75109_SGB102238 | GGB75109 |
| GGB75109_SGB102238 | GGB75109 |
| GGB81440_SGB45230 | GGB81440 |
| GGB81440_SGB45230 | GGB81440 |
| Lachnospiraceae_bacterium | Lachnospiraceae_unclassified |
| Lachnospiraceae_bacterium | Lachnospiraceae_unclassified |
| Lachnospiraceae_bacterium_A2 | Lachnospiraceae_unclassified |
| Lachnospiraceae_bacterium_A2 | Lachnospiraceae_unclassified |
| Lachnospiraceae_bacterium_MD308 | Lachnospiraceae_unclassified |
| Lachnospiraceae_bacterium_MD308 | Lachnospiraceae_unclassified |
| Lachnospiraceae_bacterium_MD329 | Lachnospiraceae_unclassified |
| Lachnospiraceae_bacterium_MD329 | Lachnospiraceae_unclassified |
| Lachnospiraceae_unclassified_SGB41414 | Lachnospiraceae_unclassified |
| Lachnospiraceae_unclassified_SGB41414 | Lachnospiraceae_unclassified |
| Lachnospiraceae_unclassified_SGB41418 | Lachnospiraceae_unclassified |
| Lachnospiraceae_unclassified_SGB41418 | Lachnospiraceae_unclassified |
| Lachnospiraceae_unclassified_SGB41424 | Lachnospiraceae_unclassified |
| Lachnospiraceae_unclassified_SGB41424 | Lachnospiraceae_unclassified |
| Lachnospiraceae_unclassified_SGB41589 | Lachnospiraceae_unclassified |
| Lachnospiraceae_unclassified_SGB41589 | Lachnospiraceae_unclassified |
| Lactobacillus_johnsonii | Lactobacillus |
| Lactobacillus_johnsonii | Lactobacillus |
| Muribaculaceae_bacterium | Muribaculaceae_unclassified |
| Muribaculaceae_bacterium | Muribaculaceae_unclassified |
| Neglectibacter_sp_X4 | Neglectibacter |
| Neglectibacter_sp_X4 | Neglectibacter |
| Oscillospiraceae_bacterium | Oscillospiraceae_unclassified |
| Oscillospiraceae_bacterium | Oscillospiraceae_unclassified |
| Oscillospiraceae_unclassified_SGB43502 | Oscillospiraceae_unclassified |
| Oscillospiraceae_unclassified_SGB43502 | Oscillospiraceae_unclassified |

| | |
|---|---|
| Oscillospiraceae_unclassified_SGB43505 | Oscillospiraceae_unclassified |
| Oscillospiraceae_unclassified_SGB43505 | Oscillospiraceae_unclassified |
| Parasutterella_excrementihominis | Parasutterella |
| Parasutterella_excrementihominis | Parasutterella |
| richness | |
| richness | |
| Romboutsia_ilealis | Romboutsia |
| Romboutsia_ilealis | Romboutsia |
| Schaedlerella_arabinosiphila | Schaedlerella |
| Schaedlerella_arabinosiphila | Schaedlerella |
| shannon | |
| shannon | |
| Turicibacter_sp_1E2 | Turicibacter |
| Turicibacter_sp_1E2 | Turicibacter |
| Acetatifactor_muris | Acetatifactor |
| Acetatifactor_muris | Acetatifactor |
| Acetatifactor_SGB41546 | Acetatifactor |
| Acetatifactor_SGB41546 | Acetatifactor |
| Acutalibacter_muris | Acutalibacter |
| Acutalibacter_muris | Acutalibacter |
| Acutalibacter_sp_1XD8_36 | Acutalibacter |
| Acutalibacter_sp_1XD8_36 | Acutalibacter |
| Adlercreutzia_caecimuris | Adlercreutzia |
| Adlercreutzia_caecimuris | Adlercreutzia |
| Adlercreutzia_mucosicola | Adlercreutzia |
| Adlercreutzia_mucosicola | Adlercreutzia |
| Adlercreutzia_muris | Adlercreutzia |
| Adlercreutzia_muris | Adlercreutzia |
| Akkermansia_muciniphila | Akkermansia |
| Akkermansia_muciniphila | Akkermansia |
| Alistipes_sp_DSM_112343 | Alistipes |
| Alistipes_sp_DSM_112343 | Alistipes |
| Anaerotruncus_sp_1XD42_93 | Anaerotruncus |
| Anaerotruncus_sp_1XD42_93 | Anaerotruncus |
| Bacteria_unclassified_SGB102200 | Bacteria_unclassified |
| Bacteria_unclassified_SGB102200 | Bacteria_unclassified |
| Bacteria_unclassified_SGB41677 | Bacteria_unclassified |
| Bacteria_unclassified_SGB41677 | Bacteria_unclassified |
| Bacteria_unclassified_SGB43546 | Bacteria_unclassified |
| Bacteria_unclassified_SGB43546 | Bacteria_unclassified |
| bacterium_1XD42_54 | Bacteria_unclassified |
| bacterium_1XD42_54 | Bacteria_unclassified |
| bacterium_1XD42_76 | Bacteria_unclassified |
| bacterium_1XD42_76 | Bacteria_unclassified |
| bacterium_1xD8_48 | Bacteria_unclassified |
| bacterium_1xD8_48 | Bacteria_unclassified |

| | |
|---|---|
| Bacteroides_thetaiotaomicron | Bacteroides |
| Bacteroides_thetaiotaomicron | Bacteroides |
| berger_parker | |
| berger_parker | |
| Bifidobacterium_pseudolongum | Bifidobacterium |
| Bifidobacterium_pseudolongum | Bifidobacterium |
| Clostridia_bacterium | Clostridia_unclassified |
| Clostridia_bacterium | Clostridia_unclassified |
| Clostridiaceae_bacterium | Clostridiaceae_unclassified |
| Clostridiaceae_bacterium | Clostridiaceae_unclassified |
| Clostridiaceae_unclassified_SGB41663 | Clostridiaceae_unclassified |
| Clostridiaceae_unclassified_SGB41663 | Clostridiaceae_unclassified |
| Clostridiales_bacterium | Eubacteriales_unclassified |
| Clostridiales_bacterium | Eubacteriales_unclassified |
| Clostridium_cocleatum | Erysipelatoclostridium |
| Clostridium_cocleatum | Erysipelatoclostridium |
| Coriobacteriaceae_bacterium | Coriobacteriaceae_unclassified |
| Coriobacteriaceae_bacterium | Coriobacteriaceae_unclassified |
| Dorea_sp_5_2 | Dorea |
| Dorea_sp_5_2 | Dorea |
| Dubosiella_newyorkensis | Dubosiella |
| Dubosiella_newyorkensis | Dubosiella |
| Erysipelotrichales_bacterium | Erysipelotrichales_unclassified |
| Erysipelotrichales_bacterium | Erysipelotrichales_unclassified |
| Eubacteriaceae_bacterium | Eubacteriaceae_unclassified |
| Eubacteriaceae_bacterium | Eubacteriaceae_unclassified |
| Eubacteriaceae_unclassified_SGB94922 | Eubacteriaceae_unclassified |
| Eubacteriaceae_unclassified_SGB94922 | Eubacteriaceae_unclassified |
| GGB20149_SGB29430 | GGB20149 |
| GGB20149_SGB29430 | GGB20149 |
| GGB22635_SGB63107 | GGB22635 |
| GGB22635_SGB63107 | GGB22635 |
| GGB25041_SGB36960 | GGB25041 |
| GGB25041_SGB36960 | GGB25041 |
| GGB27876_SGB40310 | GGB27876 |
| GGB27876_SGB40310 | GGB27876 |
| GGB27878_SGB40312 | GGB27878 |
| GGB27878_SGB40312 | GGB27878 |
| GGB27918_SGB40356 | GGB27918 |
| GGB27918_SGB40356 | GGB27918 |
| GGB28382_SGB40962 | GGB28382 |
| GGB28382_SGB40962 | GGB28382 |
| GGB28399_SGB40980 | GGB28399 |
| GGB28399_SGB40980 | GGB28399 |
| GGB28411_SGB40993 | GGB28411 |
| GGB28411_SGB40993 | GGB28411 |

| | |
|---|---|
| GGB28415_SGB40997 | GGB28415 |
| GGB28415_SGB40997 | GGB28415 |
| GGB28430_SGB41013 | GGB28430 |
| GGB28430_SGB41013 | GGB28430 |
| GGB28439_SGB41022 | GGB28439 |
| GGB28439_SGB41022 | GGB28439 |
| GGB28778_SGB41431 | GGB28778 |
| GGB28778_SGB41431 | GGB28778 |
| GGB28784_SGB41437 | GGB28784 |
| GGB28784_SGB41437 | GGB28784 |
| GGB28792_SGB41445 | GGB28792 |
| GGB28792_SGB41445 | GGB28792 |
| GGB28798_SGB41451 | GGB28798 |
| GGB28798_SGB41451 | GGB28798 |
| GGB28802_SGB41455 | GGB28802 |
| GGB28802_SGB41455 | GGB28802 |
| GGB28818_SGB41473 | GGB28818 |
| GGB28818_SGB41473 | GGB28818 |
| GGB28828_SGB41484 | GGB28828 |
| GGB28828_SGB41484 | GGB28828 |
| GGB28851_SGB41518 | GGB28851 |
| GGB28851_SGB41518 | GGB28851 |
| GGB28859_SGB41528 | GGB28859 |
| GGB28859_SGB41528 | GGB28859 |
| GGB28864_SGB41535 | GGB28864 |
| GGB28864_SGB41535 | GGB28864 |
| GGB28869_SGB41543 | GGB28869 |
| GGB28869_SGB41543 | GGB28869 |
| GGB28883_SGB41564 | GGB28883 |
| GGB28883_SGB41564 | GGB28883 |
| GGB28892_SGB41573 | GGB28892 |
| GGB28892_SGB41573 | GGB28892 |
| GGB28893_SGB41574 | GGB28893 |
| GGB28893_SGB41574 | GGB28893 |
| GGB28898_SGB41580 | GGB28898 |
| GGB28898_SGB41580 | GGB28898 |
| GGB28904_SGB41597 | GGB28904 |
| GGB28904_SGB41597 | GGB28904 |
| GGB28916_SGB41612 | GGB28916 |
| GGB28916_SGB41612 | GGB28916 |
| GGB28924_SGB41621 | GGB28924 |
| GGB28924_SGB41621 | GGB28924 |
| GGB28926_SGB41624 | GGB28926 |
| GGB28926_SGB41624 | GGB28926 |
| GGB28927_SGB41625 | GGB28927 |
| GGB28927_SGB41625 | GGB28927 |

| | |
|---|---|
| GGB28934_SGB41635 | GGB28934 |
| GGB28934_SGB41635 | GGB28934 |
| GGB28946_SGB41652 | GGB28946 |
| GGB28946_SGB41652 | GGB28946 |
| GGB28949_SGB41655 | GGB28949 |
| GGB28949_SGB41655 | GGB28949 |
| GGB28949_SGB41656 | GGB28949 |
| GGB28949_SGB41656 | GGB28949 |
| GGB28950_SGB41657 | GGB28950 |
| GGB28950_SGB41657 | GGB28950 |
| GGB28951_SGB102295 | GGB28951 |
| GGB28951_SGB102295 | GGB28951 |
| GGB28951_SGB41658 | GGB28951 |
| GGB28951_SGB41658 | GGB28951 |
| GGB28954_SGB41662 | GGB28954 |
| GGB28954_SGB41662 | GGB28954 |
| GGB28956_SGB41665 | GGB28956 |
| GGB28956_SGB41665 | GGB28956 |
| GGB28960_SGB41669 | GGB28960 |
| GGB28960_SGB41669 | GGB28960 |
| GGB28967_SGB41678 | GGB28967 |
| GGB28967_SGB41678 | GGB28967 |
| GGB28991_SGB41705 | GGB28991 |
| GGB28991_SGB41705 | GGB28991 |
| GGB29002_SGB41718 | GGB29002 |
| GGB29002_SGB41718 | GGB29002 |
| GGB29003_SGB41719 | GGB29003 |
| GGB29003_SGB41719 | GGB29003 |
| GGB29011_SGB41731 | GGB29011 |
| GGB29011_SGB41731 | GGB29011 |
| GGB29531_SGB42317 | GGB29531 |
| GGB29531_SGB42317 | GGB29531 |
| GGB29685_SGB42494 | GGB29685 |
| GGB29685_SGB42494 | GGB29685 |
| GGB30141_SGB43066 | GGB30141 |
| GGB30141_SGB43066 | GGB30141 |
| GGB30286_SGB43248 | GGB30286 |
| GGB30286_SGB43248 | GGB30286 |
| GGB30303_SGB43268 | GGB30303 |
| GGB30303_SGB43268 | GGB30303 |
| GGB30413_SGB43452 | GGB30413 |
| GGB30413_SGB43452 | GGB30413 |
| GGB30454_SGB43514 | GGB30454 |
| GGB30454_SGB43514 | GGB30454 |
| GGB30455_SGB43519 | GGB30455 |
| GGB30455_SGB43519 | GGB30455 |

| | |
|---|---|
| GGB30461_SGB43527 | GGB30461 |
| GGB30461_SGB43527 | GGB30461 |
| GGB30461_SGB43530 | GGB30461 |
| GGB30461_SGB43530 | GGB30461 |
| GGB30463_SGB43537 | GGB30463 |
| GGB30463_SGB43537 | GGB30463 |
| GGB30473_SGB43557 | GGB30473 |
| GGB30473_SGB43557 | GGB30473 |
| GGB30475_SGB63182 | GGB30475 |
| GGB30475_SGB63182 | GGB30475 |
| GGB30861_SGB44083 | GGB30861 |
| GGB30861_SGB44083 | GGB30861 |
| GGB31312_SGB44628 | GGB31312 |
| GGB31312_SGB44628 | GGB31312 |
| GGB31438_SGB44768 | GGB31438 |
| GGB31438_SGB44768 | GGB31438 |
| GGB3171_SGB4185 | GGB3171 |
| GGB3171_SGB4185 | GGB3171 |
| GGB31823_SGB45199 | GGB31823 |
| GGB31823_SGB45199 | GGB31823 |
| GGB31853_SGB45233 | GGB31853 |
| GGB31853_SGB45233 | GGB31853 |
| GGB32371_SGB41694 | GGB32371 |
| GGB32371_SGB41694 | GGB32371 |
| GGB3793_SGB5158 | GGB3793 |
| GGB3793_SGB5158 | GGB3793 |
| GGB42598_SGB59794 | GGB42598 |
| GGB42598_SGB59794 | GGB42598 |
| GGB45656_SGB63370 | GGB45656 |
| GGB45656_SGB63370 | GGB45656 |
| GGB47127_SGB65054 | GGB47127 |
| GGB47127_SGB65054 | GGB47127 |
| GGB74395_SGB43521 | GGB74395 |
| GGB74395_SGB43521 | GGB74395 |
| GGB75053_SGB43494 | GGB75053 |
| GGB75053_SGB43494 | GGB75053 |
| GGB75109_SGB102238 | GGB75109 |
| GGB75109_SGB102238 | GGB75109 |
| GGB81440_SGB45230 | GGB81440 |
| GGB81440_SGB45230 | GGB81440 |
| Lachnospiraceae_bacterium | Lachnospiraceae_unclassified |
| Lachnospiraceae_bacterium | Lachnospiraceae_unclassified |
| Lachnospiraceae_bacterium_A2 | Lachnospiraceae_unclassified |
| Lachnospiraceae_bacterium_A2 | Lachnospiraceae_unclassified |
| Lachnospiraceae_bacterium_MD308 | Lachnospiraceae_unclassified |
| Lachnospiraceae_bacterium_MD308 | Lachnospiraceae_unclassified |

| | |
|---|---|
| Lachnospiraceae_bacterium_MD329 | Lachnospiraceae_unclassified |
| Lachnospiraceae_bacterium_MD329 | Lachnospiraceae_unclassified |
| Lachnospiraceae_unclassified_SGB41414 | Lachnospiraceae_unclassified |
| Lachnospiraceae_unclassified_SGB41414 | Lachnospiraceae_unclassified |
| Lachnospiraceae_unclassified_SGB41418 | Lachnospiraceae_unclassified |
| Lachnospiraceae_unclassified_SGB41418 | Lachnospiraceae_unclassified |
| Lachnospiraceae_unclassified_SGB41424 | Lachnospiraceae_unclassified |
| Lachnospiraceae_unclassified_SGB41424 | Lachnospiraceae_unclassified |
| Lachnospiraceae_unclassified_SGB41589 | Lachnospiraceae_unclassified |
| Lachnospiraceae_unclassified_SGB41589 | Lachnospiraceae_unclassified |
| Lactobacillus_johnsonii | Lactobacillus |
| Lactobacillus_johnsonii | Lactobacillus |
| Muribaculaceae_bacterium | Muribaculaceae_unclassified |
| Muribaculaceae_bacterium | Muribaculaceae_unclassified |
| Neglectibacter_sp_X4 | Neglectibacter |
| Neglectibacter_sp_X4 | Neglectibacter |
| Oscillospiraceae_bacterium | Oscillospiraceae_unclassified |
| Oscillospiraceae_bacterium | Oscillospiraceae_unclassified |
| Oscillospiraceae_unclassified_SGB43502 | Oscillospiraceae_unclassified |
| Oscillospiraceae_unclassified_SGB43502 | Oscillospiraceae_unclassified |
| Oscillospiraceae_unclassified_SGB43505 | Oscillospiraceae_unclassified |
| Oscillospiraceae_unclassified_SGB43505 | Oscillospiraceae_unclassified |
| Parasutterella_excrementihominis | Parasutterella |
| Parasutterella_excrementihominis | Parasutterella |
| richness | |
| richness | |
| Romboutsia_ilealis | Romboutsia |
| Romboutsia_ilealis | Romboutsia |
| Schaedlerella_arabinosiphila | Schaedlerella |
| Schaedlerella_arabinosiphila | Schaedlerella |
| shannon | |
| shannon | |
| Turicibacter_sp_1E2 | Turicibacter |
| Turicibacter_sp_1E2 | Turicibacter |
| Acetatifactor_muris | Acetatifactor |
| Acetatifactor_muris | Acetatifactor |
| Acetatifactor_SGB41546 | Acetatifactor |
| Acetatifactor_SGB41546 | Acetatifactor |
| Acutalibacter_muris | Acutalibacter |
| Acutalibacter_muris | Acutalibacter |
| Acutalibacter_sp_1XD8_36 | Acutalibacter |
| Acutalibacter_sp_1XD8_36 | Acutalibacter |
| Adlercreutzia_caecimuris | Adlercreutzia |
| Adlercreutzia_caecimuris | Adlercreutzia |
| Adlercreutzia_mucosicola | Adlercreutzia |
| Adlercreutzia_mucosicola | Adlercreutzia |

| | |
|---|---|
| Adlercreutzia_muris | Adlercreutzia |
| Adlercreutzia_muris | Adlercreutzia |
| Akkermansia_muciniphila | Akkermansia |
| Akkermansia_muciniphila | Akkermansia |
| Alistipes_sp_DSM_112343 | Alistipes |
| Alistipes_sp_DSM_112343 | Alistipes |
| Anaerotruncus_sp_1XD42_93 | Anaerotruncus |
| Anaerotruncus_sp_1XD42_93 | Anaerotruncus |
| Bacteria_unclassified_SGB102200 | Bacteria_unclassified |
| Bacteria_unclassified_SGB102200 | Bacteria_unclassified |
| Bacteria_unclassified_SGB41677 | Bacteria_unclassified |
| Bacteria_unclassified_SGB41677 | Bacteria_unclassified |
| Bacteria_unclassified_SGB43546 | Bacteria_unclassified |
| Bacteria_unclassified_SGB43546 | Bacteria_unclassified |
| bacterium_1XD42_54 | Bacteria_unclassified |
| bacterium_1XD42_54 | Bacteria_unclassified |
| bacterium_1XD42_76 | Bacteria_unclassified |
| bacterium_1XD42_76 | Bacteria_unclassified |
| bacterium_1xD8_48 | Bacteria_unclassified |
| bacterium_1xD8_48 | Bacteria_unclassified |
| Bacteroides_thetaiotaomicron | Bacteroides |
| Bacteroides_thetaiotaomicron | Bacteroides |
| berger_parker | |
| berger_parker | |
| Bifidobacterium_pseudolongum | Bifidobacterium |
| Bifidobacterium_pseudolongum | Bifidobacterium |
| Clostridia_bacterium | Clostridia_unclassified |
| Clostridia_bacterium | Clostridia_unclassified |
| Clostridiaceae_bacterium | Clostridiaceae_unclassified |
| Clostridiaceae_bacterium | Clostridiaceae_unclassified |
| Clostridiaceae_unclassified_SGB41663 | Clostridiaceae_unclassified |
| Clostridiaceae_unclassified_SGB41663 | Clostridiaceae_unclassified |
| Clostridiales_bacterium | Eubacteriales_unclassified |
| Clostridiales_bacterium | Eubacteriales_unclassified |
| Clostridium_cocleatum | Erysipelatoclostridium |
| Clostridium_cocleatum | Erysipelatoclostridium |
| Coriobacteriaceae_bacterium | Coriobacteriaceae_unclassified |
| Coriobacteriaceae_bacterium | Coriobacteriaceae_unclassified |
| Dorea_sp_5_2 | Dorea |
| Dorea_sp_5_2 | Dorea |
| Dubosiella_newyorkensis | Dubosiella |
| Dubosiella_newyorkensis | Dubosiella |
| Erysipelotrichales_bacterium | Erysipelotrichales_unclassified |
| Erysipelotrichales_bacterium | Erysipelotrichales_unclassified |
| Eubacteriaceae_bacterium | Eubacteriaceae_unclassified |
| Eubacteriaceae_bacterium | Eubacteriaceae_unclassified |

| | |
|---|---|
| Eubacteriaceae_unclassified_SGB94922 | Eubacteriaceae_unclassified |
| Eubacteriaceae_unclassified_SGB94922 | Eubacteriaceae_unclassified |
| GGB20149_SGB29430 | GGB20149 |
| GGB20149_SGB29430 | GGB20149 |
| GGB22635_SGB63107 | GGB22635 |
| GGB22635_SGB63107 | GGB22635 |
| GGB25041_SGB36960 | GGB25041 |
| GGB25041_SGB36960 | GGB25041 |
| GGB27876_SGB40310 | GGB27876 |
| GGB27876_SGB40310 | GGB27876 |
| GGB27878_SGB40312 | GGB27878 |
| GGB27878_SGB40312 | GGB27878 |
| GGB27918_SGB40356 | GGB27918 |
| GGB27918_SGB40356 | GGB27918 |
| GGB28382_SGB40962 | GGB28382 |
| GGB28382_SGB40962 | GGB28382 |
| GGB28399_SGB40980 | GGB28399 |
| GGB28399_SGB40980 | GGB28399 |
| GGB28411_SGB40993 | GGB28411 |
| GGB28411_SGB40993 | GGB28411 |
| GGB28415_SGB40997 | GGB28415 |
| GGB28415_SGB40997 | GGB28415 |
| GGB28430_SGB41013 | GGB28430 |
| GGB28430_SGB41013 | GGB28430 |
| GGB28439_SGB41022 | GGB28439 |
| GGB28439_SGB41022 | GGB28439 |
| GGB28778_SGB41431 | GGB28778 |
| GGB28778_SGB41431 | GGB28778 |
| GGB28784_SGB41437 | GGB28784 |
| GGB28784_SGB41437 | GGB28784 |
| GGB28792_SGB41445 | GGB28792 |
| GGB28792_SGB41445 | GGB28792 |
| GGB28798_SGB41451 | GGB28798 |
| GGB28798_SGB41451 | GGB28798 |
| GGB28802_SGB41455 | GGB28802 |
| GGB28802_SGB41455 | GGB28802 |
| GGB28818_SGB41473 | GGB28818 |
| GGB28818_SGB41473 | GGB28818 |
| GGB28828_SGB41484 | GGB28828 |
| GGB28828_SGB41484 | GGB28828 |
| GGB28851_SGB41518 | GGB28851 |
| GGB28851_SGB41518 | GGB28851 |
| GGB28859_SGB41528 | GGB28859 |
| GGB28859_SGB41528 | GGB28859 |
| GGB28864_SGB41535 | GGB28864 |
| GGB28864_SGB41535 | GGB28864 |

| | |
|---|---|
| GGB28869_SGB41543 | GGB28869 |
| GGB28869_SGB41543 | GGB28869 |
| GGB28883_SGB41564 | GGB28883 |
| GGB28883_SGB41564 | GGB28883 |
| GGB28892_SGB41573 | GGB28892 |
| GGB28892_SGB41573 | GGB28892 |
| GGB28893_SGB41574 | GGB28893 |
| GGB28893_SGB41574 | GGB28893 |
| GGB28898_SGB41580 | GGB28898 |
| GGB28898_SGB41580 | GGB28898 |
| GGB28904_SGB41597 | GGB28904 |
| GGB28904_SGB41597 | GGB28904 |
| GGB28916_SGB41612 | GGB28916 |
| GGB28916_SGB41612 | GGB28916 |
| GGB28924_SGB41621 | GGB28924 |
| GGB28924_SGB41621 | GGB28924 |
| GGB28926_SGB41624 | GGB28926 |
| GGB28926_SGB41624 | GGB28926 |
| GGB28927_SGB41625 | GGB28927 |
| GGB28927_SGB41625 | GGB28927 |
| GGB28934_SGB41635 | GGB28934 |
| GGB28934_SGB41635 | GGB28934 |
| GGB28946_SGB41652 | GGB28946 |
| GGB28946_SGB41652 | GGB28946 |
| GGB28949_SGB41655 | GGB28949 |
| GGB28949_SGB41655 | GGB28949 |
| GGB28949_SGB41656 | GGB28949 |
| GGB28949_SGB41656 | GGB28949 |
| GGB28950_SGB41657 | GGB28950 |
| GGB28950_SGB41657 | GGB28950 |
| GGB28951_SGB102295 | GGB28951 |
| GGB28951_SGB102295 | GGB28951 |
| GGB28951_SGB41658 | GGB28951 |
| GGB28951_SGB41658 | GGB28951 |
| GGB28954_SGB41662 | GGB28954 |
| GGB28954_SGB41662 | GGB28954 |
| GGB28956_SGB41665 | GGB28956 |
| GGB28956_SGB41665 | GGB28956 |
| GGB28960_SGB41669 | GGB28960 |
| GGB28960_SGB41669 | GGB28960 |
| GGB28967_SGB41678 | GGB28967 |
| GGB28967_SGB41678 | GGB28967 |
| GGB28991_SGB41705 | GGB28991 |
| GGB28991_SGB41705 | GGB28991 |
| GGB29002_SGB41718 | GGB29002 |
| GGB29002_SGB41718 | GGB29002 |

| | |
|---|---|
| GGB29003_SGB41719 | GGB29003 |
| GGB29003_SGB41719 | GGB29003 |
| GGB29011_SGB41731 | GGB29011 |
| GGB29011_SGB41731 | GGB29011 |
| GGB29531_SGB42317 | GGB29531 |
| GGB29531_SGB42317 | GGB29531 |
| GGB29685_SGB42494 | GGB29685 |
| GGB29685_SGB42494 | GGB29685 |
| GGB30141_SGB43066 | GGB30141 |
| GGB30141_SGB43066 | GGB30141 |
| GGB30286_SGB43248 | GGB30286 |
| GGB30286_SGB43248 | GGB30286 |
| GGB30303_SGB43268 | GGB30303 |
| GGB30303_SGB43268 | GGB30303 |
| GGB30413_SGB43452 | GGB30413 |
| GGB30413_SGB43452 | GGB30413 |
| GGB30454_SGB43514 | GGB30454 |
| GGB30454_SGB43514 | GGB30454 |
| GGB30455_SGB43519 | GGB30455 |
| GGB30455_SGB43519 | GGB30455 |
| GGB30461_SGB43527 | GGB30461 |
| GGB30461_SGB43527 | GGB30461 |
| GGB30461_SGB43530 | GGB30461 |
| GGB30461_SGB43530 | GGB30461 |
| GGB30463_SGB43537 | GGB30463 |
| GGB30463_SGB43537 | GGB30463 |
| GGB30473_SGB43557 | GGB30473 |
| GGB30473_SGB43557 | GGB30473 |
| GGB30475_SGB63182 | GGB30475 |
| GGB30475_SGB63182 | GGB30475 |
| GGB30861_SGB44083 | GGB30861 |
| GGB30861_SGB44083 | GGB30861 |
| GGB31312_SGB44628 | GGB31312 |
| GGB31312_SGB44628 | GGB31312 |
| GGB31438_SGB44768 | GGB31438 |
| GGB31438_SGB44768 | GGB31438 |
| GGB3171_SGB4185 | GGB3171 |
| GGB3171_SGB4185 | GGB3171 |
| GGB31823_SGB45199 | GGB31823 |
| GGB31823_SGB45199 | GGB31823 |
| GGB31853_SGB45233 | GGB31853 |
| GGB31853_SGB45233 | GGB31853 |
| GGB32371_SGB41694 | GGB32371 |
| GGB32371_SGB41694 | GGB32371 |
| GGB3793_SGB5158 | GGB3793 |
| GGB3793_SGB5158 | GGB3793 |

| | |
|---|---|
| GGB42598_SGB59794 | GGB42598 |
| GGB42598_SGB59794 | GGB42598 |
| GGB45656_SGB63370 | GGB45656 |
| GGB45656_SGB63370 | GGB45656 |
| GGB47127_SGB65054 | GGB47127 |
| GGB47127_SGB65054 | GGB47127 |
| GGB74395_SGB43521 | GGB74395 |
| GGB74395_SGB43521 | GGB74395 |
| GGB75053_SGB43494 | GGB75053 |
| GGB75053_SGB43494 | GGB75053 |
| GGB75109_SGB102238 | GGB75109 |
| GGB75109_SGB102238 | GGB75109 |
| GGB81440_SGB45230 | GGB81440 |
| GGB81440_SGB45230 | GGB81440 |
| Lachnospiraceae_bacterium | Lachnospiraceae_unclassified |
| Lachnospiraceae_bacterium | Lachnospiraceae_unclassified |
| Lachnospiraceae_bacterium_A2 | Lachnospiraceae_unclassified |
| Lachnospiraceae_bacterium_A2 | Lachnospiraceae_unclassified |
| Lachnospiraceae_bacterium_MD308 | Lachnospiraceae_unclassified |
| Lachnospiraceae_bacterium_MD308 | Lachnospiraceae_unclassified |
| Lachnospiraceae_bacterium_MD329 | Lachnospiraceae_unclassified |
| Lachnospiraceae_bacterium_MD329 | Lachnospiraceae_unclassified |
| Lachnospiraceae_unclassified_SGB41414 | Lachnospiraceae_unclassified |
| Lachnospiraceae_unclassified_SGB41414 | Lachnospiraceae_unclassified |
| Lachnospiraceae_unclassified_SGB41418 | Lachnospiraceae_unclassified |
| Lachnospiraceae_unclassified_SGB41418 | Lachnospiraceae_unclassified |
| Lachnospiraceae_unclassified_SGB41424 | Lachnospiraceae_unclassified |
| Lachnospiraceae_unclassified_SGB41424 | Lachnospiraceae_unclassified |
| Lachnospiraceae_unclassified_SGB41589 | Lachnospiraceae_unclassified |
| Lachnospiraceae_unclassified_SGB41589 | Lachnospiraceae_unclassified |
| Lactobacillus_johnsonii | Lactobacillus |
| Lactobacillus_johnsonii | Lactobacillus |
| Muribaculaceae_bacterium | Muribaculaceae_unclassified |
| Muribaculaceae_bacterium | Muribaculaceae_unclassified |
| Neglectibacter_sp_X4 | Neglectibacter |
| Neglectibacter_sp_X4 | Neglectibacter |
| Oscillospiraceae_bacterium | Oscillospiraceae_unclassified |
| Oscillospiraceae_bacterium | Oscillospiraceae_unclassified |
| Oscillospiraceae_unclassified_SGB43502 | Oscillospiraceae_unclassified |
| Oscillospiraceae_unclassified_SGB43502 | Oscillospiraceae_unclassified |
| Oscillospiraceae_unclassified_SGB43505 | Oscillospiraceae_unclassified |
| Oscillospiraceae_unclassified_SGB43505 | Oscillospiraceae_unclassified |
| Parasutterella_excrementihominis | Parasutterella |
| Parasutterella_excrementihominis | Parasutterella |
| richness | |
| richness | |

| | |
|---|---|
| Romboutsia_ilealis | Romboutsia |
| Romboutsia_ilealis | Romboutsia |
| Schaedlerella_arabinosiphila | Schaedlerella |
| Schaedlerella_arabinosiphila | Schaedlerella |
| shannon | |
| shannon | |
| Turicibacter_sp_1E2 | Turicibacter |
| Turicibacter_sp_1E2 | Turicibacter |
| Acetatifactor_muris | Acetatifactor |
| Acetatifactor_muris | Acetatifactor |
| Acetatifactor_SGB41546 | Acetatifactor |
| Acetatifactor_SGB41546 | Acetatifactor |
| Acutalibacter_muris | Acutalibacter |
| Acutalibacter_muris | Acutalibacter |
| Acutalibacter_sp_1XD8_36 | Acutalibacter |
| Acutalibacter_sp_1XD8_36 | Acutalibacter |
| Adlercreutzia_caecimuris | Adlercreutzia |
| Adlercreutzia_caecimuris | Adlercreutzia |
| Adlercreutzia_mucosicola | Adlercreutzia |
| Adlercreutzia_mucosicola | Adlercreutzia |
| Adlercreutzia_muris | Adlercreutzia |
| Adlercreutzia_muris | Adlercreutzia |
| Akkermansia_muciniphila | Akkermansia |
| Akkermansia_muciniphila | Akkermansia |
| Alistipes_sp_DSM_112343 | Alistipes |
| Alistipes_sp_DSM_112343 | Alistipes |
| Anaerotruncus_sp_1XD42_93 | Anaerotruncus |
| Anaerotruncus_sp_1XD42_93 | Anaerotruncus |
| Bacteria_unclassified_SGB102200 | Bacteria_unclassified |
| Bacteria_unclassified_SGB102200 | Bacteria_unclassified |
| Bacteria_unclassified_SGB41677 | Bacteria_unclassified |
| Bacteria_unclassified_SGB41677 | Bacteria_unclassified |
| Bacteria_unclassified_SGB43546 | Bacteria_unclassified |
| Bacteria_unclassified_SGB43546 | Bacteria_unclassified |
| bacterium_1XD42_54 | Bacteria_unclassified |
| bacterium_1XD42_54 | Bacteria_unclassified |
| bacterium_1XD42_76 | Bacteria_unclassified |
| bacterium_1XD42_76 | Bacteria_unclassified |
| bacterium_1xD8_48 | Bacteria_unclassified |
| bacterium_1xD8_48 | Bacteria_unclassified |
| Bacteroides_thetaiotaomicron | Bacteroides |
| Bacteroides_thetaiotaomicron | Bacteroides |
| berger_parker | |
| berger_parker | |
| Bifidobacterium_pseudolongum | Bifidobacterium |
| Bifidobacterium_pseudolongum | Bifidobacterium |

| | |
|---|---|
| Clostridia_bacterium | Clostridia_unclassified |
| Clostridia_bacterium | Clostridia_unclassified |
| Clostridiaceae_bacterium | Clostridiaceae_unclassified |
| Clostridiaceae_bacterium | Clostridiaceae_unclassified |
| Clostridiaceae_unclassified_SGB41663 | Clostridiaceae_unclassified |
| Clostridiaceae_unclassified_SGB41663 | Clostridiaceae_unclassified |
| Clostridiales_bacterium | Eubacteriales_unclassified |
| Clostridiales_bacterium | Eubacteriales_unclassified |
| Clostridium_cocleatum | Erysipelatoclostridium |
| Clostridium_cocleatum | Erysipelatoclostridium |
| Coriobacteriaceae_bacterium | Coriobacteriaceae_unclassified |
| Coriobacteriaceae_bacterium | Coriobacteriaceae_unclassified |
| Dorea_sp_5_2 | Dorea |
| Dorea_sp_5_2 | Dorea |
| Dubosiella_newyorkensis | Dubosiella |
| Dubosiella_newyorkensis | Dubosiella |
| Erysipelotrichales_bacterium | Erysipelotrichales_unclassified |
| Erysipelotrichales_bacterium | Erysipelotrichales_unclassified |
| Eubacteriaceae_bacterium | Eubacteriaceae_unclassified |
| Eubacteriaceae_bacterium | Eubacteriaceae_unclassified |
| Eubacteriaceae_unclassified_SGB94922 | Eubacteriaceae_unclassified |
| Eubacteriaceae_unclassified_SGB94922 | Eubacteriaceae_unclassified |
| GGB20149_SGB29430 | GGB20149 |
| GGB20149_SGB29430 | GGB20149 |
| GGB22635_SGB63107 | GGB22635 |
| GGB22635_SGB63107 | GGB22635 |
| GGB25041_SGB36960 | GGB25041 |
| GGB25041_SGB36960 | GGB25041 |
| GGB27876_SGB40310 | GGB27876 |
| GGB27876_SGB40310 | GGB27876 |
| GGB27878_SGB40312 | GGB27878 |
| GGB27878_SGB40312 | GGB27878 |
| GGB27918_SGB40356 | GGB27918 |
| GGB27918_SGB40356 | GGB27918 |
| GGB28382_SGB40962 | GGB28382 |
| GGB28382_SGB40962 | GGB28382 |
| GGB28399_SGB40980 | GGB28399 |
| GGB28399_SGB40980 | GGB28399 |
| GGB28411_SGB40993 | GGB28411 |
| GGB28411_SGB40993 | GGB28411 |
| GGB28415_SGB40997 | GGB28415 |
| GGB28415_SGB40997 | GGB28415 |
| GGB28430_SGB41013 | GGB28430 |
| GGB28430_SGB41013 | GGB28430 |
| GGB28439_SGB41022 | GGB28439 |
| GGB28439_SGB41022 | GGB28439 |

| | |
|---|---|
| GGB28778_SGB41431 | GGB28778 |
| GGB28778_SGB41431 | GGB28778 |
| GGB28784_SGB41437 | GGB28784 |
| GGB28784_SGB41437 | GGB28784 |
| GGB28792_SGB41445 | GGB28792 |
| GGB28792_SGB41445 | GGB28792 |
| GGB28798_SGB41451 | GGB28798 |
| GGB28798_SGB41451 | GGB28798 |
| GGB28802_SGB41455 | GGB28802 |
| GGB28802_SGB41455 | GGB28802 |
| GGB28818_SGB41473 | GGB28818 |
| GGB28818_SGB41473 | GGB28818 |
| GGB28828_SGB41484 | GGB28828 |
| GGB28828_SGB41484 | GGB28828 |
| GGB28851_SGB41518 | GGB28851 |
| GGB28851_SGB41518 | GGB28851 |
| GGB28859_SGB41528 | GGB28859 |
| GGB28859_SGB41528 | GGB28859 |
| GGB28864_SGB41535 | GGB28864 |
| GGB28864_SGB41535 | GGB28864 |
| GGB28869_SGB41543 | GGB28869 |
| GGB28869_SGB41543 | GGB28869 |
| GGB28883_SGB41564 | GGB28883 |
| GGB28883_SGB41564 | GGB28883 |
| GGB28892_SGB41573 | GGB28892 |
| GGB28892_SGB41573 | GGB28892 |
| GGB28893_SGB41574 | GGB28893 |
| GGB28893_SGB41574 | GGB28893 |
| GGB28898_SGB41580 | GGB28898 |
| GGB28898_SGB41580 | GGB28898 |
| GGB28904_SGB41597 | GGB28904 |
| GGB28904_SGB41597 | GGB28904 |
| GGB28916_SGB41612 | GGB28916 |
| GGB28916_SGB41612 | GGB28916 |
| GGB28924_SGB41621 | GGB28924 |
| GGB28924_SGB41621 | GGB28924 |
| GGB28926_SGB41624 | GGB28926 |
| GGB28926_SGB41624 | GGB28926 |
| GGB28927_SGB41625 | GGB28927 |
| GGB28927_SGB41625 | GGB28927 |
| GGB28934_SGB41635 | GGB28934 |
| GGB28934_SGB41635 | GGB28934 |
| GGB28946_SGB41652 | GGB28946 |
| GGB28946_SGB41652 | GGB28946 |
| GGB28949_SGB41655 | GGB28949 |
| GGB28949_SGB41655 | GGB28949 |

| | |
|---|---|
| GGB28949_SGB41656 | GGB28949 |
| GGB28949_SGB41656 | GGB28949 |
| GGB28950_SGB41657 | GGB28950 |
| GGB28950_SGB41657 | GGB28950 |
| GGB28951_SGB102295 | GGB28951 |
| GGB28951_SGB102295 | GGB28951 |
| GGB28951_SGB41658 | GGB28951 |
| GGB28951_SGB41658 | GGB28951 |
| GGB28954_SGB41662 | GGB28954 |
| GGB28954_SGB41662 | GGB28954 |
| GGB28956_SGB41665 | GGB28956 |
| GGB28956_SGB41665 | GGB28956 |
| GGB28960_SGB41669 | GGB28960 |
| GGB28960_SGB41669 | GGB28960 |
| GGB28967_SGB41678 | GGB28967 |
| GGB28967_SGB41678 | GGB28967 |
| GGB28991_SGB41705 | GGB28991 |
| GGB28991_SGB41705 | GGB28991 |
| GGB29002_SGB41718 | GGB29002 |
| GGB29002_SGB41718 | GGB29002 |
| GGB29003_SGB41719 | GGB29003 |
| GGB29003_SGB41719 | GGB29003 |
| GGB29011_SGB41731 | GGB29011 |
| GGB29011_SGB41731 | GGB29011 |
| GGB29531_SGB42317 | GGB29531 |
| GGB29531_SGB42317 | GGB29531 |
| GGB29685_SGB42494 | GGB29685 |
| GGB29685_SGB42494 | GGB29685 |
| GGB30141_SGB43066 | GGB30141 |
| GGB30141_SGB43066 | GGB30141 |
| GGB30286_SGB43248 | GGB30286 |
| GGB30286_SGB43248 | GGB30286 |
| GGB30303_SGB43268 | GGB30303 |
| GGB30303_SGB43268 | GGB30303 |
| GGB30413_SGB43452 | GGB30413 |
| GGB30413_SGB43452 | GGB30413 |
| GGB30454_SGB43514 | GGB30454 |
| GGB30454_SGB43514 | GGB30454 |
| GGB30455_SGB43519 | GGB30455 |
| GGB30455_SGB43519 | GGB30455 |
| GGB30461_SGB43527 | GGB30461 |
| GGB30461_SGB43527 | GGB30461 |
| GGB30461_SGB43530 | GGB30461 |
| GGB30461_SGB43530 | GGB30461 |
| GGB30463_SGB43537 | GGB30463 |
| GGB30463_SGB43537 | GGB30463 |

| | |
|---|---|
| GGB30473_SGB43557 | GGB30473 |
| GGB30473_SGB43557 | GGB30473 |
| GGB30475_SGB63182 | GGB30475 |
| GGB30475_SGB63182 | GGB30475 |
| GGB30861_SGB44083 | GGB30861 |
| GGB30861_SGB44083 | GGB30861 |
| GGB31312_SGB44628 | GGB31312 |
| GGB31312_SGB44628 | GGB31312 |
| GGB31438_SGB44768 | GGB31438 |
| GGB31438_SGB44768 | GGB31438 |
| GGB3171_SGB4185 | GGB3171 |
| GGB3171_SGB4185 | GGB3171 |
| GGB31823_SGB45199 | GGB31823 |
| GGB31823_SGB45199 | GGB31823 |
| GGB31853_SGB45233 | GGB31853 |
| GGB31853_SGB45233 | GGB31853 |
| GGB32371_SGB41694 | GGB32371 |
| GGB32371_SGB41694 | GGB32371 |
| GGB3793_SGB5158 | GGB3793 |
| GGB3793_SGB5158 | GGB3793 |
| GGB42598_SGB59794 | GGB42598 |
| GGB42598_SGB59794 | GGB42598 |
| GGB45656_SGB63370 | GGB45656 |
| GGB45656_SGB63370 | GGB45656 |
| GGB47127_SGB65054 | GGB47127 |
| GGB47127_SGB65054 | GGB47127 |
| GGB74395_SGB43521 | GGB74395 |
| GGB74395_SGB43521 | GGB74395 |
| GGB75053_SGB43494 | GGB75053 |
| GGB75053_SGB43494 | GGB75053 |
| GGB75109_SGB102238 | GGB75109 |
| GGB75109_SGB102238 | GGB75109 |
| GGB81440_SGB45230 | GGB81440 |
| GGB81440_SGB45230 | GGB81440 |
| Lachnospiraceae_bacterium | Lachnospiraceae_unclassified |
| Lachnospiraceae_bacterium | Lachnospiraceae_unclassified |
| Lachnospiraceae_bacterium_A2 | Lachnospiraceae_unclassified |
| Lachnospiraceae_bacterium_A2 | Lachnospiraceae_unclassified |
| Lachnospiraceae_bacterium_MD308 | Lachnospiraceae_unclassified |
| Lachnospiraceae_bacterium_MD308 | Lachnospiraceae_unclassified |
| Lachnospiraceae_bacterium_MD329 | Lachnospiraceae_unclassified |
| Lachnospiraceae_bacterium_MD329 | Lachnospiraceae_unclassified |
| Lachnospiraceae_unclassified_SGB41414 | Lachnospiraceae_unclassified |
| Lachnospiraceae_unclassified_SGB41414 | Lachnospiraceae_unclassified |
| Lachnospiraceae_unclassified_SGB41418 | Lachnospiraceae_unclassified |
| Lachnospiraceae_unclassified_SGB41418 | Lachnospiraceae_unclassified |

| | |
|---|---|
| Lachnospiraceae_unclassified_SGB41424 | Lachnospiraceae_unclassified |
| Lachnospiraceae_unclassified_SGB41424 | Lachnospiraceae_unclassified |
| Lachnospiraceae_unclassified_SGB41589 | Lachnospiraceae_unclassified |
| Lachnospiraceae_unclassified_SGB41589 | Lachnospiraceae_unclassified |
| Lactobacillus_johnsonii | Lactobacillus |
| Lactobacillus_johnsonii | Lactobacillus |
| Muribaculaceae_bacterium | Muribaculaceae_unclassified |
| Muribaculaceae_bacterium | Muribaculaceae_unclassified |
| Neglectibacter_sp_X4 | Neglectibacter |
| Neglectibacter_sp_X4 | Neglectibacter |
| Oscillospiraceae_bacterium | Oscillospiraceae_unclassified |
| Oscillospiraceae_bacterium | Oscillospiraceae_unclassified |
| Oscillospiraceae_unclassified_SGB43502 | Oscillospiraceae_unclassified |
| Oscillospiraceae_unclassified_SGB43502 | Oscillospiraceae_unclassified |
| Oscillospiraceae_unclassified_SGB43505 | Oscillospiraceae_unclassified |
| Oscillospiraceae_unclassified_SGB43505 | Oscillospiraceae_unclassified |
| Parasutterella_excrementihominis | Parasutterella |
| Parasutterella_excrementihominis | Parasutterella |
| richness | |
| richness | |
| Romboutsia_ilealis | Romboutsia |
| Romboutsia_ilealis | Romboutsia |
| Schaedlerella_arabinosiphila | Schaedlerella |
| Schaedlerella_arabinosiphila | Schaedlerella |
| shannon | |
| shannon | |
| Turicibacter_sp_1E2 | Turicibacter |
| Turicibacter_sp_1E2 | Turicibacter |
| Acetatifactor_muris | Acetatifactor |
| Acetatifactor_muris | Acetatifactor |
| Acetatifactor_SGB41546 | Acetatifactor |
| Acetatifactor_SGB41546 | Acetatifactor |
| Acutalibacter_muris | Acutalibacter |
| Acutalibacter_muris | Acutalibacter |
| Acutalibacter_sp_1XD8_36 | Acutalibacter |
| Acutalibacter_sp_1XD8_36 | Acutalibacter |
| Adlercreutzia_caecimuris | Adlercreutzia |
| Adlercreutzia_caecimuris | Adlercreutzia |
| Adlercreutzia_mucosicola | Adlercreutzia |
| Adlercreutzia_mucosicola | Adlercreutzia |
| Adlercreutzia_muris | Adlercreutzia |
| Adlercreutzia_muris | Adlercreutzia |
| Akkermansia_muciniphila | Akkermansia |
| Akkermansia_muciniphila | Akkermansia |
| Alistipes_sp_DSM_112343 | Alistipes |
| Alistipes_sp_DSM_112343 | Alistipes |

| | |
|---|---|
| Anaerotruncus_sp_1XD42_93 | Anaerotruncus |
| Anaerotruncus_sp_1XD42_93 | Anaerotruncus |
| Bacteria_unclassified_SGB102200 | Bacteria_unclassified |
| Bacteria_unclassified_SGB102200 | Bacteria_unclassified |
| Bacteria_unclassified_SGB41677 | Bacteria_unclassified |
| Bacteria_unclassified_SGB41677 | Bacteria_unclassified |
| Bacteria_unclassified_SGB43546 | Bacteria_unclassified |
| Bacteria_unclassified_SGB43546 | Bacteria_unclassified |
| bacterium_1XD42_54 | Bacteria_unclassified |
| bacterium_1XD42_54 | Bacteria_unclassified |
| bacterium_1XD42_76 | Bacteria_unclassified |
| bacterium_1XD42_76 | Bacteria_unclassified |
| bacterium_1xD8_48 | Bacteria_unclassified |
| bacterium_1xD8_48 | Bacteria_unclassified |
| Bacteroides_thetaiotaomicron | Bacteroides |
| Bacteroides_thetaiotaomicron | Bacteroides |
| berger_parker | |
| berger_parker | |
| Bifidobacterium_pseudolongum | Bifidobacterium |
| Bifidobacterium_pseudolongum | Bifidobacterium |
| Clostridia_bacterium | Clostridia_unclassified |
| Clostridia_bacterium | Clostridia_unclassified |
| Clostridiaceae_bacterium | Clostridiaceae_unclassified |
| Clostridiaceae_bacterium | Clostridiaceae_unclassified |
| Clostridiaceae_unclassified_SGB41663 | Clostridiaceae_unclassified |
| Clostridiaceae_unclassified_SGB41663 | Clostridiaceae_unclassified |
| Clostridiales_bacterium | Eubacteriales_unclassified |
| Clostridiales_bacterium | Eubacteriales_unclassified |
| Clostridium_cocleatum | Erysipelatoclostridium |
| Clostridium_cocleatum | Erysipelatoclostridium |
| Coriobacteriaceae_bacterium | Coriobacteriaceae_unclassified |
| Coriobacteriaceae_bacterium | Coriobacteriaceae_unclassified |
| Dorea_sp_5_2 | Dorea |
| Dorea_sp_5_2 | Dorea |
| Dubosiella_newyorkensis | Dubosiella |
| Dubosiella_newyorkensis | Dubosiella |
| Erysipelotrichales_bacterium | Erysipelotrichales_unclassified |
| Erysipelotrichales_bacterium | Erysipelotrichales_unclassified |
| Eubacteriaceae_bacterium | Eubacteriaceae_unclassified |
| Eubacteriaceae_bacterium | Eubacteriaceae_unclassified |
| Eubacteriaceae_unclassified_SGB94922 | Eubacteriaceae_unclassified |
| Eubacteriaceae_unclassified_SGB94922 | Eubacteriaceae_unclassified |
| GGB20149_SGB29430 | GGB20149 |
| GGB20149_SGB29430 | GGB20149 |
| GGB22635_SGB63107 | GGB22635 |
| GGB22635_SGB63107 | GGB22635 |

| | |
|---|---|
| GGB25041_SGB36960 | GGB25041 |
| GGB25041_SGB36960 | GGB25041 |
| GGB27876_SGB40310 | GGB27876 |
| GGB27876_SGB40310 | GGB27876 |
| GGB27878_SGB40312 | GGB27878 |
| GGB27878_SGB40312 | GGB27878 |
| GGB27918_SGB40356 | GGB27918 |
| GGB27918_SGB40356 | GGB27918 |
| GGB28382_SGB40962 | GGB28382 |
| GGB28382_SGB40962 | GGB28382 |
| GGB28399_SGB40980 | GGB28399 |
| GGB28399_SGB40980 | GGB28399 |
| GGB28411_SGB40993 | GGB28411 |
| GGB28411_SGB40993 | GGB28411 |
| GGB28415_SGB40997 | GGB28415 |
| GGB28415_SGB40997 | GGB28415 |
| GGB28430_SGB41013 | GGB28430 |
| GGB28430_SGB41013 | GGB28430 |
| GGB28439_SGB41022 | GGB28439 |
| GGB28439_SGB41022 | GGB28439 |
| GGB28778_SGB41431 | GGB28778 |
| GGB28778_SGB41431 | GGB28778 |
| GGB28784_SGB41437 | GGB28784 |
| GGB28784_SGB41437 | GGB28784 |
| GGB28792_SGB41445 | GGB28792 |
| GGB28792_SGB41445 | GGB28792 |
| GGB28798_SGB41451 | GGB28798 |
| GGB28798_SGB41451 | GGB28798 |
| GGB28802_SGB41455 | GGB28802 |
| GGB28802_SGB41455 | GGB28802 |
| GGB28818_SGB41473 | GGB28818 |
| GGB28818_SGB41473 | GGB28818 |
| GGB28828_SGB41484 | GGB28828 |
| GGB28828_SGB41484 | GGB28828 |
| GGB28851_SGB41518 | GGB28851 |
| GGB28851_SGB41518 | GGB28851 |
| GGB28859_SGB41528 | GGB28859 |
| GGB28859_SGB41528 | GGB28859 |
| GGB28864_SGB41535 | GGB28864 |
| GGB28864_SGB41535 | GGB28864 |
| GGB28869_SGB41543 | GGB28869 |
| GGB28869_SGB41543 | GGB28869 |
| GGB28883_SGB41564 | GGB28883 |
| GGB28883_SGB41564 | GGB28883 |
| GGB28892_SGB41573 | GGB28892 |
| GGB28892_SGB41573 | GGB28892 |

| | |
|---|---|
| GGB28893_SGB41574 | GGB28893 |
| GGB28893_SGB41574 | GGB28893 |
| GGB28898_SGB41580 | GGB28898 |
| GGB28898_SGB41580 | GGB28898 |
| GGB28904_SGB41597 | GGB28904 |
| GGB28904_SGB41597 | GGB28904 |
| GGB28916_SGB41612 | GGB28916 |
| GGB28916_SGB41612 | GGB28916 |
| GGB28924_SGB41621 | GGB28924 |
| GGB28924_SGB41621 | GGB28924 |
| GGB28926_SGB41624 | GGB28926 |
| GGB28926_SGB41624 | GGB28926 |
| GGB28927_SGB41625 | GGB28927 |
| GGB28927_SGB41625 | GGB28927 |
| GGB28934_SGB41635 | GGB28934 |
| GGB28934_SGB41635 | GGB28934 |
| GGB28946_SGB41652 | GGB28946 |
| GGB28946_SGB41652 | GGB28946 |
| GGB28949_SGB41655 | GGB28949 |
| GGB28949_SGB41655 | GGB28949 |
| GGB28949_SGB41656 | GGB28949 |
| GGB28949_SGB41656 | GGB28949 |
| GGB28950_SGB41657 | GGB28950 |
| GGB28950_SGB41657 | GGB28950 |
| GGB28951_SGB102295 | GGB28951 |
| GGB28951_SGB102295 | GGB28951 |
| GGB28951_SGB41658 | GGB28951 |
| GGB28951_SGB41658 | GGB28951 |
| GGB28954_SGB41662 | GGB28954 |
| GGB28954_SGB41662 | GGB28954 |
| GGB28956_SGB41665 | GGB28956 |
| GGB28956_SGB41665 | GGB28956 |
| GGB28960_SGB41669 | GGB28960 |
| GGB28960_SGB41669 | GGB28960 |
| GGB28967_SGB41678 | GGB28967 |
| GGB28967_SGB41678 | GGB28967 |
| GGB28991_SGB41705 | GGB28991 |
| GGB28991_SGB41705 | GGB28991 |
| GGB29002_SGB41718 | GGB29002 |
| GGB29002_SGB41718 | GGB29002 |
| GGB29003_SGB41719 | GGB29003 |
| GGB29003_SGB41719 | GGB29003 |
| GGB29011_SGB41731 | GGB29011 |
| GGB29011_SGB41731 | GGB29011 |
| GGB29531_SGB42317 | GGB29531 |
| GGB29531_SGB42317 | GGB29531 |

| | |
|---|---|
| GGB29685_SGB42494 | GGB29685 |
| GGB29685_SGB42494 | GGB29685 |
| GGB30141_SGB43066 | GGB30141 |
| GGB30141_SGB43066 | GGB30141 |
| GGB30286_SGB43248 | GGB30286 |
| GGB30286_SGB43248 | GGB30286 |
| GGB30303_SGB43268 | GGB30303 |
| GGB30303_SGB43268 | GGB30303 |
| GGB30413_SGB43452 | GGB30413 |
| GGB30413_SGB43452 | GGB30413 |
| GGB30454_SGB43514 | GGB30454 |
| GGB30454_SGB43514 | GGB30454 |
| GGB30455_SGB43519 | GGB30455 |
| GGB30455_SGB43519 | GGB30455 |
| GGB30461_SGB43527 | GGB30461 |
| GGB30461_SGB43527 | GGB30461 |
| GGB30461_SGB43530 | GGB30461 |
| GGB30461_SGB43530 | GGB30461 |
| GGB30463_SGB43537 | GGB30463 |
| GGB30463_SGB43537 | GGB30463 |
| GGB30473_SGB43557 | GGB30473 |
| GGB30473_SGB43557 | GGB30473 |
| GGB30475_SGB63182 | GGB30475 |
| GGB30475_SGB63182 | GGB30475 |
| GGB30861_SGB44083 | GGB30861 |
| GGB30861_SGB44083 | GGB30861 |
| GGB31312_SGB44628 | GGB31312 |
| GGB31312_SGB44628 | GGB31312 |
| GGB31438_SGB44768 | GGB31438 |
| GGB31438_SGB44768 | GGB31438 |
| GGB3171_SGB4185 | GGB3171 |
| GGB3171_SGB4185 | GGB3171 |
| GGB31823_SGB45199 | GGB31823 |
| GGB31823_SGB45199 | GGB31823 |
| GGB31853_SGB45233 | GGB31853 |
| GGB31853_SGB45233 | GGB31853 |
| GGB32371_SGB41694 | GGB32371 |
| GGB32371_SGB41694 | GGB32371 |
| GGB3793_SGB5158 | GGB3793 |
| GGB3793_SGB5158 | GGB3793 |
| GGB42598_SGB59794 | GGB42598 |
| GGB42598_SGB59794 | GGB42598 |
| GGB45656_SGB63370 | GGB45656 |
| GGB45656_SGB63370 | GGB45656 |
| GGB47127_SGB65054 | GGB47127 |
| GGB47127_SGB65054 | GGB47127 |

| | |
|---|---|
| GGB74395_SGB43521 | GGB74395 |
| GGB74395_SGB43521 | GGB74395 |
| GGB75053_SGB43494 | GGB75053 |
| GGB75053_SGB43494 | GGB75053 |
| GGB75109_SGB102238 | GGB75109 |
| GGB75109_SGB102238 | GGB75109 |
| GGB81440_SGB45230 | GGB81440 |
| GGB81440_SGB45230 | GGB81440 |
| Lachnospiraceae_bacterium | Lachnospiraceae_unclassified |
| Lachnospiraceae_bacterium | Lachnospiraceae_unclassified |
| Lachnospiraceae_bacterium_A2 | Lachnospiraceae_unclassified |
| Lachnospiraceae_bacterium_A2 | Lachnospiraceae_unclassified |
| Lachnospiraceae_bacterium_MD308 | Lachnospiraceae_unclassified |
| Lachnospiraceae_bacterium_MD308 | Lachnospiraceae_unclassified |
| Lachnospiraceae_bacterium_MD329 | Lachnospiraceae_unclassified |
| Lachnospiraceae_bacterium_MD329 | Lachnospiraceae_unclassified |
| Lachnospiraceae_unclassified_SGB41414 | Lachnospiraceae_unclassified |
| Lachnospiraceae_unclassified_SGB41414 | Lachnospiraceae_unclassified |
| Lachnospiraceae_unclassified_SGB41418 | Lachnospiraceae_unclassified |
| Lachnospiraceae_unclassified_SGB41418 | Lachnospiraceae_unclassified |
| Lachnospiraceae_unclassified_SGB41424 | Lachnospiraceae_unclassified |
| Lachnospiraceae_unclassified_SGB41424 | Lachnospiraceae_unclassified |
| Lachnospiraceae_unclassified_SGB41589 | Lachnospiraceae_unclassified |
| Lachnospiraceae_unclassified_SGB41589 | Lachnospiraceae_unclassified |
| Lactobacillus_johnsonii | Lactobacillus |
| Lactobacillus_johnsonii | Lactobacillus |
| Muribaculaceae_bacterium | Muribaculaceae_unclassified |
| Muribaculaceae_bacterium | Muribaculaceae_unclassified |
| Neglectibacter_sp_X4 | Neglectibacter |
| Neglectibacter_sp_X4 | Neglectibacter |
| Oscillospiraceae_bacterium | Oscillospiraceae_unclassified |
| Oscillospiraceae_bacterium | Oscillospiraceae_unclassified |
| Oscillospiraceae_unclassified_SGB43502 | Oscillospiraceae_unclassified |
| Oscillospiraceae_unclassified_SGB43502 | Oscillospiraceae_unclassified |
| Oscillospiraceae_unclassified_SGB43505 | Oscillospiraceae_unclassified |
| Oscillospiraceae_unclassified_SGB43505 | Oscillospiraceae_unclassified |
| Parasutterella_excrementihominis | Parasutterella |
| Parasutterella_excrementihominis | Parasutterella |
| richness | |
| richness | |
| Romboutsia_ilealis | Romboutsia |
| Romboutsia_ilealis | Romboutsia |
| Schaedlerella_arabinosiphila | Schaedlerella |
| Schaedlerella_arabinosiphila | Schaedlerella |
| shannon | |
| shannon | |

| | |
|---|---|
| Turicibacter_sp_1E2 | Turicibacter |
| Turicibacter_sp_1E2 | Turicibacter |
| Acetatifactor_muris | Acetatifactor |
| Acetatifactor_muris | Acetatifactor |
| Acetatifactor_SGB41546 | Acetatifactor |
| Acetatifactor_SGB41546 | Acetatifactor |
| Acutalibacter_muris | Acutalibacter |
| Acutalibacter_muris | Acutalibacter |
| Acutalibacter_sp_1XD8_36 | Acutalibacter |
| Acutalibacter_sp_1XD8_36 | Acutalibacter |
| Adlercreutzia_caecimuris | Adlercreutzia |
| Adlercreutzia_caecimuris | Adlercreutzia |
| Adlercreutzia_mucosicola | Adlercreutzia |
| Adlercreutzia_mucosicola | Adlercreutzia |
| Adlercreutzia_muris | Adlercreutzia |
| Adlercreutzia_muris | Adlercreutzia |
| Akkermansia_muciniphila | Akkermansia |
| Akkermansia_muciniphila | Akkermansia |
| Alistipes_sp_DSM_112343 | Alistipes |
| Alistipes_sp_DSM_112343 | Alistipes |
| Anaerotruncus_sp_1XD42_93 | Anaerotruncus |
| Anaerotruncus_sp_1XD42_93 | Anaerotruncus |
| Bacteria_unclassified_SGB102200 | Bacteria_unclassified |
| Bacteria_unclassified_SGB102200 | Bacteria_unclassified |
| Bacteria_unclassified_SGB41677 | Bacteria_unclassified |
| Bacteria_unclassified_SGB41677 | Bacteria_unclassified |
| Bacteria_unclassified_SGB43546 | Bacteria_unclassified |
| Bacteria_unclassified_SGB43546 | Bacteria_unclassified |
| bacterium_1XD42_54 | Bacteria_unclassified |
| bacterium_1XD42_54 | Bacteria_unclassified |
| bacterium_1XD42_76 | Bacteria_unclassified |
| bacterium_1XD42_76 | Bacteria_unclassified |
| bacterium_1xD8_48 | Bacteria_unclassified |
| bacterium_1xD8_48 | Bacteria_unclassified |
| Bacteroides_thetaiotaomicron | Bacteroides |
| Bacteroides_thetaiotaomicron | Bacteroides |
| berger_parker | |
| berger_parker | |
| Bifidobacterium_pseudolongum | Bifidobacterium |
| Bifidobacterium_pseudolongum | Bifidobacterium |
| Clostridia_bacterium | Clostridia_unclassified |
| Clostridia_bacterium | Clostridia_unclassified |
| Clostridiaceae_bacterium | Clostridiaceae_unclassified |
| Clostridiaceae_bacterium | Clostridiaceae_unclassified |
| Clostridiaceae_unclassified_SGB41663 | Clostridiaceae_unclassified |
| Clostridiaceae_unclassified_SGB41663 | Clostridiaceae_unclassified |

| | |
|---|---|
| Clostridiales_bacterium | Eubacteriales_unclassified |
| Clostridiales_bacterium | Eubacteriales_unclassified |
| Clostridium_cocleatum | Erysipelatoclostridium |
| Clostridium_cocleatum | Erysipelatoclostridium |
| Coriobacteriaceae_bacterium | Coriobacteriaceae_unclassified |
| Coriobacteriaceae_bacterium | Coriobacteriaceae_unclassified |
| Dorea_sp_5_2 | Dorea |
| Dorea_sp_5_2 | Dorea |
| Dubosiella_newyorkensis | Dubosiella |
| Dubosiella_newyorkensis | Dubosiella |
| Erysipelotrichales_bacterium | Erysipelotrichales_unclassified |
| Erysipelotrichales_bacterium | Erysipelotrichales_unclassified |
| Eubacteriaceae_bacterium | Eubacteriaceae_unclassified |
| Eubacteriaceae_bacterium | Eubacteriaceae_unclassified |
| Eubacteriaceae_unclassified_SGB94922 | Eubacteriaceae_unclassified |
| Eubacteriaceae_unclassified_SGB94922 | Eubacteriaceae_unclassified |
| GGB20149_SGB29430 | GGB20149 |
| GGB20149_SGB29430 | GGB20149 |
| GGB22635_SGB63107 | GGB22635 |
| GGB22635_SGB63107 | GGB22635 |
| GGB25041_SGB36960 | GGB25041 |
| GGB25041_SGB36960 | GGB25041 |
| GGB27876_SGB40310 | GGB27876 |
| GGB27876_SGB40310 | GGB27876 |
| GGB27878_SGB40312 | GGB27878 |
| GGB27878_SGB40312 | GGB27878 |
| GGB27918_SGB40356 | GGB27918 |
| GGB27918_SGB40356 | GGB27918 |
| GGB28382_SGB40962 | GGB28382 |
| GGB28382_SGB40962 | GGB28382 |
| GGB28399_SGB40980 | GGB28399 |
| GGB28399_SGB40980 | GGB28399 |
| GGB28411_SGB40993 | GGB28411 |
| GGB28411_SGB40993 | GGB28411 |
| GGB28415_SGB40997 | GGB28415 |
| GGB28415_SGB40997 | GGB28415 |
| GGB28430_SGB41013 | GGB28430 |
| GGB28430_SGB41013 | GGB28430 |
| GGB28439_SGB41022 | GGB28439 |
| GGB28439_SGB41022 | GGB28439 |
| GGB28778_SGB41431 | GGB28778 |
| GGB28778_SGB41431 | GGB28778 |
| GGB28784_SGB41437 | GGB28784 |
| GGB28784_SGB41437 | GGB28784 |
| GGB28792_SGB41445 | GGB28792 |
| GGB28792_SGB41445 | GGB28792 |

| | |
|---|---|
| GGB28798_SGB41451 | GGB28798 |
| GGB28798_SGB41451 | GGB28798 |
| GGB28802_SGB41455 | GGB28802 |
| GGB28802_SGB41455 | GGB28802 |
| GGB28818_SGB41473 | GGB28818 |
| GGB28818_SGB41473 | GGB28818 |
| GGB28828_SGB41484 | GGB28828 |
| GGB28828_SGB41484 | GGB28828 |
| GGB28851_SGB41518 | GGB28851 |
| GGB28851_SGB41518 | GGB28851 |
| GGB28859_SGB41528 | GGB28859 |
| GGB28859_SGB41528 | GGB28859 |
| GGB28864_SGB41535 | GGB28864 |
| GGB28864_SGB41535 | GGB28864 |
| GGB28869_SGB41543 | GGB28869 |
| GGB28869_SGB41543 | GGB28869 |
| GGB28883_SGB41564 | GGB28883 |
| GGB28883_SGB41564 | GGB28883 |
| GGB28892_SGB41573 | GGB28892 |
| GGB28892_SGB41573 | GGB28892 |
| GGB28893_SGB41574 | GGB28893 |
| GGB28893_SGB41574 | GGB28893 |
| GGB28898_SGB41580 | GGB28898 |
| GGB28898_SGB41580 | GGB28898 |
| GGB28904_SGB41597 | GGB28904 |
| GGB28904_SGB41597 | GGB28904 |
| GGB28916_SGB41612 | GGB28916 |
| GGB28916_SGB41612 | GGB28916 |
| GGB28924_SGB41621 | GGB28924 |
| GGB28924_SGB41621 | GGB28924 |
| GGB28926_SGB41624 | GGB28926 |
| GGB28926_SGB41624 | GGB28926 |
| GGB28927_SGB41625 | GGB28927 |
| GGB28927_SGB41625 | GGB28927 |
| GGB28934_SGB41635 | GGB28934 |
| GGB28934_SGB41635 | GGB28934 |
| GGB28946_SGB41652 | GGB28946 |
| GGB28946_SGB41652 | GGB28946 |
| GGB28949_SGB41655 | GGB28949 |
| GGB28949_SGB41655 | GGB28949 |
| GGB28949_SGB41656 | GGB28949 |
| GGB28949_SGB41656 | GGB28949 |
| GGB28950_SGB41657 | GGB28950 |
| GGB28950_SGB41657 | GGB28950 |
| GGB28951_SGB102295 | GGB28951 |
| GGB28951_SGB102295 | GGB28951 |

| | |
|---|---|
| GGB28951_SGB41658 | GGB28951 |
| GGB28951_SGB41658 | GGB28951 |
| GGB28954_SGB41662 | GGB28954 |
| GGB28954_SGB41662 | GGB28954 |
| GGB28956_SGB41665 | GGB28956 |
| GGB28956_SGB41665 | GGB28956 |
| GGB28960_SGB41669 | GGB28960 |
| GGB28960_SGB41669 | GGB28960 |
| GGB28967_SGB41678 | GGB28967 |
| GGB28967_SGB41678 | GGB28967 |
| GGB28991_SGB41705 | GGB28991 |
| GGB28991_SGB41705 | GGB28991 |
| GGB29002_SGB41718 | GGB29002 |
| GGB29002_SGB41718 | GGB29002 |
| GGB29003_SGB41719 | GGB29003 |
| GGB29003_SGB41719 | GGB29003 |
| GGB29011_SGB41731 | GGB29011 |
| GGB29011_SGB41731 | GGB29011 |
| GGB29531_SGB42317 | GGB29531 |
| GGB29531_SGB42317 | GGB29531 |
| GGB29685_SGB42494 | GGB29685 |
| GGB29685_SGB42494 | GGB29685 |
| GGB30141_SGB43066 | GGB30141 |
| GGB30141_SGB43066 | GGB30141 |
| GGB30286_SGB43248 | GGB30286 |
| GGB30286_SGB43248 | GGB30286 |
| GGB30303_SGB43268 | GGB30303 |
| GGB30303_SGB43268 | GGB30303 |
| GGB30413_SGB43452 | GGB30413 |
| GGB30413_SGB43452 | GGB30413 |
| GGB30454_SGB43514 | GGB30454 |
| GGB30454_SGB43514 | GGB30454 |
| GGB30455_SGB43519 | GGB30455 |
| GGB30455_SGB43519 | GGB30455 |
| GGB30461_SGB43527 | GGB30461 |
| GGB30461_SGB43527 | GGB30461 |
| GGB30461_SGB43530 | GGB30461 |
| GGB30461_SGB43530 | GGB30461 |
| GGB30463_SGB43537 | GGB30463 |
| GGB30463_SGB43537 | GGB30463 |
| GGB30473_SGB43557 | GGB30473 |
| GGB30473_SGB43557 | GGB30473 |
| GGB30475_SGB63182 | GGB30475 |
| GGB30475_SGB63182 | GGB30475 |
| GGB30861_SGB44083 | GGB30861 |
| GGB30861_SGB44083 | GGB30861 |

| | |
|---|---|
| GGB31312_SGB44628 | GGB31312 |
| GGB31312_SGB44628 | GGB31312 |
| GGB31438_SGB44768 | GGB31438 |
| GGB31438_SGB44768 | GGB31438 |
| GGB3171_SGB4185 | GGB3171 |
| GGB3171_SGB4185 | GGB3171 |
| GGB31823_SGB45199 | GGB31823 |
| GGB31823_SGB45199 | GGB31823 |
| GGB31853_SGB45233 | GGB31853 |
| GGB31853_SGB45233 | GGB31853 |
| GGB32371_SGB41694 | GGB32371 |
| GGB32371_SGB41694 | GGB32371 |
| GGB3793_SGB5158 | GGB3793 |
| GGB3793_SGB5158 | GGB3793 |
| GGB42598_SGB59794 | GGB42598 |
| GGB42598_SGB59794 | GGB42598 |
| GGB45656_SGB63370 | GGB45656 |
| GGB45656_SGB63370 | GGB45656 |
| GGB47127_SGB65054 | GGB47127 |
| GGB47127_SGB65054 | GGB47127 |
| GGB74395_SGB43521 | GGB74395 |
| GGB74395_SGB43521 | GGB74395 |
| GGB75053_SGB43494 | GGB75053 |
| GGB75053_SGB43494 | GGB75053 |
| GGB75109_SGB102238 | GGB75109 |
| GGB75109_SGB102238 | GGB75109 |
| GGB81440_SGB45230 | GGB81440 |
| GGB81440_SGB45230 | GGB81440 |
| Lachnospiraceae_bacterium | Lachnospiraceae_unclassified |
| Lachnospiraceae_bacterium | Lachnospiraceae_unclassified |
| Lachnospiraceae_bacterium_A2 | Lachnospiraceae_unclassified |
| Lachnospiraceae_bacterium_A2 | Lachnospiraceae_unclassified |
| Lachnospiraceae_bacterium_MD308 | Lachnospiraceae_unclassified |
| Lachnospiraceae_bacterium_MD308 | Lachnospiraceae_unclassified |
| Lachnospiraceae_bacterium_MD329 | Lachnospiraceae_unclassified |
| Lachnospiraceae_bacterium_MD329 | Lachnospiraceae_unclassified |
| Lachnospiraceae_unclassified_SGB41414 | Lachnospiraceae_unclassified |
| Lachnospiraceae_unclassified_SGB41414 | Lachnospiraceae_unclassified |
| Lachnospiraceae_unclassified_SGB41418 | Lachnospiraceae_unclassified |
| Lachnospiraceae_unclassified_SGB41418 | Lachnospiraceae_unclassified |
| Lachnospiraceae_unclassified_SGB41424 | Lachnospiraceae_unclassified |
| Lachnospiraceae_unclassified_SGB41424 | Lachnospiraceae_unclassified |
| Lachnospiraceae_unclassified_SGB41589 | Lachnospiraceae_unclassified |
| Lachnospiraceae_unclassified_SGB41589 | Lachnospiraceae_unclassified |
| Lactobacillus_johnsonii | Lactobacillus |
| Lactobacillus_johnsonii | Lactobacillus |

| | |
|---|---|
| Muribaculaceae_bacterium | Muribaculaceae_unclassified |
| Muribaculaceae_bacterium | Muribaculaceae_unclassified |
| Neglectibacter_sp_X4 | Neglectibacter |
| Neglectibacter_sp_X4 | Neglectibacter |
| Oscillospiraceae_bacterium | Oscillospiraceae_unclassified |
| Oscillospiraceae_bacterium | Oscillospiraceae_unclassified |
| Oscillospiraceae_unclassified_SGB43502 | Oscillospiraceae_unclassified |
| Oscillospiraceae_unclassified_SGB43502 | Oscillospiraceae_unclassified |
| Oscillospiraceae_unclassified_SGB43505 | Oscillospiraceae_unclassified |
| Oscillospiraceae_unclassified_SGB43505 | Oscillospiraceae_unclassified |
| Parasutterella_excrementihominis | Parasutterella |
| Parasutterella_excrementihominis | Parasutterella |
| richness | |
| richness | |
| Romboutsia_ilealis | Romboutsia |
| Romboutsia_ilealis | Romboutsia |
| Schaedlerella_arabinosiphila | Schaedlerella |
| Schaedlerella_arabinosiphila | Schaedlerella |
| shannon | |
| shannon | |
| Turicibacter_sp_1E2 | Turicibacter |
| Turicibacter_sp_1E2 | Turicibacter |
| Acetatifactor_muris | Acetatifactor |
| Acetatifactor_muris | Acetatifactor |
| Acetatifactor_SGB41546 | Acetatifactor |
| Acetatifactor_SGB41546 | Acetatifactor |
| Acutalibacter_muris | Acutalibacter |
| Acutalibacter_muris | Acutalibacter |
| Acutalibacter_sp_1XD8_36 | Acutalibacter |
| Acutalibacter_sp_1XD8_36 | Acutalibacter |
| Adlercreutzia_caecimuris | Adlercreutzia |
| Adlercreutzia_caecimuris | Adlercreutzia |
| Adlercreutzia_mucosicola | Adlercreutzia |
| Adlercreutzia_mucosicola | Adlercreutzia |
| Adlercreutzia_muris | Adlercreutzia |
| Adlercreutzia_muris | Adlercreutzia |
| Akkermansia_muciniphila | Akkermansia |
| Akkermansia_muciniphila | Akkermansia |
| Alistipes_sp_DSM_112343 | Alistipes |
| Alistipes_sp_DSM_112343 | Alistipes |
| Anaerotruncus_sp_1XD42_93 | Anaerotruncus |
| Anaerotruncus_sp_1XD42_93 | Anaerotruncus |
| Bacteria_unclassified_SGB102200 | Bacteria_unclassified |
| Bacteria_unclassified_SGB102200 | Bacteria_unclassified |
| Bacteria_unclassified_SGB41677 | Bacteria_unclassified |
| Bacteria_unclassified_SGB41677 | Bacteria_unclassified |

| | |
|---|---|
| Bacteria_unclassified_SGB43546 | Bacteria_unclassified |
| Bacteria_unclassified_SGB43546 | Bacteria_unclassified |
| bacterium_1XD42_54 | Bacteria_unclassified |
| bacterium_1XD42_54 | Bacteria_unclassified |
| bacterium_1XD42_76 | Bacteria_unclassified |
| bacterium_1XD42_76 | Bacteria_unclassified |
| bacterium_1xD8_48 | Bacteria_unclassified |
| bacterium_1xD8_48 | Bacteria_unclassified |
| Bacteroides_thetaiotaomicron | Bacteroides |
| Bacteroides_thetaiotaomicron | Bacteroides |
| berger_parker | |
| berger_parker | |
| Bifidobacterium_pseudolongum | Bifidobacterium |
| Bifidobacterium_pseudolongum | Bifidobacterium |
| Clostridia_bacterium | Clostridia_unclassified |
| Clostridia_bacterium | Clostridia_unclassified |
| Clostridiaceae_bacterium | Clostridiaceae_unclassified |
| Clostridiaceae_bacterium | Clostridiaceae_unclassified |
| Clostridiaceae_unclassified_SGB41663 | Clostridiaceae_unclassified |
| Clostridiaceae_unclassified_SGB41663 | Clostridiaceae_unclassified |
| Clostridiales_bacterium | Eubacteriales_unclassified |
| Clostridiales_bacterium | Eubacteriales_unclassified |
| Clostridium_cocleatum | Erysipelatoclostridium |
| Clostridium_cocleatum | Erysipelatoclostridium |
| Coriobacteriaceae_bacterium | Coriobacteriaceae_unclassified |
| Coriobacteriaceae_bacterium | Coriobacteriaceae_unclassified |
| Dorea_sp_5_2 | Dorea |
| Dorea_sp_5_2 | Dorea |
| Dubosiella_newyorkensis | Dubosiella |
| Dubosiella_newyorkensis | Dubosiella |
| Erysipelotrichales_bacterium | Erysipelotrichales_unclassified |
| Erysipelotrichales_bacterium | Erysipelotrichales_unclassified |
| Eubacteriaceae_bacterium | Eubacteriaceae_unclassified |
| Eubacteriaceae_bacterium | Eubacteriaceae_unclassified |
| Eubacteriaceae_unclassified_SGB94922 | Eubacteriaceae_unclassified |
| Eubacteriaceae_unclassified_SGB94922 | Eubacteriaceae_unclassified |
| GGB20149_SGB29430 | GGB20149 |
| GGB20149_SGB29430 | GGB20149 |
| GGB22635_SGB63107 | GGB22635 |
| GGB22635_SGB63107 | GGB22635 |
| GGB25041_SGB36960 | GGB25041 |
| GGB25041_SGB36960 | GGB25041 |
| GGB27876_SGB40310 | GGB27876 |
| GGB27876_SGB40310 | GGB27876 |
| GGB27878_SGB40312 | GGB27878 |
| GGB27878_SGB40312 | GGB27878 |

| | |
|---|---|
| GGB27918_SGB40356 | GGB27918 |
| GGB27918_SGB40356 | GGB27918 |
| GGB28382_SGB40962 | GGB28382 |
| GGB28382_SGB40962 | GGB28382 |
| GGB28399_SGB40980 | GGB28399 |
| GGB28399_SGB40980 | GGB28399 |
| GGB28411_SGB40993 | GGB28411 |
| GGB28411_SGB40993 | GGB28411 |
| GGB28415_SGB40997 | GGB28415 |
| GGB28415_SGB40997 | GGB28415 |
| GGB28430_SGB41013 | GGB28430 |
| GGB28430_SGB41013 | GGB28430 |
| GGB28439_SGB41022 | GGB28439 |
| GGB28439_SGB41022 | GGB28439 |
| GGB28778_SGB41431 | GGB28778 |
| GGB28778_SGB41431 | GGB28778 |
| GGB28784_SGB41437 | GGB28784 |
| GGB28784_SGB41437 | GGB28784 |
| GGB28792_SGB41445 | GGB28792 |
| GGB28792_SGB41445 | GGB28792 |
| GGB28798_SGB41451 | GGB28798 |
| GGB28798_SGB41451 | GGB28798 |
| GGB28802_SGB41455 | GGB28802 |
| GGB28802_SGB41455 | GGB28802 |
| GGB28818_SGB41473 | GGB28818 |
| GGB28818_SGB41473 | GGB28818 |
| GGB28828_SGB41484 | GGB28828 |
| GGB28828_SGB41484 | GGB28828 |
| GGB28851_SGB41518 | GGB28851 |
| GGB28851_SGB41518 | GGB28851 |
| GGB28859_SGB41528 | GGB28859 |
| GGB28859_SGB41528 | GGB28859 |
| GGB28864_SGB41535 | GGB28864 |
| GGB28864_SGB41535 | GGB28864 |
| GGB28869_SGB41543 | GGB28869 |
| GGB28869_SGB41543 | GGB28869 |
| GGB28883_SGB41564 | GGB28883 |
| GGB28883_SGB41564 | GGB28883 |
| GGB28892_SGB41573 | GGB28892 |
| GGB28892_SGB41573 | GGB28892 |
| GGB28893_SGB41574 | GGB28893 |
| GGB28893_SGB41574 | GGB28893 |
| GGB28898_SGB41580 | GGB28898 |
| GGB28898_SGB41580 | GGB28898 |
| GGB28904_SGB41597 | GGB28904 |
| GGB28904_SGB41597 | GGB28904 |

| | |
|---|---|
| GGB28916_SGB41612 | GGB28916 |
| GGB28916_SGB41612 | GGB28916 |
| GGB28924_SGB41621 | GGB28924 |
| GGB28924_SGB41621 | GGB28924 |
| GGB28926_SGB41624 | GGB28926 |
| GGB28926_SGB41624 | GGB28926 |
| GGB28927_SGB41625 | GGB28927 |
| GGB28927_SGB41625 | GGB28927 |
| GGB28934_SGB41635 | GGB28934 |
| GGB28934_SGB41635 | GGB28934 |
| GGB28946_SGB41652 | GGB28946 |
| GGB28946_SGB41652 | GGB28946 |
| GGB28949_SGB41655 | GGB28949 |
| GGB28949_SGB41655 | GGB28949 |
| GGB28949_SGB41656 | GGB28949 |
| GGB28949_SGB41656 | GGB28949 |
| GGB28950_SGB41657 | GGB28950 |
| GGB28950_SGB41657 | GGB28950 |
| GGB28951_SGB102295 | GGB28951 |
| GGB28951_SGB102295 | GGB28951 |
| GGB28951_SGB41658 | GGB28951 |
| GGB28951_SGB41658 | GGB28951 |
| GGB28954_SGB41662 | GGB28954 |
| GGB28954_SGB41662 | GGB28954 |
| GGB28956_SGB41665 | GGB28956 |
| GGB28956_SGB41665 | GGB28956 |
| GGB28960_SGB41669 | GGB28960 |
| GGB28960_SGB41669 | GGB28960 |
| GGB28967_SGB41678 | GGB28967 |
| GGB28967_SGB41678 | GGB28967 |
| GGB28991_SGB41705 | GGB28991 |
| GGB28991_SGB41705 | GGB28991 |
| GGB29002_SGB41718 | GGB29002 |
| GGB29002_SGB41718 | GGB29002 |
| GGB29003_SGB41719 | GGB29003 |
| GGB29003_SGB41719 | GGB29003 |
| GGB29011_SGB41731 | GGB29011 |
| GGB29011_SGB41731 | GGB29011 |
| GGB29531_SGB42317 | GGB29531 |
| GGB29531_SGB42317 | GGB29531 |
| GGB29685_SGB42494 | GGB29685 |
| GGB29685_SGB42494 | GGB29685 |
| GGB30141_SGB43066 | GGB30141 |
| GGB30141_SGB43066 | GGB30141 |
| GGB30286_SGB43248 | GGB30286 |
| GGB30286_SGB43248 | GGB30286 |

| | |
|---|---|
| GGB30303_SGB43268 | GGB30303 |
| GGB30303_SGB43268 | GGB30303 |
| GGB30413_SGB43452 | GGB30413 |
| GGB30413_SGB43452 | GGB30413 |
| GGB30454_SGB43514 | GGB30454 |
| GGB30454_SGB43514 | GGB30454 |
| GGB30455_SGB43519 | GGB30455 |
| GGB30455_SGB43519 | GGB30455 |
| GGB30461_SGB43527 | GGB30461 |
| GGB30461_SGB43527 | GGB30461 |
| GGB30461_SGB43530 | GGB30461 |
| GGB30461_SGB43530 | GGB30461 |
| GGB30463_SGB43537 | GGB30463 |
| GGB30463_SGB43537 | GGB30463 |
| GGB30473_SGB43557 | GGB30473 |
| GGB30473_SGB43557 | GGB30473 |
| GGB30475_SGB63182 | GGB30475 |
| GGB30475_SGB63182 | GGB30475 |
| GGB30861_SGB44083 | GGB30861 |
| GGB30861_SGB44083 | GGB30861 |
| GGB31312_SGB44628 | GGB31312 |
| GGB31312_SGB44628 | GGB31312 |
| GGB31438_SGB44768 | GGB31438 |
| GGB31438_SGB44768 | GGB31438 |
| GGB3171_SGB4185 | GGB3171 |
| GGB3171_SGB4185 | GGB3171 |
| GGB31823_SGB45199 | GGB31823 |
| GGB31823_SGB45199 | GGB31823 |
| GGB31853_SGB45233 | GGB31853 |
| GGB31853_SGB45233 | GGB31853 |
| GGB32371_SGB41694 | GGB32371 |
| GGB32371_SGB41694 | GGB32371 |
| GGB3793_SGB5158 | GGB3793 |
| GGB3793_SGB5158 | GGB3793 |
| GGB42598_SGB59794 | GGB42598 |
| GGB42598_SGB59794 | GGB42598 |
| GGB45656_SGB63370 | GGB45656 |
| GGB45656_SGB63370 | GGB45656 |
| GGB47127_SGB65054 | GGB47127 |
| GGB47127_SGB65054 | GGB47127 |
| GGB74395_SGB43521 | GGB74395 |
| GGB74395_SGB43521 | GGB74395 |
| GGB75053_SGB43494 | GGB75053 |
| GGB75053_SGB43494 | GGB75053 |
| GGB75109_SGB102238 | GGB75109 |
| GGB75109_SGB102238 | GGB75109 |

| | |
|---|---|
| GGB81440_SGB45230 | GGB81440 |
| GGB81440_SGB45230 | GGB81440 |
| Lachnospiraceae_bacterium | Lachnospiraceae_unclassified |
| Lachnospiraceae_bacterium | Lachnospiraceae_unclassified |
| Lachnospiraceae_bacterium_A2 | Lachnospiraceae_unclassified |
| Lachnospiraceae_bacterium_A2 | Lachnospiraceae_unclassified |
| Lachnospiraceae_bacterium_MD308 | Lachnospiraceae_unclassified |
| Lachnospiraceae_bacterium_MD308 | Lachnospiraceae_unclassified |
| Lachnospiraceae_bacterium_MD329 | Lachnospiraceae_unclassified |
| Lachnospiraceae_bacterium_MD329 | Lachnospiraceae_unclassified |
| Lachnospiraceae_unclassified_SGB41414 | Lachnospiraceae_unclassified |
| Lachnospiraceae_unclassified_SGB41414 | Lachnospiraceae_unclassified |
| Lachnospiraceae_unclassified_SGB41418 | Lachnospiraceae_unclassified |
| Lachnospiraceae_unclassified_SGB41418 | Lachnospiraceae_unclassified |
| Lachnospiraceae_unclassified_SGB41424 | Lachnospiraceae_unclassified |
| Lachnospiraceae_unclassified_SGB41424 | Lachnospiraceae_unclassified |
| Lachnospiraceae_unclassified_SGB41589 | Lachnospiraceae_unclassified |
| Lachnospiraceae_unclassified_SGB41589 | Lachnospiraceae_unclassified |
| Lactobacillus_johnsonii | Lactobacillus |
| Lactobacillus_johnsonii | Lactobacillus |
| Muribaculaceae_bacterium | Muribaculaceae_unclassified |
| Muribaculaceae_bacterium | Muribaculaceae_unclassified |
| Neglectibacter_sp_X4 | Neglectibacter |
| Neglectibacter_sp_X4 | Neglectibacter |
| Oscillospiraceae_bacterium | Oscillospiraceae_unclassified |
| Oscillospiraceae_bacterium | Oscillospiraceae_unclassified |
| Oscillospiraceae_unclassified_SGB43502 | Oscillospiraceae_unclassified |
| Oscillospiraceae_unclassified_SGB43502 | Oscillospiraceae_unclassified |
| Oscillospiraceae_unclassified_SGB43505 | Oscillospiraceae_unclassified |
| Oscillospiraceae_unclassified_SGB43505 | Oscillospiraceae_unclassified |
| Parasutterella_excrementihominis | Parasutterella |
| Parasutterella_excrementihominis | Parasutterella |
| richness | |
| richness | |
| Romboutsia_ilealis | Romboutsia |
| Romboutsia_ilealis | Romboutsia |
| Schaedlerella_arabinosiphila | Schaedlerella |
| Schaedlerella_arabinosiphila | Schaedlerella |
| shannon | |
| shannon | |
| Turicibacter_sp_1E2 | Turicibacter |
| Turicibacter_sp_1E2 | Turicibacter |
| Acetatifactor_muris | Acetatifactor |
| Acetatifactor_muris | Acetatifactor |
| Acetatifactor_SGB41546 | Acetatifactor |
| Acetatifactor_SGB41546 | Acetatifactor |

| | |
|---|---|
| Acutalibacter_muris | Acutalibacter |
| Acutalibacter_muris | Acutalibacter |
| Acutalibacter_sp_1XD8_36 | Acutalibacter |
| Acutalibacter_sp_1XD8_36 | Acutalibacter |
| Adlercreutzia_caecimuris | Adlercreutzia |
| Adlercreutzia_caecimuris | Adlercreutzia |
| Adlercreutzia_mucosicola | Adlercreutzia |
| Adlercreutzia_mucosicola | Adlercreutzia |
| Adlercreutzia_muris | Adlercreutzia |
| Adlercreutzia_muris | Adlercreutzia |
| Akkermansia_muciniphila | Akkermansia |
| Akkermansia_muciniphila | Akkermansia |
| Alistipes_sp_DSM_112343 | Alistipes |
| Alistipes_sp_DSM_112343 | Alistipes |
| Anaerotruncus_sp_1XD42_93 | Anaerotruncus |
| Anaerotruncus_sp_1XD42_93 | Anaerotruncus |
| Bacteria_unclassified_SGB102200 | Bacteria_unclassified |
| Bacteria_unclassified_SGB102200 | Bacteria_unclassified |
| Bacteria_unclassified_SGB41677 | Bacteria_unclassified |
| Bacteria_unclassified_SGB41677 | Bacteria_unclassified |
| Bacteria_unclassified_SGB43546 | Bacteria_unclassified |
| Bacteria_unclassified_SGB43546 | Bacteria_unclassified |
| bacterium_1XD42_54 | Bacteria_unclassified |
| bacterium_1XD42_54 | Bacteria_unclassified |
| bacterium_1XD42_76 | Bacteria_unclassified |
| bacterium_1XD42_76 | Bacteria_unclassified |
| bacterium_1xD8_48 | Bacteria_unclassified |
| bacterium_1xD8_48 | Bacteria_unclassified |
| Bacteroides_thetaiotaomicron | Bacteroides |
| Bacteroides_thetaiotaomicron | Bacteroides |
| berger_parker | |
| berger_parker | |
| Bifidobacterium_pseudolongum | Bifidobacterium |
| Bifidobacterium_pseudolongum | Bifidobacterium |
| Clostridia_bacterium | Clostridia_unclassified |
| Clostridia_bacterium | Clostridia_unclassified |
| Clostridiaceae_bacterium | Clostridiaceae_unclassified |
| Clostridiaceae_bacterium | Clostridiaceae_unclassified |
| Clostridiaceae_unclassified_SGB41663 | Clostridiaceae_unclassified |
| Clostridiaceae_unclassified_SGB41663 | Clostridiaceae_unclassified |
| Clostridiales_bacterium | Eubacteriales_unclassified |
| Clostridiales_bacterium | Eubacteriales_unclassified |
| Clostridium_cocleatum | Erysipelatoclostridium |
| Clostridium_cocleatum | Erysipelatoclostridium |
| Coriobacteriaceae_bacterium | Coriobacteriaceae_unclassified |
| Coriobacteriaceae_bacterium | Coriobacteriaceae_unclassified |

| | |
|---|---|
| Dorea_sp_5_2 | Dorea |
| Dorea_sp_5_2 | Dorea |
| Dubosiella_newyorkensis | Dubosiella |
| Dubosiella_newyorkensis | Dubosiella |
| Erysipelotrichales_bacterium | Erysipelotrichales_unclassified |
| Erysipelotrichales_bacterium | Erysipelotrichales_unclassified |
| Eubacteriaceae_bacterium | Eubacteriaceae_unclassified |
| Eubacteriaceae_bacterium | Eubacteriaceae_unclassified |
| Eubacteriaceae_unclassified_SGB94922 | Eubacteriaceae_unclassified |
| Eubacteriaceae_unclassified_SGB94922 | Eubacteriaceae_unclassified |
| GGB20149_SGB29430 | GGB20149 |
| GGB20149_SGB29430 | GGB20149 |
| GGB22635_SGB63107 | GGB22635 |
| GGB22635_SGB63107 | GGB22635 |
| GGB25041_SGB36960 | GGB25041 |
| GGB25041_SGB36960 | GGB25041 |
| GGB27876_SGB40310 | GGB27876 |
| GGB27876_SGB40310 | GGB27876 |
| GGB27878_SGB40312 | GGB27878 |
| GGB27878_SGB40312 | GGB27878 |
| GGB27918_SGB40356 | GGB27918 |
| GGB27918_SGB40356 | GGB27918 |
| GGB28382_SGB40962 | GGB28382 |
| GGB28382_SGB40962 | GGB28382 |
| GGB28399_SGB40980 | GGB28399 |
| GGB28399_SGB40980 | GGB28399 |
| GGB28411_SGB40993 | GGB28411 |
| GGB28411_SGB40993 | GGB28411 |
| GGB28415_SGB40997 | GGB28415 |
| GGB28415_SGB40997 | GGB28415 |
| GGB28430_SGB41013 | GGB28430 |
| GGB28430_SGB41013 | GGB28430 |
| GGB28439_SGB41022 | GGB28439 |
| GGB28439_SGB41022 | GGB28439 |
| GGB28778_SGB41431 | GGB28778 |
| GGB28778_SGB41431 | GGB28778 |
| GGB28784_SGB41437 | GGB28784 |
| GGB28784_SGB41437 | GGB28784 |
| GGB28792_SGB41445 | GGB28792 |
| GGB28792_SGB41445 | GGB28792 |
| GGB28798_SGB41451 | GGB28798 |
| GGB28798_SGB41451 | GGB28798 |
| GGB28802_SGB41455 | GGB28802 |
| GGB28802_SGB41455 | GGB28802 |
| GGB28818_SGB41473 | GGB28818 |
| GGB28818_SGB41473 | GGB28818 |

| | |
|---|---|
| GGB28828_SGB41484 | GGB28828 |
| GGB28828_SGB41484 | GGB28828 |
| GGB28851_SGB41518 | GGB28851 |
| GGB28851_SGB41518 | GGB28851 |
| GGB28859_SGB41528 | GGB28859 |
| GGB28859_SGB41528 | GGB28859 |
| GGB28864_SGB41535 | GGB28864 |
| GGB28864_SGB41535 | GGB28864 |
| GGB28869_SGB41543 | GGB28869 |
| GGB28869_SGB41543 | GGB28869 |
| GGB28883_SGB41564 | GGB28883 |
| GGB28883_SGB41564 | GGB28883 |
| GGB28892_SGB41573 | GGB28892 |
| GGB28892_SGB41573 | GGB28892 |
| GGB28893_SGB41574 | GGB28893 |
| GGB28893_SGB41574 | GGB28893 |
| GGB28898_SGB41580 | GGB28898 |
| GGB28898_SGB41580 | GGB28898 |
| GGB28904_SGB41597 | GGB28904 |
| GGB28904_SGB41597 | GGB28904 |
| GGB28916_SGB41612 | GGB28916 |
| GGB28916_SGB41612 | GGB28916 |
| GGB28924_SGB41621 | GGB28924 |
| GGB28924_SGB41621 | GGB28924 |
| GGB28926_SGB41624 | GGB28926 |
| GGB28926_SGB41624 | GGB28926 |
| GGB28927_SGB41625 | GGB28927 |
| GGB28927_SGB41625 | GGB28927 |
| GGB28934_SGB41635 | GGB28934 |
| GGB28934_SGB41635 | GGB28934 |
| GGB28946_SGB41652 | GGB28946 |
| GGB28946_SGB41652 | GGB28946 |
| GGB28949_SGB41655 | GGB28949 |
| GGB28949_SGB41655 | GGB28949 |
| GGB28949_SGB41656 | GGB28949 |
| GGB28949_SGB41656 | GGB28949 |
| GGB28950_SGB41657 | GGB28950 |
| GGB28950_SGB41657 | GGB28950 |
| GGB28951_SGB102295 | GGB28951 |
| GGB28951_SGB102295 | GGB28951 |
| GGB28951_SGB41658 | GGB28951 |
| GGB28951_SGB41658 | GGB28951 |
| GGB28954_SGB41662 | GGB28954 |
| GGB28954_SGB41662 | GGB28954 |
| GGB28956_SGB41665 | GGB28956 |
| GGB28956_SGB41665 | GGB28956 |

| | |
|---|---|
| GGB28960_SGB41669 | GGB28960 |
| GGB28960_SGB41669 | GGB28960 |
| GGB28967_SGB41678 | GGB28967 |
| GGB28967_SGB41678 | GGB28967 |
| GGB28991_SGB41705 | GGB28991 |
| GGB28991_SGB41705 | GGB28991 |
| GGB29002_SGB41718 | GGB29002 |
| GGB29002_SGB41718 | GGB29002 |
| GGB29003_SGB41719 | GGB29003 |
| GGB29003_SGB41719 | GGB29003 |
| GGB29011_SGB41731 | GGB29011 |
| GGB29011_SGB41731 | GGB29011 |
| GGB29531_SGB42317 | GGB29531 |
| GGB29531_SGB42317 | GGB29531 |
| GGB29685_SGB42494 | GGB29685 |
| GGB29685_SGB42494 | GGB29685 |
| GGB30141_SGB43066 | GGB30141 |
| GGB30141_SGB43066 | GGB30141 |
| GGB30286_SGB43248 | GGB30286 |
| GGB30286_SGB43248 | GGB30286 |
| GGB30303_SGB43268 | GGB30303 |
| GGB30303_SGB43268 | GGB30303 |
| GGB30413_SGB43452 | GGB30413 |
| GGB30413_SGB43452 | GGB30413 |
| GGB30454_SGB43514 | GGB30454 |
| GGB30454_SGB43514 | GGB30454 |
| GGB30455_SGB43519 | GGB30455 |
| GGB30455_SGB43519 | GGB30455 |
| GGB30461_SGB43527 | GGB30461 |
| GGB30461_SGB43527 | GGB30461 |
| GGB30461_SGB43530 | GGB30461 |
| GGB30461_SGB43530 | GGB30461 |
| GGB30463_SGB43537 | GGB30463 |
| GGB30463_SGB43537 | GGB30463 |
| GGB30473_SGB43557 | GGB30473 |
| GGB30473_SGB43557 | GGB30473 |
| GGB30475_SGB63182 | GGB30475 |
| GGB30475_SGB63182 | GGB30475 |
| GGB30861_SGB44083 | GGB30861 |
| GGB30861_SGB44083 | GGB30861 |
| GGB31312_SGB44628 | GGB31312 |
| GGB31312_SGB44628 | GGB31312 |
| GGB31438_SGB44768 | GGB31438 |
| GGB31438_SGB44768 | GGB31438 |
| GGB3171_SGB4185 | GGB3171 |
| GGB3171_SGB4185 | GGB3171 |

| | |
|---|---|
| GGB31823_SGB45199 | GGB31823 |
| GGB31823_SGB45199 | GGB31823 |
| GGB31853_SGB45233 | GGB31853 |
| GGB31853_SGB45233 | GGB31853 |
| GGB32371_SGB41694 | GGB32371 |
| GGB32371_SGB41694 | GGB32371 |
| GGB3793_SGB5158 | GGB3793 |
| GGB3793_SGB5158 | GGB3793 |
| GGB42598_SGB59794 | GGB42598 |
| GGB42598_SGB59794 | GGB42598 |
| GGB45656_SGB63370 | GGB45656 |
| GGB45656_SGB63370 | GGB45656 |
| GGB47127_SGB65054 | GGB47127 |
| GGB47127_SGB65054 | GGB47127 |
| GGB74395_SGB43521 | GGB74395 |
| GGB74395_SGB43521 | GGB74395 |
| GGB75053_SGB43494 | GGB75053 |
| GGB75053_SGB43494 | GGB75053 |
| GGB75109_SGB102238 | GGB75109 |
| GGB75109_SGB102238 | GGB75109 |
| GGB81440_SGB45230 | GGB81440 |
| GGB81440_SGB45230 | GGB81440 |
| Lachnospiraceae_bacterium | Lachnospiraceae_unclassified |
| Lachnospiraceae_bacterium | Lachnospiraceae_unclassified |
| Lachnospiraceae_bacterium_A2 | Lachnospiraceae_unclassified |
| Lachnospiraceae_bacterium_A2 | Lachnospiraceae_unclassified |
| Lachnospiraceae_bacterium_MD308 | Lachnospiraceae_unclassified |
| Lachnospiraceae_bacterium_MD308 | Lachnospiraceae_unclassified |
| Lachnospiraceae_bacterium_MD329 | Lachnospiraceae_unclassified |
| Lachnospiraceae_bacterium_MD329 | Lachnospiraceae_unclassified |
| Lachnospiraceae_unclassified_SGB41414 | Lachnospiraceae_unclassified |
| Lachnospiraceae_unclassified_SGB41414 | Lachnospiraceae_unclassified |
| Lachnospiraceae_unclassified_SGB41418 | Lachnospiraceae_unclassified |
| Lachnospiraceae_unclassified_SGB41418 | Lachnospiraceae_unclassified |
| Lachnospiraceae_unclassified_SGB41424 | Lachnospiraceae_unclassified |
| Lachnospiraceae_unclassified_SGB41424 | Lachnospiraceae_unclassified |
| Lachnospiraceae_unclassified_SGB41589 | Lachnospiraceae_unclassified |
| Lachnospiraceae_unclassified_SGB41589 | Lachnospiraceae_unclassified |
| Lactobacillus_johnsonii | Lactobacillus |
| Lactobacillus_johnsonii | Lactobacillus |
| Muribaculaceae_bacterium | Muribaculaceae_unclassified |
| Muribaculaceae_bacterium | Muribaculaceae_unclassified |
| Neglectibacter_sp_X4 | Neglectibacter |
| Neglectibacter_sp_X4 | Neglectibacter |
| Oscillospiraceae_bacterium | Oscillospiraceae_unclassified |
| Oscillospiraceae_bacterium | Oscillospiraceae_unclassified |

| | |
|---|---|
| Oscillospiraceae_unclassified_SGB43502 | Oscillospiraceae_unclassified |
| Oscillospiraceae_unclassified_SGB43502 | Oscillospiraceae_unclassified |
| Oscillospiraceae_unclassified_SGB43505 | Oscillospiraceae_unclassified |
| Oscillospiraceae_unclassified_SGB43505 | Oscillospiraceae_unclassified |
| Parasutterella_excrementihominis | Parasutterella |
| Parasutterella_excrementihominis | Parasutterella |
| richness | |
| richness | |
| Romboutsia_ilealis | Romboutsia |
| Romboutsia_ilealis | Romboutsia |
| Schaedlerella_arabinosiphila | Schaedlerella |
| Schaedlerella_arabinosiphila | Schaedlerella |
| shannon | |
| shannon | |
| Turicibacter_sp_1E2 | Turicibacter |
| Turicibacter_sp_1E2 | Turicibacter |

| Family | Order | Class |
|---|---|---|
| Lachnospiraceae | Eubacteriales | Clostridia |
| Lachnospiraceae | Eubacteriales | Clostridia |
| Lachnospiraceae | Eubacteriales | Clostridia |
| Lachnospiraceae | Eubacteriales | Clostridia |
| Oscillospiraceae | Eubacteriales | Clostridia |
| Oscillospiraceae | Eubacteriales | Clostridia |
| Oscillospiraceae | Eubacteriales | Clostridia |
| Oscillospiraceae | Eubacteriales | Clostridia |
| Eggerthellaceae | Eggerthellales | Coriobacteriia |
| Eggerthellaceae | Eggerthellales | Coriobacteriia |
| Eggerthellaceae | Eggerthellales | Coriobacteriia |
| Eggerthellaceae | Eggerthellales | Coriobacteriia |
| Eggerthellaceae | Eggerthellales | Coriobacteriia |
| Eggerthellaceae | Eggerthellales | Coriobacteriia |
| Akkermansiaceae | Verrucomicrobiales | Verrucomicrobiae |
| Akkermansiaceae | Verrucomicrobiales | Verrucomicrobiae |
| Rikenellaceae | Bacteroidales | Bacteroidia |
| Rikenellaceae | Bacteroidales | Bacteroidia |
| Oscillospiraceae | Eubacteriales | Clostridia |
| Oscillospiraceae | Eubacteriales | Clostridia |
| Bacteria_unclassified | Bacteria_unclassified | Bacteria_unclassified |
| Bacteria_unclassified | Bacteria_unclassified | Bacteria_unclassified |
| Bacteria_unclassified | Bacteria_unclassified | Bacteria_unclassified |
| Bacteria_unclassified | Bacteria_unclassified | Bacteria_unclassified |
| Bacteria_unclassified | Bacteria_unclassified | Bacteria_unclassified |
| Bacteria_unclassified | Bacteria_unclassified | Bacteria_unclassified |
| Bacteria_unclassified | Bacteria_unclassified | Bacteria_unclassified |
| Bacteria_unclassified | Bacteria_unclassified | Bacteria_unclassified |
| Bacteria_unclassified | Bacteria_unclassified | Bacteria_unclassified |
| Bacteria_unclassified | Bacteria_unclassified | Bacteria_unclassified |
| Bacteria_unclassified | Bacteria_unclassified | Bacteria_unclassified |
| Bacteria_unclassified | Bacteria_unclassified | Bacteria_unclassified |
| Bacteroidaceae | Bacteroidales | Bacteroidia |
| Bacteroidaceae | Bacteroidales | Bacteroidia |
| Bifidobacteriaceae | Bifidobacteriales | Actinomycetia |
| Bifidobacteriaceae | Bifidobacteriales | Actinomycetia |
| Clostridia_unclassified | Clostridia_unclassified | Clostridia |
| Clostridia_unclassified | Clostridia_unclassified | Clostridia |
| Clostridiaceae | Eubacteriales | Clostridia |
| Clostridiaceae | Eubacteriales | Clostridia |
| Clostridiaceae | Eubacteriales | Clostridia |
| Clostridiaceae | Eubacteriales | Clostridia |
| Eubacteriales_unclassified | Eubacteriales | Clostridia |
| Eubacteriales_unclassified | Eubacteriales | Clostridia |

| | | |
|---|---|---|
| Erysipelotrichaceae | Erysipelotrichales | Erysipelotrichia |
| Erysipelotrichaceae | Erysipelotrichales | Erysipelotrichia |
| Coriobacteriaceae | Coriobacteriales | Coriobacteriia |
| Coriobacteriaceae | Coriobacteriales | Coriobacteriia |
| Lachnospiraceae | Eubacteriales | Clostridia |
| Lachnospiraceae | Eubacteriales | Clostridia |
| Erysipelotrichaceae | Erysipelotrichales | Erysipelotrichia |
| Erysipelotrichaceae | Erysipelotrichales | Erysipelotrichia |
| Erysipelotrichales_unclassified | Erysipelotrichales | Erysipelotrichia |
| Erysipelotrichales_unclassified | Erysipelotrichales | Erysipelotrichia |
| Eubacteriaceae | Eubacteriales | Clostridia |
| Eubacteriaceae | Eubacteriales | Clostridia |
| Eubacteriaceae | Eubacteriales | Clostridia |
| Eubacteriaceae | Eubacteriales | Clostridia |
| Lachnospiraceae | Eubacteriales | Clostridia |
| Lachnospiraceae | Eubacteriales | Clostridia |
| Eggerthellaceae | Eggerthellales | Coriobacteriia |
| Eggerthellaceae | Eggerthellales | Coriobacteriia |
| Lachnospiraceae | Eubacteriales | Clostridia |
| Lachnospiraceae | Eubacteriales | Clostridia |
| Muribaculaceae | Bacteroidales | Bacteroidia |
| Muribaculaceae | Bacteroidales | Bacteroidia |
| Muribaculaceae | Bacteroidales | Bacteroidia |
| Muribaculaceae | Bacteroidales | Bacteroidia |
| Muribaculaceae | Bacteroidales | Bacteroidia |
| Muribaculaceae | Bacteroidales | Bacteroidia |
| FGB9508 | OFGB9508 | CFGB9508 |
| FGB9508 | OFGB9508 | CFGB9508 |
| FGB2838 | OFGB2838 | CFGB2838 |
| FGB2838 | OFGB2838 | CFGB2838 |
| FGB2838 | OFGB2838 | CFGB2838 |
| FGB2838 | OFGB2838 | CFGB2838 |
| FGB2838 | OFGB2838 | CFGB2838 |
| FGB2838 | OFGB2838 | CFGB2838 |
| Pumilibacteraceae | Eubacteriales | Clostridia |
| Pumilibacteraceae | Eubacteriales | Clostridia |
| FGB28439 | OFGB28439 | CFGB28439 |
| FGB28439 | OFGB28439 | CFGB28439 |
| Clostridia_unclassified | Clostridia_unclassified | Clostridia |
| Clostridia_unclassified | Clostridia_unclassified | Clostridia |
| Eubacteriaceae | Eubacteriales | Clostridia |
| Eubacteriaceae | Eubacteriales | Clostridia |
| Lachnospiraceae | Eubacteriales | Clostridia |
| Lachnospiraceae | Eubacteriales | Clostridia |
| Lachnospiraceae | Eubacteriales | Clostridia |
| Lachnospiraceae | Eubacteriales | Clostridia |

| | | |
|---|---|---|
| Lachnospiraceae | Eubacteriales | Clostridia |
| Lachnospiraceae | Eubacteriales | Clostridia |
| Lachnospiraceae | Eubacteriales | Clostridia |
| Lachnospiraceae | Eubacteriales | Clostridia |
| FGB77305 | OFGB77305 | CFGB77305 |
| FGB77305 | OFGB77305 | CFGB77305 |
| Clostridiaceae | Eubacteriales | Clostridia |
| Clostridiaceae | Eubacteriales | Clostridia |
| Lachnospiraceae | Eubacteriales | Clostridia |
| Lachnospiraceae | Eubacteriales | Clostridia |
| Lachnospiraceae | Eubacteriales | Clostridia |
| Lachnospiraceae | Eubacteriales | Clostridia |
| Lachnospiraceae | Eubacteriales | Clostridia |
| Lachnospiraceae | Eubacteriales | Clostridia |
| FGB9633 | OFGB9633 | CFGB9633 |
| FGB9633 | OFGB9633 | CFGB9633 |
| Bacteria_unclassified | Bacteria_unclassified | Bacteria_unclassified |
| Bacteria_unclassified | Bacteria_unclassified | Bacteria_unclassified |
| Bacteria_unclassified | Bacteria_unclassified | Bacteria_unclassified |
| Bacteria_unclassified | Bacteria_unclassified | Bacteria_unclassified |
| Bacteria_unclassified | Bacteria_unclassified | Bacteria_unclassified |
| Bacteria_unclassified | Bacteria_unclassified | Bacteria_unclassified |
| Bacteria_unclassified | Bacteria_unclassified | Bacteria_unclassified |
| Bacteria_unclassified | Bacteria_unclassified | Bacteria_unclassified |
| Lachnospiraceae | Eubacteriales | Clostridia |
| Lachnospiraceae | Eubacteriales | Clostridia |
| Lachnospiraceae | Eubacteriales | Clostridia |
| Lachnospiraceae | Eubacteriales | Clostridia |
| Lachnospiraceae | Eubacteriales | Clostridia |
| Lachnospiraceae | Eubacteriales | Clostridia |
| FGB77359 | OFGB77359 | CFGB77359 |
| FGB77359 | OFGB77359 | CFGB77359 |
| FGB9639 | OFGB9639 | CFGB9639 |
| FGB9639 | OFGB9639 | CFGB9639 |
| Lachnospiraceae | Eubacteriales | Clostridia |
| Lachnospiraceae | Eubacteriales | Clostridia |
| Lachnospiraceae | Eubacteriales | Clostridia |
| Lachnospiraceae | Eubacteriales | Clostridia |
| Lachnospiraceae | Eubacteriales | Clostridia |
| Lachnospiraceae | Eubacteriales | Clostridia |
| Clostridiaceae | Eubacteriales | Clostridia |
| Clostridiaceae | Eubacteriales | Clostridia |
| Clostridiaceae | Eubacteriales | Clostridia |
| Clostridiaceae | Eubacteriales | Clostridia |
| Clostridiaceae | Eubacteriales | Clostridia |
| Clostridiaceae | Eubacteriales | Clostridia |

| | | |
|---|---|---|
| Clostridiaceae | Eubacteriales | Clostridia |
| Clostridiaceae | Eubacteriales | Clostridia |
| Clostridiaceae | Eubacteriales | Clostridia |
| Clostridiaceae | Eubacteriales | Clostridia |
| Clostridiaceae | Eubacteriales | Clostridia |
| Clostridiaceae | Eubacteriales | Clostridia |
| Clostridiaceae | Eubacteriales | Clostridia |
| Clostridiaceae | Eubacteriales | Clostridia |
| Eubacteriaceae | Eubacteriales | Clostridia |
| Eubacteriaceae | Eubacteriales | Clostridia |
| FGB9658 | OFGB9658 | CFGB9658 |
| FGB9658 | OFGB9658 | CFGB9658 |
| FGB9659 | OFGB9659 | CFGB9659 |
| FGB9659 | OFGB9659 | CFGB9659 |
| Bacteria_unclassified | Bacteria_unclassified | Bacteria_unclassified |
| Bacteria_unclassified | Bacteria_unclassified | Bacteria_unclassified |
| FGB9827 | OFGB9827 | CFGB9827 |
| FGB9827 | OFGB9827 | CFGB9827 |
| Eubacteriaceae | Eubacteriales | Clostridia |
| Eubacteriaceae | Eubacteriales | Clostridia |
| FGB77303 | OFGB77303 | CFGB77303 |
| FGB77303 | OFGB77303 | CFGB77303 |
| Eubacteriales_unclassified | Eubacteriales | Clostridia |
| Eubacteriales_unclassified | Eubacteriales | Clostridia |
| Oscillospiraceae | Eubacteriales | Clostridia |
| Oscillospiraceae | Eubacteriales | Clostridia |
| FGB30328 | OFGB30328 | CFGB30328 |
| FGB30328 | OFGB30328 | CFGB30328 |
| Oscillospiraceae | Eubacteriales | Clostridia |
| Oscillospiraceae | Eubacteriales | Clostridia |
| Oscillospiraceae | Eubacteriales | Clostridia |
| Oscillospiraceae | Eubacteriales | Clostridia |
| Oscillospiraceae | Eubacteriales | Clostridia |
| Oscillospiraceae | Eubacteriales | Clostridia |
| Oscillospiraceae | Eubacteriales | Clostridia |
| Oscillospiraceae | Eubacteriales | Clostridia |
| Oscillospiraceae | Eubacteriales | Clostridia |
| Oscillospiraceae | Eubacteriales | Clostridia |
| Oscillospiraceae | Eubacteriales | Clostridia |
| Oscillospiraceae | Eubacteriales | Clostridia |
| Oscillospiraceae | Eubacteriales | Clostridia |
| Oscillospiraceae | Eubacteriales | Clostridia |
| FGB77153 | OFGB77153 | CFGB77153 |
| FGB77153 | OFGB77153 | CFGB77153 |
| FGB1791 | OFGB1791 | CFGB1791 |
| FGB1791 | OFGB1791 | CFGB1791 |

| | | |
|---|---|---|
| FGB10290 | OFGB10290 | CFGB10290 |
| FGB10290 | OFGB10290 | CFGB10290 |
| Oscillospiraceae | Eubacteriales | Clostridia |
| Oscillospiraceae | Eubacteriales | Clostridia |
| FGB1765 | OFGB1765 | CFGB1765 |
| FGB1765 | OFGB1765 | CFGB1765 |
| FGB10349 | OFGB10349 | CFGB10349 |
| FGB10349 | OFGB10349 | CFGB10349 |
| FGB10667 | OFGB10667 | CFGB10667 |
| FGB10667 | OFGB10667 | CFGB10667 |
| Lachnospiraceae | Eubacteriales | Clostridia |
| Lachnospiraceae | Eubacteriales | Clostridia |
| Lachnospiraceae | Eubacteriales | Clostridia |
| Lachnospiraceae | Eubacteriales | Clostridia |
| Christensenellaceae | Eubacteriales | Clostridia |
| Christensenellaceae | Eubacteriales | Clostridia |
| FGB10299 | OFGB10299 | CFGB10299 |
| FGB10299 | OFGB10299 | CFGB10299 |
| Oscillospiraceae | Eubacteriales | Clostridia |
| Oscillospiraceae | Eubacteriales | Clostridia |
| Oscillospiraceae | Eubacteriales | Clostridia |
| Oscillospiraceae | Eubacteriales | Clostridia |
| Lachnospiraceae | Eubacteriales | Clostridia |
| Lachnospiraceae | Eubacteriales | Clostridia |
| Clostridia_unclassified | Clostridia_unclassified | Clostridia |
| Clostridia_unclassified | Clostridia_unclassified | Clostridia |
| Lachnospiraceae | Eubacteriales | Clostridia |
| Lachnospiraceae | Eubacteriales | Clostridia |
| Lachnospiraceae | Eubacteriales | Clostridia |
| Lachnospiraceae | Eubacteriales | Clostridia |
| Lachnospiraceae | Eubacteriales | Clostridia |
| Lachnospiraceae | Eubacteriales | Clostridia |
| Lachnospiraceae | Eubacteriales | Clostridia |
| Lachnospiraceae | Eubacteriales | Clostridia |
| Lachnospiraceae | Eubacteriales | Clostridia |
| Lachnospiraceae | Eubacteriales | Clostridia |
| Lachnospiraceae | Eubacteriales | Clostridia |
| Lachnospiraceae | Eubacteriales | Clostridia |
| Lachnospiraceae | Eubacteriales | Clostridia |
| Lachnospiraceae | Eubacteriales | Clostridia |
| Lachnospiraceae | Eubacteriales | Clostridia |
| Lachnospiraceae | Eubacteriales | Clostridia |
| Lactobacillaceae | Lactobacillales | Bacilli |
| Lactobacillaceae | Lactobacillales | Bacilli |
| Muribaculaceae | Bacteroidales | Bacteroidia |
| Muribaculaceae | Bacteroidales | Bacteroidia |

| | | |
|---|---|---|
| Oscillospiraceae | Eubacteriales | Clostridia |
| Oscillospiraceae | Eubacteriales | Clostridia |
| Oscillospiraceae | Eubacteriales | Clostridia |
| Oscillospiraceae | Eubacteriales | Clostridia |
| Oscillospiraceae | Eubacteriales | Clostridia |
| Oscillospiraceae | Eubacteriales | Clostridia |
| Oscillospiraceae | Eubacteriales | Clostridia |
| Oscillospiraceae | Eubacteriales | Clostridia |
| Sutterellaceae | Burkholderiales | Betaproteobacteria |
| Sutterellaceae | Burkholderiales | Betaproteobacteria |
| Peptostreptococcaceae | Eubacteriales | Clostridia |
| Peptostreptococcaceae | Eubacteriales | Clostridia |
| Lachnospiraceae | Eubacteriales | Clostridia |
| Lachnospiraceae | Eubacteriales | Clostridia |
| Turicibacteraceae | Erysipelotrichales | Erysipelotrichia |
| Turicibacteraceae | Erysipelotrichales | Erysipelotrichia |
| Lachnospiraceae | Eubacteriales | Clostridia |
| Lachnospiraceae | Eubacteriales | Clostridia |
| Lachnospiraceae | Eubacteriales | Clostridia |
| Lachnospiraceae | Eubacteriales | Clostridia |
| Oscillospiraceae | Eubacteriales | Clostridia |
| Oscillospiraceae | Eubacteriales | Clostridia |
| Oscillospiraceae | Eubacteriales | Clostridia |
| Oscillospiraceae | Eubacteriales | Clostridia |
| Eggerthellaceae | Eggerthellales | Coriobacteriia |
| Eggerthellaceae | Eggerthellales | Coriobacteriia |
| Eggerthellaceae | Eggerthellales | Coriobacteriia |
| Eggerthellaceae | Eggerthellales | Coriobacteriia |
| Eggerthellaceae | Eggerthellales | Coriobacteriia |
| Eggerthellaceae | Eggerthellales | Coriobacteriia |
| Akkermansiaceae | Verrucomicrobiales | Verrucomicrobiae |
| Akkermansiaceae | Verrucomicrobiales | Verrucomicrobiae |
| Rikenellaceae | Bacteroidales | Bacteroidia |
| Rikenellaceae | Bacteroidales | Bacteroidia |
| Oscillospiraceae | Eubacteriales | Clostridia |
| Oscillospiraceae | Eubacteriales | Clostridia |
| Bacteria_unclassified | Bacteria_unclassified | Bacteria_unclassified |
| Bacteria_unclassified | Bacteria_unclassified | Bacteria_unclassified |
| Bacteria_unclassified | Bacteria_unclassified | Bacteria_unclassified |
| Bacteria_unclassified | Bacteria_unclassified | Bacteria_unclassified |
| Bacteria_unclassified | Bacteria_unclassified | Bacteria_unclassified |
| Bacteria_unclassified | Bacteria_unclassified | Bacteria_unclassified |

| | | |
|---|---|---|
| Bacteria_unclassified | Bacteria_unclassified | Bacteria_unclassified |
| Bacteria_unclassified | Bacteria_unclassified | Bacteria_unclassified |
| Bacteria_unclassified | Bacteria_unclassified | Bacteria_unclassified |
| Bacteria_unclassified | Bacteria_unclassified | Bacteria_unclassified |
| Bacteria_unclassified | Bacteria_unclassified | Bacteria_unclassified |
| Bacteria_unclassified | Bacteria_unclassified | Bacteria_unclassified |
| Bacteroidaceae | Bacteroidales | Bacteroidia |
| Bacteroidaceae | Bacteroidales | Bacteroidia |
| Bifidobacteriaceae | Bifidobacteriales | Actinomycetia |
| Bifidobacteriaceae | Bifidobacteriales | Actinomycetia |
| Clostridia_unclassified | Clostridia_unclassified | Clostridia |
| Clostridia_unclassified | Clostridia_unclassified | Clostridia |
| Clostridiaceae | Eubacteriales | Clostridia |
| Clostridiaceae | Eubacteriales | Clostridia |
| Clostridiaceae | Eubacteriales | Clostridia |
| Clostridiaceae | Eubacteriales | Clostridia |
| Eubacteriales_unclassified | Eubacteriales | Clostridia |
| Eubacteriales_unclassified | Eubacteriales | Clostridia |
| Erysipelotrichaceae | Erysipelotrichales | Erysipelotrichia |
| Erysipelotrichaceae | Erysipelotrichales | Erysipelotrichia |
| Coriobacteriaceae | Coriobacteriales | Coriobacteriia |
| Coriobacteriaceae | Coriobacteriales | Coriobacteriia |
| Lachnospiraceae | Eubacteriales | Clostridia |
| Lachnospiraceae | Eubacteriales | Clostridia |
| Erysipelotrichaceae | Erysipelotrichales | Erysipelotrichia |
| Erysipelotrichaceae | Erysipelotrichales | Erysipelotrichia |
| Erysipelotrichales_unclassified | Erysipelotrichales | Erysipelotrichia |
| Erysipelotrichales_unclassified | Erysipelotrichales | Erysipelotrichia |
| Eubacteriaceae | Eubacteriales | Clostridia |
| Eubacteriaceae | Eubacteriales | Clostridia |
| Eubacteriaceae | Eubacteriales | Clostridia |
| Eubacteriaceae | Eubacteriales | Clostridia |
| Lachnospiraceae | Eubacteriales | Clostridia |
| Lachnospiraceae | Eubacteriales | Clostridia |
| Eggerthellaceae | Eggerthellales | Coriobacteriia |
| Eggerthellaceae | Eggerthellales | Coriobacteriia |
| Lachnospiraceae | Eubacteriales | Clostridia |
| Lachnospiraceae | Eubacteriales | Clostridia |
| Muribaculaceae | Bacteroidales | Bacteroidia |
| Muribaculaceae | Bacteroidales | Bacteroidia |
| Muribaculaceae | Bacteroidales | Bacteroidia |
| Muribaculaceae | Bacteroidales | Bacteroidia |
| Muribaculaceae | Bacteroidales | Bacteroidia |
| Muribaculaceae | Bacteroidales | Bacteroidia |

| | | |
|---|---|---|
| FGB9508 | OFGB9508 | CFGB9508 |
| FGB9508 | OFGB9508 | CFGB9508 |
| FGB2838 | OFGB2838 | CFGB2838 |
| FGB2838 | OFGB2838 | CFGB2838 |
| FGB2838 | OFGB2838 | CFGB2838 |
| FGB2838 | OFGB2838 | CFGB2838 |
| FGB2838 | OFGB2838 | CFGB2838 |
| FGB2838 | OFGB2838 | CFGB2838 |
| Pumilibacteraceae | Eubacteriales | Clostridia |
| Pumilibacteraceae | Eubacteriales | Clostridia |
| FGB28439 | OFGB28439 | CFGB28439 |
| FGB28439 | OFGB28439 | CFGB28439 |
| Clostridia_unclassified | Clostridia_unclassified | Clostridia |
| Clostridia_unclassified | Clostridia_unclassified | Clostridia |
| Eubacteriaceae | Eubacteriales | Clostridia |
| Eubacteriaceae | Eubacteriales | Clostridia |
| Lachnospiraceae | Eubacteriales | Clostridia |
| Lachnospiraceae | Eubacteriales | Clostridia |
| Lachnospiraceae | Eubacteriales | Clostridia |
| Lachnospiraceae | Eubacteriales | Clostridia |
| Lachnospiraceae | Eubacteriales | Clostridia |
| Lachnospiraceae | Eubacteriales | Clostridia |
| Lachnospiraceae | Eubacteriales | Clostridia |
| Lachnospiraceae | Eubacteriales | Clostridia |
| FGB77305 | OFGB77305 | CFGB77305 |
| FGB77305 | OFGB77305 | CFGB77305 |
| Clostridiaceae | Eubacteriales | Clostridia |
| Clostridiaceae | Eubacteriales | Clostridia |
| Lachnospiraceae | Eubacteriales | Clostridia |
| Lachnospiraceae | Eubacteriales | Clostridia |
| Lachnospiraceae | Eubacteriales | Clostridia |
| Lachnospiraceae | Eubacteriales | Clostridia |
| Lachnospiraceae | Eubacteriales | Clostridia |
| Lachnospiraceae | Eubacteriales | Clostridia |
| FGB9633 | OFGB9633 | CFGB9633 |
| FGB9633 | OFGB9633 | CFGB9633 |
| Bacteria_unclassified | Bacteria_unclassified | Bacteria_unclassified |
| Bacteria_unclassified | Bacteria_unclassified | Bacteria_unclassified |
| Bacteria_unclassified | Bacteria_unclassified | Bacteria_unclassified |
| Bacteria_unclassified | Bacteria_unclassified | Bacteria_unclassified |
| Bacteria_unclassified | Bacteria_unclassified | Bacteria_unclassified |
| Bacteria_unclassified | Bacteria_unclassified | Bacteria_unclassified |
| Bacteria_unclassified | Bacteria_unclassified | Bacteria_unclassified |
| Bacteria_unclassified | Bacteria_unclassified | Bacteria_unclassified |
| Lachnospiraceae | Eubacteriales | Clostridia |
| Lachnospiraceae | Eubacteriales | Clostridia |

| | | |
|---|---|---|
| Lachnospiraceae | Eubacteriales | Clostridia |
| Lachnospiraceae | Eubacteriales | Clostridia |
| Lachnospiraceae | Eubacteriales | Clostridia |
| Lachnospiraceae | Eubacteriales | Clostridia |
| FGB77359 | OFGB77359 | CFGB77359 |
| FGB77359 | OFGB77359 | CFGB77359 |
| FGB9639 | OFGB9639 | CFGB9639 |
| FGB9639 | OFGB9639 | CFGB9639 |
| Lachnospiraceae | Eubacteriales | Clostridia |
| Lachnospiraceae | Eubacteriales | Clostridia |
| Lachnospiraceae | Eubacteriales | Clostridia |
| Lachnospiraceae | Eubacteriales | Clostridia |
| Lachnospiraceae | Eubacteriales | Clostridia |
| Lachnospiraceae | Eubacteriales | Clostridia |
| Clostridiaceae | Eubacteriales | Clostridia |
| Clostridiaceae | Eubacteriales | Clostridia |
| Clostridiaceae | Eubacteriales | Clostridia |
| Clostridiaceae | Eubacteriales | Clostridia |
| Clostridiaceae | Eubacteriales | Clostridia |
| Clostridiaceae | Eubacteriales | Clostridia |
| Clostridiaceae | Eubacteriales | Clostridia |
| Clostridiaceae | Eubacteriales | Clostridia |
| Clostridiaceae | Eubacteriales | Clostridia |
| Clostridiaceae | Eubacteriales | Clostridia |
| Clostridiaceae | Eubacteriales | Clostridia |
| Clostridiaceae | Eubacteriales | Clostridia |
| Clostridiaceae | Eubacteriales | Clostridia |
| Clostridiaceae | Eubacteriales | Clostridia |
| Eubacteriaceae | Eubacteriales | Clostridia |
| Eubacteriaceae | Eubacteriales | Clostridia |
| FGB9658 | OFGB9658 | CFGB9658 |
| FGB9658 | OFGB9658 | CFGB9658 |
| FGB9659 | OFGB9659 | CFGB9659 |
| FGB9659 | OFGB9659 | CFGB9659 |
| Bacteria_unclassified | Bacteria_unclassified | Bacteria_unclassified |
| Bacteria_unclassified | Bacteria_unclassified | Bacteria_unclassified |
| FGB9827 | OFGB9827 | CFGB9827 |
| FGB9827 | OFGB9827 | CFGB9827 |
| Eubacteriaceae | Eubacteriales | Clostridia |
| Eubacteriaceae | Eubacteriales | Clostridia |
| FGB77303 | OFGB77303 | CFGB77303 |
| FGB77303 | OFGB77303 | CFGB77303 |
| Eubacteriales_unclassified | Eubacteriales | Clostridia |
| Eubacteriales_unclassified | Eubacteriales | Clostridia |
| Oscillospiraceae | Eubacteriales | Clostridia |
| Oscillospiraceae | Eubacteriales | Clostridia |

| | | |
|---|---|---|
| FGB30328 | OFGB30328 | CFGB30328 |
| FGB30328 | OFGB30328 | CFGB30328 |
| Oscillospiraceae | Eubacteriales | Clostridia |
| Oscillospiraceae | Eubacteriales | Clostridia |
| Oscillospiraceae | Eubacteriales | Clostridia |
| Oscillospiraceae | Eubacteriales | Clostridia |
| Oscillospiraceae | Eubacteriales | Clostridia |
| Oscillospiraceae | Eubacteriales | Clostridia |
| Oscillospiraceae | Eubacteriales | Clostridia |
| Oscillospiraceae | Eubacteriales | Clostridia |
| Oscillospiraceae | Eubacteriales | Clostridia |
| Oscillospiraceae | Eubacteriales | Clostridia |
| Oscillospiraceae | Eubacteriales | Clostridia |
| Oscillospiraceae | Eubacteriales | Clostridia |
| Oscillospiraceae | Eubacteriales | Clostridia |
| Oscillospiraceae | Eubacteriales | Clostridia |
| FGB77153 | OFGB77153 | CFGB77153 |
| FGB77153 | OFGB77153 | CFGB77153 |
| FGB1791 | OFGB1791 | CFGB1791 |
| FGB1791 | OFGB1791 | CFGB1791 |
| FGB10290 | OFGB10290 | CFGB10290 |
| FGB10290 | OFGB10290 | CFGB10290 |
| Oscillospiraceae | Eubacteriales | Clostridia |
| Oscillospiraceae | Eubacteriales | Clostridia |
| FGB1765 | OFGB1765 | CFGB1765 |
| FGB1765 | OFGB1765 | CFGB1765 |
| FGB10349 | OFGB10349 | CFGB10349 |
| FGB10349 | OFGB10349 | CFGB10349 |
| FGB10667 | OFGB10667 | CFGB10667 |
| FGB10667 | OFGB10667 | CFGB10667 |
| Lachnospiraceae | Eubacteriales | Clostridia |
| Lachnospiraceae | Eubacteriales | Clostridia |
| Lachnospiraceae | Eubacteriales | Clostridia |
| Lachnospiraceae | Eubacteriales | Clostridia |
| Christensenellaceae | Eubacteriales | Clostridia |
| Christensenellaceae | Eubacteriales | Clostridia |
| FGB10299 | OFGB10299 | CFGB10299 |
| FGB10299 | OFGB10299 | CFGB10299 |
| Oscillospiraceae | Eubacteriales | Clostridia |
| Oscillospiraceae | Eubacteriales | Clostridia |
| Oscillospiraceae | Eubacteriales | Clostridia |
| Oscillospiraceae | Eubacteriales | Clostridia |
| Lachnospiraceae | Eubacteriales | Clostridia |
| Lachnospiraceae | Eubacteriales | Clostridia |
| Clostridia_unclassified | Clostridia_unclassified | Clostridia |
| Clostridia_unclassified | Clostridia_unclassified | Clostridia |

| | | |
|---|---|---|
| Lachnospiraceae | Eubacteriales | Clostridia |
| Lachnospiraceae | Eubacteriales | Clostridia |
| Lachnospiraceae | Eubacteriales | Clostridia |
| Lachnospiraceae | Eubacteriales | Clostridia |
| Lachnospiraceae | Eubacteriales | Clostridia |
| Lachnospiraceae | Eubacteriales | Clostridia |
| Lachnospiraceae | Eubacteriales | Clostridia |
| Lachnospiraceae | Eubacteriales | Clostridia |
| Lachnospiraceae | Eubacteriales | Clostridia |
| Lachnospiraceae | Eubacteriales | Clostridia |
| Lachnospiraceae | Eubacteriales | Clostridia |
| Lachnospiraceae | Eubacteriales | Clostridia |
| Lachnospiraceae | Eubacteriales | Clostridia |
| Lachnospiraceae | Eubacteriales | Clostridia |
| Lachnospiraceae | Eubacteriales | Clostridia |
| Lachnospiraceae | Eubacteriales | Clostridia |
| Lactobacillaceae | Lactobacillales | Bacilli |
| Lactobacillaceae | Lactobacillales | Bacilli |
| Muribaculaceae | Bacteroidales | Bacteroidia |
| Muribaculaceae | Bacteroidales | Bacteroidia |
| Oscillospiraceae | Eubacteriales | Clostridia |
| Oscillospiraceae | Eubacteriales | Clostridia |
| Oscillospiraceae | Eubacteriales | Clostridia |
| Oscillospiraceae | Eubacteriales | Clostridia |
| Oscillospiraceae | Eubacteriales | Clostridia |
| Oscillospiraceae | Eubacteriales | Clostridia |
| Oscillospiraceae | Eubacteriales | Clostridia |
| Oscillospiraceae | Eubacteriales | Clostridia |
| Sutterellaceae | Burkholderiales | Betaproteobacteria |
| Sutterellaceae | Burkholderiales | Betaproteobacteria |
| Peptostreptococcaceae | Eubacteriales | Clostridia |
| Peptostreptococcaceae | Eubacteriales | Clostridia |
| Lachnospiraceae | Eubacteriales | Clostridia |
| Lachnospiraceae | Eubacteriales | Clostridia |
| Turicibacteraceae | Erysipelotrichales | Erysipelotrichia |
| Turicibacteraceae | Erysipelotrichales | Erysipelotrichia |
| Lachnospiraceae | Eubacteriales | Clostridia |
| Lachnospiraceae | Eubacteriales | Clostridia |
| Lachnospiraceae | Eubacteriales | Clostridia |
| Lachnospiraceae | Eubacteriales | Clostridia |
| Oscillospiraceae | Eubacteriales | Clostridia |
| Oscillospiraceae | Eubacteriales | Clostridia |

| | | |
|---|---|---|
| Oscillospiraceae | Eubacteriales | Clostridia |
| Oscillospiraceae | Eubacteriales | Clostridia |
| Eggerthellaceae | Eggerthellales | Coriobacteriia |
| Eggerthellaceae | Eggerthellales | Coriobacteriia |
| Eggerthellaceae | Eggerthellales | Coriobacteriia |
| Eggerthellaceae | Eggerthellales | Coriobacteriia |
| Eggerthellaceae | Eggerthellales | Coriobacteriia |
| Eggerthellaceae | Eggerthellales | Coriobacteriia |
| Akkermansiaceae | Verrucomicrobiales | Verrucomicrobiae |
| Akkermansiaceae | Verrucomicrobiales | Verrucomicrobiae |
| Rikenellaceae | Bacteroidales | Bacteroidia |
| Rikenellaceae | Bacteroidales | Bacteroidia |
| Oscillospiraceae | Eubacteriales | Clostridia |
| Oscillospiraceae | Eubacteriales | Clostridia |
| Bacteria_unclassified | Bacteria_unclassified | Bacteria_unclassified |
| Bacteria_unclassified | Bacteria_unclassified | Bacteria_unclassified |
| Bacteria_unclassified | Bacteria_unclassified | Bacteria_unclassified |
| Bacteria_unclassified | Bacteria_unclassified | Bacteria_unclassified |
| Bacteria_unclassified | Bacteria_unclassified | Bacteria_unclassified |
| Bacteria_unclassified | Bacteria_unclassified | Bacteria_unclassified |
| Bacteria_unclassified | Bacteria_unclassified | Bacteria_unclassified |
| Bacteria_unclassified | Bacteria_unclassified | Bacteria_unclassified |
| Bacteria_unclassified | Bacteria_unclassified | Bacteria_unclassified |
| Bacteria_unclassified | Bacteria_unclassified | Bacteria_unclassified |
| Bacteria_unclassified | Bacteria_unclassified | Bacteria_unclassified |
| Bacteria_unclassified | Bacteria_unclassified | Bacteria_unclassified |
| Bacteroidaceae | Bacteroidales | Bacteroidia |
| Bacteroidaceae | Bacteroidales | Bacteroidia |

| | | |
|---|---|---|
| Bifidobacteriaceae | Bifidobacteriales | Actinomycetia |
| Bifidobacteriaceae | Bifidobacteriales | Actinomycetia |
| Clostridia_unclassified | Clostridia_unclassified | Clostridia |
| Clostridia_unclassified | Clostridia_unclassified | Clostridia |
| Clostridiaceae | Eubacteriales | Clostridia |
| Clostridiaceae | Eubacteriales | Clostridia |
| Clostridiaceae | Eubacteriales | Clostridia |
| Clostridiaceae | Eubacteriales | Clostridia |
| Eubacteriales_unclassified | Eubacteriales | Clostridia |
| Eubacteriales_unclassified | Eubacteriales | Clostridia |
| Erysipelotrichaceae | Erysipelotrichales | Erysipelotrichia |
| Erysipelotrichaceae | Erysipelotrichales | Erysipelotrichia |
| Coriobacteriaceae | Coriobacteriales | Coriobacteriia |
| Coriobacteriaceae | Coriobacteriales | Coriobacteriia |
| Lachnospiraceae | Eubacteriales | Clostridia |
| Lachnospiraceae | Eubacteriales | Clostridia |

| | | |
|---|---|---|
| Erysipelotrichaceae | Erysipelotrichales | Erysipelotrichia |
| Erysipelotrichaceae | Erysipelotrichales | Erysipelotrichia |
| Erysipelotrichales_unclassified | Erysipelotrichales | Erysipelotrichia |
| Erysipelotrichales_unclassified | Erysipelotrichales | Erysipelotrichia |
| Eubacteriaceae | Eubacteriales | Clostridia |
| Eubacteriaceae | Eubacteriales | Clostridia |
| Eubacteriaceae | Eubacteriales | Clostridia |
| Eubacteriaceae | Eubacteriales | Clostridia |
| Lachnospiraceae | Eubacteriales | Clostridia |
| Lachnospiraceae | Eubacteriales | Clostridia |
| Eggerthellaceae | Eggerthellales | Coriobacteriia |
| Eggerthellaceae | Eggerthellales | Coriobacteriia |
| Lachnospiraceae | Eubacteriales | Clostridia |
| Lachnospiraceae | Eubacteriales | Clostridia |
| Muribaculaceae | Bacteroidales | Bacteroidia |
| Muribaculaceae | Bacteroidales | Bacteroidia |
| Muribaculaceae | Bacteroidales | Bacteroidia |
| Muribaculaceae | Bacteroidales | Bacteroidia |
| Muribaculaceae | Bacteroidales | Bacteroidia |
| Muribaculaceae | Bacteroidales | Bacteroidia |
| FGB9508 | OFGB9508 | CFGB9508 |
| FGB9508 | OFGB9508 | CFGB9508 |
| FGB2838 | OFGB2838 | CFGB2838 |
| FGB2838 | OFGB2838 | CFGB2838 |
| FGB2838 | OFGB2838 | CFGB2838 |
| FGB2838 | OFGB2838 | CFGB2838 |
| FGB2838 | OFGB2838 | CFGB2838 |
| FGB2838 | OFGB2838 | CFGB2838 |
| Pumilibacteraceae | Eubacteriales | Clostridia |
| Pumilibacteraceae | Eubacteriales | Clostridia |
| FGB28439 | OFGB28439 | CFGB28439 |
| FGB28439 | OFGB28439 | CFGB28439 |
| Clostridia_unclassified | Clostridia_unclassified | Clostridia |
| Clostridia_unclassified | Clostridia_unclassified | Clostridia |
| Eubacteriaceae | Eubacteriales | Clostridia |
| Eubacteriaceae | Eubacteriales | Clostridia |
| Lachnospiraceae | Eubacteriales | Clostridia |
| Lachnospiraceae | Eubacteriales | Clostridia |
| Lachnospiraceae | Eubacteriales | Clostridia |
| Lachnospiraceae | Eubacteriales | Clostridia |
| Lachnospiraceae | Eubacteriales | Clostridia |
| Lachnospiraceae | Eubacteriales | Clostridia |
| Lachnospiraceae | Eubacteriales | Clostridia |
| Lachnospiraceae | Eubacteriales | Clostridia |
| FGB77305 | OFGB77305 | CFGB77305 |
| FGB77305 | OFGB77305 | CFGB77305 |

| | | |
|---|---|---|
| Clostridiaceae | Eubacteriales | Clostridia |
| Clostridiaceae | Eubacteriales | Clostridia |
| Lachnospiraceae | Eubacteriales | Clostridia |
| Lachnospiraceae | Eubacteriales | Clostridia |
| Lachnospiraceae | Eubacteriales | Clostridia |
| Lachnospiraceae | Eubacteriales | Clostridia |
| Lachnospiraceae | Eubacteriales | Clostridia |
| Lachnospiraceae | Eubacteriales | Clostridia |
| FGB9633 | OFGB9633 | CFGB9633 |
| FGB9633 | OFGB9633 | CFGB9633 |
| Bacteria_unclassified | Bacteria_unclassified | Bacteria_unclassified |
| Bacteria_unclassified | Bacteria_unclassified | Bacteria_unclassified |
| Bacteria_unclassified | Bacteria_unclassified | Bacteria_unclassified |
| Bacteria_unclassified | Bacteria_unclassified | Bacteria_unclassified |
| Bacteria_unclassified | Bacteria_unclassified | Bacteria_unclassified |
| Bacteria_unclassified | Bacteria_unclassified | Bacteria_unclassified |
| Bacteria_unclassified | Bacteria_unclassified | Bacteria_unclassified |
| Bacteria_unclassified | Bacteria_unclassified | Bacteria_unclassified |
| Lachnospiraceae | Eubacteriales | Clostridia |
| Lachnospiraceae | Eubacteriales | Clostridia |
| Lachnospiraceae | Eubacteriales | Clostridia |
| Lachnospiraceae | Eubacteriales | Clostridia |
| Lachnospiraceae | Eubacteriales | Clostridia |
| Lachnospiraceae | Eubacteriales | Clostridia |
| FGB77359 | OFGB77359 | CFGB77359 |
| FGB77359 | OFGB77359 | CFGB77359 |
| FGB9639 | OFGB9639 | CFGB9639 |
| FGB9639 | OFGB9639 | CFGB9639 |
| Lachnospiraceae | Eubacteriales | Clostridia |
| Lachnospiraceae | Eubacteriales | Clostridia |
| Lachnospiraceae | Eubacteriales | Clostridia |
| Lachnospiraceae | Eubacteriales | Clostridia |
| Lachnospiraceae | Eubacteriales | Clostridia |
| Lachnospiraceae | Eubacteriales | Clostridia |
| Clostridiaceae | Eubacteriales | Clostridia |
| Clostridiaceae | Eubacteriales | Clostridia |
| Clostridiaceae | Eubacteriales | Clostridia |
| Clostridiaceae | Eubacteriales | Clostridia |
| Clostridiaceae | Eubacteriales | Clostridia |
| Clostridiaceae | Eubacteriales | Clostridia |
| Clostridiaceae | Eubacteriales | Clostridia |
| Clostridiaceae | Eubacteriales | Clostridia |
| Clostridiaceae | Eubacteriales | Clostridia |
| Clostridiaceae | Eubacteriales | Clostridia |
| Clostridiaceae | Eubacteriales | Clostridia |
| Clostridiaceae | Eubacteriales | Clostridia |

| | | |
|---|---|---|
| Clostridiaceae | Eubacteriales | Clostridia |
| Clostridiaceae | Eubacteriales | Clostridia |
| Eubacteriaceae | Eubacteriales | Clostridia |
| Eubacteriaceae | Eubacteriales | Clostridia |
| FGB9658 | OFGB9658 | CFGB9658 |
| FGB9658 | OFGB9658 | CFGB9658 |
| FGB9659 | OFGB9659 | CFGB9659 |
| FGB9659 | OFGB9659 | CFGB9659 |
| Bacteria_unclassified | Bacteria_unclassified | Bacteria_unclassified |
| Bacteria_unclassified | Bacteria_unclassified | Bacteria_unclassified |
| FGB9827 | OFGB9827 | CFGB9827 |
| FGB9827 | OFGB9827 | CFGB9827 |
| Eubacteriaceae | Eubacteriales | Clostridia |
| Eubacteriaceae | Eubacteriales | Clostridia |
| FGB77303 | OFGB77303 | CFGB77303 |
| FGB77303 | OFGB77303 | CFGB77303 |
| Eubacteriales_unclassified | Eubacteriales | Clostridia |
| Eubacteriales_unclassified | Eubacteriales | Clostridia |
| Oscillospiraceae | Eubacteriales | Clostridia |
| Oscillospiraceae | Eubacteriales | Clostridia |
| FGB30328 | OFGB30328 | CFGB30328 |
| FGB30328 | OFGB30328 | CFGB30328 |
| Oscillospiraceae | Eubacteriales | Clostridia |
| Oscillospiraceae | Eubacteriales | Clostridia |
| Oscillospiraceae | Eubacteriales | Clostridia |
| Oscillospiraceae | Eubacteriales | Clostridia |
| Oscillospiraceae | Eubacteriales | Clostridia |
| Oscillospiraceae | Eubacteriales | Clostridia |
| Oscillospiraceae | Eubacteriales | Clostridia |
| Oscillospiraceae | Eubacteriales | Clostridia |
| Oscillospiraceae | Eubacteriales | Clostridia |
| Oscillospiraceae | Eubacteriales | Clostridia |
| Oscillospiraceae | Eubacteriales | Clostridia |
| Oscillospiraceae | Eubacteriales | Clostridia |
| Oscillospiraceae | Eubacteriales | Clostridia |
| Oscillospiraceae | Eubacteriales | Clostridia |
| FGB77153 | OFGB77153 | CFGB77153 |
| FGB77153 | OFGB77153 | CFGB77153 |
| FGB1791 | OFGB1791 | CFGB1791 |
| FGB1791 | OFGB1791 | CFGB1791 |
| FGB10290 | OFGB10290 | CFGB10290 |
| FGB10290 | OFGB10290 | CFGB10290 |
| Oscillospiraceae | Eubacteriales | Clostridia |
| Oscillospiraceae | Eubacteriales | Clostridia |
| FGB1765 | OFGB1765 | CFGB1765 |
| FGB1765 | OFGB1765 | CFGB1765 |

| | | |
|---|---|---|
| FGB10349 | OFGB10349 | CFGB10349 |
| FGB10349 | OFGB10349 | CFGB10349 |
| FGB10667 | OFGB10667 | CFGB10667 |
| FGB10667 | OFGB10667 | CFGB10667 |
| Lachnospiraceae | Eubacteriales | Clostridia |
| Lachnospiraceae | Eubacteriales | Clostridia |
| Lachnospiraceae | Eubacteriales | Clostridia |
| Lachnospiraceae | Eubacteriales | Clostridia |
| Christensenellaceae | Eubacteriales | Clostridia |
| Christensenellaceae | Eubacteriales | Clostridia |
| FGB10299 | OFGB10299 | CFGB10299 |
| FGB10299 | OFGB10299 | CFGB10299 |
| Oscillospiraceae | Eubacteriales | Clostridia |
| Oscillospiraceae | Eubacteriales | Clostridia |
| Oscillospiraceae | Eubacteriales | Clostridia |
| Oscillospiraceae | Eubacteriales | Clostridia |
| Lachnospiraceae | Eubacteriales | Clostridia |
| Lachnospiraceae | Eubacteriales | Clostridia |
| Clostridia_unclassified | Clostridia_unclassified | Clostridia |
| Clostridia_unclassified | Clostridia_unclassified | Clostridia |
| Lachnospiraceae | Eubacteriales | Clostridia |
| Lachnospiraceae | Eubacteriales | Clostridia |
| Lachnospiraceae | Eubacteriales | Clostridia |
| Lachnospiraceae | Eubacteriales | Clostridia |
| Lachnospiraceae | Eubacteriales | Clostridia |
| Lachnospiraceae | Eubacteriales | Clostridia |
| Lachnospiraceae | Eubacteriales | Clostridia |
| Lachnospiraceae | Eubacteriales | Clostridia |
| Lachnospiraceae | Eubacteriales | Clostridia |
| Lachnospiraceae | Eubacteriales | Clostridia |
| Lachnospiraceae | Eubacteriales | Clostridia |
| Lachnospiraceae | Eubacteriales | Clostridia |
| Lachnospiraceae | Eubacteriales | Clostridia |
| Lachnospiraceae | Eubacteriales | Clostridia |
| Lachnospiraceae | Eubacteriales | Clostridia |
| Lachnospiraceae | Eubacteriales | Clostridia |
| Lactobacillaceae | Lactobacillales | Bacilli |
| Lactobacillaceae | Lactobacillales | Bacilli |
| Muribaculaceae | Bacteroidales | Bacteroidia |
| Muribaculaceae | Bacteroidales | Bacteroidia |
| Oscillospiraceae | Eubacteriales | Clostridia |
| Oscillospiraceae | Eubacteriales | Clostridia |
| Oscillospiraceae | Eubacteriales | Clostridia |
| Oscillospiraceae | Eubacteriales | Clostridia |
| Oscillospiraceae | Eubacteriales | Clostridia |
| Oscillospiraceae | Eubacteriales | Clostridia |

| | | |
|---|---|---|
| Oscillospiraceae | Eubacteriales | Clostridia |
| Oscillospiraceae | Eubacteriales | Clostridia |
| Sutterellaceae | Burkholderiales | Betaproteobacteria |
| Sutterellaceae | Burkholderiales | Betaproteobacteria |
| Peptostreptococcaceae | Eubacteriales | Clostridia |
| Peptostreptococcaceae | Eubacteriales | Clostridia |
| Lachnospiraceae | Eubacteriales | Clostridia |
| Lachnospiraceae | Eubacteriales | Clostridia |
| Turicibacteraceae | Erysipelotrichales | Erysipelotrichia |
| Turicibacteraceae | Erysipelotrichales | Erysipelotrichia |
| Lachnospiraceae | Eubacteriales | Clostridia |
| Lachnospiraceae | Eubacteriales | Clostridia |
| Lachnospiraceae | Eubacteriales | Clostridia |
| Lachnospiraceae | Eubacteriales | Clostridia |
| Oscillospiraceae | Eubacteriales | Clostridia |
| Oscillospiraceae | Eubacteriales | Clostridia |
| Oscillospiraceae | Eubacteriales | Clostridia |
| Oscillospiraceae | Eubacteriales | Clostridia |
| Eggerthellaceae | Eggerthellales | Coriobacteriia |
| Eggerthellaceae | Eggerthellales | Coriobacteriia |
| Eggerthellaceae | Eggerthellales | Coriobacteriia |
| Eggerthellaceae | Eggerthellales | Coriobacteriia |
| Eggerthellaceae | Eggerthellales | Coriobacteriia |
| Eggerthellaceae | Eggerthellales | Coriobacteriia |
| Akkermansiaceae | Verrucomicrobiales | Verrucomicrobiae |
| Akkermansiaceae | Verrucomicrobiales | Verrucomicrobiae |
| Rikenellaceae | Bacteroidales | Bacteroidia |
| Rikenellaceae | Bacteroidales | Bacteroidia |
| Oscillospiraceae | Eubacteriales | Clostridia |
| Oscillospiraceae | Eubacteriales | Clostridia |
| Bacteria_unclassified | Bacteria_unclassified | Bacteria_unclassified |
| Bacteria_unclassified | Bacteria_unclassified | Bacteria_unclassified |
| Bacteria_unclassified | Bacteria_unclassified | Bacteria_unclassified |
| Bacteria_unclassified | Bacteria_unclassified | Bacteria_unclassified |
| Bacteria_unclassified | Bacteria_unclassified | Bacteria_unclassified |
| Bacteria_unclassified | Bacteria_unclassified | Bacteria_unclassified |
| Bacteria_unclassified | Bacteria_unclassified | Bacteria_unclassified |
| Bacteria_unclassified | Bacteria_unclassified | Bacteria_unclassified |
| Bacteria_unclassified | Bacteria_unclassified | Bacteria_unclassified |
| Bacteria_unclassified | Bacteria_unclassified | Bacteria_unclassified |
| Bacteria_unclassified | Bacteria_unclassified | Bacteria_unclassified |
| Bacteria_unclassified | Bacteria_unclassified | Bacteria_unclassified |

| | | |
|---|---|---|
| Bacteroidaceae | Bacteroidales | Bacteroidia |
| Bacteroidaceae | Bacteroidales | Bacteroidia |
| Bifidobacteriaceae | Bifidobacteriales | Actinomycetia |
| Bifidobacteriaceae | Bifidobacteriales | Actinomycetia |
| Clostridia_unclassified | Clostridia_unclassified | Clostridia |
| Clostridia_unclassified | Clostridia_unclassified | Clostridia |
| Clostridiaceae | Eubacteriales | Clostridia |
| Clostridiaceae | Eubacteriales | Clostridia |
| Clostridiaceae | Eubacteriales | Clostridia |
| Clostridiaceae | Eubacteriales | Clostridia |
| Eubacteriales_unclassified | Eubacteriales | Clostridia |
| Eubacteriales_unclassified | Eubacteriales | Clostridia |
| Erysipelotrichaceae | Erysipelotrichales | Erysipelotrichia |
| Erysipelotrichaceae | Erysipelotrichales | Erysipelotrichia |
| Coriobacteriaceae | Coriobacteriales | Coriobacteriia |
| Coriobacteriaceae | Coriobacteriales | Coriobacteriia |
| Lachnospiraceae | Eubacteriales | Clostridia |
| Lachnospiraceae | Eubacteriales | Clostridia |
| Erysipelotrichaceae | Erysipelotrichales | Erysipelotrichia |
| Erysipelotrichaceae | Erysipelotrichales | Erysipelotrichia |
| Erysipelotrichales_unclassified | Erysipelotrichales | Erysipelotrichia |
| Erysipelotrichales_unclassified | Erysipelotrichales | Erysipelotrichia |
| Eubacteriaceae | Eubacteriales | Clostridia |
| Eubacteriaceae | Eubacteriales | Clostridia |
| Eubacteriaceae | Eubacteriales | Clostridia |
| Eubacteriaceae | Eubacteriales | Clostridia |
| Lachnospiraceae | Eubacteriales | Clostridia |
| Lachnospiraceae | Eubacteriales | Clostridia |
| Eggerthellaceae | Eggerthellales | Coriobacteriia |
| Eggerthellaceae | Eggerthellales | Coriobacteriia |
| Lachnospiraceae | Eubacteriales | Clostridia |
| Lachnospiraceae | Eubacteriales | Clostridia |
| Muribaculaceae | Bacteroidales | Bacteroidia |
| Muribaculaceae | Bacteroidales | Bacteroidia |
| Muribaculaceae | Bacteroidales | Bacteroidia |
| Muribaculaceae | Bacteroidales | Bacteroidia |
| Muribaculaceae | Bacteroidales | Bacteroidia |
| Muribaculaceae | Bacteroidales | Bacteroidia |
| FGB9508 | OFGB9508 | CFGB9508 |
| FGB9508 | OFGB9508 | CFGB9508 |
| FGB2838 | OFGB2838 | CFGB2838 |
| FGB2838 | OFGB2838 | CFGB2838 |
| FGB2838 | OFGB2838 | CFGB2838 |
| FGB2838 | OFGB2838 | CFGB2838 |

| | | |
|---|---|---|
| FGB2838 | OFGB2838 | CFGB2838 |
| FGB2838 | OFGB2838 | CFGB2838 |
| Pumilibacteraceae | Eubacteriales | Clostridia |
| Pumilibacteraceae | Eubacteriales | Clostridia |
| FGB28439 | OFGB28439 | CFGB28439 |
| FGB28439 | OFGB28439 | CFGB28439 |
| Clostridia_unclassified | Clostridia_unclassified | Clostridia |
| Clostridia_unclassified | Clostridia_unclassified | Clostridia |
| Eubacteriaceae | Eubacteriales | Clostridia |
| Eubacteriaceae | Eubacteriales | Clostridia |
| Lachnospiraceae | Eubacteriales | Clostridia |
| Lachnospiraceae | Eubacteriales | Clostridia |
| Lachnospiraceae | Eubacteriales | Clostridia |
| Lachnospiraceae | Eubacteriales | Clostridia |
| Lachnospiraceae | Eubacteriales | Clostridia |
| Lachnospiraceae | Eubacteriales | Clostridia |
| Lachnospiraceae | Eubacteriales | Clostridia |
| Lachnospiraceae | Eubacteriales | Clostridia |
| FGB77305 | OFGB77305 | CFGB77305 |
| FGB77305 | OFGB77305 | CFGB77305 |
| Clostridiaceae | Eubacteriales | Clostridia |
| Clostridiaceae | Eubacteriales | Clostridia |
| Lachnospiraceae | Eubacteriales | Clostridia |
| Lachnospiraceae | Eubacteriales | Clostridia |
| Lachnospiraceae | Eubacteriales | Clostridia |
| Lachnospiraceae | Eubacteriales | Clostridia |
| Lachnospiraceae | Eubacteriales | Clostridia |
| Lachnospiraceae | Eubacteriales | Clostridia |
| FGB9633 | OFGB9633 | CFGB9633 |
| FGB9633 | OFGB9633 | CFGB9633 |
| Bacteria_unclassified | Bacteria_unclassified | Bacteria_unclassified |
| Bacteria_unclassified | Bacteria_unclassified | Bacteria_unclassified |
| Bacteria_unclassified | Bacteria_unclassified | Bacteria_unclassified |
| Bacteria_unclassified | Bacteria_unclassified | Bacteria_unclassified |
| Bacteria_unclassified | Bacteria_unclassified | Bacteria_unclassified |
| Bacteria_unclassified | Bacteria_unclassified | Bacteria_unclassified |
| Bacteria_unclassified | Bacteria_unclassified | Bacteria_unclassified |
| Bacteria_unclassified | Bacteria_unclassified | Bacteria_unclassified |
| Lachnospiraceae | Eubacteriales | Clostridia |
| Lachnospiraceae | Eubacteriales | Clostridia |
| Lachnospiraceae | Eubacteriales | Clostridia |
| Lachnospiraceae | Eubacteriales | Clostridia |
| Lachnospiraceae | Eubacteriales | Clostridia |
| Lachnospiraceae | Eubacteriales | Clostridia |
| FGB77359 | OFGB77359 | CFGB77359 |
| FGB77359 | OFGB77359 | CFGB77359 |

| | | |
|---|---|---|
| FGB9639 | OFGB9639 | CFGB9639 |
| FGB9639 | OFGB9639 | CFGB9639 |
| Lachnospiraceae | Eubacteriales | Clostridia |
| Lachnospiraceae | Eubacteriales | Clostridia |
| Lachnospiraceae | Eubacteriales | Clostridia |
| Lachnospiraceae | Eubacteriales | Clostridia |
| Lachnospiraceae | Eubacteriales | Clostridia |
| Lachnospiraceae | Eubacteriales | Clostridia |
| Clostridiaceae | Eubacteriales | Clostridia |
| Clostridiaceae | Eubacteriales | Clostridia |
| Clostridiaceae | Eubacteriales | Clostridia |
| Clostridiaceae | Eubacteriales | Clostridia |
| Clostridiaceae | Eubacteriales | Clostridia |
| Clostridiaceae | Eubacteriales | Clostridia |
| Clostridiaceae | Eubacteriales | Clostridia |
| Clostridiaceae | Eubacteriales | Clostridia |
| Clostridiaceae | Eubacteriales | Clostridia |
| Clostridiaceae | Eubacteriales | Clostridia |
| Clostridiaceae | Eubacteriales | Clostridia |
| Clostridiaceae | Eubacteriales | Clostridia |
| Clostridiaceae | Eubacteriales | Clostridia |
| Clostridiaceae | Eubacteriales | Clostridia |
| Eubacteriaceae | Eubacteriales | Clostridia |
| Eubacteriaceae | Eubacteriales | Clostridia |
| FGB9658 | OFGB9658 | CFGB9658 |
| FGB9658 | OFGB9658 | CFGB9658 |
| FGB9659 | OFGB9659 | CFGB9659 |
| FGB9659 | OFGB9659 | CFGB9659 |
| Bacteria_unclassified | Bacteria_unclassified | Bacteria_unclassified |
| Bacteria_unclassified | Bacteria_unclassified | Bacteria_unclassified |
| FGB9827 | OFGB9827 | CFGB9827 |
| FGB9827 | OFGB9827 | CFGB9827 |
| Eubacteriaceae | Eubacteriales | Clostridia |
| Eubacteriaceae | Eubacteriales | Clostridia |
| FGB77303 | OFGB77303 | CFGB77303 |
| FGB77303 | OFGB77303 | CFGB77303 |
| Eubacteriales_unclassified | Eubacteriales | Clostridia |
| Eubacteriales_unclassified | Eubacteriales | Clostridia |
| Oscillospiraceae | Eubacteriales | Clostridia |
| Oscillospiraceae | Eubacteriales | Clostridia |
| FGB30328 | OFGB30328 | CFGB30328 |
| FGB30328 | OFGB30328 | CFGB30328 |
| Oscillospiraceae | Eubacteriales | Clostridia |
| Oscillospiraceae | Eubacteriales | Clostridia |
| Oscillospiraceae | Eubacteriales | Clostridia |
| Oscillospiraceae | Eubacteriales | Clostridia |

| | | |
|---|---|---|
| Oscillospiraceae | Eubacteriales | Clostridia |
| Oscillospiraceae | Eubacteriales | Clostridia |
| Oscillospiraceae | Eubacteriales | Clostridia |
| Oscillospiraceae | Eubacteriales | Clostridia |
| Oscillospiraceae | Eubacteriales | Clostridia |
| Oscillospiraceae | Eubacteriales | Clostridia |
| Oscillospiraceae | Eubacteriales | Clostridia |
| Oscillospiraceae | Eubacteriales | Clostridia |
| Oscillospiraceae | Eubacteriales | Clostridia |
| Oscillospiraceae | Eubacteriales | Clostridia |
| FGB77153 | OFGB77153 | CFGB77153 |
| FGB77153 | OFGB77153 | CFGB77153 |
| FGB1791 | OFGB1791 | CFGB1791 |
| FGB1791 | OFGB1791 | CFGB1791 |
| FGB10290 | OFGB10290 | CFGB10290 |
| FGB10290 | OFGB10290 | CFGB10290 |
| Oscillospiraceae | Eubacteriales | Clostridia |
| Oscillospiraceae | Eubacteriales | Clostridia |
| FGB1765 | OFGB1765 | CFGB1765 |
| FGB1765 | OFGB1765 | CFGB1765 |
| FGB10349 | OFGB10349 | CFGB10349 |
| FGB10349 | OFGB10349 | CFGB10349 |
| FGB10667 | OFGB10667 | CFGB10667 |
| FGB10667 | OFGB10667 | CFGB10667 |
| Lachnospiraceae | Eubacteriales | Clostridia |
| Lachnospiraceae | Eubacteriales | Clostridia |
| Lachnospiraceae | Eubacteriales | Clostridia |
| Lachnospiraceae | Eubacteriales | Clostridia |
| Christensenellaceae | Eubacteriales | Clostridia |
| Christensenellaceae | Eubacteriales | Clostridia |
| FGB10299 | OFGB10299 | CFGB10299 |
| FGB10299 | OFGB10299 | CFGB10299 |
| Oscillospiraceae | Eubacteriales | Clostridia |
| Oscillospiraceae | Eubacteriales | Clostridia |
| Oscillospiraceae | Eubacteriales | Clostridia |
| Oscillospiraceae | Eubacteriales | Clostridia |
| Lachnospiraceae | Eubacteriales | Clostridia |
| Lachnospiraceae | Eubacteriales | Clostridia |
| Clostridia_unclassified | Clostridia_unclassified | Clostridia |
| Clostridia_unclassified | Clostridia_unclassified | Clostridia |
| Lachnospiraceae | Eubacteriales | Clostridia |
| Lachnospiraceae | Eubacteriales | Clostridia |
| Lachnospiraceae | Eubacteriales | Clostridia |
| Lachnospiraceae | Eubacteriales | Clostridia |
| Lachnospiraceae | Eubacteriales | Clostridia |
| Lachnospiraceae | Eubacteriales | Clostridia |

| | | |
|---|---|---|
| Lachnospiraceae | Eubacteriales | Clostridia |
| Lachnospiraceae | Eubacteriales | Clostridia |
| Lachnospiraceae | Eubacteriales | Clostridia |
| Lachnospiraceae | Eubacteriales | Clostridia |
| Lachnospiraceae | Eubacteriales | Clostridia |
| Lachnospiraceae | Eubacteriales | Clostridia |
| Lachnospiraceae | Eubacteriales | Clostridia |
| Lachnospiraceae | Eubacteriales | Clostridia |
| Lachnospiraceae | Eubacteriales | Clostridia |
| Lachnospiraceae | Eubacteriales | Clostridia |
| Lactobacillaceae | Lactobacillales | Bacilli |
| Lactobacillaceae | Lactobacillales | Bacilli |
| Muribaculaceae | Bacteroidales | Bacteroidia |
| Muribaculaceae | Bacteroidales | Bacteroidia |
| Oscillospiraceae | Eubacteriales | Clostridia |
| Oscillospiraceae | Eubacteriales | Clostridia |
| Oscillospiraceae | Eubacteriales | Clostridia |
| Oscillospiraceae | Eubacteriales | Clostridia |
| Oscillospiraceae | Eubacteriales | Clostridia |
| Oscillospiraceae | Eubacteriales | Clostridia |
| Oscillospiraceae | Eubacteriales | Clostridia |
| Oscillospiraceae | Eubacteriales | Clostridia |
| Sutterellaceae | Burkholderiales | Betaproteobacteria |
| Sutterellaceae | Burkholderiales | Betaproteobacteria |
| Peptostreptococcaceae | Eubacteriales | Clostridia |
| Peptostreptococcaceae | Eubacteriales | Clostridia |
| Lachnospiraceae | Eubacteriales | Clostridia |
| Lachnospiraceae | Eubacteriales | Clostridia |
| Turicibacteraceae | Erysipelotrichales | Erysipelotrichia |
| Turicibacteraceae | Erysipelotrichales | Erysipelotrichia |
| Lachnospiraceae | Eubacteriales | Clostridia |
| Lachnospiraceae | Eubacteriales | Clostridia |
| Lachnospiraceae | Eubacteriales | Clostridia |
| Lachnospiraceae | Eubacteriales | Clostridia |
| Oscillospiraceae | Eubacteriales | Clostridia |
| Oscillospiraceae | Eubacteriales | Clostridia |
| Oscillospiraceae | Eubacteriales | Clostridia |
| Oscillospiraceae | Eubacteriales | Clostridia |
| Eggerthellaceae | Eggerthellales | Coriobacteriia |
| Eggerthellaceae | Eggerthellales | Coriobacteriia |
| Eggerthellaceae | Eggerthellales | Coriobacteriia |
| Eggerthellaceae | Eggerthellales | Coriobacteriia |

| | | |
|---|---|---|
| Eggerthellaceae | Eggerthellales | Coriobacteriia |
| Eggerthellaceae | Eggerthellales | Coriobacteriia |
| Akkermansiaceae | Verrucomicrobiales | Verrucomicrobiae |
| Akkermansiaceae | Verrucomicrobiales | Verrucomicrobiae |
| Rikenellaceae | Bacteroidales | Bacteroidia |
| Rikenellaceae | Bacteroidales | Bacteroidia |
| Oscillospiraceae | Eubacteriales | Clostridia |
| Oscillospiraceae | Eubacteriales | Clostridia |
| Bacteria_unclassified | Bacteria_unclassified | Bacteria_unclassified |
| Bacteria_unclassified | Bacteria_unclassified | Bacteria_unclassified |
| Bacteria_unclassified | Bacteria_unclassified | Bacteria_unclassified |
| Bacteria_unclassified | Bacteria_unclassified | Bacteria_unclassified |
| Bacteria_unclassified | Bacteria_unclassified | Bacteria_unclassified |
| Bacteria_unclassified | Bacteria_unclassified | Bacteria_unclassified |
| Bacteria_unclassified | Bacteria_unclassified | Bacteria_unclassified |
| Bacteria_unclassified | Bacteria_unclassified | Bacteria_unclassified |
| Bacteria_unclassified | Bacteria_unclassified | Bacteria_unclassified |
| Bacteria_unclassified | Bacteria_unclassified | Bacteria_unclassified |
| Bacteria_unclassified | Bacteria_unclassified | Bacteria_unclassified |
| Bacteria_unclassified | Bacteria_unclassified | Bacteria_unclassified |
| Bacteroidaceae | Bacteroidales | Bacteroidia |
| Bacteroidaceae | Bacteroidales | Bacteroidia |

| | | |
|---|---|---|
| Bifidobacteriaceae | Bifidobacteriales | Actinomycetia |
| Bifidobacteriaceae | Bifidobacteriales | Actinomycetia |
| Clostridia_unclassified | Clostridia_unclassified | Clostridia |
| Clostridia_unclassified | Clostridia_unclassified | Clostridia |
| Clostridiaceae | Eubacteriales | Clostridia |
| Clostridiaceae | Eubacteriales | Clostridia |
| Clostridiaceae | Eubacteriales | Clostridia |
| Clostridiaceae | Eubacteriales | Clostridia |
| Eubacteriales_unclassified | Eubacteriales | Clostridia |
| Eubacteriales_unclassified | Eubacteriales | Clostridia |
| Erysipelotrichaceae | Erysipelotrichales | Erysipelotrichia |
| Erysipelotrichaceae | Erysipelotrichales | Erysipelotrichia |
| Coriobacteriaceae | Coriobacteriales | Coriobacteriia |
| Coriobacteriaceae | Coriobacteriales | Coriobacteriia |
| Lachnospiraceae | Eubacteriales | Clostridia |
| Lachnospiraceae | Eubacteriales | Clostridia |
| Erysipelotrichaceae | Erysipelotrichales | Erysipelotrichia |
| Erysipelotrichaceae | Erysipelotrichales | Erysipelotrichia |
| Erysipelotrichales_unclassified | Erysipelotrichales | Erysipelotrichia |
| Erysipelotrichales_unclassified | Erysipelotrichales | Erysipelotrichia |
| Eubacteriaceae | Eubacteriales | Clostridia |
| Eubacteriaceae | Eubacteriales | Clostridia |

| | | |
|---|---|---|
| Eubacteriaceae | Eubacteriales | Clostridia |
| Eubacteriaceae | Eubacteriales | Clostridia |
| Lachnospiraceae | Eubacteriales | Clostridia |
| Lachnospiraceae | Eubacteriales | Clostridia |
| Eggerthellaceae | Eggerthellales | Coriobacteriia |
| Eggerthellaceae | Eggerthellales | Coriobacteriia |
| Lachnospiraceae | Eubacteriales | Clostridia |
| Lachnospiraceae | Eubacteriales | Clostridia |
| Muribaculaceae | Bacteroidales | Bacteroidia |
| Muribaculaceae | Bacteroidales | Bacteroidia |
| Muribaculaceae | Bacteroidales | Bacteroidia |
| Muribaculaceae | Bacteroidales | Bacteroidia |
| Muribaculaceae | Bacteroidales | Bacteroidia |
| Muribaculaceae | Bacteroidales | Bacteroidia |
| FGB9508 | OFGB9508 | CFGB9508 |
| FGB9508 | OFGB9508 | CFGB9508 |
| FGB2838 | OFGB2838 | CFGB2838 |
| FGB2838 | OFGB2838 | CFGB2838 |
| FGB2838 | OFGB2838 | CFGB2838 |
| FGB2838 | OFGB2838 | CFGB2838 |
| FGB2838 | OFGB2838 | CFGB2838 |
| FGB2838 | OFGB2838 | CFGB2838 |
| Pumilibacteraceae | Eubacteriales | Clostridia |
| Pumilibacteraceae | Eubacteriales | Clostridia |
| FGB28439 | OFGB28439 | CFGB28439 |
| FGB28439 | OFGB28439 | CFGB28439 |
| Clostridia_unclassified | Clostridia_unclassified | Clostridia |
| Clostridia_unclassified | Clostridia_unclassified | Clostridia |
| Eubacteriaceae | Eubacteriales | Clostridia |
| Eubacteriaceae | Eubacteriales | Clostridia |
| Lachnospiraceae | Eubacteriales | Clostridia |
| Lachnospiraceae | Eubacteriales | Clostridia |
| Lachnospiraceae | Eubacteriales | Clostridia |
| Lachnospiraceae | Eubacteriales | Clostridia |
| Lachnospiraceae | Eubacteriales | Clostridia |
| Lachnospiraceae | Eubacteriales | Clostridia |
| Lachnospiraceae | Eubacteriales | Clostridia |
| Lachnospiraceae | Eubacteriales | Clostridia |
| FGB77305 | OFGB77305 | CFGB77305 |
| FGB77305 | OFGB77305 | CFGB77305 |
| Clostridiaceae | Eubacteriales | Clostridia |
| Clostridiaceae | Eubacteriales | Clostridia |
| Lachnospiraceae | Eubacteriales | Clostridia |
| Lachnospiraceae | Eubacteriales | Clostridia |
| Lachnospiraceae | Eubacteriales | Clostridia |
| Lachnospiraceae | Eubacteriales | Clostridia |

| | | |
|---|---|---|
| Lachnospiraceae | Eubacteriales | Clostridia |
| Lachnospiraceae | Eubacteriales | Clostridia |
| FGB9633 | OFGB9633 | CFGB9633 |
| FGB9633 | OFGB9633 | CFGB9633 |
| Bacteria_unclassified | Bacteria_unclassified | Bacteria_unclassified |
| Bacteria_unclassified | Bacteria_unclassified | Bacteria_unclassified |
| Bacteria_unclassified | Bacteria_unclassified | Bacteria_unclassified |
| Bacteria_unclassified | Bacteria_unclassified | Bacteria_unclassified |
| Bacteria_unclassified | Bacteria_unclassified | Bacteria_unclassified |
| Bacteria_unclassified | Bacteria_unclassified | Bacteria_unclassified |
| Bacteria_unclassified | Bacteria_unclassified | Bacteria_unclassified |
| Bacteria_unclassified | Bacteria_unclassified | Bacteria_unclassified |
| Lachnospiraceae | Eubacteriales | Clostridia |
| Lachnospiraceae | Eubacteriales | Clostridia |
| Lachnospiraceae | Eubacteriales | Clostridia |
| Lachnospiraceae | Eubacteriales | Clostridia |
| Lachnospiraceae | Eubacteriales | Clostridia |
| Lachnospiraceae | Eubacteriales | Clostridia |
| FGB77359 | OFGB77359 | CFGB77359 |
| FGB77359 | OFGB77359 | CFGB77359 |
| FGB9639 | OFGB9639 | CFGB9639 |
| FGB9639 | OFGB9639 | CFGB9639 |
| Lachnospiraceae | Eubacteriales | Clostridia |
| Lachnospiraceae | Eubacteriales | Clostridia |
| Lachnospiraceae | Eubacteriales | Clostridia |
| Lachnospiraceae | Eubacteriales | Clostridia |
| Lachnospiraceae | Eubacteriales | Clostridia |
| Lachnospiraceae | Eubacteriales | Clostridia |
| Clostridiaceae | Eubacteriales | Clostridia |
| Clostridiaceae | Eubacteriales | Clostridia |
| Clostridiaceae | Eubacteriales | Clostridia |
| Clostridiaceae | Eubacteriales | Clostridia |
| Clostridiaceae | Eubacteriales | Clostridia |
| Clostridiaceae | Eubacteriales | Clostridia |
| Clostridiaceae | Eubacteriales | Clostridia |
| Clostridiaceae | Eubacteriales | Clostridia |
| Clostridiaceae | Eubacteriales | Clostridia |
| Clostridiaceae | Eubacteriales | Clostridia |
| Clostridiaceae | Eubacteriales | Clostridia |
| Clostridiaceae | Eubacteriales | Clostridia |
| Clostridiaceae | Eubacteriales | Clostridia |
| Clostridiaceae | Eubacteriales | Clostridia |
| Eubacteriaceae | Eubacteriales | Clostridia |
| Eubacteriaceae | Eubacteriales | Clostridia |
| FGB9658 | OFGB9658 | CFGB9658 |
| FGB9658 | OFGB9658 | CFGB9658 |

| | | |
|---|---|---|
| FGB9659 | OFGB9659 | CFGB9659 |
| FGB9659 | OFGB9659 | CFGB9659 |
| Bacteria_unclassified | Bacteria_unclassified | Bacteria_unclassified |
| Bacteria_unclassified | Bacteria_unclassified | Bacteria_unclassified |
| FGB9827 | OFGB9827 | CFGB9827 |
| FGB9827 | OFGB9827 | CFGB9827 |
| Eubacteriaceae | Eubacteriales | Clostridia |
| Eubacteriaceae | Eubacteriales | Clostridia |
| FGB77303 | OFGB77303 | CFGB77303 |
| FGB77303 | OFGB77303 | CFGB77303 |
| Eubacteriales_unclassified | Eubacteriales | Clostridia |
| Eubacteriales_unclassified | Eubacteriales | Clostridia |
| Oscillospiraceae | Eubacteriales | Clostridia |
| Oscillospiraceae | Eubacteriales | Clostridia |
| FGB30328 | OFGB30328 | CFGB30328 |
| FGB30328 | OFGB30328 | CFGB30328 |
| Oscillospiraceae | Eubacteriales | Clostridia |
| Oscillospiraceae | Eubacteriales | Clostridia |
| Oscillospiraceae | Eubacteriales | Clostridia |
| Oscillospiraceae | Eubacteriales | Clostridia |
| Oscillospiraceae | Eubacteriales | Clostridia |
| Oscillospiraceae | Eubacteriales | Clostridia |
| Oscillospiraceae | Eubacteriales | Clostridia |
| Oscillospiraceae | Eubacteriales | Clostridia |
| Oscillospiraceae | Eubacteriales | Clostridia |
| Oscillospiraceae | Eubacteriales | Clostridia |
| Oscillospiraceae | Eubacteriales | Clostridia |
| Oscillospiraceae | Eubacteriales | Clostridia |
| Oscillospiraceae | Eubacteriales | Clostridia |
| Oscillospiraceae | Eubacteriales | Clostridia |
| FGB77153 | OFGB77153 | CFGB77153 |
| FGB77153 | OFGB77153 | CFGB77153 |
| FGB1791 | OFGB1791 | CFGB1791 |
| FGB1791 | OFGB1791 | CFGB1791 |
| FGB10290 | OFGB10290 | CFGB10290 |
| FGB10290 | OFGB10290 | CFGB10290 |
| Oscillospiraceae | Eubacteriales | Clostridia |
| Oscillospiraceae | Eubacteriales | Clostridia |
| FGB1765 | OFGB1765 | CFGB1765 |
| FGB1765 | OFGB1765 | CFGB1765 |
| FGB10349 | OFGB10349 | CFGB10349 |
| FGB10349 | OFGB10349 | CFGB10349 |
| FGB10667 | OFGB10667 | CFGB10667 |
| FGB10667 | OFGB10667 | CFGB10667 |
| Lachnospiraceae | Eubacteriales | Clostridia |
| Lachnospiraceae | Eubacteriales | Clostridia |

| | | |
|---|---|---|
| Lachnospiraceae | Eubacteriales | Clostridia |
| Lachnospiraceae | Eubacteriales | Clostridia |
| Christensenellaceae | Eubacteriales | Clostridia |
| Christensenellaceae | Eubacteriales | Clostridia |
| FGB10299 | OFGB10299 | CFGB10299 |
| FGB10299 | OFGB10299 | CFGB10299 |
| Oscillospiraceae | Eubacteriales | Clostridia |
| Oscillospiraceae | Eubacteriales | Clostridia |
| Oscillospiraceae | Eubacteriales | Clostridia |
| Oscillospiraceae | Eubacteriales | Clostridia |
| Lachnospiraceae | Eubacteriales | Clostridia |
| Lachnospiraceae | Eubacteriales | Clostridia |
| Clostridia_unclassified | Clostridia_unclassified | Clostridia |
| Clostridia_unclassified | Clostridia_unclassified | Clostridia |
| Lachnospiraceae | Eubacteriales | Clostridia |
| Lachnospiraceae | Eubacteriales | Clostridia |
| Lachnospiraceae | Eubacteriales | Clostridia |
| Lachnospiraceae | Eubacteriales | Clostridia |
| Lachnospiraceae | Eubacteriales | Clostridia |
| Lachnospiraceae | Eubacteriales | Clostridia |
| Lachnospiraceae | Eubacteriales | Clostridia |
| Lachnospiraceae | Eubacteriales | Clostridia |
| Lachnospiraceae | Eubacteriales | Clostridia |
| Lachnospiraceae | Eubacteriales | Clostridia |
| Lachnospiraceae | Eubacteriales | Clostridia |
| Lachnospiraceae | Eubacteriales | Clostridia |
| Lachnospiraceae | Eubacteriales | Clostridia |
| Lachnospiraceae | Eubacteriales | Clostridia |
| Lachnospiraceae | Eubacteriales | Clostridia |
| Lachnospiraceae | Eubacteriales | Clostridia |
| Lactobacillaceae | Lactobacillales | Bacilli |
| Lactobacillaceae | Lactobacillales | Bacilli |
| Muribaculaceae | Bacteroidales | Bacteroidia |
| Muribaculaceae | Bacteroidales | Bacteroidia |
| Oscillospiraceae | Eubacteriales | Clostridia |
| Oscillospiraceae | Eubacteriales | Clostridia |
| Oscillospiraceae | Eubacteriales | Clostridia |
| Oscillospiraceae | Eubacteriales | Clostridia |
| Oscillospiraceae | Eubacteriales | Clostridia |
| Oscillospiraceae | Eubacteriales | Clostridia |
| Oscillospiraceae | Eubacteriales | Clostridia |
| Oscillospiraceae | Eubacteriales | Clostridia |
| Sutterellaceae | Burkholderiales | Betaproteobacteria |
| Sutterellaceae | Burkholderiales | Betaproteobacteria |

| | | |
|---|---|---|
| Peptostreptococcaceae | Eubacteriales | Clostridia |
| Peptostreptococcaceae | Eubacteriales | Clostridia |
| Lachnospiraceae | Eubacteriales | Clostridia |
| Lachnospiraceae | Eubacteriales | Clostridia |
| Turicibacteraceae | Erysipelotrichales | Erysipelotrichia |
| Turicibacteraceae | Erysipelotrichales | Erysipelotrichia |
| Lachnospiraceae | Eubacteriales | Clostridia |
| Lachnospiraceae | Eubacteriales | Clostridia |
| Lachnospiraceae | Eubacteriales | Clostridia |
| Lachnospiraceae | Eubacteriales | Clostridia |
| Oscillospiraceae | Eubacteriales | Clostridia |
| Oscillospiraceae | Eubacteriales | Clostridia |
| Oscillospiraceae | Eubacteriales | Clostridia |
| Oscillospiraceae | Eubacteriales | Clostridia |
| Eggerthellaceae | Eggerthellales | Coriobacteriia |
| Eggerthellaceae | Eggerthellales | Coriobacteriia |
| Eggerthellaceae | Eggerthellales | Coriobacteriia |
| Eggerthellaceae | Eggerthellales | Coriobacteriia |
| Eggerthellaceae | Eggerthellales | Coriobacteriia |
| Eggerthellaceae | Eggerthellales | Coriobacteriia |
| Akkermansiaceae | Verrucomicrobiales | Verrucomicrobiae |
| Akkermansiaceae | Verrucomicrobiales | Verrucomicrobiae |
| Rikenellaceae | Bacteroidales | Bacteroidia |
| Rikenellaceae | Bacteroidales | Bacteroidia |
| Oscillospiraceae | Eubacteriales | Clostridia |
| Oscillospiraceae | Eubacteriales | Clostridia |
| Bacteria_unclassified | Bacteria_unclassified | Bacteria_unclassified |
| Bacteria_unclassified | Bacteria_unclassified | Bacteria_unclassified |
| Bacteria_unclassified | Bacteria_unclassified | Bacteria_unclassified |
| Bacteria_unclassified | Bacteria_unclassified | Bacteria_unclassified |
| Bacteria_unclassified | Bacteria_unclassified | Bacteria_unclassified |
| Bacteria_unclassified | Bacteria_unclassified | Bacteria_unclassified |
| Bacteria_unclassified | Bacteria_unclassified | Bacteria_unclassified |
| Bacteria_unclassified | Bacteria_unclassified | Bacteria_unclassified |
| Bacteria_unclassified | Bacteria_unclassified | Bacteria_unclassified |
| Bacteria_unclassified | Bacteria_unclassified | Bacteria_unclassified |
| Bacteria_unclassified | Bacteria_unclassified | Bacteria_unclassified |
| Bacteria_unclassified | Bacteria_unclassified | Bacteria_unclassified |
| Bacteroidaceae | Bacteroidales | Bacteroidia |
| Bacteroidaceae | Bacteroidales | Bacteroidia |
| Bifidobacteriaceae | Bifidobacteriales | Actinomycetia |
| Bifidobacteriaceae | Bifidobacteriales | Actinomycetia |

| | | |
|---|---|---|
| Clostridia_unclassified | Clostridia_unclassified | Clostridia |
| Clostridia_unclassified | Clostridia_unclassified | Clostridia |
| Clostridiaceae | Eubacteriales | Clostridia |
| Clostridiaceae | Eubacteriales | Clostridia |
| Clostridiaceae | Eubacteriales | Clostridia |
| Clostridiaceae | Eubacteriales | Clostridia |
| Eubacteriales_unclassified | Eubacteriales | Clostridia |
| Eubacteriales_unclassified | Eubacteriales | Clostridia |
| Erysipelotrichaceae | Erysipelotrichales | Erysipelotrichia |
| Erysipelotrichaceae | Erysipelotrichales | Erysipelotrichia |
| Coriobacteriaceae | Coriobacteriales | Coriobacteriia |
| Coriobacteriaceae | Coriobacteriales | Coriobacteriia |
| Lachnospiraceae | Eubacteriales | Clostridia |
| Lachnospiraceae | Eubacteriales | Clostridia |
| Erysipelotrichaceae | Erysipelotrichales | Erysipelotrichia |
| Erysipelotrichaceae | Erysipelotrichales | Erysipelotrichia |
| Erysipelotrichales_unclassified | Erysipelotrichales | Erysipelotrichia |
| Erysipelotrichales_unclassified | Erysipelotrichales | Erysipelotrichia |
| Eubacteriaceae | Eubacteriales | Clostridia |
| Eubacteriaceae | Eubacteriales | Clostridia |
| Eubacteriaceae | Eubacteriales | Clostridia |
| Eubacteriaceae | Eubacteriales | Clostridia |
| Lachnospiraceae | Eubacteriales | Clostridia |
| Lachnospiraceae | Eubacteriales | Clostridia |
| Eggerthellaceae | Eggerthellales | Coriobacteriia |
| Eggerthellaceae | Eggerthellales | Coriobacteriia |
| Lachnospiraceae | Eubacteriales | Clostridia |
| Lachnospiraceae | Eubacteriales | Clostridia |
| Muribaculaceae | Bacteroidales | Bacteroidia |
| Muribaculaceae | Bacteroidales | Bacteroidia |
| Muribaculaceae | Bacteroidales | Bacteroidia |
| Muribaculaceae | Bacteroidales | Bacteroidia |
| Muribaculaceae | Bacteroidales | Bacteroidia |
| Muribaculaceae | Bacteroidales | Bacteroidia |
| FGB9508 | OFGB9508 | CFGB9508 |
| FGB9508 | OFGB9508 | CFGB9508 |
| FGB2838 | OFGB2838 | CFGB2838 |
| FGB2838 | OFGB2838 | CFGB2838 |
| FGB2838 | OFGB2838 | CFGB2838 |
| FGB2838 | OFGB2838 | CFGB2838 |
| FGB2838 | OFGB2838 | CFGB2838 |
| FGB2838 | OFGB2838 | CFGB2838 |
| Pumilibacteraceae | Eubacteriales | Clostridia |
| Pumilibacteraceae | Eubacteriales | Clostridia |
| FGB28439 | OFGB28439 | CFGB28439 |
| FGB28439 | OFGB28439 | CFGB28439 |

| | | |
|---|---|---|
| Clostridia_unclassified | Clostridia_unclassified | Clostridia |
| Clostridia_unclassified | Clostridia_unclassified | Clostridia |
| Eubacteriaceae | Eubacteriales | Clostridia |
| Eubacteriaceae | Eubacteriales | Clostridia |
| Lachnospiraceae | Eubacteriales | Clostridia |
| Lachnospiraceae | Eubacteriales | Clostridia |
| Lachnospiraceae | Eubacteriales | Clostridia |
| Lachnospiraceae | Eubacteriales | Clostridia |
| Lachnospiraceae | Eubacteriales | Clostridia |
| Lachnospiraceae | Eubacteriales | Clostridia |
| Lachnospiraceae | Eubacteriales | Clostridia |
| Lachnospiraceae | Eubacteriales | Clostridia |
| FGB77305 | OFGB77305 | CFGB77305 |
| FGB77305 | OFGB77305 | CFGB77305 |
| Clostridiaceae | Eubacteriales | Clostridia |
| Clostridiaceae | Eubacteriales | Clostridia |
| Lachnospiraceae | Eubacteriales | Clostridia |
| Lachnospiraceae | Eubacteriales | Clostridia |
| Lachnospiraceae | Eubacteriales | Clostridia |
| Lachnospiraceae | Eubacteriales | Clostridia |
| Lachnospiraceae | Eubacteriales | Clostridia |
| Lachnospiraceae | Eubacteriales | Clostridia |
| FGB9633 | OFGB9633 | CFGB9633 |
| FGB9633 | OFGB9633 | CFGB9633 |
| Bacteria_unclassified | Bacteria_unclassified | Bacteria_unclassified |
| Bacteria_unclassified | Bacteria_unclassified | Bacteria_unclassified |
| Bacteria_unclassified | Bacteria_unclassified | Bacteria_unclassified |
| Bacteria_unclassified | Bacteria_unclassified | Bacteria_unclassified |
| Bacteria_unclassified | Bacteria_unclassified | Bacteria_unclassified |
| Bacteria_unclassified | Bacteria_unclassified | Bacteria_unclassified |
| Bacteria_unclassified | Bacteria_unclassified | Bacteria_unclassified |
| Bacteria_unclassified | Bacteria_unclassified | Bacteria_unclassified |
| Lachnospiraceae | Eubacteriales | Clostridia |
| Lachnospiraceae | Eubacteriales | Clostridia |
| Lachnospiraceae | Eubacteriales | Clostridia |
| Lachnospiraceae | Eubacteriales | Clostridia |
| Lachnospiraceae | Eubacteriales | Clostridia |
| Lachnospiraceae | Eubacteriales | Clostridia |
| FGB77359 | OFGB77359 | CFGB77359 |
| FGB77359 | OFGB77359 | CFGB77359 |
| FGB9639 | OFGB9639 | CFGB9639 |
| FGB9639 | OFGB9639 | CFGB9639 |
| Lachnospiraceae | Eubacteriales | Clostridia |
| Lachnospiraceae | Eubacteriales | Clostridia |
| Lachnospiraceae | Eubacteriales | Clostridia |
| Lachnospiraceae | Eubacteriales | Clostridia |

| | | |
|---|---|---|
| Lachnospiraceae | Eubacteriales | Clostridia |
| Lachnospiraceae | Eubacteriales | Clostridia |
| Clostridiaceae | Eubacteriales | Clostridia |
| Clostridiaceae | Eubacteriales | Clostridia |
| Clostridiaceae | Eubacteriales | Clostridia |
| Clostridiaceae | Eubacteriales | Clostridia |
| Clostridiaceae | Eubacteriales | Clostridia |
| Clostridiaceae | Eubacteriales | Clostridia |
| Clostridiaceae | Eubacteriales | Clostridia |
| Clostridiaceae | Eubacteriales | Clostridia |
| Clostridiaceae | Eubacteriales | Clostridia |
| Clostridiaceae | Eubacteriales | Clostridia |
| Clostridiaceae | Eubacteriales | Clostridia |
| Clostridiaceae | Eubacteriales | Clostridia |
| Clostridiaceae | Eubacteriales | Clostridia |
| Clostridiaceae | Eubacteriales | Clostridia |
| Eubacteriaceae | Eubacteriales | Clostridia |
| Eubacteriaceae | Eubacteriales | Clostridia |
| FGB9658 | OFGB9658 | CFGB9658 |
| FGB9658 | OFGB9658 | CFGB9658 |
| FGB9659 | OFGB9659 | CFGB9659 |
| FGB9659 | OFGB9659 | CFGB9659 |
| Bacteria_unclassified | Bacteria_unclassified | Bacteria_unclassified |
| Bacteria_unclassified | Bacteria_unclassified | Bacteria_unclassified |
| FGB9827 | OFGB9827 | CFGB9827 |
| FGB9827 | OFGB9827 | CFGB9827 |
| Eubacteriaceae | Eubacteriales | Clostridia |
| Eubacteriaceae | Eubacteriales | Clostridia |
| FGB77303 | OFGB77303 | CFGB77303 |
| FGB77303 | OFGB77303 | CFGB77303 |
| Eubacteriales_unclassified | Eubacteriales | Clostridia |
| Eubacteriales_unclassified | Eubacteriales | Clostridia |
| Oscillospiraceae | Eubacteriales | Clostridia |
| Oscillospiraceae | Eubacteriales | Clostridia |
| FGB30328 | OFGB30328 | CFGB30328 |
| FGB30328 | OFGB30328 | CFGB30328 |
| Oscillospiraceae | Eubacteriales | Clostridia |
| Oscillospiraceae | Eubacteriales | Clostridia |
| Oscillospiraceae | Eubacteriales | Clostridia |
| Oscillospiraceae | Eubacteriales | Clostridia |
| Oscillospiraceae | Eubacteriales | Clostridia |
| Oscillospiraceae | Eubacteriales | Clostridia |
| Oscillospiraceae | Eubacteriales | Clostridia |
| Oscillospiraceae | Eubacteriales | Clostridia |
| Oscillospiraceae | Eubacteriales | Clostridia |
| Oscillospiraceae | Eubacteriales | Clostridia |

| | | |
|---|---|---|
| Oscillospiraceae | Eubacteriales | Clostridia |
| Oscillospiraceae | Eubacteriales | Clostridia |
| Oscillospiraceae | Eubacteriales | Clostridia |
| Oscillospiraceae | Eubacteriales | Clostridia |
| FGB77153 | OFGB77153 | CFGB77153 |
| FGB77153 | OFGB77153 | CFGB77153 |
| FGB1791 | OFGB1791 | CFGB1791 |
| FGB1791 | OFGB1791 | CFGB1791 |
| FGB10290 | OFGB10290 | CFGB10290 |
| FGB10290 | OFGB10290 | CFGB10290 |
| Oscillospiraceae | Eubacteriales | Clostridia |
| Oscillospiraceae | Eubacteriales | Clostridia |
| FGB1765 | OFGB1765 | CFGB1765 |
| FGB1765 | OFGB1765 | CFGB1765 |
| FGB10349 | OFGB10349 | CFGB10349 |
| FGB10349 | OFGB10349 | CFGB10349 |
| FGB10667 | OFGB10667 | CFGB10667 |
| FGB10667 | OFGB10667 | CFGB10667 |
| Lachnospiraceae | Eubacteriales | Clostridia |
| Lachnospiraceae | Eubacteriales | Clostridia |
| Lachnospiraceae | Eubacteriales | Clostridia |
| Lachnospiraceae | Eubacteriales | Clostridia |
| Christensenellaceae | Eubacteriales | Clostridia |
| Christensenellaceae | Eubacteriales | Clostridia |
| FGB10299 | OFGB10299 | CFGB10299 |
| FGB10299 | OFGB10299 | CFGB10299 |
| Oscillospiraceae | Eubacteriales | Clostridia |
| Oscillospiraceae | Eubacteriales | Clostridia |
| Oscillospiraceae | Eubacteriales | Clostridia |
| Oscillospiraceae | Eubacteriales | Clostridia |
| Lachnospiraceae | Eubacteriales | Clostridia |
| Lachnospiraceae | Eubacteriales | Clostridia |
| Clostridia_unclassified | Clostridia_unclassified | Clostridia |
| Clostridia_unclassified | Clostridia_unclassified | Clostridia |
| Lachnospiraceae | Eubacteriales | Clostridia |
| Lachnospiraceae | Eubacteriales | Clostridia |
| Lachnospiraceae | Eubacteriales | Clostridia |
| Lachnospiraceae | Eubacteriales | Clostridia |
| Lachnospiraceae | Eubacteriales | Clostridia |
| Lachnospiraceae | Eubacteriales | Clostridia |
| Lachnospiraceae | Eubacteriales | Clostridia |
| Lachnospiraceae | Eubacteriales | Clostridia |
| Lachnospiraceae | Eubacteriales | Clostridia |
| Lachnospiraceae | Eubacteriales | Clostridia |
| Lachnospiraceae | Eubacteriales | Clostridia |
| Lachnospiraceae | Eubacteriales | Clostridia |

| | | |
|---|---|---|
| Lachnospiraceae | Eubacteriales | Clostridia |
| Lachnospiraceae | Eubacteriales | Clostridia |
| Lachnospiraceae | Eubacteriales | Clostridia |
| Lachnospiraceae | Eubacteriales | Clostridia |
| Lactobacillaceae | Lactobacillales | Bacilli |
| Lactobacillaceae | Lactobacillales | Bacilli |
| Muribaculaceae | Bacteroidales | Bacteroidia |
| Muribaculaceae | Bacteroidales | Bacteroidia |
| Oscillospiraceae | Eubacteriales | Clostridia |
| Oscillospiraceae | Eubacteriales | Clostridia |
| Oscillospiraceae | Eubacteriales | Clostridia |
| Oscillospiraceae | Eubacteriales | Clostridia |
| Oscillospiraceae | Eubacteriales | Clostridia |
| Oscillospiraceae | Eubacteriales | Clostridia |
| Oscillospiraceae | Eubacteriales | Clostridia |
| Oscillospiraceae | Eubacteriales | Clostridia |
| Sutterellaceae | Burkholderiales | Betaproteobacteria |
| Sutterellaceae | Burkholderiales | Betaproteobacteria |
| Peptostreptococcaceae | Eubacteriales | Clostridia |
| Peptostreptococcaceae | Eubacteriales | Clostridia |
| Lachnospiraceae | Eubacteriales | Clostridia |
| Lachnospiraceae | Eubacteriales | Clostridia |
| Turicibacteraceae | Erysipelotrichales | Erysipelotrichia |
| Turicibacteraceae | Erysipelotrichales | Erysipelotrichia |
| Lachnospiraceae | Eubacteriales | Clostridia |
| Lachnospiraceae | Eubacteriales | Clostridia |
| Lachnospiraceae | Eubacteriales | Clostridia |
| Lachnospiraceae | Eubacteriales | Clostridia |
| Oscillospiraceae | Eubacteriales | Clostridia |
| Oscillospiraceae | Eubacteriales | Clostridia |
| Oscillospiraceae | Eubacteriales | Clostridia |
| Oscillospiraceae | Eubacteriales | Clostridia |
| Eggerthellaceae | Eggerthellales | Coriobacteriia |
| Eggerthellaceae | Eggerthellales | Coriobacteriia |
| Eggerthellaceae | Eggerthellales | Coriobacteriia |
| Eggerthellaceae | Eggerthellales | Coriobacteriia |
| Eggerthellaceae | Eggerthellales | Coriobacteriia |
| Eggerthellaceae | Eggerthellales | Coriobacteriia |
| Akkermansiaceae | Verrucomicrobiales | Verrucomicrobiae |
| Akkermansiaceae | Verrucomicrobiales | Verrucomicrobiae |
| Rikenellaceae | Bacteroidales | Bacteroidia |
| Rikenellaceae | Bacteroidales | Bacteroidia |

| | | |
|---|---|---|
| Oscillospiraceae | Eubacteriales | Clostridia |
| Oscillospiraceae | Eubacteriales | Clostridia |
| Bacteria_unclassified | Bacteria_unclassified | Bacteria_unclassified |
| Bacteria_unclassified | Bacteria_unclassified | Bacteria_unclassified |
| Bacteria_unclassified | Bacteria_unclassified | Bacteria_unclassified |
| Bacteria_unclassified | Bacteria_unclassified | Bacteria_unclassified |
| Bacteria_unclassified | Bacteria_unclassified | Bacteria_unclassified |
| Bacteria_unclassified | Bacteria_unclassified | Bacteria_unclassified |
| Bacteria_unclassified | Bacteria_unclassified | Bacteria_unclassified |
| Bacteria_unclassified | Bacteria_unclassified | Bacteria_unclassified |
| Bacteria_unclassified | Bacteria_unclassified | Bacteria_unclassified |
| Bacteria_unclassified | Bacteria_unclassified | Bacteria_unclassified |
| Bacteria_unclassified | Bacteria_unclassified | Bacteria_unclassified |
| Bacteria_unclassified | Bacteria_unclassified | Bacteria_unclassified |
| Bacteroidaceae | Bacteroidales | Bacteroidia |
| Bacteroidaceae | Bacteroidales | Bacteroidia |
| Bifidobacteriaceae | Bifidobacteriales | Actinomycetia |
| Bifidobacteriaceae | Bifidobacteriales | Actinomycetia |
| Clostridia_unclassified | Clostridia_unclassified | Clostridia |
| Clostridia_unclassified | Clostridia_unclassified | Clostridia |
| Clostridiaceae | Eubacteriales | Clostridia |
| Clostridiaceae | Eubacteriales | Clostridia |
| Clostridiaceae | Eubacteriales | Clostridia |
| Clostridiaceae | Eubacteriales | Clostridia |
| Eubacteriales_unclassified | Eubacteriales | Clostridia |
| Eubacteriales_unclassified | Eubacteriales | Clostridia |
| Erysipelotrichaceae | Erysipelotrichales | Erysipelotrichia |
| Erysipelotrichaceae | Erysipelotrichales | Erysipelotrichia |
| Coriobacteriaceae | Coriobacteriales | Coriobacteriia |
| Coriobacteriaceae | Coriobacteriales | Coriobacteriia |
| Lachnospiraceae | Eubacteriales | Clostridia |
| Lachnospiraceae | Eubacteriales | Clostridia |
| Erysipelotrichaceae | Erysipelotrichales | Erysipelotrichia |
| Erysipelotrichaceae | Erysipelotrichales | Erysipelotrichia |
| Erysipelotrichales_unclassified | Erysipelotrichales | Erysipelotrichia |
| Erysipelotrichales_unclassified | Erysipelotrichales | Erysipelotrichia |
| Eubacteriaceae | Eubacteriales | Clostridia |
| Eubacteriaceae | Eubacteriales | Clostridia |
| Eubacteriaceae | Eubacteriales | Clostridia |
| Eubacteriaceae | Eubacteriales | Clostridia |
| Lachnospiraceae | Eubacteriales | Clostridia |
| Lachnospiraceae | Eubacteriales | Clostridia |
| Eggerthellaceae | Eggerthellales | Coriobacteriia |
| Eggerthellaceae | Eggerthellales | Coriobacteriia |

| | | |
|---|---|---|
| Lachnospiraceae | Eubacteriales | Clostridia |
| Lachnospiraceae | Eubacteriales | Clostridia |
| Muribaculaceae | Bacteroidales | Bacteroidia |
| Muribaculaceae | Bacteroidales | Bacteroidia |
| Muribaculaceae | Bacteroidales | Bacteroidia |
| Muribaculaceae | Bacteroidales | Bacteroidia |
| Muribaculaceae | Bacteroidales | Bacteroidia |
| Muribaculaceae | Bacteroidales | Bacteroidia |
| FGB9508 | OFGB9508 | CFGB9508 |
| FGB9508 | OFGB9508 | CFGB9508 |
| FGB2838 | OFGB2838 | CFGB2838 |
| FGB2838 | OFGB2838 | CFGB2838 |
| FGB2838 | OFGB2838 | CFGB2838 |
| FGB2838 | OFGB2838 | CFGB2838 |
| FGB2838 | OFGB2838 | CFGB2838 |
| FGB2838 | OFGB2838 | CFGB2838 |
| Pumilibacteraceae | Eubacteriales | Clostridia |
| Pumilibacteraceae | Eubacteriales | Clostridia |
| FGB28439 | OFGB28439 | CFGB28439 |
| FGB28439 | OFGB28439 | CFGB28439 |
| Clostridia_unclassified | Clostridia_unclassified | Clostridia |
| Clostridia_unclassified | Clostridia_unclassified | Clostridia |
| Eubacteriaceae | Eubacteriales | Clostridia |
| Eubacteriaceae | Eubacteriales | Clostridia |
| Lachnospiraceae | Eubacteriales | Clostridia |
| Lachnospiraceae | Eubacteriales | Clostridia |
| Lachnospiraceae | Eubacteriales | Clostridia |
| Lachnospiraceae | Eubacteriales | Clostridia |
| Lachnospiraceae | Eubacteriales | Clostridia |
| Lachnospiraceae | Eubacteriales | Clostridia |
| Lachnospiraceae | Eubacteriales | Clostridia |
| Lachnospiraceae | Eubacteriales | Clostridia |
| FGB77305 | OFGB77305 | CFGB77305 |
| FGB77305 | OFGB77305 | CFGB77305 |
| Clostridiaceae | Eubacteriales | Clostridia |
| Clostridiaceae | Eubacteriales | Clostridia |
| Lachnospiraceae | Eubacteriales | Clostridia |
| Lachnospiraceae | Eubacteriales | Clostridia |
| Lachnospiraceae | Eubacteriales | Clostridia |
| Lachnospiraceae | Eubacteriales | Clostridia |
| Lachnospiraceae | Eubacteriales | Clostridia |
| Lachnospiraceae | Eubacteriales | Clostridia |
| FGB9633 | OFGB9633 | CFGB9633 |
| FGB9633 | OFGB9633 | CFGB9633 |
| Bacteria_unclassified | Bacteria_unclassified | Bacteria_unclassified |
| Bacteria_unclassified | Bacteria_unclassified | Bacteria_unclassified |

| | | |
|---|---|---|
| Bacteria_unclassified | Bacteria_unclassified | Bacteria_unclassified |
| Bacteria_unclassified | Bacteria_unclassified | Bacteria_unclassified |
| Bacteria_unclassified | Bacteria_unclassified | Bacteria_unclassified |
| Bacteria_unclassified | Bacteria_unclassified | Bacteria_unclassified |
| Bacteria_unclassified | Bacteria_unclassified | Bacteria_unclassified |
| Bacteria_unclassified | Bacteria_unclassified | Bacteria_unclassified |
| Lachnospiraceae | Eubacteriales | Clostridia |
| Lachnospiraceae | Eubacteriales | Clostridia |
| Lachnospiraceae | Eubacteriales | Clostridia |
| Lachnospiraceae | Eubacteriales | Clostridia |
| Lachnospiraceae | Eubacteriales | Clostridia |
| Lachnospiraceae | Eubacteriales | Clostridia |
| FGB77359 | OFGB77359 | CFGB77359 |
| FGB77359 | OFGB77359 | CFGB77359 |
| FGB9639 | OFGB9639 | CFGB9639 |
| FGB9639 | OFGB9639 | CFGB9639 |
| Lachnospiraceae | Eubacteriales | Clostridia |
| Lachnospiraceae | Eubacteriales | Clostridia |
| Lachnospiraceae | Eubacteriales | Clostridia |
| Lachnospiraceae | Eubacteriales | Clostridia |
| Lachnospiraceae | Eubacteriales | Clostridia |
| Lachnospiraceae | Eubacteriales | Clostridia |
| Clostridiaceae | Eubacteriales | Clostridia |
| Clostridiaceae | Eubacteriales | Clostridia |
| Clostridiaceae | Eubacteriales | Clostridia |
| Clostridiaceae | Eubacteriales | Clostridia |
| Clostridiaceae | Eubacteriales | Clostridia |
| Clostridiaceae | Eubacteriales | Clostridia |
| Clostridiaceae | Eubacteriales | Clostridia |
| Clostridiaceae | Eubacteriales | Clostridia |
| Clostridiaceae | Eubacteriales | Clostridia |
| Clostridiaceae | Eubacteriales | Clostridia |
| Clostridiaceae | Eubacteriales | Clostridia |
| Clostridiaceae | Eubacteriales | Clostridia |
| Clostridiaceae | Eubacteriales | Clostridia |
| Clostridiaceae | Eubacteriales | Clostridia |
| Eubacteriaceae | Eubacteriales | Clostridia |
| Eubacteriaceae | Eubacteriales | Clostridia |
| FGB9658 | OFGB9658 | CFGB9658 |
| FGB9658 | OFGB9658 | CFGB9658 |
| FGB9659 | OFGB9659 | CFGB9659 |
| FGB9659 | OFGB9659 | CFGB9659 |
| Bacteria_unclassified | Bacteria_unclassified | Bacteria_unclassified |
| Bacteria_unclassified | Bacteria_unclassified | Bacteria_unclassified |
| FGB9827 | OFGB9827 | CFGB9827 |
| FGB9827 | OFGB9827 | CFGB9827 |

| | | |
|---|---|---|
| Eubacteriaceae | Eubacteriales | Clostridia |
| Eubacteriaceae | Eubacteriales | Clostridia |
| FGB77303 | OFGB77303 | CFGB77303 |
| FGB77303 | OFGB77303 | CFGB77303 |
| Eubacteriales_unclassified | Eubacteriales | Clostridia |
| Eubacteriales_unclassified | Eubacteriales | Clostridia |
| Oscillospiraceae | Eubacteriales | Clostridia |
| Oscillospiraceae | Eubacteriales | Clostridia |
| FGB30328 | OFGB30328 | CFGB30328 |
| FGB30328 | OFGB30328 | CFGB30328 |
| Oscillospiraceae | Eubacteriales | Clostridia |
| Oscillospiraceae | Eubacteriales | Clostridia |
| Oscillospiraceae | Eubacteriales | Clostridia |
| Oscillospiraceae | Eubacteriales | Clostridia |
| Oscillospiraceae | Eubacteriales | Clostridia |
| Oscillospiraceae | Eubacteriales | Clostridia |
| Oscillospiraceae | Eubacteriales | Clostridia |
| Oscillospiraceae | Eubacteriales | Clostridia |
| Oscillospiraceae | Eubacteriales | Clostridia |
| Oscillospiraceae | Eubacteriales | Clostridia |
| Oscillospiraceae | Eubacteriales | Clostridia |
| Oscillospiraceae | Eubacteriales | Clostridia |
| Oscillospiraceae | Eubacteriales | Clostridia |
| Oscillospiraceae | Eubacteriales | Clostridia |
| FGB77153 | OFGB77153 | CFGB77153 |
| FGB77153 | OFGB77153 | CFGB77153 |
| FGB1791 | OFGB1791 | CFGB1791 |
| FGB1791 | OFGB1791 | CFGB1791 |
| FGB10290 | OFGB10290 | CFGB10290 |
| FGB10290 | OFGB10290 | CFGB10290 |
| Oscillospiraceae | Eubacteriales | Clostridia |
| Oscillospiraceae | Eubacteriales | Clostridia |
| FGB1765 | OFGB1765 | CFGB1765 |
| FGB1765 | OFGB1765 | CFGB1765 |
| FGB10349 | OFGB10349 | CFGB10349 |
| FGB10349 | OFGB10349 | CFGB10349 |
| FGB10667 | OFGB10667 | CFGB10667 |
| FGB10667 | OFGB10667 | CFGB10667 |
| Lachnospiraceae | Eubacteriales | Clostridia |
| Lachnospiraceae | Eubacteriales | Clostridia |
| Lachnospiraceae | Eubacteriales | Clostridia |
| Lachnospiraceae | Eubacteriales | Clostridia |
| Christensenellaceae | Eubacteriales | Clostridia |
| Christensenellaceae | Eubacteriales | Clostridia |
| FGB10299 | OFGB10299 | CFGB10299 |
| FGB10299 | OFGB10299 | CFGB10299 |

| | | |
|---|---|---|
| Oscillospiraceae | Eubacteriales | Clostridia |
| Oscillospiraceae | Eubacteriales | Clostridia |
| Oscillospiraceae | Eubacteriales | Clostridia |
| Oscillospiraceae | Eubacteriales | Clostridia |
| Lachnospiraceae | Eubacteriales | Clostridia |
| Lachnospiraceae | Eubacteriales | Clostridia |
| Clostridia_unclassified | Clostridia_unclassified | Clostridia |
| Clostridia_unclassified | Clostridia_unclassified | Clostridia |
| Lachnospiraceae | Eubacteriales | Clostridia |
| Lachnospiraceae | Eubacteriales | Clostridia |
| Lachnospiraceae | Eubacteriales | Clostridia |
| Lachnospiraceae | Eubacteriales | Clostridia |
| Lachnospiraceae | Eubacteriales | Clostridia |
| Lachnospiraceae | Eubacteriales | Clostridia |
| Lachnospiraceae | Eubacteriales | Clostridia |
| Lachnospiraceae | Eubacteriales | Clostridia |
| Lachnospiraceae | Eubacteriales | Clostridia |
| Lachnospiraceae | Eubacteriales | Clostridia |
| Lachnospiraceae | Eubacteriales | Clostridia |
| Lachnospiraceae | Eubacteriales | Clostridia |
| Lachnospiraceae | Eubacteriales | Clostridia |
| Lachnospiraceae | Eubacteriales | Clostridia |
| Lachnospiraceae | Eubacteriales | Clostridia |
| Lachnospiraceae | Eubacteriales | Clostridia |
| Lactobacillaceae | Lactobacillales | Bacilli |
| Lactobacillaceae | Lactobacillales | Bacilli |
| Muribaculaceae | Bacteroidales | Bacteroidia |
| Muribaculaceae | Bacteroidales | Bacteroidia |
| Oscillospiraceae | Eubacteriales | Clostridia |
| Oscillospiraceae | Eubacteriales | Clostridia |
| Oscillospiraceae | Eubacteriales | Clostridia |
| Oscillospiraceae | Eubacteriales | Clostridia |
| Oscillospiraceae | Eubacteriales | Clostridia |
| Oscillospiraceae | Eubacteriales | Clostridia |
| Oscillospiraceae | Eubacteriales | Clostridia |
| Oscillospiraceae | Eubacteriales | Clostridia |
| Sutterellaceae | Burkholderiales | Betaproteobacteria |
| Sutterellaceae | Burkholderiales | Betaproteobacteria |
| Peptostreptococcaceae | Eubacteriales | Clostridia |
| Peptostreptococcaceae | Eubacteriales | Clostridia |
| Lachnospiraceae | Eubacteriales | Clostridia |
| Lachnospiraceae | Eubacteriales | Clostridia |

| | | |
|---|---|---|
| Turicibacteraceae | Erysipelotrichales | Erysipelotrichia |
| Turicibacteraceae | Erysipelotrichales | Erysipelotrichia |
| Lachnospiraceae | Eubacteriales | Clostridia |
| Lachnospiraceae | Eubacteriales | Clostridia |
| Lachnospiraceae | Eubacteriales | Clostridia |
| Lachnospiraceae | Eubacteriales | Clostridia |
| Oscillospiraceae | Eubacteriales | Clostridia |
| Oscillospiraceae | Eubacteriales | Clostridia |
| Oscillospiraceae | Eubacteriales | Clostridia |
| Oscillospiraceae | Eubacteriales | Clostridia |
| Eggerthellaceae | Eggerthellales | Coriobacteriia |
| Eggerthellaceae | Eggerthellales | Coriobacteriia |
| Eggerthellaceae | Eggerthellales | Coriobacteriia |
| Eggerthellaceae | Eggerthellales | Coriobacteriia |
| Eggerthellaceae | Eggerthellales | Coriobacteriia |
| Eggerthellaceae | Eggerthellales | Coriobacteriia |
| Akkermansiaceae | Verrucomicrobiales | Verrucomicrobiae |
| Akkermansiaceae | Verrucomicrobiales | Verrucomicrobiae |
| Rikenellaceae | Bacteroidales | Bacteroidia |
| Rikenellaceae | Bacteroidales | Bacteroidia |
| Oscillospiraceae | Eubacteriales | Clostridia |
| Oscillospiraceae | Eubacteriales | Clostridia |
| Bacteria_unclassified | Bacteria_unclassified | Bacteria_unclassified |
| Bacteria_unclassified | Bacteria_unclassified | Bacteria_unclassified |
| Bacteria_unclassified | Bacteria_unclassified | Bacteria_unclassified |
| Bacteria_unclassified | Bacteria_unclassified | Bacteria_unclassified |
| Bacteria_unclassified | Bacteria_unclassified | Bacteria_unclassified |
| Bacteria_unclassified | Bacteria_unclassified | Bacteria_unclassified |
| Bacteria_unclassified | Bacteria_unclassified | Bacteria_unclassified |
| Bacteria_unclassified | Bacteria_unclassified | Bacteria_unclassified |
| Bacteria_unclassified | Bacteria_unclassified | Bacteria_unclassified |
| Bacteria_unclassified | Bacteria_unclassified | Bacteria_unclassified |
| Bacteria_unclassified | Bacteria_unclassified | Bacteria_unclassified |
| Bacteria_unclassified | Bacteria_unclassified | Bacteria_unclassified |
| Bacteroidaceae | Bacteroidales | Bacteroidia |
| Bacteroidaceae | Bacteroidales | Bacteroidia |

| | | |
|---|---|---|
| Bifidobacteriaceae | Bifidobacteriales | Actinomycetia |
| Bifidobacteriaceae | Bifidobacteriales | Actinomycetia |
| Clostridia_unclassified | Clostridia_unclassified | Clostridia |
| Clostridia_unclassified | Clostridia_unclassified | Clostridia |
| Clostridiaceae | Eubacteriales | Clostridia |
| Clostridiaceae | Eubacteriales | Clostridia |
| Clostridiaceae | Eubacteriales | Clostridia |
| Clostridiaceae | Eubacteriales | Clostridia |

| | | |
|---|---|---|
| Eubacteriales_unclassified | Eubacteriales | Clostridia |
| Eubacteriales_unclassified | Eubacteriales | Clostridia |
| Erysipelotrichaceae | Erysipelotrichales | Erysipelotrichia |
| Erysipelotrichaceae | Erysipelotrichales | Erysipelotrichia |
| Coriobacteriaceae | Coriobacteriales | Coriobacteriia |
| Coriobacteriaceae | Coriobacteriales | Coriobacteriia |
| Lachnospiraceae | Eubacteriales | Clostridia |
| Lachnospiraceae | Eubacteriales | Clostridia |
| Erysipelotrichaceae | Erysipelotrichales | Erysipelotrichia |
| Erysipelotrichaceae | Erysipelotrichales | Erysipelotrichia |
| Erysipelotrichales_unclassified | Erysipelotrichales | Erysipelotrichia |
| Erysipelotrichales_unclassified | Erysipelotrichales | Erysipelotrichia |
| Eubacteriaceae | Eubacteriales | Clostridia |
| Eubacteriaceae | Eubacteriales | Clostridia |
| Eubacteriaceae | Eubacteriales | Clostridia |
| Eubacteriaceae | Eubacteriales | Clostridia |
| Lachnospiraceae | Eubacteriales | Clostridia |
| Lachnospiraceae | Eubacteriales | Clostridia |
| Eggerthellaceae | Eggerthellales | Coriobacteriia |
| Eggerthellaceae | Eggerthellales | Coriobacteriia |
| Lachnospiraceae | Eubacteriales | Clostridia |
| Lachnospiraceae | Eubacteriales | Clostridia |
| Muribaculaceae | Bacteroidales | Bacteroidia |
| Muribaculaceae | Bacteroidales | Bacteroidia |
| Muribaculaceae | Bacteroidales | Bacteroidia |
| Muribaculaceae | Bacteroidales | Bacteroidia |
| Muribaculaceae | Bacteroidales | Bacteroidia |
| Muribaculaceae | Bacteroidales | Bacteroidia |
| FGB9508 | OFGB9508 | CFGB9508 |
| FGB9508 | OFGB9508 | CFGB9508 |
| FGB2838 | OFGB2838 | CFGB2838 |
| FGB2838 | OFGB2838 | CFGB2838 |
| FGB2838 | OFGB2838 | CFGB2838 |
| FGB2838 | OFGB2838 | CFGB2838 |
| FGB2838 | OFGB2838 | CFGB2838 |
| FGB2838 | OFGB2838 | CFGB2838 |
| Pumilibacteraceae | Eubacteriales | Clostridia |
| Pumilibacteraceae | Eubacteriales | Clostridia |
| FGB28439 | OFGB28439 | CFGB28439 |
| FGB28439 | OFGB28439 | CFGB28439 |
| Clostridia_unclassified | Clostridia_unclassified | Clostridia |
| Clostridia_unclassified | Clostridia_unclassified | Clostridia |
| Eubacteriaceae | Eubacteriales | Clostridia |
| Eubacteriaceae | Eubacteriales | Clostridia |
| Lachnospiraceae | Eubacteriales | Clostridia |
| Lachnospiraceae | Eubacteriales | Clostridia |

| | | |
|---|---|---|
| Lachnospiraceae | Eubacteriales | Clostridia |
| Lachnospiraceae | Eubacteriales | Clostridia |
| Lachnospiraceae | Eubacteriales | Clostridia |
| Lachnospiraceae | Eubacteriales | Clostridia |
| Lachnospiraceae | Eubacteriales | Clostridia |
| Lachnospiraceae | Eubacteriales | Clostridia |
| FGB77305 | OFGB77305 | CFGB77305 |
| FGB77305 | OFGB77305 | CFGB77305 |
| Clostridiaceae | Eubacteriales | Clostridia |
| Clostridiaceae | Eubacteriales | Clostridia |
| Lachnospiraceae | Eubacteriales | Clostridia |
| Lachnospiraceae | Eubacteriales | Clostridia |
| Lachnospiraceae | Eubacteriales | Clostridia |
| Lachnospiraceae | Eubacteriales | Clostridia |
| Lachnospiraceae | Eubacteriales | Clostridia |
| Lachnospiraceae | Eubacteriales | Clostridia |
| FGB9633 | OFGB9633 | CFGB9633 |
| FGB9633 | OFGB9633 | CFGB9633 |
| Bacteria_unclassified | Bacteria_unclassified | Bacteria_unclassified |
| Bacteria_unclassified | Bacteria_unclassified | Bacteria_unclassified |
| Bacteria_unclassified | Bacteria_unclassified | Bacteria_unclassified |
| Bacteria_unclassified | Bacteria_unclassified | Bacteria_unclassified |
| Bacteria_unclassified | Bacteria_unclassified | Bacteria_unclassified |
| Bacteria_unclassified | Bacteria_unclassified | Bacteria_unclassified |
| Bacteria_unclassified | Bacteria_unclassified | Bacteria_unclassified |
| Bacteria_unclassified | Bacteria_unclassified | Bacteria_unclassified |
| Lachnospiraceae | Eubacteriales | Clostridia |
| Lachnospiraceae | Eubacteriales | Clostridia |
| Lachnospiraceae | Eubacteriales | Clostridia |
| Lachnospiraceae | Eubacteriales | Clostridia |
| Lachnospiraceae | Eubacteriales | Clostridia |
| Lachnospiraceae | Eubacteriales | Clostridia |
| FGB77359 | OFGB77359 | CFGB77359 |
| FGB77359 | OFGB77359 | CFGB77359 |
| FGB9639 | OFGB9639 | CFGB9639 |
| FGB9639 | OFGB9639 | CFGB9639 |
| Lachnospiraceae | Eubacteriales | Clostridia |
| Lachnospiraceae | Eubacteriales | Clostridia |
| Lachnospiraceae | Eubacteriales | Clostridia |
| Lachnospiraceae | Eubacteriales | Clostridia |
| Lachnospiraceae | Eubacteriales | Clostridia |
| Lachnospiraceae | Eubacteriales | Clostridia |
| Clostridiaceae | Eubacteriales | Clostridia |
| Clostridiaceae | Eubacteriales | Clostridia |
| Clostridiaceae | Eubacteriales | Clostridia |
| Clostridiaceae | Eubacteriales | Clostridia |

| | | |
|---|---|---|
| Clostridiaceae | Eubacteriales | Clostridia |
| Clostridiaceae | Eubacteriales | Clostridia |
| Clostridiaceae | Eubacteriales | Clostridia |
| Clostridiaceae | Eubacteriales | Clostridia |
| Clostridiaceae | Eubacteriales | Clostridia |
| Clostridiaceae | Eubacteriales | Clostridia |
| Clostridiaceae | Eubacteriales | Clostridia |
| Clostridiaceae | Eubacteriales | Clostridia |
| Clostridiaceae | Eubacteriales | Clostridia |
| Clostridiaceae | Eubacteriales | Clostridia |
| Eubacteriaceae | Eubacteriales | Clostridia |
| Eubacteriaceae | Eubacteriales | Clostridia |
| FGB9658 | OFGB9658 | CFGB9658 |
| FGB9658 | OFGB9658 | CFGB9658 |
| FGB9659 | OFGB9659 | CFGB9659 |
| FGB9659 | OFGB9659 | CFGB9659 |
| Bacteria_unclassified | Bacteria_unclassified | Bacteria_unclassified |
| Bacteria_unclassified | Bacteria_unclassified | Bacteria_unclassified |
| FGB9827 | OFGB9827 | CFGB9827 |
| FGB9827 | OFGB9827 | CFGB9827 |
| Eubacteriaceae | Eubacteriales | Clostridia |
| Eubacteriaceae | Eubacteriales | Clostridia |
| FGB77303 | OFGB77303 | CFGB77303 |
| FGB77303 | OFGB77303 | CFGB77303 |
| Eubacteriales_unclassified | Eubacteriales | Clostridia |
| Eubacteriales_unclassified | Eubacteriales | Clostridia |
| Oscillospiraceae | Eubacteriales | Clostridia |
| Oscillospiraceae | Eubacteriales | Clostridia |
| FGB30328 | OFGB30328 | CFGB30328 |
| FGB30328 | OFGB30328 | CFGB30328 |
| Oscillospiraceae | Eubacteriales | Clostridia |
| Oscillospiraceae | Eubacteriales | Clostridia |
| Oscillospiraceae | Eubacteriales | Clostridia |
| Oscillospiraceae | Eubacteriales | Clostridia |
| Oscillospiraceae | Eubacteriales | Clostridia |
| Oscillospiraceae | Eubacteriales | Clostridia |
| Oscillospiraceae | Eubacteriales | Clostridia |
| Oscillospiraceae | Eubacteriales | Clostridia |
| Oscillospiraceae | Eubacteriales | Clostridia |
| Oscillospiraceae | Eubacteriales | Clostridia |
| Oscillospiraceae | Eubacteriales | Clostridia |
| Oscillospiraceae | Eubacteriales | Clostridia |
| Oscillospiraceae | Eubacteriales | Clostridia |
| Oscillospiraceae | Eubacteriales | Clostridia |
| FGB77153 | OFGB77153 | CFGB77153 |
| FGB77153 | OFGB77153 | CFGB77153 |

| | | |
|---|---|---|
| FGB1791 | OFGB1791 | CFGB1791 |
| FGB1791 | OFGB1791 | CFGB1791 |
| FGB10290 | OFGB10290 | CFGB10290 |
| FGB10290 | OFGB10290 | CFGB10290 |
| Oscillospiraceae | Eubacteriales | Clostridia |
| Oscillospiraceae | Eubacteriales | Clostridia |
| FGB1765 | OFGB1765 | CFGB1765 |
| FGB1765 | OFGB1765 | CFGB1765 |
| FGB10349 | OFGB10349 | CFGB10349 |
| FGB10349 | OFGB10349 | CFGB10349 |
| FGB10667 | OFGB10667 | CFGB10667 |
| FGB10667 | OFGB10667 | CFGB10667 |
| Lachnospiraceae | Eubacteriales | Clostridia |
| Lachnospiraceae | Eubacteriales | Clostridia |
| Lachnospiraceae | Eubacteriales | Clostridia |
| Lachnospiraceae | Eubacteriales | Clostridia |
| Christensenellaceae | Eubacteriales | Clostridia |
| Christensenellaceae | Eubacteriales | Clostridia |
| FGB10299 | OFGB10299 | CFGB10299 |
| FGB10299 | OFGB10299 | CFGB10299 |
| Oscillospiraceae | Eubacteriales | Clostridia |
| Oscillospiraceae | Eubacteriales | Clostridia |
| Oscillospiraceae | Eubacteriales | Clostridia |
| Oscillospiraceae | Eubacteriales | Clostridia |
| Lachnospiraceae | Eubacteriales | Clostridia |
| Lachnospiraceae | Eubacteriales | Clostridia |
| Clostridia_unclassified | Clostridia_unclassified | Clostridia |
| Clostridia_unclassified | Clostridia_unclassified | Clostridia |
| Lachnospiraceae | Eubacteriales | Clostridia |
| Lachnospiraceae | Eubacteriales | Clostridia |
| Lachnospiraceae | Eubacteriales | Clostridia |
| Lachnospiraceae | Eubacteriales | Clostridia |
| Lachnospiraceae | Eubacteriales | Clostridia |
| Lachnospiraceae | Eubacteriales | Clostridia |
| Lachnospiraceae | Eubacteriales | Clostridia |
| Lachnospiraceae | Eubacteriales | Clostridia |
| Lachnospiraceae | Eubacteriales | Clostridia |
| Lachnospiraceae | Eubacteriales | Clostridia |
| Lachnospiraceae | Eubacteriales | Clostridia |
| Lachnospiraceae | Eubacteriales | Clostridia |
| Lachnospiraceae | Eubacteriales | Clostridia |
| Lachnospiraceae | Eubacteriales | Clostridia |
| Lachnospiraceae | Eubacteriales | Clostridia |
| Lachnospiraceae | Eubacteriales | Clostridia |
| Lactobacillaceae | Lactobacillales | Bacilli |
| Lactobacillaceae | Lactobacillales | Bacilli |

| | | |
|---|---|---|
| Muribaculaceae | Bacteroidales | Bacteroidia |
| Muribaculaceae | Bacteroidales | Bacteroidia |
| Oscillospiraceae | Eubacteriales | Clostridia |
| Oscillospiraceae | Eubacteriales | Clostridia |
| Oscillospiraceae | Eubacteriales | Clostridia |
| Oscillospiraceae | Eubacteriales | Clostridia |
| Oscillospiraceae | Eubacteriales | Clostridia |
| Oscillospiraceae | Eubacteriales | Clostridia |
| Oscillospiraceae | Eubacteriales | Clostridia |
| Oscillospiraceae | Eubacteriales | Clostridia |
| Sutterellaceae | Burkholderiales | Betaproteobacteria |
| Sutterellaceae | Burkholderiales | Betaproteobacteria |
| Peptostreptococcaceae | Eubacteriales | Clostridia |
| Peptostreptococcaceae | Eubacteriales | Clostridia |
| Lachnospiraceae | Eubacteriales | Clostridia |
| Lachnospiraceae | Eubacteriales | Clostridia |
| Turicibacteraceae | Erysipelotrichales | Erysipelotrichia |
| Turicibacteraceae | Erysipelotrichales | Erysipelotrichia |
| Lachnospiraceae | Eubacteriales | Clostridia |
| Lachnospiraceae | Eubacteriales | Clostridia |
| Lachnospiraceae | Eubacteriales | Clostridia |
| Lachnospiraceae | Eubacteriales | Clostridia |
| Oscillospiraceae | Eubacteriales | Clostridia |
| Oscillospiraceae | Eubacteriales | Clostridia |
| Oscillospiraceae | Eubacteriales | Clostridia |
| Oscillospiraceae | Eubacteriales | Clostridia |
| Eggerthellaceae | Eggerthellales | Coriobacteriia |
| Eggerthellaceae | Eggerthellales | Coriobacteriia |
| Eggerthellaceae | Eggerthellales | Coriobacteriia |
| Eggerthellaceae | Eggerthellales | Coriobacteriia |
| Eggerthellaceae | Eggerthellales | Coriobacteriia |
| Eggerthellaceae | Eggerthellales | Coriobacteriia |
| Akkermansiaceae | Verrucomicrobiales | Verrucomicrobiae |
| Akkermansiaceae | Verrucomicrobiales | Verrucomicrobiae |
| Rikenellaceae | Bacteroidales | Bacteroidia |
| Rikenellaceae | Bacteroidales | Bacteroidia |
| Oscillospiraceae | Eubacteriales | Clostridia |
| Oscillospiraceae | Eubacteriales | Clostridia |
| Bacteria_unclassified | Bacteria_unclassified | Bacteria_unclassified |
| Bacteria_unclassified | Bacteria_unclassified | Bacteria_unclassified |
| Bacteria_unclassified | Bacteria_unclassified | Bacteria_unclassified |
| Bacteria_unclassified | Bacteria_unclassified | Bacteria_unclassified |

| | | |
|---|---|---|
| Bacteria_unclassified | Bacteria_unclassified | Bacteria_unclassified |
| Bacteria_unclassified | Bacteria_unclassified | Bacteria_unclassified |
| Bacteria_unclassified | Bacteria_unclassified | Bacteria_unclassified |
| Bacteria_unclassified | Bacteria_unclassified | Bacteria_unclassified |
| Bacteria_unclassified | Bacteria_unclassified | Bacteria_unclassified |
| Bacteria_unclassified | Bacteria_unclassified | Bacteria_unclassified |
| Bacteria_unclassified | Bacteria_unclassified | Bacteria_unclassified |
| Bacteria_unclassified | Bacteria_unclassified | Bacteria_unclassified |
| Bacteroidaceae | Bacteroidales | Bacteroidia |
| Bacteroidaceae | Bacteroidales | Bacteroidia |
| Bifidobacteriaceae | Bifidobacteriales | Actinomycetia |
| Bifidobacteriaceae | Bifidobacteriales | Actinomycetia |
| Clostridia_unclassified | Clostridia_unclassified | Clostridia |
| Clostridia_unclassified | Clostridia_unclassified | Clostridia |
| Clostridiaceae | Eubacteriales | Clostridia |
| Clostridiaceae | Eubacteriales | Clostridia |
| Clostridiaceae | Eubacteriales | Clostridia |
| Clostridiaceae | Eubacteriales | Clostridia |
| Eubacteriales_unclassified | Eubacteriales | Clostridia |
| Eubacteriales_unclassified | Eubacteriales | Clostridia |
| Erysipelotrichaceae | Erysipelotrichales | Erysipelotrichia |
| Erysipelotrichaceae | Erysipelotrichales | Erysipelotrichia |
| Coriobacteriaceae | Coriobacteriales | Coriobacteriia |
| Coriobacteriaceae | Coriobacteriales | Coriobacteriia |
| Lachnospiraceae | Eubacteriales | Clostridia |
| Lachnospiraceae | Eubacteriales | Clostridia |
| Erysipelotrichaceae | Erysipelotrichales | Erysipelotrichia |
| Erysipelotrichaceae | Erysipelotrichales | Erysipelotrichia |
| Erysipelotrichales_unclassified | Erysipelotrichales | Erysipelotrichia |
| Erysipelotrichales_unclassified | Erysipelotrichales | Erysipelotrichia |
| Eubacteriaceae | Eubacteriales | Clostridia |
| Eubacteriaceae | Eubacteriales | Clostridia |
| Eubacteriaceae | Eubacteriales | Clostridia |
| Eubacteriaceae | Eubacteriales | Clostridia |
| Lachnospiraceae | Eubacteriales | Clostridia |
| Lachnospiraceae | Eubacteriales | Clostridia |
| Eggerthellaceae | Eggerthellales | Coriobacteriia |
| Eggerthellaceae | Eggerthellales | Coriobacteriia |
| Lachnospiraceae | Eubacteriales | Clostridia |
| Lachnospiraceae | Eubacteriales | Clostridia |
| Muribaculaceae | Bacteroidales | Bacteroidia |
| Muribaculaceae | Bacteroidales | Bacteroidia |
| Muribaculaceae | Bacteroidales | Bacteroidia |
| Muribaculaceae | Bacteroidales | Bacteroidia |

| | | |
|---|---|---|
| Muribaculaceae | Bacteroidales | Bacteroidia |
| Muribaculaceae | Bacteroidales | Bacteroidia |
| FGB9508 | OFGB9508 | CFGB9508 |
| FGB9508 | OFGB9508 | CFGB9508 |
| FGB2838 | OFGB2838 | CFGB2838 |
| FGB2838 | OFGB2838 | CFGB2838 |
| FGB2838 | OFGB2838 | CFGB2838 |
| FGB2838 | OFGB2838 | CFGB2838 |
| FGB2838 | OFGB2838 | CFGB2838 |
| FGB2838 | OFGB2838 | CFGB2838 |
| Pumilibacteraceae | Eubacteriales | Clostridia |
| Pumilibacteraceae | Eubacteriales | Clostridia |
| FGB28439 | OFGB28439 | CFGB28439 |
| FGB28439 | OFGB28439 | CFGB28439 |
| Clostridia_unclassified | Clostridia_unclassified | Clostridia |
| Clostridia_unclassified | Clostridia_unclassified | Clostridia |
| Eubacteriaceae | Eubacteriales | Clostridia |
| Eubacteriaceae | Eubacteriales | Clostridia |
| Lachnospiraceae | Eubacteriales | Clostridia |
| Lachnospiraceae | Eubacteriales | Clostridia |
| Lachnospiraceae | Eubacteriales | Clostridia |
| Lachnospiraceae | Eubacteriales | Clostridia |
| Lachnospiraceae | Eubacteriales | Clostridia |
| Lachnospiraceae | Eubacteriales | Clostridia |
| Lachnospiraceae | Eubacteriales | Clostridia |
| Lachnospiraceae | Eubacteriales | Clostridia |
| FGB77305 | OFGB77305 | CFGB77305 |
| FGB77305 | OFGB77305 | CFGB77305 |
| Clostridiaceae | Eubacteriales | Clostridia |
| Clostridiaceae | Eubacteriales | Clostridia |
| Lachnospiraceae | Eubacteriales | Clostridia |
| Lachnospiraceae | Eubacteriales | Clostridia |
| Lachnospiraceae | Eubacteriales | Clostridia |
| Lachnospiraceae | Eubacteriales | Clostridia |
| Lachnospiraceae | Eubacteriales | Clostridia |
| Lachnospiraceae | Eubacteriales | Clostridia |
| FGB9633 | OFGB9633 | CFGB9633 |
| FGB9633 | OFGB9633 | CFGB9633 |
| Bacteria_unclassified | Bacteria_unclassified | Bacteria_unclassified |
| Bacteria_unclassified | Bacteria_unclassified | Bacteria_unclassified |
| Bacteria_unclassified | Bacteria_unclassified | Bacteria_unclassified |
| Bacteria_unclassified | Bacteria_unclassified | Bacteria_unclassified |
| Bacteria_unclassified | Bacteria_unclassified | Bacteria_unclassified |
| Bacteria_unclassified | Bacteria_unclassified | Bacteria_unclassified |
| Bacteria_unclassified | Bacteria_unclassified | Bacteria_unclassified |
| Bacteria_unclassified | Bacteria_unclassified | Bacteria_unclassified |

| | | |
|---|---|---|
| Lachnospiraceae | Eubacteriales | Clostridia |
| Lachnospiraceae | Eubacteriales | Clostridia |
| Lachnospiraceae | Eubacteriales | Clostridia |
| Lachnospiraceae | Eubacteriales | Clostridia |
| Lachnospiraceae | Eubacteriales | Clostridia |
| Lachnospiraceae | Eubacteriales | Clostridia |
| FGB77359 | OFGB77359 | CFGB77359 |
| FGB77359 | OFGB77359 | CFGB77359 |
| FGB9639 | OFGB9639 | CFGB9639 |
| FGB9639 | OFGB9639 | CFGB9639 |
| Lachnospiraceae | Eubacteriales | Clostridia |
| Lachnospiraceae | Eubacteriales | Clostridia |
| Lachnospiraceae | Eubacteriales | Clostridia |
| Lachnospiraceae | Eubacteriales | Clostridia |
| Lachnospiraceae | Eubacteriales | Clostridia |
| Lachnospiraceae | Eubacteriales | Clostridia |
| Clostridiaceae | Eubacteriales | Clostridia |
| Clostridiaceae | Eubacteriales | Clostridia |
| Clostridiaceae | Eubacteriales | Clostridia |
| Clostridiaceae | Eubacteriales | Clostridia |
| Clostridiaceae | Eubacteriales | Clostridia |
| Clostridiaceae | Eubacteriales | Clostridia |
| Clostridiaceae | Eubacteriales | Clostridia |
| Clostridiaceae | Eubacteriales | Clostridia |
| Clostridiaceae | Eubacteriales | Clostridia |
| Clostridiaceae | Eubacteriales | Clostridia |
| Clostridiaceae | Eubacteriales | Clostridia |
| Clostridiaceae | Eubacteriales | Clostridia |
| Clostridiaceae | Eubacteriales | Clostridia |
| Clostridiaceae | Eubacteriales | Clostridia |
| Eubacteriaceae | Eubacteriales | Clostridia |
| Eubacteriaceae | Eubacteriales | Clostridia |
| FGB9658 | OFGB9658 | CFGB9658 |
| FGB9658 | OFGB9658 | CFGB9658 |
| FGB9659 | OFGB9659 | CFGB9659 |
| FGB9659 | OFGB9659 | CFGB9659 |
| Bacteria_unclassified | Bacteria_unclassified | Bacteria_unclassified |
| Bacteria_unclassified | Bacteria_unclassified | Bacteria_unclassified |
| FGB9827 | OFGB9827 | CFGB9827 |
| FGB9827 | OFGB9827 | CFGB9827 |
| Eubacteriaceae | Eubacteriales | Clostridia |
| Eubacteriaceae | Eubacteriales | Clostridia |
| FGB77303 | OFGB77303 | CFGB77303 |
| FGB77303 | OFGB77303 | CFGB77303 |
| Eubacteriales_unclassified | Eubacteriales | Clostridia |
| Eubacteriales_unclassified | Eubacteriales | Clostridia |

| | | |
|---|---|---|
| Oscillospiraceae | Eubacteriales | Clostridia |
| Oscillospiraceae | Eubacteriales | Clostridia |
| FGB30328 | OFGB30328 | CFGB30328 |
| FGB30328 | OFGB30328 | CFGB30328 |
| Oscillospiraceae | Eubacteriales | Clostridia |
| Oscillospiraceae | Eubacteriales | Clostridia |
| Oscillospiraceae | Eubacteriales | Clostridia |
| Oscillospiraceae | Eubacteriales | Clostridia |
| Oscillospiraceae | Eubacteriales | Clostridia |
| Oscillospiraceae | Eubacteriales | Clostridia |
| Oscillospiraceae | Eubacteriales | Clostridia |
| Oscillospiraceae | Eubacteriales | Clostridia |
| Oscillospiraceae | Eubacteriales | Clostridia |
| Oscillospiraceae | Eubacteriales | Clostridia |
| Oscillospiraceae | Eubacteriales | Clostridia |
| Oscillospiraceae | Eubacteriales | Clostridia |
| Oscillospiraceae | Eubacteriales | Clostridia |
| Oscillospiraceae | Eubacteriales | Clostridia |
| FGB77153 | OFGB77153 | CFGB77153 |
| FGB77153 | OFGB77153 | CFGB77153 |
| FGB1791 | OFGB1791 | CFGB1791 |
| FGB1791 | OFGB1791 | CFGB1791 |
| FGB10290 | OFGB10290 | CFGB10290 |
| FGB10290 | OFGB10290 | CFGB10290 |
| Oscillospiraceae | Eubacteriales | Clostridia |
| Oscillospiraceae | Eubacteriales | Clostridia |
| FGB1765 | OFGB1765 | CFGB1765 |
| FGB1765 | OFGB1765 | CFGB1765 |
| FGB10349 | OFGB10349 | CFGB10349 |
| FGB10349 | OFGB10349 | CFGB10349 |
| FGB10667 | OFGB10667 | CFGB10667 |
| FGB10667 | OFGB10667 | CFGB10667 |
| Lachnospiraceae | Eubacteriales | Clostridia |
| Lachnospiraceae | Eubacteriales | Clostridia |
| Lachnospiraceae | Eubacteriales | Clostridia |
| Lachnospiraceae | Eubacteriales | Clostridia |
| Christensenellaceae | Eubacteriales | Clostridia |
| Christensenellaceae | Eubacteriales | Clostridia |
| FGB10299 | OFGB10299 | CFGB10299 |
| FGB10299 | OFGB10299 | CFGB10299 |
| Oscillospiraceae | Eubacteriales | Clostridia |
| Oscillospiraceae | Eubacteriales | Clostridia |
| Oscillospiraceae | Eubacteriales | Clostridia |
| Oscillospiraceae | Eubacteriales | Clostridia |
| Lachnospiraceae | Eubacteriales | Clostridia |
| Lachnospiraceae | Eubacteriales | Clostridia |

| | | |
|---|---|---|
| Clostridia_unclassified | Clostridia_unclassified | Clostridia |
| Clostridia_unclassified | Clostridia_unclassified | Clostridia |
| Lachnospiraceae | Eubacteriales | Clostridia |
| Lachnospiraceae | Eubacteriales | Clostridia |
| Lachnospiraceae | Eubacteriales | Clostridia |
| Lachnospiraceae | Eubacteriales | Clostridia |
| Lachnospiraceae | Eubacteriales | Clostridia |
| Lachnospiraceae | Eubacteriales | Clostridia |
| Lachnospiraceae | Eubacteriales | Clostridia |
| Lachnospiraceae | Eubacteriales | Clostridia |
| Lachnospiraceae | Eubacteriales | Clostridia |
| Lachnospiraceae | Eubacteriales | Clostridia |
| Lachnospiraceae | Eubacteriales | Clostridia |
| Lachnospiraceae | Eubacteriales | Clostridia |
| Lachnospiraceae | Eubacteriales | Clostridia |
| Lachnospiraceae | Eubacteriales | Clostridia |
| Lachnospiraceae | Eubacteriales | Clostridia |
| Lachnospiraceae | Eubacteriales | Clostridia |
| Lactobacillaceae | Lactobacillales | Bacilli |
| Lactobacillaceae | Lactobacillales | Bacilli |
| Muribaculaceae | Bacteroidales | Bacteroidia |
| Muribaculaceae | Bacteroidales | Bacteroidia |
| Oscillospiraceae | Eubacteriales | Clostridia |
| Oscillospiraceae | Eubacteriales | Clostridia |
| Oscillospiraceae | Eubacteriales | Clostridia |
| Oscillospiraceae | Eubacteriales | Clostridia |
| Oscillospiraceae | Eubacteriales | Clostridia |
| Oscillospiraceae | Eubacteriales | Clostridia |
| Oscillospiraceae | Eubacteriales | Clostridia |
| Oscillospiraceae | Eubacteriales | Clostridia |
| Sutterellaceae | Burkholderiales | Betaproteobacteria |
| Sutterellaceae | Burkholderiales | Betaproteobacteria |
| Peptostreptococcaceae | Eubacteriales | Clostridia |
| Peptostreptococcaceae | Eubacteriales | Clostridia |
| Lachnospiraceae | Eubacteriales | Clostridia |
| Lachnospiraceae | Eubacteriales | Clostridia |
| Turicibacteraceae | Erysipelotrichales | Erysipelotrichia |
| Turicibacteraceae | Erysipelotrichales | Erysipelotrichia |
| Lachnospiraceae | Eubacteriales | Clostridia |
| Lachnospiraceae | Eubacteriales | Clostridia |
| Lachnospiraceae | Eubacteriales | Clostridia |
| Lachnospiraceae | Eubacteriales | Clostridia |

| | | |
|---|---|---|
| Oscillospiraceae | Eubacteriales | Clostridia |
| Oscillospiraceae | Eubacteriales | Clostridia |
| Oscillospiraceae | Eubacteriales | Clostridia |
| Oscillospiraceae | Eubacteriales | Clostridia |
| Eggerthellaceae | Eggerthellales | Coriobacteriia |
| Eggerthellaceae | Eggerthellales | Coriobacteriia |
| Eggerthellaceae | Eggerthellales | Coriobacteriia |
| Eggerthellaceae | Eggerthellales | Coriobacteriia |
| Eggerthellaceae | Eggerthellales | Coriobacteriia |
| Eggerthellaceae | Eggerthellales | Coriobacteriia |
| Akkermansiaceae | Verrucomicrobiales | Verrucomicrobiae |
| Akkermansiaceae | Verrucomicrobiales | Verrucomicrobiae |
| Rikenellaceae | Bacteroidales | Bacteroidia |
| Rikenellaceae | Bacteroidales | Bacteroidia |
| Oscillospiraceae | Eubacteriales | Clostridia |
| Oscillospiraceae | Eubacteriales | Clostridia |
| Bacteria_unclassified | Bacteria_unclassified | Bacteria_unclassified |
| Bacteria_unclassified | Bacteria_unclassified | Bacteria_unclassified |
| Bacteria_unclassified | Bacteria_unclassified | Bacteria_unclassified |
| Bacteria_unclassified | Bacteria_unclassified | Bacteria_unclassified |
| Bacteria_unclassified | Bacteria_unclassified | Bacteria_unclassified |
| Bacteria_unclassified | Bacteria_unclassified | Bacteria_unclassified |
| Bacteria_unclassified | Bacteria_unclassified | Bacteria_unclassified |
| Bacteria_unclassified | Bacteria_unclassified | Bacteria_unclassified |
| Bacteria_unclassified | Bacteria_unclassified | Bacteria_unclassified |
| Bacteria_unclassified | Bacteria_unclassified | Bacteria_unclassified |
| Bacteria_unclassified | Bacteria_unclassified | Bacteria_unclassified |
| Bacteria_unclassified | Bacteria_unclassified | Bacteria_unclassified |
| Bacteroidaceae | Bacteroidales | Bacteroidia |
| Bacteroidaceae | Bacteroidales | Bacteroidia |

| | | |
|---|---|---|
| Bifidobacteriaceae | Bifidobacteriales | Actinomycetia |
| Bifidobacteriaceae | Bifidobacteriales | Actinomycetia |
| Clostridia_unclassified | Clostridia_unclassified | Clostridia |
| Clostridia_unclassified | Clostridia_unclassified | Clostridia |
| Clostridiaceae | Eubacteriales | Clostridia |
| Clostridiaceae | Eubacteriales | Clostridia |
| Clostridiaceae | Eubacteriales | Clostridia |
| Clostridiaceae | Eubacteriales | Clostridia |
| Eubacteriales_unclassified | Eubacteriales | Clostridia |
| Eubacteriales_unclassified | Eubacteriales | Clostridia |
| Erysipelotrichaceae | Erysipelotrichales | Erysipelotrichia |
| Erysipelotrichaceae | Erysipelotrichales | Erysipelotrichia |
| Coriobacteriaceae | Coriobacteriales | Coriobacteriia |
| Coriobacteriaceae | Coriobacteriales | Coriobacteriia |

| | | |
|---|---|---|
| Lachnospiraceae | Eubacteriales | Clostridia |
| Lachnospiraceae | Eubacteriales | Clostridia |
| Erysipelotrichaceae | Erysipelotrichales | Erysipelotrichia |
| Erysipelotrichaceae | Erysipelotrichales | Erysipelotrichia |
| Erysipelotrichales_unclassified | Erysipelotrichales | Erysipelotrichia |
| Erysipelotrichales_unclassified | Erysipelotrichales | Erysipelotrichia |
| Eubacteriaceae | Eubacteriales | Clostridia |
| Eubacteriaceae | Eubacteriales | Clostridia |
| Eubacteriaceae | Eubacteriales | Clostridia |
| Eubacteriaceae | Eubacteriales | Clostridia |
| Lachnospiraceae | Eubacteriales | Clostridia |
| Lachnospiraceae | Eubacteriales | Clostridia |
| Eggerthellaceae | Eggerthellales | Coriobacteriia |
| Eggerthellaceae | Eggerthellales | Coriobacteriia |
| Lachnospiraceae | Eubacteriales | Clostridia |
| Lachnospiraceae | Eubacteriales | Clostridia |
| Muribaculaceae | Bacteroidales | Bacteroidia |
| Muribaculaceae | Bacteroidales | Bacteroidia |
| Muribaculaceae | Bacteroidales | Bacteroidia |
| Muribaculaceae | Bacteroidales | Bacteroidia |
| Muribaculaceae | Bacteroidales | Bacteroidia |
| Muribaculaceae | Bacteroidales | Bacteroidia |
| FGB9508 | OFGB9508 | CFGB9508 |
| FGB9508 | OFGB9508 | CFGB9508 |
| FGB2838 | OFGB2838 | CFGB2838 |
| FGB2838 | OFGB2838 | CFGB2838 |
| FGB2838 | OFGB2838 | CFGB2838 |
| FGB2838 | OFGB2838 | CFGB2838 |
| FGB2838 | OFGB2838 | CFGB2838 |
| FGB2838 | OFGB2838 | CFGB2838 |
| Pumilibacteraceae | Eubacteriales | Clostridia |
| Pumilibacteraceae | Eubacteriales | Clostridia |
| FGB28439 | OFGB28439 | CFGB28439 |
| FGB28439 | OFGB28439 | CFGB28439 |
| Clostridia_unclassified | Clostridia_unclassified | Clostridia |
| Clostridia_unclassified | Clostridia_unclassified | Clostridia |
| Eubacteriaceae | Eubacteriales | Clostridia |
| Eubacteriaceae | Eubacteriales | Clostridia |
| Lachnospiraceae | Eubacteriales | Clostridia |
| Lachnospiraceae | Eubacteriales | Clostridia |
| Lachnospiraceae | Eubacteriales | Clostridia |
| Lachnospiraceae | Eubacteriales | Clostridia |
| Lachnospiraceae | Eubacteriales | Clostridia |
| Lachnospiraceae | Eubacteriales | Clostridia |
| Lachnospiraceae | Eubacteriales | Clostridia |
| Lachnospiraceae | Eubacteriales | Clostridia |

| | | |
|---|---|---|
| FGB77305 | OFGB77305 | CFGB77305 |
| FGB77305 | OFGB77305 | CFGB77305 |
| Clostridiaceae | Eubacteriales | Clostridia |
| Clostridiaceae | Eubacteriales | Clostridia |
| Lachnospiraceae | Eubacteriales | Clostridia |
| Lachnospiraceae | Eubacteriales | Clostridia |
| Lachnospiraceae | Eubacteriales | Clostridia |
| Lachnospiraceae | Eubacteriales | Clostridia |
| Lachnospiraceae | Eubacteriales | Clostridia |
| Lachnospiraceae | Eubacteriales | Clostridia |
| FGB9633 | OFGB9633 | CFGB9633 |
| FGB9633 | OFGB9633 | CFGB9633 |
| Bacteria_unclassified | Bacteria_unclassified | Bacteria_unclassified |
| Bacteria_unclassified | Bacteria_unclassified | Bacteria_unclassified |
| Bacteria_unclassified | Bacteria_unclassified | Bacteria_unclassified |
| Bacteria_unclassified | Bacteria_unclassified | Bacteria_unclassified |
| Bacteria_unclassified | Bacteria_unclassified | Bacteria_unclassified |
| Bacteria_unclassified | Bacteria_unclassified | Bacteria_unclassified |
| Bacteria_unclassified | Bacteria_unclassified | Bacteria_unclassified |
| Bacteria_unclassified | Bacteria_unclassified | Bacteria_unclassified |
| Lachnospiraceae | Eubacteriales | Clostridia |
| Lachnospiraceae | Eubacteriales | Clostridia |
| Lachnospiraceae | Eubacteriales | Clostridia |
| Lachnospiraceae | Eubacteriales | Clostridia |
| Lachnospiraceae | Eubacteriales | Clostridia |
| Lachnospiraceae | Eubacteriales | Clostridia |
| FGB77359 | OFGB77359 | CFGB77359 |
| FGB77359 | OFGB77359 | CFGB77359 |
| FGB9639 | OFGB9639 | CFGB9639 |
| FGB9639 | OFGB9639 | CFGB9639 |
| Lachnospiraceae | Eubacteriales | Clostridia |
| Lachnospiraceae | Eubacteriales | Clostridia |
| Lachnospiraceae | Eubacteriales | Clostridia |
| Lachnospiraceae | Eubacteriales | Clostridia |
| Lachnospiraceae | Eubacteriales | Clostridia |
| Lachnospiraceae | Eubacteriales | Clostridia |
| Clostridiaceae | Eubacteriales | Clostridia |
| Clostridiaceae | Eubacteriales | Clostridia |
| Clostridiaceae | Eubacteriales | Clostridia |
| Clostridiaceae | Eubacteriales | Clostridia |
| Clostridiaceae | Eubacteriales | Clostridia |
| Clostridiaceae | Eubacteriales | Clostridia |
| Clostridiaceae | Eubacteriales | Clostridia |
| Clostridiaceae | Eubacteriales | Clostridia |
| Clostridiaceae | Eubacteriales | Clostridia |
| Clostridiaceae | Eubacteriales | Clostridia |

| | | |
|---|---|---|
| Clostridiaceae | Eubacteriales | Clostridia |
| Clostridiaceae | Eubacteriales | Clostridia |
| Clostridiaceae | Eubacteriales | Clostridia |
| Clostridiaceae | Eubacteriales | Clostridia |
| Eubacteriaceae | Eubacteriales | Clostridia |
| Eubacteriaceae | Eubacteriales | Clostridia |
| FGB9658 | OFGB9658 | CFGB9658 |
| FGB9658 | OFGB9658 | CFGB9658 |
| FGB9659 | OFGB9659 | CFGB9659 |
| FGB9659 | OFGB9659 | CFGB9659 |
| Bacteria_unclassified | Bacteria_unclassified | Bacteria_unclassified |
| Bacteria_unclassified | Bacteria_unclassified | Bacteria_unclassified |
| FGB9827 | OFGB9827 | CFGB9827 |
| FGB9827 | OFGB9827 | CFGB9827 |
| Eubacteriaceae | Eubacteriales | Clostridia |
| Eubacteriaceae | Eubacteriales | Clostridia |
| FGB77303 | OFGB77303 | CFGB77303 |
| FGB77303 | OFGB77303 | CFGB77303 |
| Eubacteriales_unclassified | Eubacteriales | Clostridia |
| Eubacteriales_unclassified | Eubacteriales | Clostridia |
| Oscillospiraceae | Eubacteriales | Clostridia |
| Oscillospiraceae | Eubacteriales | Clostridia |
| FGB30328 | OFGB30328 | CFGB30328 |
| FGB30328 | OFGB30328 | CFGB30328 |
| Oscillospiraceae | Eubacteriales | Clostridia |
| Oscillospiraceae | Eubacteriales | Clostridia |
| Oscillospiraceae | Eubacteriales | Clostridia |
| Oscillospiraceae | Eubacteriales | Clostridia |
| Oscillospiraceae | Eubacteriales | Clostridia |
| Oscillospiraceae | Eubacteriales | Clostridia |
| Oscillospiraceae | Eubacteriales | Clostridia |
| Oscillospiraceae | Eubacteriales | Clostridia |
| Oscillospiraceae | Eubacteriales | Clostridia |
| Oscillospiraceae | Eubacteriales | Clostridia |
| Oscillospiraceae | Eubacteriales | Clostridia |
| Oscillospiraceae | Eubacteriales | Clostridia |
| Oscillospiraceae | Eubacteriales | Clostridia |
| Oscillospiraceae | Eubacteriales | Clostridia |
| FGB77153 | OFGB77153 | CFGB77153 |
| FGB77153 | OFGB77153 | CFGB77153 |
| FGB1791 | OFGB1791 | CFGB1791 |
| FGB1791 | OFGB1791 | CFGB1791 |
| FGB10290 | OFGB10290 | CFGB10290 |
| FGB10290 | OFGB10290 | CFGB10290 |
| Oscillospiraceae | Eubacteriales | Clostridia |
| Oscillospiraceae | Eubacteriales | Clostridia |

| | | |
|---|---|---|
| FGB1765 | OFGB1765 | CFGB1765 |
| FGB1765 | OFGB1765 | CFGB1765 |
| FGB10349 | OFGB10349 | CFGB10349 |
| FGB10349 | OFGB10349 | CFGB10349 |
| FGB10667 | OFGB10667 | CFGB10667 |
| FGB10667 | OFGB10667 | CFGB10667 |
| Lachnospiraceae | Eubacteriales | Clostridia |
| Lachnospiraceae | Eubacteriales | Clostridia |
| Lachnospiraceae | Eubacteriales | Clostridia |
| Lachnospiraceae | Eubacteriales | Clostridia |
| Christensenellaceae | Eubacteriales | Clostridia |
| Christensenellaceae | Eubacteriales | Clostridia |
| FGB10299 | OFGB10299 | CFGB10299 |
| FGB10299 | OFGB10299 | CFGB10299 |
| Oscillospiraceae | Eubacteriales | Clostridia |
| Oscillospiraceae | Eubacteriales | Clostridia |
| Oscillospiraceae | Eubacteriales | Clostridia |
| Oscillospiraceae | Eubacteriales | Clostridia |
| Lachnospiraceae | Eubacteriales | Clostridia |
| Lachnospiraceae | Eubacteriales | Clostridia |
| Clostridia_unclassified | Clostridia_unclassified | Clostridia |
| Clostridia_unclassified | Clostridia_unclassified | Clostridia |
| Lachnospiraceae | Eubacteriales | Clostridia |
| Lachnospiraceae | Eubacteriales | Clostridia |
| Lachnospiraceae | Eubacteriales | Clostridia |
| Lachnospiraceae | Eubacteriales | Clostridia |
| Lachnospiraceae | Eubacteriales | Clostridia |
| Lachnospiraceae | Eubacteriales | Clostridia |
| Lachnospiraceae | Eubacteriales | Clostridia |
| Lachnospiraceae | Eubacteriales | Clostridia |
| Lachnospiraceae | Eubacteriales | Clostridia |
| Lachnospiraceae | Eubacteriales | Clostridia |
| Lachnospiraceae | Eubacteriales | Clostridia |
| Lachnospiraceae | Eubacteriales | Clostridia |
| Lachnospiraceae | Eubacteriales | Clostridia |
| Lachnospiraceae | Eubacteriales | Clostridia |
| Lachnospiraceae | Eubacteriales | Clostridia |
| Lachnospiraceae | Eubacteriales | Clostridia |
| Lactobacillaceae | Lactobacillales | Bacilli |
| Lactobacillaceae | Lactobacillales | Bacilli |
| Muribaculaceae | Bacteroidales | Bacteroidia |
| Muribaculaceae | Bacteroidales | Bacteroidia |
| Oscillospiraceae | Eubacteriales | Clostridia |
| Oscillospiraceae | Eubacteriales | Clostridia |
| Oscillospiraceae | Eubacteriales | Clostridia |
| Oscillospiraceae | Eubacteriales | Clostridia |

| | | |
|---|---|---|
| Oscillospiraceae | Eubacteriales | Clostridia |
| Oscillospiraceae | Eubacteriales | Clostridia |
| Oscillospiraceae | Eubacteriales | Clostridia |
| Oscillospiraceae | Eubacteriales | Clostridia |
| Sutterellaceae | Burkholderiales | Betaproteobacteria |
| Sutterellaceae | Burkholderiales | Betaproteobacteria |
| Peptostreptococcaceae | Eubacteriales | Clostridia |
| Peptostreptococcaceae | Eubacteriales | Clostridia |
| Lachnospiraceae | Eubacteriales | Clostridia |
| Lachnospiraceae | Eubacteriales | Clostridia |
| Turicibacteraceae | Erysipelotrichales | Erysipelotrichia |
| Turicibacteraceae | Erysipelotrichales | Erysipelotrichia |

| Phylum | Kingdom |
|---|---|
| Firmicutes | Bacteria |
| Firmicutes | Bacteria |
| Firmicutes | Bacteria |
| Firmicutes | Bacteria |
| Firmicutes | Bacteria |
| Firmicutes | Bacteria |
| Firmicutes | Bacteria |
| Firmicutes | Bacteria |
| Actinobacteria | Bacteria |
| Actinobacteria | Bacteria |
| Actinobacteria | Bacteria |
| Actinobacteria | Bacteria |
| Actinobacteria | Bacteria |
| Actinobacteria | Bacteria |
| Verrucomicrobia | Bacteria |
| Verrucomicrobia | Bacteria |
| Bacteroidota | Bacteria |
| Bacteroidota | Bacteria |
| Firmicutes | Bacteria |
| Firmicutes | Bacteria |
| Bacteria_unclassified | Bacteria |
| Bacteria_unclassified | Bacteria |
| Bacteria_unclassified | Bacteria |
| Bacteria_unclassified | Bacteria |
| Bacteria_unclassified | Bacteria |
| Bacteria_unclassified | Bacteria |
| Bacteria_unclassified | Bacteria |
| Bacteria_unclassified | Bacteria |
| Bacteria_unclassified | Bacteria |
| Bacteria_unclassified | Bacteria |
| Bacteria_unclassified | Bacteria |
| Bacteria_unclassified | Bacteria |
| Bacteroidota | Bacteria |
| Bacteroidota | Bacteria |
| Actinobacteria | Bacteria |
| Actinobacteria | Bacteria |
| Firmicutes | Bacteria |
| Firmicutes | Bacteria |
| Firmicutes | Bacteria |
| Firmicutes | Bacteria |
| Firmicutes | Bacteria |
| Firmicutes | Bacteria |
| Firmicutes | Bacteria |
| Firmicutes | Bacteria |

| | |
|---|---|
| Firmicutes | Bacteria |
| Firmicutes | Bacteria |
| Actinobacteria | Bacteria |
| Actinobacteria | Bacteria |
| Firmicutes | Bacteria |
| Firmicutes | Bacteria |
| Firmicutes | Bacteria |
| Firmicutes | Bacteria |
| Firmicutes | Bacteria |
| Firmicutes | Bacteria |
| Firmicutes | Bacteria |
| Firmicutes | Bacteria |
| Firmicutes | Bacteria |
| Firmicutes | Bacteria |
| Firmicutes | Bacteria |
| Firmicutes | Bacteria |
| Actinobacteria | Bacteria |
| Actinobacteria | Bacteria |
| Firmicutes | Bacteria |
| Firmicutes | Bacteria |
| Bacteroidota | Bacteria |
| Bacteroidota | Bacteria |
| Bacteroidota | Bacteria |
| Bacteroidota | Bacteria |
| Bacteroidota | Bacteria |
| Bacteroidota | Bacteria |
| Firmicutes | Bacteria |
| Firmicutes | Bacteria |
| Firmicutes | Bacteria |
| Firmicutes | Bacteria |
| Firmicutes | Bacteria |
| Firmicutes | Bacteria |
| Firmicutes | Bacteria |
| Firmicutes | Bacteria |
| Firmicutes | Bacteria |
| Firmicutes | Bacteria |
| Firmicutes | Bacteria |
| Firmicutes | Bacteria |
| Firmicutes | Bacteria |
| Firmicutes | Bacteria |
| Firmicutes | Bacteria |
| Firmicutes | Bacteria |
| Firmicutes | Bacteria |
| Firmicutes | Bacteria |
| Firmicutes | Bacteria |
| Firmicutes | Bacteria |

| | |
|---|---|
| Firmicutes | Bacteria |
| Firmicutes | Bacteria |
| Firmicutes | Bacteria |
| Firmicutes | Bacteria |
| Firmicutes | Bacteria |
| Firmicutes | Bacteria |
| Firmicutes | Bacteria |
| Firmicutes | Bacteria |
| Firmicutes | Bacteria |
| Firmicutes | Bacteria |
| Firmicutes | Bacteria |
| Firmicutes | Bacteria |
| Firmicutes | Bacteria |
| Firmicutes | Bacteria |
| Firmicutes | Bacteria |
| Firmicutes | Bacteria |
| Bacteria_unclassified | Bacteria |
| Bacteria_unclassified | Bacteria |
| Bacteria_unclassified | Bacteria |
| Bacteria_unclassified | Bacteria |
| Bacteria_unclassified | Bacteria |
| Bacteria_unclassified | Bacteria |
| Bacteria_unclassified | Bacteria |
| Bacteria_unclassified | Bacteria |
| Firmicutes | Bacteria |
| Firmicutes | Bacteria |
| Firmicutes | Bacteria |
| Firmicutes | Bacteria |
| Firmicutes | Bacteria |
| Firmicutes | Bacteria |
| Bacteria_unclassified | Bacteria |
| Bacteria_unclassified | Bacteria |
| Firmicutes | Bacteria |
| Firmicutes | Bacteria |
| Firmicutes | Bacteria |
| Firmicutes | Bacteria |
| Firmicutes | Bacteria |
| Firmicutes | Bacteria |
| Firmicutes | Bacteria |
| Firmicutes | Bacteria |
| Firmicutes | Bacteria |
| Firmicutes | Bacteria |
| Firmicutes | Bacteria |
| Firmicutes | Bacteria |
| Firmicutes | Bacteria |
| Firmicutes | Bacteria |

| | |
|---|---|
| Firmicutes | Bacteria |
| Firmicutes | Bacteria |
| Firmicutes | Bacteria |
| Firmicutes | Bacteria |
| Firmicutes | Bacteria |
| Firmicutes | Bacteria |
| Firmicutes | Bacteria |
| Firmicutes | Bacteria |
| Firmicutes | Bacteria |
| Firmicutes | Bacteria |
| Firmicutes | Bacteria |
| Firmicutes | Bacteria |
| Firmicutes | Bacteria |
| Firmicutes | Bacteria |
| Bacteria_unclassified | Bacteria |
| Bacteria_unclassified | Bacteria |
| Firmicutes | Bacteria |
| Firmicutes | Bacteria |
| Firmicutes | Bacteria |
| Firmicutes | Bacteria |
| Bacteria_unclassified | Bacteria |
| Bacteria_unclassified | Bacteria |
| Firmicutes | Bacteria |
| Firmicutes | Bacteria |
| Firmicutes | Bacteria |
| Firmicutes | Bacteria |
| Firmicutes | Bacteria |
| Firmicutes | Bacteria |
| Firmicutes | Bacteria |
| Firmicutes | Bacteria |
| Firmicutes | Bacteria |
| Firmicutes | Bacteria |
| Firmicutes | Bacteria |
| Firmicutes | Bacteria |
| Firmicutes | Bacteria |
| Firmicutes | Bacteria |
| Firmicutes | Bacteria |
| Firmicutes | Bacteria |
| Firmicutes | Bacteria |
| Firmicutes | Bacteria |
| Firmicutes | Bacteria |
| Firmicutes | Bacteria |
| Actinobacteria | Bacteria |
| Actinobacteria | Bacteria |
| Tenericutes | Bacteria |
| Tenericutes | Bacteria |

| | |
|---|---|
| Firmicutes | Bacteria |
| Firmicutes | Bacteria |
| Firmicutes | Bacteria |
| Firmicutes | Bacteria |
| Firmicutes | Bacteria |
| Firmicutes | Bacteria |
| Firmicutes | Bacteria |
| Firmicutes | Bacteria |
| Firmicutes | Bacteria |
| Firmicutes | Bacteria |
| Firmicutes | Bacteria |
| Firmicutes | Bacteria |
| Firmicutes | Bacteria |
| Firmicutes | Bacteria |
| Firmicutes | Bacteria |
| Firmicutes | Bacteria |
| Firmicutes | Bacteria |
| Firmicutes | Bacteria |
| Firmicutes | Bacteria |
| Firmicutes | Bacteria |
| Firmicutes | Bacteria |
| Firmicutes | Bacteria |
| Firmicutes | Bacteria |
| Firmicutes | Bacteria |
| Firmicutes | Bacteria |
| Firmicutes | Bacteria |
| Firmicutes | Bacteria |
| Firmicutes | Bacteria |
| Firmicutes | Bacteria |
| Firmicutes | Bacteria |
| Firmicutes | Bacteria |
| Firmicutes | Bacteria |
| Firmicutes | Bacteria |
| Firmicutes | Bacteria |
| Firmicutes | Bacteria |
| Firmicutes | Bacteria |
| Firmicutes | Bacteria |
| Firmicutes | Bacteria |
| Firmicutes | Bacteria |
| Firmicutes | Bacteria |
| Firmicutes | Bacteria |
| Firmicutes | Bacteria |
| Firmicutes | Bacteria |
| Firmicutes | Bacteria |
| Bacteroidota | Bacteria |
| Bacteroidota | Bacteria |

| | |
|---|---|
| Firmicutes | Bacteria |
| Firmicutes | Bacteria |
| Firmicutes | Bacteria |
| Firmicutes | Bacteria |
| Firmicutes | Bacteria |
| Firmicutes | Bacteria |
| Firmicutes | Bacteria |
| Firmicutes | Bacteria |
| Proteobacteria | Bacteria |
| Proteobacteria | Bacteria |
| Firmicutes | Bacteria |
| Firmicutes | Bacteria |
| Firmicutes | Bacteria |
| Firmicutes | Bacteria |
| Firmicutes | Bacteria |
| Firmicutes | Bacteria |
| Firmicutes | Bacteria |
| Firmicutes | Bacteria |
| Firmicutes | Bacteria |
| Firmicutes | Bacteria |
| Firmicutes | Bacteria |
| Firmicutes | Bacteria |
| Firmicutes | Bacteria |
| Firmicutes | Bacteria |
| Actinobacteria | Bacteria |
| Actinobacteria | Bacteria |
| Actinobacteria | Bacteria |
| Actinobacteria | Bacteria |
| Actinobacteria | Bacteria |
| Actinobacteria | Bacteria |
| Verrucomicrobia | Bacteria |
| Verrucomicrobia | Bacteria |
| Bacteroidota | Bacteria |
| Bacteroidota | Bacteria |
| Firmicutes | Bacteria |
| Firmicutes | Bacteria |
| Bacteria_unclassified | Bacteria |
| Bacteria_unclassified | Bacteria |
| Bacteria_unclassified | Bacteria |
| Bacteria_unclassified | Bacteria |
| Bacteria_unclassified | Bacteria |
| Bacteria_unclassified | Bacteria |

| | |
|---|---|
| Bacteria_unclassified | Bacteria |
| Bacteria_unclassified | Bacteria |
| Bacteria_unclassified | Bacteria |
| Bacteria_unclassified | Bacteria |
| Bacteria_unclassified | Bacteria |
| Bacteria_unclassified | Bacteria |
| Bacteroidota | Bacteria |
| Bacteroidota | Bacteria |

| | |
|---|---|
| Actinobacteria | Bacteria |
| Actinobacteria | Bacteria |
| Firmicutes | Bacteria |
| Firmicutes | Bacteria |
| Firmicutes | Bacteria |
| Firmicutes | Bacteria |
| Firmicutes | Bacteria |
| Firmicutes | Bacteria |
| Firmicutes | Bacteria |
| Firmicutes | Bacteria |
| Firmicutes | Bacteria |
| Firmicutes | Bacteria |
| Actinobacteria | Bacteria |
| Actinobacteria | Bacteria |
| Firmicutes | Bacteria |
| Firmicutes | Bacteria |
| Firmicutes | Bacteria |
| Firmicutes | Bacteria |
| Firmicutes | Bacteria |
| Firmicutes | Bacteria |
| Firmicutes | Bacteria |
| Firmicutes | Bacteria |
| Firmicutes | Bacteria |
| Firmicutes | Bacteria |
| Firmicutes | Bacteria |
| Firmicutes | Bacteria |
| Actinobacteria | Bacteria |
| Actinobacteria | Bacteria |
| Firmicutes | Bacteria |
| Firmicutes | Bacteria |
| Bacteroidota | Bacteria |
| Bacteroidota | Bacteria |
| Bacteroidota | Bacteria |
| Bacteroidota | Bacteria |
| Bacteroidota | Bacteria |
| Bacteroidota | Bacteria |

| | |
|---|---|
| Firmicutes | Bacteria |
| Firmicutes | Bacteria |
| Firmicutes | Bacteria |
| Firmicutes | Bacteria |
| Firmicutes | Bacteria |
| Firmicutes | Bacteria |
| Firmicutes | Bacteria |
| Firmicutes | Bacteria |
| Firmicutes | Bacteria |
| Firmicutes | Bacteria |
| Firmicutes | Bacteria |
| Firmicutes | Bacteria |
| Firmicutes | Bacteria |
| Firmicutes | Bacteria |
| Firmicutes | Bacteria |
| Firmicutes | Bacteria |
| Firmicutes | Bacteria |
| Firmicutes | Bacteria |
| Firmicutes | Bacteria |
| Firmicutes | Bacteria |
| Firmicutes | Bacteria |
| Firmicutes | Bacteria |
| Firmicutes | Bacteria |
| Firmicutes | Bacteria |
| Firmicutes | Bacteria |
| Firmicutes | Bacteria |
| Firmicutes | Bacteria |
| Firmicutes | Bacteria |
| Firmicutes | Bacteria |
| Firmicutes | Bacteria |
| Firmicutes | Bacteria |
| Firmicutes | Bacteria |
| Firmicutes | Bacteria |
| Firmicutes | Bacteria |
| Firmicutes | Bacteria |
| Firmicutes | Bacteria |
| Bacteria_unclassified | Bacteria |
| Bacteria_unclassified | Bacteria |
| Bacteria_unclassified | Bacteria |
| Bacteria_unclassified | Bacteria |
| Bacteria_unclassified | Bacteria |
| Bacteria_unclassified | Bacteria |
| Bacteria_unclassified | Bacteria |
| Bacteria_unclassified | Bacteria |
| Firmicutes | Bacteria |
| Firmicutes | Bacteria |

| | |
|---|---|
| Firmicutes | Bacteria |
| Firmicutes | Bacteria |
| Firmicutes | Bacteria |
| Firmicutes | Bacteria |
| Bacteria_unclassified | Bacteria |
| Bacteria_unclassified | Bacteria |
| Firmicutes | Bacteria |
| Firmicutes | Bacteria |
| Firmicutes | Bacteria |
| Firmicutes | Bacteria |
| Firmicutes | Bacteria |
| Firmicutes | Bacteria |
| Firmicutes | Bacteria |
| Firmicutes | Bacteria |
| Firmicutes | Bacteria |
| Firmicutes | Bacteria |
| Firmicutes | Bacteria |
| Firmicutes | Bacteria |
| Firmicutes | Bacteria |
| Firmicutes | Bacteria |
| Firmicutes | Bacteria |
| Firmicutes | Bacteria |
| Firmicutes | Bacteria |
| Firmicutes | Bacteria |
| Firmicutes | Bacteria |
| Firmicutes | Bacteria |
| Firmicutes | Bacteria |
| Firmicutes | Bacteria |
| Firmicutes | Bacteria |
| Firmicutes | Bacteria |
| Firmicutes | Bacteria |
| Firmicutes | Bacteria |
| Firmicutes | Bacteria |
| Firmicutes | Bacteria |
| Bacteria_unclassified | Bacteria |
| Bacteria_unclassified | Bacteria |
| Firmicutes | Bacteria |
| Firmicutes | Bacteria |
| Firmicutes | Bacteria |
| Firmicutes | Bacteria |
| Bacteria_unclassified | Bacteria |
| Bacteria_unclassified | Bacteria |
| Firmicutes | Bacteria |
| Firmicutes | Bacteria |
| Firmicutes | Bacteria |
| Firmicutes | Bacteria |

| | |
|---|---|
| Firmicutes | Bacteria |
| Firmicutes | Bacteria |
| Firmicutes | Bacteria |
| Firmicutes | Bacteria |
| Firmicutes | Bacteria |
| Firmicutes | Bacteria |
| Firmicutes | Bacteria |
| Firmicutes | Bacteria |
| Firmicutes | Bacteria |
| Firmicutes | Bacteria |
| Firmicutes | Bacteria |
| Firmicutes | Bacteria |
| Firmicutes | Bacteria |
| Firmicutes | Bacteria |
| Firmicutes | Bacteria |
| Firmicutes | Bacteria |
| Actinobacteria | Bacteria |
| Actinobacteria | Bacteria |
| Tenericutes | Bacteria |
| Tenericutes | Bacteria |
| Firmicutes | Bacteria |
| Firmicutes | Bacteria |
| Firmicutes | Bacteria |
| Firmicutes | Bacteria |
| Firmicutes | Bacteria |
| Firmicutes | Bacteria |
| Firmicutes | Bacteria |
| Firmicutes | Bacteria |
| Firmicutes | Bacteria |
| Firmicutes | Bacteria |
| Firmicutes | Bacteria |
| Firmicutes | Bacteria |
| Firmicutes | Bacteria |
| Firmicutes | Bacteria |
| Firmicutes | Bacteria |
| Firmicutes | Bacteria |
| Firmicutes | Bacteria |
| Firmicutes | Bacteria |
| Firmicutes | Bacteria |
| Firmicutes | Bacteria |
| Firmicutes | Bacteria |
| Firmicutes | Bacteria |
| Firmicutes | Bacteria |
| Firmicutes | Bacteria |
| Firmicutes | Bacteria |
| Firmicutes | Bacteria |

| | |
|---|---|
| Firmicutes | Bacteria |
| Firmicutes | Bacteria |
| Firmicutes | Bacteria |
| Firmicutes | Bacteria |
| Firmicutes | Bacteria |
| Firmicutes | Bacteria |
| Firmicutes | Bacteria |
| Firmicutes | Bacteria |
| Firmicutes | Bacteria |
| Firmicutes | Bacteria |
| Firmicutes | Bacteria |
| Firmicutes | Bacteria |
| Firmicutes | Bacteria |
| Firmicutes | Bacteria |
| Firmicutes | Bacteria |
| Firmicutes | Bacteria |
| Firmicutes | Bacteria |
| Firmicutes | Bacteria |
| Bacteroidota | Bacteria |
| Bacteroidota | Bacteria |
| Firmicutes | Bacteria |
| Firmicutes | Bacteria |
| Firmicutes | Bacteria |
| Firmicutes | Bacteria |
| Firmicutes | Bacteria |
| Firmicutes | Bacteria |
| Firmicutes | Bacteria |
| Firmicutes | Bacteria |
| Proteobacteria | Bacteria |
| Proteobacteria | Bacteria |

| | |
|---|---|
| Firmicutes | Bacteria |
| Firmicutes | Bacteria |
| Firmicutes | Bacteria |
| Firmicutes | Bacteria |

| | |
|---|---|
| Firmicutes | Bacteria |
| Firmicutes | Bacteria |
| Firmicutes | Bacteria |
| Firmicutes | Bacteria |
| Firmicutes | Bacteria |
| Firmicutes | Bacteria |
| Firmicutes | Bacteria |
| Firmicutes | Bacteria |

| | |
|---|---|
| Firmicutes | Bacteria |
| Firmicutes | Bacteria |
| Actinobacteria | Bacteria |
| Actinobacteria | Bacteria |
| Actinobacteria | Bacteria |
| Actinobacteria | Bacteria |
| Actinobacteria | Bacteria |
| Actinobacteria | Bacteria |
| Verrucomicrobia | Bacteria |
| Verrucomicrobia | Bacteria |
| Bacteroidota | Bacteria |
| Bacteroidota | Bacteria |
| Firmicutes | Bacteria |
| Firmicutes | Bacteria |
| Bacteria_unclassified | Bacteria |
| Bacteria_unclassified | Bacteria |
| Bacteria_unclassified | Bacteria |
| Bacteria_unclassified | Bacteria |
| Bacteria_unclassified | Bacteria |
| Bacteria_unclassified | Bacteria |
| Bacteria_unclassified | Bacteria |
| Bacteria_unclassified | Bacteria |
| Bacteria_unclassified | Bacteria |
| Bacteria_unclassified | Bacteria |
| Bacteria_unclassified | Bacteria |
| Bacteria_unclassified | Bacteria |
| Bacteroidota | Bacteria |
| Bacteroidota | Bacteria |
| Actinobacteria | Bacteria |
| Actinobacteria | Bacteria |
| Firmicutes | Bacteria |
| Firmicutes | Bacteria |
| Firmicutes | Bacteria |
| Firmicutes | Bacteria |
| Firmicutes | Bacteria |
| Firmicutes | Bacteria |
| Firmicutes | Bacteria |
| Firmicutes | Bacteria |
| Firmicutes | Bacteria |
| Firmicutes | Bacteria |
| Actinobacteria | Bacteria |
| Actinobacteria | Bacteria |
| Firmicutes | Bacteria |
| Firmicutes | Bacteria |

| | |
|---|---|
| Firmicutes | Bacteria |
| Firmicutes | Bacteria |
| Firmicutes | Bacteria |
| Firmicutes | Bacteria |
| Firmicutes | Bacteria |
| Firmicutes | Bacteria |
| Firmicutes | Bacteria |
| Firmicutes | Bacteria |
| Firmicutes | Bacteria |
| Firmicutes | Bacteria |
| Actinobacteria | Bacteria |
| Actinobacteria | Bacteria |
| Firmicutes | Bacteria |
| Firmicutes | Bacteria |
| Bacteroidota | Bacteria |
| Bacteroidota | Bacteria |
| Bacteroidota | Bacteria |
| Bacteroidota | Bacteria |
| Bacteroidota | Bacteria |
| Bacteroidota | Bacteria |
| Firmicutes | Bacteria |
| Firmicutes | Bacteria |
| Firmicutes | Bacteria |
| Firmicutes | Bacteria |
| Firmicutes | Bacteria |
| Firmicutes | Bacteria |
| Firmicutes | Bacteria |
| Firmicutes | Bacteria |
| Firmicutes | Bacteria |
| Firmicutes | Bacteria |
| Firmicutes | Bacteria |
| Firmicutes | Bacteria |
| Firmicutes | Bacteria |
| Firmicutes | Bacteria |
| Firmicutes | Bacteria |
| Firmicutes | Bacteria |
| Firmicutes | Bacteria |
| Firmicutes | Bacteria |
| Firmicutes | Bacteria |
| Firmicutes | Bacteria |
| Firmicutes | Bacteria |
| Firmicutes | Bacteria |
| Firmicutes | Bacteria |
| Firmicutes | Bacteria |
| Firmicutes | Bacteria |
| Firmicutes | Bacteria |

| | |
|---|---|
| Firmicutes | Bacteria |
| Firmicutes | Bacteria |
| Firmicutes | Bacteria |
| Firmicutes | Bacteria |
| Firmicutes | Bacteria |
| Firmicutes | Bacteria |
| Firmicutes | Bacteria |
| Firmicutes | Bacteria |
| Firmicutes | Bacteria |
| Firmicutes | Bacteria |
| Bacteria_unclassified | Bacteria |
| Bacteria_unclassified | Bacteria |
| Bacteria_unclassified | Bacteria |
| Bacteria_unclassified | Bacteria |
| Bacteria_unclassified | Bacteria |
| Bacteria_unclassified | Bacteria |
| Bacteria_unclassified | Bacteria |
| Bacteria_unclassified | Bacteria |
| Firmicutes | Bacteria |
| Firmicutes | Bacteria |
| Firmicutes | Bacteria |
| Firmicutes | Bacteria |
| Firmicutes | Bacteria |
| Firmicutes | Bacteria |
| Bacteria_unclassified | Bacteria |
| Bacteria_unclassified | Bacteria |
| Firmicutes | Bacteria |
| Firmicutes | Bacteria |
| Firmicutes | Bacteria |
| Firmicutes | Bacteria |
| Firmicutes | Bacteria |
| Firmicutes | Bacteria |
| Firmicutes | Bacteria |
| Firmicutes | Bacteria |
| Firmicutes | Bacteria |
| Firmicutes | Bacteria |
| Firmicutes | Bacteria |
| Firmicutes | Bacteria |
| Firmicutes | Bacteria |
| Firmicutes | Bacteria |
| Firmicutes | Bacteria |
| Firmicutes | Bacteria |
| Firmicutes | Bacteria |
| Firmicutes | Bacteria |
| Firmicutes | Bacteria |
| Firmicutes | Bacteria |

| | |
|---|---|
| Firmicutes | Bacteria |
| Firmicutes | Bacteria |
| Firmicutes | Bacteria |
| Firmicutes | Bacteria |
| Firmicutes | Bacteria |
| Firmicutes | Bacteria |
| Firmicutes | Bacteria |
| Firmicutes | Bacteria |
| Bacteria_unclassified | Bacteria |
| Bacteria_unclassified | Bacteria |
| Firmicutes | Bacteria |
| Firmicutes | Bacteria |
| Firmicutes | Bacteria |
| Firmicutes | Bacteria |
| Bacteria_unclassified | Bacteria |
| Bacteria_unclassified | Bacteria |
| Firmicutes | Bacteria |
| Firmicutes | Bacteria |
| Firmicutes | Bacteria |
| Firmicutes | Bacteria |
| Firmicutes | Bacteria |
| Firmicutes | Bacteria |
| Firmicutes | Bacteria |
| Firmicutes | Bacteria |
| Firmicutes | Bacteria |
| Firmicutes | Bacteria |
| Firmicutes | Bacteria |
| Firmicutes | Bacteria |
| Firmicutes | Bacteria |
| Firmicutes | Bacteria |
| Firmicutes | Bacteria |
| Firmicutes | Bacteria |
| Firmicutes | Bacteria |
| Firmicutes | Bacteria |
| Firmicutes | Bacteria |
| Firmicutes | Bacteria |
| Actinobacteria | Bacteria |
| Actinobacteria | Bacteria |
| Tenericutes | Bacteria |
| Tenericutes | Bacteria |
| Firmicutes | Bacteria |
| Firmicutes | Bacteria |
| Firmicutes | Bacteria |
| Firmicutes | Bacteria |
| Firmicutes | Bacteria |
| Firmicutes | Bacteria |

| | |
|---|---|
| Firmicutes | Bacteria |
| Firmicutes | Bacteria |
| Firmicutes | Bacteria |
| Firmicutes | Bacteria |
| Firmicutes | Bacteria |
| Firmicutes | Bacteria |
| Firmicutes | Bacteria |
| Firmicutes | Bacteria |
| Firmicutes | Bacteria |
| Firmicutes | Bacteria |
| Firmicutes | Bacteria |
| Firmicutes | Bacteria |
| Firmicutes | Bacteria |
| Firmicutes | Bacteria |
| Firmicutes | Bacteria |
| Firmicutes | Bacteria |
| Firmicutes | Bacteria |
| Firmicutes | Bacteria |
| Firmicutes | Bacteria |
| Firmicutes | Bacteria |
| Firmicutes | Bacteria |
| Firmicutes | Bacteria |
| Firmicutes | Bacteria |
| Firmicutes | Bacteria |
| Firmicutes | Bacteria |
| Firmicutes | Bacteria |
| Firmicutes | Bacteria |
| Firmicutes | Bacteria |
| Firmicutes | Bacteria |
| Firmicutes | Bacteria |
| Firmicutes | Bacteria |
| Firmicutes | Bacteria |
| Firmicutes | Bacteria |
| Firmicutes | Bacteria |
| Firmicutes | Bacteria |
| Firmicutes | Bacteria |
| Firmicutes | Bacteria |
| Firmicutes | Bacteria |
| Bacteroidota | Bacteria |
| Bacteroidota | Bacteria |
| Firmicutes | Bacteria |
| Firmicutes | Bacteria |
| Firmicutes | Bacteria |
| Firmicutes | Bacteria |
| Firmicutes | Bacteria |
| Firmicutes | Bacteria |

| | |
|---|---|
| Firmicutes | Bacteria |
| Firmicutes | Bacteria |
| Proteobacteria | Bacteria |
| Proteobacteria | Bacteria |
| Firmicutes | Bacteria |
| Firmicutes | Bacteria |
| Firmicutes | Bacteria |
| Firmicutes | Bacteria |
| Firmicutes | Bacteria |
| Firmicutes | Bacteria |
| Firmicutes | Bacteria |
| Firmicutes | Bacteria |
| Firmicutes | Bacteria |
| Firmicutes | Bacteria |
| Firmicutes | Bacteria |
| Firmicutes | Bacteria |
| Firmicutes | Bacteria |
| Firmicutes | Bacteria |
| Actinobacteria | Bacteria |
| Actinobacteria | Bacteria |
| Actinobacteria | Bacteria |
| Actinobacteria | Bacteria |
| Actinobacteria | Bacteria |
| Actinobacteria | Bacteria |
| Verrucomicrobia | Bacteria |
| Verrucomicrobia | Bacteria |
| Bacteroidota | Bacteria |
| Bacteroidota | Bacteria |
| Firmicutes | Bacteria |
| Firmicutes | Bacteria |
| Bacteria_unclassified | Bacteria |
| Bacteria_unclassified | Bacteria |
| Bacteria_unclassified | Bacteria |
| Bacteria_unclassified | Bacteria |
| Bacteria_unclassified | Bacteria |
| Bacteria_unclassified | Bacteria |
| Bacteria_unclassified | Bacteria |
| Bacteria_unclassified | Bacteria |
| Bacteria_unclassified | Bacteria |
| Bacteria_unclassified | Bacteria |
| Bacteria_unclassified | Bacteria |
| Bacteria_unclassified | Bacteria |

| | |
|---|---|
| Bacteroidota | Bacteria |
| Bacteroidota | Bacteria |
| Actinobacteria | Bacteria |
| Actinobacteria | Bacteria |
| Firmicutes | Bacteria |
| Firmicutes | Bacteria |
| Firmicutes | Bacteria |
| Firmicutes | Bacteria |
| Firmicutes | Bacteria |
| Firmicutes | Bacteria |
| Firmicutes | Bacteria |
| Firmicutes | Bacteria |
| Firmicutes | Bacteria |
| Firmicutes | Bacteria |
| Actinobacteria | Bacteria |
| Actinobacteria | Bacteria |
| Firmicutes | Bacteria |
| Firmicutes | Bacteria |
| Firmicutes | Bacteria |
| Firmicutes | Bacteria |
| Firmicutes | Bacteria |
| Firmicutes | Bacteria |
| Firmicutes | Bacteria |
| Firmicutes | Bacteria |
| Firmicutes | Bacteria |
| Firmicutes | Bacteria |
| Firmicutes | Bacteria |
| Firmicutes | Bacteria |
| Actinobacteria | Bacteria |
| Actinobacteria | Bacteria |
| Firmicutes | Bacteria |
| Firmicutes | Bacteria |
| Bacteroidota | Bacteria |
| Bacteroidota | Bacteria |
| Bacteroidota | Bacteria |
| Bacteroidota | Bacteria |
| Bacteroidota | Bacteria |
| Bacteroidota | Bacteria |
| Firmicutes | Bacteria |
| Firmicutes | Bacteria |
| Firmicutes | Bacteria |
| Firmicutes | Bacteria |
| Firmicutes | Bacteria |
| Firmicutes | Bacteria |

| | |
|---|---|
| Firmicutes | Bacteria |
| Firmicutes | Bacteria |
| Firmicutes | Bacteria |
| Firmicutes | Bacteria |
| Firmicutes | Bacteria |
| Firmicutes | Bacteria |
| Firmicutes | Bacteria |
| Firmicutes | Bacteria |
| Firmicutes | Bacteria |
| Firmicutes | Bacteria |
| Firmicutes | Bacteria |
| Firmicutes | Bacteria |
| Firmicutes | Bacteria |
| Firmicutes | Bacteria |
| Firmicutes | Bacteria |
| Firmicutes | Bacteria |
| Firmicutes | Bacteria |
| Firmicutes | Bacteria |
| Firmicutes | Bacteria |
| Firmicutes | Bacteria |
| Firmicutes | Bacteria |
| Firmicutes | Bacteria |
| Firmicutes | Bacteria |
| Firmicutes | Bacteria |
| Firmicutes | Bacteria |
| Firmicutes | Bacteria |
| Firmicutes | Bacteria |
| Firmicutes | Bacteria |
| Firmicutes | Bacteria |
| Firmicutes | Bacteria |
| Bacteria_unclassified | Bacteria |
| Bacteria_unclassified | Bacteria |
| Bacteria_unclassified | Bacteria |
| Bacteria_unclassified | Bacteria |
| Bacteria_unclassified | Bacteria |
| Bacteria_unclassified | Bacteria |
| Bacteria_unclassified | Bacteria |
| Bacteria_unclassified | Bacteria |
| Firmicutes | Bacteria |
| Firmicutes | Bacteria |
| Firmicutes | Bacteria |
| Firmicutes | Bacteria |
| Firmicutes | Bacteria |
| Firmicutes | Bacteria |
| Bacteria_unclassified | Bacteria |
| Bacteria_unclassified | Bacteria |

| | |
|---|---|
| Firmicutes | Bacteria |
| Firmicutes | Bacteria |
| Firmicutes | Bacteria |
| Firmicutes | Bacteria |
| Firmicutes | Bacteria |
| Firmicutes | Bacteria |
| Firmicutes | Bacteria |
| Firmicutes | Bacteria |
| Firmicutes | Bacteria |
| Firmicutes | Bacteria |
| Firmicutes | Bacteria |
| Firmicutes | Bacteria |
| Firmicutes | Bacteria |
| Firmicutes | Bacteria |
| Firmicutes | Bacteria |
| Firmicutes | Bacteria |
| Firmicutes | Bacteria |
| Firmicutes | Bacteria |
| Firmicutes | Bacteria |
| Firmicutes | Bacteria |
| Firmicutes | Bacteria |
| Firmicutes | Bacteria |
| Firmicutes | Bacteria |
| Firmicutes | Bacteria |
| Firmicutes | Bacteria |
| Firmicutes | Bacteria |
| Firmicutes | Bacteria |
| Firmicutes | Bacteria |
| Bacteria_unclassified | Bacteria |
| Bacteria_unclassified | Bacteria |
| Firmicutes | Bacteria |
| Firmicutes | Bacteria |
| Firmicutes | Bacteria |
| Firmicutes | Bacteria |
| Bacteria_unclassified | Bacteria |
| Bacteria_unclassified | Bacteria |
| Firmicutes | Bacteria |
| Firmicutes | Bacteria |
| Firmicutes | Bacteria |
| Firmicutes | Bacteria |
| Firmicutes | Bacteria |
| Firmicutes | Bacteria |
| Firmicutes | Bacteria |
| Firmicutes | Bacteria |
| Firmicutes | Bacteria |
| Firmicutes | Bacteria |

| | |
|---|---|
| Firmicutes | Bacteria |
| Firmicutes | Bacteria |
| Firmicutes | Bacteria |
| Firmicutes | Bacteria |
| Firmicutes | Bacteria |
| Firmicutes | Bacteria |
| Firmicutes | Bacteria |
| Firmicutes | Bacteria |
| Firmicutes | Bacteria |
| Firmicutes | Bacteria |
| Actinobacteria | Bacteria |
| Actinobacteria | Bacteria |
| Tenericutes | Bacteria |
| Tenericutes | Bacteria |
| Firmicutes | Bacteria |
| Firmicutes | Bacteria |
| Firmicutes | Bacteria |
| Firmicutes | Bacteria |
| Firmicutes | Bacteria |
| Firmicutes | Bacteria |
| Firmicutes | Bacteria |
| Firmicutes | Bacteria |
| Firmicutes | Bacteria |
| Firmicutes | Bacteria |
| Firmicutes | Bacteria |
| Firmicutes | Bacteria |
| Firmicutes | Bacteria |
| Firmicutes | Bacteria |
| Firmicutes | Bacteria |
| Firmicutes | Bacteria |
| Firmicutes | Bacteria |
| Firmicutes | Bacteria |
| Firmicutes | Bacteria |
| Firmicutes | Bacteria |
| Firmicutes | Bacteria |
| Firmicutes | Bacteria |
| Firmicutes | Bacteria |
| Firmicutes | Bacteria |
| Firmicutes | Bacteria |
| Firmicutes | Bacteria |
| Firmicutes | Bacteria |
| Firmicutes | Bacteria |
| Firmicutes | Bacteria |
| Firmicutes | Bacteria |
| Firmicutes | Bacteria |
| Firmicutes | Bacteria |

| | |
|---|---|
| Firmicutes | Bacteria |
| Firmicutes | Bacteria |
| Firmicutes | Bacteria |
| Firmicutes | Bacteria |
| Firmicutes | Bacteria |
| Firmicutes | Bacteria |
| Firmicutes | Bacteria |
| Firmicutes | Bacteria |
| Firmicutes | Bacteria |
| Firmicutes | Bacteria |
| Firmicutes | Bacteria |
| Firmicutes | Bacteria |
| Bacteroidota | Bacteria |
| Bacteroidota | Bacteria |
| Firmicutes | Bacteria |
| Firmicutes | Bacteria |
| Firmicutes | Bacteria |
| Firmicutes | Bacteria |
| Firmicutes | Bacteria |
| Firmicutes | Bacteria |
| Firmicutes | Bacteria |
| Firmicutes | Bacteria |
| Proteobacteria | Bacteria |
| Proteobacteria | Bacteria |
| Firmicutes | Bacteria |
| Firmicutes | Bacteria |
| Firmicutes | Bacteria |
| Firmicutes | Bacteria |
| Firmicutes | Bacteria |
| Firmicutes | Bacteria |
| Firmicutes | Bacteria |
| Firmicutes | Bacteria |
| Firmicutes | Bacteria |
| Firmicutes | Bacteria |
| Firmicutes | Bacteria |
| Firmicutes | Bacteria |
| Firmicutes | Bacteria |
| Firmicutes | Bacteria |
| Actinobacteria | Bacteria |
| Actinobacteria | Bacteria |
| Actinobacteria | Bacteria |
| Actinobacteria | Bacteria |

| | |
|---|---|
| Actinobacteria | Bacteria |
| Actinobacteria | Bacteria |
| Verrucomicrobia | Bacteria |
| Verrucomicrobia | Bacteria |
| Bacteroidota | Bacteria |
| Bacteroidota | Bacteria |
| Firmicutes | Bacteria |
| Firmicutes | Bacteria |
| Bacteria_unclassified | Bacteria |
| Bacteria_unclassified | Bacteria |
| Bacteria_unclassified | Bacteria |
| Bacteria_unclassified | Bacteria |
| Bacteria_unclassified | Bacteria |
| Bacteria_unclassified | Bacteria |
| Bacteria_unclassified | Bacteria |
| Bacteria_unclassified | Bacteria |
| Bacteria_unclassified | Bacteria |
| Bacteria_unclassified | Bacteria |
| Bacteria_unclassified | Bacteria |
| Bacteria_unclassified | Bacteria |
| Bacteroidota | Bacteria |
| Bacteroidota | Bacteria |

| | |
|---|---|
| Actinobacteria | Bacteria |
| Actinobacteria | Bacteria |
| Firmicutes | Bacteria |
| Firmicutes | Bacteria |
| Firmicutes | Bacteria |
| Firmicutes | Bacteria |
| Firmicutes | Bacteria |
| Firmicutes | Bacteria |
| Firmicutes | Bacteria |
| Firmicutes | Bacteria |
| Firmicutes | Bacteria |
| Firmicutes | Bacteria |
| Actinobacteria | Bacteria |
| Actinobacteria | Bacteria |
| Firmicutes | Bacteria |
| Firmicutes | Bacteria |
| Firmicutes | Bacteria |
| Firmicutes | Bacteria |
| Firmicutes | Bacteria |
| Firmicutes | Bacteria |
| Firmicutes | Bacteria |
| Firmicutes | Bacteria |

| | |
|---|---|
| Firmicutes | Bacteria |
| Firmicutes | Bacteria |
| Firmicutes | Bacteria |
| Firmicutes | Bacteria |
| Actinobacteria | Bacteria |
| Actinobacteria | Bacteria |
| Firmicutes | Bacteria |
| Firmicutes | Bacteria |
| Bacteroidota | Bacteria |
| Bacteroidota | Bacteria |
| Bacteroidota | Bacteria |
| Bacteroidota | Bacteria |
| Bacteroidota | Bacteria |
| Bacteroidota | Bacteria |
| Firmicutes | Bacteria |
| Firmicutes | Bacteria |
| Firmicutes | Bacteria |
| Firmicutes | Bacteria |
| Firmicutes | Bacteria |
| Firmicutes | Bacteria |
| Firmicutes | Bacteria |
| Firmicutes | Bacteria |
| Firmicutes | Bacteria |
| Firmicutes | Bacteria |
| Firmicutes | Bacteria |
| Firmicutes | Bacteria |
| Firmicutes | Bacteria |
| Firmicutes | Bacteria |
| Firmicutes | Bacteria |
| Firmicutes | Bacteria |
| Firmicutes | Bacteria |
| Firmicutes | Bacteria |
| Firmicutes | Bacteria |
| Firmicutes | Bacteria |
| Firmicutes | Bacteria |
| Firmicutes | Bacteria |
| Firmicutes | Bacteria |
| Firmicutes | Bacteria |
| Firmicutes | Bacteria |
| Firmicutes | Bacteria |
| Firmicutes | Bacteria |
| Firmicutes | Bacteria |
| Firmicutes | Bacteria |
| Firmicutes | Bacteria |
| Firmicutes | Bacteria |
| Firmicutes | Bacteria |

| | |
|---|---|
| Firmicutes | Bacteria |
| Firmicutes | Bacteria |
| Firmicutes | Bacteria |
| Firmicutes | Bacteria |
| Bacteria_unclassified | Bacteria |
| Bacteria_unclassified | Bacteria |
| Bacteria_unclassified | Bacteria |
| Bacteria_unclassified | Bacteria |
| Bacteria_unclassified | Bacteria |
| Bacteria_unclassified | Bacteria |
| Bacteria_unclassified | Bacteria |
| Bacteria_unclassified | Bacteria |
| Firmicutes | Bacteria |
| Firmicutes | Bacteria |
| Firmicutes | Bacteria |
| Firmicutes | Bacteria |
| Firmicutes | Bacteria |
| Firmicutes | Bacteria |
| Bacteria_unclassified | Bacteria |
| Bacteria_unclassified | Bacteria |
| Firmicutes | Bacteria |
| Firmicutes | Bacteria |
| Firmicutes | Bacteria |
| Firmicutes | Bacteria |
| Firmicutes | Bacteria |
| Firmicutes | Bacteria |
| Firmicutes | Bacteria |
| Firmicutes | Bacteria |
| Firmicutes | Bacteria |
| Firmicutes | Bacteria |
| Firmicutes | Bacteria |
| Firmicutes | Bacteria |
| Firmicutes | Bacteria |
| Firmicutes | Bacteria |
| Firmicutes | Bacteria |
| Firmicutes | Bacteria |
| Firmicutes | Bacteria |
| Firmicutes | Bacteria |
| Firmicutes | Bacteria |
| Firmicutes | Bacteria |
| Firmicutes | Bacteria |
| Firmicutes | Bacteria |
| Firmicutes | Bacteria |
| Firmicutes | Bacteria |
| Firmicutes | Bacteria |
| Firmicutes | Bacteria |

| | |
|---|---|
| Firmicutes | Bacteria |
| Firmicutes | Bacteria |
| Bacteria_unclassified | Bacteria |
| Bacteria_unclassified | Bacteria |
| Firmicutes | Bacteria |
| Firmicutes | Bacteria |
| Firmicutes | Bacteria |
| Firmicutes | Bacteria |
| Bacteria_unclassified | Bacteria |
| Bacteria_unclassified | Bacteria |
| Firmicutes | Bacteria |
| Firmicutes | Bacteria |
| Firmicutes | Bacteria |
| Firmicutes | Bacteria |
| Firmicutes | Bacteria |
| Firmicutes | Bacteria |
| Firmicutes | Bacteria |
| Firmicutes | Bacteria |
| Firmicutes | Bacteria |
| Firmicutes | Bacteria |
| Firmicutes | Bacteria |
| Firmicutes | Bacteria |
| Firmicutes | Bacteria |
| Firmicutes | Bacteria |
| Firmicutes | Bacteria |
| Firmicutes | Bacteria |
| Firmicutes | Bacteria |
| Firmicutes | Bacteria |
| Firmicutes | Bacteria |
| Firmicutes | Bacteria |
| Actinobacteria | Bacteria |
| Actinobacteria | Bacteria |
| Tenericutes | Bacteria |
| Tenericutes | Bacteria |
| Firmicutes | Bacteria |
| Firmicutes | Bacteria |
| Firmicutes | Bacteria |
| Firmicutes | Bacteria |
| Firmicutes | Bacteria |
| Firmicutes | Bacteria |
| Firmicutes | Bacteria |
| Firmicutes | Bacteria |
| Firmicutes | Bacteria |
| Firmicutes | Bacteria |
| Firmicutes | Bacteria |
| Firmicutes | Bacteria |

| | |
|---|---|
| Firmicutes | Bacteria |
| Firmicutes | Bacteria |
| Firmicutes | Bacteria |
| Firmicutes | Bacteria |
| Firmicutes | Bacteria |
| Firmicutes | Bacteria |
| Firmicutes | Bacteria |
| Firmicutes | Bacteria |
| Firmicutes | Bacteria |
| Firmicutes | Bacteria |
| Firmicutes | Bacteria |
| Firmicutes | Bacteria |
| Firmicutes | Bacteria |
| Firmicutes | Bacteria |
| Firmicutes | Bacteria |
| Firmicutes | Bacteria |
| Firmicutes | Bacteria |
| Firmicutes | Bacteria |
| Firmicutes | Bacteria |
| Firmicutes | Bacteria |
| Firmicutes | Bacteria |
| Firmicutes | Bacteria |
| Firmicutes | Bacteria |
| Firmicutes | Bacteria |
| Firmicutes | Bacteria |
| Firmicutes | Bacteria |
| Firmicutes | Bacteria |
| Firmicutes | Bacteria |
| Firmicutes | Bacteria |
| Firmicutes | Bacteria |
| Firmicutes | Bacteria |
| Firmicutes | Bacteria |
| Bacteroidota | Bacteria |
| Bacteroidota | Bacteria |
| Firmicutes | Bacteria |
| Firmicutes | Bacteria |
| Firmicutes | Bacteria |
| Firmicutes | Bacteria |
| Firmicutes | Bacteria |
| Firmicutes | Bacteria |
| Firmicutes | Bacteria |
| Firmicutes | Bacteria |
| Proteobacteria | Bacteria |
| Proteobacteria | Bacteria |

| | |
|---|---|
| Firmicutes | Bacteria |
| Firmicutes | Bacteria |
| Firmicutes | Bacteria |
| Firmicutes | Bacteria |
| Firmicutes | Bacteria |
| Firmicutes | Bacteria |
| Firmicutes | Bacteria |
| Firmicutes | Bacteria |
| Firmicutes | Bacteria |
| Firmicutes | Bacteria |
| Firmicutes | Bacteria |
| Firmicutes | Bacteria |
| Firmicutes | Bacteria |
| Firmicutes | Bacteria |
| Actinobacteria | Bacteria |
| Actinobacteria | Bacteria |
| Actinobacteria | Bacteria |
| Actinobacteria | Bacteria |
| Actinobacteria | Bacteria |
| Actinobacteria | Bacteria |
| Verrucomicrobia | Bacteria |
| Verrucomicrobia | Bacteria |
| Bacteroidota | Bacteria |
| Bacteroidota | Bacteria |
| Firmicutes | Bacteria |
| Firmicutes | Bacteria |
| Bacteria_unclassified | Bacteria |
| Bacteria_unclassified | Bacteria |
| Bacteria_unclassified | Bacteria |
| Bacteria_unclassified | Bacteria |
| Bacteria_unclassified | Bacteria |
| Bacteria_unclassified | Bacteria |
| Bacteria_unclassified | Bacteria |
| Bacteria_unclassified | Bacteria |
| Bacteria_unclassified | Bacteria |
| Bacteria_unclassified | Bacteria |
| Bacteria_unclassified | Bacteria |
| Bacteria_unclassified | Bacteria |
| Bacteroidota | Bacteria |
| Bacteroidota | Bacteria |
| Actinobacteria | Bacteria |
| Actinobacteria | Bacteria |

| | |
|---|---|
| Firmicutes | Bacteria |
| Firmicutes | Bacteria |
| Firmicutes | Bacteria |
| Firmicutes | Bacteria |
| Firmicutes | Bacteria |
| Firmicutes | Bacteria |
| Firmicutes | Bacteria |
| Firmicutes | Bacteria |
| Firmicutes | Bacteria |
| Firmicutes | Bacteria |
| Actinobacteria | Bacteria |
| Actinobacteria | Bacteria |
| Firmicutes | Bacteria |
| Firmicutes | Bacteria |
| Firmicutes | Bacteria |
| Firmicutes | Bacteria |
| Firmicutes | Bacteria |
| Firmicutes | Bacteria |
| Firmicutes | Bacteria |
| Firmicutes | Bacteria |
| Firmicutes | Bacteria |
| Firmicutes | Bacteria |
| Firmicutes | Bacteria |
| Firmicutes | Bacteria |
| Actinobacteria | Bacteria |
| Actinobacteria | Bacteria |
| Firmicutes | Bacteria |
| Firmicutes | Bacteria |
| Bacteroidota | Bacteria |
| Bacteroidota | Bacteria |
| Bacteroidota | Bacteria |
| Bacteroidota | Bacteria |
| Bacteroidota | Bacteria |
| Bacteroidota | Bacteria |
| Firmicutes | Bacteria |
| Firmicutes | Bacteria |
| Firmicutes | Bacteria |
| Firmicutes | Bacteria |
| Firmicutes | Bacteria |
| Firmicutes | Bacteria |
| Firmicutes | Bacteria |
| Firmicutes | Bacteria |
| Firmicutes | Bacteria |
| Firmicutes | Bacteria |
| Firmicutes | Bacteria |
| Firmicutes | Bacteria |

| | |
|---|---|
| Firmicutes | Bacteria |
| Firmicutes | Bacteria |
| Firmicutes | Bacteria |
| Firmicutes | Bacteria |
| Firmicutes | Bacteria |
| Firmicutes | Bacteria |
| Firmicutes | Bacteria |
| Firmicutes | Bacteria |
| Firmicutes | Bacteria |
| Firmicutes | Bacteria |
| Firmicutes | Bacteria |
| Firmicutes | Bacteria |
| Firmicutes | Bacteria |
| Firmicutes | Bacteria |
| Firmicutes | Bacteria |
| Firmicutes | Bacteria |
| Firmicutes | Bacteria |
| Firmicutes | Bacteria |
| Firmicutes | Bacteria |
| Firmicutes | Bacteria |
| Firmicutes | Bacteria |
| Firmicutes | Bacteria |
| Firmicutes | Bacteria |
| Firmicutes | Bacteria |
| Bacteria_unclassified | Bacteria |
| Bacteria_unclassified | Bacteria |
| Bacteria_unclassified | Bacteria |
| Bacteria_unclassified | Bacteria |
| Bacteria_unclassified | Bacteria |
| Bacteria_unclassified | Bacteria |
| Bacteria_unclassified | Bacteria |
| Bacteria_unclassified | Bacteria |
| Firmicutes | Bacteria |
| Firmicutes | Bacteria |
| Firmicutes | Bacteria |
| Firmicutes | Bacteria |
| Firmicutes | Bacteria |
| Firmicutes | Bacteria |
| Bacteria_unclassified | Bacteria |
| Bacteria_unclassified | Bacteria |
| Firmicutes | Bacteria |
| Firmicutes | Bacteria |
| Firmicutes | Bacteria |
| Firmicutes | Bacteria |
| Firmicutes | Bacteria |
| Firmicutes | Bacteria |

| | |
|---|---|
| Firmicutes | Bacteria |
| Firmicutes | Bacteria |
| Firmicutes | Bacteria |
| Firmicutes | Bacteria |
| Firmicutes | Bacteria |
| Firmicutes | Bacteria |
| Firmicutes | Bacteria |
| Firmicutes | Bacteria |
| Firmicutes | Bacteria |
| Firmicutes | Bacteria |
| Firmicutes | Bacteria |
| Firmicutes | Bacteria |
| Firmicutes | Bacteria |
| Firmicutes | Bacteria |
| Firmicutes | Bacteria |
| Firmicutes | Bacteria |
| Firmicutes | Bacteria |
| Firmicutes | Bacteria |
| Firmicutes | Bacteria |
| Firmicutes | Bacteria |
| Firmicutes | Bacteria |
| Firmicutes | Bacteria |
| Bacteria_unclassified | Bacteria |
| Bacteria_unclassified | Bacteria |
| Firmicutes | Bacteria |
| Firmicutes | Bacteria |
| Firmicutes | Bacteria |
| Firmicutes | Bacteria |
| Bacteria_unclassified | Bacteria |
| Bacteria_unclassified | Bacteria |
| Firmicutes | Bacteria |
| Firmicutes | Bacteria |
| Firmicutes | Bacteria |
| Firmicutes | Bacteria |
| Firmicutes | Bacteria |
| Firmicutes | Bacteria |
| Firmicutes | Bacteria |
| Firmicutes | Bacteria |
| Firmicutes | Bacteria |
| Firmicutes | Bacteria |
| Firmicutes | Bacteria |
| Firmicutes | Bacteria |
| Firmicutes | Bacteria |
| Firmicutes | Bacteria |
| Firmicutes | Bacteria |
| Firmicutes | Bacteria |

| | |
|---|---|
| Firmicutes | Bacteria |
| Firmicutes | Bacteria |
| Firmicutes | Bacteria |
| Firmicutes | Bacteria |
| Actinobacteria | Bacteria |
| Actinobacteria | Bacteria |
| Tenericutes | Bacteria |
| Tenericutes | Bacteria |
| Firmicutes | Bacteria |
| Firmicutes | Bacteria |
| Firmicutes | Bacteria |
| Firmicutes | Bacteria |
| Firmicutes | Bacteria |
| Firmicutes | Bacteria |
| Firmicutes | Bacteria |
| Firmicutes | Bacteria |
| Firmicutes | Bacteria |
| Firmicutes | Bacteria |
| Firmicutes | Bacteria |
| Firmicutes | Bacteria |
| Firmicutes | Bacteria |
| Firmicutes | Bacteria |
| Firmicutes | Bacteria |
| Firmicutes | Bacteria |
| Firmicutes | Bacteria |
| Firmicutes | Bacteria |
| Firmicutes | Bacteria |
| Firmicutes | Bacteria |
| Firmicutes | Bacteria |
| Firmicutes | Bacteria |
| Firmicutes | Bacteria |
| Firmicutes | Bacteria |
| Firmicutes | Bacteria |
| Firmicutes | Bacteria |
| Firmicutes | Bacteria |
| Firmicutes | Bacteria |
| Firmicutes | Bacteria |
| Firmicutes | Bacteria |
| Firmicutes | Bacteria |
| Firmicutes | Bacteria |
| Firmicutes | Bacteria |
| Firmicutes | Bacteria |
| Firmicutes | Bacteria |
| Firmicutes | Bacteria |
| Firmicutes | Bacteria |
| Firmicutes | Bacteria |

| | |
|---|---|
| Firmicutes | Bacteria |
| Firmicutes | Bacteria |
| Firmicutes | Bacteria |
| Firmicutes | Bacteria |
| Firmicutes | Bacteria |
| Firmicutes | Bacteria |
| Bacteroidota | Bacteria |
| Bacteroidota | Bacteria |
| Firmicutes | Bacteria |
| Firmicutes | Bacteria |
| Firmicutes | Bacteria |
| Firmicutes | Bacteria |
| Firmicutes | Bacteria |
| Firmicutes | Bacteria |
| Firmicutes | Bacteria |
| Firmicutes | Bacteria |
| Proteobacteria | Bacteria |
| Proteobacteria | Bacteria |
| Firmicutes | Bacteria |
| Firmicutes | Bacteria |
| Firmicutes | Bacteria |
| Firmicutes | Bacteria |
| Firmicutes | Bacteria |
| Firmicutes | Bacteria |
| Firmicutes | Bacteria |
| Firmicutes | Bacteria |
| Firmicutes | Bacteria |
| Firmicutes | Bacteria |
| Firmicutes | Bacteria |
| Firmicutes | Bacteria |
| Firmicutes | Bacteria |
| Firmicutes | Bacteria |
| Actinobacteria | Bacteria |
| Actinobacteria | Bacteria |
| Actinobacteria | Bacteria |
| Actinobacteria | Bacteria |
| Actinobacteria | Bacteria |
| Actinobacteria | Bacteria |
| Verrucomicrobia | Bacteria |
| Verrucomicrobia | Bacteria |
| Bacteroidota | Bacteria |
| Bacteroidota | Bacteria |

| | |
|---|---|
| Firmicutes | Bacteria |
| Firmicutes | Bacteria |
| Bacteria_unclassified | Bacteria |
| Bacteria_unclassified | Bacteria |
| Bacteria_unclassified | Bacteria |
| Bacteria_unclassified | Bacteria |
| Bacteria_unclassified | Bacteria |
| Bacteria_unclassified | Bacteria |
| Bacteria_unclassified | Bacteria |
| Bacteria_unclassified | Bacteria |
| Bacteria_unclassified | Bacteria |
| Bacteria_unclassified | Bacteria |
| Bacteria_unclassified | Bacteria |
| Bacteria_unclassified | Bacteria |
| Bacteroidota | Bacteria |
| Bacteroidota | Bacteria |

| | |
|---|---|
| Actinobacteria | Bacteria |
| Actinobacteria | Bacteria |
| Firmicutes | Bacteria |
| Firmicutes | Bacteria |
| Firmicutes | Bacteria |
| Firmicutes | Bacteria |
| Firmicutes | Bacteria |
| Firmicutes | Bacteria |
| Firmicutes | Bacteria |
| Firmicutes | Bacteria |
| Firmicutes | Bacteria |
| Firmicutes | Bacteria |
| Actinobacteria | Bacteria |
| Actinobacteria | Bacteria |
| Firmicutes | Bacteria |
| Firmicutes | Bacteria |
| Firmicutes | Bacteria |
| Firmicutes | Bacteria |
| Firmicutes | Bacteria |
| Firmicutes | Bacteria |
| Firmicutes | Bacteria |
| Firmicutes | Bacteria |
| Firmicutes | Bacteria |
| Firmicutes | Bacteria |
| Firmicutes | Bacteria |
| Firmicutes | Bacteria |
| Actinobacteria | Bacteria |
| Actinobacteria | Bacteria |

| | |
|---|---|
| Firmicutes | Bacteria |
| Firmicutes | Bacteria |
| Bacteroidota | Bacteria |
| Bacteroidota | Bacteria |
| Bacteroidota | Bacteria |
| Bacteroidota | Bacteria |
| Bacteroidota | Bacteria |
| Bacteroidota | Bacteria |
| Firmicutes | Bacteria |
| Firmicutes | Bacteria |
| Firmicutes | Bacteria |
| Firmicutes | Bacteria |
| Firmicutes | Bacteria |
| Firmicutes | Bacteria |
| Firmicutes | Bacteria |
| Firmicutes | Bacteria |
| Firmicutes | Bacteria |
| Firmicutes | Bacteria |
| Firmicutes | Bacteria |
| Firmicutes | Bacteria |
| Firmicutes | Bacteria |
| Firmicutes | Bacteria |
| Firmicutes | Bacteria |
| Firmicutes | Bacteria |
| Firmicutes | Bacteria |
| Firmicutes | Bacteria |
| Firmicutes | Bacteria |
| Firmicutes | Bacteria |
| Firmicutes | Bacteria |
| Firmicutes | Bacteria |
| Firmicutes | Bacteria |
| Firmicutes | Bacteria |
| Firmicutes | Bacteria |
| Firmicutes | Bacteria |
| Firmicutes | Bacteria |
| Firmicutes | Bacteria |
| Firmicutes | Bacteria |
| Firmicutes | Bacteria |
| Firmicutes | Bacteria |
| Firmicutes | Bacteria |
| Firmicutes | Bacteria |
| Firmicutes | Bacteria |
| Firmicutes | Bacteria |
| Firmicutes | Bacteria |
| Bacteria_unclassified | Bacteria |
| Bacteria_unclassified | Bacteria |

| | |
|---|---|
| Bacteria_unclassified | Bacteria |
| Bacteria_unclassified | Bacteria |
| Bacteria_unclassified | Bacteria |
| Bacteria_unclassified | Bacteria |
| Bacteria_unclassified | Bacteria |
| Bacteria_unclassified | Bacteria |
| Firmicutes | Bacteria |
| Firmicutes | Bacteria |
| Firmicutes | Bacteria |
| Firmicutes | Bacteria |
| Firmicutes | Bacteria |
| Firmicutes | Bacteria |
| Bacteria_unclassified | Bacteria |
| Bacteria_unclassified | Bacteria |
| Firmicutes | Bacteria |
| Firmicutes | Bacteria |
| Firmicutes | Bacteria |
| Firmicutes | Bacteria |
| Firmicutes | Bacteria |
| Firmicutes | Bacteria |
| Firmicutes | Bacteria |
| Firmicutes | Bacteria |
| Firmicutes | Bacteria |
| Firmicutes | Bacteria |
| Firmicutes | Bacteria |
| Firmicutes | Bacteria |
| Firmicutes | Bacteria |
| Firmicutes | Bacteria |
| Firmicutes | Bacteria |
| Firmicutes | Bacteria |
| Firmicutes | Bacteria |
| Firmicutes | Bacteria |
| Firmicutes | Bacteria |
| Firmicutes | Bacteria |
| Firmicutes | Bacteria |
| Firmicutes | Bacteria |
| Firmicutes | Bacteria |
| Firmicutes | Bacteria |
| Firmicutes | Bacteria |
| Firmicutes | Bacteria |
| Firmicutes | Bacteria |
| Firmicutes | Bacteria |
| Bacteria_unclassified | Bacteria |
| Bacteria_unclassified | Bacteria |
| Firmicutes | Bacteria |
| Firmicutes | Bacteria |

| | |
|---|---|
| Firmicutes | Bacteria |
| Firmicutes | Bacteria |
| Bacteria_unclassified | Bacteria |
| Bacteria_unclassified | Bacteria |
| Firmicutes | Bacteria |
| Firmicutes | Bacteria |
| Firmicutes | Bacteria |
| Firmicutes | Bacteria |
| Firmicutes | Bacteria |
| Firmicutes | Bacteria |
| Firmicutes | Bacteria |
| Firmicutes | Bacteria |
| Firmicutes | Bacteria |
| Firmicutes | Bacteria |
| Firmicutes | Bacteria |
| Firmicutes | Bacteria |
| Firmicutes | Bacteria |
| Firmicutes | Bacteria |
| Firmicutes | Bacteria |
| Firmicutes | Bacteria |
| Firmicutes | Bacteria |
| Firmicutes | Bacteria |
| Firmicutes | Bacteria |
| Firmicutes | Bacteria |
| Actinobacteria | Bacteria |
| Actinobacteria | Bacteria |
| Tenericutes | Bacteria |
| Tenericutes | Bacteria |
| Firmicutes | Bacteria |
| Firmicutes | Bacteria |
| Firmicutes | Bacteria |
| Firmicutes | Bacteria |
| Firmicutes | Bacteria |
| Firmicutes | Bacteria |
| Firmicutes | Bacteria |
| Firmicutes | Bacteria |
| Firmicutes | Bacteria |
| Firmicutes | Bacteria |
| Firmicutes | Bacteria |
| Firmicutes | Bacteria |
| Firmicutes | Bacteria |
| Firmicutes | Bacteria |
| Firmicutes | Bacteria |
| Firmicutes | Bacteria |
| Firmicutes | Bacteria |
| Firmicutes | Bacteria |

| | |
|---|---|
| Firmicutes | Bacteria |
| Firmicutes | Bacteria |
| Firmicutes | Bacteria |
| Firmicutes | Bacteria |
| Firmicutes | Bacteria |
| Firmicutes | Bacteria |
| Firmicutes | Bacteria |
| Firmicutes | Bacteria |
| Firmicutes | Bacteria |
| Firmicutes | Bacteria |
| Firmicutes | Bacteria |
| Firmicutes | Bacteria |
| Firmicutes | Bacteria |
| Firmicutes | Bacteria |
| Firmicutes | Bacteria |
| Firmicutes | Bacteria |
| Firmicutes | Bacteria |
| Firmicutes | Bacteria |
| Firmicutes | Bacteria |
| Firmicutes | Bacteria |
| Firmicutes | Bacteria |
| Firmicutes | Bacteria |
| Firmicutes | Bacteria |
| Firmicutes | Bacteria |
| Firmicutes | Bacteria |
| Firmicutes | Bacteria |
| Bacteroidota | Bacteria |
| Bacteroidota | Bacteria |
| Firmicutes | Bacteria |
| Firmicutes | Bacteria |
| Firmicutes | Bacteria |
| Firmicutes | Bacteria |
| Firmicutes | Bacteria |
| Firmicutes | Bacteria |
| Firmicutes | Bacteria |
| Firmicutes | Bacteria |
| Proteobacteria | Bacteria |
| Proteobacteria | Bacteria |

| | |
|---|---|
| Firmicutes | Bacteria |
| Firmicutes | Bacteria |
| Firmicutes | Bacteria |
| Firmicutes | Bacteria |

| | |
|---|---|
| Firmicutes | Bacteria |
| Firmicutes | Bacteria |
| Firmicutes | Bacteria |
| Firmicutes | Bacteria |
| Firmicutes | Bacteria |
| Firmicutes | Bacteria |
| Firmicutes | Bacteria |
| Firmicutes | Bacteria |
| Firmicutes | Bacteria |
| Firmicutes | Bacteria |
| Actinobacteria | Bacteria |
| Actinobacteria | Bacteria |
| Actinobacteria | Bacteria |
| Actinobacteria | Bacteria |
| Actinobacteria | Bacteria |
| Actinobacteria | Bacteria |
| Verrucomicrobia | Bacteria |
| Verrucomicrobia | Bacteria |
| Bacteroidota | Bacteria |
| Bacteroidota | Bacteria |
| Firmicutes | Bacteria |
| Firmicutes | Bacteria |
| Bacteria_unclassified | Bacteria |
| Bacteria_unclassified | Bacteria |
| Bacteria_unclassified | Bacteria |
| Bacteria_unclassified | Bacteria |
| Bacteria_unclassified | Bacteria |
| Bacteria_unclassified | Bacteria |
| Bacteria_unclassified | Bacteria |
| Bacteria_unclassified | Bacteria |
| Bacteria_unclassified | Bacteria |
| Bacteria_unclassified | Bacteria |
| Bacteria_unclassified | Bacteria |
| Bacteria_unclassified | Bacteria |
| Bacteroidota | Bacteria |
| Bacteroidota | Bacteria |
| Actinobacteria | Bacteria |
| Actinobacteria | Bacteria |
| Firmicutes | Bacteria |
| Firmicutes | Bacteria |
| Firmicutes | Bacteria |
| Firmicutes | Bacteria |
| Firmicutes | Bacteria |
| Firmicutes | Bacteria |

| | |
|---|---|
| Firmicutes | Bacteria |
| Firmicutes | Bacteria |
| Firmicutes | Bacteria |
| Firmicutes | Bacteria |
| Actinobacteria | Bacteria |
| Actinobacteria | Bacteria |
| Firmicutes | Bacteria |
| Firmicutes | Bacteria |
| Firmicutes | Bacteria |
| Firmicutes | Bacteria |
| Firmicutes | Bacteria |
| Firmicutes | Bacteria |
| Firmicutes | Bacteria |
| Firmicutes | Bacteria |
| Firmicutes | Bacteria |
| Firmicutes | Bacteria |
| Firmicutes | Bacteria |
| Firmicutes | Bacteria |
| Actinobacteria | Bacteria |
| Actinobacteria | Bacteria |
| Firmicutes | Bacteria |
| Firmicutes | Bacteria |
| Bacteroidota | Bacteria |
| Bacteroidota | Bacteria |
| Bacteroidota | Bacteria |
| Bacteroidota | Bacteria |
| Bacteroidota | Bacteria |
| Bacteroidota | Bacteria |
| Firmicutes | Bacteria |
| Firmicutes | Bacteria |
| Firmicutes | Bacteria |
| Firmicutes | Bacteria |
| Firmicutes | Bacteria |
| Firmicutes | Bacteria |
| Firmicutes | Bacteria |
| Firmicutes | Bacteria |
| Firmicutes | Bacteria |
| Firmicutes | Bacteria |
| Firmicutes | Bacteria |
| Firmicutes | Bacteria |
| Firmicutes | Bacteria |
| Firmicutes | Bacteria |
| Firmicutes | Bacteria |
| Firmicutes | Bacteria |
| Firmicutes | Bacteria |
| Firmicutes | Bacteria |

| | |
|---|---|
| Firmicutes | Bacteria |
| Firmicutes | Bacteria |
| Firmicutes | Bacteria |
| Firmicutes | Bacteria |
| Firmicutes | Bacteria |
| Firmicutes | Bacteria |
| Firmicutes | Bacteria |
| Firmicutes | Bacteria |
| Firmicutes | Bacteria |
| Firmicutes | Bacteria |
| Firmicutes | Bacteria |
| Firmicutes | Bacteria |
| Firmicutes | Bacteria |
| Firmicutes | Bacteria |
| Firmicutes | Bacteria |
| Firmicutes | Bacteria |
| Firmicutes | Bacteria |
| Firmicutes | Bacteria |
| Bacteria_unclassified | Bacteria |
| Bacteria_unclassified | Bacteria |
| Bacteria_unclassified | Bacteria |
| Bacteria_unclassified | Bacteria |
| Bacteria_unclassified | Bacteria |
| Bacteria_unclassified | Bacteria |
| Bacteria_unclassified | Bacteria |
| Bacteria_unclassified | Bacteria |
| Firmicutes | Bacteria |
| Firmicutes | Bacteria |
| Firmicutes | Bacteria |
| Firmicutes | Bacteria |
| Firmicutes | Bacteria |
| Firmicutes | Bacteria |
| Bacteria_unclassified | Bacteria |
| Bacteria_unclassified | Bacteria |
| Firmicutes | Bacteria |
| Firmicutes | Bacteria |
| Firmicutes | Bacteria |
| Firmicutes | Bacteria |
| Firmicutes | Bacteria |
| Firmicutes | Bacteria |
| Firmicutes | Bacteria |
| Firmicutes | Bacteria |
| Firmicutes | Bacteria |
| Firmicutes | Bacteria |
| Firmicutes | Bacteria |
| Firmicutes | Bacteria |

| | |
|---|---|
| Firmicutes | Bacteria |
| Firmicutes | Bacteria |
| Firmicutes | Bacteria |
| Firmicutes | Bacteria |
| Firmicutes | Bacteria |
| Firmicutes | Bacteria |
| Firmicutes | Bacteria |
| Firmicutes | Bacteria |
| Firmicutes | Bacteria |
| Firmicutes | Bacteria |
| Firmicutes | Bacteria |
| Firmicutes | Bacteria |
| Firmicutes | Bacteria |
| Firmicutes | Bacteria |
| Firmicutes | Bacteria |
| Firmicutes | Bacteria |
| Bacteria_unclassified | Bacteria |
| Bacteria_unclassified | Bacteria |
| Firmicutes | Bacteria |
| Firmicutes | Bacteria |
| Firmicutes | Bacteria |
| Firmicutes | Bacteria |
| Bacteria_unclassified | Bacteria |
| Bacteria_unclassified | Bacteria |
| Firmicutes | Bacteria |
| Firmicutes | Bacteria |
| Firmicutes | Bacteria |
| Firmicutes | Bacteria |
| Firmicutes | Bacteria |
| Firmicutes | Bacteria |
| Firmicutes | Bacteria |
| Firmicutes | Bacteria |
| Firmicutes | Bacteria |
| Firmicutes | Bacteria |
| Firmicutes | Bacteria |
| Firmicutes | Bacteria |
| Firmicutes | Bacteria |
| Firmicutes | Bacteria |
| Firmicutes | Bacteria |
| Firmicutes | Bacteria |
| Firmicutes | Bacteria |
| Firmicutes | Bacteria |
| Firmicutes | Bacteria |
| Firmicutes | Bacteria |
| Actinobacteria | Bacteria |
| Actinobacteria | Bacteria |

| | |
|---|---|
| Tenericutes | Bacteria |
| Tenericutes | Bacteria |
| Firmicutes | Bacteria |
| Firmicutes | Bacteria |
| Firmicutes | Bacteria |
| Firmicutes | Bacteria |
| Firmicutes | Bacteria |
| Firmicutes | Bacteria |
| Firmicutes | Bacteria |
| Firmicutes | Bacteria |
| Firmicutes | Bacteria |
| Firmicutes | Bacteria |
| Firmicutes | Bacteria |
| Firmicutes | Bacteria |
| Firmicutes | Bacteria |
| Firmicutes | Bacteria |
| Firmicutes | Bacteria |
| Firmicutes | Bacteria |
| Firmicutes | Bacteria |
| Firmicutes | Bacteria |
| Firmicutes | Bacteria |
| Firmicutes | Bacteria |
| Firmicutes | Bacteria |
| Firmicutes | Bacteria |
| Firmicutes | Bacteria |
| Firmicutes | Bacteria |
| Firmicutes | Bacteria |
| Firmicutes | Bacteria |
| Firmicutes | Bacteria |
| Firmicutes | Bacteria |
| Firmicutes | Bacteria |
| Firmicutes | Bacteria |
| Firmicutes | Bacteria |
| Firmicutes | Bacteria |
| Firmicutes | Bacteria |
| Firmicutes | Bacteria |
| Firmicutes | Bacteria |
| Firmicutes | Bacteria |
| Firmicutes | Bacteria |
| Firmicutes | Bacteria |
| Firmicutes | Bacteria |
| Firmicutes | Bacteria |
| Firmicutes | Bacteria |
| Firmicutes | Bacteria |
| Firmicutes | Bacteria |
| Firmicutes | Bacteria |

| | |
|---|---|
| Bacteroidota | Bacteria |
| Bacteroidota | Bacteria |
| Firmicutes | Bacteria |
| Firmicutes | Bacteria |
| Firmicutes | Bacteria |
| Firmicutes | Bacteria |
| Firmicutes | Bacteria |
| Firmicutes | Bacteria |
| Firmicutes | Bacteria |
| Firmicutes | Bacteria |
| Proteobacteria | Bacteria |
| Proteobacteria | Bacteria |
| Firmicutes | Bacteria |
| Firmicutes | Bacteria |
| Firmicutes | Bacteria |
| Firmicutes | Bacteria |
| Firmicutes | Bacteria |
| Firmicutes | Bacteria |
| Firmicutes | Bacteria |
| Firmicutes | Bacteria |
| Firmicutes | Bacteria |
| Firmicutes | Bacteria |
| Firmicutes | Bacteria |
| Firmicutes | Bacteria |
| Firmicutes | Bacteria |
| Firmicutes | Bacteria |
| Actinobacteria | Bacteria |
| Actinobacteria | Bacteria |
| Actinobacteria | Bacteria |
| Actinobacteria | Bacteria |
| Actinobacteria | Bacteria |
| Actinobacteria | Bacteria |
| Verrucomicrobia | Bacteria |
| Verrucomicrobia | Bacteria |
| Bacteroidota | Bacteria |
| Bacteroidota | Bacteria |
| Firmicutes | Bacteria |
| Firmicutes | Bacteria |
| Bacteria_unclassified | Bacteria |
| Bacteria_unclassified | Bacteria |
| Bacteria_unclassified | Bacteria |
| Bacteria_unclassified | Bacteria |

| | |
|---|---|
| Bacteria_unclassified | Bacteria |
| Bacteria_unclassified | Bacteria |
| Bacteria_unclassified | Bacteria |
| Bacteria_unclassified | Bacteria |
| Bacteria_unclassified | Bacteria |
| Bacteria_unclassified | Bacteria |
| Bacteria_unclassified | Bacteria |
| Bacteria_unclassified | Bacteria |
| Bacteroidota | Bacteria |
| Bacteroidota | Bacteria |

| | |
|---|---|
| Actinobacteria | Bacteria |
| Actinobacteria | Bacteria |
| Firmicutes | Bacteria |
| Firmicutes | Bacteria |
| Firmicutes | Bacteria |
| Firmicutes | Bacteria |
| Firmicutes | Bacteria |
| Firmicutes | Bacteria |
| Firmicutes | Bacteria |
| Firmicutes | Bacteria |
| Firmicutes | Bacteria |
| Firmicutes | Bacteria |
| Actinobacteria | Bacteria |
| Actinobacteria | Bacteria |
| Firmicutes | Bacteria |
| Firmicutes | Bacteria |
| Firmicutes | Bacteria |
| Firmicutes | Bacteria |
| Firmicutes | Bacteria |
| Firmicutes | Bacteria |
| Firmicutes | Bacteria |
| Firmicutes | Bacteria |
| Firmicutes | Bacteria |
| Firmicutes | Bacteria |
| Firmicutes | Bacteria |
| Firmicutes | Bacteria |
| Actinobacteria | Bacteria |
| Actinobacteria | Bacteria |
| Firmicutes | Bacteria |
| Firmicutes | Bacteria |
| Bacteroidota | Bacteria |
| Bacteroidota | Bacteria |
| Bacteroidota | Bacteria |
| Bacteroidota | Bacteria |

| | |
|---|---|
| Bacteroidota | Bacteria |
| Bacteroidota | Bacteria |
| Firmicutes | Bacteria |
| Firmicutes | Bacteria |
| Firmicutes | Bacteria |
| Firmicutes | Bacteria |
| Firmicutes | Bacteria |
| Firmicutes | Bacteria |
| Firmicutes | Bacteria |
| Firmicutes | Bacteria |
| Firmicutes | Bacteria |
| Firmicutes | Bacteria |
| Firmicutes | Bacteria |
| Firmicutes | Bacteria |
| Firmicutes | Bacteria |
| Firmicutes | Bacteria |
| Firmicutes | Bacteria |
| Firmicutes | Bacteria |
| Firmicutes | Bacteria |
| Firmicutes | Bacteria |
| Firmicutes | Bacteria |
| Firmicutes | Bacteria |
| Firmicutes | Bacteria |
| Firmicutes | Bacteria |
| Firmicutes | Bacteria |
| Firmicutes | Bacteria |
| Firmicutes | Bacteria |
| Firmicutes | Bacteria |
| Firmicutes | Bacteria |
| Firmicutes | Bacteria |
| Firmicutes | Bacteria |
| Firmicutes | Bacteria |
| Firmicutes | Bacteria |
| Firmicutes | Bacteria |
| Firmicutes | Bacteria |
| Firmicutes | Bacteria |
| Firmicutes | Bacteria |
| Firmicutes | Bacteria |
| Bacteria_unclassified | Bacteria |
| Bacteria_unclassified | Bacteria |
| Bacteria_unclassified | Bacteria |
| Bacteria_unclassified | Bacteria |
| Bacteria_unclassified | Bacteria |
| Bacteria_unclassified | Bacteria |
| Bacteria_unclassified | Bacteria |
| Bacteria_unclassified | Bacteria |

| | |
|---|---|
| Firmicutes | Bacteria |
| Firmicutes | Bacteria |
| Firmicutes | Bacteria |
| Firmicutes | Bacteria |
| Firmicutes | Bacteria |
| Firmicutes | Bacteria |
| Bacteria_unclassified | Bacteria |
| Bacteria_unclassified | Bacteria |
| Firmicutes | Bacteria |
| Firmicutes | Bacteria |
| Firmicutes | Bacteria |
| Firmicutes | Bacteria |
| Firmicutes | Bacteria |
| Firmicutes | Bacteria |
| Firmicutes | Bacteria |
| Firmicutes | Bacteria |
| Firmicutes | Bacteria |
| Firmicutes | Bacteria |
| Firmicutes | Bacteria |
| Firmicutes | Bacteria |
| Firmicutes | Bacteria |
| Firmicutes | Bacteria |
| Firmicutes | Bacteria |
| Firmicutes | Bacteria |
| Firmicutes | Bacteria |
| Firmicutes | Bacteria |
| Firmicutes | Bacteria |
| Firmicutes | Bacteria |
| Firmicutes | Bacteria |
| Firmicutes | Bacteria |
| Firmicutes | Bacteria |
| Firmicutes | Bacteria |
| Firmicutes | Bacteria |
| Firmicutes | Bacteria |
| Firmicutes | Bacteria |
| Firmicutes | Bacteria |
| Bacteria_unclassified | Bacteria |
| Bacteria_unclassified | Bacteria |
| Firmicutes | Bacteria |
| Firmicutes | Bacteria |
| Firmicutes | Bacteria |
| Firmicutes | Bacteria |
| Bacteria_unclassified | Bacteria |
| Bacteria_unclassified | Bacteria |
| Firmicutes | Bacteria |
| Firmicutes | Bacteria |

| | |
|---|---|
| Firmicutes | Bacteria |
| Firmicutes | Bacteria |
| Firmicutes | Bacteria |
| Firmicutes | Bacteria |
| Firmicutes | Bacteria |
| Firmicutes | Bacteria |
| Firmicutes | Bacteria |
| Firmicutes | Bacteria |
| Firmicutes | Bacteria |
| Firmicutes | Bacteria |
| Firmicutes | Bacteria |
| Firmicutes | Bacteria |
| Firmicutes | Bacteria |
| Firmicutes | Bacteria |
| Firmicutes | Bacteria |
| Firmicutes | Bacteria |
| Firmicutes | Bacteria |
| Firmicutes | Bacteria |
| Actinobacteria | Bacteria |
| Actinobacteria | Bacteria |
| Tenericutes | Bacteria |
| Tenericutes | Bacteria |
| Firmicutes | Bacteria |
| Firmicutes | Bacteria |
| Firmicutes | Bacteria |
| Firmicutes | Bacteria |
| Firmicutes | Bacteria |
| Firmicutes | Bacteria |
| Firmicutes | Bacteria |
| Firmicutes | Bacteria |
| Firmicutes | Bacteria |
| Firmicutes | Bacteria |
| Firmicutes | Bacteria |
| Firmicutes | Bacteria |
| Firmicutes | Bacteria |
| Firmicutes | Bacteria |
| Firmicutes | Bacteria |
| Firmicutes | Bacteria |
| Firmicutes | Bacteria |
| Firmicutes | Bacteria |
| Firmicutes | Bacteria |
| Firmicutes | Bacteria |
| Firmicutes | Bacteria |
| Firmicutes | Bacteria |
| Firmicutes | Bacteria |
| Firmicutes | Bacteria |

| | |
|---|---|
| Firmicutes | Bacteria |
| Firmicutes | Bacteria |
| Firmicutes | Bacteria |
| Firmicutes | Bacteria |
| Firmicutes | Bacteria |
| Firmicutes | Bacteria |
| Firmicutes | Bacteria |
| Firmicutes | Bacteria |
| Firmicutes | Bacteria |
| Firmicutes | Bacteria |
| Firmicutes | Bacteria |
| Firmicutes | Bacteria |
| Firmicutes | Bacteria |
| Firmicutes | Bacteria |
| Firmicutes | Bacteria |
| Firmicutes | Bacteria |
| Firmicutes | Bacteria |
| Firmicutes | Bacteria |
| Firmicutes | Bacteria |
| Firmicutes | Bacteria |
| Bacteroidota | Bacteria |
| Bacteroidota | Bacteria |
| Firmicutes | Bacteria |
| Firmicutes | Bacteria |
| Firmicutes | Bacteria |
| Firmicutes | Bacteria |
| Firmicutes | Bacteria |
| Firmicutes | Bacteria |
| Firmicutes | Bacteria |
| Firmicutes | Bacteria |
| Proteobacteria | Bacteria |
| Proteobacteria | Bacteria |
| Firmicutes | Bacteria |
| Firmicutes | Bacteria |
| Firmicutes | Bacteria |
| Firmicutes | Bacteria |
| Firmicutes | Bacteria |
| Firmicutes | Bacteria |
| Firmicutes | Bacteria |
| Firmicutes | Bacteria |
| Firmicutes | Bacteria |
| Firmicutes | Bacteria |

| | |
|---|---|
| Firmicutes | Bacteria |
| Firmicutes | Bacteria |
| Firmicutes | Bacteria |
| Firmicutes | Bacteria |
| Actinobacteria | Bacteria |
| Actinobacteria | Bacteria |
| Actinobacteria | Bacteria |
| Actinobacteria | Bacteria |
| Actinobacteria | Bacteria |
| Actinobacteria | Bacteria |
| Verrucomicrobia | Bacteria |
| Verrucomicrobia | Bacteria |
| Bacteroidota | Bacteria |
| Bacteroidota | Bacteria |
| Firmicutes | Bacteria |
| Firmicutes | Bacteria |
| Bacteria_unclassified | Bacteria |
| Bacteria_unclassified | Bacteria |
| Bacteria_unclassified | Bacteria |
| Bacteria_unclassified | Bacteria |
| Bacteria_unclassified | Bacteria |
| Bacteria_unclassified | Bacteria |
| Bacteria_unclassified | Bacteria |
| Bacteria_unclassified | Bacteria |
| Bacteria_unclassified | Bacteria |
| Bacteria_unclassified | Bacteria |
| Bacteria_unclassified | Bacteria |
| Bacteria_unclassified | Bacteria |
| Bacteroidota | Bacteria |
| Bacteroidota | Bacteria |

| | |
|---|---|
| Actinobacteria | Bacteria |
| Actinobacteria | Bacteria |
| Firmicutes | Bacteria |
| Firmicutes | Bacteria |
| Firmicutes | Bacteria |
| Firmicutes | Bacteria |
| Firmicutes | Bacteria |
| Firmicutes | Bacteria |
| Firmicutes | Bacteria |
| Firmicutes | Bacteria |
| Firmicutes | Bacteria |
| Firmicutes | Bacteria |
| Actinobacteria | Bacteria |
| Actinobacteria | Bacteria |

| | |
|---|---|
| Firmicutes | Bacteria |
| Firmicutes | Bacteria |
| Firmicutes | Bacteria |
| Firmicutes | Bacteria |
| Firmicutes | Bacteria |
| Firmicutes | Bacteria |
| Firmicutes | Bacteria |
| Firmicutes | Bacteria |
| Firmicutes | Bacteria |
| Firmicutes | Bacteria |
| Firmicutes | Bacteria |
| Firmicutes | Bacteria |
| Actinobacteria | Bacteria |
| Actinobacteria | Bacteria |
| Firmicutes | Bacteria |
| Firmicutes | Bacteria |
| Bacteroidota | Bacteria |
| Bacteroidota | Bacteria |
| Bacteroidota | Bacteria |
| Bacteroidota | Bacteria |
| Bacteroidota | Bacteria |
| Bacteroidota | Bacteria |
| Firmicutes | Bacteria |
| Firmicutes | Bacteria |
| Firmicutes | Bacteria |
| Firmicutes | Bacteria |
| Firmicutes | Bacteria |
| Firmicutes | Bacteria |
| Firmicutes | Bacteria |
| Firmicutes | Bacteria |
| Firmicutes | Bacteria |
| Firmicutes | Bacteria |
| Firmicutes | Bacteria |
| Firmicutes | Bacteria |
| Firmicutes | Bacteria |
| Firmicutes | Bacteria |
| Firmicutes | Bacteria |
| Firmicutes | Bacteria |
| Firmicutes | Bacteria |
| Firmicutes | Bacteria |
| Firmicutes | Bacteria |
| Firmicutes | Bacteria |
| Firmicutes | Bacteria |
| Firmicutes | Bacteria |
| Firmicutes | Bacteria |
| Firmicutes | Bacteria |

| | |
|---|---|
| Firmicutes | Bacteria |
| Firmicutes | Bacteria |
| Firmicutes | Bacteria |
| Firmicutes | Bacteria |
| Firmicutes | Bacteria |
| Firmicutes | Bacteria |
| Firmicutes | Bacteria |
| Firmicutes | Bacteria |
| Firmicutes | Bacteria |
| Firmicutes | Bacteria |
| Firmicutes | Bacteria |
| Firmicutes | Bacteria |
| Bacteria_unclassified | Bacteria |
| Bacteria_unclassified | Bacteria |
| Bacteria_unclassified | Bacteria |
| Bacteria_unclassified | Bacteria |
| Bacteria_unclassified | Bacteria |
| Bacteria_unclassified | Bacteria |
| Bacteria_unclassified | Bacteria |
| Bacteria_unclassified | Bacteria |
| Firmicutes | Bacteria |
| Firmicutes | Bacteria |
| Firmicutes | Bacteria |
| Firmicutes | Bacteria |
| Firmicutes | Bacteria |
| Firmicutes | Bacteria |
| Bacteria_unclassified | Bacteria |
| Bacteria_unclassified | Bacteria |
| Firmicutes | Bacteria |
| Firmicutes | Bacteria |
| Firmicutes | Bacteria |
| Firmicutes | Bacteria |
| Firmicutes | Bacteria |
| Firmicutes | Bacteria |
| Firmicutes | Bacteria |
| Firmicutes | Bacteria |
| Firmicutes | Bacteria |
| Firmicutes | Bacteria |
| Firmicutes | Bacteria |
| Firmicutes | Bacteria |
| Firmicutes | Bacteria |
| Firmicutes | Bacteria |
| Firmicutes | Bacteria |
| Firmicutes | Bacteria |
| Firmicutes | Bacteria |
| Firmicutes | Bacteria |

| | |
|---|---|
| Firmicutes | Bacteria |
| Firmicutes | Bacteria |
| Firmicutes | Bacteria |
| Firmicutes | Bacteria |
| Firmicutes | Bacteria |
| Firmicutes | Bacteria |
| Firmicutes | Bacteria |
| Firmicutes | Bacteria |
| Firmicutes | Bacteria |
| Firmicutes | Bacteria |
| Bacteria_unclassified | Bacteria |
| Bacteria_unclassified | Bacteria |
| Firmicutes | Bacteria |
| Firmicutes | Bacteria |
| Firmicutes | Bacteria |
| Firmicutes | Bacteria |
| Bacteria_unclassified | Bacteria |
| Bacteria_unclassified | Bacteria |
| Firmicutes | Bacteria |
| Firmicutes | Bacteria |
| Firmicutes | Bacteria |
| Firmicutes | Bacteria |
| Firmicutes | Bacteria |
| Firmicutes | Bacteria |
| Firmicutes | Bacteria |
| Firmicutes | Bacteria |
| Firmicutes | Bacteria |
| Firmicutes | Bacteria |
| Firmicutes | Bacteria |
| Firmicutes | Bacteria |
| Firmicutes | Bacteria |
| Firmicutes | Bacteria |
| Firmicutes | Bacteria |
| Firmicutes | Bacteria |
| Firmicutes | Bacteria |
| Firmicutes | Bacteria |
| Firmicutes | Bacteria |
| Firmicutes | Bacteria |
| Actinobacteria | Bacteria |
| Actinobacteria | Bacteria |
| Tenericutes | Bacteria |
| Tenericutes | Bacteria |
| Firmicutes | Bacteria |
| Firmicutes | Bacteria |
| Firmicutes | Bacteria |
| Firmicutes | Bacteria |

| | |
|---|---|
| Firmicutes | Bacteria |
| Firmicutes | Bacteria |
| Firmicutes | Bacteria |
| Firmicutes | Bacteria |
| Firmicutes | Bacteria |
| Firmicutes | Bacteria |
| Firmicutes | Bacteria |
| Firmicutes | Bacteria |
| Firmicutes | Bacteria |
| Firmicutes | Bacteria |
| Firmicutes | Bacteria |
| Firmicutes | Bacteria |
| Firmicutes | Bacteria |
| Firmicutes | Bacteria |
| Firmicutes | Bacteria |
| Firmicutes | Bacteria |
| Firmicutes | Bacteria |
| Firmicutes | Bacteria |
| Firmicutes | Bacteria |
| Firmicutes | Bacteria |
| Firmicutes | Bacteria |
| Firmicutes | Bacteria |
| Firmicutes | Bacteria |
| Firmicutes | Bacteria |
| Firmicutes | Bacteria |
| Firmicutes | Bacteria |
| Firmicutes | Bacteria |
| Firmicutes | Bacteria |
| Firmicutes | Bacteria |
| Firmicutes | Bacteria |
| Firmicutes | Bacteria |
| Firmicutes | Bacteria |
| Firmicutes | Bacteria |
| Firmicutes | Bacteria |
| Firmicutes | Bacteria |
| Firmicutes | Bacteria |
| Firmicutes | Bacteria |
| Firmicutes | Bacteria |
| Firmicutes | Bacteria |
| Firmicutes | Bacteria |
| Bacteroidota | Bacteria |
| Bacteroidota | Bacteria |
| Firmicutes | Bacteria |
| Firmicutes | Bacteria |
| Firmicutes | Bacteria |
| Firmicutes | Bacteria |

| | |
|---|---|
| Firmicutes | Bacteria |
| Firmicutes | Bacteria |
| Firmicutes | Bacteria |
| Firmicutes | Bacteria |
| Proteobacteria | Bacteria |
| Proteobacteria | Bacteria |
| Firmicutes | Bacteria |
| Firmicutes | Bacteria |
| Firmicutes | Bacteria |
| Firmicutes | Bacteria |
| Firmicutes | Bacteria |
| Firmicutes | Bacteria |

k__Bacteria|p__Firmicutes|c__Clostridia|o__Eubacteriales|f__Peptostreptococcaceae|g__Romboutsia|s__Romboutsia_ilea[illegible]
k__Bacteria|p__Firmicutes|c__Clostridia|o__Eubacteriales|f__Peptostreptococcaceae|g__Romboutsia|s__Romboutsia_ilea[illegible]
k__Bacteria|p__Firmicutes|c__Clostridia|o__Eubacteriales|f__Lachnospiraceae|g__Schaedlerella|s__Schaedlerella_arabino[illegible]
k__Bacteria|p__Firmicutes|c__Clostridia|o__Eubacteriales|f__Lachnospiraceae|g__Schaedlerella|s__Schaedlerella_arabino[illegible]

k__Bacteria|p__Firmicutes|c__Clostridia|o__Eubacteriales|f__Peptostreptococcaceae|g__Romboutsia|s__Romboutsia_ilea[illegible]
k__Bacteria|p__Firmicutes|c__Clostridia|o__Eubacteriales|f__Peptostreptococcaceae|g__Romboutsia|s__Romboutsia_ilea[illegible]
k__Bacteria|p__Firmicutes|c__Clostridia|o__Eubacteriales|f__Lachnospiraceae|g__Schaedlerella|s__Schaedlerella_arabino[illegible]
k__Bacteria|p__Firmicutes|c__Clostridia|o__Eubacteriales|f__Lachnospiraceae|g__Schaedlerella|s__Schaedlerella_arabino[illegible]

k__Bacteria|p__Firmicutes|c__Erysipelotrichia|o__Erysipelotrichales|f__Turicibacteraceae|g__Turicibacter|s__Turicibacter[illegible]
k__Bacteria|p__Firmicutes|c__Erysipelotrichia|o__Erysipelotrichales|f__Turicibacteraceae|g__Turicibacter|s__Turicibacter[illegible]
k__Bacteria|p__Firmicutes|c__Clostridia|o__Eubacteriales|f__Lachnospiraceae|g__Acetatifactor|s__Acetatifactor_muris
k__Bacteria|p__Firmicutes|c__Clostridia|o__Eubacteriales|f__Lachnospiraceae|g__Acetatifactor|s__Acetatifactor_muris
k__Bacteria|p__Firmicutes|c__Clostridia|o__Eubacteriales|f__Lachnospiraceae|g__Acetatifactor|s__Acetatifactor_SGB415[illegible]
k__Bacteria|p__Firmicutes|c__Clostridia|o__Eubacteriales|f__Lachnospiraceae|g__Acetatifactor|s__Acetatifactor_SGB415[illegible]
