## Supplementary material for "Non-invasive Vagal Nerve Stimulation as a Potential Treatment for Repetitive Blast Trauma": st9

| | | | 95% Confidence Intervals of Blast effect on outcome (at specified VNS conditio | | | | |
|---|---|---|---|---|---|---|---|
| | | | VNS (–) | | | VNS (+) | |
| Exposure | Mediator (Species-level featur | Outcome | ACME | ADE | Total | ACME | ADE |
| Blast | Clostridiales_bacterium | EtOH_pref_24h_final | [-0.039, 0.418] | [0.008, 1.327] | [0.147, 1.516] | [0.068, 0.758] | [-1.109, 0.053] |
| Blast | bacterium_1XD42_54 | EtOH_pref_24h_final | [-0.582, -0.066] | [0.356, 1.818] | [0.149, 1.477] | [-0.263, 0.019] | [-0.585, 0.48] |
| Blast | GGB29002_SGB41718 | OFB_Clatency | [-0.064, 0.188] | [0.208, 0.851] | [0.228, 0.948] | [0.124, 0.497] | [0.41, 1.204] |
| Blast | GGB3171_SGB4185 | PD_choice_filt | [-0.434, -0.021] | [0.594, 1.863] | [0.377, 1.66] | [-0.5, -0.012] | [-0.454, 0.877] |

| | | | 95% Confidence Intervals of VNS effect on outcome (at specified Blast conditio | | | | |
|---|---|---|---|---|---|---|---|
| | | | Blast (–) | | | Blast (+) | |
| Exposure | Mediator (Species-level featur | Outcome | ACME | ADE | Total | ACME | ADE |
| VNS | GGB28883_SGB41564 | OFB_Cdistance | [-0.028, 0.236] | [-0.475, 0.565] | [-0.412, 0.675] | [-0.024, 0.307] | [-0.681, 0.272] |

| n) | | | | | |
|---|---|---|---|---|---|
| | Taxonomic Information | | | | |
| **Total** | **Genus** | **Family** | **Order** | **Class** | **Phylum** |
| [-0.74, 0.321] | Eubacteriales_unclassified | Eubacteriales_unclassified | Eubacteriales | Clostridia | Firmicutes |
| [-0.67, 0.375] | Bacteria_unclassified | Bacteria_unclassified | Bacteria_unclassified | Bacteria_unclassified | Bacteria_unclassified |
| [0.605, 1.52] | GGB29002 | FGB9658 | OFGB9658 | CFGB9658 | Firmicutes |
| [-0.769, 0.777] | GGB3171 | Oscillospiraceae | Eubacteriales | Clostridia | Firmicutes |

| n) | | | | | |
|---|---|---|---|---|---|
| | Taxonomic Information | | | | |
| **Total** | **Genus** | **Family** | **Order** | **Class** | **Phylum** |
| [-0.56, 0.356] | GGB28883 | FGB9633 | OFGB9633 | CFGB9633 | Firmicutes |

| Kingdom | MetaPhlan Annotation |
|---|---|
| Bacteria | k__Bacteria\|p__Firmicutes\|c__Clostridia\|o__Eubacteriales\|f__Eubacteriales_unclassified\|g__Eubacteriales_unclassified\|s__Clostridiales_bacterium |
| Bacteria | k__Bacteria\|p__Bacteria_unclassified\|c__Bacteria_unclassified\|o__Bacteria_unclassified\|f__Bacteria_unclassified\|g__Bacteria_unclassified\|s__bacter |
| Bacteria | k__Bacteria\|p__Firmicutes\|c__CFGB9658\|o__OFGB9658\|f__FGB9658\|g__GGB29002\|s__GGB29002_SGB41718 |
| Bacteria | k__Bacteria\|p__Firmicutes\|c__Clostridia\|o__Eubacteriales\|f__Oscillospiraceae\|g__GGB3171\|s__GGB3171_SGB4185 |

| Kingdom | MetaPhlan Annotation |
|---|---|
| Bacteria | k__Bacteria\|p__Firmicutes\|c__CFGB9633\|o__OFGB9633\|f__FGB9633\|g__GGB28883\|s__GGB28883_SGB41564 |

rium_1XD42_54
